# Heterotrophic prokaryotes internal carbon recycling compensates mismatches between phytoplankton production and heterotrophic prokaryotic consumption

**DOI:** 10.1101/2024.01.10.574976

**Authors:** Falk Eigemann, Karen Tait, Ben Temperton, Ferdi L. Hellweger

## Abstract

Molecular observational tools are great for characterizing the composition and genetic endowment of microbial communities, but cannot measure fluxes, which are critical for the understanding of ecosystems. To overcome these limitations, we use a mechanistic inference approach to estimate dissolved organic carbon (DOC) production and consumption by phytoplankton operational taxonomic units (OTUs) and heterotrophic prokaryotic amplicon sequences variants (ASVs), and infer carbon fluxes between members of this microbial community from Western English Channel (WEC) time-series data. Our analyses focus on phytoplankton spring and summer blooms, as well as bacteria summer blooms. In spring blooms, phytoplankton DOC production exceeds heterotrophic prokaryotic consumption, but in bacterial summer blooms heterotrophic prokaryotes consume almost 3 times more DOC than produced by the phytoplankton. This mismatch is compensated by heterotrophic prokaryotic DOC release by death, presumably viral lysis. In both types of summer blooms, large amounts of the DOC liberated by heterotrophic prokaryotes are reused, i.e. internally recycled, and fluxes between different heterotrophic prokaryotes are at the same level as fluxes between phytoplankton and heterotrophic prokaryotes. Contextualized, internal recycling accounts for approximately 75% and 30% of the estimated net primary production (0.16 vs 0.22 and 0.08 vs 0.29 µmol l^-1^ d^-1^) in bacteria and phytoplankton summer blooms, respectively, and thus represents a major component of the WEC carbon cycle. We conclude that internal recycling compensates mismatches between phytoplankton DOC production and heterotrophic prokaryotic consumption, and encourage future analyses on aquatic carbon cycles to consider fluxes between heterotrophic prokaryotes, i.e. internal recycling.

## Introduction

Phytoplankton-bacteria interactions have global consequences on carbon and nutrient cycling (Azam and Malfatti 2007, Cole 1982, York 2018). Briefly, inorganic carbon is photosynthetically fixed by phytoplankton, and a substantial fraction subsequently released in the form of dissolved organic matter (DOM), which is recycled/consumed by heterotrophic bacteria (Azam and Malfatti 2007). The microbial recycling of photosynthates, in turn, drives the microbial loop (Azam et al 1983), as well as the microbial carbon pump (Jiao and Azam 2011). By upscaling observations of participants, next generation sequencing (NGS) advances understanding of microbial ecosystems. For example, high taxonomical resolution enabled the identification of pronounced seasonal differences in composition and richness of microbial communities in the North Atlantic (Bolaños et al 2021), and temporal distributions of closely related bacteria in the Mediterranean (Auladell et al 2021). However, understanding these ecosystems also requires quantification of interactions between individual components, which cannot be so readily observed. Most interaction analyses are based on empirical measures such as species co-occurrences or expression of specific genes (Ahlgren et al 2019, Faust et al 2015), but co-occurrence of species does not necessarily imply interaction (Blanchet et al 2020), and the expression of genes e.g. for the degradation of specific polysaccharides by a defined bacterium may indeed target different producers/phytoplankton species (Teeling et al 2012). Mechanistic inference, which is based on mass balances, offers the potential to overcome these limitations. We enhanced the mass-balancing, mechanistically constrained inference approach FluxNet (Mayerhofer et al 2021) to 158 heterotrophic, prokaryotic amplicon sequence variants (ASVs, derived from 16S rRNA), 135 phototrophic, eukaryotic operational taxonomic units (OTUs, derived from 18S rRNA), *Synechococcus* (derived from flow-cytometric counts), and 162 hypothetical dissolved organic matter (DOM) species. The model consists of a set of differential mass balance equations and a customized optimization/calibration routine. Organic carbon is represented in phytoplankton, DOM, POM and heterotrophic prokaryotes compartments, each with many “species”. It is a high-resolution biogeochemical model, automatically calibrated to time-series data. The resulting flux network includes quantitative carbon fluxes between all members in microbial communities for any time point, enabling insights in the functioning of the respective ecosystem. In the present study, we applied the enhanced inference approach to a seven-year time-series (2012 – 2018) of the Western English Channel (WEC) station L4.

WEC station L4 depicts a temperate coastal-ocean site off Plymouth, UK, with well-mixed waters in autumn and winter months, but weak stratification and declining nutrient concentrations from spring into summer (www.westernchannelobservatory.org.uk/data). It’s phytoplankton community is dominated by phytoflagellates, diatoms, *Phaeocystis*, coccolithophorids and dinoflagellates (Widdicombe et al 2010), and pronounced phytoplankton blooms occur in spring as well as in summer/autumn (Irigoien et al 2000, Smyth et al 2009), which vary in their timing, intensity and key taxa present (Widdicombe et al 2010). The bacterial community is dominated by Alphaproteobacteria and Bacteroidetes (Gilbert et al 2009), with occasional peaks of Gammaproteobacteria (Gilbert et al 2012), and reveals strong seasonal patterns (Gilbert et al 2009, Gilbert et al 2012). However, despite a few single-year analyses on seasonal primary and/or bacterial production in the English Channel (Lamy et al 2006, Napoléon et al 2014), a transformation of phytoplankton and bacteria seasonality into season-specific estimates of carbon production, consumption and fluxes has not been undertaken. Furthermore, correlations between bacteria and phytoplankton were markedly low (Gilbert et al 2009, Gilbert et al 2012), posing the question of the bacterial carbon source in periods with high bacterial but low phytoplankton concentrations. Interestingly, different bacterial taxa revealed the highest correlations in association network analyses (Gilbert et al 2012), but the mechanisms underlying these putative bacteria-bacteria interactions were not explored.

Here, we aimed to answer the following questions: i) which periods/seasons of the year have the highest carbon fluxes at station L4? ii) how do heterotrophic prokaryotes (i.e. heterotrophic archaea and heterotrophic bacteria) meet their carbon demands in periods with high heterotrophic prokaryote but low phytoplankton abundances? iii) how do fluxes between phytoplankton and heterotrophic prokaryotes as well as between different heterotrophic prokaryotes develop during the course of microbial blooms? and iv) which resources limit phytoplankton and heterotrophic prokaryotes at different seasons of the year?

## Results

### Model performance

The model provides a continuous picture of fluxes over the 7-year period, with distinct periods of high growth, i.e. blooms, and we focus our analyses on those periods. Specifically, we defined phytoplankton spring and summer blooms, as well as bacterial summer blooms for each year (Fig. 1A, Supplementary table 1). The model reproduces the main patterns of the blooms (Fig. 1A), as well as concentration patterns of single phytoplankton OTUs and heterotrophic prokaryotic ASVs (Fig. 1B-D left panel and Supplementary Data 1).

**Fig. 1:**
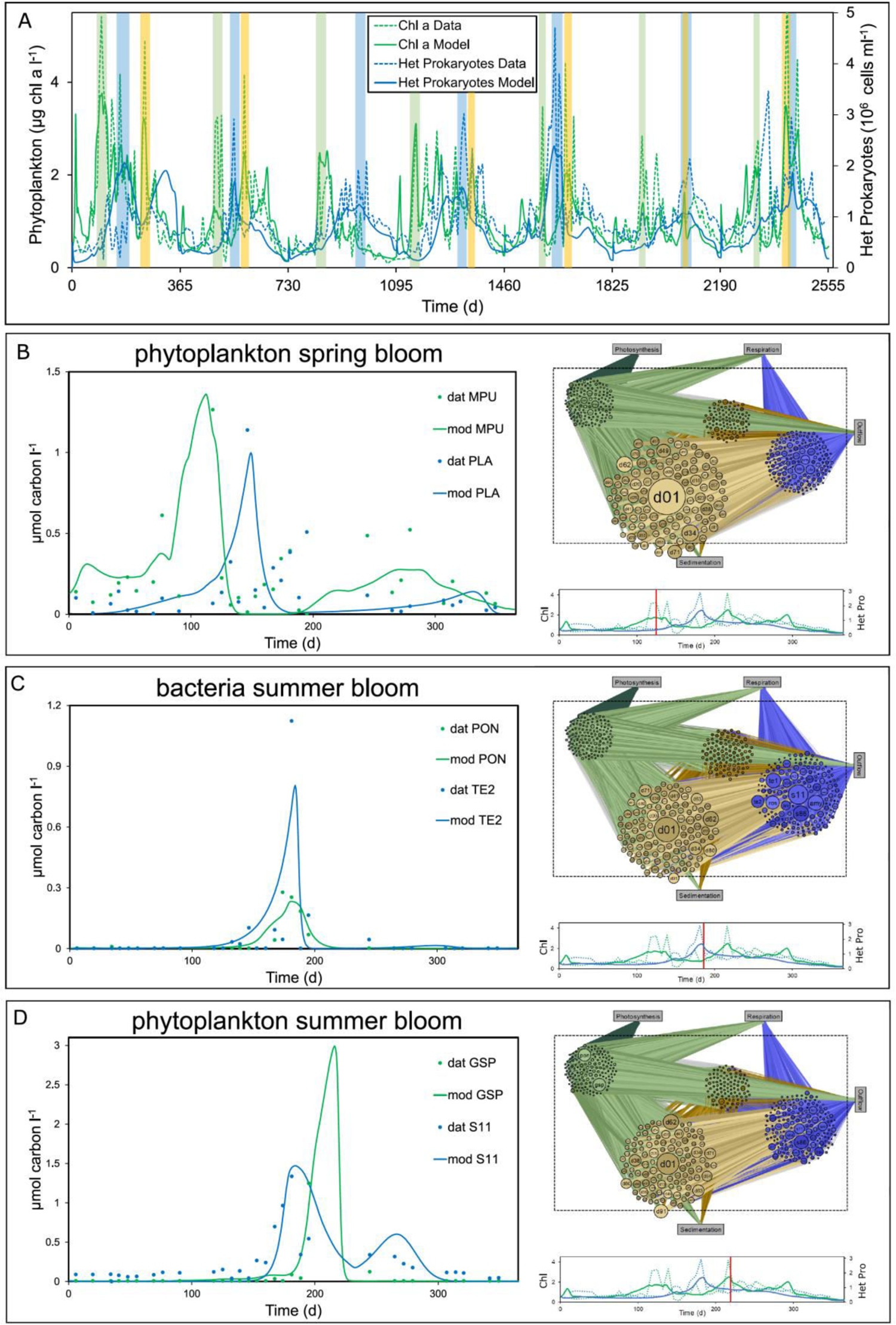
Model-data comparisons for the 7-year time-series and illustration of the inferred flux network (A) Model-data comparison for total phytoplankton (green) and heterotrophic prokaryotes (blue) concentrations with t0 = 01.01.2012. Phytoplankton spring blooms are shaded green, bacteria summer blooms blue and phytoplankton summer blooms yellow. (B-D) Model-data comparisons for phytoplankton OTUs (green) and heterotrophic prokaryotic ASVs (blue) with the highest carbon flux for each bloom type in the example year 2013 with t0 = 01.01.2013. MPU = Micromonas pusila, PON = Prorocentrum donghaiense, GSP = Gonyaulax spinifera, PLA = Planktomarina, TE2 = Tenacibaculum, S11 = SAR11. The right panel displays the network for the median day in each bloom period. Phytoplankton is depicted in green, DOM in light brown, POM in dark brown, and heterotrophic prokaryotes in blue. The size of the species reflects the carbon outflow (phytoplankton), throughflow (average of inflow and outflow, DOM and POM) and inflow (heterotrophic prokaryotes), and the size of the lines between the compartments the magnitude of fluxes. Below each network a timeline with the corresponding chl. a and heterotrophic prokaryotes concentrations is given (green: phytoplankton, blue: heterotrophic prokaryotes, solid lines: model, dashed lines: data). The red bar indicates the day for the network. (B) phytoplankton spring bloom, (C) bacteria summer bloom, (D) phytoplankton summer bloom.

The networks show the direct flux of organic carbon from phytoplankton to heterotrophic prokaryotes via DOM, as well as the indirect transfer via POM (i.e. dead phytoplankton cells that partly dissolve) to DOM before uptake by heterotrophic prokaryotes, but also a pronounced flux from heterotrophic prokaryotes back to the DOM pool in bacterial summer blooms. Varying sizes of phytoplankton and heterotrophic prokaryotic OTUs/ASVs illustrate the changing carbon inflows and outflows for the different bloom types (Fig. 1B-D). For instance, *Micromonas pusila* (MPU) plays a major role in the 2013 phytoplankton spring bloom, but vanishes towards the bacterial summer bloom, whereas SAR11 (S11) plays a minor role in the phytoplankton spring bloom, but has the highest DOC influx in bacterial summer blooms. Detailed analyses on important phytoplankton producer OTUs and heterotrophic prokaryotic consumer ASVs and their recurrences are given in the SI and in Supplementary Table 2.

Besides the individual microbe concentrations time-series given in Supplementary Data 1, the model results can be compared to observations of bulk parameters. We calculated gross and net primary production, as well as gross and net heterotrophic prokaryotic production as integrated values for each complete year, but also for the selected bloom types of each year (Supplementary Table 3). Whole year gross primary production ranged between 32 (year 2013) and 18 (year 2016) mmol C m^-2^ d^-1^, whereas net primary production was highest in 2012 and 2013 (each 23 mmol C m^-2^ d^-1^) and lowest in 2014 (13 mmol C m^-2^ d^-1^, Supplementary Table 3). For the different bloom types, phytoplankton spring blooms revealed the highest, and bacterial summer blooms the lowest primary production, with the highest value for gross primary production achieved in the phytoplankton spring bloom in 2014 (91 mmol C m^-2^ d^-1^), and the lowest in bacterial summer blooms in the same year (8 mmol C m^-2^ d^-1^). These estimated primary production values are in the range of measured primary production rates in the English Channel (e.g. 8.3 - 217 mmol C m^-2^ d^-1^ ca. 20 km south of Plymouth (Boalch et al 1978), 0.42 – 2.4 mmol C m^-2^ d^-1^, measured at a transect between Portsmouth and Ouistreham (Napoléon et al 2014), or 6 – 60 µmol C l^-1^ d^-1^ at Southampton surface waters (Derenbach and Williams 1974)). The whole year gross heterotrophic prokaryotic production inferred with the model revealed contrasting patterns to the primary production with the highest value obtained in 2018 (0.24 µmol l^-1^ d^-1^), and lowest in 2017 (0.13 µmol l^-1^ d^-1^), whereas net heterotrophic prokaryotic production ranged between 0.12 µmol l^-1^ d^-1^ (2012) and 0.07 µmol l^-1^ d^-1^ (2017, Supplementary Table 3).

Heterotrophic prokaryotic production was highest in bacterial summer blooms (max value 0.73 µmol C l^-1^ d^-1^ in 2013), and mostly lowest in phytoplankton spring blooms (min value 0.04 µmol C l^-1^ d^-1^ in 2015). Similar to the phytoplankton, the estimated values for heterotrophic prokaryotic production are in the range of measurements from the English Channel, with e.g. 0.01 – 0.16 µmol C l^-1^ d^-1^ after a *Phaeocystis* bloom (Lamy et al 2006), or 0.5 µmol C l^-1^ d^-1^ measured by Derenbach and Williams (1974).

### Heterotrophic prokaryotes dominate DOC production in summer blooms

In order to explore the importance of the different bloom types for the WEC carbon cycle, we estimated the DOC concentration, the DOC consumption and the DOC production (Fig. 2). We note that DOC in the model is solely from microbes in the system, i.e. it does not include allochthonous input or recalcitrant background DOC.

**Fig. 2:**
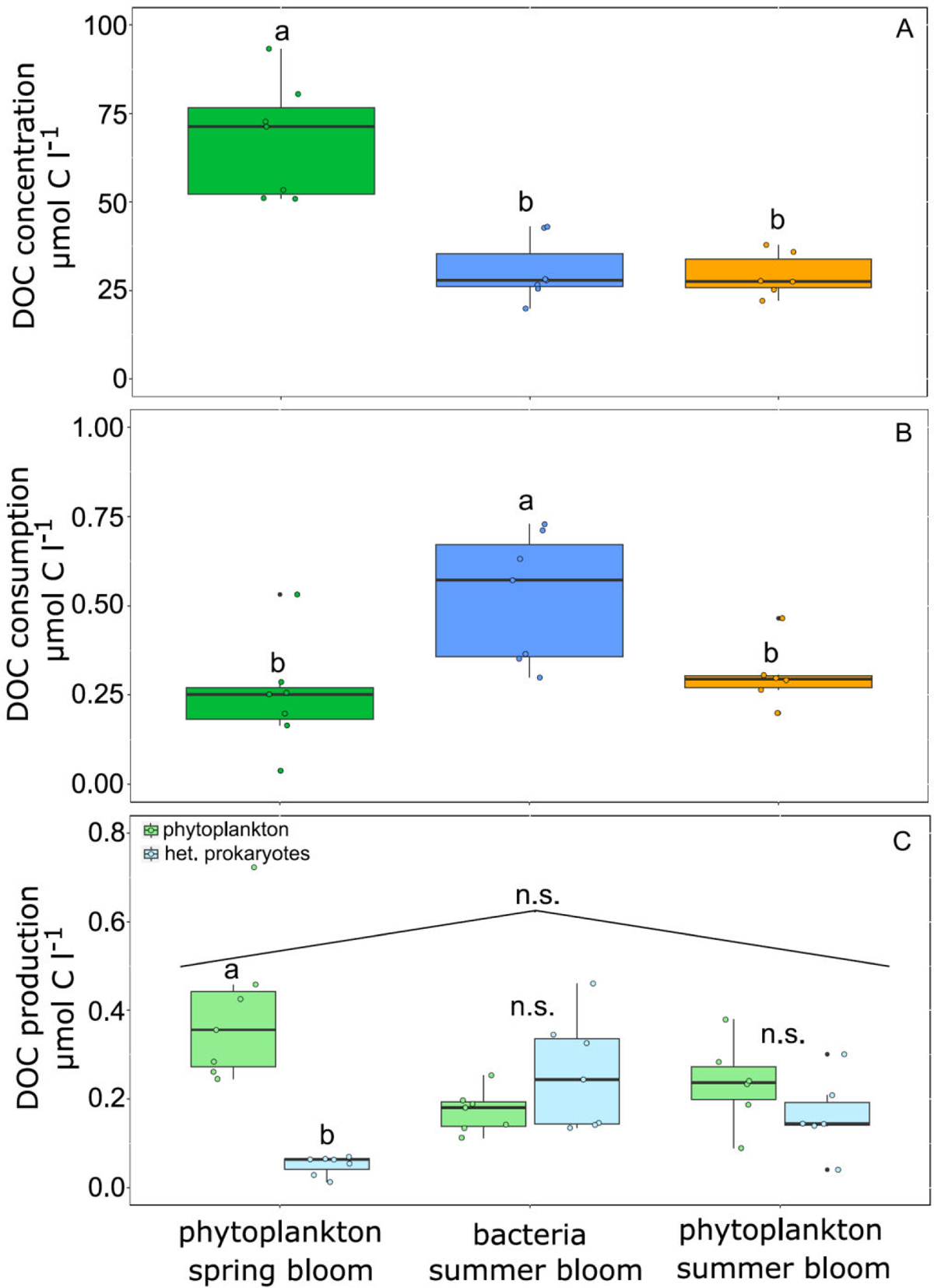
DOC concentration (A), heterotrophic prokaryotes DOC consumption (B), and DOC production of phytoplankton and heterotrophic prokaryotes (C) for the three different bloom types. Letters on the box-plots refer to Tukey post-hoc tests between the different bloom types (combined production of heterotrophic prokaryotes and phytoplankton for panel C), and to T-tests for differences in DOC production of phytoplankton and heterotrophic prokaryotes in the same bloom type (panel C). n.s. = not significant.

DOC concentrations were highest in phytoplankton spring blooms, whereas the DOC uptake by heterotrophic prokaryotes showed the opposite pattern, with highest uptake rates in bacterial summer blooms, and lowest uptake in phytoplankton spring blooms (Fig. 2A-B). We next asked how much DOC is produced and who (phytoplankton or heterotrophic prokaryotes) are the producers. We hypothesized that in periods of high DOC consumption (bacterial summer blooms) the mostly low phytoplankton abundances (Fig. 1A) were balanced with a high phytoplankton DOC production, i.e. a disconnect between phytoplankton abundance and productivity. However, this was not the case, with lowest production in bacterial summer blooms, and highest production in phytoplankton spring blooms (Fig. 2C). Despite higher phytoplankton than heterotrophic prokaryotes DOC production in spring blooms, we indeed found the heterotrophic prokaryotes DOC production exceeding that of the phytoplankton in bacterial summer blooms, leading to overall similar combined production rates in the different bloom types (Fig. 2C). Taken together the outcomes for phytoplankton DOC production and heterotrophic prokaryotes consumption, two points become obvious: First, summer blooms are important to the WEC carbon cycle, with higher carbon fluxes compared to spring blooms (Fig. 2B-C). Second, the heterotrophic prokaryotes carbon consumption in bacterial summer blooms cannot be satisfied by the phytoplankton DOC production, as almost three times the amount of the phytoplankton produced DOC is taken up (Fig. 2B-C).

### Fluxes between heterotrophic prokaryotes substantially contribute to the WEC carbon cycle

High shares of heterotrophic prokaryotes produced DOC in combination with high uptake rates in bacterial summer blooms pose the question whether heterotrophic prokaryotes may consume significant amounts of heterotrophic prokaryotes derived DOC, and hence we quantified carbon fluxes between heterotrophic prokaryotes as well as between phytoplankton and heterotrophic prokaryotes (Fig. 3).

**Fig. 3:**
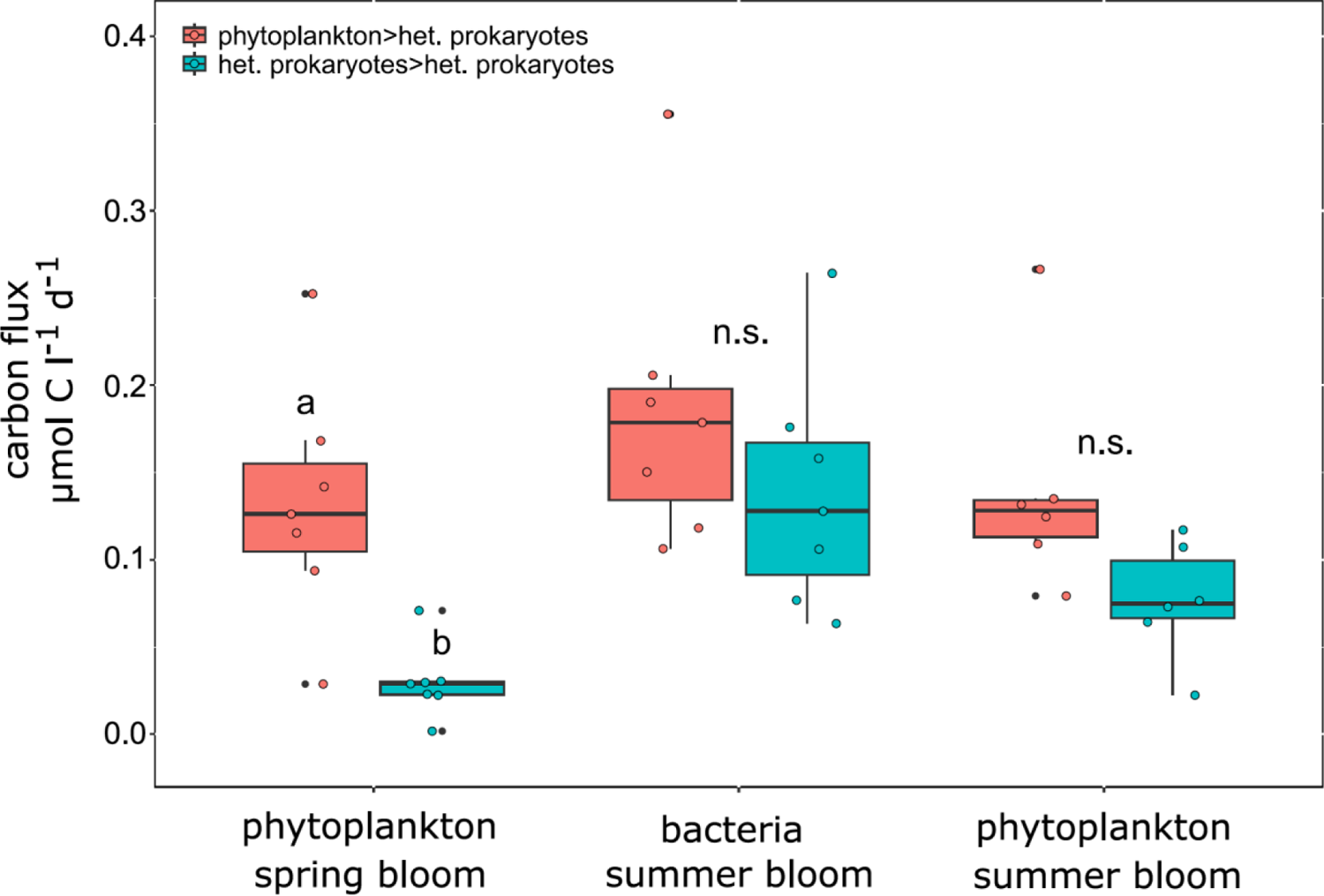
Quantitative bloom averaged carbon fluxes from phytoplankton to heterotrophic prokaryotes (mandarin) and heterotrophic prokaryotes to heterotrophic prokaryotes (turquoise) for the three bloom types for WEC station L4. Letters on the box-plots refer to T-tests for differences between phytoplankton>heterotrophic prokaryotes and heterotrophic prokaryotes>heterotrophic prokaryotes fluxes. n.s. = not significant.

In phytoplankton spring blooms, fluxes from phytoplankton to heterotrophic prokaryotes dominated, whereas fluxes between heterotrophic prokaryotes were negligible. However, in bacteria summer blooms heterotrophic prokaryotes>heterotrophic prokaryotes fluxes were in the same range as phytoplankton>heterotrophic prokaryotes fluxes, and also in phytoplankton summer blooms both flux types did not differ significantly (Fig. 3). These results imply that the mostly neglected fluxes between heterotrophic prokaryotes are of major importance for summer blooms in the WEC, and should be considered in ecosystem analyses. It is possible that allochthonous sources supply some of the carbon and that fluxes between heterotrophic prokaryotes estimated with the model, which assumes a closed system of production and consumption (see model description), are thus overestimated. However, a plot of Tamar river (the major source of allochthonous input for L4) flow rate and bacterial bloom starts did not reveal a relation (Supplementary Fig. 1).

### Bacteria summer blooms are initiated by the recycling of photosynthetically fixed carbon and transition into an internal recycling between heterotrophic prokaryotes

In order to get insights into temporal developments of phytoplankton>heterotrophic prokaryotes and heterotrophic prokaryotes>heterotrophic prokaryotes fluxes, we examined the evolution of fluxes over time. Phytoplankton>heterotrophic prokaryotes fluxes were consistently higher than heterotrophic prokaryotes>heterotrophic prokaryotes fluxes in spring blooms, whereas in bacterial summer blooms phytoplankton>heterotrophic prokaryotes fluxes decreased and heterotrophic prokaryotes>heterotrophic prokaryotes fluxes significantly increased during the course of the bloom (Fig. 4, Supplementary Table 4). These opposed trends suggest that bacterial summer blooms are primarily initiated by the recycling of photosynthetically fixed carbon, and then transition into a DOC re-recycling between different heterotrophic prokaryotes, a process for which the term “internal recycling” may be most fitting (to distinguish from “recycling” for consumption of phytoplankton-derived DOM, sensu Buchan et al (2014)).

**Fig. 4:**
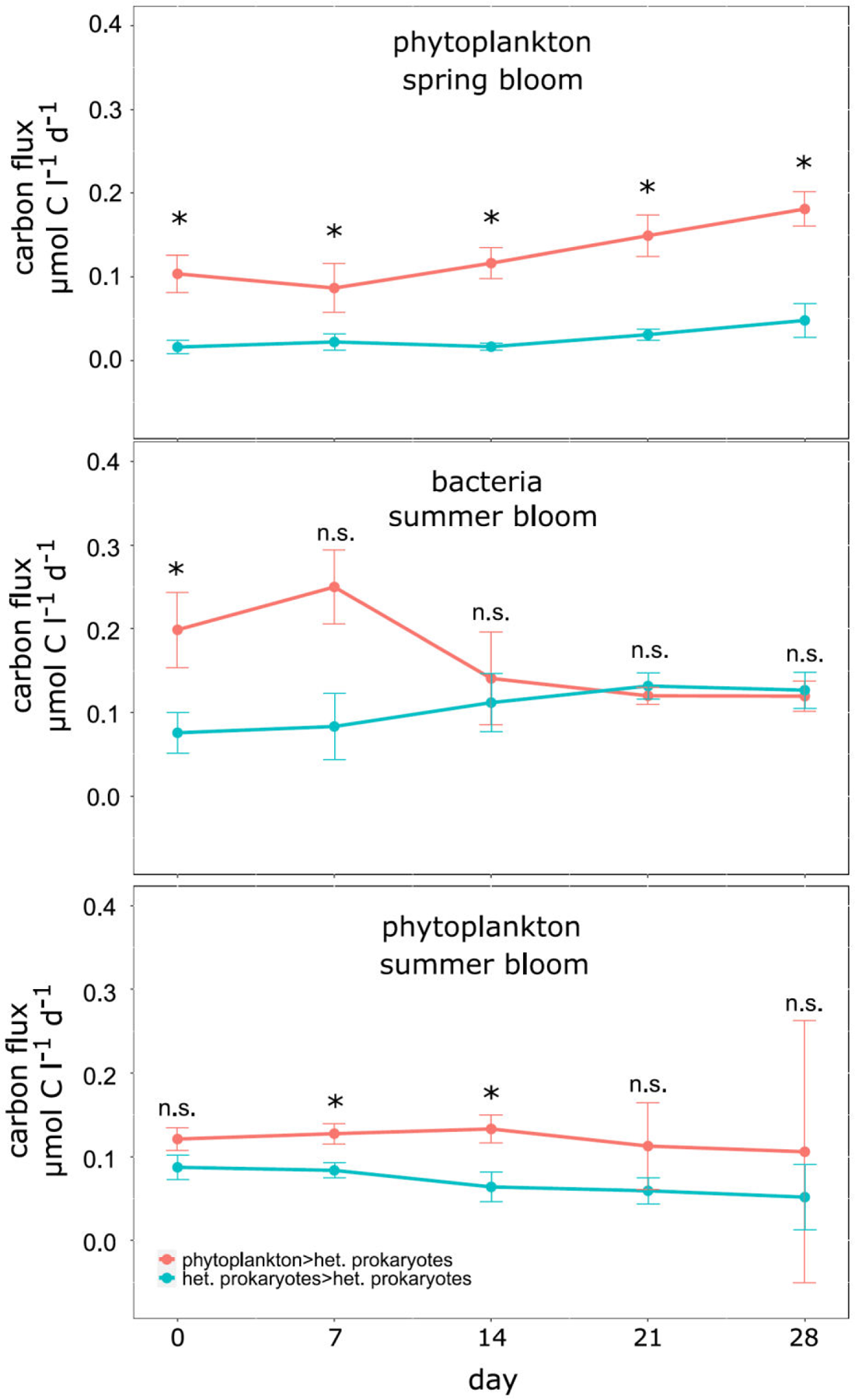
Dynamic carbon fluxes ± standard errors of phytoplankton>heterotrophic prokaryotes (mandarin) and heterotrophic prokaryotes>heterotrophic prokaryotes (turquoise) from bloom-day 0 until 28, at weekly intervals for the three bloom types phytoplankton spring blooms, bacteria summer blooms and phytoplankton summer blooms. Asterix indicate significant differences between the flux types, n.s. = not significant.

### Phages may cause fluxes between heterotrophic prokaryotes

In our analyses carbon fluxes from phytoplankton to heterotrophic prokaryotes are based on a combination of phytoplankton DOM exudation and DOM release due to phytoplankton death (directly or via POM). However, for heterotrophic prokaryotes, we did not include DOM exudation in the model, and consequently all carbon fluxes between heterotrophic prokaryotes are based on DOM liberation at death, i.e. viral predation or protist grazing.

Fast and specific loss processes as drivers for the internal carbon recycling between heterotrophic prokaryotes are corroborated by high fluxes in bacterial summer blooms in years with pronounced successions of abundant ASVs in periods of days (SI, Supplementary Fig. 2, Supplementary Table 5). However, our model does not explicitly resolve the death mechanism, rather a general death process is simulated. Nevertheless, the model calibrates the proportion of DOM and POM released, and viral lysis predominantly liberates DOM and protist grazing POM. The estimated DOM/POM ratio of 1.14±0.12 (Supplementary Table 6) supports viral lysis as predominant cause of death, because dominating protist grazing would result in a ratio far below 1.

### Bacteria summer blooms are initiated by increasing temperatures and may recycle nitrogen for subsequent phytoplankton summer blooms

Bacteria summer blooms occur at times with comparable low phytoplankton DOC production (Fig. 2C) and decreasing estimated DOC concentrations (Supplementary Data 1), raising the question why they occur in summer and not after phytoplankton spring blooms, i.e. periods with high phytoplankton DOC production and concentrations. DOC concentration normalized heterotrophy rates (consumed DOC (mol C consumed d^-1^)/heterotrophic prokaryotes concentration (mol C l^-1^)/DOC concentration (mol C l^-1^)) revealed significantly lower rates in spring blooms compared to summer blooms, suggesting inhibition in the former (Fig. 5). We analyzed the relative strength of substrate and temperature inhibition for the most abundant heterotrophic prokaryotic ASV for each year, and found that the onsets of bacterial summer blooms are caused by increases in water temperature (i.e. the temperature limitation decreases considerably, Supplementary Fig. 3).

**Fig. 5:**
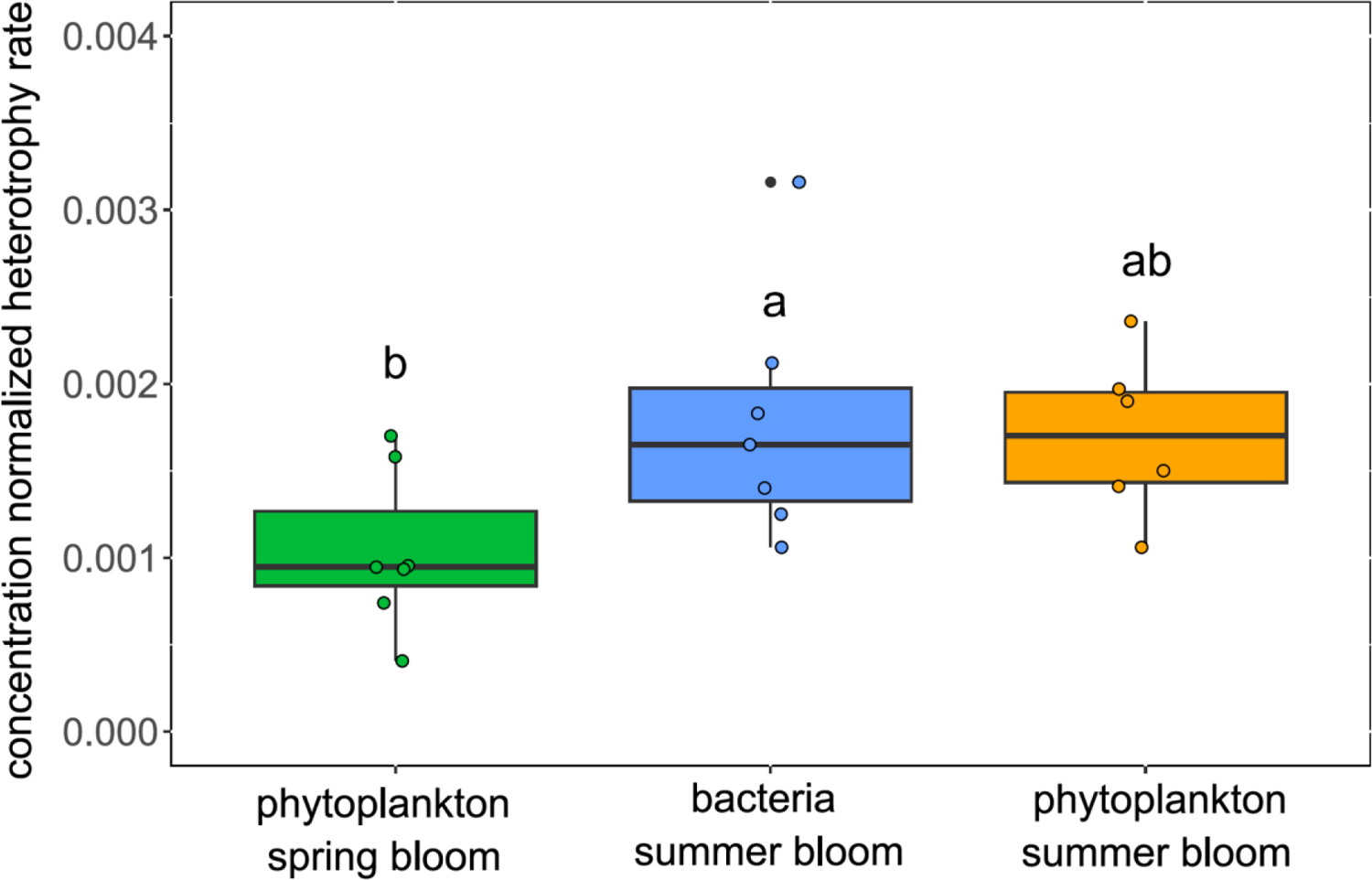
Concentration normalized heterotrophy rates (mol C consumed d^-1^/mol C bacteria l^-1^/mol DOC l^-1^). Letters on top of the box plots refer to Tukey post-hoc tests between the different bloom types.

Our data series depicts several phytoplankton summer blooms following bacterial summer blooms with a delay of ∼15-30 days (Fig. 1A). To test whether we can mechanistically explain summer bloom sequences, we examined the growth inhibition due to PO_4_ and dissolved inorganic nitrogen (DIN) concentrations, as well as due to light and temperature for the most abundant phytoplankton OTU for each year. Phytoplankton was predominantly limited by nitrogen (N) in summer, and several onsets of phytoplankton summer blooms followed a decrease of N limitation accompanying bacteria summer blooms (e.g. in 2012, Supplementary Fig. 3). Thus, reductions of N limitation for the phytoplankton might be partly caused by previous occurring bacterial blooms. Ultimately, the mechanistic inhibition analyses suggest that the sequences of phytoplankton and bacteria blooms at station L4 not only interdepend from each other due to the liberation of organic carbon by the phytoplankton, but also due to the liberation of nitrogen during bacterial blooms.

## Discussion

By analyzing a seven-year time-series of WEC station L4 with a mechanistically constrained inference approach, we identified carbon fluxes between heterotrophic prokaryotes as important components of L4’s carbon cycle (Fig. 3), which are most likely mediated by viruses, and suggested temperature and nitrogen limitations as drivers for microbial summer bloom sequences. A summary of these outcomes is presented as Fig. 6.

**Fig. 6:**
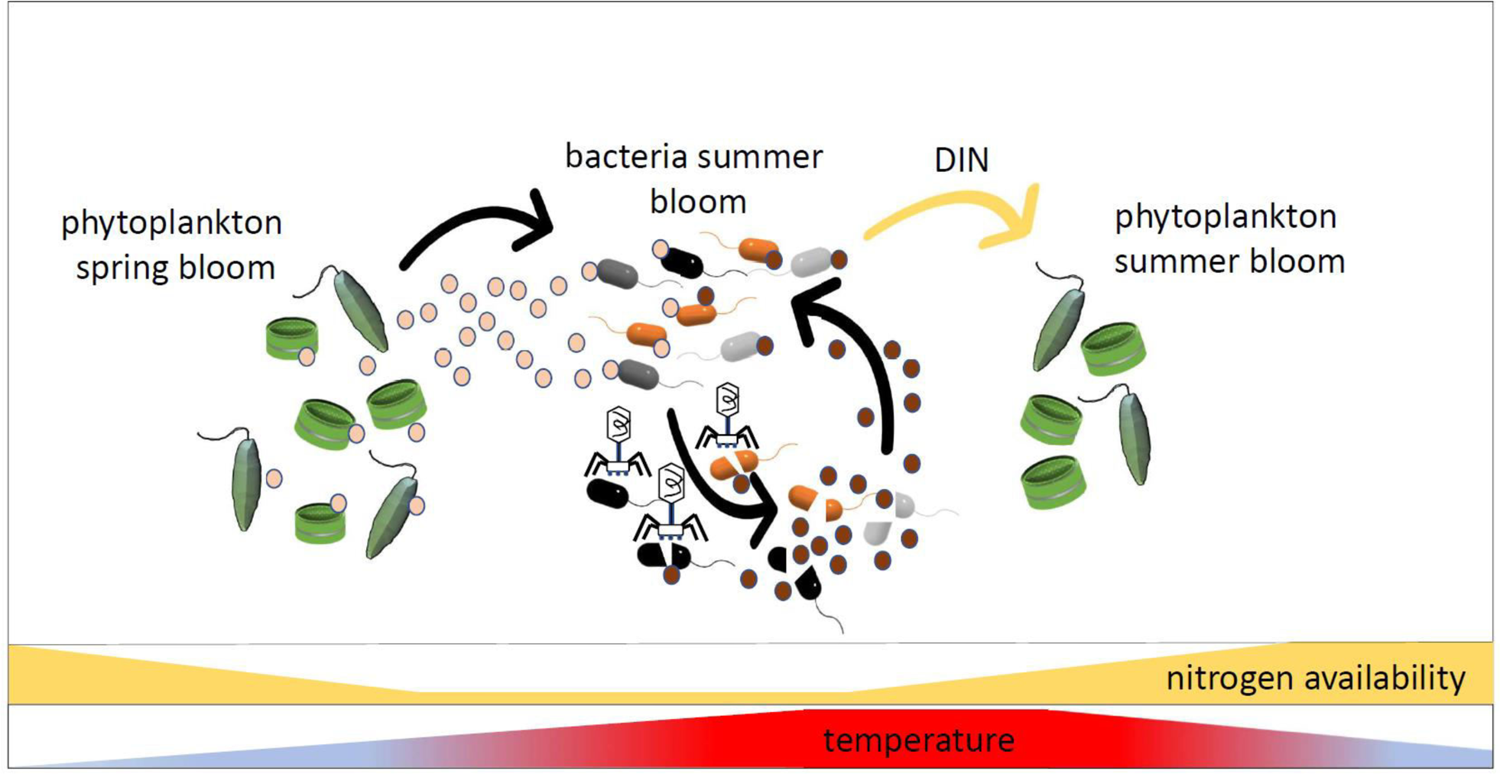
Summary figure for the major outcomes of the present study. Phytoplankton produces large amounts of dissolved organic matter (DOM, light-brown spheres) in spring blooms, whose instantaneous utilization is partly inhibited by low water temperatures. With increasing water temperatures, heterotrophic prokaryotes consume higher amounts of DOM, leading to bacteria summer blooms. Viruses infect abundant heterotrophic prokaryotes, and the lysed cells liberate heterotrophic prokaryotic DOM (dark-brown spheres) that is internally recycled. Bacteria summer blooms also liberate nitrogen, which may foster phytoplankton summer booms.

Although heterotrophic prokaryotes utilization of carbon allocated from other heterotrophic prokaryotes has been observed (Kawasaki and Benner 2006, Lønborg et al 2009, Ortega-Retuerta et al 2021, Zhang et al 2015), this process is not considered in most studies on aquatic carbon cycles, because heterotrophic prokaryotes derived organic carbon is thought to be mostly refractory (Kawasaki et al 2013, Kramer and Herndl 2004, Ogawa et al 2001), and additionally many studies are only focused on the net flux (e.g. Baetge et al (2021)).

To the best of our knowledge, this is the first study that explicitly quantifies the importance of carbon fluxes between heterotrophic prokaryotes in relation to that of phytoplankton to heterotrophic prokaryotes, and hence it is difficult to contextualize our outcomes. Nevertheless, our findings are supported by studies with different scopes. First, microbial association network analysis at station L4 showed that abundance correlations were more pronounced within bacterial taxa than between bacteria and phytoplankton or between bacteria and environmental variables (Gilbert et al 2012). The rationale for the strong correlations within the bacterial compartment, however, were not addressed in that study. Second, several studies showed that heterotrophic bacteria exude large fractions/quantities of DOM, with e.g. 14-31% of the before ingested DOM being extracellularly released by an ocean beach side bacterial community (Kawasaki and Benner 2006), or exudation efficiencies of 11% for DOC, 18% for DON and 17% for DOP by bacterial communities of Scottish coastal waters (Lønborg et al 2009). Third, despite the assumption that heterotrophic bacteria are responsible for a high share of recalcitrant oceanic DOM (Kramer and Herndl 2004, Ogawa et al 2001), the released DOM indeed consists of many bioavailable compounds such as amino acids, amino sugars (Kawasaki and Benner 2006), vitamins, polysaccharides (Wienhausen et al 2017), peptides, and saturated fatty acids (Romano et al 2014). Fourth, recent results from the Helgoland-roads time-series suggests that alpha-glucans from bacterial necromass (i.e. lysed bacteria) are used as carbon source by other bacteria, and that specific PULs (polysaccharide utilization-loci) for this recycling exist (Beidler et al (2023), also see SI for overlaps in the suggested taxonomy of important ASVs between their and our study). The authors of that study conclude that large amounts of carbon may get redirected via an intrapopulation loop, which corroborates the outcomes of our mechanistic inference approach by meta-transcriptomic analyses. Fifth, incubation experiments confirmed that bacteria can grow on DOM exclusively derived from other heterotrophic bacteria (Jørgensen et al 2003, Ortega-Retuerta et al 2021), yet yielding the same cell numbers as if grown on glucose (Zhang et al 2015). Thus, fluxes between heterotrophic prokaryotes indeed reflect well-known processes, whose importance for marine carbon cycles, however, may have been drastically underestimated. A short classification of fluxes between heterotrophic prokaryotes in relation to primary production for bacterial summer blooms (different blooms treated pari passu, then means got calculated), revealed that overall fluxes between heterotrophic prokaryotes accounted for 50% of gross primary production (0.16 vs 0.32 µmol C l^-1^ d^-1^), and were in the same range as net primary production (0.22 µmol l^-1^ d^-1^), as well as the flux from phytoplankton to heterotrophic prokaryotes (0.2 µmol l^-1^ d^-1^, Supplementary Table 3).

The starting point for the quantitative analyses of fluxes between heterotrophic prokaryotes, however, were temporal mismatches between phytoplankton DOC production and DOC consumption by heterotrophic prokaryotes (Fig. 2). Observations that heterotrophic prokaryotes consume more DOC than it is allocated by the phytoplankton also exist from other environments (Fouilland and Mostajir 2010, Fouilland et al 2014, Morán et al 2002). Fouilland and Mostajir (2010) reviewed numerous studies on aquatic primary vs bacterial production (among them 20 studies from marine environments), and concluded that the bacterial carbon utilization significantly exceeds the corresponding total primary production, raising the question whether fluxes between heterotrophic prokaryotes are important components of the carbon cycle also in other systems. Indeed, the study by Beidler et al (2023) suggests that the proposed internal recycling also takes place at other places in significant amounts, and we encourage future analyses of aquatic microbial communities to also resolve fluxes between heterotrophic prokaryotes. The so far applied assumption that heterotrophic prokaryotes just consume DOM and do not also produce it for further consumption by others would be a gross oversimplification.

All fluxes between heterotrophic prokaryotes are a result of DOM liberation upon death (see above), and viral predation and grazing by protists are the major death causes for heterotrophic prokaryotes (e.g. Breitbart et al. 2018). Both processes (due to cell lysis and sloppy feeding, respectively) liberate bioavailable DOC, but although sloppy feeding liberates significant amounts of DOC for e.g. copepod grazing on phytoplankton (up to 17% of the prey’s carbon content, Lampert (1978)), protist grazing on bacteria only liberates minor amounts of DOC (because of phagotrophy and whole prey engulfment, Weisse et al (2016)). Thus, the major part of DOC liberation can be attributed to viral lysis. Estimated DOM/POM release ratios >1 furthermore indicate viral lysis as predominant death mechanism during bacterial blooms. As consequence, the inferred fluxes between heterotrophic prokaryotes may be largely attributed to viral infections and considered as a component of the “viral shunt” (Wilhelm and Suttle 1999), i.e. the retainment of carbon for higher trophic levels due to the recycling of DOM derived from viral lysis of fish, zooplankton, phytoplankton, archaea and bacteria by heterotrophic prokaryotes.

The growth of marine heterotrophic prokaryotes predominantly depends on the availability of organic carbon (Christie-Oleza et al 2017, Hellweger et al 2018, Liu et al 2012), and the question arises why bacteria blooms occur at times with comparably low DOC concentration and production. Analyses of growth limiting factors in the model, however, suggest that the onsets of bacteria summer blooms are caused by increased water temperatures (Supplementary Fig. 3). Higher temperatures enable the utilization of less bioavailable DOM (Lønborg et al 2020), including less bioavailable compounds of phytoplankton exudates (Zhang et al 2022). The occurrence of the latter is indicated by high phytoplankton>heterotrophic prokaryotes fluxes despite low phytoplankton DOC production in periods with higher water temperatures (as the start of bacterial summer blooms, Figs 2-3). However, increased temperatures may also enable the internal recycling occurring at later phases in bacterial summer blooms, i.e. the utilization of less bioavailable DOM derived from heterotrophic prokaryotes (Kawasaki et al 2013, Kramer and Herndl 2004, Ogawa et al 2001). On the phytoplankton side, mechanistic analyses suggested nitrogen deficiency as major inhibitor in summer (Supplementary Fig. 3), and that limitation decreases following bacterial blooms led to phytoplankton summer blooms in several years. We speculate that the bacteria blooms recycled nitrogen and thus partly initialized phytoplankton summer blooms. Our speculation that bacterial blooms foster phytoplankton summer blooms is supported by studies showing that a net production of DIN took place during bacterial utilization of glucose (Lønborg et al 2009), and that the exploitation of organic material by coastal bacterioplankton communities led to increased NH_4_ levels (Kawasaki and Benner 2006). Bloom sequences with distinct phytoplankton spring blooms, and bacterial summer blooms preceding additional phytoplankton blooms also occur in other than the used years at station L4 (Gilbert et al 2009), as well as at other places (Helgoland roads time-series, Teeling et al (2016)). Yet, due to lacking analyses and availability of data, we may not speculate on the underlying processes. Nevertheless, modeling efforts and/or experimental approaches with different time-series/at different places may help to understand the sequences of phytoplankton and bacteria blooms, and would help to support or refute the suggested mechanisms.

Decreasing costs and efforts of modern molecular tools (e.g. proteomics, next generation sequencing) enable high resolutions of observations and increase e.g. the number of heterotrophic prokaryotic ASVs or DOM species in ecosystem analyses. Increased numbers of analyzed components in microbial communities, however, are accompanied by exponential increases of their interactions, which makes quantitative measurements of multi-species interactions impossible. As consequence, such analyses require the deployment of models. In the present study, we used a mechanistic inference approach with hundreds of phytoplankton and heterotrophic prokaryotic species and were able to quantitatively estimate interactions (i.e. carbon fluxes) between phytoplankton OTUs and heterotrophic prokaryotic ASVs, as well as between the latter. The code for the inference approach is open-source, and readily extendable in terms of dimensions (e.g. zooplankton and viruses), model agents/species as well as constrains (e.g. the implementation of transcriptomic data), and thus may allow mechanistic analyses to keep track with progressions/outcomes made by modern molecular tools.

## Materials and Methods

### Data acquisition and selection

All data, except where noted otherwise, come from WEC station L4, and were obtained from the Western Channel Observatory (westernchannelobservatory.org.uk), with sampling performed by the Plymouth Marine Laboratory. Station L4 depicts a northern-temperate coastal-ocean side ca. 50 km off Plymouth, UK, with occasional pulses of increased nitrate concentrations due to riverine input (Smyth et al 2009). Data reach from the beginning of 2012 until the end of 2018 with a weekly sampling period. Eukaryotic phytoplankton concentrations were derived from 18S rRNA sequence data (full ASV Table given as Supplementary Data 2) after conversion of individual read counts into carbon concentrations (for details see SI), and prokaryotic phytoplankton (*Synechococcus*) concentrations from flow-cytometer cell counts (Supplementary Table 7). The full 18S table contained 69,207 amplicon sequence variants (ASVs), but in order to emerge manageable data loads, only the 10,000 most abundant ASVs were considered for further analyses. Heterotrophic as well as unassigned ASVs were filtered out, and the remaining phytoplankton ASVs combined into 135 operational taxonomic units (OTUs, for details see SI). Heterotrophic prokaryotes concentrations were derived from 16S rRNA sequence data after conversion of relative read abundances into carbon concentrations (for details see SI). The 16S dataset contained the 200 most abundant ASVs (Supplementary Table 8), and after filtering out chloroplast, unassigned, cyanobacterial and ammonia-oxidizing archaeal ASVs, 157 ASVs were used for elaborated analyses. Additional data that were incorporated into the model were photosynthetic active radiation (obtained from MODIS satellite, https://modis.gsfc.nasa.gov/data/), daylength (obtained from https://timeanddate.com for Plymouth), temperature, NO_2_ + NO_3_, NH_4_, PO_4_, silicate, salinity, oxygen concentrations, particulate organic carbon (POC), and chlorophyll (Supplementary Data 3).

### FluxNet method - Mechanistic microbial ecosystem model inference

A complete description and explanation of the inference approach is available in Mayerhofer et al (2021) and details on the English Channel application are provided in the SI. Major differences (compared to Mayerhofer et al (2021)) are the upscaling of components and separate model runs for separate years. The latter was established in order to improve the fit between data and model. Briefly, modeling concepts and equations are based on past models of phytoplankton and bacteria (Chapra 1997, Hellweger and Lall 2004, Mieleitner and Reichert 2008, Pinto et al 2017, Weitz et al 2015), where fluxes between ecological compartments, e.g. phytoplankton to DOM to heterotrophic prokaryotes, are explicitly simulated. For example, phytoplankton growth rates are a function of light, temperature and inorganic nutrients. They exude DOM in a basal and growth rate-proportional manner. Heterotrophic prokaryotes grow by heterotrophy with different affinities for each hypothetical DOM species. The model parameters are automatically calibrated to time series observations. Key features of the approach are: an optimization routine customized for microbial ecosystems; inclusion of dormancy to avoid extinction and support co-existence; gradual de-lumping/diversification to increase the resolution (i.e. number of species).

### Definition and duration of blooms

Phytoplankton spring blooms were defined as the highest chlorophyll concentration between February and May, phytoplankton summer blooms as the highest chlorophyll concentration between July and August. In the year 2014, no pronounced phytoplankton summer bloom was observed, and consequently no bloom assigned. Bacterial summer blooms were defined as the highest bacterial concentrations between June and August. To ensure model/data overlaps, bloom starting days were not defined on strict parameters, and assigned manually. Bloom durations were set to 28 days, except when concentrations fell below the starting concentration during that period. In this case blooms ended at the day concentrations fell below the bloom starting concentration. However, in the years 2012 and 2017, bacterial summer bloom duration was increased to 34 and 42 days, respectively, because concentrations remained high. An overview on bloom durations, start- and end-days is given as Supplementary Table 1.

### Statistics

All applied statistical tests and statistical outcomes are provided as Supplementary Table 4. Statistical tests and graphics were executed with the free software R (R Development Team 2020) and R studio (RStudio Team 2020).

## Data availability

The open-source model code is provided at https://github.com/fhellweger, and all input data for the model (except of 16S and 18S sequences) given in Supplementary Data 3. 16S and 18S raw data have been uploaded to NCBI with the Submission ID of SUB13994125 and BioProject ID of PRJNA1045854. The top 200 16S ASVs are given as Supplementary Table 8, and the full 18S ASV table is given as Supplementary Data 2.

## Acknowledgements

We thank Carsten Lackner, Jutta Hoffmann, Behnam Zamani und Charlotte Schampera for discussions on the manuscript. Funding was provided by the BIOS-SCOPE program (Award number 409923FY20 to FLH), and the ECOHAB project (Award number NA18NOS4780175 to FLH).

## References

Ahlgren NA, Perelman JN, Yeh YC, Fuhrman JA (2019). Multi-year dynamics of fine-scale marine cyanobacterial populations are more strongly explained by phage interactions than abiotic, bottom-up factors. Environ Microbiol 21: 2948–2963.

Auladell A, Barberán A, Logares R, Garcés E, Gasol JM, Ferrera I (2021). Seasonal niche differentiation among closely related marine bacteria. The ISME Journal.

Azam F, Fenchel T, Field J, Gray JS, Meyer-Reil LA, Thingstad F (1983). The Ecological Role of Water-Column Microbes in the Sea. Marine Ecology Progress Series 10: 257–263.

Azam F, Malfatti F (2007). Microbial structuring of marine ecosystems. Nat Rev Microbiol 5: 782–791.

Baetge N, Behrenfeld MJ, Fox J, Halsey KH, Mojica KDA, Novoa A et al (2021). The Seasonal Flux and Fate of Dissolved Organic Carbon Through Bacterioplankton in the Western North Atlantic. Frontiers in Microbiology 12.

Beidler I, Steinke N, Schulze T, Sidhu C, Bartosik D, Krull J et al (2023). Alpha-glucans from bacterial necromass indicate an intra-population loop within the marine carbon cycle. PREPRINT Research Square Version 1.

Blanchet FG, Cazelles K, Gravel D (2020). Co-occurrence is not evidence of ecological interactions. Ecology Letters 23: 1050–1063.

Boalch GT, Harbour DS, Butler EI (1978). Seasonal phytoplankton production in the western English Channel 1964–1974. Journal of the Marine Biological Association of the United Kingdom 58: 943–953.

Bolaños LM, Choi CJ, Worden AZ, Baetge N, Carlson CA, Giovannoni S (2021). Seasonality of the Microbial Community Composition in the North Atlantic. Frontiers in Marine Science 8.

Buchan A, LeCleir GR, Gulvik CA, Gonzalez JM (2014). Master recyclers: features and functions of bacteria associated with phytoplankton blooms. Nat Rev Microbiol 12: 686–698.

Chapra SC (1997). Surface Water-Quality Modeling. McGraw-Hill: Boston.

Christie-Oleza JA, Sousoni D, Lloyd M, Armengaud J, Scanlan DJ (2017). Nutrient recycling facilitates long-term stability of marine microbial phototroph–heterotroph interactions. Nature Microbiology 2: 17100.

Cole JJ (1982). Interactions between bacteria and algae in aquatic ecosystems. Annual Review of Ecology and Systematics 13: 291–314.

Derenbach JB, Williams PJLB (1974). Autotrophic and bacterial production: Fractionation of plankton populations by differential filtration of samples from the English channel. Marine Biology 25: 263–269.

Faust K, Lahti L, Gonze D, de Vos WM, Raes J (2015). Metagenomics meets time series analysis: unraveling microbial community dynamics. Current opinion in microbiology 25: 56–66.

Fouilland E, Mostajir B (2010). Revisited phytoplanktonic carbon dependency of heterotrophic bacteria in freshwaters, transitional, coastal and oceanic waters. FEMS Microbiology Ecology 73: 419–429.

Fouilland E, Tolosa I, Bonnet D, Bouvier C, Bouvier T, Bouvy M et al (2014). Bacterial carbon dependence on freshly produced phytoplankton exudates under different nutrient availability and grazing pressure conditions in coastal marine waters. FEMS Microbiol Ecol 87: 757–769.

Gilbert JA, Field D, Swift P, Newbold L, Oliver A, Smyth T et al (2009). The seasonal structure of microbial communities in the Western English Channel. Environ Microbiol 11: 3132–3139.

Gilbert JA, Steele JA, Caporaso JG, Steinbruck L, Reeder J, Temperton B et al (2012). Defining seasonal marine microbial community dynamics. ISME J 6: 298–308.

Hellweger FL, Lall U (2004). Modeling the Effect of Algal Dynamics on Arsenic Speciation in Lake Biwa. Environmental Science & Technology 38: 6716–6723.

Hellweger FL, Huang Y, Luo H (2018). Carbon limitation drives GC content evolution of a marine bacterium in an individual-based genome-scale model. The ISME Journal 12: 1180–1187.

Irigoien X, Harris RP, Head RN, Harbour D (2000). North Atlantic Oscillation and spring bloom phytoplankton composition in the English Channel. Journal of Plankton Research 22: 2367–2371.

Jiao N, Azam F (2011). Microbial carbon pump and its significance for carbon sequestration in the ocean. Microbial Carbon Pump in the Ocean 10: 43–45.

Jørgensen NOG, Stepanaukas R, Pedersen A-GU, Hansen M, Nybroe O (2003). Occurrence and degradation of peptidoglycan in aquatic environments. FEMS Microbiology Ecology 46: 269–280.

Kawasaki N, Benner R (2006). Bacterial release of dissolved organic matter during cell growth and decline: Molecular origin and composition. Limnology and Oceanography 51: 2170–2180.

Kawasaki N, Komatsu K, Kohzu A, Tomioka N, Shinohara R, Satou T et al (2013). Bacterial contribution to dissolved organic matter in eutrophic Lake Kasumigaura, Japan. Appl Environ Microbiol 79: 7160–7168.

Kramer G, D, Herndl G, J (2004). Photo- and bioreactivity of chromophoric dissolved organic matter produced by marine bacterioplankton. Aquatic Microbial Ecology 36: 239–246.

Lampert W (1978). Release of dissolved organic carbon by grazing zooplankton. Limnology and Oceanography 23: 831–834.

Lamy D, Artigas L, Jauzein C, Lizon F, Cornille V (2006). Coastal bacterial viability and production in the eastern English Channel: A case study during a Phaeocystis globosa bloom. Journal of Sea Research 56: 227–238.

Liu H, Zhou Y, Xiao W, Ji L, Cao X, Song C (2012). Shifting nutrient-mediated interactions between algae and bacteria in a microcosm: evidence from alkaline phosphatase assay. Microbiol Res 167: 292–298.

Lønborg C, Álvarez-Salgado XA, Davidson K, Miller AEJ (2009). Production of bioavailable and refractory dissolved organic matter by coastal heterotrophic microbial populations. Estuarine, Coastal and Shelf Science 82: 682–688.

Lønborg C, Carreira C, Jickells T, Álvarez-Salgado XA (2020). Impacts of Global Change on Ocean Dissolved Organic Carbon (DOC) Cycling. Frontiers in Marine Science 7.

Mayerhofer MM, Eigemann F, Lackner C, Hoffmann J, Hellweger FL (2021). Dynamic carbon flux network of a diverse marine microbial community. ISME Communications 1: 50.

Mieleitner J, Reichert P (2008). Modelling functional groups of phytoplankton in three lakes of different trophic state. Ecological Modelling 211: 279–291.

Morán XA, G., Josep MG, Pedros-Alio C, Marta E (2002). Partitioning of phytoplanktonic organic carbon production and bacterial production along a coastal-offshore gradient in the NE Atlantic during different hydrographic regimes. Aquatic Microbial Ecology 29: 239–252.

Napoléon C, Fiant L, Raimbault V, Riou P, Claquin P (2014). Dynamics of phytoplankton diversity structure and primary productivity in the English Channel. Marine Ecology Progress Series 505: 49–64.

Ogawa H, Amagai Y, Koike I, Kaiser K, Benner R (2001). Production of Refractory Dissolved Organic Matter by Bacteria. Science 292: 917–920.

Ortega-Retuerta E, Devresse Q, Caparros J, Marie B, Crispi O, Catala P et al (2021). Dissolved organic matter released by two marine heterotrophic bacterial strains and its bioavailability for natural prokaryotic communities. Environmental Microbiology 23: 1363–1378.

Pinto F, Medina DA, Pérez-Correa JR, Garrido D (2017). Modeling Metabolic Interactions in a Consortium of the Infant Gut Microbiome. Frontiers in Microbiology 8.

R Development Team (2020). R: A language and environment for statistical computing. R Foundation for Statistical Computing, Vienna, Austria. URL https://www.R-project.org/.

Romano S, Dittmar T, Bondarev V, Weber RJM, Viant MR, Schulz-Vogt HN (2014). Exo-Metabolome of Pseudovibrio sp. FO-BEG1 Analyzed by Ultra-High Resolution Mass Spectrometry and the Effect of Phosphate Limitation. PLOS ONE 9: e96038.

RStudio Team (2020). RStudio: Integrated Development for R. RStudio, PBC, Boston, MA. URL https://www.rstudio.com/.

Smyth TJ, Fishwick JR, Al-Moosawi L, Cummings DG, Harris C, Kitidis V et al (2009). A broad spatio-temporal view of the Western English Channel observatory. Journal of Plankton Research 32: 585–601.

Teeling H, Fuchs BM, Becher D, Klockow C, Gardebrecht A, Bennke CM et al (2012). Substrate-controlled succession of marine bacterioplankton populations induced by a phytoplankton bloom. Science 336: 608–611.

Teeling H, Fuchs BM, Bennke CM, Kruger K, Chafee M, Kappelmann L et al (2016). Recurring patterns in bacterioplankton dynamics during coastal spring algae blooms. Elife 5: e11888.

Weisse T, Anderson R, Arndt H, Calbet A, Hansen PJ, Montagnes DJS (2016). Functional ecology of aquatic phagotrophic protists – Concepts, limitations, and perspectives. European Journal of Protistology 55: 50–74.

Weitz JS, Stock CA, Wilhelm SW, Bourouiba L, Coleman ML, Buchan A et al (2015). A multitrophic model to quantify the effects of marine viruses on microbial food webs and ecosystem processes. The Isme Journal 9: 1352.

Widdicombe CE, Eloire D, Harbour D, Harris RP, Somerfield PJ (2010). Long-term phytoplankton community dynamics in the Western English Channel. Journal of Plankton Research 32: 643–655.

Wienhausen G, Noriega-Ortega BE, Niggemann J, Dittmar T, Simon M (2017). The Exometabolome of Two Model Strains of the Roseobacter Group: A Marketplace of Microbial Metabolites. Front Microbiol 8: 1985.

Wilhelm SW, Suttle CA (1999). Viruses and Nutrient Cycles in the Sea: Viruses play critical roles in the structure and function of aquatic food webs. BioScience 49: 781–788.

York A (2018). Marine biogeochemical cycles in a changing world. Nature Reviews Microbiology 16: 259–259.

Zhang J, Liu J, Liu D, Chen X, Shi Q, He C et al (2022). Temperature Rise Increases the Bioavailability of Marine Synechococcus-Derived Dissolved Organic Matter. Frontiers in Microbiology 13.

Zhang Z, Chen Y, Wang R, Cai R, Fu Y, Jiao N (2015). The Fate of Marine Bacterial Exopolysaccharide in Natural Marine Microbial Communities. PLoS One 10: e0142690.

