## Supplementary for "Heterotrophic prokaryotes internal carbon recycling compensates mismatches between phytoplankton production and heterotrophic prokaryotic consumption"

**This file includes:**

S1. Introduction
S2. Details of the mechanistic microbial ecosystem model FluxNet
S3. Optimization routine
S4. Data acquisition and processing
S5. Additional results and discussion
S6. References

#### S1: Introduction

The purpose of this section is twofold: (1) document the modeling framework with examples for the English Channel application, and (2) present additional analyses for the English Channel application. General information on model details can be found in the SI in Mayerhofer et al (2021) from the original Helgoland-roads application. In the paragraphs S1-S2 the most important model features are summarized.

#### S2: Details of the mechanistic microbial ecosystem model

##### S2.1: Concepts, components and processes

The modeling concepts and equations are generally based on past models of phytoplankton, bacteria, zooplankton and viruses (e.g. Chapra (1997), Weitz et al (2015)). Novel aspects (compared to Mayerhofer et al (2021)) include up-scaling of components, and separate model runs for separate years.
The model consists of a number of components (concentrations, state variables) that interact via a number of processes (functions) (see Supplementary Table 9 for components used in the English Channel application). The model structure is flexible and puts no constraints on the number and types of processes that connect components, so the model can, for example, simulate mixotrophy. However, it helps to think in terms of and organize components into

traditional ecological compartments. In that case, the processes corresponding to each component depend on the ecological compartment it is in.
Model components are generally selected based on observations. However, for a mechanistic model, it is important to close the mass balance and that puts some constraints on the definition of components. First, state variables cannot overlap. Second, the state variables have to cover the trophic levels. To cover the trophic level, "hypothetical" species are added (conceptually similar to "others" in Hellweger et al (2008)).
For processes, the model includes loading [w], flow [q], settling [s], first-order [f], respiration [r], photosynthesis [p], grazing [z], heterotrophy [h], viral [v], exudation [e], death [y], loss [u] and inhibition [i].
To illustrate the approach, we present mass balance equations for a reduced system consisting of one phytoplankton OTU (mpu), two DOM species (d01, d02) and one heterotrophic prokaryotic ASV (s11).
Processes affecting phytoplankton concentration include photosynthesis, respiration, exudation, inhibition, death, settling and outflow. The mass balance equation for mpu is:

$$\frac{d}{dt} C_{mpu} = kp_{mpu}\, Lp_{mpu}\, C_{mpu} - kr_{mpu}\, Lr_{mpu}\, C_{mpu} - \left(ke_{mpu} + ef_{mpu}\, kp_{mpu}\, Lp_{mpu}\right) C_{mpu} - ku_{mpu}\, Lu_{mpu}\, C_{mpu} - \frac{vs_{mpu}}{H} C_{mpu} - \frac{Q}{V} C_{mpu} \tag{1}$$

t (d) = time, C (mmolC/L) = concentration, kp (1/d) = max. photosynthesis rate constant, Lp = limitation/modification factor for photosynthesis (light, nutrients, temperature, salinity), kr (1/d) = max. respiration rate constant, Lr = limitation/modification factor for respiration (temperature), ke (1/d) = basal exudation rate constant, ef (1/d) = exudation/photosynthesis fraction, ku (1/d) = maximum death rate, Lu = limitation/modification factor for death (time-of-year, salinity), vs (m/d) = settling velocity, H (m) = water column depth, Q (m3/d) = flow rate, V (m3) = volume.

Processes affecting DOM include microbial exudation and death, POM dissolution, heterotrophy and outflow. The mass balance equation for d01, considering only mpu and s11 as source, s11 as sink, and p01 as corresponding POM species, is:

$$\begin{aligned}\frac{d}{dt} C_{d01} = {} & Fe_{mpu,d01} \left(ke_{mpu} + ef_{mpu}\, kp_{mpu}\, Lp_{mpu}\right) C_{mpu} + Fx_{mpu,d01}\, ku_{mpu}\, Lu_{mpu}\, C_{mpu} \\ & + Fx_{s11,d01}\, ku_{s11}\, Lu_{s11} C_{s11} + kf_{p01}\, Lf_{p01}\, C_{p01} \\ & - kh_{s11} \frac{C_{d01}/Ksh_{s11,d01}}{1 + C_{d01}/Ksh_{s11,d01} + C_{d02}/Ksh_{s11,d02}}\, Lh_{s11}\, C_{s11} - \frac{Q}{V} C_{d01}\end{aligned} \tag{2}$$

Fe = exudation fraction, Fx = composition fraction, kf (1/d) = dissolution rate constant, Lf = limitation/modification factor for dissolution (temperature), kh (1/d) = max. heterotrophy rate, Ksh (mmolC/L) = half-saturation constant, Lh = limitation/modification factor for heterotrophy (temperature, salinity, light).

Processes affecting heterotrophic prokaryotes concentration include heterotrophy, death and outflow. The mass balance equation for s11, considering only growth on d01 and d02, is:

$$\frac{d}{dt}C_{s11} = Yh_{s11}\, kh_{s11}\, \frac{C_{d01}/Ksh_{s11,d01} + C_{d02}/Ksh_{s11,d02}}{1 + C_{d01}/Ksh_{s11,d01} + C_{d02}/Ksh_{s11,d02}}\, Lh_{s11}\, C_{s11} - ku_{s11}\, Lu_{s11}\, C_{s11} - \frac{Q}{V} C_{s11} \quad (3)$$

Yh = yield coefficient.

##### S2.2: Dormancy

Dormancy is modeled by specifying a floor concentration (Cflr) and reducing any loss rate that would result in concentration below this value, an approach similar to previous models (Daines et al 2014). Specifically, a floor concentration (*Cflr*) of ¼ of the lowest, measured concentration is given, and any loss process that would reduce the concentration below this value is blocked.

##### S2.3: Environment and boundary conditions

The environment is a completely-mixed reactor with specified area (*A*) and depth (*H*). Boundary conditions include light intensity (*IT*), photoperiod (*f*), temperature (*T*) and nutrient loadings (*WNOX*, *WNH4*, *WPO4*, *WSIL*).

For the English Channel application, external nutrient inputs (loadings) were calculated from observations *a priori* using a simplified approach. For nitrogen, it was assumed total nitrogen (TN) is made up of NOX, NH4 and phytoplankton nitrogen (PN), which was calculated from Chlorophyll *a* using conversions (see SI in Mayerhofer et al (2021)). It was further assumed the only loss processes are settling of PN and outflow of TN. These assumptions result in the following mass balance equation.

$$V\, \frac{dTN}{dt} = W_{NOX} - vs\, A\, PN - Q\, TN \quad (4)$$

This equation can be discretized and solved for the nitrogen input.

$$W_{NOX} = V \frac{\Delta TN}{\Delta t} + vs\ A\ \overline{PN} + Q\ \overline{TN} \quad (5)$$

When this calculation is applied to a data set with high variability, it can lead to negative values, which does not agree with the concept of external input. Therefore, the load is kept positive, but a deficit is tracked so that the resulting cumulative input is consistent with the mass balance. Input of PO4 and SIL are handled equivalently.

#### S3: Optimization routine

The optimization method adjusts parameter values within literature ranges to minimize the disagreement i.e. error between model and data. The method generally follows previous numerical optimization approaches for microbial ecosystem models (Mieleitner and Reichert 2008, Pinto et al 2017, Weitz et al 2015). Novel aspects include a two-dimensional (concentration and time) quantification of model-data disagreement, numerical optimization methods customized for microbial ecosystems and gradual increase in model complexity (de-lumping).

##### S3.1: De-lumping

Species are introduced at specified de-lump levels, by de-lumping from an existing species. The new species generally inherits the parameter values (i.e. the genome, sensu Daines et al (2014)) from the old species. Subsequent optimization then diversifies the population. This is illustrated in Fig. 1C of Mayerhofer et al (2021), which shows the affinity of bacteria species for *mpu*. However, if parameter values are specified for the new species, they are adopted and overwrite those inherited from the old species. This is used, for example, to assign species-specific cell sizes (*MC*).

#### S4. Data acquisition and processing

##### S4.1: 18S Illumina sequencing and phototroph/heterotroph factor

18S rRNA sequences were derived from Illumina sequencing and analyzed with the dada2 pipeline (Callahan et al 2016). The full 18S dataset comprised 69,207 amplicon sequence variants (ASVs, Supplementary Data 2). Absolute read counts were converted into relative abundances, and the 10,000 most abundant ASVs used for elaborated analyses (all dates/samples contained ~20,000 reads, and thus no subsampling was necessary, Supplementary Data 2), which covered 61-98% of all reads. In order to account for 18S reads that do not derive from phototrophic organisms, and are as such not in the scope of our study, all ASVs (10,000 most abundant) were manually classified into phototroph/not phototroph, whereby mixotrophic ASVs were treated as phototrophs. From these phototroph/not phototroph ratios, a factor for each sampling date was calculated, and each phototrophic ASV multiplied with the factor at the respective sampling point. This factor ensured that we received the true share of each ASV in the phototrophic community.

##### S4.2: Assignment of OTUs

Because most 18S ASVs only occurred at one or few sampling dates, ASVs were pooled to OTUs. For all OTUs, we set the minimum occurrence to 4 sampling dates in the entire time-series. If less than 4 sampling dates were obtained, ASVs were pooled to the next lower taxonomic level until 4 or more occurrences in the data set were reached. This means, at first different ASVs of the same species were combined to OTUs, then all species with less than 4 occurrences were pooled to genera a.s.o. This was done successively for each taxonomic level (species/genera/family/order/class/phyla). If several ASVs of the same OTU occurred at the same sampling date(s), relative abundances of these ASVs were totalized. If ASVs were originally assigned to taxonomic levels lower than species, the highest resolved taxonomic level was taken and treated as described above. Altogether, we obtained 138 OTUs, consisting of 120 species, 7 genera (undefined family), 1 family (undefined species and family), 2 orders, 5 classes, and 3 phyla. In order to account for the share of the phytoplankton not covered by the 10,000 most abundant OTUs (see above), we also added 24 hypothetical phytoplankton OTUs.

##### S4.3: Transformation of 18S data in carbon concentrations

Relative abundances of OTUs were transferred into carbon concentrations based on chlorophyll concentrations. Illumina read abundances of OTUs depend on their cell numbers per volume as well as 18S gene copy numbers (GCN) per cell. 18S GCN, for their part, depend on the cell volume as follows:

$$\log (\text{cell volume } \mu m^3) = -0.61 + 1.22 \log GCN \qquad (6)$$

(Godhe et al 2008).

The carbon concentration of an OTU depends on the OTU cell numbers per volume as well as the cell volume, with carbon content per cell calculated as follows:

$$\text{pg carbon cell}^{-1} = 0.76 \text{ cell volume}^{0.819} \text{ for dinoflagellates}$$

$$\text{pg carbon cell}^{-1} = 0.288 \text{ cell volume}^{0.811} \text{ for diatoms, and}$$

$$\text{pg carbon cell}^{-1} = 0.216 \text{ cell volume}^{0.939} \text{ for other protists} \qquad (7)$$

*with cell volume given in* $\mu m^3$.
(Menden-Deuer and Lessard 2000).

By opposing 18S read abundance and carbon contents of OTUs, the cell numbers per volume can be eliminated, and consequently GCN per cell can be compared with carbon content per cell. By combining the equations for GCN and carbon content per cell, and solving it for log(carbon cell$^{-1}$), one gets:

$$-0.61*0.819 + \log(0.76) + 1.22*0.819 * \log(GCN) = \log(\text{carbon cell}^{-1}) \text{ for dinoflagellates}$$

$$-0.61*0.811 + \log(0.288) + 1.22*0.811 * \log(GCN) = \log(\text{carbon cell}^{-1}) \text{ for diatoms}$$

$$-0.61*0.939 + \log(0.216) + 1.22*0.939 * \log(GCN) = \log(\text{carbon cell}^{-1}) \text{ for other protists} \quad (8)$$

Thus, almost linear relations between GCN and carbon contents exist (0.999 log (GCN) = log (carbon cell$^{-1}$) for dinoflagellates, (0.989 log (GCN) = log (carbon cell$^{-1}$) for diatoms, and 1.14 log (GCN) = log (carbon cell$^{-1}$) for diverse protists).
In other words, the relative share of read abundances of a specific phytoplankton OTU corresponds to its relative share of carbon concentration. We then applied a fixed conversion of chlorophyll:carbon of 1:40, and calculated the concentration of each phytoplankton OTU as mmol carbon l$^{-1}$ for each sampling point. As example, if 20% of the phototrophic 18S reads (cleaned and filtered, see above) would come from OTU A at 01.01.2012, and the chlorophyll concentration at this date is 5 µg l$^{-1}$, the carbon concentration of OTU A at 01.01.2012 would be 0.2*5 µg l$^{-1}$ * 40 = 40 µg carbon l$^{-1}$.

##### S4.4: 16S rRNA Illumina sequencing and factor for relative abundances

16S sequencing was performed with primers 515F–806R targeting the V4 region of the 16S SSU rRNA), and analyzed with the dada2 pipeline (Callahan et al 2016). The current primers have been modified from the original 515F–806R primer pair (Caporaso et al 2012) in the following ways: Barcodes are now on the forward primer 515F (Parada et al., 2016). This enables the usage of various reverse primer constructs to obtain longer amplicons, for example the V4–V5 region using reverse primer 926R (Quince et al., 2011; Parada et al., 2016). Degeneracy was added to both the forward and reverse primers to remove known biases against Crenarachaeota/Thaumarchaeota (515F, also called 515F-Y, Parada et al., 2016) and the marine and freshwater Alphaproteobacterial clade SAR11 (806R, Apprill et al., 2015).
In order to reduce the full 16S dataset into manageable amounts of data, the 200 most abundant ASVs were used for elaborated analyses. After filtering out chloroplast, unassigned, cyanobacterial and ammonia-oxidizing archaeal ASVs, 157 ASVs were used for elaborated analyses, which covered between 18 and 94% of the reads at different sampling dates (average 82%). For each sampling date, a factor for heterotrophic 16S ASVs was introduced, considering the share of non-heterotrophic (i.e. cyanobacteria) and non-bacterial cell ASVs (i.e. chloroplasts).

##### S4.5: Transformation of 16S data in carbon concentrations

Transformations of relative read abundances into carbon concentrations were based on flow

cytometer counts of heterotrophic prokaryotic cells, and individual cell volumes.
The corrected relative abundance of each heterotrophic 16S ASV was multiplied with the overall heterotrophic prokaryotic cell number (heterotrophic prokaryotic cell numbers derived from flow cytometry, Supplementary Table 7), yielding cell numbers per volume for each ASV. Next, the carbon content of each 16S ASV cell was calculated with the formula

$$fgC\ cell^{-1} = 133.754 \times V^{0.438} \qquad (9)$$

*(Romanova and Sazhin 2010)*

Cell volumes were looked up from the literature (Avci et al 2017, Buchan et al 2014, Cho and Giovannoni 2004, Cottrell and Kirchman 2016, Duhaime et al 2016, Ho et al 2017, Kirchman 2016, Kruger et al 2019, Lauro et al 2009, Mann et al 2013, Morris et al 2006, Pelve et al 2017, Seo et al 2009, Sheik 2012, Sheik et al 2014, Voget et al 2015, Yan et al 2009, Yao et al 2017). If cell volumes were not known, Gammaproteobacterial ASVs were set as 13 fmol carbon $cell^{-1}$, and all other ASVs with unknown cell volume as 5 fmol carbon $cell^{-1}$ (all given carbon contents and sources are given in Supplementary Table 10).

#### S5: Additional results and discussion

##### S5.1: Important phytoplankton producer OTUs and heterotrophic prokaryotic consumer ASVs

In order to identify phytoplankton OTUs and heterotrophic prokaryotic ASVs that are important for the carbon production and consumption at station L4, we calculated for each year and bloom type: the ten primary phytoplankton DOM producers; the ten primary heterotrophic prokaryotic DOM consumers along with their top phytoplankton donor and top DOM species used; and the ten phytoplankton>heterotrophic prokaryotes as well as heterotrophic prokaryote>heterotrophic prokaryote pairs with the highest carbon fluxes (Supplementary Table 3). Primary phytoplankton DOM producers mostly differed between years as well as bloom types, with few exceptions: In phytoplankton spring blooms *Bathycoccus prasinos* (BPR), in bacteria summer blooms *Gonyaulax spinifera* (GSP), and in phytoplankton summer blooms *Pterocystis* sp. (PTE) were found in the top 10 of DOM producers, 71%, 57% and 83% of the years, respectively (Supplementary Table 3). On the consumer side, phytoplankton spring blooms revealed frequent occurrences of SAR86 (S86) and *Amylibacter* (AMY), as primary heterotrophic prokaryotes (top 10 rank in 100% and 86% of the years), whereas SAR11 (S11) showed top 10 occurrences in phytoplankton and bacteria summer blooms in 86% and 57% of the years, respectively (Supplementary Table 3). The highest obtained carbon flux for phytoplankton>heterotrophic prokaryote pairs was in the 2018 bacteria summer bloom with 0.082 µmol C $l^{-1}$ $d^{-1}$ for *Dinophysis acuminata*>Marine group II archaea (DAC>M22). In contrast to the paradigm of carbon flux being dominated by flow from phototrophs to heterotrophs, fluxes between heterotrophic prokaryotic pairs were in the same range as fluxes for phytoplankton heterotrophic prokaryote pairs, with a maximum of 0.035 C $l^{-1}$ $d^{-1}$ for

*Amylibacter*>SAR86 (AMY>S86) in the bacteria summer bloom of 2016 (Supplementary Table 3).

**S5.2: Taxonomy of important heterotrophic prokaryotes DOM consumers:**

In all bloom events, FLUXNET predicted that most primary DOC consumers belong to Gammaproteobacteria, the Rhodobacterales clade of Alphaproteobacteria or the Flavobacteriales clade of Bacteroidetes (Supplementary Table 3). This outcome is supported by numerous studies, that defined these three bacterial groups as principle utilizers of DOM (Becker et al 2019, Buchan et al 2014, Eigemann et al 2022, Eigemann et al 2023, Kieft et al 2021, Pinhassi et al 2004, Riemann et al 2000, Sarmento and Gasol 2012, Teeling et al 2012, Teeling et al 2016). However, in bacterial summer blooms, we found especially many Flavobacteriales such as Aquibacter (AQU), Cryomorphaceae (CR1), Flavicella (FAC)), Tenacibaculum (TE2), NS5 marine group (N51), and Fluviicola (FU2) as major carbon consumers (Supplementary Table 3). The (predicted) pivotal role of Flavobacteriales in periods with high heterotrophic prokaryotes>heterotrophic prokaryotes fluxes is supported by their genetic capacity for using manifold different sources of high-molecular-weight (HMW) DOM (Lombard et al 2014, Teeling et al 2012), as well as experiments: Flavobacteriales were the principle degraders of bacterial derived polysaccharides (Zhang et al 2015), and also dominated bacterial communities enriched with exudates of the copiotrophic Gammaproteobacterium Photobacterium angustum (Ortega-Retuerta et al 2021). A recent preview additionally showed that 75% from 53 sequenced Bacteroidetes/Flavobacteriia strains had specific polysaccharide utilization loci (PULs) for α-glucans (which is the bacterial storage polysaccharide), and that the Bacteroidetes strain *Polaribacter* was able to grew solely on bacterial lysates (Beidler et al 2023).

**S5.3: Lower recurrences of producers compared to consumers:**

In order to quantify the recurrences of producers, consumers and phytoplankton>heterotrophic prokaryotes and heterotrophic prokaryote>heterotrophic prokaryote interactions (fluxes), we calculated for each bloom type Bray-Curtis similarities of phytoplankton producer, heterotrophic prokaryotic consumers, phytoplankton>heterotrophic prokaryote and heterotrophic prokaryote>heterotrophic prokaryote pairs between all years. The highest recurrences were obtained for heterotrophic prokaryotic consumers with Bray-Curtis similarities of ~0.32 in spring blooms, whereas phytoplankton DOM producer revealed similarities between ~0.1 (phytoplankton summer blooms) and ~0.17 (phytoplankton spring and bacteria summer blooms). This decoupling of producers and consumers (same consumers supplied from different producers) is presumably driven by the production of similar DOM species by different phytoplankton species (Supplementary Table 3), which was also found at Helgoland Island (Mayerhofer et al 2021, Teeling et al 2016), and is further supported by low recurrences of phytoplankton>heterotrophic prokaryote and heterotrophic prokaryote>heterotrophic prokaryote pairs (SI Figure 1).

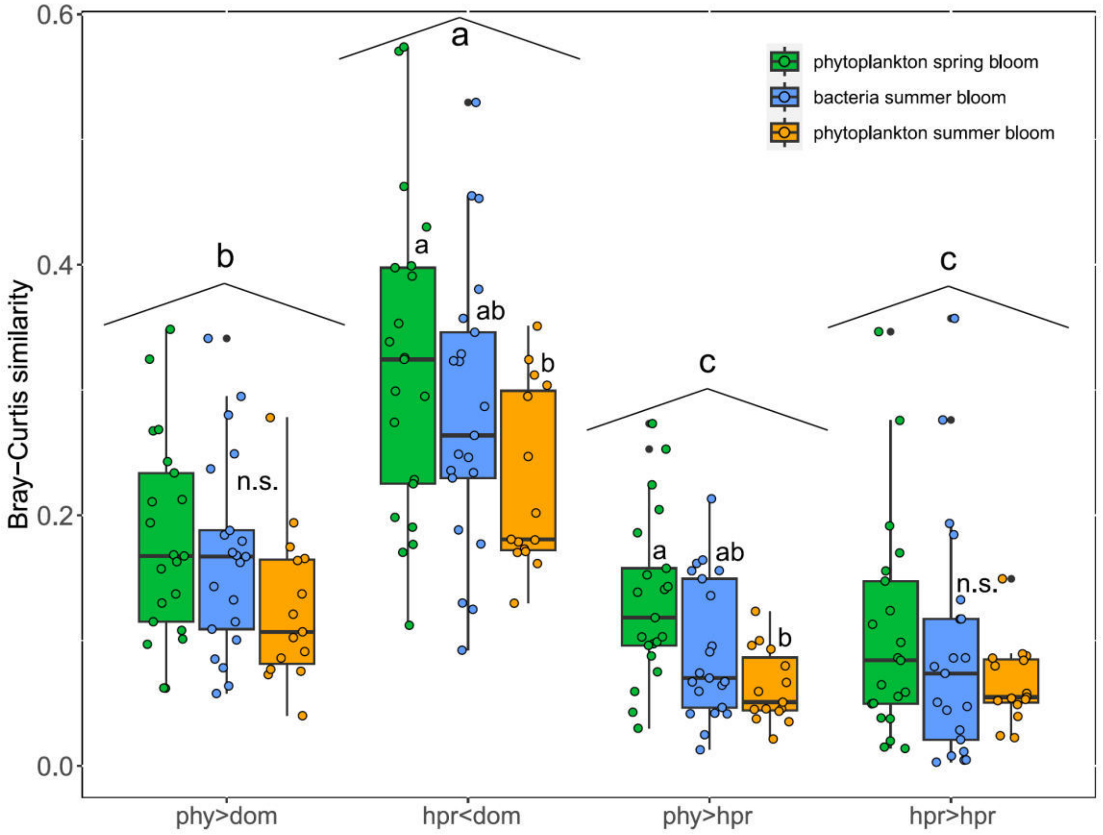


SI Figure 1: Recurrence of phytoplankton DOM producers (phy>dom), heterotrophic prokaryotic DOM consumers (hpr<dom), phytoplankton>heterotrophic prokaryote pairs (phy>hpr), and heterotrophic prokaryote>heterotrophic prokaryote pairs (hpr>hpr) in the years 2012-2018. Letters on top of the box plots refer to outcomes of Tukey post-hoc tests between the different bloom types, letters on top of the roofs to differences between variables (combined bloom types, Supplementary Table 4). n.s. = not significant.

##### S5.4: Succession patterns of abundant phytoplankton OTUs and heterotrophic prokaryotic ASVs differ between years and bloom types:

To get a deeper insight into the dynamics of abundant phytoplankton OTUs and heterotrophic prokaryotic ASVs, we chose the OTUs/ASVs with the highest concentrations in each of the three different bloom types, and plotted their concentrations throughout bloom events for the example year 2013 (SI Fig. 2), and all other years (Supplementary Fig. 2). 2013's phytoplankton spring bloom revealed a succession of phytoplankton OTUs to consecutive heterotrophic prokaryotic ASVs (SI Fig. 2). Accordingly, carbon concentrations of the phytoplankton OTUs *Bathycoccus prasinos* (bpr) and *Micromonas pusilla* (mpu) exceeded that of SAR86 (s86) and *Planktomarina* (pla) at the bloom start, but decreased below that of the heterotrophic prokaryotes at later phases. In contrast, the bacterial summer bloom in 2013 showed distinct successions of the most abundant heterotrophic prokaryotic ASVs without preceding abundant phytoplankton OTUs (SI Fig. 2). The increase of the most abundant phytoplankton OTU

*Gonyaulax spinifera* (gsp) towards the end of the bacterial bloom ultimately led to the onset of the phytoplankton summer bloom with consecutive phytoplankton OTUs *Gonyaulax spinifera* (gsp) to Dinophyceae (din)), in which, however, no successions of the three most abundant heterotrophic prokaryotic ASVs were obvious (SI Fig. 2). Successions from phytoplankton to consecutive heterotrophic prokaryotic ASVs in phytoplankton spring blooms also occurred in other years (Supplementary Fig. 2), and display a well-known process for marine phytoplankton blooms (Luria et al 2017, Sison-Mangus et al 2016), representing an initial DOM liberation by the phytoplankton and a subsequent exploitation of highly towards less bioavailable phytoplankton DOM by different heterotrophic prokaryotes (Buchan et al 2014, Eigemann et al 2023, Teeling et al 2012, Teeling et al 2016). Phytoplankton summer blooms also revealed consistent patterns with consecutive abundant phytoplankton OTUs in all years, but without subsequent successions of the abundant heterotrophic prokaryotic ASVs (SI Fig. 2 and Supplementary Fig. 2). However, the latter may have been masked by high concentrations of heterotrophic prokaryotic ASVs remaining from forgoing bacteria summer blooms (SI Fig. 2 and Supplementary Fig. 2). In contrast, bacterial summer blooms did not show consistent patterns, with some years revealing successions of abundant heterotrophic prokaryotes (2013, 2014, 2018), whereas others did not (2015, 2017, SI Fig. 2 and Supplementary Fig. 2). If successions occurred, succession periods in the range of days (i.e. DOM liberation by one ASV followed by the uptake/growth of another ASV, SI Fig. 2 and Supplementary Fig. 2) indicate the occurrence of fast and specific loss processes, such as protist grazing or infections with phages, where the latter may cause up to 61% of bacterial mortality after 24 h (Fouilland et al 2014). Interestingly, bacterial summer blooms in years displaying successions of abundant heterotrophic prokaryotes showed higher fluxes between heterotrophic prokaryotes than those in years without successions (Supplementary Table 5), highlighting the importance of specific loss processes (and the pivotal role of phages, see main paper) as drivers of fluxes between heterotrophic prokaryotes.

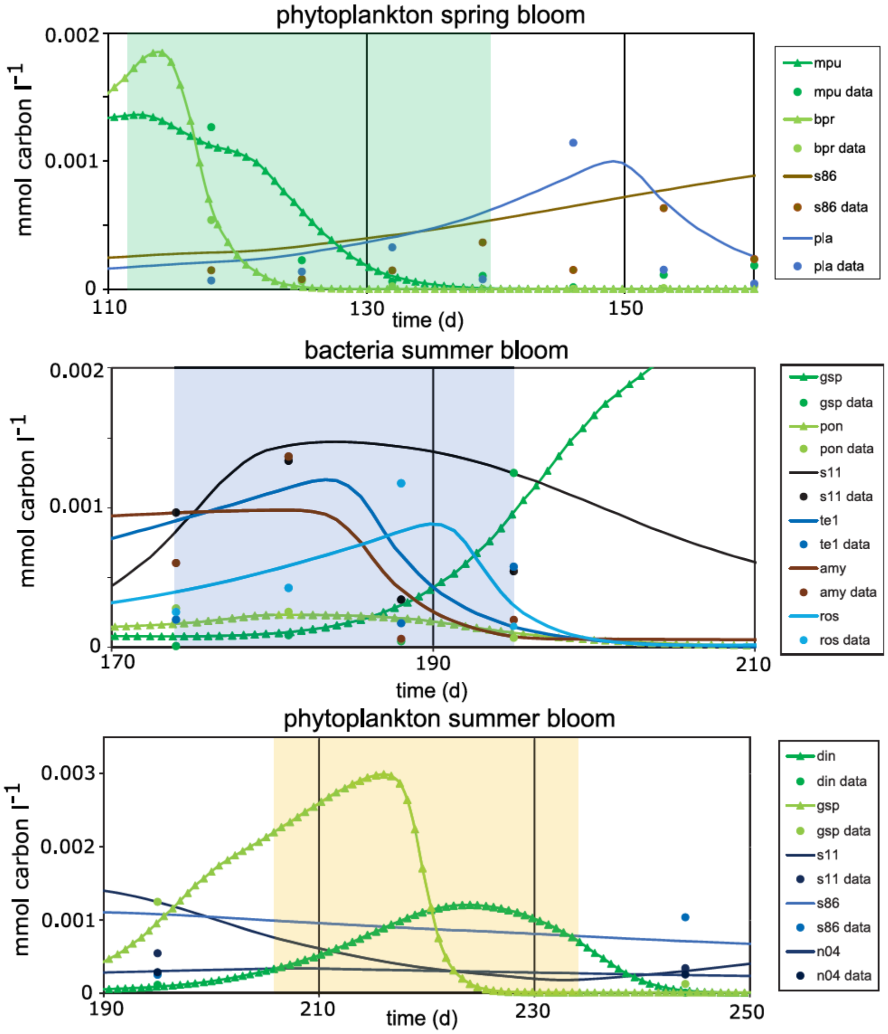


SI Fig. 2: Temporal development of phytoplankton OTUs and heterotrophic prokaryotic ASVs with the highest concentrations in the three different bloom types in the example year 2013. Model outputs are indicated by lines (phytoplankton: lines including triangles, heterotrophic prokaryotes: smooth lines), data by dots. The day of the year 2013 is assigned to the x-axis, and the concentrations on the y-axis. The shaded areas correspond to the periods that were defined as bloom periods. mpu = Micromonas pusilla, bpr = Bathycoccus prasinos, s86 = SAR86, pla = Planktomarina, din = Dinophyceae, gsp = Gonyaulax spinifera, s11 = SAR11, n04 = NS4 marine group, pon = Prorocentrum donghaiense, te1 = Tenacibaculum, amy = Amylibacter, ros = Roseobacter.

#### S6: References

Avci B, Hahnke RL, Chafee M, Fischer T, Gruber-Vodicka H, Tegetmeyer HE *et al* (2017). Genomic and physiological analyses of 'Reinekea forsetii' reveal a versatile opportunistic lifestyle during spring algae blooms. *Environ Microbiol* **19:** 1209-1221.

Becker JW, Hogle SL, Rosendo K, Chisholm SW (2019). Co-culture and biogeography of Prochlorococcus and SAR11. *The ISME Journal* **13:** 1506-1519.

Beidler I, Steinke N, Schulze T, Sidhu C, Bartosik D, Krull J *et al* (2023). Alpha-glucans from bacterial necromass indicate an intra-population loop within the marine carbon cycle. *PREPRINT Research Square* **Version 1**.

Buchan A, LeCleir GR, Gulvik CA, Gonzalez JM (2014). Master recyclers: features and functions of bacteria associated with phytoplankton blooms. *Nat Rev Microbiol* **12:** 686-698.

Callahan BJ, McMurdie PJ, Rosen MJ, Han AW, Johnson AJA, Holmes SP (2016). DADA2: High-resolution sample inference from Illumina amplicon data. *Nat Methods* **13:** 581-583.

Caporaso JG, Lauber CL, Walters WA, Berg-Lyons D, Huntley J, Fierer N *et al* (2012). Ultra-high-throughput microbial community analysis on the Illumina HiSeq and MiSeq platforms. *ISME J* **6:** 1621-1624.

Chapra SC (1997). *Surface Water-Quality Modeling*. McGraw-Hill: Boston.

Cho JC, Giovannoni SJ (2004). Cultivation and growth characteristics of a diverse group of oligotrophic marine Gammaproteobacteria. *Appl Environ Microbiol* **70:** 432-440.

Cottrell MT, Kirchman DL (2016). Transcriptional Control in Marine Copiotrophic and Oligotrophic Bacteria with Streamlined Genomes. *Appl Environ Microbiol* **82:** 6010-6018.

Daines SJ, Clark JR, Lenton TM (2014). Multiple environmental controls on phytoplankton growth strategies determine adaptive responses of the N : P ratio. *Ecology Letters* **17:** 414-425.

Duhaime MB, Wichels A, Sullivan MB (2016). Six Pseudoalteromonas Strains Isolated from Surface Waters of Kabeltonne, Offshore Helgoland, North Sea. *Genome Announc* **4**.

Eigemann F, Rahav E, Grossart H-P, Aharonovich D, Sher D, Vogts A *et al* (2022). Phytoplankton exudates provide full nutrition to a subset of accompanying heterotrophic bacteria via carbon, nitrogen and phosphorus allocation. *Environmental Microbiology* **24:** 2467-2483.

Eigemann F, Rahav E, Grossart H-P, Aharonovich D, Voss M, Sher D (2023). Phytoplankton Producer Species and Transformation of Released Compounds over Time Define Bacterial Communities following Phytoplankton Dissolved Organic Matter Pulses. *Applied and Environmental Microbiology* **89:** e00539-00523.

Fouilland E, Tolosa I, Bonnet D, Bouvier C, Bouvier T, Bouvy M *et al* (2014). Bacterial carbon dependence on freshly produced phytoplankton exudates under different nutrient availability and grazing pressure conditions in coastal marine waters. *FEMS Microbiol Ecol* **87:** 757-769.

Godhe A, Asplund ME, Härnström K, Saravanan V, Tyagi A, Karunasagar I (2008). Quantification of diatom and dinoflagellate biomasses in coastal marine seawater samples by real-time PCR. *Appl Environ Microbiol* **74:** 7174-7182.

Hellweger FL, Kravchuk ES, Novotny V, Gladyshev MI (2008). Agent-Based Modeling of the Complex Life Cycle of a Cyanobacterium (Anabaena) in a Shallow Reservoir. *Limnology and Oceanography* **53:** 1227-1241.

Ho A, Di Lonardo DP, Bodelier PL (2017). Revisiting life strategy concepts in environmental microbial ecology. *FEMS Microbiol Ecol* **93**.

Kieft B, Li Z, Bryson S, Hettich RL, Pan C, Mayali X *et al* (2021). Phytoplankton exudates and lysates support distinct microbial consortia with specialized metabolic and ecophysiological traits. *Proceedings of the National Academy of Sciences* **118:** e2101178118.

Kirchman DL (2016). Growth Rates of Microbes in the Oceans. *Ann Rev Mar Sci* **8:** 285-309.

Kruger K, Chafee M, Ben Francis T, Glavina Del Rio T, Becher D, Schweder T *et al* (2019). In marine Bacteroidetes the bulk of glycan degradation during algae blooms is mediated by few clades using a restricted set of genes. *ISME J* **13:** 2800-2816.

Lauro FM, McDougald D, Thomas T, Williams TJ, Egan S, Rice S *et al* (2009). The genomic basis of trophic strategy in marine bacteria. *Proc Natl Acad Sci U S A* **106:** 15527-15533.

Lombard V, Golaconda Ramulu H, Drula E, Coutinho PM, Henrissat B (2014). The carbohydrate-active enzymes database (CAZy) in 2013. *Nucleic Acids Res* **42:** D490-495.

Luria CM, Amaral-Zettler LA, Ducklow HW, Repeta DJ, Rhyne AL, Rich JJ (2017). Seasonal Shifts in Bacterial Community Responses to Phytoplankton-Derived Dissolved Organic Matter in the Western Antarctic Peninsula. *Frontiers in Microbiology* **8**.

Mann AJ, Hahnke RL, Huang S, Werner J, Xing P, Barbeyron T *et al* (2013). The genome of the alga-associated marine flavobacterium Formosa agariphila KMM 3901T reveals a broad potential for degradation of algal polysaccharides. *Appl Environ Microbiol* **79:** 6813-6822.

Mayerhofer MM, Eigemann F, Lackner C, Hoffmann J, Hellweger FL (2021). Dynamic carbon flux network of a diverse marine microbial community. *ISME Communications* **1:** 50.

Menden-Deuer S, Lessard EJ (2000). Carbon to volume relationships for dinoflagellates, diatoms, and other protist plankton. *Limnology and Oceanography* **45:** 569-579.

Mieleitner J, Reichert P (2008). Modelling functional groups of phytoplankton in three lakes of different trophic state. *Ecological Modelling* **211:** 279-291.

Morris RM, Longnecker K, Giovannoni SJ (2006). Pirellula and OM43 are among the dominant lineages identified in an Oregon coast diatom bloom. *Environ Microbiol* **8:** 1361-1370.

Ortega-Retuerta E, Devresse Q, Caparros J, Marie B, Crispi O, Catala P *et al* (2021). Dissolved organic matter released by two marine heterotrophic bacterial strains and its bioavailability for natural prokaryotic communities. *Environmental Microbiology* **23:** 1363-1378.

Pelve EA, Fontanez KM, DeLong EF (2017). Bacterial Succession on Sinking Particles in the Ocean's Interior. *Front Microbiol* **8:** 2269.

Pinhassi J, Sala MM, Havskum H, Peters F, Guadayol O, Malits A *et al* (2004). Changes in bacterioplankton composition under different phytoplankton regimens. *Appl Environ Microbiol* **70:** 6753-6766.

Pinto F, Medina DA, Pérez-Correa JR, Garrido D (2017). Modeling Metabolic Interactions in a Consortium of the Infant Gut Microbiome. *Frontiers in Microbiology* **8**.

Riemann L, Steward GF, Azam F (2000). Dynamics of bacterial community composition and activity during a mesocosm diatom bloom. *Appl Environ Microbiol* **66:** 578-587.

Romanova ND, Sazhin AF (2010). Relationships between the cell volume and the carbon content of bacteria. *Oceanology* **50:** 522-530.

Sarmento H, Gasol JM (2012). Use of phytoplankton-derived dissolved organic carbon by different types of bacterioplankton. *Environ Microbiol* **14:** 2348-2360.

Seo HS, Kwon KK, Yang SH, Lee HS, Bae SS, Lee JH *et al* (2009). Marinoscillum gen. nov., a member of the family 'Flexibacteraceae', with Marinoscillum pacificum sp. nov. from a marine sponge and Marinoscillum furvescens nom. rev., comb. nov. *Int J Syst Evol Microbiol* **59:** 1204-1208.

Sheik AR (2012). Viral regulation of nutrient assimilation by algae and prokaryotes. PhD thesis, University of Bremen, Bremen.

Sheik CS, Jain S, Dick GJ (2014). Metabolic flexibility of enigmatic SAR324 revealed through metagenomics and metatranscriptomics. *Environ Microbiol* **16:** 304-317.

Sison-Mangus MP, Jiang S, Kudela RM, Mehic S (2016). Phytoplankton-Associated Bacterial Community Composition and Succession during Toxic Diatom Bloom and Non-Bloom Events. *Front Microbiol* **7:** 1433.

Teeling H, Fuchs BM, Becher D, Klockow C, Gardebrecht A, Bennke CM *et al* (2012). Substrate-controlled succession of marine bacterioplankton populations induced by a phytoplankton bloom. *Science* **336:** 608-611.

Teeling H, Fuchs BM, Bennke CM, Kruger K, Chafee M, Kappelmann L *et al* (2016). Recurring patterns in bacterioplankton dynamics during coastal spring algae blooms. *Elife* **5:** e11888.

Voget S, Wemheuer B, Brinkhoff T, Vollmers J, Dietrich S, Giebel HA *et al* (2015). Adaptation of an abundant Roseobacter RCA organism to pelagic systems revealed by genomic and transcriptomic analyses. *ISME J* **9:** 371-384.

Weitz JS, Stock CA, Wilhelm SW, Bourouiba L, Coleman ML, Buchan A *et al* (2015). A multitrophic model to quantify the effects of marine viruses on microbial food webs and ecosystem processes. *The Isme Journal* **9:** 1352.

Yan S, Fuchs BM, Lenk S, Harder J, Wulf J, Jiao NZ *et al* (2009). Biogeography and phylogeny of the NOR5/OM60 clade of Gammaproteobacteria. *Syst Appl Microbiol* **32:** 124-139.

Yao F, Yang S, Wang Z, Wang X, Ye J, Wang X *et al* (2017). Microbial Taxa Distribution Is Associated with Ecological Trophic Cascades along an Elevation Gradient. *Front Microbiol* **8:** 2071.

Zhang Z, Chen Y, Wang R, Cai R, Fu Y, Jiao N (2015). The Fate of Marine Bacterial Exopolysaccharide in Natural Marine Microbial Communities. *PLoS One* **10:** e0142690.

Supplementary Figure 1: Tamar river run-off

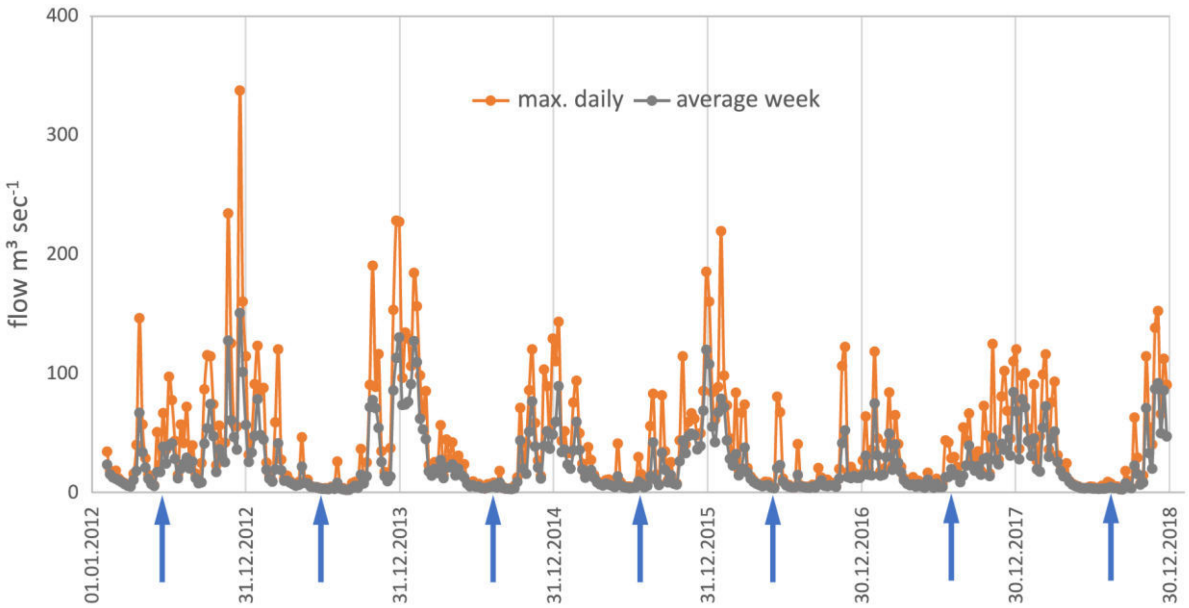


Maximum daily (orange) and weekly average (grey) Tamar river run-off. The starting dates for bacterial summer blooms are indicated by blue arrows.

### Supplementary Fig. 2

#### succession of phytoplankton and heterotrophic prokaryotes

### 2012

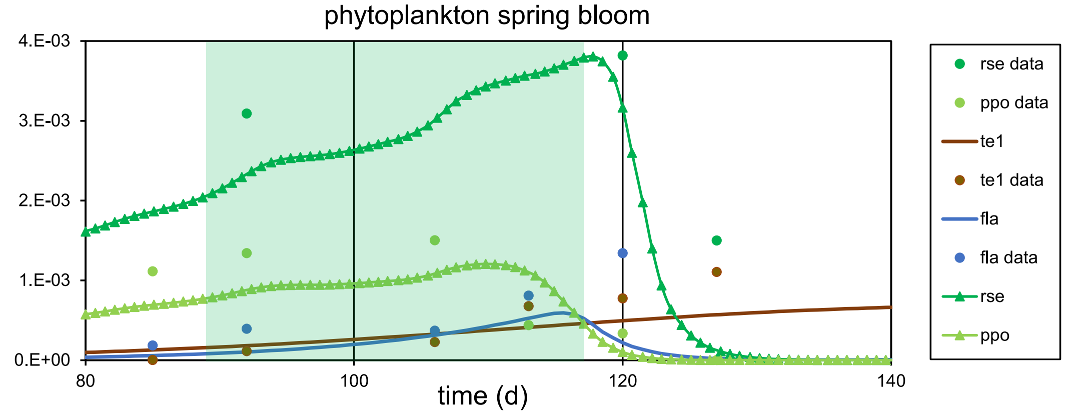


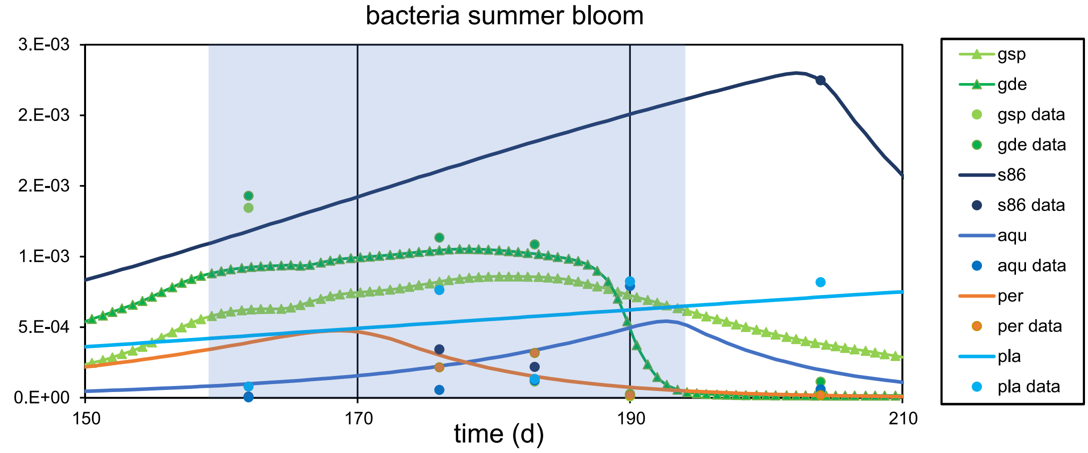


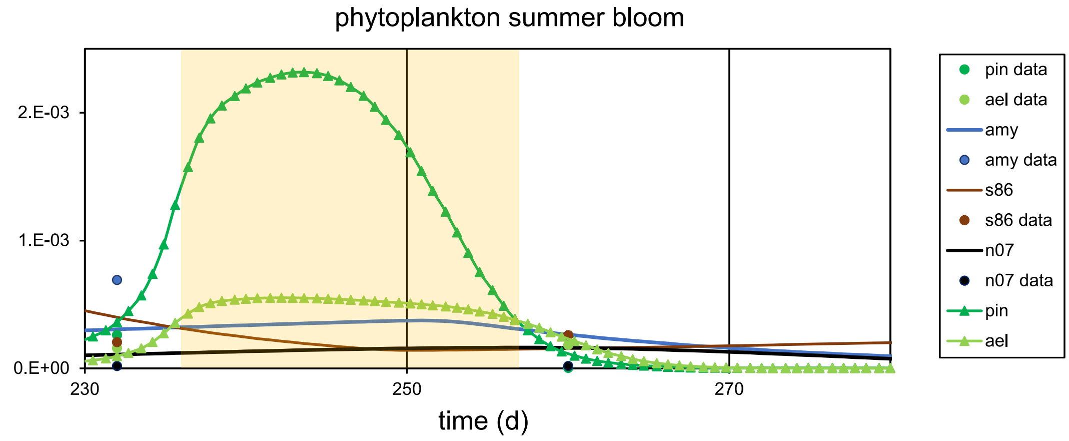

# 2014

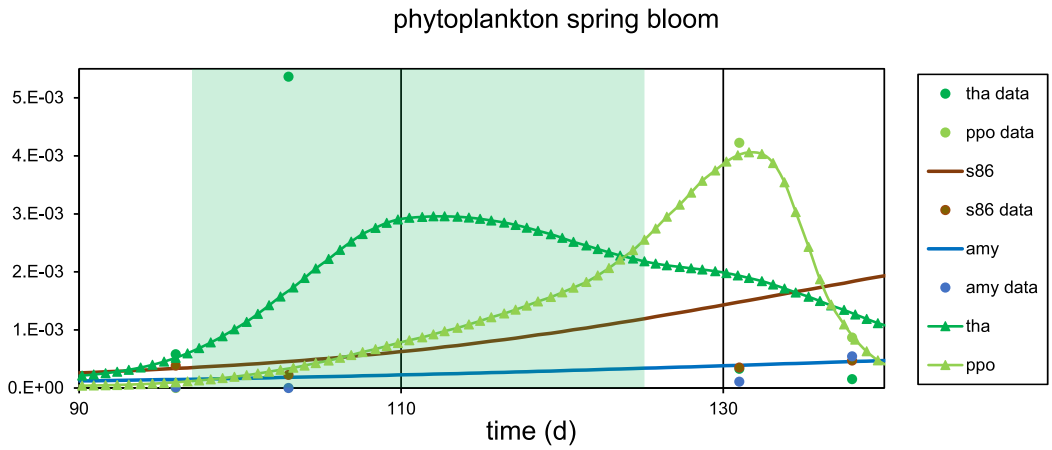
phytoplankton spring bloom
5.E-03
4.E-03
3.E-03
2.E-03
1.E-03
0.E+00
90
110
130
time (d)
tha data
ppo data
s86
s86 data
amy
amy data
tha
ppo

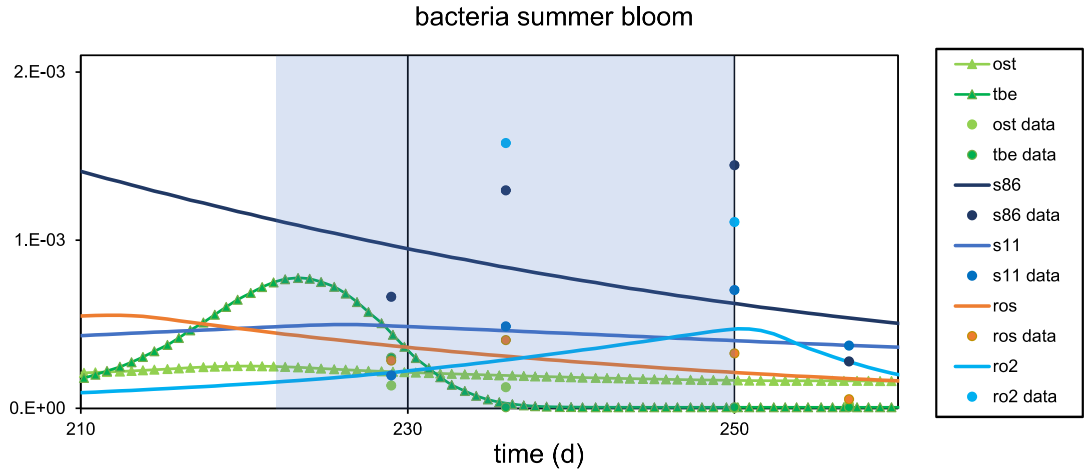
bacteria summer bloom
2.E-03
1.E-03
0.E+00
210
230
250
time (d)
ost
tbe
ost data
tbe data
s86
s86 data
s11
s11 data
ros
ros data
ro2
ro2 data

# 2015

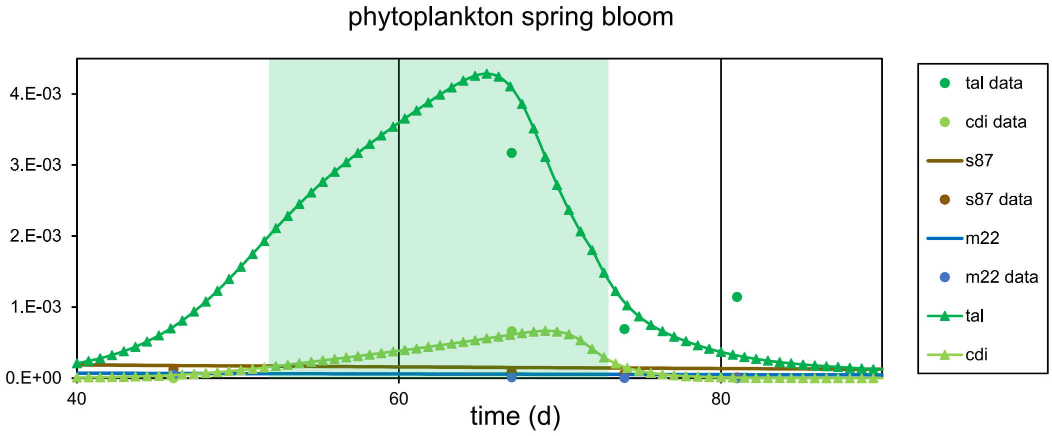

phytoplankton spring bloom
4.E-03
3.E-03
2.E-03
1.E-03
0.E+00
40
60
80
time (d)
tal data
cdi data
s87
s87 data
m22
m22 data
tal
cdi


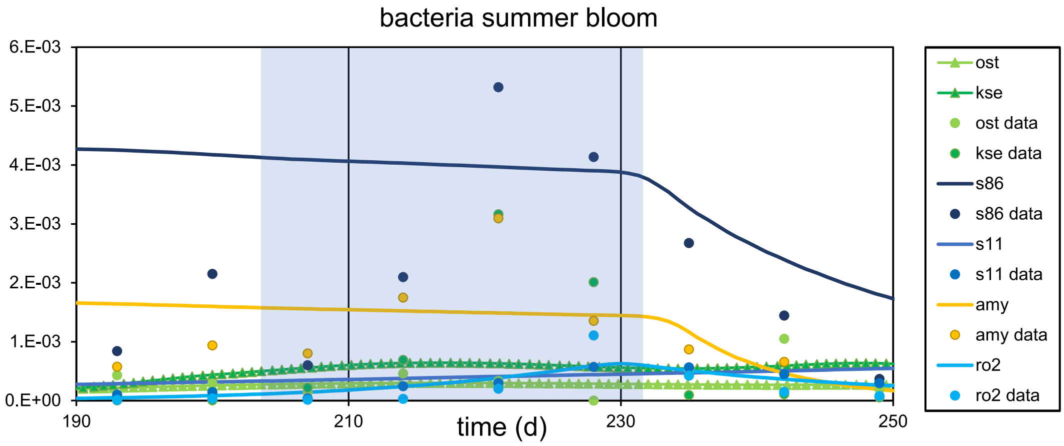

bacteria summer bloom
6.E-03
5.E-03
4.E-03
3.E-03
2.E-03
1.E-03
0.E+00
190
210
230
250
time (d)
ost
kse
ost data
kse data
s86
s86 data
s11
s11 data
amy
amy data
ro2
ro2 data


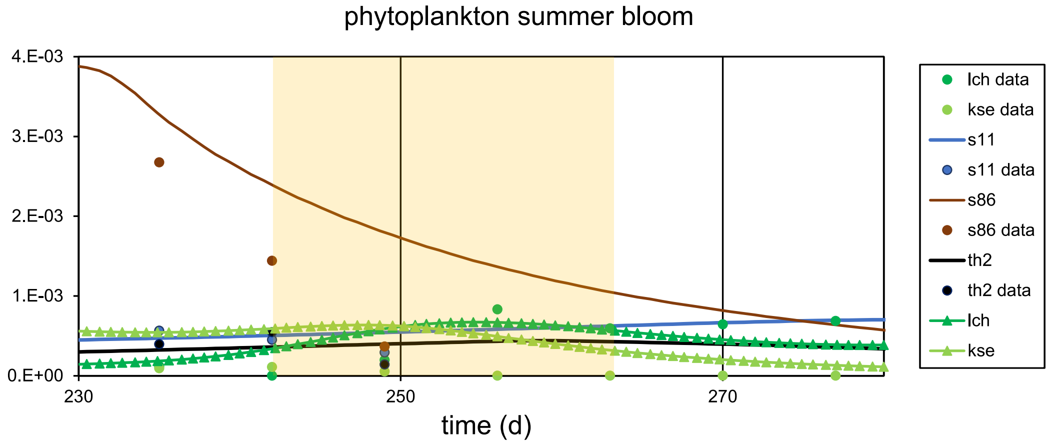

phytoplankton summer bloom
4.E-03
3.E-03
2.E-03
1.E-03
0.E+00
230
250
270
time (d)
lch data
kse data
s11
s11 data
s86
s86 data
th2
th2 data
lch
kse

# 2016

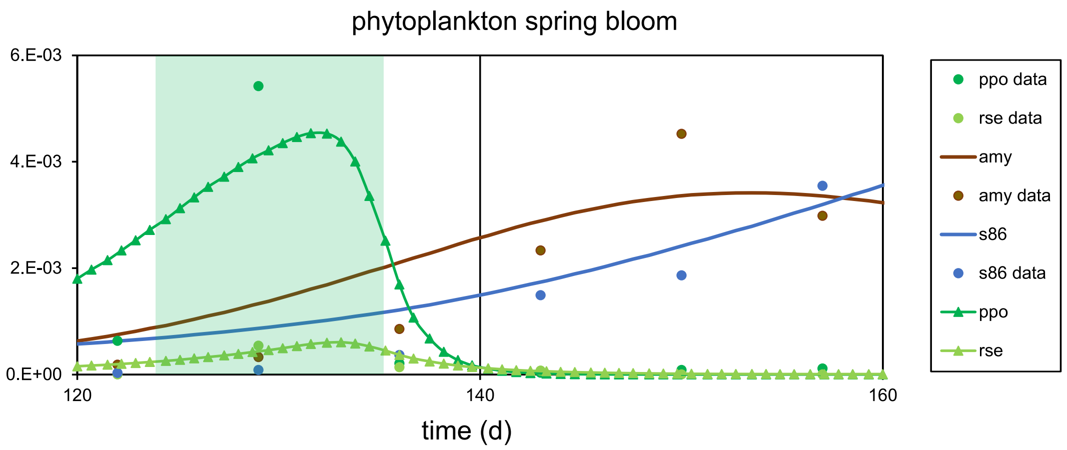
phytoplankton spring bloom
6.E-03
4.E-03
2.E-03
0.E+00
120
140
160
time (d)
ppo data
rse data
amy
amy data
s86
s86 data
ppo
rse

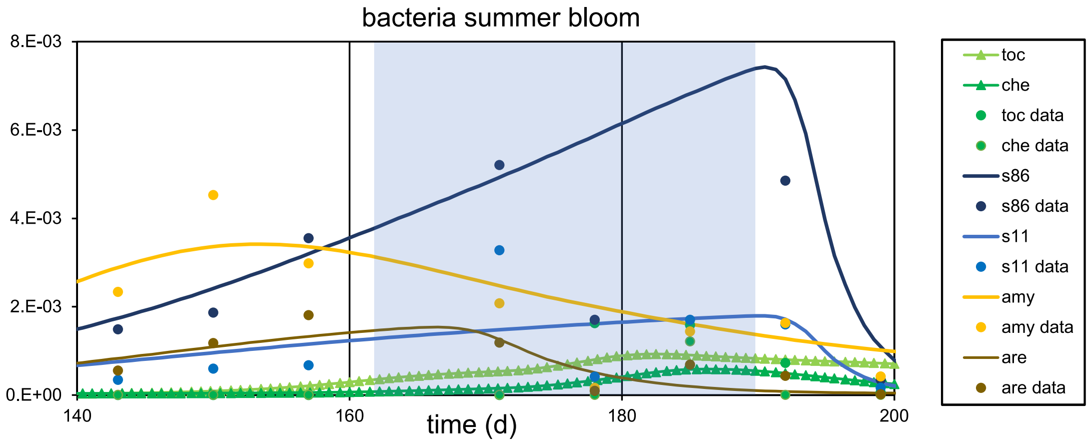
bacteria summer bloom
8.E-03
6.E-03
4.E-03
2.E-03
0.E+00
140
160
180
200
time (d)
toc
che
toc data
che data
s86
s86 data
s11
s11 data
amy
amy data
are
are data

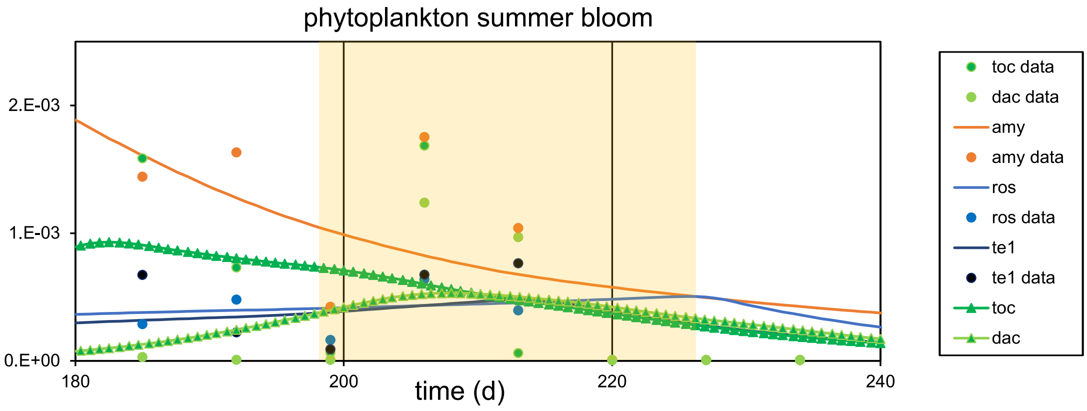
phytoplankton summer bloom
2.E-03
1.E-03
0.E+00
180
200
220
240
time (d)
toc data
dac data
amy
amy data
ros
ros data
te1
te1 data
toc
dac

# 2017

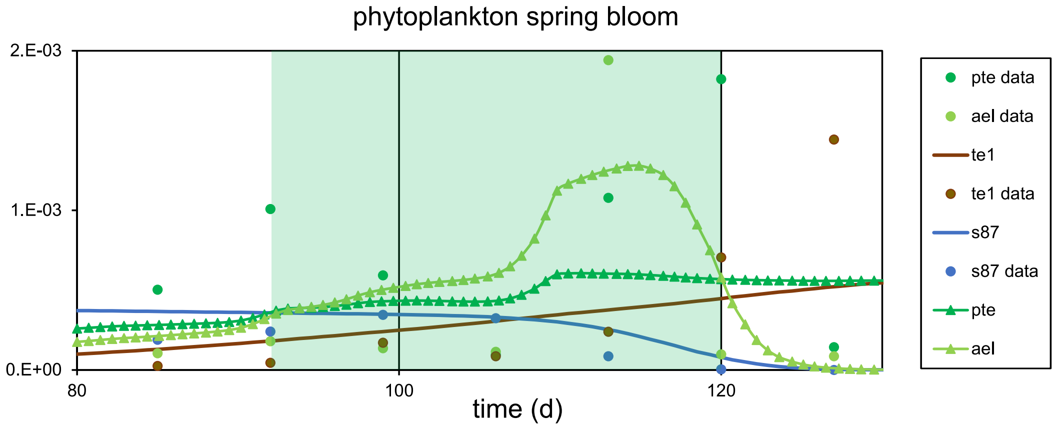
phytoplankton spring bloom
2.E-03
1.E-03
0.E+00
80
100
120
time (d)
pte data
ael data
te1
te1 data
s87
s87 data
pte
ael

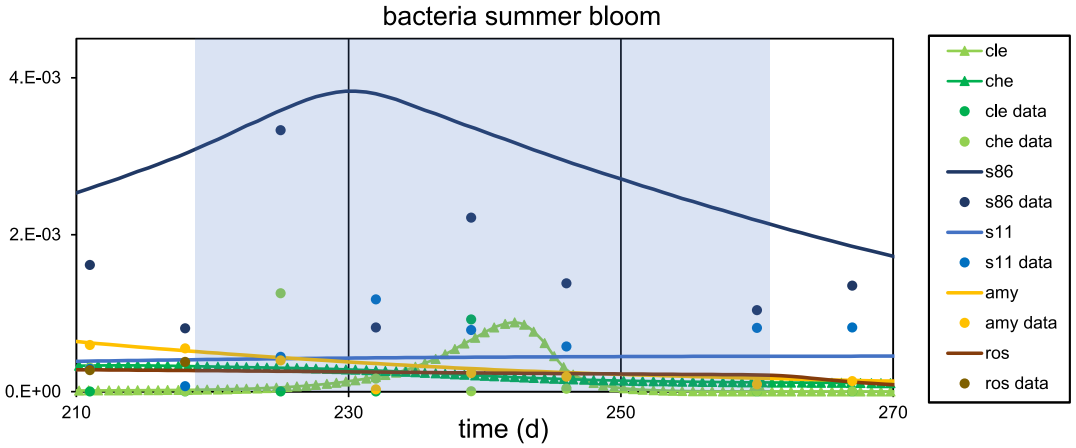
bacteria summer bloom
4.E-03
2.E-03
0.E+00
210
230
250
270
time (d)
cle
che
cle data
che data
s86
s86 data
s11
s11 data
amy
amy data
ros
ros data

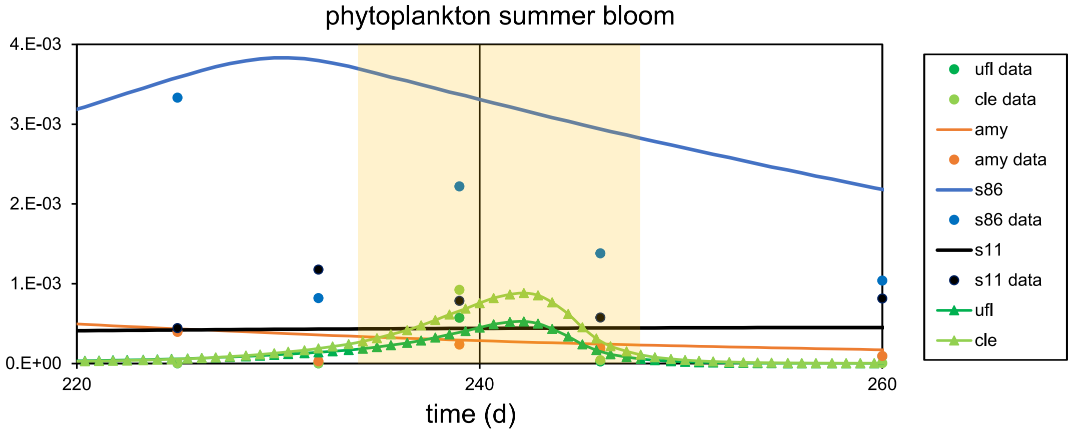
phytoplankton summer bloom
4.E-03
3.E-03
2.E-03
1.E-03
0.E+00
220
240
260
time (d)
ufl data
cle data
amy
amy data
s86
s86 data
s11
s11 data
ufl
cle

# 2018

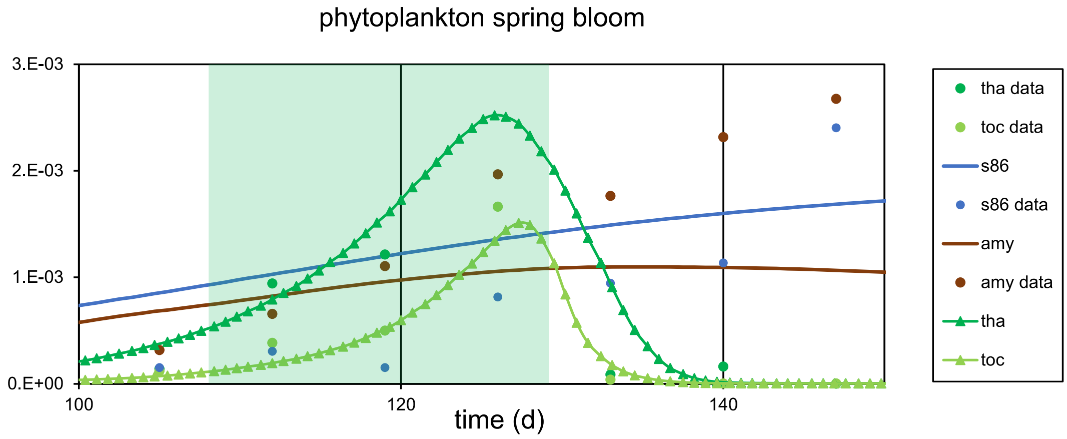

phytoplankton spring bloom
3.E-03
2.E-03
1.E-03
0.E+00
100
120
140
time (d)
tha data
toc data
s86
s86 data
amy
amy data
tha
toc


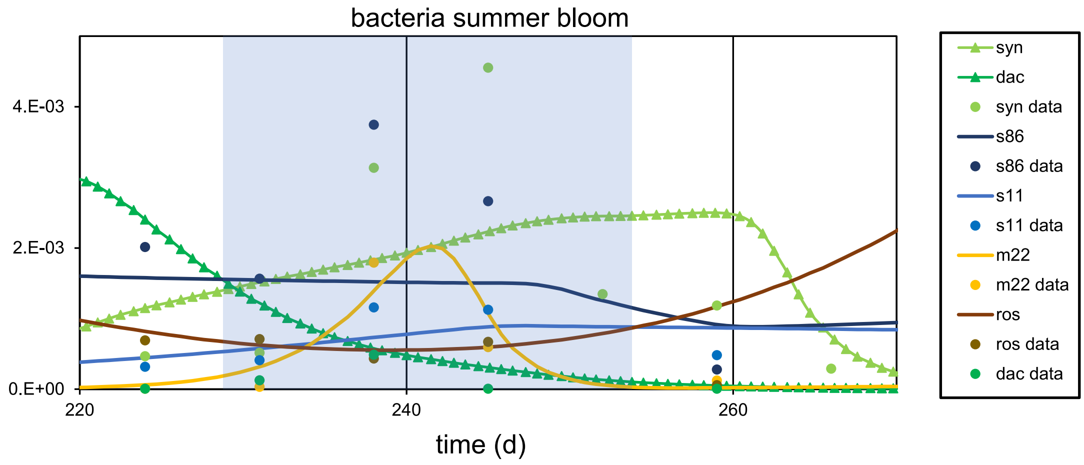

bacteria summer bloom
4.E-03
2.E-03
0.E+00
220
240
260
time (d)
syn
dac
syn data
s86
s86 data
s11
s11 data
m22
m22 data
ros
ros data
dac data


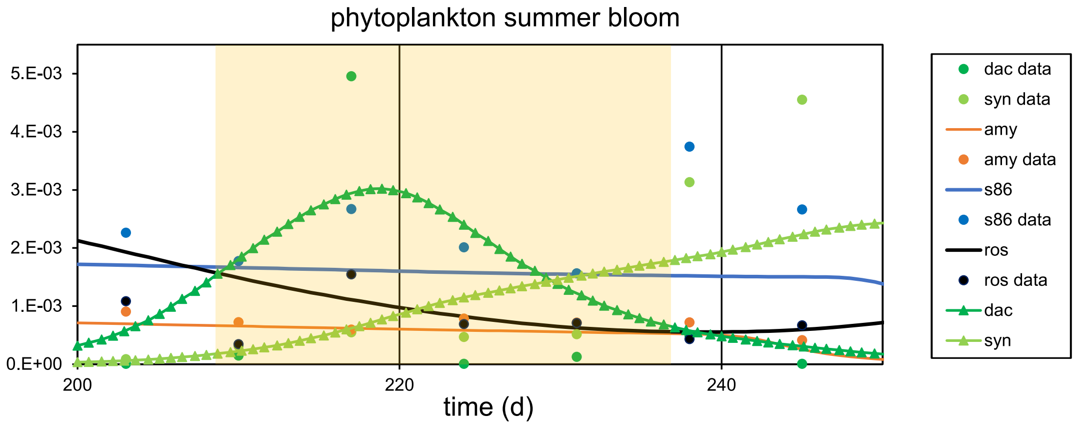

phytoplankton summer bloom
5.E-03
4.E-03
3.E-03
2.E-03
1.E-03
0.E+00
200
220
240
time (d)
dac data
syn data
amy
amy data
s86
s86 data
ros
ros data
dac
syn

### Supplementary Fig. 3: Inhibition analyses

2012

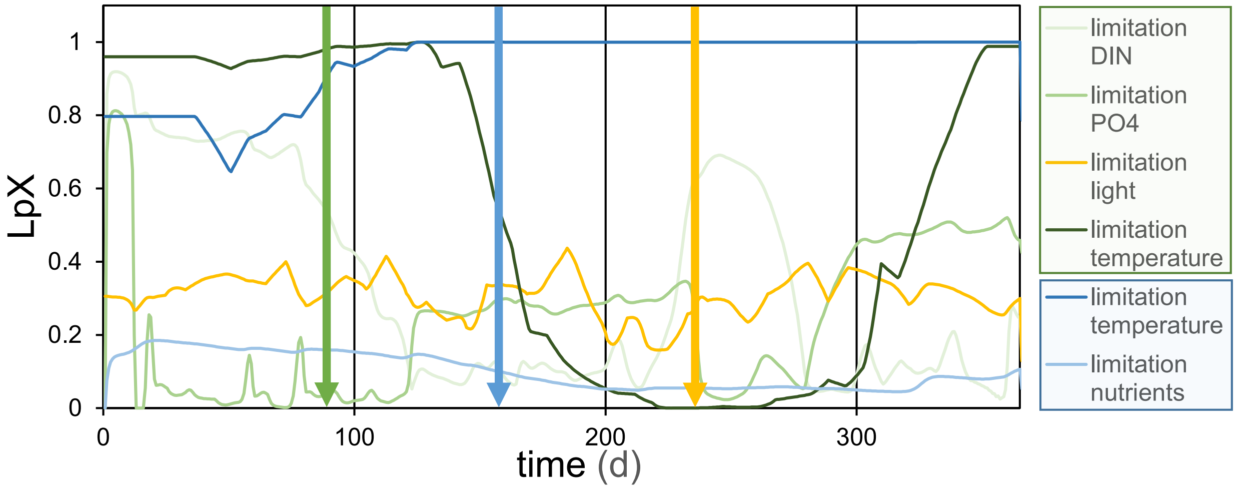


2013

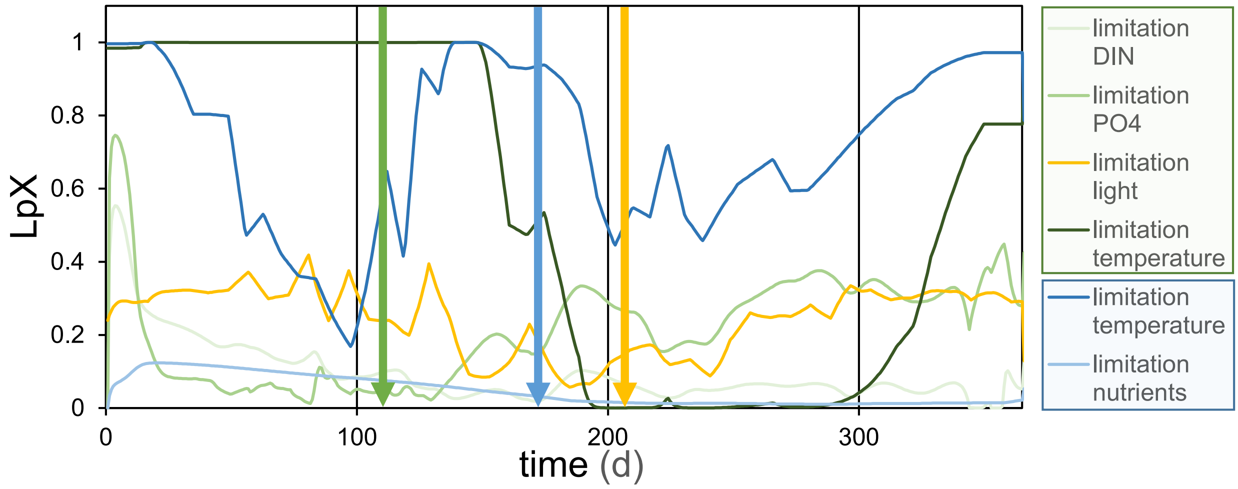


2014

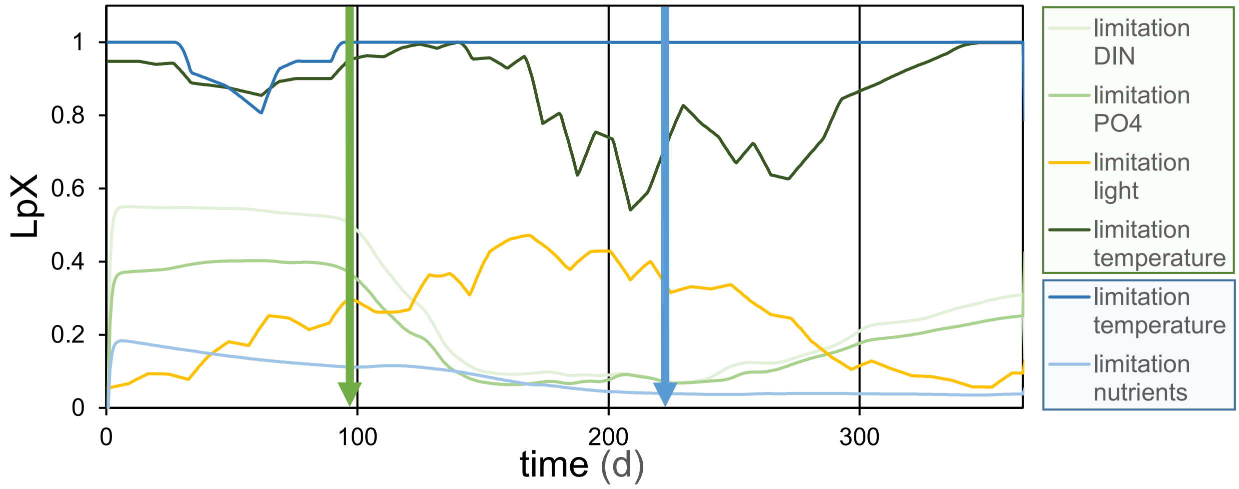


2015

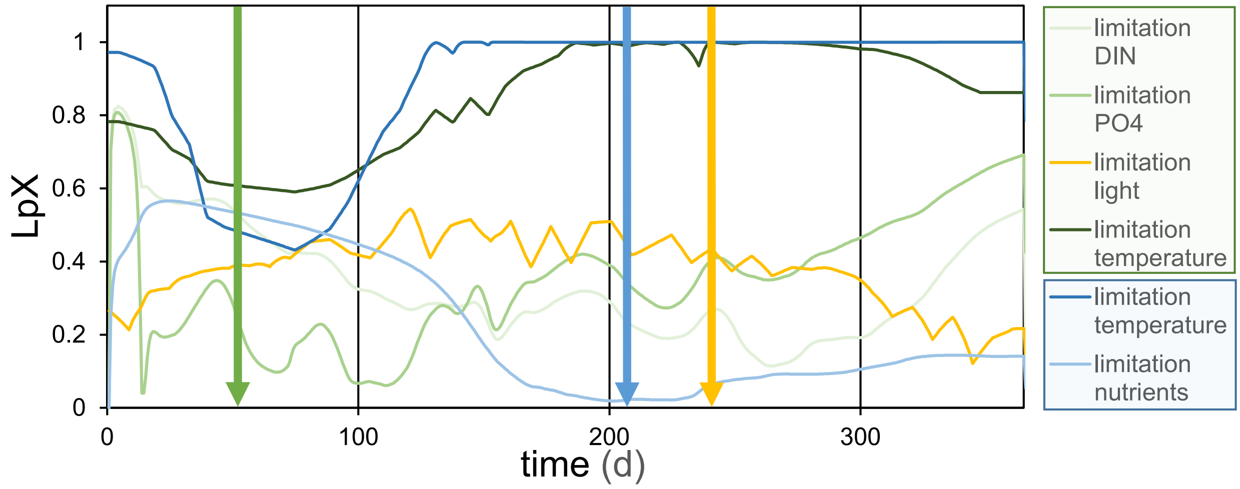

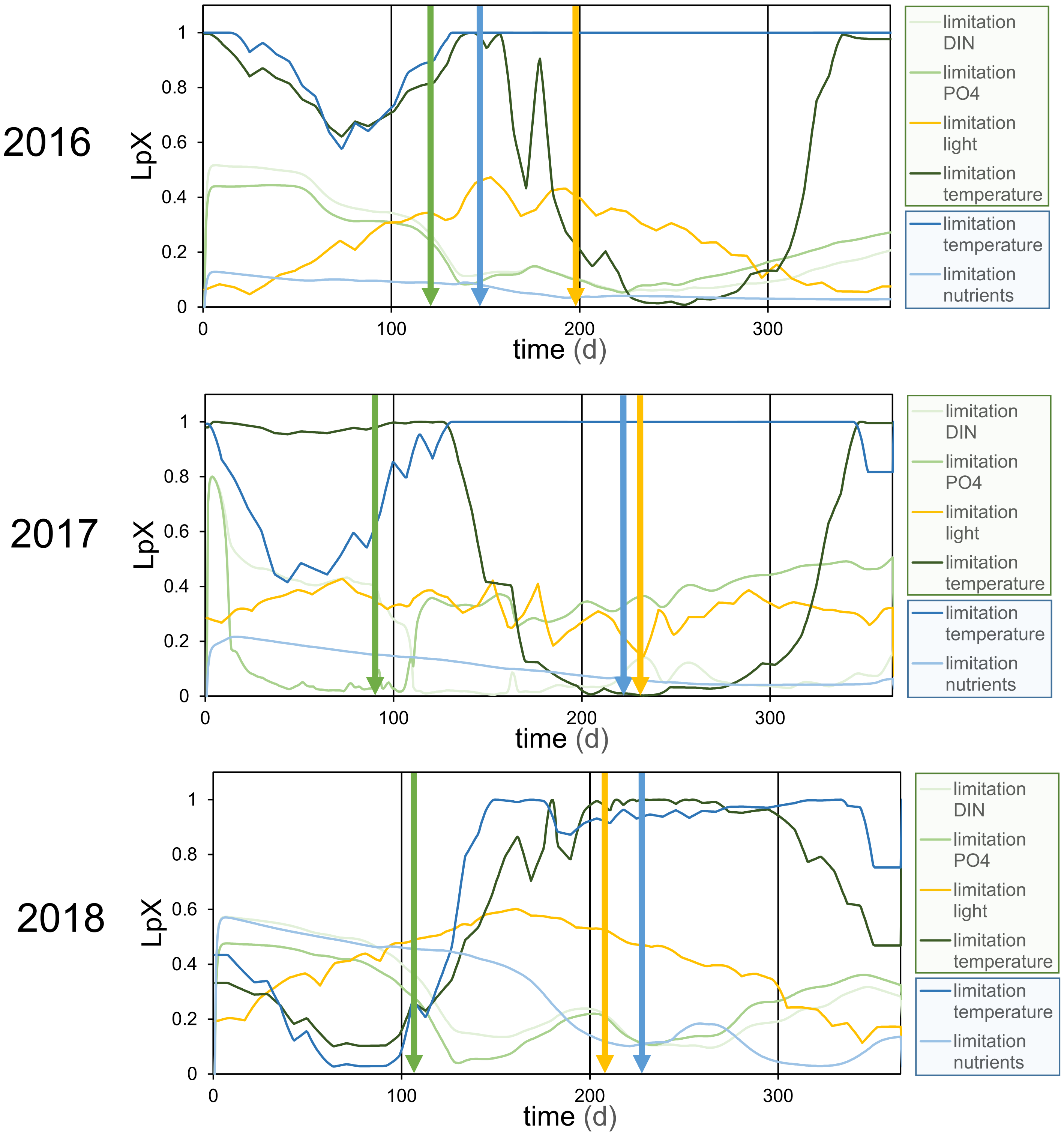


The y-axis refers to the relative inhibition, where a higher value stands for lower inhibition. The green shaded legend refers to phytoplankton inhibition, the blue shaded legend to heterotrophic prokaryotic inhibition. DIN = dissolved inoranic nitrogen, PO4 = phosphate. The arrows indicate the start of the blooms: green: phytoplankton spring bloom, blue: bacteria summer bloom, orange: phytoplankton summer bloom.

Supplemental Table 1:

Definition of bloom periods that were chosen for elaborated analyses. Phytoplankton spring blooms were defined as the highest chlorophyll concentration between February and May, phytoplankton summer blooms as the highest chlorophyll concentration between July and August. Bacterial summer blooms were defined as the highest bacterial concentrations between June and August. Model-data overlaps, estimated peak chl. a or bacterial cell number concentrations, and estimated growth rates (positive/negative) were considered for the assignments of bloom periods without applying strict cut-offs to ensure reasonable definitions (for general rules see main text).

| Year | bloom | Days of the year | Comments | Duration (d) |
|---|---|---|---|---|
| 2012 | Phytoplankton spring bloom | 89-117 | Start: 3.1 µg Chl. a $l^{-1}$, end: 3.32 µg Chl. a $l^{-1}$. First overlap between model and data with positive growth rate. | 28 |
| 2013 | Phytoplankton spring bloom | 112-140 | First day above 2.5 µg Chl. a $l^{-1}$. | 28 |
| 2014 | Phytoplankton spring bloom | 97-125 | First day above 1.5 µg Chl. a $l^{-1}$ that fits the data. | 28 |
| 2015 | Phytoplankton spring bloom | 52-73 | First bloom of the year. First day above 1.7 µg Chl. a $l^{-1,}$ last day above 1.3 µg Chl. a $l^{-1}$. Only 21-day period applied. | 21 |
| 2016 | Phytoplankton spring bloom | 124-135 | Short, small spring bloom. Starting day: first day above 1.4 µg Chl. a $l^{-1}$, last day: last day above 1.4 µg Chl. a $l^{-1}$. | 11 |
| 2017 | Phytoplankton spring bloom | 92-120 | Double peak, first day above 2.8 µg Chl. a $l^{-1}$ with data match. Full period, last day above 2 µg µg Chl. a $l^{-1}$. | 28 |
| 2018 | Phytoplankton spring bloom | 108-129 | First day that fits the data and is above 2.5 µg Chl. a $l^{-1}$. Last day above 2 µg Chl. a $l^{-1}$. Sharp decrease. | 21 |
| 2012 | Phytoplankton summer bloom | 234-255 | First day 1.34 µg Chl. a $l^{-1}$, last day 1.87 µg Chl. a $l^{-1}$. | 21 |
| 2013 | Phytoplankton summer bloom | 206-234 | Fist day above 1.8 µg Chl. a $l^{-1}$. | 28 |
| 2015 | Phytoplankton summer bloom | 242-265 | First day above 1.5 µg Chl. a $l^{-1}$, last day with 1.5 µg Chl. a $l^{-1}$. | 23 |
| 2016 | Phytoplankton summer bloom | 198-226 | Pronounced long bloom. First day that fits the data, and above 1.6 µg Chl. a $l^{-1}$. | 28 |
| 2017 | Phytoplankton summer bloom | 231-246 | Small peak that overlaps with the data. First day above 0.9 µg Chl. a $l^{-1}$ | 15 |

| | | | | |
|---|---|---|---|---|
| 2018 | Phytoplankton summer bloom | 209-237 | Strong double peak. First day above 2 µg Chl. a $l^{-1}$. | 28 |
| 2012 | Bacteria summer bloom | 159-193 | First day above 1.8 million cells $ml^{-1}$ that matches the data, afterwards a long plateau. Last day above 1.8 million cells $ml^{-1}$. | 34 |
| 2013 | Bacteria summer bloom | 174-200 | First day above 1.3 million cells $ml^{-1}$ with positive growth rate that fits the data, last day above 1 million cells $ml^{-1}$. | 26 |
| 2014 | Bacteria summer bloom | 222-250 | Second increase of a long plateau that reflects the data. | 28 |
| 2015 | Bacteria summer bloom | 208-236 | Chosen period reflects the data. Start: first day above 1.4 million cells $ml^{-1}$. | 28 |
| 2016 | Bacteria summer bloom | 148-194 | Long, pronounced plateau. First day above 1.7 million cells $ml^{-1}$ until last day above 1.7 million cells $ml^{-1}$. | 46 |
| 2017 | Bacteria summer bloom | 221-263 | Long plateau. First day: positive growth rate that matches the data. Last day: first day with negative growth rates, i.e. decline. | 42 |
| 2018 | Bacteria summer bloom | 229-254 | Peak is reflected by the data (other peaks not). First day above 1.2 million cells $ml^{-1}$. | 25 |

Supplementary Table 2: (A) Top ten phytoplankton DOM provider, (B) top ten heterotrophic prokaryotic DOM consumers with the corresponding largest phytoplankton OTU provider, and consumed DOM species, (C) top ten phytoplankton>heterotrophic prokaryotes and heterotrophic prokaryotes>heterotrophic prokaryotes pairs with the fluxes between them (µmol C l$^{-1}$ d$^{-1}$). Analyses were done for separate years and the three different bloom types. For taxonomies/names of phytoplankton OTUs and heterotrophic prokaryotic ASVs see Supplementary Table 8. Bloom types: Psb = phytoplankton spring bloom, Bsu = bacteria summer bloom, Psu = phytoplankton summer bloom.

(A) Top ten phytoplankton DOM provider

| year | 2012 | | | 2013 | | | 2014 | | 2015 | | | 2016 | | | 2017 | | | 2018 | | |
|---|---|---|---|---|---|---|---|---|---|---|---|---|---|---|---|---|---|---|---|---|
| Bloom /rank | Psb | Bsu | Psu | Psb | Bsu | Psu | Psb | Bsu | Psb | Bsu | Psu | Psb | Bsu | Psu | Psb | Bsu | Psu | Psb | Bsu | Psu |
| 1 | ppo | lmi | pin | mpu | h37 | gsp | tha | tbe | tal | kse | kse | rse | che | toc | ael | cle | cle | tha | dac | dac |
| 2 | toc | ezo | dio | tro | pon | din | h39 | ost | bpr | syn | syn | ppo | ns1 | pte | tam | h44 | pcu | fna | gsp | gsp |
| 3 | rse | gde | ael | bpr | dsp | pha | pcu | dia | syn | dac | lch | cro | pte | dia | h50 | pcu | ufl | tam | syn | syn |
| 4 | kve | hap | acu | h37 | gsp | pin | bpr | che | cdi | pte | tte | csi | cwe | che | h44 | ufl | cwi | toc | ale | ale |
| 5 | lbo | lgr | tac | tam | stu | pte | h36 | pha | lch | che | pte | csp | cs1 | dac | bpr | cwi | che | csi | ael | ael |
| 6 | gcr | csi | ske | cra | pve | mpu | ppo | dno | ezo | gsm | rse | kse | toc | pre | mpu | che | lco | cca | chs | chs |
| 7 | tam | tre | lvi | tha | pha | gcr | h40 | rat | csp | lch | pon | stu | cro | ezo | pte | lco | gos | isp | rsh | ns1 |
| 8 | mpu | och | lda | ha2 | psp | ael | h35 | gsp | toc | lgr | yye | ns1 | pha | dno | aan | pte | pon | mpu | pre | pre |
| 9 | da2 | gsp | pon | cs1 | psh | str | tco | hap | tco | lco | gsm | kve | gde | gsm | cca | pve | pte | bpr | cci | lmi |
| 10 | tro | gsm | pte | h56 | pte | che | syn | gca | pin | str | ost | pte | pve | cle | h39 | toc | pve | sha | cwe | cwe |

(B): Top ten bacterial carbon consumers (con), corresponding top phytoplankton provider (prov) and corresponding top DOM species consumed for (B1) phytoplankton spring blooms, (B2) bacteria summer blooms, and (B3) phytoplankton summer blooms.

(B1) phytoplankton spring blooms

| year | 2012 | | | 2013 | | | 2014 | | | 2015 | | | 2016 | | | 2017 | | | 2018 | | |
|---|---|---|---|---|---|---|---|---|---|---|---|---|---|---|---|---|---|---|---|---|---|
| rank | con | prov | DOM | con | prov | DOM | con | prov | DOM | con | prov | DOM | con | prov | DOM | con | prov | DOM | con | prov | DOM |
| 1 | b03 | ppo | d01 | pla | mpu | d01 | s86 | h39 | d01 | s86 | tal | d60 | amy | rse | d01 | amy | ael | d01 | s86 | fna | d01 |
| 2 | fla | ppo | d01 | amy | bpr | d01 | s88 | tha | d69 | pla | tal | d01 | s86 | rse | d01 | te2 | ael | d01 | fo1 | tha | d46 |
| 3 | ulv | ppo | d05 | s86 | mpu | f31 | s87 | h39 | d01 | ro4 | tal | d05 | are | csi | d34 | pla | ael | d01 | amy | fna | d01 |
| 4 | b02 | ppo | d01 | cr1 | mpu | d45 | amy | h39 | d01 | n04 | tal | d01 | s11 | rse | d01 | te1 | h50 | d12 | m27 | tam | d44 |
| 5 | s86 | ppo | f14 | pl1 | mpu | d49 | s89 | tha | d01 | pmy | tal | d15 | pu1 | rse | d01 | s86 | ael | d01 | cr1 | fna | d01 |
| 6 | te1 | ppo | d06 | te1 | tro | d01 | b05 | tha | d01 | te1 | tal | d01 | ro1 | rse | d01 | n04 | ael | d01 | pla | fna | d01 |
| 7 | b05 | ppo | d01 | sul | mpu | d62 | m21 | h39 | d31 | cr1 | tal | d01 | n55 | ppo | d21 | hyp | ael | d01 | pl1 | fna | d01 |
| 8 | psm | da2 | d01 | hyp | mpu | d01 | are | pcu | d01 | amy | tal | d47 | sul | ppo | d34 | s87 | h44 | d76 | pl2 | tha | d01 |
| 9 | sa2 | ppo | d01 | n55 | mpu | d01 | ro4 | tha | d73 | te2 | tal | d01 | m21 | rse | d34 | ros | ael | d01 | cr3 | tam | d97 |
| 10 | pla | ppo | d01 | te2 | h37 | d69 | at2 | tha | d01 | n94 | tal | d01 | nmo | ppo | d01 | pl1 | tam | d01 | te1 | tha | d01 |

(B2) bacteria summer blooms

| year | 2012 | | | 2013 | | | 2014 | | | 2015 | | | 2016 | | | 2017 | | | 2018 | | |
|---|---|---|---|---|---|---|---|---|---|---|---|---|---|---|---|---|---|---|---|---|---|
| rank | con | prov | DOM | con | prov | DOM | con | prov | DOM | con | prov | DOM | con | prov | DOM | con | prov | DOM | con | prov | DOM |
| 1 | b02 | lmi | d01 | s11 | h37 | d01 | ro1 | ost | d61 | s86 | syn | d01 | s86 | che | d21 | s86 | cle | d01 | m22 | dac | d42 |
| 2 | s86 | lmi | d13 | te2 | h37 | d63 | ro2 | ost | d71 | ro2 | kse | d01 | amy | che | d01 | n91 | cle | d72 | th2 | dac | d42 |
| 3 | b05 | lmi | d01 | ros | h37 | d80 | b05 | tbe | d01 | ro3 | kse | d84 | pu1 | che | d01 | ma1 | cle | d01 | ro2 | dac | d42 |
| 4 | b03 | lmi | d01 | te1 | h37 | d01 | s86 | tbe | d01 | amy | syn | d01 | s11 | che | d01 | ect | cle | d34 | s11 | syn | d94 |
| 5 | aqu | gde | d82 | m21 | h37 | d01 | s89 | tbe | d01 | aqu | kse | d74 | are | ns1 | d34 | s11 | cle | d01 | s90 | dac | d42 |
| 6 | per | ezo | d01 | s86 | h37 | d01 | n51 | ost | d07 | n91 | kse | d74 | nmo | ns1 | d21 | s13 | cle | d08 | ro4 | dac | d42 |
| 7 | cr1 | gde | d01 | sa2 | h37 | d62 | pi2 | ost | d68 | fu2 | kse | d84 | fo1 | che | d21 | ro2 | cle | d81 | ros | syn | d01 |
| 8 | pla | gde | d01 | are | h37 | d01 | s11 | tbe | d01 | th2 | kse | d39 | po9 | che | d21 | m22 | cle | d01 | are | dac | d42 |
| 9 | om7 | gde | d01 | rub | h37 | d34 | ro3 | ost | d62 | s11 | kse | d01 | ro3 | che | d21 | amy | cle | d01 | oce | dac | d42 |
| 10 | fac | ezo | d01 | pi1 | gsp | d63 | s88 | tbe | d69 | pu2 | kse | d01 | ros | che | d01 | s14 | cle | d01 | ae2 | dac | d42 |

(B3) phytoplankton summer blooms

| year | 2012 | | | 2013 | | | 2015 | | | 2016 | | | 2017 | | | 2018 | | |
|---|---|---|---|---|---|---|---|---|---|---|---|---|---|---|---|---|---|---|
| rank | con | prov | DOM | con | prov | DOM | con | prov | DOM | con | prov | DOM | con | prov | DOM | con | prov | DOM |
| 1 | b02 | pin | d01 | s11 | din | d01 | s86 | syn | d47 | aqu | toc | d75 | s86 | cle | d01 | m22 | dac | d42 |
| 2 | b05 | pin | d01 | aqu | gsp | d62 | ro2 | kse | d01 | amy | pte | d01 | ect | cle | d34 | ros | syn | d01 |
| 3 | s86 | pin | d13 | n05 | gsp | d62 | th2 | kse | d39 | pu1 | toc | d75 | n91 | cle | d72 | th2 | dac | d42 |
| 4 | amy | pin | f12 | lum | gsp | d34 | s11 | kse | d01 | pl1 | toc | d75 | ma1 | cle | d01 | ro2 | dac | d42 |
| 5 | n51 | pin | d79 | fo1 | din | d62 | ro3 | kse | d01 | cr1 | toc | d75 | s13 | cle | d08 | s90 | dac | d42 |
| 6 | n07 | pin | f28 | pi1 | gsp | d63 | s12 | kse | d70 | lem | toc | d79 | s11 | cle | d01 | are | dac | d42 |
| 7 | cr1 | pin | d68 | om7 | gsp | d62 | ect | kse | d74 | te1 | pte | d76 | ro2 | cle | d81 | s11 | syn | d94 |
| 8 | fu1 | pin | f17 | m22 | gsp | d38 | pu2 | kse | d01 | fu1 | toc | d79 | m22 | cle | d01 | n91 | dac | d42 |
| 9 | ub1 | pin | d01 | n04 | din | d01 | n91 | kse | d74 | ns2 | toc | d75 | fo2 | cle | d03 | ro3 | dac | d42 |
| 10 | s87 | pin | d01 | fla | gsp | d63 | ae1 | kse | d01 | te2 | toc | d97 | amy | cle | d01 | ro4 | dac | d42 |

(C) Top ten phytoplankton>heterotrophic prokaryotes and heterotrophic prokaryotes>heterotrophic prokaryotes carbon flux pairs for the respective bloom types and years. Fluxes are given in µmol C $l^{-1}$ $d^{-1}$.

(C1) Phytoplankton spring blooms

| year | 2012 | | 2013 | | 2014 | | 2015 | | 2016 | | 2017 | | 2018 | |
|---|---|---|---|---|---|---|---|---|---|---|---|---|---|---|
| | pair | flux | pair | flux | pair | flux | pair | flux | pair | flux | pair | flux | pair | flux |
| rank | phytoplankton>heterotrophic prokaryotes | | | | | | | | | | | | | |
| 1 | ppo>b03 | 0.0060 | mpu>pla | 0.0018 | tha>s88 | 0.0066 | tal>s86 | 0.00137 | rse>amy | 0.0156 | ael>te2 | 0.0021 | tha>fo1 | 0.0047 |
| 2 | ppo>fla | 0.0037 | bpr>amy | 0.0018 | h39>s86 | 0.0052 | tal>pla | 0.00082 | ppo>am | 0.0068 | ael>amy | 0.0016 | fna>s86 | 0.0025 |
| 3 | toc>b03 | 0.0030 | tro>pla | 0.0016 | tha>s86 | 0.0043 | tal>ro4 | 0.00070 | cro>amy | 0.0066 | tam>amy | 0.0011 | tam>m27 | 0.0023 |
| 4 | ppo>b02 | 0.0027 | mpu>am | 0.0014 | h39>s88 | 0.0040 | tal>n04 | 0.00052 | rse>s86 | 0.0063 | h50>amy | 0.0008 | tam>fo1 | 0.0020 |
| 5 | rse>b03 | 0.0023 | tro>amy | 0.0011 | tha>s89 | 0.0021 | bpr>s86 | 0.00051 | ppo>s86 | 0.0058 | tam>te2 | 0.0008 | tam>s86 | 0.0018 |
| 6 | ppo>s86 | 0.0021 | mpu>cr1 | 0.0011 | pcu>s86 | 0.0020 | tal>pmy | 0.00047 | csi>amy | 0.0044 | h44>te2 | 0.0008 | csi>s86 | 0.0017 |
| 7 | rse>fla | 0.0021 | mpu>sul | 0.0010 | tha>ro4 | 0.0012 | syn>s86 | 0.00041 | csp>am | 0.0033 | ael>pla | 0.0007 | toc>s86 | 0.0017 |
| 8 | kve>b03 | 0.0017 | h37>sul | 0.0009 | h39>s87 | 0.0012 | tal>te1 | 0.00039 | cro>s86 | 0.0033 | h44>amy | 0.0007 | tha>s86 | 0.0017 |
| 9 | toc>fla | 0.0017 | tam>am | 0.0009 | tha>s87 | 0.0011 | tal>amy | 0.00035 | kse>am | 0.0032 | h50>te1 | 0.0007 | fna>fo1 | 0.0014 |
| 10 | ppo>ulv | 0.0016 | bpr>pla | 0.0008 | bpr>s86 | 0.0011 | syn>pmy | 0.00035 | stu>amy | 0.0028 | ael>s86 | 0.0006 | tha>n56 | 0.0013 |
| rank | heterotrophic prokaryotes>heterotrophic prokaryotes | | | | | | | | | | | | | |
| 1 | psm>ps | 0.0017 | s12>pla | 0.0011 | s87>s86 | 0.0020 | pmy>s86 | 0.00002 | sul>amy | 0.0120 | s87>amy | 0.0005 | m27>s86 | 0.0011 |
| 2 | fla>b03 | 0.0013 | s12>amy | 0.0007 | s88>s86 | 0.0019 | pmy>pla | 0.00001 | n55>am | 0.0096 | s88>amy | 0.0004 | at1>s86 | 0.0011 |
| 3 | psm>b03 | 0.0011 | s92>amy | 0.0007 | ma1>s86 | 0.0010 | pmy>ro4 | 0.00001 | sul>s86 | 0.0028 | s87>pla | 0.0003 | m27>amy | 0.0006 |
| 4 | ths>psm | 0.0009 | s87>pla | 0.0007 | pmy>s86 | 0.0008 | pmy>te1 | 0.00001 | s88>am | 0.0028 | s87>te1 | 0.0002 | at1>amy | 0.0005 |
| 5 | fla>ulv | 0.0008 | s12>s86 | 0.0005 | s89>s86 | 0.0005 | m23>s86 | 0.00001 | n55>s86 | 0.0021 | s87>te2 | 0.0002 | m25>s86 | 0.0005 |
| 6 | fla>fla | 0.0007 | s92>pla | 0.0004 | n51>s86 | 0.0005 | pmy>n04 | 0.00001 | m25>a | 0.0017 | s92>amy | 0.0002 | psa>s86 | 0.0005 |
| 7 | fla>b02 | 0.0006 | s87>amy | 0.0004 | s87>amy | 0.0005 | pmy>cr1 | 0.00001 | s93>am | 0.0012 | s13>amy | 0.0002 | cr3>s86 | 0.0004 |
| 8 | psm>ths | 0.0006 | s92>s86 | 0.0003 | sul>s86 | 0.0005 | m04>s86 | 0.00001 | amy>s8 | 0.0011 | s13>te1 | 0.0002 | m27>te1 | 0.0003 |
| 9 | psm>b02 | 0.0005 | s12>cr1 | 0.0003 | s92>s86 | 0.0003 | n51>s86 | 0.00001 | mas>am | 0.0011 | s12>amy | 0.0002 | m27>pl1 | 0.0003 |
| 10 | psm>ulv | 0.0005 | pav>amy | 0.0003 | s88>s89 | 0.0003 | hoc>s86 | 0.00001 | pav>am | 0.0011 | leb>amy | 0.0001 | m27>pla | 0.0003 |

(C2) Bacteria summer bloom

| year | 2012 | | 2013 | | 2014 | | 2015 | | 2016 | | 2017 | | 2018 | |
|---|---|---|---|---|---|---|---|---|---|---|---|---|---|---|
| | pair | flux | pair | flux | pair | flux | pair | flux | pair | flux | pair | flux | pair | flux |
| rank | phytoplankton>heterotrophic prokaryotes | | | | | | | | | | | | | |
| 1 | lmi>b02 | 0.0067 | h37>s11 | 0.0104 | ost>ro1 | 0.0047 | syn>s86 | 0.0281 | che>s86 | 0.008 | cle>s86 | 0.0120 | dac>m22 | 0.082 |
| 2 | ezo>b02 | 0.0050 | pon>s11 | 0.0073 | ost>ro2 | 0.0028 | kse>ro2 | 0.0126 | che>pu1 | 0.002 | h44>s86 | 0.0063 | gsp>m22 | 0.0273 |
| 3 | gde>b02 | 0.0050 | dsp>s11 | 0.0062 | tbe>s86 | 0.0024 | kse>ro3 | 0.0042 | che>amy | 0.002 | pcu>s86 | 0.0052 | dac>th2 | 0.0221 |
| 4 | lmi>s86 | 0.0048 | h37>te2 | 0.0053 | tbe>b05 | 0.0022 | kse>n91 | 0.0041 | pte>s86 | 0.002 | cwi>s86 | 0.0039 | dac>ro2 | 0.0175 |
| 5 | hap>b02 | 0.0045 | pve>s11 | 0.0032 | dia>ro1 | 0.0016 | kse>aqu | 0.0040 | ns1>s86 | 0.001 | ufl>s86 | 0.0035 | syn>m22 | 0.0150 |
| 6 | hap>s86 | 0.0039 | pha>s11 | 0.0029 | tbe>ro2 | 0.0014 | dac>ro2 | 0.0038 | pte>amy | 0.001 | che>s86 | 0.0032 | syn>s11 | 0.0124 |
| 7 | lmi>b03 | 0.0036 | stu>s11 | 0.0029 | tbe>s11 | 0.0012 | syn>amy | 0.0033 | cwe>s86 | 0.001 | lco>s86 | 0.0016 | dac>s90 | 0.0115 |
| 8 | lmi>b05 | 0.0034 | psp>s11 | 0.0025 | tbe>s89 | 0.0012 | kse>fu2 | 0.0032 | pha>s86 | 0.001 | pte>s86 | 0.0014 | dac>ro4 | 0.0103 |
| 9 | gde>s86 | 0.0030 | h37>te1 | 0.0024 | tbe>ro1 | 0.0011 | dac>s86 | 0.0031 | gde>s86 | 0.001 | toc>s86 | 0.0012 | dac>are | 0.0080 |
| 10 | csi>b02 | 0.0029 | h37>ros | 0.0023 | dia>ro2 | 0.0011 | kse>th2 | 0.0029 | ns1>amy | 0.001 | pve>s86 | 0.0009 | gsp>th2 | 0.0072 |
| rank | heterotrophic prokaryotes>heterotrophic prokaryotes | | | | | | | | | | | | | |
| 1 | b03>b02 | 0.0205 | ros>s11 | 0.0170 | s86>ro1 | 0.0059 | s86>s86 | 0.0038 | ros>s11 | 0.027 | s86>s86 | 0.0176 | ros>s11 | 0.0115 |
| 2 | b02>b02 | 0.0129 | rub>s11 | 0.0153 | s86>ro2 | 0.0041 | amy>s86 | 0.0023 | ros>th2 | 0.013 | amy>s86 | 0.0064 | ros>m22 | 0.0103 |
| 3 | b03>s86 | 0.0125 | lem>s11 | 0.0131 | s86>n51 | 0.0011 | aqu>s86 | 0.0021 | ros>n07 | 0.009 | te1>s86 | 0.0054 | m22>ros | 0.0084 |
| 4 | b05>b02 | 0.0109 | te1>s11 | 0.0123 | s86>om | 0.0011 | aqu>ro2 | 0.0021 | ros>n91 | 0.006 | ect>s86 | 0.0023 | m22>s11 | 0.0061 |
| 5 | b03>b03 | 0.0103 | te2>s11 | 0.0118 | pla>s86 | 0.0010 | ro2>s86 | 0.0020 | ros>are | 0.005 | s86>n91 | 0.0023 | ros>th2 | 0.0045 |
| 6 | b03>b05 | 0.0098 | amy>s1 | 0.0086 | n51>s86 | 0.0010 | fu2>s86 | 0.0016 | ros>ro2 | 0.005 | s86>ma1 | 0.0015 | m22>m22 | 0.0041 |
| 7 | b02>s86 | 0.0074 | m21>s1 | 0.0077 | s86>s89 | 0.0009 | fu2>ro2 | 0.0015 | n07>s11 | 0.004 | s19>s86 | 0.0011 | pla>ros | 0.0030 |
| 8 | b05>s86 | 0.0062 | s11>s11 | 0.0074 | s86>ro3 | 0.0009 | te1>s86 | 0.0014 | n07>ros | 0.004 | s86>ect | 0.0010 | ros>ro2 | 0.0029 |
| 9 | b02>b05 | 0.0047 | s11>te2 | 0.0054 | s86>b05 | 0.0008 | cr1>s86 | 0.0013 | ros>ros | 0.003 | ae2>s86 | 0.0009 | m22>s86 | 0.0025 |
| 10 | per>b02 | 0.0046 | ro1>s11 | 0.0052 | amy>s86 | 0.0008 | s86>ro2 | 0.0012 | n91>s11 | 0.003 | fu2>s86 | 0.0009 | ros>ros | 0.0023 |

(C3) Phytoplankton summer blooms

| year | 2012 | | 2013 | | 2015 | | 2016 | | 2017 | | 2018 | |
|---|---|---|---|---|---|---|---|---|---|---|---|---|
| | pair | flux | pair | flux | pair | flux | pair | flux | Pair | Flux | pair | |
| rank | phytoplankton>heterotrophic prokaryotes | | | | | | | | | | | |
| 1 | pin>b02 | 0.01612 | din>s11 | 0.007575 | syn>s86 | 0.02362 | toc>aqu | 0.004074 | cle>s86 | 0.02606 | dac>m22 | 0.03109 |
| 2 | pin>b05 | 0.009112 | gsp>s11 | 0.005128 | lch>s86 | 0.005699 | toc>lem | 0.001895 | pcu>s86 | 0.01484 | dac>th2 | 0.01448 |
| 3 | dio>b02 | 0.003694 | pha>s11 | 0.003383 | kse>ro2 | 0.005616 | pte>amy | 0.001858 | ufl>s86 | 0.007476 | dac>ro2 | 0.01378 |
| 4 | ael>b02 | 0.003689 | pin>s11 | 0.002761 | tte>s86 | 0.003823 | toc>pl1 | 0.001734 | cwi>s86 | 0.00408 | gsp>m22 | 0.00978 |
| 5 | acu>b02 | 0.003469 | gsp>aqu | 0.00256 | kse>th2 | 0.003379 | che>aqu | 0.001497 | che>s86 | 0.0033 | dac>s90 | 0.008944 |
| 6 | pin>amy | 0.002478 | gsp>lum | 0.002326 | kse>ro3 | 0.002091 | toc>cr1 | 0.001414 | lco>s86 | 0.001947 | dac>are | 0.008385 |
| 7 | ael>b05 | 0.002176 | gsp>pi1 | 0.002319 | kse>s11 | 0.002057 | toc>pu1 | 0.00117 | cle>ect | 0.001572 | dac>n91 | 0.006132 |
| 8 | pin>s86 | 0.00205 | gsp>om7 | 0.00225 | kse>s12 | 0.002028 | pte>aqu | 0.001032 | pte>s86 | 0.001507 | dac>ro3 | 0.00611 |
| 9 | acu>b05 | 0.002027 | gsp>n05 | 0.002216 | lch>ro2 | 0.001797 | toc>fu1 | 0.00103 | cle>n91 | 0.001498 | dac>ro4 | 0.00573 |
| 10 | dio>b05 | 0.002025 | pte>s11 | 0.001793 | rse>s86 | 0.001788 | toc>n05 | 0.001007 | cle>ma1 | 0.001395 | syn>s11 | 0.0053 |
| rank | heterotrophic prokaryotes>heterotrophic prokaryotes | | | | | | | | | | | |
| 1 | s86>b02 | 0.001864 | pi1>s11 | 0.003938 | s86>s86 | 0.01585 | s86>aqu | 0.005872 | s86>s86 | 0.0255 | ros>s11 | 0.01522 |
| 2 | cr1>b02 | 0.0009606 | s11>n05 | 0.002879 | ro2>s86 | 0.004515 | amy>aqu | 0.004685 | amy>s86 | 0.006559 | ros>ros | 0.008367 |
| 3 | s86>b05 | 0.0009451 | aqu>s11 | 0.002877 | amy>s86 | 0.004429 | s86>pl1 | 0.004224 | ect>s86 | 0.004169 | pla>ros | 0.004953 |
| 4 | amy>b02 | 0.0007938 | s11>pi1 | 0.002838 | s86>ro2 | 0.002252 | s86>lem | 0.002324 | s19>s86 | 0.001947 | te1>ros | 0.003863 |
| 5 | cr1>b05 | 0.000556 | s11>aqu | 0.002229 | ro3>s86 | 0.001624 | aqu>amy | 0.001823 | s86>n91 | 0.00187 | ros>m22 | 0.003848 |
| 6 | n07>b02 | 0.000554 | s11>lum | 0.001868 | m21>s86 | 0.001375 | s86>ns2 | 0.001652 | s86>ect | 0.001555 | ros>th2 | 0.003116 |
| 7 | s86>amy | 0.0004857 | n58>s11 | 0.001835 | ro2>ro2 | 0.001164 | s86>n05 | 0.00163 | ae2>s86 | 0.001524 | n05>ros | 0.002609 |
| 8 | amy>b05 | 0.0004601 | lum>s11 | 0.001706 | s86>s11 | 0.0009686 | amy>pl1 | 0.001575 | fo2>s86 | 0.001342 | ros>ro2 | 0.002472 |
| 9 | ps1>b02 | 0.0003854 | om7>s11 | 0.001706 | n51>s86 | 0.0008687 | amy>lem | 0.001481 | fu2>s86 | 0.00109 | n07>ros | 0.002432 |
| 10 | s11>b02 | 0.0003813 | om7>s11 | 0.001639 | pu2>s86 | 0.0007934 | amy>cr1 | 0.001406 | s86>ma1 | 0.001066 | ros>fu2 | 0.002402 |

Supplementary Table 3:

Primary and heterotrophic prokaryotic production.

Net primary production was calculated as gross primary production – respiration. Gross heterotrophic prokaryotic production: all dissolved organic carbon (DOC) fluxes into heterotrophic prokaryotes, net heterotrophic prokaryotic production: gross production – heterotrophic respiration. Bloom types: Psp: phytoplankton spring bloom, Bsu: bacteria summer bloom, Psu: phytoplankton summer bloom.

| year | 2012 | | | 2013 | | | 2014 | | 2015 | | | 2016 | | | 2017 | | | 2018 | | |
|---|---|---|---|---|---|---|---|---|---|---|---|---|---|---|---|---|---|---|---|---|
| Primary production gross mmol C $m^{-2}$ $d^{-1}$ | 25 | | | 32 | | | 20 | | 22 | | | 18 | | | 22 | | | 25 | | |
| Primary production net mmol C $m^{-2}$ $d^{-1}$ | 23 | | | 23 | | | 13 | | 20 | | | 13 | | | 19 | | | 15 | | |
| Heterotrophic prokaryotic production gross µmol C $l^{-1}$ $d^{-1}$ | 0.2 | | | 0.14 | | | 0.18 | | 0.21 | | | 0.2 | | | 0.13 | | | 0.24 | | |
| Heterotrophic prokaryotic production net µmol C $l^{-1}$ $d^{-1}$ | 0.12 | | | 0.07 | | | 0.08 | | 0.08 | | | 0.11 | | | 0.07 | | | 0.1 | | |
| Bloom type | Psp | Bsu | Psu | Psp | Bsu | Psu | Psp | Bsu | Psp | Bsu | Psu | Psp | Bsu | Psu | Psp | Bsu | Psu | Psp | Bsu | Psu |
| Primary production gross mmol C $m^{-2}$ $d^{-1}$ | 16 | 17 | 19 | 25 | 16 | 18 | 91 | 8 | 22 | 14 | 19 | 55 | 19 | 19 | 21 | 11 | 13 | 71 | 31 | 52 |
| Primary production net mmol C | 15 | 12 | 16 | 12 | 13 | 14 | 61 | 5 | 19 | 12 | 16 | 47 | 13 | 11 | 16 | 10 | 12 | 39 | 14 | 24 |

| | | | | | | | | | | | | | | | | | | | | |
|---|---|---|---|---|---|---|---|---|---|---|---|---|---|---|---|---|---|---|---|---|
| $m^{-2}$ $d^{-1}$ | | | | | | | | | | | | | | | | | | | | |
| Heterotrophic prokaryotic production gross µmol C $l^{-1}$ $d^{-1}$ | 0.26 | 0.57 | 0.19 | 0.2 | 0.73 | 0.26 | 0.25 | 0.35 | 0.04 | 0.37 | 0.29 | 0.53 | 0.65 | 0.3 | 0.17 | 0.3 | 0.31 | 0.29 | 0.62 | 0.47 |
| Heterotrophic prokaryotic production net µmol C $l^{-1}$ $d^{-1}$ | 0.25 | 0.55 | 0.19 | 0.19 | 0.4 | 0.15 | 0.13 | 0.19 | 0.02 | 0.18 | 0.13 | 0.36 | 0.45 | 0.22 | 0.12 | 0.18 | 0.18 | 0.2 | 0.42 | 0.3 |

Supplementary Table 4

Statistical tests and statistical outcomes.

(A) Tests for differences of recurrences between different variables, and test for differences of recurrences in the tested variables between the different bloom types.
(B) Tests for differences in DOM concentration, DOM consumption and DOM production between the three different bloom types
(C) Differences in averaged fluxes of phytoplankton>heterotrophic prokaryotes and heterotrophic prokaryotes>heterotrophic prokaryotes
(D) Differences in instantaneous fluxes between phytoplankton>heterotrophic prokaryotes and heterotrophic prokaryotes>heterotrophic prokaryotes between bloom types and trends over time
(E) Differences in concentration normalized heterotrophy rates between bloom types

#### (A) Tests for differences of recurrences between different variables, and test for differences of recurrences in the tested variables between the different bloom types.

Test for homogeneity of variances, Levene test

| Tested variables | F | p |
|---|---|---|
| Variables (phyto>dom, pro. het<dom, phyto>pro. het, pro. het>pro. het) | 7.5 | <0.001 |
| Phyto>dom (different bloom types) | 0.64 | 0.53 |
| Pro. het<dom (different bloom types) | 1.56 | 0.22 |
| Phyto>pro. het (different bloom types) | 2.72 | 0.07 |
| pro. het>pro. het (different bloom types) | 3 | 0.06 |

Kruskal-Wallis and Tukey posthoc test for differences between variables

| Tested variables | $Chi^2$ | p | Tukey |
|---|---|---|---|
| Variables (phyto>dom, pro. het<dom, phyto>pro. | 104.68 | <0.0001 | Phyto>dom = b, pro. het<dom = a, |

| | | | |
|---|---|---|---|
| het, pro. het>pro. het) | | | phyto>pro. het = c, pro. het>pro. het = c |

ANOVA and posthoc test for differences in the recurrence in the different variables between the three bloom types

| Tested variables | F | p | TukeyHSD |
|---|---|---|---|
| Phyto>dom | 2.37 | 0.1 | - |
| Pro. het<dom | 3.32 | 0.04 | Phytoplankton spring bloom = a, bacteria summer bloom = ab, phytoplankton summer bloom = b |
| Phyto>pro. het | 7.42 | 0.001 | Phytoplankton spring bloom = a, bacteria summer bloom = ab, phytoplankton summer bloom = b |
| Pro. het>pro. het | 1.06 | 0.35 | - |

#### (B) Tests for differences in DOM concentration, DOM consumption and DOM production between the three different bloom types

Test for homogeneity of variances, Levene test

| Tested variables | F | p |
|---|---|---|
| concentration~ bloom types | 2.68 | 0.1 |
| production combined~ bloom types | 0.61 | 0.56 |
| consumption ~ bloom types | 1.83 | 0.19 |

ANOVA and Tukey posthoc to test for differences between blooms

| Tested | F | p | TukeyHSD |
|---|---|---|---|

| | | | |
|---|---|---|---|
| variables | | | |
| DOC concentration | 23.98 | <0.0001 | Phytoplankton spring bloom = a, bacteria summer bloom and phytoplankton summer bloom = b |
| DOC production combined | 0.18 | 0.84 | - |
| DOC consumption | 6.73 | 0.007 | bacteria summer bloom = a, phytoplankton spring and summer bloom = b |

Test for differences of phytoplankton and heterotrophic prokaryotic DOC production

Test for homogeneity of variances, Levene test

| Tested variables | F | p |
|---|---|---|
| Phytoplankton spring bloom: Phytoplankton production~ Heterotrophic prokaryotes production | 5.22 | 0.04 |
| Bacteria summer bloom: Phytoplankton production~ Heterotrophic prokaryotes production | 6.27 | 0.03 |
| Phytoplankton summer bloom: Phytoplankton production~ Heterotrophic prokaryotes production | 0.08 | 0.78 |

Mann-Whitney-U and T-Tests for differences between phytoplankton and pheterotrophic prokaryotes production

| Tested variables | W or T value | p |
|---|---|---|
| Phytoplankton spring bloom | W: 49 | <0.001 |
| Bacteria summer bloom | W: 15 | 0.26 |
| Phytoplankton summer bloom | T: 1.37 | 0.2 |

#### (C) Differences in averaged fluxes of phytoplankton>pro. hetteria and pro. hetteria>pro. hetteria

Test for homogeneity of variances, Levene test

| Tested variables | F | p |
|---|---|---|
| Phytoplankton spring bloom | 3.28 | 0.1 |
| Bacteria summer bloom | 0.02 | 0.9 |
| Phytoplankton summer bloom | 0.32 | 0.58 |

T-tests for differences between phytoplankton>heterotrophic prokaryotes and heterotrophic prokaryotes>heterotrophic prokaryotes fluxes for bloom averaged fluxes

| Tested variables | t | p | average µmol C $l^{-1}$ $d^{-1}$ |
|---|---|---|---|
| Phytoplankton spring bloom | 3.79 | 0.0026 | phytoplankton> heterotrophic prokaryotes 0.13, heterotrophic prokaryotes > heterotrophic prokaryotes 0.03 |
| Bacteria summer bloom | 1.17 | 0.27 | phytoplankton> heterotrophic prokaryotes 0.18, heterotrophic prokaryotes>heterotrophic prokaryotes 0.15 |
| Phytoplankton summer bloom | 2.16 | 0.056 | phytoplankton>heterotrophic prokaryotes 0.14, heterotrophic prokaryotes>heterotrophic prokaryotes 0.08 |

#### (D) Differences in instantaneous fluxes between phytoplankton>heterotrophic prokaryotes and heterotrophic prokaryotes>heterotrophic prokaryotes

Test for homogeneity of variances, Levene test

| Tested variables | F | p |
|---|---|---|
| Phytoplankton spring bloom day 0 | 3.32 | 0.09 |
| Phytoplankton spring bloom day 7 | 2.21 | 0.16 |
| Phytoplankton spring bloom day 14 | 2.77 | 0.13 |
| Phytoplankton spring bloom day 21 | 3.11 | 0.11 |
| Phytoplankton spring bloom day 28 | 0 | 0.99 |
| Bacteria summer bloom day 0 | 1.64 | 0.22 |
| Bacteria summer bloom day 7 | 0.07 | 0.79 |
| Bacteria summer bloom day 14 | 0.3 | 0.59 |
| Bacteria summer bloom day 21 | 0.6 | 0.45 |
| Bacteria summer bloom day 28 | 0.31 | 0.6 |
| Phytoplankton summer bloom day 0 | 0.23 | 0.64 |
| Phytoplankton summer bloom day 7 | 0.77 | 0.4 |
| Phytoplankton summer bloom day 14 | 0.1 | 0.75 |
| Phytoplankton summer bloom day 21 | 0.62 | 0.45 |
| Phytoplankton summer bloom day 28 | 0.73 | 0.44 |

T-tests for differences between phytoplankton>heterotrophic prokaryotes and heterotrophic prokaryotes>heterotrophic prokaryotes fluxes for bloom averaged fluxes

| Tested | t | p |
|---|---|---|

| | | |
|---|---|---|
| variables | | |
| Phytoplankton spring bloom day 0 | 3.52 | 0.004 |
| Phytoplankton spring bloom day 7 | 2.87 | 0.01 |
| Phytoplankton spring bloom day 14 | 5 | 0.0005 |
| Phytoplankton spring bloom day 21 | 3.95 | 0.003 |
| Phytoplankton spring bloom day 28 | 4.18 | 0.006 |
| Bacteria summer bloom day 0 | 2.92 | 0.01 |
| Bacteria summer bloom day 7 | 1.94 | 0.08 |
| Bacteria summer bloom day 14 | 0.95 | 0.36 |
| Bacteria summer bloom day 21 | 0.03 | 0.97 |
| Bacteria summer bloom day 28 | 0.79 | 0.45 |
| Phytoplankton summer bloom day 0 | 1.66 | 0.13 |
| Phytoplankton summer bloom day 7 | 3.99 | 0.003 |
| Phytoplankton summer bloom day 14 | 2.69 | 0.02 |
| Phytoplankton summer bloom day 21 | 1.59 | 0.15 |
| Phytoplankton summer bloom day 28 | 0.95 | 0.39 |

Differences in trends of instantaneous fluxes between bloom types

Linear models to test for increases/decreases of phytoplankton>heterotrophic prokaryotes or heterotrophic prokaryotes>heterotrophic prokaryotes fluxes over time

| Tested variables | Formula | $R^2$ | p |
|---|---|---|---|
| Bacteria summer bloom heterotrophic prokaryotes>heterotrophic prokaryotes | Y=0.11+0.00081x | 0.011 | 0.56 |
| Bacteria summer bloom phytoplankton>heterotrophic prokaryotes | Y=0.25-0.005x | 0.15 | 0.016 |
| Phytoplankton summer bloom heterotrophic prokaryotes>heterotrophic prokaryotes | Y=0.07-0.00016x | 0.002 | 0.84 |
| Phytoplankton summer bloom phytoplankton>heterotrophic prokaryotes | Y=0.1+0.003x | 0.09 | 0.13 |
| Phytoplankton spring bloom heterotrophic prokaryotes>heterotrophic prokaryotes | Y=0.02+0.0005x | 0.05 | 0.23 |
| Phytoplankton spring bloom phytoplankton>heterotrophic prokaryotes | Y=0.1+0.0017x | 0.08 | 0.13 |

**(E) Differences in concentration normalized heterotrophy rates:**

Test for homogeneity of variances of the concentration normalized heterotrophy rates: Levene test

| Tested variables | F | p |
|---|---|---|
| Concentration normalized heterotrophy rate~ bloom types | 0.41 | 0.67 |

ANOVA and posthoc test for differences in the concentration normalized heterotrophy rate (heterotrophy rate: mol C consumed per day by mol C of pro. hetteria)

| Tested variables | F | p | TukeyHSD |
|---|---|---|---|
| concentration | 3.67 | 0.047 | Phytoplankton spring bloom = b, bacteria summer bloom = a, phytoplankton summer bloom = ab |

Supplementary Table 5: Correlations between heterotrophic prokaryotes>heterotrophic prokaryotes fluxes and occurrence of successions of abundant 16S ASVs.

| Year | Flux het. prokaryotes>het. prokaryotes µmol C l-1 d-1 | Successions of abundant 16S ASVs |
|---|---|---|
| 2012 | 0.176007 | partly |
| 2013 | 0.264127 | yes |
| 2014 | 0.105966 | partly |
| 2015 | 0.063408 | no |
| 2016 | 0.127775 | partly/yes |
| 2017 | 0.076782 | no/partly |
| 2018 | 0.158104 | yes/partly |

Supplementary Table 9: Components with associated names, groups and compartments used in the English Chanel application.

| icom | input | name | genus/group | compartment |
|---|---|---|---|---|
| 1 | nox | reactive nitrogen oxide | $NO_2 + NO_3$ | nutrients |
| 2 | nh4 | ammonium | - | nutrients |
| 3 | po4 | phosphate | - | nutrients |
| 4 | sil | silicate | - | nutrients |
| 11 | p01 | - | hypothetical phytoplankton | 18S/phytoplankton |
| 12 | syn | Synechococcus | phototrophic cyanobacteria | 18S/phytoplankton |
| 13 | pte | Prorocentrum texanum | Dinophyceae | 18S/phytoplankton |
| 14 | mpu | Micromonas pusilla | Chlorophyta | 18S/phytoplankton |
| 15 | bpr | Bathycoccus prasinos | Chlorophyta | 18S/phytoplankton |
| 16 | lch | Lepidodinium chlorophorum | Dinophyceae | 18S/phytoplankton |
| 17 | ppo | Phaeocystis pouchetii | haptophyceae | 18S/phytoplankton |
| 18 | tam | Teleaulax amphioxeia | cryptophyta | 18S/phytoplankton |
| 19 | kve | Karlodinium veneficum | Dinophyceae | 18S/phytoplankton |
| 20 | kse | Karenia selliformis | Dinophyceae | 18S/phytoplankton |
| 21 | gcr | Geminigera cryophila | cryptophyta | 18S/phytoplankton |
| 22 | str | Scrippsiella trochoidea | Dinophyceae | 18S/phytoplankton |
| 23 | tal | Thalassiosira allenii | Bacillariophyta | 18S/phytoplankton |
| 24 | rse | Rhizosolenia setigera | Bacillariophyta | 18S/phytoplankton |
| 25 | toc | Thalassiosira oceanica | Bacillariophyta | 18S/phytoplankton |
| 26 | pin | Parvodinium inconspicuum | Dinophyceae | 18S/phytoplankton |
| 27 | aan | Aureococcus anophagefferens | Pelagophyceae | 18S/phytoplankton |
| 28 | ael | Adenoides eludens | Dinophyceae | 18S/phytoplankton |
| 29 | pha | Pheopolykrikos hartmannii | Dinophyceae | 18S/phytoplankton |
| 30 | che | Chaetoceros tenuissimus | Bacillariophyta | 18S/phytoplankton |
| 31 | tcf | Takayama cf. Pulchellum | Dinophyceae | 18S/phytoplankton |
| 32 | gsp | Gonyaulax spinifera | Dinophyceae | 18S/phytoplankton |
| 33 | yye | Yihiella yeosuensis | Dinophyceae | 18S/phytoplankton |
| 34 | pon | Prorocentrum donghaiense | Dinophyceae | 18S/phytoplankton |
| 35 | ezo | Eucampia zodiacus | Bacillariophyta | 18S/phytoplankton |
| 36 | lmi | Leptocylindrus minimus | Bacillariophyta | 18S/phytoplankton |
| 37 | dac | Dinophysis acuminata | Dinophyceae | 18S/phytoplankton |
| 38 | tte | Thalassiosira tenera | Bacillariophyta | 18S/phytoplankton |
| 39 | pdi | Pyramimonas disomata | Chlorophyta | 18S/phytoplankton |
| 40 | cbe | Cymatosira belgica | Bacillariophyta | 18S/phytoplankton |

| | | | | |
|---|---|---|---|---|
| 41 | tfu | Tripos fusus | Dinophyceae | 18S/phytoplankton |
| 42 | pve | Pseudochattonella verruculosa | Dictyochophyceae | 18S/phytoplankton |
| 43 | dia | Dinophysis acuta | Dinophyceae | 18S/phytoplankton |
| 44 | cca | Chrysochromulina campanulifera | Haptophyceae | 18S/phytoplankton |
| 45 | lda | Leptocylindrus danicus | Bacillariophyta | 18S/phytoplankton |
| 46 | lvi | Lepidodinium viride | Dinophyceae | 18S/phytoplankton |
| 47 | fna | Fragilaria nanana | Bacillariophyta | 18S/phytoplankton |
| 48 | isp | Imantonia_cf. Imantonia sp. CCMP1404 | Haptophyceae | 18S/phytoplankton |
| 49 | nar | Navicula arenaria | Bacillariophyta | 18S/phytoplankton |
| 50 | sgr | Skeletonema grevillei | Bacillariophyta | 18S/phytoplankton |
| 51 | csp | Chrysoxys sp. CCMP591 | Chrysophyceae | 18S/phytoplankton |
| 52 | gde | Guinardia delicatula | Bacillariophyta | 18S/phytoplankton |
| 53 | csi | Chrysochromulina simplex | Haptophyceae | 18S/phytoplankton |
| 54 | ppa | Pyramimonas parkeae | Chlorophyta | 18S/phytoplankton |
| 55 | tco | Thalassiosira concaviuscula | Bacillariophyta | 18S/phytoplankton |
| 56 | tpa | Triparma pacifica | Bolidophyceae | 18S/phytoplankton |
| 57 | pcu | Pseudo-nitzschia cuspidata | Bacillariophyta | 18S/phytoplankton |
| 58 | gos | Goniomonas sp. SH-8 | Cryptophyta | 18S/phytoplankton |
| 59 | pel | Pseudopedinella elastica | Dictyochophyceae | 18S/phytoplankton |
| 60 | trt | Tripos tenuis | Dinophyceae | 18S/phytoplankton |
| 61 | psp | Proboscia sp. | Bacillariophyta | 18S/phytoplankton |
| 62 | stu | Stephanopyxis turris | Bacillariophyta | 18S/phytoplankton |
| 63 | lbo | Lauderia borealis | Bacillariophyta | 18S/phytoplankton |
| 64 | pol | Pyramimonas olivacea | Chlorophyta | 18S/phytoplankton |
| 65 | cle | Chrysochromulina leadbeateri | Haptophyceae | 18S/phytoplankton |
| 66 | gsm | Gymnodinium smaydae | Dinophyceae | 18S/phytoplankton |
| 67 | dsp | Dictyocha speculum | Dictyochophyceae | 18S/phytoplankton |
| 68 | ehu | Emiliania huxleyi | Haptophyceae | 18S/phytoplankton |
| 69 | chs | Chaetoceros sp. UNC1415 | Bacillariophyta | 18S/phytoplankton |
| 70 | pau | Pseudo-nitzschia australis | Bacillariophyta | 18S/phytoplankton |
| 71 | pls | Pleurosigma sp. 102 | Bacillariophyta | 18S/phytoplankton |
| 72 | cry | Cryptophyta sp. CCMP2293 | Cryptophyta | 18S/phytoplankton |
| 73 | esp | Esoptrodinium sp. RP | Dinophyceae | 18S/phytoplankton |
| 74 | fdo | Fragilariopsis doliolus | Bacillariophyta | 18S/phytoplankton |
| 75 | tbe | Tenuicylindrus belgicus | Bacillariophyta | 18S/phytoplankton |
| 76 | tro | Thalassiosira rotula | Bacillariophyta | 18S/phytoplankton |
| 77 | cdi | Chaetoceros diadema | Bacillariophyta | 18S/phytoplankton |

| | | | | |
|---|---|---|---|---|
| 78 | ufl | Ulva flexuosa | Chlorophyta | 18S/phytoplankton |
| 79 | php | Phaeomonas parva | Pinguiophyceae | 18S/phytoplankton |
| 80 | pre | Protoceratium reticulatum | Dinophyceae | 18S/phytoplankton |
| 81 | hak | Heterosigma akashiwo | Raphidophyceae | 18S/phytoplankton |
| 82 | lpo | Lingulodinium polyedra | Dinophyceae | 18S/phytoplankton |
| 83 | cs1 | Chaetoceros sp. CCAP 1010/16 | Bacillariophyta | 18S/phytoplankton |
| 84 | tac | Teleaulax acuta | Cryptophyta | 18S/phytoplankton |
| 85 | cro | Chaetoceros rostratus | Bacillariophyta | 18S/phytoplankton |
| 86 | ssp | Spumella sp. GOT220 | Chrysophyceae | 18S/phytoplankton |
| 87 | cde | Chaetoceros debilis | Bacillariophyta | 18S/phytoplankton |
| 88 | phe | Pseliodinium helix | Dinophyceae | 18S/phytoplankton |
| 89 | lco | Leptocylindrus convexus | Bacillariophyta | 18S/phytoplankton |
| 90 | psi | Protodinium simplex | Dinophyceae | 18S/phytoplankton |
| 91 | pne | Pseudo-nitzschia heimii | Bacillariophyta | 18S/phytoplankton |
| 92 | lgr | Leonella granifera | Dinophyceae | 18S/phytoplankton |
| 93 | ctt | Chlorellidium tetrabotrys | Xanthophyceae | 18S/phytoplankton |
| 94 | mco | Micromonas commoda | Chlorophyta | 18S/phytoplankton |
| 95 | gca | Gymnodinium catenatum | Dinophyceae | 18S/phytoplankton |
| 96 | sha | Scrippsiella hangoei | Dinophyceae | 18S/phytoplankton |
| 97 | mpo | Margalefidinium polykrikoides | Dinophyceae | 18S/phytoplankton |
| 98 | vac | Vacuolaria | Raphidophyceae | 18S/phytoplankton |
| 99 | tcu | Thalassiosira curviseriata | Bacillariophyta | 18S/phytoplankton |
| 100 | cwi | Chaetoceros cf. Wighamii | Bacillariophyta | 18S/phytoplankton |
| 101 | pge | Polykrikos geminatum | Dinophyceae | 18S/phytoplankton |
| 102 | ha2 | Haptophyceae sp. EE-2014 | Haptophyceae | 18S/phytoplankton |
| 103 | han | Hemiselmis andersenii | Cryptophyta | 18S/phytoplankton |
| 104 | cs2 | Coscinodiscus sp. GGM-2004 | Bacillariophyta | 18S/phytoplankton |
| 105 | acu | Azadinium cuneatum | Dinophyceae | 18S/phytoplankton |
| 106 | rsh | Rhizosolenia shrubsolei | Bacillariophyta | 18S/phytoplankton |
| 107 | ps2 | Plagiolemma sp. NC-2018a | Bacillariophyta | 18S/phytoplankton |
| 108 | cc2 | Crypthecodinium sp. CAAE-CL2 | Dinophyceae | 18S/phytoplankton |
| 109 | pya | Prymnesium parvum | Haptophyceae | 18S/phytoplankton |
| 110 | cte | Cymbomonas tetramitiformis | Chlorophyta | 18S/phytoplankton |
| 111 | mgi | Mamiella gilva | Chlorophyta | 18S/phytoplankton |
| 112 | cci | Chaetoceros cinctus | Bacillariophyta | 18S/phytoplankton |
| 113 | tch | Trieres chinensis | Bacillariophyta | 18S/phytoplankton |
| 114 | pan | Phaeocystis | Haptophyceae | 18S/phytoplankton |

| | | | | |
|---|---|---|---|---|
| | | antarctica | | |
| 115 | rat | Rhodomonas atrorosea | Cryptophyta | 18S/phytoplankton |
| 116 | ns1 | Nannochloris sp. MBIC10055 | Chlorophyta | 18S/phytoplankton |
| 117 | dno | Dinophysis norvegica | Dinophyceae | 18S/phytoplankton |
| 118 | lfi | Levanderina fissa | Dinophyceae | 18S/phytoplankton |
| 119 | cra | Chaetoceros radicans | Bacillariophyta | 18S/phytoplankton |
| 120 | cwe | Conticribra weissflogii | Bacillariophyta | 18S/phytoplankton |
| 121 | chy | Corethron hystrix | Bacillariophyta | 18S/phytoplankton |
| 122 | mlo | Madanidinium loirii | Dinophyceae | 18S/phytoplankton |
| 123 | rsi | Rhizosolenia similoides | Bacillariophyta | 18S/phytoplankton |
| 124 | ppi | Prymnesium pigrum | Haptophyceae | 18S/phytoplankton |
| 125 | psh | Paragymnodinium shiwhaense | Dinophyceae | 18S/phytoplankton |
| 126 | pts | Pterocystis sp. | Acanthocystidae | 18S/phytoplankton |
| 127 | cpu | Ceratodon purpureus | Streptophyta | 18S/phytoplankton |
| 128 | pbl | Prototheca blaschkeae | Chlorophyta | 18S/phytoplankton |
| 129 | asp | Amphora sp. 38 | Bacillariophyta | 18S/phytoplankton |
| 130 | cpa | Cyanophora paradoxa | Glaucocystophyceae | 18S/phytoplankton |
| 131 | pio | Prototheca eriobotryae | Chlorophyta | 18S/phytoplankton |
| 132 | ale | Alexandrium | Dinophyceae | 18S/phytoplankton |
| 133 | och | Ochromonas | Chrysophyceae | 18S/phytoplankton |
| 134 | ost | Ostreococcus | Chlorophyta | 18S/phytoplankton |
| 135 | da2 | Dactylodinium | Dinophyceae | 18S/phytoplankton |
| 136 | tre | Trebouxiophyceae | Chlorophyta | 18S/phytoplankton |
| 137 | pry | Prymnesiales | Haptophyceae | 18S/phytoplankton |
| 138 | pep | Pelagophyceae | Pelagophyceae | 18S/phytoplankton |
| 139 | cym | Cymatosiraceae | Bacillariophyta | 18S/phytoplankton |
| 140 | dit | Ditylum | Bacillariophyta | 18S/phytoplankton |
| 141 | cha | Chaetoceros | Bacillariophyta | 18S/phytoplankton |
| 142 | tha | Thalassiosirales | Bacillariophyta | 18S/phytoplankton |
| 143 | ske | Skeletonema | Bacillariophyta | 18S/phytoplankton |
| 144 | dio | Bacillariophyta | Bacillariophyta | 18S/phytoplankton |
| 145 | cph | Chlorophyta | Chlorophyta | 18S/phytoplankton |
| 146 | cy2 | Cryptophyta | Cryptophyta | 18S/phytoplankton |
| 147 | crs | Chrysophyceae | Chrysophyceae | 18S/phytoplankton |
| 148 | din | Dinophyceae | Dinophyceae | 18S/phytoplankton |
| 149 | hap | Haptophyceae | Haptophyceae | 18S/phytoplankton |
| 150-172 | h35-h57 | - | hypothetical phytoplankton species | 18S/phytoplankton |
| 174 | b01 | - | hypothetical 16S ASV | 16S/heterotrophic prokaryotes |
| 175,178,180,181,206,211,250,257,269,279,282,305,322,325,329 | s11-s23 | SAR11 | Alphaproteo-bacteriaa | 16S/heterotrophic prokaryotes |
| 176,182,215,2 | s86-s93 | SAR86 | Gammaproteo- | 16S/heterotrophic |

| | | | | |
|---|---|---|---|---|
| 31,232,245,255,272 | | | bacteria | prokaryotes |
| 177 | amy | Amylibacter | Alphaproteo-bacteriaa | 16S/heterotrophic prokaryotes |
| 179 | act | Actinomarina | Actinobacteria | 16S/heterotrophic prokaryotes |
| 183,284,288,289 | n04-n07 | NS4 marine group | Bacteroidetes | 16S/heterotrophic prokaryotes |
| 184 | pla | Planktomarina | Alphaproteo-bacteriaa | 16S/heterotrophic prokaryotes |
| 185,233,236,237,256,286,298,301,336 | n51-n59 | NS5 marine group | Bacteroidetes | 16S/heterotrophic prokaryotes |
| 186 | are | Arenicellaceae | Gammaproteo-bacteria | 16S/heterotrophic prokaryotes |
| 187,190,203,235,238,247,251,264,281,294,297,317,326 | m21-m33 | Marine group II archaea | Euryarchaeota | 16S/heterotrophic prokaryotes |
| 188,202,290 | pu1-pu3 | SAR116 clade | Alphaproteo-bacteriaa | 16S/heterotrophic prokaryotes |
| 189,222,310 | cr1-cr3 | Cryomorphoceae | Bacteroidetes | 16S/heterotrophic prokaryotes |
| 191,266,306 | o43-o45 | OM43 clade | Betaproteobacteriales | 16S/heterotrophic prokaryotes |
| 192 | ros | Roseobacter | Alphaproteo-bacteriaa | 16S/heterotrophic prokaryotes |
| 193 | ns2 | NS2b marine group | Bacteroidetes | 16S/heterotrophic prokaryotes |
| 194,260 | te1, te2 | Tenacibaculum | Bacteroidetes | 16S/heterotrophic prokaryotes |
| 195,242,263,278,291,299,304,318 | ma1-ma8 | Marinimicrobia | SAR406 clade | 16S/heterotrophic prokaryotes |
| 196,204 | th1, th2 | Thioglobaceae | Gammaproteo-bacteria | 16S/heterotrophic prokaryotes |
| 197 | sa2 | SAR92 clade | Gammaproteo-bacteria | 16S/heterotrophic prokaryotes |
| 198 | pav | Parvibaculales | Alphaproteo-bacteriaa | 16S/heterotrophic prokaryotes |
| 199,225,270 | m03-m05 | Marine group III archaea | Euryarchaeota | 16S/heterotrophic prokaryotes |
| 200,221 | pl1, pl2 | Polaribacter | Bacteroidetes | 16S/heterotrophic prokaryotes |
| 201 | fla | Flavobacteriaceae | Bacteroidetes | 16S/heterotrophic prokaryotes |
| 205 | ulv | Ulvibacter | Bacteroidetes | 16S/heterotrophic prokaryotes |
| 207,218,311,315 | om6-om9 | OM60(NOR5) clade | Gammaproteo-bacteria | 16S/heterotrophic prokaryotes |
| 208 | psa | Pseudoalteromonas | Gammaproteo-bacteria | 16S/heterotrophic prokaryotes |
| 209 | lum | Luminiphilus | Gammaproteo-bacteria | 16S/heterotrophic prokaryotes |
| 210,234 | fo1, fo2 | Formosa | Bacteroidetes | 16S/heterotrophic prokaryotes |
| 212,273 | fu1, fu2 | Fluviicola | Bacteroidetes | 16S/heterotrophic prokaryotes |
| 213,216,220,312 | ae1-ae4 | AEGEAN-169 | Alphaproteo-bacteria | 16S/heterotrophic prokaryotes |

| | | | | |
|---|---|---|---|---|
| 214 | sa3 | SAR92 clade | Gammaproteo-bacteria | 16S/heterotrophic prokaryotes |
| 217 | dad | Dadabacteriales | Dadabacteria | 16S/heterotrophic prokaryotes |
| 219 | pmy | Pla3 lineage | Planctomycetes | 16S/heterotrophic prokaryotes |
| 223,226,229,244 | ro1-ro4 | Rhodobacteraceae | Alphaproteo-bacteria | 16S/heterotrophic prokaryotes |
| 224 | o75 | OM75 clade | Alphaproteo-bacteria | 16S/heterotrophic prokaryotes |
| 227 | rub | Rubritaleaceae | Verrucomicrobia | 16S/heterotrophic prokaryotes |
| 228 | hoc | HOC36 | Gammaproteo-bacteria | 16S/heterotrophic prokaryotes |
| 230 | leb | Lentibacter | Alphaproteo-bacteria | 16S/heterotrophic prokaryotes |
| 239 | n3a | NS3a marine group | Bacteroidetes | 16S/heterotrophic prokaryotes |
| 240,287,293,296 | n91-n94 | NS9 marine group | Bacteroidetes | 16S/heterotrophic prokaryotes |
| 241,246,248,259,327 | po1-po3, po9, po5 | Pseudohongiella | Gammaproteo-bacteria | 16S/heterotrophic prokaryotes |
| 243 | aqu | Aquibacter | Bacteroidetes | 16S/heterotrophic prokaryotes |
| 249 | per | Persicirhabdus | Verrucomicrobia | 16S/heterotrophic prokaryotes |
| 252 | oce | Nitrincolaceae | Gammaproteo-bacteria | 16S/heterotrophic prokaryotes |
| 253 | flc | Flavicella | Bacteroidetes | 16S/heterotrophic prokaryotes |
| 254 | hyp | Hellea | Alphaproteo-bacteria | 16S/heterotrophic prokaryotes |
| 258 | n7m | NS7 marine group | Bacteroidetes | 16S/heterotrophic prokaryotes |
| 261,328 | ub1, ub2 | UBA10353 marine group | Gammaproteo-bacteria | 16S/heterotrophic prokaryotes |
| 262 | mas | Marinoscillum | Bacteroidetes | 16S/heterotrophic prokaryotes |
| 265,276 | o18, o19 | OM182 clade | Gammaproteo-bacteria | 16S/heterotrophic prokaryotes |
| 267 | vib | Vibrio sp. hMe3-9 | Gammaproteo-bacteria | 16S/heterotrophic prokaryotes |
| 268,316 | s32, del | SAR324 | Deltaproteobacteria | 16S/heterotrophic prokaryotes |
| 271 | ect | Ectothiorhodospiraceae | Gammaproteo-bacteria | 16S/heterotrophic prokaryotes |
| 274,277,331 | pi1-pi3 | Pirrelulaceae | Planctomycetes | 16S/heterotrophic prokaryotes |
| 275 | sul | Sulfitobacter sp. | Alphaproteo-bacteria | 16S/heterotrophic prokaryotes |
| 280,308,313 | al1-al3 | Alphaproteo-bacteria | Alphaproteo-bacteria | 16S/heterotrophic prokaryotes |
| 283 | nmo | Nitrosomonadaceae | Gammaproteo-bacteria | 16S/heterotrophic prokaryotes |
| 285 | pga | Paraglaciecola | Gammaproteo-bacteria | 16S/heterotrophic prokaryotes |
| 292,314 | at1, at2 | Alteromonas sp. | Gammaproteo-bacteria | 16S/heterotrophic prokaryotes |
| 295 | ki1 | KI89A clade | Gammaproteo-bacteria | 16S/heterotrophic prokaryotes |

| | | | | |
|---|---|---|---|---|
| 300 | fac | Flavicella | Bacteroidetes | 16S/heterotrophic prokaryotes |
| 302 | wos | Woeseia | Gammaproteo-bacteria | 16S/heterotrophic prokaryotes |
| 303 | ps1 | PS1 clade | Alphaproteo-bacteria | 16S/heterotrophic prokaryotes |
| 307 | ga1 | Gammaproteo-bacteria | Gammaproteo-bacteria | 16S/heterotrophic prokaryotes |
| 309 | clo | SAR202 clade | Chloroflexi | 16S/heterotrophic prokaryotes |
| 319 | lem | Lentimonas | Verrucomicrobia | 16S/heterotrophic prokaryotes |
| 320 | sap | Saprospiraceae | Bacteroidetes | 16S/heterotrophic prokaryotes |
| 321 | psm | Pseudomonas | Gammaproteo-bacteria | 16S/heterotrophic prokaryotes |
| 323 | slp | Salinisphaera | Gammaproteo-bacteria | 16S/heterotrophic prokaryotes |
| 324 | ths | Thalassospira sp. | Alphaproteo-bacteria | 16S/heterotrophic prokaryotes |
| 332 | b02-b05 | - | hypothetical 16S ASV | 16S/heterotrophic prokaryotes |
| 336-497 | a01-a99, c01-c63 | - | hypothetical POM species | POM |
| 503-664 | d01-d99, f01-f63 | - | hypothetical DOM species | DOM |
