## Supplementary Data 1 for "Heterotrophic prokaryotes internal carbon recycling compensates mismatches between phytoplankton production and heterotrophic prokaryotic consumption"

Supplementary Data 1: Model-data comparsions for nutrients, sum-parameters (phytoplankton, heterotrophic prokaryotes, etc), and each individual phytoplankton OTU and heterotrophic prokaryotic ASV for the years 2012-2018. Each year has 303 pages, i.e. year 2012: page 2-304, year 2013: page 305-607, year 2014: page 608-910, year 2015: page 911-1213, year 2016: page 1214-1516, year 2017: page 1517-1819, and year 2018: page 1820-2122. The first page for each year is an overview of the most important parameters (nutrients, chlorophyll, heterotrophic prokaryotes, POM, DOM), pages 2-5 are nutrients, page 6: summed up phytoplankton OTUs, pages 7-144: single phytoplankton OTUs, page 145: summed up Bacillariophyceae (BAP), pages 146-302: single heterotrophic prokaryotic ASVs, page 303: summed up heterotrophic prokaryotic ASVs.

324_Calib_S_A

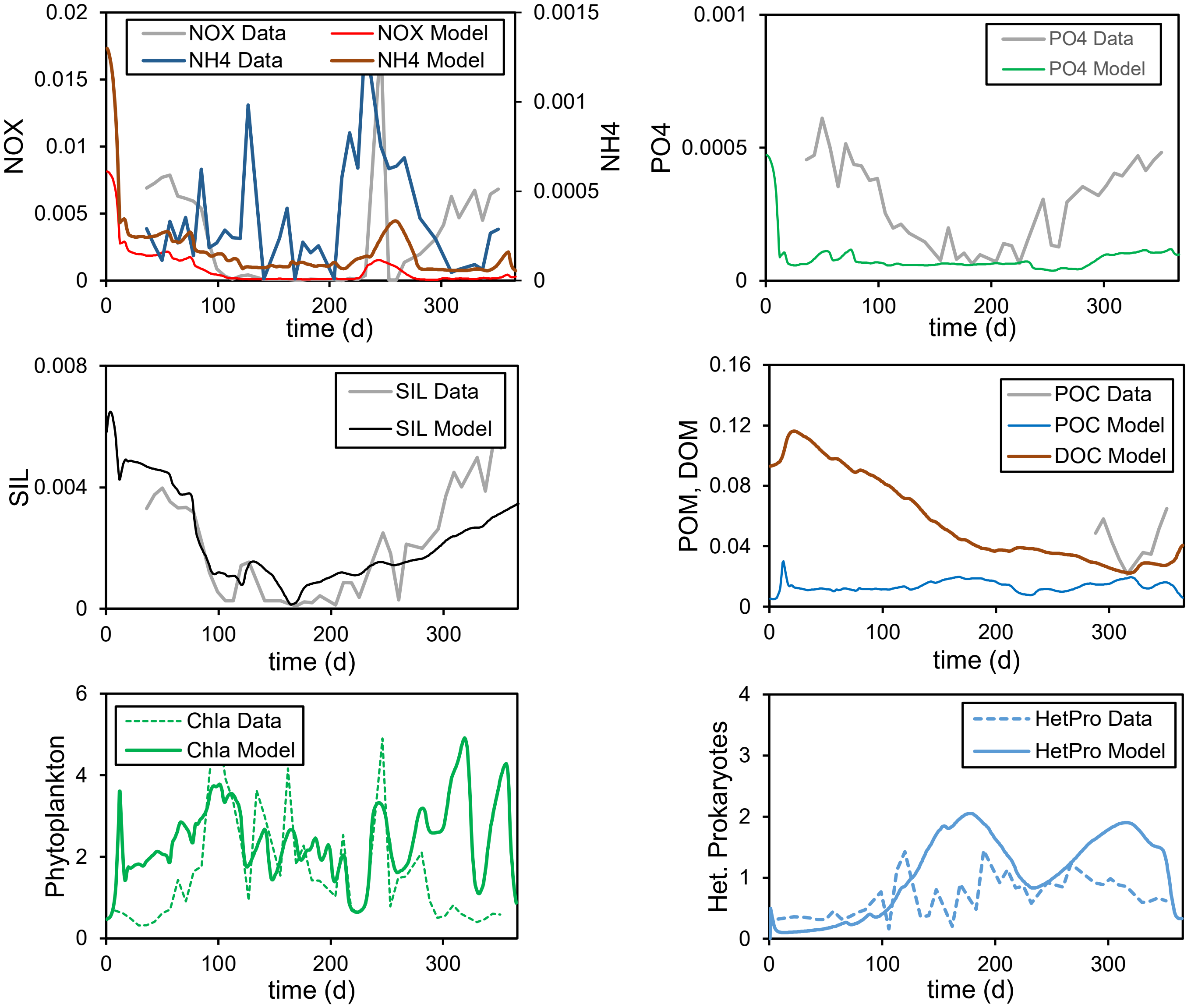
NOX Data
NOX Model
NH4 Data
NH4 Model
NOX
NH4
time (d)
0.02
0.015
0.01
0.005
0
0.0015
0.001
0.0005
0
100
200
300
PO4 Data
PO4 Model
PO4
0.001
0.0005
time (d)
SIL Data
SIL Model
SIL
0.008
0.004
time (d)
POC Data
POC Model
DOC Model
POM, DOM
0.16
0.12
0.08
0.04
time (d)
Chla Data
Chla Model
Phytoplankton
6
4
2
time (d)
HetPro Data
HetPro Model
Het. Prokaryotes
4
3
2
1
time (d)

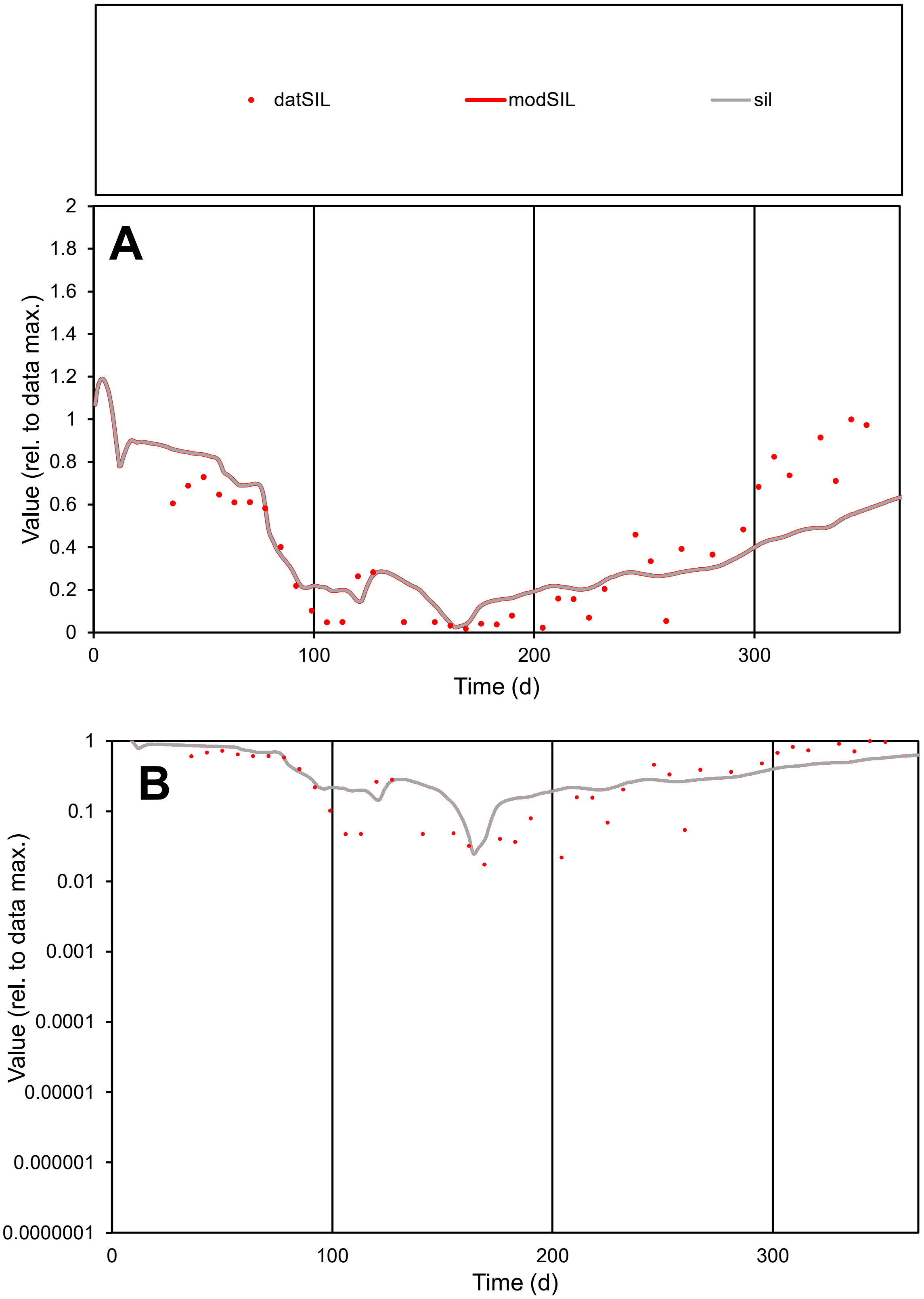


**C1**

Variables
SIL Provide jtd or Dst
CalibOnly?
2 1 to reproduce of
oNormalize?
2 1=no, 2=yes

ofxn calc*
1.45675
ofxn mod
1.4996

sRun
RunA1
zout
1

Ave
Data
0.00205
Model
0.00195

data/vMax
0.00545

ofxng
321.551

Min d/m
0.66387

ave/min d
21.5731
ave/min m
13.643

324_Calib_S_A

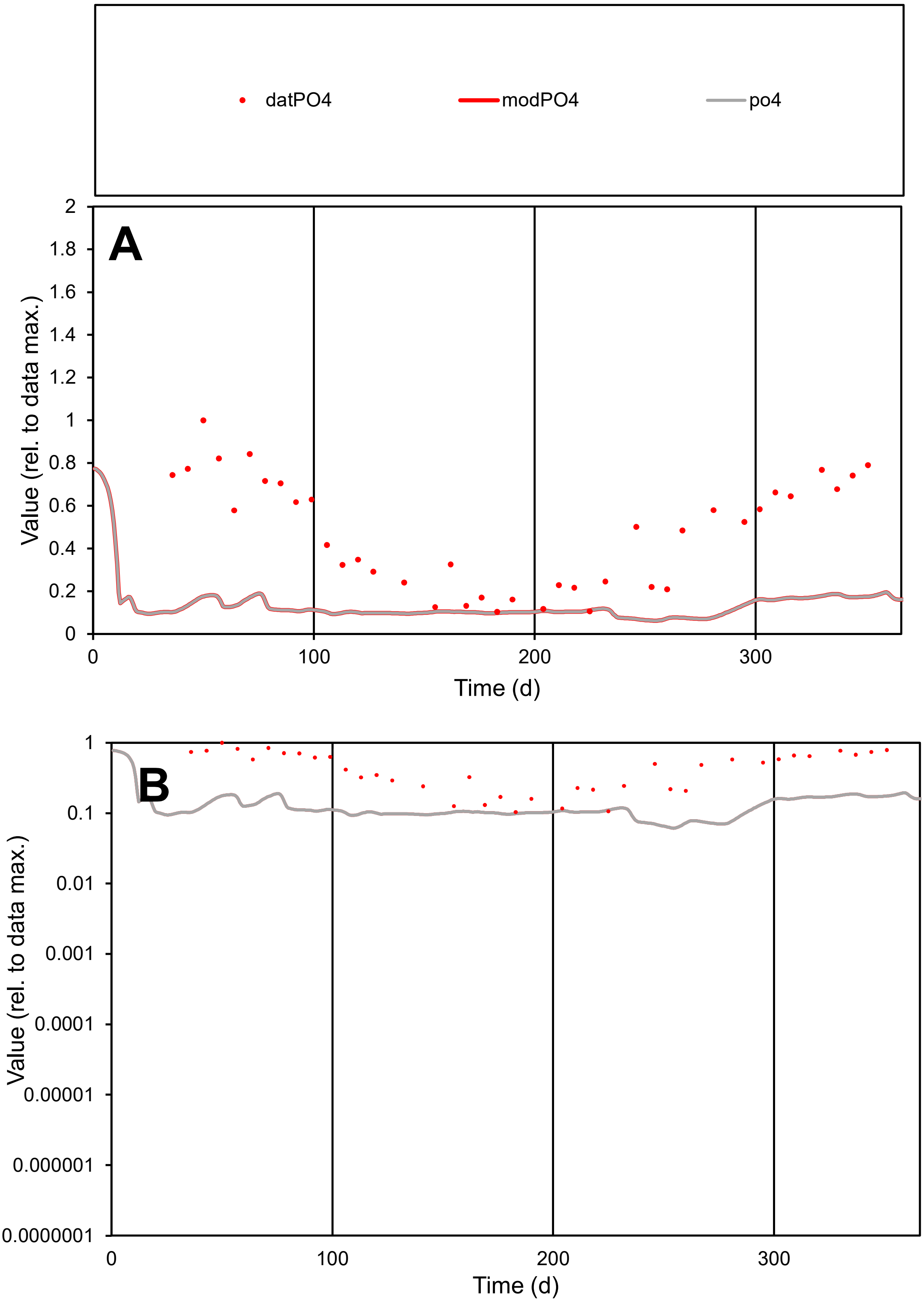


**C1**

Variables
PO4 Provide jtd or Dst
CalibOnly?
2 1 to reproduce of
oNormalize?
2 1=no, 2=yes

ofxn calc*
2.67649
ofxn mod
2.68748

sRun
RunA1
zout
1

Ave
Data
0.00029
Model
7.4E-05

data/vMax
0.00061

ofxng
321.551

Min d/m
1.68738

ave/min d
4.54642
ave/min m
1.97555

324_Calib_S_A

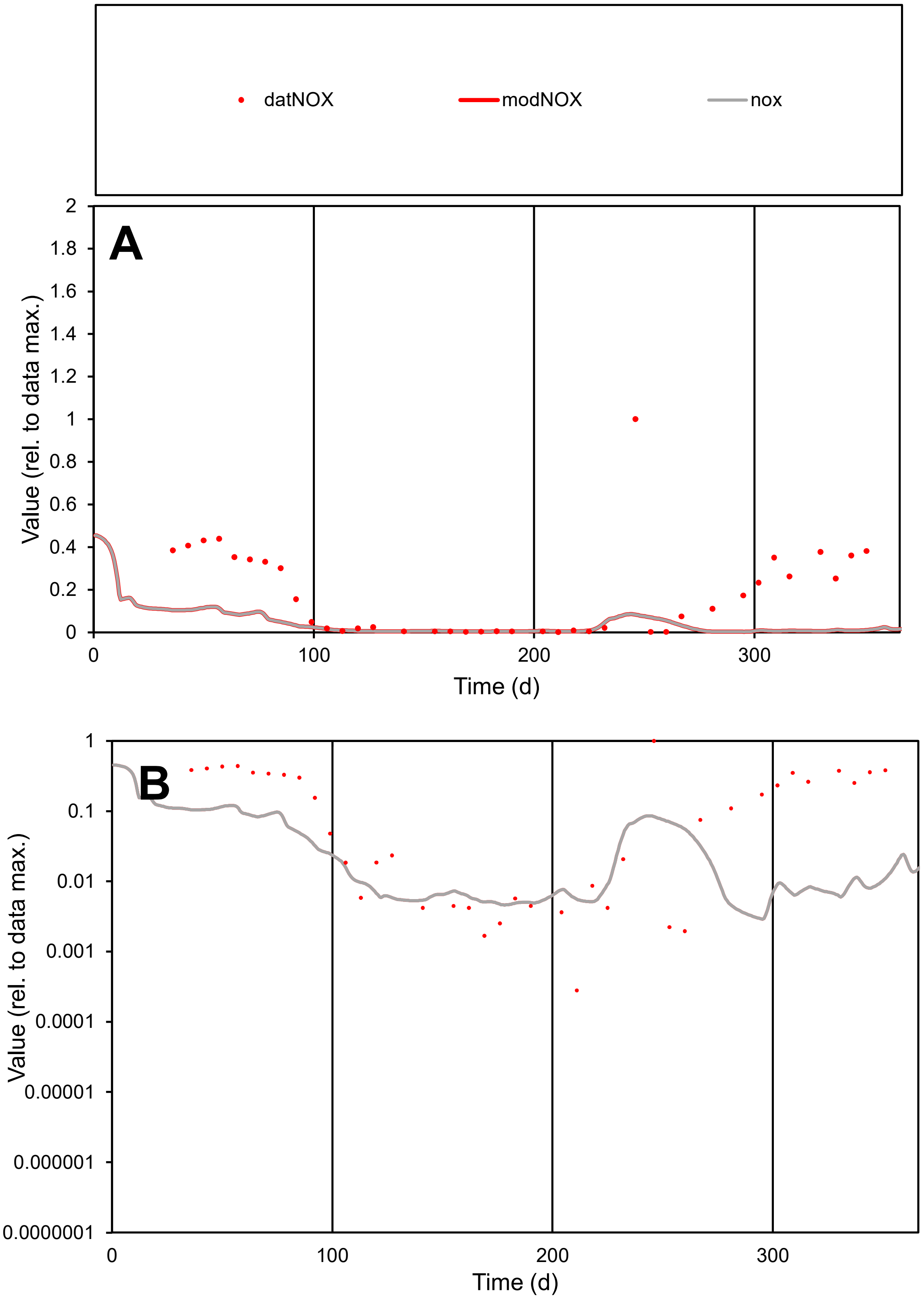


**C1**

Variables
NOX Provide jtd or Dst
CalibOnly?
2 1 to reproduce of
oNormalize?
2 1=no, 2=yes

ofxn calc*
4.06308
ofxn mod
4.07732

sRun
RunA1
zout
1

Ave
Data
0.00316
Model
0.00056

data/vMax
0.01789

ofxng
321.551

Min d/m
0.09463

ave/min d
632.862
ave/min m
10.6686

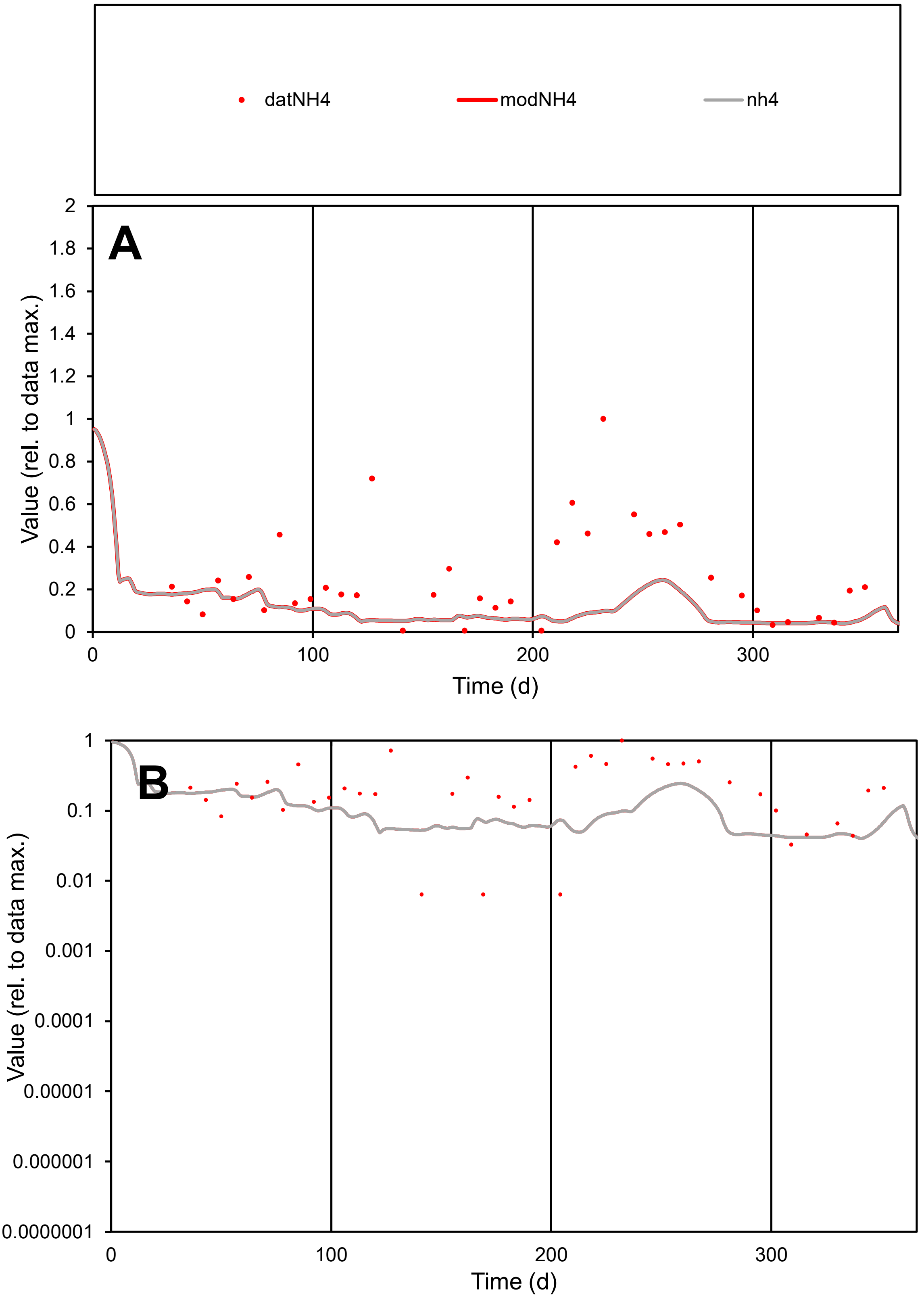


**C1**

Variables
NH4 Provide jtd or Dst
CalibOnly?
2 1 to reproduce of
oNormalize?
2 1=no, 2=yes

ofxn calc*
3.06601
ofxn mod
3.07966

sRun
RunA1
zout
1

Ave
Data
0.00034
Model
0.00014

data/vMax
0.00137

ofxng
321.551

Min d/m
0.15443

ave/min d
38.8368
ave/min m
2.43053

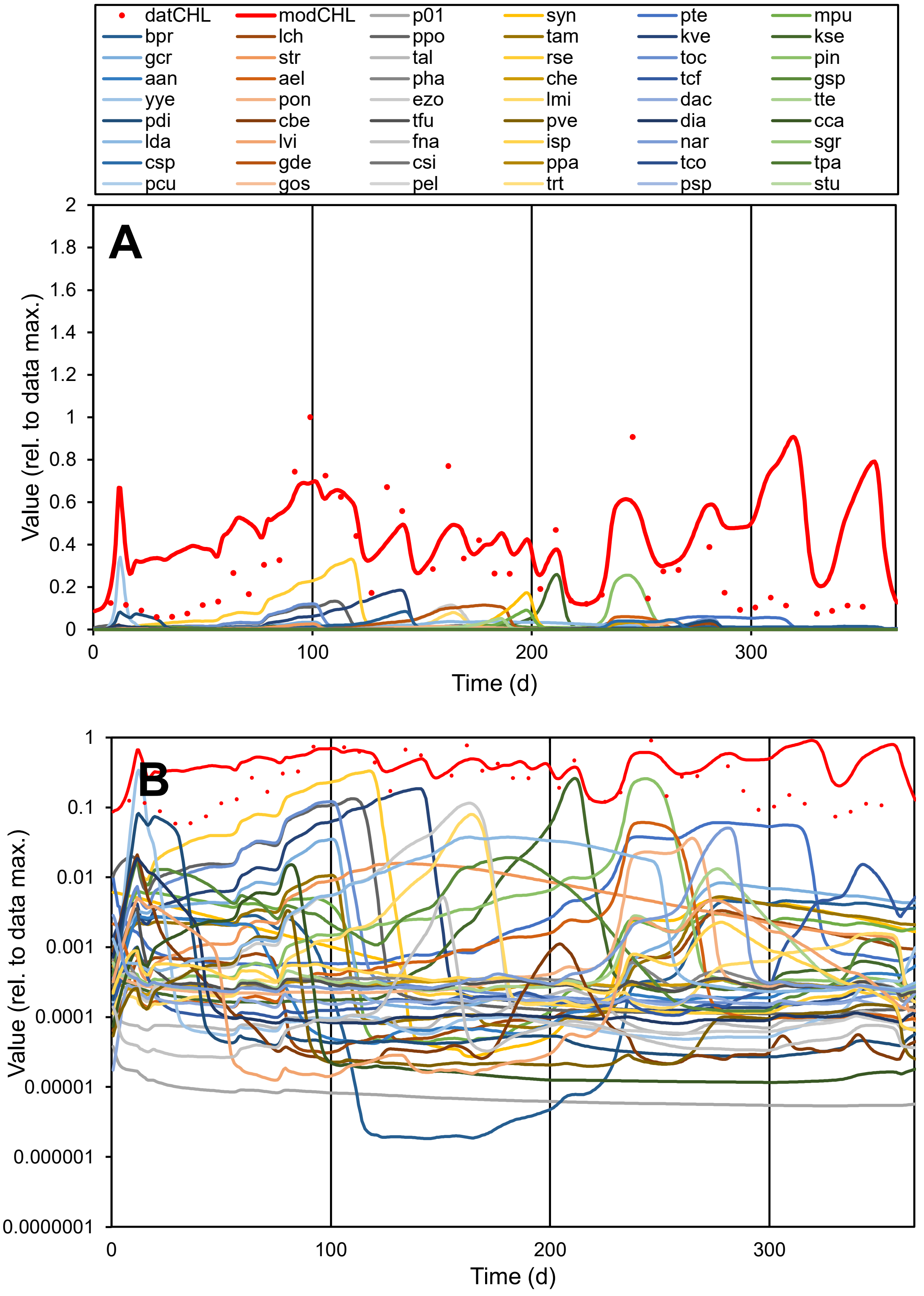


**C1**

Variables

CHL Provide jtd or Dst

CalibOnly?

2 1 to reproduce of

oNormalize?

2 1=no, 2=yes

ofxn calc*

4.05998

ofxn mod

4.31784

sRun

RunA1

zout

1

Ave

Data

1.58547

Model

1.59705

data/vMax

5.413

ofxng

321.551

Min d/m

0.795

ave/min d

4.98575

ave/min m

3.99263

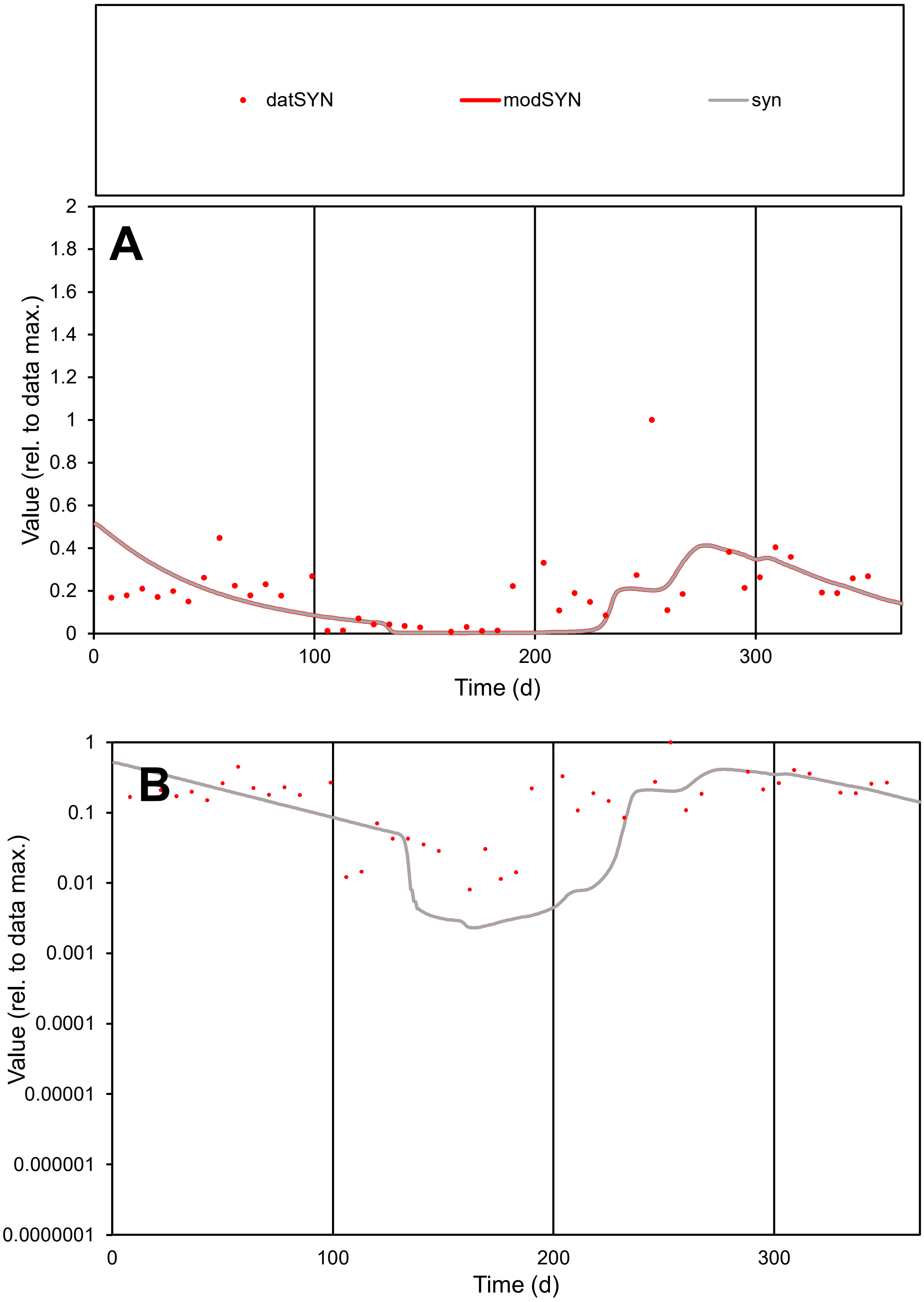


**C1**

Variables
SYN Provide jtd or Dst
CalibOnly?
2 1 to reproduce of
oNormalize?
2 1=no, 2=yes

ofxn calc*
0.87922
ofxn mod
0.88836

sRun
RunA1
zout
1

Ave
Data
7.37757
Model
6.20448

data/vMax
38

ofxng
321.551

Min d/m
3.48877

ave/min d
24.1097
ave/min m
70.7386

324_Calib_S_A

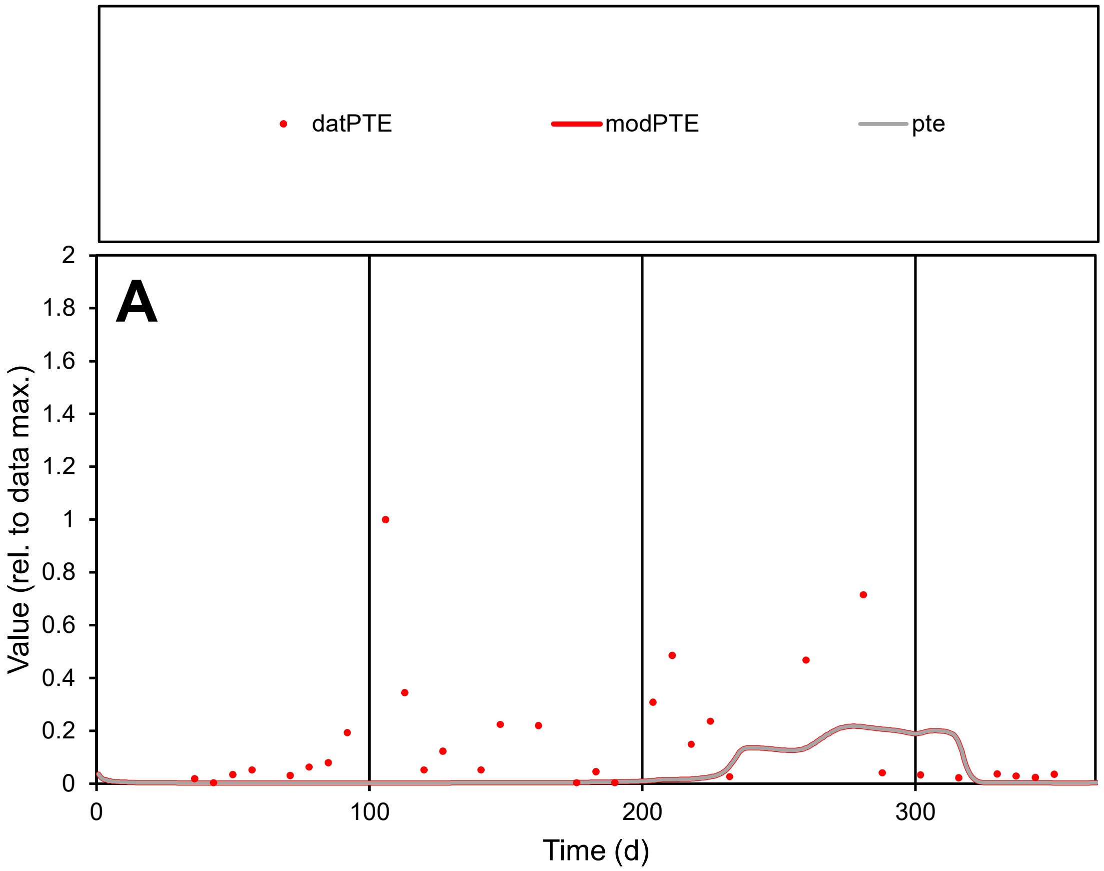


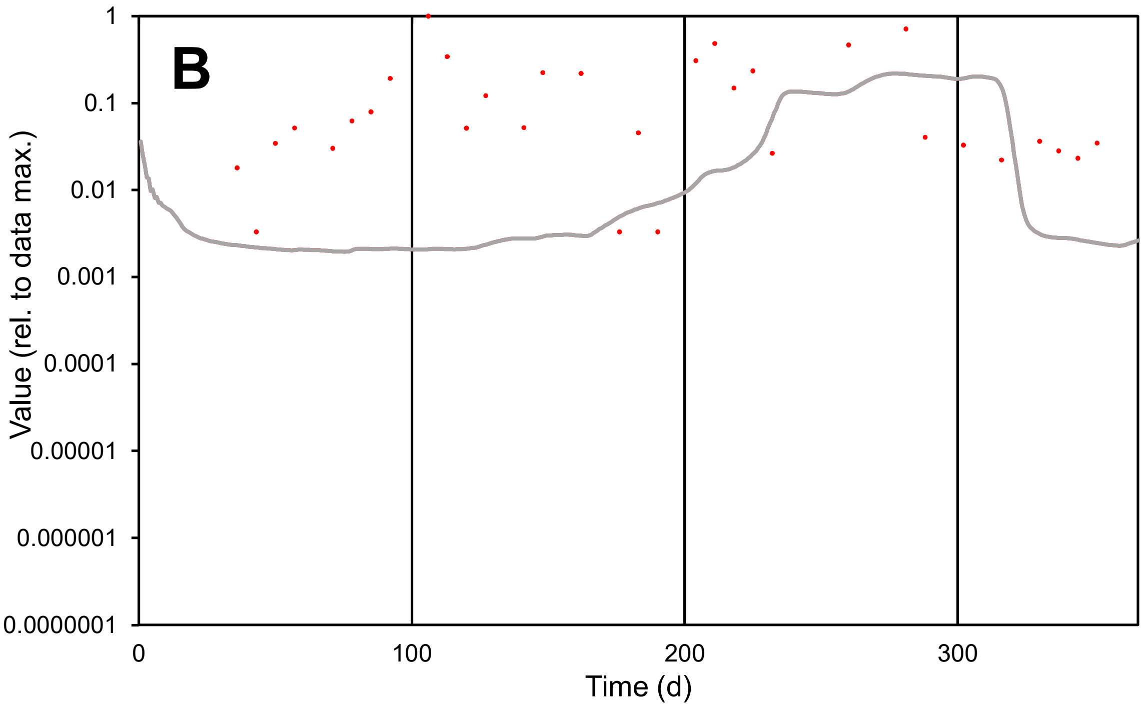


**C1**

Variables
PTE Provide jtd or Dst
CalibOnly?
2 1 to reproduce of
oNormalize?
2 1=no, 2=yes

ofxn calc*
1.58023
ofxn mod
1.58419

sRun
RunA1
zout
1

Ave
Data
0.00041
Model
8.1E-05

data/vMax
0.00252

ofxng
321.551

Min d/m
1.67602

ave/min d
48.4738
ave/min m
16.1986

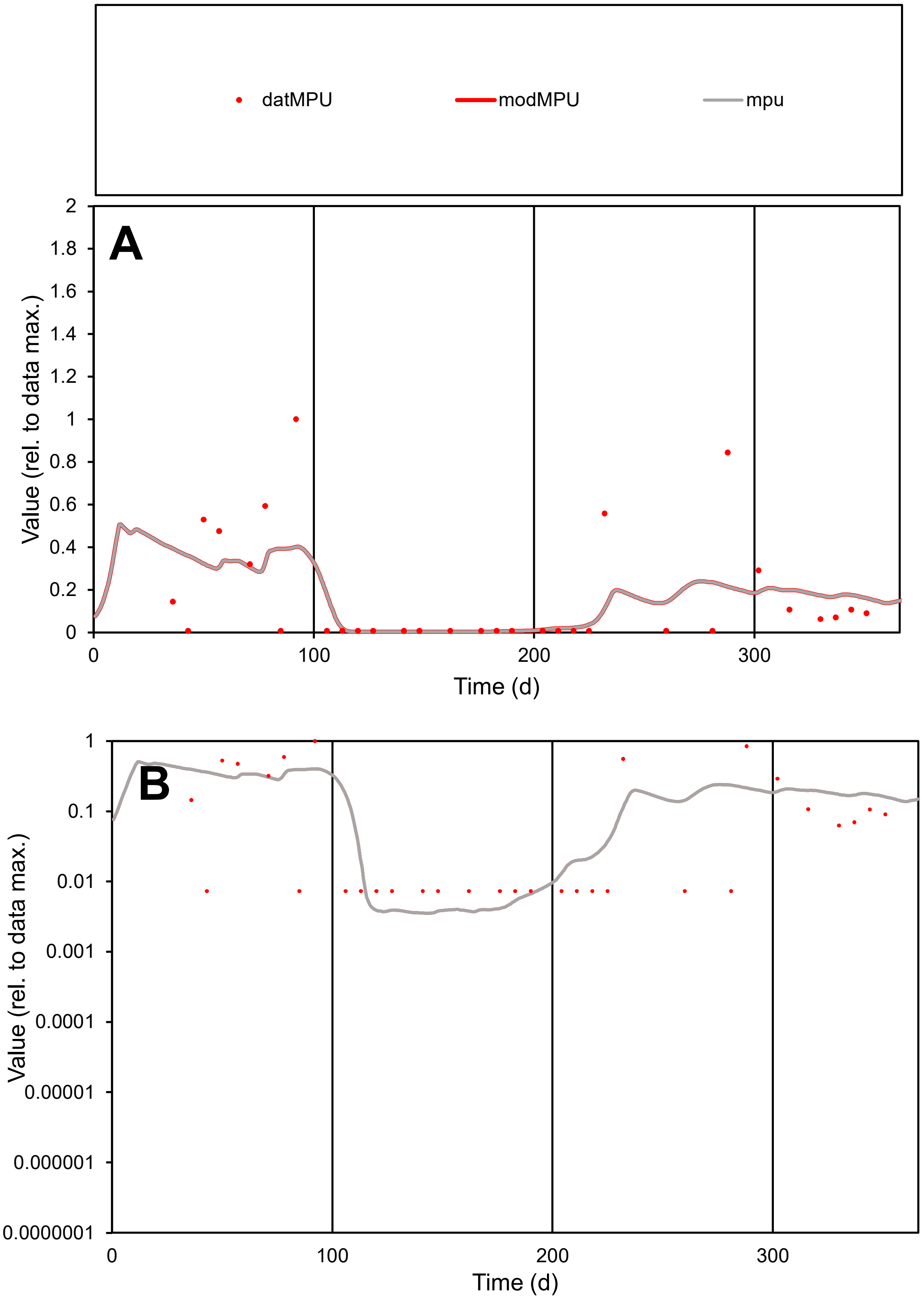


**C1**

Variables
MPU Provide jtd or Dst
CalibOnly?
2 1 to reproduce of
oNormalize?
2 1=no, 2=yes

ofxn calc*
1.30223
ofxn mod
1.31043

sRun
RunA1
zout
1

Ave
Data
9.3E-05
Model
8.5E-05

data/vMax
0.00056

ofxng
321.551

Min d/m
2.07306

ave/min d
22.7615
ave/min m
43.1917

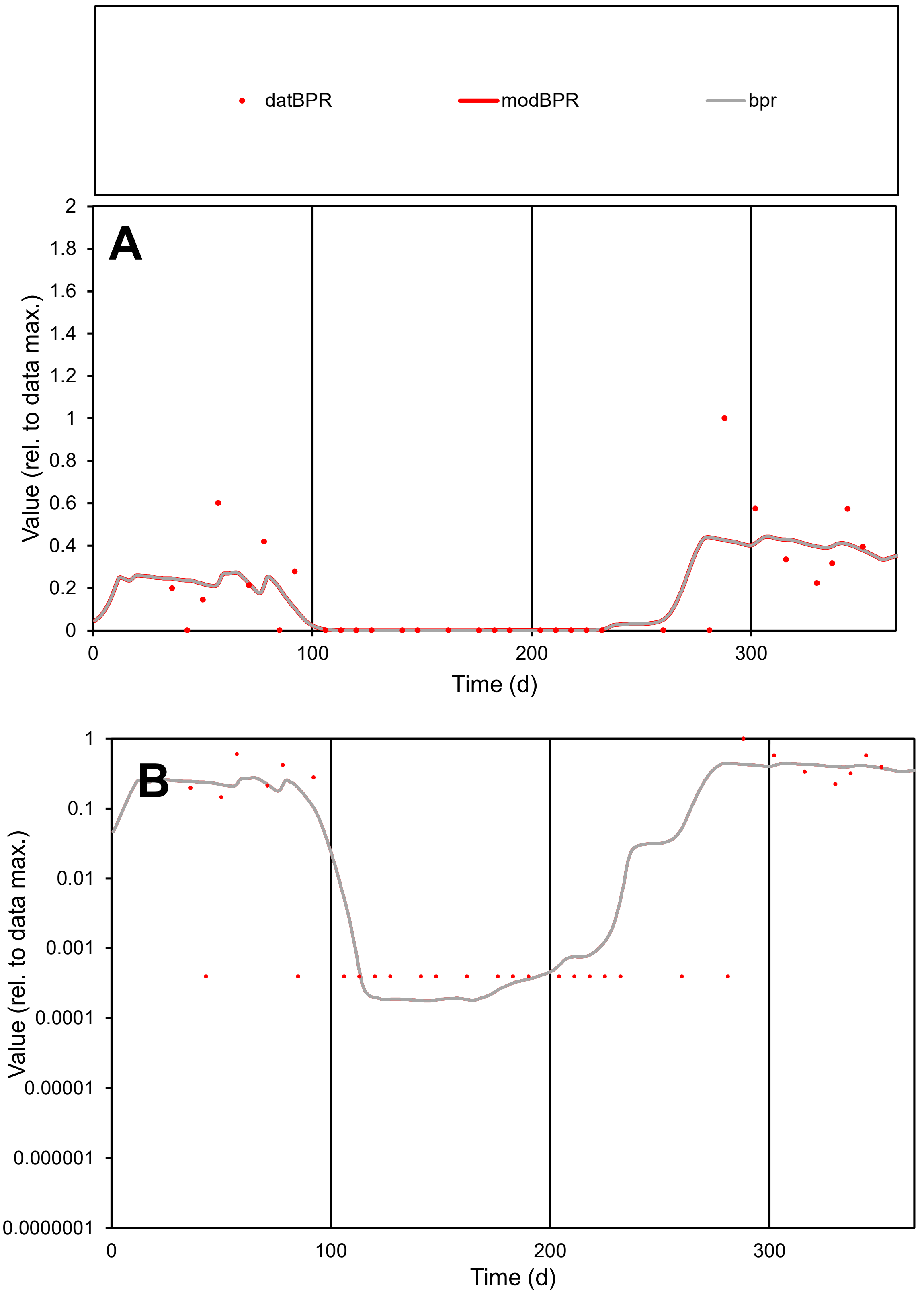


**C1**

Variables

BPR Provide jtd or Dst

CalibOnly?

2 1 to reproduce of

oNormalize?

2 1=no, 2=yes

ofxn calc*

1.00944

ofxn mod

1.02476

sRun

RunA1

zout

1

Ave

Data

7.7E-05

Model

7.3E-05

data/vMax

0.00047

ofxng

321.551

Min d/m

2.24497

ave/min d

417.634

ave/min m

883.266

324_Calib_S_A

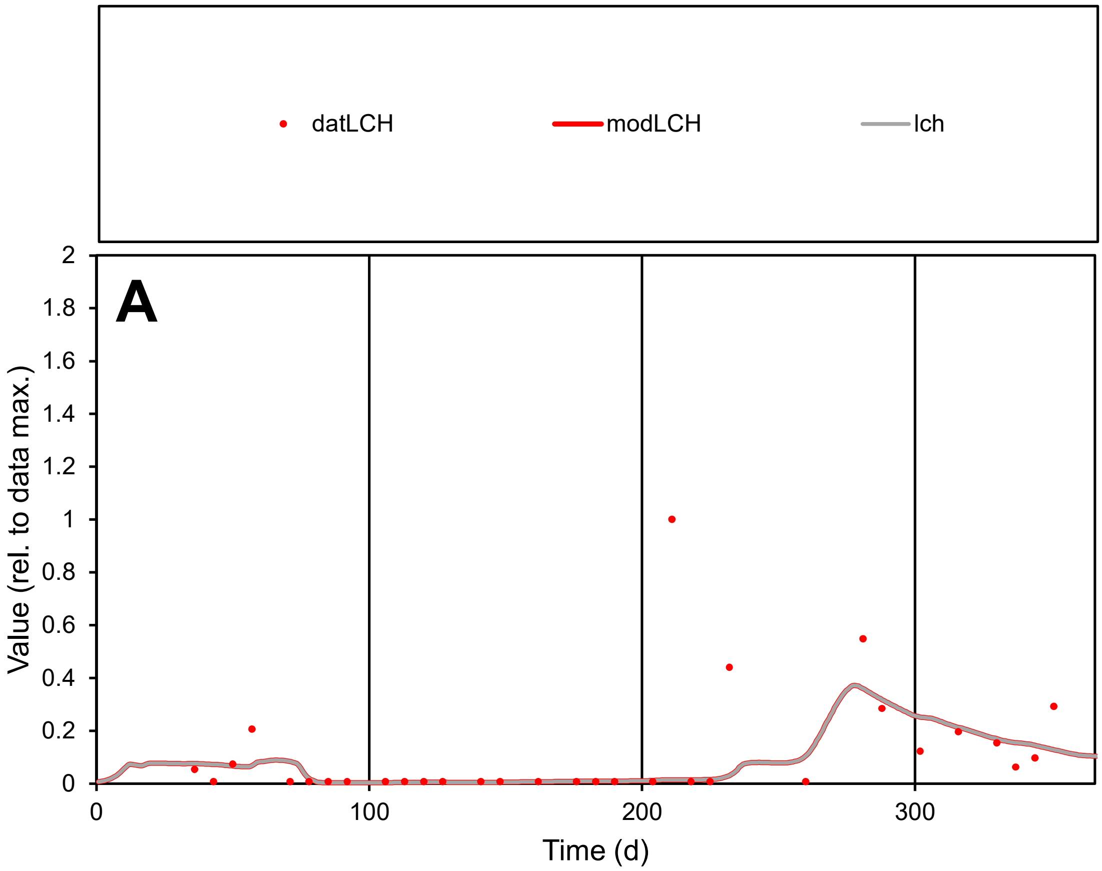


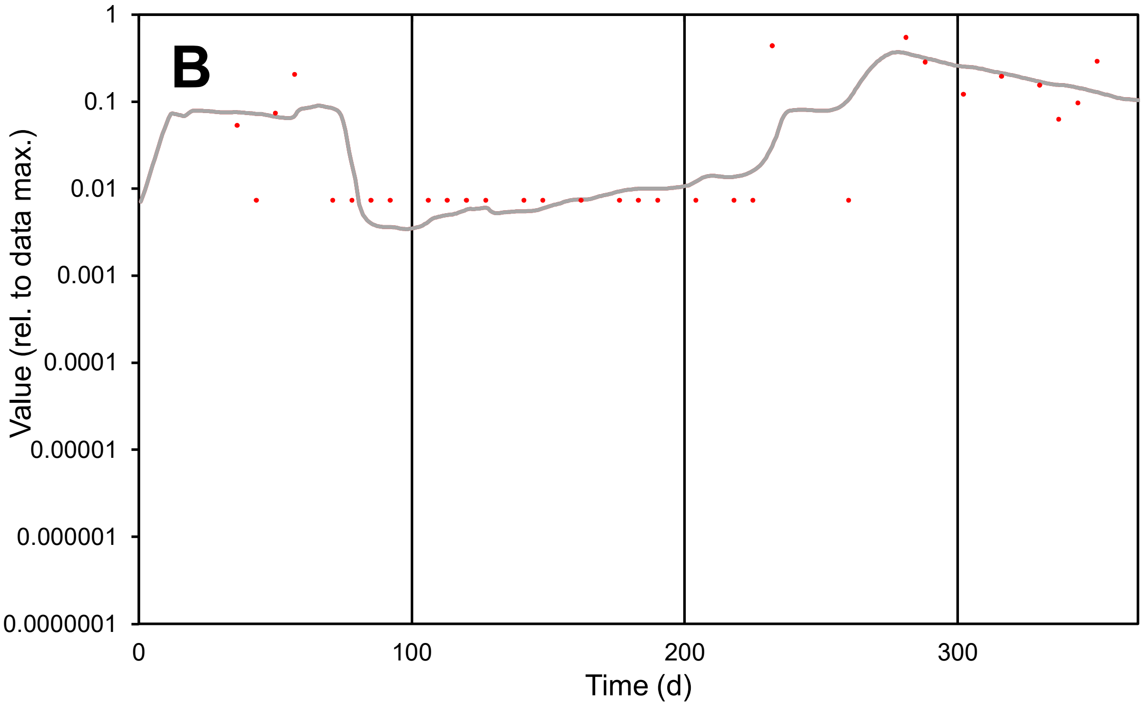


**C1**

Variables

LCH Provide jtd or Dst

CalibOnly?

2 1 to reproduce of

oNormalize?

2 1=no, 2=yes

ofxn calc*

1.72811

ofxn mod

1.7369

sRun

RunA1

zout

1

Ave

Data

3.8E-05

Model

2.4E-05

data/vMax

0.00033

ofxng

321.551

Min d/m

2.03833

ave/min d

15.5889

ave/min m

20.105

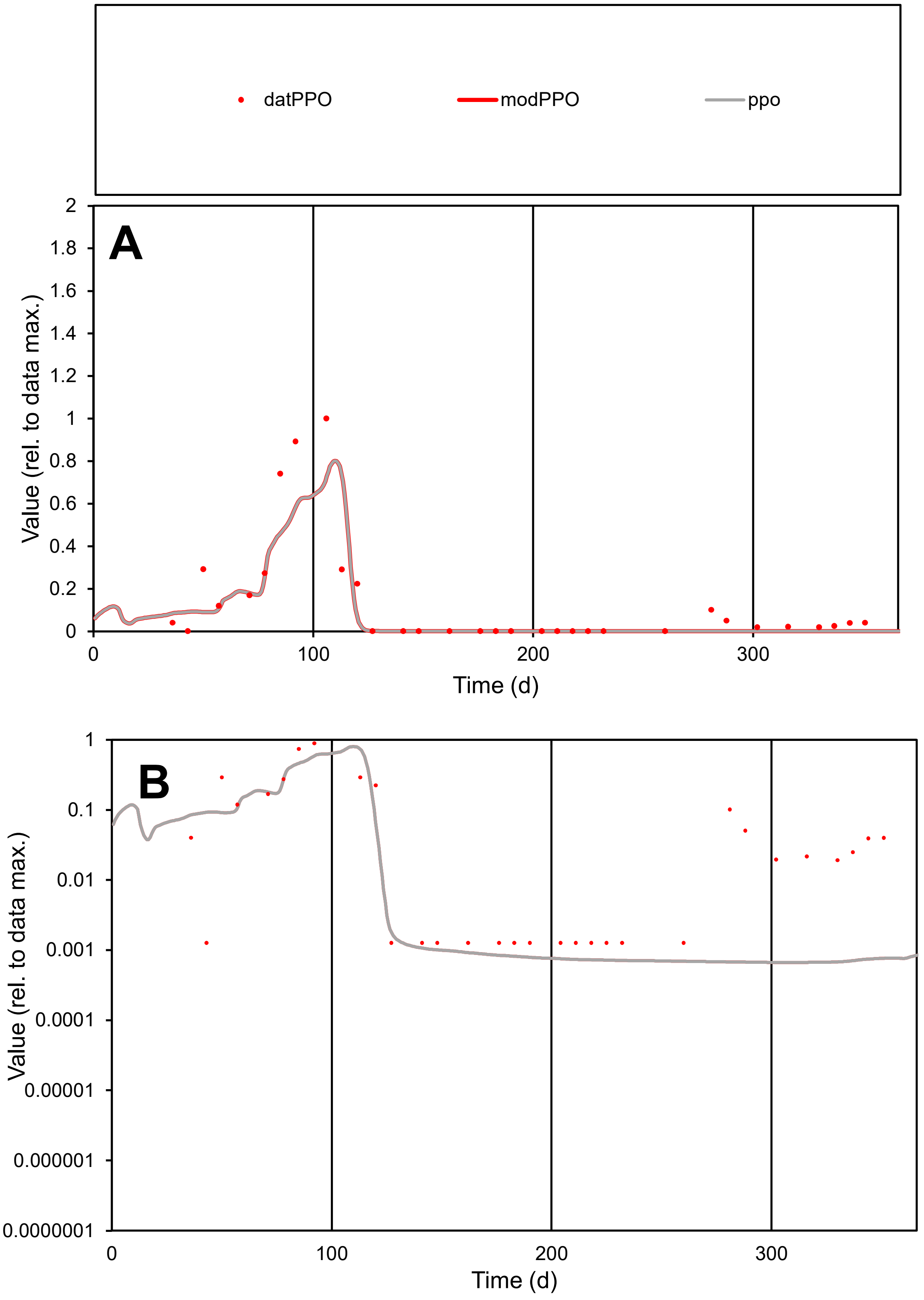


**C1**

Variables
PPO Provide jtd or Dst
CalibOnly?
2 1 to reproduce of
oNormalize?
2 1=no, 2=yes

ofxn calc*
0.72354
ofxn mod
0.82074

sRun
RunA1
zout
1

Ave
Data
0.00021
Model
0.00016

data/vMax
0.0015

ofxng
321.551

Min d/m
1.90348

ave/min d
107.521
ave/min m
158.968

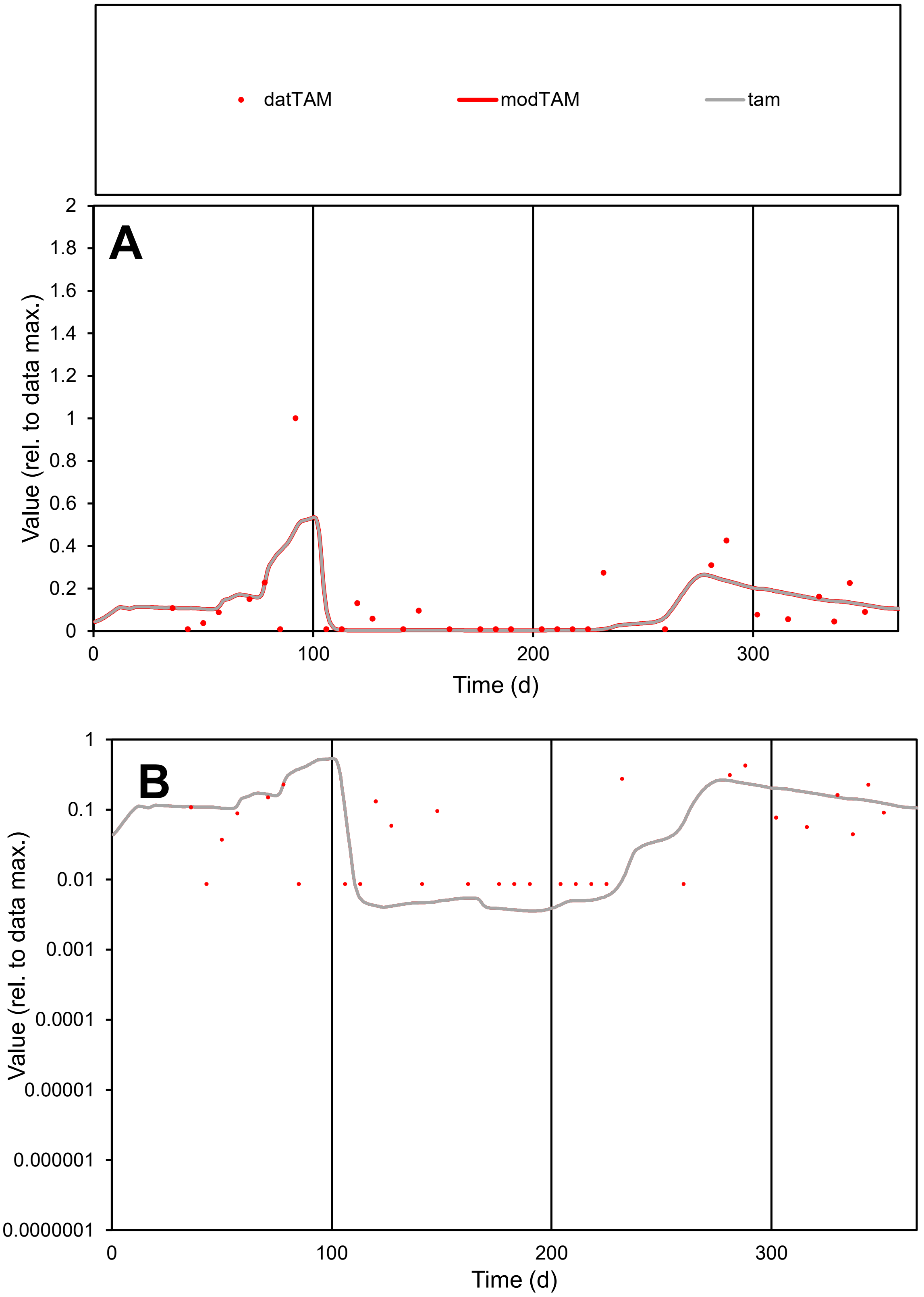


**C1**

Variables
TAM Provide jtd or Dst
CalibOnly?
2 1 to reproduce of
oNormalize?
2 1=no, 2=yes

ofxn calc*
1.17895
ofxn mod
1.19103

sRun
RunA1
zout
1

Ave
Data
7.7E-05
Model
6.8E-05

data/vMax
0.00066

ofxng
321.551

Min d/m
2.43407

ave/min d
13.2841
ave/min m
28.8877

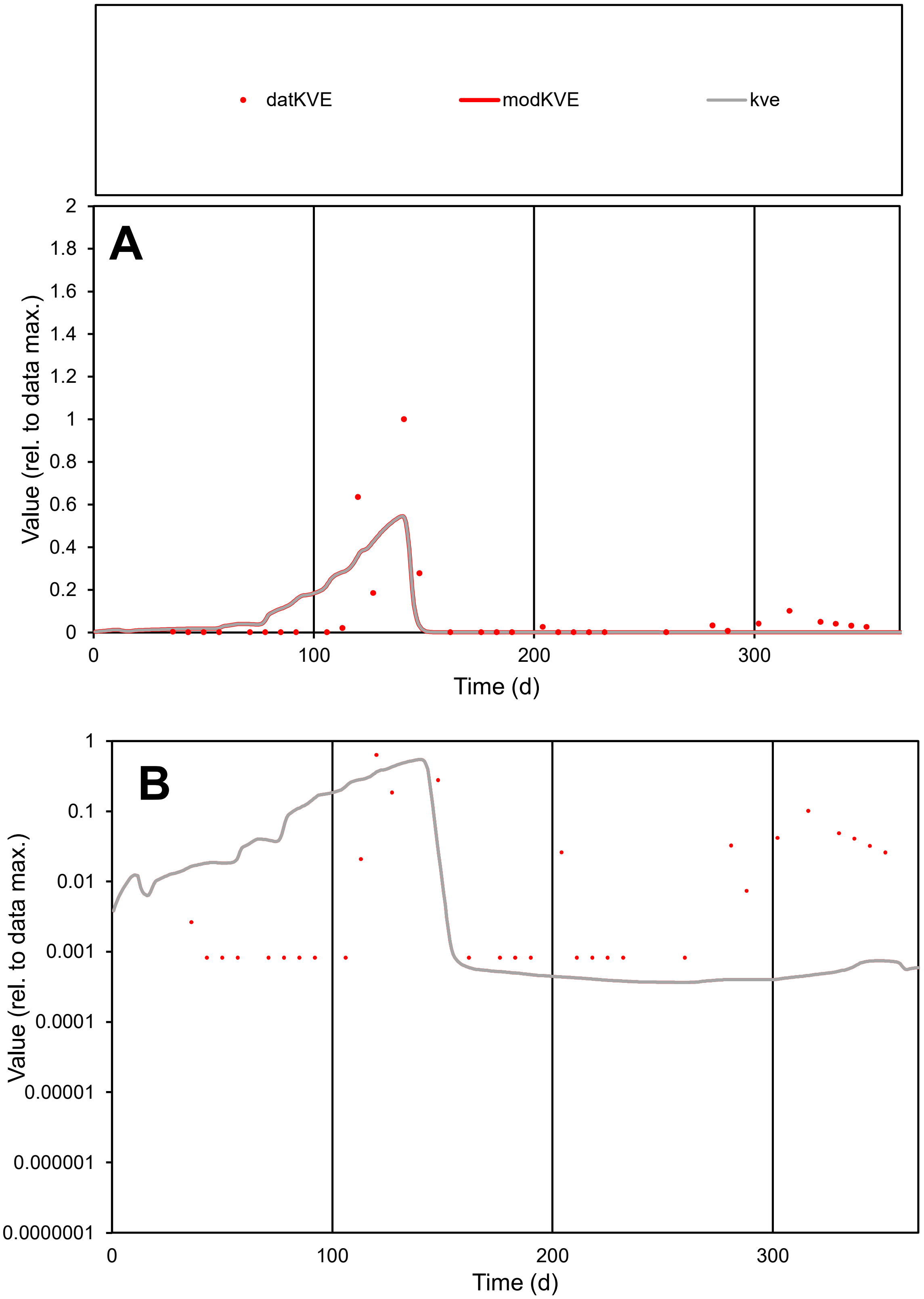


**C1**

Variables
KVE Provide jtd or Dst
CalibOnly?
2 1 to reproduce of
oNormalize?
2 1=no, 2=yes

ofxn calc*
1.60784
ofxn mod
1.76006

sRun
RunA1
zout
1

Ave
Data
0.0003
Model
0.0003

data/vMax
0.00381

ofxng
321.551

Min d/m
2.24284

ave/min d
94.7286
ave/min m
216.604

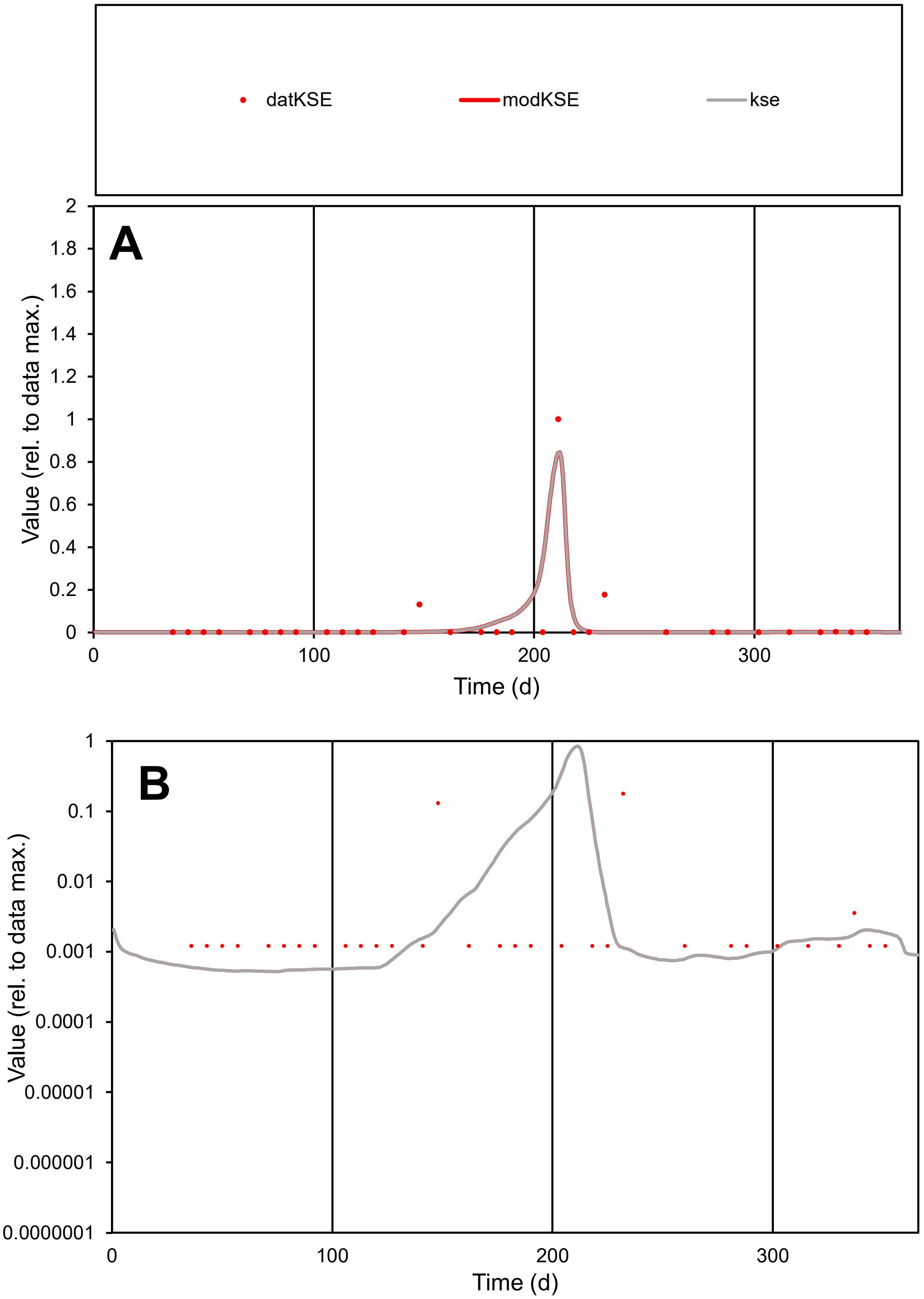


**C1**

Variables
KSE Provide jtd or Dst
CalibOnly?
2 1 to reproduce of
oNormalize?
2 1=no, 2=yes

ofxn calc*
1.57806
ofxn mod
1.83255

sRun
RunA1
zout
1

Ave
Data
0.00013
Model
0.00012

data/vMax
0.00309

ofxng
321.551

Min d/m
2.31215

ave/min d
34.7001
ave/min m
76.682

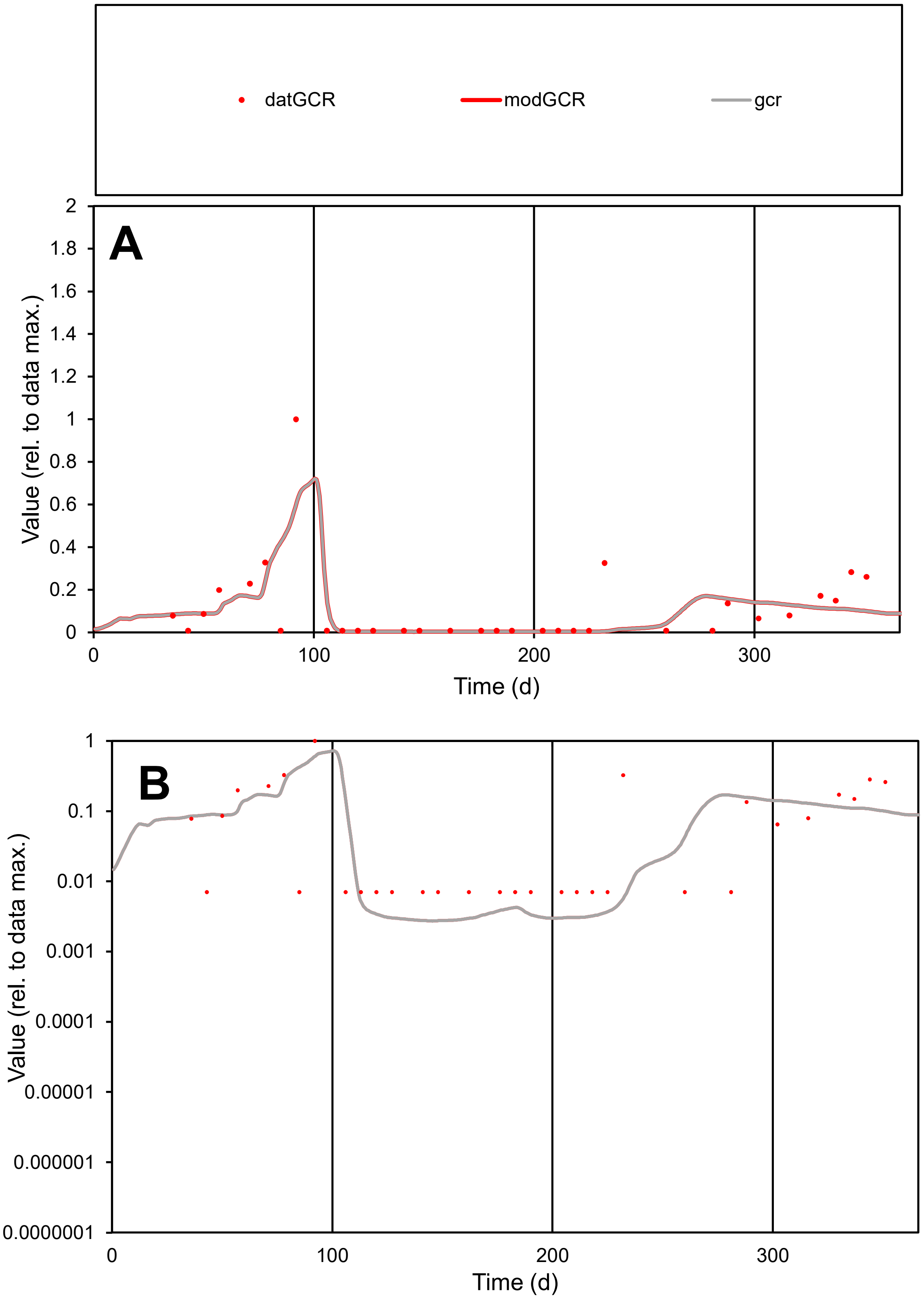


**C1**

Variables
GCR Provide jtd or Dst
CalibOnly?
2 1 to reproduce of
oNormalize?
2 1=no, 2=yes

ofxn calc*
1.15427
ofxn mod
1.17513

sRun
RunA1
zout
1

Ave
Data
4.9E-05
Model
4.2E-05

data/vMax
0.00044

ofxng
321.551

Min d/m
2.56255

ave/min d
15.5658
ave/min m
34.6041

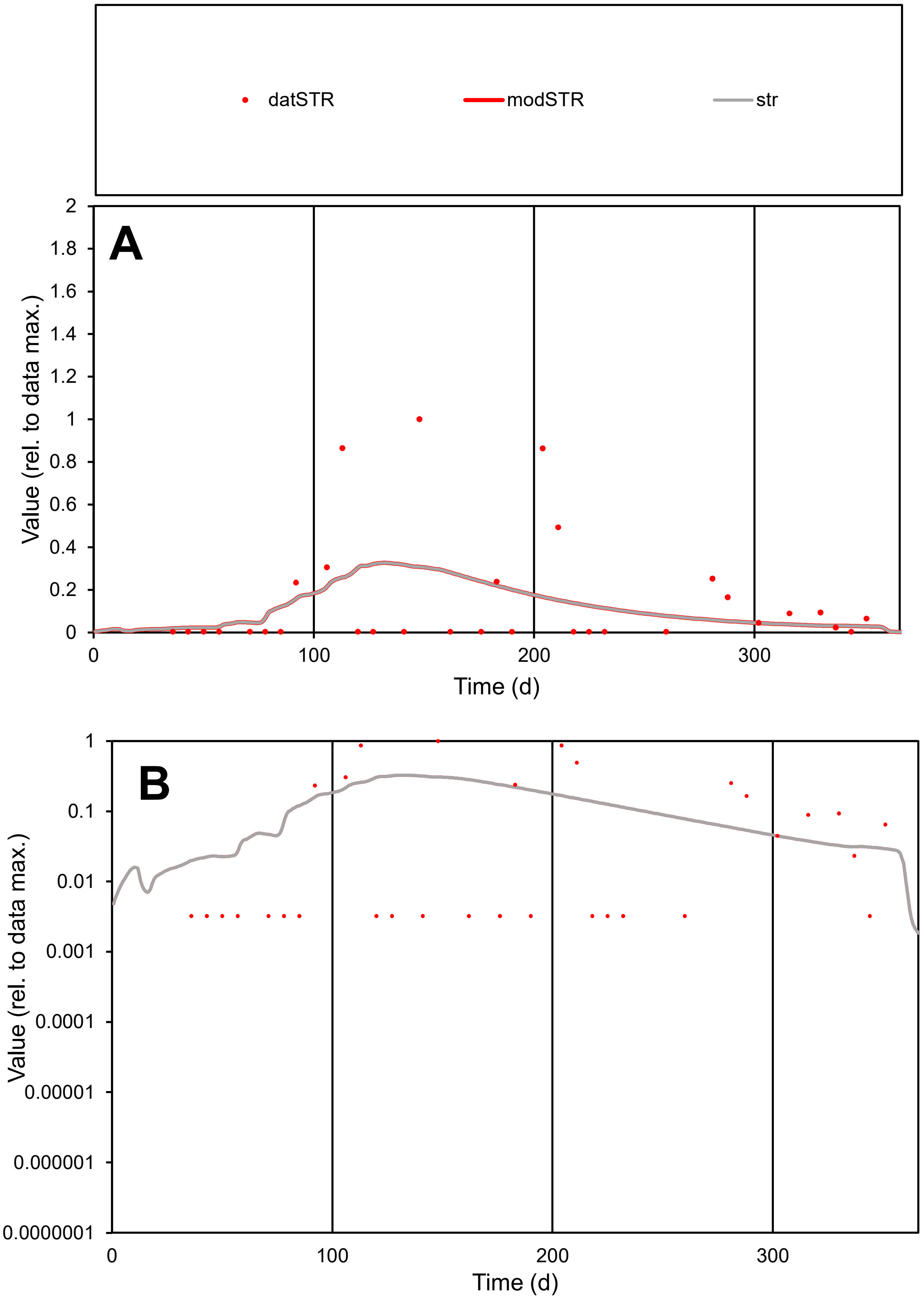


**C1**

Variables
STR Provide jtd or Dst
CalibOnly?
2 1 to reproduce of
oNormalize?
2 1=no, 2=yes

ofxn calc*
1.68925
ofxn mod
1.69541

sRun
RunA1
zout
1

Ave
Data
7.6E-05
Model
6.7E-05

data/vMax
0.00051

ofxng
321.551

Min d/m
0.16049

ave/min d
46.4835
ave/min m
6.57411

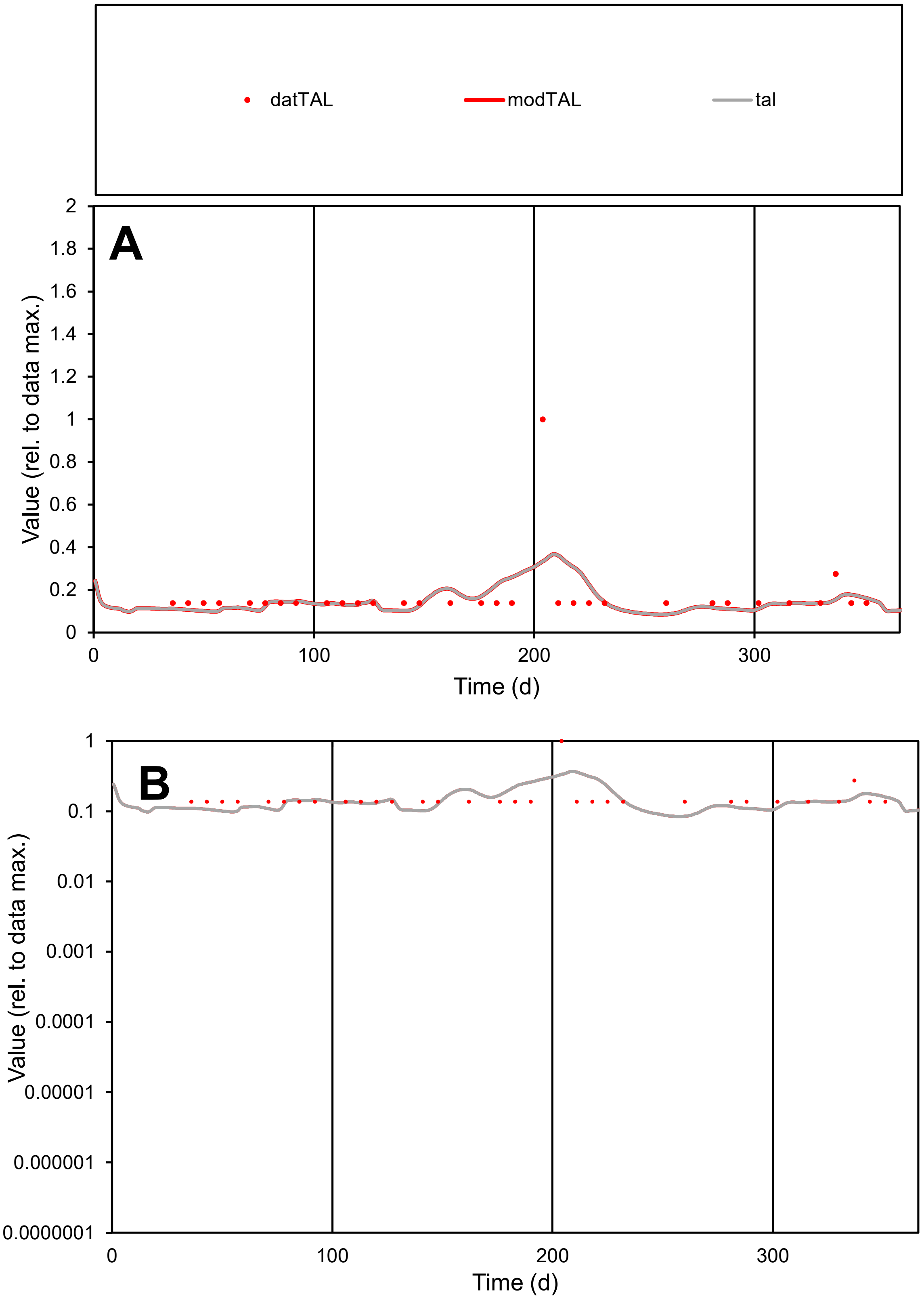


**C1**

Variables
TAL Provide jtd or Dst
CalibOnly?
2 1 to reproduce of
oNormalize?
2 1=no, 2=yes

ofxn calc*
0.78646
ofxn mod
0.79419

sRun
RunA1
zout
1

Ave
Data
5.1E-06
Model
4.8E-06

data/vMax
3E-05

ofxng
321.551

Min d/m
1.60309

ave/min d
1.22743
ave/min m
1.87513

324_Calib_S_A

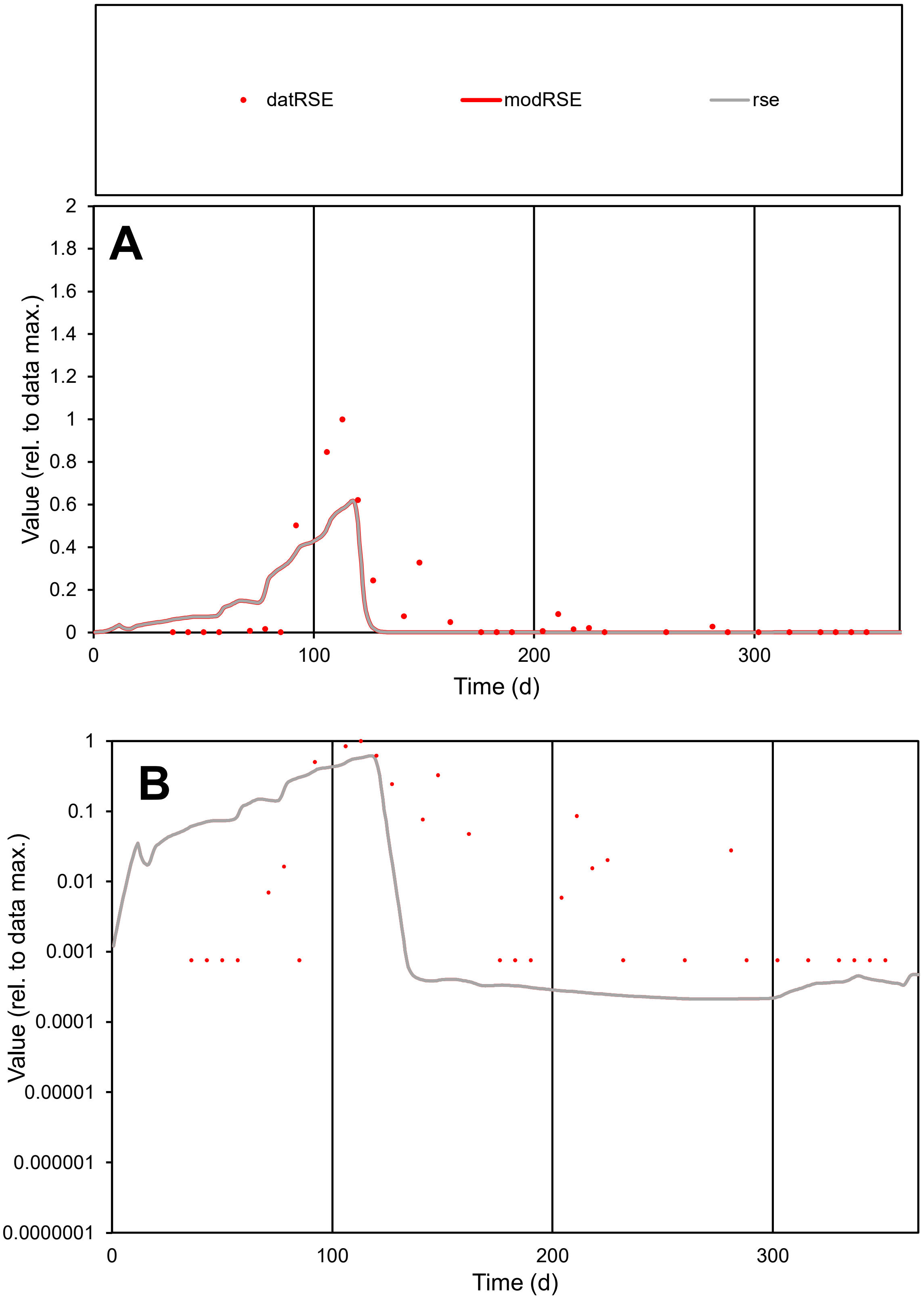


**C1**

Variables
RSE Provide jtd or Dst
CalibOnly?
2 1 to reproduce of
oNormalize?
2 1=no, 2=yes

ofxn calc*
1.14398
ofxn mod
1.18813

sRun
RunA1
zout
1

Ave
Data
0.00074
Model
0.00057

data/vMax
0.00616

ofxng
321.551

Min d/m
3.54062

ave/min d
159.036
ave/min m
434.881

324_Calib_S_A

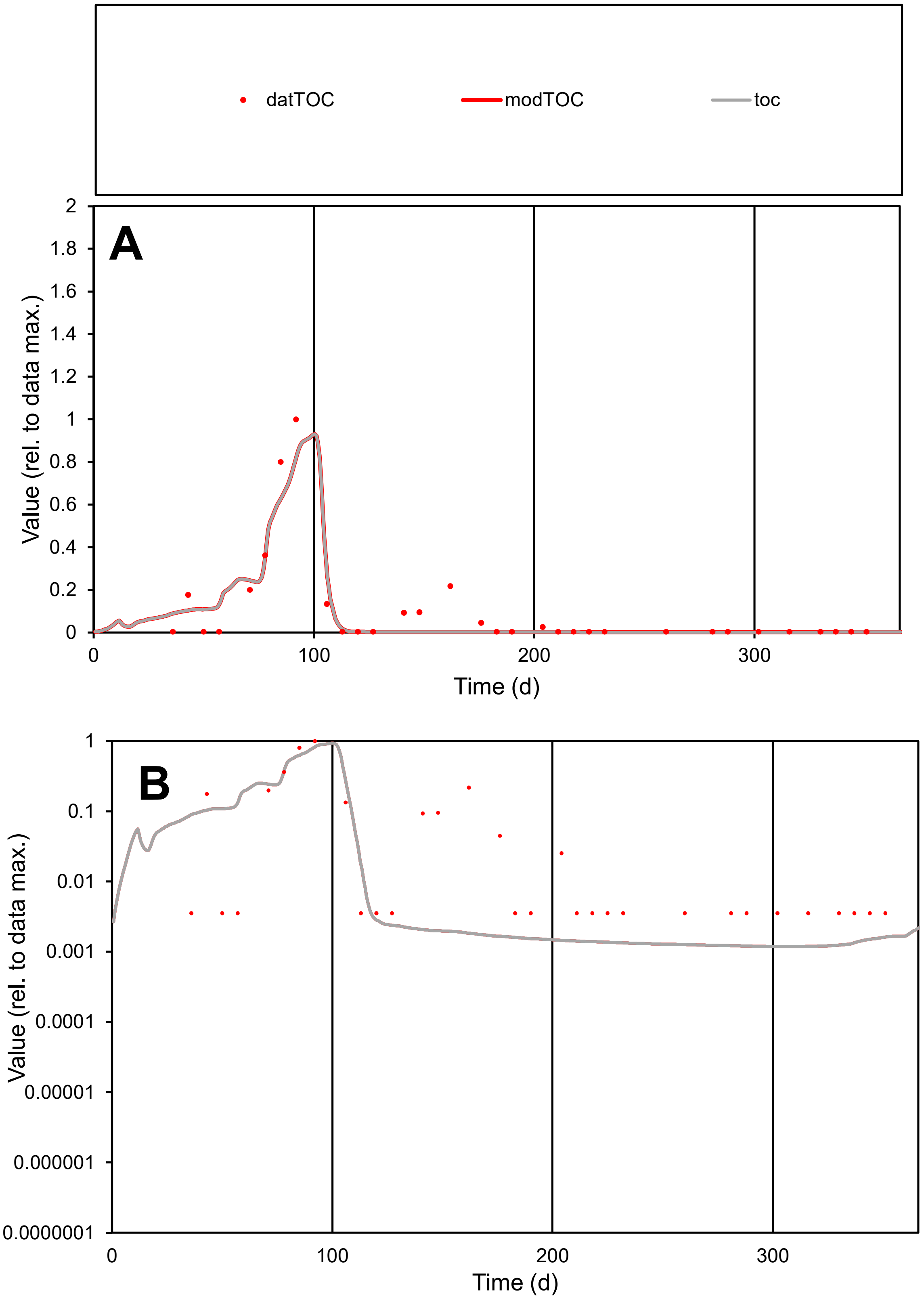


**C1**

Variables
TOC Provide jtd or Dst
CalibOnly?
2 1 to reproduce of
oNormalize?
2 1=no, 2=yes

ofxn calc*
0.6728
ofxn mod
0.71936

sRun
RunA1
zout
1

Ave
Data
0.00012
Model
0.00011

data/vMax
0.00122

ofxng
321.551

Min d/m
2.97658

ave/min d
28.4892
ave/min m
73.3878

324_Calib_S_A

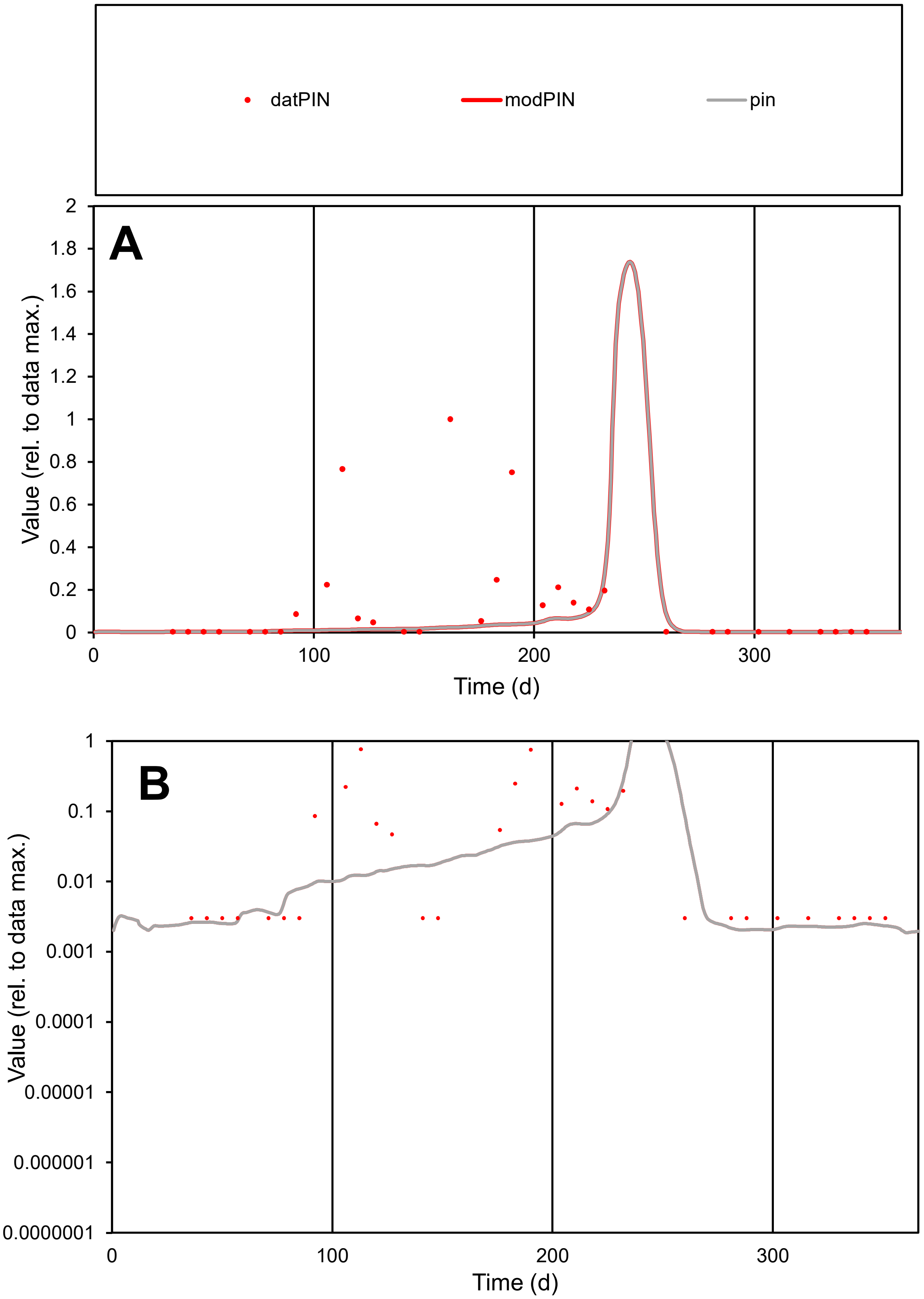


**C1**

Variables
PIN Provide jtd or Dst
CalibOnly?
2 1 to reproduce of
oNormalize?
2 1=no, 2=yes

ofxn calc*
2.04013
ofxn mod
2.04817

sRun
RunA1
zout
1

Ave
Data
0.00017
Model
3.4E-05

data/vMax
0.00133

ofxng
321.551

Min d/m
1.45924

ave/min d
42.3529
ave/min m
12.5292

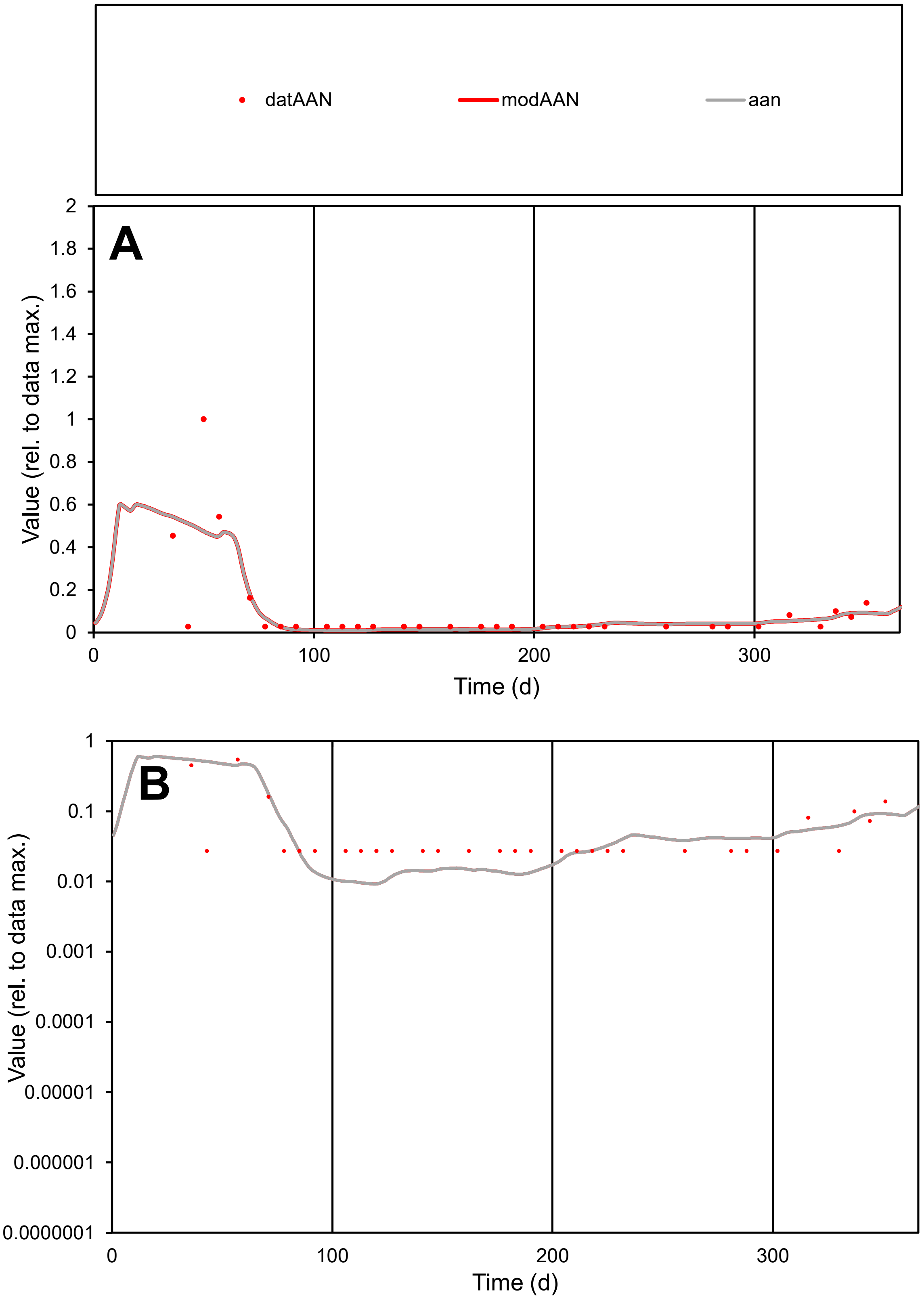


**C1**

Variables
AAN Provide jtd or Dst
CalibOnly?
2 1 to reproduce of
oNormalize?
2 1=no, 2=yes

ofxn calc*
1.29237
ofxn mod
1.3096

sRun
RunA1
zout
1

Ave
Data
2.1E-05
Model
1.9E-05

data/vMax
0.0002

ofxng
321.551

Min d/m
2.88269

ave/min d
3.67539
ave/min m
10.0694

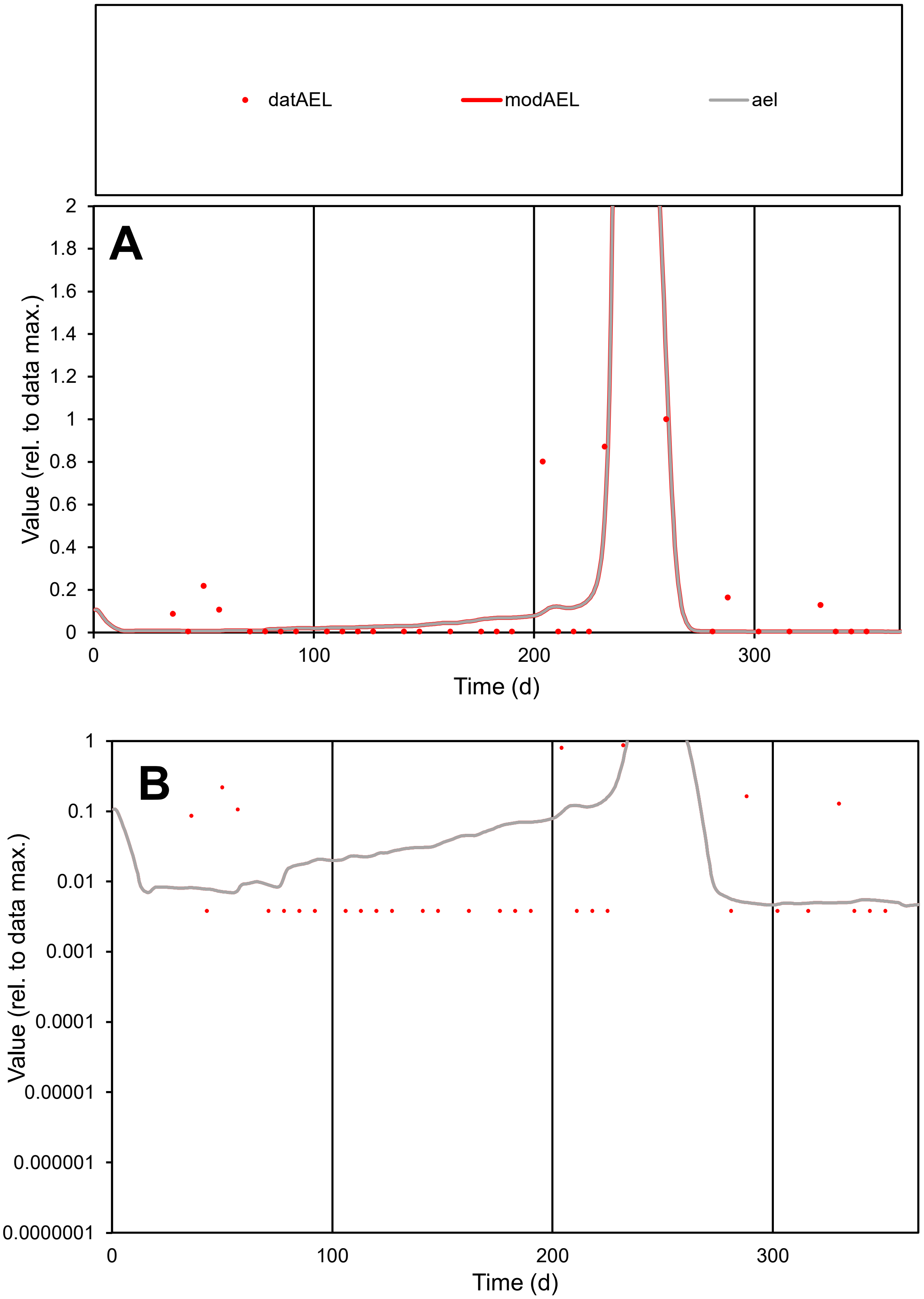


**C1**

Variables
AEL Provide jtd or Dst
CalibOnly?
2 1 to reproduce of
oNormalize?
2 1=no, 2=yes

ofxn calc*
1.31735
ofxn mod
1.33467

sRun
RunA1
zout
1

Ave
Data
2E-05
Model
1.6E-05

data/vMax
0.00018

ofxng
321.551

Min d/m
0.79621
ave/min d
28.2983
ave/min m
18.6393

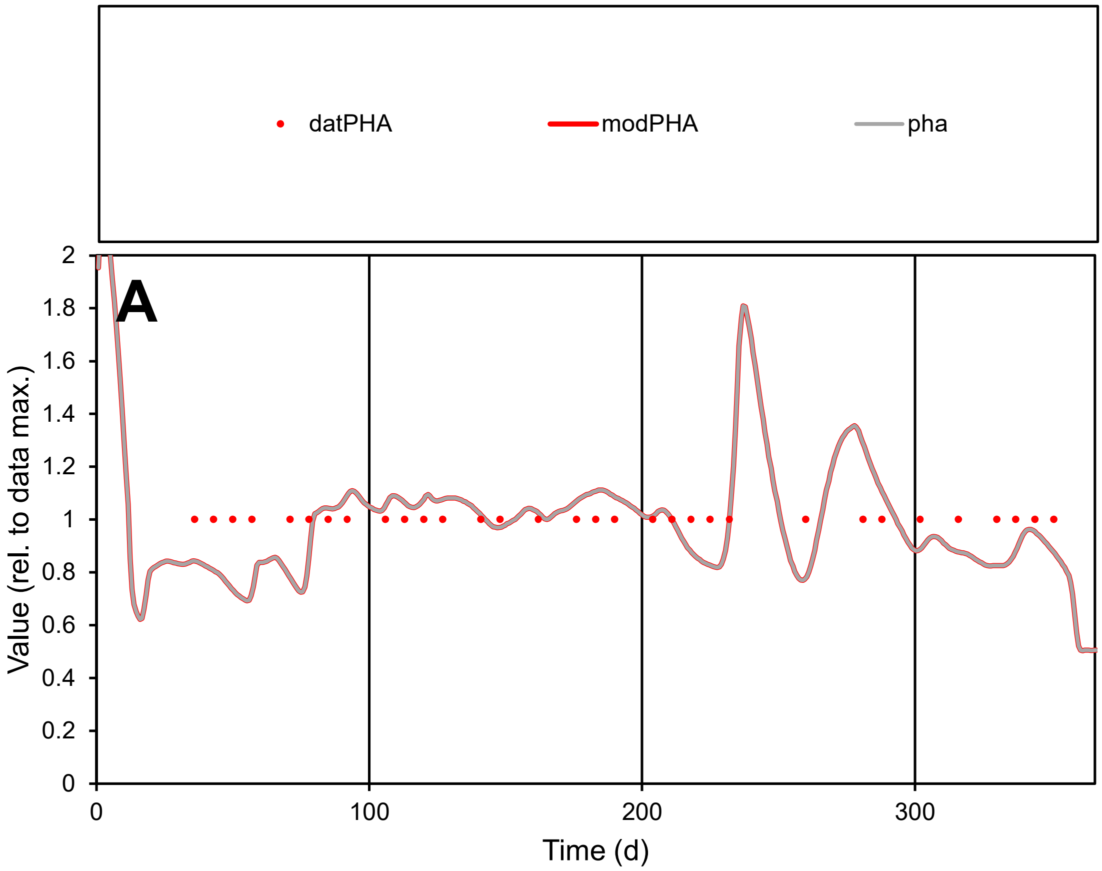


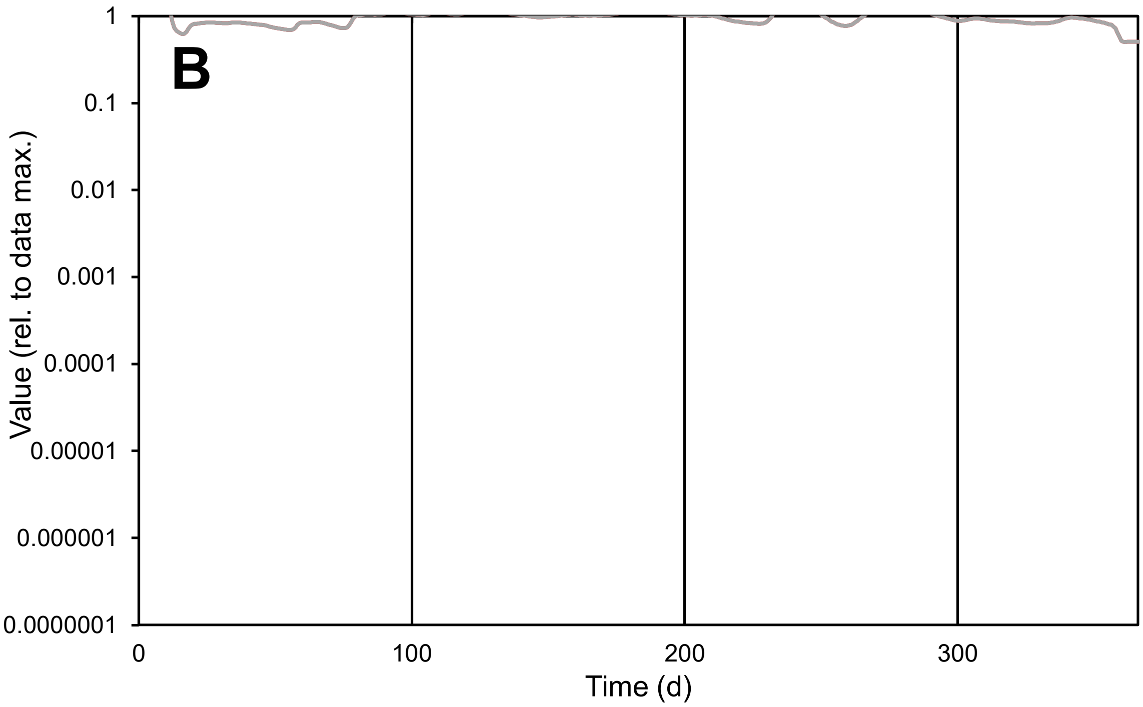


**C1**

Variables
PHA Provide jtd or Dst
CalibOnly?
2 1 to reproduce of
oNormalize?
2 1=no, 2=yes

ofxn calc*
0.13312
ofxn mod
0.16657

sRun
RunA1
zout
1

Ave
Data
1E-05
Model
1E-05

data/vMax
1E-05

ofxng
321.551

Min d/m
1.39488

ave/min d
1
ave/min m
1.34369

324_Calib_S_A

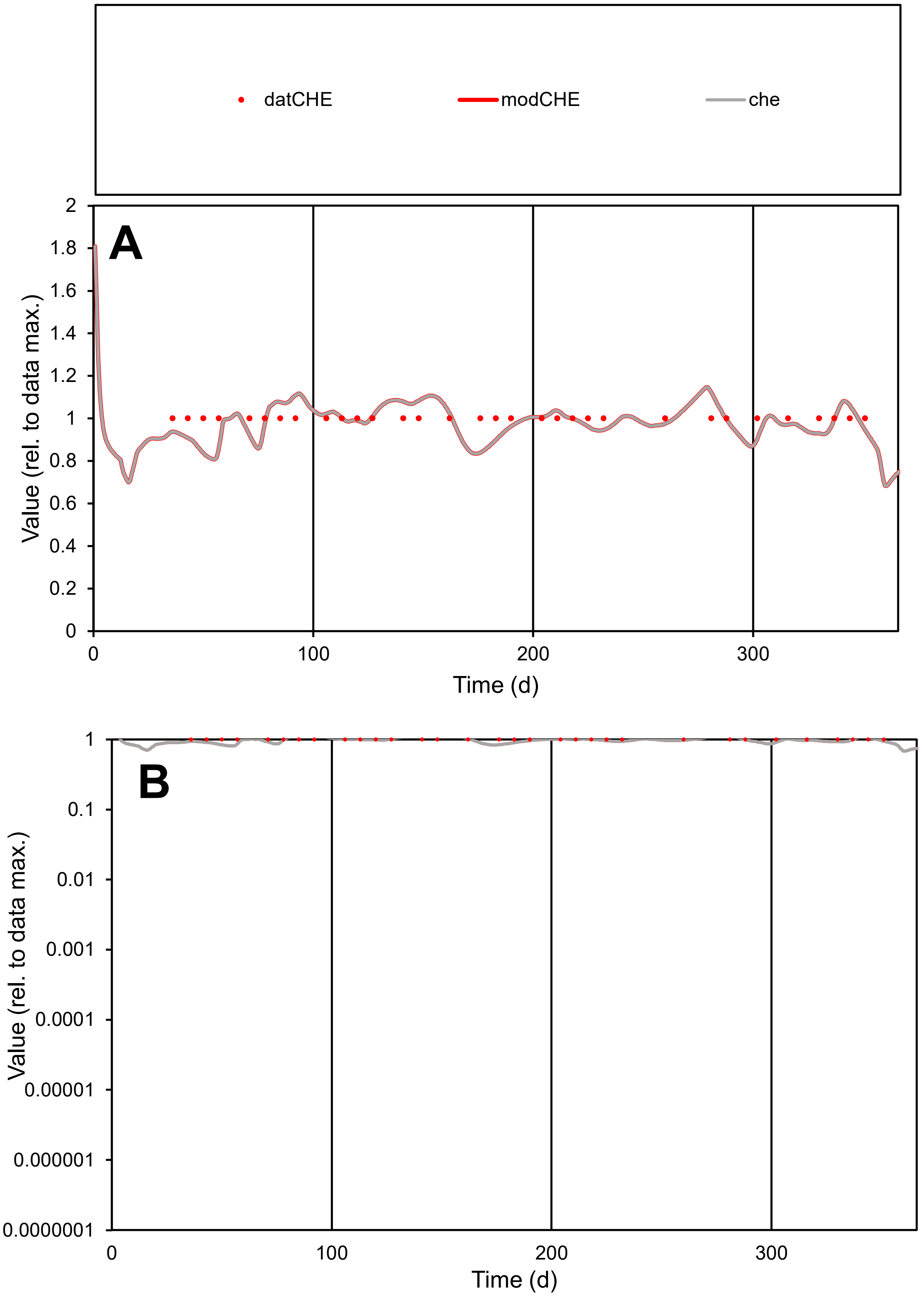


**C1**

Variables

CHE Provide jtd or Dst

CalibOnly?

2 1 to reproduce of

oNormalize?

2 1=no, 2=yes

ofxn calc*

0.07053

ofxn mod

0.11943

sRun

RunA1

zout

1

Ave

Data

7E-06

Model

6.9E-06

data/vMax

7E-06

ofxng

321.551

Min d/m

1.21579

ave/min d

1

ave/min m

1.19499

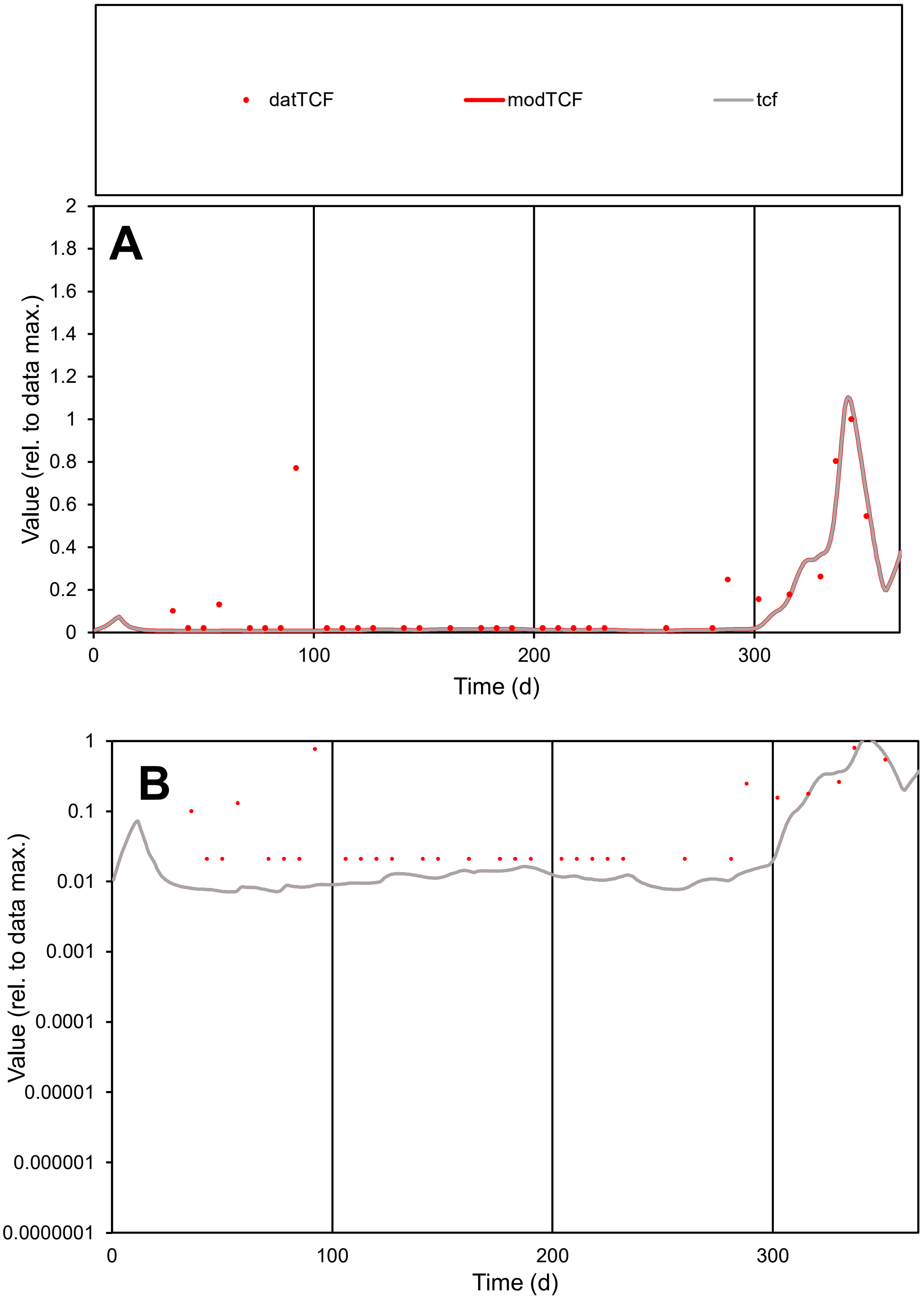


**C1**

Variables
TCF Provide jtd or Dst
CalibOnly?
2 1 to reproduce of
oNormalize?
2 1=no, 2=yes

ofxn calc*
1.00885
ofxn mod
1.02452

sRun
RunA1
zout
1

Ave
Data
2.2E-05
Model
1.5E-05

data/vMax
0.00015

ofxng
321.551

Min d/m
2.92864

ave/min d
6.91606
ave/min m
13.9276

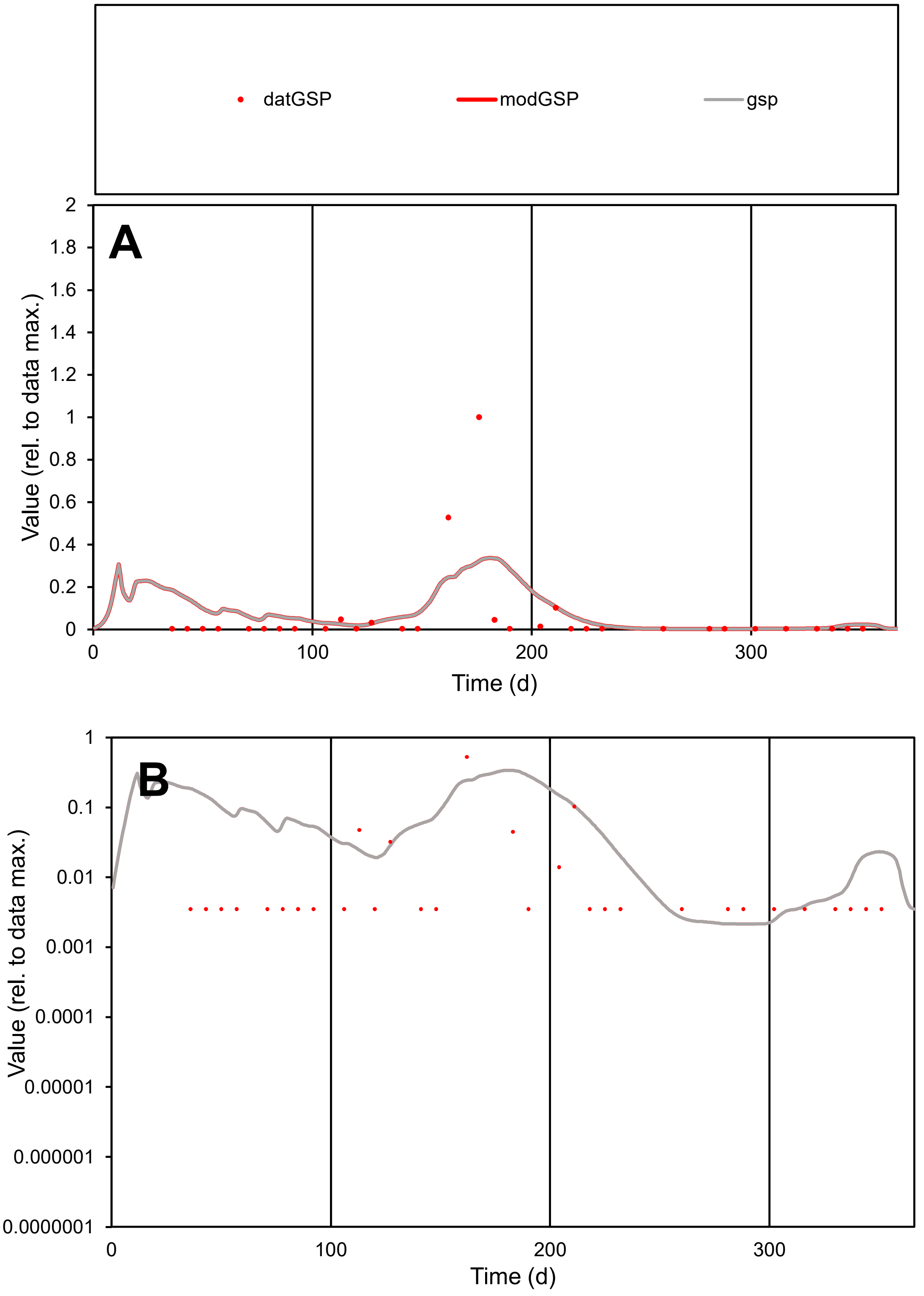


**C1**

Variables
GSP Provide jtd or Dst
CalibOnly?
2 1 to reproduce of
oNormalize?
2 1=no, 2=yes

ofxn calc*
2.66445
ofxn mod
2.70518

sRun
RunA1
zout
1

Ave
Data
0.00015
Model
0.0002

data/vMax
0.00255

ofxng
321.551

Min d/m
1.63641

ave/min d
16.4528
ave/min m
35.7555

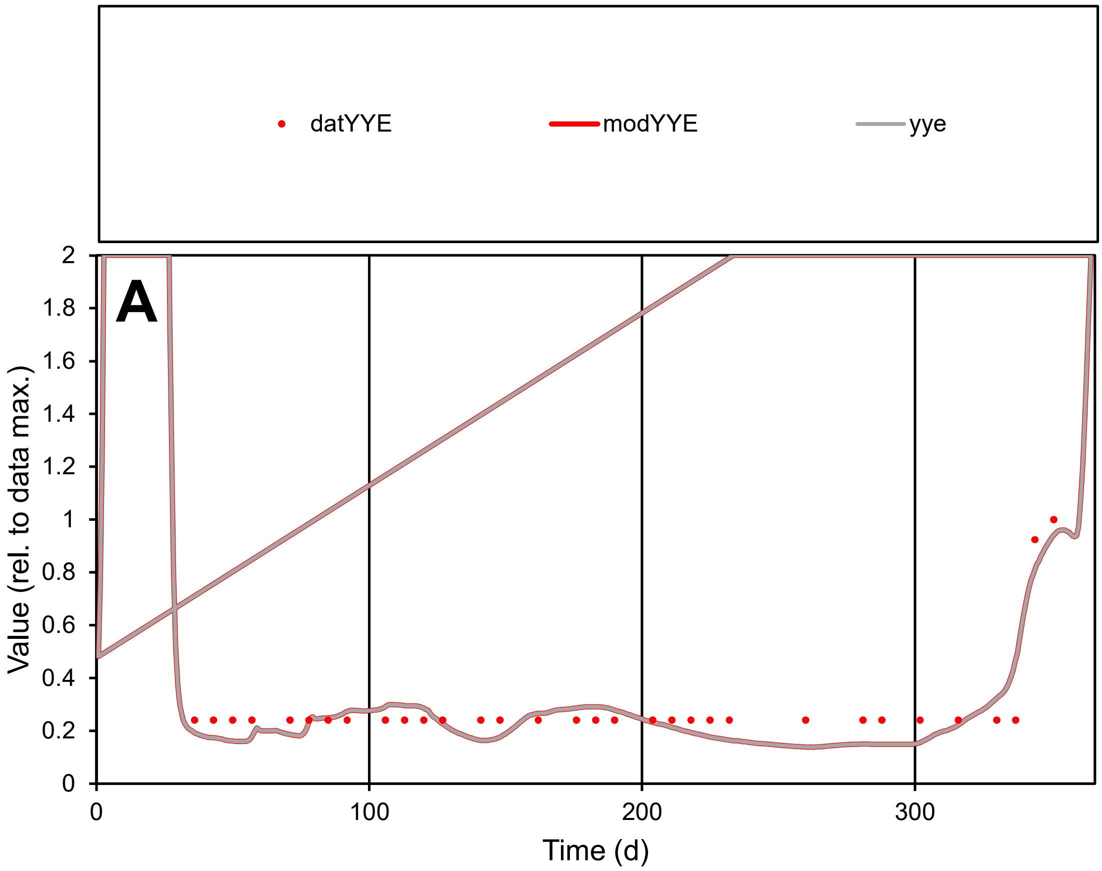


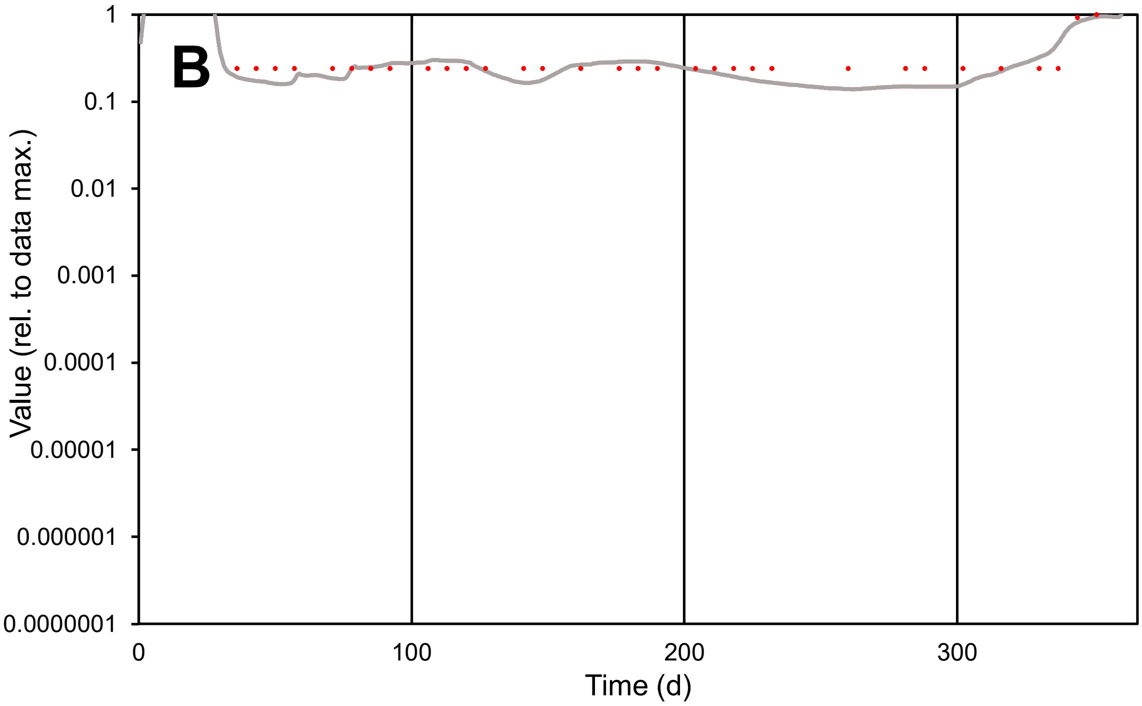


**C1**

Variables

YYE Provide jtd or Dst

CalibOnly?

2 1 to reproduce of

oNormalize?

2 1=no, 2=yes

ofxn calc*

0.25828

ofxn mod

0.28045

sRun

RunA1

zout

1

Ave

Data

4.5E-06

Model

4.2E-06

data/vMax

1.6E-05

ofxng

321.551

Min d/m

1.73011

ave/min d

1.18738

ave/min m

1.94948

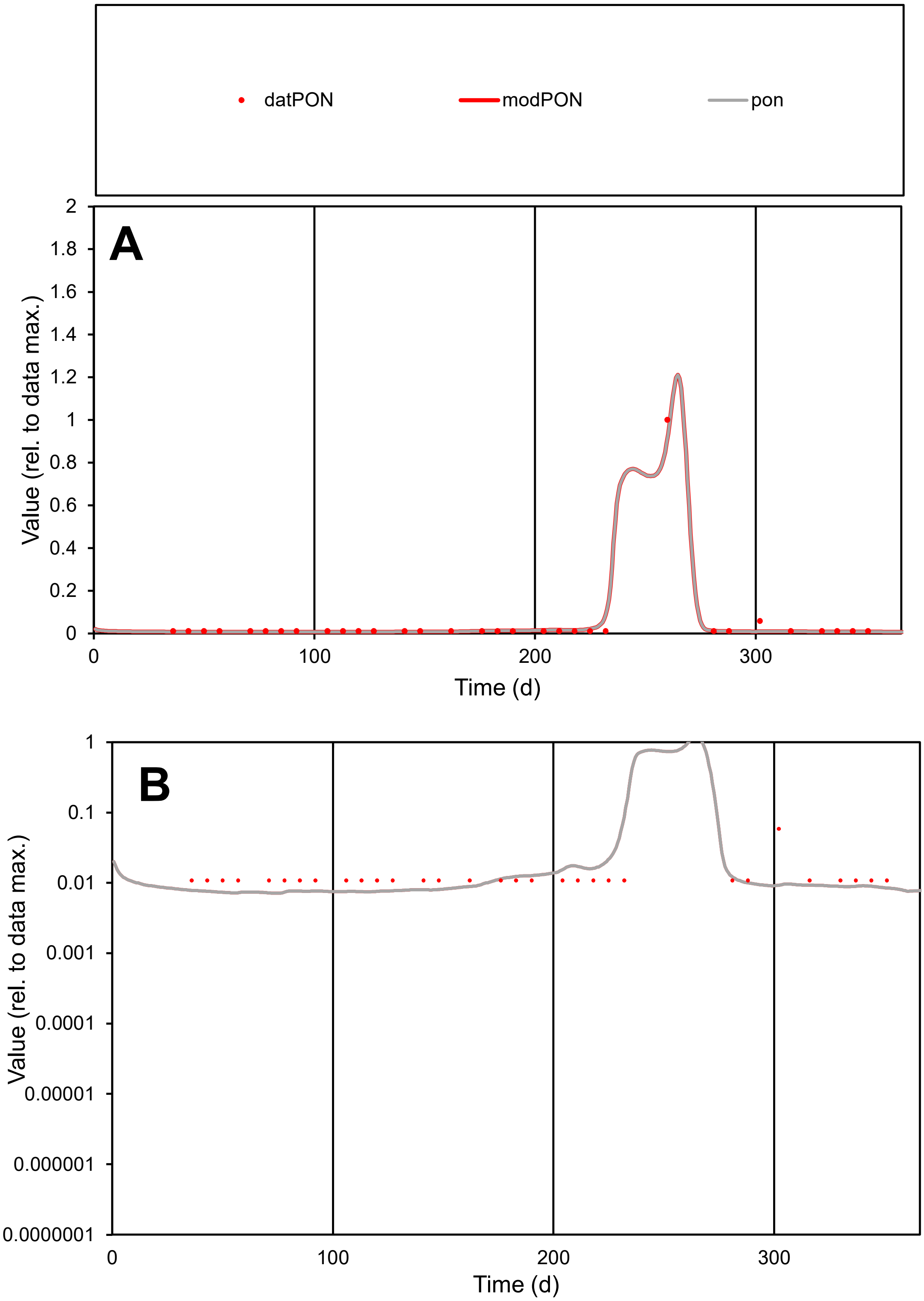


**C1**

Variables
PON Provide jtd or Dst
CalibOnly?
2 1 to reproduce of
oNormalize?
2 1=no, 2=yes

ofxn calc*
0.31705
ofxn mod
0.44233

sRun
RunA1
zout
1

Ave
Data
1.2E-05
Model
1.1E-05

data/vMax
0.00027

ofxng
321.551

Min d/m
1.51004

ave/min d
3.98209
ave/min m
5.89705

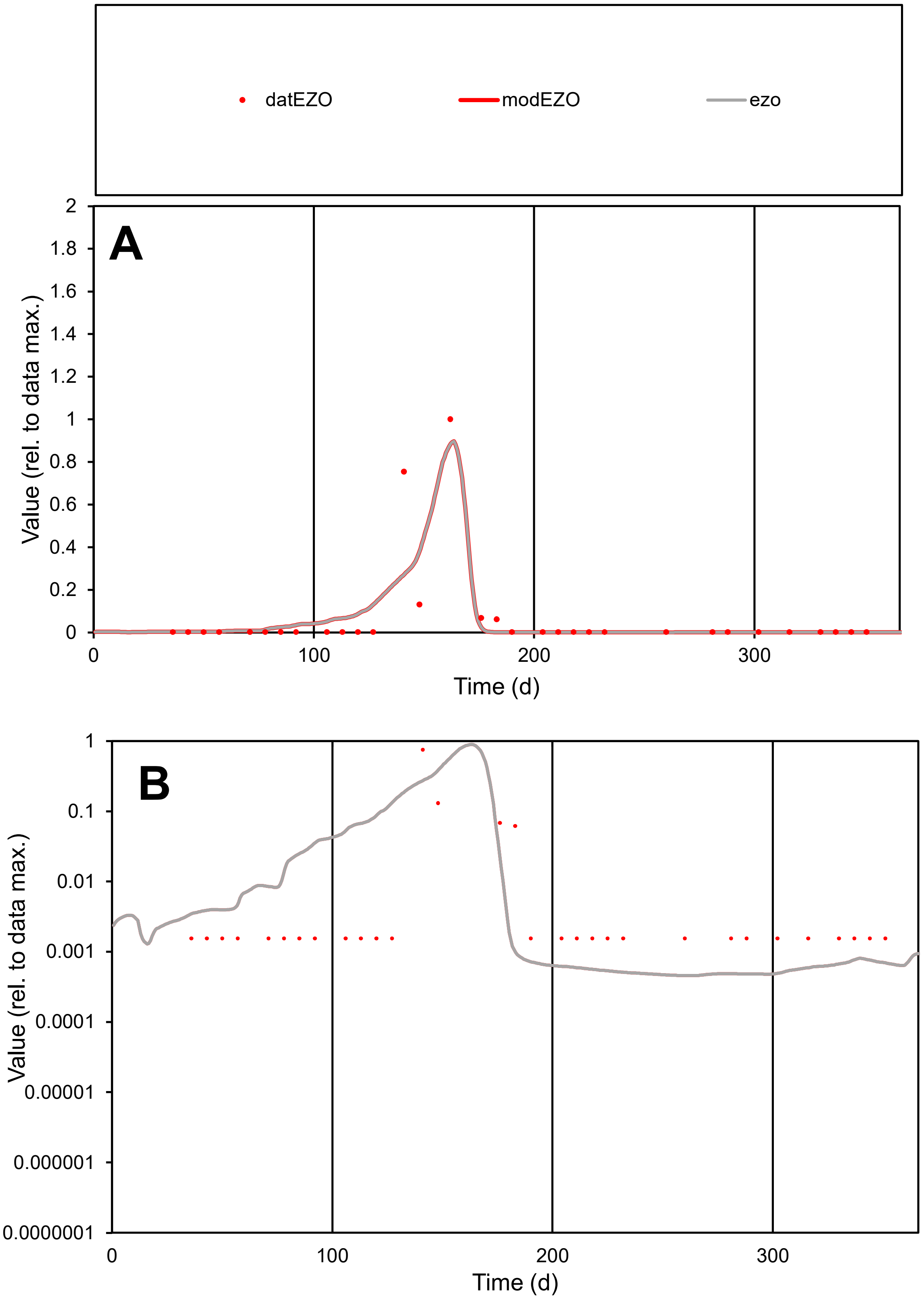


**C1**

Variables

| | |
|---|---|
| EZO | Provide jtd or Dst |

CalibOnly?

| | |
|---|---|
| 2 | 1 to reproduce of |

oNormalize?

| | |
|---|---|
| 2 | 1=no, 2=yes |

ofxn calc*
1.5755
ofxn mod
1.63906

sRun
RunA1
zout
1

Ave
Data
0.00016
Model
0.00016

data/vMax
0.00254

ofxng
321.551

Min d/m
3.39914

ave/min d
41.3618
ave/min m
140.117

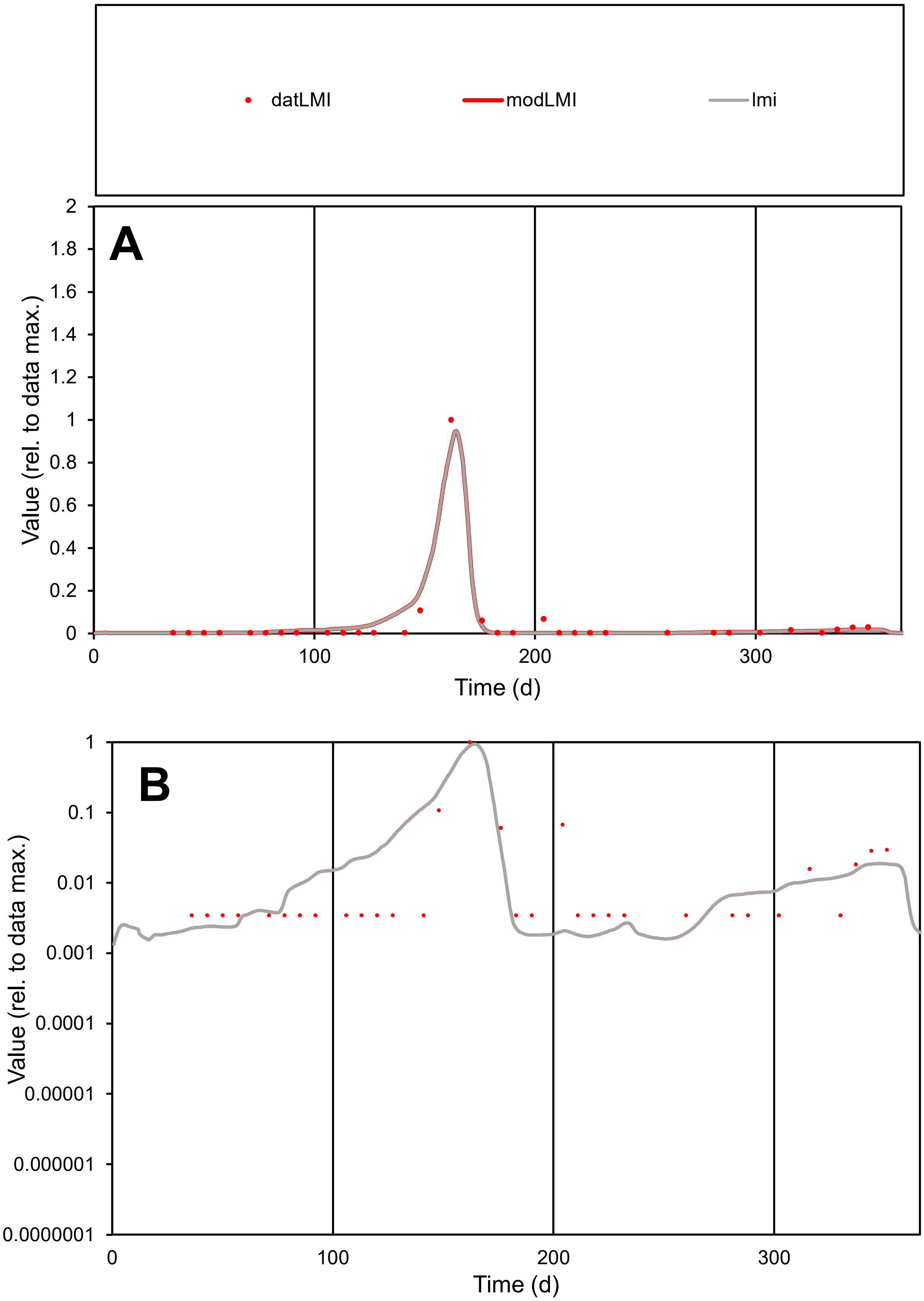


**C1**

Variables
LMI Provide jtd or Dst
CalibOnly?
2 1 to reproduce of
oNormalize?
2 1=no, 2=yes

ofxn calc*
0.72893
ofxn mod
0.9116

sRun
RunA1
zout
1

Ave
Data
3.8E-05
Model
4.3E-05

data/vMax
0.00087

ofxng
321.551

Min d/m
1.92401
ave/min d
12.7161
ave/min m
27.3531

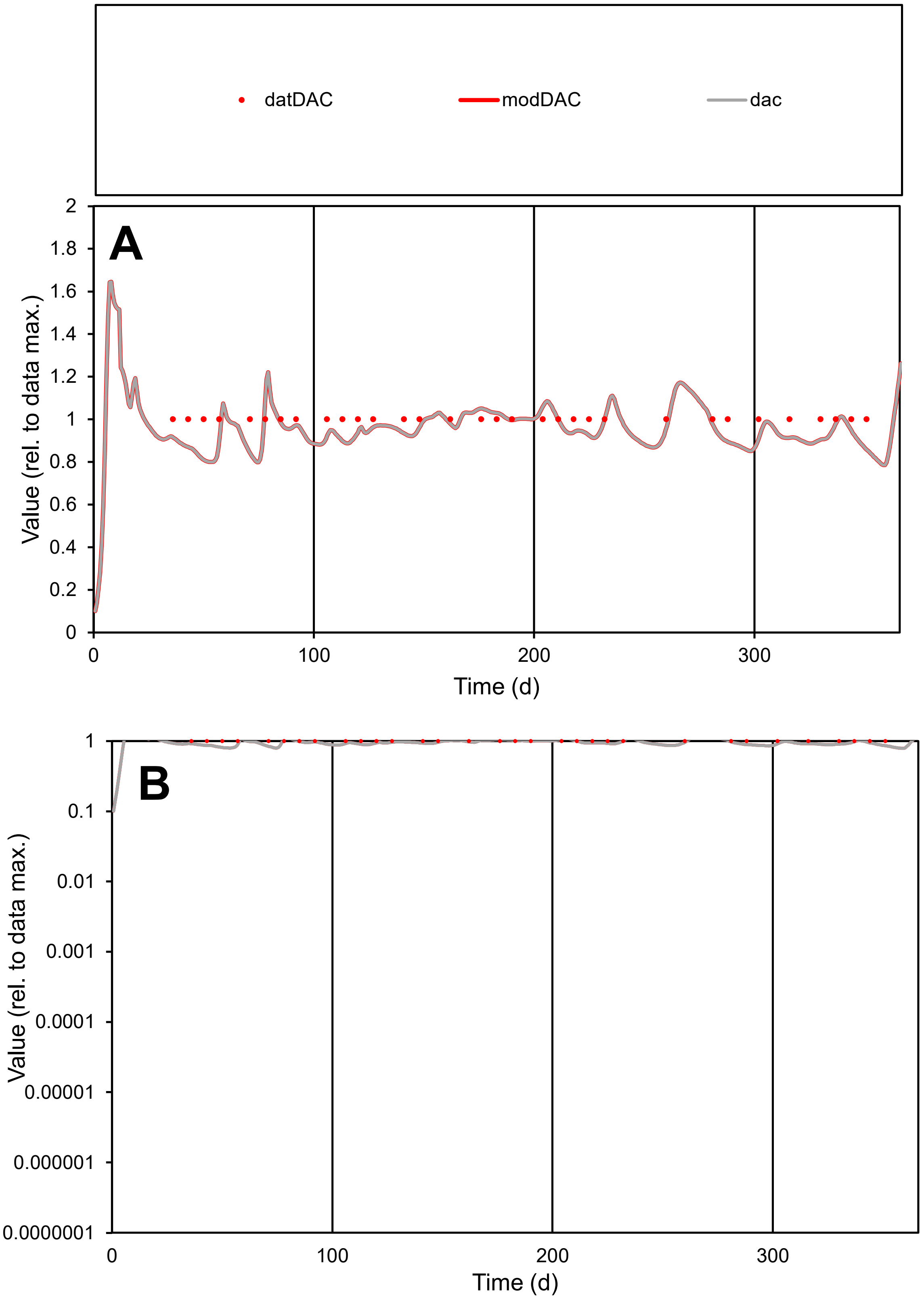


C1

Variables
DAC Provide jtd or Dst
CalibOnly?
2 1 to reproduce of
oNormalize?
2 1=no, 2=yes

ofxn calc*
0.08921
ofxn mod
0.13038

sRun
RunA1
zout
1

Ave
Data
7.7E-06
Model
7.4E-06

data/vMax
7.7E-06

ofxng
321.551

Min d/m
1.24612

ave/min d
1
ave/min m
1.19275

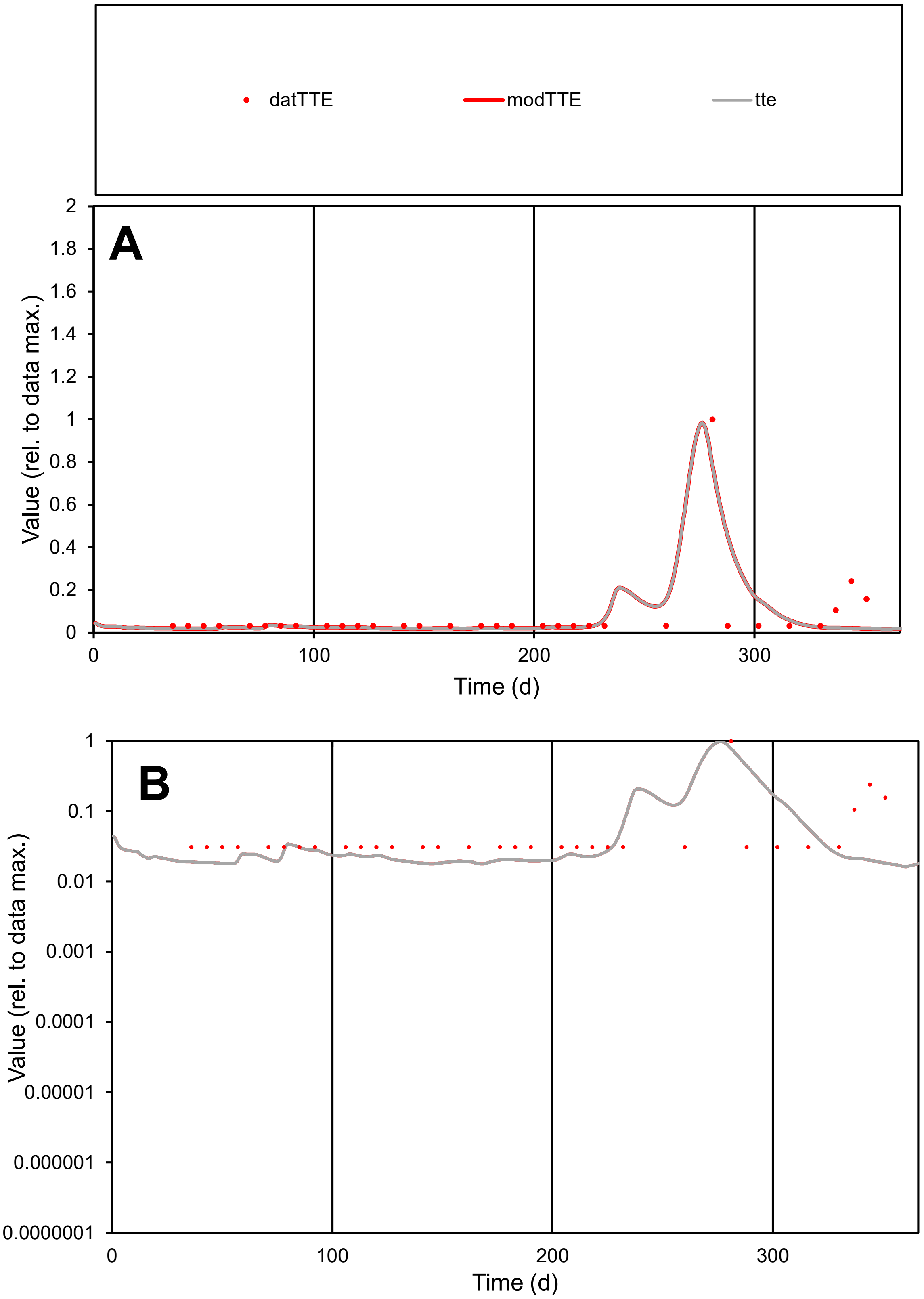


**C1**

Variables
TTE Provide jtd or Dst
CalibOnly?
2 1 to reproduce of
oNormalize?
2 1=no, 2=yes

ofxn calc*
1.20965
ofxn mod
1.34693

sRun
RunA1
zout
1

Ave
Data
9E-06
Model
7.8E-06

data/vMax
0.00012

ofxng
321.551

Min d/m
1.72761

ave/min d
2.3923
ave/min m
3.60523

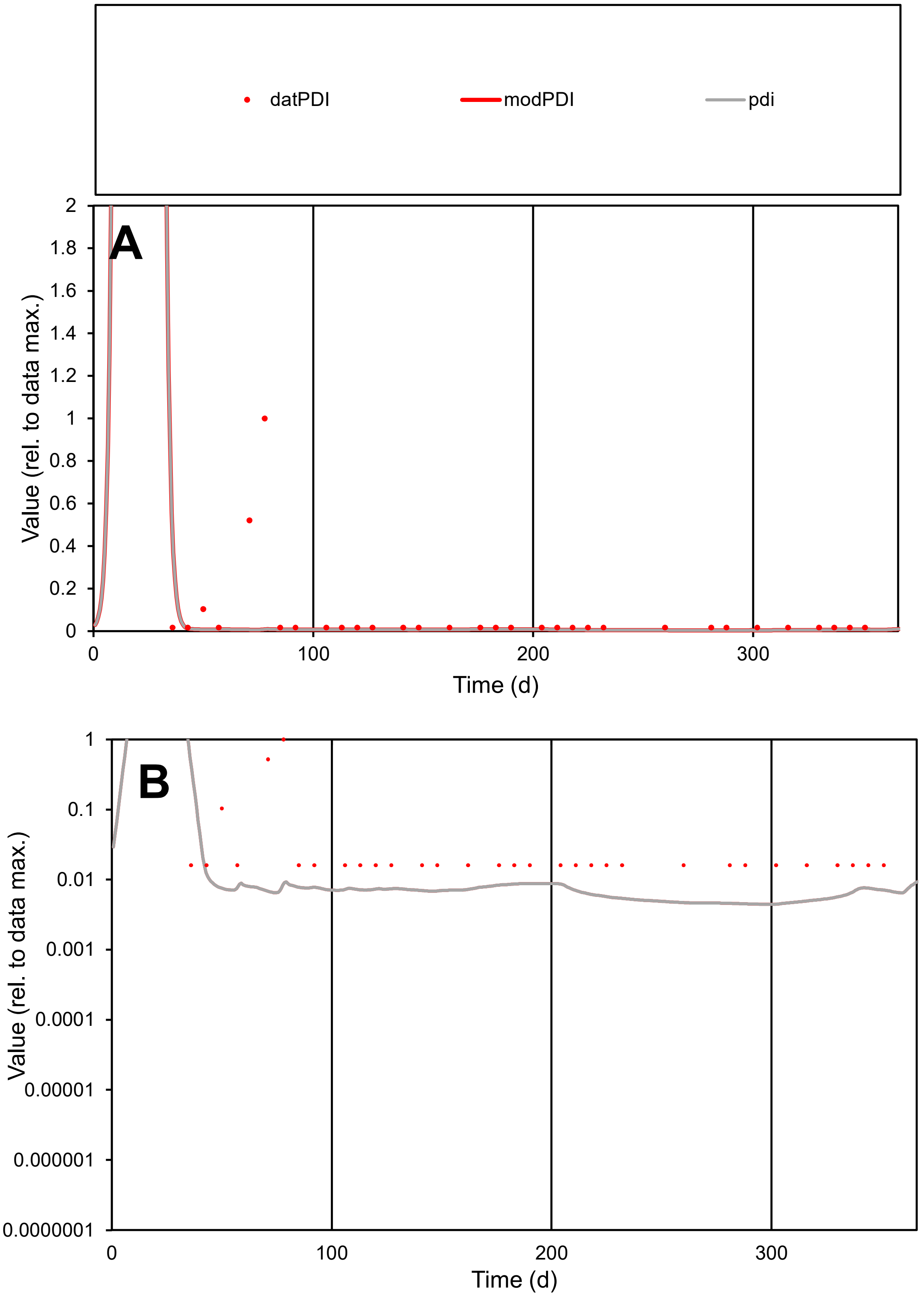


**C1**

Variables

PDI Provide jtd or Dst

CalibOnly?

2 1 to reproduce of

oNormalize?

2 1=no, 2=yes

ofxn calc*

3.04485

ofxn mod

3.05582

sRun

RunA1

zout

1

Ave

Data

1.8E-05

Model

2.8E-06

data/vMax

0.00027

ofxng

321.551

Min d/m

3.5798

ave/min d

4.06551

ave/min m

2.25662

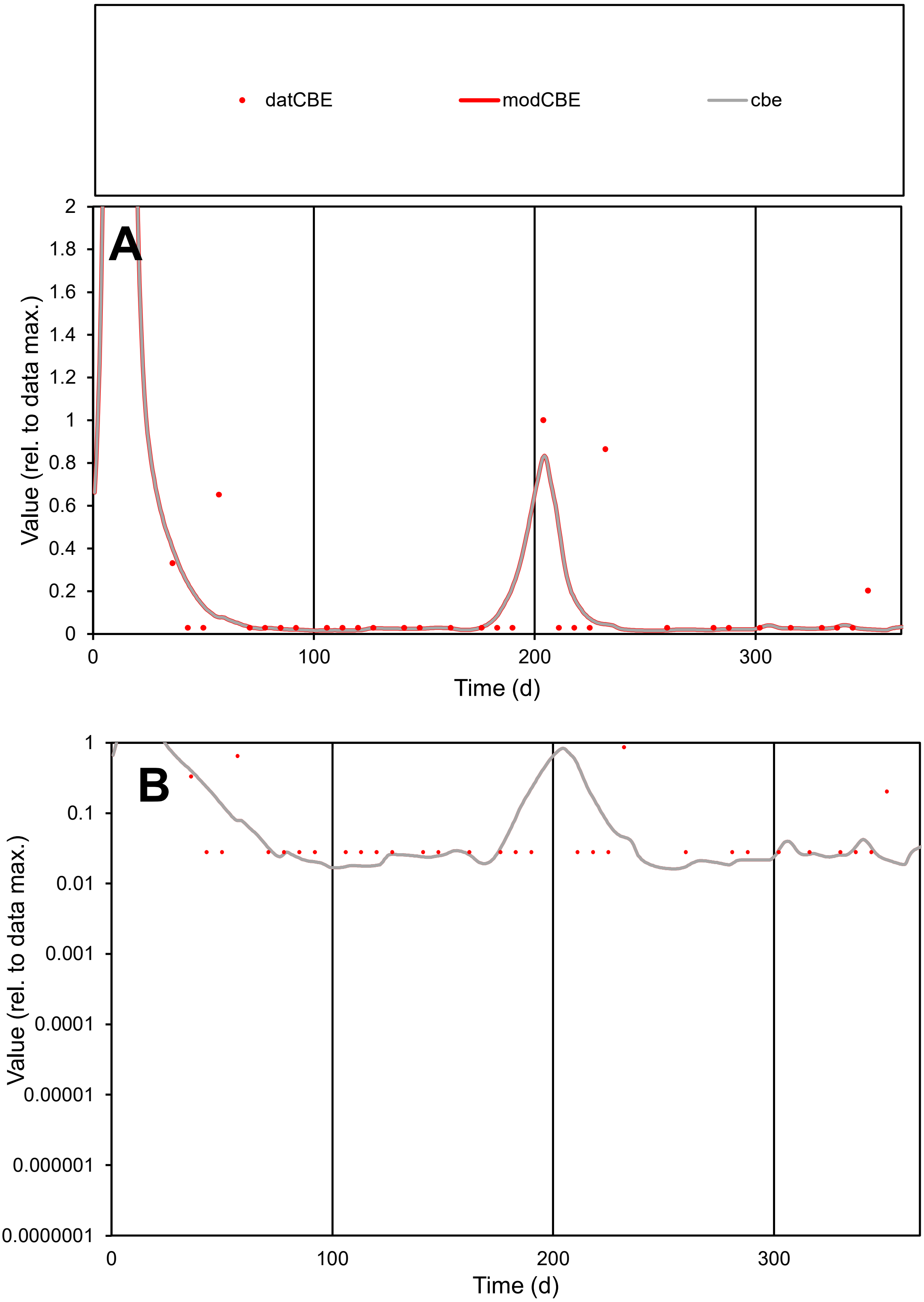


**C1**

Variables
CBE Provide jtd or Dst
CalibOnly?
2 1 to reproduce of
oNormalize?
2 1=no, 2=yes

ofxn calc*
1.62501
ofxn mod
1.66294

sRun
RunA1
zout
1

Ave
Data
7.1E-06
Model
5.3E-06

data/vMax
6E-05

ofxng
321.551

Min d/m
1.58254

ave/min d
4.25689
ave/min m
5.03128

324_Calib_S_A

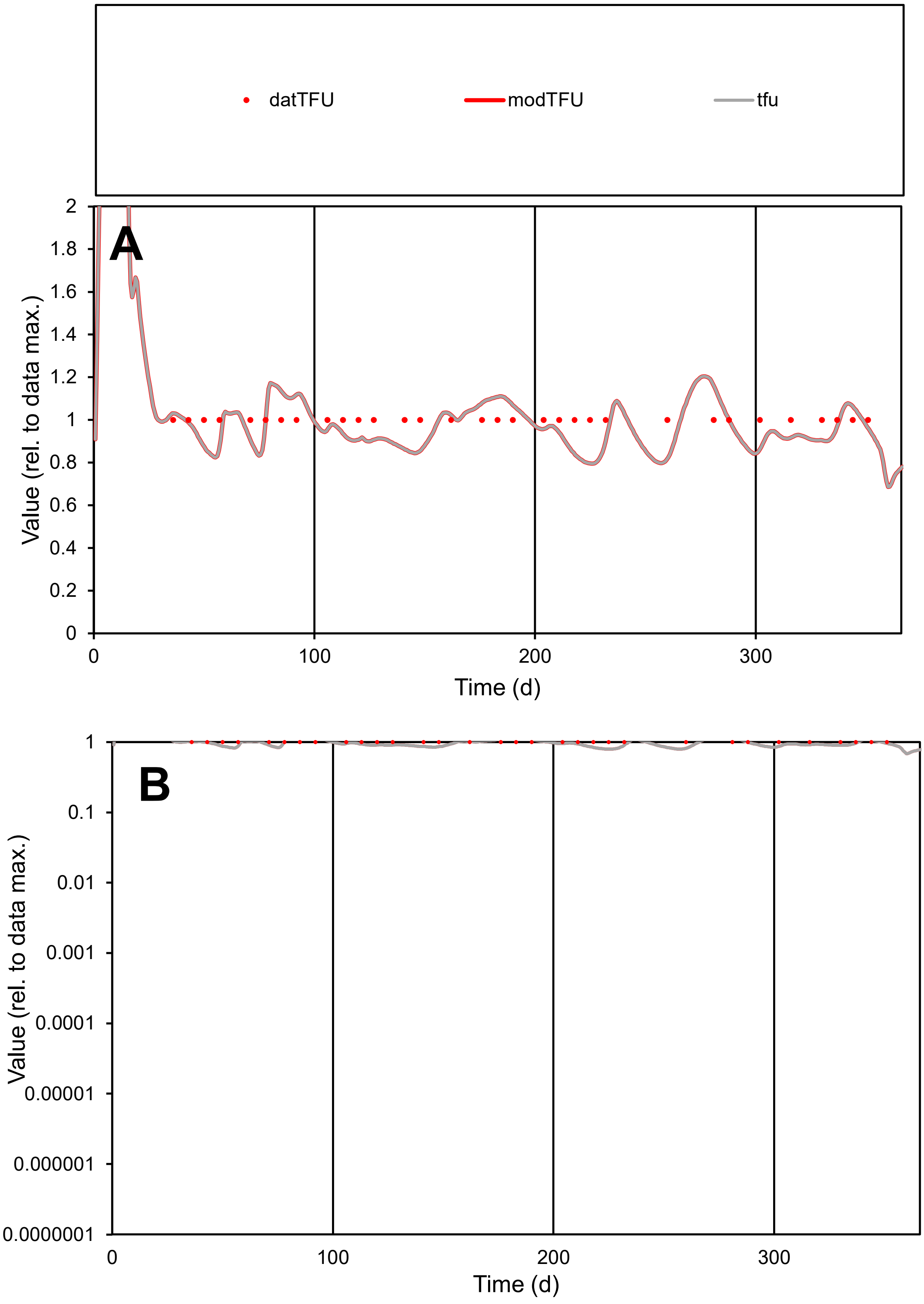


**C1**

Variables
TFU Provide jtd or Dst
CalibOnly?
2 1 to reproduce of
oNormalize?
2 1=no, 2=yes

ofxn calc*
0.10174
ofxn mod
0.13933

sRun
RunA1
zout
1

Ave
Data
1.3E-05
Model
1.2E-05

data/vMax
1.3E-05

ofxng
321.551

Min d/m
1.2574

ave/min d
1
ave/min m
1.21521

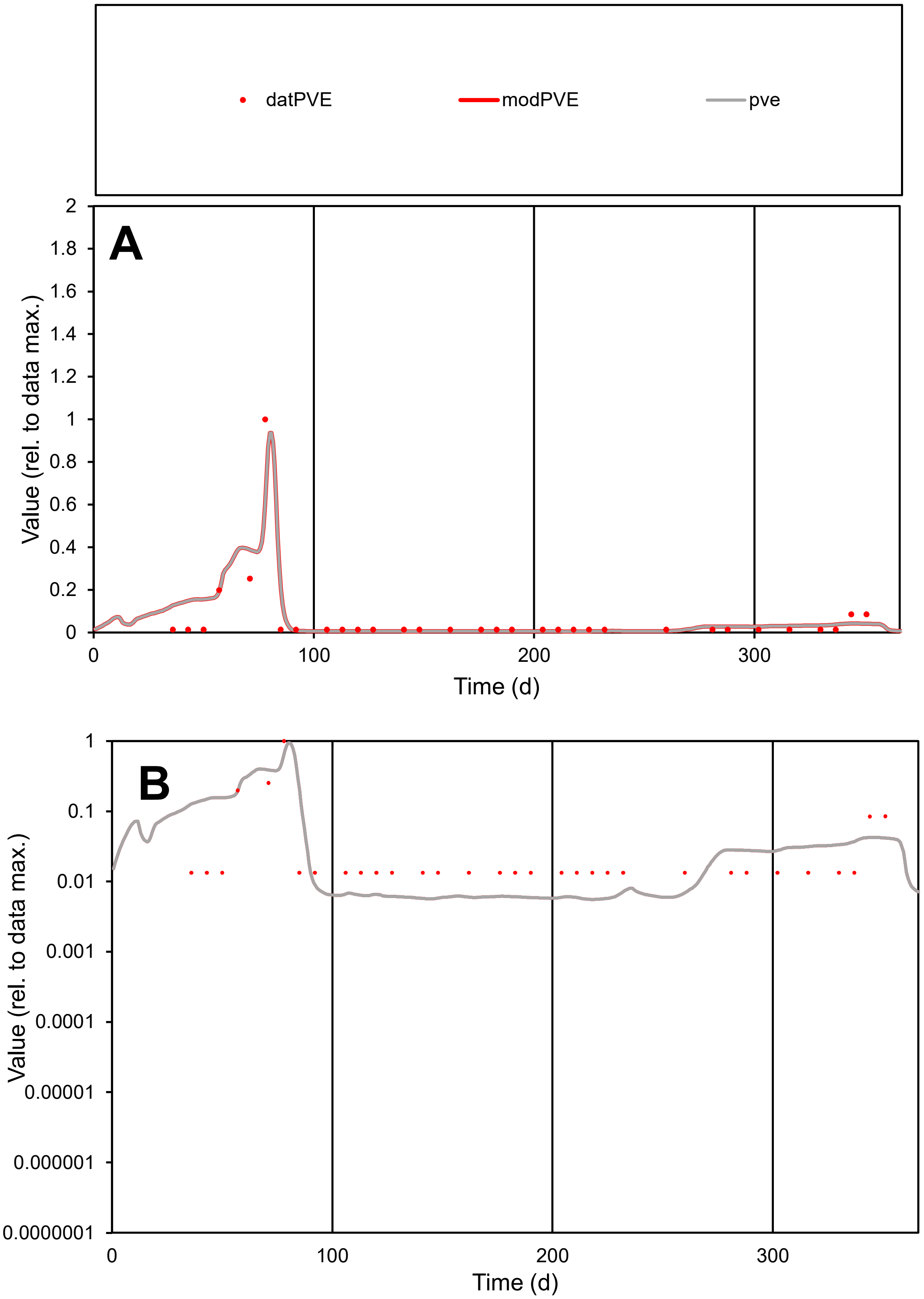


**C1**

Variables
PVE Provide jtd or Dst
CalibOnly?
2 1 to reproduce of
oNormalize?
2 1=no, 2=yes

ofxn calc*
0.88725
ofxn mod
0.94148

sRun
RunA1
zout
1

Ave
Data
9.7E-06
Model
1.1E-05

data/vMax
0.00016

ofxng
321.551

Min d/m
2.38052
ave/min d
4.65051
ave/min m
13.0562

324_Calib_S_A

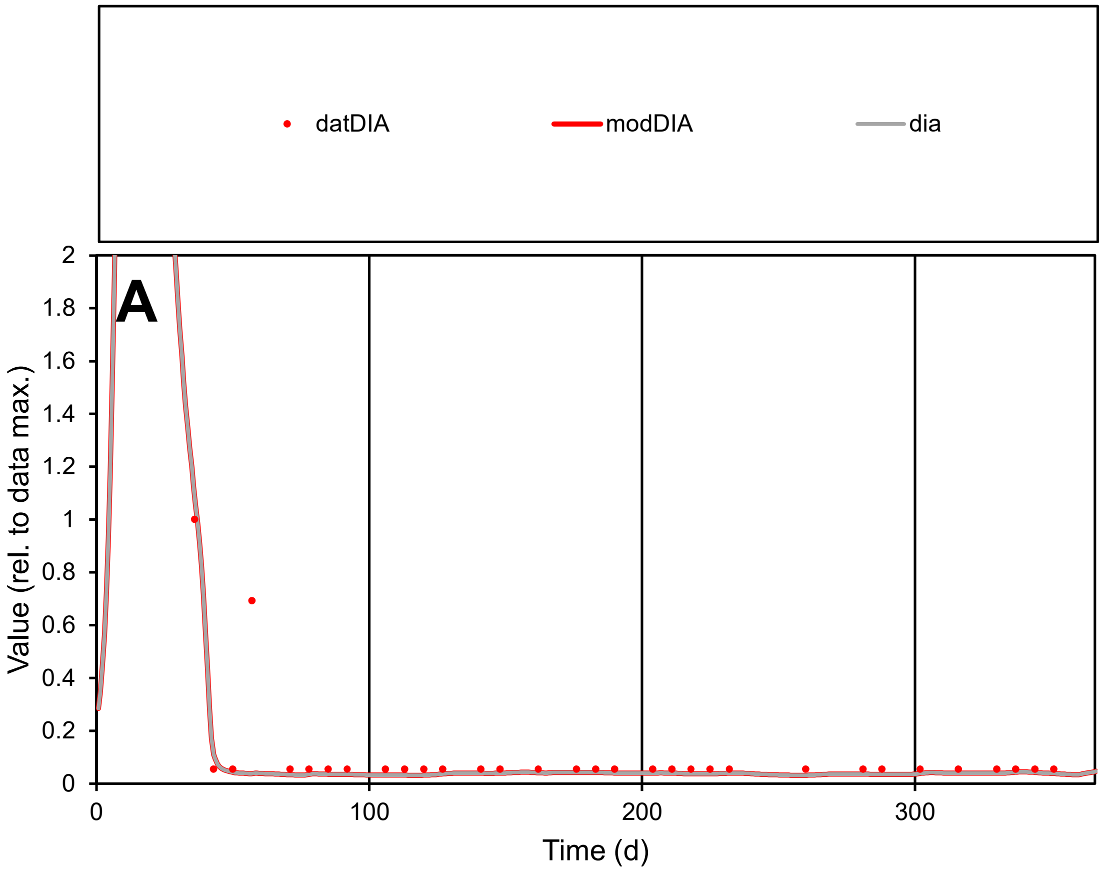


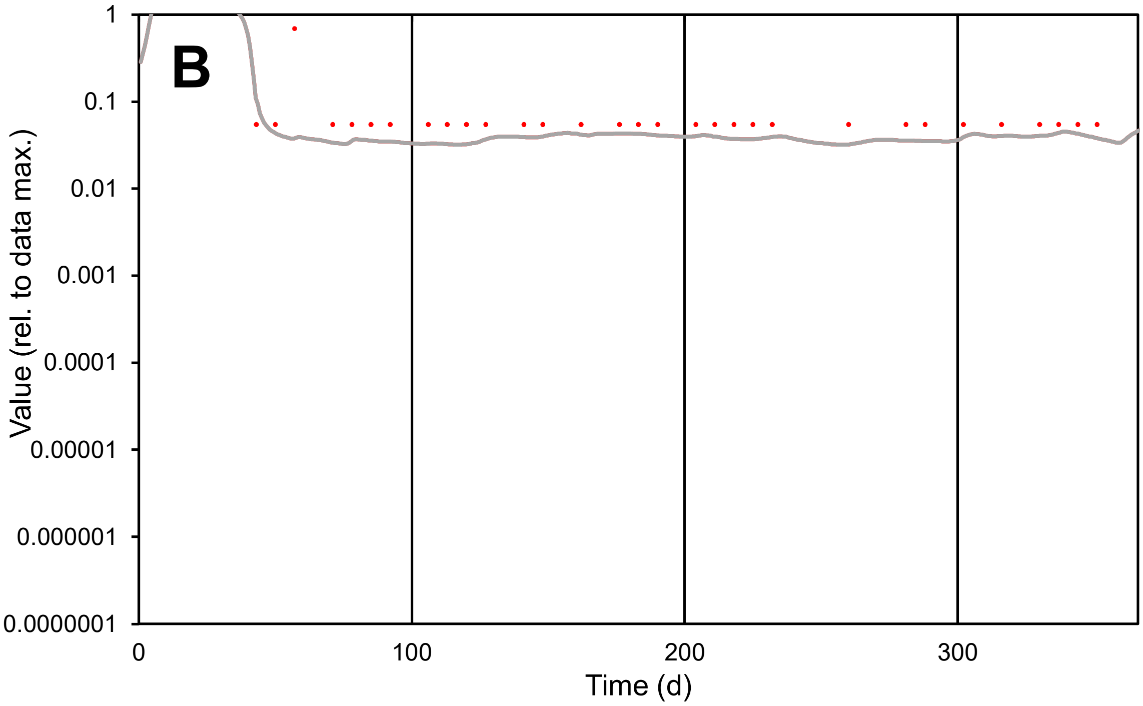


**C1**

Variables
DIA Provide jtd or Dst
CalibOnly?
2 1 to reproduce of
oNormalize?
2 1=no, 2=yes

ofxn calc*
1.11884
ofxn mod
1.12821

sRun
RunA1
zout
1

Ave
Data
1.2E-05
Model
7.8E-06

data/vMax
0.00011

ofxng
321.551

Min d/m
1.69997

ave/min d
1.90215
ave/min m
2.13966

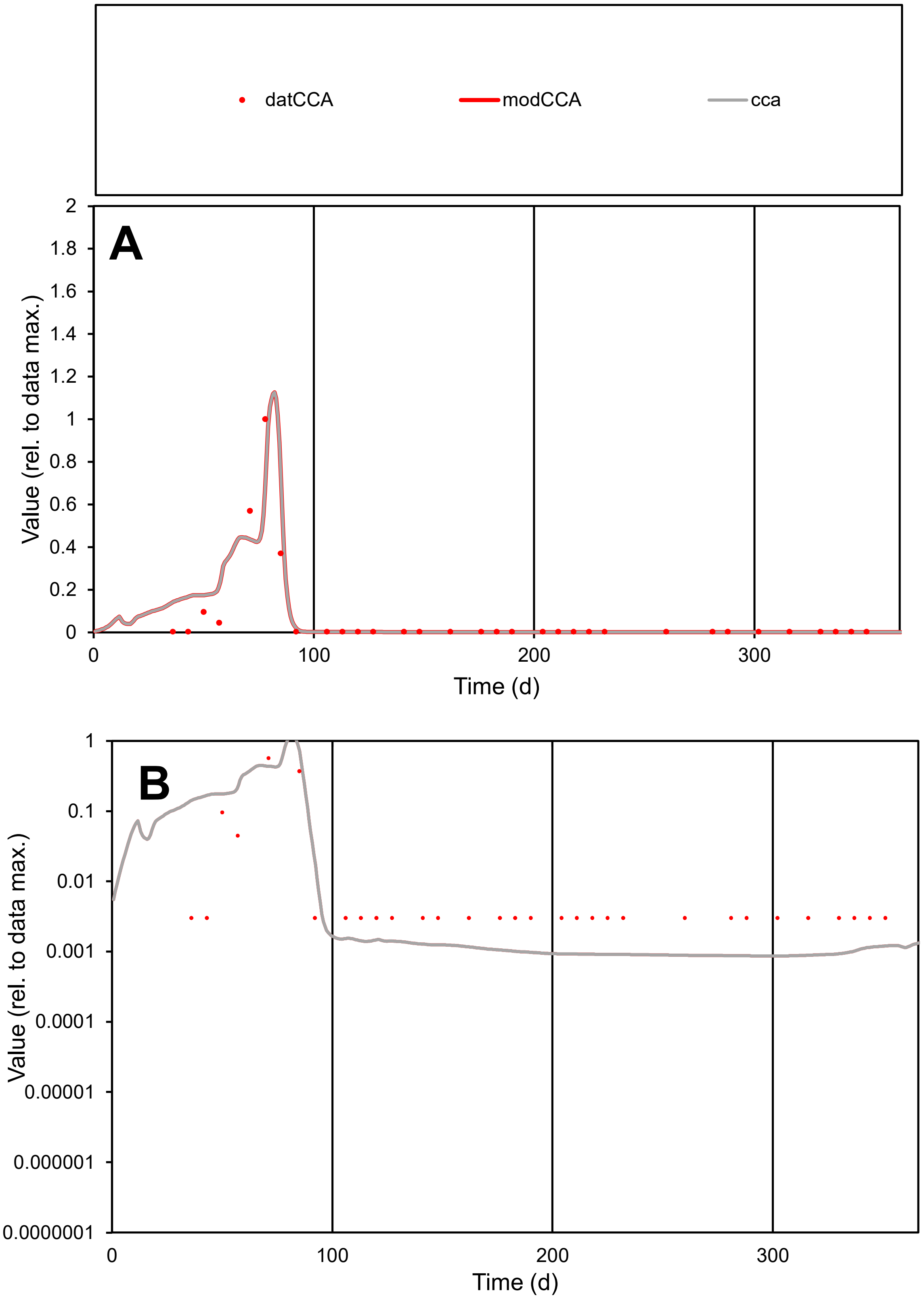


**C1**

Variables
CCA Provide jtd or Dst
CalibOnly?
2 1 to reproduce of
oNormalize?
2 1=no, 2=yes

ofxn calc*
0.81691
ofxn mod
0.89189

sRun
RunA1
zout
1

Ave
Data
4.1E-05
Model
4.7E-05

data/vMax
0.00061

ofxng
321.551

Min d/m
3.47067

ave/min d
22.482
ave/min m
89.8714

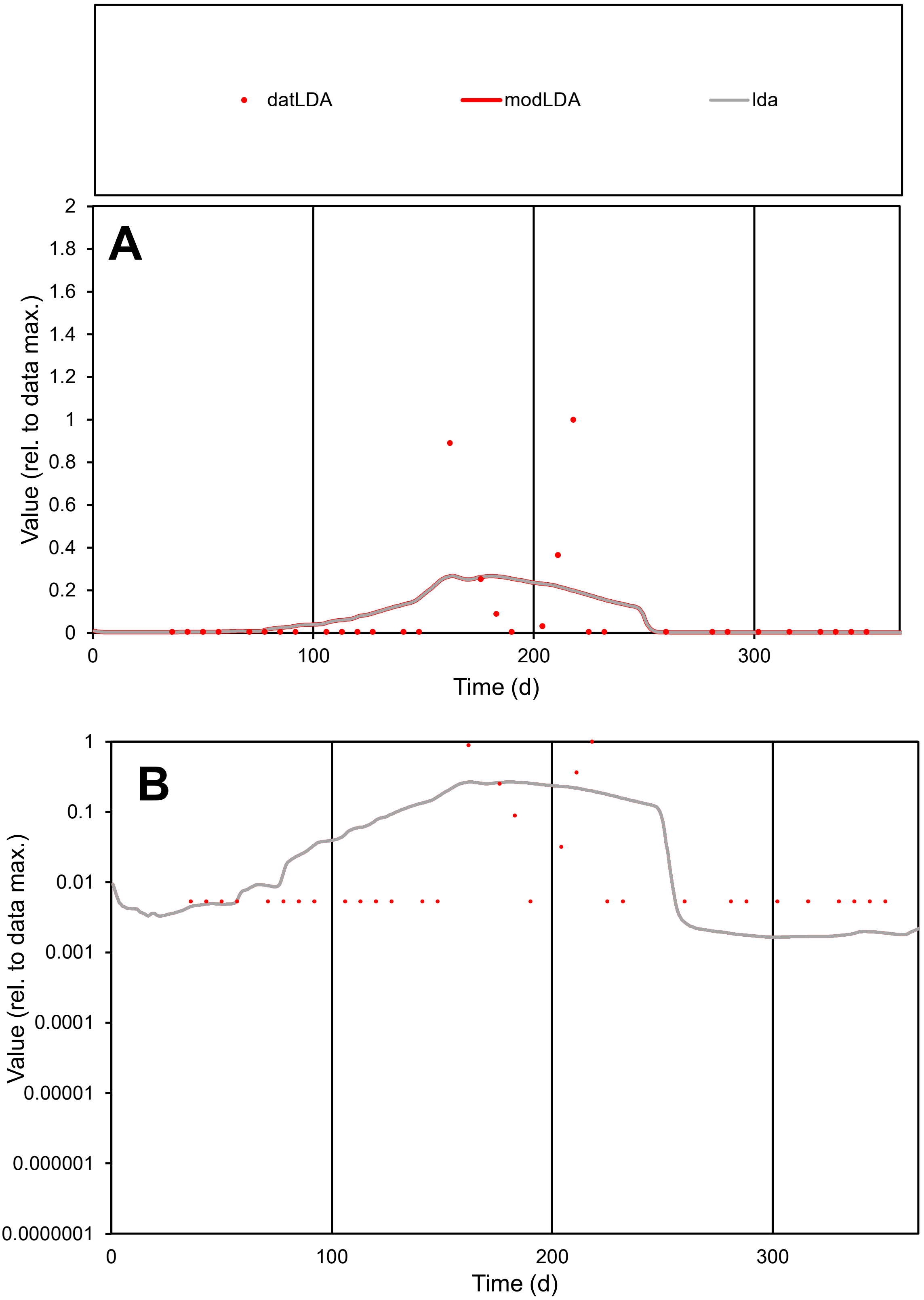


**C1**

Variables
LDA Provide jtd or Dst
CalibOnly?
2 1 to reproduce of
oNormalize?
2 1=no, 2=yes

ofxn calc*
2.32637
ofxn mod
2.34841

sRun
RunA1
zout
1

Ave
Data
0.00011
Model
0.00011

data/vMax
0.0013

ofxng
321.551

Min d/m
3.22041
ave/min d
16.2699
ave/min m
51.1145

324_Calib_S_A

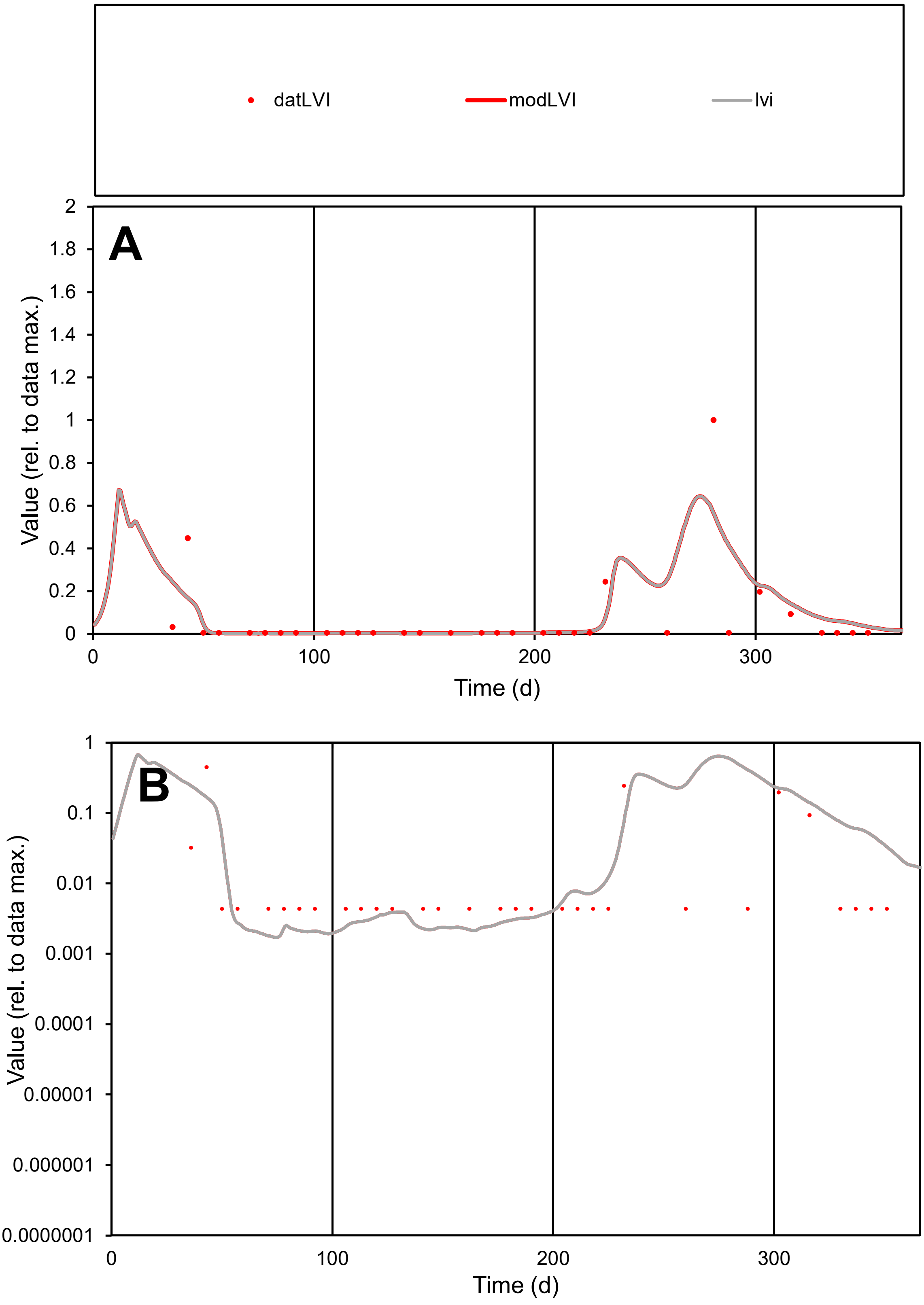


**C1**

Variables

LVI Provide jtd or Dst

CalibOnly?

2 1 to reproduce of

oNormalize?

2 1=no, 2=yes

ofxn calc*

1.93438

ofxn mod

2.00786

sRun

RunA1

zout

1

Ave

Data

2.2E-05

Model

2.4E-05

data/vMax

0.00033

ofxng

321.551

Min d/m

2.46021

ave/min d

15.2416

ave/min m

41.8323

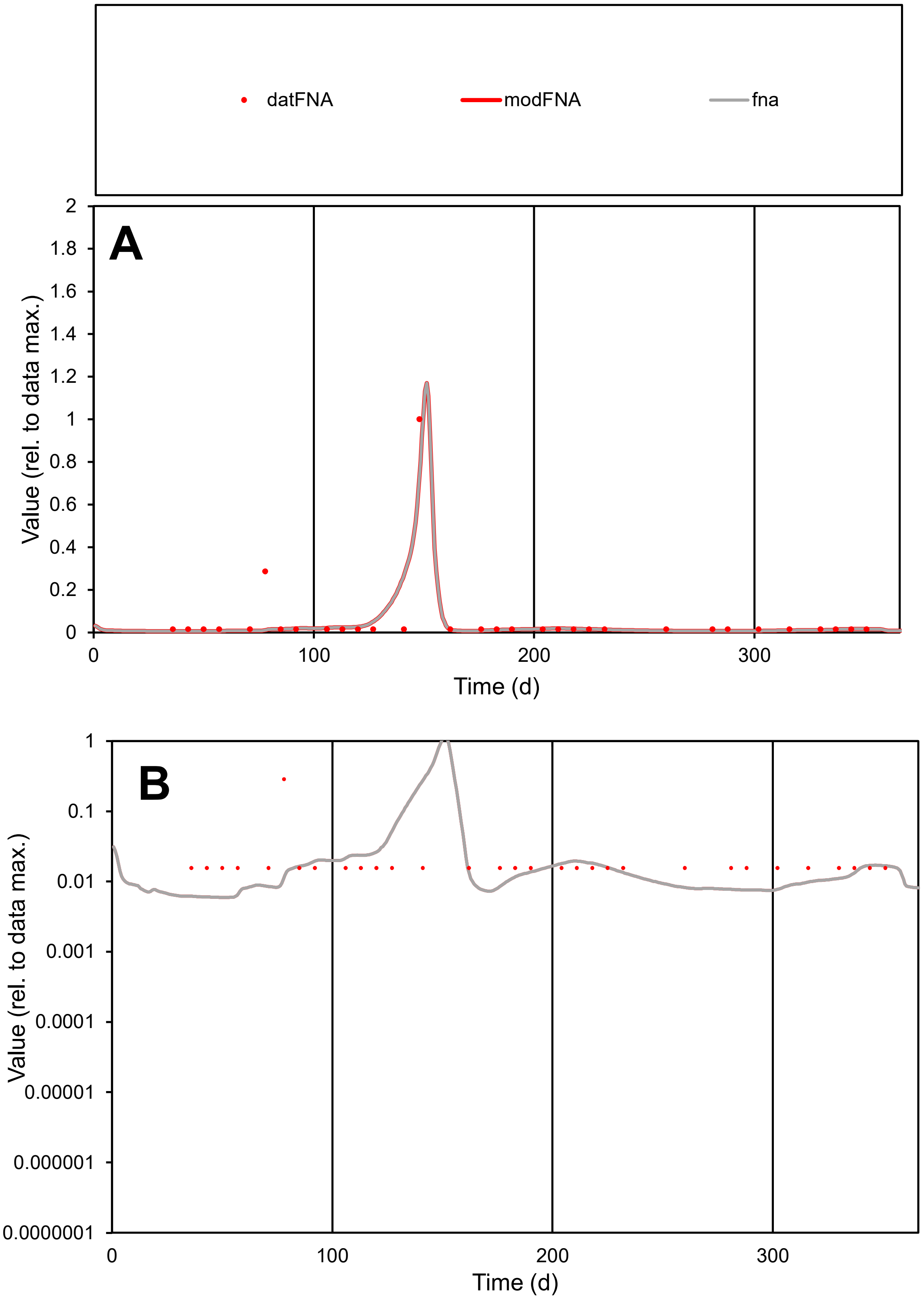


**C1**

Variables
FNA Provide jtd or Dst
CalibOnly?
2 1 to reproduce of
oNormalize?
2 1=no, 2=yes

ofxn calc*
1.16804
ofxn mod
1.24352

sRun
RunA1
zout
1

Ave
Data
1.1E-05
Model
9.9E-06

data/vMax
0.0002

ofxng
321.551

Min d/m
2.64461

ave/min d
3.51307
ave/min m
8.42364

324_Calib_S_A

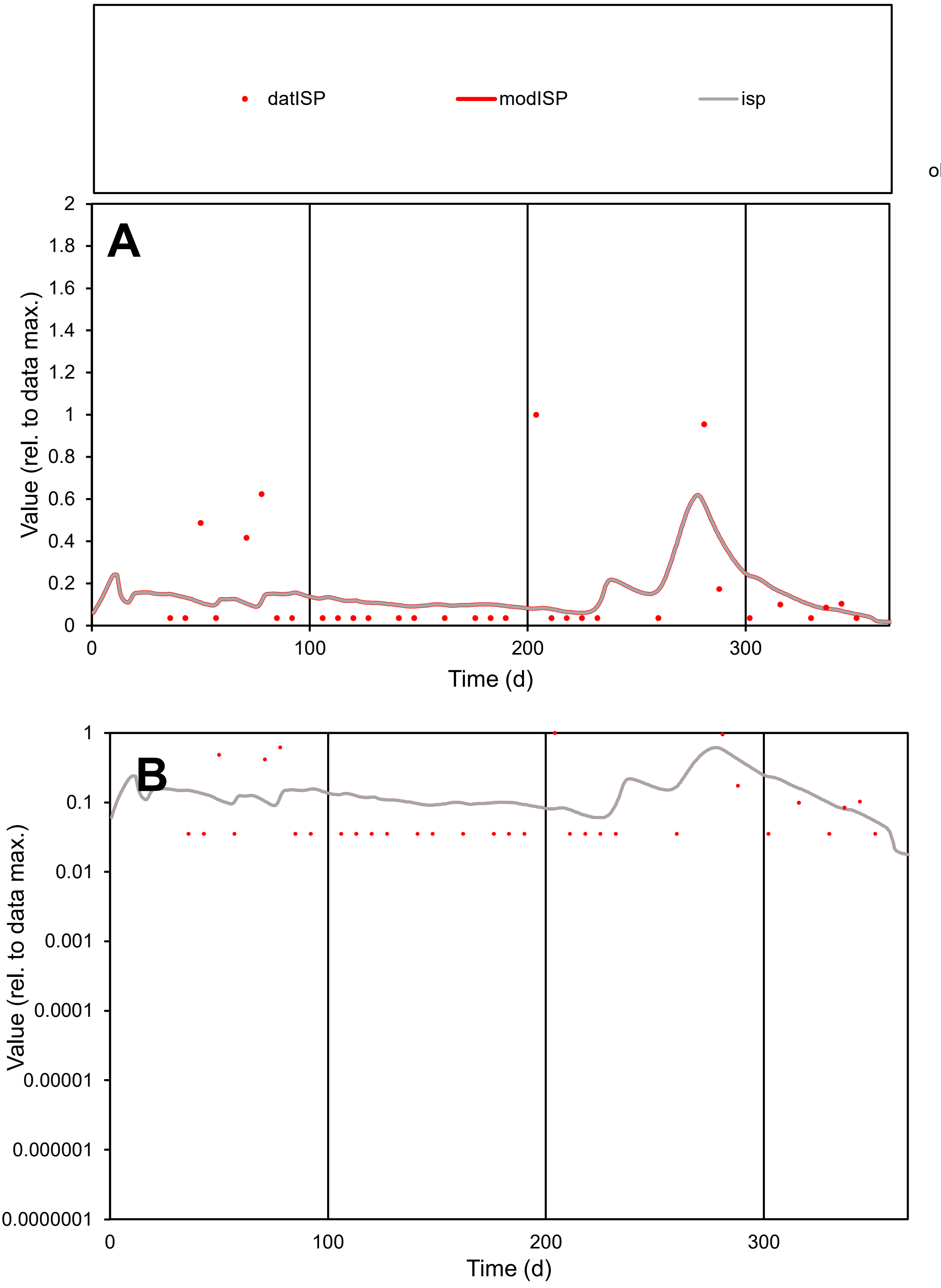


**C1**

Variables
ISP Provide jtd or Dst
CalibOnly?
2 1 to reproduce of
oNormalize?
2 1=no, 2=yes

ofxn calc*
1.54635
ofxn mod
1.56045

sRun
RunA1
zout
1

Ave
Data
2.1E-05
Model
1.9E-05

data/vMax
0.00014

ofxng
321.551

Min d/m
0.6786

ave/min d
4.21023
ave/min m
2.55958

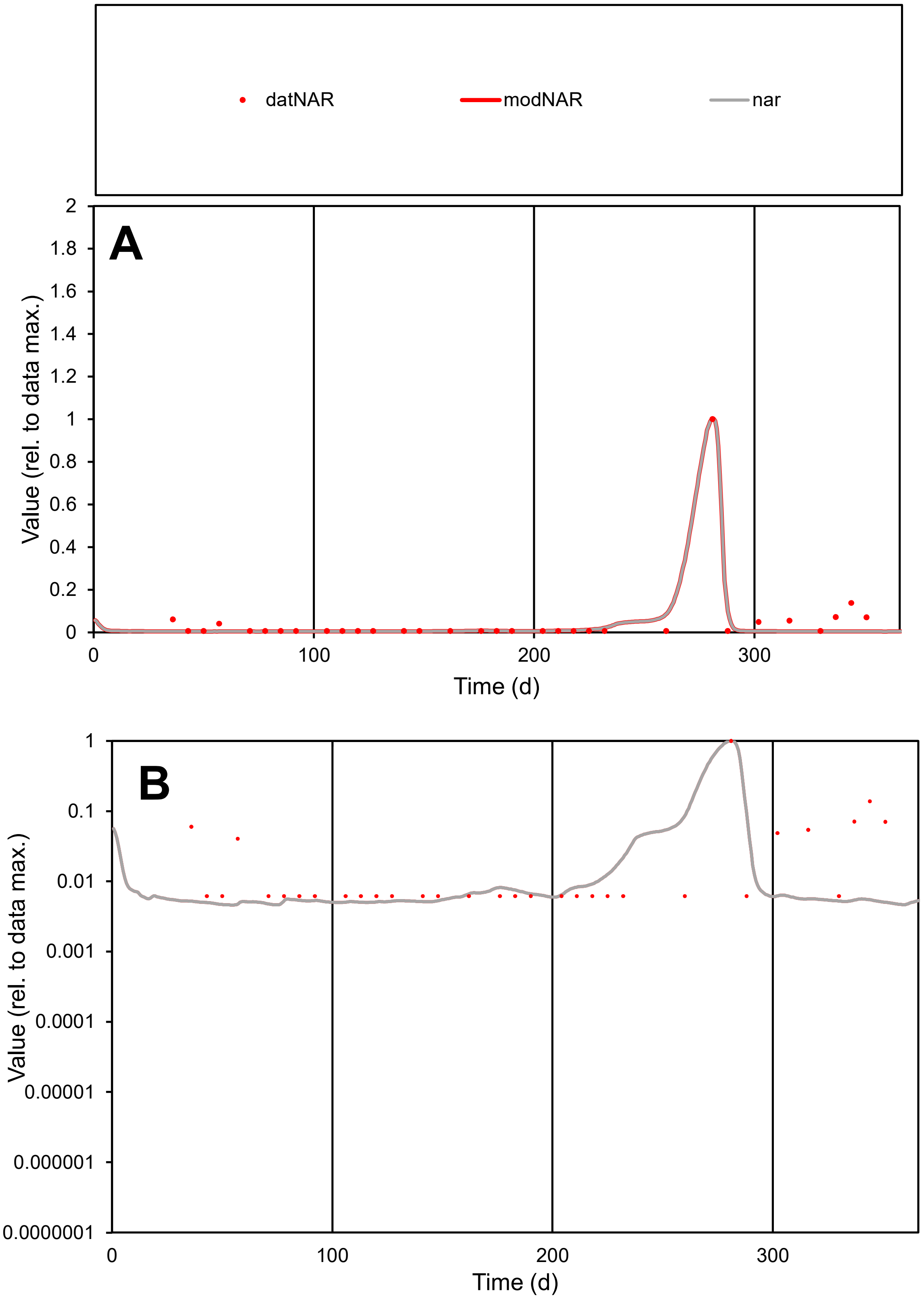


**C1**

Variables

NAR Provide jtd or Dst

CalibOnly?

2 1 to reproduce of

oNormalize?

2 1=no, 2=yes

ofxn calc*

0.68982

ofxn mod

0.75715

sRun

RunA1

zout

1

Ave

Data

2.3E-05

Model

1.8E-05

data/vMax

0.00046

ofxng

321.551

Min d/m

1.33318

ave/min d

8.24347

ave/min m

8.62367

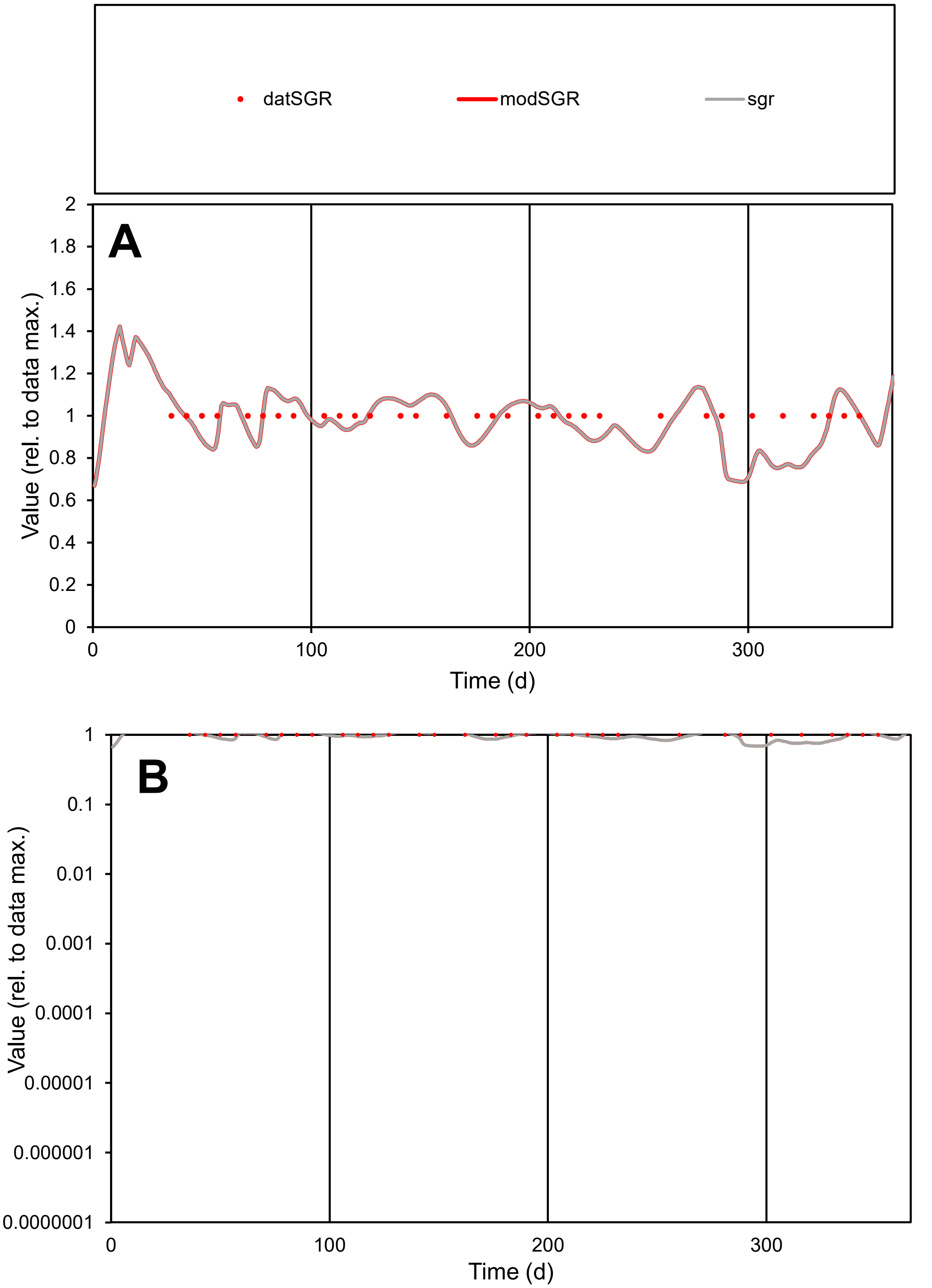


**C1**

Variables

| | |
|---|---|
| SGR | Provide jtd or Dst |
| CalibOnly? | |
| 2 | 1 to reproduce of |
| oNormalize? | |
| 2 | 1=no, 2=yes |

ofxn calc*
0.09217
ofxn mod
0.13797

sRun
RunA1
zout
1

Ave
Data
4.5E-06
Model
4.4E-06

data/vMax
4.5E-06

ofxng
321.551

Min d/m
1.30166

ave/min d
1
ave/min m
1.2702

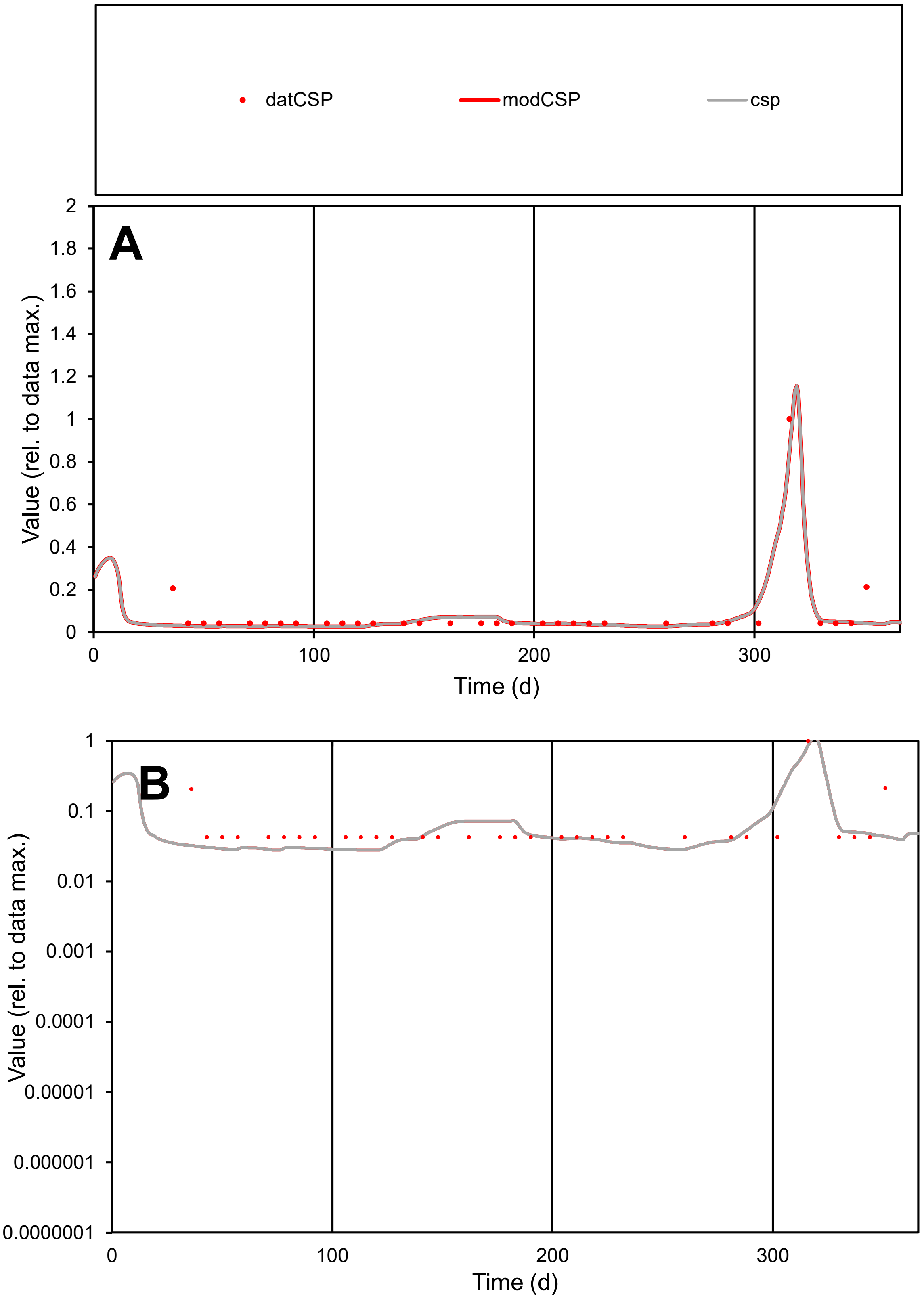


**C1**

Variables
CSP Provide jtd or Dst
CalibOnly?
2 1 to reproduce of
oNormalize?
2 1=no, 2=yes

ofxn calc*
0.56932
ofxn mod
0.60432

sRun
RunA1
zout
1

Ave
Data
4.6E-06
Model
4.1E-06

data/vMax
5.6E-05

ofxng
321.551

Min d/m
1.53013

ave/min d
1.94416
ave/min m
2.62316

324_Calib_S_A

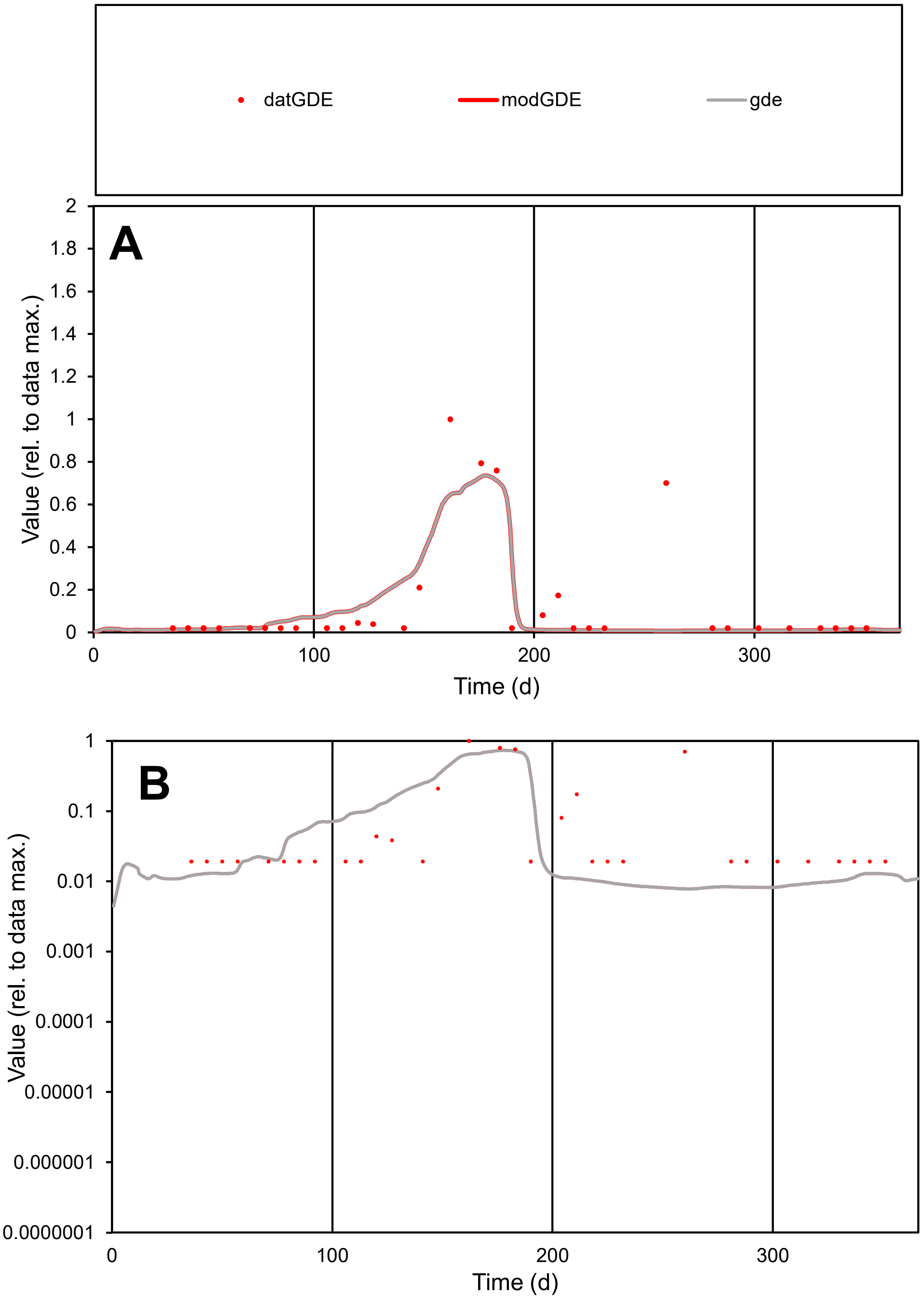


**C1**

Variables

GDE Provide jtd or Dst

CalibOnly?

2 1 to reproduce of

oNormalize?

2 1=no, 2=yes

ofxn calc*

1.14895

ofxn mod

1.17258

sRun

RunA1

zout

1

Ave

Data

0.00019

Model

0.00016

data/vMax

0.00143

ofxng

321.551

Min d/m

2.45138

ave/min d

6.8984

ave/min m

14.1142

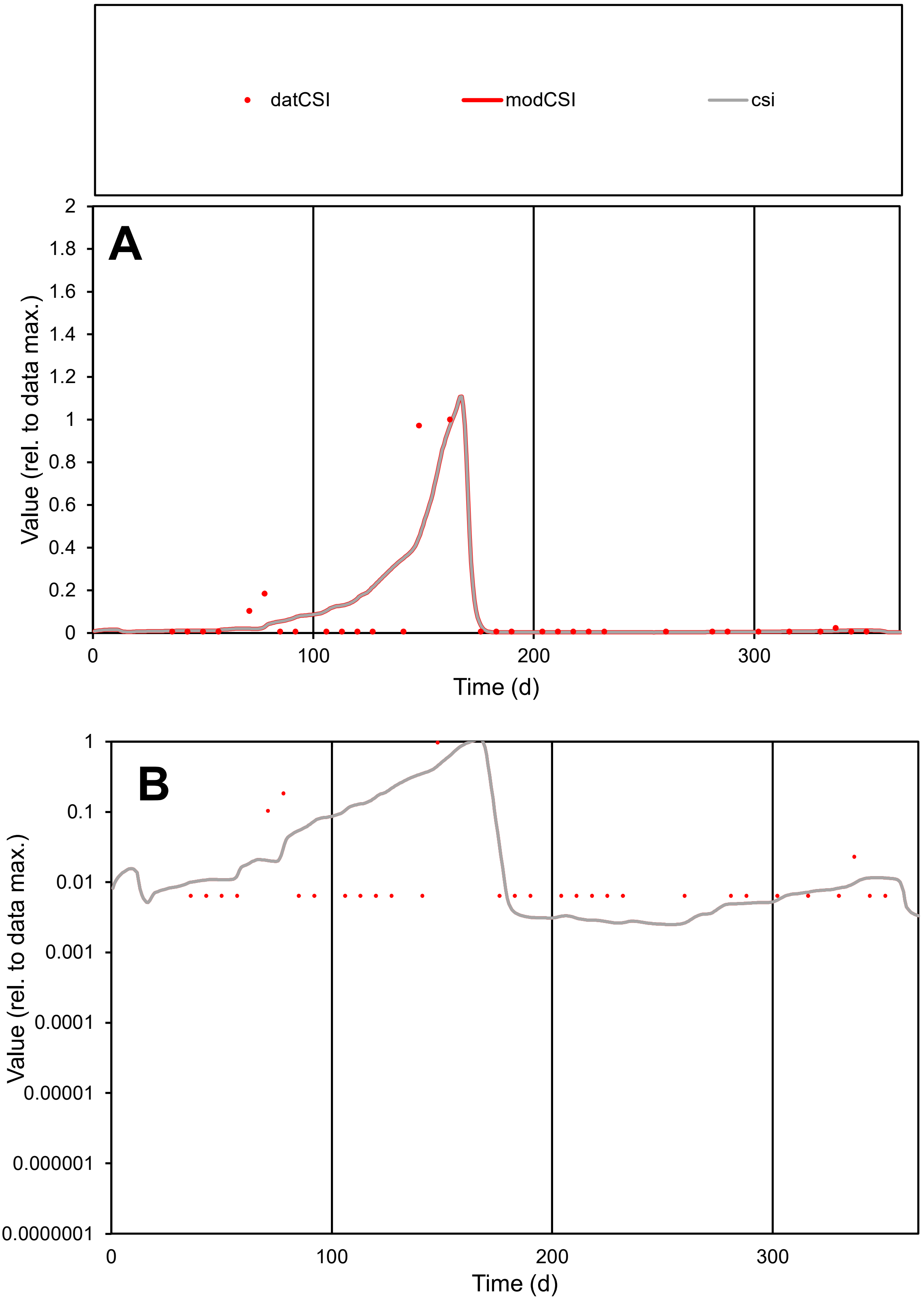


**C1**

Variables
CSI Provide jtd or Dst
CalibOnly?
2 1 to reproduce of
oNormalize?
2 1=no, 2=yes

ofxn calc*
1.59901
ofxn mod
1.6453

sRun
RunA1
zout
1

Ave
Data
2.8E-05
Model
3.1E-05

data/vMax
0.00036

ofxng
321.551

Min d/m
2.38046

ave/min d
11.9807
ave/min m
32.2635

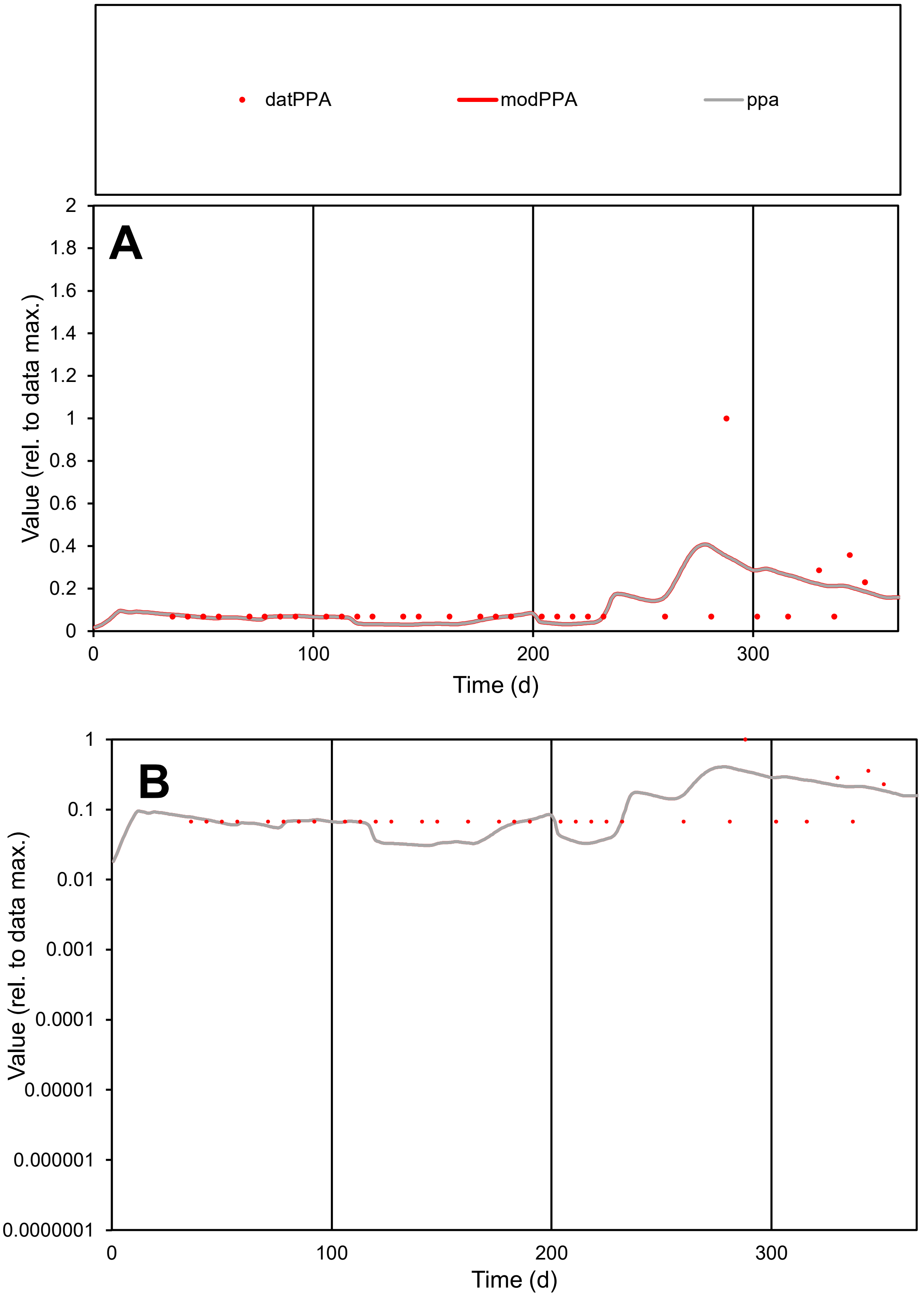


**C1**

Variables
PPA Provide jtd or Dst
CalibOnly?
2 1 to reproduce of
oNormalize?
2 1=no, 2=yes

ofxn calc*
1.2489
ofxn mod
1.25728

sRun
RunA1
zout
1

Ave
Data
5.6E-06
Model
5.2E-06

data/vMax
4.8E-05

ofxng
321.551

Min d/m
2.19631

ave/min d
1.74029
ave/min m
3.57644

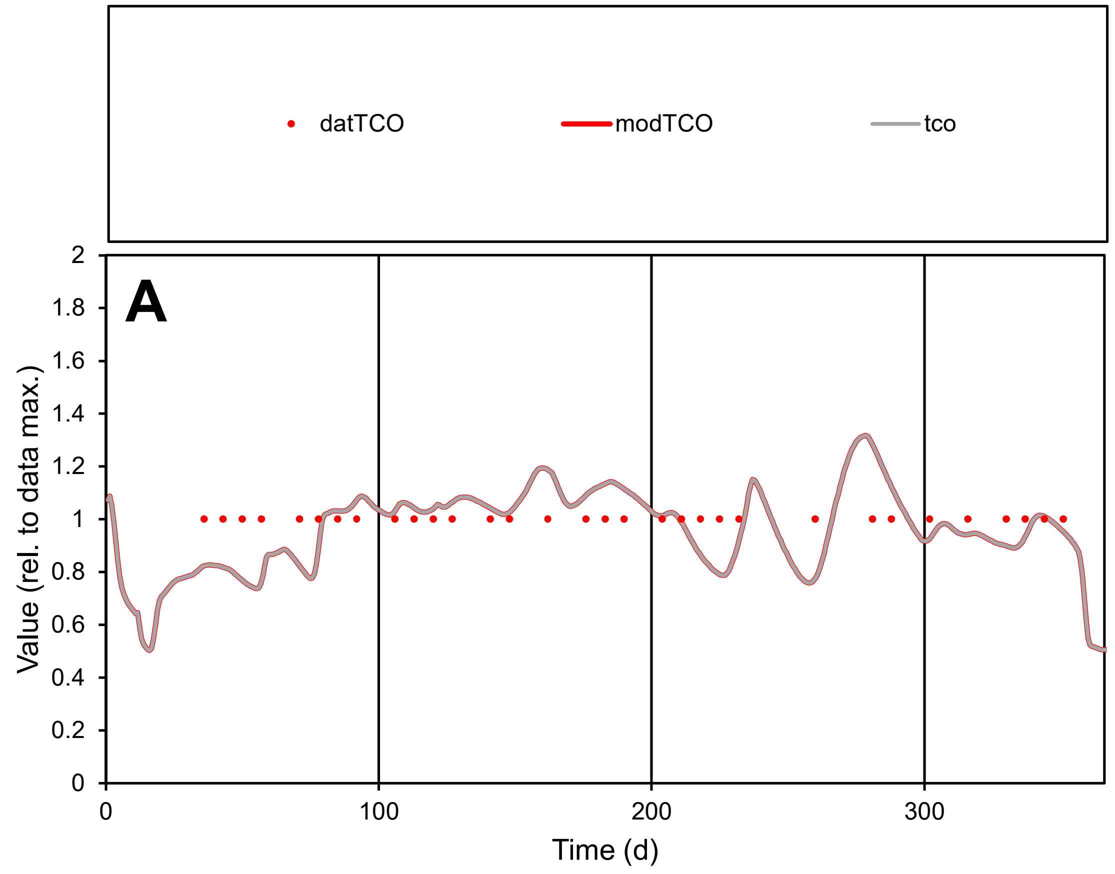


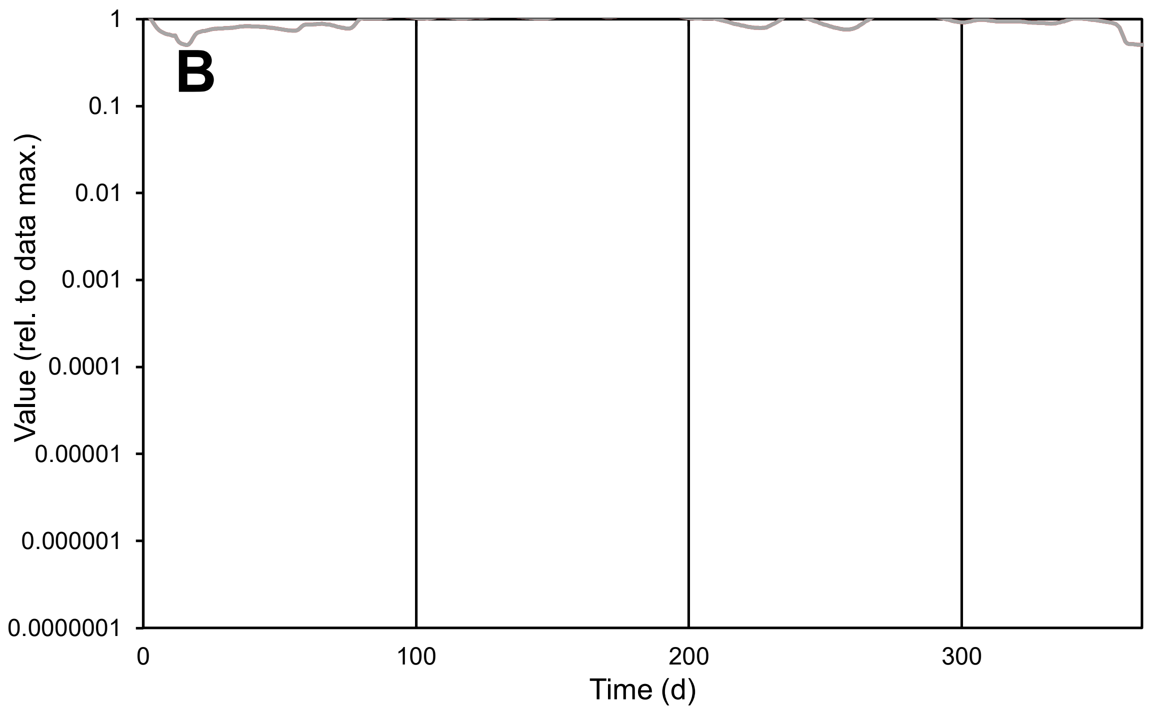


**C1**

Variables
TCO Provide jtd or Dst
CalibOnly?
2 1 to reproduce of
oNormalize?
2 1=no, 2=yes

ofxn calc*
0.12467
ofxn mod
0.15715

sRun
RunA1
zout
1

Ave
Data
8.6E-06
Model
8.5E-06

data/vMax
8.6E-06

ofxng
321.551

Min d/m
1.32376

ave/min d
1
ave/min m
1.29799

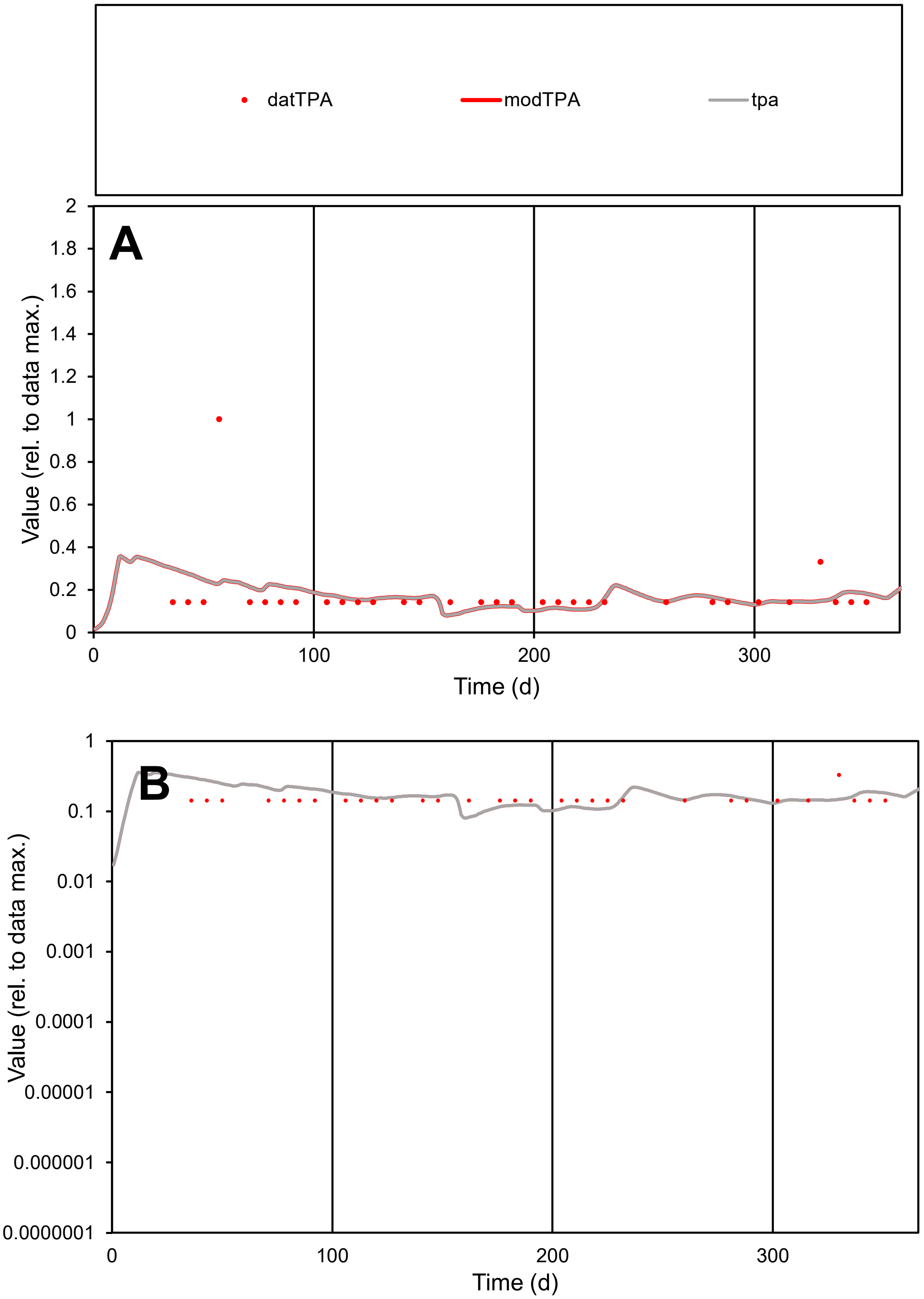


**C1**

Variables
TPA Provide jtd or Dst
CalibOnly?
2 1 to reproduce of
oNormalize?
2 1=no, 2=yes

ofxn calc*
0.84314
ofxn mod
0.85017

sRun
RunA1
zout
1

Ave
Data
5.8E-06
Model
5.6E-06

data/vMax
3.3E-05

ofxng
321.551

Min d/m
1.68534

ave/min d
1.22943
ave/min m
1.98561

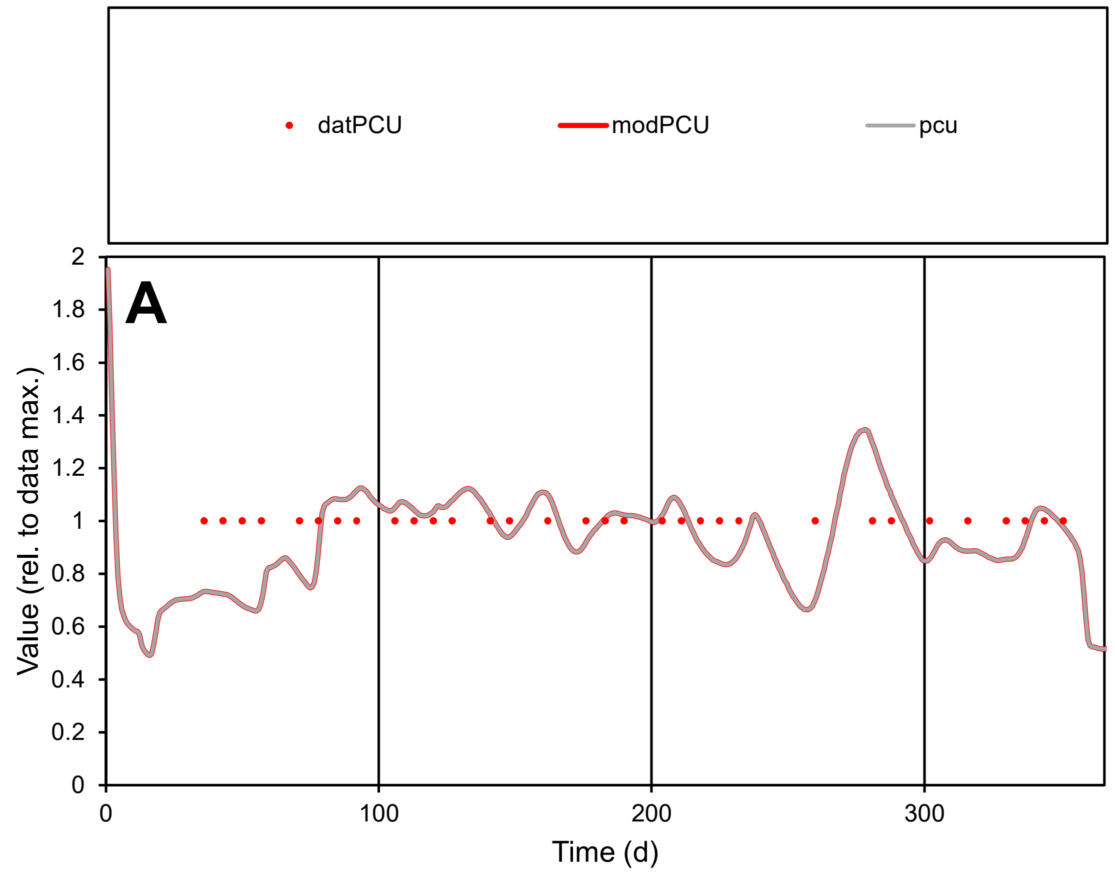


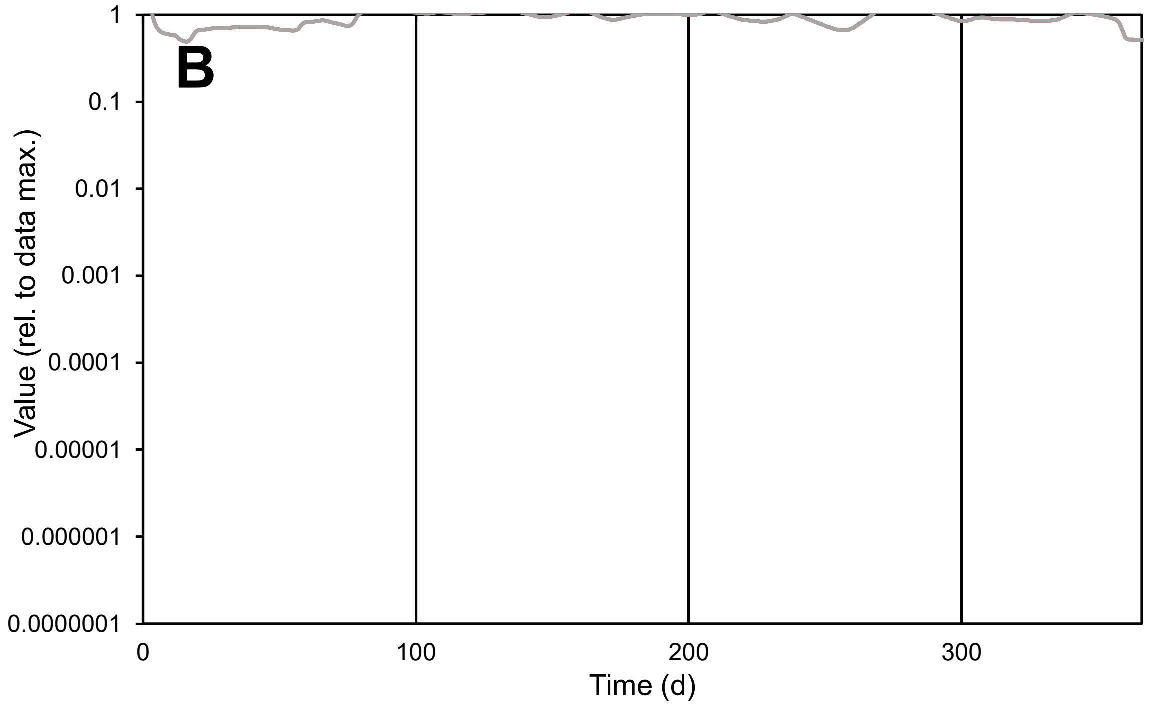


**C1**

Variables
PCU Provide jtd or Dst
CalibOnly?
2 1 to reproduce of
oNormalize?
2 1=no, 2=yes

ofxn calc*
0.14311
ofxn mod
0.17156

sRun
RunA1
zout
1

Ave
Data
6.2E-06
Model
5.9E-06

data/vMax
6.2E-06

ofxng
321.551

Min d/m
1.48966

ave/min d
1
ave/min m
1.41997

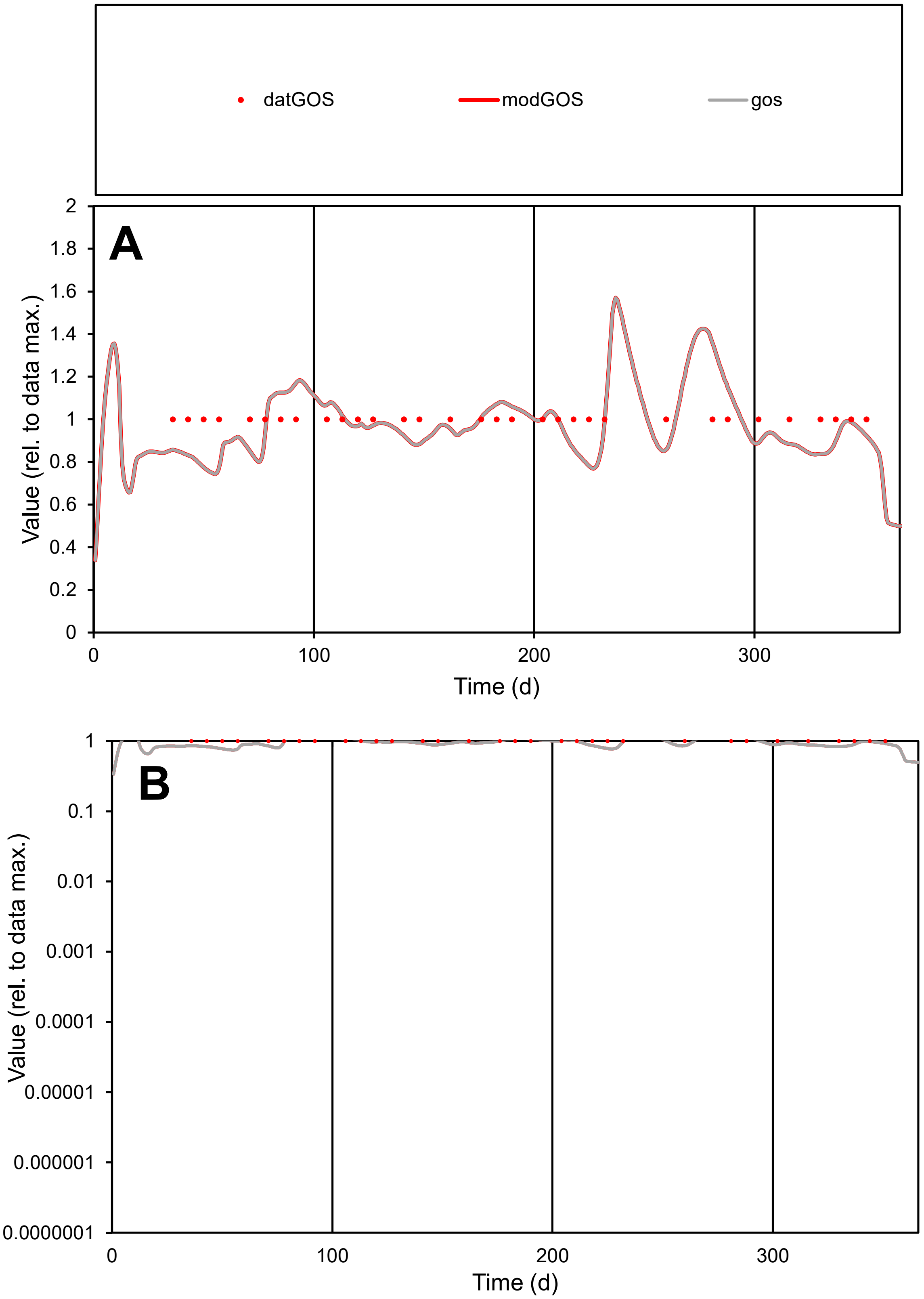


**C1**

Variables
GOS Provide jtd or Dst
CalibOnly?
2 1 to reproduce of
oNormalize?
2 1=no, 2=yes

ofxn calc*
0.12915
ofxn mod
0.16307

sRun
RunA1
zout
1

Ave
Data
1.8E-06
Model
1.7E-06

data/vMax
1.8E-06

ofxng
321.551

Min d/m
1.31209

ave/min d
1
ave/min m
1.26521

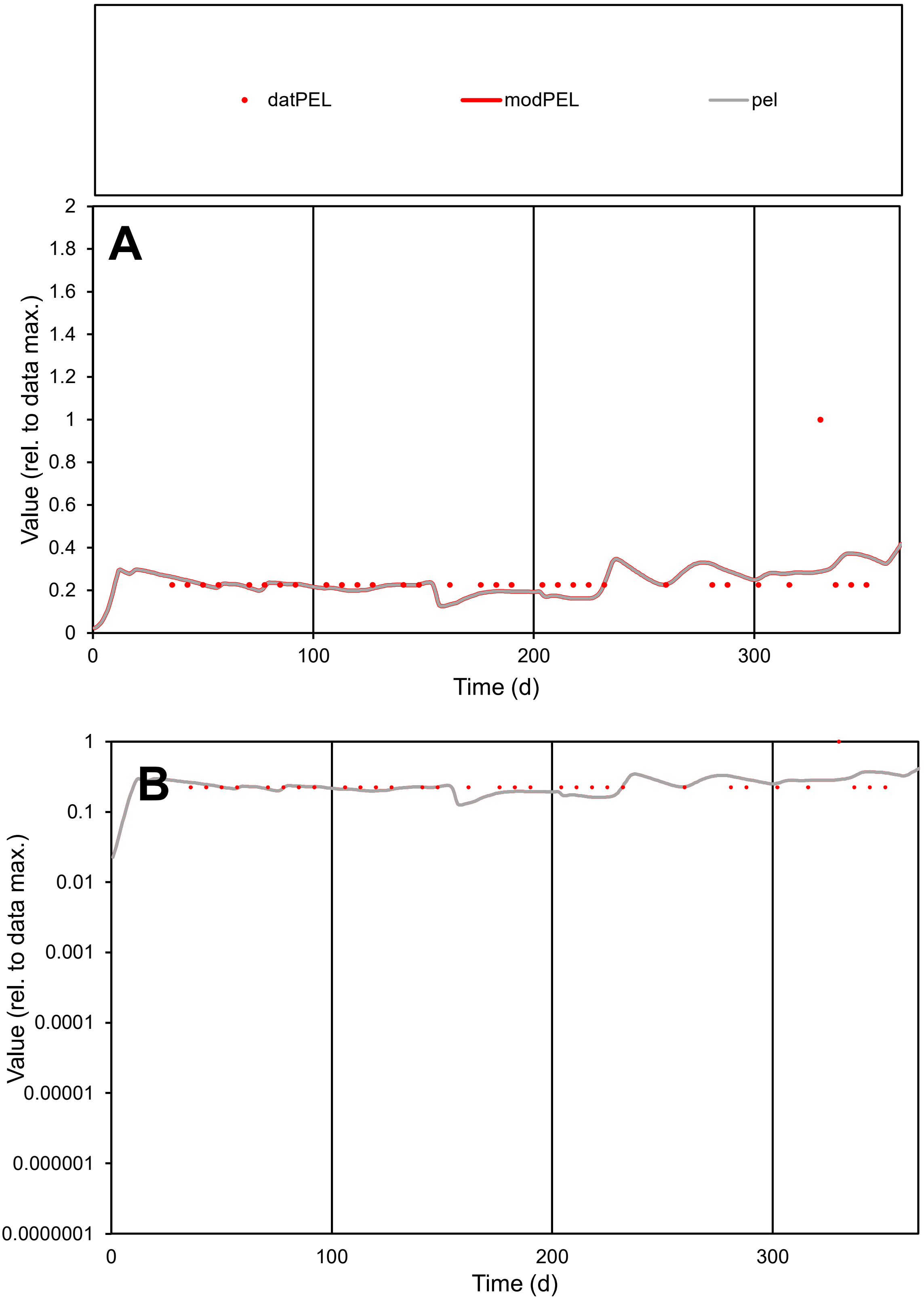


**C1**

Variables
PEL Provide jtd or Dst
CalibOnly?
2 1 to reproduce of
oNormalize?
2 1=no, 2=yes

ofxn calc*
0.55041
ofxn mod
0.56131

sRun
RunA1
zout
1

Ave
Data
3.6E-06
Model
3.4E-06

data/vMax
1.5E-05

ofxng
321.551

Min d/m
1.62959

ave/min d
1.10805
ave/min m
1.70294

324_Calib_S_A

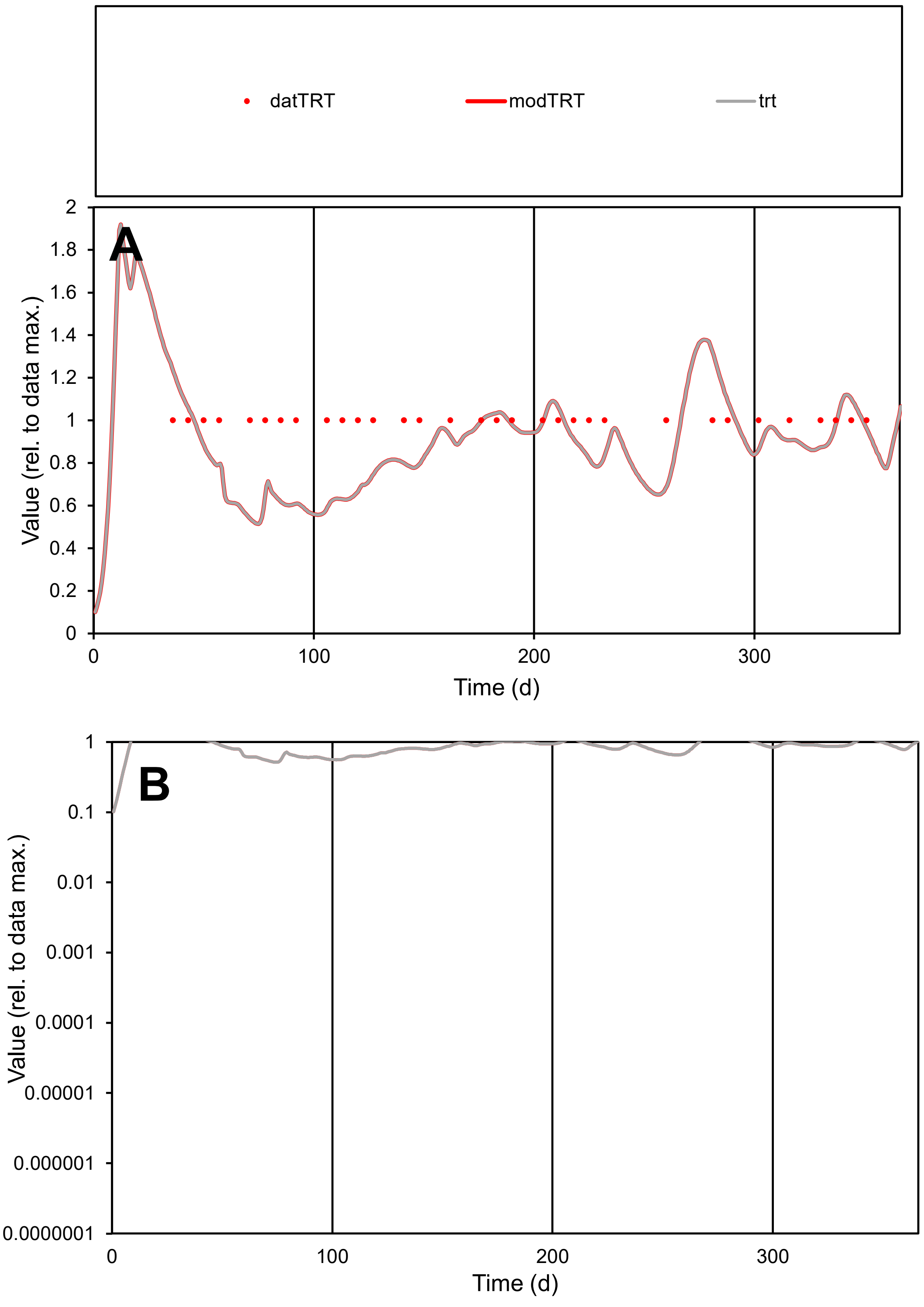


**C1**

Variables
TRT Provide jtd or Dst
CalibOnly?
2 1 to reproduce of
oNormalize?
2 1=no, 2=yes

ofxn calc*
0.21838
ofxn mod
0.23666

sRun
RunA1
zout
1

Ave
Data
2.3E-05
Model
2E-05

data/vMax
2.3E-05

ofxng
321.551

Min d/m
1.88374

ave/min d
1
ave/min m
1.64499

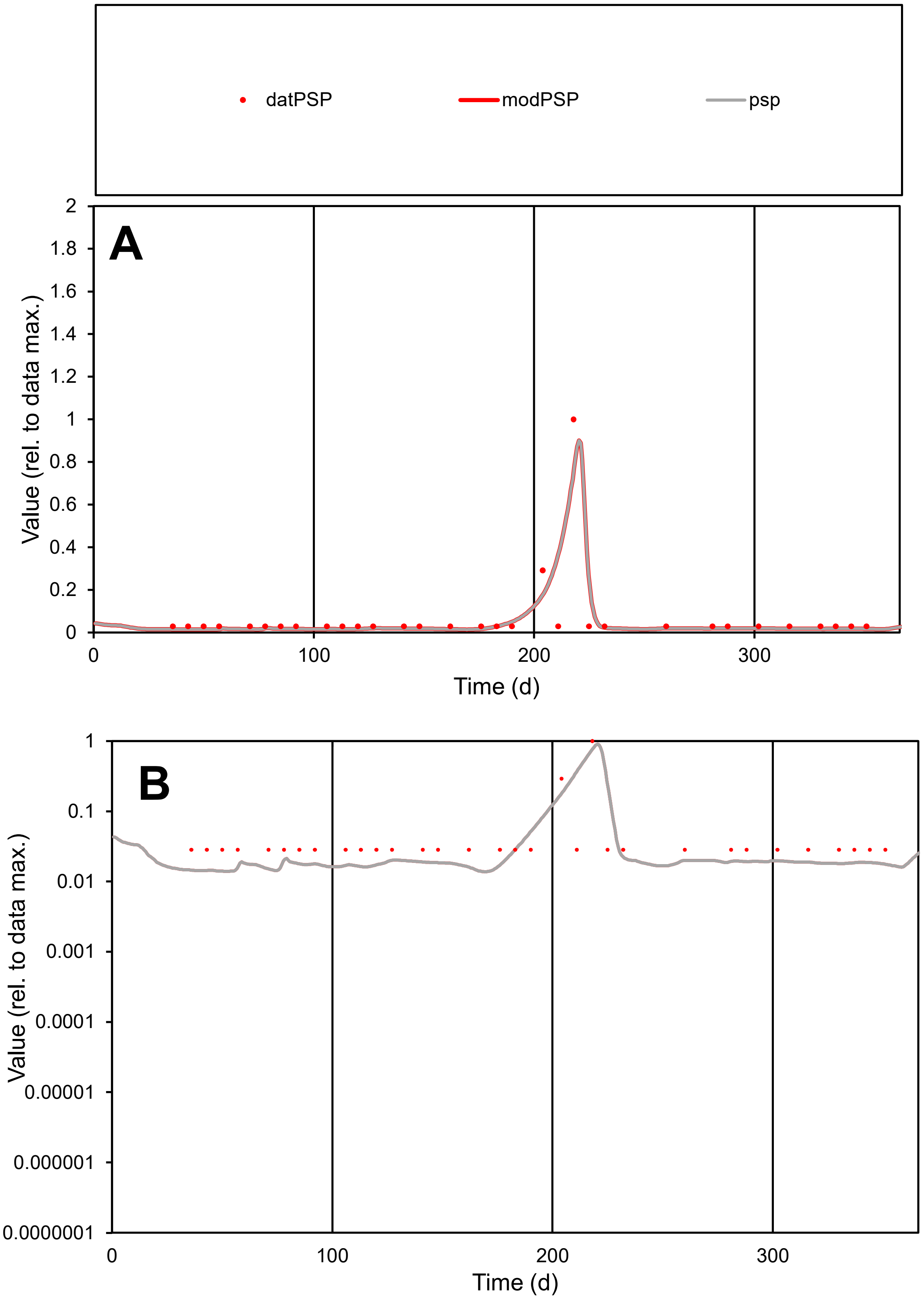


**C1**

Variables
PSP Provide jtd or Dst
CalibOnly?
2 1 to reproduce of
oNormalize?
2 1=no, 2=yes

ofxn calc*
0.87984
ofxn mod
1.04027

sRun
RunA1
zout
1

Ave
Data
1E-05
Model
9.5E-06

data/vMax
0.00015

ofxng
321.551

Min d/m
2.03151
ave/min d
2.36173
ave/min m
4.48881

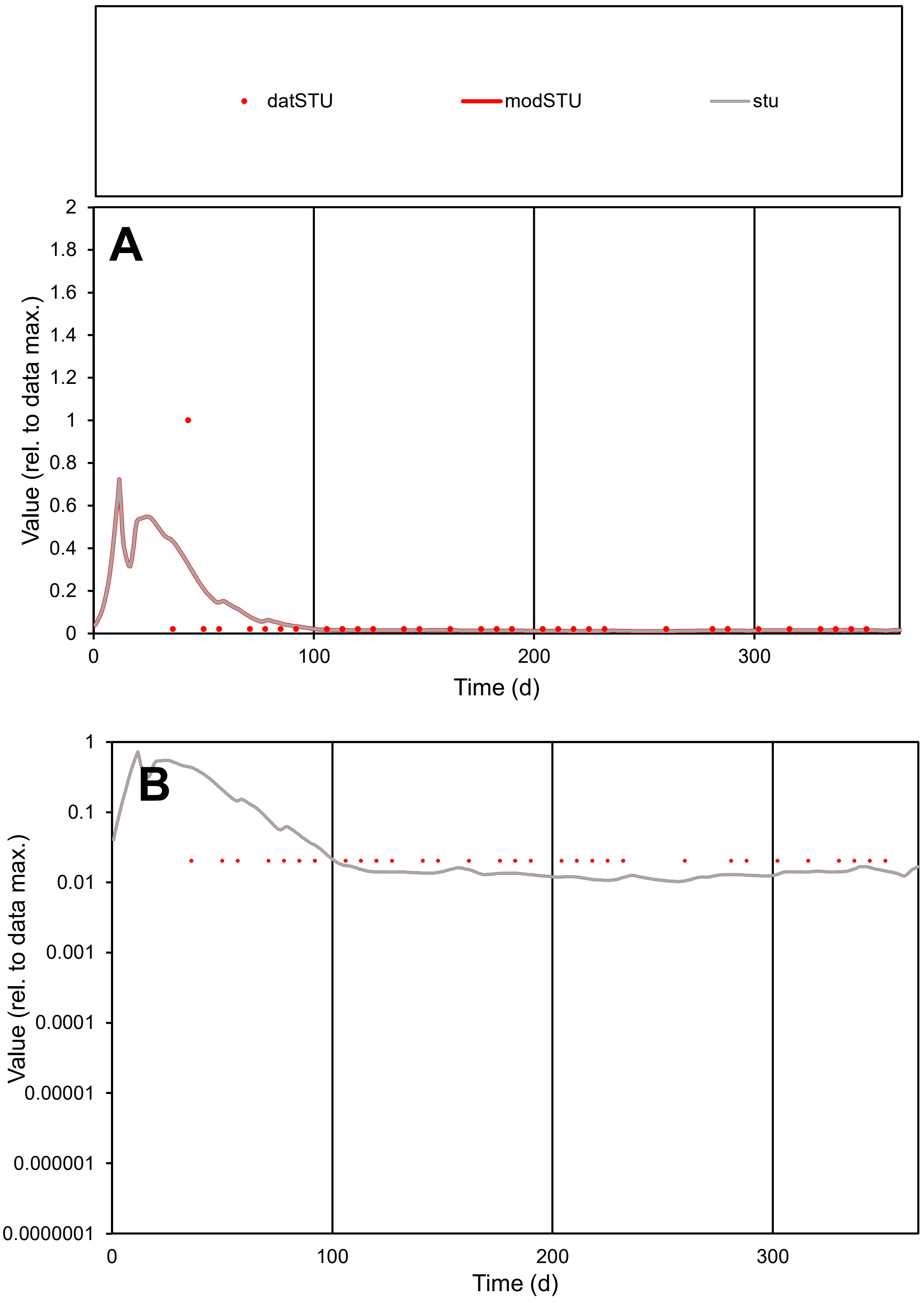


**C1**

Variables
STU Provide jtd or Dst
CalibOnly?
2 1 to reproduce of
oNormalize?
2 1=no, 2=yes

ofxn calc*
2.81773
ofxn mod
2.86889

sRun
RunA1
zout
1

Ave
Data
6.2E-06
Model
6E-06

data/vMax
0.00012

ofxng
321.551

Min d/m
1.91181

ave/min d
2.50054
ave/min m
4.58675

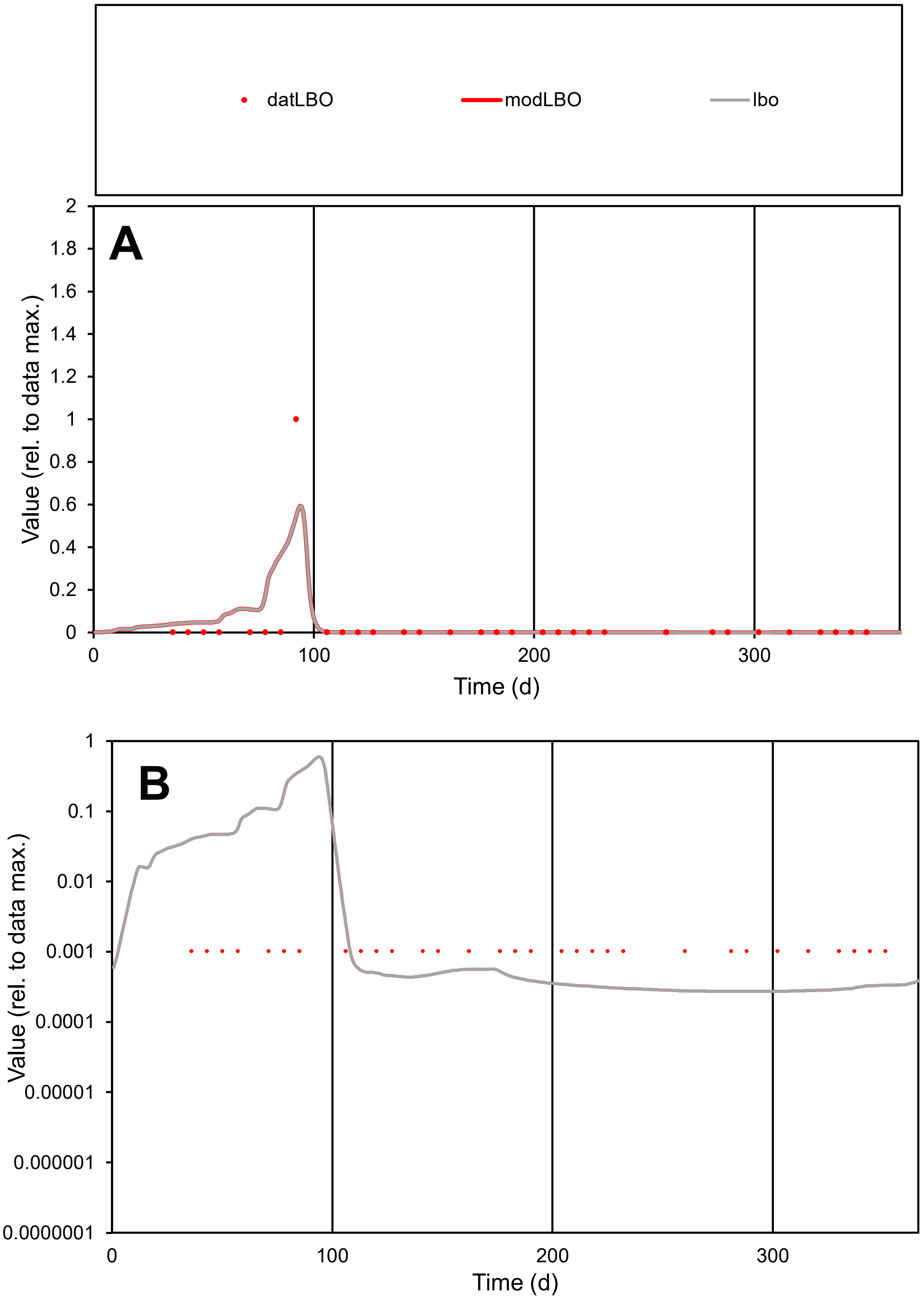


**C1**

Variables
LBO Provide jtd or Dst
CalibOnly?
2 1 to reproduce of
oNormalize?
2 1=no, 2=yes

ofxn calc*
3.16426
ofxn mod
3.24972

sRun
RunA1
zout
1

Ave
Data
4.4E-05
Model
5.7E-05

data/vMax
0.00135

ofxng
321.551

Min d/m
3.74258

ave/min d
31.3851
ave/min m
154.296

324_Calib_S_A

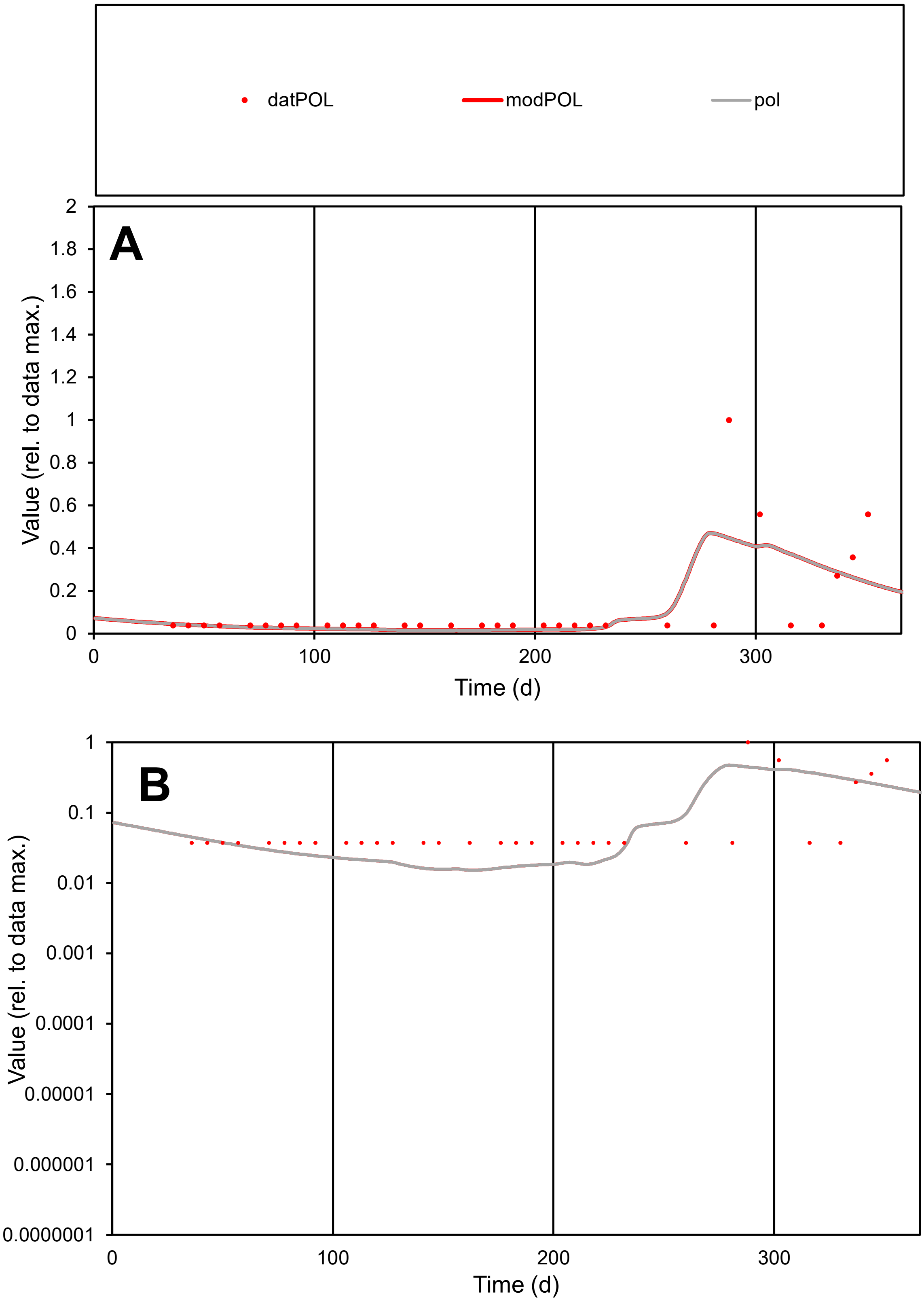


**C1**

Variables

| | |
|---|---|
| POL | Provide jtd or Dst |

CalibOnly?

| | |
|---|---|
| 2 | 1 to reproduce of |

oNormalize?

| | |
|---|---|
| 2 | 1=no, 2=yes |

ofxn calc*
1.36996

ofxn mod
1.37903

sRun
RunA1

zout
1

Ave
Data
5E-06

Model
4.6E-06

data/vMax
4.3E-05

ofxng
321.551

Min d/m
2.45199

ave/min d
3.1548

ave/min m
7.09757

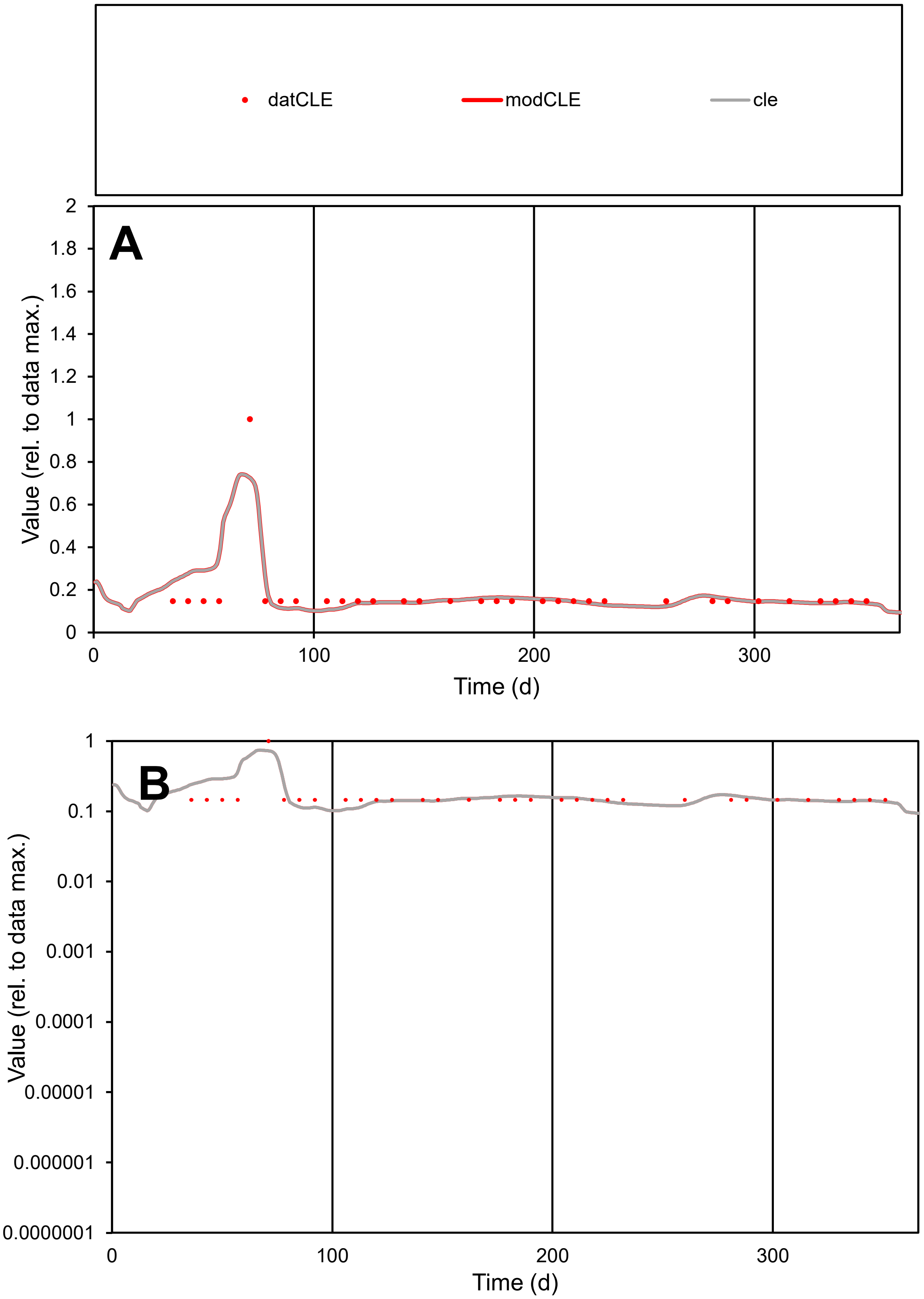


**C1**

Variables
CLE Provide jtd or Dst
CalibOnly?
2 1 to reproduce of
oNormalize?
2 1=no, 2=yes

ofxn calc*
0.43238
ofxn mod
0.45831

sRun
RunA1
zout
1

Ave
Data
5.6E-06
Model
5.7E-06

data/vMax
3.2E-05

ofxng
321.551

Min d/m
1.35137

ave/min d
1.1825
ave/min m
1.65397

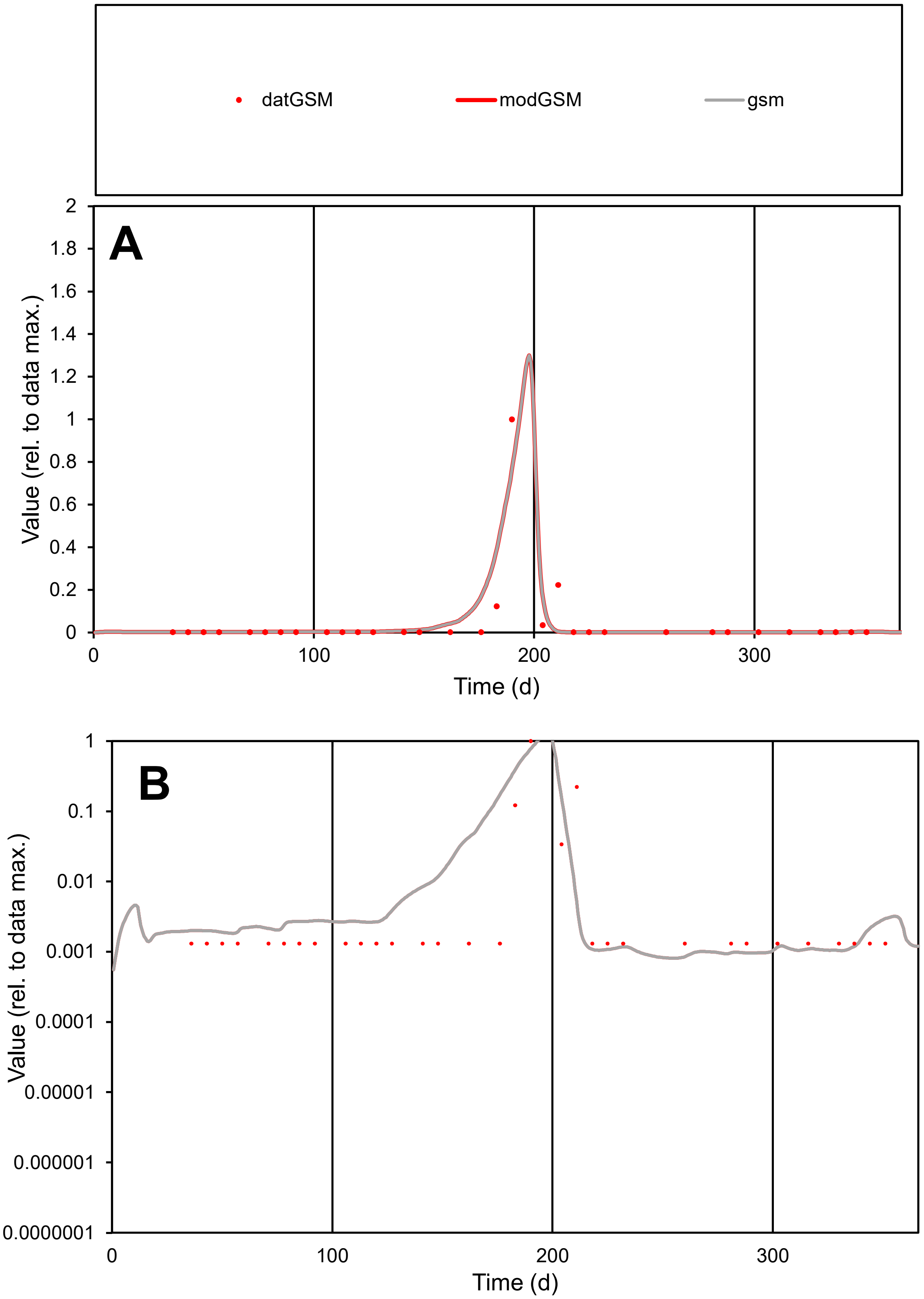


**C1**

Variables

GSM Provide jtd or Dst

CalibOnly?

2 1 to reproduce of

oNormalize?

2 1=no, 2=yes

ofxn calc*

1.52797

ofxn mod

1.70759

sRun

RunA1

zout

1

Ave

Data

5.3E-05

Model

5.7E-05

data/vMax

0.00121

ofxng

321.551

Min d/m

1.48682

ave/min d

33.8575

ave/min m

53.2512

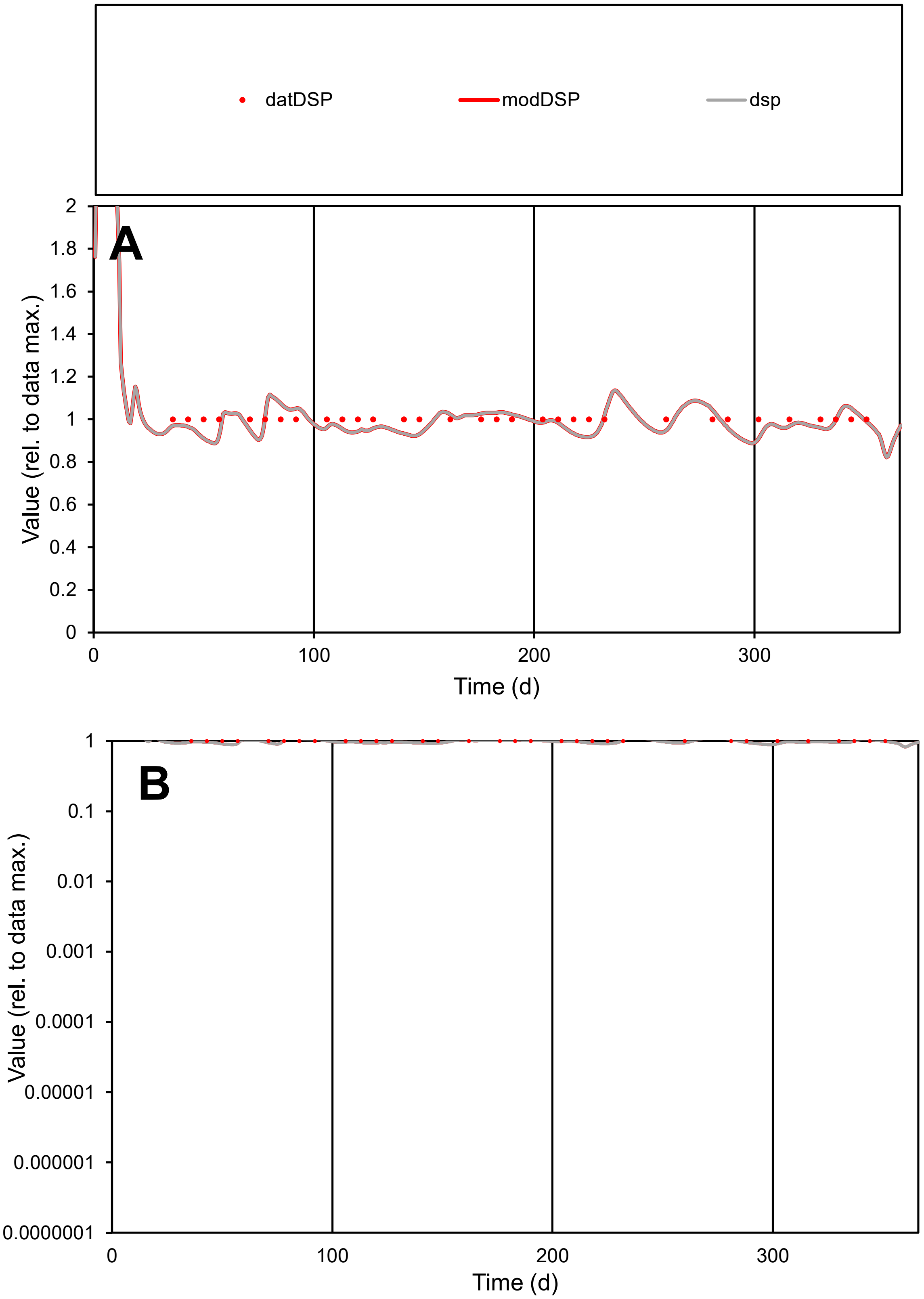


**C1**

Variables
DSP Provide jtd or Dst
CalibOnly?
2 1 to reproduce of
oNormalize?
2 1=no, 2=yes

ofxn calc*
0.05055
ofxn mod
0.10892

sRun
RunA1
zout
1

Ave
Data
5E-06
Model
4.9E-06

data/vMax
5E-06

ofxng
321.551

Min d/m
1.11123

ave/min d
1
ave/min m
1.09244

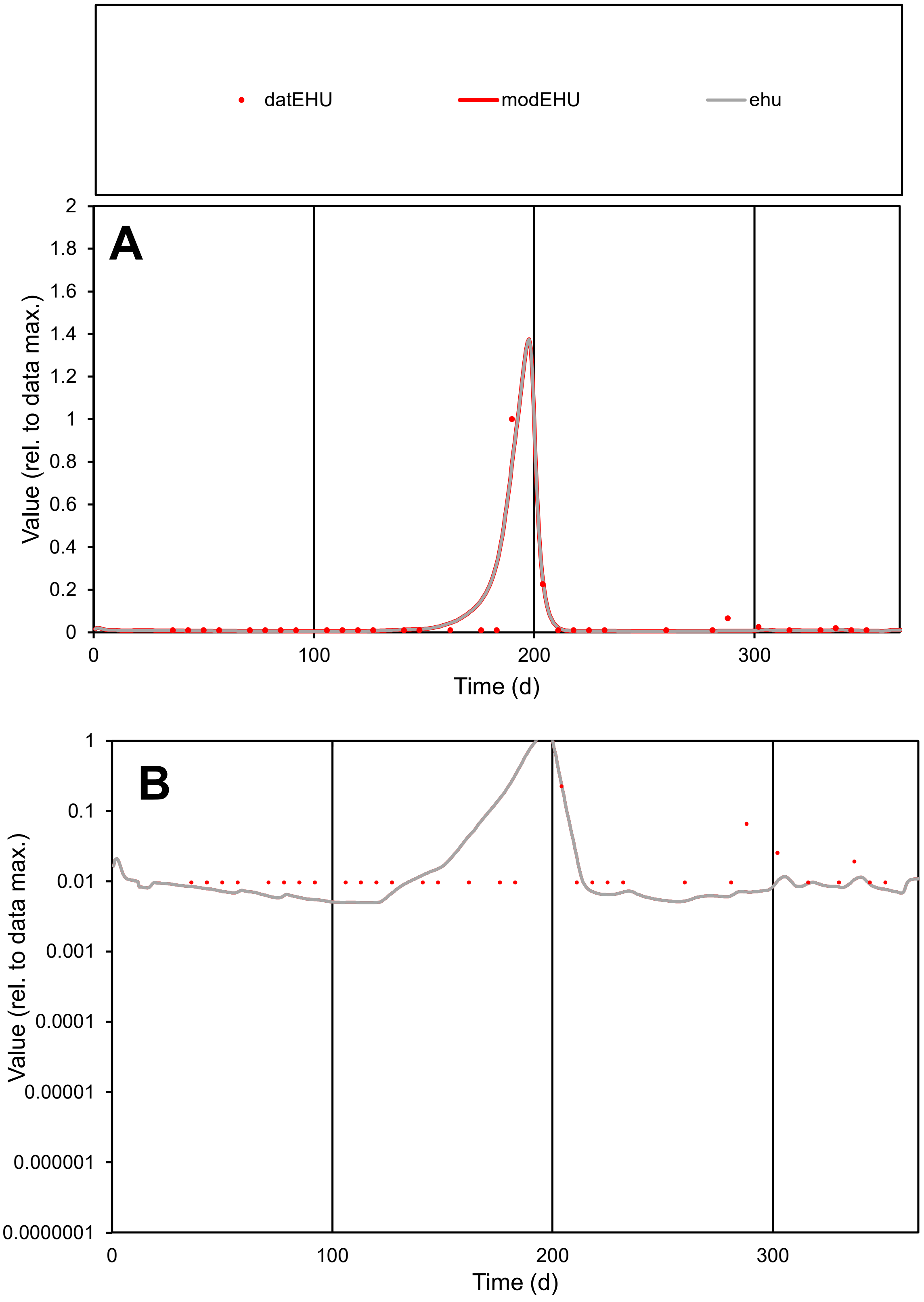


**C1**

Variables

EHU Provide jtd or Dst

CalibOnly?

2 1 to reproduce of

oNormalize?

2 1=no, 2=yes

ofxn calc*

1.08225

ofxn mod

1.25522

sRun

RunA1

zout

1

Ave

Data

3E-05

Model

3.3E-05

data/vMax

0.0006

ofxng

321.551

Min d/m

1.9436

ave/min d

5.18704

ave/min m

11.1308

324_Calib_S_A

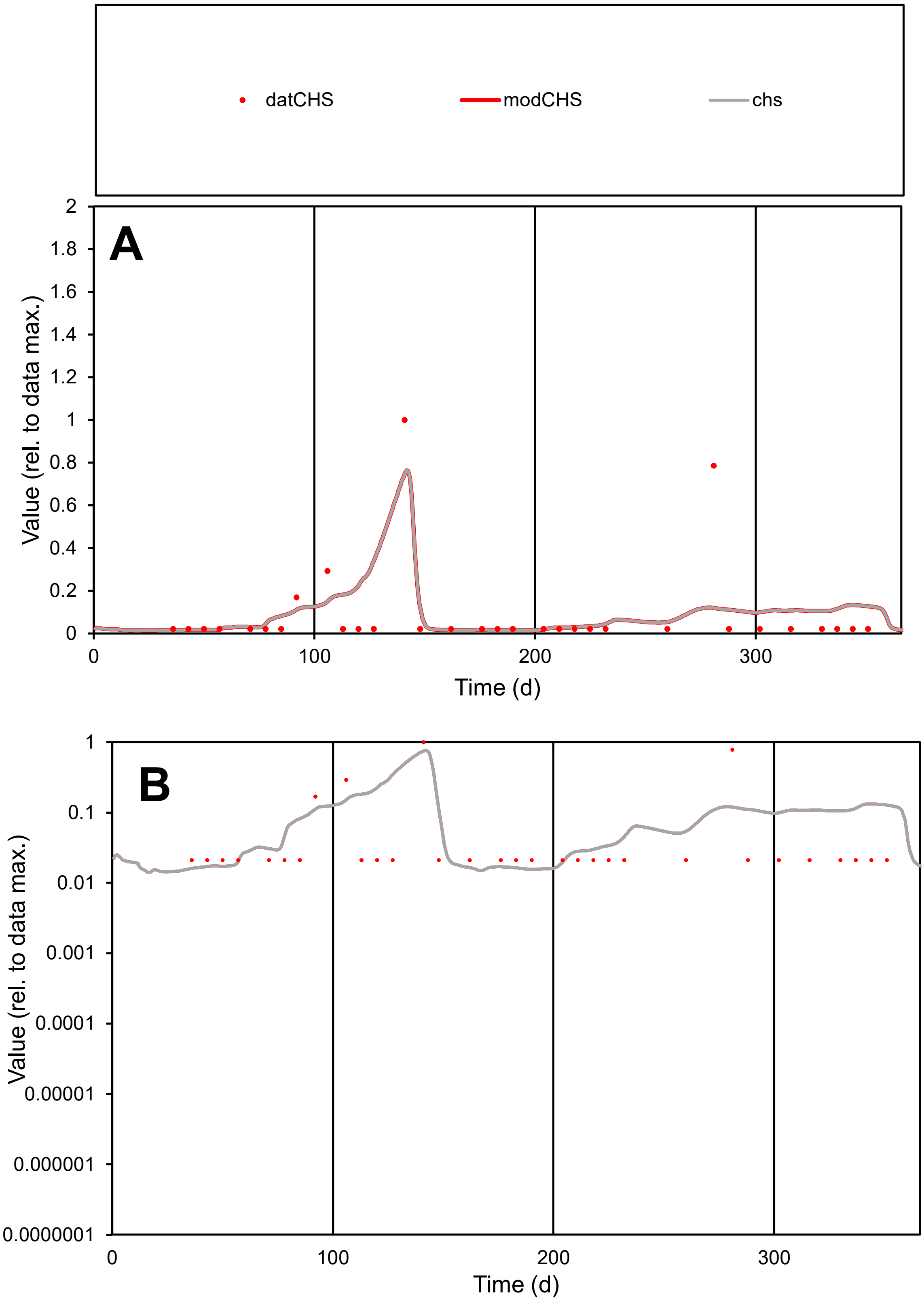


**C1**

Variables
CHS Provide jtd or Dst
CalibOnly?
2 1 to reproduce of
oNormalize?
2 1=no, 2=yes

ofxn calc*
1.70163
ofxn mod
1.7502

sRun
RunA1
zout
1

Ave
Data
9E-05
Model
0.0001

data/vMax
0.00101

ofxng
321.551

Min d/m
1.35123
ave/min d
4.20322
ave/min m
6.38884

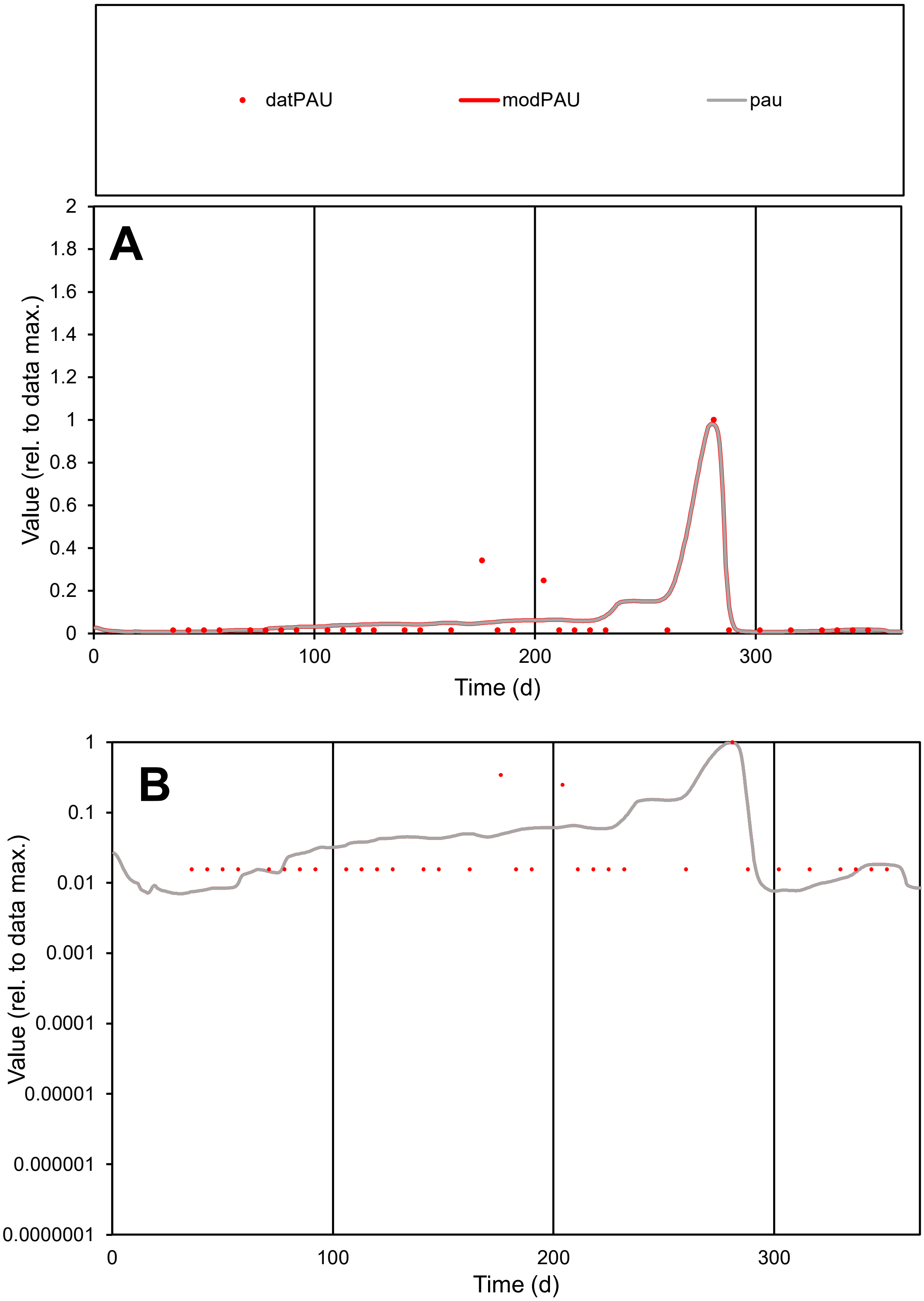


**C1**

Variables
PAU Provide jtd or Dst
CalibOnly?
2 1 to reproduce of
oNormalize?
2 1=no, 2=yes

ofxn calc*
1.12671
ofxn mod
1.16106

sRun
RunA1
zout
1

Ave
Data
1.6E-05
Model
1.7E-05

data/vMax
0.00025

ofxng
321.551

Min d/m
2.06332
ave/min d
4.10036
ave/min m
8.97115

324_Calib_S_A

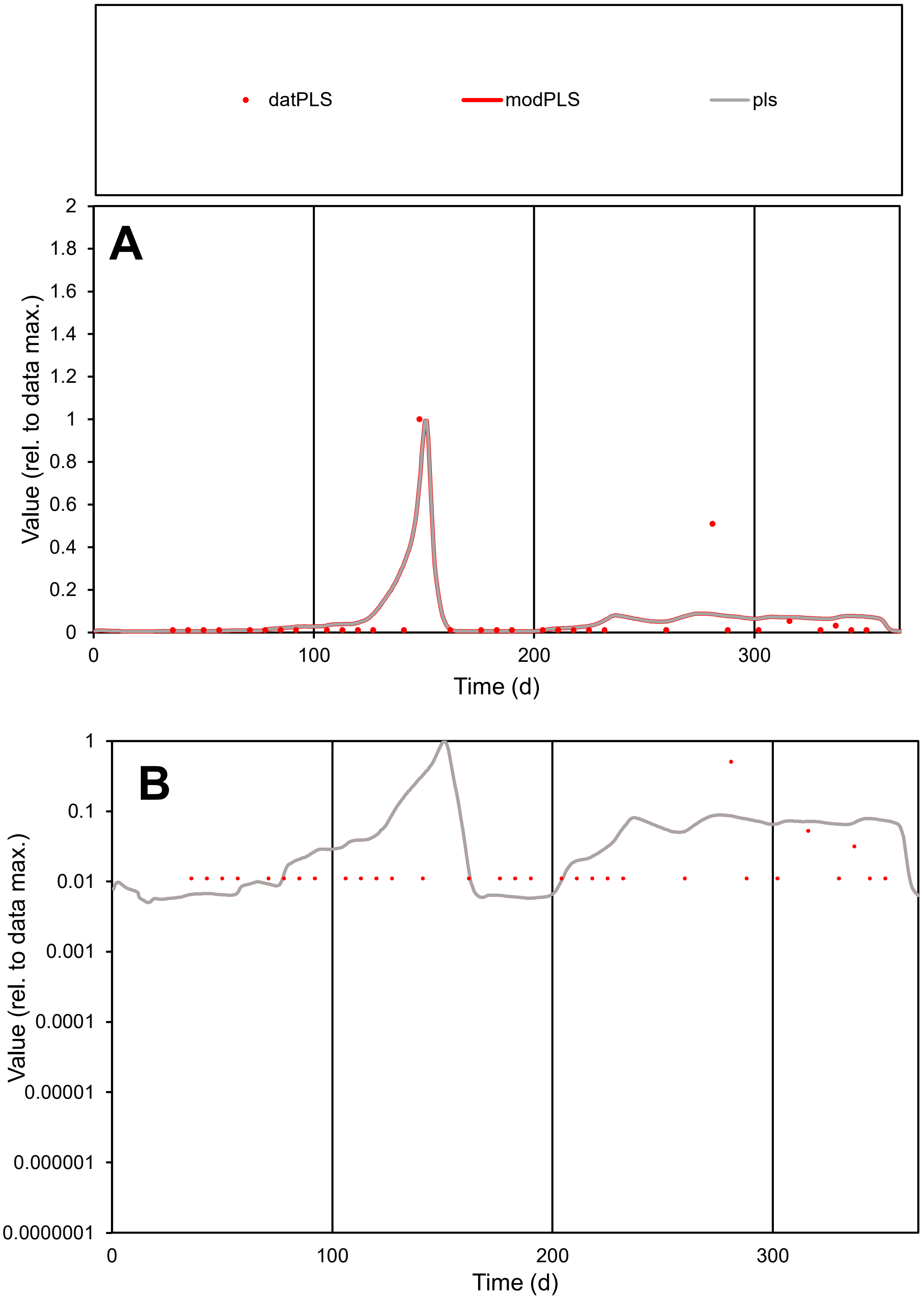


**C1**

Variables
PLS Provide jtd or Dst
CalibOnly?
2 1 to reproduce of
oNormalize?
2 1=no, 2=yes

ofxn calc*
1.69337
ofxn mod
1.72935

sRun
RunA1
zout
1

Ave
Data
2.1E-05
Model
2.4E-05

data/vMax
0.00035

ofxng
321.551

Min d/m
1.90715

ave/min d
5.3661
ave/min m
12.0676

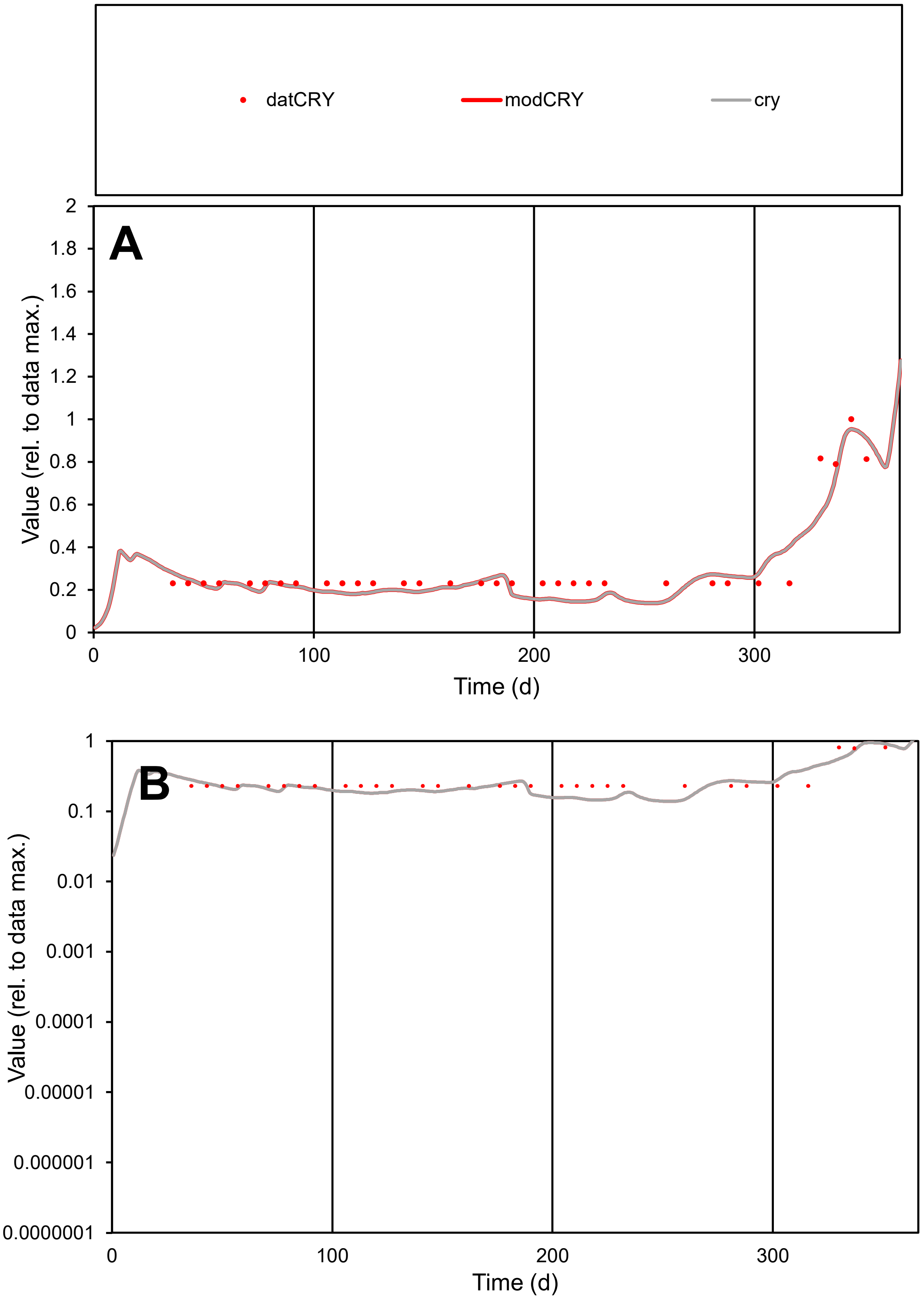


**C1**

Variables
CRY Provide jtd or Dst
CalibOnly?
2 1 to reproduce of
oNormalize?
2 1=no, 2=yes

ofxn calc*
0.22648
ofxn mod
0.25324

sRun
RunA1
zout
1

Ave
Data
4E-06
Model
3.8E-06

data/vMax
1.3E-05

ofxng
321.551

Min d/m
1.58914

ave/min d
1.33894
ave/min m
2.00138

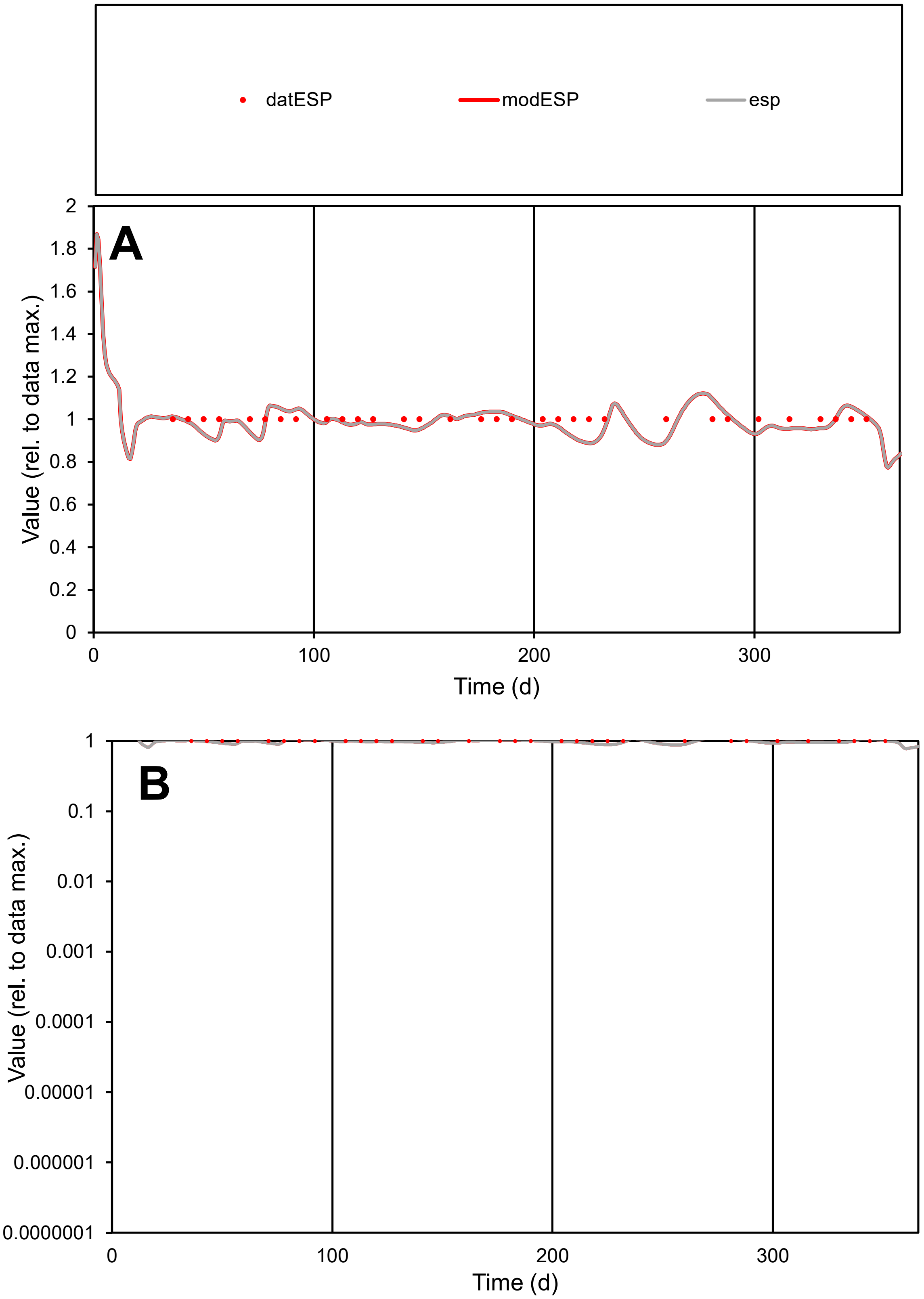


**C1**

Variables
ESP Provide jtd or Dst
CalibOnly?
2 1 to reproduce of
oNormalize?
2 1=no, 2=yes

ofxn calc*
0.04888
ofxn mod
0.10782

sRun
RunA1
zout
1

Ave
Data
8E-06
Model
7.9E-06

data/vMax
8E-06

ofxng
321.551

Min d/m
1.12507

ave/min d
1
ave/min m
1.10926

324_Calib_S_A

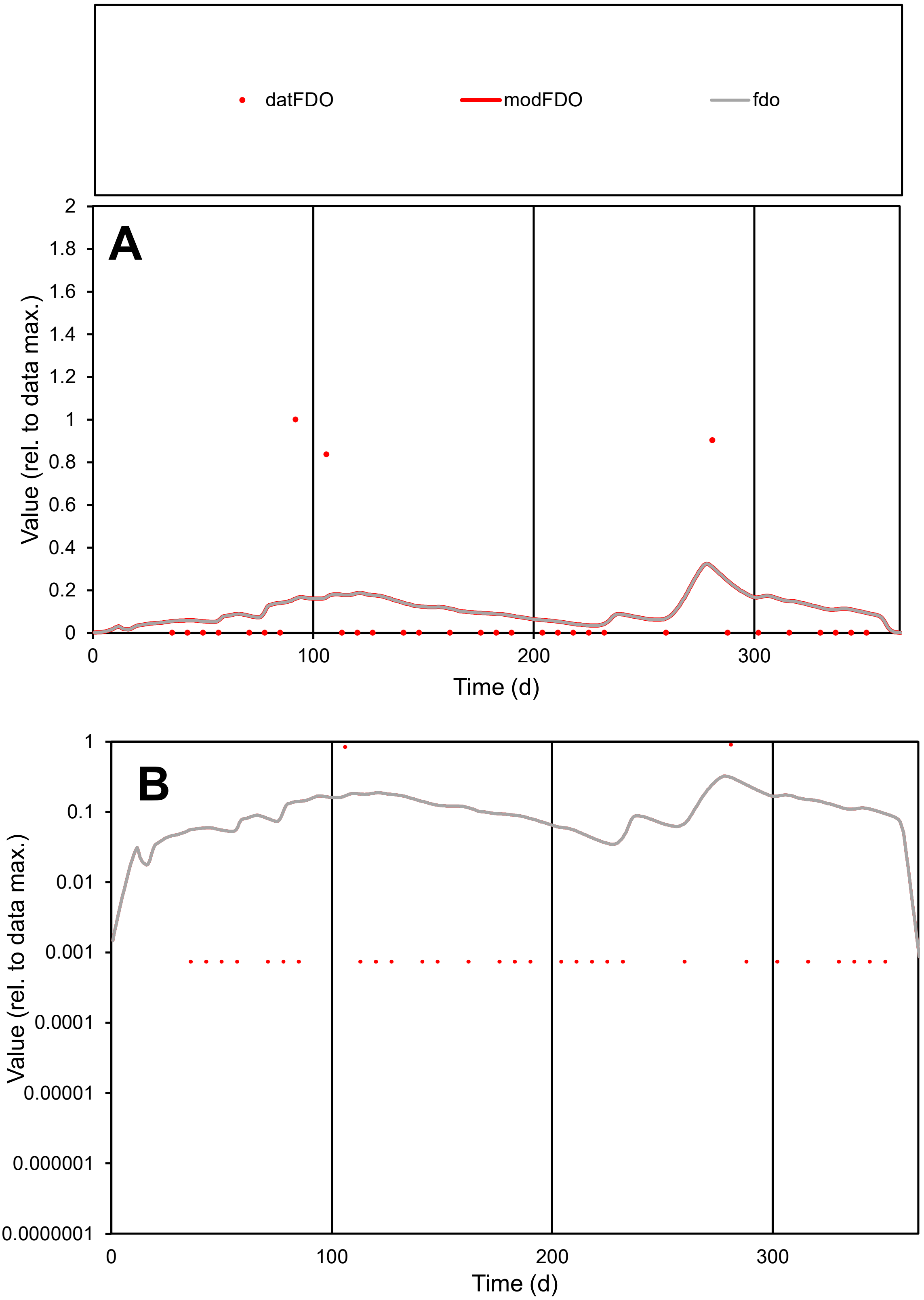


**C1**

Variables

FDO Provide jtd or Dst

CalibOnly?

2 1 to reproduce of

oNormalize?

2 1=no, 2=yes

ofxn calc*

2.78999

ofxn mod

2.80797

sRun

RunA1

zout

1

Ave

Data

1.2E-05

Model

1.5E-05

data/vMax

0.00013

ofxng

321.551

Min d/m

0.02118

ave/min d

116.584

ave/min m

3.21851

324_Calib_S_A

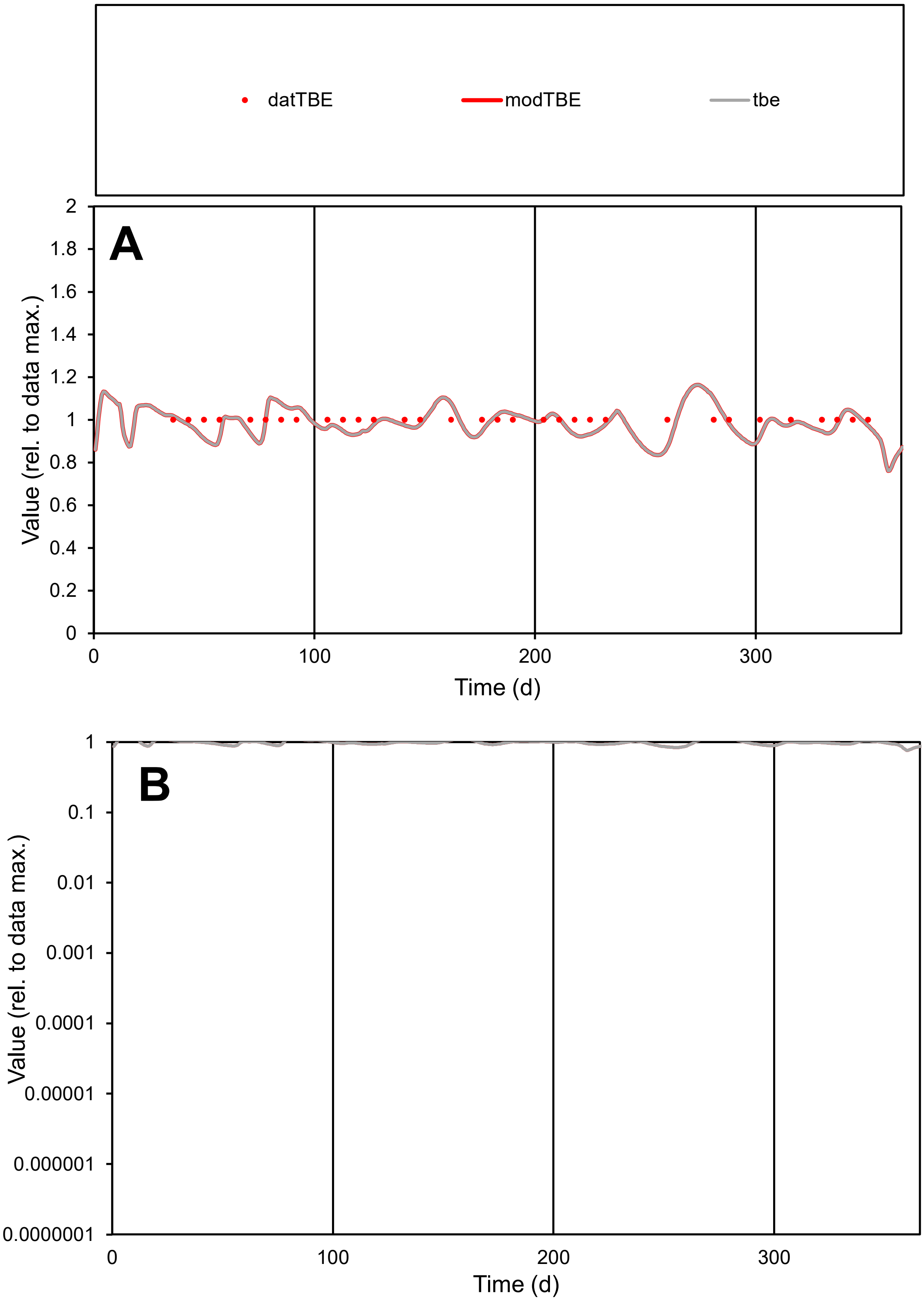


**C1**

Variables
TBE Provide jtd or Dst
CalibOnly?
2 1 to reproduce of
oNormalize?
2 1=no, 2=yes

ofxn calc*
0.05111
ofxn mod
0.10887

sRun
RunA1
zout
1

Ave
Data
6.7E-06
Model
6.6E-06

data/vMax
6.7E-06

ofxng
321.551

Min d/m
1.11176

ave/min d
1
ave/min m
1.09584

324_Calib_S_A

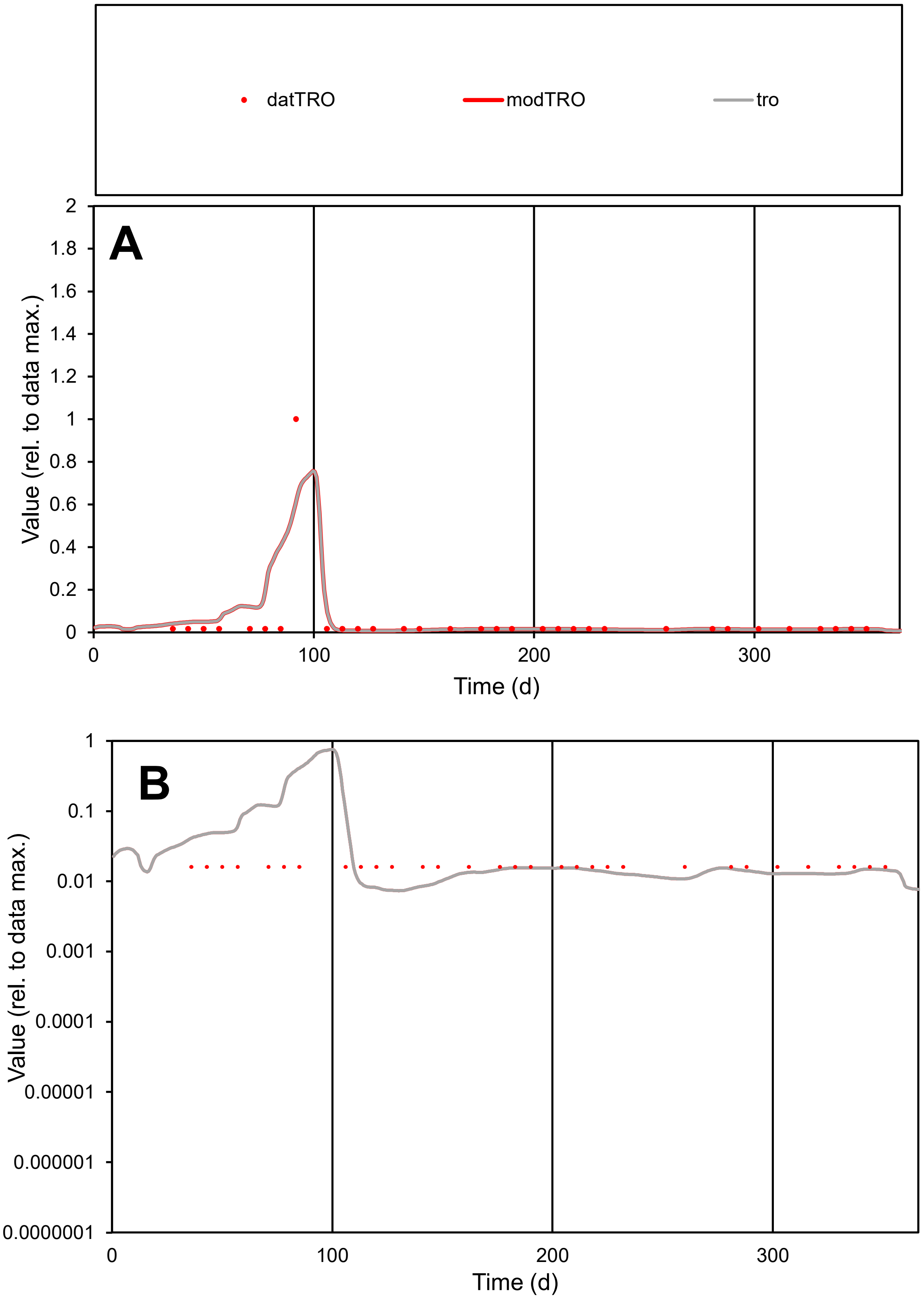


**C1**

Variables
TRO Provide jtd or Dst
CalibOnly?
2 1 to reproduce of
oNormalize?
2 1=no, 2=yes

ofxn calc*
2.03579
ofxn mod
2.11912

sRun
RunA1
zout
1

Ave
Data
1.5E-05
Model
1.8E-05

data/vMax
0.00031

ofxng
321.551

Min d/m
2.15979

ave/min d
2.92094
ave/min m
7.72715

324_Calib_S_A

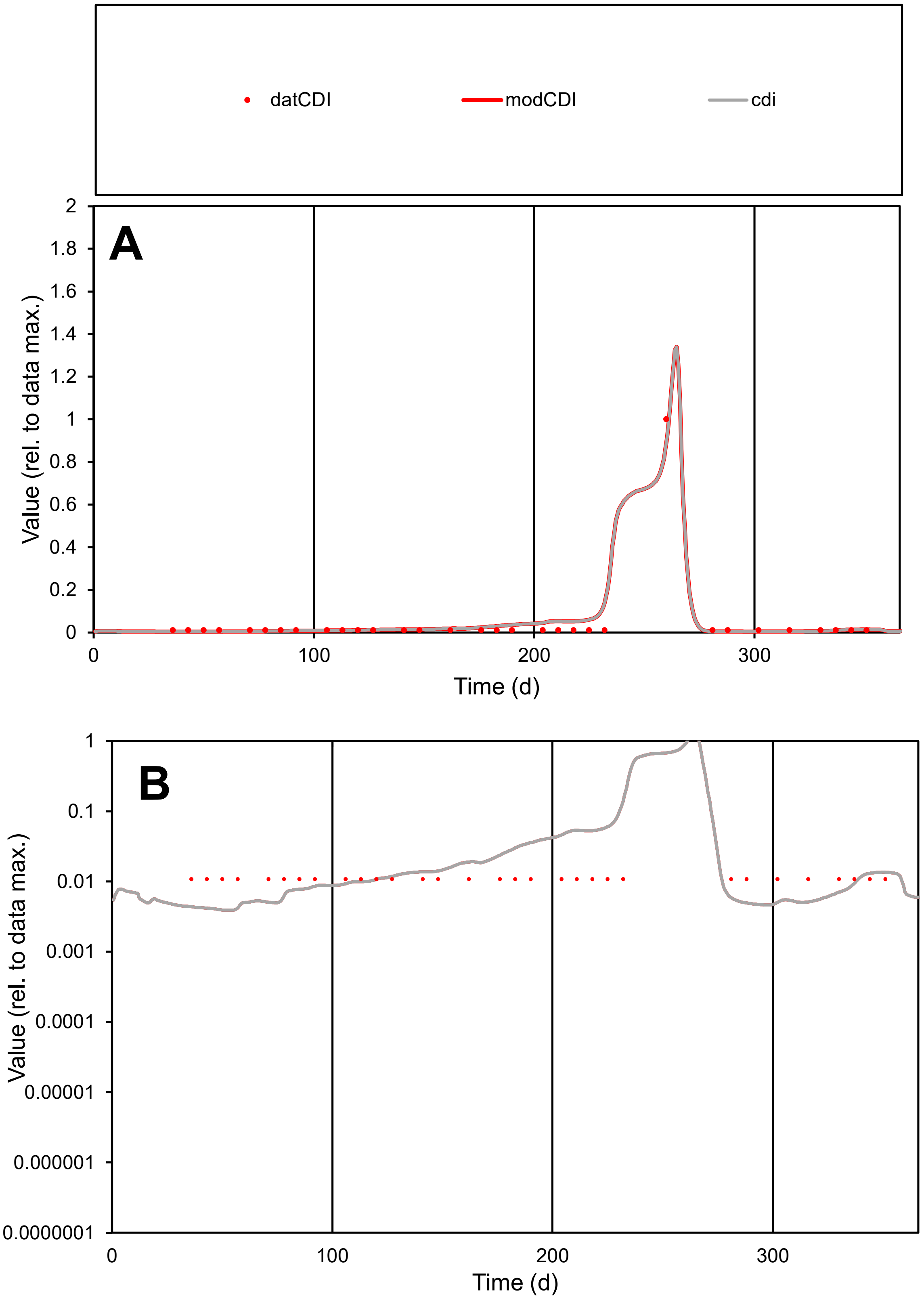


**C1**

Variables
CDI Provide jtd or Dst
CalibOnly?
2 1 to reproduce of
oNormalize?
2 1=no, 2=yes

ofxn calc*
0.55836
ofxn mod
0.68649

sRun
RunA1
zout
1

Ave
Data
6.7E-07
Model
8E-07

data/vMax
1.6E-05

ofxng
321.551

Min d/m
2.76324

ave/min d
3.85559
ave/min m
12.6881

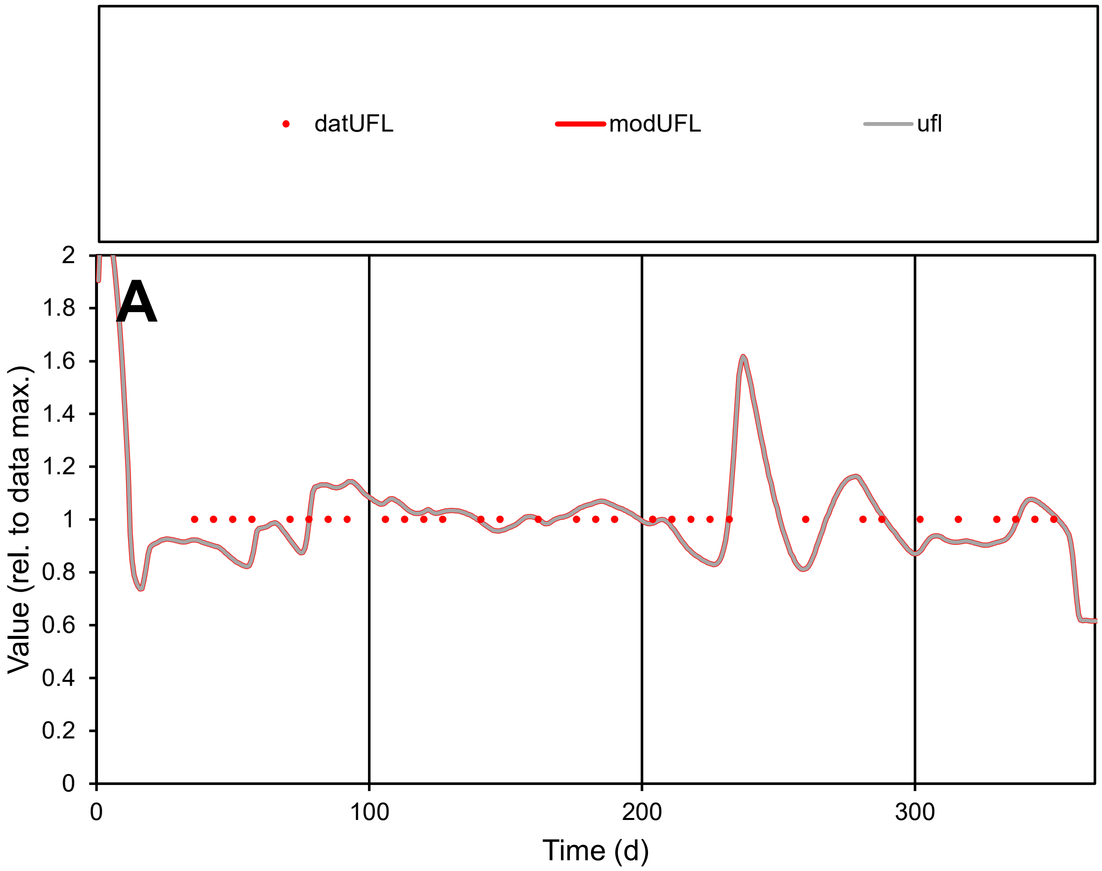


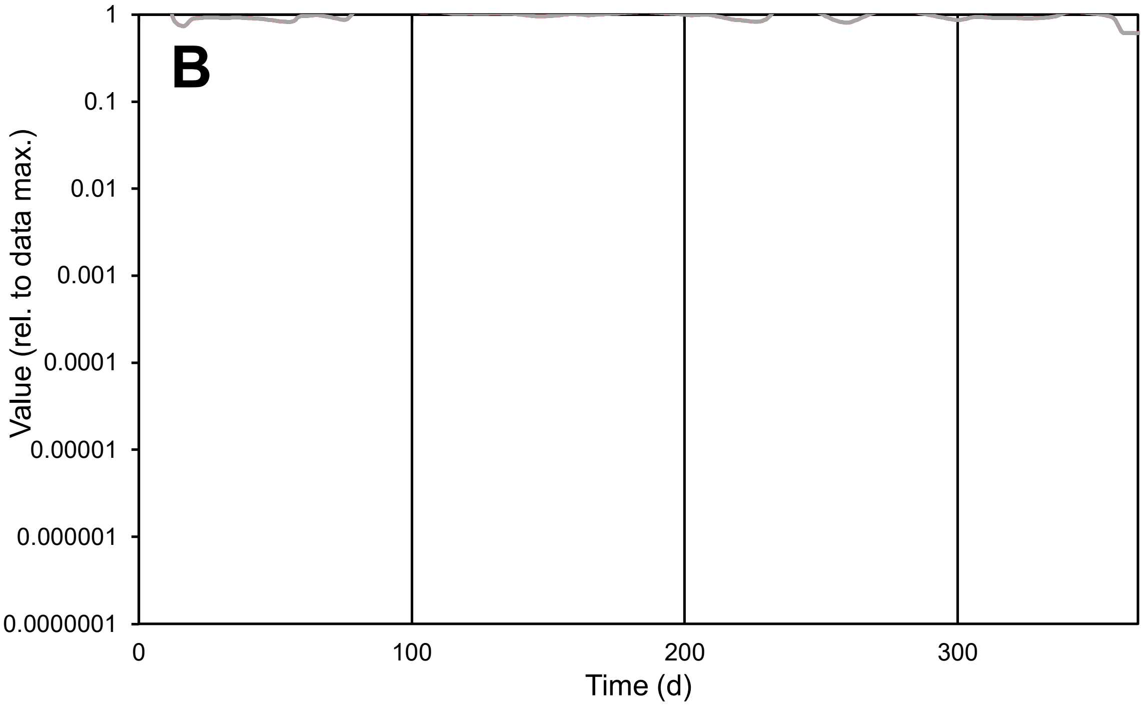


**C1**

Variables
UFL Provide jtd or Dst
CalibOnly?
2 1 to reproduce of
oNormalize?
2 1=no, 2=yes

ofxn calc*
0.09045
ofxn mod
0.13663

sRun
RunA1
zout
1

Ave
Data
3.1E-06
Model
3E-06

data/vMax
3.1E-06

ofxng
321.551

Min d/m
1.21171

ave/min d
1
ave/min m
1.19797

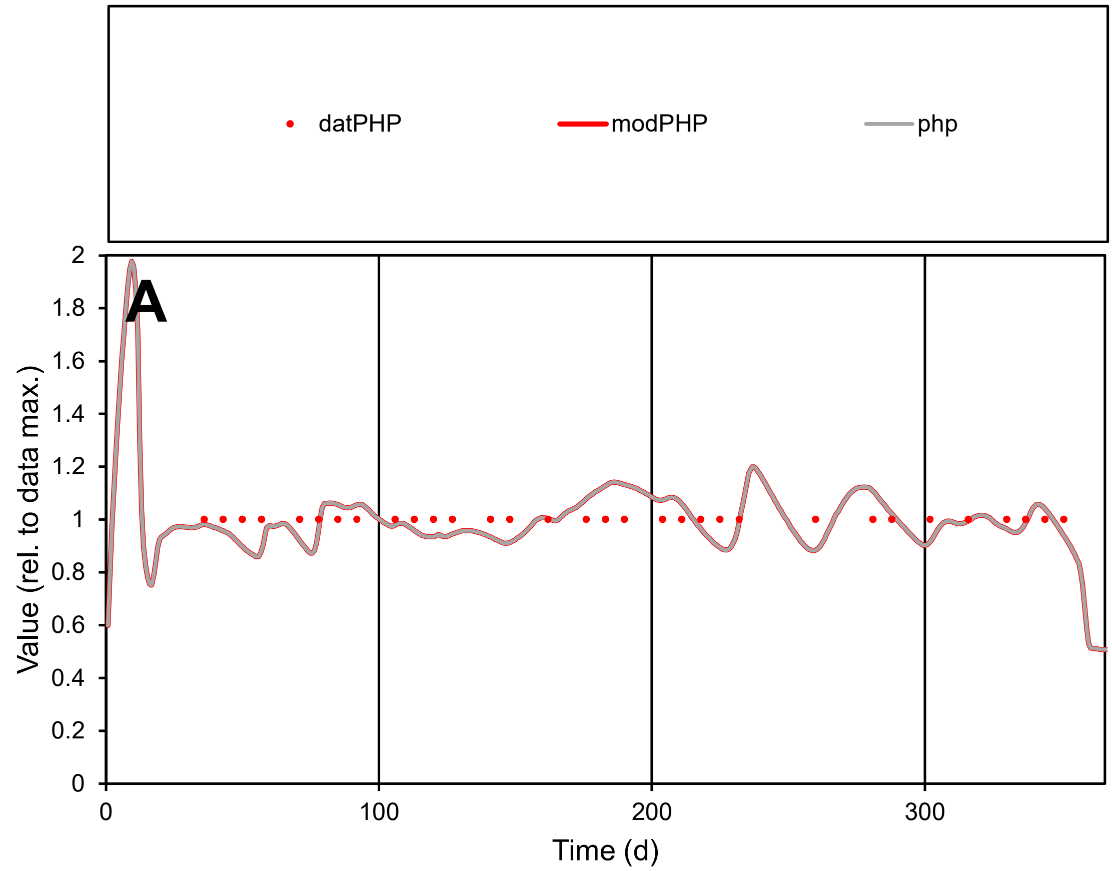


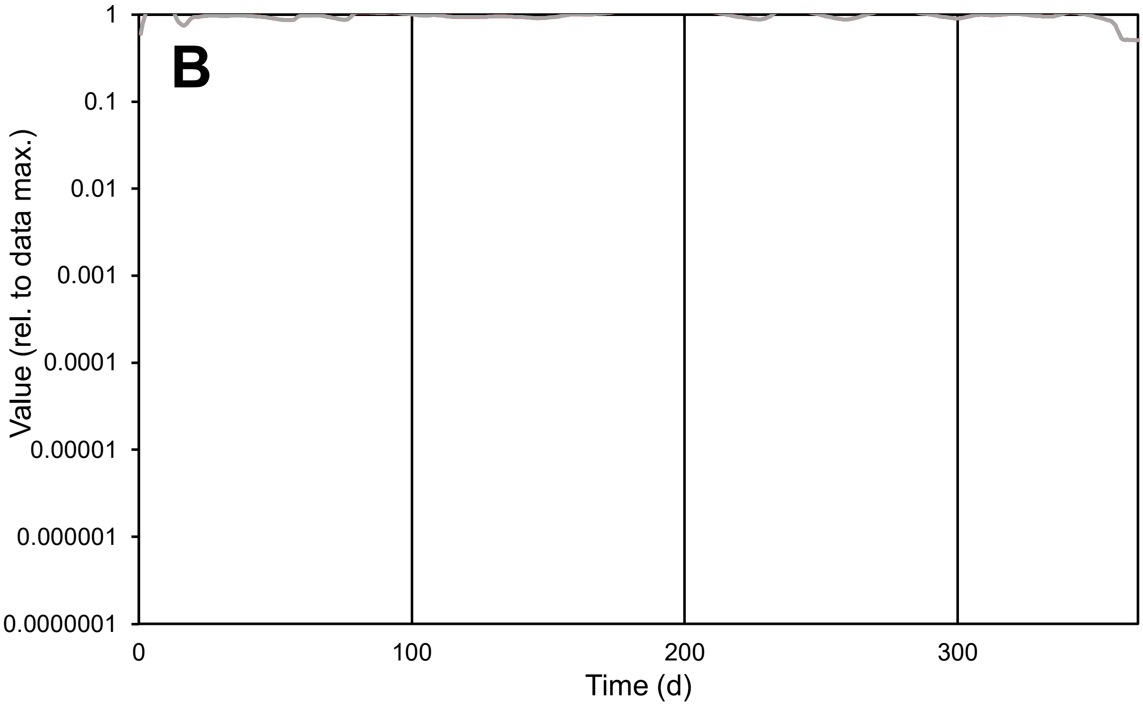


**C1**

Variables
PHP Provide jtd or Dst
CalibOnly?
2 1 to reproduce of
oNormalize?
2 1=no, 2=yes

ofxn calc*
0.07065
ofxn mod
0.11998

sRun
RunA1
zout
1

Ave
Data
6.9E-06
Model
6.9E-06

data/vMax
6.9E-06

ofxng
321.551

Min d/m
1.13304

ave/min d
1
ave/min m
1.12104

324_Calib_S_A

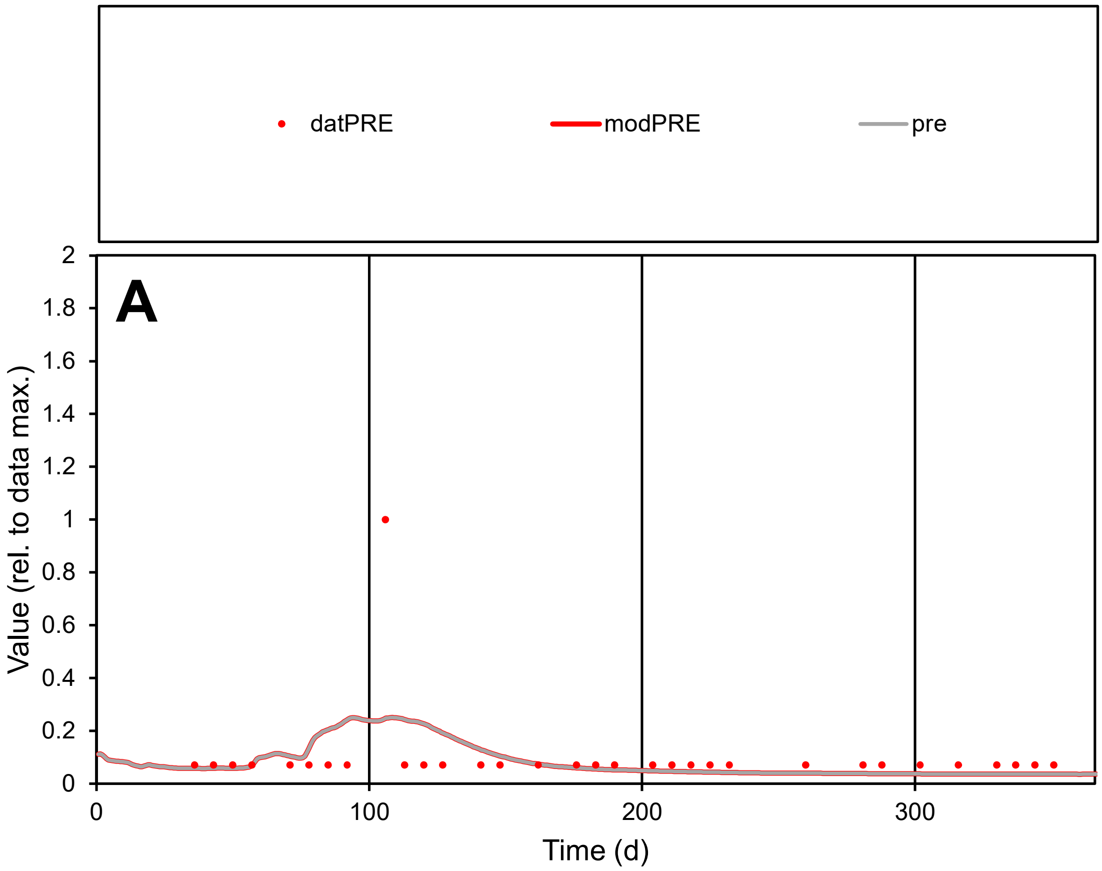


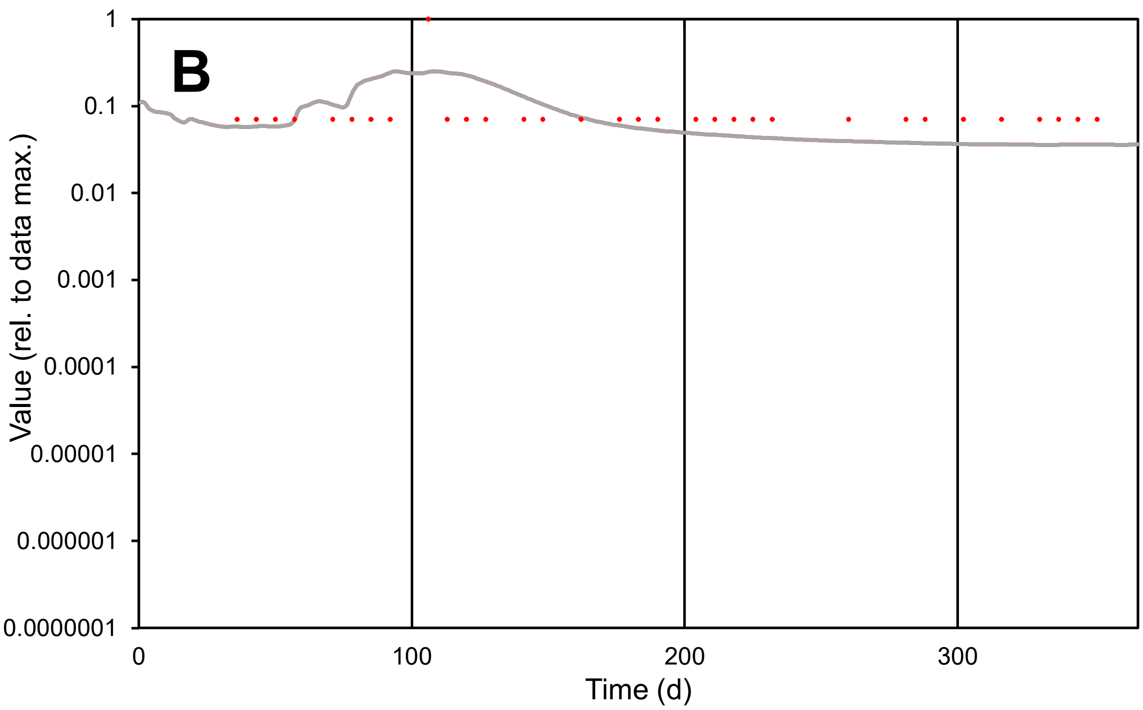


**C1**

Variables
PRE Provide jtd or Dst
CalibOnly?
2 1 to reproduce of
oNormalize?
2 1=no, 2=yes

ofxn calc*
1.47965
ofxn mod
1.49411

sRun
RunA1
zout
1

Ave
Data
1.9E-05
Model
1.7E-05

data/vMax
0.00019

ofxng
321.551

Min d/m
1.97733

ave/min d
1.41086
ave/min m
2.47894

324_Calib_S_A

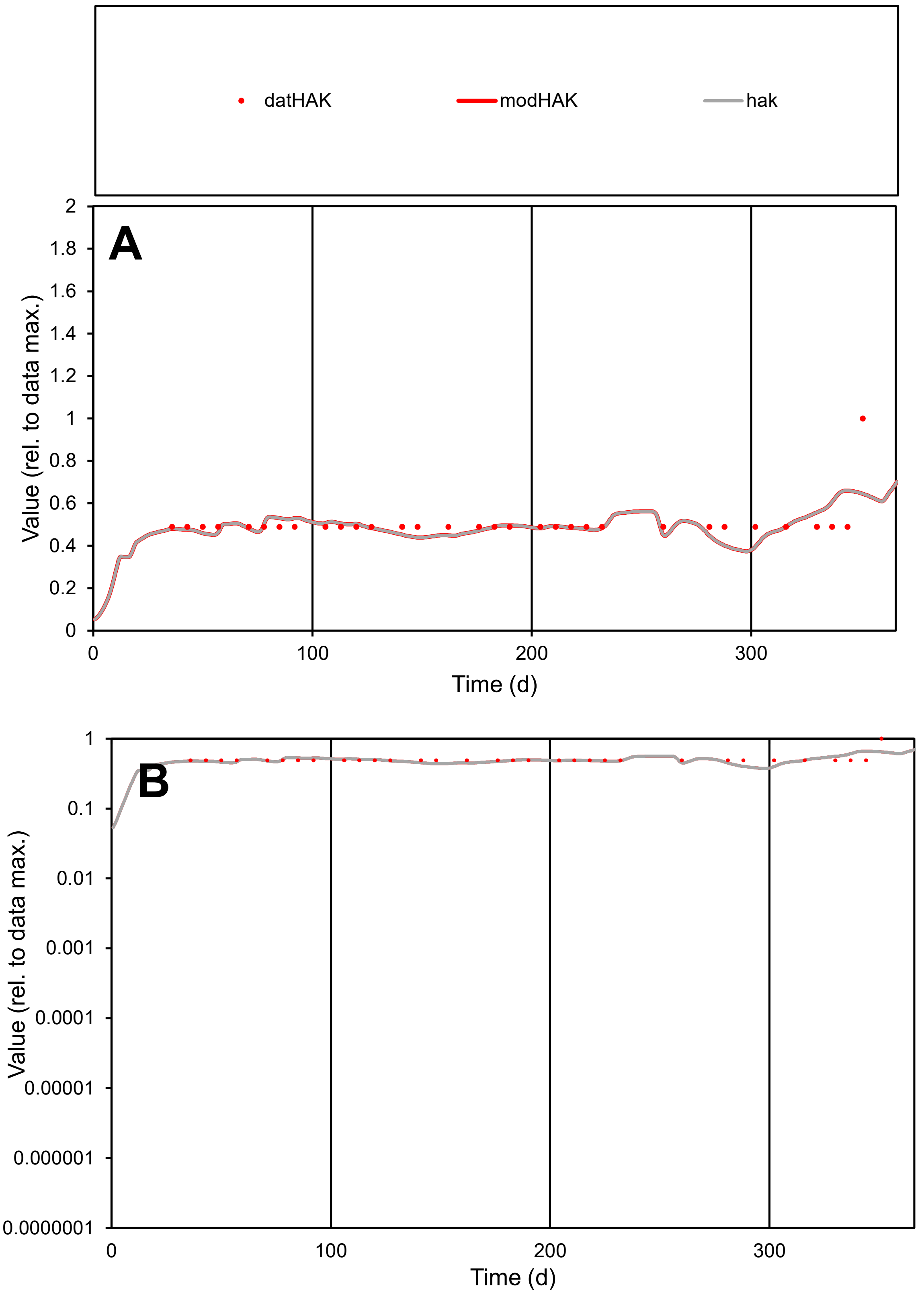


**C1**

Variables
HAK Provide jtd or Dst
CalibOnly?
2 1 to reproduce of
oNormalize?
2 1=no, 2=yes

ofxn calc*
0.16197
ofxn mod
0.18767

sRun
RunA1
zout
1

Ave
Data
5.3E-06
Model
5.2E-06

data/vMax
1E-05

ofxng
321.551

Min d/m
1.23901

ave/min d
1.03265
ave/min m
1.25539

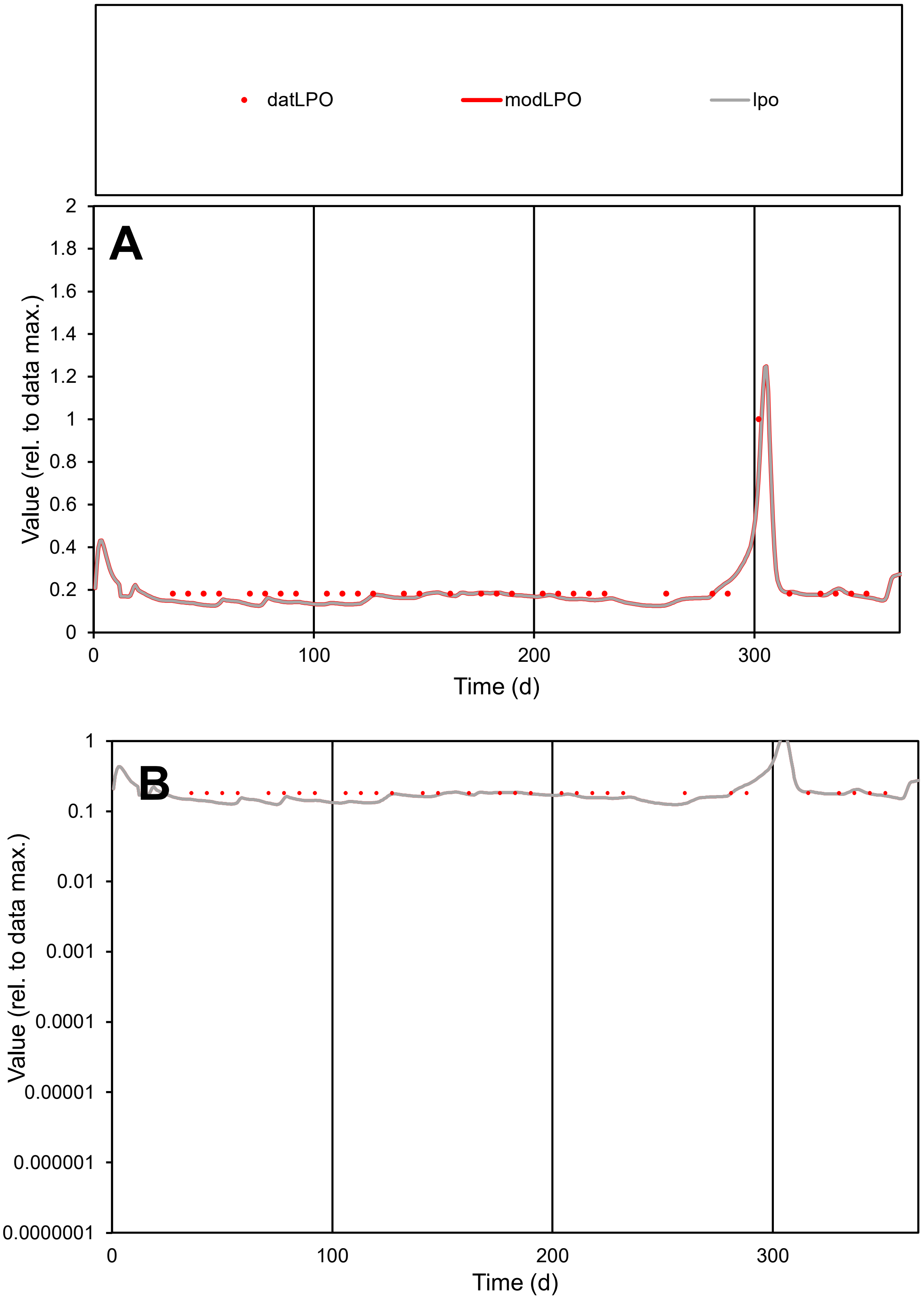


**C1**

Variables
LPO Provide jtd or Dst
CalibOnly?
2 1 to reproduce of
oNormalize?
2 1=no, 2=yes

ofxn calc*
0.15064
ofxn mod
0.18836

sRun
RunA1
zout
1

Ave
Data
6.7E-06
Model
6.1E-06

data/vMax
3.2E-05

ofxng
321.551

Min d/m
1.42274

ave/min d
1.14058
ave/min m
1.48056

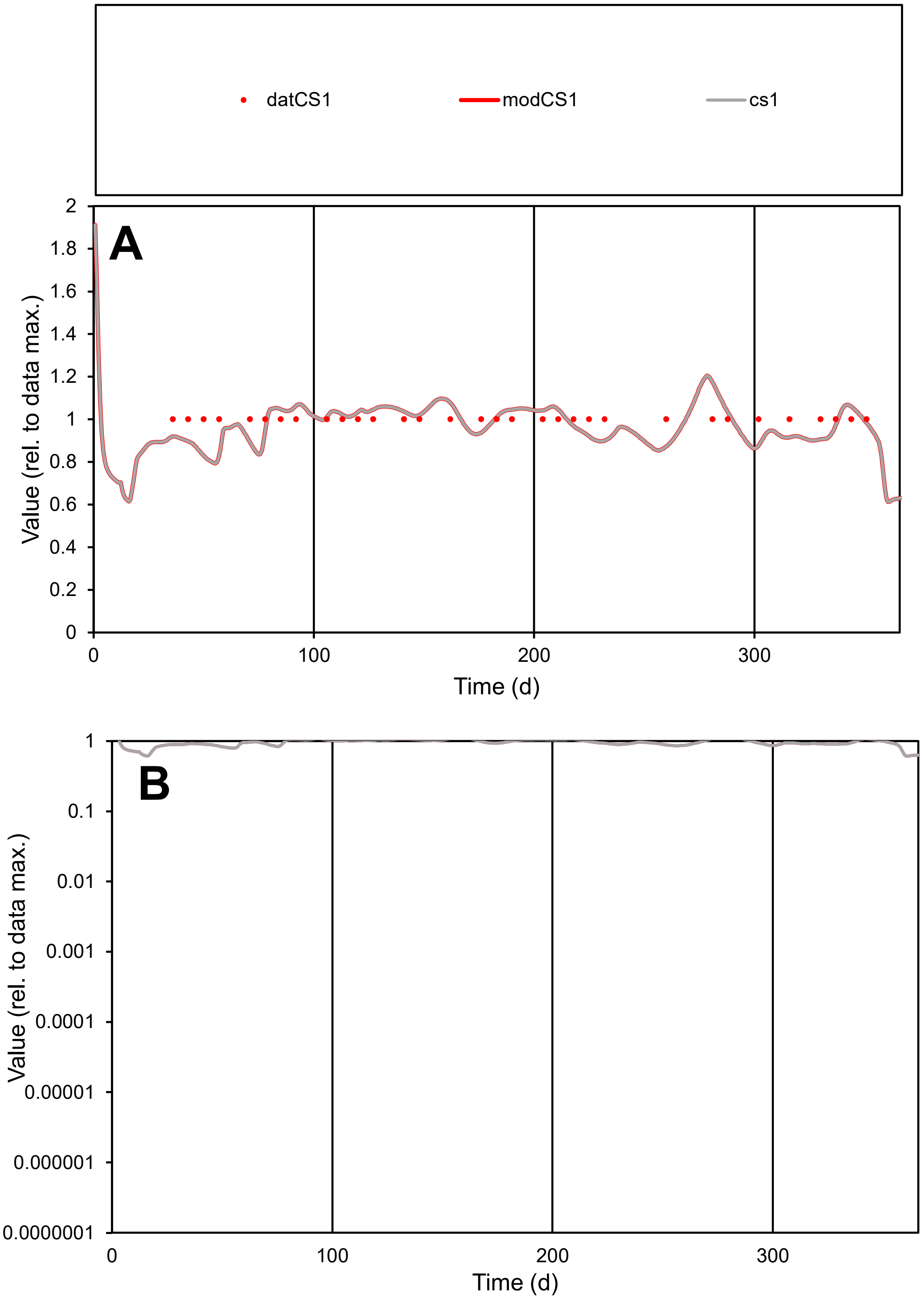


**C1**

Variables
CS1 Provide jtd or Dst
CalibOnly?
2 1 to reproduce of
oNormalize?
2 1=no, 2=yes

ofxn calc*
0.07668
ofxn mod
0.12281

sRun
RunA1
zout
1

Ave
Data
1.8E-05
Model
1.7E-05

data/vMax
1.8E-05

ofxng
321.551

Min d/m
1.22775

ave/min d
1
ave/min m
1.20692

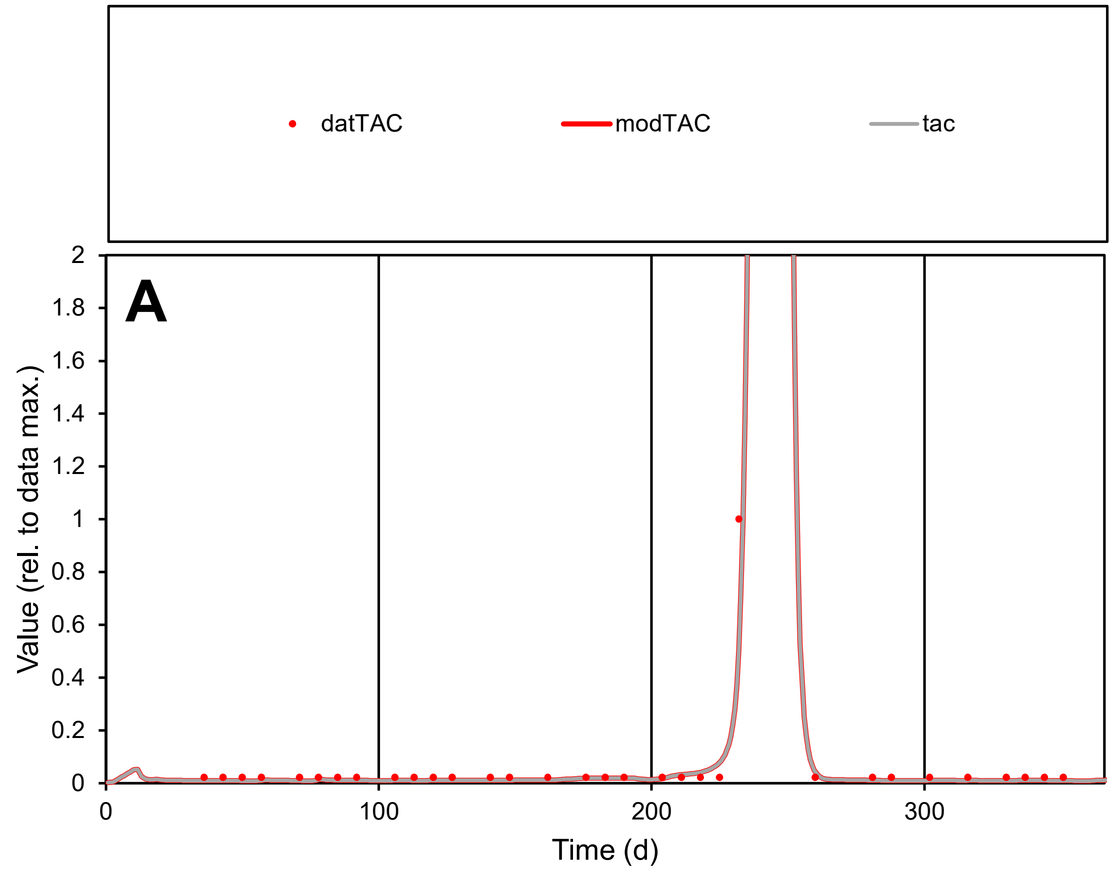


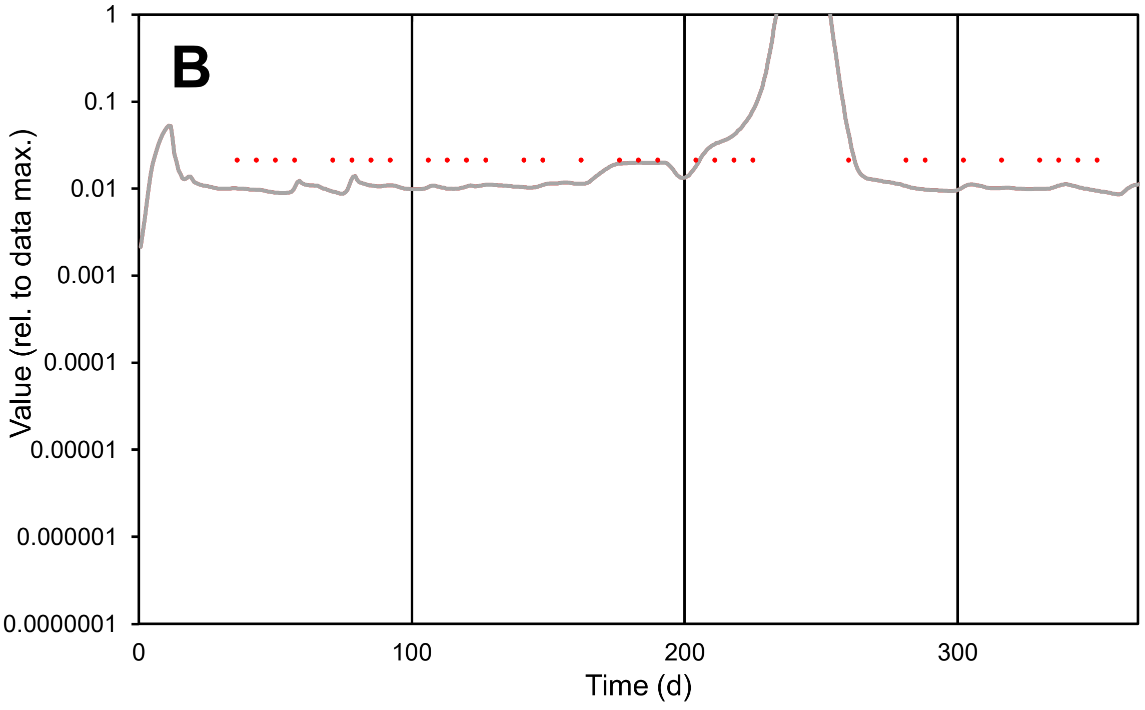


**C1**

Variables
TAC Provide jtd or Dst
CalibOnly?
2 1 to reproduce of
oNormalize?
2 1=no, 2=yes

ofxn calc*
0.27286
ofxn mod
0.37123

sRun
RunA1
zout
1

Ave
Data
3.2E-06
Model
2.8E-06

data/vMax
6.1E-05

ofxng
321.551

Min d/m
2.38666

ave/min d
2.43424
ave/min m
5.24003

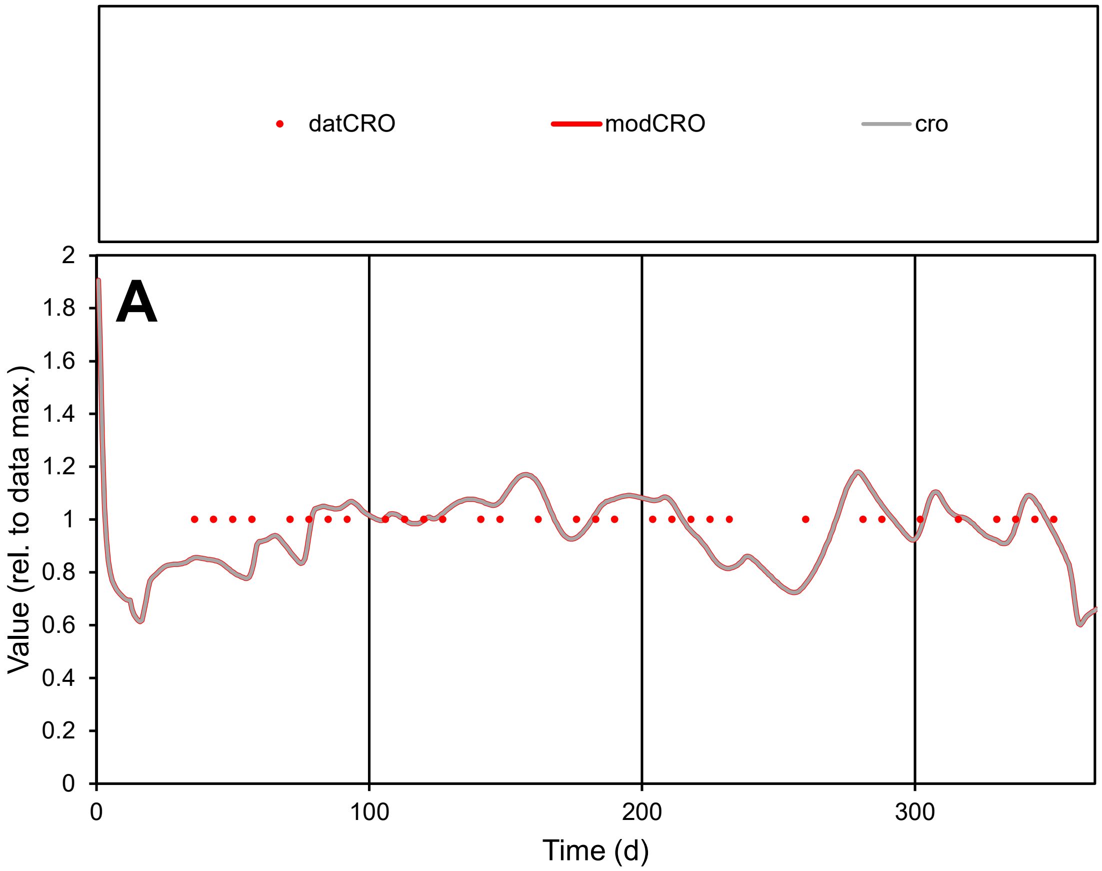


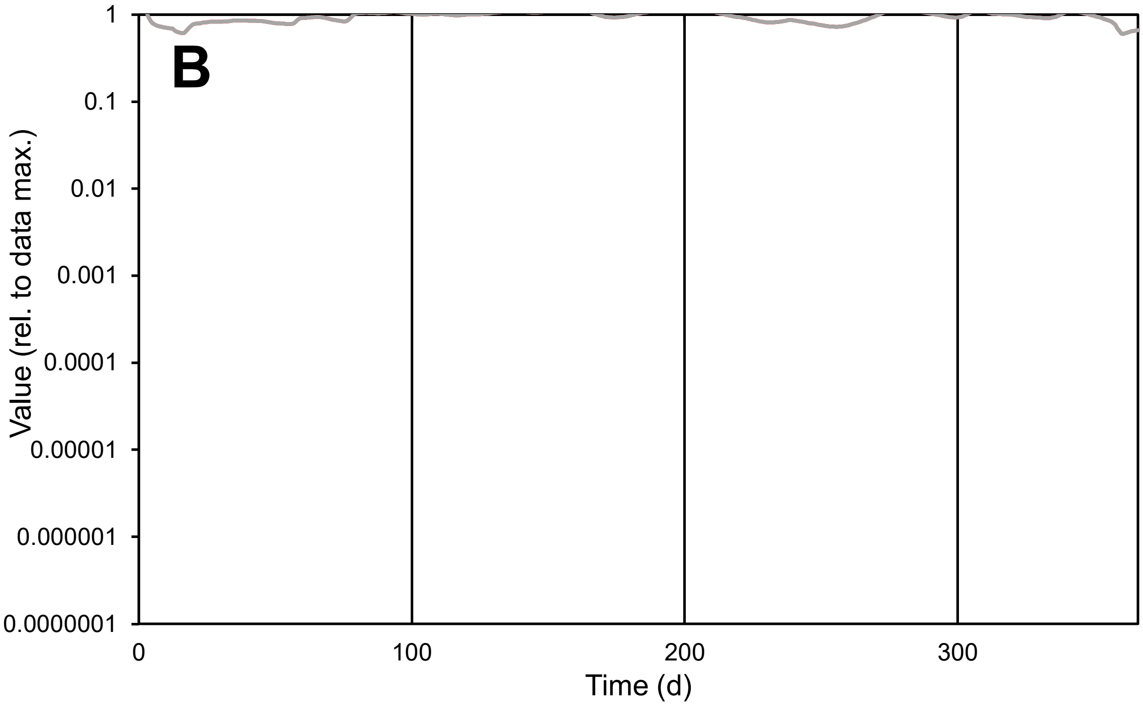


**C1**

Variables
CRO Provide jtd or Dst
CalibOnly?
2 1 to reproduce of
oNormalize?
2 1=no, 2=yes

ofxn calc*
0.10042
ofxn mod
0.13856

sRun
RunA1
zout
1

Ave
Data
1E-05
Model
1E-05

data/vMax
1E-05

ofxng
321.551

Min d/m
1.29555

ave/min d
1
ave/min m
1.26811

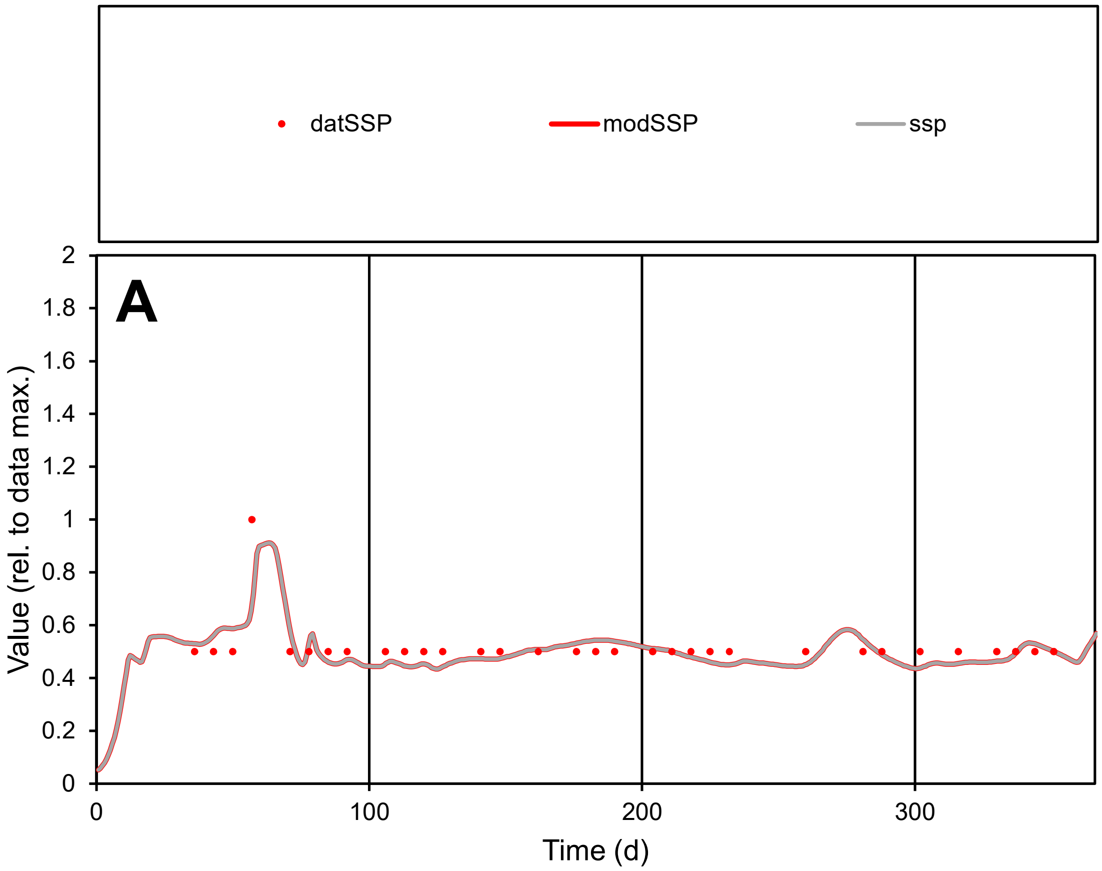


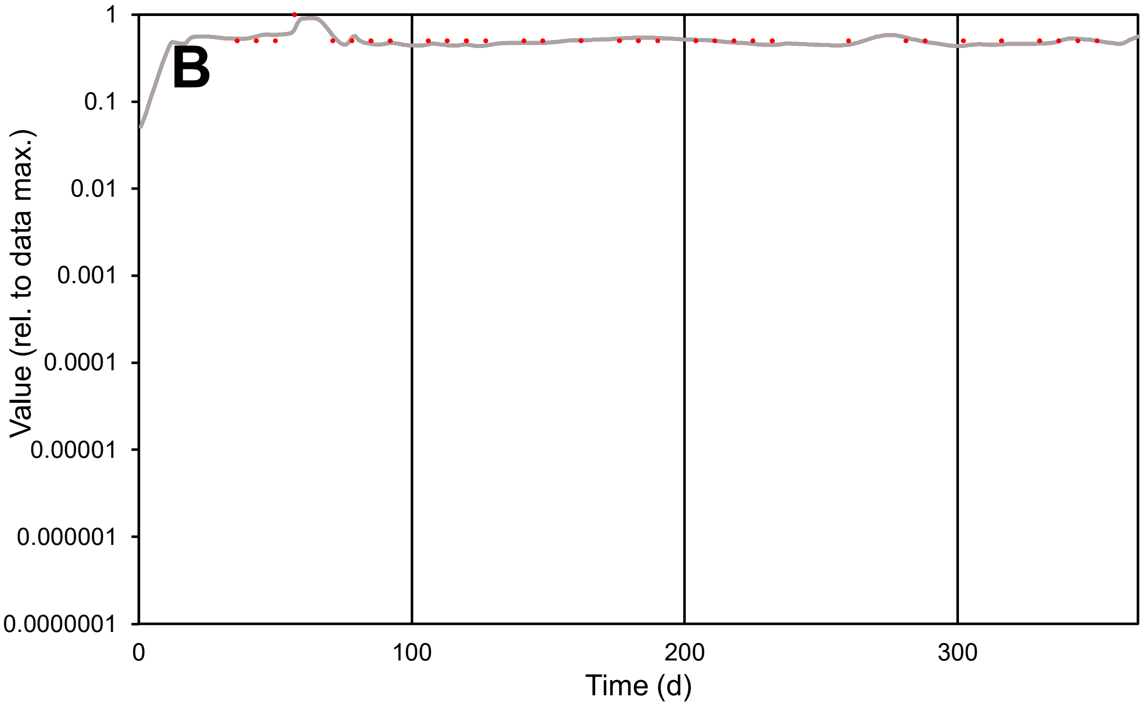


**C1**

Variables
SSP Provide jtd or Dst
CalibOnly?
2 1 to reproduce of
oNormalize?
2 1=no, 2=yes

ofxn calc*
0.10609
ofxn mod
0.14259

sRun
RunA1
zout
1

Ave
Data
1.4E-05
Model
1.3E-05

data/vMax
2.6E-05

ofxng
321.551

Min d/m
1.12884

ave/min d
1.03125
ave/min m
1.13929

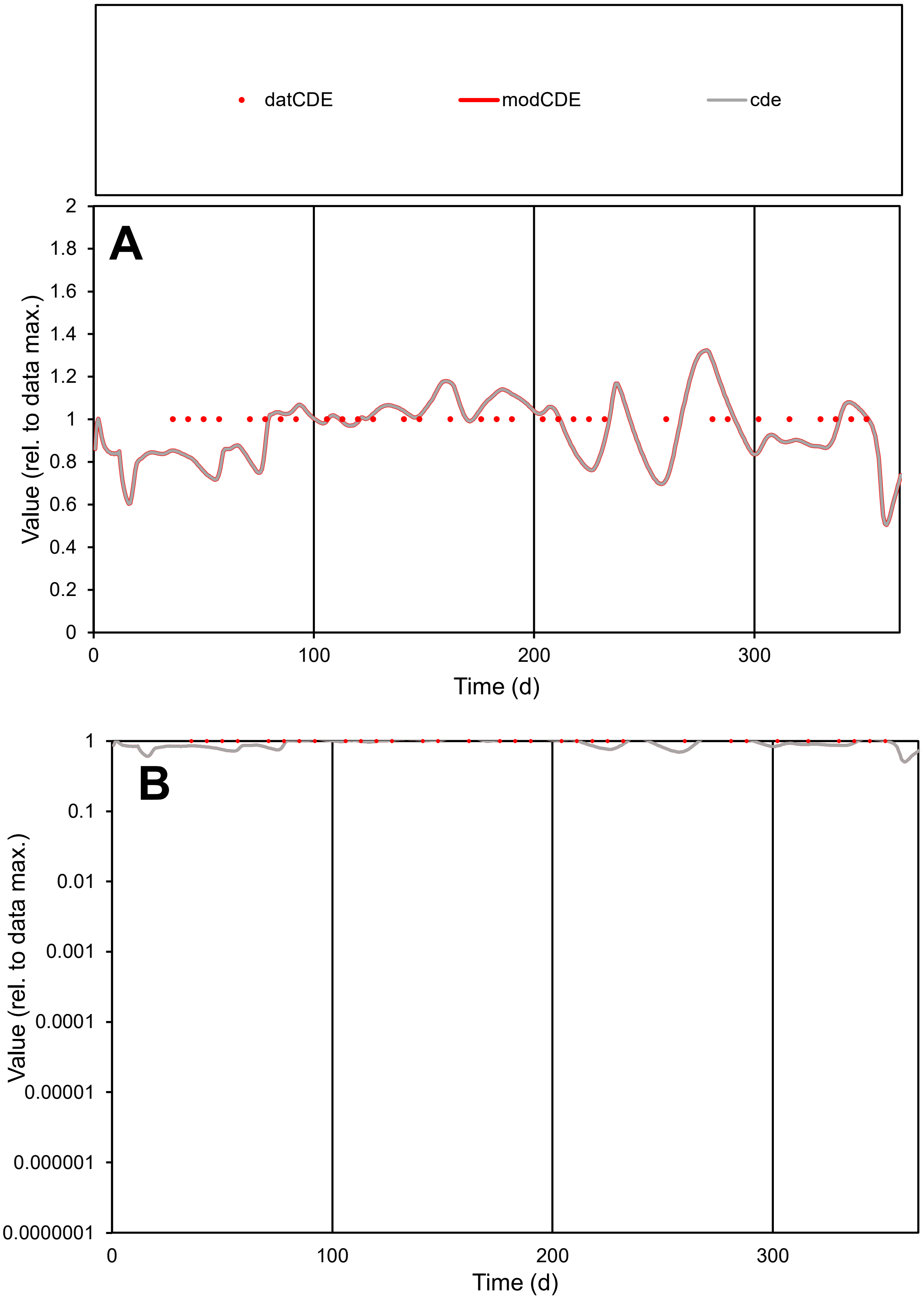


**C1**

Variables
CDE Provide jtd or Dst
CalibOnly?
2 1 to reproduce of
oNormalize?
2 1=no, 2=yes

ofxn calc*
0.12952
ofxn mod
0.16086

sRun
RunA1
zout
1

Ave
Data
1E-05
Model
9.7E-06

data/vMax
1E-05

ofxng
321.551

Min d/m
1.35069

ave/min d
1
ave/min m
1.30671

324_Calib_S_A

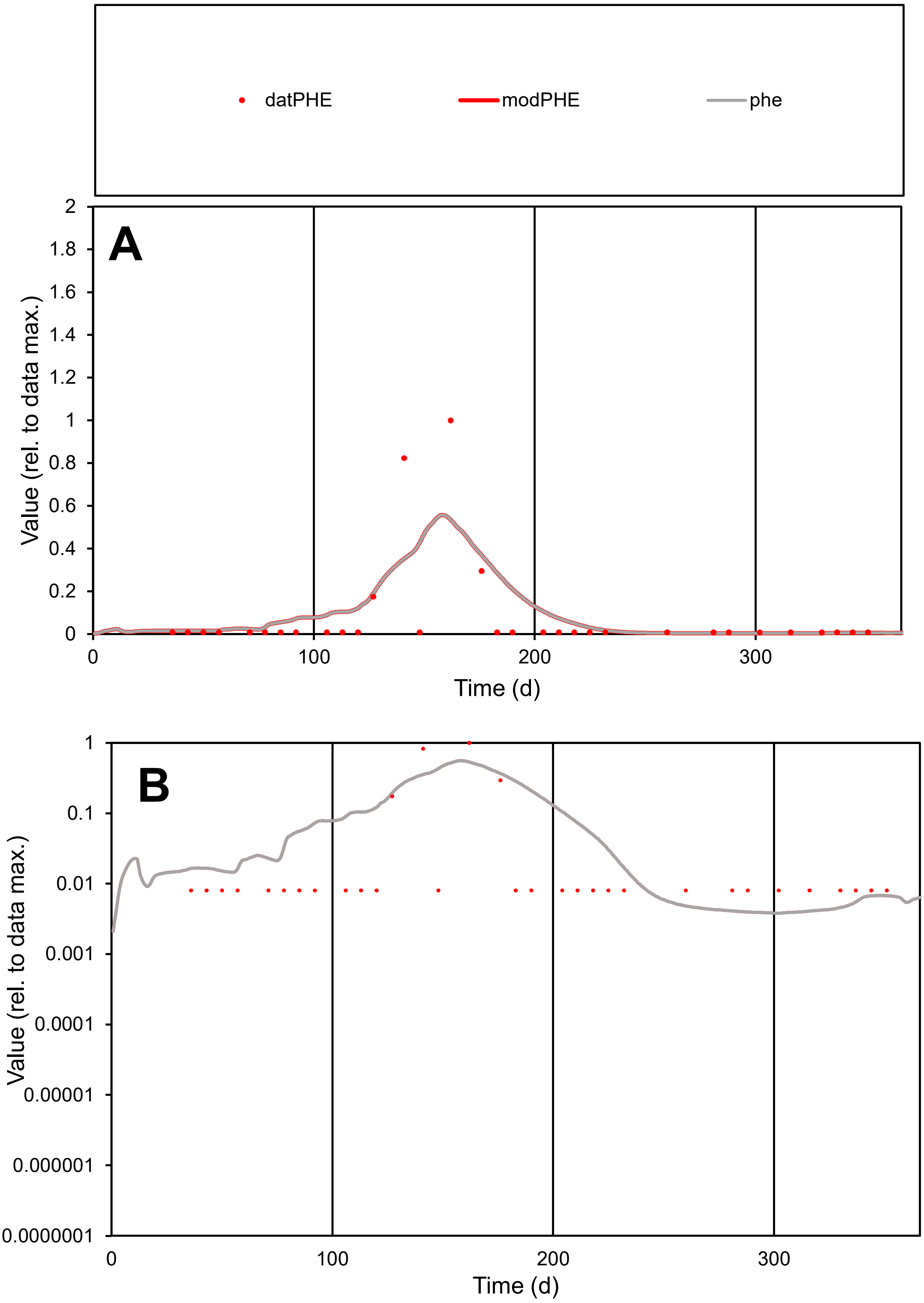


**C1**

Variables
PHE Provide jtd or Dst
CalibOnly?
2 1 to reproduce of
oNormalize?
2 1=no, 2=yes

ofxn calc*
1.96282
ofxn mod
2.0103

sRun
RunA1
zout
1

Ave
Data
3.9E-05
Model
4.9E-05

data/vMax
0.0005

ofxng
321.551

Min d/m
2.0923

ave/min d
9.82722
ave/min m
25.6765

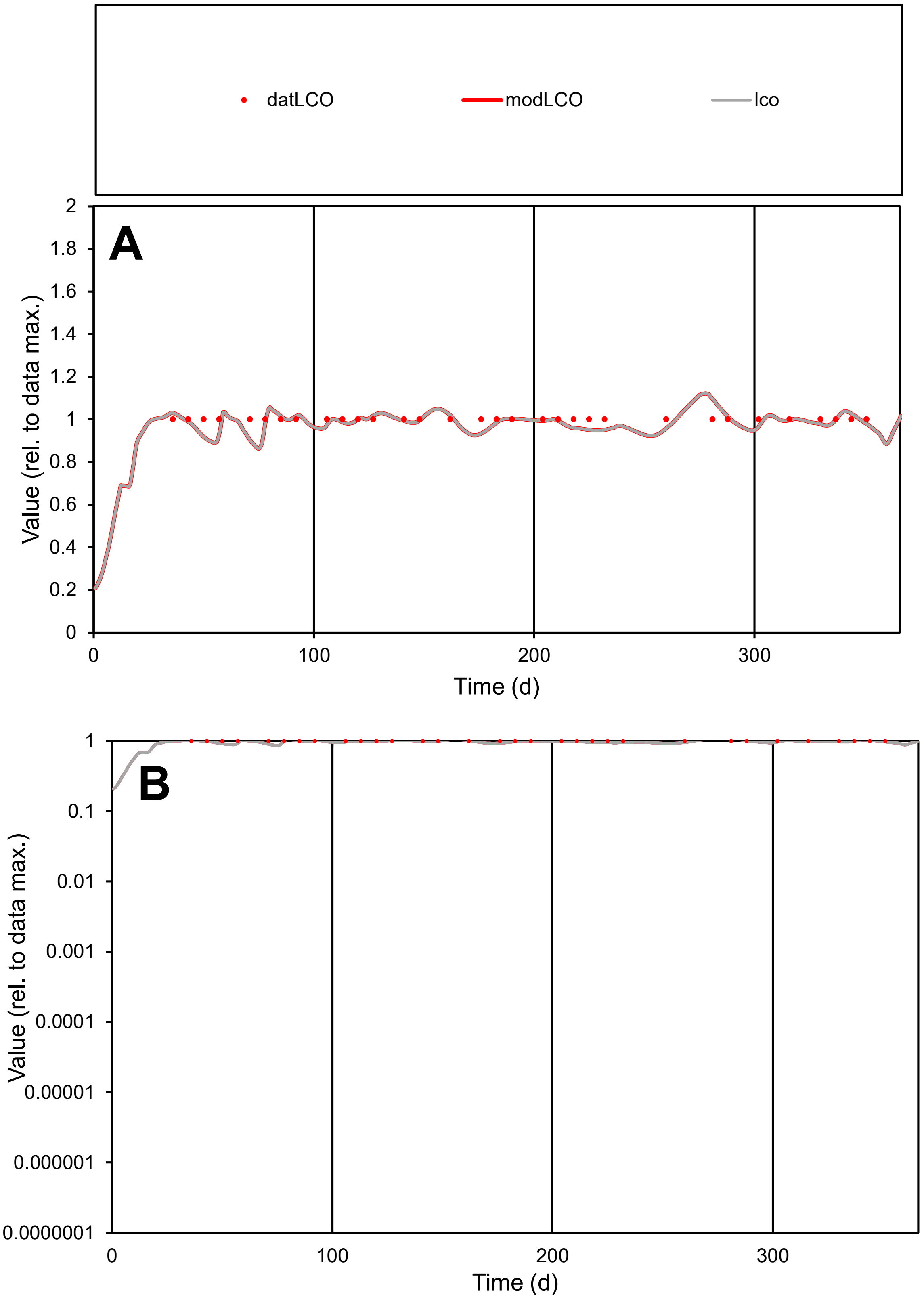


**C1**

Variables
LCO Provide jtd or Dst
CalibOnly?
2 1 to reproduce of
oNormalize?
2 1=no, 2=yes

ofxn calc*
0.03775
ofxn mod
0.10344

sRun
RunA1
zout
1

Ave
Data
7.4E-06
Model
7.3E-06

data/vMax
7.4E-06

ofxng
321.551

Min d/m
1.12329

ave/min d
1
ave/min m
1.11117

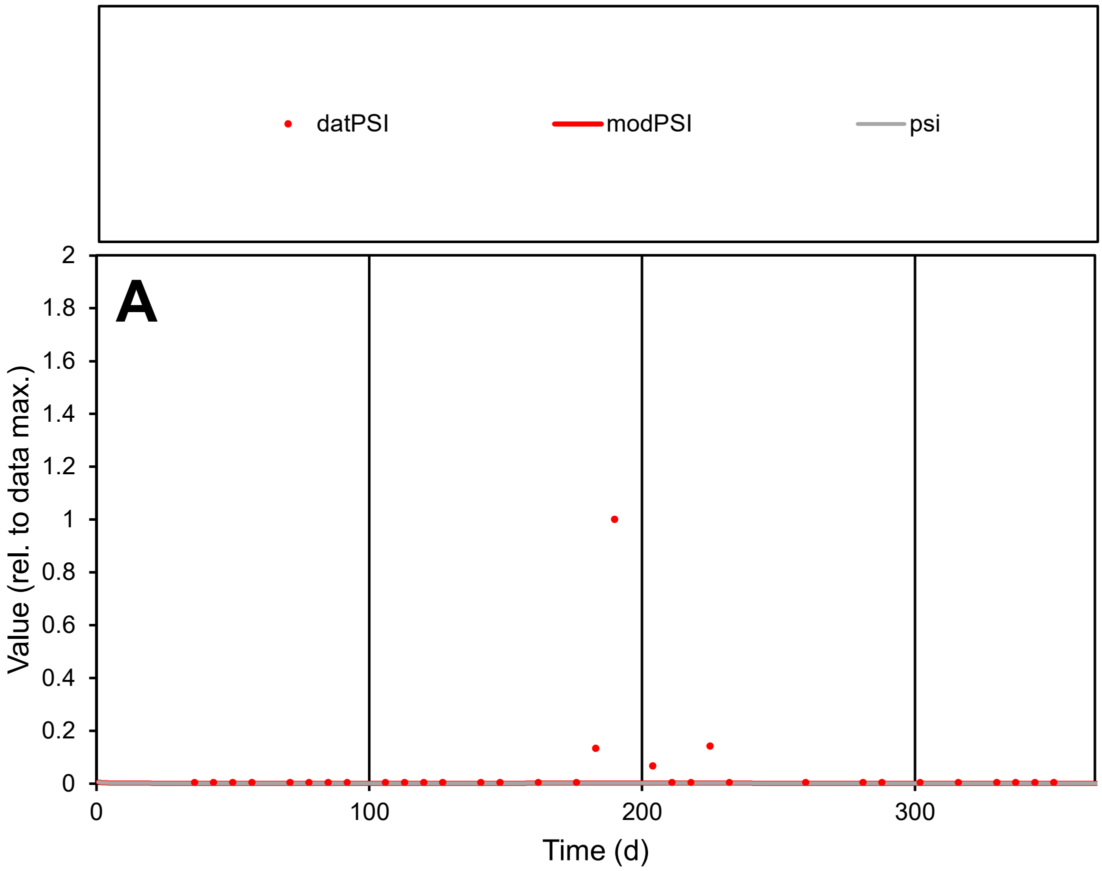


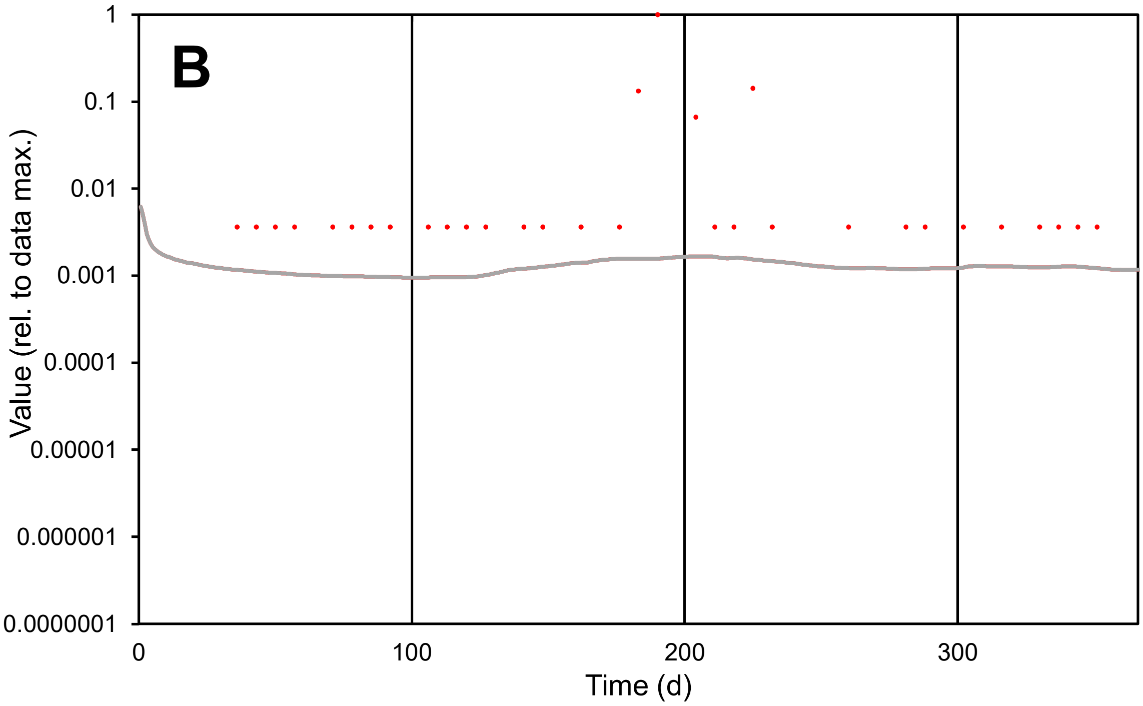


**C1**

Variables

| | |
|---|---|
| PSI | Provide jtd or Dst |
| CalibOnly? | |
| 2 | 1 to reproduce of |
| oNormalize? | |
| 2 | 1=no, 2=yes |

ofxn calc*
3.9932
ofxn mod
3.99829

sRun
RunA1
zout
1

Ave
Data
4.4E-05
Model
1.2E-06

data/vMax
0.00098

ofxng
321.551

Min d/m
3.80428

ave/min d
12.4077
ave/min m
1.30324

324_Calib_S_A

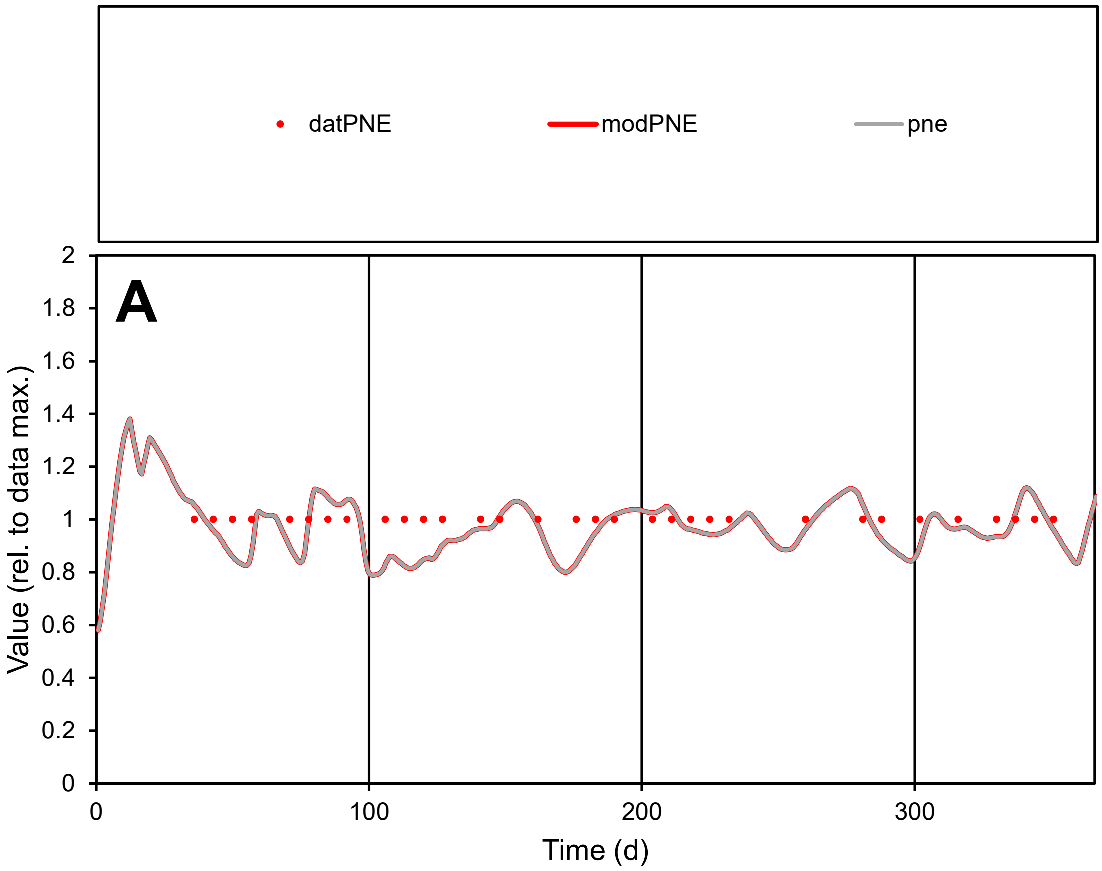


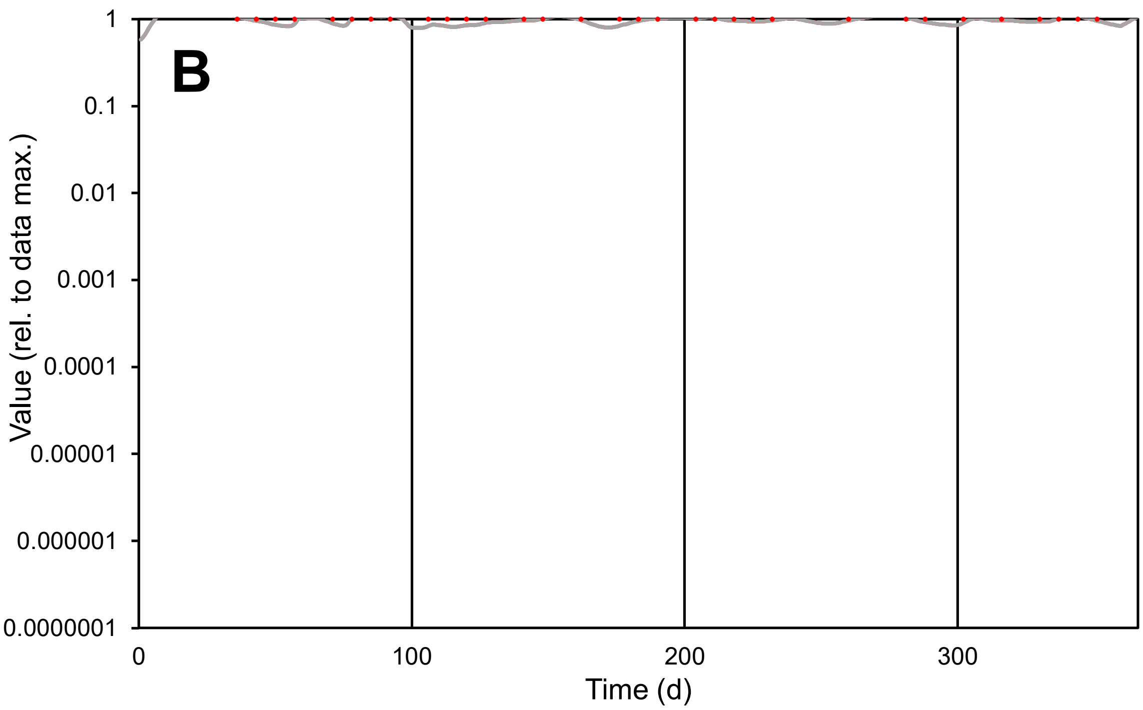


**C1**

Variables

| | |
|---|---|
| PNE | Provide jtd or Dst |
| CalibOnly? | |
| 2 | 1 to reproduce of |
| oNormalize? | |
| 2 | 1=no, 2=yes |

ofxn calc*
0.08092
ofxn mod
0.12499

sRun
RunA1
zout
1

Ave
Data
1.5E-05
Model
1.4E-05

data/vMax
1.5E-05

ofxng
321.551

Min d/m
1.22532

ave/min d
1
ave/min m
1.18241

324_Calib_S_A

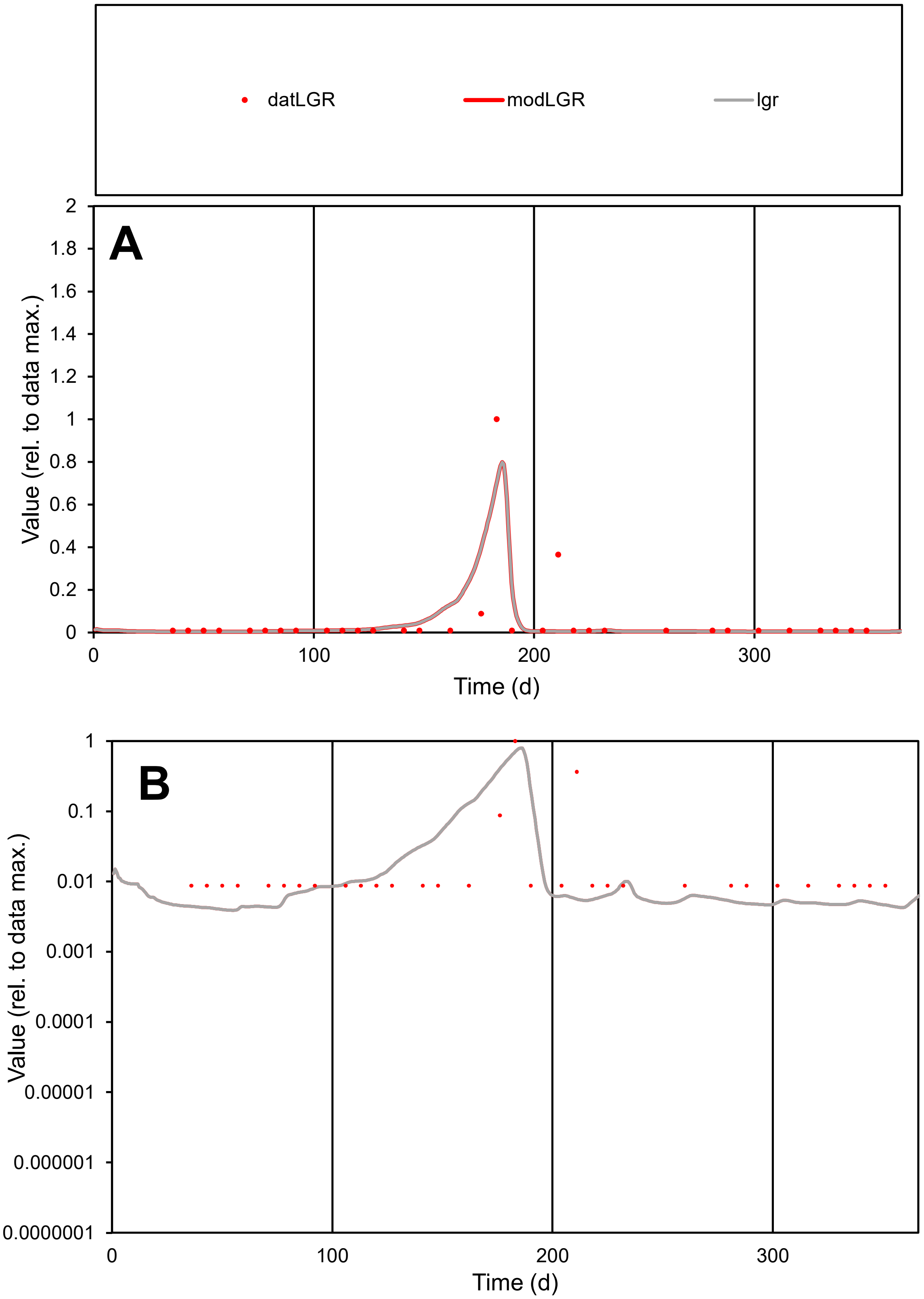


**C1**

Variables
LGR Provide jtd or Dst
CalibOnly?
2 1 to reproduce of
oNormalize?
2 1=no, 2=yes

ofxn calc*
1.74974
ofxn mod
1.85605

sRun
RunA1
zout
1

Ave
Data
3.2E-05
Model
2.9E-05

data/vMax
0.00059

ofxng
321.551

Min d/m
2.20787

ave/min d
6.10911
ave/min m
12.5738

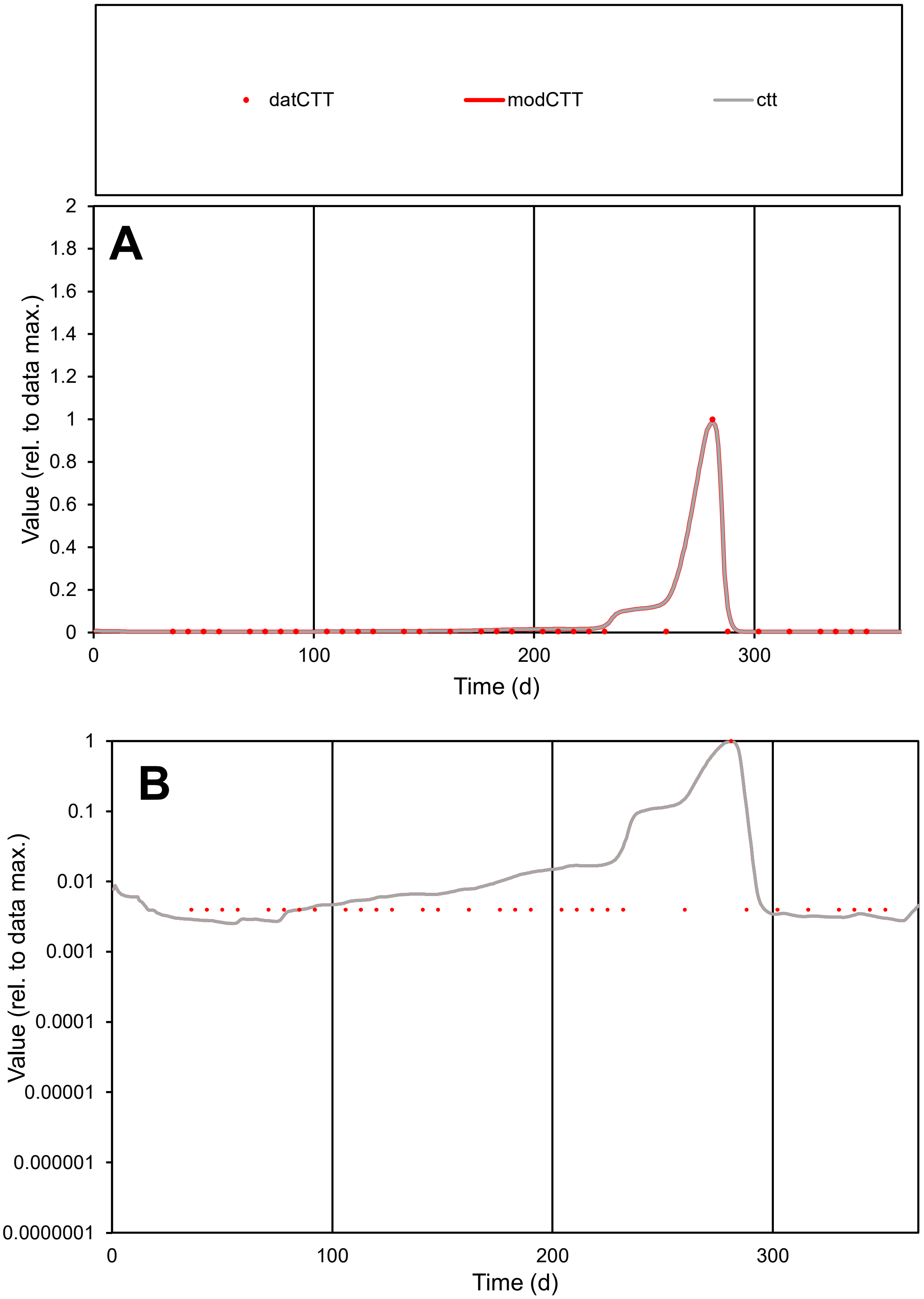


**C1**

Variables
CTT Provide jtd or Dst
CalibOnly?
2 1 to reproduce of
oNormalize?
2 1=no, 2=yes

ofxn calc*
0.70902
ofxn mod
0.88118

sRun
RunA1
zout
1

Ave
Data
1.3E-05
Model
1.6E-05

data/vMax
0.00037

ofxng
321.551

Min d/m
1.54705

ave/min d
8.84131
ave/min m
16.5407

324_Calib_S_A

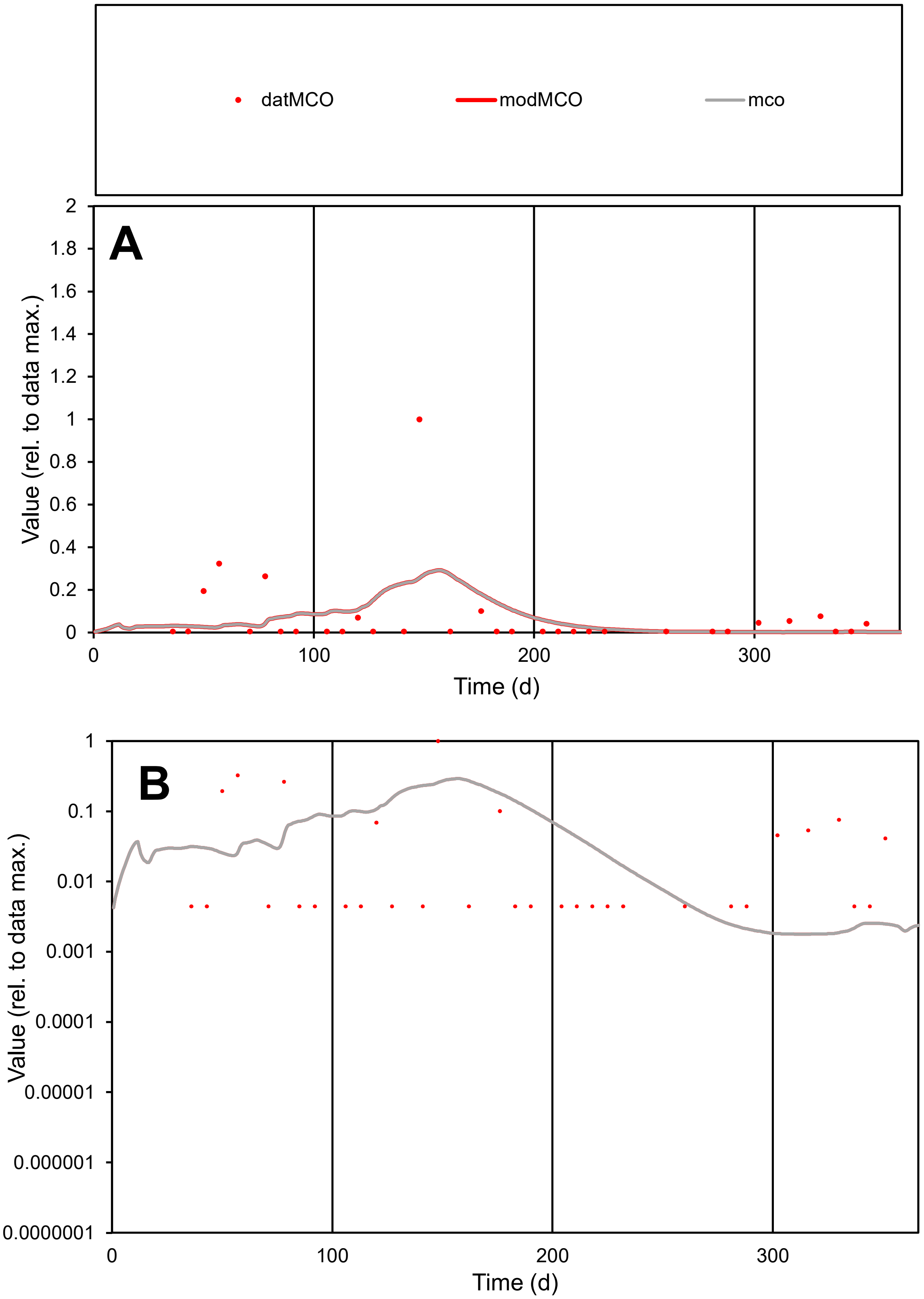


**C1**

Variables

MCO Provide jtd or Dst

CalibOnly?

2 1 to reproduce of

oNormalize?

2 1=no, 2=yes

ofxn calc*

2.37978

ofxn mod

2.39527

sRun

RunA1

zout

1

Ave

Data

9.8E-07

Model

9.2E-07

data/vMax

1.4E-05

ofxng

321.551

Min d/m

2.48274

ave/min d

15.9851

ave/min m

37.2463

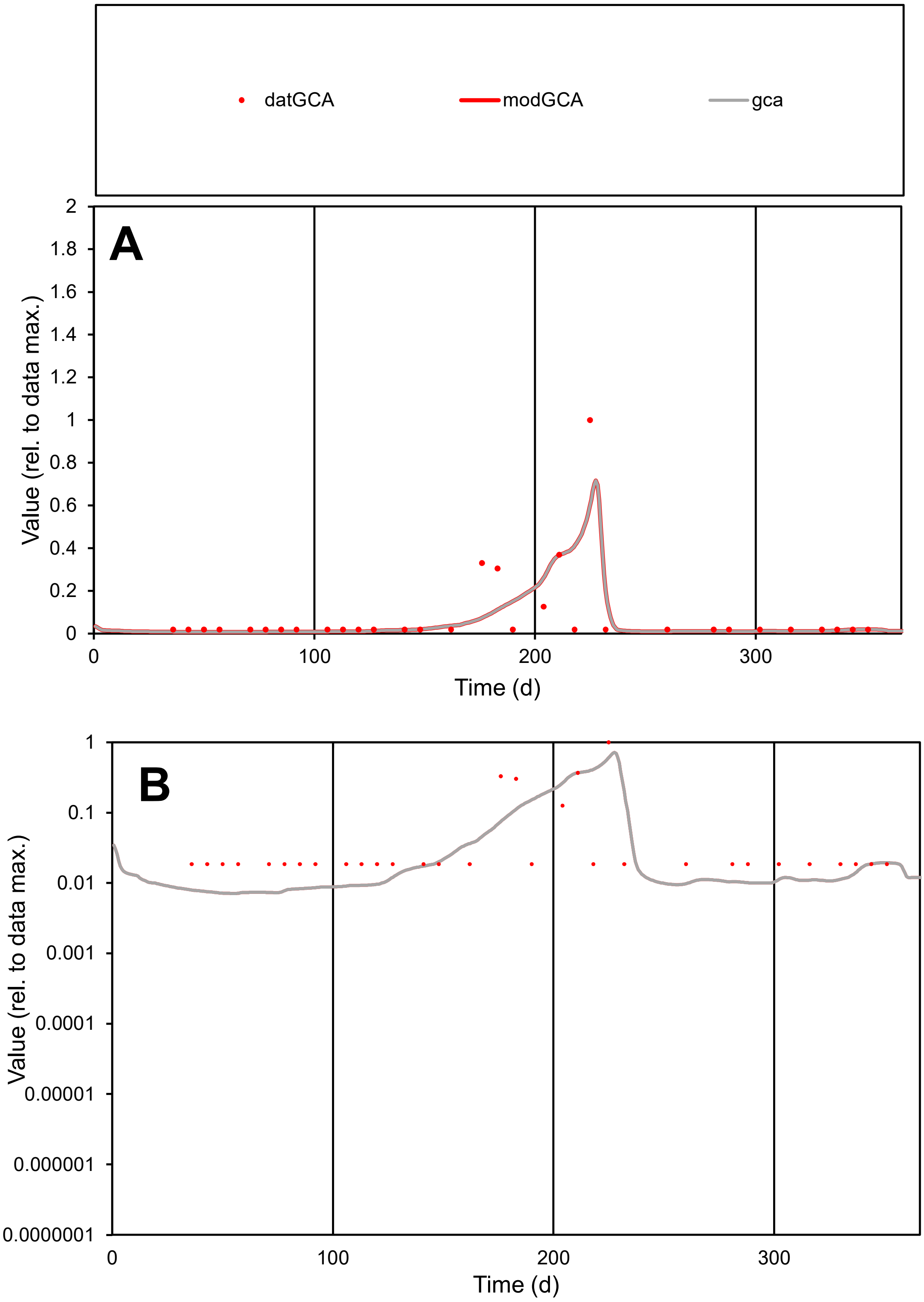


**C1**

Variables

GCA Provide jtd or Dst

CalibOnly?

2 1 to reproduce of

oNormalize?

2 1=no, 2=yes

ofxn calc*

1.36205

ofxn mod

1.39329

sRun

RunA1

zout

1

Ave

Data

3.7E-05

Model

3.4E-05

data/vMax

0.00045

ofxng

321.551

Min d/m

2.60174

ave/min d

4.42368

ave/min m

10.548

324_Calib_S_A

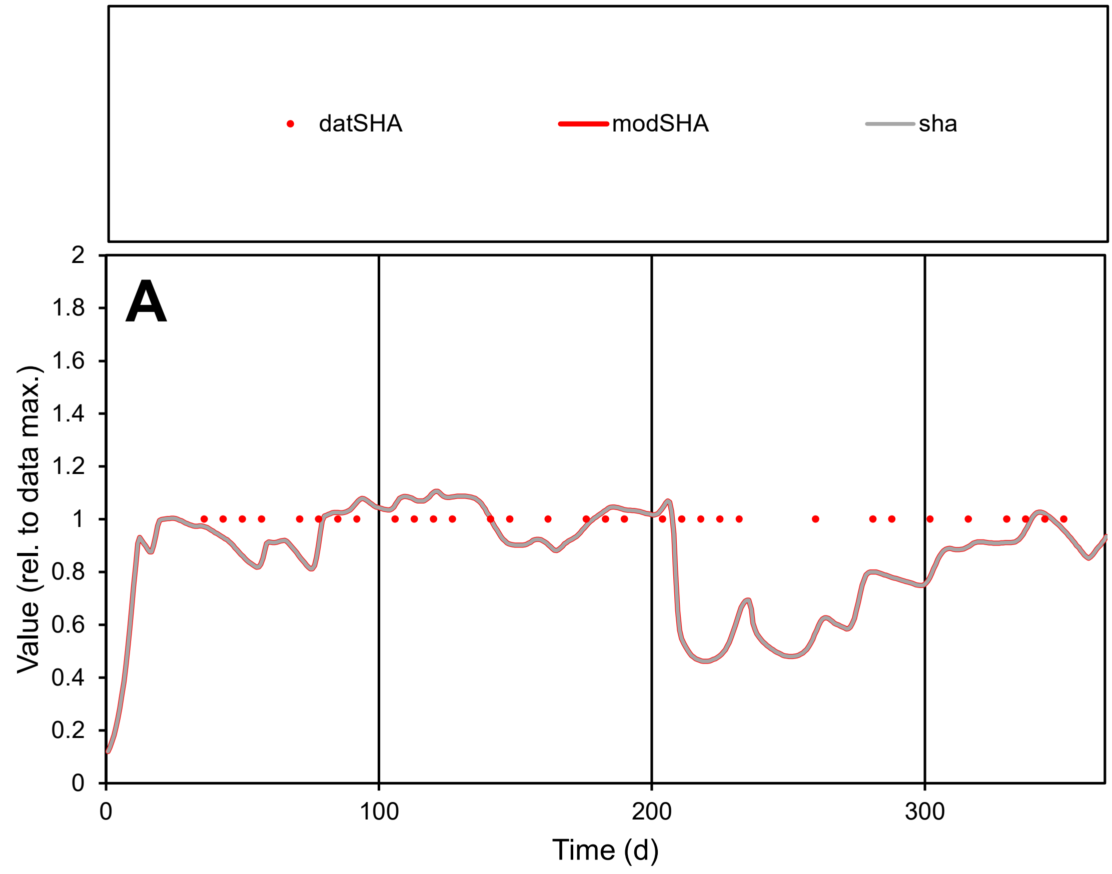


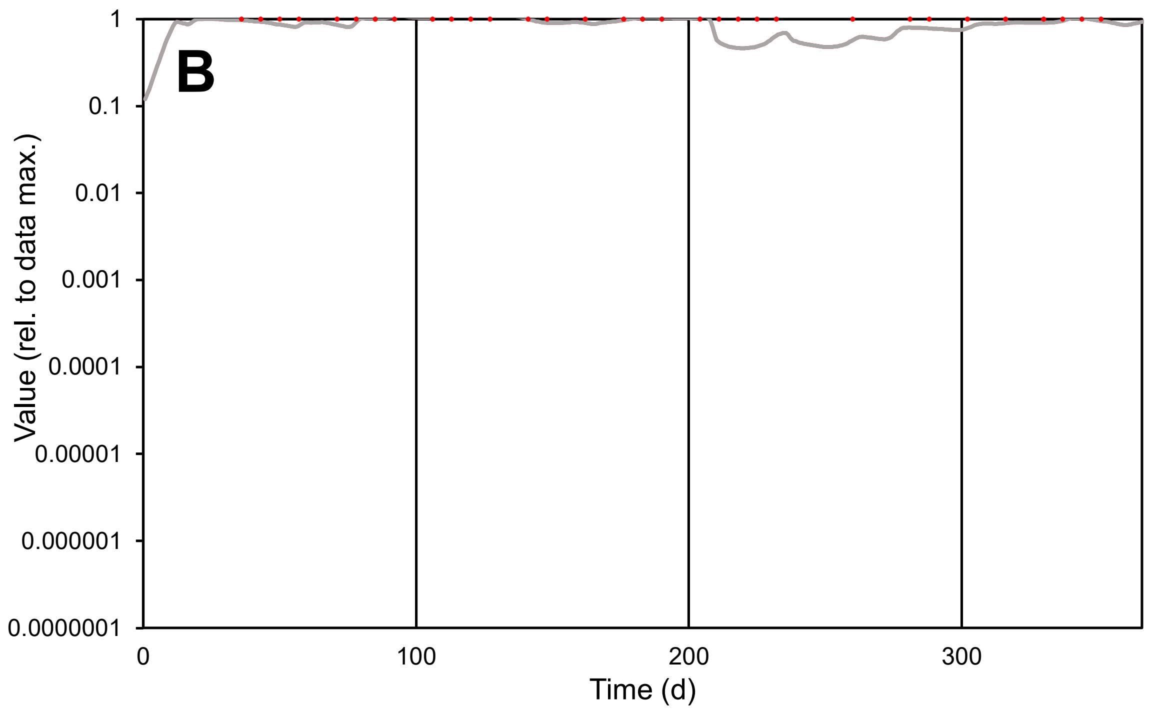


**C1**

Variables

SHA Provide jtd or Dst

CalibOnly?

2 1 to reproduce of

oNormalize?

2 1=no, 2=yes

ofxn calc*

0.20225

ofxn mod

0.22248

sRun

RunA1

zout

1

Ave

Data

7E-07

Model

6.3E-07

data/vMax

7E-07

ofxng

321.551

Min d/m

2.16517

ave/min d

1

ave/min m

1.94224

324_Calib_S_A

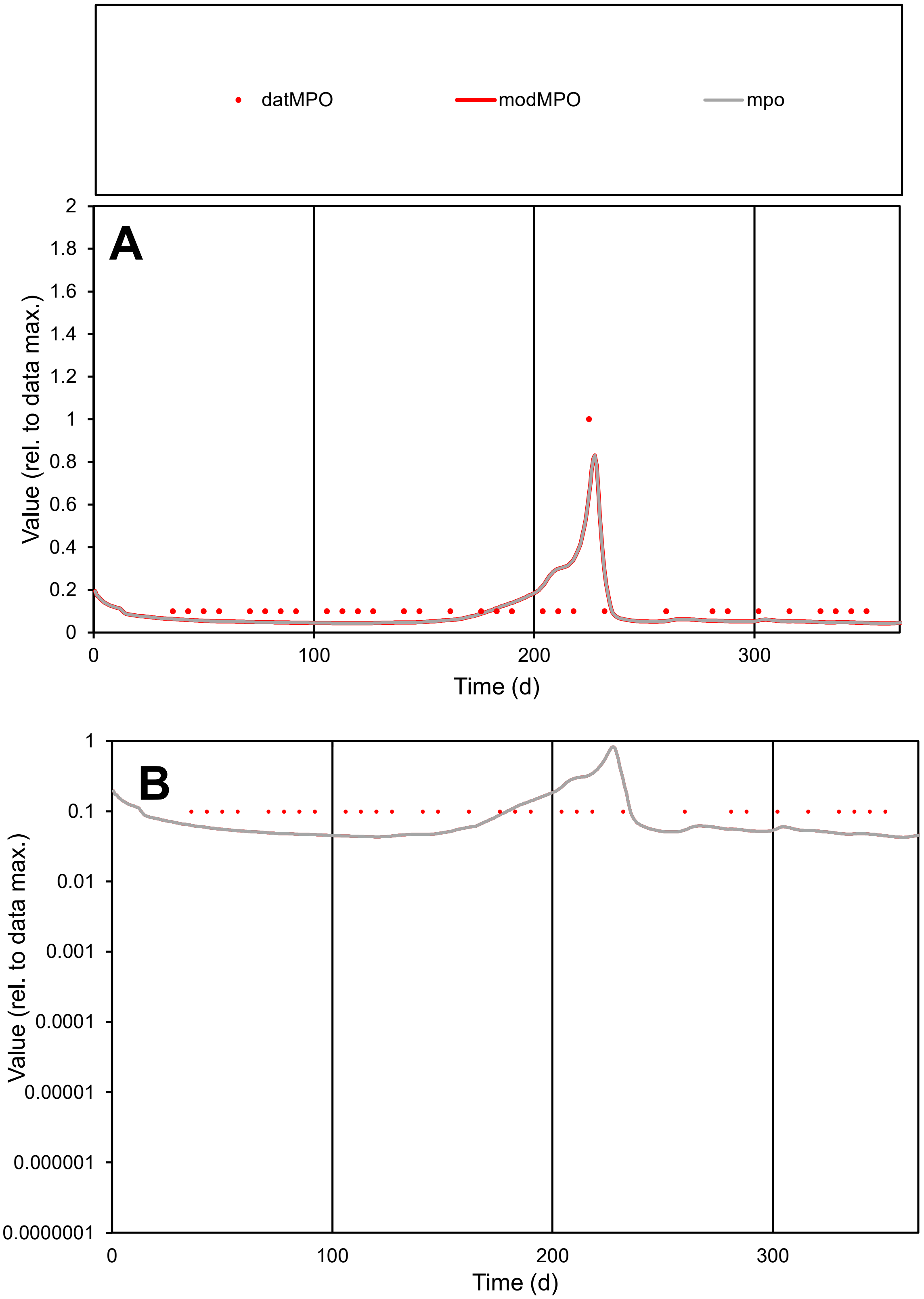


**C1**

Variables
MPO Provide jtd or Dst
CalibOnly?
2 1 to reproduce of
oNormalize?
2 1=no, 2=yes

ofxn calc*
0.67401
ofxn mod
0.6962

sRun
RunA1
zout
1

Ave
Data
1.6E-05
Model
1.3E-05

data/vMax
0.00012

ofxng
321.551

Min d/m
2.29305

ave/min d
1.28326
ave/min m
2.41483

324_Calib_S_A

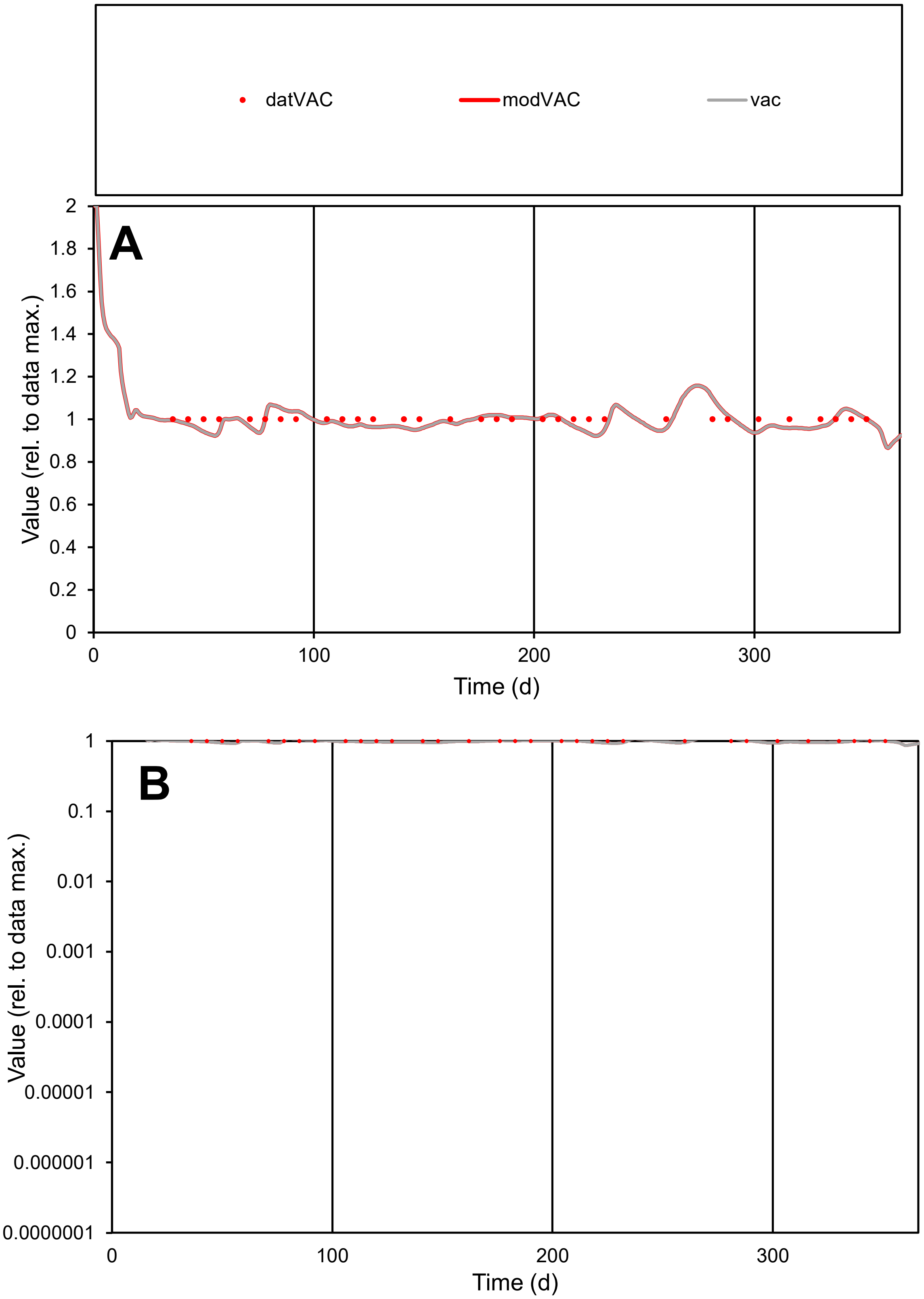


**C1**

Variables
VAC Provide jtd or Dst
CalibOnly?
2 1 to reproduce of
oNormalize?
2 1=no, 2=yes

ofxn calc*
0.03758
ofxn mod
0.10334

sRun
RunA1
zout
1

Ave
Data
9.5E-06
Model
9.4E-06

data/vMax
9.5E-06

ofxng
321.551

Min d/m
1.07832

ave/min d
1
ave/min m
1.06772

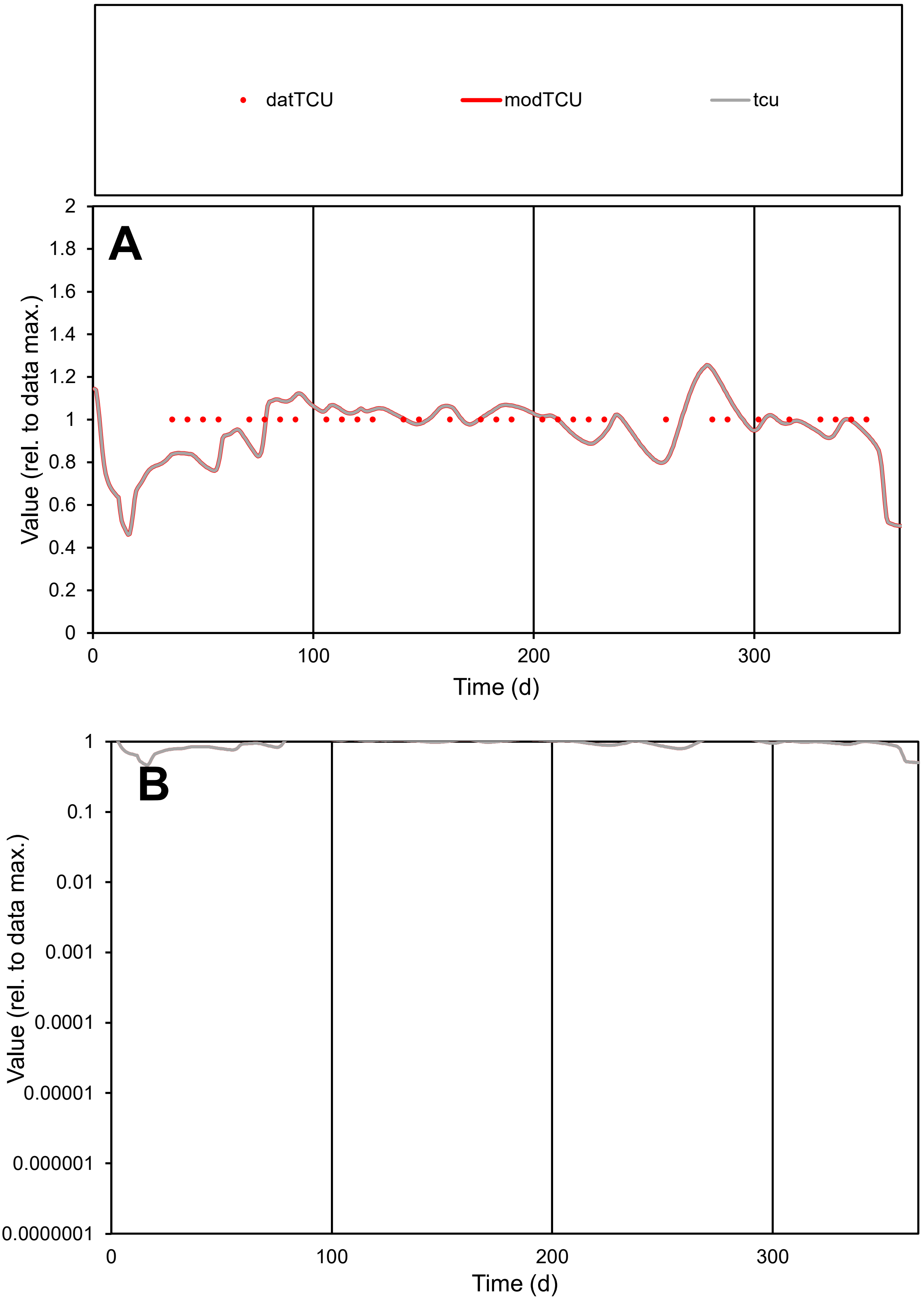


**C1**

Variables
TCU Provide jtd or Dst
CalibOnly?
2 1 to reproduce of
oNormalize?
2 1=no, 2=yes

ofxn calc*
0.09801
ofxn mod
0.13703

sRun
RunA1
zout
1

Ave
Data
9.8E-06
Model
9.6E-06

data/vMax
9.8E-06

ofxng
321.551

Min d/m
1.28609

ave/min d
1
ave/min m
1.26542

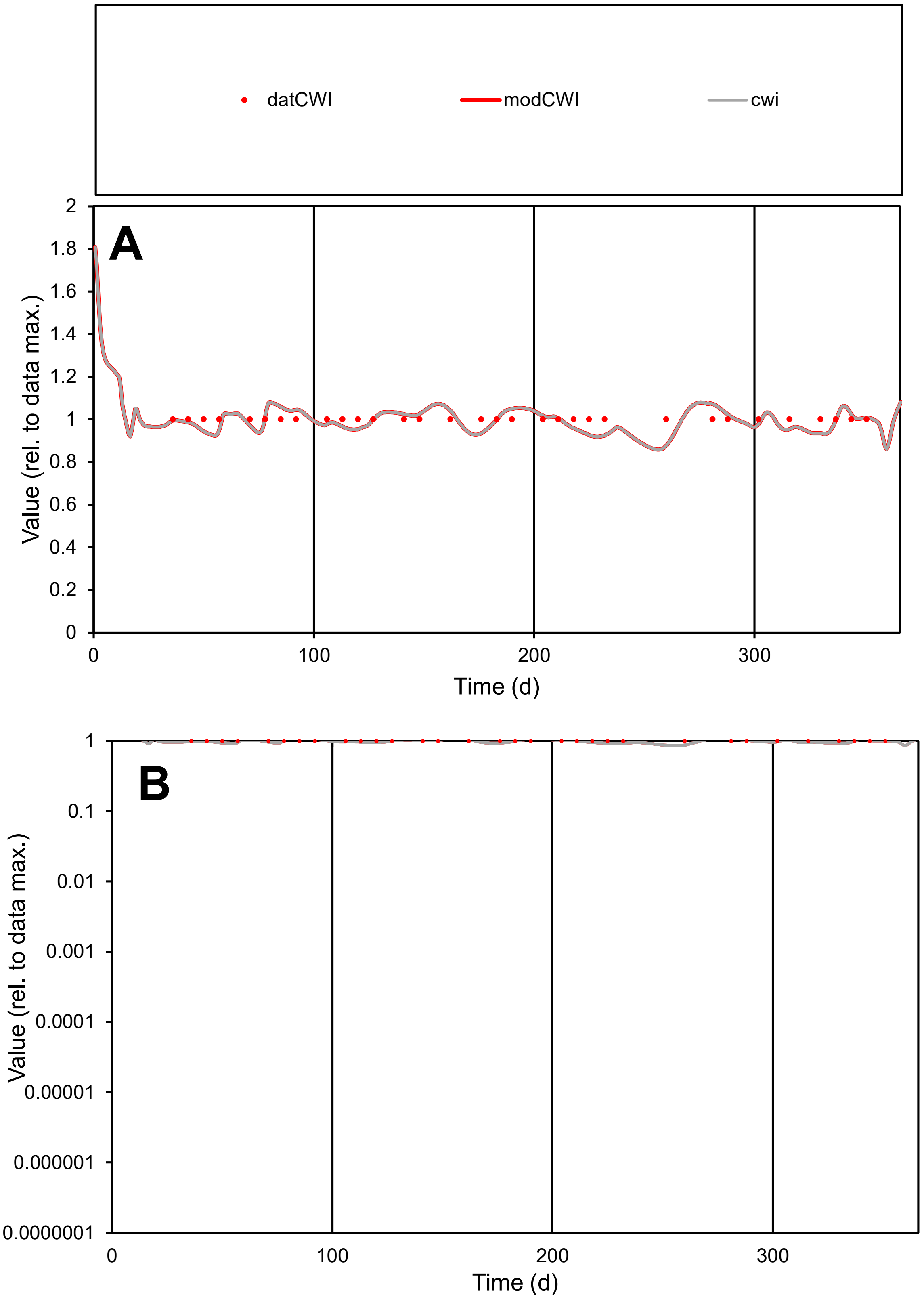


**C1**

Variables
CWI Provide jtd or Dst
CalibOnly?
2 1 to reproduce of
oNormalize?
2 1=no, 2=yes

ofxn calc*
0.04501
ofxn mod
0.1064

sRun
RunA1
zout
1

Ave
Data
1.3E-05
Model
1.3E-05

data/vMax
1.3E-05

ofxng
321.551

Min d/m
1.12247

ave/min d
1
ave/min m
1.11386

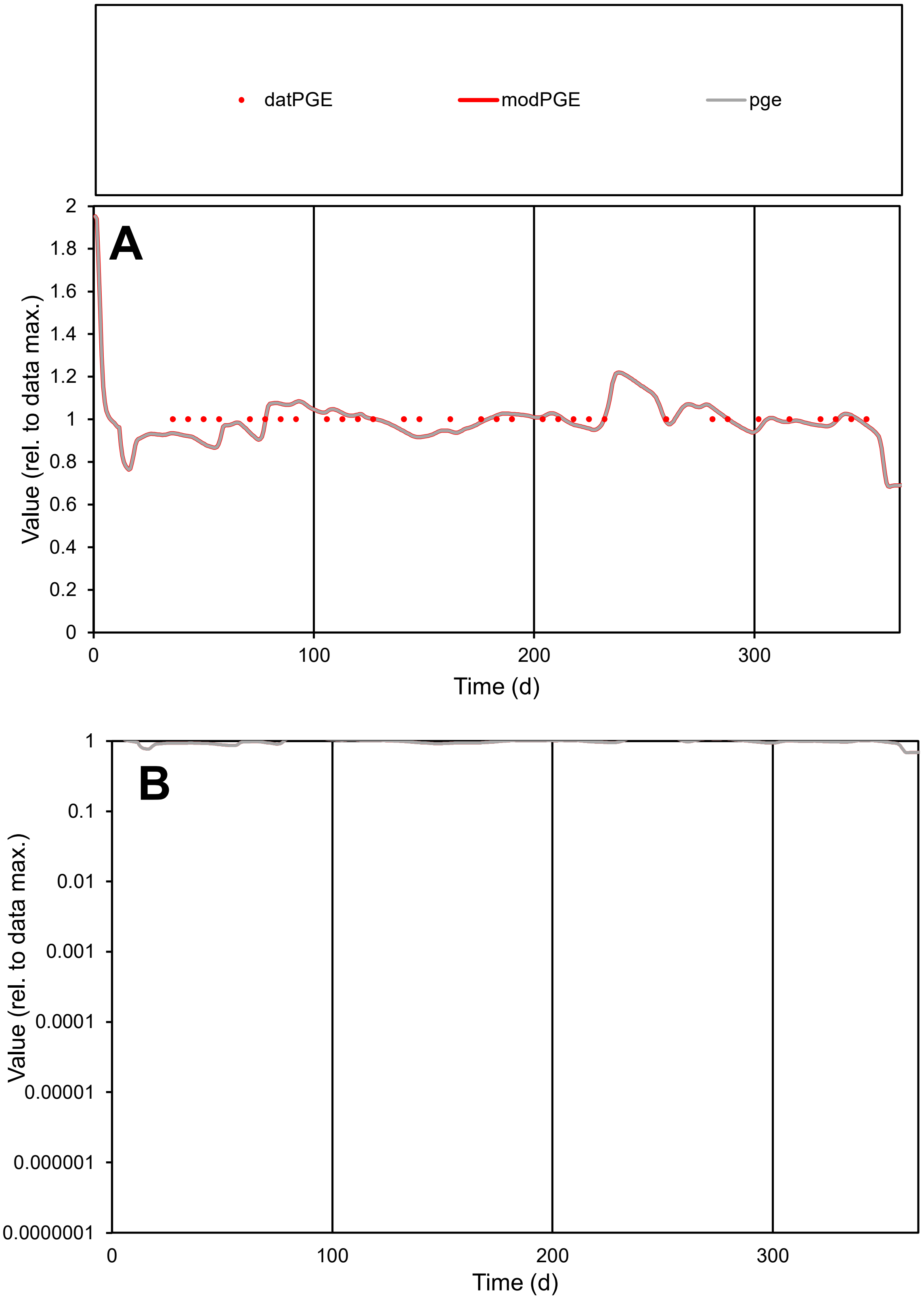


**C1**

Variables
PGE Provide jtd or Dst
CalibOnly?
2 1 to reproduce of
oNormalize?
2 1=no, 2=yes

ofxn calc*
0.0501
ofxn mod
0.10887

sRun
RunA1
zout
1

Ave
Data
5.9E-06
Model
5.8E-06

data/vMax
5.9E-06

ofxng
321.551

Min d/m
1.14006

ave/min d
1
ave/min m
1.12738

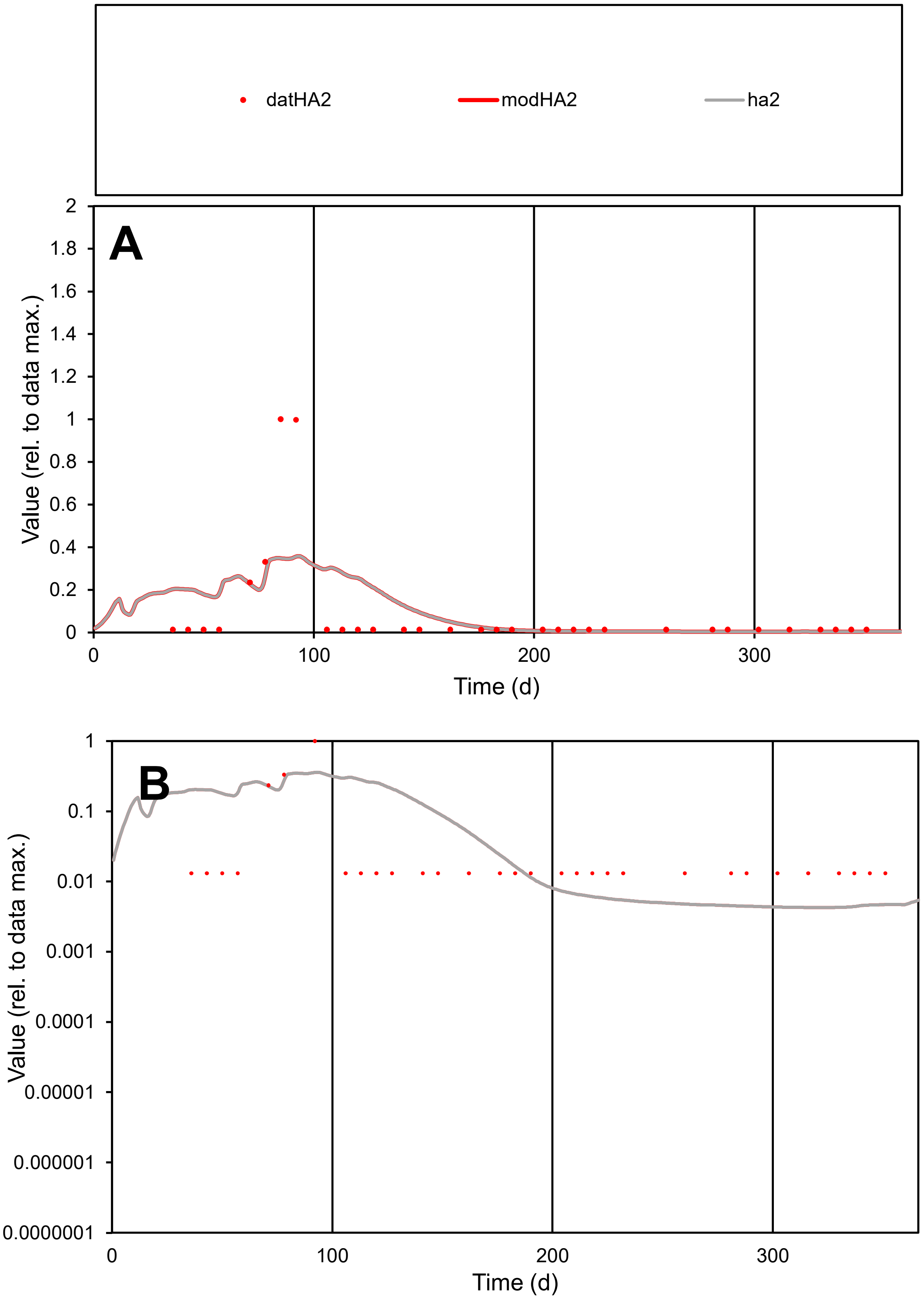


**C1**

Variables
HA2 Provide jtd or Dst
CalibOnly?
2 1 to reproduce of
oNormalize?
2 1=no, 2=yes

ofxn calc*
2.12622
ofxn mod
2.14191

sRun
RunA1
zout
1

Ave
Data
1.8E-05
Model
2.1E-05

data/vMax
0.0002

ofxng
321.551

Min d/m
3.07317

ave/min d
6.95053
ave/min m
24.6361

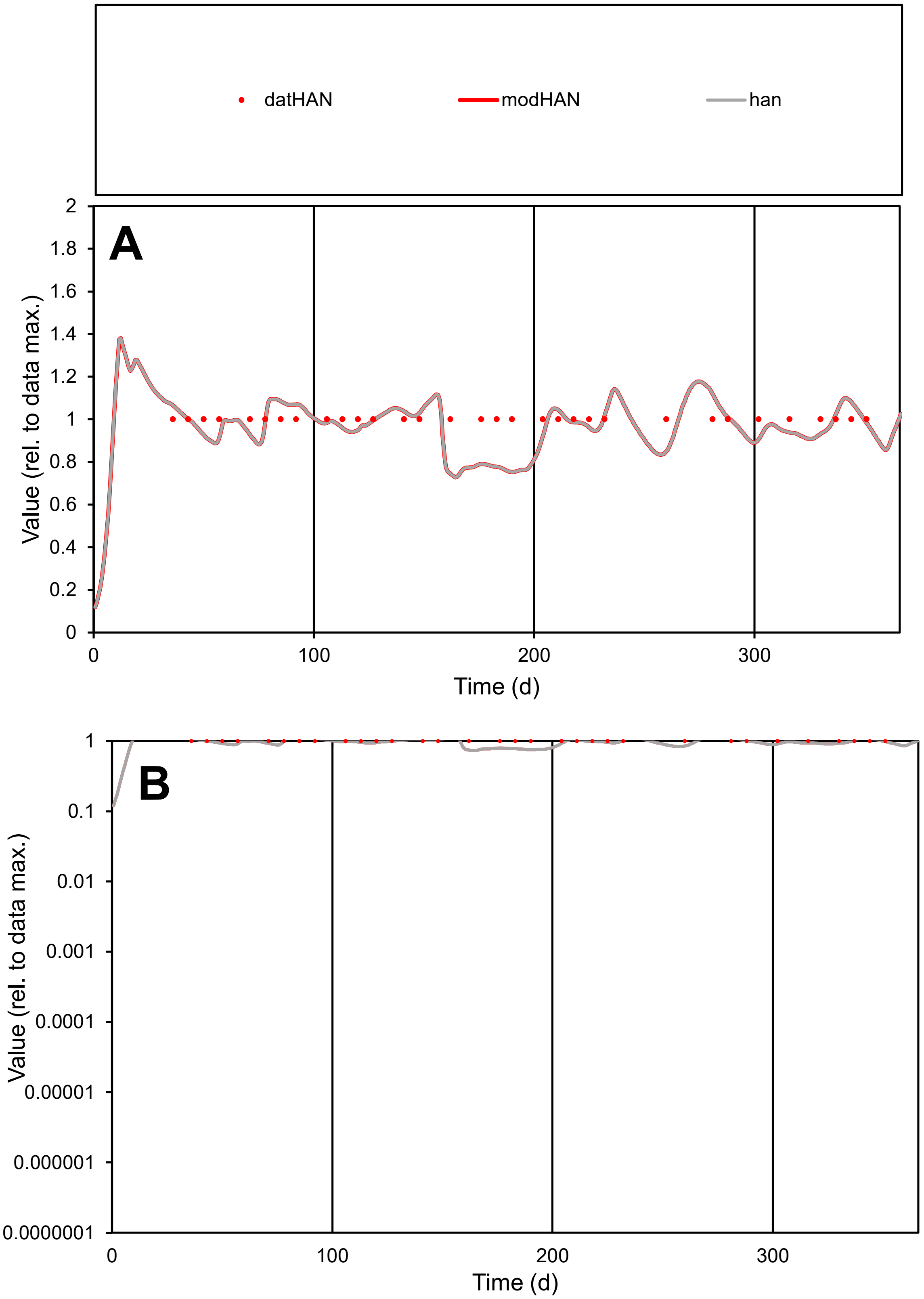


**C1**

Variables
HAN Provide jtd or Dst
CalibOnly?
2 1 to reproduce of
oNormalize?
2 1=no, 2=yes

ofxn calc*
0.10068
ofxn mod
0.13889

sRun
RunA1
zout
1

Ave
Data
5.1E-06
Model
5E-06

data/vMax
5.1E-06

ofxng
321.551

Min d/m
1.36604

ave/min d
1
ave/min m
1.31968

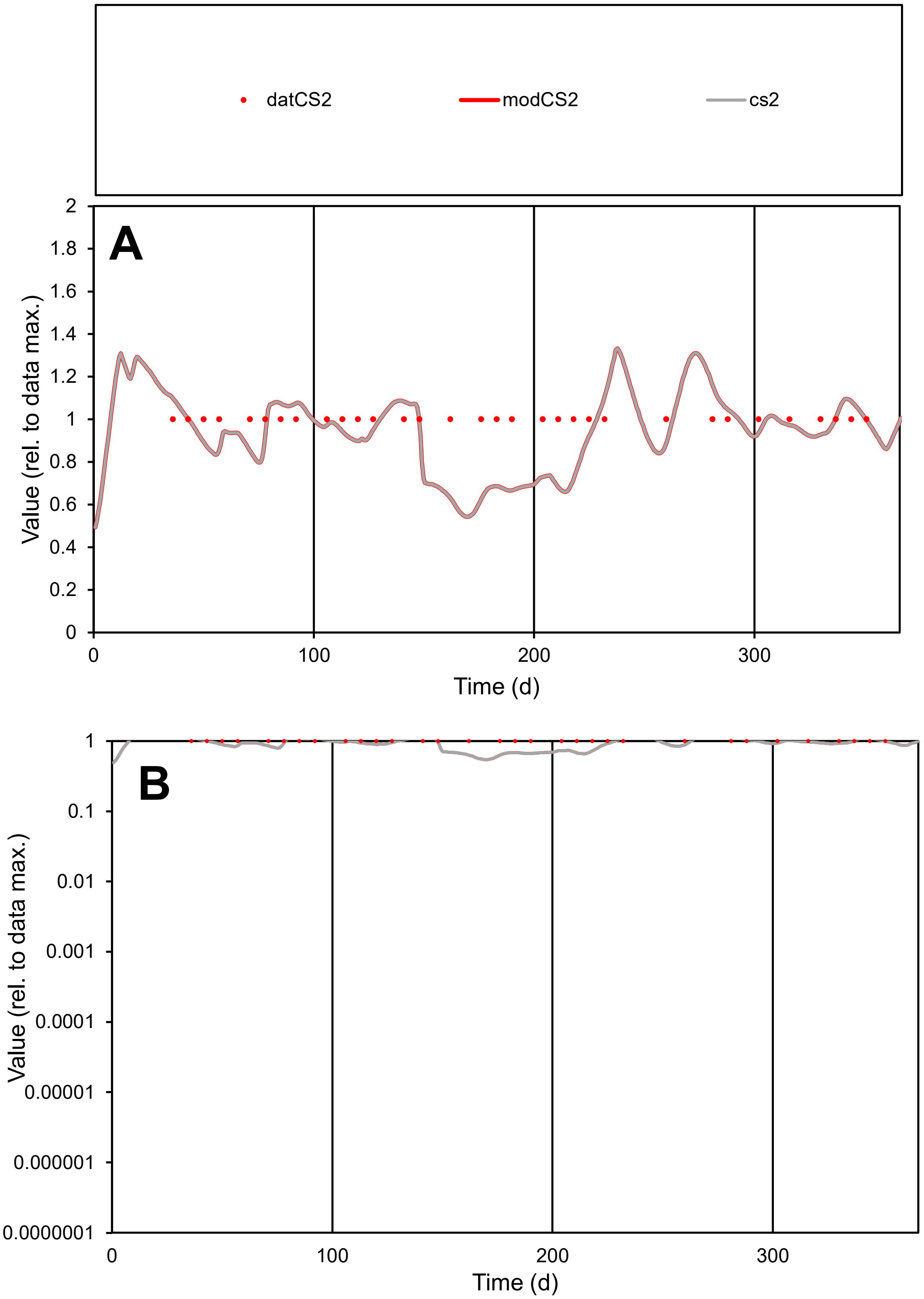


**C1**

Variables
CS2 Provide jtd or Dst
CalibOnly?
2 1 to reproduce of
oNormalize?
2 1=no, 2=yes

ofxn calc*
0.16523
ofxn mod
0.19211

sRun
RunA1
zout
1

Ave
Data
7.6E-06
Model
7E-06

data/vMax
7.6E-06

ofxng
321.551

Min d/m
1.64525

ave/min d
1
ave/min m
1.52056

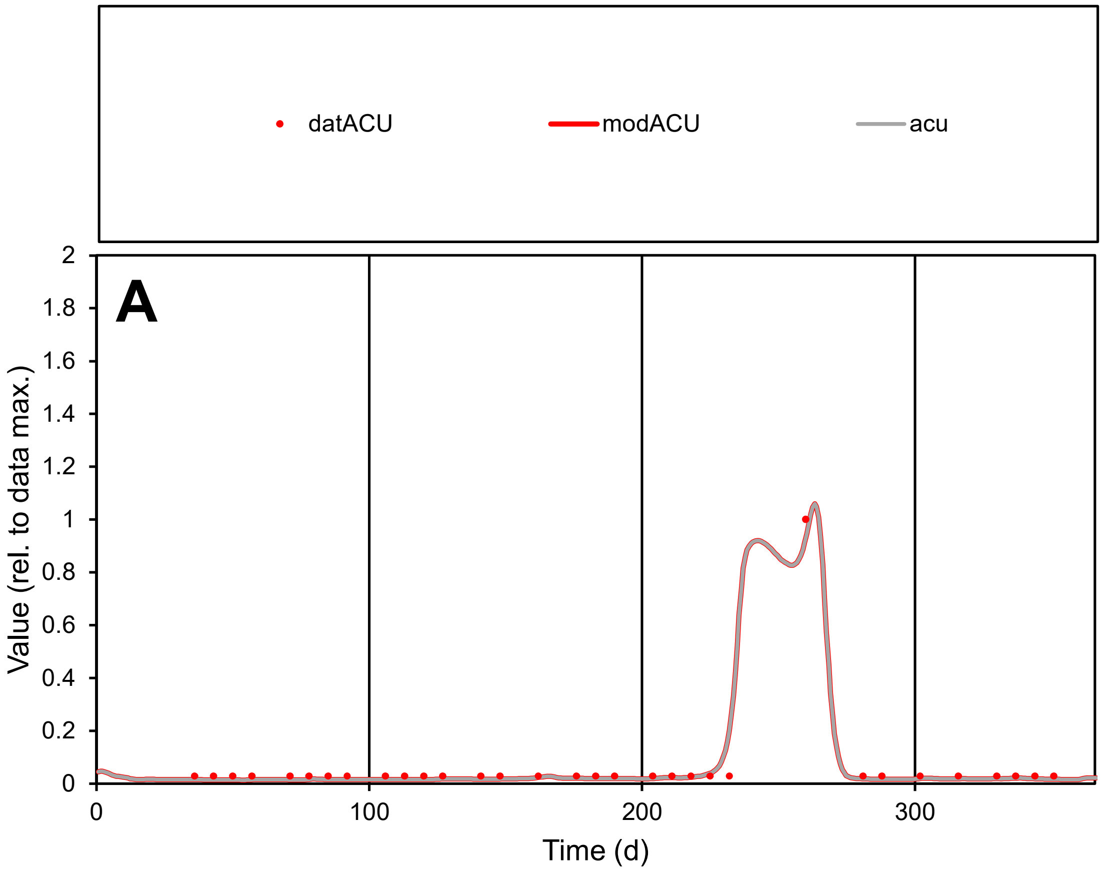


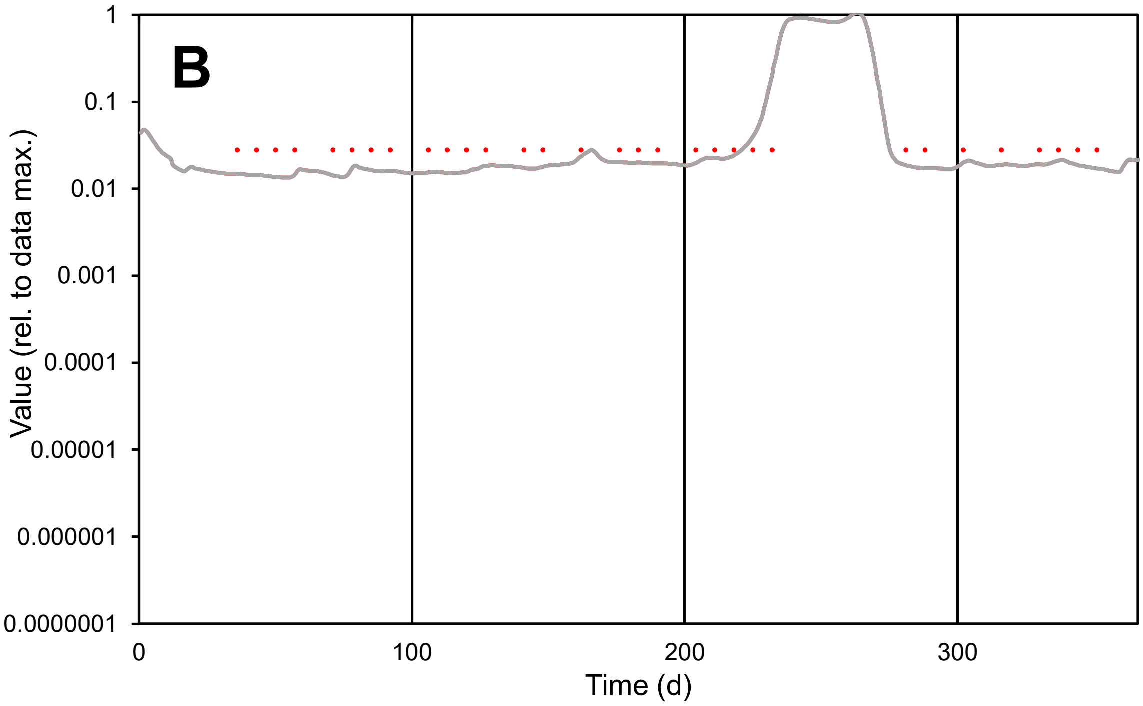


**C1**

Variables
ACU Provide jtd or Dst
CalibOnly?
2 1 to reproduce of
oNormalize?
2 1=no, 2=yes

ofxn calc*
0.34741
ofxn mod
0.49726

sRun
RunA1
zout
1

Ave
Data
2.3E-05
Model
2.1E-05

data/vMax
0.0004

ofxng
321.551

Min d/m
2.05817

ave/min d
2.08824
ave/min m
3.87083

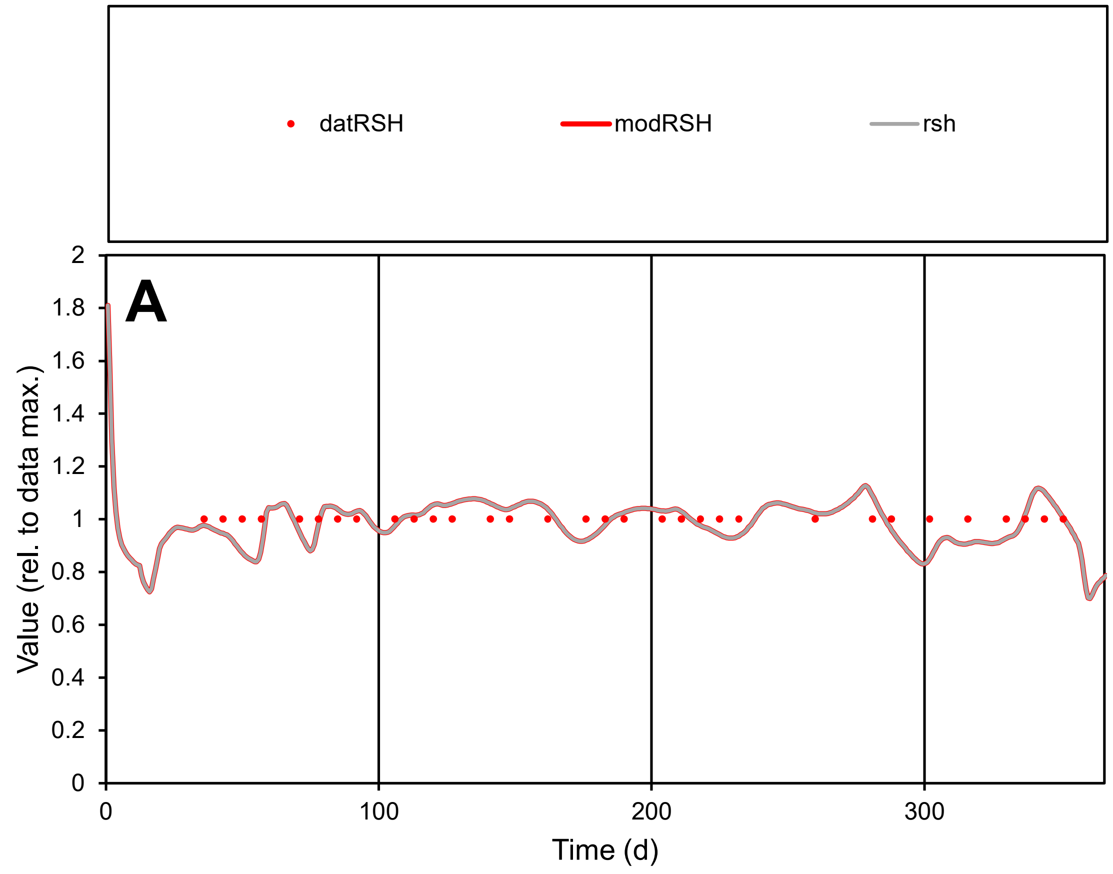


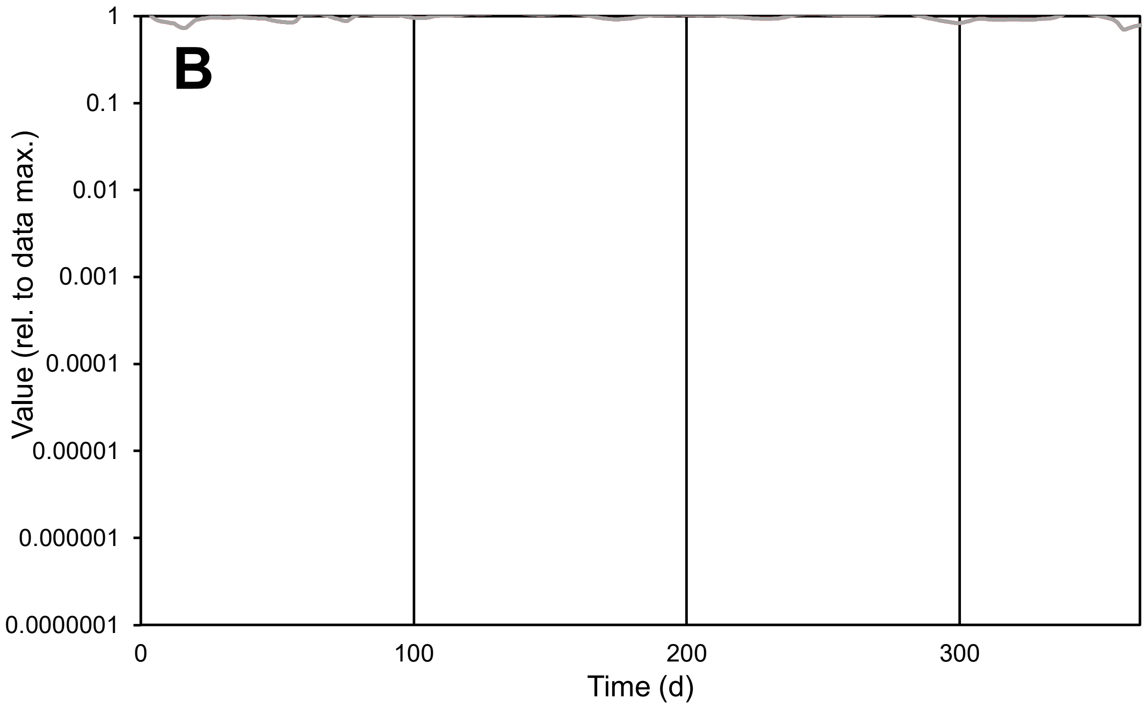


**C1**

Variables
RSH Provide jtd or Dst
CalibOnly?
2 1 to reproduce of
oNormalize?
2 1=no, 2=yes

ofxn calc*
0.05832
ofxn mod
0.11254

sRun
RunA1
zout
1

Ave
Data
1.8E-05
Model
1.8E-05

data/vMax
1.8E-05

ofxng
321.551

Min d/m
1.16656

ave/min d
1
ave/min m
1.15678

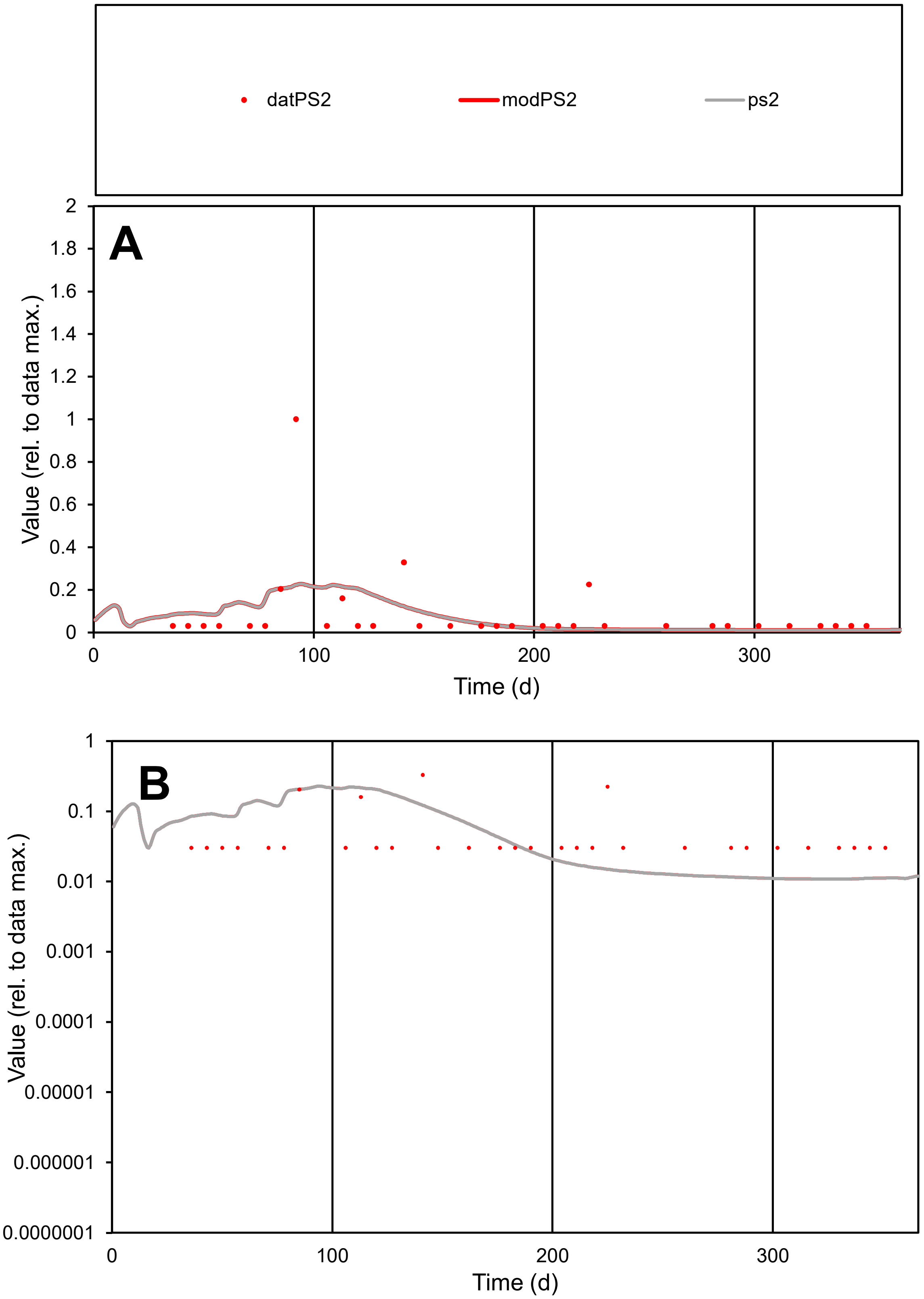


**C1**

Variables
PS2 Provide jtd or Dst
CalibOnly?
2 1 to reproduce of
oNormalize?
2 1=no, 2=yes

ofxn calc*
1.86721
ofxn mod
1.88123

sRun
RunA1
zout
1

Ave
Data
5.4E-05
Model
4.8E-05

data/vMax
0.00063

ofxng
321.551

Min d/m
2.79736

ave/min d
2.8326
ave/min m
6.97819

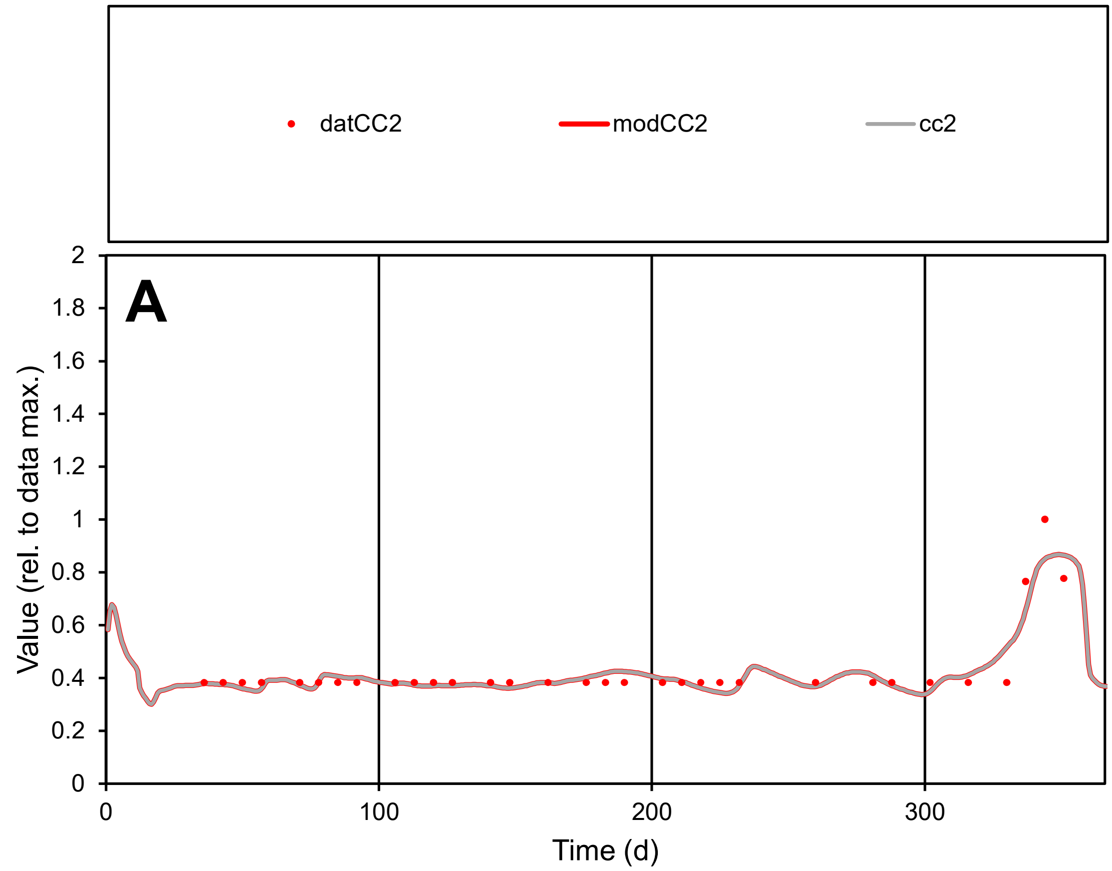


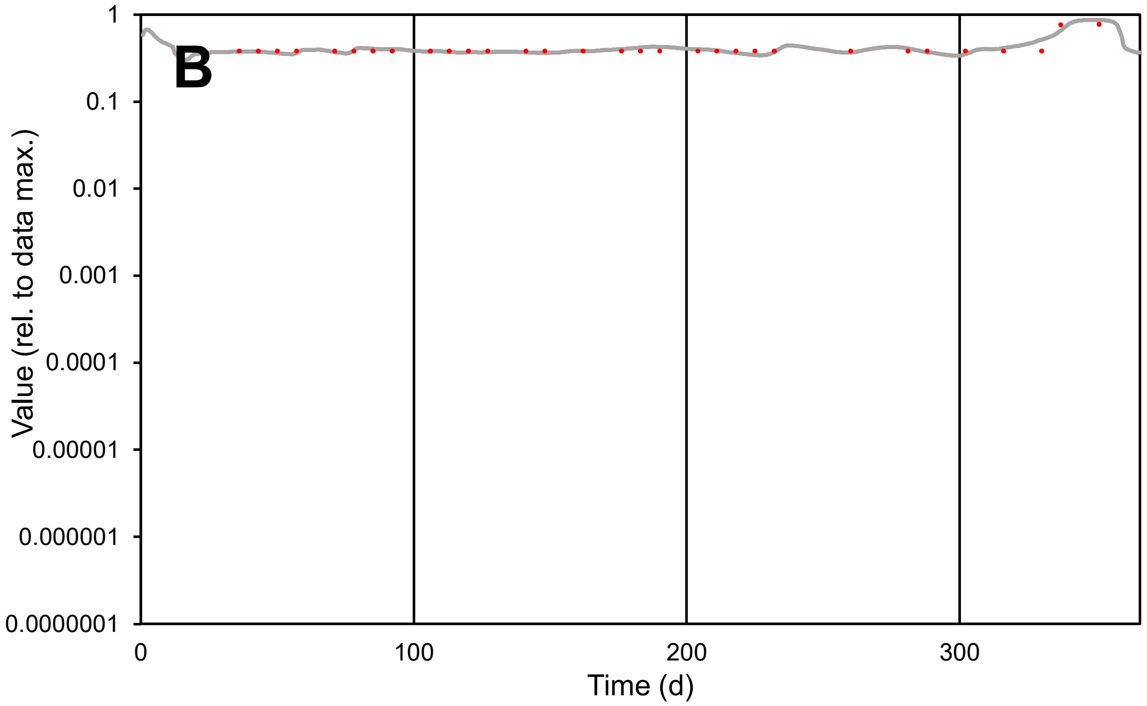


**C1**

Variables

CC2 Provide jtd or Dst

CalibOnly?

2 1 to reproduce of

oNormalize?

2 1=no, 2=yes

ofxn calc*

0.10713

ofxn mod

0.14481

sRun

RunA1

zout

1

Ave

Data

6.5E-06

Model

6.5E-06

data/vMax

1.5E-05

ofxng

321.551

Min d/m

1.11767

ave/min d

1.11371

ave/min m

1.24695

324_Calib_S_A

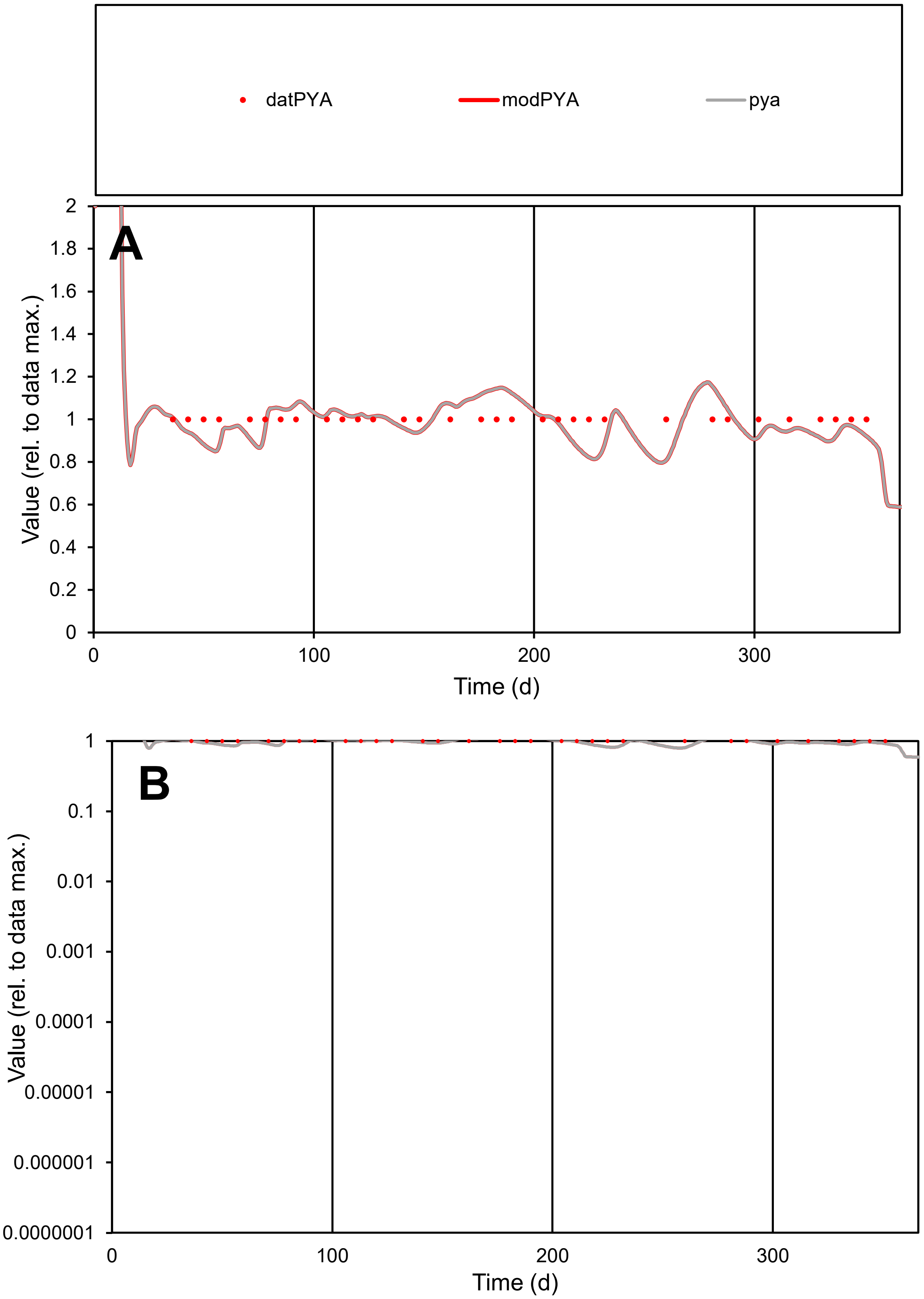


**C1**

Variables
PYA Provide jtd or Dst
CalibOnly?
2 1 to reproduce of
oNormalize?
2 1=no, 2=yes

ofxn calc*
0.08781
ofxn mod
0.13018

sRun
RunA1
zout
1

Ave
Data
1.8E-06
Model
1.8E-06

data/vMax
1.8E-06

ofxng
321.551

Min d/m
1.22515

ave/min d
1
ave/min m
1.20305

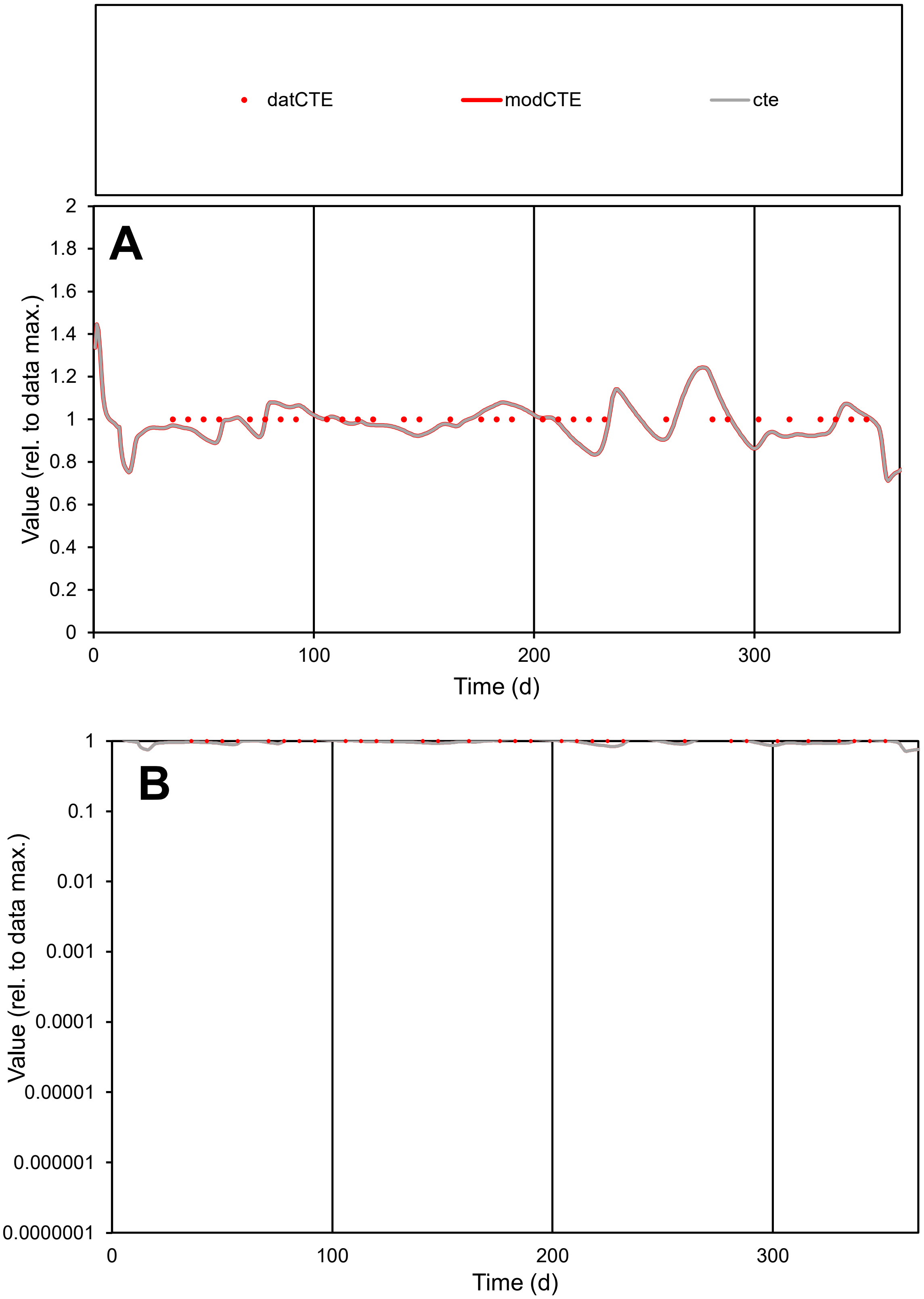


**C1**

Variables

CTE Provide jtd or Dst

CalibOnly?

2 1 to reproduce of

oNormalize?

2 1=no, 2=yes

ofxn calc*

0.068

ofxn mod

0.11752

sRun

RunA1

zout

1

Ave

Data

6.1E-06

Model

6E-06

data/vMax

6.1E-06

ofxng

321.551

Min d/m

1.19299

ave/min d

1

ave/min m

1.17406

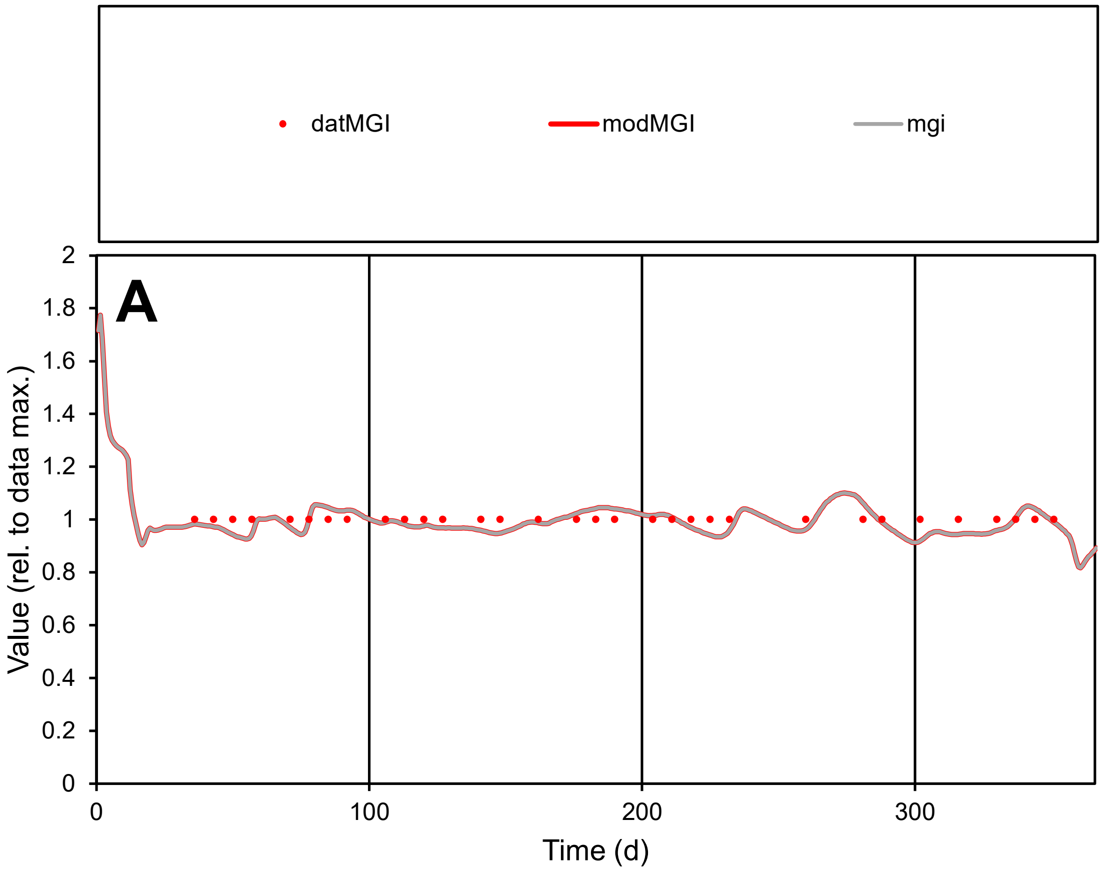


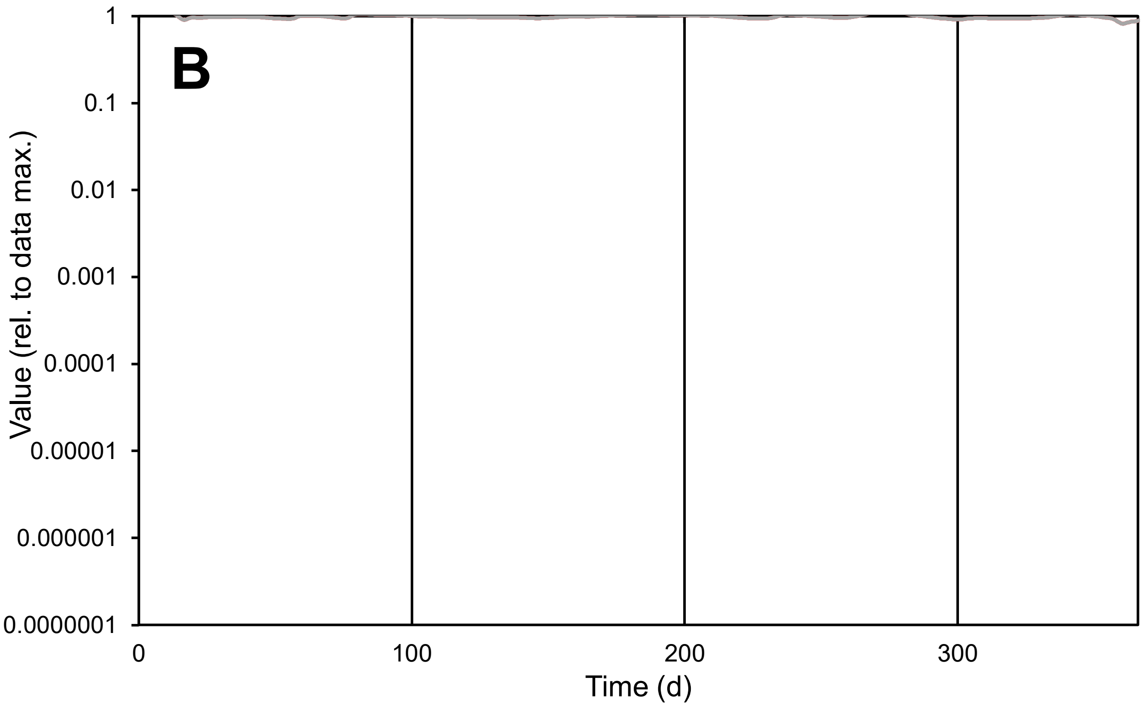


**C1**

Variables
MGI Provide jtd or Dst
CalibOnly?
2 1 to reproduce of
oNormalize?
2 1=no, 2=yes

ofxn calc*
0.03757
ofxn mod
0.10331

sRun
RunA1
zout
1

Ave
Data
1E-05
Model
1E-05

data/vMax
1E-05

ofxng
321.551

Min d/m
1.07792

ave/min d
1
ave/min m
1.06557

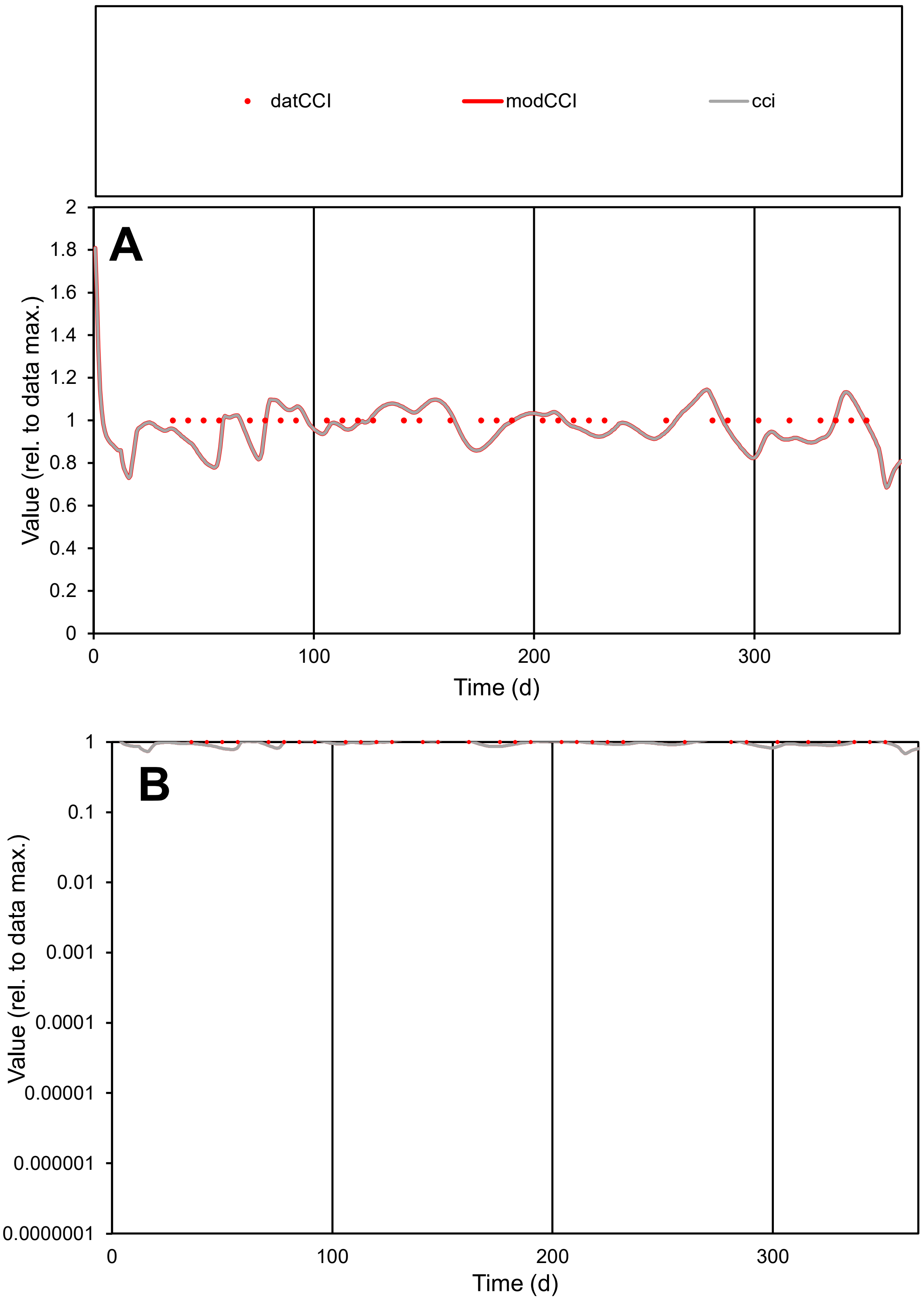


**C1**

Variables
CCI Provide jtd or Dst
CalibOnly?
2 1 to reproduce of
oNormalize?
2 1=no, 2=yes

ofxn calc*
0.0749
ofxn mod
0.12156

sRun
RunA1
zout
1

Ave
Data
1.1E-05
Model
1.1E-05

data/vMax
1.1E-05

ofxng
321.551

Min d/m
1.25602

ave/min d
1
ave/min m
1.226

324_Calib_S_A

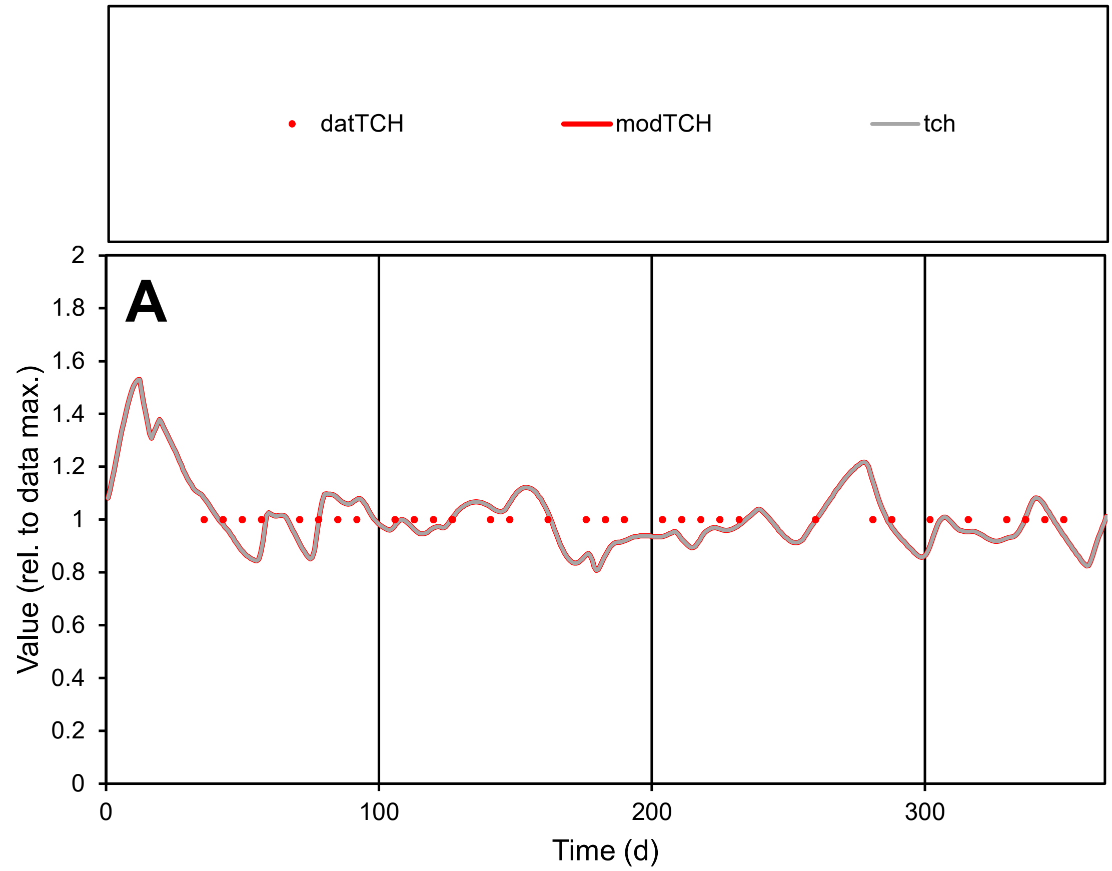


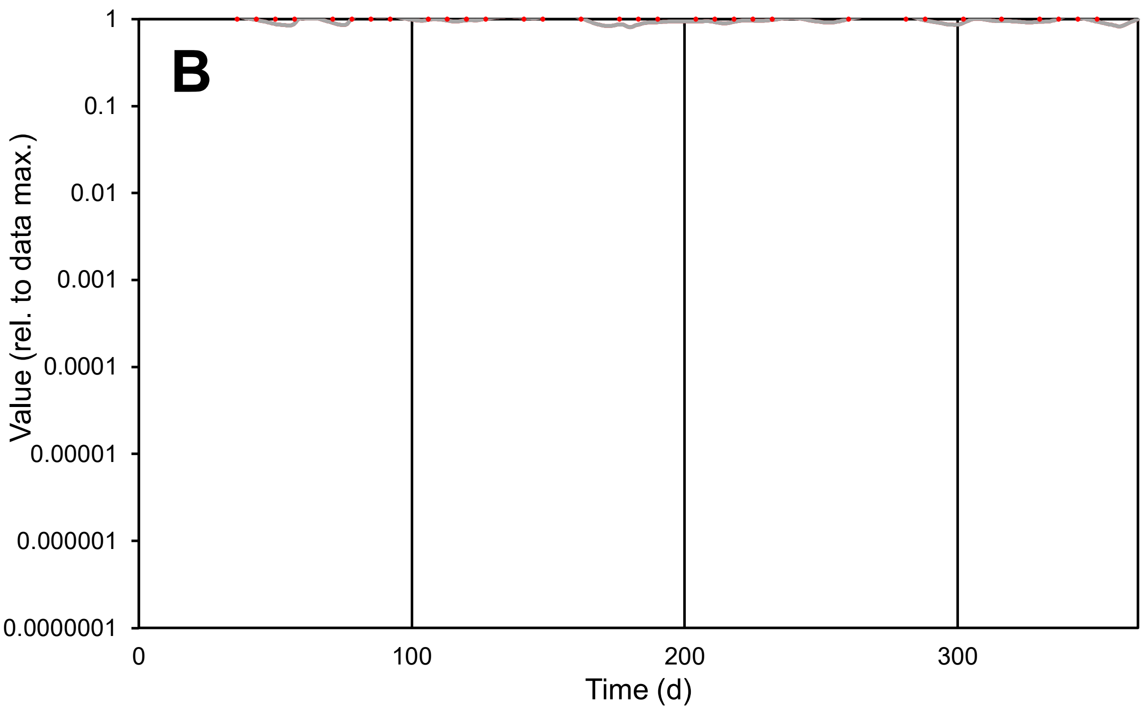


**C1**

Variables

TCH Provide jtd or Dst

CalibOnly?

2 1 to reproduce of

oNormalize?

2 1=no, 2=yes

ofxn calc*

0.06787

ofxn mod

0.1178

sRun

RunA1

zout

1

Ave

Data

1.7E-06

Model

1.6E-06

data/vMax

1.7E-06

ofxng

321.551

Min d/m

1.1576

ave/min d

1

ave/min m

1.13367

324_Calib_S_A

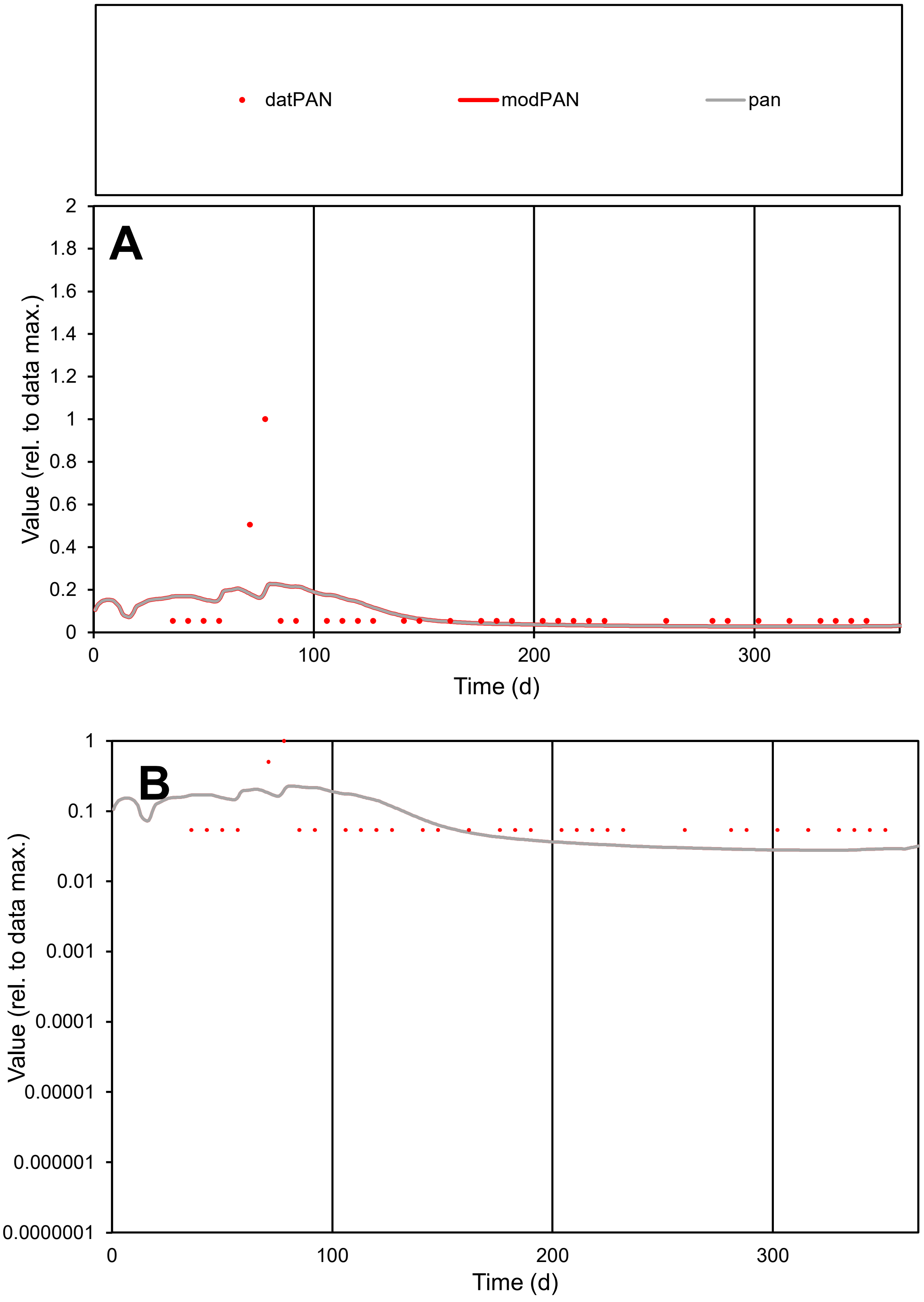


**C1**

Variables
PAN Provide jtd or Dst
CalibOnly?
2 1 to reproduce of
oNormalize?
2 1=no, 2=yes

ofxn calc*
1.67748
ofxn mod
1.68538

sRun
RunA1
zout
1

Ave
Data
1.7E-05
Model
1.5E-05

data/vMax
0.00017

ofxng
321.551

Min d/m
1.94872

ave/min d
1.80931
ave/min m
3.15023

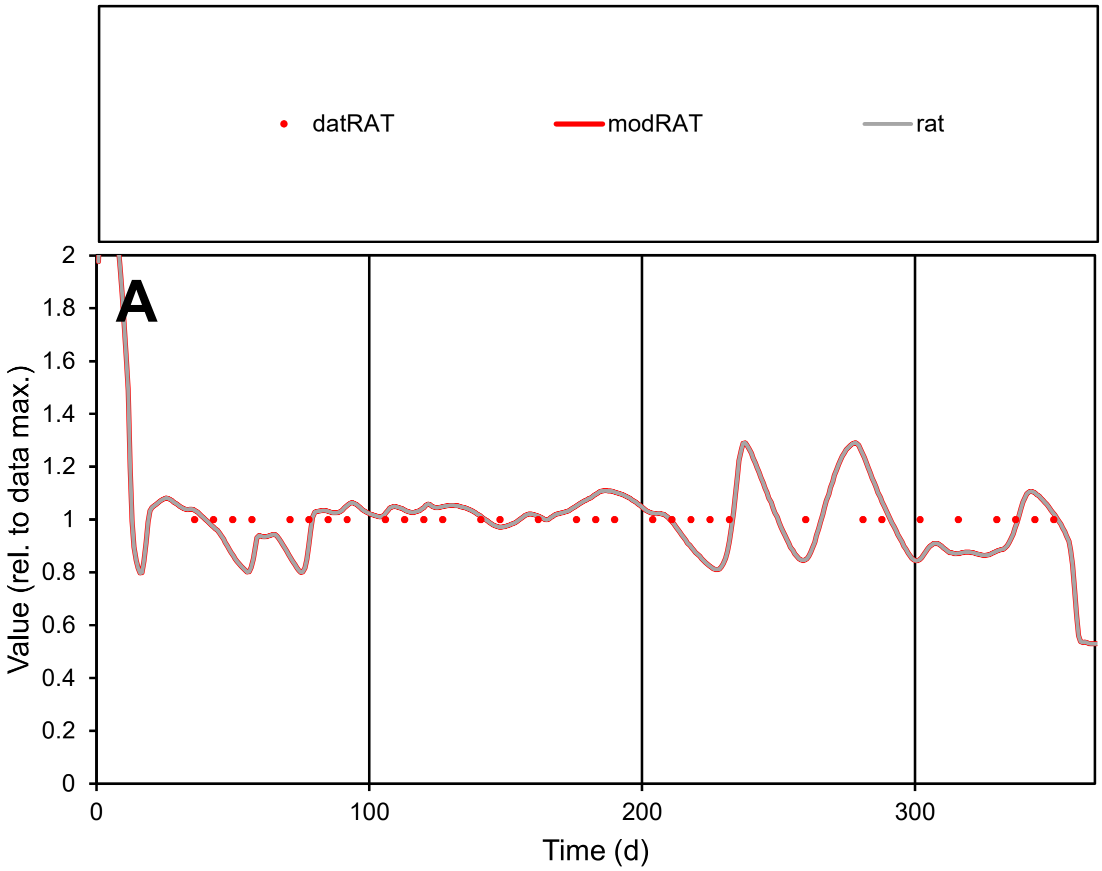


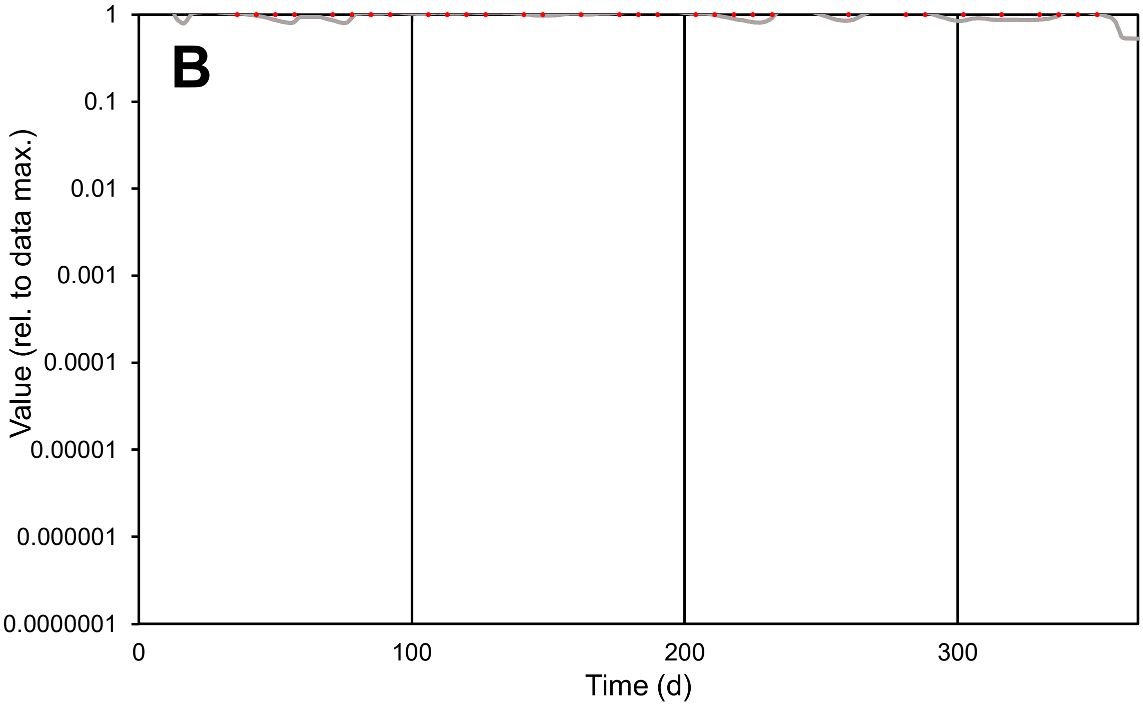


**C1**

Variables
RAT Provide jtd or Dst
CalibOnly?
2 1 to reproduce of
oNormalize?
2 1=no, 2=yes

ofxn calc*
0.09314
ofxn mod
0.13359

sRun
RunA1
zout
1

Ave
Data
1.1E-05
Model
1.1E-05

data/vMax
1.1E-05

ofxng
321.551

Min d/m
1.22919

ave/min d
1
ave/min m
1.21336

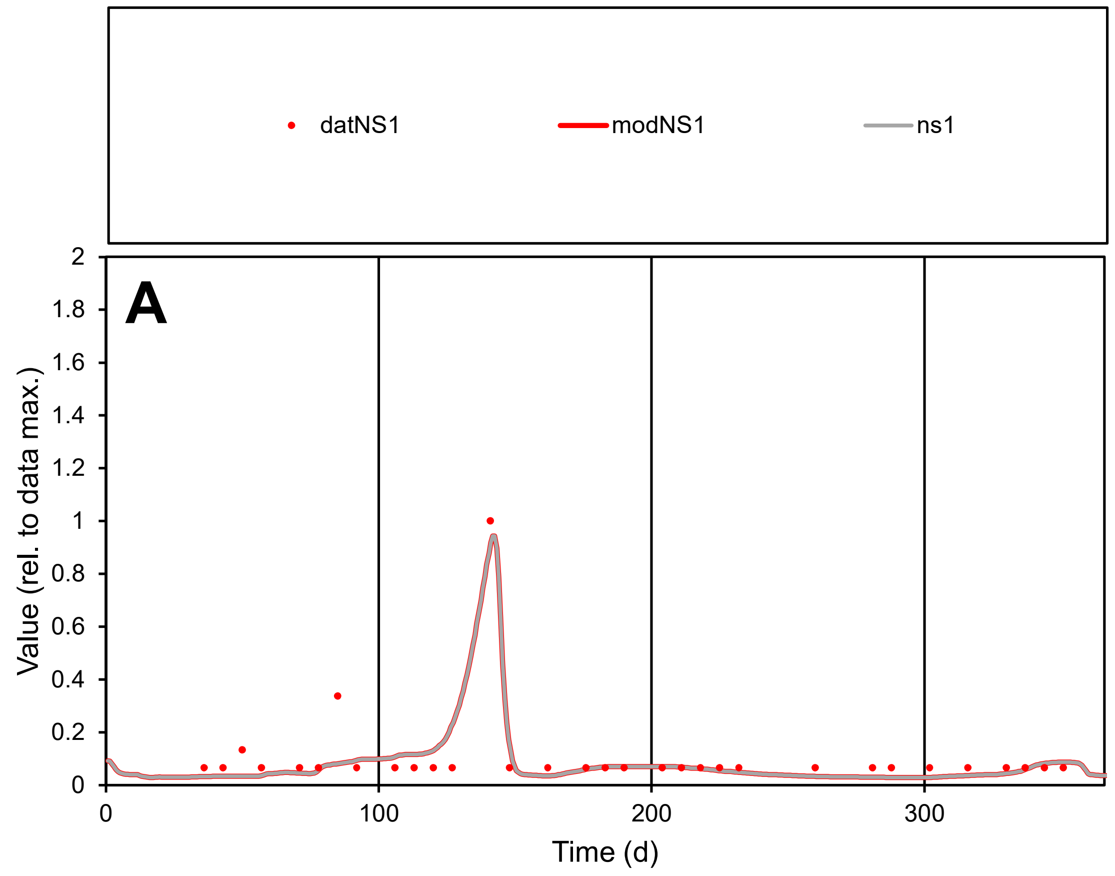


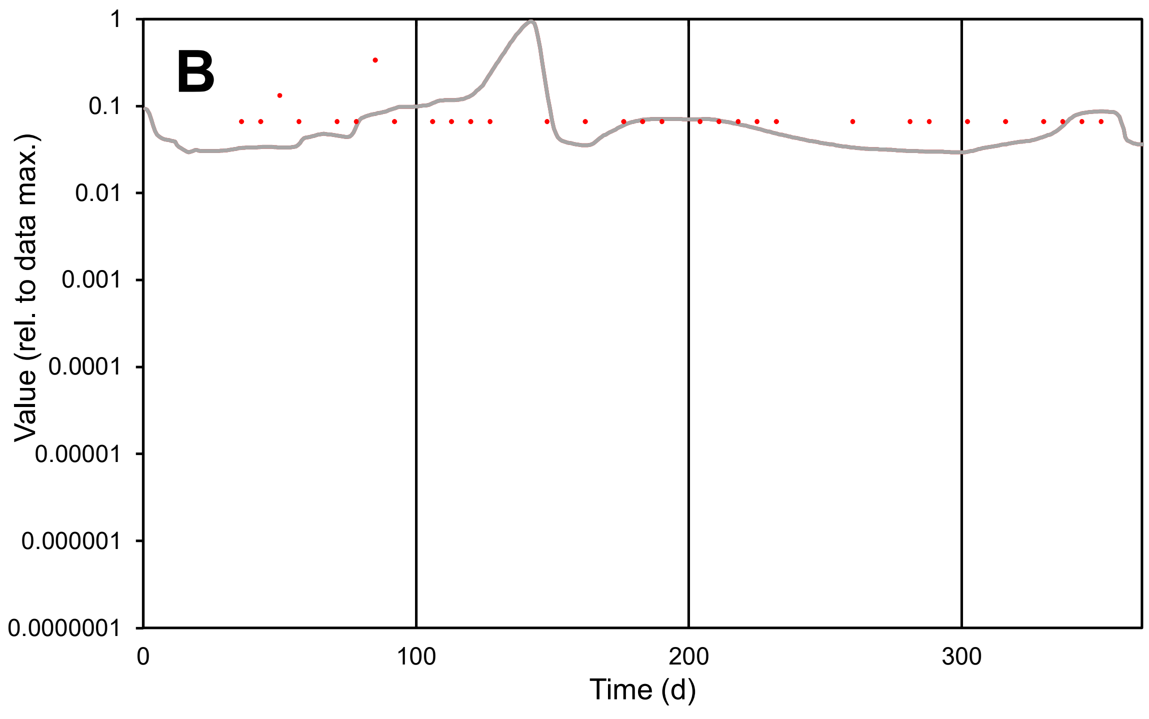


**C1**

Variables
NS1 Provide jtd or Dst
CalibOnly?
2 1 to reproduce of
oNormalize?
2 1=no, 2=yes

ofxn calc*
0.58358
ofxn mod
0.63191

sRun
RunA1
zout
1

Ave
Data
4.4E-05
Model
3.9E-05

data/vMax
0.00041

ofxng
321.551

Min d/m
2.21989

ave/min d
1.59763
ave/min m
3.16241

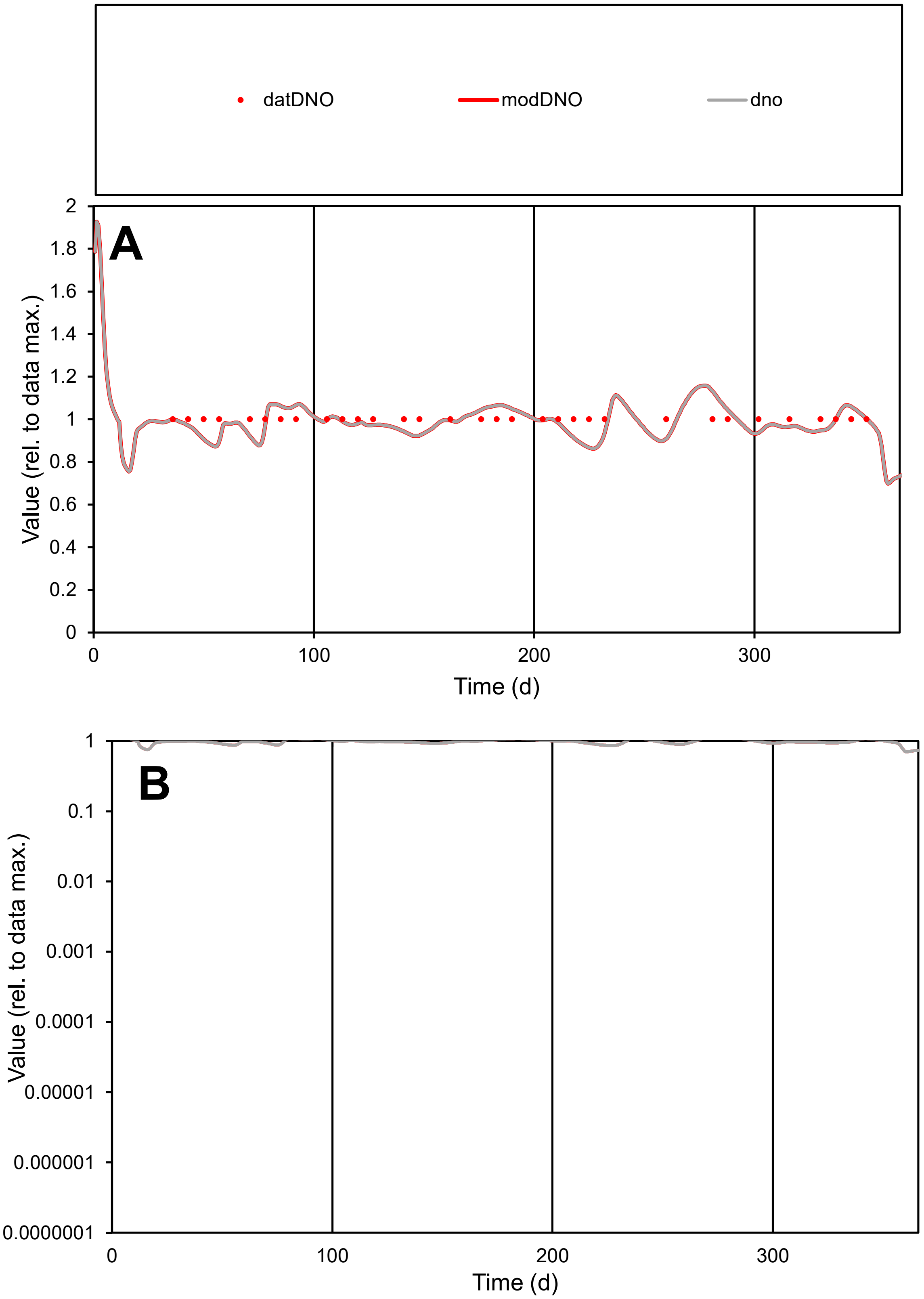


C1

Variables
DNO Provide jtd or Dst
CalibOnly?
2 1 to reproduce of
oNormalize?
2 1=no, 2=yes

ofxn calc*
0.06004
ofxn mod
0.11318

sRun
RunA1
zout
1

Ave
Data
1.8E-05
Model
1.8E-05

data/vMax
1.8E-05

ofxng
321.551

Min d/m
1.15803

ave/min d
1

ave/min m
1.14207

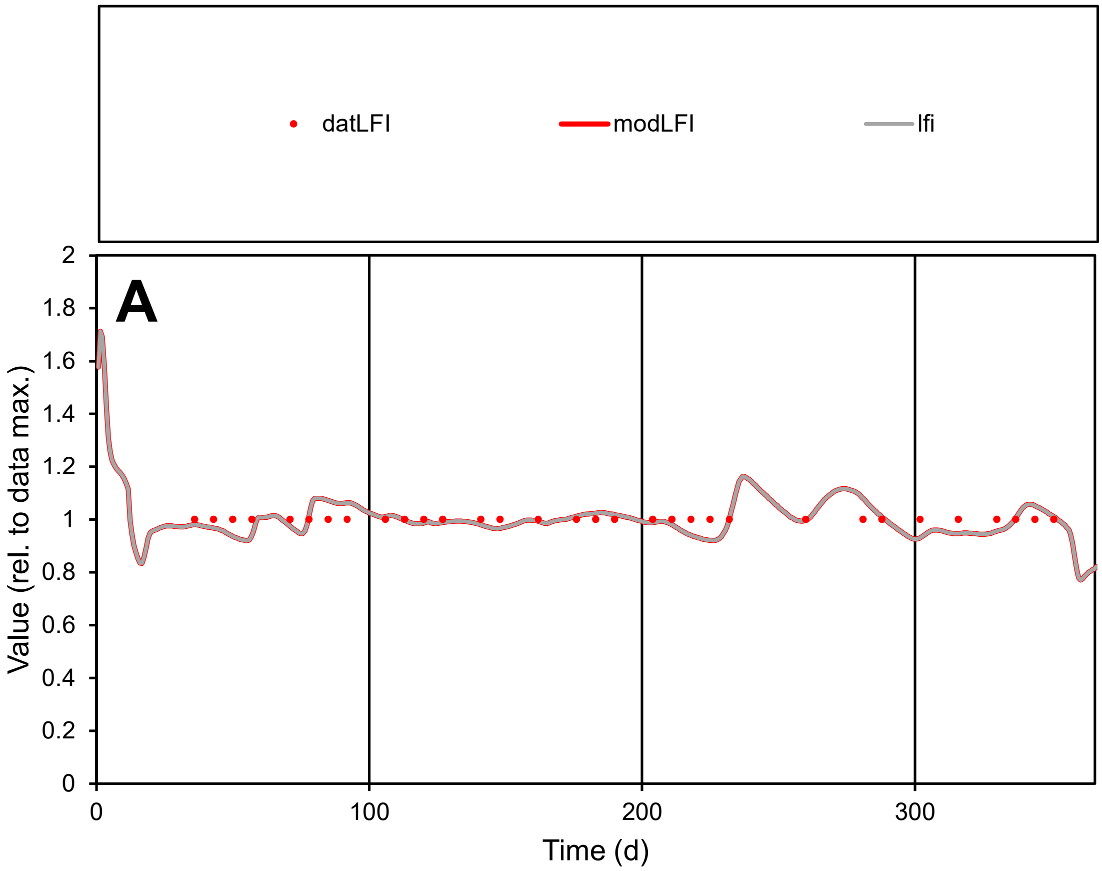


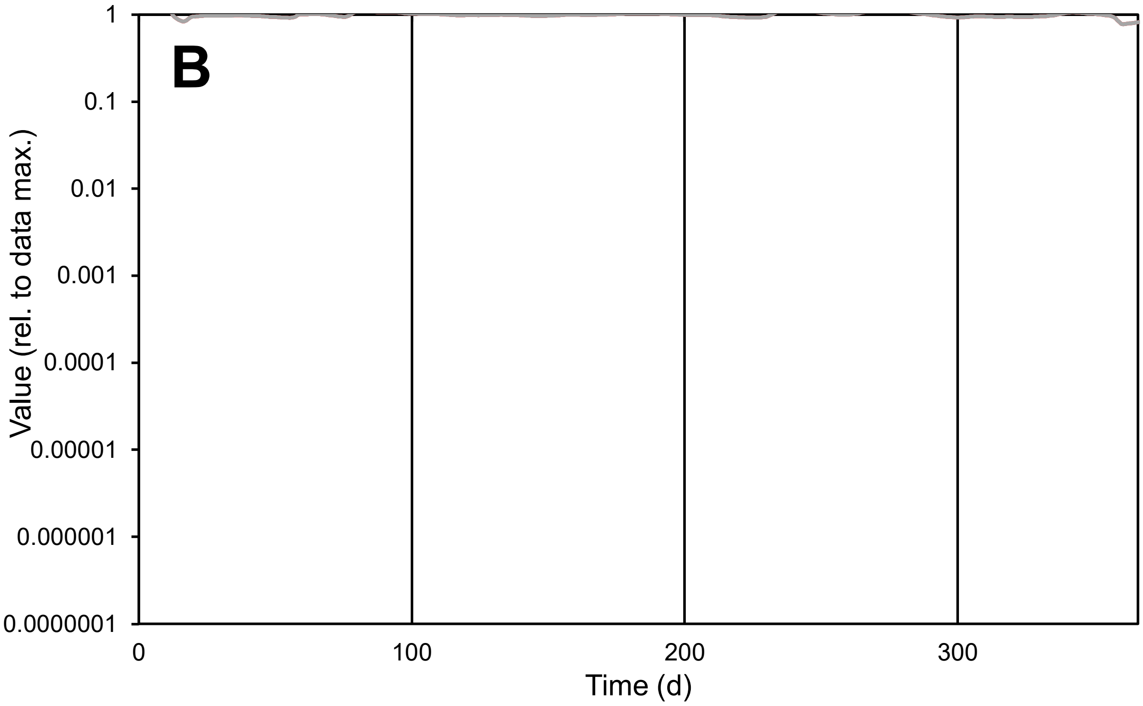


**C1**

Variables
LFI Provide jtd or Dst
CalibOnly?
2 1 to reproduce of
oNormalize?
2 1=no, 2=yes

ofxn calc*
0.04055
ofxn mod
0.10517

sRun
RunA1
zout
1

Ave
Data
8.3E-06
Model
8.3E-06

data/vMax
8.3E-06

ofxng
321.551

Min d/m
1.08643

ave/min d
1
ave/min m
1.08103

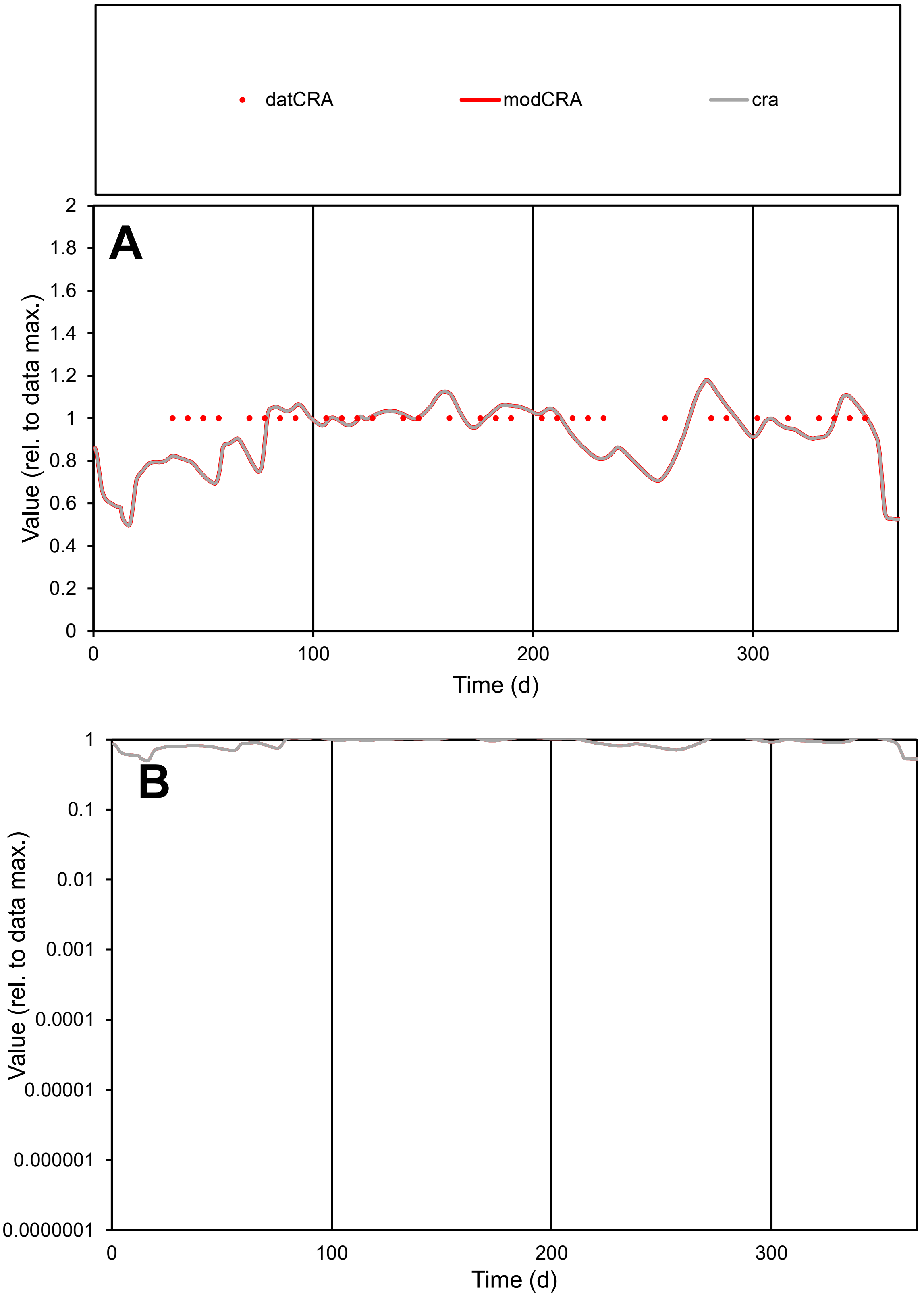


**C1**

Variables
CRA Provide jtd or Dst
CalibOnly?
2 1 to reproduce of
oNormalize?
2 1=no, 2=yes

ofxn calc*
0.11631
ofxn mod
0.15026

sRun
RunA1
zout
1

Ave
Data
4.9E-06
Model
4.7E-06

data/vMax
4.9E-06

ofxng
321.551

Min d/m
1.3969

ave/min d
1
ave/min m
1.34163

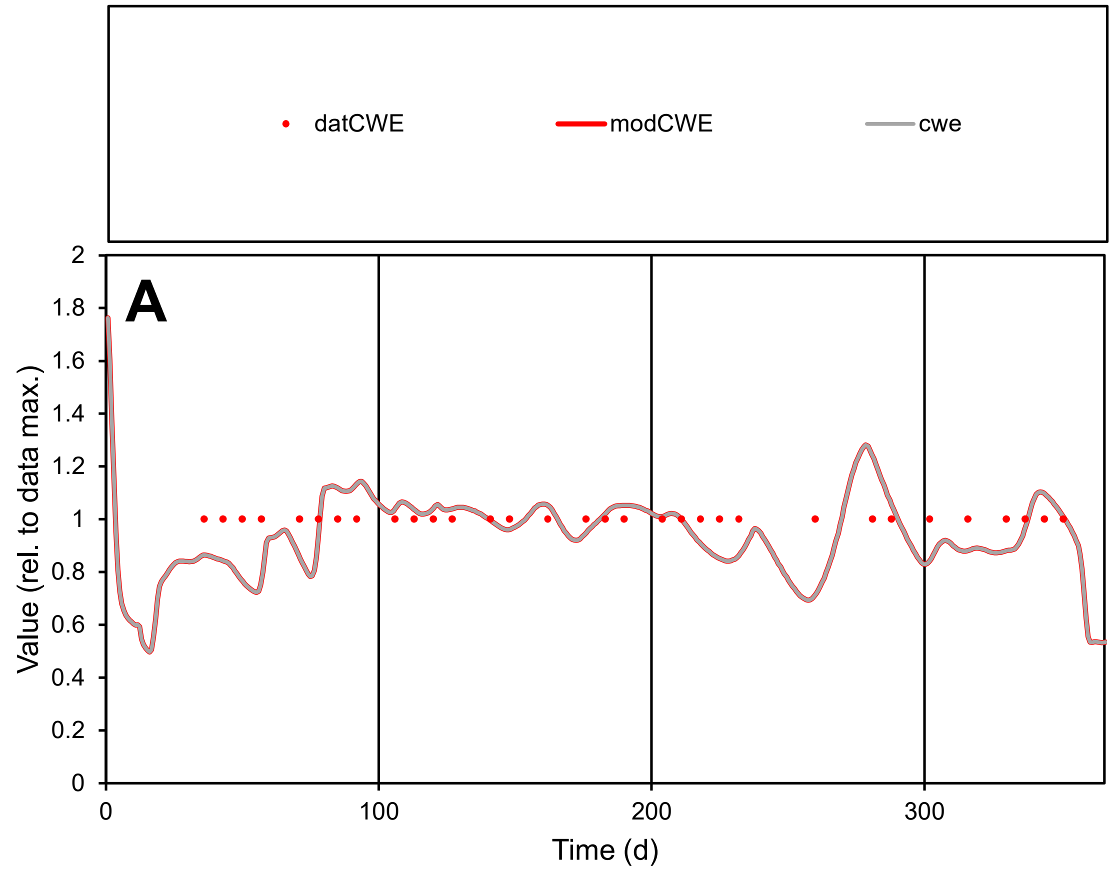


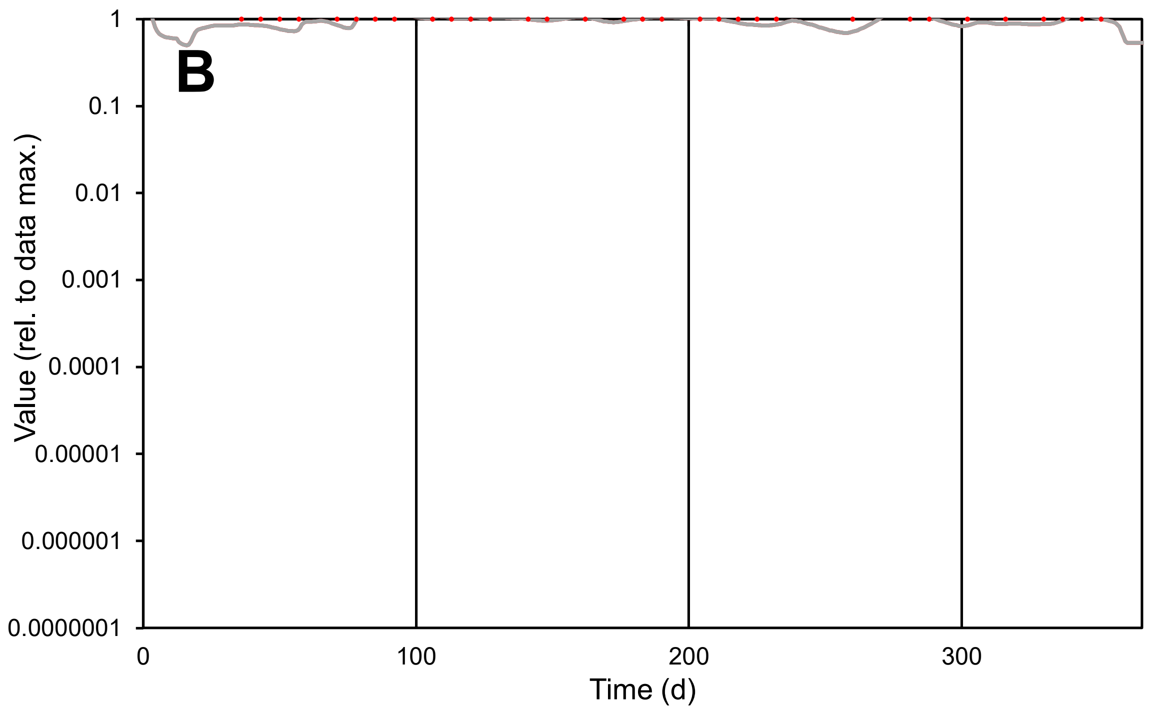


**C1**

Variables
CWE Provide jtd or Dst
CalibOnly?
2 1 to reproduce of
oNormalize?
2 1=no, 2=yes

ofxn calc*
0.1161
ofxn mod
0.15038

sRun
RunA1
zout
1

Ave
Data
2.6E-05
Model
2.5E-05

data/vMax
2.6E-05

ofxng
321.551

Min d/m
1.36656

ave/min d
1
ave/min m
1.32412

324_Calib_S_A

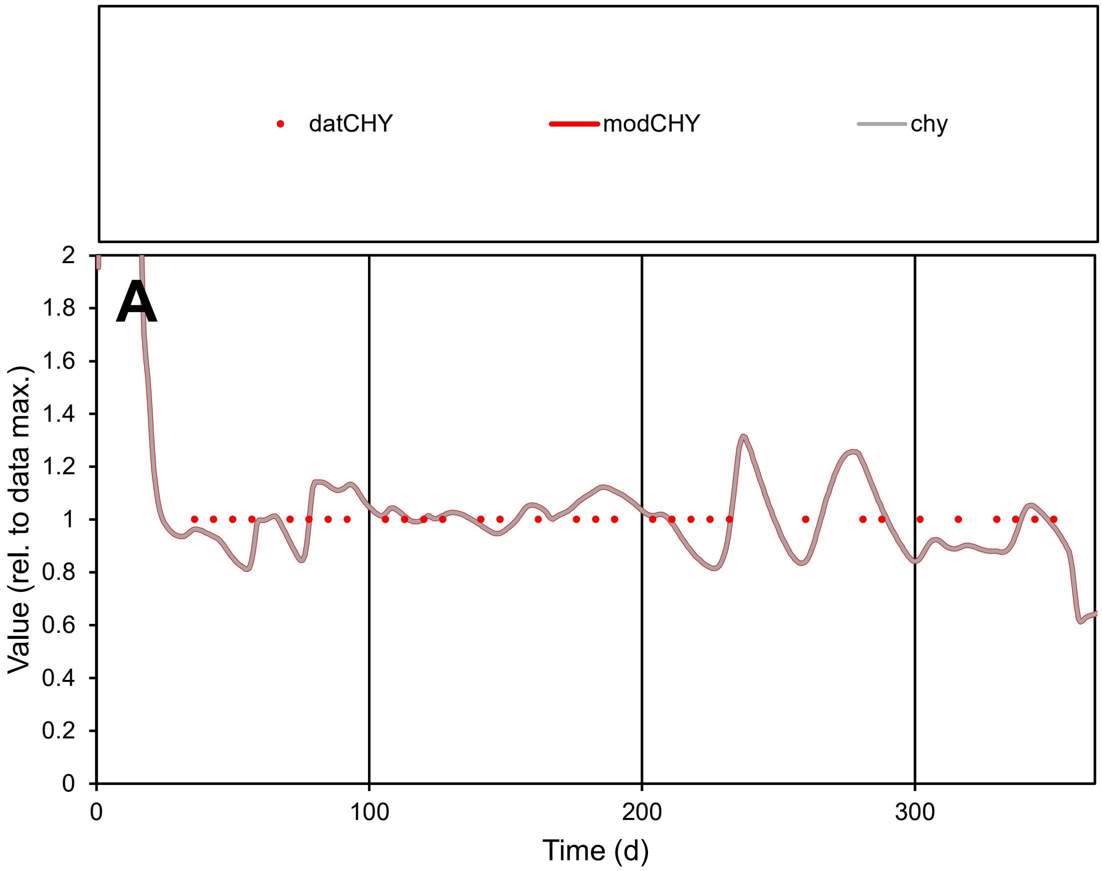


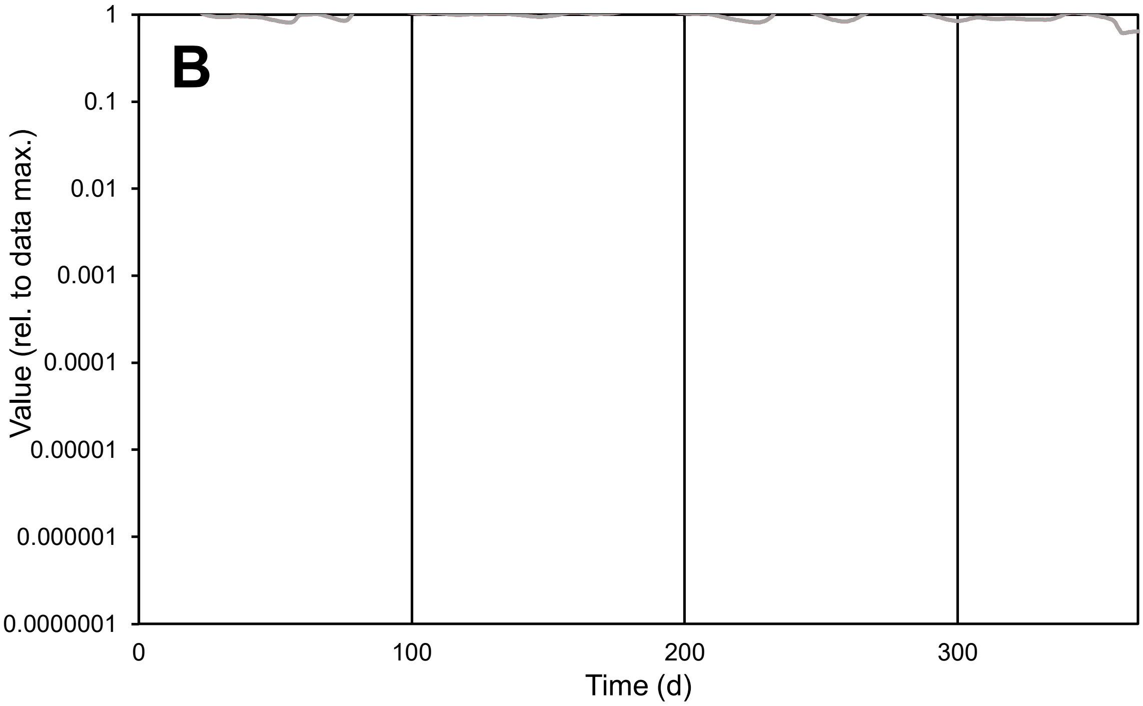


**C1**

Variables
CHY Provide jtd or Dst
CalibOnly?
2 1 to reproduce of
oNormalize?
2 1=no, 2=yes

ofxn calc*
0.09454
ofxn mod
0.13556

sRun
RunA1
zout
1

Ave
Data
6.7E-06
Model
6.6E-06

data/vMax
6.7E-06

ofxng
321.551

Min d/m
1.22723

ave/min d
1
ave/min m
1.21239

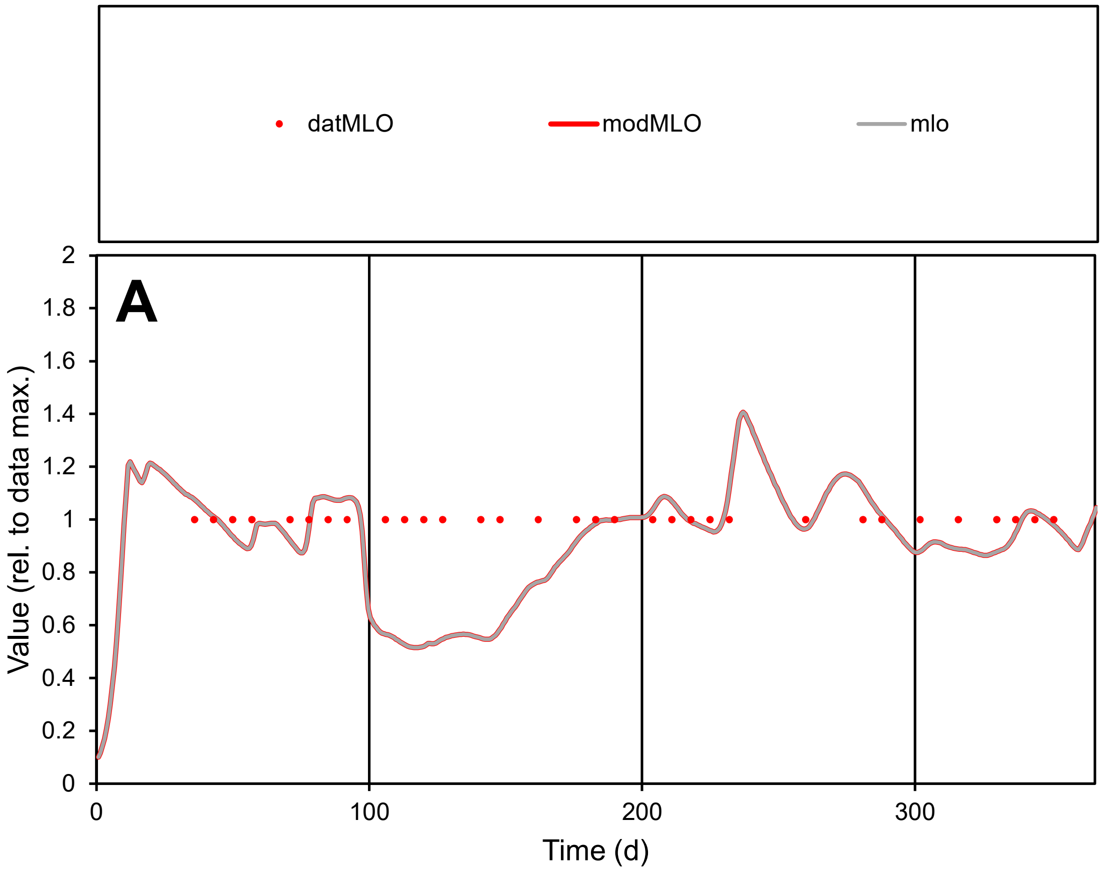


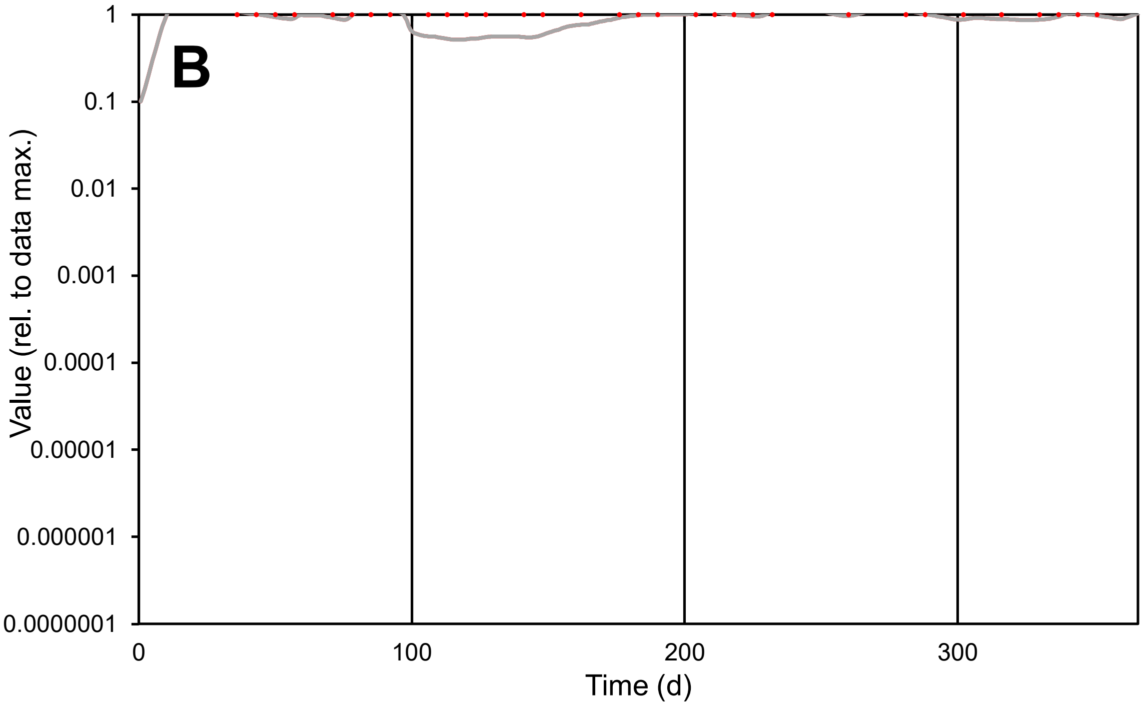


**C1**

Variables
MLO Provide jtd or Dst
CalibOnly?
2 1 to reproduce of
oNormalize?
2 1=no, 2=yes

ofxn calc*
0.20829
ofxn mod
0.22932

sRun
RunA1
zout
1

Ave
Data
1E-05
Model
9.5E-06

data/vMax
1E-05

ofxng
321.551

Min d/m
1.92081

ave/min d
1
ave/min m
1.74088

324_Calib_S_A

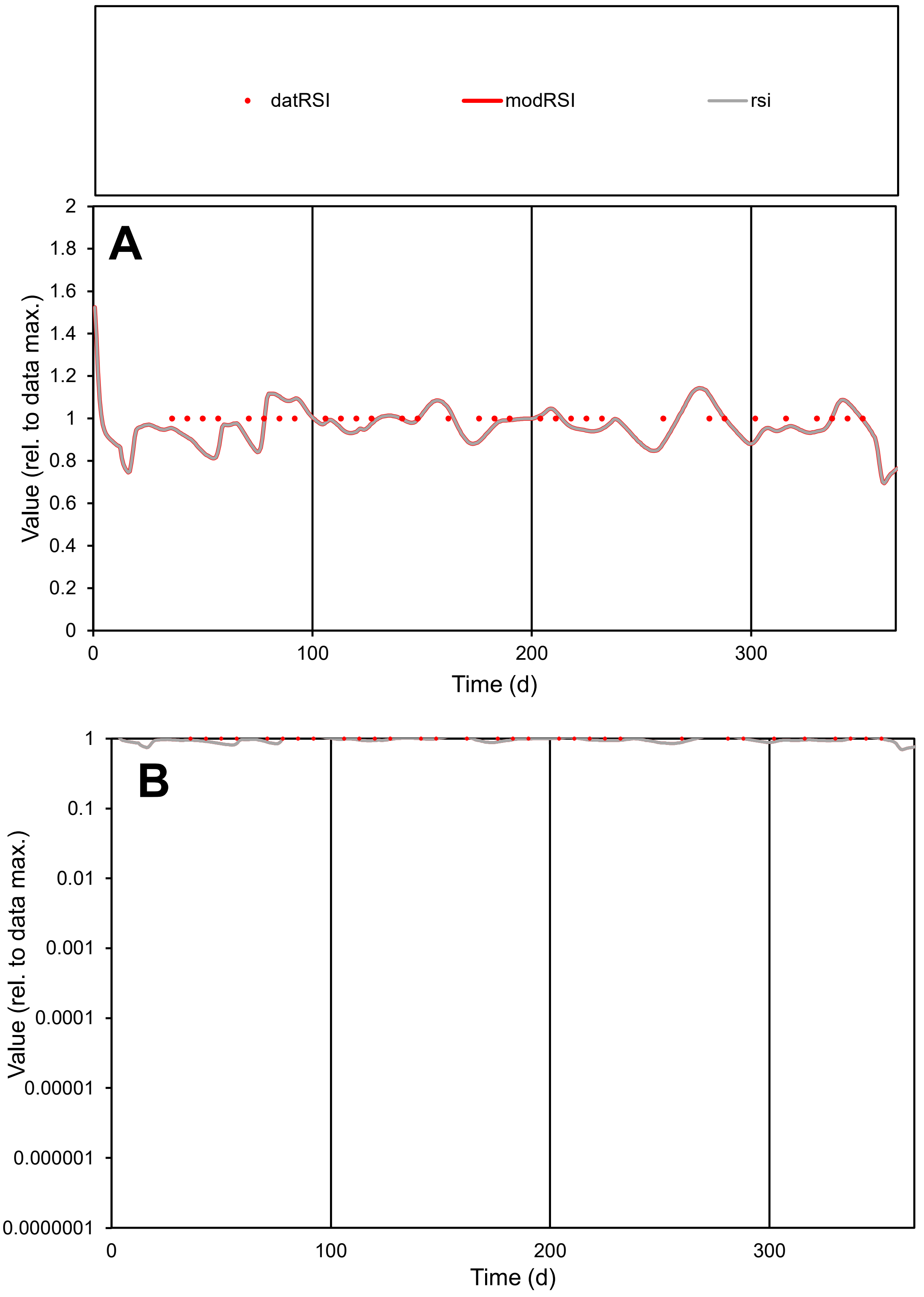


**C1**

Variables

RSI Provide jtd or Dst

CalibOnly?

2 1 to reproduce of

oNormalize?

2 1=no, 2=yes

ofxn calc*

0.06802

ofxn mod

0.11754

sRun

RunA1

zout

1

Ave

Data

1E-05

Model

1E-05

data/vMax

1E-05

ofxng

321.551

Min d/m

1.20649

ave/min d

1

ave/min m

1.17959

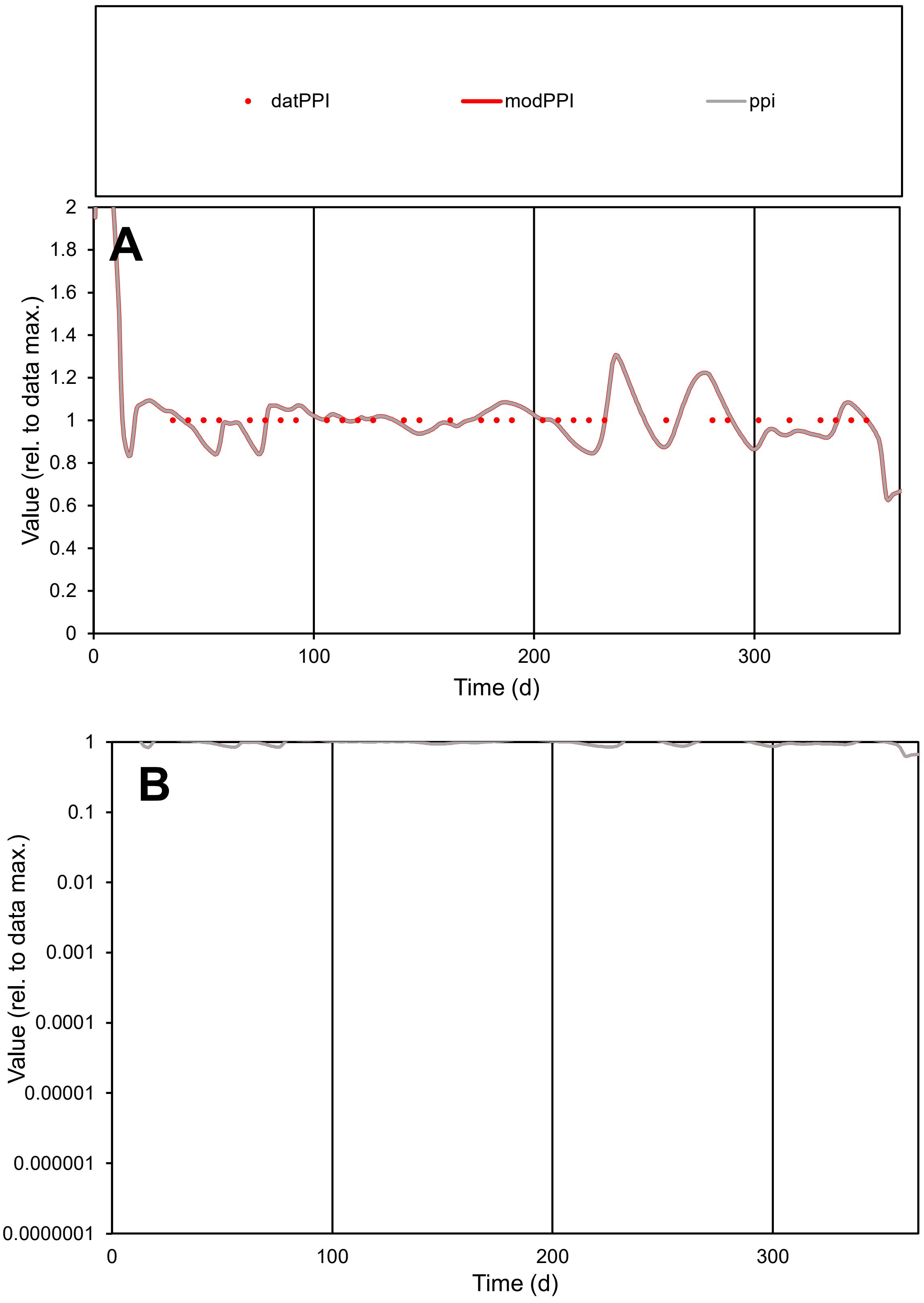


**C1**

Variables
PPI Provide jtd or Dst
CalibOnly?
2 1 to reproduce of
oNormalize?
2 1=no, 2=yes

ofxn calc*
0.07309
ofxn mod
0.12179

sRun
RunA1
zout
1

Ave
Data
1.4E-06
Model
1.4E-06

data/vMax
1.4E-06

ofxng
321.551

Min d/m
1.18331

ave/min d
1
ave/min m
1.16991

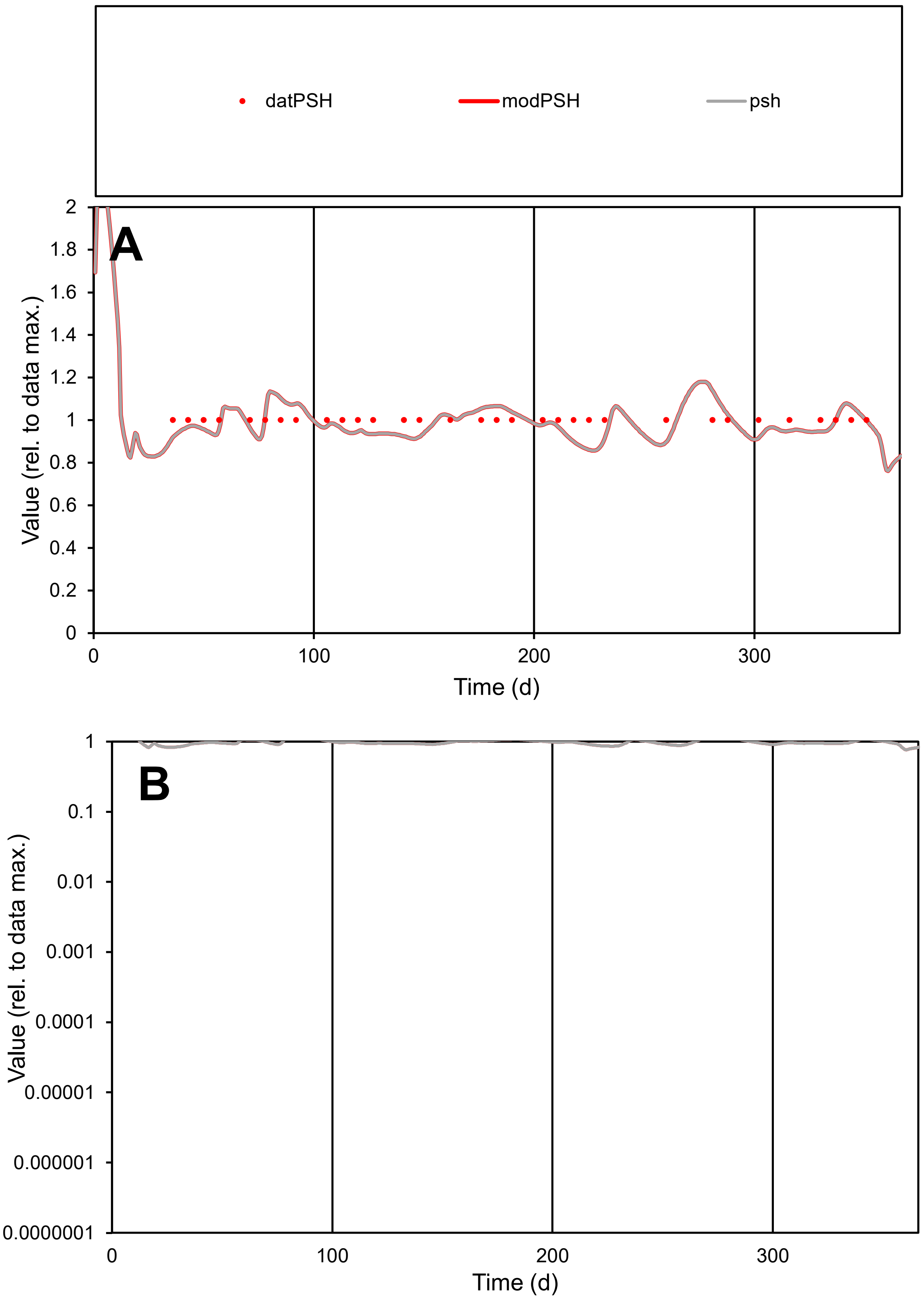


**C1**

Variables
PSH Provide jtd or Dst
CalibOnly?
2 1 to reproduce of
oNormalize?
2 1=no, 2=yes

ofxn calc*
0.0665
ofxn mod
0.11696

sRun
RunA1
zout
1

Ave
Data
1.3E-05
Model
1.3E-05

data/vMax
1.3E-05

ofxng
321.551

Min d/m
1.16637

ave/min d
1
ave/min m
1.14913

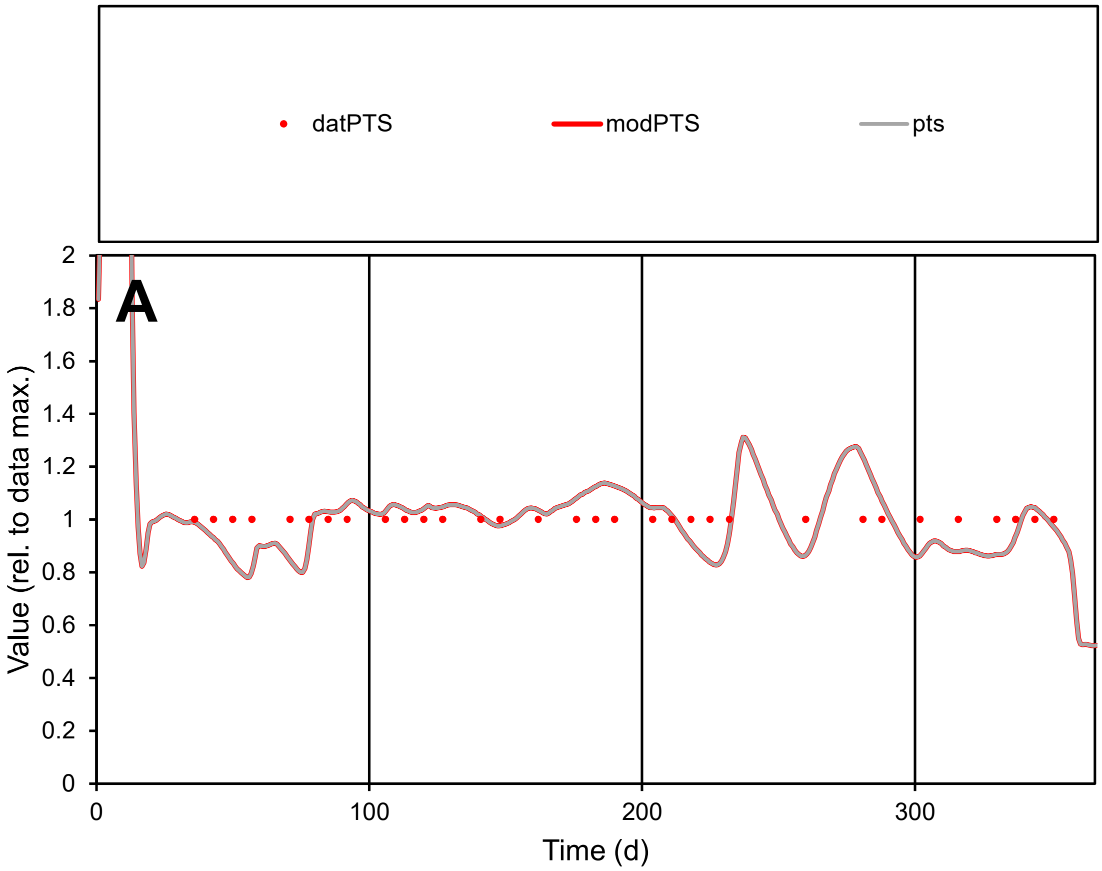


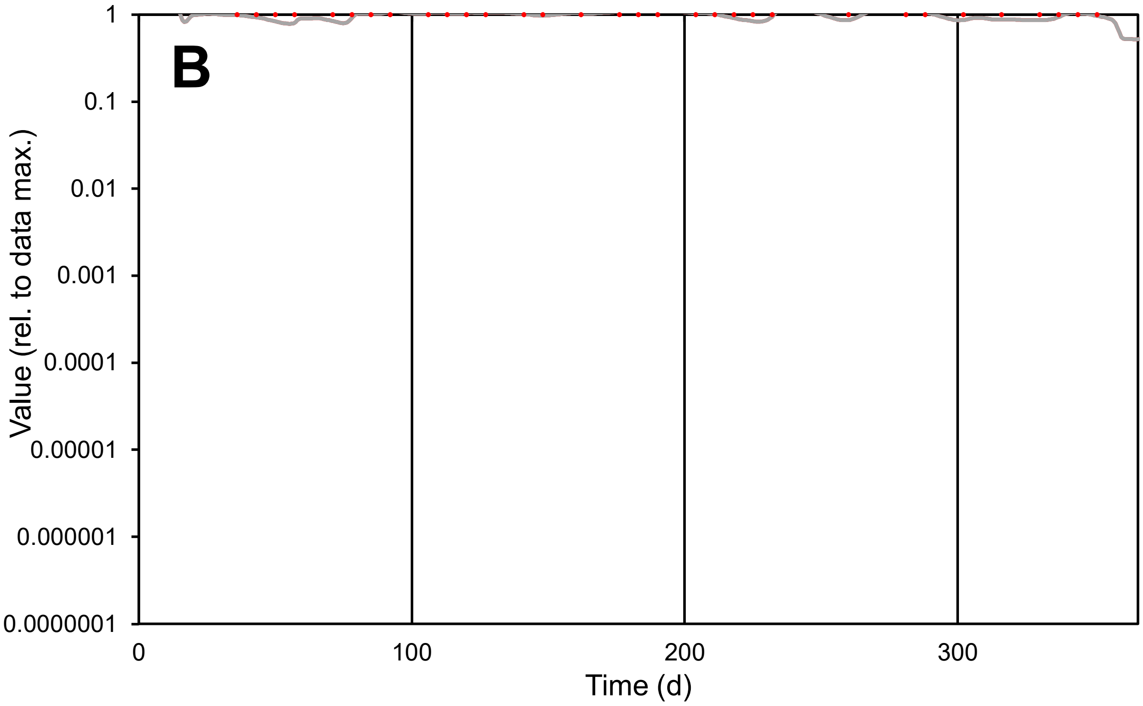


**C1**

Variables

| | |
|---|---|
| PTS | Provide jtd or Dst |
| CalibOnly? | |
| 2 | 1 to reproduce of |
| oNormalize? | |
| 2 | 1=no, 2=yes |

ofxn calc*
0.09652
ofxn mod
0.13674

sRun
RunA1
zout
1

Ave
Data
9.7E-06
Model
9.5E-06

data/vMax
9.7E-06

ofxng
321.551

Min d/m
1.22787

ave/min d
1
ave/min m
1.20972

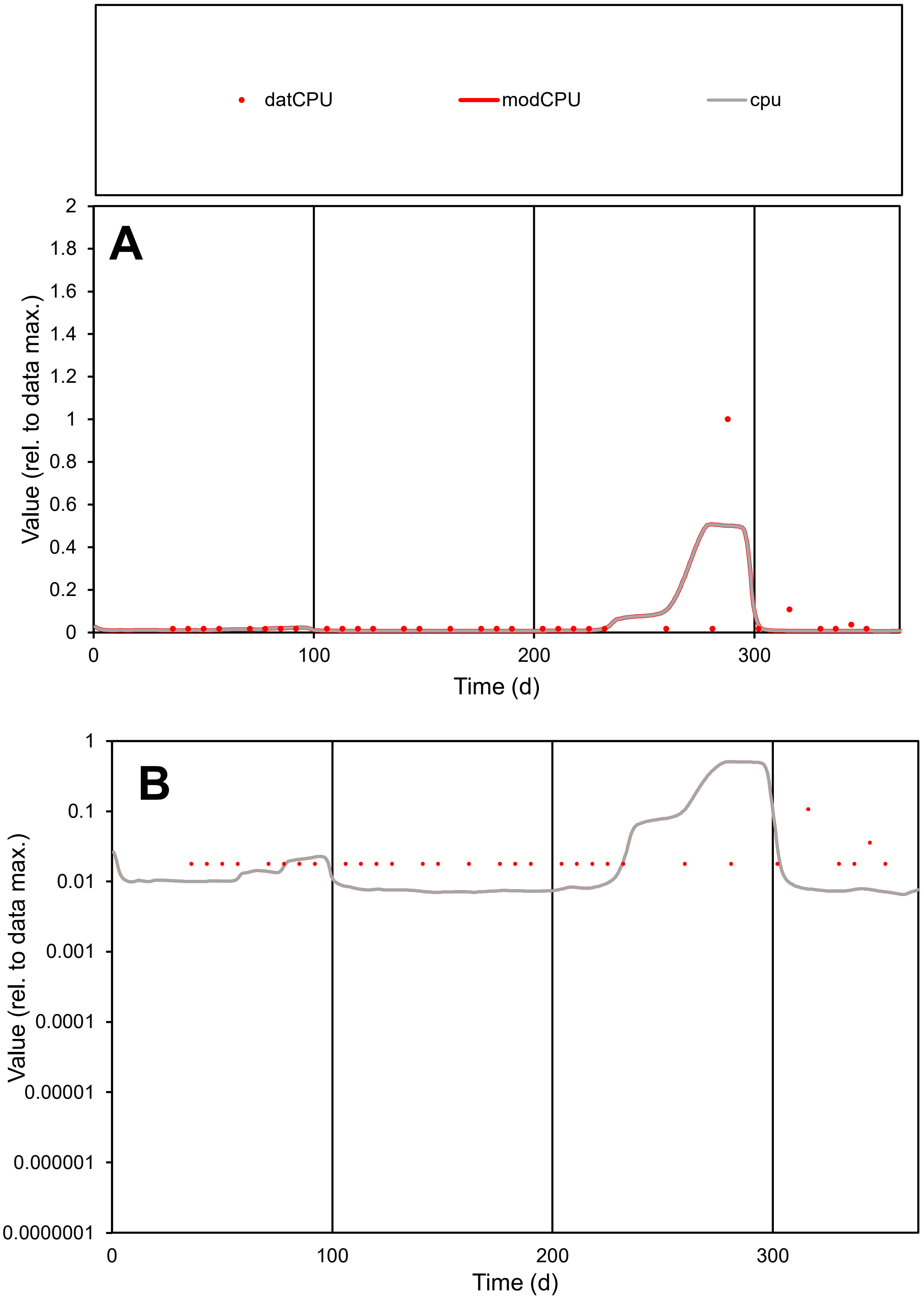


**C1**

Variables
CPU Provide jtd or Dst
CalibOnly?
2 1 to reproduce of
oNormalize?
2 1=no, 2=yes

ofxn calc*
2.42994
ofxn mod
2.44574

sRun
RunA1
zout
1

Ave
Data
1.3E-06
Model
1.1E-06

data/vMax
2.5E-05

ofxng
321.551

Min d/m
2.54284

ave/min d
2.9087
ave/min m
6.30267

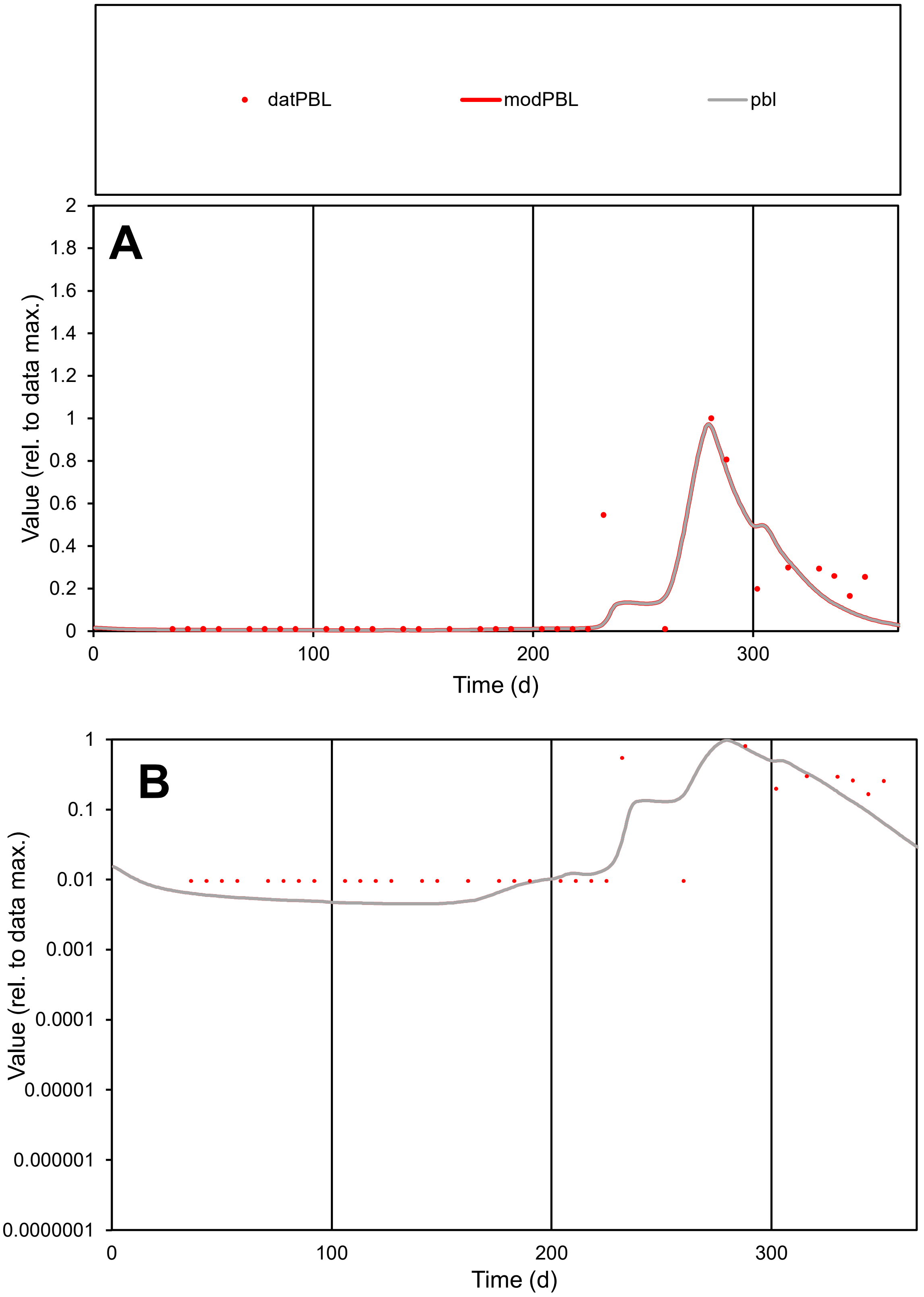


**C1**

Variables

PBL Provide jtd or Dst

CalibOnly?

2 1 to reproduce of

oNormalize?

2 1=no, 2=yes

ofxn calc*

0.9137

ofxn mod

0.95044

sRun

RunA1

zout

1

Ave

Data

5.6E-07

Model

4.6E-07

data/vMax

4.4E-06

ofxng

321.551

Min d/m

2.12481

ave/min d

13.2001

ave/min m

22.7212

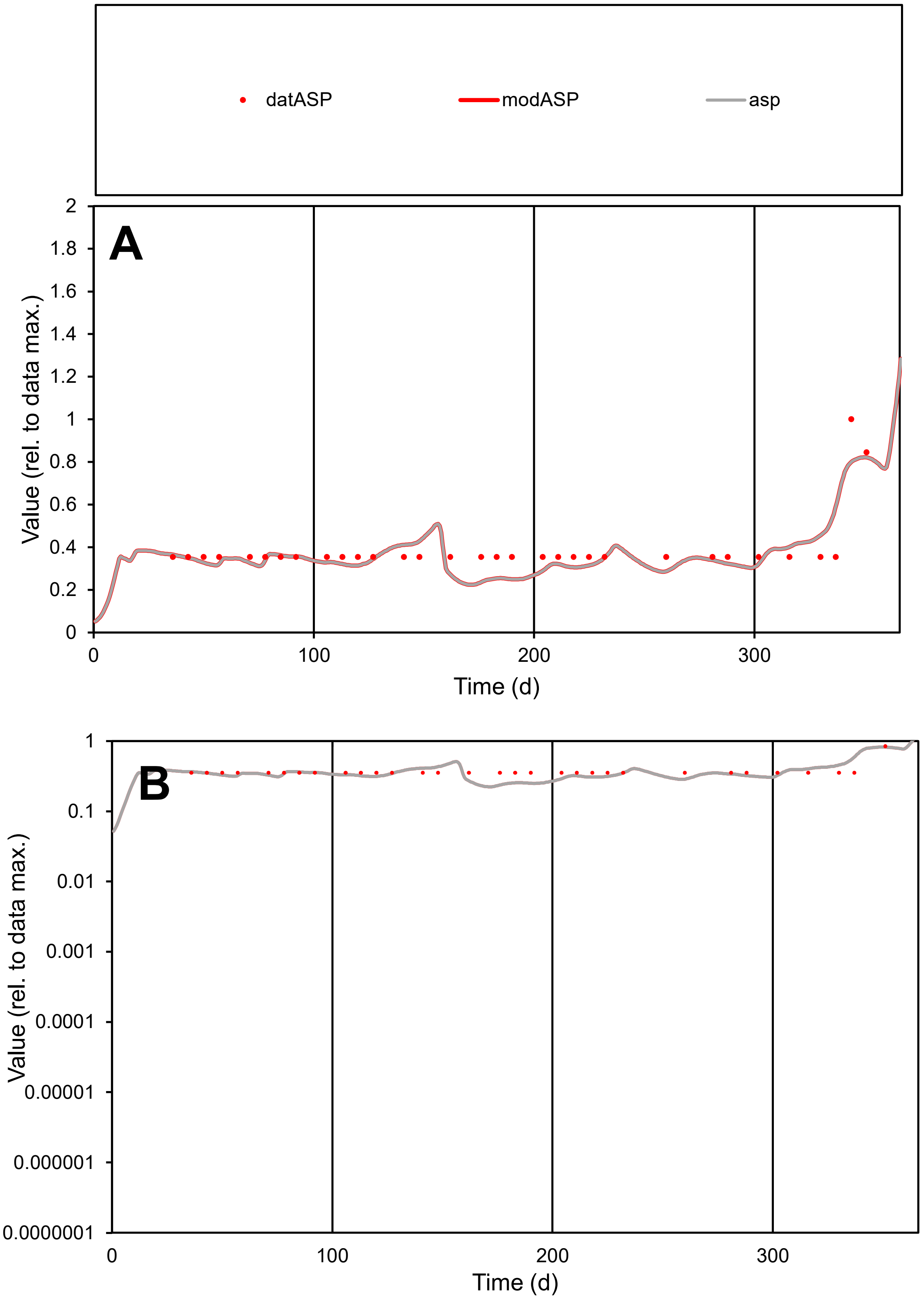


**C1**

Variables
ASP Provide jtd or Dst
CalibOnly?
2 1 to reproduce of
oNormalize?
2 1=no, 2=yes

ofxn calc*
0.19807
ofxn mod
0.22402

sRun
RunA1
zout
1

Ave
Data
6.4E-06
Model
6.1E-06

data/vMax
1.6E-05

ofxng
321.551

Min d/m
1.45972

ave/min d
1.10016
ave/min m
1.53802

324_Calib_S_A

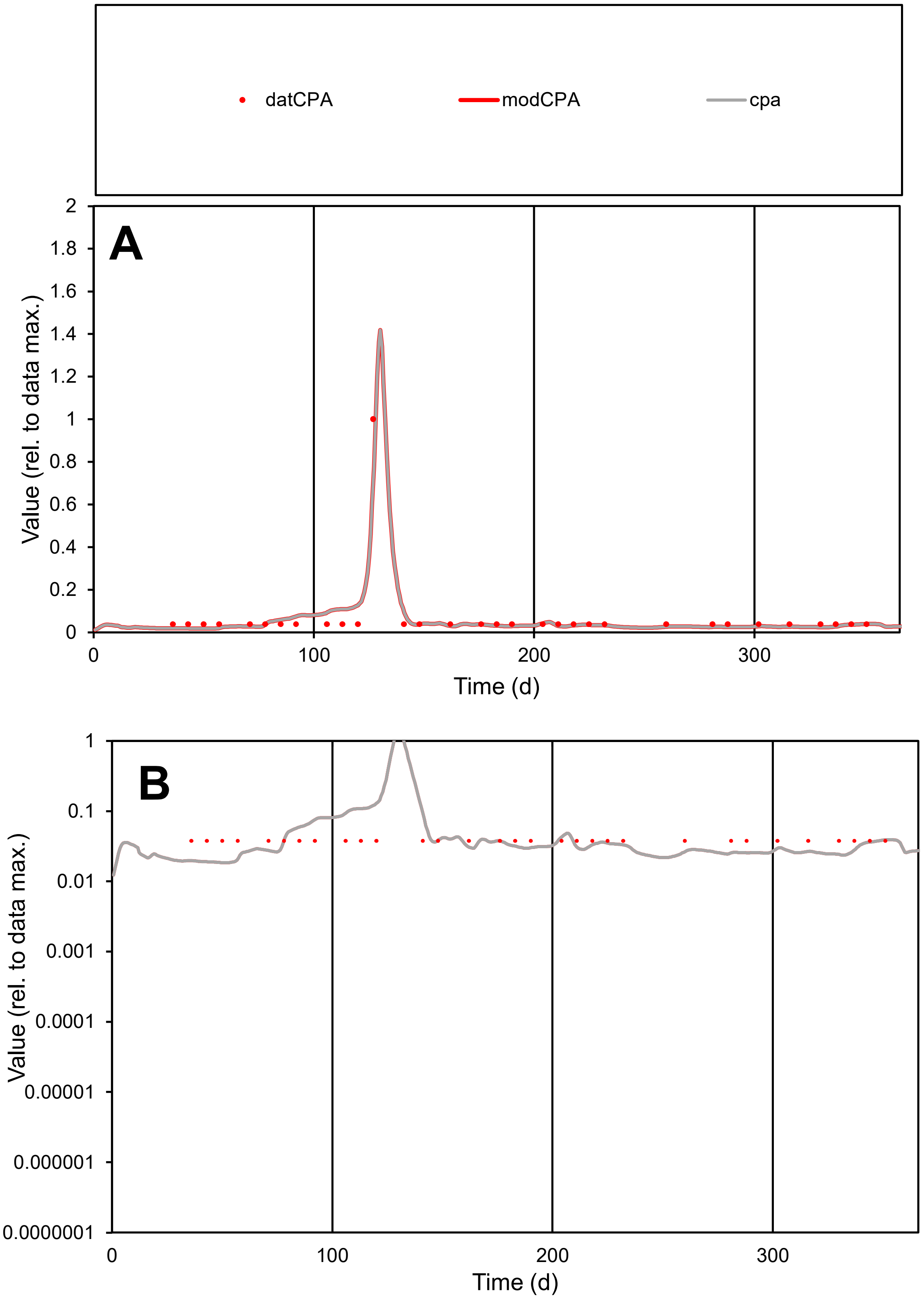


**C1**

Variables

CPA Provide jtd or Dst

CalibOnly?

2 1 to reproduce of

oNormalize?

2 1=no, 2=yes

ofxn calc*

0.38335

ofxn mod

0.41247

sRun

RunA1

zout

1

Ave

Data

1.2E-07

Model

1.2E-07

data/vMax

1.8E-06

ofxng

321.551

Min d/m

2.04621

ave/min d

1.79399

ave/min m

3.84579

324_Calib_S_A

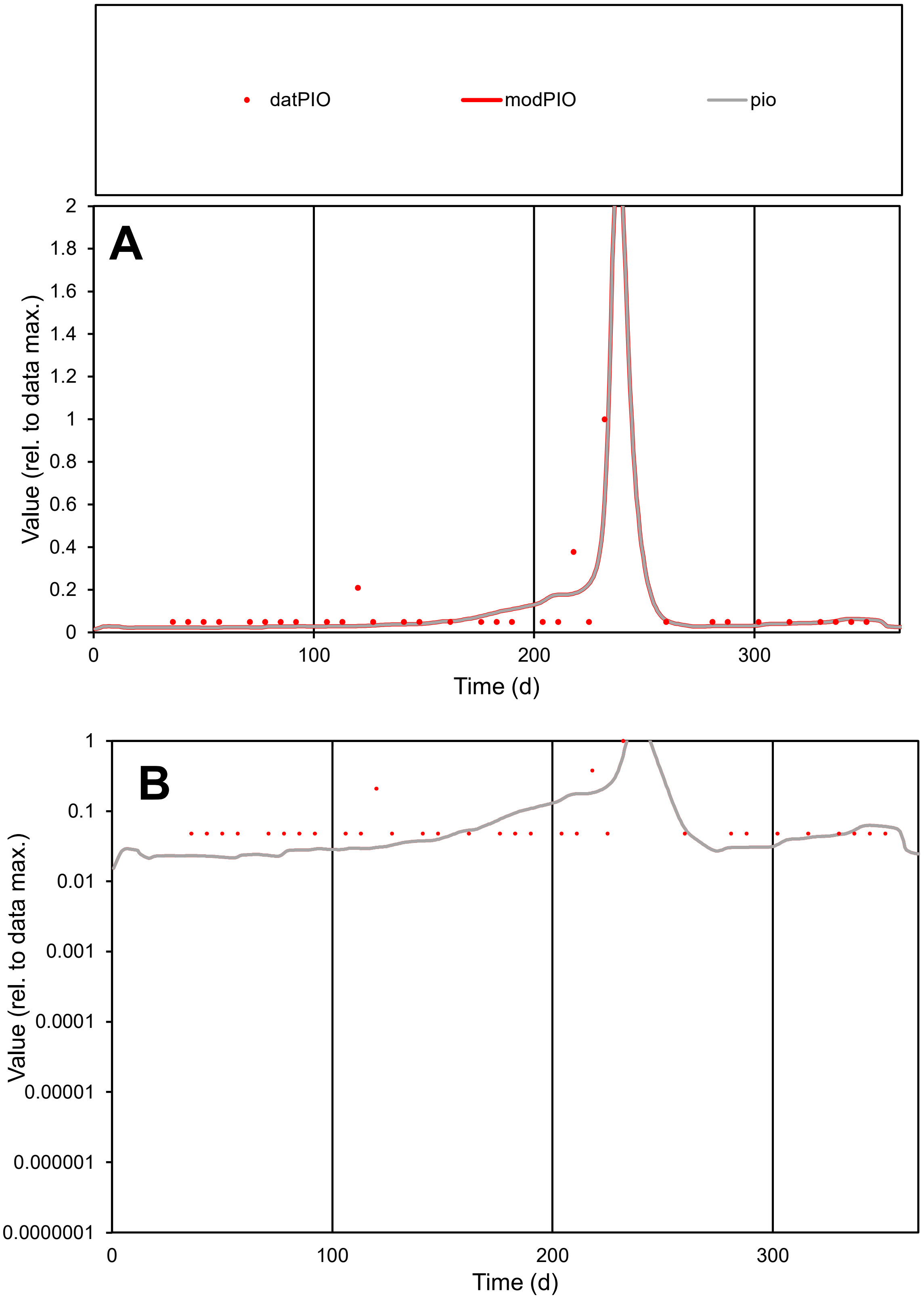


**C1**

Variables
PIO Provide jtd or Dst
CalibOnly?
2 1 to reproduce of
oNormalize?
2 1=no, 2=yes

ofxn calc*
0.71466
ofxn mod
0.74422

sRun
RunA1
zout
1

Ave
Data
2.8E-07
Model
2.7E-07

data/vMax
3E-06

ofxng
321.551

Min d/m
2.17659

ave/min d
1.94301
ave/min m
4.05151

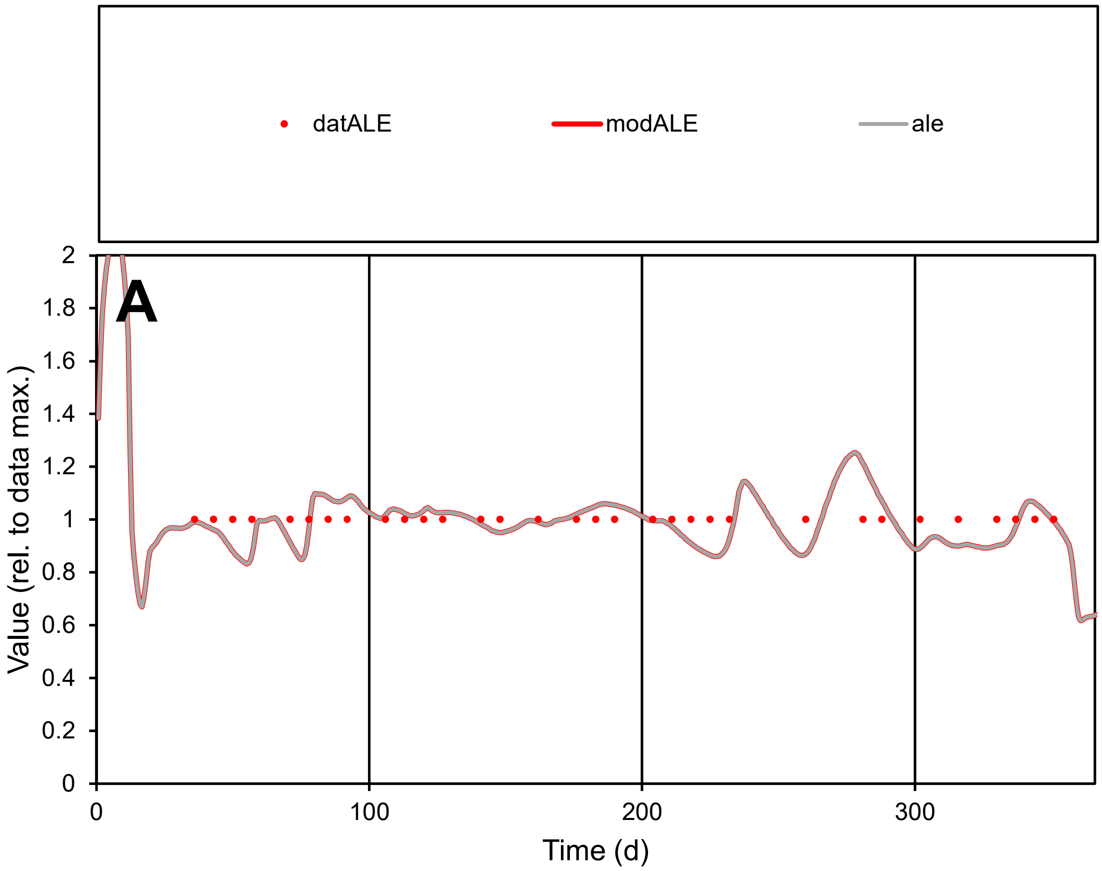


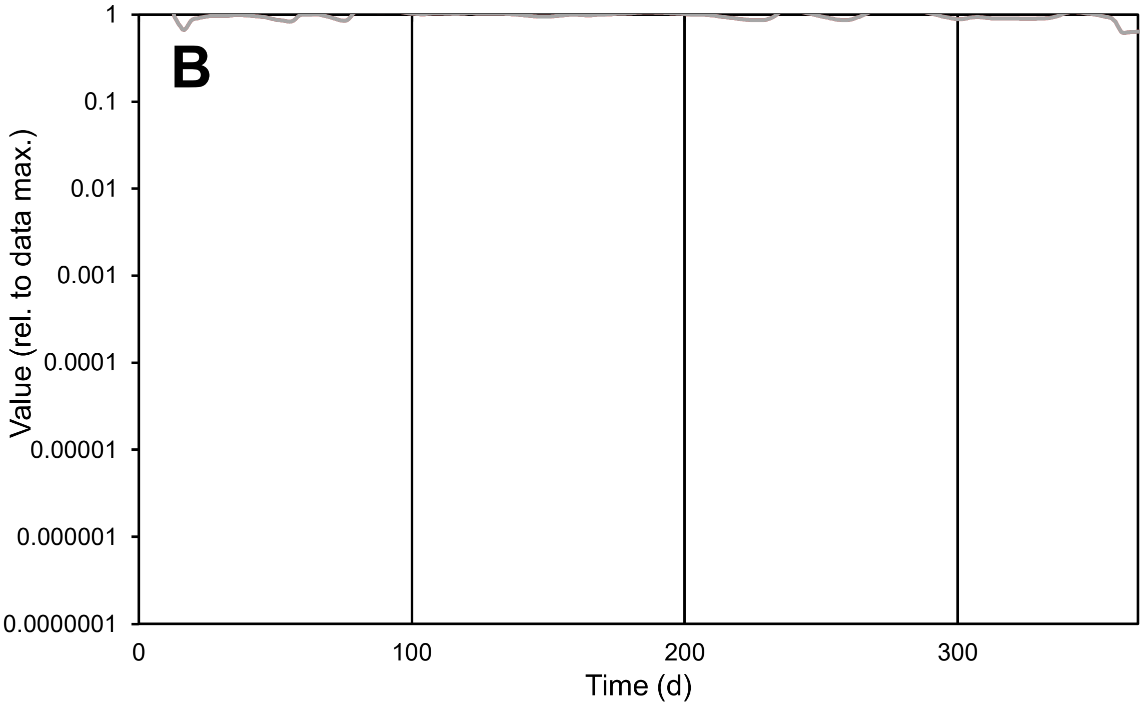


**C1**

Variables
ALE Provide jtd or Dst
CalibOnly?
2 1 to reproduce of
oNormalize?
2 1=no, 2=yes

ofxn calc*
0.0762
ofxn mod
0.12233

sRun
RunA1
zout
1

Ave
Data
1.1E-05
Model
1.1E-05

data/vMax
1.1E-05

ofxng
321.551

Min d/m
1.16201

ave/min d
1
ave/min m
1.14666

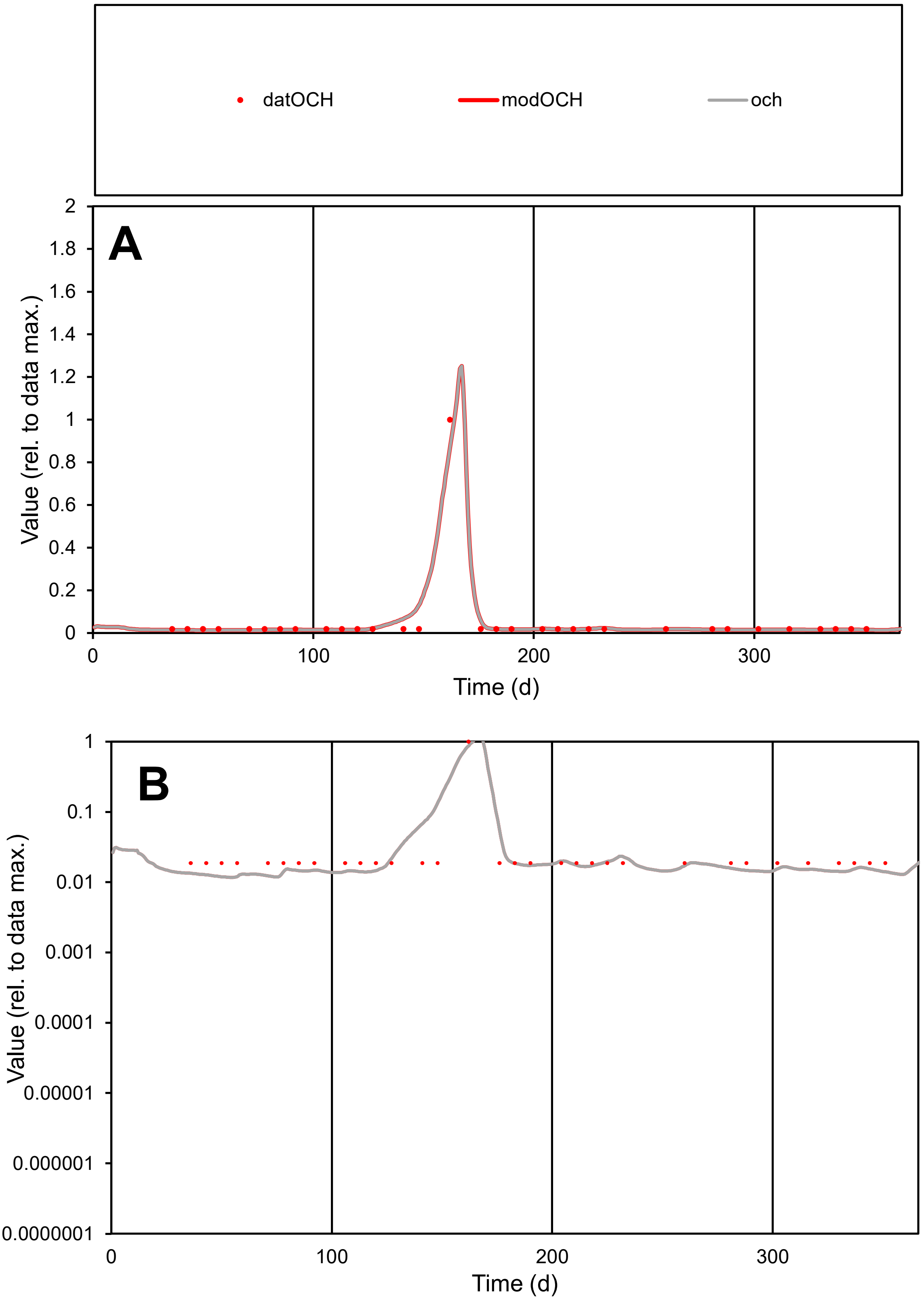


**C1**

Variables
OCH Provide jtd or Dst
CalibOnly?
2 1 to reproduce of
oNormalize?
2 1=no, 2=yes

ofxn calc*
0.37428
ofxn mod
0.63279

sRun
RunA1
zout
1

Ave
Data
1.2E-05
Model
1.2E-05

data/vMax
0.00023

ofxng
321.551

Min d/m
1.58279

ave/min d
2.63821
ave/min m
4.29487

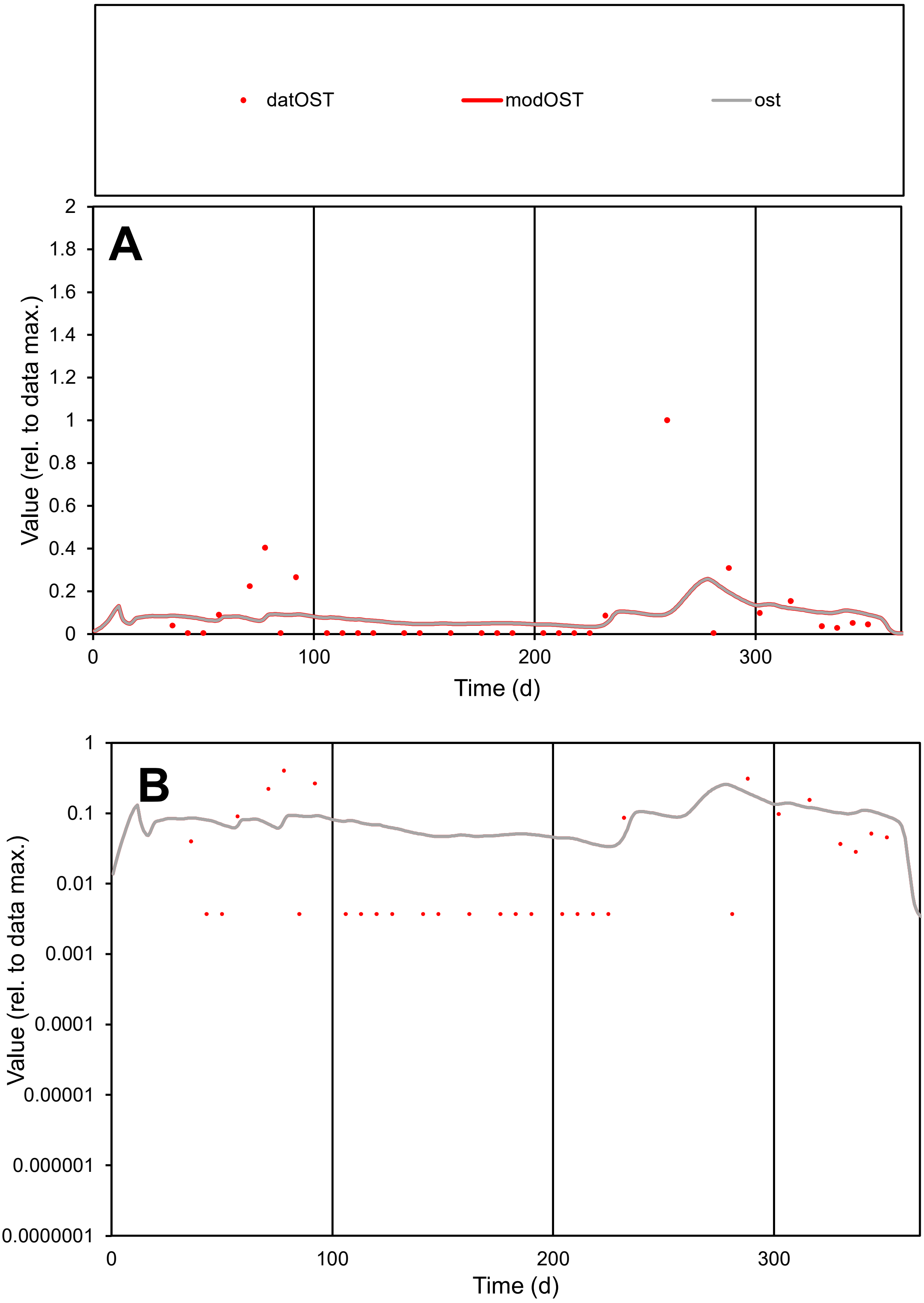


**C1**

Variables
OST Provide jtd or Dst
CalibOnly?
2 1 to reproduce of
oNormalize?
2 1=no, 2=yes

ofxn calc*
2.0452
ofxn mod
2.05117

sRun
RunA1
zout
1

Ave
Data
5.3E-05
Model
4.7E-05

data/vMax
0.00058

ofxng
321.551

Min d/m
0.11

ave/min d
24.4059
ave/min m
2.40376

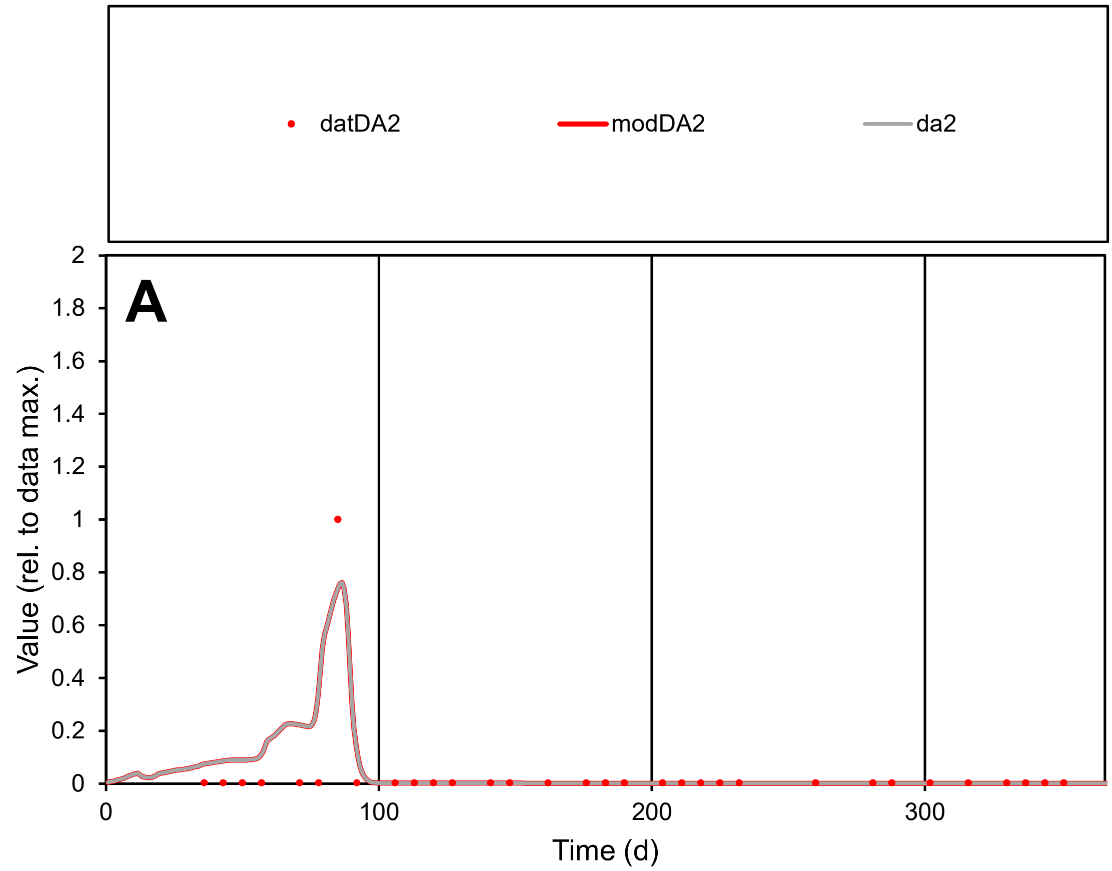


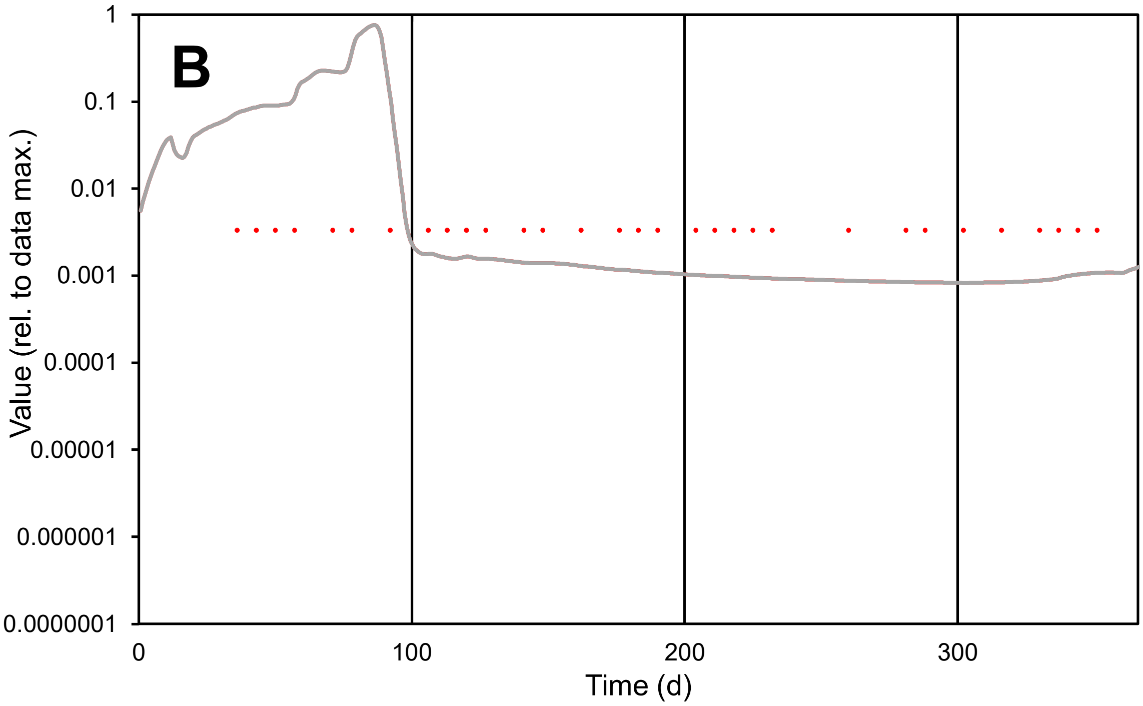


**C1**

| | |
|---|---|
| Variables | |
| DA2 | Provide jtd or Dst |
| CalibOnly? | |
| 2 | 1 to reproduce of |
| oNormalize? | |
| 2 | 1=no, 2=yes |
| ofxn calc* | |
| 2.23639 | |
| ofxn mod | |
| 2.46729 | |
| sRun | |
| RunA1 | |
| zout | |
| 1 | |
| Ave | |
| Data | |
| 3.2E-05 | |
| Model | |
| 4.8E-05 | |
| data/vMax | |
| 0.00093 | |
| ofxng | |
| 321.551 | |
| Min d/m | |
| 4.05037 | |
| ave/min d | |
| 10.3257 | |
| ave/min m | |
| 61.7059 | |

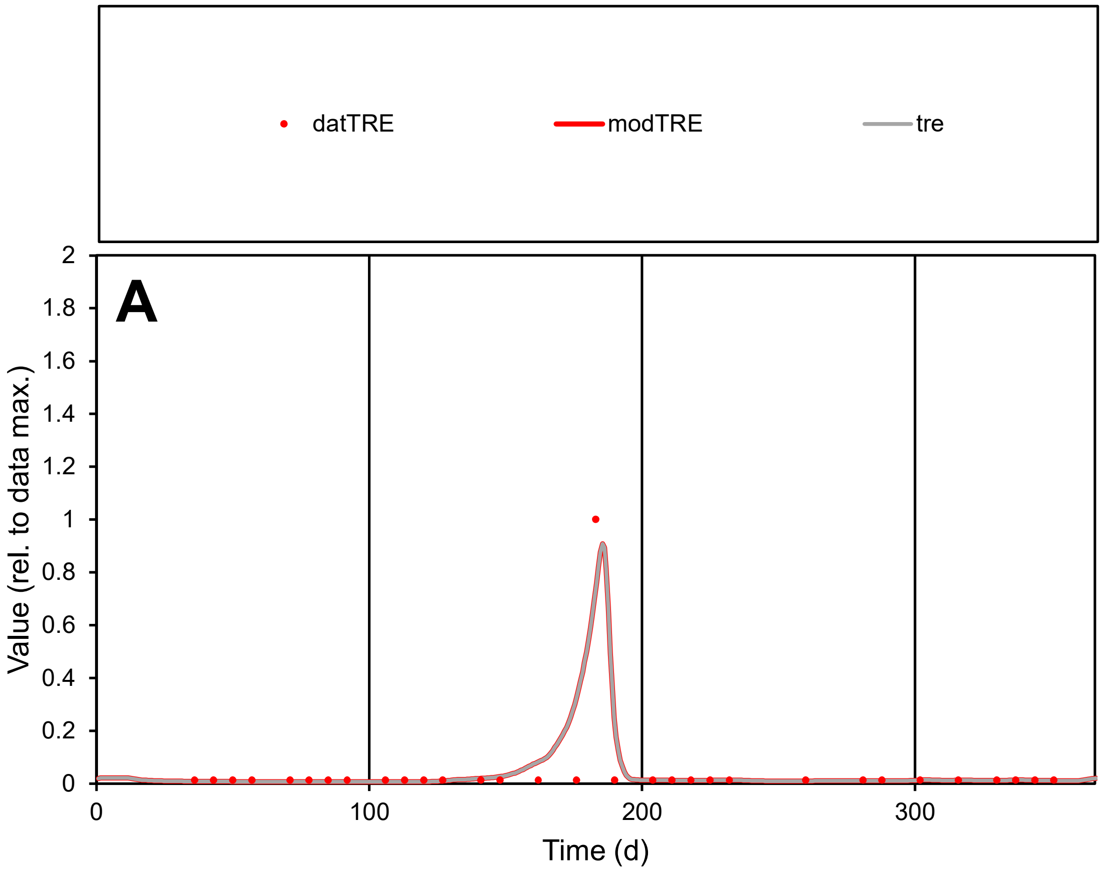


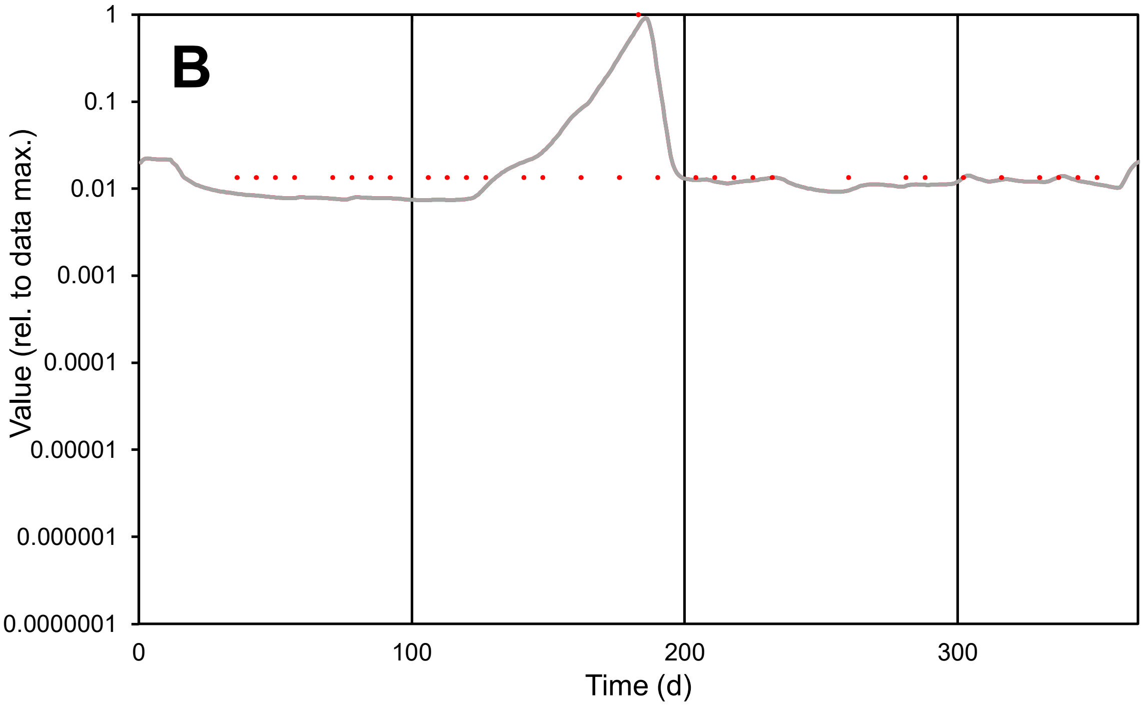


**C1**

Variables
TRE Provide jtd or Dst
CalibOnly?
2 1 to reproduce of
oNormalize?
2 1=no, 2=yes

ofxn calc*
1.06229
ofxn mod
1.43589

sRun
RunA1
zout
1

Ave
Data
2.3E-05
Model
2.5E-05

data/vMax
0.00052

ofxng
321.551

Min d/m
1.80946

ave/min d
3.29839
ave/min m
6.62061

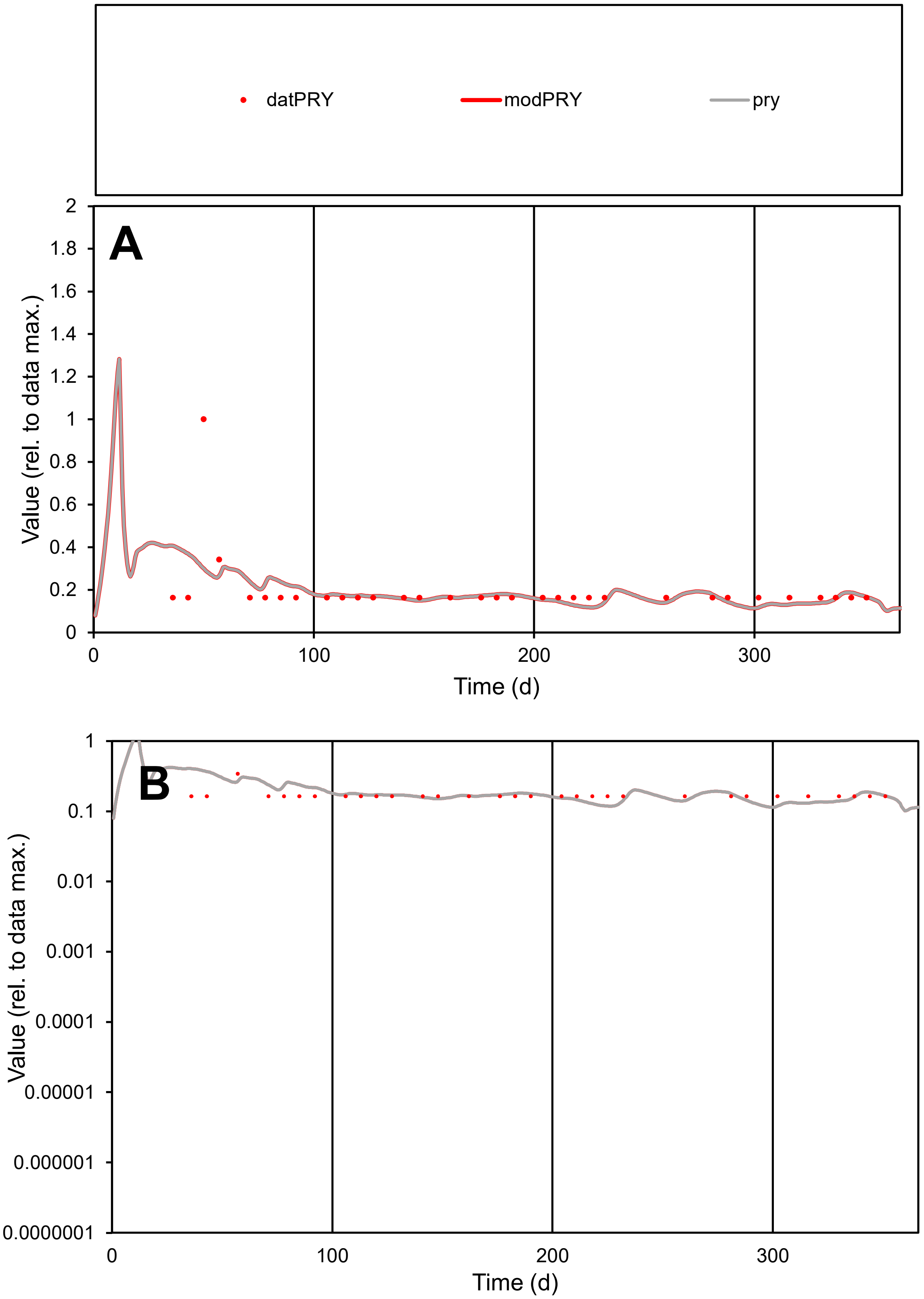


**C1**

| | |
|---|---|
| Variables | |
| PRY | Provide jtd or Dst |
| CalibOnly? | |
| 2 | 1 to reproduce of |
| oNormalize? | |
| 2 | 1=no, 2=yes |
| ofxn calc* | |
| 0.7138 | |
| ofxn mod | |
| 0.73175 | |
| sRun | |
| RunA1 | |
| zout | |
| 1 | |
| Ave | |
| Data | |
| 1.1E-05 | |
| Model | |
| 1E-05 | |
| data/vMax | |
| 5.4E-05 | |
| ofxng | |
| 321.551 | |
| Min d/m | |
| 1.38172 | |
| ave/min d | |
| 1.19472 | |
| ave/min m | |
| 1.59737 | |

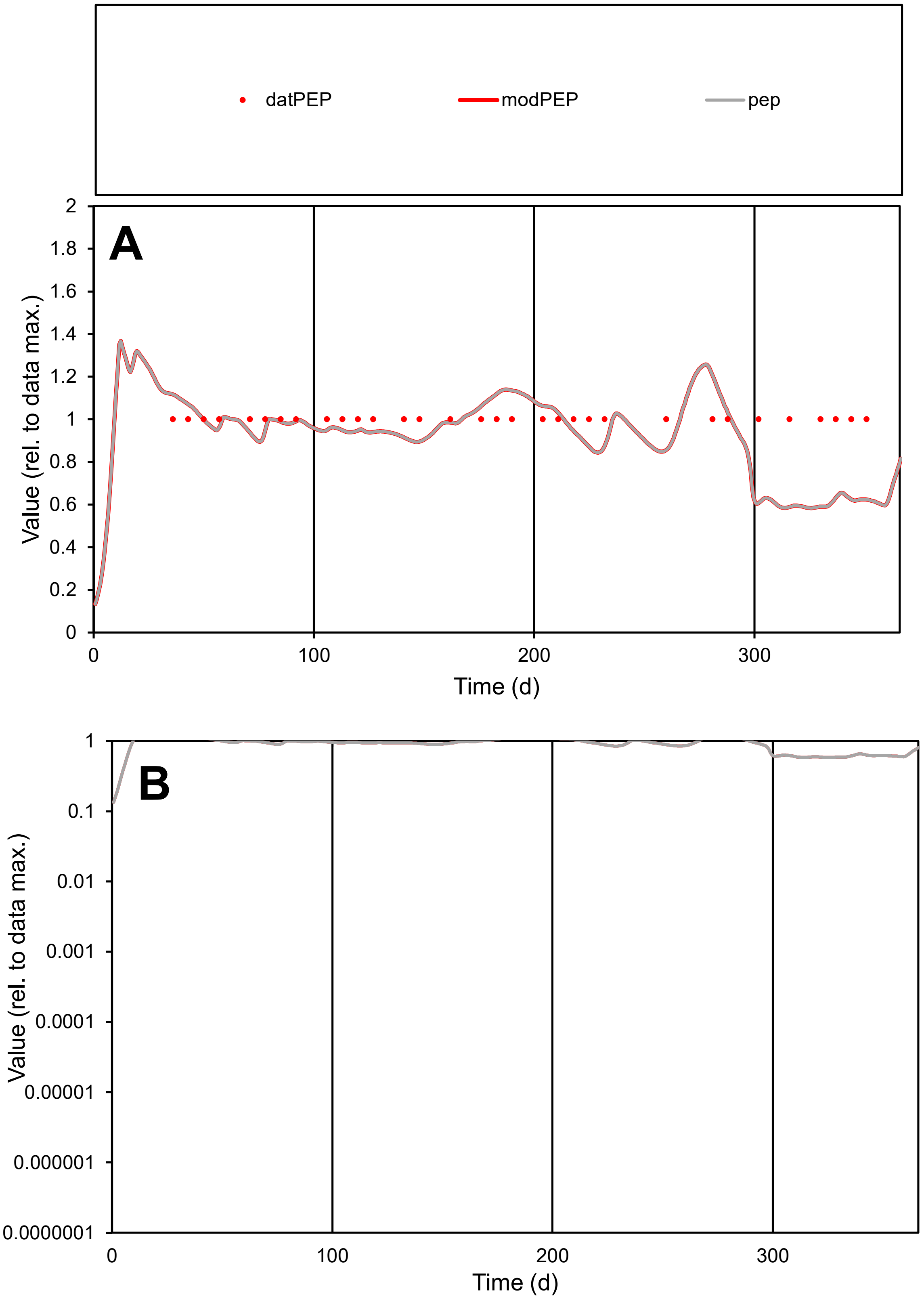


**C1**

Variables

PEP Provide jtd or Dst

CalibOnly?

2 1 to reproduce of

oNormalize?

2 1=no, 2=yes

ofxn calc*

0.18285

ofxn mod

0.20521

sRun

RunA1

zout

1

Ave

Data

4.9E-06

Model

4.5E-06

data/vMax

4.9E-06

ofxng

321.551

Min d/m

1.69553

ave/min d

1

ave/min m

1.56226

324_Calib_S_A

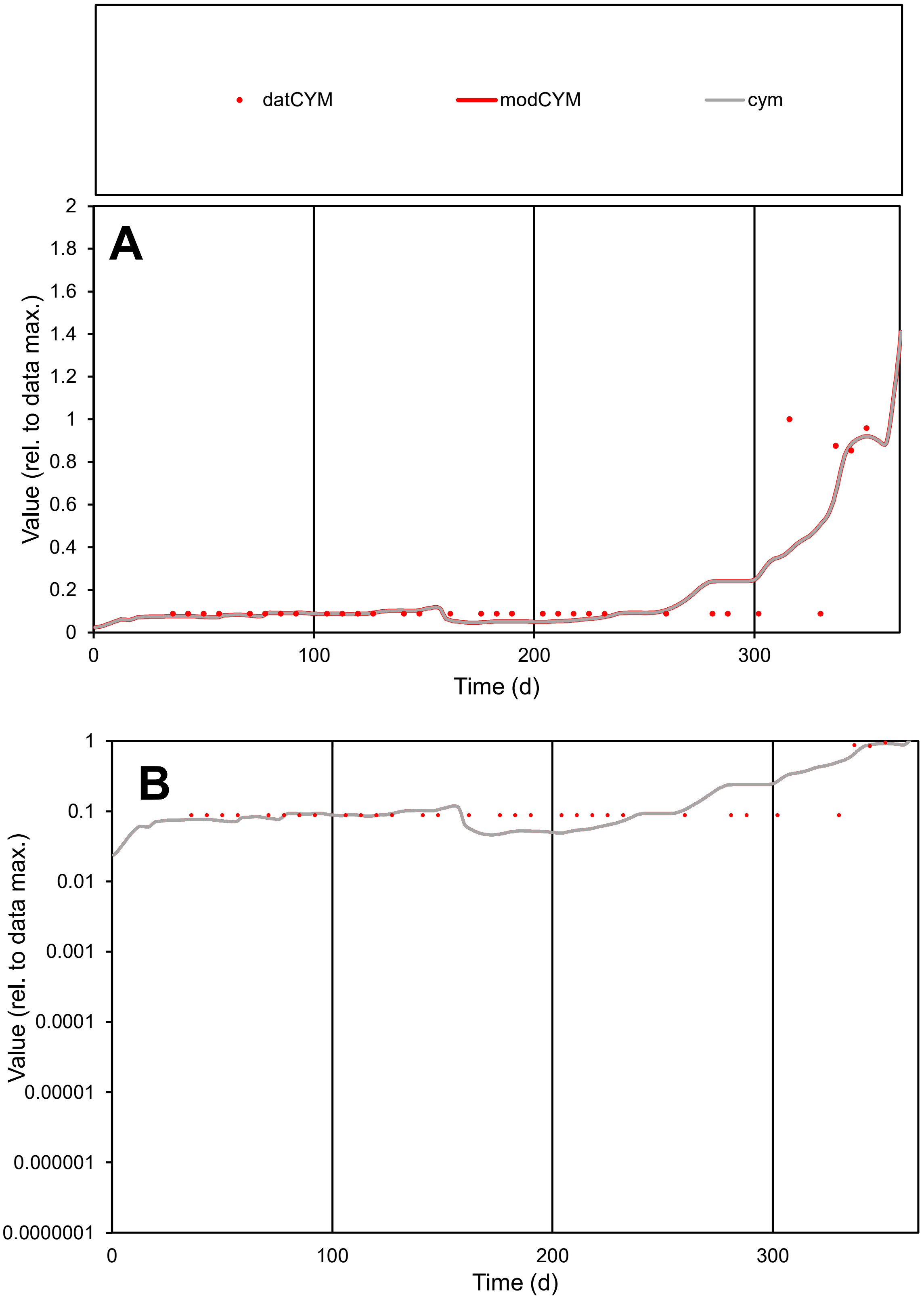


**C1**

Variables
CYM Provide jtd or Dst
CalibOnly?
2 1 to reproduce of
oNormalize?
2 1=no, 2=yes

ofxn calc*
0.74811
ofxn mod
0.76144

sRun
RunA1
zout
1

Ave
Data
3E-06
Model
2.9E-06

data/vMax
1.5E-05

ofxng
321.551

Min d/m
1.83374

ave/min d
2.17666
ave/min m
3.90541

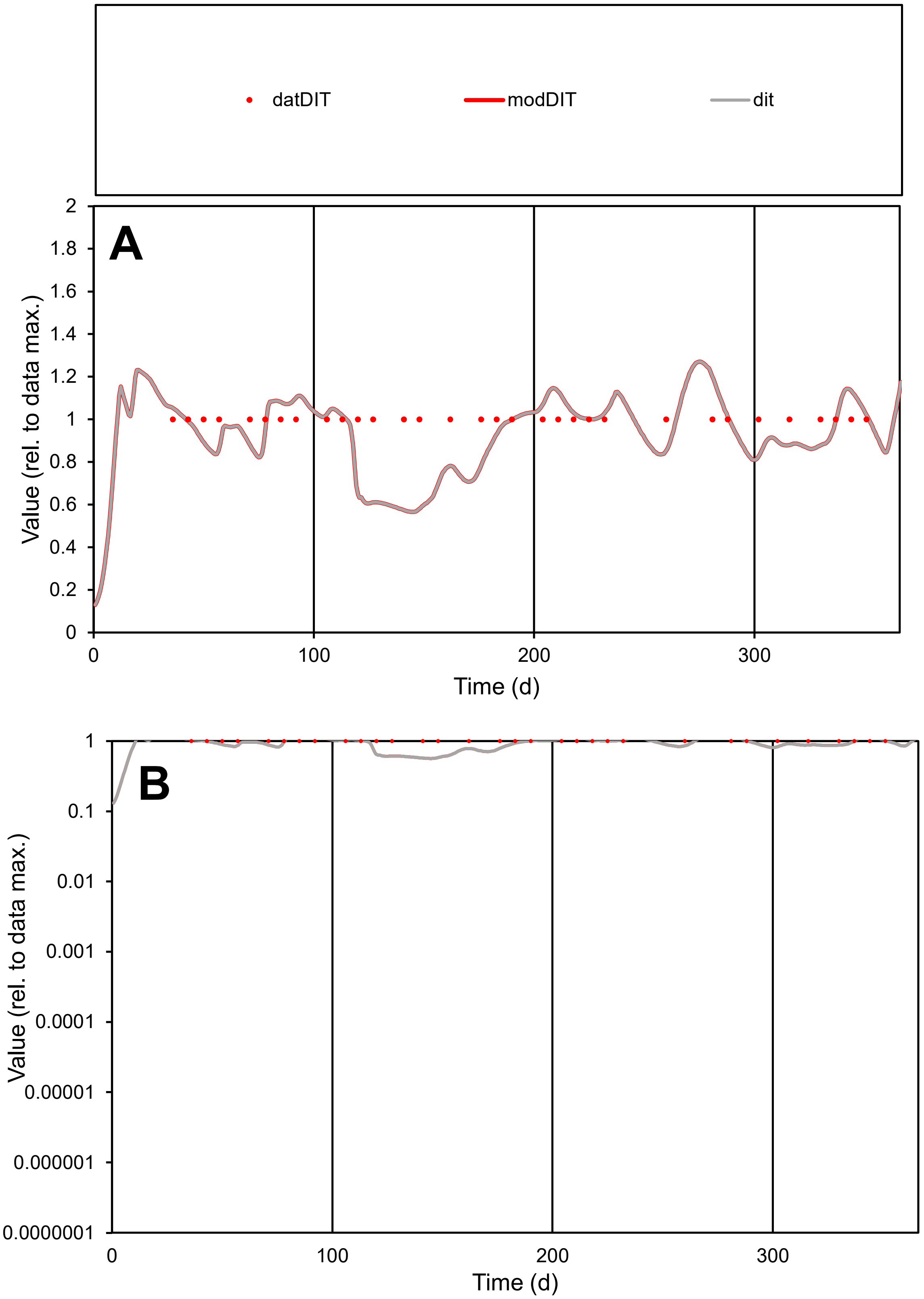


**C1**

Variables
DIT Provide jtd or Dst
CalibOnly?
2 1 to reproduce of
oNormalize?
2 1=no, 2=yes

ofxn calc*
0.16955
ofxn mod
0.19402

sRun
RunA1
zout
1

Ave
Data
5.1E-06
Model
4.8E-06

data/vMax
5.1E-06

ofxng
321.551

Min d/m
1.7567

ave/min d
1
ave/min m
1.64732

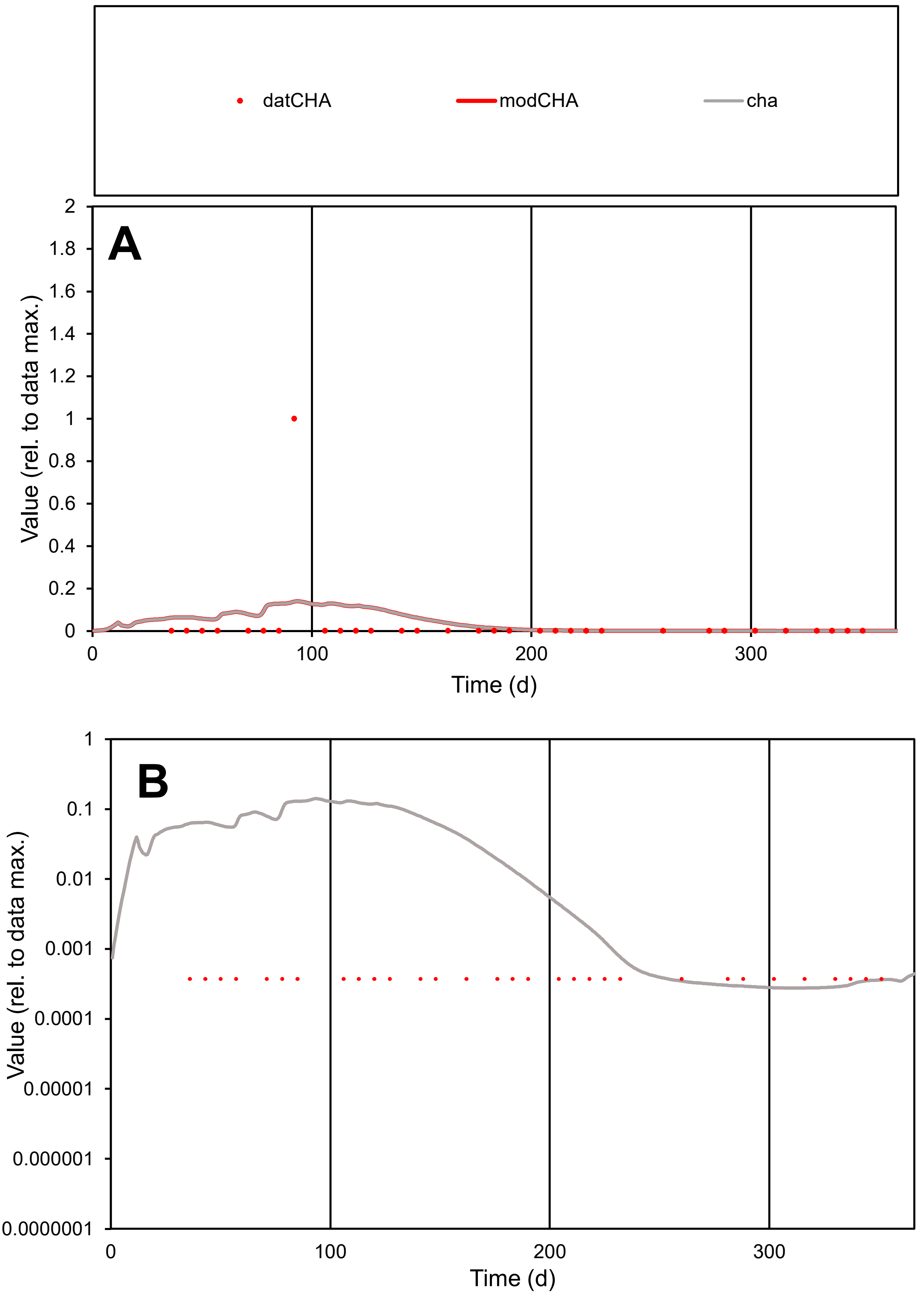


**C1**

Variables
CHA Provide jtd or Dst
CalibOnly?
2 1 to reproduce of
oNormalize?
2 1=no, 2=yes

ofxn calc*
5.14584
ofxn mod
5.16177

sRun
RunA1
zout
1

Ave
Data
7.3E-06
Model
9.8E-06

data/vMax
0.00023

ofxng
321.551

Min d/m
1.33792

ave/min d
84.9159
ave/min m
151.347

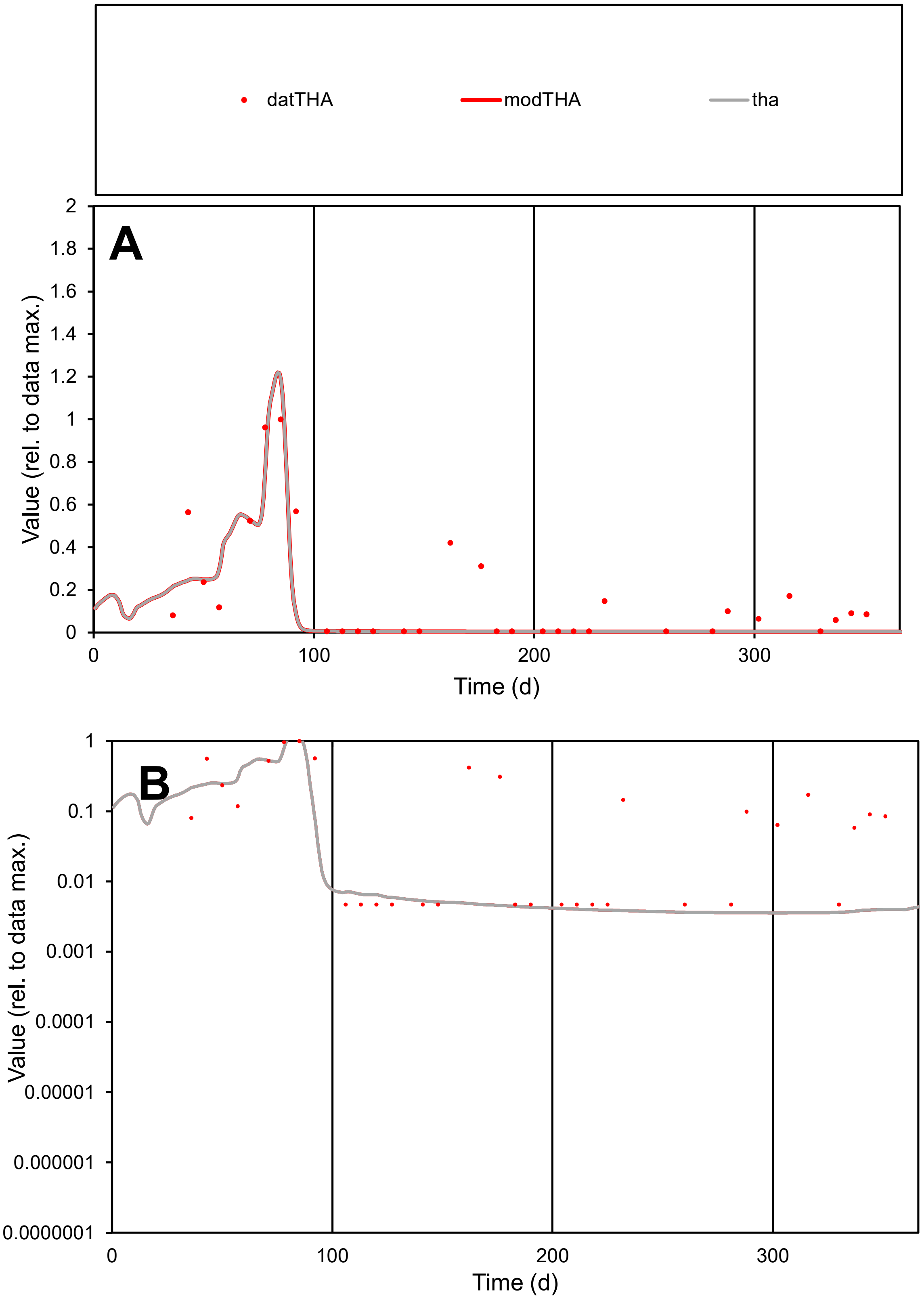


**C1**

Variables
THA Provide jtd or Dst
CalibOnly?
2 1 to reproduce of
oNormalize?
2 1=no, 2=yes

ofxn calc*
0.75122
ofxn mod
0.85652

sRun
RunA1
zout
1

Ave
Data
0.0001
Model
7.2E-05

data/vMax
0.00058

ofxng
321.551

Min d/m
1.3269

ave/min d
36.7898
ave/min m
34.6756

324_Calib_S_A

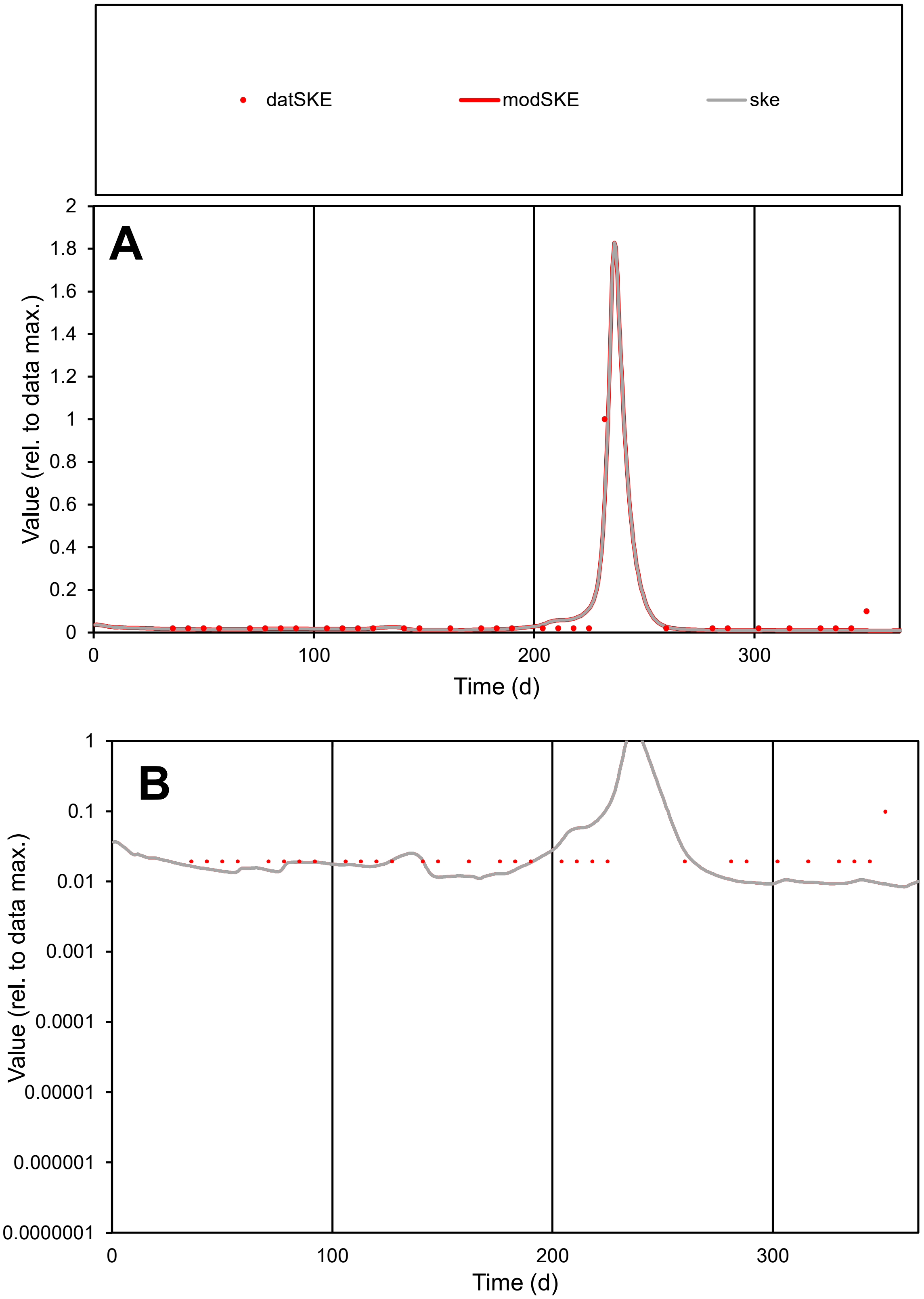


**C1**

Variables
SKE Provide jtd or Dst
CalibOnly?
2 1 to reproduce of
oNormalize?
2 1=no, 2=yes

ofxn calc*
0.4618
ofxn mod
0.55787

sRun
RunA1
zout
1

Ave
Data
6.2E-06
Model
6.1E-06

data/vMax
0.00012

ofxng
321.551

Min d/m
2.13056

ave/min d
2.71377
ave/min m
5.6364

324_Calib_S_A

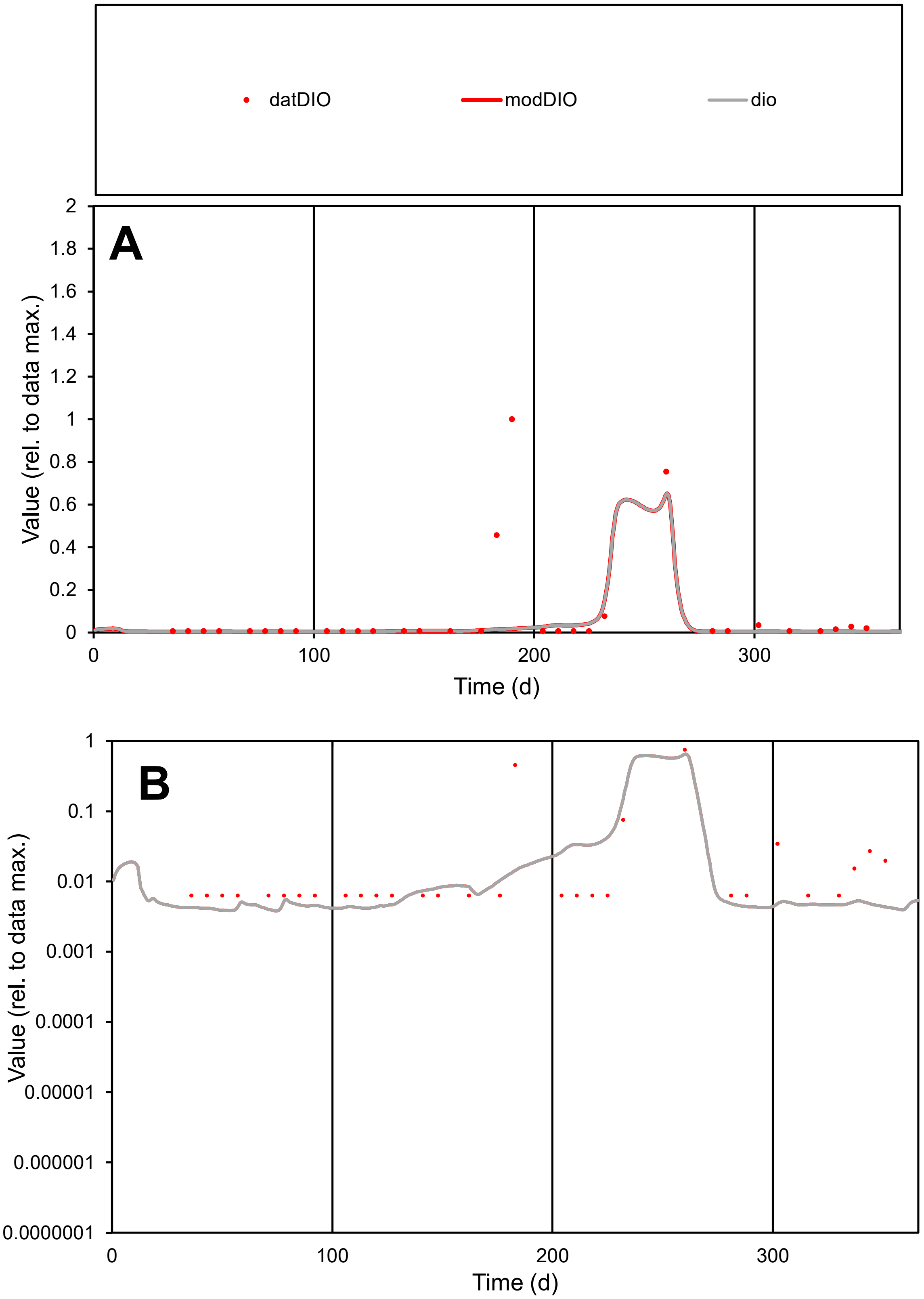


**C1**

Variables
DIO Provide jtd or Dst
CalibOnly?
2 1 to reproduce of
oNormalize?
2 1=no, 2=yes

ofxn calc*
2.41988
ofxn mod
2.43078

sRun
RunA1
zout
1

Ave
Data
5.8E-05
Model
2.4E-05

data/vMax
0.00074

ofxng
321.551

Min d/m
1.64055

ave/min d
12.5052
ave/min m
8.53849

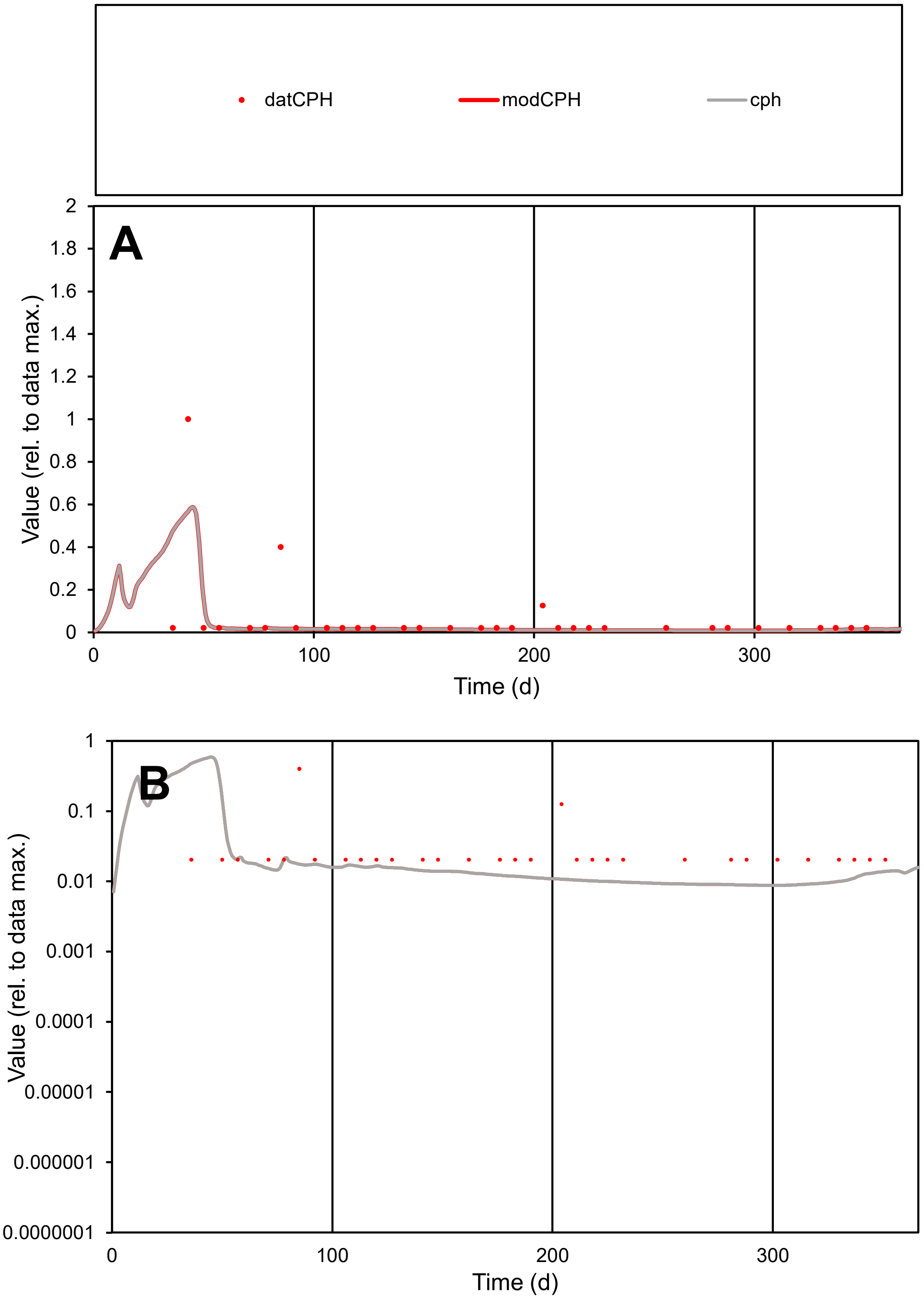


**C1**

Variables
CPH Provide jtd or Dst
CalibOnly?
2 1 to reproduce of
oNormalize?
2 1=no, 2=yes

ofxn calc*
1.90414
ofxn mod
1.99904

sRun
RunA1
zout
1

Ave
Data
3.1E-05
Model
2.1E-05

data/vMax
0.00047

ofxng
321.551

Min d/m
2.33423

ave/min d
3.24139
ave/min m
5.20484

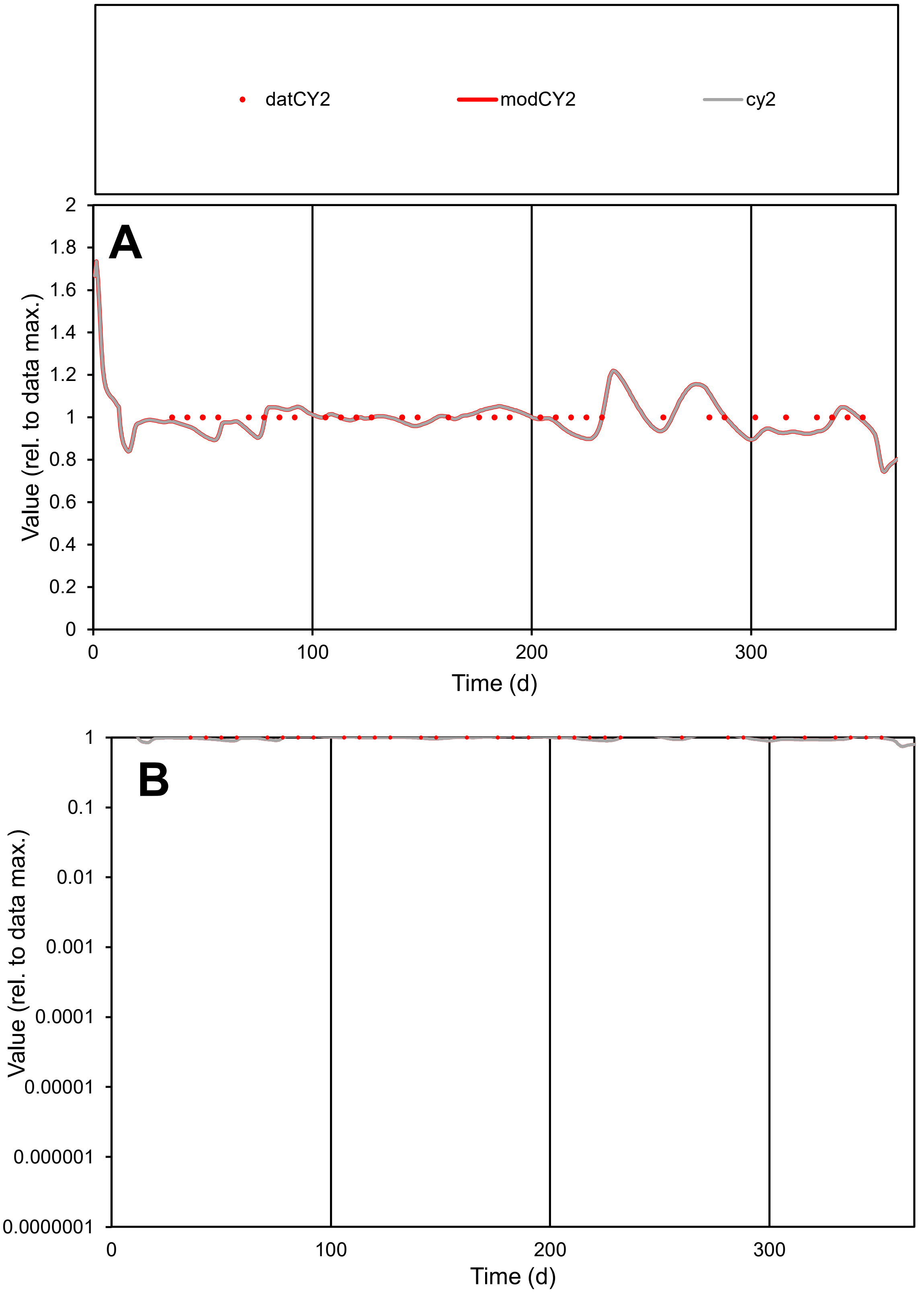


**C1**

Variables
CY2 Provide jtd or Dst
CalibOnly?
2 1 to reproduce of
oNormalize?
2 1=no, 2=yes

ofxn calc*
0.05029
ofxn mod
0.10911

sRun
RunA1
zout
1

Ave
Data
1.1E-05
Model
1.1E-05

data/vMax
1.1E-05

ofxng
321.551

Min d/m
1.11289

ave/min d
1
ave/min m
1.09672

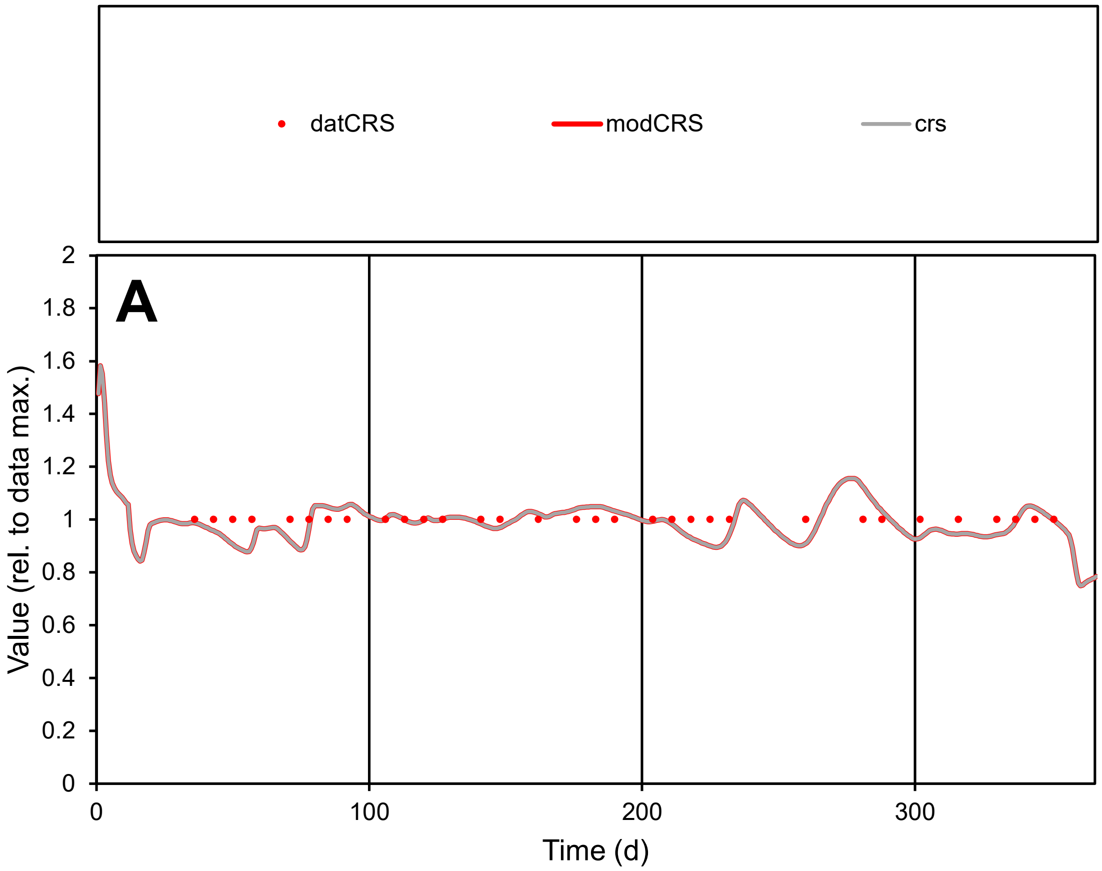


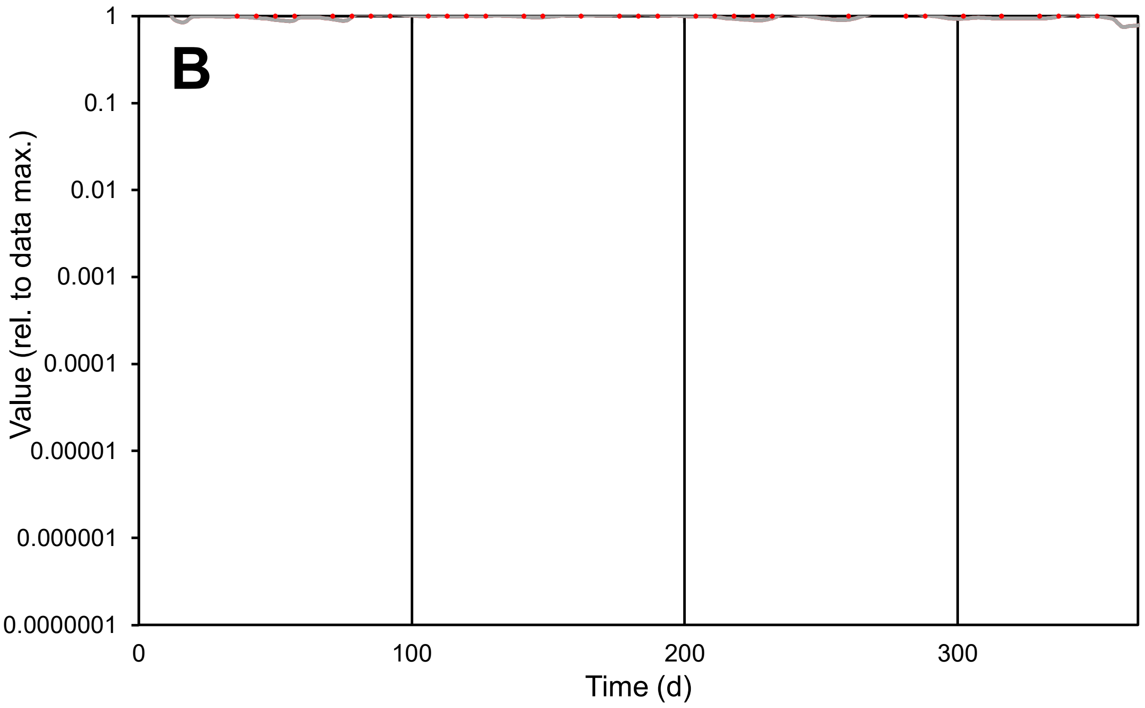


**C1**

Variables

CRS Provide jtd or Dst

CalibOnly?

2 1 to reproduce of

oNormalize?

2 1=no, 2=yes

ofxn calc*

0.05233

ofxn mod

0.10937

sRun

RunA1

zout

1

Ave

Data

5.4E-06

Model

5.3E-06

data/vMax

5.4E-06

ofxng

321.551

Min d/m

1.11823

ave/min d

1

ave/min m

1.10383

324_Calib_S_A

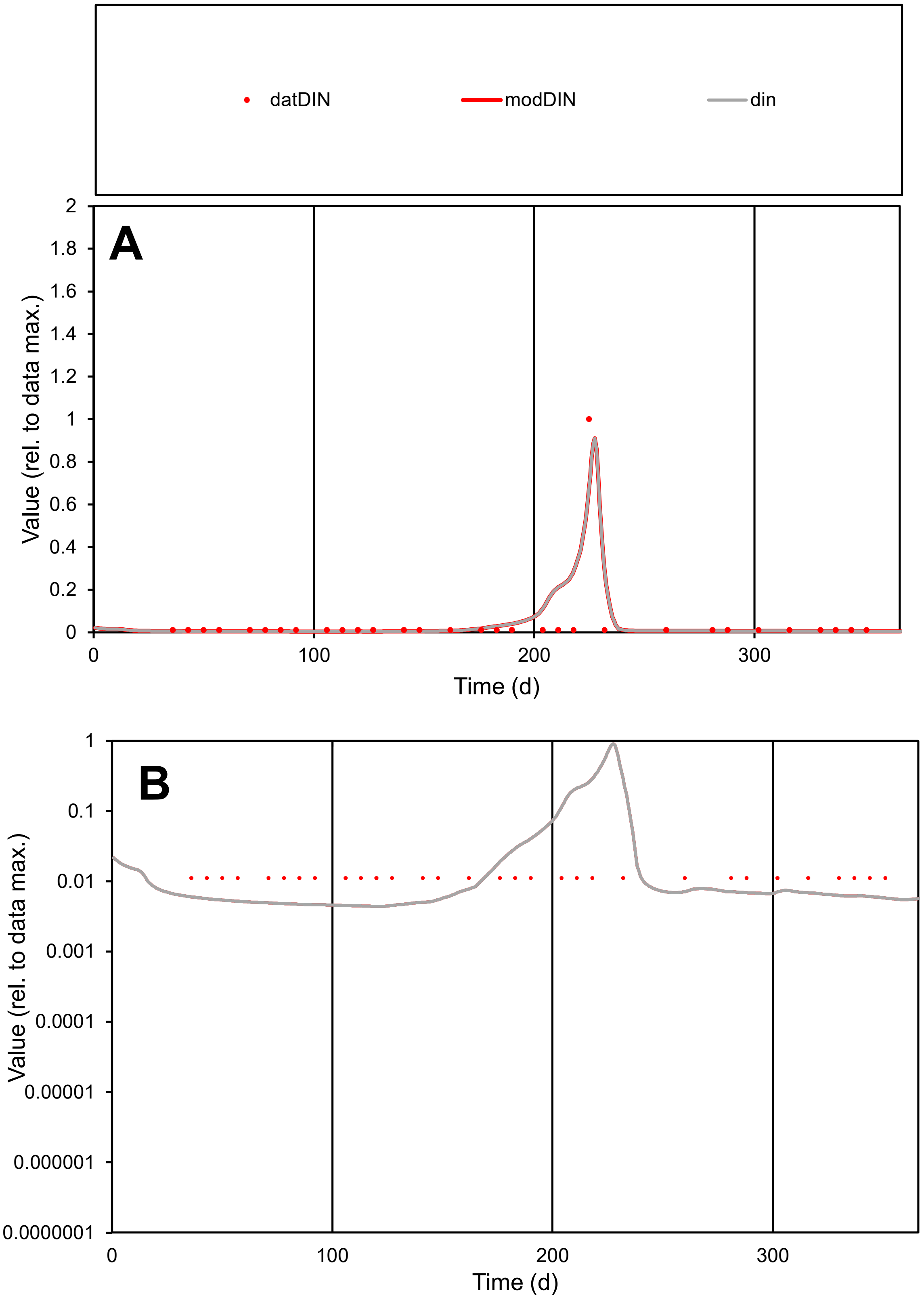


**C1**

Variables

DIN Provide jtd or Dst

CalibOnly?

2 1 to reproduce of

oNormalize?

2 1=no, 2=yes

ofxn calc*

1.6368

ofxn mod

1.7763

sRun

RunA1

zout

1

Ave

Data

1.1E-05

Model

1.4E-05

data/vMax

0.00026

ofxng

321.551

Min d/m

2.52761

ave/min d

3.76976

ave/min m

11.994

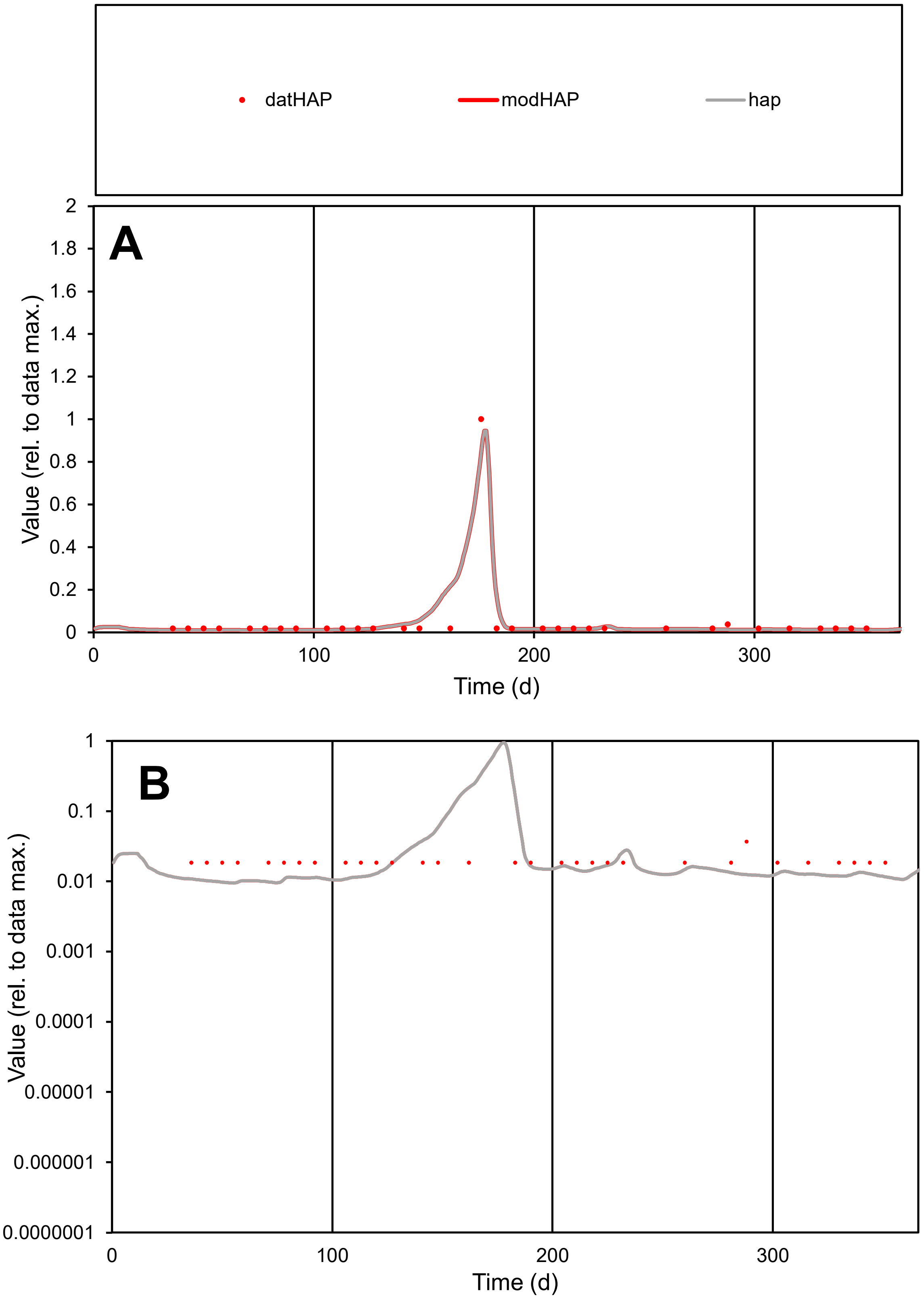


**C1**

Variables
HAP Provide jtd or Dst
CalibOnly?
2 1 to reproduce of
oNormalize?
2 1=no, 2=yes

ofxn calc*
0.69123
ofxn mod
0.84944

sRun
RunA1
zout
1

Ave
Data
3.1E-05
Model
3.2E-05

data/vMax
0.00062

ofxng
321.551

Min d/m
1.91432

ave/min d
2.69411
ave/min m
5.34485

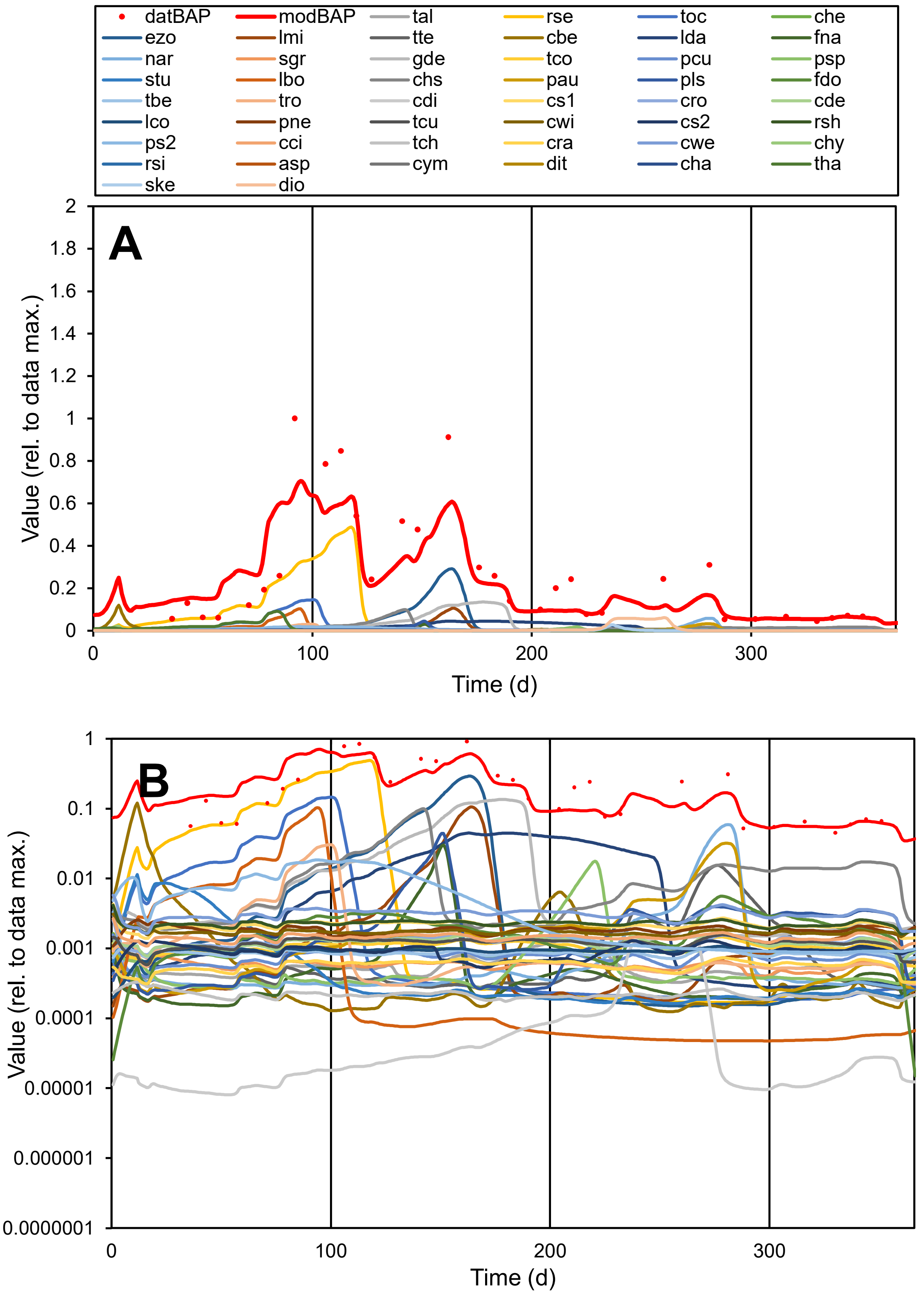


**C1**

Variables

BAP Provide jtd or Dst

CalibOnly?

2 1 to reproduce of

oNormalize?

2 1=no, 2=yes

ofxn calc*

0.51612

ofxn mod

0.54321

sRun

RunA1

zout

1

Ave

Data

0.00208

Model

0.00179

data/vMax

0.00777

ofxng

321.551

Min d/m

0.8331

ave/min d

5.92081

ave/min m

4.24395

324_Calib_S_A

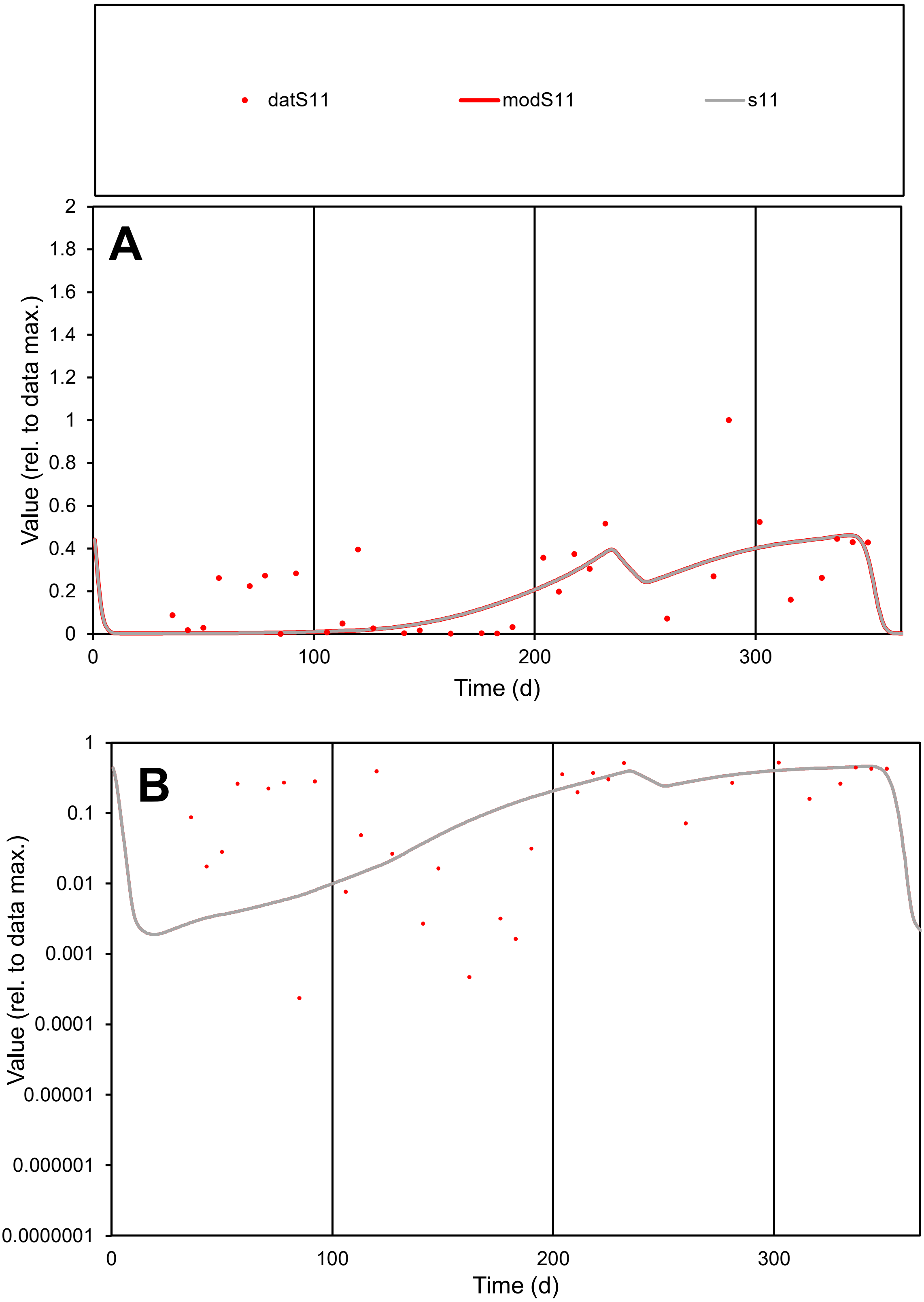


**C1**

Variables
S11 Provide jtd or Dst
CalibOnly?
2 1 to reproduce of
oNormalize?
2 1=no, 2=yes

ofxn calc*
0.83583
ofxn mod
0.84652

sRun
RunA1
zout
1

Ave
Data
0.01629
Model
0.01335

data/vMax
0.07402

ofxng
321.551

Min d/m
0.08224

ave/min d
933.164
ave/min m
62.8988

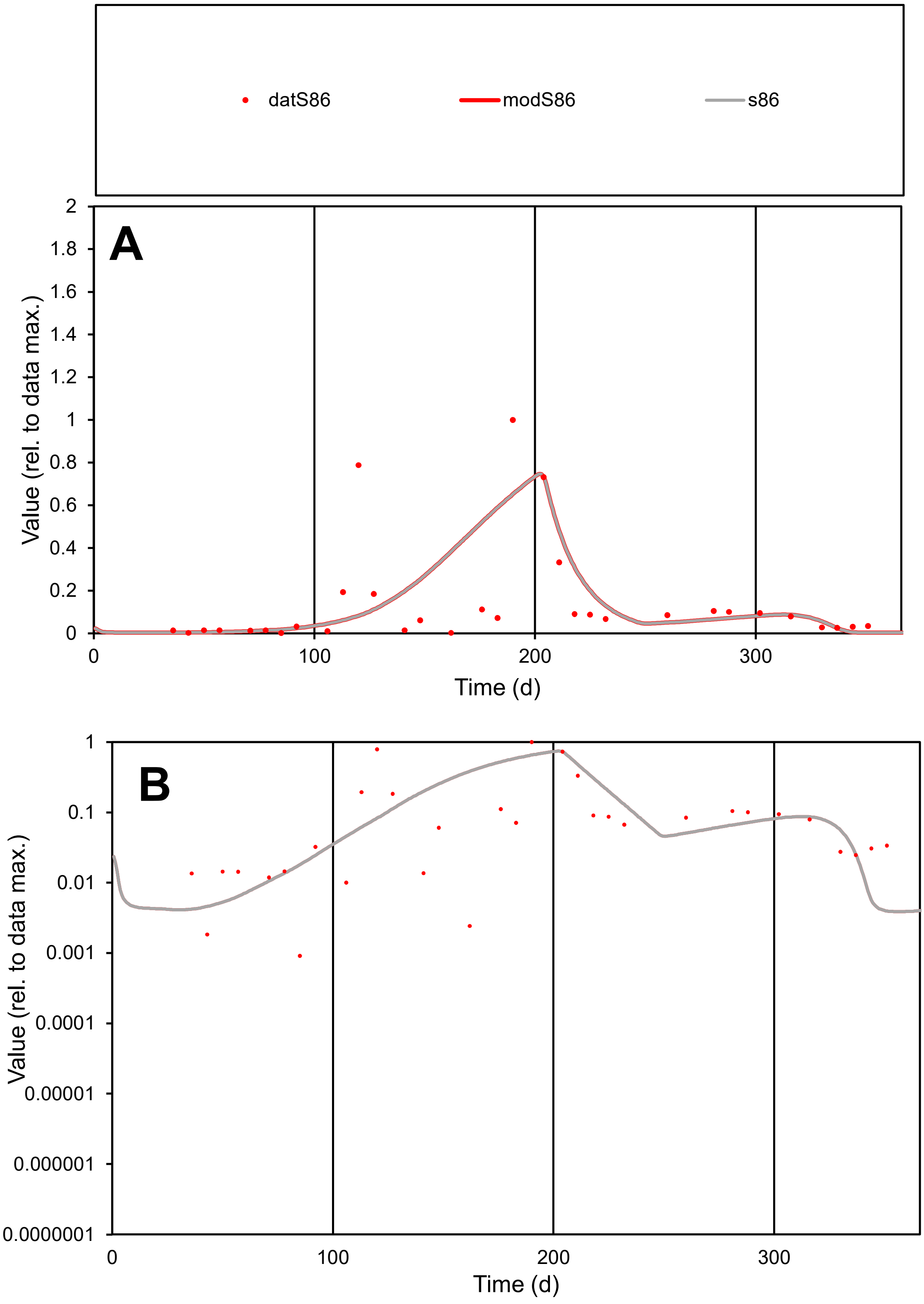


**C1**

Variables
S86 Provide jtd or Dst
CalibOnly?
2 1 to reproduce of
oNormalize?
2 1=no, 2=yes

ofxn calc*
1.48706
ofxn mod
1.50293

sRun
RunA1
zout
1

Ave
Data
0.0327
Model
0.03833

data/vMax
0.2366

ofxng
321.551

Min d/m
0.23211
ave/min d
151.616
ave/min m
41.2502

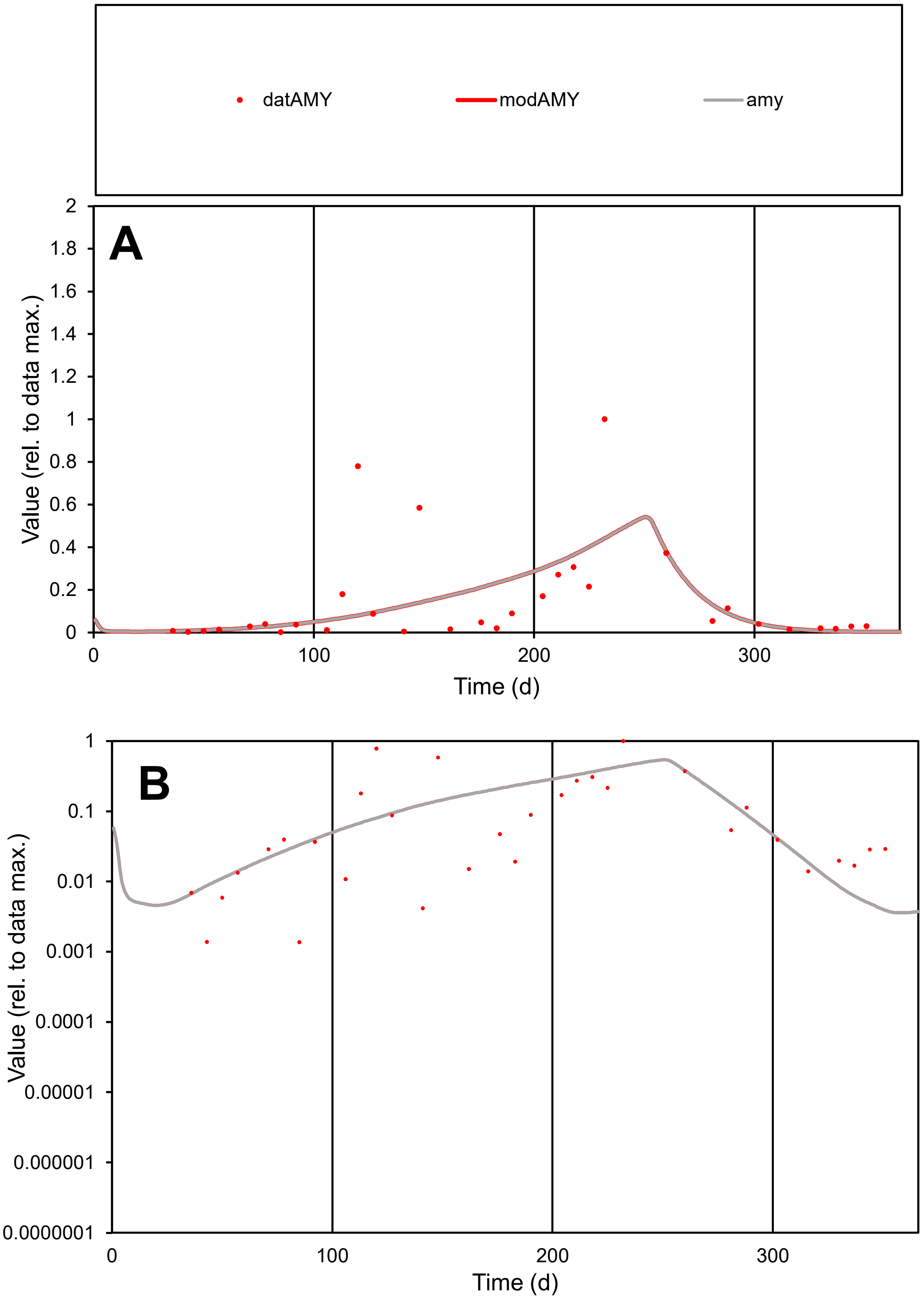


**C1**

Variables
AMY Provide jtd or Dst
CalibOnly?
2 1 to reproduce of
oNormalize?
2 1=no, 2=yes

ofxn calc*
1.35268
ofxn mod
1.3624

sRun
RunA1
zout
1

Ave
Data
0.01057
Model
0.00951

data/vMax
0.07351

ofxng
321.551

Min d/m
0.35913

ave/min d
104.953
ave/min m
33.9029

324_Calib_S_A

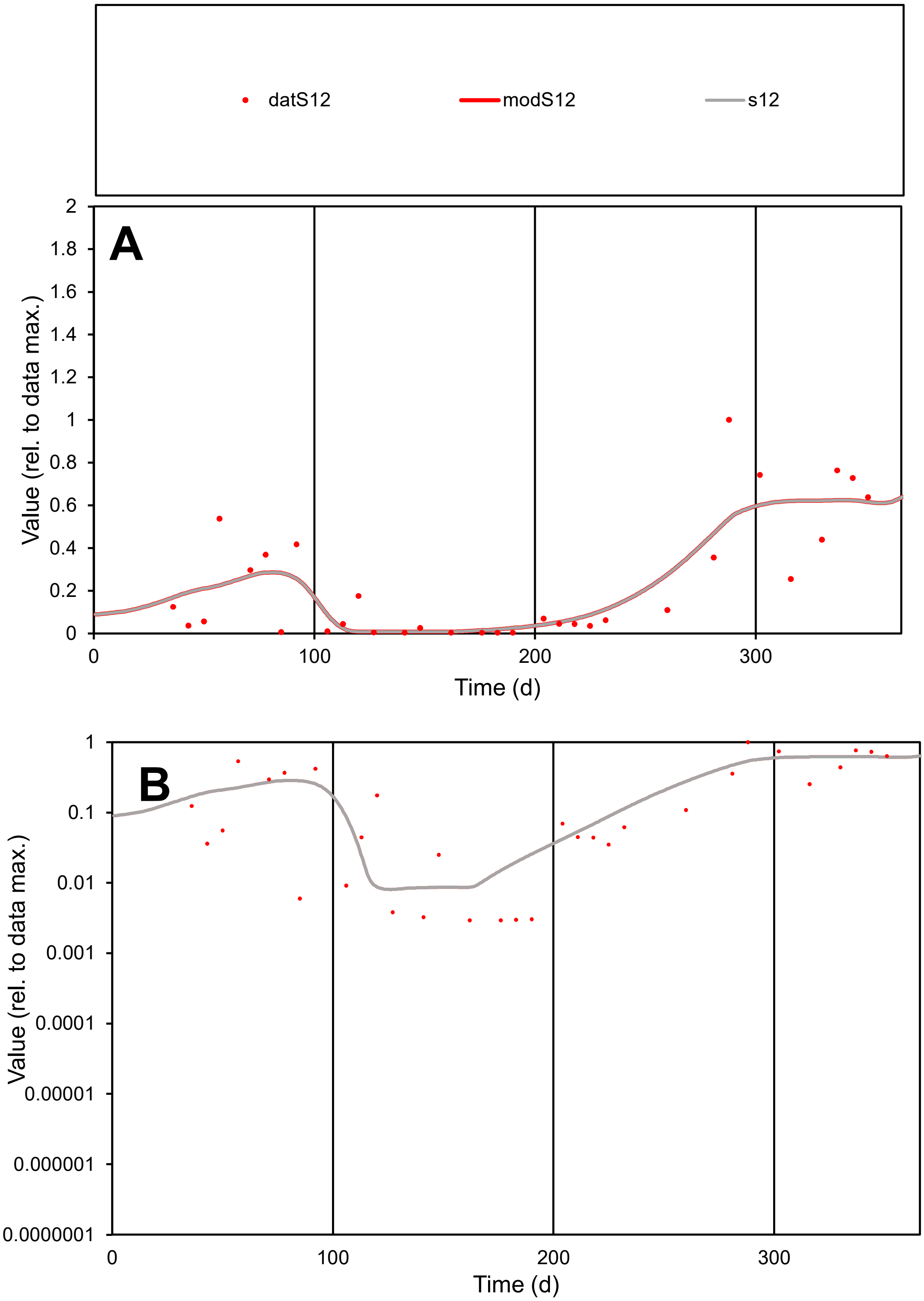


**C1**

Variables
S12 Provide jtd or Dst
CalibOnly?
2 1 to reproduce of
oNormalize?
2 1=no, 2=yes

ofxn calc*
0.66723
ofxn mod
0.68189

sRun
RunA1
zout
1

Ave
Data
0.0108
Model
0.01091

data/vMax
0.04671

ofxng
321.551

Min d/m
0.36229

ave/min d
78.6873
ave/min m
28.8172

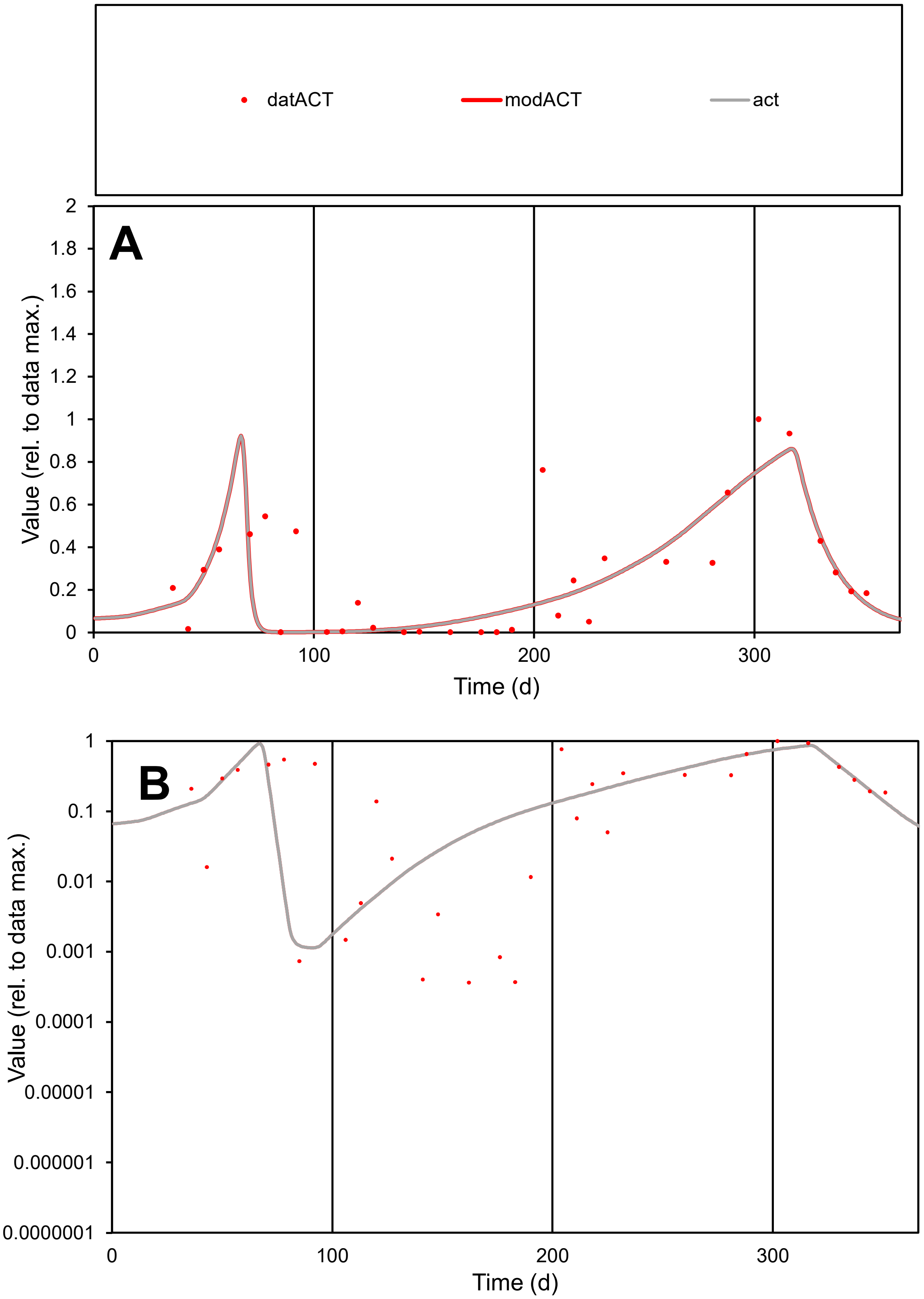


**C1**

Variables
ACT Provide jtd or Dst
CalibOnly?
2 1 to reproduce of
oNormalize?
2 1=no, 2=yes

ofxn calc*
0.72847
ofxn mod
0.75112

sRun
RunA1
zout
1

Ave
Data
0.01836
Model
0.01556

data/vMax
0.07003

ofxng
321.551

Min d/m
0.31294

ave/min d
722.16
ave/min m
191.604

324_Calib_S_A

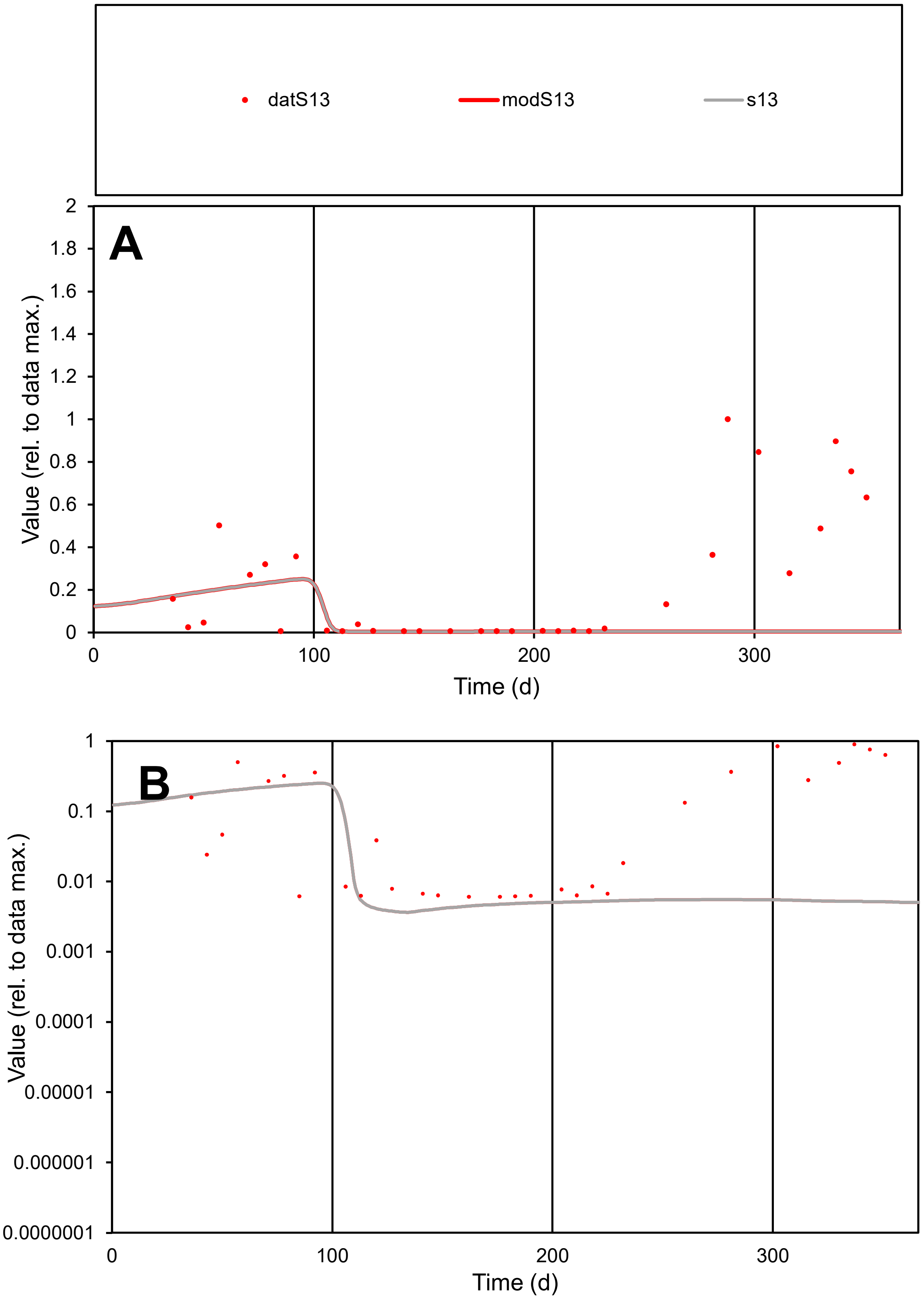


**C1**

Variables

S13 Provide jtd or Dst

CalibOnly?

2 1 to reproduce of

oNormalize?

2 1=no, 2=yes

ofxn calc*

1.58842

ofxn mod

1.59094

sRun

RunA1

zout

1

Ave

Data

0.01016

Model

0.00259

data/vMax

0.04502

ofxng

321.551

Min d/m

1.61775

ave/min d

37.1655

ave/min m

15.3113

324_Calib_S_A

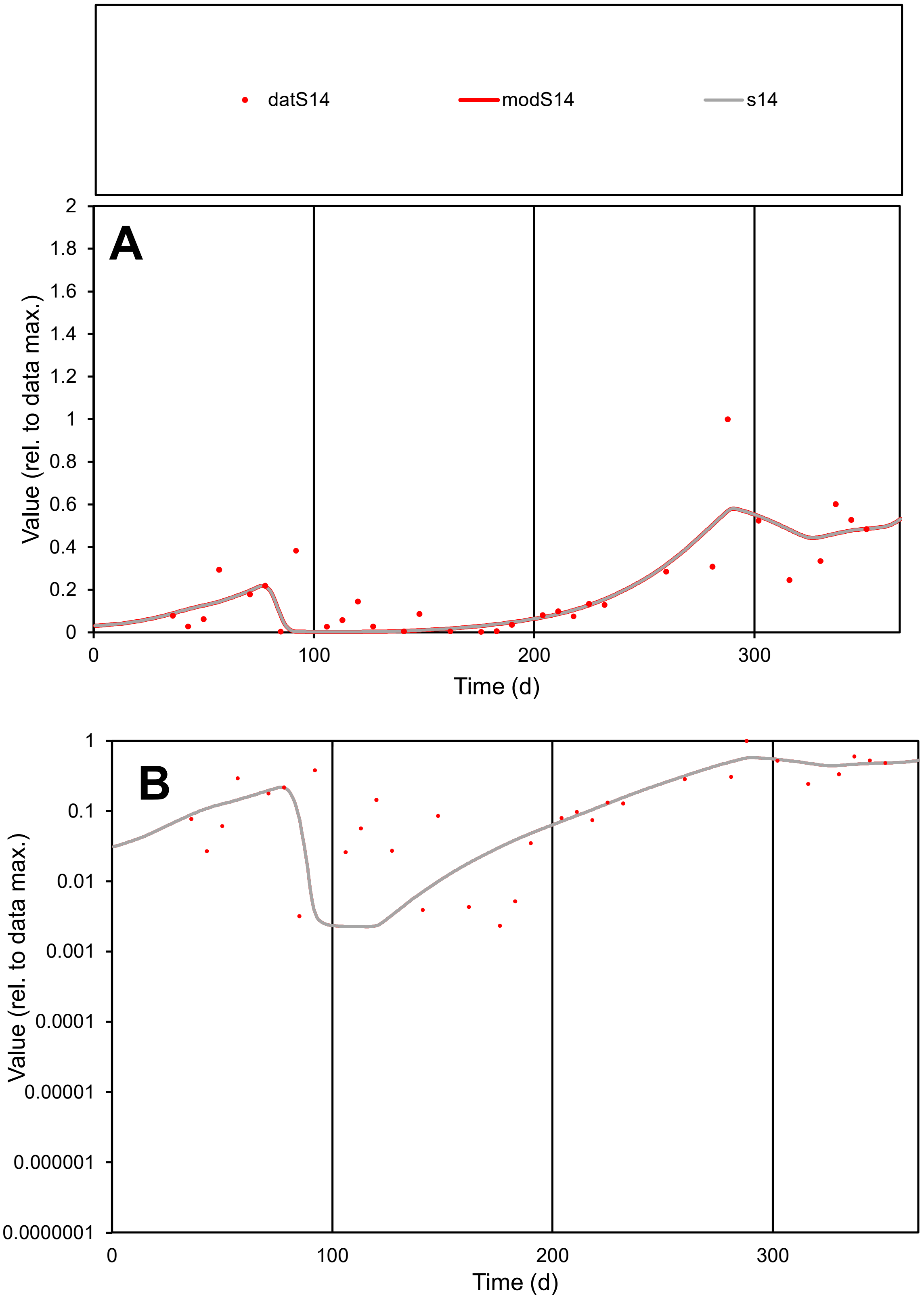


**C1**

Variables
S14 Provide jtd or Dst
CalibOnly?
2 1 to reproduce of
oNormalize?
2 1=no, 2=yes

ofxn calc*
0.63145
ofxn mod
0.64642

sRun
RunA1
zout
1

Ave
Data
0.00847
Model
0.00781

data/vMax
0.04201

ofxng
321.551

Min d/m
1.03068

ave/min d
86.3407
ave/min m
82.0862

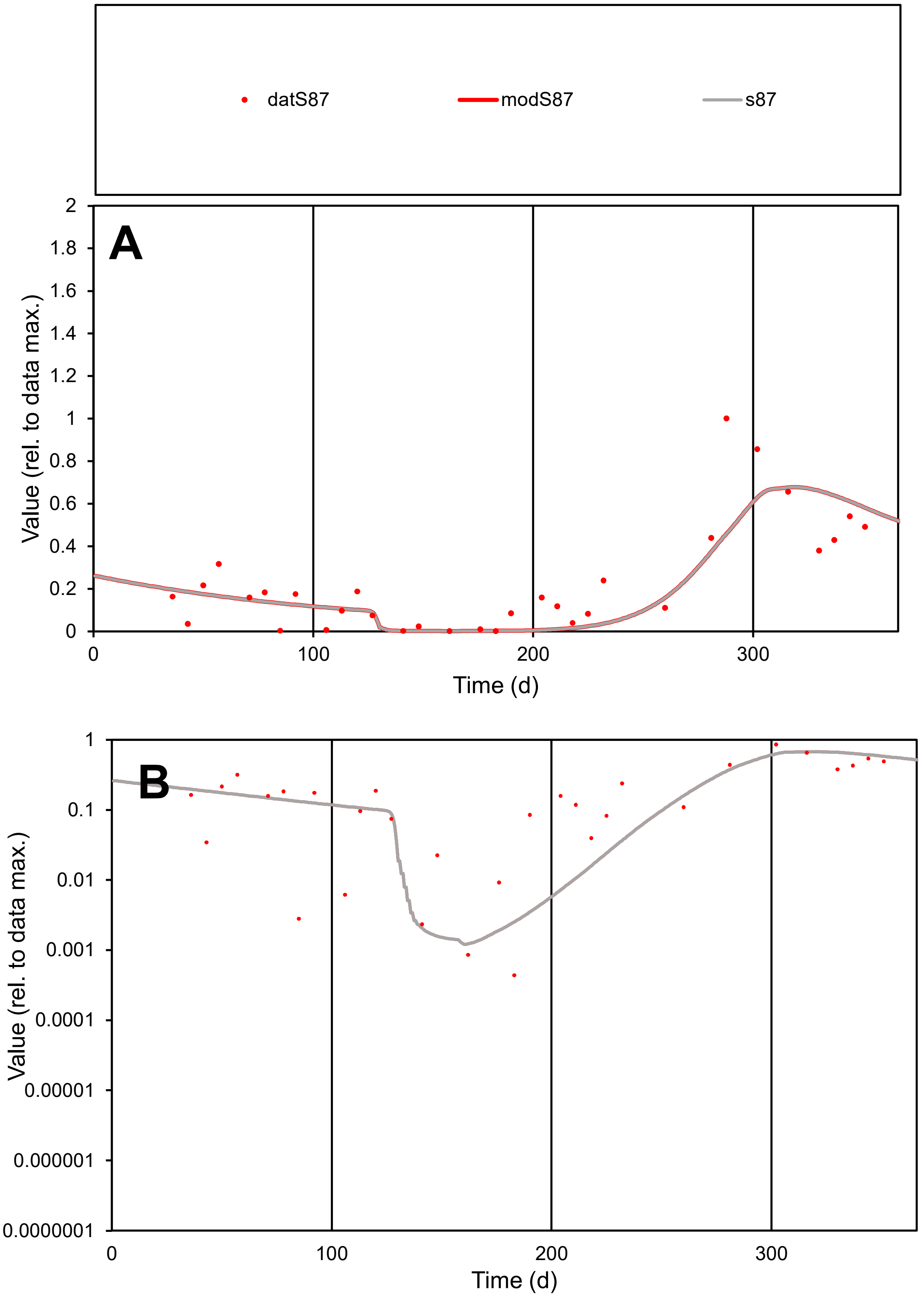


**C1**

Variables
S87 Provide jtd or Dst
CalibOnly?
2 1 to reproduce of
oNormalize?
2 1=no, 2=yes

ofxn calc*
0.6161
ofxn mod
0.62986

sRun
RunA1
zout
1

Ave
Data
0.00917
Model
0.0083

data/vMax
0.04035

ofxng
321.551

Min d/m
0.34521

ave/min d
519.757
ave/min m
162.462

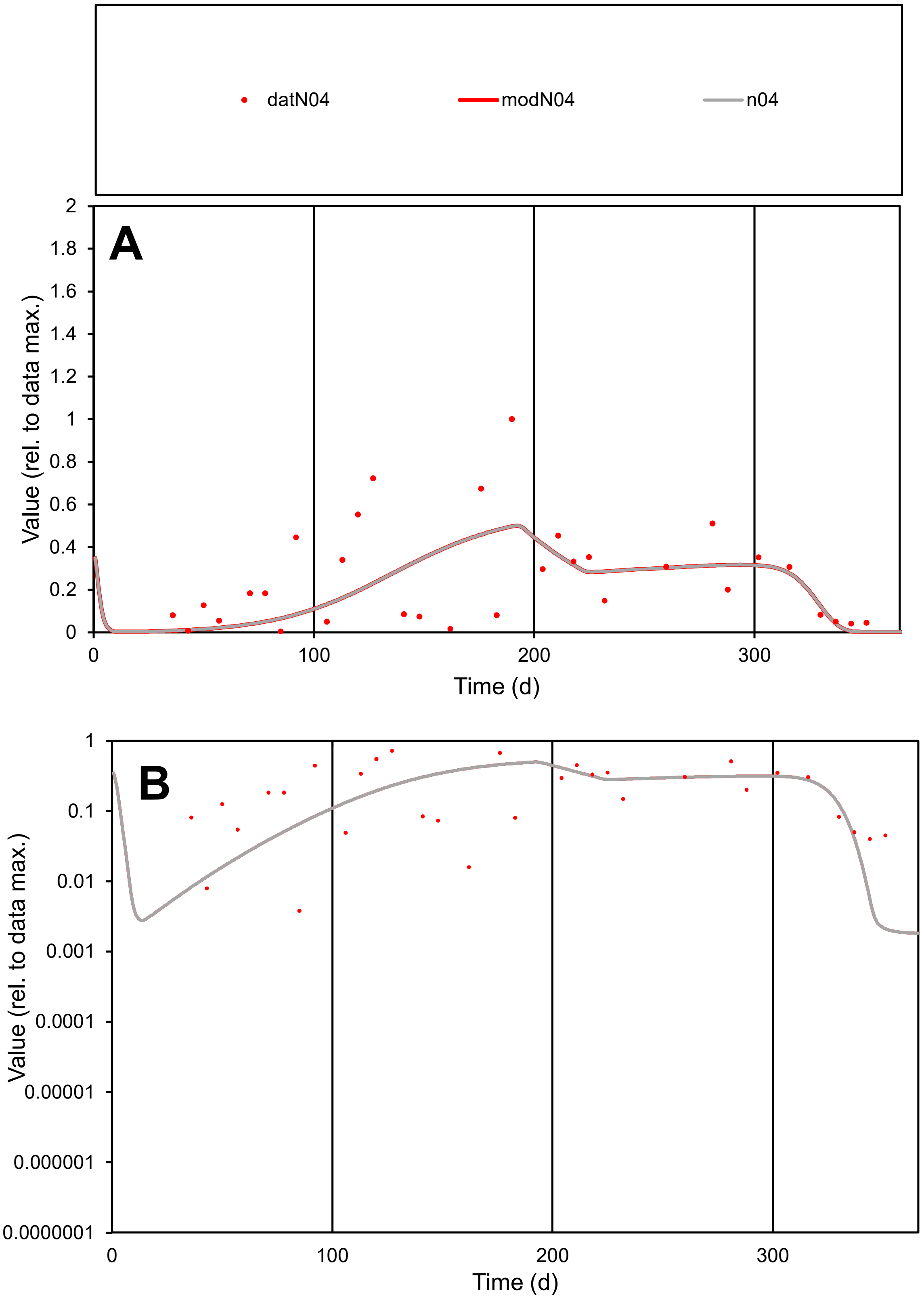


**C1**

Variables
N04 Provide jtd or Dst
CalibOnly?
2 1 to reproduce of
oNormalize?
2 1=no, 2=yes

ofxn calc*
0.82597
ofxn mod
0.83471

sRun
RunA1
zout
1

Ave
Data
0.01589
Model
0.01335

data/vMax
0.06235

ofxng
321.551

Min d/m
1.84825

ave/min d
66.9171
ave/min m
103.859

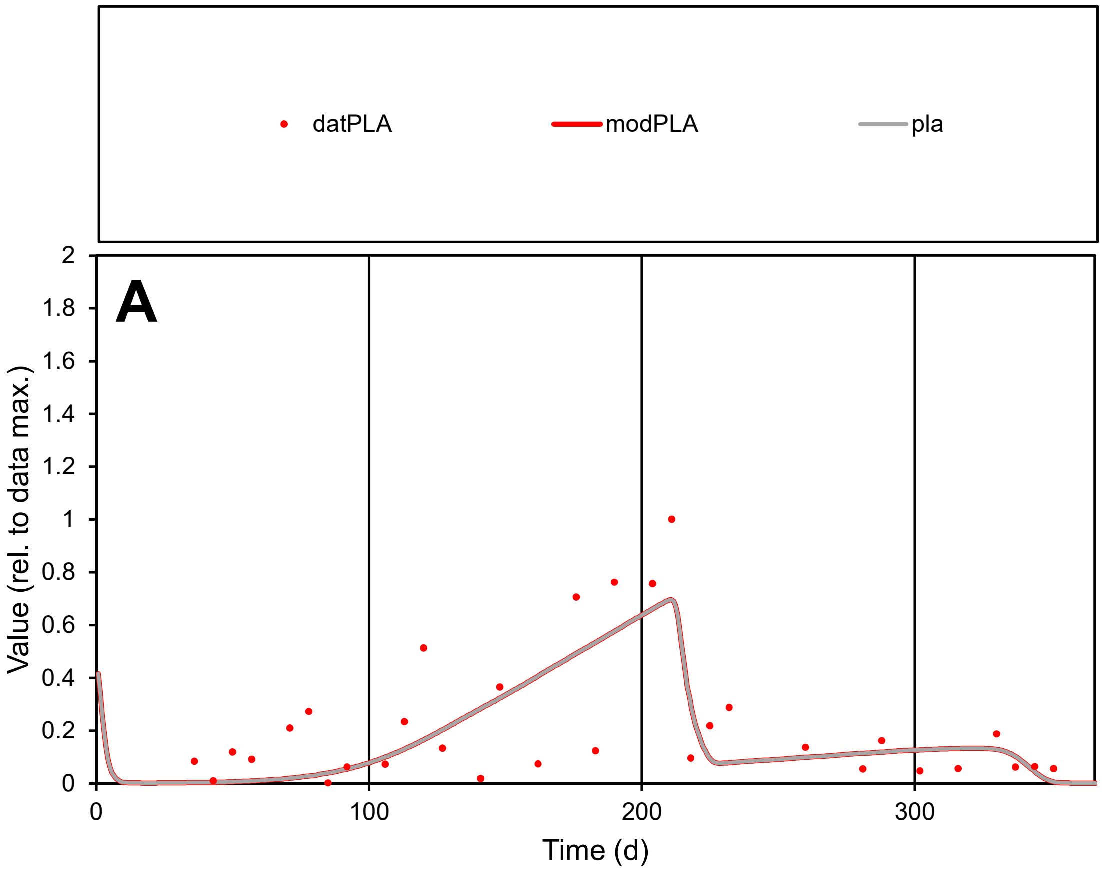


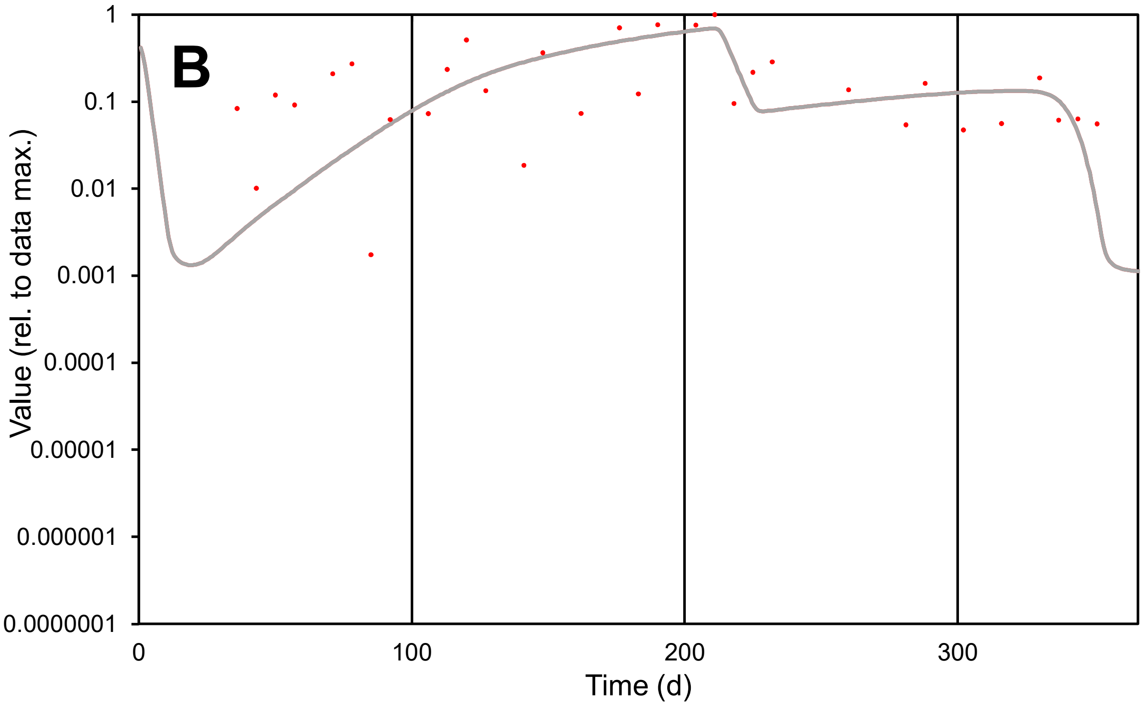


**C1**

Variables

| | |
|---|---|
| PLA | Provide jtd or Dst |
| CalibOnly? | |
| 2 | 1 to reproduce of |
| oNormalize? | |
| 2 | 1=no, 2=yes |
| ofxn calc* | |
| 0.75531 | |
| ofxn mod | |
| 0.77674 | |
| sRun | |
| RunA1 | |
| zout | |
| 1 | |
| Ave | |
| Data | |
| 0.01949 | |
| Model | |
| 0.01673 | |
| data/vMax | |
| 0.08867 | |
| ofxng | |
| 321.551 | |
| Min d/m | |
| 0.55396 | |
| ave/min d | |
| 126.128 | |
| ave/min m | |
| 59.971 | |

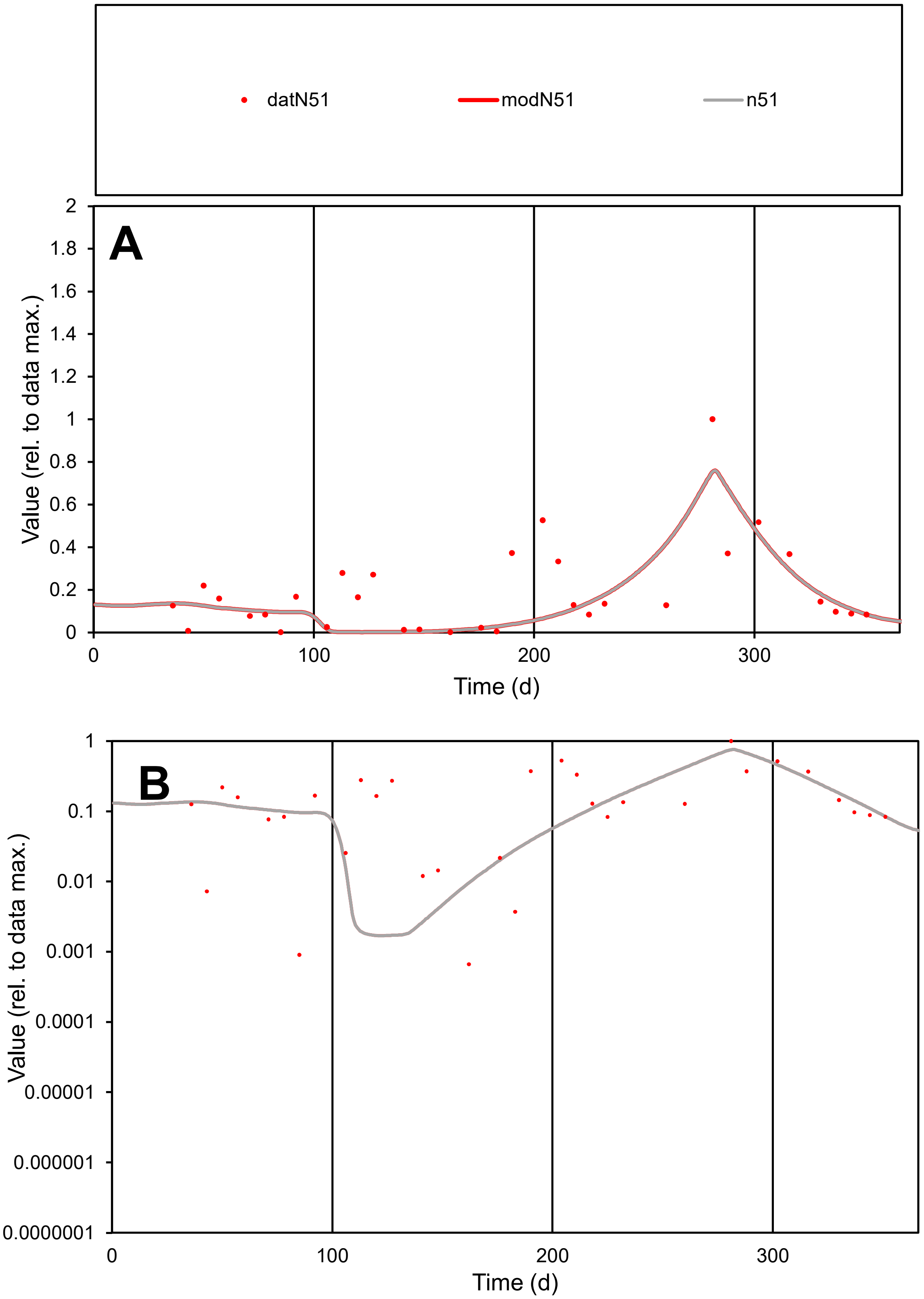


**C1**

Variables
N51 Provide jtd or Dst
CalibOnly?
2 1 to reproduce of
oNormalize?
2 1=no, 2=yes

ofxn calc*
0.86421
ofxn mod
0.87322

sRun
RunA1
zout
1

Ave
Data
0.01715
Model
0.01338

data/vMax
0.09136

ofxng
321.551

Min d/m
0.39187

ave/min d
282.544
ave/min m
86.3488

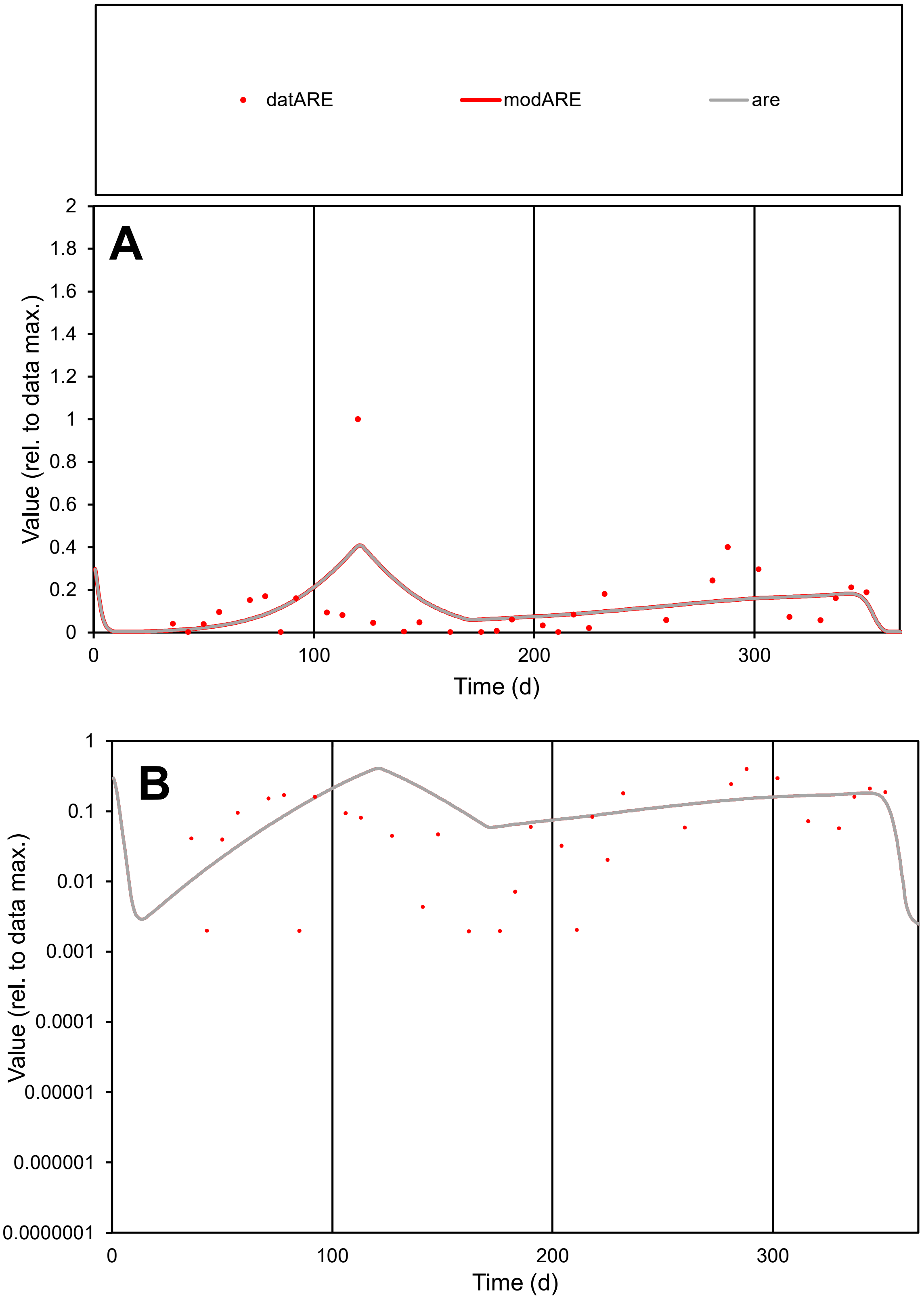


**C1**

Variables
ARE Provide jtd or Dst
CalibOnly?
2 1 to reproduce of
oNormalize?
2 1=no, 2=yes

ofxn calc*
1.22132
ofxn mod
1.24007

sRun
RunA1
zout
1

Ave
Data
0.0034
Model
0.00366

data/vMax
0.02708

ofxng
321.551

Min d/m
0.17782

ave/min d
63.9911
ave/min m
12.2537

324_Calib_S_A

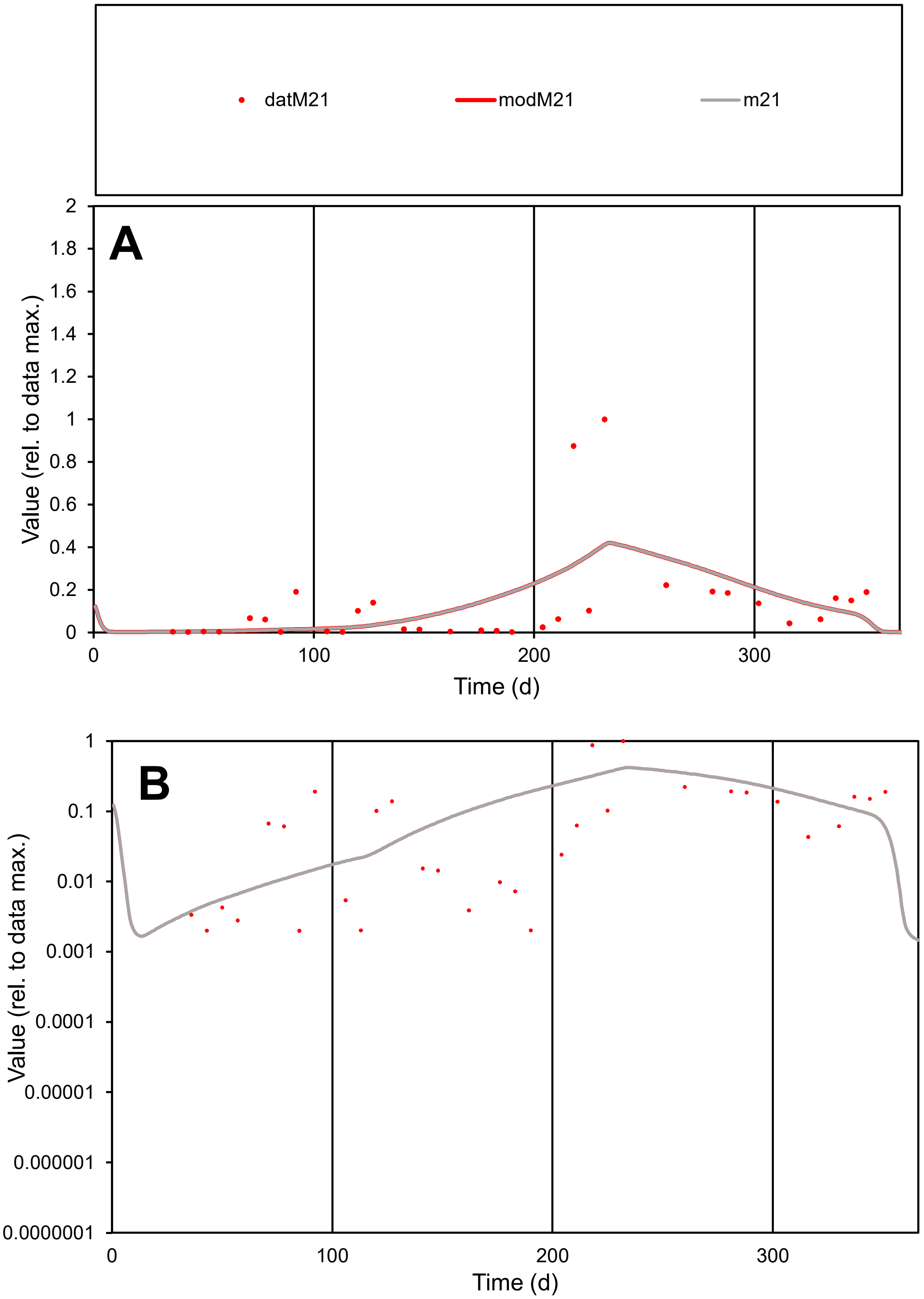


**C1**

Variables
M21 Provide jtd or Dst
CalibOnly?
2 1 to reproduce of
oNormalize?
2 1=no, 2=yes

ofxn calc*
1.41845
ofxn mod
1.43123

sRun
RunA1
zout
1

Ave
Data
0.00126
Model
0.0013

data/vMax
0.01001

ofxng
321.551

Min d/m
0.50873

ave/min d
63.7315
ave/min m
33.3484

324_Calib_S_A

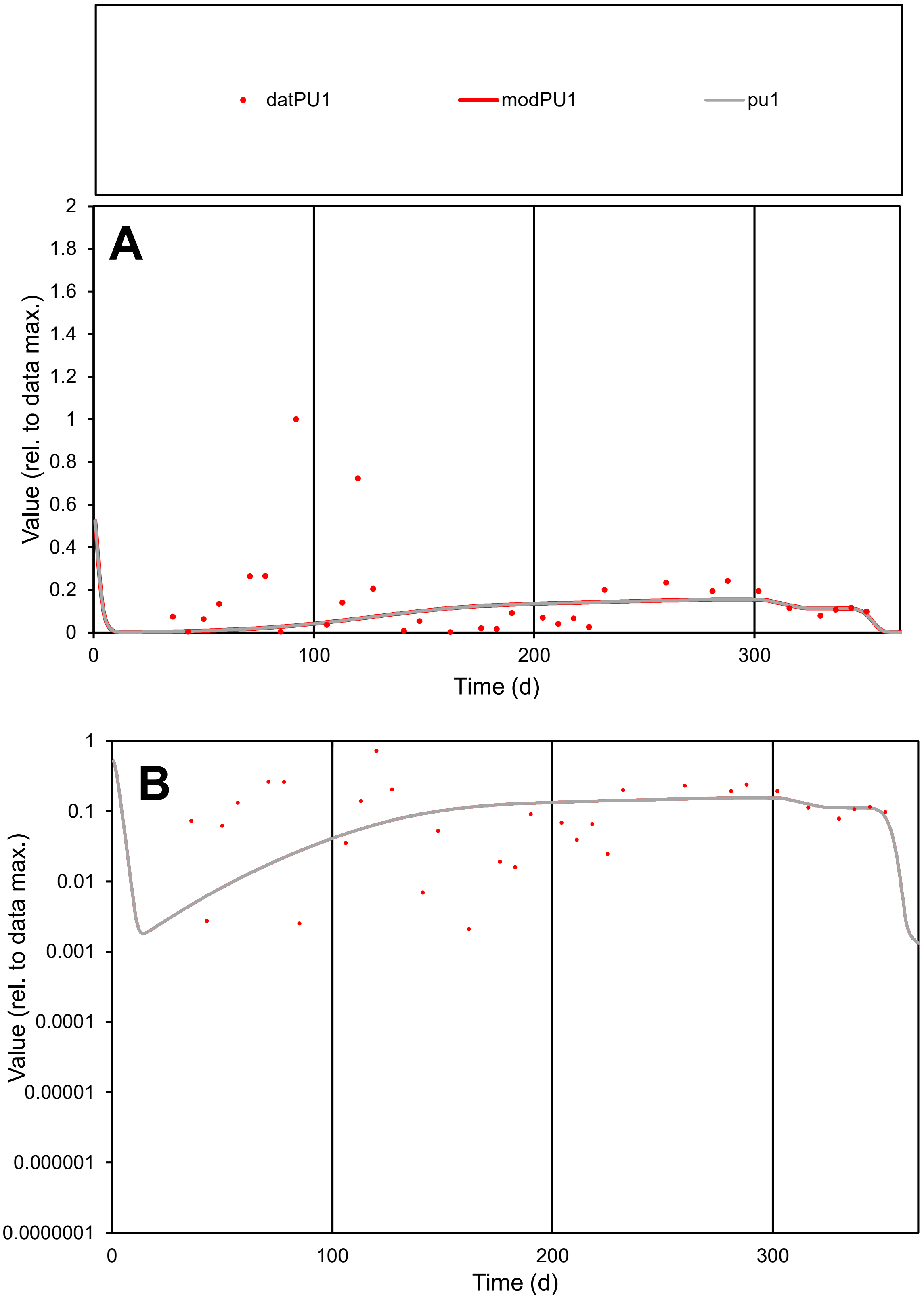


**C1**

Variables
PU1 Provide jtd or Dst
CalibOnly?
2 1 to reproduce of
oNormalize?
2 1=no, 2=yes

ofxn calc*
1.48493
ofxn mod
1.48794

sRun
RunA1
zout
1

Ave
Data
0.00435
Model
0.00262

data/vMax
0.02863

ofxng
321.551

Min d/m
0.43388

ave/min d
71.694
ave/min m
18.7336

324_Calib_S_A

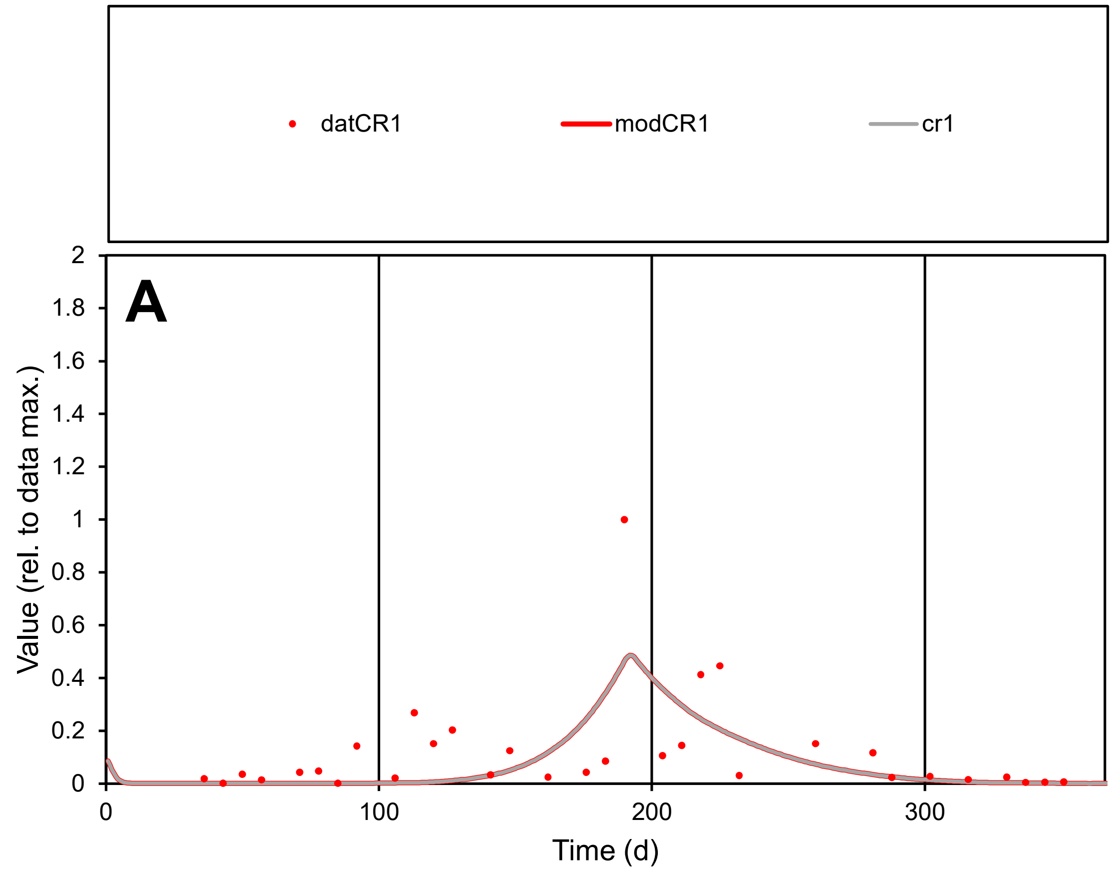


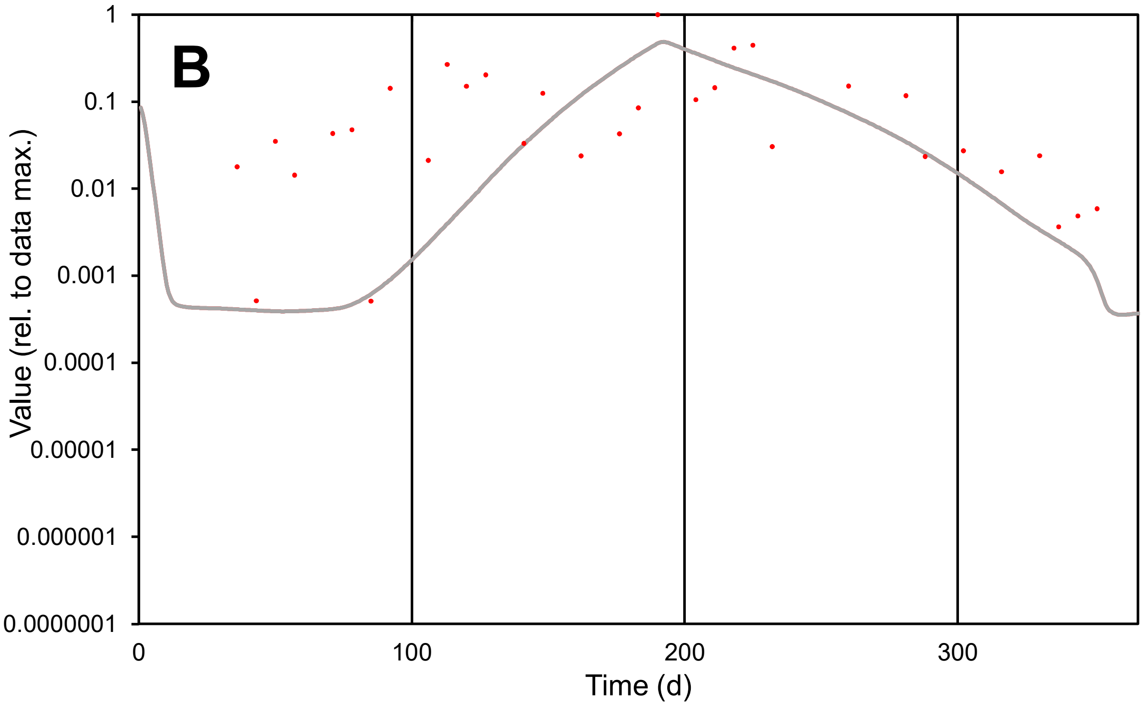


**C1**

Variables
CR1 Provide jtd or Dst
CalibOnly?
2 1 to reproduce of
oNormalize?
2 1=no, 2=yes

ofxn calc*
1.2958
ofxn mod
1.31119

sRun
RunA1
zout
1

Ave
Data
0.01501
Model
0.01084

data/vMax
0.1276

ofxng
321.551

Min d/m
1.30584

ave/min d
230.64
ave/min m
217.624

324_Calib_S_A

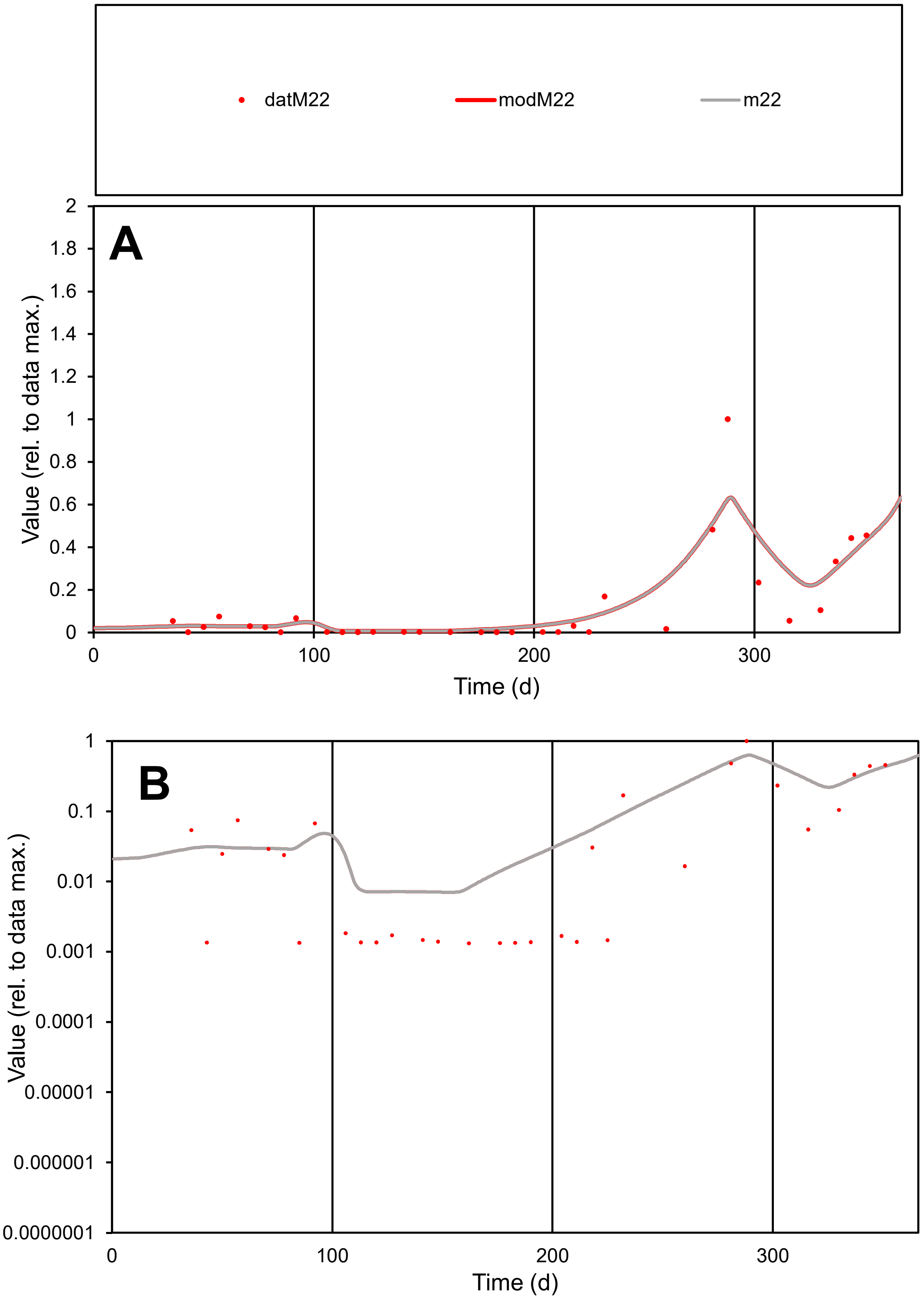


**C1**

Variables

M22 Provide jtd or Dst

CalibOnly?

2 1 to reproduce of

oNormalize?

2 1=no, 2=yes

ofxn calc*

0.87817

ofxn mod

0.90441

sRun

RunA1

zout

1

Ave

Data

0.0018

Model

0.00207

data/vMax

0.01596

ofxng

321.551

Min d/m

0.18798

ave/min d

85.4301

ave/min m

18.4327

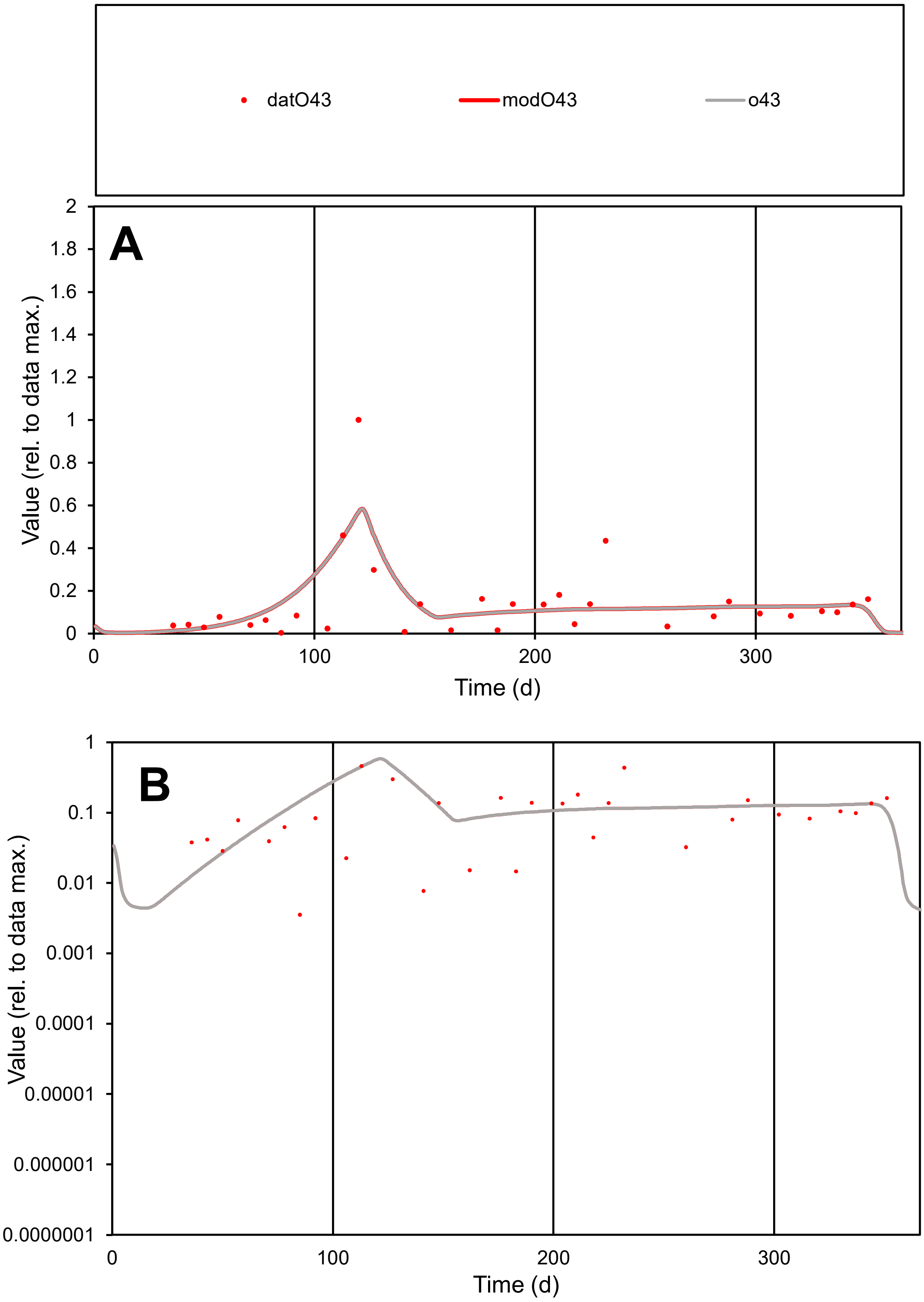


**C1**

Variables
O43 Provide jtd or Dst
CalibOnly?
2 1 to reproduce of
oNormalize?
2 1=no, 2=yes

ofxn calc*
0.901
ofxn mod
0.91521

sRun
RunA1
zout
1

Ave
Data
0.00557
Model
0.00597

data/vMax
0.03966

ofxng
321.551

Min d/m
0.26289

ave/min d
39.7632
ave/min m
11.1859

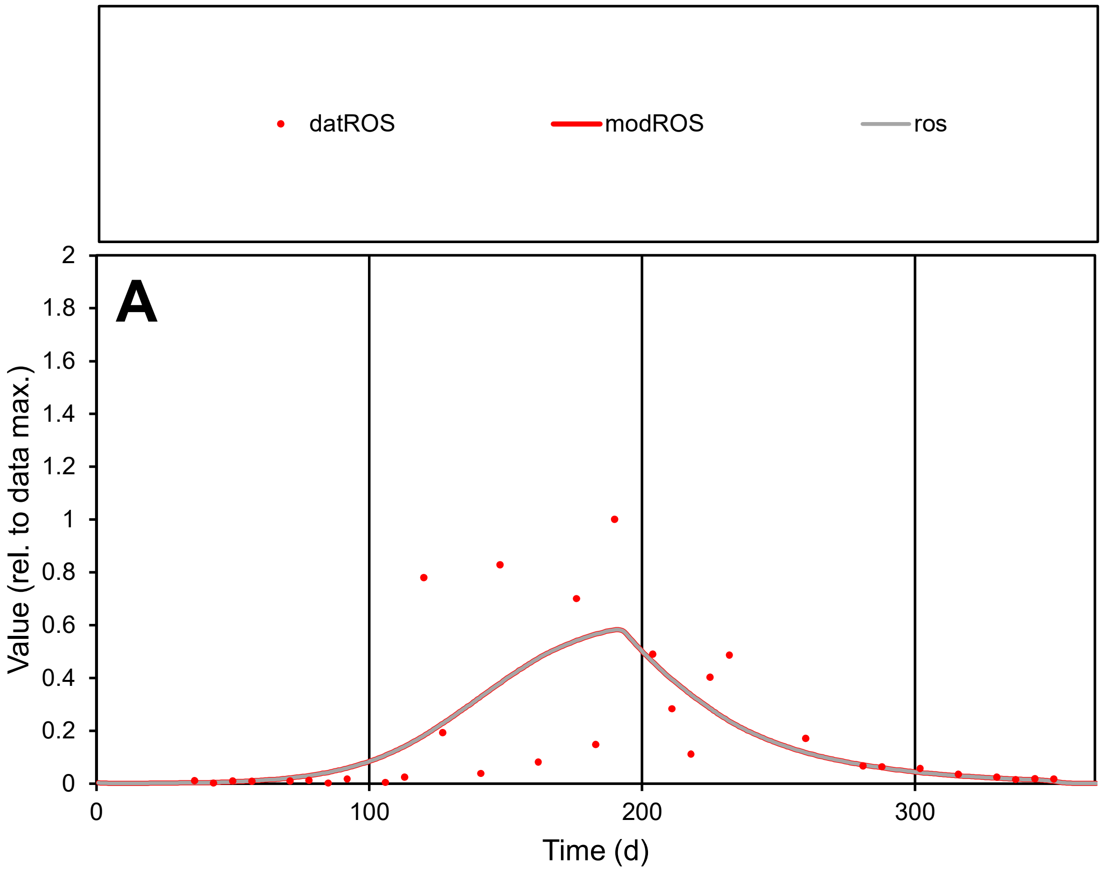


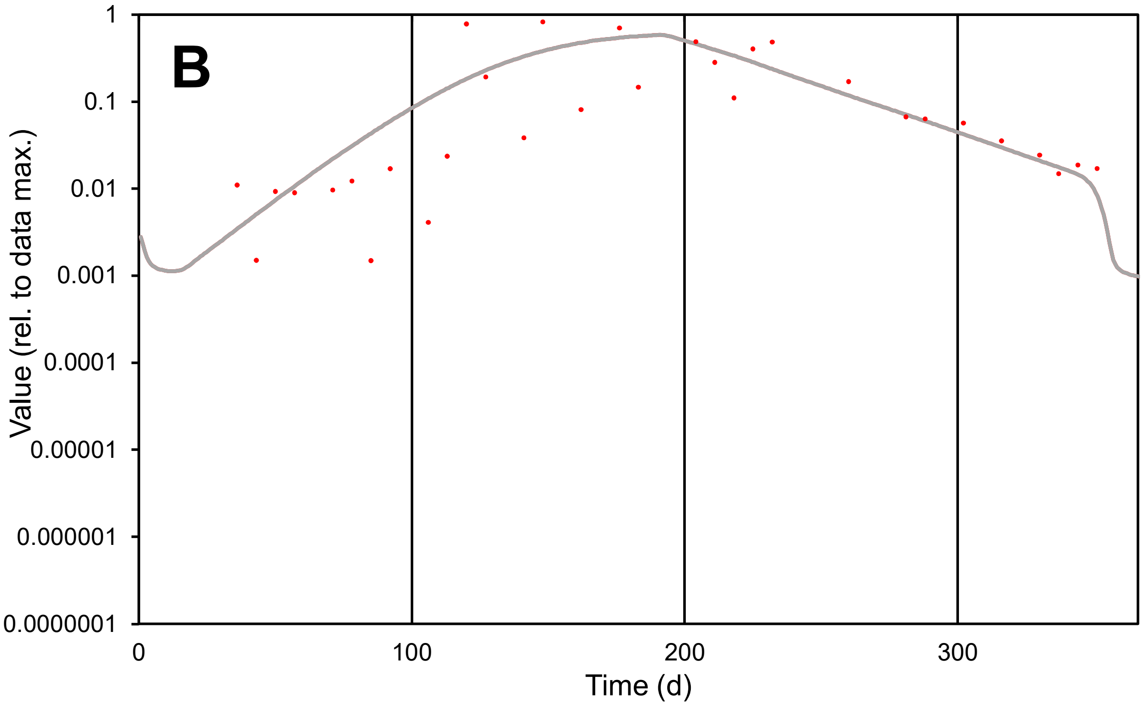


**C1**

Variables
ROS Provide jtd or Dst
CalibOnly?
2 1 to reproduce of
oNormalize?
2 1=no, 2=yes

ofxn calc*
1.06655
ofxn mod
1.07898

sRun
RunA1
zout
1

Ave
Data
0.00633
Model
0.00601

data/vMax
0.03317

ofxng
321.551

Min d/m
0.40923

ave/min d
127.557
ave/min m
49.5101

324_Calib_S_A

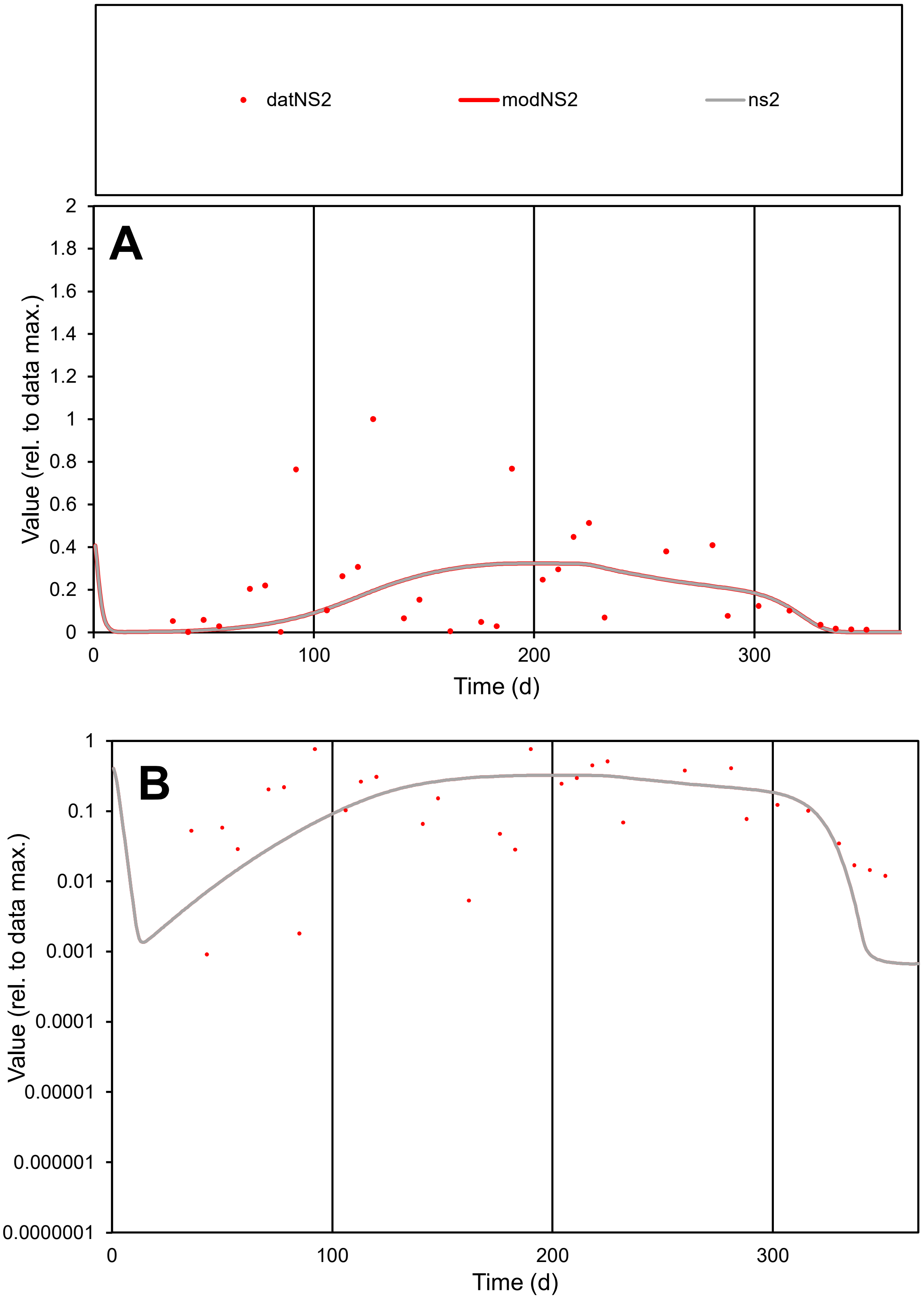


**C1**

Variables

NS2 Provide jtd or Dst

CalibOnly?

2 1 to reproduce of

oNormalize?

2 1=no, 2=yes

ofxn calc*

1.14439

ofxn mod

1.14947

sRun

RunA1

zout

1

Ave

Data

0.01338

Model

0.01022

data/vMax

0.06286

ofxng

321.551

Min d/m

1.27904

ave/min d

233.709

ave/min m

228.41

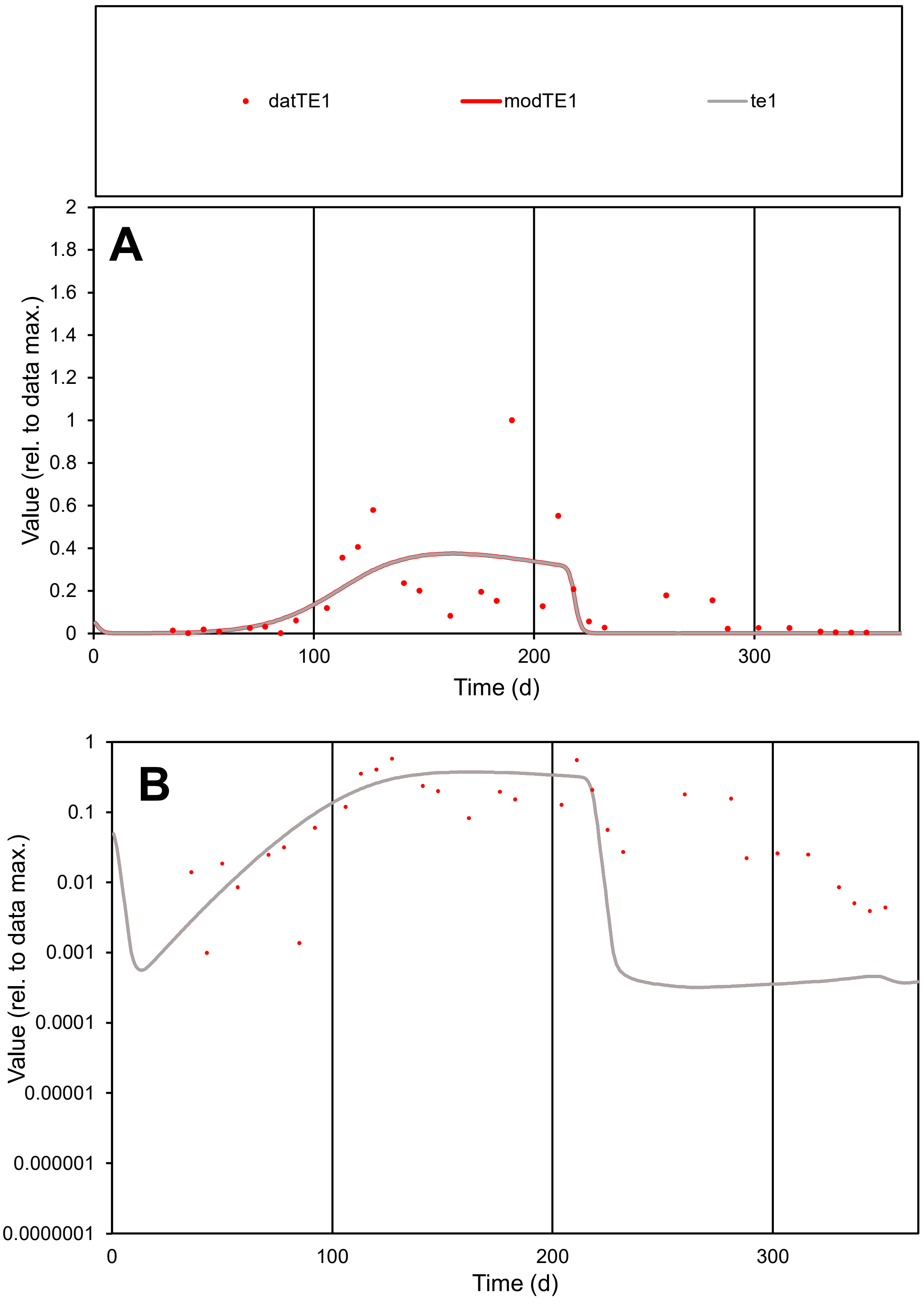


**C1**

Variables

TE1 Provide jtd or Dst

CalibOnly?

2 1 to reproduce of

oNormalize?

2 1=no, 2=yes

ofxn calc*

1.09705

ofxn mod

1.10877

sRun

RunA1

zout

1

Ave

Data

0.01612

Model

0.01413

data/vMax

0.1056

ofxng

321.551

Min d/m

3.07919

ave/min d

154.111

ave/min m

416.086

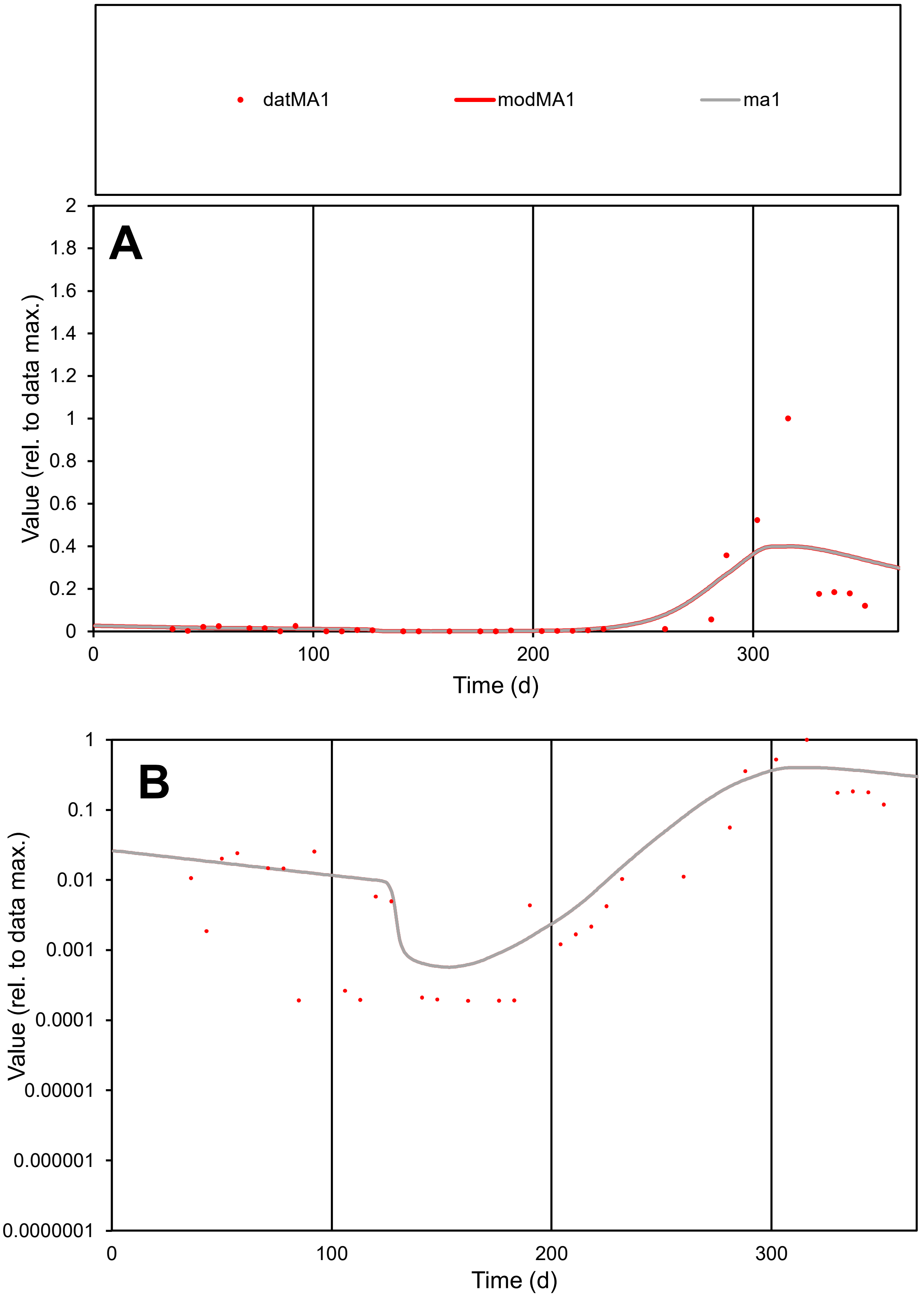


**C1**

Variables
MA1 Provide jtd or Dst
CalibOnly?
2 1 to reproduce of
oNormalize?
2 1=no, 2=yes

ofxn calc*
1.55026
ofxn mod
1.56452

sRun
RunA1
zout
1

Ave
Data
0.00643
Model
0.00701

data/vMax
0.07472

ofxng
321.551

Min d/m
0.32588

ave/min d
454.56
ave/min m
161.554

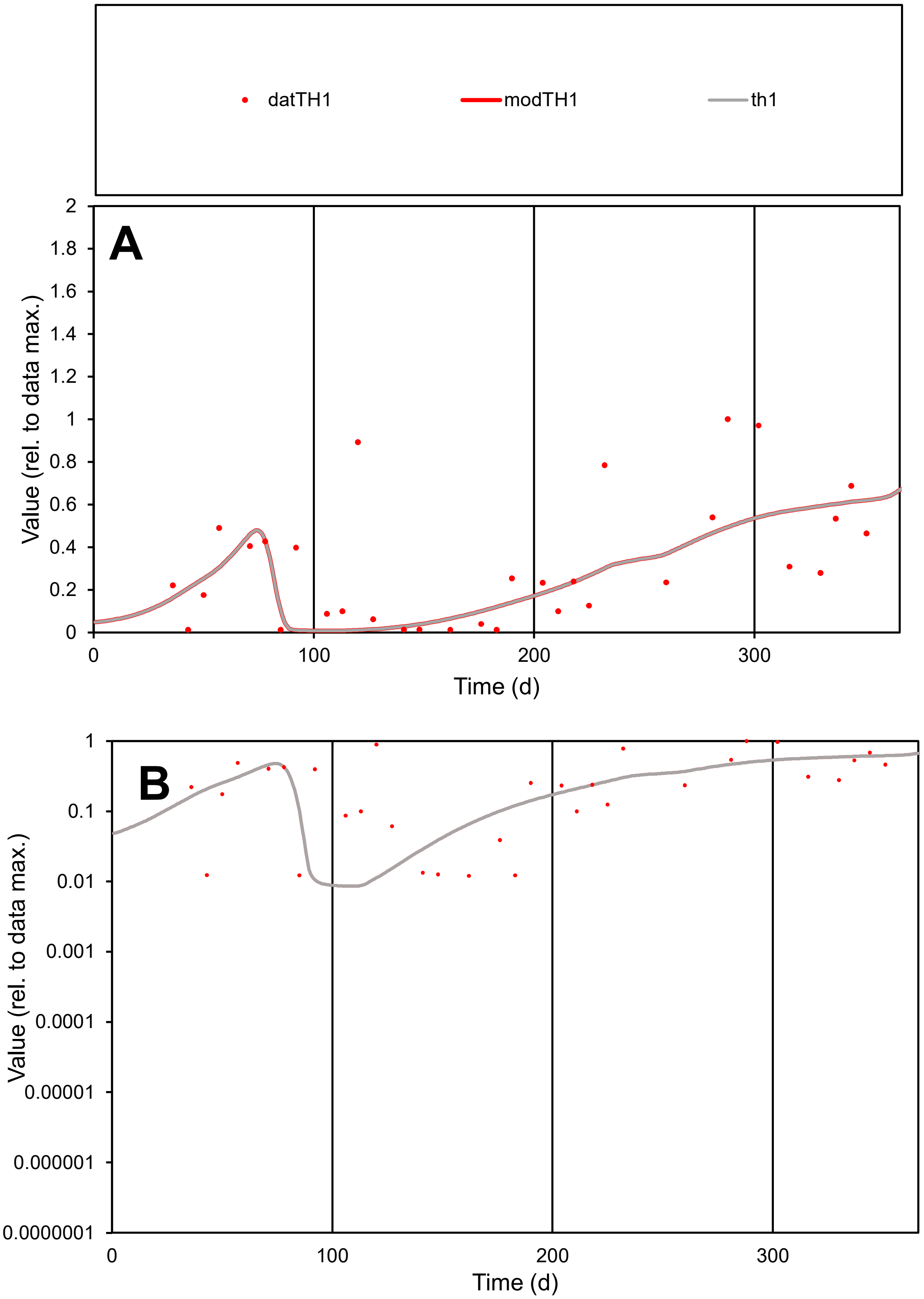


**C1**

Variables

TH1 Provide jtd or Dst

CalibOnly?

2 1 to reproduce of

oNormalize?

2 1=no, 2=yes

ofxn calc*

0.78604

ofxn mod

0.79468

sRun

RunA1

zout

1

Ave

Data

0.00239

Model

0.00201

data/vMax

0.00756

ofxng

321.551

Min d/m

1.39755

ave/min d

26.233

ave/min m

30.8464

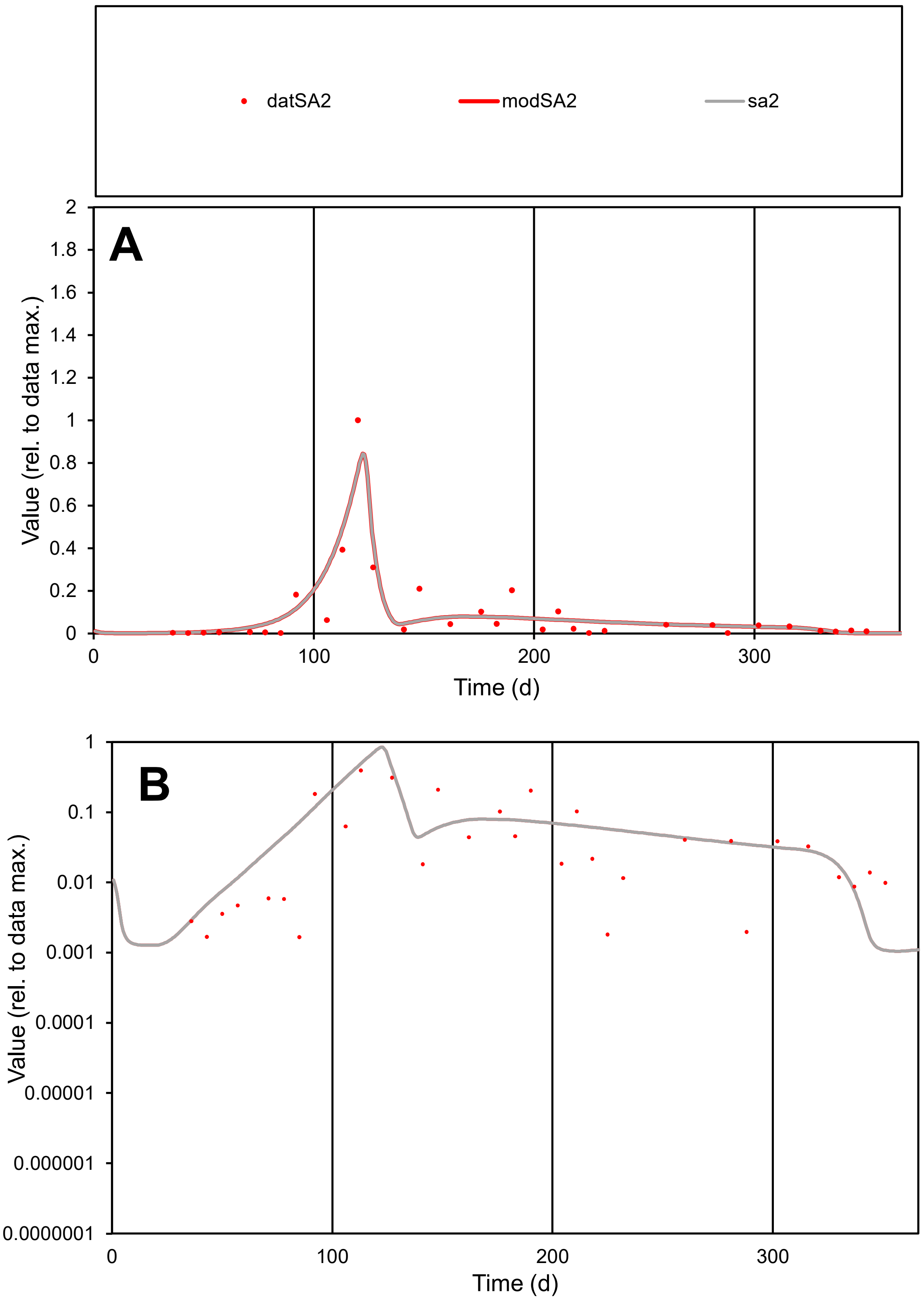


**C1**

Variables

SA2 Provide jtd or Dst

CalibOnly?

2 1 to reproduce of

oNormalize?

2 1=no, 2=yes

ofxn calc*

0.79051

ofxn mod

0.83309

sRun

RunA1

zout

1

Ave

Data

0.00561

Model

0.00612

data/vMax

0.06091

ofxng

321.551

Min d/m

1.55682

ave/min d

55.5189

ave/min m

94.3152

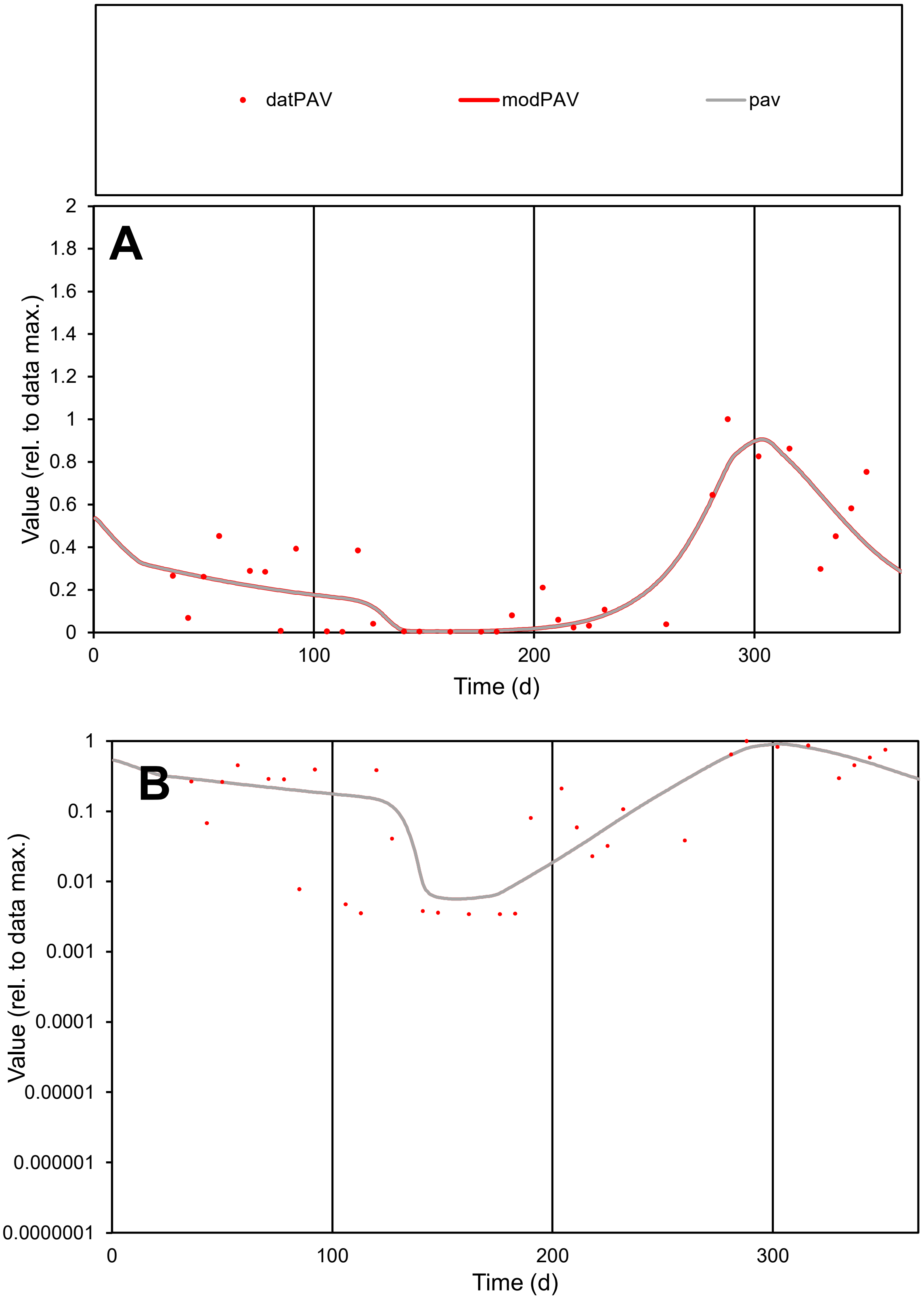


**C1**

Variables
PAV Provide jtd or Dst
CalibOnly?
2 1 to reproduce of
oNormalize?
2 1=no, 2=yes

ofxn calc*
0.55687
ofxn mod
0.57221

sRun
RunA1
zout
1

Ave
Data
0.00281
Model
0.00274

data/vMax
0.01064

ofxng
321.551

Min d/m
0.59374

ave/min d
76.9879
ave/min m
44.6126

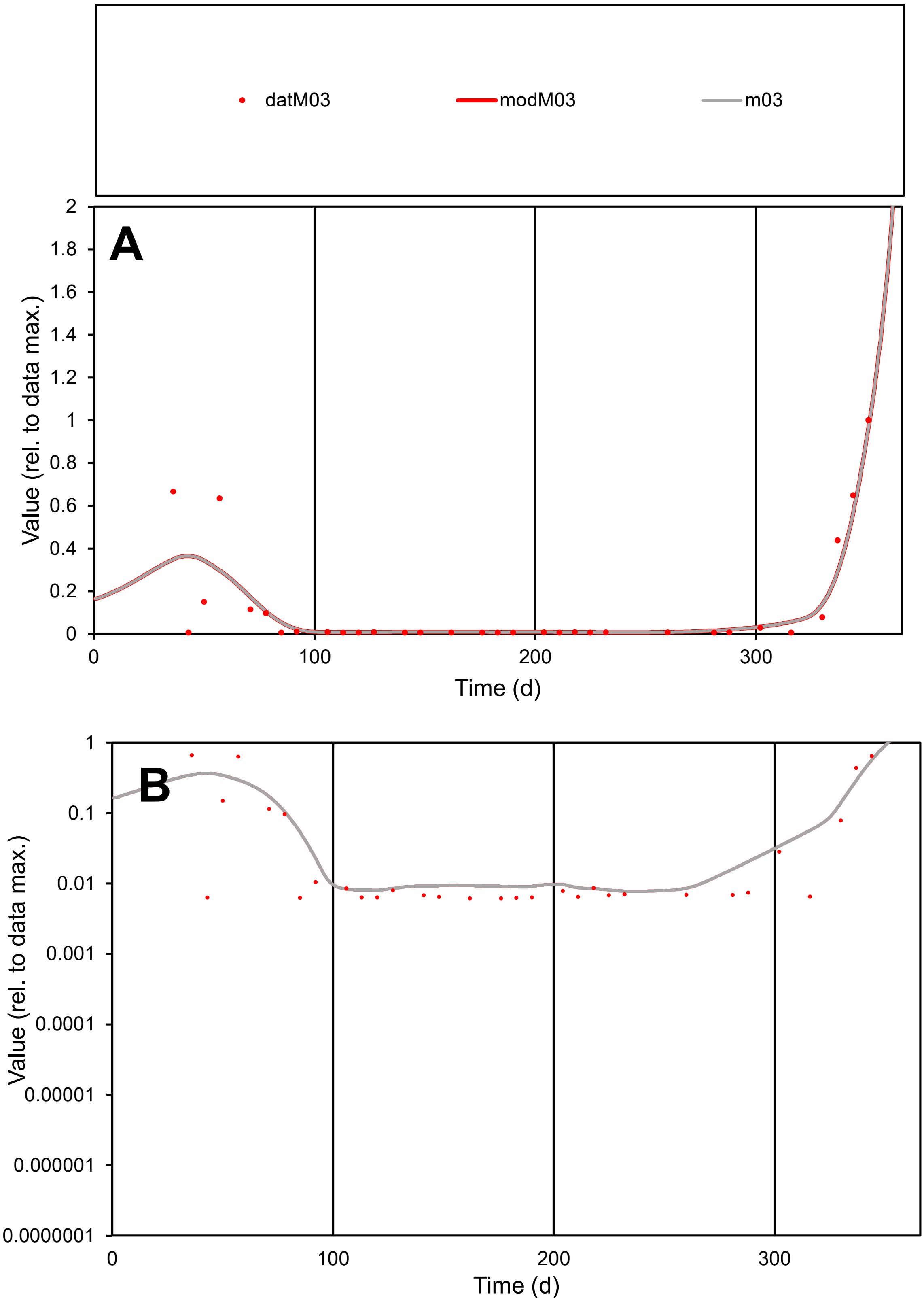


**C1**

Variables
M03 Provide jtd or Dst
CalibOnly?
2 1 to reproduce of
oNormalize?
2 1=no, 2=yes

ofxn calc*
0.89015
ofxn mod
0.90805

sRun
RunA1
zout
1

Ave
Data
0.00035
Model
0.00036

data/vMax
0.00279

ofxng
321.551

Min d/m
0.78836

ave/min d
20.3356
ave/min m
16.2996

324_Calib_S_A

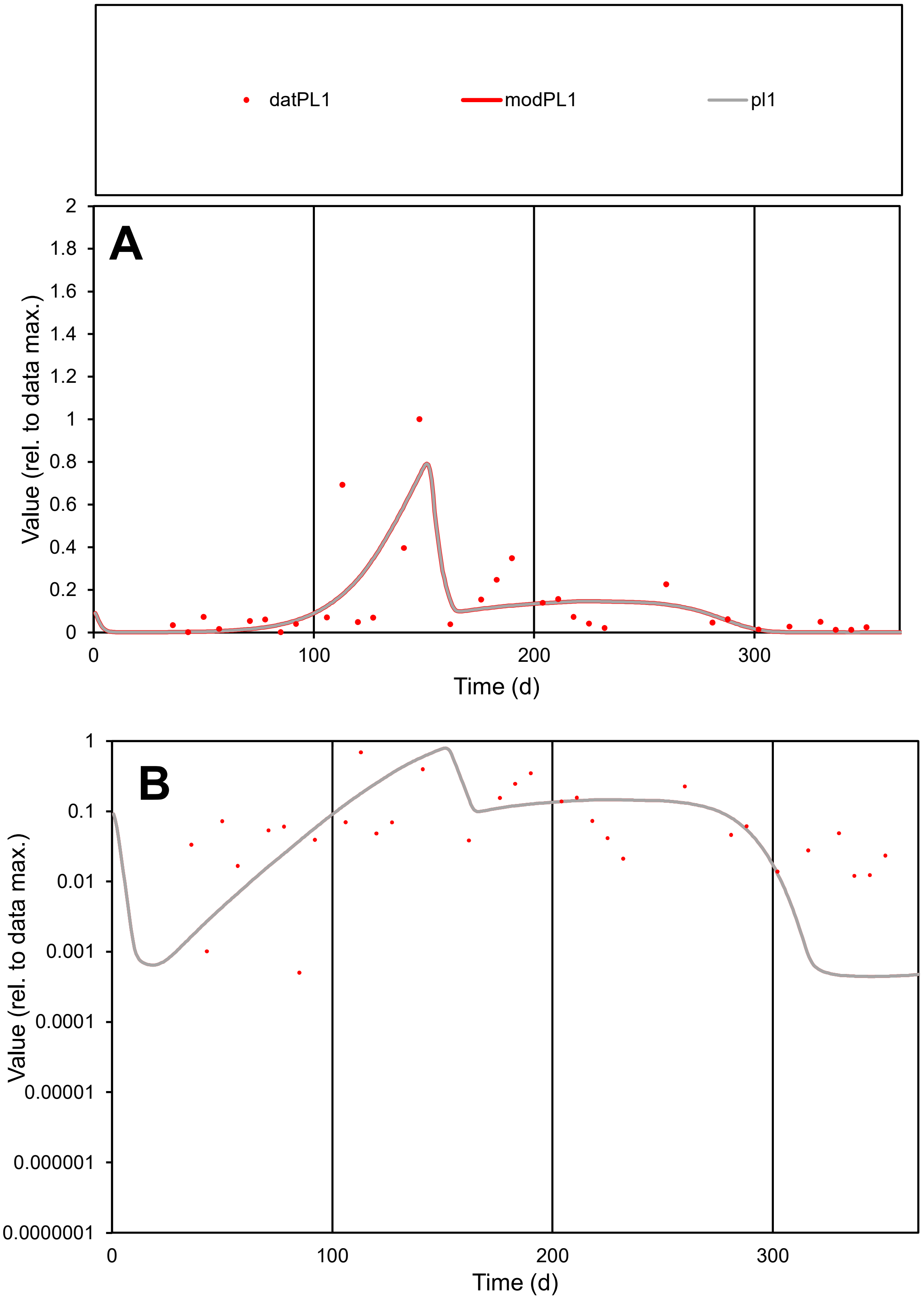


**C1**

Variables

| PL1 | Provide jtd or Dst |
|---|---|
| CalibOnly? | |
| 2 | 1 to reproduce of |
| oNormalize? | |
| 2 | 1=no, 2=yes |

ofxn calc*
1.0344
ofxn mod
1.05488

sRun
RunA1
zout
1

Ave
Data
0.00909
Model
0.00834

data/vMax
0.06855

ofxng
321.551

Min d/m
1.13377

ave/min d
262.93
ave/min m
273.538

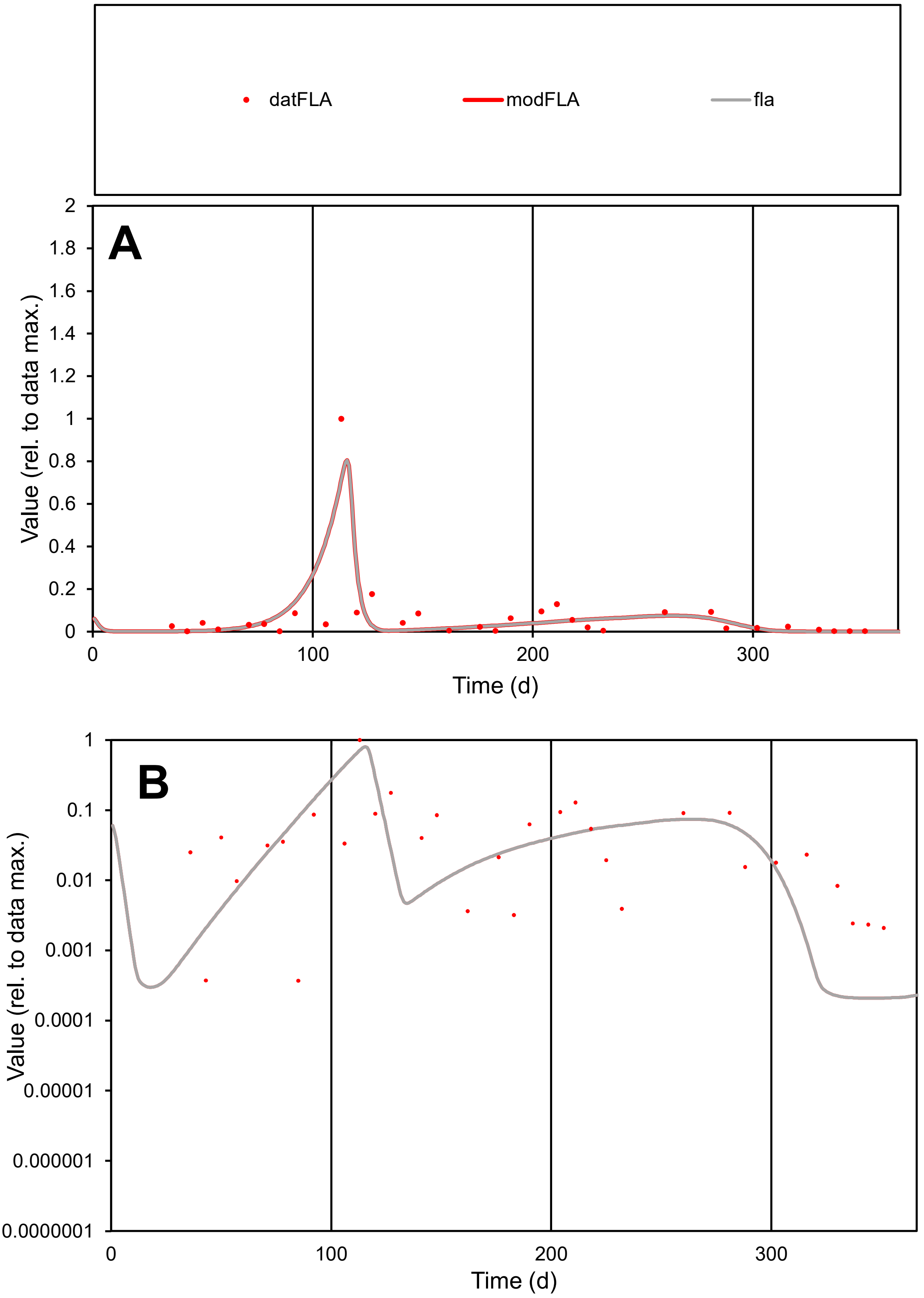


**C1**

Variables
FLA Provide jtd or Dst
CalibOnly?
2 1 to reproduce of
oNormalize?
2 1=no, 2=yes

ofxn calc*
1.14451
ofxn mod
1.28731

sRun
RunA1
zout
1

Ave
Data
0.0106
Model
0.01022

data/vMax
0.1475

ofxng
321.551

Min d/m
1.7846

ave/min d
193.835
ave/min m
333.422

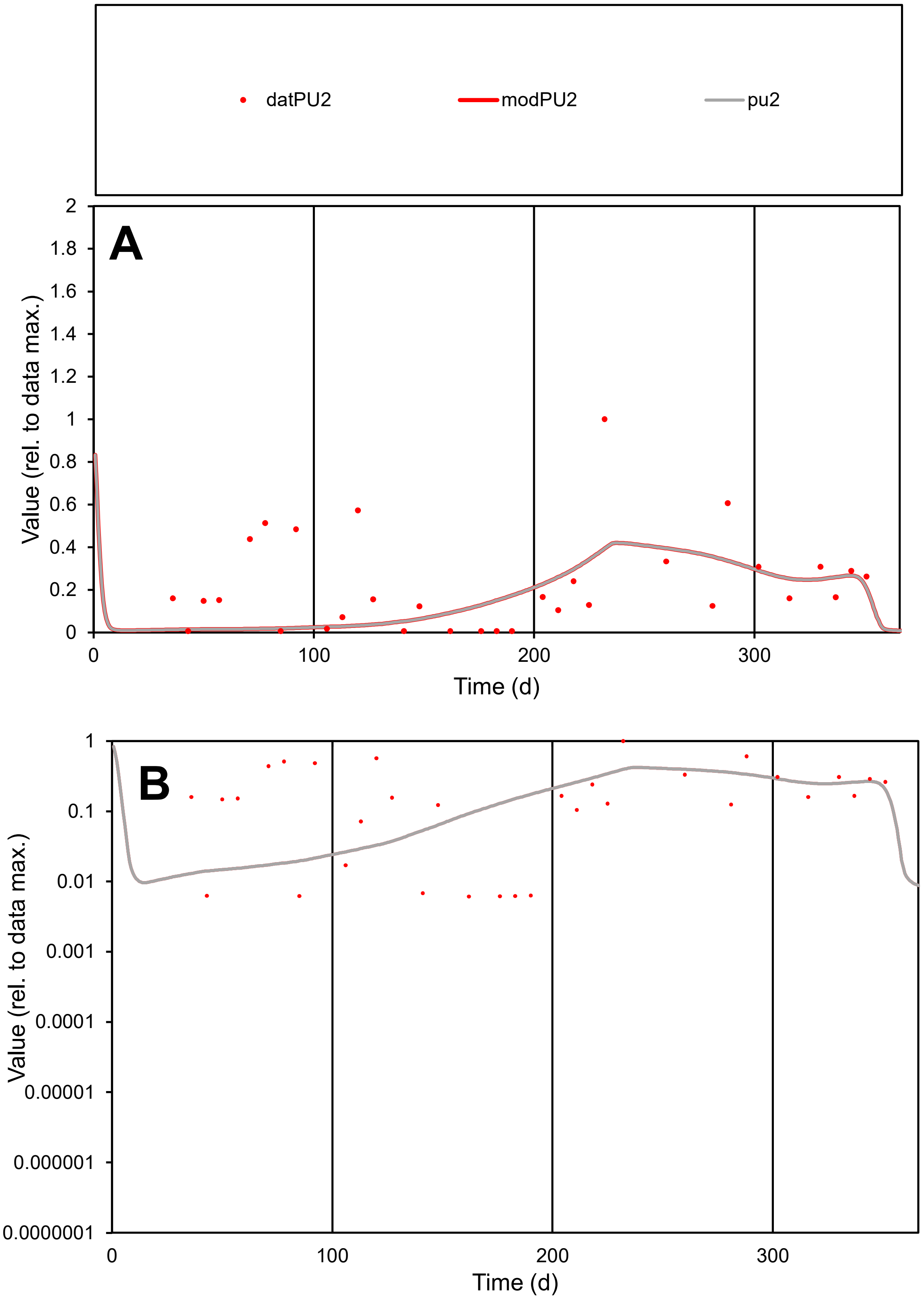


**C1**

Variables
PU2 Provide jtd or Dst
CalibOnly?
2 1 to reproduce of
oNormalize?
2 1=no, 2=yes

ofxn calc*
1.03295
ofxn mod
1.03953

sRun
RunA1
zout
1

Ave
Data
0.00194
Model
0.00139

data/vMax
0.00876

ofxng
321.551

Min d/m
0.46385

ave/min d
36.0381
ave/min m
12.0117

324_Calib_S_A

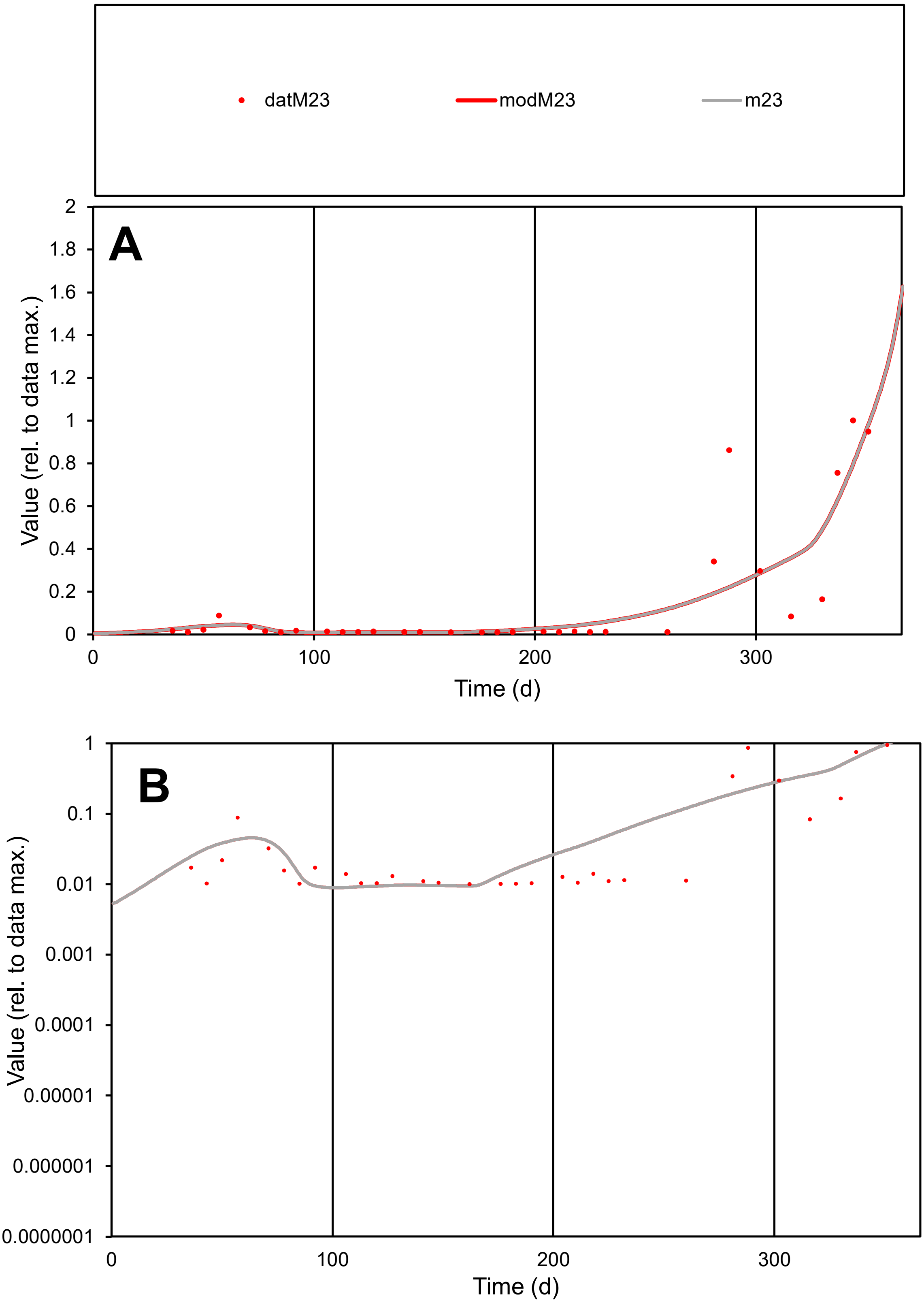


**C1**

Variables
M23 Provide jtd or Dst
CalibOnly?
2 1 to reproduce of
oNormalize?
2 1=no, 2=yes

ofxn calc*
0.95533
ofxn mod
0.97269

sRun
RunA1
zout
1

Ave
Data
0.00159
Model
0.00156

data/vMax
0.0105

ofxng
321.551

Min d/m
1.11381

ave/min d
15.0836
ave/min m
16.4779

324_Calib_S_A

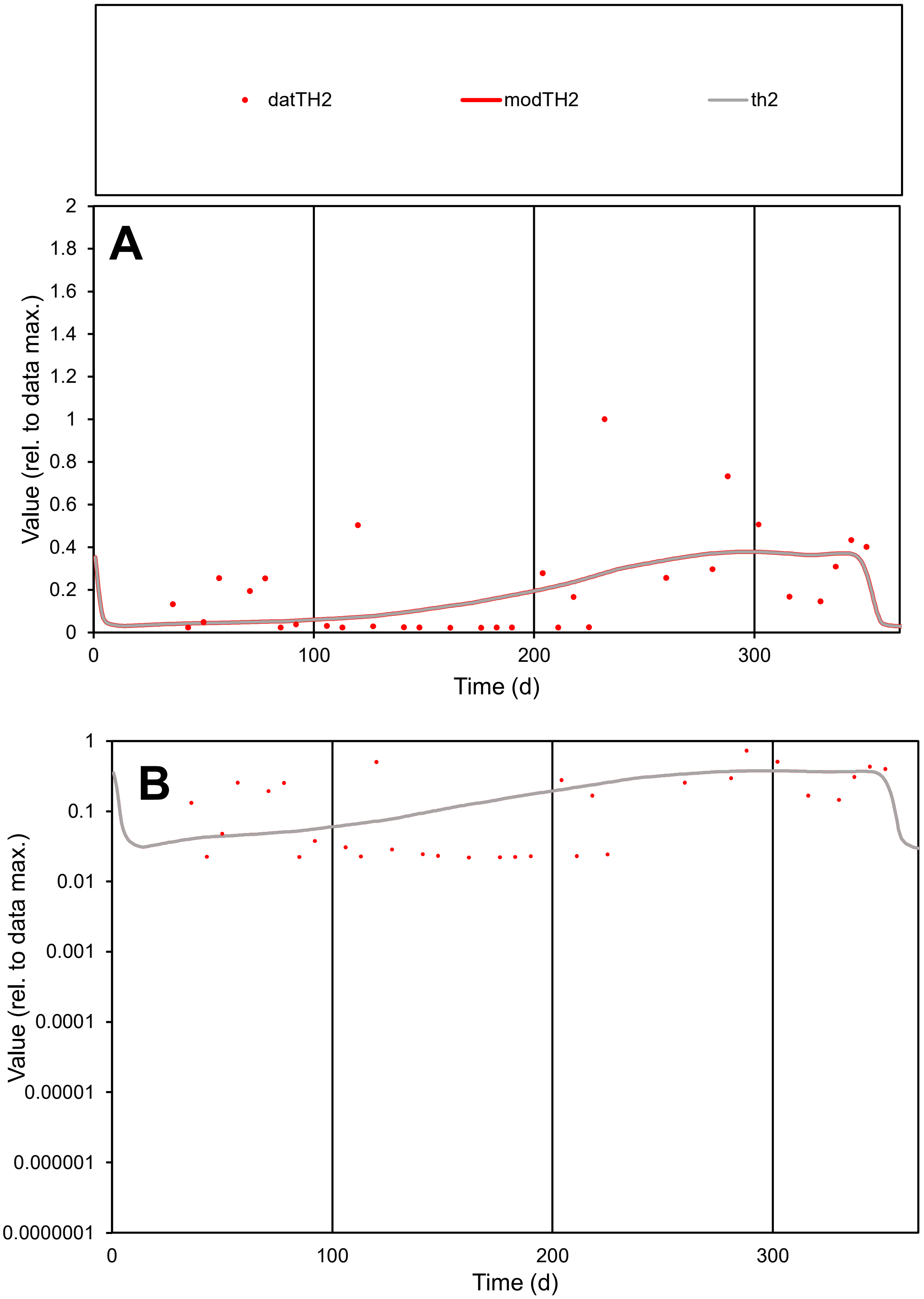


**C1**

Variables
TH2 Provide jtd or Dst
CalibOnly?
2 1 to reproduce of
oNormalize?
2 1=no, 2=yes

ofxn calc*
0.99476
ofxn mod
1.00382

sRun
RunA1
zout
1

Ave
Data
0.00094
Model
0.00086

data/vMax
0.0047

ofxng
321.551

Min d/m
0.53982

ave/min d
9.09682
ave/min m
4.50171

324_Calib_S_A

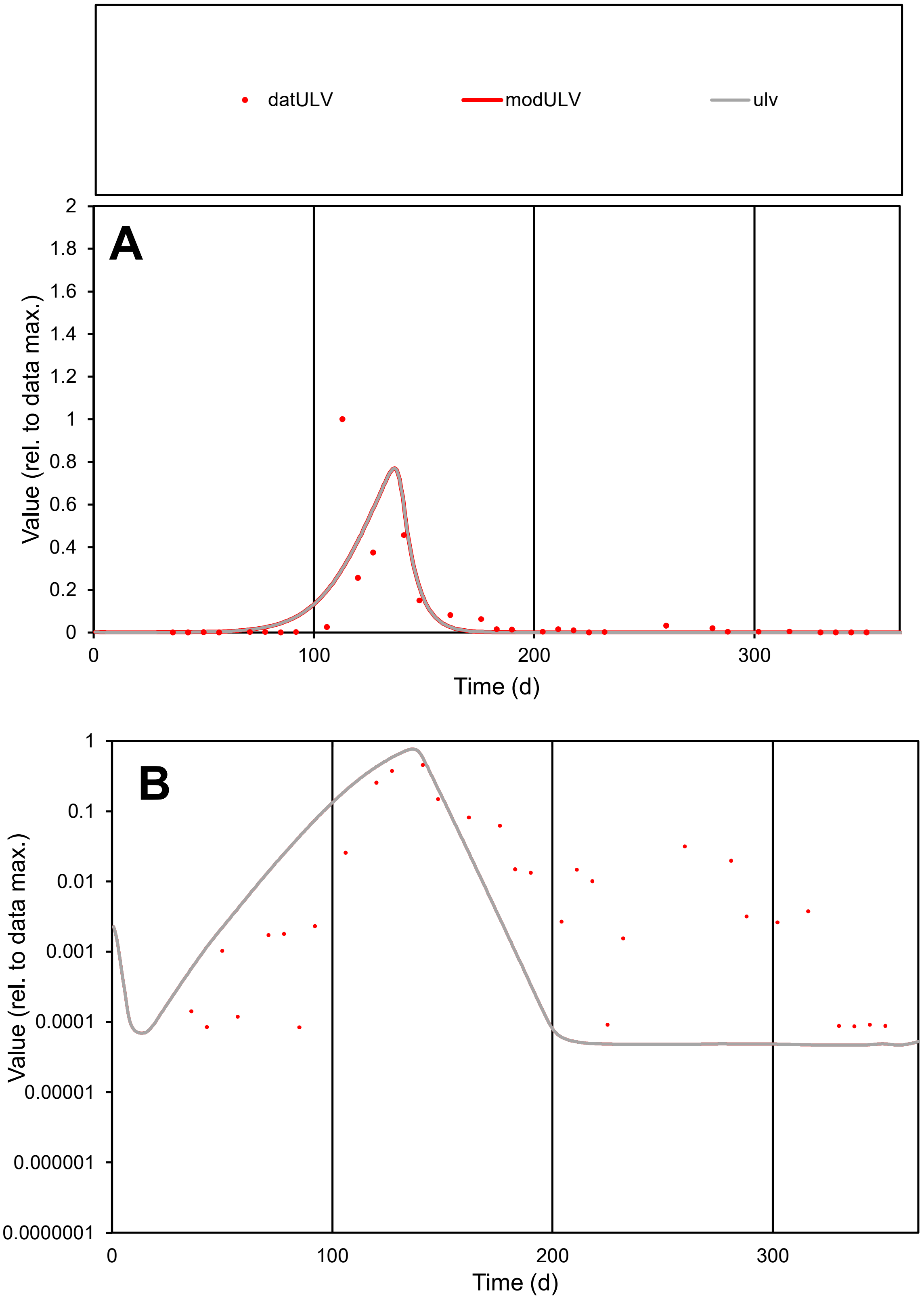


**C1**

Variables
ULV Provide jtd or Dst
CalibOnly?
2 1 to reproduce of
oNormalize?
2 1=no, 2=yes

ofxn calc*
1.71419
ofxn mod
1.74207

sRun
RunA1
zout
1

Ave
Data
0.01612
Model
0.01508

data/vMax
0.2037

ofxng
321.551

Min d/m
1.79005

ave/min d
941.729
ave/min m
1576.99

324_Calib_S_A

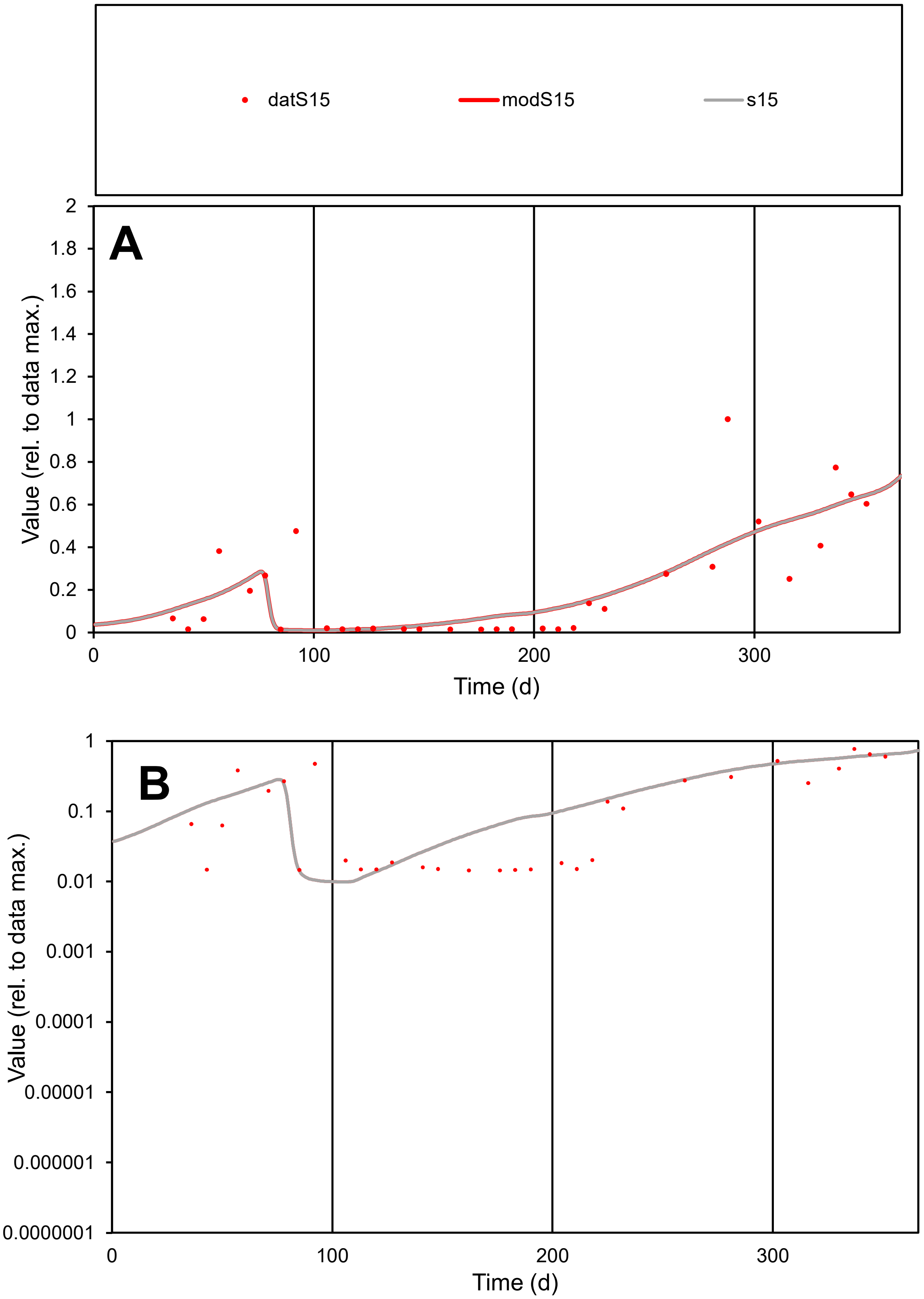


**C1**

Variables
S15 Provide jtd or Dst
CalibOnly?
2 1 to reproduce of
oNormalize?
2 1=no, 2=yes

ofxn calc*
0.75716
ofxn mod
0.76975

sRun
RunA1
zout
1

Ave
Data
0.00178
Model
0.00178

data/vMax
0.00847

ofxng
321.551

Min d/m
1.454

ave/min d
14.6064
ave/min m
21.2892

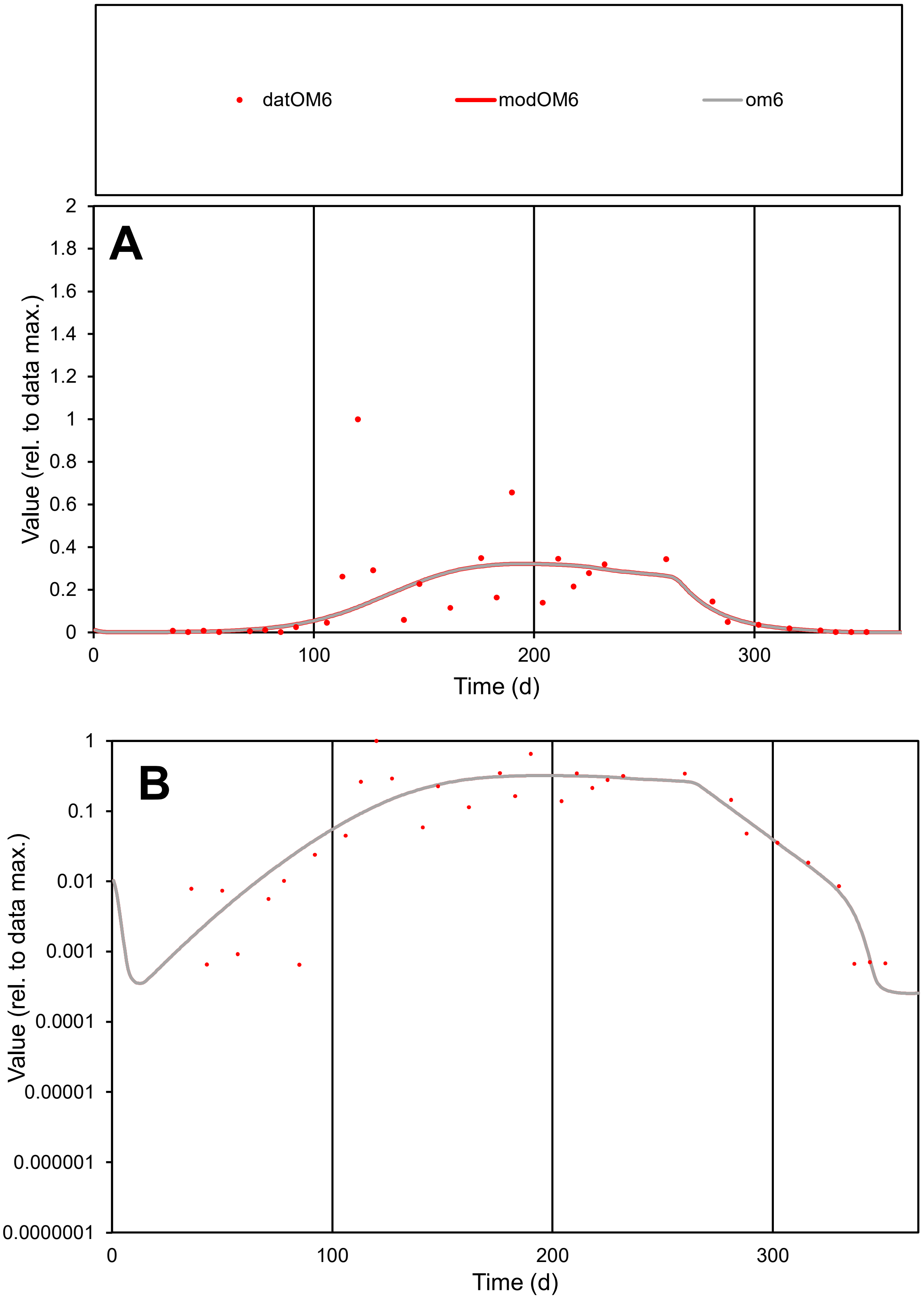


**C1**

Variables
OM6 Provide jtd or Dst
CalibOnly?
2 1 to reproduce of
oNormalize?
2 1=no, 2=yes

ofxn calc*
1.13717
ofxn mod
1.1448

sRun
RunA1
zout
1

Ave
Data
0.00615
Model
0.00517

data/vMax
0.03846

ofxng
321.551

Min d/m
2.31628

ave/min d
246.963
ave/min m
480.612

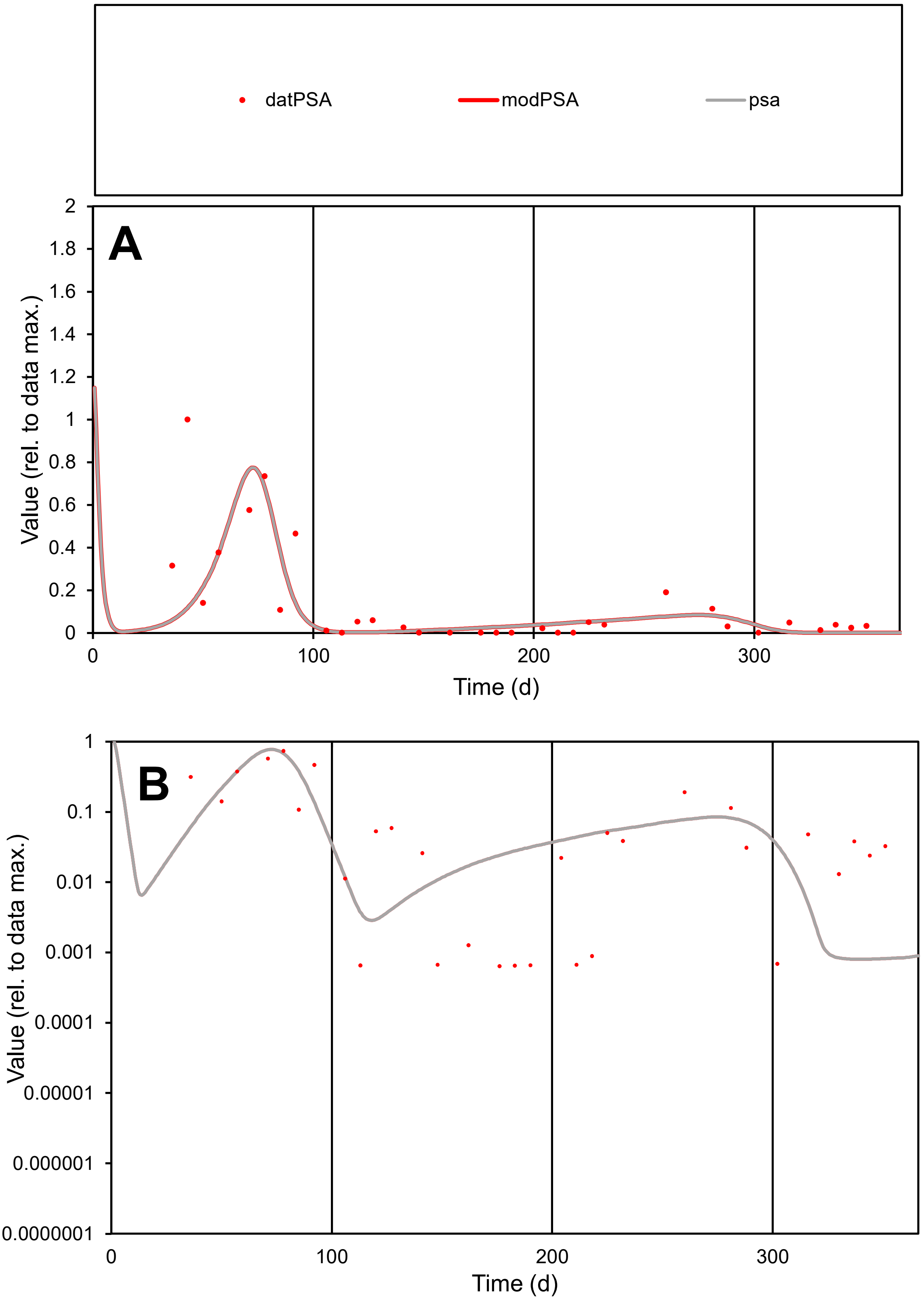


**C1**

Variables

PSA Provide jtd or Dst

CalibOnly?

2 1 to reproduce of

oNormalize?

2 1=no, 2=yes

ofxn calc*

1.29206

ofxn mod

1.30818

sRun

RunA1

zout

1

Ave

Data

0.0019

Model

0.00141

data/vMax

0.01362

ofxng

321.551

Min d/m

0.79817

ave/min d

218.654

ave/min m

128.847

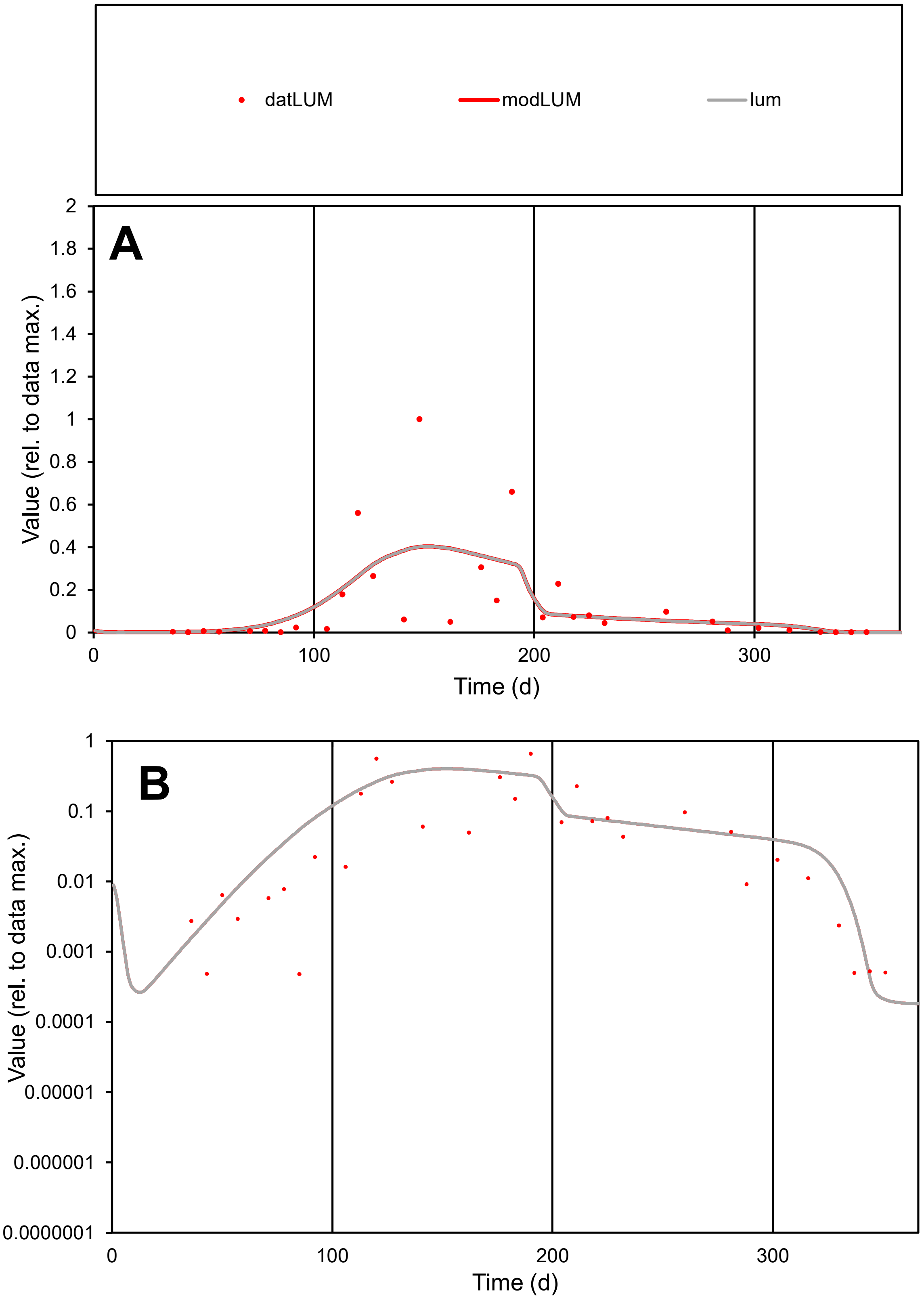


**C1**

Variables
LUM Provide jtd or Dst
CalibOnly?
2 1 to reproduce of
oNormalize?
2 1=no, 2=yes

ofxn calc*
1.32592
ofxn mod
1.33658

sRun
RunA1
zout
1

Ave
Data
0.01148
Model
0.01153

data/vMax
0.09231

ofxng
321.551

Min d/m
2.37567

ave/min d
257.957
ave/min m
615.358

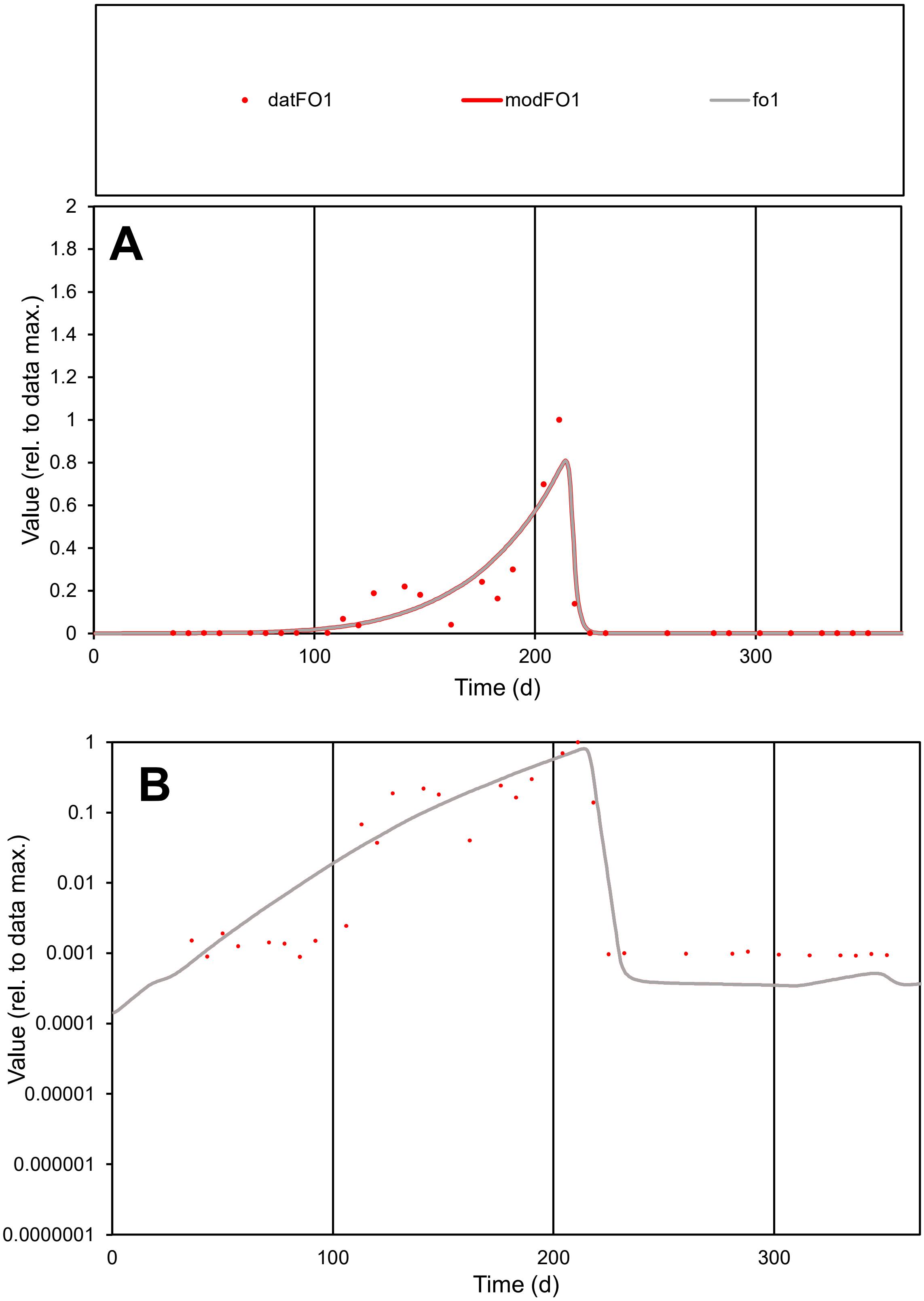


**C1**

Variables
FO1 Provide jtd or Dst
CalibOnly?
2 1 to reproduce of
oNormalize?
2 1=no, 2=yes

ofxn calc*
0.71603
ofxn mod
0.75912

sRun
RunA1
zout
1

Ave
Data
0.00669
Model
0.00684

data/vMax
0.06485

ofxng
321.551

Min d/m
2.57035

ave/min d
115.439
ave/min m
303.625

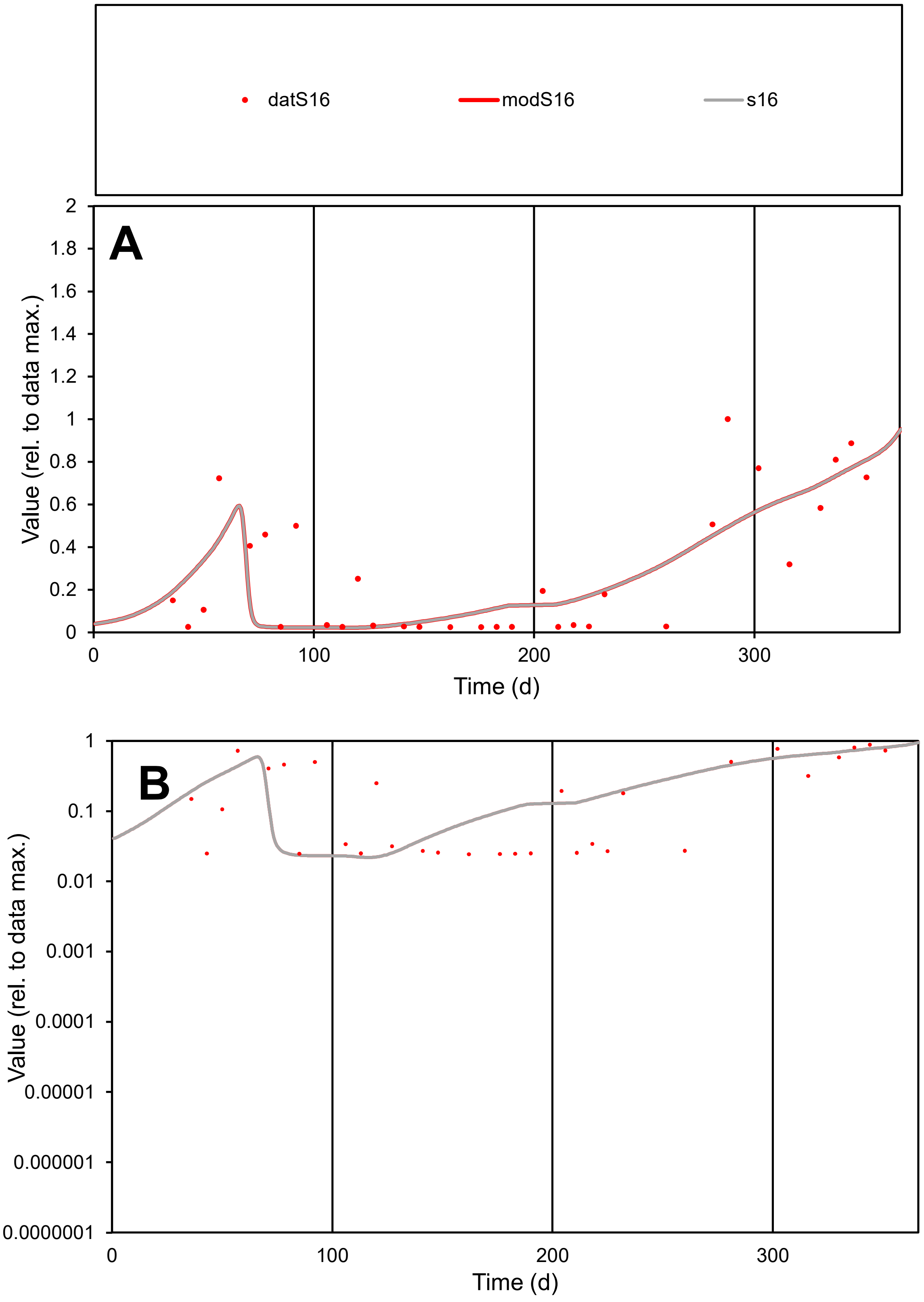


**C1**

Variables

S16 Provide jtd or Dst

CalibOnly?

2 1 to reproduce of

oNormalize?

2 1=no, 2=yes

ofxn calc*

0.70324

ofxn mod

0.72923

sRun

RunA1

zout

1

Ave

Data

0.0014

Model

0.00135

data/vMax

0.005

ofxng

321.551

Min d/m

1.10536

ave/min d

11.5091

ave/min m

12.2305

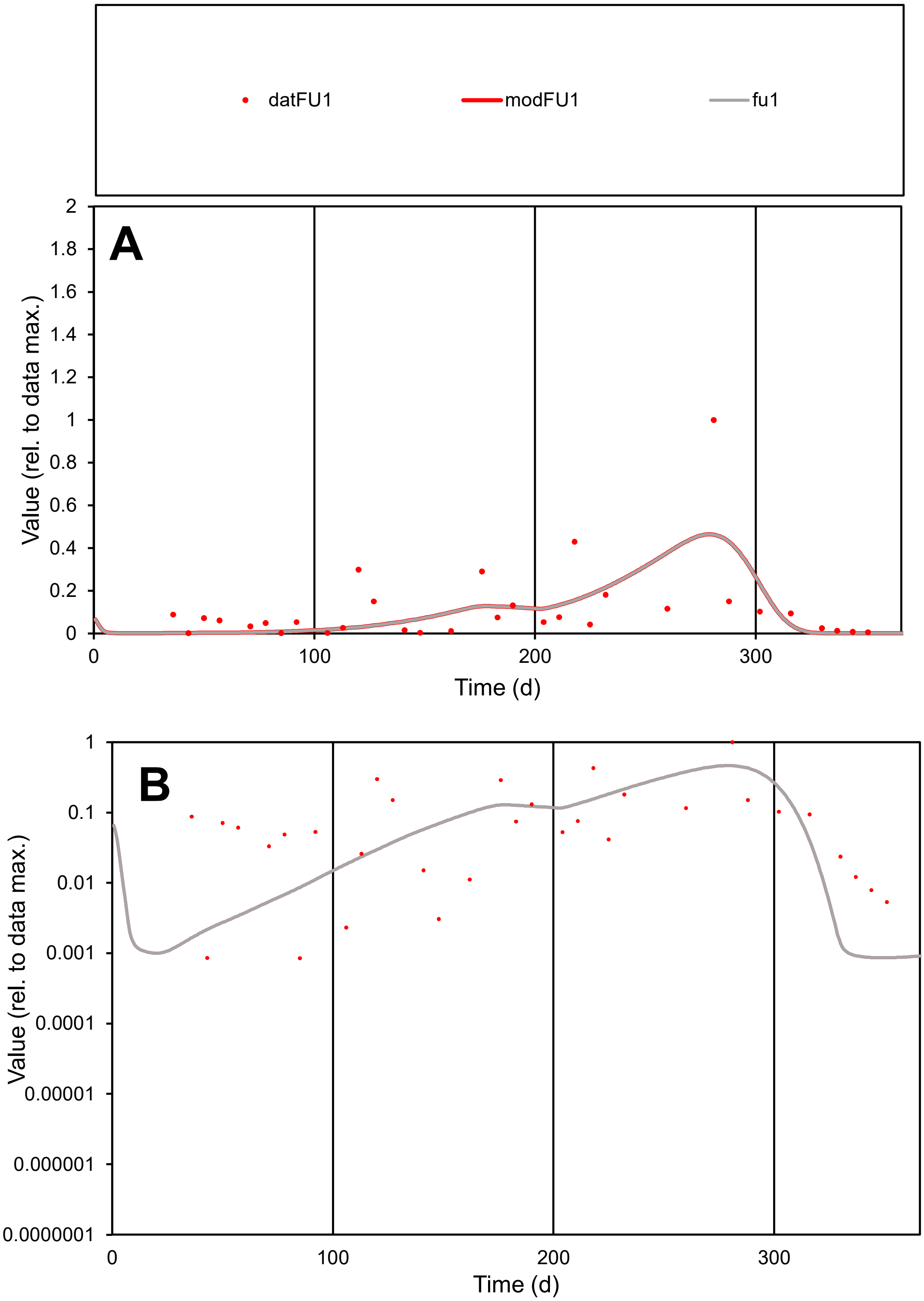


**C1**

Variables
FU1 Provide jtd or Dst
CalibOnly?
2 1 to reproduce of
oNormalize?
2 1=no, 2=yes

ofxn calc*
1.29296
ofxn mod
1.30261

sRun
RunA1
zout
1

Ave
Data
0.00622
Model
0.00527

data/vMax
0.0546

ofxng
321.551

Min d/m
0.98011

ave/min d
134.333
ave/min m
111.434

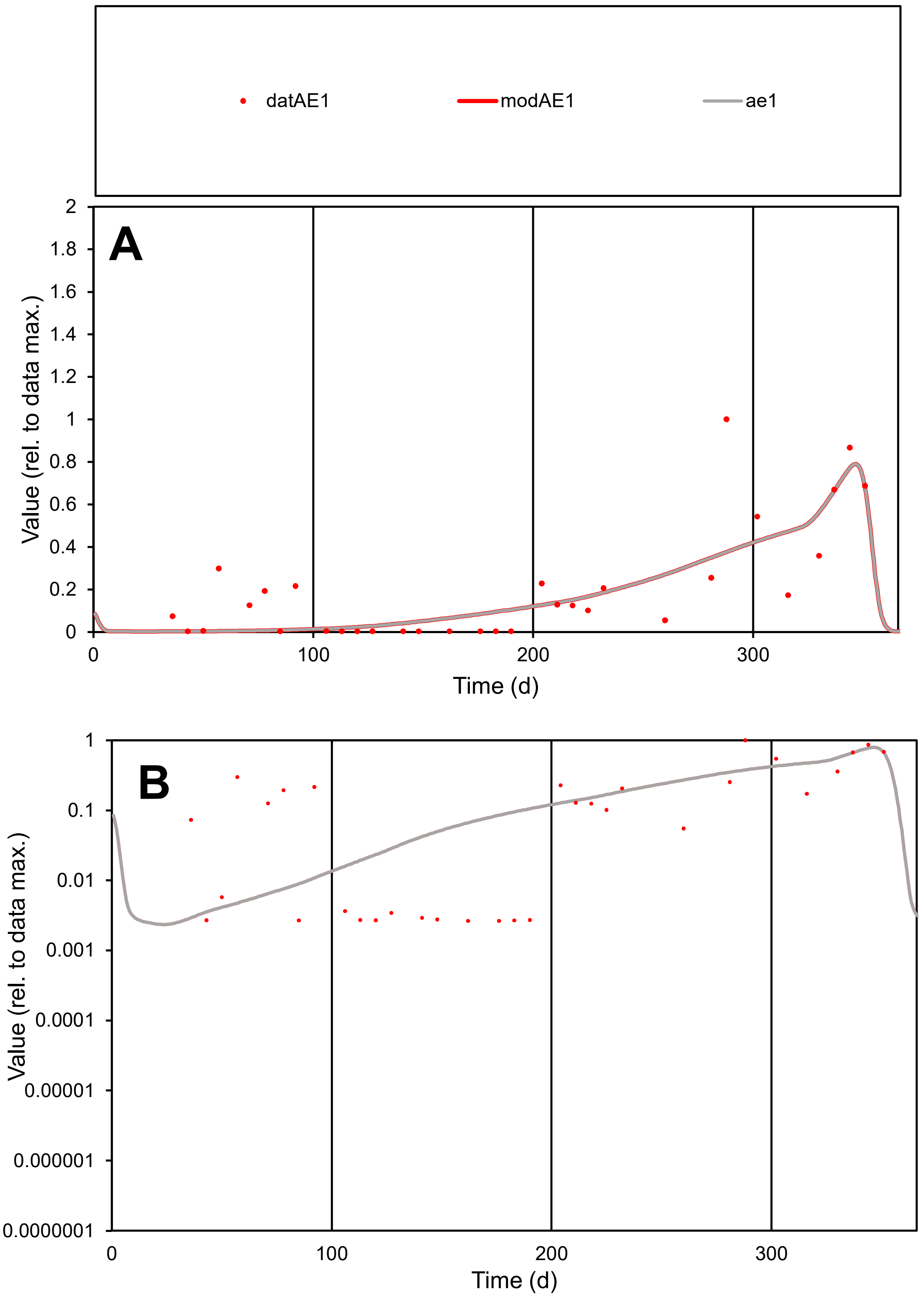


**C1**

Variables

AE1 Provide jtd or Dst

CalibOnly?

2 1 to reproduce of

oNormalize?

2 1=no, 2=yes

ofxn calc*

0.81403

ofxn mod

0.82961

sRun

RunA1

zout

1

Ave

Data

0.00103

Model

0.00096

data/vMax

0.00519

ofxng

321.551

Min d/m

0.87173

ave/min d

75.307

ave/min m

61.3139

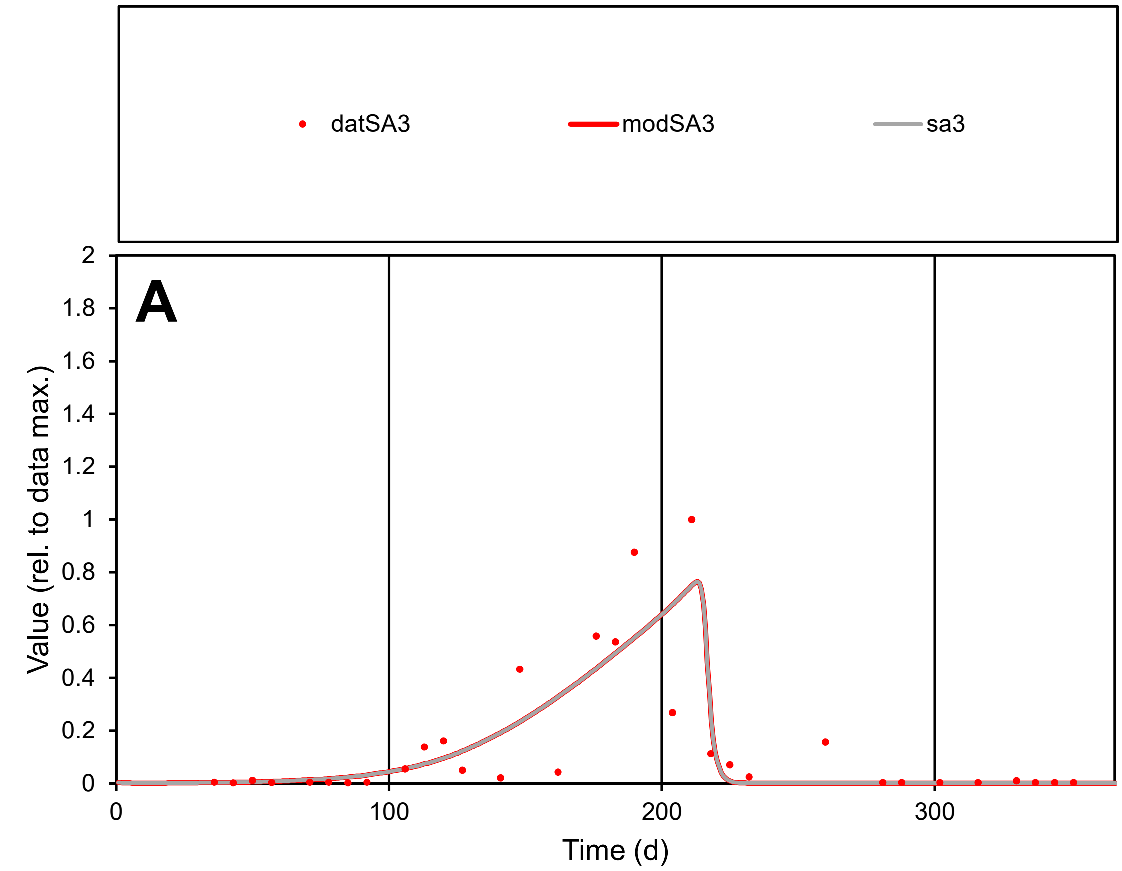


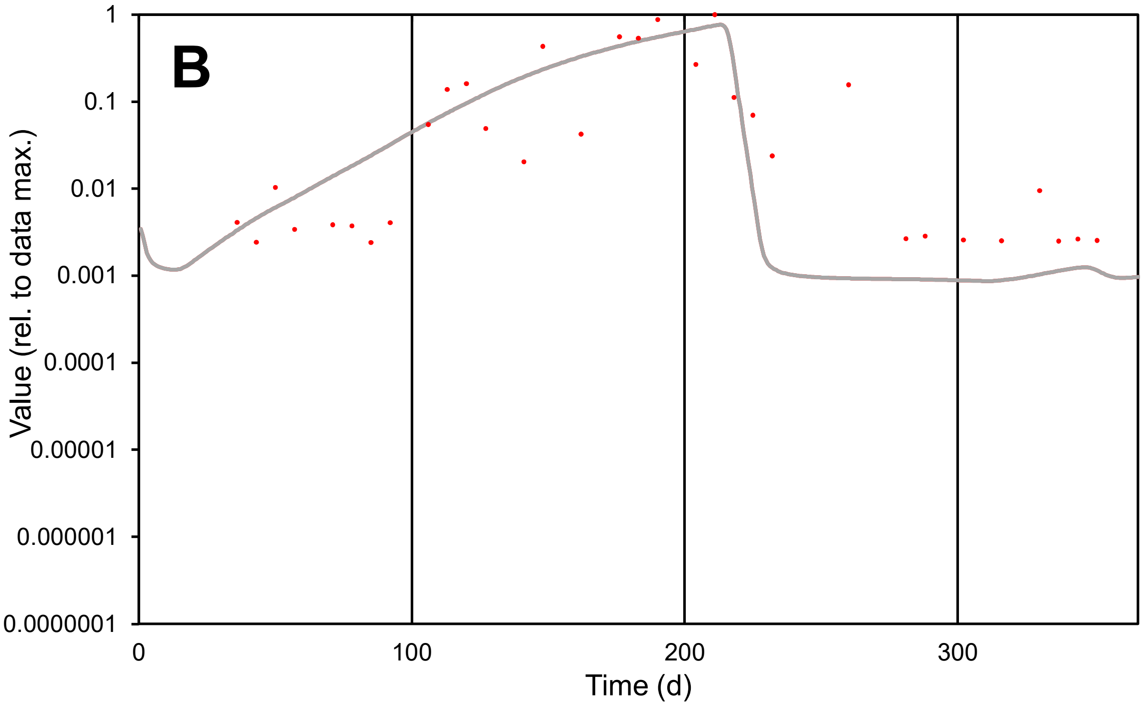


**C1**

Variables

SA3 Provide jtd or Dst

CalibOnly?

2 1 to reproduce of

oNormalize?

2 1=no, 2=yes

ofxn calc*

0.90053

ofxn mod

0.92959

sRun

RunA1

zout

1

Ave

Data

0.00576

Model

0.00555

data/vMax

0.04045

ofxng

321.551

Min d/m

2.74866

ave/min d

58.9588

ave/min m

155.977

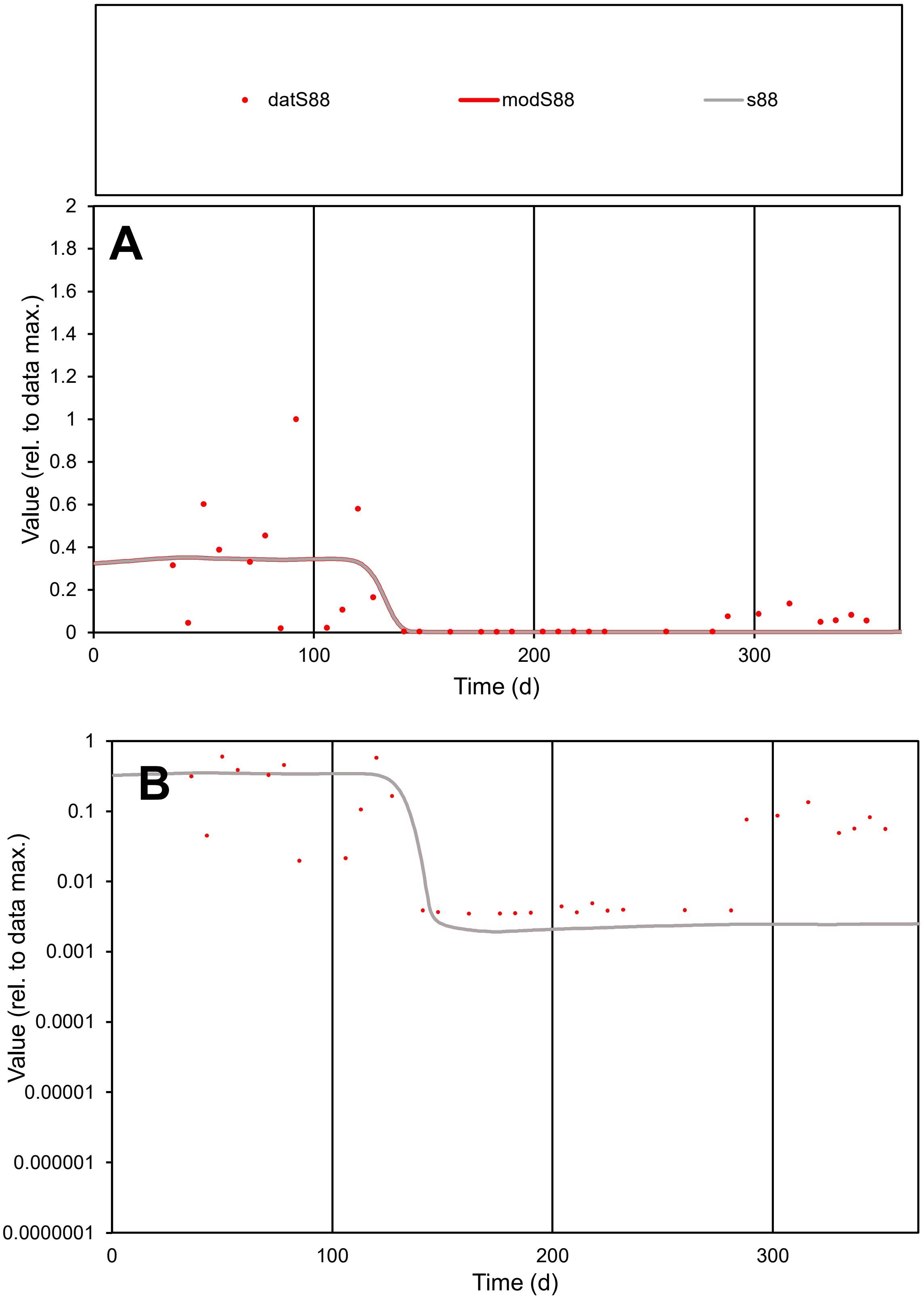


C1

Variables
S88 Provide jtd or Dst
CalibOnly?
2 1 to reproduce of
oNormalize?
2 1=no, 2=yes

ofxn calc*
1.21683
ofxn mod
1.22536

sRun
RunA1
zout
1

Ave
Data
0.00234
Model
0.00206

data/vMax
0.01619

ofxng
321.551

Min d/m
1.81166

ave/min d
41.3364
ave/min m
66.0209

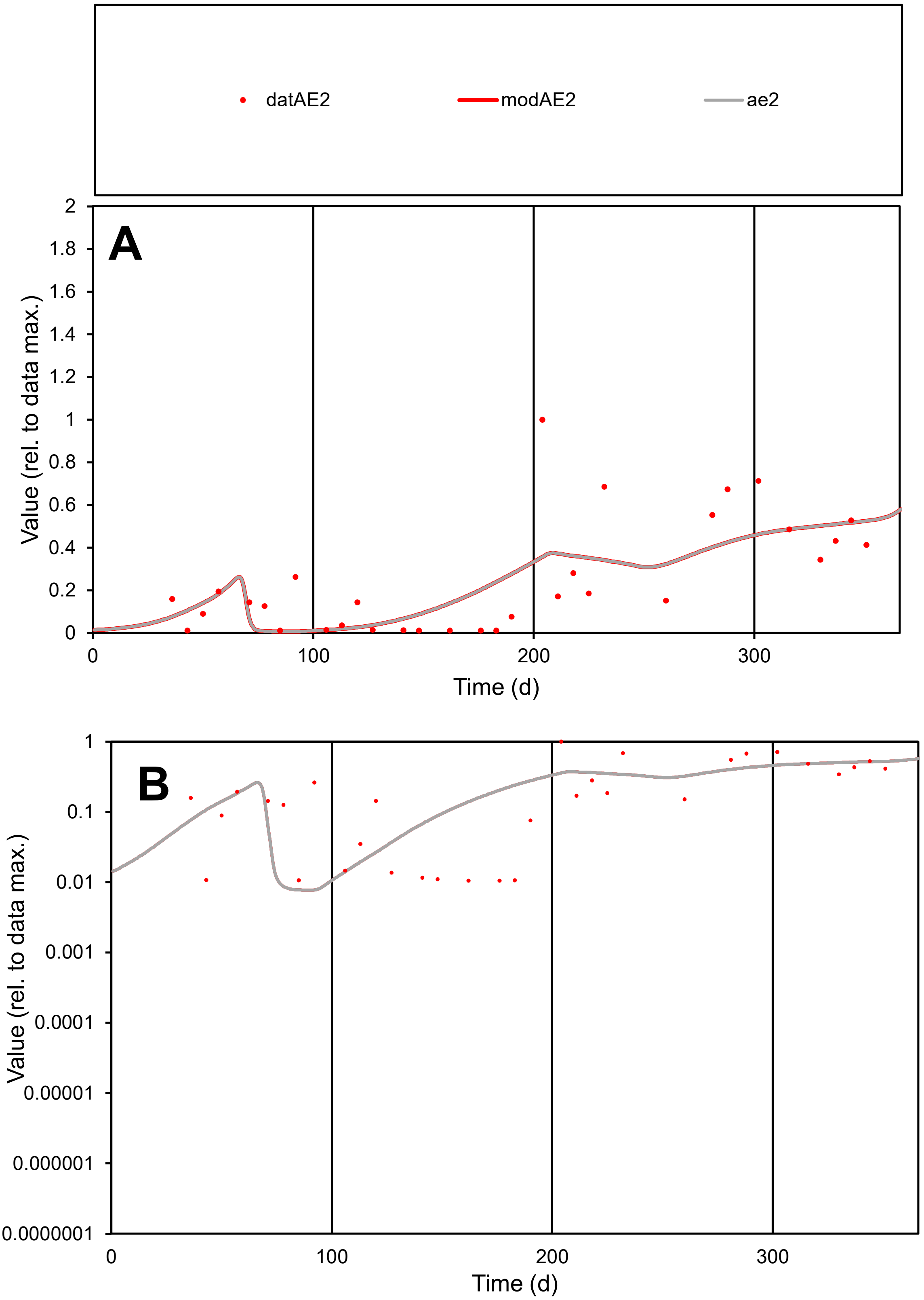


**C1**

Variables

| | |
|---|---|
| AE2 | Provide jtd or Dst |
| CalibOnly? | |
| 2 | 1 to reproduce of |
| oNormalize? | |
| 2 | 1=no, 2=yes |

ofxn calc*
0.74254
ofxn mod
0.75404

sRun
RunA1
zout
1

Ave
Data
0.00134
Model
0.0013

data/vMax
0.0054

ofxng
321.551

Min d/m
1.36276

ave/min d
23.637
ave/min m
31.2867

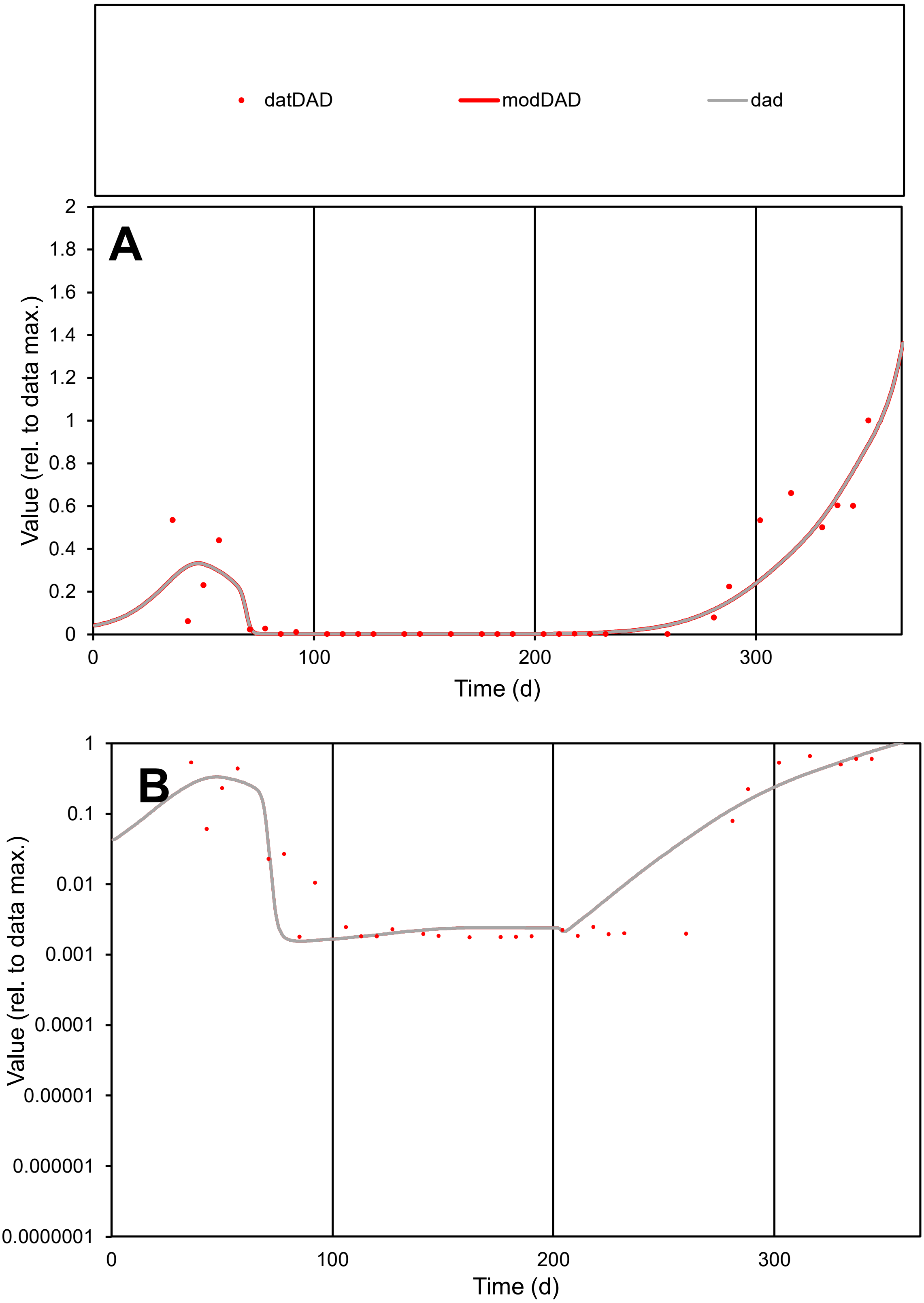


**C1**

Variables

DAD Provide jtd or Dst

CalibOnly?

2 1 to reproduce of

oNormalize?

2 1=no, 2=yes

ofxn calc*

0.61992

ofxn mod

0.63865

sRun

RunA1

zout

1

Ave

Data

0.00149

Model

0.00139

data/vMax

0.00855

ofxng

321.551

Min d/m

1.13835

ave/min d

98.1937

ave/min m

104.328

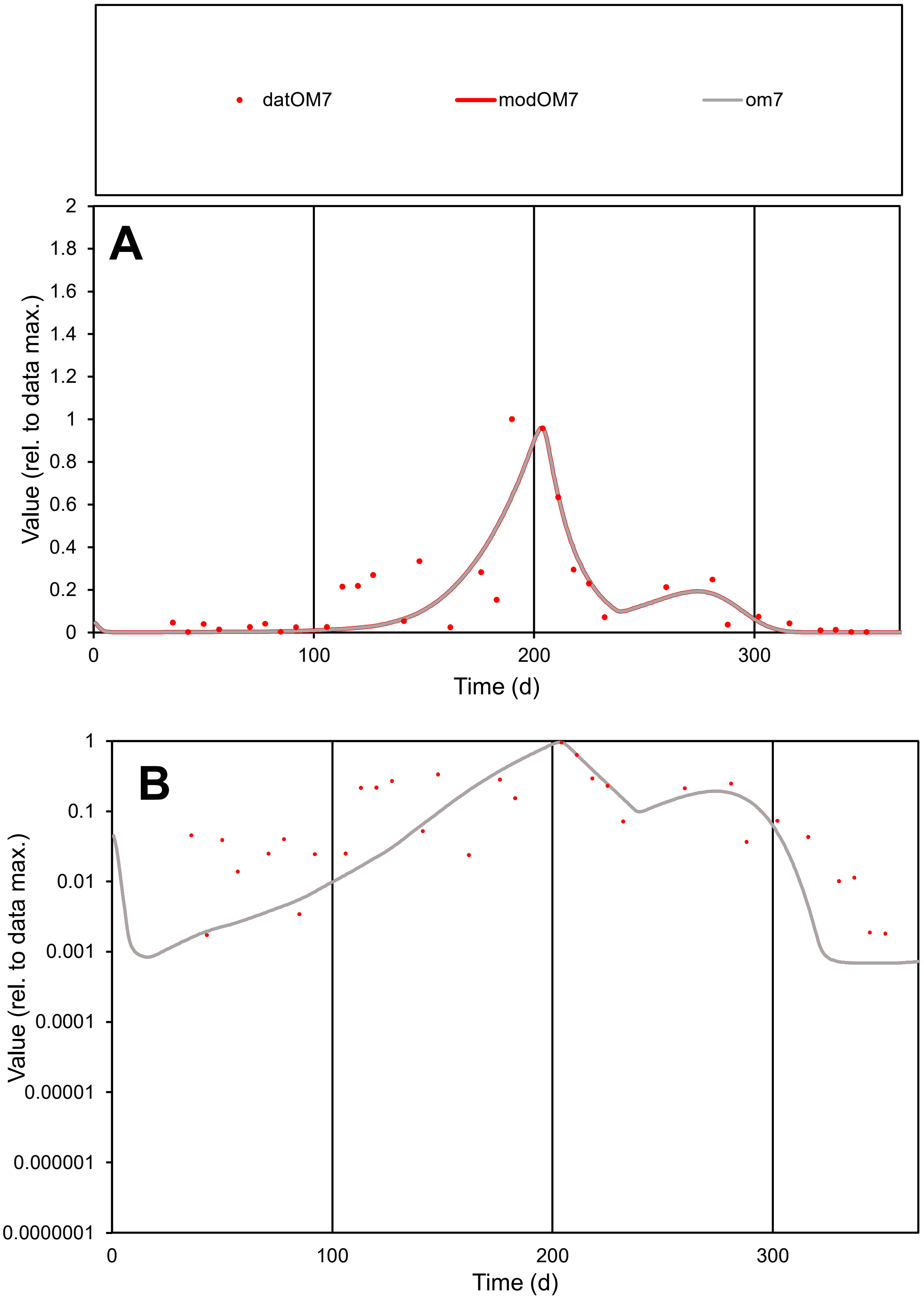


**C1**

Variables
OM7 Provide jtd or Dst
CalibOnly?
2 1 to reproduce of
oNormalize?
2 1=no, 2=yes

ofxn calc*
0.71317
ofxn mod
0.733

sRun
RunA1
zout
1

Ave
Data
0.00945
Model
0.00814

data/vMax
0.05409

ofxng
321.551

Min d/m
2.52616
ave/min d
100.91
ave/min m
219.453

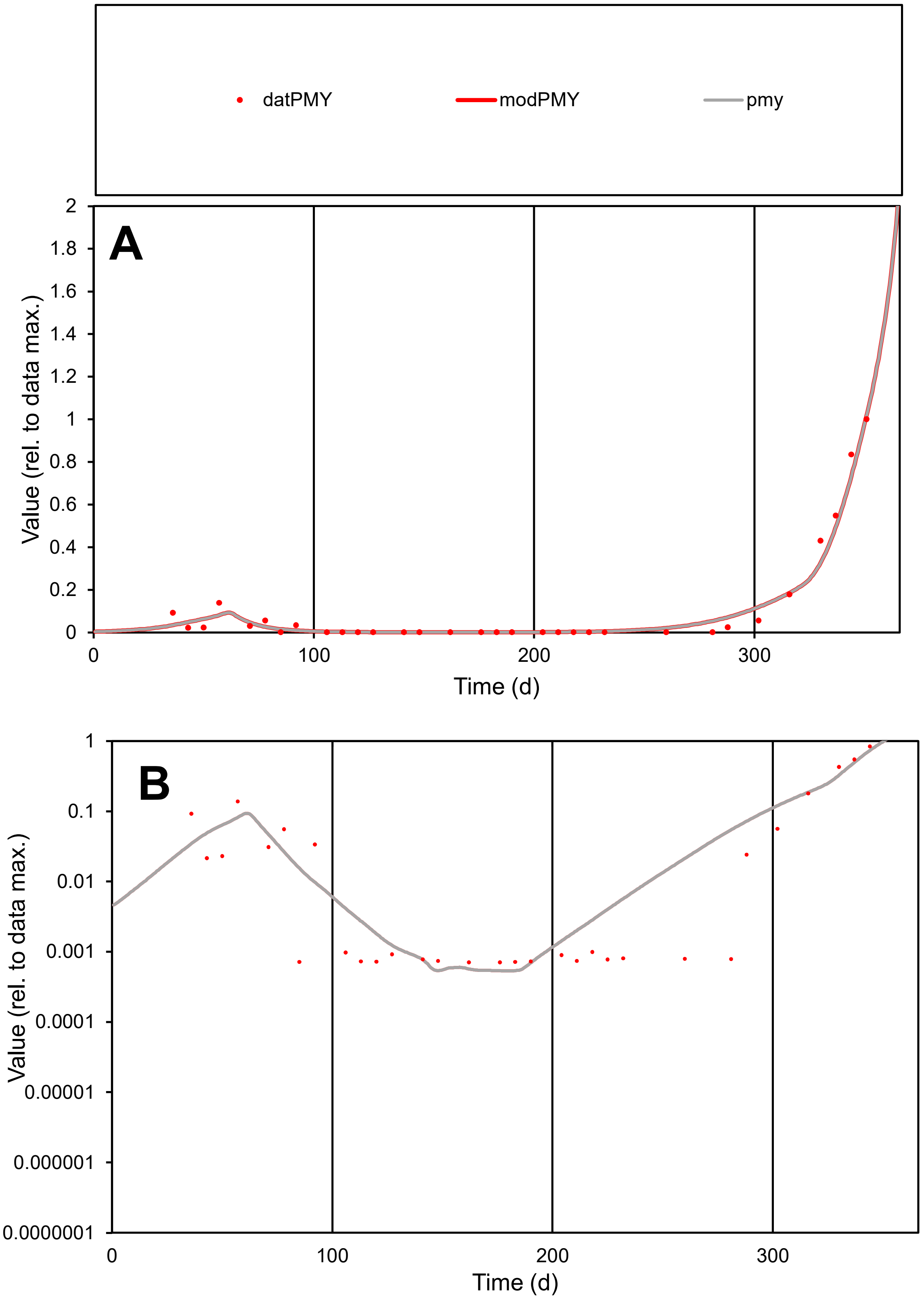


**C1**

Variables
PMY Provide jtd or Dst
CalibOnly?
2 1 to reproduce of
oNormalize?
2 1=no, 2=yes

ofxn calc*
0.29002
ofxn mod
0.34859

sRun
RunA1
zout
1

Ave
Data
0.00146
Model
0.00149

data/vMax
0.01338

ofxng
321.551

Min d/m
1.31699

ave/min d
154.456
ave/min m
208.027

324_Calib_S_A

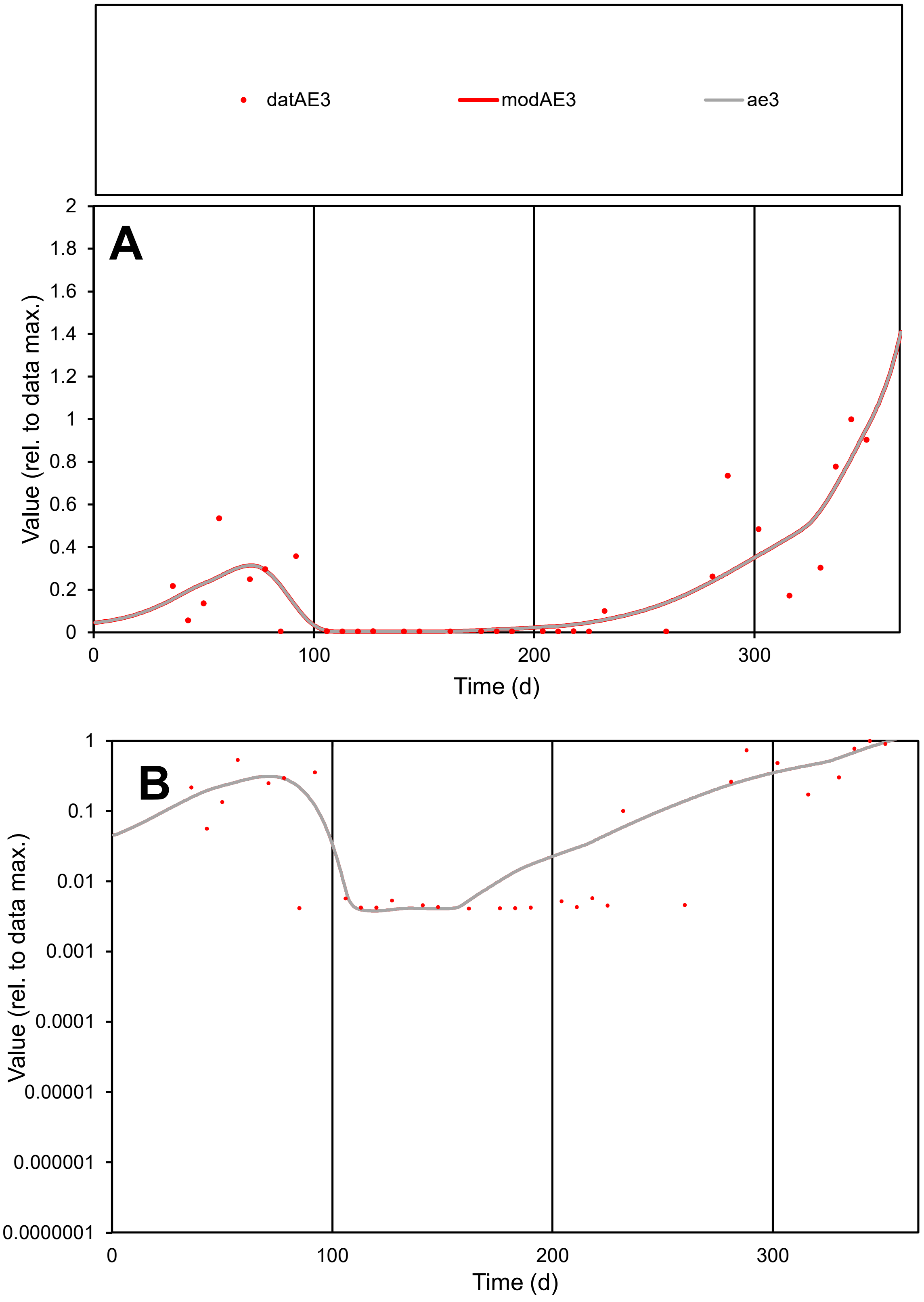


C1

Variables
AE3 Provide jtd or Dst
CalibOnly?
2 1 to reproduce of
oNormalize?
2 1=no, 2=yes

ofxn calc*
0.66888
ofxn mod
0.69003

sRun
RunA1
zout
1

Ave
Data
0.00084
Model
0.00083

data/vMax
0.00402

ofxng
321.551

Min d/m
1.08055

ave/min d
50.6894
ave/min m
54.5212

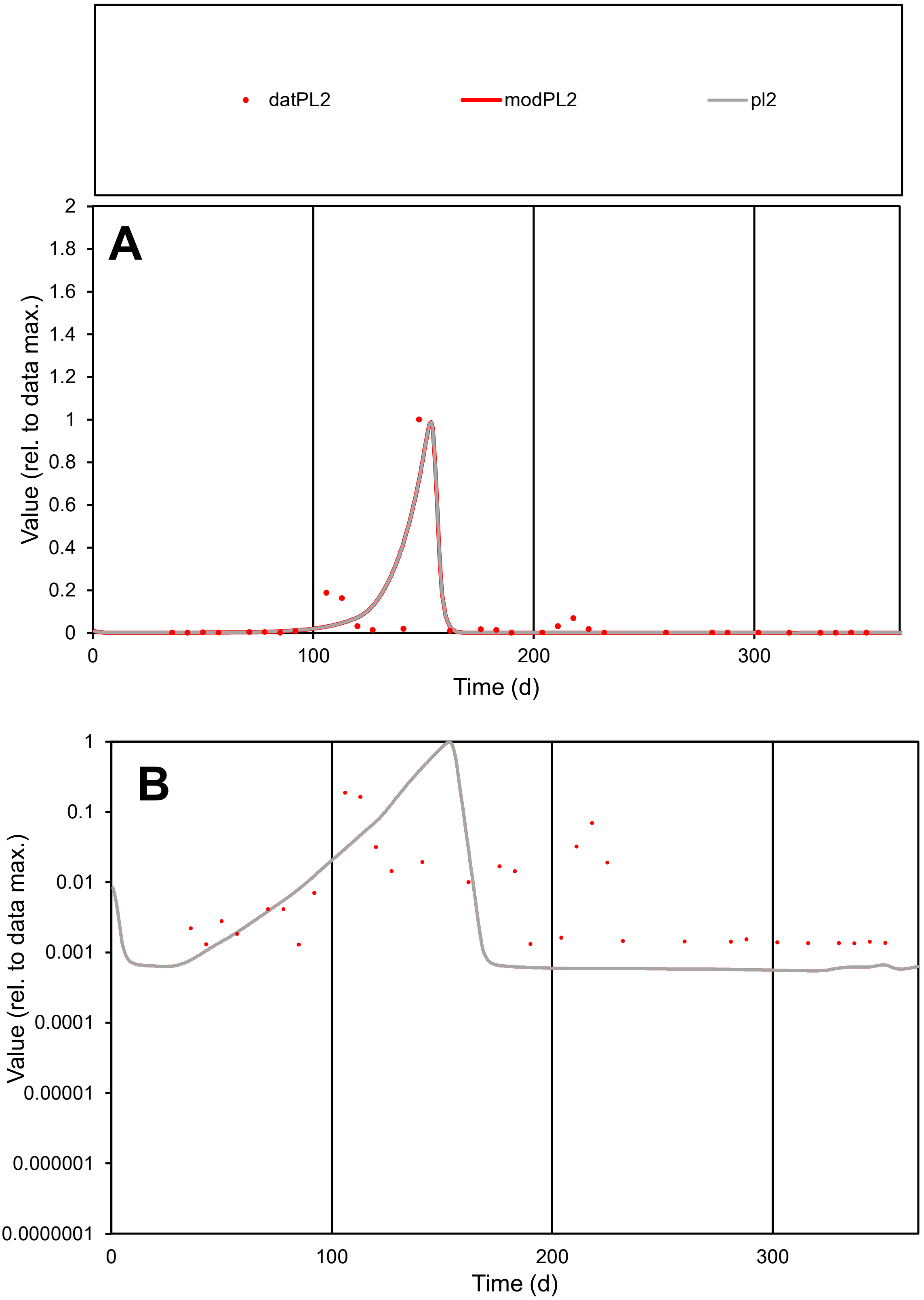


**C1**

Variables
PL2 Provide jtd or Dst
CalibOnly?
2 1 to reproduce of
oNormalize?
2 1=no, 2=yes

ofxn calc*
1.75122
ofxn mod
1.8192

sRun
RunA1
zout
1

Ave
Data
0.0063
Model
0.00587

data/vMax
0.1245

ofxng
321.551

Min d/m
2.35953

ave/min d
38.8893
ave/min m
85.3875

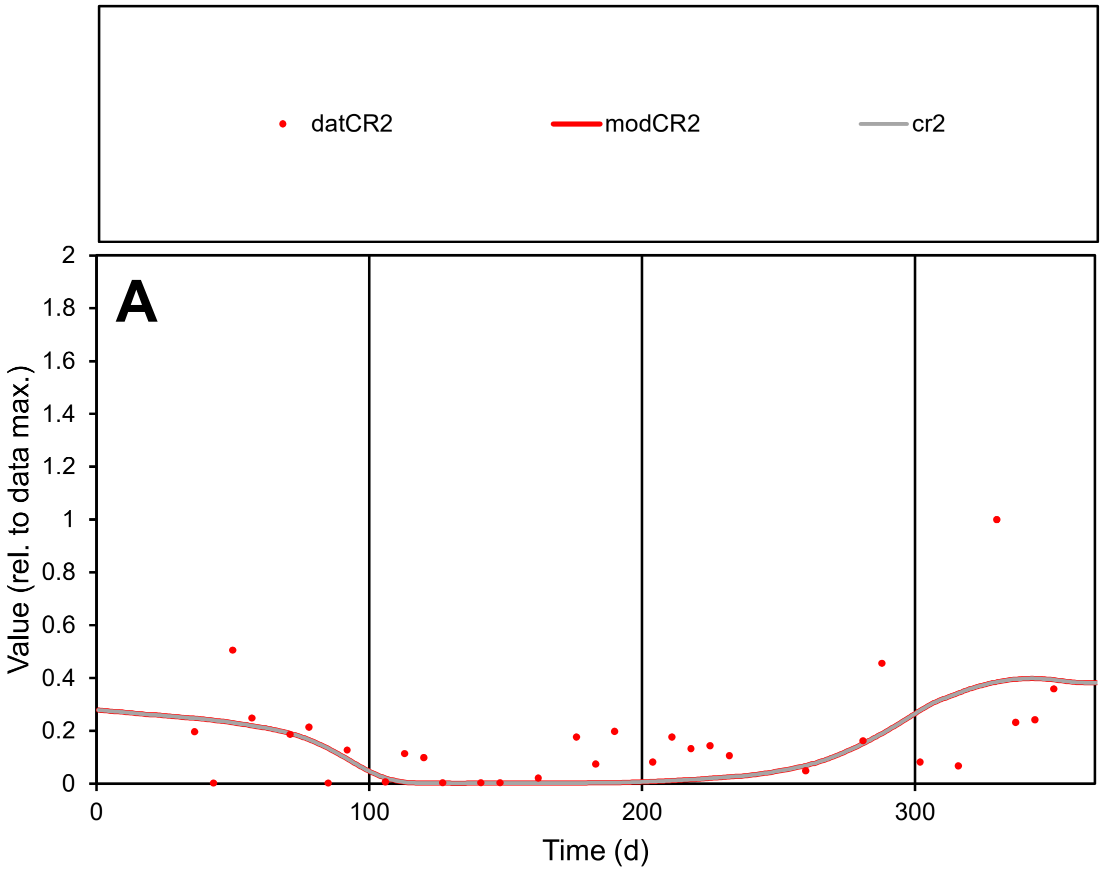


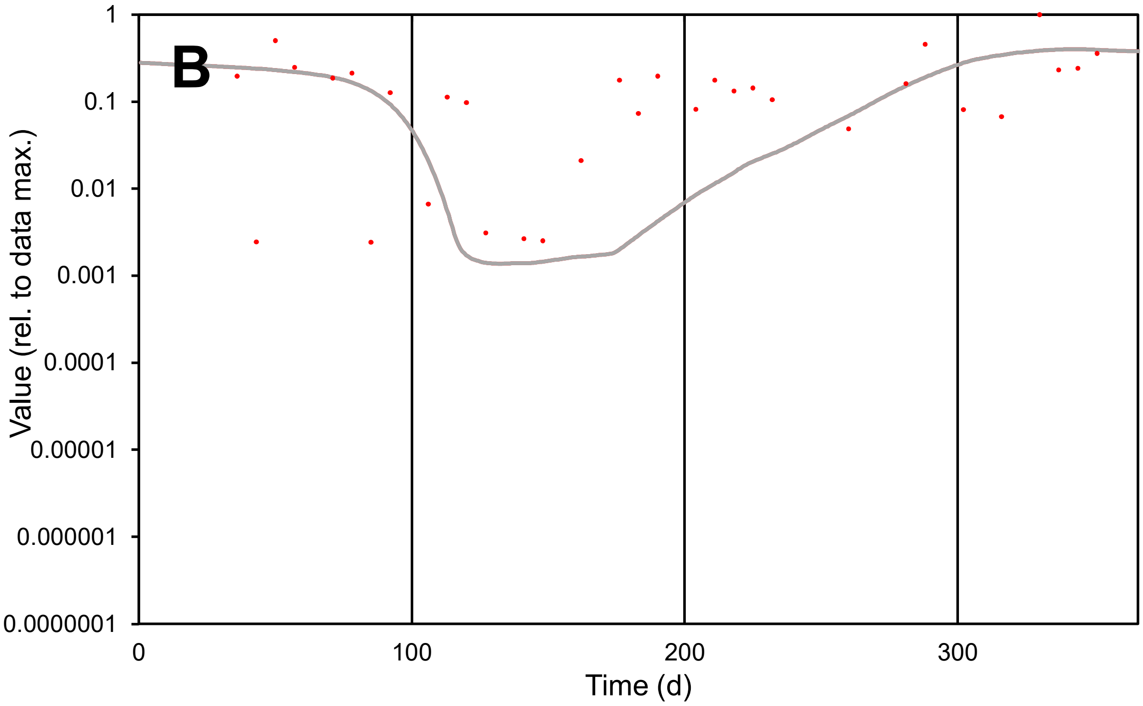


**C1**

Variables
CR2 Provide jtd or Dst
CalibOnly?
2 1 to reproduce of
oNormalize?
2 1=no, 2=yes

ofxn calc*
0.99631
ofxn mod
1.00682

sRun
RunA1
zout
1

Ave
Data
0.00279
Model
0.00216

data/vMax
0.01633

ofxng
321.551

Min d/m
1.7409

ave/min d
70.1737
ave/min m
94.8661

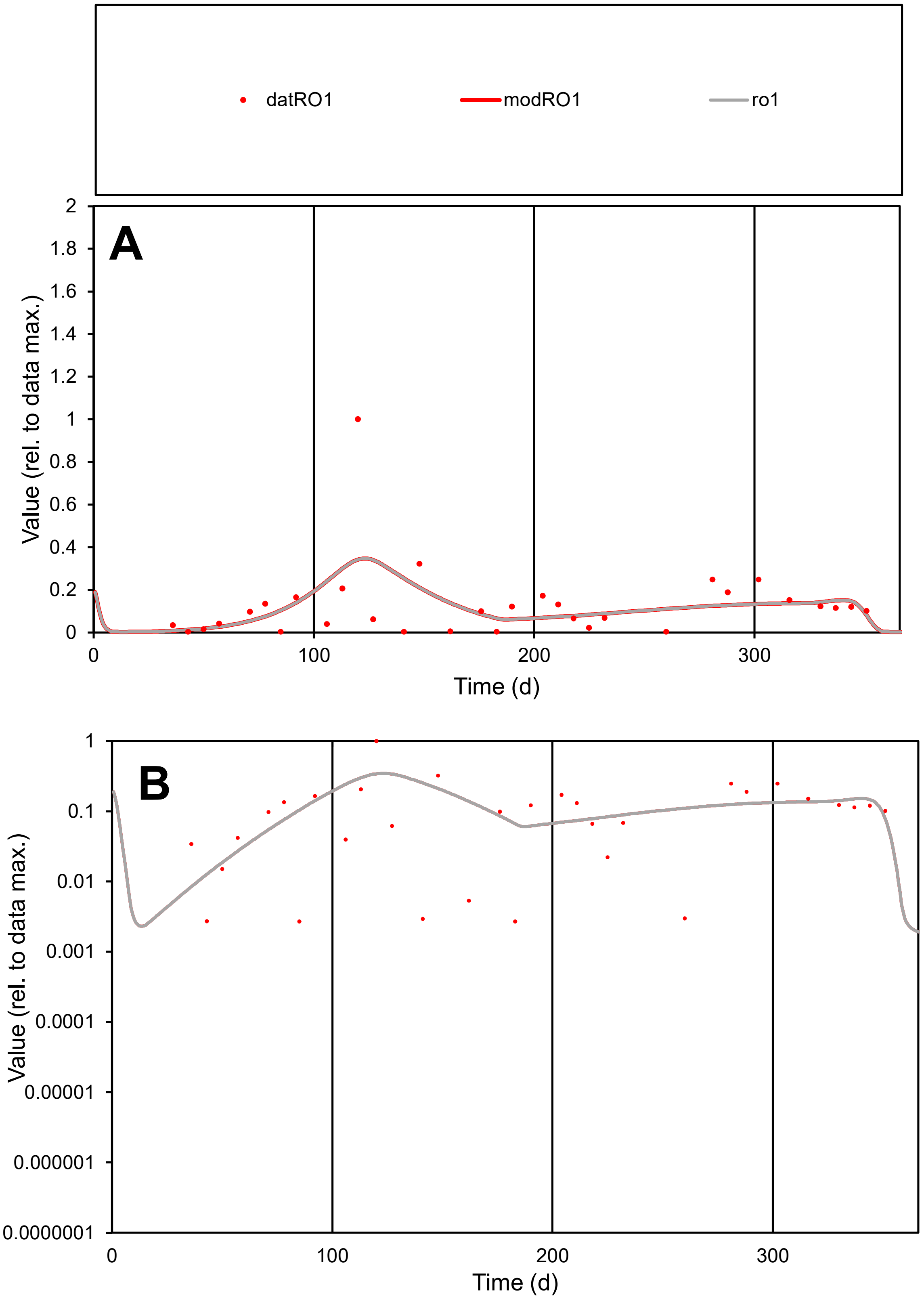


**C1**

Variables
RO1 Provide jtd or Dst
CalibOnly?
2 1 to reproduce of
oNormalize?
2 1=no, 2=yes

ofxn calc*
1.17585
ofxn mod
1.18367

sRun
RunA1
zout
1

Ave
Data
0.00168
Model
0.00164

data/vMax
0.01306

ofxng
321.551

Min d/m
0.30009

ave/min d
47.6311
ave/min m
13.9746

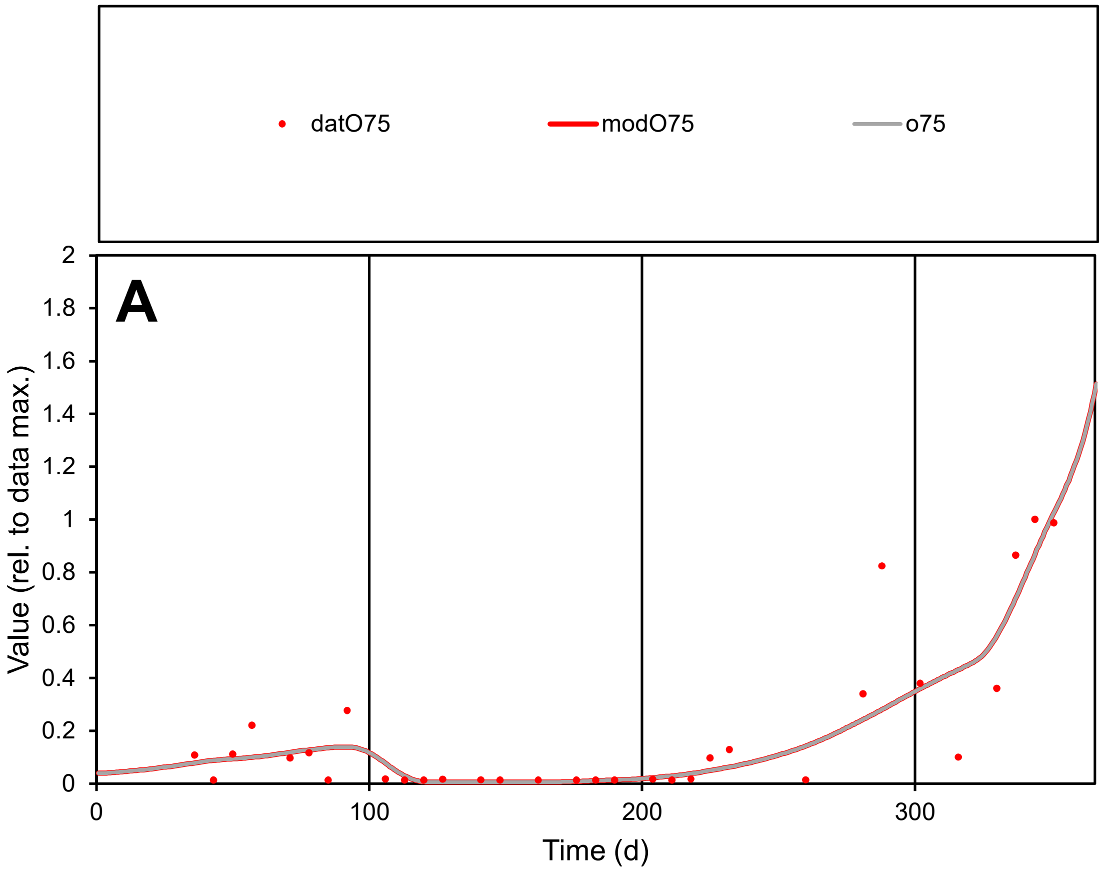


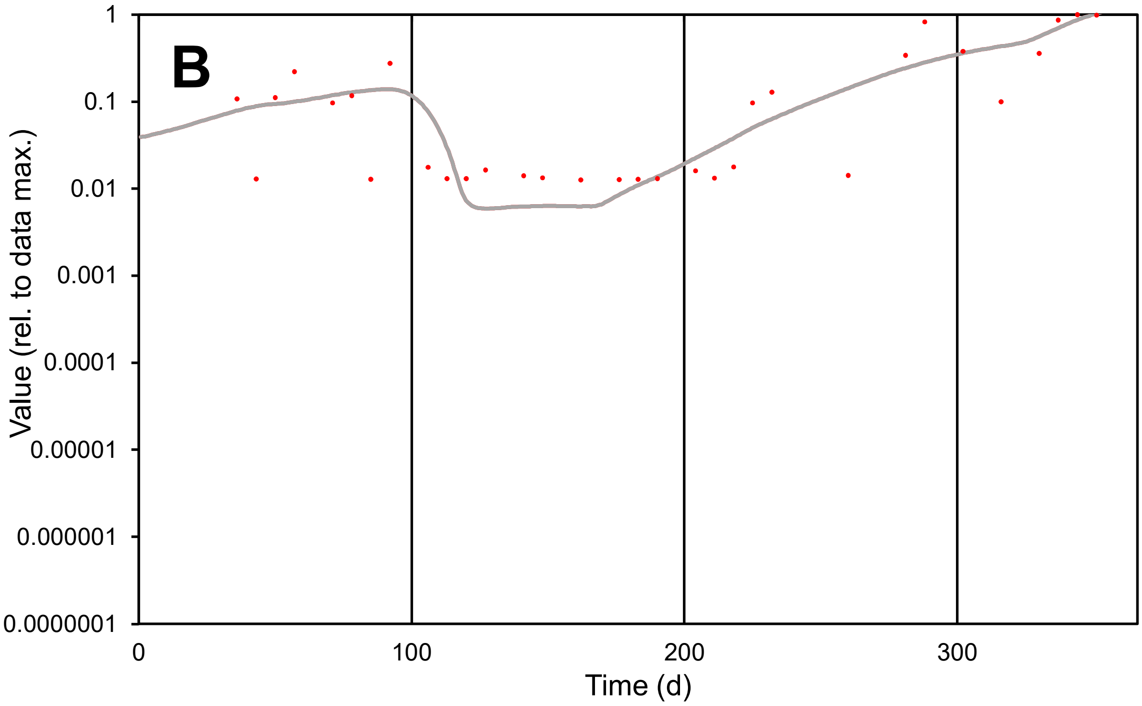


C1

Variables
O75 Provide jtd or Dst
CalibOnly?
2 1 to reproduce of
oNormalize?
2 1=no, 2=yes

ofxn calc*
0.67809
ofxn mod
0.69878

sRun
RunA1
zout
1

Ave
Data
0.00106
Model
0.00101

data/vMax
0.00543

ofxng
321.551

Min d/m
2.15

ave/min d
15.3945
ave/min m
31.6589

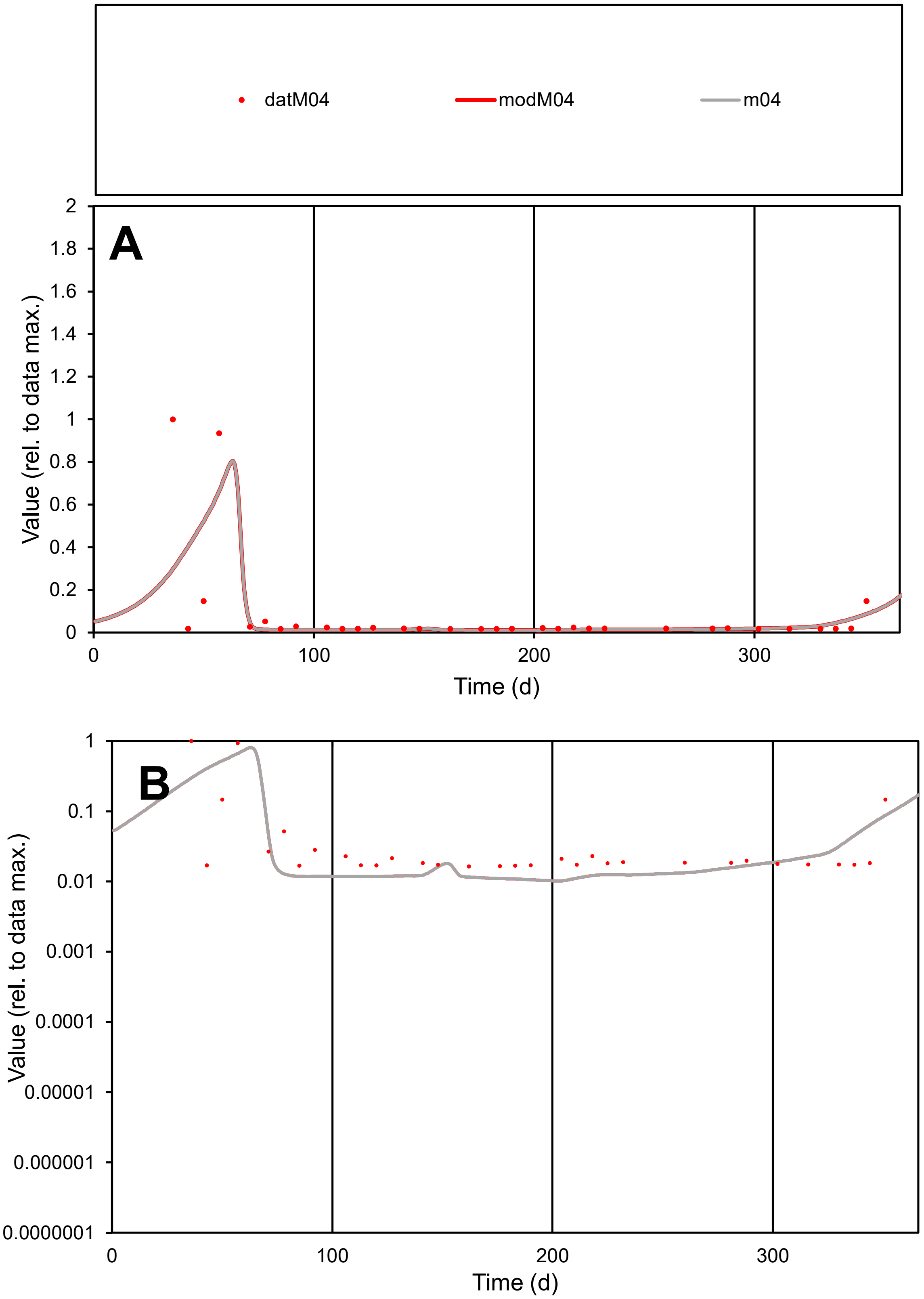


C1

Variables
M04 Provide jtd or Dst
CalibOnly?
2 1 to reproduce of
oNormalize?
2 1=no, 2=yes

ofxn calc*
1.8358
ofxn mod
1.86333

sRun
RunA1
zout
1

Ave
Data
0.00018
Model
0.00016

data/vMax
0.0021

ofxng
321.551

Min d/m
1.60259

ave/min d
5.29342
ave/min m
7.55823

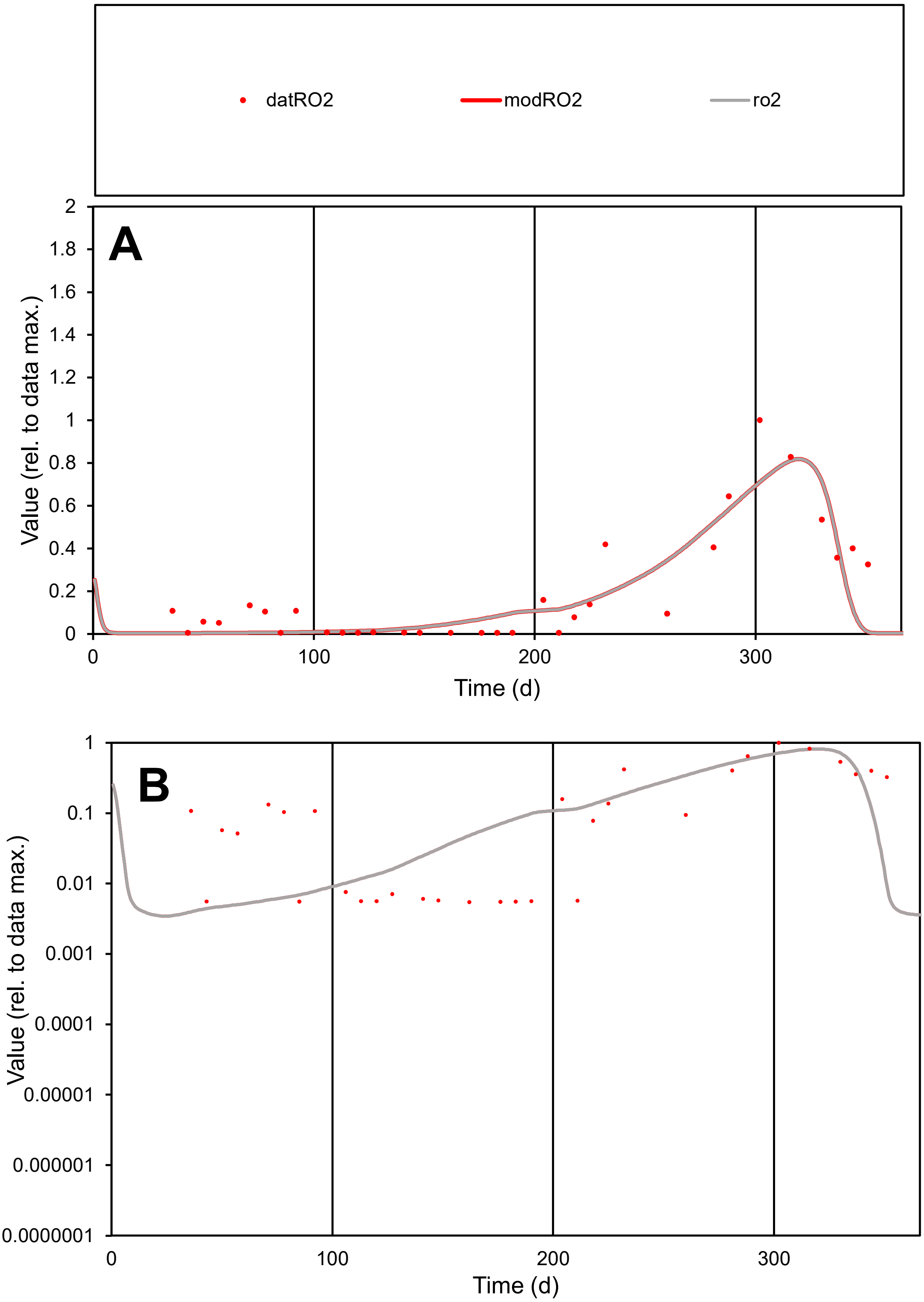


C1

Variables

RO2 Provide jtd or Dst

CalibOnly?

2 1 to reproduce of

oNormalize?

2 1=no, 2=yes

ofxn calc*

0.65504

ofxn mod

0.68549

sRun

RunA1

zout

1

Ave

Data

0.0013

Model

0.00116

data/vMax

0.0069

ofxng

321.551

Min d/m

1.35113

ave/min d

34.4269

ave/min m

41.5831

324_Calib_S_A

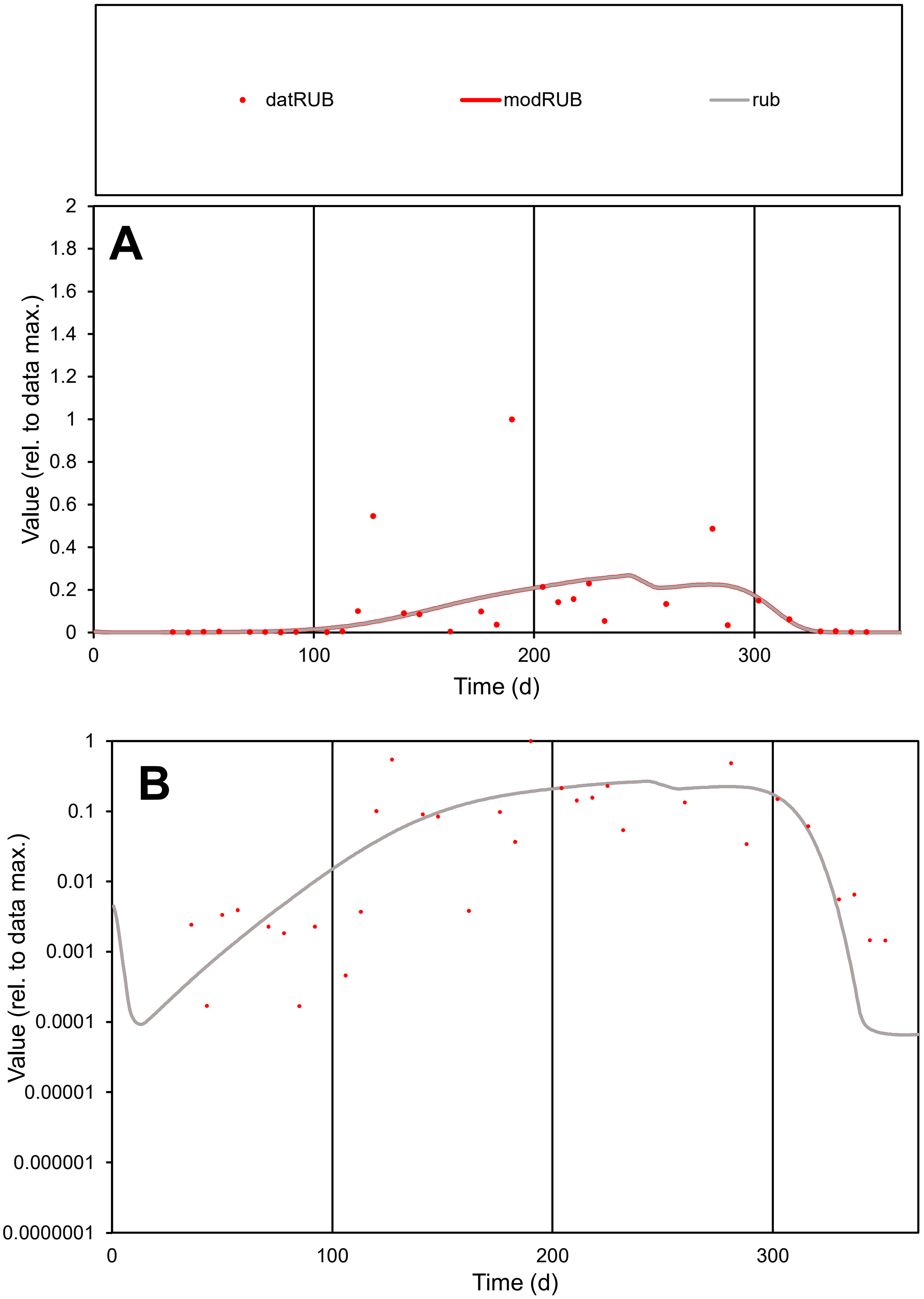


**C1**

Variables
RUB Provide jtd or Dst
CalibOnly?
2 1 to reproduce of
oNormalize?
2 1=no, 2=yes

ofxn calc*
1.62803
ofxn mod
1.63447

sRun
RunA1
zout
1

Ave
Data
0.00674
Model
0.00567

data/vMax
0.05902

ofxng
321.551

Min d/m
2.48597

ave/min d
679.131
ave/min m
1419.78

324_Calib_S_A

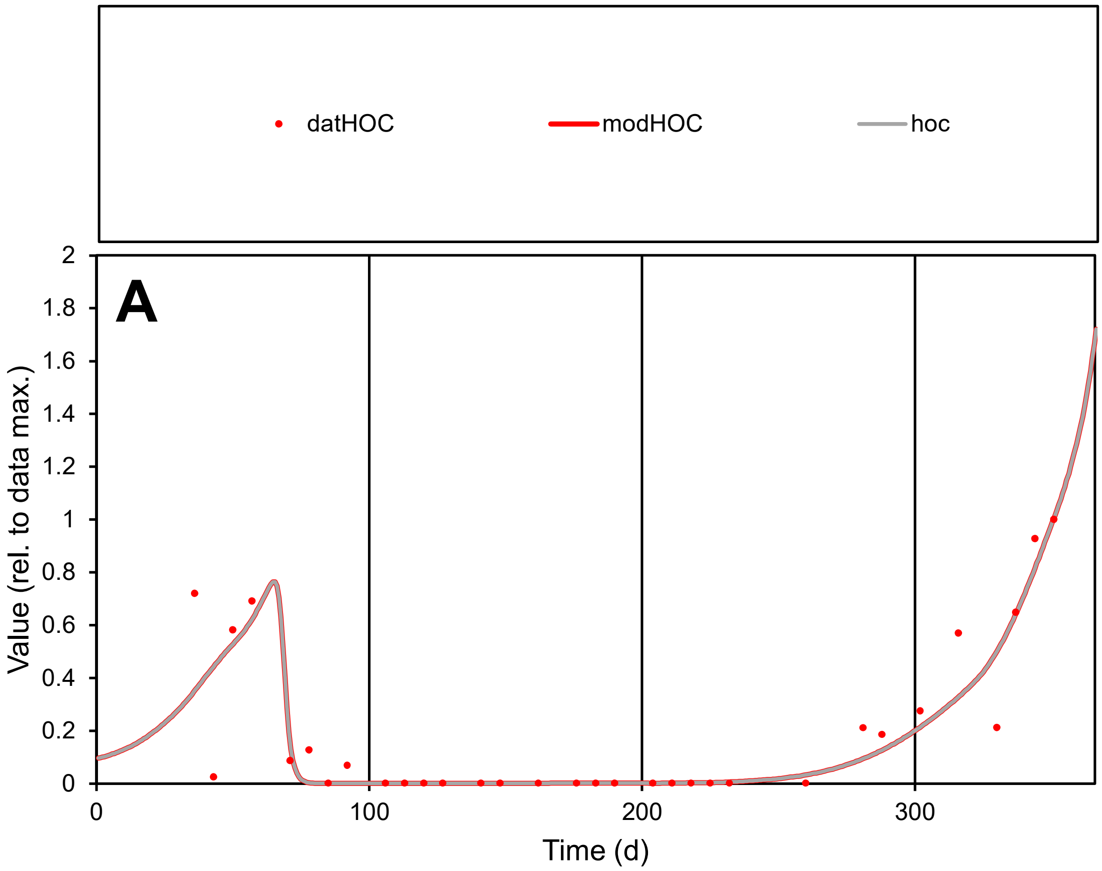


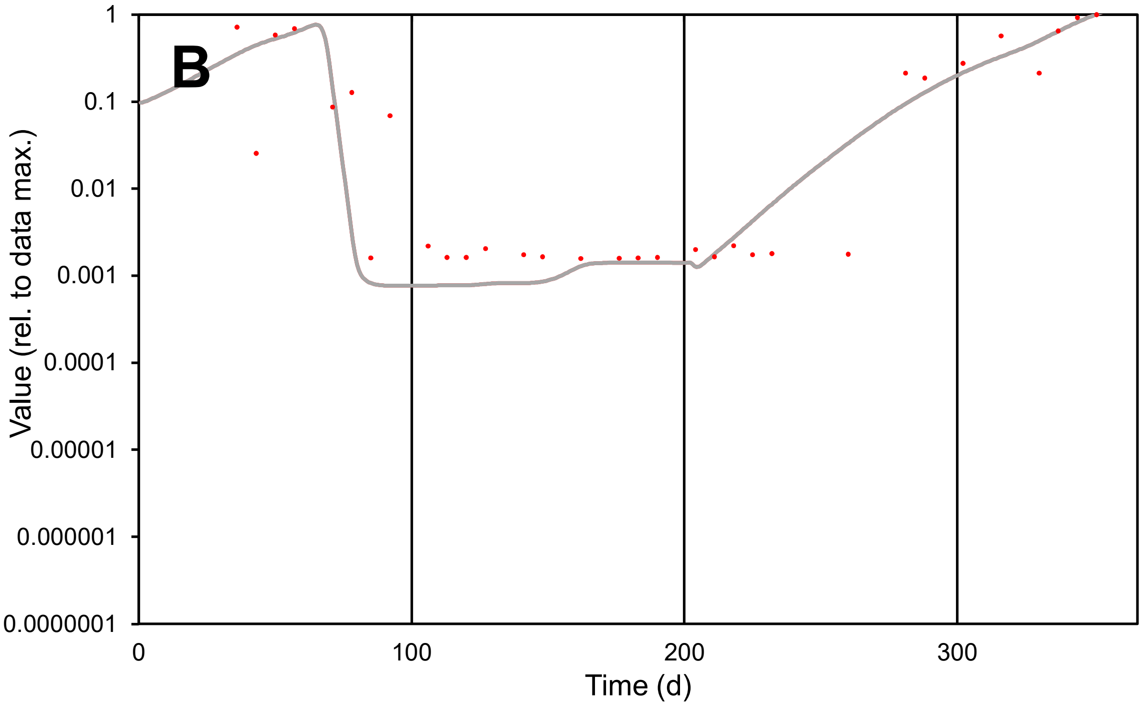


**C1**

Variables
HOC Provide jtd or Dst
CalibOnly?
2 1 to reproduce of
oNormalize?
2 1=no, 2=yes

ofxn calc*
0.61514
ofxn mod
0.64457

sRun
RunA1
zout
1

Ave
Data
0.00156
Model
0.00147

data/vMax
0.00783

ofxng
321.551

Min d/m
2.03967

ave/min d
126.306
ave/min m
242.483

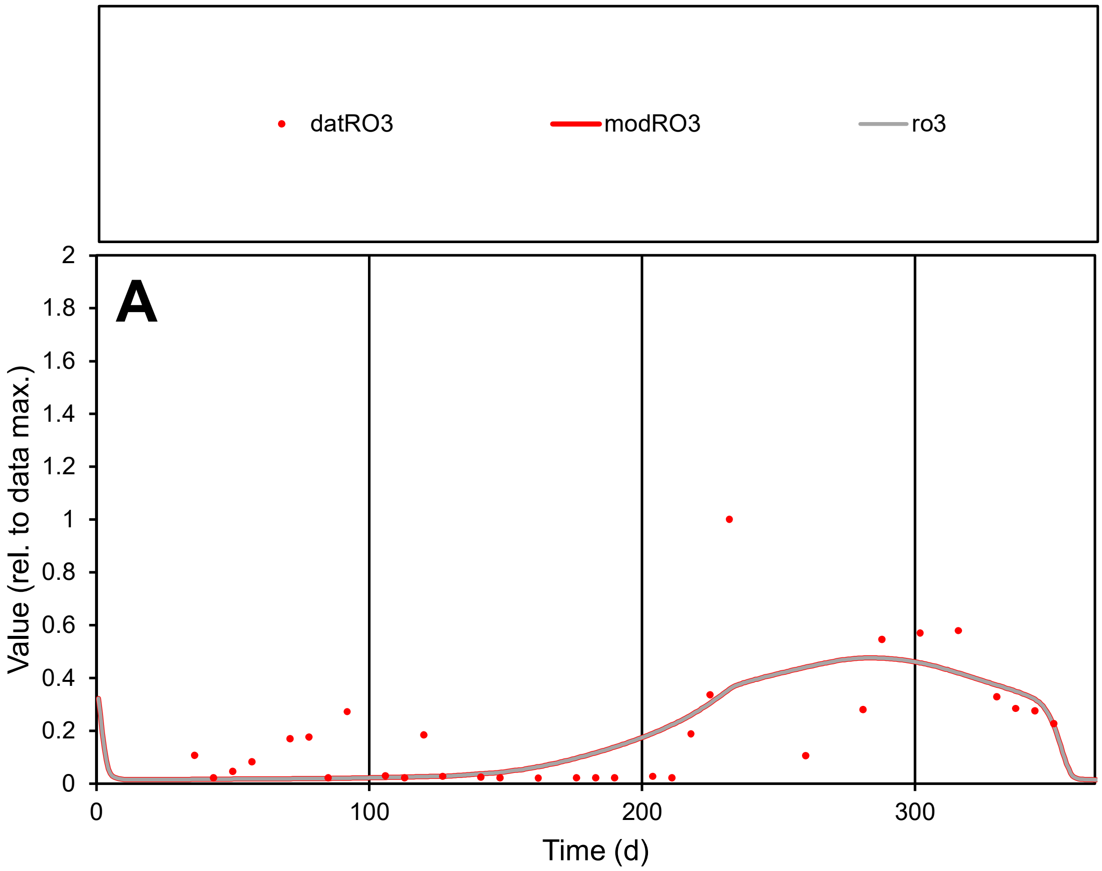


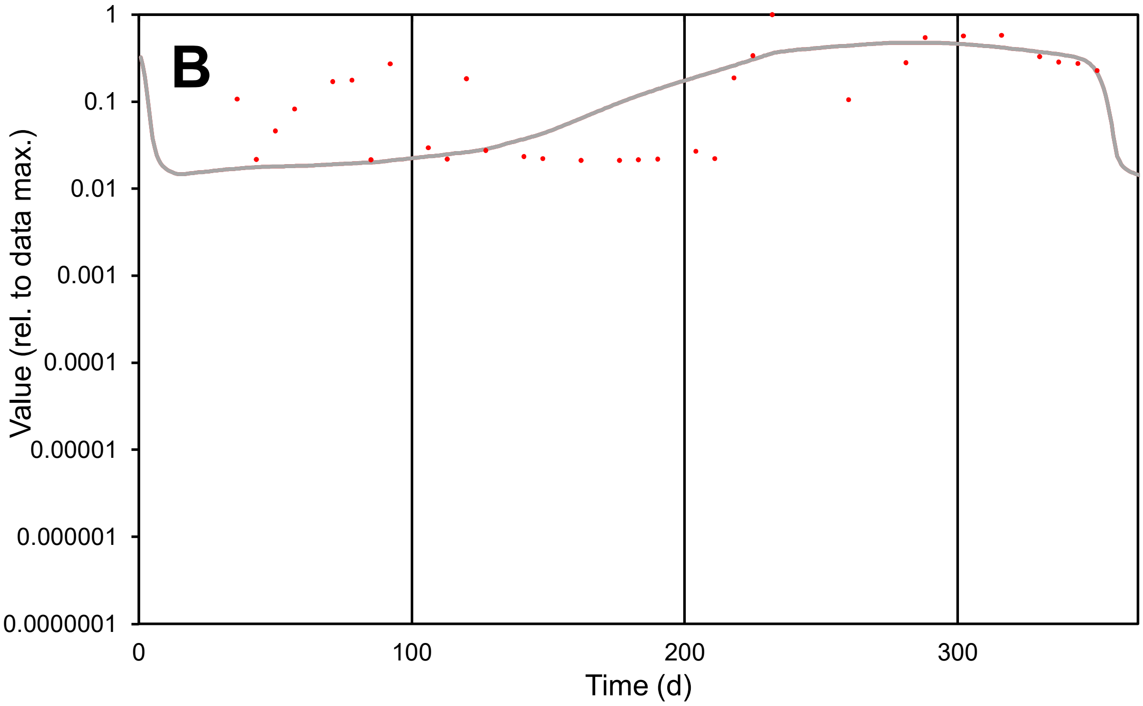


**C1**

Variables
RO3 Provide jtd or Dst
CalibOnly?
2 1 to reproduce of
oNormalize?
2 1=no, 2=yes

ofxn calc*
0.86622
ofxn mod
0.87671

sRun
RunA1
zout
1

Ave
Data
0.00084
Model
0.00078

data/vMax
0.00445

ofxng
321.551

Min d/m
1.23581

ave/min d
8.9451
ave/min m
10.2029

324_Calib_S_A

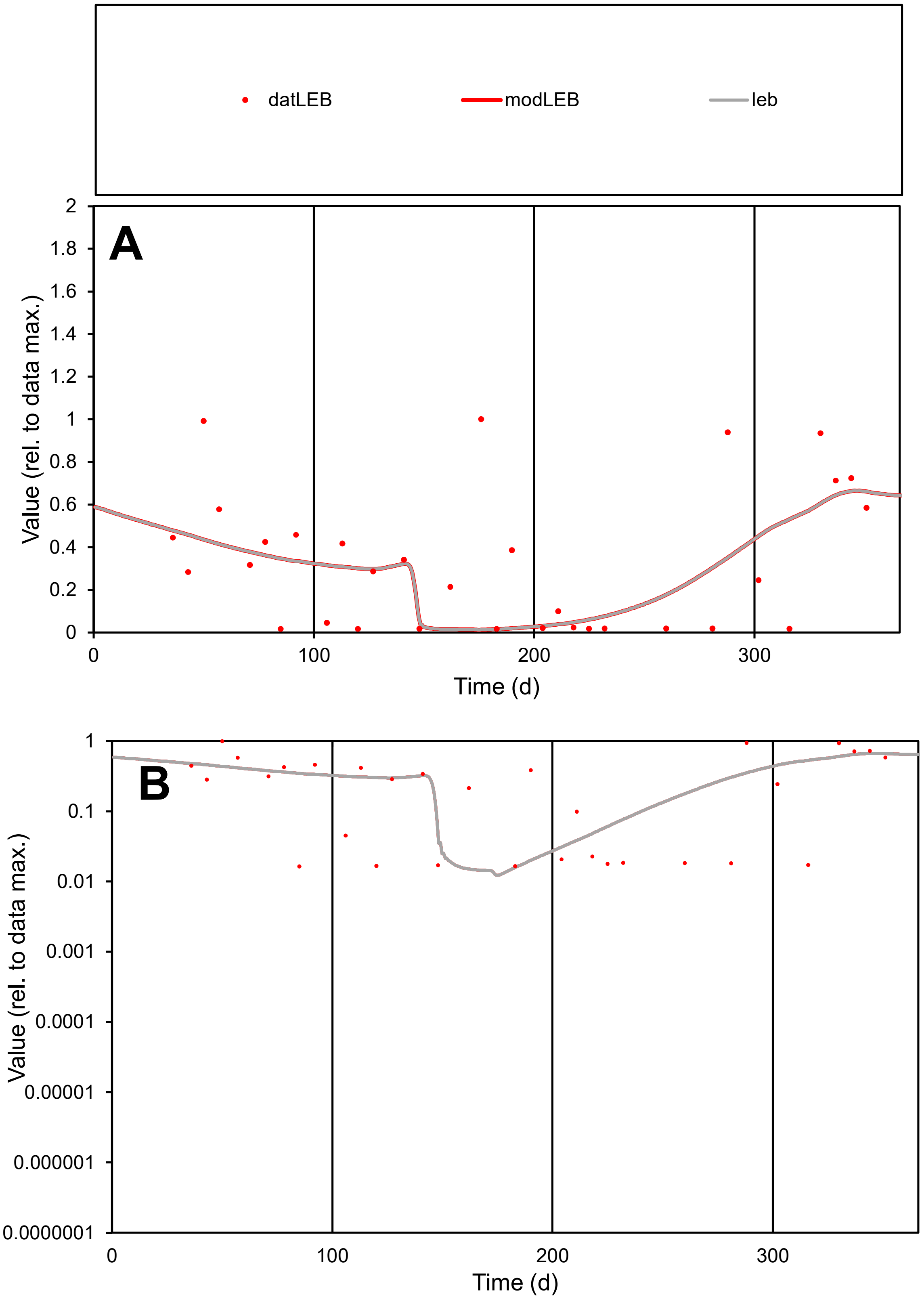


**C1**

Variables
LEB Provide jtd or Dst
CalibOnly?
2 1 to reproduce of
oNormalize?
2 1=no, 2=yes

ofxn calc*
0.87421
ofxn mod
0.88256

sRun
RunA1
zout
1

Ave
Data
0.0014
Model
0.00125

data/vMax
0.00422

ofxng
321.551

Min d/m
1.26318

ave/min d
20.1429
ave/min m
22.7786

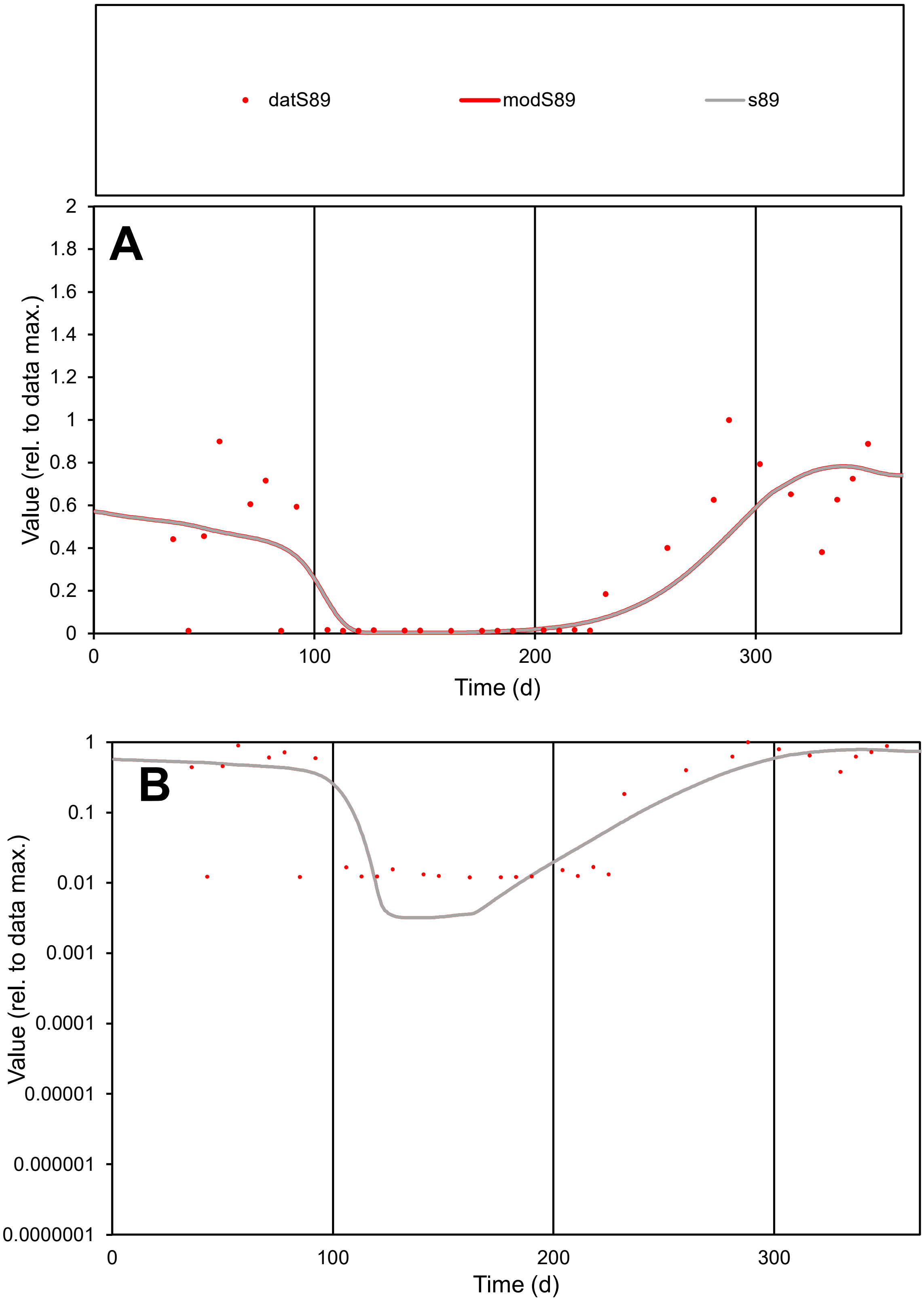


**C1**

Variables
S89 Provide jtd or Dst
CalibOnly?
2 1 to reproduce of
oNormalize?
2 1=no, 2=yes

ofxn calc*
0.64501
ofxn mod
0.65686

sRun
RunA1
zout
1

Ave
Data
0.00094
Model
0.00088

data/vMax
0.00295

ofxng
321.551

Min d/m
3.75704

ave/min d
26.6158
ave/min m
93.9281

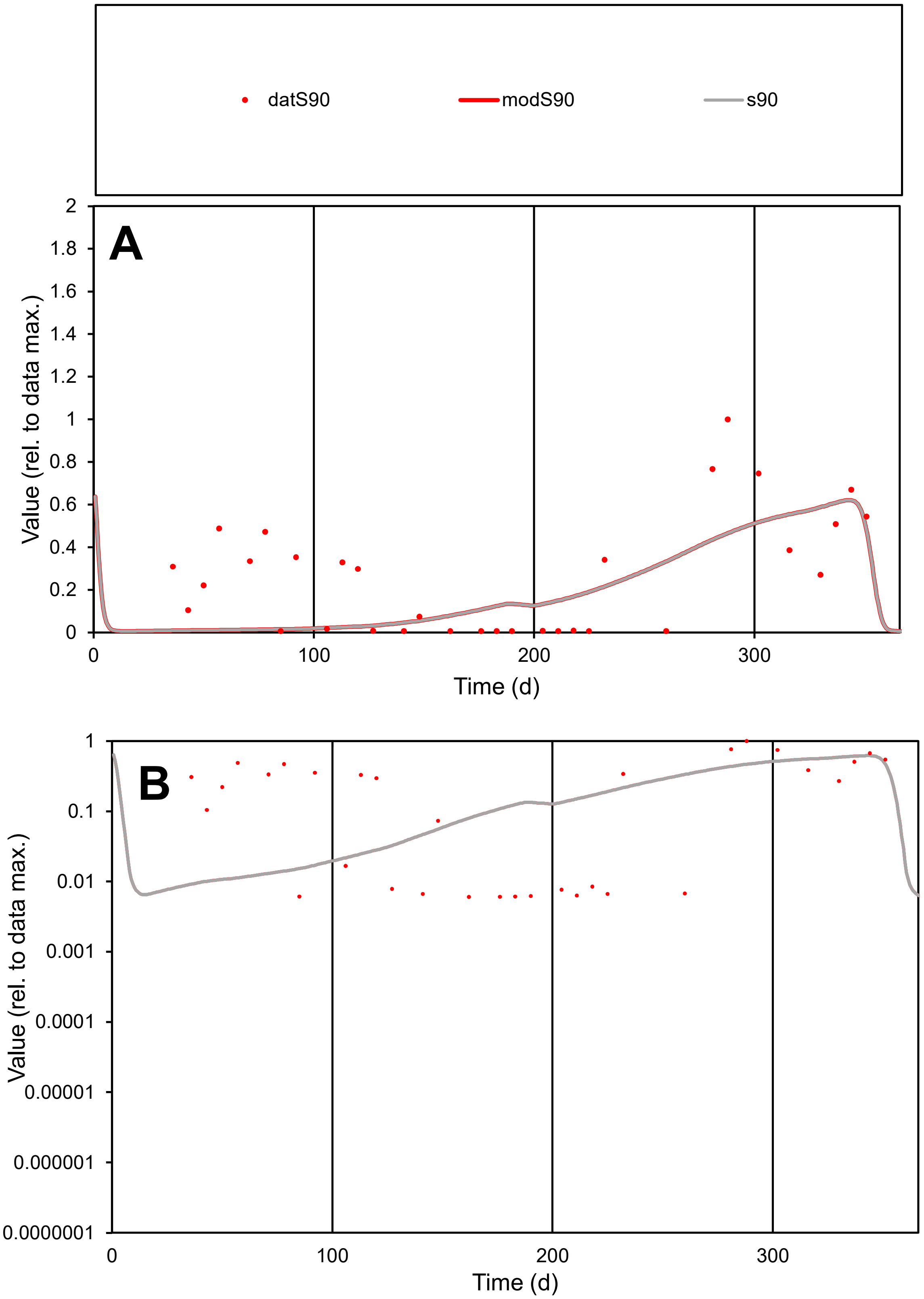


C1

Variables
S90 Provide jtd or Dst
CalibOnly?
2 1 to reproduce of
oNormalize?
2 1=no, 2=yes

ofxn calc*
0.91971
ofxn mod
0.92672

sRun
RunA1
zout
1

Ave
Data
0.00084
Model
0.00063

data/vMax
0.00324

ofxng
321.551

Min d/m
0.64585

ave/min d
43.0877
ave/min m
20.7562

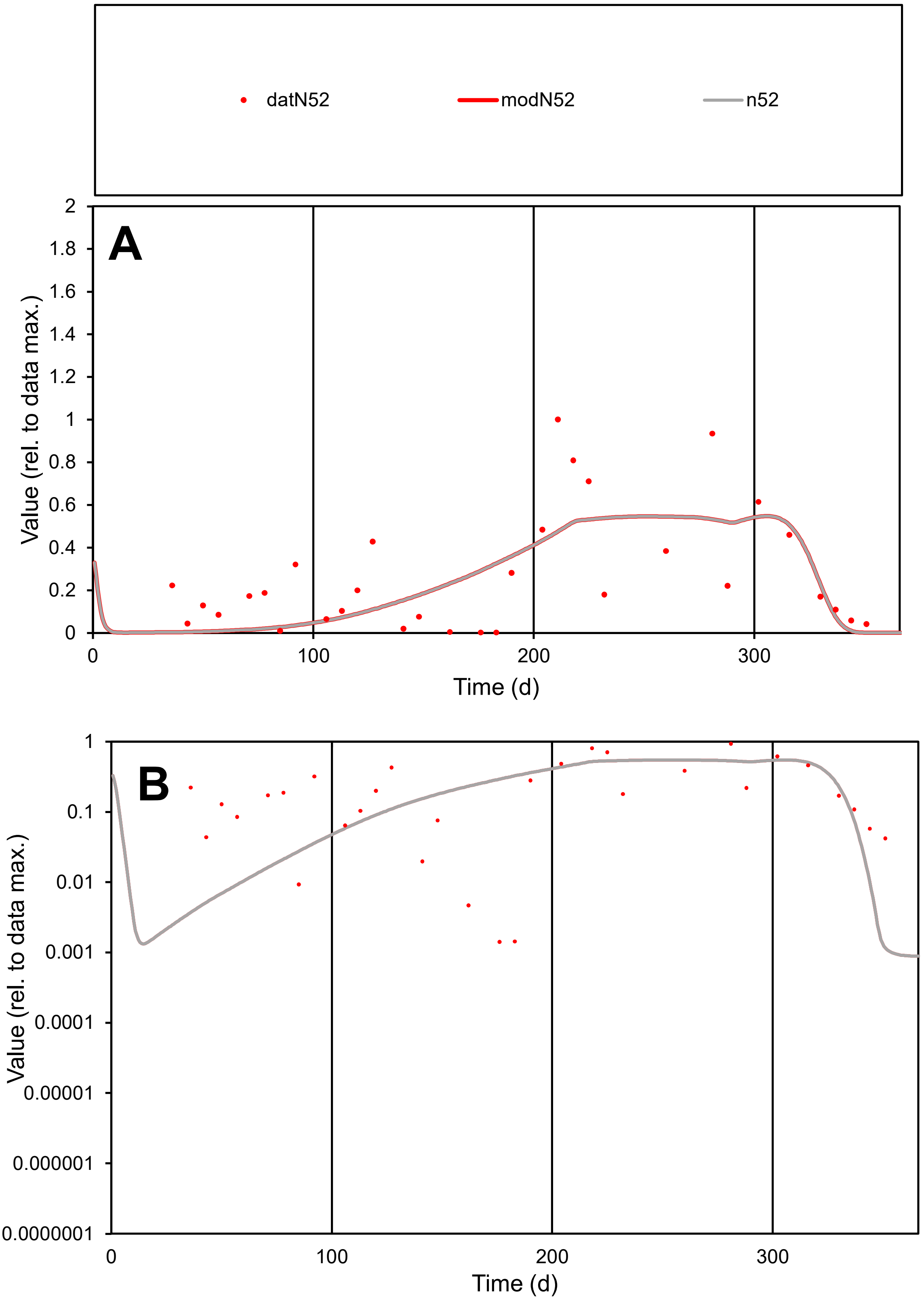


**C1**

Variables
N52 Provide jtd or Dst
CalibOnly?
2 1 to reproduce of
oNormalize?
2 1=no, 2=yes

ofxn calc*
0.78216
ofxn mod
0.79324

sRun
RunA1
zout
1

Ave
Data
0.00444
Model
0.00389

data/vMax
0.01667

ofxng
321.551

Min d/m
1.29841

ave/min d
187.552
ave/min m
213.153

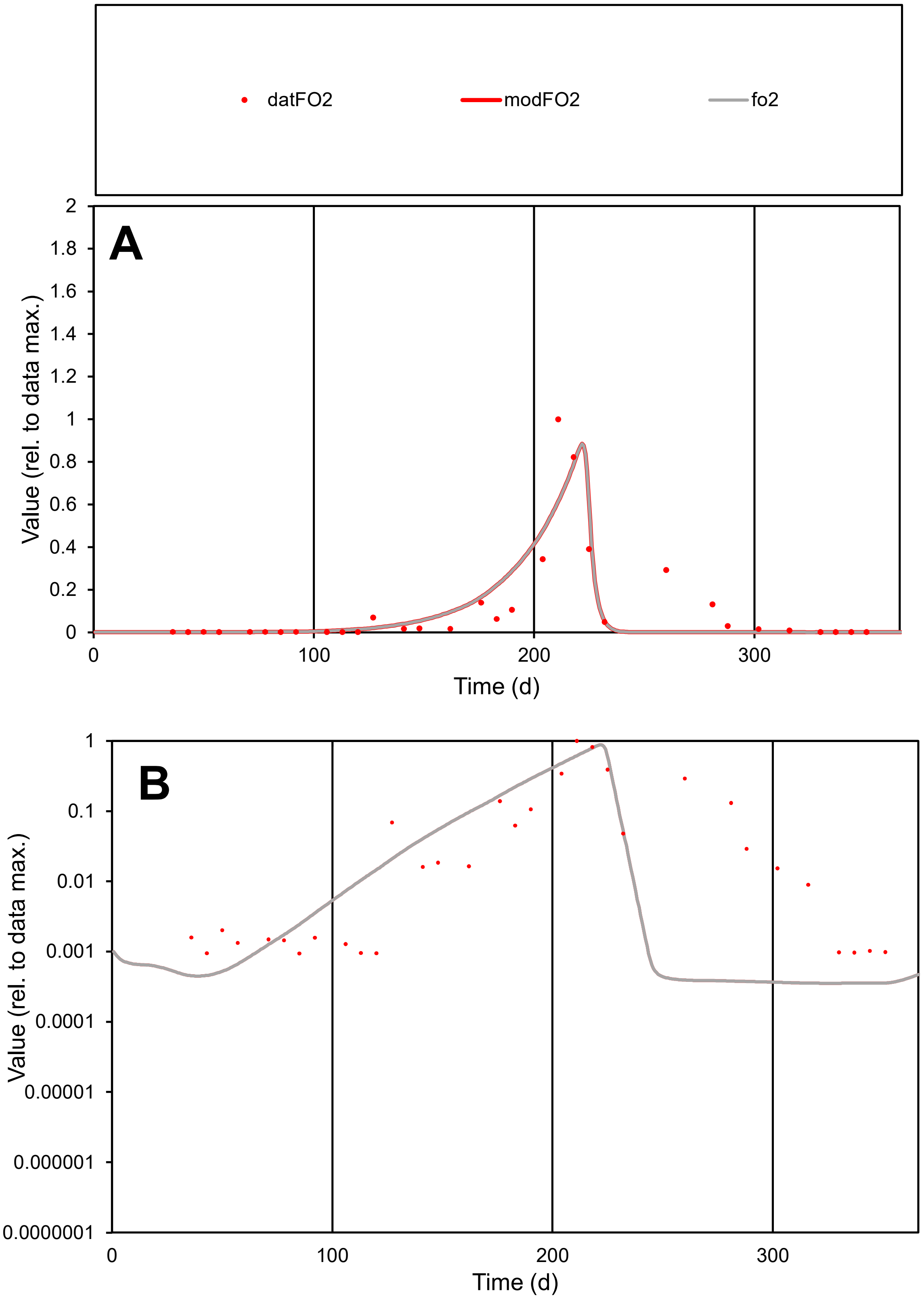


**C1**

Variables
FO2 Provide jtd or Dst
CalibOnly?
2 1 to reproduce of
oNormalize?
2 1=no, 2=yes

ofxn calc*
0.91725
ofxn mod
0.95431

sRun
RunA1
zout
1

Ave
Data
0.00786
Model
0.00753

data/vMax
0.07133

ofxng
321.551

Min d/m
2.65317

ave/min d
117.545
ave/min m
298.795

324_Calib_S_A

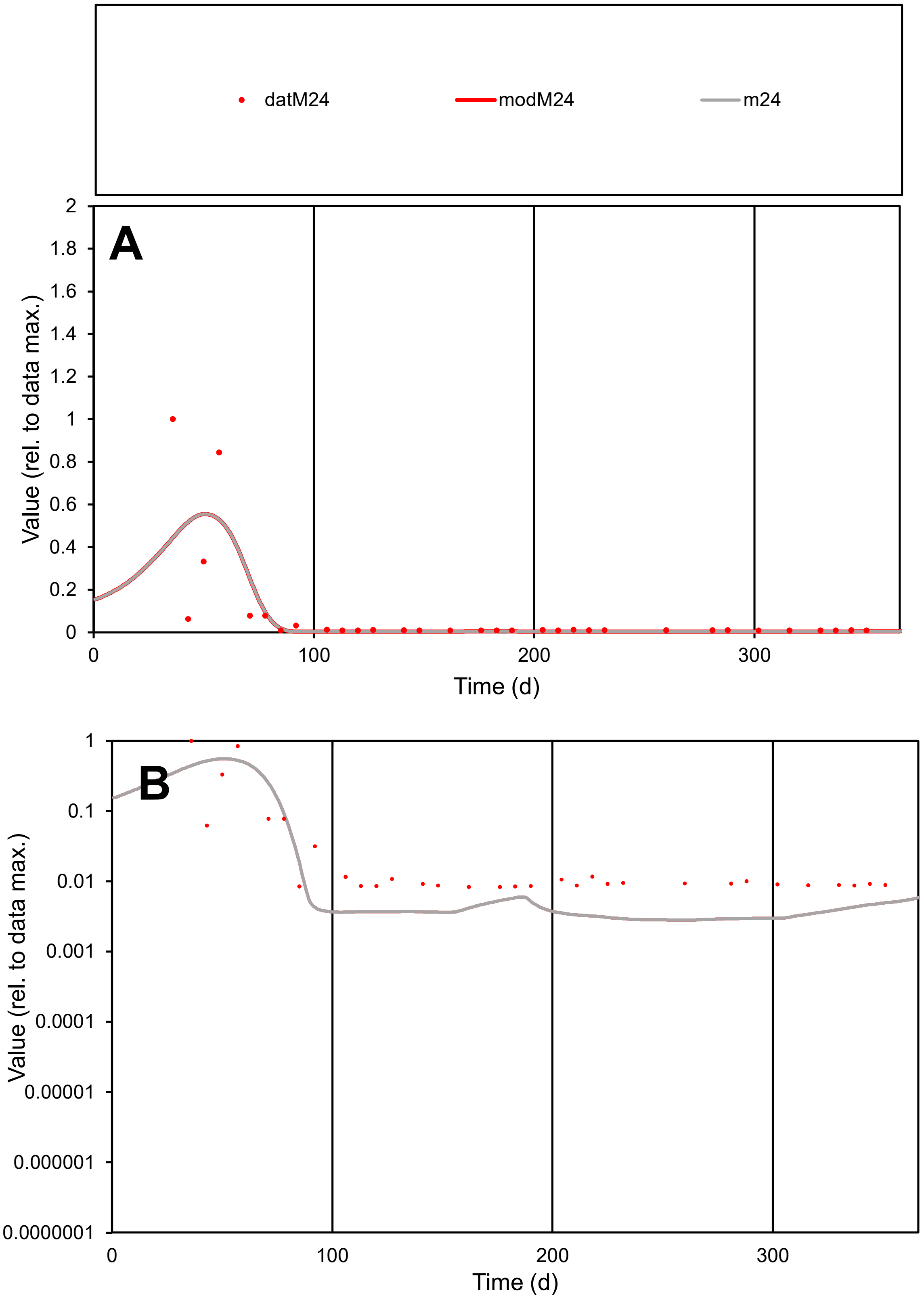


**C1**

Variables
M24 Provide jtd or Dst
CalibOnly?
2 1 to reproduce of
oNormalize?
2 1=no, 2=yes

ofxn calc*
1.75155
ofxn mod
1.77155

sRun
RunA1
zout
1

Ave
Data
0.00071
Model
0.00064

data/vMax
0.0085

ofxng
321.551

Min d/m
2.95411
ave/min d
9.97386
ave/min m
26.8905

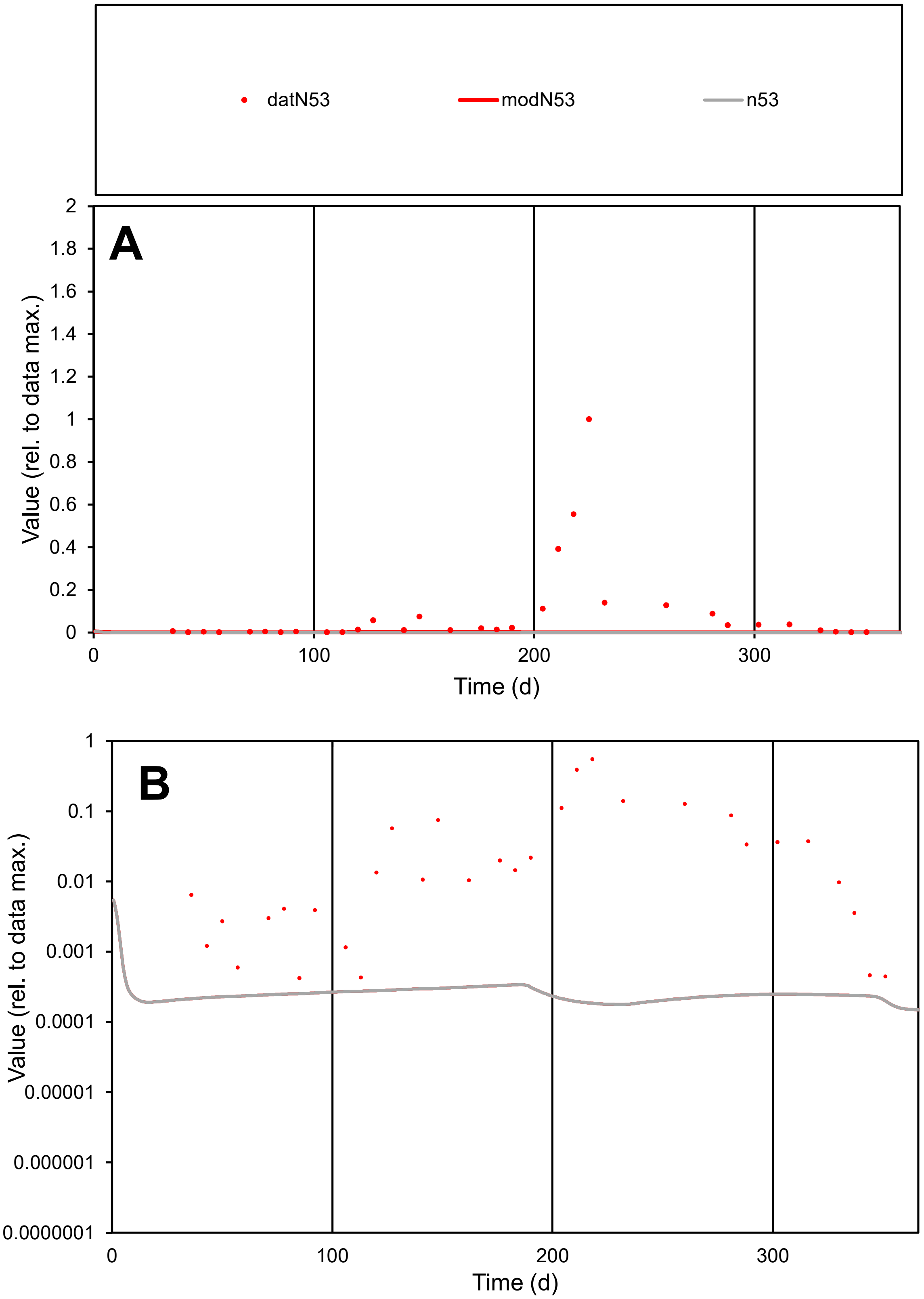


**C1**

Variables
N53 Provide jtd or Dst
CalibOnly?
2 1 to reproduce of
oNormalize?
2 1=no, 2=yes

ofxn calc*
2.5159
ofxn mod
2.51886

sRun
RunA1
zout
1

Ave
Data
0.00747
Model
2.1E-05

data/vMax
0.08598

ofxng
321.551

Min d/m
2.38376

ave/min d
205.257
ave/min m
1.4014

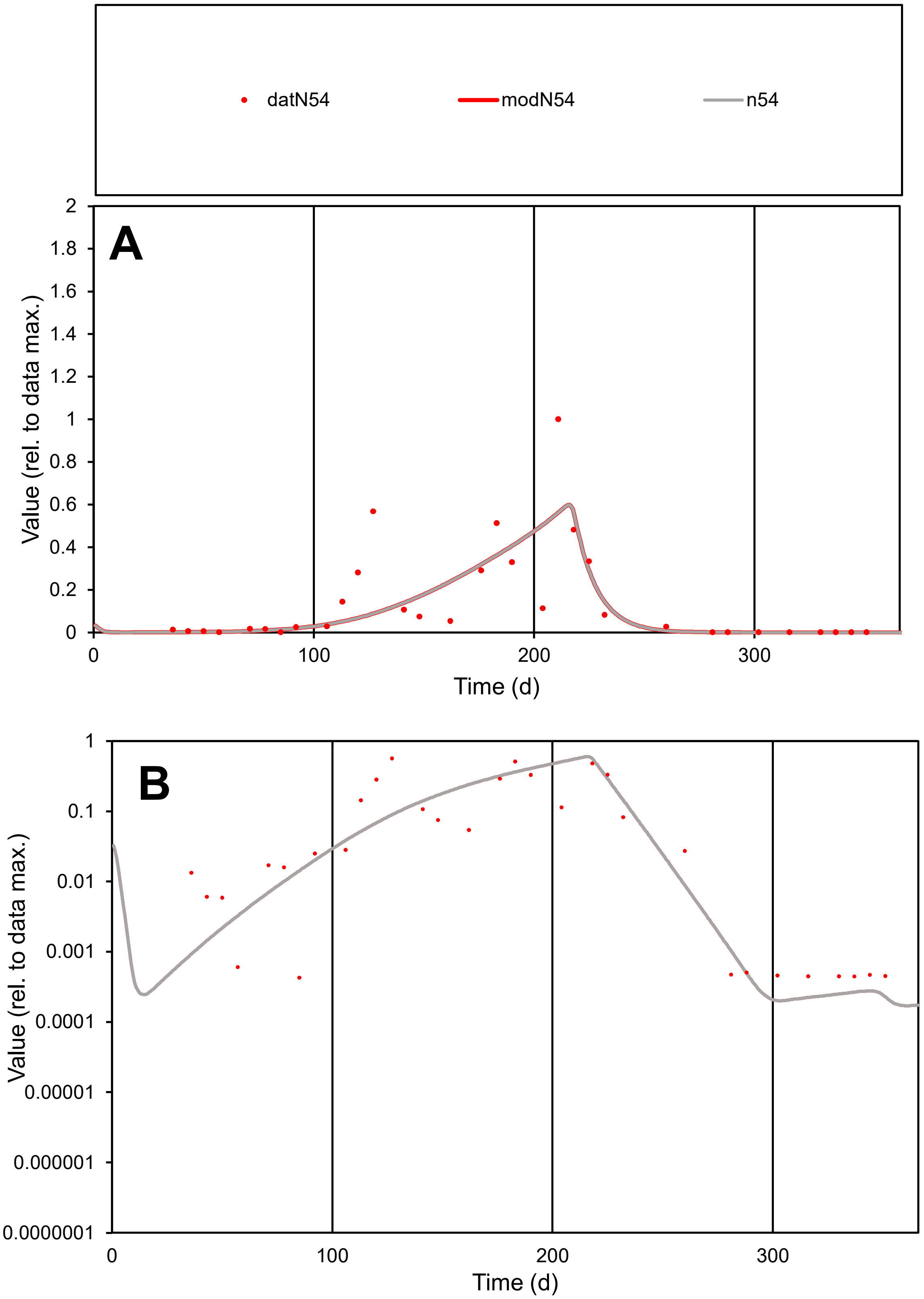


C1

| | |
|---|---|
| Variables | |
| N54 | Provide jtd or Dst |
| CalibOnly? | |
| 2 | 1 to reproduce of |
| oNormalize? | |
| 2 | 1=no, 2=yes |
| ofxn calc* | |
| 1.04284 | |
| ofxn mod | |
| 1.06303 | |
| sRun | |
| RunA1 | |
| zout | |
| 1 | |
| Ave | |
| Data | |
| 0.00708 | |
| Model | |
| 0.00623 | |
| data/vMax | |
| 0.05014 | |
| ofxng | |
| 321.551 | |
| Min d/m | |
| 2.14756 | |
| ave/min d | |
| 328.705 | |
| ave/min m | |
| 621.6 | |

324_Calib_S_A

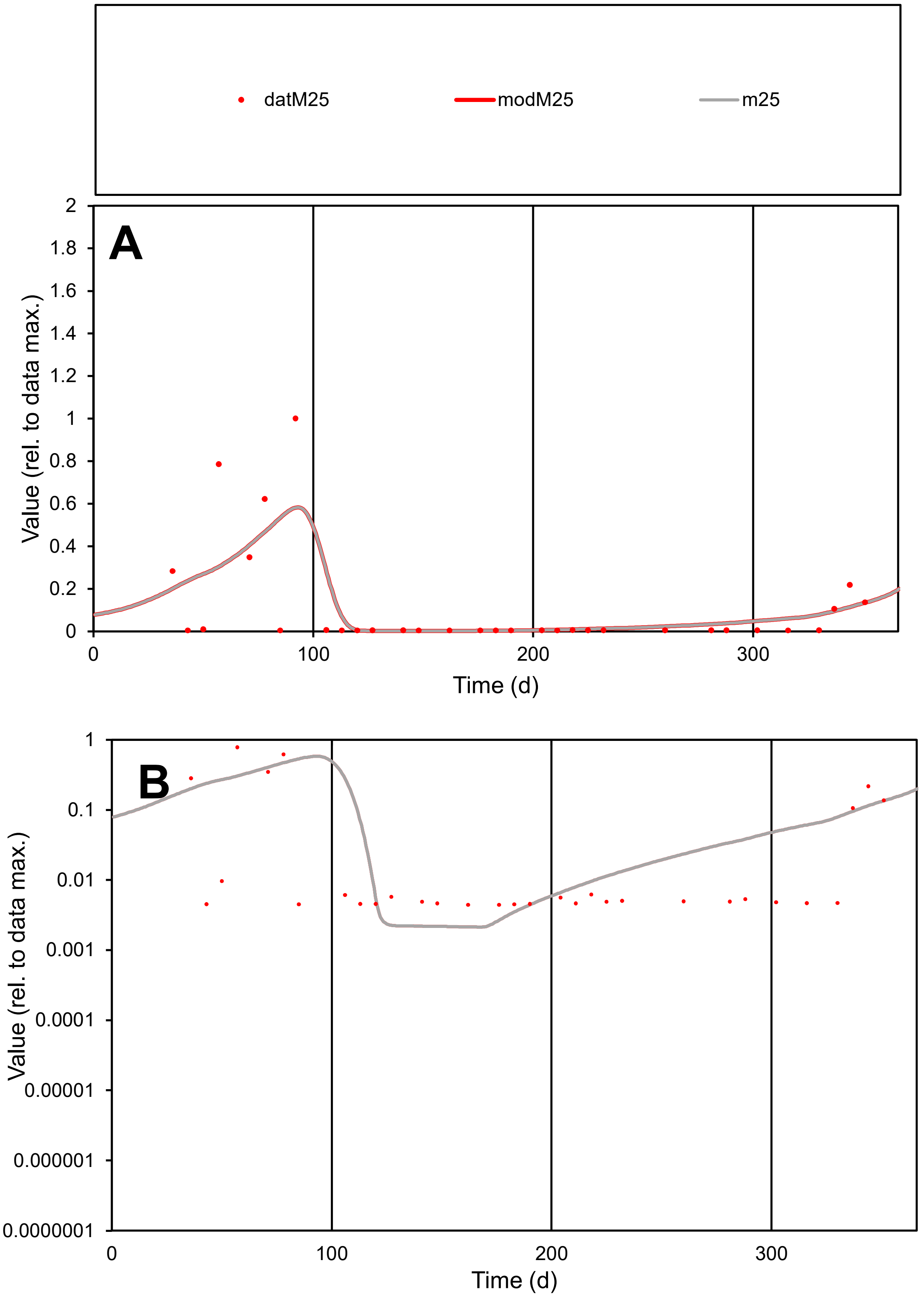


**C1**

Variables
M25 Provide jtd or Dst
CalibOnly?
2 1 to reproduce of
oNormalize?
2 1=no, 2=yes

ofxn calc*
1.45607
ofxn mod
1.48844

sRun
RunA1
zout
1

Ave
Data
0.00024
Model
0.00026

data/vMax
0.00213

ofxng
321.551

Min d/m
2.0574

ave/min d
25.6214
ave/min m
56.8642

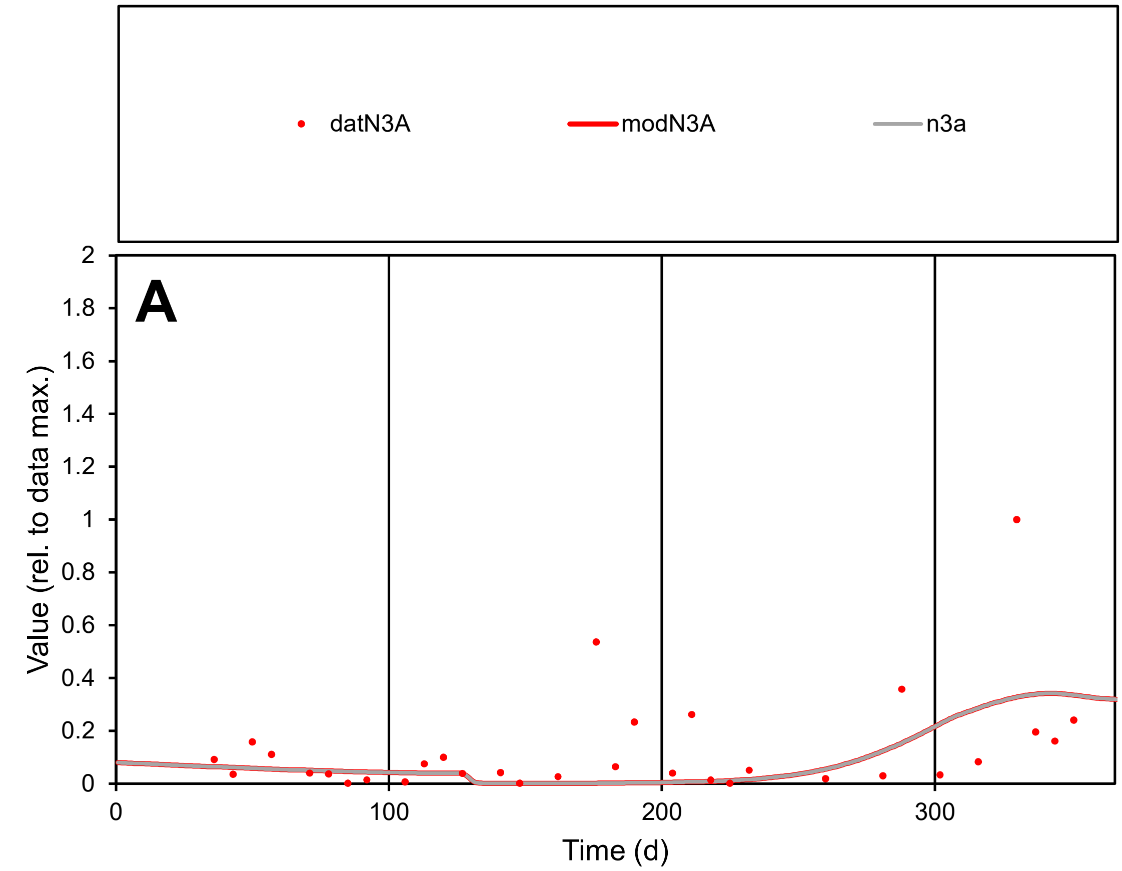


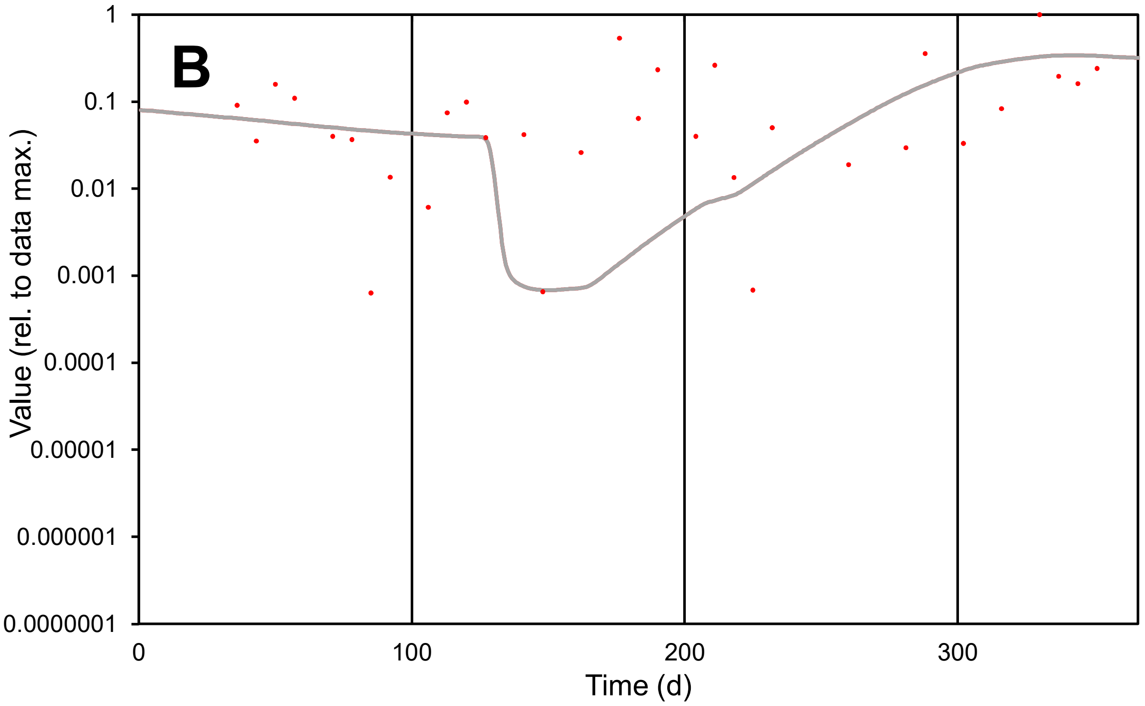


**C1**

Variables

N3A Provide jtd or Dst

CalibOnly?

2 1 to reproduce of

oNormalize?

2 1=no, 2=yes

ofxn calc*

1.43433

ofxn mod

1.44323

sRun

RunA1

zout

1

Ave

Data

0.0036

Model

0.00251

data/vMax

0.02818

ofxng

321.551

Min d/m

0.92909

ave/min d

201.995

ave/min m

130.697

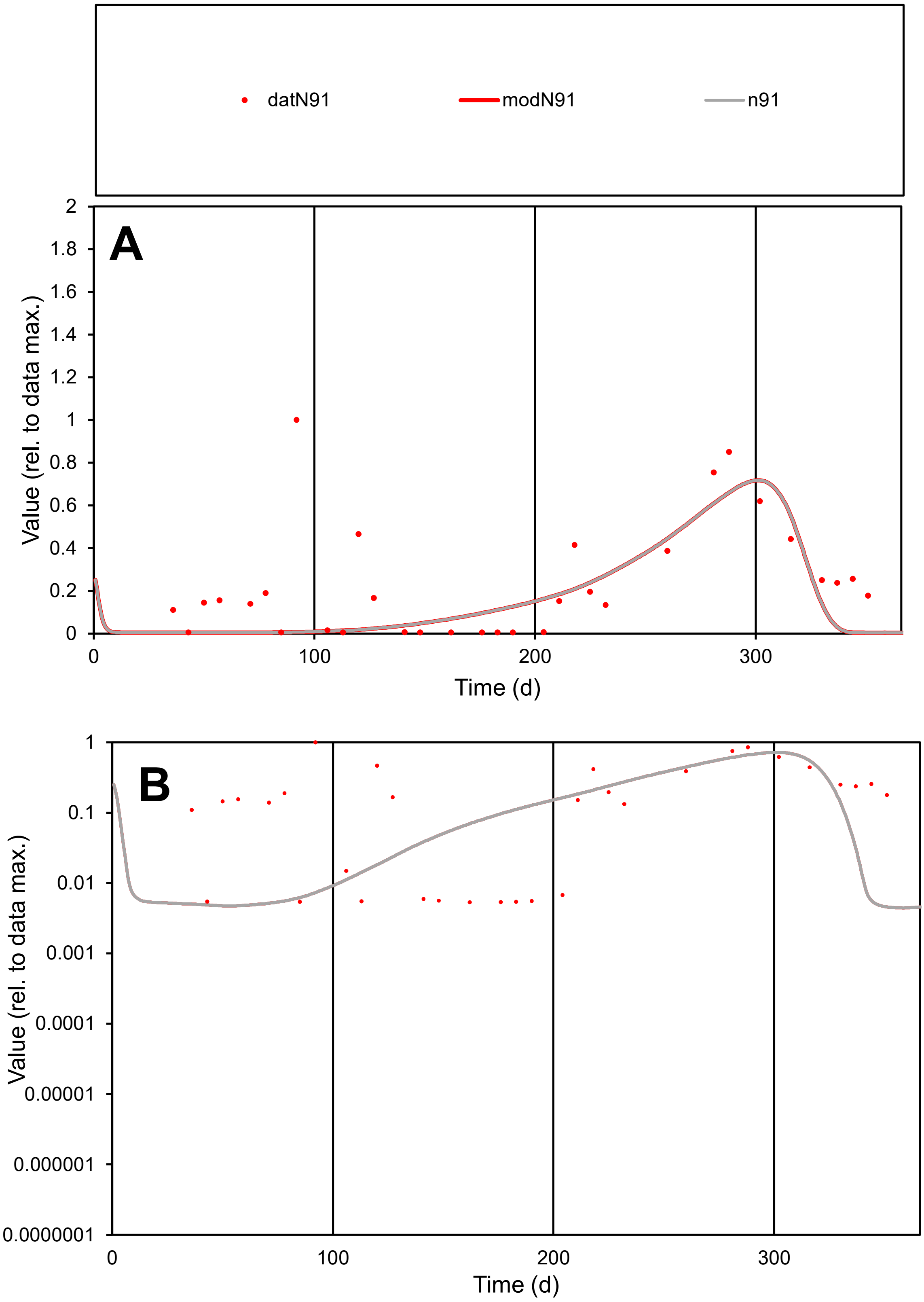


**C1**

Variables
N91 Provide jtd or Dst
CalibOnly?
2 1 to reproduce of
oNormalize?
2 1=no, 2=yes

ofxn calc*
0.9955
ofxn mod
1.00435

sRun
RunA1
zout
1

Ave
Data
0.00132
Model
0.00088

data/vMax
0.00579

ofxng
321.551

Min d/m
1.1804

ave/min d
42.7666
ave/min m
33.5727

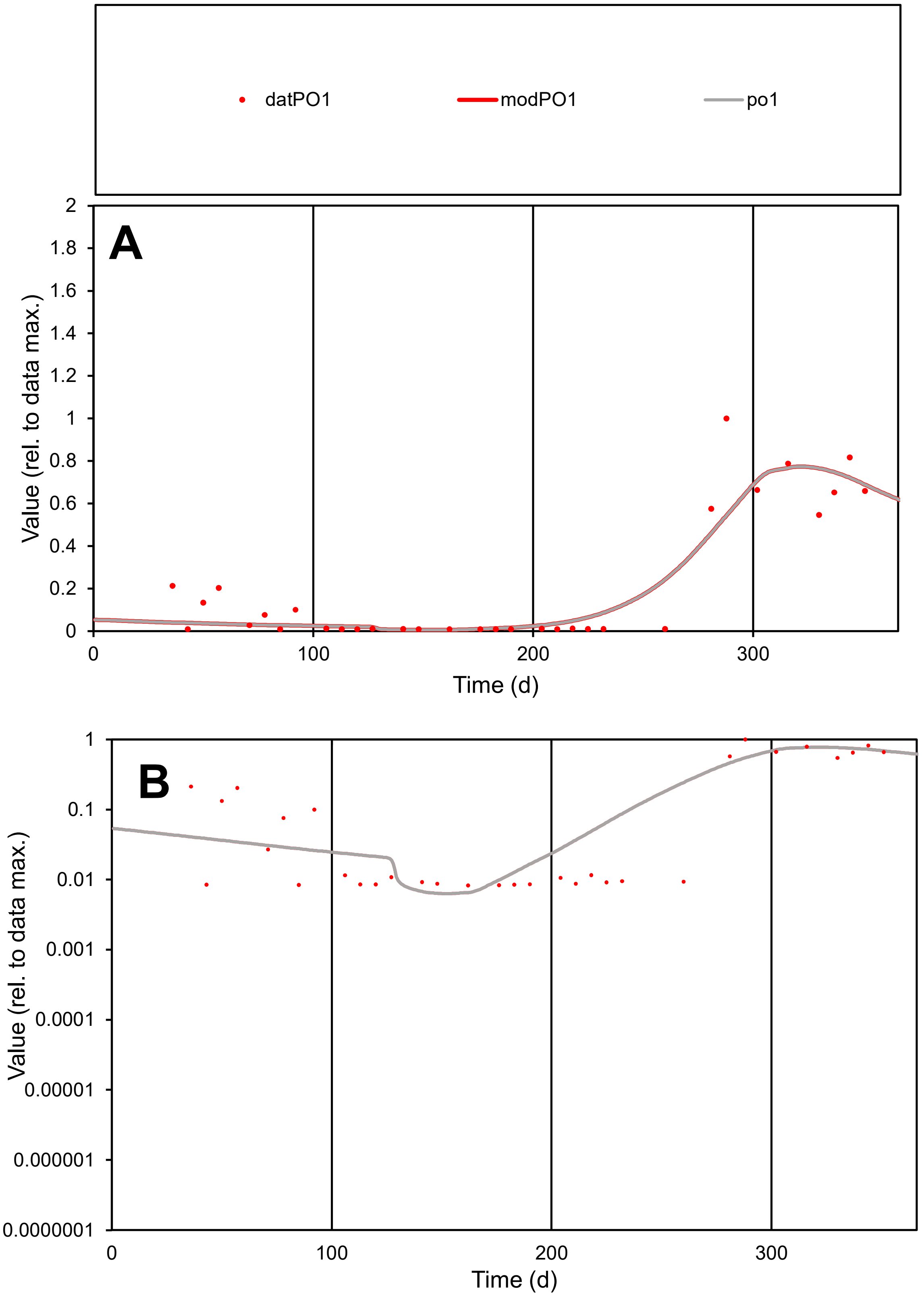


**C1**

Variables
PO1 Provide jtd or Dst
CalibOnly?
2 1 to reproduce of
oNormalize?
2 1=no, 2=yes

ofxn calc*
0.551
ofxn mod
0.57368

sRun
RunA1
zout
1

Ave
Data
0.00111
Model
0.00107

data/vMax
0.00539

ofxng
321.551

Min d/m
1.31085

ave/min d
24.907
ave/min m
31.303

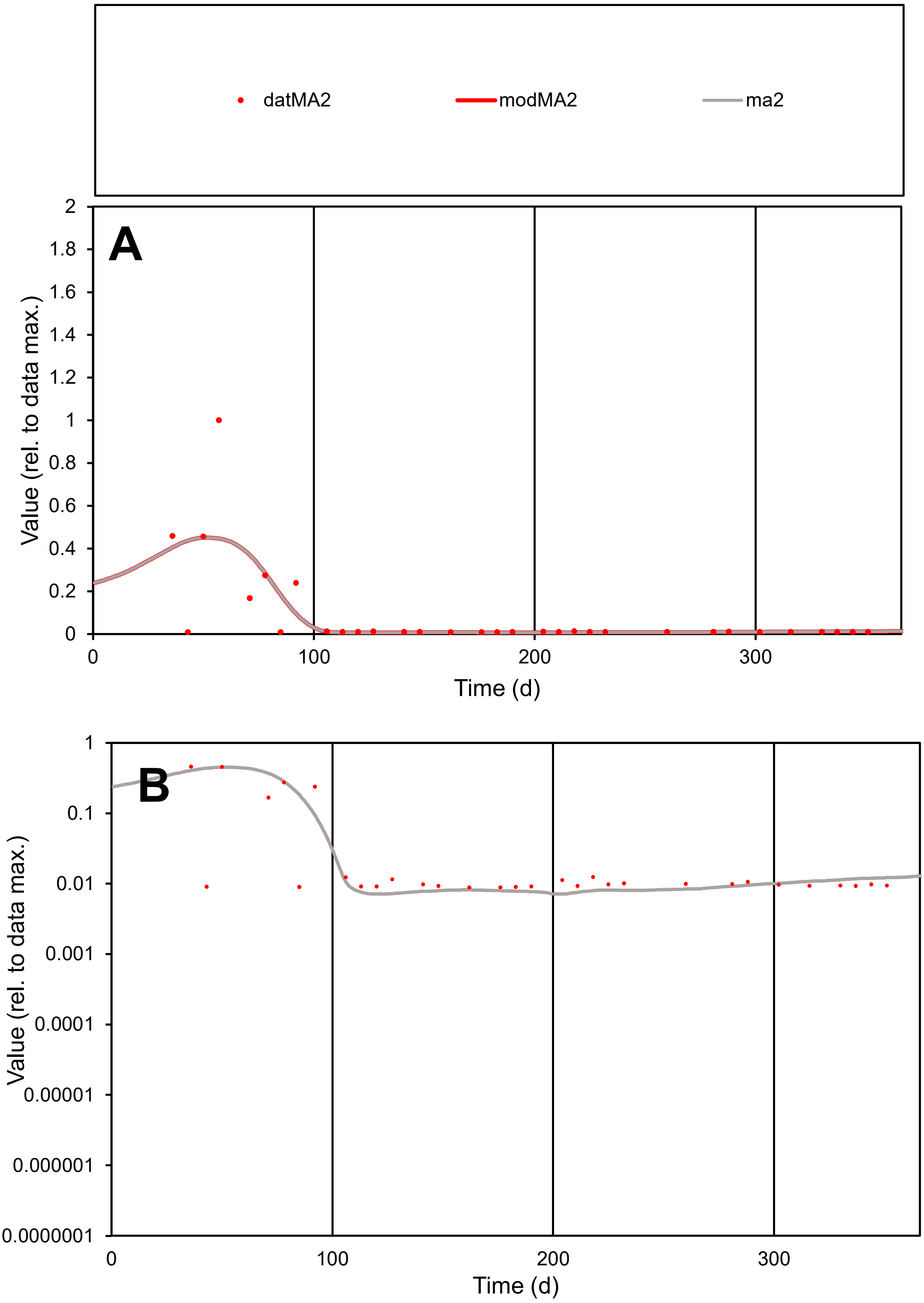


**C1**

Variables
MA2 Provide jtd or Dst
CalibOnly?
2 1 to reproduce of
oNormalize?
2 1=no, 2=yes

ofxn calc*
1.50081
ofxn mod
1.52029

sRun
RunA1
zout
1

Ave
Data
0.00018
Model
0.00017

data/vMax
0.00197

ofxng
321.551

Min d/m
1.23955

ave/min d
10.0576
ave/min m
12.3286

324_Calib_S_A

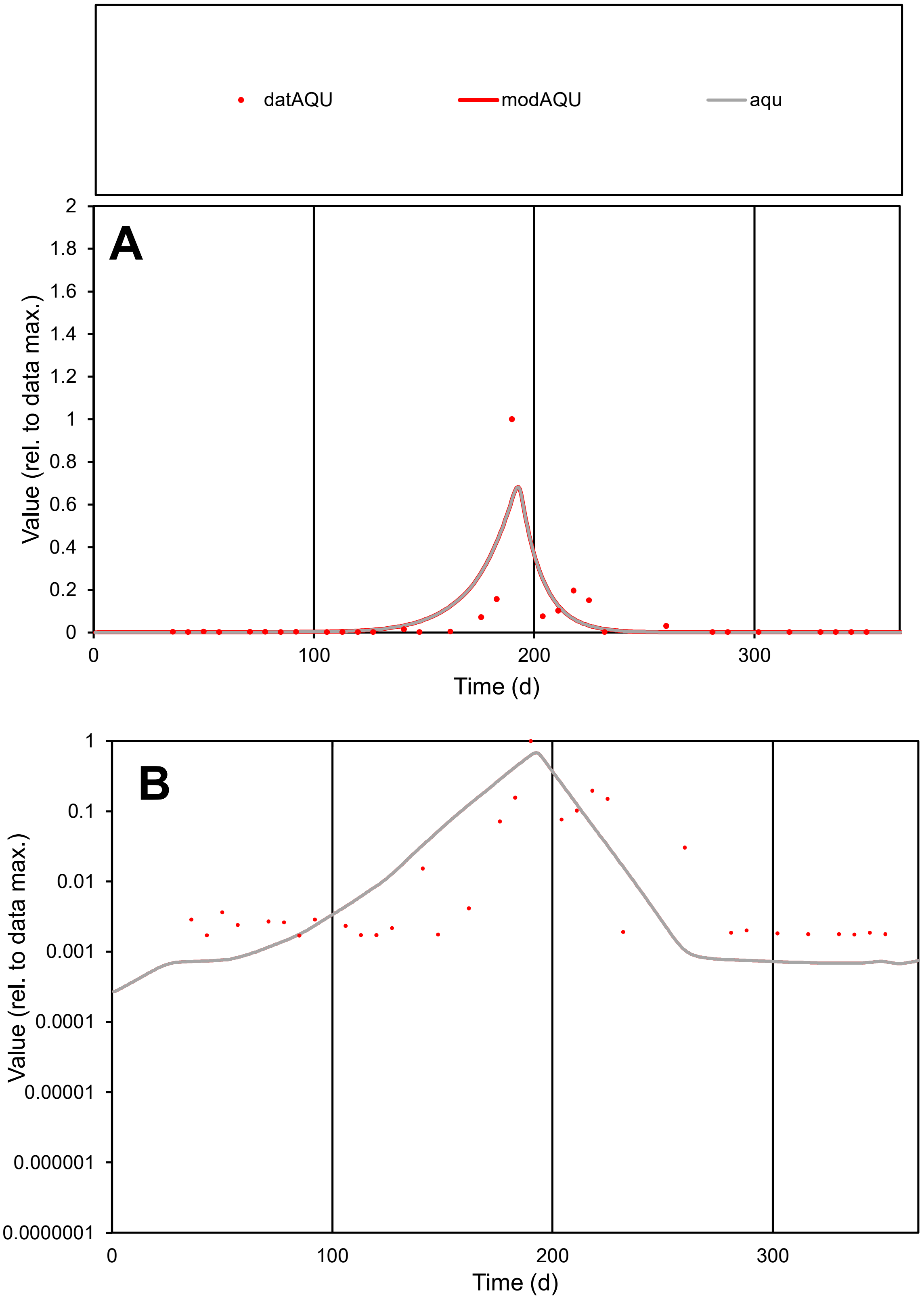


**C1**

Variables
AQU Provide jtd or Dst
CalibOnly?
2 1 to reproduce of
oNormalize?
2 1=no, 2=yes

ofxn calc*
1.65995
ofxn mod
1.72021

sRun
RunA1
zout
1

Ave
Data
0.0044
Model
0.00481

data/vMax
0.07607

ofxng
321.551

Min d/m
2.47801

ave/min d
33.939
ave/min m
91.8741

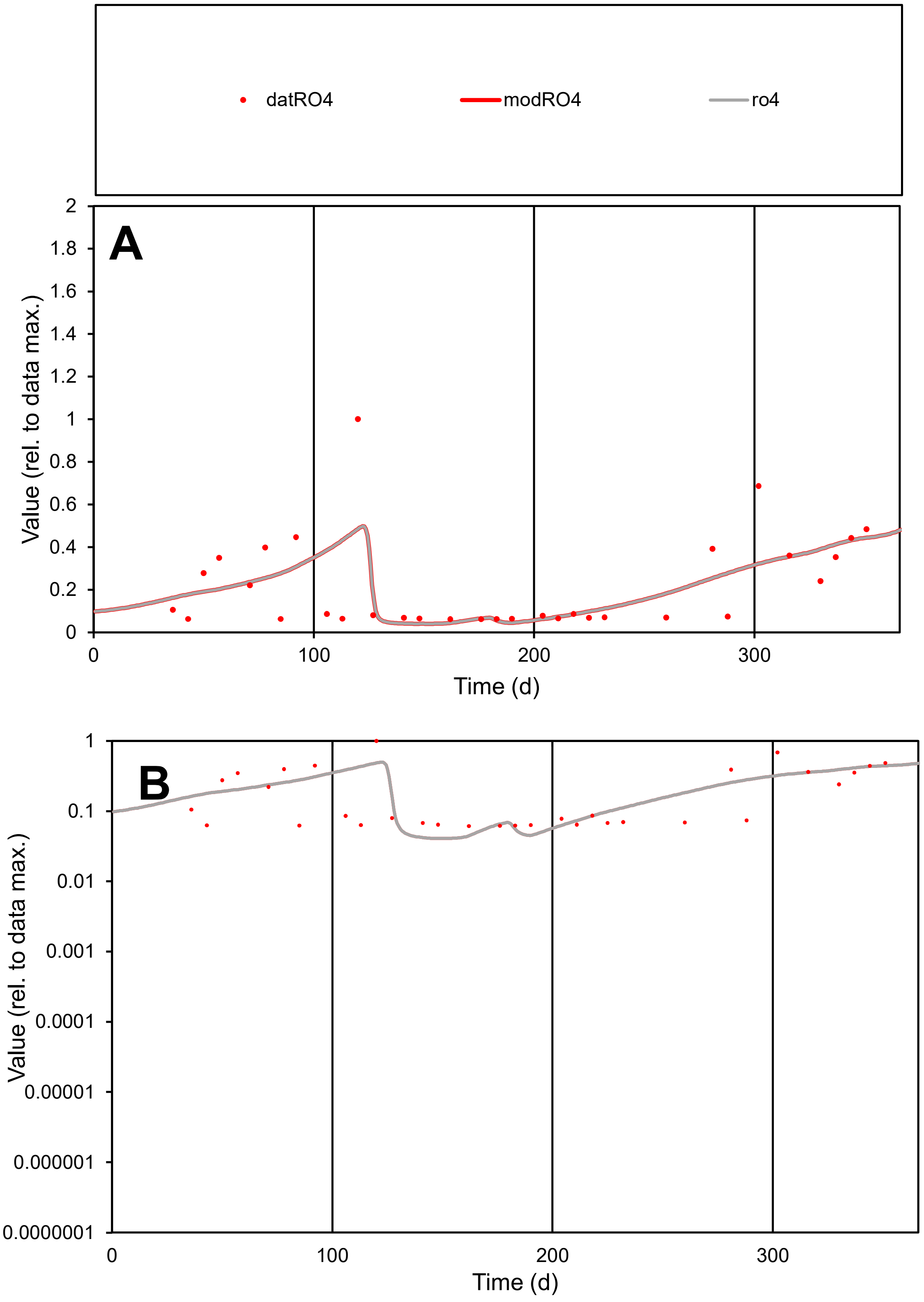


**C1**

Variables
RO4 Provide jtd or Dst
CalibOnly?
2 1 to reproduce of
oNormalize?
2 1=no, 2=yes

ofxn calc*
0.74542
ofxn mod
0.75777

sRun
RunA1
zout
1

Ave
Data
0.00076
Model
0.00077

data/vMax
0.00347

ofxng
321.551

Min d/m
1.5007

ave/min d
3.56181
ave/min m
5.41761

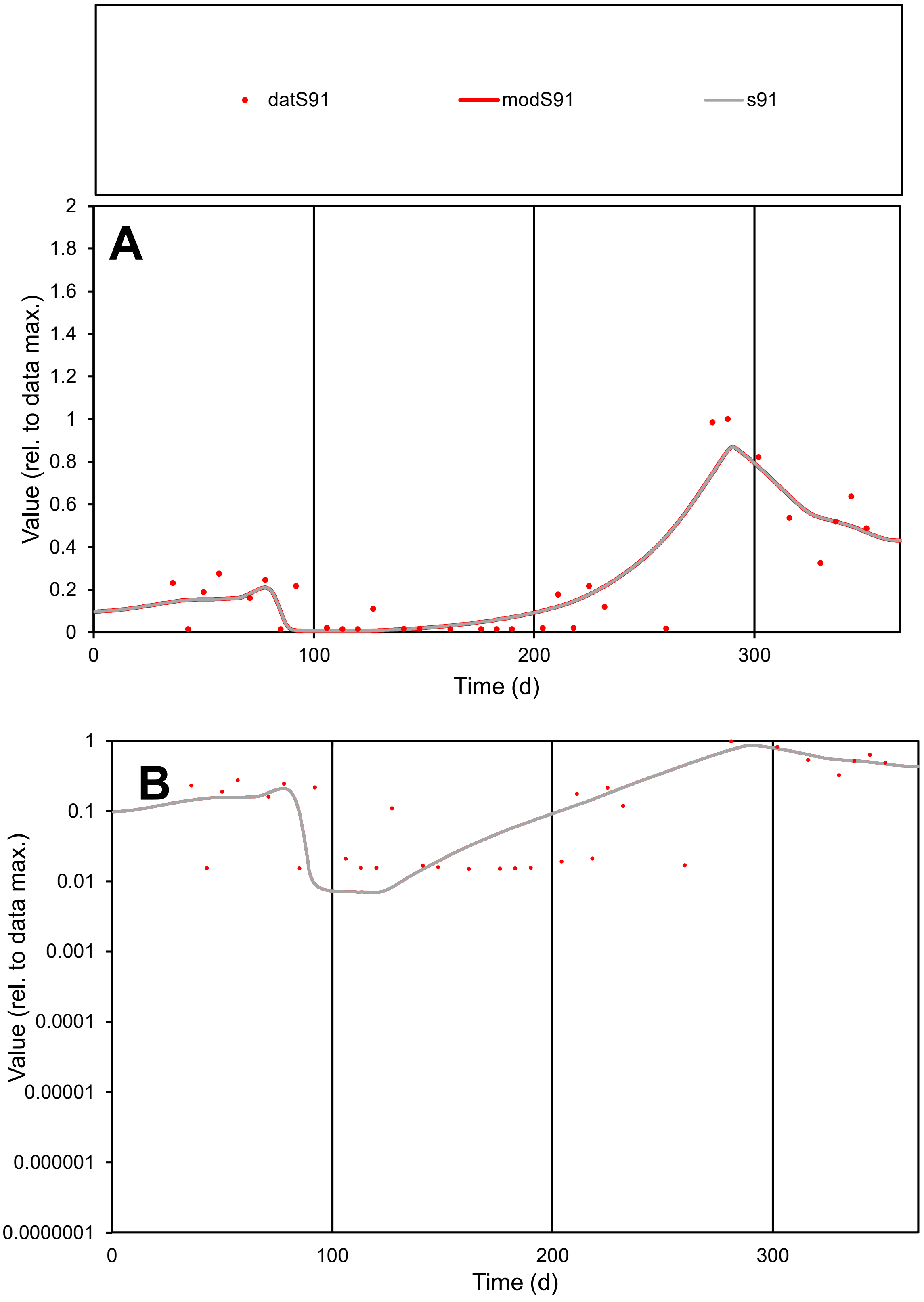


**C1**

Variables
S91 Provide jtd or Dst
CalibOnly?
2 1 to reproduce of
oNormalize?
2 1=no, 2=yes

ofxn calc*
0.52135
ofxn mod
0.5409

sRun
RunA1
zout
1

Ave
Data
0.00099
Model
0.00101

data/vMax
0.00425

ofxng
321.551

Min d/m
2.15542

ave/min d
15.4874
ave/min m
33.8976

324_Calib_S_A

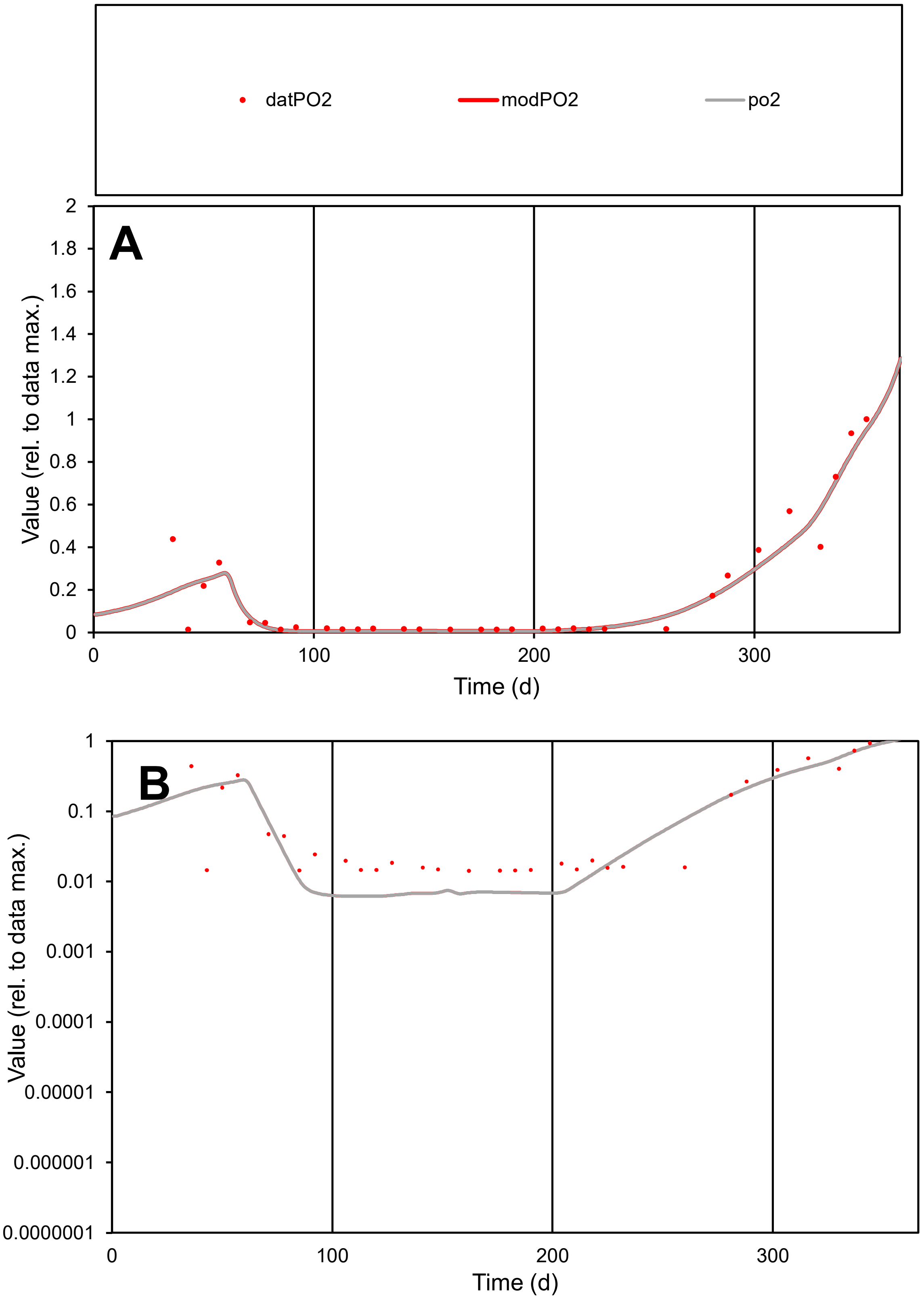


**C1**

Variables

PO2 Provide jtd or Dst

CalibOnly?

2 1 to reproduce of

oNormalize?

2 1=no, 2=yes

ofxn calc*

0.40633

ofxn mod

0.44106

sRun

RunA1

zout

1

Ave

Data

0.001

Model

0.00095

data/vMax

0.00548

ofxng

321.551

Min d/m

2.29743

ave/min d

12.8647

ave/min m

28.1401

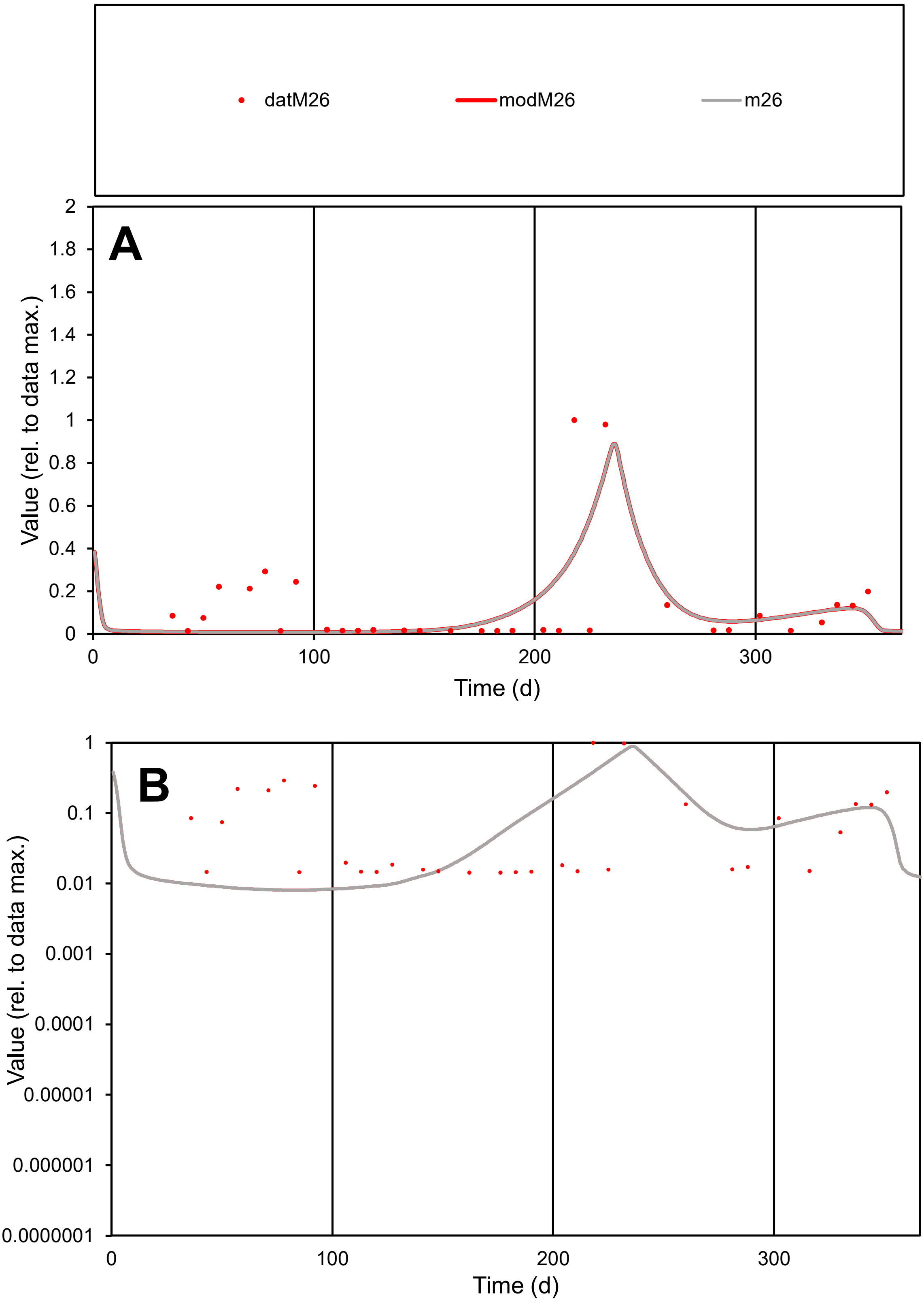


**C1**

Variables
M26 Provide jtd or Dst
CalibOnly?
2 1 to reproduce of
oNormalize?
2 1=no, 2=yes

ofxn calc*
1.34362
ofxn mod
1.39181

sRun
RunA1
zout
1

Ave
Data
0.00072
Model
0.00061

data/vMax
0.00562

ofxng
321.551

Min d/m
1.77458

ave/min d
9.04196
ave/min m
13.4113

324_Calib_S_A

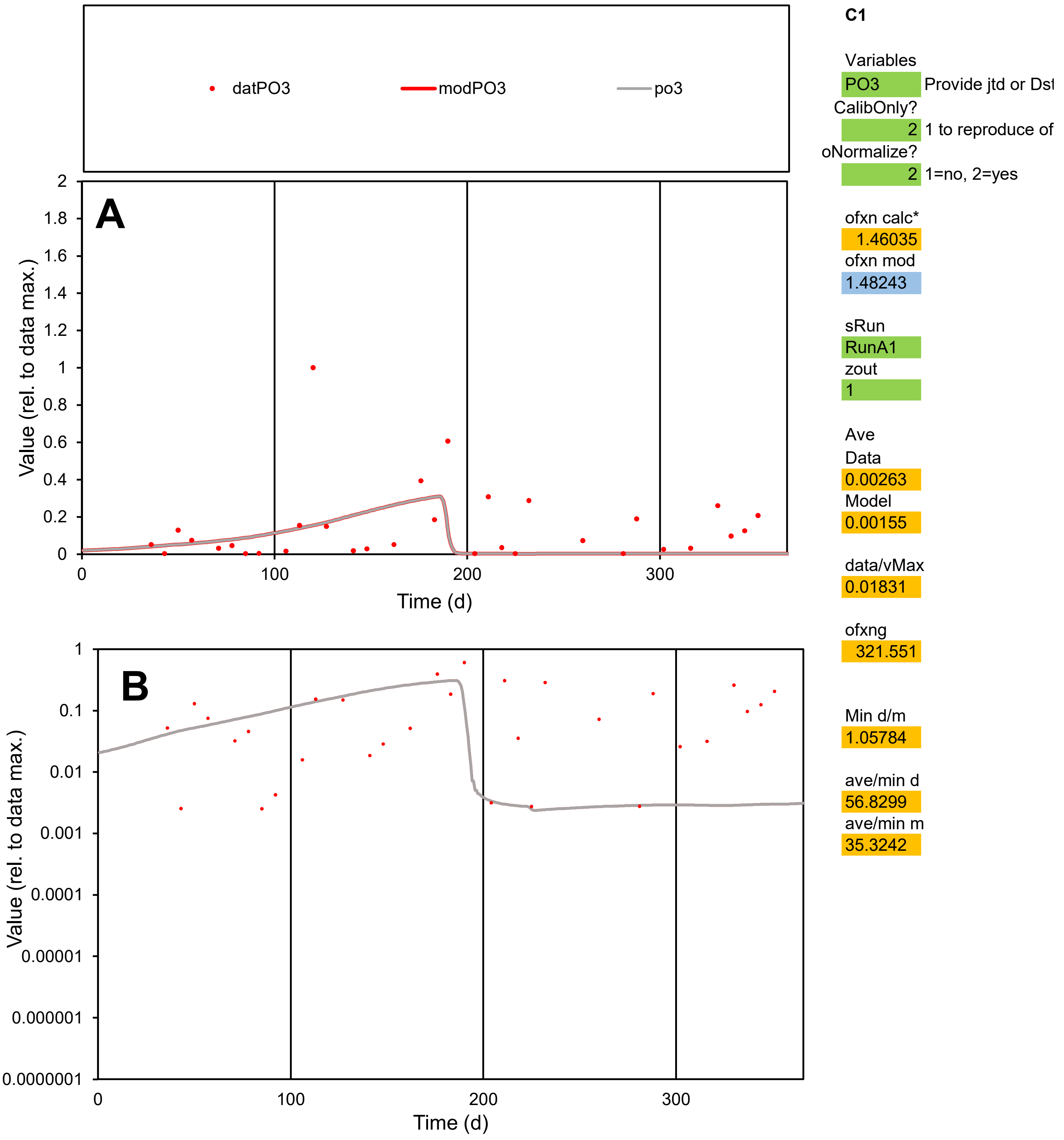

datPO3
modPO3
po3
A
Value (rel. to data max.)
Time (d)
B
Value (rel. to data max.)
Time (d)
C1
Variables
PO3
Provide jtd or Dst
CalibOnly?
2
1 to reproduce of
oNormalize?
2
1=no, 2=yes
ofxn calc*
1.46035
ofxn mod
1.48243
sRun
RunA1
zout
1
Ave
Data
0.00263
Model
0.00155
data/vMax
0.01831
ofxng
321.551
Min d/m
1.05784
ave/min d
56.8299
ave/min m
35.3242

324_Calib_S_A

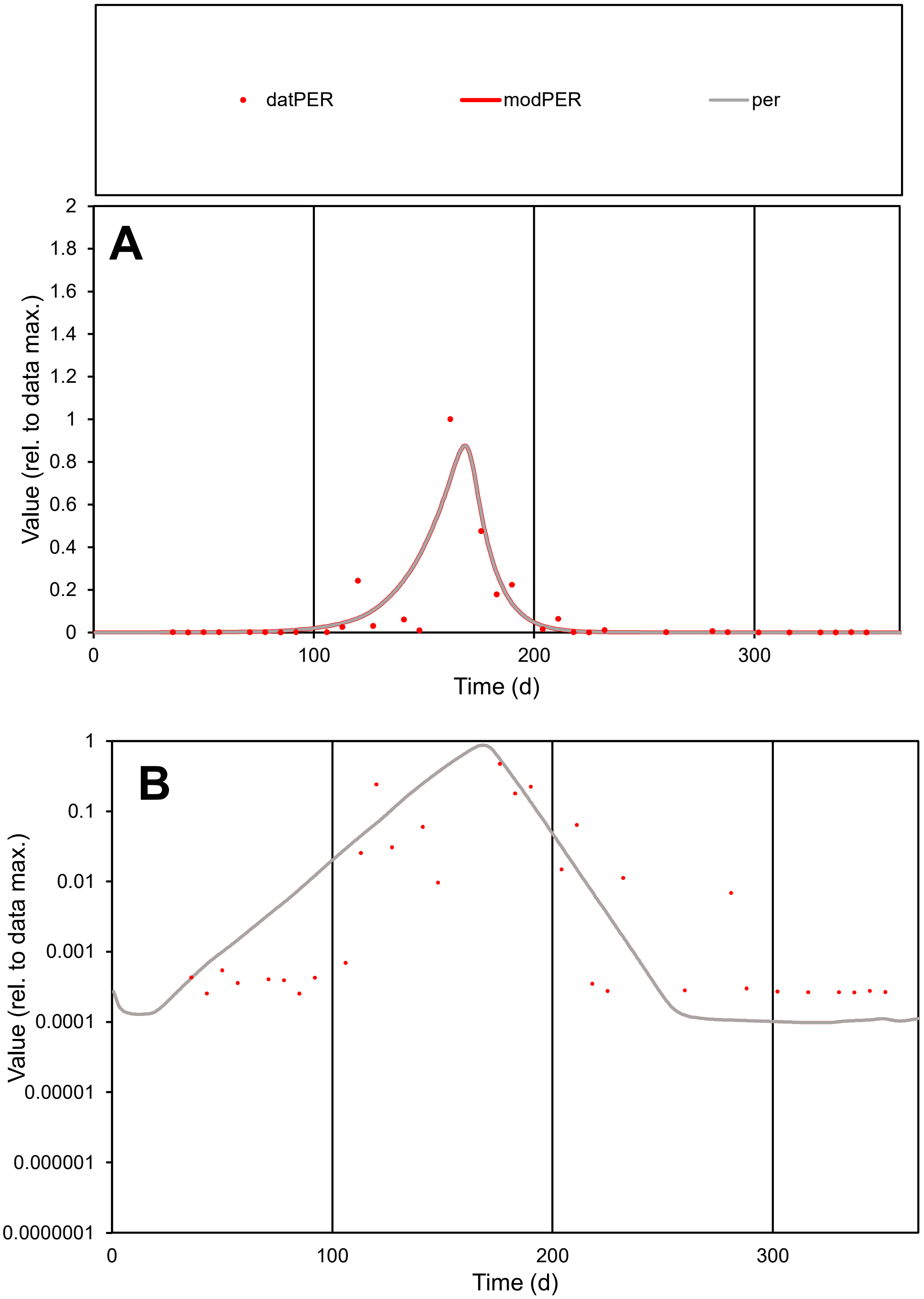


**C1**

Variables
PER Provide jtd or Dst
CalibOnly?
2 1 to reproduce of
oNormalize?
2 1=no, 2=yes

ofxn calc*
1.17676
ofxn mod
1.28516

sRun
RunA1
zout
1

Ave
Data
0.00525
Model
0.00566

data/vMax
0.07156

ofxng
321.551

Min d/m
2.59444

ave/min d
288.729
ave/min m
806.607

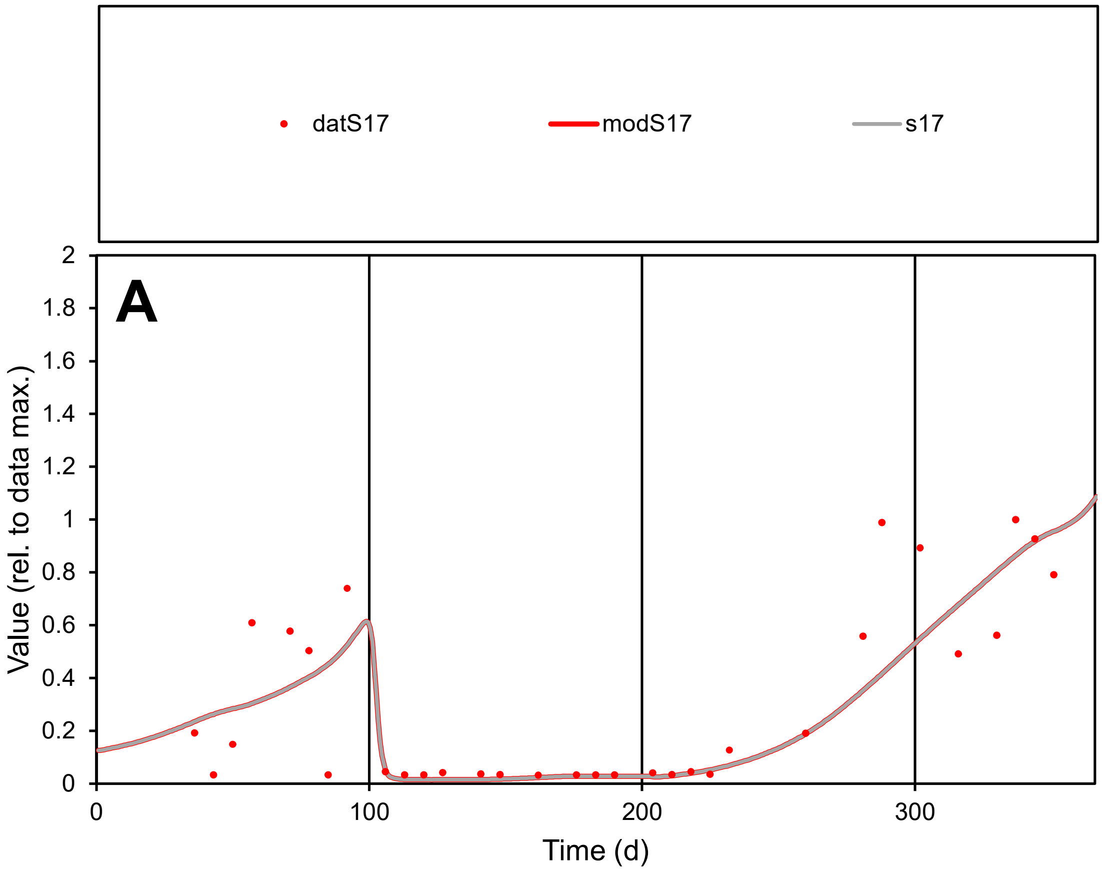


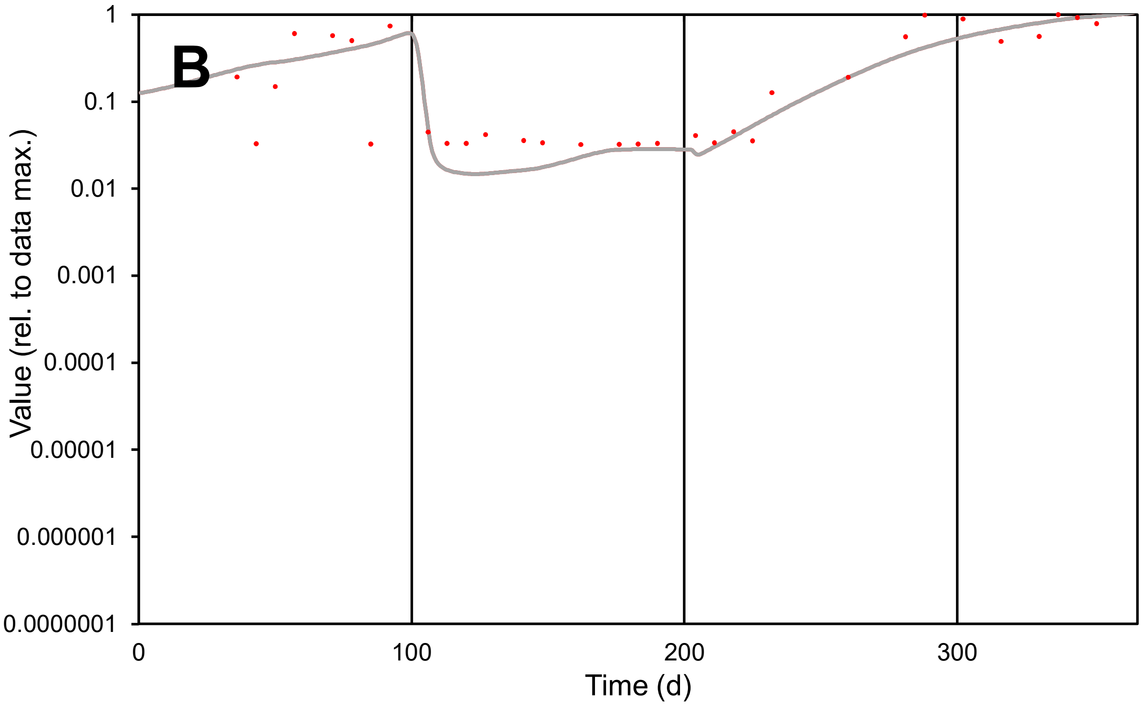


**C1**

| | |
|---|---|
| Variables | |
| S17 | Provide jtd or Dst |
| CalibOnly? | |
| 2 | 1 to reproduce of |
| oNormalize? | |
| 2 | 1=no, 2=yes |
| ofxn calc* | |
| 0.58566 | |
| ofxn mod | |
| 0.59947 | |
| sRun | |
| RunA1 | |
| zout | |
| 1 | |
| Ave | |
| Data | |
| 0.00102 | |
| Model | |
| 0.00095 | |
| data/vMax | |
| 0.00331 | |
| ofxng | |
| 321.551 | |
| Min d/m | |
| 2.17702 | |
| ave/min d | |
| 9.59316 | |
| ave/min m | |
| 19.3758 | |

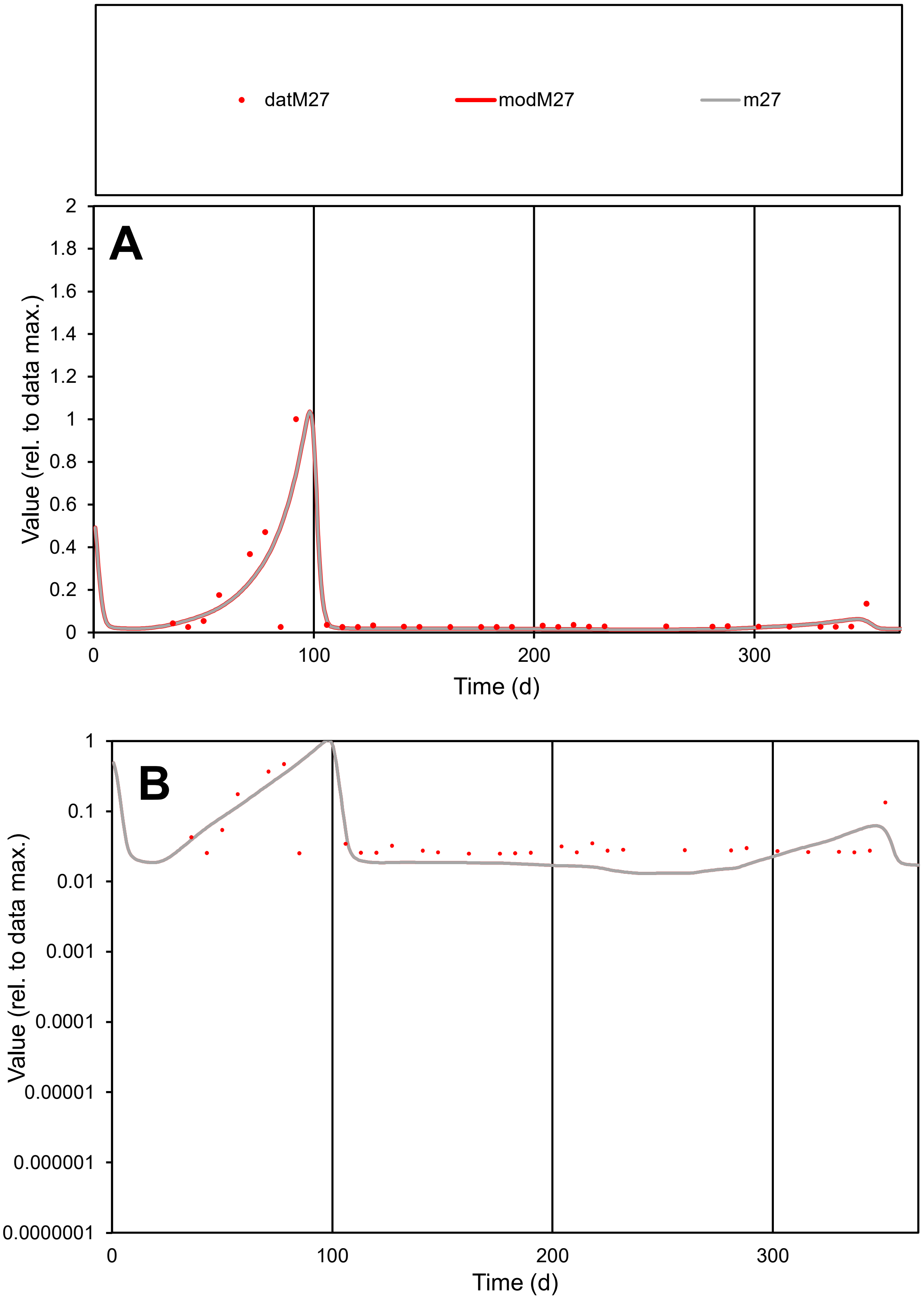


**C1**

Variables
M27 Provide jtd or Dst
CalibOnly?
2 1 to reproduce of
oNormalize?
2 1=no, 2=yes

ofxn calc*
1.03311
ofxn mod
1.07427

sRun
RunA1
zout
1

Ave
Data
0.00023
Model
0.00022

data/vMax
0.00256

ofxng
321.551

Min d/m
1.88619

ave/min d
3.69002
ave/min m
6.61932

324_Calib_S_A

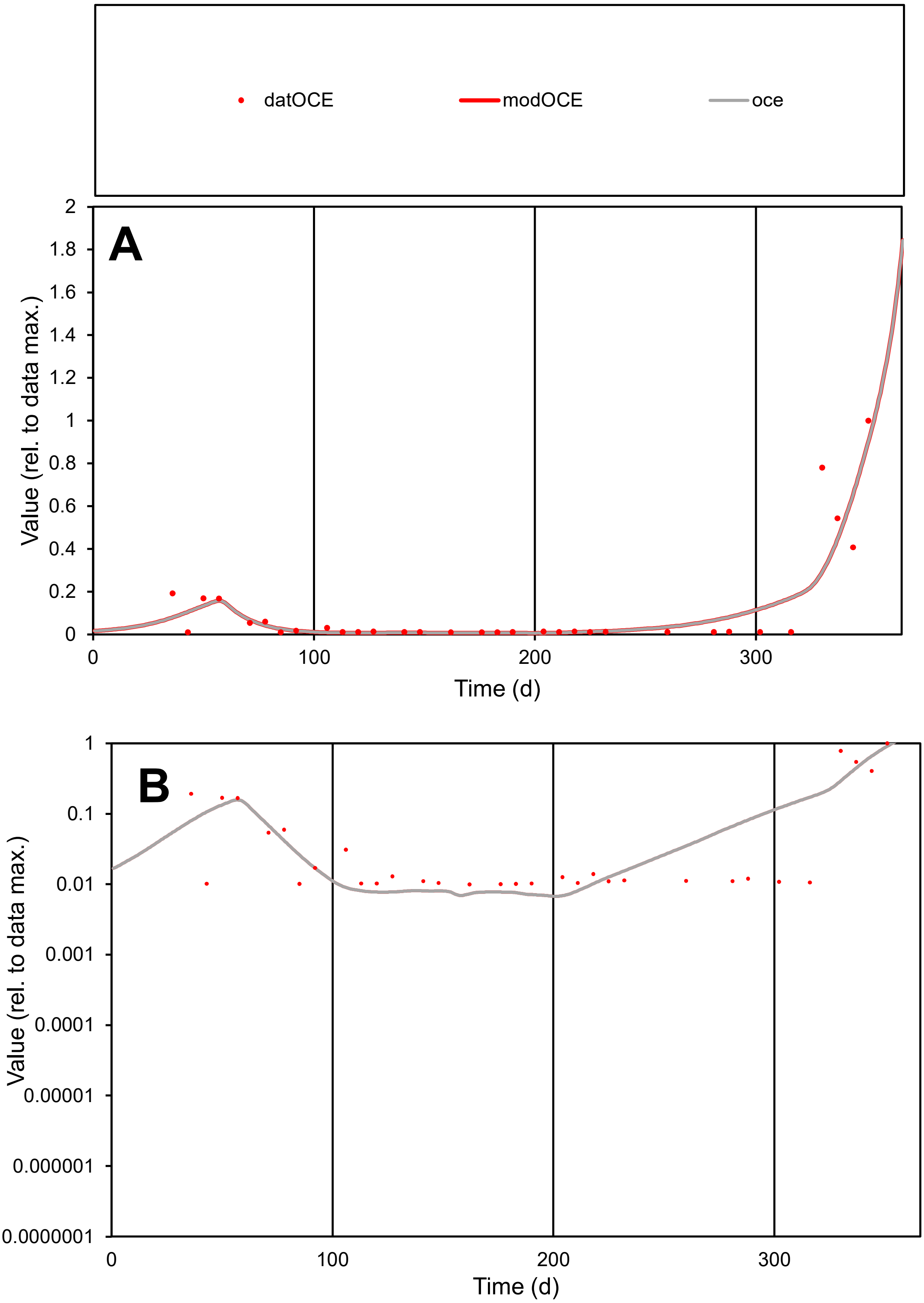


**C1**

Variables
OCE Provide jtd or Dst
CalibOnly?
2 1 to reproduce of
oNormalize?
2 1=no, 2=yes

ofxn calc*
0.91499
ofxn mod
0.95208

sRun
RunA1
zout
1

Ave
Data
0.00042
Model
0.00041

data/vMax
0.00365

ofxng
321.551

Min d/m
1.42885

ave/min d
11.4516
ave/min m
16.3075

324_Calib_S_A

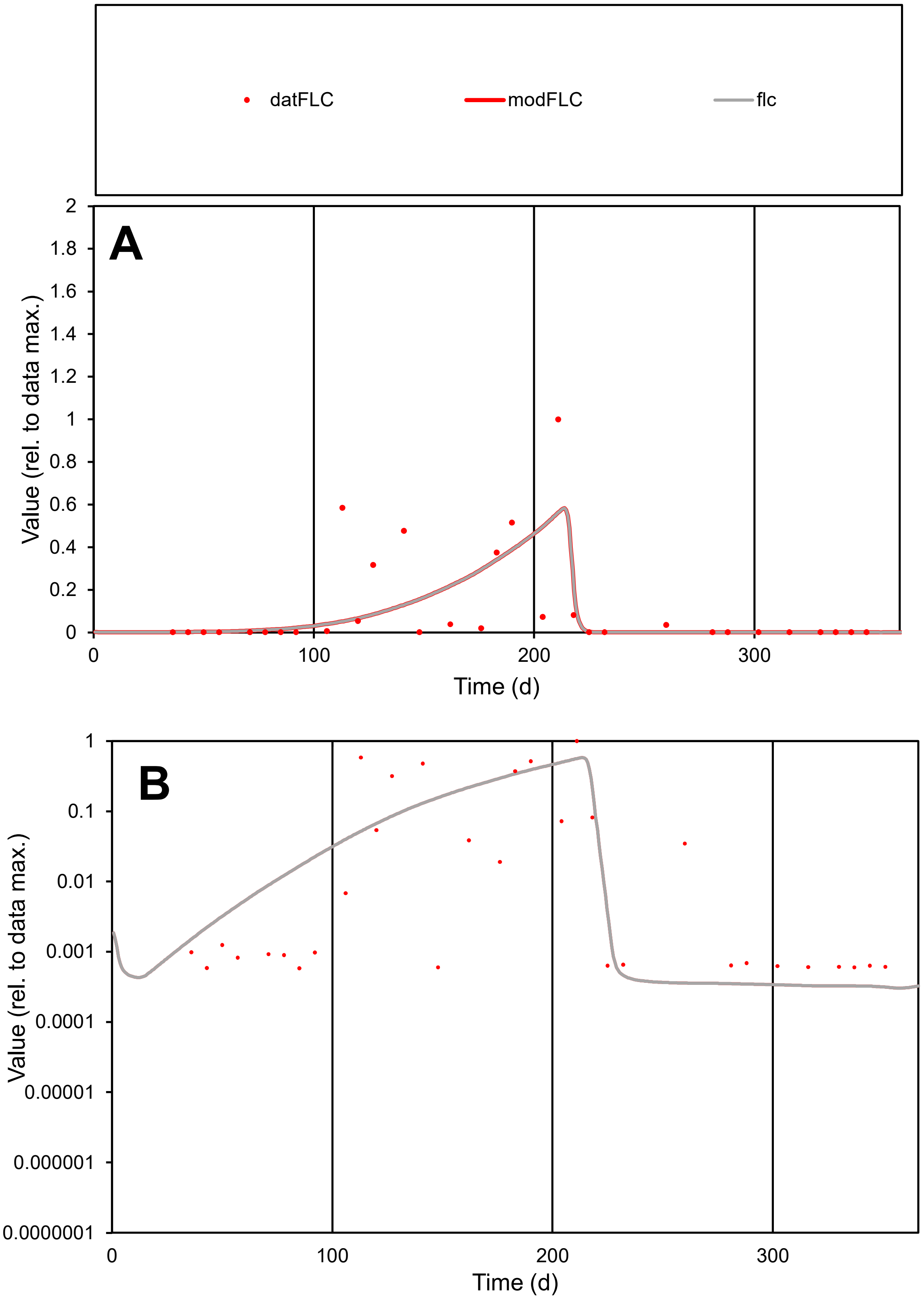


**C1**

Variables
FLC Provide jtd or Dst
CalibOnly?
2 1 to reproduce of
oNormalize?
2 1=no, 2=yes

ofxn calc*
1.54884
ofxn mod
1.56435

sRun
RunA1
zout
1

Ave
Data
0.0023
Model
0.00196

data/vMax
0.02056

ofxng
321.551

Min d/m
1.8642

ave/min d
192.527
ave/min m
304.919

324_Calib_S_A

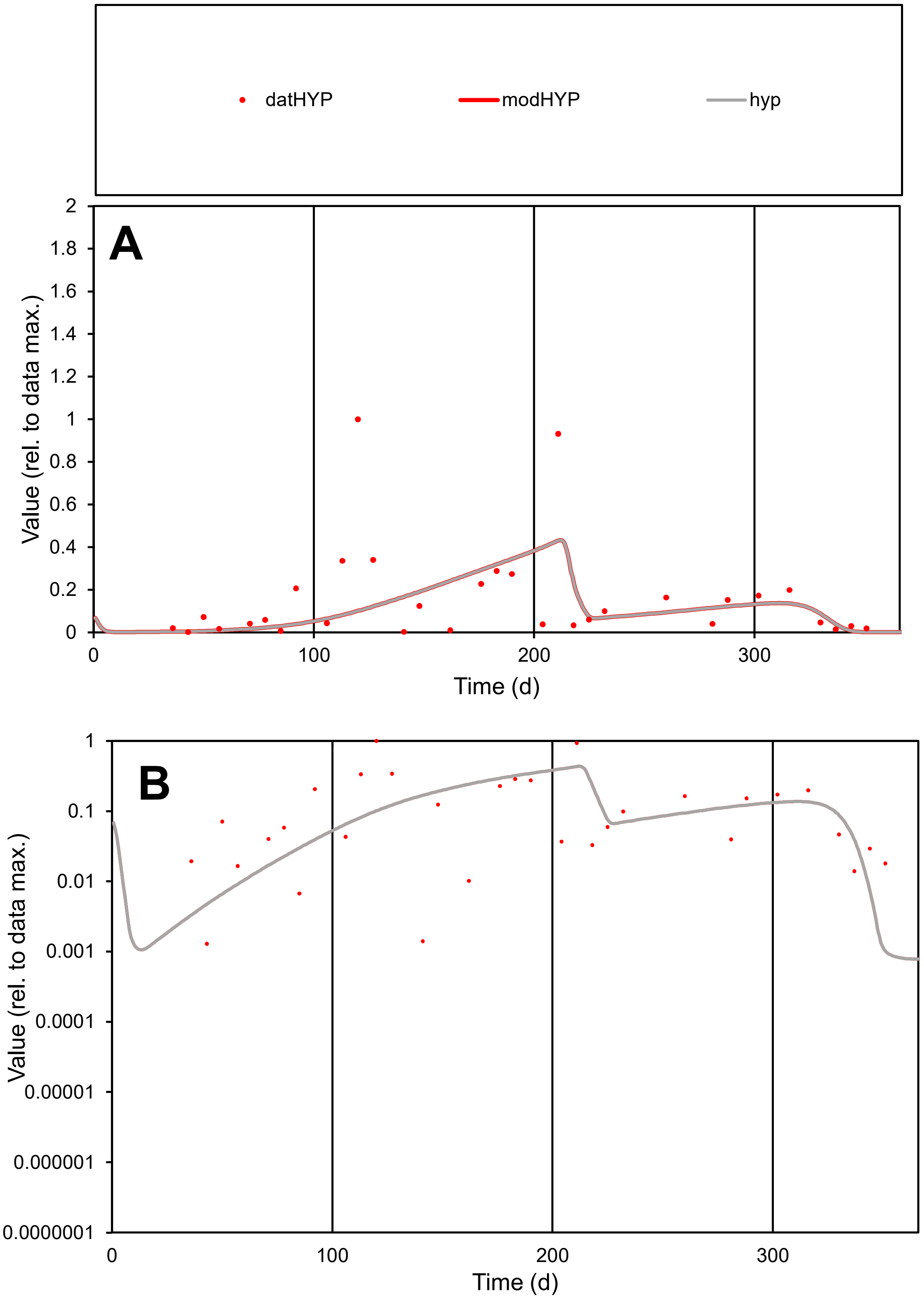


**C1**

Variables
HYP Provide jtd or Dst
CalibOnly?
2 1 to reproduce of
oNormalize?
2 1=no, 2=yes

ofxn calc*
1.35197
ofxn mod
1.36067

sRun
RunA1
zout
1

Ave
Data
0.00195
Model
0.00152

data/vMax
0.01234

ofxng
321.551

Min d/m
1.34599

ave/min d
122.204
ave/min m
128.298

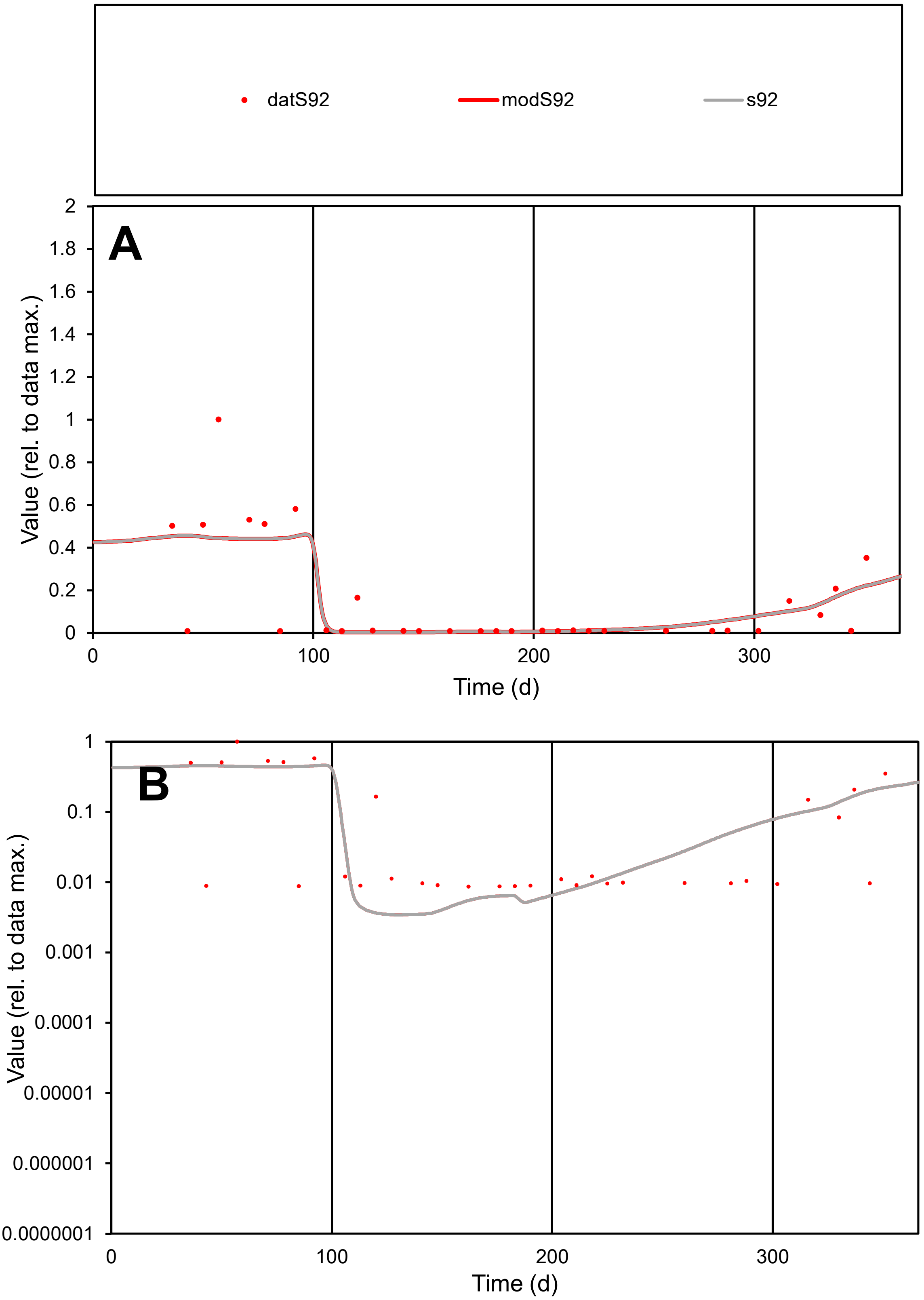


**C1**

Variables

S92 Provide jtd or Dst

CalibOnly?

2 1 to reproduce of

oNormalize?

2 1=no, 2=yes

ofxn calc*

1.07338

ofxn mod

1.0839

sRun

RunA1

zout

1

Ave

Data

0.00039

Model

0.00039

data/vMax

0.00263

ofxng

321.551

Min d/m

2.52186

ave/min d

17.2784

ave/min m

43.2474

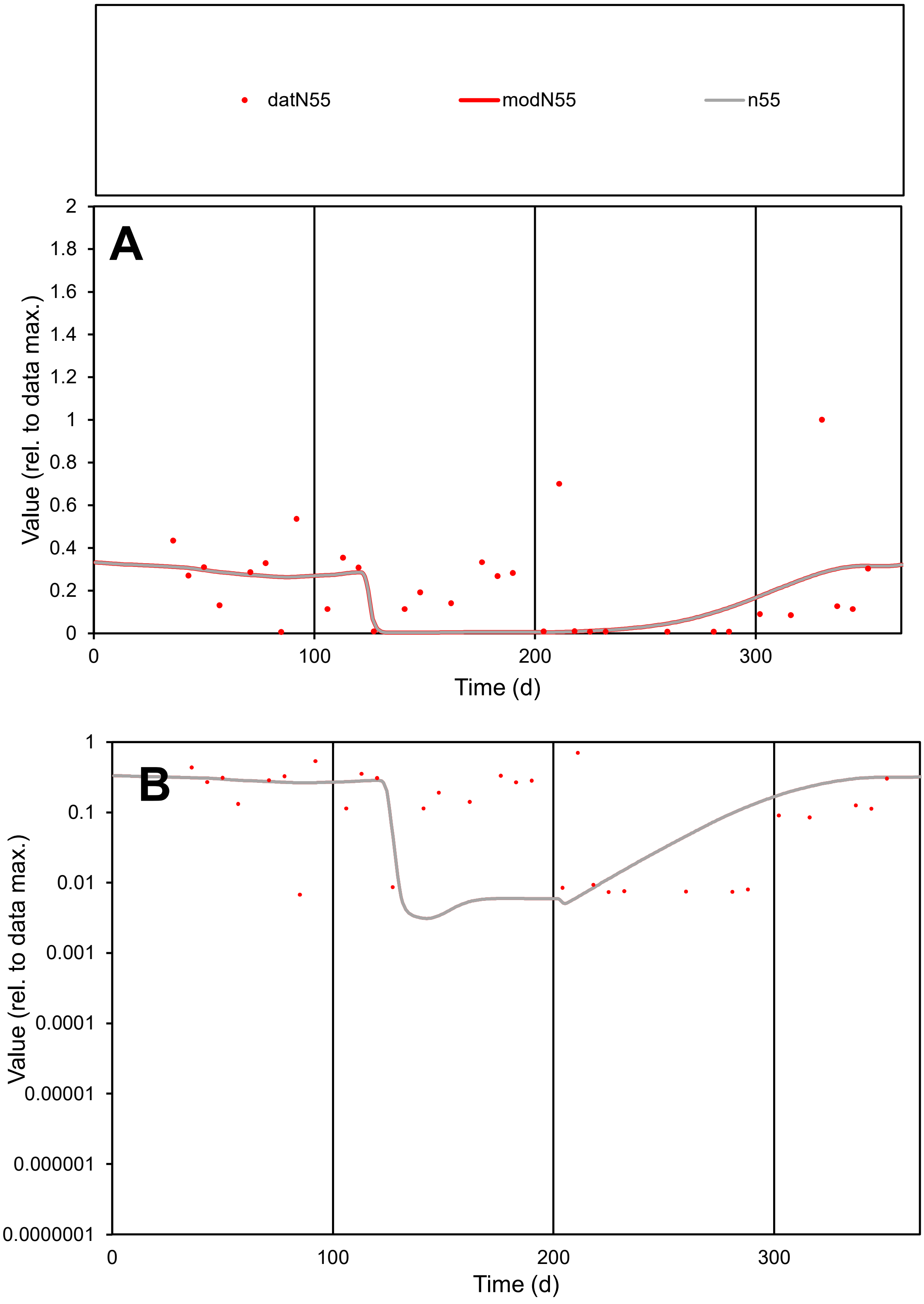


**C1**

Variables

N55 Provide jtd or Dst

CalibOnly?

2 1 to reproduce of

oNormalize?

2 1=no, 2=yes

ofxn calc*

1.05301

ofxn mod

1.05983

sRun

RunA1

zout

1

Ave

Data

0.00118

Model

0.00087

data/vMax

0.00549

ofxng

321.551

Min d/m

2.17616

ave/min d

31.8827

ave/min m

51.341

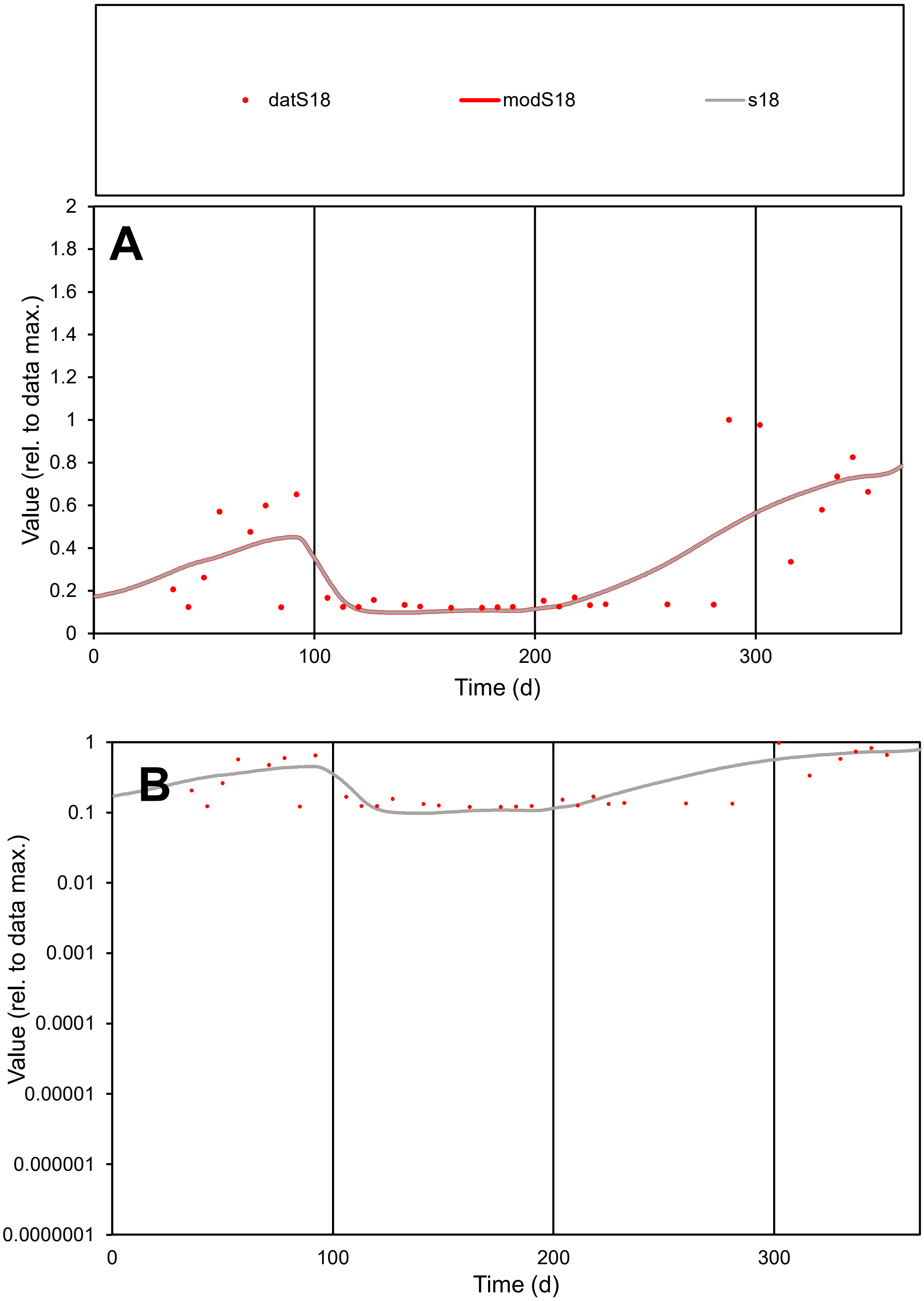


**C1**

Variables
S18 Provide jtd or Dst
CalibOnly?
2 1 to reproduce of
oNormalize?
2 1=no, 2=yes

ofxn calc*
0.53128
ofxn mod
0.54504

sRun
RunA1
zout
1

Ave
Data
0.00071
Model
0.00071

data/vMax
0.00217

ofxng
321.551

Min d/m
1.22787

ave/min d
2.71389
ave/min m
3.34618

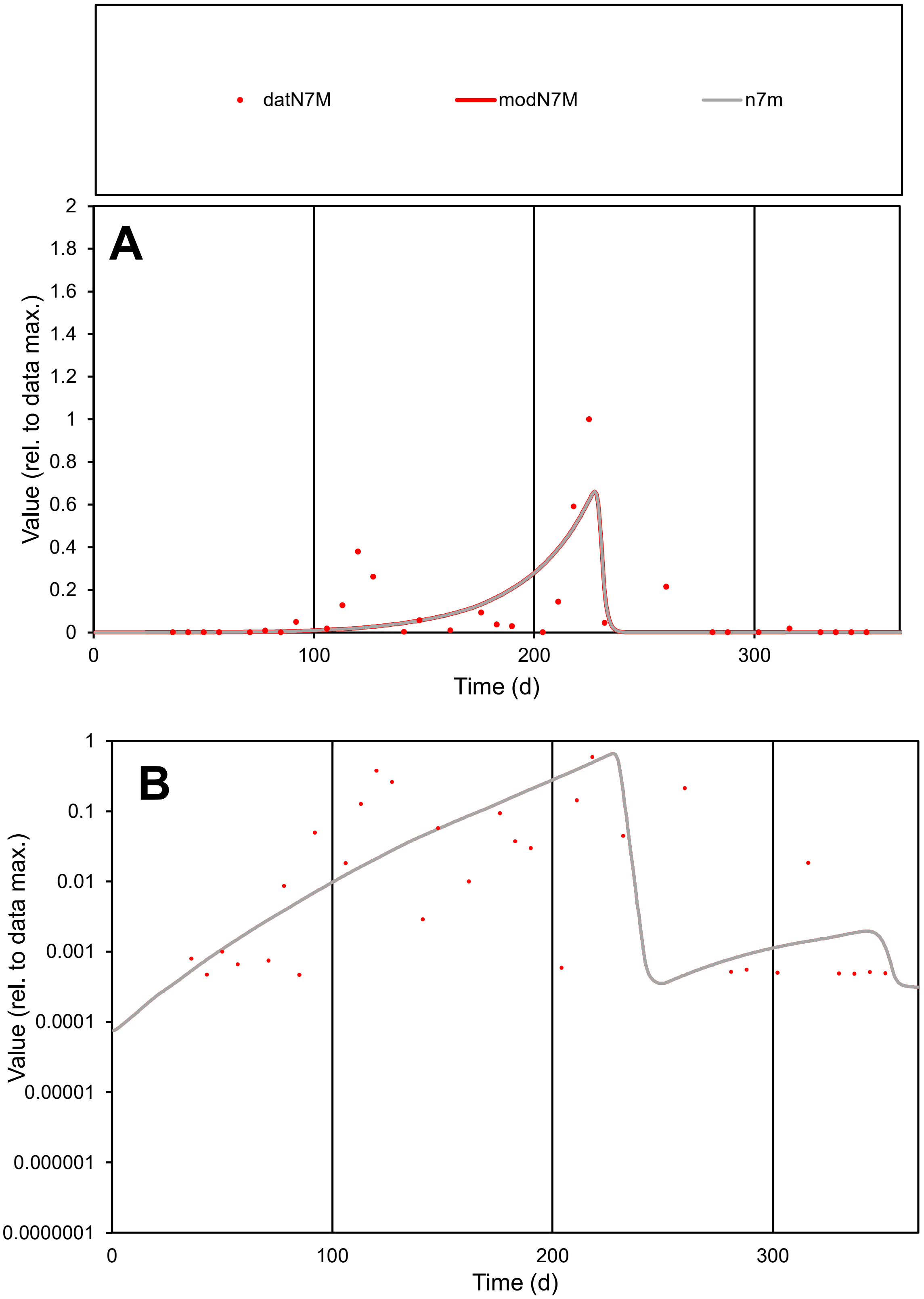


**C1**

Variables
N7M Provide jtd or Dst
CalibOnly?
2 1 to reproduce of
oNormalize?
2 1=no, 2=yes

ofxn calc*
1.40954
ofxn mod
1.43285

sRun
RunA1
zout
1

Ave
Data
0.00197
Model
0.00175

data/vMax
0.02033

ofxng
321.551

Min d/m
0.97784

ave/min d
205.57
ave/min m
178.973

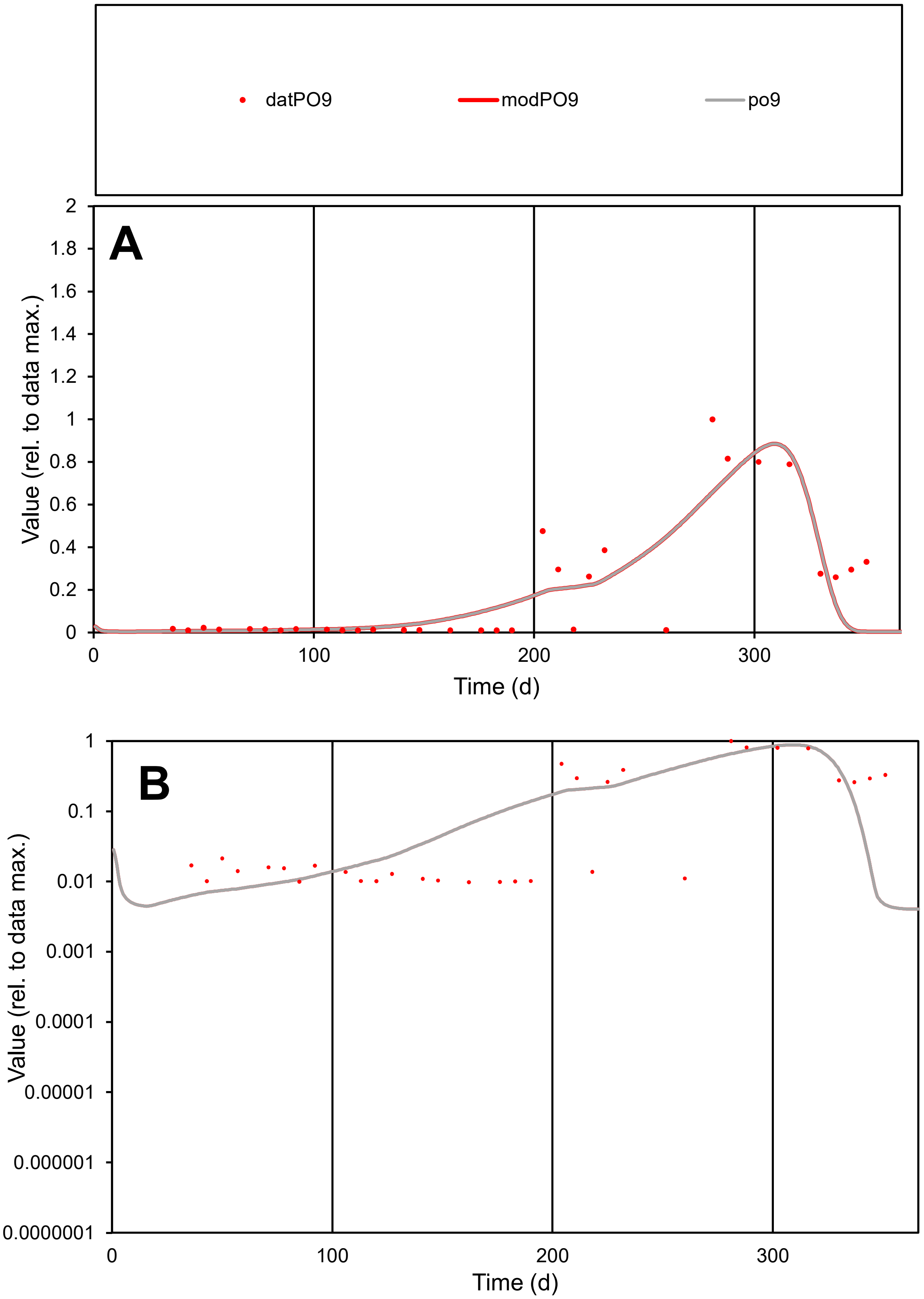


**C1**

Variables
PO9 Provide jtd or Dst
CalibOnly?
2 1 to reproduce of
oNormalize?
2 1=no, 2=yes

ofxn calc*
0.75644
ofxn mod
0.77849

sRun
RunA1
zout
1

Ave
Data
0.00113
Model
0.00105

data/vMax
0.00581

ofxng
321.551

Min d/m
2.16484

ave/min d
19.8044
ave/min m
39.966

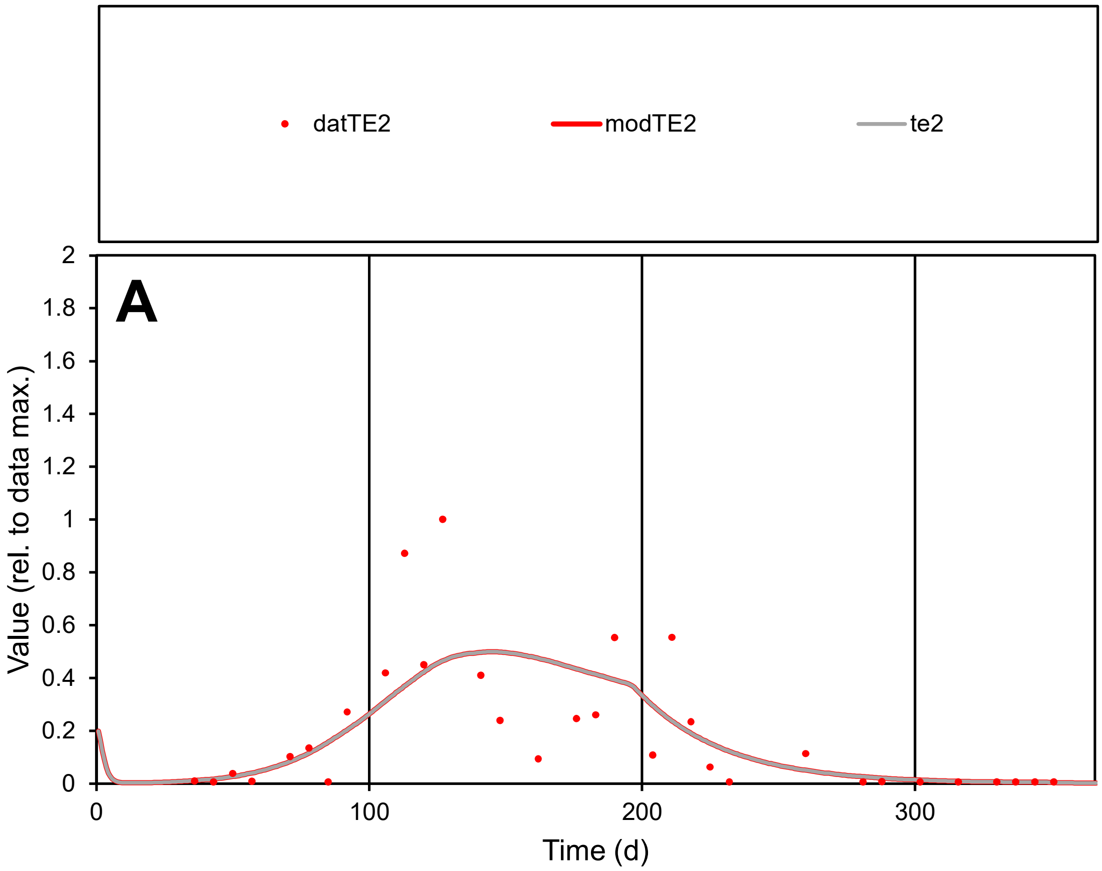


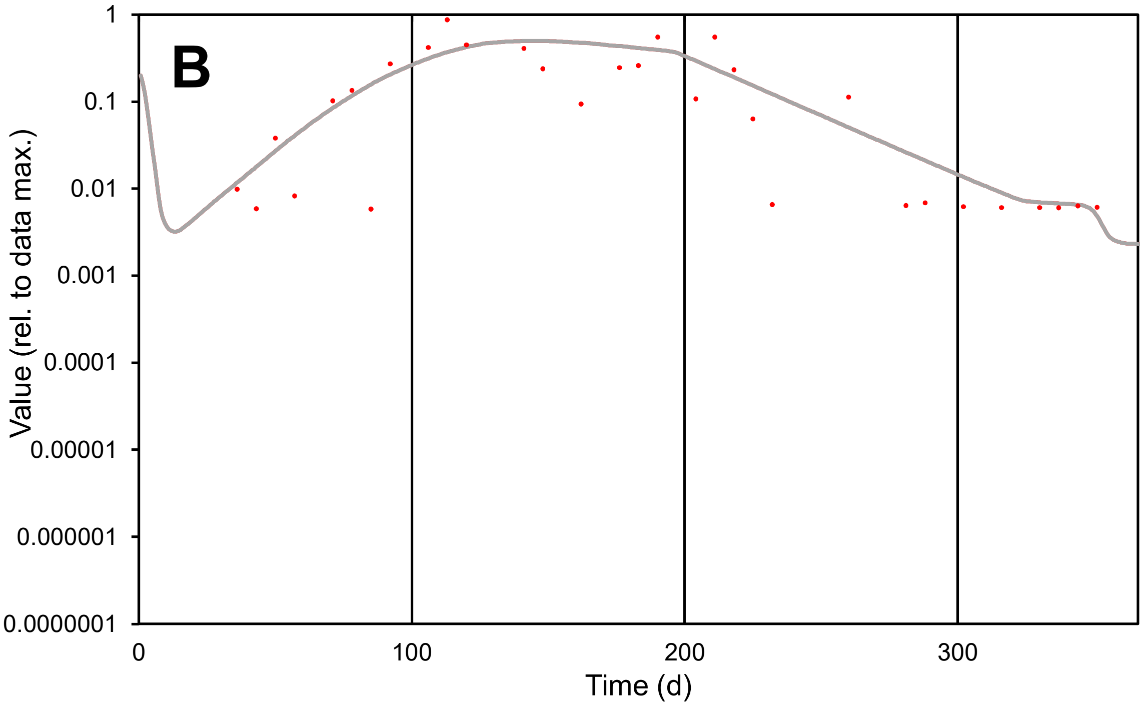


**C1**

Variables
TE2 Provide jtd or Dst
CalibOnly?
2 1 to reproduce of
oNormalize?
2 1=no, 2=yes

ofxn calc*
0.9137
ofxn mod
0.92537

sRun
RunA1
zout
1

Ave
Data
0.00285
Model
0.00278

data/vMax
0.01458

ofxng
321.551

Min d/m
1.21978

ave/min d
33.4876
ave/min m
39.8375

324_Calib_S_A

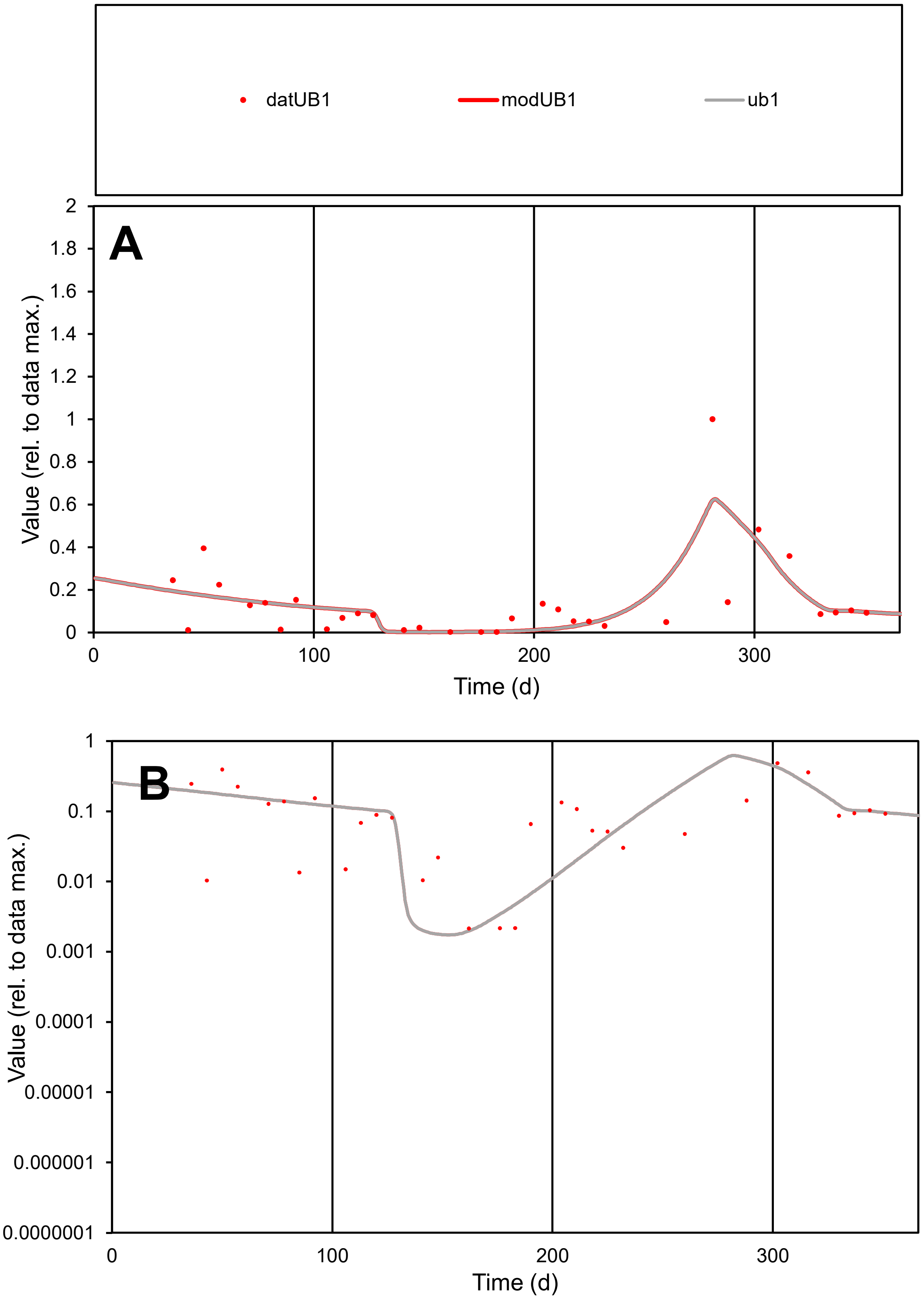


**C1**

Variables
UB1 Provide jtd or Dst
CalibOnly?
2 1 to reproduce of
oNormalize?
2 1=no, 2=yes

ofxn calc*
0.91795
ofxn mod
0.92817

sRun
RunA1
zout
1

Ave
Data
0.00362
Model
0.00354

data/vMax
0.02605

ofxng
321.551

Min d/m
1.21813

ave/min d
64.7792
ave/min m
77.0325

324_Calib_S_A

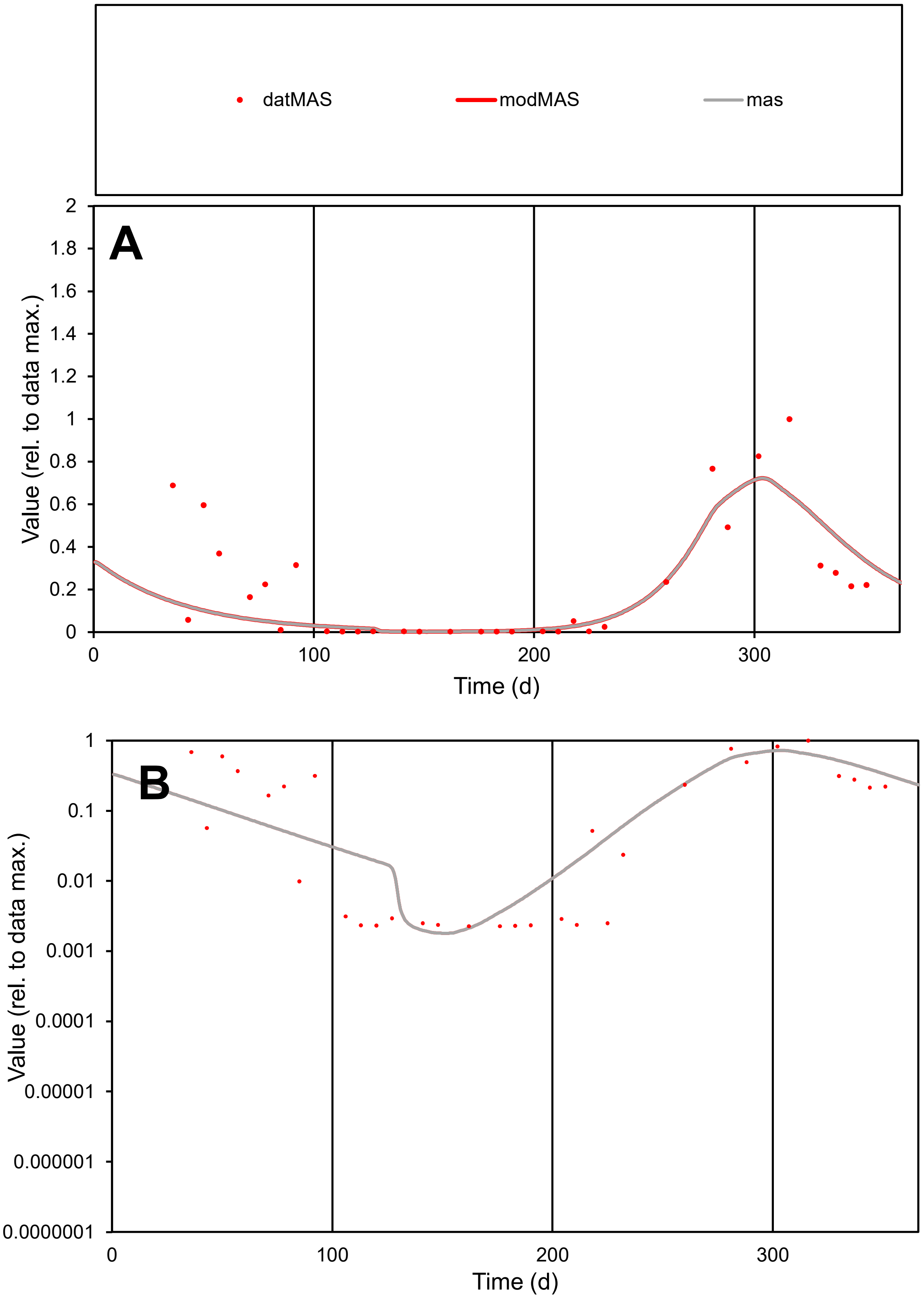


**C1**

Variables
MAS Provide jtd or Dst
CalibOnly?
2 1 to reproduce of
oNormalize?
2 1=no, 2=yes

ofxn calc*
0.84629
ofxn mod
0.85823

sRun
RunA1
zout
1

Ave
Data
0.00112
Model
0.00088

data/vMax
0.00522

ofxng
321.551

Min d/m
1.25386

ave/min d
95.0783
ave/min m
93.2118

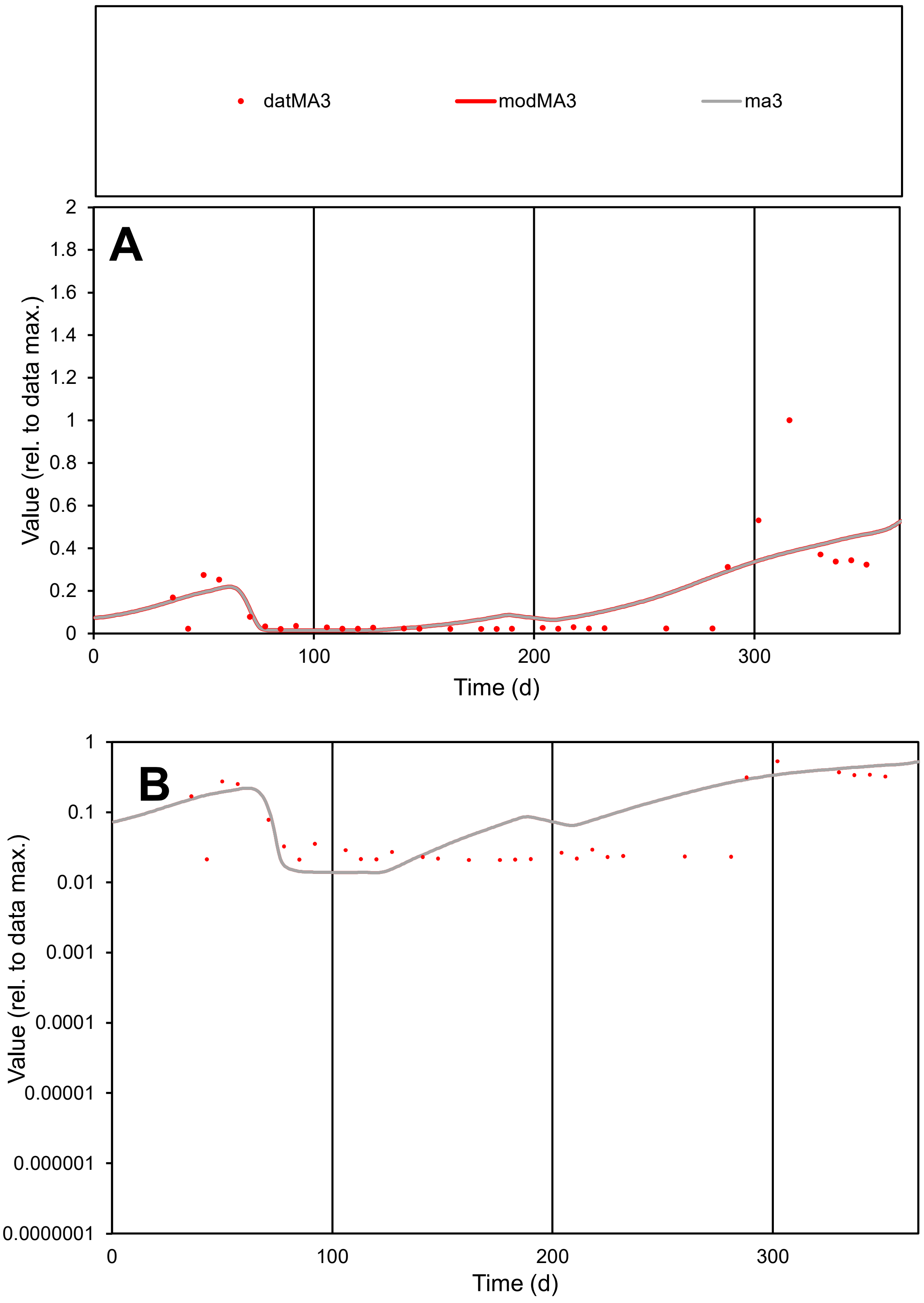


**C1**

Variables
MA3 Provide jtd or Dst
CalibOnly?
2 1 to reproduce of
oNormalize?
2 1=no, 2=yes

ofxn calc*
0.97792
ofxn mod
0.99001

sRun
RunA1
zout
1

Ave
Data
0.00041
Model
0.00045

data/vMax
0.00292

ofxng
321.551

Min d/m
1.51145

ave/min d
6.75587
ave/min m
11.2275

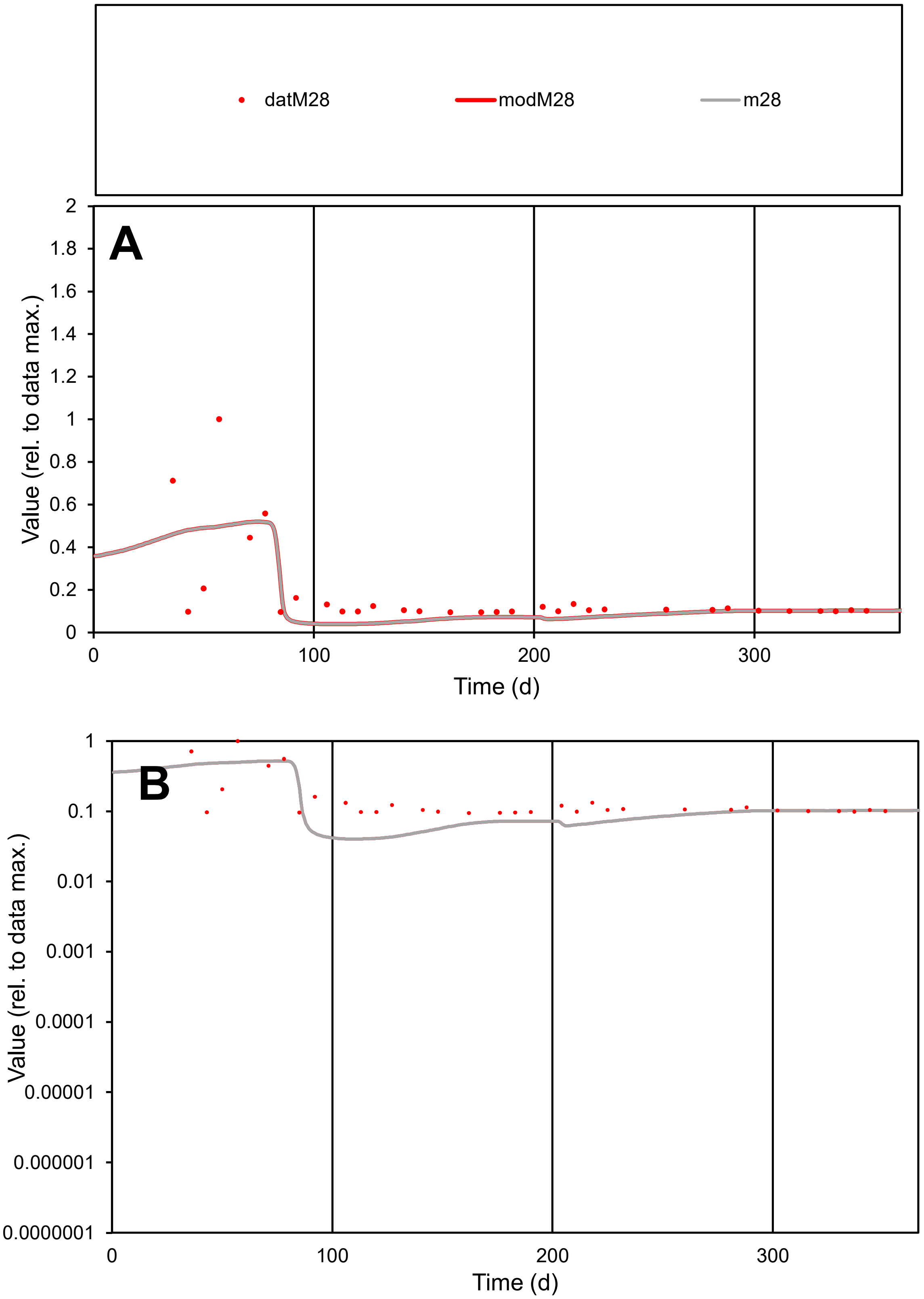


C1

Variables
M28 Provide jtd or Dst
CalibOnly?
2 1 to reproduce of
oNormalize?
2 1=no, 2=yes

ofxn calc*
0.75463
ofxn mod
0.7643

sRun
RunA1
zout
1

Ave
Data
0.0001
Model
8.6E-05

data/vMax
0.00056

ofxng
321.551

Min d/m
2.3431

ave/min d
1.92008
ave/min m
3.81661

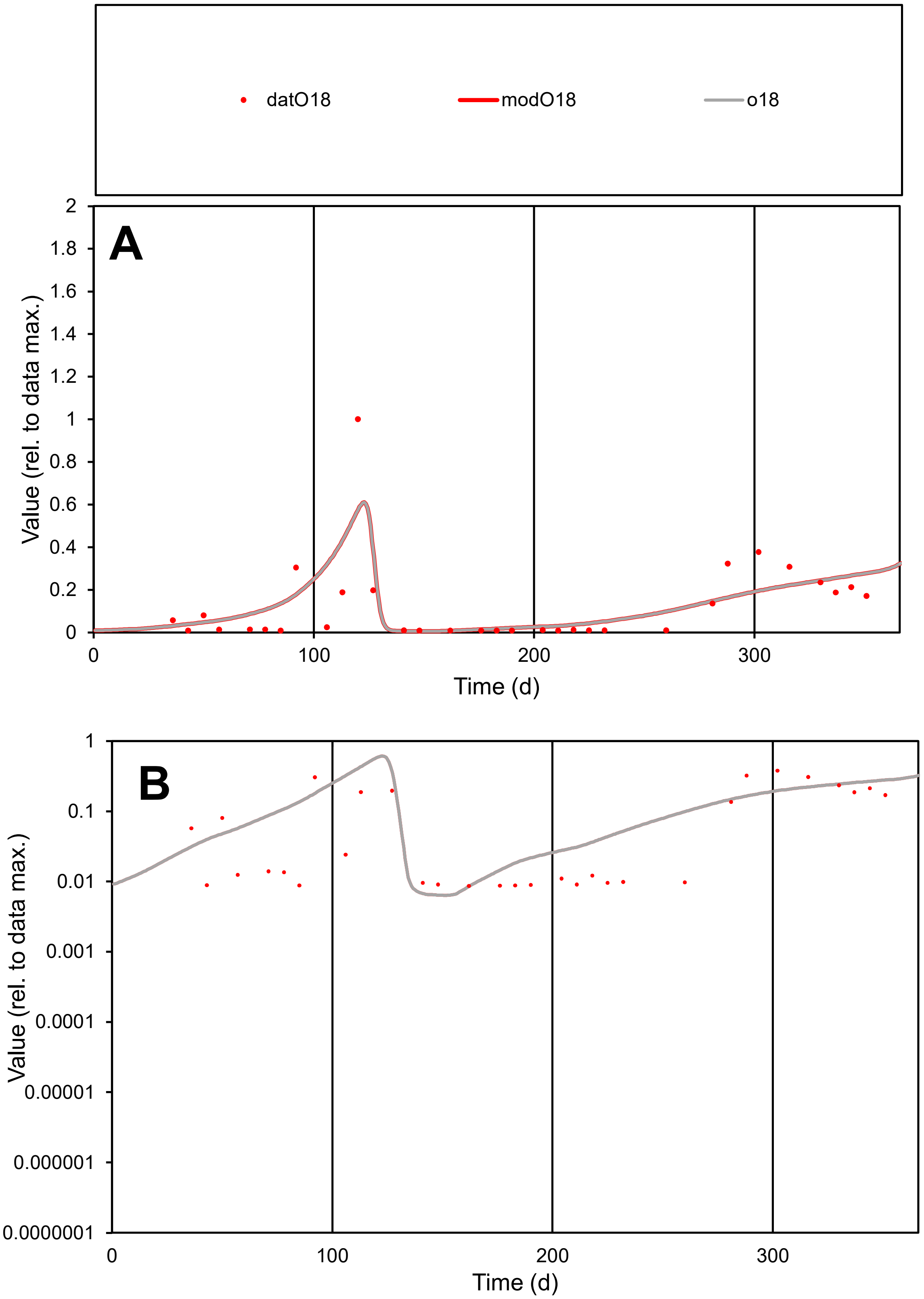


**C1**

Variables
O18 Provide jtd or Dst
CalibOnly?
2 1 to reproduce of
oNormalize?
2 1=no, 2=yes

ofxn calc*
0.97069
ofxn mod
0.98787

sRun
RunA1
zout
1

Ave
Data
0.00072
Model
0.00082

data/vMax
0.00583

ofxng
321.551

Min d/m
1.35643

ave/min d
14.3347
ave/min m
22.0606

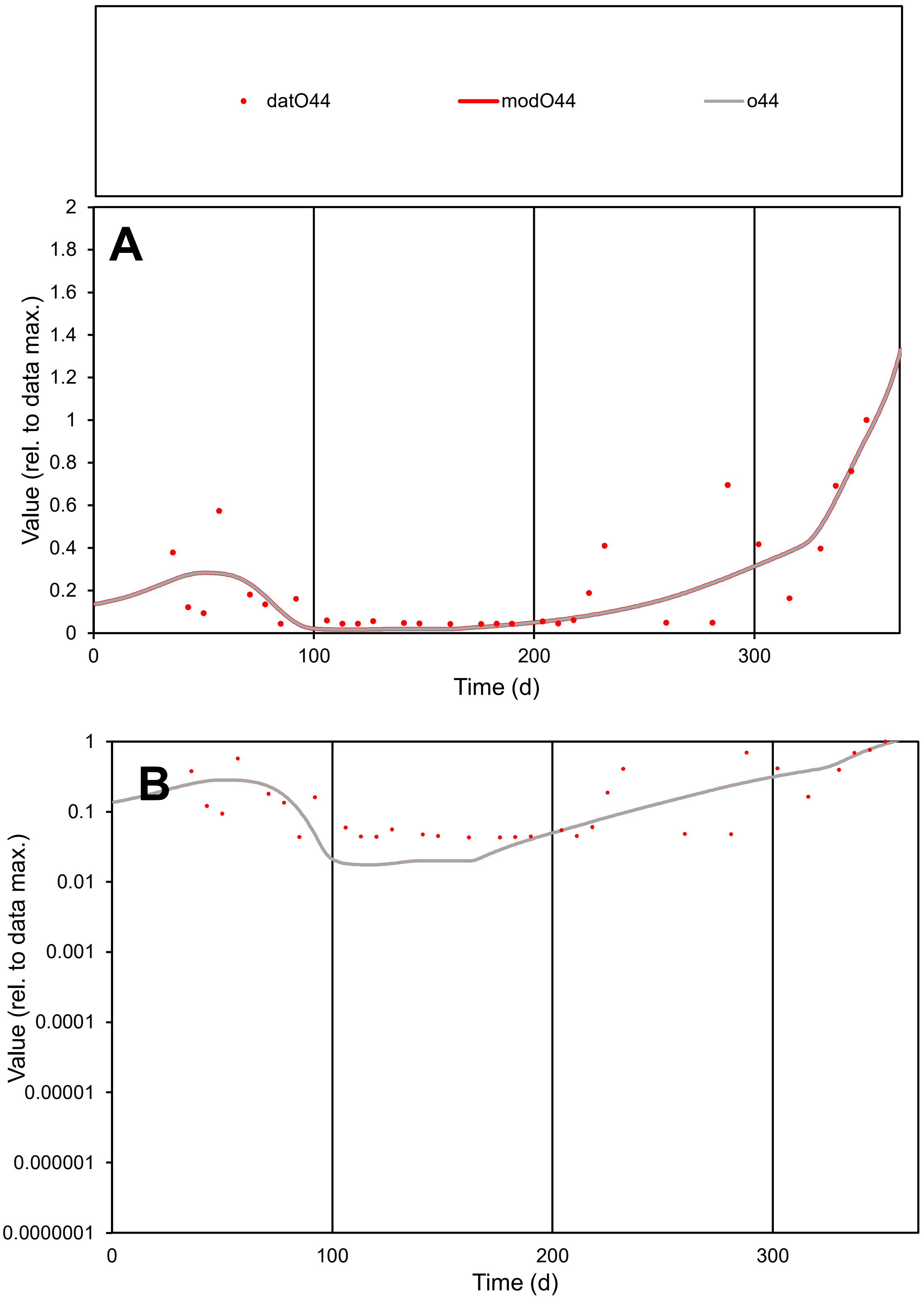


**C1**

Variables

O44 Provide jtd or Dst

CalibOnly?

2 1 to reproduce of

oNormalize?

2 1=no, 2=yes

ofxn calc*

0.61811

ofxn mod

0.63066

sRun

RunA1

zout

1

Ave

Data

0.00056

Model

0.00051

data/vMax

0.00252

ofxng

321.551

Min d/m

2.44128

ave/min d

5.19536

ave/min m

11.6251

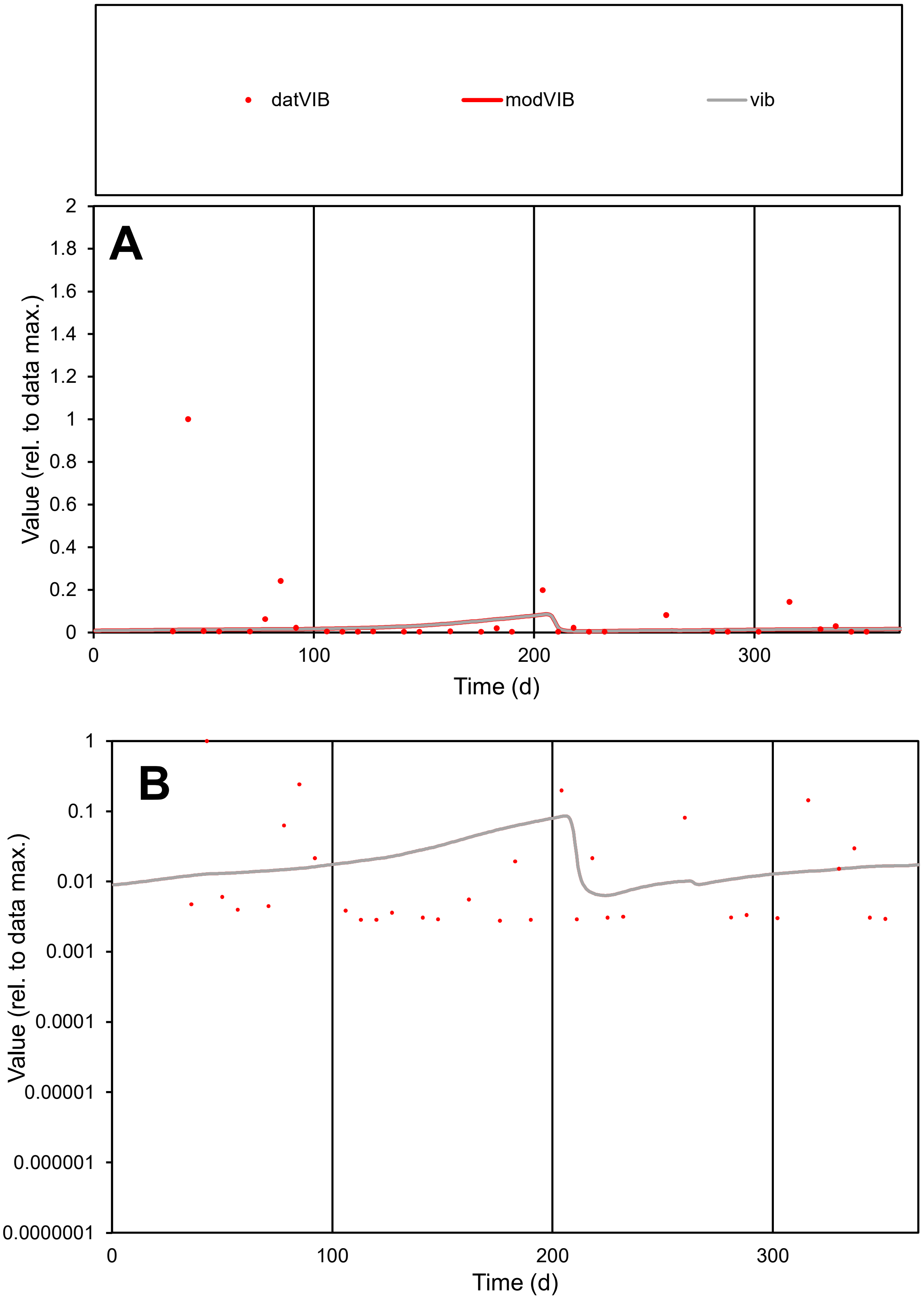


**C1**

Variables
VIB Provide jtd or Dst
CalibOnly?
2 1 to reproduce of
oNormalize?
2 1=no, 2=yes

ofxn calc*
3.07657
ofxn mod
3.08298

sRun
RunA1
zout
1

Ave
Data
0.00037
Model
0.00014

data/vMax
0.00628

ofxng
321.551

Min d/m
0.43488

ave/min d
21.4673
ave/min m
3.54932

324_Calib_S_A

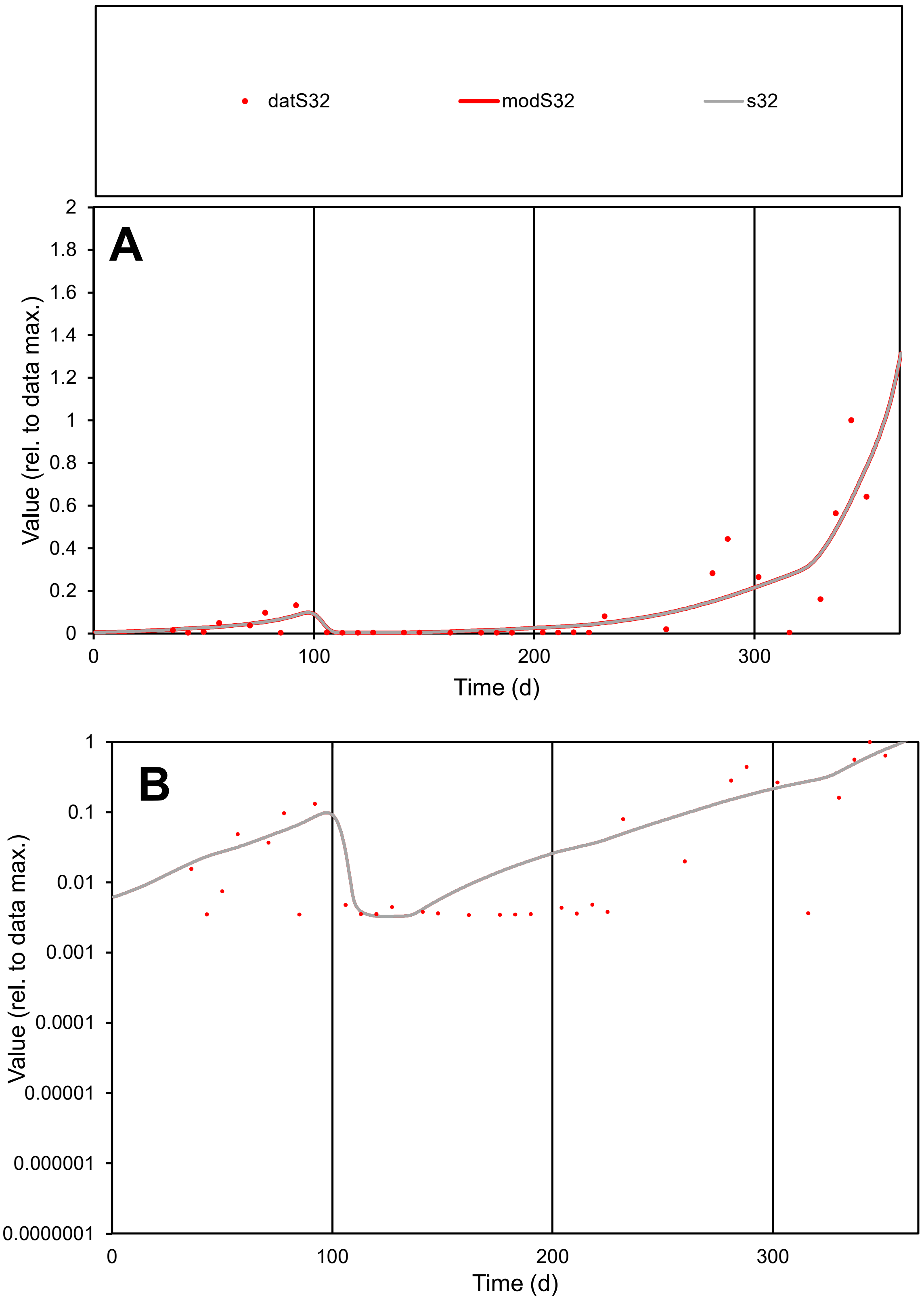


**C1**

Variables
S32 Provide jtd or Dst
CalibOnly?
2 1 to reproduce of
oNormalize?
2 1=no, 2=yes

ofxn calc*
0.89922
ofxn mod
0.9181

sRun
RunA1
zout
1

Ave
Data
0.00037
Model
0.00038

data/vMax
0.00307

ofxng
321.551

Min d/m
1.04866

ave/min d
35.0626
ave/min m
37.4801

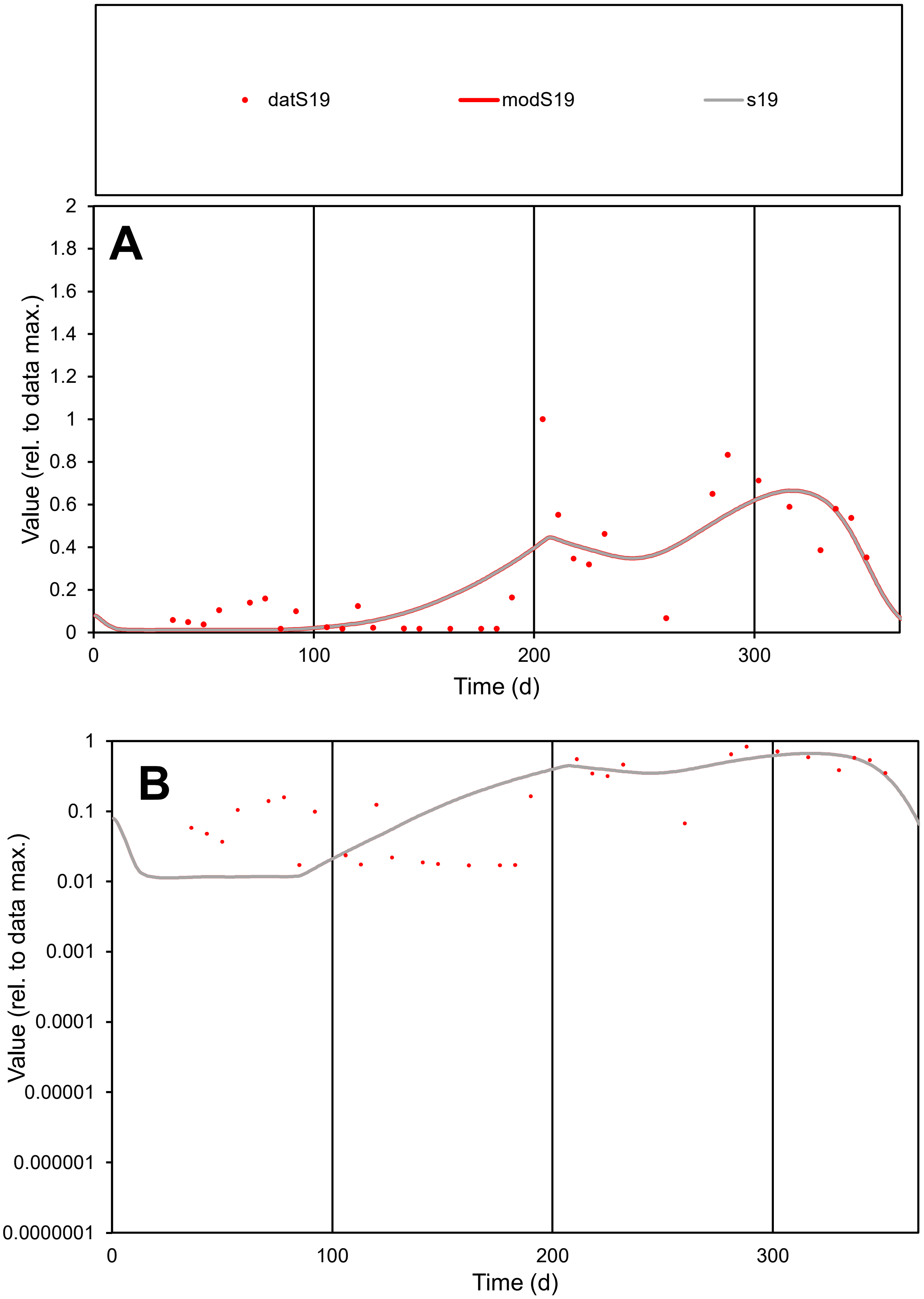


**C1**

Variables
S19 Provide jtd or Dst
CalibOnly?
2 1 to reproduce of
oNormalize?
2 1=no, 2=yes

ofxn calc*
0.62375
ofxn mod
0.63845

sRun
RunA1
zout
1

Ave
Data
0.00072
Model
0.0007

data/vMax
0.00273

ofxng
321.551

Min d/m
1.46713

ave/min d
15.661
ave/min m
22.2377

324_Calib_S_A

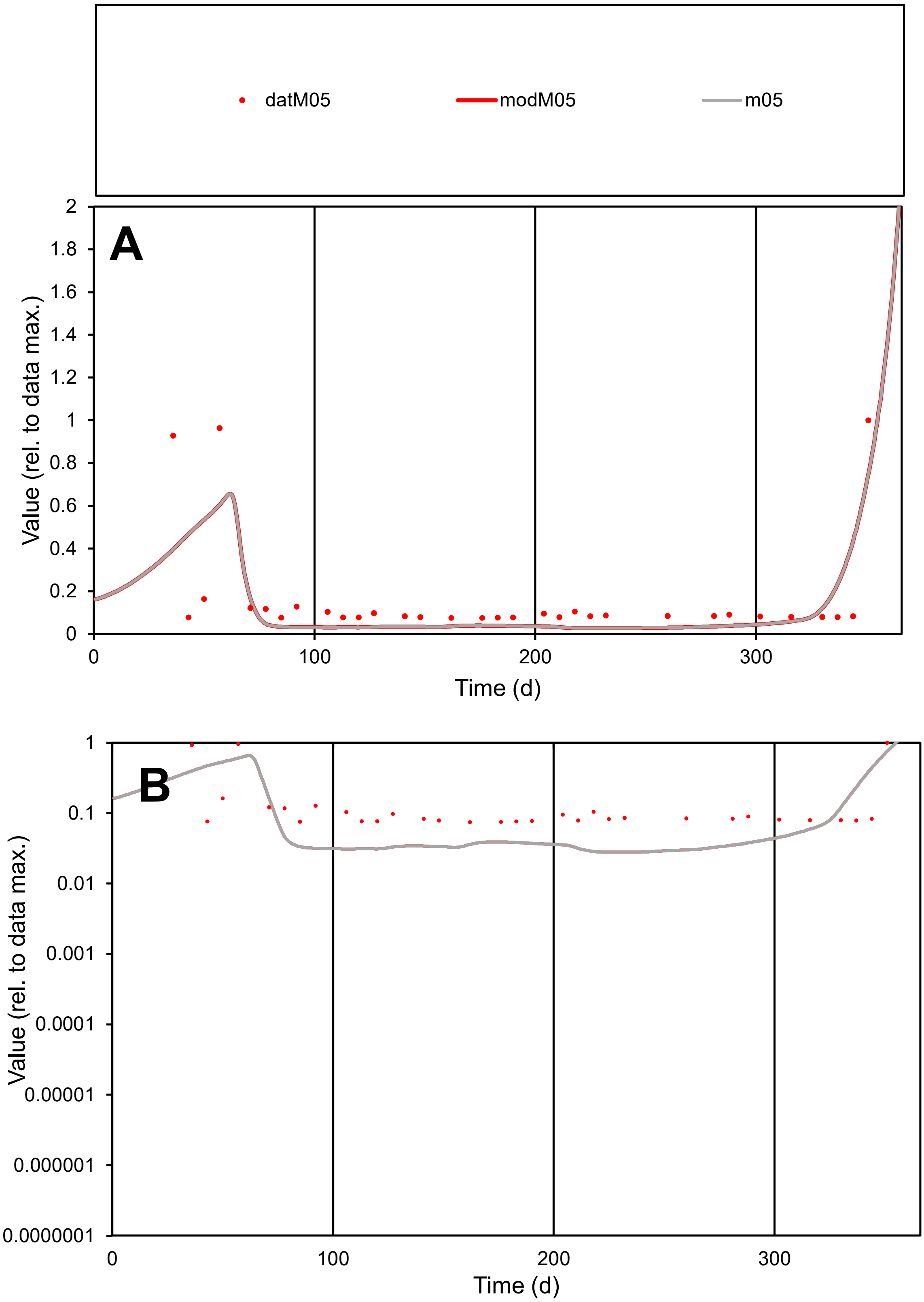


**C1**

Variables
M05 Provide jtd or Dst
CalibOnly?
2 1 to reproduce of
oNormalize?
2 1=no, 2=yes

ofxn calc*
0.98523
ofxn mod
1.00766

sRun
RunA1
zout
1

Ave
Data
9.7E-05
Model
8E-05

data/vMax
0.00057

ofxng
321.551

Min d/m
2.67914

ave/min d
2.29837
ave/min m
5.08935

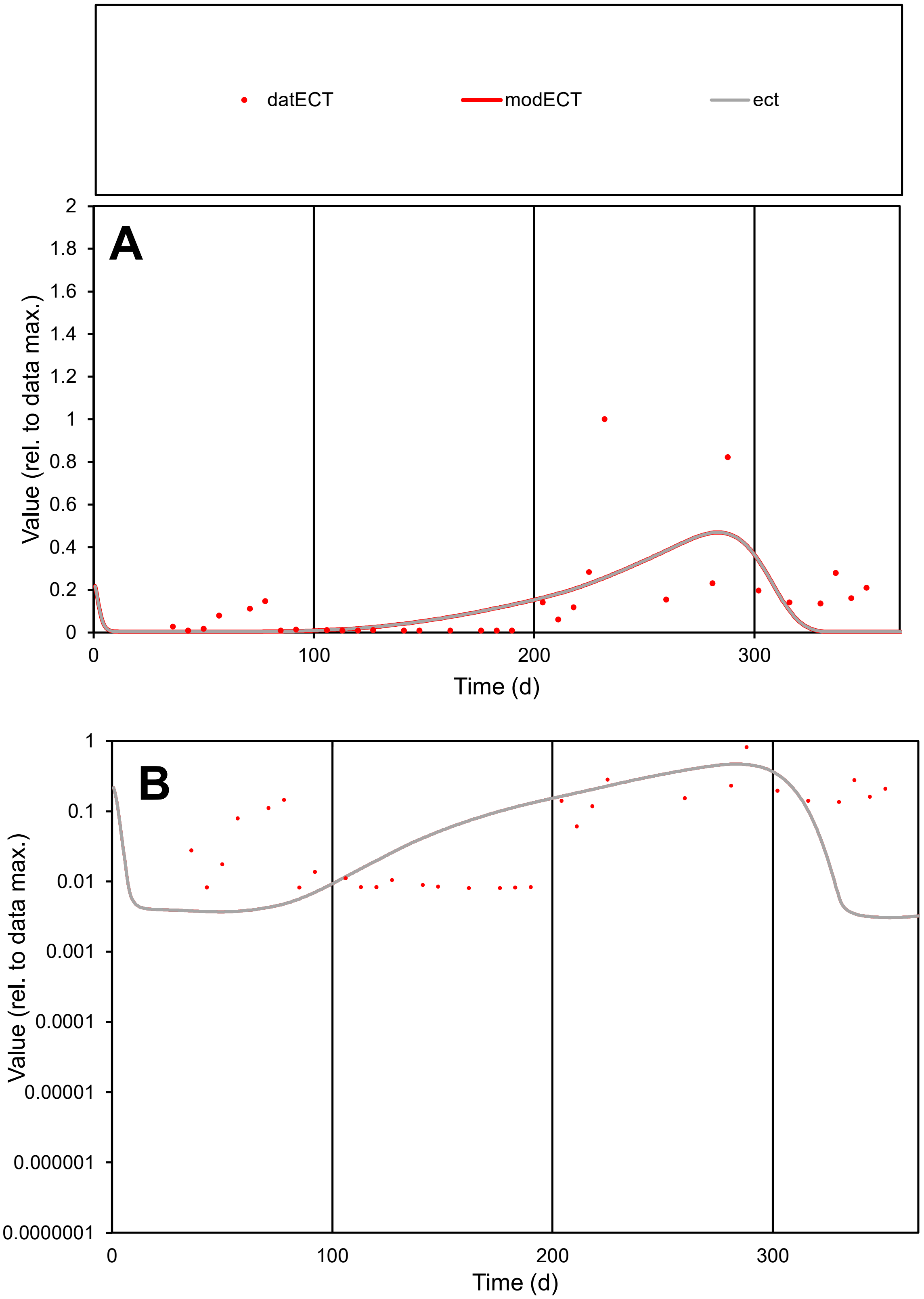


**C1**

Variables
ECT Provide jtd or Dst
CalibOnly?
2 1 to reproduce of
oNormalize?
2 1=no, 2=yes

ofxn calc*
1.32029
ofxn mod
1.32845

sRun
RunA1
zout
1

Ave
Data
0.00048
Model
0.00037

data/vMax
0.00344

ofxng
321.551

Min d/m
2.61509

ave/min d
17.1646
ave/min m
34.6573

324_Calib_S_A

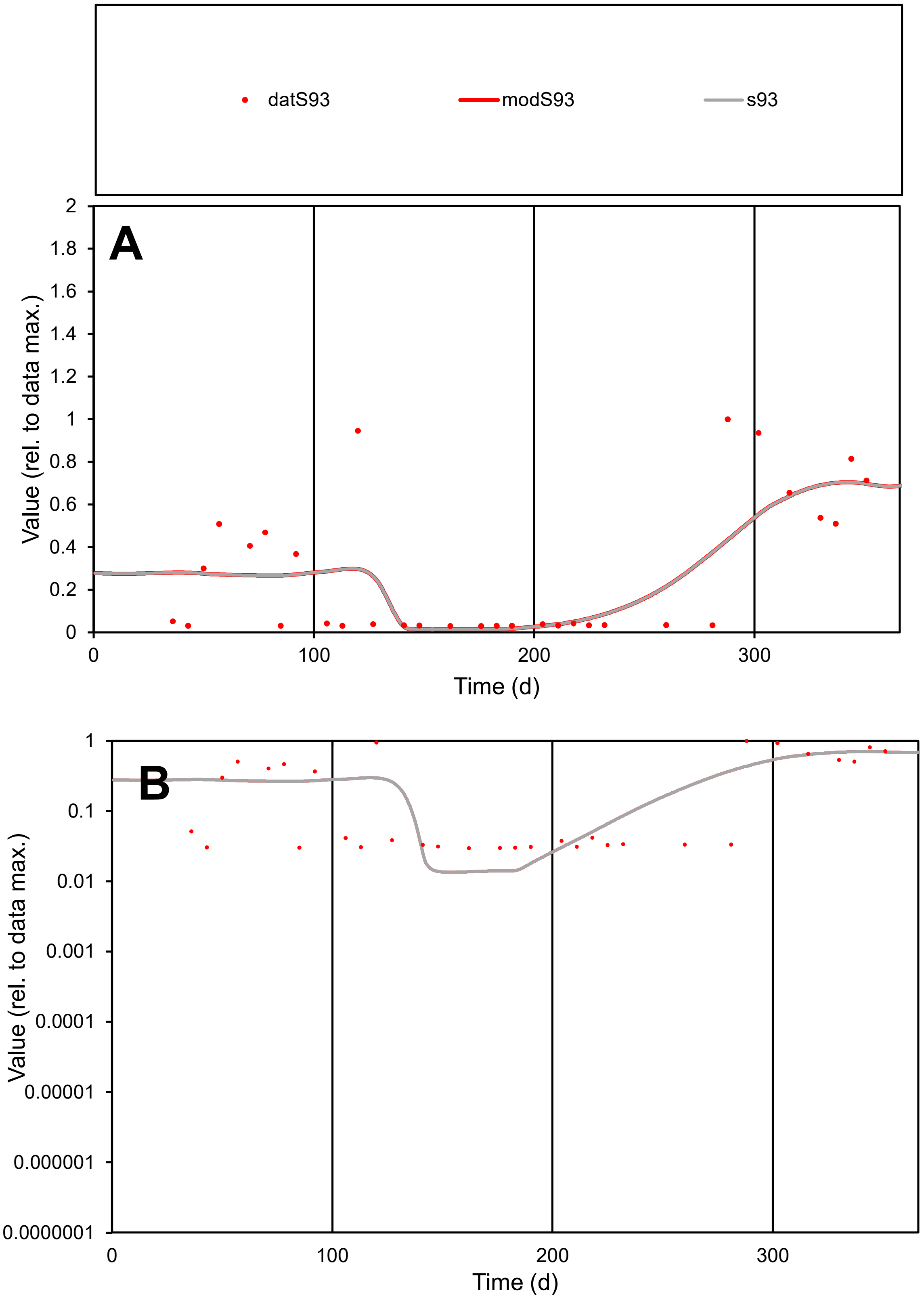


**C1**

Variables

S93 Provide jtd or Dst

CalibOnly?

2 1 to reproduce of

oNormalize?

2 1=no, 2=yes

ofxn calc*

0.80365

ofxn mod

0.81503

sRun

RunA1

zout

1

Ave

Data

0.00059

Model

0.00058

data/vMax

0.00214

ofxng

321.551

Min d/m

2.17337

ave/min d

9.23739

ave/min m

19.8746

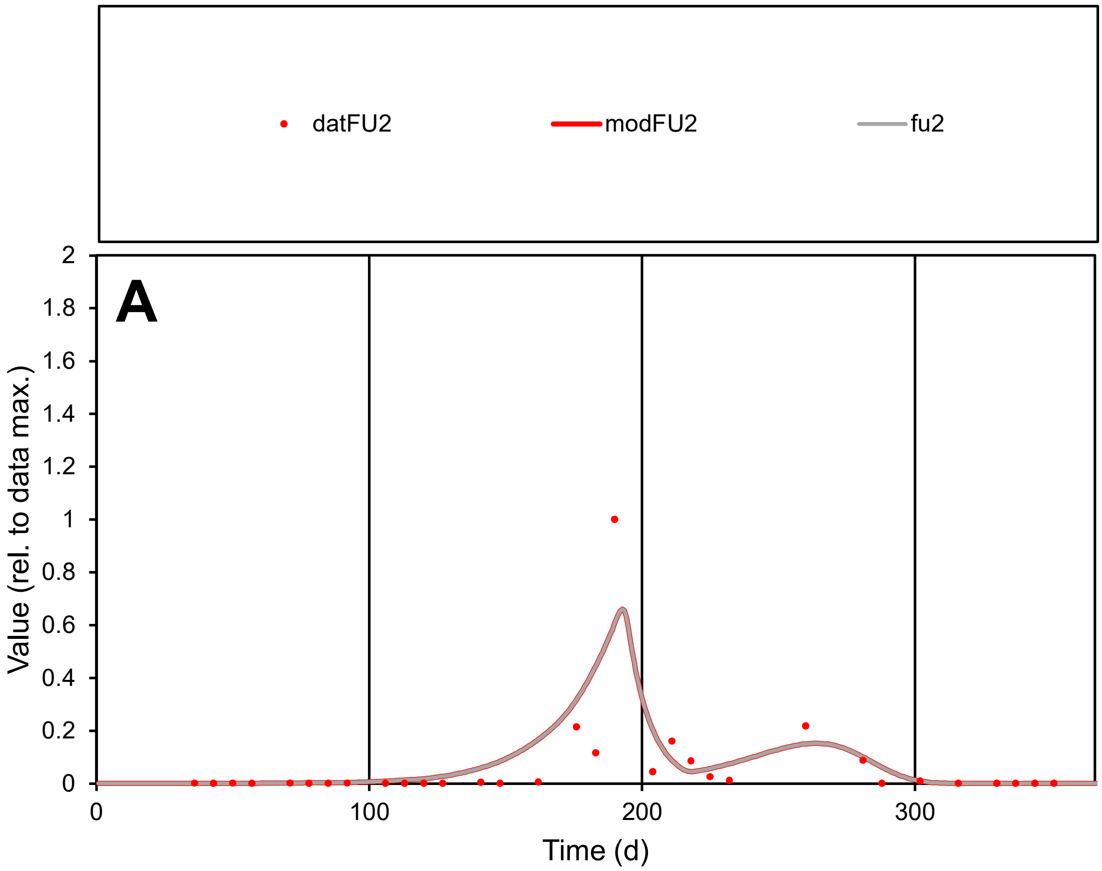


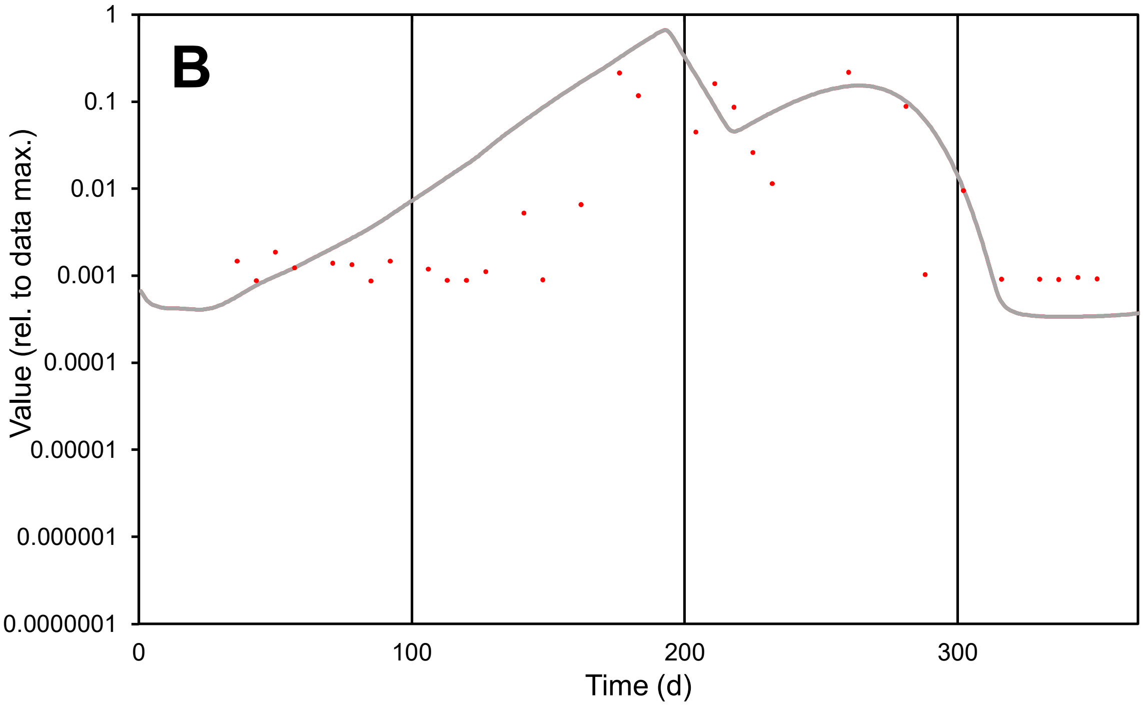


**C1**

Variables
FU2 Provide jtd or Dst
CalibOnly?
2 1 to reproduce of
oNormalize?
2 1=no, 2=yes

ofxn calc*
1.58268
ofxn mod
1.6381

sRun
RunA1
zout
1

Ave
Data
0.00183
Model
0.00228

data/vMax
0.02914

ofxng
321.551

Min d/m
2.58055

ave/min d
72.0769
ave/min m
231.443

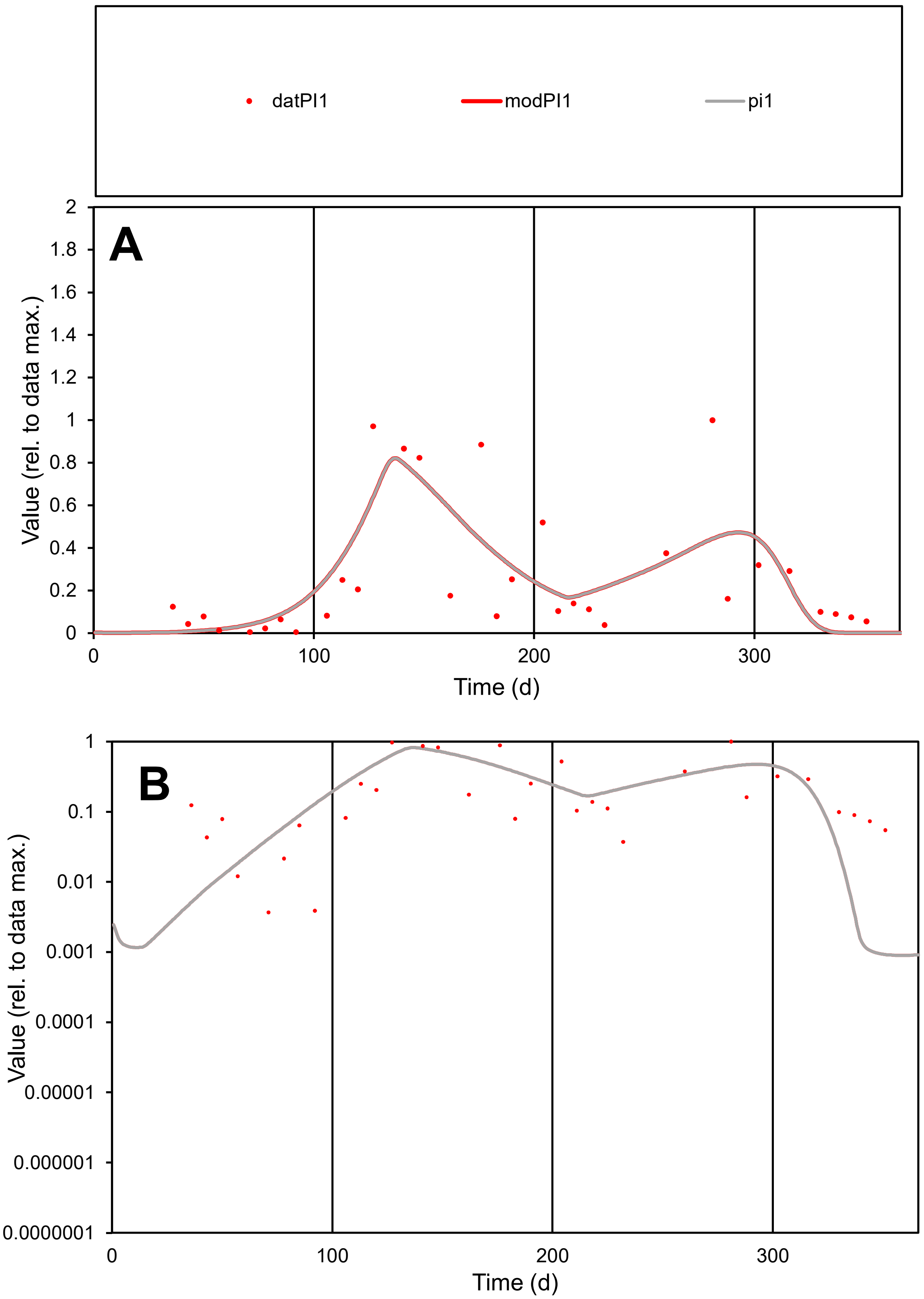


**C1**

Variables

PI1 Provide jtd or Dst

CalibOnly?

2 1 to reproduce of

oNormalize?

2 1=no, 2=yes

ofxn calc*

0.77625

ofxn mod

0.791

sRun

RunA1

zout

1

Ave

Data

0.00147

Model

0.00145

data/vMax

0.00566

ofxng

321.551

Min d/m

4.02203

ave/min d

70.581

ave/min m

280.227

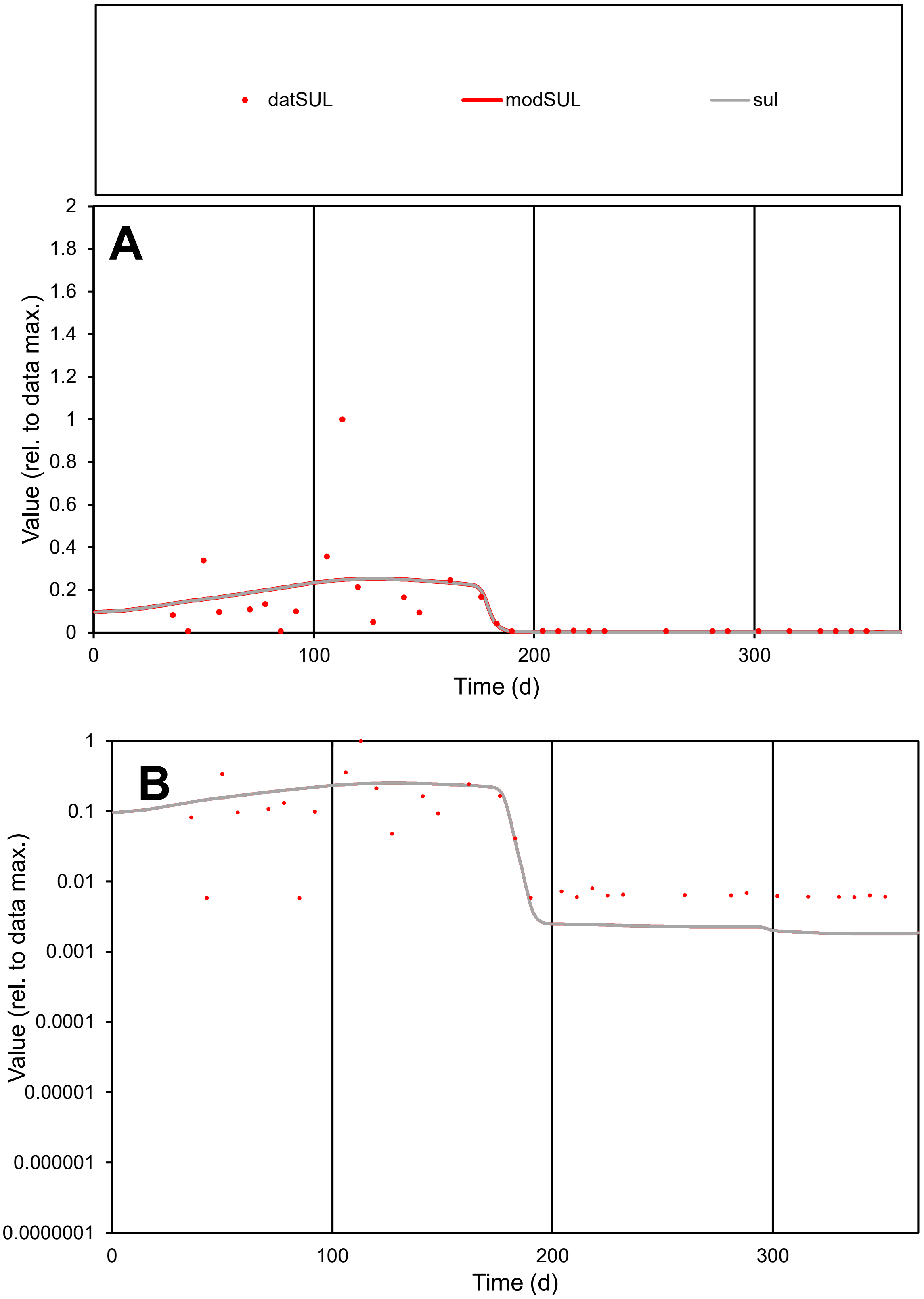


**C1**

Variables

SUL Provide jtd or Dst

CalibOnly?

2 1 to reproduce of

oNormalize?

2 1=no, 2=yes

ofxn calc*

1.51902

ofxn mod

1.52747

sRun

RunA1

zout

1

Ave

Data

0.0014

Model

0.00144

data/vMax

0.01358

ofxng

321.551

Min d/m

3.20838

ave/min d

17.703

ave/min m

58.4516

324_Calib_S_A

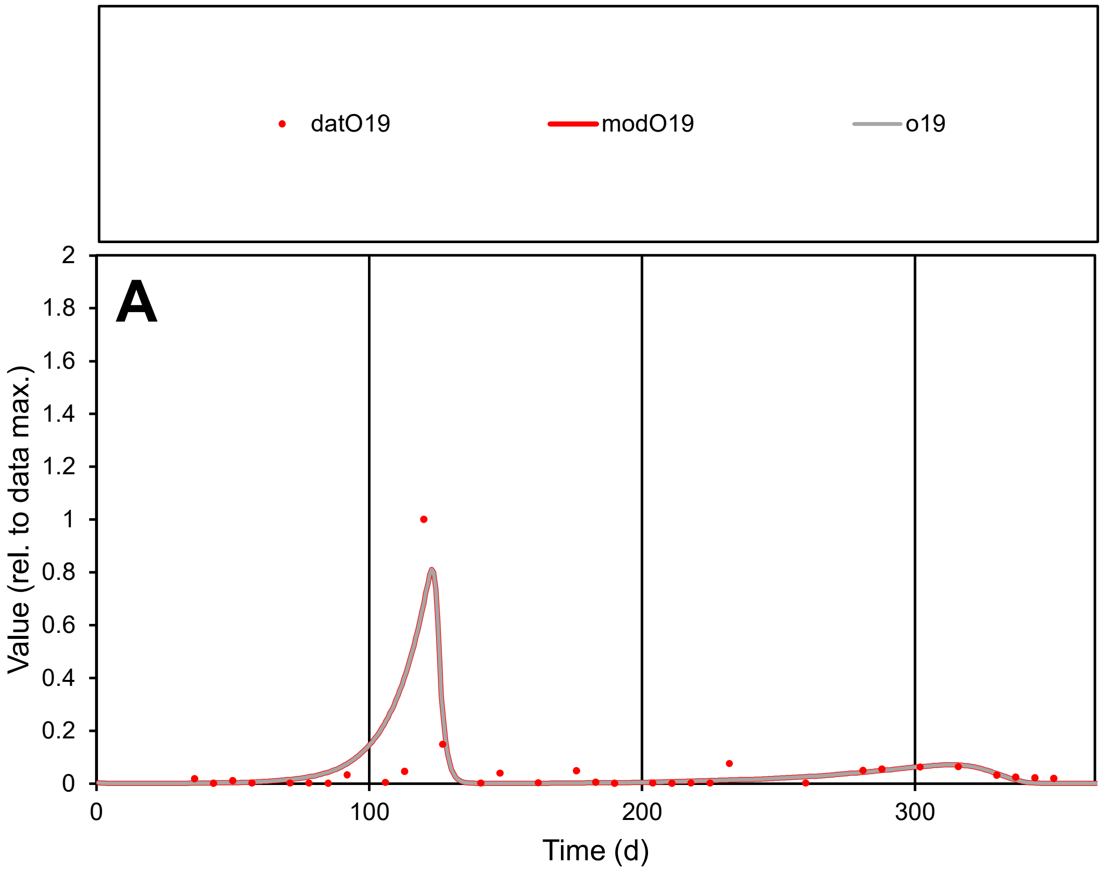


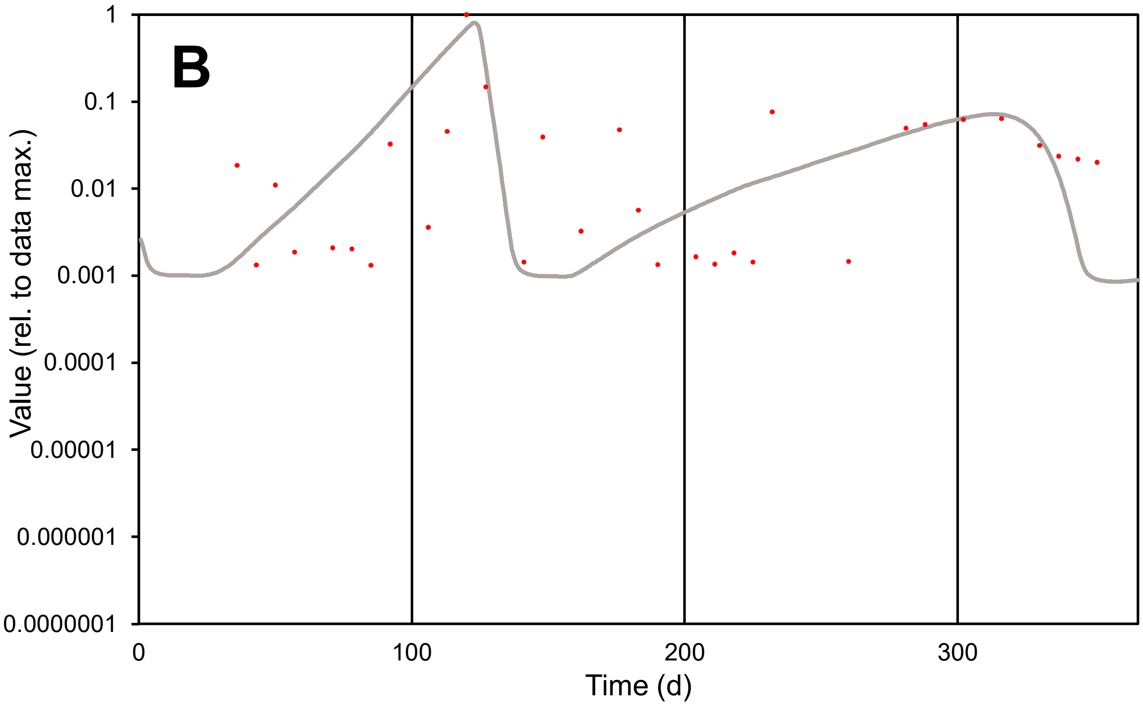


**C1**

Variables

| | |
|---|---|
| O19 | Provide jtd or Dst |
| CalibOnly? | |
| 2 | 1 to reproduce of |
| oNormalize? | |
| 2 | 1=no, 2=yes |

ofxn calc*
1.49104
ofxn mod
1.60151

sRun
RunA1
zout
1

Ave
Data
0.00119
Model
0.00136

data/vMax
0.02135

ofxng
321.551

Min d/m
1.47897

ave/min d
42.176
ave/min m
71.3825

324_Calib_S_A

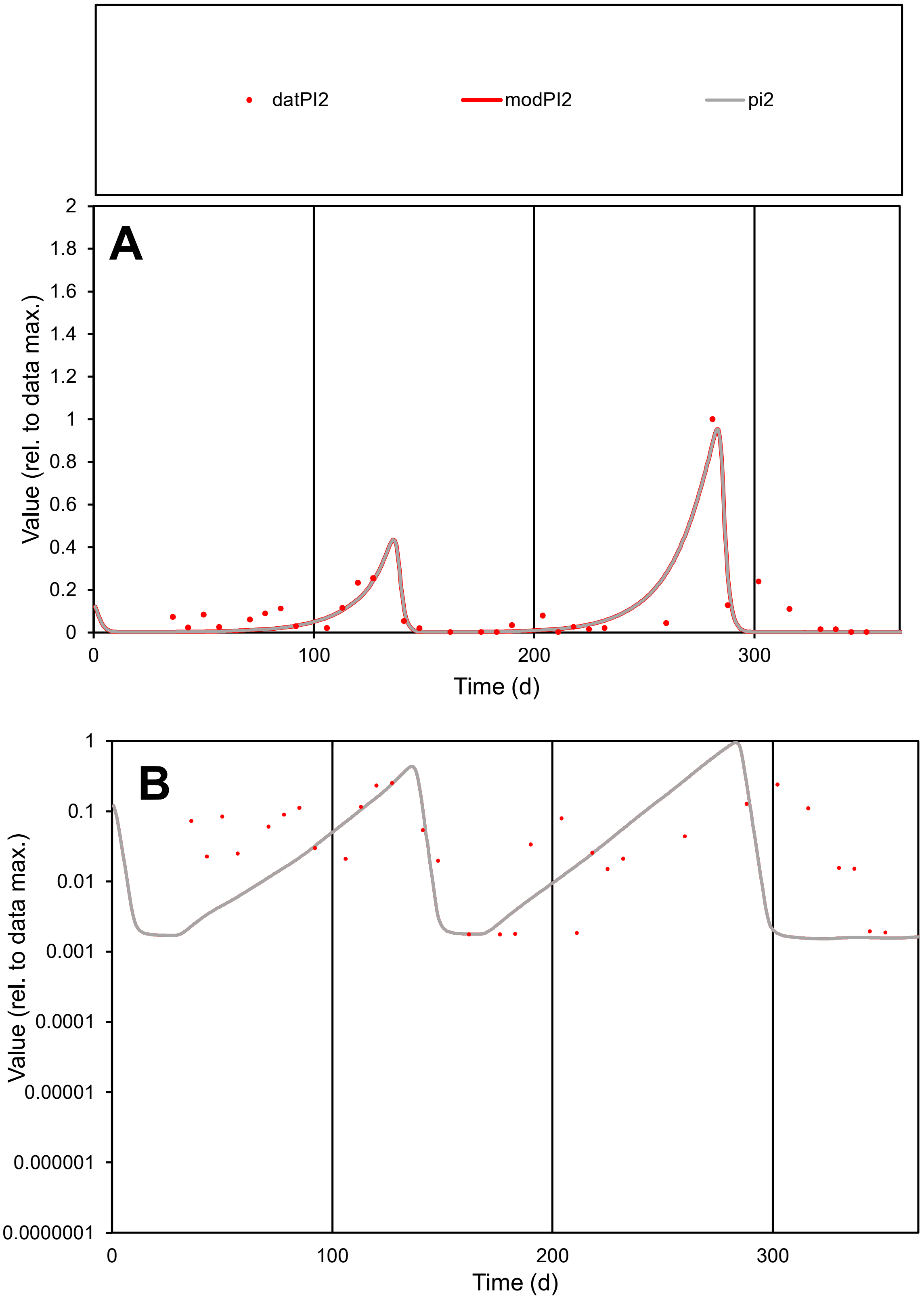


**C1**

Variables
PI2 Provide jtd or Dst
CalibOnly?
2 1 to reproduce of
oNormalize?
2 1=no, 2=yes

ofxn calc*
0.78687
ofxn mod
0.8167

sRun
RunA1
zout
1

Ave
Data
0.00144
Model
0.00113

data/vMax
0.01572

ofxng
321.551

Min d/m
1.13519

ave/min d
52.0151
ave/min m
46.1126

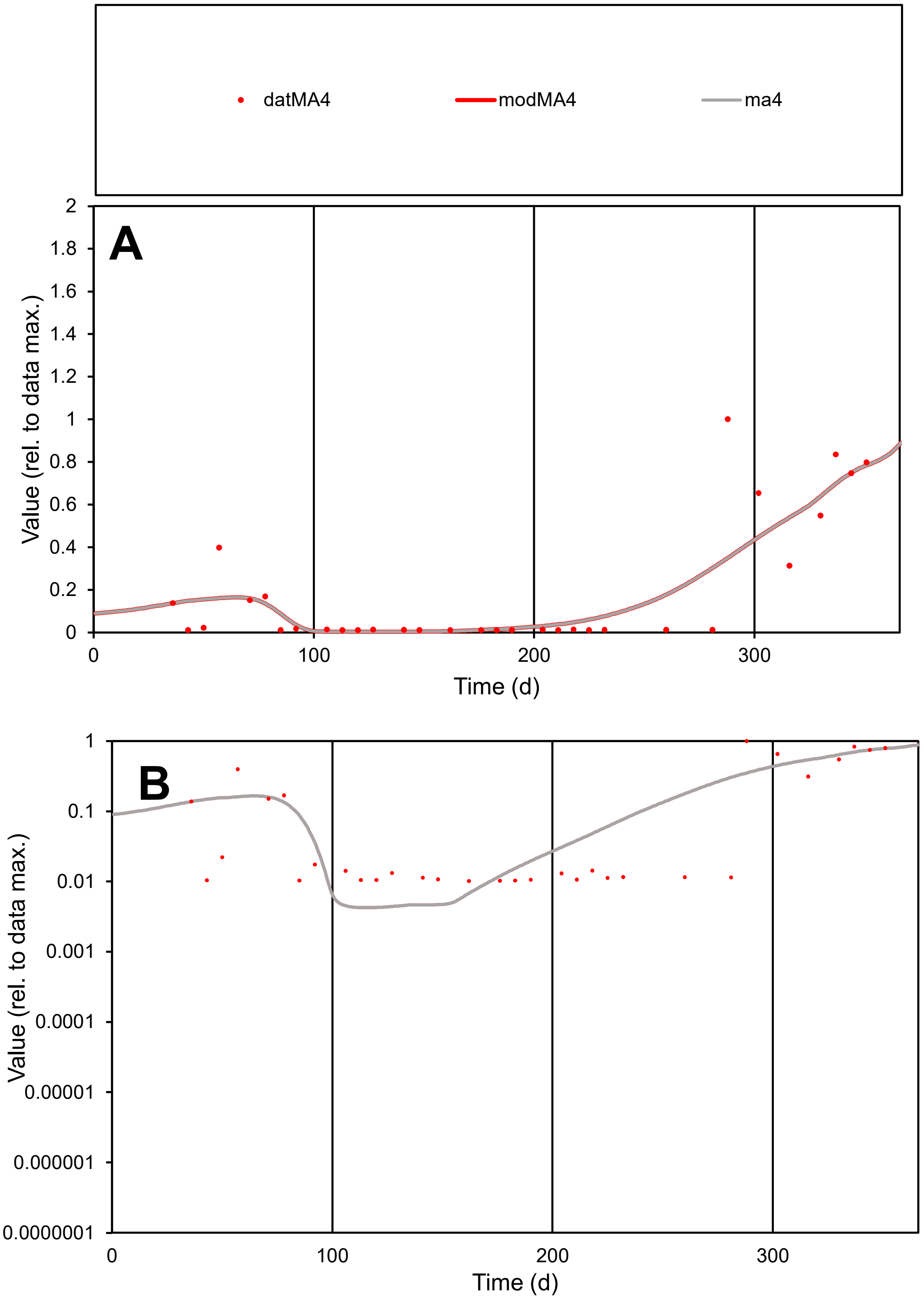


**C1**

| | |
|---|---|
| Variables | |
| MA4 | Provide jtd or Dst |
| CalibOnly? | |
| 2 | 1 to reproduce of |
| oNormalize? | |
| 2 | 1=no, 2=yes |
| ofxn calc* | |
| 0.81644 | |
| ofxn mod | |
| 0.83273 | |
| sRun | |
| RunA1 | |
| zout | |
| 1 | |
| Ave | |
| Data | |
| 0.00035 | |
| Model | |
| 0.00035 | |
| data/vMax | |
| 0.00185 | |
| ofxng | |
| 321.551 | |
| Min d/m | |
| 2.38746 | |
| ave/min d | |
| 18.377 | |
| ave/min m | |
| 44.3696 | |

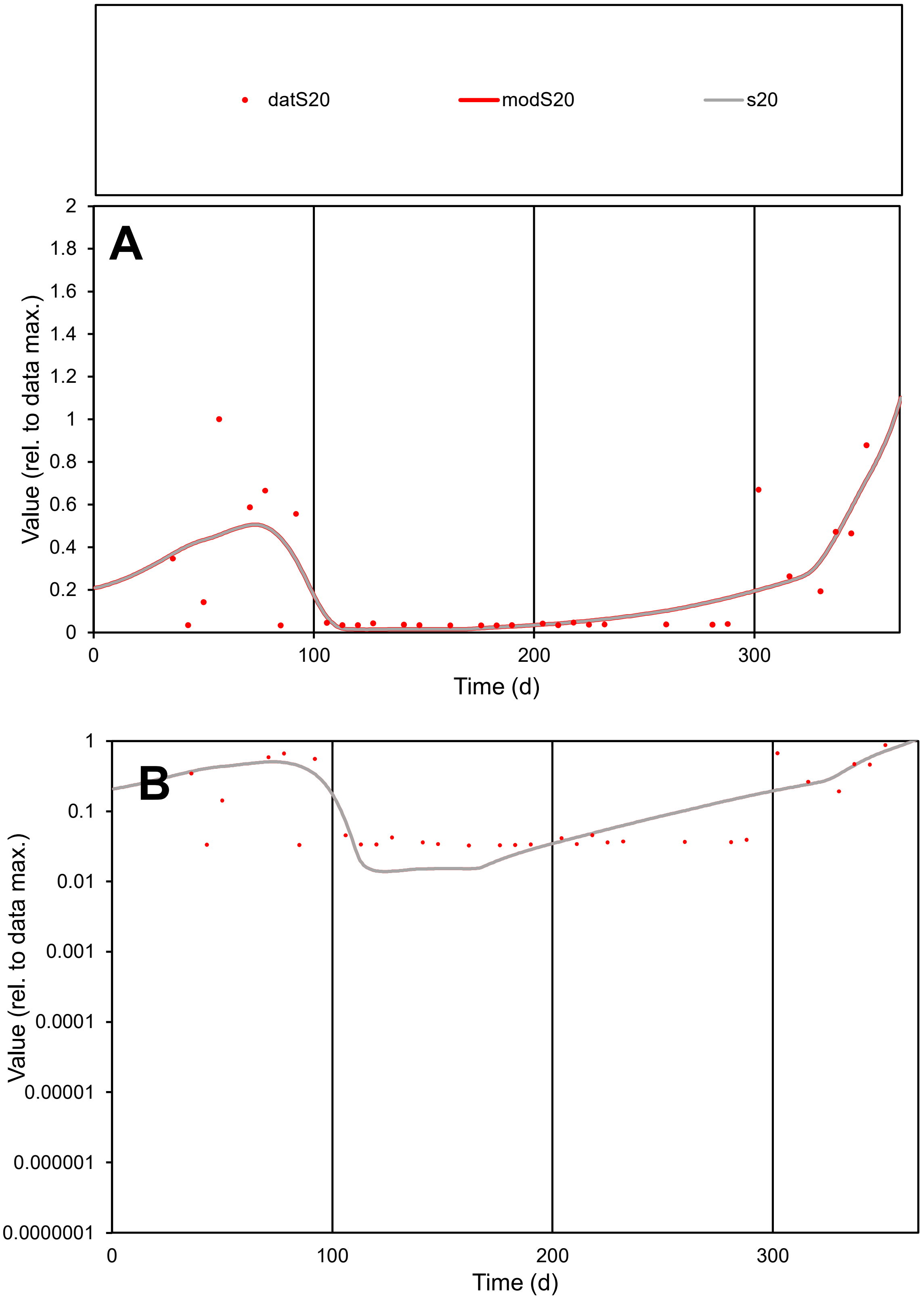


**C1**

Variables
S20 Provide jtd or Dst
CalibOnly?
2 1 to reproduce of
oNormalize?
2 1=no, 2=yes

ofxn calc*
0.84124
ofxn mod
0.85412

sRun
RunA1
zout
1

Ave
Data
0.00051
Model
0.00051

data/vMax
0.00234

ofxng
321.551

Min d/m
2.33761

ave/min d
6.67049
ave/min m
15.4998

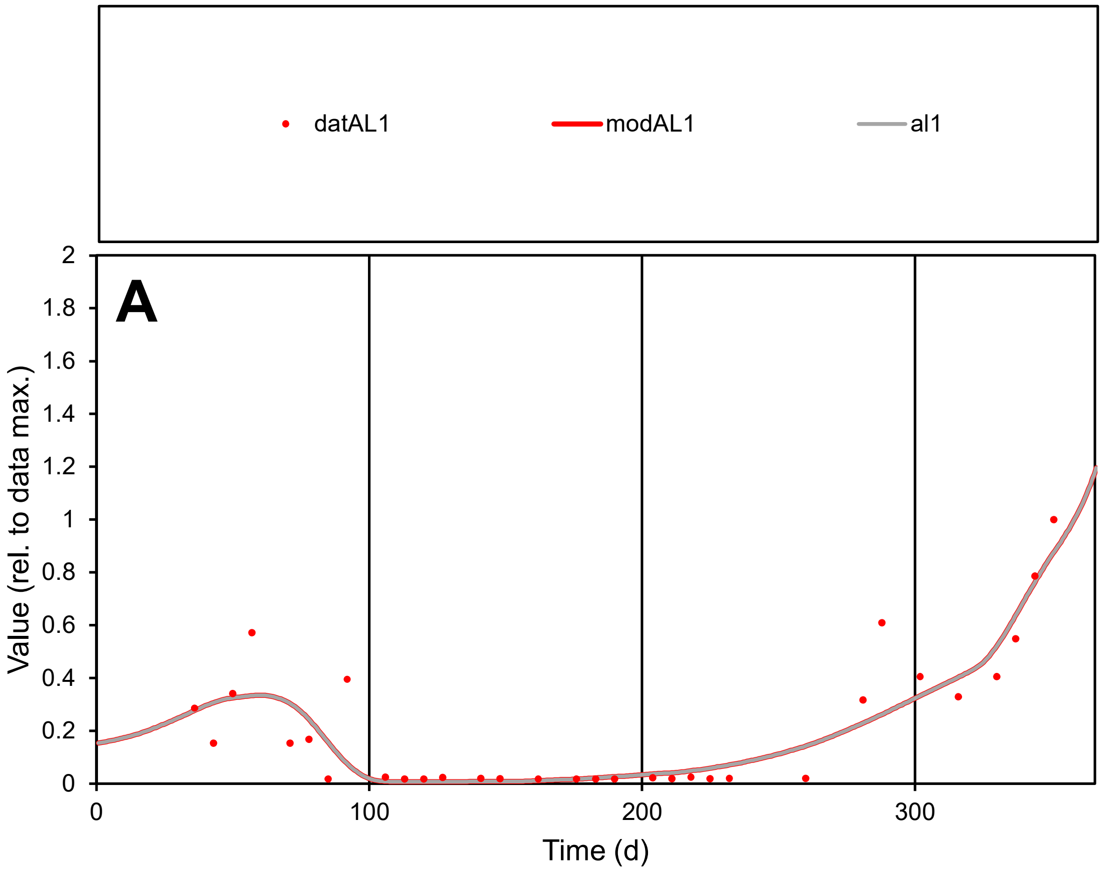


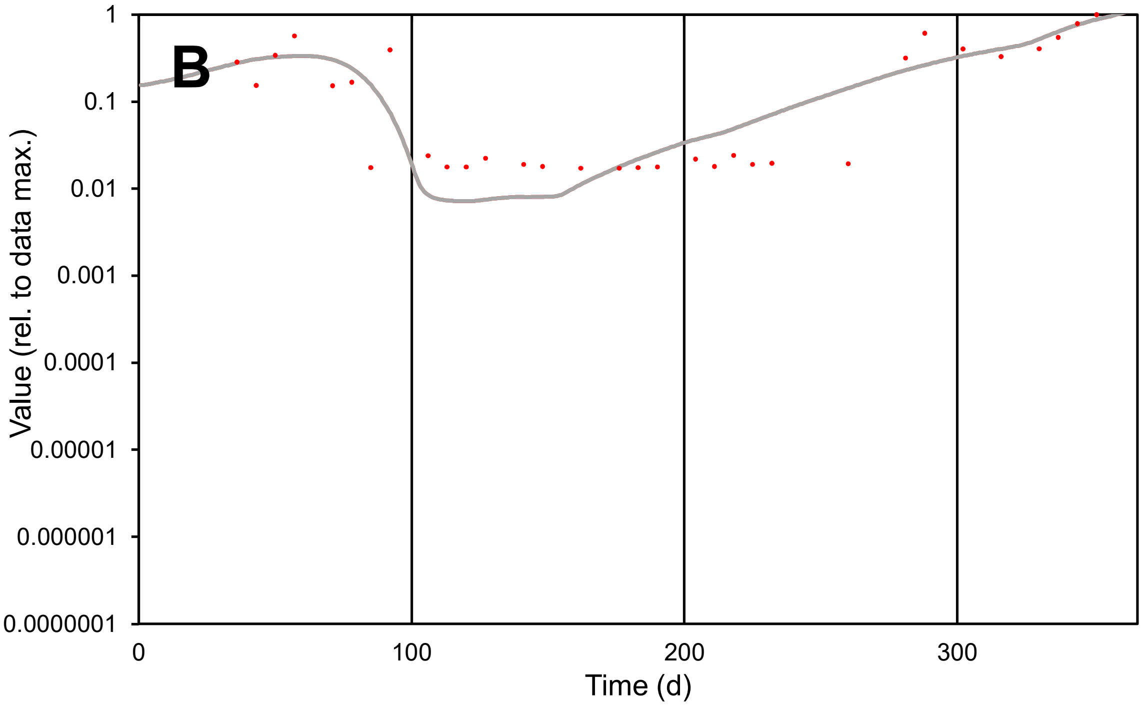


**C1**

Variables
AL1 Provide jtd or Dst
CalibOnly?
2 1 to reproduce of
oNormalize?
2 1=no, 2=yes

ofxn calc*
0.54067
ofxn mod
0.55814

sRun
RunA1
zout
1

Ave
Data
0.00053
Model
0.00052

data/vMax
0.0025

ofxng
321.551

Min d/m
2.38227

ave/min d
12.3485
ave/min m
28.7864

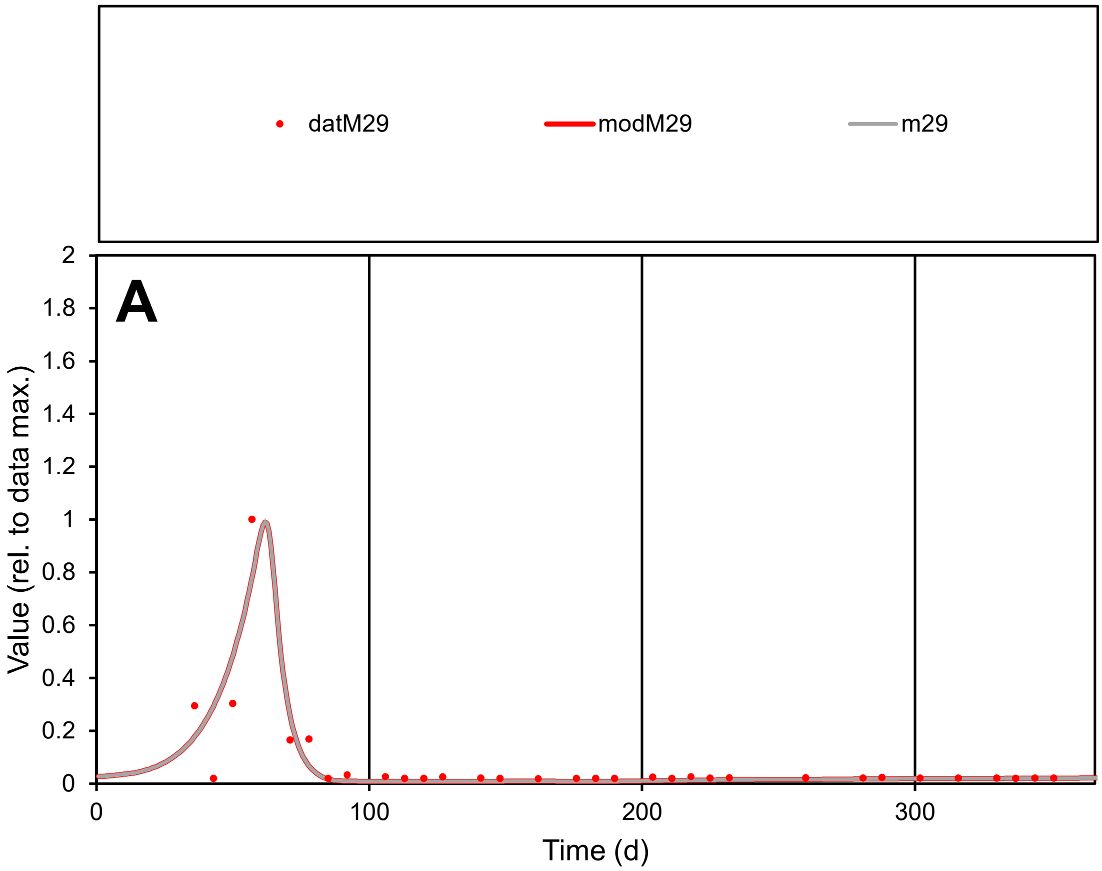


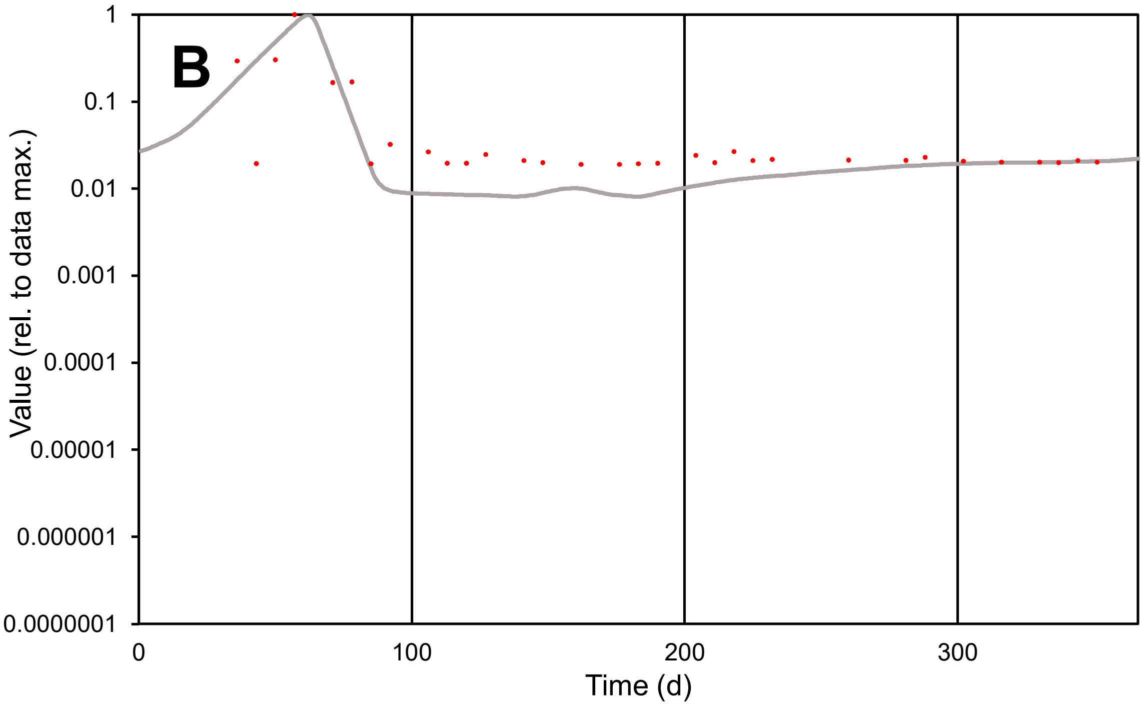


**C1**

| | |
|---|---|
| Variables | |
| M29 | Provide jtd or Dst |
| CalibOnly? | |
| 2 | 1 to reproduce of |
| oNormalize? | |
| 2 | 1=no, 2=yes |
| ofxn calc* | |
| 0.873 | |
| ofxn mod | |
| 0.94052 | |
| sRun | |
| RunA1 | |
| zout | |
| 1 | |
| Ave | |
| Data | |
| 0.00022 | |
| Model | |
| 0.00021 | |
| data/vMax | |
| 0.00281 | |
| ofxng | |
| 321.551 | |
| Min d/m | |
| 2.32737 | |
| ave/min d | |
| 4.13262 | |
| ave/min m | |
| 9.35388 | |

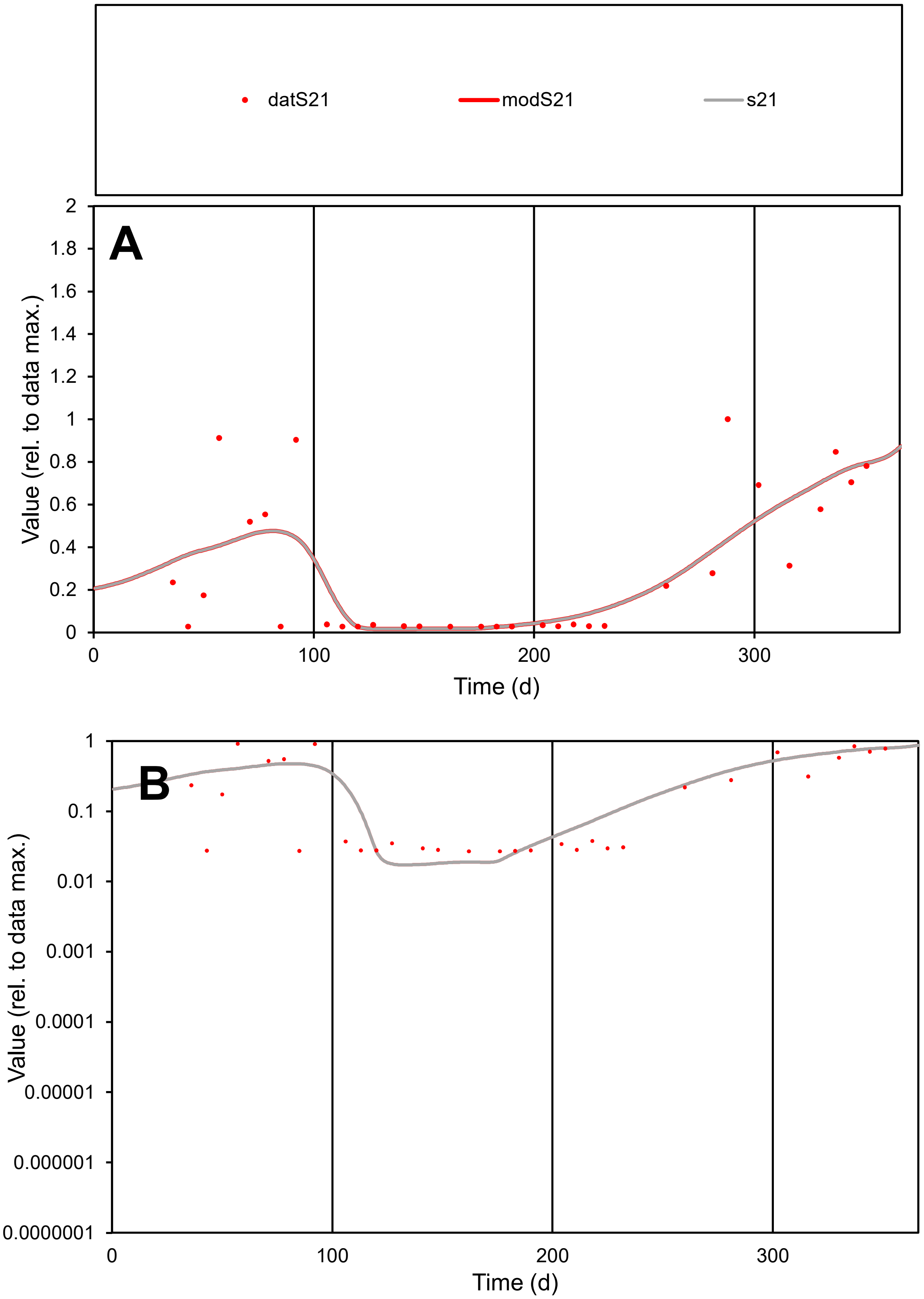


**C1**

Variables

S21 Provide jtd or Dst

CalibOnly?

2 1 to reproduce of

oNormalize?

2 1=no, 2=yes

ofxn calc*

0.71545

ofxn mod

0.72779

sRun

RunA1

zout

1

Ave

Data

0.00043

Model

0.00044

data/vMax

0.00149

ofxng

321.551

Min d/m

1.5261

ave/min d

10.7288

ave/min m

16.7928

324_Calib_S_A

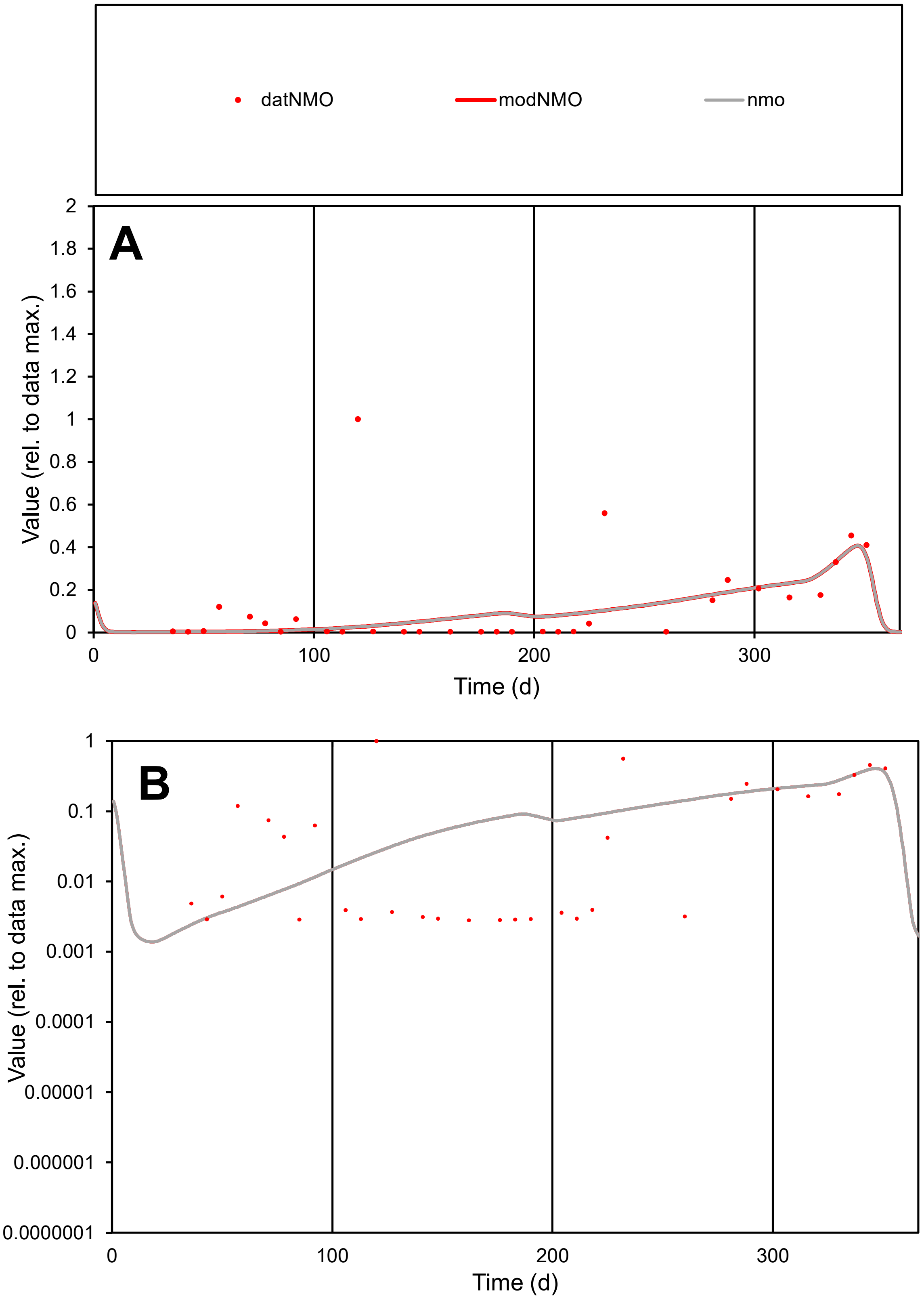


**C1**

Variables
NMO Provide jtd or Dst
CalibOnly?
2 1 to reproduce of
oNormalize?
2 1=no, 2=yes

ofxn calc*
1.56046
ofxn mod
1.56711

sRun
RunA1
zout
1

Ave
Data
0.00032
Model
0.00026

data/vMax
0.00251

ofxng
321.551

Min d/m
1.11761

ave/min d
45.3497
ave/min m
41.0456

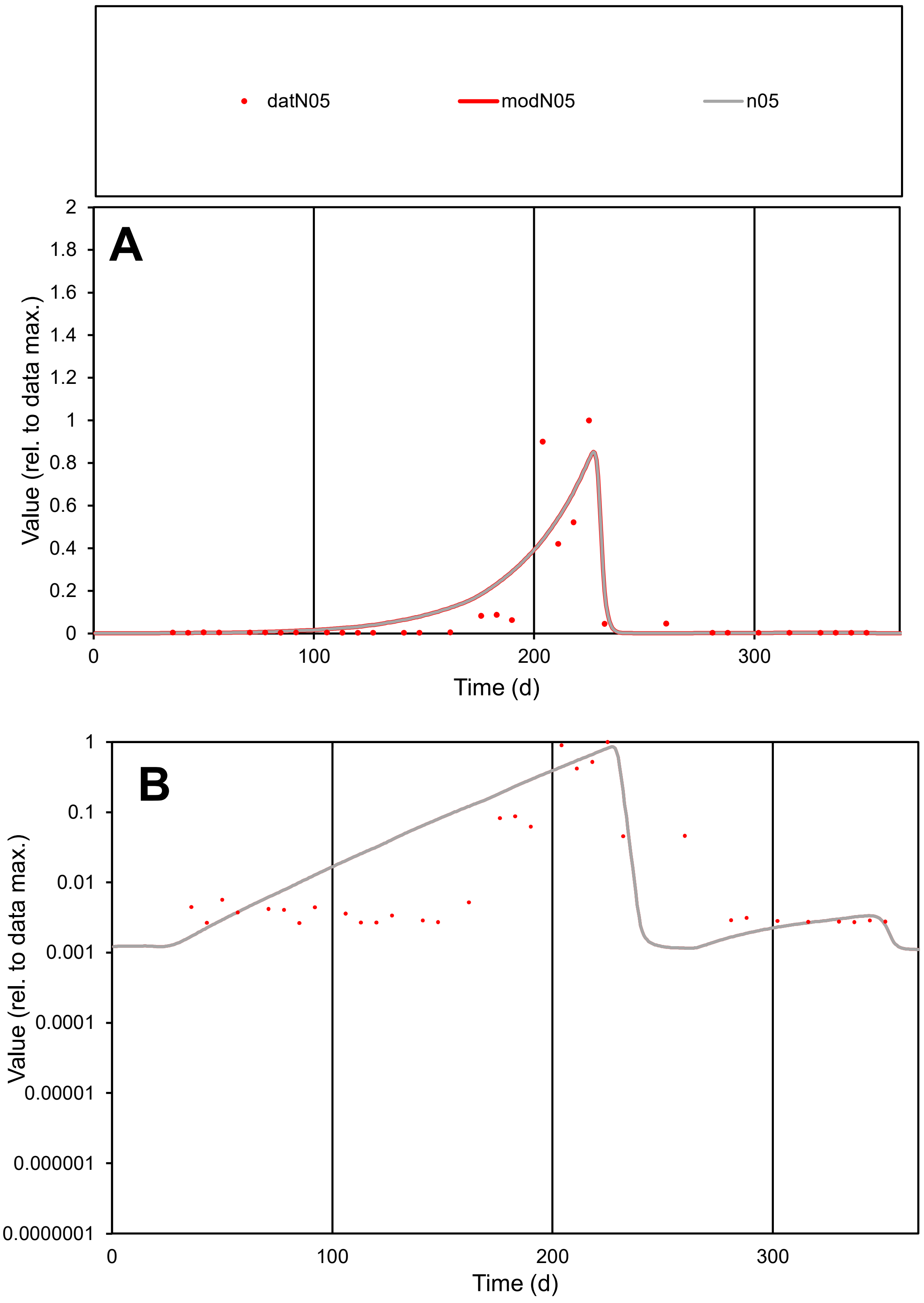


**C1**

Variables
N05 Provide jtd or Dst
CalibOnly?
2 1 to reproduce of
oNormalize?
2 1=no, 2=yes

ofxn calc*
1.07118
ofxn mod
1.12005

sRun
RunA1
zout
1

Ave
Data
0.00152
Model
0.00176

data/vMax
0.01502

ofxng
321.551

Min d/m
2.29075

ave/min d
38.4106
ave/min m
101.723

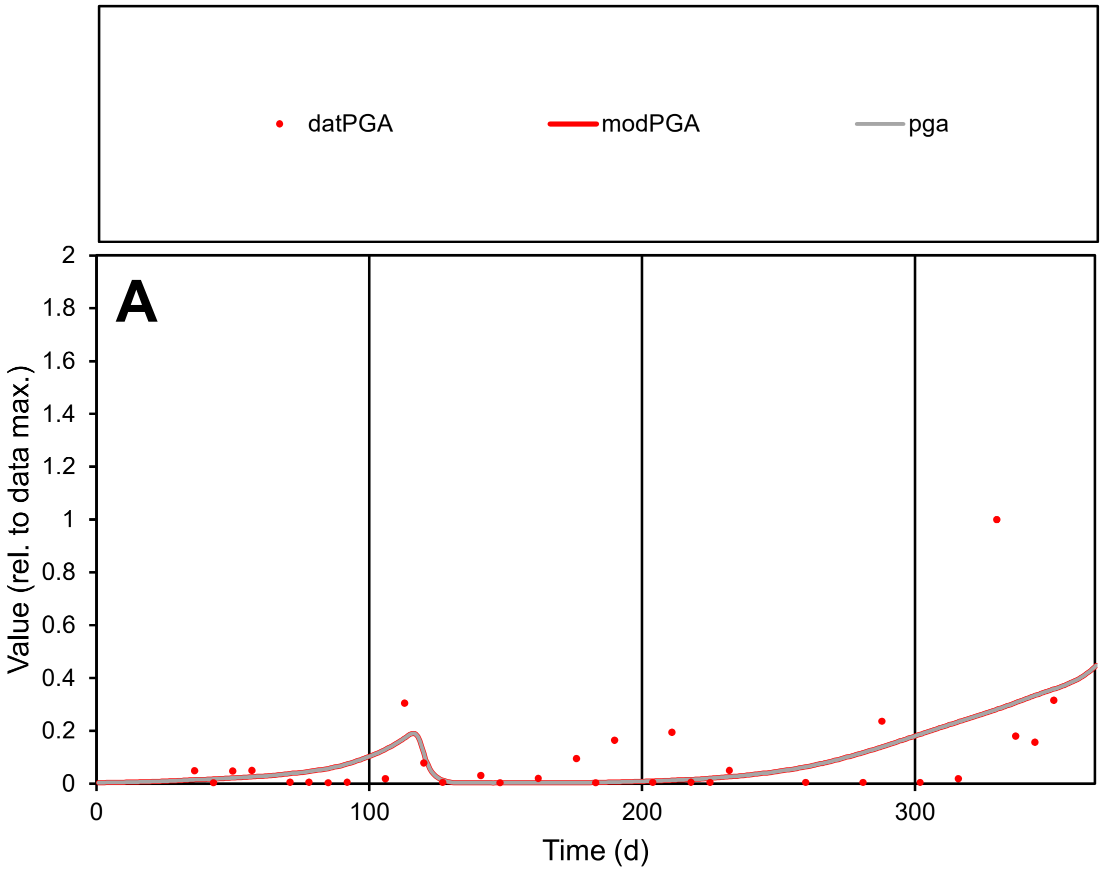


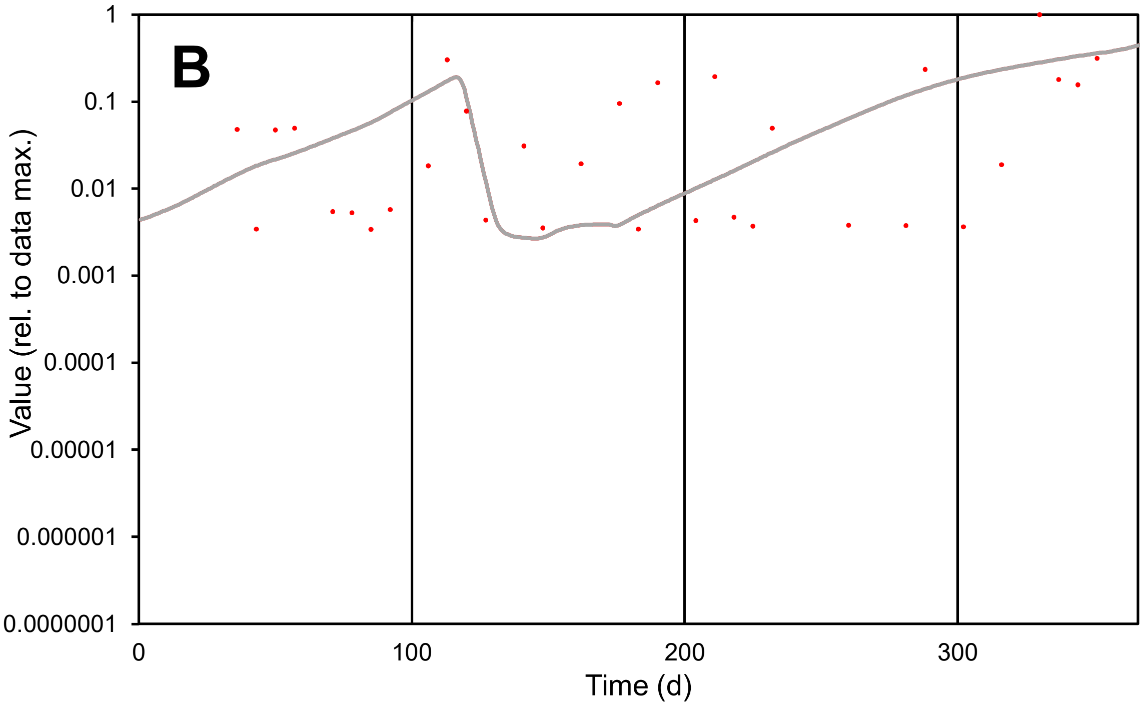


**C1**

Variables
PGA Provide jtd or Dst
CalibOnly?
2 1 to reproduce of
oNormalize?
2 1=no, 2=yes

ofxn calc*
1.62714
ofxn mod
1.63912

sRun
RunA1
zout
1

Ave
Data
0.00047
Model
0.00044

data/vMax
0.00495

ofxng
321.551

Min d/m
1.26816

ave/min d
28.003
ave/min m
33.0875

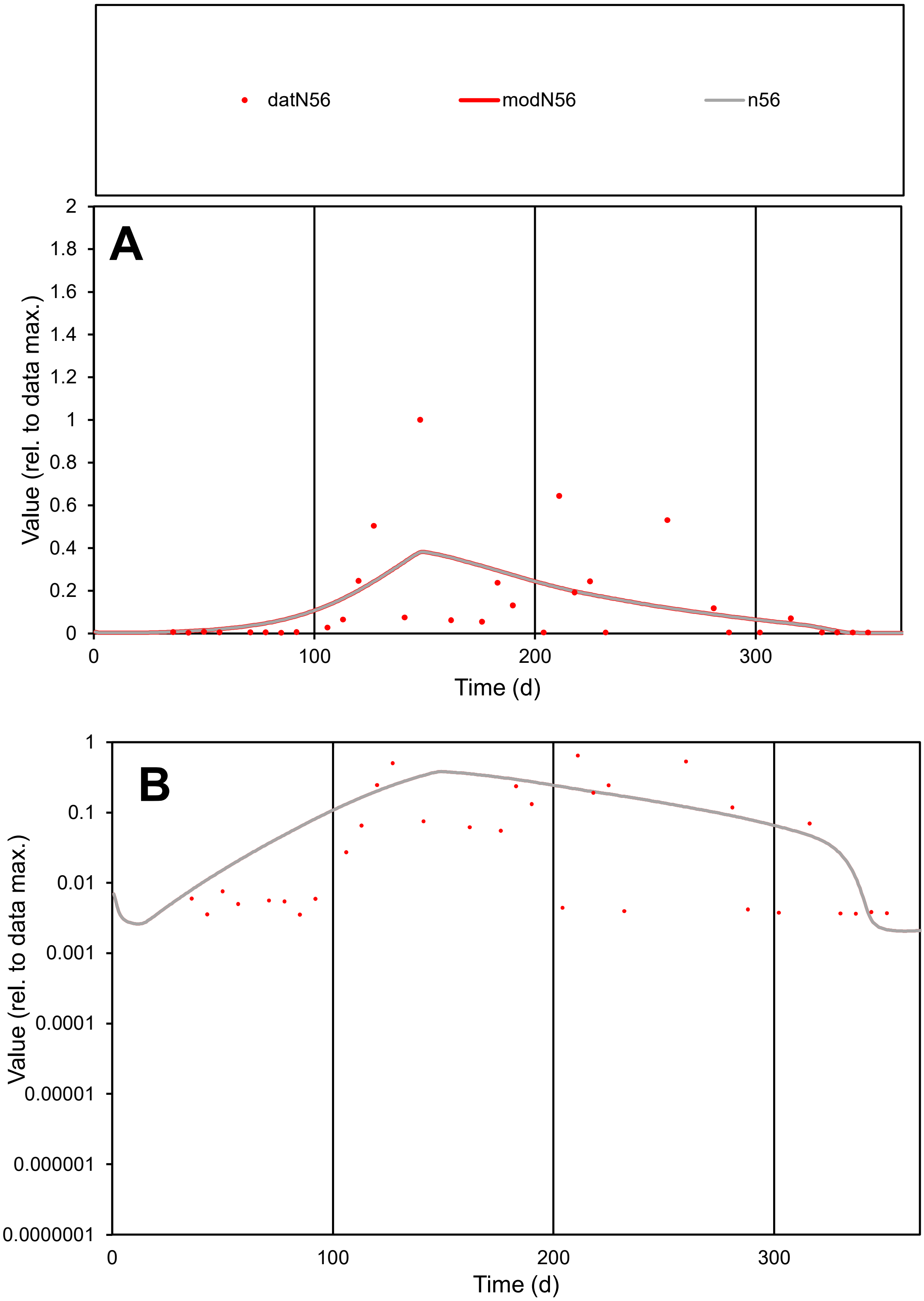


**C1**

Variables
N56 Provide jtd or Dst
CalibOnly?
2 1 to reproduce of
oNormalize?
2 1=no, 2=yes

ofxn calc*
1.43131
ofxn mod
1.44066

sRun
RunA1
zout
1

Ave
Data
0.00128
Model
0.00132

data/vMax
0.0096

ofxng
321.551

Min d/m
1.65205

ave/min d
37.8275
ave/min m
64.2839

324_Calib_S_A

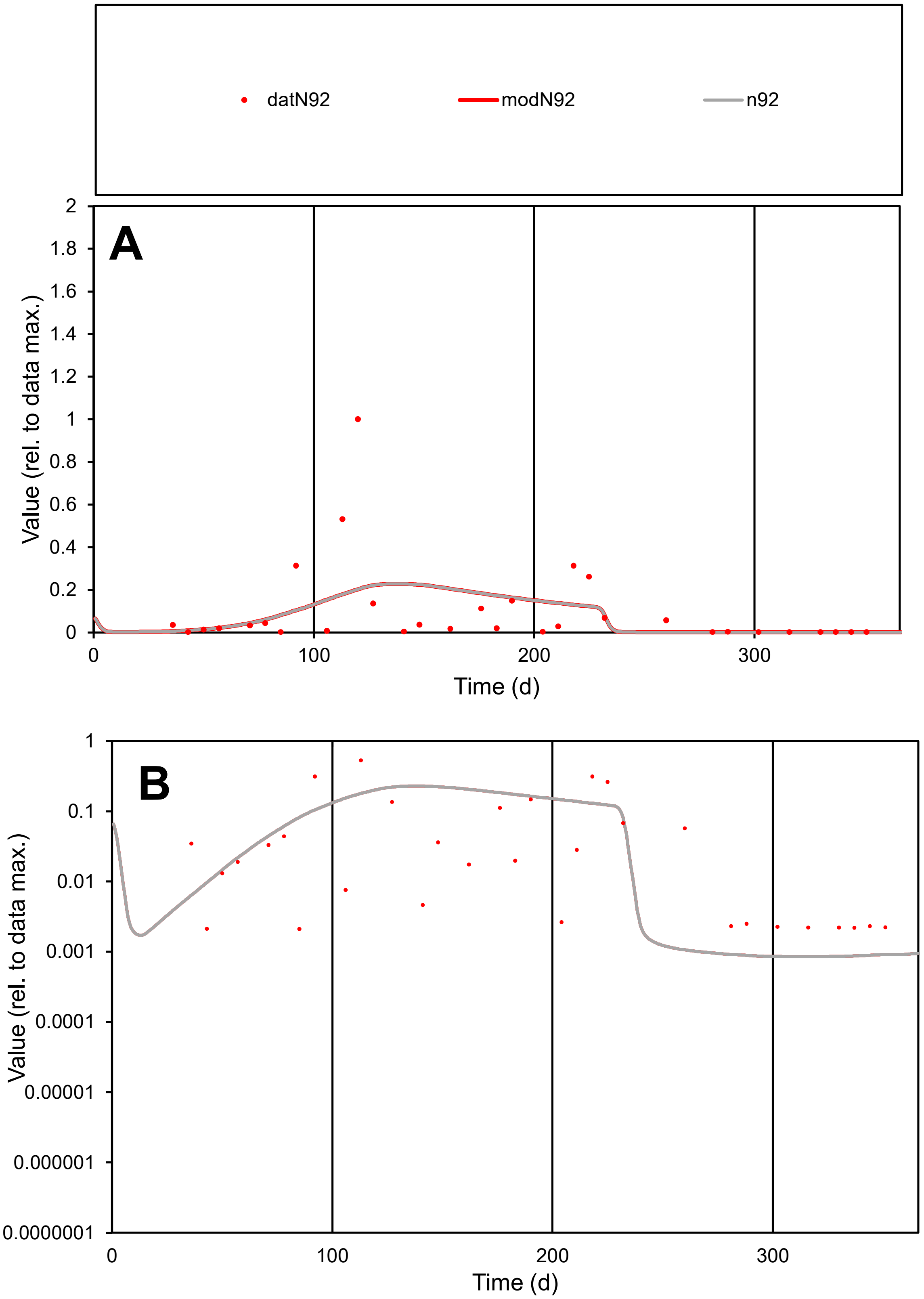


**C1**

Variables
N92 Provide jtd or Dst
CalibOnly?
2 1 to reproduce of
oNormalize?
2 1=no, 2=yes

ofxn calc*
1.80999
ofxn mod
1.81687

sRun
RunA1
zout
1

Ave
Data
0.0018
Model
0.00161

data/vMax
0.01783

ofxng
321.551

Min d/m
2.47248

ave/min d
47.6071
ave/min m
105.553

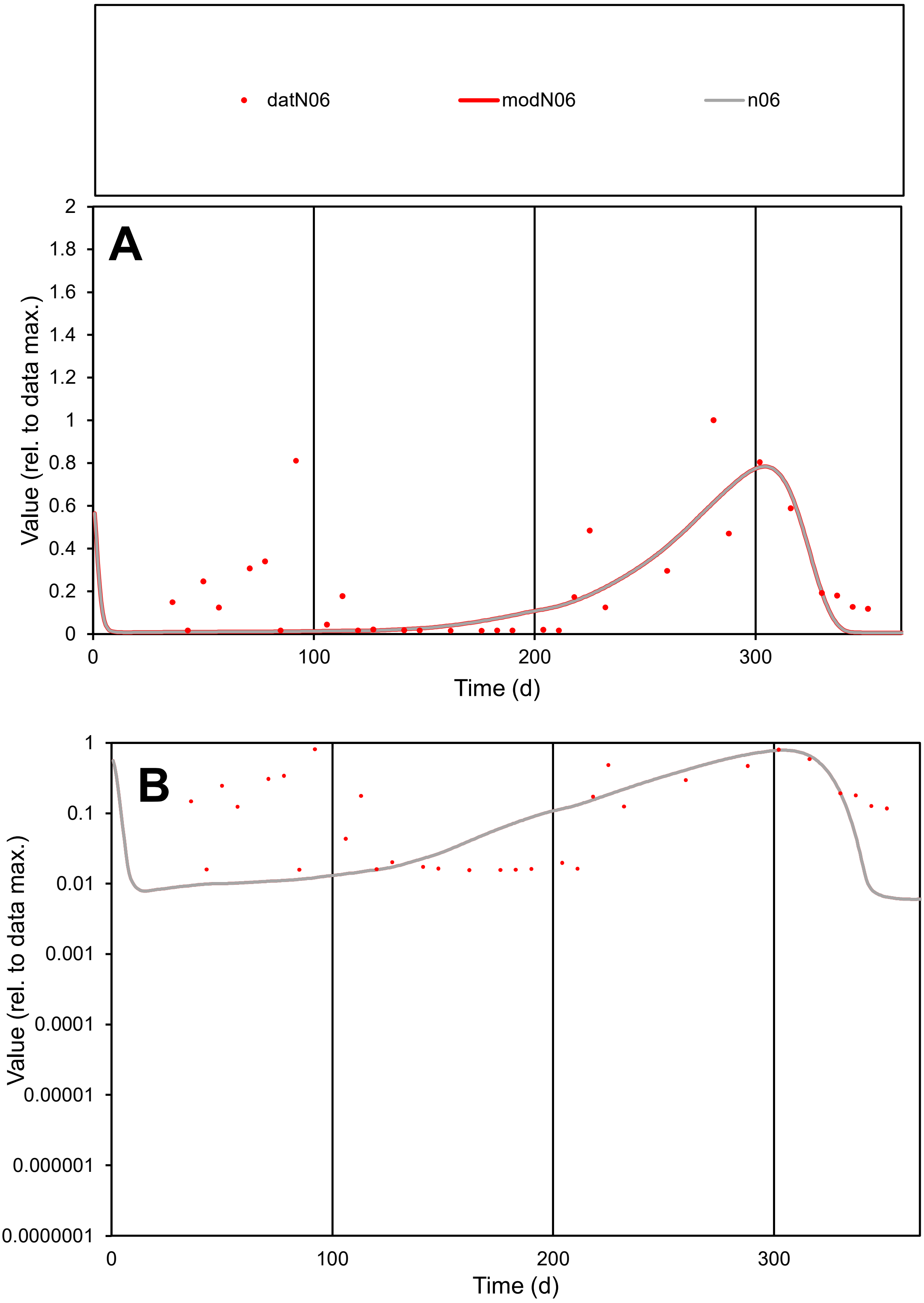


**C1**

Variables
N06 Provide jtd or Dst
CalibOnly?
2 1 to reproduce of
oNormalize?
2 1=no, 2=yes

ofxn calc*
0.9376
ofxn mod
0.94713

sRun
RunA1
zout
1

Ave
Data
0.00088
Model
0.0006

data/vMax
0.00407

ofxng
321.551

Min d/m
2.44534

ave/min d
13.9226
ave/min m
23.0837

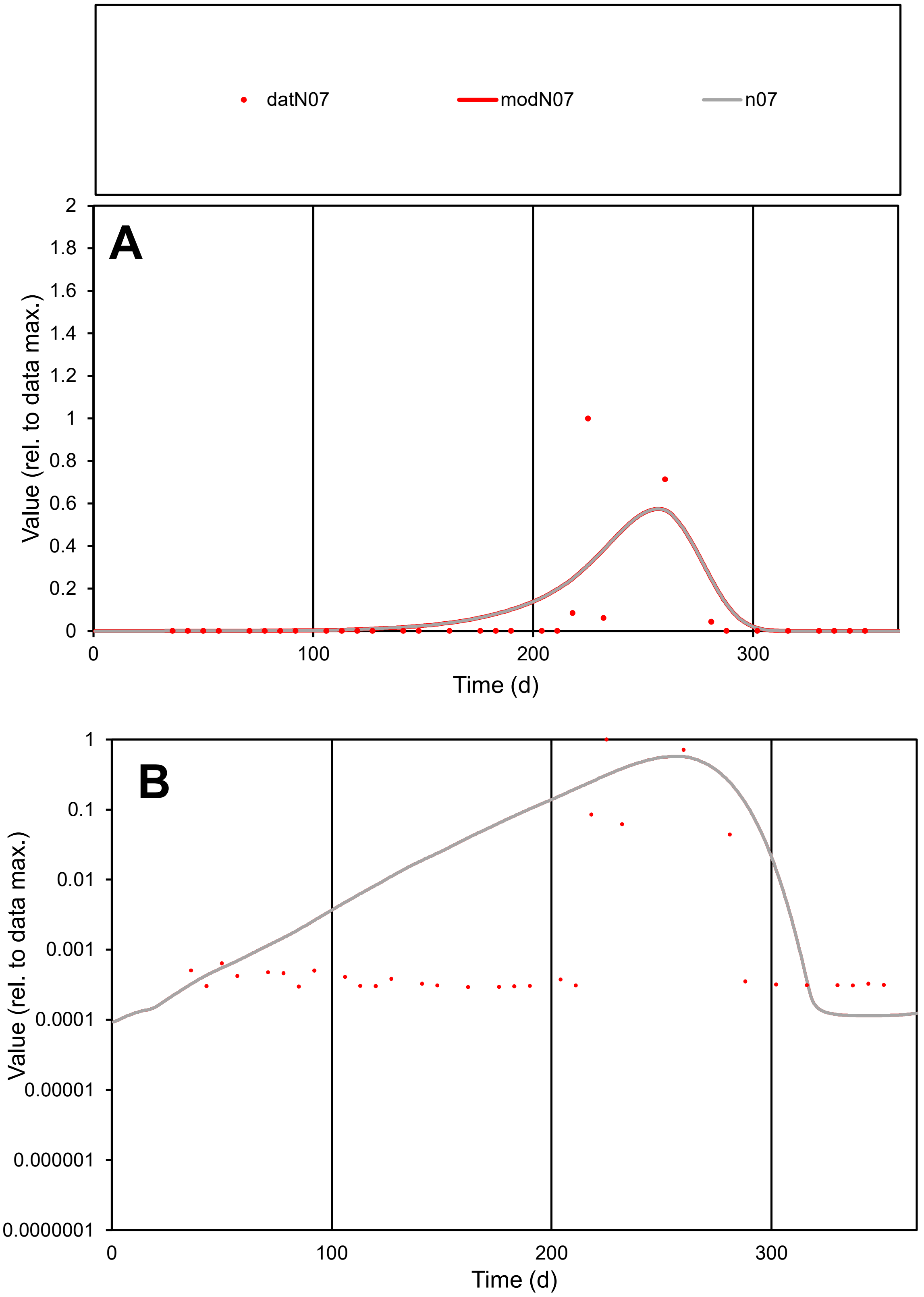


**C1**

Variables
N07 Provide jtd or Dst
CalibOnly?
2 1 to reproduce of
oNormalize?
2 1=no, 2=yes

ofxn calc*
2.52478
ofxn mod
2.55134

sRun
RunA1
zout
1

Ave
Data
0.00338
Model
0.00446

data/vMax
0.05658

ofxng
321.551

Min d/m
2.5797

ave/min d
203.052
ave/min m
690.409

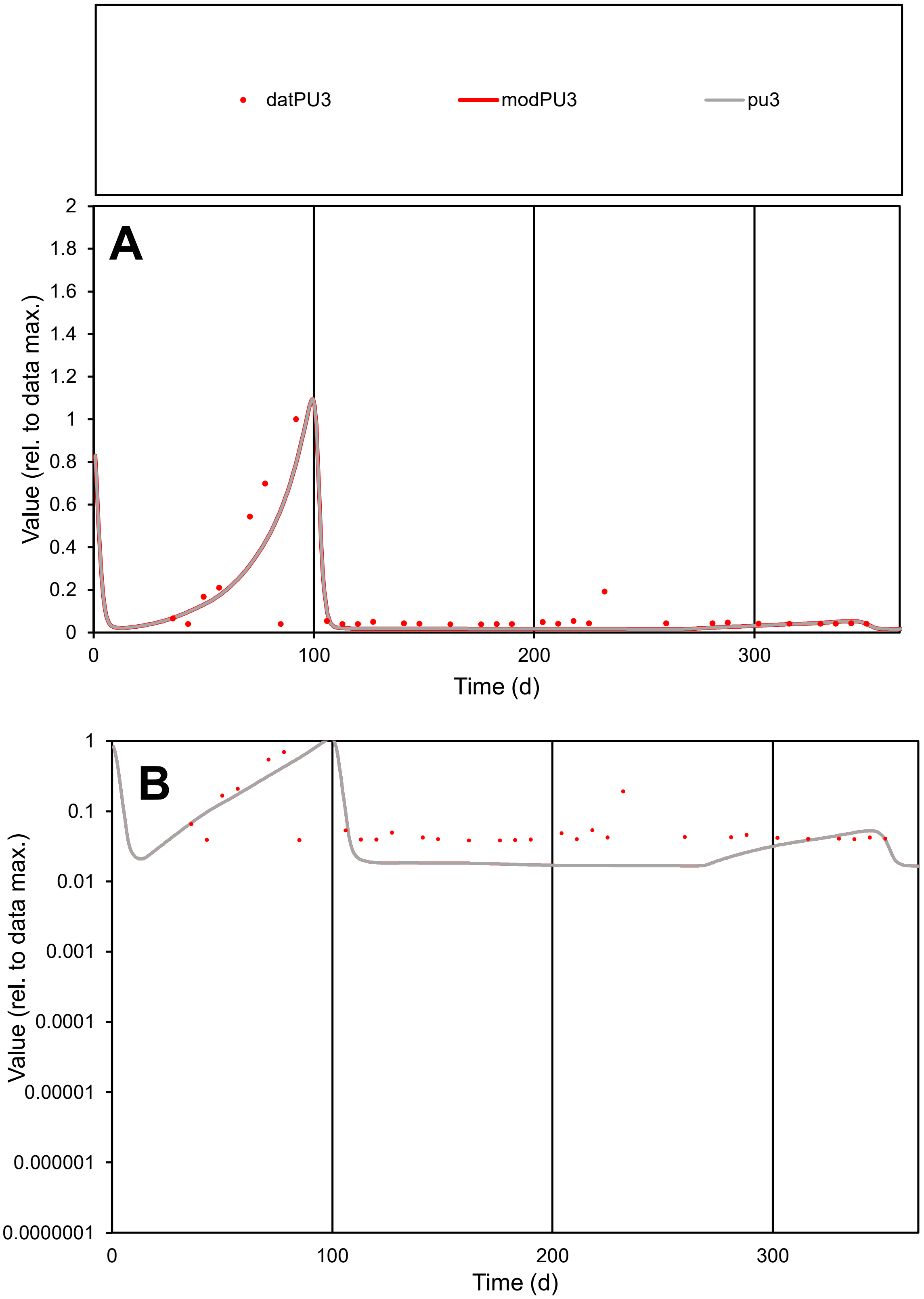


**C1**

Variables
PU3 Provide jtd or Dst
CalibOnly?
2 1 to reproduce of
oNormalize?
2 1=no, 2=yes

ofxn calc*
0.96284
ofxn mod
0.99587

sRun
RunA1
zout
1

Ave
Data
0.00047
Model
0.00039

data/vMax
0.00384

ofxng
321.551

Min d/m
2.30072

ave/min d
3.21081
ave/min m
6.17192

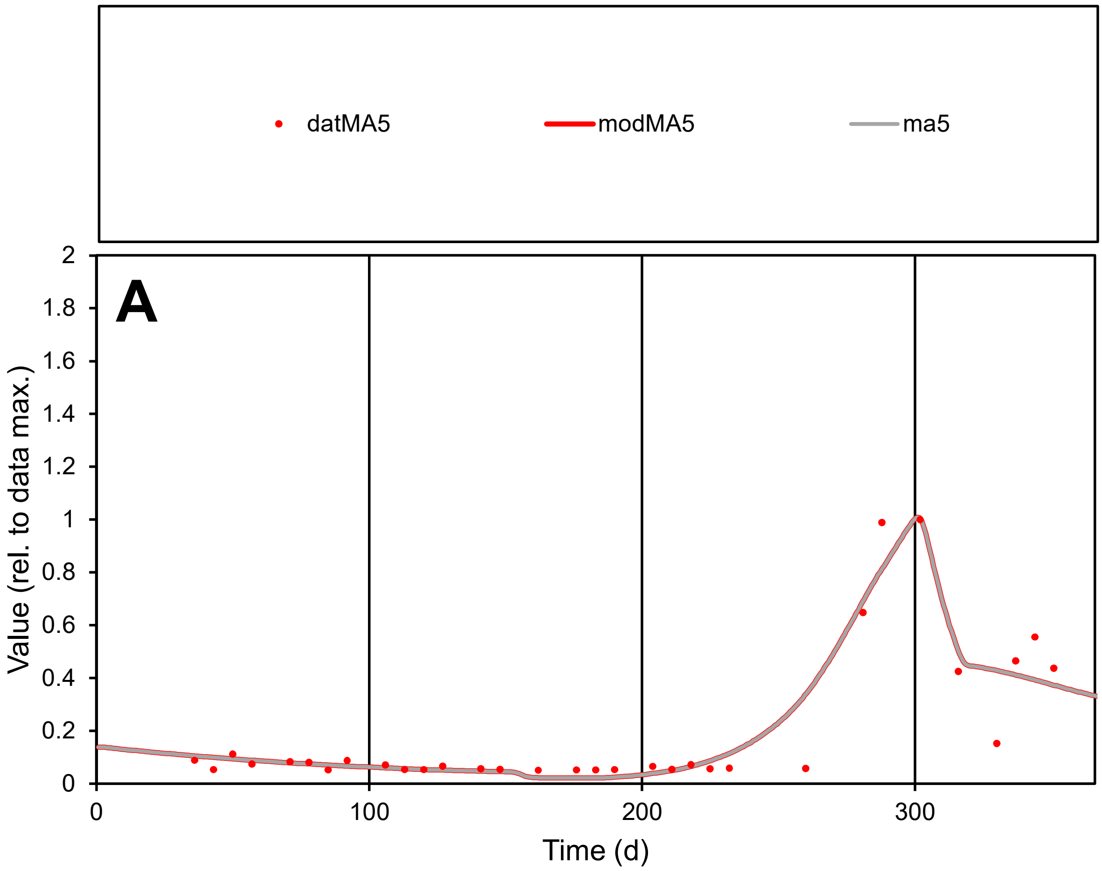


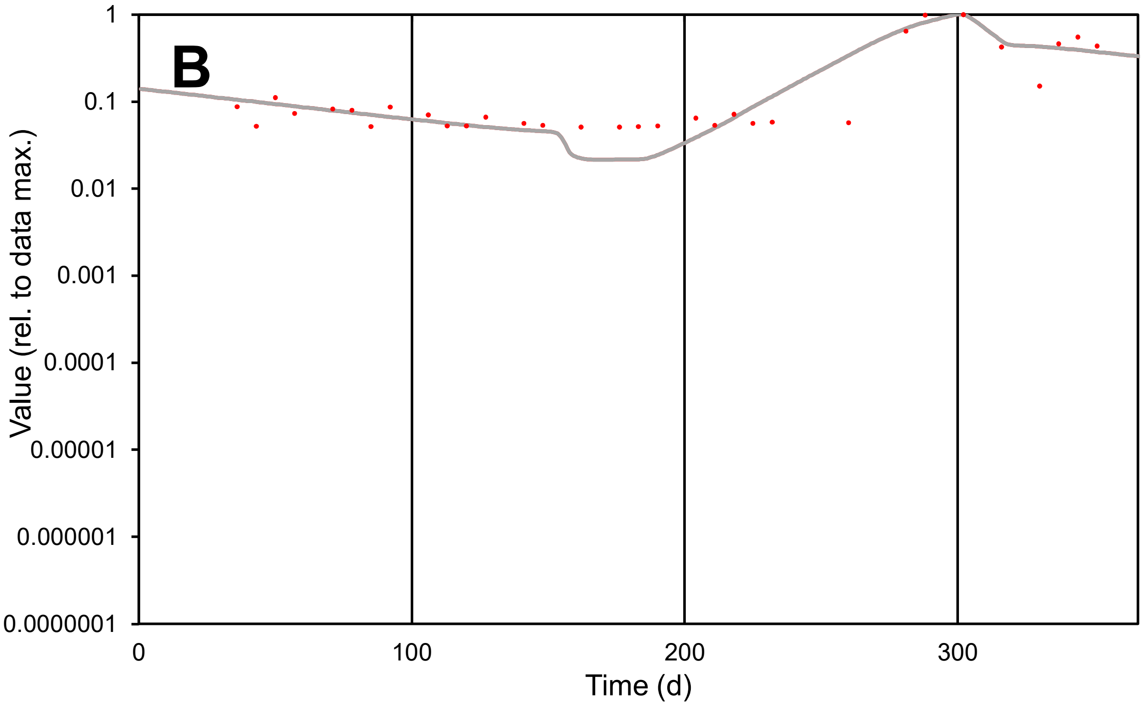


**C1**

Variables
MA5 Provide jtd or Dst
CalibOnly?
2 1 to reproduce of
oNormalize?
2 1=no, 2=yes

ofxn calc*
0.44088
ofxn mod
0.46843

sRun
RunA1
zout
1

Ave
Data
0.00035
Model
0.00036

data/vMax
0.00181

ofxng
321.551

Min d/m
2.35356

ave/min d
3.81158
ave/min m
9.20905

324_Calib_S_A

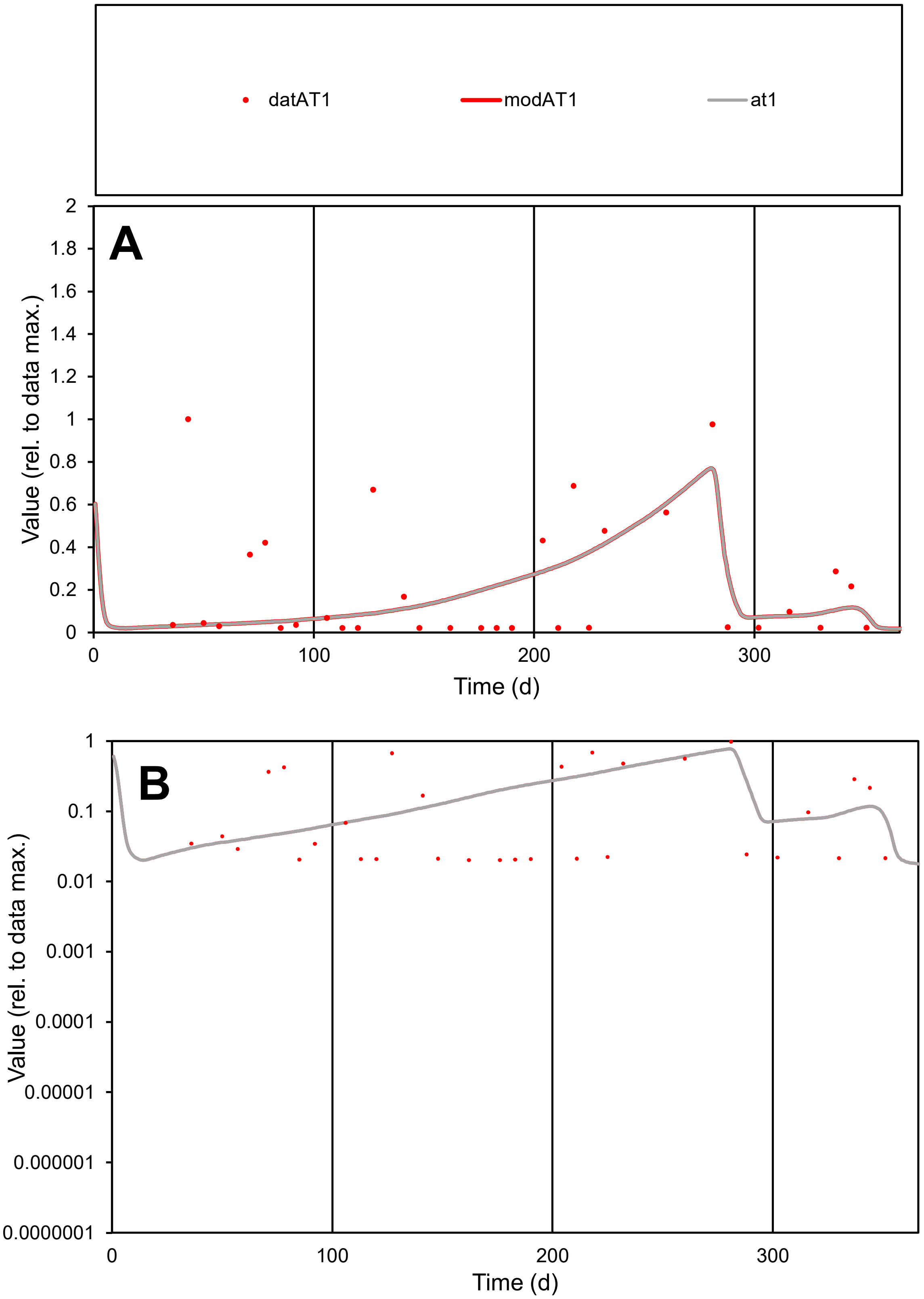


**C1**

Variables
AT1 Provide jtd or Dst
CalibOnly?
2 1 to reproduce of
oNormalize?
2 1=no, 2=yes

ofxn calc*
1.20345
ofxn mod
1.2171

sRun
RunA1
zout
1

Ave
Data
0.0005
Model
0.0004

data/vMax
0.00232

ofxng
321.551

Min d/m
0.66139

ave/min d
10.6218
ave/min m
5.6539

324_Calib_S_A

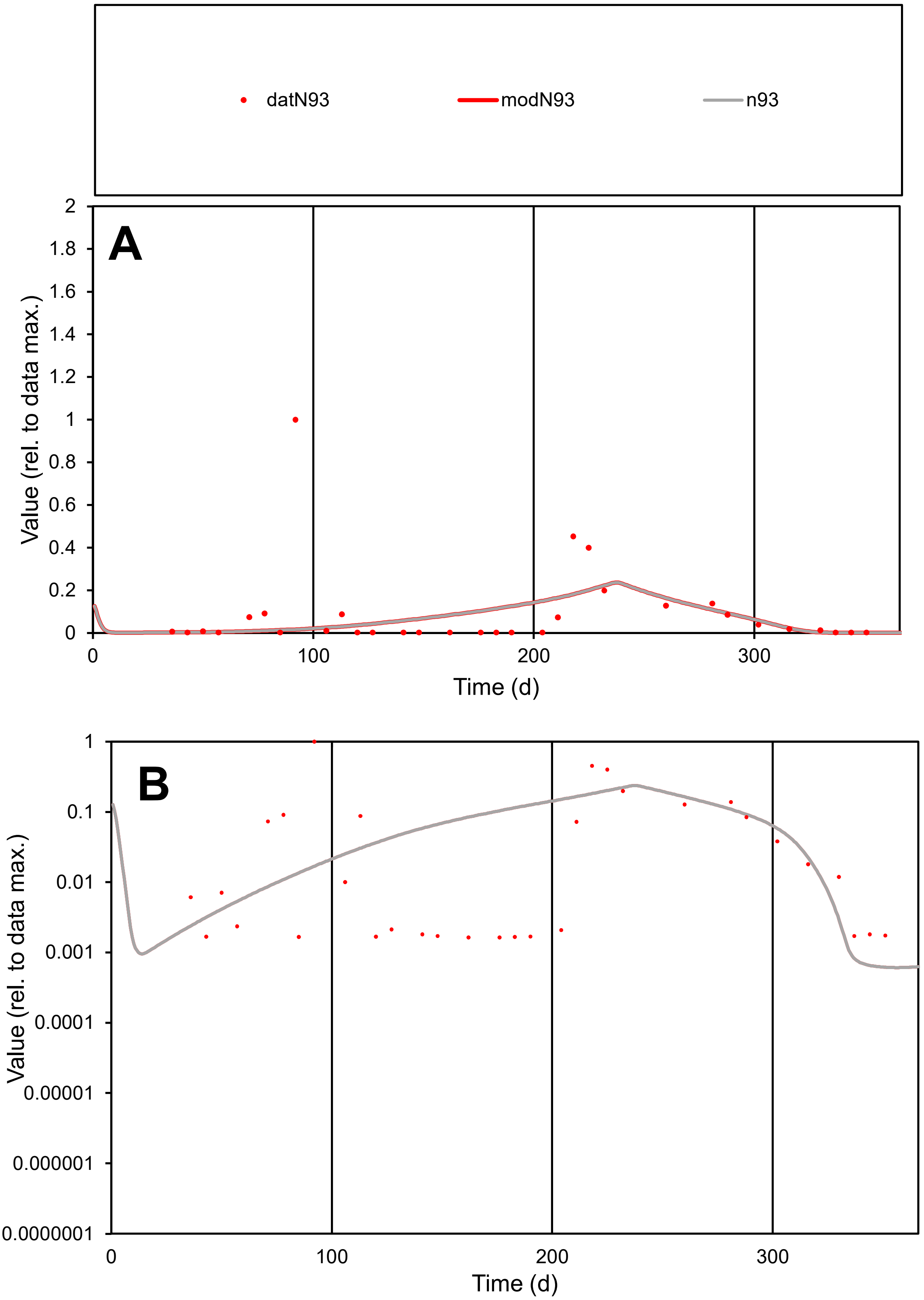


**C1**

Variables
N93 Provide jtd or Dst
CalibOnly?
2 1 to reproduce of
oNormalize?
2 1=no, 2=yes

ofxn calc*
2.16057
ofxn mod
2.16577

sRun
RunA1
zout
1

Ave
Data
0.00092
Model
0.00069

data/vMax
0.0104

ofxng
321.551

Min d/m
2.65686

ave/min d
54.4042
ave/min m
108.244

324_Calib_S_A

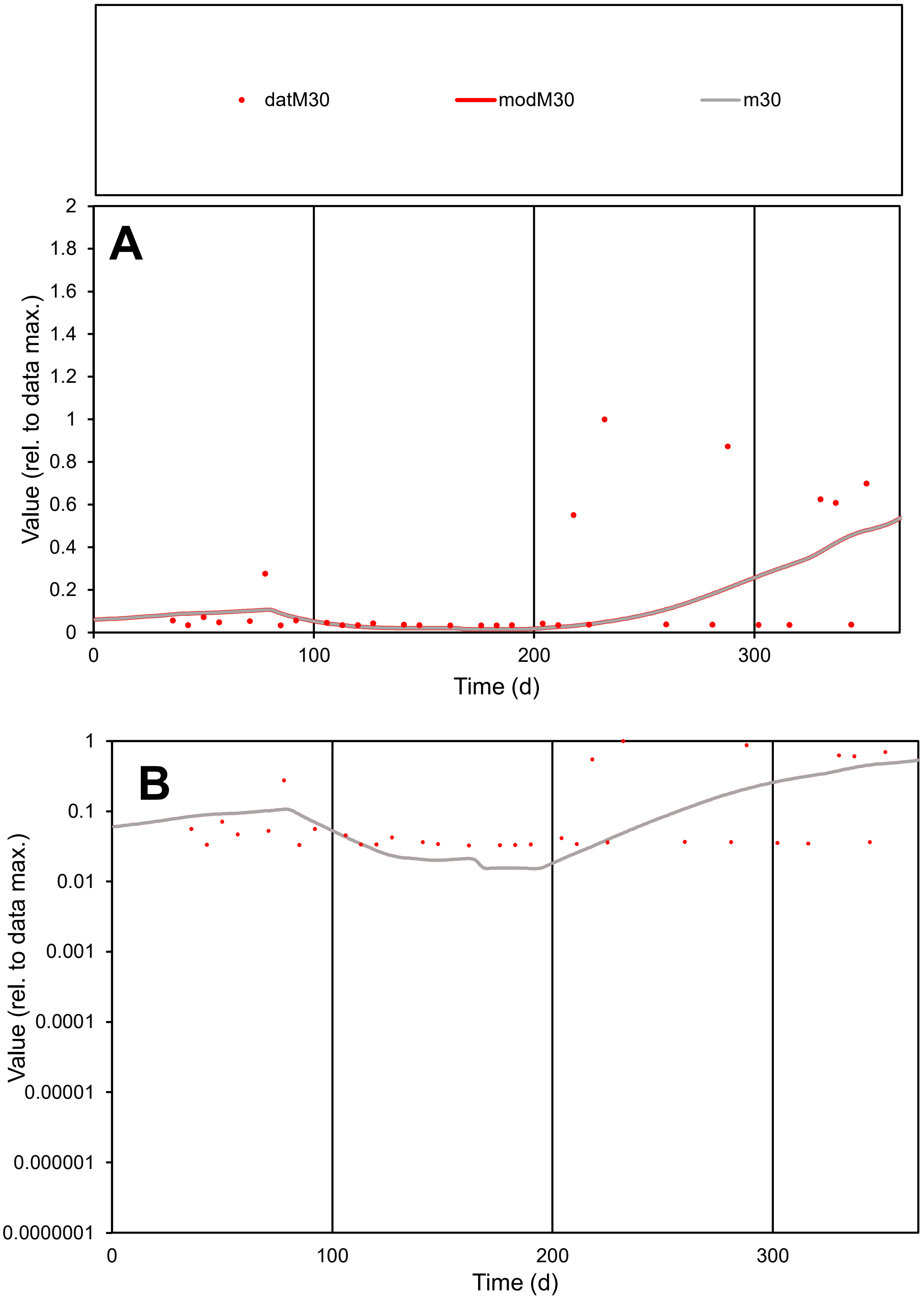


| | |
|---|---|
| **C1** | |
| Variables | |
| M30 | Provide jtd or Dst |
| CalibOnly? | |
| 2 | 1 to reproduce of |
| oNormalize? | |
| 2 | 1=no, 2=yes |
| ofxn calc* | |
| 1.46188 | |
| ofxn mod | |
| 1.46911 | |
| sRun | |
| RunA1 | |
| zout | |
| 1 | |
| Ave | |
| Data | |
| 0.00022 | |
| Model | |
| 0.00015 | |
| data/vMax | |
| 0.00123 | |
| ofxng | |
| 321.551 | |
| Min d/m | |
| 2.15145 | |
| ave/min d | |
| 5.38627 | |
| ave/min m | |
| 8.18287 | |

324_Calib_S_A

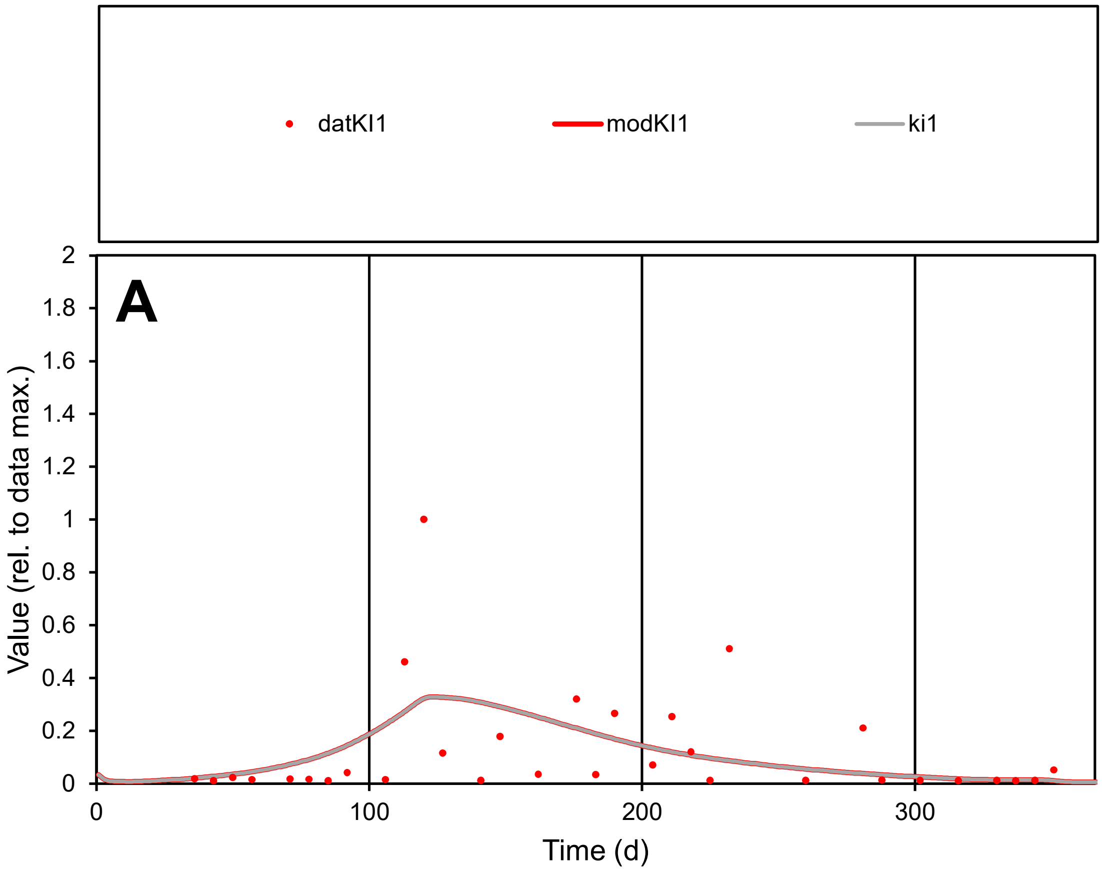


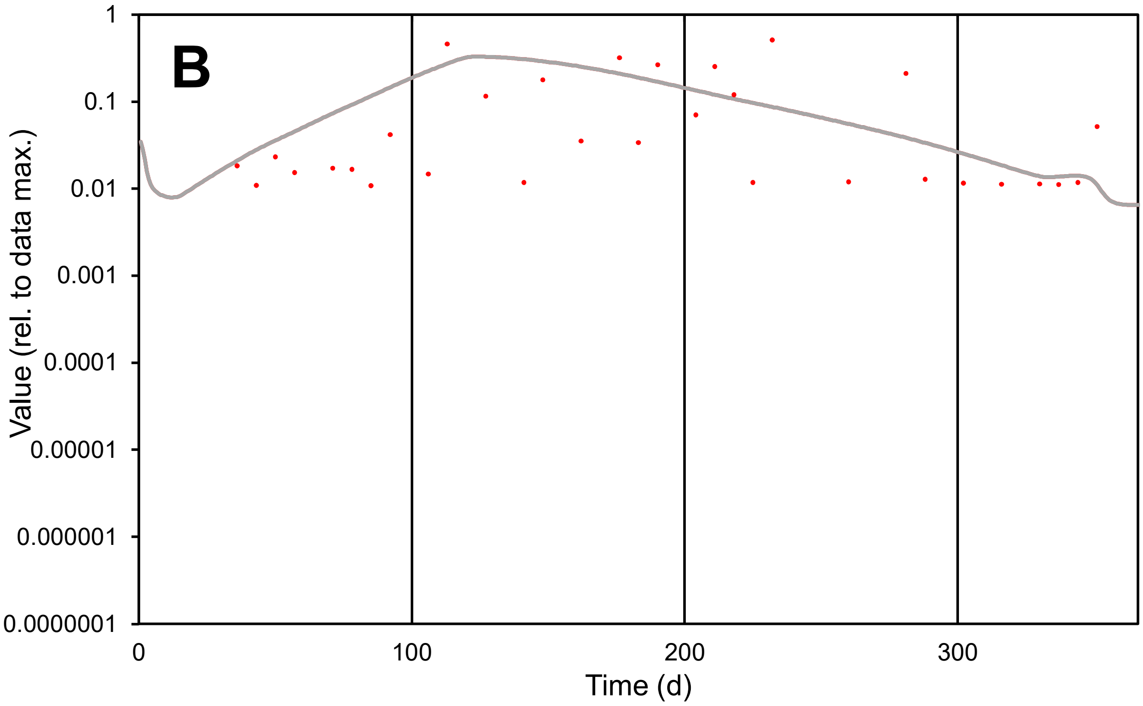


**C1**

Variables

KI1 Provide jtd or Dst

CalibOnly?

2 1 to reproduce of

oNormalize?

2 1=no, 2=yes

ofxn calc*

1.47719

ofxn mod

1.48573

sRun

RunA1

zout

1

Ave

Data

0.00052

Model

0.00052

data/vMax

0.00429

ofxng

321.551

Min d/m

0.98934

ave/min d

11.2727

ave/min m

11.0871

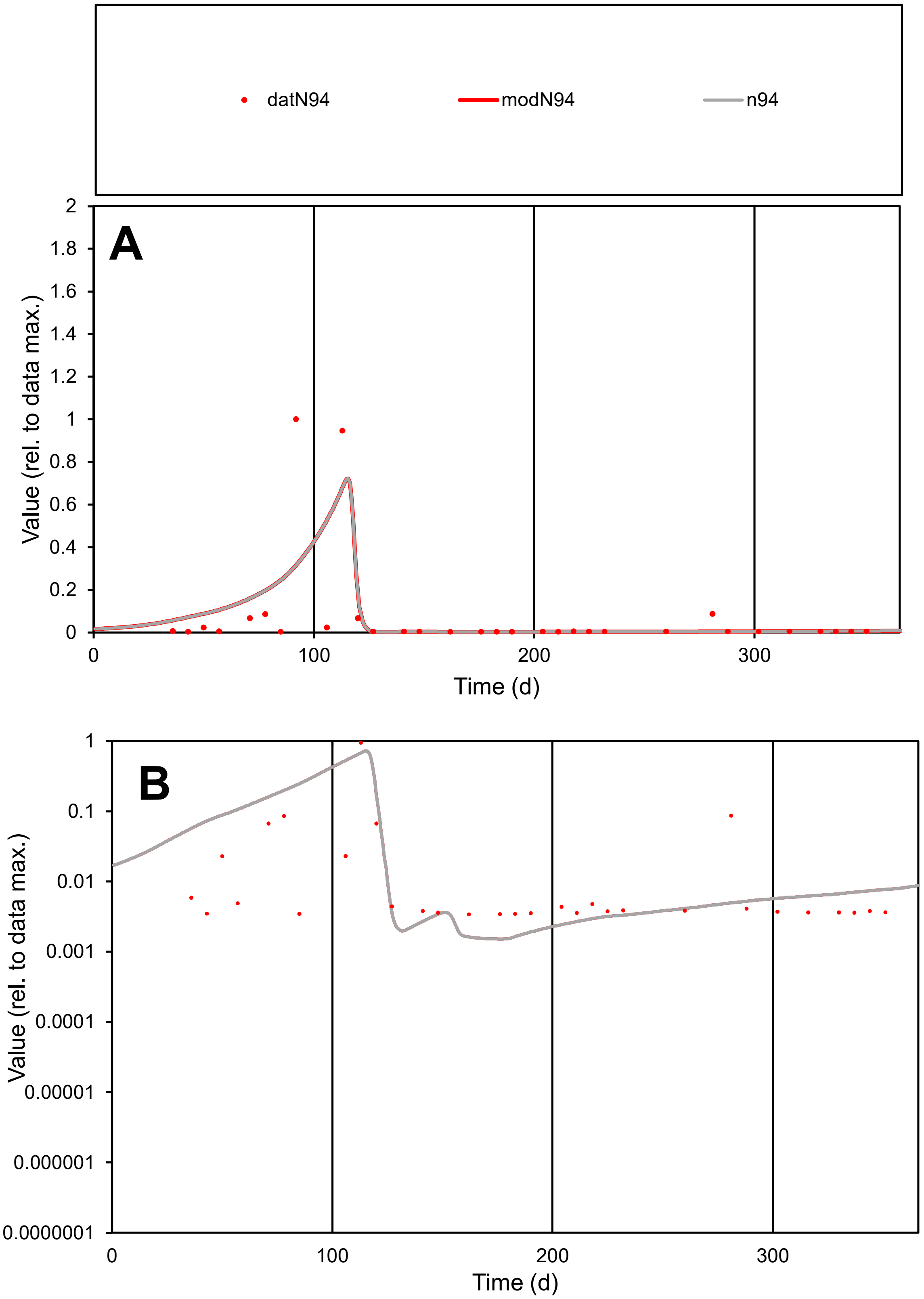


**C1**

Variables
N94 Provide jtd or Dst
CalibOnly?
2 1 to reproduce of
oNormalize?
2 1=no, 2=yes

ofxn calc*
2.20812
ofxn mod
2.22799

sRun
RunA1
zout
1

Ave
Data
0.00097
Model
0.00108

data/vMax
0.01298

ofxng
321.551

Min d/m
2.257

ave/min d
21.885
ave/min m
54.9631

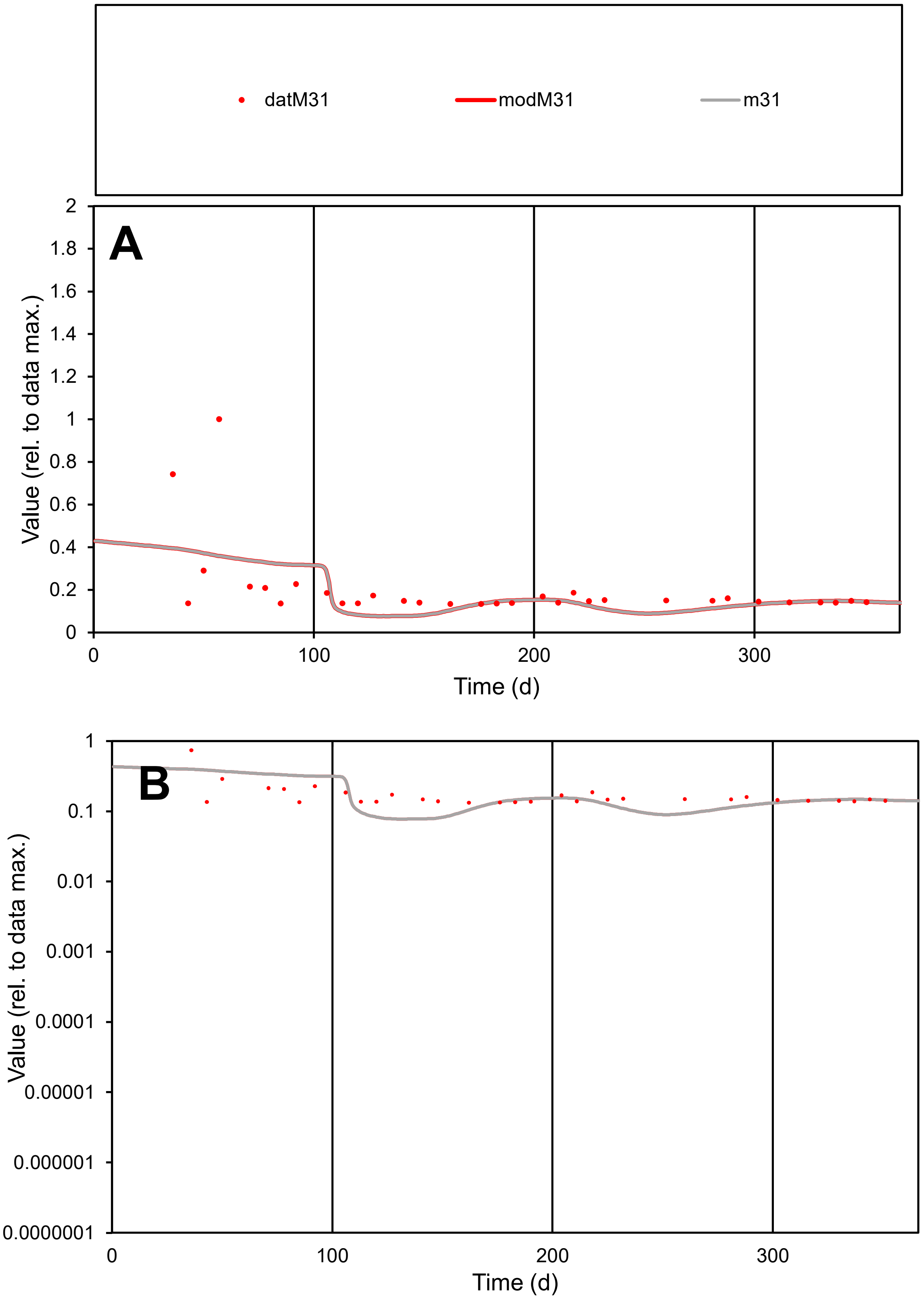


**C1**

Variables

M31 Provide jtd or Dst

CalibOnly?

2 1 to reproduce of

oNormalize?

2 1=no, 2=yes

ofxn calc*

0.73047

ofxn mod

0.73793

sRun

RunA1

zout

1

Ave

Data

0.00012

Model

0.00011

data/vMax

0.0006

ofxng

321.551

Min d/m

1.71198

ave/min d

1.53269

ave/min m

2.34189

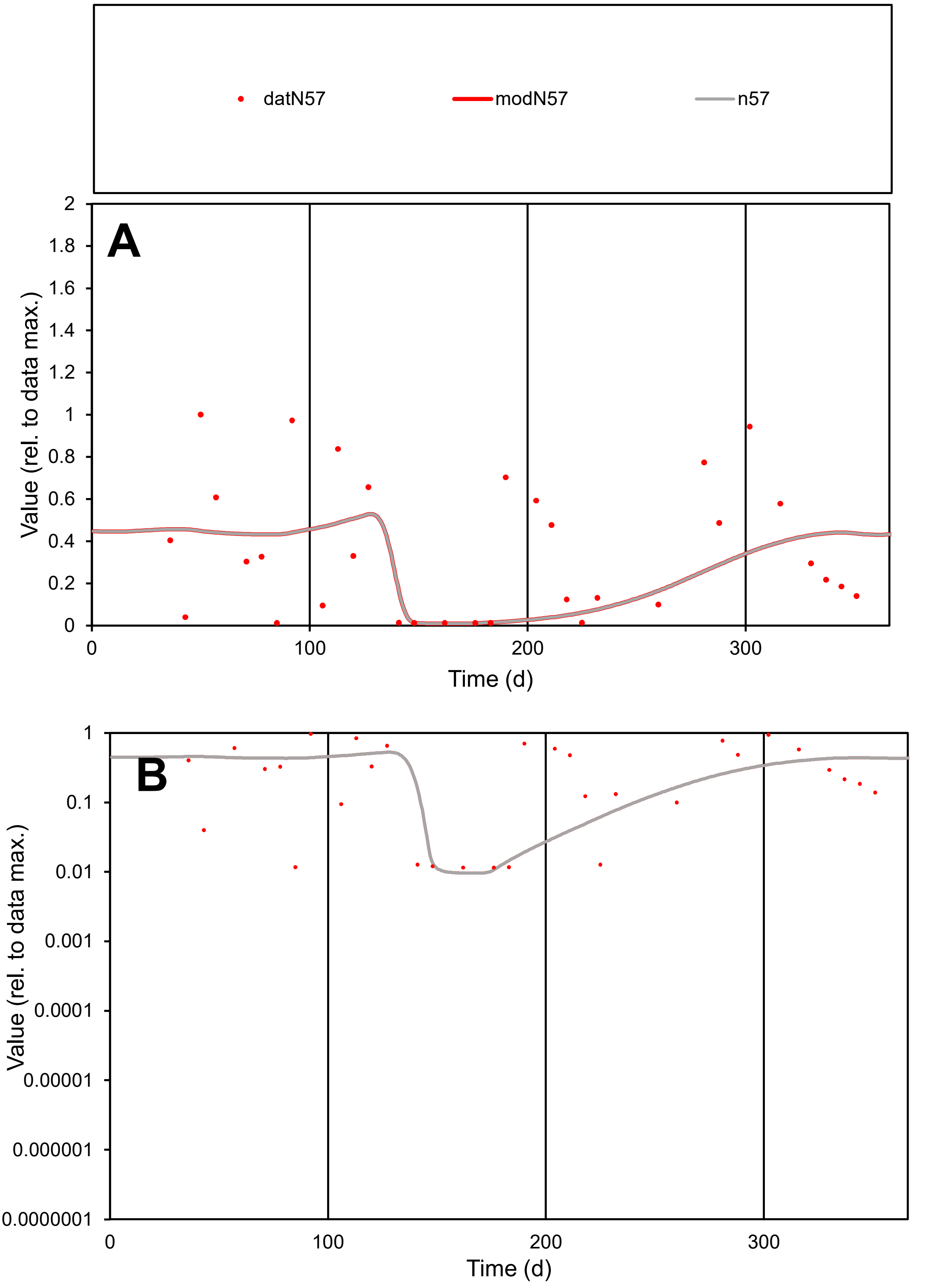


**C1**

Variables
N57 Provide jtd or Dst
CalibOnly?
2 1 to reproduce of
oNormalize?
2 1=no, 2=yes

ofxn calc*
0.8866
ofxn mod
0.89274

sRun
RunA1
zout
1

Ave
Data
0.00109
Model
0.00088

data/vMax
0.00307

ofxng
321.551

Min d/m
1.19871

ave/min d
30.9358
ave/min m
29.8447

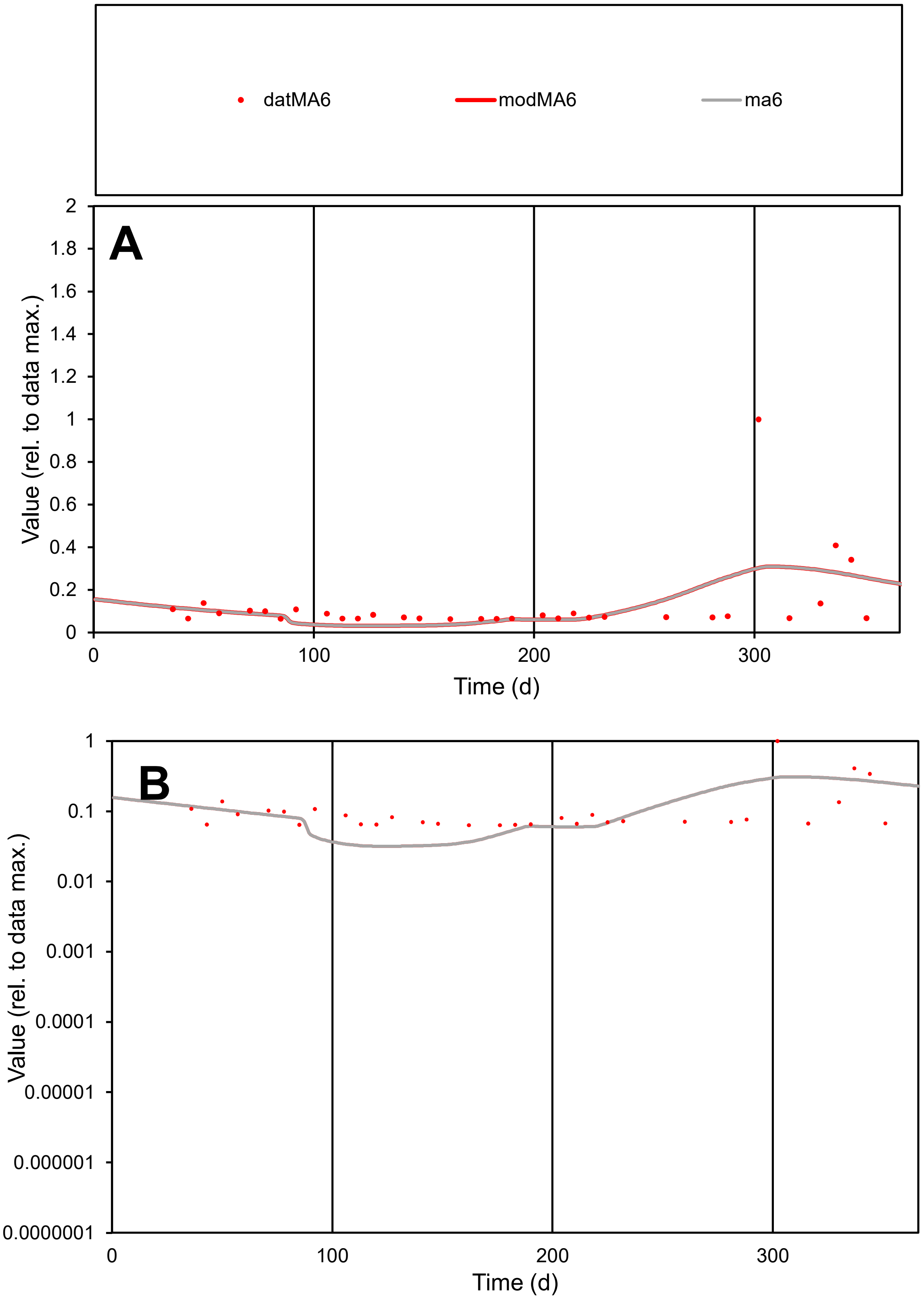


C1

Variables
MA6 Provide jtd or Dst
CalibOnly?
2 1 to reproduce of
oNormalize?
2 1=no, 2=yes

ofxn calc*
1.1685
ofxn mod
1.17792

sRun
RunA1
zout
1

Ave
Data
0.00023
Model
0.00022

data/vMax
0.00183

ofxng
321.551

Min d/m
1.98347

ave/min d
2.01953
ave/min m
3.74089

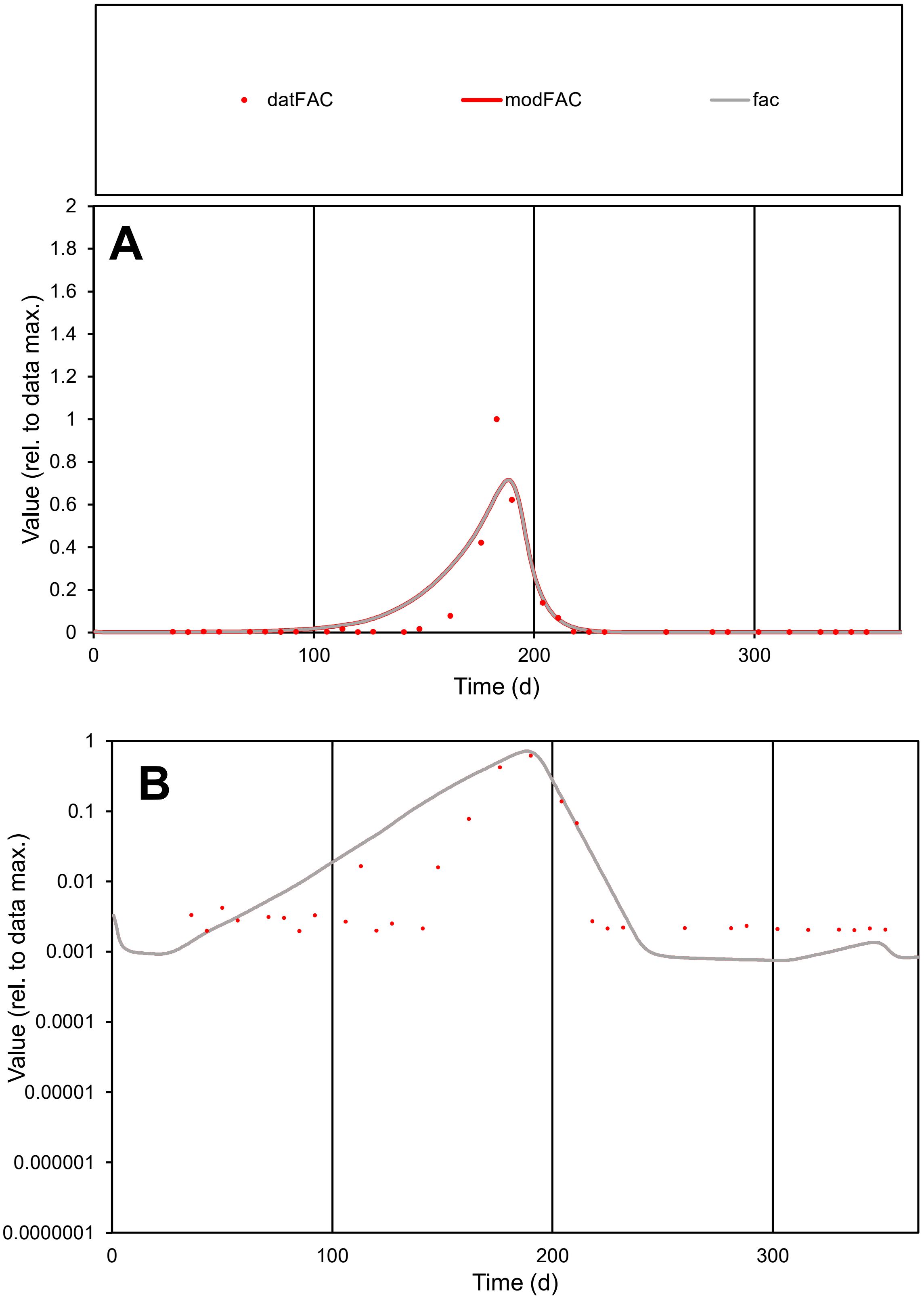


**C1**

Variables
FAC Provide jtd or Dst
CalibOnly?
2 1 to reproduce of
oNormalize?
2 1=no, 2=yes

ofxn calc*
1.0971
ofxn mod
1.15282

sRun
RunA1
zout
1

Ave
Data
0.0033
Model
0.00401

data/vMax
0.04359

ofxng
321.551

Min d/m
2.63583

ave/min d
38.4265
ave/min m
123.224

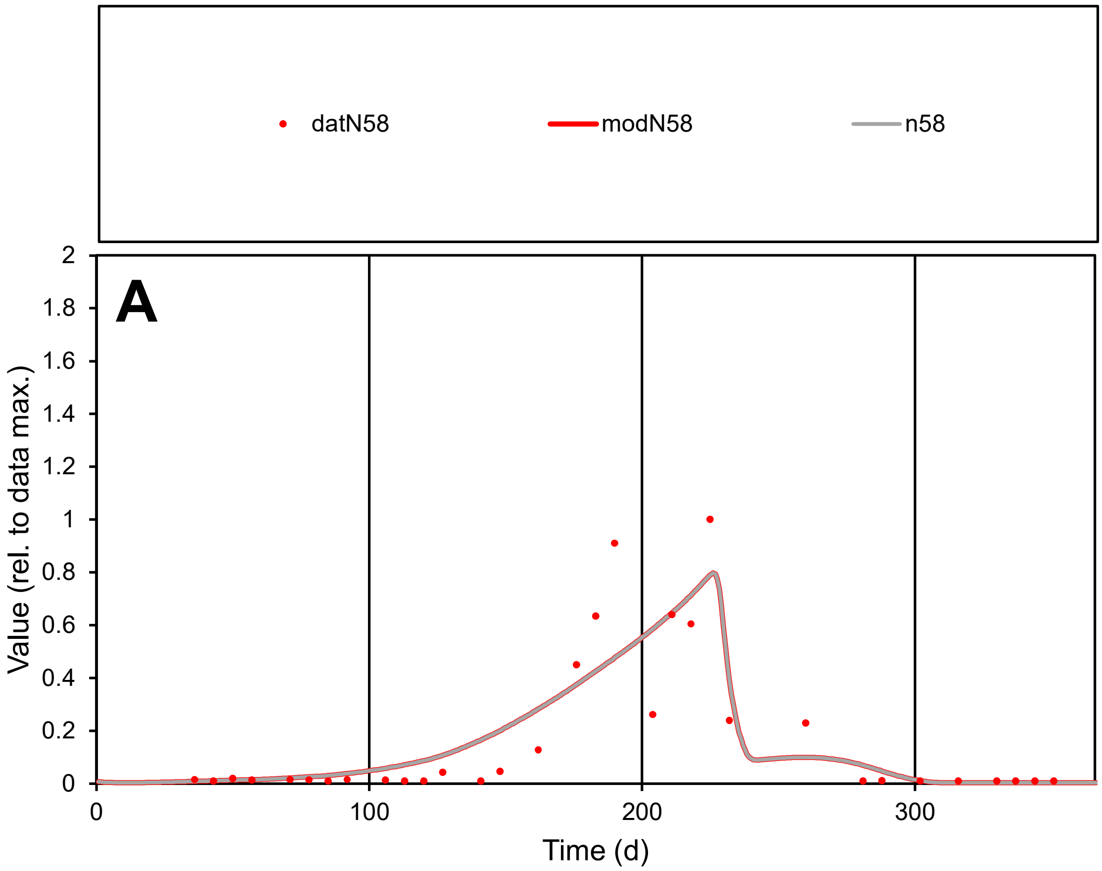


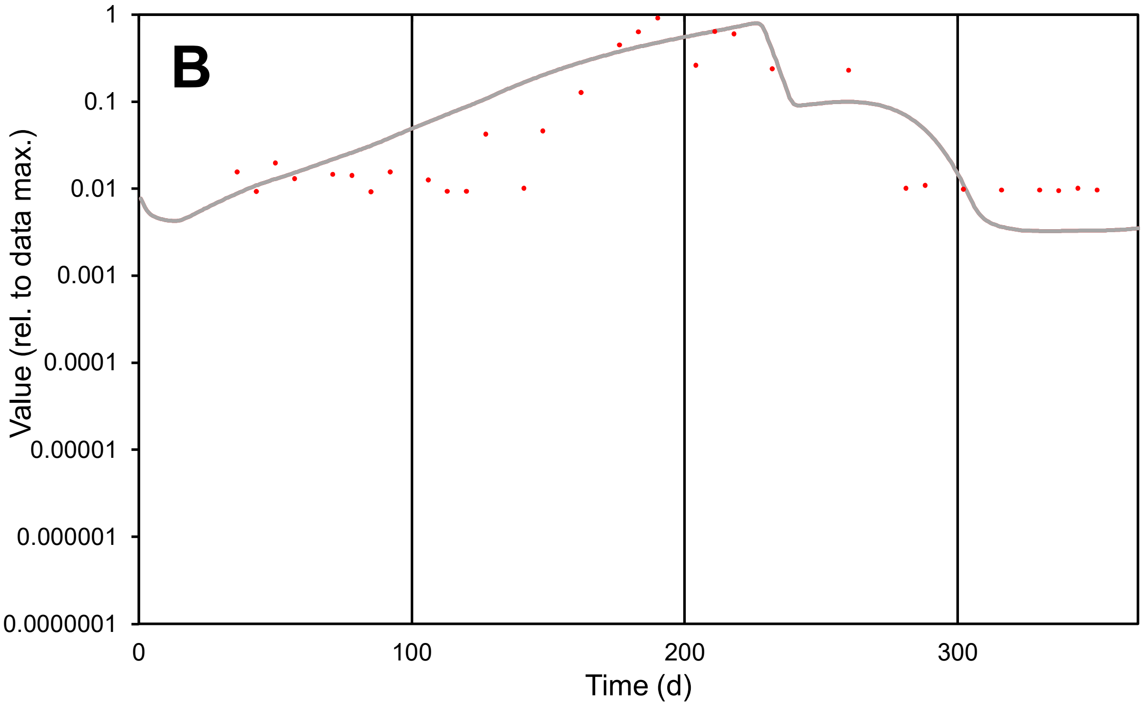


**C1**

Variables
N58 Provide jtd or Dst
CalibOnly?
2 1 to reproduce of
oNormalize?
2 1=no, 2=yes

ofxn calc*
0.7435
ofxn mod
0.76841

sRun
RunA1
zout
1

Ave
Data
0.00149
Model
0.00159

data/vMax
0.00881

ofxng
321.551

Min d/m
2.83659

ave/min d
18.3529
ave/min m
55.4445

324_Calib_S_A

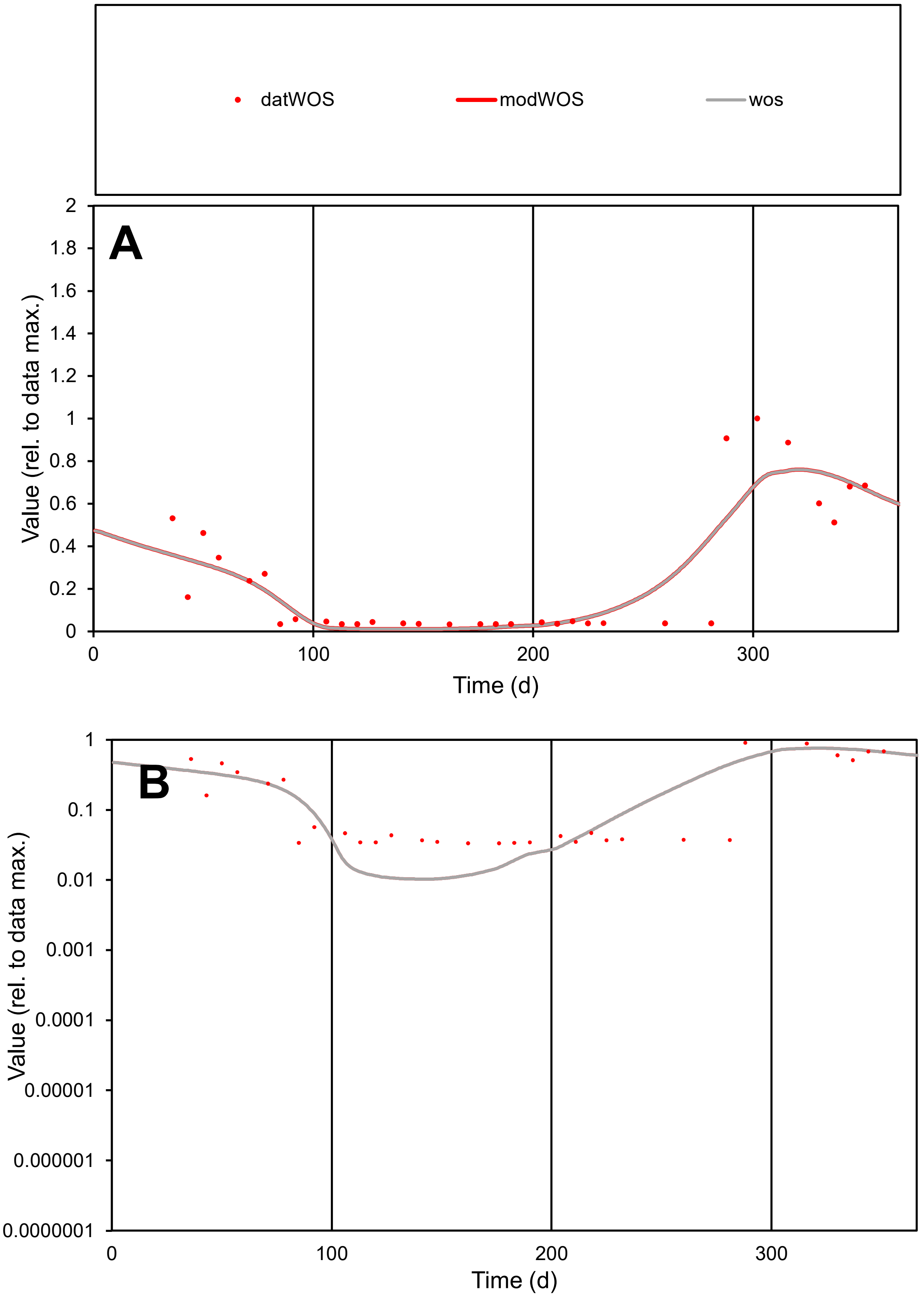


**C1**

Variables
WOS Provide jtd or Dst
CalibOnly?
2 1 to reproduce of
oNormalize?
2 1=no, 2=yes

ofxn calc*
0.55658
ofxn mod
0.57532

sRun
RunA1
zout
1

Ave
Data
0.0005
Model
0.00049

data/vMax
0.002

ofxng
321.551

Min d/m
3.2374

ave/min d
7.50367
ave/min m
23.9106

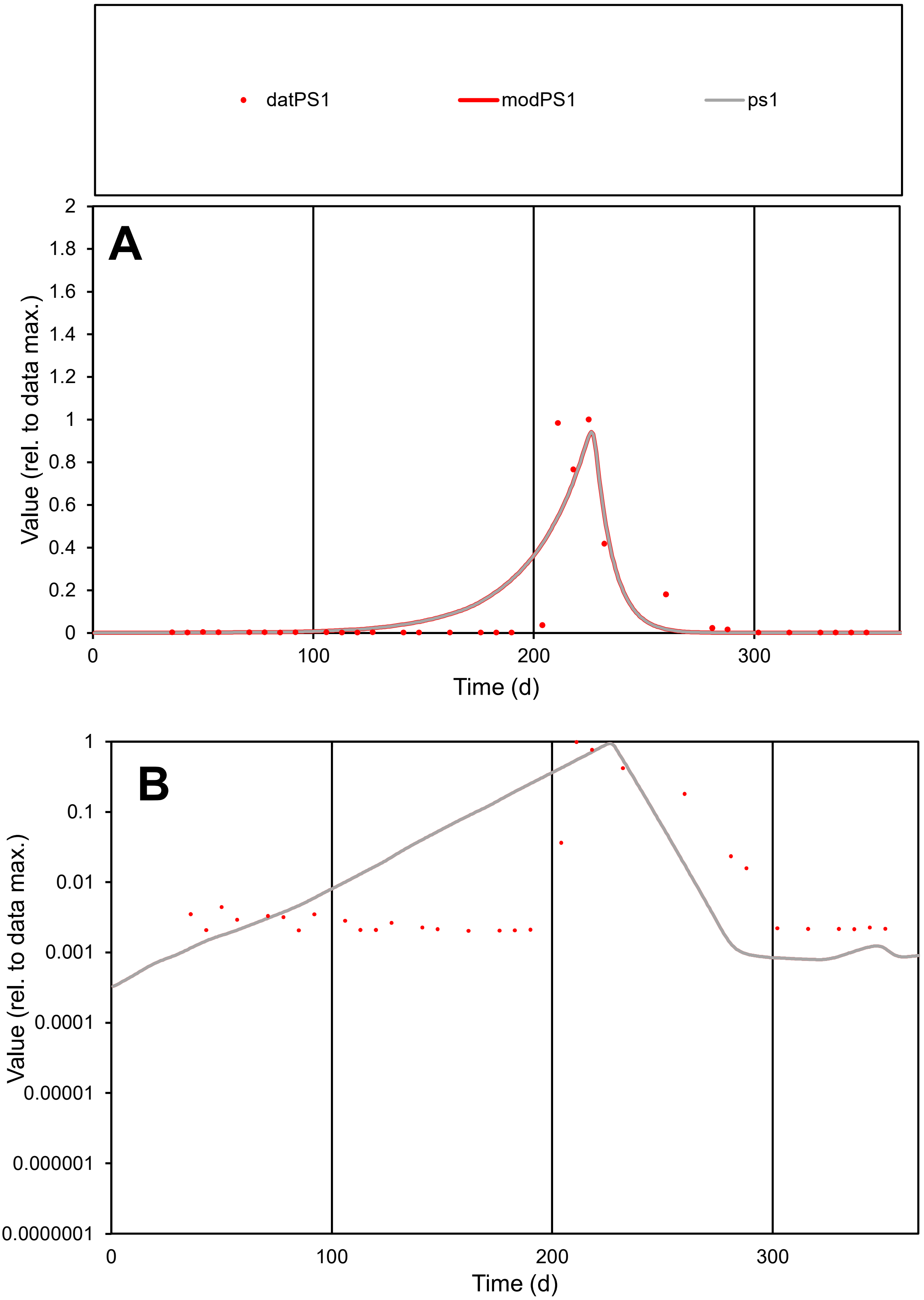


**C1**

Variables

PS1 Provide jtd or Dst

CalibOnly?

2 1 to reproduce of

oNormalize?

2 1=no, 2=yes

ofxn calc*

1.12211

ofxn mod

1.16503

sRun

RunA1

zout

1

Ave

Data

0.00246

Model

0.00285

data/vMax

0.02259

ofxng

321.551

Min d/m

2.55438

ave/min d

53.4522

ave/min m

157.946

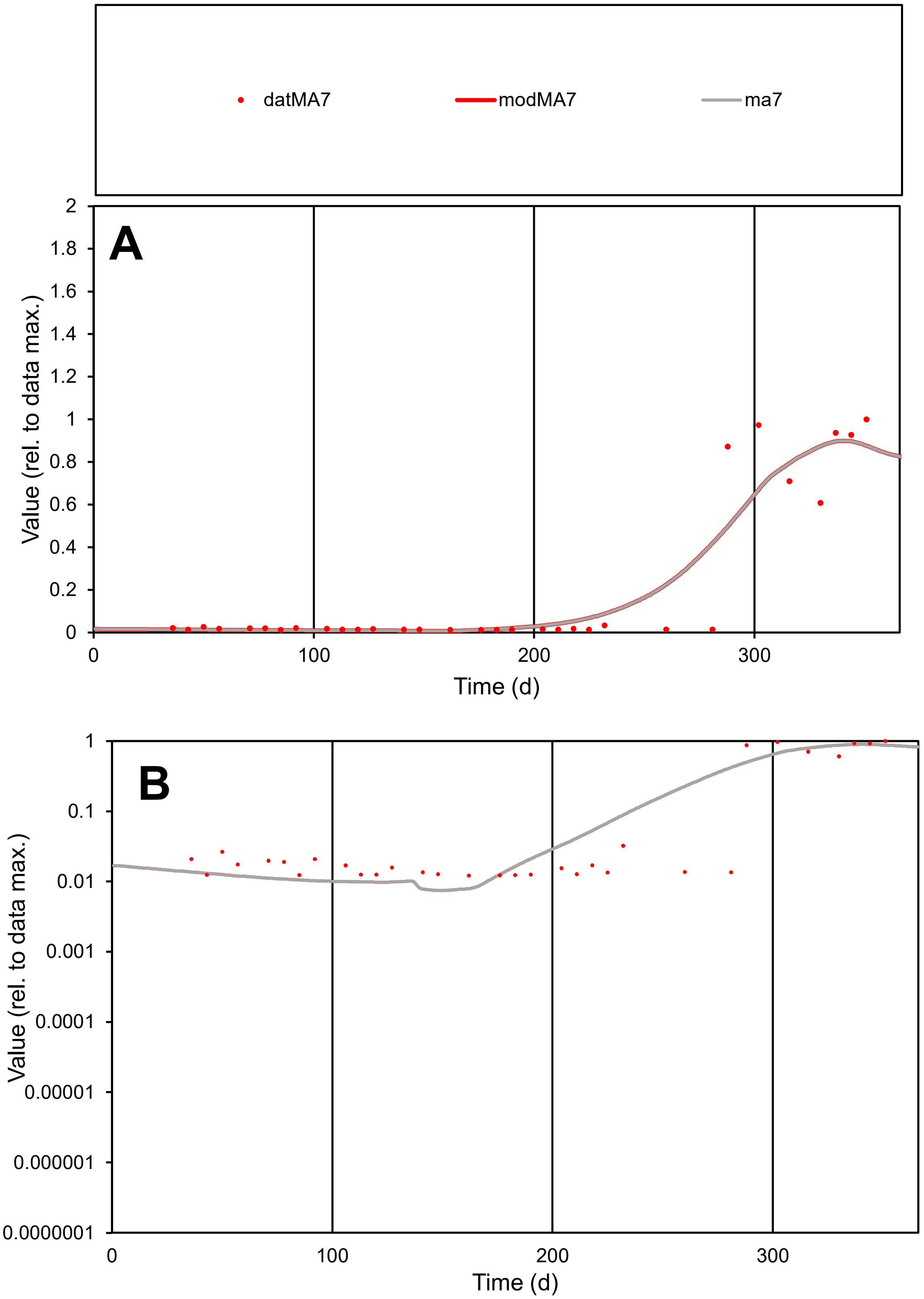


**C1**

Variables
MA7 Provide jtd or Dst
CalibOnly?
2 1 to reproduce of
oNormalize?
2 1=no, 2=yes

ofxn calc*
0.64382
ofxn mod
0.66922

sRun
RunA1
zout
1

Ave
Data
0.00022
Model
0.00023

data/vMax
0.00108

ofxng
321.551

Min d/m
1.63293

ave/min d
16.4912
ave/min m
28.0767

324_Calib_S_A

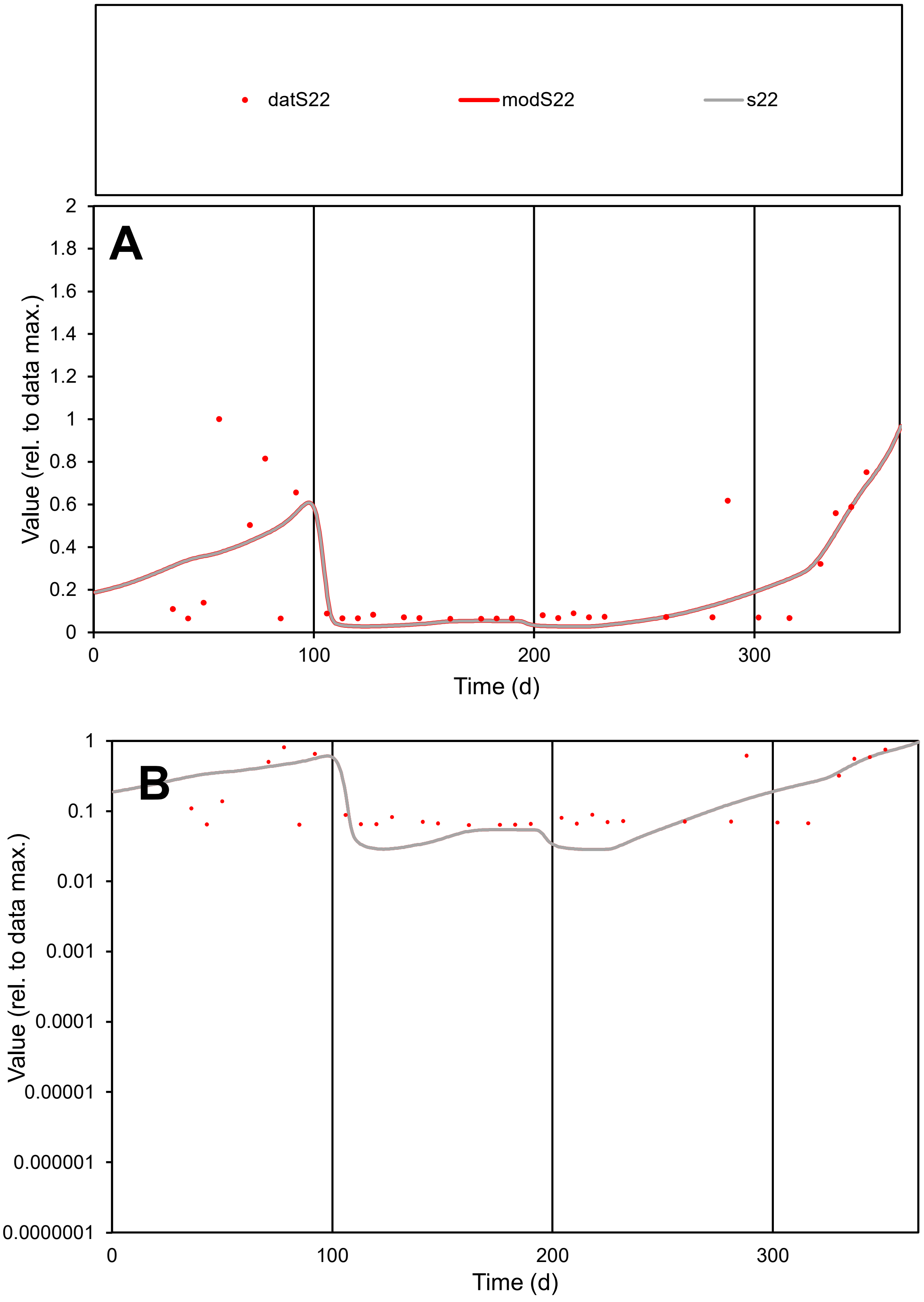


**C1**

Variables
S22 Provide jtd or Dst
CalibOnly?
2 1 to reproduce of
oNormalize?
2 1=no, 2=yes

ofxn calc*
0.81383
ofxn mod
0.82444

sRun
RunA1
zout
1

Ave
Data
0.00053
Model
0.00049

data/vMax
0.00226

ofxng
321.551

Min d/m
2.2293

ave/min d
3.71095
ave/min m
7.663

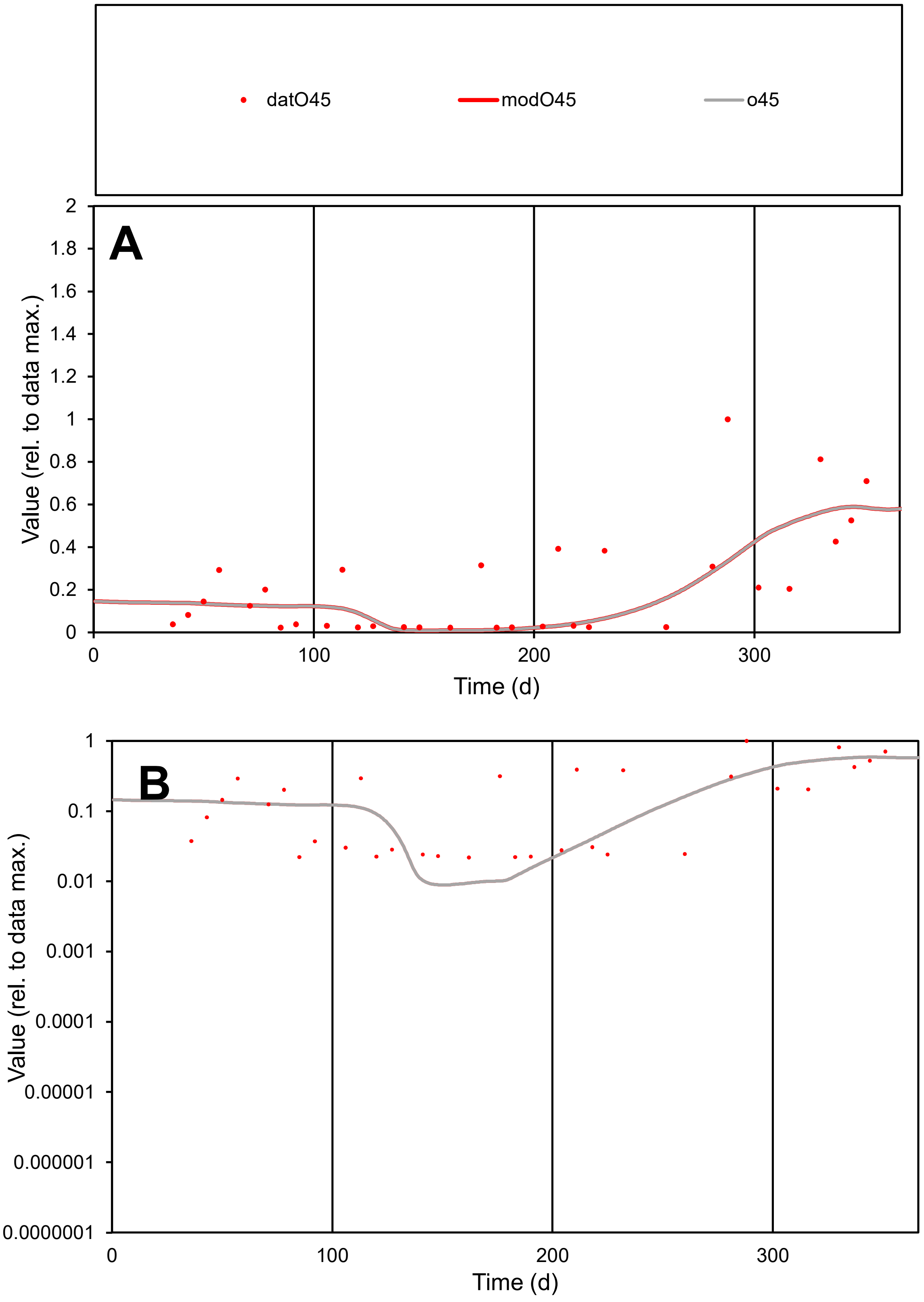


**C1**

Variables
O45 Provide jtd or Dst
CalibOnly?
2 1 to reproduce of
oNormalize?
2 1=no, 2=yes

ofxn calc*
0.88362
ofxn mod
0.8939

sRun
RunA1
zout
1

Ave
Data
0.00055
Model
0.00046

data/vMax
0.00256

ofxng
321.551

Min d/m
2.45391

ave/min d
9.76754
ave/min m
20.274

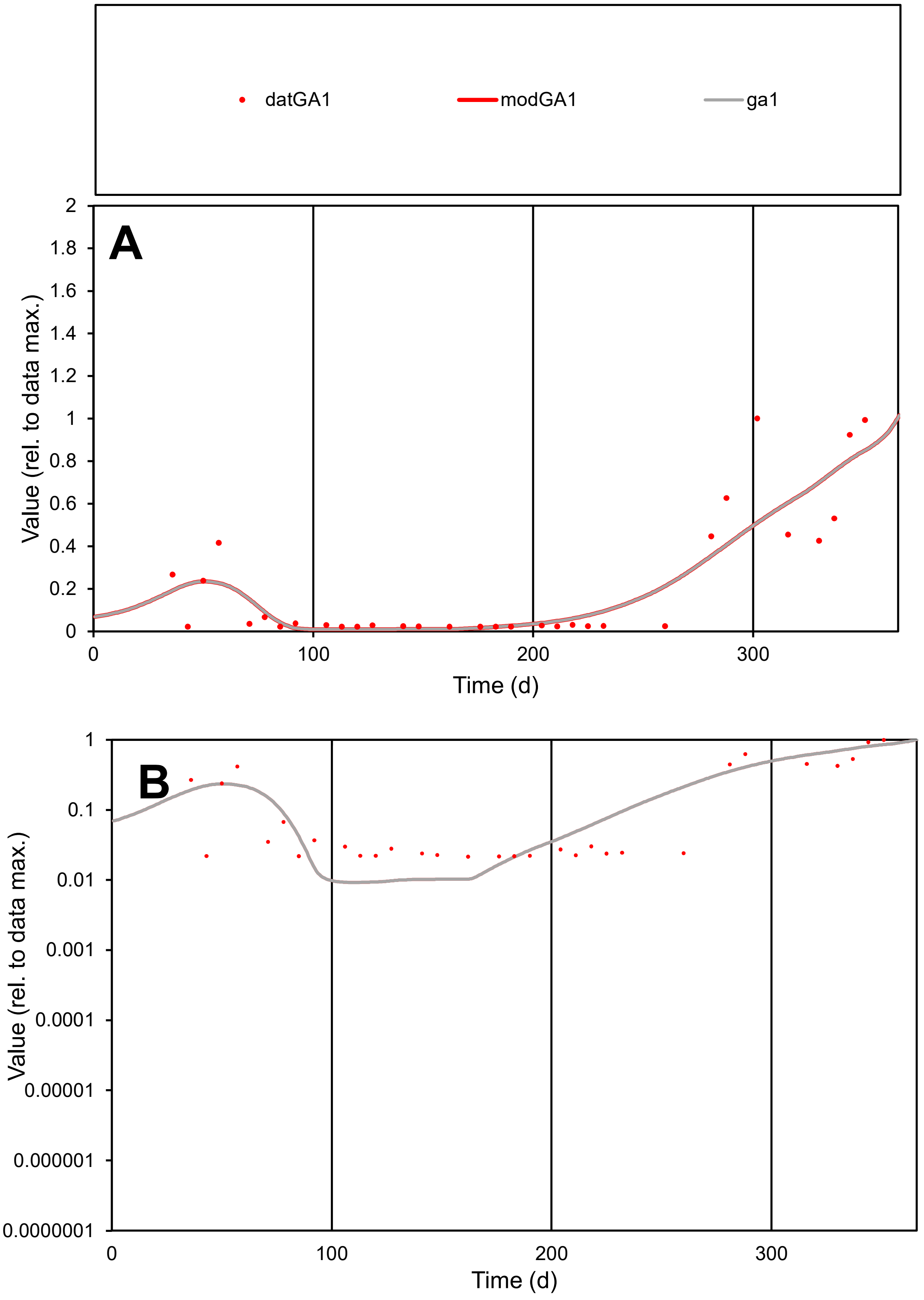


**C1**

Variables
GA1 Provide jtd or Dst
CalibOnly?
2 1 to reproduce of
oNormalize?
2 1=no, 2=yes

ofxn calc*
0.64227
ofxn mod
0.65961

sRun
RunA1
zout
1

Ave
Data
0.00035
Model
0.00035

data/vMax
0.00161

ofxng
321.551

Min d/m
2.32215

ave/min d
9.99632
ave/min m
23.1841

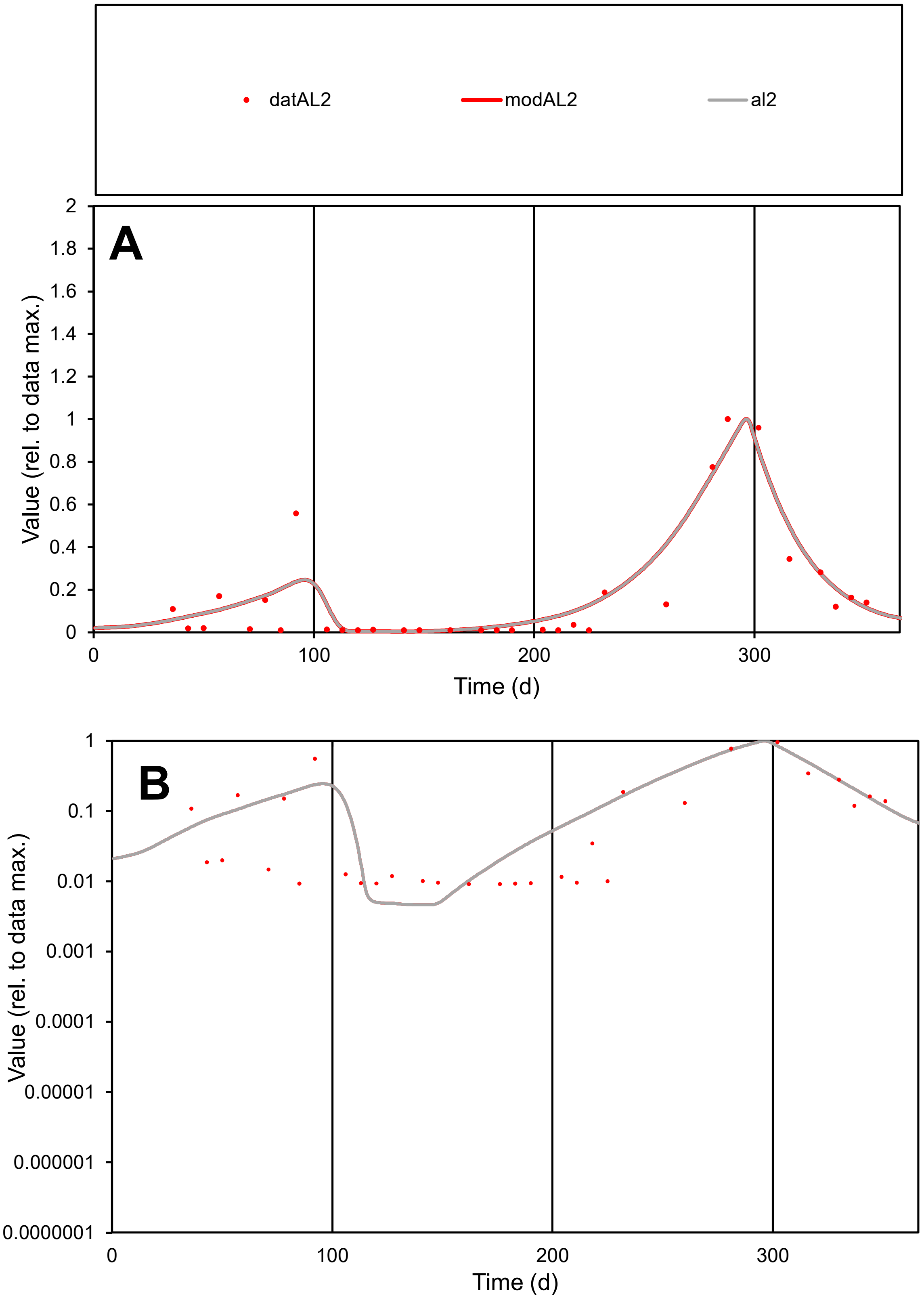


**C1**

Variables
AL2 Provide jtd or Dst
CalibOnly?
2 1 to reproduce of
oNormalize?
2 1=no, 2=yes

ofxn calc*
0.62282
ofxn mod
0.65048

sRun
RunA1
zout
1

Ave
Data
0.00039
Model
0.00043

data/vMax
0.00234

ofxng
321.551

Min d/m
1.96491

ave/min d
18.2282
ave/min m
39.9228

324_Calib_S_A

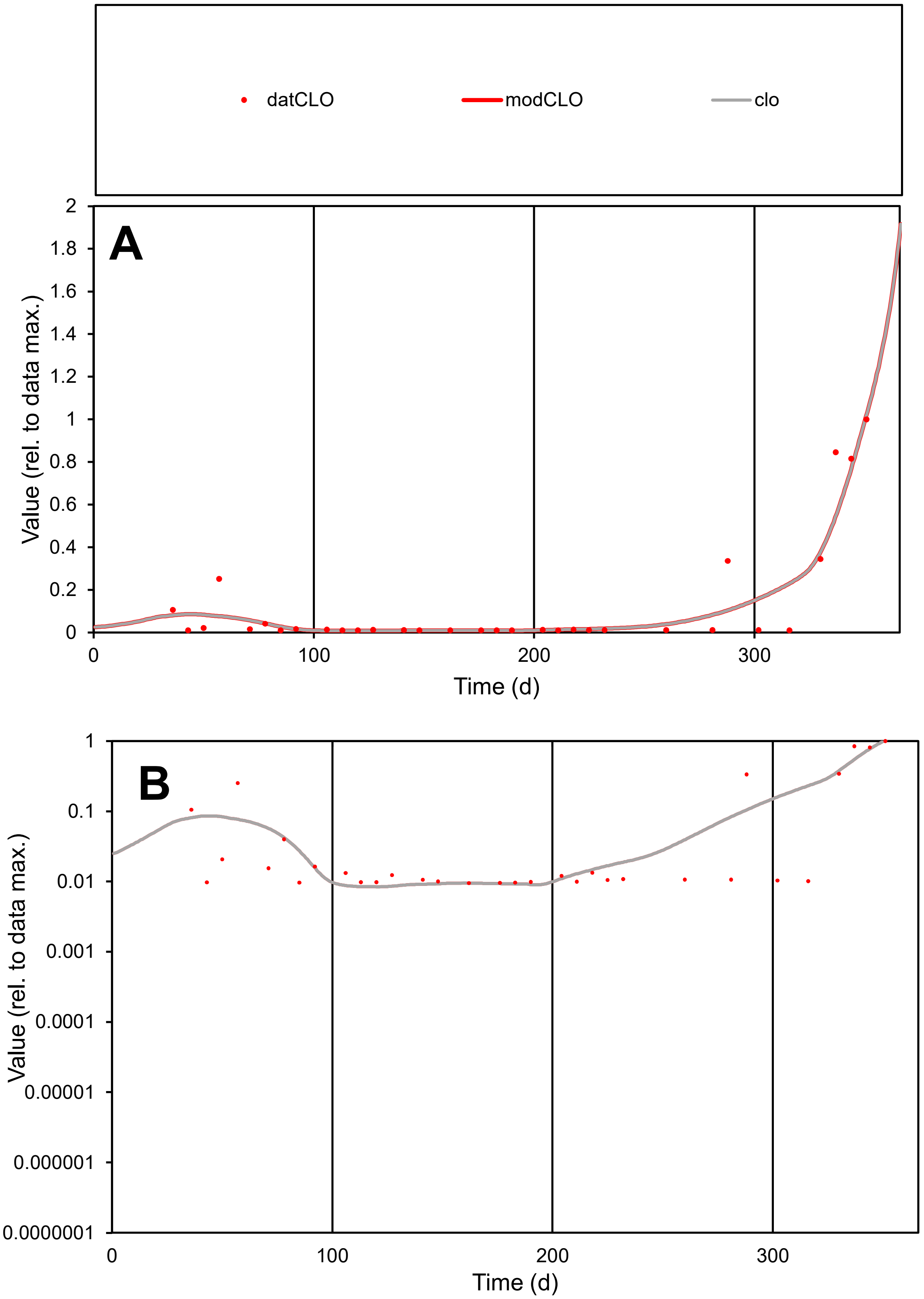


**C1**

Variables
CLO Provide jtd or Dst
CalibOnly?
2 1 to reproduce of
oNormalize?
2 1=no, 2=yes

ofxn calc*
0.70344
ofxn mod
0.72847

sRun
RunA1
zout
1

Ave
Data
0.00017
Model
0.00017

data/vMax
0.00135

ofxng
321.551

Min d/m
1.13556

ave/min d
13.1572
ave/min m
15.3354

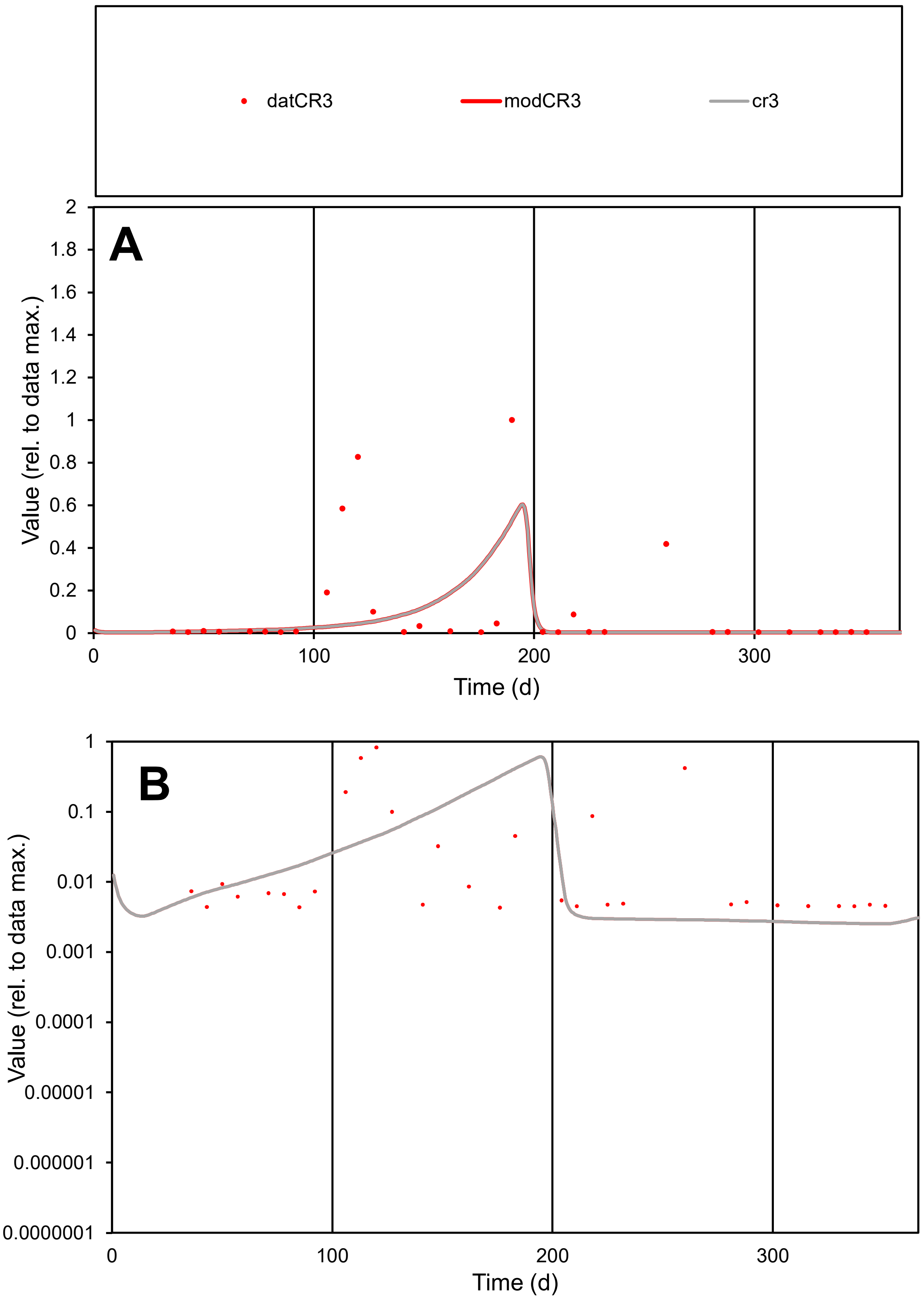


**C1**

Variables
CR3 Provide jtd or Dst
CalibOnly?
2 1 to reproduce of
oNormalize?
2 1=no, 2=yes

ofxn calc*
2.09294
ofxn mod
2.10479

sRun
RunA1
zout
1

Ave
Data
0.00086
Model
0.0005

data/vMax
0.0081

ofxng
321.551

Min d/m
1.69455

ave/min d
24.7795
ave/min m
24.4165

324_Calib_S_A

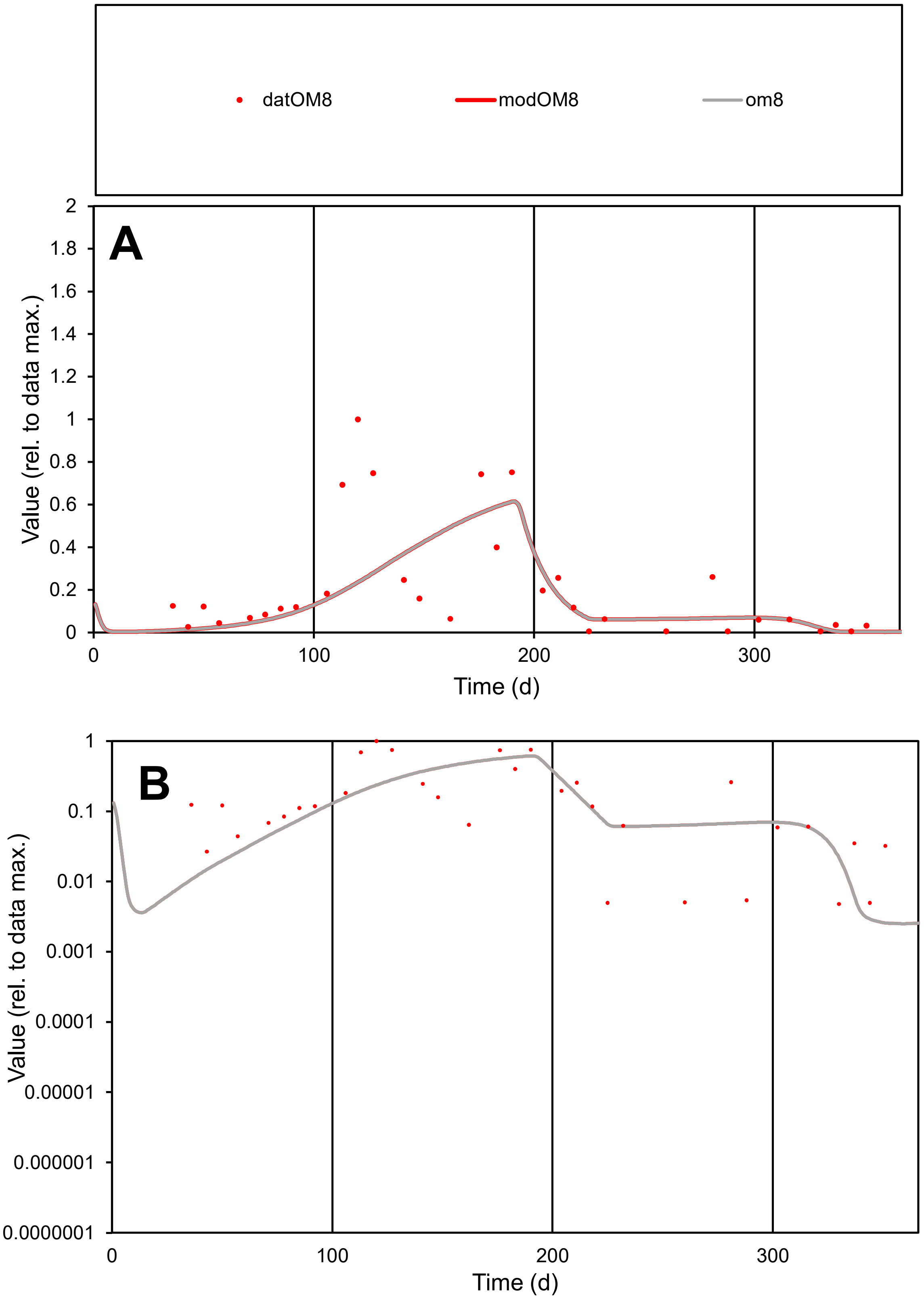


**C1**

Variables
OM8 Provide jtd or Dst
CalibOnly?
2 1 to reproduce of
oNormalize?
2 1=no, 2=yes

ofxn calc*
1.01111
ofxn mod
1.01939

sRun
RunA1
zout
1

Ave
Data
0.00249
Model
0.00197

data/vMax
0.01176

ofxng
321.551

Min d/m
1.85937

ave/min d
44.4775
ave/min m
65.2142

324_Calib_S_A

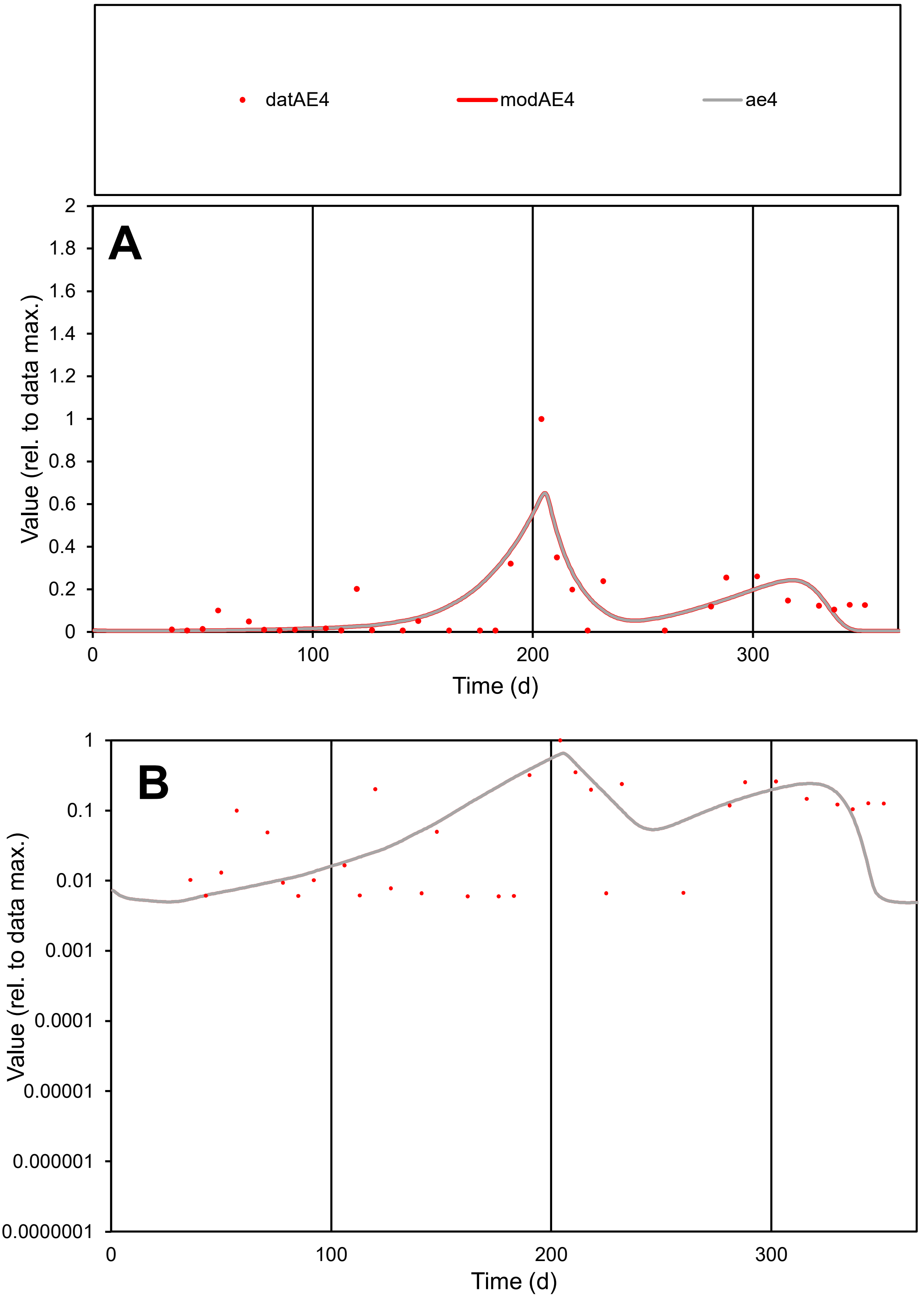


**C1**

Variables
AE4 Provide jtd or Dst
CalibOnly?
2 1 to reproduce of
oNormalize?
2 1=no, 2=yes

ofxn calc*
0.93064
ofxn mod
0.9562

sRun
RunA1
zout
1

Ave
Data
0.00023
Model
0.00024

data/vMax
0.00191

ofxng
321.551

Min d/m
1.11984

ave/min d
20.356
ave/min m
23.1273

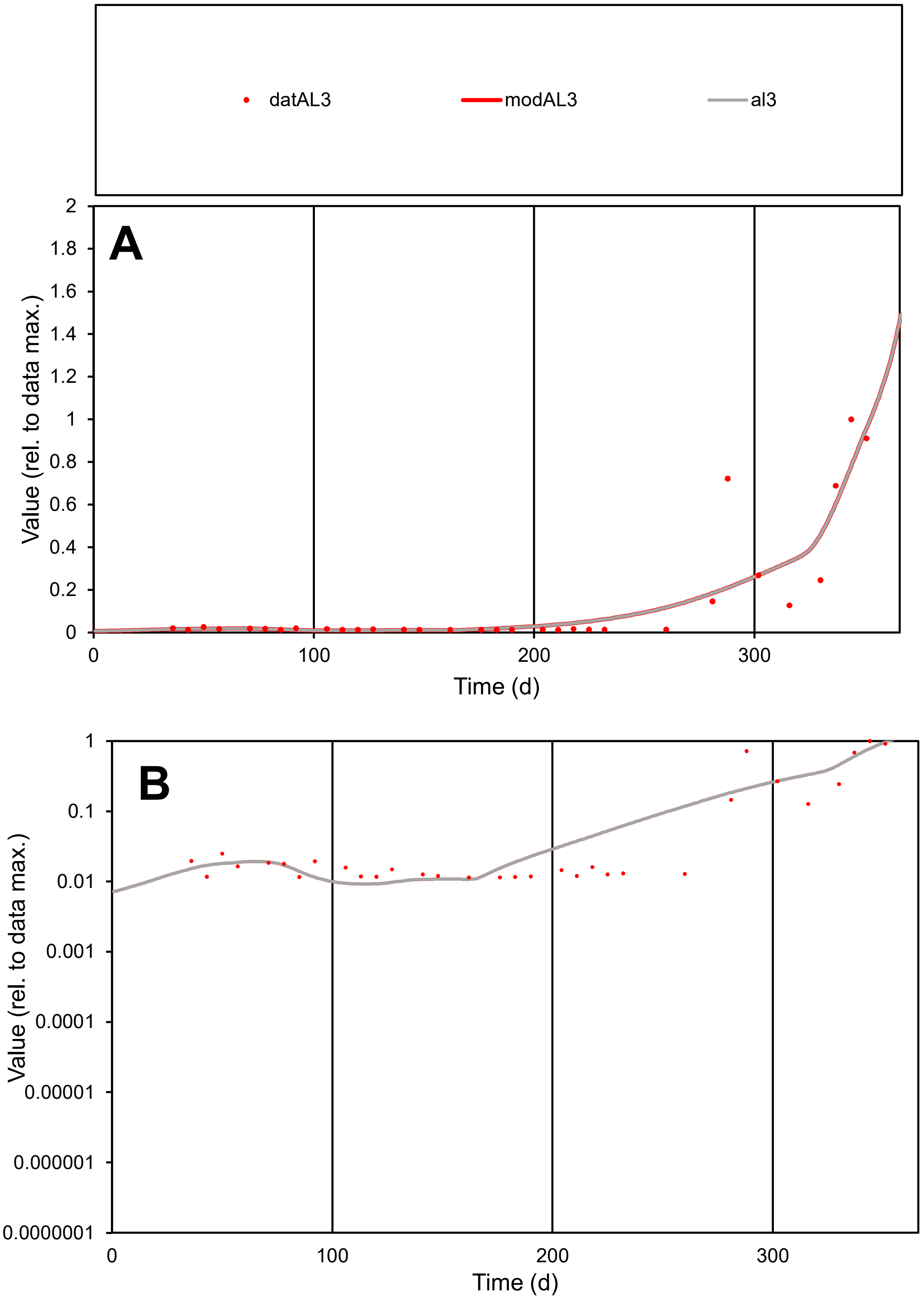


**C1**

Variables
AL3 Provide jtd or Dst
CalibOnly?
2 1 to reproduce of
oNormalize?
2 1=no, 2=yes

ofxn calc*
0.80891
ofxn mod
0.83225

sRun
RunA1
zout
1

Ave
Data
0.00029
Model
0.00029

data/vMax
0.00207

ofxng
321.551

Min d/m
1.22905

ave/min d
12.1928
ave/min m
15.2231

324_Calib_S_A

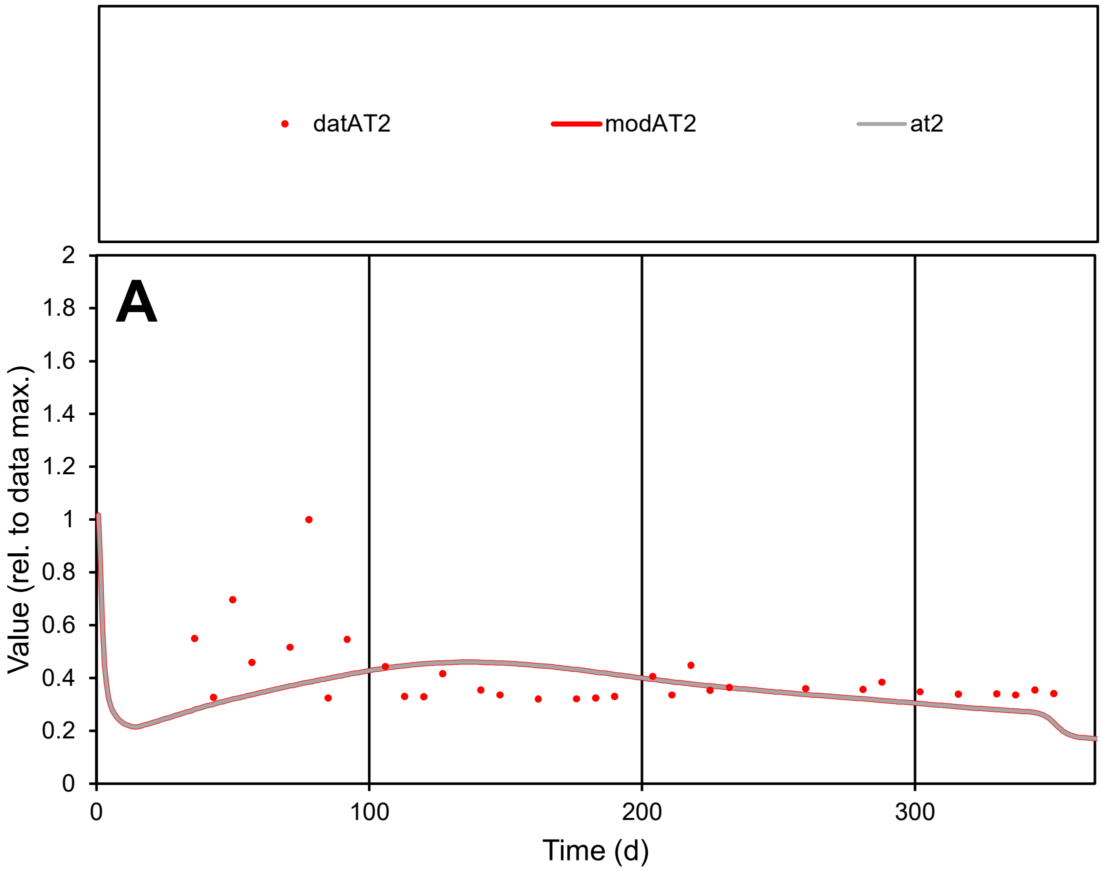


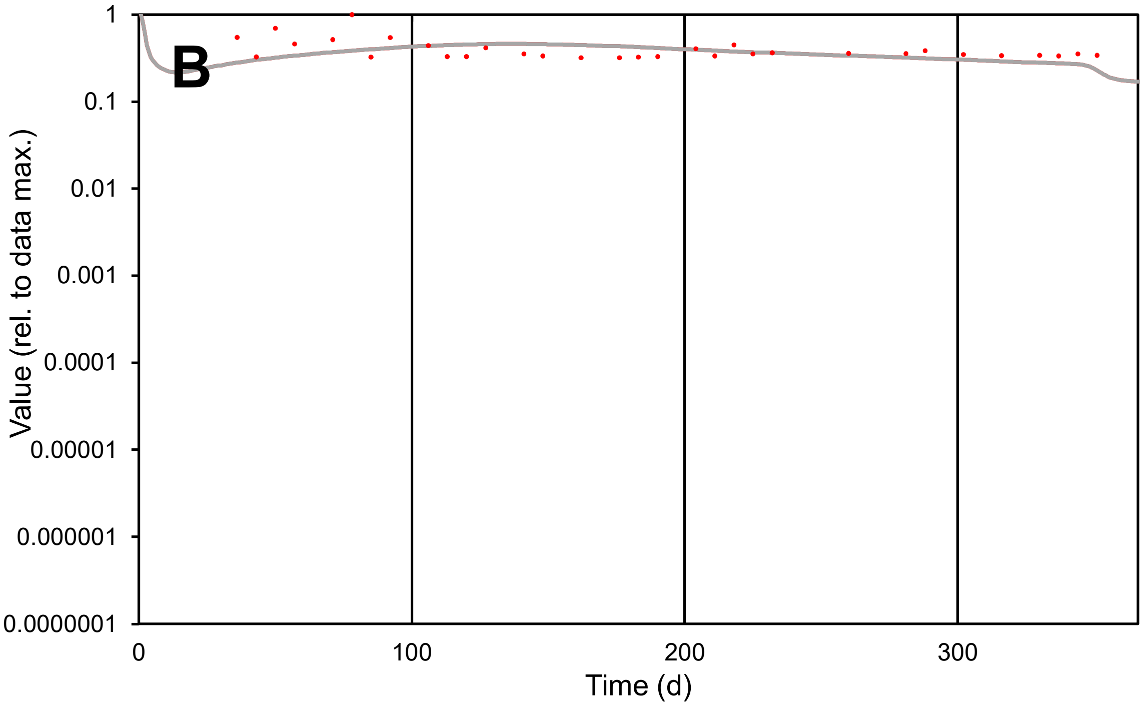


**C1**

Variables
AT2 Provide jtd or Dst
CalibOnly?
2 1 to reproduce of
oNormalize?
2 1=no, 2=yes

ofxn calc*
0.38801
ofxn mod
0.39953

sRun
RunA1
zout
1

Ave
Data
9.9E-05
Model
8.9E-05

data/vMax
0.00024

ofxng
321.551

Min d/m
1.45261

ave/min d
1.2703
ave/min m
1.66964

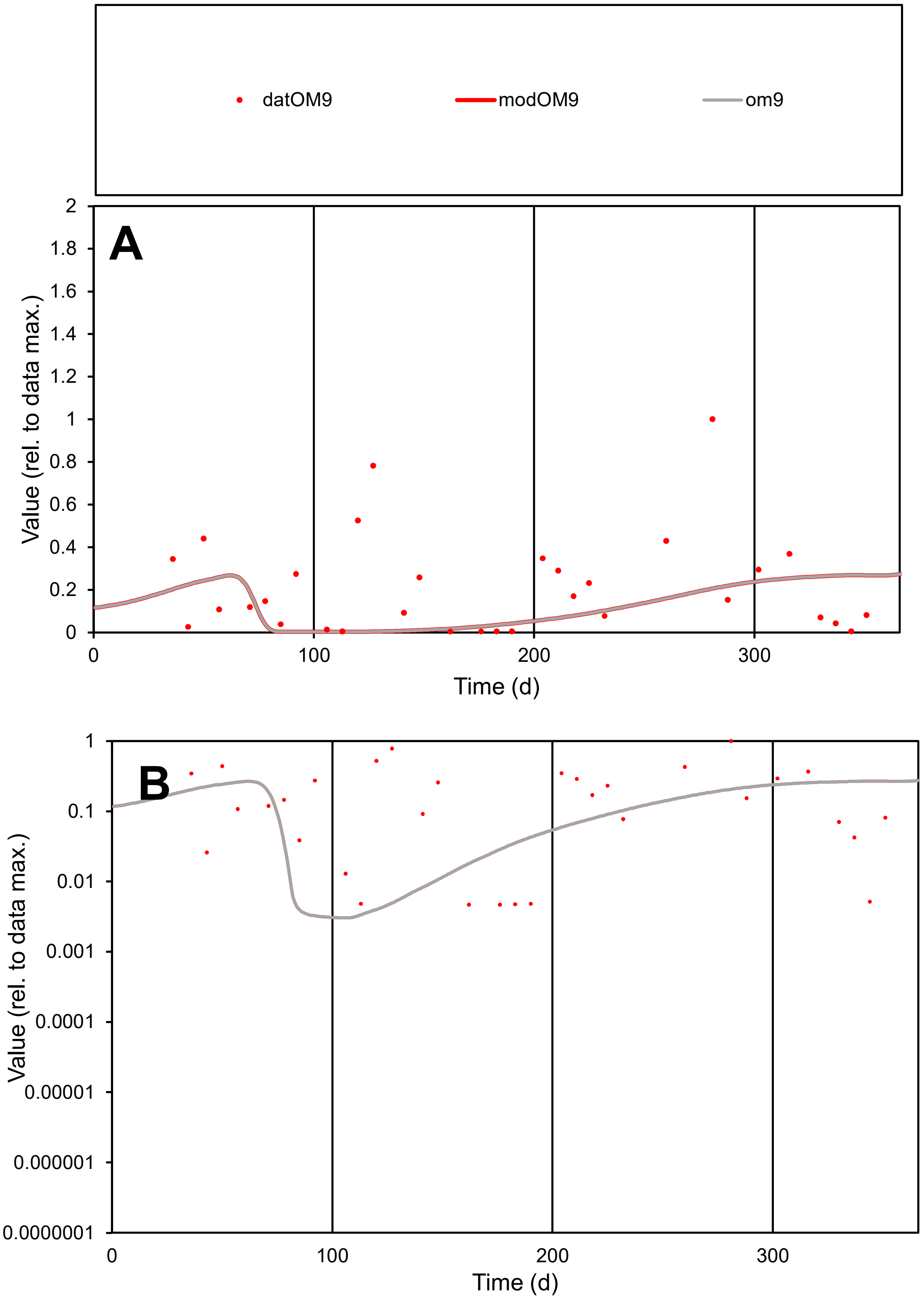


**C1**

Variables
OM9 Provide jtd or Dst
CalibOnly?
2 1 to reproduce of
oNormalize?
2 1=no, 2=yes

ofxn calc*
1.24869
ofxn mod
1.25257

sRun
RunA1
zout
1

Ave
Data
0.00192
Model
0.00111

data/vMax
0.00909

ofxng
321.551

Min d/m
1.52429

ave/min d
45.2401
ave/min m
39.8169

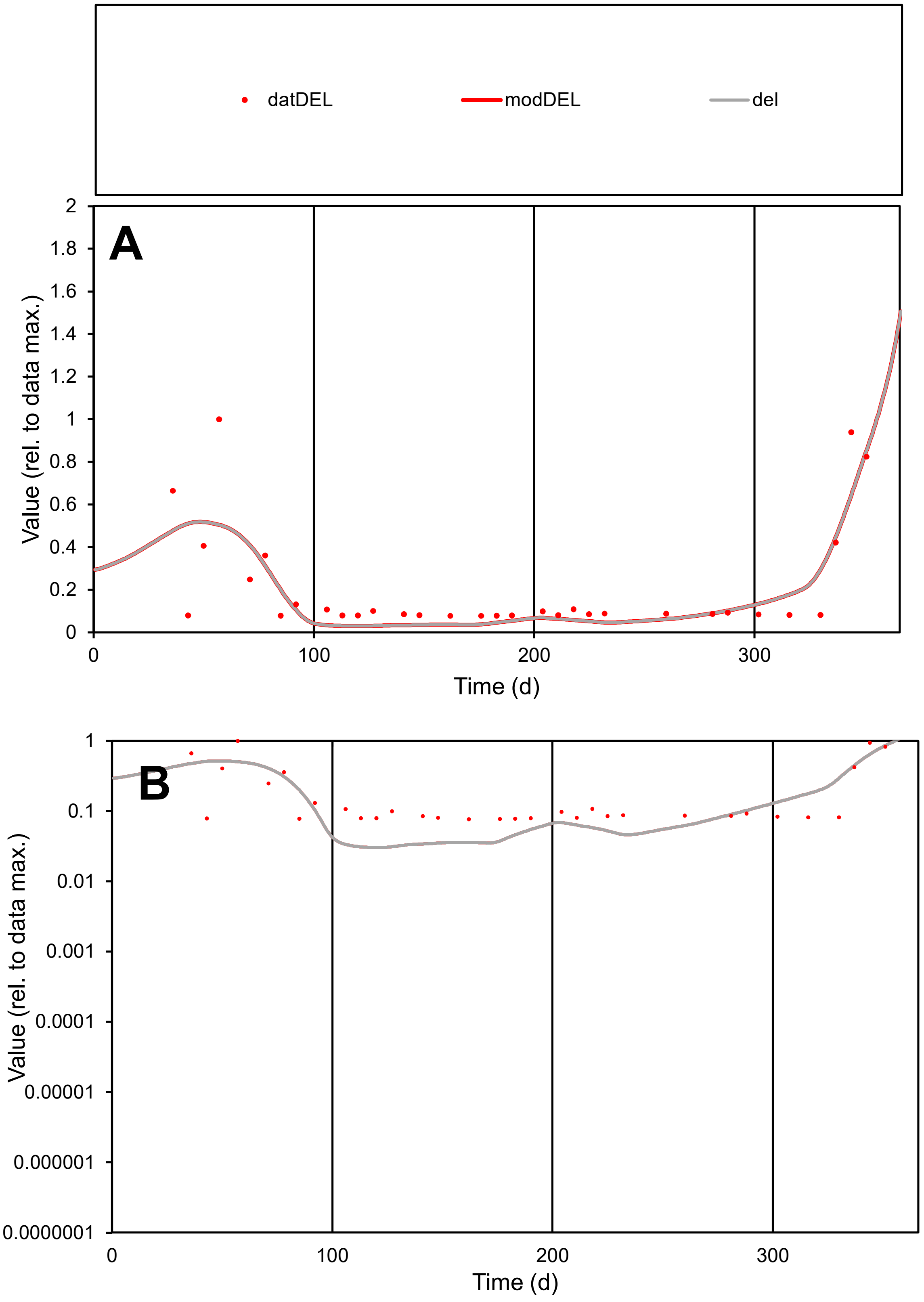


**C1**

Variables

DEL Provide jtd or Dst

CalibOnly?

2 1 to reproduce of

oNormalize?

2 1=no, 2=yes

ofxn calc*

0.66952

ofxn mod

0.68204

sRun

RunA1

zout

1

Ave

Data

0.00017

Model

0.00016

data/vMax

0.00079

ofxng

321.551

Min d/m

2.52961

ave/min d

2.83301

ave/min m

6.75571

324_Calib_S_A

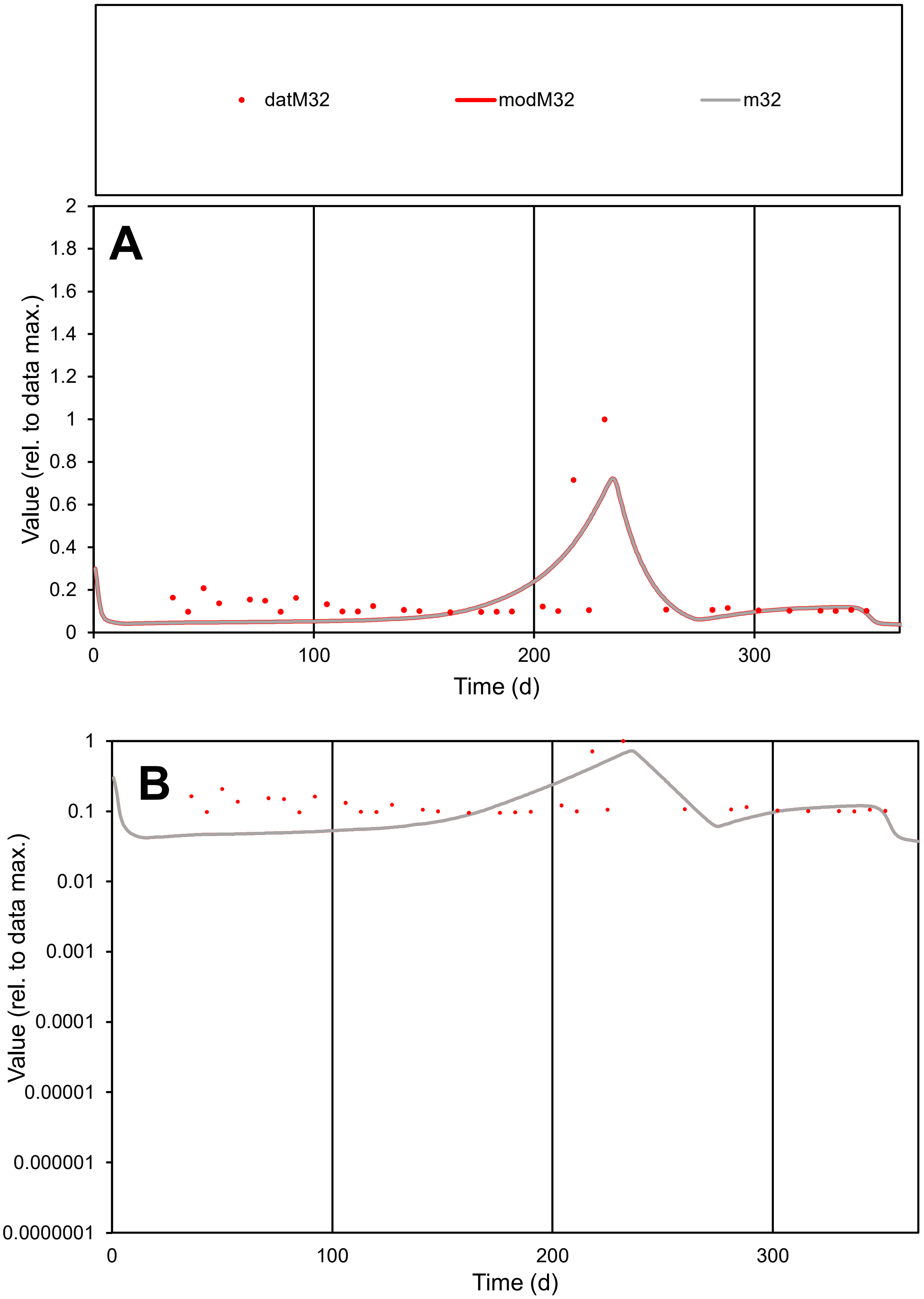


**C1**

Variables

M32 Provide jtd or Dst

CalibOnly?

2 1 to reproduce of

oNormalize?

2 1=no, 2=yes

ofxn calc*

0.81124

ofxn mod

0.8273

sRun

RunA1

zout

1

Ave

Data

0.00035

Model

0.0003

data/vMax

0.00216

ofxng

321.551

Min d/m

2.06365

ave/min d

1.70522

ave/min m

3.07008

324_Calib_S_A

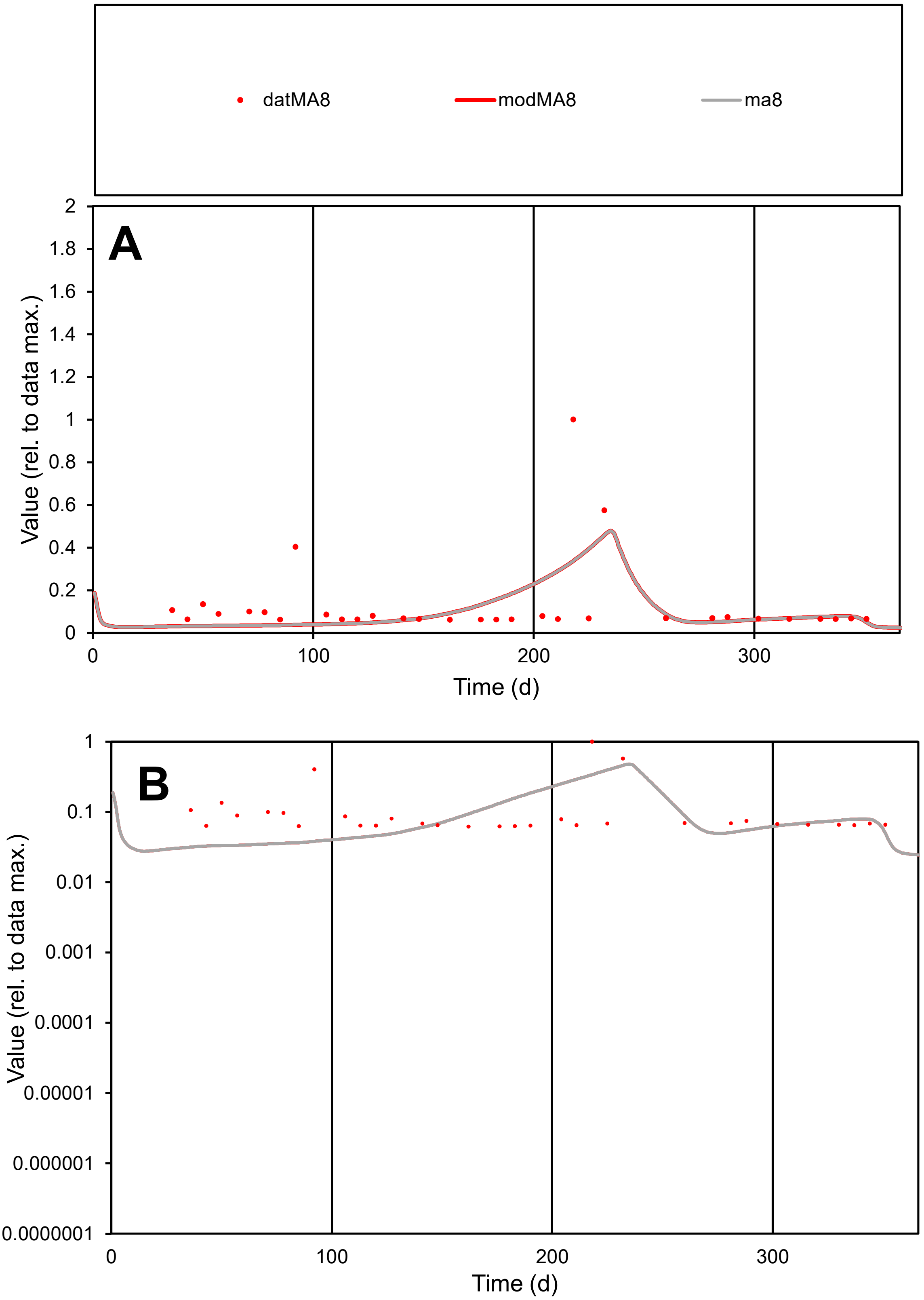


**C1**

Variables
MA8 Provide jtd or Dst
CalibOnly?
2 1 to reproduce of
oNormalize?
2 1=no, 2=yes

ofxn calc*
1.24054
ofxn mod
1.25391

sRun
RunA1
zout
1

Ave
Data
0.00012
Model
0.0001

data/vMax
0.00095

ofxng
321.551

Min d/m
1.94578

ave/min d
2.0879
ave/min m
3.47373

324_Calib_S_A

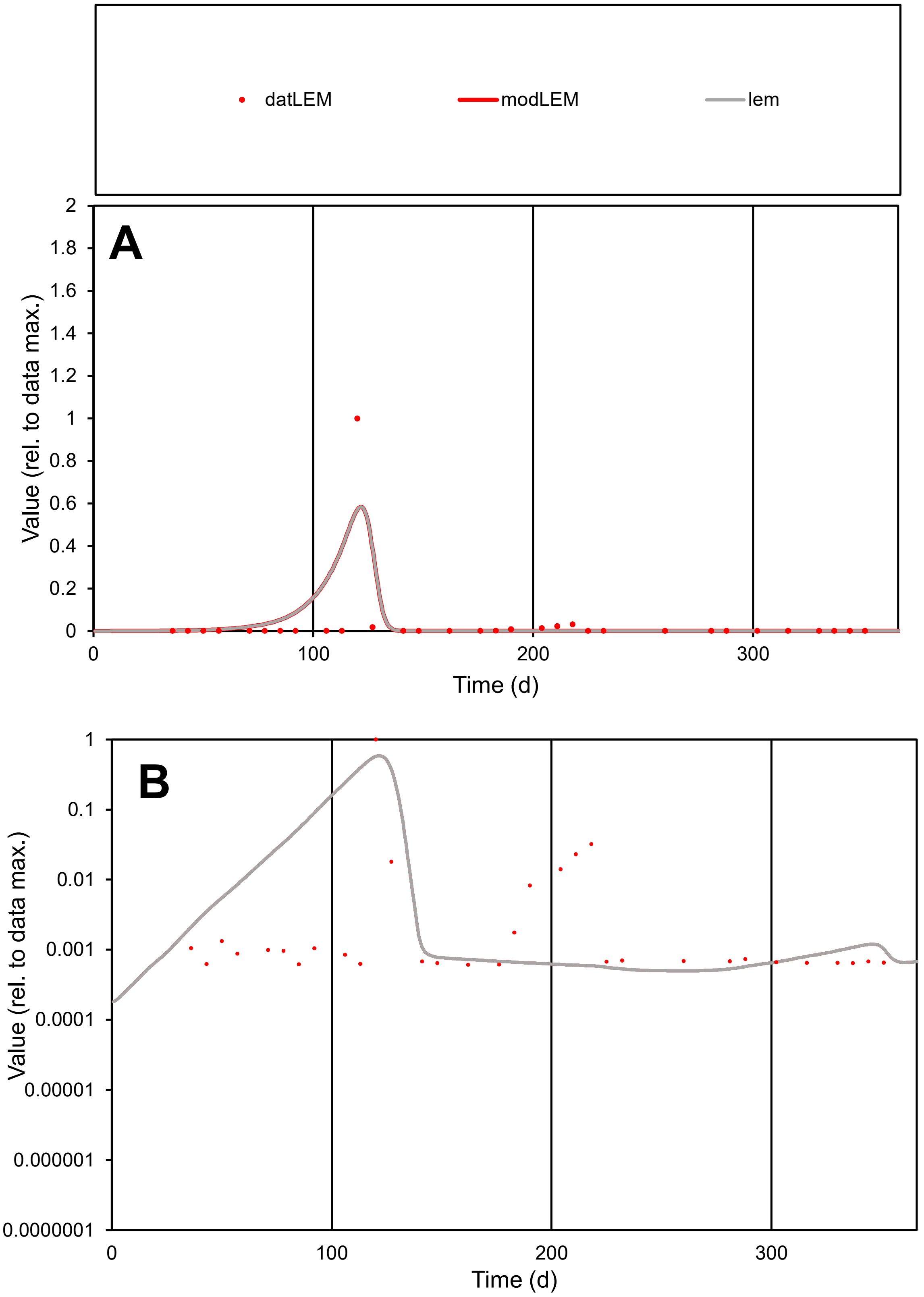


**C1**

Variables

LEM Provide jtd or Dst

CalibOnly?

2 1 to reproduce of

oNormalize?

2 1=no, 2=yes

ofxn calc*

3.1334

ofxn mod

3.32198

sRun

RunA1

zout

1

Ave

Data

0.0012

Model

0.00166

data/vMax

0.03451

ofxng

321.551

Min d/m

1.23122

ave/min d

56.9388

ave/min m

96.4306

324_Calib_S_A

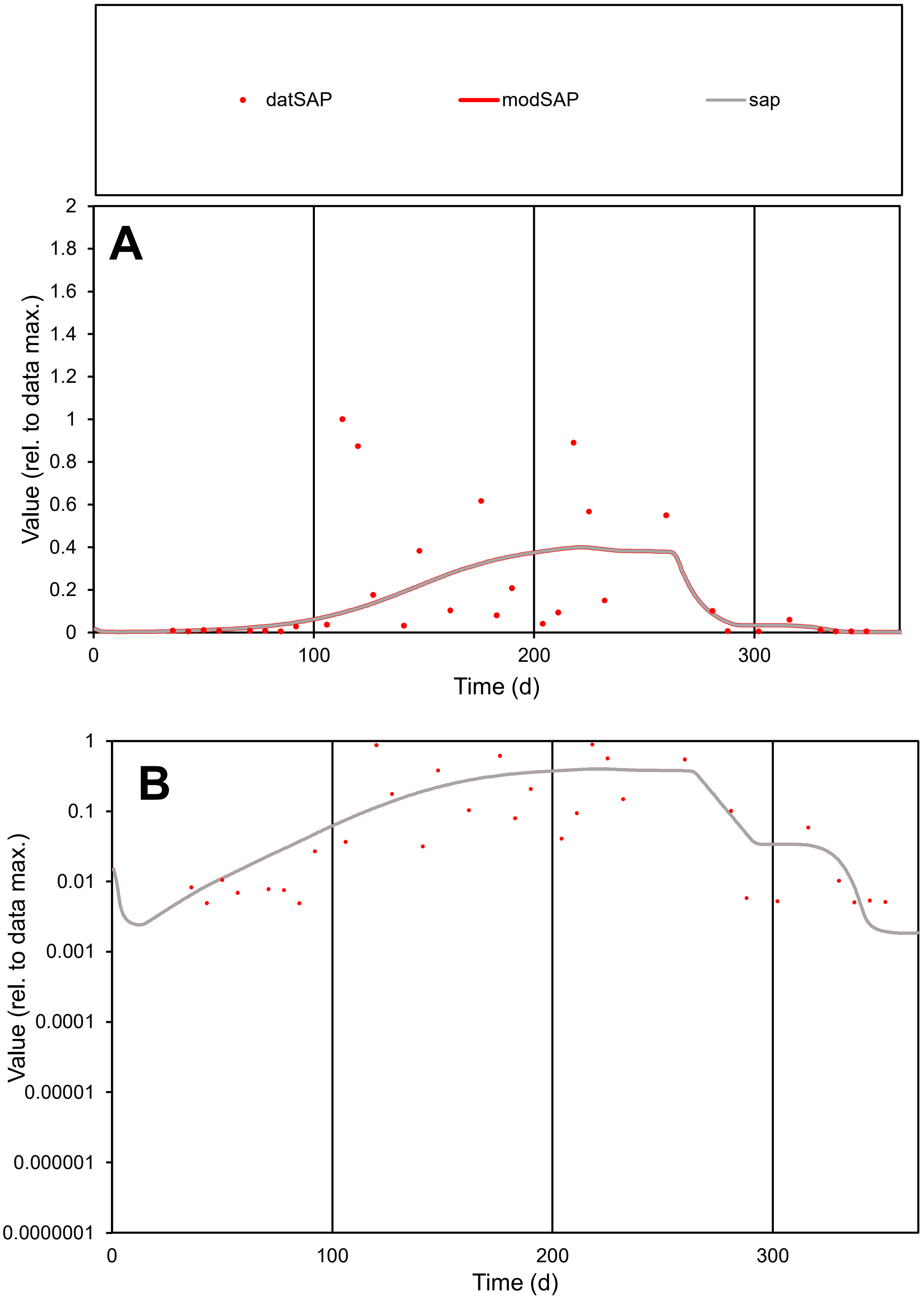


**C1**

Variables

SAP Provide jtd or Dst

CalibOnly?

2 1 to reproduce of

oNormalize?

2 1=no, 2=yes

ofxn calc*

1.38494

ofxn mod

1.39259

sRun

RunA1

zout

1

Ave

Data

0.00127

Model

0.00102

data/vMax

0.00668

ofxng

321.551

Min d/m

2.55747

ave/min d

38.7563

ave/min m

79.8779

324_Calib_S_A

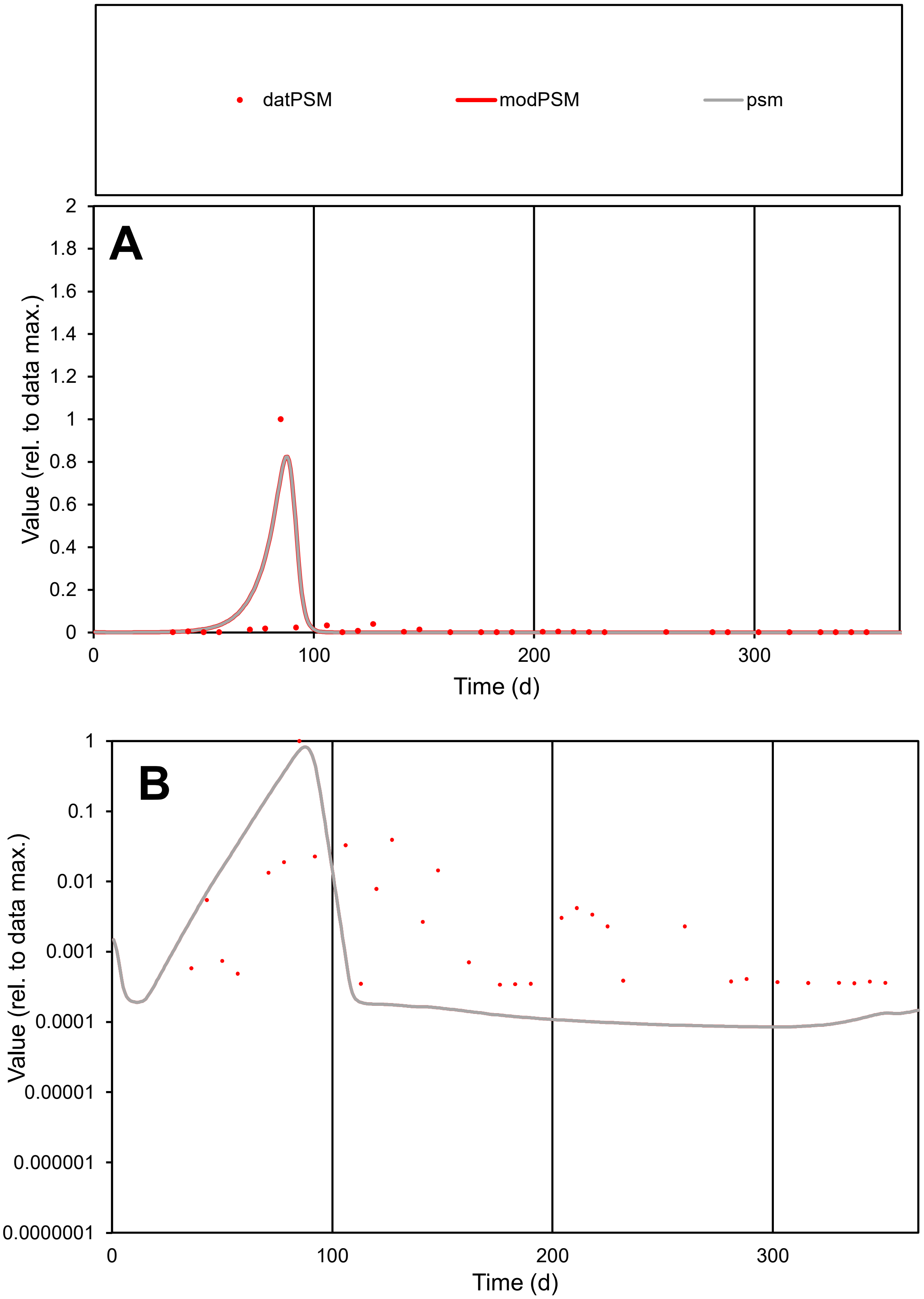


**C1**

Variables
PSM Provide jtd or Dst
CalibOnly?
2 1 to reproduce of
oNormalize?
2 1=no, 2=yes

ofxn calc*
1.92607
ofxn mod
2.20262

sRun
RunA1
zout
1

Ave
Data
0.00294
Model
0.00368

data/vMax
0.07978

ofxng
321.551

Min d/m
4.01624

ave/min d
108.125
ave/min m
543.024

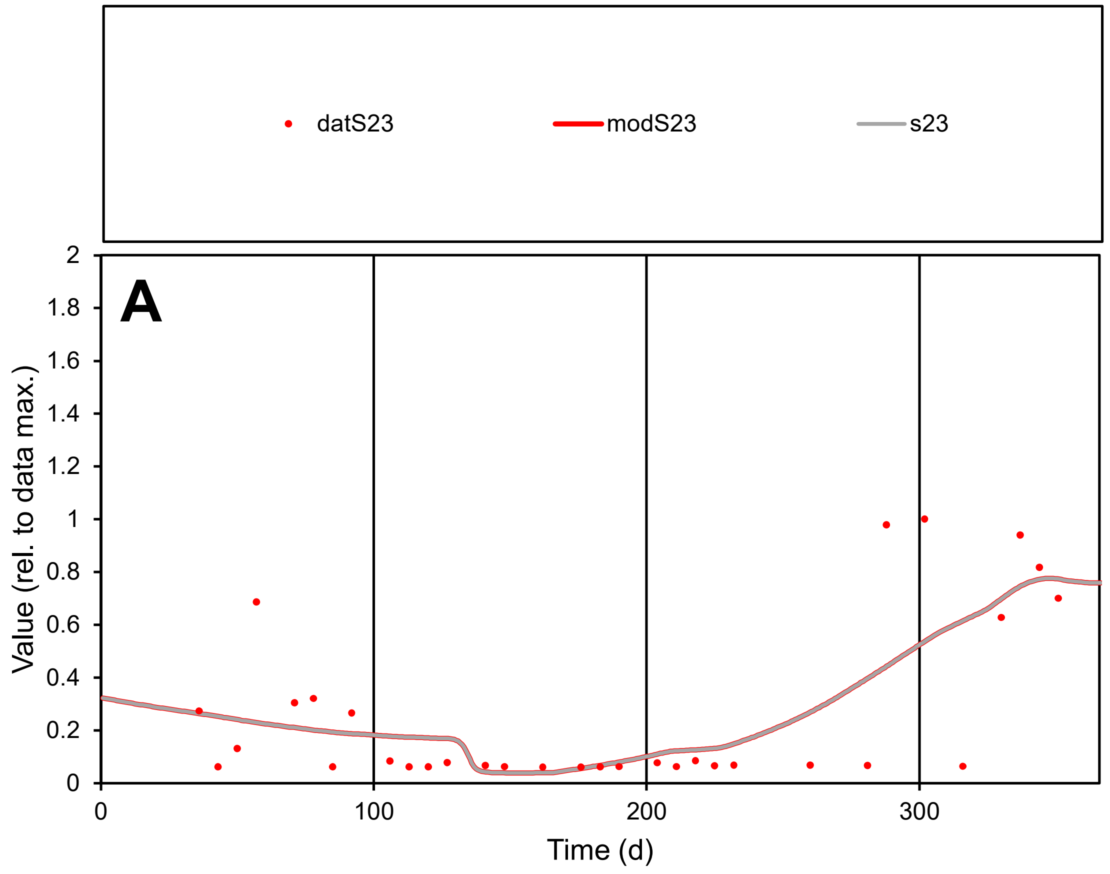


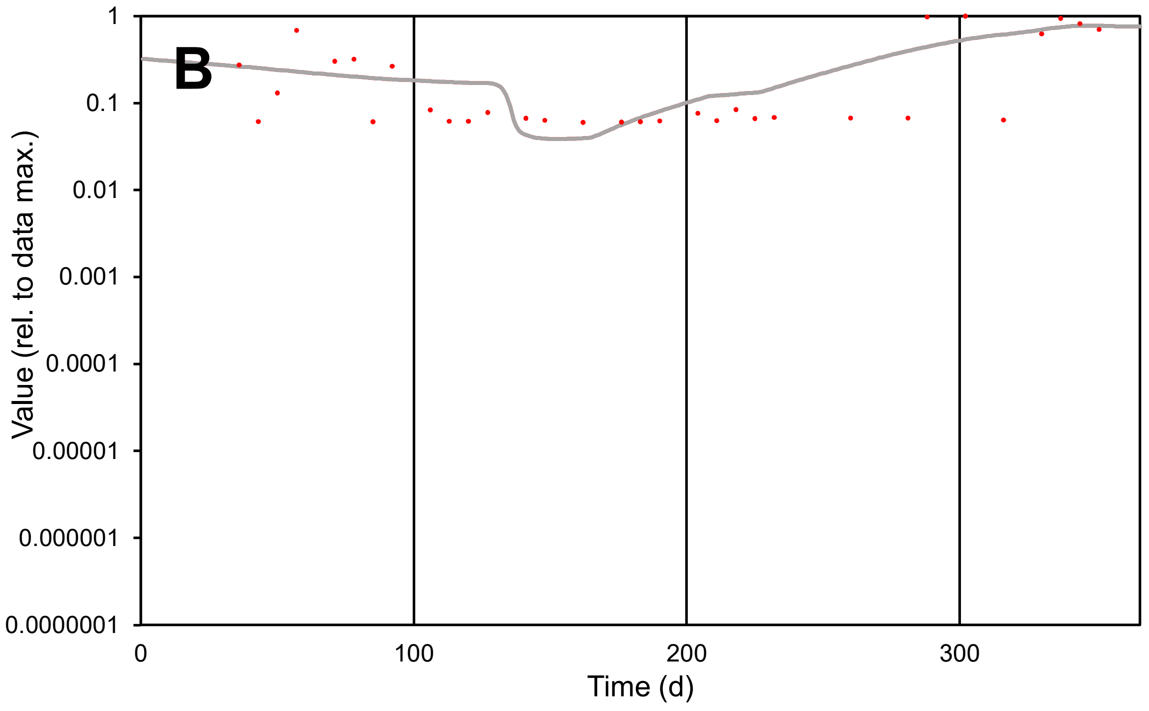


**C1**

Variables

S23 Provide jtd or Dst

CalibOnly?

2 1 to reproduce of

oNormalize?

2 1=no, 2=yes

ofxn calc*

0.78585

ofxn mod

0.79837

sRun

RunA1

zout

1

Ave

Data

0.00069

Model

0.00071

data/vMax

0.00262

ofxng

321.551

Min d/m

1.55227

ave/min d

4.35777

ave/min m

7.04888

324_Calib_S_A

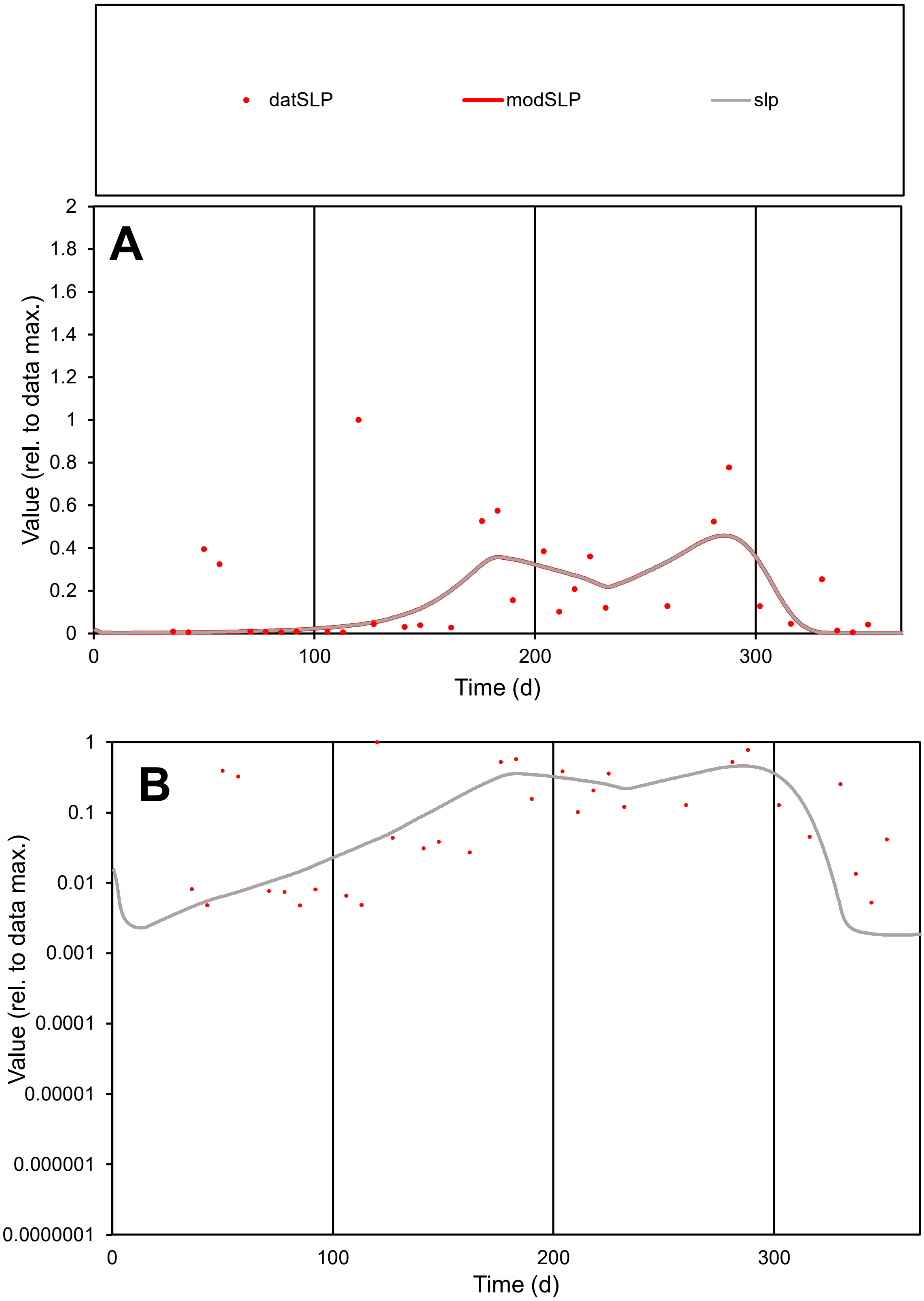


**C1**

Variables
SLP Provide jtd or Dst
CalibOnly?
2 1 to reproduce of
oNormalize?
2 1=no, 2=yes

ofxn calc*
1.16081
ofxn mod
1.16754

sRun
RunA1
zout
1

Ave
Data
0.00097
Model
0.00072

data/vMax
0.00496

ofxng
321.551

Min d/m
2.64138

ave/min d
40.7431
ave/min m
79.8586

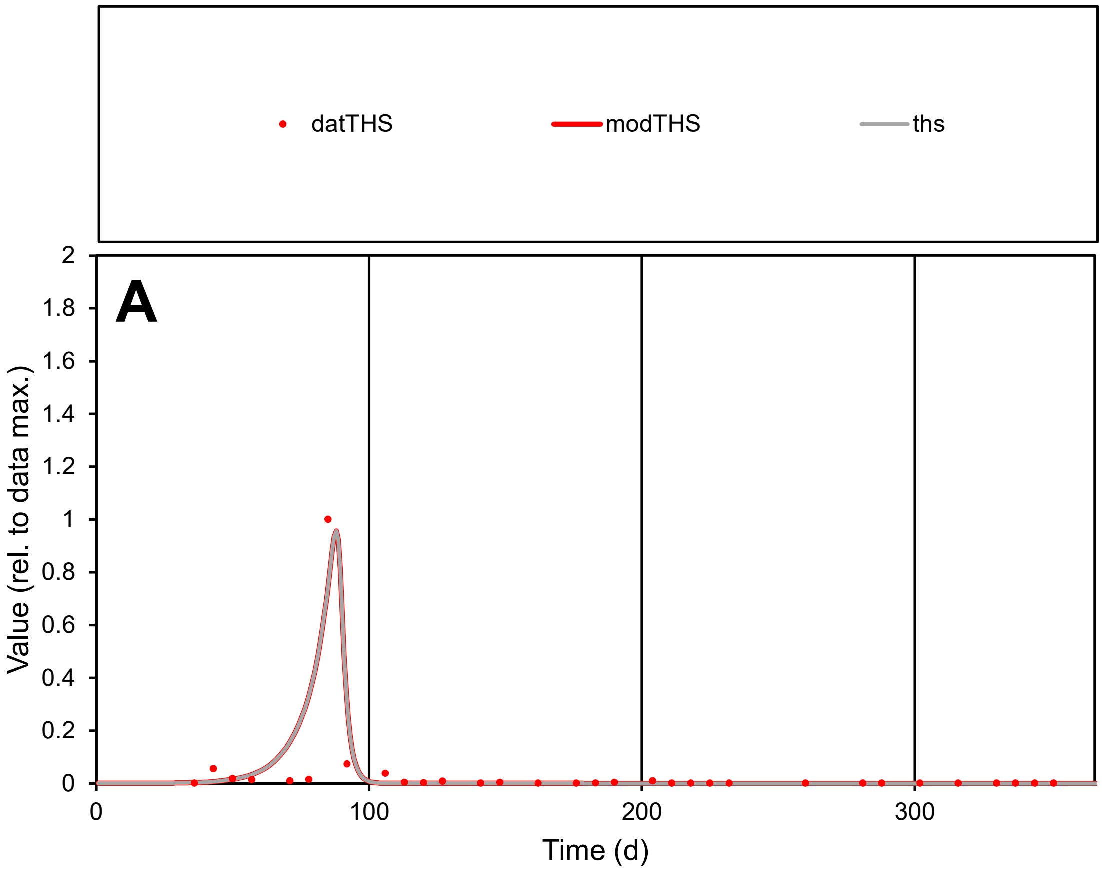


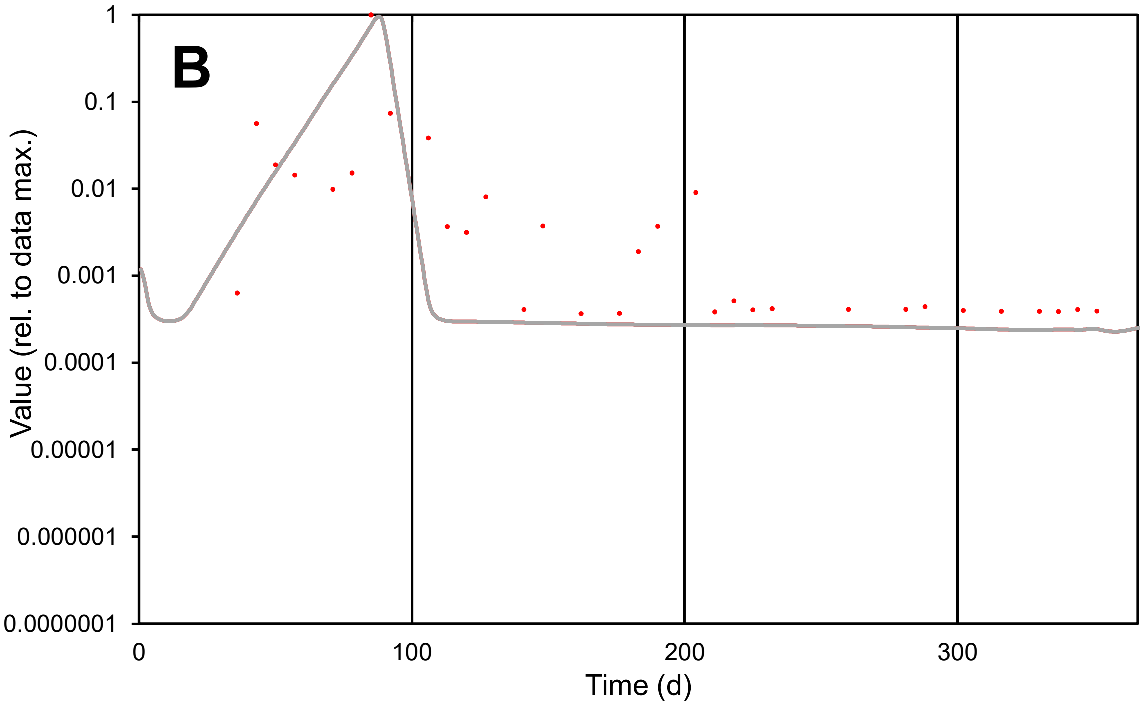


**C1**

Variables

| | |
|---|---|
| THS | Provide jtd or Dst |
| CalibOnly? | |
| 2 | 1 to reproduce of |
| oNormalize? | |
| 2 | 1=no, 2=yes |

ofxn calc*
1.42946
ofxn mod
1.69417

sRun
RunA1
zout
1

Ave
Data
0.00303
Model
0.00348

data/vMax
0.07656

ofxng
321.551

Min d/m
1.53377

ave/min d
107.646
ave/min m
189.734

324_Calib_S_A

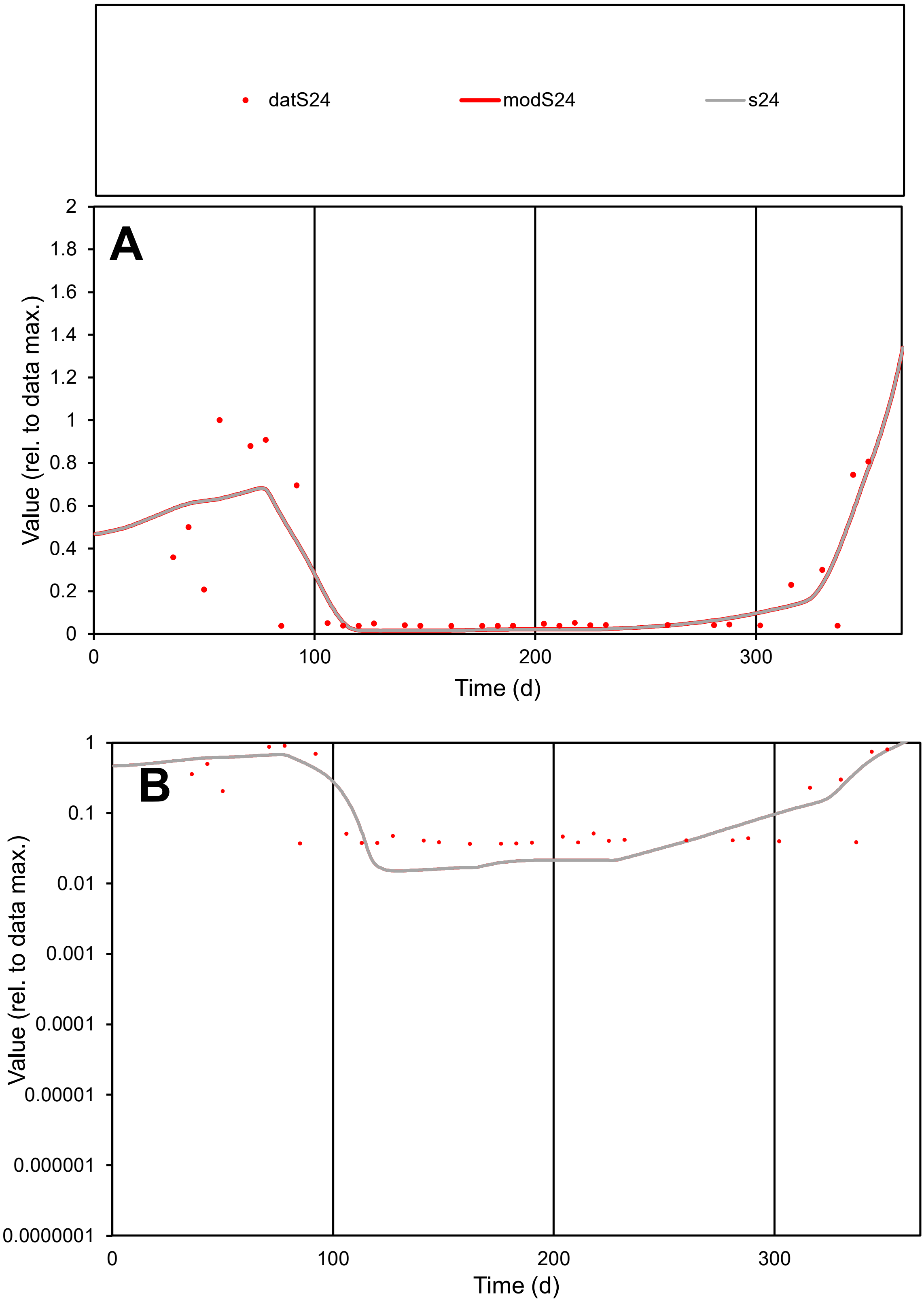


**C1**

Variables
S24 Provide jtd or Dst
CalibOnly?
2 1 to reproduce of
oNormalize?
2 1=no, 2=yes

ofxn calc*
0.73186
ofxn mod
0.75014

sRun
RunA1
zout
1

Ave
Data
0.00036
Model
0.00037

data/vMax
0.00155

ofxng
321.551

Min d/m
2.43566

ave/min d
6.37593
ave/min m
15.7759

324_Calib_S_A

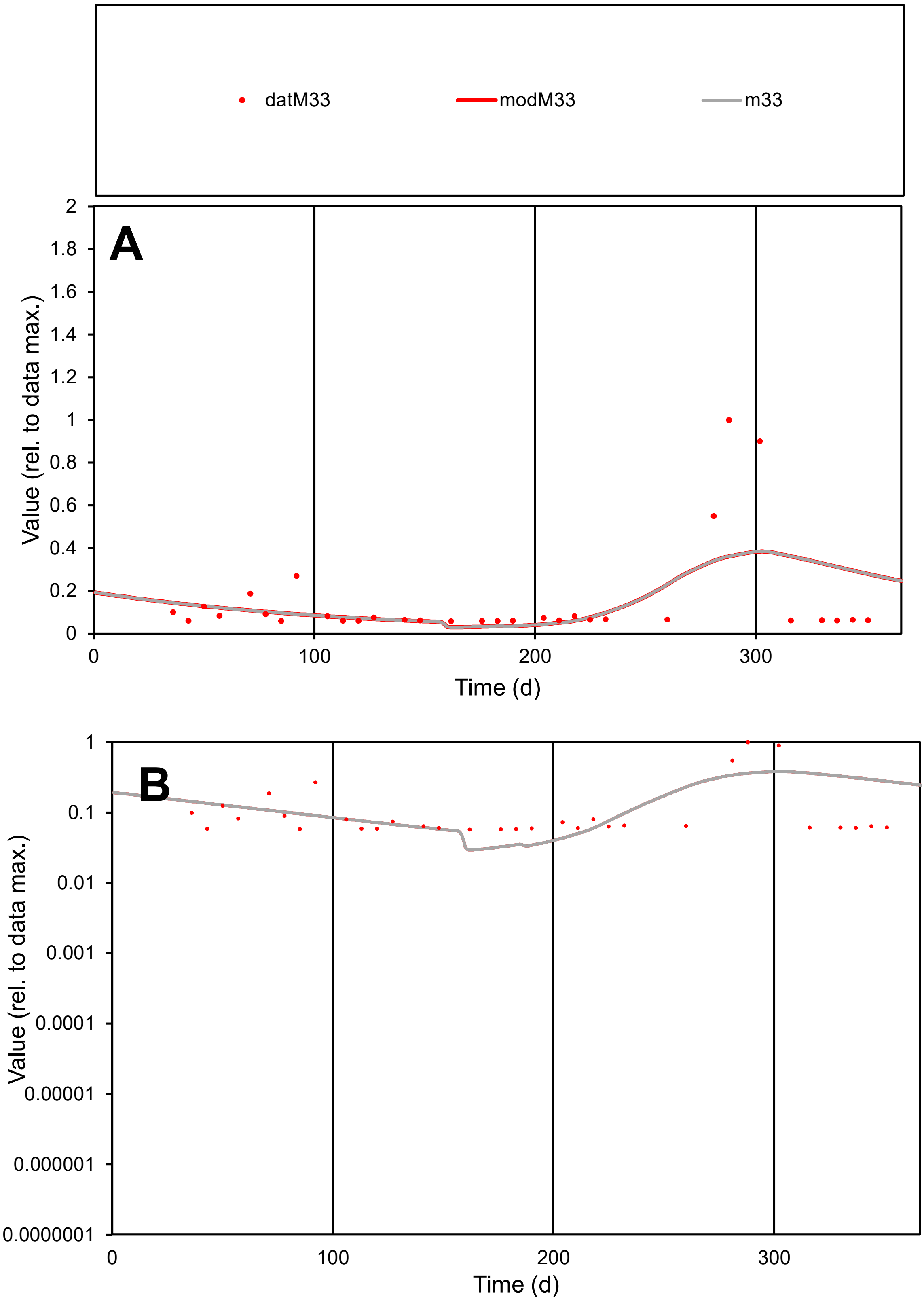


**C1**

Variables

M33 Provide jtd or Dst

CalibOnly?

2 1 to reproduce of

oNormalize?

2 1=no, 2=yes

ofxn calc*

1.25378

ofxn mod

1.26146

sRun

RunA1

zout

1

Ave

Data

0.00017

Model

0.00017

data/vMax

0.00117

ofxng

321.551

Min d/m

1.94686

ave/min d

2.59155

ave/min m

4.96204

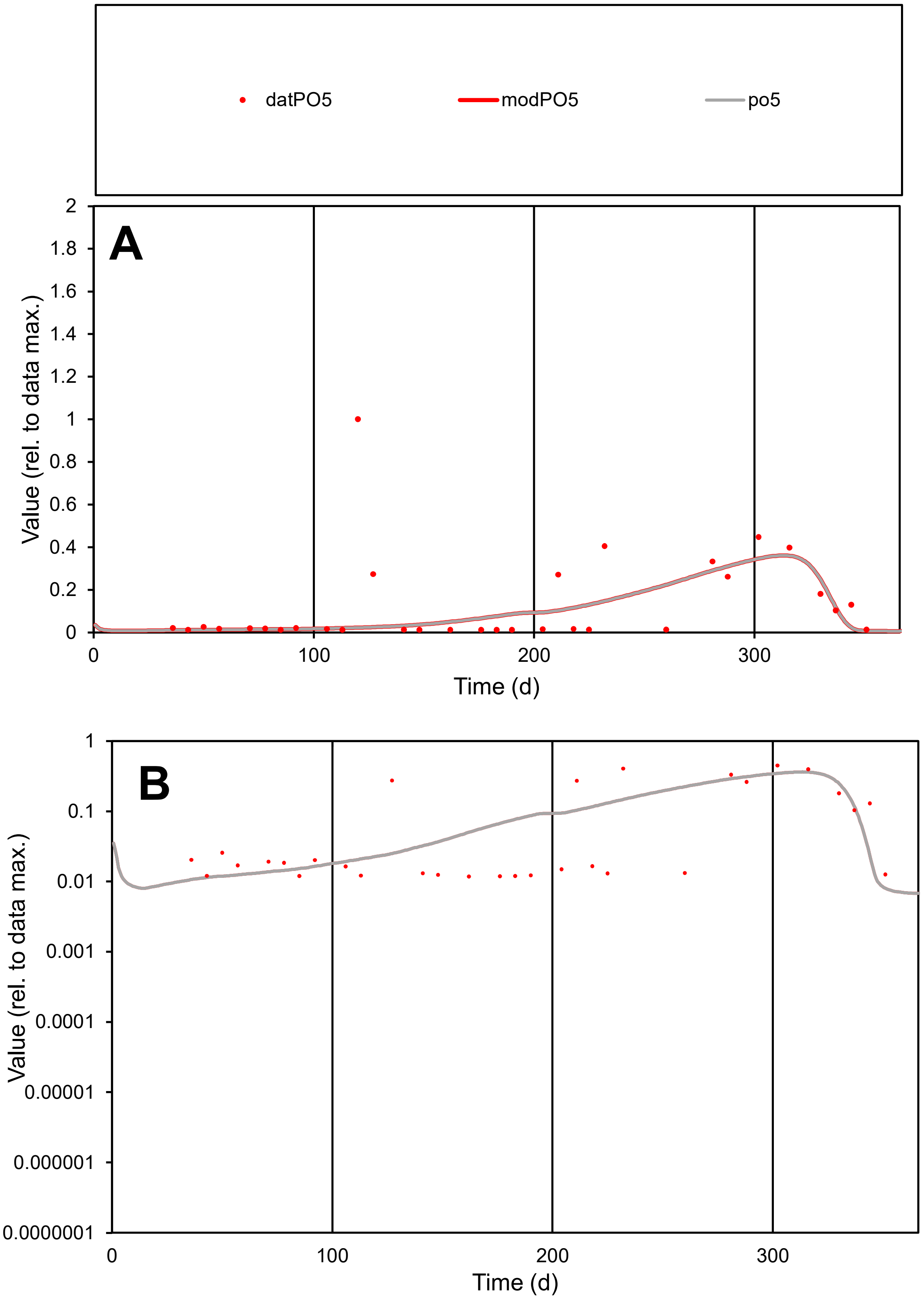


**C1**

Variables
PO5 Provide jtd or Dst
CalibOnly?
2 1 to reproduce of
oNormalize?
2 1=no, 2=yes

ofxn calc*
1.52037
ofxn mod
1.52741

sRun
RunA1
zout
1

Ave
Data
0.0005
Model
0.00037

data/vMax
0.00386

ofxng
321.551

Min d/m
1.51968

ave/min d
10.8939
ave/min m
12.2333

324_Calib_S_A

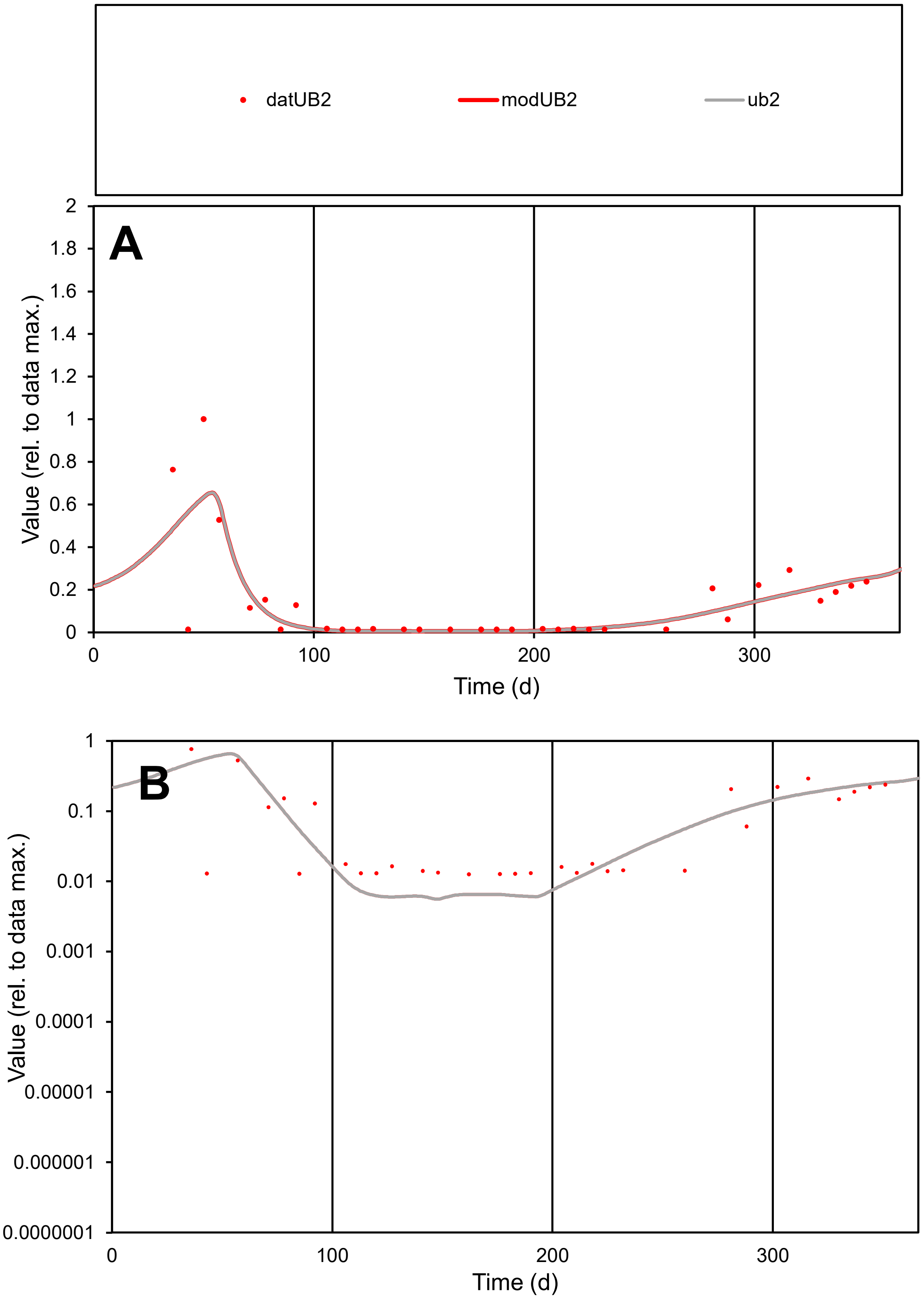


**C1**

Variables

UB2 Provide jtd or Dst

CalibOnly?

2 1 to reproduce of

oNormalize?

2 1=no, 2=yes

ofxn calc*

0.94359

ofxn mod

0.96145

sRun

RunA1

zout

1

Ave

Data

0.00114

Model

0.00109

data/vMax

0.00805

ofxng

321.551

Min d/m

2.23635

ave/min d

11.1436

ave/min m

23.8679

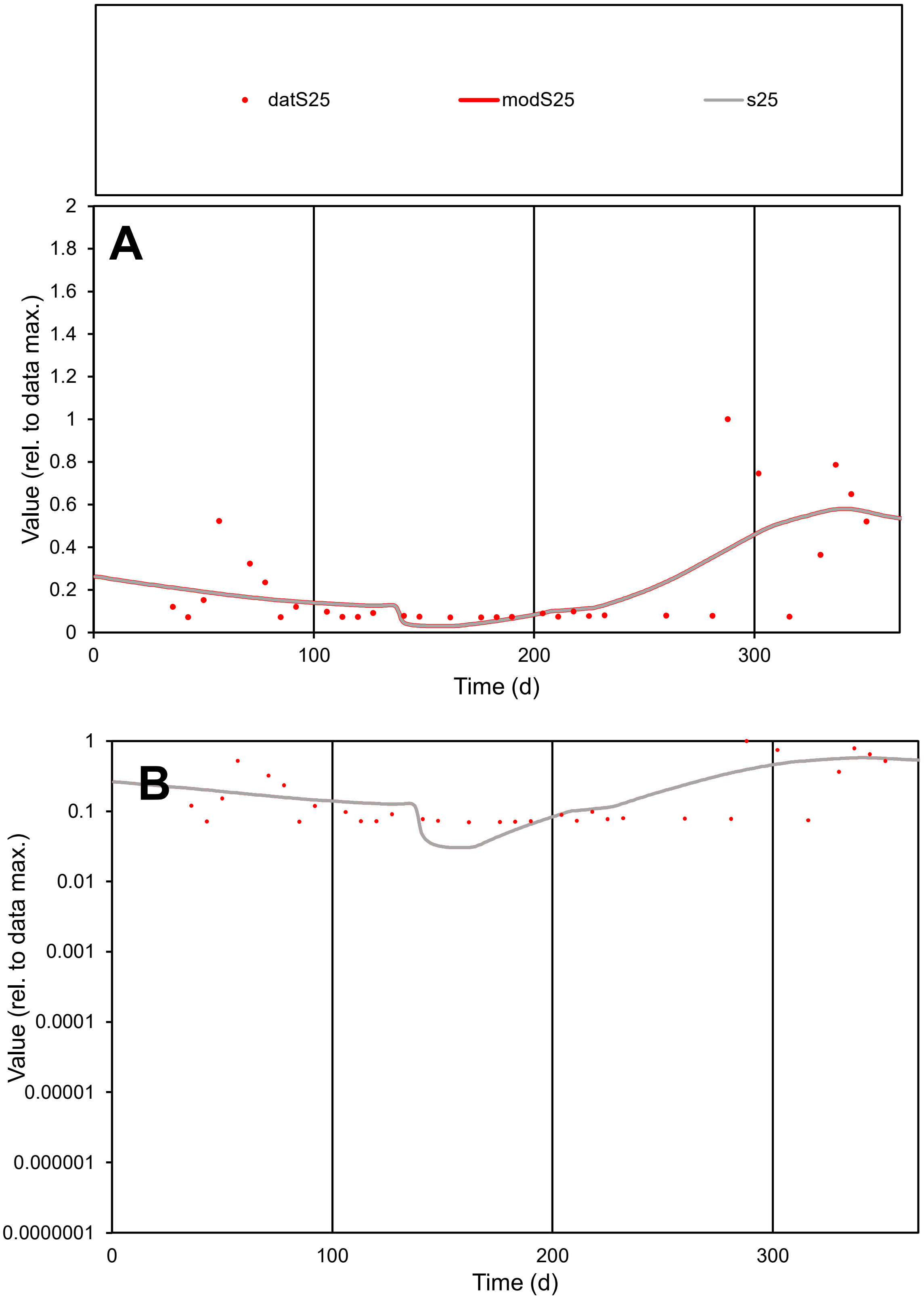


**C1**

Variables
S25 Provide jtd or Dst
CalibOnly?
2 1 to reproduce of
oNormalize?
2 1=no, 2=yes

ofxn calc*
0.8104
ofxn mod
0.82093

sRun
RunA1
zout
1

Ave
Data
0.00053
Model
0.00052

data/vMax
0.00237

ofxng
321.551

Min d/m
2.26273

ave/min d
3.1696
ave/min m
7.10553

324_Calib_S_A

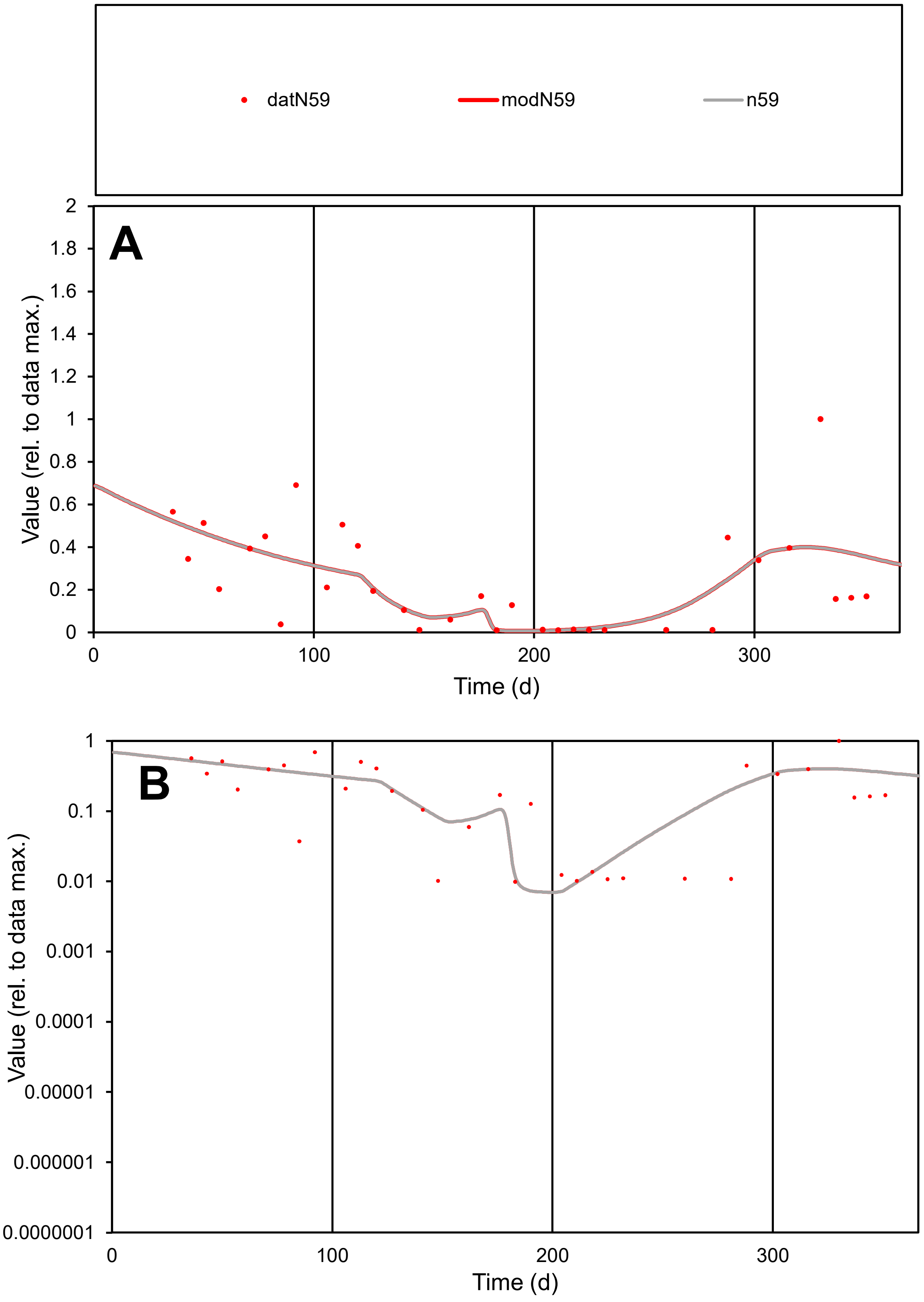


**C1**

Variables

N59 Provide jtd or Dst

CalibOnly?

2 1 to reproduce of

oNormalize?

2 1=no, 2=yes

ofxn calc*

0.7265

ofxn mod

0.7377

sRun

RunA1

zout

1

Ave

Data

0.00101

Model

0.00099

data/vMax

0.00416

ofxng

321.551

Min d/m

1.37212

ave/min d

24.4394

ave/min m

33.1222

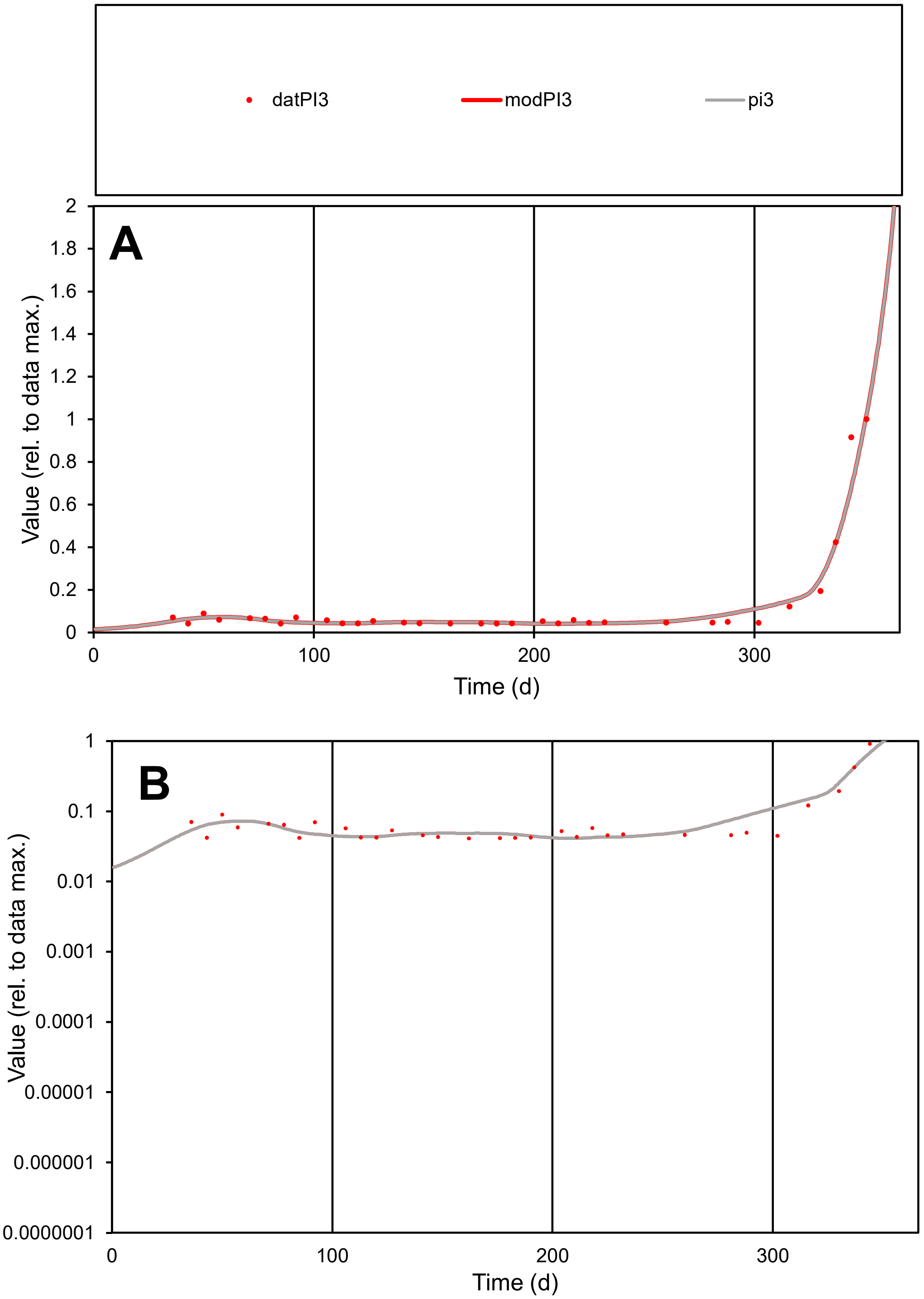


**C1**

Variables
PI3 Provide jtd or Dst
CalibOnly?
2 1 to reproduce of
oNormalize?
2 1=no, 2=yes

ofxn calc*
0.31817
ofxn mod
0.35792

sRun
RunA1
zout
1

Ave
Data
0.00021
Model
0.00022

data/vMax
0.0017

ofxng
321.551

Min d/m
0.99671

ave/min d
3.07047
ave/min m
3.167

324_Calib_S_A

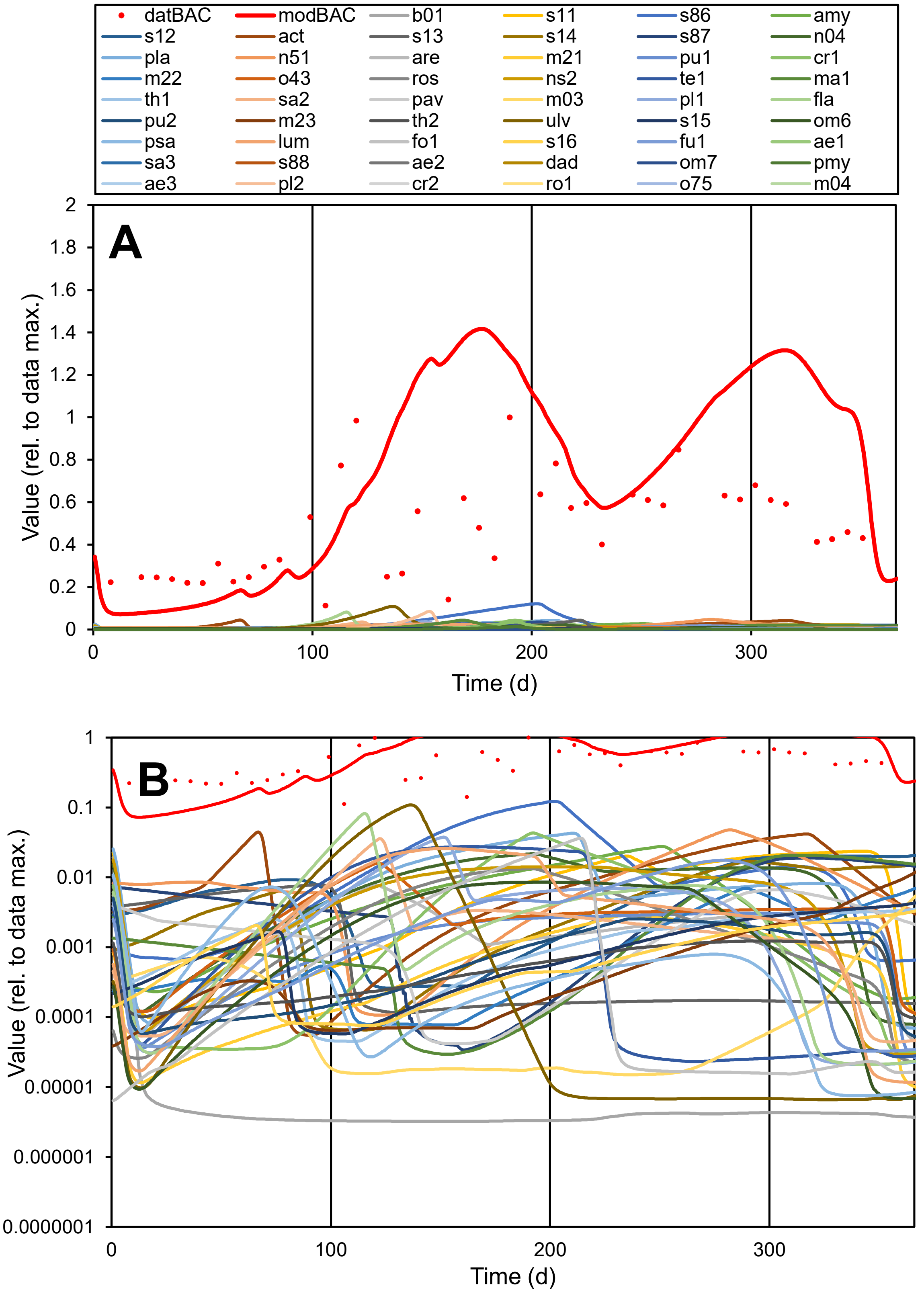


**C1**

Variables

BAC Provide jtd or Dst

CalibOnly?

2 1 to reproduce of

oNormalize?

2 1=no, 2=yes

ofxn calc*

1.01337

ofxn mod

1.02928

sRun

RunA1

zout

1

Ave

Data

0.68283

Model

1.06607

data/vMax

1.447

ofxng

321.551

Min d/m

1.51115

ave/min d

4.19944

ave/min m

9.90774

325_Calib_S_A

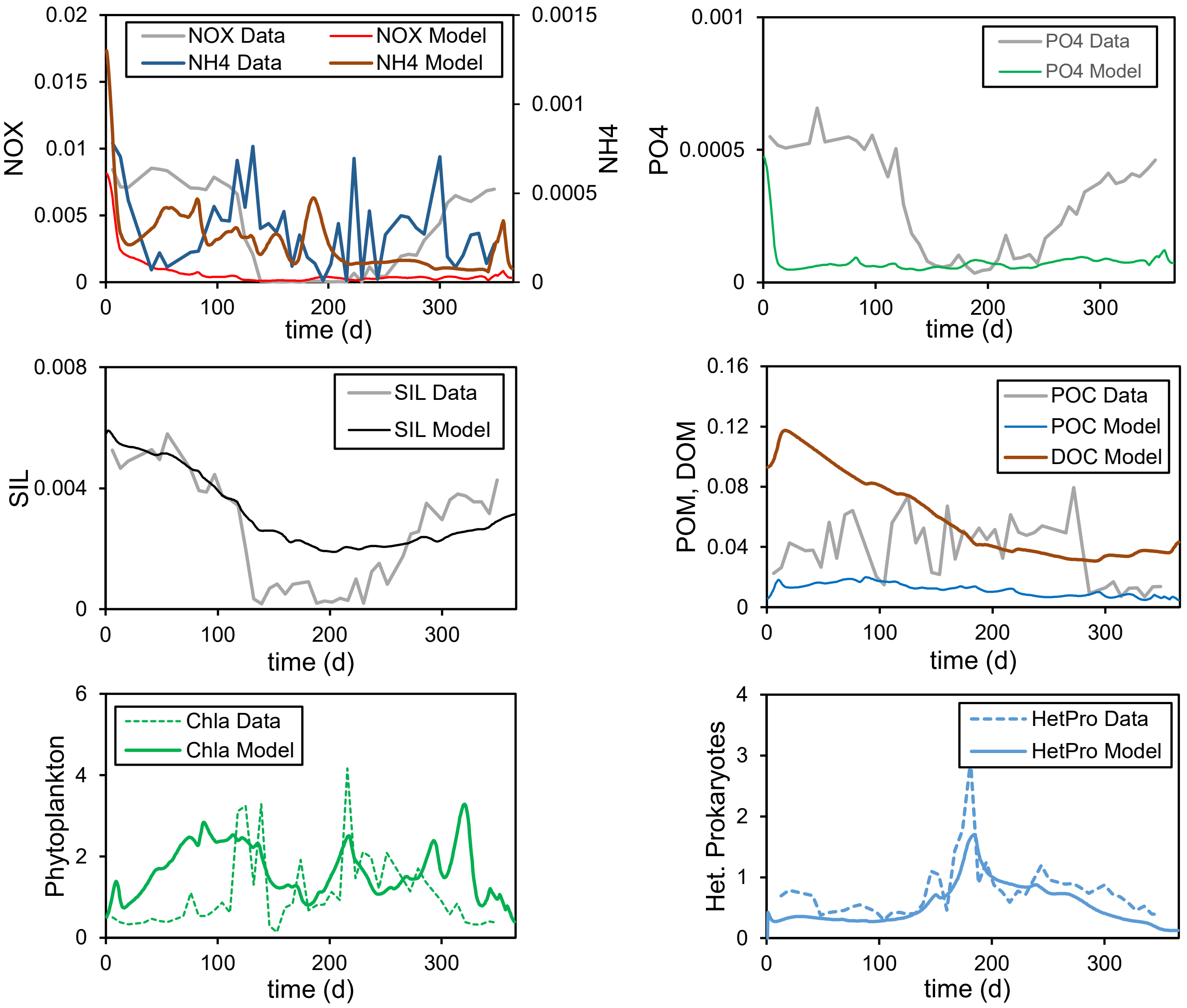

NOX Data
NOX Model
NH4 Data
NH4 Model
NOX
NH4
time (d)
PO4 Data
PO4 Model
PO4
time (d)
SIL Data
SIL Model
SIL
time (d)
POC Data
POC Model
DOC Model
POM, DOM
time (d)
Chla Data
Chla Model
Phytoplankton
time (d)
HetPro Data
HetPro Model
Het. Prokaryotes
time (d)

325_Calib_S_A

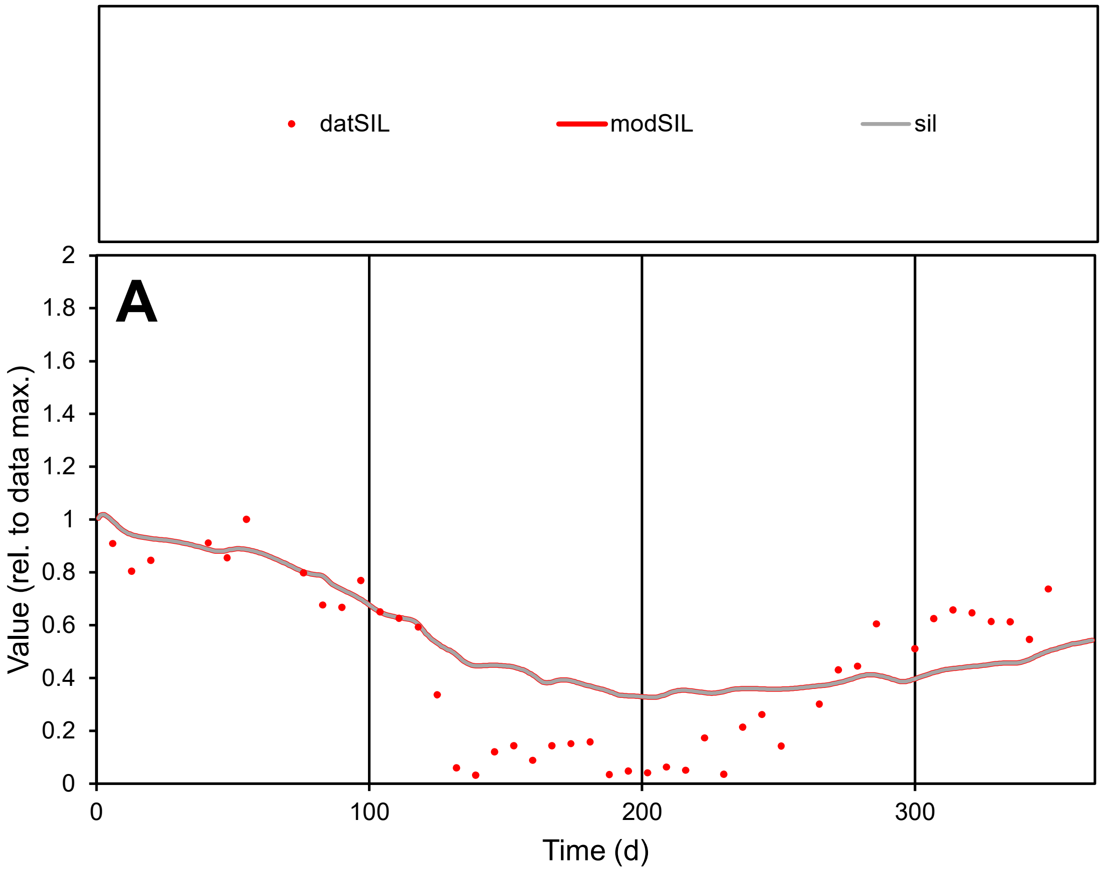


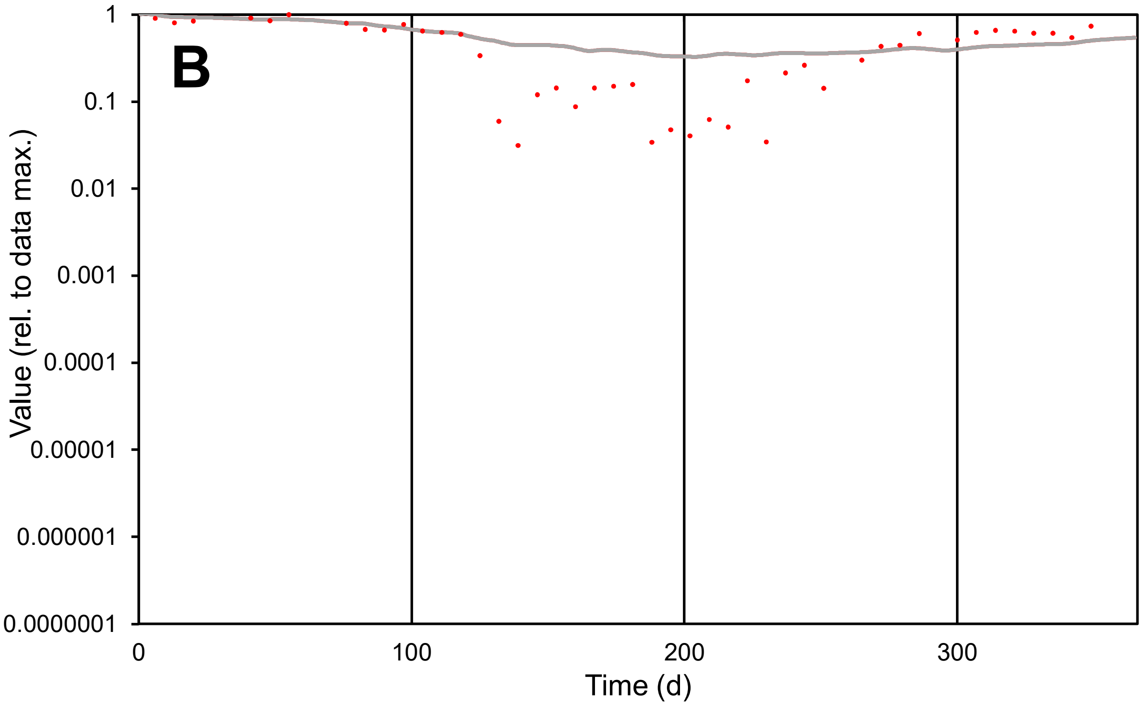


**C1**

Variables
SIL Provide jtd or Dst
CalibOnly?
2 1 to reproduce of
oNormalize?
2 1=no, 2=yes

ofxn calc*
1.40634
ofxn mod
1.46444

sRun
RunA1
zout
1

Ave
Data
0.00252
Model
0.00301

data/vMax
0.0058

ofxng
275.646

Min d/m
0.09636

ave/min d
13.7979
ave/min m
1.58969

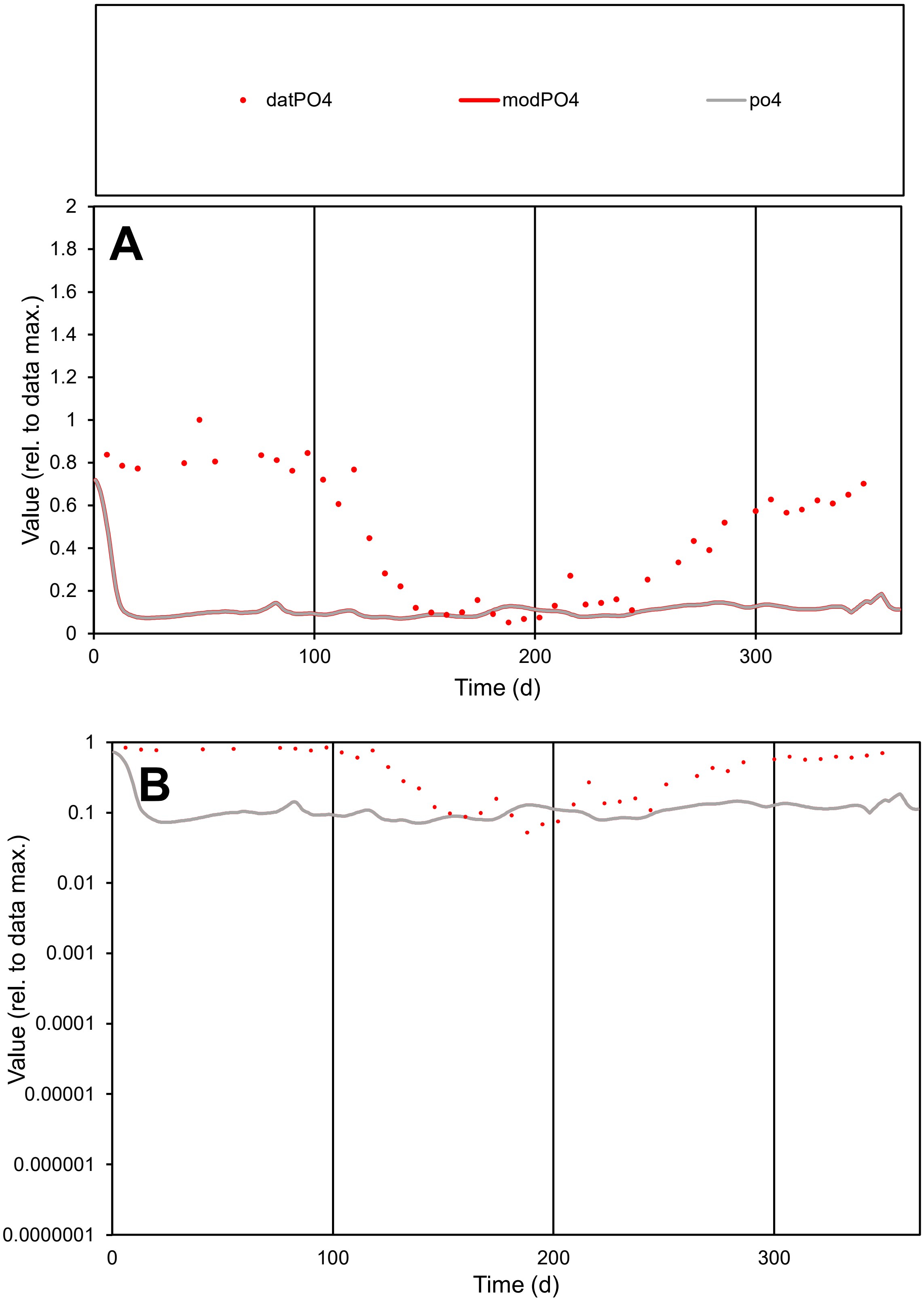


**C1**

Variables

PO4 Provide jtd or Dst

CalibOnly?

2 1 to reproduce of

oNormalize?

2 1=no, 2=yes

ofxn calc*

2.91573

ofxn mod

2.92973

sRun

RunA1

zout

1

Ave

Data

0.0003

Model

7.4E-05

data/vMax

0.00066

ofxng

275.646

Min d/m

0.73204

ave/min d

8.70723

ave/min m

1.59127

325_Calib_S_A

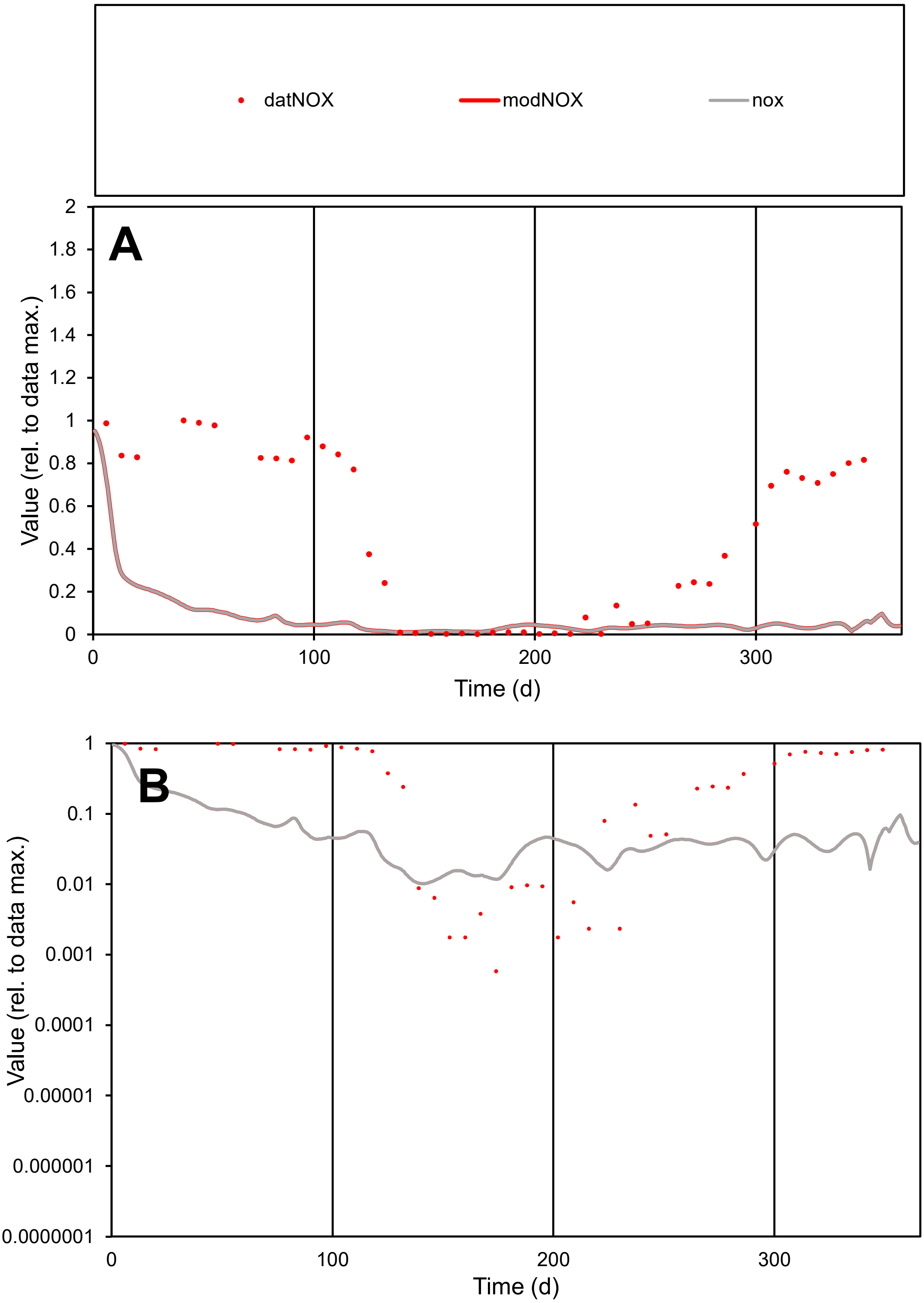


**C1**

Variables
NOX Provide jtd or Dst
CalibOnly?
2 1 to reproduce of
oNormalize?
2 1=no, 2=yes

ofxn calc*
3.49493
ofxn mod
3.51061

sRun
RunA1
zout
1

Ave
Data
0.00375
Model
0.00056

data/vMax
0.00855

ofxng
275.646

Min d/m
0.05711

ave/min d
750.993
ave/min m
6.39278

325_Calib_S_A

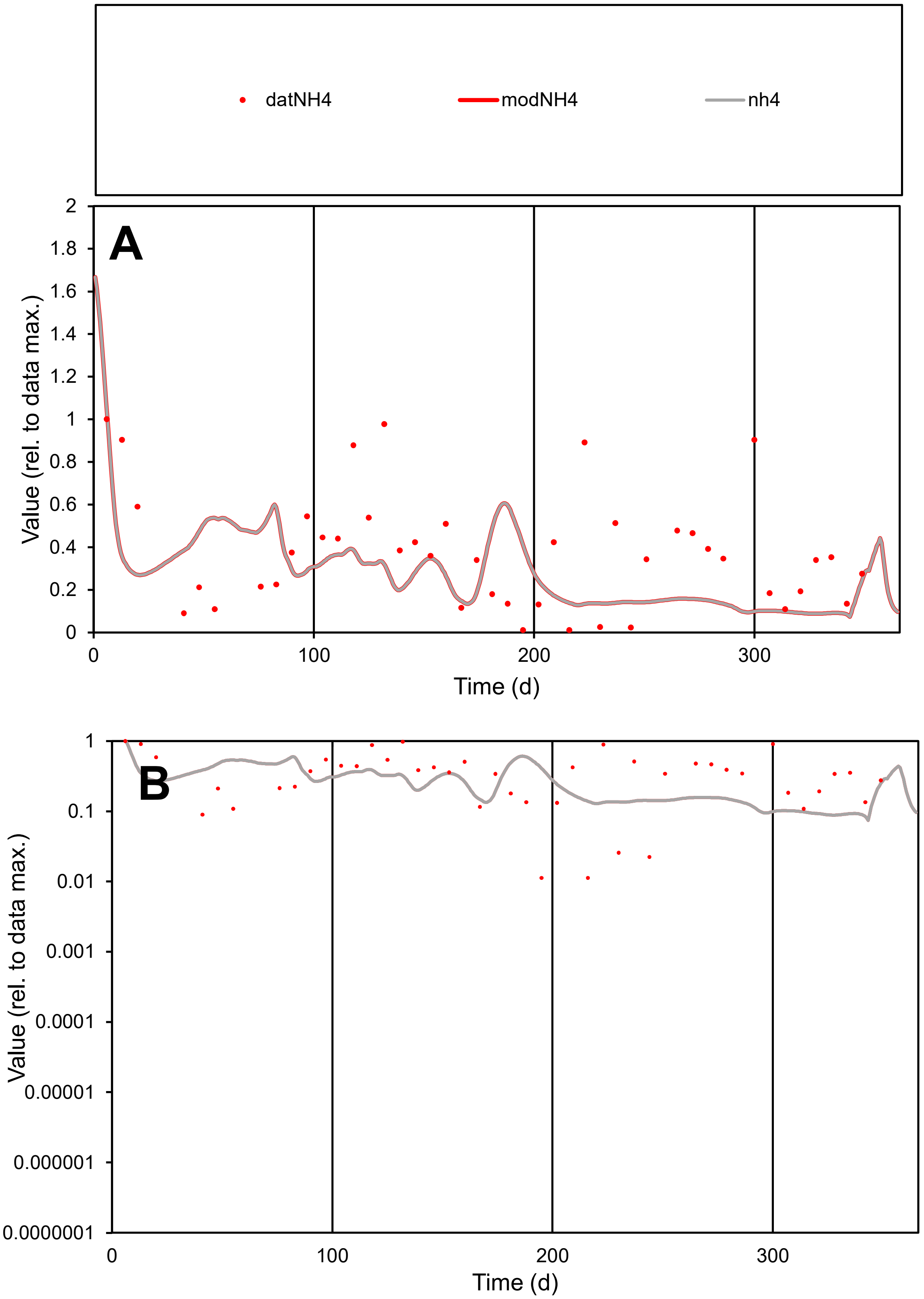


**C1**

Variables
NH4 Provide jtd or Dst
CalibOnly?
2 1 to reproduce of
oNormalize?
2 1=no, 2=yes

ofxn calc*
2.45165
ofxn mod
2.50635

sRun
RunA1
zout
1

Ave
Data
0.00029
Model
0.00021

data/vMax
0.00078

ofxng
275.646

Min d/m
0.13732

ave/min d
33.4921
ave/min m
3.2728

325_Calib_S_A

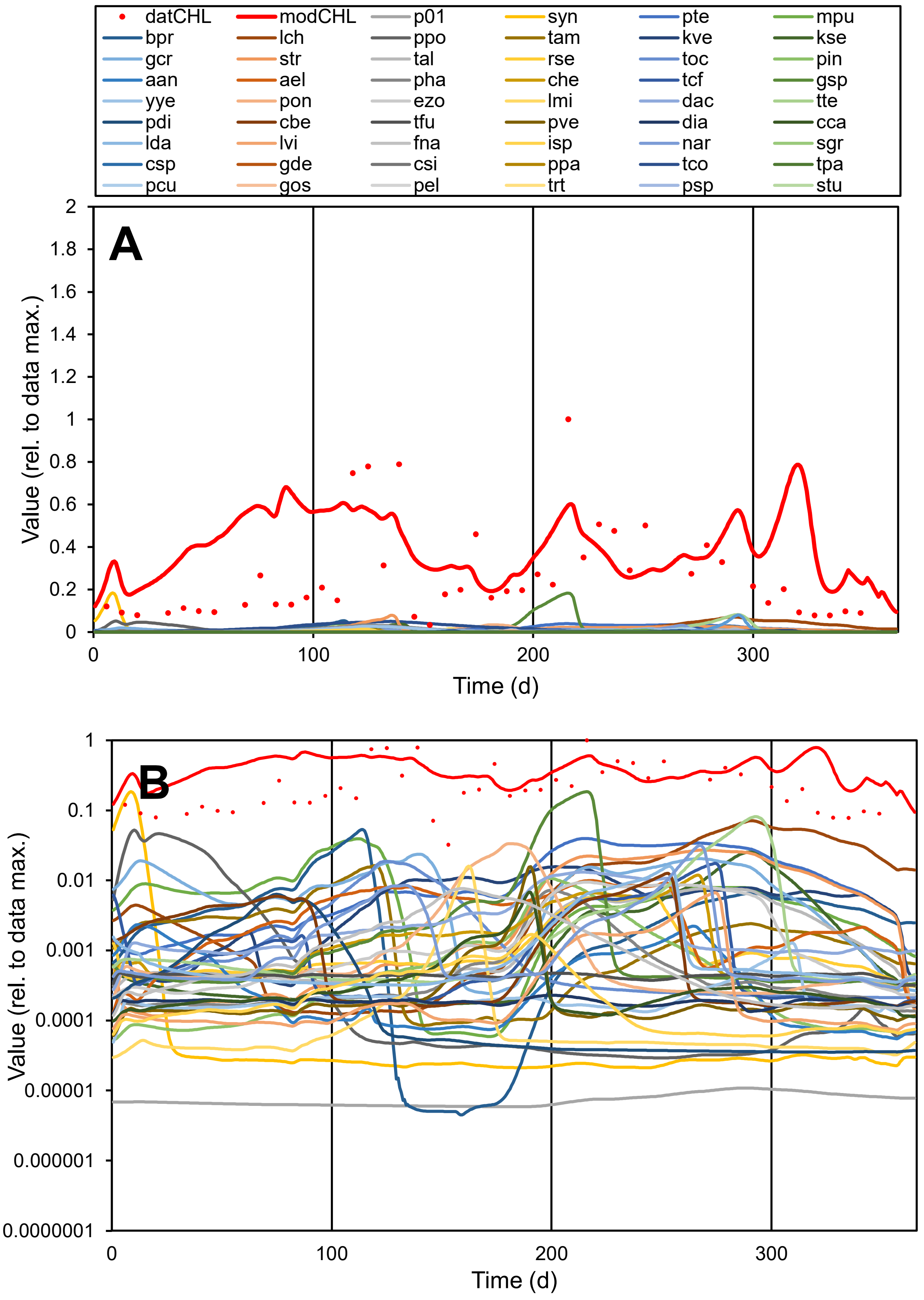


**C1**

Variables

CHL Provide jtd or Dst

CalibOnly?

2 1 to reproduce of

oNormalize?

2 1=no, 2=yes

ofxn calc*

6.05275

ofxn mod

6.27912

sRun

RunA1

zout

1

Ave

Data

1.07345

Model

1.03876

data/vMax

4.17

ofxng

275.646

Min d/m

0.42006

ave/min d

7.93388

ave/min m

3.22497

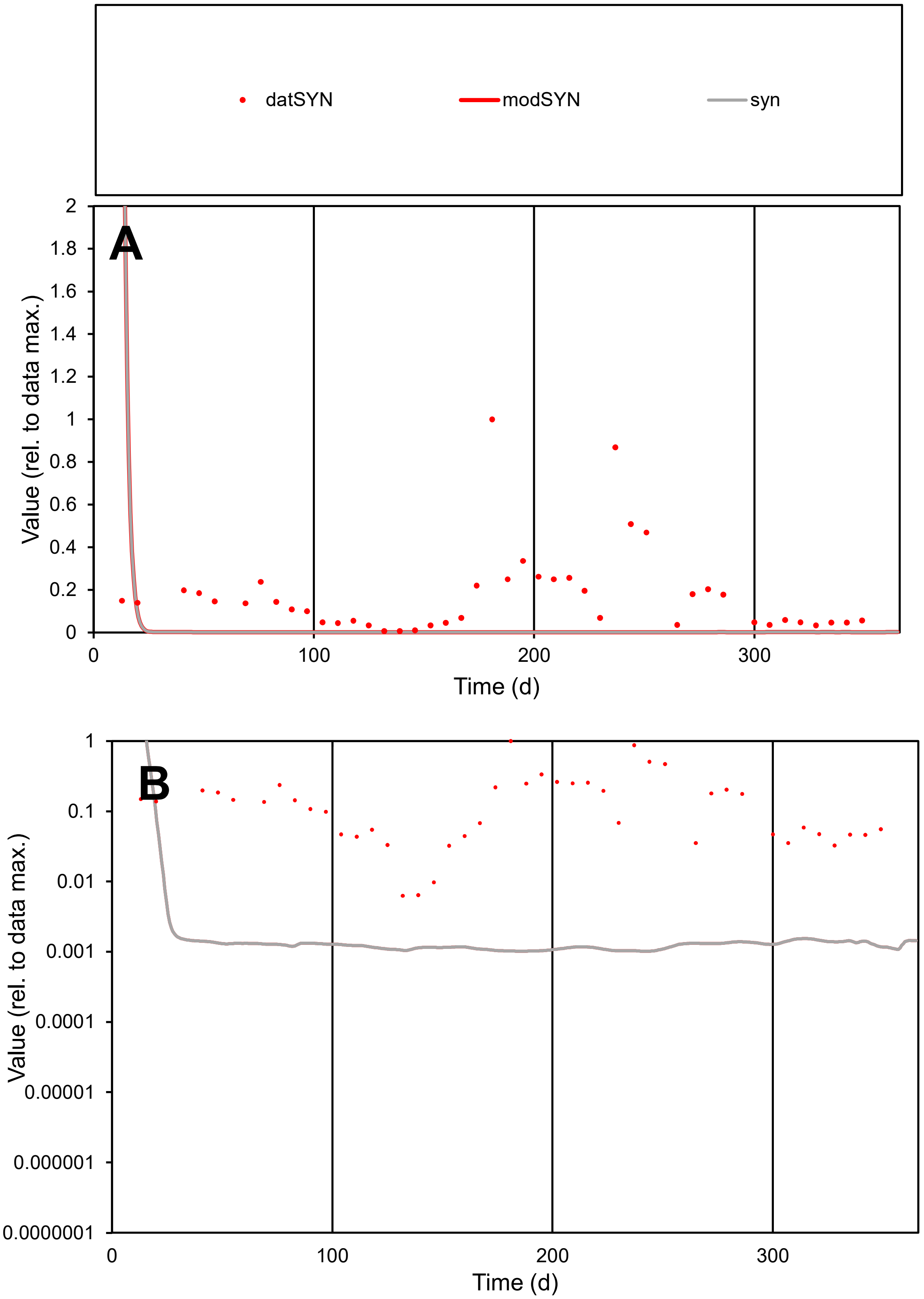


**C1**

Variables
SYN Provide jtd or Dst
CalibOnly?
2 1 to reproduce of
oNormalize?
2 1=no, 2=yes

ofxn calc*
1.78988
ofxn mod
1.89714

sRun
RunA1
zout
1

Ave
Data
8.84665
Model
1.59559

data/vMax
51.72

ofxng
275.646

Min d/m
6.11331

ave/min d
27.3298
ave/min m
30.134

325_Calib_S_A

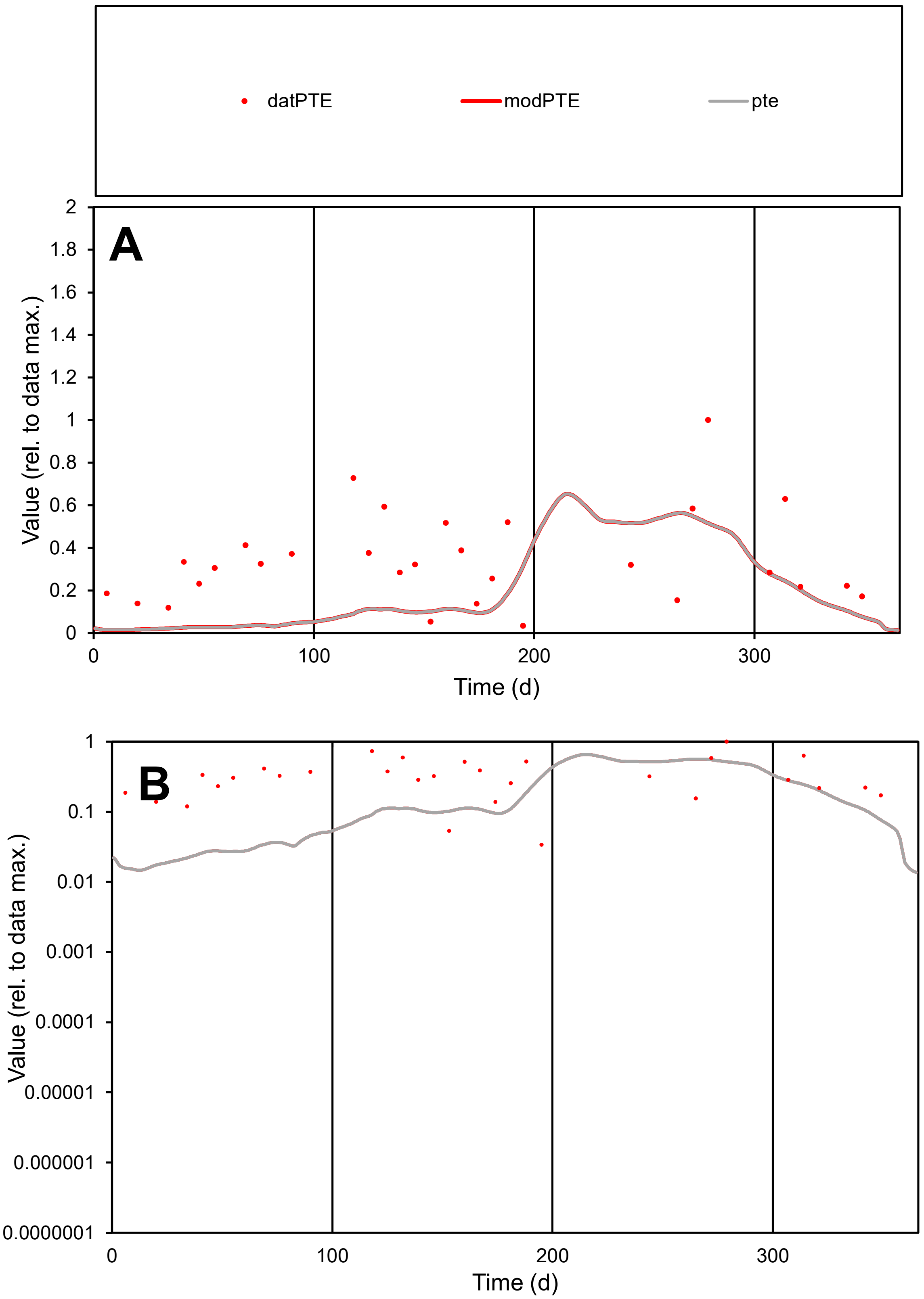


**C1**

Variables
PTE Provide jtd or Dst
CalibOnly?
2 1 to reproduce of
oNormalize?
2 1=no, 2=yes

ofxn calc*
0.83794
ofxn mod
0.84617

sRun
RunA1
zout
1

Ave
Data
0.00022
Model
0.0001

data/vMax
0.00063

ofxng
275.646

Min d/m
2.19739

ave/min d
10.0057
ave/min m
10.4027

325_Calib_S_A

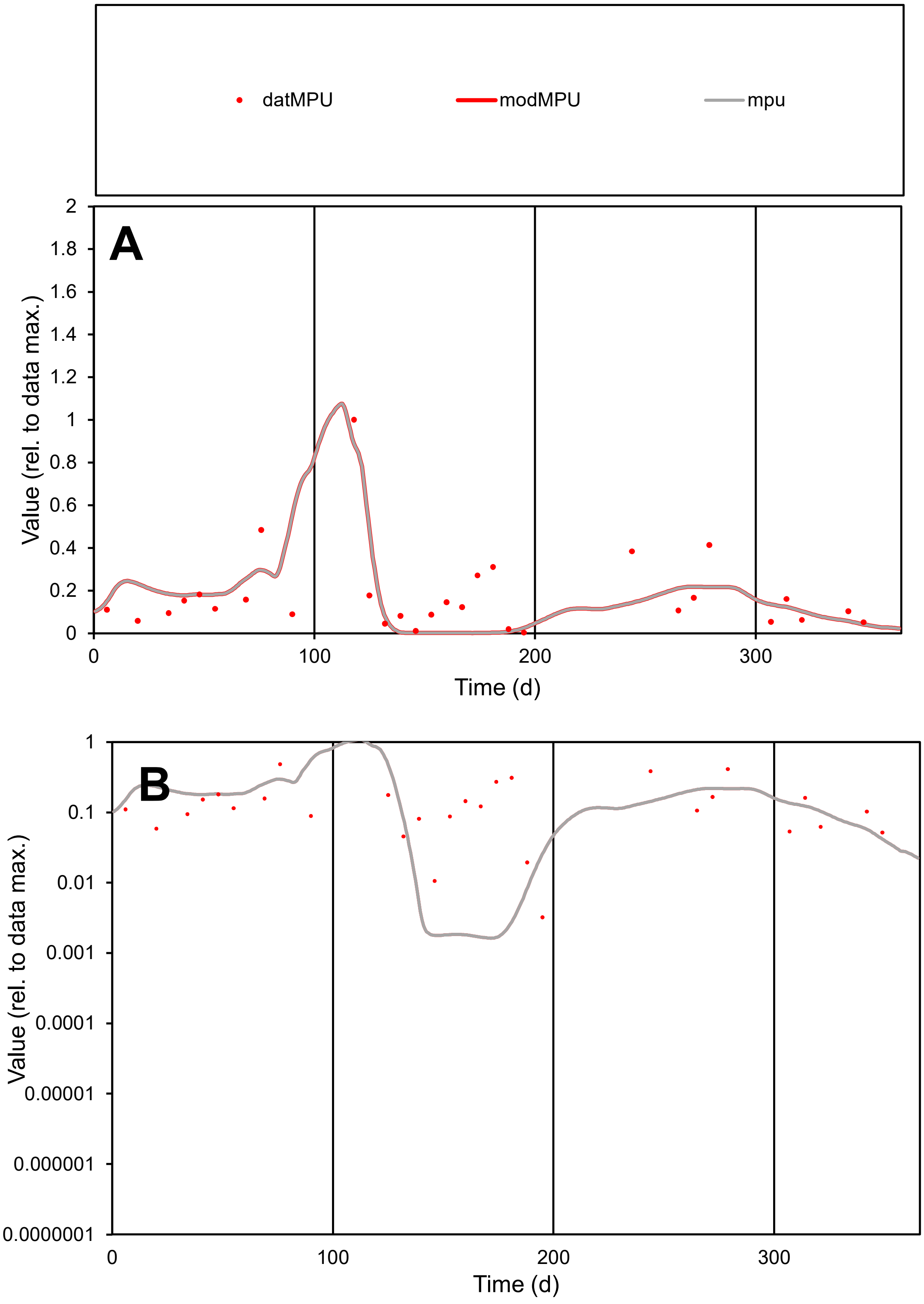


**C1**

Variables
MPU Provide jtd or Dst
CalibOnly?
2 1 to reproduce of
oNormalize?
2 1=no, 2=yes

ofxn calc*
0.81064
ofxn mod
0.88469

sRun
RunA1
zout
1

Ave
Data
0.00022
Model
0.0002

data/vMax
0.00127

ofxng
275.646

Min d/m
1.93191

ave/min d
53.8371
ave/min m
92.437

325_Calib_S_A

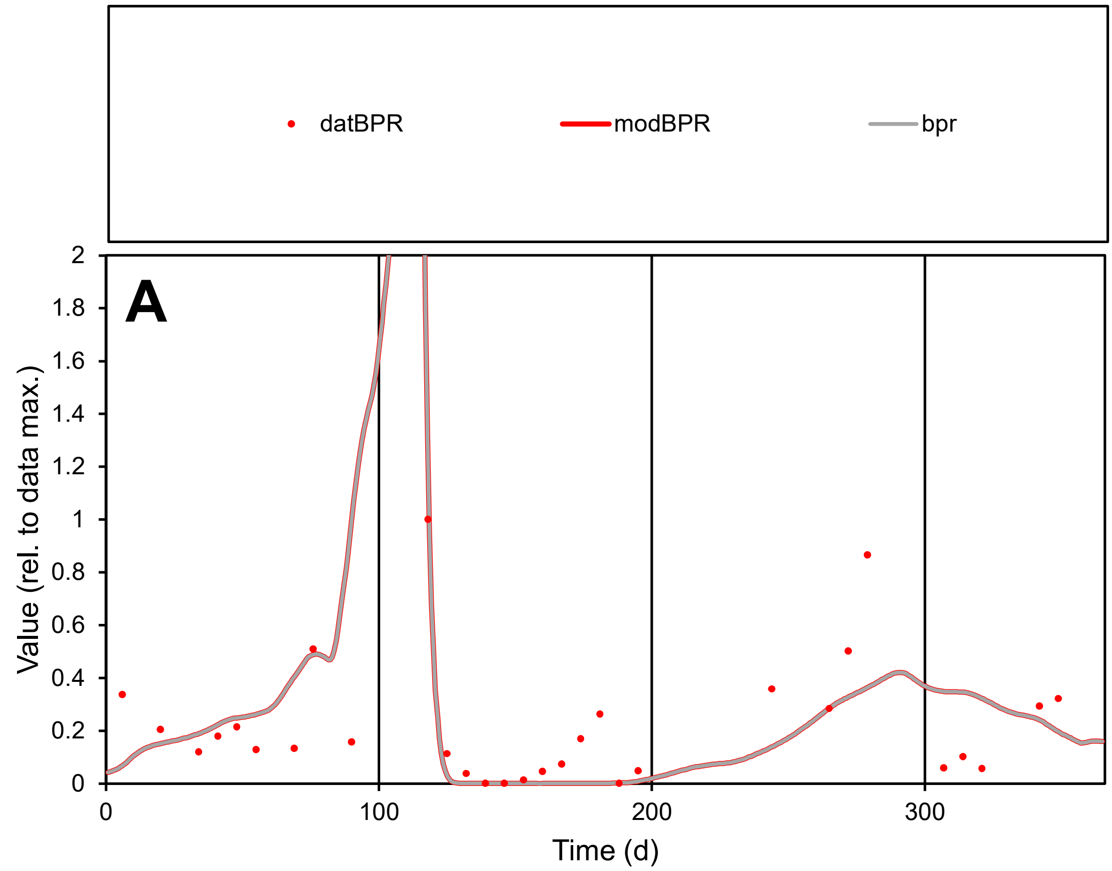


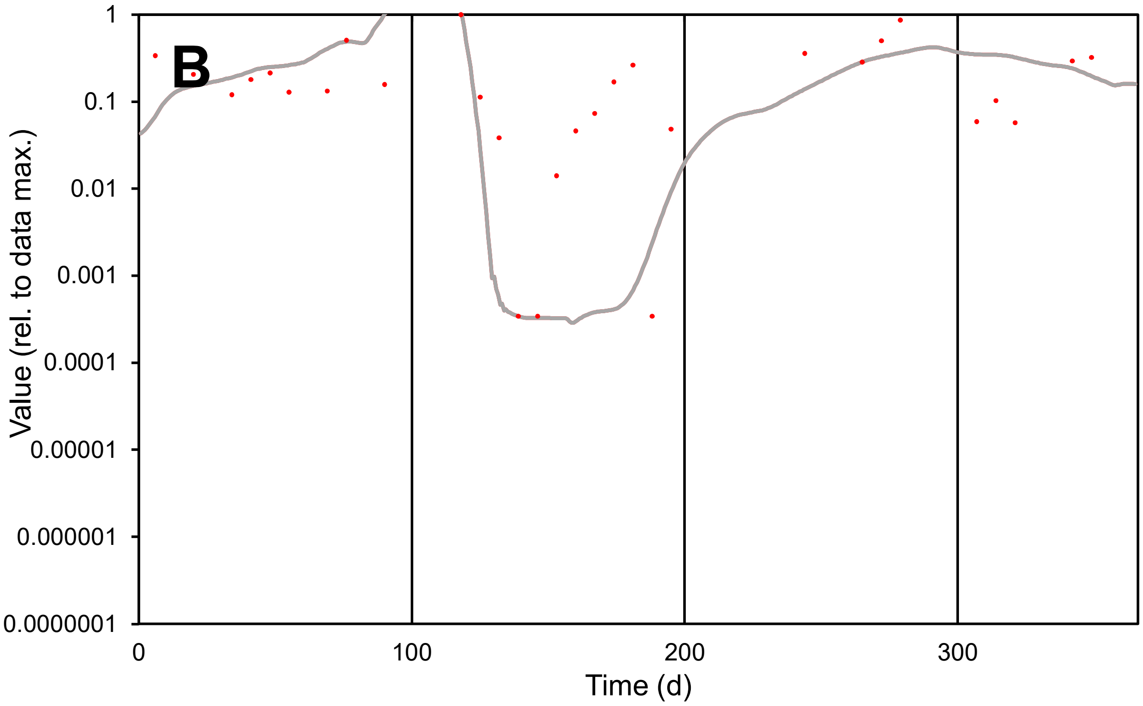


**C1**

Variables
BPR Provide jtd or Dst
CalibOnly?
2 1 to reproduce of
oNormalize?
2 1=no, 2=yes

ofxn calc*
0.9313
ofxn mod
1.03933

sRun
RunA1
zout
1

Ave
Data
0.00012
Model
0.00012

data/vMax
0.00054

ofxng
275.646

Min d/m
1.07483

ave/min d
640.405
ave/min m
675.893

325_Calib_S_A

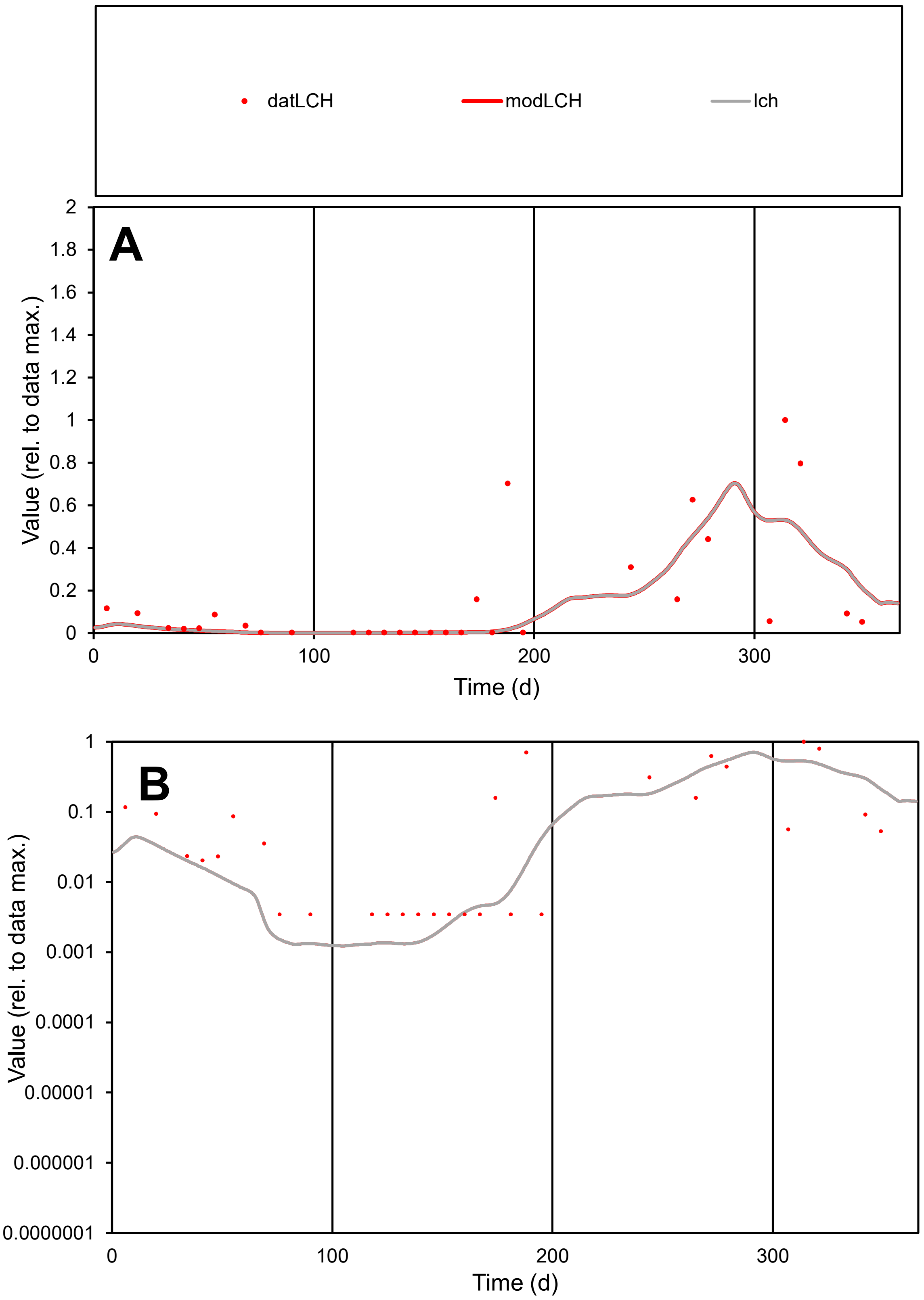


**C1**

Variables

LCH Provide jtd or Dst

CalibOnly?

2 1 to reproduce of

oNormalize?

2 1=no, 2=yes

ofxn calc*

1.24903

ofxn mod

1.2663

sRun

RunA1

zout

1

Ave

Data

0.00011

Model

8.9E-05

data/vMax

0.00071

ofxng

275.646

Min d/m

2.62503

ave/min d

46.5353

ave/min m

95.9419

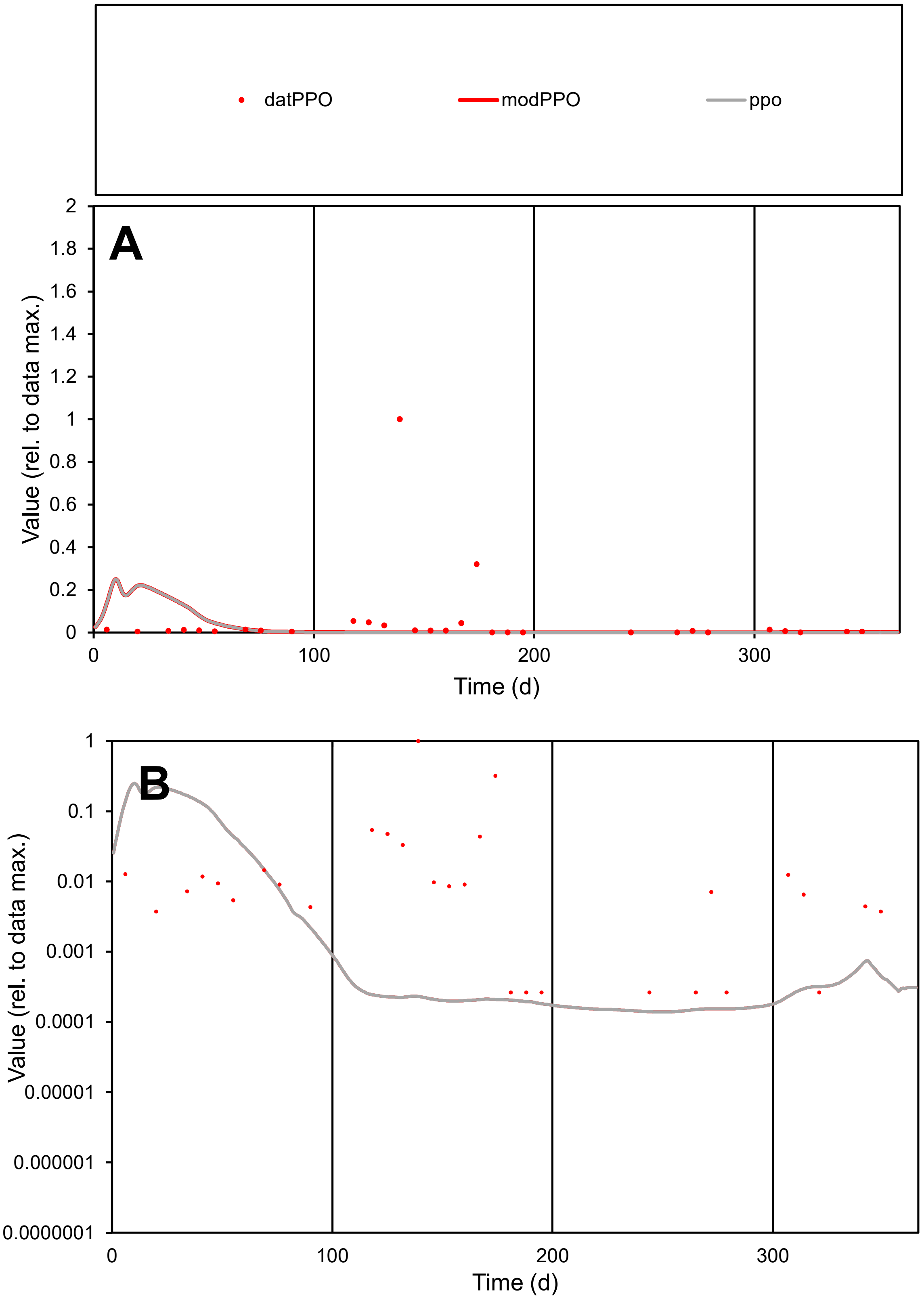


**C1**

Variables
PPO Provide jtd or Dst
CalibOnly?
2 1 to reproduce of
oNormalize?
2 1=no, 2=yes

ofxn calc*
3.65785
ofxn mod
3.67409

sRun
RunA1
zout
1

Ave
Data
0.0004
Model
0.00017

data/vMax
0.00728

ofxng
275.646

Min d/m
1.87549

ave/min d
207.953
ave/min m
167.288

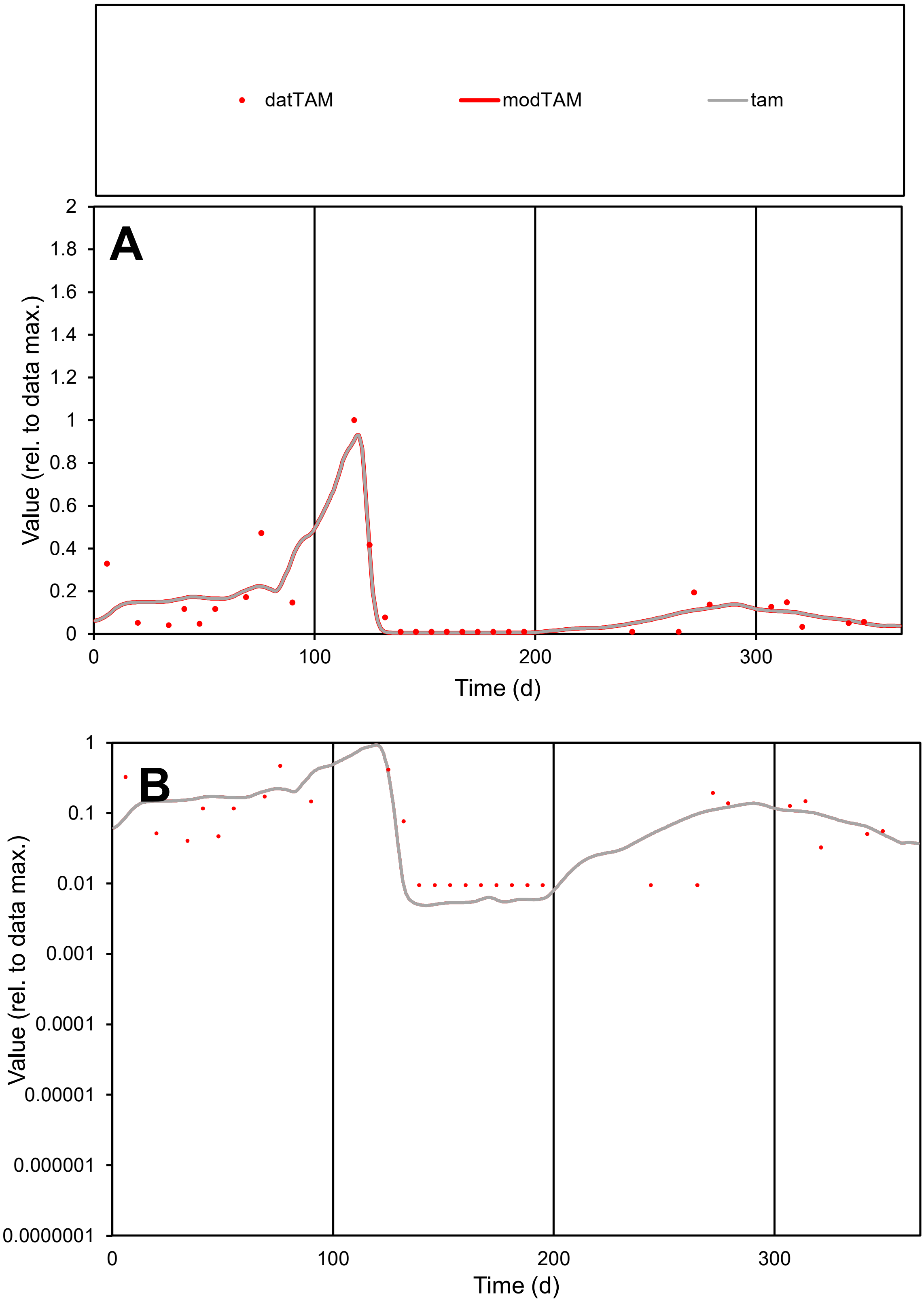


**C1**

Variables
TAM Provide jtd or Dst
CalibOnly?
2 1 to reproduce of
oNormalize?
2 1=no, 2=yes

ofxn calc*
0.69746
ofxn mod
0.73643

sRun
RunA1
zout
1

Ave
Data
7.7E-05
Model
7.8E-05

data/vMax
0.0006

ofxng
275.646

Min d/m
1.9384

ave/min d
13.3954
ave/min m
26.3242

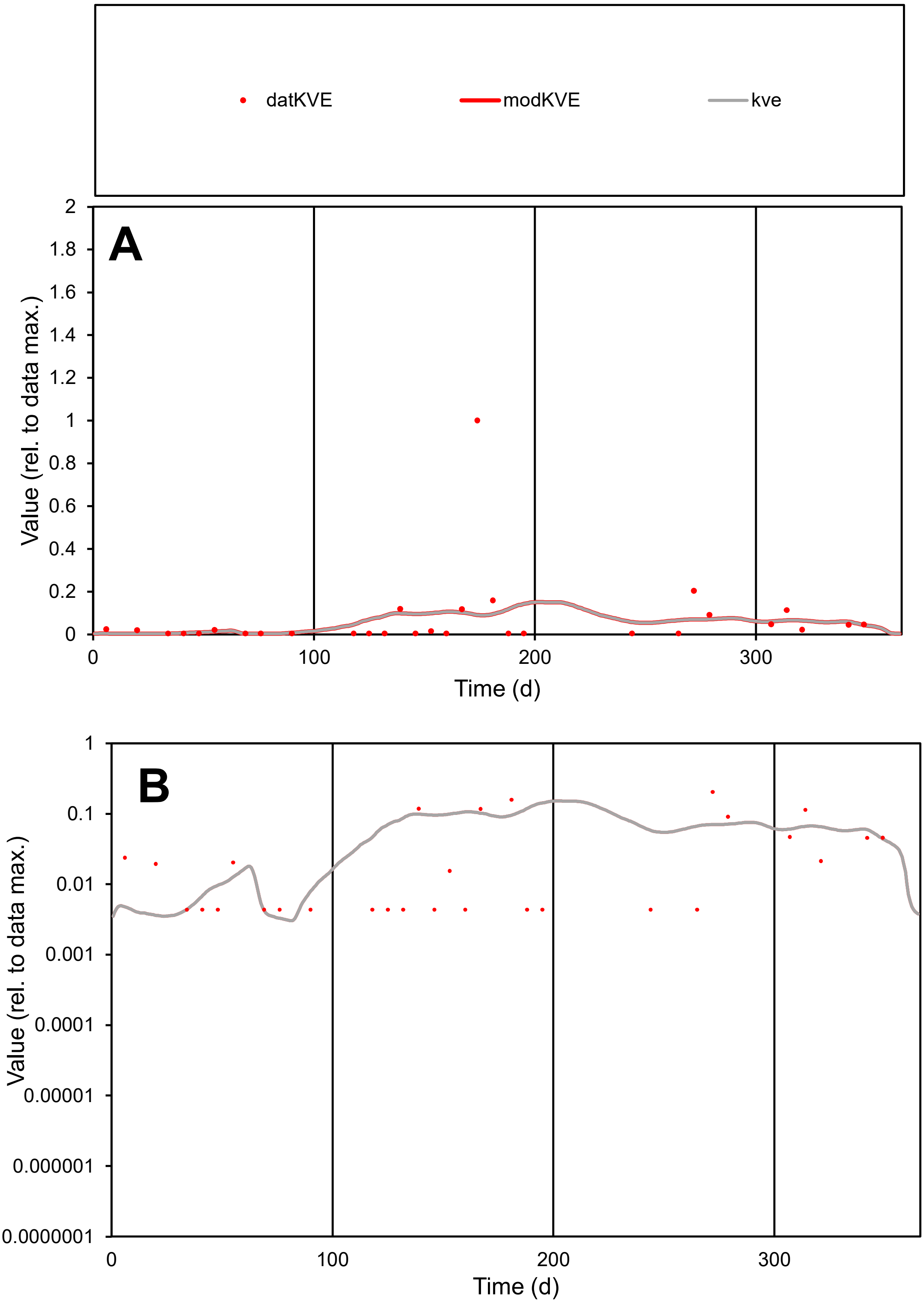


**C1**

Variables
KVE Provide jtd or Dst
CalibOnly?
2 1 to reproduce of
oNormalize?
2 1=no, 2=yes

ofxn calc*
2.50893
ofxn mod
2.51755

sRun
RunA1
zout
1

Ave
Data
5E-05
Model
4.2E-05

data/vMax
0.00072

ofxng
275.646

Min d/m
1.34841

ave/min d
16.1194
ave/min m
18.2245

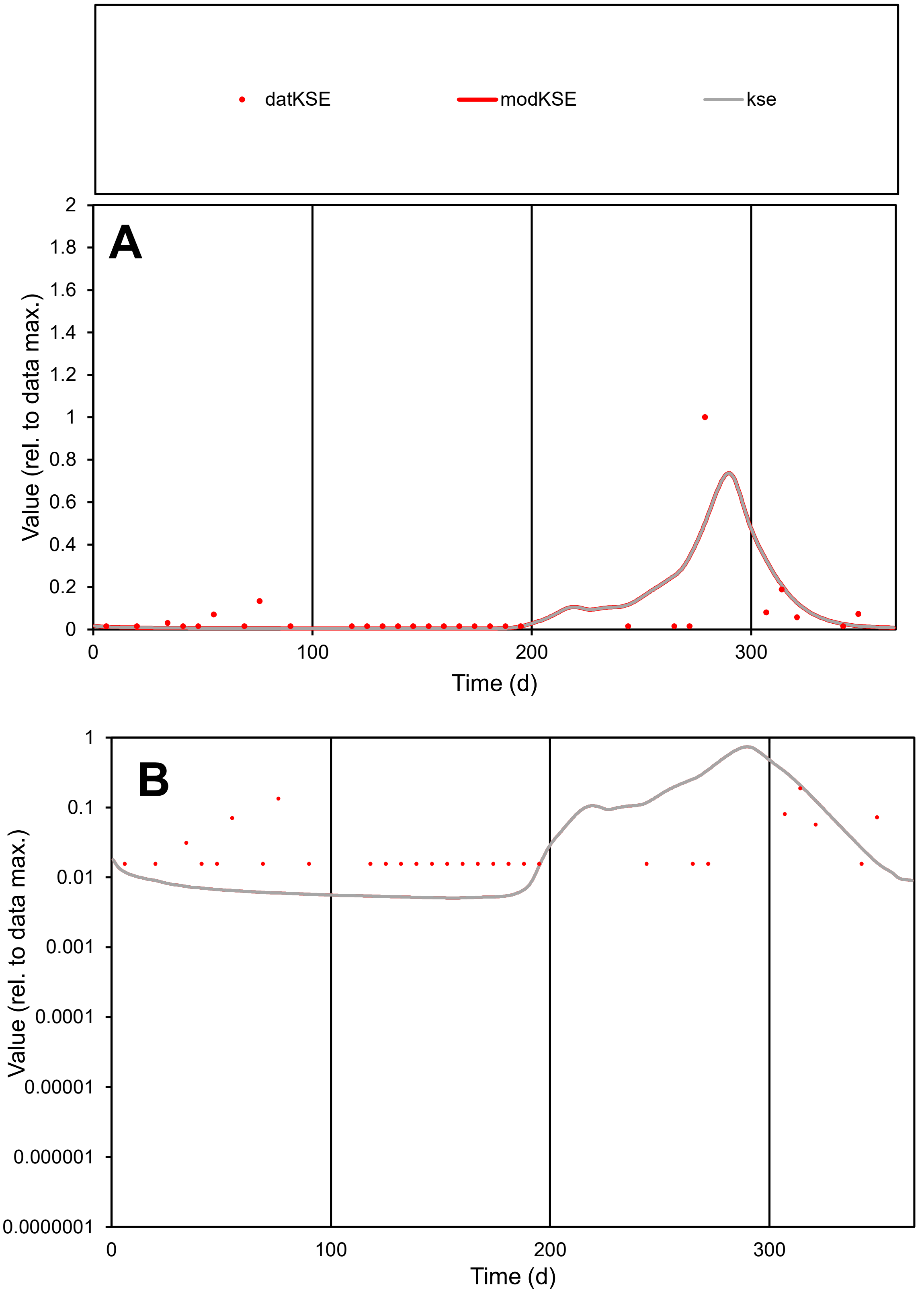


**C1**

Variables
KSE Provide jtd or Dst
CalibOnly?
2 1 to reproduce of
oNormalize?
2 1=no, 2=yes

ofxn calc*
1.84488
ofxn mod
1.90638

sRun
RunA1
zout
1

Ave
Data
1.6E-05
Model
1.6E-05

data/vMax
0.00024

ofxng
275.646

Min d/m
3.0571

ave/min d
4.23445
ave/min m
13.2685

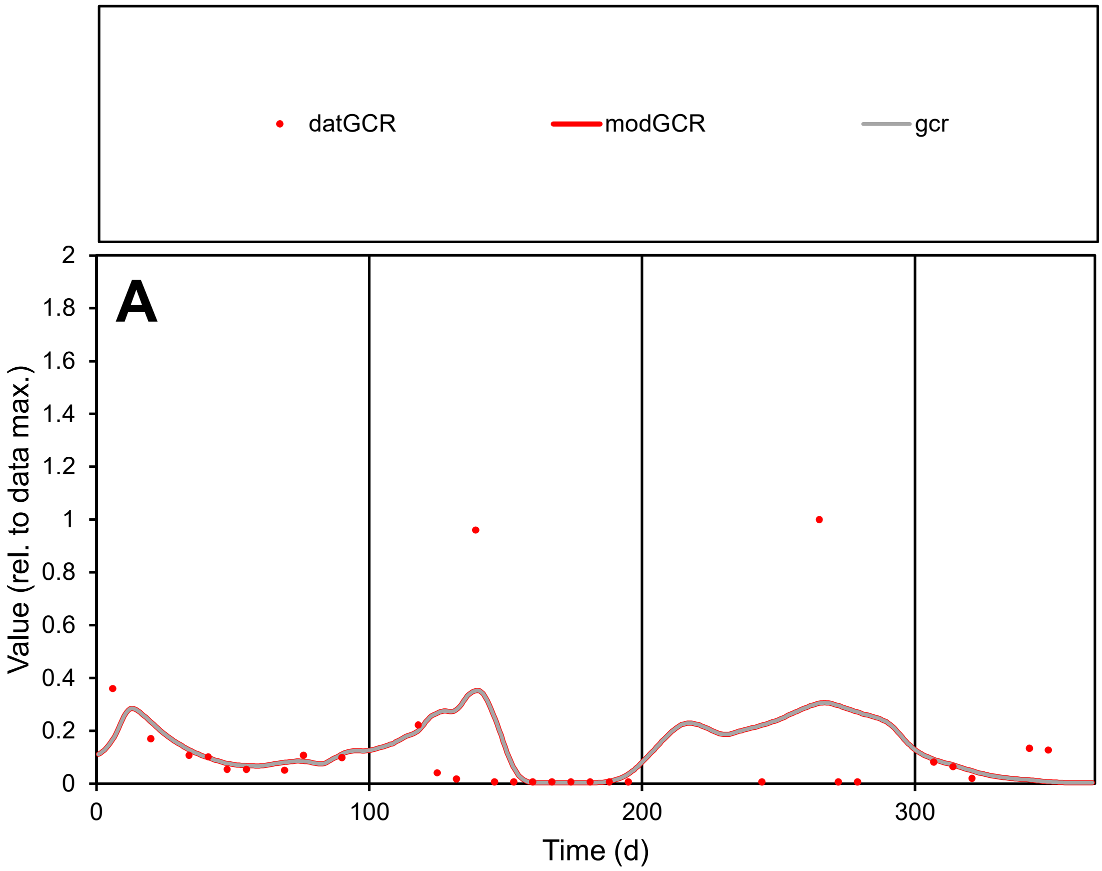


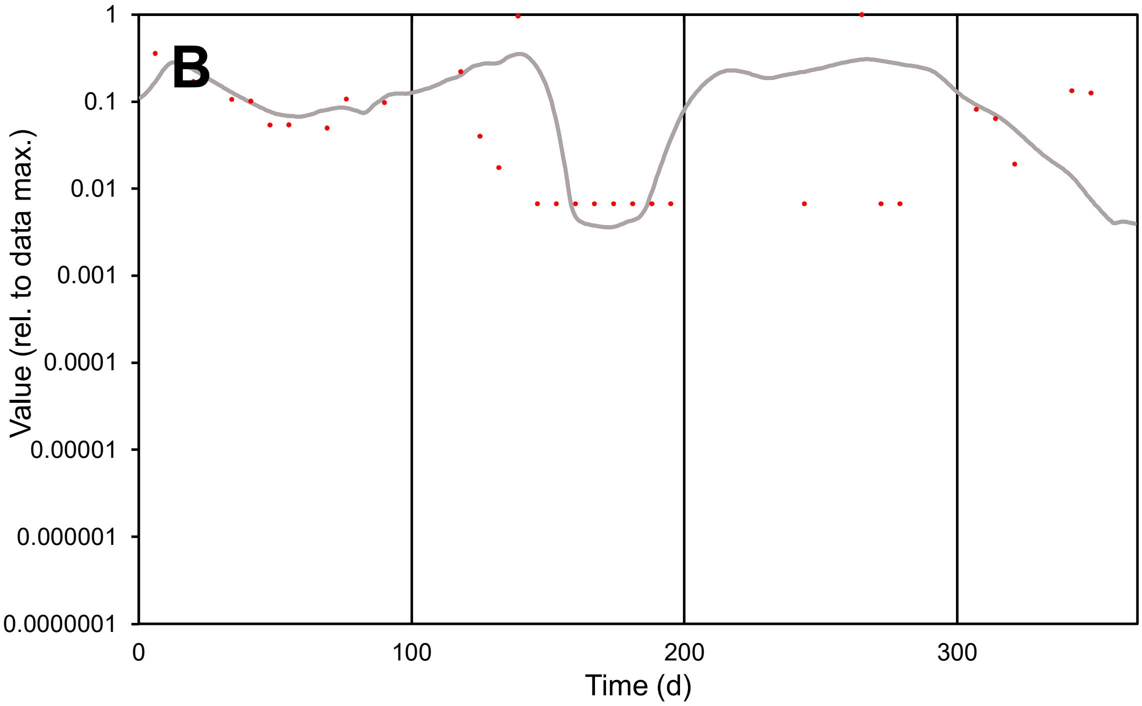


**C1**

Variables
GCR Provide jtd or Dst
CalibOnly?
2 1 to reproduce of
oNormalize?
2 1=no, 2=yes

ofxn calc*
1.59285
ofxn mod
1.61409

sRun
RunA1
zout
1

Ave
Data
6E-05
Model
5.7E-05

data/vMax
0.00047

ofxng
275.646

Min d/m
1.80634

ave/min d
19.0111
ave/min m
32.8563

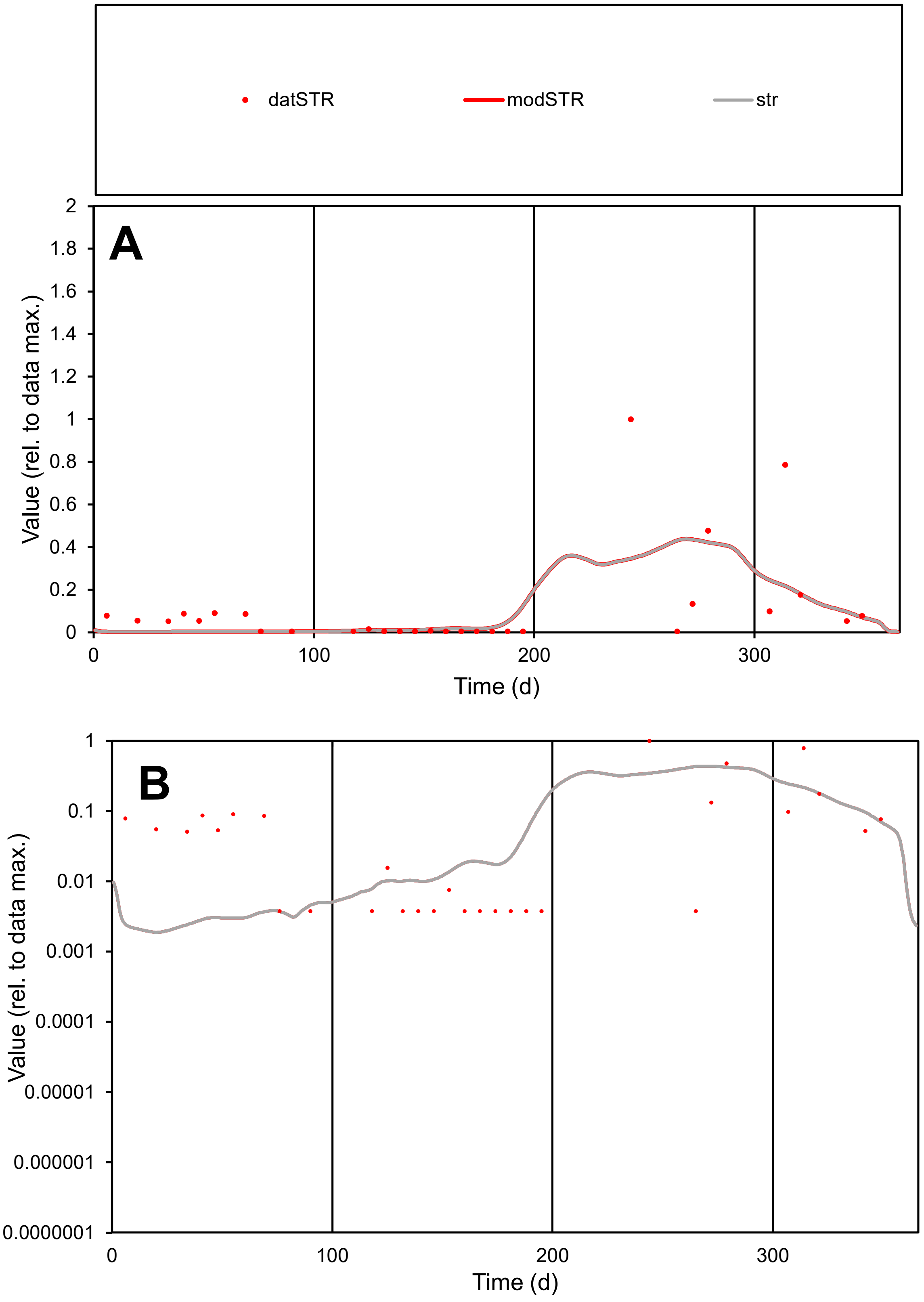


**C1**

Variables
STR Provide jtd or Dst
CalibOnly?
2 1 to reproduce of
oNormalize?
2 1=no, 2=yes

ofxn calc*
1.70596
ofxn mod
1.72061

sRun
RunA1
zout
1

Ave
Data
4.9E-05
Model
3.9E-05

data/vMax
0.00043

ofxng
275.646

Min d/m
2.00443

ave/min d
29.7746
ave/min m
48.2752

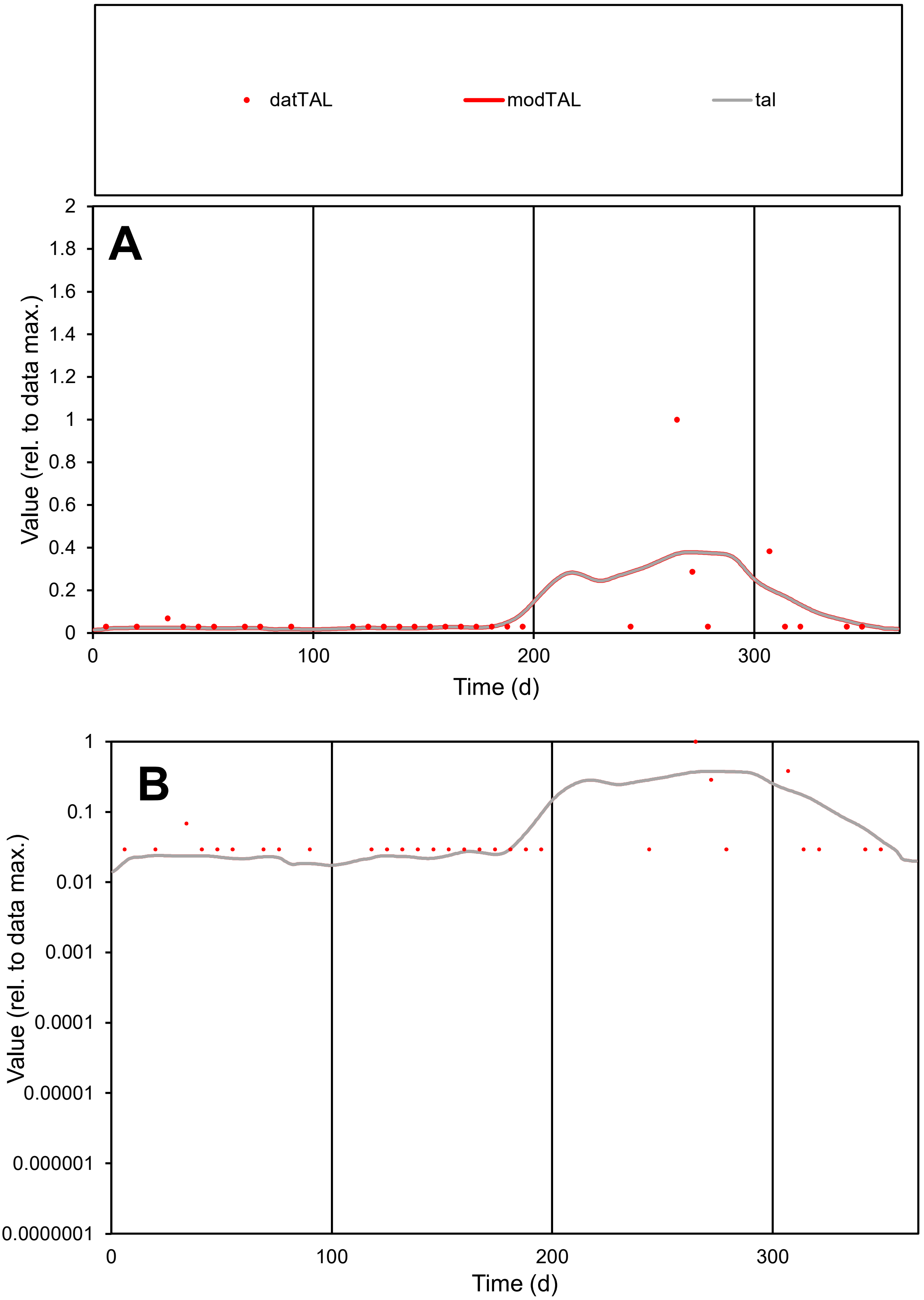


**C1**

Variables
TAL Provide jtd or Dst
CalibOnly?
2 1 to reproduce of
oNormalize?
2 1=no, 2=yes

ofxn calc*
1.76604
ofxn mod
1.78468

sRun
RunA1
zout
1

Ave
Data
1.2E-05
Model
1.2E-05

data/vMax
0.00014

ofxng
275.646

Min d/m
1.60682

ave/min d
2.83606
ave/min m
4.69131

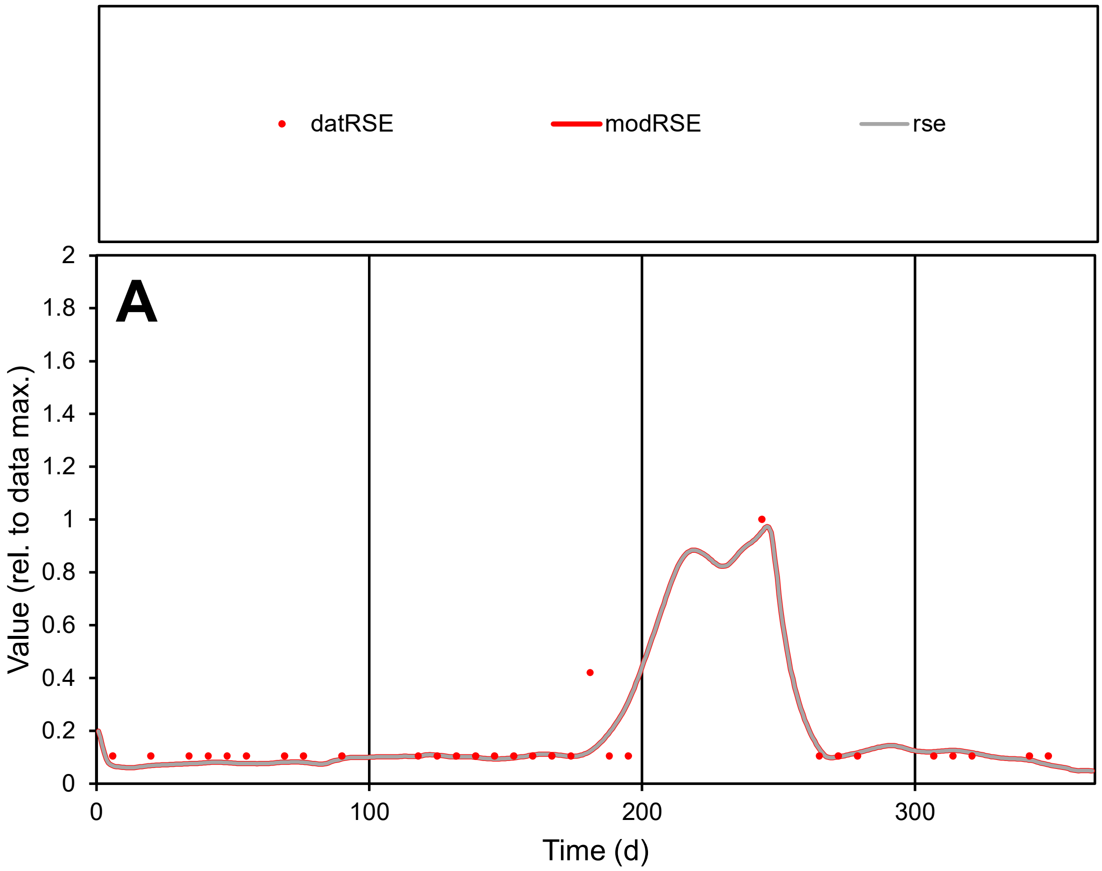


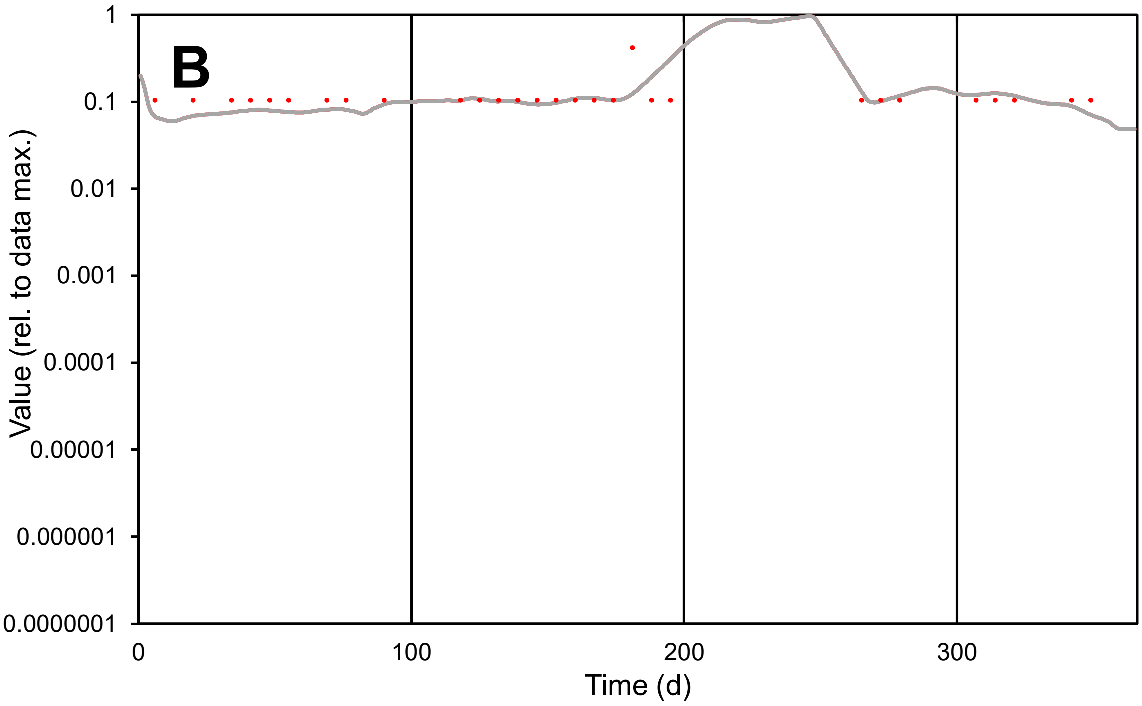


**C1**

Variables
RSE Provide jtd or Dst
CalibOnly?
2 1 to reproduce of
oNormalize?
2 1=no, 2=yes

ofxn calc*
0.46758
ofxn mod
0.50424

sRun
RunA1
zout
1

Ave
Data
6.5E-06
Model
6.1E-06

data/vMax
4.5E-05

ofxng
275.646

Min d/m
1.60904

ave/min d
1.38712
ave/min m
2.10023

325_Calib_S_A

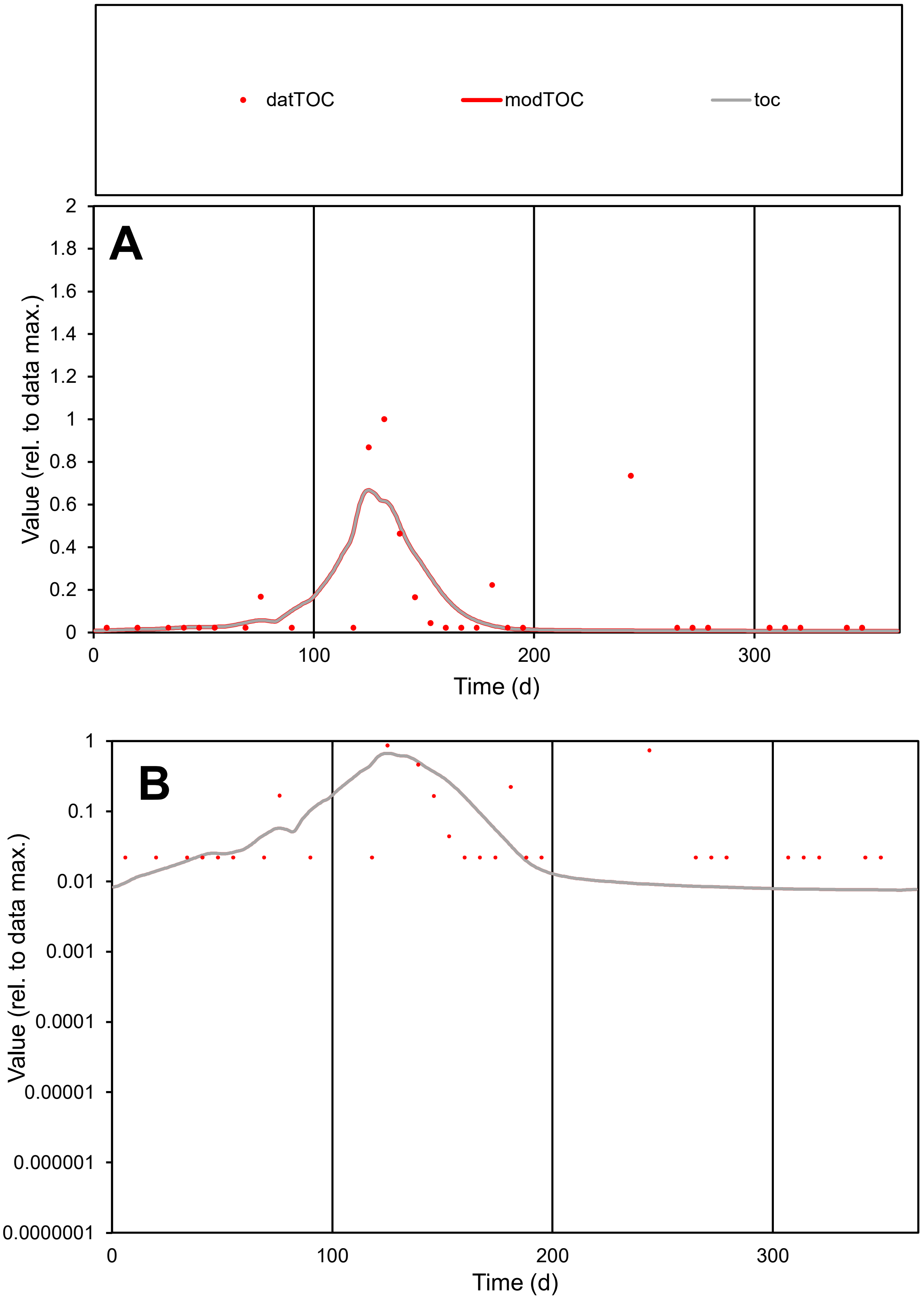


**C1**

Variables

TOC Provide jtd or Dst

CalibOnly?

2 1 to reproduce of

oNormalize?

2 1=no, 2=yes

ofxn calc*

1.34145

ofxn mod

1.36939

sRun

RunA1

zout

1

Ave

Data

2.7E-05

Model

2.3E-05

data/vMax

0.0002

ofxng

275.646

Min d/m

2.90262

ave/min d

6.28357

ave/min m

15.3168

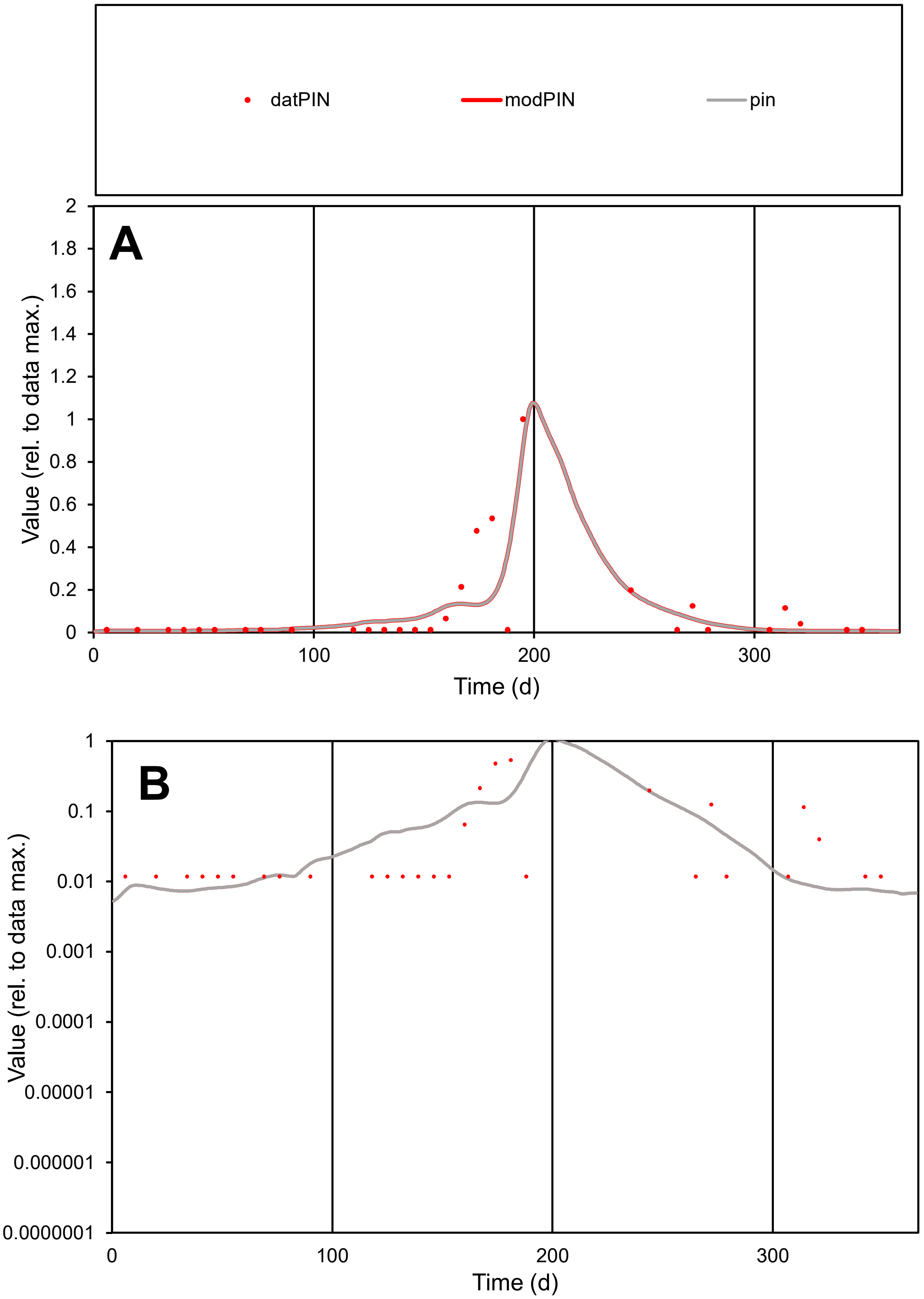


**C1**

Variables
PIN Provide jtd or Dst
CalibOnly?
2 1 to reproduce of
oNormalize?
2 1=no, 2=yes

ofxn calc*
1.0934
ofxn mod
1.16036

sRun
RunA1
zout
1

Ave
Data
3.4E-05
Model
3E-05

data/vMax
0.00034

ofxng
275.646

Min d/m
1.61955

ave/min d
8.52934
ave/min m
12.2901

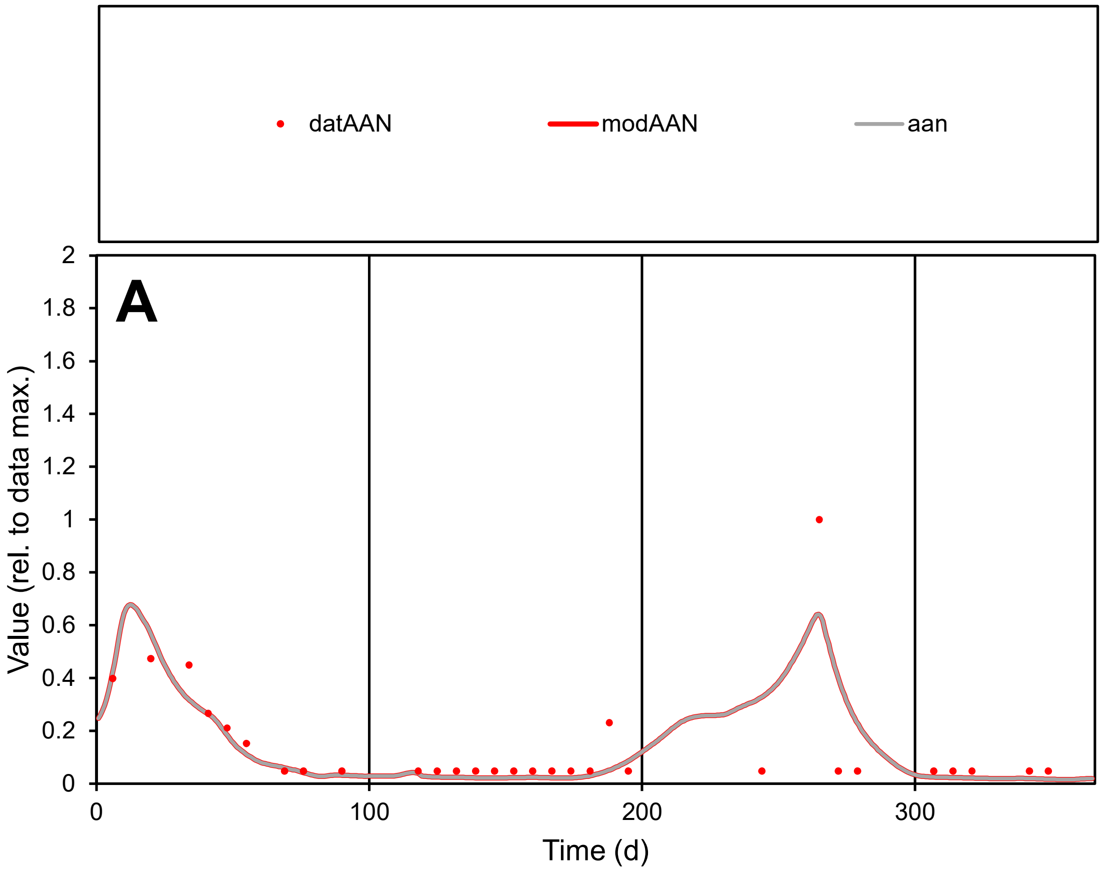


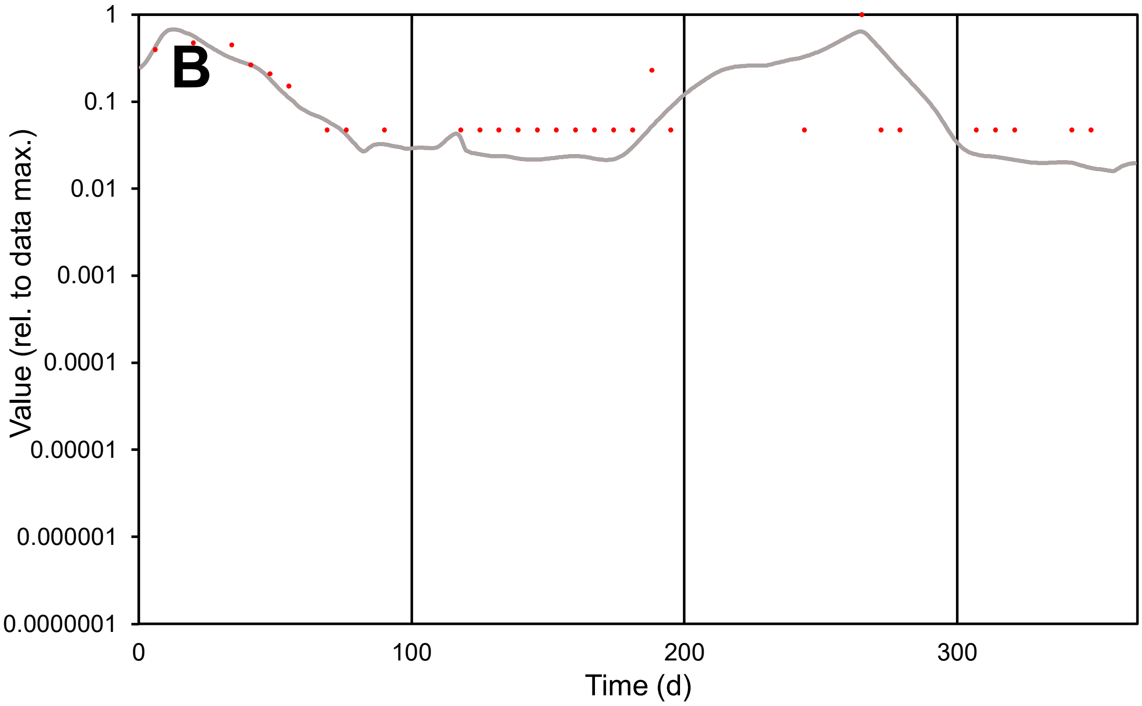


**C1**

Variables
AAN Provide jtd or Dst
CalibOnly?
2 1 to reproduce of
oNormalize?
2 1=no, 2=yes

ofxn calc*
0.78978
ofxn mod
0.84897

sRun
RunA1
zout
1

Ave
Data
1.7E-05
Model
1.6E-05

data/vMax
0.00012

ofxng
275.646

Min d/m
2.76824

ave/min d
2.98312
ave/min m
7.72073

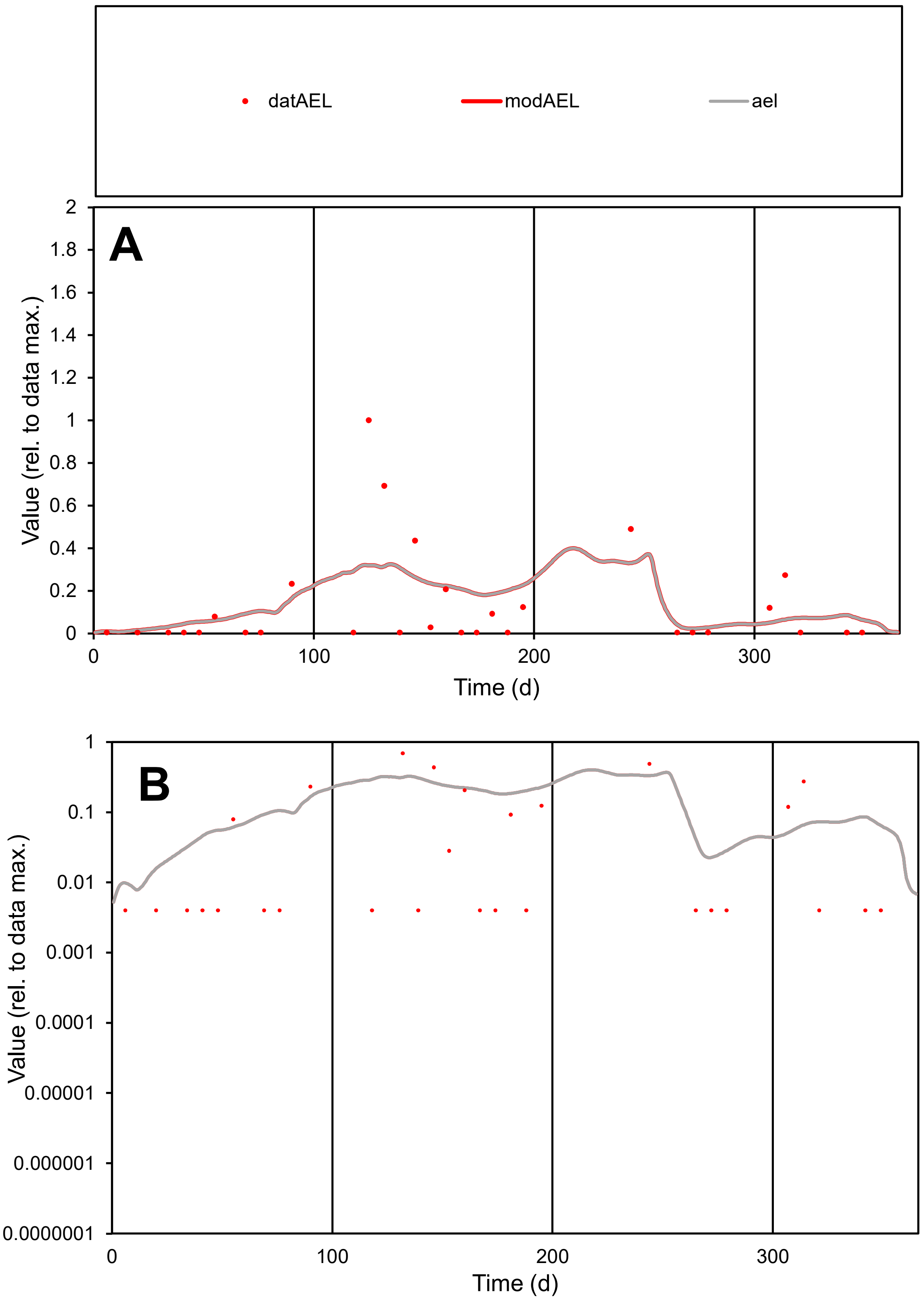


**C1**

Variables
AEL Provide jtd or Dst
CalibOnly?
2 1 to reproduce of
oNormalize?
2 1=no, 2=yes

ofxn calc*
1.48923
ofxn mod
1.50066

sRun
RunA1
zout
1

Ave
Data
2.2E-05
Model
2.5E-05

data/vMax
0.00017

ofxng
275.646

Min d/m
0.41987

ave/min d
32.0439
ave/min m
15.0475

325_Calib_S_A

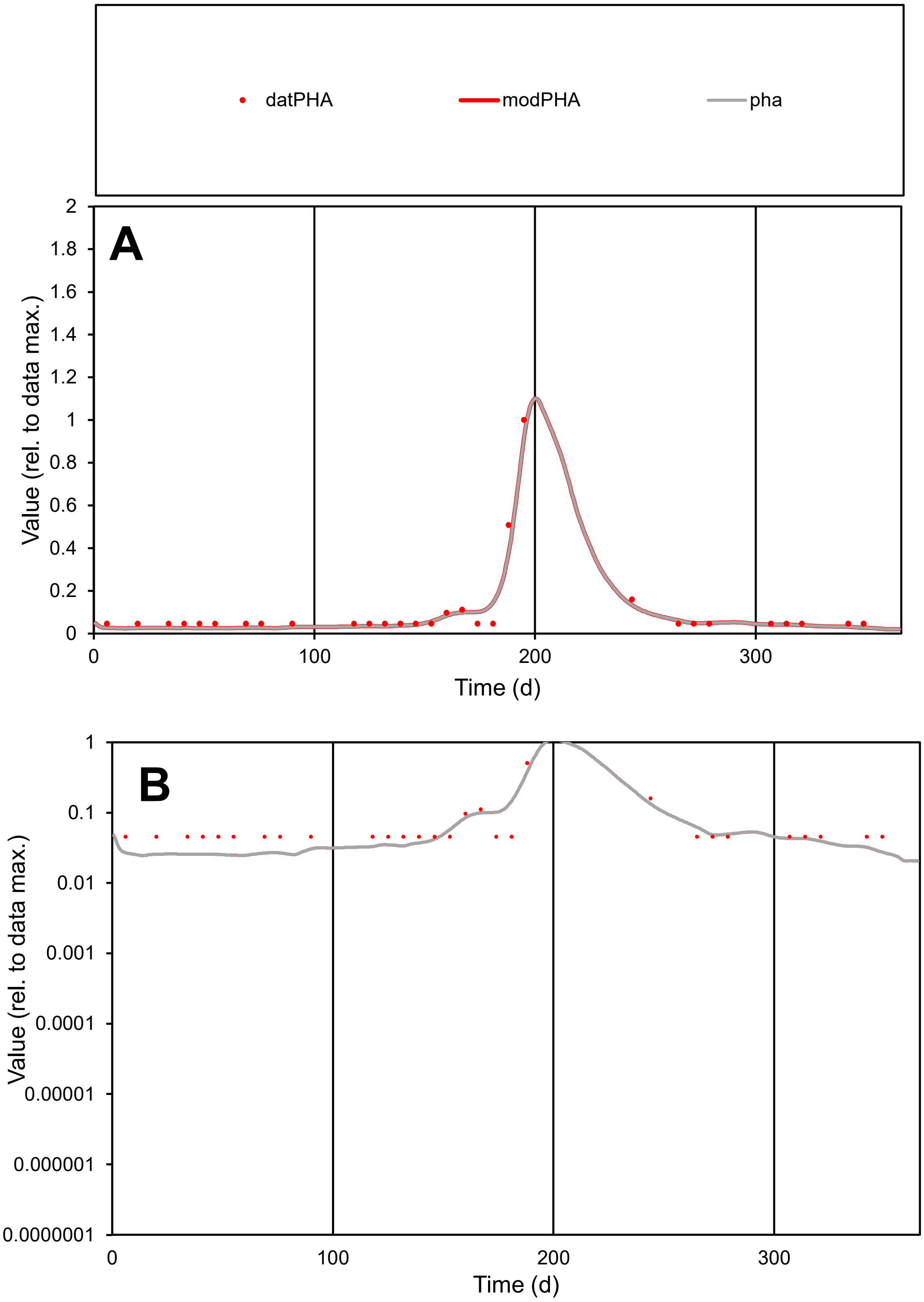


**C1**

Variables
PHA Provide jtd or Dst
CalibOnly?
2 1 to reproduce of
oNormalize?
2 1=no, 2=yes

ofxn calc*
0.26914
ofxn mod
0.37184

sRun
RunA1
zout
1

Ave
Data
2.3E-05
Model
2.2E-05

data/vMax
0.00023

ofxng
275.646

Min d/m
1.83987

ave/min d
2.2074
ave/min m
3.80596

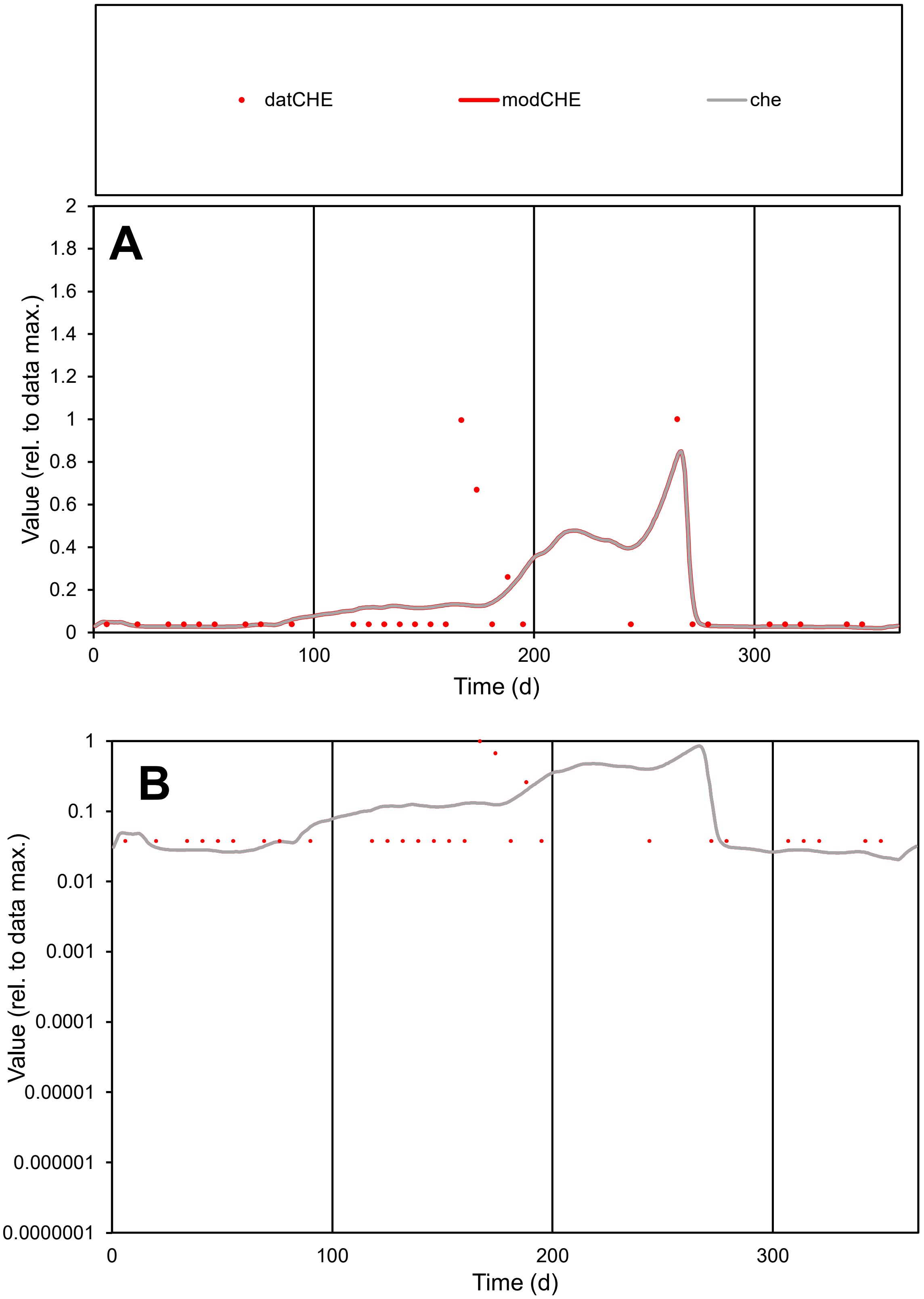


C1

Variables
CHE Provide jtd or Dst
CalibOnly?
2 1 to reproduce of
oNormalize?
2 1=no, 2=yes

ofxn calc*
1.60793
ofxn mod
1.62126

sRun
RunA1
zout
1

Ave
Data
2.4E-05
Model
2.2E-05

data/vMax
0.00019

ofxng
275.646

Min d/m
1.71932

ave/min d
3.44606
ave/min m
5.29625

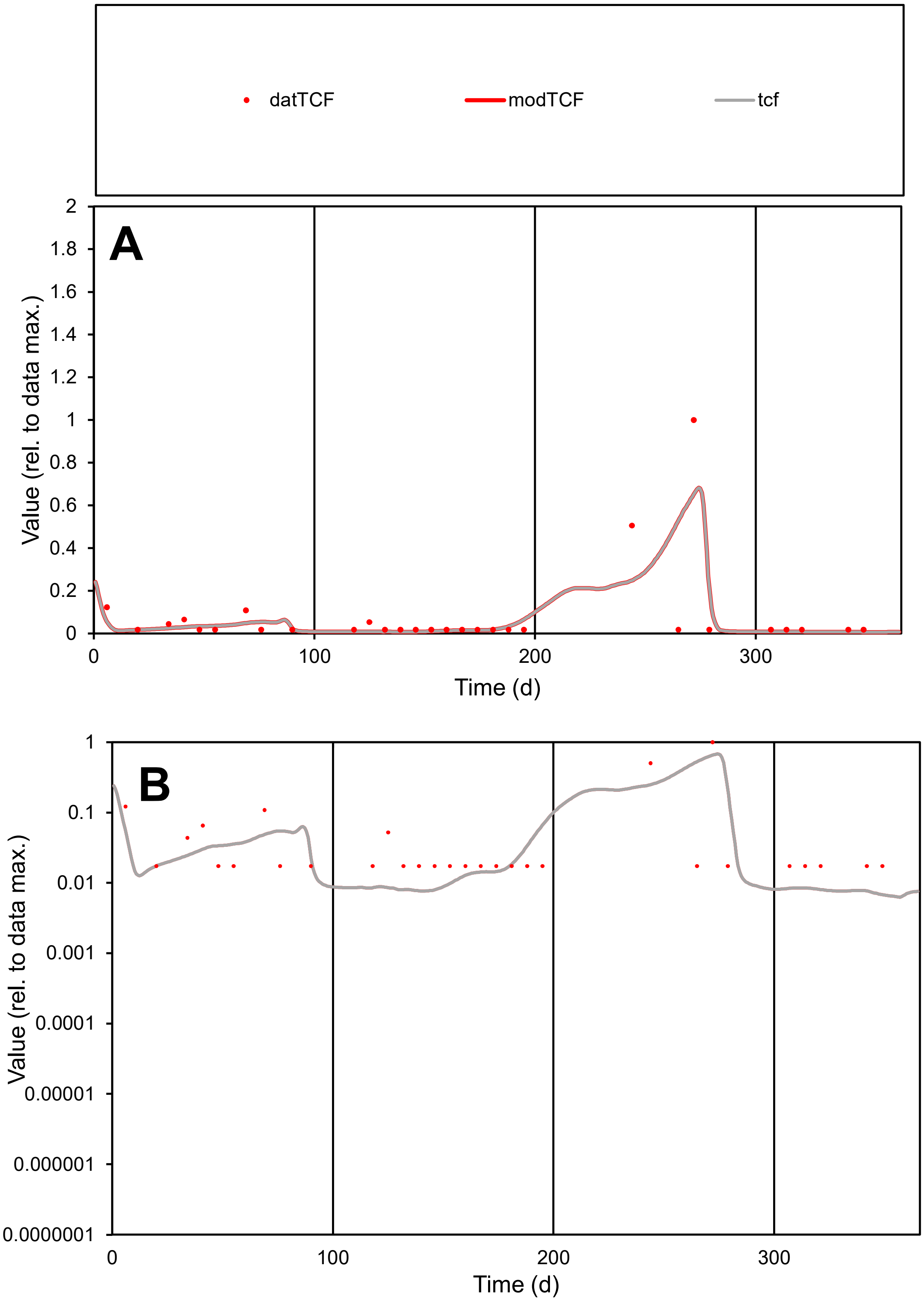


**C1**

Variables
TCF Provide jtd or Dst
CalibOnly?
2 1 to reproduce of
oNormalize?
2 1=no, 2=yes

ofxn calc*
1.58362
ofxn mod
1.64615

sRun
RunA1
zout
1

Ave
Data
1.4E-05
Model
1.3E-05

data/vMax
0.00018

ofxng
275.646

Min d/m
2.59917

ave/min d
4.41291
ave/min m
10.4548

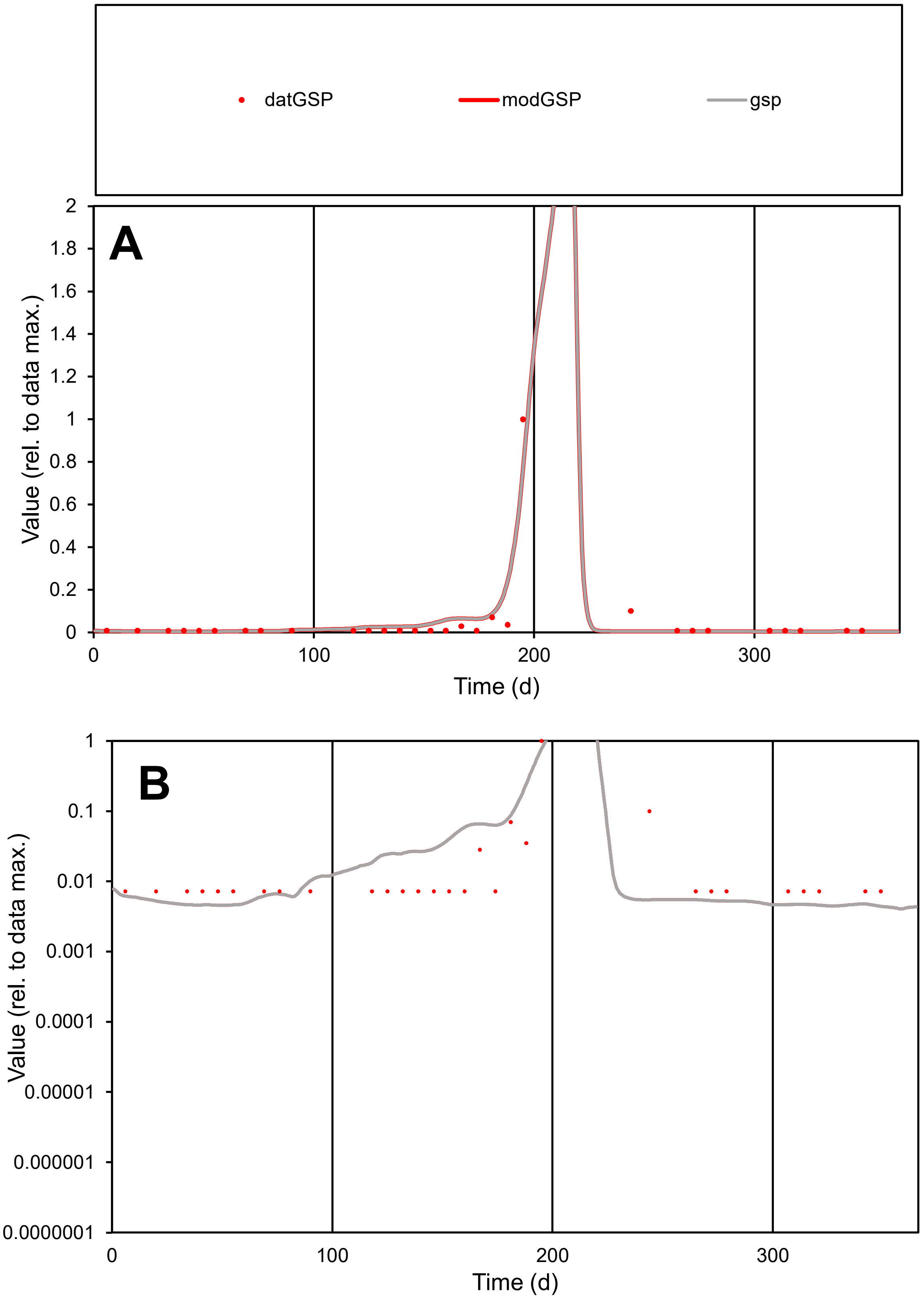


**C1**

Variables
GSP Provide jtd or Dst
CalibOnly?
2 1 to reproduce of
oNormalize?
2 1=no, 2=yes

ofxn calc*
0.81977
ofxn mod
1.04652

sRun
RunA1
zout
1

Ave
Data
5.9E-05
Model
6.8E-05

data/vMax
0.00125

ofxng
275.646

Min d/m
1.64961
ave/min d
6.53853
ave/min m
12.5244

325_Calib_S_A

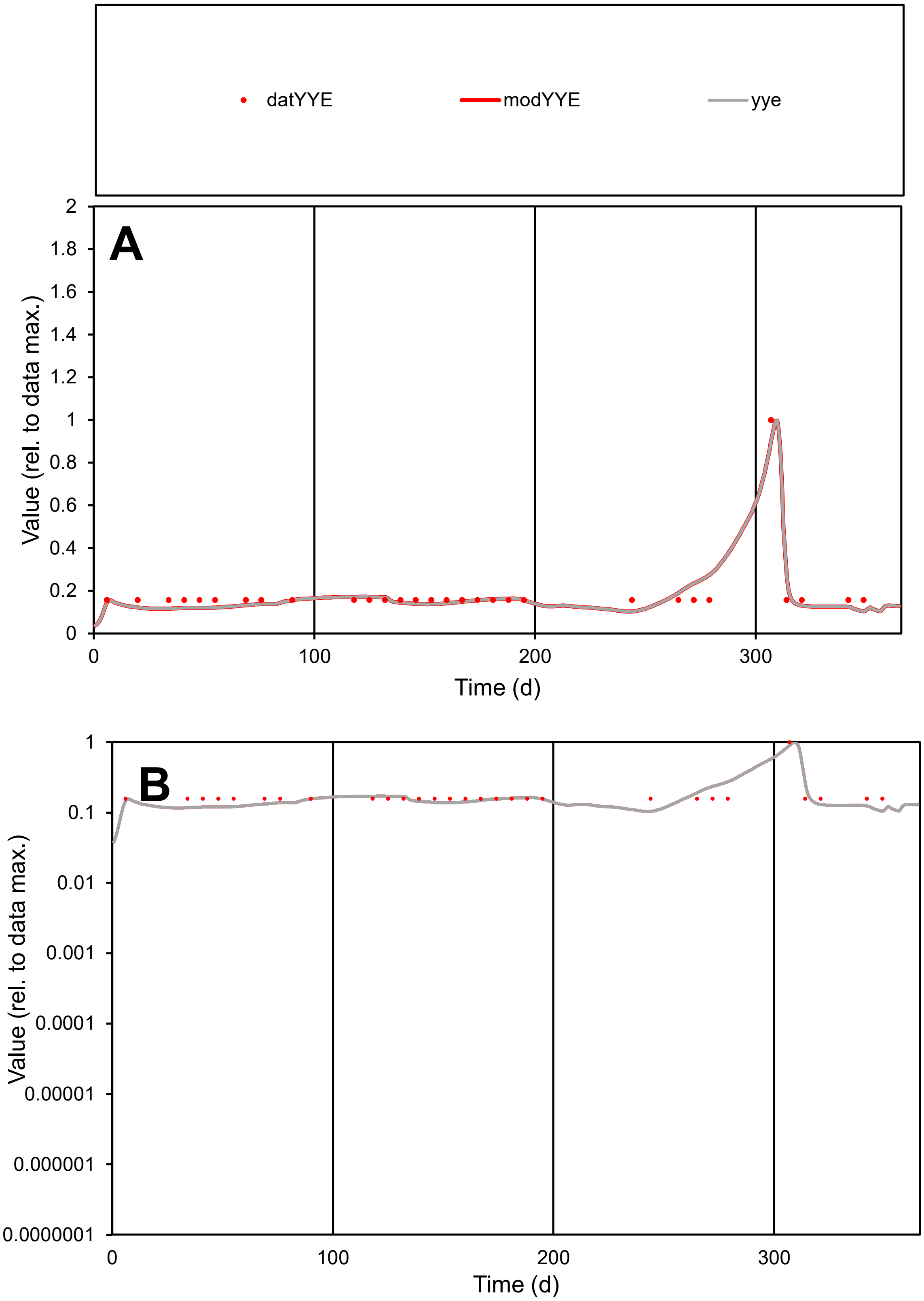


**C1**

Variables
YYE Provide jtd or Dst
CalibOnly?
2 1 to reproduce of
oNormalize?
2 1=no, 2=yes

ofxn calc*
0.19548
ofxn mod
0.24834

sRun
RunA1
zout
1

Ave
Data
4.4E-06
Model
4.3E-06

data/vMax
2.4E-05

ofxng
275.646

Min d/m
1.49325

ave/min d
1.17909
ave/min m
1.70087

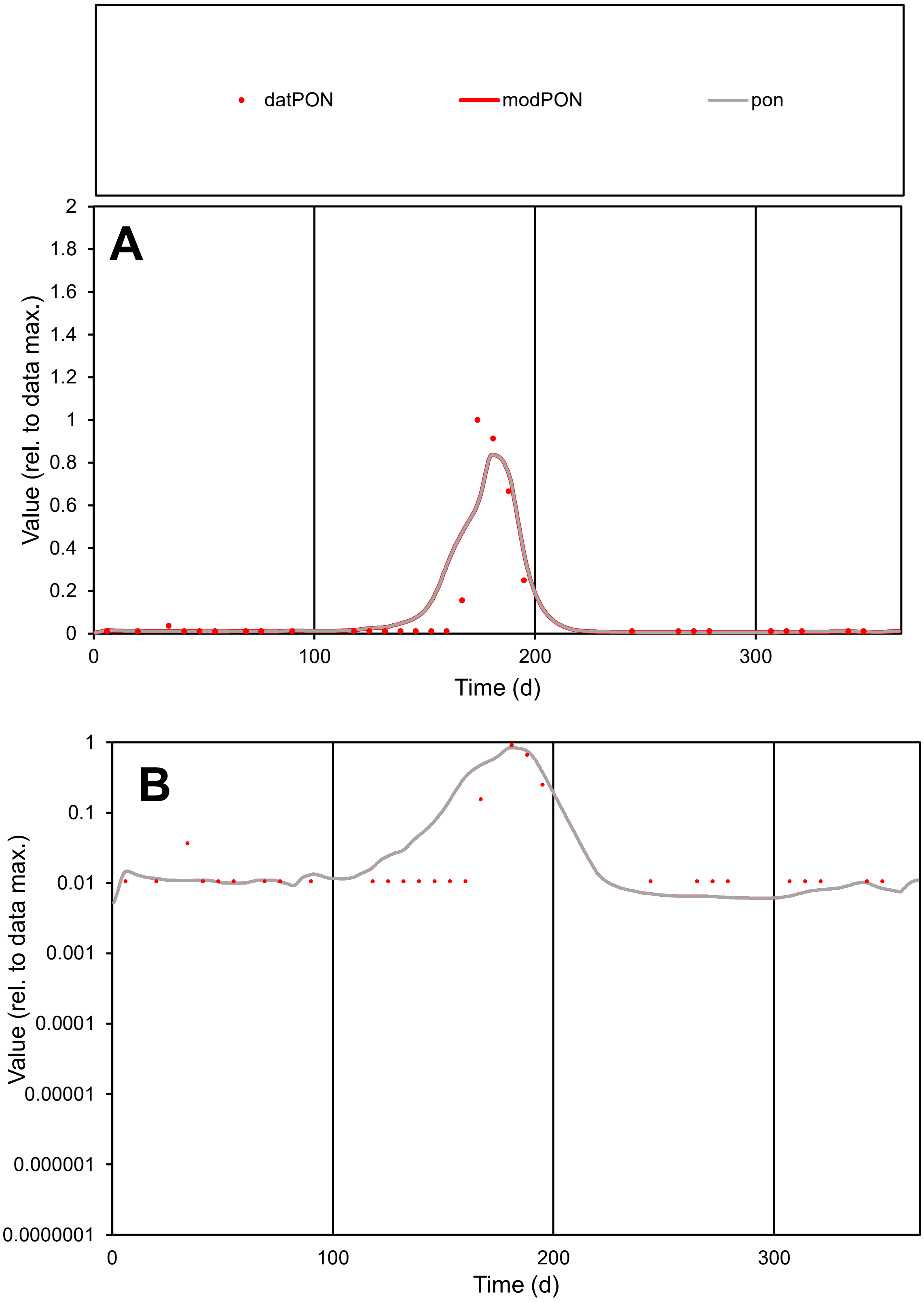


**C1**

Variables

PON Provide jtd or Dst

CalibOnly?

2 1 to reproduce of

oNormalize?

2 1=no, 2=yes

ofxn calc*

0.93724

ofxn mod

1.03099

sRun

RunA1

zout

1

Ave

Data

3E-05

Model

3.4E-05

data/vMax

0.00028

ofxng

275.646

Min d/m

1.70087

ave/min d

10.3424

ave/min m

19.9324

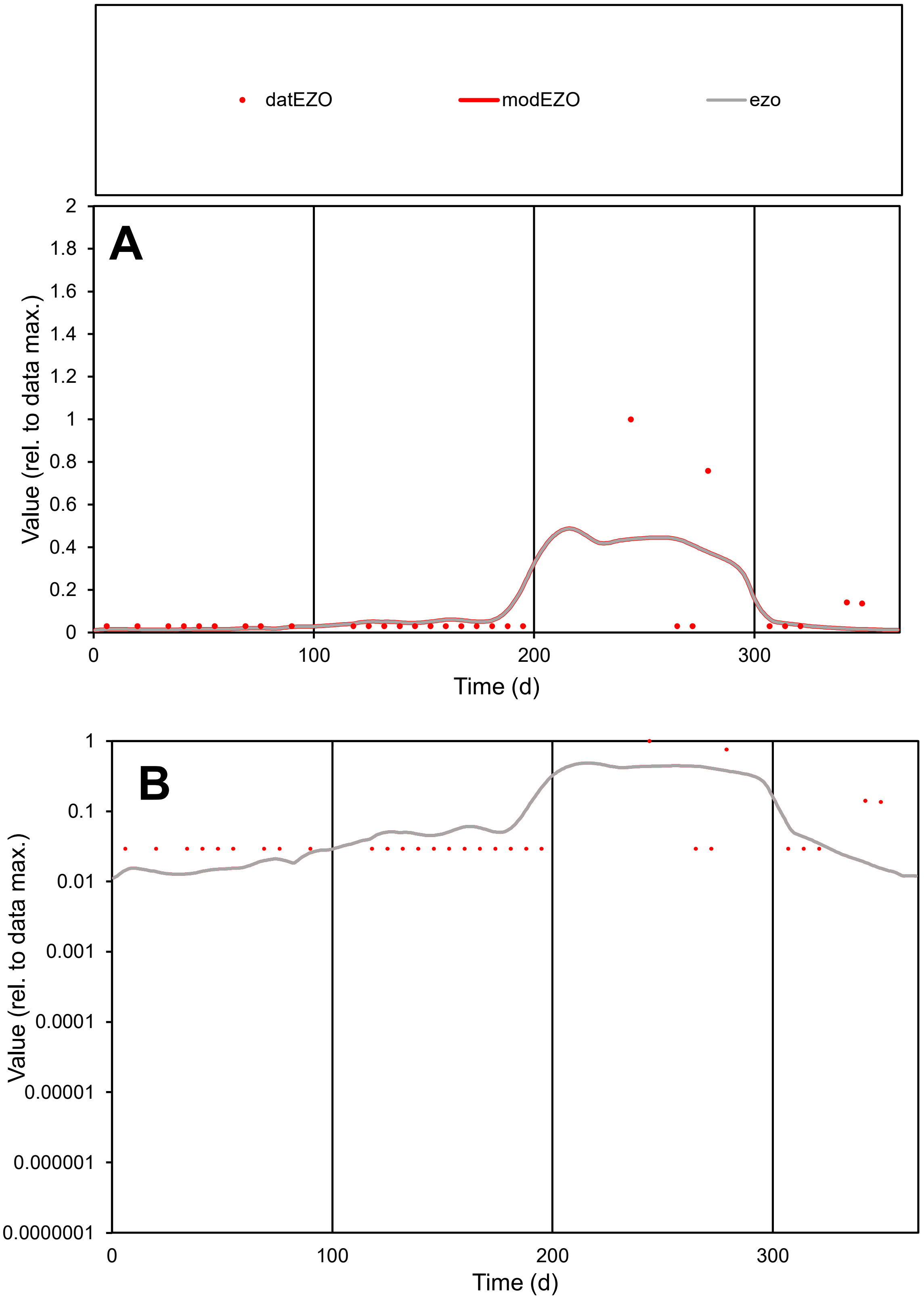


**C1**

Variables
EZO Provide jtd or Dst
CalibOnly?
2 1 to reproduce of
oNormalize?
2 1=no, 2=yes

ofxn calc*
1.79345
ofxn mod
1.81787

sRun
RunA1
zout
1

Ave
Data
1.3E-05
Model
1.3E-05

data/vMax
0.00014

ofxng
275.646

Min d/m
2.25057

ave/min d
3.19045
ave/min m
7.15732

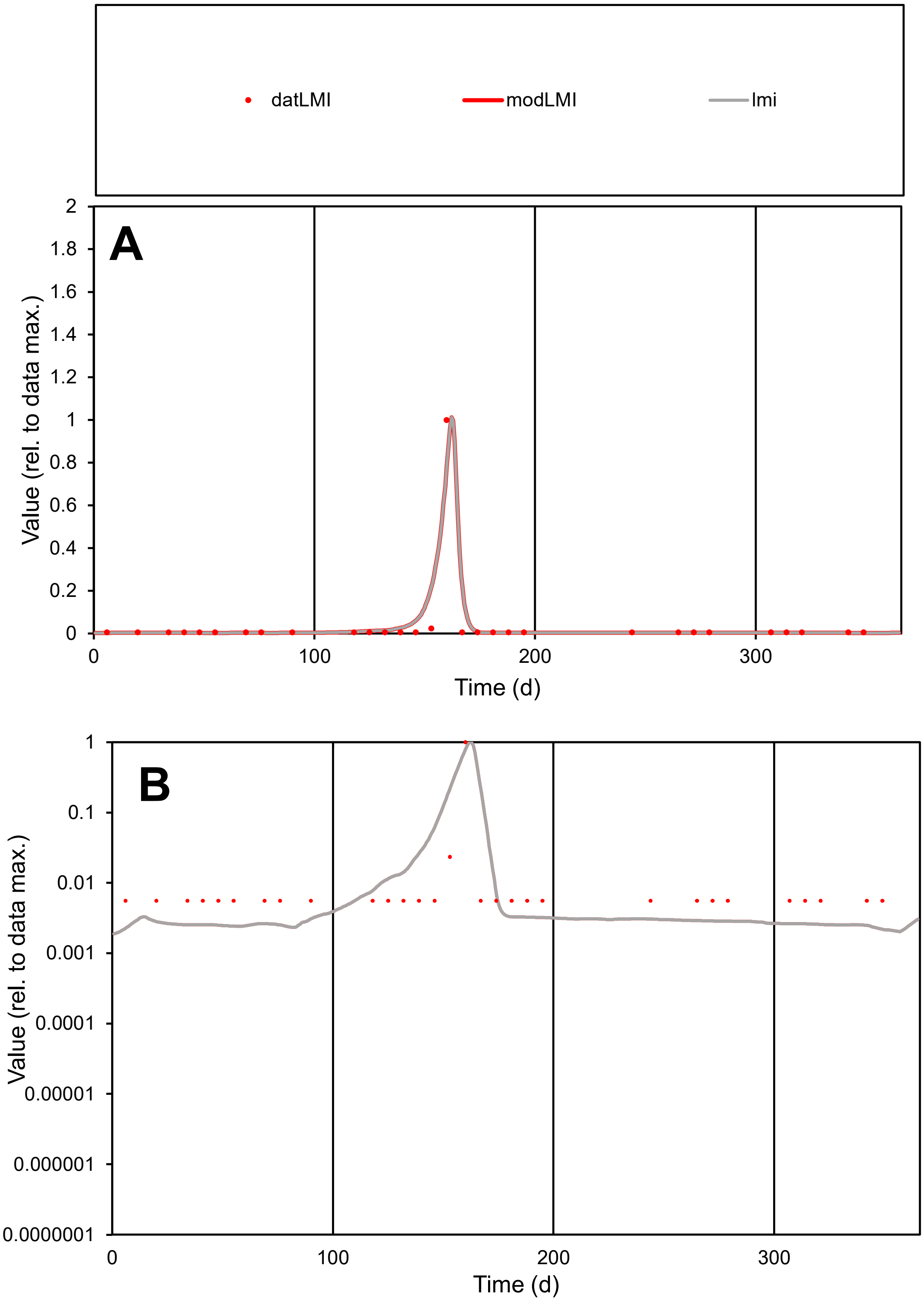


**C1**

Variables
LMI Provide jtd or Dst
CalibOnly?
2 1 to reproduce of
oNormalize?
2 1=no, 2=yes

ofxn calc*
0.73561
ofxn mod
0.98737

sRun
RunA1
zout
1

Ave
Data
2.1E-05
Model
2.4E-05

data/vMax
0.00054

ofxng
275.646

Min d/m
2.59071

ave/min d
7.035
ave/min m
20.8853

325_Calib_S_A

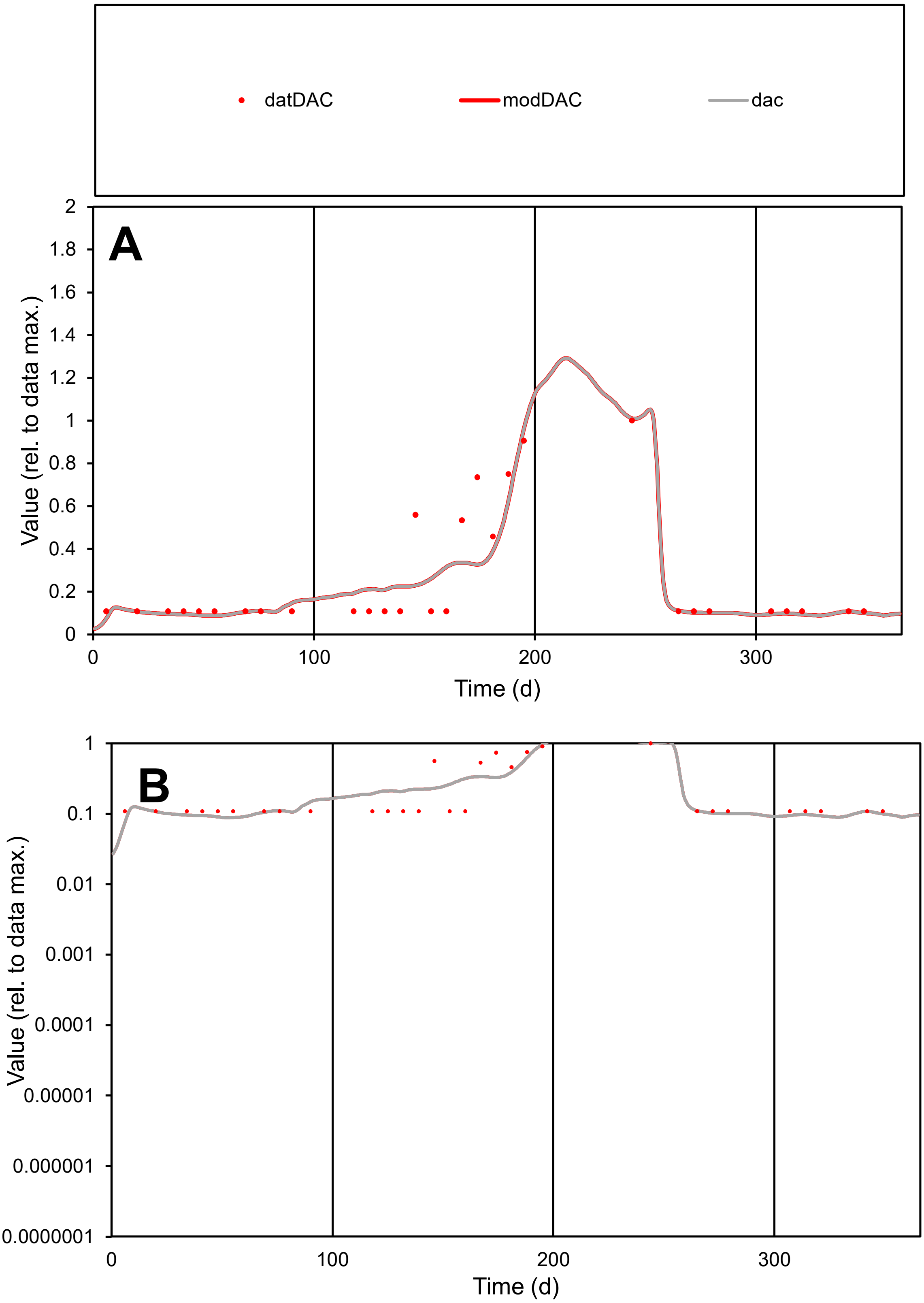


**C1**

Variables
DAC Provide jtd or Dst
CalibOnly?
2 1 to reproduce of
oNormalize?
2 1=no, 2=yes

ofxn calc*
0.48986
ofxn mod
0.51197

sRun
RunA1
zout
1

Ave
Data
1.8E-05
Model
1.7E-05

data/vMax
7.1E-05

ofxng
275.646

Min d/m
1.21469

ave/min d
2.28648
ave/min m
2.68643

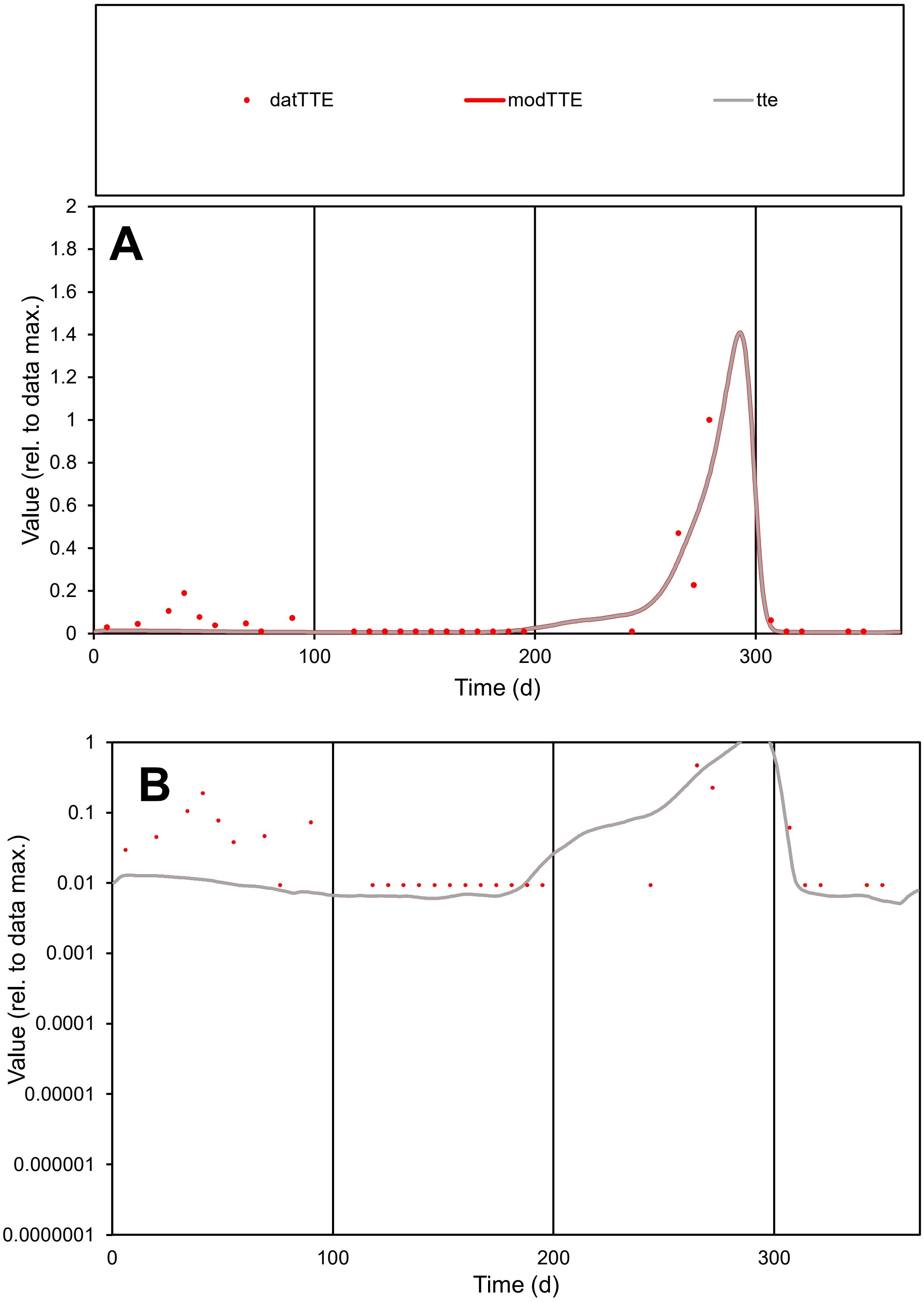


**C1**

Variables
TTE Provide jtd or Dst
CalibOnly?
2 1 to reproduce of
oNormalize?
2 1=no, 2=yes

ofxn calc*
0.97439
ofxn mod
1.03319

sRun
RunA1
zout
1

Ave
Data
3.4E-05
Model
2.8E-05

data/vMax
0.0004

ofxng
275.646

Min d/m
1.70258

ave/min d
9.01647
ave/min m
12.654

325_Calib_S_A

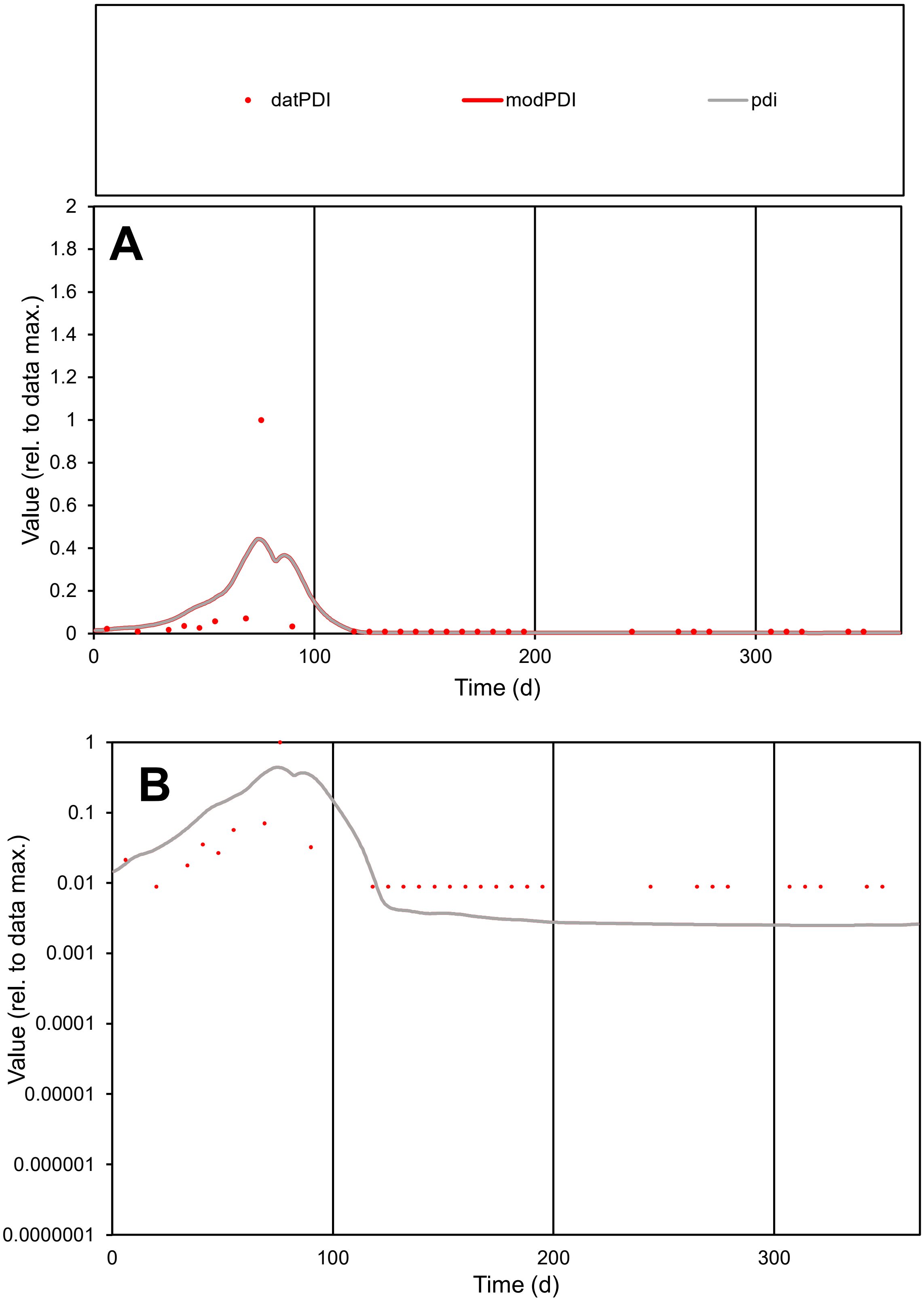


**C1**

Variables
PDI Provide jtd or Dst
CalibOnly?
2 1 to reproduce of
oNormalize?
2 1=no, 2=yes

ofxn calc*
2.69335
ofxn mod
2.72471

sRun
RunA1
zout
1

Ave
Data
2.4E-05
Model
2.8E-05

data/vMax
0.00049

ofxng
275.646

Min d/m
3.57398

ave/min d
5.45598
ave/min m
22.4614

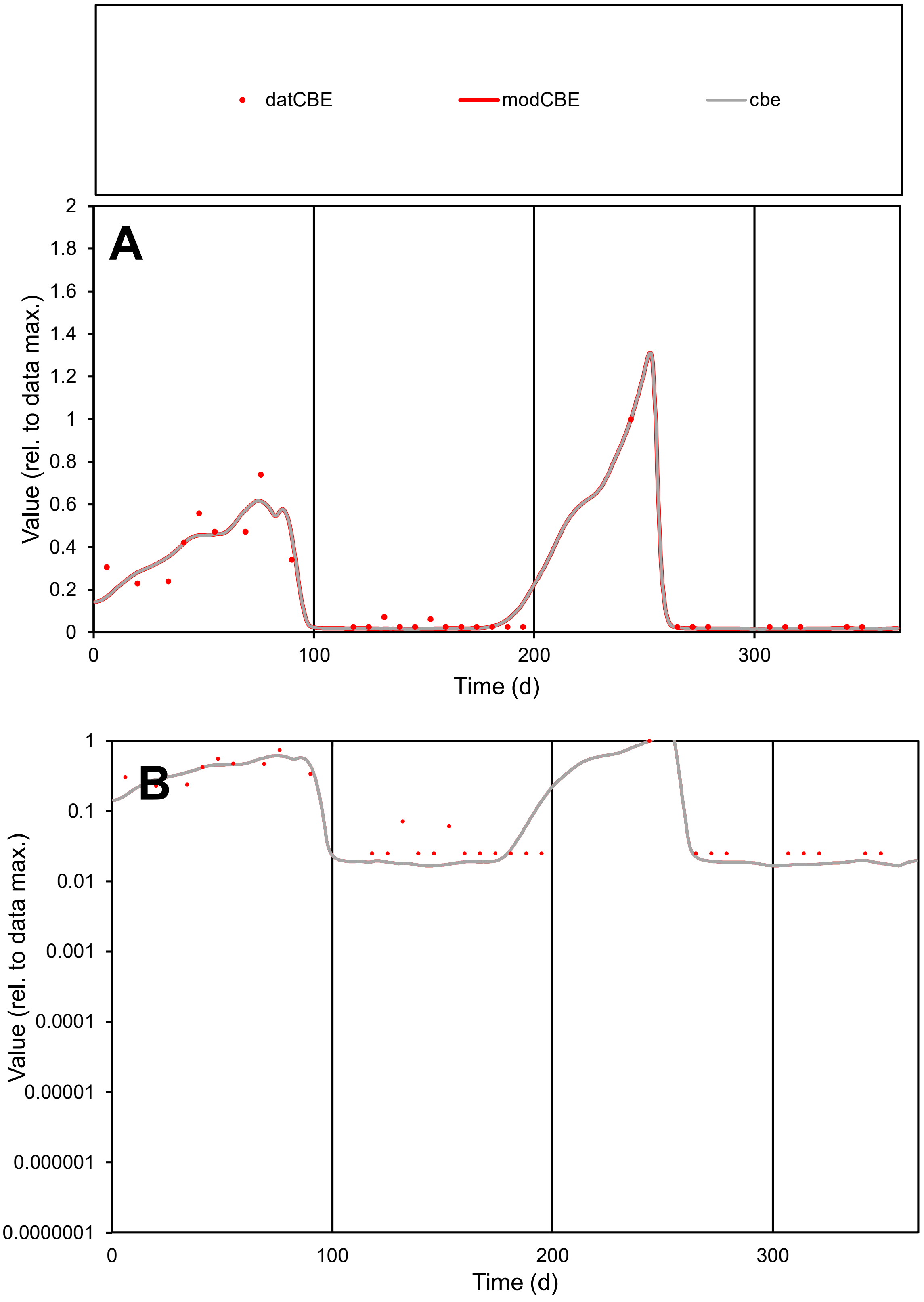


**C1**

Variables
CBE Provide jtd or Dst
CalibOnly?
2 1 to reproduce of
oNormalize?
2 1=no, 2=yes

ofxn calc*
0.31269
ofxn mod
0.36596

sRun
RunA1
zout
1

Ave
Data
1.2E-05
Model
1.2E-05

data/vMax
6.7E-05

ofxng
275.646

Min d/m
1.47872

ave/min d
7.17866
ave/min m
10.4539

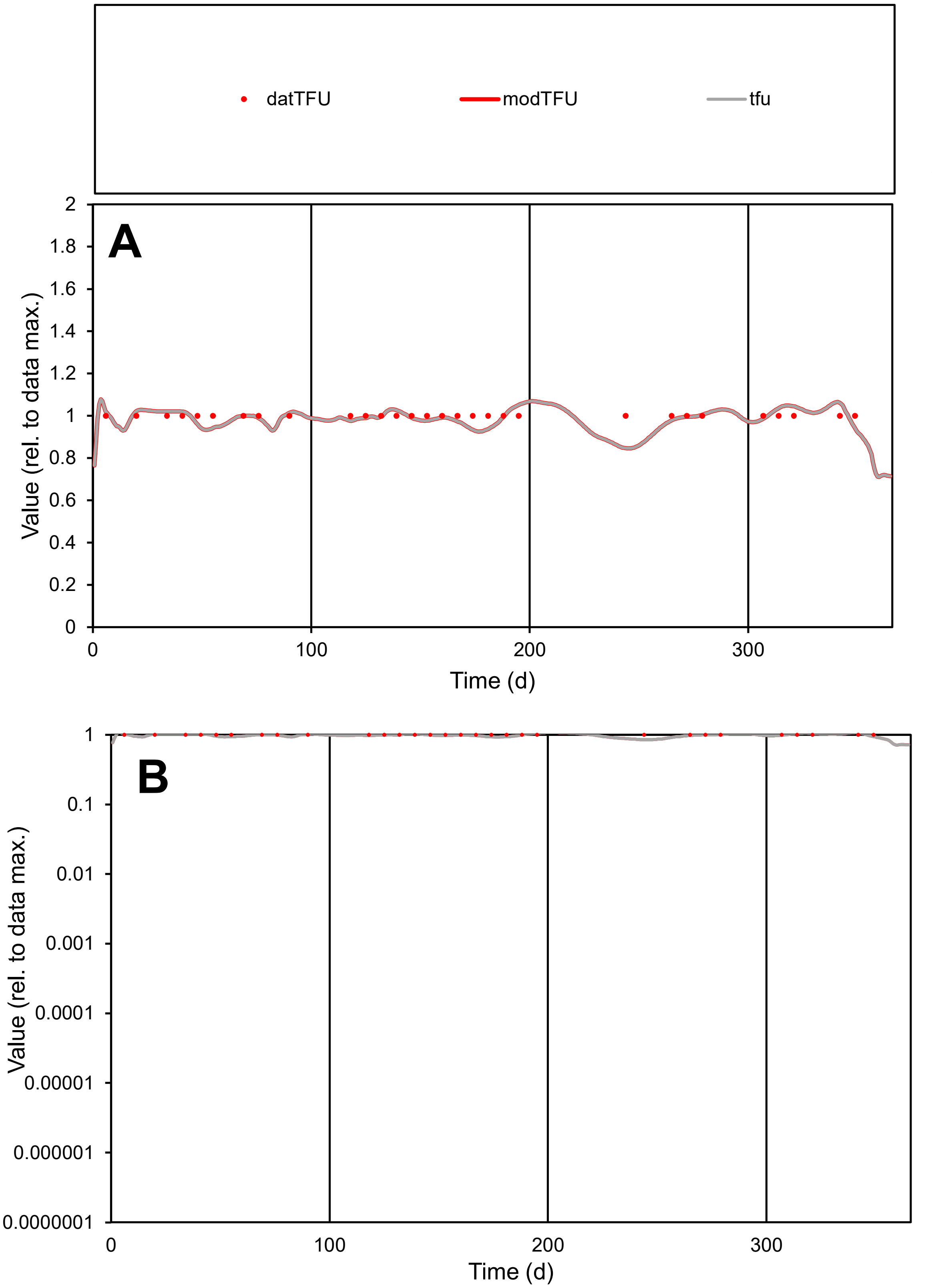


**C1**

Variables
TFU Provide jtd or Dst
CalibOnly?
2 1 to reproduce of
oNormalize?
2 1=no, 2=yes

ofxn calc*
0.04413
ofxn mod
0.12033

sRun
RunA1
zout
1

Ave
Data
1.3E-05
Model
1.3E-05

data/vMax
1.3E-05

ofxng
275.646

Min d/m
1.18275

ave/min d
1
ave/min m
1.17124

325_Calib_S_A

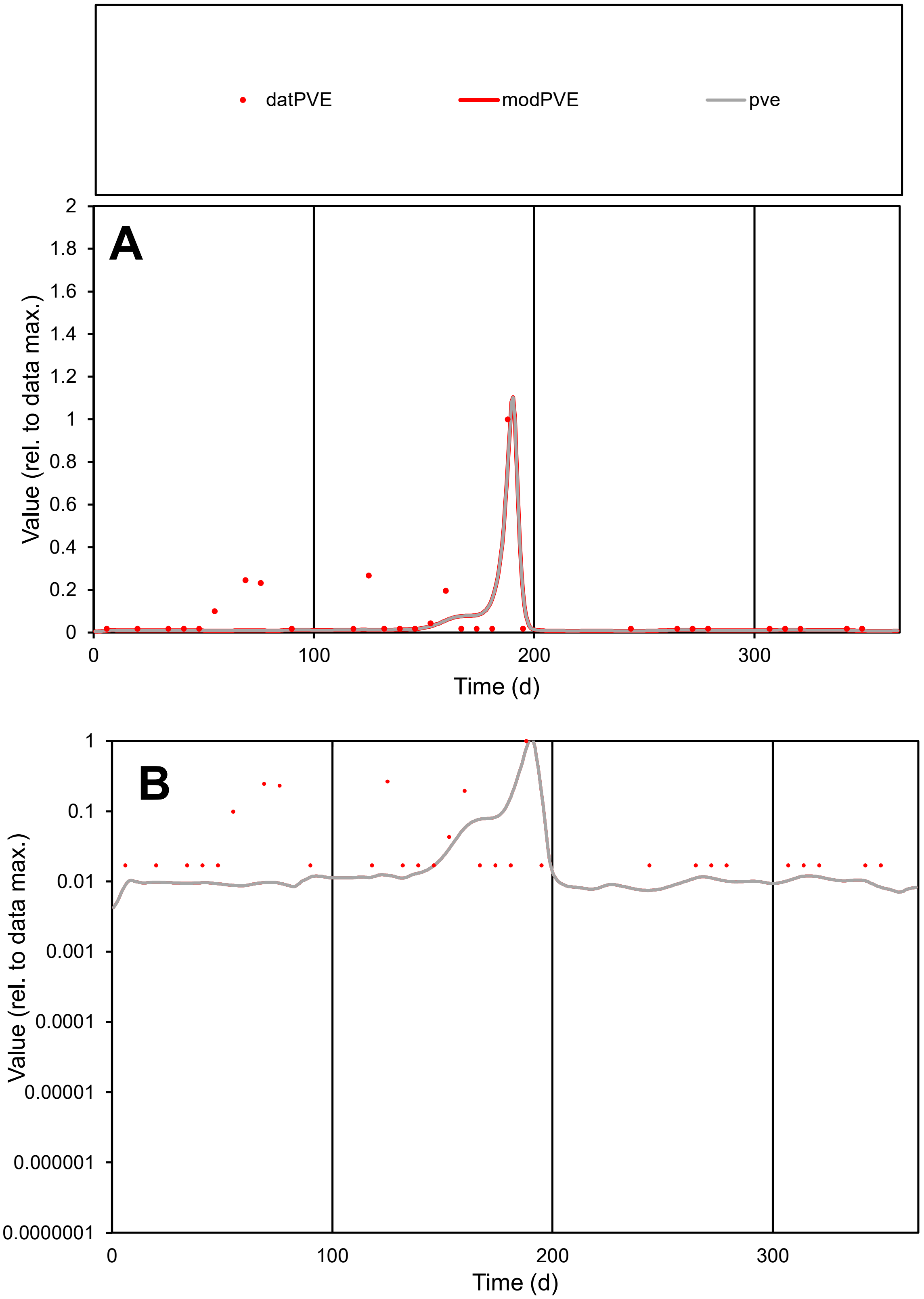


**C1**

Variables

PVE Provide jtd or Dst

CalibOnly?

2 1 to reproduce of

oNormalize?

2 1=no, 2=yes

ofxn calc*

1.03424

ofxn mod

1.06122

sRun

RunA1

zout

1

Ave

Data

1E-05

Model

6.8E-06

data/vMax

0.00012

ofxng

275.646

Min d/m

2.25577

ave/min d

4.85699

ave/min m

7.36408

325_Calib_S_A

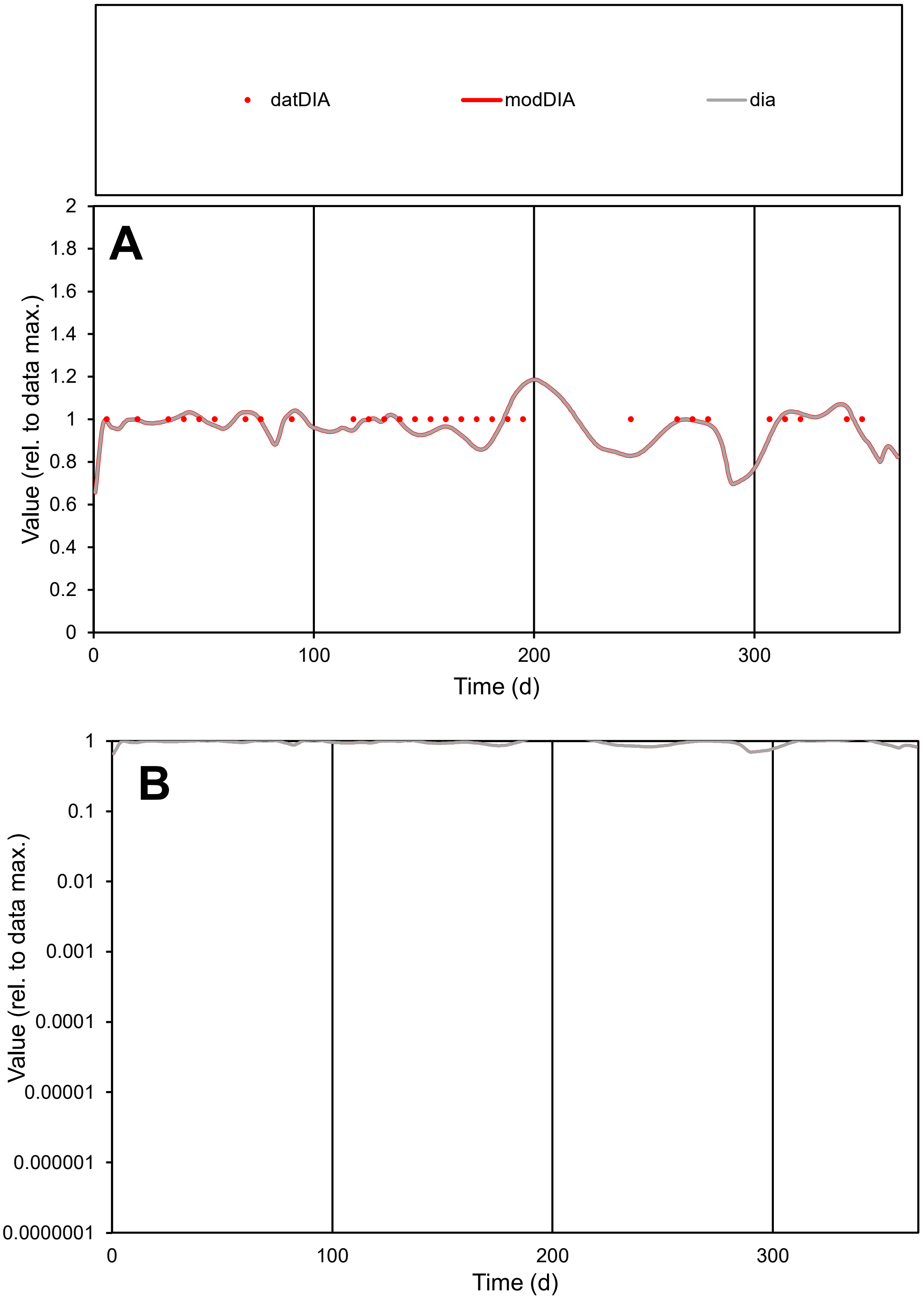


**C1**

Variables
DIA Provide jtd or Dst
CalibOnly?
2 1 to reproduce of
oNormalize?
2 1=no, 2=yes

ofxn calc*
0.06615
ofxn mod
0.12984

sRun
RunA1
zout
1

Ave
Data
6.2E-06
Model
6.1E-06

data/vMax
6.2E-06

ofxng
275.646

Min d/m
1.20445

ave/min d
1
ave/min m
1.18369

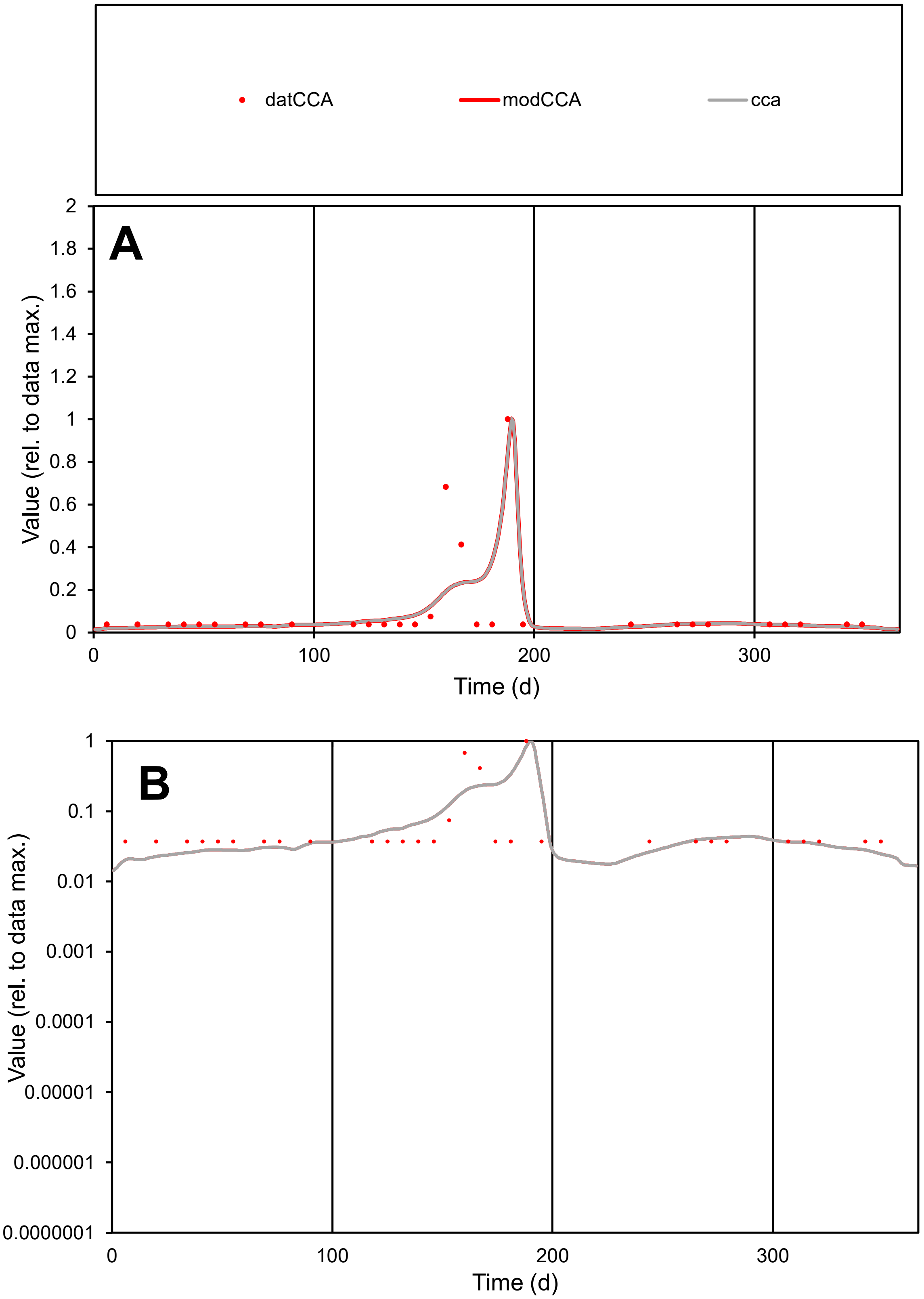


**C1**

Variables
CCA Provide jtd or Dst
CalibOnly?
2 1 to reproduce of
oNormalize?
2 1=no, 2=yes

ofxn calc*
1.09469
ofxn mod
1.11991

sRun
RunA1
zout
1

Ave
Data
5.1E-06
Model
5E-06

data/vMax
4.9E-05

ofxng
275.646

Min d/m
1.77994

ave/min d
2.80753
ave/min m
4.84106

325_Calib_S_A

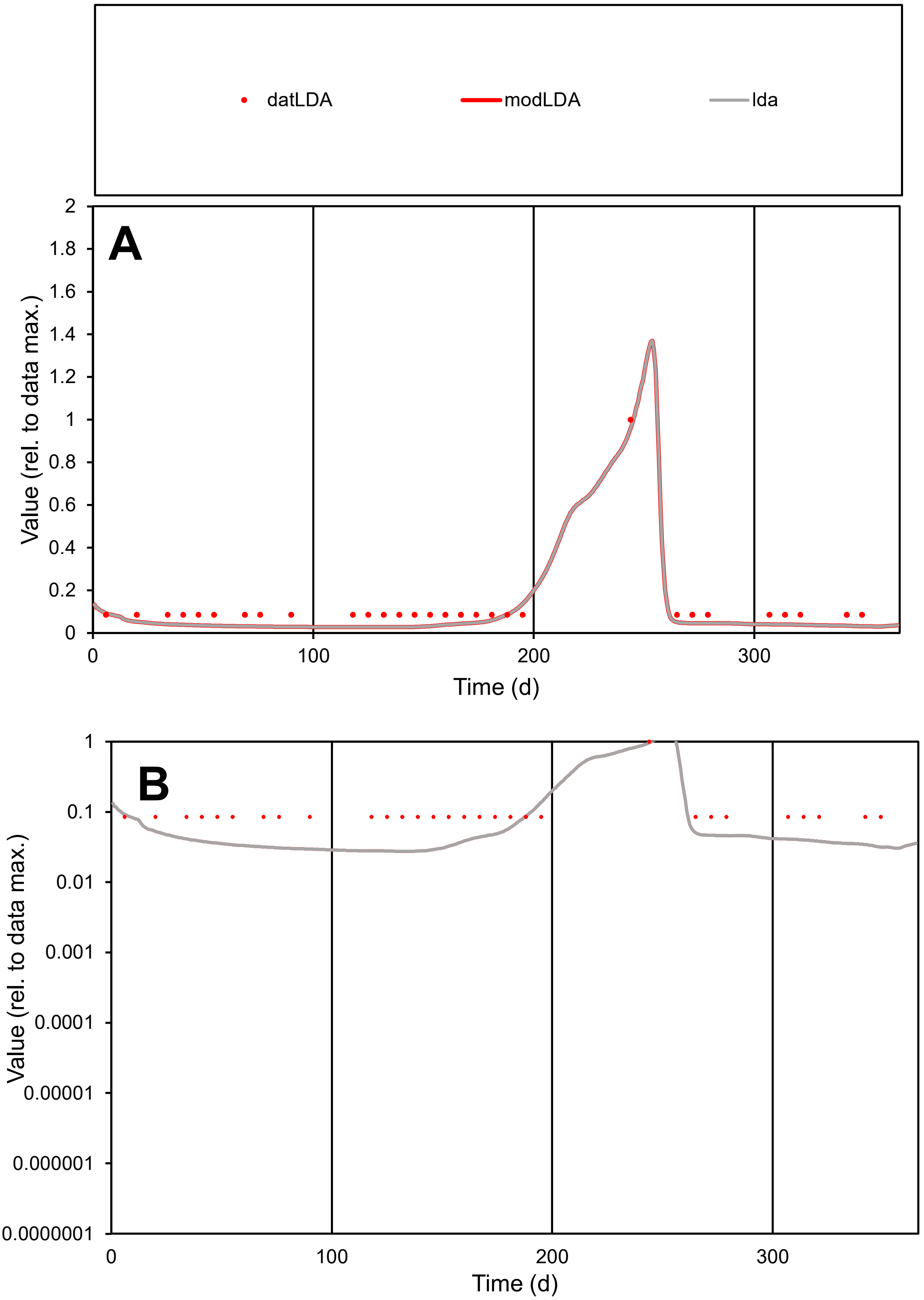


**C1**

Variables
LDA Provide jtd or Dst
CalibOnly?
2 1 to reproduce of
oNormalize?
2 1=no, 2=yes

ofxn calc*
0.39717
ofxn mod
0.42045

sRun
RunA1
zout
1

Ave
Data
9.4E-06
Model
6.3E-06

data/vMax
8.1E-05

ofxng
275.646

Min d/m
3.10467

ave/min d
1.3586
ave/min m
2.81294

325_Calib_S_A

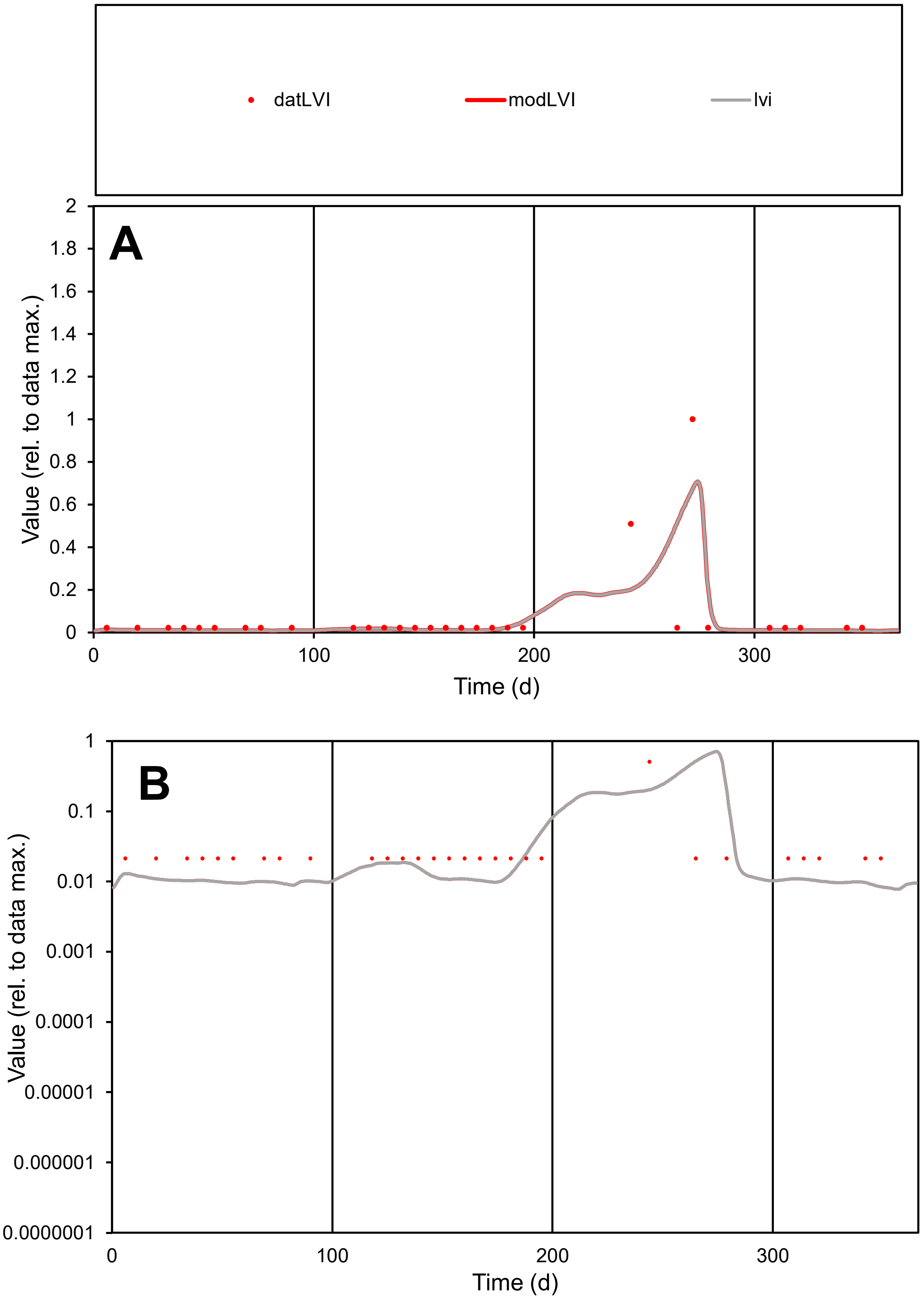


**C1**

Variables
LVI Provide jtd or Dst
CalibOnly?
2 1 to reproduce of
oNormalize?
2 1=no, 2=yes

ofxn calc*
1.69415
ofxn mod
1.7596

sRun
RunA1
zout
1

Ave
Data
4.7E-06
Model
4.1E-06

data/vMax
6.7E-05

ofxng
275.646

Min d/m
2.57515

ave/min d
3.29644
ave/min m
7.37031

325_Calib_S_A

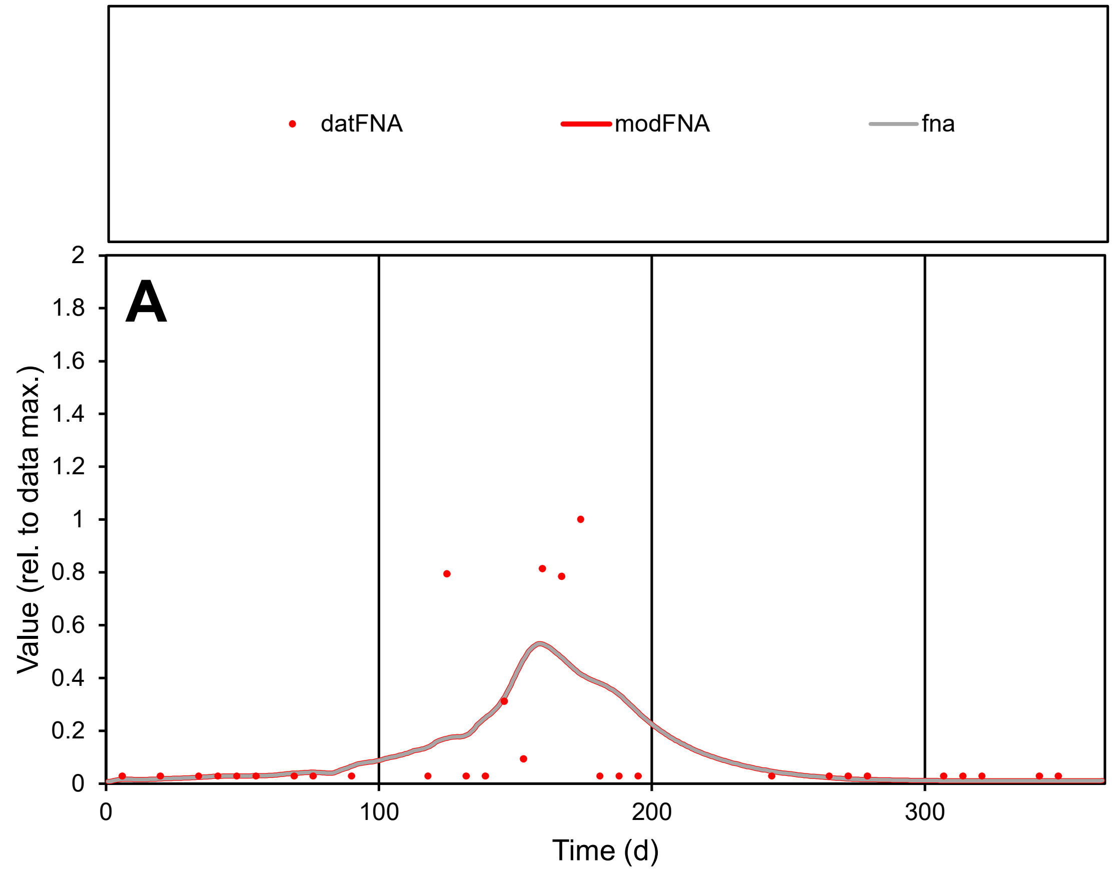


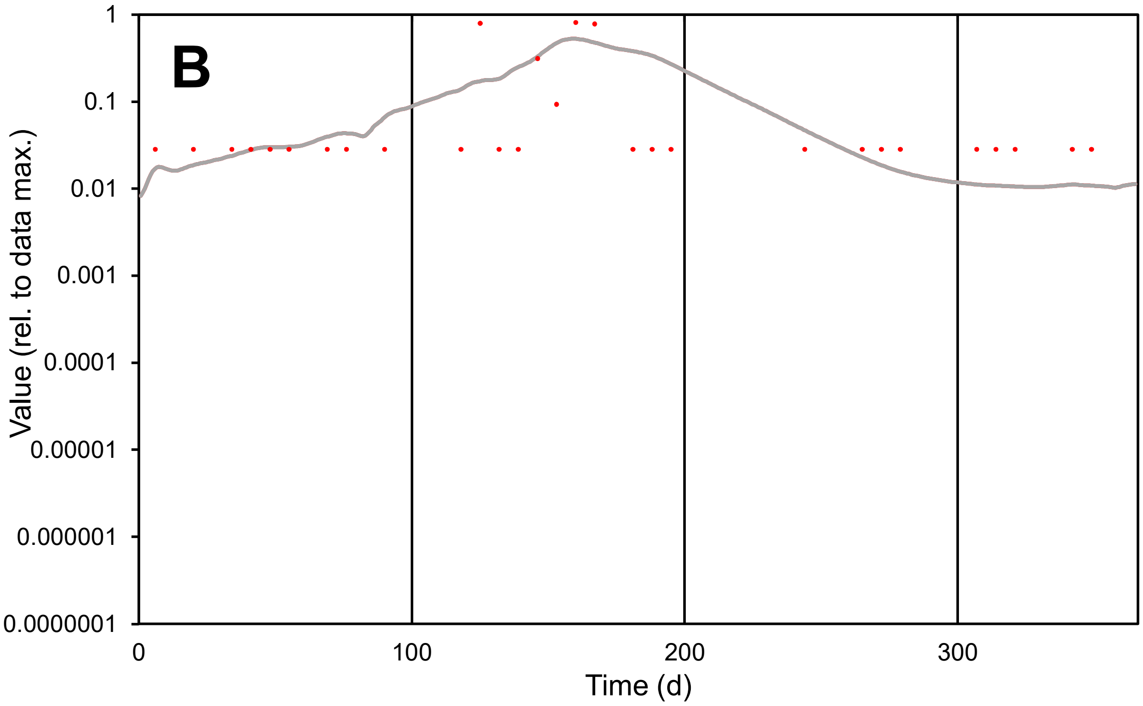


**C1**

Variables

FNA Provide jtd or Dst

CalibOnly?

2 1 to reproduce of

oNormalize?

2 1=no, 2=yes

ofxn calc*

1.45063

ofxn mod

1.46839

sRun

RunA1

zout

1

Ave

Data

1.6E-05

Model

1.6E-05

data/vMax

0.00011

ofxng

275.646

Min d/m

2.69025

ave/min d

5.26077

ave/min m

13.8244

325_Calib_S_A

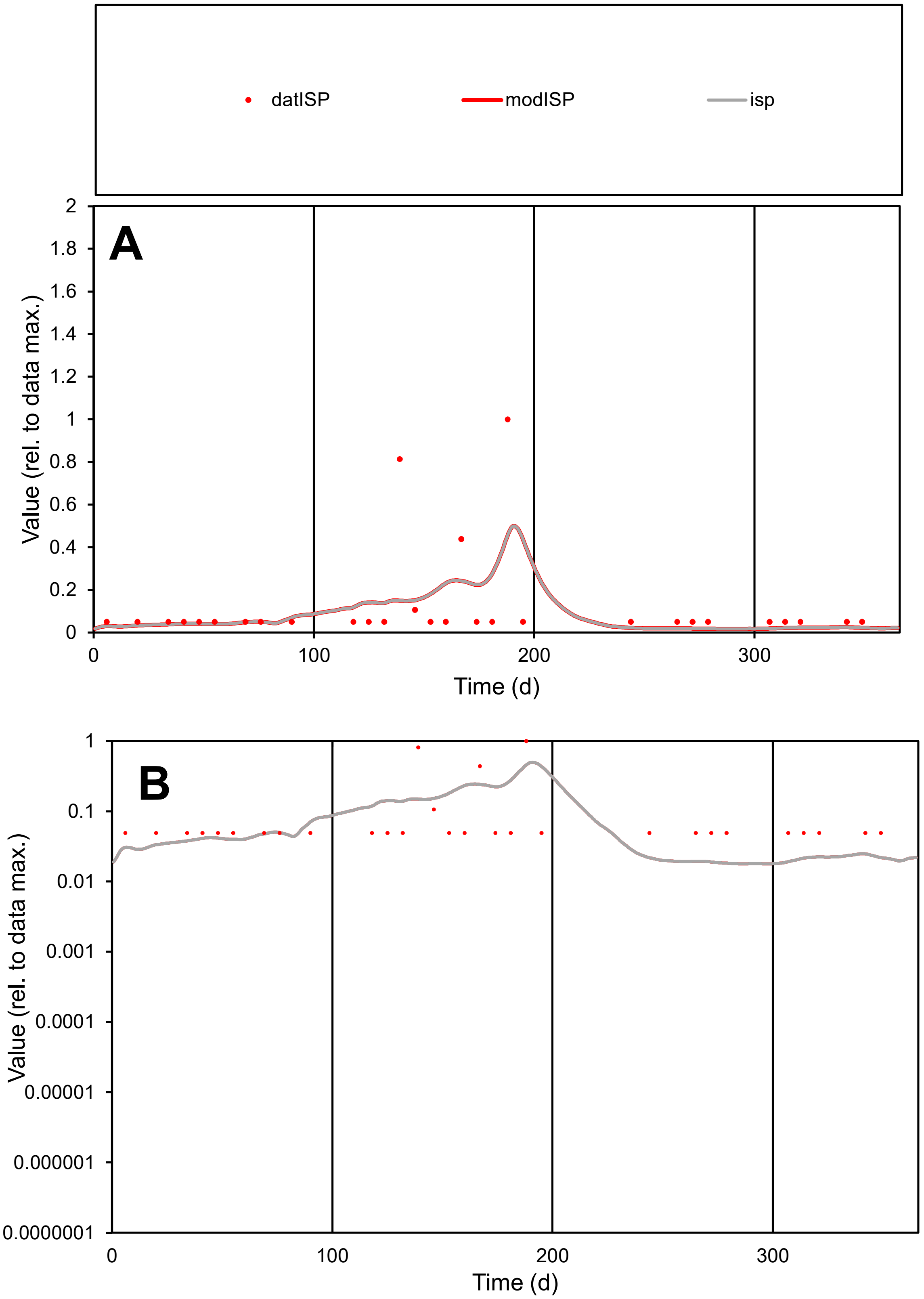


**C1**

Variables
ISP Provide jtd or Dst
CalibOnly?
2 1 to reproduce of
oNormalize?
2 1=no, 2=yes

ofxn calc*
1.50333
ofxn mod
1.53688

sRun
RunA1
zout
1

Ave
Data
1.2E-05
Model
1.1E-05

data/vMax
0.0001

ofxng
275.646

Min d/m
2.72688

ave/min d
2.46213
ave/min m
6.01741

325_Calib_S_A

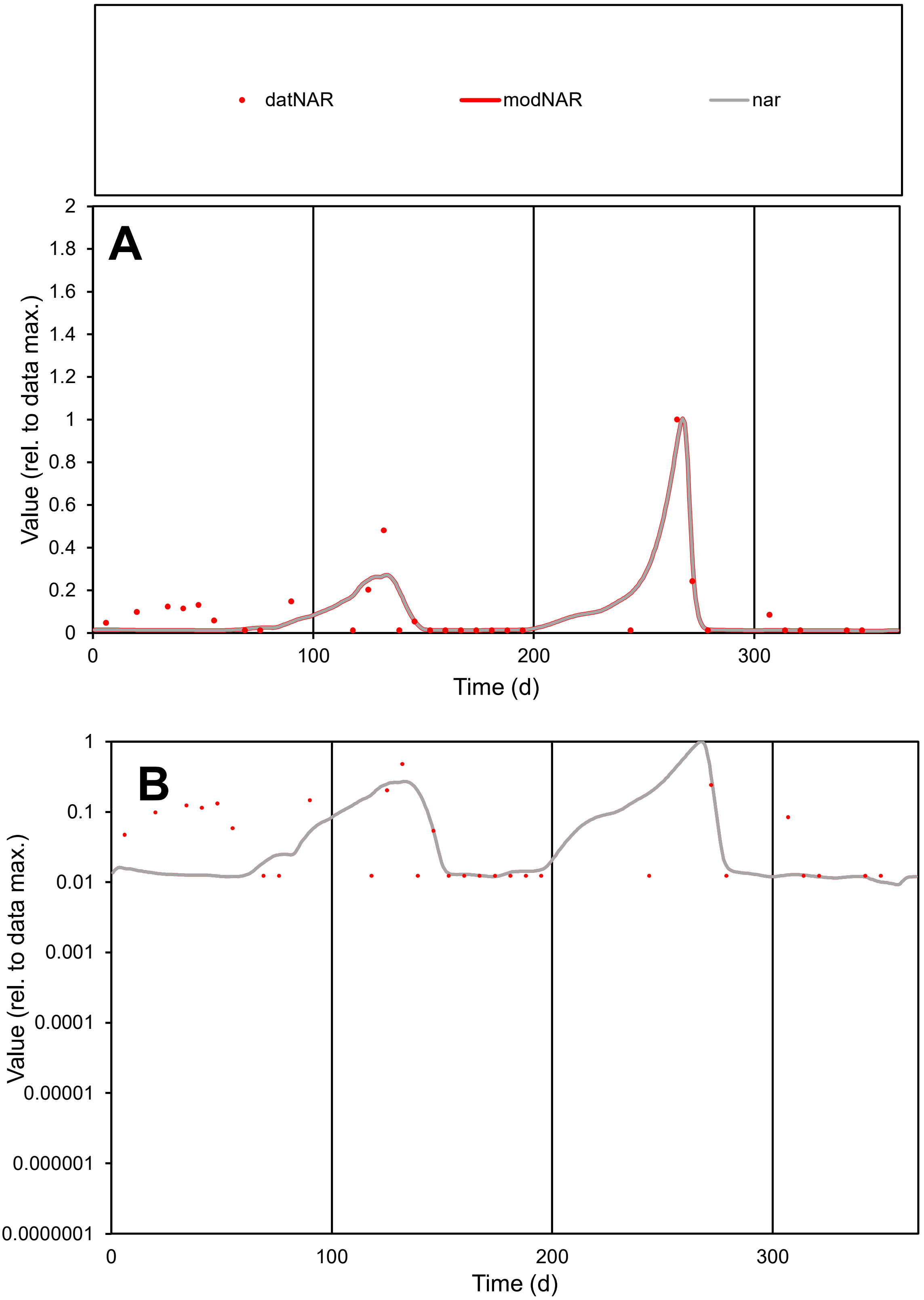


**C1**

Variables
NAR Provide jtd or Dst
CalibOnly?
2 1 to reproduce of
oNormalize?
2 1=no, 2=yes

ofxn calc*
0.76218
ofxn mod
0.80966

sRun
RunA1
zout
1

Ave
Data
2.3E-05
Model
2E-05

data/vMax
0.00023

ofxng
275.646

Min d/m
1.24669

ave/min d
8.08973
ave/min m
8.73236

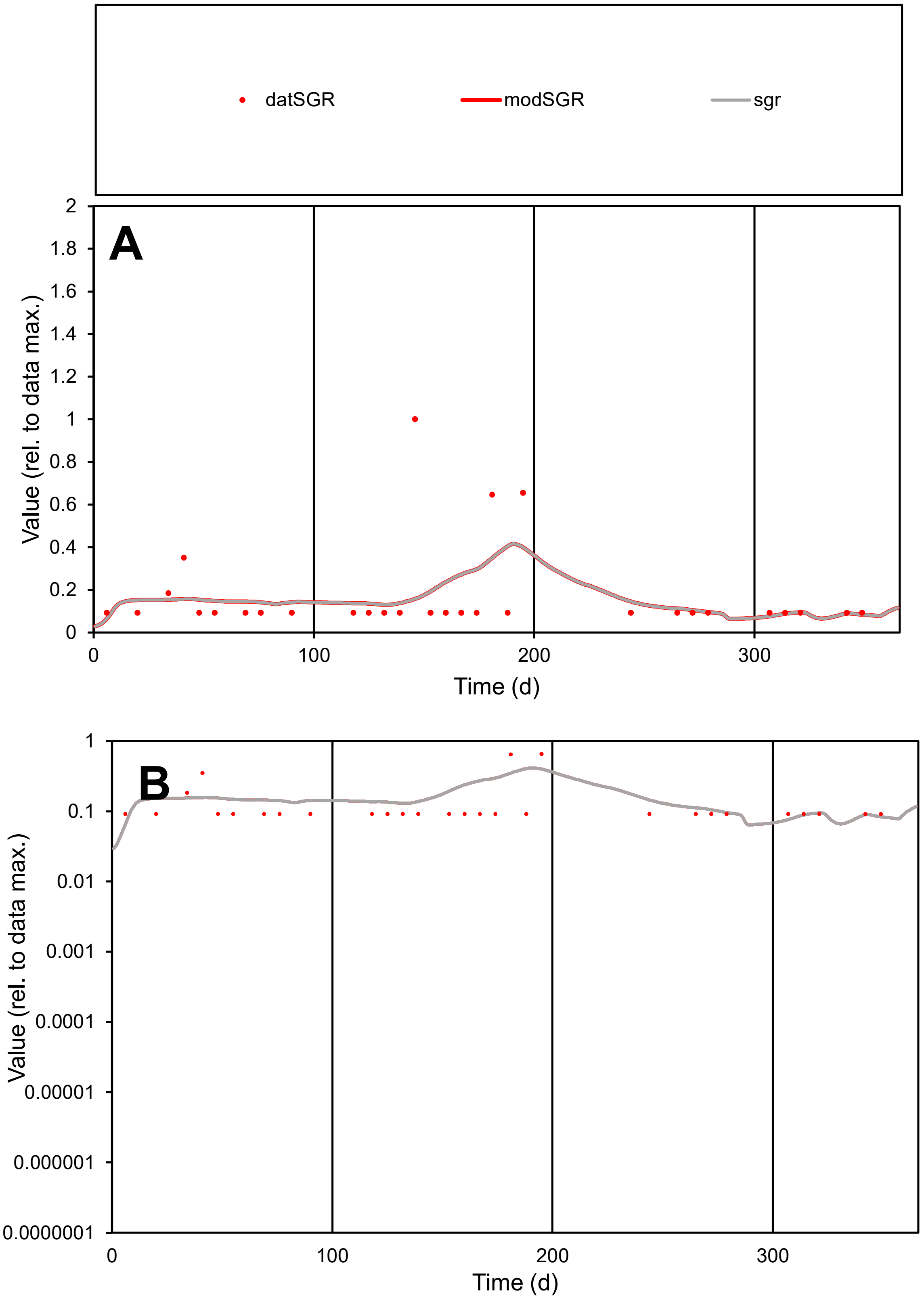


**C1**

Variables
SGR Provide jtd or Dst
CalibOnly?
2 1 to reproduce of
oNormalize?
2 1=no, 2=yes

ofxn calc*
1.13041
ofxn mod
1.14355

sRun
RunA1
zout
1

Ave
Data
8.4E-06
Model
8.3E-06

data/vMax
4.9E-05

ofxng
275.646

Min d/m
1.17123

ave/min d
1.86118
ave/min m
2.14281

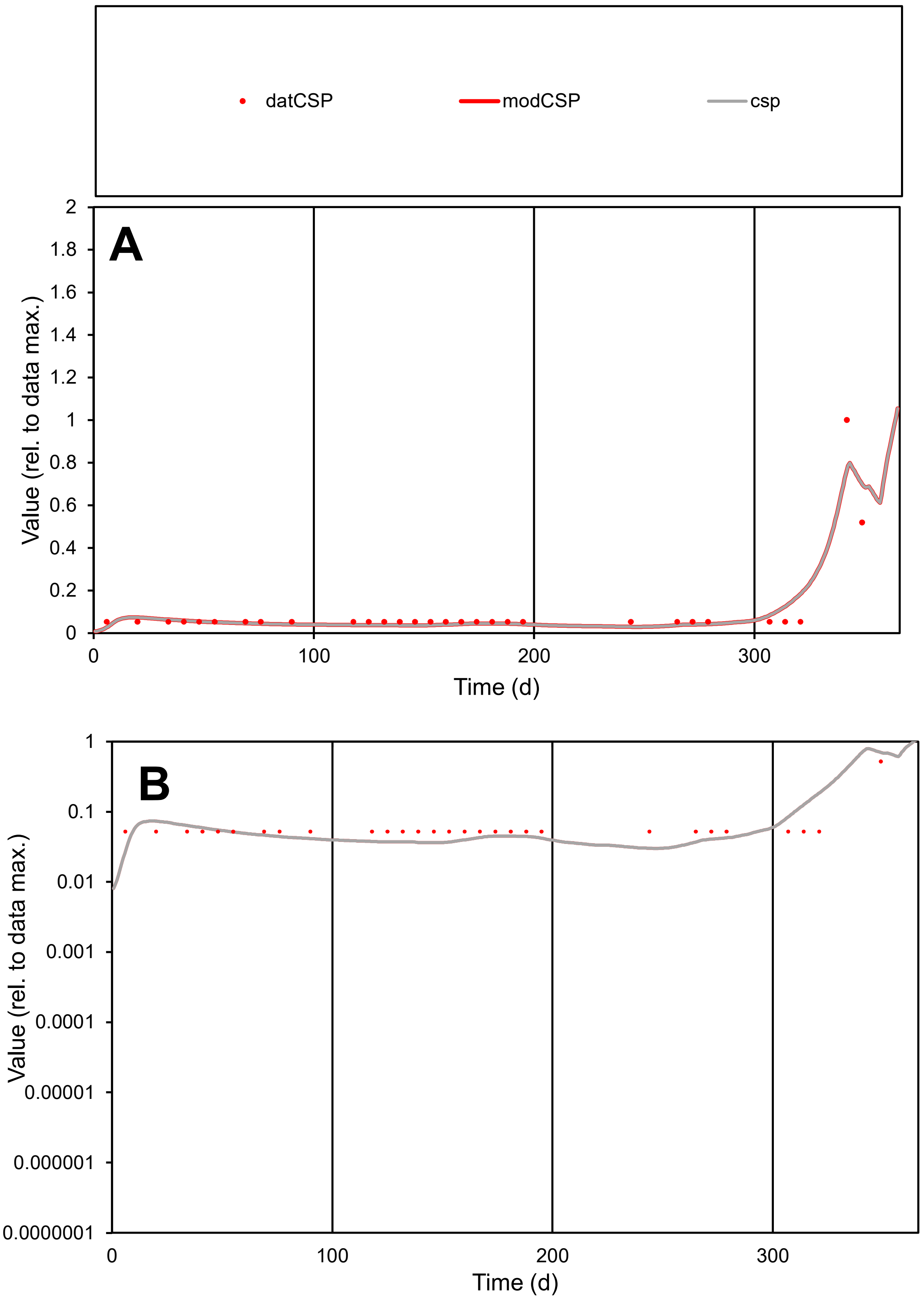


**C1**

Variables
CSP Provide jtd or Dst
CalibOnly?
2 1 to reproduce of
oNormalize?
2 1=no, 2=yes

ofxn calc*
0.57194
ofxn mod
0.61491

sRun
RunA1
zout
1

Ave
Data
4.5E-06
Model
4.5E-06

data/vMax
4.6E-05

ofxng
275.646

Min d/m
1.74488

ave/min d
1.90066
ave/min m
3.31674

325_Calib_S_A

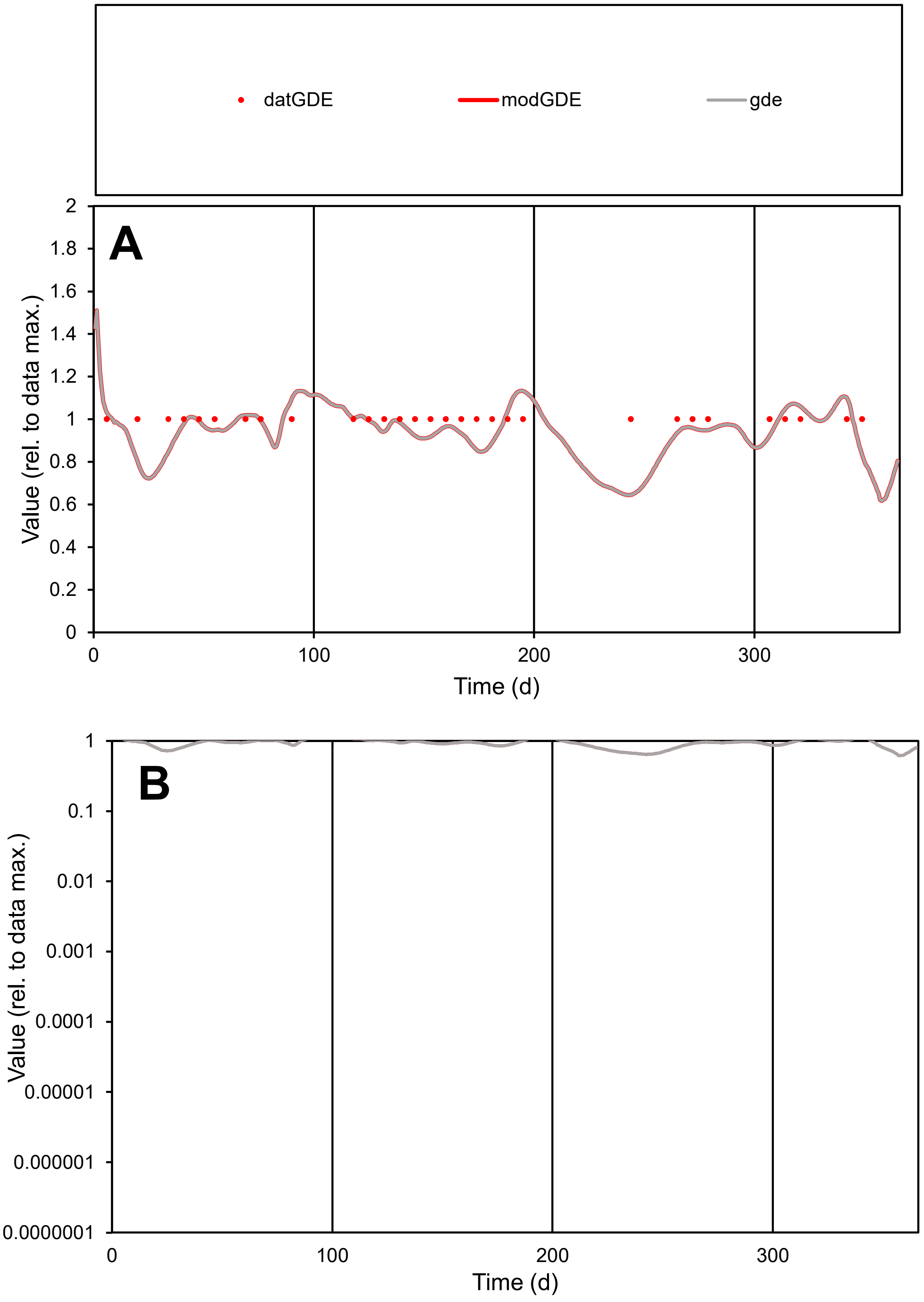


**C1**

Variables
GDE Provide jtd or Dst
CalibOnly?
2 1 to reproduce of
oNormalize?
2 1=no, 2=yes

ofxn calc*
0.10755
ofxn mod
0.15419

sRun
RunA1
zout
1

Ave
Data
2.7E-05
Model
2.6E-05

data/vMax
2.7E-05

ofxng
275.646

Min d/m
1.5395

ave/min d
1
ave/min m
1.47585

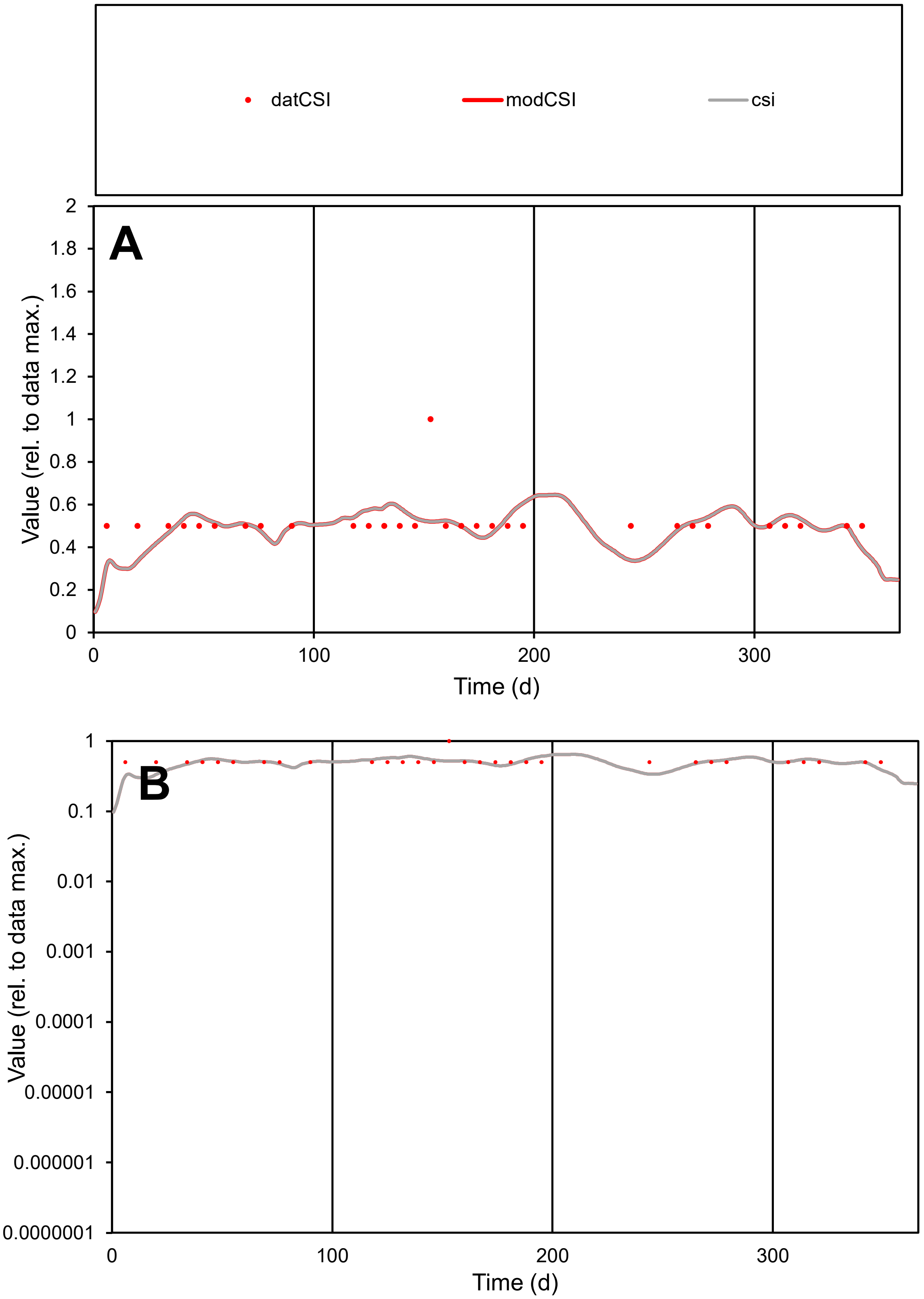


**C1**

Variables
CSI Provide jtd or Dst
CalibOnly?
2 1 to reproduce of
oNormalize?
2 1=no, 2=yes

ofxn calc*
0.21792
ofxn mod
0.24449

sRun
RunA1
zout
1

Ave
Data
2.4E-06
Model
2.3E-06

data/vMax
4.6E-06

ofxng
275.646

Min d/m
1.48362

ave/min d
1.03335
ave/min m
1.48917

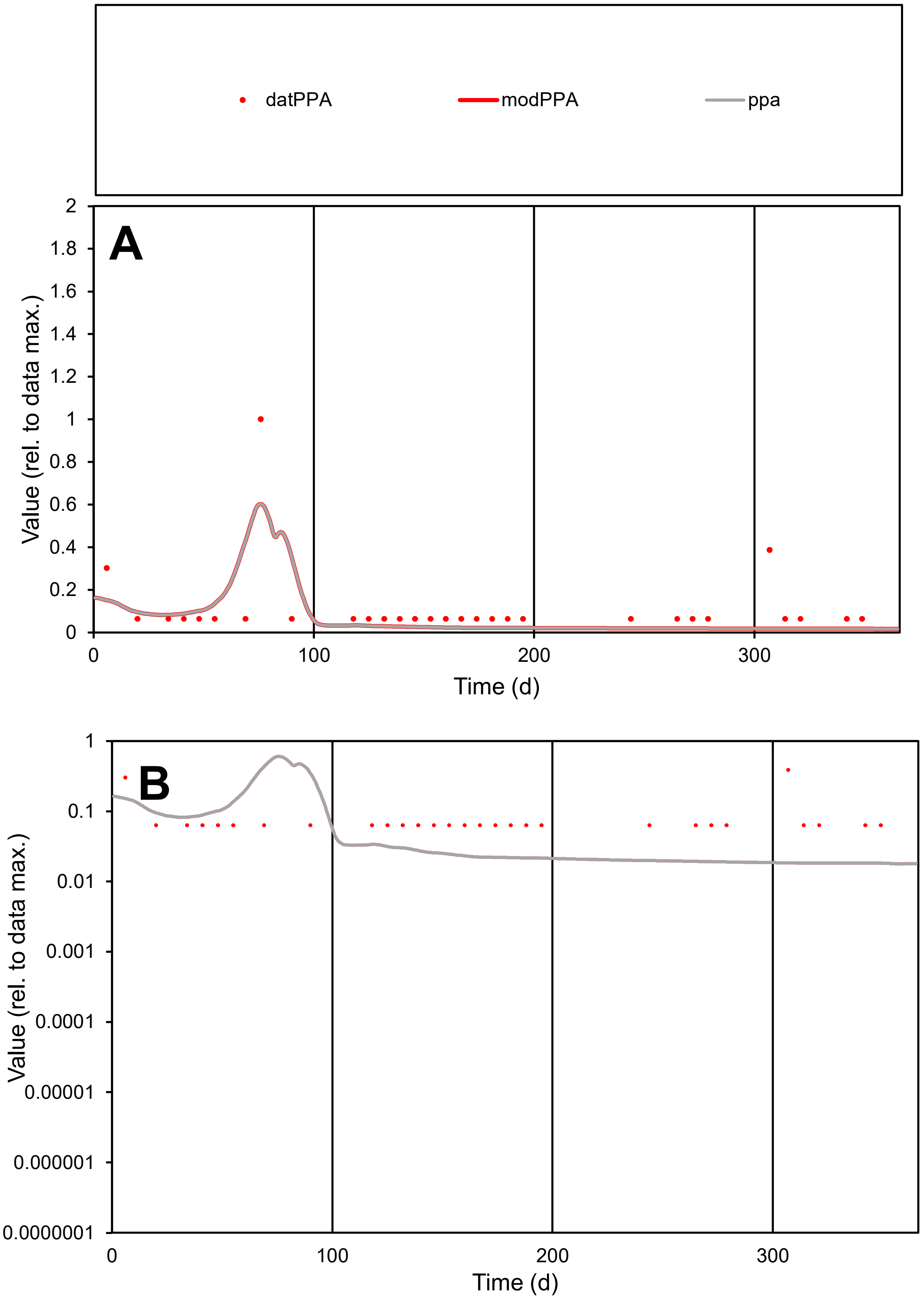


**C1**

Variables
PPA Provide jtd or Dst
CalibOnly?
2 1 to reproduce of
oNormalize?
2 1=no, 2=yes

ofxn calc*
1.16971
ofxn mod
1.19962

sRun
RunA1
zout
1

Ave
Data
5.7E-06
Model
4.1E-06

data/vMax
5.1E-05

ofxng
275.646

Min d/m
3.48454

ave/min d
1.79069
ave/min m
4.402

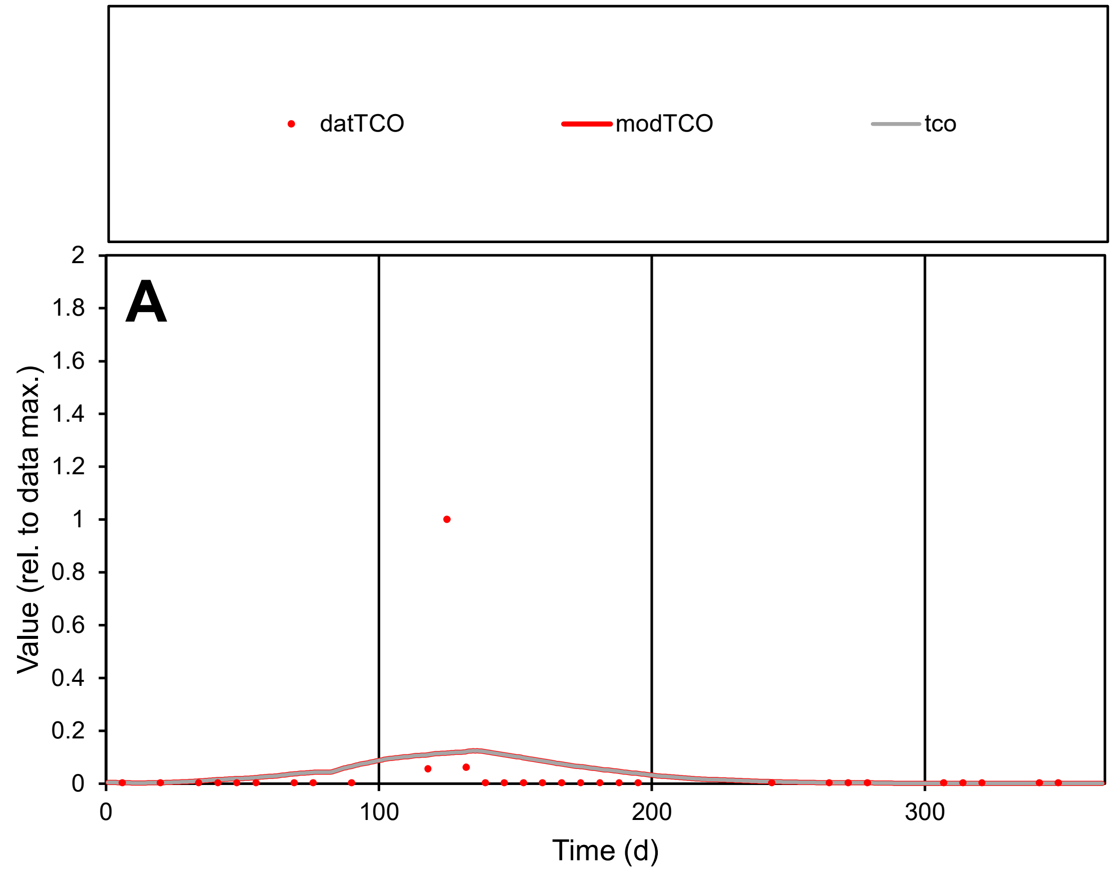


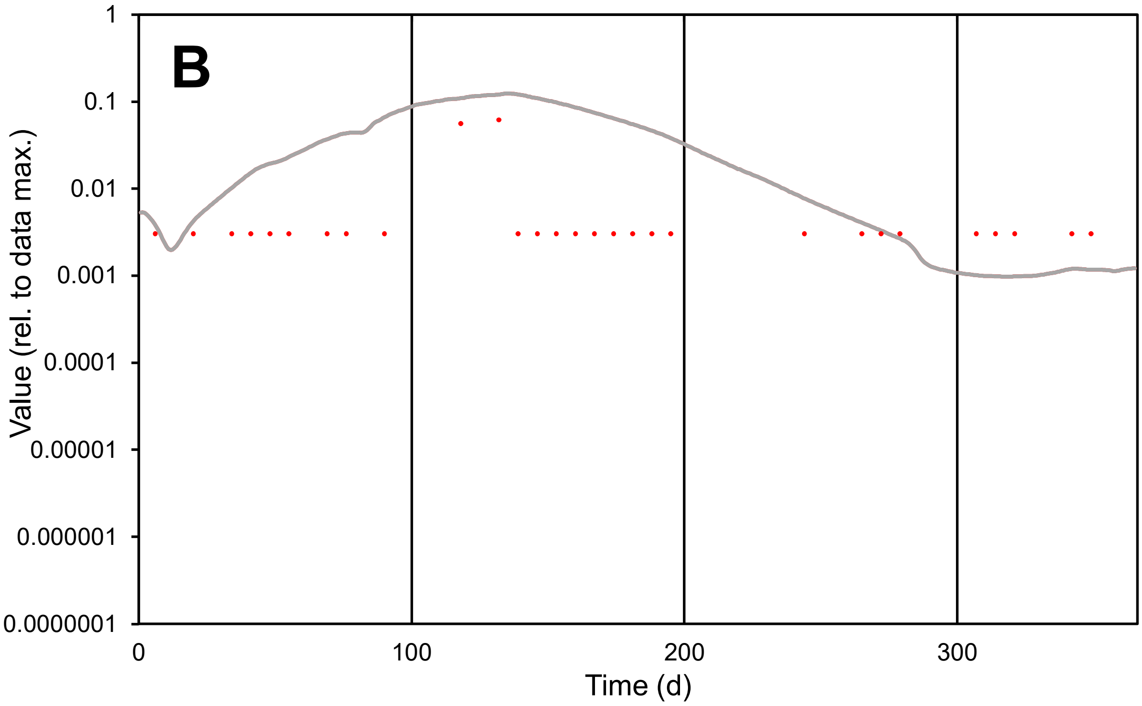


**C1**

Variables
TCO Provide jtd or Dst
CalibOnly?
2 1 to reproduce of
oNormalize?
2 1=no, 2=yes

ofxn calc*
4.19707
ofxn mod
4.20999

sRun
RunA1
zout
1

Ave
Data
0.00011
Model
0.00012

data/vMax
0.00283

ofxng
275.646

Min d/m
3.11525

ave/min d
13.1405
ave/min m
43.2451

325_Calib_S_A

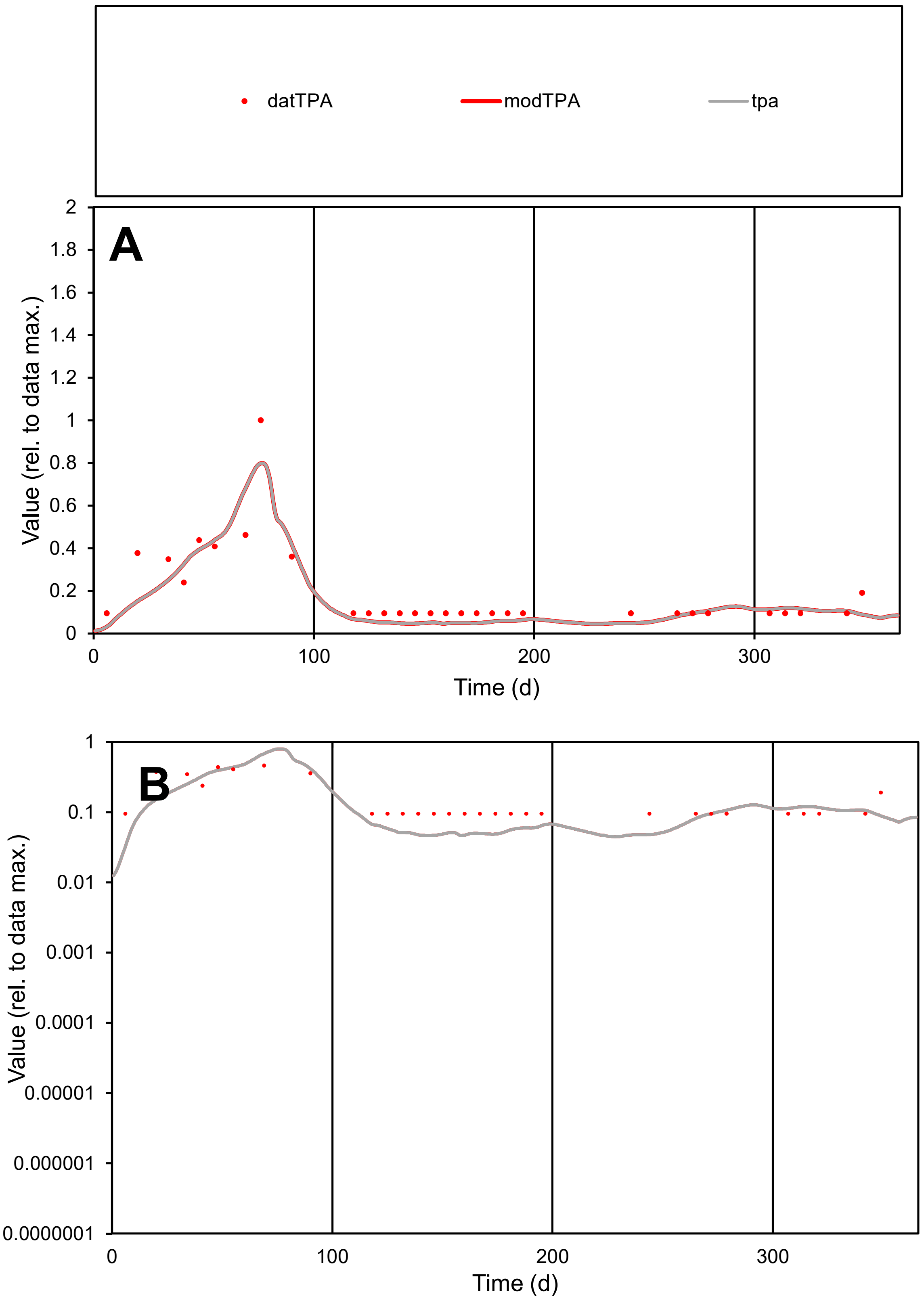


**C1**

Variables

TPA Provide jtd or Dst

CalibOnly?

2 1 to reproduce of

oNormalize?

2 1=no, 2=yes

ofxn calc*

0.4143

ofxn mod

0.45318

sRun

RunA1

zout

1

Ave

Data

9.6E-06

Model

8.4E-06

data/vMax

5E-05

ofxng

275.646

Min d/m

2.01967

ave/min d

2.03938

ave/min m

3.5713

325_Calib_S_A

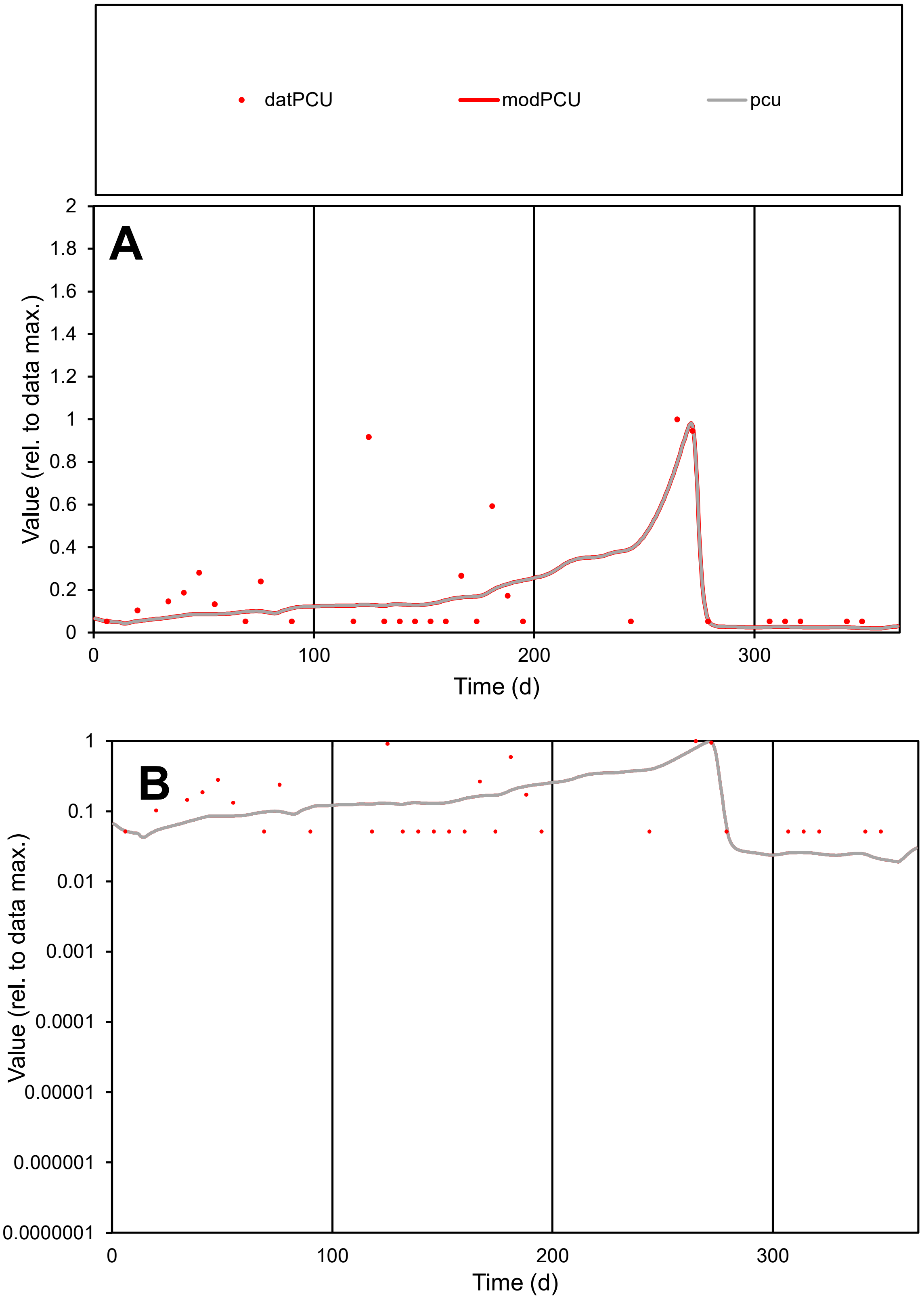


**C1**

Variables
PCU Provide jtd or Dst
CalibOnly?
2 1 to reproduce of
oNormalize?
2 1=no, 2=yes

ofxn calc*
0.9762
ofxn mod
0.99283

sRun
RunA1
zout
1

Ave
Data
2.4E-05
Model
2E-05

data/vMax
0.00012

ofxng
275.646

Min d/m
2.52816

ave/min d
3.81779
ave/min m
8.1977

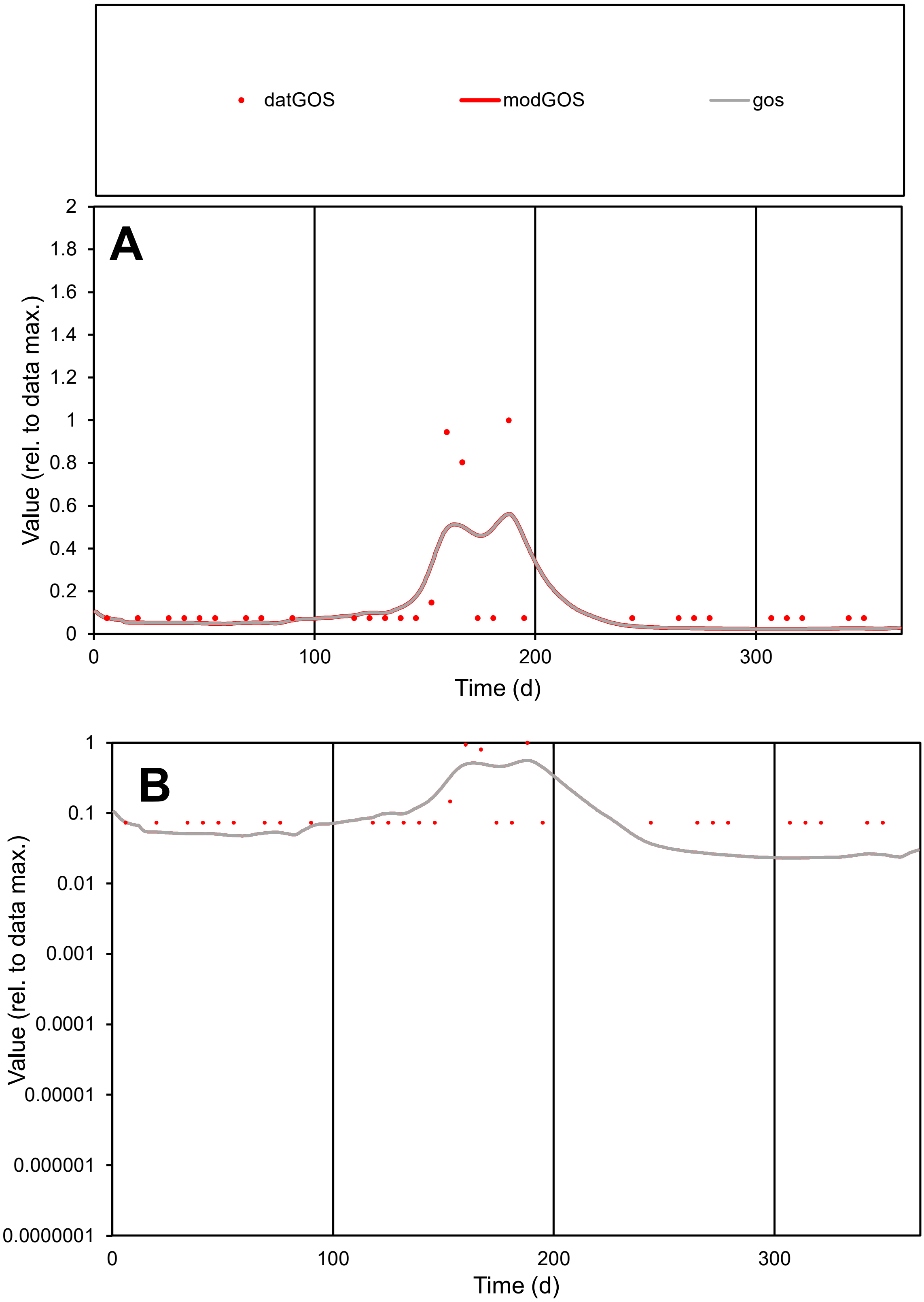


**C1**

Variables

| | |
|---|---|
| GOS | Provide jtd or Dst |
| CalibOnly? | |
| 2 | 1 to reproduce of |
| oNormalize? | |
| 2 | 1=no, 2=yes |
| ofxn calc* | |
| 1.13526 | |
| ofxn mod | |
| 1.15108 | |
| sRun | |
| RunA1 | |
| zout | |
| 1 | |
| Ave | |
| Data | |
| 4E-06 | |
| Model | |
| 3.8E-06 | |
| data/vMax | |
| 2.5E-05 | |
| ofxng | |
| 275.646 | |
| Min d/m | |
| 3.19126 | |
| ave/min d | |
| 2.18076 | |
| ave/min m | |
| 6.64833 | |

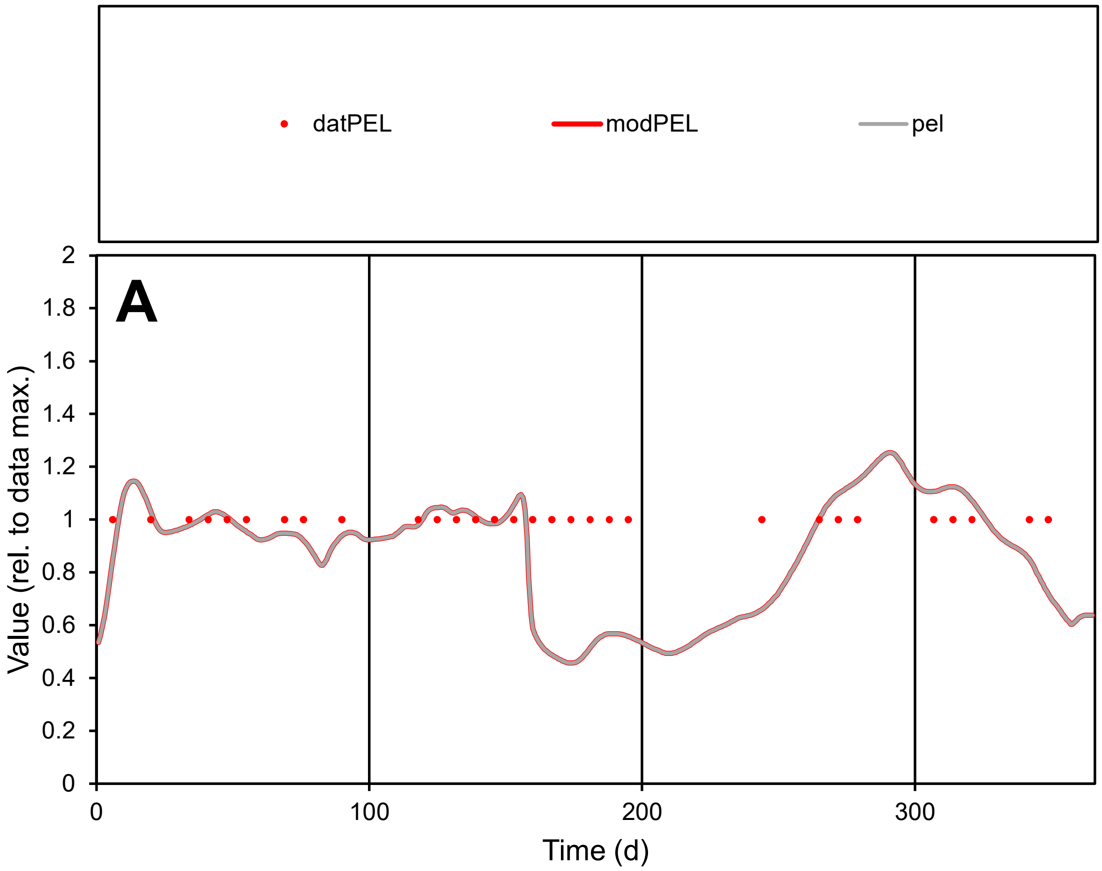


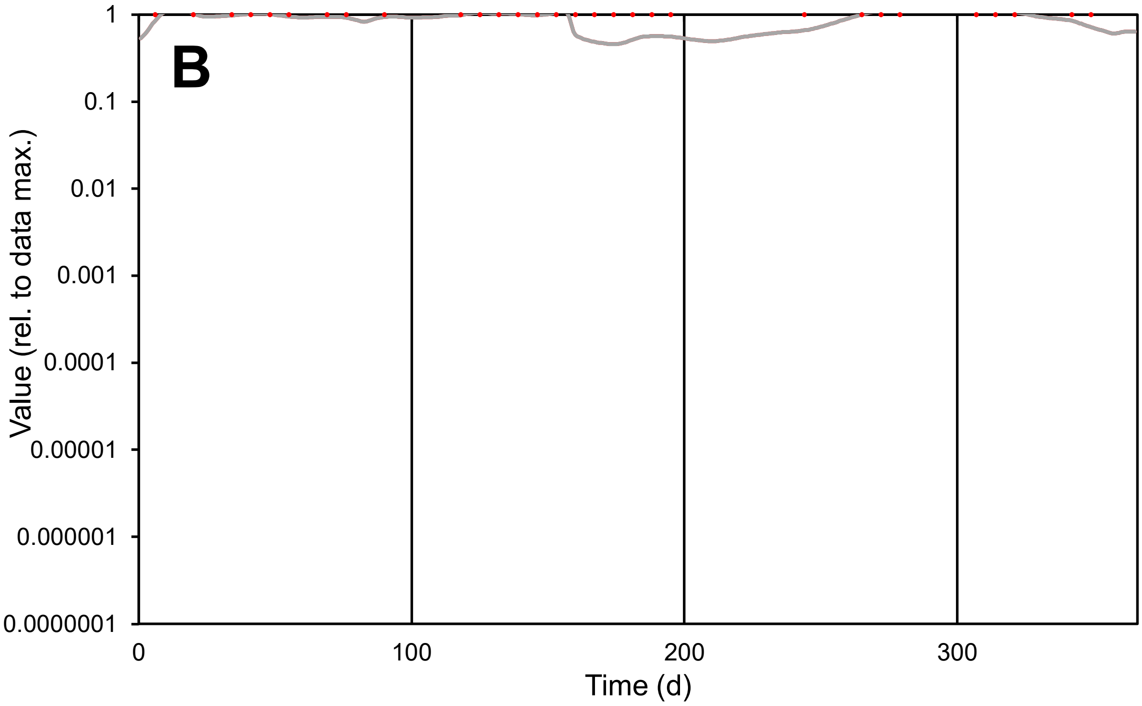


**C1**

Variables

PEL Provide jtd or Dst

CalibOnly?

2 1 to reproduce of

oNormalize?

2 1=no, 2=yes

ofxn calc*

0.23832

ofxn mod

0.26191

sRun

RunA1

zout

1

Ave

Data

3.3E-06

Model

2.9E-06

data/vMax

3.3E-06

ofxng

275.646

Min d/m

2.18218

ave/min d

1

ave/min m

1.94648

325_Calib_S_A

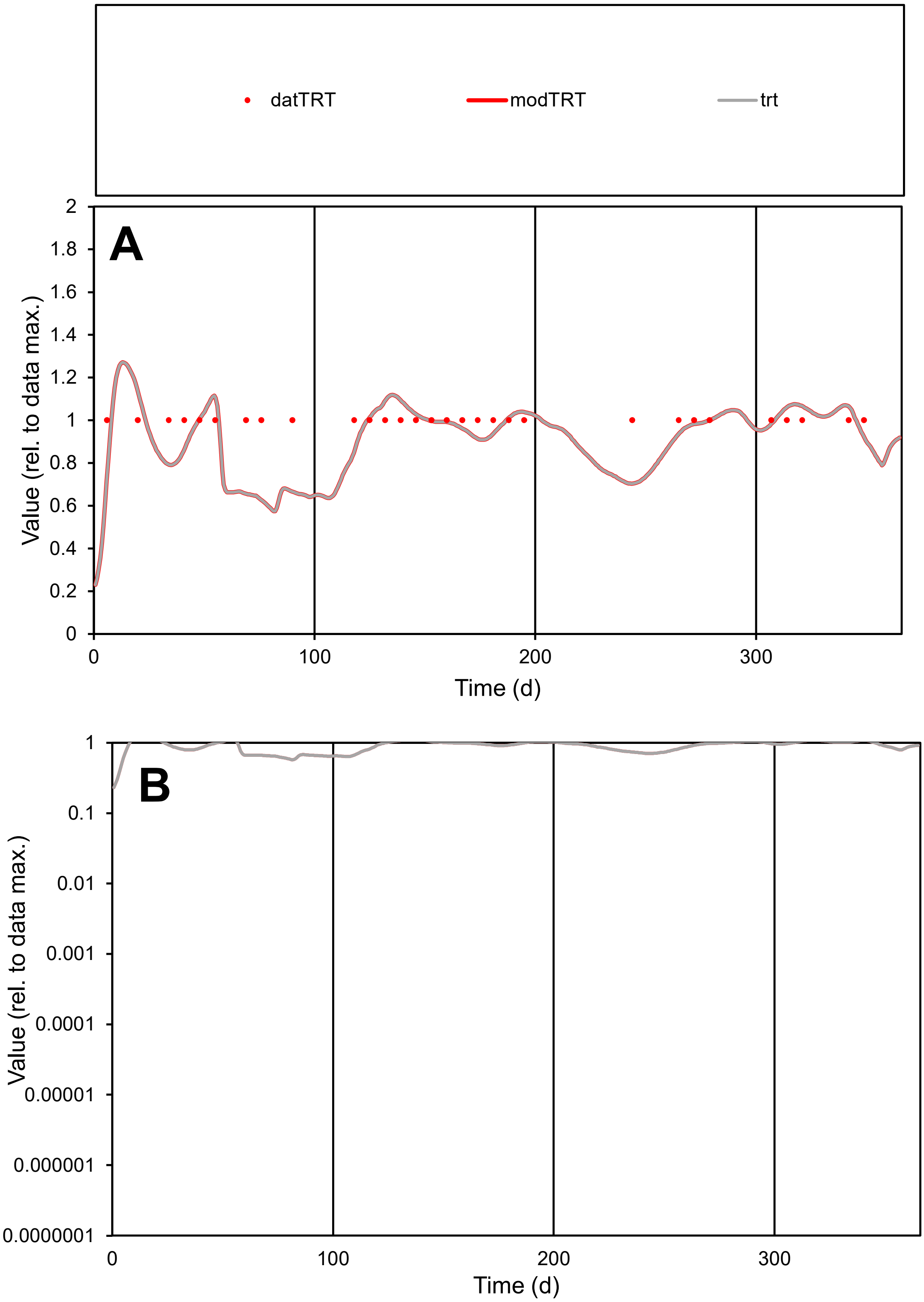


**C1**

Variables
TRT Provide jtd or Dsl
CalibOnly?
2 1 to reproduce of
oNormalize?
2 1=no, 2=yes

ofxn calc*
0.1425
ofxn mod
0.17994

sRun
RunA1
zout
1

Ave
Data
2.3E-05
Model
2.2E-05

data/vMax
2.3E-05

ofxng
275.646

Min d/m
1.61915

ave/min d
1
ave/min m
1.52986

325_Calib_S_A

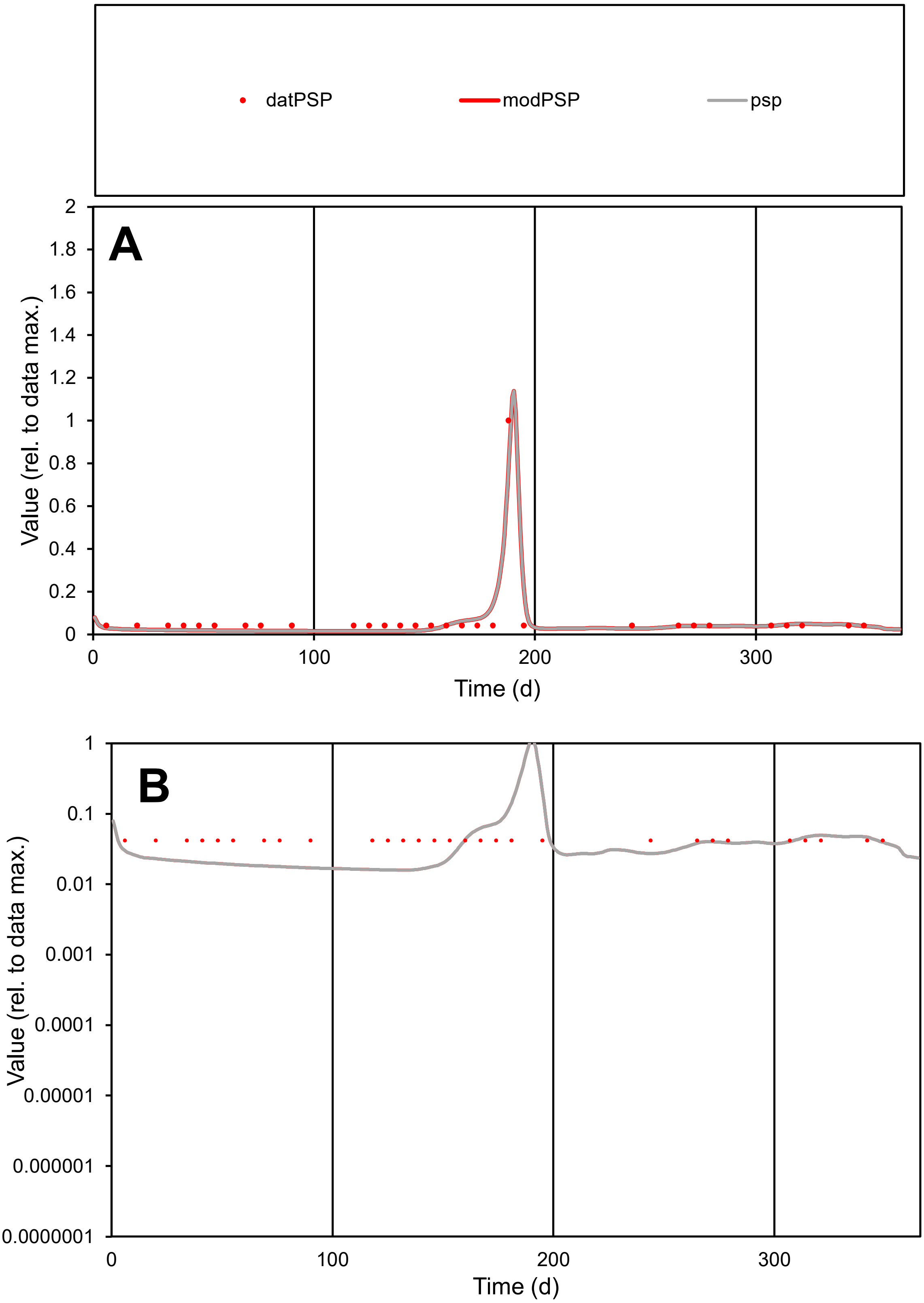


**C1**

Variables

| | |
|---|---|
| PSP | Provide jtd or Dst |
| CalibOnly? | |
| 2 | 1 to reproduce of |
| oNormalize? | |
| 2 | 1=no, 2=yes |

ofxn calc*
0.32818
ofxn mod
0.40258

sRun
RunA1
zout
1

Ave
Data
7.6E-06
Model
7E-06

data/vMax
0.0001

ofxng
275.646

Min d/m
2.63354

ave/min d
1.76315
ave/min m
4.26321

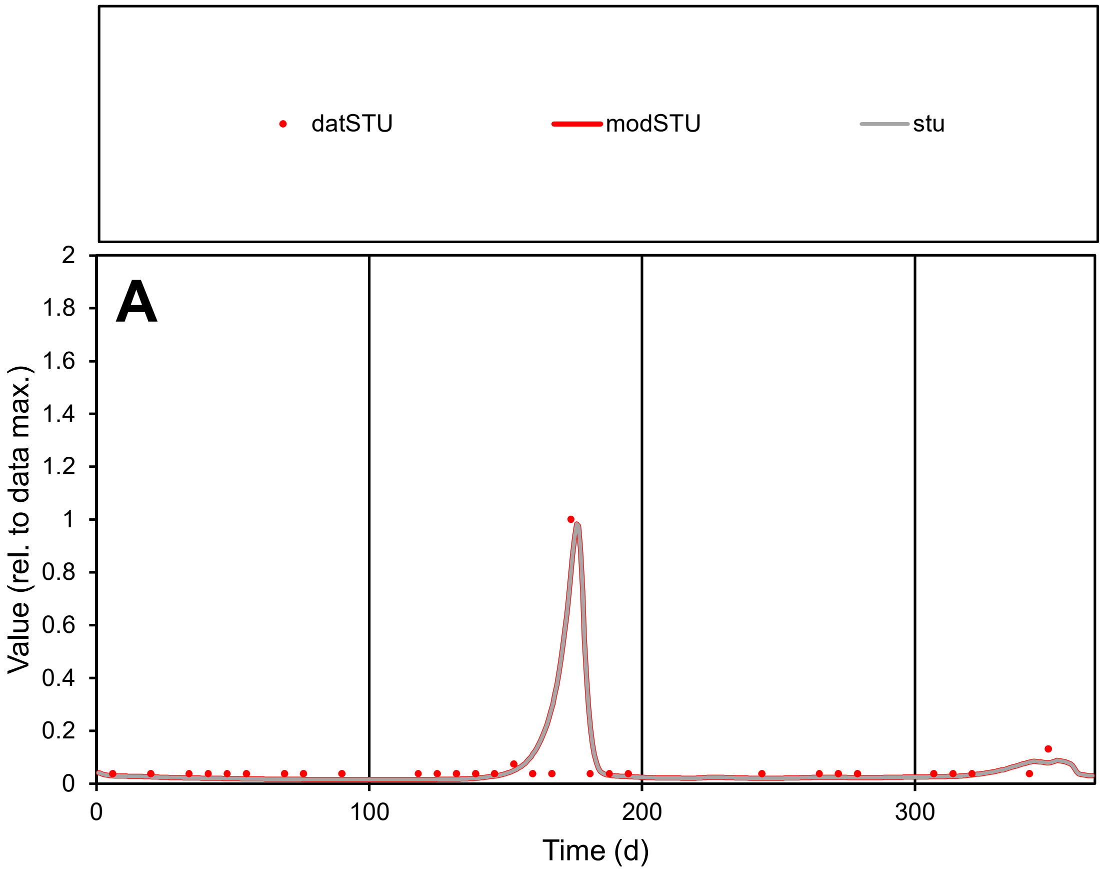


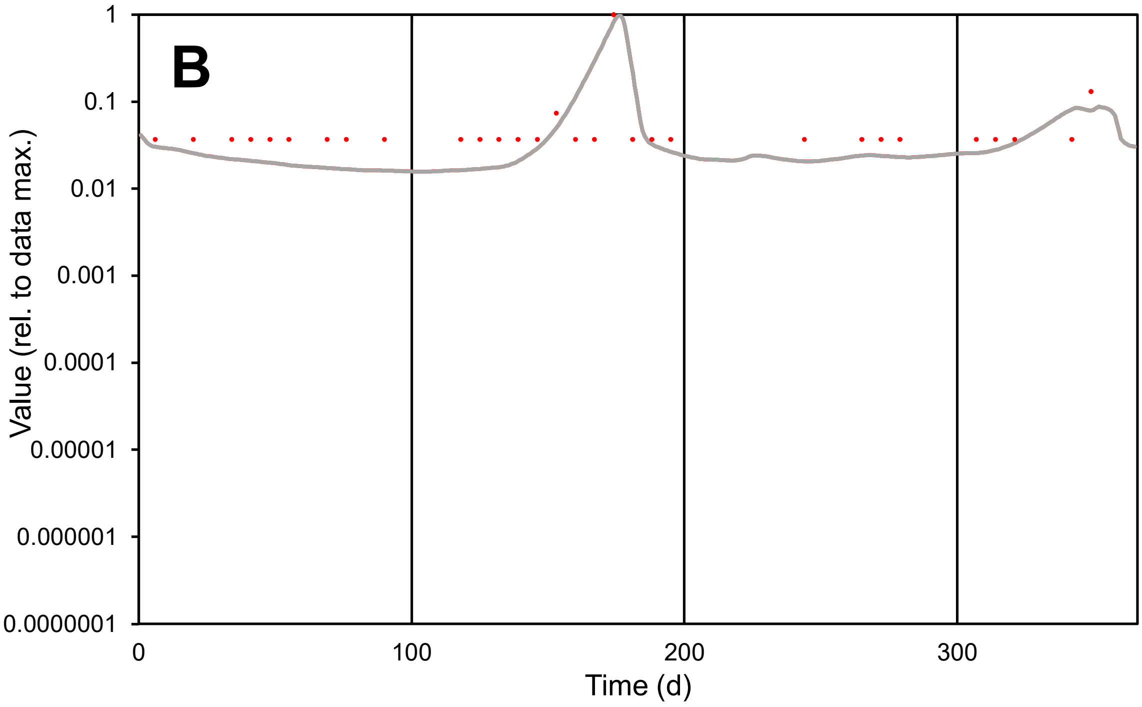


**C1**

Variables
STU Provide jtd or Dst
CalibOnly?
2 1 to reproduce of
oNormalize?
2 1=no, 2=yes

ofxn calc*
0.57746
ofxn mod
0.73179

sRun
RunA1
zout
1

Ave
Data
4.9E-06
Model
4.7E-06

data/vMax
6.7E-05

ofxng
275.646

Min d/m
2.2977

ave/min d
1.98387
ave/min m
4.32037

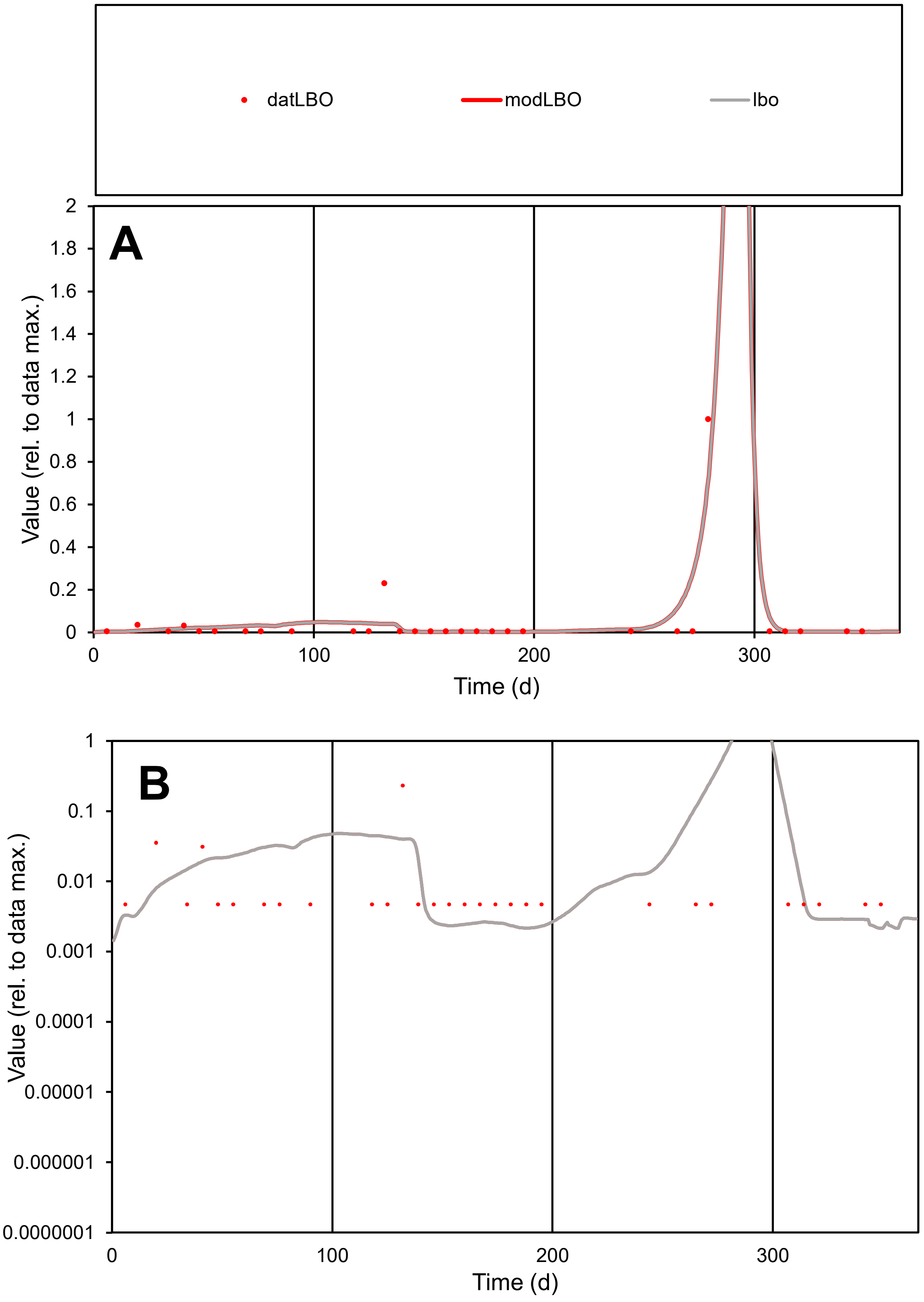


**C1**

Variables

LBO Provide jtd or Dst

CalibOnly?

2 1 to reproduce of

oNormalize?

2 1=no, 2=yes

ofxn calc*

1.17096

ofxn mod

1.46428

sRun

RunA1

zout

1

Ave

Data

1.4E-05

Model

1.6E-05

data/vMax

0.00029

ofxng

275.646

Min d/m

2.17671

ave/min d

10.0306

ave/min m

24.5555

325_Calib_S_A

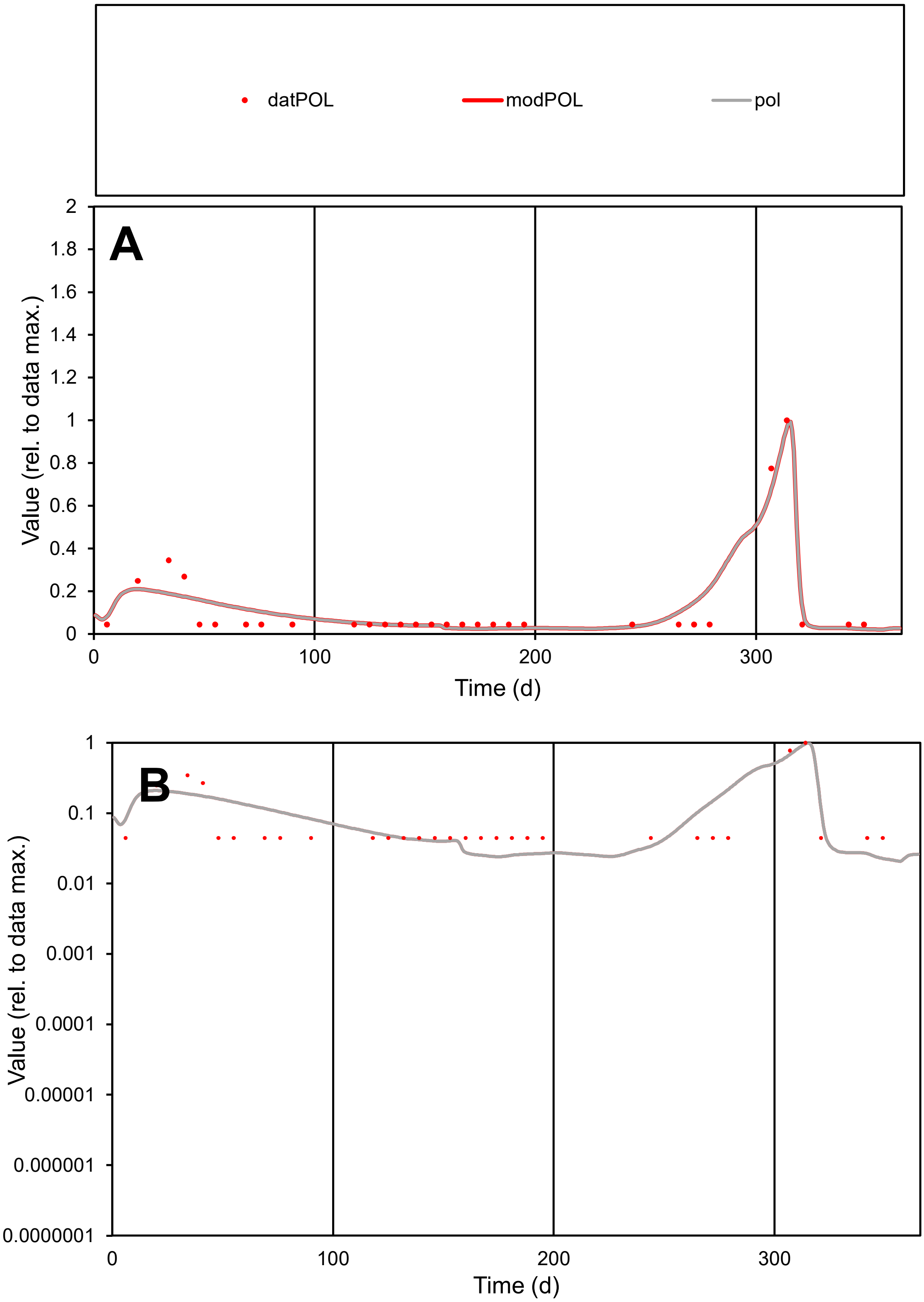


**C1**

Variables
POL Provide jtd or Dst
CalibOnly?
2 1 to reproduce of
oNormalize?
2 1=no, 2=yes

ofxn calc*
0.5024
ofxn mod
0.5679

sRun
RunA1
zout
1

Ave
Data
4.5E-06
Model
4.8E-06

data/vMax
3.6E-05

ofxng
275.646

Min d/m
1.99525

ave/min d
2.81571
ave/min m
5.9933

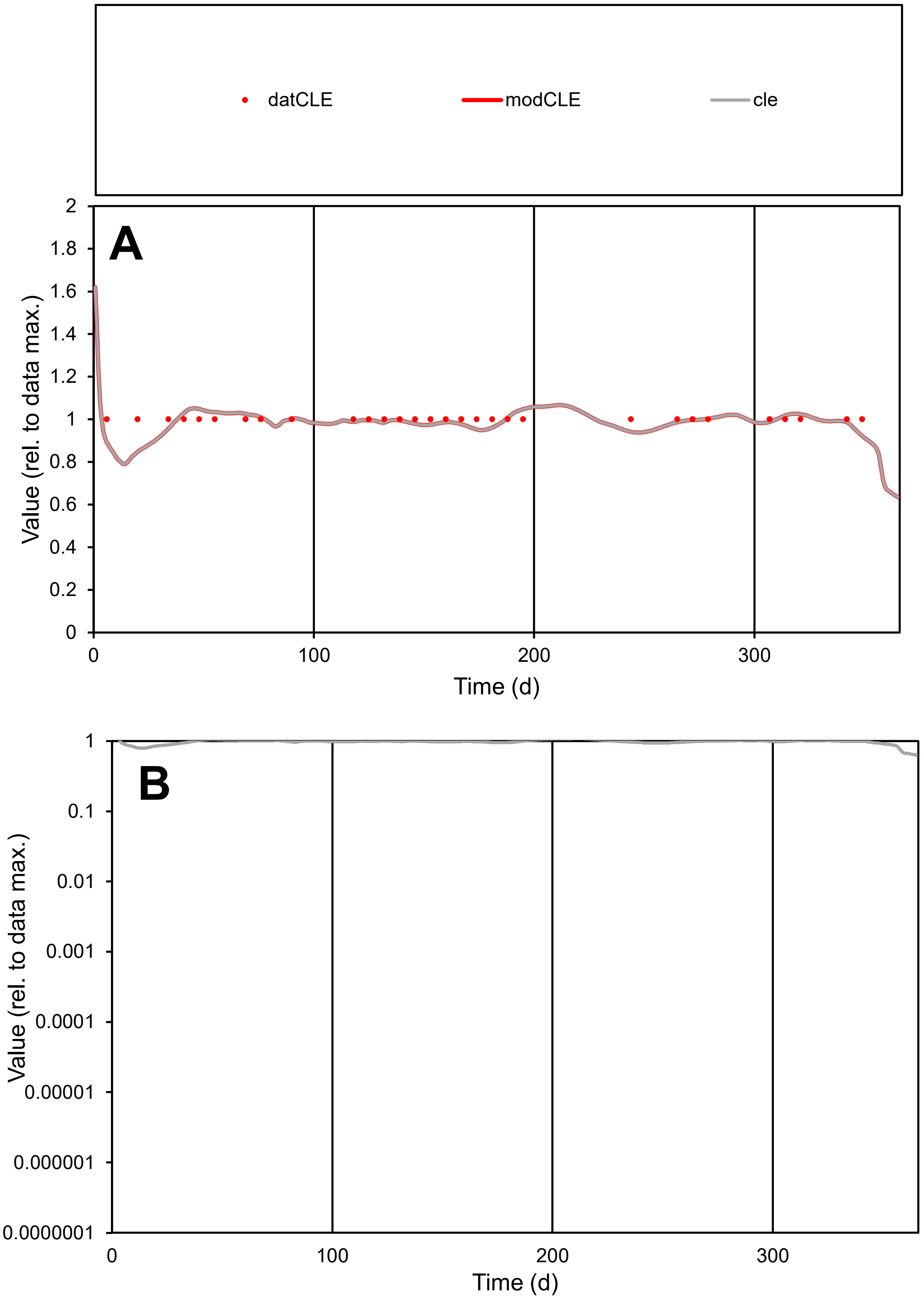


**C1**

Variables
CLE Provide jtd or Dst
CalibOnly?
2 1 to reproduce of
oNormalize?
2 1=no, 2=yes

ofxn calc*
0.04704
ofxn mod
0.12144

sRun
RunA1
zout
1

Ave
Data
4.7E-06
Model
4.6E-06

data/vMax
4.7E-06

ofxng
275.646

Min d/m
1.16845

ave/min d
1
ave/min m
1.15185

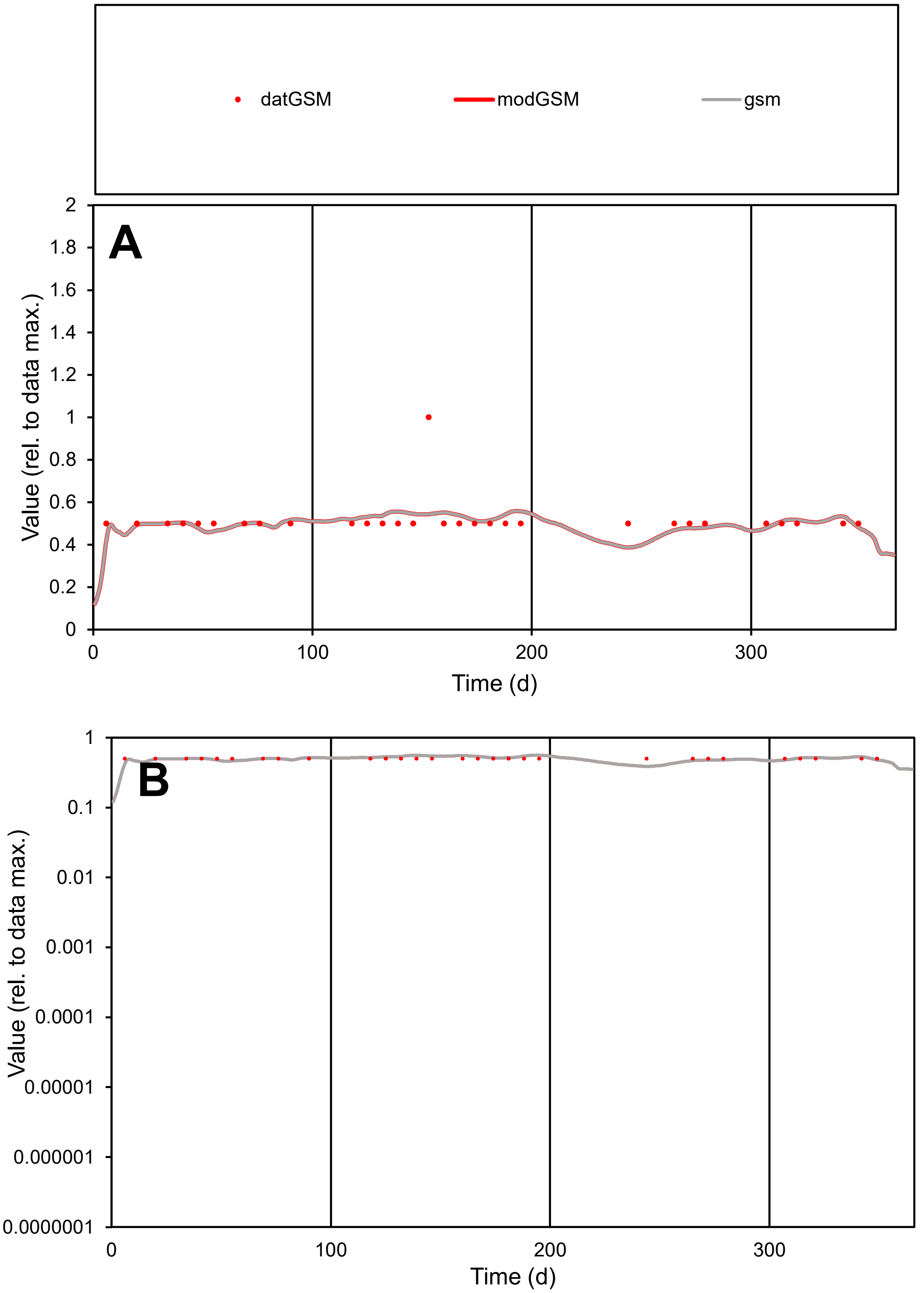


**C1**

Variables
GSM Provide jtd or Dst
CalibOnly?
2 1 to reproduce of
oNormalize?
2 1=no, 2=yes

ofxn calc*
0.17436
ofxn mod
0.20628

sRun
RunA1
zout
1

Ave
Data
1.6E-06
Model
1.6E-06

data/vMax
3.2E-06

ofxng
275.646

Min d/m
1.29003

ave/min d
1.03333
ave/min m
1.31122

325_Calib_S_A

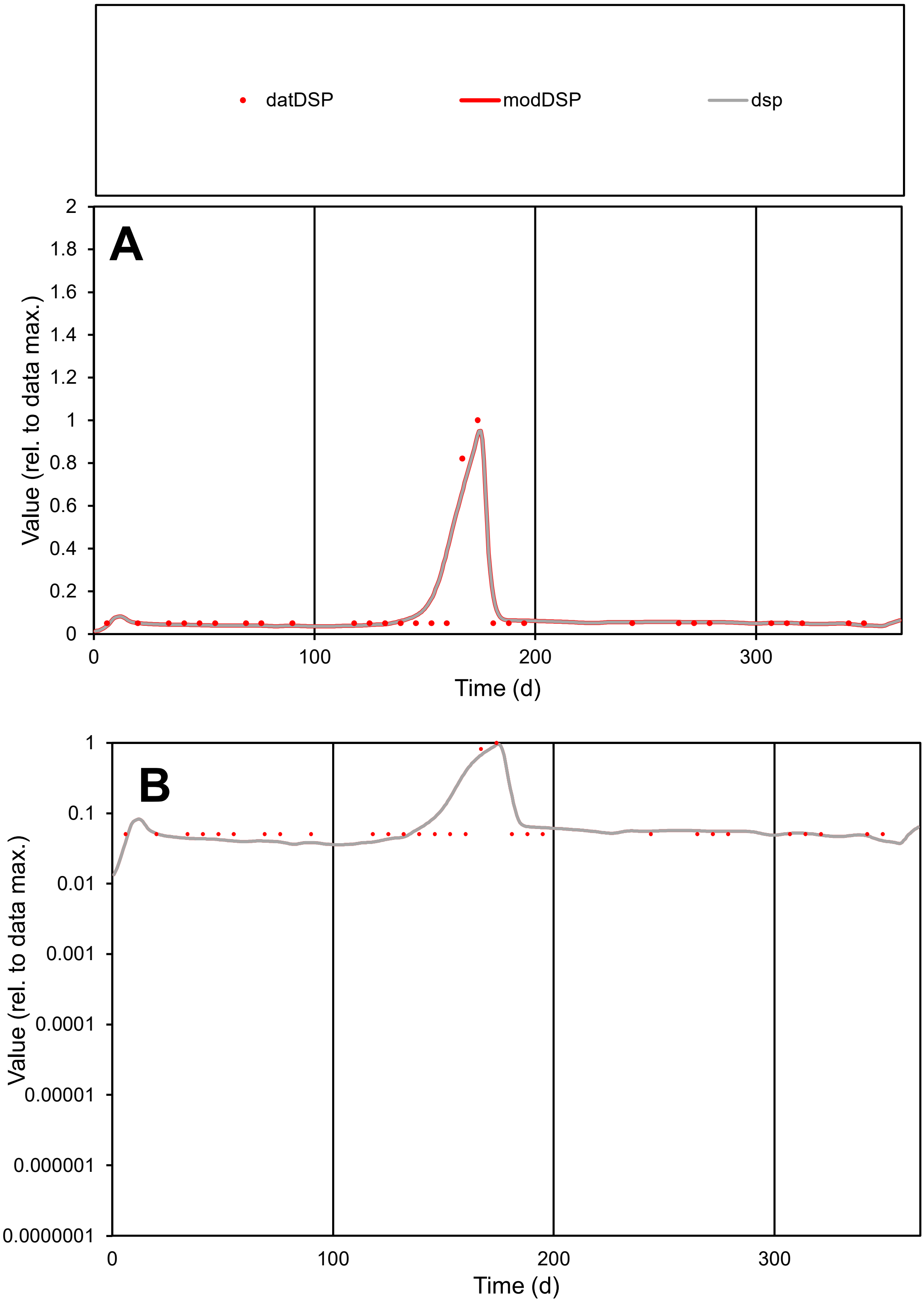


**C1**

Variables
DSP Provide jtd or Dst
CalibOnly?
2 1 to reproduce of
oNormalize?
2 1=no, 2=yes

ofxn calc*
0.59779
ofxn mod
0.66594

sRun
RunA1
zout
1

Ave
Data
1.1E-05
Model
1.2E-05

data/vMax
9.9E-05

ofxng
275.646

Min d/m
1.32198

ave/min d
2.13513
ave/min m
3.07126

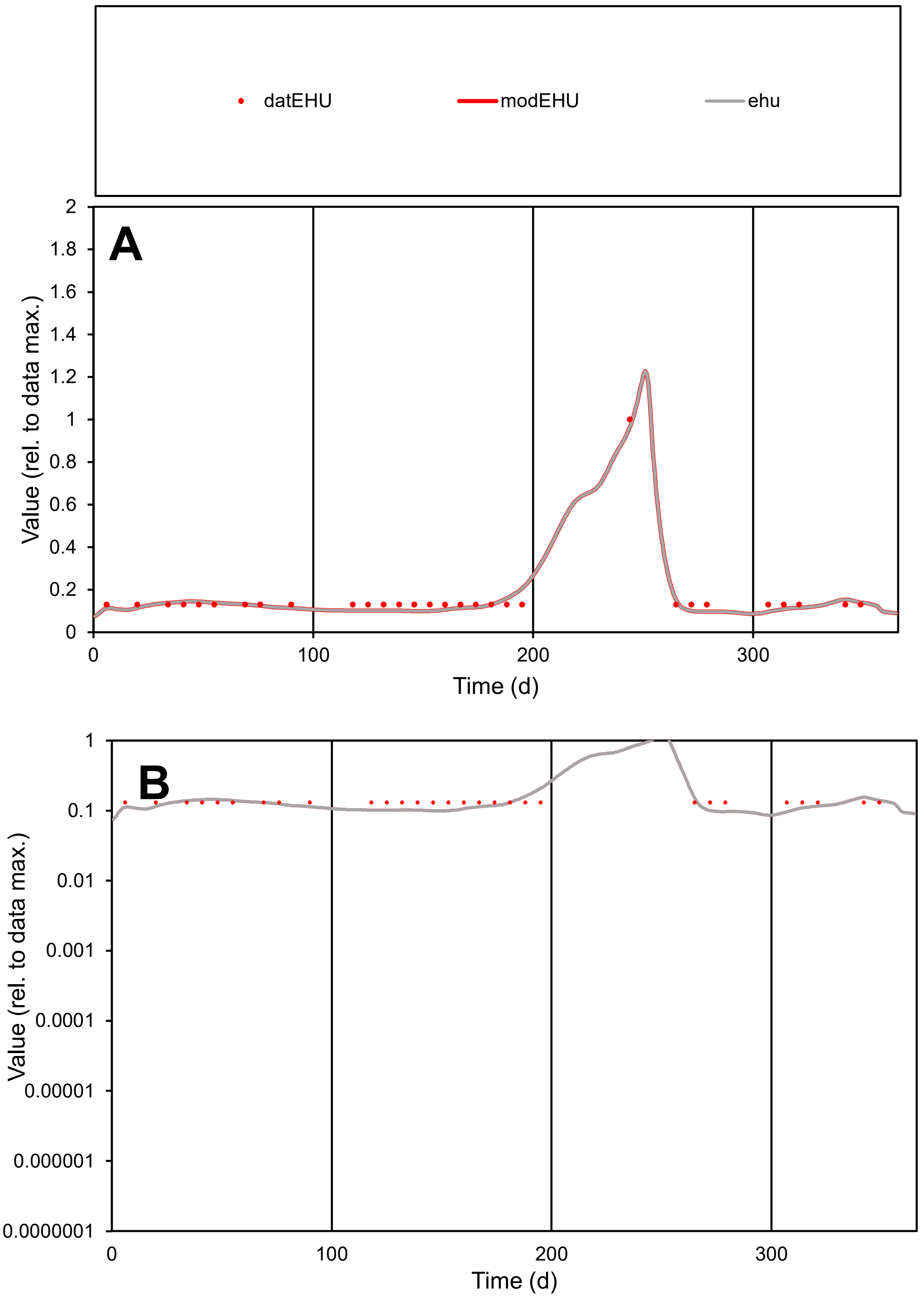


**C1**

Variables
EHU Provide jtd or Dst
CalibOnly?
2 1 to reproduce of
oNormalize?
2 1=no, 2=yes

ofxn calc*
0.16094
ofxn mod
0.20462

sRun
RunA1
zout
1

Ave
Data
7E-06
Model
6.7E-06

data/vMax
4.4E-05

ofxng
275.646

Min d/m
1.34118

ave/min d
1.22256
ave/min m
1.56747

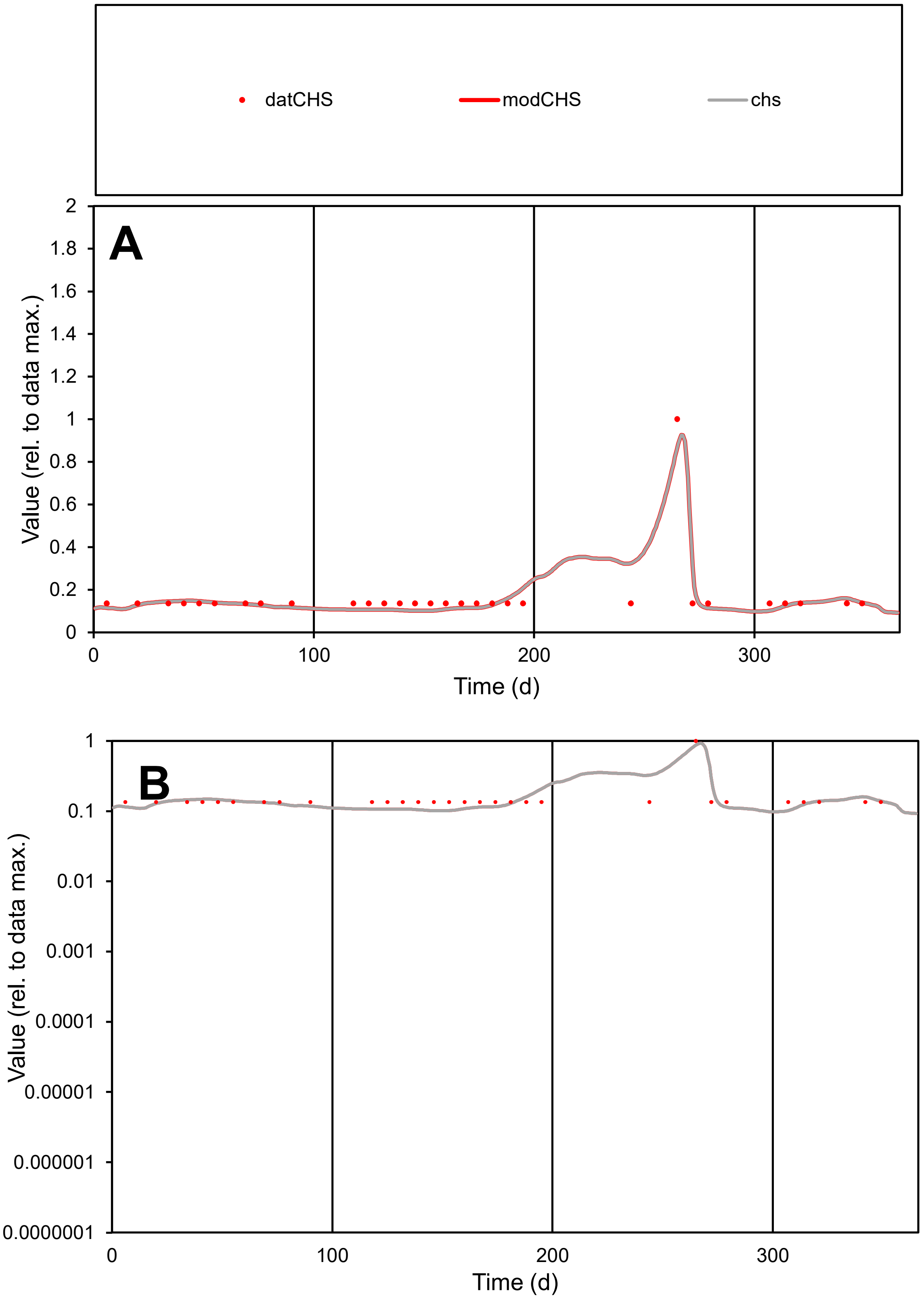


**C1**

Variables
CHS Provide jtd or Dst
CalibOnly?
2 1 to reproduce of
oNormalize?
2 1=no, 2=yes

ofxn calc*
0.28054
ofxn mod
0.3172

sRun
RunA1
zout
1

Ave
Data
2.6E-05
Model
2.6E-05

data/vMax
0.00016

ofxng
275.646

Min d/m
1.33271

ave/min d
1.21367
ave/min m
1.60399

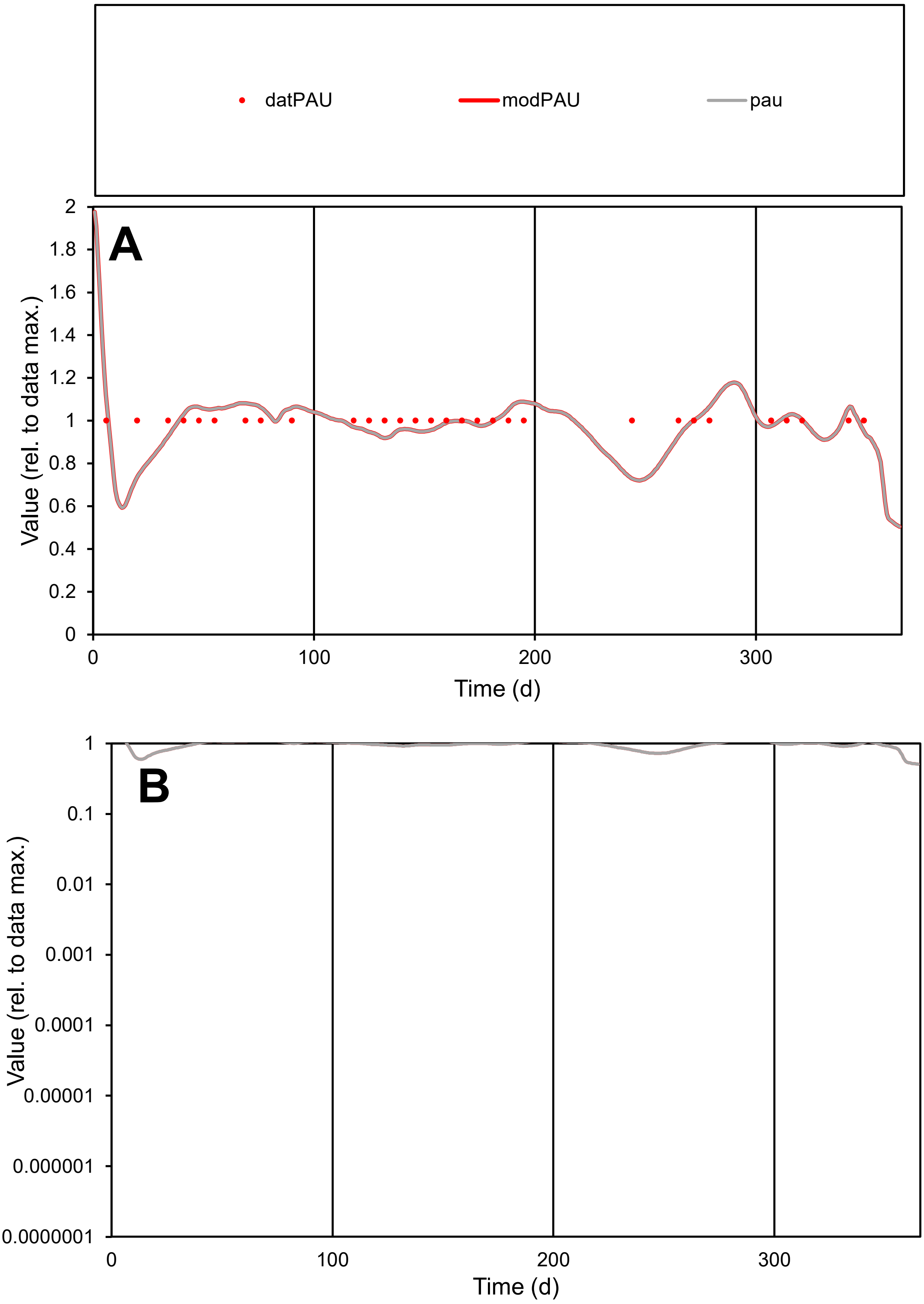


**C1**

Variables
PAU Provide jtd or Dst
CalibOnly?
2 1 to reproduce of
oNormalize?
2 1=no, 2=yes

ofxn calc*
0.08427
ofxn mod
0.1406

sRun
RunA1
zout
1

Ave
Data
3.9E-06
Model
3.9E-06

data/vMax
3.9E-06

ofxng
275.646

Min d/m
1.38012

ave/min d
1
ave/min m
1.36248

325_Calib_S_A

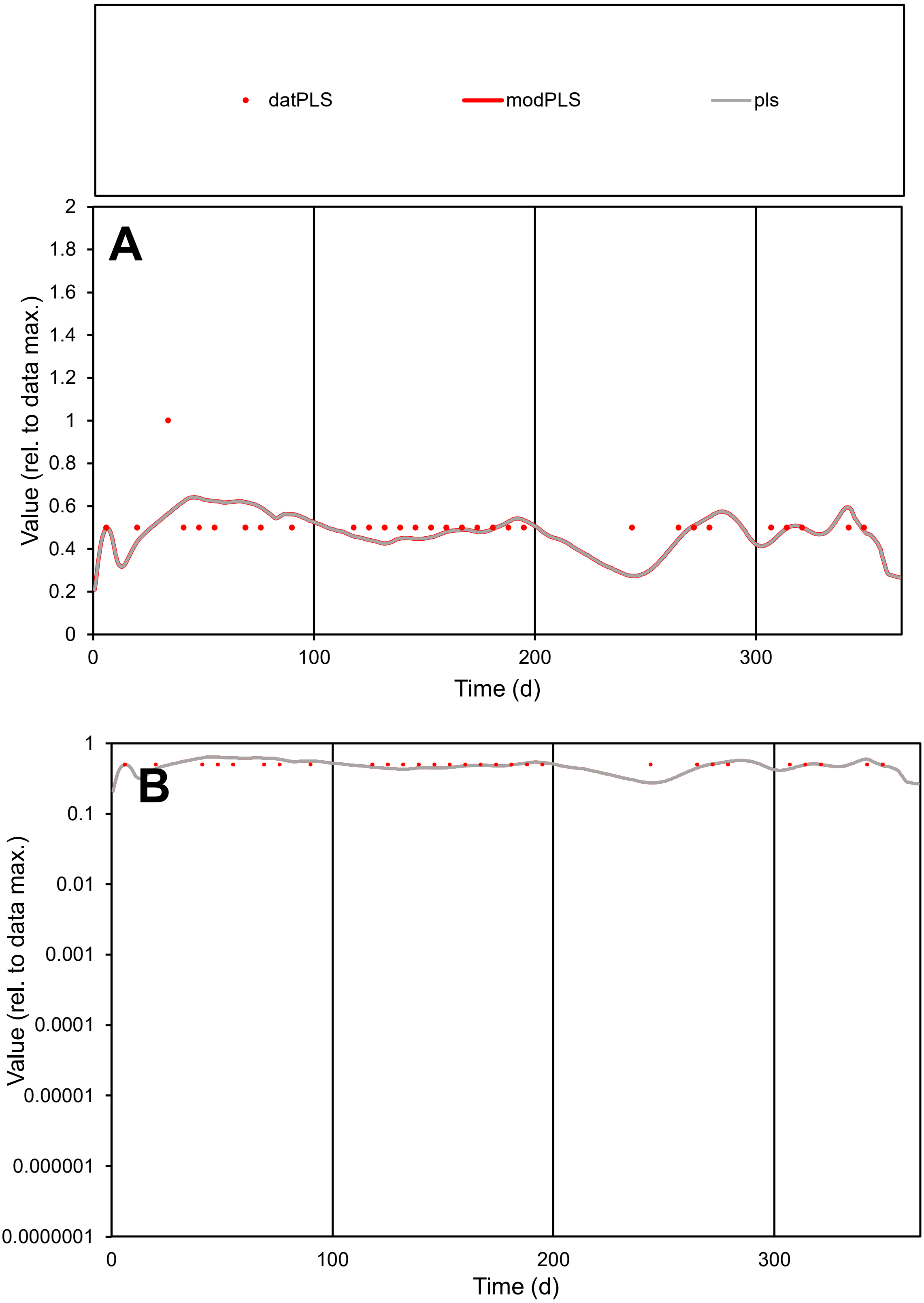


**C1**

Variables
PLS Provide jtd or Dst
CalibOnly?
2 1 to reproduce of
oNormalize?
2 1=no, 2=yes

ofxn calc*
0.20699
ofxn mod
0.23684

sRun
RunA1
zout
1

Ave
Data
4E-06
Model
3.9E-06

data/vMax
7.7E-06

ofxng
275.646

Min d/m
1.82644

ave/min d
1.03332
ave/min m
1.84603

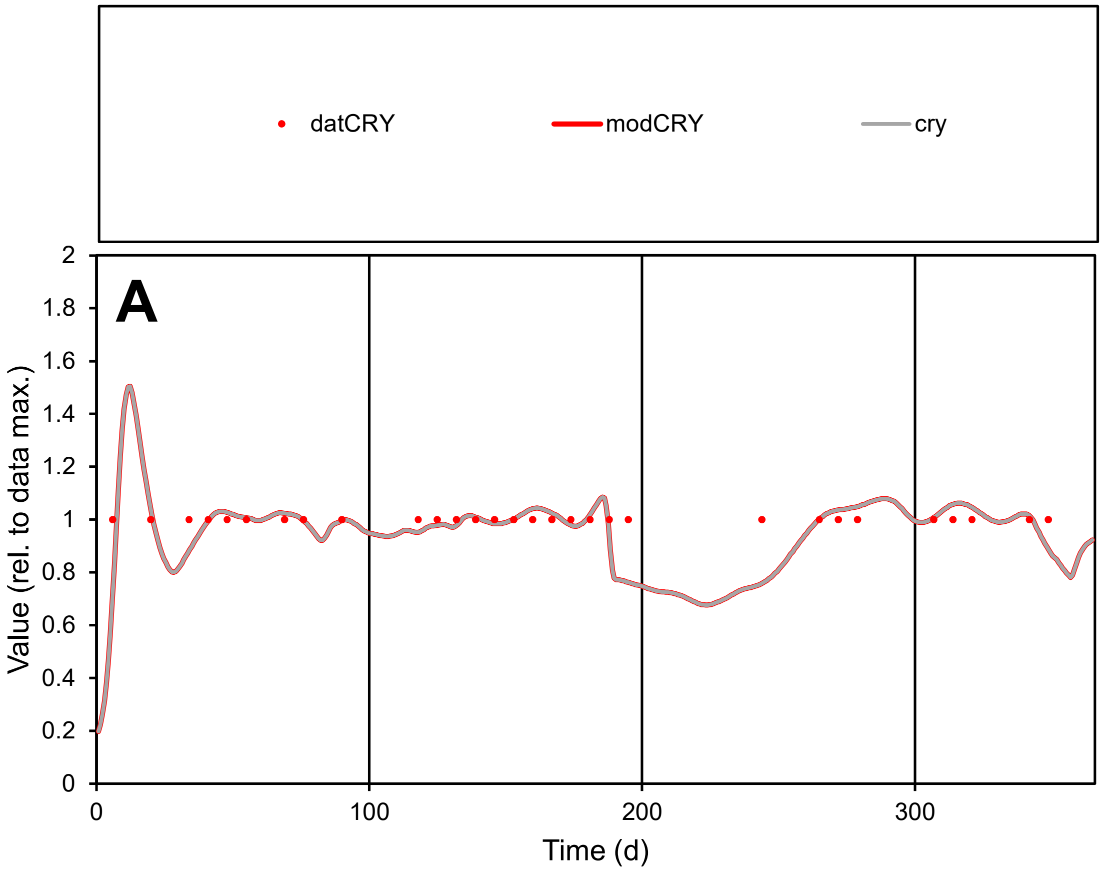


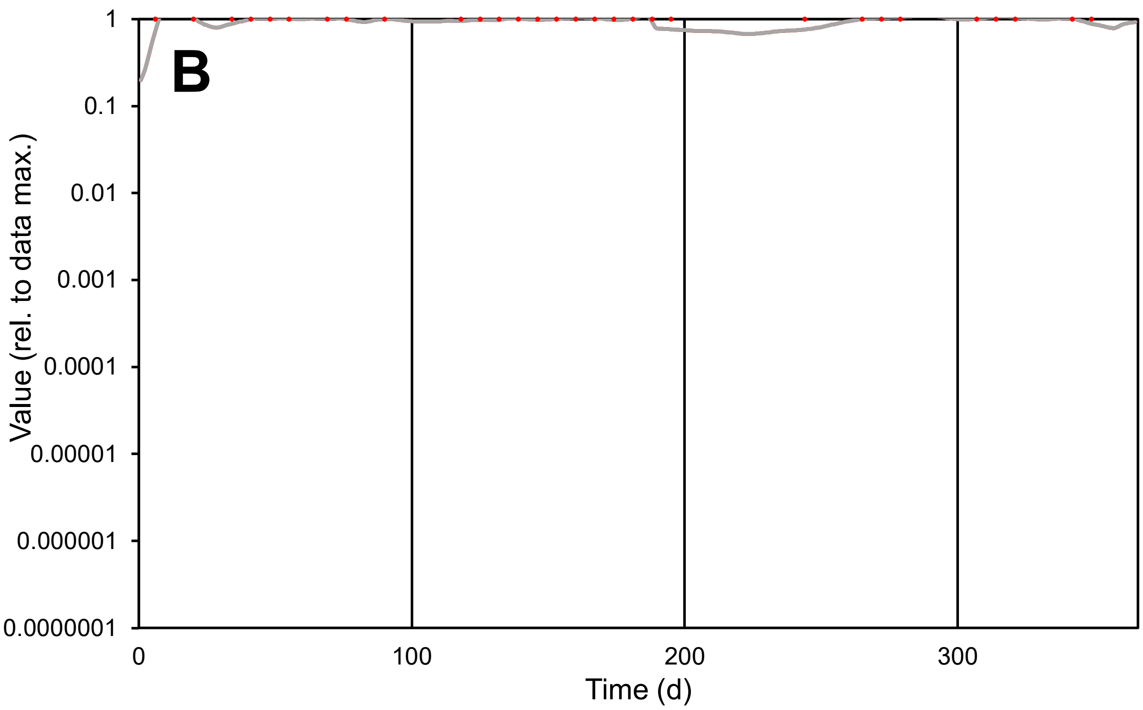


**C1**

Variables
CRY Provide jtd or Dst
CalibOnly?
2 1 to reproduce of
oNormalize?
2 1=no, 2=yes

ofxn calc*
0.07612
ofxn mod
0.13742

sRun
RunA1
zout
1

Ave
Data
3E-06
Model
2.9E-06

data/vMax
3E-06

ofxng
275.646

Min d/m
1.31949

ave/min d
1
ave/min m
1.29333

325_Calib_S_A

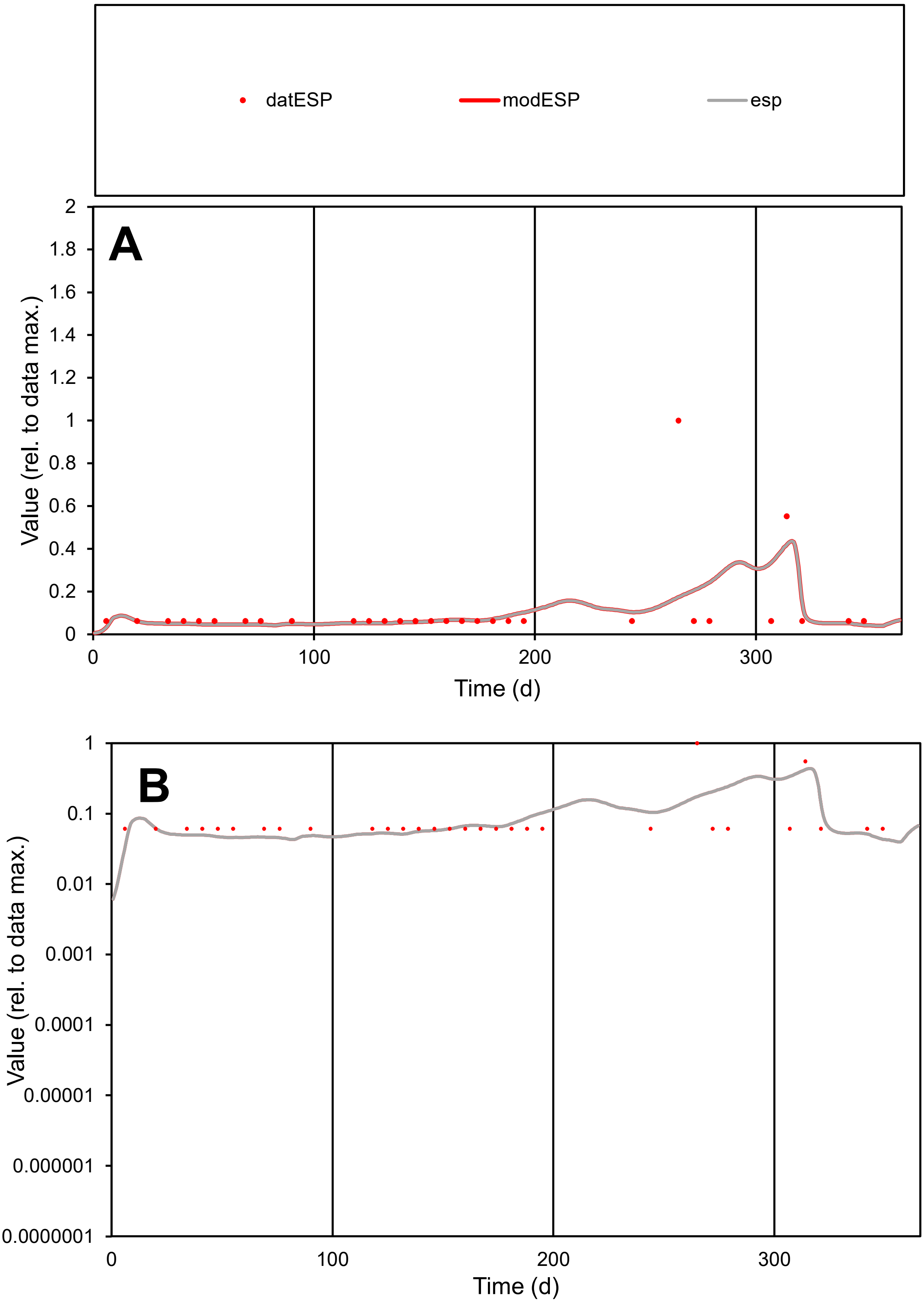


**C1**

Variables
ESP Provide jtd or Dst
CalibOnly?
2 1 to reproduce of
oNormalize?
2 1=no, 2=yes

ofxn calc*
1.5308
ofxn mod
1.54097

sRun
RunA1
zout
1

Ave
Data
1.4E-05
Model
1.3E-05

data/vMax
0.00013

ofxng
275.646

Min d/m
1.434

ave/min d
1.78153
ave/min m
2.30663

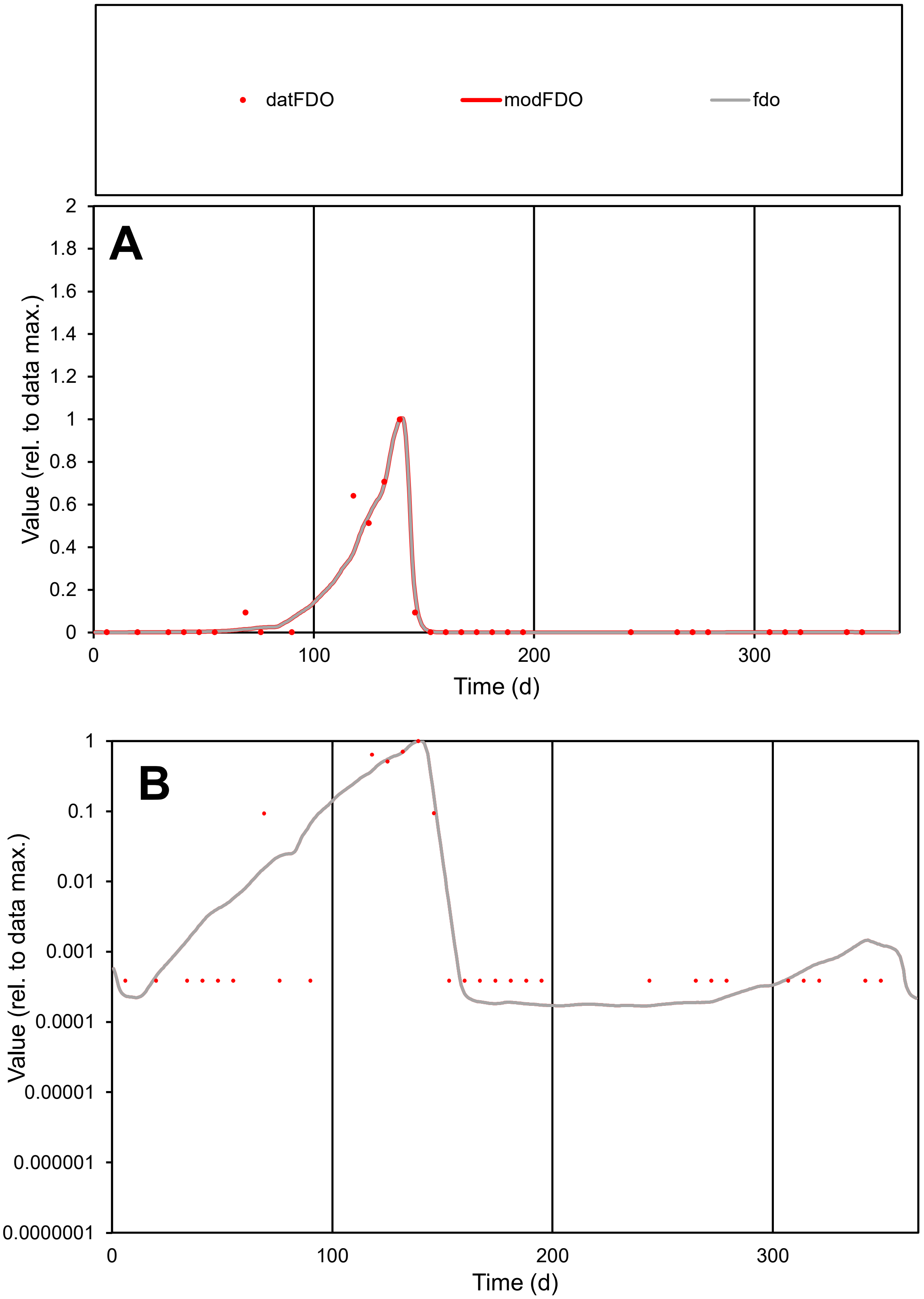


**C1**

Variables
FDO Provide jtd or Dst
CalibOnly?
2 1 to reproduce of
oNormalize?
2 1=no, 2=yes

ofxn calc*
0.46984
ofxn mod
0.54964

sRun
RunA1
zout
1

Ave
Data
2.6E-05
Model
2.5E-05

data/vMax
0.00026

ofxng
275.646

Min d/m
2.25765

ave/min d
263.408
ave/min m
568.579

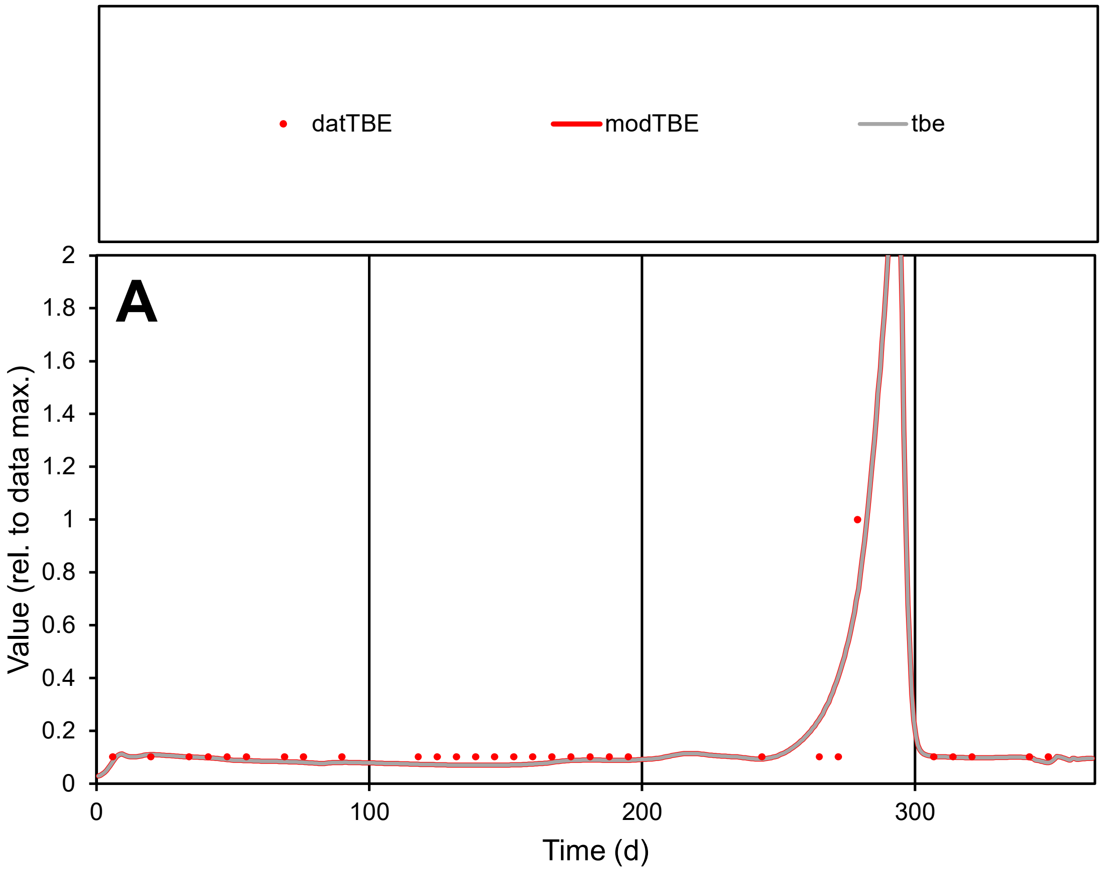


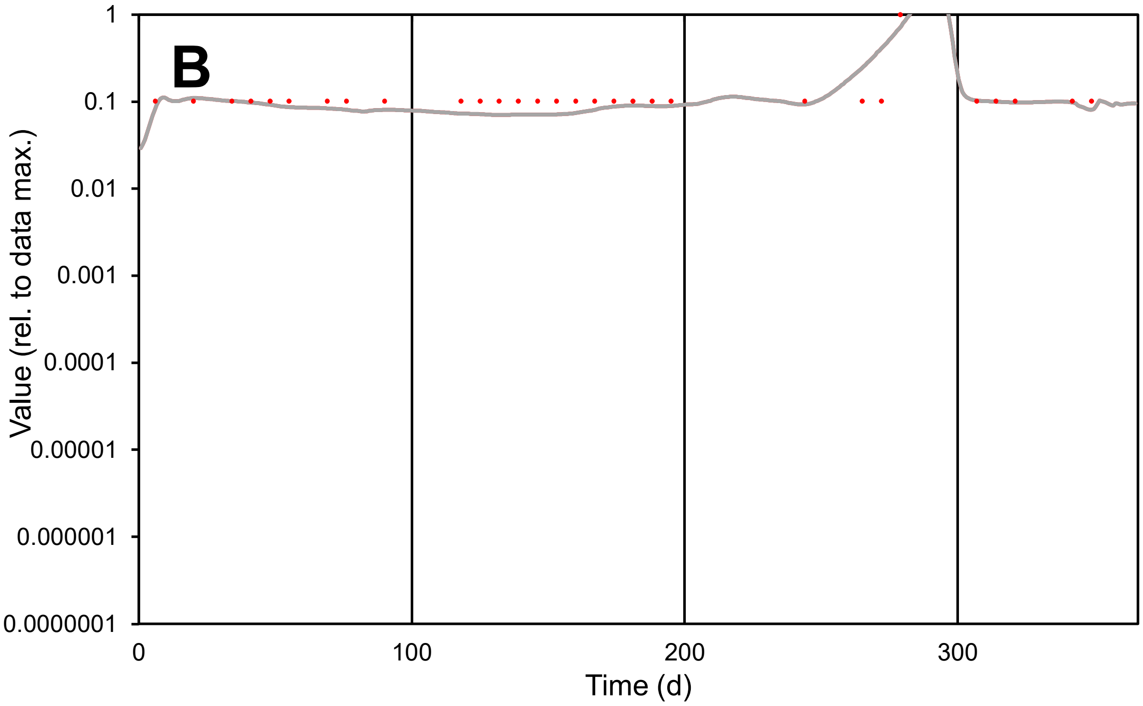


**C1**

Variables

| | |
|---|---|
| TBE | Provide jtd or Dst |
| CalibOnly? | |
| 2 | 1 to reproduce of |
| oNormalize? | |
| 2 | 1=no, 2=yes |

ofxn calc*
0.51181
ofxn mod
0.5835

sRun
RunA1
zout
1

Ave
Data
8.7E-06
Model
8.4E-06

data/vMax
6.6E-05

ofxng
275.646

Min d/m
1.44013

ave/min d
1.29655
ave/min m
1.8224

325_Calib_S_A

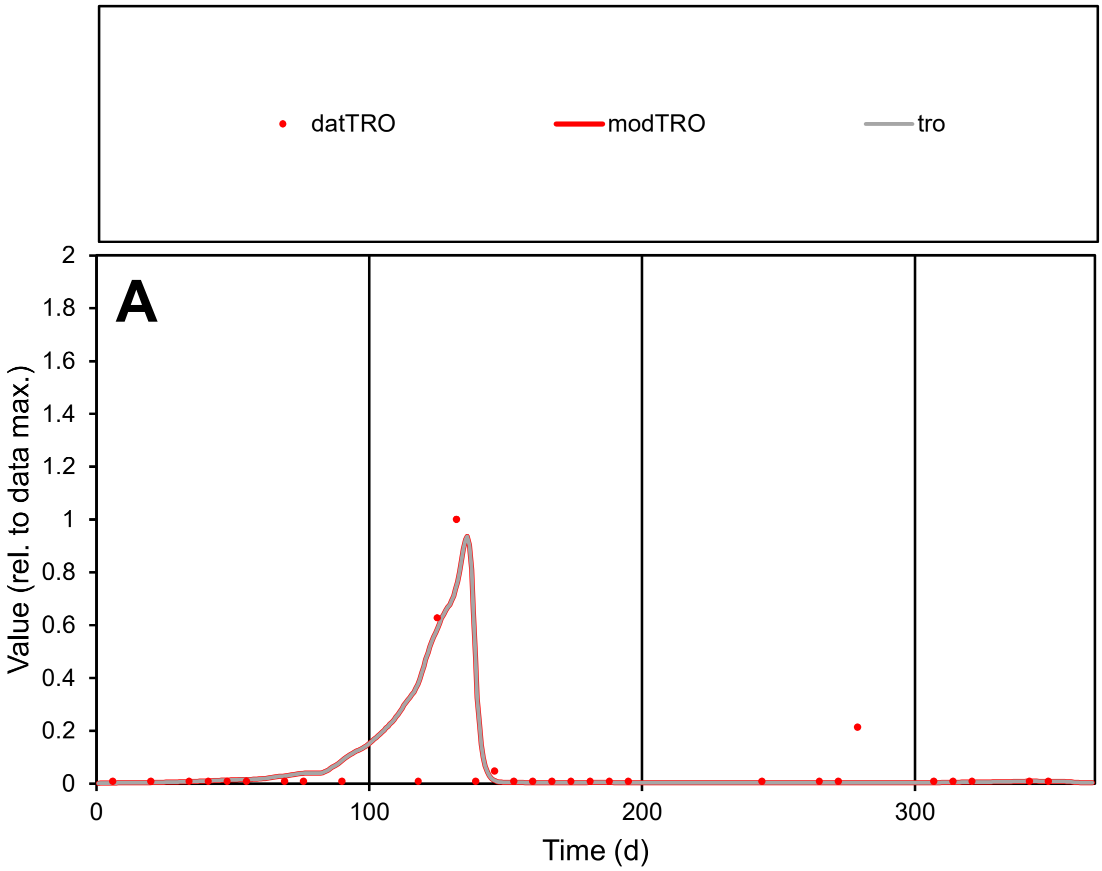


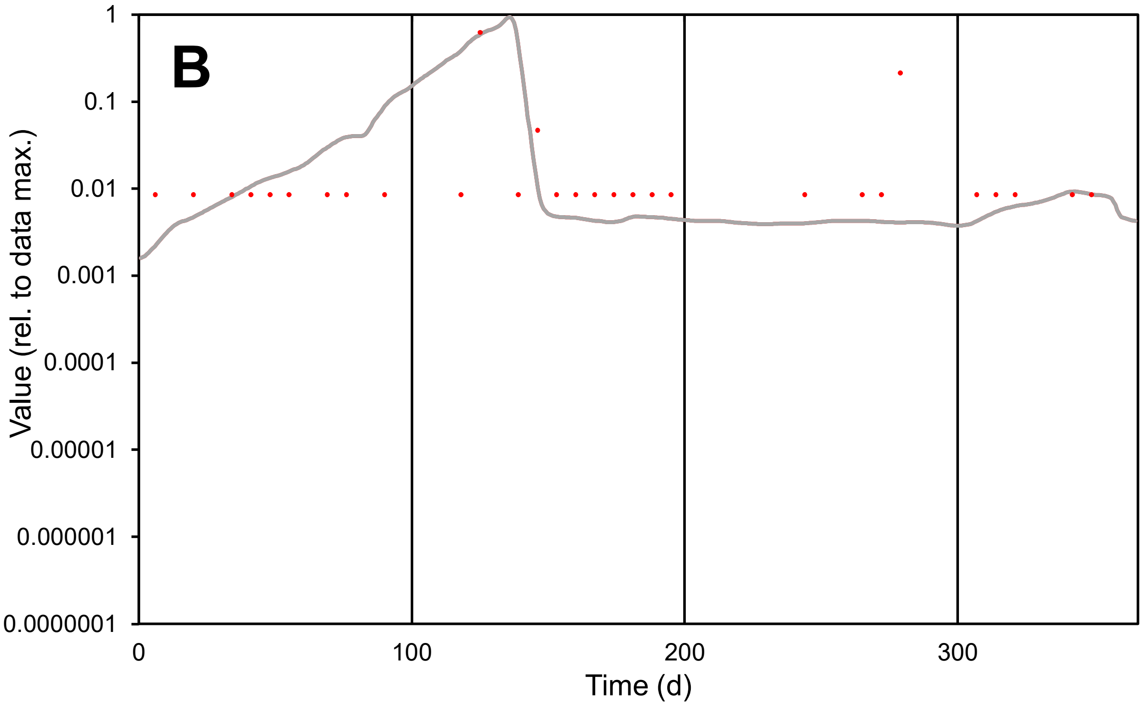


**C1**

Variables

TRO Provide jtd or Dst

CalibOnly?

2 1 to reproduce of

oNormalize?

2 1=no, 2=yes

ofxn calc*

1.20885

ofxn mod

1.31034

sRun

RunA1

zout

1

Ave

Data

4.1E-05

Model

4.2E-05

data/vMax

0.00059

ofxng

275.646

Min d/m

3.49409

ave/min d

8.2232

ave/min m

29.4583

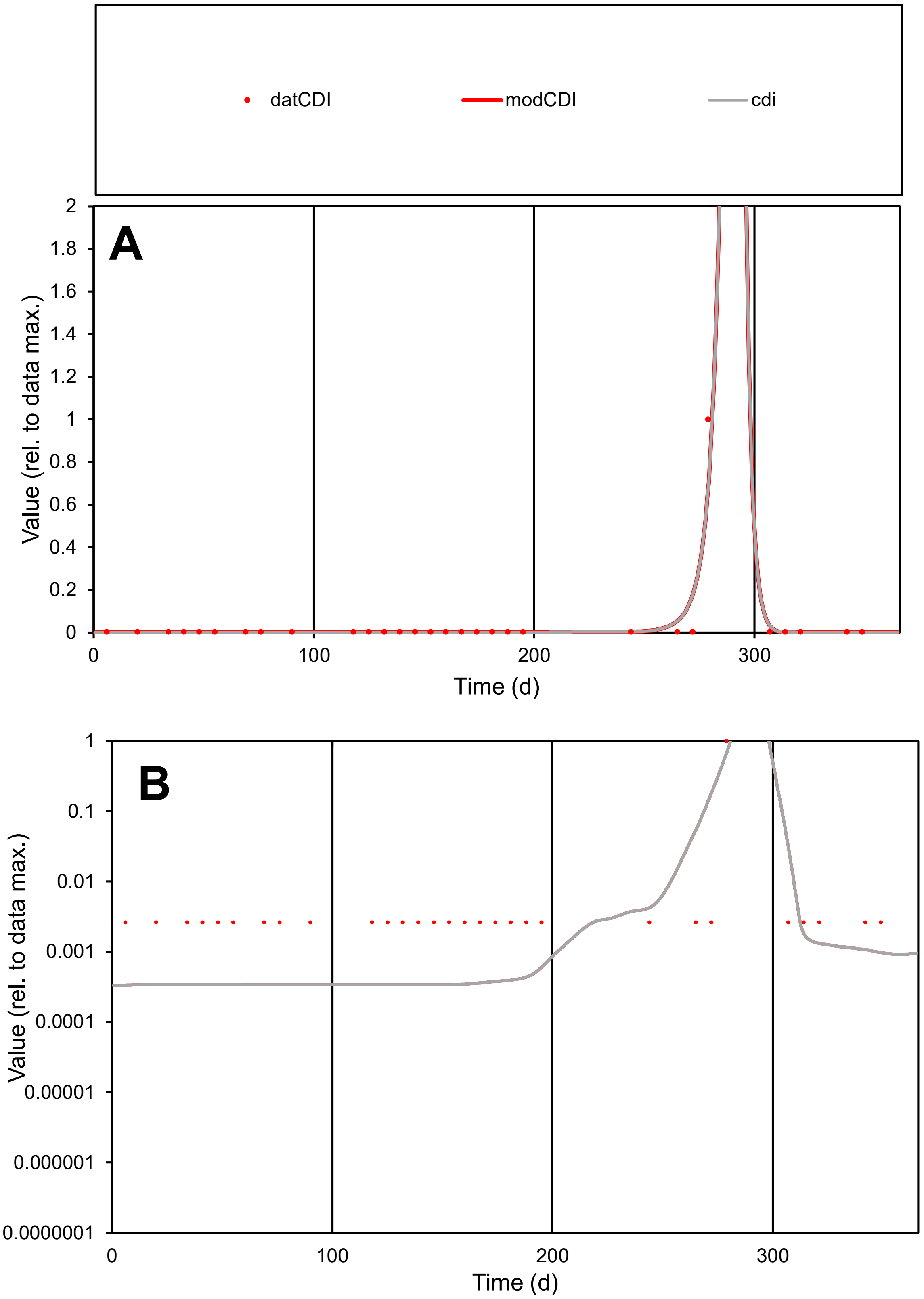


**C1**

Variables
CDI Provide jtd or Dst
CalibOnly?
2 1 to reproduce of
oNormalize?
2 1=no, 2=yes

ofxn calc*
0.58131
ofxn mod
1.08907

sRun
RunA1
zout
1

Ave
Data
2.4E-06
Model
2.5E-06

data/vMax
6.6E-05

ofxng
275.646

Min d/m
7.81588

ave/min d
13.6803
ave/min m
113.759

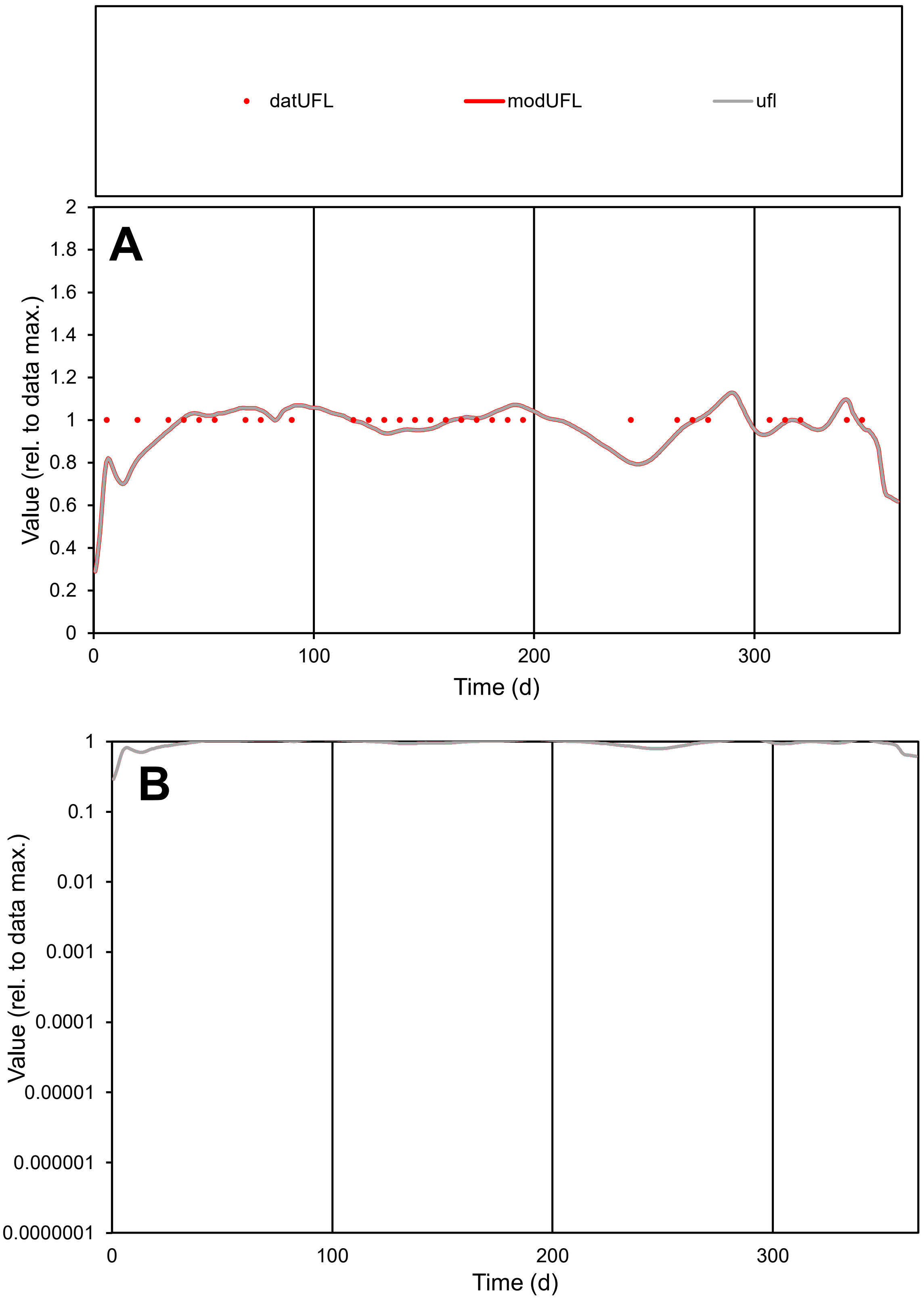


**C1**

Variables
UFL Provide jtd or Dst
CalibOnly?
2 1 to reproduce of
oNormalize?
2 1=no, 2=yes

ofxn calc*
0.07173
ofxn mod
0.13305

sRun
RunA1
zout
1

Ave
Data
3.1E-06
Model
3E-06

data/vMax
3.1E-06

ofxng
275.646

Min d/m
1.25646

ave/min d
1
ave/min m
1.23813

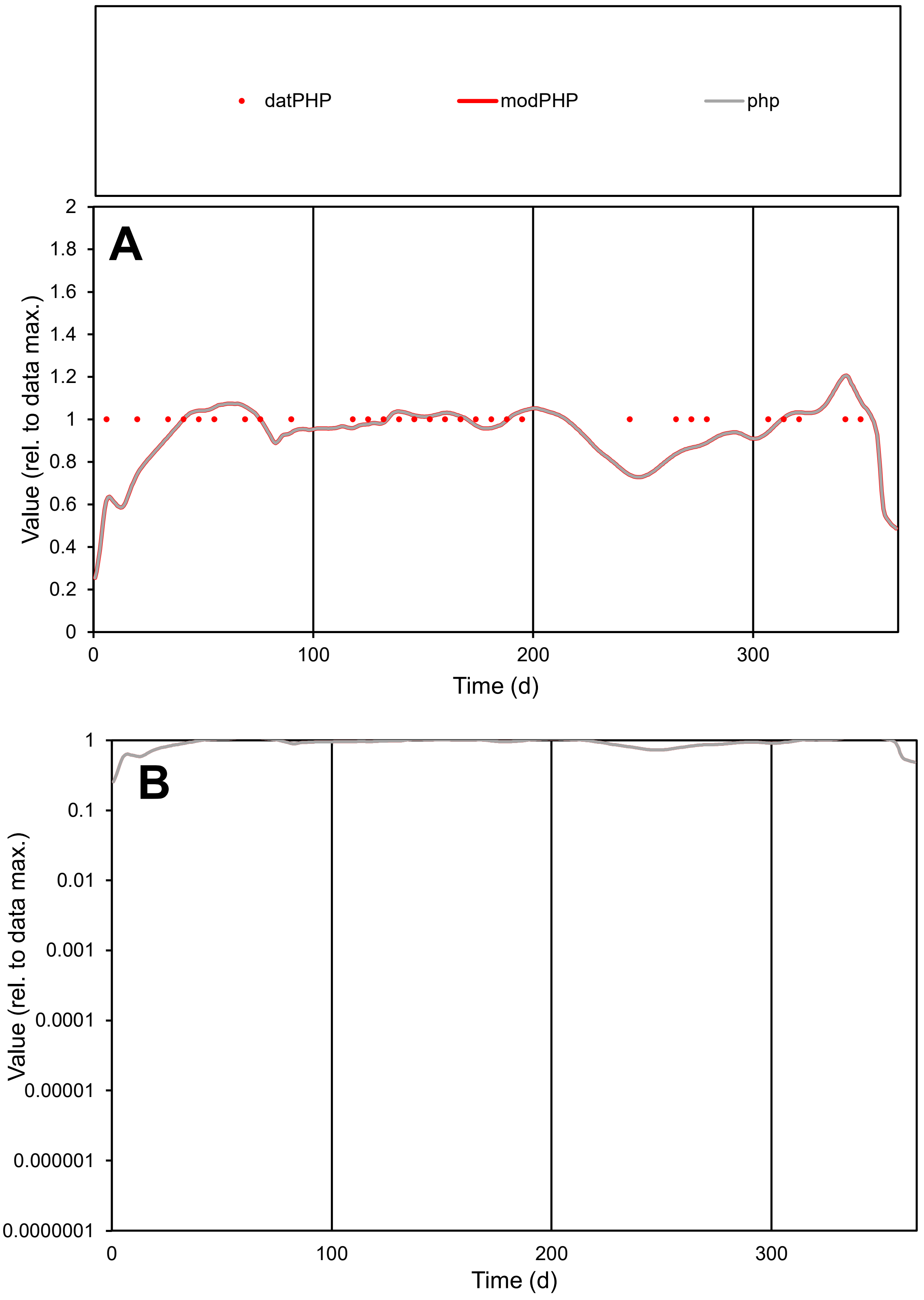


**C1**

Variables

PHP Provide jtd or Dst

CalibOnly?

2 1 to reproduce of

oNormalize?

2 1=no, 2=yes

ofxn calc*

0.11295

ofxn mod

0.15835

sRun

RunA1

zout

1

Ave

Data

6.9E-06

Model

6.7E-06

data/vMax

6.9E-06

ofxng

275.646

Min d/m

1.57143

ave/min d

1

ave/min m

1.52338

325_Calib_S_A

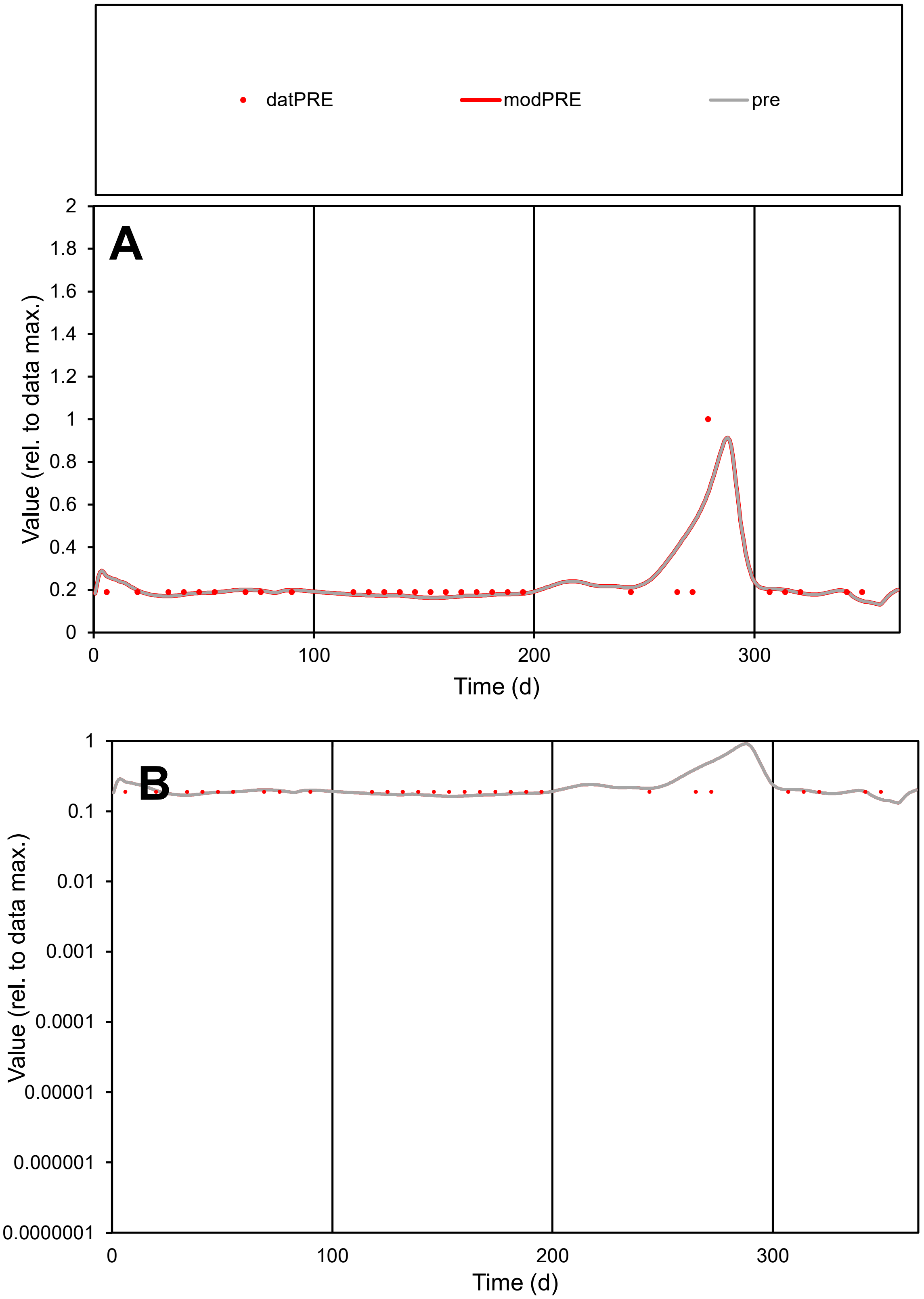


**C1**

Variables
PRE Provide jtd or Dst
CalibOnly?
2 1 to reproduce of
oNormalize?
2 1=no, 2=yes

ofxn calc*
0.42411
ofxn mod
0.45145

sRun
RunA1
zout
1

Ave
Data
1.5E-05
Model
1.6E-05

data/vMax
7.1E-05

ofxng
275.646

Min d/m
1.31024

ave/min d
1.14291
ave/min m
1.52608

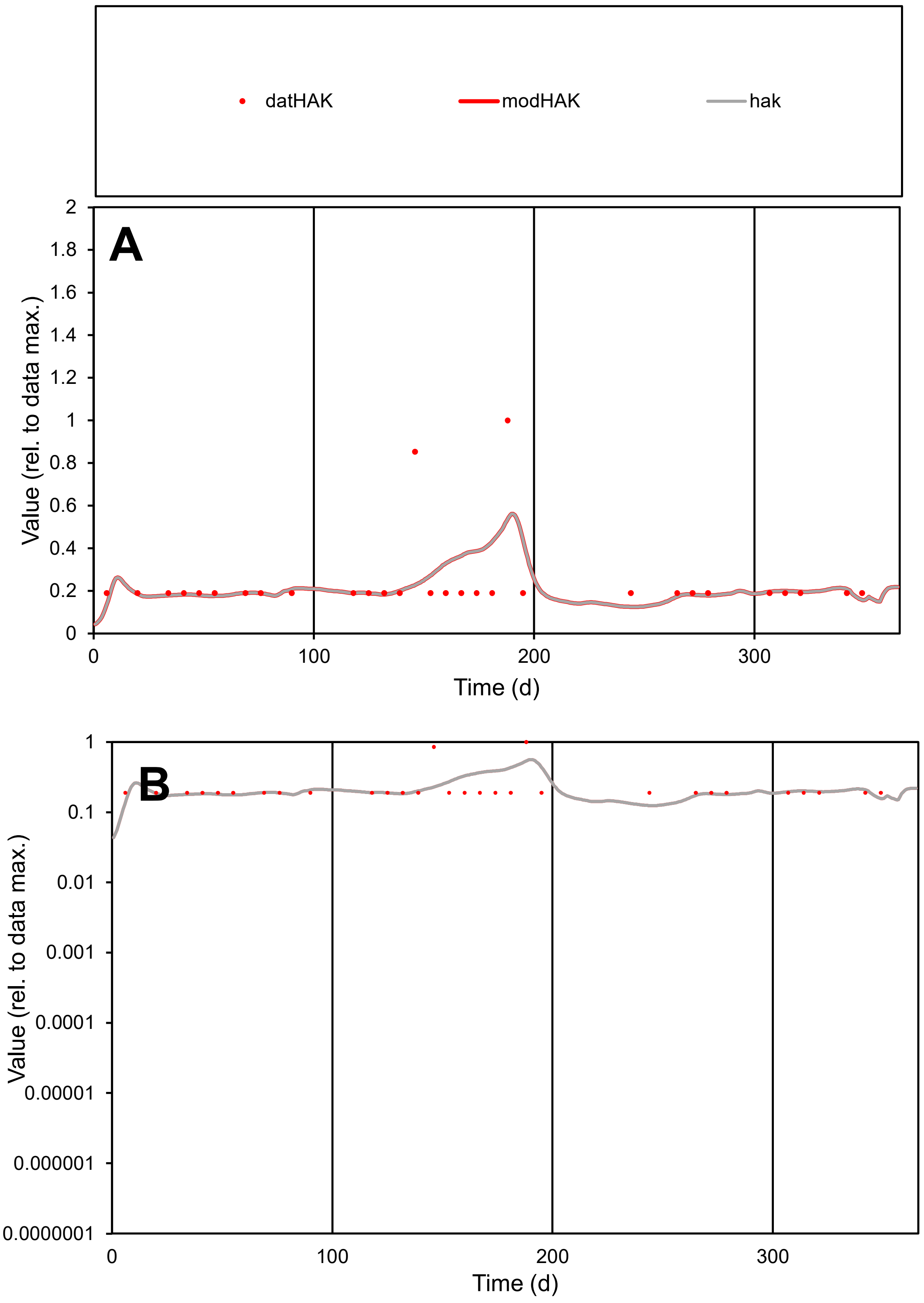


**C1**

Variables
HAK Provide jtd or Dst
CalibOnly?
2 1 to reproduce of
oNormalize?
2 1=no, 2=yes

ofxn calc*
0.67869
ofxn mod
0.69034

sRun
RunA1
zout
1

Ave
Data
6.4E-06
Model
6.3E-06

data/vMax
2.7E-05

ofxng
275.646

Min d/m
1.52269

ave/min d
1.25967
ave/min m
1.88818

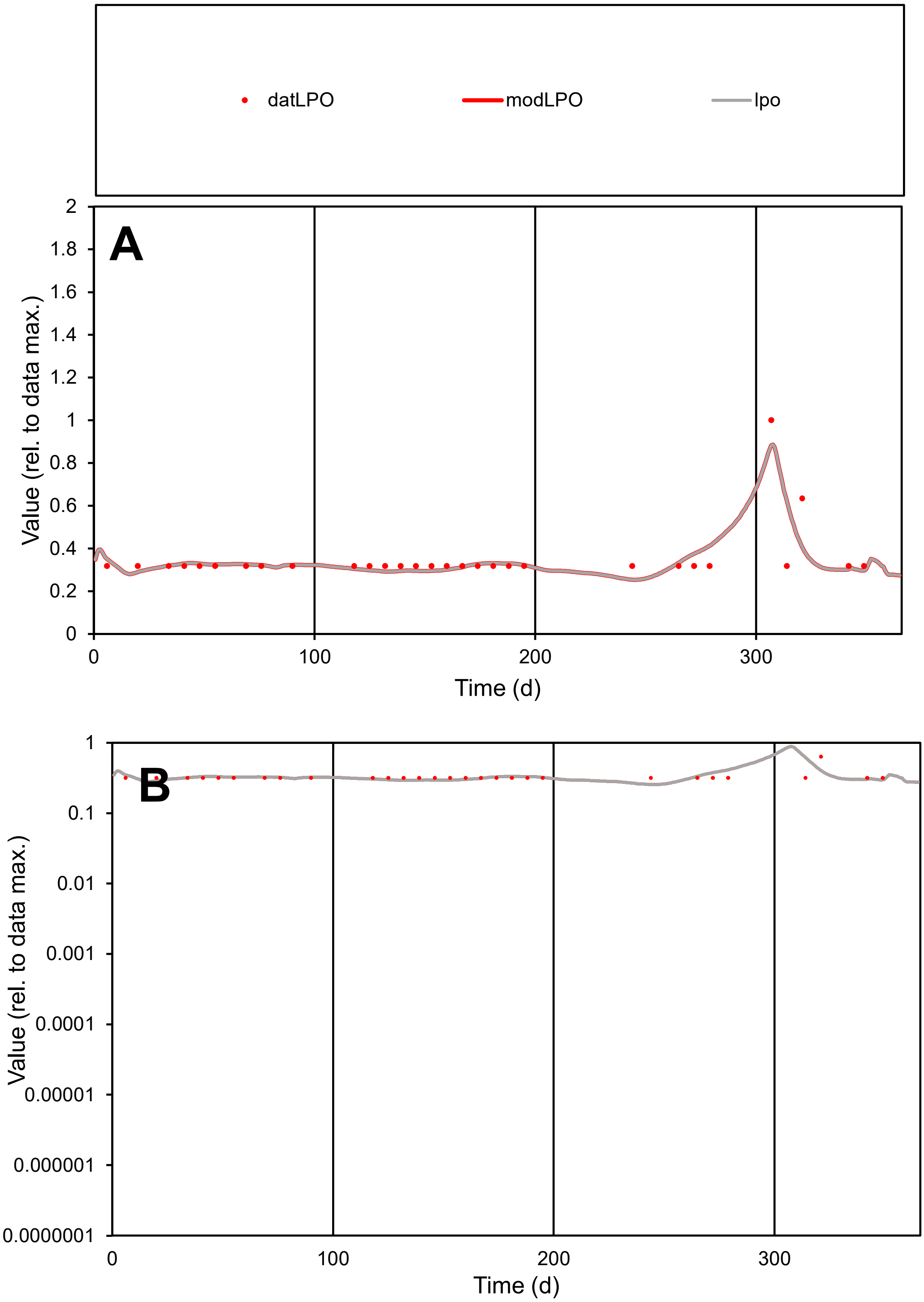


**C1**

Variables
LPO Provide jtd or Dst
CalibOnly?
2 1 to reproduce of
oNormalize?
2 1=no, 2=yes

ofxn calc*
0.20785
ofxn mod
0.24146

sRun
RunA1
zout
1

Ave
Data
6.5E-06
Model
6.4E-06

data/vMax
1.8E-05

ofxng
275.646

Min d/m
1.24915

ave/min d
1.10512
ave/min m
1.37213

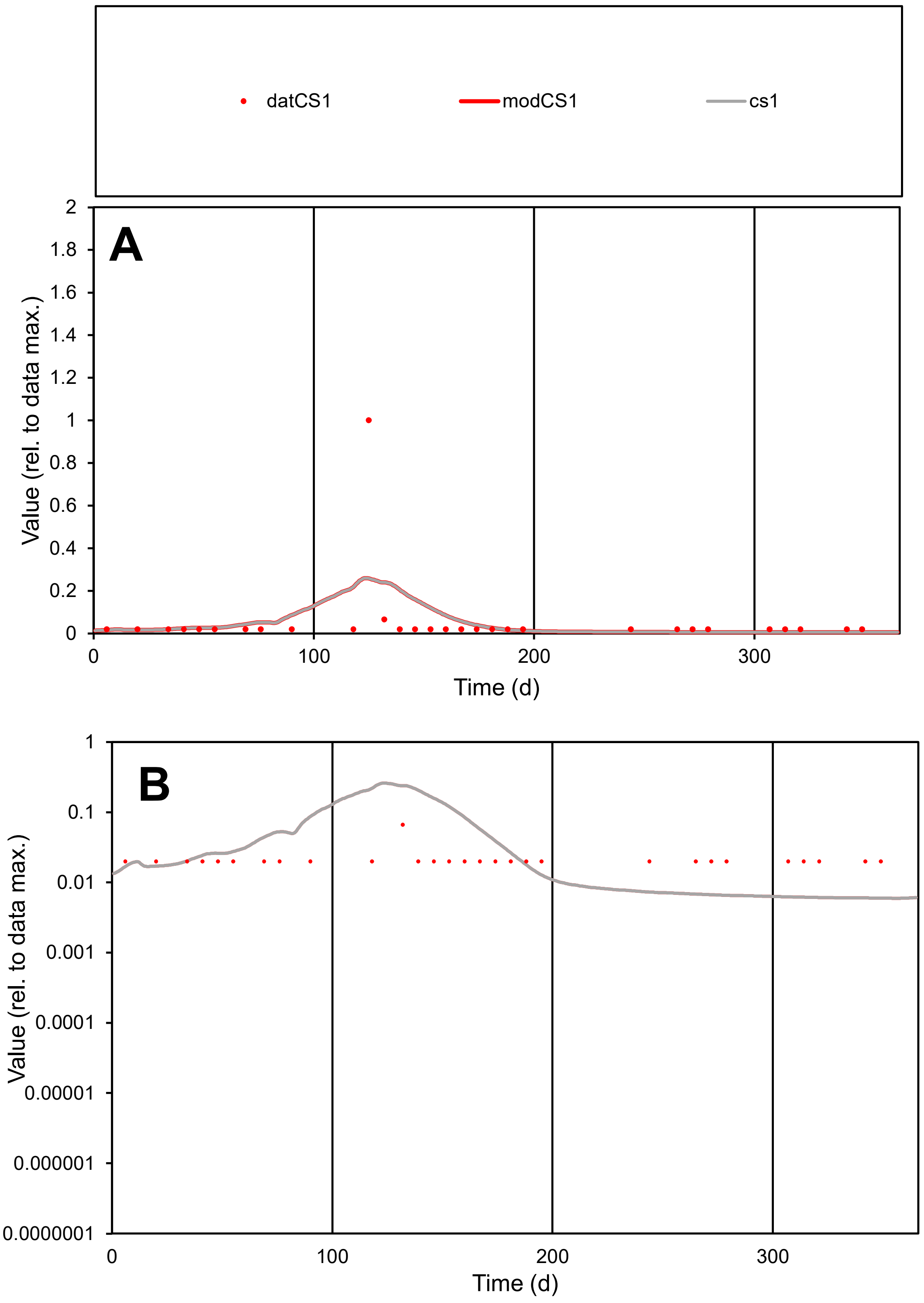


**C1**

Variables
CS1 Provide jtd or Dst
CalibOnly?
2 1 to reproduce of
oNormalize?
2 1=no, 2=yes

ofxn calc*
2.79929
ofxn mod
2.82573

sRun
RunA1
zout
1

Ave
Data
4.8E-05
Model
5.1E-05

data/vMax
0.00088

ofxng
275.646

Min d/m
3.34607

ave/min d
2.71621
ave/min m
9.77403

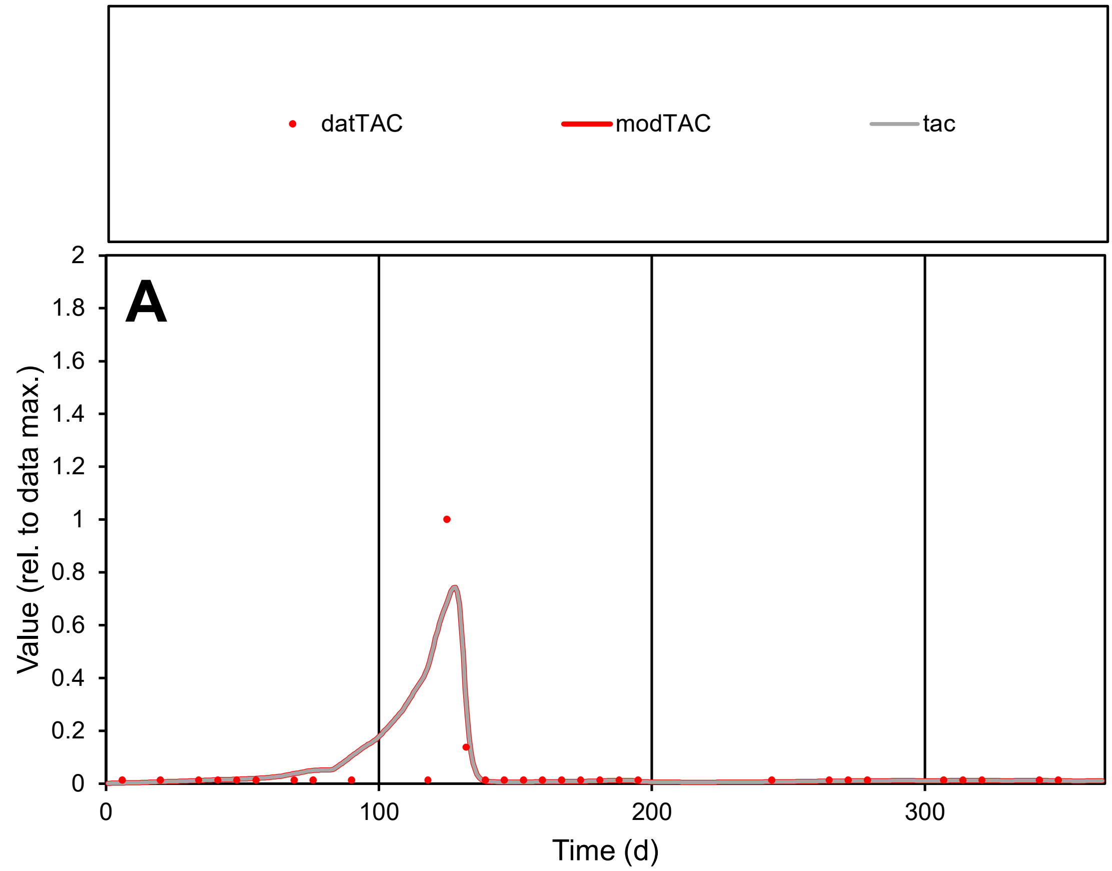


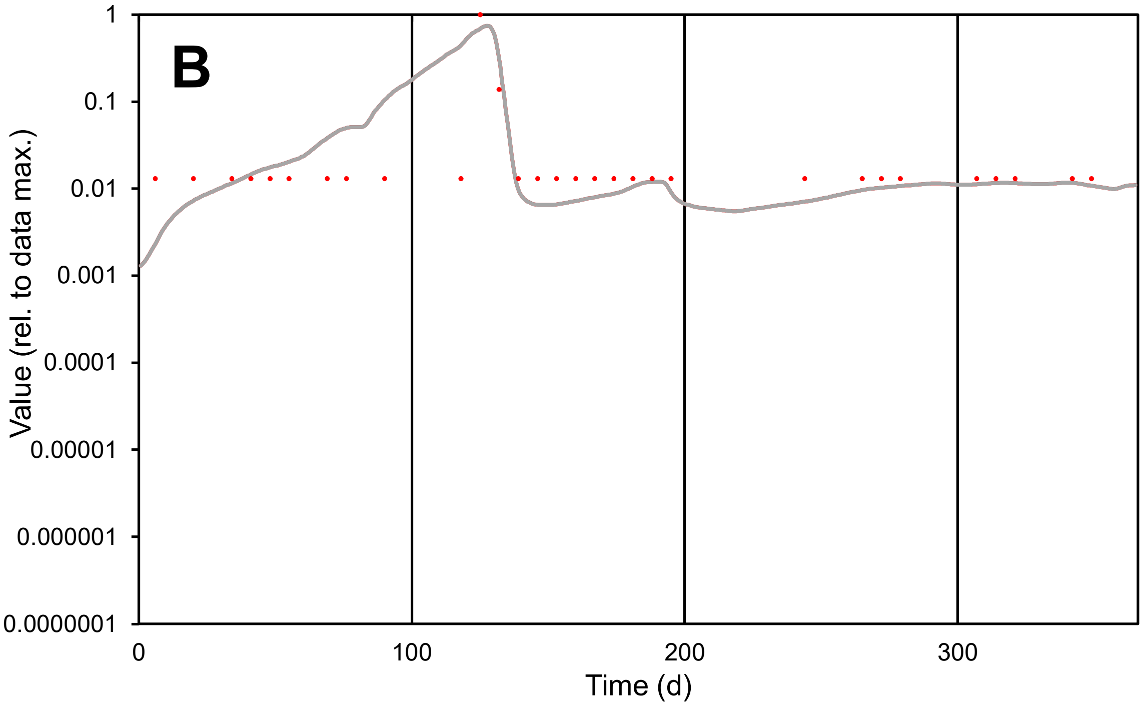


**C1**

Variables

TAC | Provide jtd or Dst

CalibOnly?

2 | 1 to reproduce of

oNormalize?

2 | 1=no, 2=yes

ofxn calc*

1.81025

ofxn mod

1.91186

sRun

RunA1

zout

1

Ave

Data

5E-06

Model

5.8E-06

data/vMax

9.9E-05

ofxng

275.646

Min d/m

4.78213

ave/min d

3.84221

ave/min m

21.3338

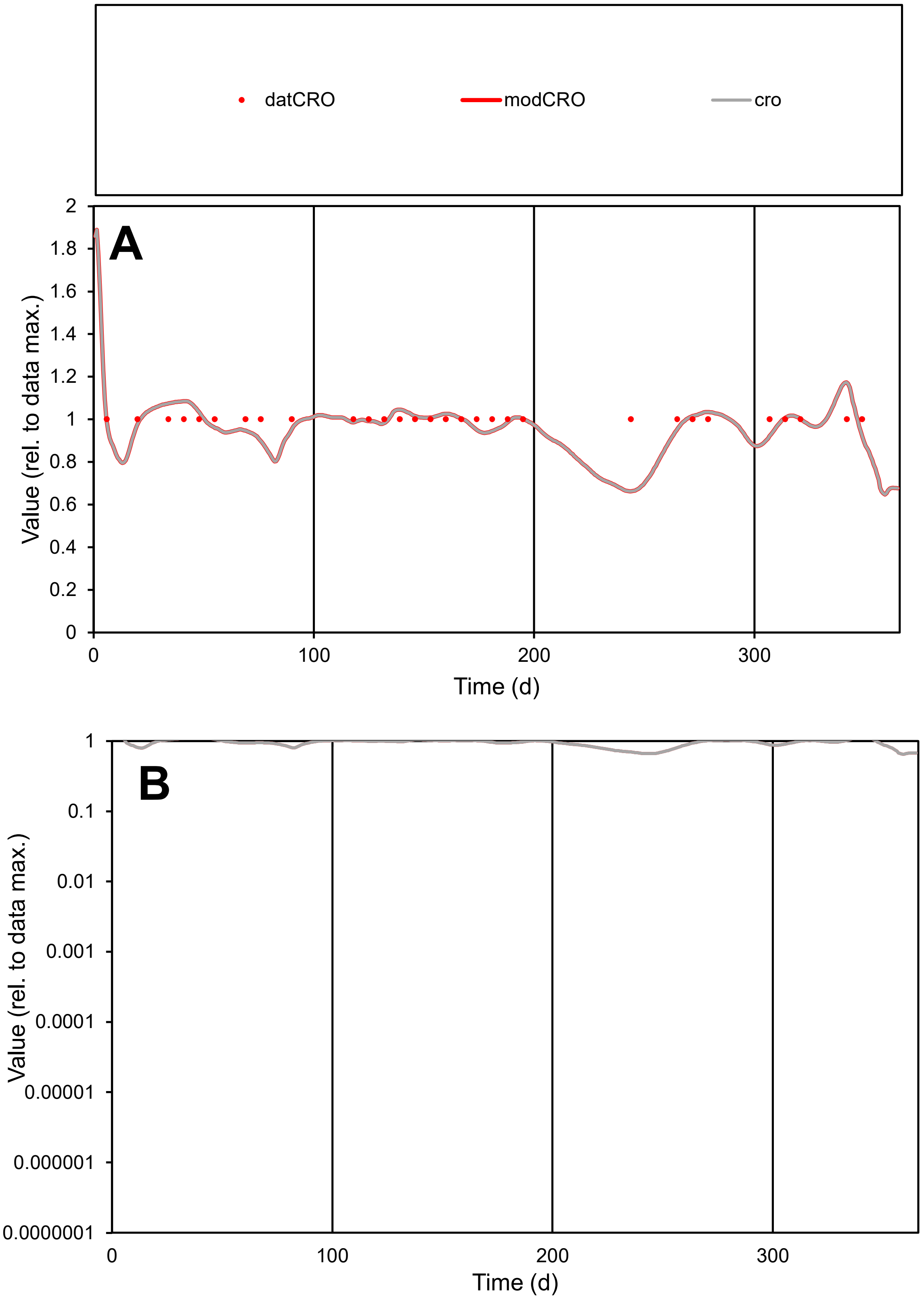


**C1**

Variables
CRO Provide jtd or Dst
CalibOnly?
2 1 to reproduce of
oNormalize?
2 1=no, 2=yes

ofxn calc*
0.08299
ofxn mod
0.13941

sRun
RunA1
zout
1

Ave
Data
1E-05
Model
1E-05

data/vMax
1E-05

ofxng
275.646

Min d/m
1.50633

ave/min d
1
ave/min m
1.47915

325_Calib_S_A

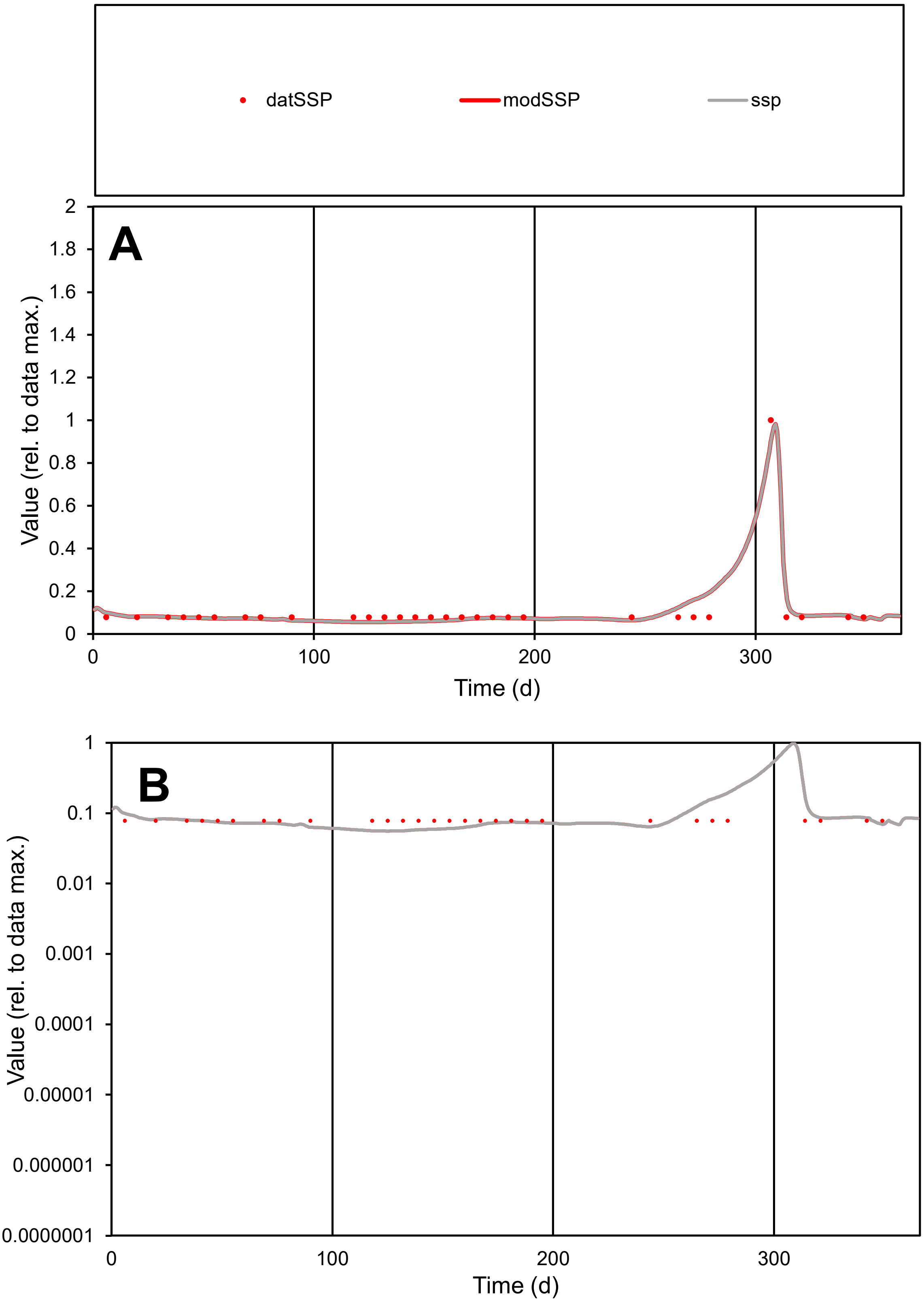


**C1**

Variables
SSP Provide jtd or Dst
CalibOnly?
2 1 to reproduce of
oNormalize?
2 1=no, 2=yes

ofxn calc*
0.28676
ofxn mod
0.36029

sRun
RunA1
zout
1

Ave
Data
1.8E-05
Model
1.9E-05

data/vMax
0.00017

ofxng
275.646

Min d/m
1.39117

ave/min d
1.39448
ave/min m
1.98691

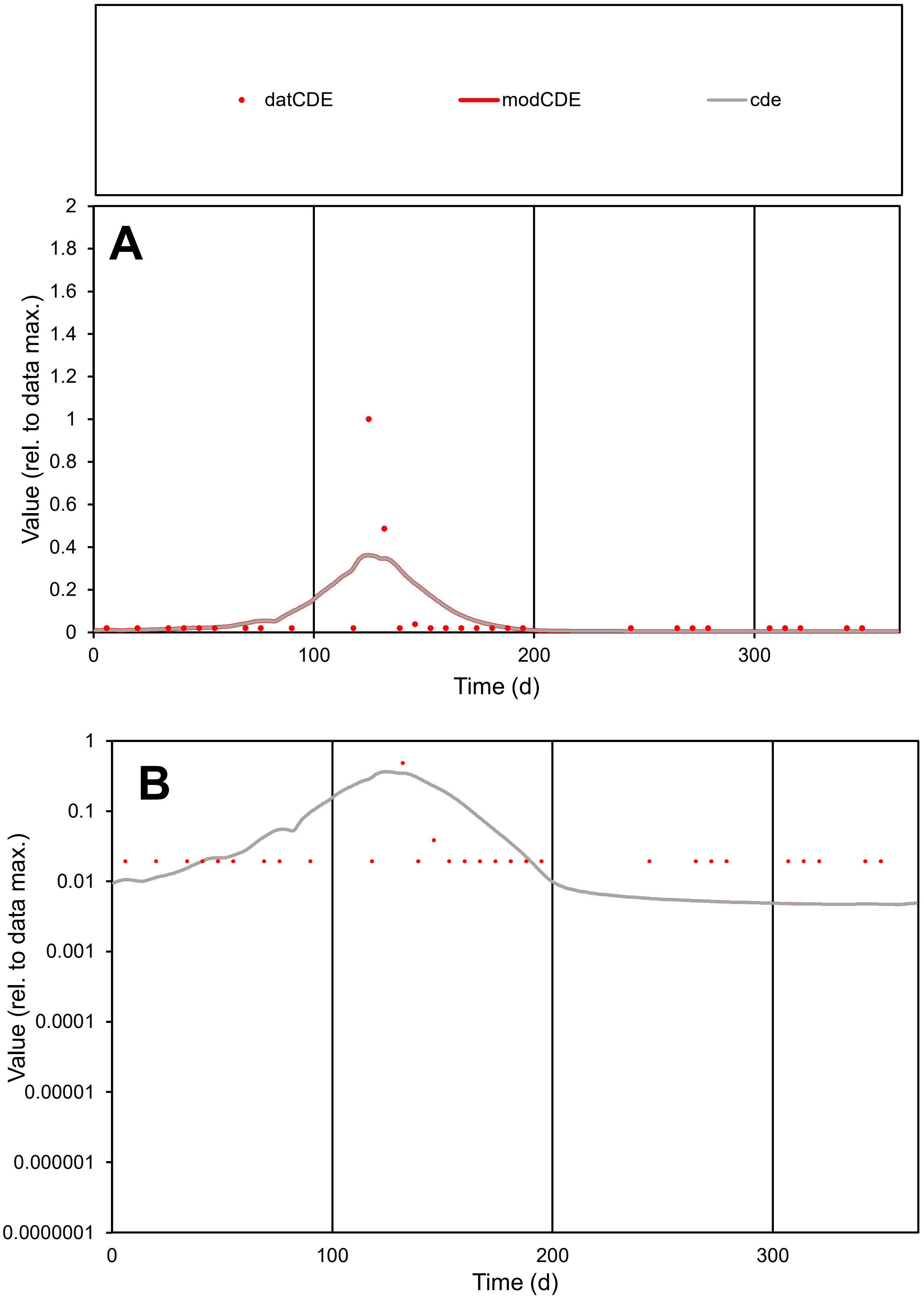


C1

Variables
CDE Provide jtd or Dst
CalibOnly?
2 1 to reproduce of
oNormalize?
2 1=no, 2=yes

ofxn calc*
2.13671
ofxn mod
2.17299

sRun
RunA1
zout
1

Ave
Data
3.5E-05
Model
3.9E-05

data/vMax
0.00052

ofxng
275.646

Min d/m
4.08299

ave/min d
3.52932
ave/min m
16.0003

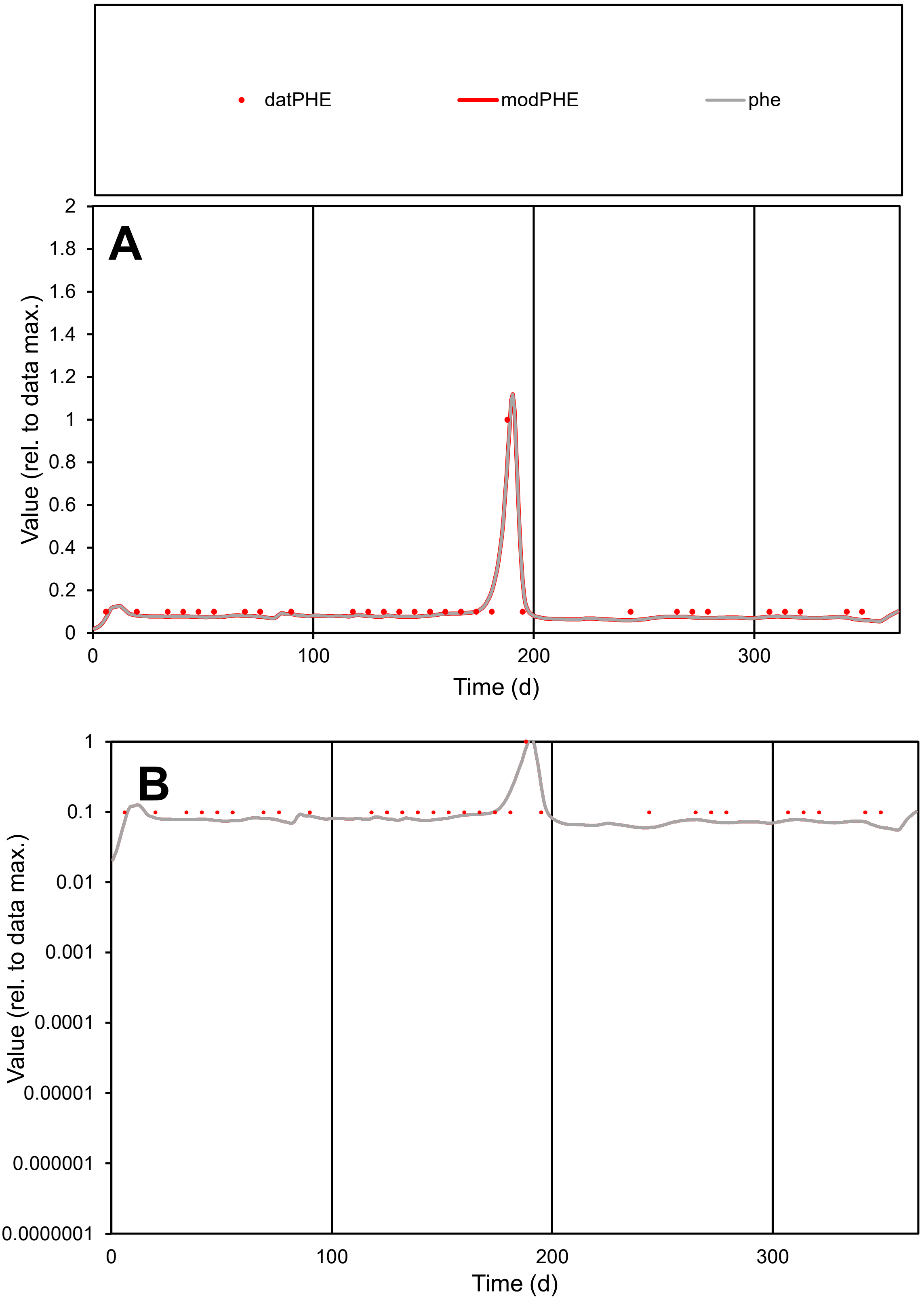


**C1**

Variables
PHE Provide jtd or Dst
CalibOnly?
2 1 to reproduce of
oNormalize?
2 1=no, 2=yes

ofxn calc*
0.22321
ofxn mod
0.26176

sRun
RunA1
zout
1

Ave
Data
5.2E-06
Model
4.6E-06

data/vMax
4E-05

ofxng
275.646

Min d/m
1.66331

ave/min d
1.30513
ave/min m
1.94261

325_Calib_S_A

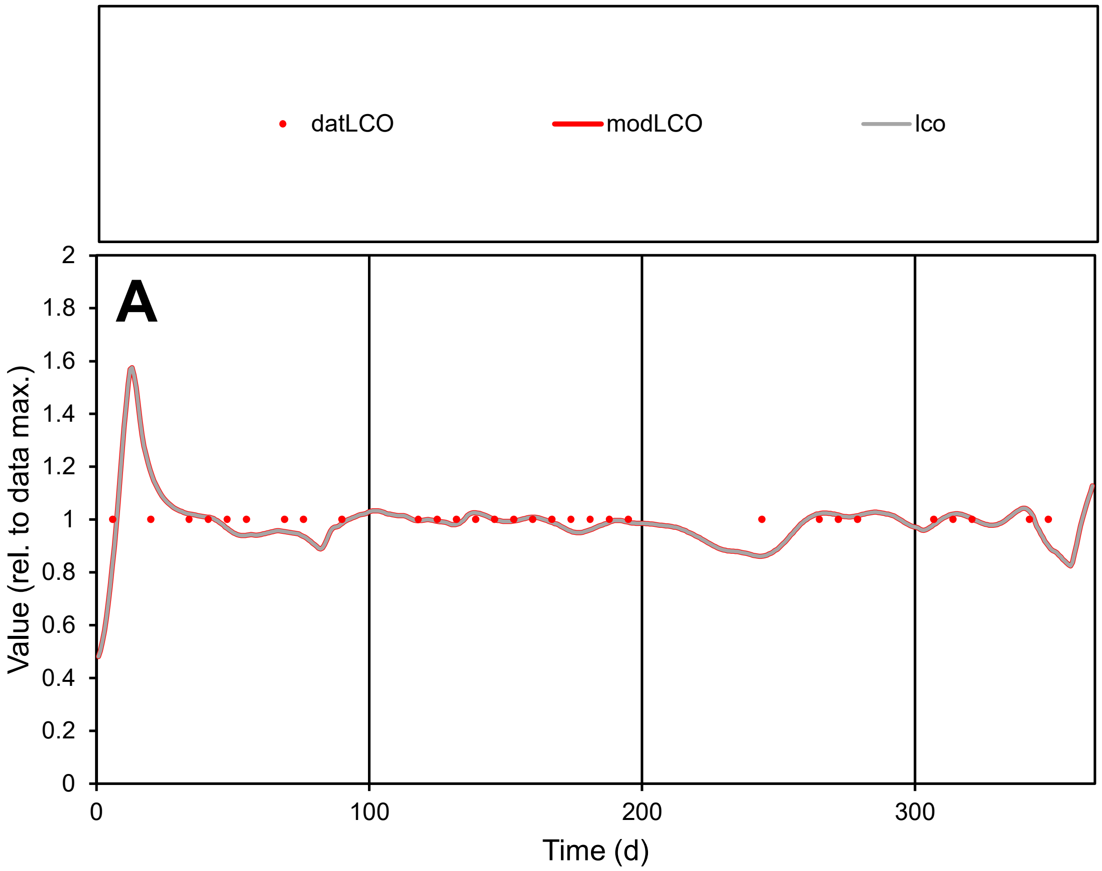


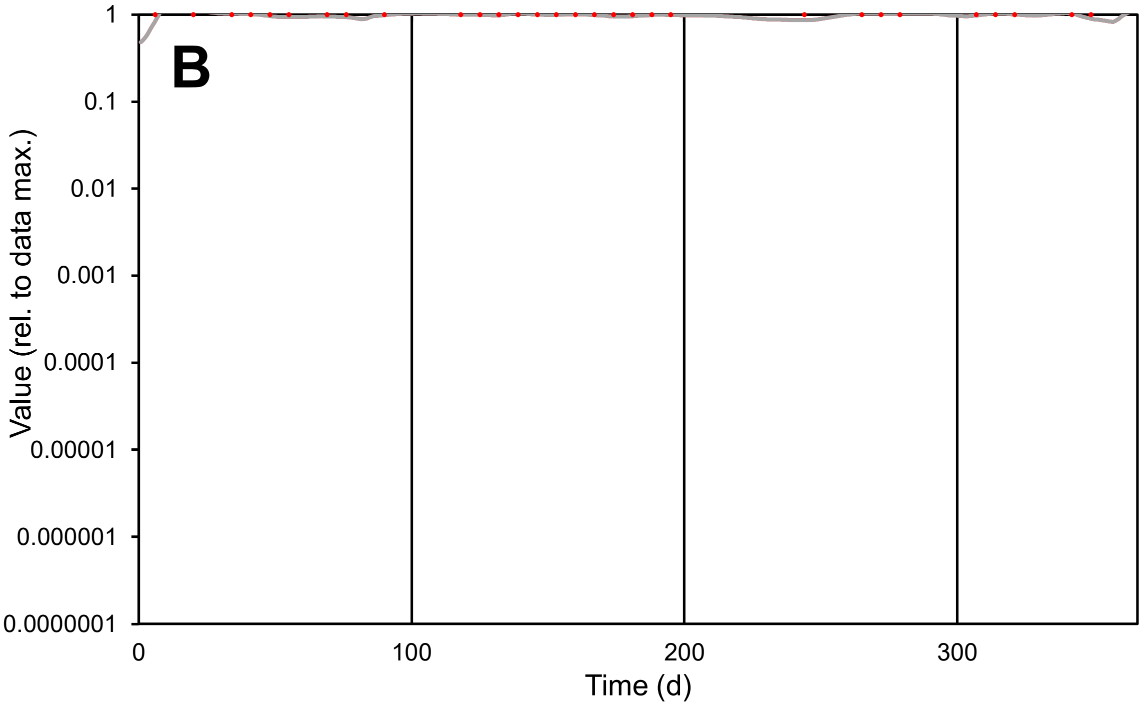


**C1**

Variables
LCO Provide jtd or Dst
CalibOnly?
2 1 to reproduce of
oNormalize?
2 1=no, 2=yes

ofxn calc*
0.04949
ofxn mod
0.12213

sRun
RunA1
zout
1

Ave
Data
7.4E-06
Model
7.3E-06

data/vMax
7.4E-06

ofxng
275.646

Min d/m
1.15734

ave/min d
1
ave/min m
1.14377

325_Calib_S_A

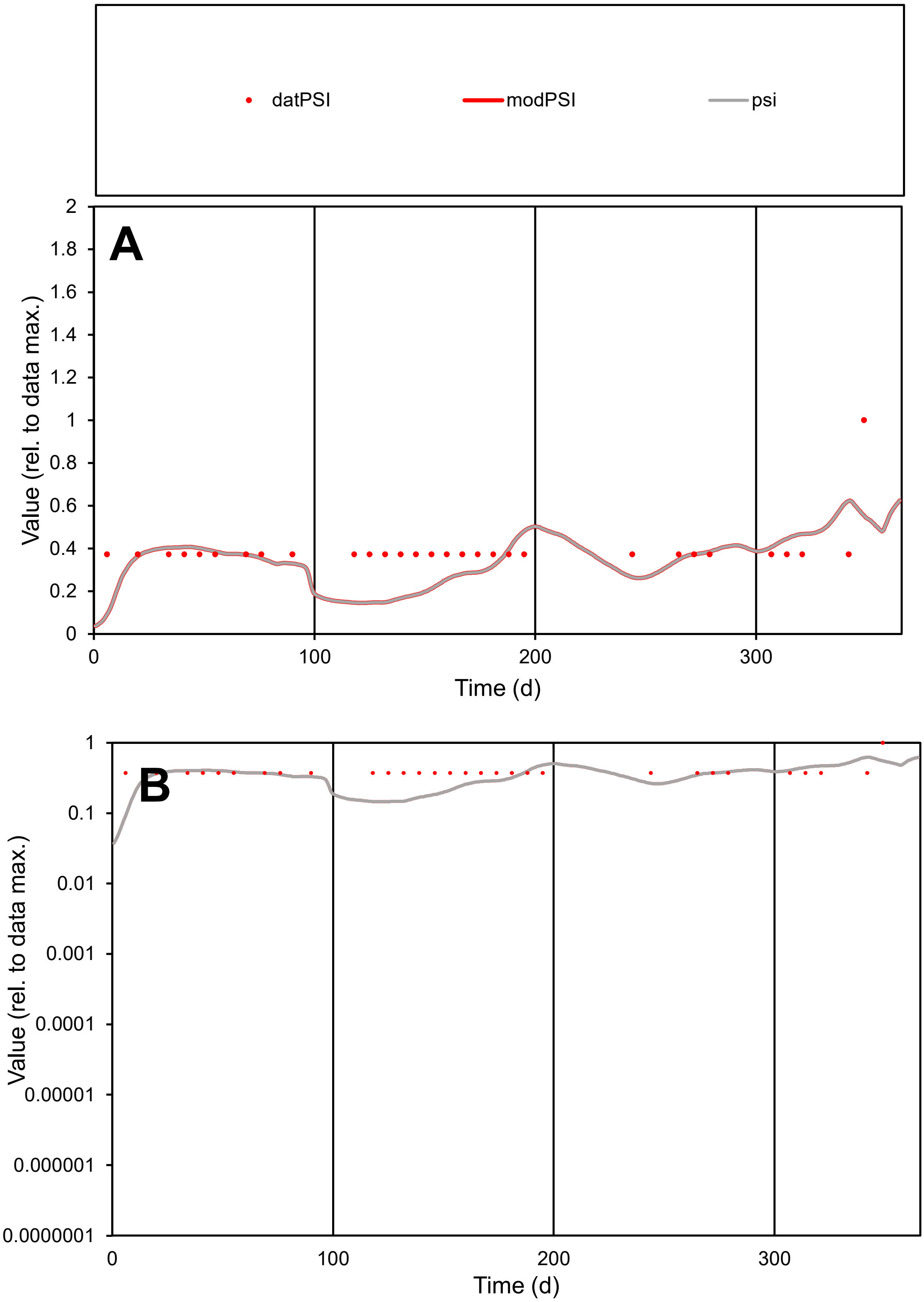


**C1**

Variables
PSI Provide jtd or Dst
CalibOnly?
2 1 to reproduce of
oNormalize?
2 1=no, 2=yes

ofxn calc*
0.37722
ofxn mod
0.39459

sRun
RunA1
zout
1

Ave
Data
3.8E-06
Model
3.2E-06

data/vMax
9.6E-06

ofxng
275.646

Min d/m
2.7898

ave/min d
1.05622
ave/min m
2.52847

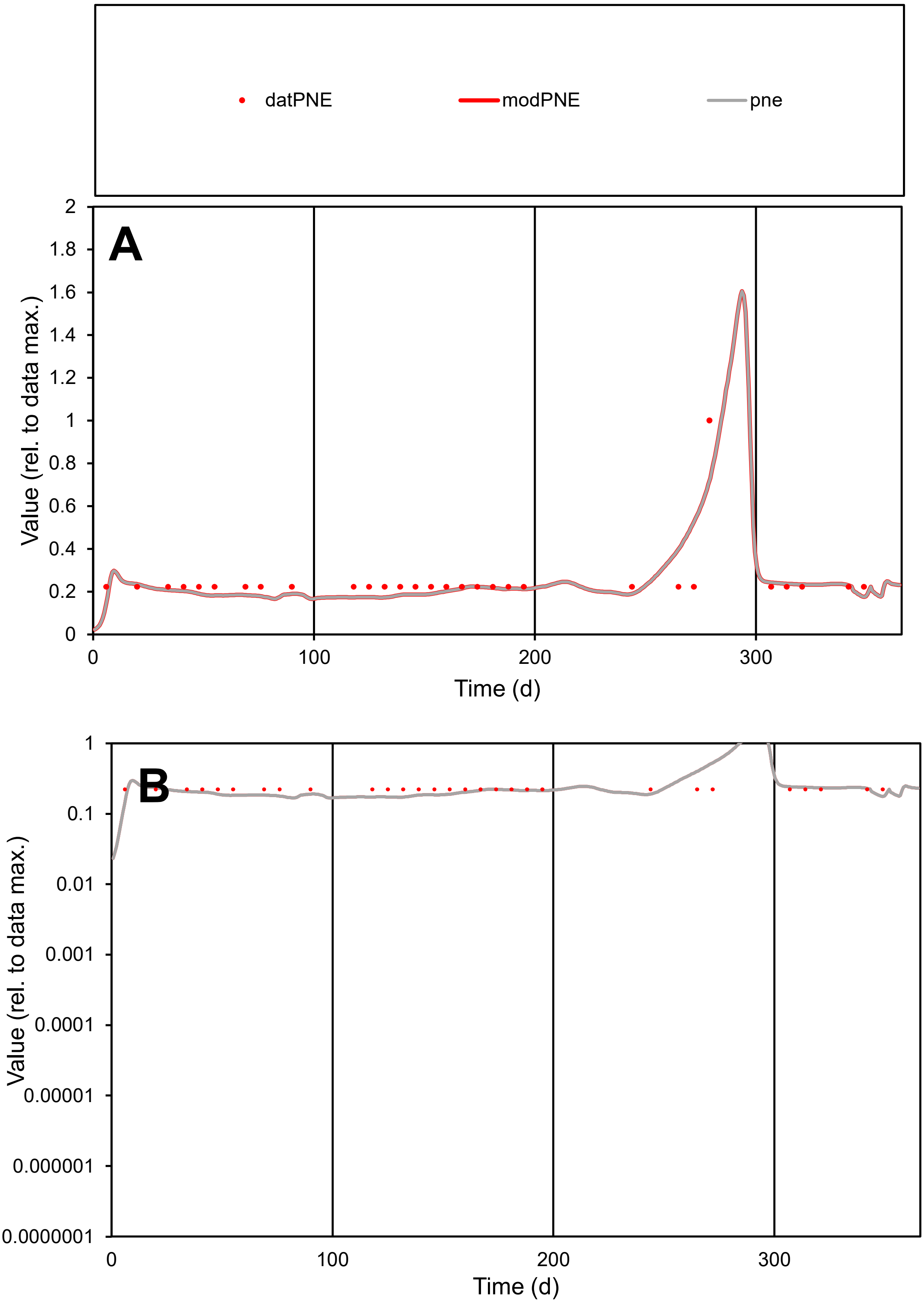


**C1**

Variables
PNE Provide jtd or Dst
CalibOnly?
2 1 to reproduce of
oNormalize?
2 1=no, 2=yes

ofxn calc*
0.32761
ofxn mod
0.35812

sRun
RunA1
zout
1

Ave
Data
1.6E-05
Model
1.6E-05

data/vMax
6.5E-05

ofxng
275.646

Min d/m
1.28155

ave/min d
1.11641
ave/min m
1.38676

325_Calib_S_A

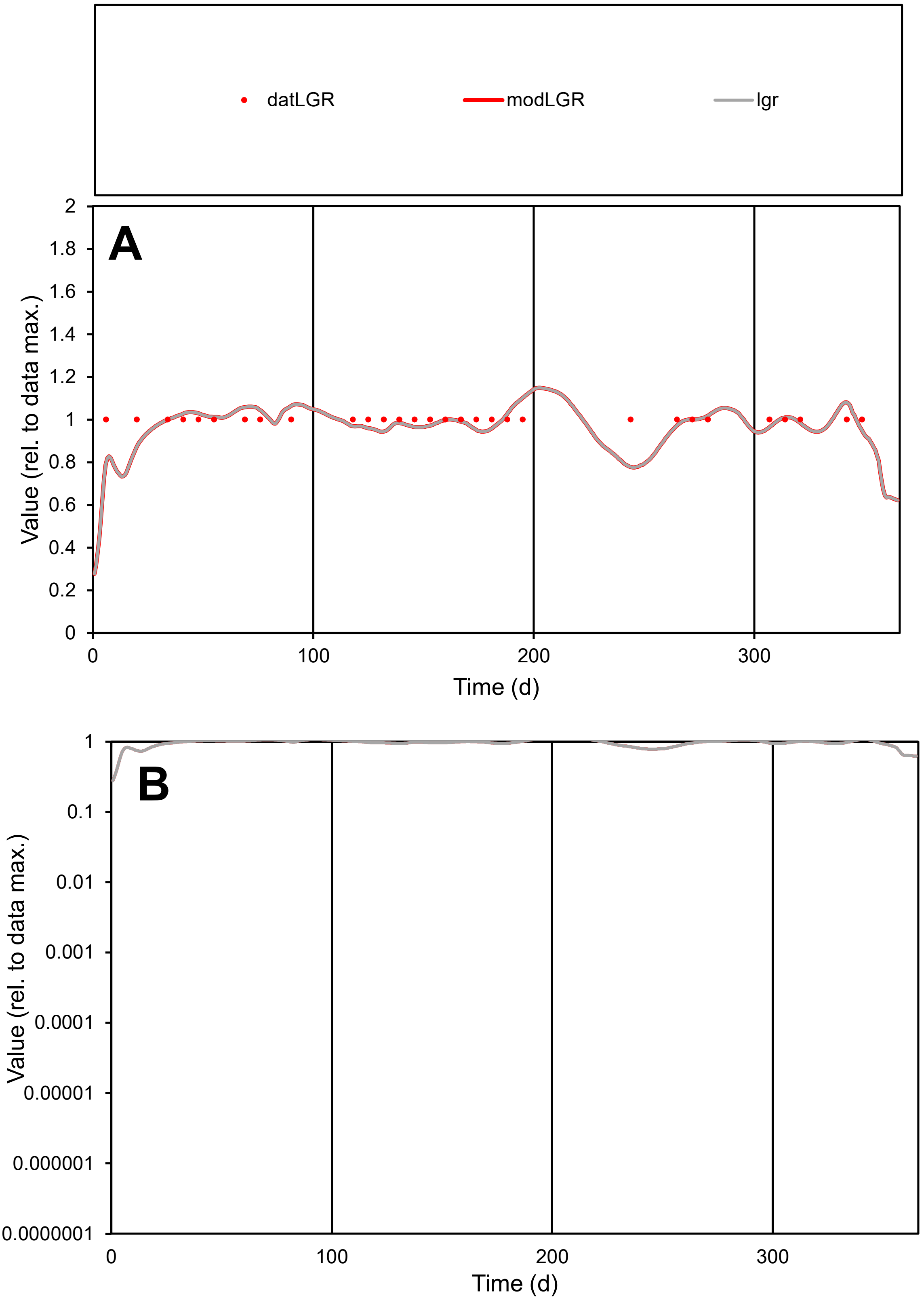


**C1**

Variables

LGR Provide jtd or Dst

CalibOnly?

2 1 to reproduce of

oNormalize?

2 1=no, 2=yes

ofxn calc*

0.06889

ofxn mod

0.13162

sRun

RunA1

zout

1

Ave

Data

5.2E-06

Model

5.1E-06

data/vMax

5.2E-06

ofxng

275.646

Min d/m

1.28696

ave/min d

1

ave/min m

1.2687

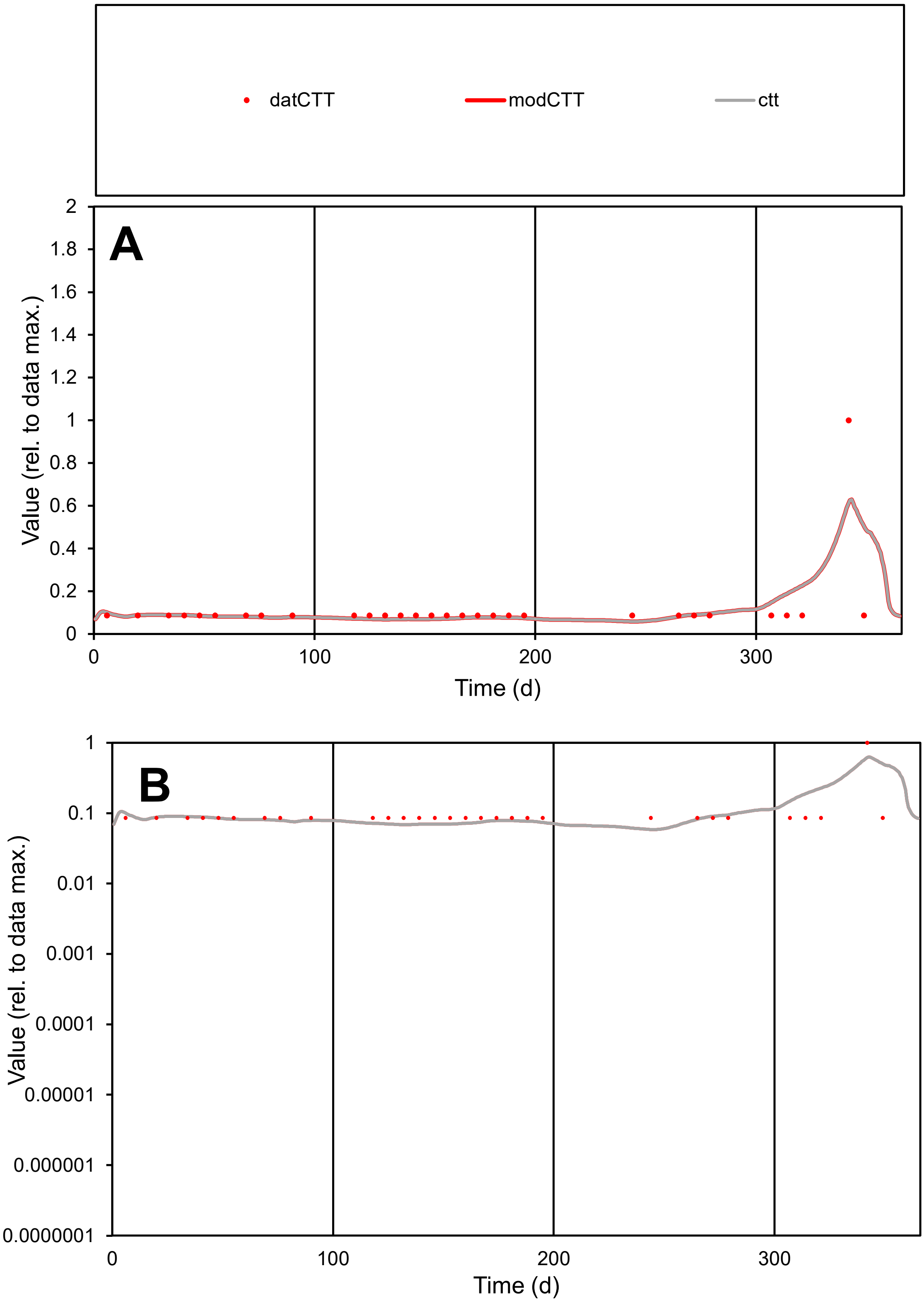


**C1**

Variables
CTT Provide jtd or Dst
CalibOnly?
2 1 to reproduce of
oNormalize?
2 1=no, 2=yes

ofxn calc*
0.9089
ofxn mod
0.92594

sRun
RunA1
zout
1

Ave
Data
2E-06
Model
2.1E-06

data/vMax
1.7E-05

ofxng
275.646

Min d/m
1.45965

ave/min d
1.35754
ave/min m
2.08163

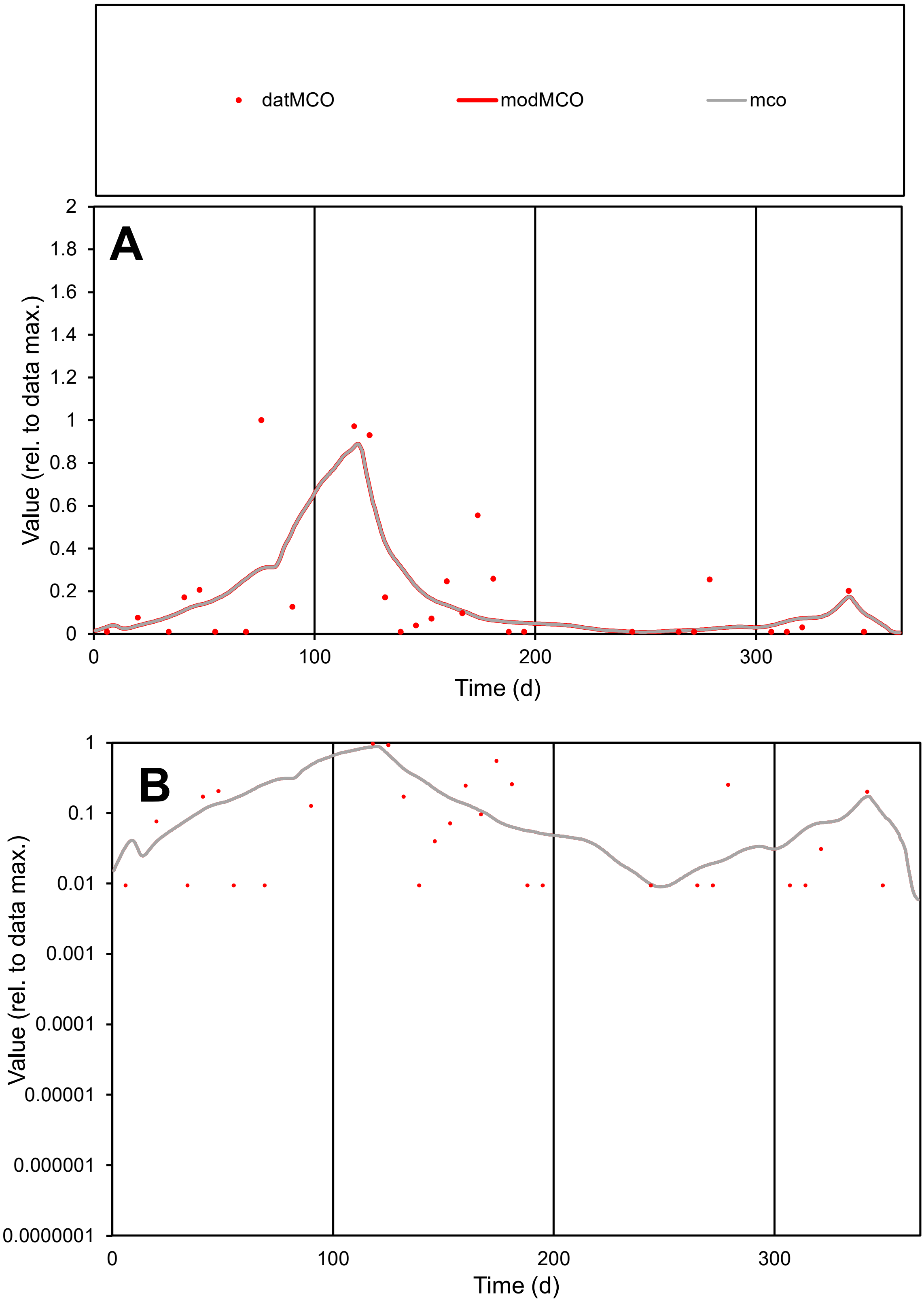


**C1**

Variables

MCO Provide jtd or Dst

CalibOnly?

2 1 to reproduce of

oNormalize?

2 1=no, 2=yes

ofxn calc*

1.1223

ofxn mod

1.15192

sRun

RunA1

zout

1

Ave

Data

1.2E-06

Model

1.1E-06

data/vMax

6.5E-06

ofxng

275.646

Min d/m

0.99512

ave/min d

19.5697

ave/min m

18.311

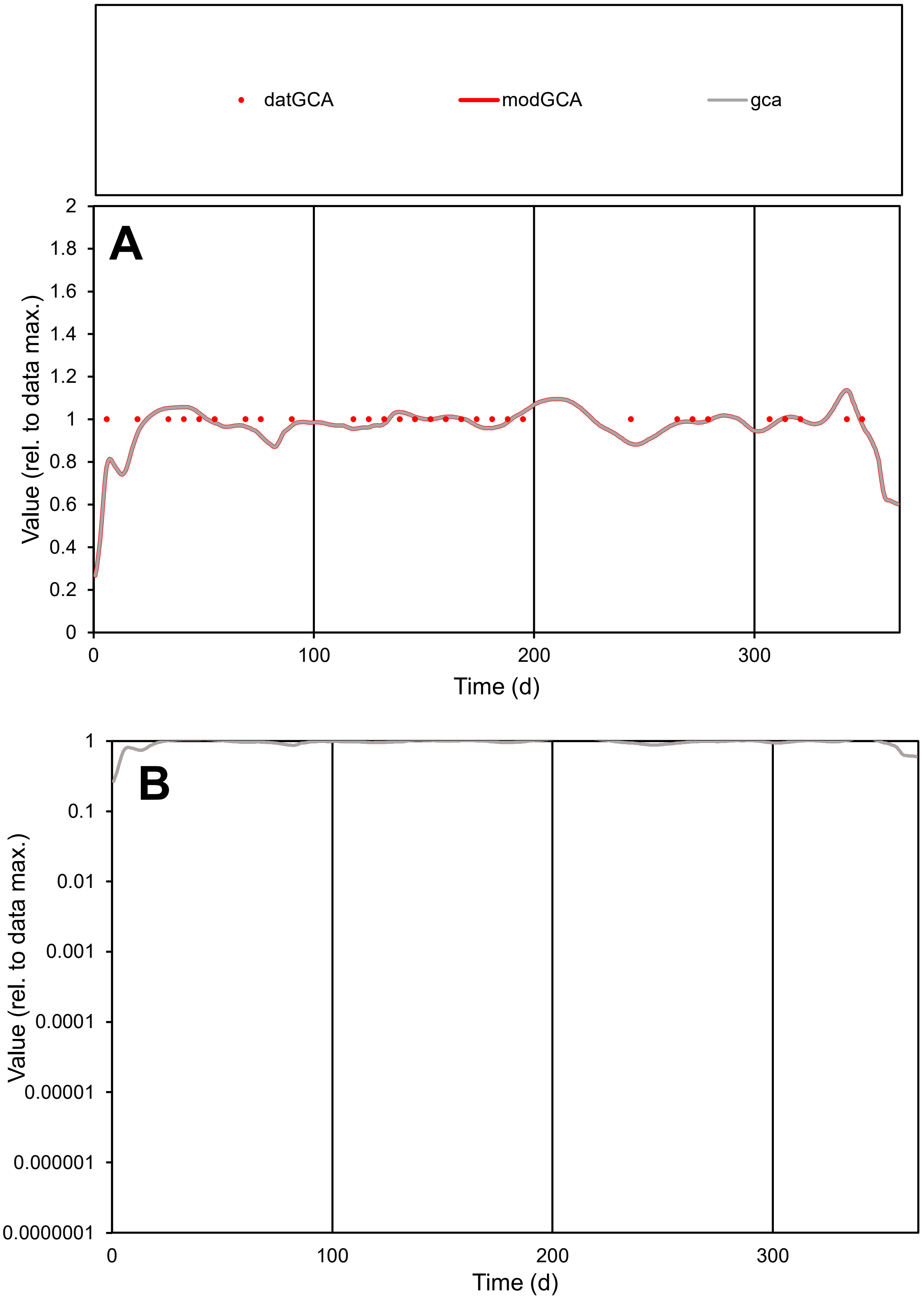


**C1**

Variables
GCA Provide jtd or Dst
CalibOnly?
2 1 to reproduce of
oNormalize?
2 1=no, 2=yes

ofxn calc*
0.05653
ofxn mod
0.1254

sRun
RunA1
zout
1

Ave
Data
8.4E-06
Model
8.3E-06

data/vMax
8.4E-06

ofxng
275.646

Min d/m
1.22693

ave/min d
1
ave/min m
1.20976

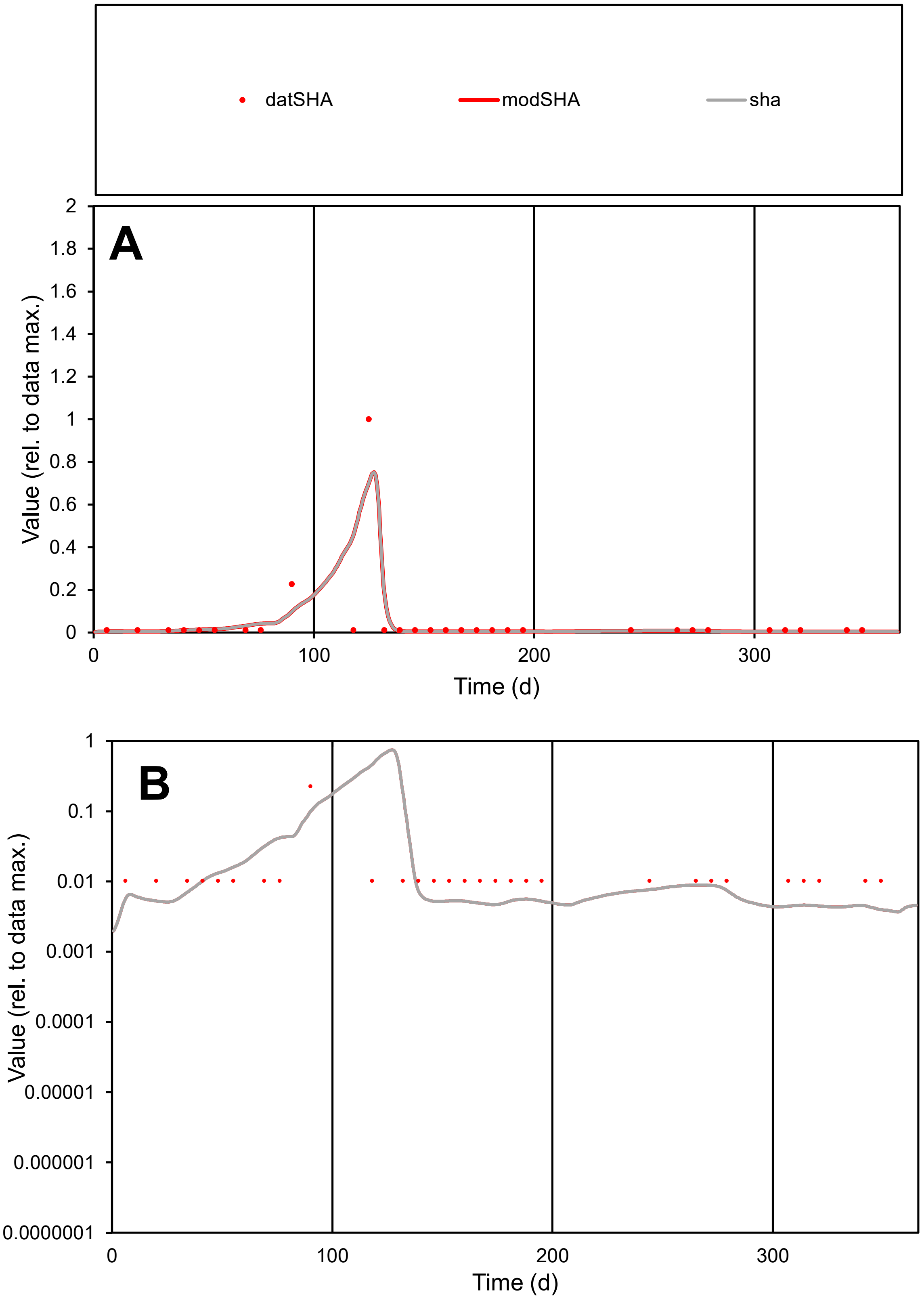


| | |
|---|---|
| **C1** | |
| Variables | |
| SHA | Provide jtd or Dst |
| CalibOnly? | |
| 2 | 1 to reproduce of |
| oNormalize? | |
| 2 | 1=no, 2=yes |
| ofxn calc* | |
| 1.84622 | |
| ofxn mod | |
| 1.94428 | |
| sRun | |
| RunA1 | |
| zout | |
| 1 | |
| Ave | |
| Data | |
| 3.4E-06 | |
| Model | |
| 3.6E-06 | |
| data/vMax | |
| 6.8E-05 | |
| ofxng | |
| 275.646 | |
| Min d/m | |
| 2.62575 | |
| ave/min d | |
| 4.90857 | |
| ave/min m | |
| 13.4191 | |

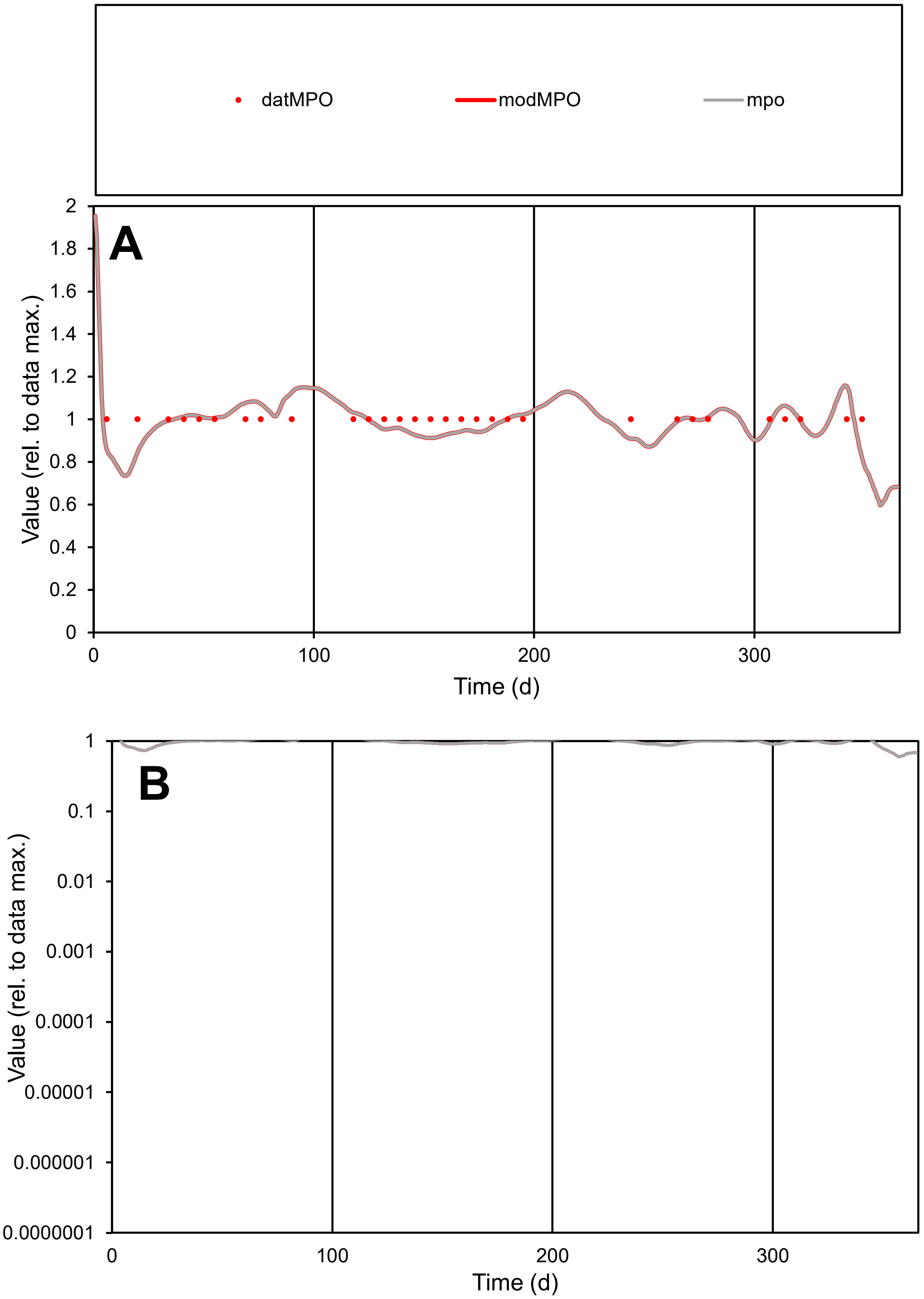


C1

Variables
MPO Provide jtd or Dst
CalibOnly?
2 1 to reproduce of
oNormalize?
2 1=no, 2=yes

ofxn calc*
0.07945
ofxn mod
0.13737

sRun
RunA1
zout
1

Ave
Data
1.2E-05
Model
1.2E-05

data/vMax
1.2E-05

ofxng
275.646

Min d/m
1.3033

ave/min d
1
ave/min m
1.27996

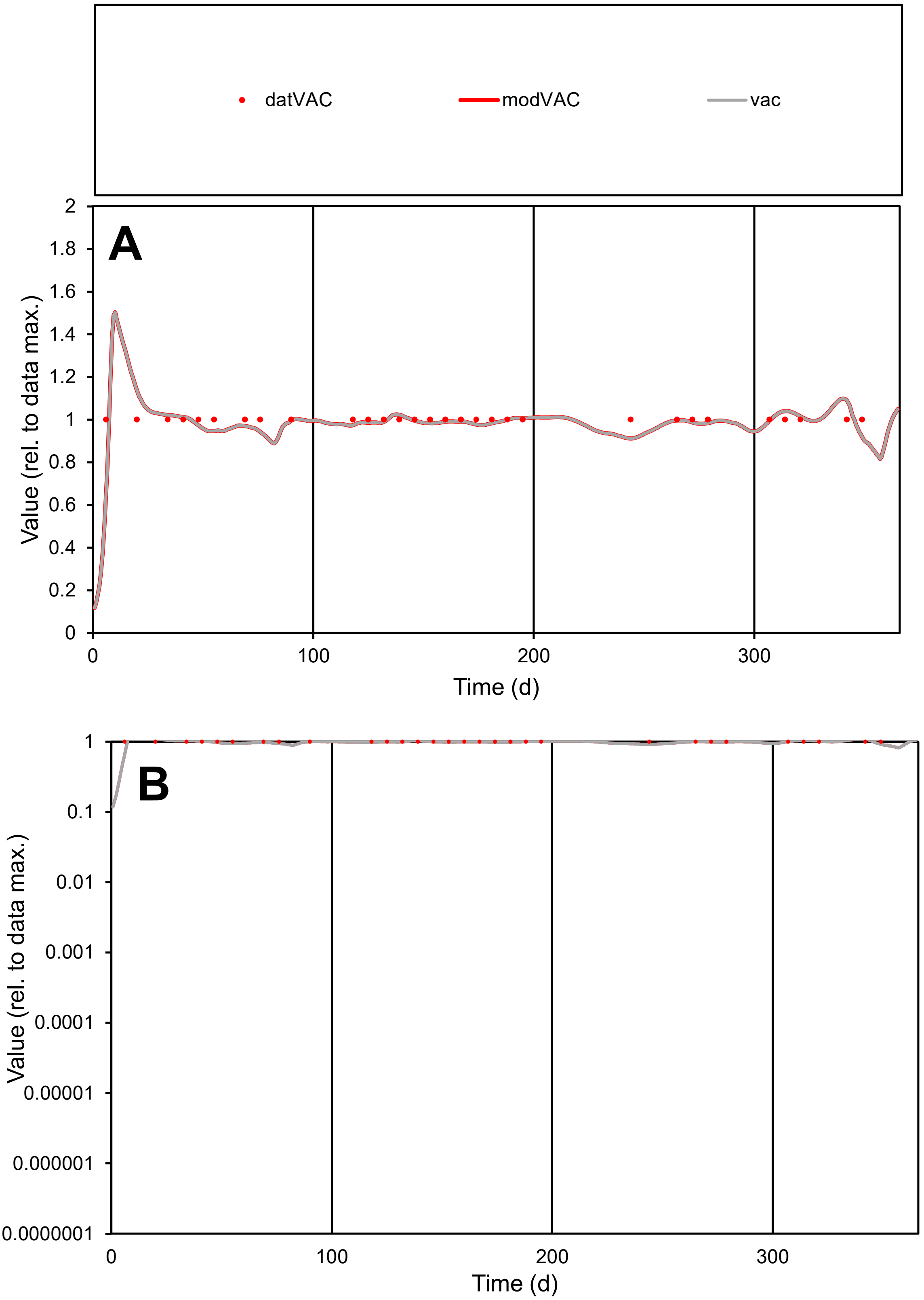


**C1**

Variables
VAC Provide jtd or Dst
CalibOnly?
2 1 to reproduce of
oNormalize?
2 1=no, 2=yes

ofxn calc*
0.04019
ofxn mod
0.11864

sRun
RunA1
zout
1

Ave
Data
9.5E-06
Model
9.4E-06

data/vMax
9.5E-06

ofxng
275.646

Min d/m
1.11529

ave/min d
1
ave/min m
1.10615

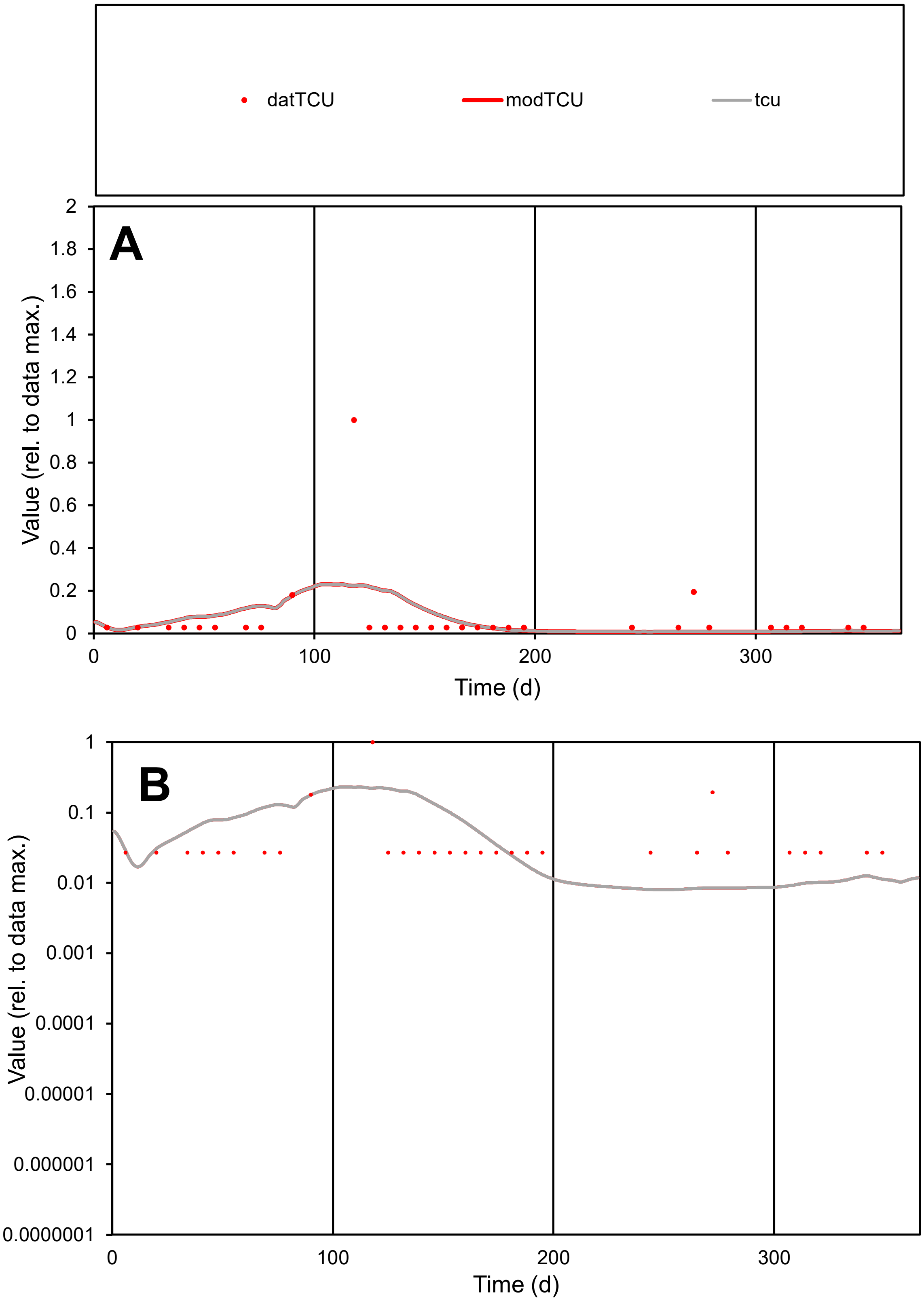


**C1**

Variables
TCU Provide jtd or Dst
CalibOnly?
2 1 to reproduce of
oNormalize?
2 1=no, 2=yes

ofxn calc*
2.27901
ofxn mod
2.29033

sRun
RunA1
zout
1

Ave
Data
2.5E-05
Model
2.5E-05

data/vMax
0.00036

ofxng
275.646

Min d/m
3.37038

ave/min d
2.59712
ave/min m
8.66882

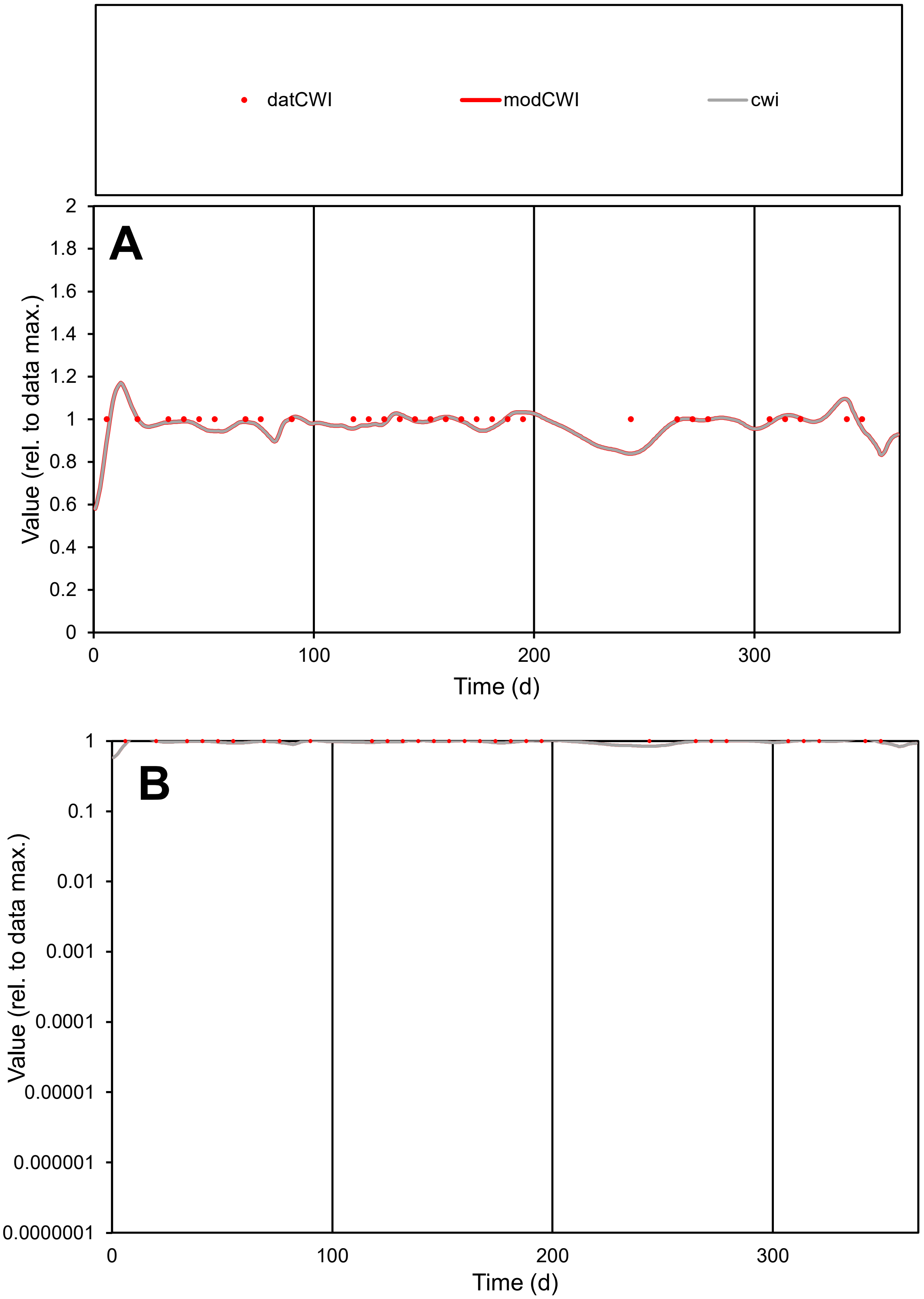


**C1**

Variables

CWI Provide jtd or Dst

CalibOnly?

2 1 to reproduce of

oNormalize?

2 1=no, 2=yes

ofxn calc*

0.04174

ofxn mod

0.11917

sRun

RunA1

zout

1

Ave

Data

1.3E-05

Model

1.3E-05

data/vMax

1.3E-05

ofxng

275.646

Min d/m

1.18979

ave/min d

1

ave/min m

1.17302

325_Calib_S_A

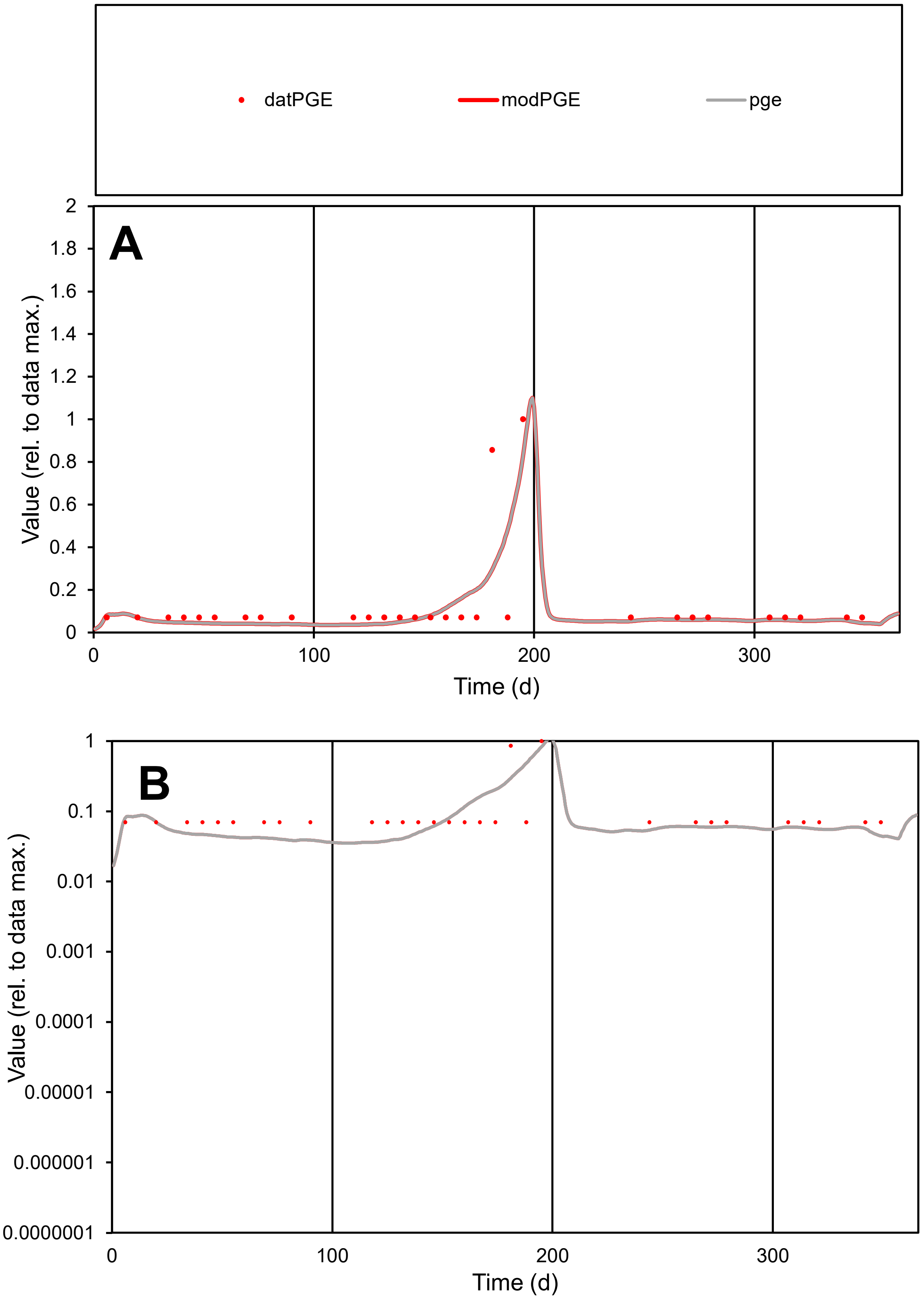


**C1**

Variables
PGE Provide jtd or Dst
CalibOnly?
2 1 to reproduce of
oNormalize?
2 1=no, 2=yes

ofxn calc*
0.95893
ofxn mod
1.03438

sRun
RunA1
zout
1

Ave
Data
1.1E-05
Model
9.7E-06

data/vMax
8.4E-05

ofxng
275.646

Min d/m
1.93204

ave/min d
1.81943
ave/min m
3.17737

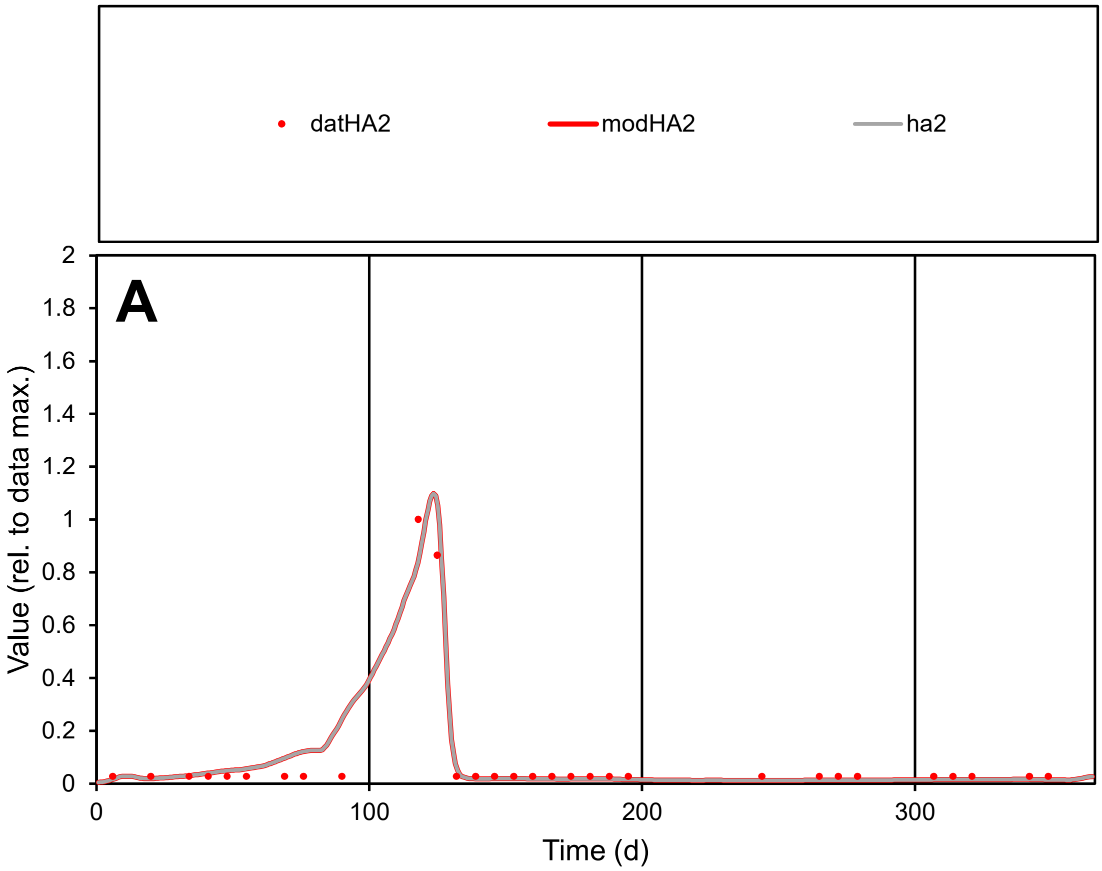


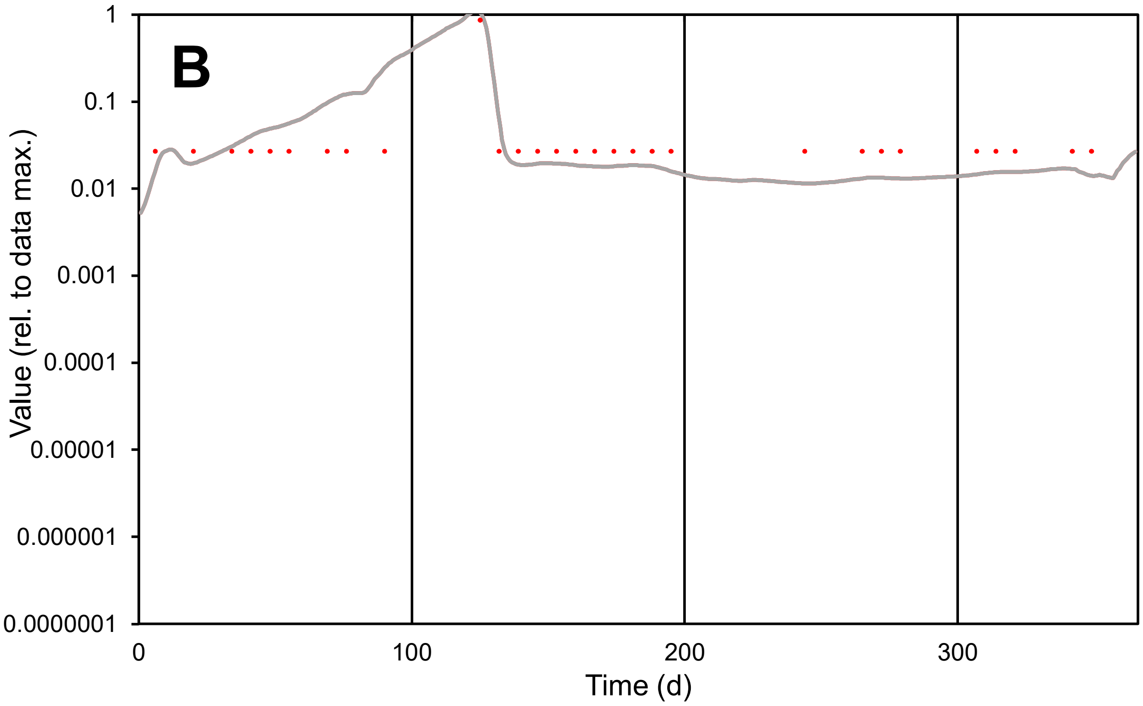


**C1**

Variables

| | |
|---|---|
| HA2 | Provide jtd or Dst |
| CalibOnly? | |
| 2 | 1 to reproduce of |
| oNormalize? | |
| 2 | 1=no, 2=yes |
| ofxn calc* | |
| 0.50272 | |
| ofxn mod | |
| 0.64009 | |
| sRun | |
| RunA1 | |
| zout | |
| 1 | |
| Ave | |
| Data | |
| 8.5E-06 | |
| Model | |
| 9E-06 | |
| data/vMax | |
| 9.8E-05 | |
| ofxng | |
| 275.646 | |
| Min d/m | |
| 2.35263 | |
| ave/min d | |
| 3.23386 | |
| ave/min m | |
| 7.9748 | |

325_Calib_S_A

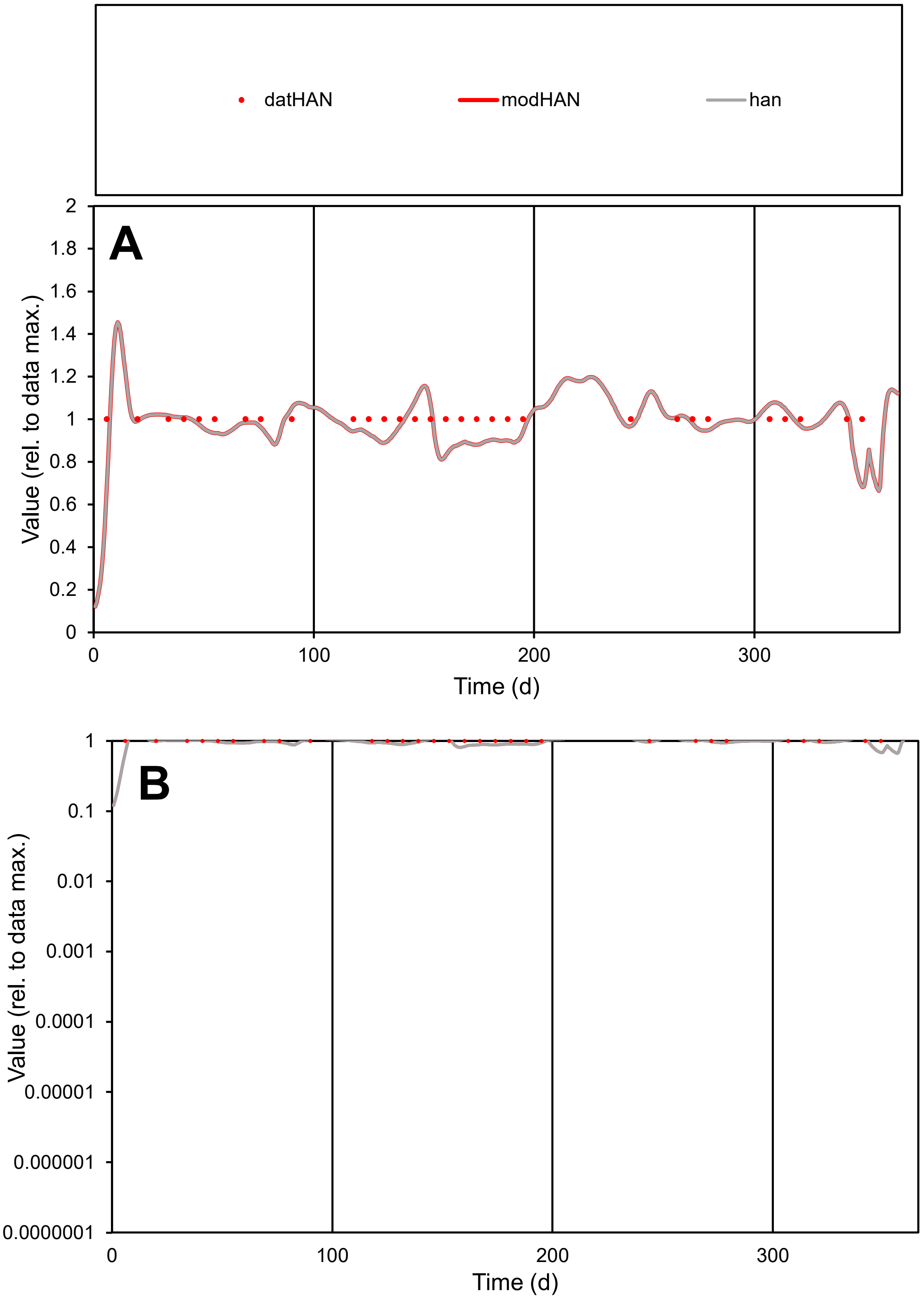


**C1**

Variables
HAN Provide jtd or Dst
CalibOnly?
2 1 to reproduce of
oNormalize?
2 1=no, 2=yes

ofxn calc*
0.08329
ofxn mod
0.13827

sRun
RunA1
zout
1

Ave
Data
5.1E-06
Model
4.9E-06

data/vMax
5.1E-06

ofxng
275.646

Min d/m
1.39088

ave/min d
1
ave/min m
1.33826

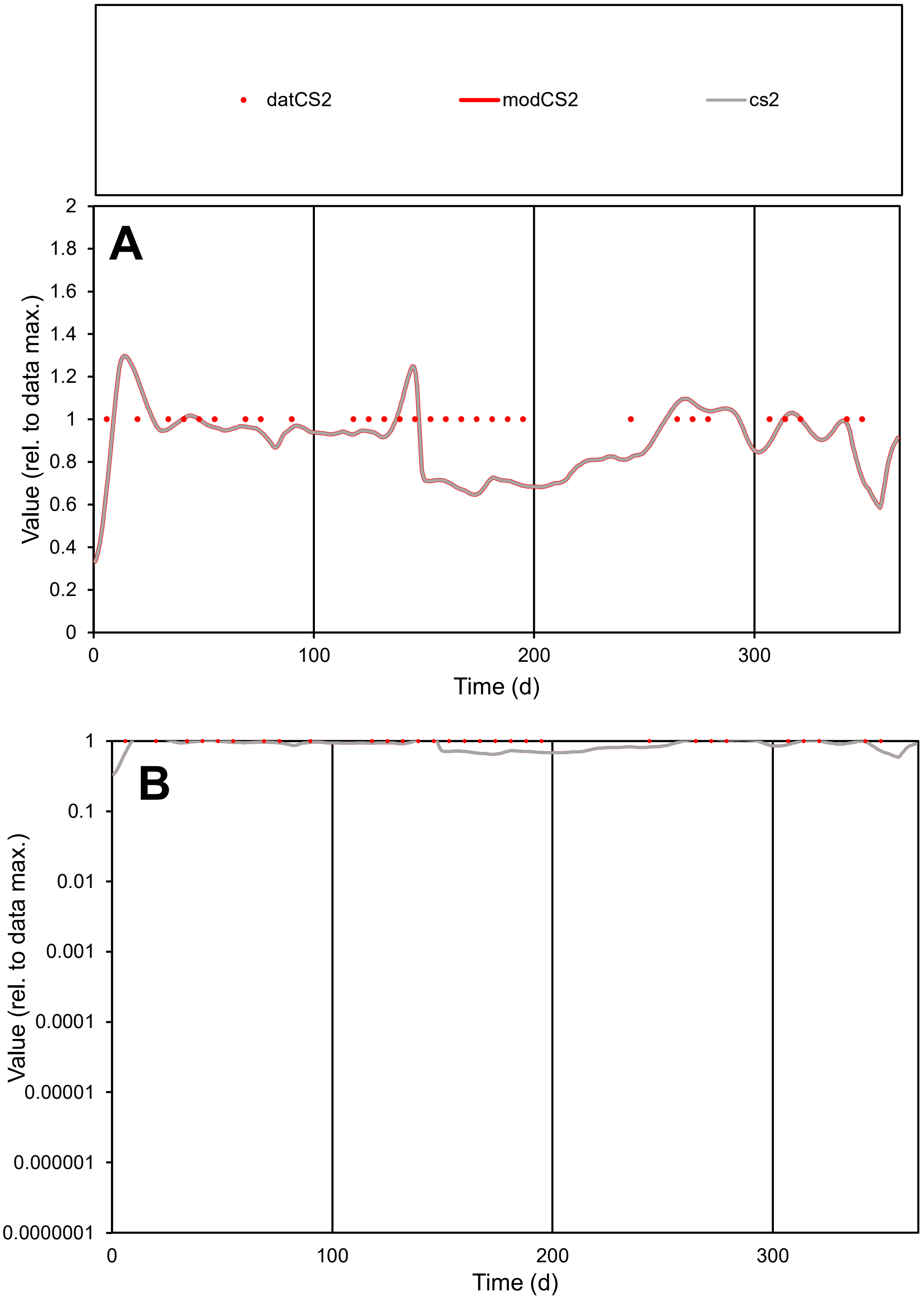


**C1**

Variables
CS2 Provide jtd or Dst
CalibOnly?
2 1 to reproduce of
oNormalize?
2 1=no, 2=yes

ofxn calc*
0.17395
ofxn mod
0.20529

sRun
RunA1
zout
1

Ave
Data
7.6E-06
Model
6.9E-06

data/vMax
7.6E-06

ofxng
275.646

Min d/m
1.52567

ave/min d
1
ave/min m
1.37948

325_Calib_S_A

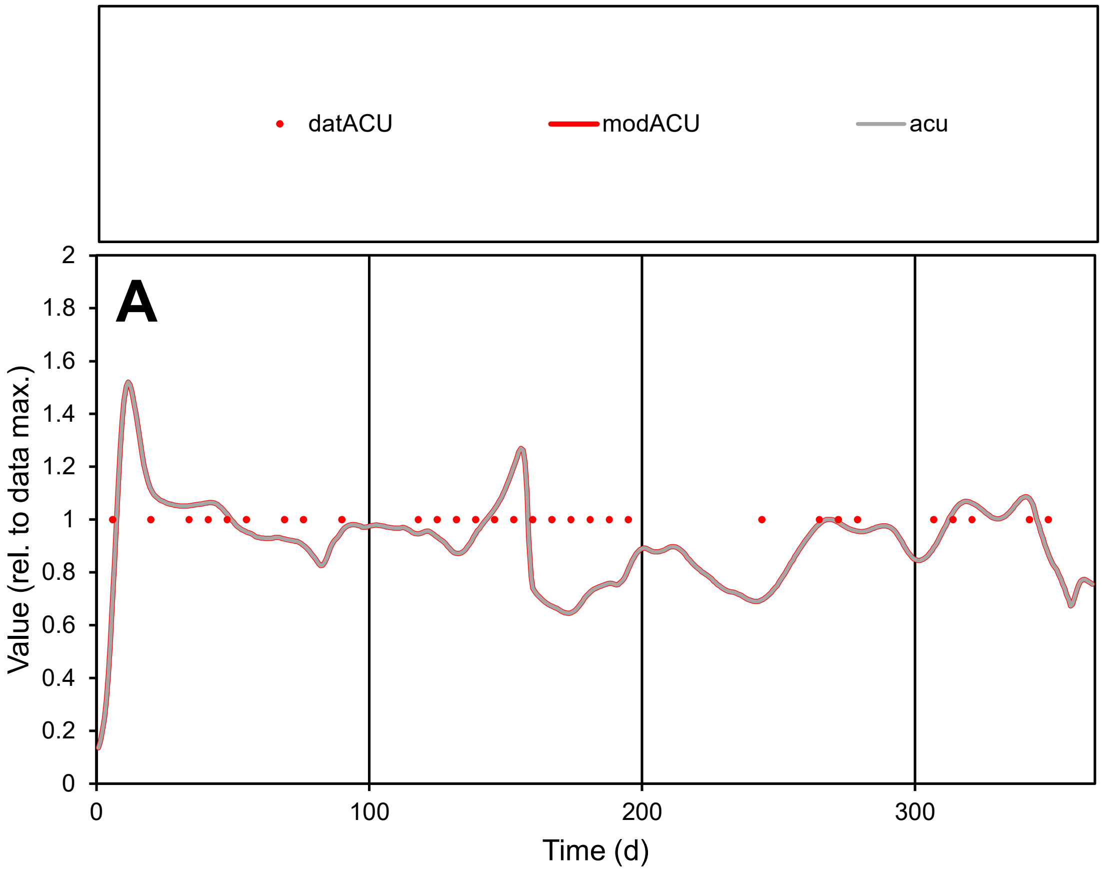


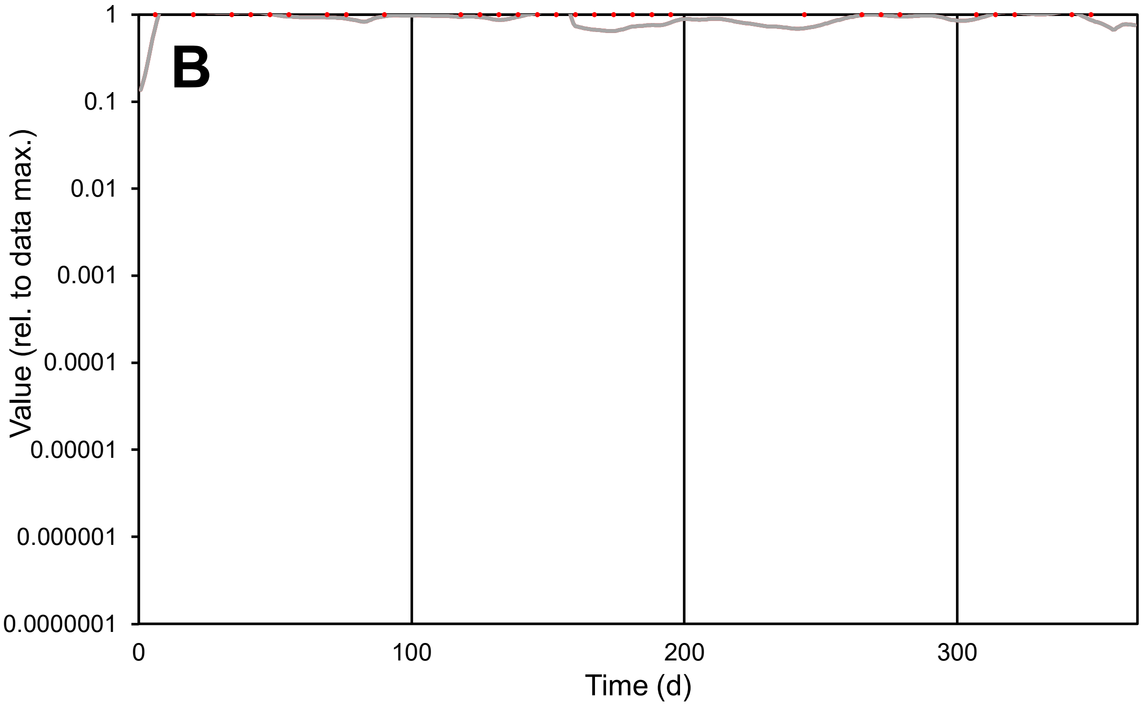


**C1**

Variables

ACU Provide jtd or Dst

CalibOnly?

2 1 to reproduce of

oNormalize?

2 1=no, 2=yes

ofxn calc*

0.155

ofxn mod

0.18948

sRun

RunA1

zout

1

Ave

Data

1.1E-05

Model

1E-05

data/vMax

1.1E-05

ofxng

275.646

Min d/m

1.52757

ave/min d

1

ave/min m

1.41972

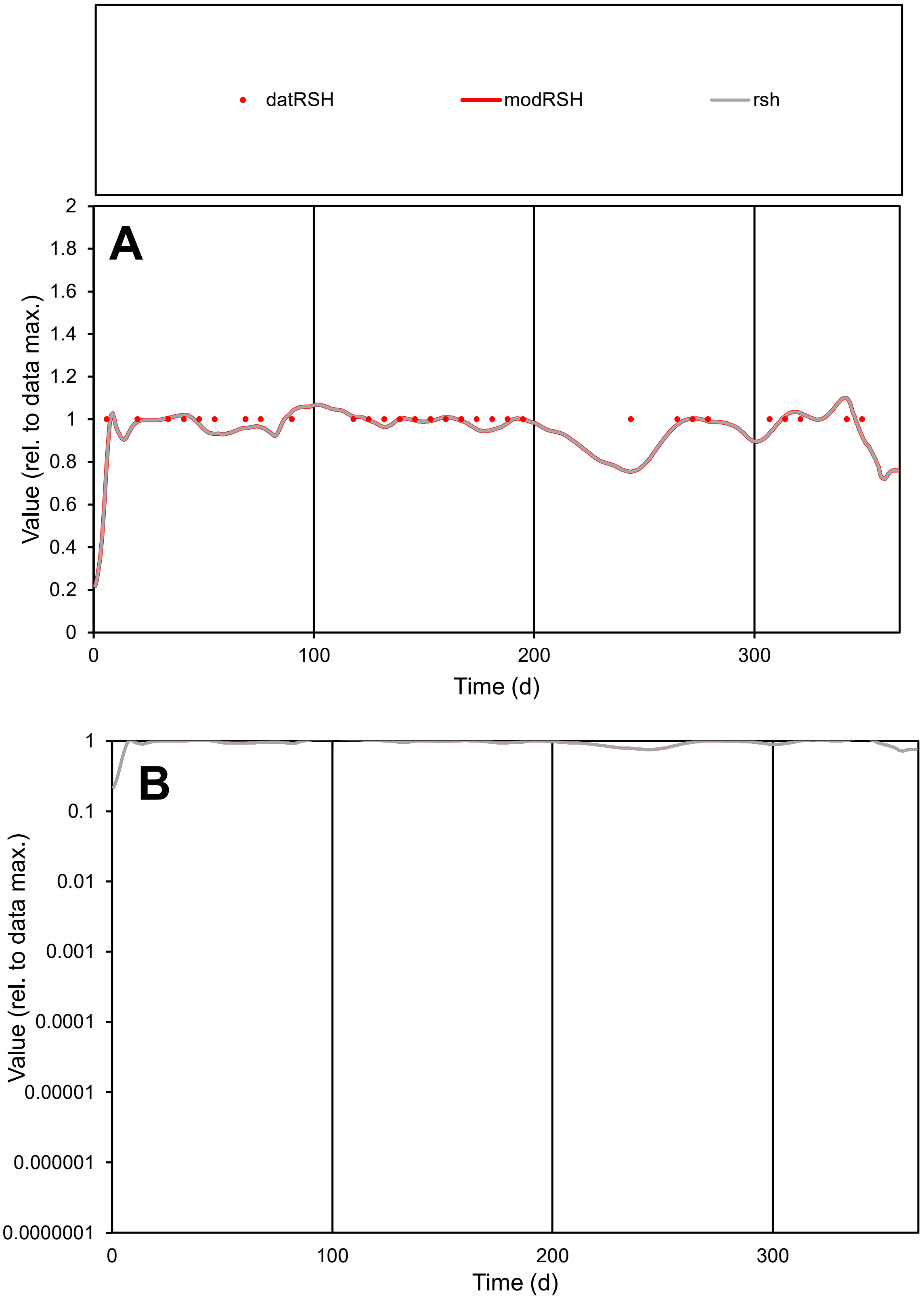


**C1**

Variables

RSH Provide jtd or Dst

CalibOnly?

2 1 to reproduce of

oNormalize?

2 1=no, 2=yes

ofxn calc*

0.05796

ofxn mod

0.12551

sRun

RunA1

zout

1

Ave

Data

1.8E-05

Model

1.8E-05

data/vMax

1.8E-05

ofxng

275.646

Min d/m

1.32198

ave/min d

1

ave/min m

1.29674

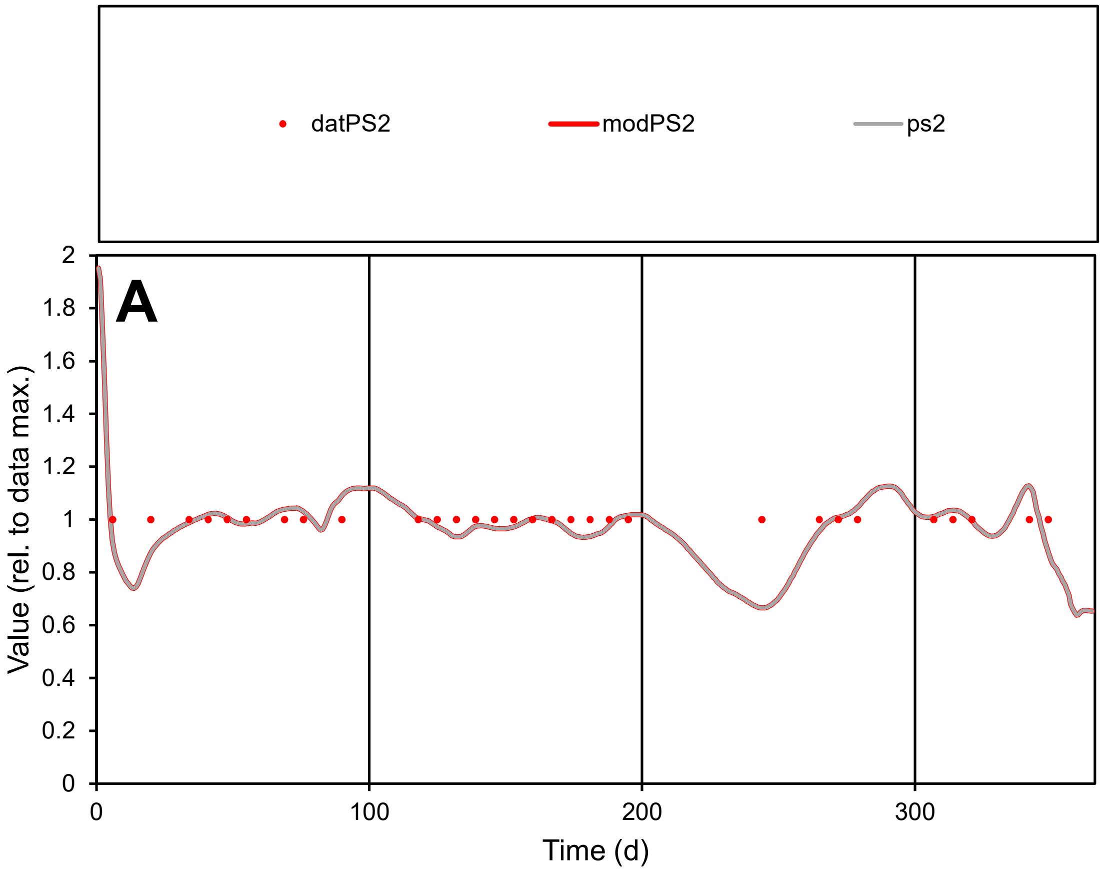


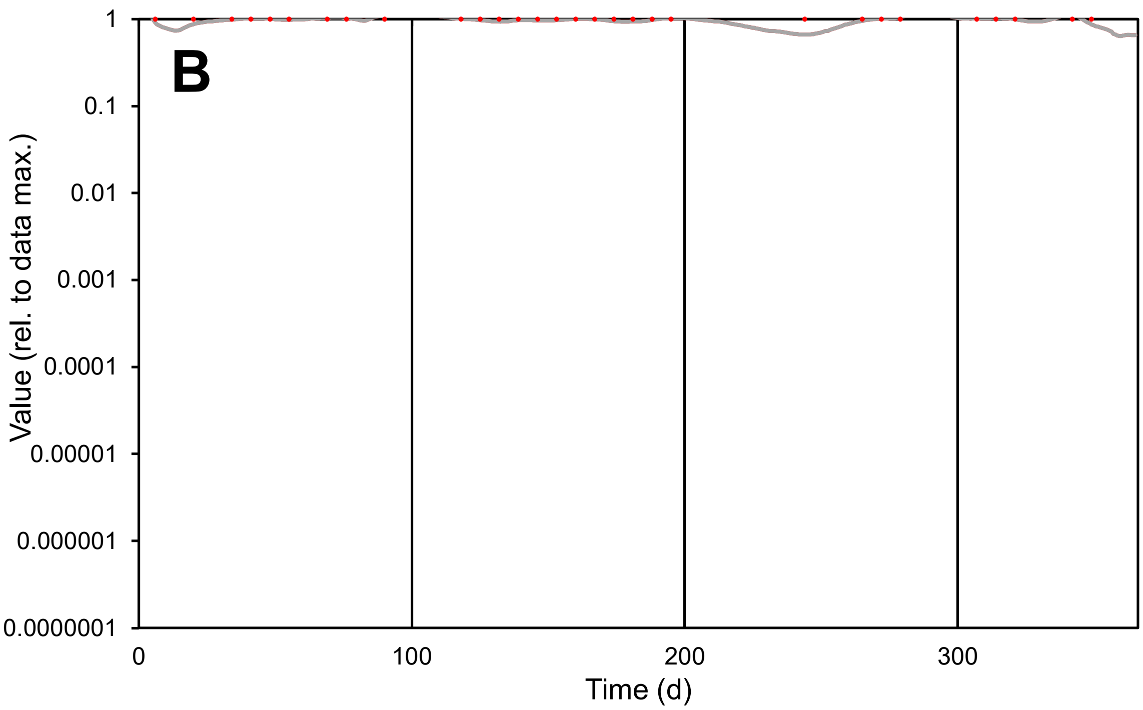


**C1**

Variables

PS2 Provide jtd or Dst

CalibOnly?

2 1 to reproduce of

oNormalize?

2 1=no, 2=yes

ofxn calc*

0.08608

ofxn mod

0.14115

sRun

RunA1

zout

1

Ave

Data

1.9E-05

Model

1.9E-05

data/vMax

1.9E-05

ofxng

275.646

Min d/m

1.5

ave/min d

1

ave/min m

1.46507

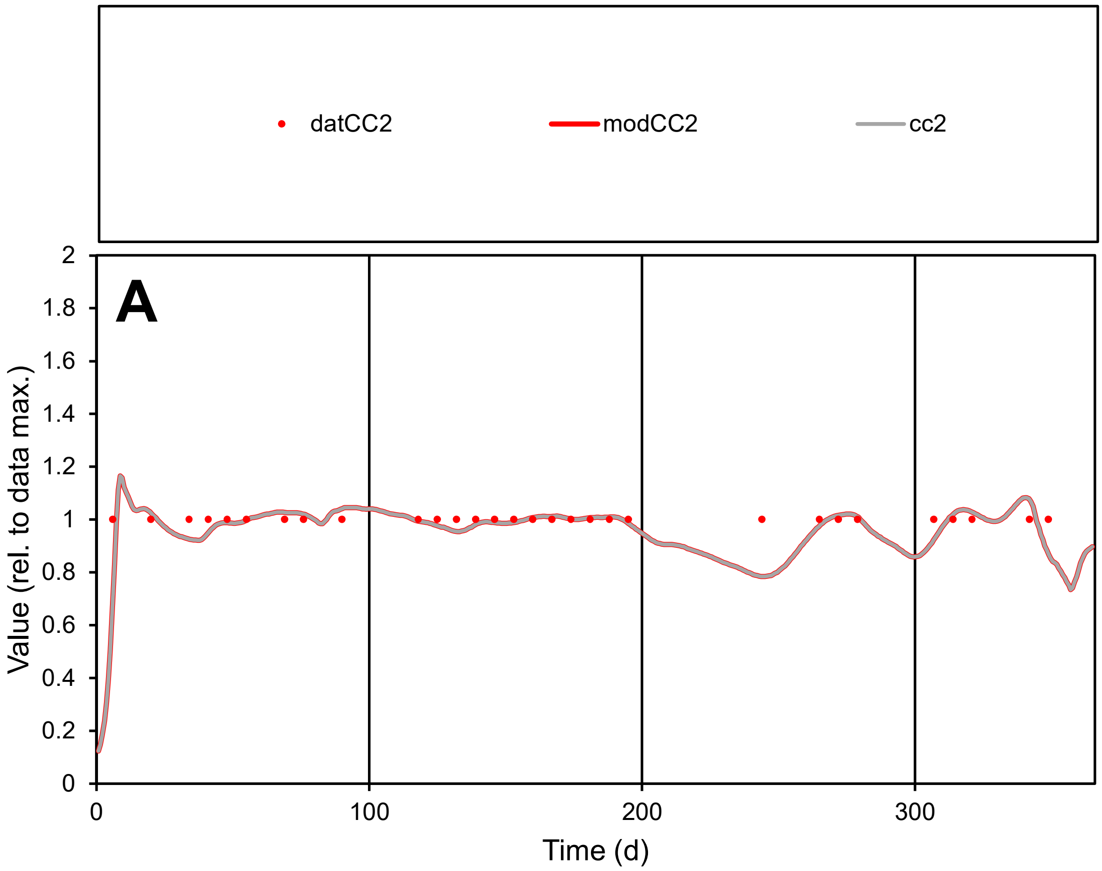


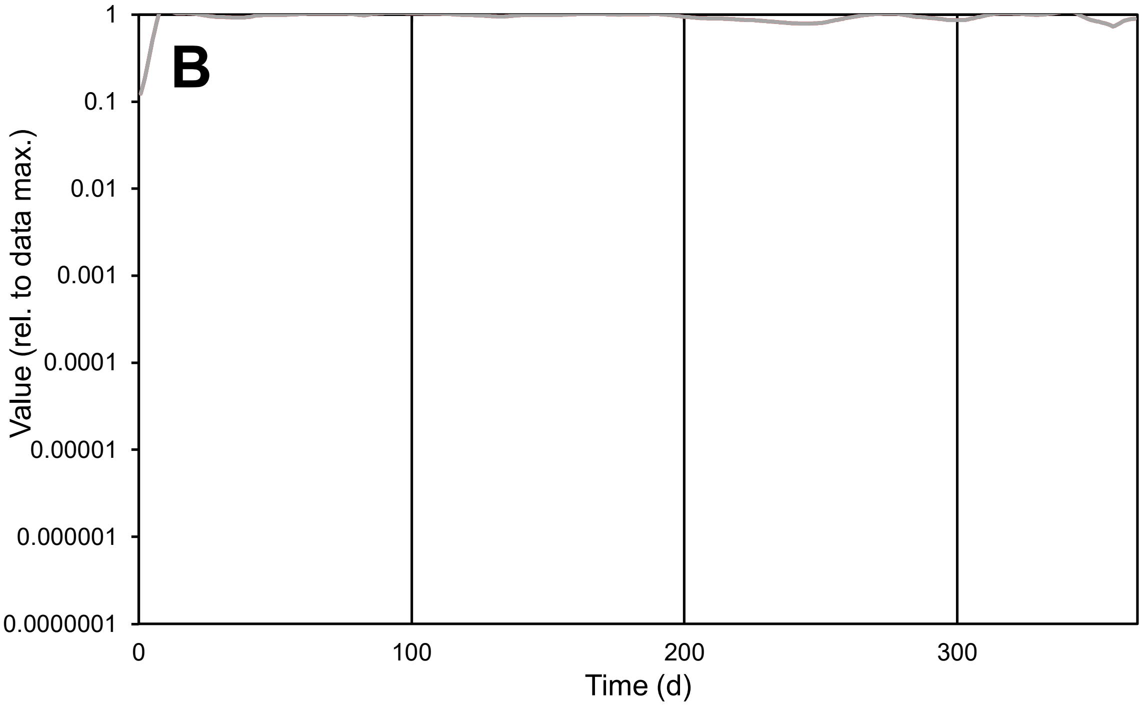


**C1**

Variables
CC2 Provide jtd or Dst
CalibOnly?
2 1 to reproduce of
oNormalize?
2 1=no, 2=yes

ofxn calc*
0.05603
ofxn mod
0.12473

sRun
RunA1
zout
1

Ave
Data
5.9E-06
Model
5.8E-06

data/vMax
5.9E-06

ofxng
275.646

Min d/m
1.27498

ave/min d
1
ave/min m
1.25573

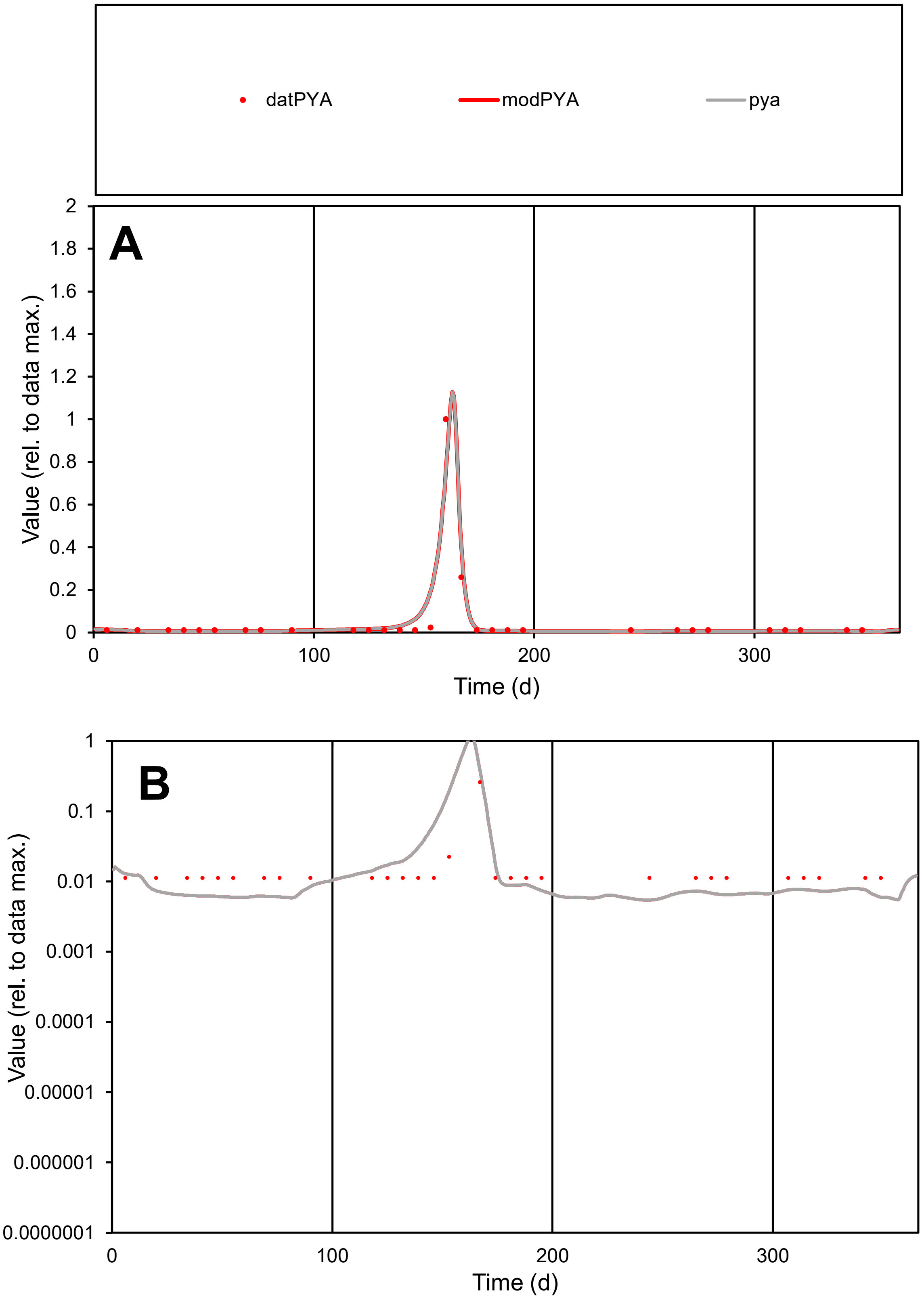


**C1**

Variables
PYA Provide jtd or Dst
CalibOnly?
2 1 to reproduce of
oNormalize?
2 1=no, 2=yes

ofxn calc*
0.4468
ofxn mod
0.63297

sRun
RunA1
zout
1

Ave
Data
8.5E-06
Model
9E-06

data/vMax
0.00016

ofxng
275.646

Min d/m
2.06543

ave/min d
4.69202
ave/min m
10.2634

325_Calib_S_A

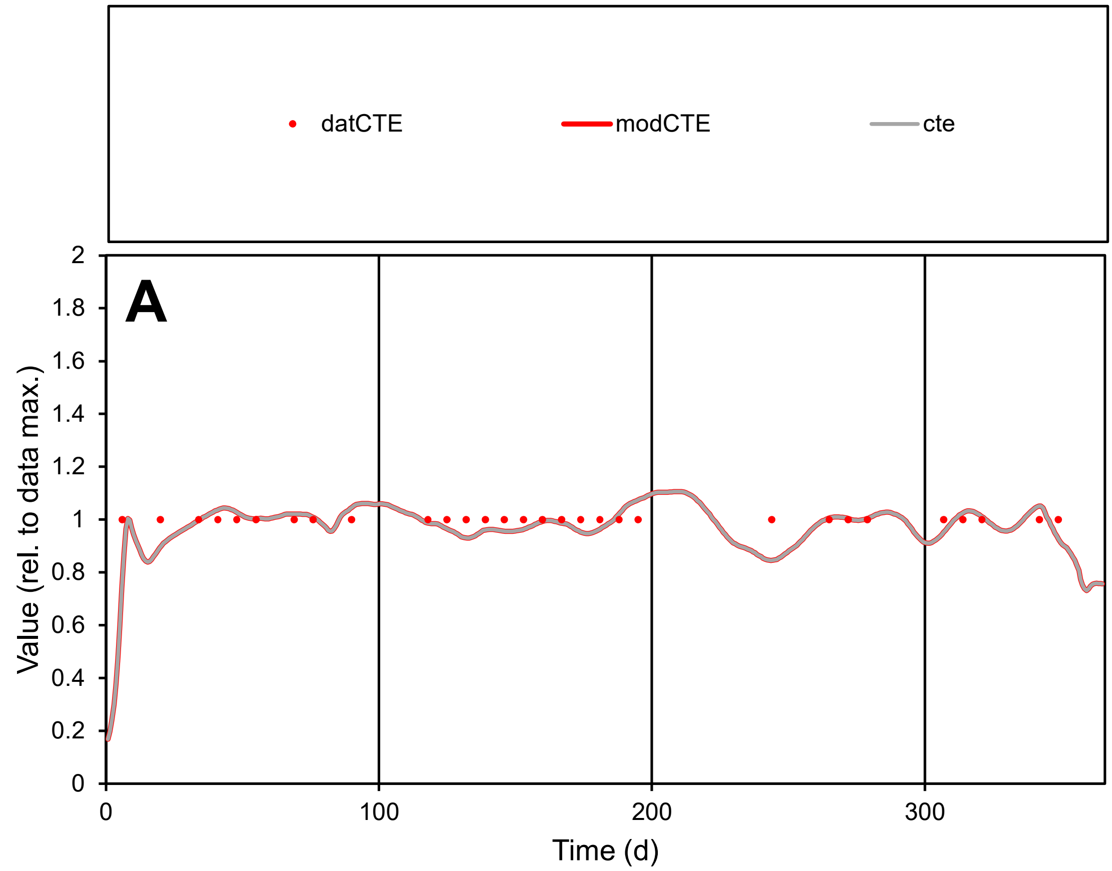


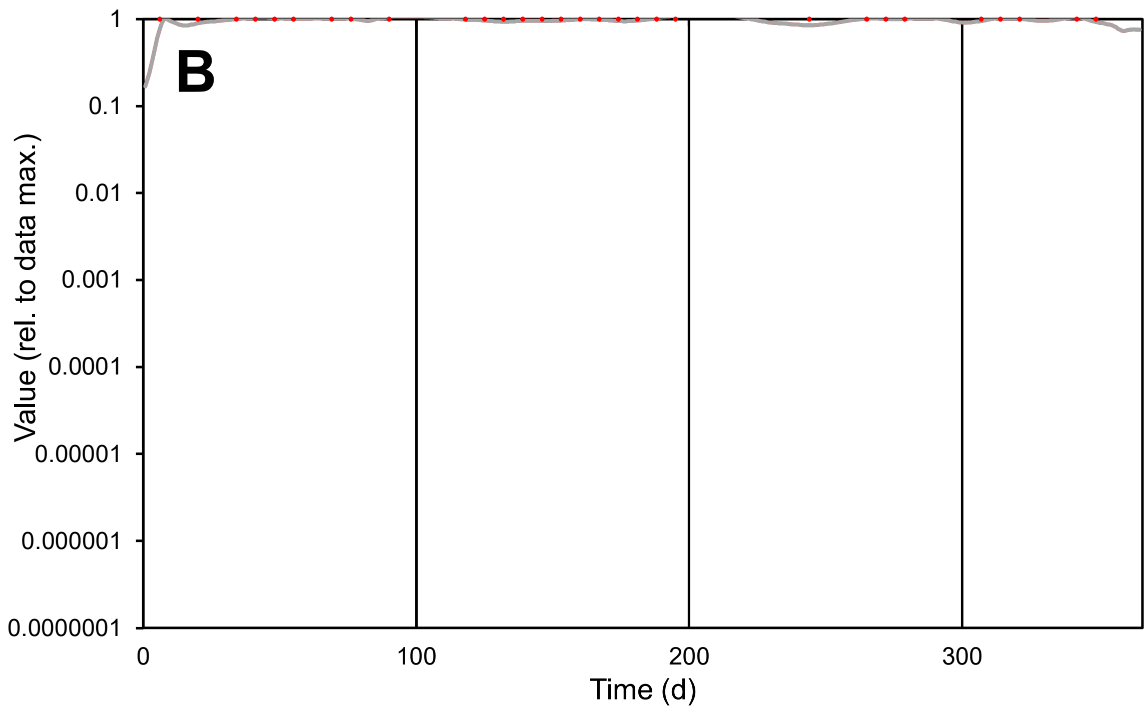


**C1**

Variables
CTE Provide jtd or Dst
CalibOnly?
2 1 to reproduce of
oNormalize?
2 1=no, 2=yes

ofxn calc*
0.04992
ofxn mod
0.12253

sRun
RunA1
zout
1

Ave
Data
6.1E-06
Model
6E-06

data/vMax
6.1E-06

ofxng
275.646

Min d/m
1.18057

ave/min d
1
ave/min m
1.16583

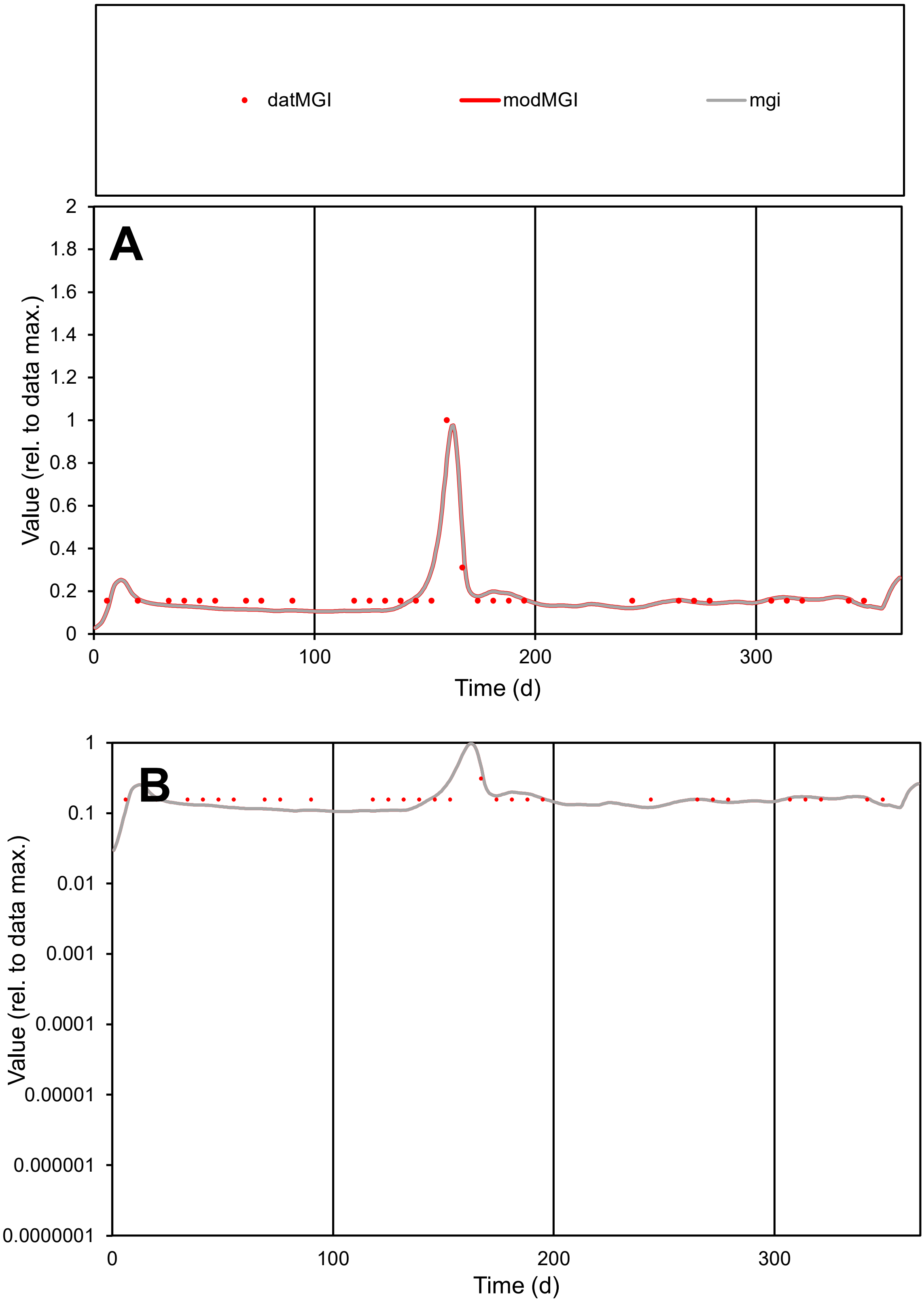


**C1**

Variables

MGI Provide jtd or Dst

CalibOnly?

2 1 to reproduce of

oNormalize?

2 1=no, 2=yes

ofxn calc*

0.21974

ofxn mod

0.25508

sRun

RunA1

zout

1

Ave

Data

1.3E-05

Model

1.2E-05

data/vMax

6.6E-05

ofxng

275.646

Min d/m

1.44255

ave/min d

1.21437

ave/min m

1.69176

325_Calib_S_A

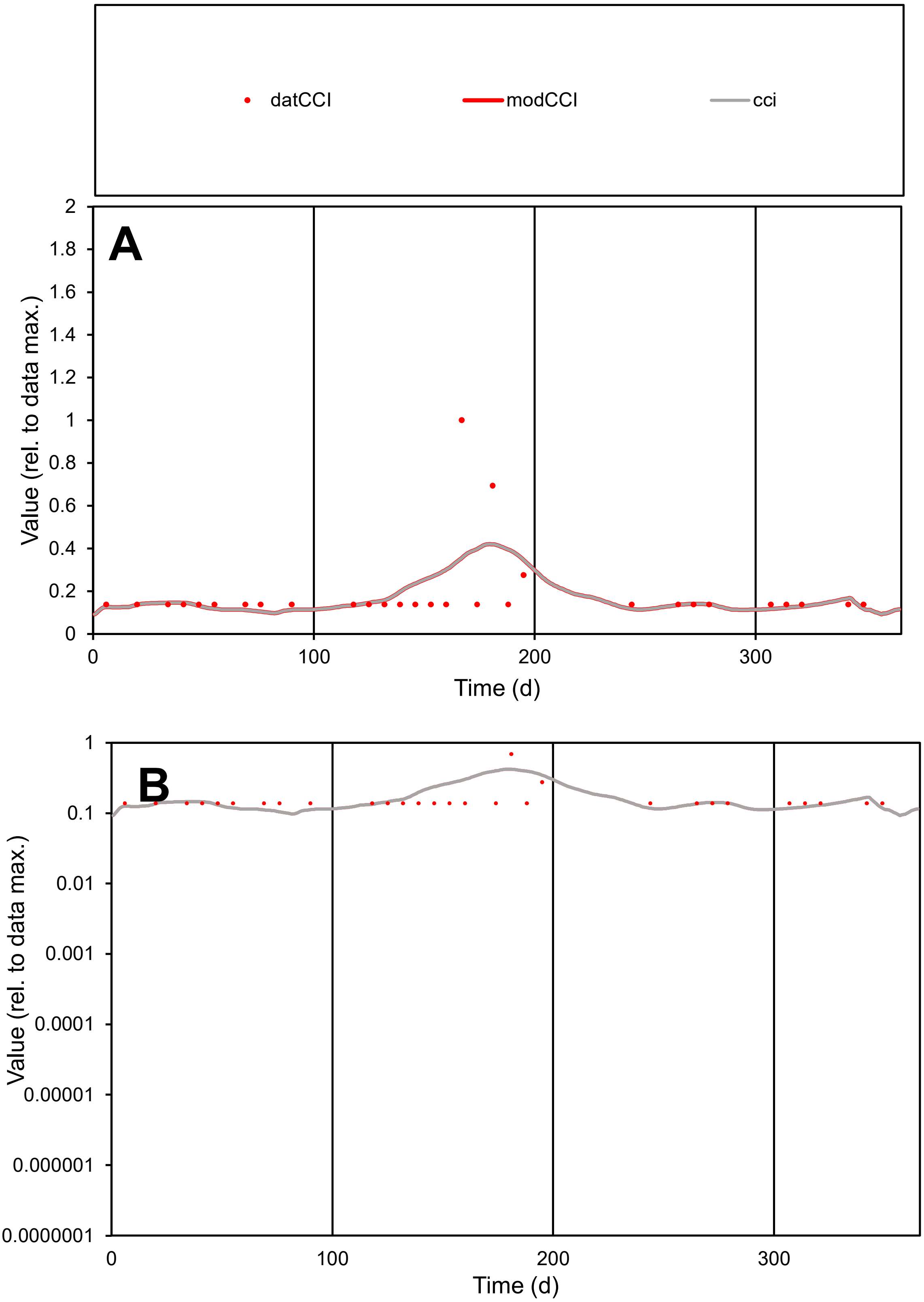


**C1**

Variables
CCI Provide jtd or Dst
CalibOnly?
2 1 to reproduce of
oNormalize?
2 1=no, 2=yes

ofxn calc*
0.79397
ofxn mod
0.8062

sRun
RunA1
zout
1

Ave
Data
1.6E-05
Model
1.6E-05

data/vMax
8.3E-05

ofxng
275.646

Min d/m
1.3355

ave/min d
1.37581
ave/min m
1.832

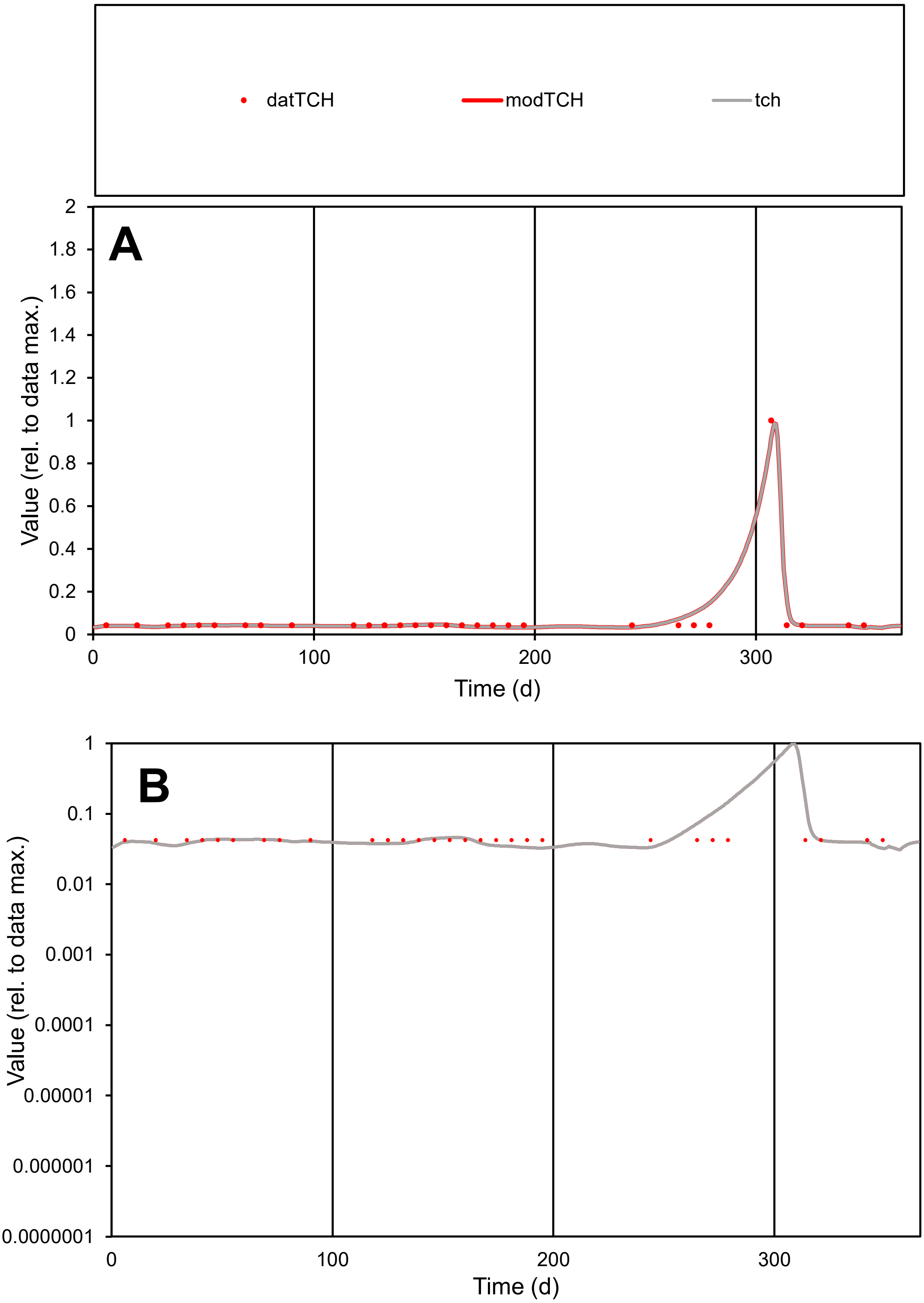


**C1**

Variables
TCH Provide jtd or Dst
CalibOnly?
2 1 to reproduce of
oNormalize?
2 1=no, 2=yes

ofxn calc*
0.32932
ofxn mod
0.44669

sRun
RunA1
zout
1

Ave
Data
2.9E-06
Model
3.1E-06

data/vMax
4E-05

ofxng
275.646

Min d/m
1.31201

ave/min d
1.74946
ave/min m
2.43593

325_Calib_S_A

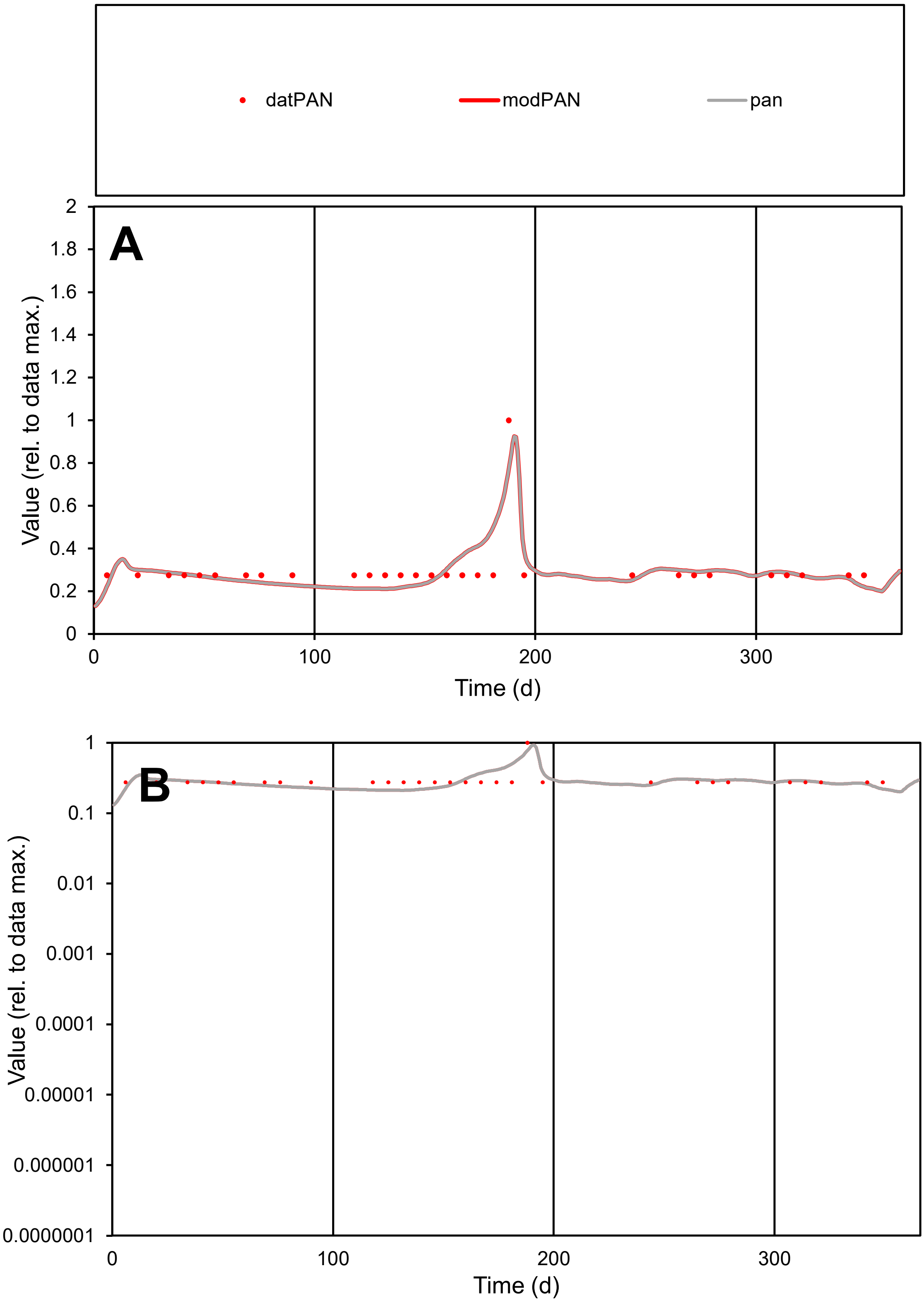


**C1**

Variables
PAN Provide jtd or Dst
CalibOnly?
2 1 to reproduce of
oNormalize?
2 1=no, 2=yes

ofxn calc*
0.23051
ofxn mod
0.26045

sRun
RunA1
zout
1

Ave
Data
9.9E-06
Model
9.9E-06

data/vMax
3.3E-05

ofxng
275.646

Min d/m
1.3023

ave/min d
1.08798
ave/min m
1.41573

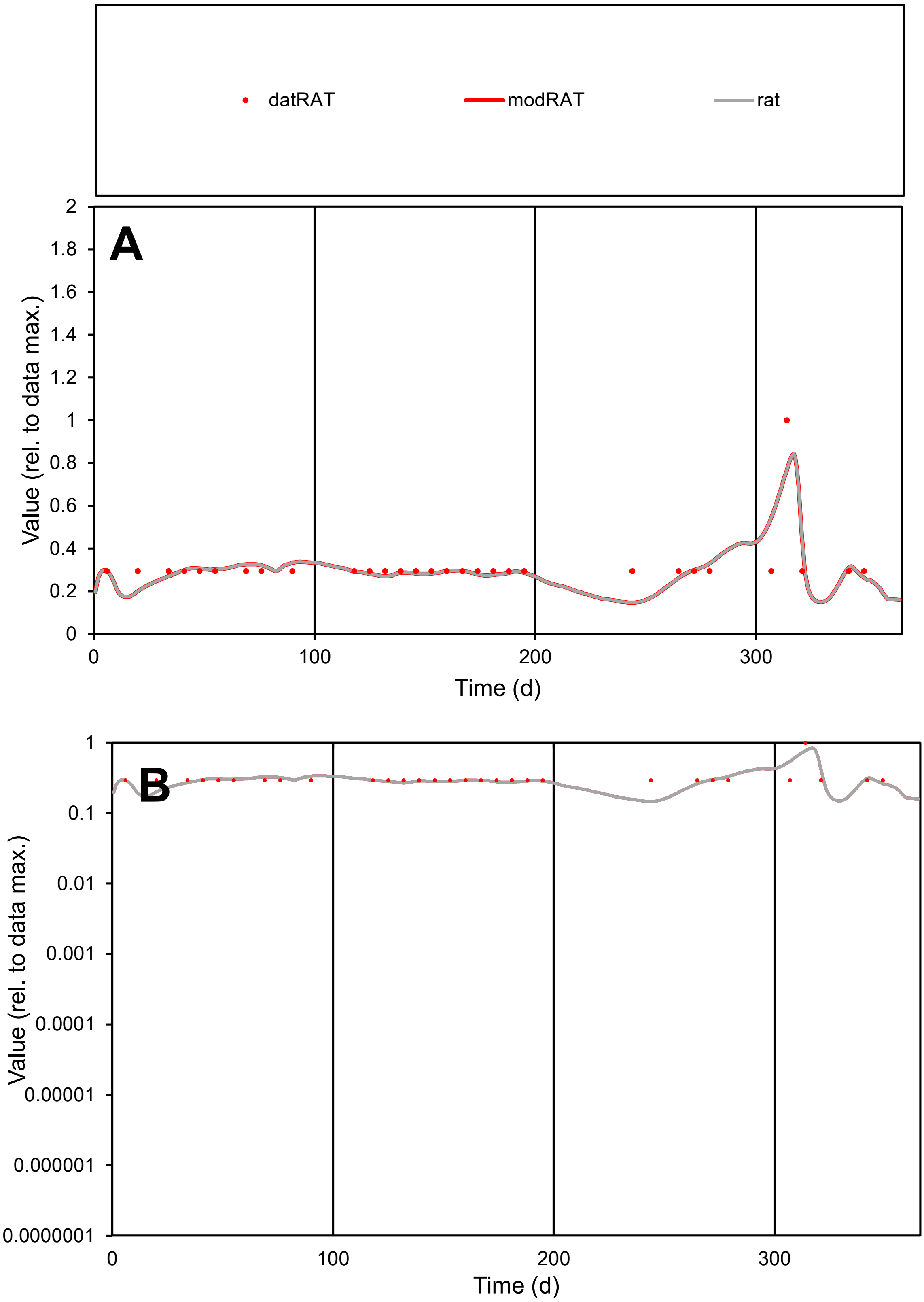


**C1**

Variables
RAT Provide jtd or Dst
CalibOnly?
2 1 to reproduce of
oNormalize?
2 1=no, 2=yes

ofxn calc*
0.21946
ofxn mod
0.25374

sRun
RunA1
zout
1

Ave
Data
1.2E-05
Model
1.2E-05

data/vMax
3.9E-05

ofxng
275.646

Min d/m
1.99754

ave/min d
1.08021
ave/min m
2.12815

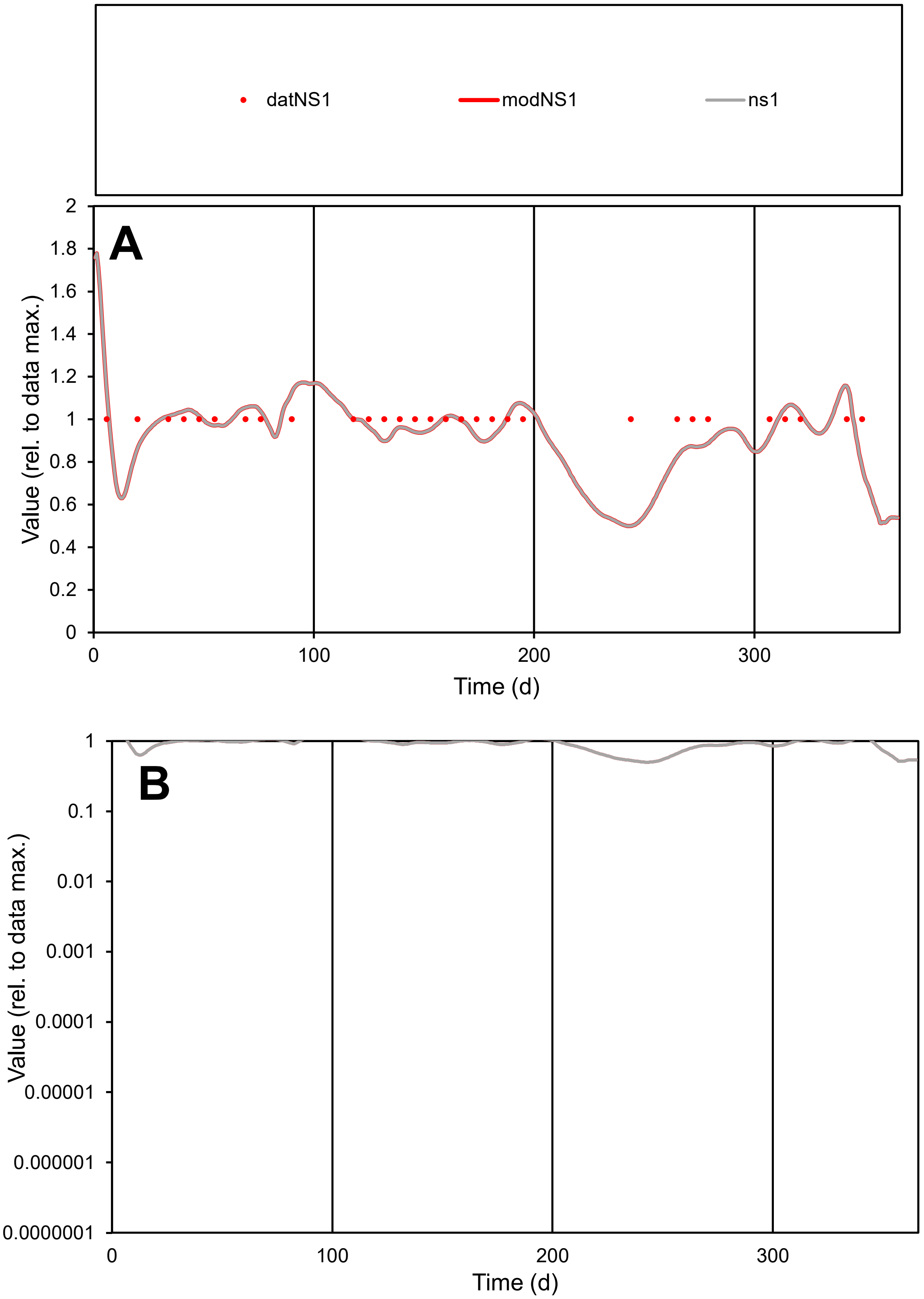


**C1**

Variables
NS1 Provide jtd or Dst
CalibOnly?
2 1 to reproduce of
oNormalize?
2 1=no, 2=yes

ofxn calc*
0.12693
ofxn mod
0.16846

sRun
RunA1
zout
1

Ave
Data
2.7E-05
Model
2.6E-05

data/vMax
2.7E-05

ofxng
275.646

Min d/m
1.98698

ave/min d
1
ave/min m
1.90523

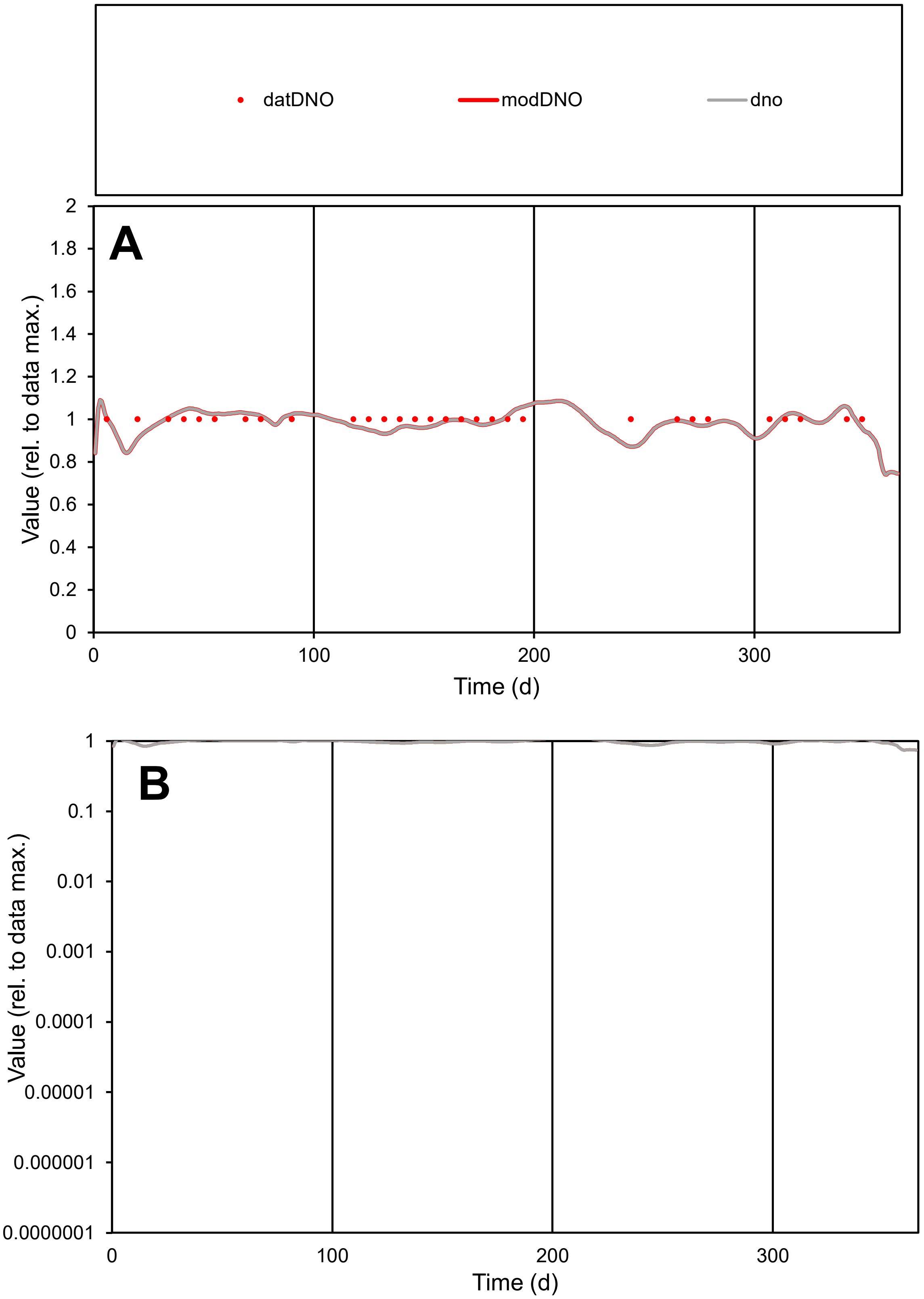


**C1**

Variables
DNO Provide jtd or Dst
CalibOnly?
2 1 to reproduce of
oNormalize?
2 1=no, 2=yes

ofxn calc*
0.04318
ofxn mod
0.12014

sRun
RunA1
zout
1

Ave
Data
1.8E-05
Model
1.8E-05

data/vMax
1.8E-05

ofxng
275.646

Min d/m
1.14775

ave/min d
1
ave/min m
1.13619

325_Calib_S_A

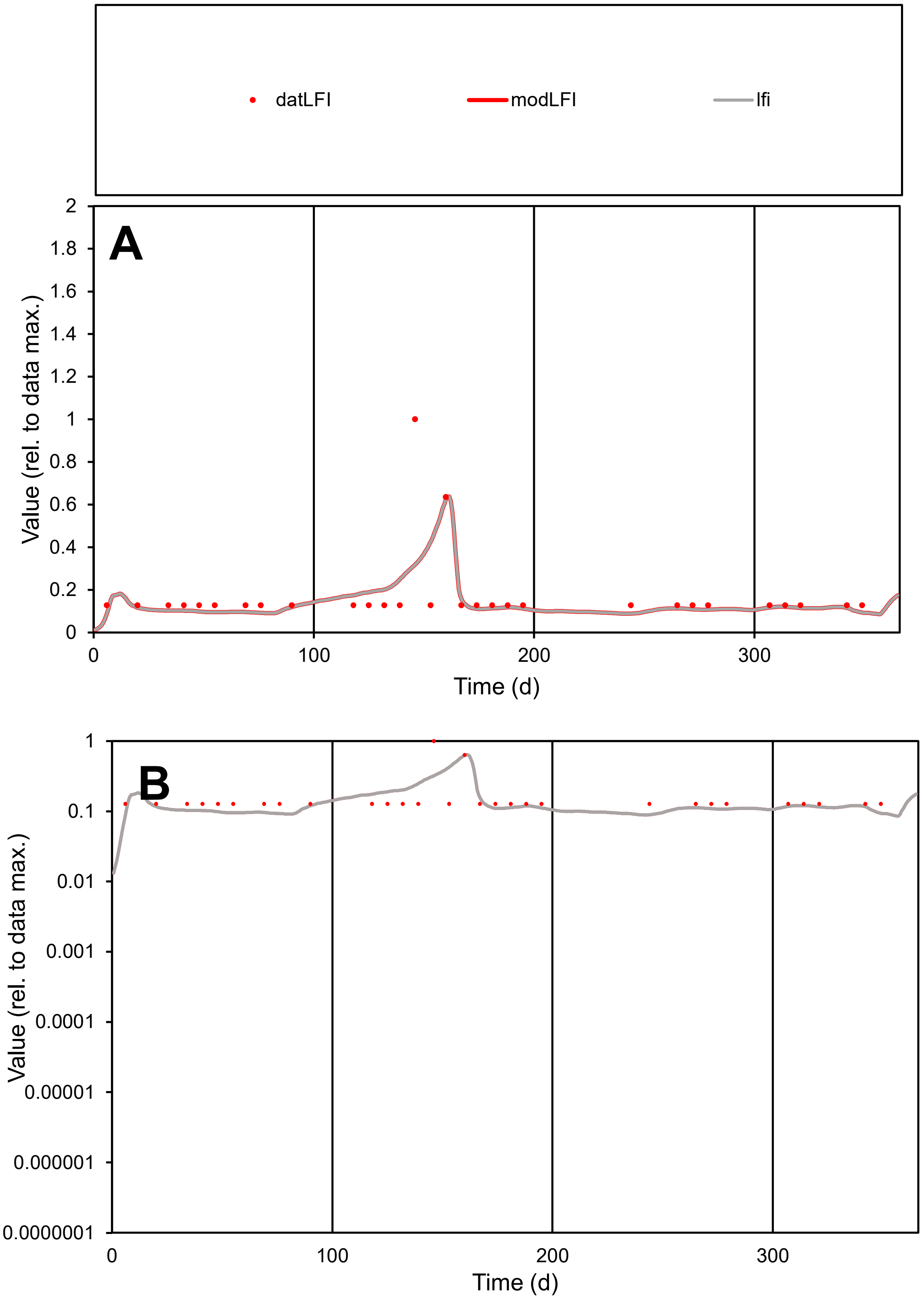


**C1**

Variables
LFI Provide jtd or Dst
CalibOnly?
2 1 to reproduce of
oNormalize?
2 1=no, 2=yes

ofxn calc*
0.79789
ofxn mod
0.81186

sRun
RunA1
zout
1

Ave
Data
1.1E-05
Model
1E-05

data/vMax
6.5E-05

ofxng
275.646

Min d/m
1.41816

ave/min d
1.36142
ave/min m
1.77051

325_Calib_S_A

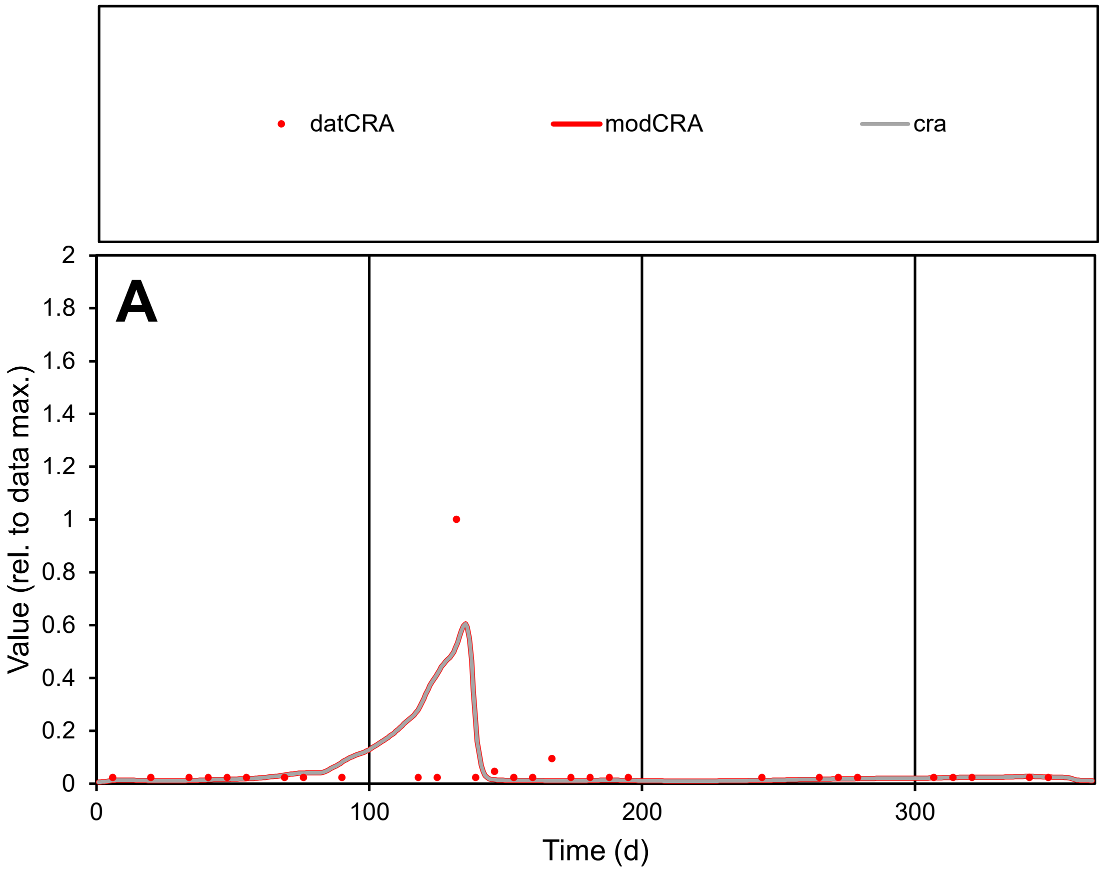


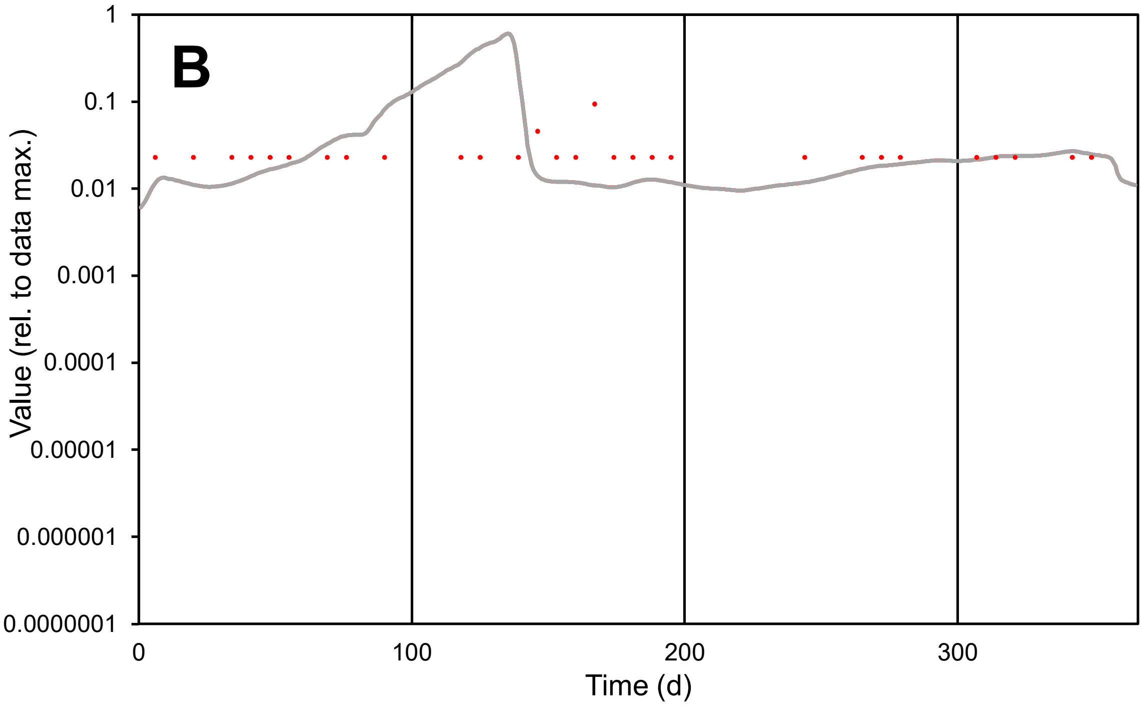


**C1**

Variables
CRA Provide jtd or Dst
CalibOnly?
2 1 to reproduce of
oNormalize?
2 1=no, 2=yes

ofxn calc*
2.01471
ofxn mod
2.06874

sRun
RunA1
zout
1

Ave
Data
1.2E-05
Model
1.3E-05

data/vMax
0.00021

ofxng
275.646

Min d/m
2.19198

ave/min d
2.56151
ave/min m
5.83484

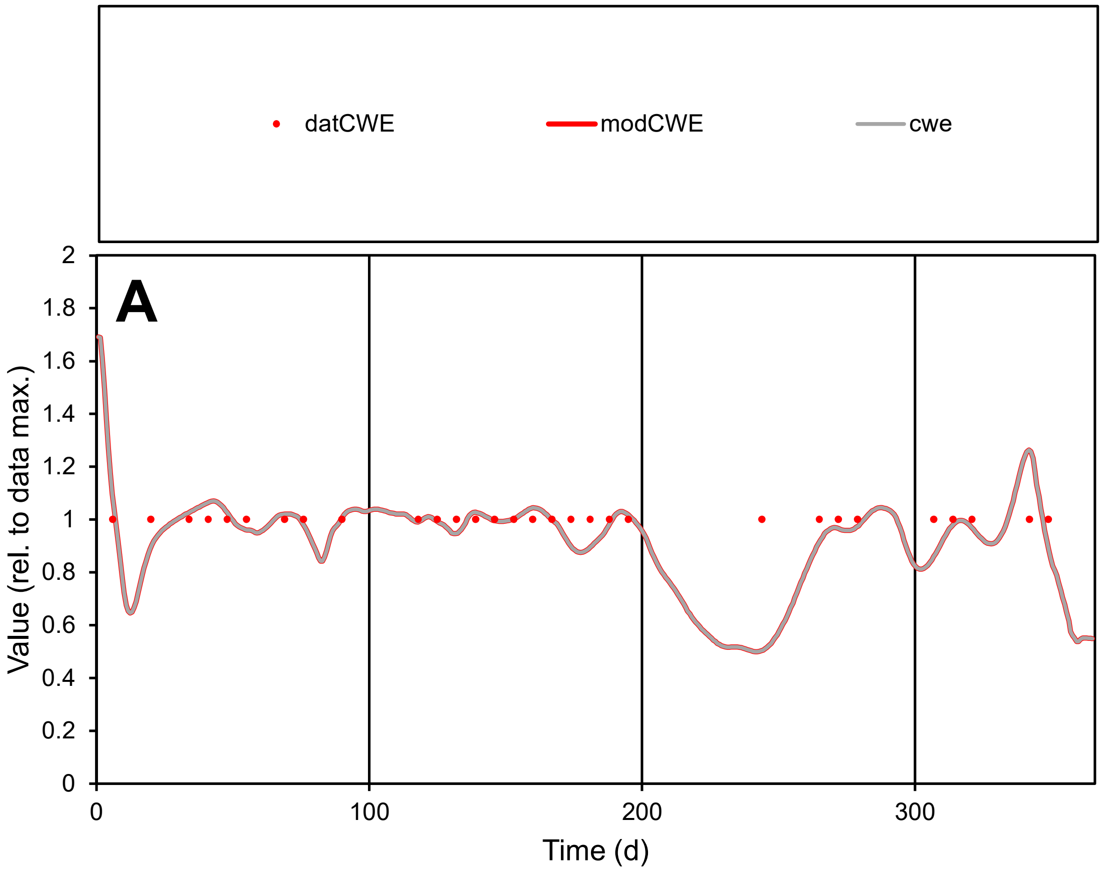


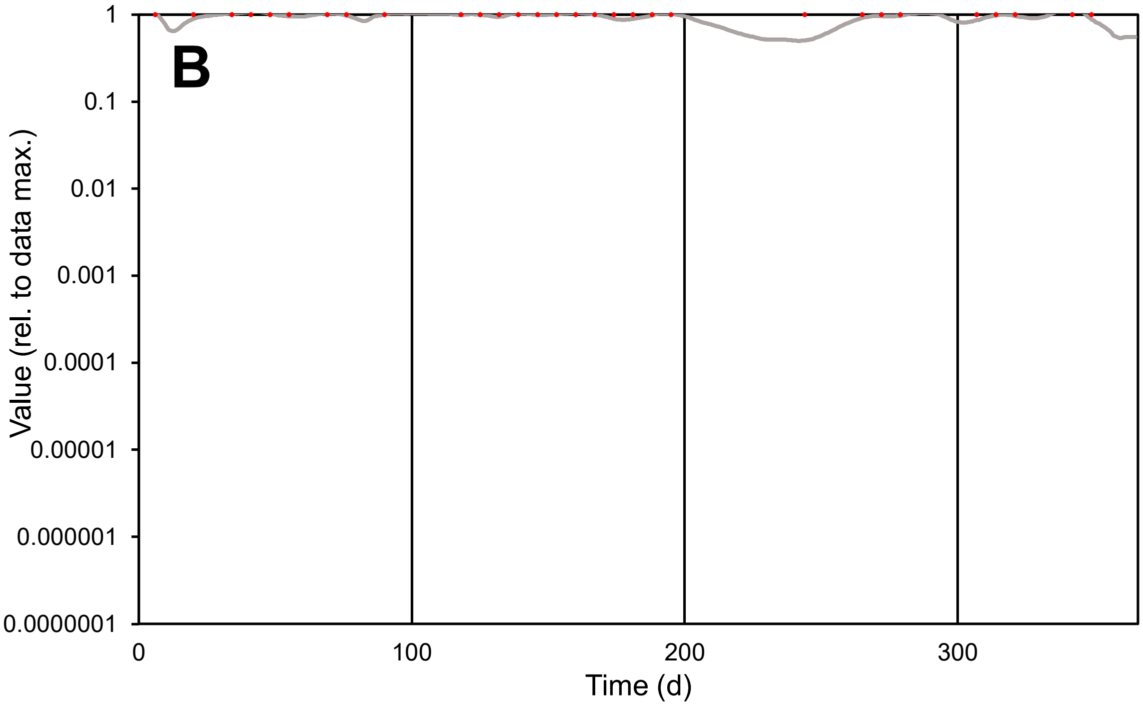


**C1**

Variables
CWE Provide jtd or Dst
CalibOnly?
2 1 to reproduce of
oNormalize?
2 1=no, 2=yes

ofxn calc*
0.11399
ofxn mod
0.15929

sRun
RunA1
zout
1

Ave
Data
2.6E-05
Model
2.5E-05

data/vMax
2.6E-05

ofxng
275.646

Min d/m
1.95853

ave/min d
1
ave/min m
1.89736

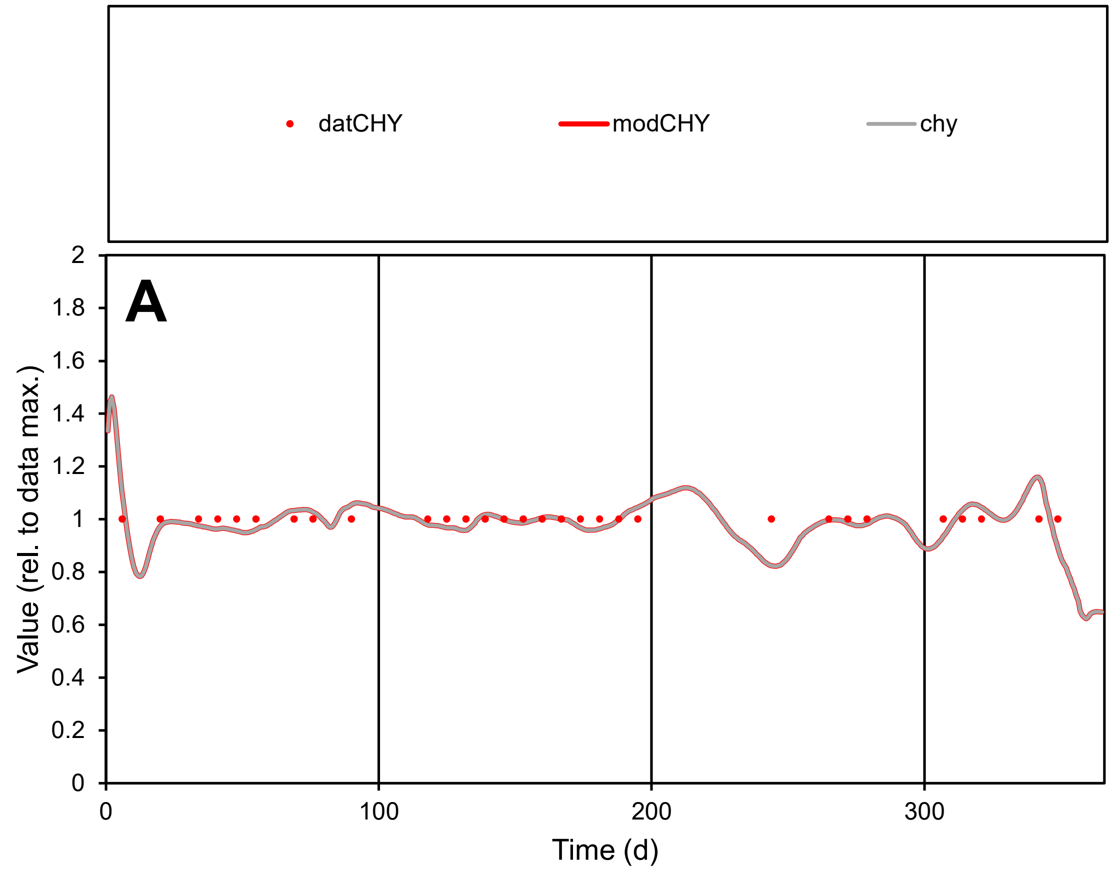


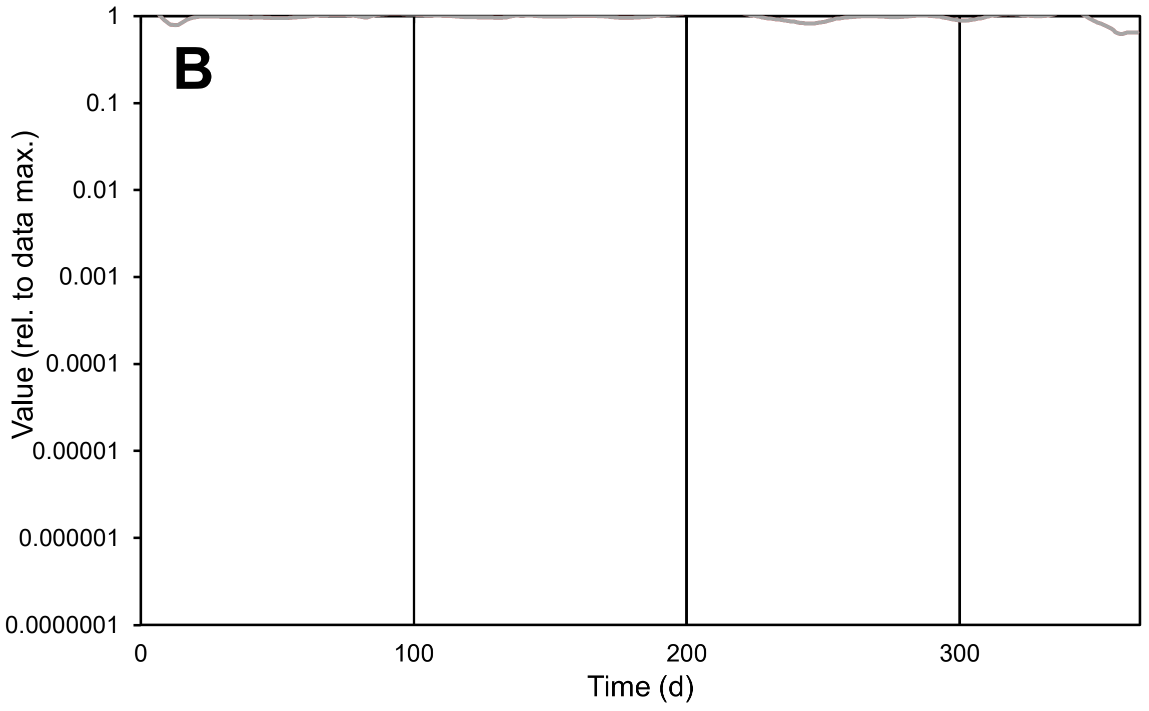


**C1**

Variables

CHY Provide jtd or Dst

CalibOnly?

2 1 to reproduce of

oNormalize?

2 1=no, 2=yes

ofxn calc*

0.05752

ofxn mod

0.12582

sRun

RunA1

zout

1

Ave

Data

6.7E-06

Model

6.6E-06

data/vMax

6.7E-06

ofxng

275.646

Min d/m

1.21719

ave/min d

1

ave/min m

1.20387

325_Calib_S_A

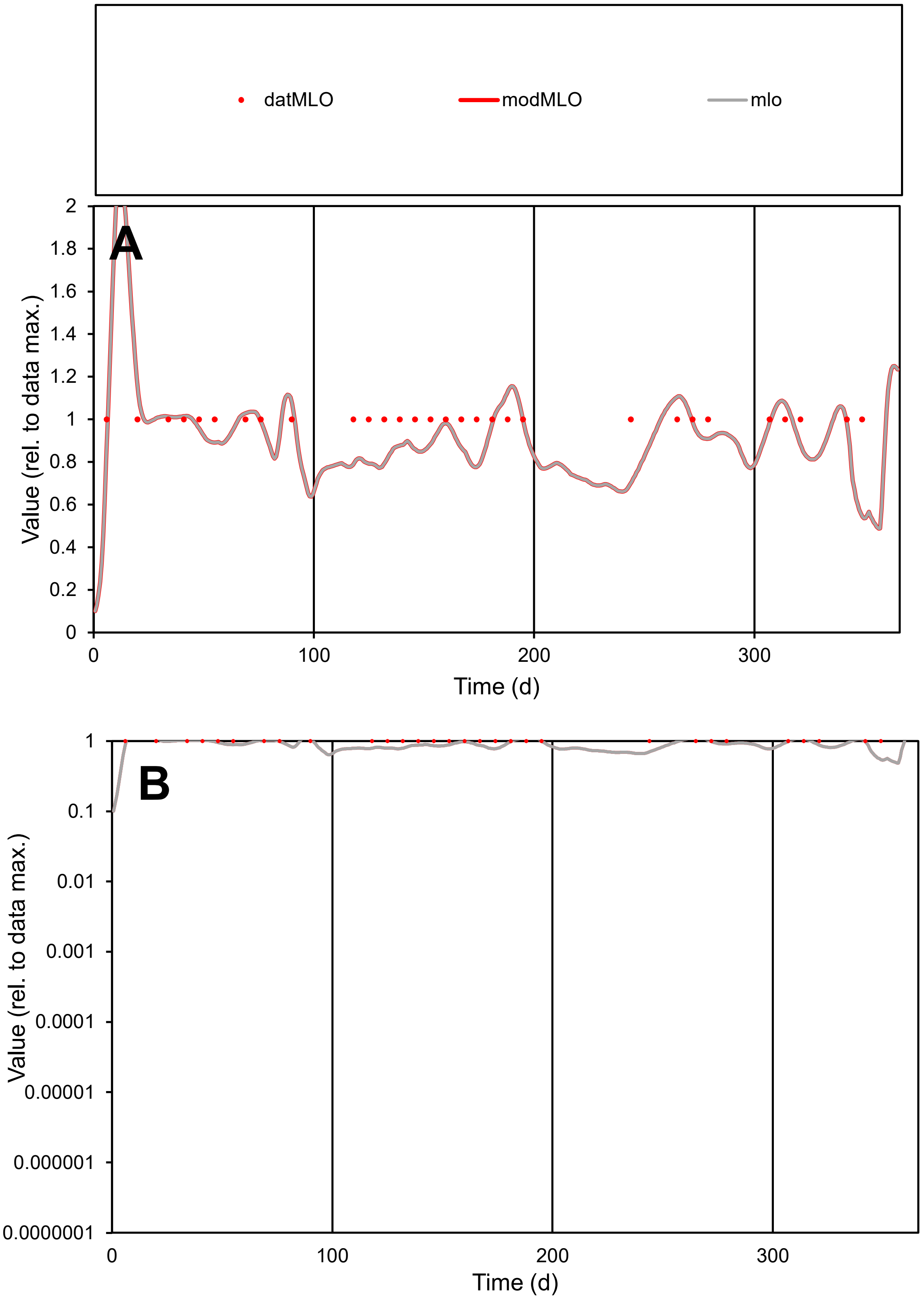


**C1**

Variables
MLO Provide jtd or Dst
CalibOnly?
2 1 to reproduce of
oNormalize?
2 1=no, 2=yes

ofxn calc*
0.14497
ofxn mod
0.18329

sRun
RunA1
zout
1

Ave
Data
1E-05
Model
9.6E-06

data/vMax
1E-05

ofxng
275.646

Min d/m
1.86017

ave/min d
1
ave/min m
1.72098

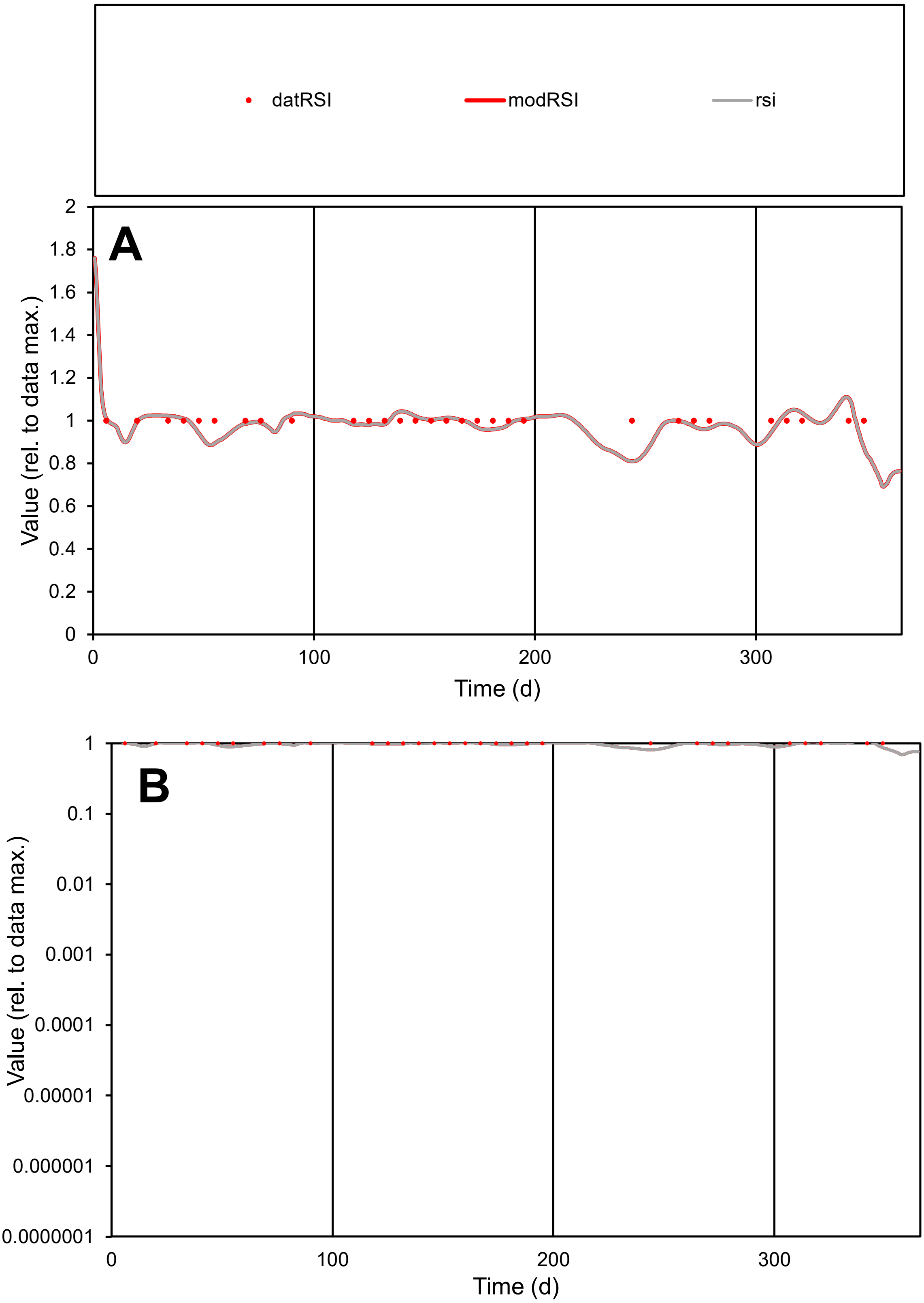


**C1**

Variables
RSI Provide jtd or Dst
CalibOnly?
2 1 to reproduce of
oNormalize?
2 1=no, 2=yes

ofxn calc*
0.05696
ofxn mod
0.12507

sRun
RunA1
zout
1

Ave
Data
1E-05
Model
1E-05

data/vMax
1E-05

ofxng
275.646

Min d/m
1.23358

ave/min d
1
ave/min m
1.2139

325_Calib_S_A

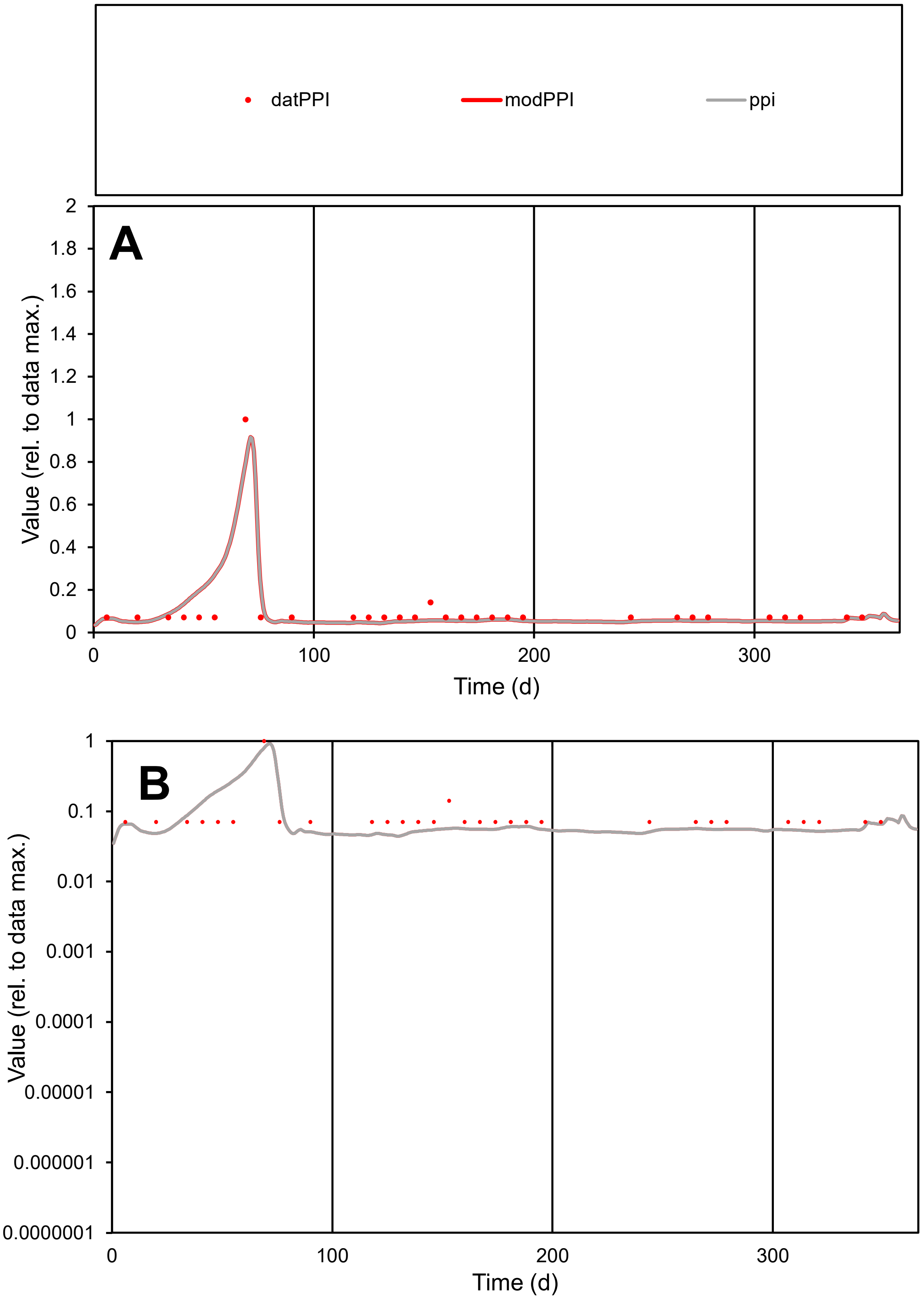


| | |
|---|---|
| **C1** | |
| Variables | |
| PPI | Provide jtd or Dst |
| CalibOnly? | |
| 2 | 1 to reproduce of |
| oNormalize? | |
| 2 | 1=no, 2=yes |
| ofxn calc* | |
| 0.53165 | |
| ofxn mod | |
| 0.57367 | |
| sRun | |
| RunA1 | |
| zout | |
| 1 | |
| Ave | |
| Data | |
| 2.1E-06 | |
| Model | |
| 2.1E-06 | |
| data/vMax | |
| 2.1E-05 | |
| ofxng | |
| 275.646 | |
| Min d/m | |
| 1.53797 | |
| ave/min d | |
| 1.47303 | |
| ave/min m | |
| 2.2088 | |

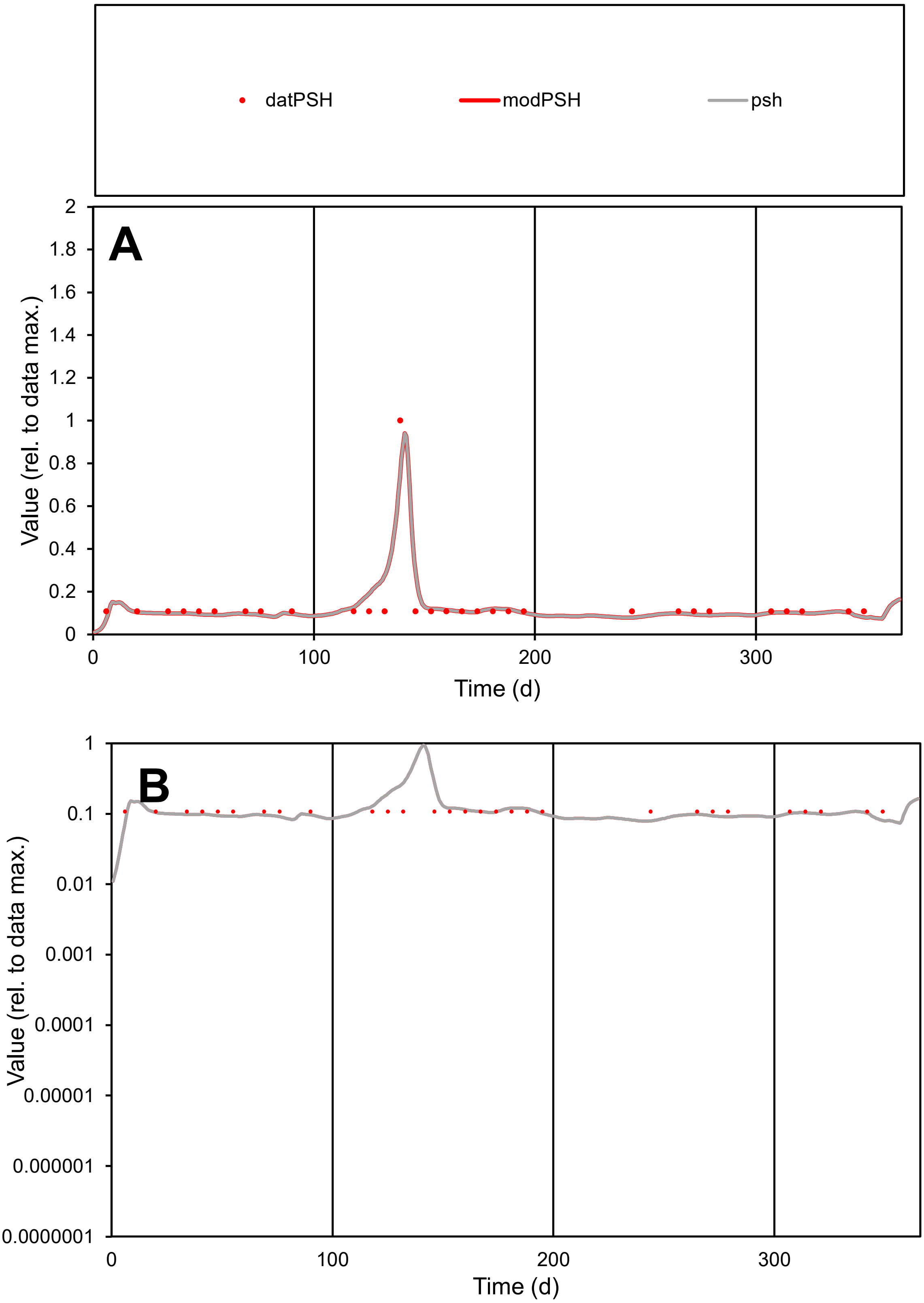


**C1**

Variables
PSH Provide jtd or Dst
CalibOnly?
2 1 to reproduce of
oNormalize?
2 1=no, 2=yes

ofxn calc*
0.32568
ofxn mod
0.37276

sRun
RunA1
zout
1

Ave
Data
1.6E-05
Model
1.7E-05

data/vMax
0.00012

ofxng
275.646

Min d/m
1.35704

ave/min d
1.27674
ave/min m
1.76729

325_Calib_S_A

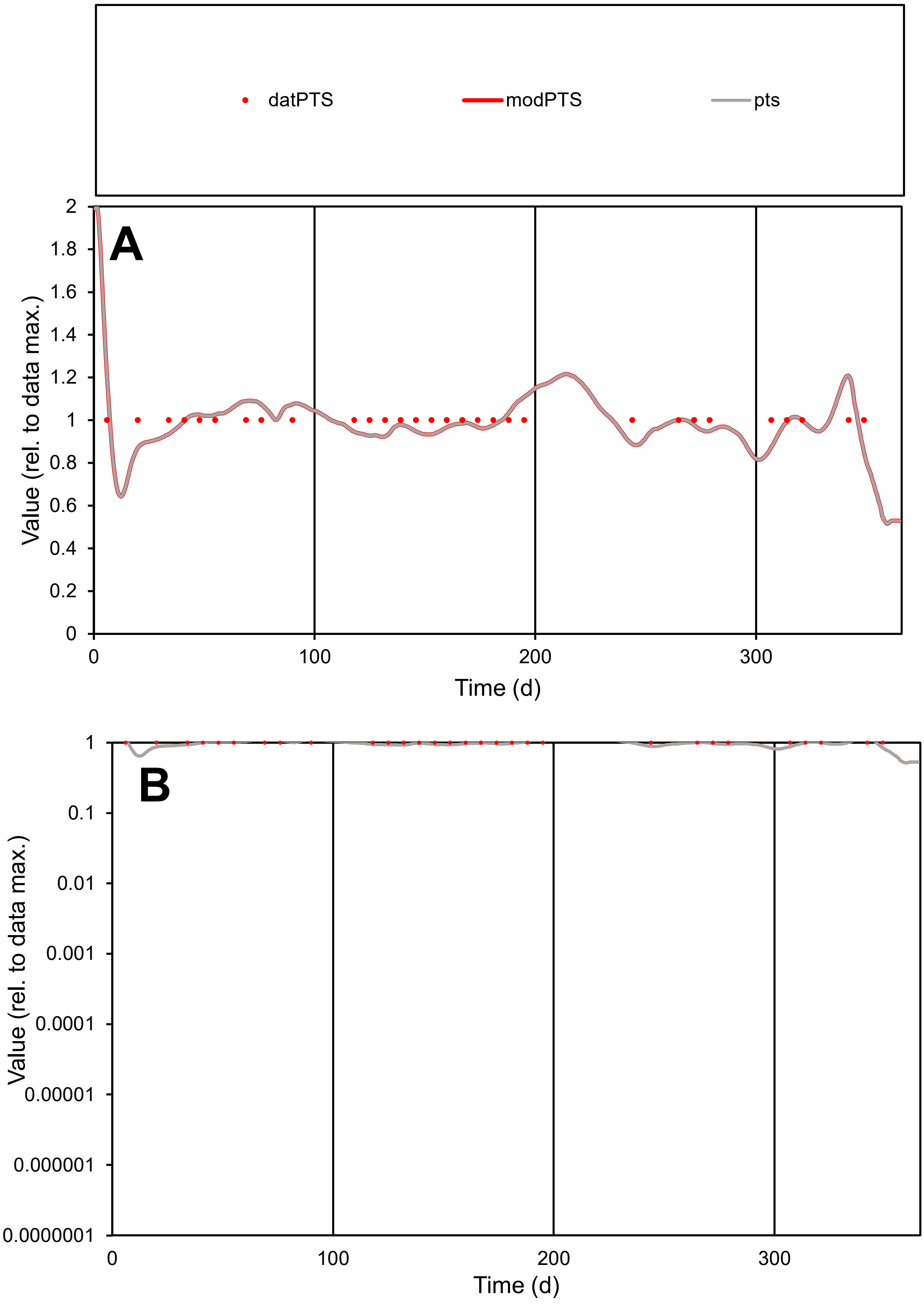


**C1**

Variables
PTS Provide jtd or Dst
CalibOnly?
2 1 to reproduce of
oNormalize?
2 1=no, 2=yes

ofxn calc*
0.07724
ofxn mod
0.136

sRun
RunA1
zout
1

Ave
Data
9.7E-06
Model
9.5E-06

data/vMax
9.7E-06

ofxng
275.646

Min d/m
1.2714

ave/min d
1
ave/min m
1.25165

325_Calib_S_A

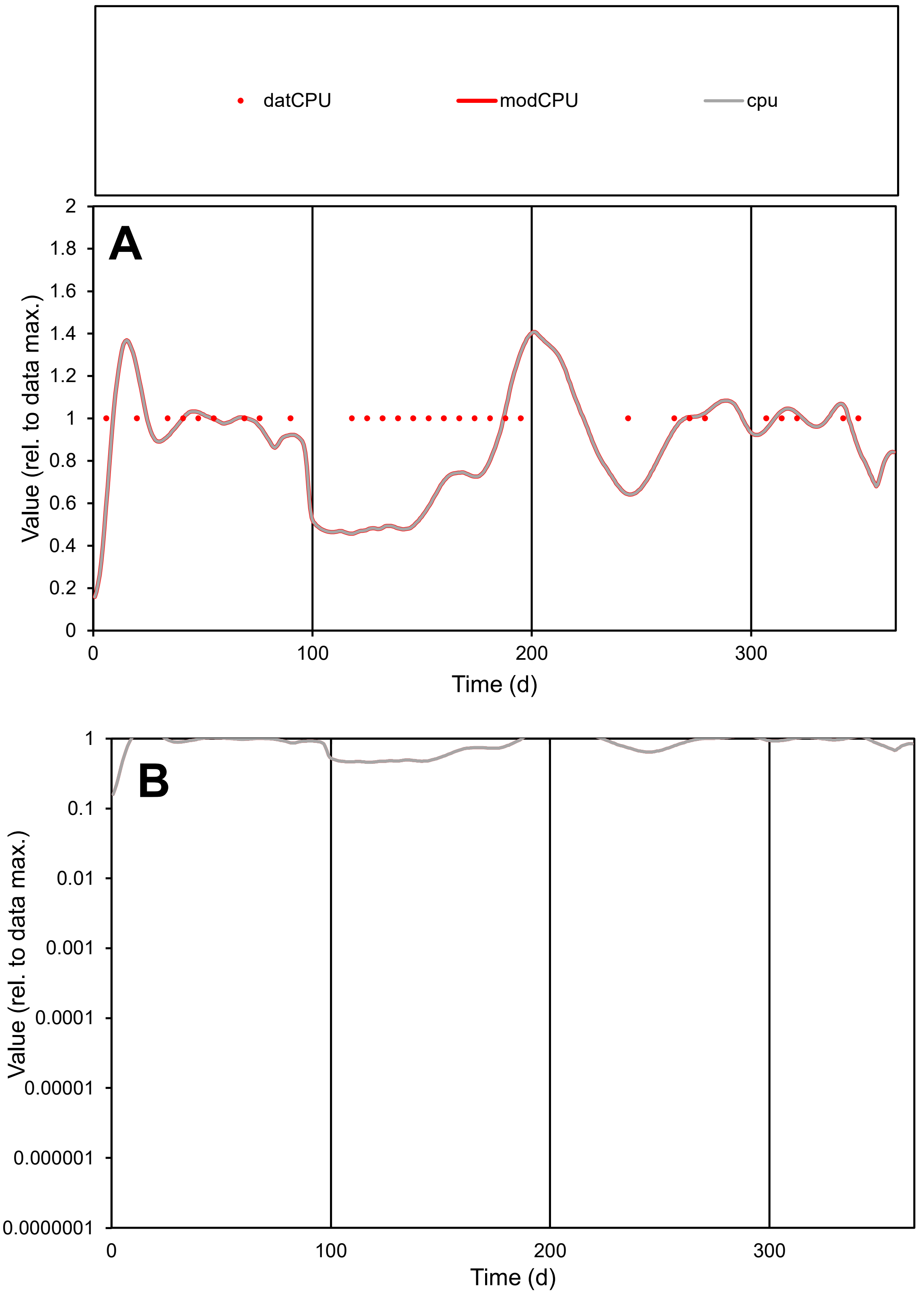


**C1**

Variables

CPU Provide jtd or Dst

CalibOnly?

2 1 to reproduce of

oNormalize?

2 1=no, 2=yes

ofxn calc*

0.26508

ofxn mod

0.28622

sRun

RunA1

zout

1

Ave

Data

4.4E-07

Model

3.8E-07

data/vMax

4.4E-07

ofxng

275.646

Min d/m

2.17511

ave/min d

1

ave/min m

1.87167

325_Calib_S_A

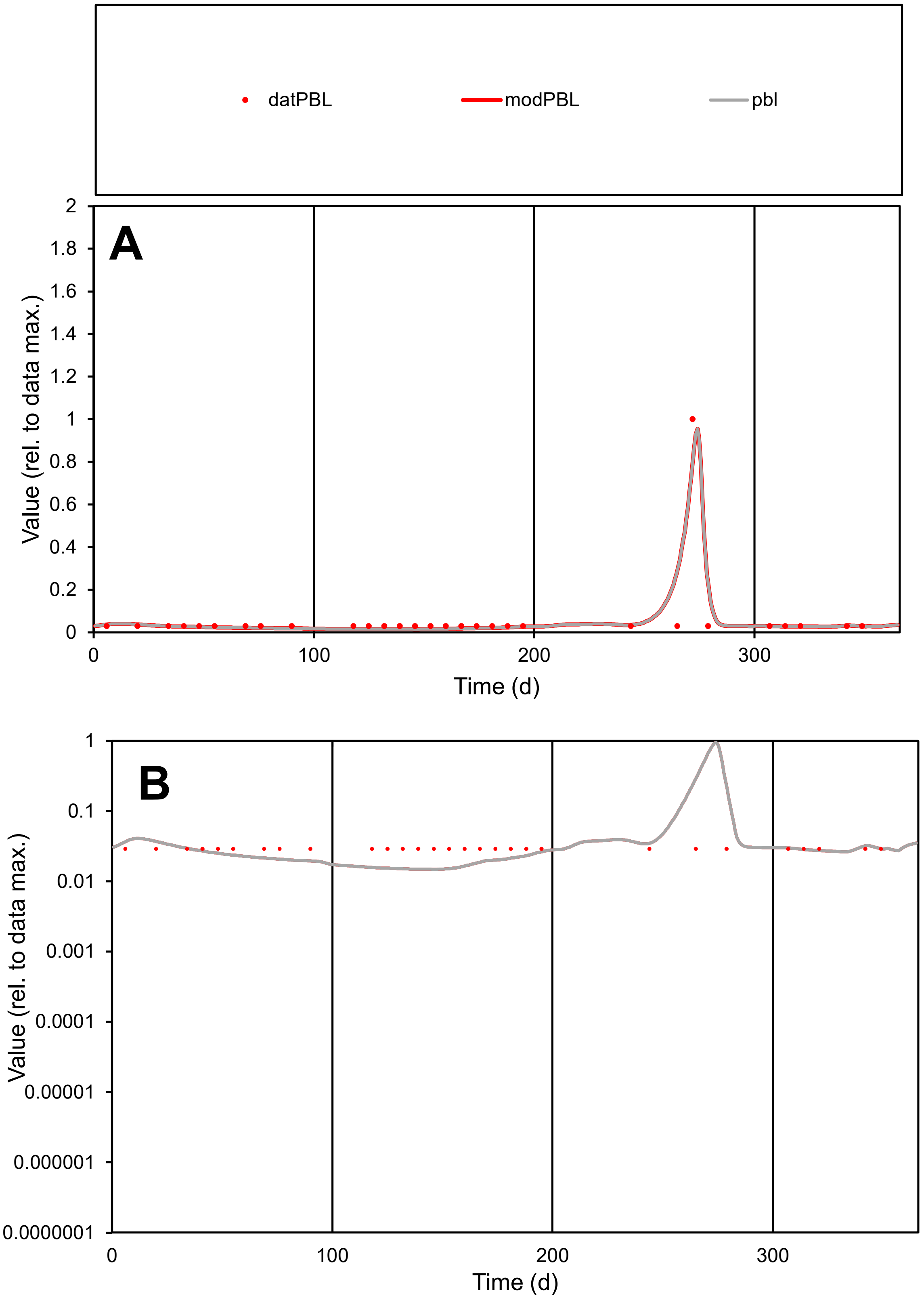


**C1**

Variables

PBL Provide jtd or Dst

CalibOnly?

2 1 to reproduce of

oNormalize?

2 1=no, 2=yes

ofxn calc*

0.6562

ofxn mod

0.88652

sRun

RunA1

zout

1

Ave

Data

9E-08

Model

9.2E-08

data/vMax

1.5E-06

ofxng

275.646

Min d/m

1.95858

ave/min d

2.11172

ave/min m

4.23214

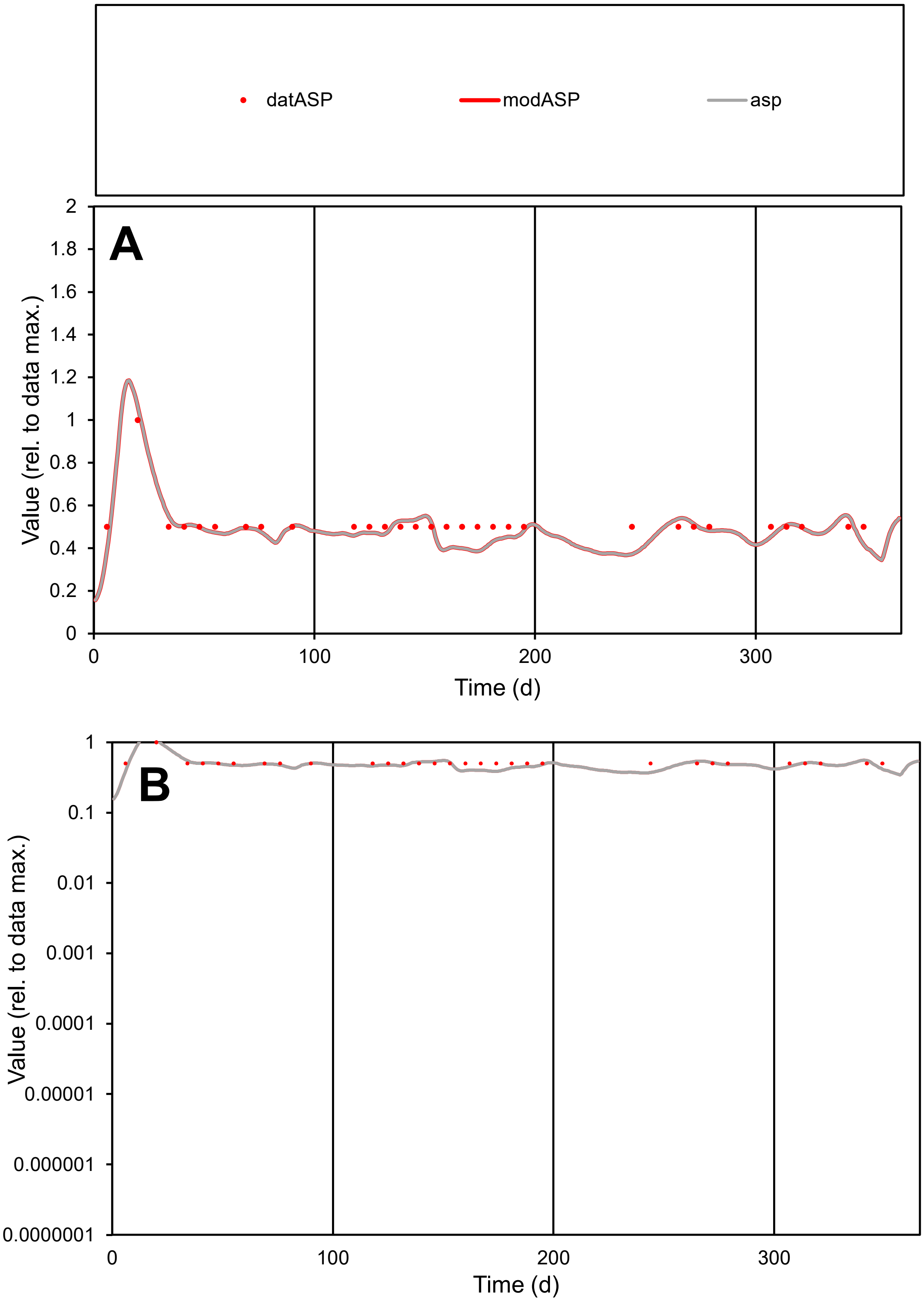


**C1**

Variables
ASP Provide jtd or Dst
CalibOnly?
2 1 to reproduce of
oNormalize?
2 1=no, 2=yes

ofxn calc*
0.09753
ofxn mod
0.14691

sRun
RunA1
zout
1

Ave
Data
6E-06
Model
5.7E-06

data/vMax
1.2E-05

ofxng
275.646

Min d/m
1.32556

ave/min d
1.03336
ave/min m
1.3098

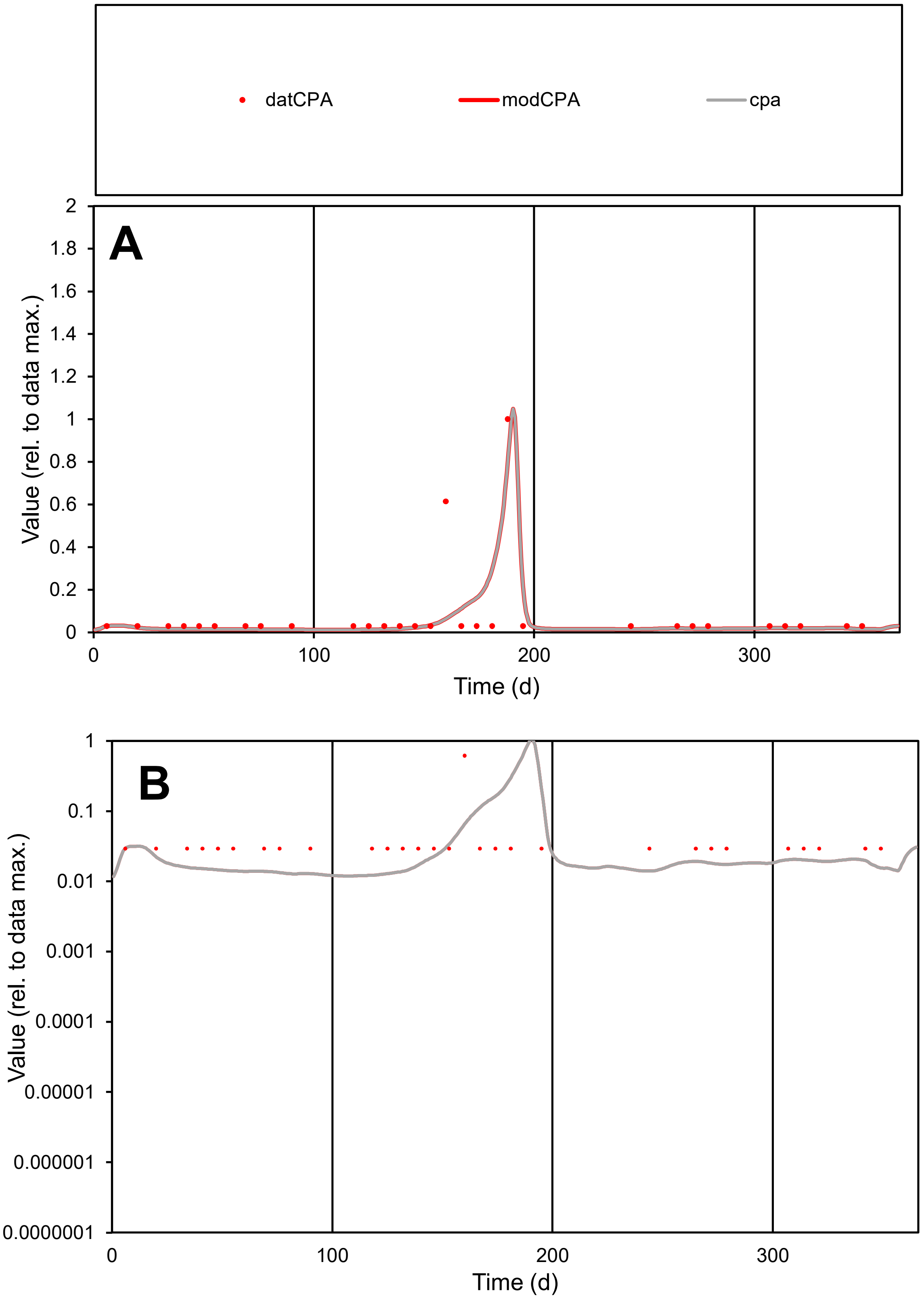


**C1**

Variables
CPA Provide jtd or Dst
CalibOnly?
2 1 to reproduce of
oNormalize?
2 1=no, 2=yes

ofxn calc*
1.37844
ofxn mod
1.42582

sRun
RunA1
zout
1

Ave
Data
1.8E-07
Model
1.5E-07

data/vMax
2.3E-06

ofxng
275.646

Min d/m
2.38406

ave/min d
2.77214
ave/min m
5.50526

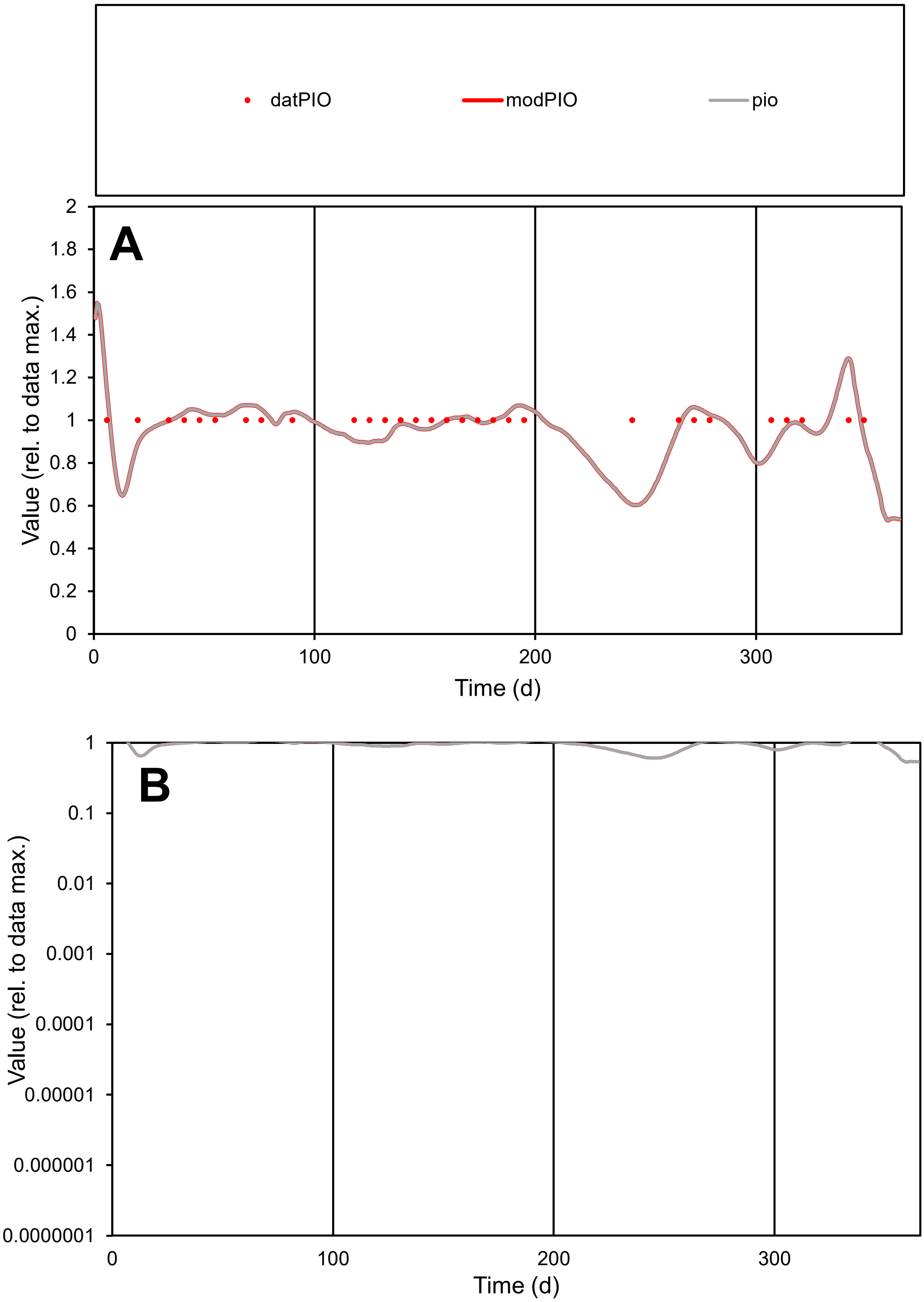


**C1**

Variables
PIO Provide jtd or Dst
CalibOnly?
2 1 to reproduce of
oNormalize?
2 1=no, 2=yes

ofxn calc*
0.10403
ofxn mod
0.15315

sRun
RunA1
zout
1

Ave
Data
1.5E-07
Model
1.4E-07

data/vMax
1.5E-07

ofxng
275.646

Min d/m
1.65848

ave/min d
1
ave/min m
1.63387

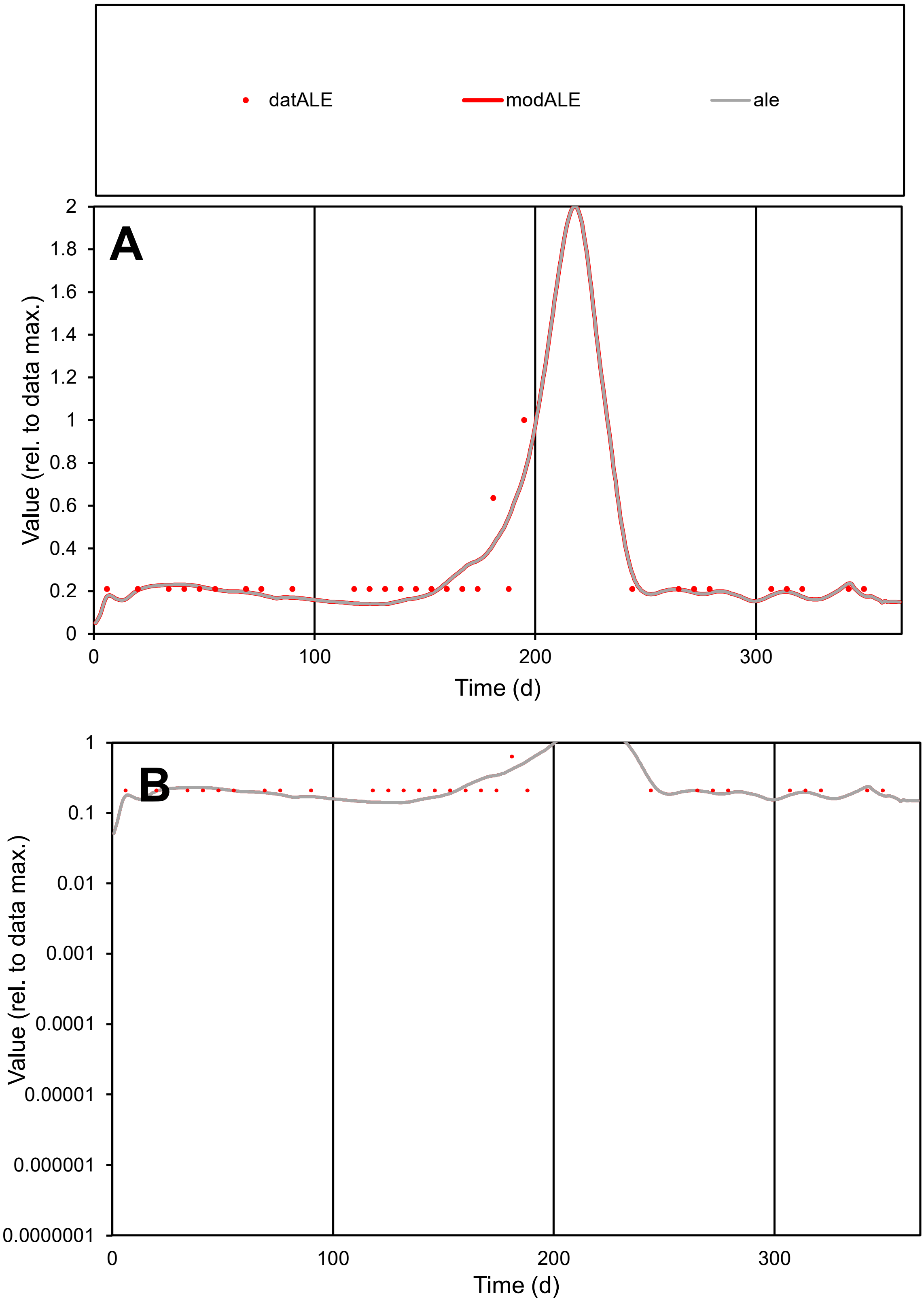


**C1**

Variables

ALE Provide jtd or Dst

CalibOnly?

2 1 to reproduce of

oNormalize?

2 1=no, 2=yes

ofxn calc*

0.37209

ofxn mod

0.39654

sRun

RunA1

zout

1

Ave

Data

1.3E-05

Model

1.3E-05

data/vMax

5.3E-05

ofxng

275.646

Min d/m

1.48956

ave/min d

1.19361

ave/min m

1.72519

325_Calib_S_A

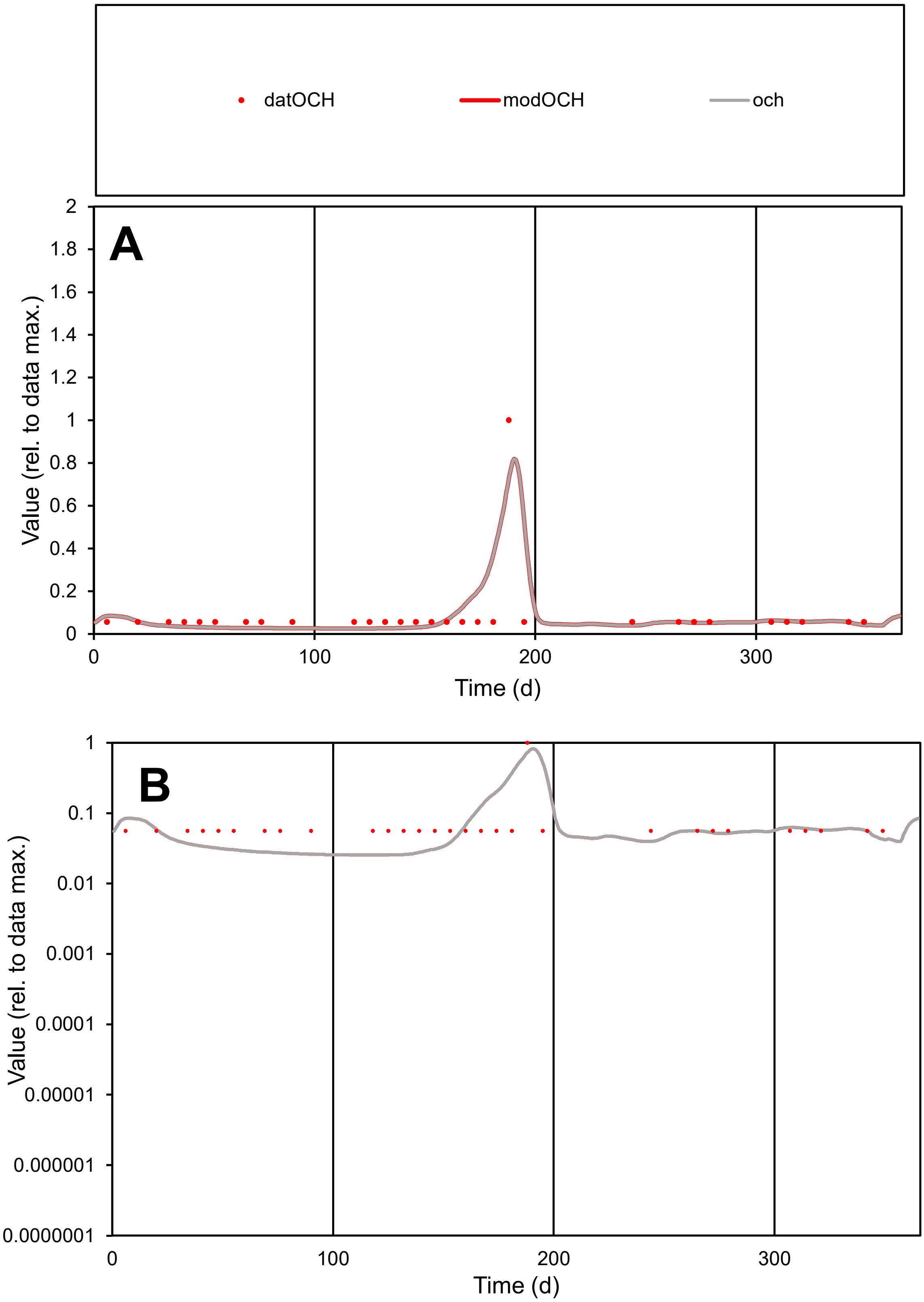


**C1**

Variables
OCH Provide jtd or Dst
CalibOnly?
2 1 to reproduce of
oNormalize?
2 1=no, 2=yes

ofxn calc*
0.89752
ofxn mod
1.02325

sRun
RunA1
zout
1

Ave
Data
6.9E-06
Model
7.1E-06

data/vMax
7.9E-05

ofxng
275.646

Min d/m
2.20844

ave/min d
1.5623
ave/min m
3.56456

325_Calib_S_A

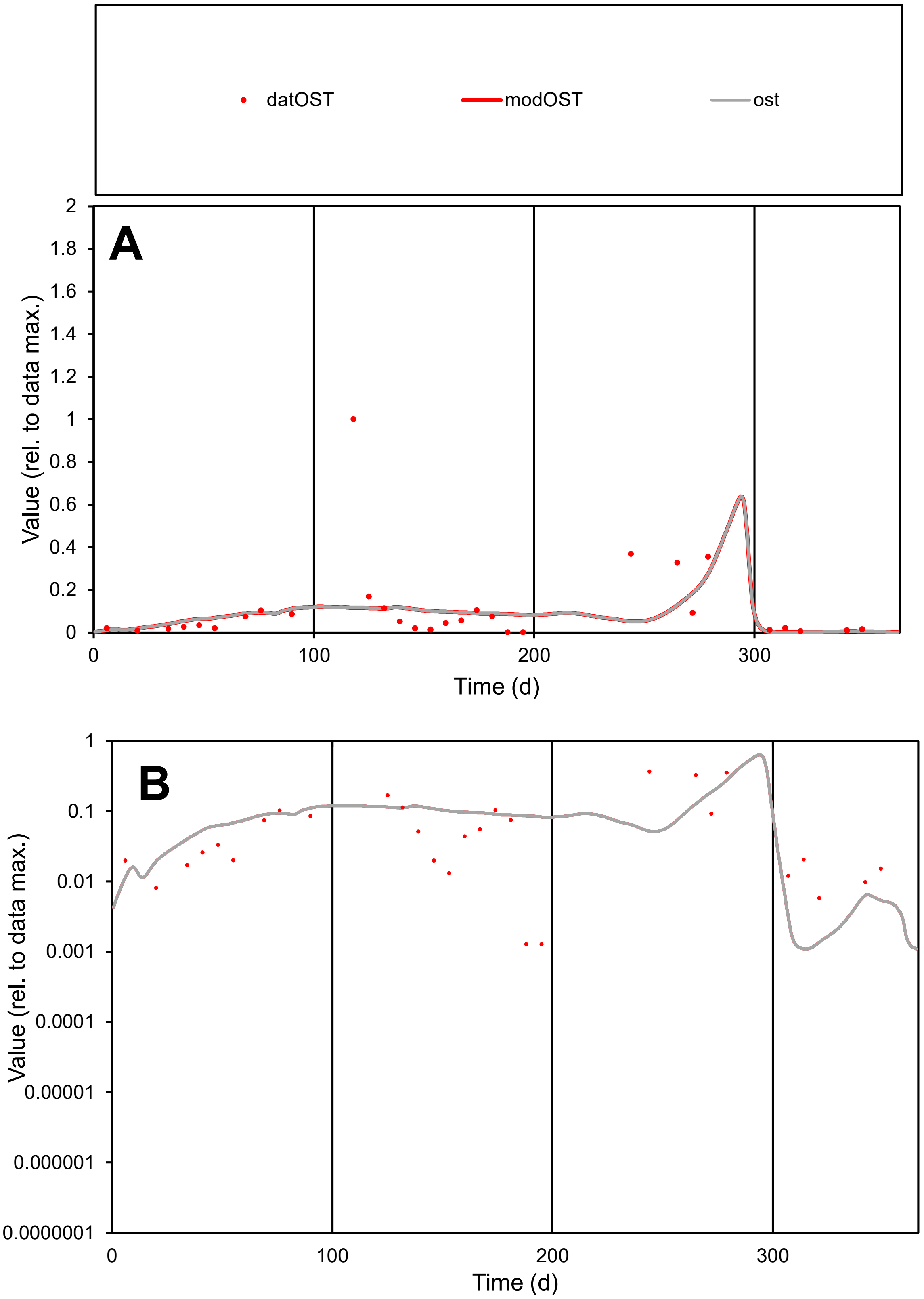


**C1**

Variables
OST Provide jtd or Dst
CalibOnly?
2 1 to reproduce of
oNormalize?
2 1=no, 2=yes

ofxn calc*
1.66859
ofxn mod
1.67564

sRun
RunA1
zout
1

Ave
Data
0.00018
Model
0.00014

data/vMax
0.00169

ofxng
275.646

Min d/m
1.16568

ave/min d
84.3763
ave/min m
75.0903

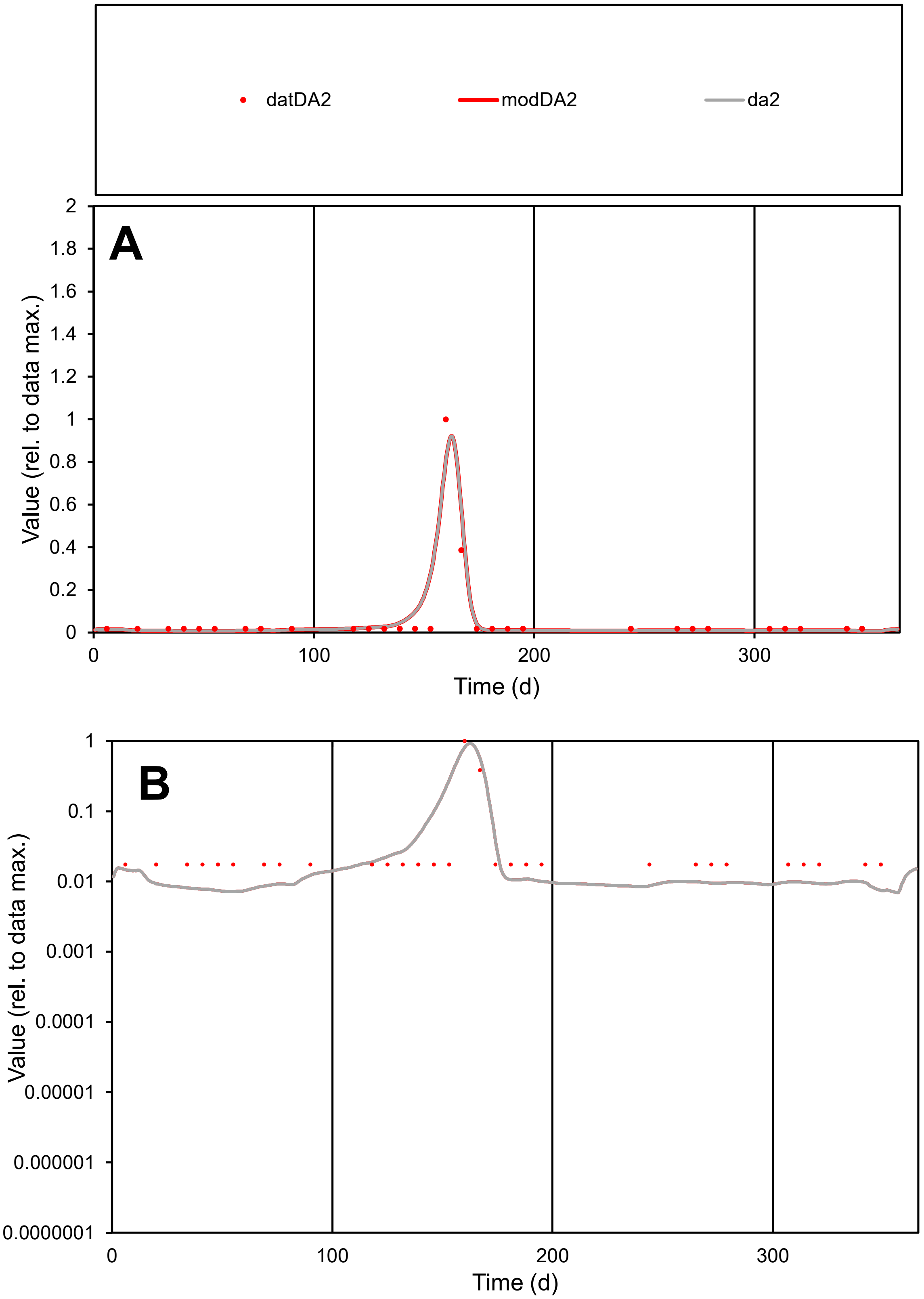


C1

Variables
DA2 Provide jtd or Dst
CalibOnly?
2 1 to reproduce of
oNormalize?
2 1=no, 2=yes

ofxn calc*
0.69207
ofxn mod
0.83872

sRun
RunA1
zout
1

Ave
Data
1.1E-05
Model
1.2E-05

data/vMax
0.00018

ofxng
275.646

Min d/m
2.40185

ave/min d
3.58323
ave/min m
9.05607

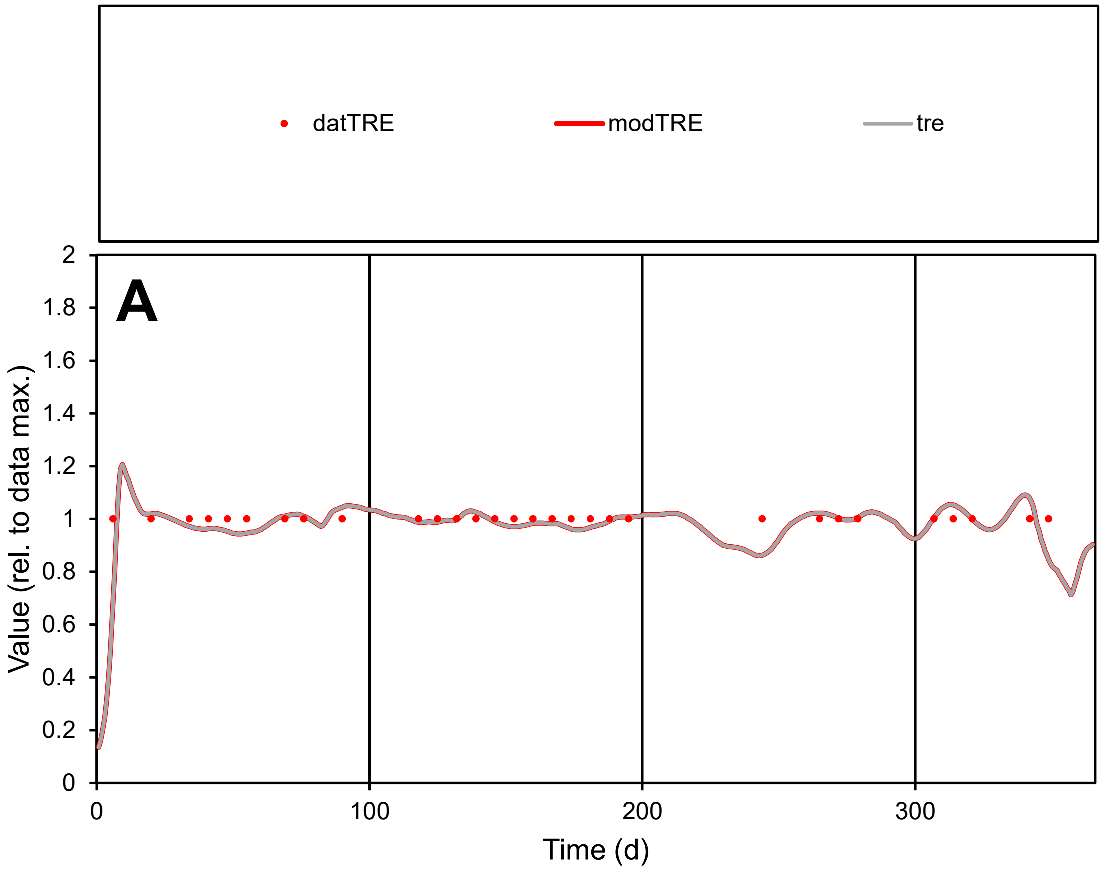


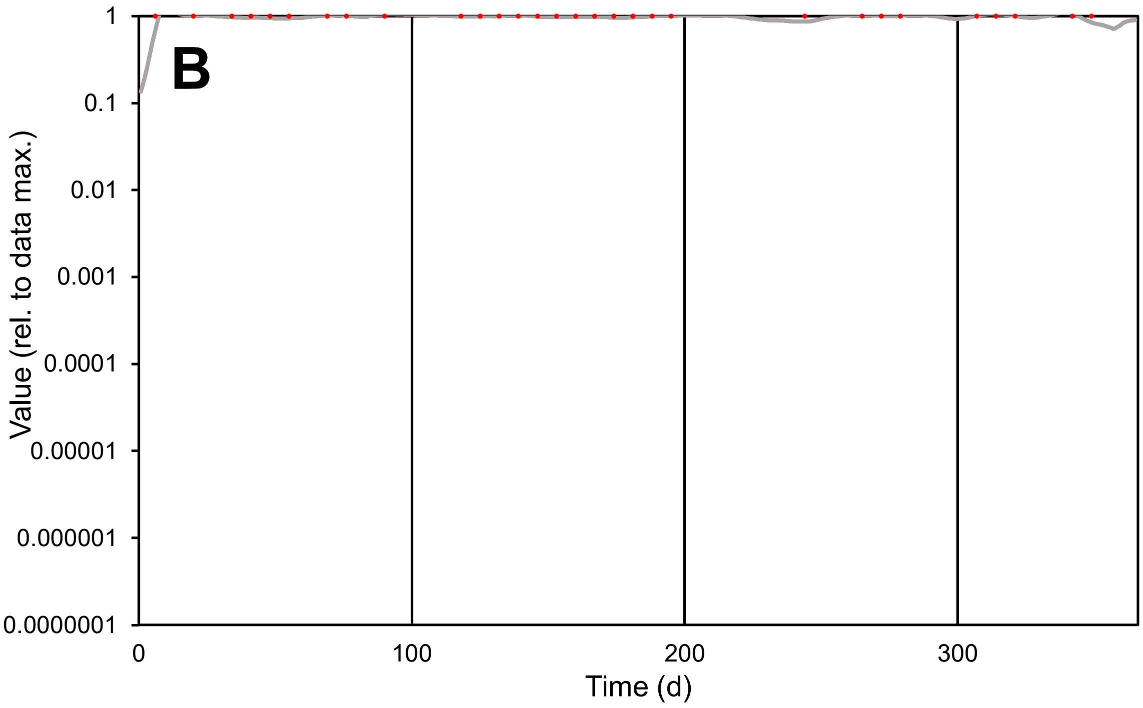


**C1**

Variables
TRE Provide jtd or Dst
CalibOnly?
2 1 to reproduce of
oNormalize?
2 1=no, 2=yes

ofxn calc*
0.04923
ofxn mod
0.12188

sRun
RunA1
zout
1

Ave
Data
6.9E-06
Model
6.8E-06

data/vMax
6.9E-06

ofxng
275.646

Min d/m
1.22379

ave/min d
1
ave/min m
1.20687

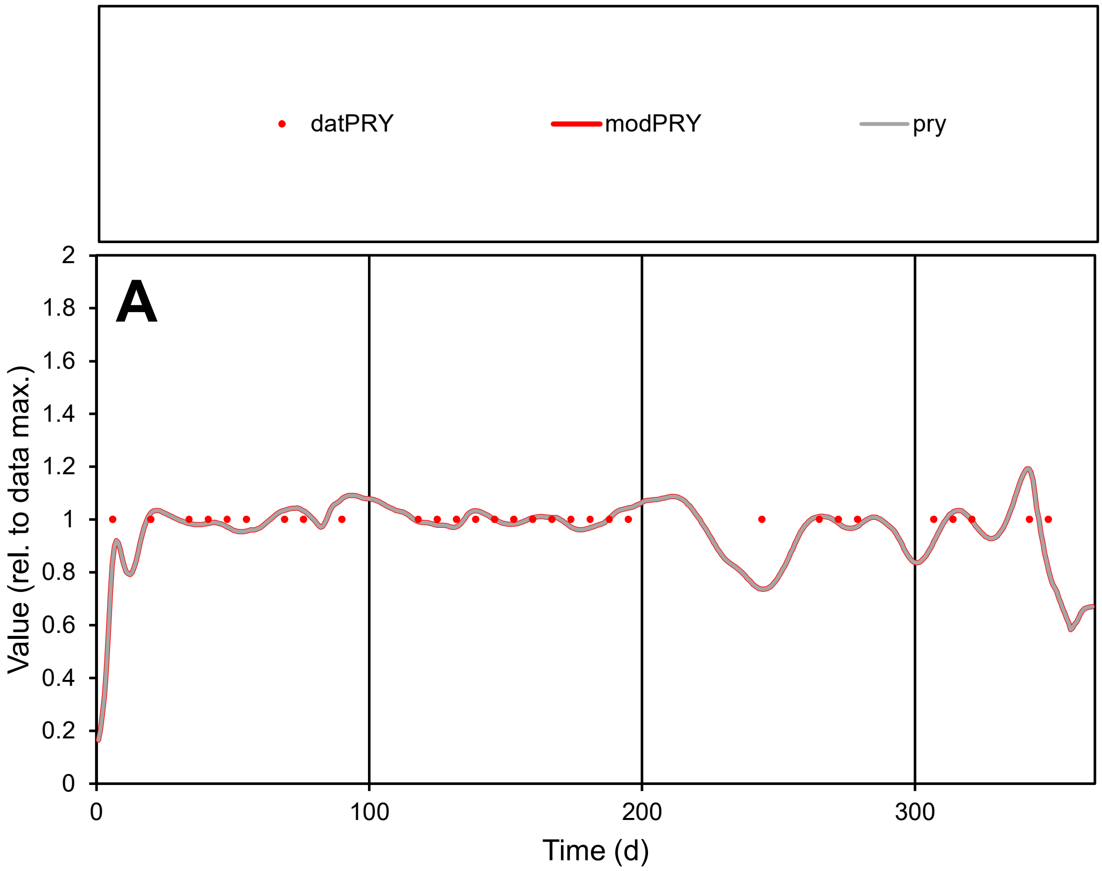


C1

Variables

| PRY | Provide jtd or Dst |
| --- | --- |

CalibOnly?

| 2 | 1 to reproduce of |
| --- | --- |

oNormalize?

| 2 | 1=no, 2=yes |
| --- | --- |

ofxn calc*
0.07827
ofxn mod
0.13638

sRun
RunA1
zout
1

Ave
Data
8.8E-06
Model
8.7E-06

data/vMax
8.8E-06

ofxng
275.646

Min d/m
1.35804

ave/min d
1
ave/min m
1.33812

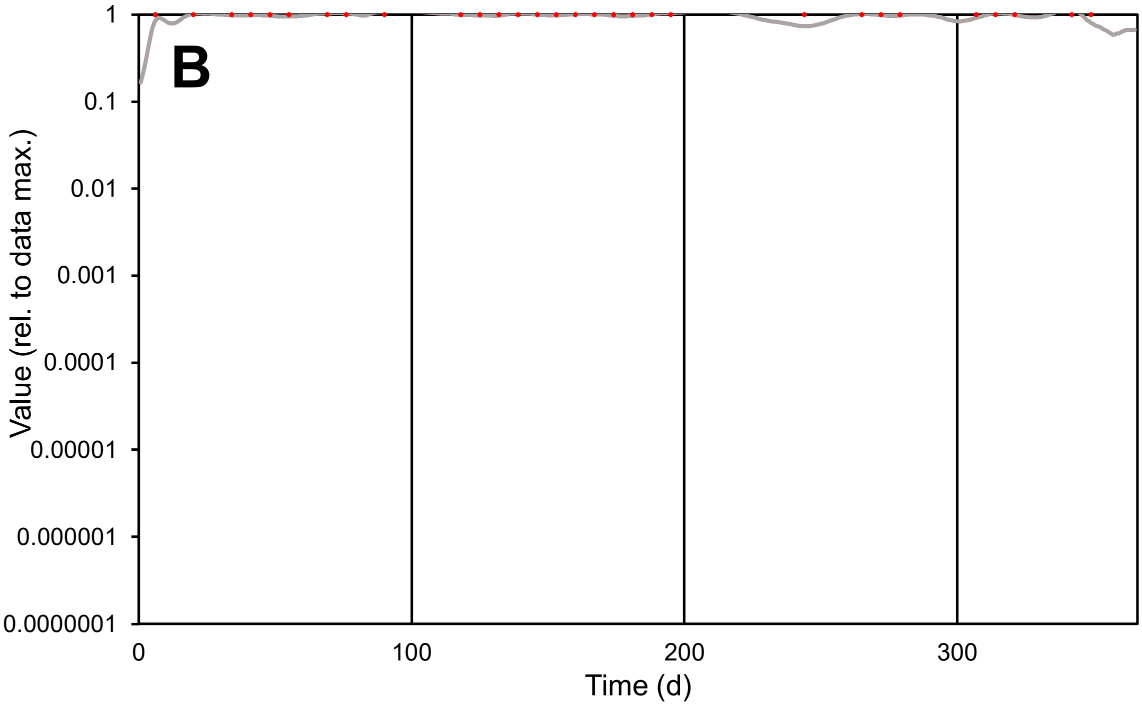

325_Calib_S_A

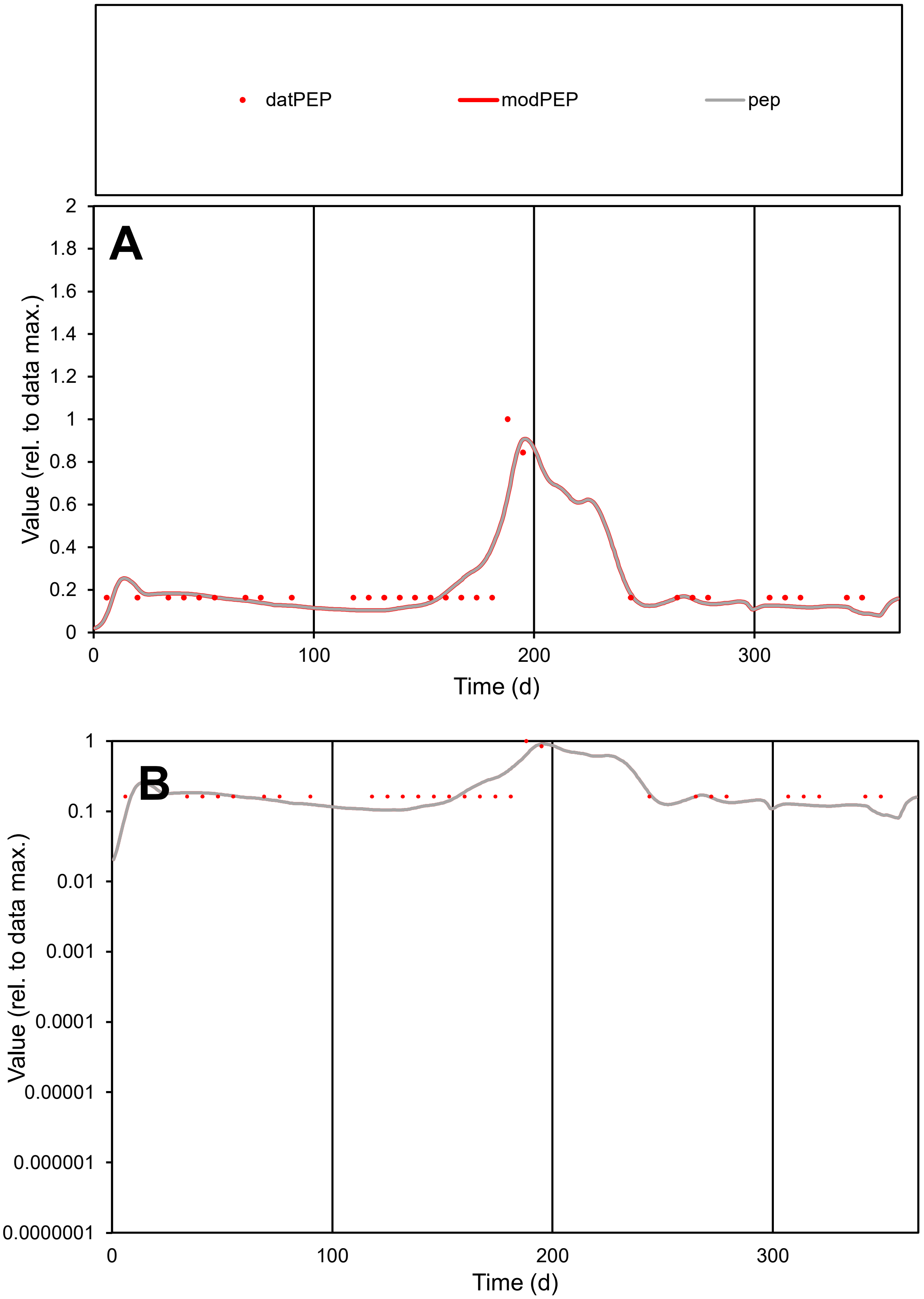


**C1**

Variables

PEP Provide jtd or Dst

CalibOnly?

2 1 to reproduce of

oNormalize?

2 1=no, 2=yes

ofxn calc*

0.40405

ofxn mod

0.43072

sRun

RunA1

zout

1

Ave

Data

6.5E-06

Model

6.2E-06

data/vMax

3E-05

ofxng

275.646

Min d/m

1.84449

ave/min d

1.31126

ave/min m

2.30041

325_Calib_S_A

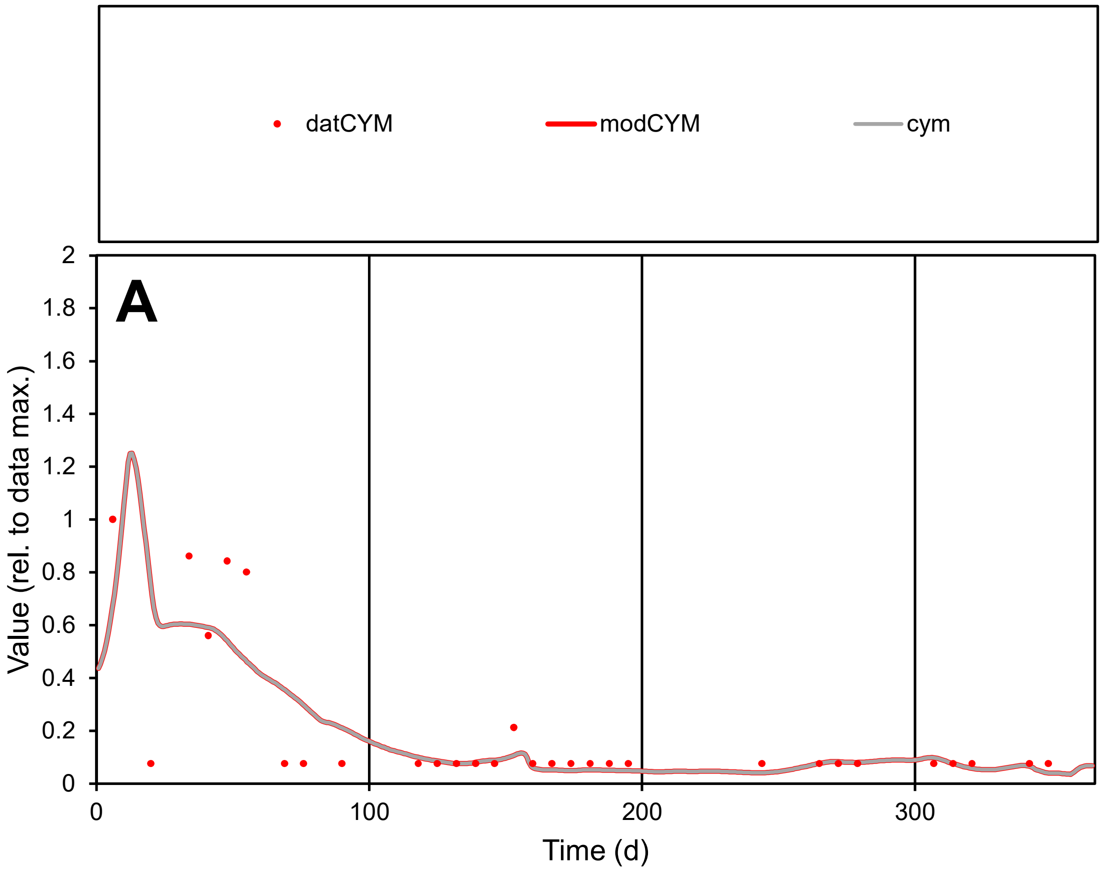


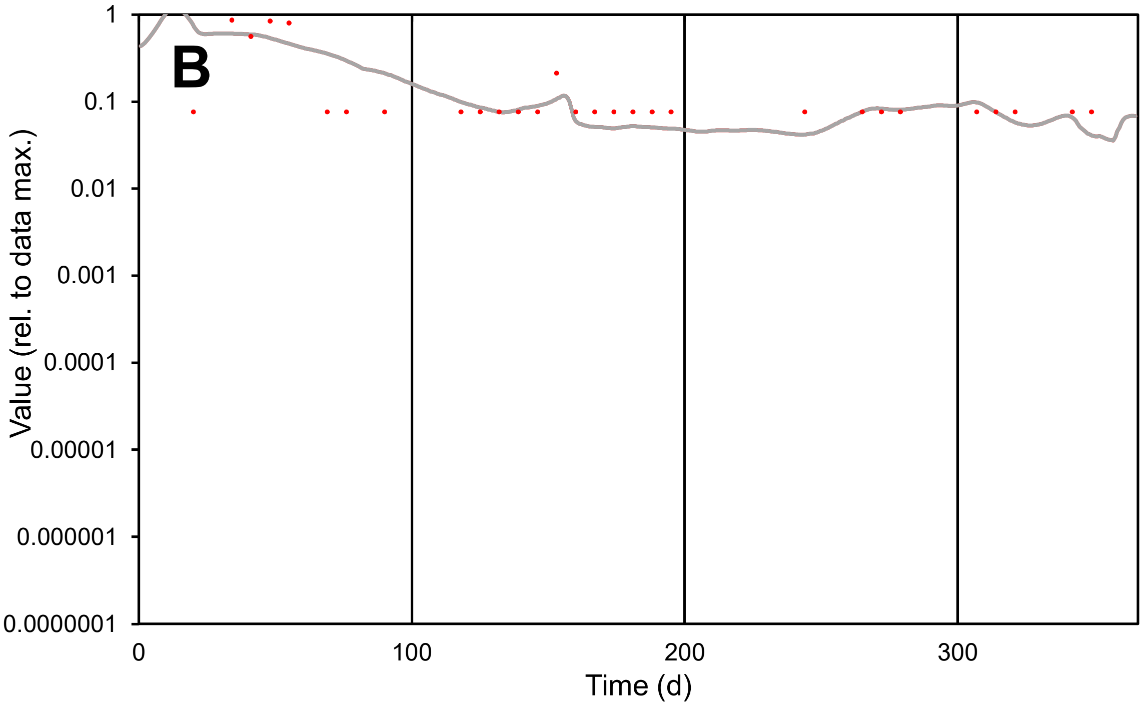


**C1**

Variables
CYM Provide jtd or Dst
CalibOnly?
2 1 to reproduce of
oNormalize?
2 1=no, 2=yes

ofxn calc*
0.7839
ofxn mod
0.82316

sRun
RunA1
zout
1

Ave
Data
3.6E-06
Model
3.5E-06

data/vMax
1.8E-05

ofxng
275.646

Min d/m
1.91367

ave/min d
2.66806
ave/min m
4.89547

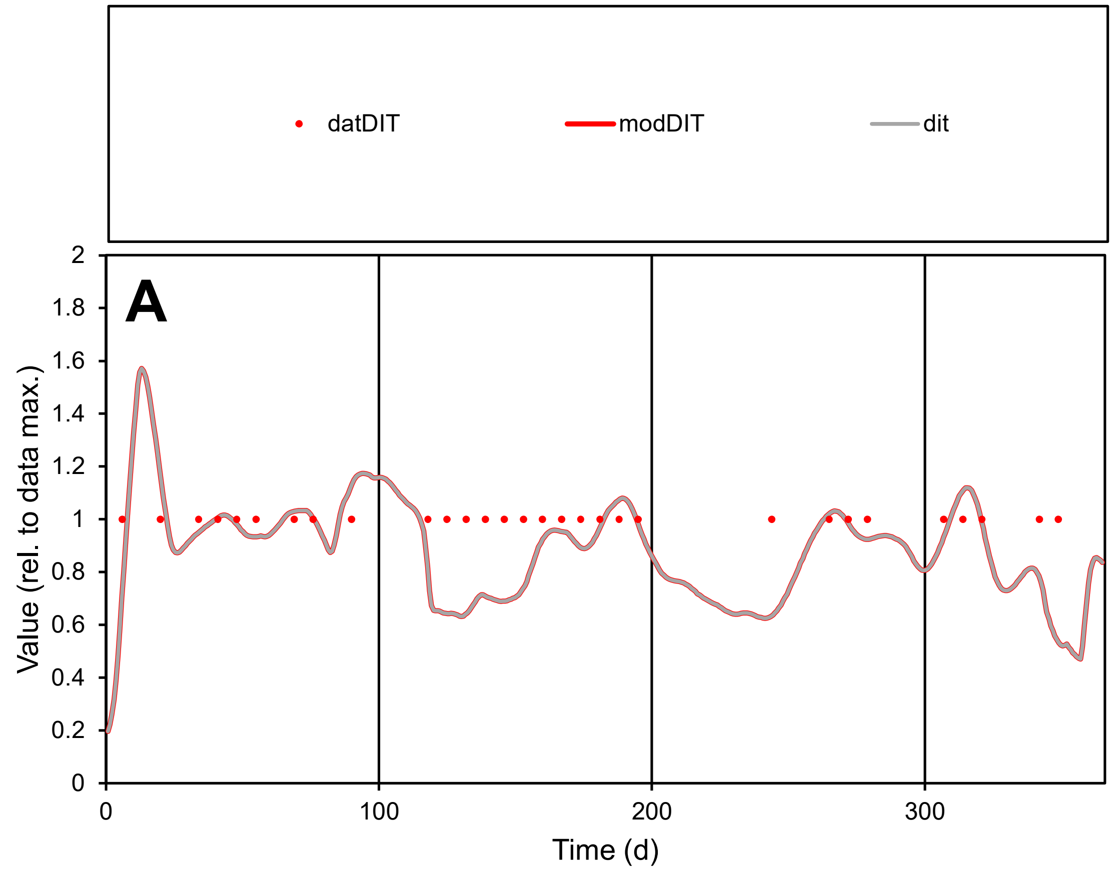


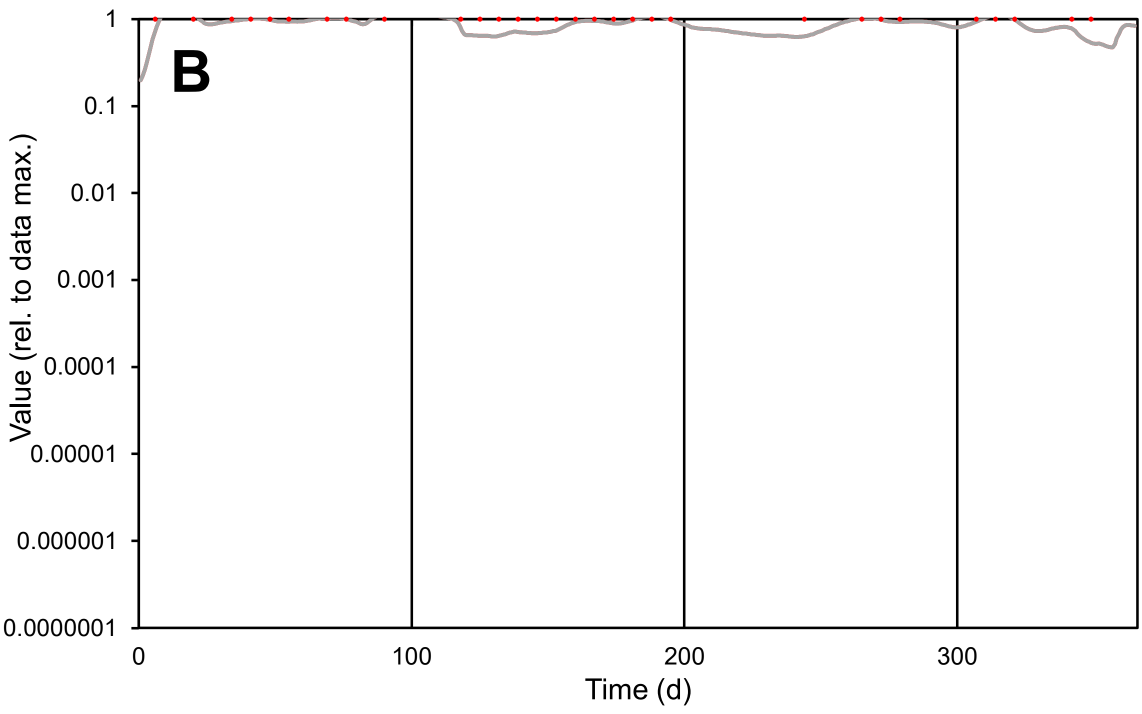


**C1**

Variables

DIT Provide jtd or Dst

CalibOnly?

2 1 to reproduce of

oNormalize?

2 1=no, 2=yes

ofxn calc*

0.19019

ofxn mod

0.21962

sRun

RunA1

zout

1

Ave

Data

5.1E-06

Model

4.6E-06

data/vMax

5.1E-06

ofxng

275.646

Min d/m

1.9218

ave/min d

1

ave/min m

1.7245

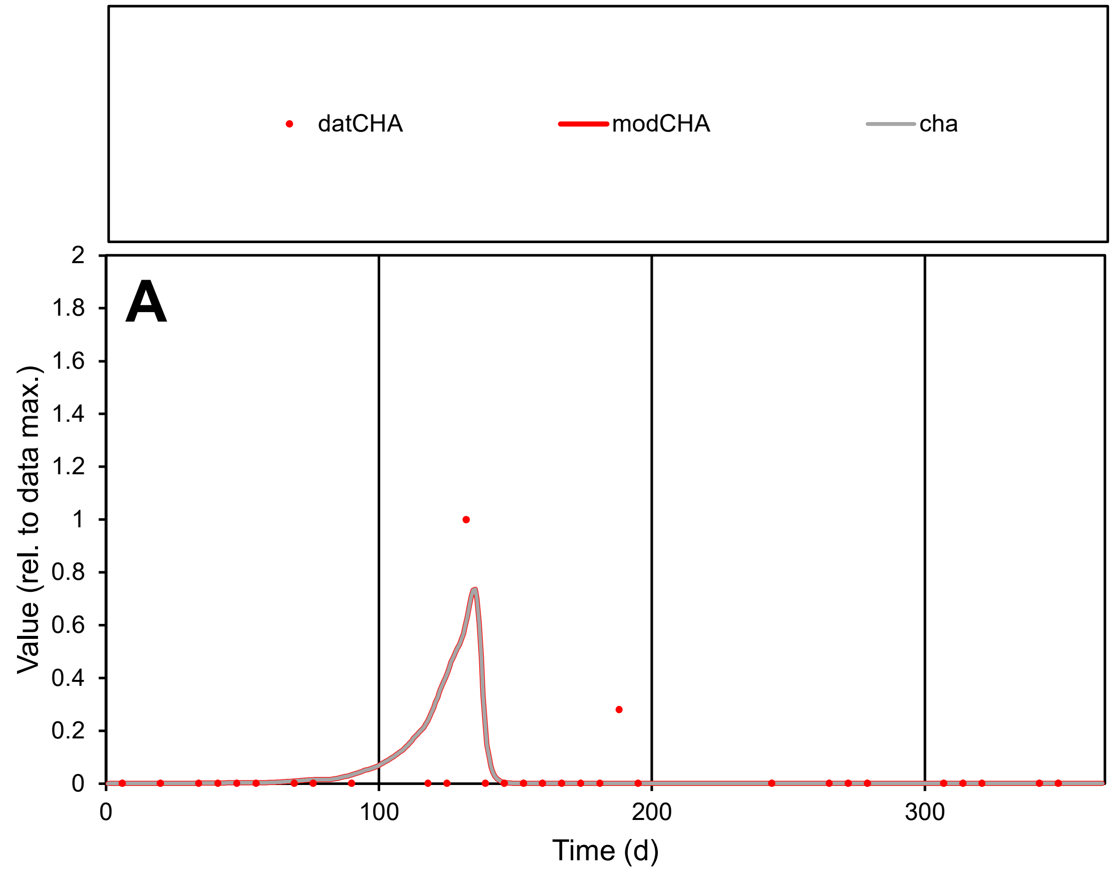


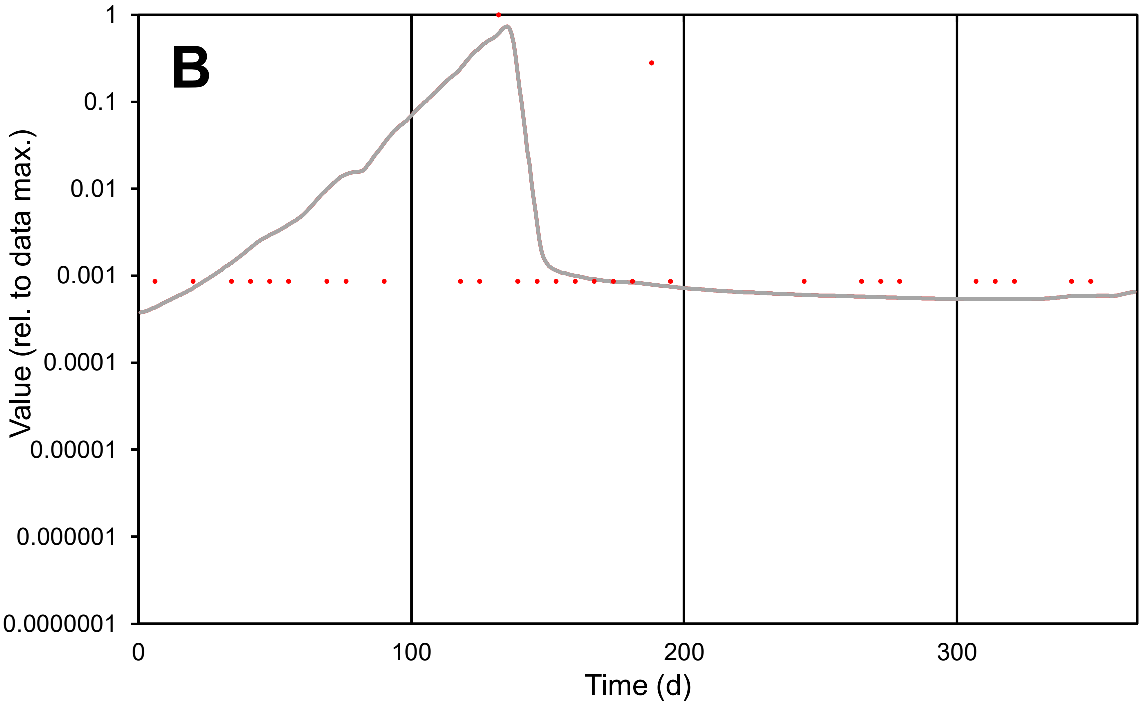


**C1**

Variables

CHA Provide jtd or Dst

CalibOnly?

2 1 to reproduce of

oNormalize?

2 1=no, 2=yes

ofxn calc*

2.6494

ofxn mod

2.71964

sRun

RunA1

zout

1

Ave

Data

4.4E-06

Model

4.7E-06

data/vMax

0.0001

ofxng

275.646

Min d/m

1.91313

ave/min d

50.2998

ave/min m

103.277

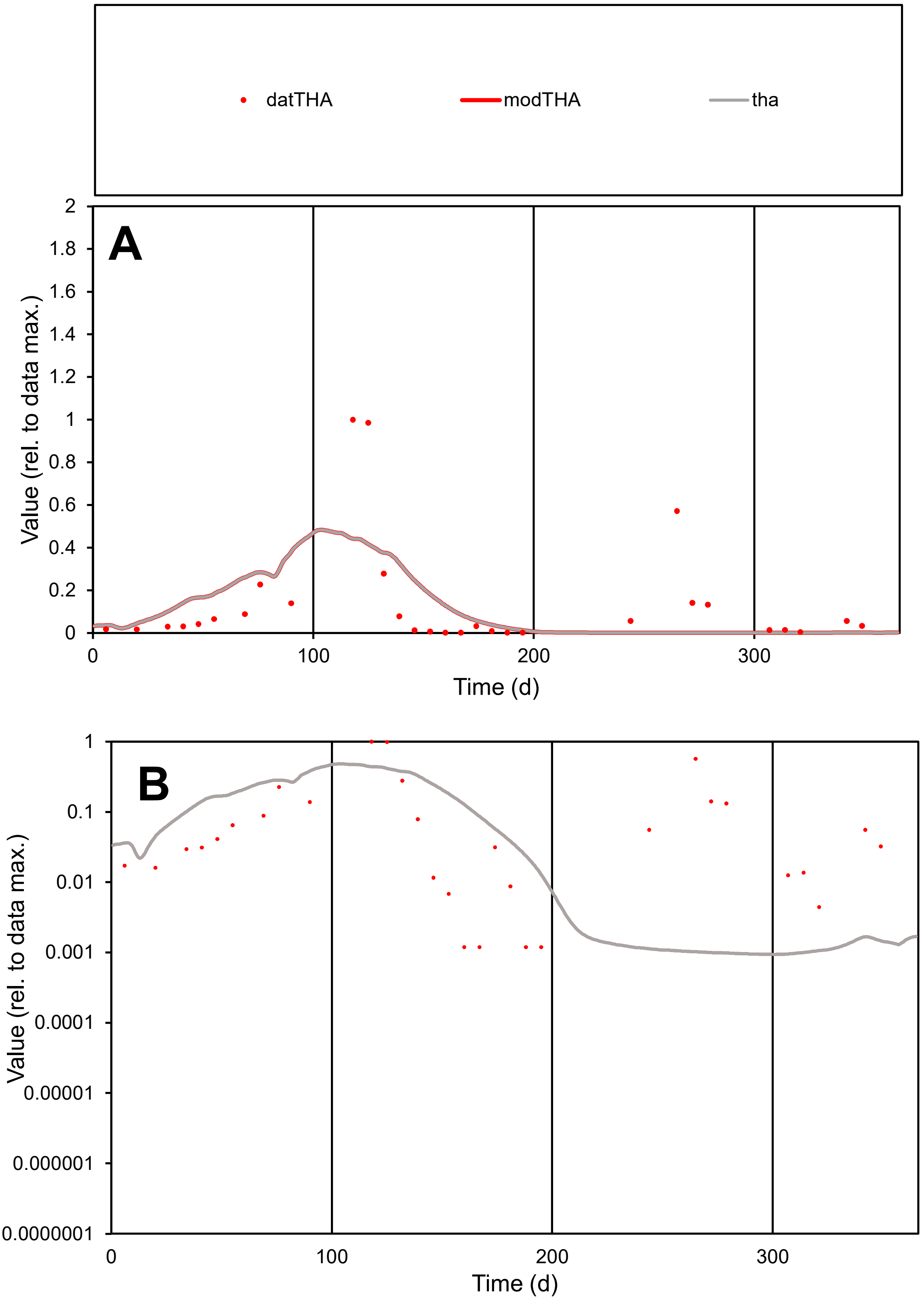


**C1**

Variables
THA Provide jtd or Dst
CalibOnly?
2 1 to reproduce of
oNormalize?
2 1=no, 2=yes

ofxn calc*
1.53644
ofxn mod
1.55279

sRun
RunA1
zout
1

Ave
Data
0.00031
Model
0.00029

data/vMax
0.0023

ofxng
275.646

Min d/m
1.24751

ave/min d
113.513
ave/min m
133.134

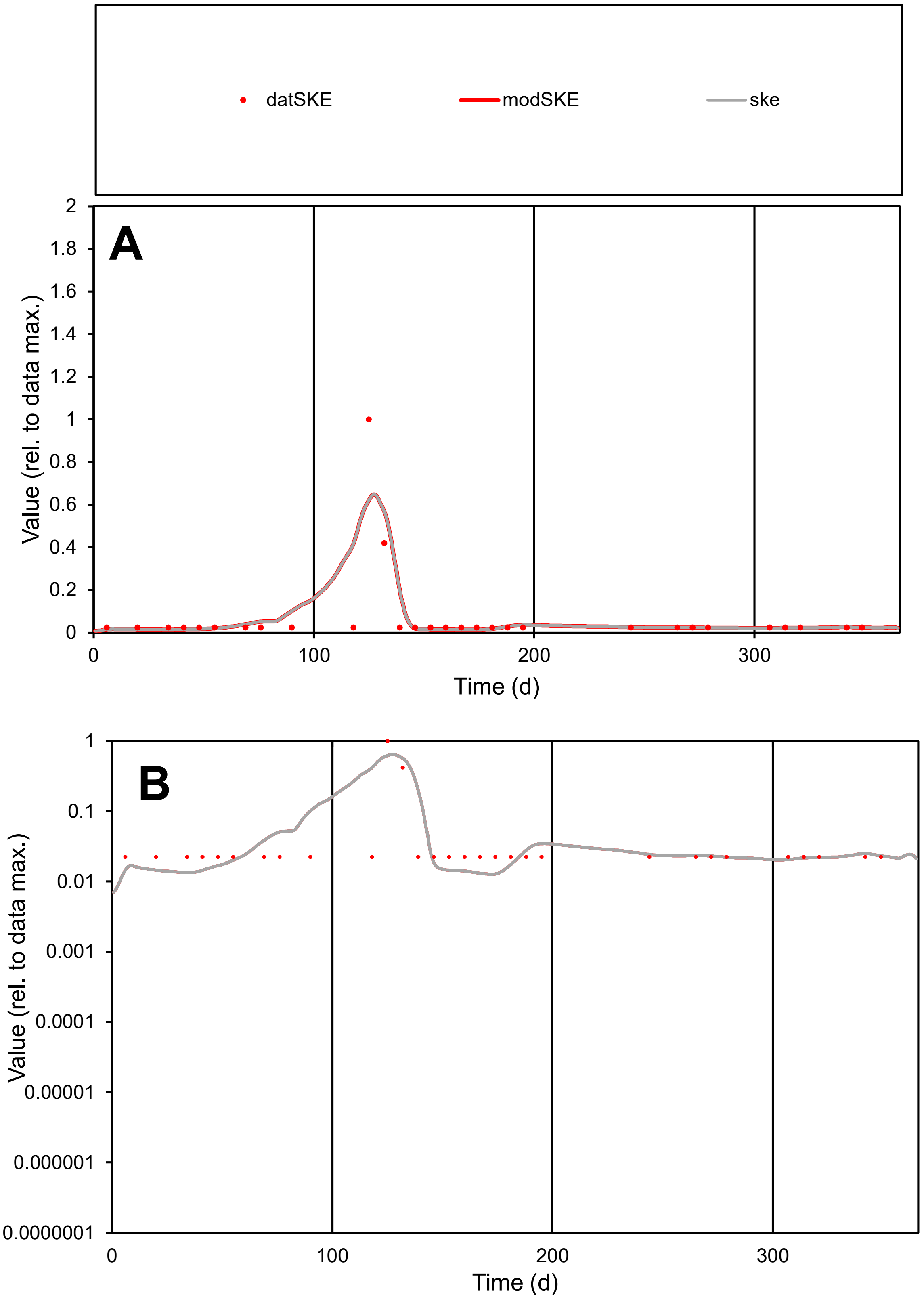


**C1**

Variables

SKE Provide jtd or Dst

CalibOnly?

2 1 to reproduce of

oNormalize?

2 1=no, 2=yes

ofxn calc*

1.43863

ofxn mod

1.51105

sRun

RunA1

zout

1

Ave

Data

7E-06

Model

7.8E-06

data/vMax

0.0001

ofxng

275.646

Min d/m

1.72102

ave/min d

3.04386

ave/min m

5.84341

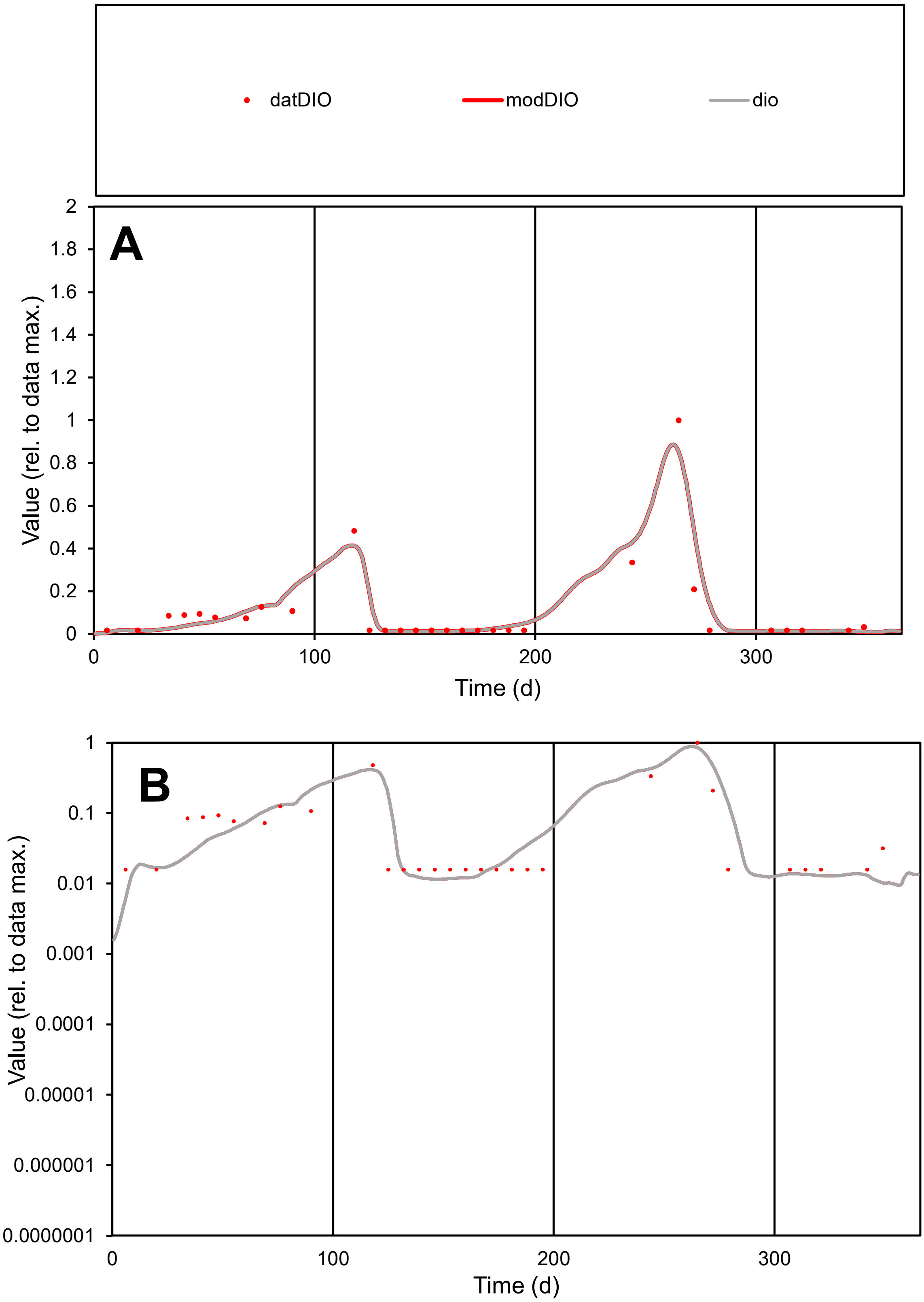


**C1**

Variables
DIO Provide jtd or Dst
CalibOnly?
2 1 to reproduce of
oNormalize?
2 1=no, 2=yes

ofxn calc*
0.62228
ofxn mod
0.67056

sRun
RunA1
zout
1

Ave
Data
2.9E-05
Model
3.1E-05

data/vMax
0.00029

ofxng
275.646

Min d/m
1.56573

ave/min d
6.28265
ave/min m
10.3259

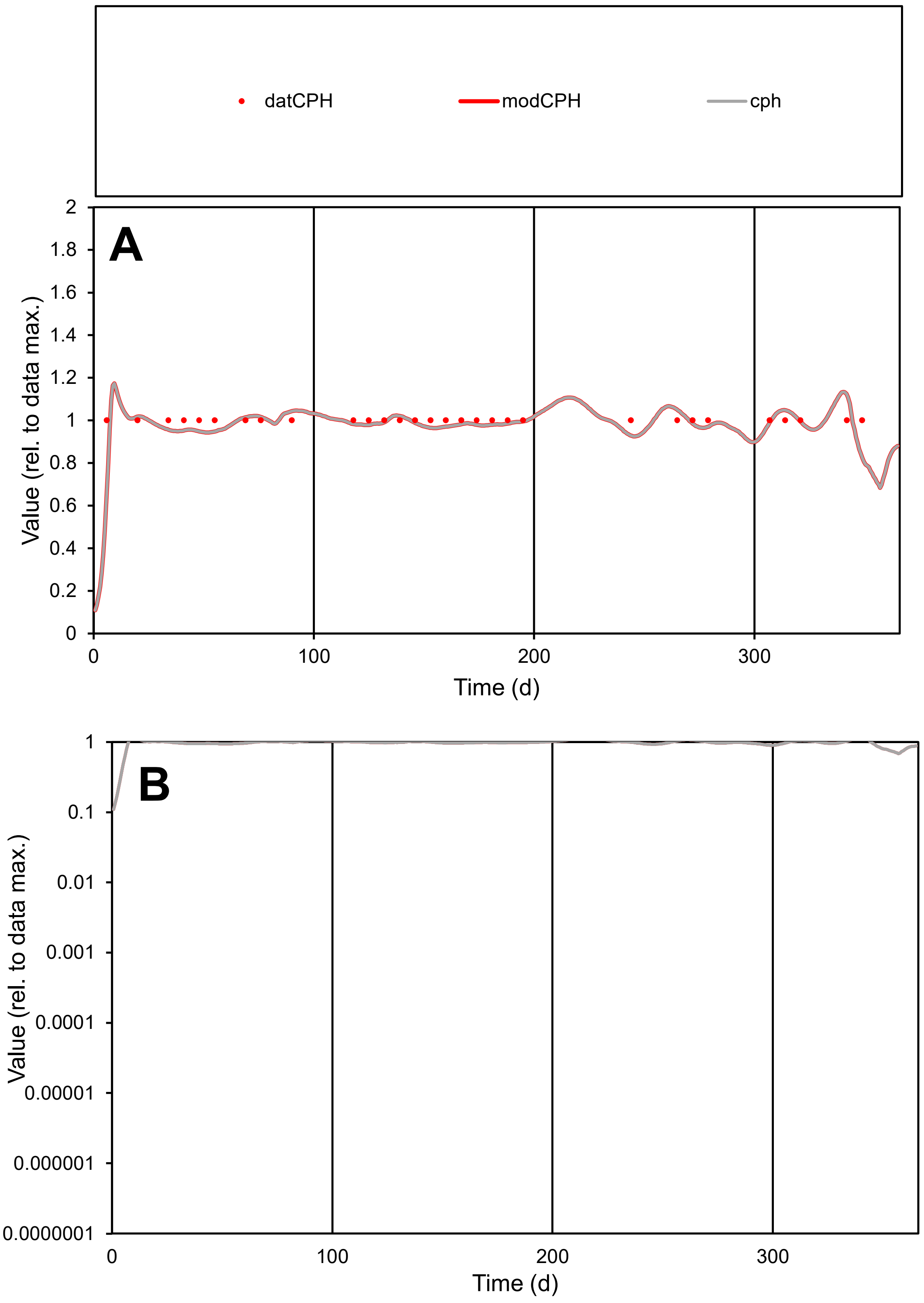


**C1**

Variables
CPH Provide jtd or Dst
CalibOnly?
2 1 to reproduce of
oNormalize?
2 1=no, 2=yes

ofxn calc*
0.05089
ofxn mod
0.12232

sRun
RunA1
zout
1

Ave
Data
9.6E-06
Model
9.4E-06

data/vMax
9.6E-06

ofxng
275.646

Min d/m
1.26096

ave/min d
1
ave/min m
1.24094

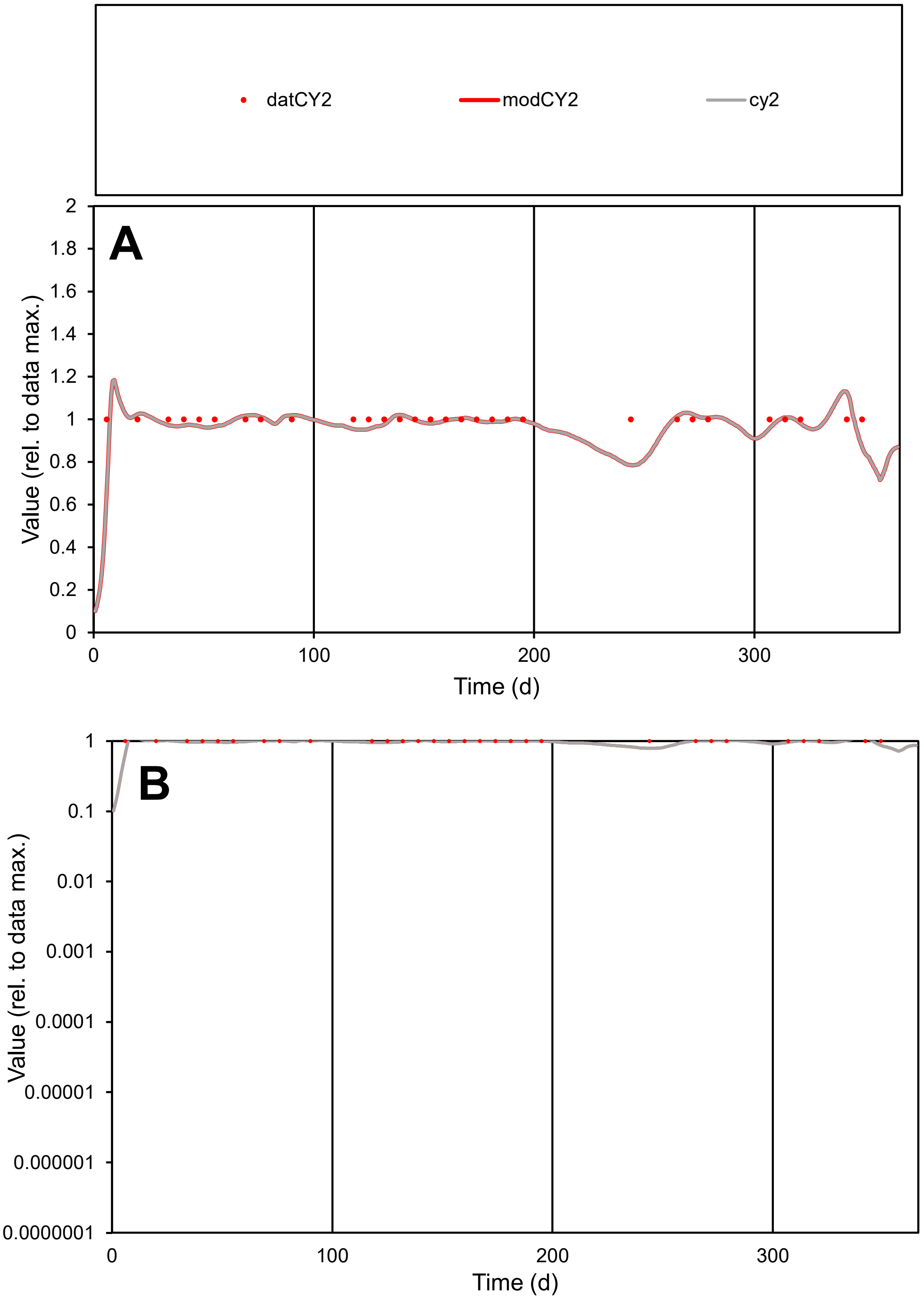


**C1**

Variables
CY2 Provide jtd or Dst
CalibOnly?
2 1 to reproduce of
oNormalize?
2 1=no, 2=yes

ofxn calc*
0.0566
ofxn mod
0.12494

sRun
RunA1
zout
1

Ave
Data
1.1E-05
Model
1.1E-05

data/vMax
1.1E-05

ofxng
275.646

Min d/m
1.27518

ave/min d
1
ave/min m
1.25437

325_Calib_S_A

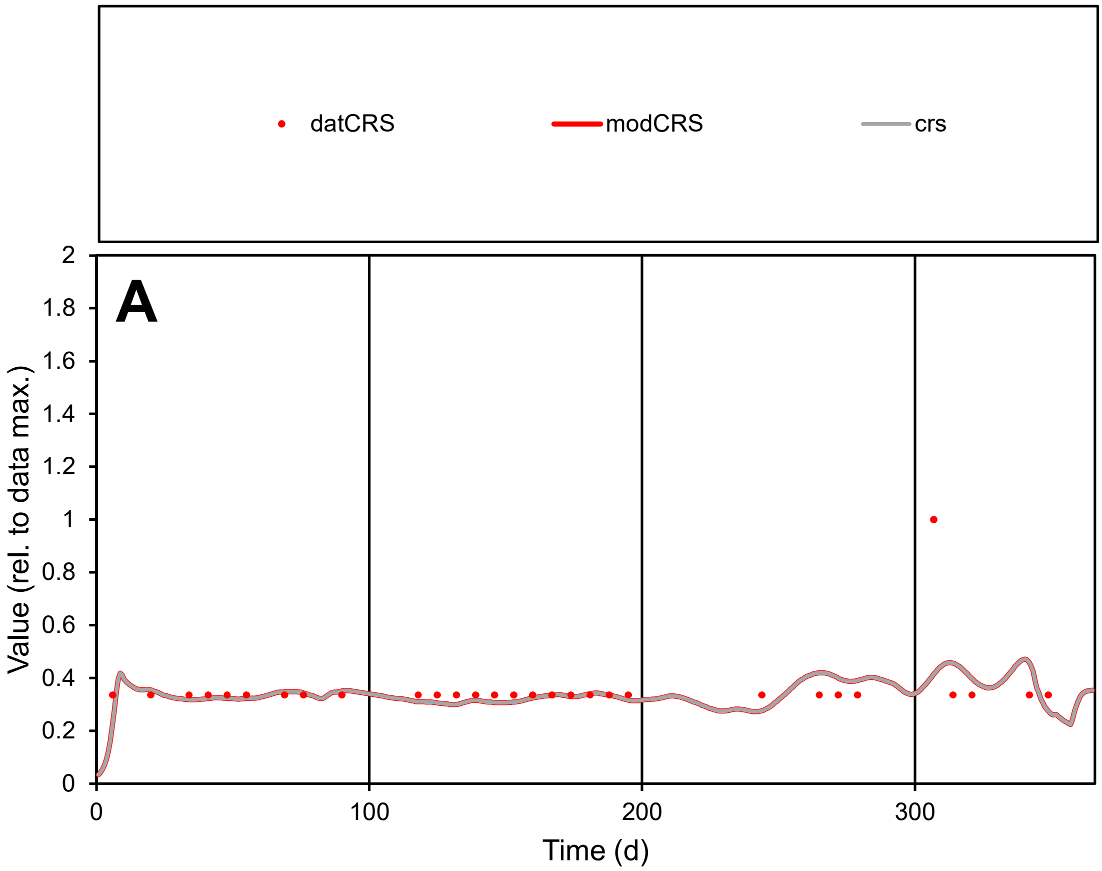


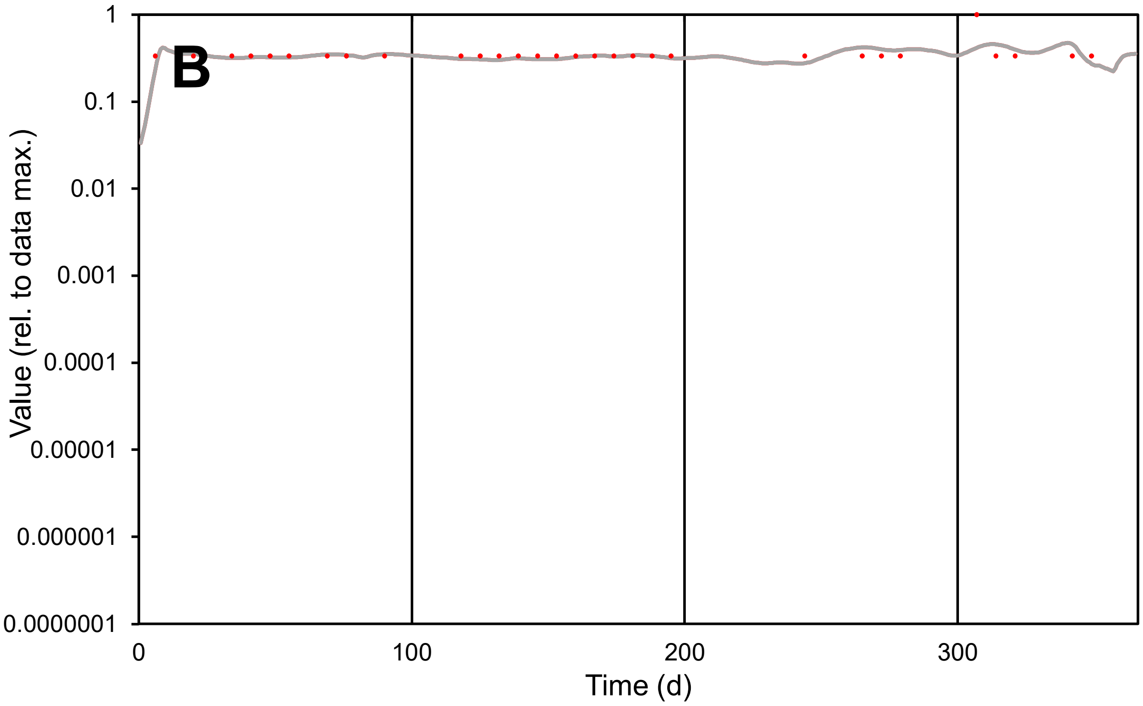


**C1**

Variables

| | |
|---|---|
| CRS | Provide jtd or Dst |
| CalibOnly? | |
| 2 | 1 to reproduce of |
| oNormalize? | |
| 2 | 1=no, 2=yes |
| ofxn calc* | |
| 0.31272 | |
| ofxn mod | |
| 0.33269 | |
| sRun | |
| RunA1 | |
| zout | |
| 1 | |
| Ave Data | |
| 5.8E-06 | |
| Model | |
| 5.5E-06 | |
| data/vMax | |
| 1.6E-05 | |
| ofxng | |
| 275.646 | |
| Min d/m | |
| 1.28382 | |
| ave/min d | |
| 1.06599 | |
| ave/min m | |
| 1.31366 | |

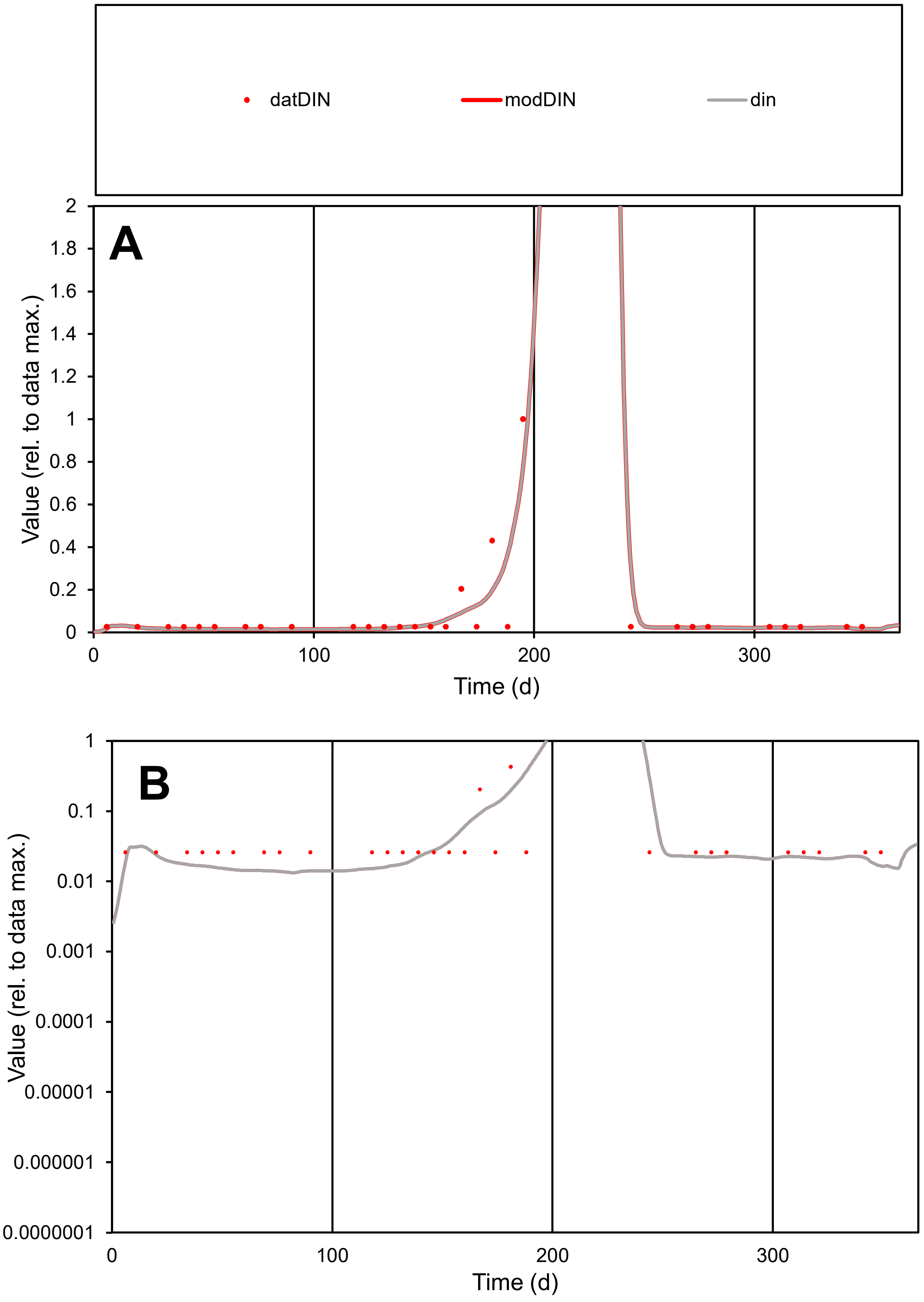


**C1**

Variables
DIN Provide jtd or Dst
CalibOnly?
2 1 to reproduce of
oNormalize?
2 1=no, 2=yes

ofxn calc*
0.931
ofxn mod
1.06159

sRun
RunA1
zout
1

Ave
Data
8.7E-06
Model
8.5E-06

data/vMax
0.00011

ofxng
275.646

Min d/m
1.88374

ave/min d
3.00599
ave/min m
5.54986

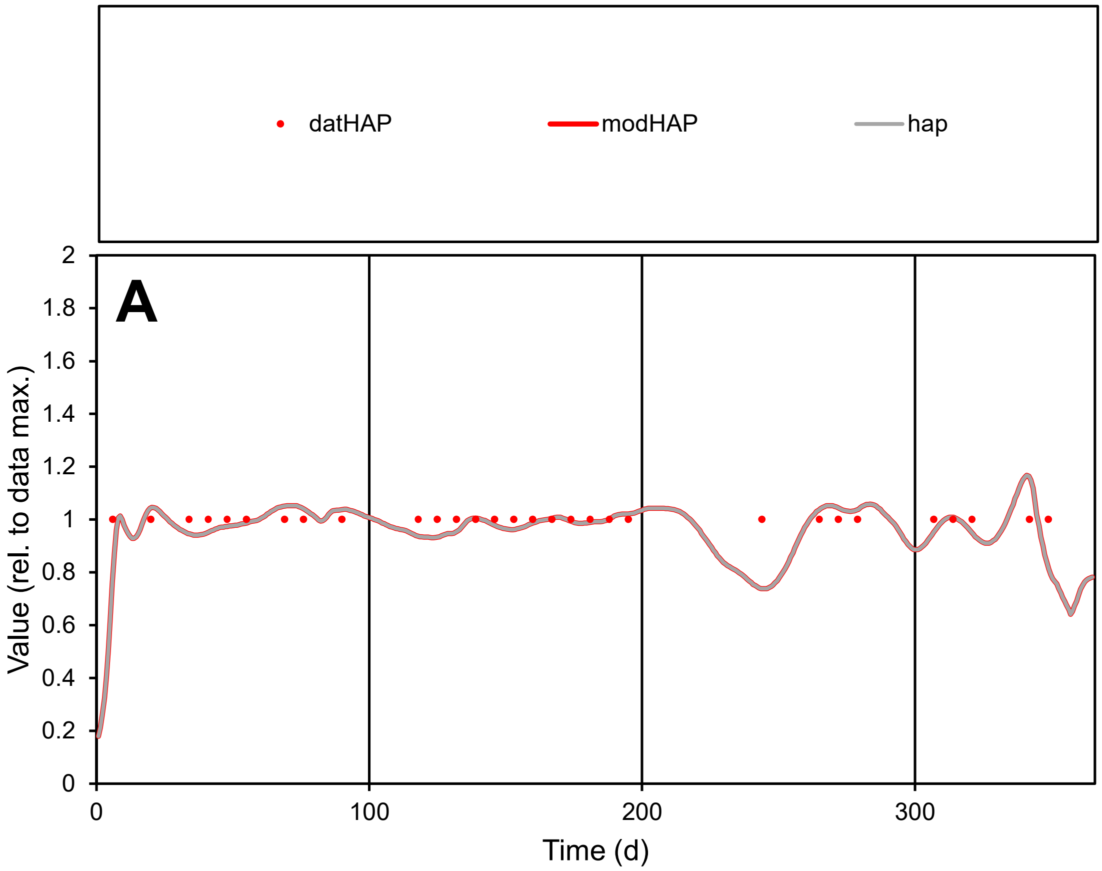


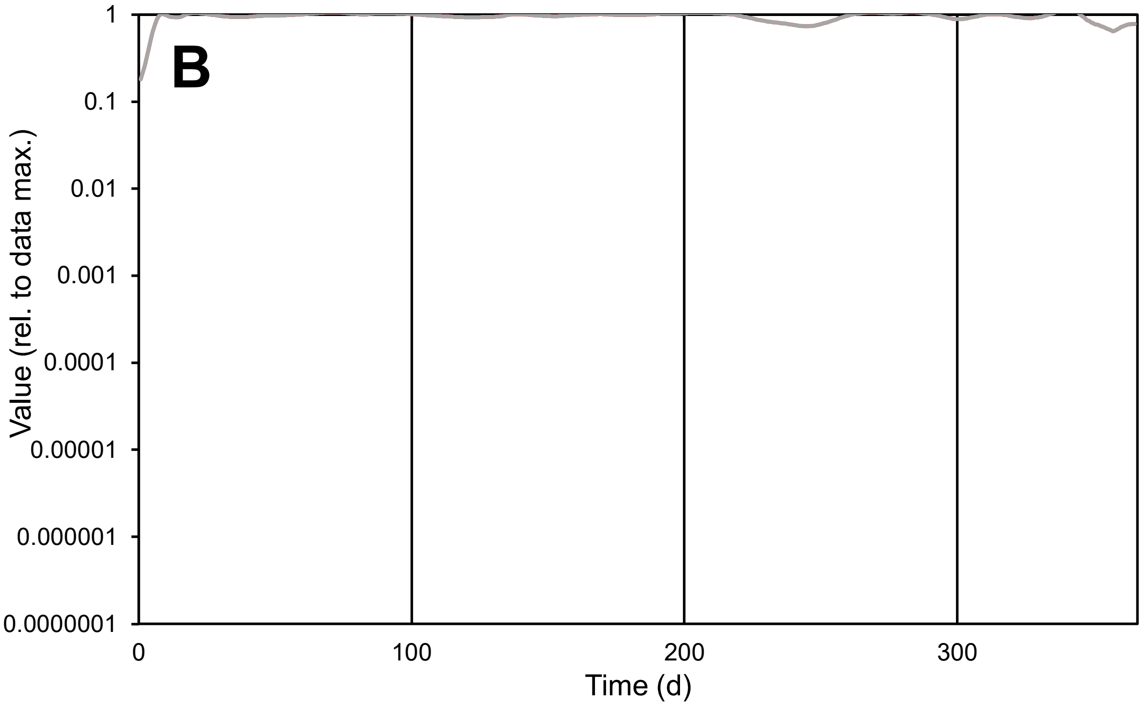


**C1**

| | |
|---|---|
| Variables | |
| HAP | Provide jtd or Dst |
| CalibOnly? | |
| 2 | 1 to reproduce of |
| oNormalize? | |
| 2 | 1=no, 2=yes |
| ofxn calc* | |
| 0.07564 | |
| ofxn mod | |
| 0.13487 | |
| sRun | |
| RunA1 | |
| zout | |
| 1 | |
| Ave | |
| Data | |
| 1.1E-05 | |
| Model | |
| 1.1E-05 | |
| data/vMax | |
| 1.1E-05 | |
| ofxng | |
| 275.646 | |
| Min d/m | |
| 1.35474 | |
| ave/min d | |
| 1 | |
| ave/min m | |
| 1.33064 | |

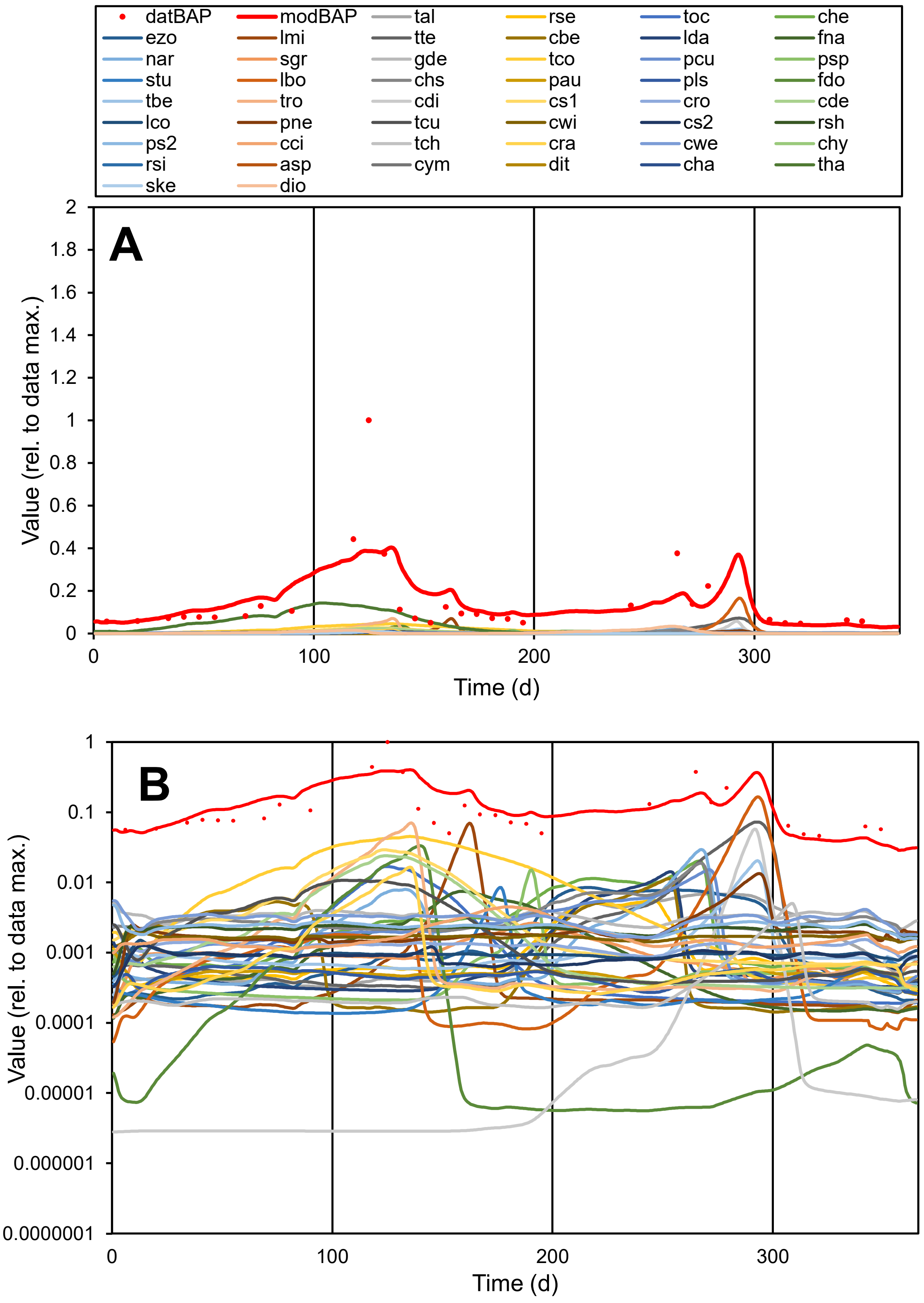


**C1**

Variables
BAP Provide jtd or Dst
CalibOnly?
2 1 to reproduce of
oNormalize?
2 1=no, 2=yes

ofxn calc*
0.90262
ofxn mod
0.91619

sRun
RunA1
zout
1

Ave
Data
0.00115
Model
0.00115

data/vMax
0.0078

ofxng
275.646

Min d/m
1.35639

ave/min d
3.20741
ave/min m
4.33886

325_Calib_S_A

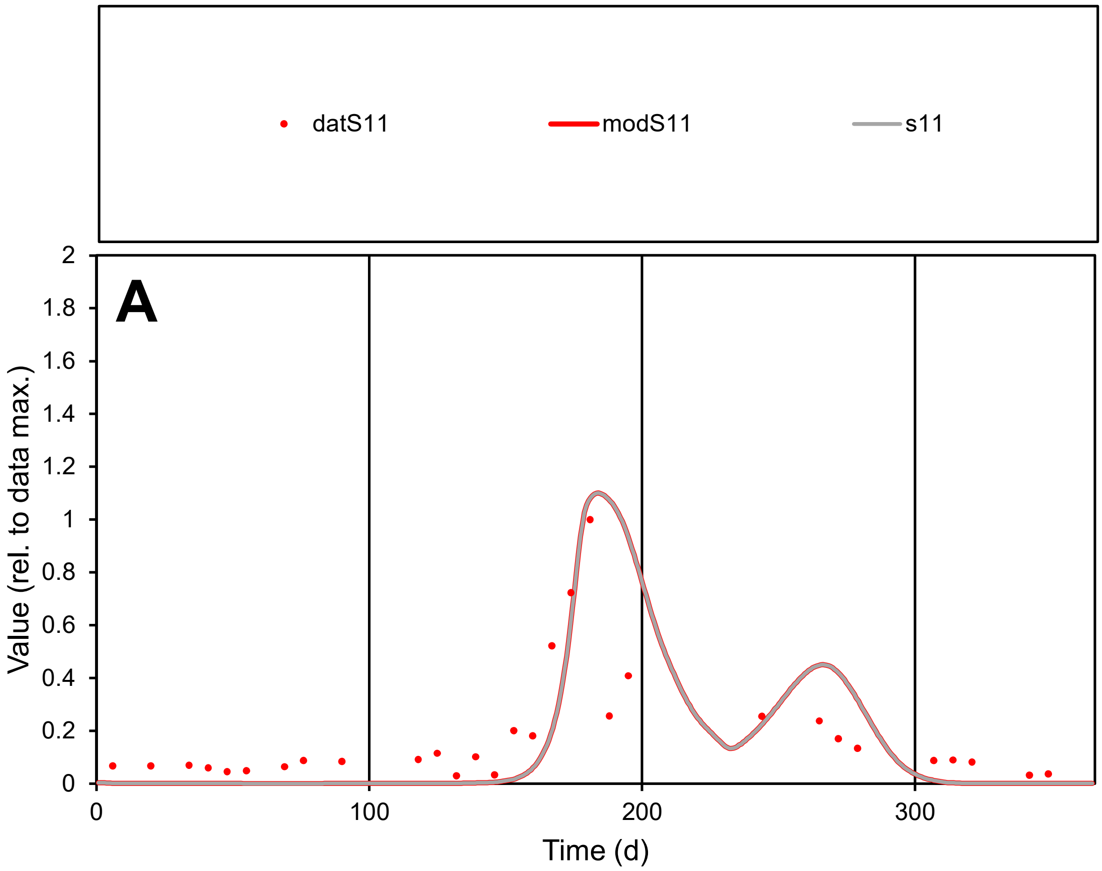


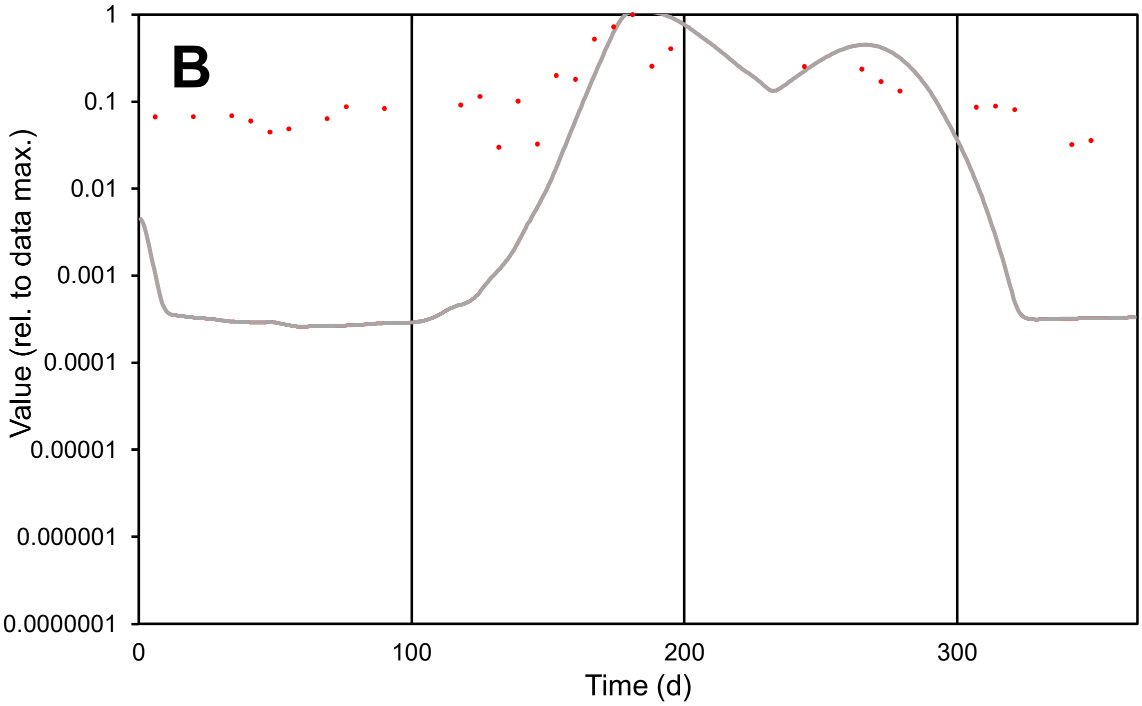


**C1**

Variables
S11 Provide jtd or Dst
CalibOnly?
2 1 to reproduce of
oNormalize?
2 1=no, 2=yes

ofxn calc*
1.11532
ofxn mod
1.14298

sRun
RunA1
zout
1

Ave
Data
0.09093
Model
0.09505

data/vMax
0.5081

ofxng
275.646

Min d/m
113.647

ave/min d
5.96636
ave/min m
708.823

325_Calib_S_A

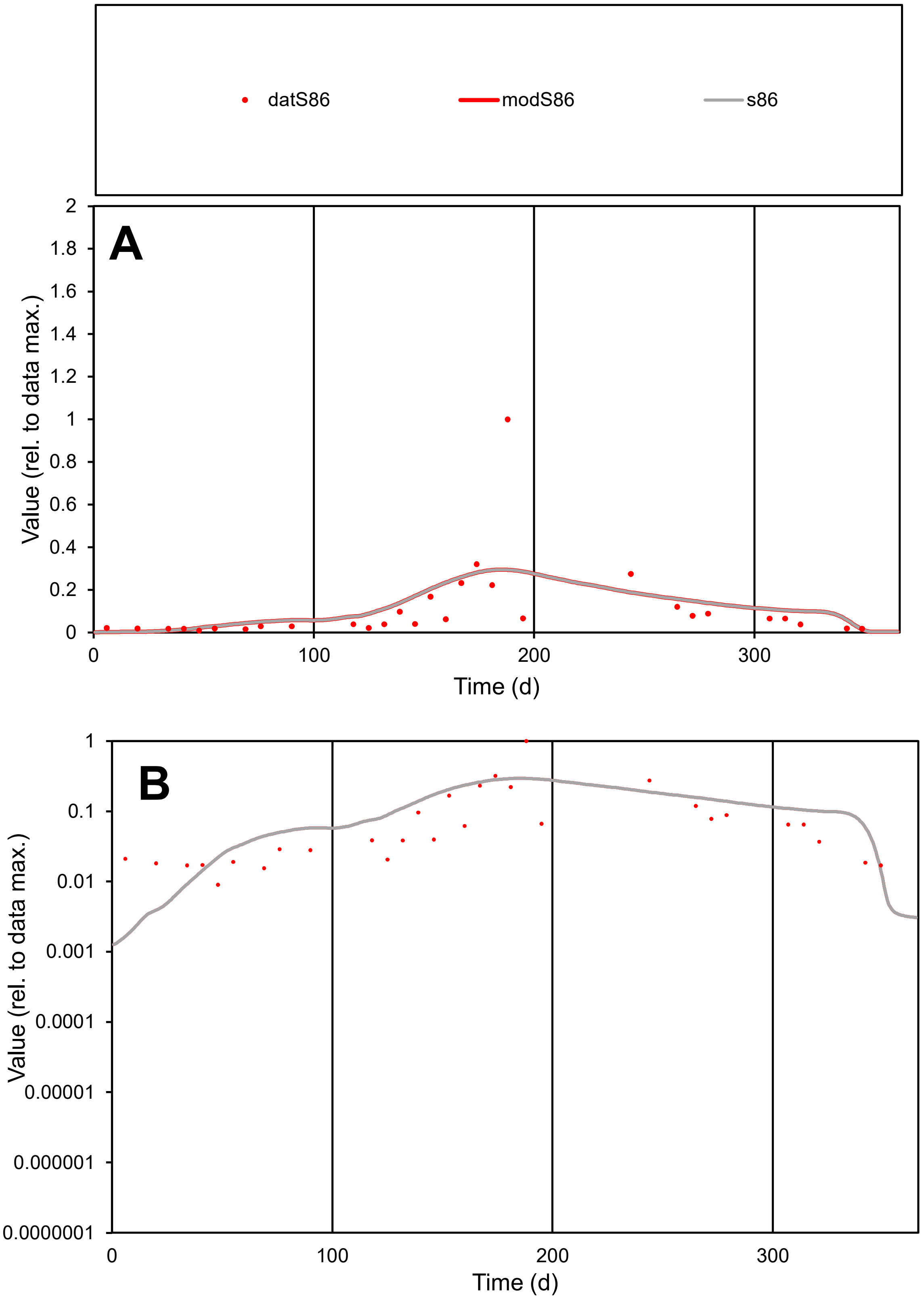


**C1**

Variables
S86 Provide jtd or Dst
CalibOnly?
2 1 to reproduce of
oNormalize?
2 1=no, 2=yes

ofxn calc*
1.35801
ofxn mod
1.36789

sRun
RunA1
zout
1

Ave
Data
0.03132
Model
0.03569

data/vMax
0.2902

ofxng
275.646

Min d/m
5.01155

ave/min d
12.0295
ave/min m
68.6913

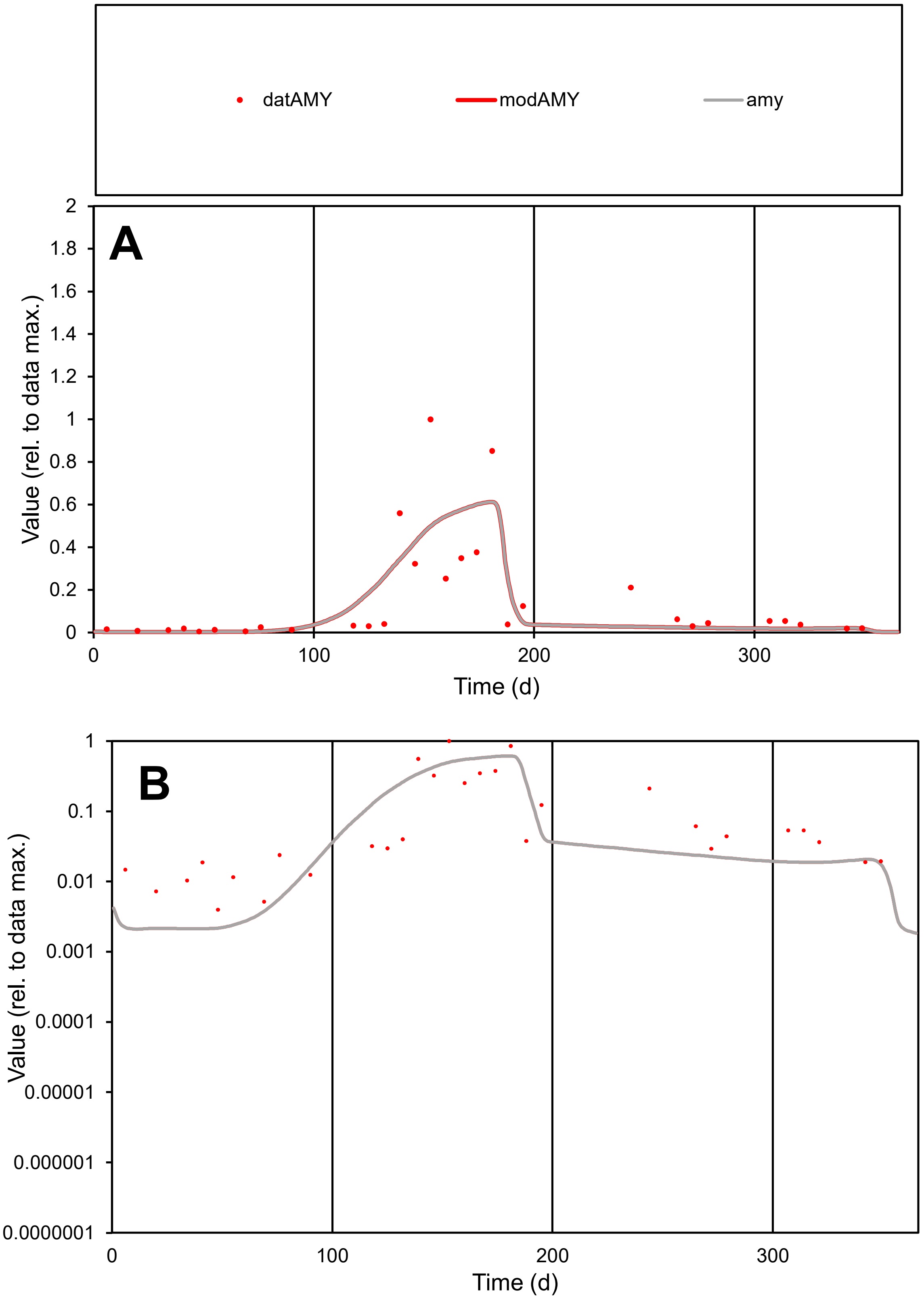


**C1**

Variables
AMY Provide jtd or Dst
CalibOnly?
2 1 to reproduce of
oNormalize?
2 1=no, 2=yes

ofxn calc*
0.98759
ofxn mod
1.00805

sRun
RunA1
zout
1

Ave
Data
0.02619
Model
0.02644

data/vMax
0.1706

ofxng
275.646

Min d/m
1.87514

ave/min d
38.6695
ave/min m
73.1898

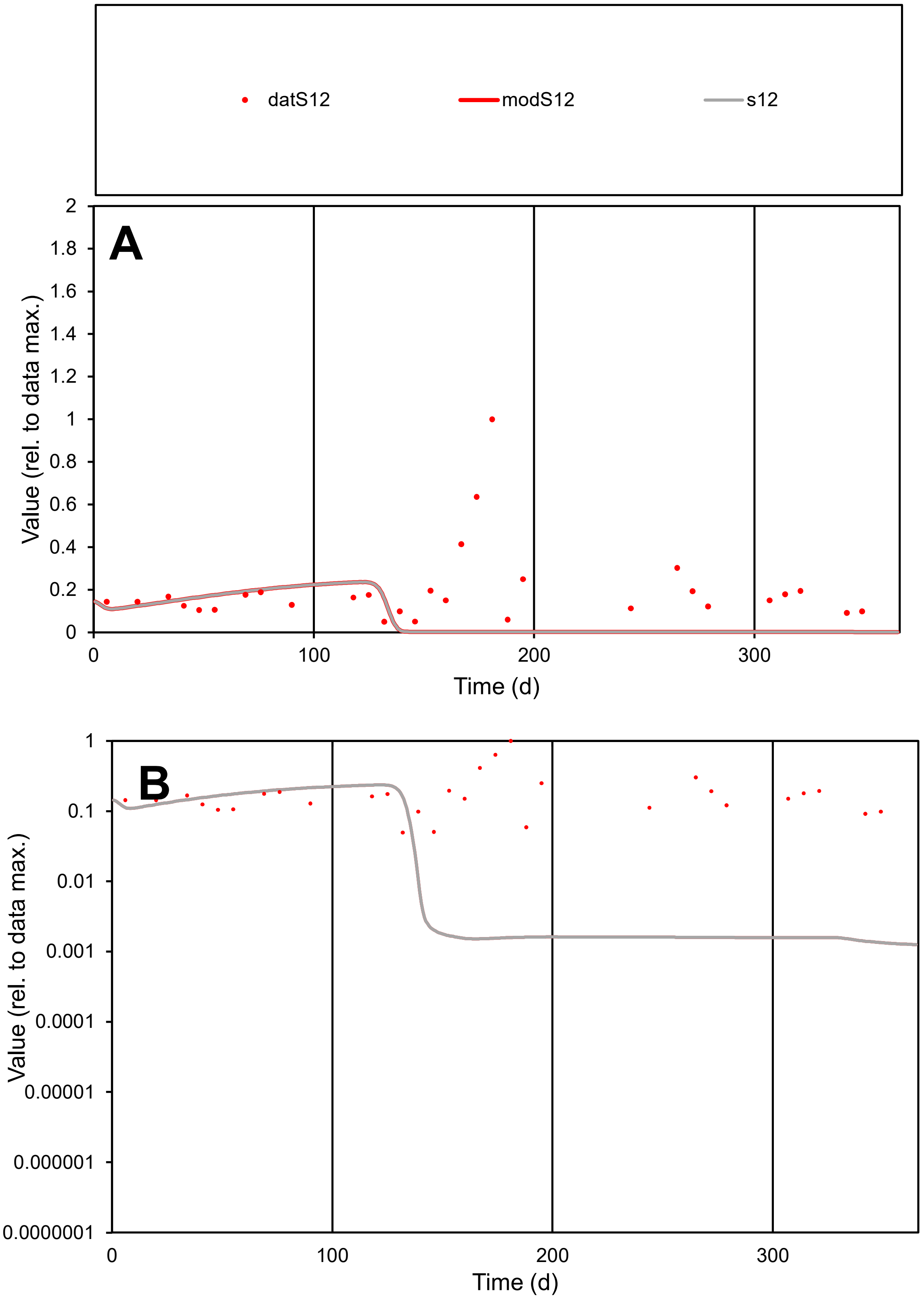


**C1**

Variables

| | |
|---|---|
| S12 | Provide jtd or Dst |
| CalibOnly? | |
| 2 | 1 to reproduce of |
| oNormalize? | |
| 2 | 1=no, 2=yes |

ofxn calc*
1.29536
ofxn mod
1.30032

sRun
RunA1
zout
1

Ave
Data
0.04112
Model
0.0147

data/vMax
0.2069

ofxng
275.646

Min d/m
37.1511

ave/min d
4.01174
ave/min m
53.2806

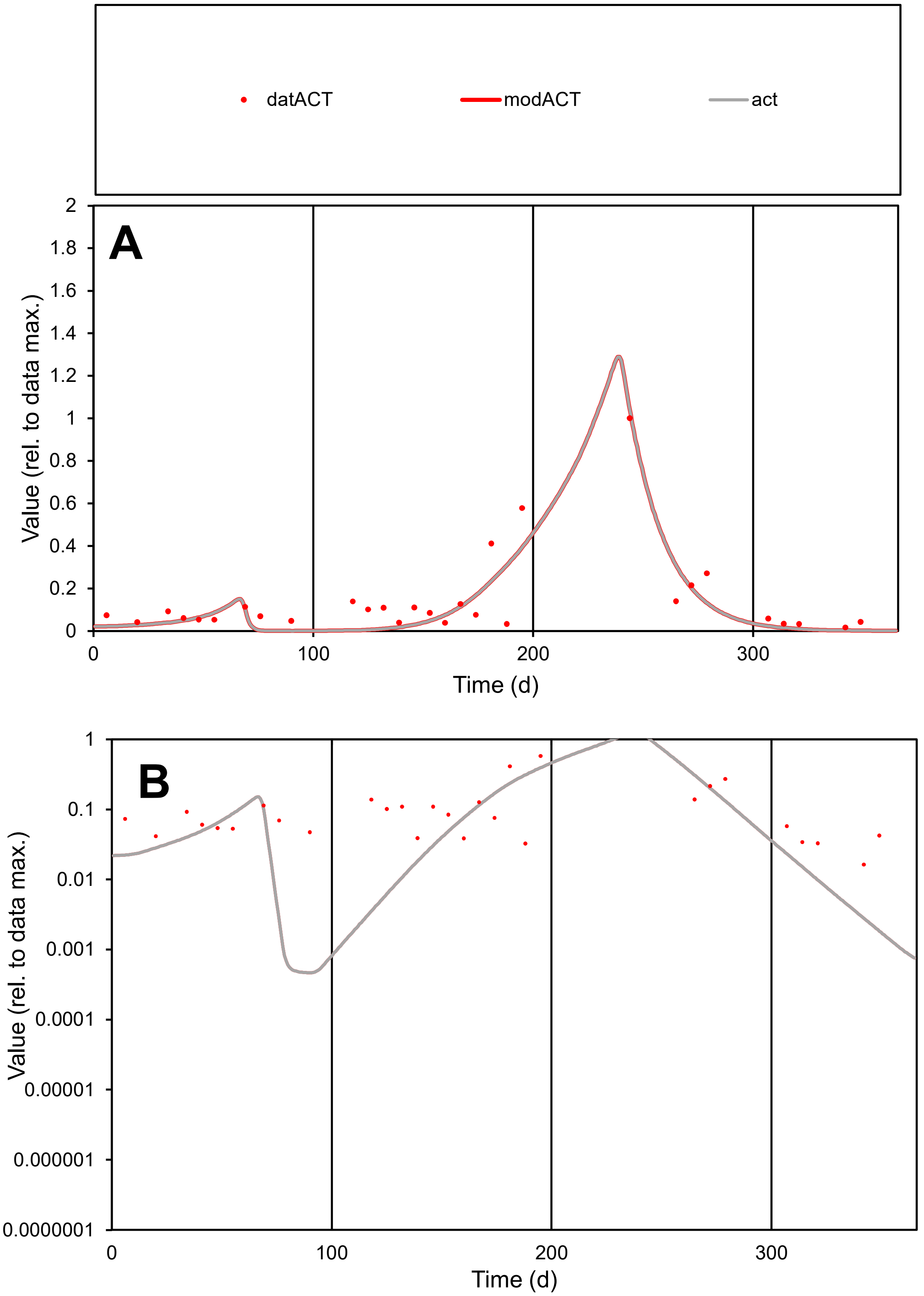


**C1**

Variables

ACT Provide jtd or Dst

CalibOnly?

2 1 to reproduce of

oNormalize?

2 1=no, 2=yes

ofxn calc*

0.63974

ofxn mod

0.66182

sRun

RunA1

zout

1

Ave

Data

0.0246

Model

0.02032

data/vMax

0.1733

ofxng

275.646

Min d/m

34.541

ave/min d

8.7186

ave/min m

248.682

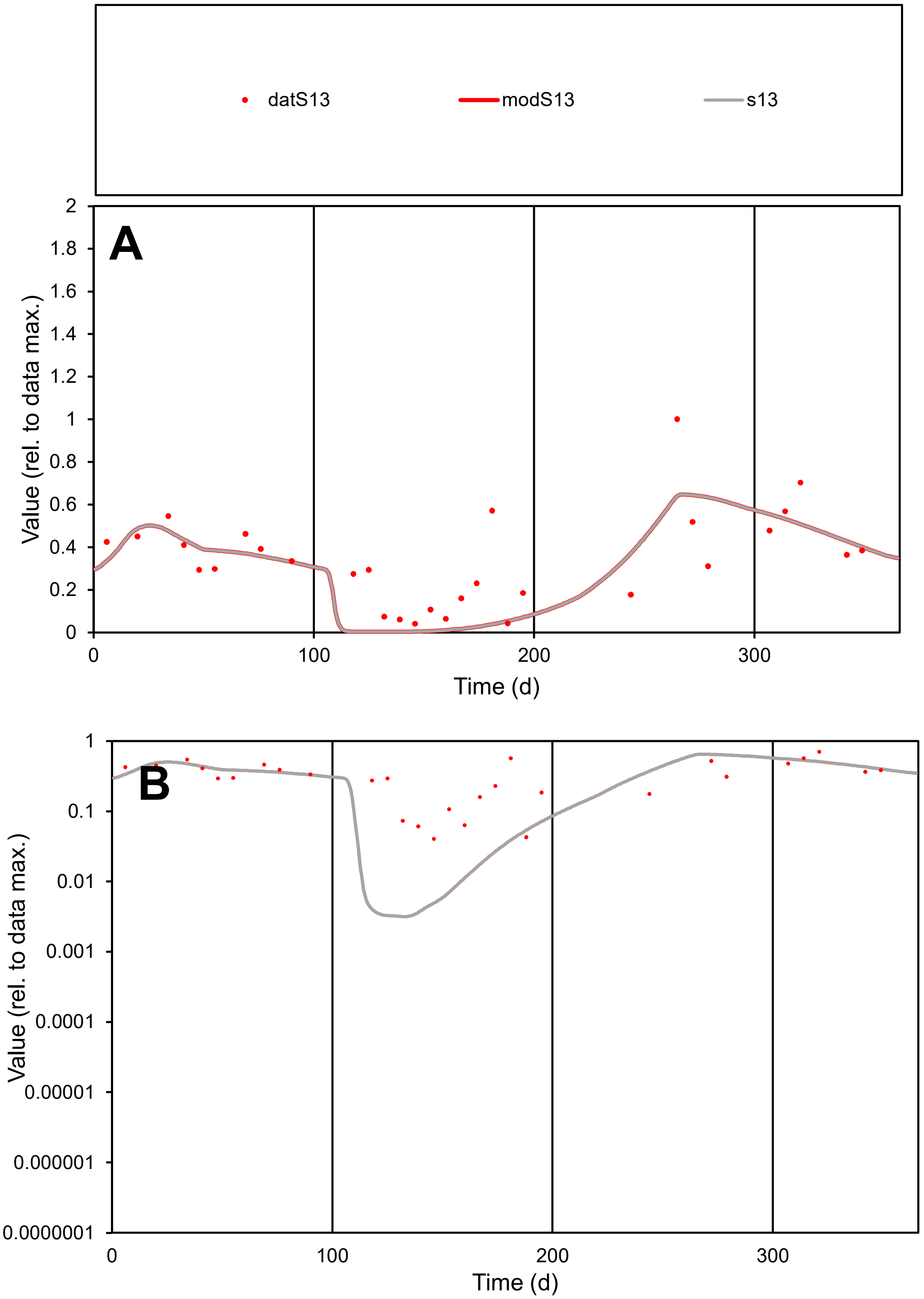


**C1**

| | |
|---|---|
| Variables | |
| S13 | Provide jtd or Dst |
| CalibOnly? | |
| 2 | 1 to reproduce of |
| oNormalize? | |
| 2 | 1=no, 2=yes |
| ofxn calc* | |
| 0.51078 | |
| ofxn mod | |
| 0.52738 | |
| sRun | |
| RunA1 | |
| zout | |
| 1 | |
| Ave | |
| Data | |
| 0.01909 | |
| Model | |
| 0.01592 | |
| data/vMax | |
| 0.05605 | |
| ofxng | |
| 275.646 | |
| Min d/m | |
| 12.7872 | |
| ave/min d | |
| 8.4038 | |
| ave/min m | |
| 89.6293 | |

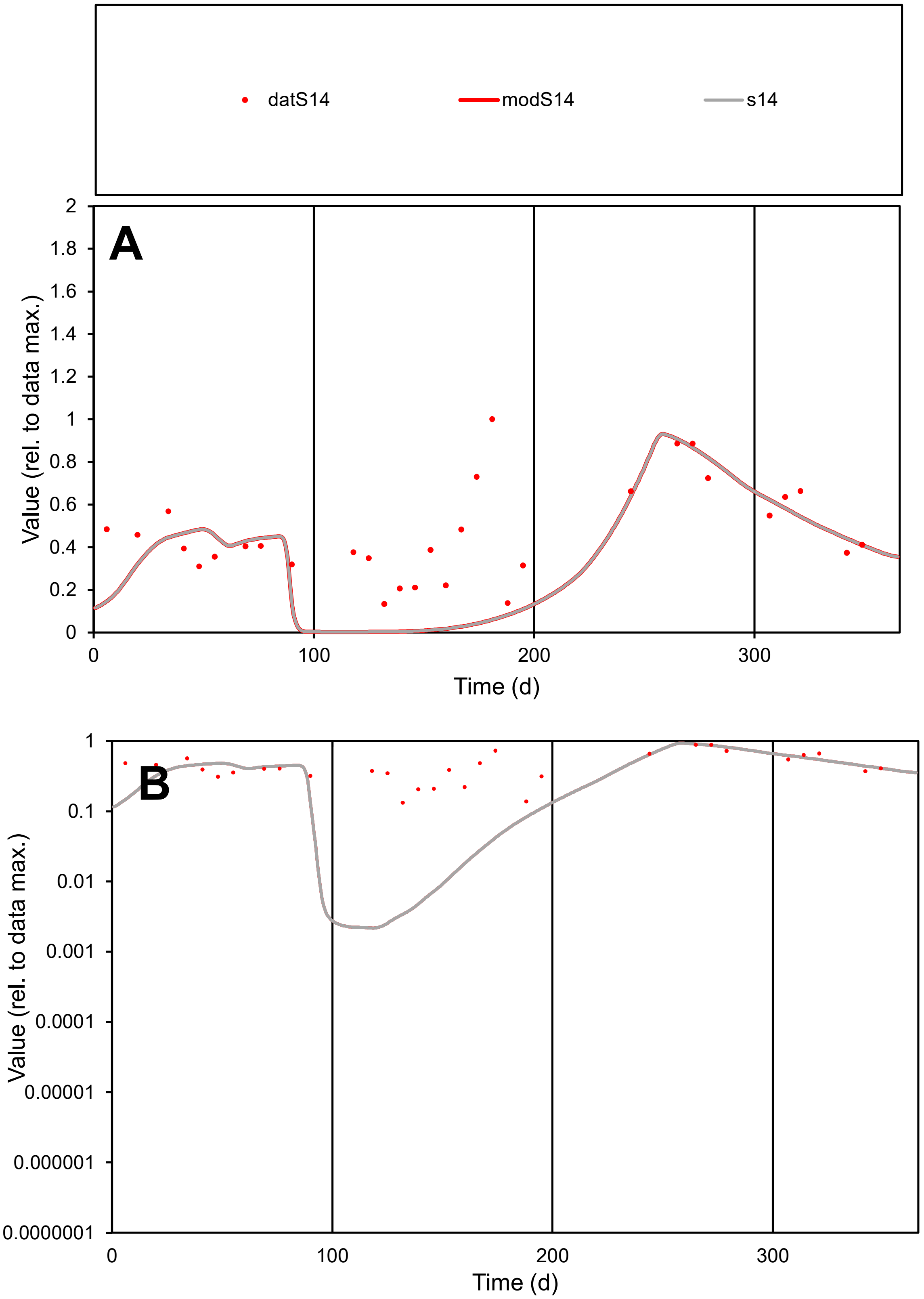


**C1**

Variables
S14 Provide jtd or Dst
CalibOnly?
2 1 to reproduce of
oNormalize?
2 1=no, 2=yes

ofxn calc*
0.59999
ofxn mod
0.61251

sRun
RunA1
zout
1

Ave
Data
0.01746
Model
0.01196

data/vMax
0.03734

ofxng
275.646

Min d/m
61.1179

ave/min d
3.50986
ave/min m
146.88

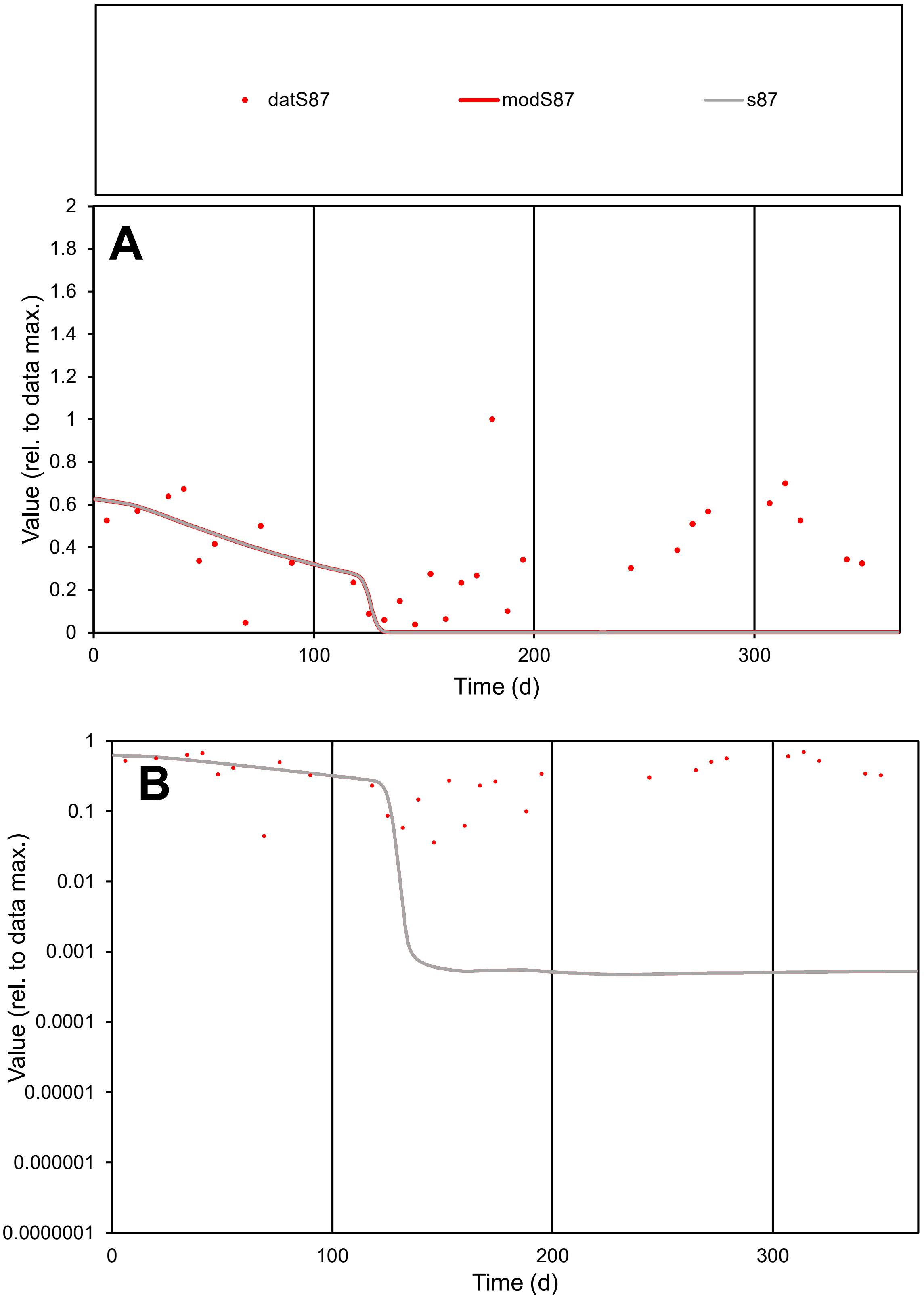


**C1**

Variables
S87 Provide jtd or Dst
CalibOnly?
2 1 to reproduce of
oNormalize?
2 1=no, 2=yes

ofxn calc*
0.95194
ofxn mod
0.95762

sRun
RunA1
zout
1

Ave
Data
0.01745
Model
0.00747

data/vMax
0.04707

ofxng
275.646

Min d/m
75.2983

ave/min d
10.2425
ave/min m
329.898

325_Calib_S_A

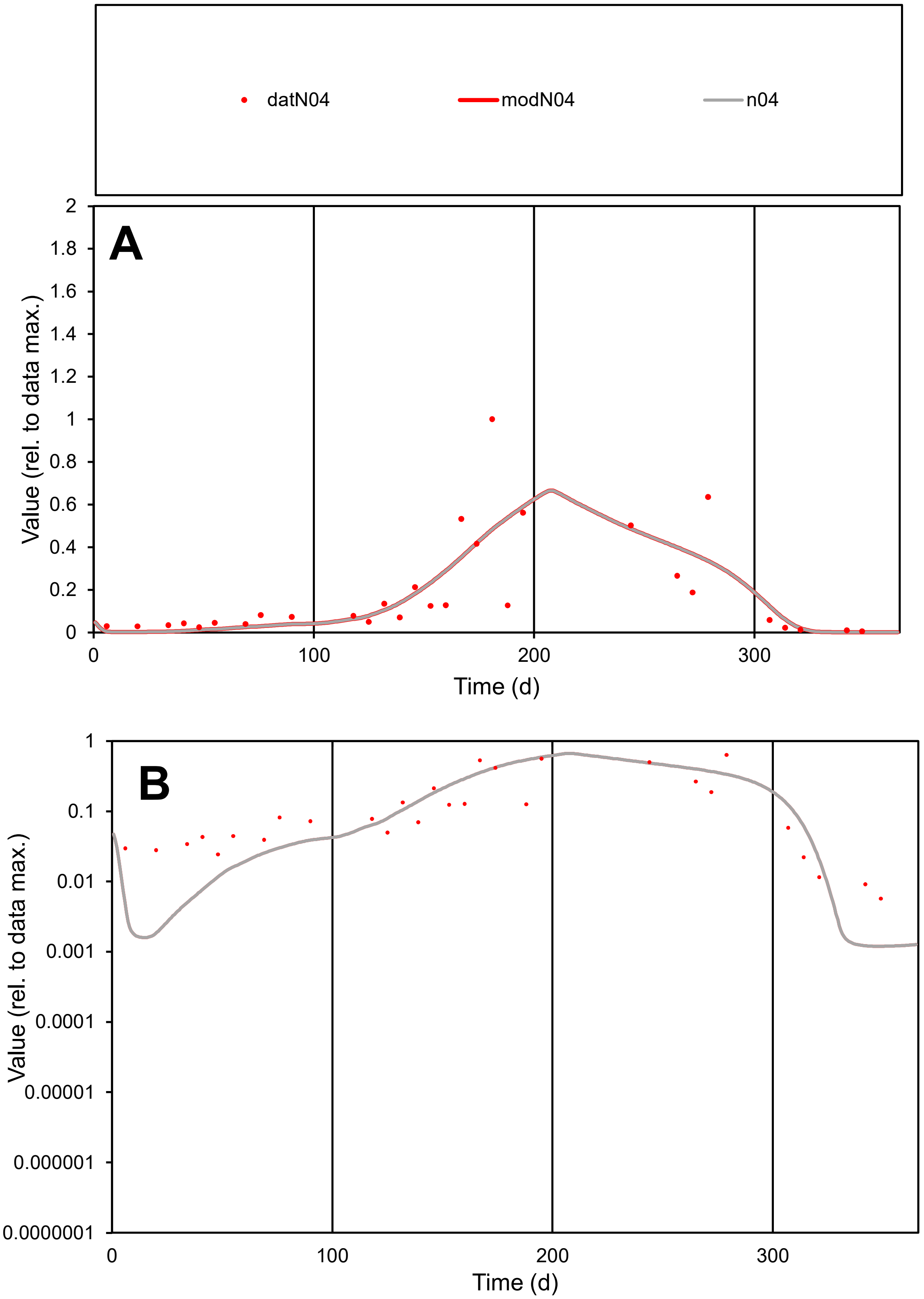


**C1**

Variables
N04 Provide jtd or Dst
CalibOnly?
2 1 to reproduce of
oNormalize?
2 1=no, 2=yes

ofxn calc*
0.80039
ofxn mod
0.82373

sRun
RunA1
zout
1

Ave
Data
0.01862
Model
0.01841

data/vMax
0.1011

ofxng
275.646

Min d/m
4.74589

ave/min d
32.2651
ave/min m
151.379

325_Calib_S_A

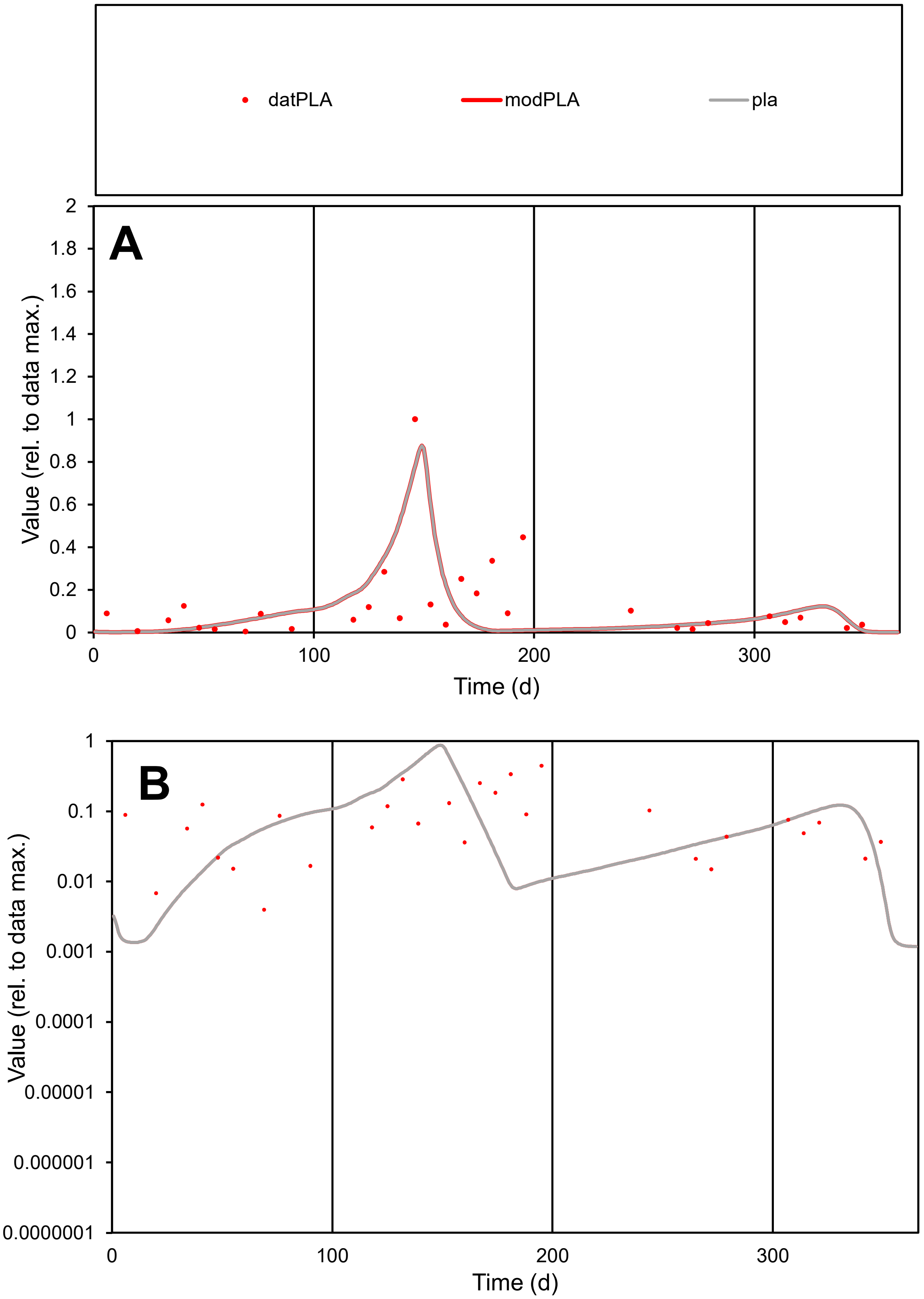


C1

Variables
PLA Provide jtd or Dst
CalibOnly?
2 1 to reproduce of
oNormalize?
2 1=no, 2=yes

ofxn calc*
1.22314
ofxn mod
1.28584

sRun
RunA1
zout
1

Ave
Data
0.01205
Model
0.01125

data/vMax
0.09369

ofxng
275.646

Min d/m
2.87693

ave/min d
32.421
ave/min m
87.0882

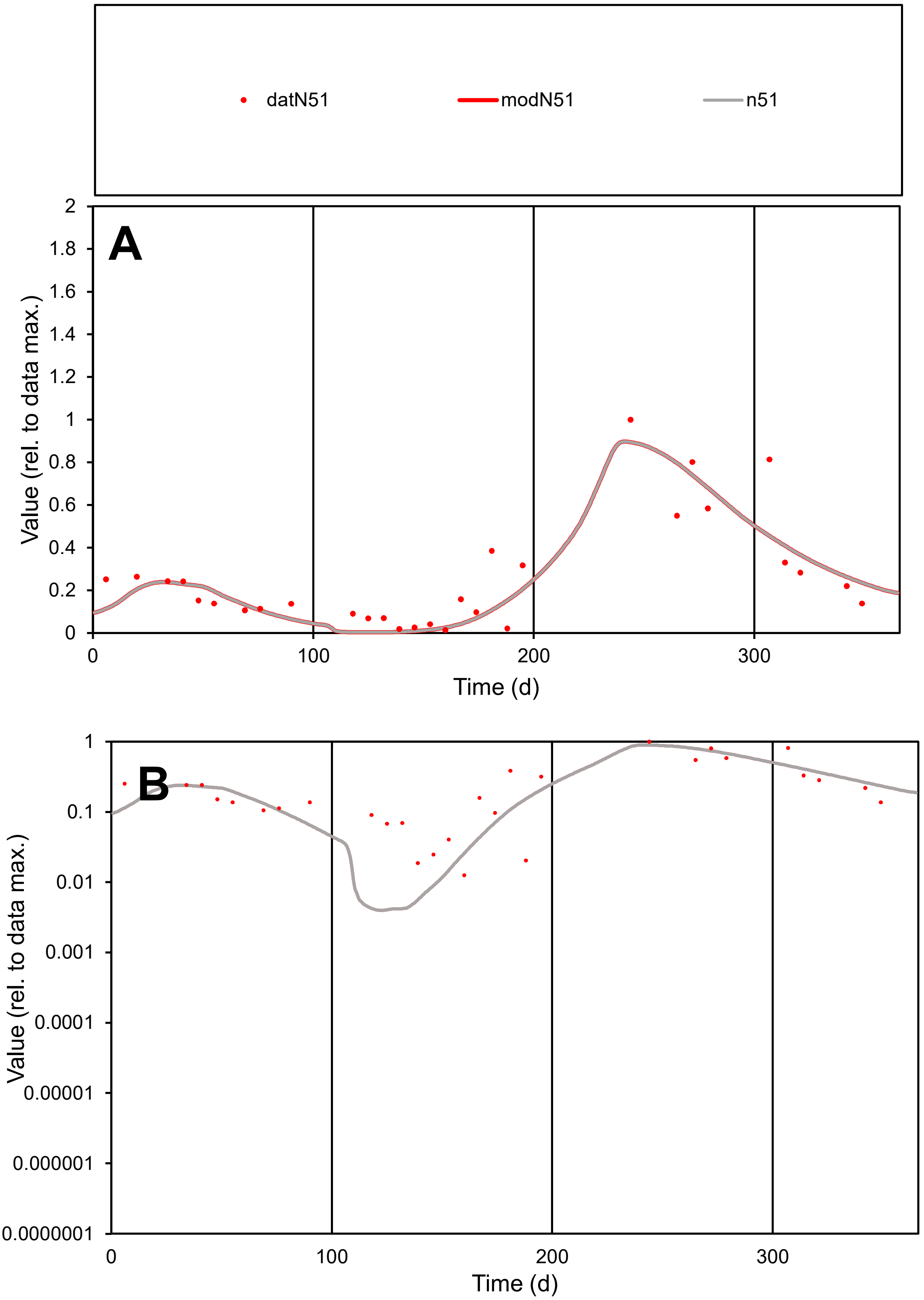


**C1**

Variables
N51 Provide jtd or Dst
CalibOnly?
2 1 to reproduce of
oNormalize?
2 1=no, 2=yes

ofxn calc*
0.45041
ofxn mod
0.47709

sRun
RunA1
zout
1

Ave
Data
0.00881
Model
0.00799

data/vMax
0.03451

ofxng
275.646

Min d/m
3.04645

ave/min d
20.3493
ave/min m
56.214

325_Calib_S_A

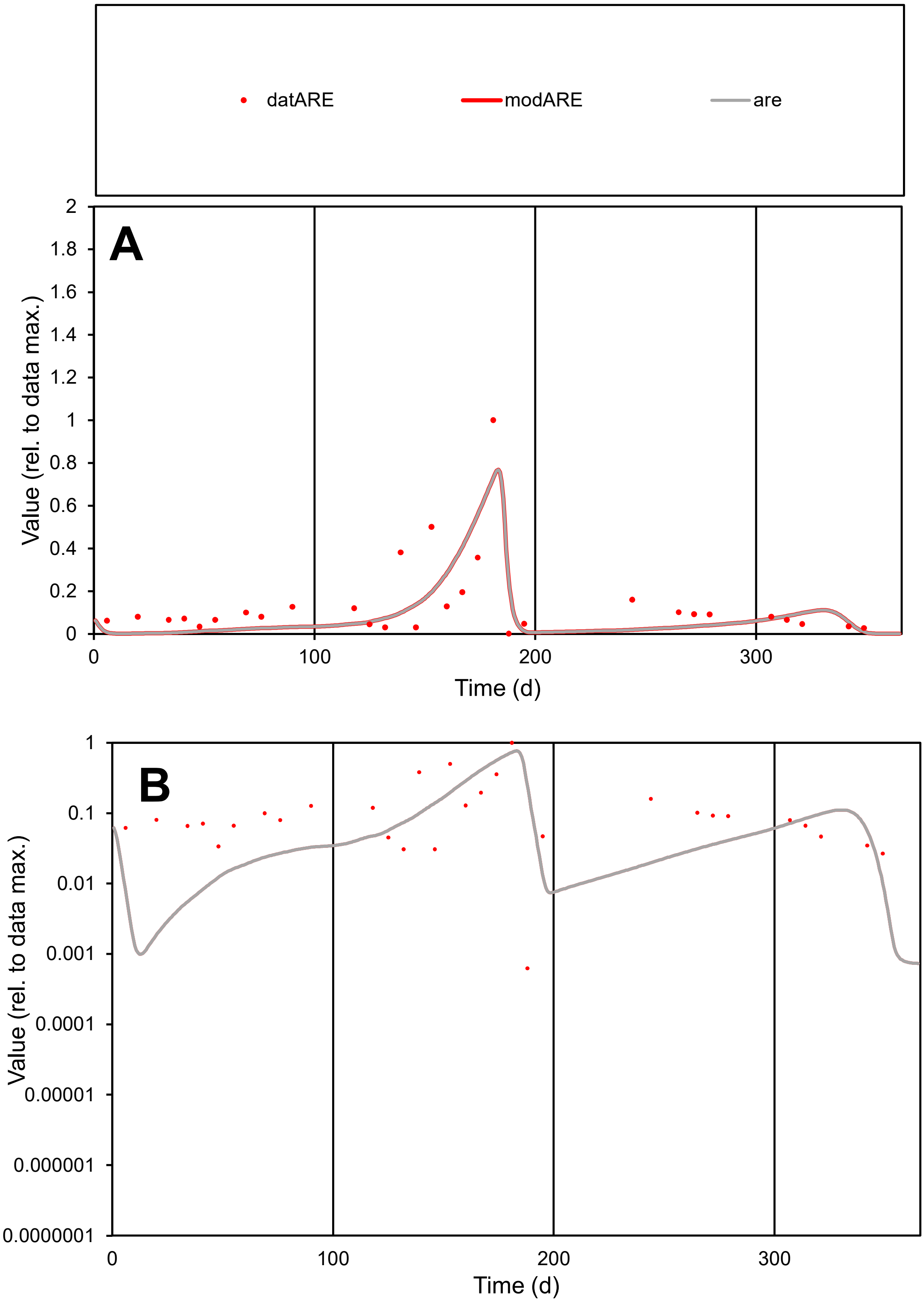


**C1**

Variables

ARE Provide jtd or Dst

CalibOnly?

2 1 to reproduce of

oNormalize?

2 1=no, 2=yes

ofxn calc*

0.85191

ofxn mod

0.88287

sRun

RunA1

zout

1

Ave

Data

0.01225

Model

0.00989

data/vMax

0.08707

ofxng

275.646

Min d/m

0.30334

ave/min d

224.498

ave/min m

54.975

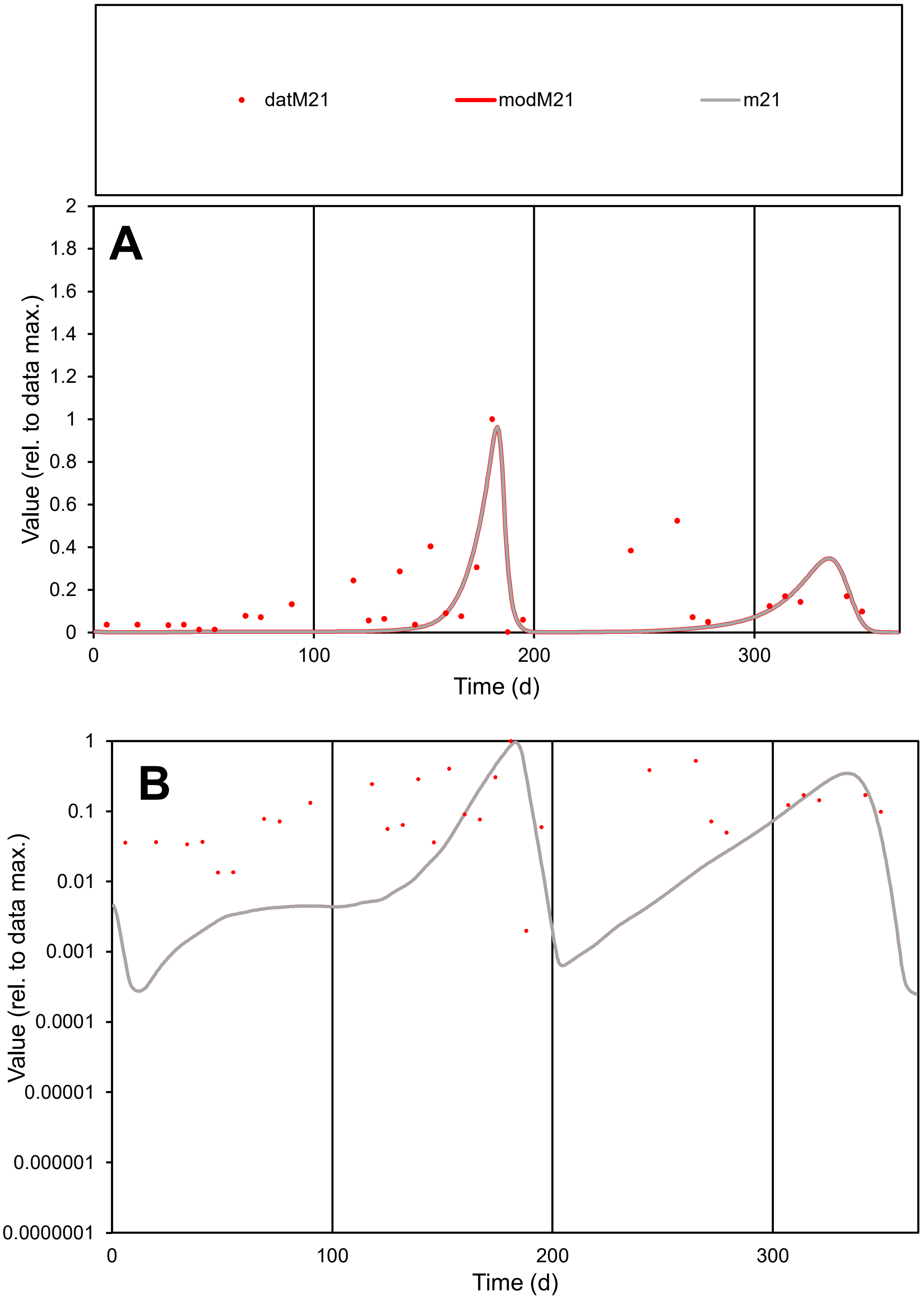


**C1**

Variables

M21 Provide jtd or Dst

CalibOnly?

2 1 to reproduce of

oNormalize?

2 1=no, 2=yes

ofxn calc*

0.99993

ofxn mod

1.02064

sRun

RunA1

zout

1

Ave

Data

0.01013

Model

0.00595

data/vMax

0.06319

ofxng

275.646

Min d/m

3.54674

ave/min d

80.9014

ave/min m

168.53

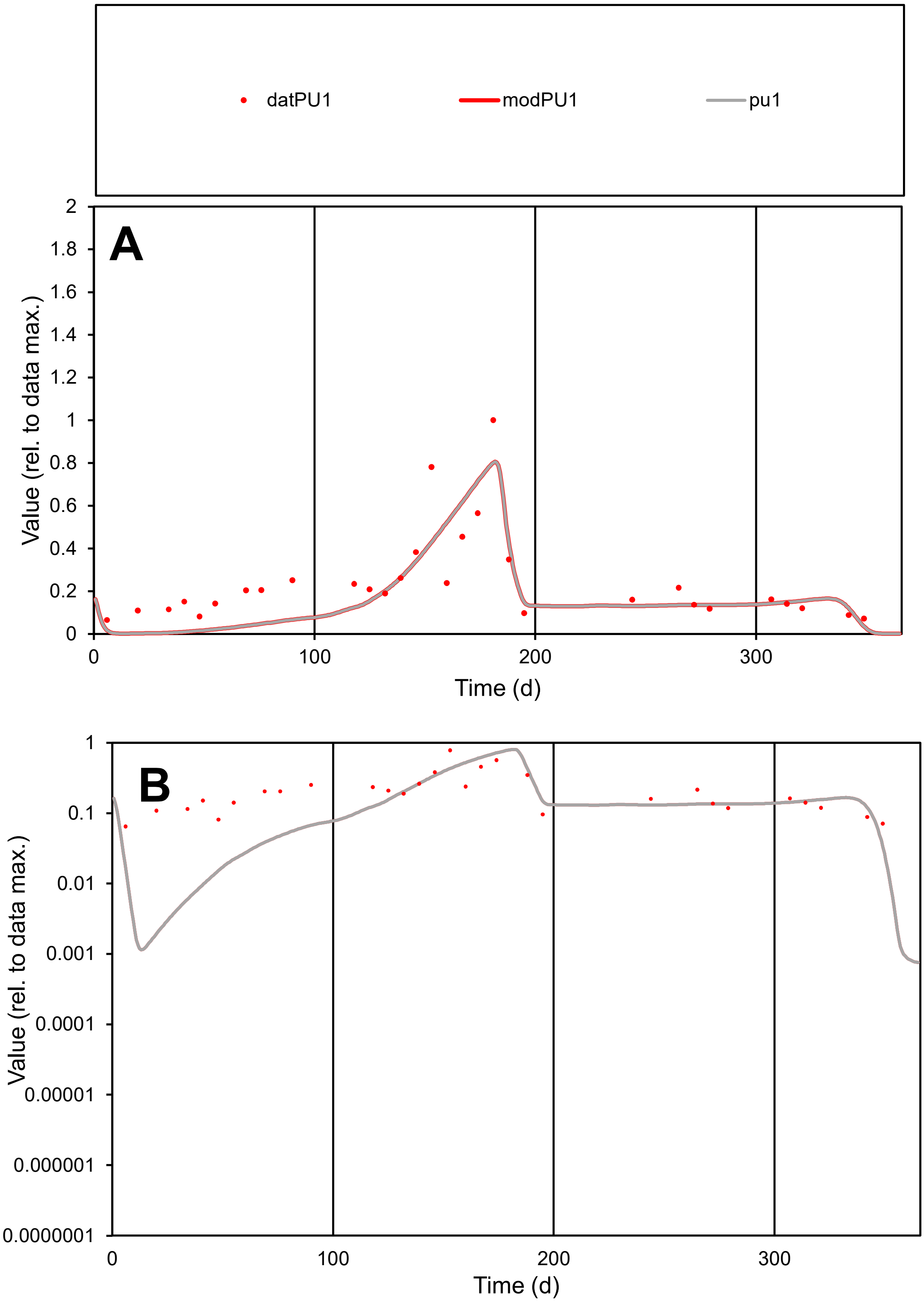


**C1**

Variables
PU1 Provide jtd or Dst
CalibOnly?
2 1 to reproduce of
oNormalize?
2 1=no, 2=yes

ofxn calc*
0.51264
ofxn mod
0.53281

sRun
RunA1
zout
1

Ave
Data
0.01157
Model
0.00978

data/vMax
0.04756

ofxng
275.646

Min d/m
30.3846

ave/min d
3.75419
ave/min m
96.4702

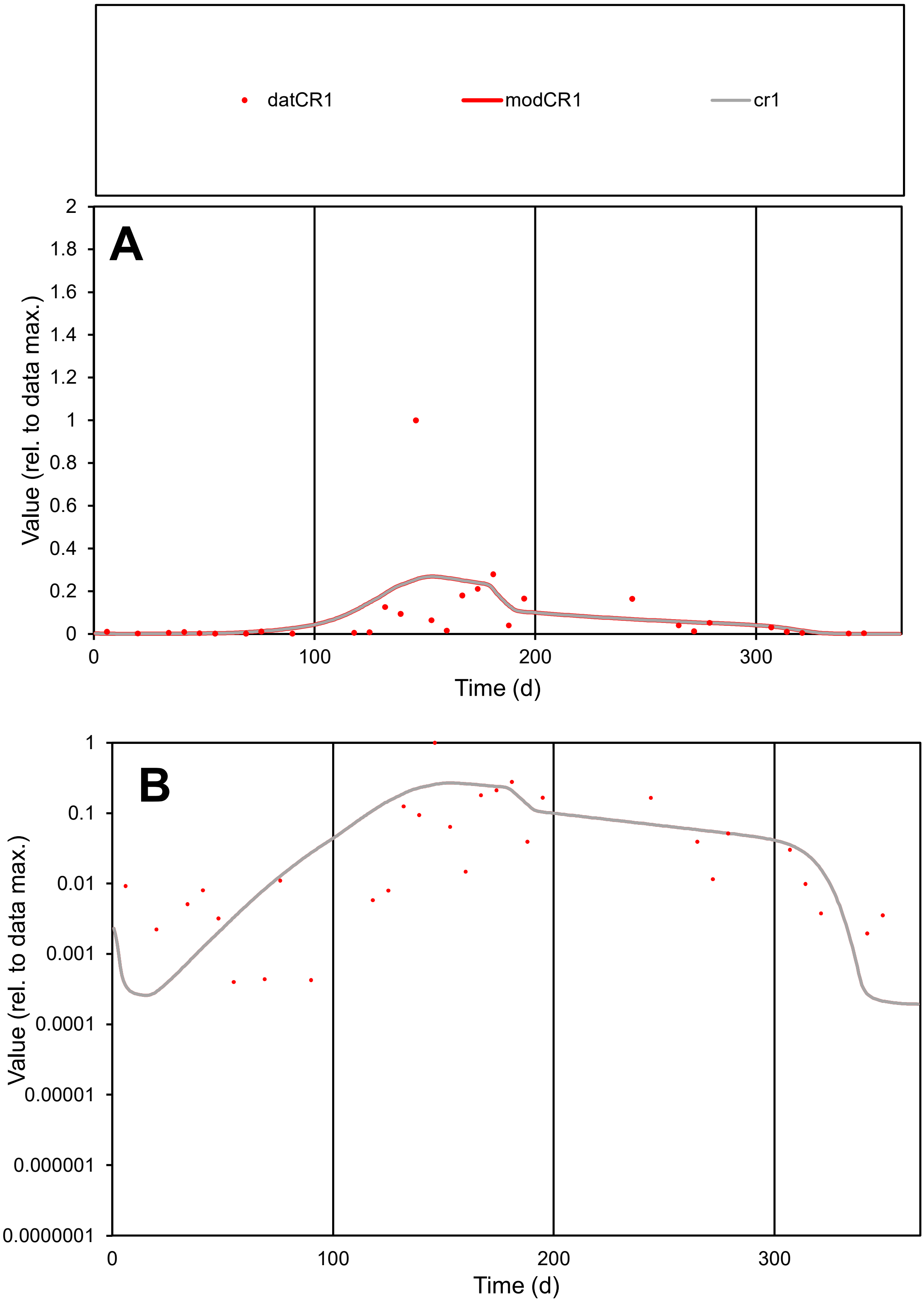


**C1**

Variables
CR1 Provide jtd or Dst
CalibOnly?
2 1 to reproduce of
oNormalize?
2 1=no, 2=yes

ofxn calc*
1.84449
ofxn mod
1.86527

sRun
RunA1
zout
1

Ave
Data
0.01516
Model
0.01629

data/vMax
0.179

ofxng
275.646

Min d/m
1.9118

ave/min d
211.313
ave/min m
434.182

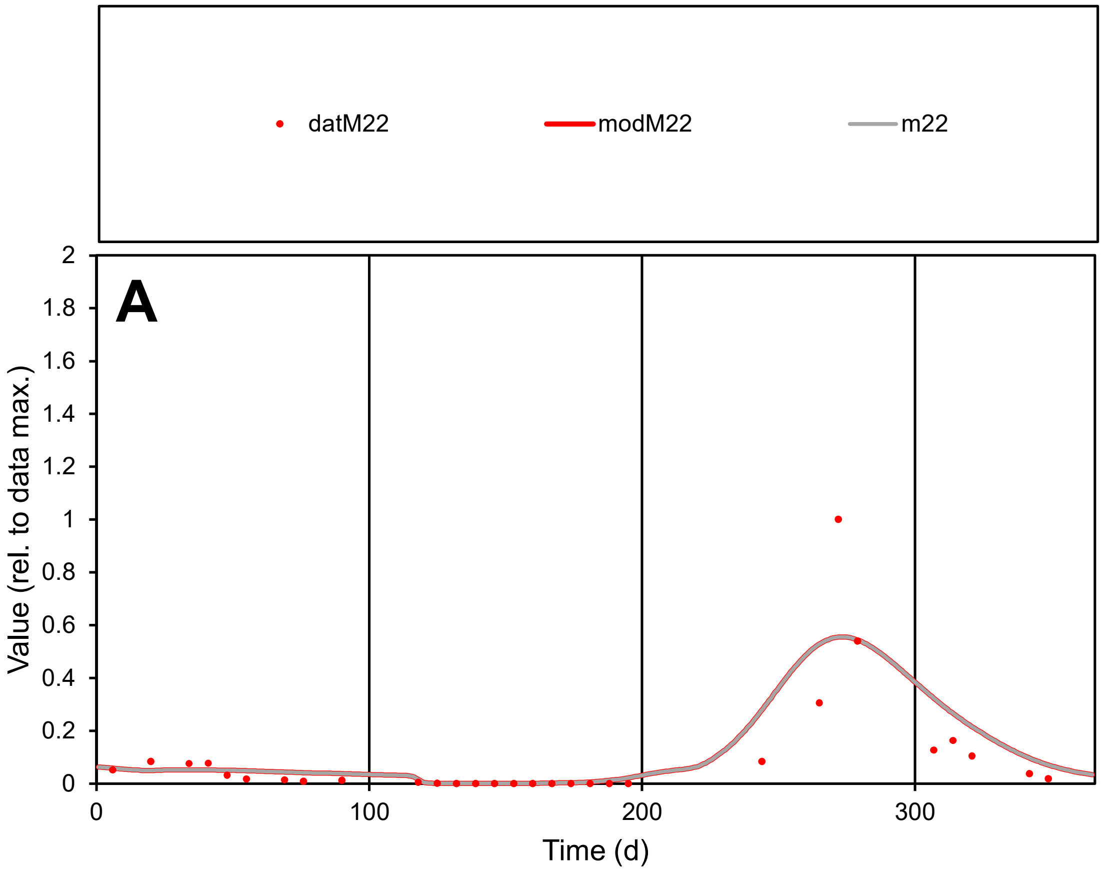


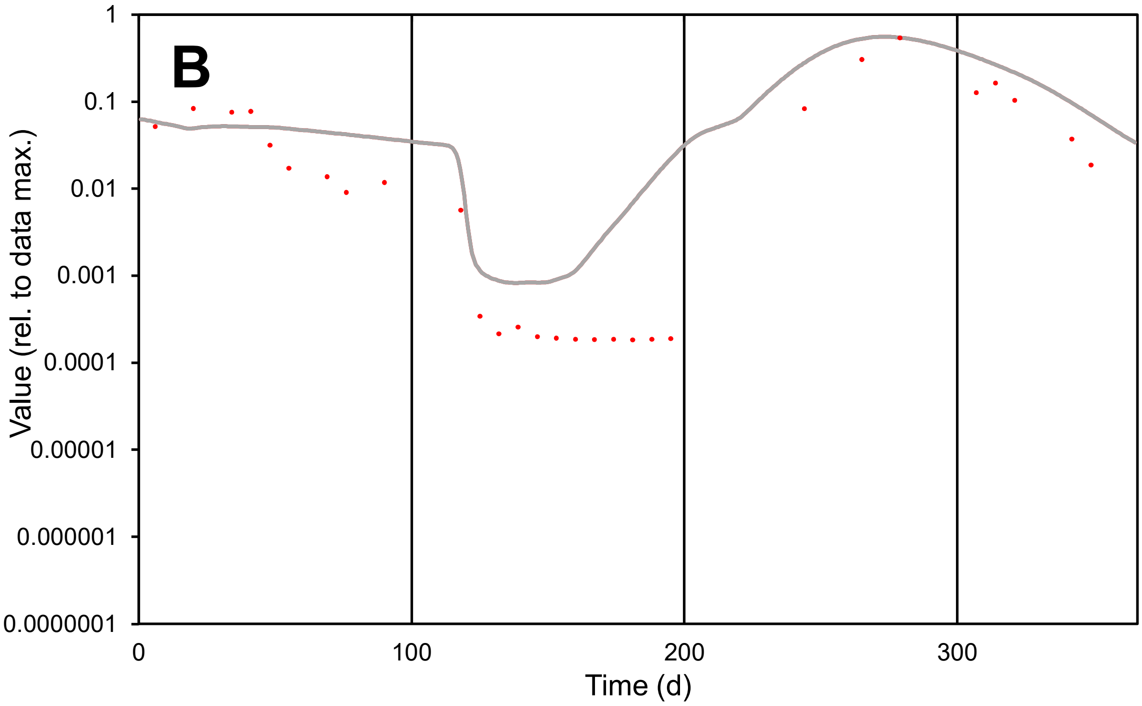


**C1**

Variables
M22 Provide jtd or Dst
CalibOnly?
2 1 to reproduce of
oNormalize?
2 1=no, 2=yes

ofxn calc*
1.18546
ofxn mod
1.22549

sRun
RunA1
zout
1

Ave
Data
0.01075
Model
0.01298

data/vMax
0.1169

ofxng
275.646

Min d/m
0.22094

ave/min d
502.18
ave/min m
134.002

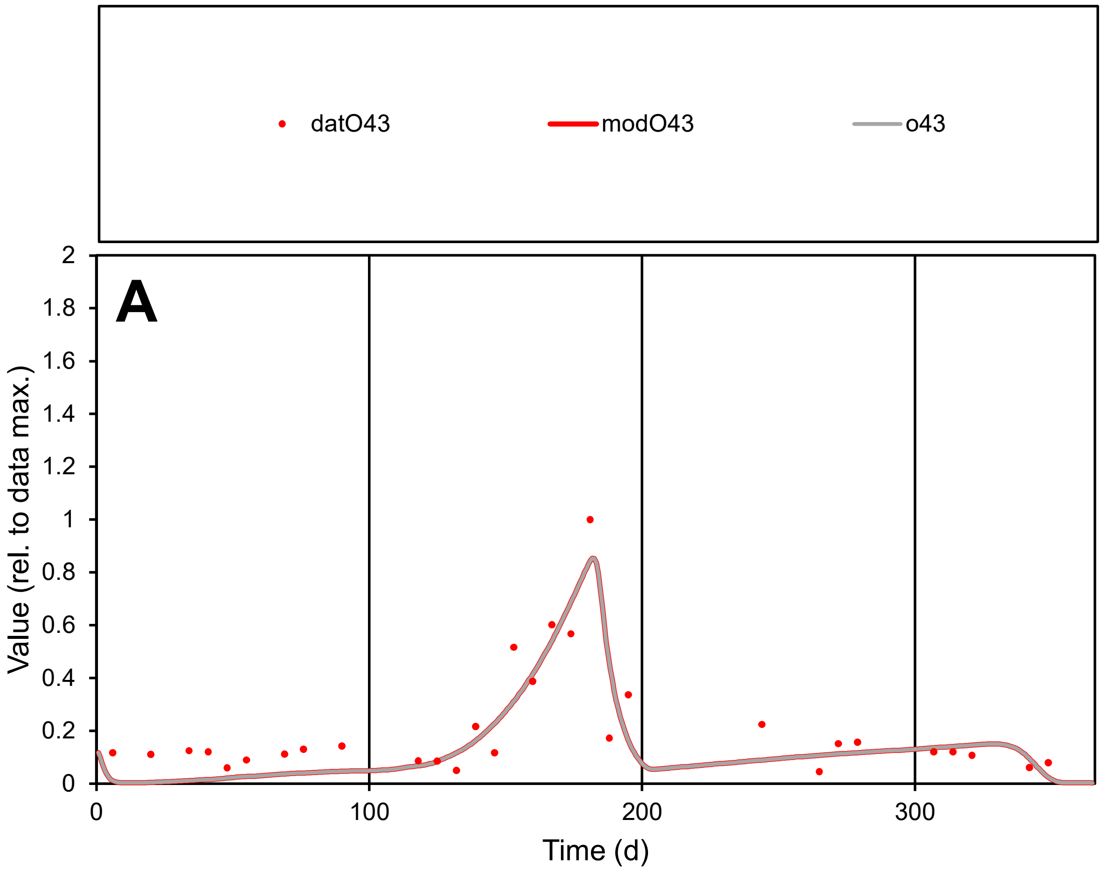


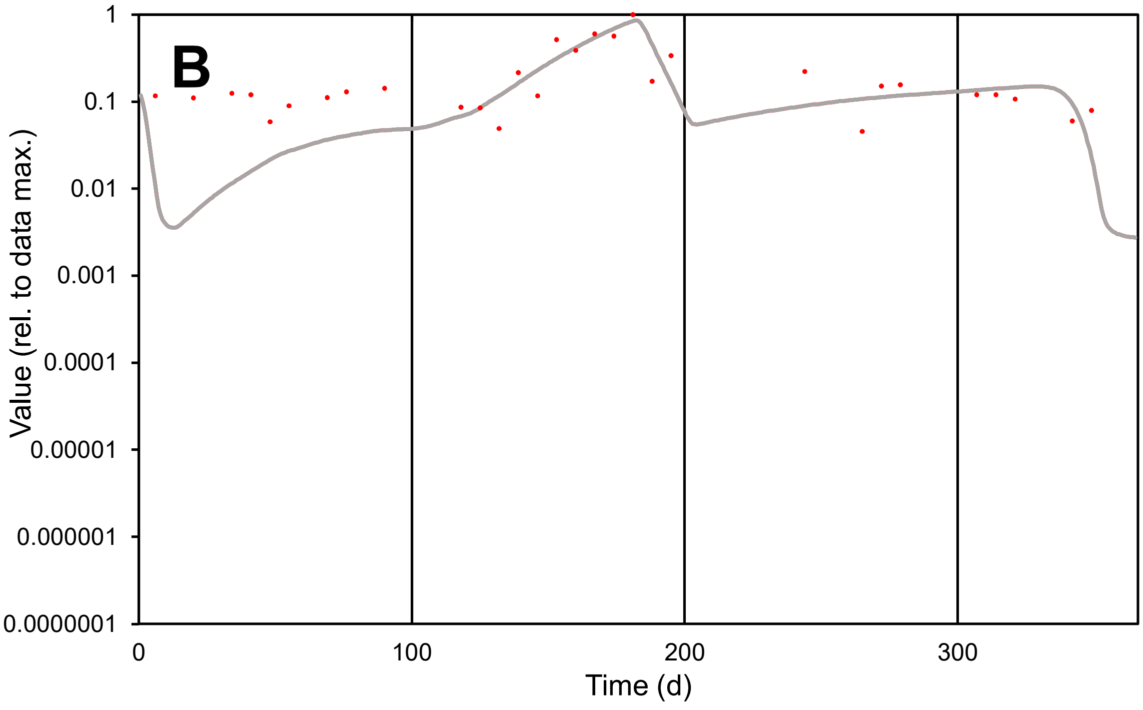


**C1**

Variables
O43 Provide jtd or Dst
CalibOnly?
2 1 to reproduce of
oNormalize?
2 1=no, 2=yes

ofxn calc*
0.44474
ofxn mod
0.48625

sRun
RunA1
zout
1

Ave
Data
0.01121
Model
0.00958

data/vMax
0.05425

ofxng
275.646

Min d/m
8.08134

ave/min d
4.54786
ave/min m
31.4306

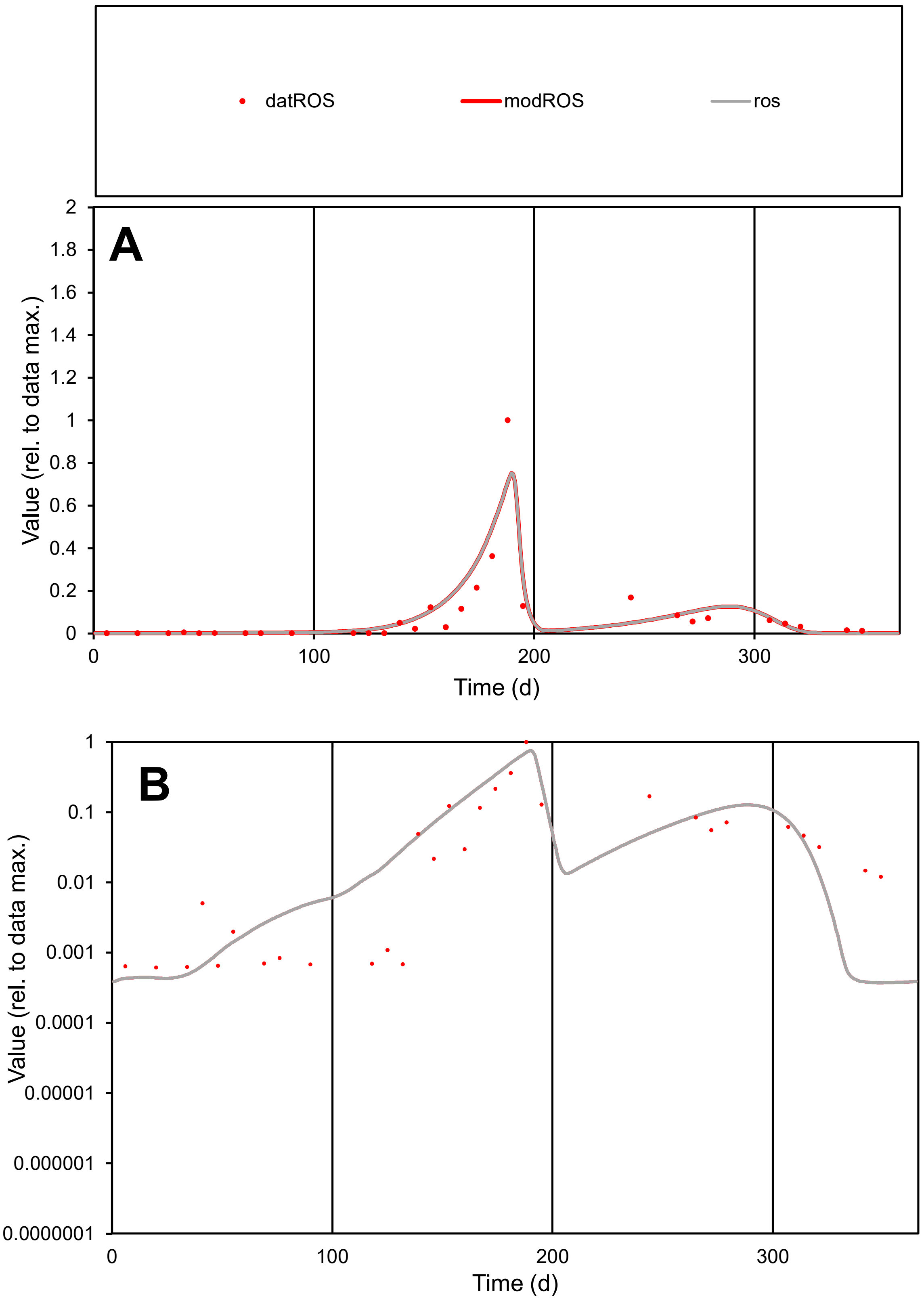


**C1**

Variables
ROS Provide jtd or Dst
CalibOnly?
2 1 to reproduce of
oNormalize?
2 1=no, 2=yes

ofxn calc*
0.83717
ofxn mod
0.88875

sRun
RunA1
zout
1

Ave
Data
0.0074
Model
0.00834

data/vMax
0.08513

ofxng
275.646

Min d/m
1.6485

ave/min d
141.311
ave/min m
262.788

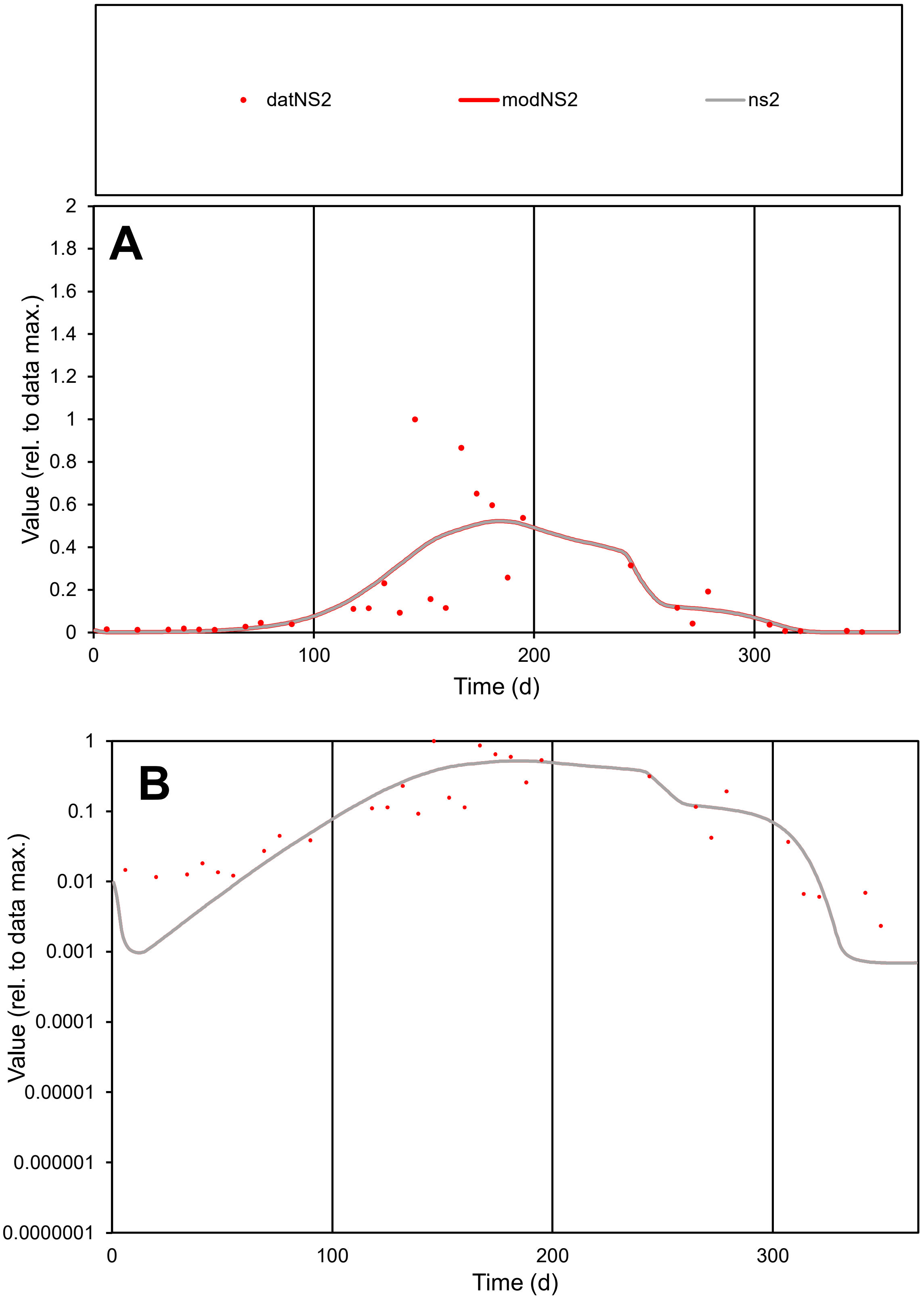


**C1**

Variables
NS2 Provide jtd or Dst
CalibOnly?
2 1 to reproduce of
oNormalize?
2 1=no, 2=yes

ofxn calc*
0.91864
ofxn mod
0.93339

sRun
RunA1
zout
1

Ave
Data
0.01118
Model
0.01119

data/vMax
0.05945

ofxng
275.646

Min d/m
3.36247

ave/min d
80.7126
ave/min m
271.742

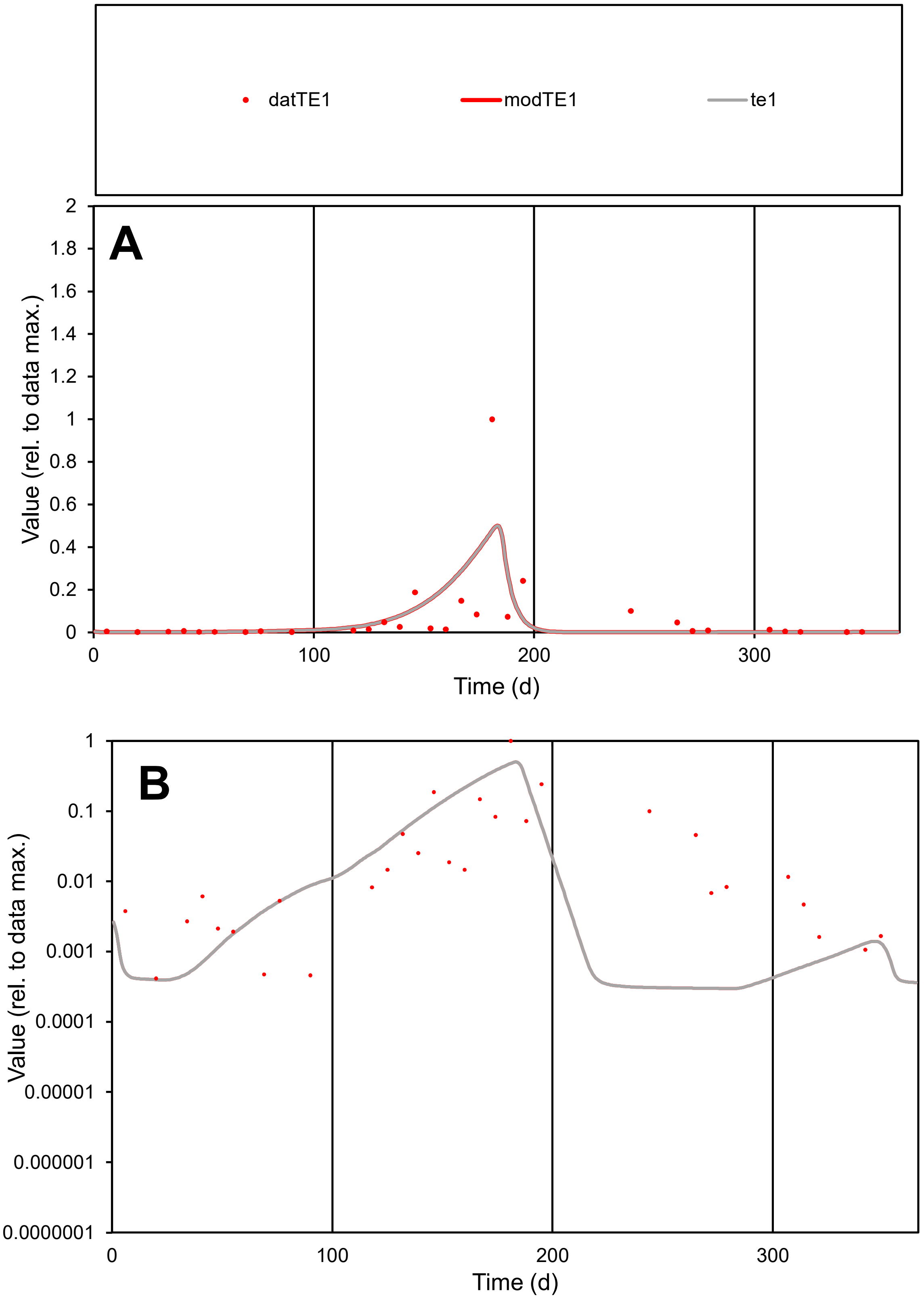


**C1**

Variables
TE1 Provide jtd or Dst
CalibOnly?
2 1 to reproduce of
oNormalize?
2 1=no, 2=yes

ofxn calc*
1.83217
ofxn mod
1.88488

sRun
RunA1
zout
1

Ave
Data
0.00908
Model
0.00931

data/vMax
0.1319

ofxng
275.646

Min d/m
1.38246

ave/min d
166.033
ave/min m
235.263

325_Calib_S_A

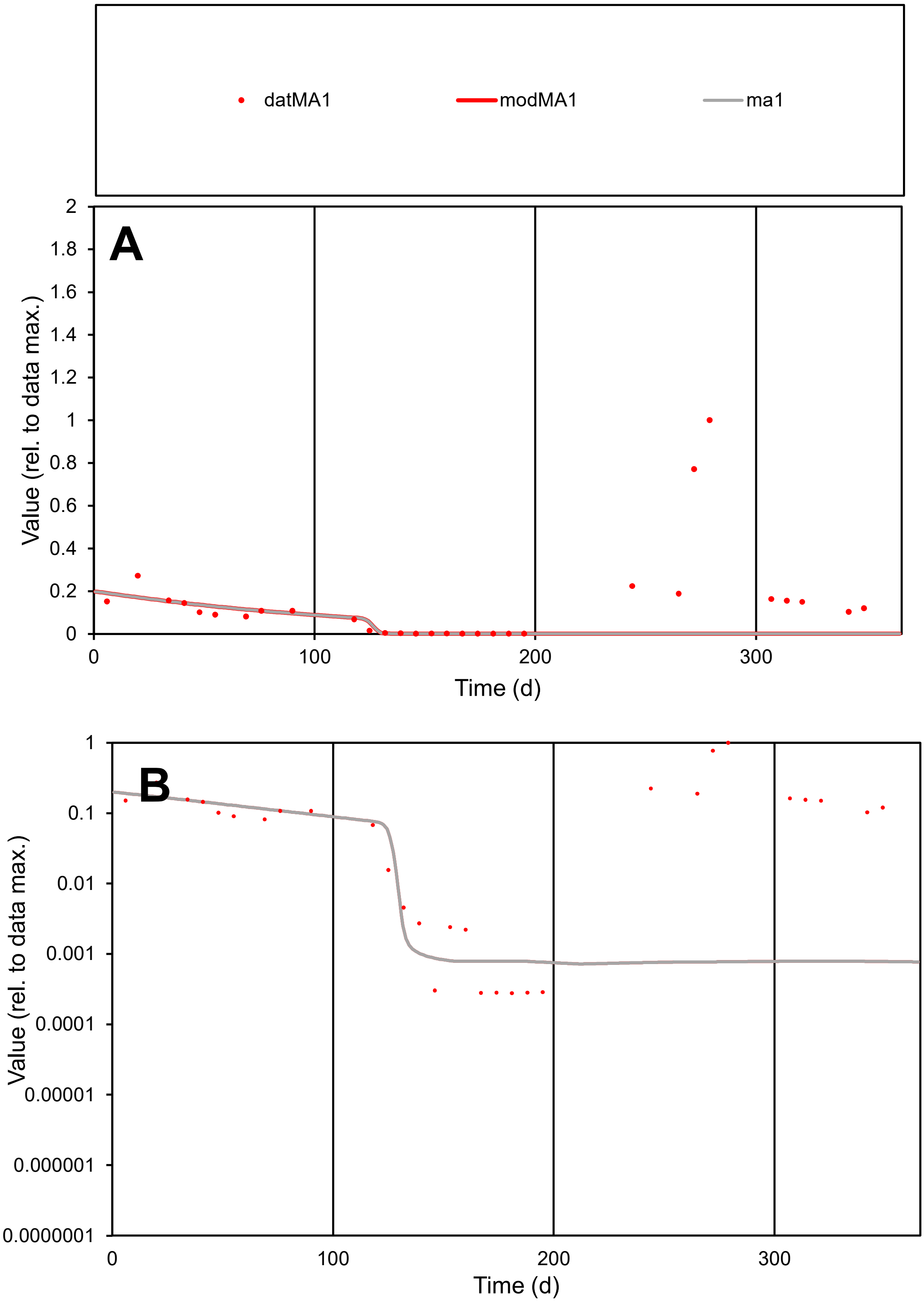


**C1**

Variables
MA1 Provide jtd or Dst
CalibOnly?
2 1 to reproduce of
oNormalize?
2 1=no, 2=yes

ofxn calc*
1.75129
ofxn mod
1.75732

sRun
RunA1
zout
1

Ave
Data
0.00718
Model
0.00232

data/vMax
0.05151

ofxng
275.646

Min d/m
0.36807

ave/min d
500.99
ave/min m
59.5934

325_Calib_S_A

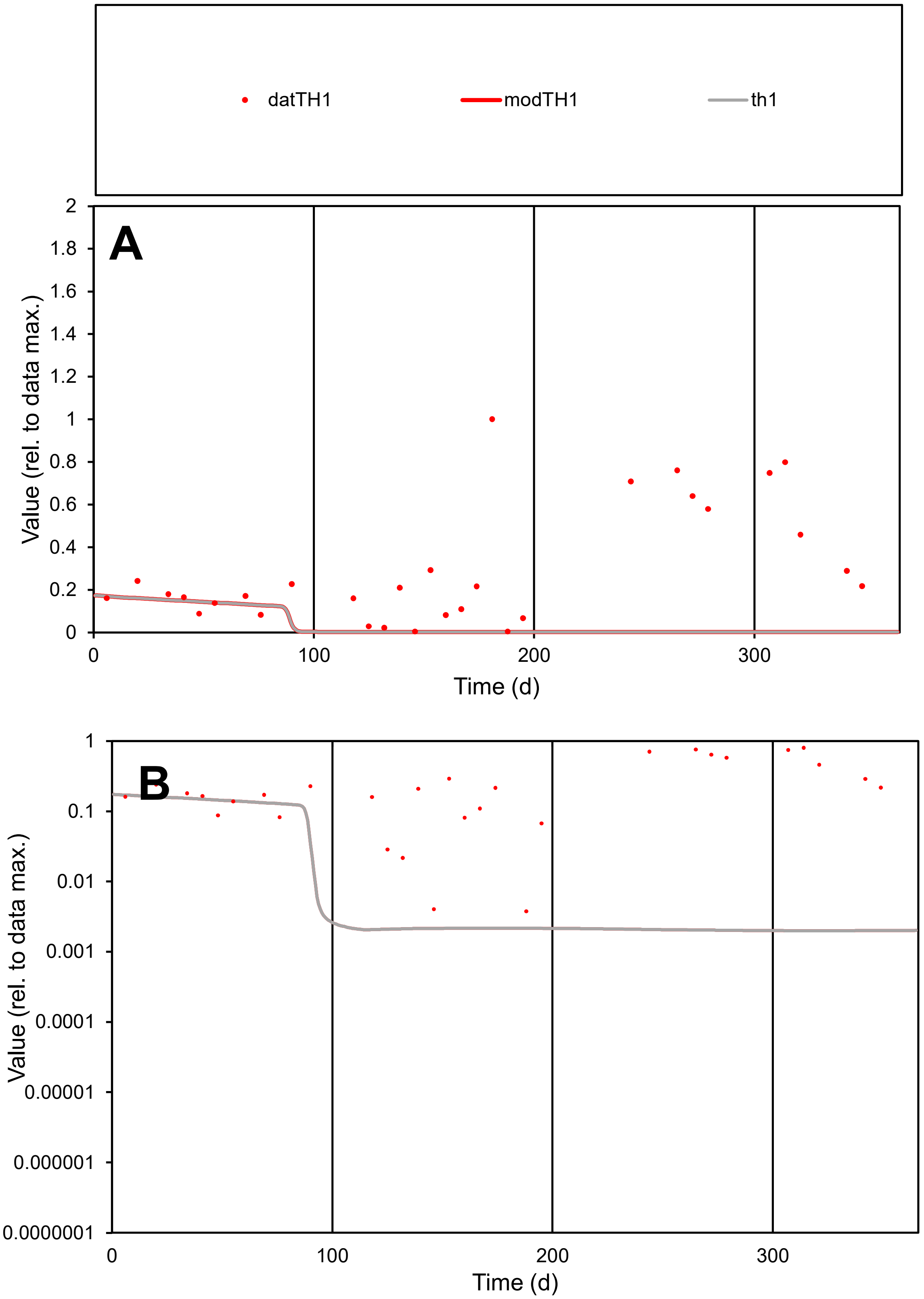


**C1**

| | |
|---|---|
| Variables | |
| TH1 | Provide jtd or Dst |
| CalibOnly? | |
| 2 | 1 to reproduce of |
| oNormalize? | |
| 2 | 1=no, 2=yes |
| ofxn calc* | |
| 1.32664 | |
| ofxn mod | |
| 1.33108 | |
| sRun | |
| RunA1 | |
| zout | |
| 1 | |
| Ave | |
| Data | |
| 0.00736 | |
| Model | |
| 0.00109 | |
| data/vMax | |
| 0.02496 | |
| ofxng | |
| 275.646 | |
| Min d/m | |
| 1.87848 | |
| ave/min d | |
| 78.5634 | |
| ave/min m | |
| 21.8767 | |

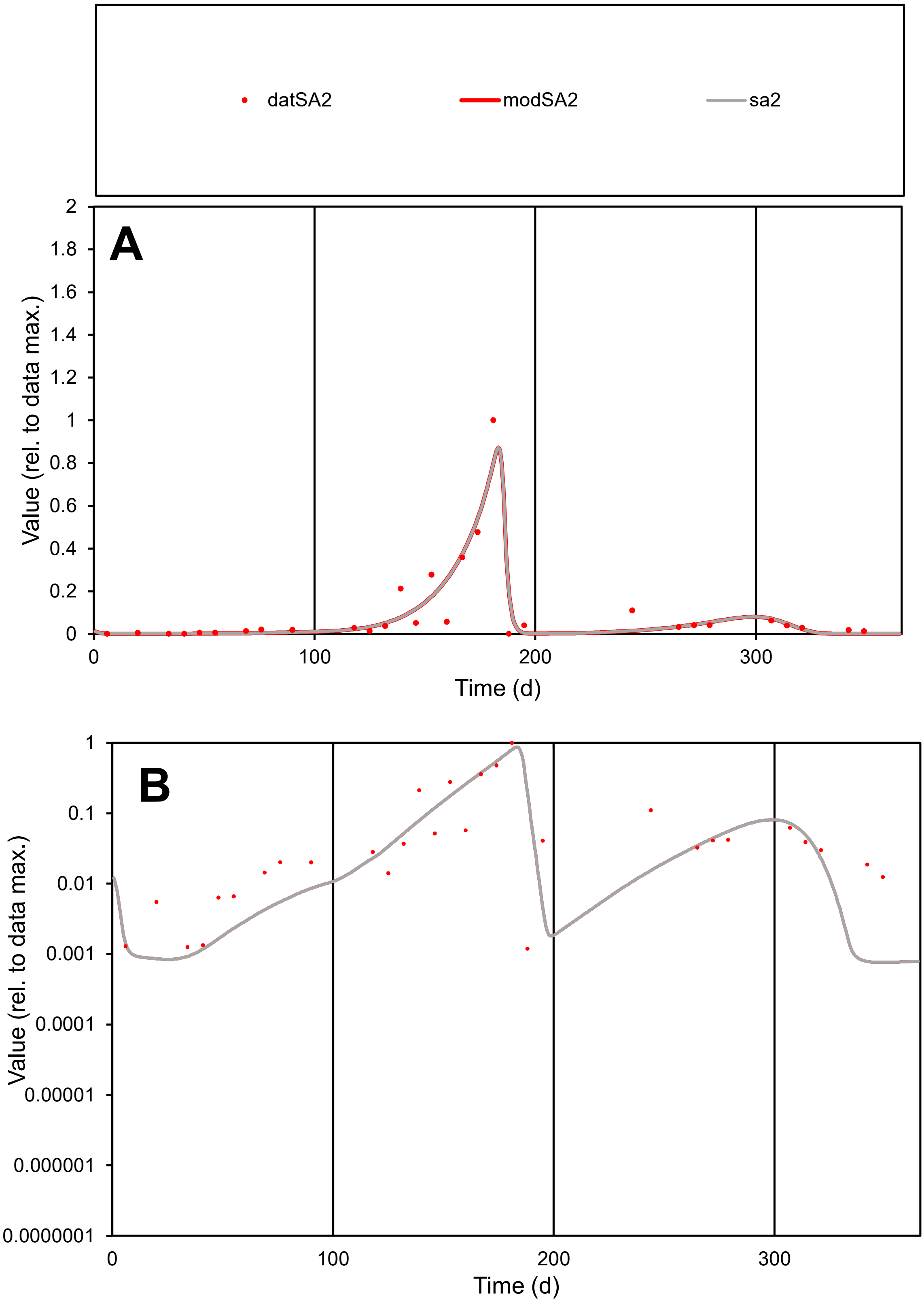


**C1**

Variables
SA2 Provide jtd or Dst
CalibOnly?
2 1 to reproduce of
oNormalize?
2 1=no, 2=yes

ofxn calc*
0.61662
ofxn mod
0.6669

sRun
RunA1
zout
1

Ave
Data
0.00866
Model
0.0086

data/vMax
0.08599

ofxng
275.646

Min d/m
1.55165

ave/min d
84.6655
ave/min m
130.45

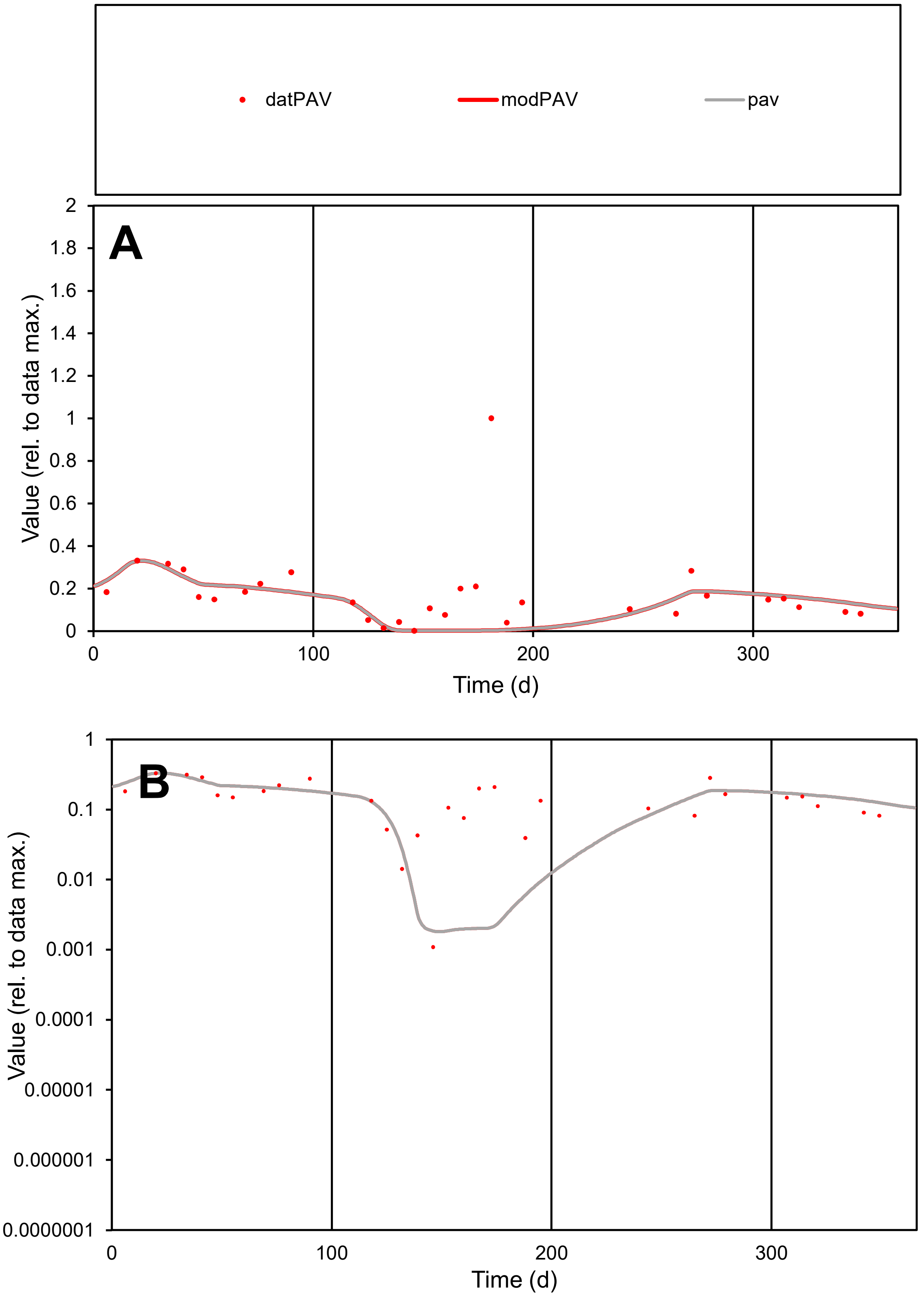


**C1**

Variables
PAV Provide jtd or Dst
CalibOnly?
2 1 to reproduce of
oNormalize?
2 1=no, 2=yes

ofxn calc*
1.10279
ofxn mod
1.10984

sRun
RunA1
zout
1

Ave
Data
0.00656
Model
0.0046

data/vMax
0.03685

ofxng
275.646

Min d/m
0.59857

ave/min d
163.088
ave/min m
68.4424

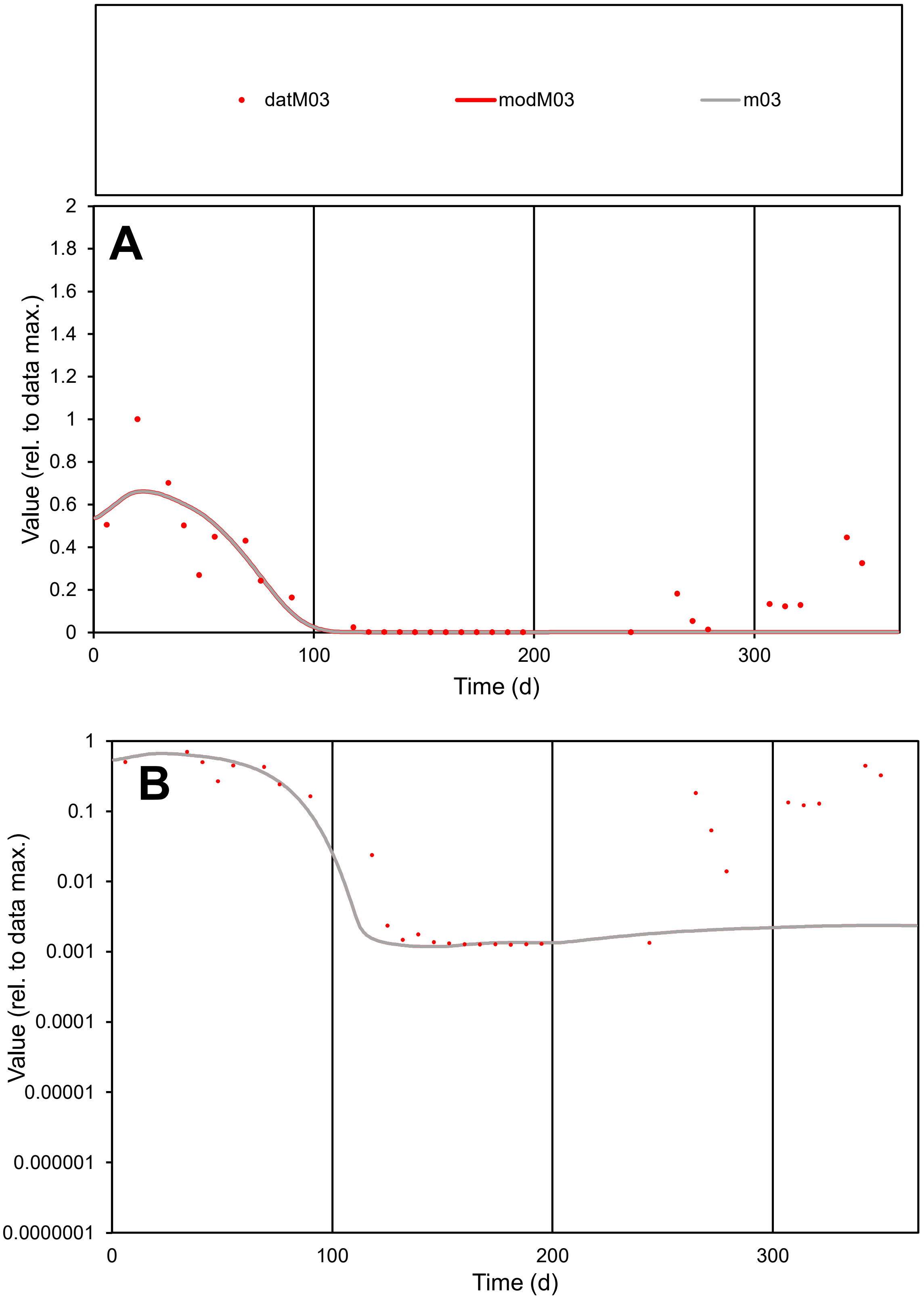


C1

Variables
M03 Provide jtd or Dst
CalibOnly?
2 1 to reproduce of
oNormalize?
2 1=no, 2=yes

ofxn calc*
0.7535
ofxn mod
0.77403

sRun
RunA1
zout
1

Ave
Data
0.00264
Model
0.00195

data/vMax
0.01386

ofxng
275.646

Min d/m
1.06147

ave/min d
151.262
ave/min m
118.65

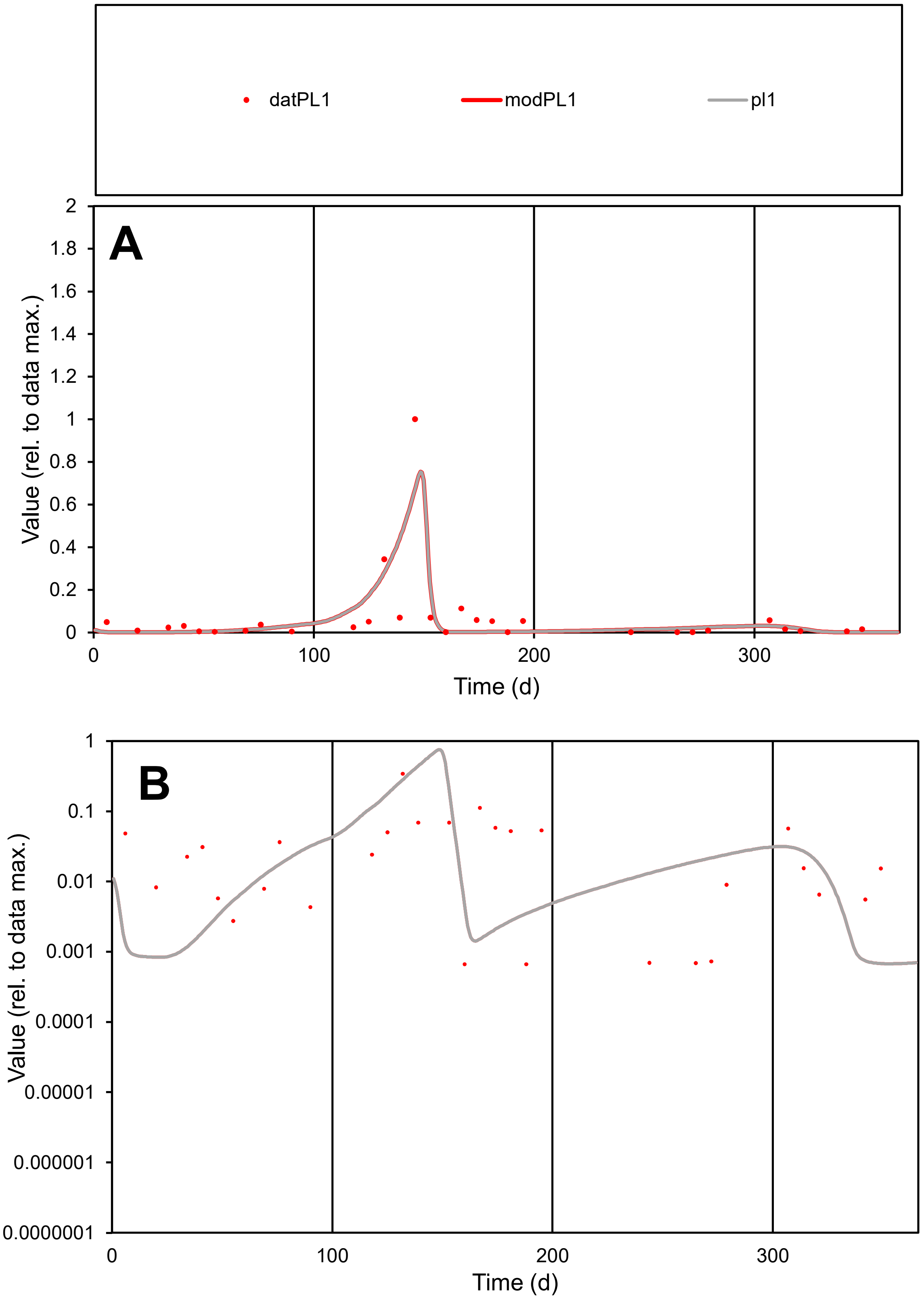


**C1**

Variables

PL1 Provide jtd or Dst

CalibOnly?

2 1 to reproduce of

oNormalize?

2 1=no, 2=yes

ofxn calc*

1.2881

ofxn mod

1.37554

sRun

RunA1

zout

1

Ave

Data

0.00371

Model

0.00369

data/vMax

0.05277

ofxng

275.646

Min d/m

0.98149

ave/min d

106.084

ave/min m

103.459

325_Calib_S_A

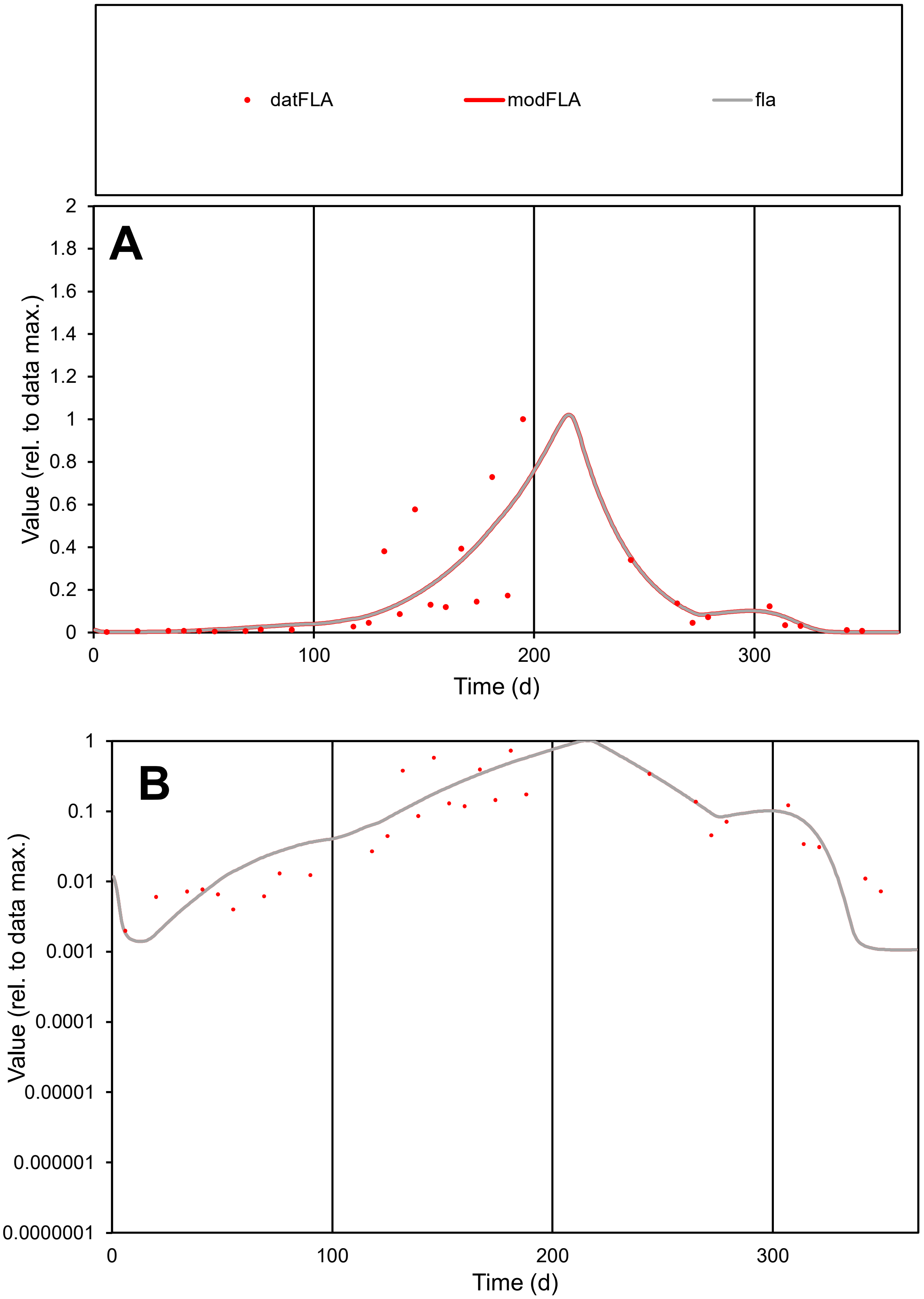


**C1**

Variables

FLA Provide jtd or Dst

CalibOnly?

2 1 to reproduce of

oNormalize?

2 1=no, 2=yes

ofxn calc*

0.9498

ofxn mod

0.97238

sRun

RunA1

zout

1

Ave

Data

0.0047

Model

0.00463

data/vMax

0.03019

ofxng

275.646

Min d/m

1.85185

ave/min d

78.2627

ave/min m

142.944

325_Calib_S_A

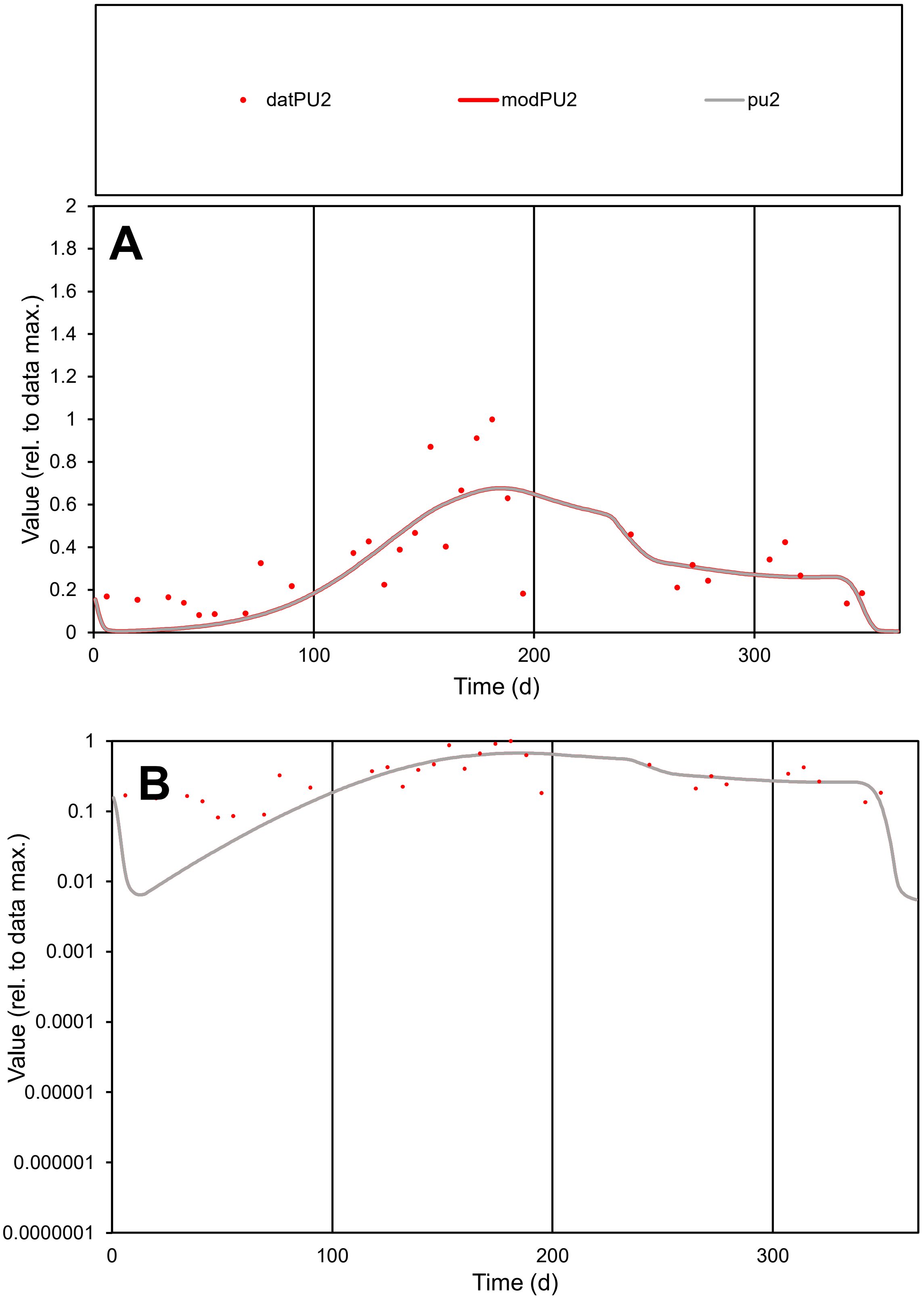


**C1**

Variables
PU2 Provide jtd or Dst
CalibOnly?
2 1 to reproduce of
oNormalize?
2 1=no, 2=yes

ofxn calc*
0.45735
ofxn mod
0.47637

sRun
RunA1
zout
1

Ave
Data
0.00463
Model
0.00421

data/vMax
0.01317

ofxng
275.646

Min d/m
9.25065

ave/min d
4.30999
ave/min m
36.2835

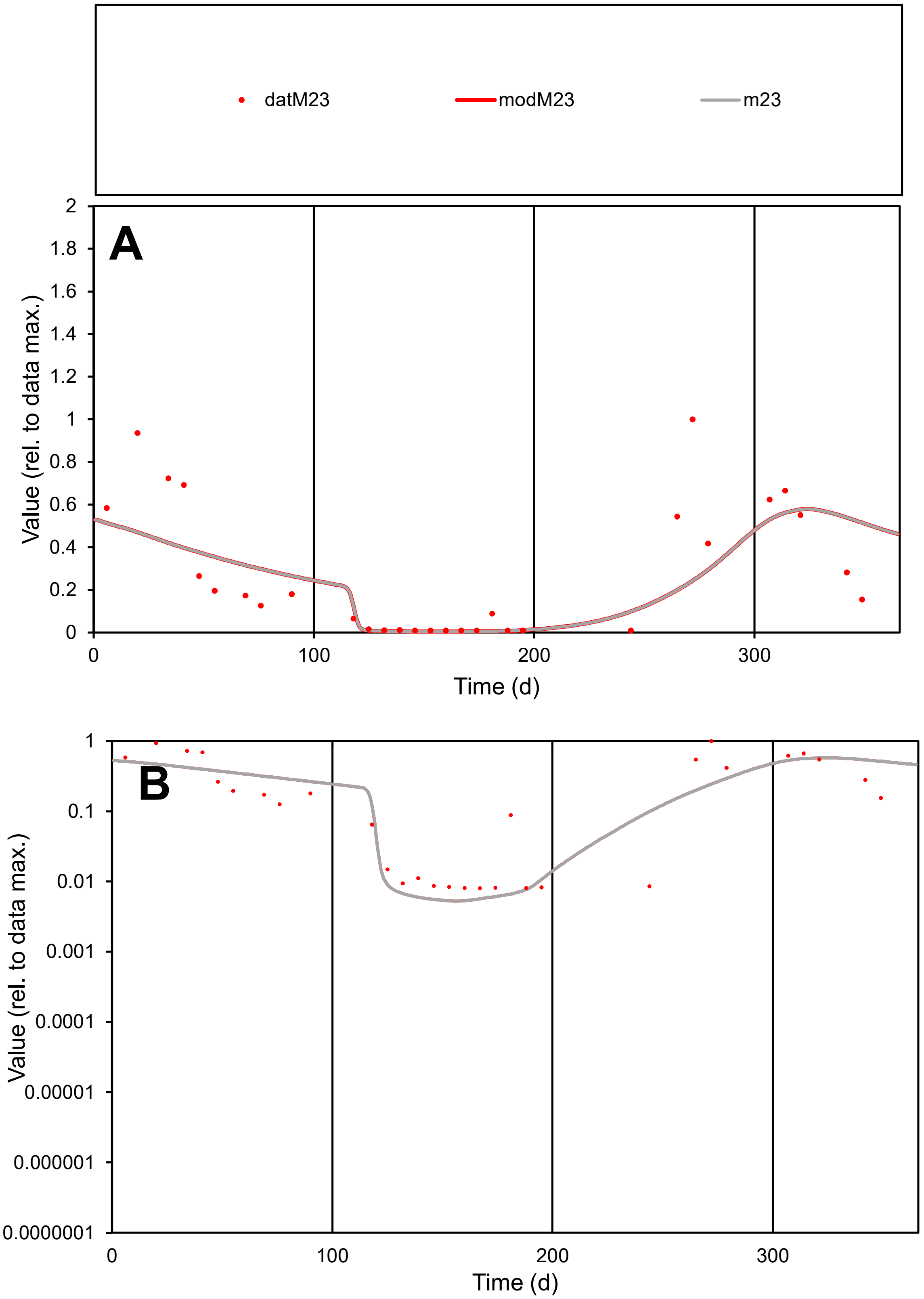


**C1**

Variables
M23 Provide jtd or Dst
CalibOnly?
2 1 to reproduce of
oNormalize?
2 1=no, 2=yes

ofxn calc*
0.77677
ofxn mod
0.79092

sRun
RunA1
zout
1

Ave
Data
0.00373
Model
0.00317

data/vMax
0.01338

ofxng
275.646

Min d/m
1.51849

ave/min d
34.6434
ave/min m
44.7632

325_Calib_S_A

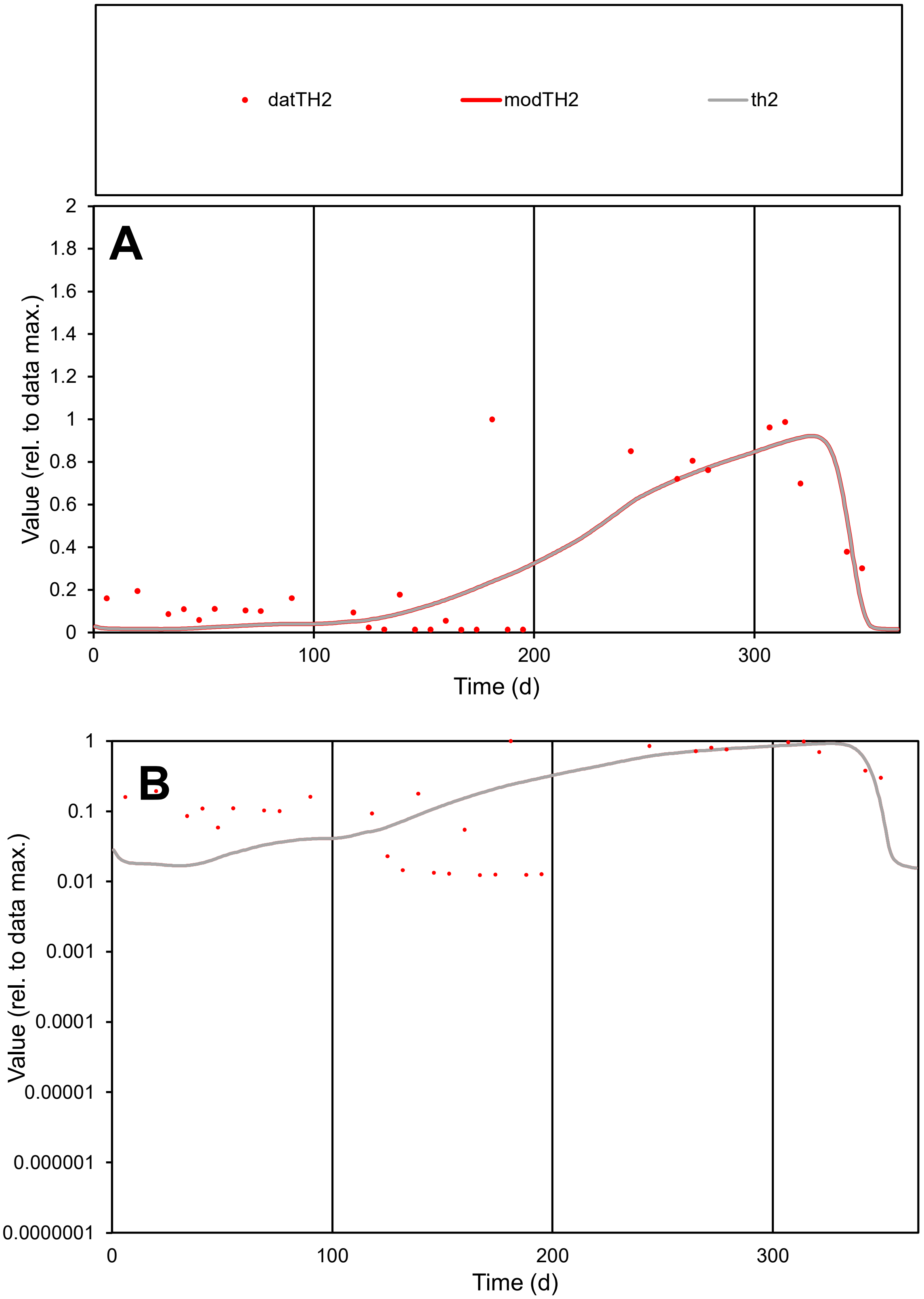


**C1**

Variables
TH2 Provide jtd or Dst
CalibOnly?
2 1 to reproduce of
oNormalize?
2 1=no, 2=yes

ofxn calc*
0.63331
ofxn mod
0.65883

sRun
RunA1
zout
1

Ave
Data
0.00256
Model
0.00236

data/vMax
0.00855

ofxng
275.646

Min d/m
0.72287

ave/min d
24.1806
ave/min m
16.1204

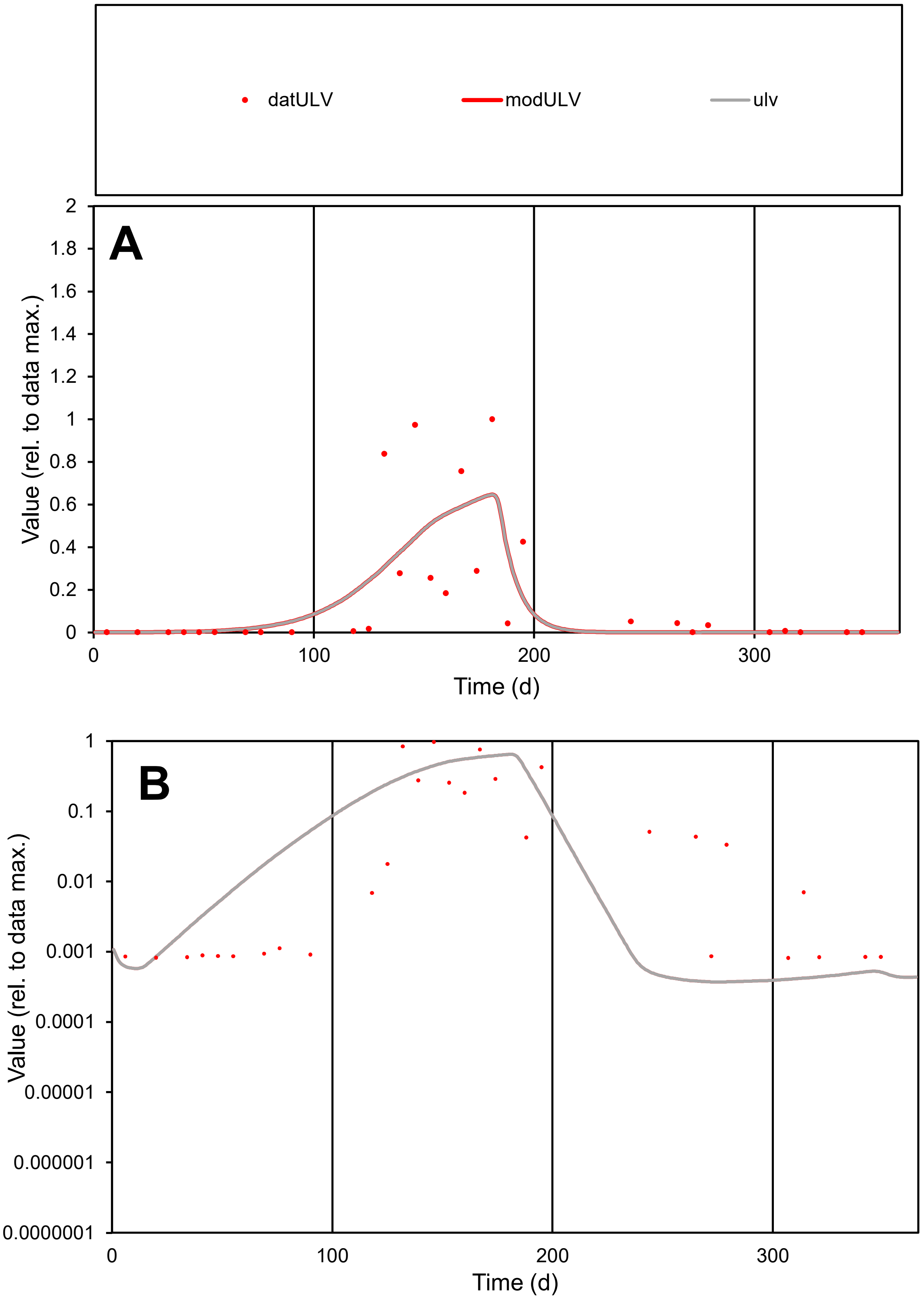


**C1**

Variables
ULV Provide jtd or Dst
CalibOnly?
2 1 to reproduce of
oNormalize?
2 1=no, 2=yes

ofxn calc*
1.17527
ofxn mod
1.19511

sRun
RunA1
zout
1

Ave
Data
0.00381
Model
0.00371

data/vMax
0.02191

ofxng
275.646

Min d/m
2.2041

ave/min d
211.96
ave/min m
454.905

325_Calib_S_A

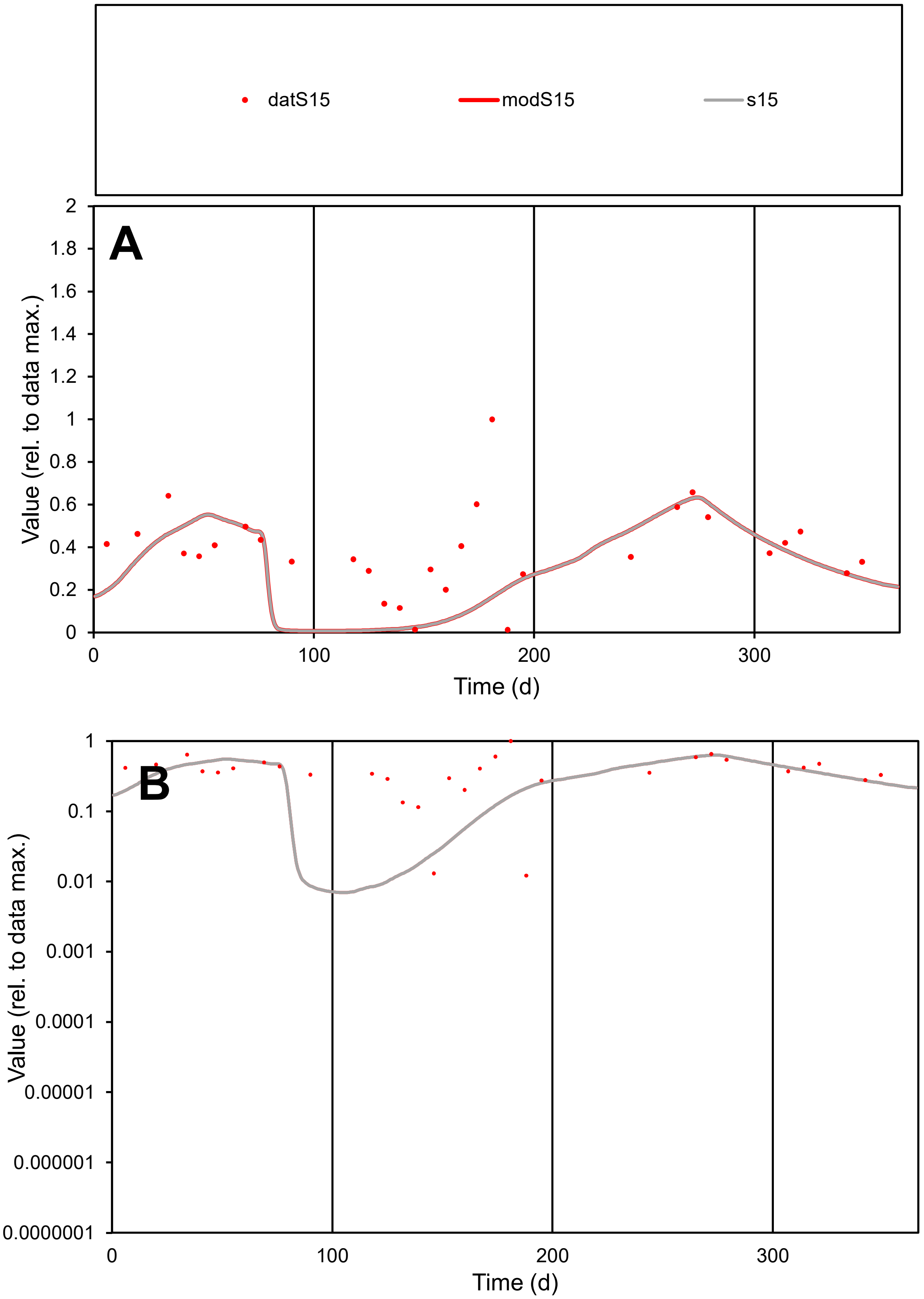


**C1**

Variables
S15 Provide jtd or Dst
CalibOnly?
2 1 to reproduce of
oNormalize?
2 1=no, 2=yes

ofxn calc*
0.60542
ofxn mod
0.61695

sRun
RunA1
zout
1

Ave
Data
0.00399
Model
0.00295

data/vMax
0.01032

ofxng
275.646

Min d/m
1.47485

ave/min d
31.9072
ave/min m
34.7637

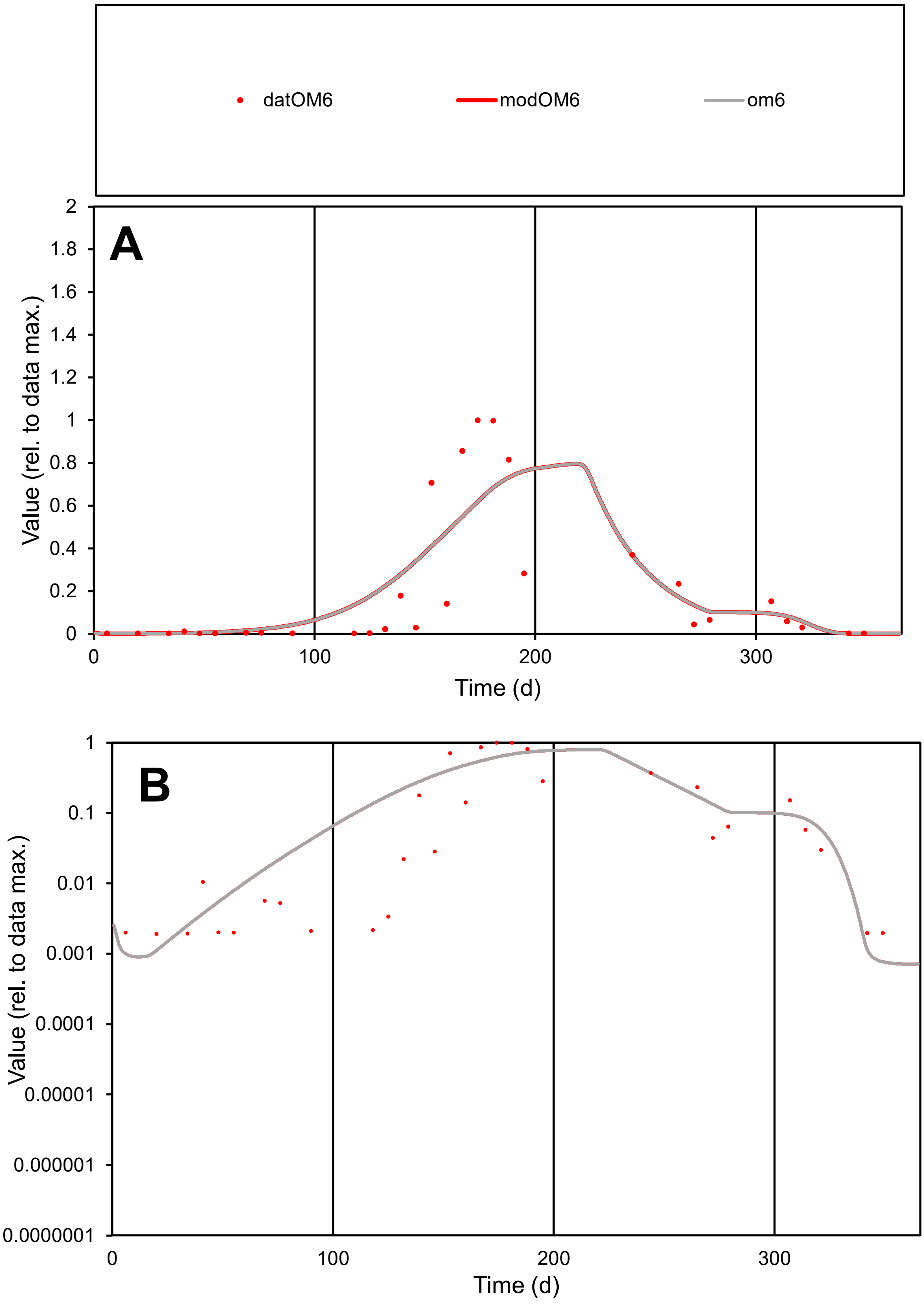


**C1**

Variables
OM6 Provide jtd or Dst
CalibOnly?
2 1 to reproduce of
oNormalize?
2 1=no, 2=yes

ofxn calc*
0.8937
ofxn mod
0.91337

sRun
RunA1
zout
1

Ave
Data
0.00275
Model
0.00298

data/vMax
0.0137

ofxng
275.646

Min d/m
2.54607

ave/min d
104.781
ave/min m
289.366

325_Calib_S_A

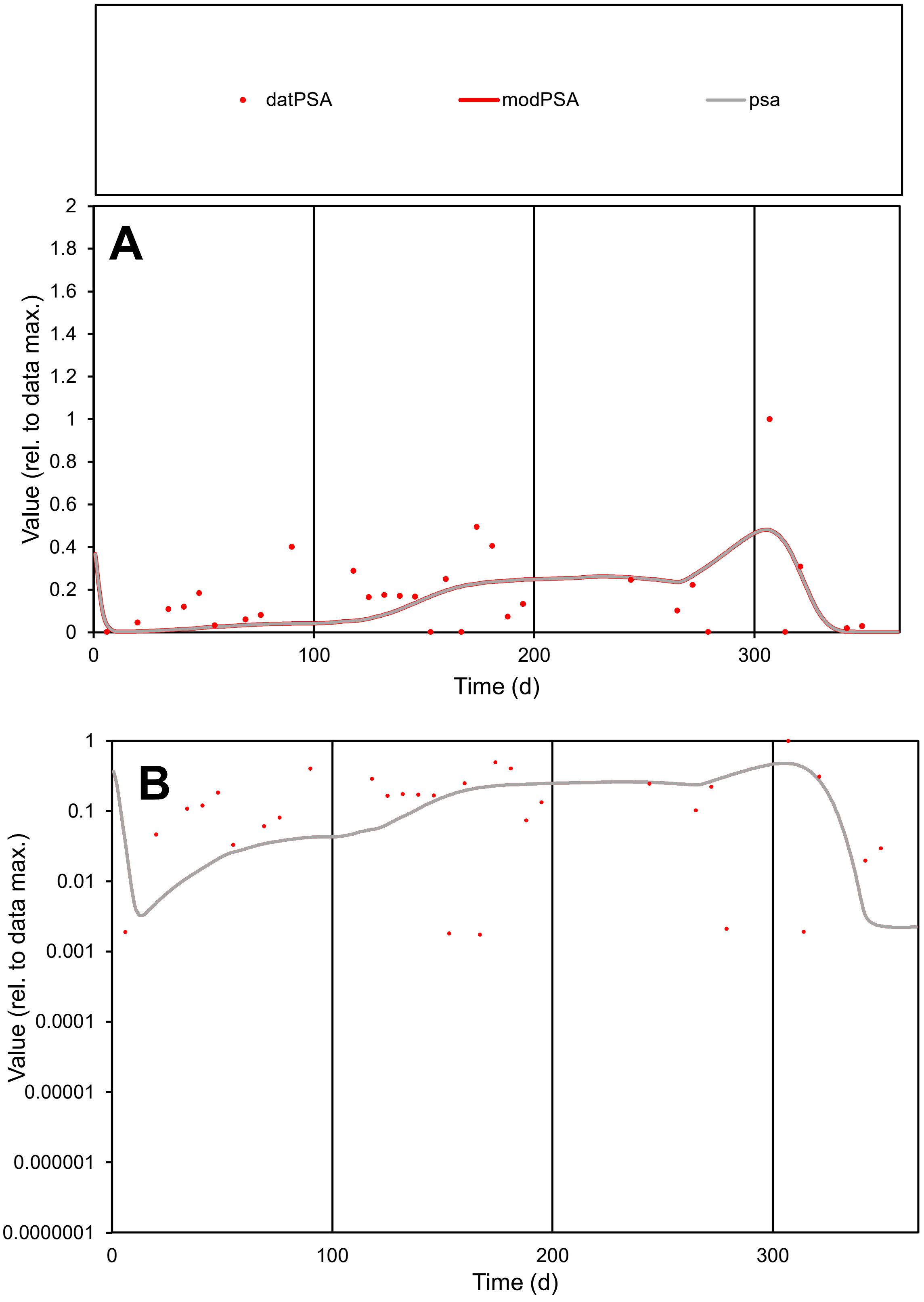


**C1**

Variables
PSA Provide jtd or Dst
CalibOnly?
2 1 to reproduce of
oNormalize?
2 1=no, 2=yes

ofxn calc*
1.03872
ofxn mod
1.05321

sRun
RunA1
zout
1

Ave
Data
0.0009
Model
0.00075

data/vMax
0.00509

ofxng
275.646

Min d/m
0.76534

ave/min d
101.611
ave/min m
64.5401

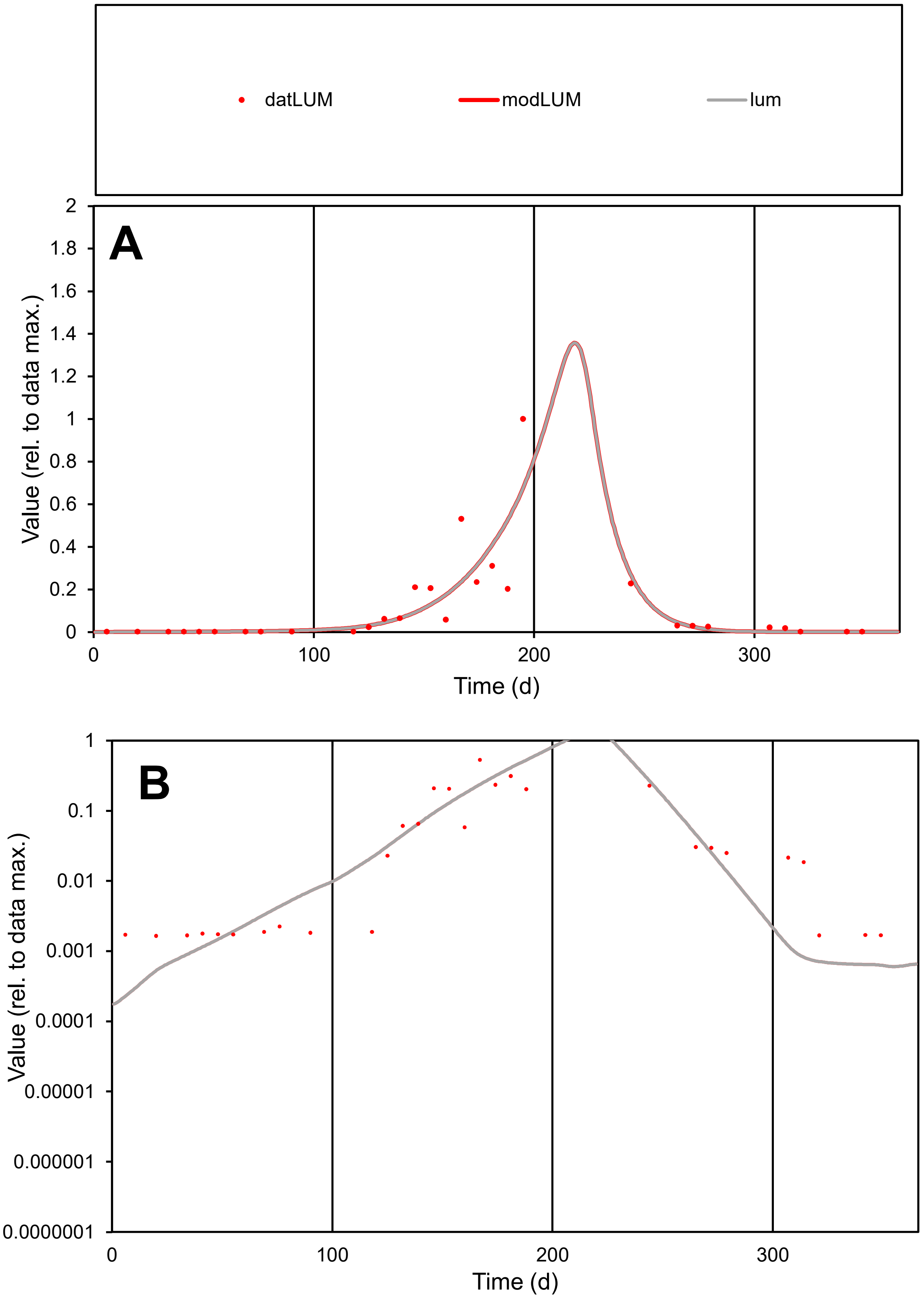


**C1**

Variables
LUM Provide jtd or Dst
CalibOnly?
2 1 to reproduce of
oNormalize?
2 1=no, 2=yes

ofxn calc*
0.95272
ofxn mod
0.98934

sRun
RunA1
zout
1

Ave
Data
0.00311
Model
0.003

data/vMax
0.0284

ofxng
275.646

Min d/m
6.74232

ave/min d
66.1651
ave/min m
430.271

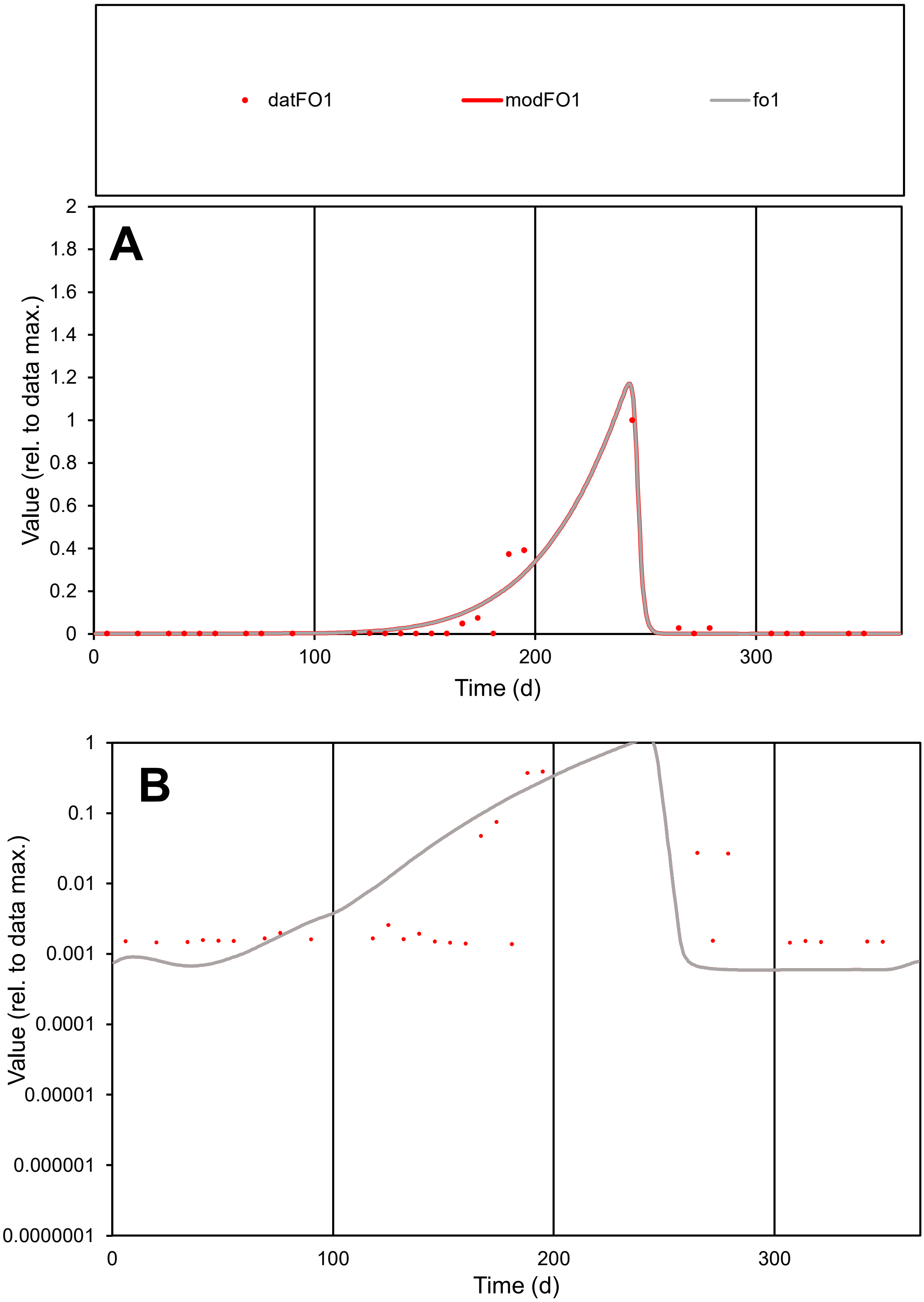


**C1**

Variables
FO1 Provide jtd or Dst
CalibOnly?
2 1 to reproduce of
oNormalize?
2 1=no, 2=yes

ofxn calc*
0.74716
ofxn mod
0.80683

sRun
RunA1
zout
1

Ave
Data
0.00276
Model
0.00302

data/vMax
0.04184

ofxng
275.646

Min d/m
2.30963

ave/min d
47.7364
ave/min m
120.632

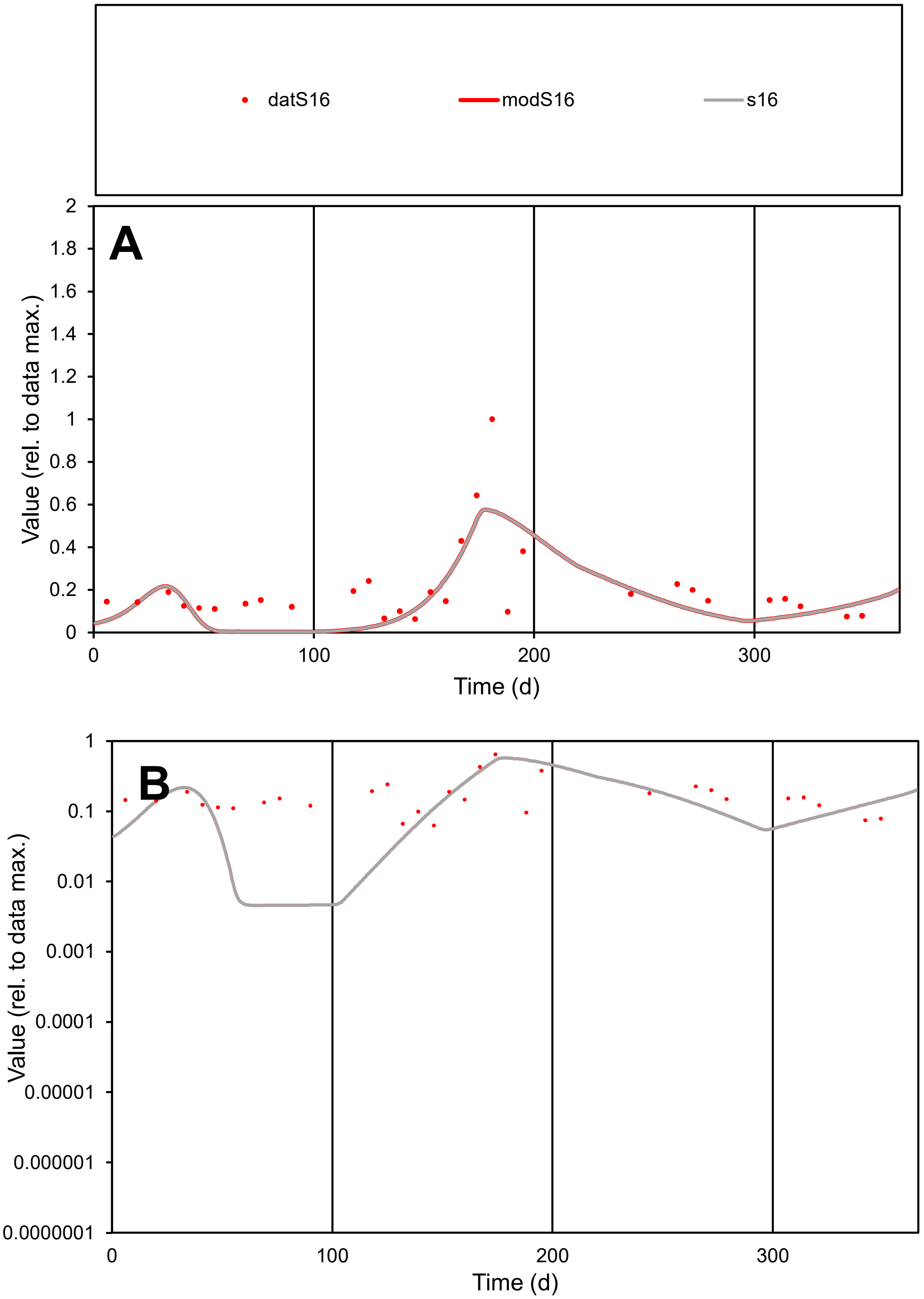


C1

Variables
S16 Provide jtd or Dst
CalibOnly?
2 1 to reproduce of
oNormalize?
2 1=no, 2=yes

ofxn calc*
0.69785
ofxn mod
0.71284

sRun
RunA1
zout
1

Ave
Data
0.00551
Model
0.00442

data/vMax
0.02702

ofxng
275.646

Min d/m
13.753

ave/min d
3.234
ave/min m
35.6592

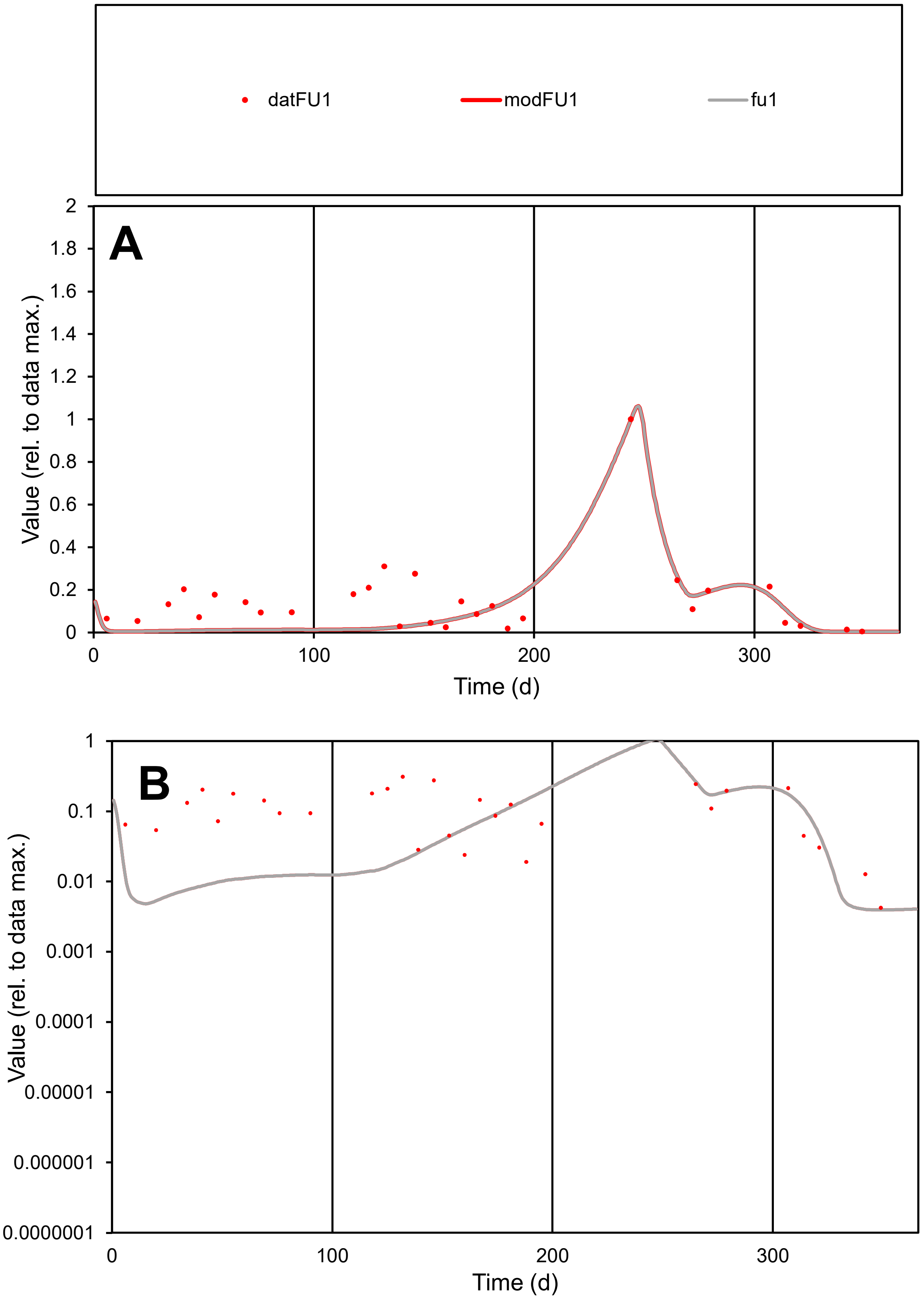


**C1**

Variables
FU1 Provide jtd or Dst
CalibOnly?
2 1 to reproduce of
oNormalize?
2 1=no, 2=yes

ofxn calc*
0.75841
ofxn mod
0.77168

sRun
RunA1
zout
1

Ave
Data
0.00174
Model
0.00115

data/vMax
0.01185

ofxng
275.646

Min d/m
1.07048

ave/min d
34.7189
ave/min m
24.5625

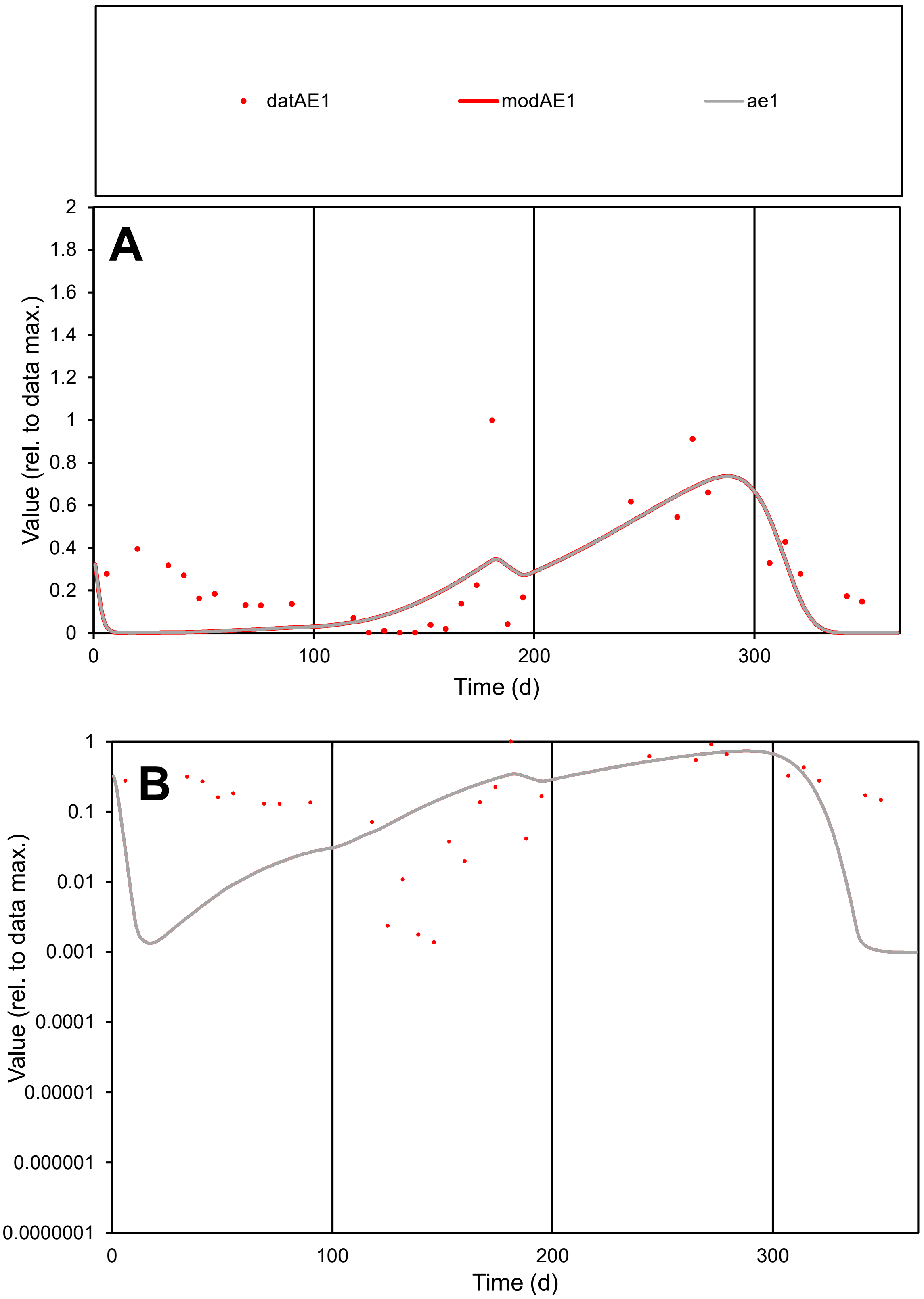


C1

Variables
AE1 Provide jtd or Dst
CalibOnly?
2 1 to reproduce of
oNormalize?
2 1=no, 2=yes

ofxn calc*
0.77346
ofxn mod
0.80567

sRun
RunA1
zout
1

Ave
Data
0.00284
Model
0.00225

data/vMax
0.01091

ofxng
275.646

Min d/m
1.35036

ave/min d
188.409
ave/min m
201.631

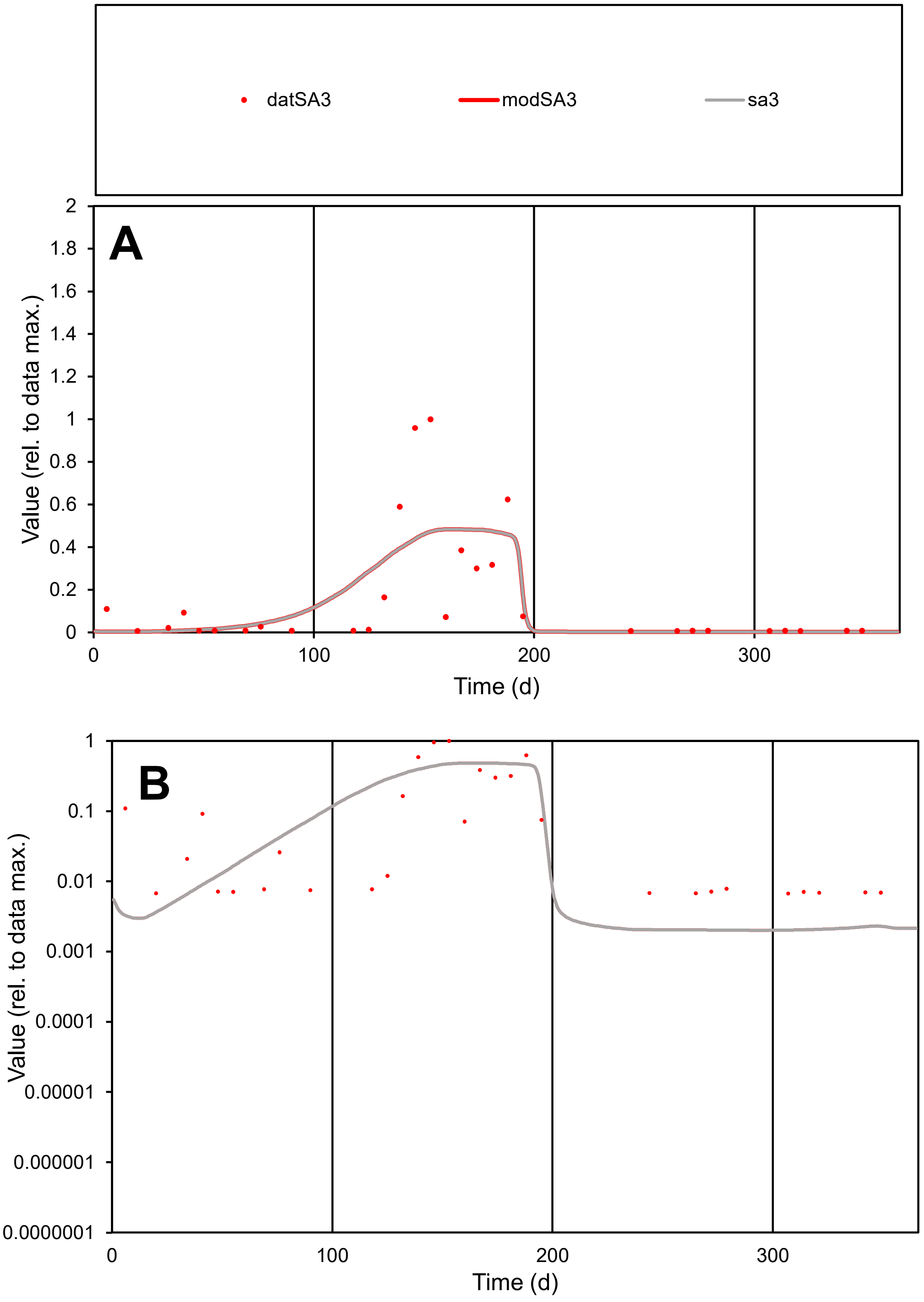


**C1**

Variables

SA3 Provide jtd or Dst

CalibOnly?

2 1 to reproduce of

oNormalize?

2 1=no, 2=yes

ofxn calc*

1.13692

ofxn mod

1.15314

sRun

RunA1

zout

1

Ave

Data

0.00246

Model

0.00246

data/vMax

0.01522

ofxng

275.646

Min d/m

3.33225

ave/min d

23.9845

ave/min m

79.954

325_Calib_S_A

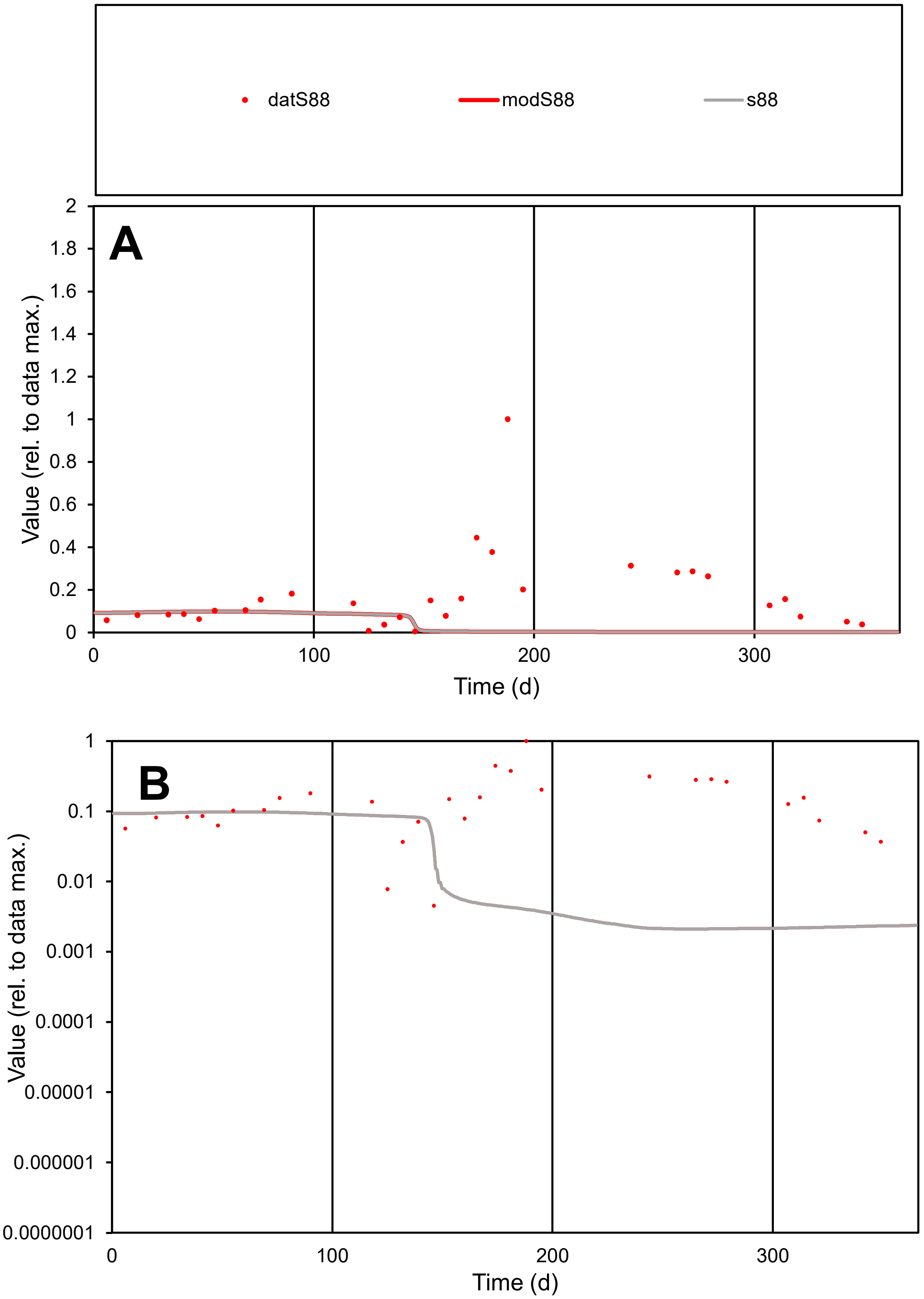


**C1**

Variables

S88 Provide jtd or Dst

CalibOnly?

2 1 to reproduce of

oNormalize?

2 1=no, 2=yes

ofxn calc*

1.42568

ofxn mod

1.42797

sRun

RunA1

zout

1

Ave

Data

0.00238

Model

0.00058

data/vMax

0.0138

ofxng

275.646

Min d/m

2.14364

ave/min d

38.1233

ave/min m

19.9979

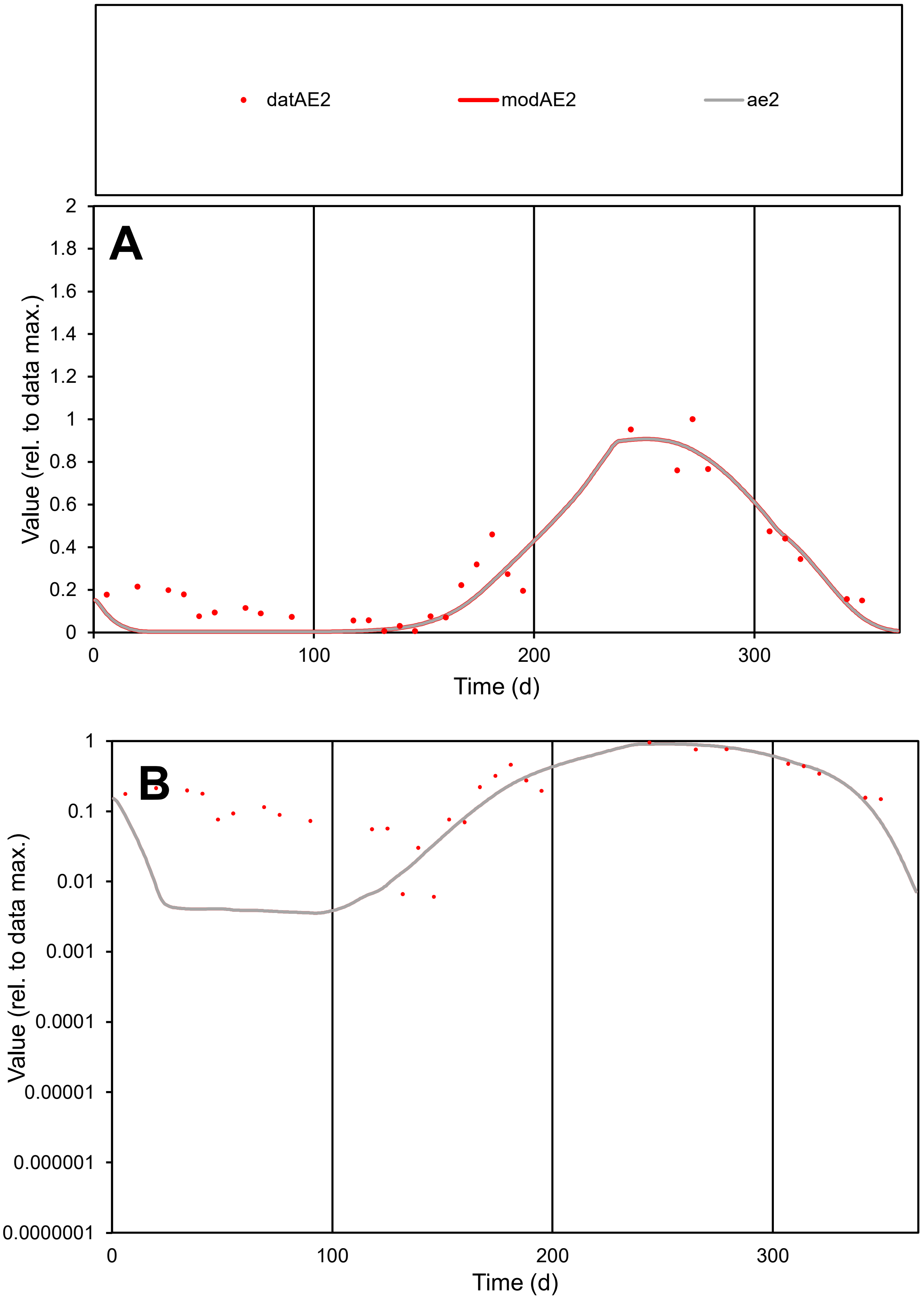


**C1**

Variables
AE2 Provide jtd or Dst
CalibOnly?
2 1 to reproduce of
oNormalize?
2 1=no, 2=yes

ofxn calc*
0.38653
ofxn mod
0.42132

sRun
RunA1
zout
1

Ave
Data
0.00275
Model
0.00225

data/vMax
0.01026

ofxng
275.646

Min d/m
1.71979

ave/min d
43.9394
ave/min m
61.854

325_Calib_S_A

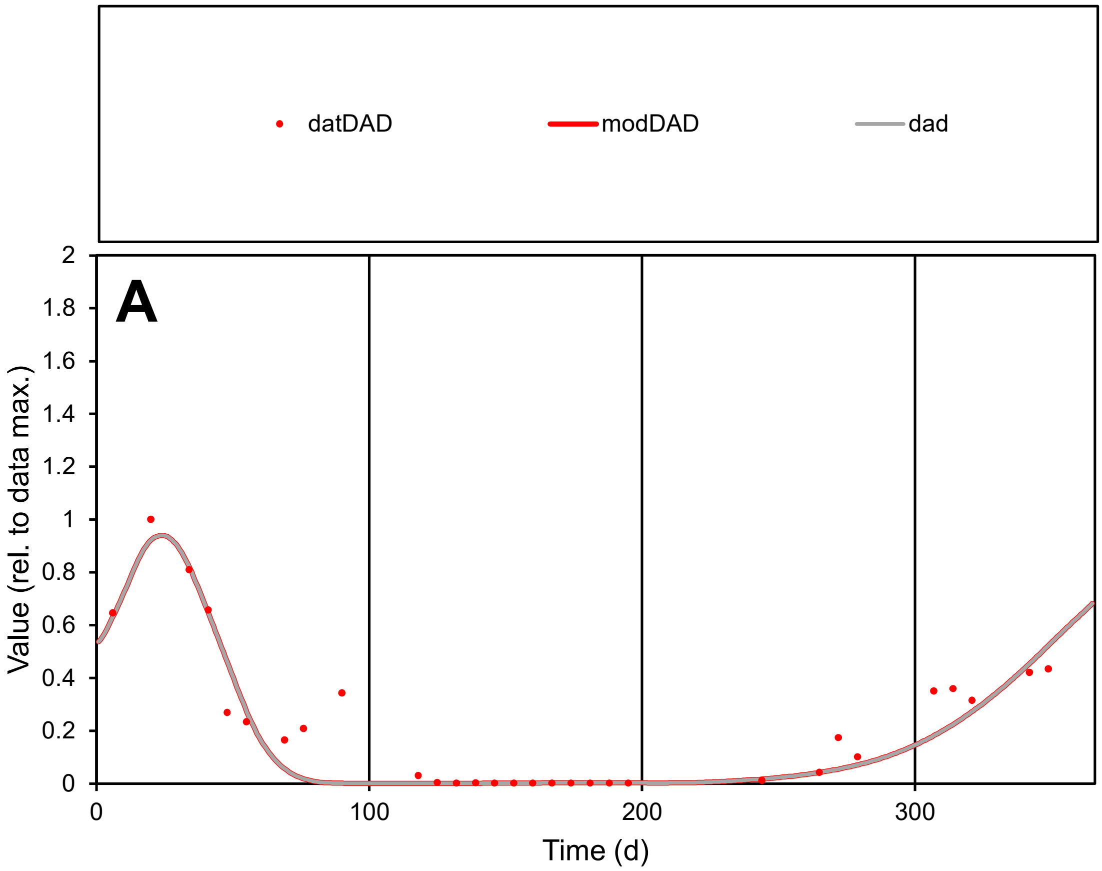


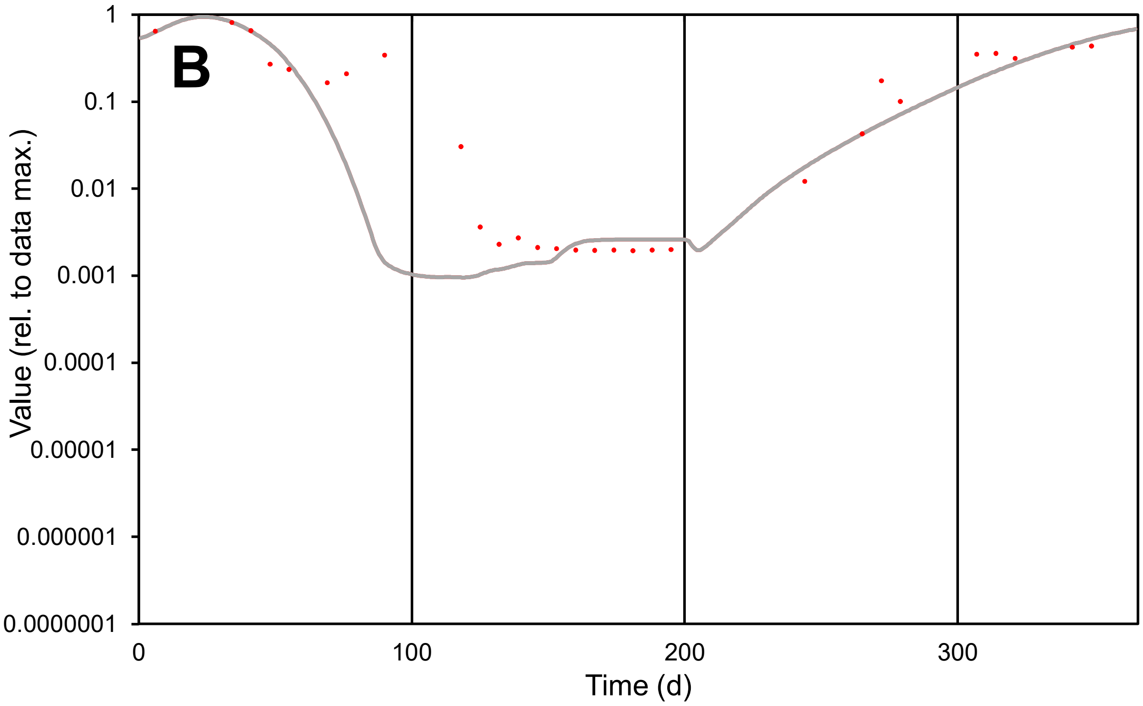


**C1**

Variables
DAD Provide jtd or Dst
CalibOnly?
2 1 to reproduce of
oNormalize?
2 1=no, 2=yes

ofxn calc*
0.42823
ofxn mod
0.46148

sRun
RunA1
zout
1

Ave
Data
0.00174
Model
0.00149

data/vMax
0.0079

ofxng
275.646

Min d/m
2.03797

ave/min d
113.144
ave/min m
197.549

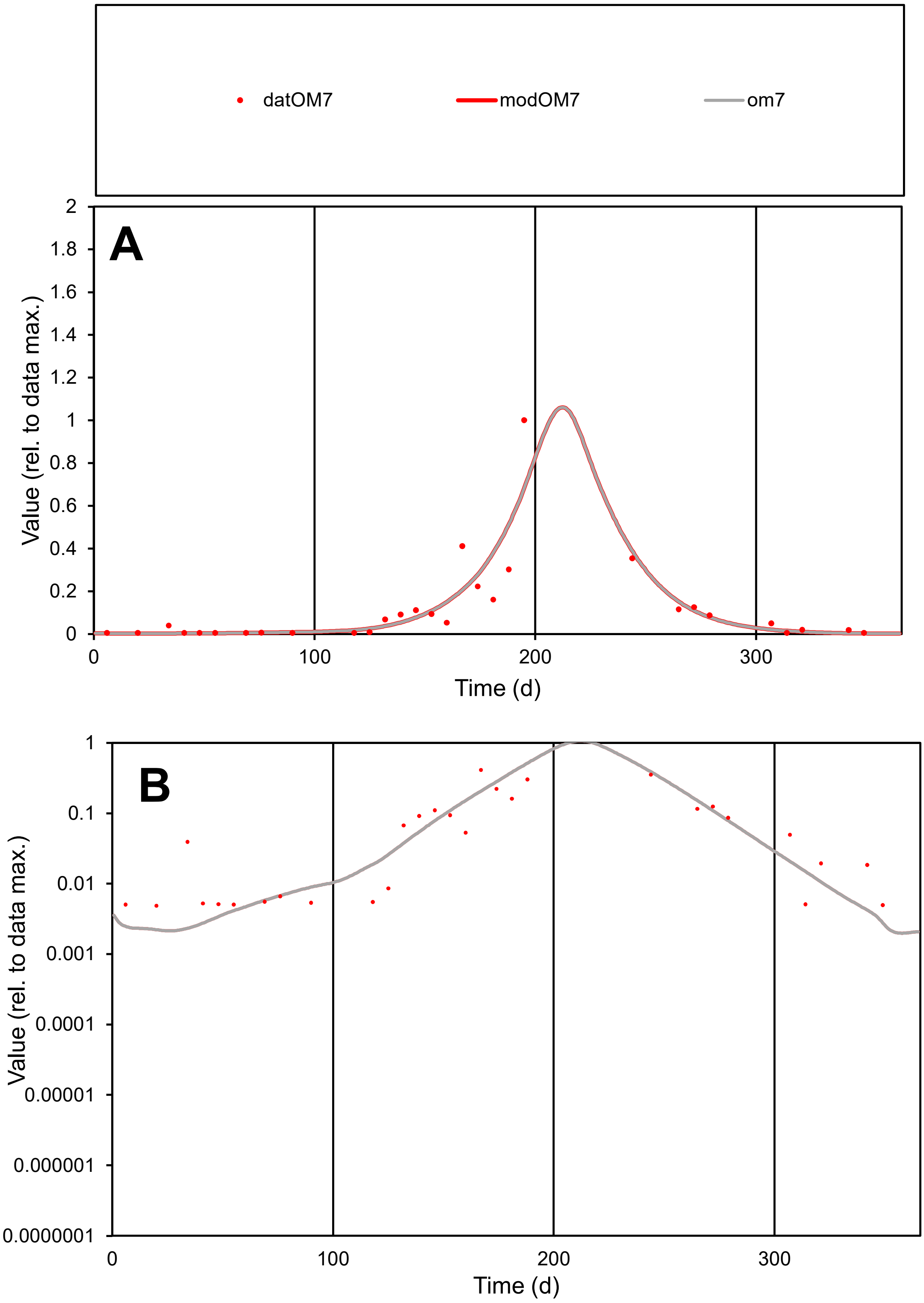


**C1**

Variables
OM7 Provide jtd or Dst
CalibOnly?
2 1 to reproduce of
oNormalize?
2 1=no, 2=yes

ofxn calc*
0.78481
ofxn mod
0.82372

sRun
RunA1
zout
1

Ave
Data
0.00228
Model
0.0023

data/vMax
0.02021

ofxng
275.646

Min d/m
2.20108

ave/min d
23.2988
ave/min m
51.7521

325_Calib_S_A

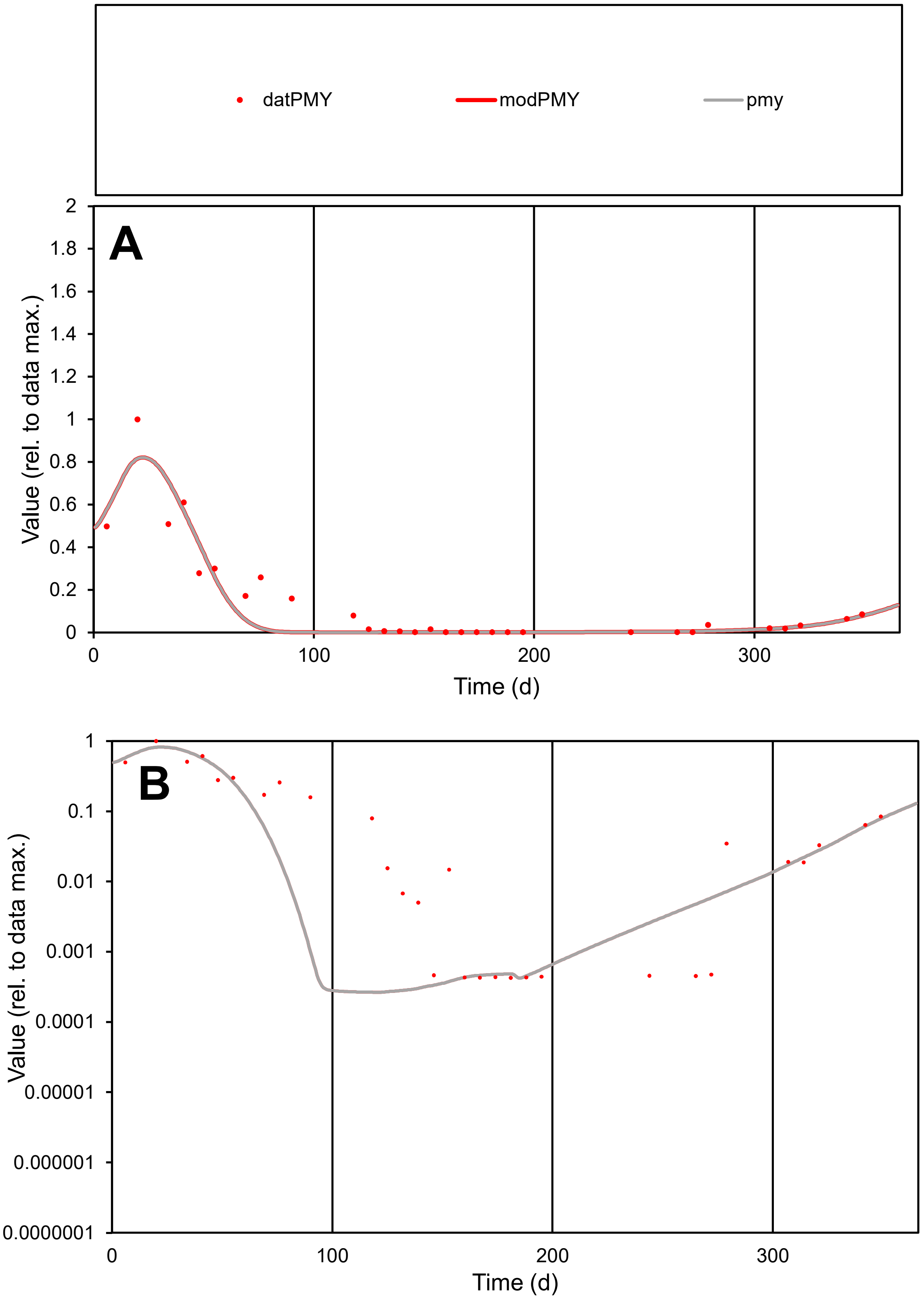


**C1**

Variables

PMY Provide jtd or Dst

CalibOnly?

2 1 to reproduce of

oNormalize?

2 1=no, 2=yes

ofxn calc*

0.57677

ofxn mod

0.61536

sRun

RunA1

zout

1

Ave

Data

0.00311

Model

0.00269

data/vMax

0.02242

ofxng

275.646

Min d/m

1.60828

ave/min d

325.368

ave/min m

453.189

325_Calib_S_A

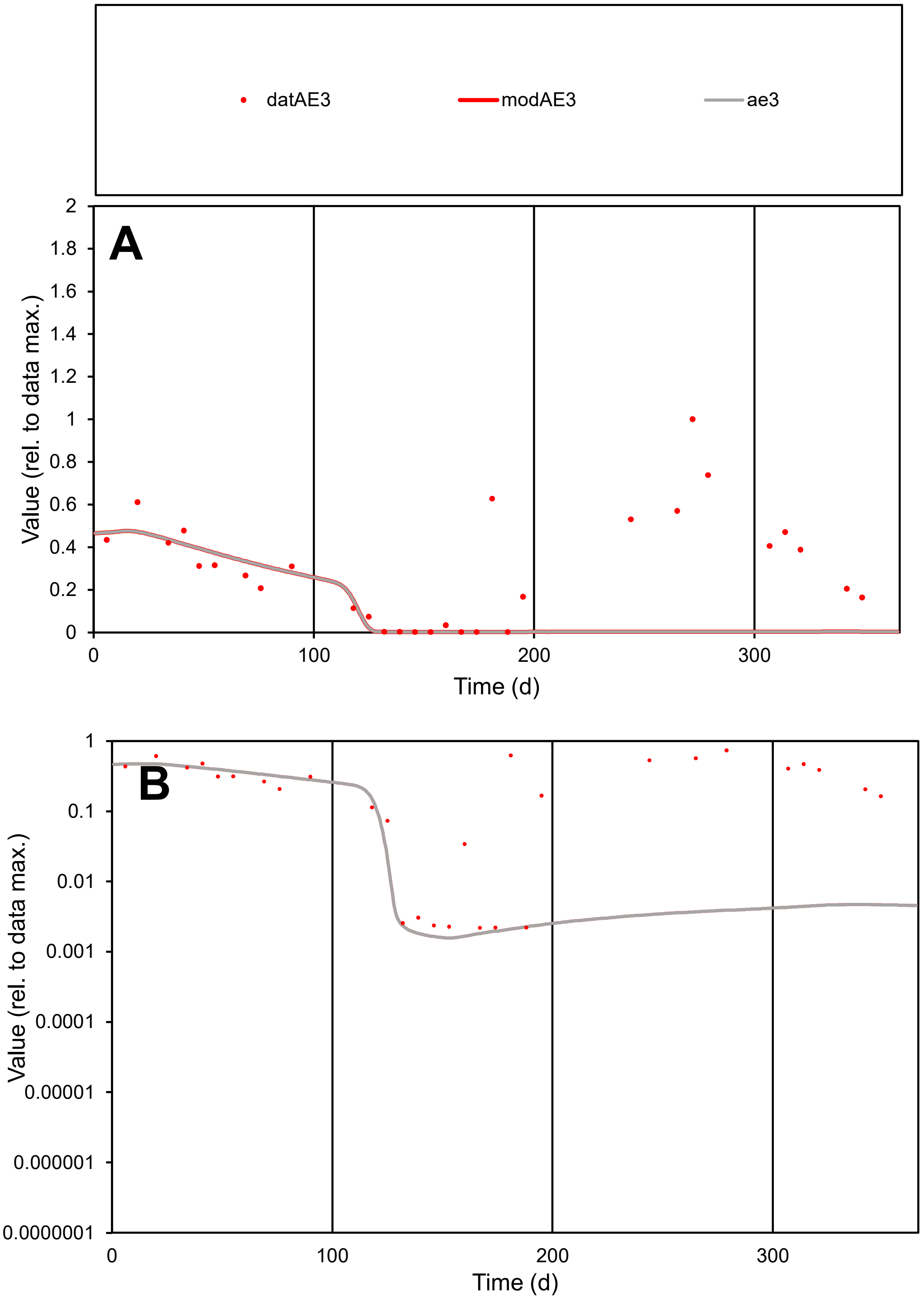


**C1**

Variables

AE3 Provide jtd or Dst

CalibOnly?

2 1 to reproduce of

oNormalize?

2 1=no, 2=yes

ofxn calc*

1.10474

ofxn mod

1.1104

sRun

RunA1

zout

1

Ave

Data

0.00227

Model

0.00094

data/vMax

0.00768

ofxng

275.646

Min d/m

1.39371

ave/min d

134.534

ave/min m

77.7127

325_Calib_S_A

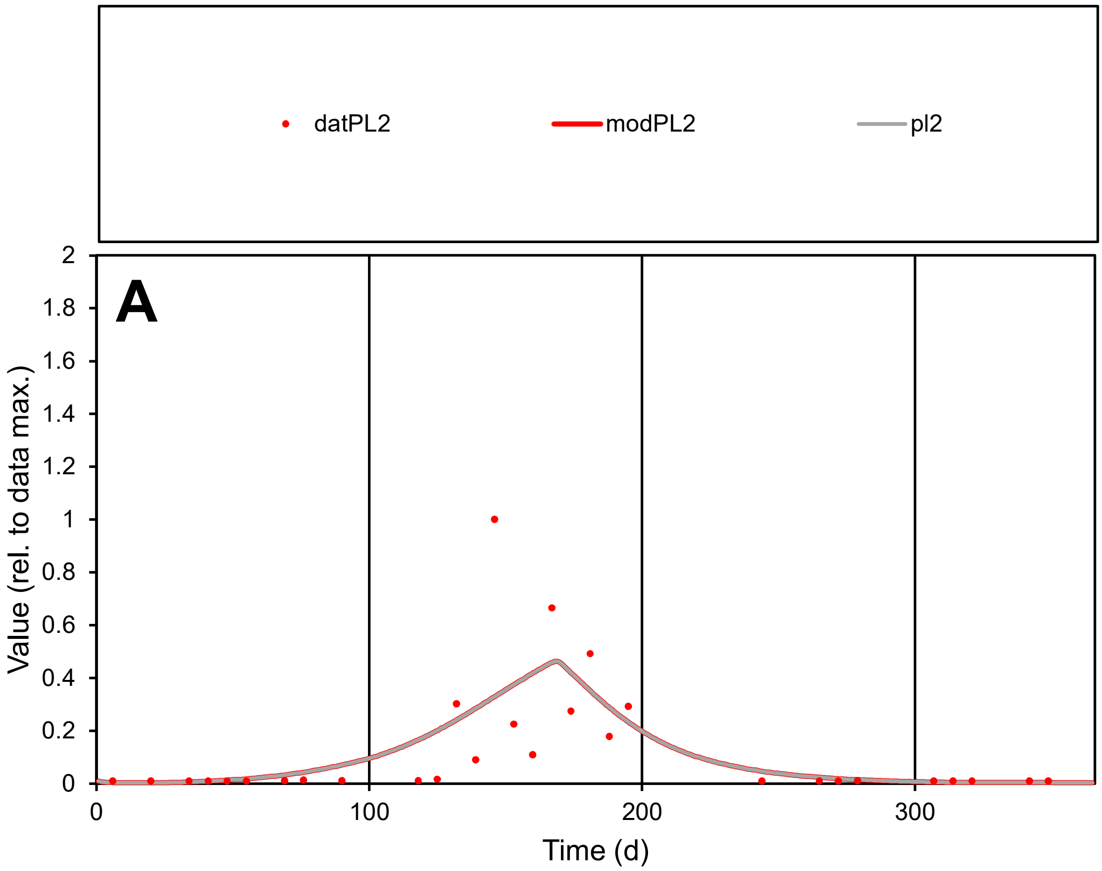


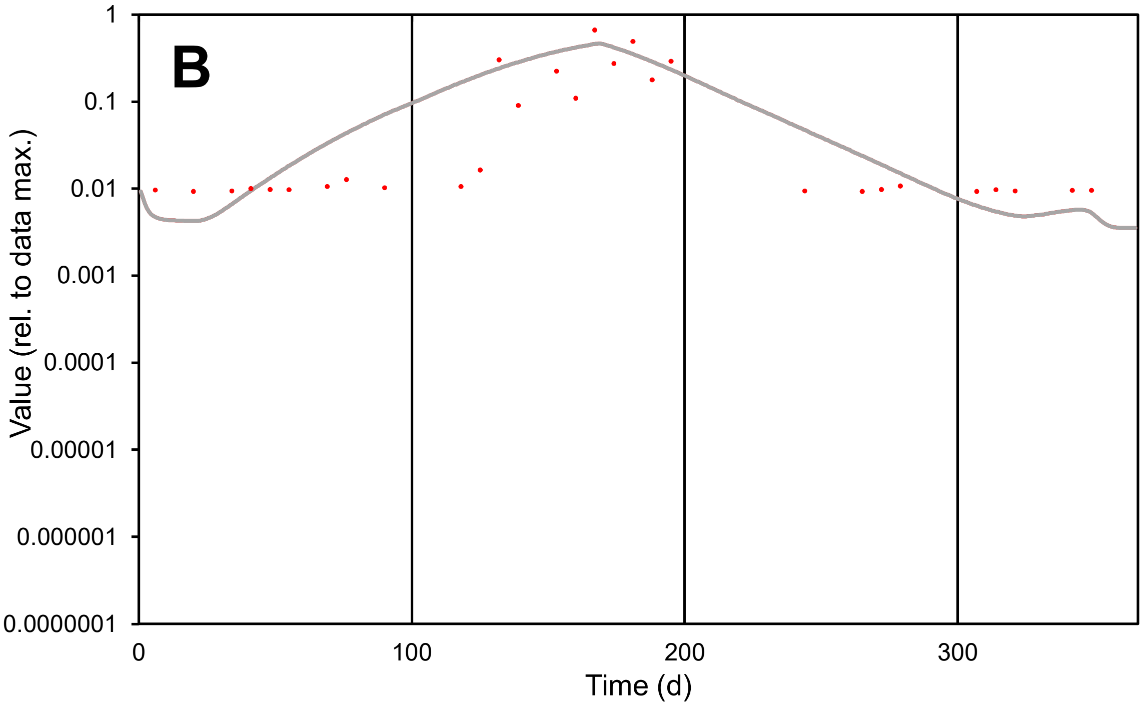


**C1**

Variables
PL2 Provide jtd or Dst
CalibOnly?
2 1 to reproduce of
oNormalize?
2 1=no, 2=yes

ofxn calc*
1.24492
ofxn mod
1.26424

sRun
RunA1
zout
1

Ave
Data
0.00235
Model
0.00251

data/vMax
0.01838

ofxng
275.646

Min d/m
2.17258

ave/min d
13.7978
ave/min m
31.9958

325_Calib_S_A

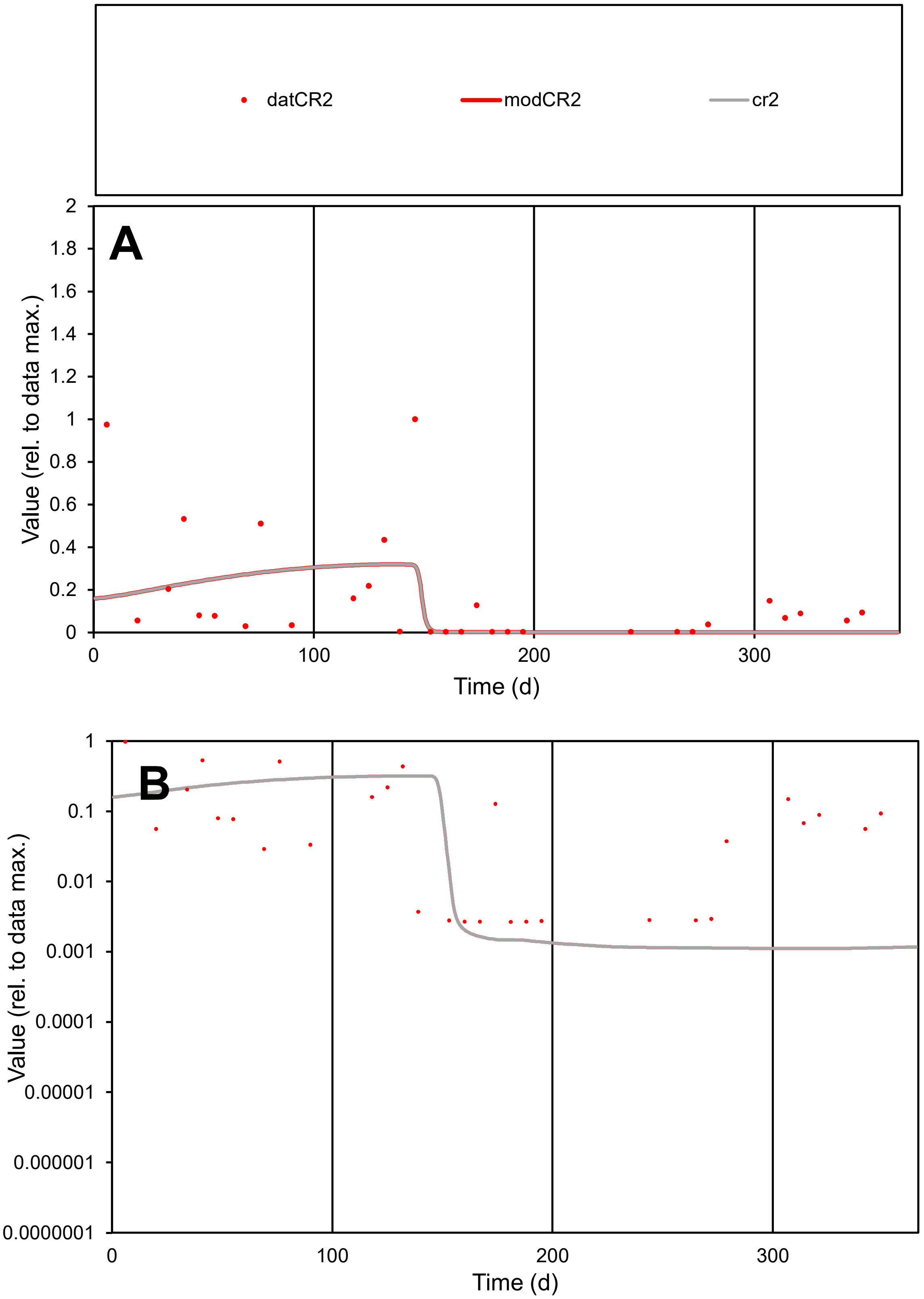


**C1**

Variables
CR2 Provide jtd or Dst
CalibOnly?
2 1 to reproduce of
oNormalize?
2 1=no, 2=yes

ofxn calc*
1.43438
ofxn mod
1.44922

sRun
RunA1
zout
1

Ave
Data
0.00247
Model
0.00187

data/vMax
0.01494

ofxng
275.646

Min d/m
2.38138

ave/min d
62.2938
ave/min m
112.478

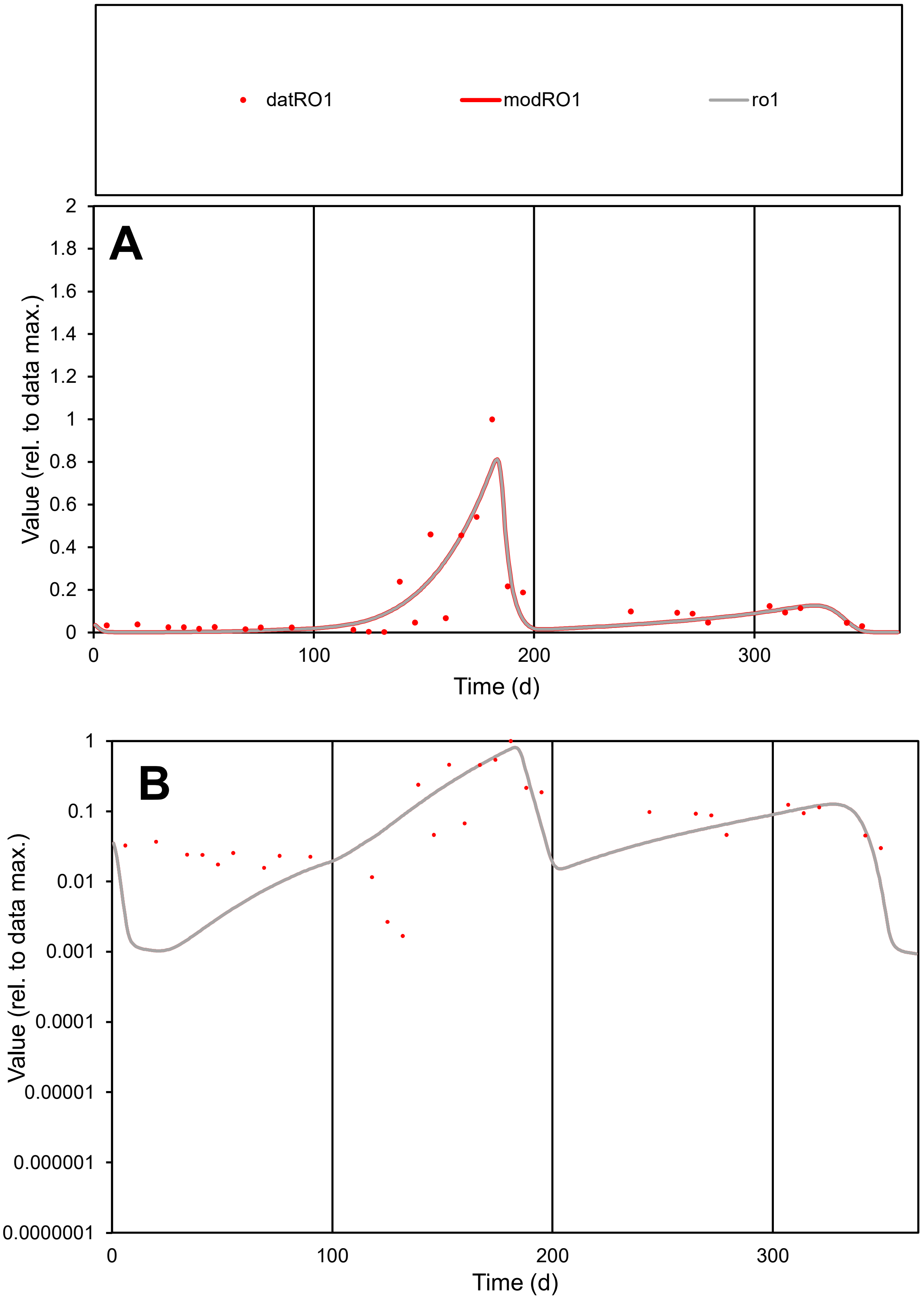


**C1**

Variables
RO1 Provide jtd or Dst
CalibOnly?
2 1 to reproduce of
oNormalize?
2 1=no, 2=yes

ofxn calc*
0.61974
ofxn mod
0.65546

sRun
RunA1
zout
1

Ave
Data
0.00344
Model
0.00332

data/vMax
0.02467

ofxng
275.646

Min d/m
1.63515

ave/min d
82.8191
ave/min m
130.796

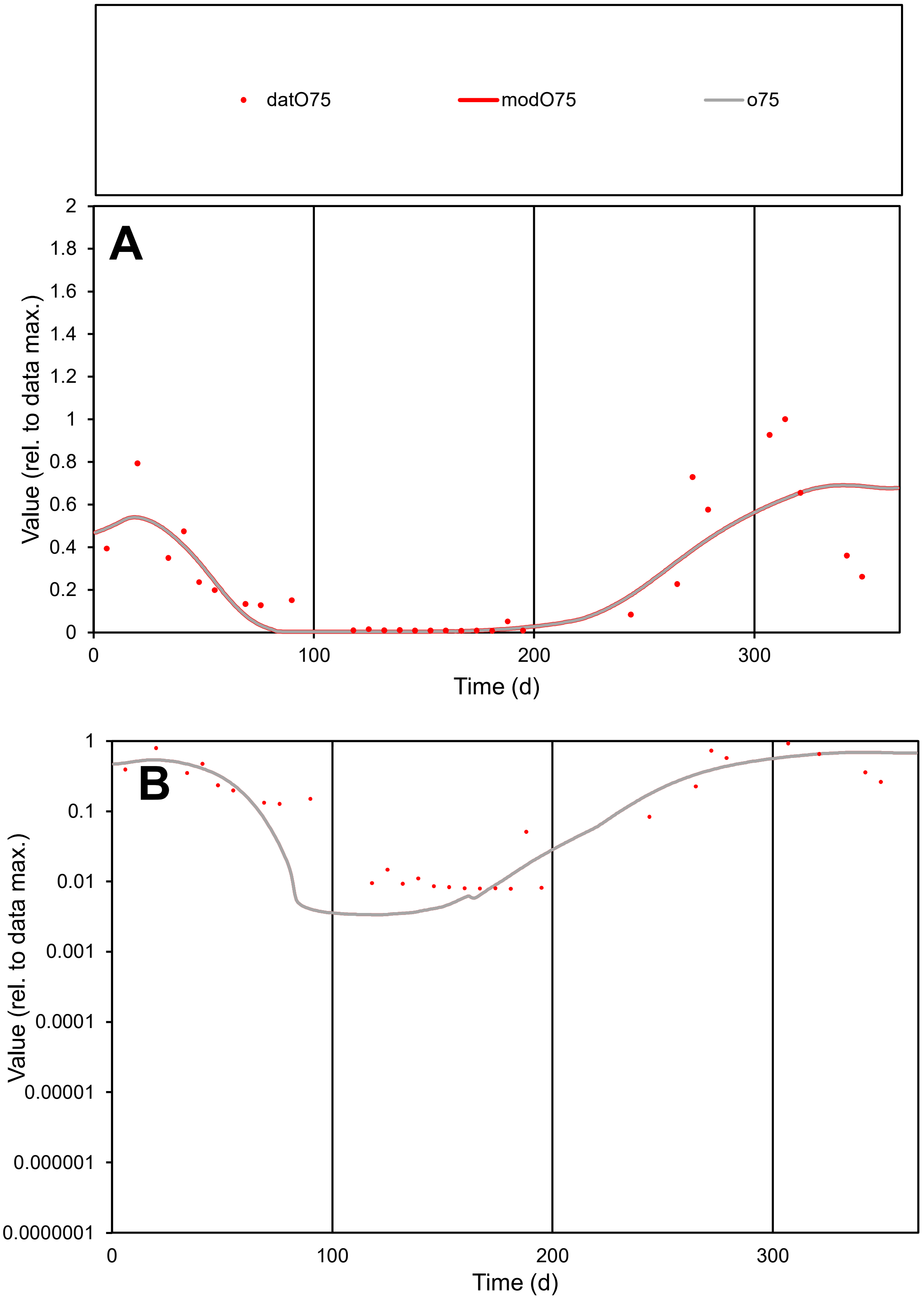


**C1**

Variables
O75 Provide jtd or Dst
CalibOnly?
2 1 to reproduce of
oNormalize?
2 1=no, 2=yes

ofxn calc*
0.62916
ofxn mod
0.65139

sRun
RunA1
zout
1

Ave
Data
0.0023
Model
0.00214

data/vMax
0.00883

ofxng
275.646

Min d/m
2.3406

ave/min d
33.0355
ave/min m
71.7486

325_Calib_S_A

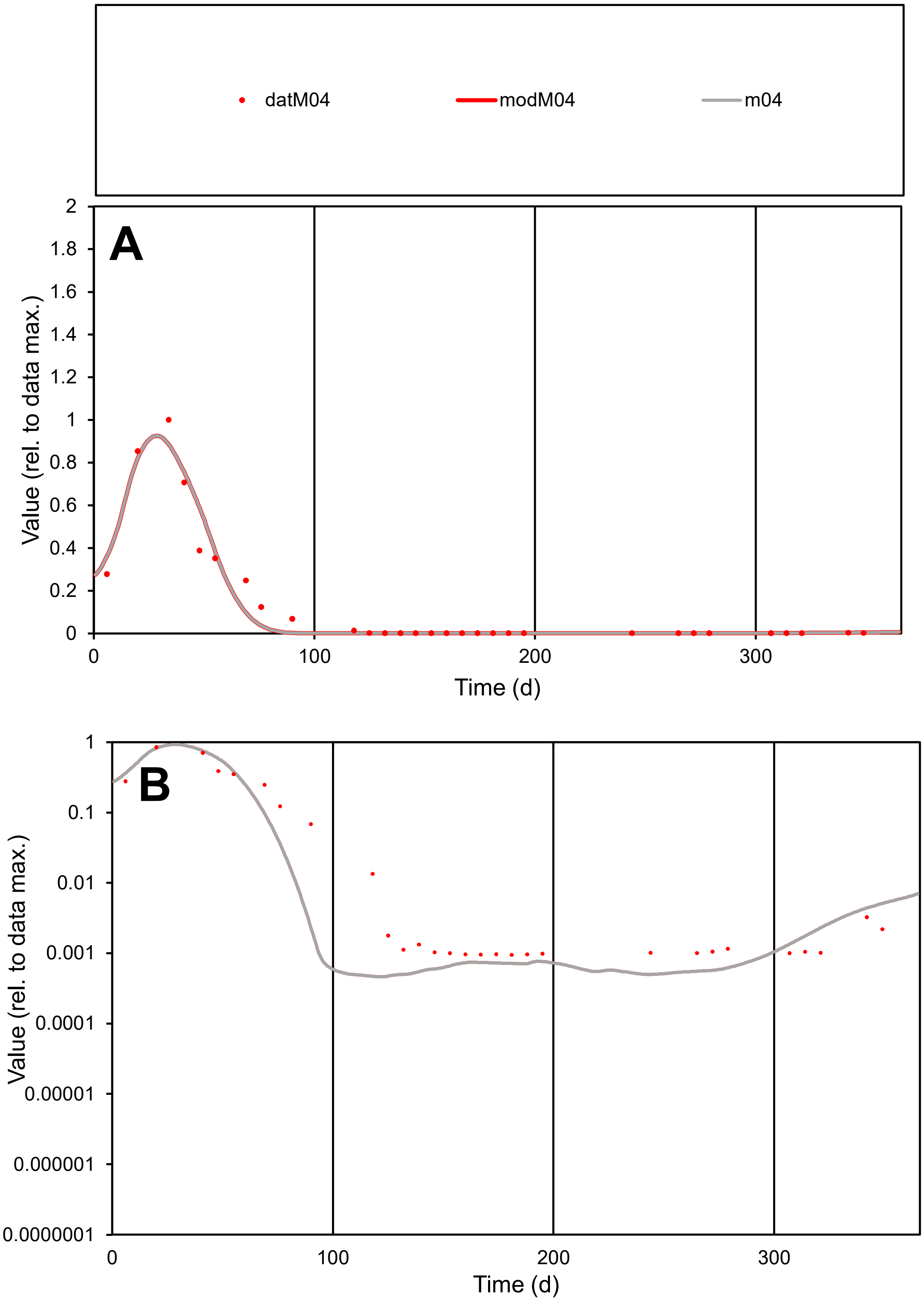


**C1**

Variables
M04 Provide jtd or Dst
CalibOnly?
2 1 to reproduce of
oNormalize?
2 1=no, 2=yes

ofxn calc*
0.40787
ofxn mod
0.49305

sRun
RunA1
zout
1

Ave
Data
0.00499
Model
0.00474

data/vMax
0.03689

ofxng
275.646

Min d/m
2.0348

ave/min d
142.168
ave/min m
274.953

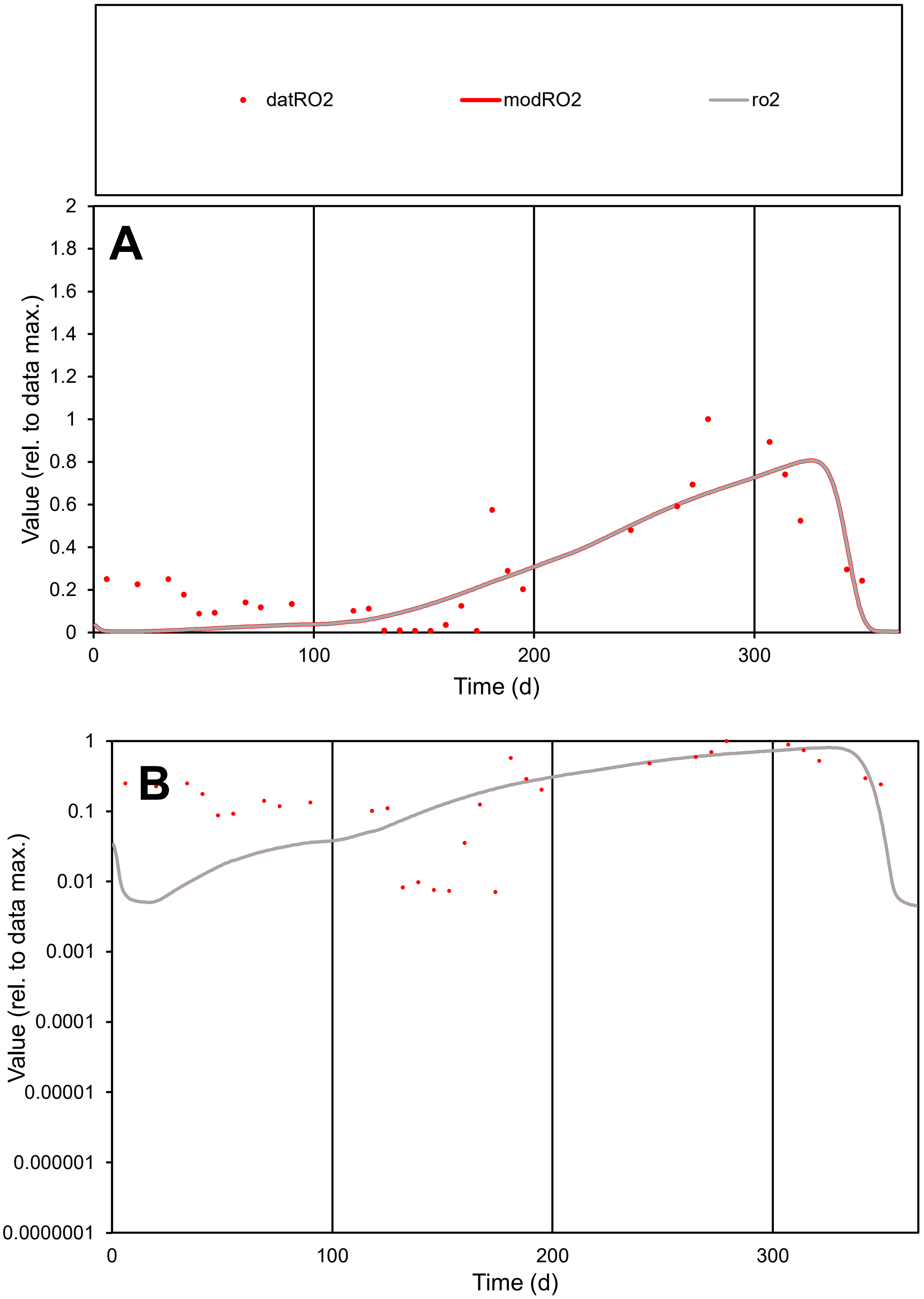


**C1**

Variables
RO2 Provide jtd or Dst
CalibOnly?
2 1 to reproduce of
oNormalize?
2 1=no, 2=yes

ofxn calc*
0.54267
ofxn mod
0.56856

sRun
RunA1
zout
1

Ave
Data
0.00153
Model
0.00132

data/vMax
0.00547

ofxng
275.646

Min d/m
1.31451

ave/min d
39.4904
ave/min m
44.7074

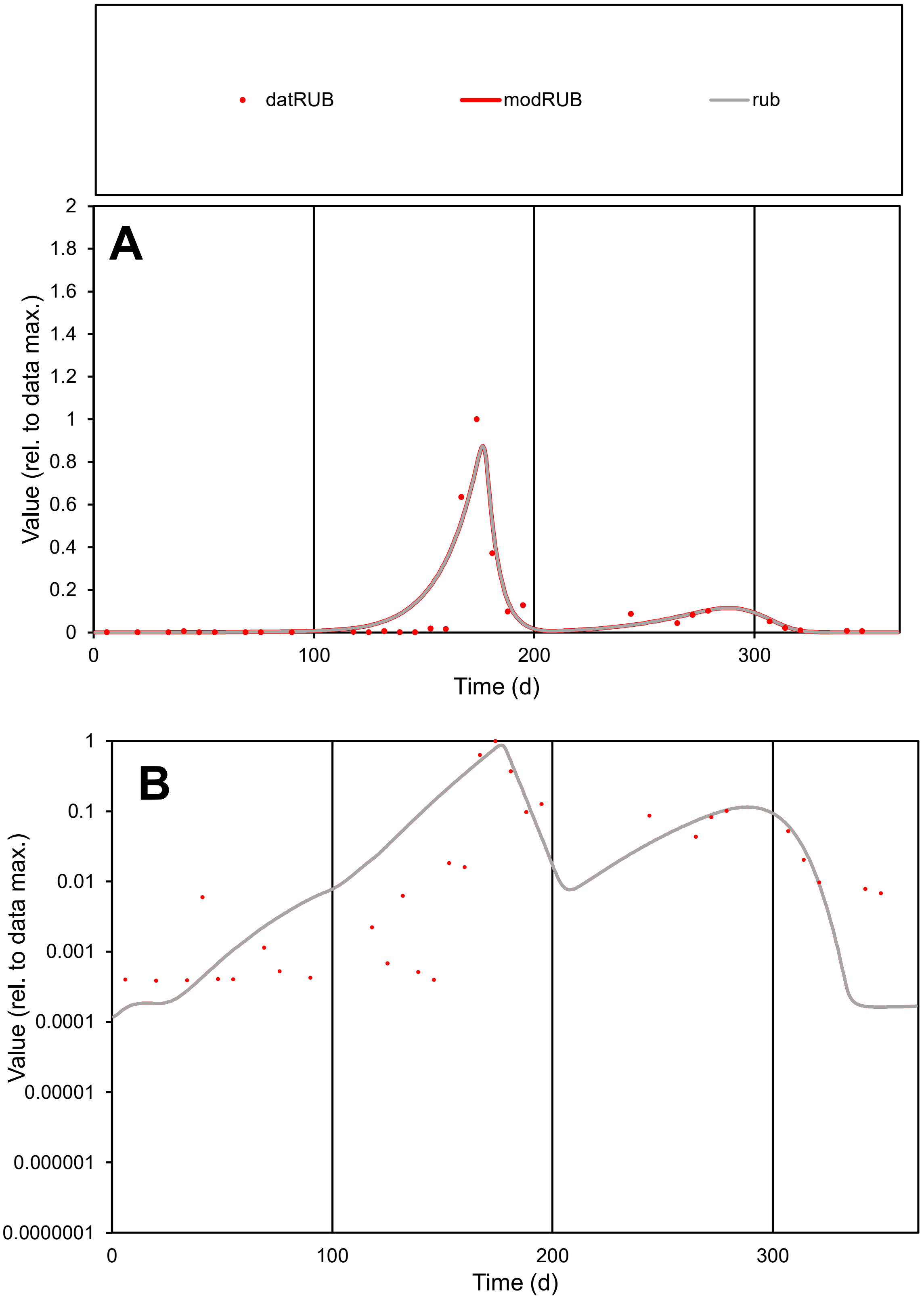


**C1**

Variables

RUB Provide jtd or Dst

CalibOnly?

2 1 to reproduce of

oNormalize?

2 1=no, 2=yes

ofxn calc*

0.902

ofxn mod

0.9714

sRun

RunA1

zout

1

Ave

Data

0.00243

Model

0.00288

data/vMax

0.02699

ofxng

275.646

Min d/m

2.37457

ave/min d

231.837

ave/min m

654.842

325_Calib_S_A

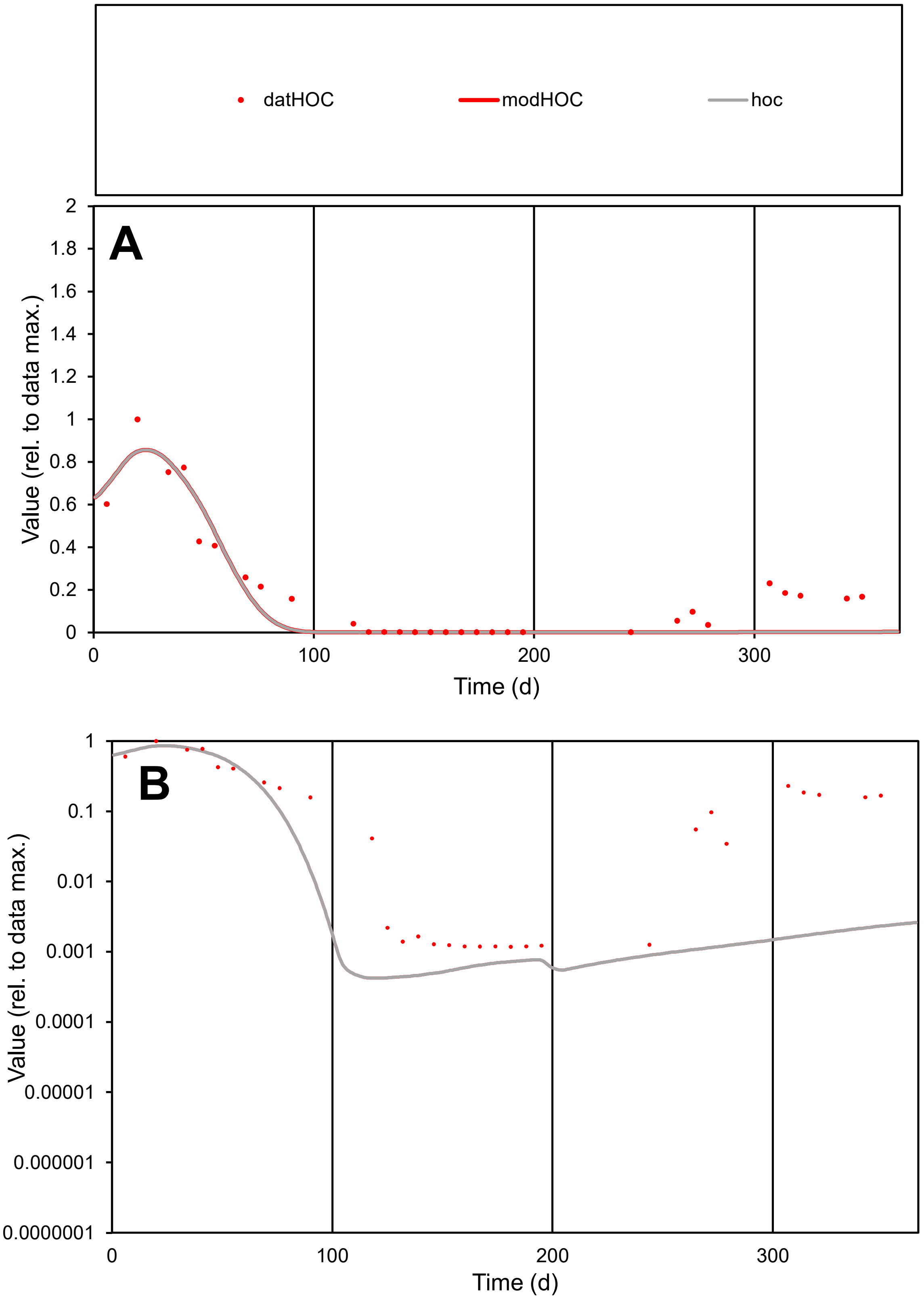


**C1**

Variables
HOC Provide jtd or Dst
CalibOnly?
2 1 to reproduce of
oNormalize?
2 1=no, 2=yes

ofxn calc*
0.50703
ofxn mod
0.54414

sRun
RunA1
zout
1

Ave
Data
0.00203
Model
0.00156

data/vMax
0.01061

ofxng
275.646

Min d/m
2.78495

ave/min d
162.512
ave/min m
346.239

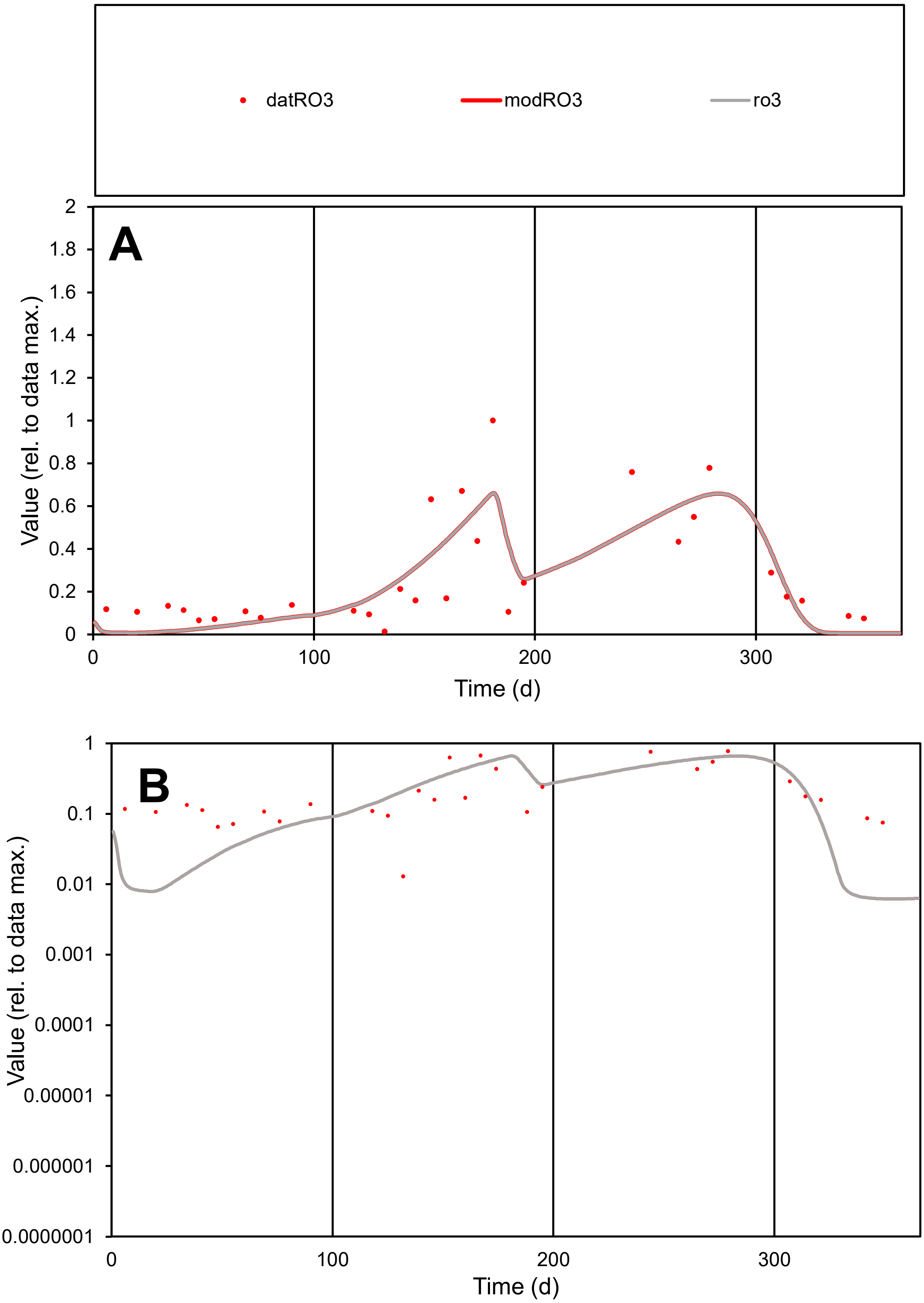


**C1**

Variables

RO3 Provide jtd or Dst

CalibOnly?

2 1 to reproduce of

oNormalize?

2 1=no, 2=yes

ofxn calc*

0.53955

ofxn mod

0.56746

sRun

RunA1

zout

1

Ave

Data

0.00235

Model

0.00223

data/vMax

0.00875

ofxng

275.646

Min d/m

2.07811

ave/min d

20.8684

ave/min m

40.9932

325_Calib_S_A

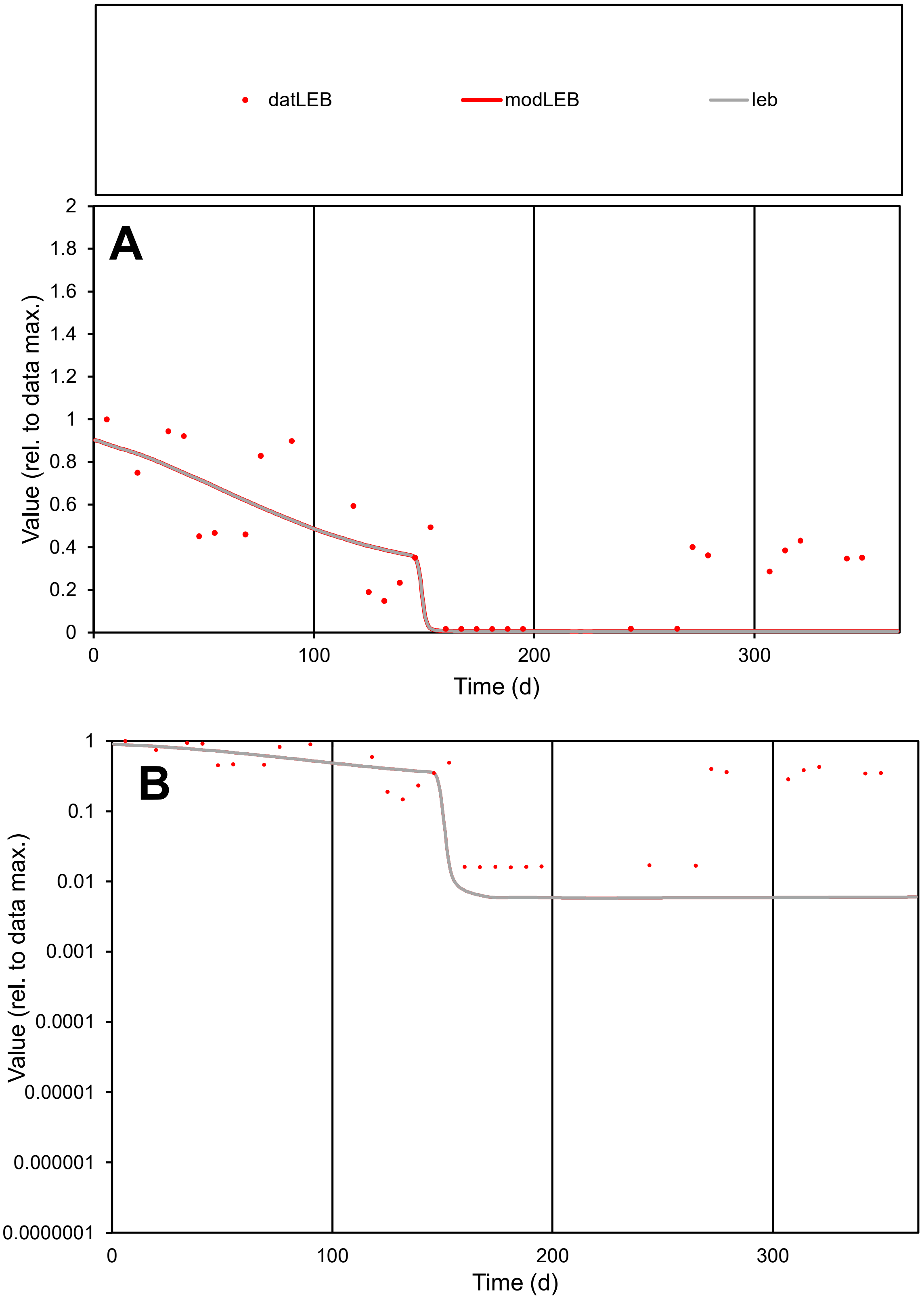


**C1**

Variables

LEB Provide jtd or Dst

CalibOnly?

2 1 to reproduce of

oNormalize?

2 1=no, 2=yes

ofxn calc*

0.60505

ofxn mod

0.6439

sRun

RunA1

zout

1

Ave

Data

0.00165

Model

0.00123

data/vMax

0.00435

ofxng

275.646

Min d/m

2.73218

ave/min d

23.8481

ave/min m

48.4188

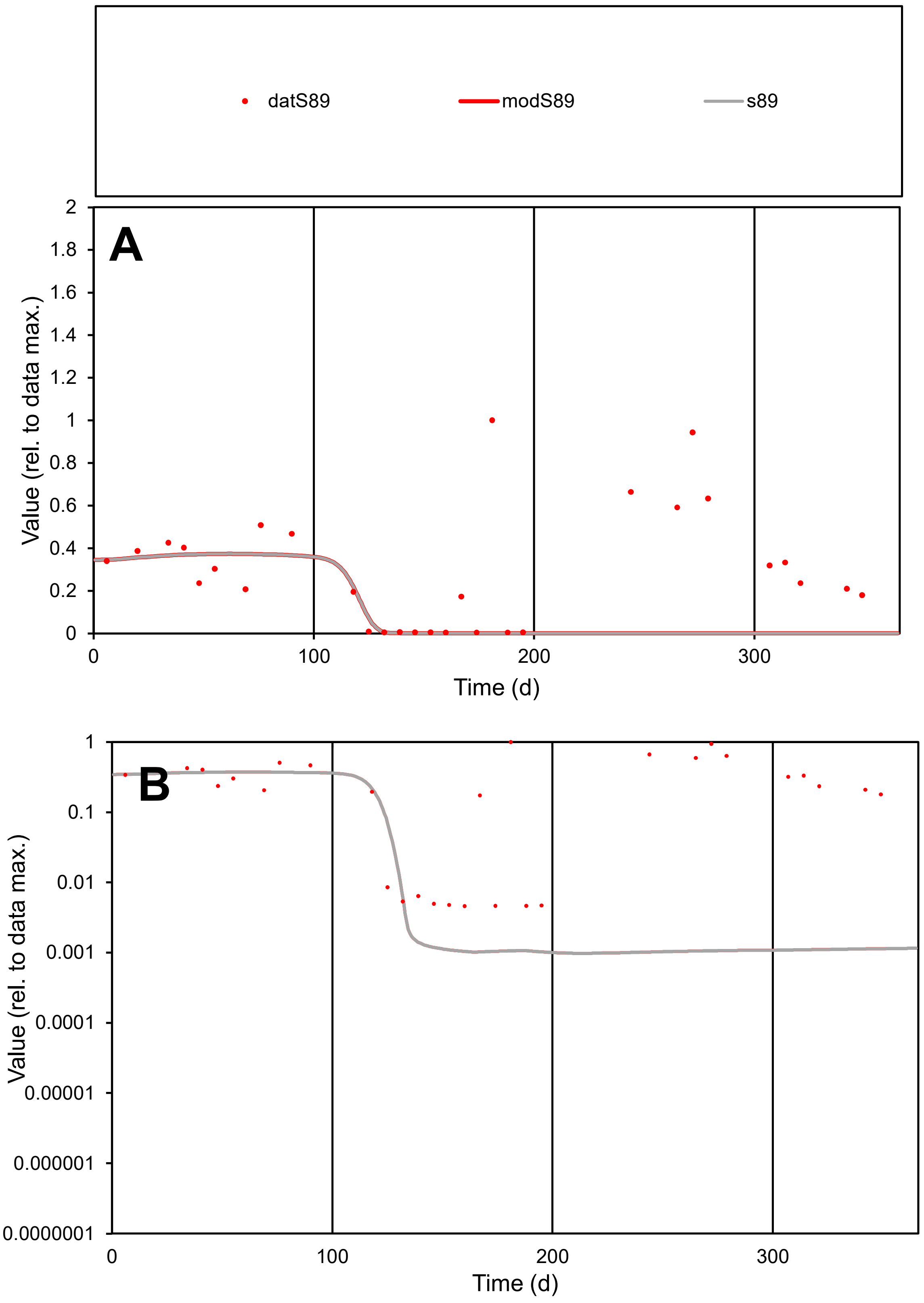


**C1**

Variables

S89 Provide jtd or Dst

CalibOnly?

2 1 to reproduce of

oNormalize?

2 1=no, 2=yes

ofxn calc*

1.16941

ofxn mod

1.17458

sRun

RunA1

zout

1

Ave

Data

0.00231

Model

0.00093

data/vMax

0.00786

ofxng

275.646

Min d/m

4.52316

ave/min d

63.4786

ave/min m

115.747

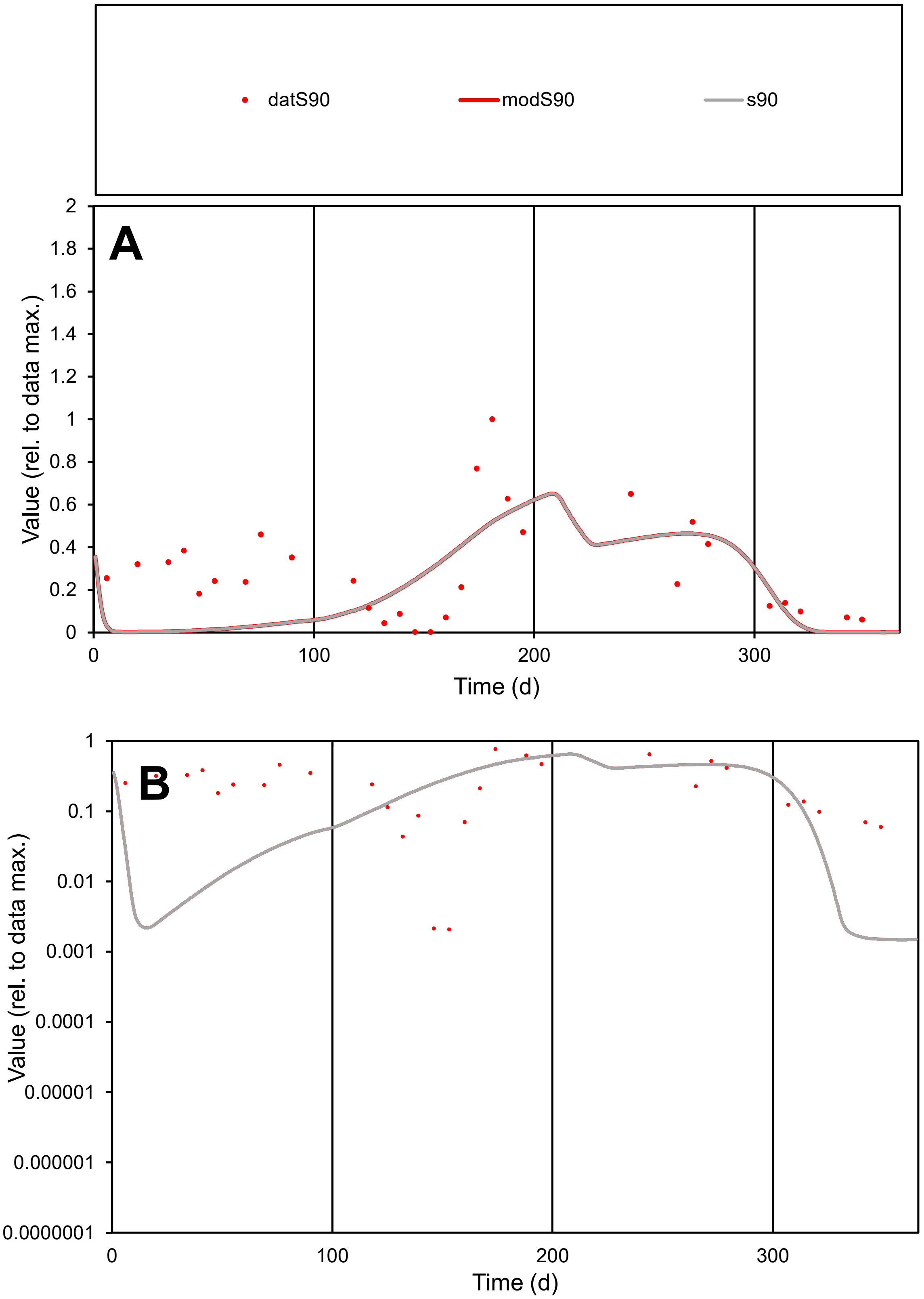


**C1**

Variables
S90 Provide jtd or Dst
CalibOnly?
2 1 to reproduce of
oNormalize?
2 1=no, 2=yes

ofxn calc*
0.77235
ofxn mod
0.79381

sRun
RunA1
zout
1

Ave
Data
0.00291
Model
0.00221

data/vMax
0.01002

ofxng
275.646

Min d/m
1.39093

ave/min d
139.393
ave/min m
147.479

325_Calib_S_A

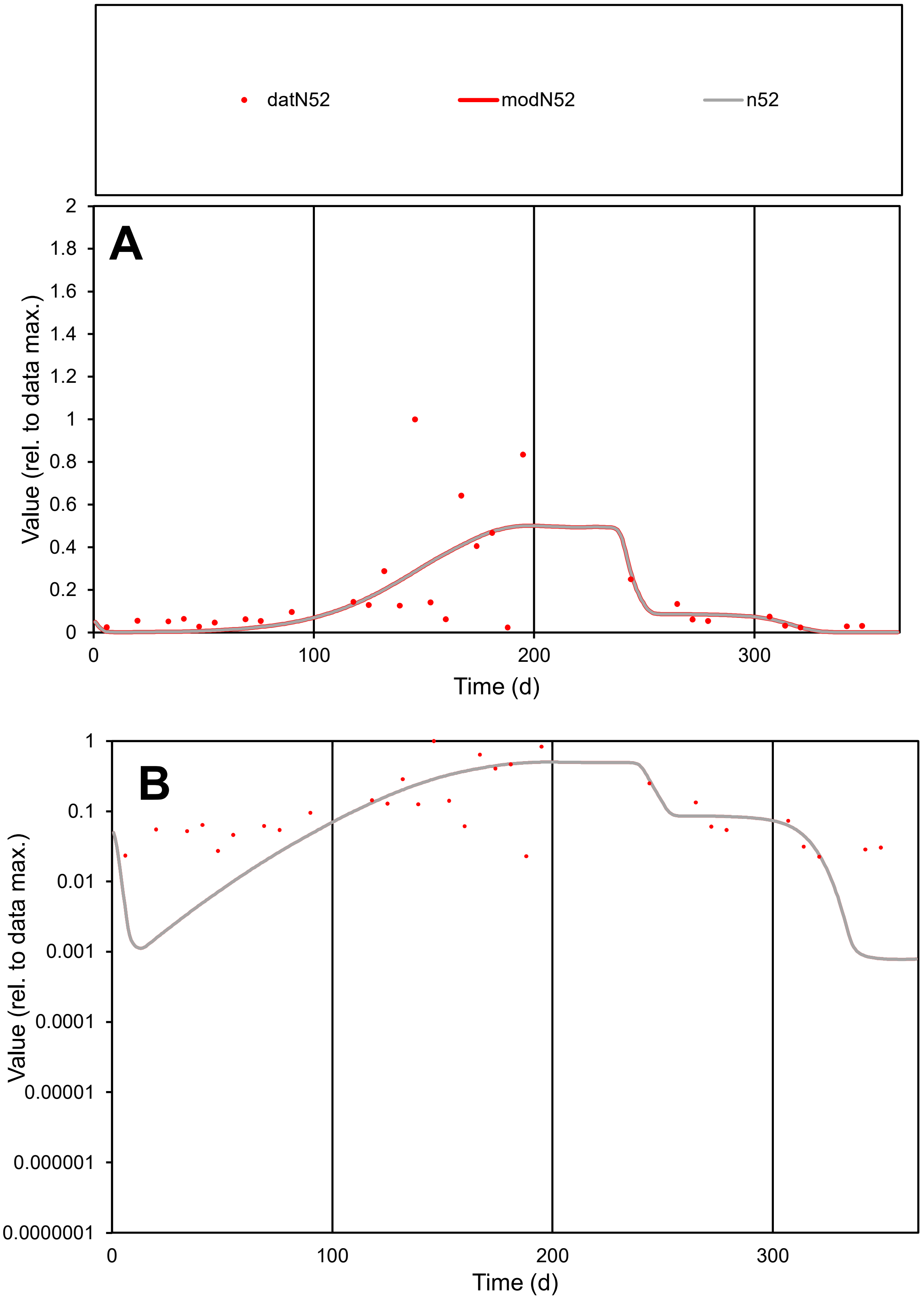


**C1**

Variables
N52 Provide jtd or Dst
CalibOnly?
2 1 to reproduce of
oNormalize?
2 1=no, 2=yes

ofxn calc*
1.03968
ofxn mod
1.0527

sRun
RunA1
zout
1

Ave
Data
0.00316
Model
0.00284

data/vMax
0.0175

ofxng
275.646

Min d/m
28.4043

ave/min d
8.03759
ave/min m
205.235

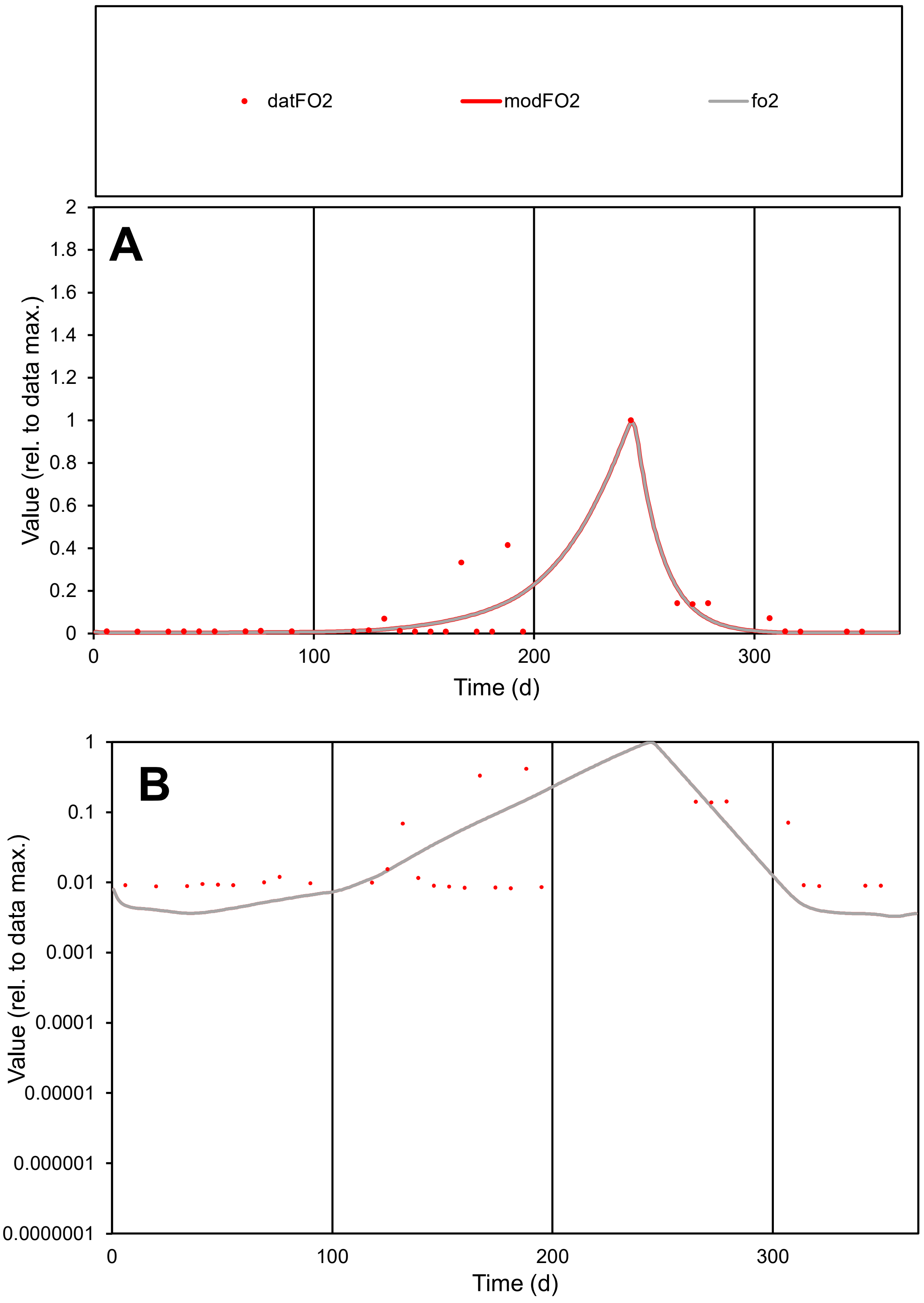


**C1**

Variables
FO2 Provide jtd or Dst
CalibOnly?
2 1 to reproduce of
oNormalize?
2 1=no, 2=yes

ofxn calc*
0.98187
ofxn mod
1.00876

sRun
RunA1
zout
1

Ave
Data
0.00068
Model
0.00061

data/vMax
0.00804

ofxng
275.646

Min d/m
2.45007

ave/min d
10.1226
ave/min m
22.3047

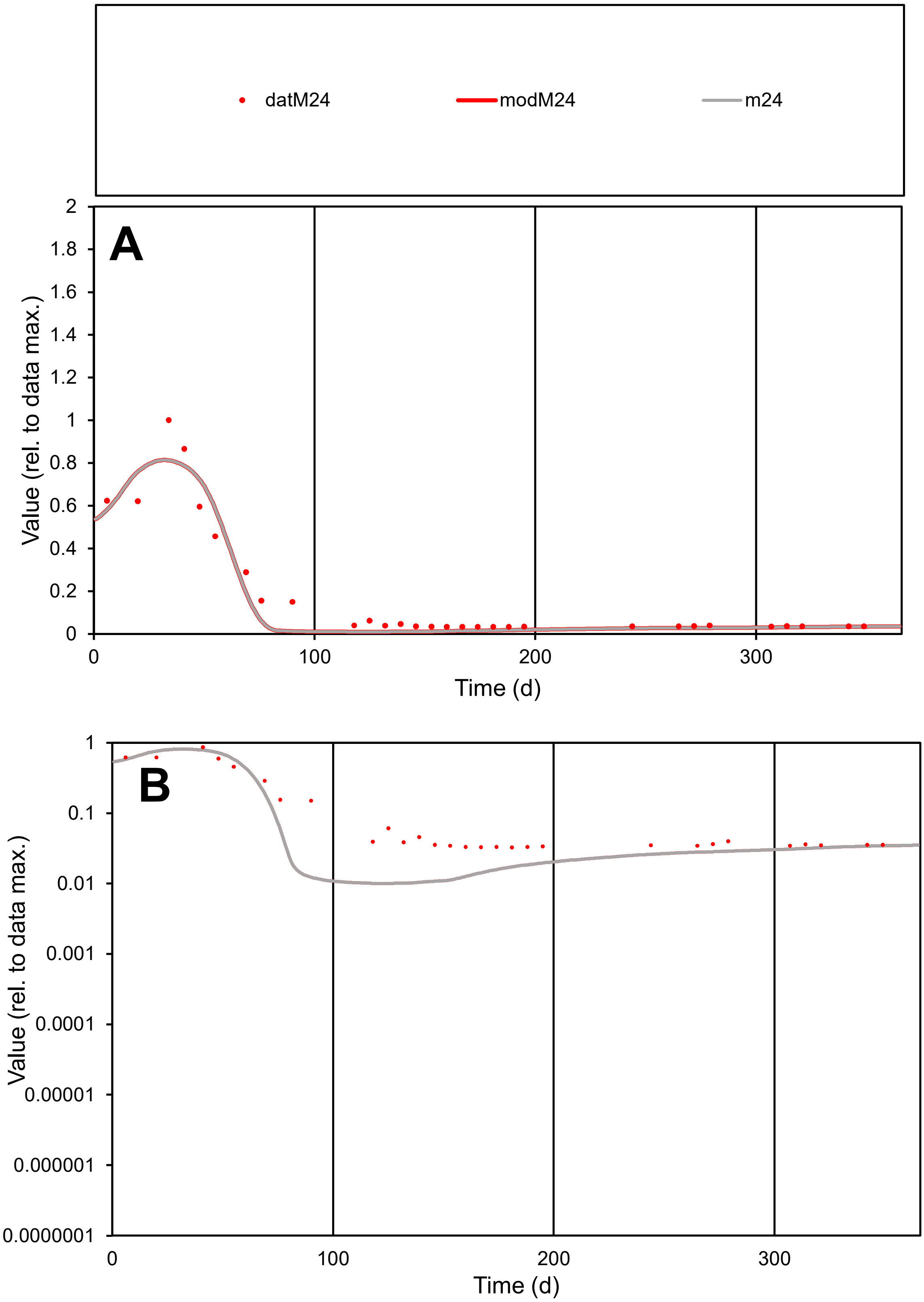


**C1**

Variables

M24 Provide jtd or Dst

CalibOnly?

2 1 to reproduce of

oNormalize?

2 1=no, 2=yes

ofxn calc*

0.35983

ofxn mod

0.42383

sRun

RunA1

zout

1

Ave

Data

0.00041

Model

0.00036

data/vMax

0.0022

ofxng

275.646

Min d/m

3.24695

ave/min d

5.64328

ave/min m

16.2067

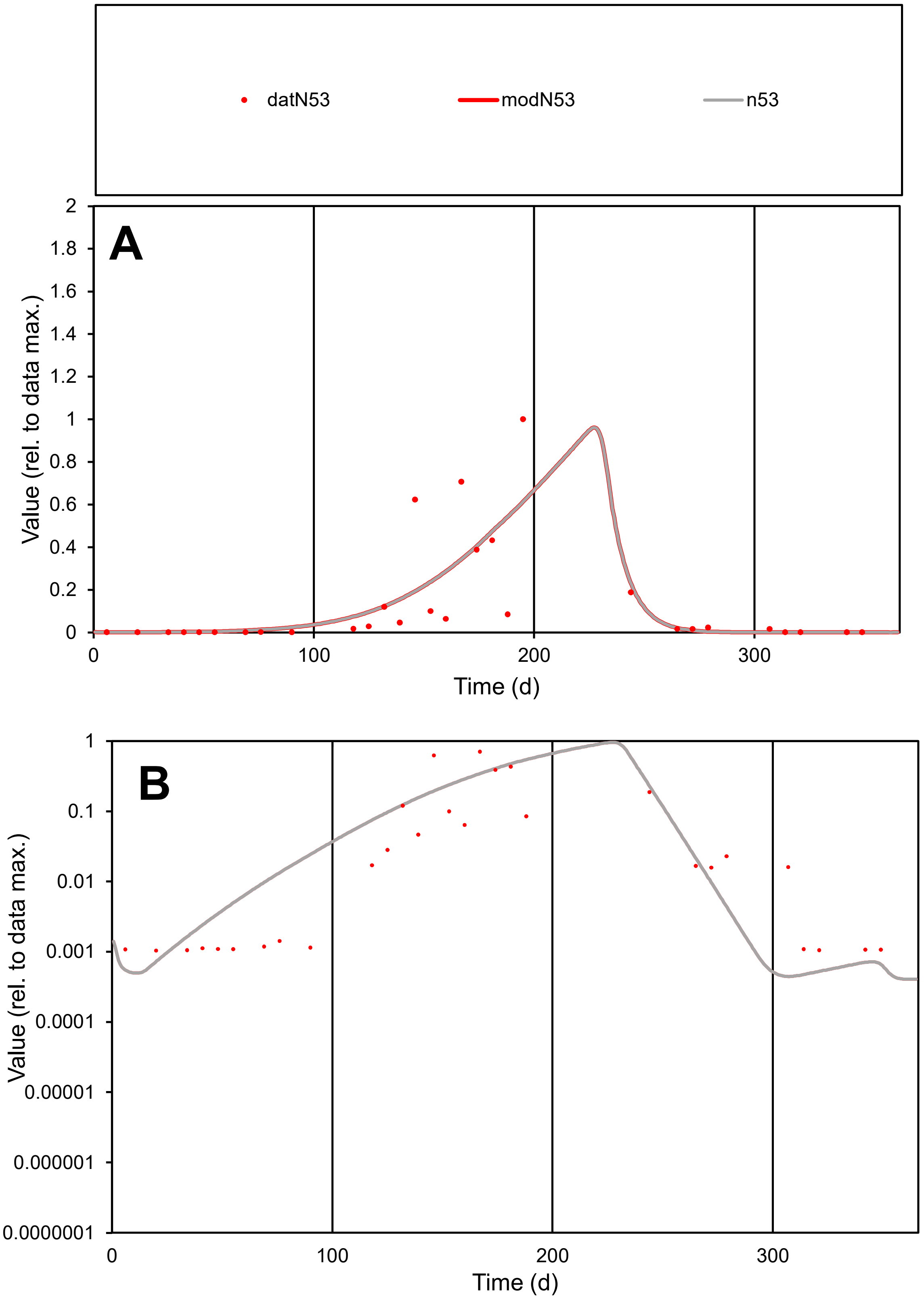


**C1**

Variables
N53 Provide jtd or Dst
CalibOnly?
2 1 to reproduce of
oNormalize?
2 1=no, 2=yes

ofxn calc*
1.22056
ofxn mod
1.24142

sRun
RunA1
zout
1

Ave
Data
0.00477
Model
0.00482

data/vMax
0.03684

ofxng
275.646

Min d/m
2.33536

ave/min d
124.344
ave/min m
293.502

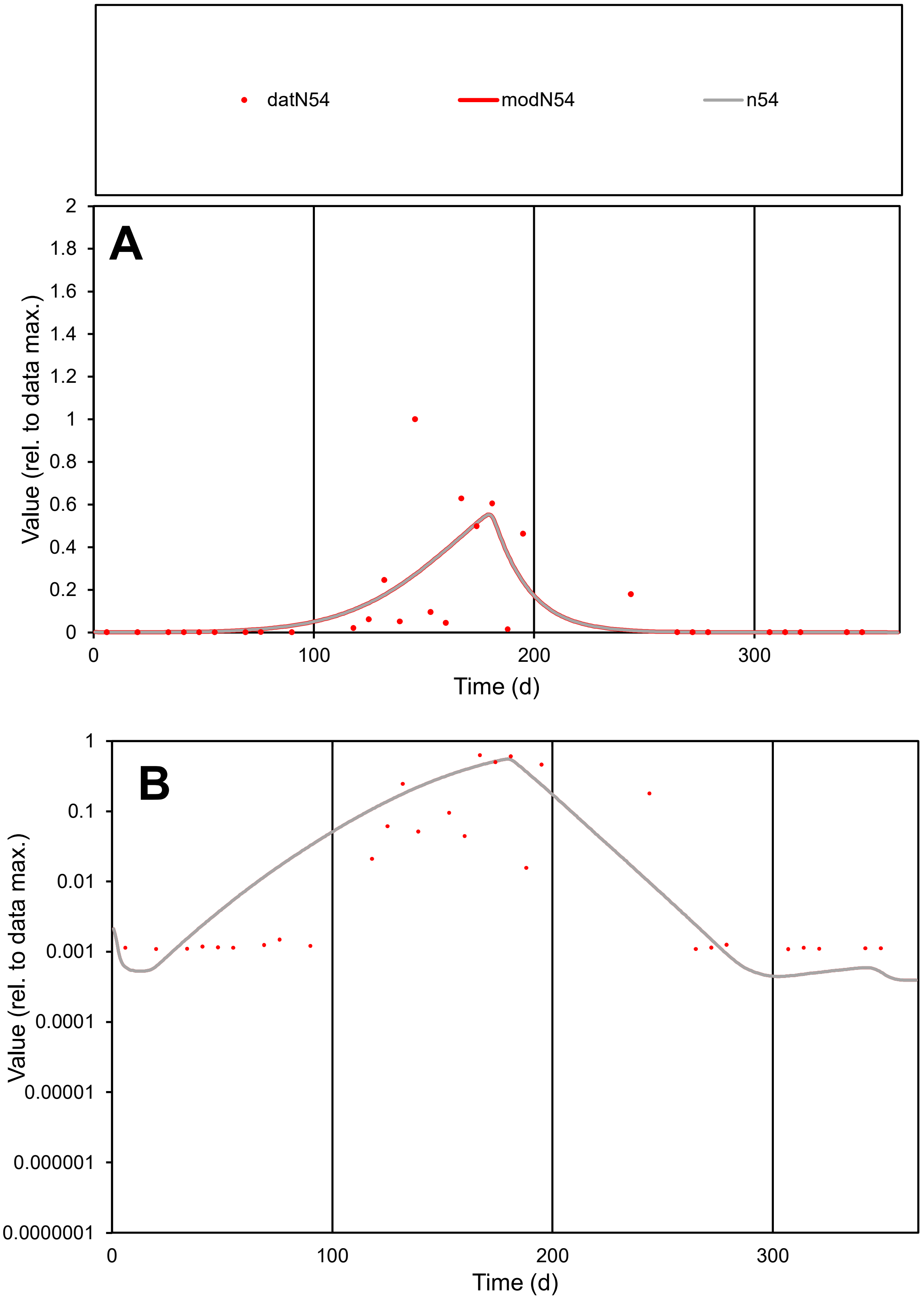


**C1**

Variables
N54 Provide jtd or Dst
CalibOnly?
2 1 to reproduce of
oNormalize?
2 1=no, 2=yes

ofxn calc*
1.35559
ofxn mod
1.37748

sRun
RunA1
zout
1

Ave
Data
0.00272
Model
0.00262

data/vMax
0.02076

ofxng
275.646

Min d/m
2.38578

ave/min d
120.291
ave/min m
276.966

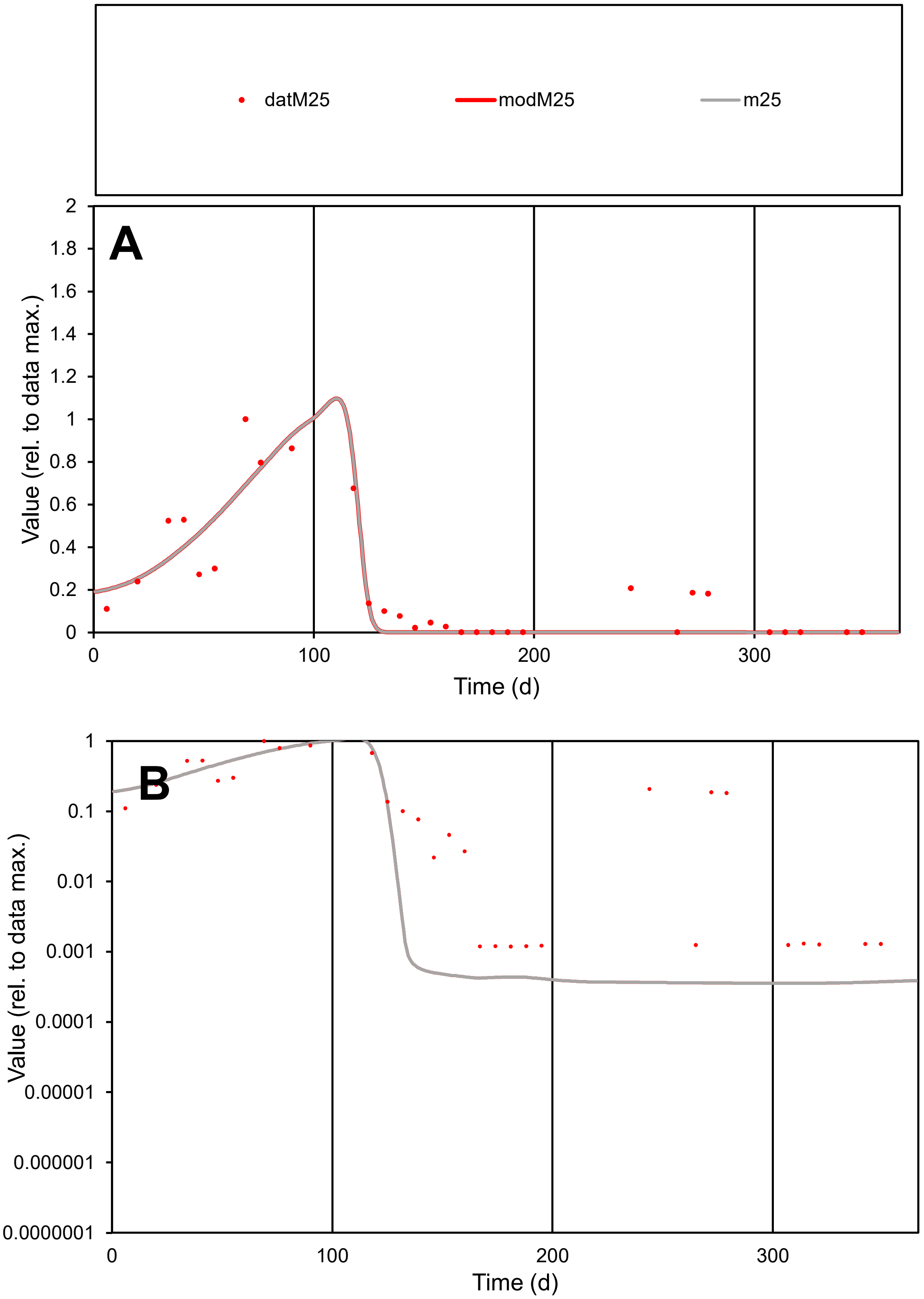


**C1**

Variables
M25 Provide jtd or Dst
CalibOnly?
2 1 to reproduce of
oNormalize?
2 1=no, 2=yes

ofxn calc*
0.52569
ofxn mod
0.56106

sRun
RunA1
zout
1

Ave
Data
0.00169
Model
0.00147

data/vMax
0.00805

ofxng
275.646

Min d/m
3.33147

ave/min d
177.215
ave/min m
513.902

325_Calib_S_A

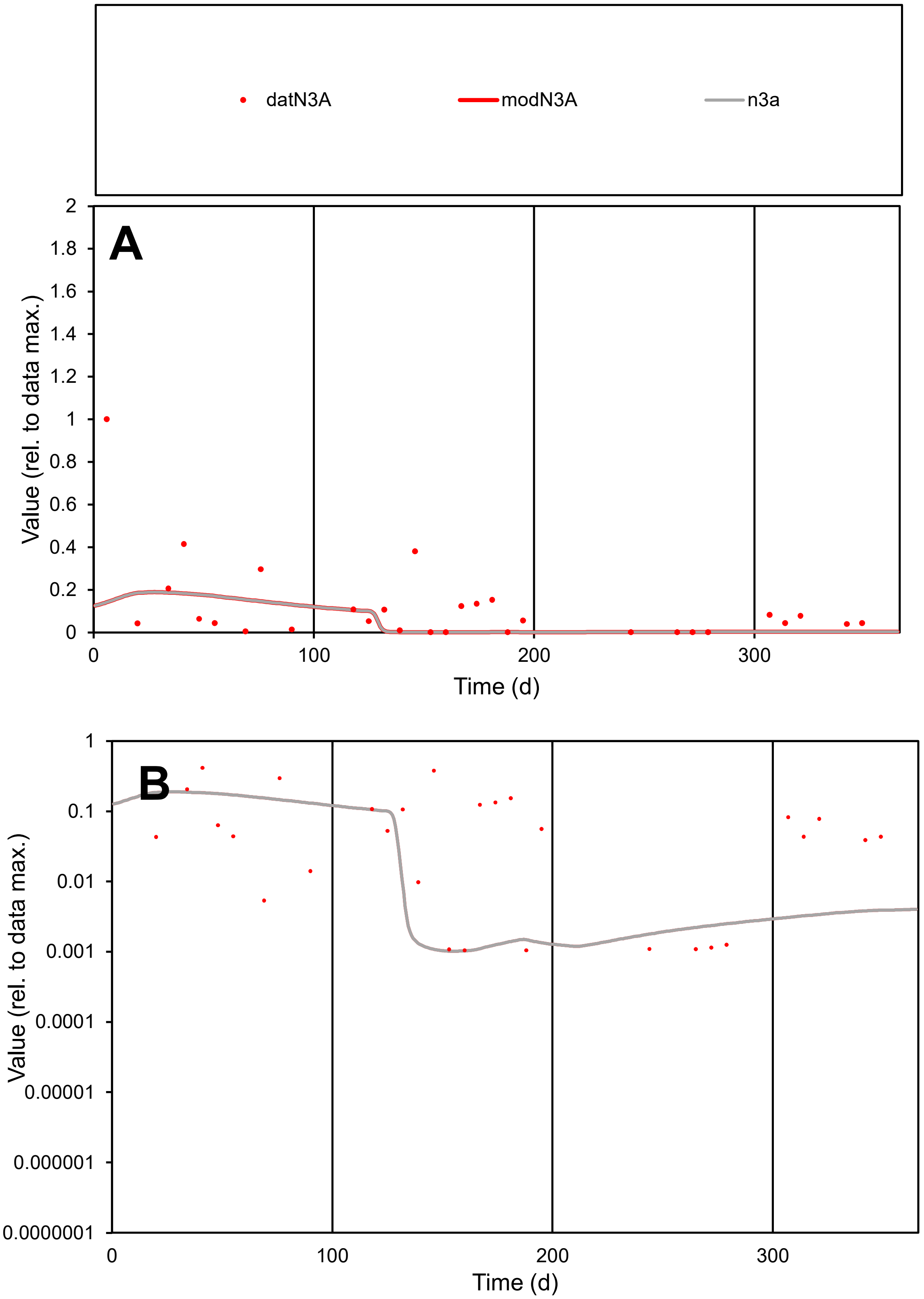


**C1**

| | | |
|---|---|---|
| | Variables | |
| | N3A | Provide jtd or Dst |
| | CalibOnly? | |
| | 2 | 1 to reproduce of |
| | oNormalize? | |
| | 2 | 1=no, 2=yes |
| | ofxn calc* | |
| | 1.64671 | |
| | ofxn mod | |
| | 1.65715 | |
| | sRun | |
| | RunA1 | |
| | zout | |
| | 1 | |
| | Ave | |
| | Data | |
| | 0.00202 | |
| | Model | |
| | 0.00103 | |
| | data/vMax | |
| | 0.01727 | |
| | ofxng | |
| | 275.646 | |
| | Min d/m | |
| | 1.02269 | |
| | ave/min d | |
| | 111.948 | |
| | ave/min m | |
| | 58.4537 | |

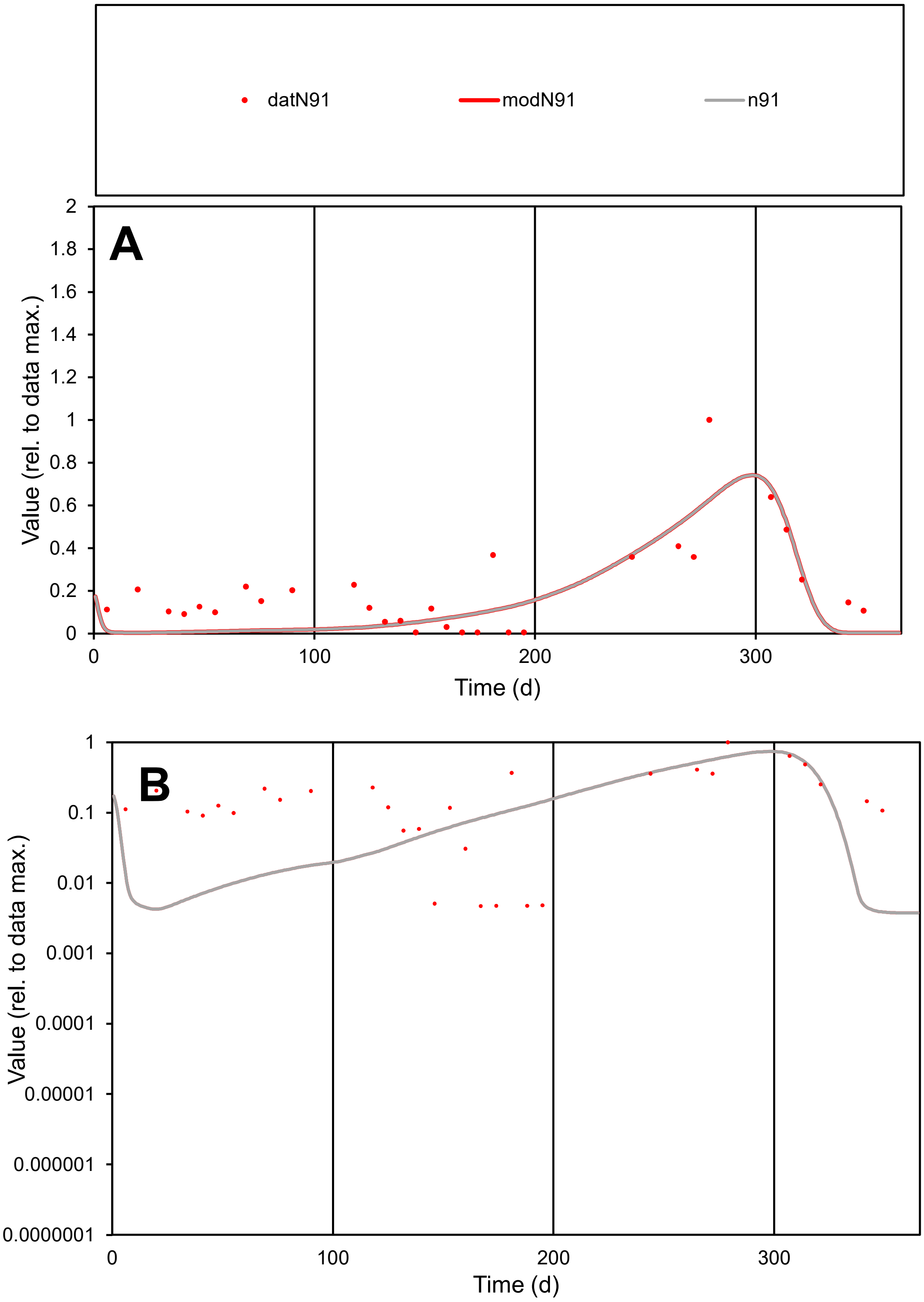


**C1**

Variables
N91 Provide jtd or Dst
CalibOnly?
2 1 to reproduce of
oNormalize?
2 1=no, 2=yes

ofxn calc*
0.67706
ofxn mod
0.70353

sRun
RunA1
zout
1

Ave
Data
0.00136
Model
0.00103

data/vMax
0.0067

ofxng
275.646

Min d/m
1.23083

ave/min d
42.8626
ave/min m
39.9224

325_Calib_S_A

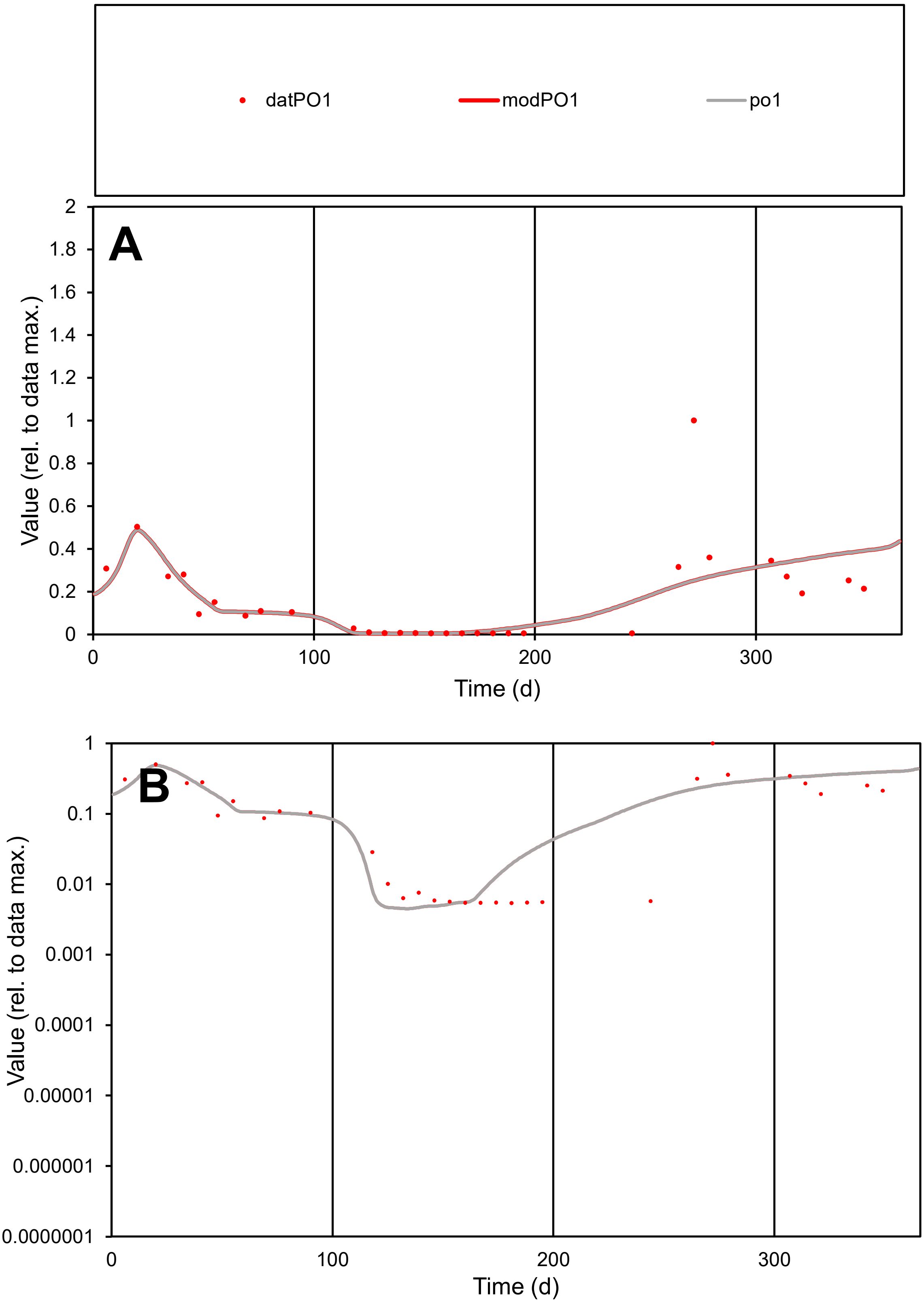


**C1**

Variables

PO1 Provide jtd or Dst

CalibOnly?

2 1 to reproduce of

oNormalize?

2 1=no, 2=yes

ofxn calc*

0.91541

ofxn mod

0.93162

sRun

RunA1

zout

1

Ave

Data

0.00139

Model

0.00132

data/vMax

0.00839

ofxng

275.646

Min d/m

1.20276

ave/min d

30.6011

ave/min m

35.0175

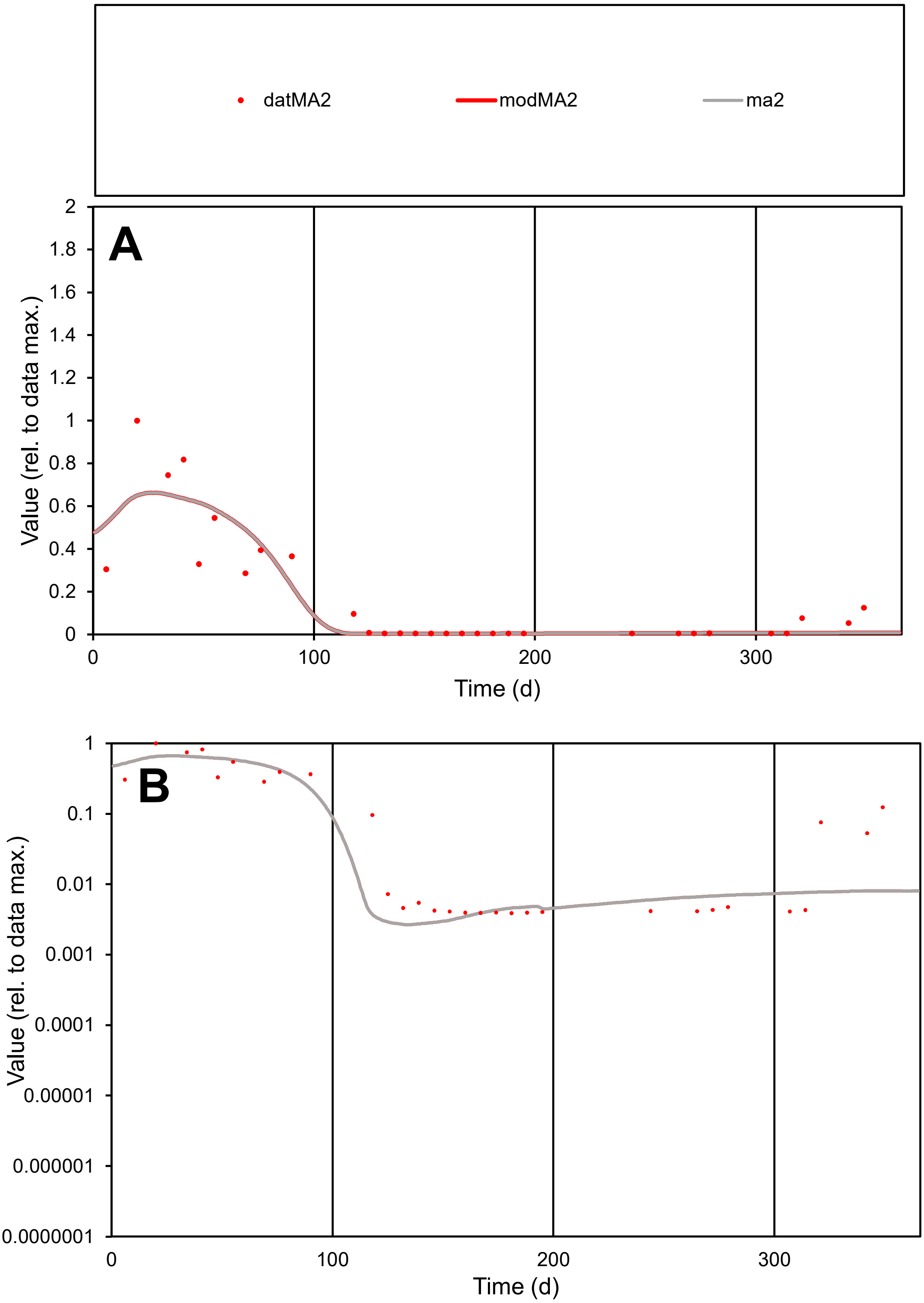


**C1**

Variables

| | |
|---|---|
| MA2 | Provide jtd or Dst |
| CalibOnly? | |
| 2 | 1 to reproduce of |
| oNormalize? | |
| 2 | 1=no, 2=yes |

ofxn calc*
0.64926
ofxn mod
0.68421

sRun
RunA1
zout
1

Ave
Data
0.00079
Model
0.00074

data/vMax
0.00455

ofxng
275.646

Min d/m
1.46287

ave/min d
44.5386
ave/min m
60.9769

325_Calib_S_A

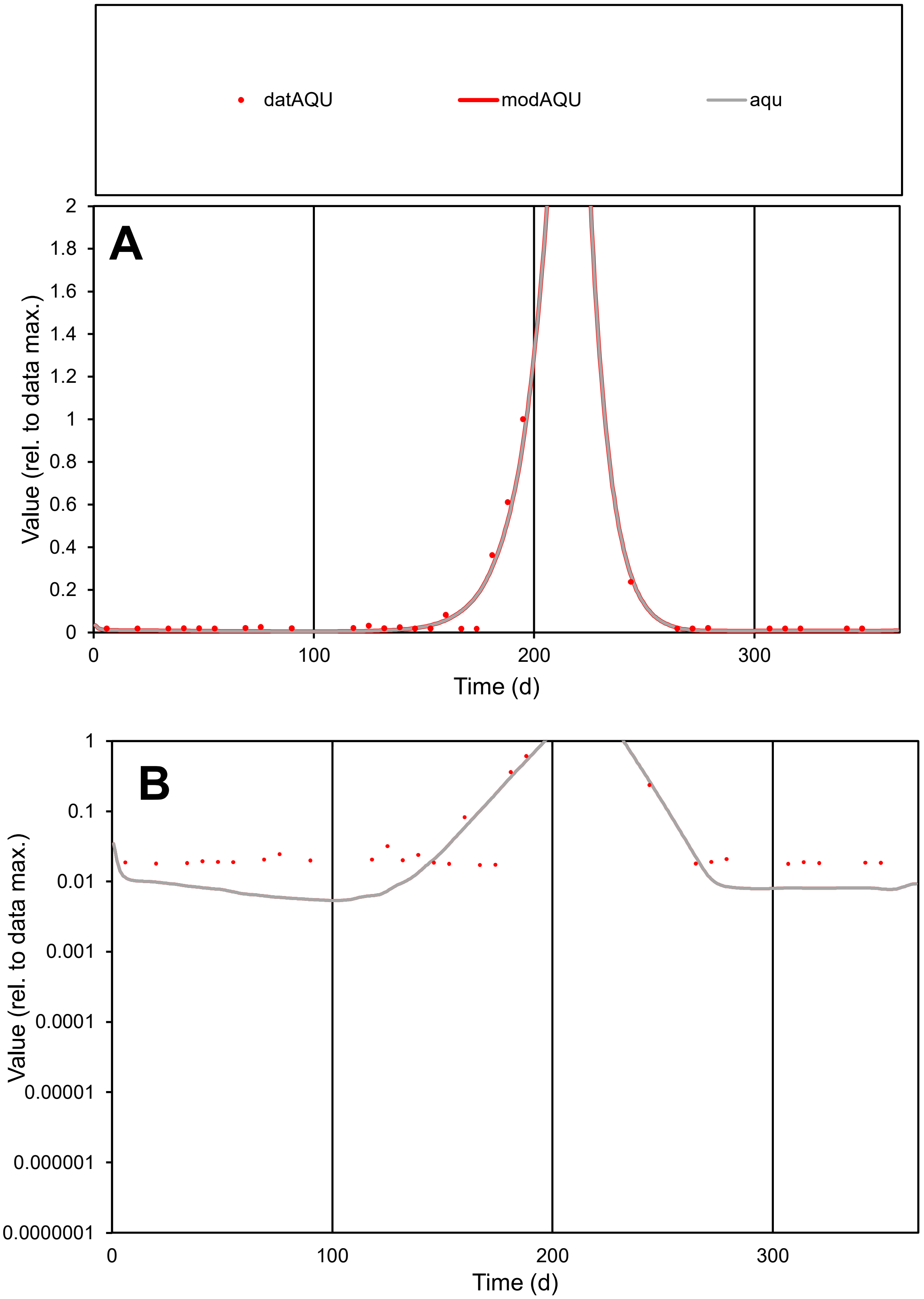


**C1**

Variables
AQU Provide jtd or Dst
CalibOnly?
2 1 to reproduce of
oNormalize?
2 1=no, 2=yes

ofxn calc*
0.35959
ofxn mod
0.46665

sRun
RunA1
zout
1

Ave
Data
0.0007
Model
0.00068

data/vMax
0.00758

ofxng
275.646

Min d/m
3.11291

ave/min d
5.40427
ave/min m
16.333

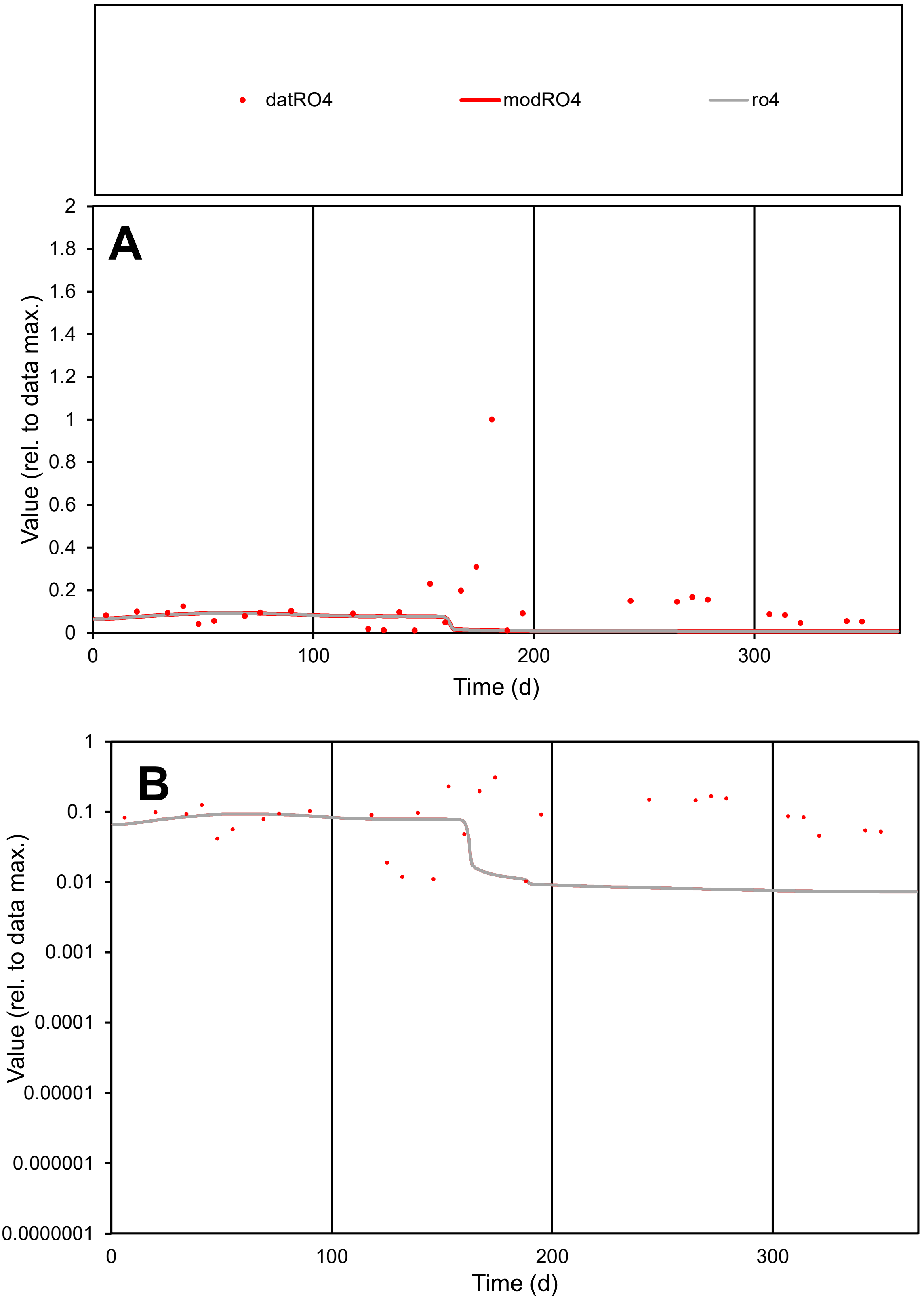


**C1**

Variables
RO4 Provide jtd or Dst
CalibOnly?
2 1 to reproduce of
oNormalize?
2 1=no, 2=yes

ofxn calc*
1.60127
ofxn mod
1.60506

sRun
RunA1
zout
1

Ave
Data
0.00273
Model
0.00102

data/vMax
0.02142

ofxng
275.646

Min d/m
1.4

ave/min d
12.4694
ave/min m
6.51042

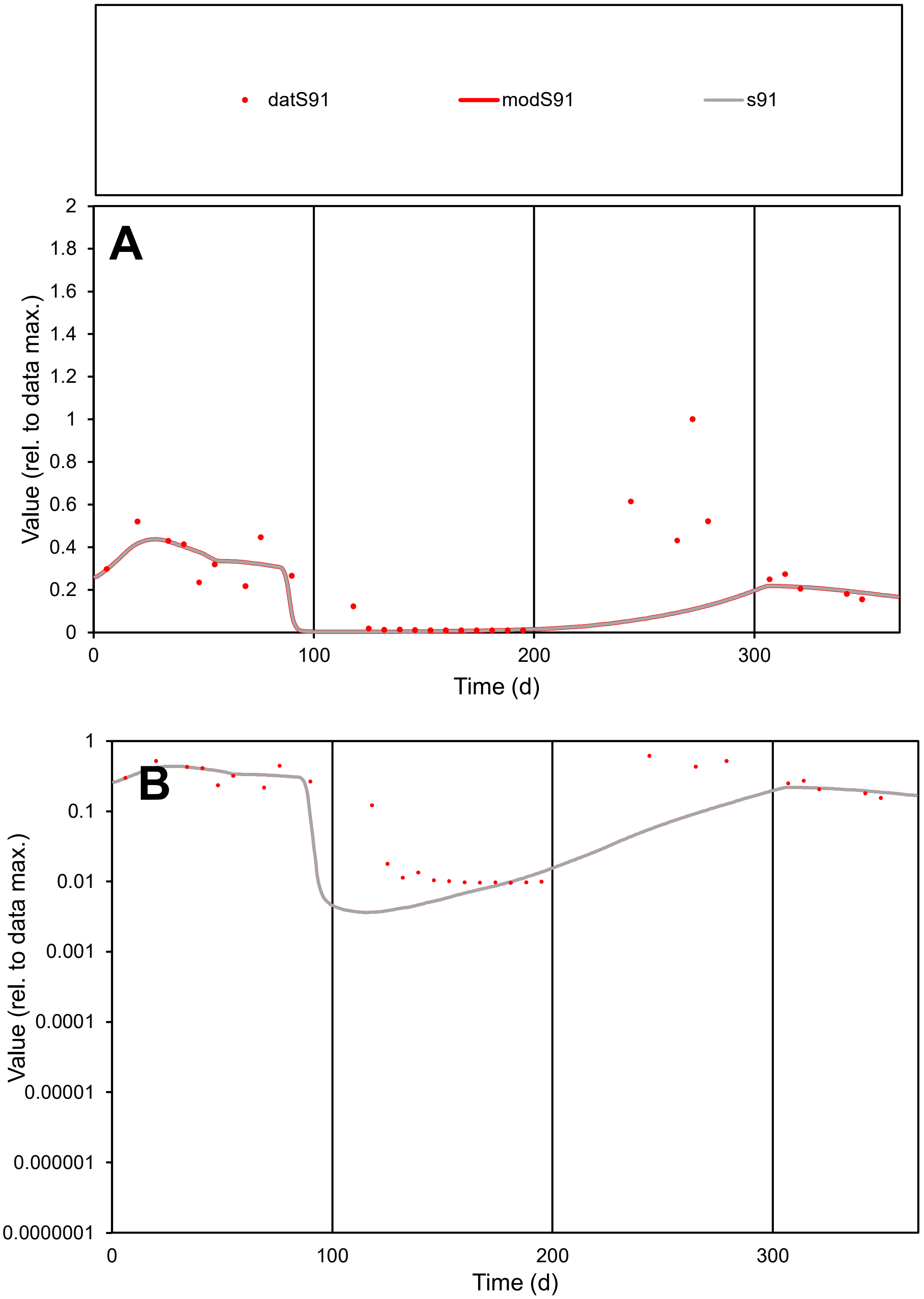


**C1**

Variables
S91 Provide jtd or Dst
CalibOnly?
2 1 to reproduce of
oNormalize?
2 1=no, 2=yes

ofxn calc*
0.93734
ofxn mod
0.95334

sRun
RunA1
zout
1

Ave
Data
0.00159
Model
0.00105

data/vMax
0.00678

ofxng
275.646

Min d/m
2.61236

ave/min d
24.3609
ave/min m
42.0682

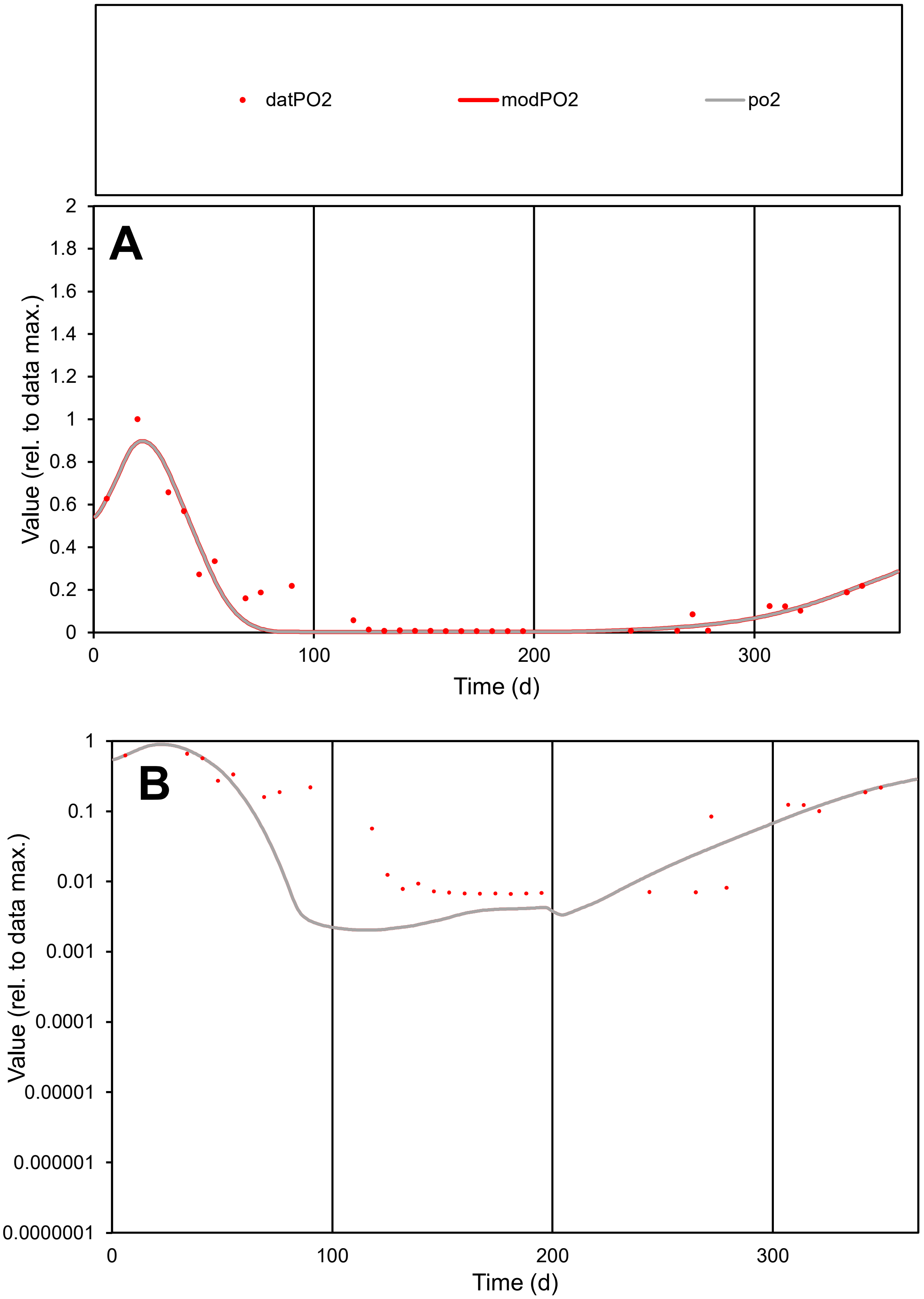


**C1**

Variables

PO2 Provide jtd or Dst

CalibOnly?

2 1 to reproduce of

oNormalize?

2 1=no, 2=yes

ofxn calc*

0.39505

ofxn mod

0.44042

sRun

RunA1

zout

1

Ave

Data

0.00199

Model

0.00173

data/vMax

0.01186

ofxng

275.646

Min d/m

3.2566

ave/min d

25.1707

ave/min m

71.5713

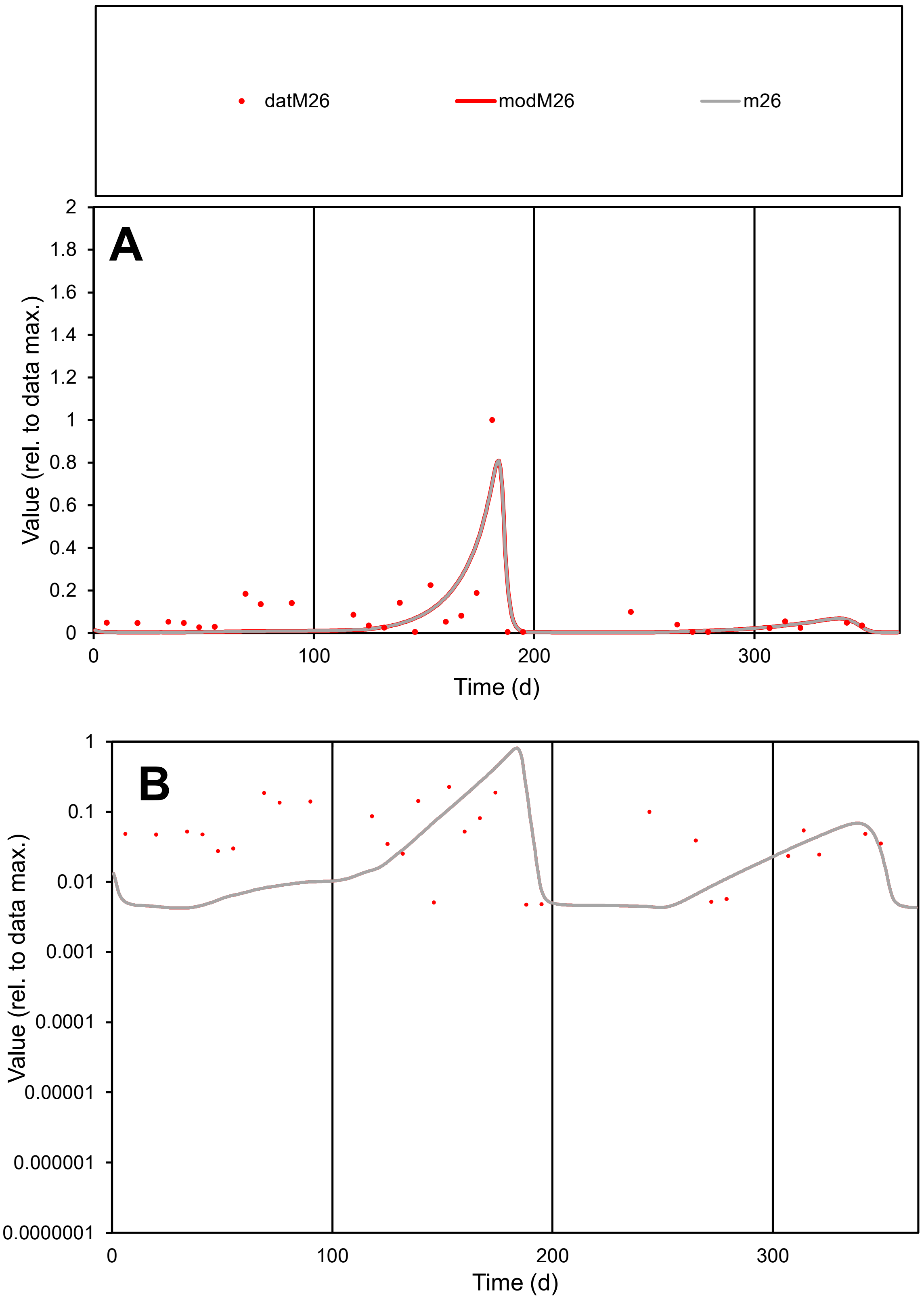


**C1**

Variables
M26 Provide jtd or Dst
CalibOnly?
2 1 to reproduce of
oNormalize?
2 1=no, 2=yes

ofxn calc*
0.99476
ofxn mod
1.02816

sRun
RunA1
zout
1

Ave
Data
0.00168
Model
0.00134

data/vMax
0.01739

ofxng
275.646

Min d/m
1.09912

ave/min d
20.3908
ave/min m
17.8783

325_Calib_S_A

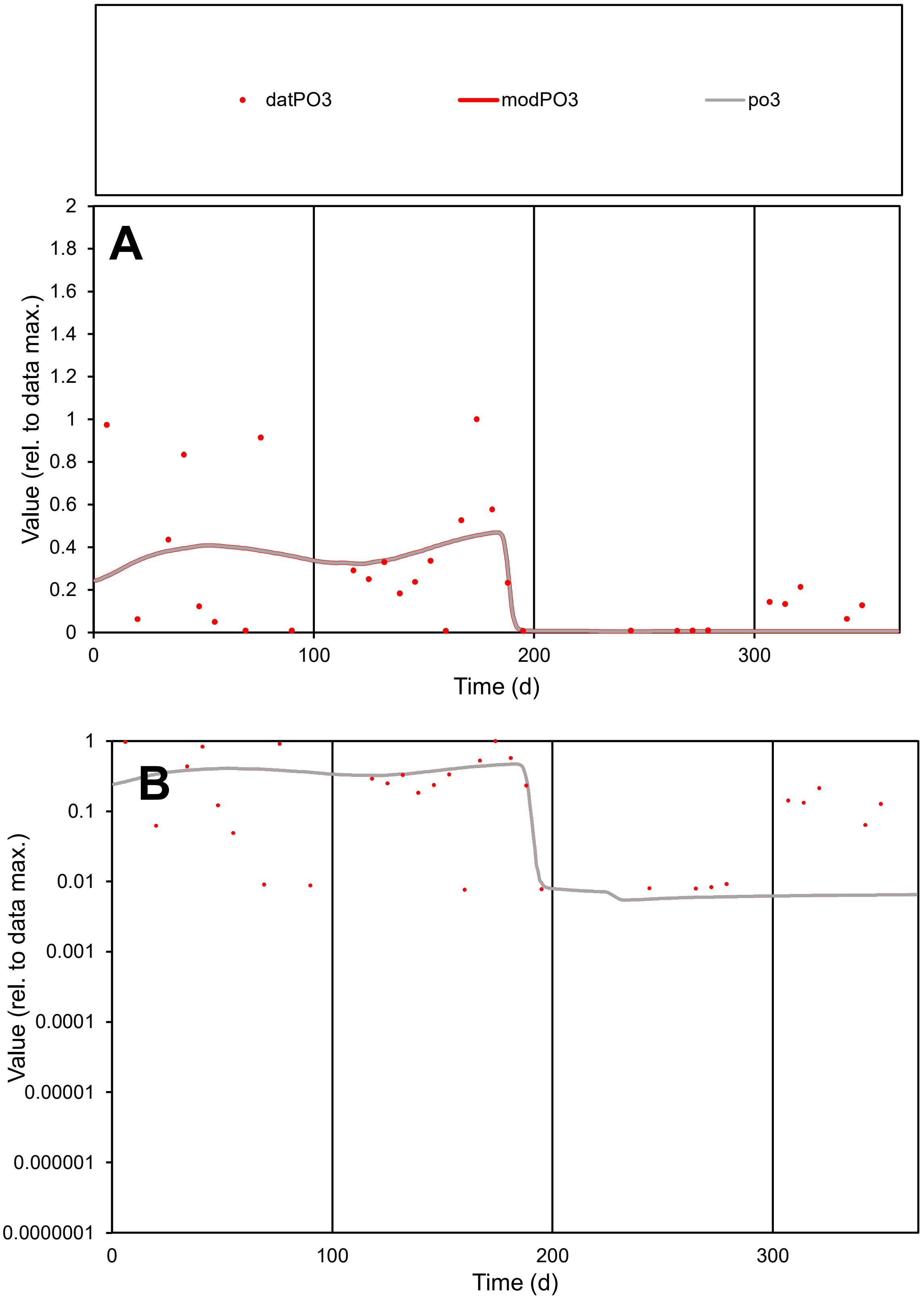


**C1**

Variables
PO3 Provide jtd or Dst
CalibOnly?
2 1 to reproduce of
oNormalize?
2 1=no, 2=yes

ofxn calc*
0.995
ofxn mod
1.00377

sRun
RunA1
zout
1

Ave
Data
0.00166
Model
0.00153

data/vMax
0.00613

ofxng
275.646

Min d/m
1.35123

ave/min d
35.3879
ave/min m
44.2909

325_Calib_S_A

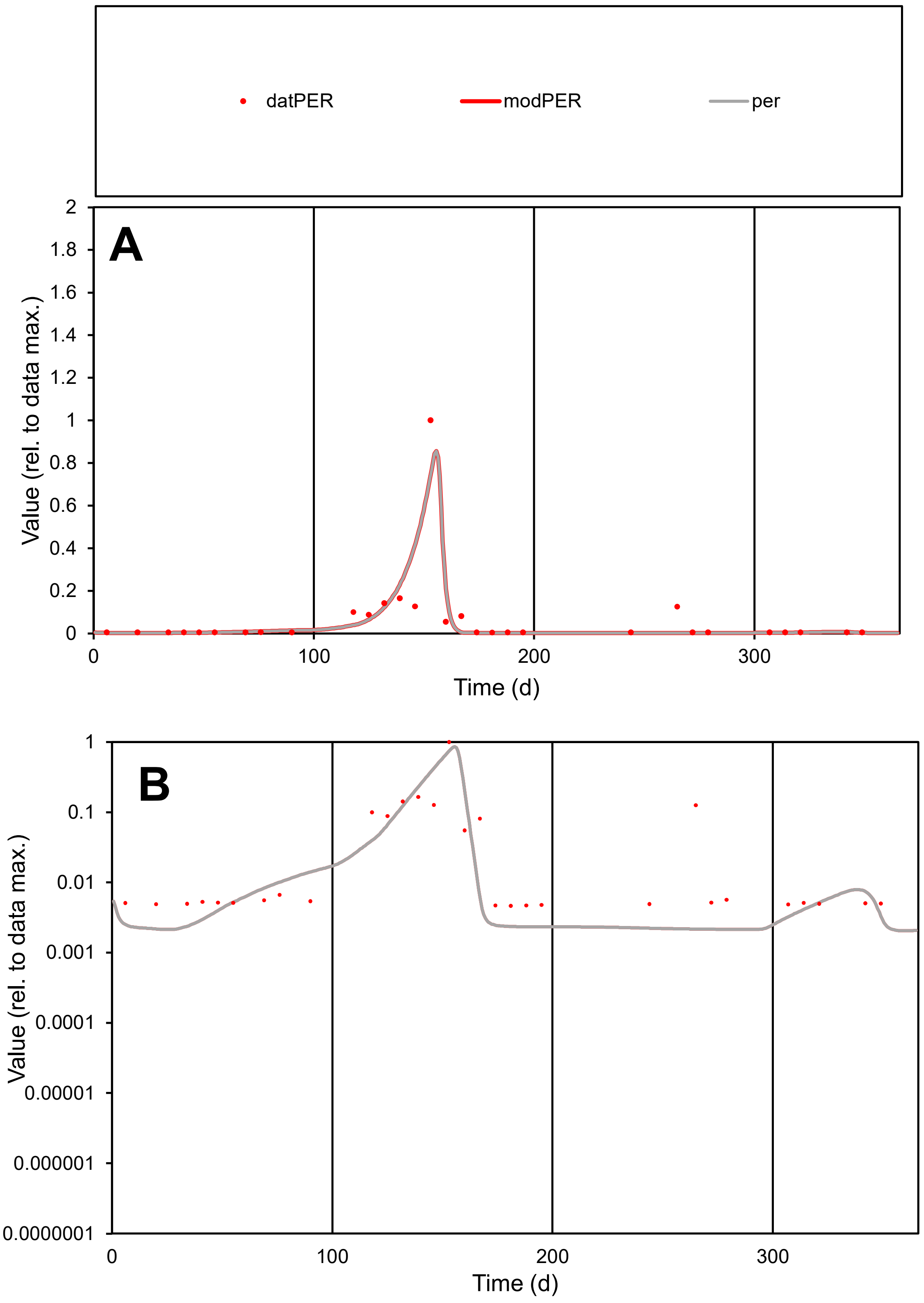


**C1**

Variables

PER Provide jtd or Dst

CalibOnly?

2 1 to reproduce of

oNormalize?

2 1=no, 2=yes

ofxn calc*

1.08695

ofxn mod

1.12492

sRun

RunA1

zout

1

Ave

Data

0.00026

Model

0.00025

data/vMax

0.00392

ofxng

275.646

Min d/m

2.15131

ave/min d

14.3179

ave/min m

30.0345

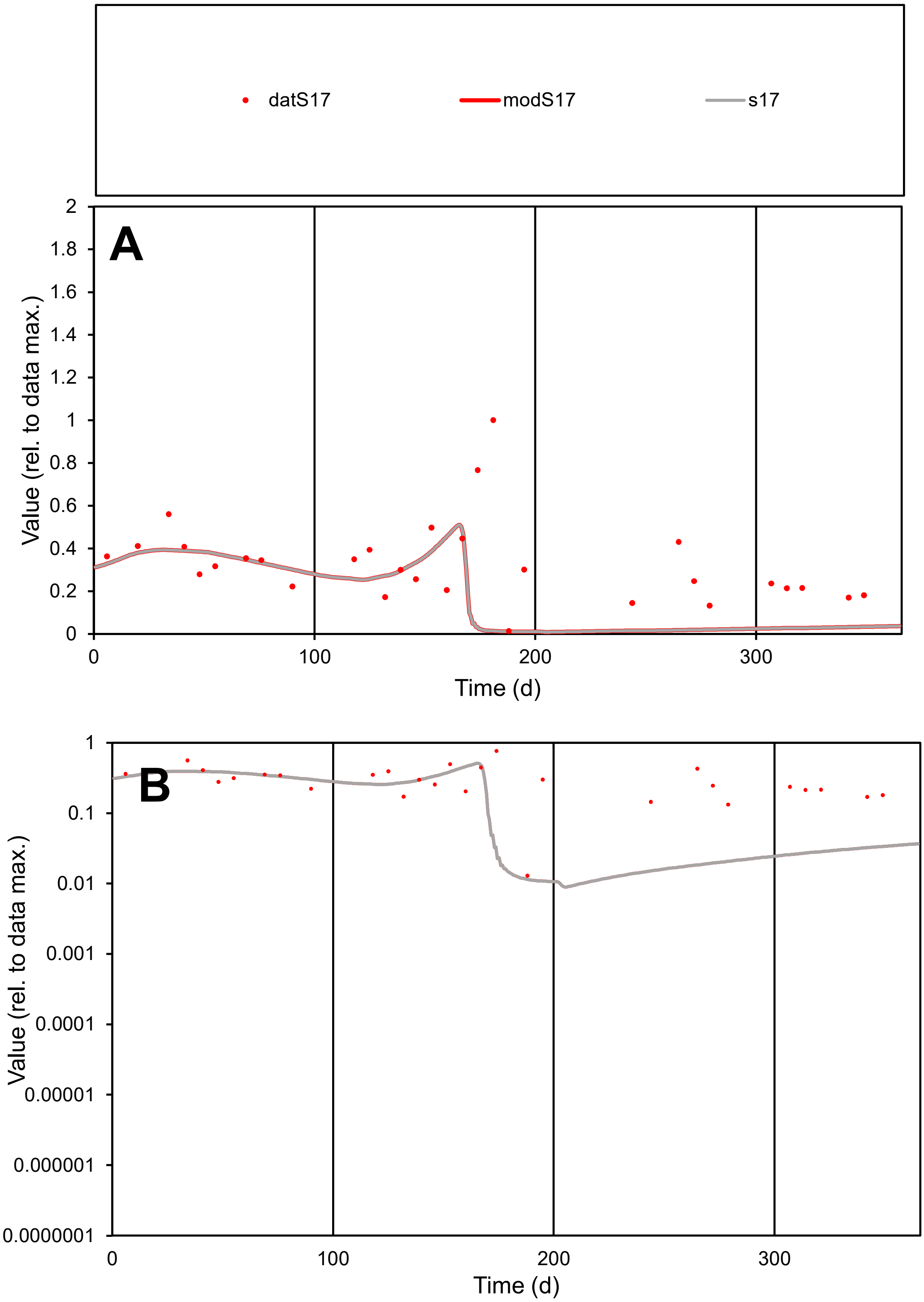


| | |
|---|---|
| **C1** | |
| Variables | |
| S17 | Provide jtd or Dst |
| CalibOnly? | |
| 2 | 1 to reproduce of |
| oNormalize? | |
| 2 | 1=no, 2=yes |
| ofxn calc* | |
| 0.81573 | |
| ofxn mod | |
| 0.823 | |
| sRun | |
| RunA1 | |
| zout | |
| 1 | |
| Ave Data | |
| 0.0028 | |
| Model | |
| 0.00175 | |
| data/vMax | |
| 0.00845 | |
| ofxng | |
| 275.646 | |
| Min d/m | |
| 1.20396 | |
| ave/min d | |
| 25.5374 | |
| ave/min m | |
| 19.22 | |

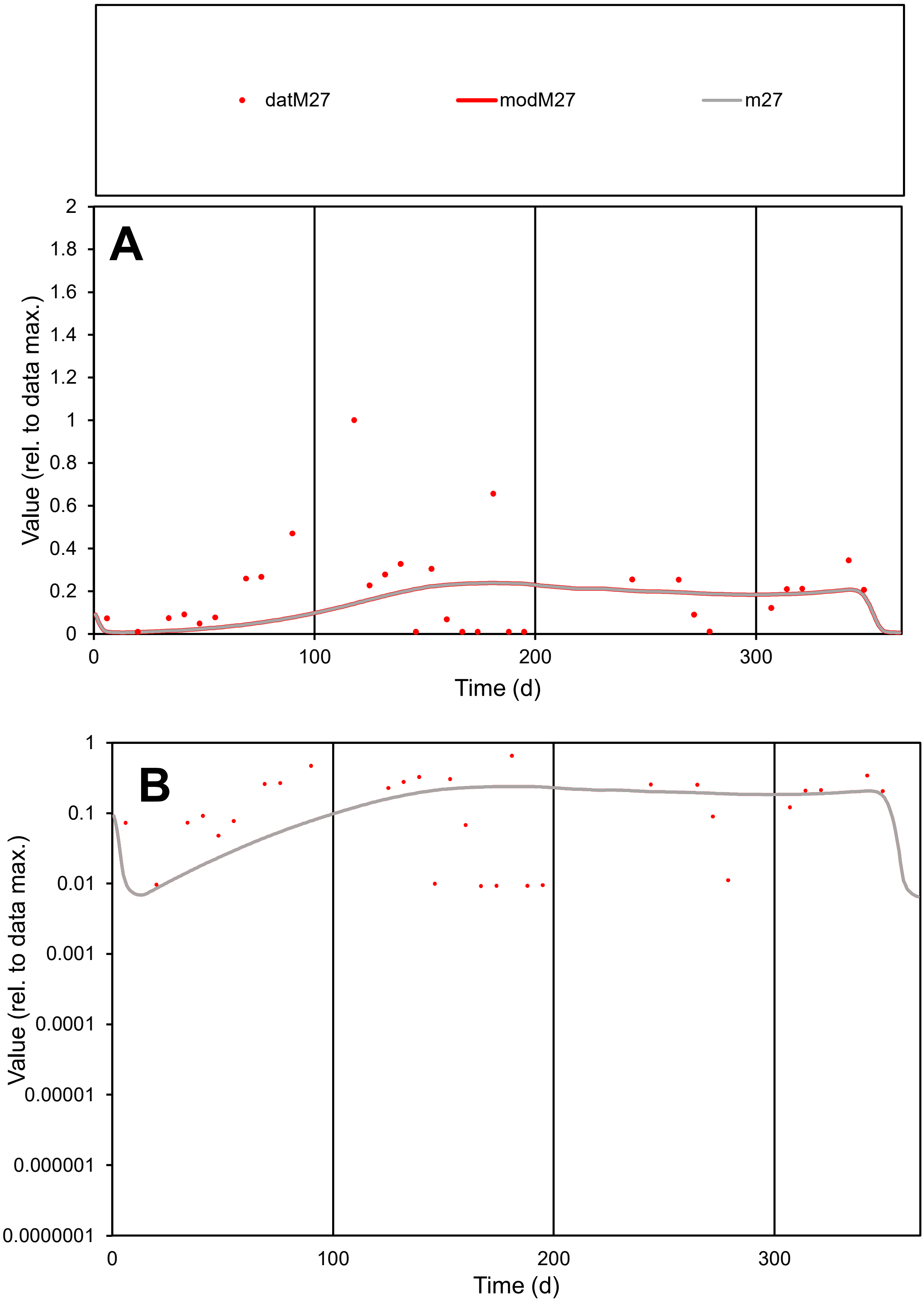


**C1**

Variables

M27 Provide jtd or Dst

CalibOnly?

2 1 to reproduce of

oNormalize?

2 1=no, 2=yes

ofxn calc*

1.13594

ofxn mod

1.14331

sRun

RunA1

zout

1

Ave

Data

0.0014

Model

0.00108

data/vMax

0.00704

ofxng

275.646

Min d/m

1.03784

ave/min d

21.5891

ave/min m

17.2209

325_Calib_S_A

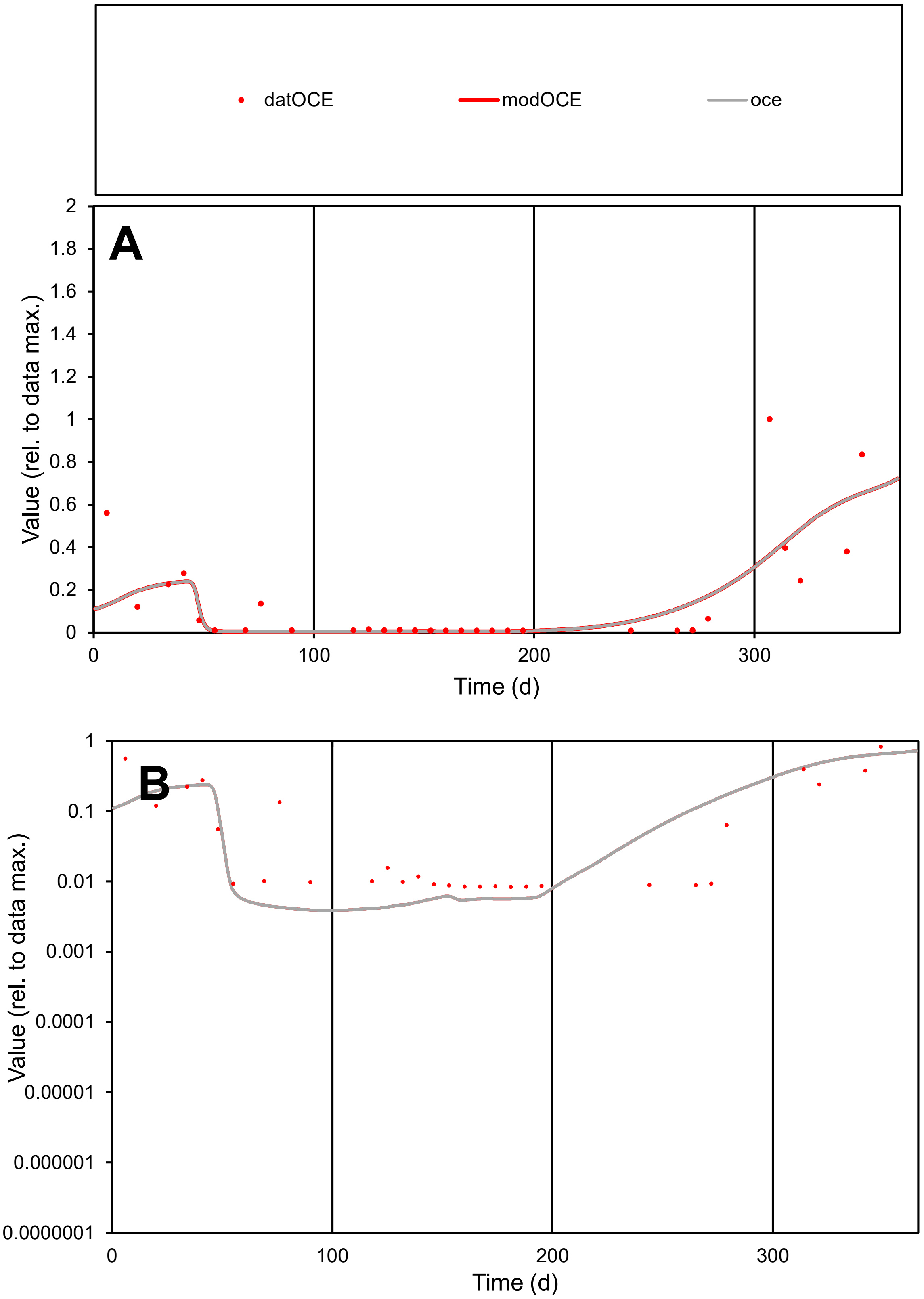


**C1**

Variables
OCE Provide jtd or Dst
CalibOnly?
2 1 to reproduce of
oNormalize?
2 1=no, 2=yes

ofxn calc*
1.09509
ofxn mod
1.11313

sRun
RunA1
zout
1

Ave
Data
0.00065
Model
0.00059

data/vMax
0.0044

ofxng
275.646

Min d/m
2.14302

ave/min d
17.7428
ave/min m
34.2867

325_Calib_S_A

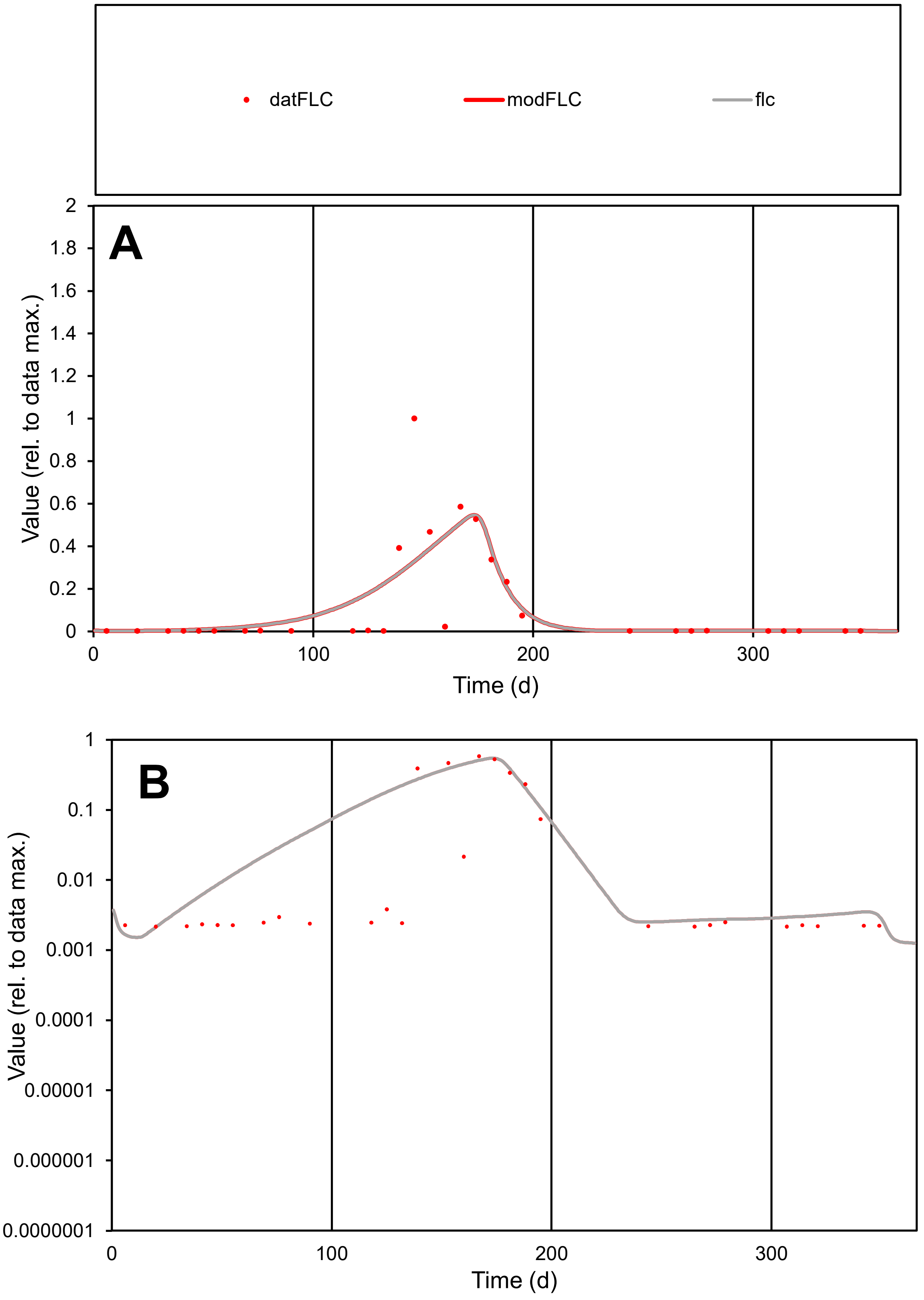


**C1**

Variables
FLC Provide jtd or Dst
CalibOnly?
2 1 to reproduce of
oNormalize?
2 1=no, 2=yes

ofxn calc*
1.27841
ofxn mod
1.29742

sRun
RunA1
zout
1

Ave
Data
0.00072
Model
0.00075

data/vMax
0.00582

ofxng
275.646

Min d/m
1.34923

ave/min d
56.9542
ave/min m
80.2935

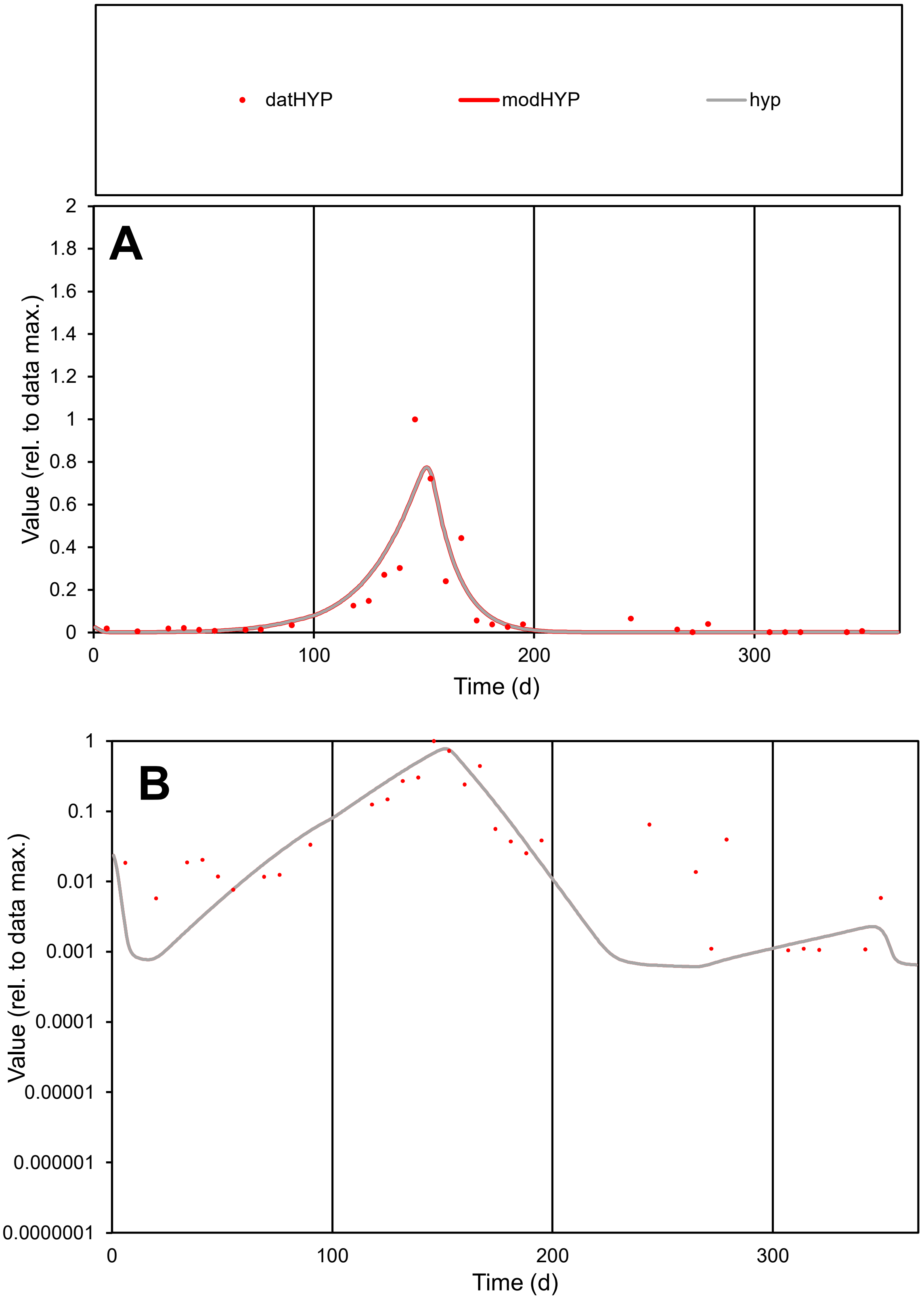


C1

Variables

HYP Provide jtd or Dst

CalibOnly?

2 1 to reproduce of

oNormalize?

2 1=no, 2=yes

ofxn calc*

0.72433

ofxn mod

0.77251

sRun

RunA1

zout

1

Ave

Data

0.00194

Model

0.002

data/vMax

0.01583

ofxng

275.646

Min d/m

1.70887

ave/min d

116.803

ave/min m

205.703

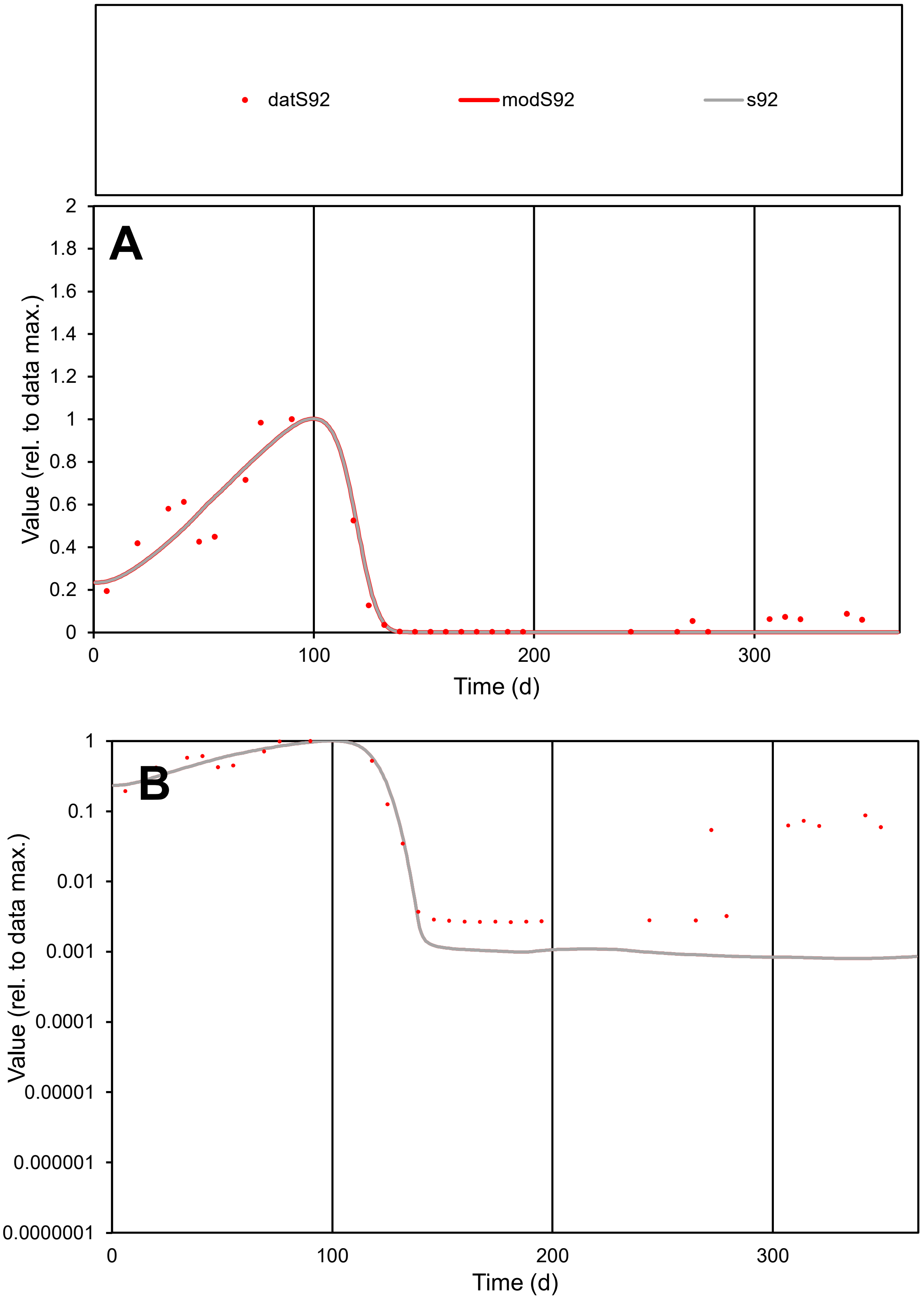


**C1**

Variables
S92 Provide jtd or Dst
CalibOnly?
2 1 to reproduce of
oNormalize?
2 1=no, 2=yes

ofxn calc*
0.33024
ofxn mod
0.38909

sRun
RunA1
zout
1

Ave
Data
0.00189
Model
0.00177

data/vMax
0.00874

ofxng
275.646

Min d/m
3.28543

ave/min d
82.0084
ave/min m
251.983

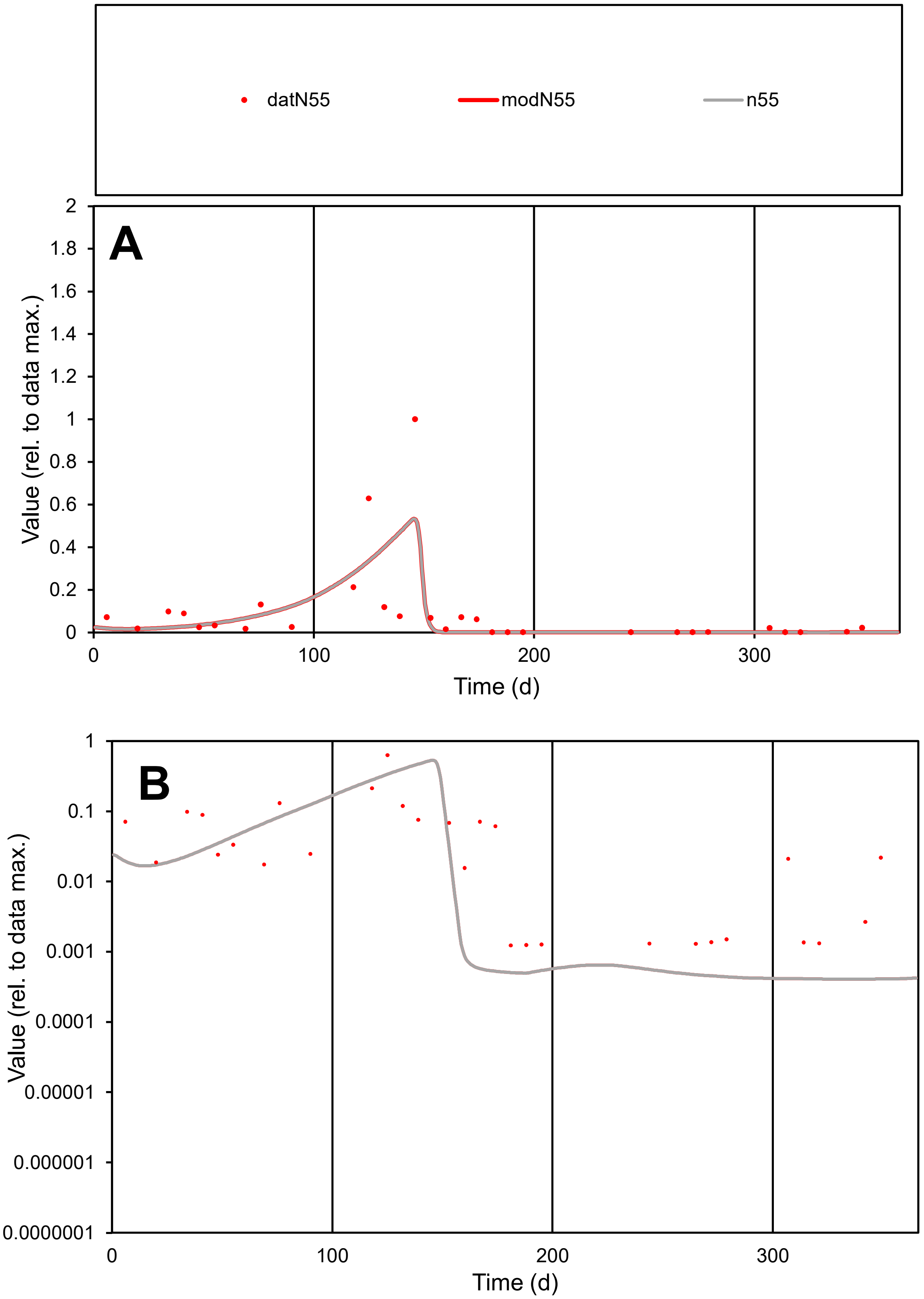


**C1**

Variables

N55 Provide jtd or Dst

CalibOnly?

2 1 to reproduce of

oNormalize?

2 1=no, 2=yes

ofxn calc*

1.48282

ofxn mod

1.53235

sRun

RunA1

zout

1

Ave

Data

0.00282

Model

0.00257

data/vMax

0.03006

ofxng

275.646

Min d/m

3.01058

ave/min d

76.294

ave/min m

208.886

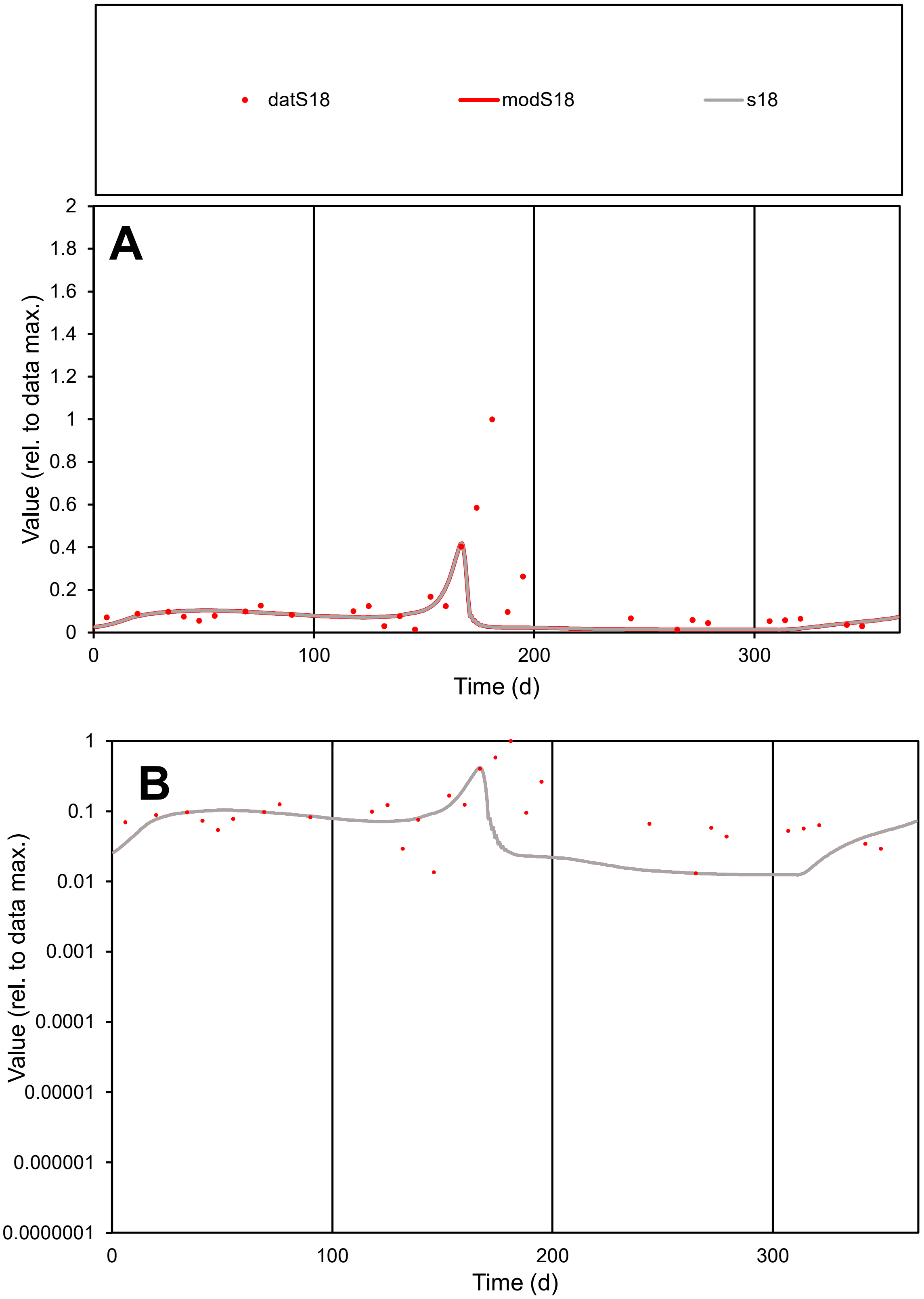


**C1**

Variables
S18 Provide jtd or Dst
CalibOnly?
2 1 to reproduce of
oNormalize?
2 1=no, 2=yes

ofxn calc*
1.52588
ofxn mod
1.53219

sRun
RunA1
zout
1

Ave
Data
0.00295
Model
0.00158

data/vMax
0.02123

ofxng
275.646

Min d/m
1.05356

ave/min d
10.5451
ave/min m
5.97211

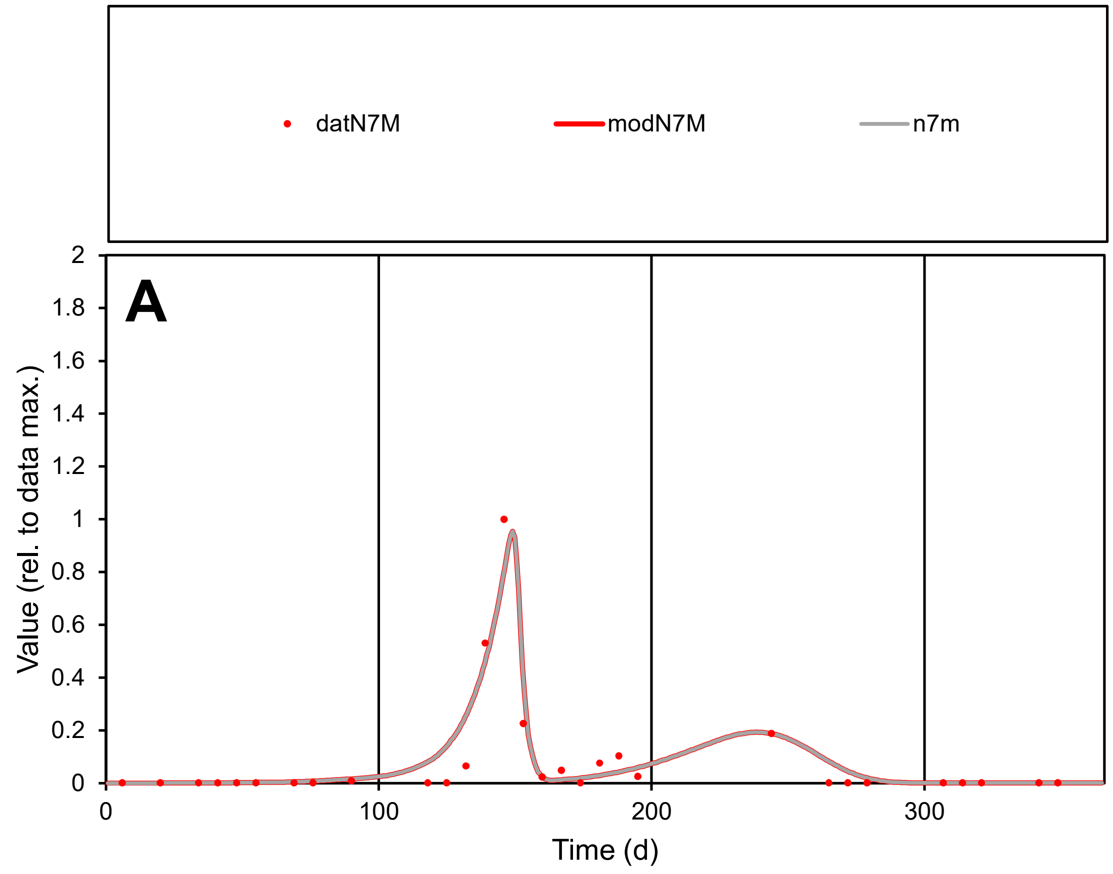


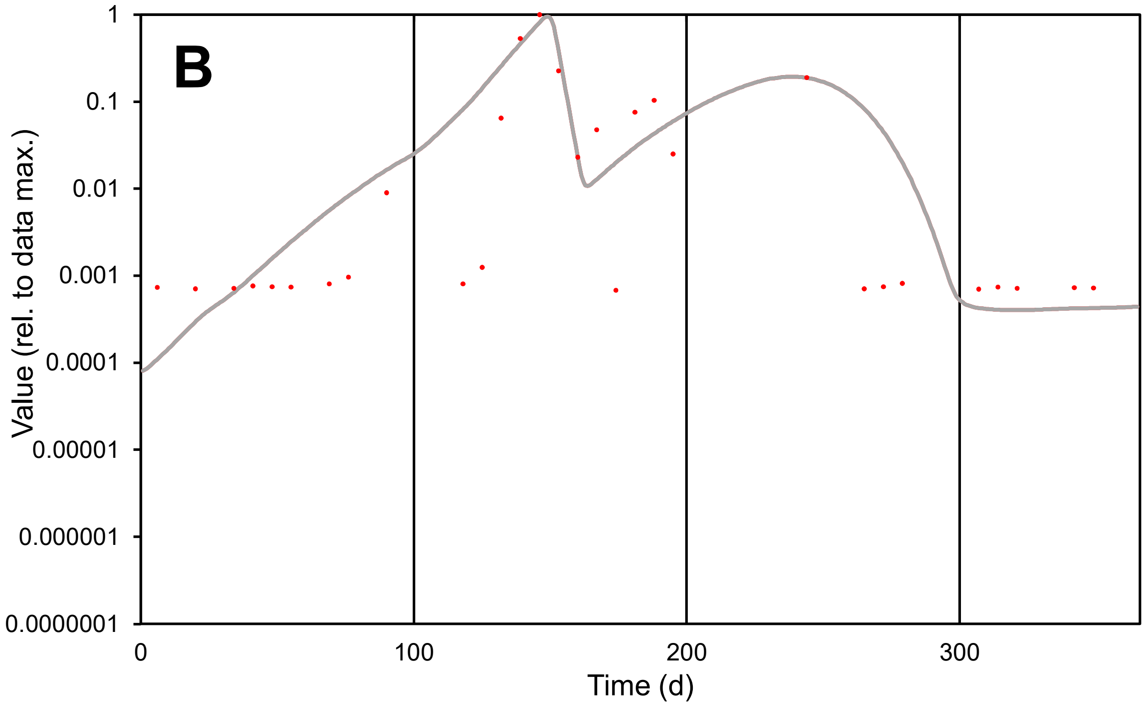


**C1**

Variables
N7M Provide jtd or Dst
CalibOnly?
2 1 to reproduce of
oNormalize?
2 1=no, 2=yes

ofxn calc*
0.66476
ofxn mod
0.77879

sRun
RunA1
zout
1

Ave
Data
0.0011
Model
0.00126

data/vMax
0.01427

ofxng
275.646

Min d/m
5.71252

ave/min d
112.996
ave/min m
740.646

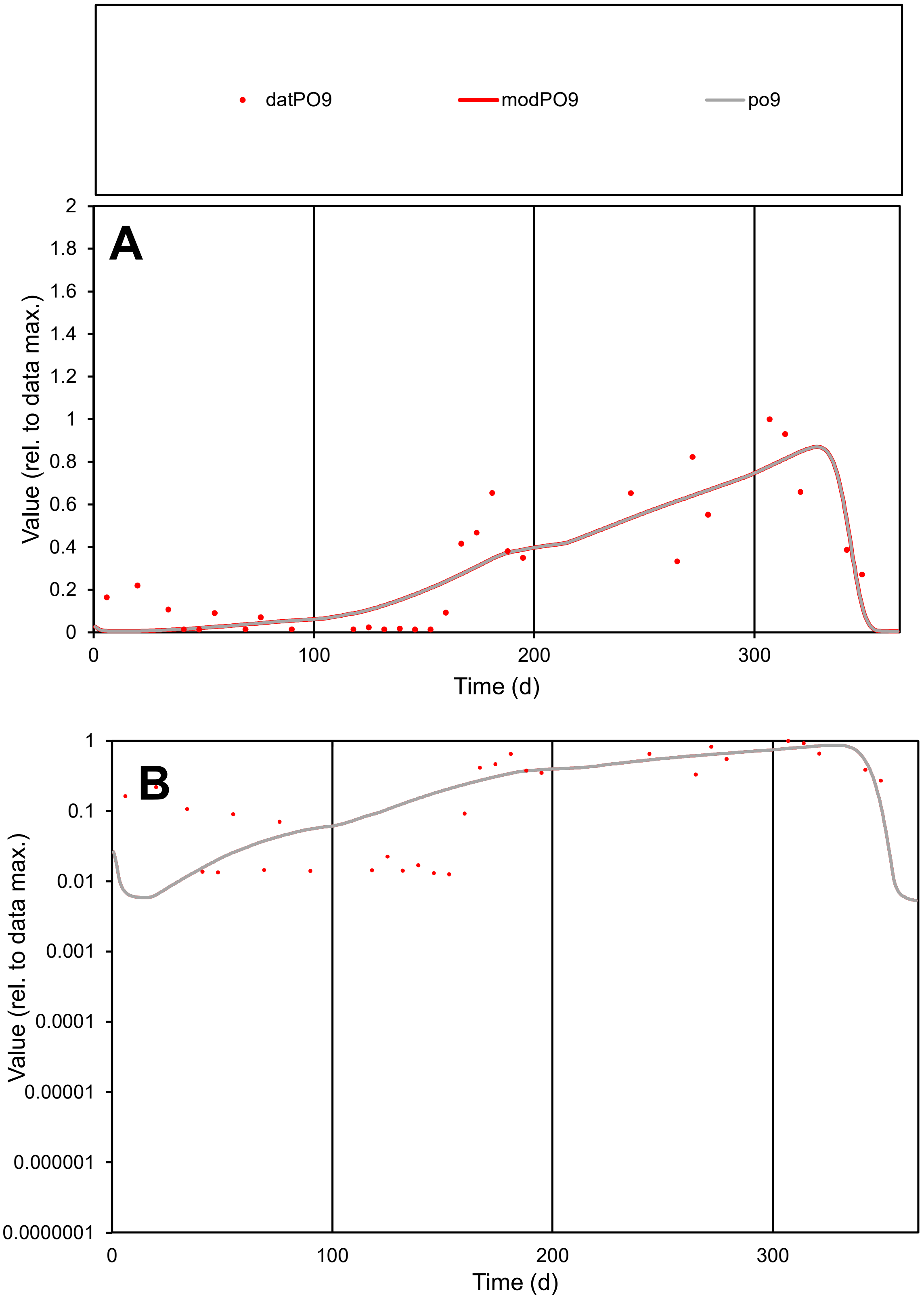


**C1**

Variables
PO9 Provide jtd or Dst
CalibOnly?
2 1 to reproduce of
oNormalize?
2 1=no, 2=yes

ofxn calc*
0.48765
ofxn mod
0.51696

sRun
RunA1
zout
1

Ave
Data
0.00141
Model
0.00138

data/vMax
0.00481

ofxng
275.646

Min d/m
1.93611

ave/min d
23.0681
ave/min m
43.7348

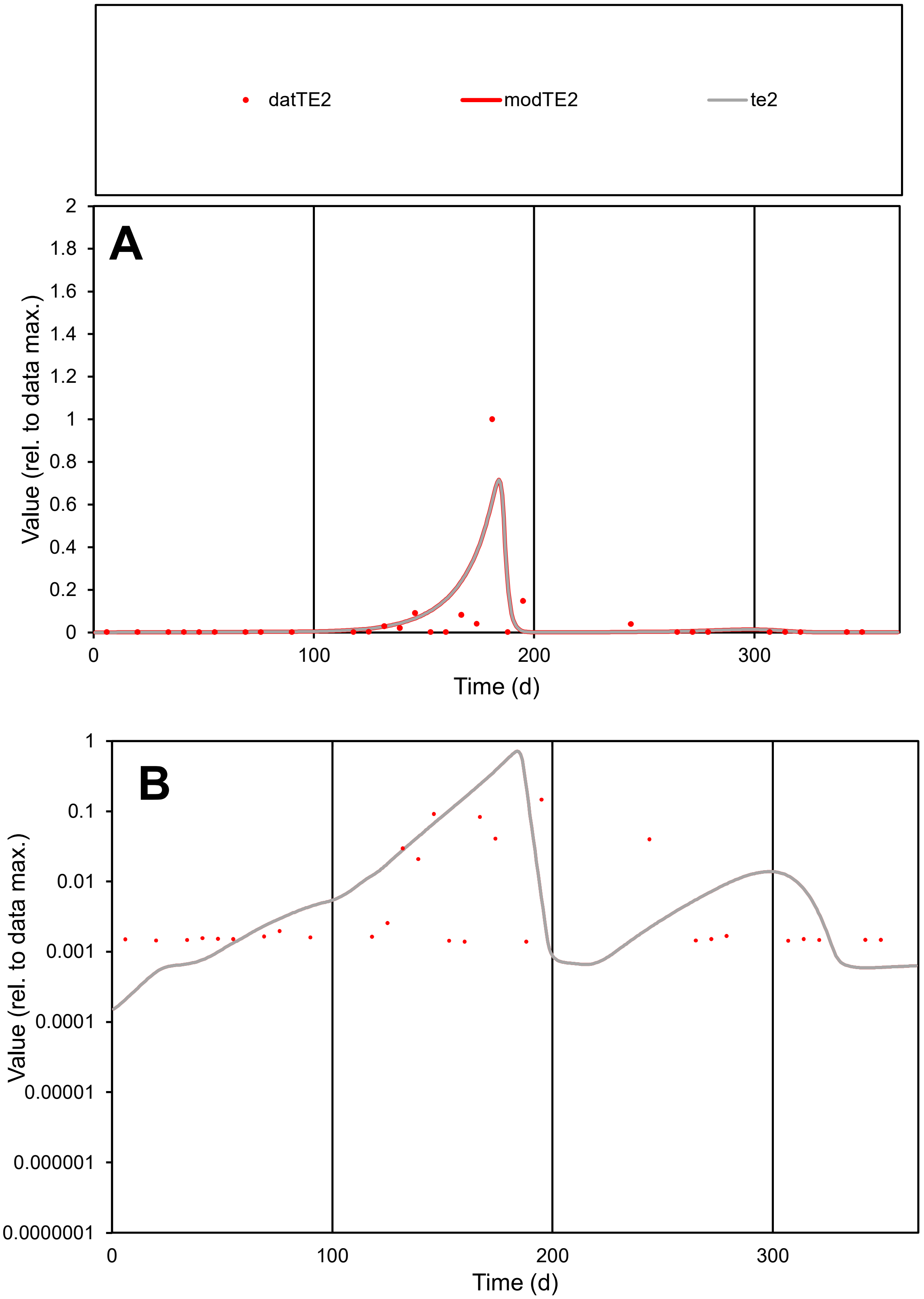


**C1**

Variables
TE2 Provide jtd or Dst
CalibOnly?
2 1 to reproduce of
oNormalize?
2 1=no, 2=yes

ofxn calc*
1.96538
ofxn mod
2.06282

sRun
RunA1
zout
1

Ave
Data
0.00307
Model
0.00378

data/vMax
0.06191

ofxng
275.646

Min d/m
6.13195

ave/min d
35.7163
ave/min m
269.853

325_Calib_S_A

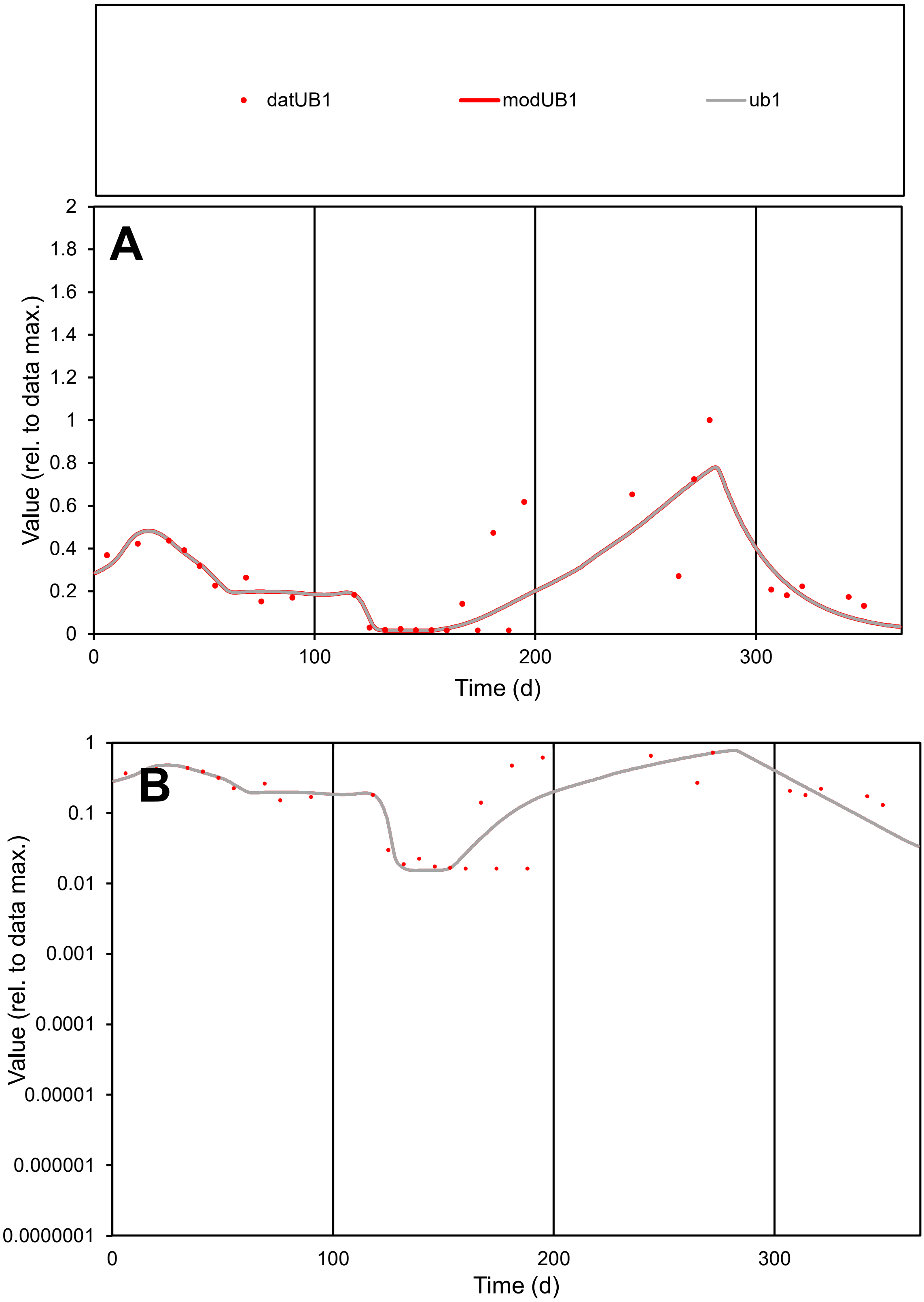


**C1**

Variables
UB1 Provide jtd or Dst
CalibOnly?
2 1 to reproduce of
oNormalize?
2 1=no, 2=yes

ofxn calc*
0.54464
ofxn mod
0.5697

sRun
RunA1
zout
1

Ave
Data
0.00093
Model
0.00083

data/vMax
0.00353

ofxng
275.646

Min d/m
1.05452

ave/min d
16.1523
ave/min m
15.2729

325_Calib_S_A

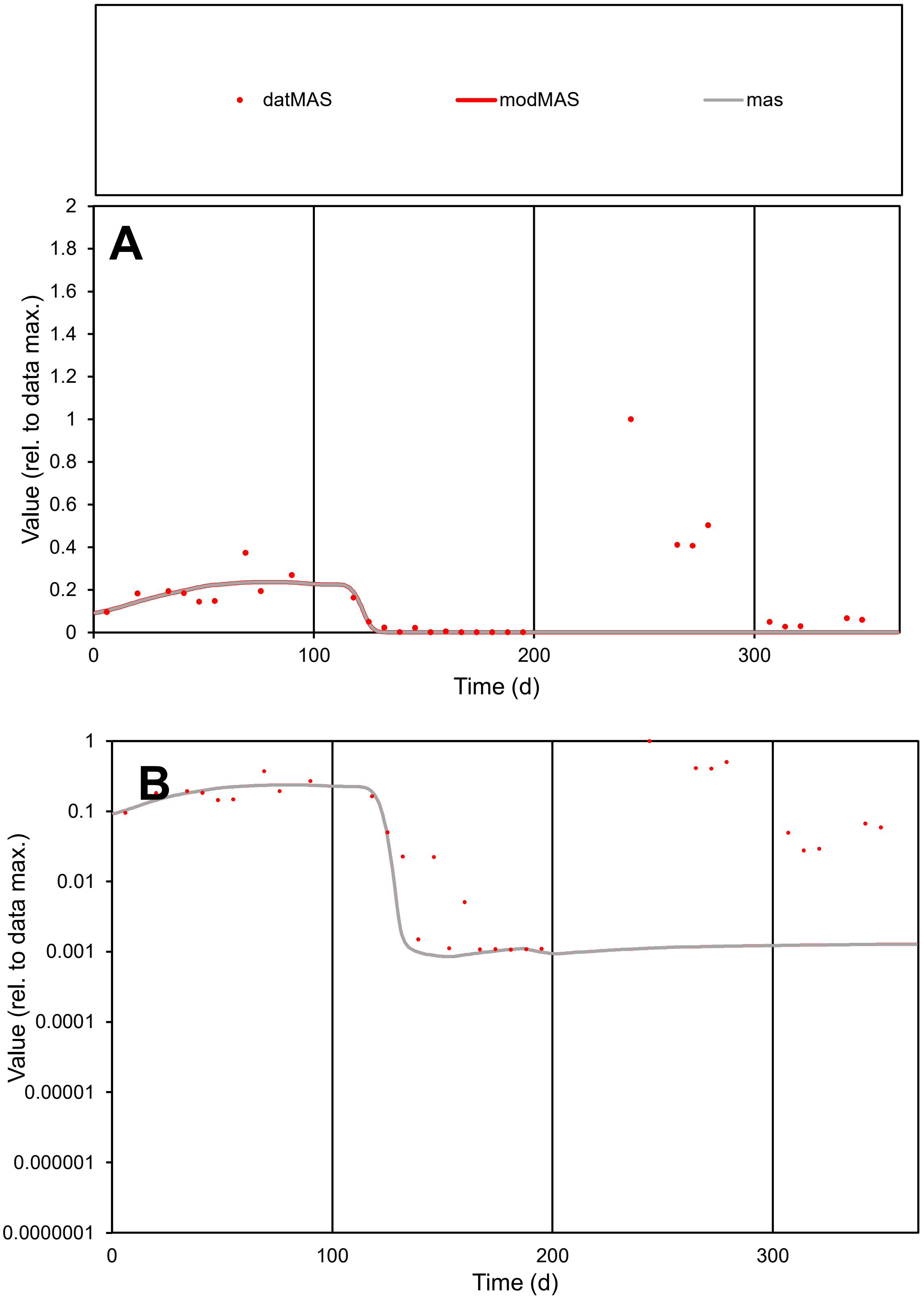


**C1**

Variables

MAS Provide jtd or Dst

CalibOnly?

2 1 to reproduce of

oNormalize?

2 1=no, 2=yes

ofxn calc*

1.51685

ofxn mod

1.52174

sRun

RunA1

zout

1

Ave

Data

0.00171

Model

0.00075

data/vMax

0.01115

ofxng

275.646

Min d/m

1.25829

ave/min d

143.349

ave/min m

78.9169

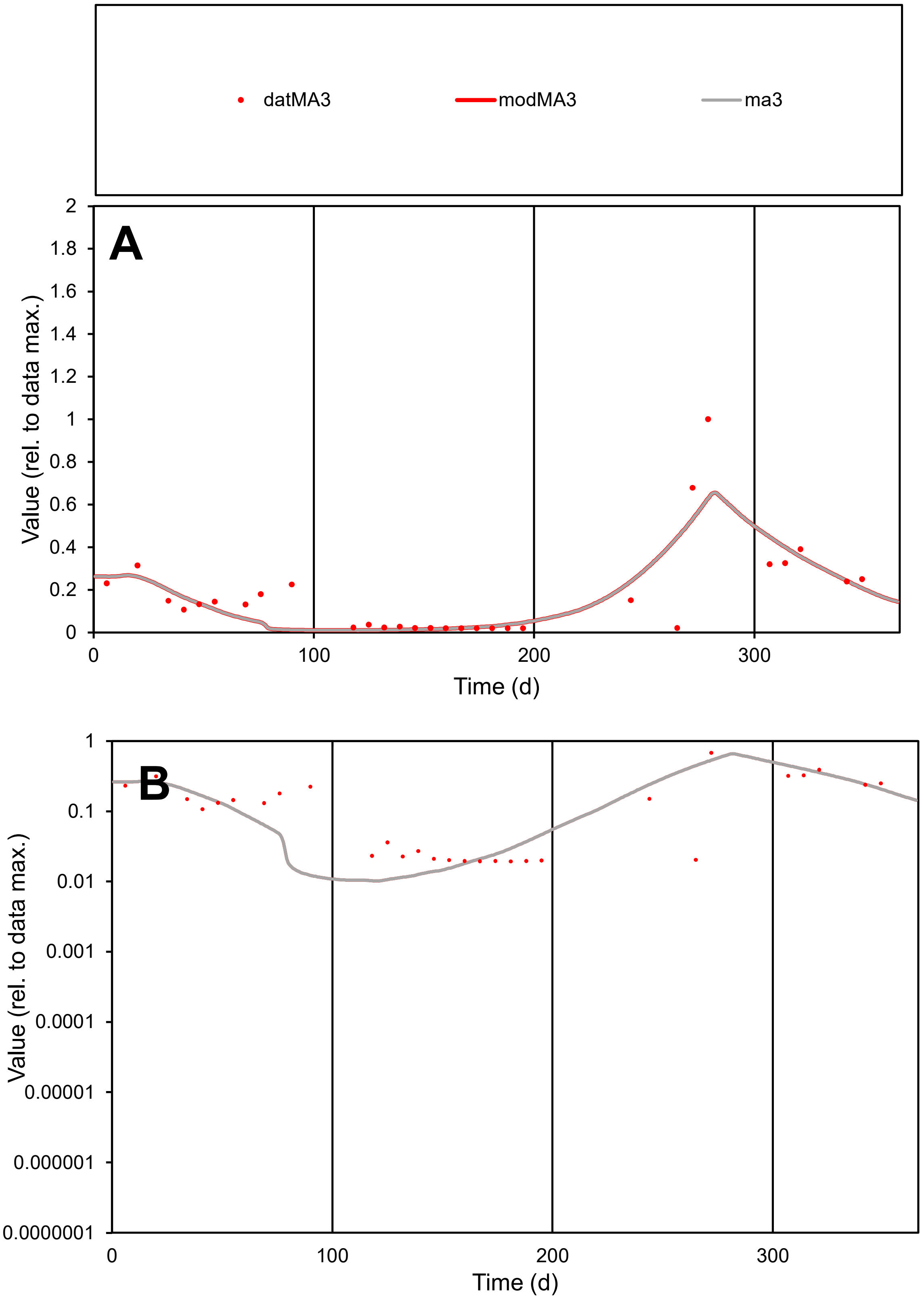


**C1**

Variables
MA3 Provide jtd or Dst
CalibOnly?
2 1 to reproduce of
oNormalize?
2 1=no, 2=yes

ofxn calc*
0.68798
ofxn mod
0.71757

sRun
RunA1
zout
1

Ave
Data
0.00056
Model
0.00053

data/vMax
0.00319

ofxng
275.646

Min d/m
1.88657

ave/min d
9.07213
ave/min m
16.2951

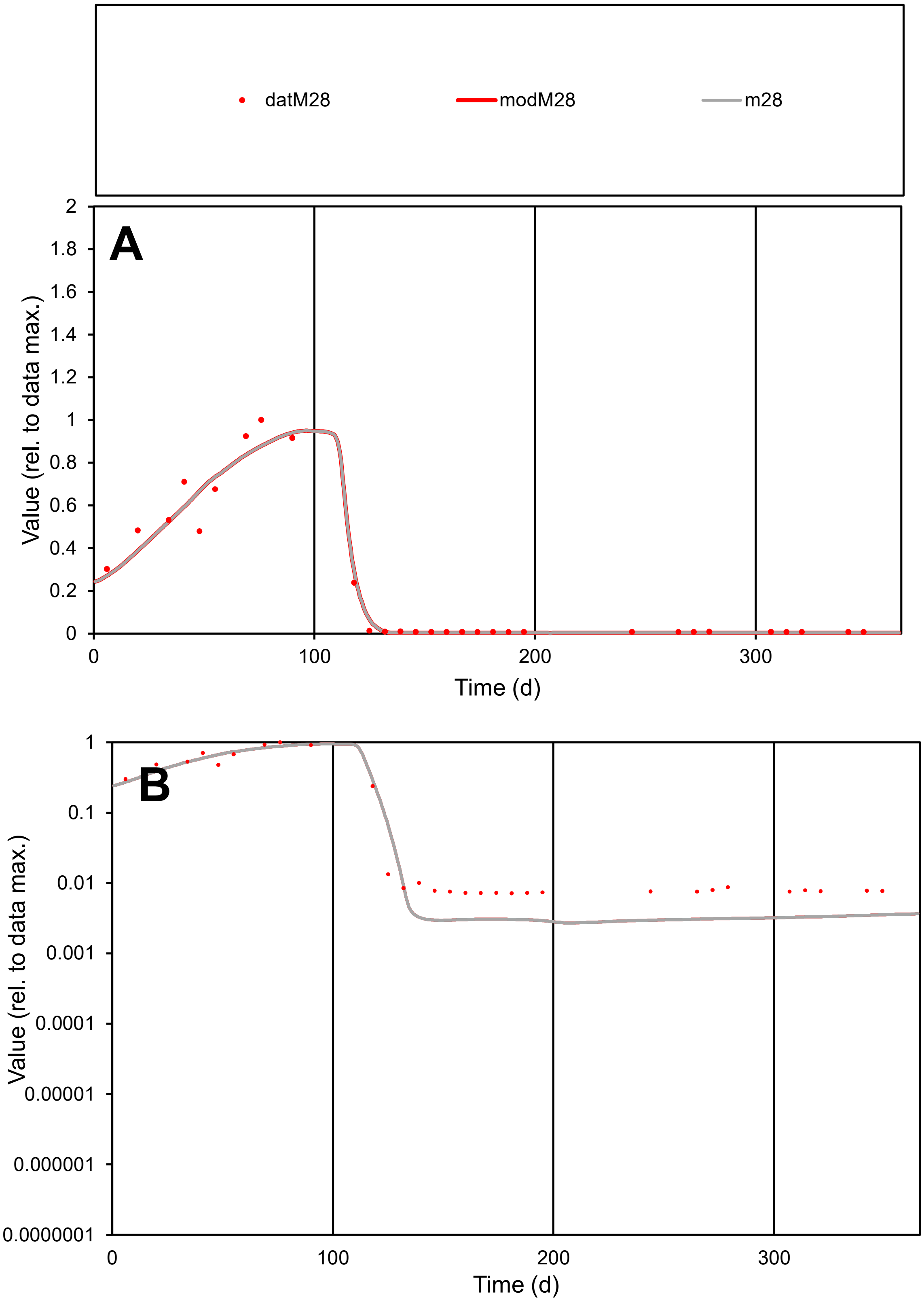


**C1**

Variables

M28 Provide jtd or Dst

CalibOnly?

2 1 to reproduce of

oNormalize?

2 1=no, 2=yes

ofxn calc*

0.24656

ofxn mod

0.32737

sRun

RunA1

zout

1

Ave

Data

0.0016

Model

0.00156

data/vMax

0.00749

ofxng

275.646

Min d/m

2.47091

ave/min d

29.9437

ave/min m

72.0714

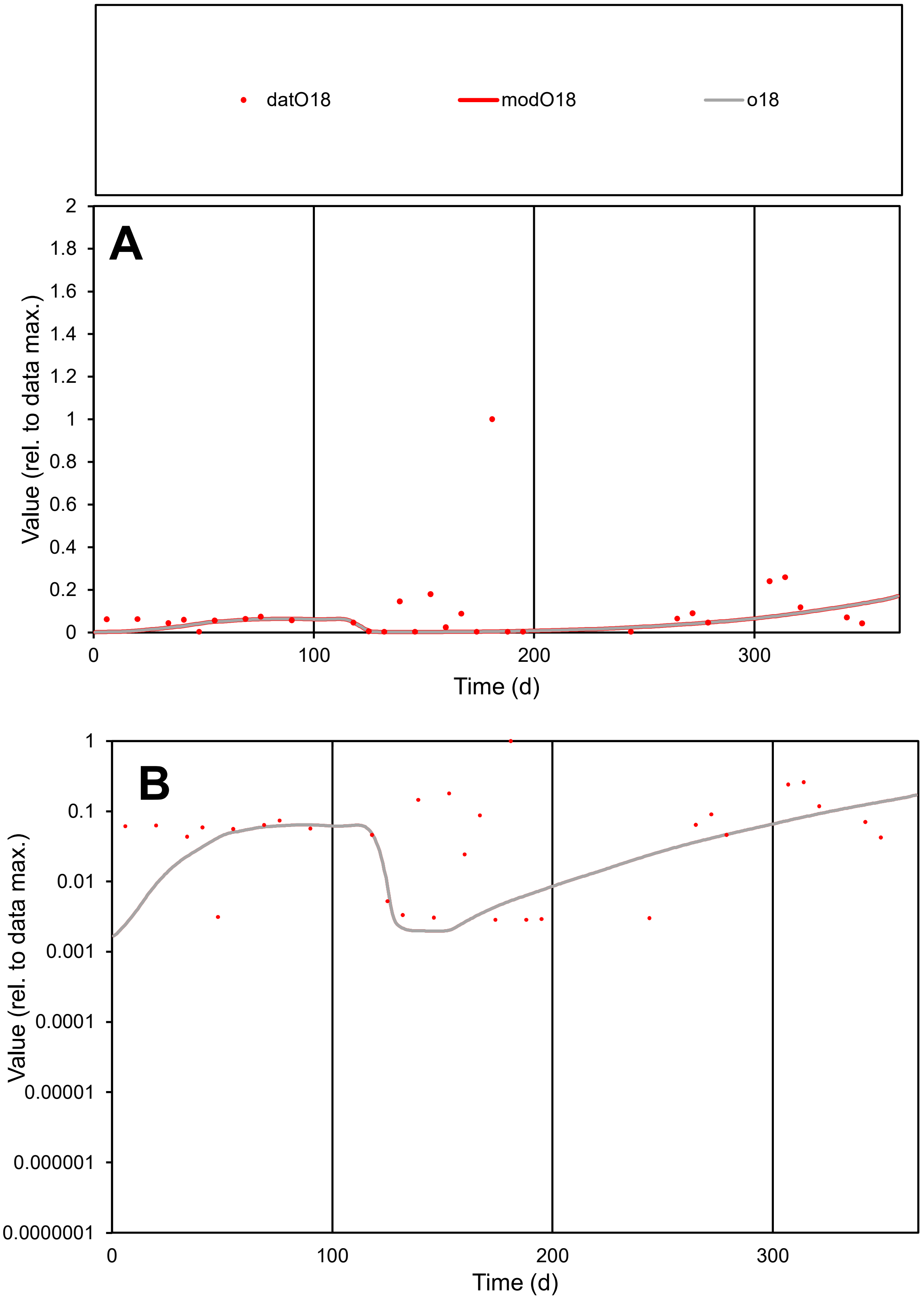


**C1**

Variables
O18 Provide jtd or Dst
CalibOnly?
2 1 to reproduce of
oNormalize?
2 1=no, 2=yes

ofxn calc*
1.99619
ofxn mod
2.00125

sRun
RunA1
zout
1

Ave
Data
0.00177
Model
0.00067

data/vMax
0.01818

ofxng
275.646

Min d/m
1.44866

ave/min d
34.053
ave/min m
18.6163

325_Calib_S_A

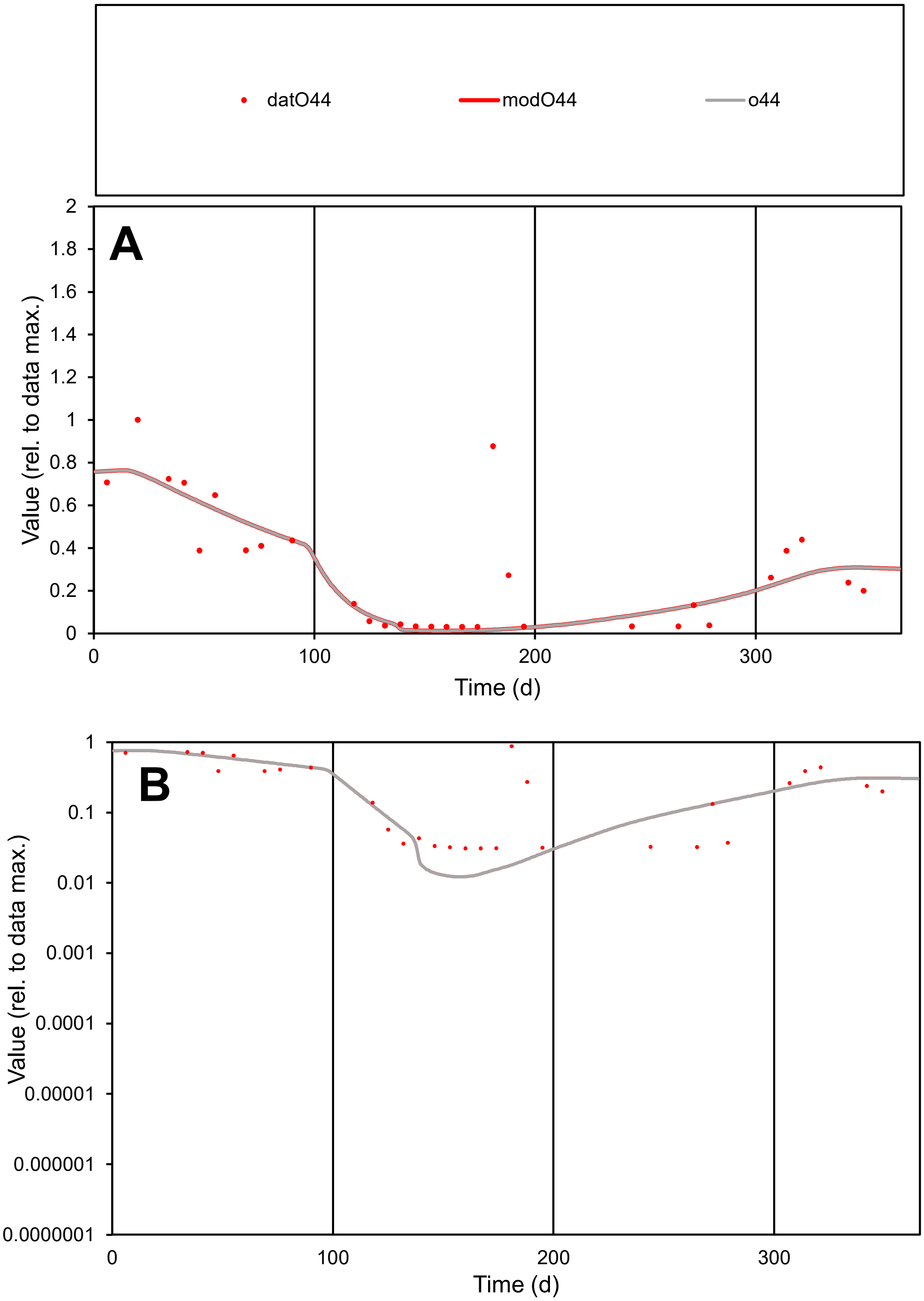


**C1**

Variables
O44 Provide jtd or Dst
CalibOnly?
2 1 to reproduce of
oNormalize?
2 1=no, 2=yes

ofxn calc*
0.63535
ofxn mod
0.65272

sRun
RunA1
zout
1

Ave
Data
0.00105
Model
0.00092

data/vMax
0.00358

ofxng
275.646

Min d/m
2.50283

ave/min d
9.49348
ave/min m
20.8861

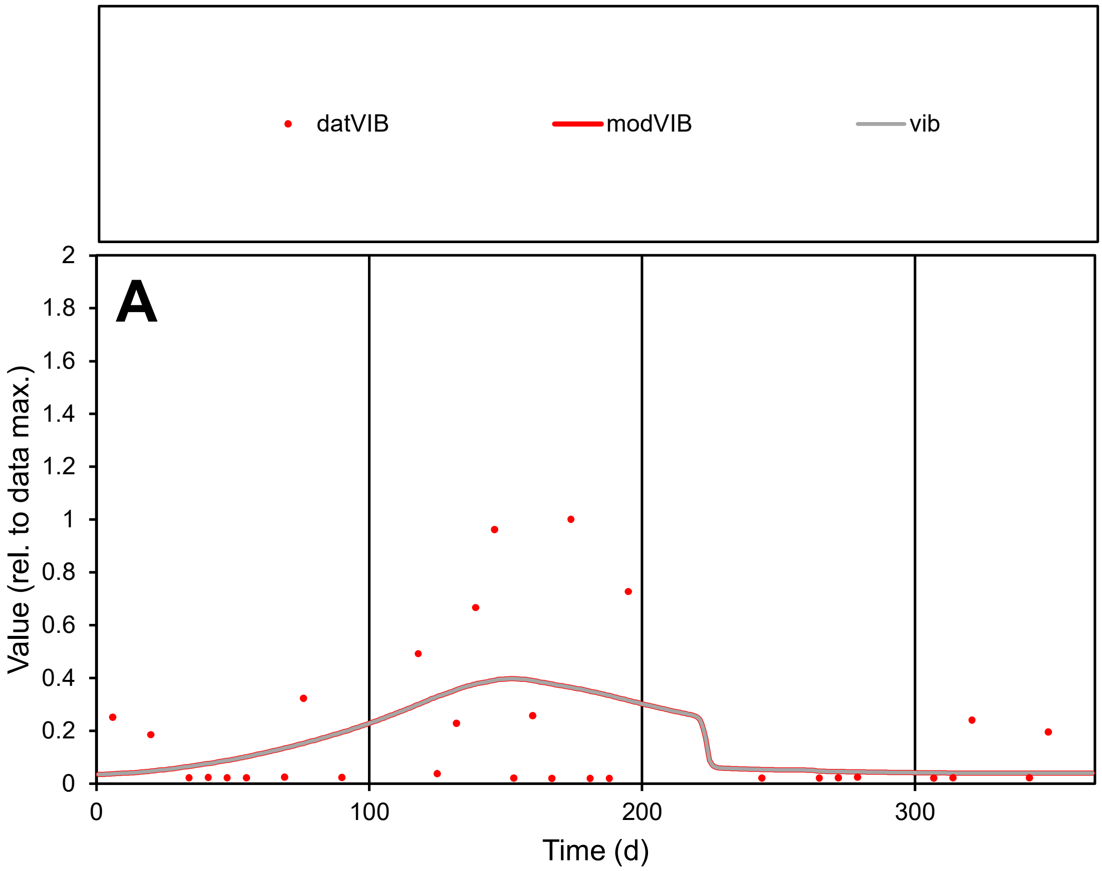


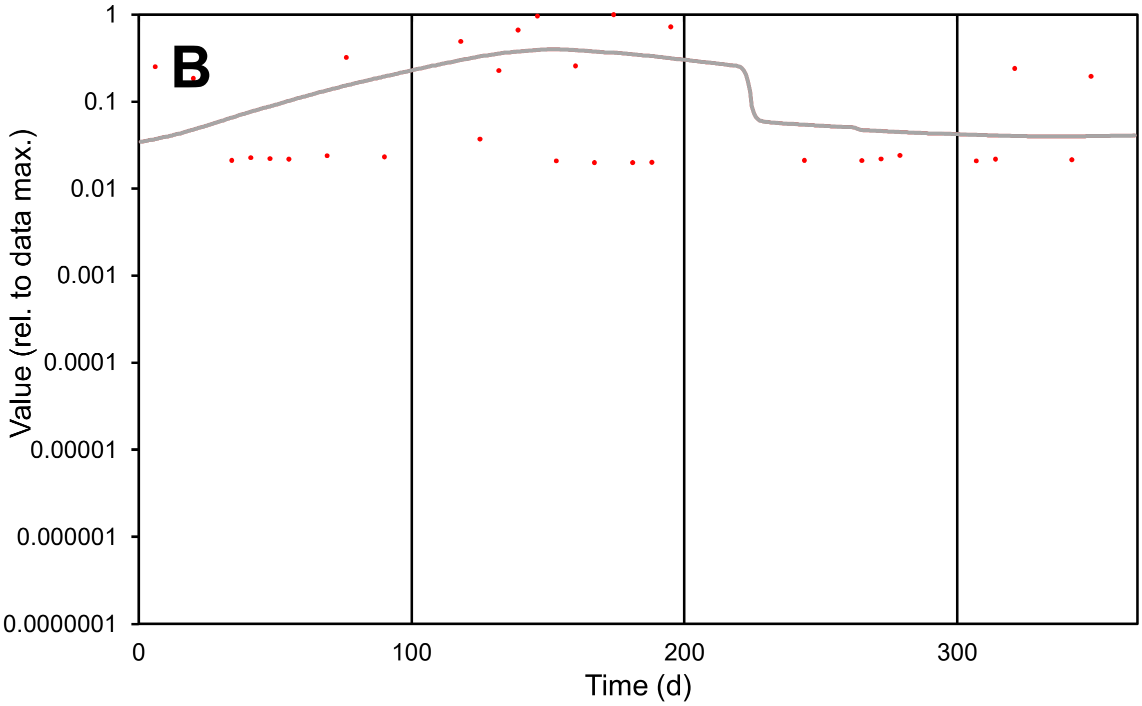


**C1**

Variables
VIB Provide jtd or Dst
CalibOnly?
2 1 to reproduce of
oNormalize?
2 1=no, 2=yes

ofxn calc*
1.24755
ofxn mod
1.25748

sRun
RunA1
zout
1

Ave
Data
0.00018
Model
0.00016

data/vMax
0.00089

ofxng
275.646

Min d/m
0.52429

ave/min d
9.96123
ave/min m
4.91536

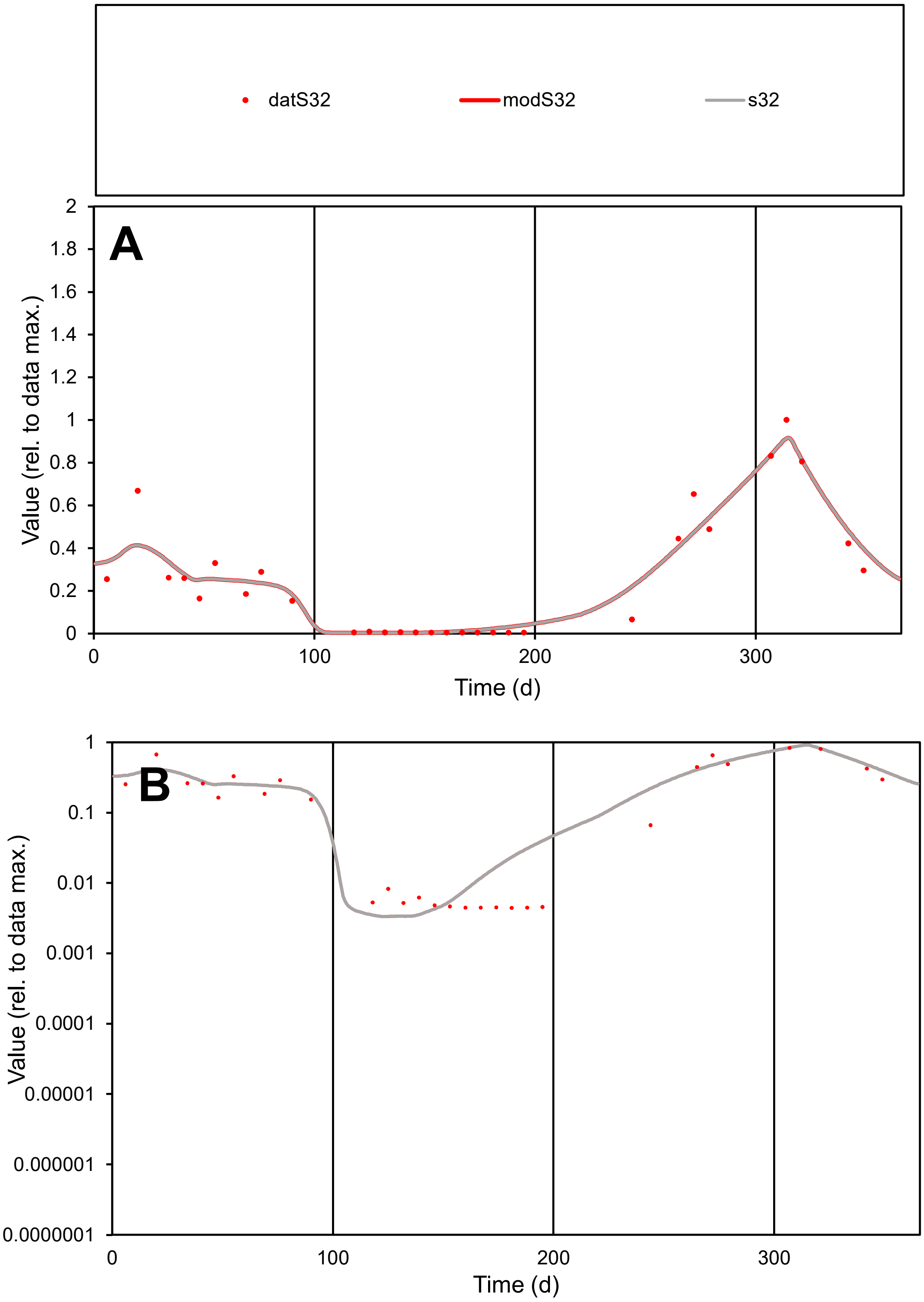


**C1**

Variables
S32 Provide jtd or Dst
CalibOnly?
2 1 to reproduce of
oNormalize?
2 1=no, 2=yes

ofxn calc*
0.29975
ofxn mod
0.34746

sRun
RunA1
zout
1

Ave
Data
0.00062
Model
0.00063

data/vMax
0.00242

ofxng
275.646

Min d/m
1.3133

ave/min d
57.5632
ave/min m
76.6899

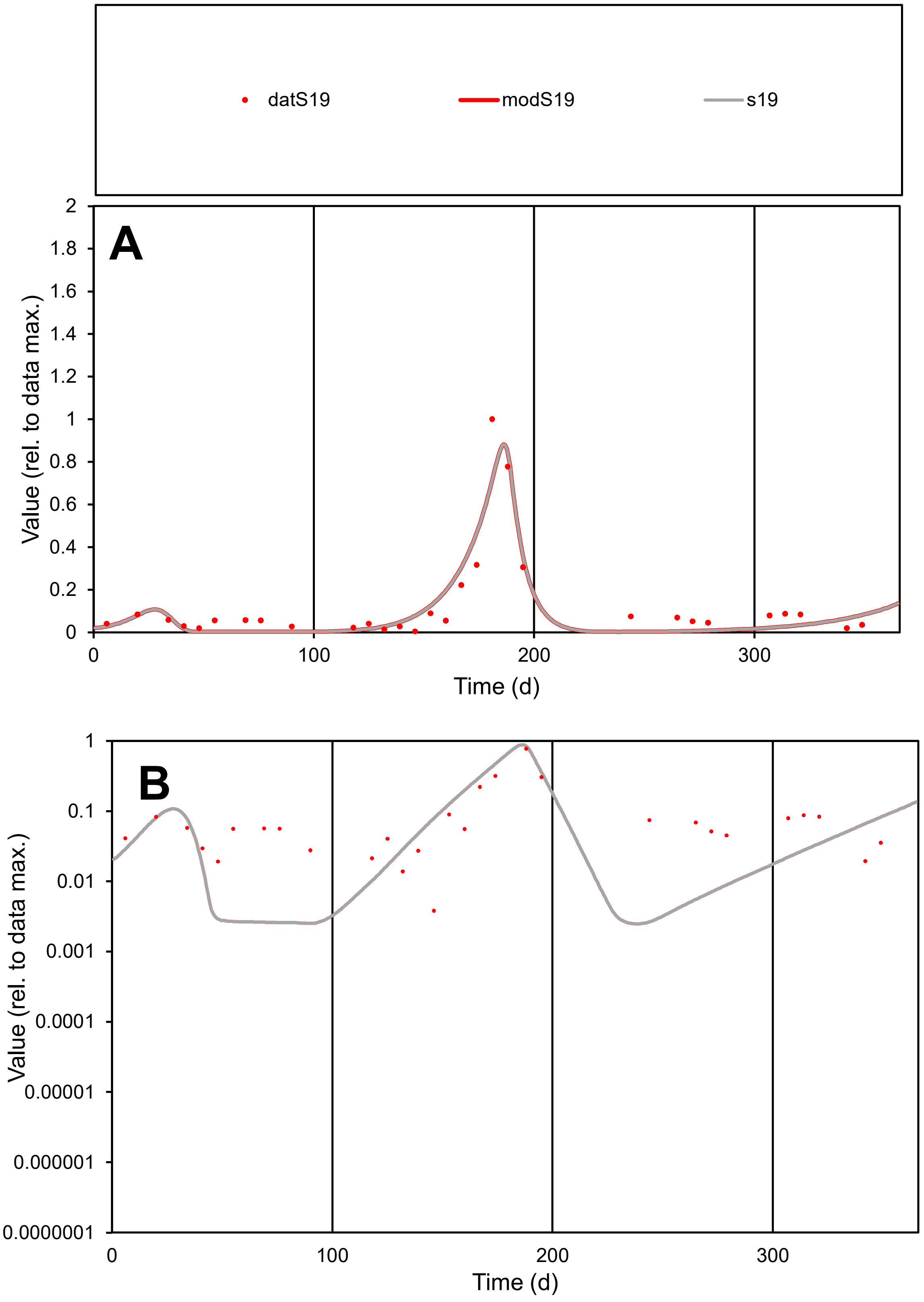


**C1**

Variables
S19 Provide jtd or Dst
CalibOnly?
2 1 to reproduce of
oNormalize?
2 1=no, 2=yes

ofxn calc*
0.54848
ofxn mod
0.59728

sRun
RunA1
zout
1

Ave
Data
0.00171
Model
0.00162

data/vMax
0.01336

ofxng
275.646

Min d/m
1.50473

ave/min d
33.6065
ave/min m
47.7043

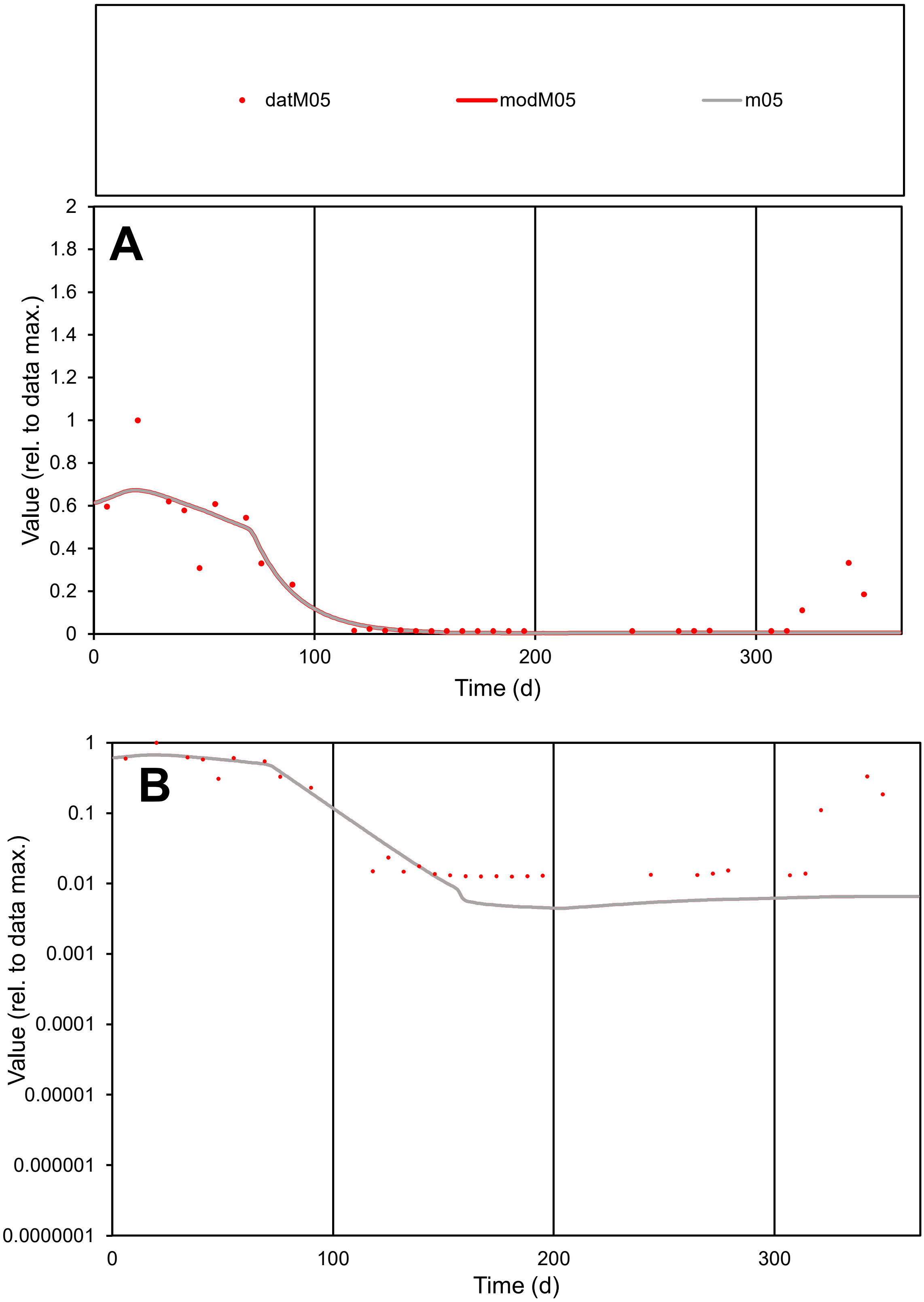


**C1**

Variables

M05 Provide jtd or Dst

CalibOnly?

2 1 to reproduce of

oNormalize?

2 1=no, 2=yes

ofxn calc*

0.56549

ofxn mod

0.59648

sRun

RunA1

zout

1

Ave

Data

0.00065

Model

0.00056

data/vMax

0.00342

ofxng

275.646

Min d/m

2.77038

ave/min d

15.1609

ave/min m

36.3731

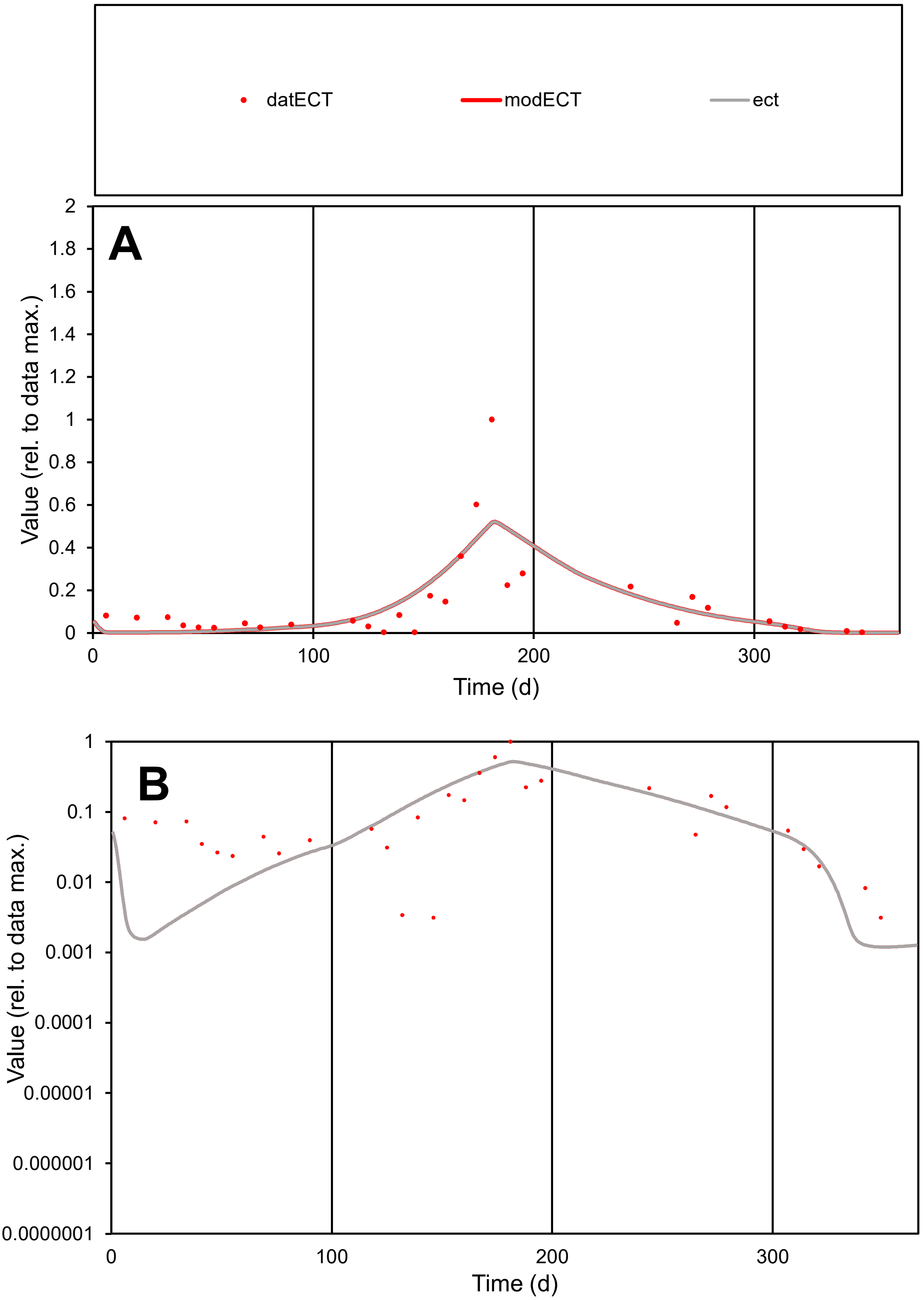


**C1**

Variables
ECT Provide jtd or Dst
CalibOnly?
2 1 to reproduce of
oNormalize?
2 1=no, 2=yes

ofxn calc*
0.90695
ofxn mod
0.93057

sRun
RunA1
zout
1

Ave
Data
0.00131
Model
0.00133

data/vMax
0.00973

ofxng
275.646

Min d/m
2.61235

ave/min d
43.0908
ave/min m
114.19

325_Calib_S_A

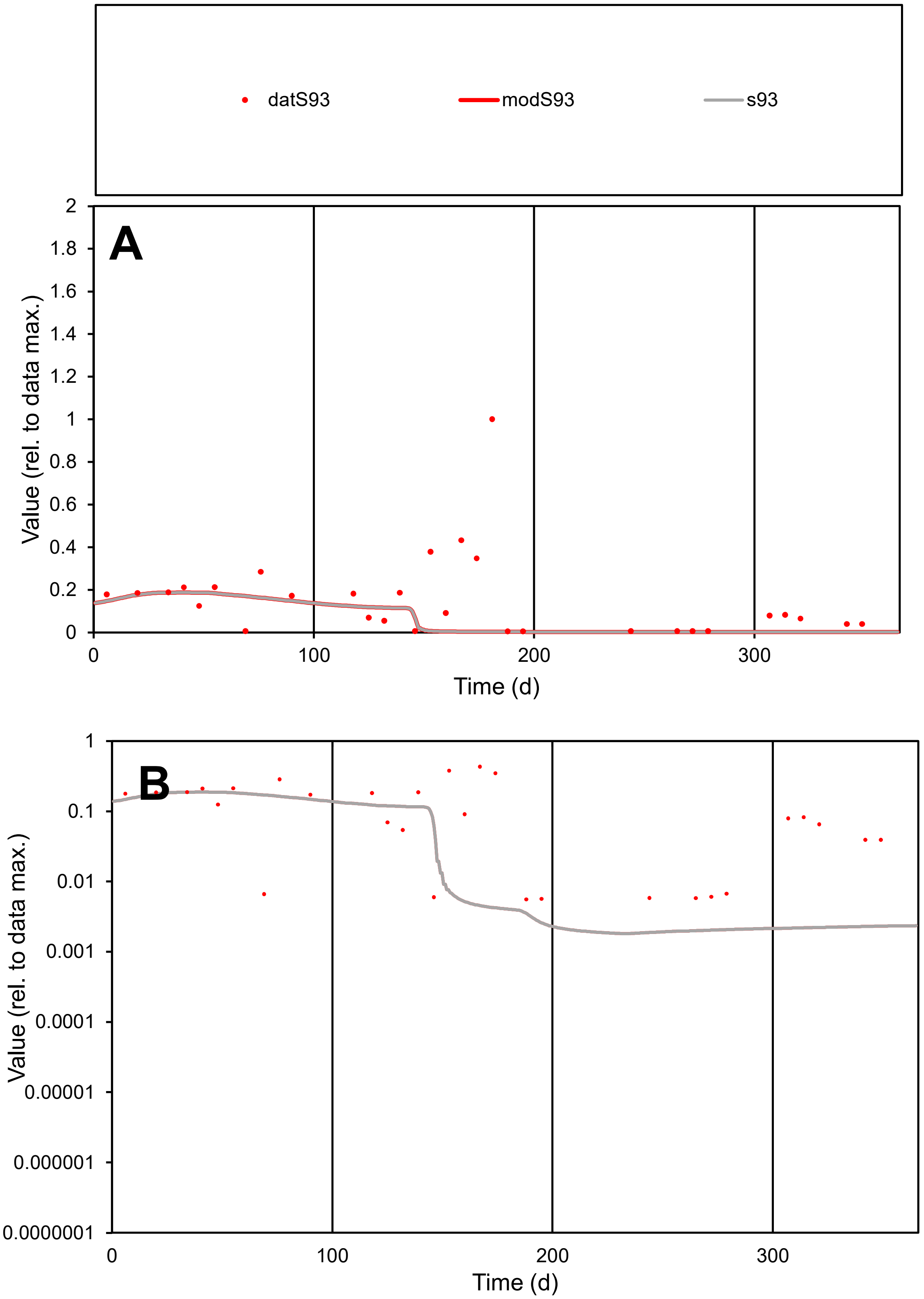


**C1**

Variables

S93 Provide jtd or Dst

CalibOnly?

2 1 to reproduce of

oNormalize?

2 1=no, 2=yes

ofxn calc*

1.45162

ofxn mod

1.45605

sRun

RunA1

zout

1

Ave

Data

0.00183

Model

0.00082

data/vMax

0.01179

ofxng

275.646

Min d/m

2.94831

ave/min d

27.8637

ave/min m

36.9832

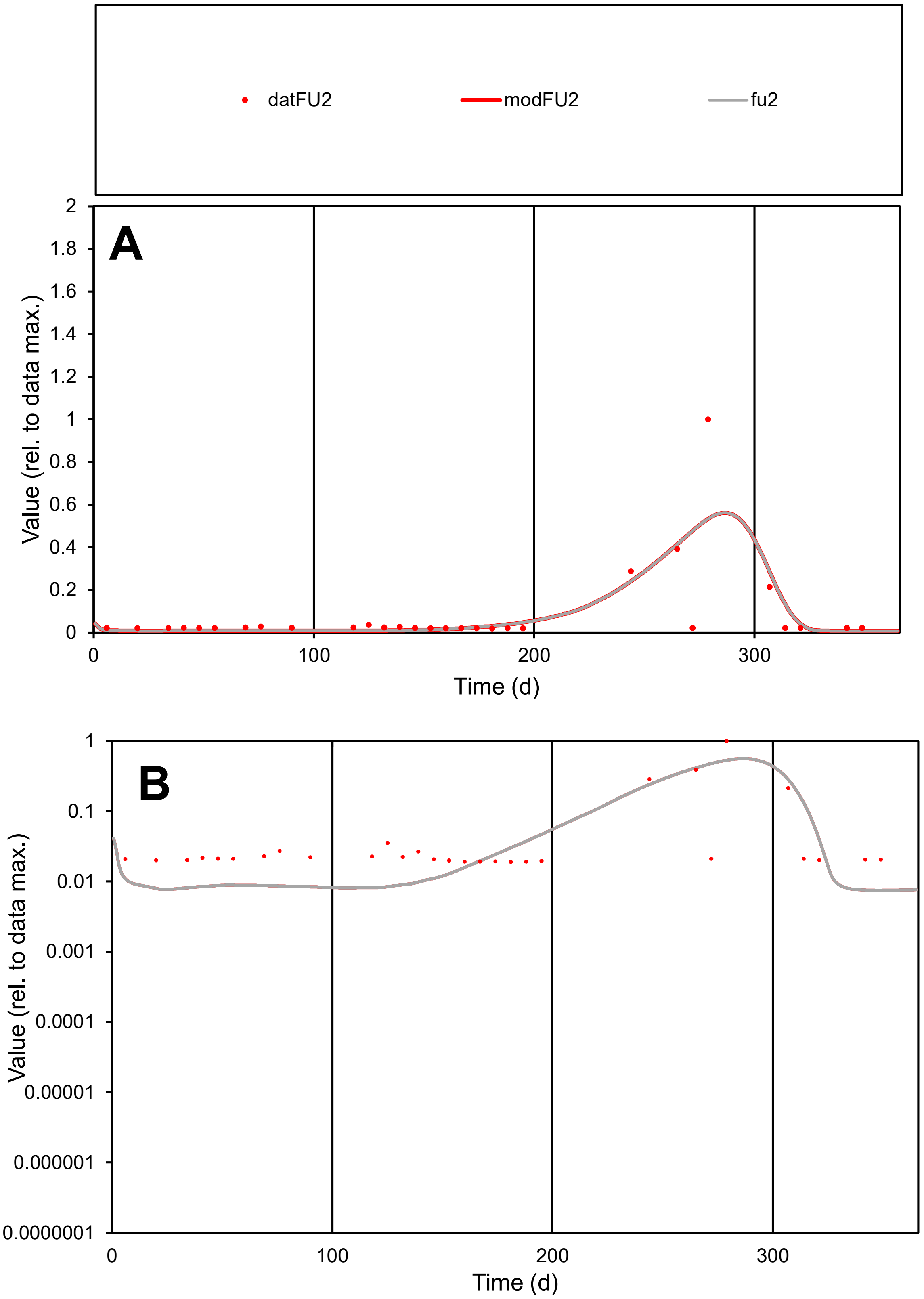


**C1**

Variables
FU2 Provide jtd or Dst
CalibOnly?
2 1 to reproduce of
oNormalize?
2 1=no, 2=yes

ofxn calc*
1.47343
ofxn mod
1.50716

sRun
RunA1
zout
1

Ave
Data
0.00011
Model
0.00011

data/vMax
0.00134

ofxng
275.646

Min d/m
2.535

ave/min d
4.31682
ave/min m
10.7374

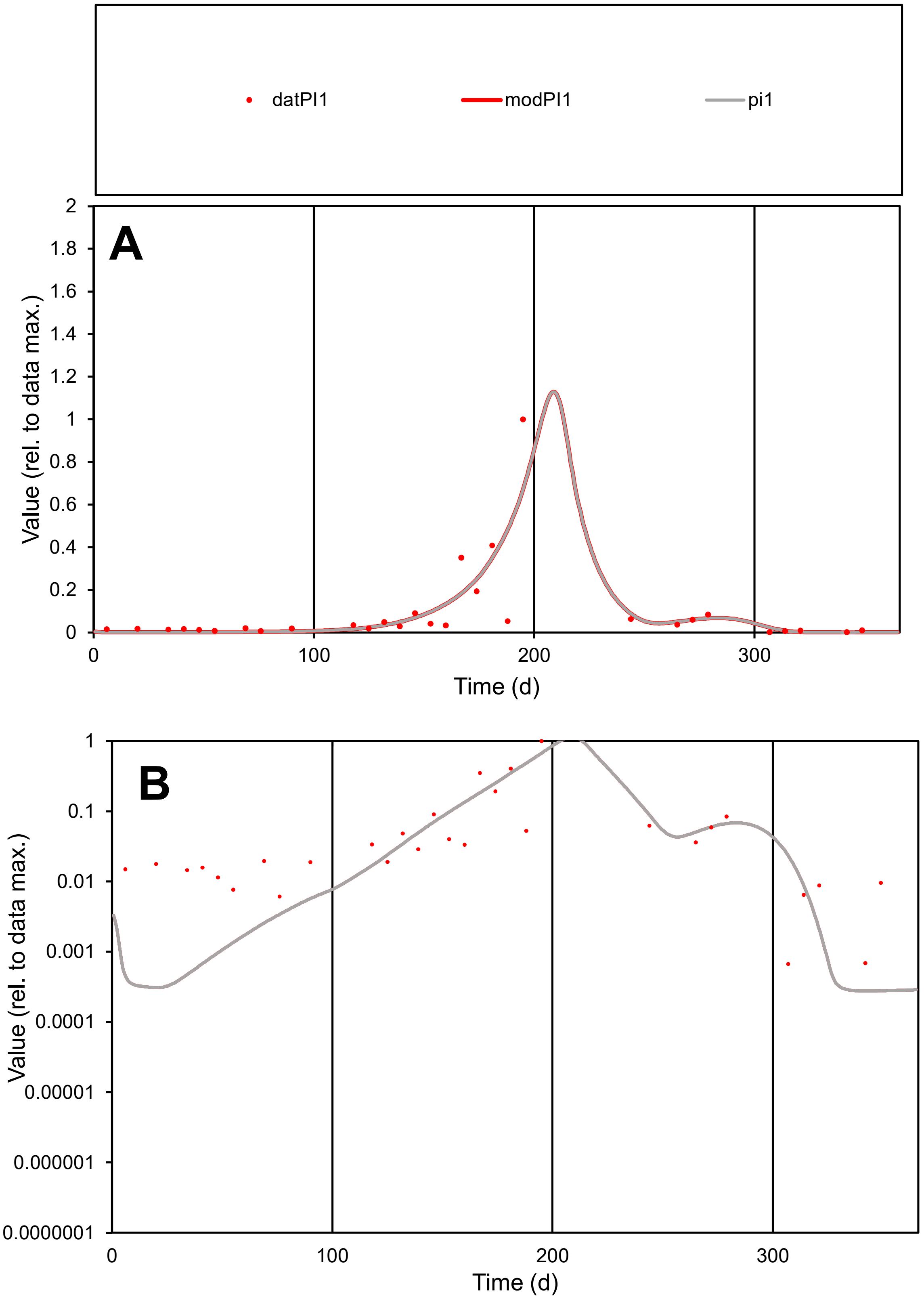


**C1**

Variables
PI1 Provide jtd or Dst
CalibOnly?
2 1 to reproduce of
oNormalize?
2 1=no, 2=yes

ofxn calc*
1.12367
ofxn mod
1.18133

sRun
RunA1
zout
1

Ave
Data
0.00184
Model
0.00189

data/vMax
0.0205

ofxng
275.646

Min d/m
2.42274

ave/min d
134.052
ave/min m
333.187

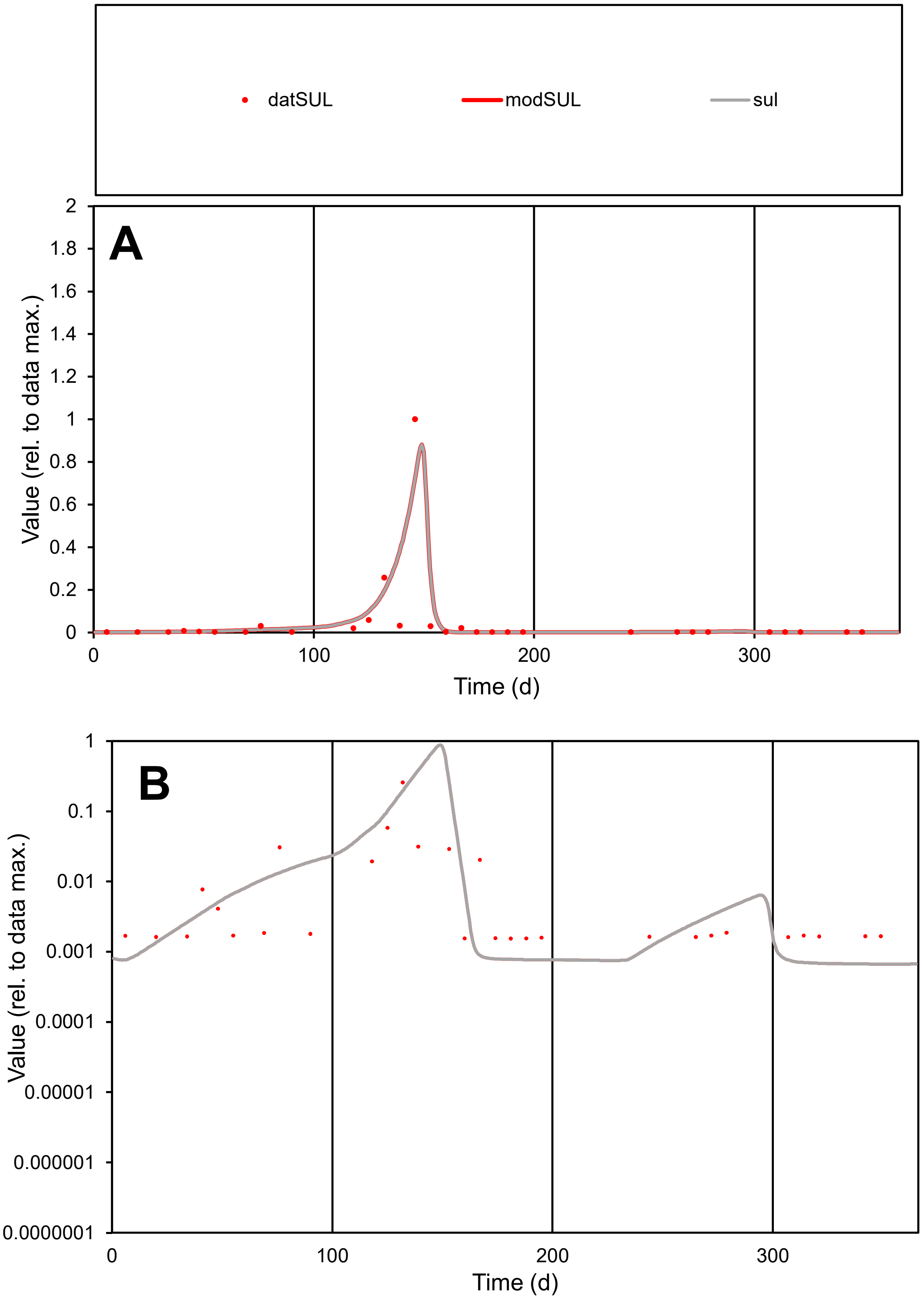


**C1**

Variables
SUL Provide jtd or Dst
CalibOnly?
2 1 to reproduce of
oNormalize?
2 1=no, 2=yes

ofxn calc*
1.37778
ofxn mod
1.51718

sRun
RunA1
zout
1

Ave
Data
0.00255
Model
0.00294

data/vMax
0.05121

ofxng
275.646

Min d/m
2.3005

ave/min d
32.3475
ave/min m
86.0317

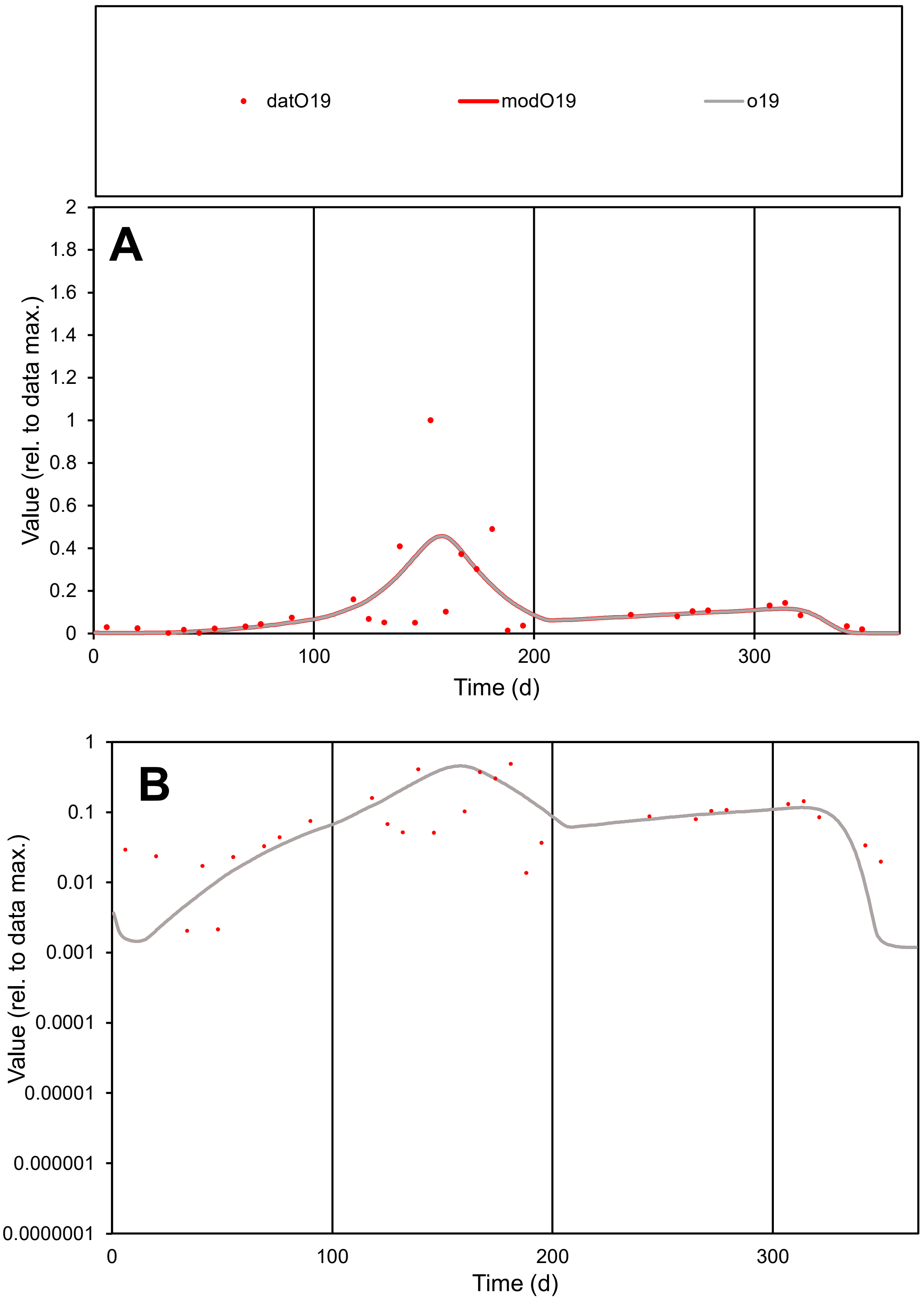


**C1**

Variables
O19 Provide jtd or Dst
CalibOnly?
2 1 to reproduce of
oNormalize?
2 1=no, 2=yes

ofxn calc*
1.09444
ofxn mod
1.11026

sRun
RunA1
zout
1

Ave
Data
0.002
Model
0.00198

data/vMax
0.01464

ofxng
275.646

Min d/m
1.36653

ave/min d
66.5687
ave/min m
90.1318

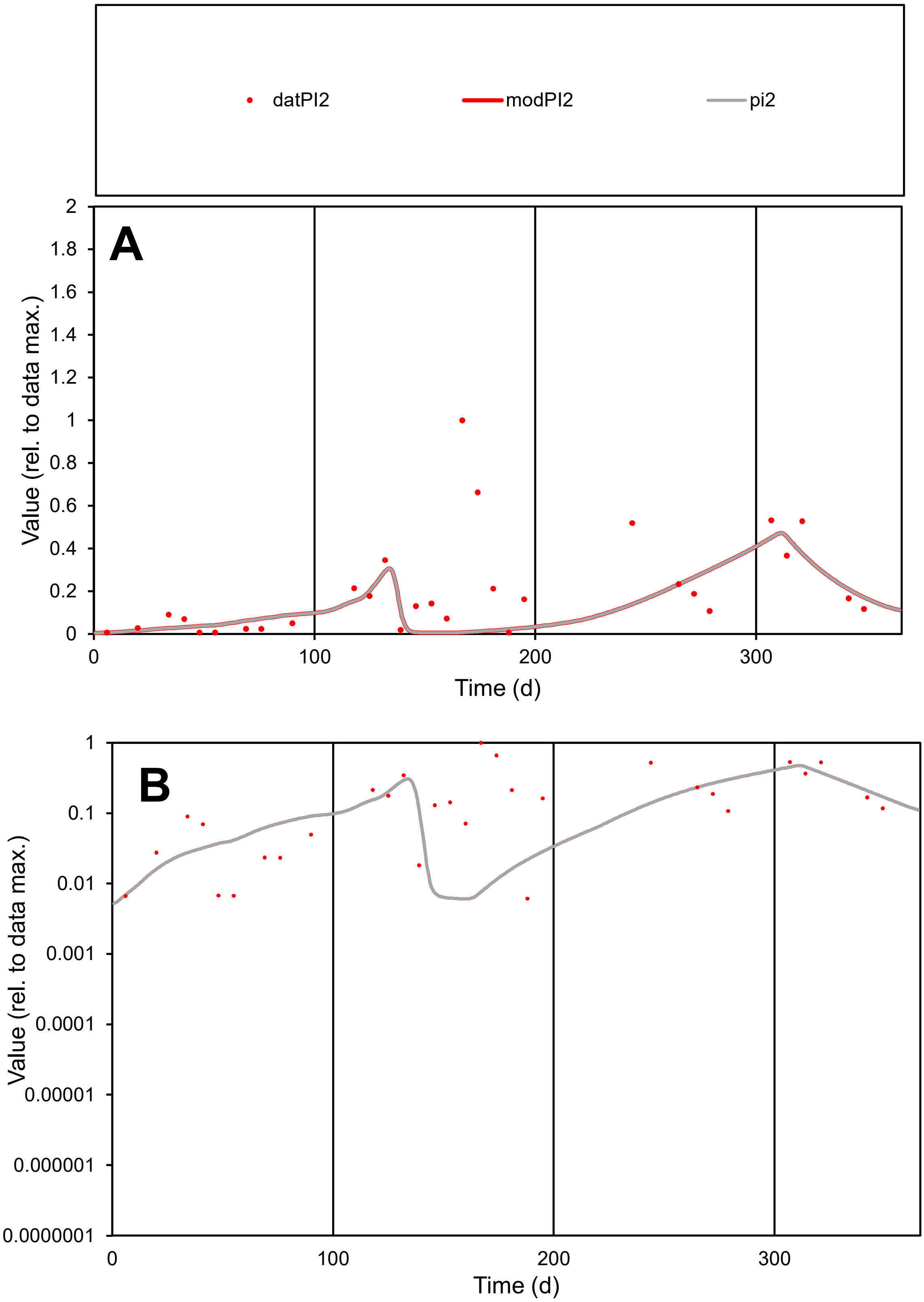


**C1**

Variables
PI2 Provide jtd or Dst
CalibOnly?
2 1 to reproduce of
oNormalize?
2 1=no, 2=yes

ofxn calc*
1.16726
ofxn mod
1.17275

sRun
RunA1
zout
1

Ave
Data
0.00096
Model
0.00059

data/vMax
0.00464

ofxng
275.646

Min d/m
1.00743

ave/min d
33.6113
ave/min m
20.7677

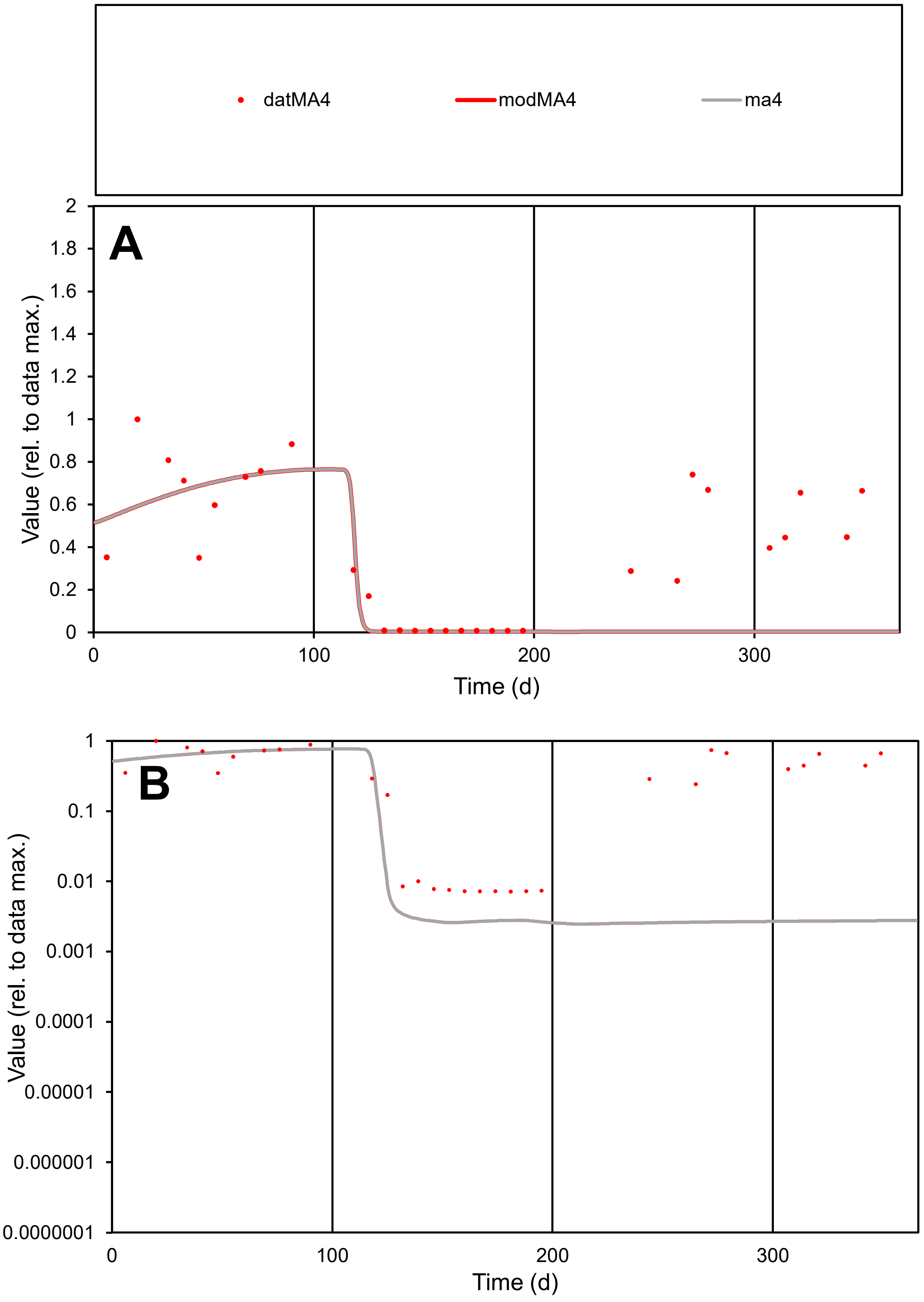


**C1**

Variables

MA4 Provide jtd or Dst

CalibOnly?

2 1 to reproduce of

oNormalize?

2 1=no, 2=yes

ofxn calc*

0.82959

ofxn mod

0.84199

sRun

RunA1

zout

1

Ave

Data

0.001

Model

0.00058

data/vMax

0.00268

ofxng

275.646

Min d/m

2.78181

ave/min d

52.4667

ave/min m

84.5546

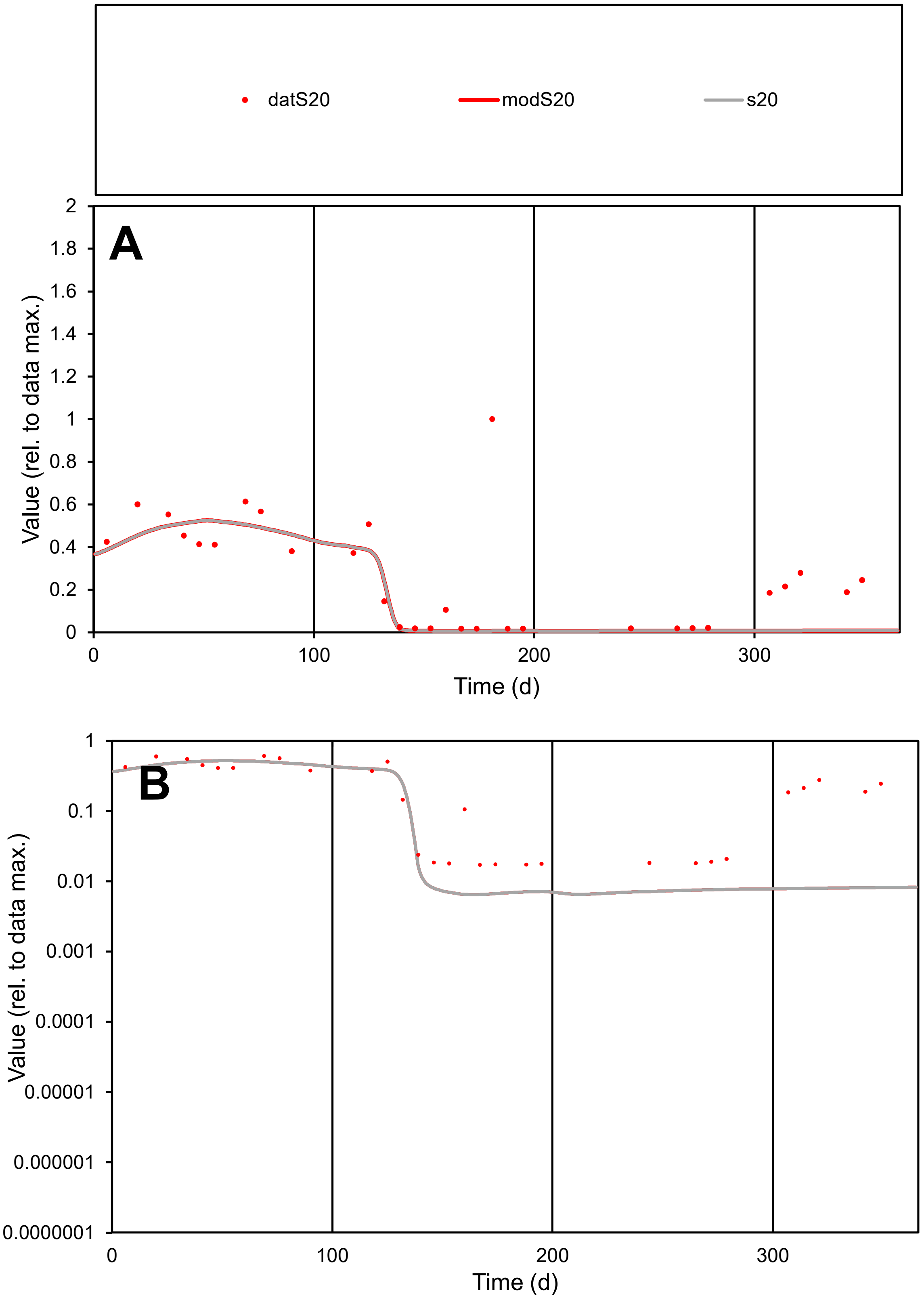


**C1**

Variables
S20 Provide jtd or Dst
CalibOnly?
2 1 to reproduce of
oNormalize?
2 1=no, 2=yes

ofxn calc*
0.80287
ofxn mod
0.81483

sRun
RunA1
zout
1

Ave
Data
0.00118
Model
0.00083

data/vMax
0.00452

ofxng
275.646

Min d/m
2.66849

ave/min d
15.174
ave/min m
28.2104

325_Calib_S_A

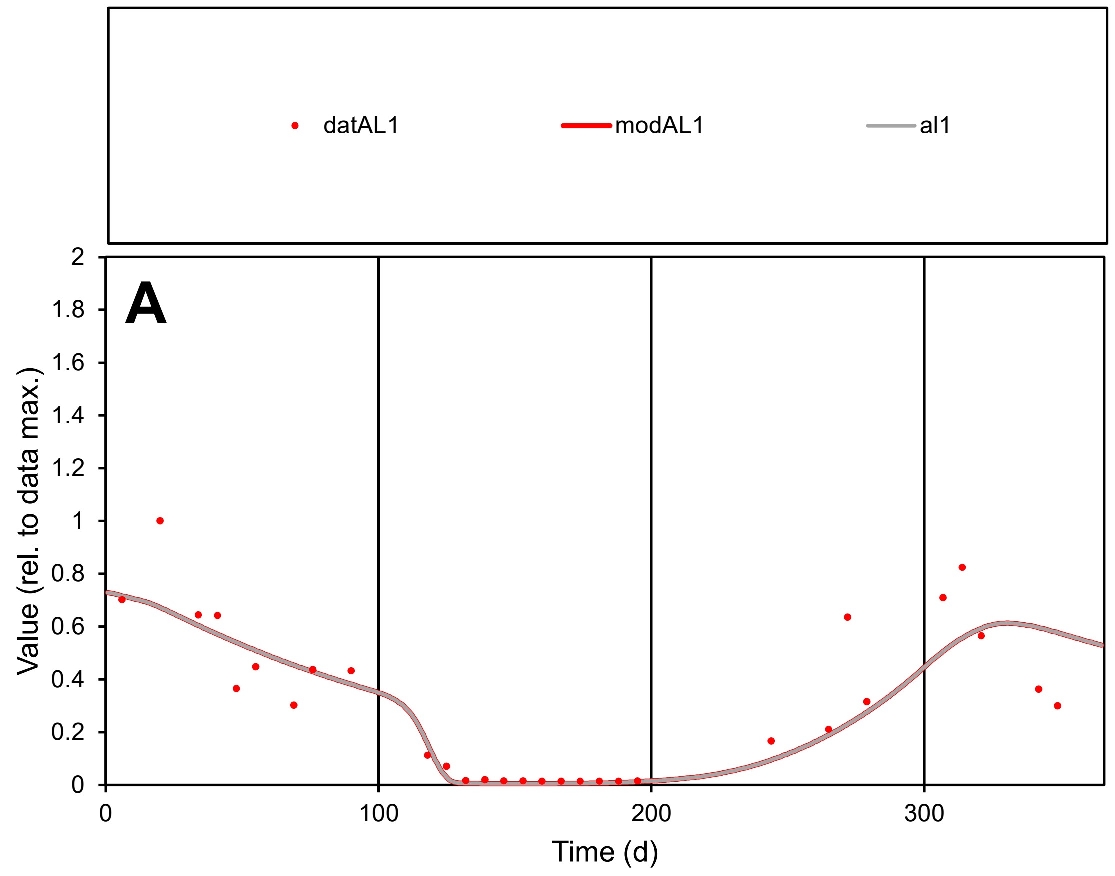


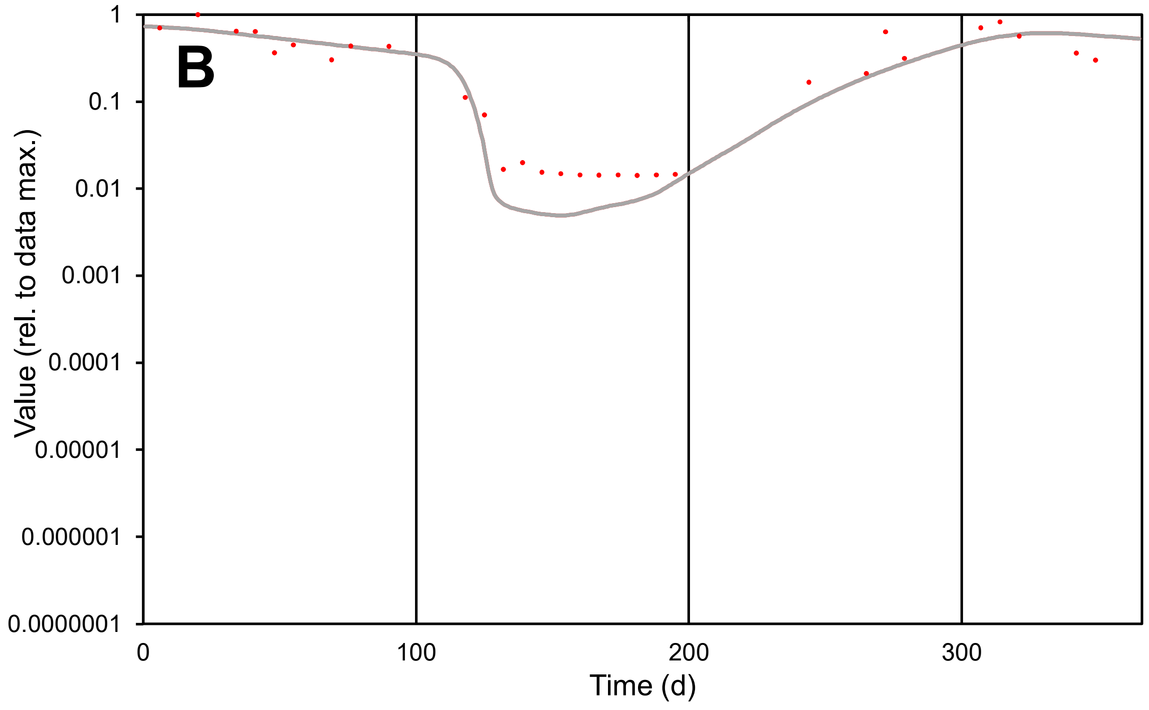


**C1**

Variables

AL1 Provide jtd or Dst

CalibOnly?

2 1 to reproduce of

oNormalize?

2 1=no, 2=yes

ofxn calc*

0.44015

ofxn mod

0.46496

sRun

RunA1

zout

1

Ave

Data

0.00096

Model

0.00089

data/vMax

0.00308

ofxng

275.646

Min d/m

2.88484

ave/min d

22.0931

ave/min m

59.2193

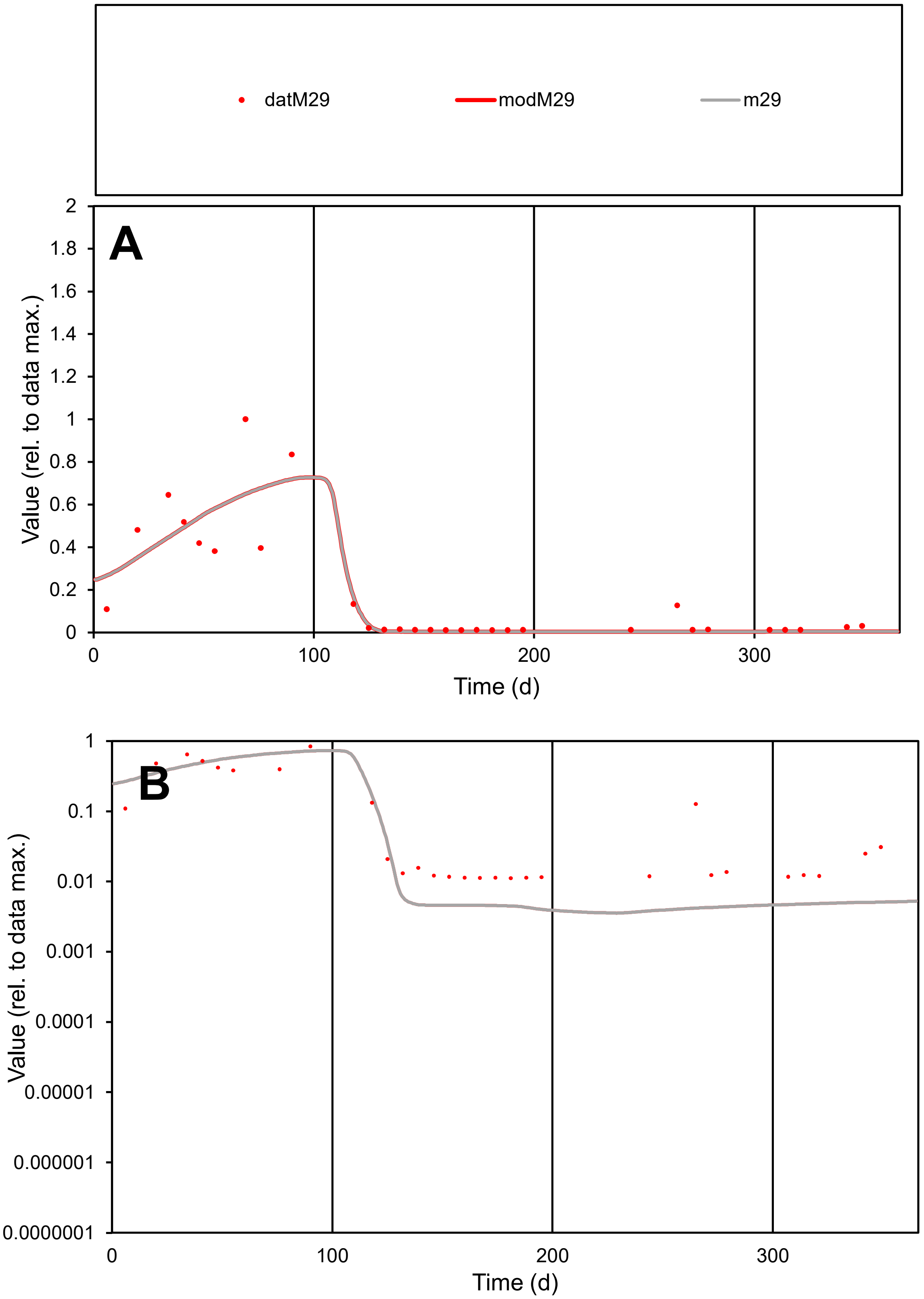


**C1**

Variables
M29 Provide jtd or Dst
CalibOnly?
2 1 to reproduce of
oNormalize?
2 1=no, 2=yes

ofxn calc*
0.62252
ofxn mod
0.6591

sRun
RunA1
zout
1

Ave
Data
0.00086
Model
0.00081

data/vMax
0.00485

ofxng
275.646

Min d/m
2.91272

ave/min d
15.891
ave/min m
43.7459

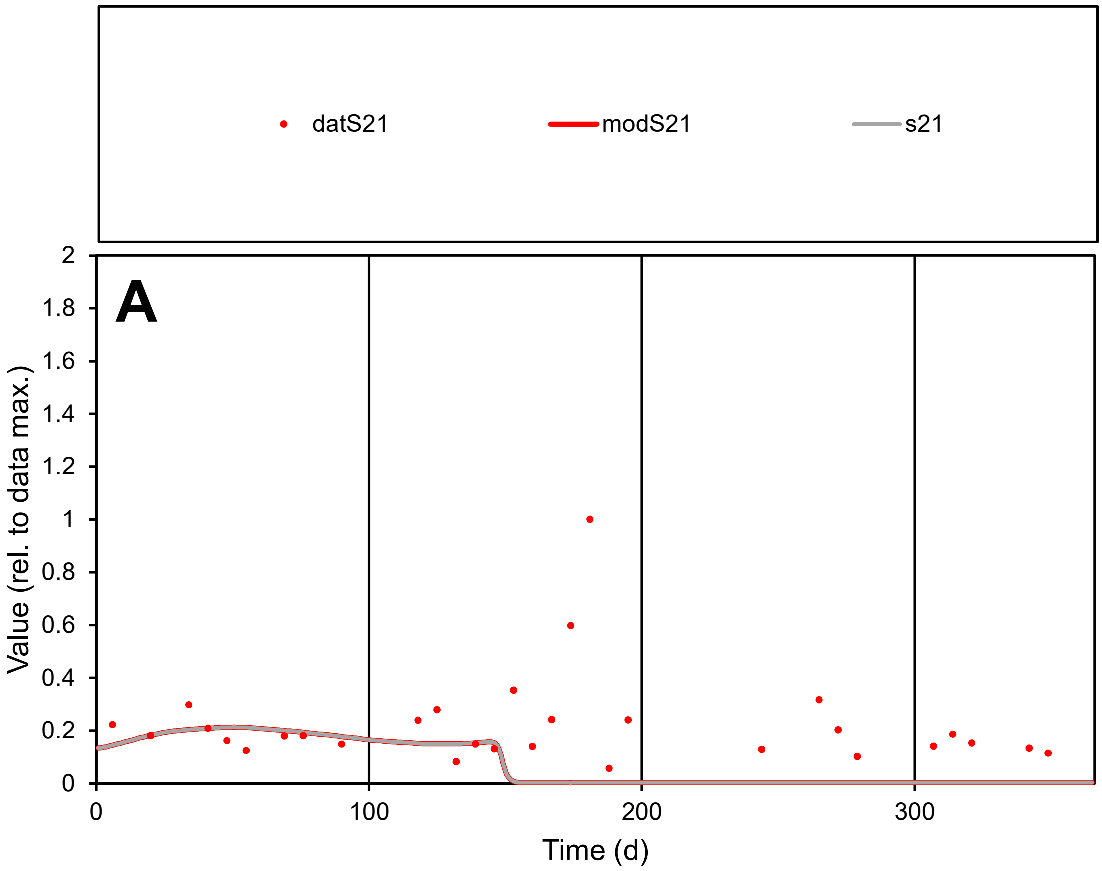


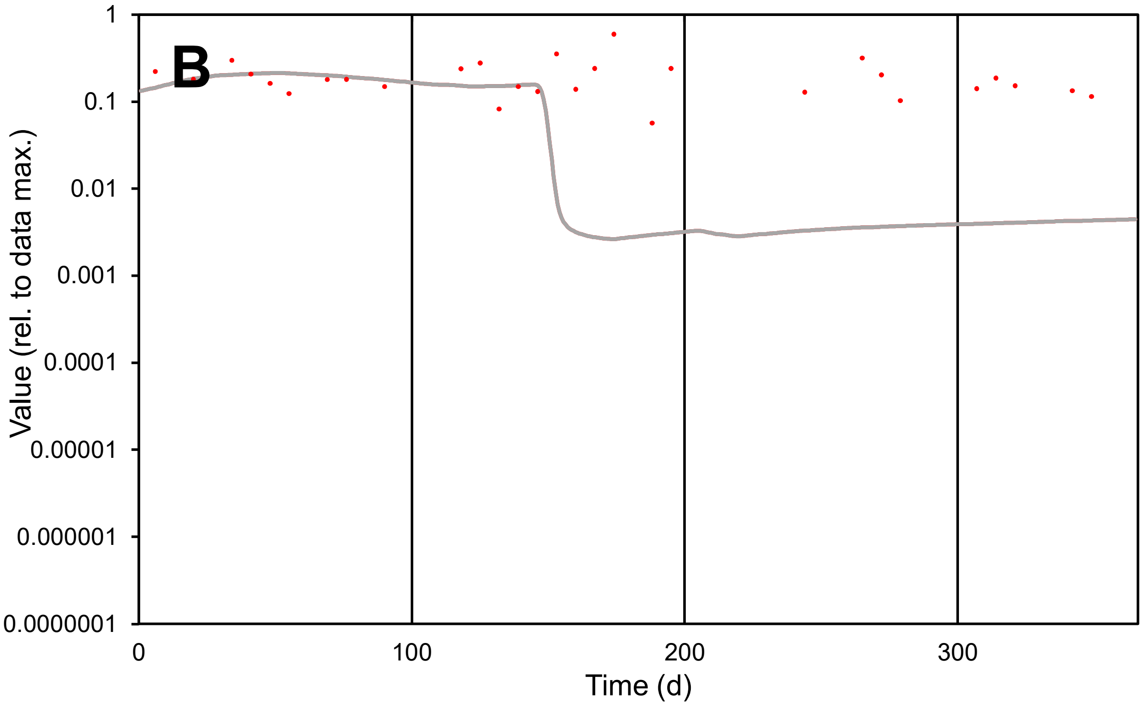


**C1**

Variables

S21 Provide jtd or Dst

CalibOnly?

2 1 to reproduce of

oNormalize?

2 1=no, 2=yes

ofxn calc*

1.13552

ofxn mod

1.1393

sRun

RunA1

zout

1

Ave

Data

0.00206

Model

0.00078

data/vMax

0.00924

ofxng

275.646

Min d/m

21.4869

ave/min d

3.9164

ave/min m

31.8597

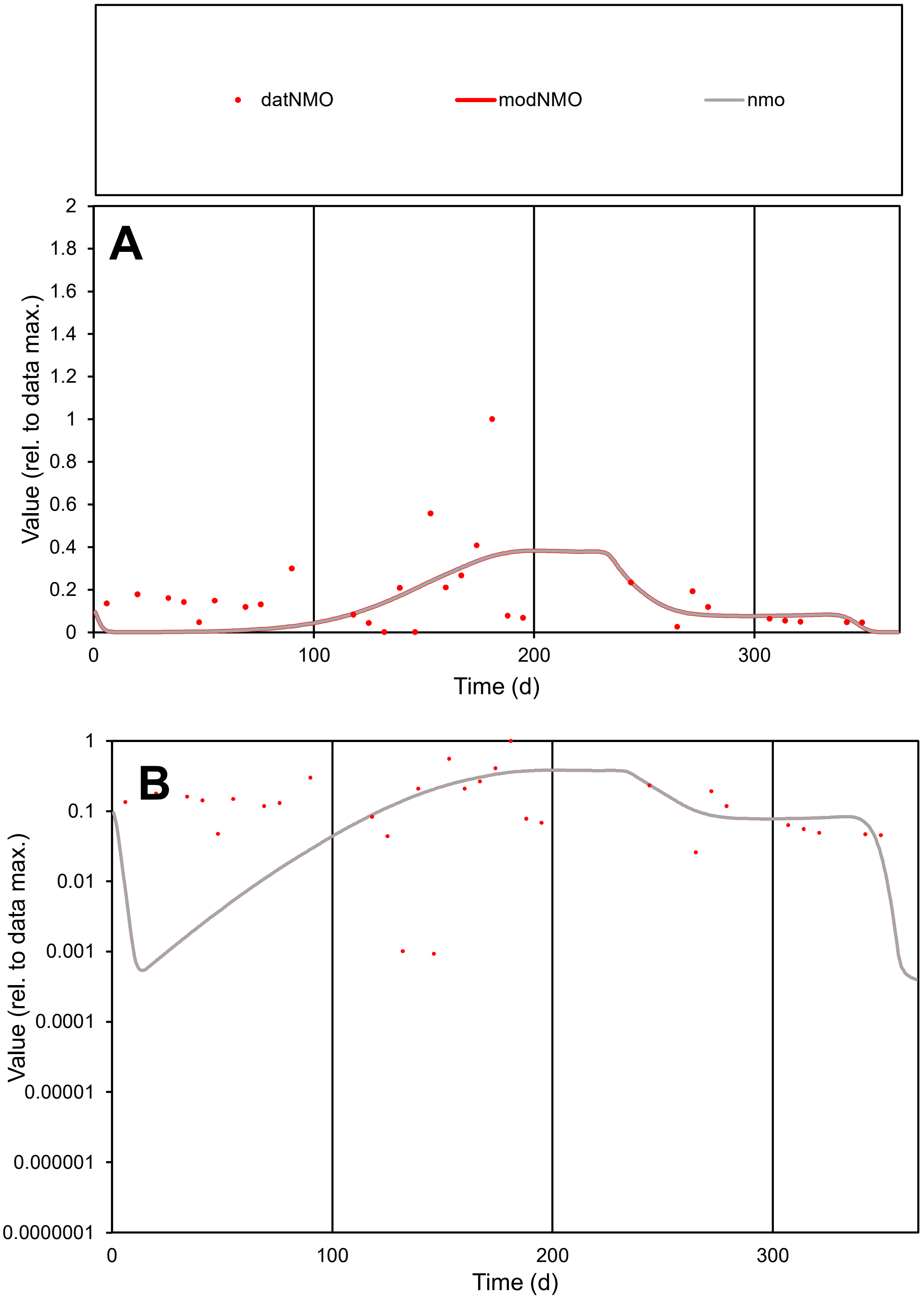


C1

Variables
NMO Provide jtd or Dst
CalibOnly?
2 1 to reproduce of
oNormalize?
2 1=no, 2=yes

ofxn calc*
1.06567
ofxn mod
1.07834

sRun
RunA1
zout
1

Ave
Data
0.00143
Model
0.00111

data/vMax
0.00835

ofxng
275.646

Min d/m
1.18748

ave/min d
182.857
ave/min m
168.347

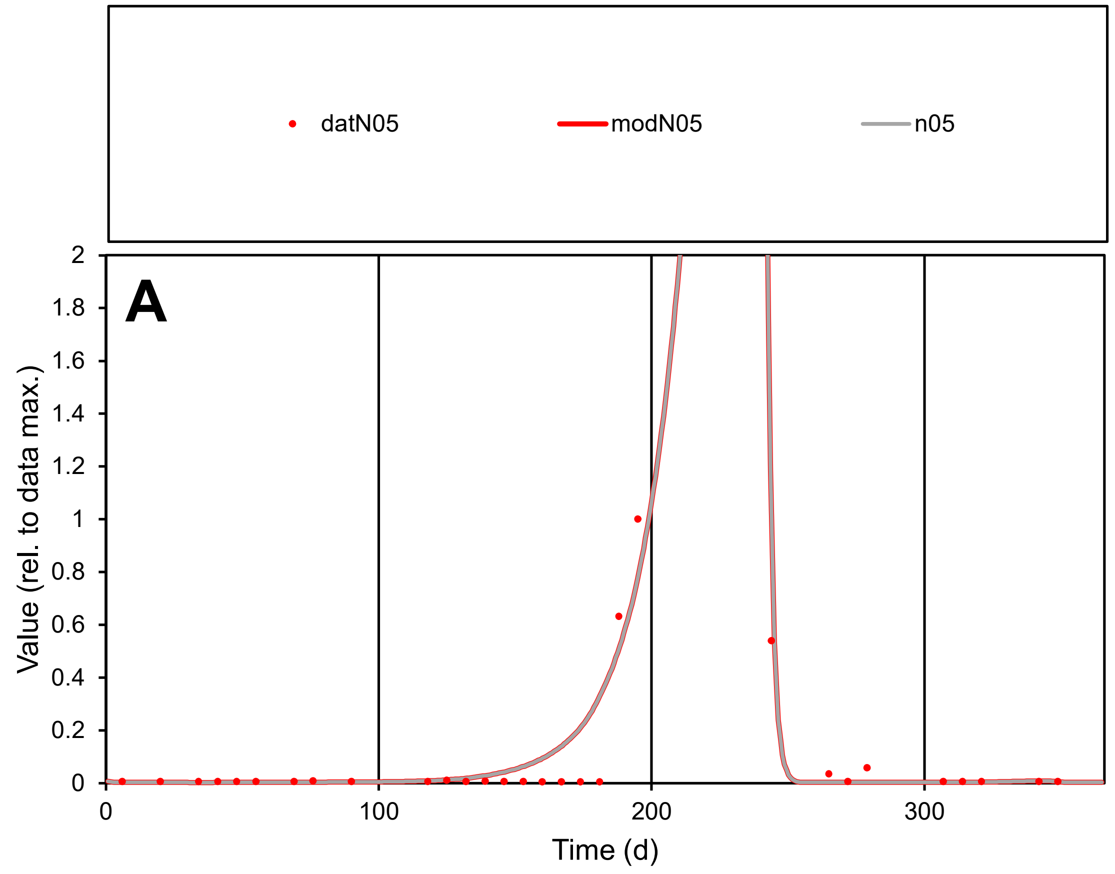


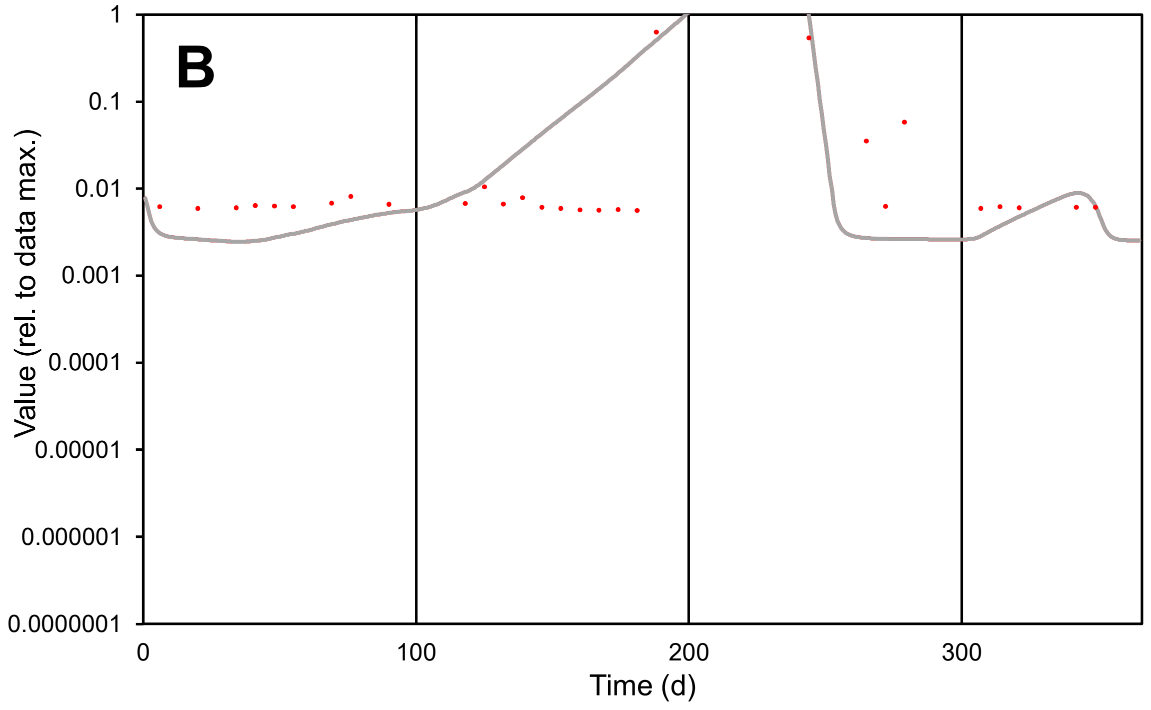


**C1**

Variables

N05 Provide jtd or Dst

CalibOnly?

2 1 to reproduce of

oNormalize?

2 1=no, 2=yes

ofxn calc*

0.99747

ofxn mod

1.07862

sRun

RunA1

zout

1

Ave

Data

0.00057

Model

0.00068

data/vMax

0.00701

ofxng

275.646

Min d/m

2.29866

ave/min d

14.3391

ave/min m

39.3017

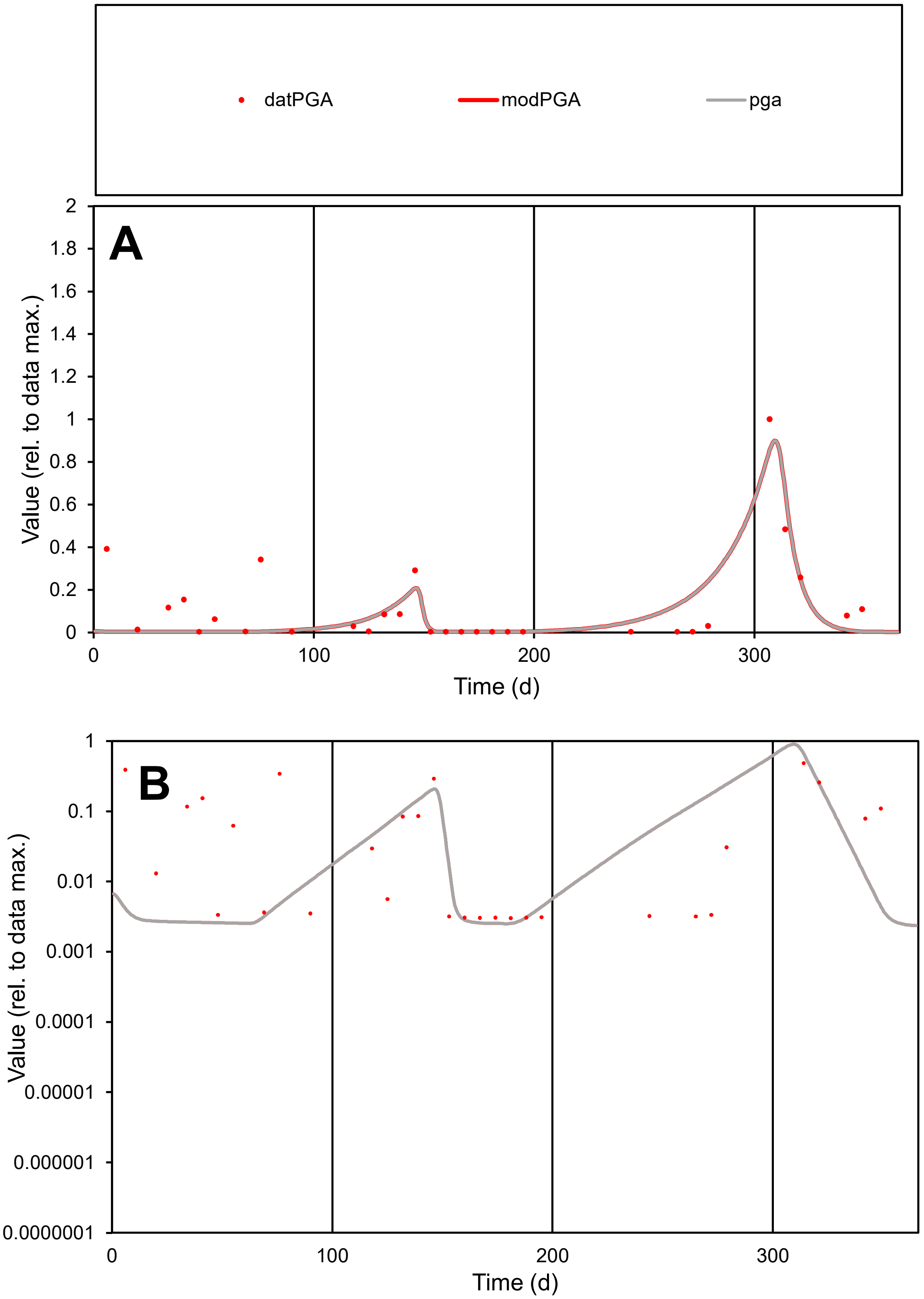


**C1**

Variables
PGA Provide jtd or Dst
CalibOnly?
2 1 to reproduce of
oNormalize?
2 1=no, 2=yes

ofxn calc*
1.01845
ofxn mod
1.04977

sRun
RunA1
zout
1

Ave
Data
0.00067
Model
0.00054

data/vMax
0.0056

ofxng
275.646

Min d/m
1.19182
ave/min d
39.5243
ave/min m
37.8345

325_Calib_S_A

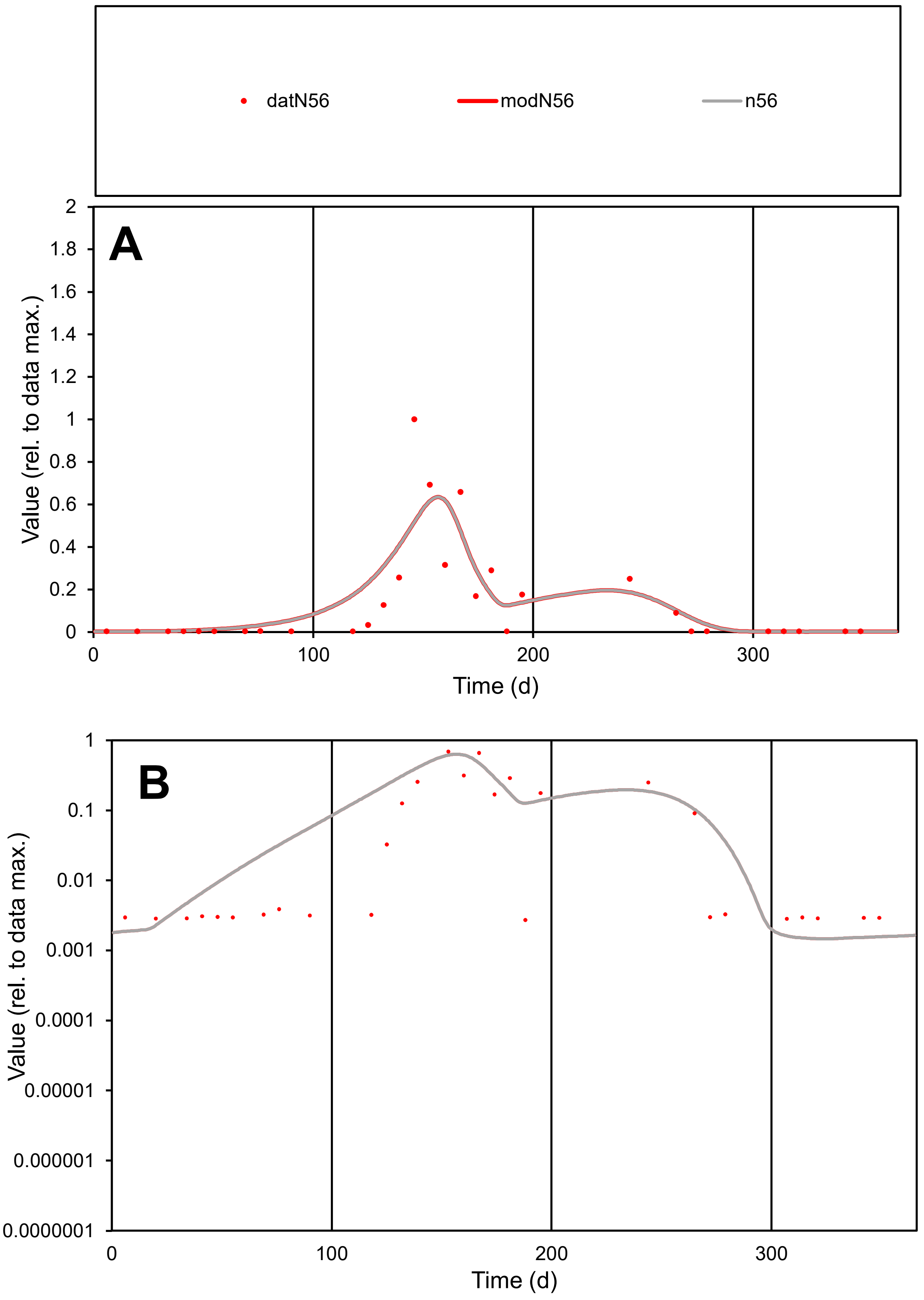


**C1**

Variables
N56 Provide jtd or Dst
CalibOnly?
2 1 to reproduce of
oNormalize?
2 1=no, 2=yes

ofxn calc*
0.9587
ofxn mod
0.98442

sRun
RunA1
zout
1

Ave
Data
0.00172
Model
0.00192

data/vMax
0.01256

ofxng
275.646

Min d/m
1.86366

ave/min d
50.1549
ave/min m
104.124

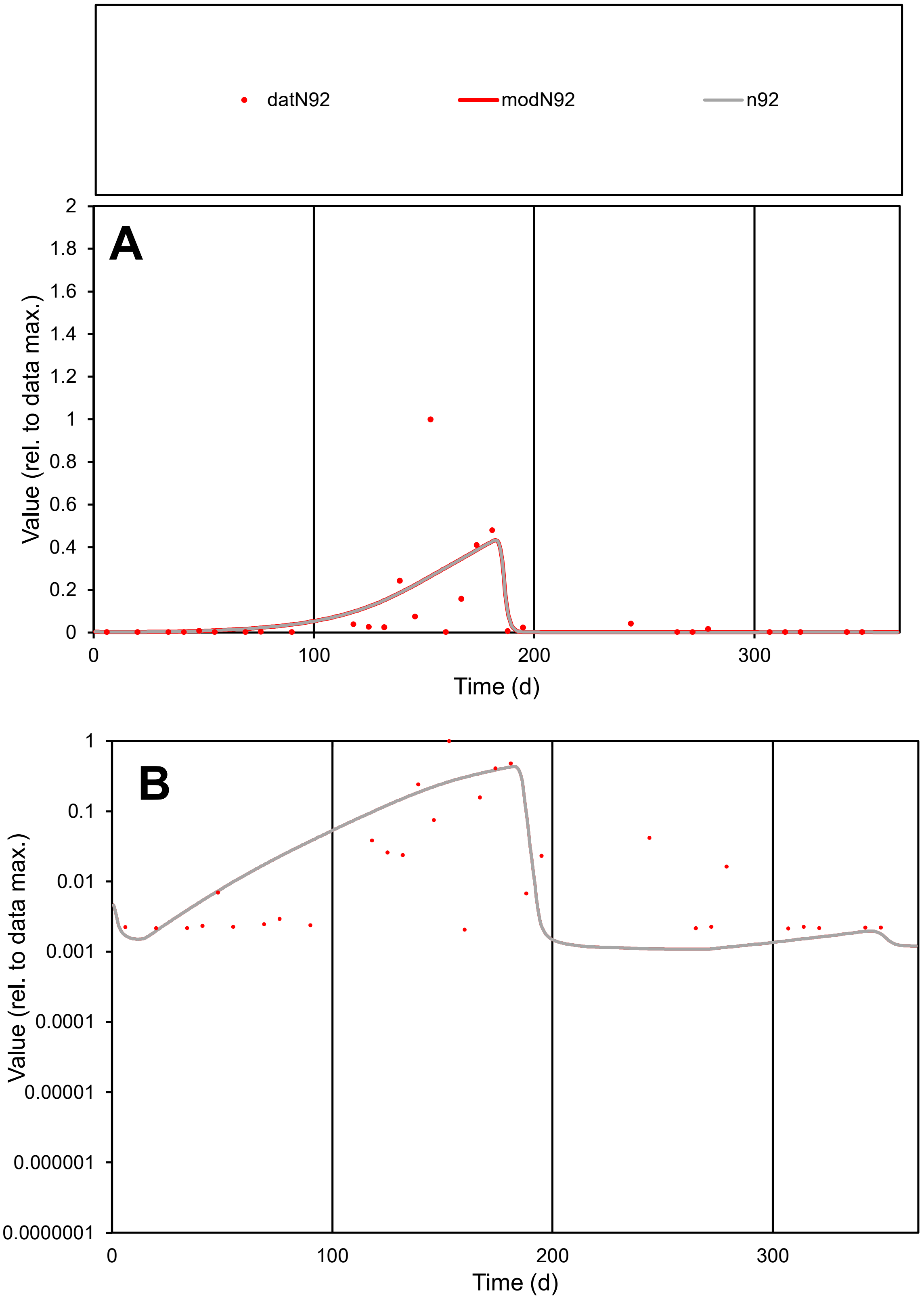


**C1**

Variables
N92 Provide jtd or Dst
CalibOnly?
2 1 to reproduce of
oNormalize?
2 1=no, 2=yes

ofxn calc*
1.80055
ofxn mod
1.81463

sRun
RunA1
zout
1

Ave
Data
0.00159
Model
0.00166

data/vMax
0.01844

ofxng
275.646

Min d/m
1.90045

ave/min d
41.6094
ave/min m
82.7084

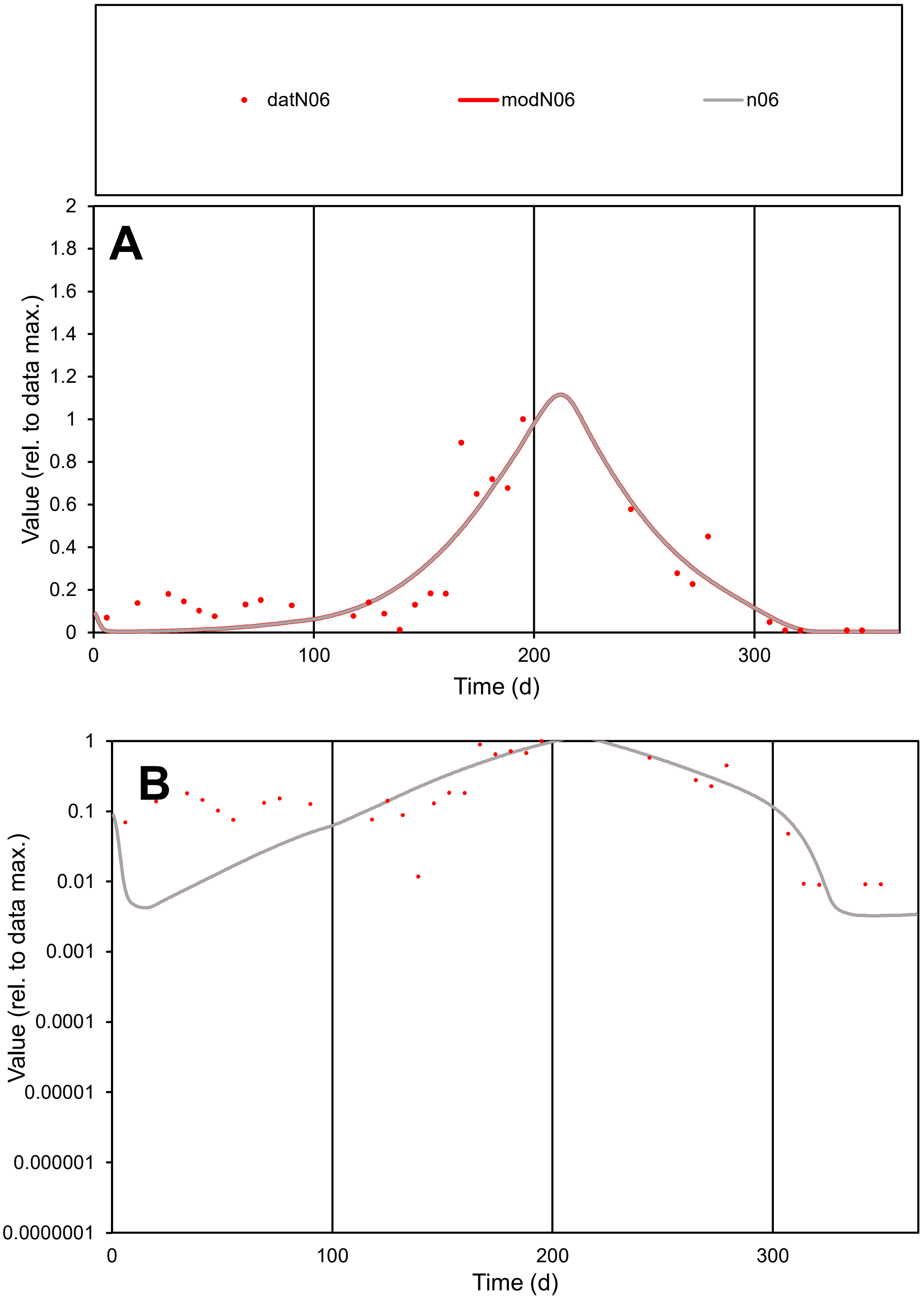


**C1**

Variables
N06 Provide jtd or Dst
CalibOnly?
2 1 to reproduce of
oNormalize?
2 1=no, 2=yes

ofxn calc*
0.5104
ofxn mod
0.5387

sRun
RunA1
zout
1

Ave
Data
0.00191
Model
0.00179

data/vMax
0.00766

ofxng
275.646

Min d/m
2.76796

ave/min d
27.713
ave/min m
71.8835

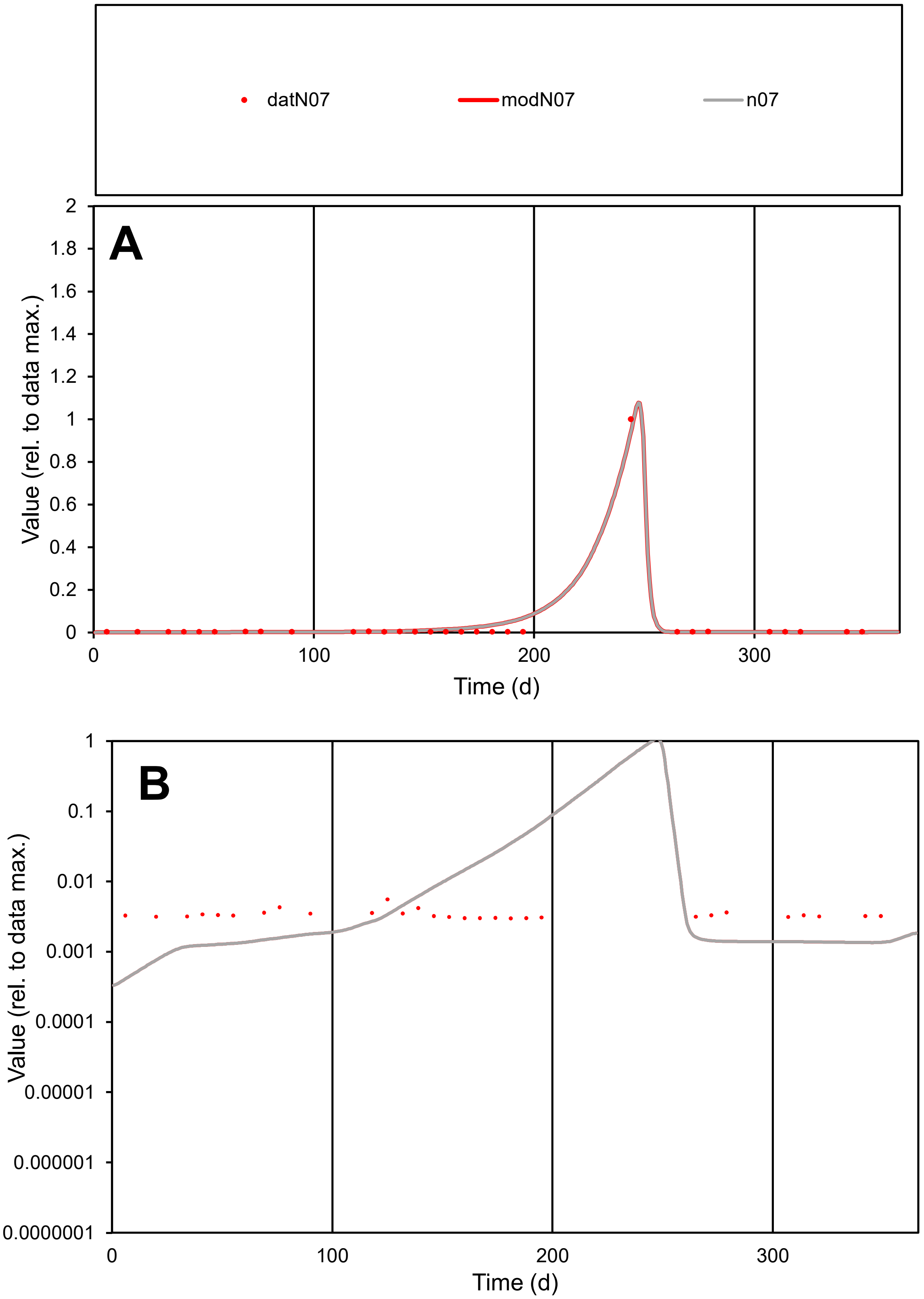


**C1**

Variables
N07 Provide jtd or Dst
CalibOnly?
2 1 to reproduce of
oNormalize?
2 1=no, 2=yes

ofxn calc*
0.44244
ofxn mod
0.59543

sRun
RunA1
zout
1

Ave
Data
0.00021
Model
0.00024

data/vMax
0.00567

ofxng
275.646

Min d/m
6.87551

ave/min d
12.2875
ave/min m
96.7448

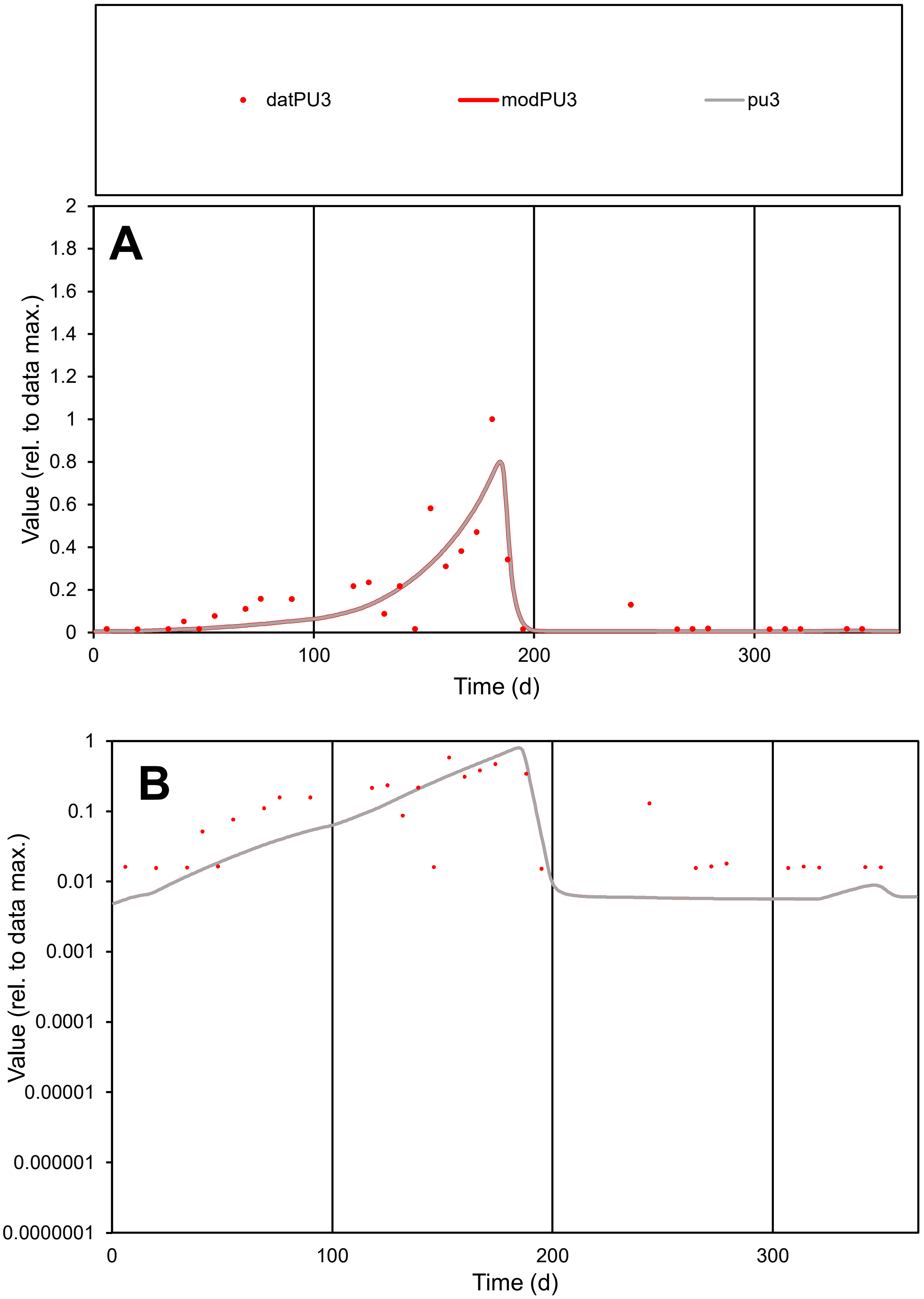


**C1**

Variables
PU3 Provide jtd or Dst
CalibOnly?
2 1 to reproduce of
oNormalize?
2 1=no, 2=yes

ofxn calc*
0.63255
ofxn mod
0.65395

sRun
RunA1
zout
1

Ave
Data
0.0016
Model
0.00141

data/vMax
0.01012

ofxng
275.646

Min d/m
2.69755

ave/min d
10.3832
ave/min m
24.5837

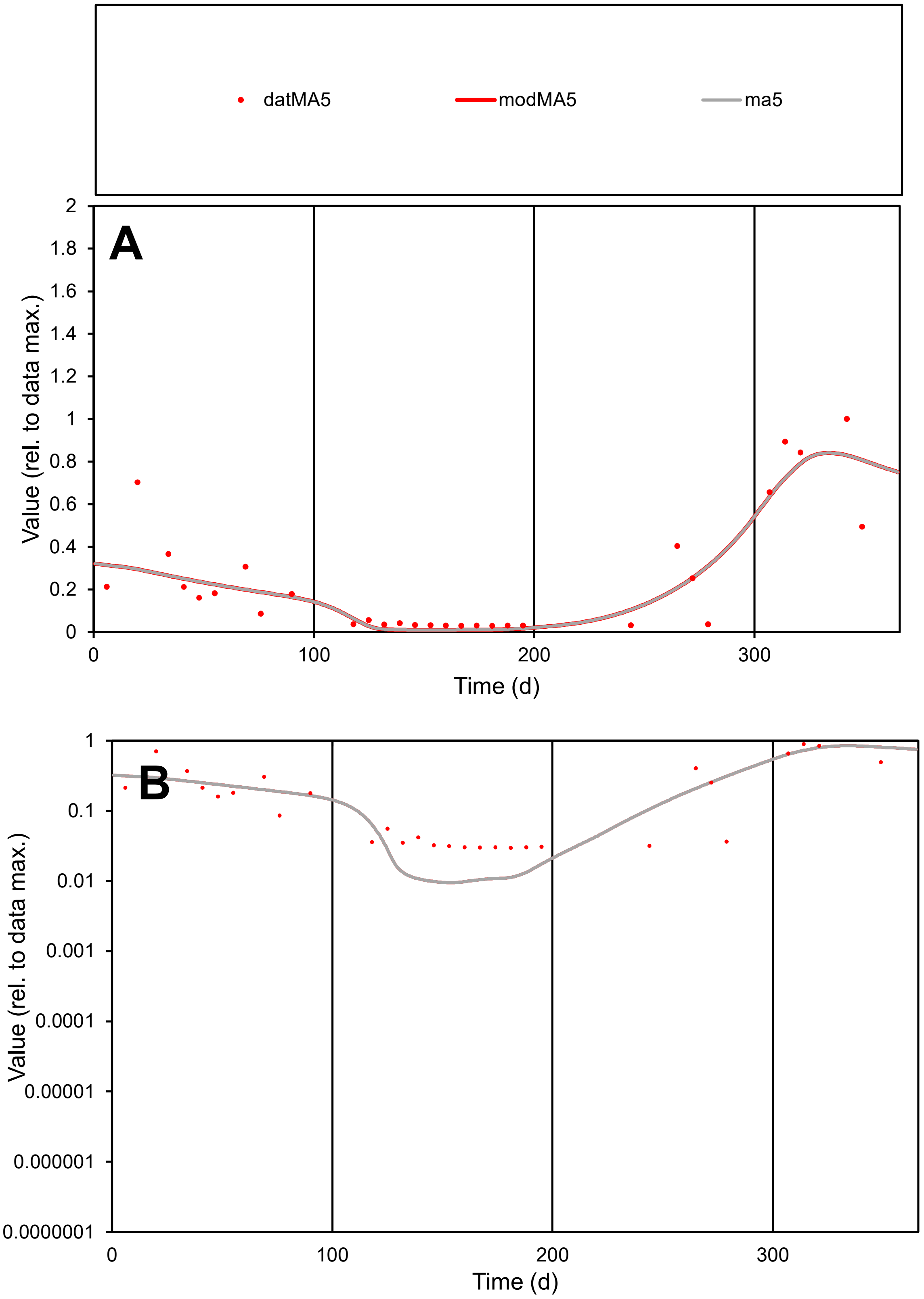


**C1**

Variables

MA5 Provide jtd or Dst

CalibOnly?

2 1 to reproduce of

oNormalize?

2 1=no, 2=yes

ofxn calc*

0.51768

ofxn mod

0.54218

sRun

RunA1

zout

1

Ave

Data

0.00078

Model

0.00074

data/vMax

0.00313

ofxng

275.646

Min d/m

3.1405

ave/min d

8.32342

ave/min m

24.7681

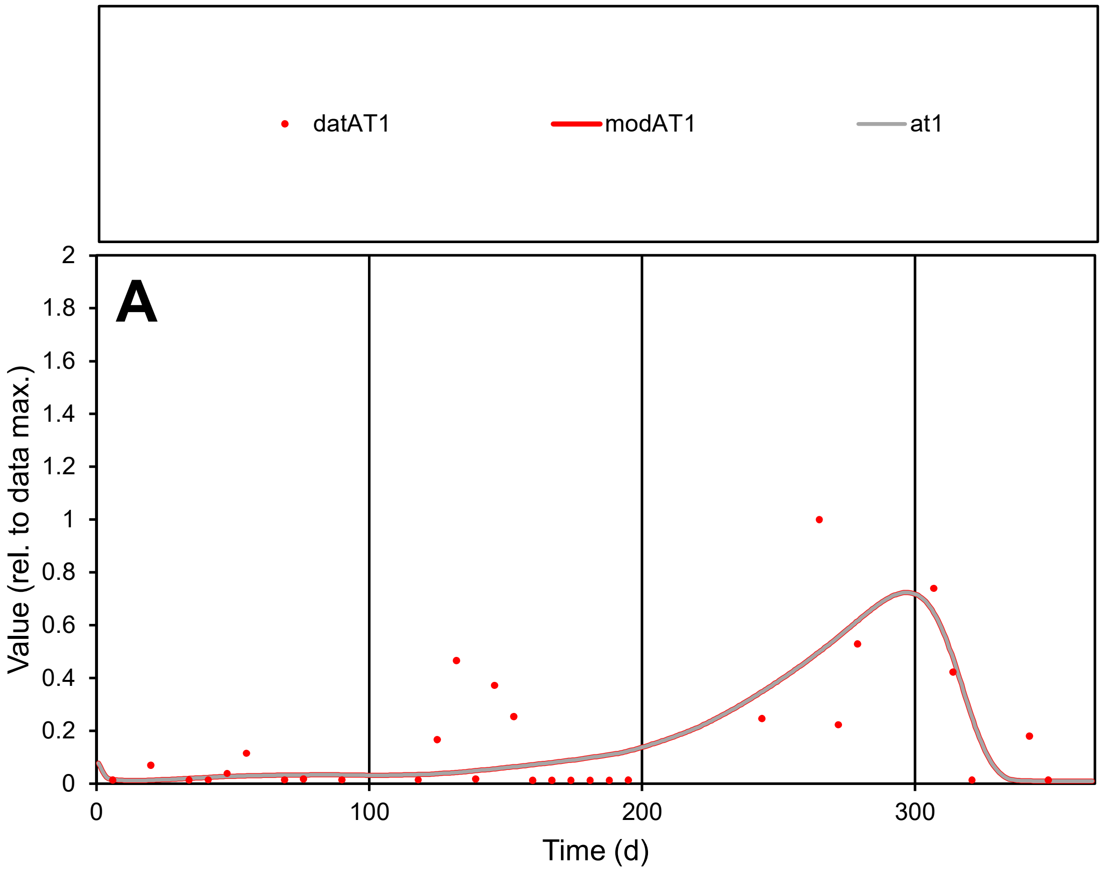


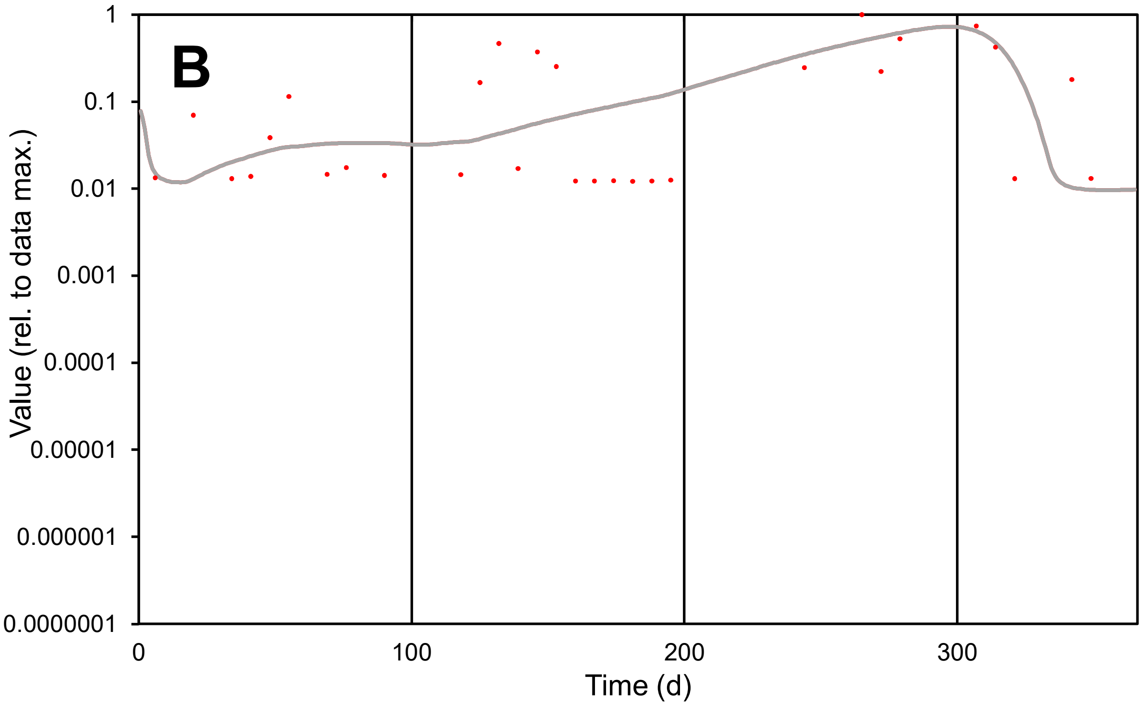


**C1**

Variables
AT1 Provide jtd or Dst
CalibOnly?
2 1 to reproduce of
oNormalize?
2 1=no, 2=yes

ofxn calc*
0.98989
ofxn mod
1.01264

sRun
RunA1
zout
1

Ave
Data
0.00066
Model
0.00057

data/vMax
0.00391

ofxng
275.646

Min d/m
1.25648

ave/min d
13.8321
ave/min m
15.0941

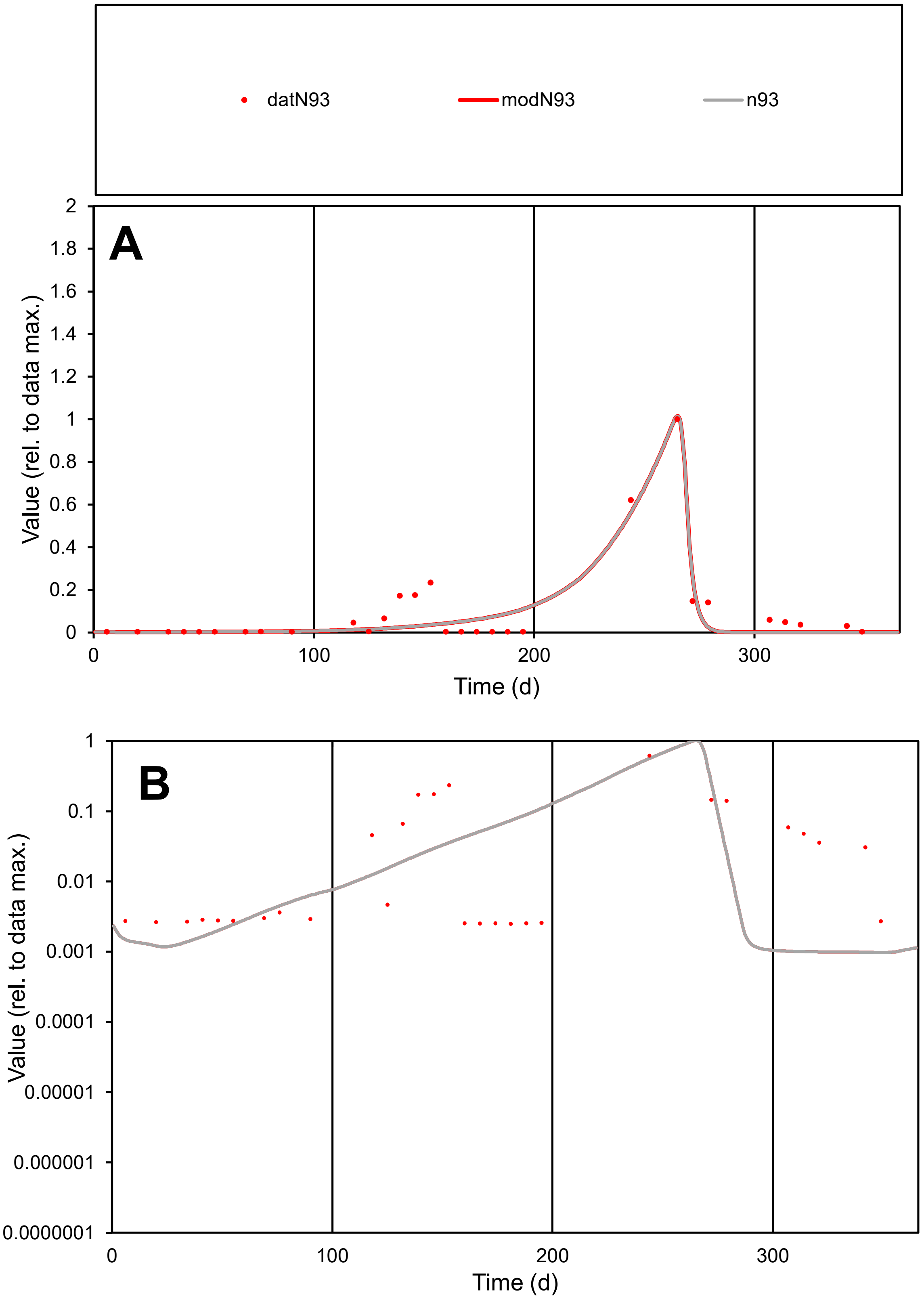


**C1**

Variables
N93 Provide jtd or Dst
CalibOnly?
2 1 to reproduce of
oNormalize?
2 1=no, 2=yes

ofxn calc*
0.7098
ofxn mod
0.75993

sRun
RunA1
zout
1

Ave
Data
0.00065
Model
0.00055

data/vMax
0.00687

ofxng
275.646

Min d/m
2.5763

ave/min d
37.5474
ave/min m
81.7758

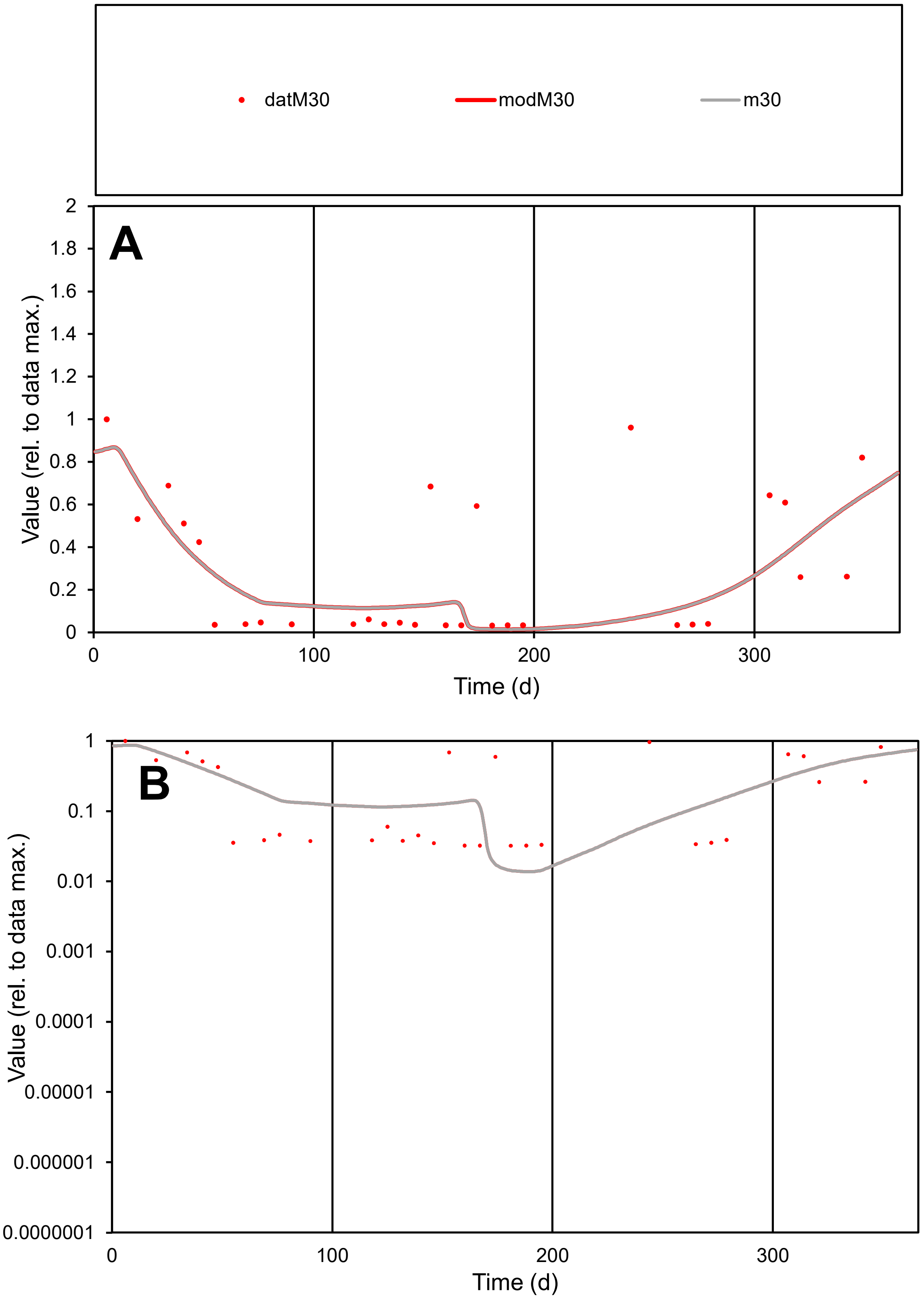


**C1**

Variables
M30 Provide jtd or Dst
CalibOnly?
2 1 to reproduce of
oNormalize?
2 1=no, 2=yes

ofxn calc*
0.90391
ofxn mod
0.91581

sRun
RunA1
zout
1

Ave
Data
0.00036
Model
0.00031

data/vMax
0.00127

ofxng
275.646

Min d/m
2.33851

ave/min d
8.96921
ave/min m
17.7067

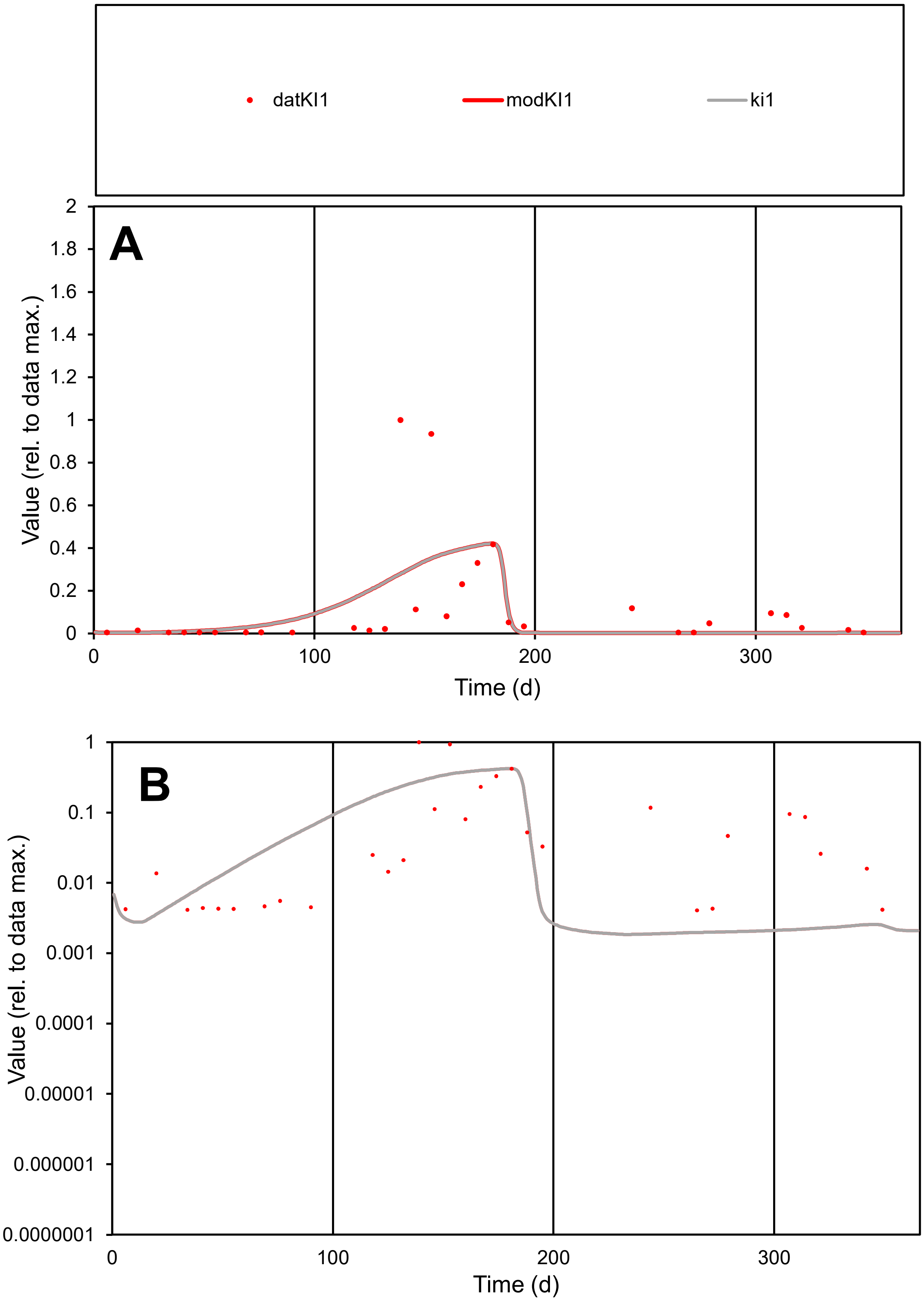


**C1**

Variables
KI1 Provide jtd or Dst
CalibOnly?
2 1 to reproduce of
oNormalize?
2 1=no, 2=yes

ofxn calc*
1.59033
ofxn mod
1.60185

sRun
RunA1
zout
1

Ave
Data
0.00148
Model
0.00138

data/vMax
0.01201

ofxng
275.646

Min d/m
2.16313

ave/min d
30.2648
ave/min m
61.0384

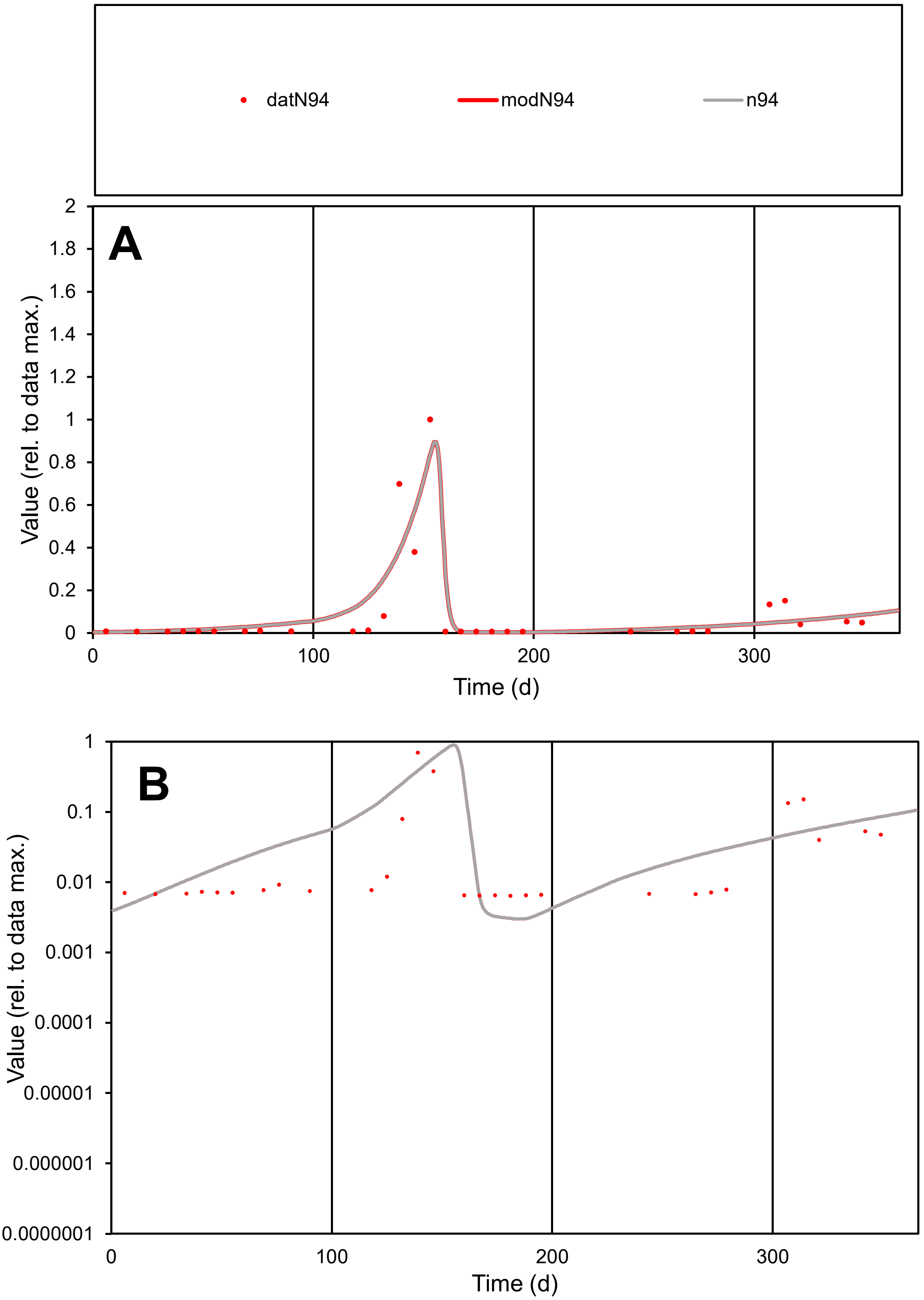


**C1**

Variables

N94 Provide jtd or Dst

CalibOnly?

2 1 to reproduce of

oNormalize?

2 1=no, 2=yes

ofxn calc*

0.95798

ofxn mod

1.01491

sRun

RunA1

zout

1

Ave

Data

0.00064

Model

0.00073

data/vMax

0.00701

ofxng

275.646

Min d/m

2.11325

ave/min d

14.2176

ave/min m

34.2187

325_Calib_S_A

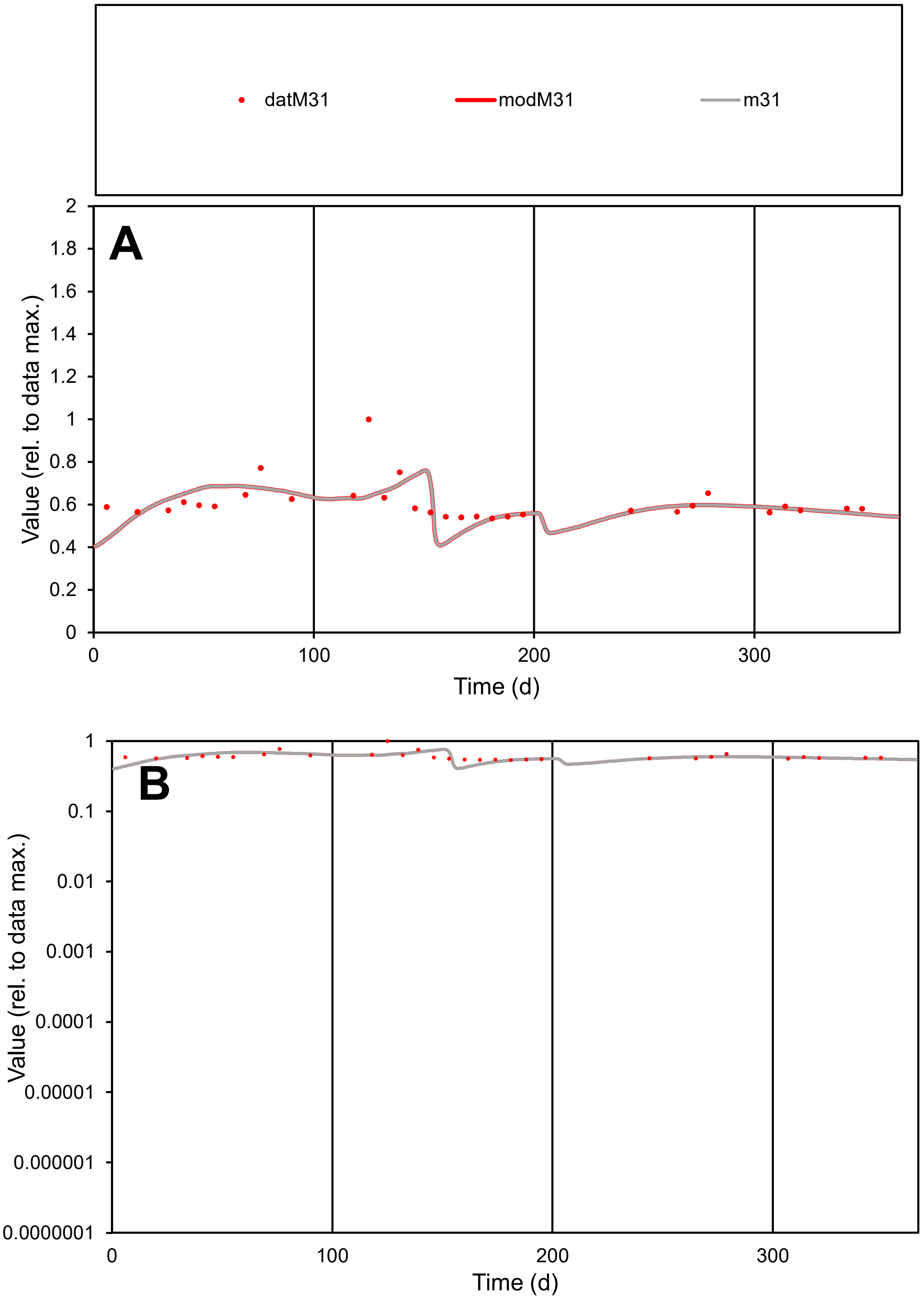


**C1**

Variables
M31 Provide jtd or Dst
CalibOnly?
2 1 to reproduce of
oNormalize?
2 1=no, 2=yes

ofxn calc*
0.14448
ofxn mod
0.18382

sRun
RunA1
zout
1

Ave
Data
9.2E-05
Model
9E-05

data/vMax
0.00015

ofxng
275.646

Min d/m
1.24422

ave/min d
1.1376
ave/min m
1.38866

325_Calib_S_A

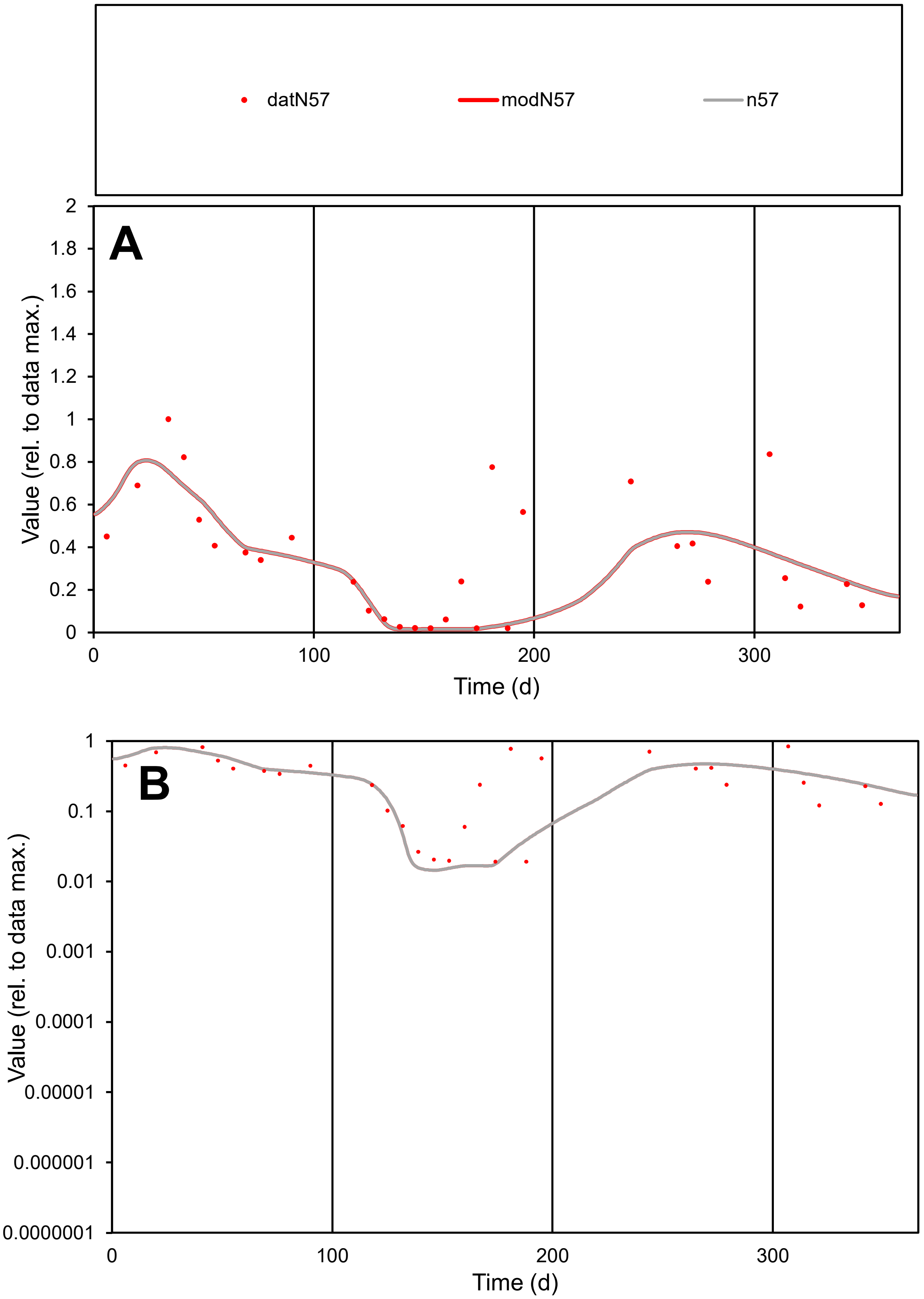


**C1**

Variables
N57 Provide jtd or Dst
CalibOnly?
2 1 to reproduce of
oNormalize?
2 1=no, 2=yes

ofxn calc*
0.62642
ofxn mod
0.64146

sRun
RunA1
zout
1

Ave
Data
0.00067
Model
0.00057

data/vMax
0.0019

ofxng
275.646

Min d/m
1.31858

ave/min d
18.3663
ave/min m
20.6275

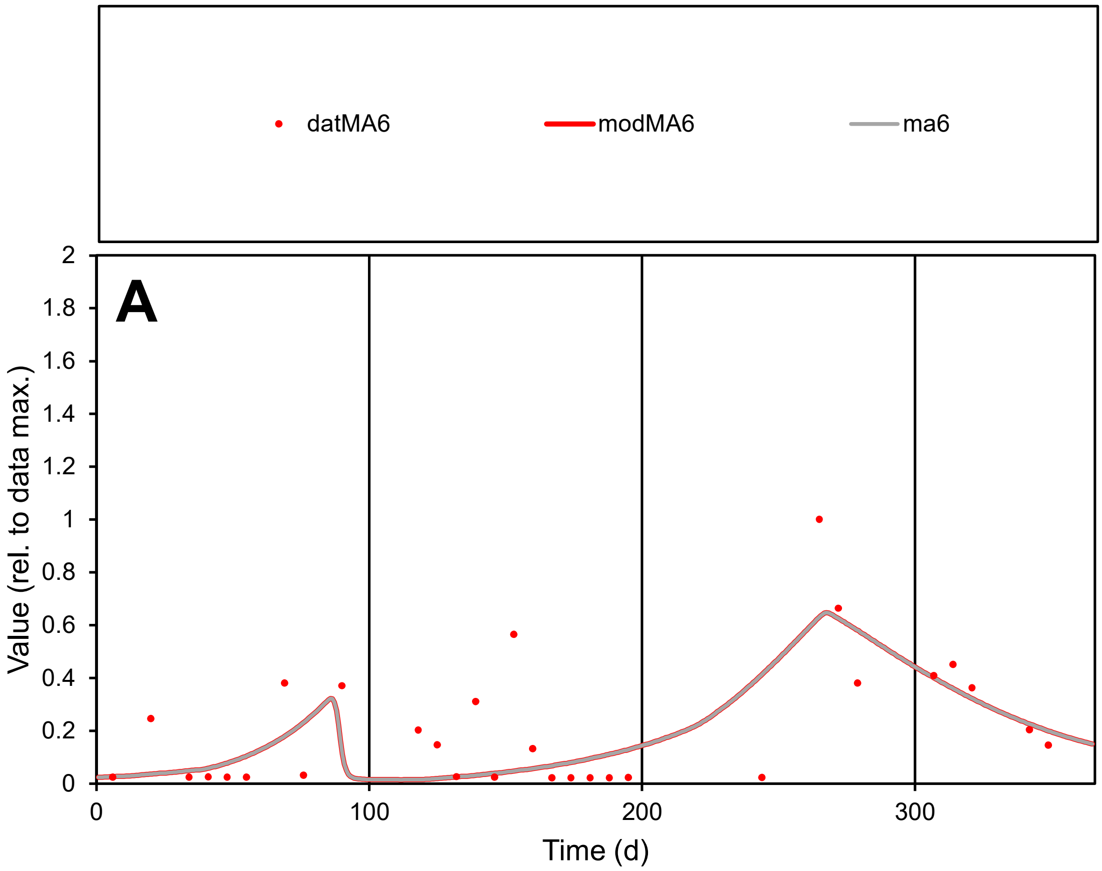


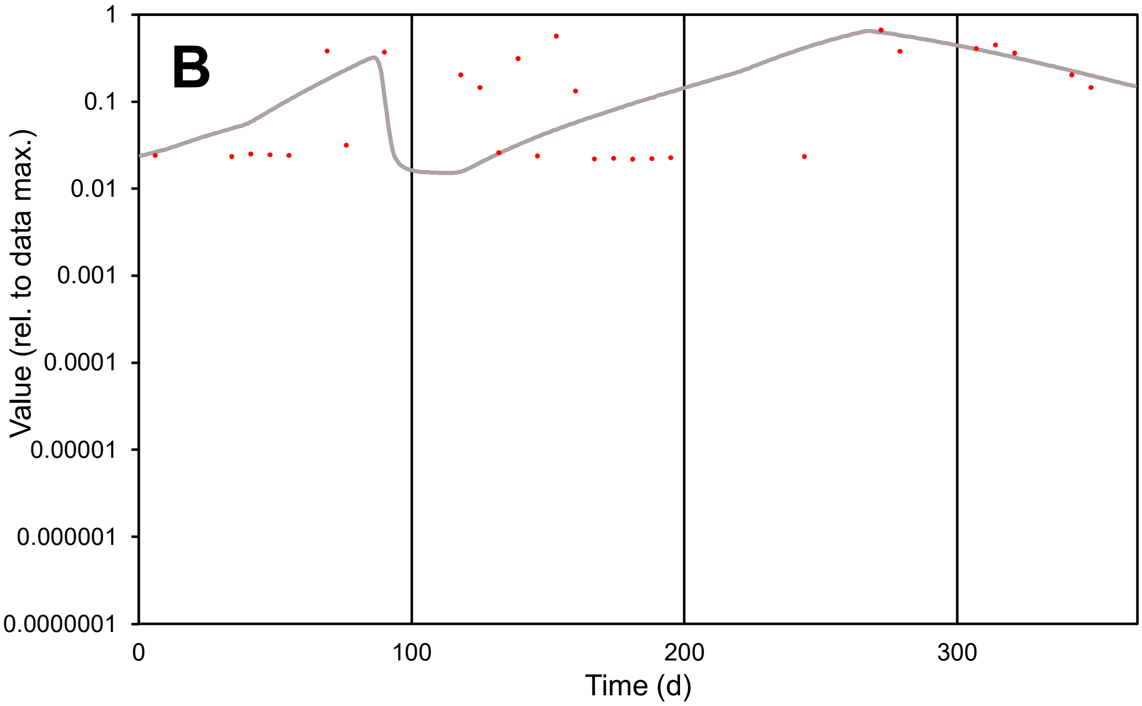


**C1**

Variables

MA6 Provide jtd or Dst

CalibOnly?

2 1 to reproduce of

oNormalize?

2 1=no, 2=yes

ofxn calc*

0.82765

ofxn mod

0.86112

sRun

RunA1

zout

1

Ave

Data

0.00112

Model

0.00098

data/vMax

0.00533

ofxng

275.646

Min d/m

1.3553

ave/min d

9.59084

ave/min m

11.375

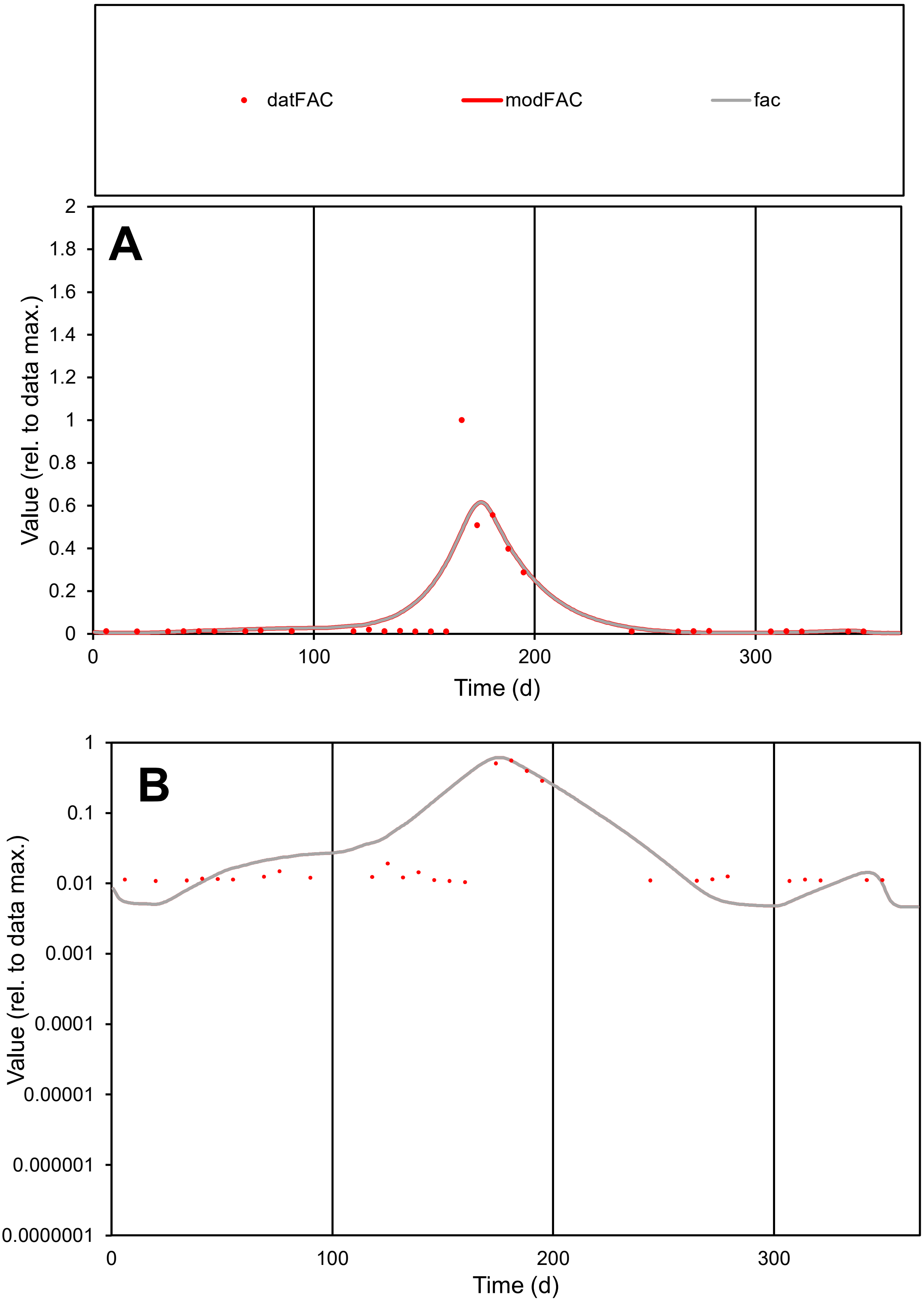


**C1**

Variables

FAC Provide jtd or Dst

CalibOnly?

2 1 to reproduce of

oNormalize?

2 1=no, 2=yes

ofxn calc*

1.16446

ofxn mod

1.1997

sRun

RunA1

zout

1

Ave

Data

0.00085

Model

0.00097

data/vMax

0.00832

ofxng

275.646

Min d/m

2.03805

ave/min d

9.738

ave/min m

22.6855

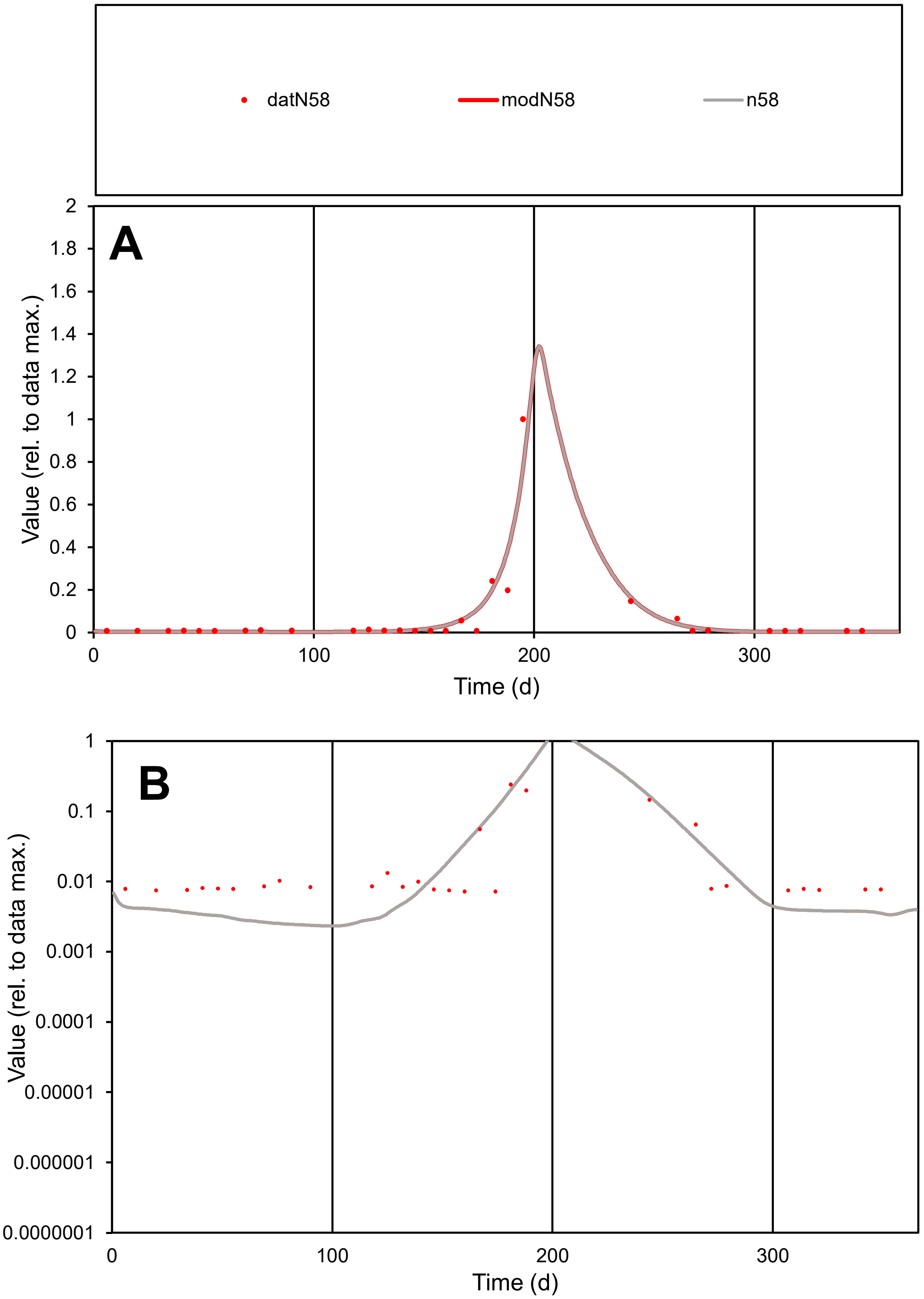


**C1**

Variables
N58 Provide jtd or Dst
CalibOnly?
2 1 to reproduce of
oNormalize?
2 1=no, 2=yes

ofxn calc*
0.72579
ofxn mod
0.82343

sRun
RunA1
zout
1

Ave
Data
0.00072
Model
0.00078

data/vMax
0.01141

ofxng
275.646

Min d/m
3.03209
ave/min d
8.80948
ave/min m
28.6504

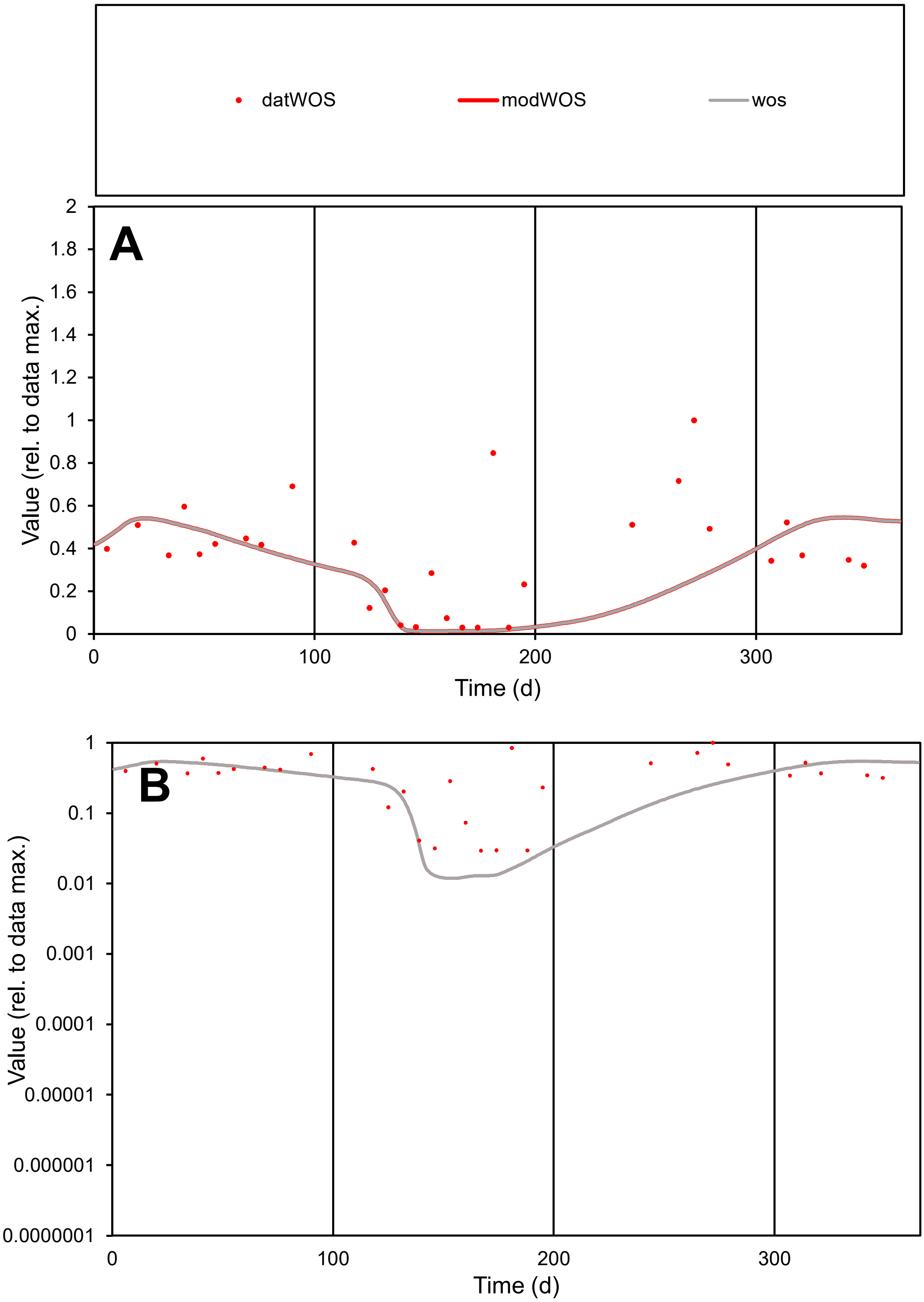


**C1**

Variables

WOS Provide jtd or Dst

CalibOnly?

2 1 to reproduce of

oNormalize?

2 1=no, 2=yes

ofxn calc*

0.70942

ofxn mod

0.7199

sRun

RunA1

zout

1

Ave

Data

0.00087

Model

0.00065

data/vMax

0.00232

ofxng

275.646

Min d/m

2.47588

ave/min d

12.675

ave/min m

23.5509

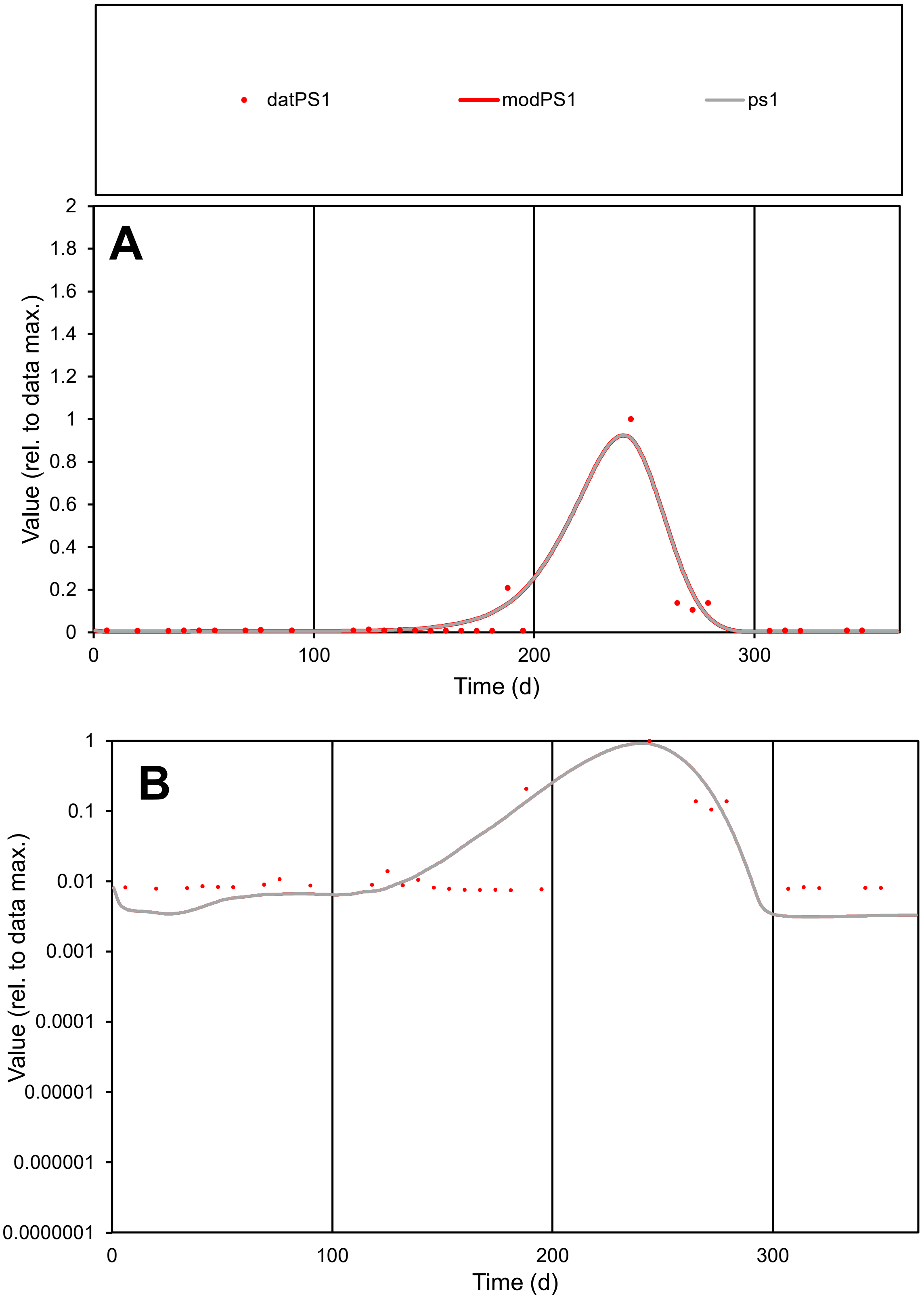


**C1**

Variables

PS1 Provide jtd or Dst

CalibOnly?

2 1 to reproduce of

oNormalize?

2 1=no, 2=yes

ofxn calc*

0.89975

ofxn mod

1.00597

sRun

RunA1

zout

1

Ave

Data

0.00038

Model

0.00044

data/vMax

0.00625

ofxng

275.646

Min d/m

2.38843

ave/min d

8.05032

ave/min m

22.5503

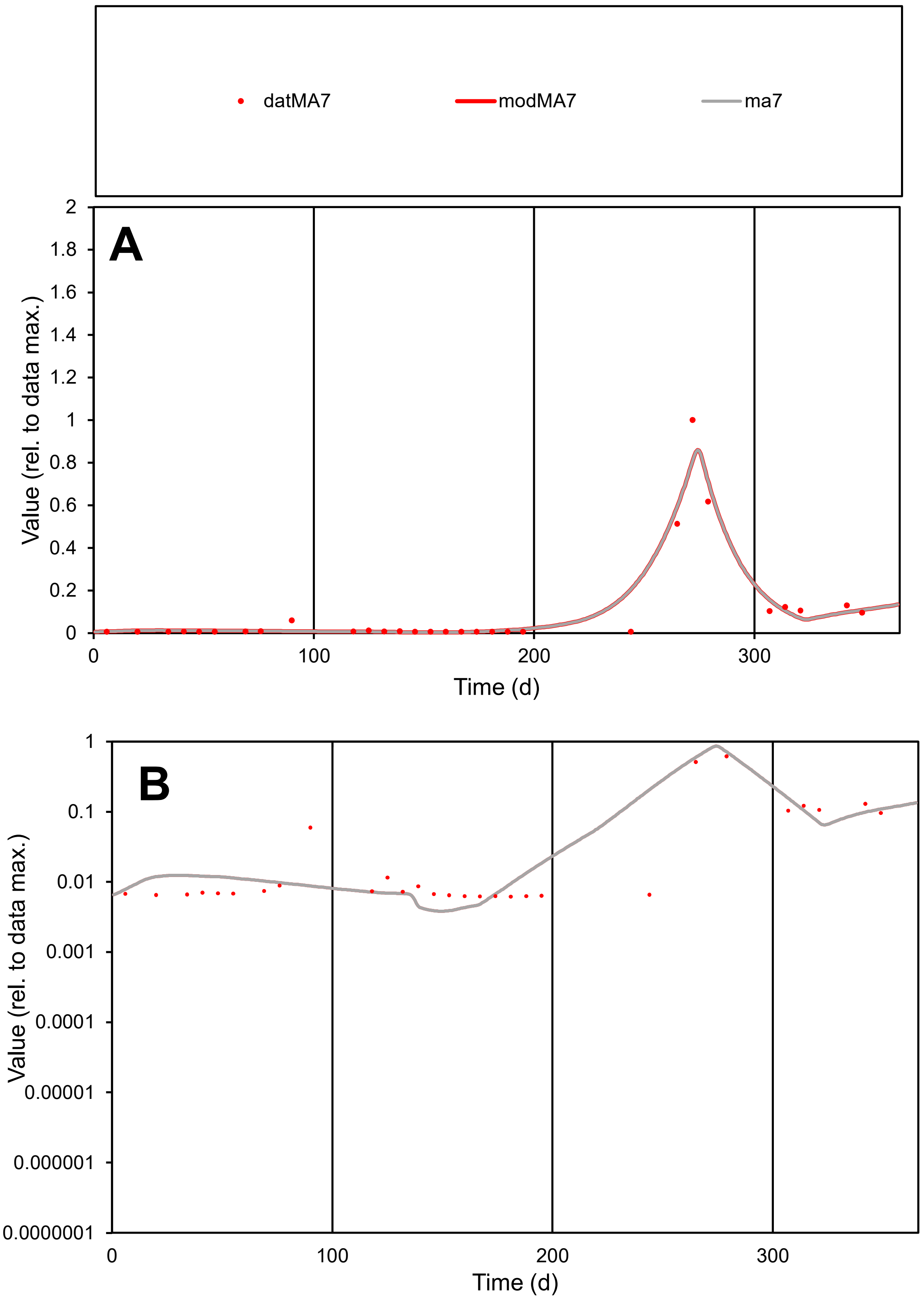


**C1**

Variables
MA7 Provide jtd or Dst
CalibOnly?
2 1 to reproduce of
oNormalize?
2 1=no, 2=yes

ofxn calc*
0.51083
ofxn mod
0.65018

sRun
RunA1
zout
1

Ave
Data
0.00021
Model
0.00022

data/vMax
0.00216

ofxng
275.646

Min d/m
1.60757

ave/min d
15.6687
ave/min m
26.0979

325_Calib_S_A

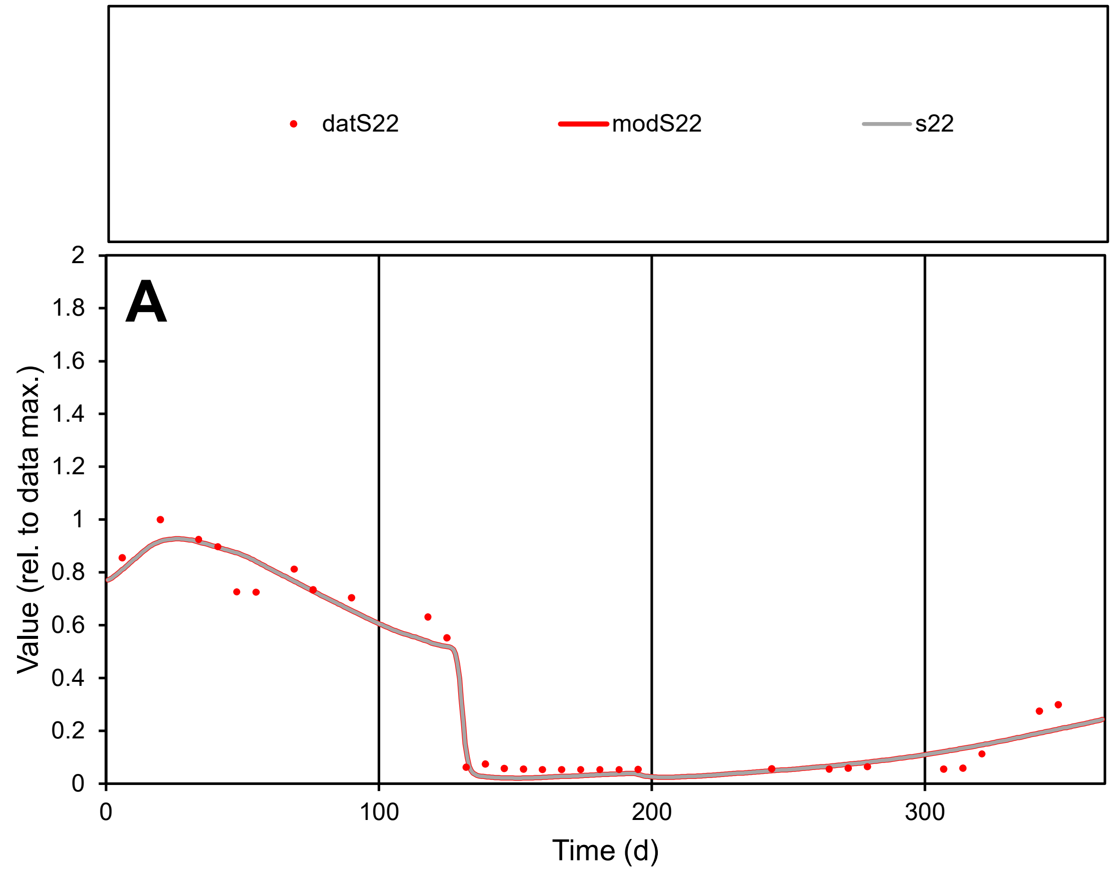


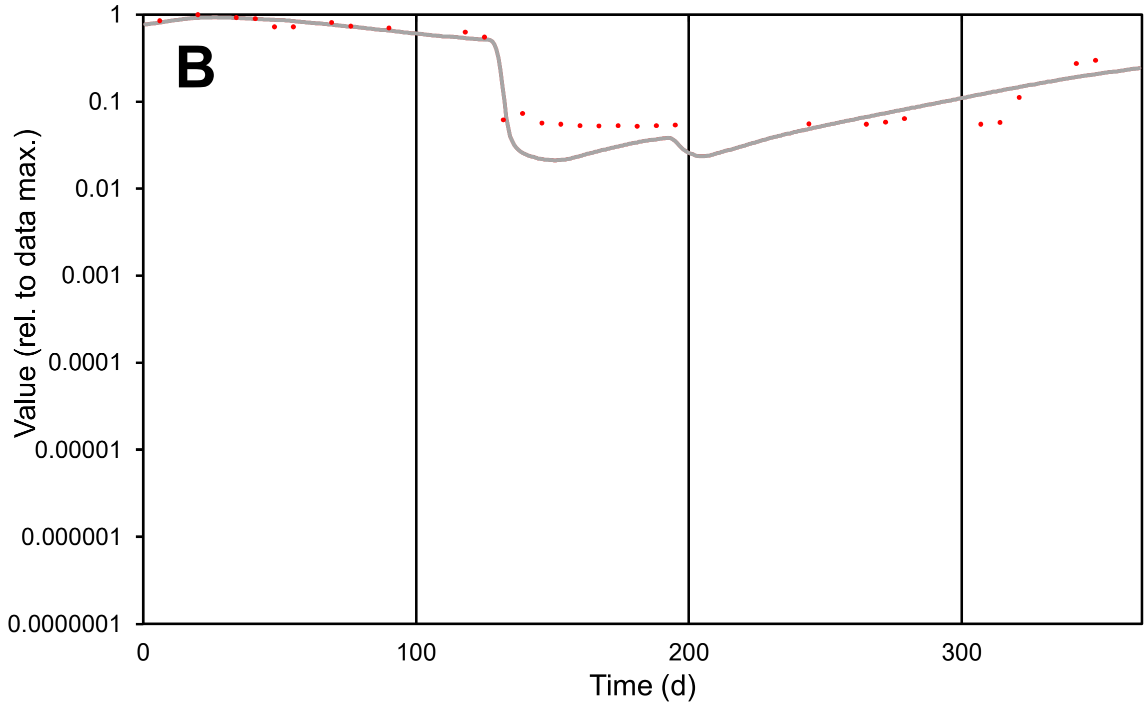


**C1**

Variables
S22 Provide jtd or Dst
CalibOnly?
2 1 to reproduce of
oNormalize?
2 1=no, 2=yes

ofxn calc*
0.16343
ofxn mod
0.23466

sRun
RunA1
zout
1

Ave
Data
0.00094
Model
0.00091

data/vMax
0.00278

ofxng
275.646

Min d/m
2.43027

ave/min d
6.47544
ave/min m
15.2408

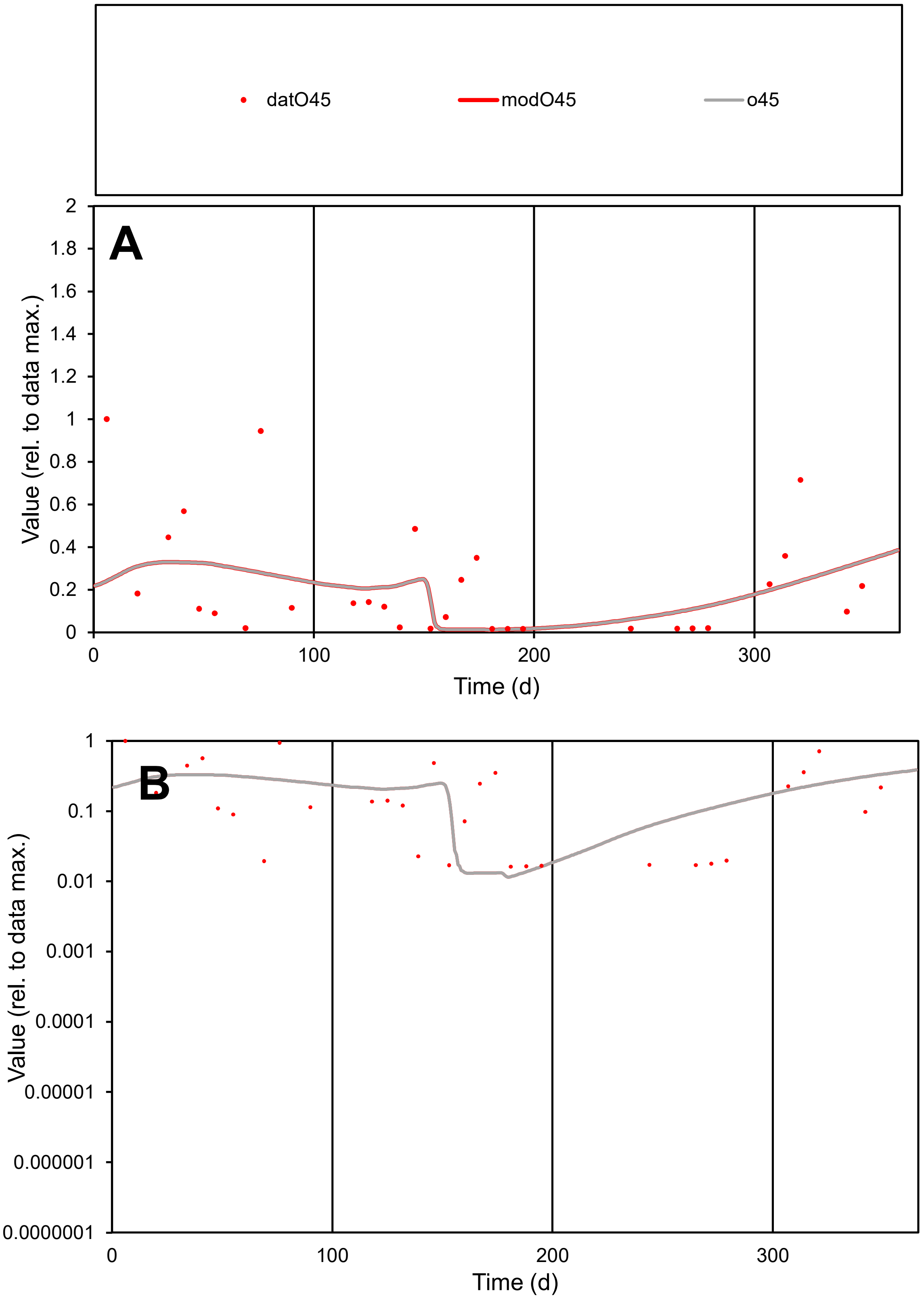


**C1**

Variables
O45 Provide jtd or Dst
CalibOnly?
2 1 to reproduce of
oNormalize?
2 1=no, 2=yes

ofxn calc*
1.11079
ofxn mod
1.11835

sRun
RunA1
zout
1

Ave
Data
0.0008
Model
0.00066

data/vMax
0.00351

ofxng
275.646

Min d/m
1.34568

ave/min d
14.0295
ave/min m
15.5649

325_Calib_S_A

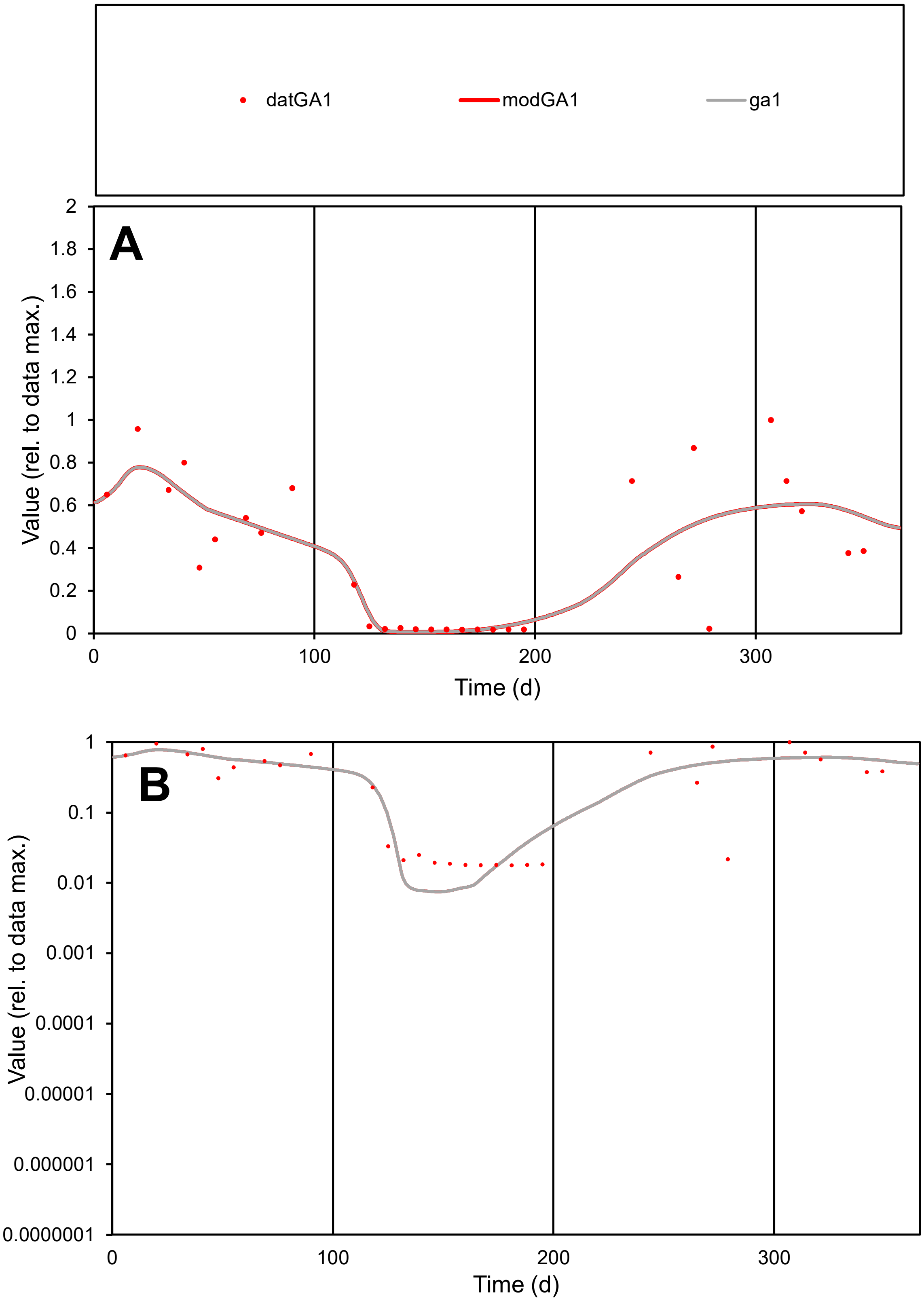


**C1**

Variables

GA1 Provide jtd or Dst

CalibOnly?

2 1 to reproduce of

oNormalize?

2 1=no, 2=yes

ofxn calc*

0.51048

ofxn mod

0.5318

sRun

RunA1

zout

1

Ave

Data

0.00072

Model

0.0007

data/vMax

0.00198

ofxng

275.646

Min d/m

2.38802

ave/min d

20.4429

ave/min m

47.7723

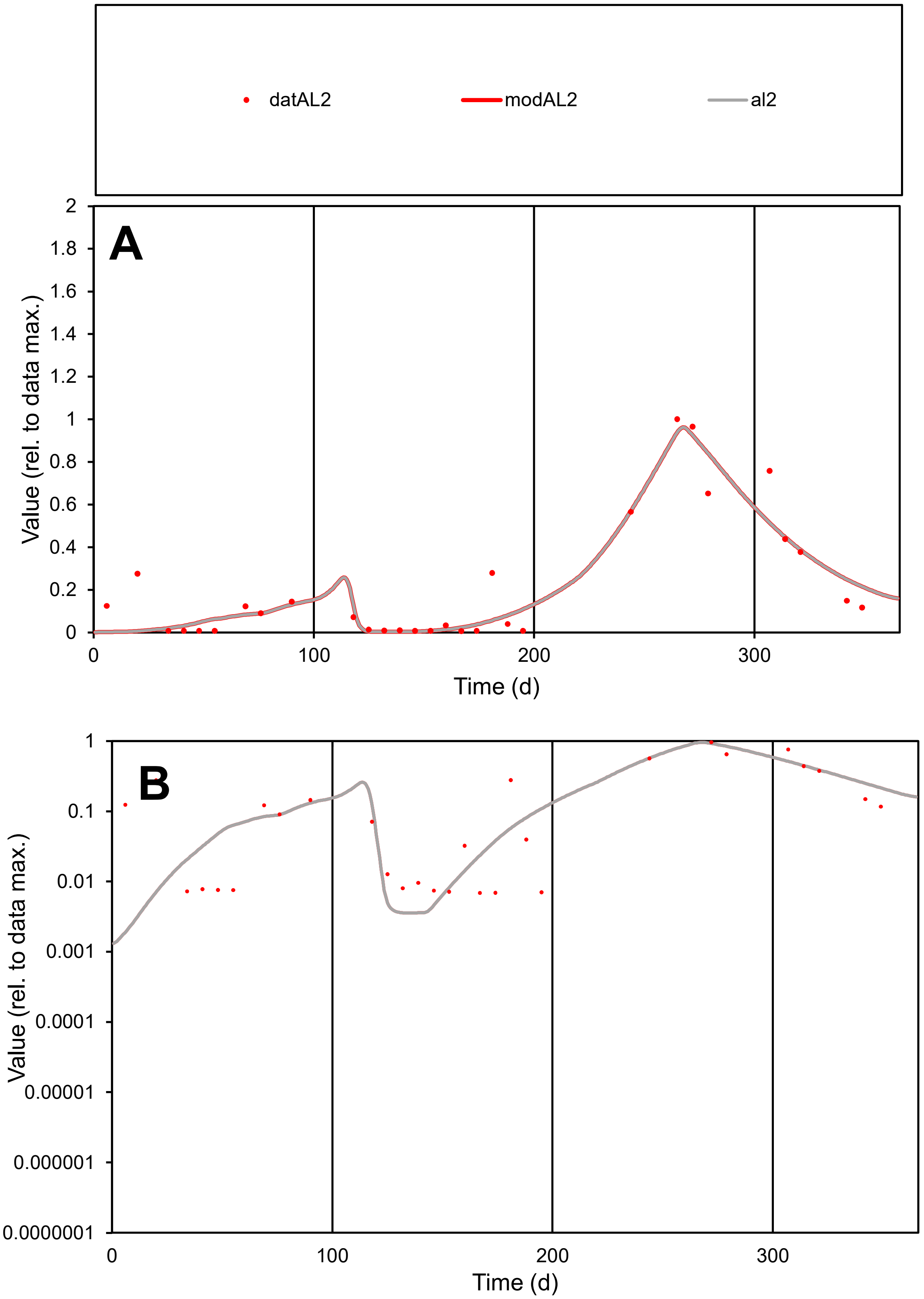


**C1**

Variables
AL2 Provide jtd or Dst
CalibOnly?
2 1 to reproduce of
oNormalize?
2 1=no, 2=yes

ofxn calc*
0.4518
ofxn mod
0.49704

sRun
RunA1
zout
1

Ave
Data
0.00067
Model
0.00063

data/vMax
0.00317

ofxng
275.646

Min d/m
3.25108

ave/min d
30.6072
ave/min m
94.0272

325_Calib_S_A

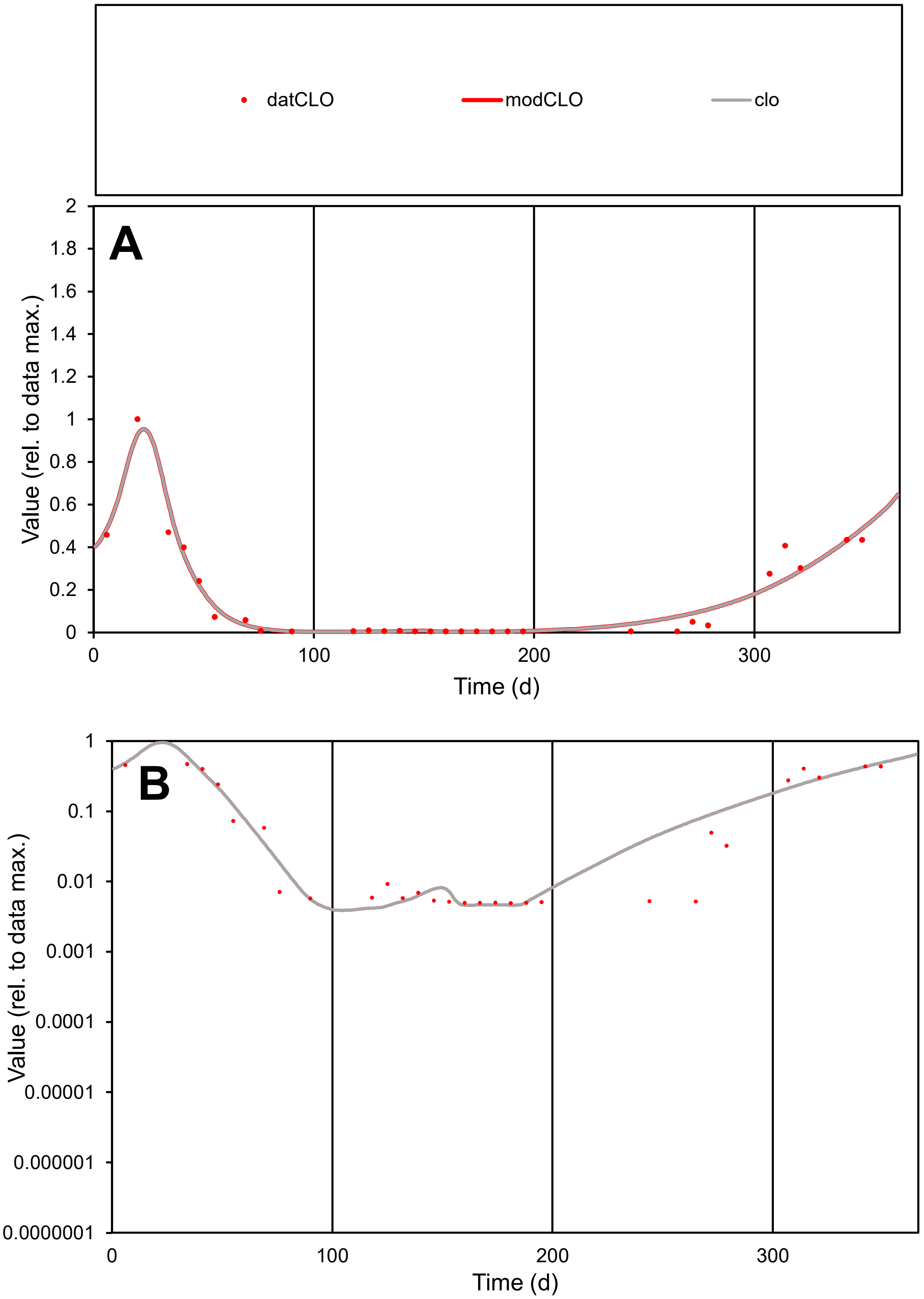


**C1**

Variables
CLO Provide jtd or Dst
CalibOnly?
2 1 to reproduce of
oNormalize?
2 1=no, 2=yes

ofxn calc*
0.28338
ofxn mod
0.34194

sRun
RunA1
zout
1

Ave
Data
0.00042
Model
0.00043

data/vMax
0.00266

ofxng
275.646

Min d/m
1.1752

ave/min d
31.9936
ave/min m
38.3191

325_Calib_S_A

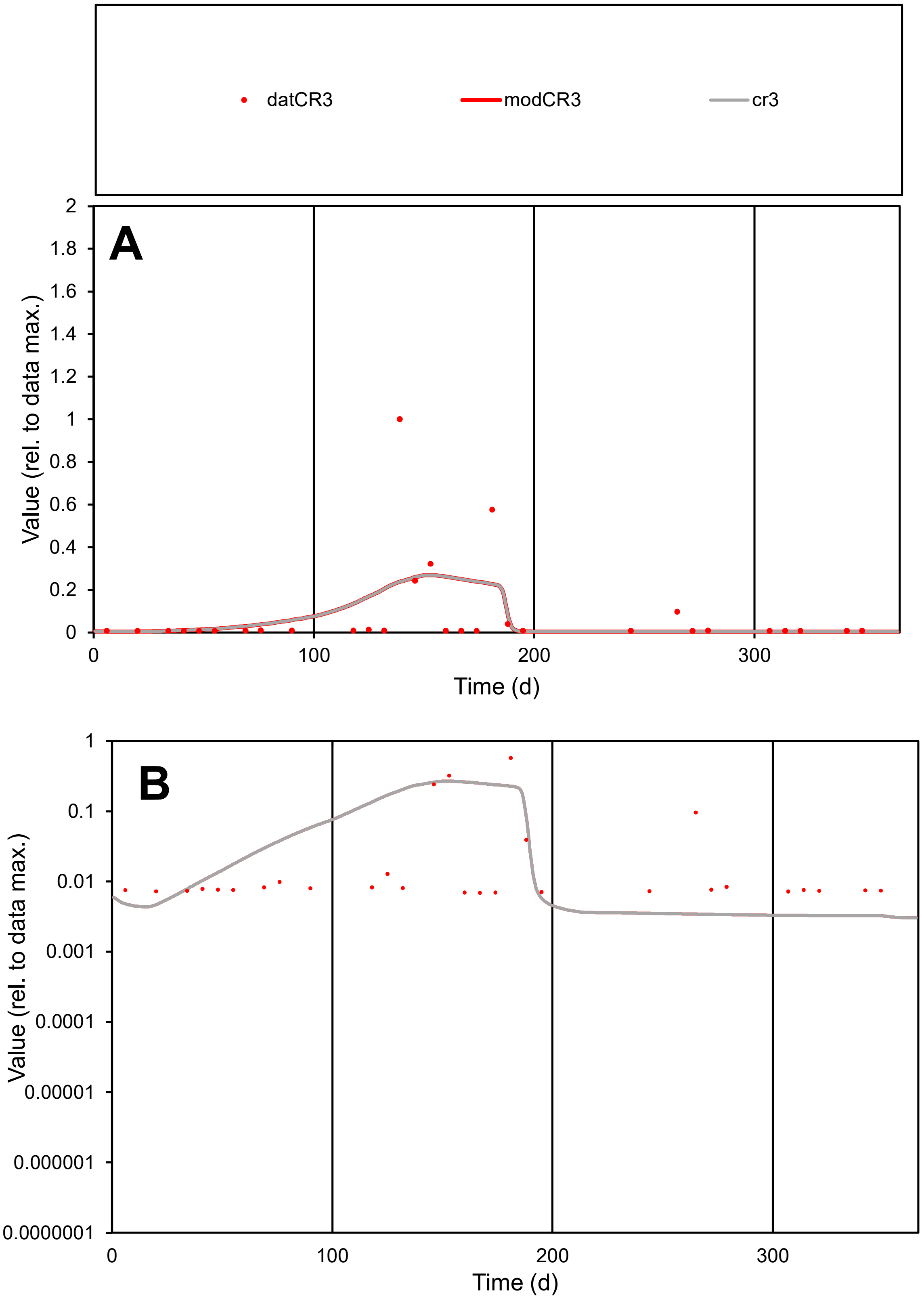


**C1**

Variables
CR3 Provide jtd or Dst
CalibOnly?
2 1 to reproduce of
oNormalize?
2 1=no, 2=yes

ofxn calc*
2.18683
ofxn mod
2.20062

sRun
RunA1
zout
1

Ave
Data
0.00042
Model
0.00043

data/vMax
0.00512

ofxng
275.646

Min d/m
2.14988
ave/min d
11.872
ave/min m
26.0837

325_Calib_S_A

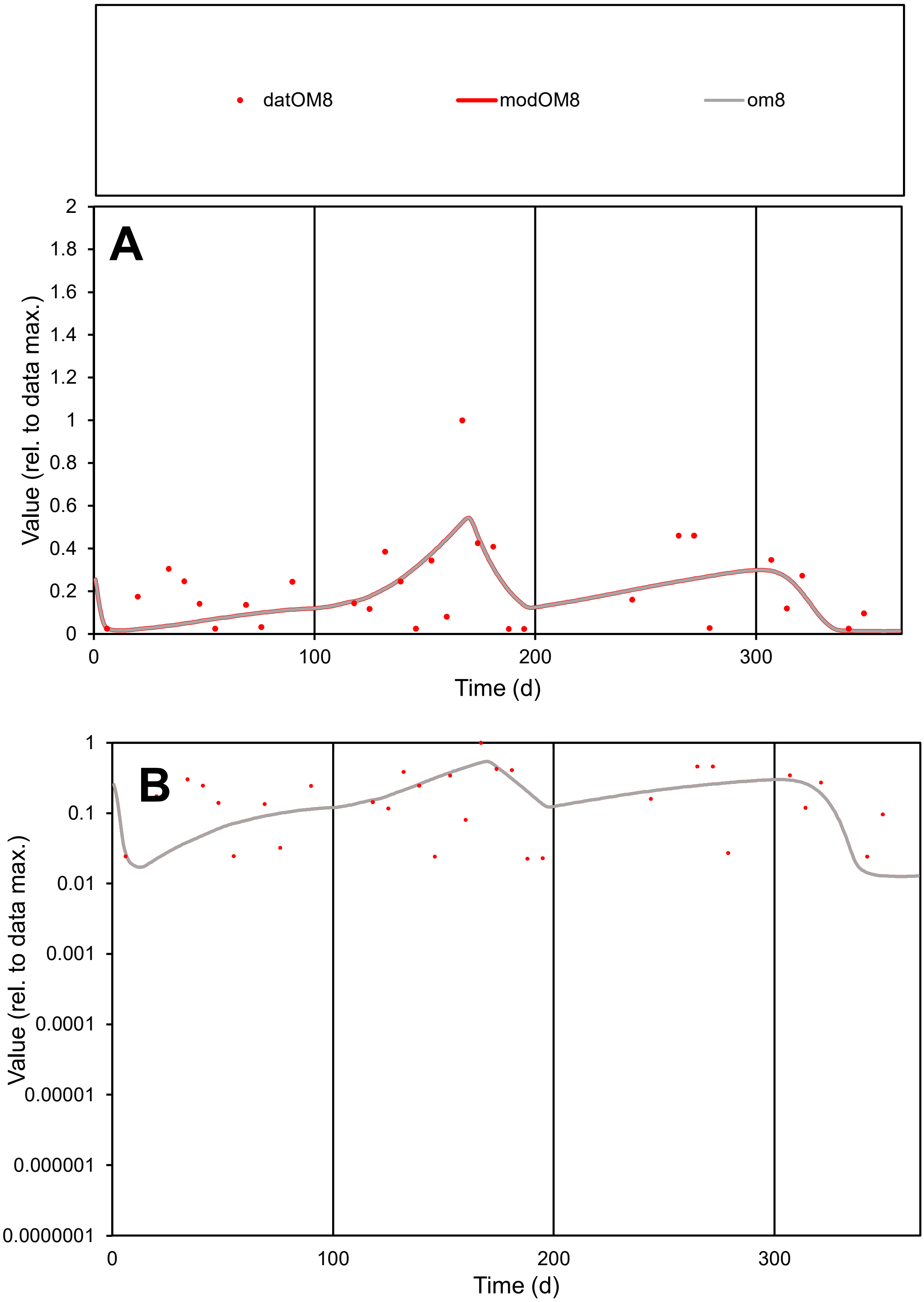


| | |
|---|---|
| **C1** | |
| Variables | |
| OM8 | Provide jtd or Dst |
| CalibOnly? | |
| 2 | 1 to reproduce of |
| oNormalize? | |
| 2 | 1=no, 2=yes |
| ofxn calc* | |
| 0.78049 | |
| ofxn mod | |
| 0.79448 | |
| sRun | |
| RunA1 | |
| zout | |
| 1 | |
| Ave | |
| Data | |
| 0.00052 | |
| Model | |
| 0.00047 | |
| data/vMax | |
| 0.00241 | |
| ofxng | |
| 275.646 | |
| Min d/m | |
| 1.76112 | |
| ave/min d | |
| 9.62441 | |
| ave/min m | |
| 15.2456 | |

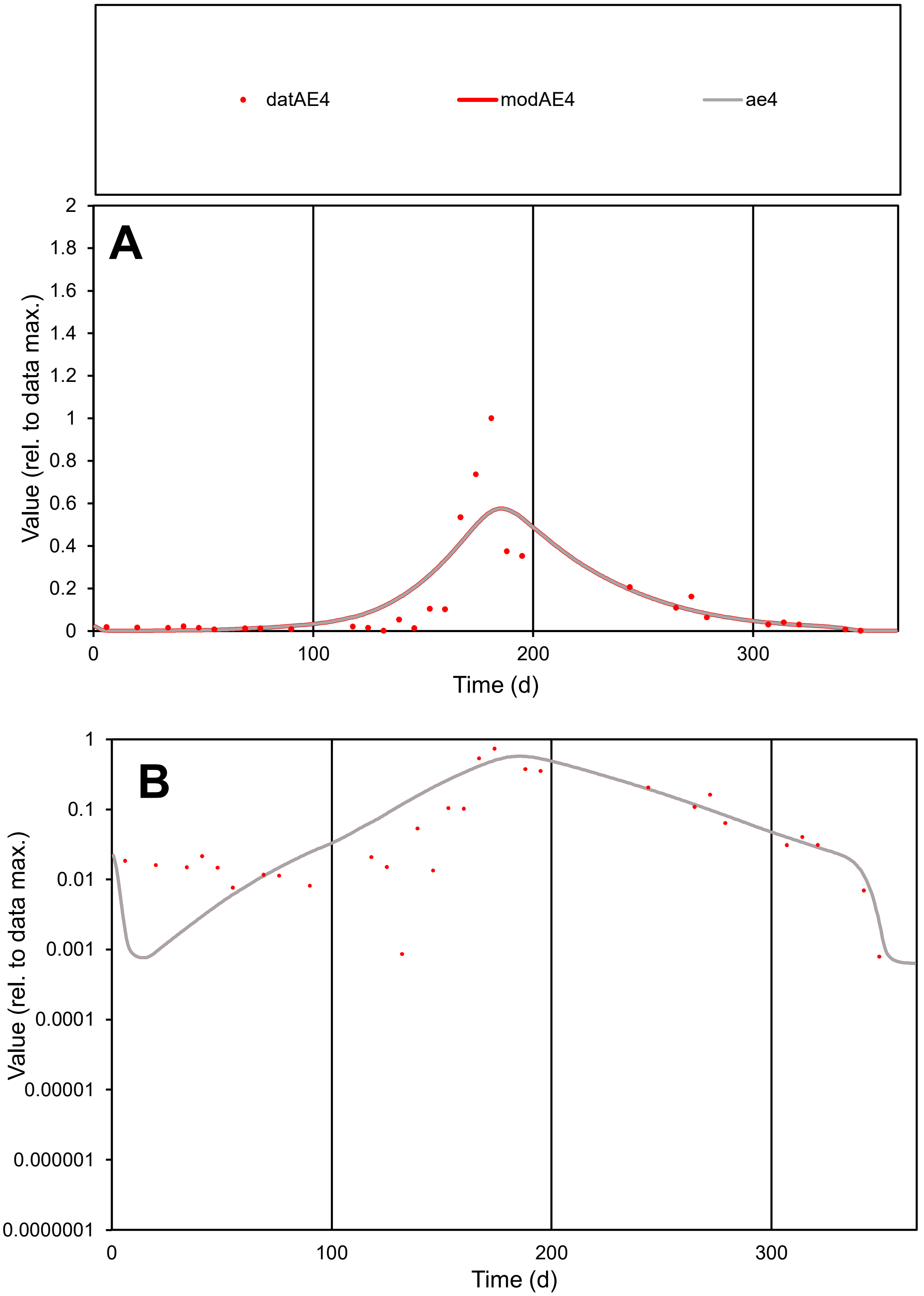


**C1**

Variables
AE4 Provide jtd or Dst
CalibOnly?
2 1 to reproduce of
oNormalize?
2 1=no, 2=yes

ofxn calc*
0.9174
ofxn mod
0.94175

sRun
RunA1
zout
1

Ave
Data
0.00215
Model
0.00238

data/vMax
0.01582

ofxng
275.646

Min d/m
0.80566

ave/min d
172.059
ave/min m
153.103

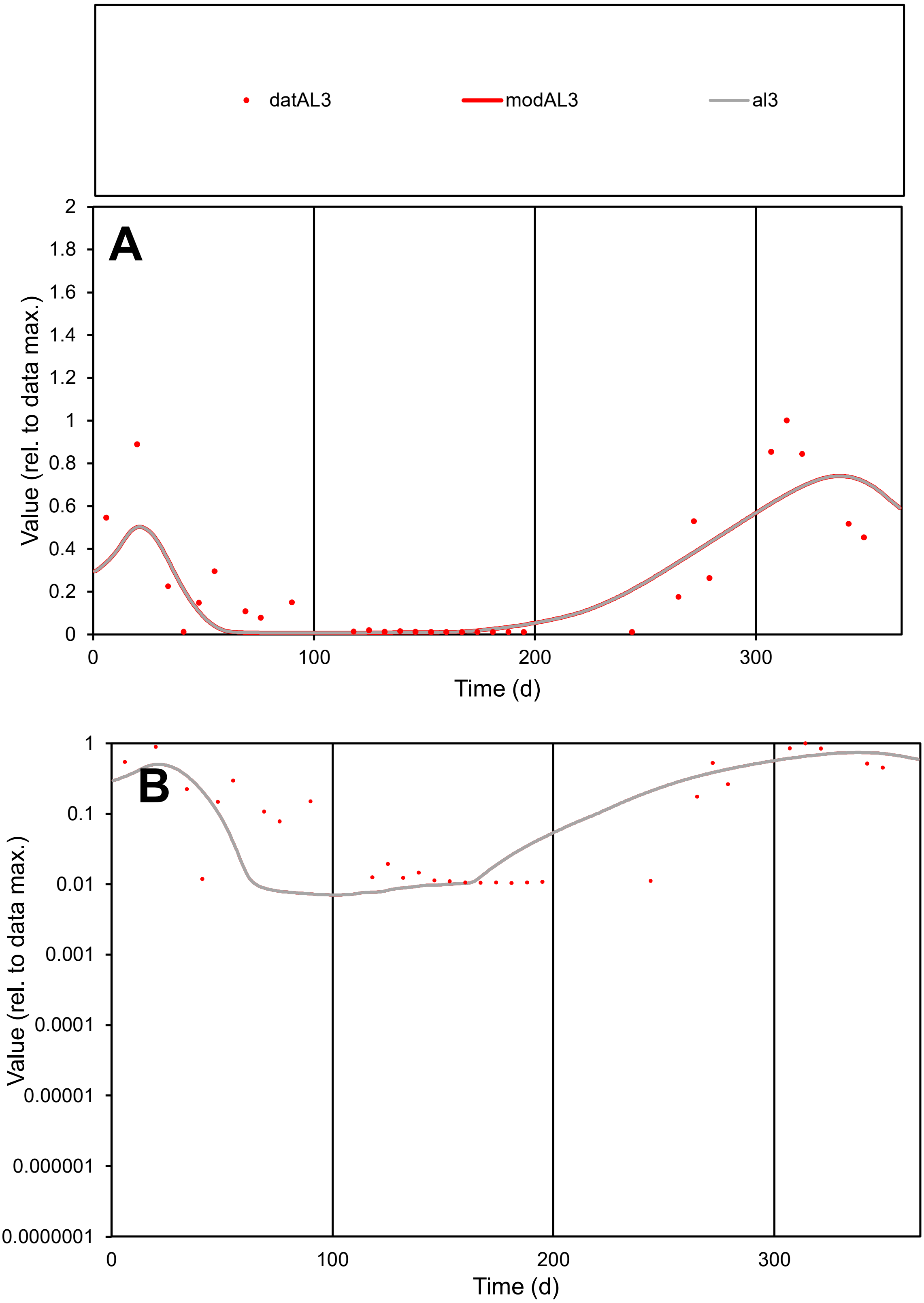


**C1**

Variables
AL3 Provide jtd or Dst
CalibOnly?
2 1 to reproduce of
oNormalize?
2 1=no, 2=yes

ofxn calc*
0.65067
ofxn mod
0.67392

sRun
RunA1
zout
1

Ave
Data
0.00055
Model
0.0005

data/vMax
0.00229

ofxng
275.646

Min d/m
1.4407

ave/min d
23.1163
ave/min m
29.898

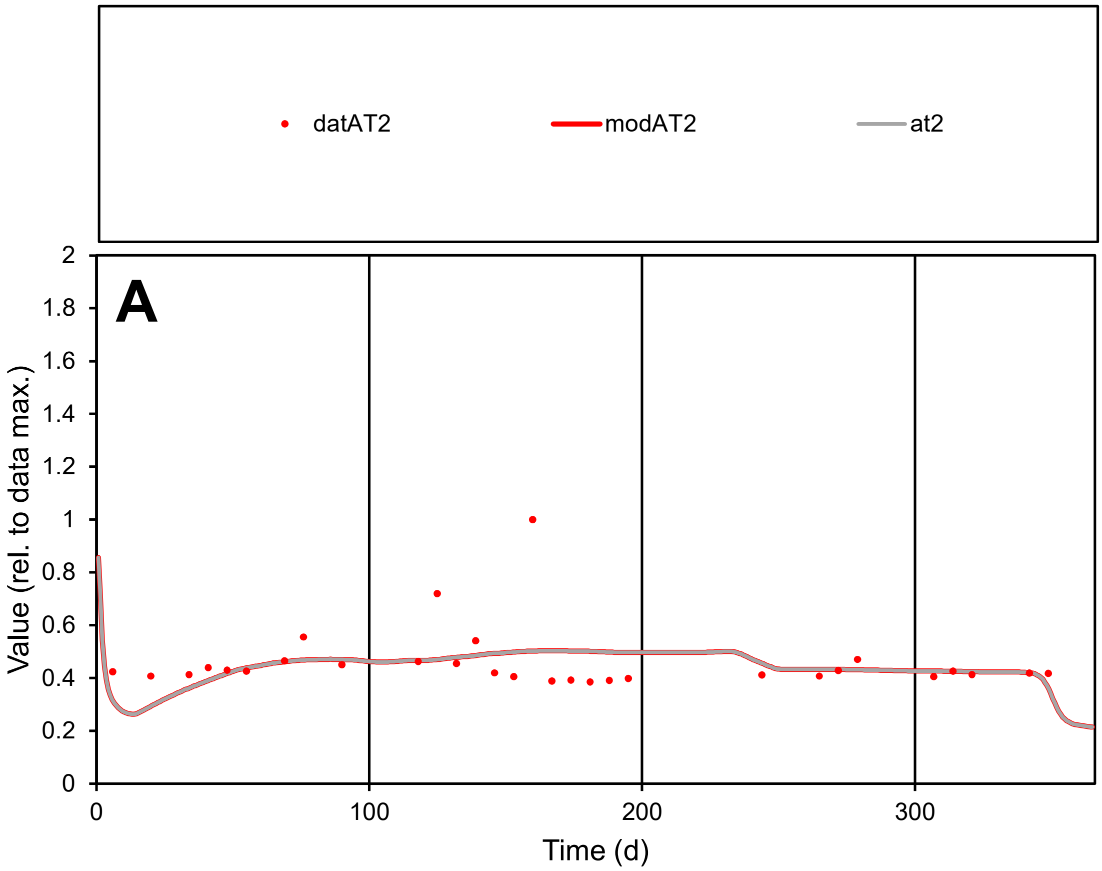


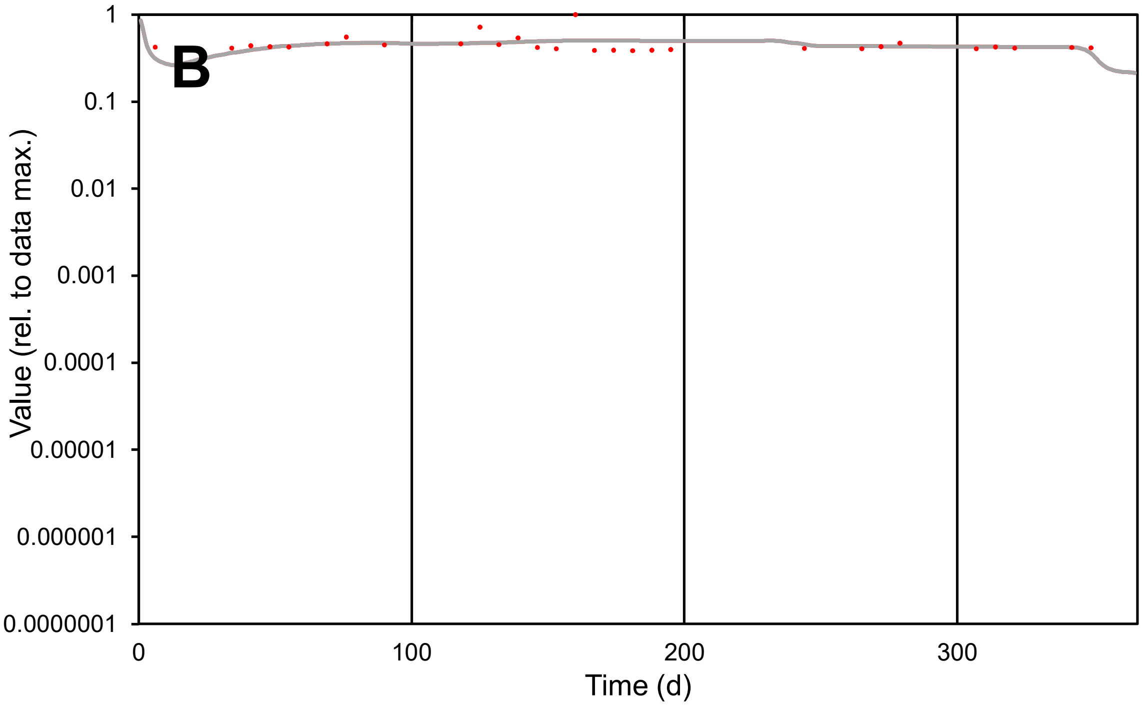


**C1**

Variables

AT2 Provide jtd or Dst

CalibOnly?

2 1 to reproduce of

oNormalize?

2 1=no, 2=yes

ofxn calc*

0.26439

ofxn mod

0.28663

sRun

RunA1

zout

1

Ave

Data

9.4E-05

Model

9.1E-05

data/vMax

0.00021

ofxng

275.646

Min d/m

1.30479

ave/min d

1.19052

ave/min m

1.50159

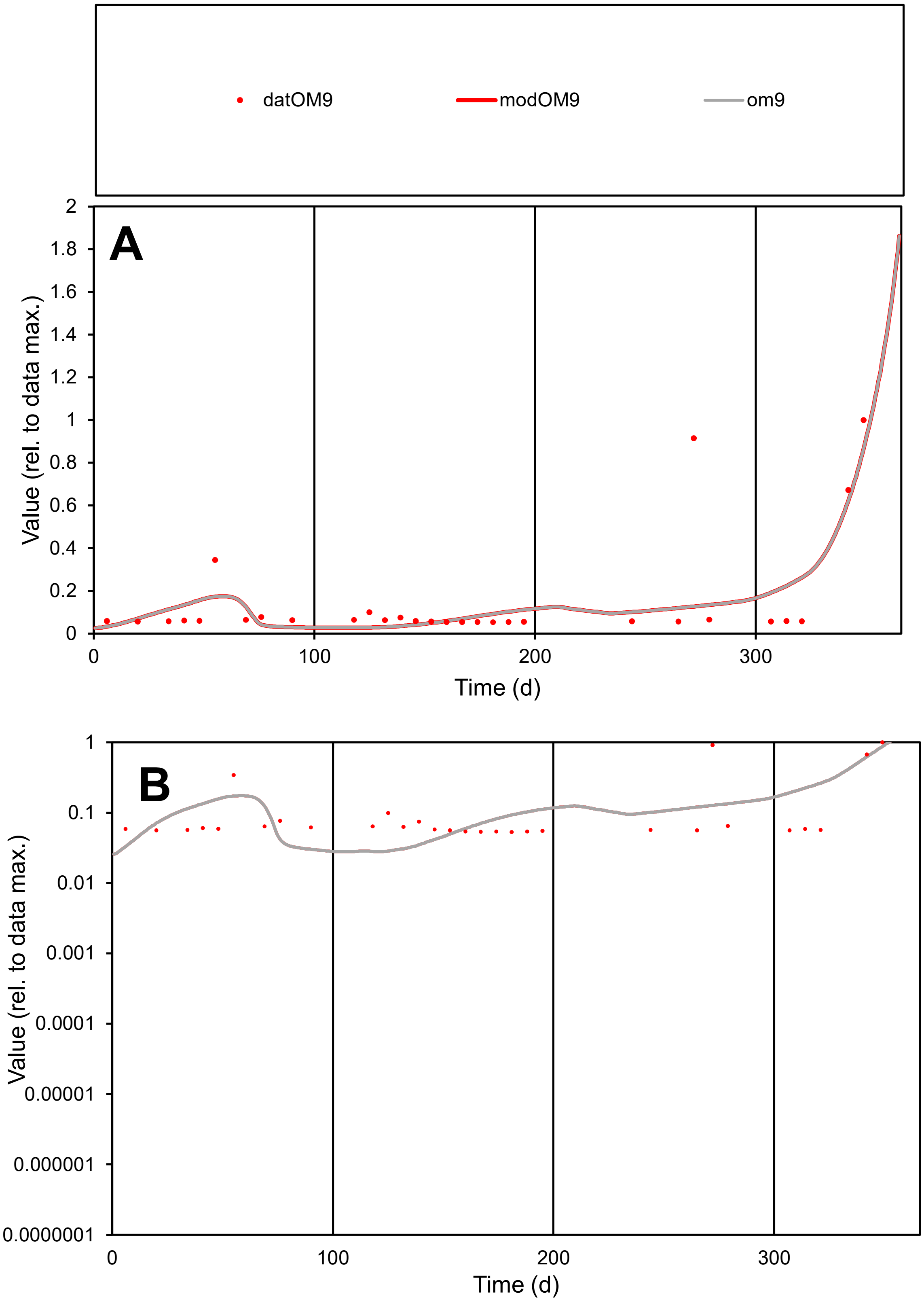


**C1**

Variables
OM9 Provide jtd or Dst
CalibOnly?
2 1 to reproduce of
oNormalize?
2 1=no, 2=yes

ofxn calc*
1.08191
ofxn mod
1.10156

sRun
RunA1
zout
1

Ave
Data
0.00012
Model
0.00012

data/vMax
0.00081

ofxng
275.646

Min d/m
1.89374

ave/min d
2.82836
ave/min m
5.20097

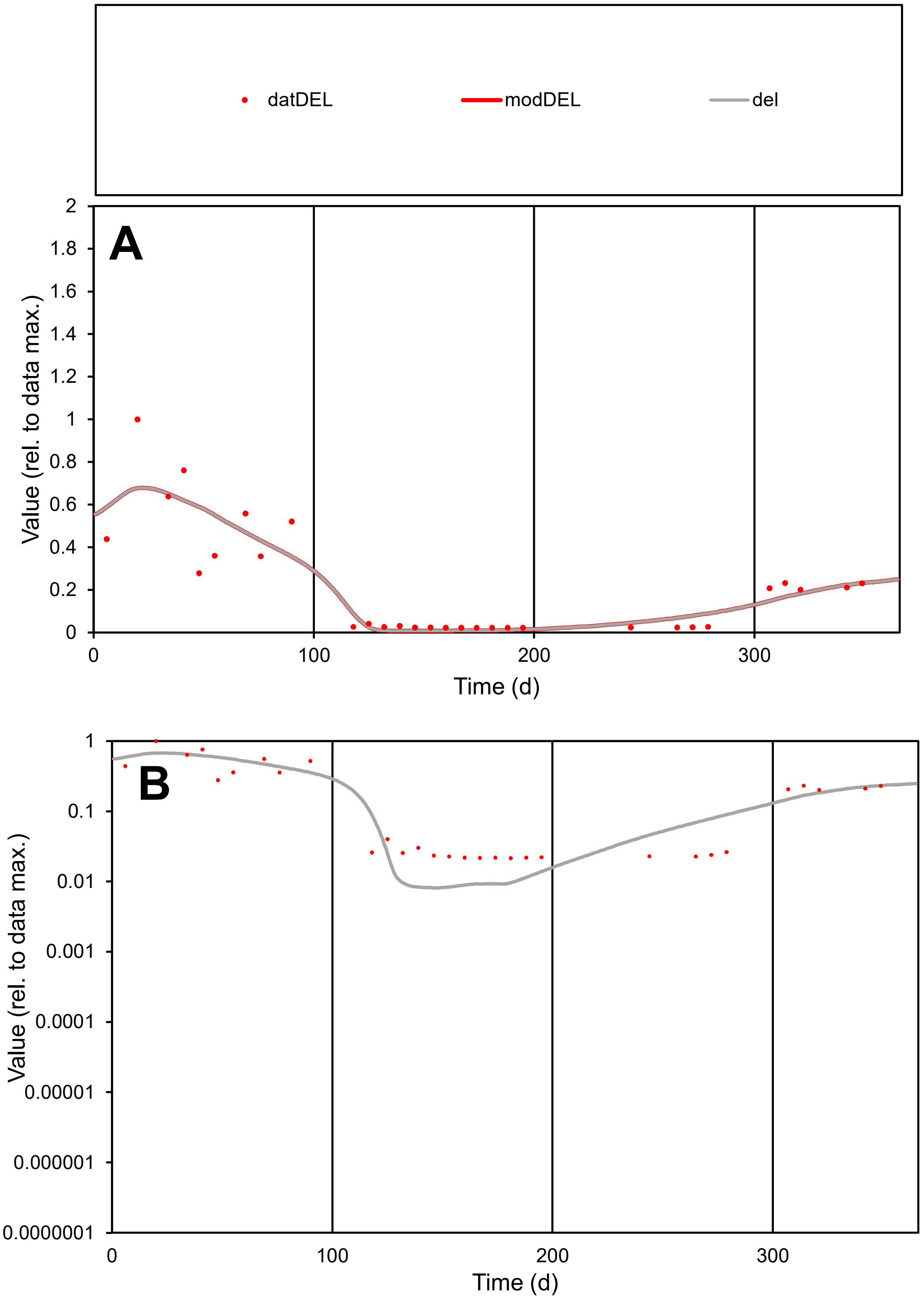


**C1**

Variables

DEL Provide jtd or Dst

CalibOnly?

2 1 to reproduce of

oNormalize?

2 1=no, 2=yes

ofxn calc*

0.50152

ofxn mod

0.53099

sRun

RunA1

zout

1

Ave

Data

0.00061

Model

0.0006

data/vMax

0.00286

ofxng

275.646

Min d/m

2.64858

ave/min d

9.88615

ave/min m

25.9798

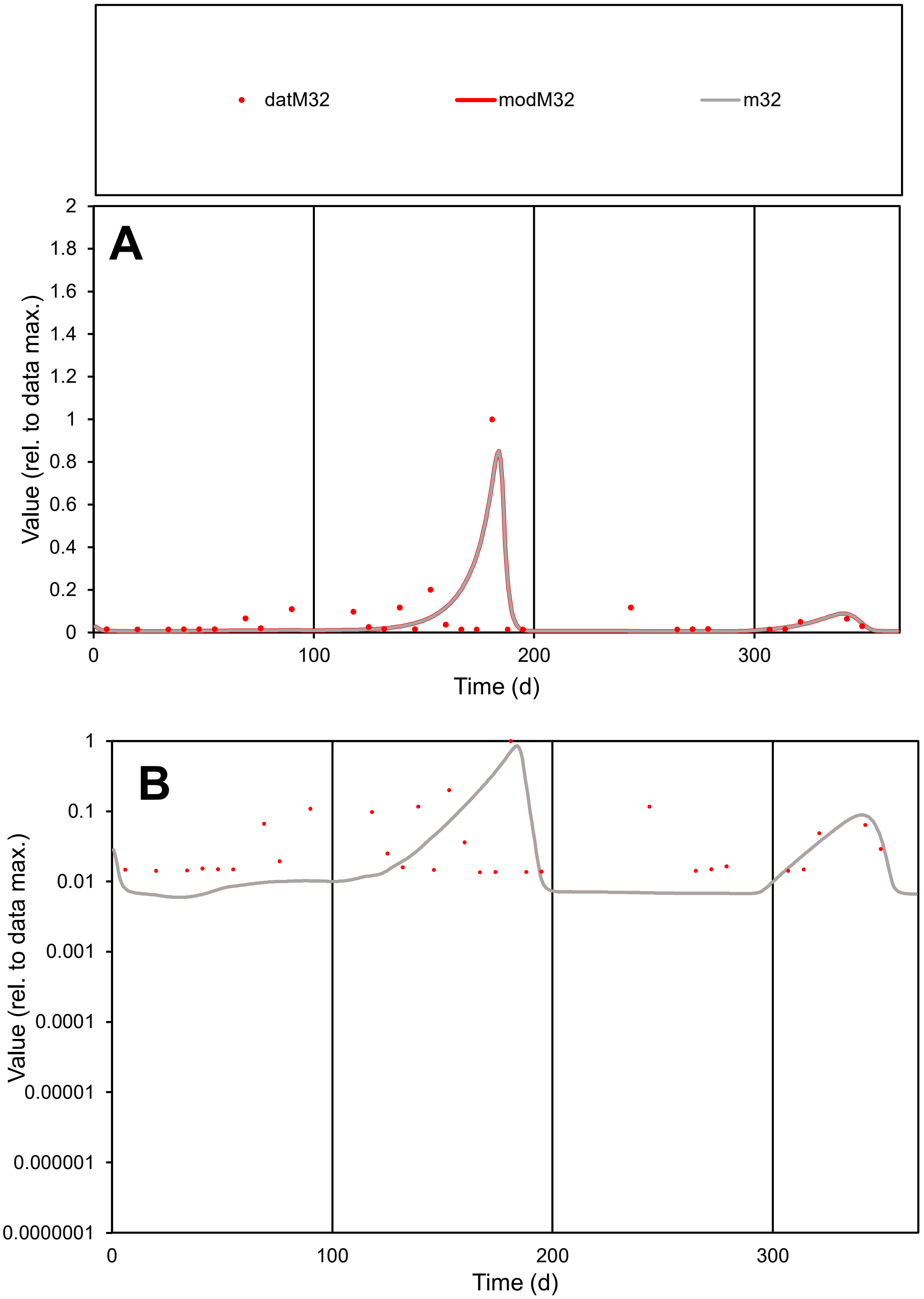


C1

Variables
M32 Provide jtd or Dst
CalibOnly?
2 1 to reproduce of
oNormalize?
2 1=no, 2=yes

ofxn calc*
1.21911
ofxn mod
1.30028

sRun
RunA1
zout
1

Ave
Data
0.00112
Model
0.00103

data/vMax
0.01543

ofxng
275.646

Min d/m
2.22376

ave/min d
5.34861
ave/min m
10.9669

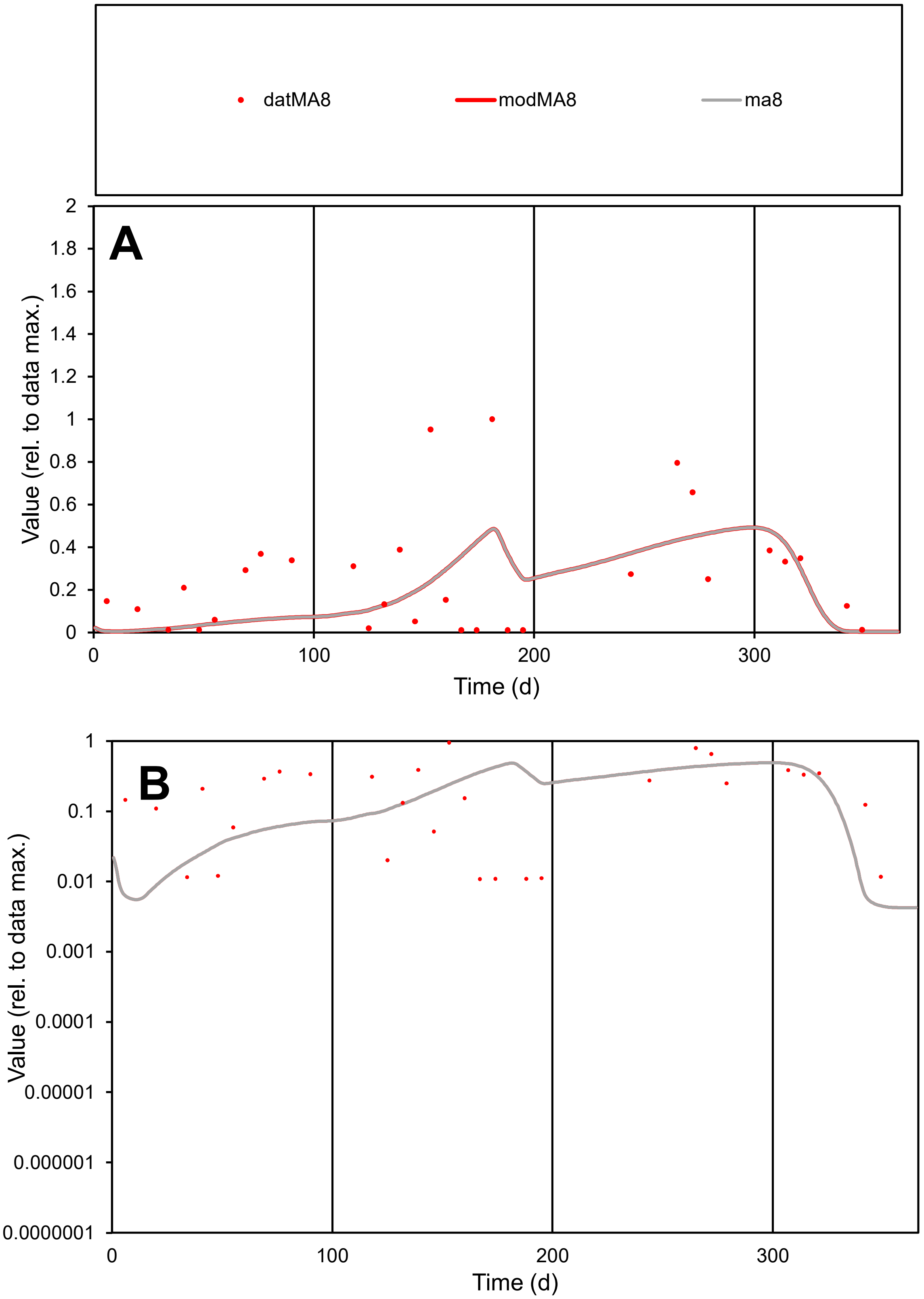


**C1**

Variables
MA8 Provide jtd or Dst
CalibOnly?
2 1 to reproduce of
oNormalize?
2 1=no, 2=yes

ofxn calc*
0.9718
ofxn mod
0.98356

sRun
RunA1
zout
1

Ave
Data
0.00143
Model
0.00115

data/vMax
0.0055

ofxng
275.646

Min d/m
2.45986

ave/min d
23.8669
ave/min m
47.472

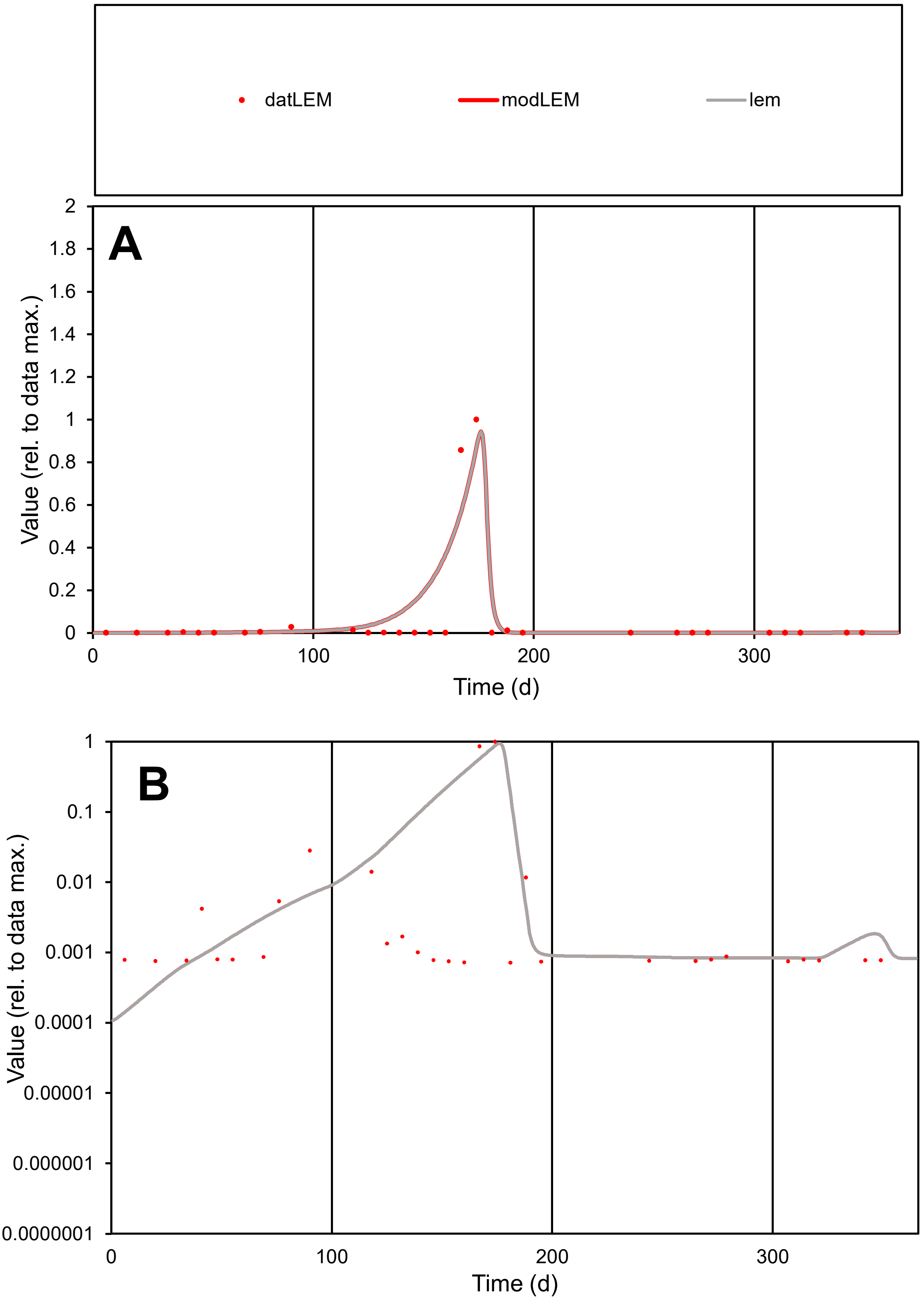


**C1**

Variables
LEM Provide jtd or Dst
CalibOnly?
2 1 to reproduce of
oNormalize?
2 1=no, 2=yes

ofxn calc*
1.47383
ofxn mod
1.55353

sRun
RunA1
zout
1

Ave
Data
0.00193
Model
0.00252

data/vMax
0.02992

ofxng
275.646

Min d/m
4.71819

ave/min d
90.2745
ave/min m
555.168

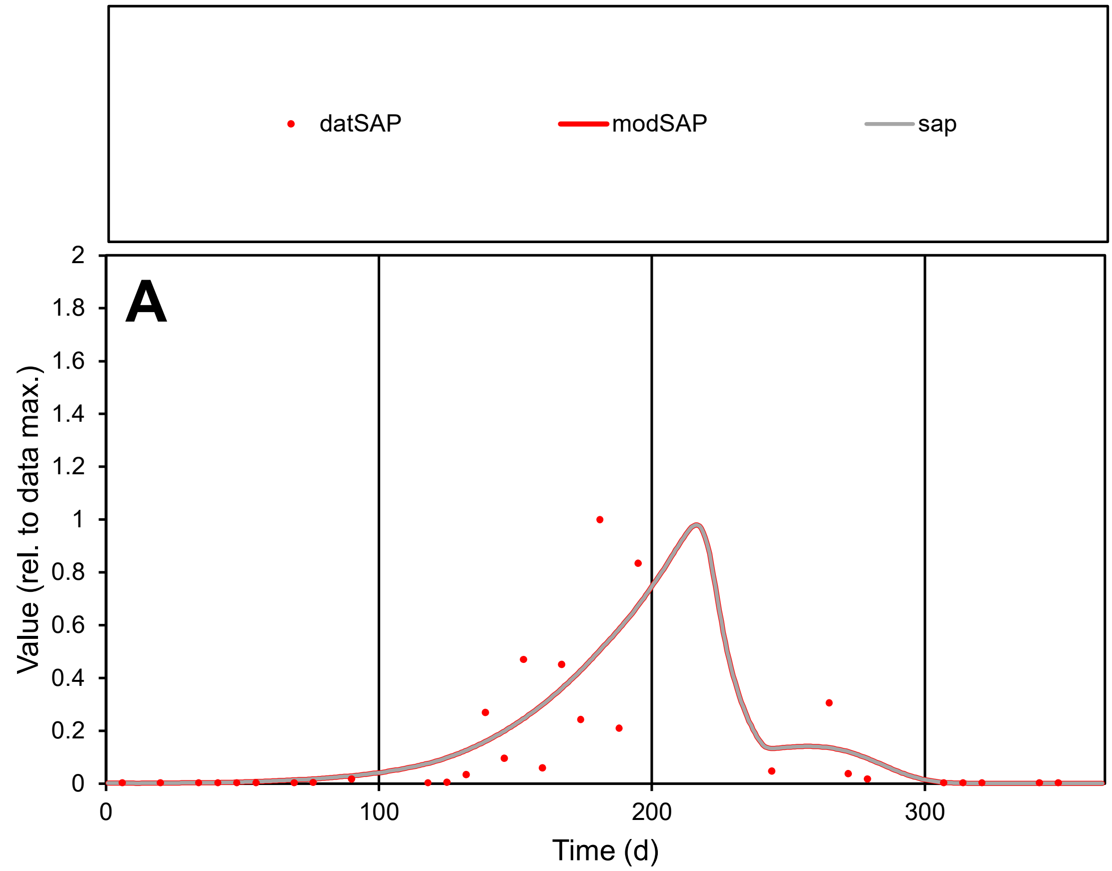


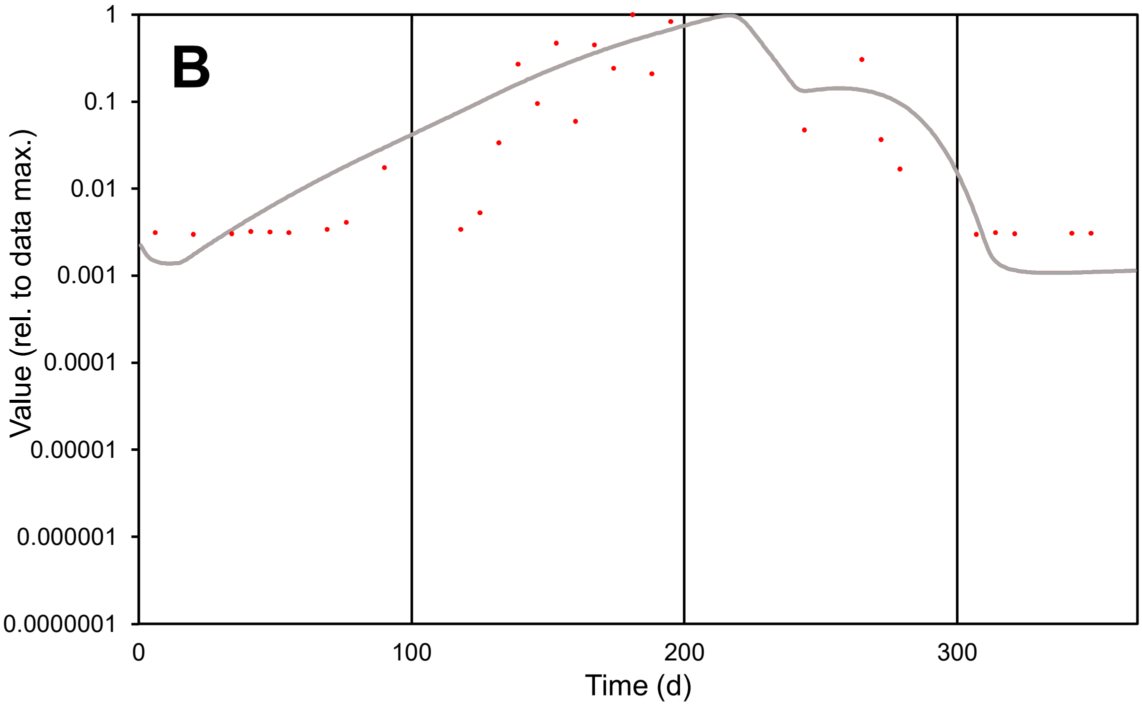


**C1**

Variables
SAP Provide jtd or Dst
CalibOnly?
2 1 to reproduce of
oNormalize?
2 1=no, 2=yes

ofxn calc*
1.05204
ofxn mod
1.07863

sRun
RunA1
zout
1

Ave
Data
0.00159
Model
0.00169

data/vMax
0.0115

ofxng
275.646

Min d/m
2.73328

ave/min d
46.2019
ave/min m
134.428

325_Calib_S_A

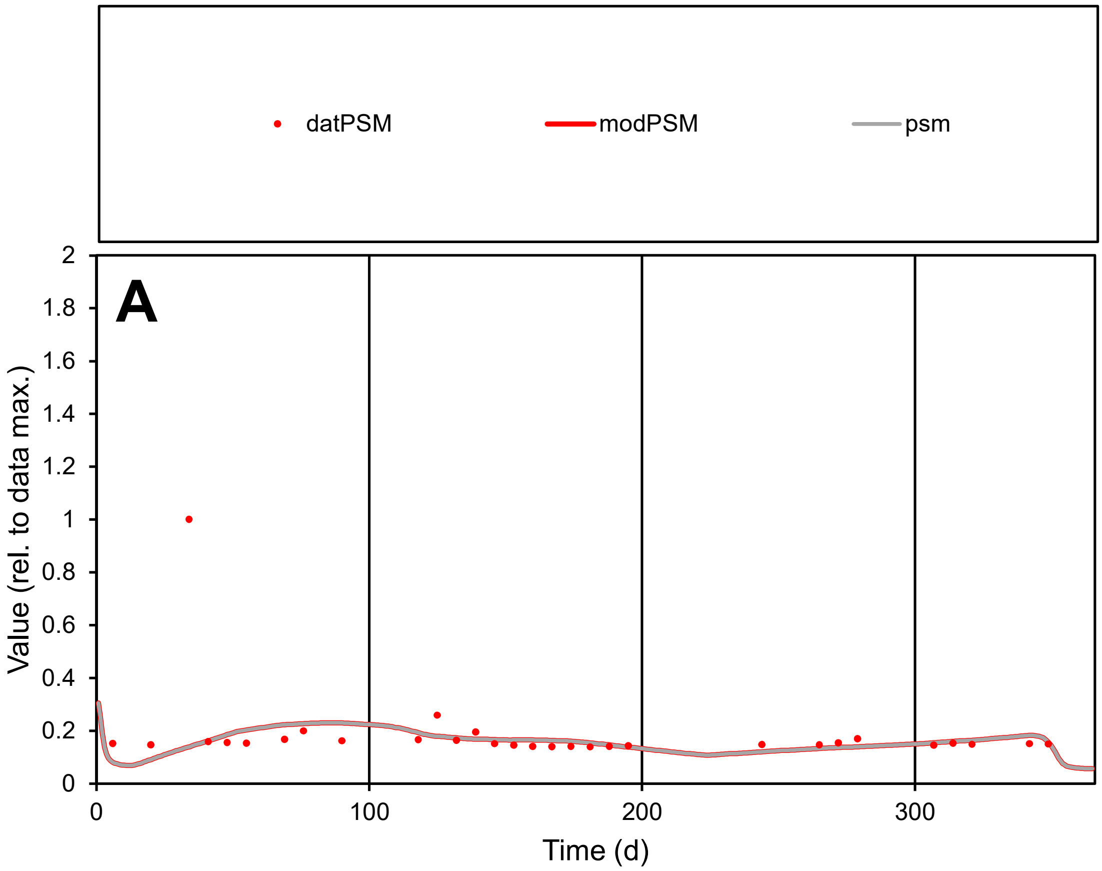


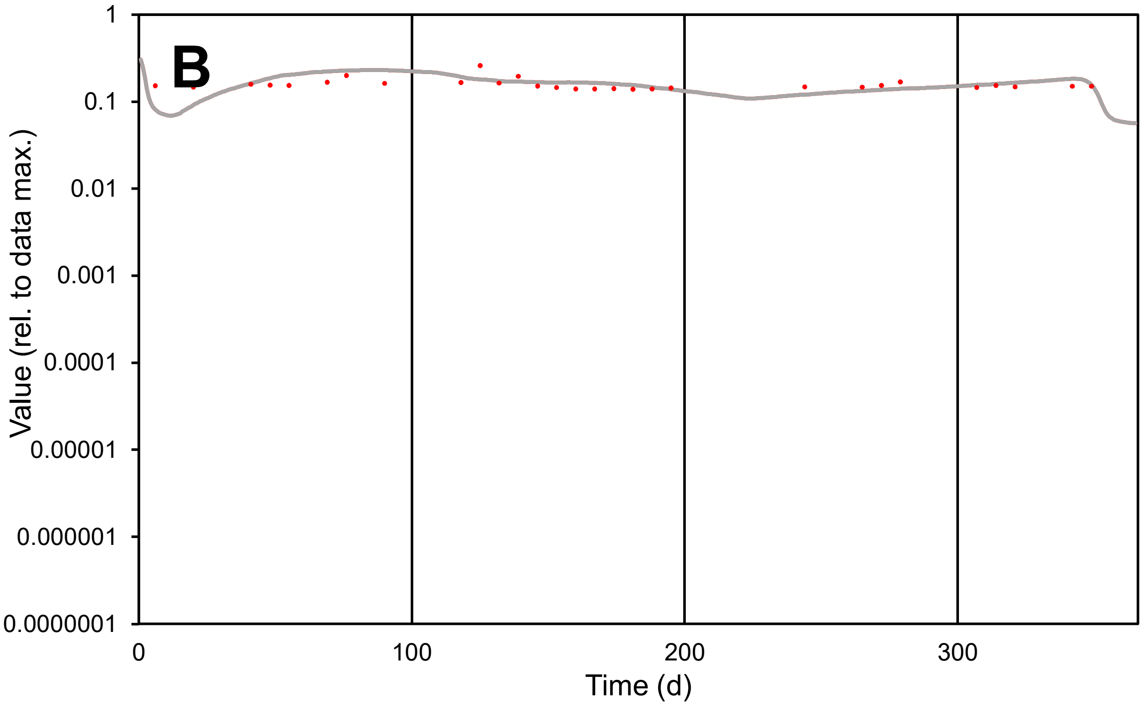


**C1**

Variables
PSM Provide jtd or Dst
CalibOnly?
2 1 to reproduce of
oNormalize?
2 1=no, 2=yes

ofxn calc*
0.85815
ofxn mod
0.86767

sRun
RunA1
zout
1

Ave
Data
3.7E-05
Model
3.2E-05

data/vMax
0.0002

ofxng
275.646

Min d/m
1.77161

ave/min d
1.34244
ave/min m
2.06923

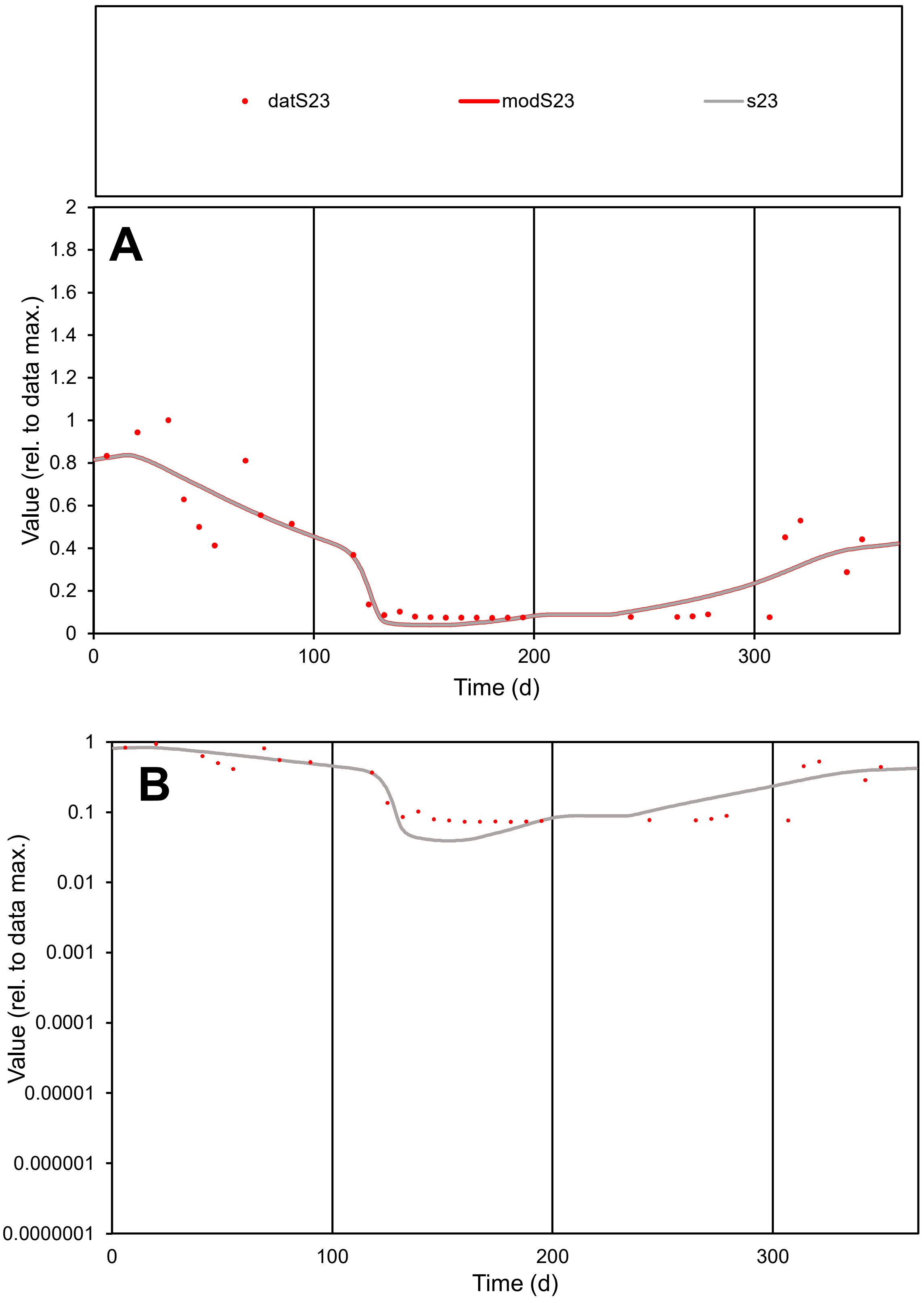


**C1**

Variables
S23 Provide jtd or Dst
CalibOnly?
2 1 to reproduce of
oNormalize?
2 1=no, 2=yes

ofxn calc*
0.3474
ofxn mod
0.38005

sRun
RunA1
zout
1

Ave
Data
0.0007
Model
0.00069

data/vMax
0.0022

ofxng
275.646

Min d/m
1.85884

ave/min d
4.40457
ave/min m
7.99294

325_Calib_S_A

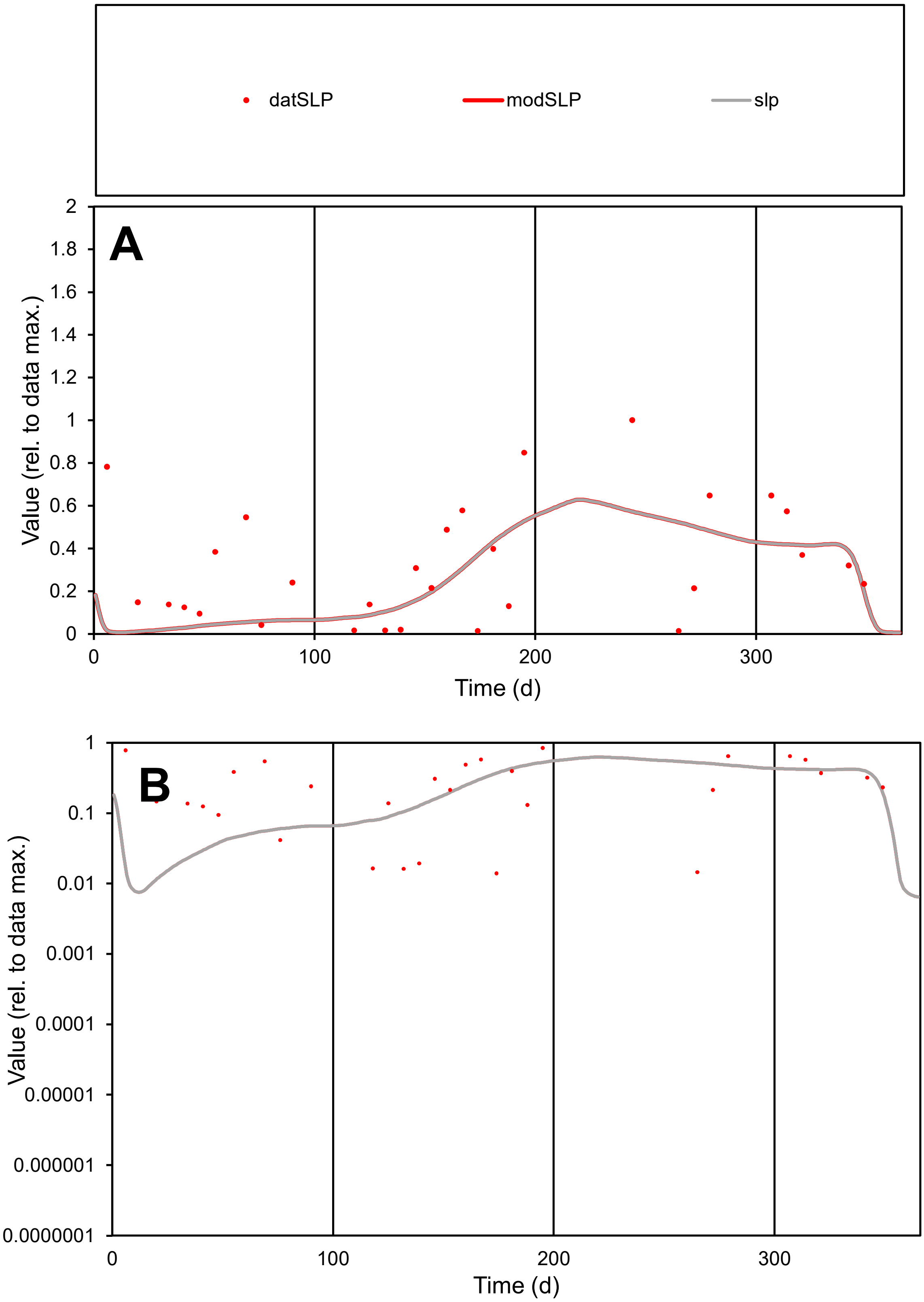


**C1**

Variables

SLP Provide jtd or Dst

CalibOnly?

2 1 to reproduce of

oNormalize?

2 1=no, 2=yes

ofxn calc*

0.83194

ofxn mod

0.84406

sRun

RunA1

zout

1

Ave

Data

0.00056

Model

0.00043

data/vMax

0.00173

ofxng

275.646

Min d/m

1.1438

ave/min d

23.09

ave/min m

20.2228

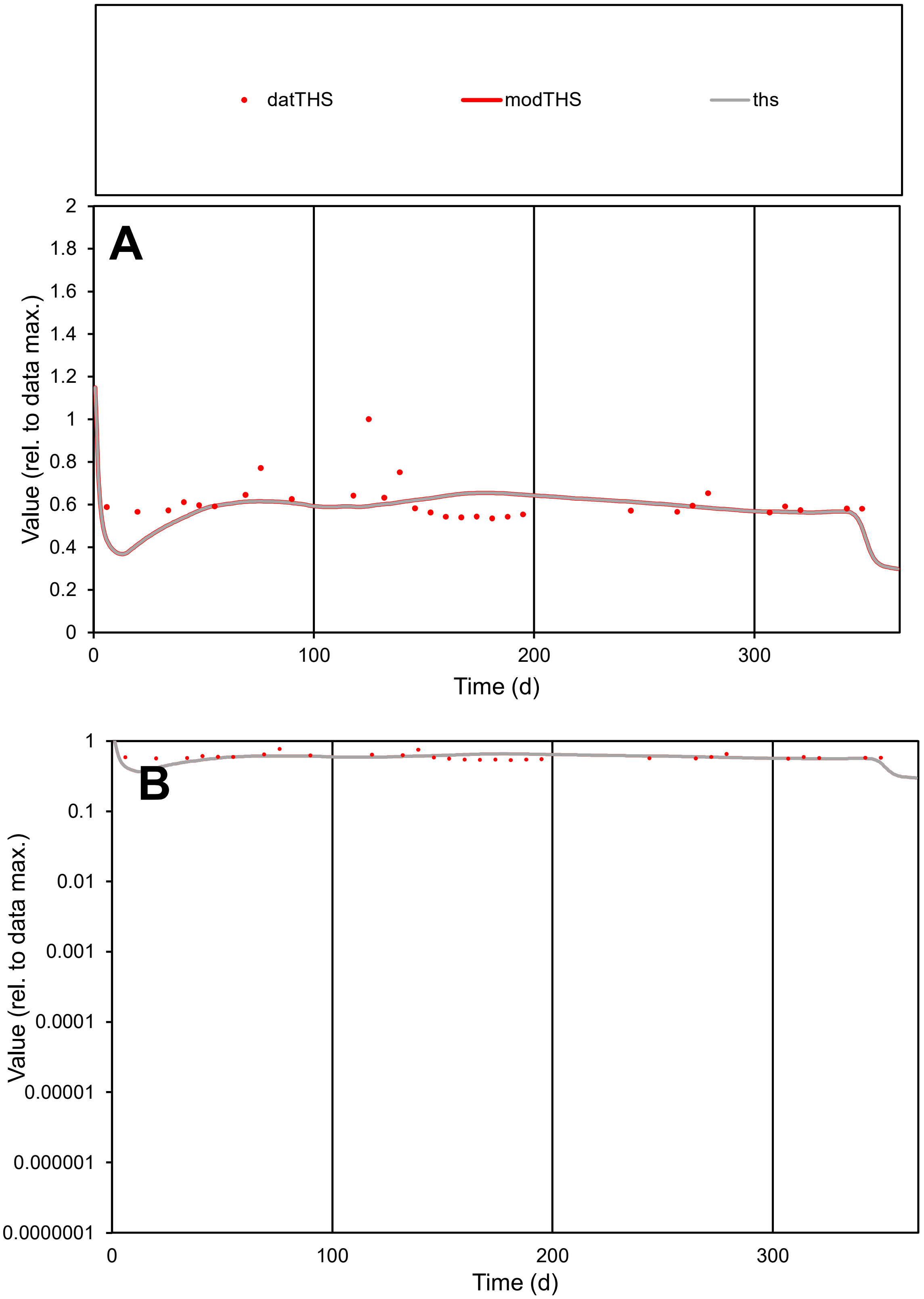


**C1**

Variables
THS Provide jtd or Dst
CalibOnly?
2 1 to reproduce of
oNormalize?
2 1=no, 2=yes

ofxn calc*
0.1841
ofxn mod
0.21493

sRun
RunA1
zout
1

Ave
Data
3.2E-05
Model
3.1E-05

data/vMax
5.3E-05

ofxng
275.646

Min d/m
1.31023

ave/min d
1.13763
ave/min m
1.4307

325_Calib_S_A

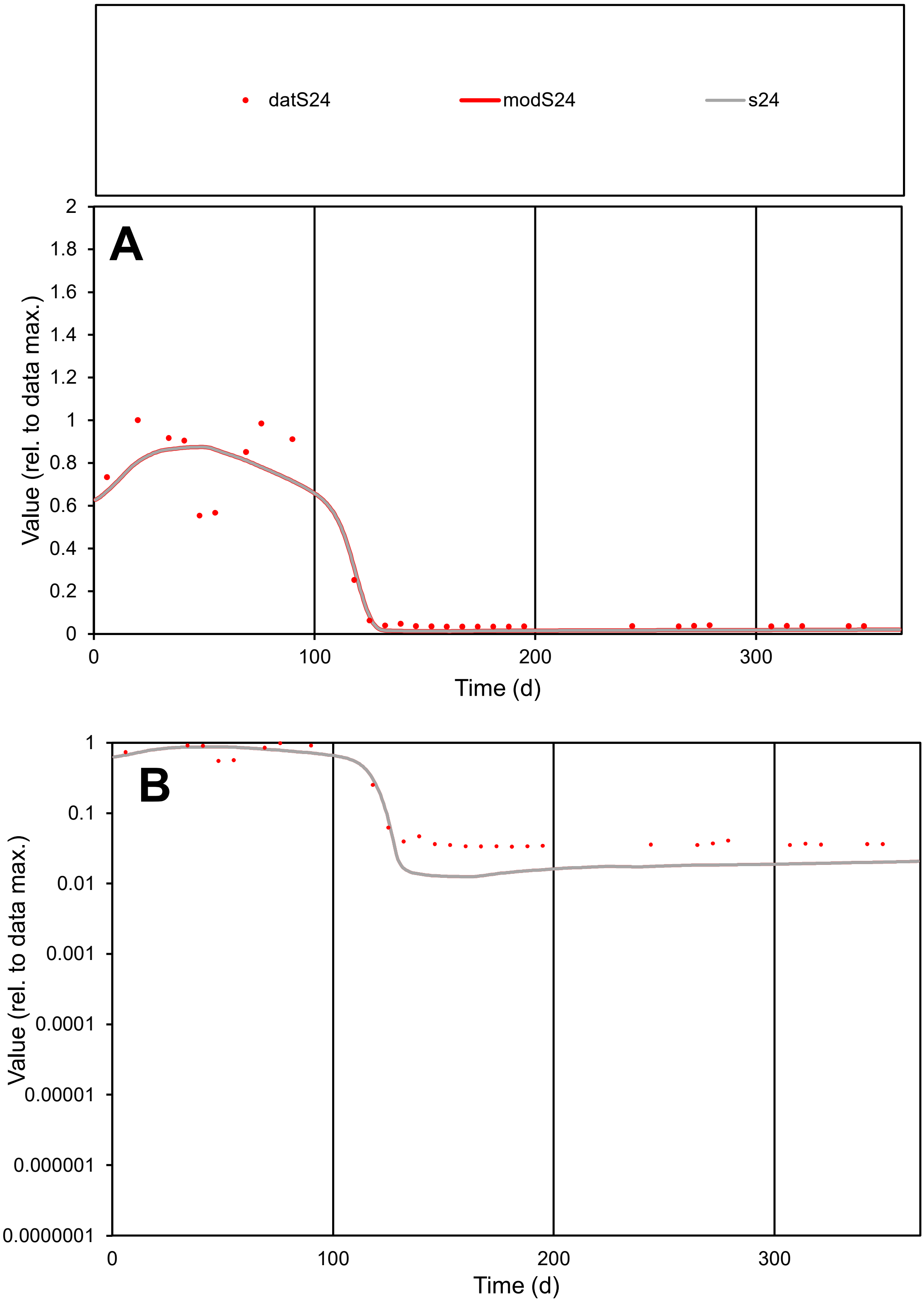


**C1**

Variables

S24 Provide jtd or Dst

CalibOnly?

2 1 to reproduce of

oNormalize?

2 1=no, 2=yes

ofxn calc*

0.37007

ofxn mod

0.41574

sRun

RunA1

zout

1

Ave

Data

0.00049

Model

0.00045

data/vMax

0.00173

ofxng

275.646

Min d/m

2.67531

ave/min d

8.4031

ave/min m

21.018

325_Calib_S_A

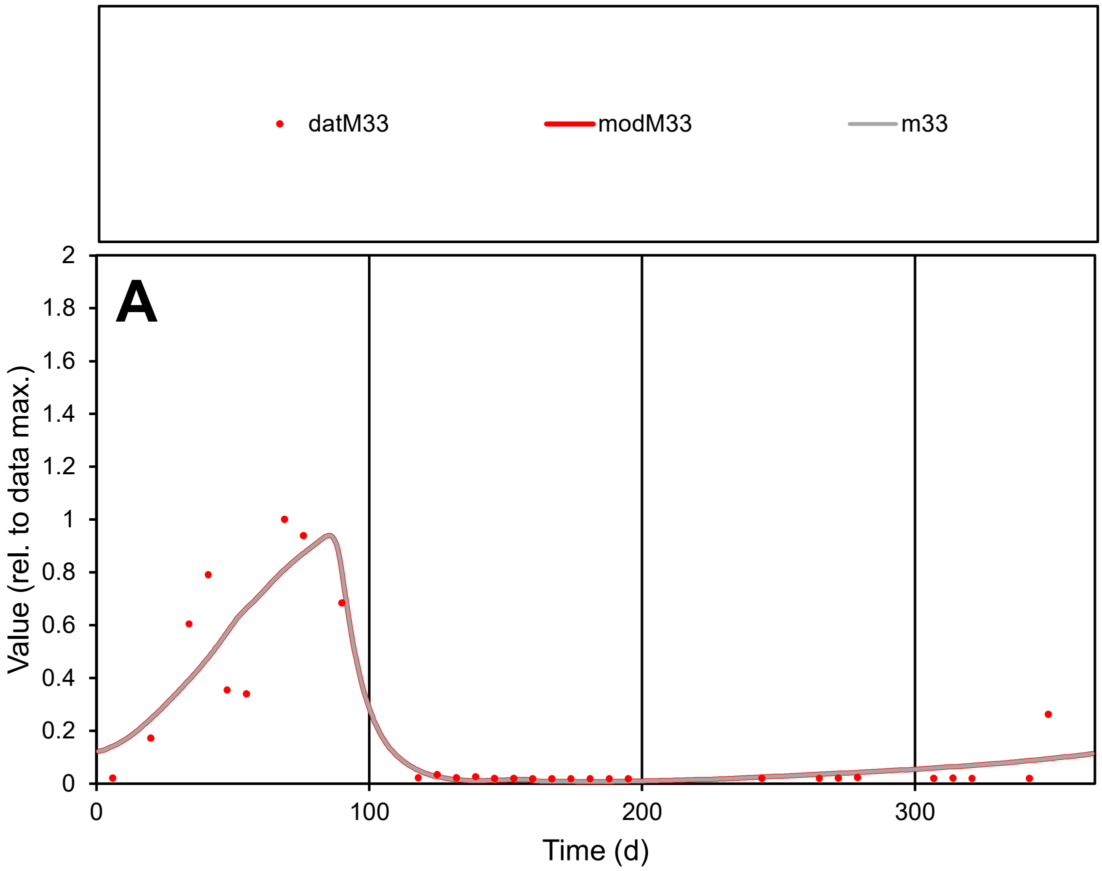


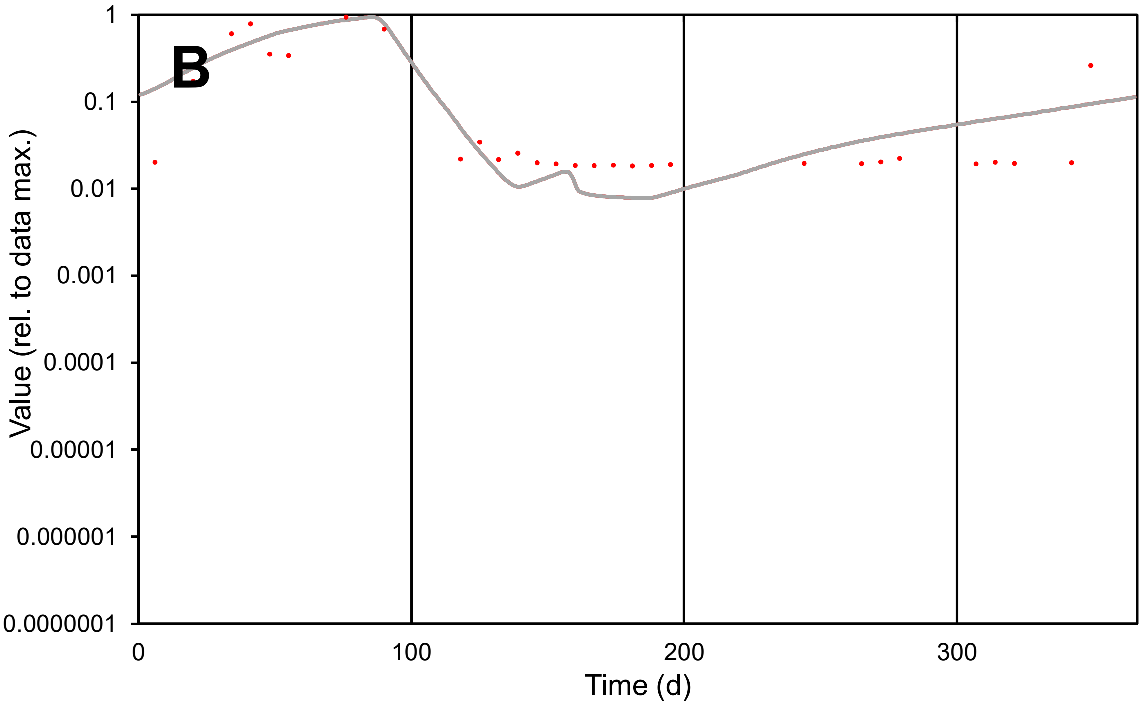


**C1**

Variables
M33 Provide jtd or Dst
CalibOnly?
2 1 to reproduce of
oNormalize?
2 1=no, 2=yes

ofxn calc*
0.60579
ofxn mod
0.64726

sRun
RunA1
zout
1

Ave
Data
0.00069
Model
0.0007

data/vMax
0.0037

ofxng
275.646

Min d/m
2.33608

ave/min d
10.1296
ave/min m
23.9021

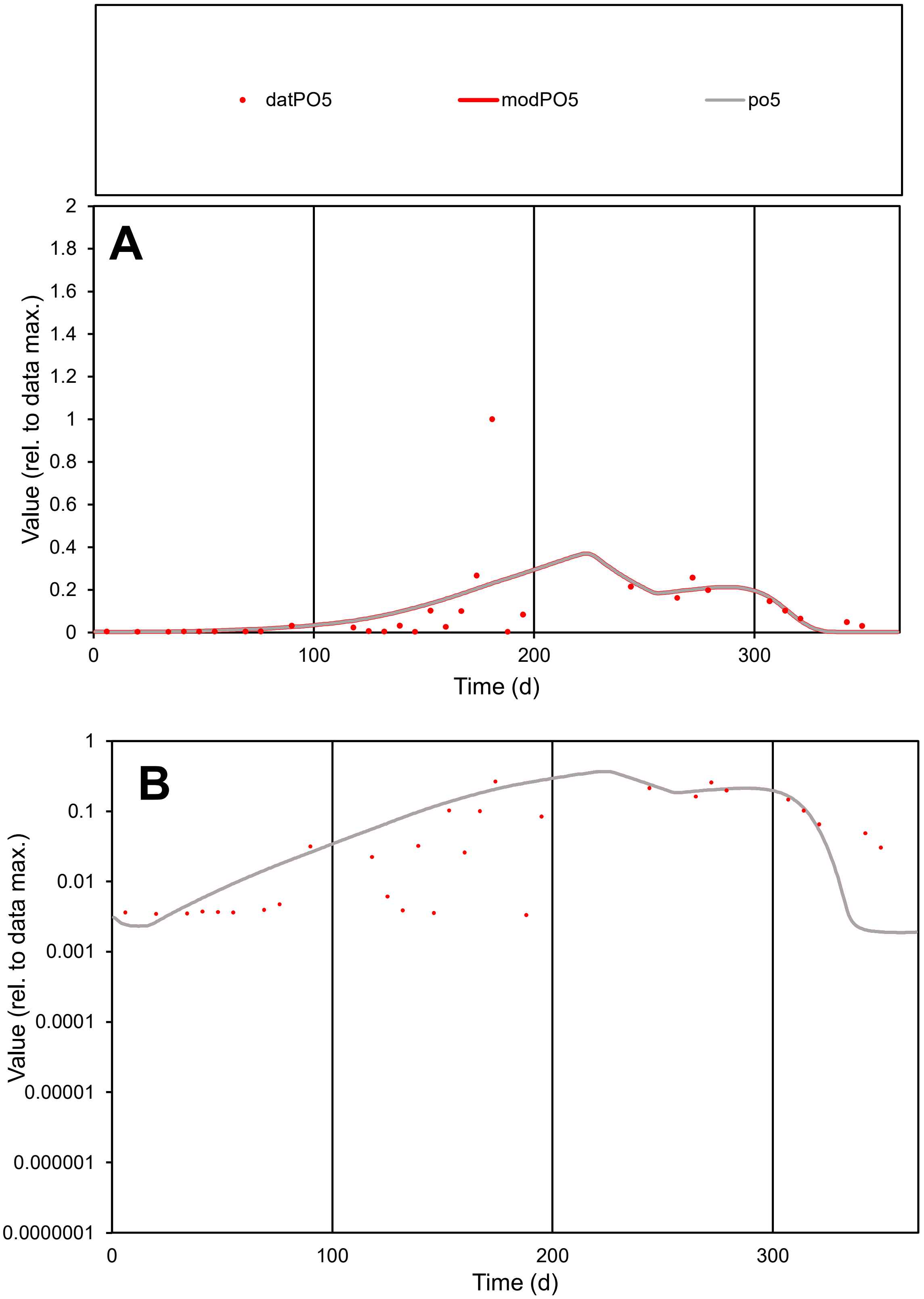


**C1**

Variables
PO5 Provide jtd or Dst
CalibOnly?
2 1 to reproduce of
oNormalize?
2 1=no, 2=yes

ofxn calc*
1.61613
ofxn mod
1.62893

sRun
RunA1
zout
1

Ave
Data
0.00138
Model
0.00147

data/vMax
0.01405

ofxng
275.646

Min d/m
1.75393

ave/min d
29.3747
ave/min m
54.8936

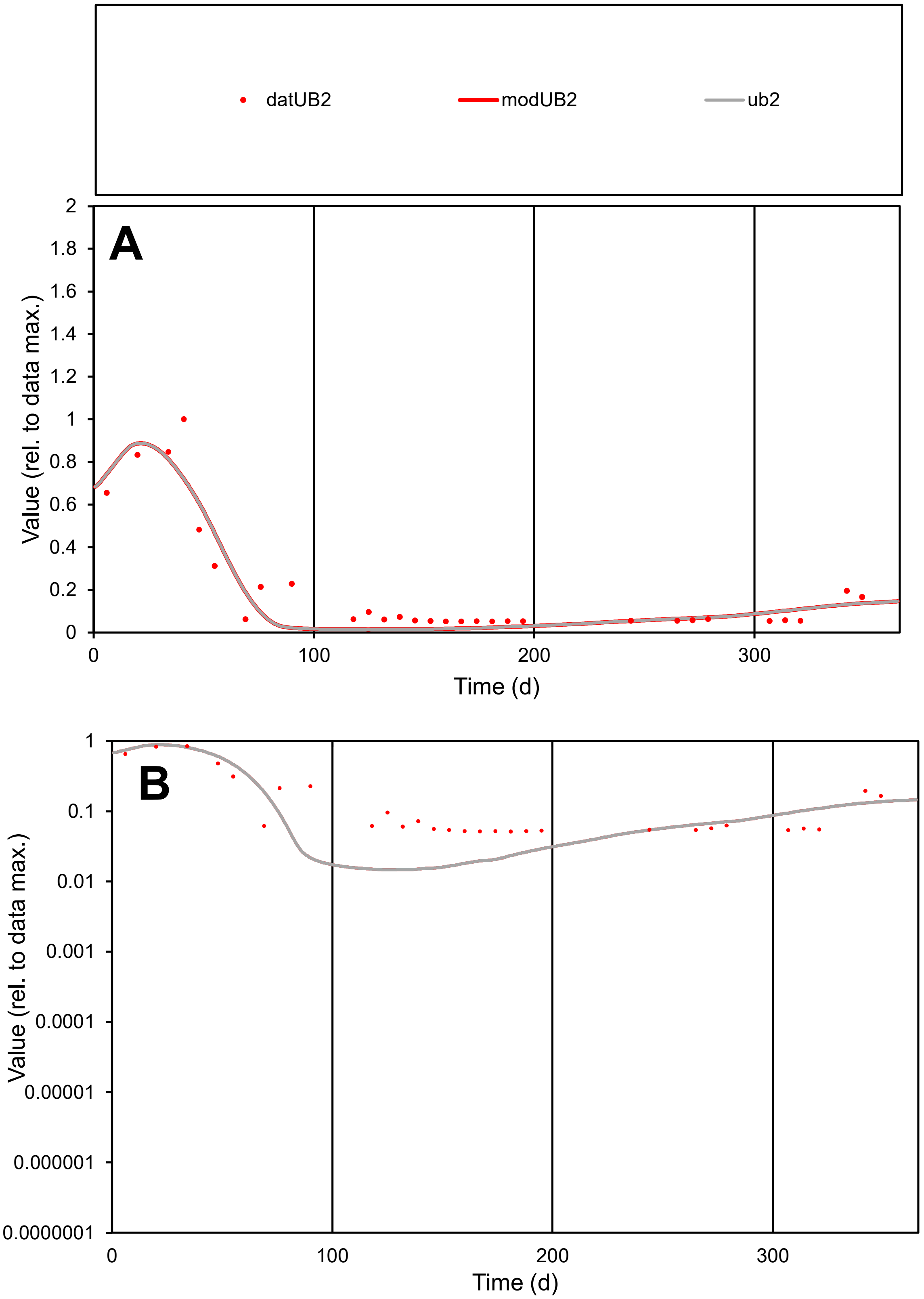


**C1**

Variables
UB2 Provide jtd or Dst
CalibOnly?
2 1 to reproduce of
oNormalize?
2 1=no, 2=yes

ofxn calc*
0.43181
ofxn mod
0.47132

sRun
RunA1
zout
1

Ave
Data
0.00041
Model
0.00037

data/vMax
0.00201

ofxng
275.646

Min d/m
3.48971

ave/min d
3.95616
ave/min m
12.4433

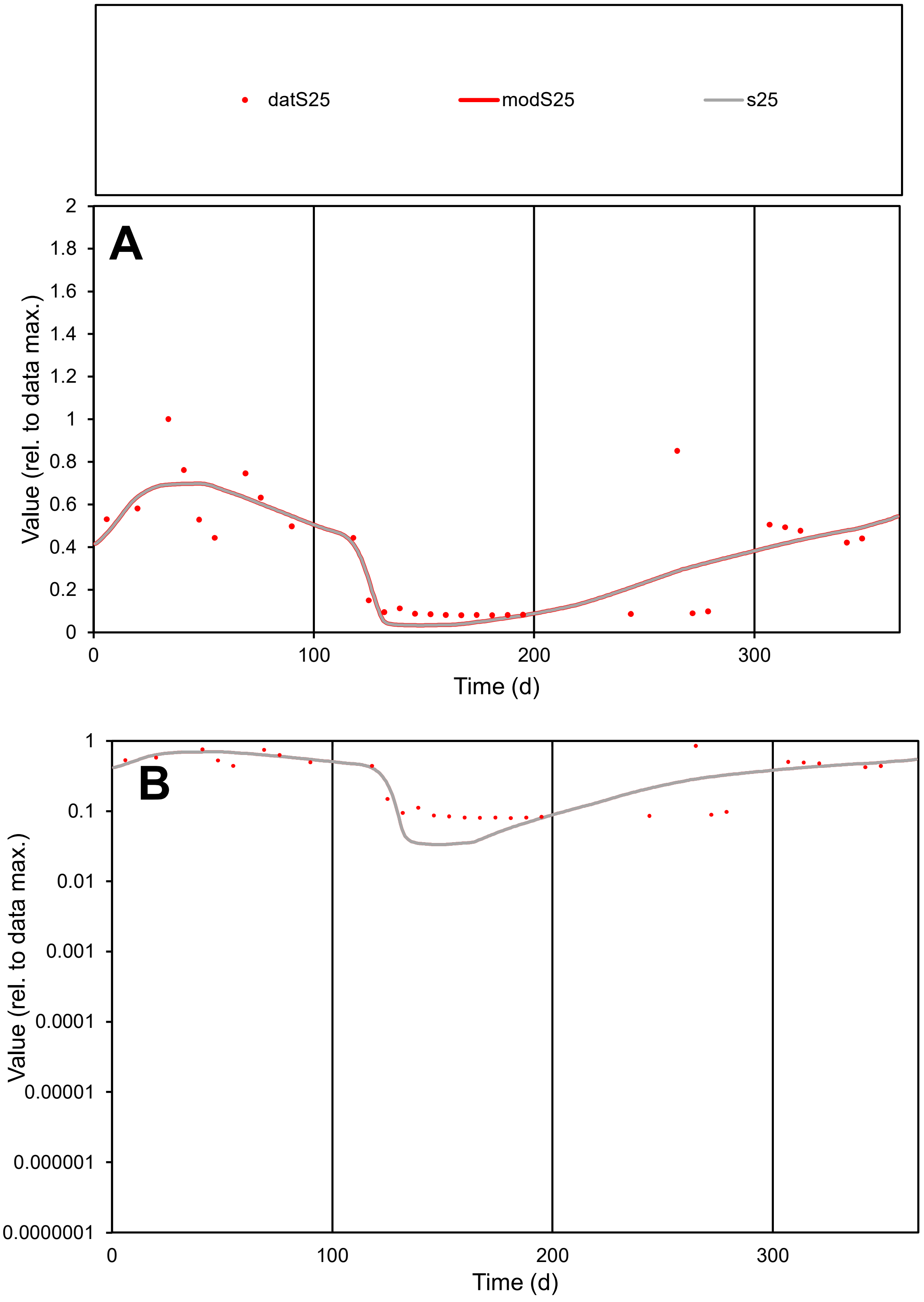


**C1**

Variables
S25 Provide jtd or Dst
CalibOnly?
2 1 to reproduce of
oNormalize?
2 1=no, 2=yes

ofxn calc*
0.4255
ofxn mod
0.44903

sRun
RunA1
zout
1

Ave
Data
0.00074
Model
0.00071

data/vMax
0.0021

ofxng
275.646

Min d/m
2.4024

ave/min d
4.42871
ave/min m
10.1158

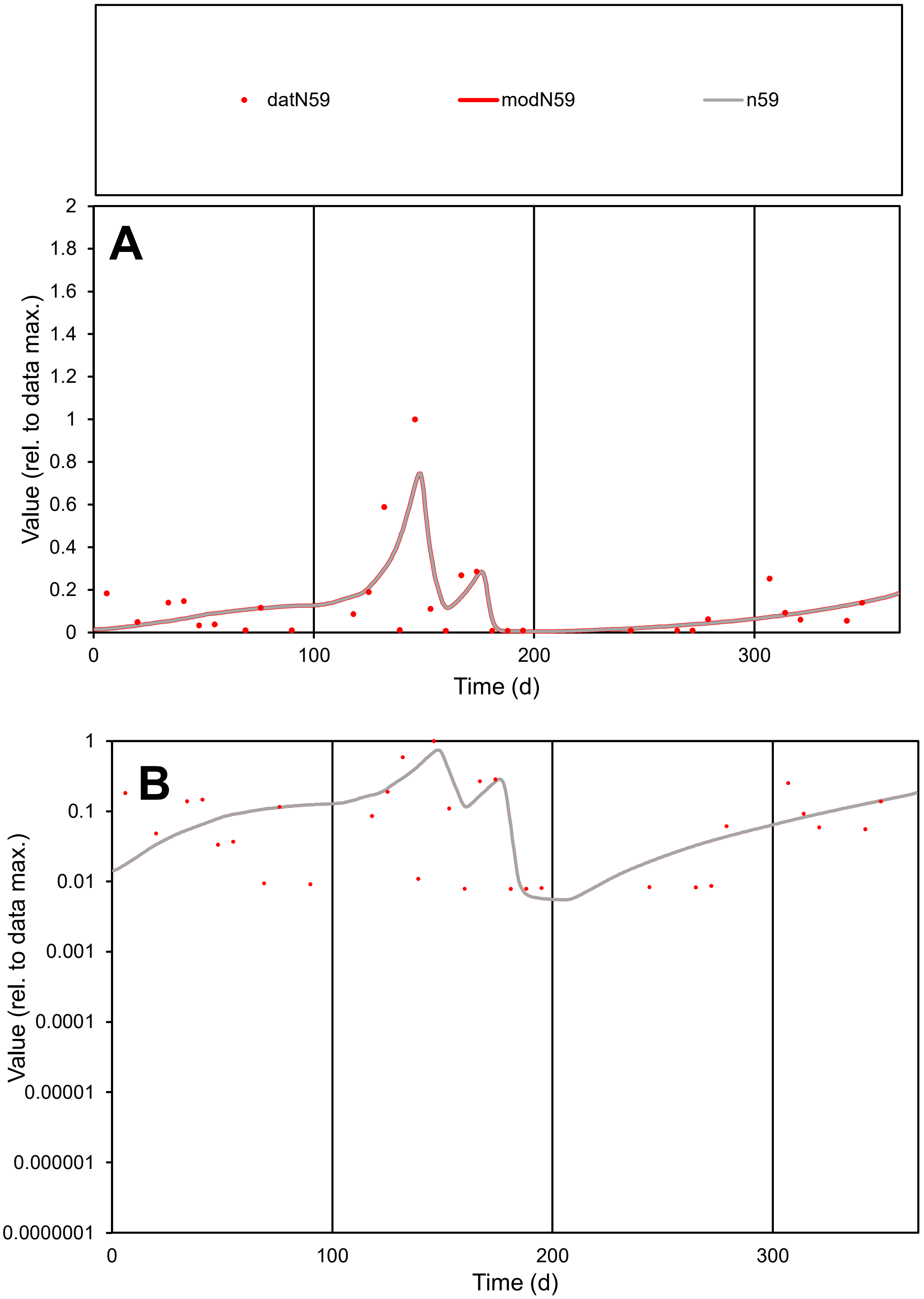


**C1**

Variables
N59 Provide jtd or Dst
CalibOnly?
2 1 to reproduce of
oNormalize?
2 1=no, 2=yes

ofxn calc*
0.93384
ofxn mod
0.97679

sRun
RunA1
zout
1

Ave
Data
0.0007
Model
0.00071

data/vMax
0.00525

ofxng
275.646

Min d/m
1.37638

ave/min d
16.9642
ave/min m
23.9445

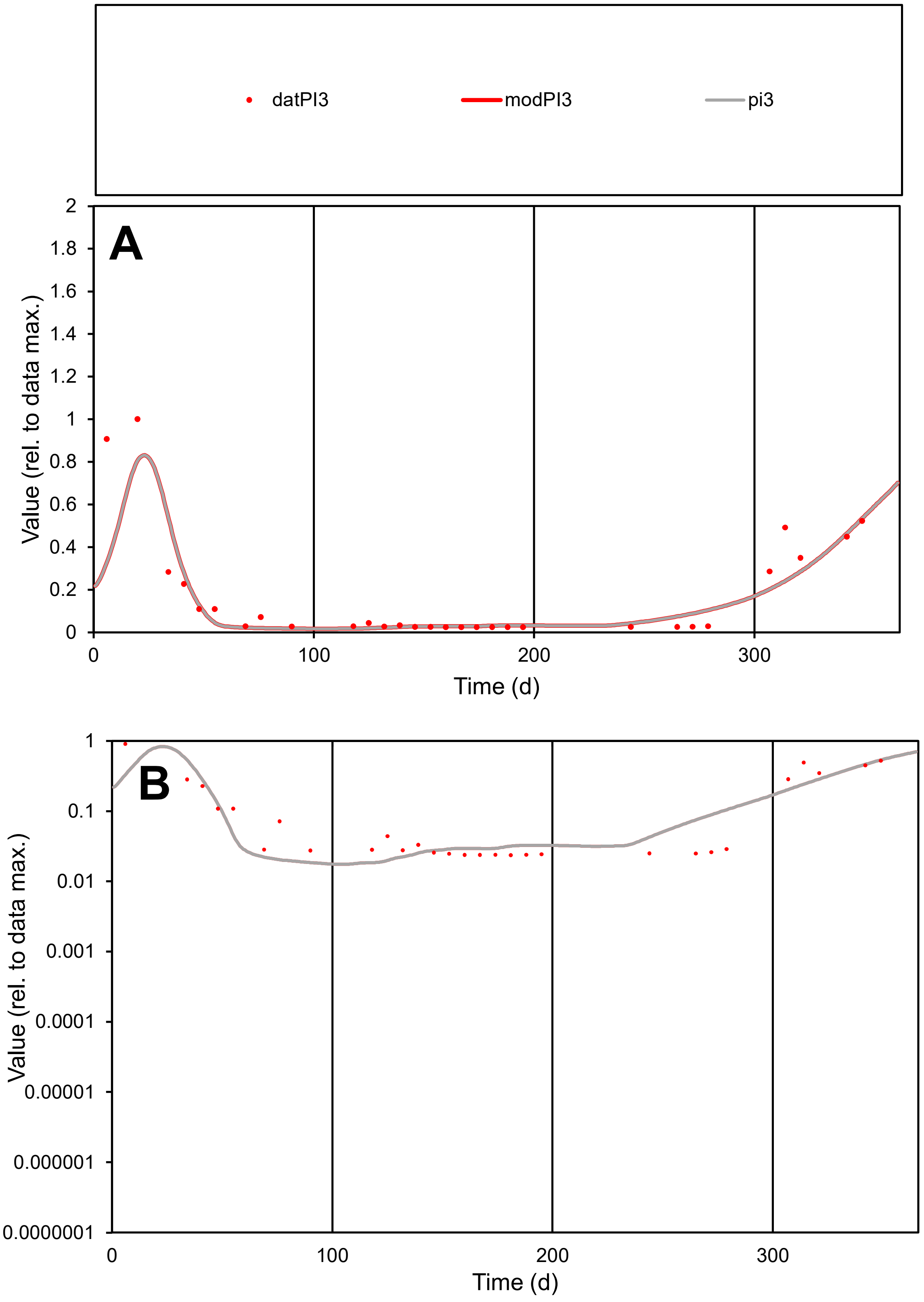


**C1**

Variables
PI3 Provide jtd or Dst
CalibOnly?
2 1 to reproduce of
oNormalize?
2 1=no, 2=yes

ofxn calc*
0.69344
ofxn mod
0.71165

sRun
RunA1
zout
1

Ave
Data
0.00053
Model
0.00045

data/vMax
0.003

ofxng
275.646

Min d/m
1.28186

ave/min d
7.48807
ave/min m
8.24096

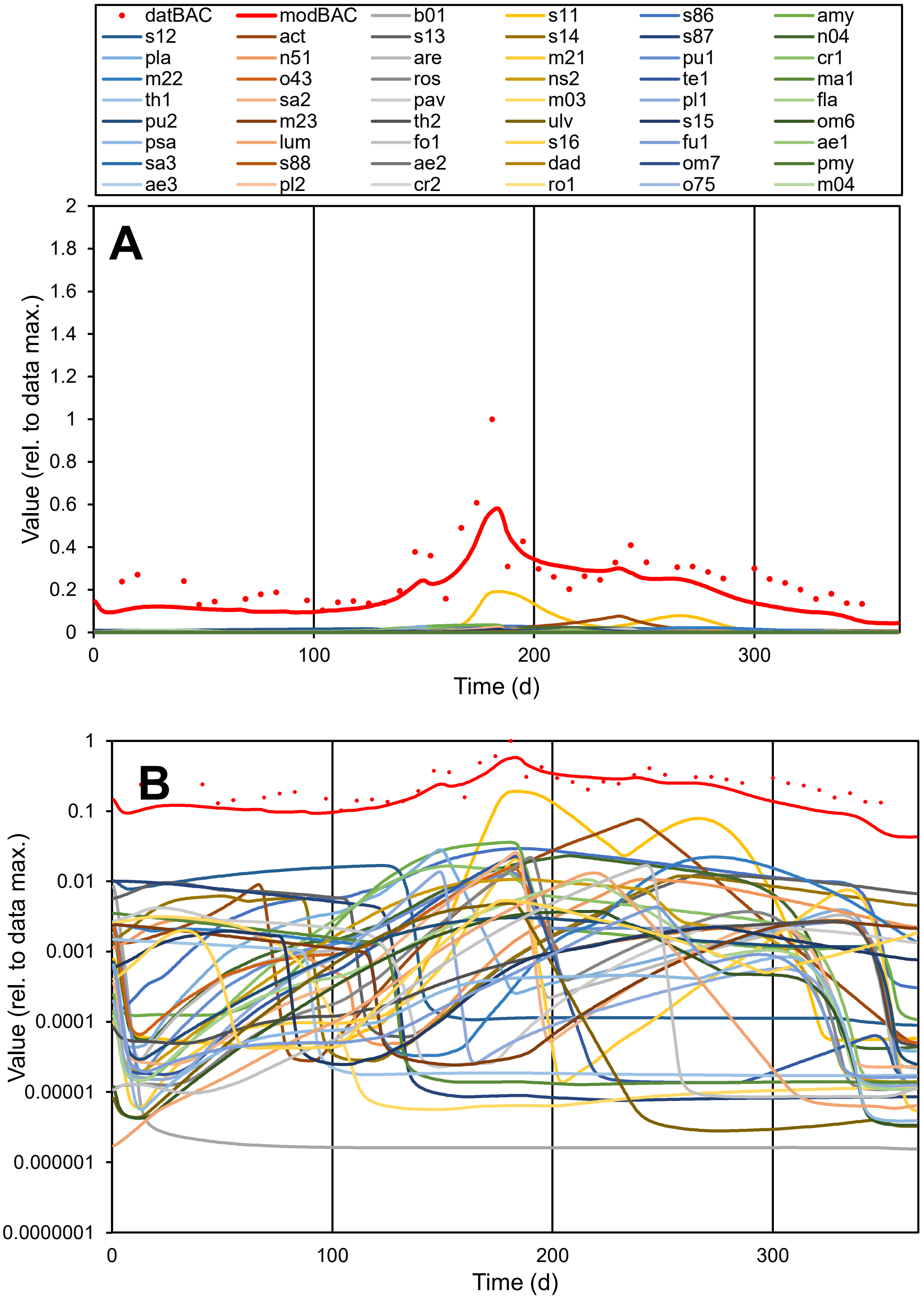


**C1**

Variables

BAC Provide jtd or Dst

CalibOnly?

2 1 to reproduce of

oNormalize?

2 1=no, 2=yes

ofxn calc*

0.41963

ofxn mod

0.43501

sRun

RunA1

zout

1

Ave

Data

0.76978

Model

0.58298

data/vMax

2.916

ofxng

275.646

Min d/m

2.00066

ave/min d

2.55657

ave/min m

3.87365

326_Calib_S_A

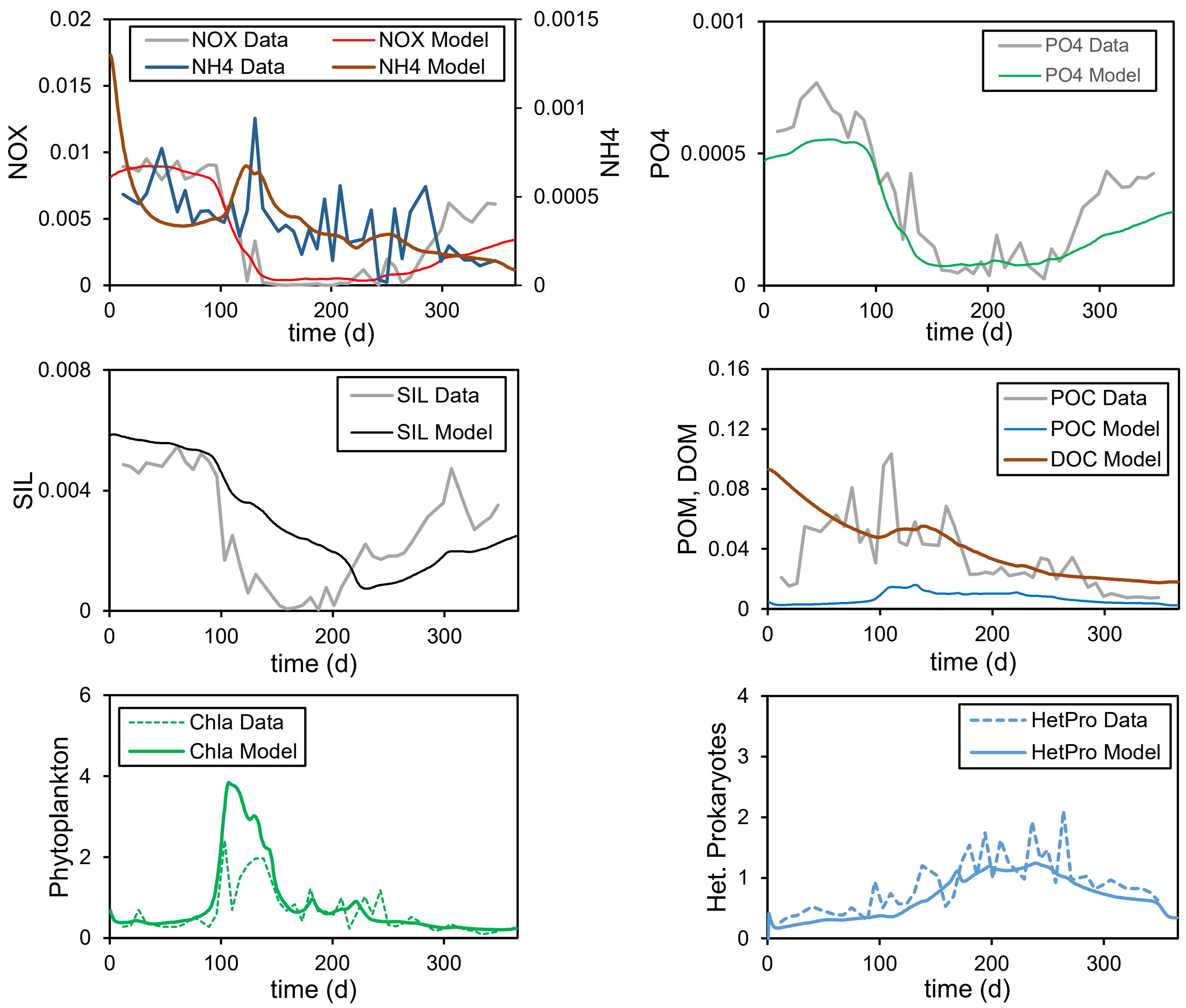

NOX Data
NOX Model
NH4 Data
NH4 Model
NOX
NH4
time (d)
PO4 Data
PO4 Model
PO4
time (d)
SIL Data
SIL Model
SIL
time (d)
POC Data
POC Model
DOC Model
POM, DOM
time (d)
Chla Data
Chla Model
Phytoplankton
time (d)
HetPro Data
HetPro Model
Het. Prokaryotes
time (d)

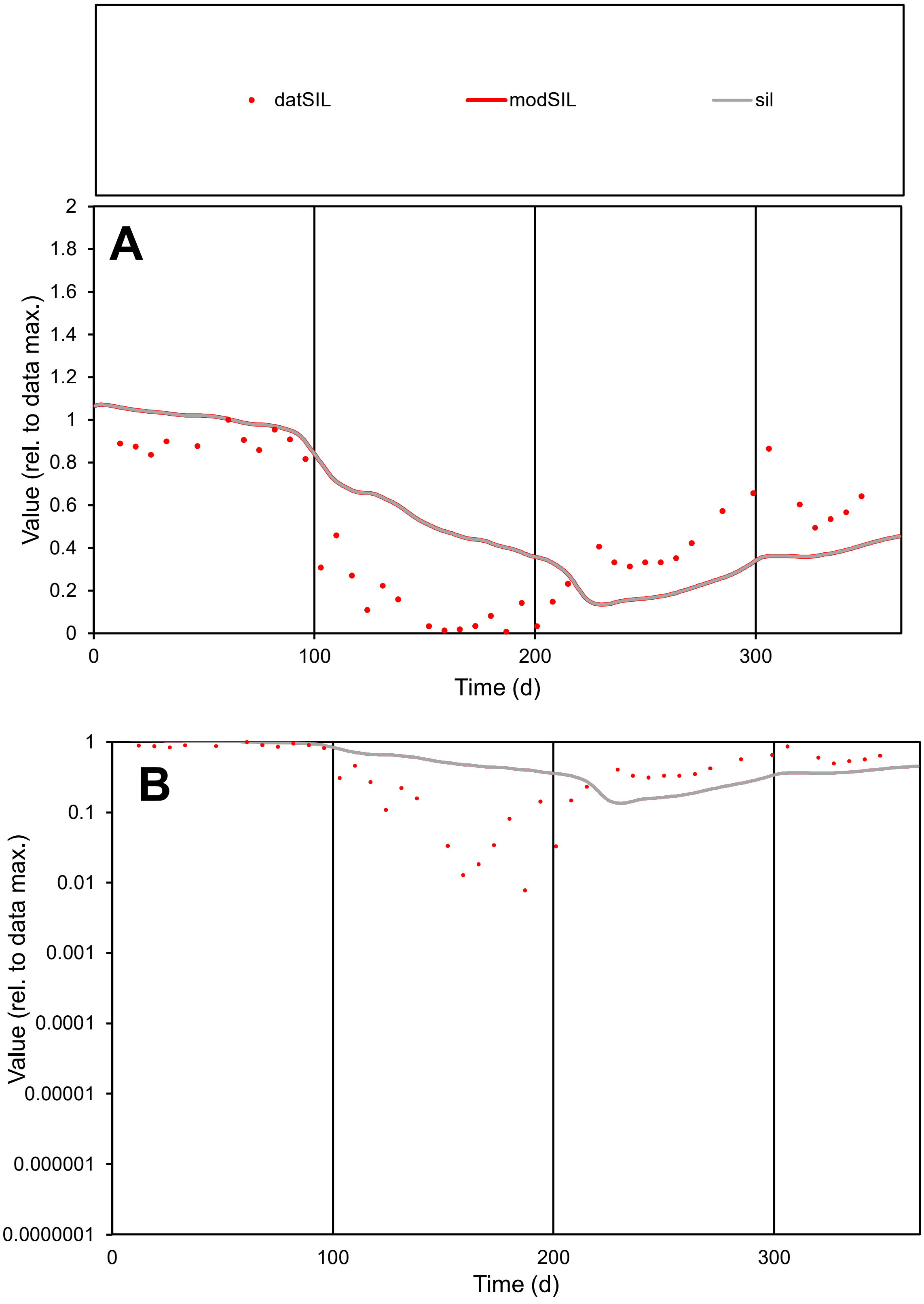


**C1**

Variables
SIL Provide jtd or Dst
CalibOnly?
2 1 to reproduce of
oNormalize?
2 1=no, 2=yes

ofxn calc*
1.82787
ofxn mod
1.8767

sRun
RunA1
zout
1

Ave
Data
0.00254
Model
0.003

data/vMax
0.00547

ofxng
263.459

Min d/m
0.05764

ave/min d
59.7881
ave/min m
4.06705

326_Calib_S_A

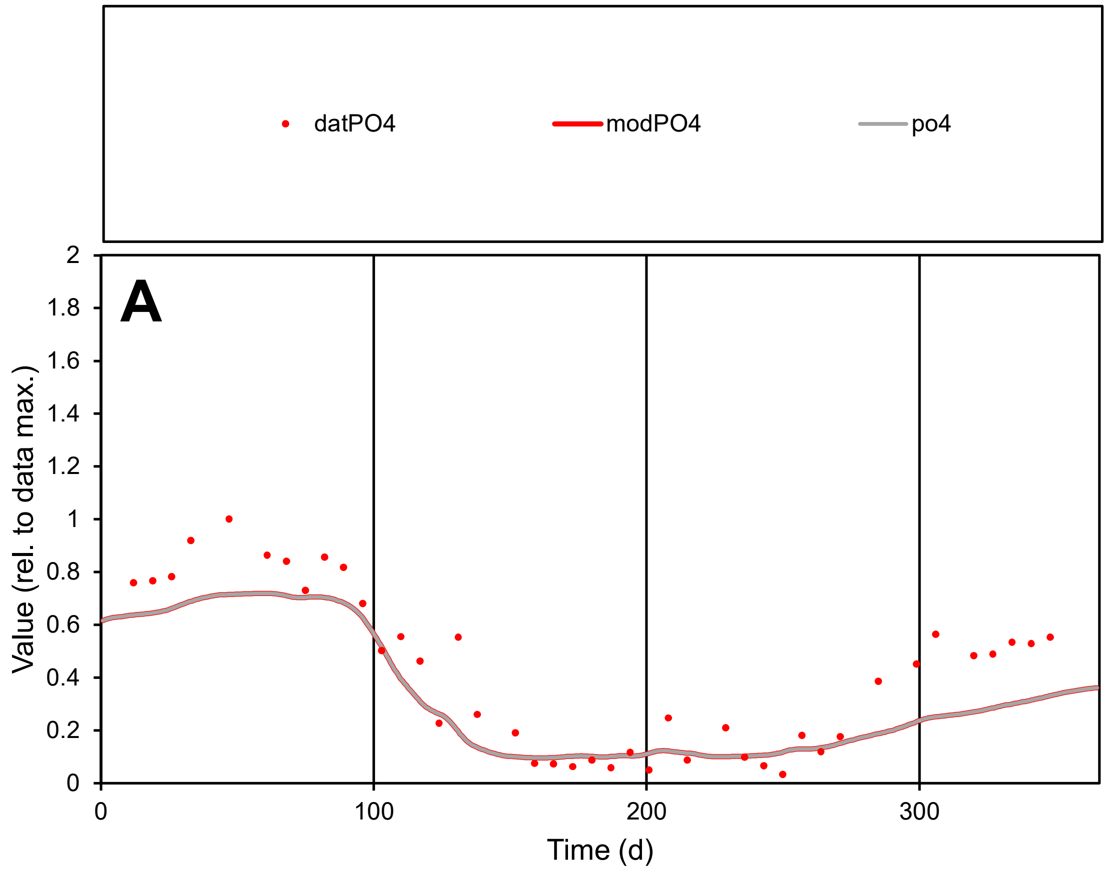


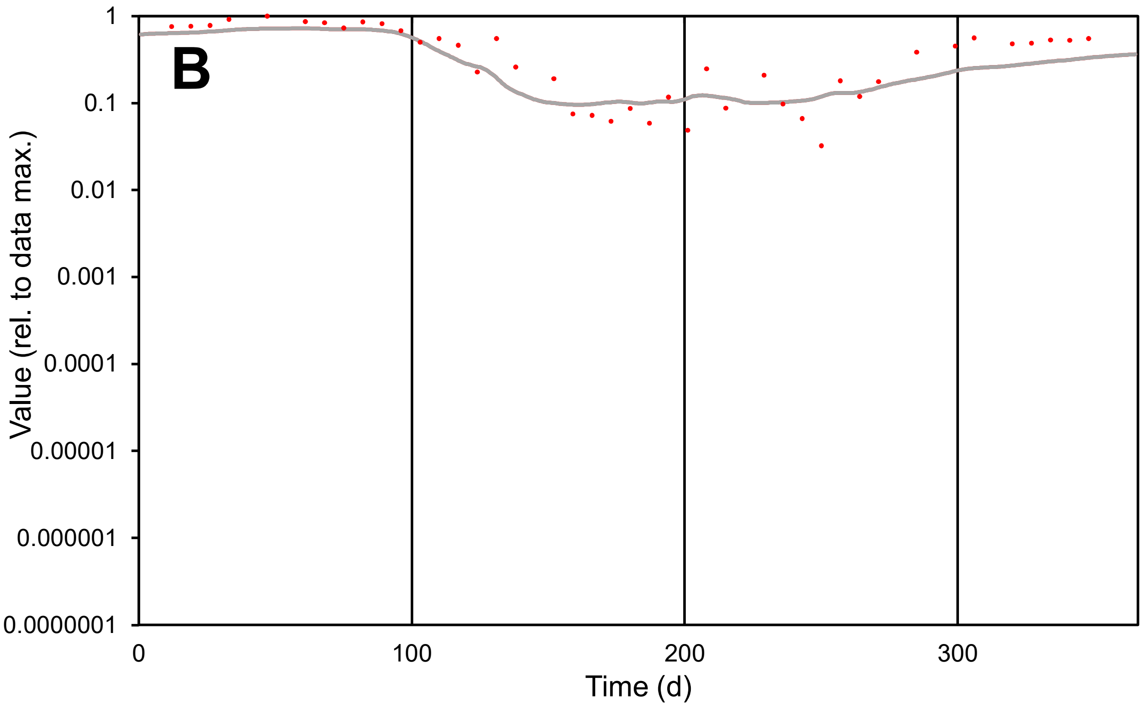


**C1**

Variables

PO4 Provide jtd or Dst

CalibOnly?

2 1 to reproduce of

oNormalize?

2 1=no, 2=yes

ofxn calc*

1.07645

ofxn mod

1.1394

sRun

RunA1

zout

1

Ave

Data

0.00032

Model

0.00024

data/vMax

0.00077

ofxng

263.459

Min d/m

0.33696

ave/min d

12.9022

ave/min m

3.31481

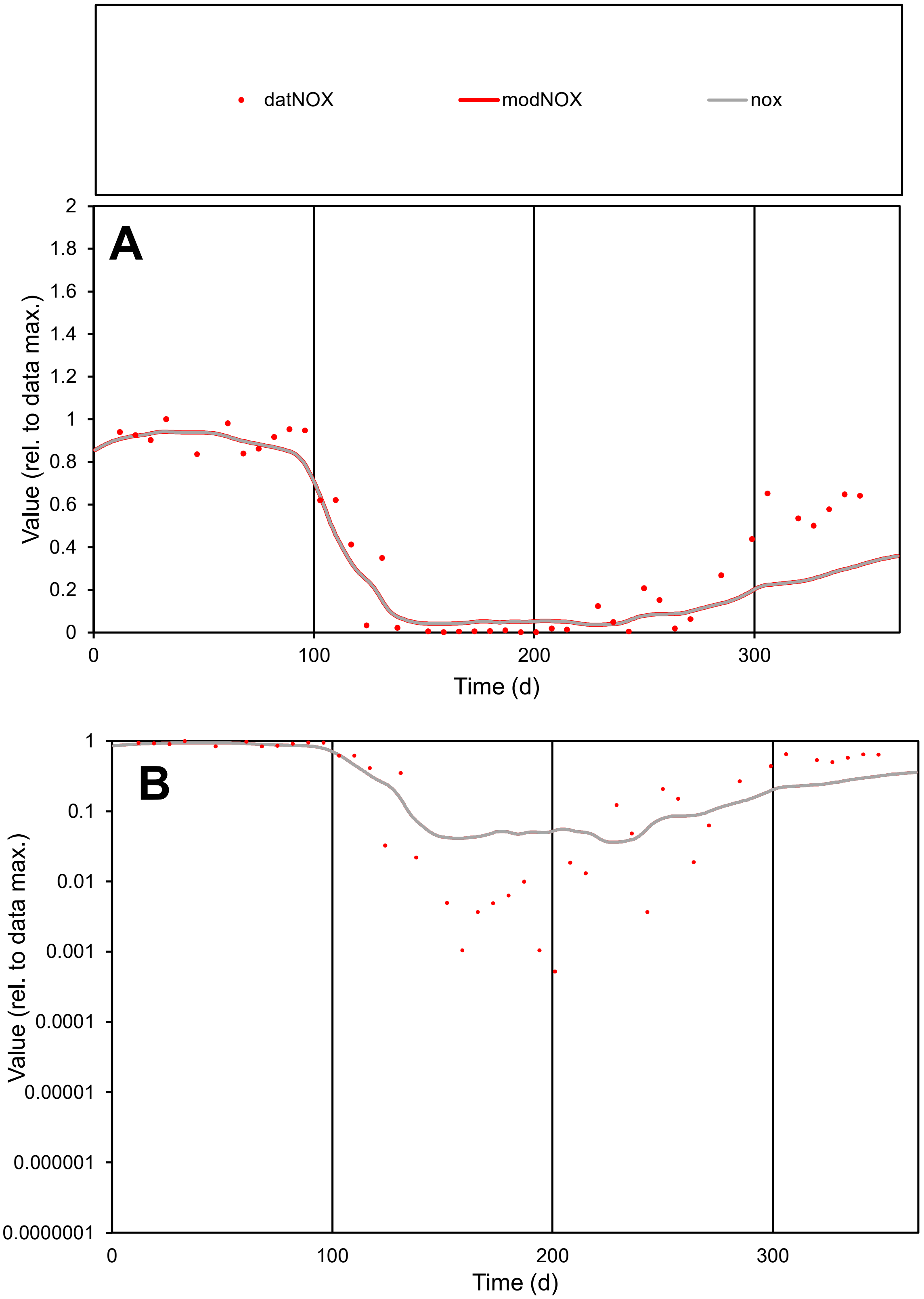


C1

Variables
NOX Provide jtd or Dst
CalibOnly?
2 1 to reproduce of
oNormalize?
2 1=no, 2=yes

ofxn calc*
1.11743
ofxn mod
1.19869

sRun
RunA1
zout
1

Ave
Data
0.00388
Model
0.00329

data/vMax
0.00952

ofxng
263.459

Min d/m
0.01433

ave/min d
775.146
ave/min m
9.43392

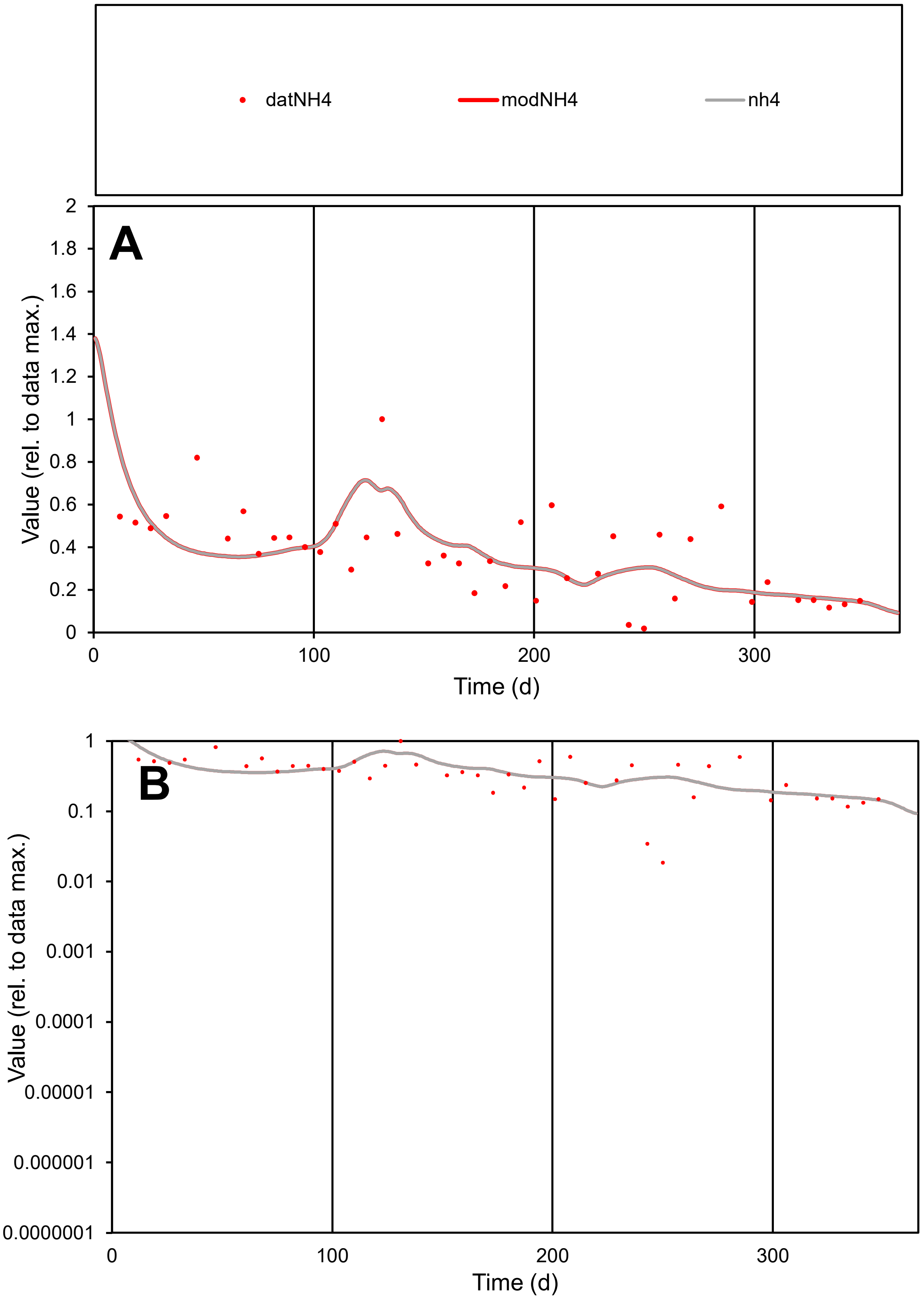


**C1**

Variables
NH4 Provide jtd or Dst
CalibOnly?
2 1 to reproduce of
oNormalize?
2 1=no, 2=yes

ofxn calc*
1.45721
ofxn mod
1.50925

sRun
RunA1
zout
1

Ave
Data
0.00035
Model
0.00034

data/vMax
0.00094

ofxng
263.459

Min d/m
0.1305

ave/min d
19.7924
ave/min m
2.56278

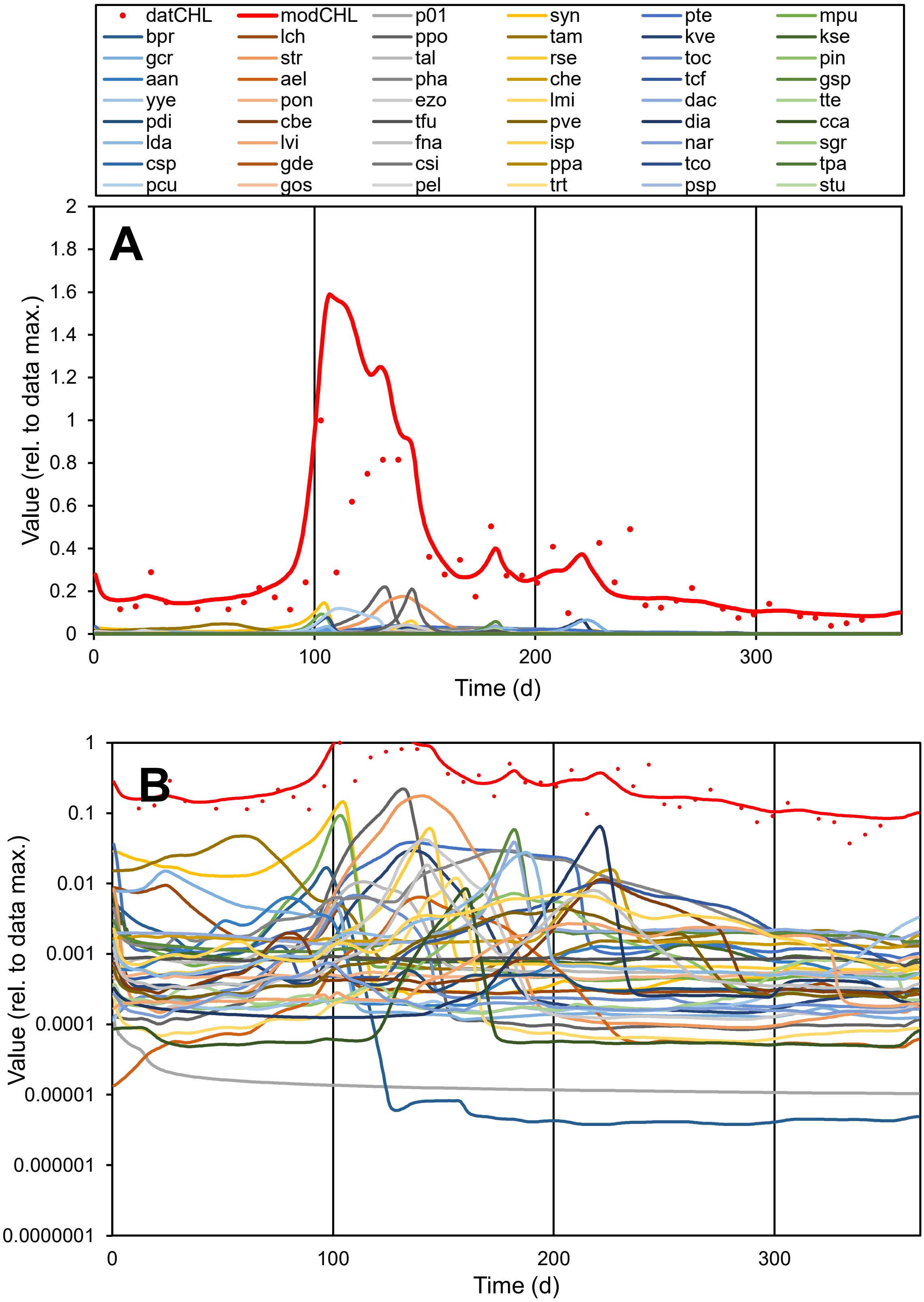


**C1**

Variables

CHL Provide jtd or Dst

CalibOnly?

2 1 to reproduce of

oNormalize?

2 1=no, 2=yes

ofxn calc*

3.8297

ofxn mod

4.1006

sRun

RunA1

zout

1

Ave

Data

0.64951

Model

0.63412

data/vMax

2.417

ofxng

263.459

Min d/m

0.45478

ave/min d

7.2168

ave/min m

3.20423

326_Calib_S_A

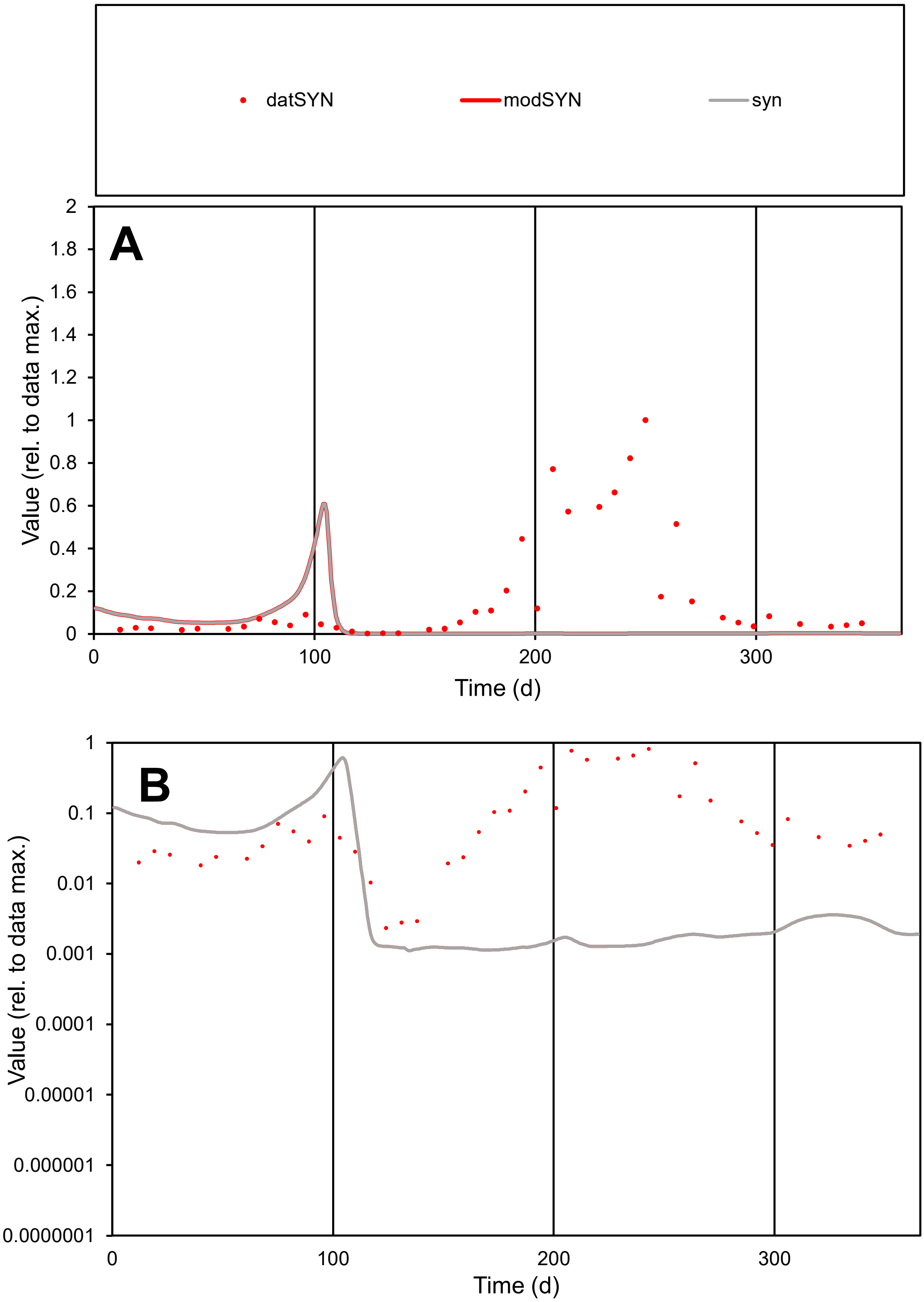


**C1**

Variables
SYN Provide jtd or Dst
CalibOnly?
2 1 to reproduce of
oNormalize?
2 1=no, 2=yes

ofxn calc*
1.80271
ofxn mod
1.83168

sRun
RunA1
zout
1

Ave
Data
11.9291
Model
2.34417

data/vMax
68.87

ofxng
263.459

Min d/m
2.05905

ave/min d
74.048
ave/min m
29.9613

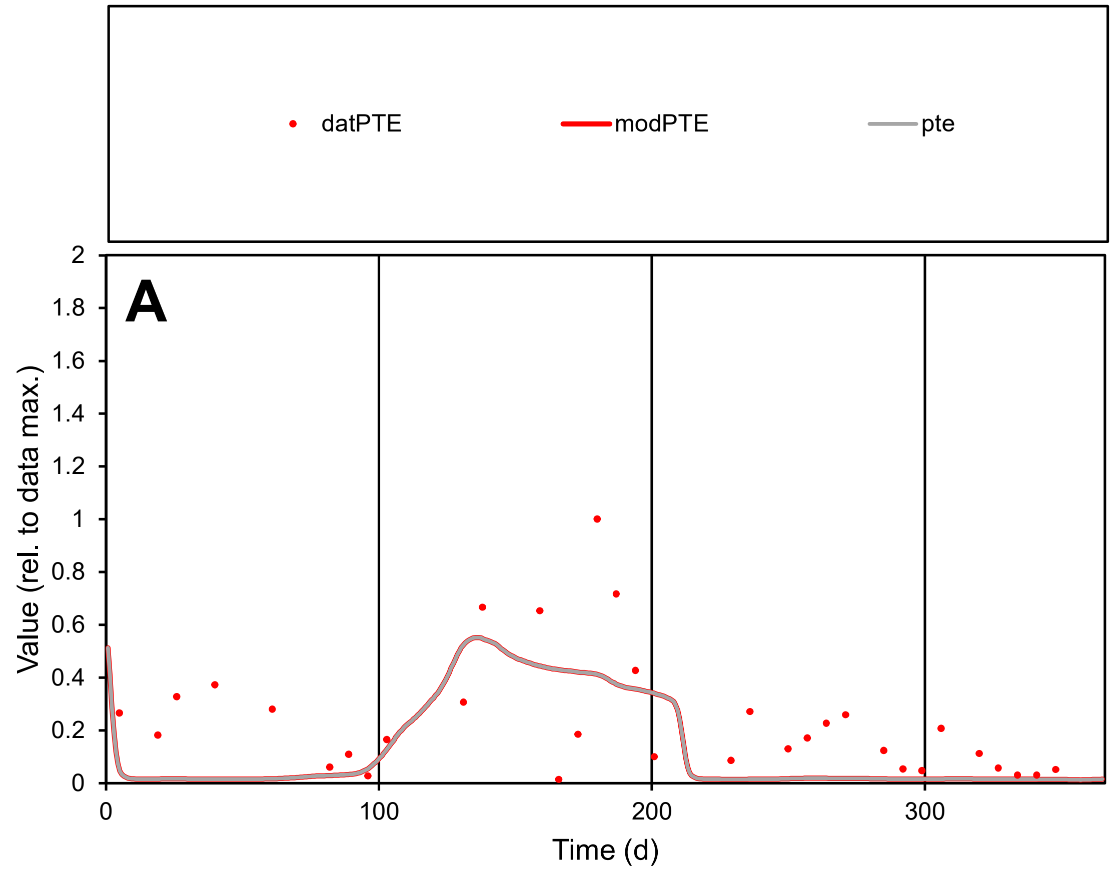


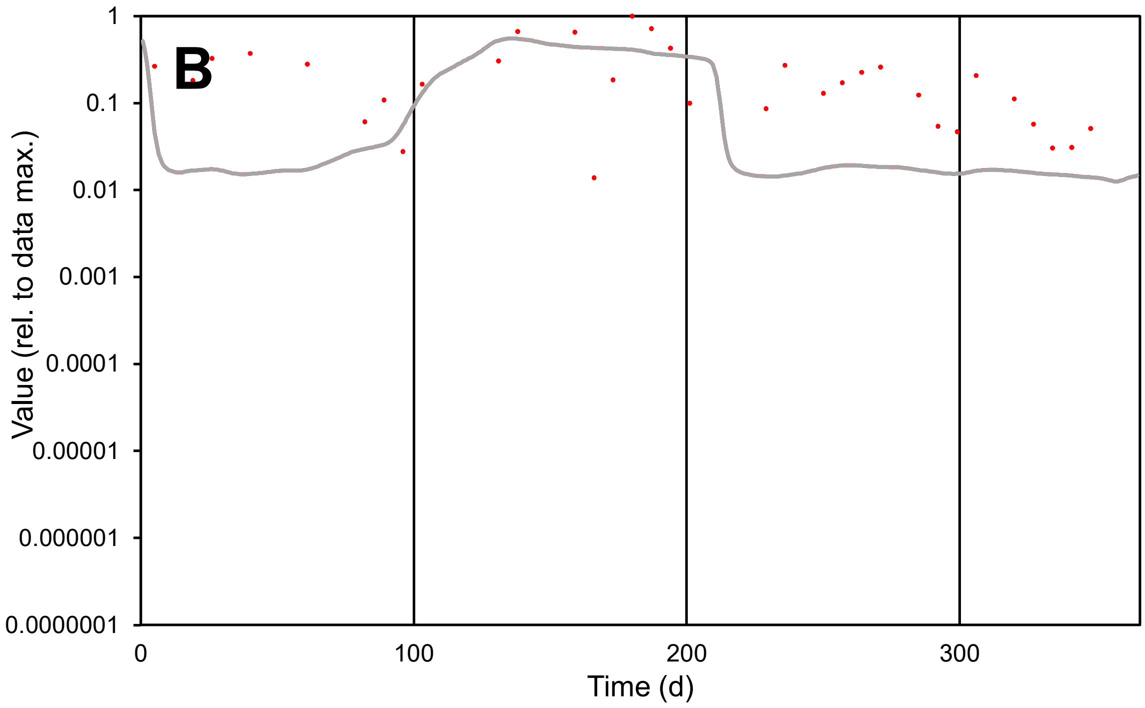


**C1**

Variables
PTE Provide jtd or Dst
CalibOnly?
2 1 to reproduce of
oNormalize?
2 1=no, 2=yes

ofxn calc*
0.90207
ofxn mod
0.91918

sRun
RunA1
zout
1

Ave
Data
0.00014
Model
8.6E-05

data/vMax
0.0006

ofxng
263.459

Min d/m
0.99749

ave/min d
16.8781
ave/min m
10.2483

326_Calib_S_A

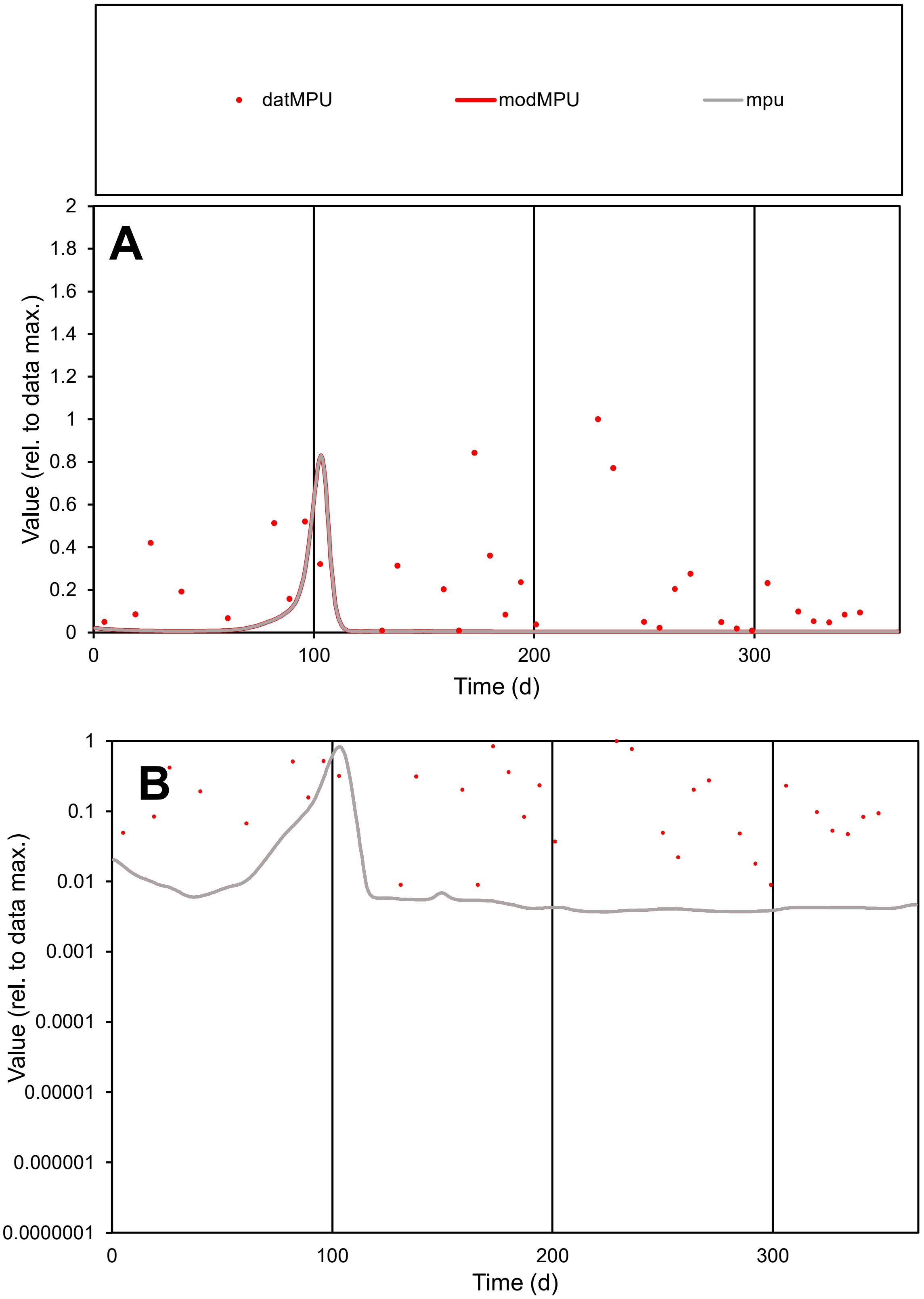


**C1**

Variables
MPU Provide jtd or Dst
CalibOnly?
2 1 to reproduce of
oNormalize?
2 1=no, 2=yes

ofxn calc*
1.41856
ofxn mod
1.43216

sRun
RunA1
zout
1

Ave
Data
0.0001
Model
1.9E-05

data/vMax
0.00045

ofxng
263.459

Min d/m
2.42349
ave/min d
24.9568
ave/min m
11.5431

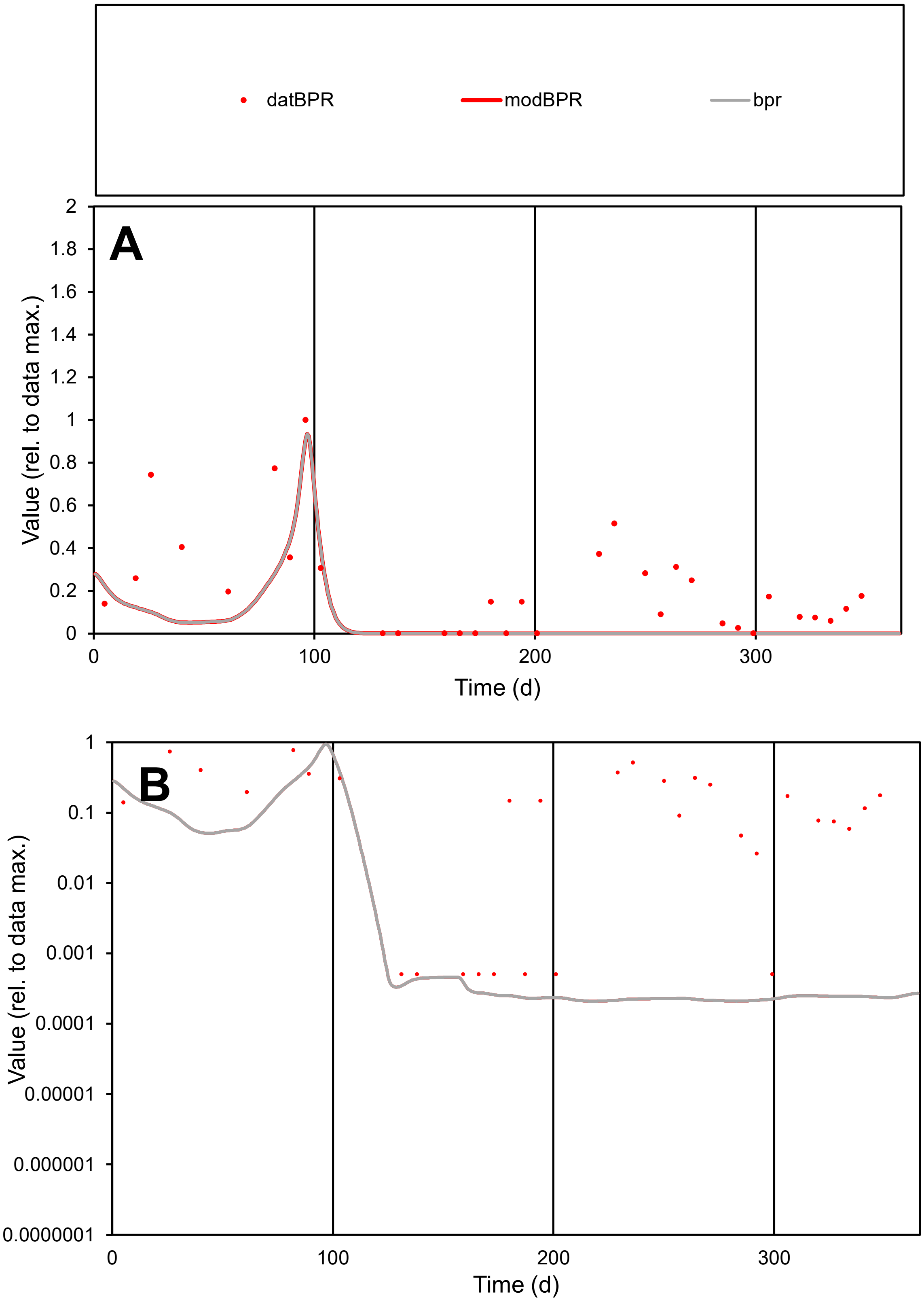


**C1**

Variables
BPR Provide jtd or Dst
CalibOnly?
2 1 to reproduce of
oNormalize?
2 1=no, 2=yes

ofxn calc*
1.02908
ofxn mod
1.03945

sRun
RunA1
zout
1

Ave
Data
7.8E-05
Model
2.8E-05

data/vMax
0.00036

ofxng
263.459

Min d/m
2.42857

ave/min d
419.475
ave/min m
368.701

326_Calib_S_A

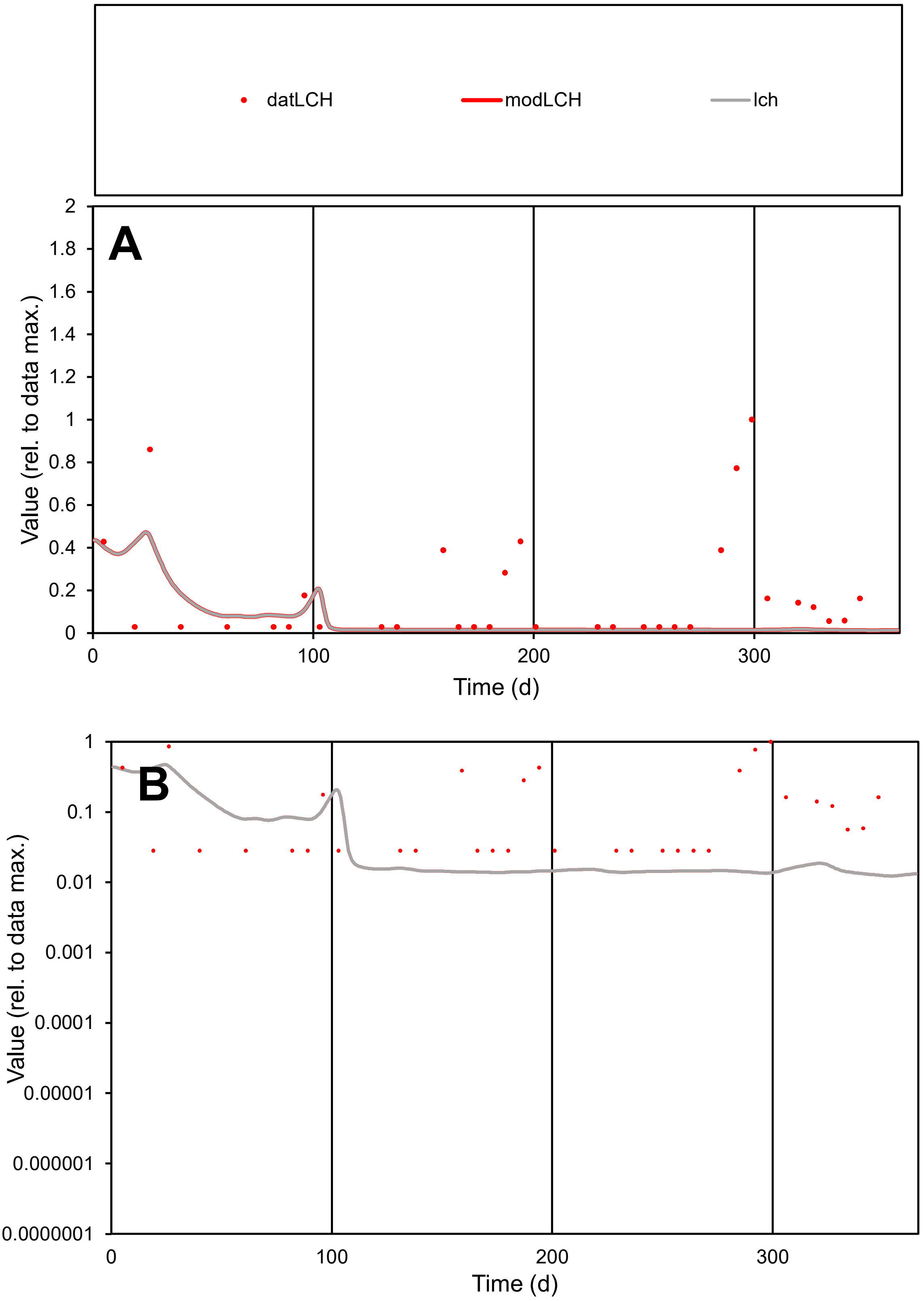


**C1**

Variables
LCH Provide jtd or Dst
CalibOnly?
2 1 to reproduce of
oNormalize?
2 1=no, 2=yes

ofxn calc*
1.53552
ofxn mod
1.54513

sRun
RunA1
zout
1

Ave
Data
1.6E-05
Model
5.9E-06

data/vMax
8.7E-05

ofxng
263.459

Min d/m
2.24404

ave/min d
6.39238
ave/min m
5.4109

326_Calib_S_A

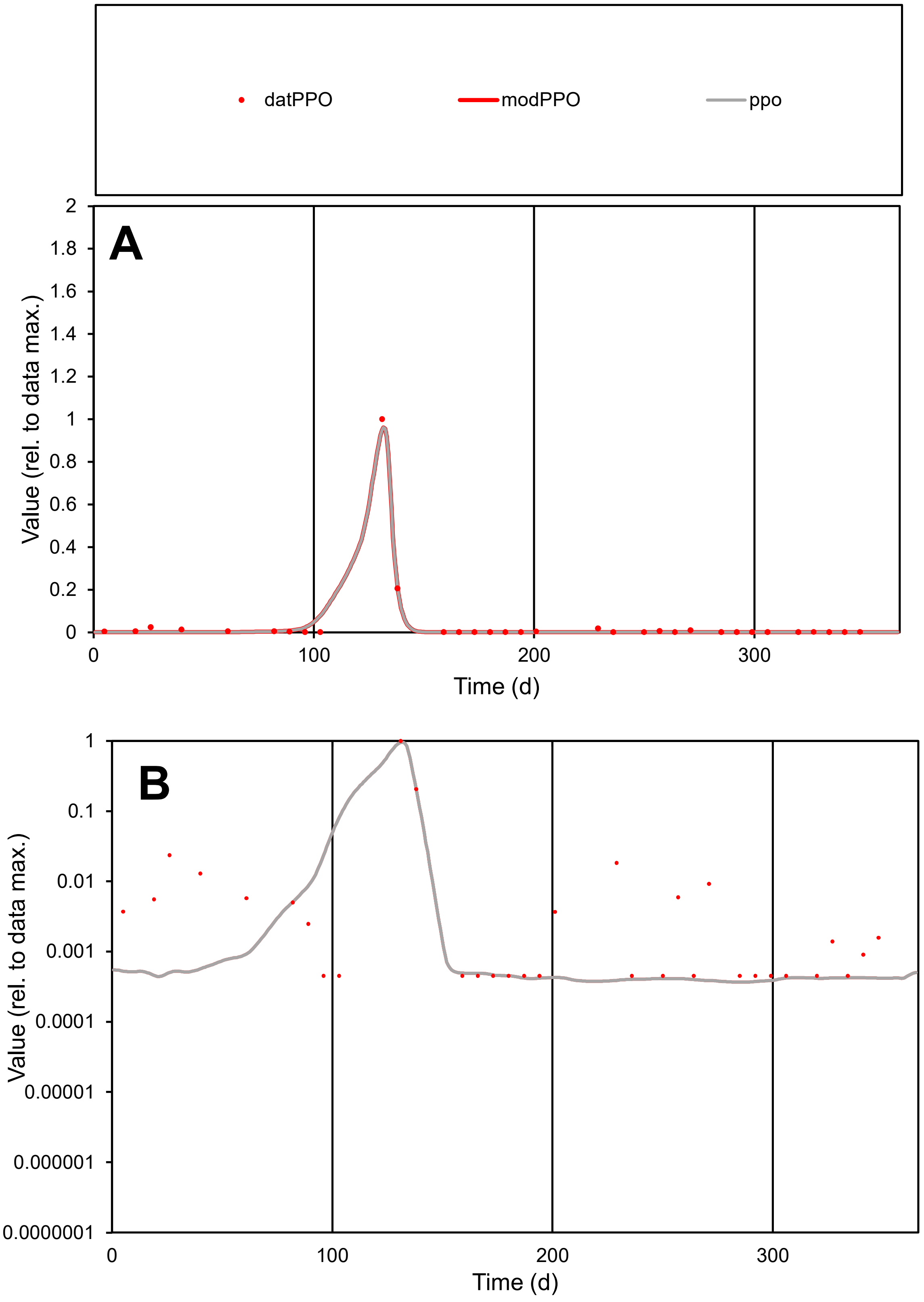


**C1**

Variables
PPO Provide jtd or Dst
CalibOnly?
2 1 to reproduce of
oNormalize?
2 1=no, 2=yes

ofxn calc*
0.36485
ofxn mod
0.6836

sRun
RunA1
zout
1

Ave
Data
0.00017
Model
0.00016

data/vMax
0.00422

ofxng
263.459

Min d/m
1.23181

ave/min d
87.9118
ave/min m
103.126

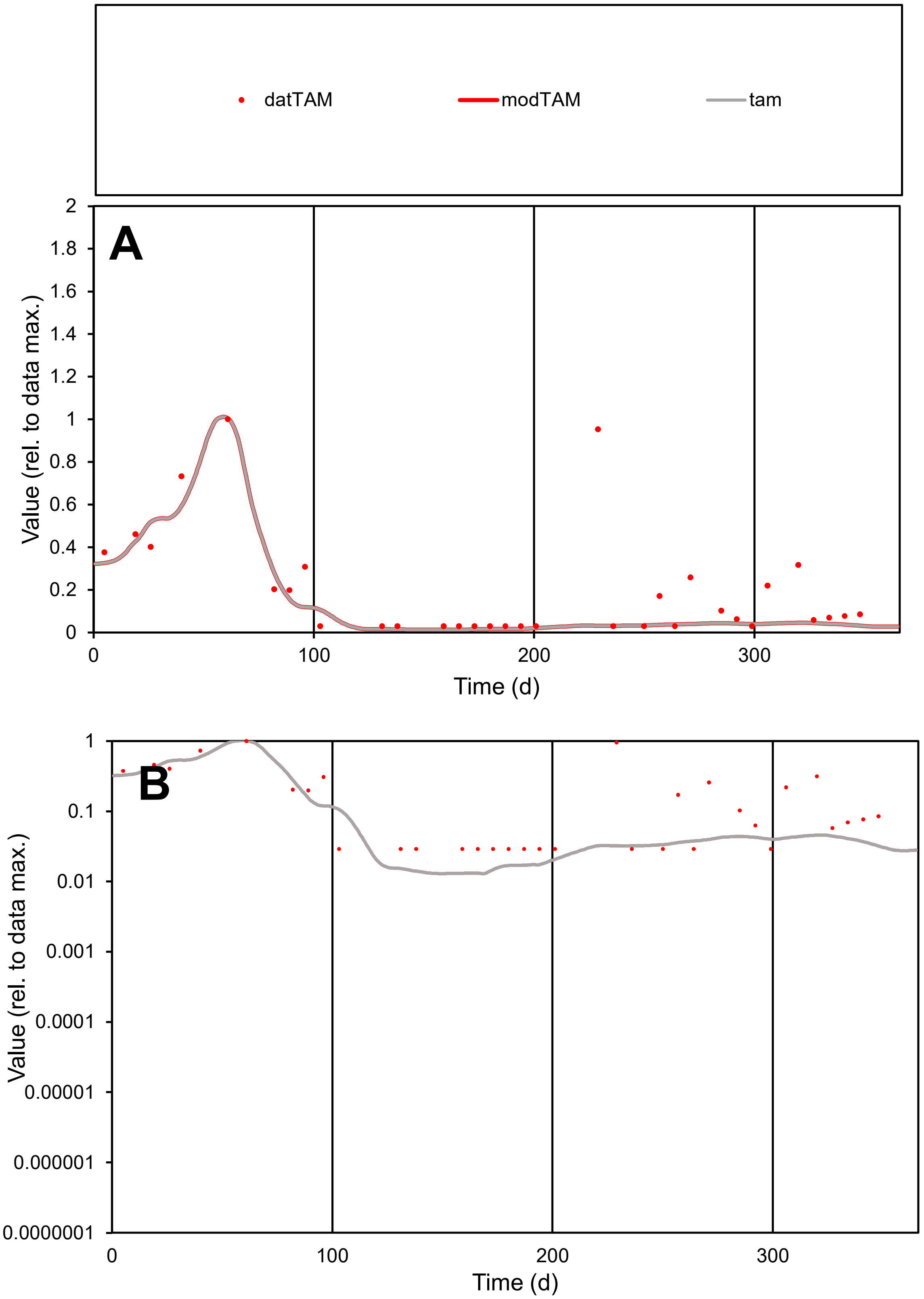


C1

Variables
TAM Provide jtd or Dst
CalibOnly?
2 1 to reproduce of
oNormalize?
2 1=no, 2=yes

ofxn calc*
0.93473
ofxn mod
0.95216

sRun
RunA1
zout
1

Ave
Data
3.9E-05
Model
2.5E-05

data/vMax
0.0002

ofxng
263.459

Min d/m
2.21585

ave/min d
6.75562
ave/min m
9.79765

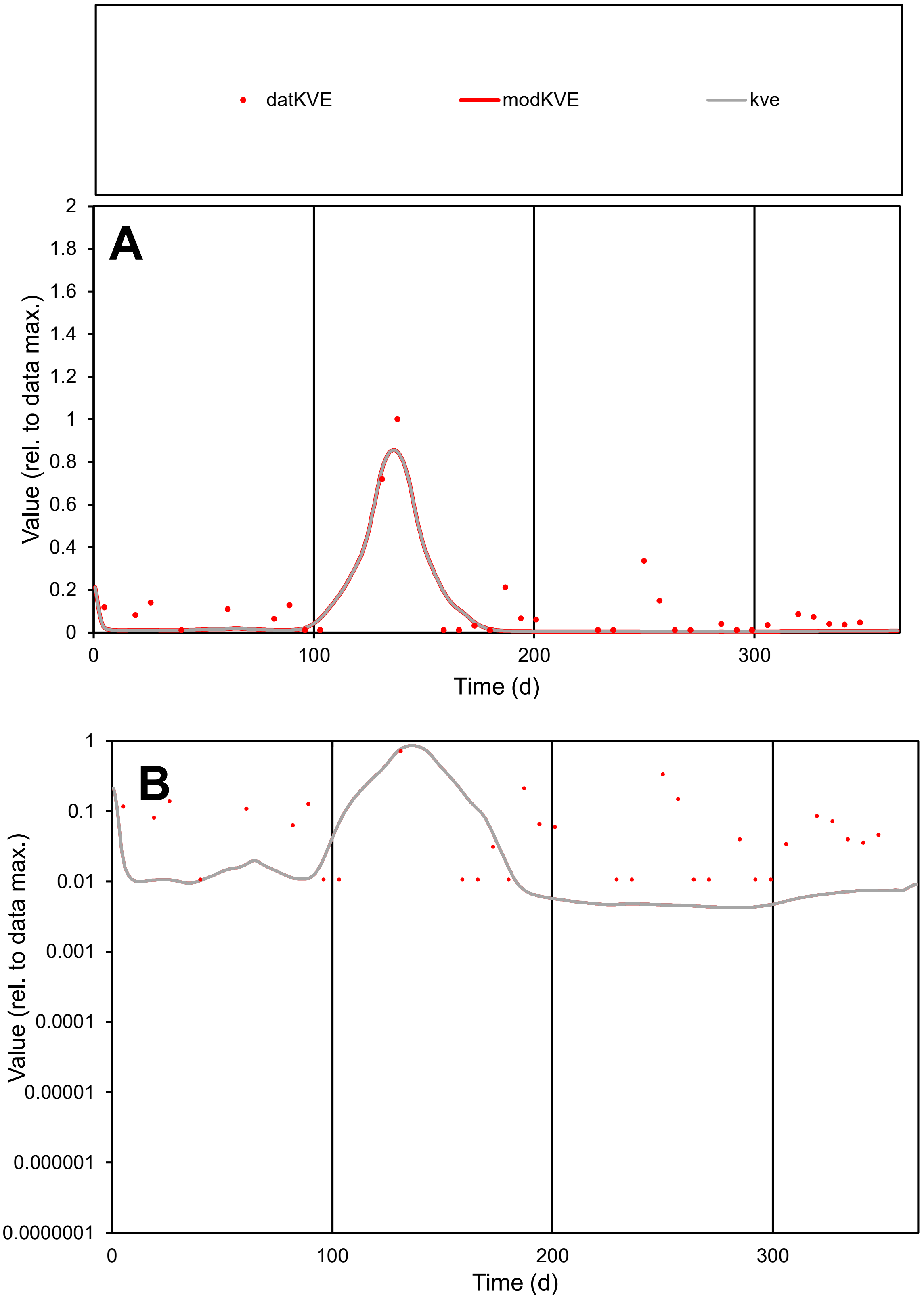


**C1**

Variables

KVE Provide jtd or Dst

CalibOnly?

2 1 to reproduce of

oNormalize?

2 1=no, 2=yes

ofxn calc*

0.8625

ofxn mod

0.9092

sRun

RunA1

zout

1

Ave

Data

3.3E-05

Model

2E-05

data/vMax

0.00029

ofxng

263.459

Min d/m

2.5008

ave/min d

10.5202

ave/min m

16.2034

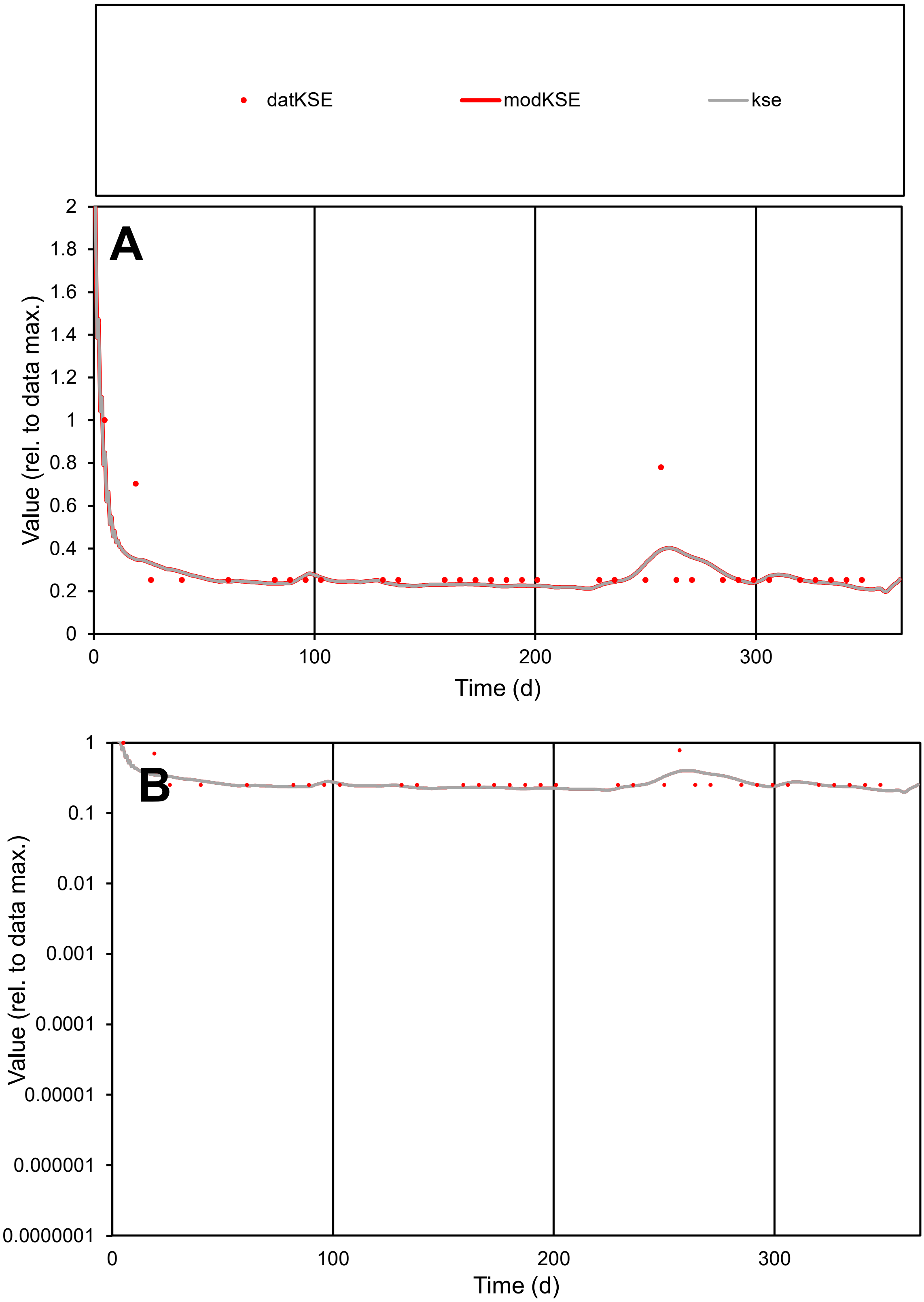


**C1**

Variables

KSE Provide jtd or Dst

CalibOnly?

2 1 to reproduce of

oNormalize?

2 1=no, 2=yes

ofxn calc*

0.33705

ofxn mod

0.37247

sRun

RunA1

zout

1

Ave

Data

4.5E-06

Model

4.2E-06

data/vMax

1.5E-05

ofxng

263.459

Min d/m

1.19706

ave/min d

1.20767

ave/min m

1.3387

326_Calib_S_A

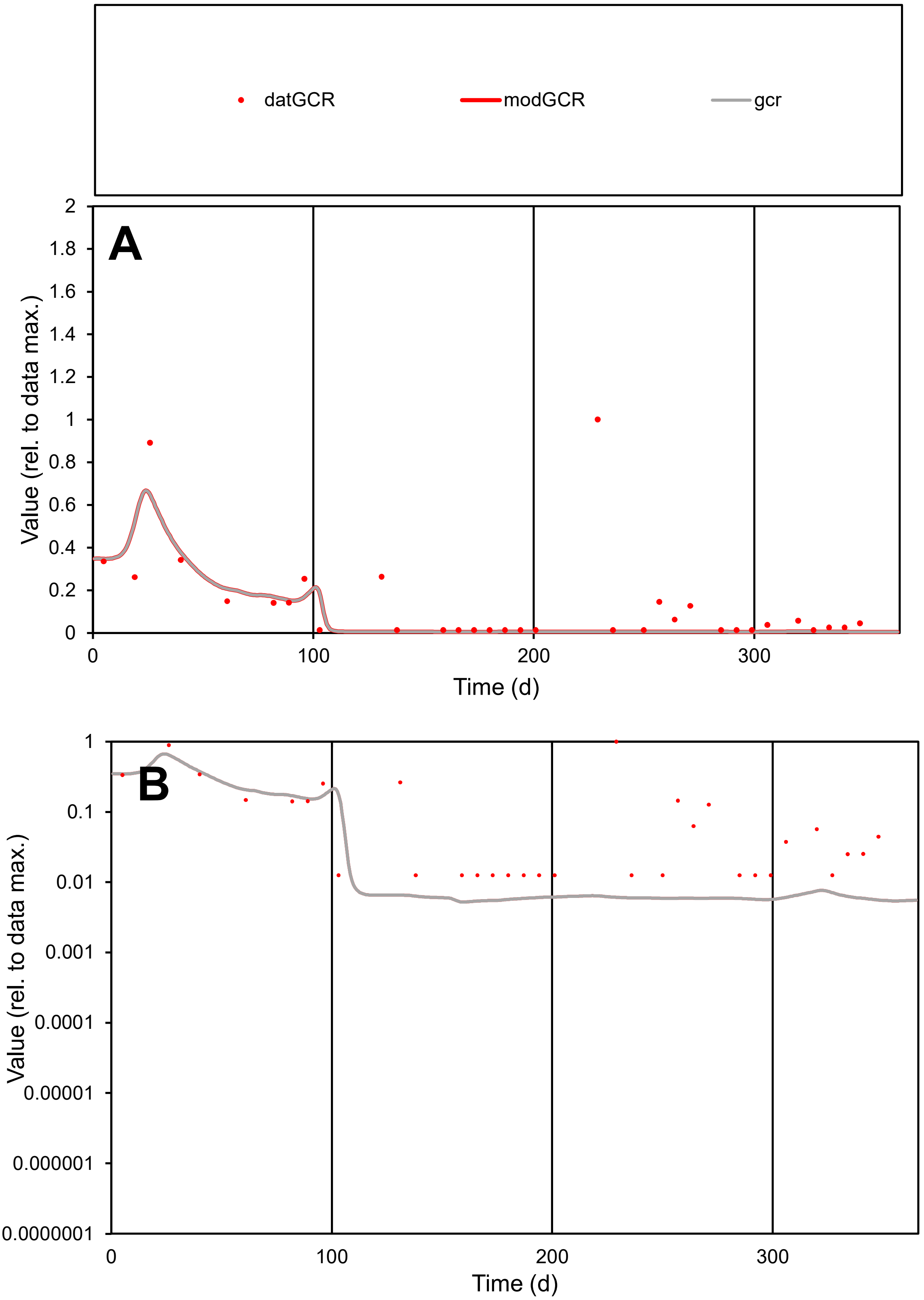


**C1**

Variables

GCR Provide jtd or Dst

CalibOnly?

2 1 to reproduce of

oNormalize?

2 1=no, 2=yes

ofxn calc*

1.43222

ofxn mod

1.45401

sRun

RunA1

zout

1

Ave

Data

3.4E-05

Model

2.1E-05

data/vMax

0.00025

ofxng

263.459

Min d/m

2.40707

ave/min d

10.821

ave/min m

16.3114

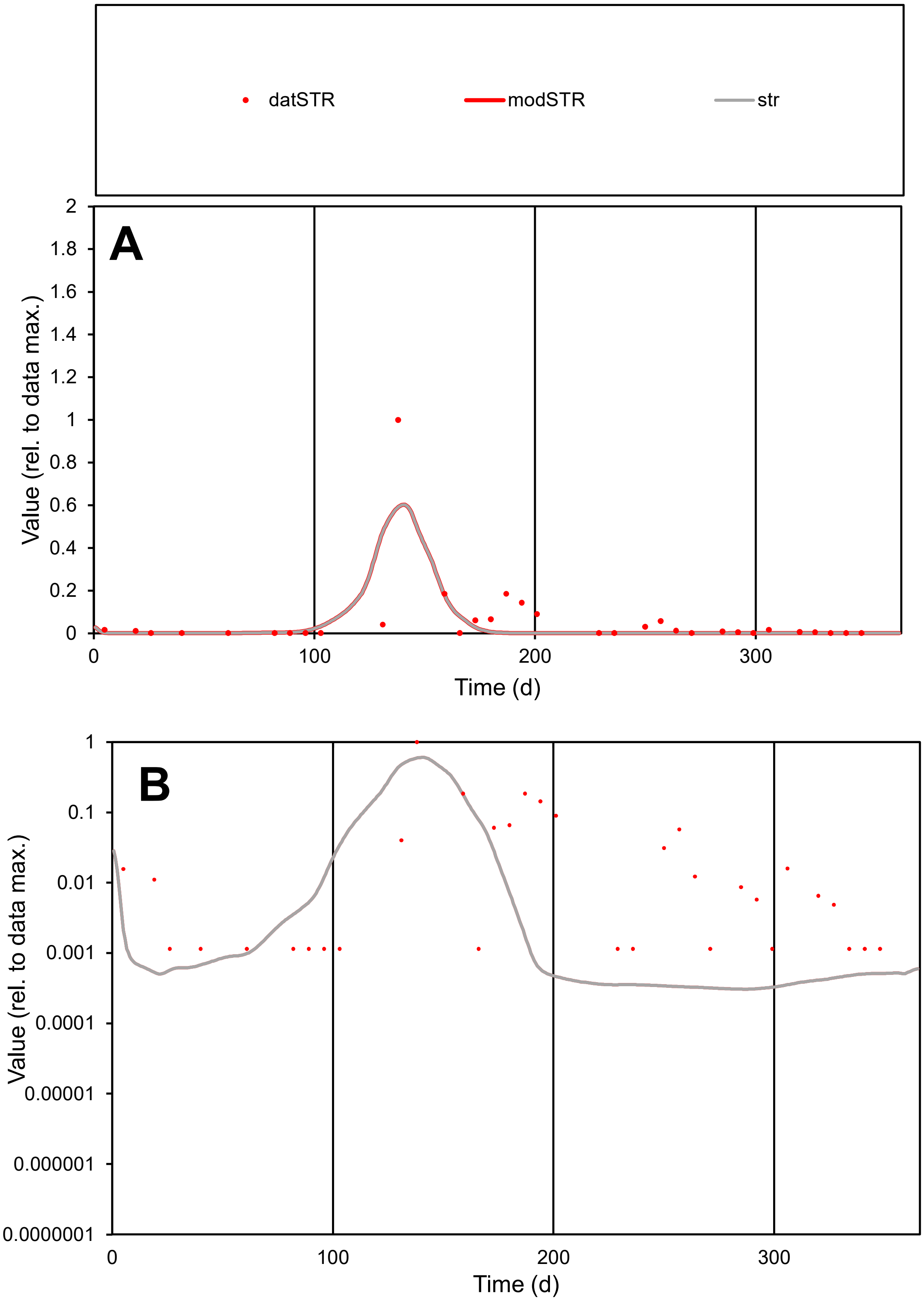


**C1**

Variables

STR Provide jtd or Dst

CalibOnly?

2 1 to reproduce of

oNormalize?

2 1=no, 2=yes

ofxn calc*

1.87155

ofxn mod

1.94416

sRun

RunA1

zout

1

Ave

Data

8.4E-05

Model

5.9E-05

data/vMax

0.00142

ofxng

263.459

Min d/m

3.72769

ave/min d

51.7566

ave/min m

135.142

326_Calib_S_A

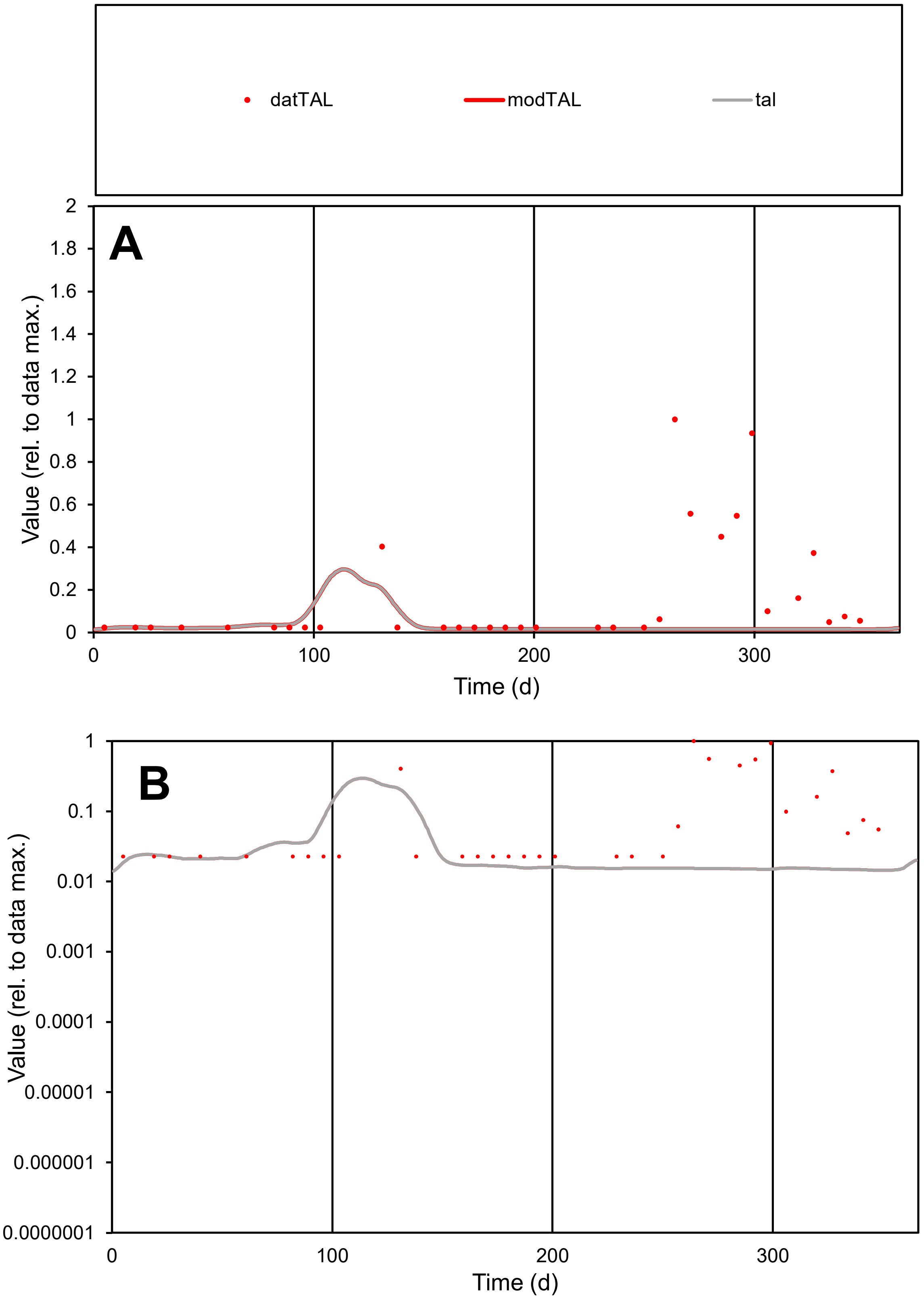


**C1**

Variables
TAL Provide jtd or Dst
CalibOnly?
2 1 to reproduce of
oNormalize?
2 1=no, 2=yes

ofxn calc*
1.84669
ofxn mod
1.85452

sRun
RunA1
zout
1

Ave
Data
2.9E-05
Model
5.7E-06

data/vMax
0.00018

ofxng
263.459

Min d/m
1.56614

ave/min d
6.98226
ave/min m
2.16961

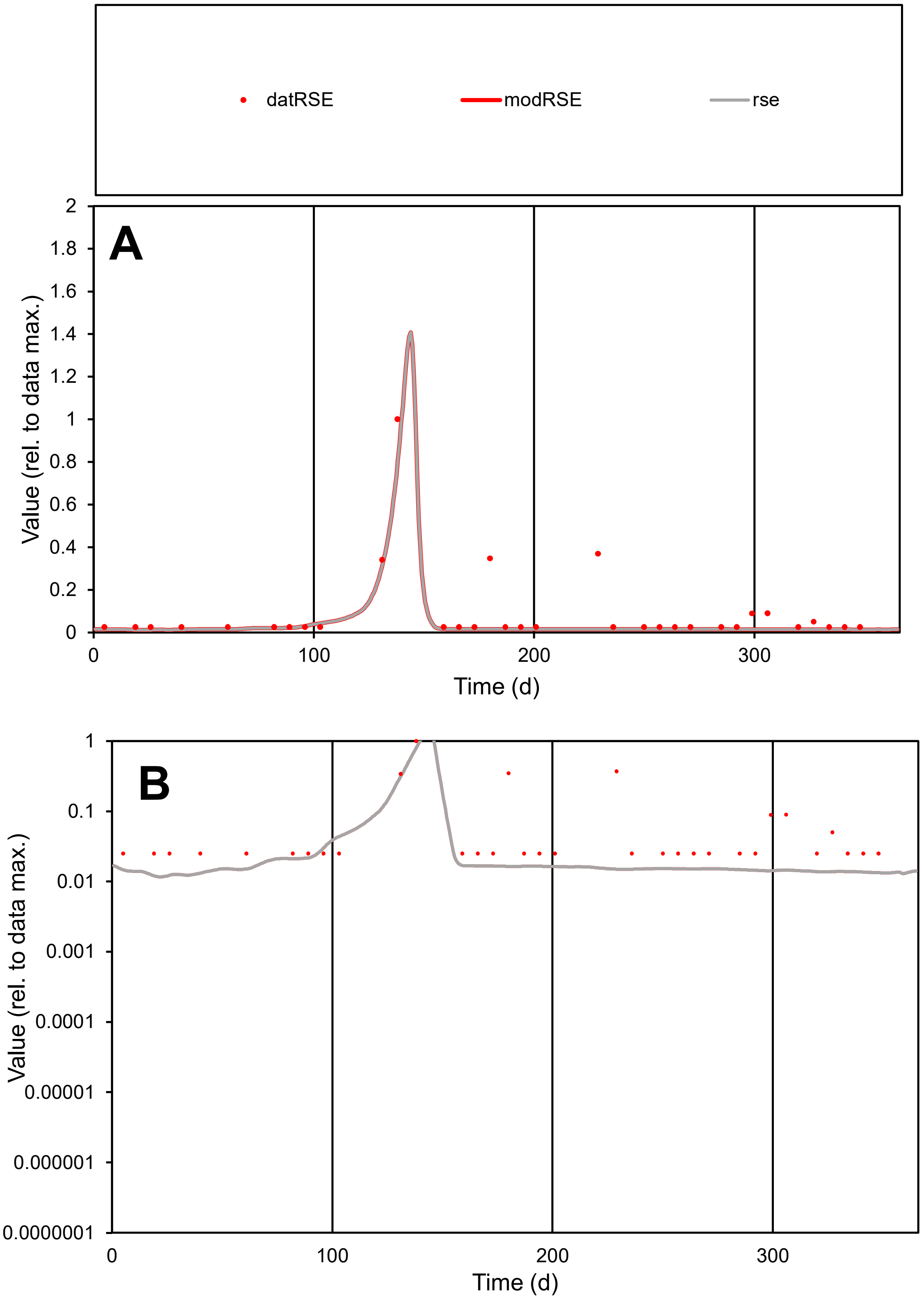


**C1**

Variables

RSE Provide jtd or Dst

CalibOnly?

2 1 to reproduce of

oNormalize?

2 1=no, 2=yes

ofxn calc*

0.98264

ofxn mod

1.01528

sRun

RunA1

zout

1

Ave

Data

1.7E-05

Model

1.1E-05

data/vMax

0.00019

ofxng

263.459

Min d/m

2.11378

ave/min d

3.55009

ave/min m

4.76236

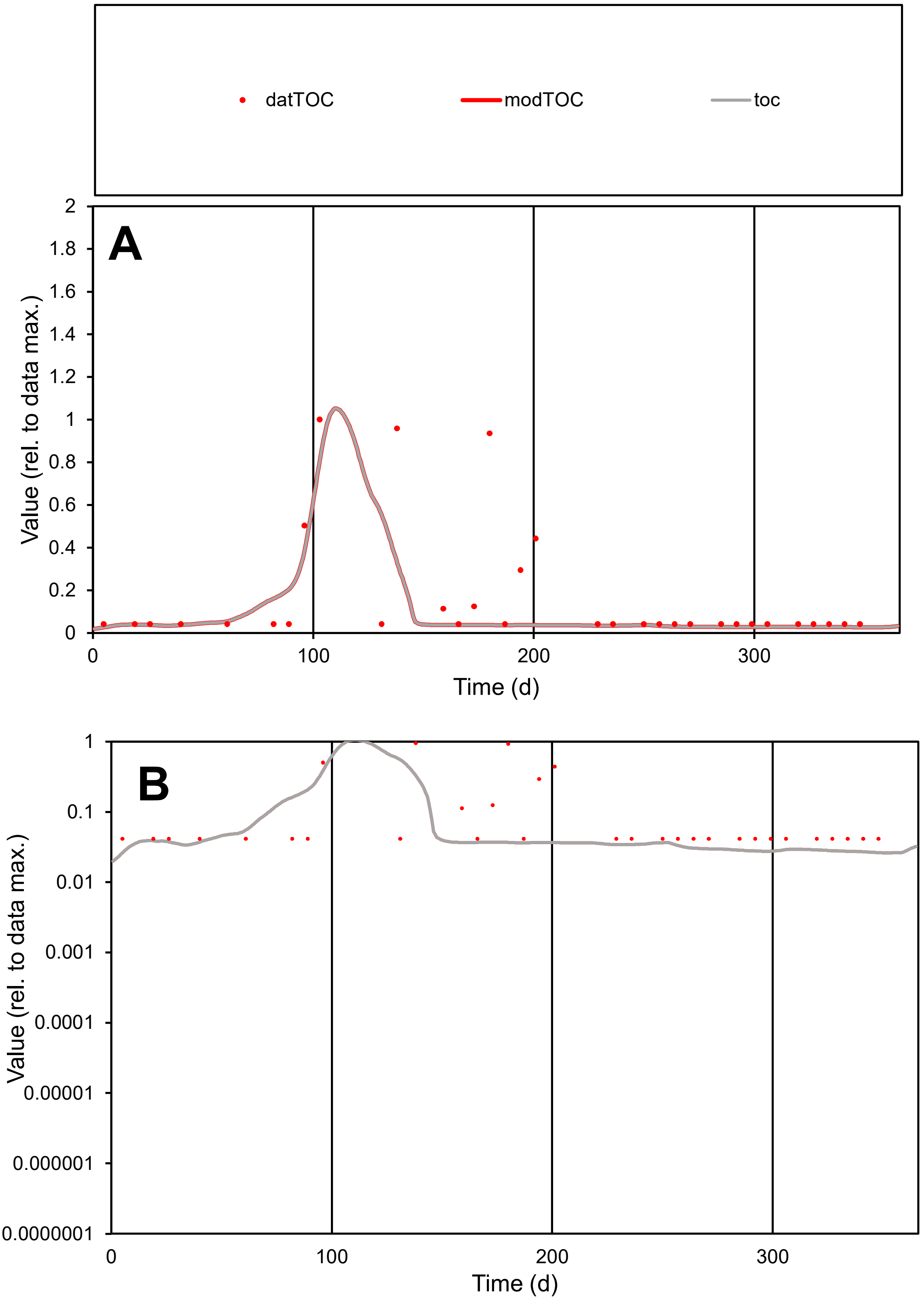


**C1**

Variables
TOC Provide jtd or Dst
CalibOnly?
2 1 to reproduce of
oNormalize?
2 1=no, 2=yes

ofxn calc*
1.37739
ofxn mod
1.40672

sRun
RunA1
zout
1

Ave
Data
1.7E-05
Model
1.1E-05

data/vMax
0.0001

ofxng
263.459

Min d/m
1.5745

ave/min d
3.95314
ave/min m
3.89494

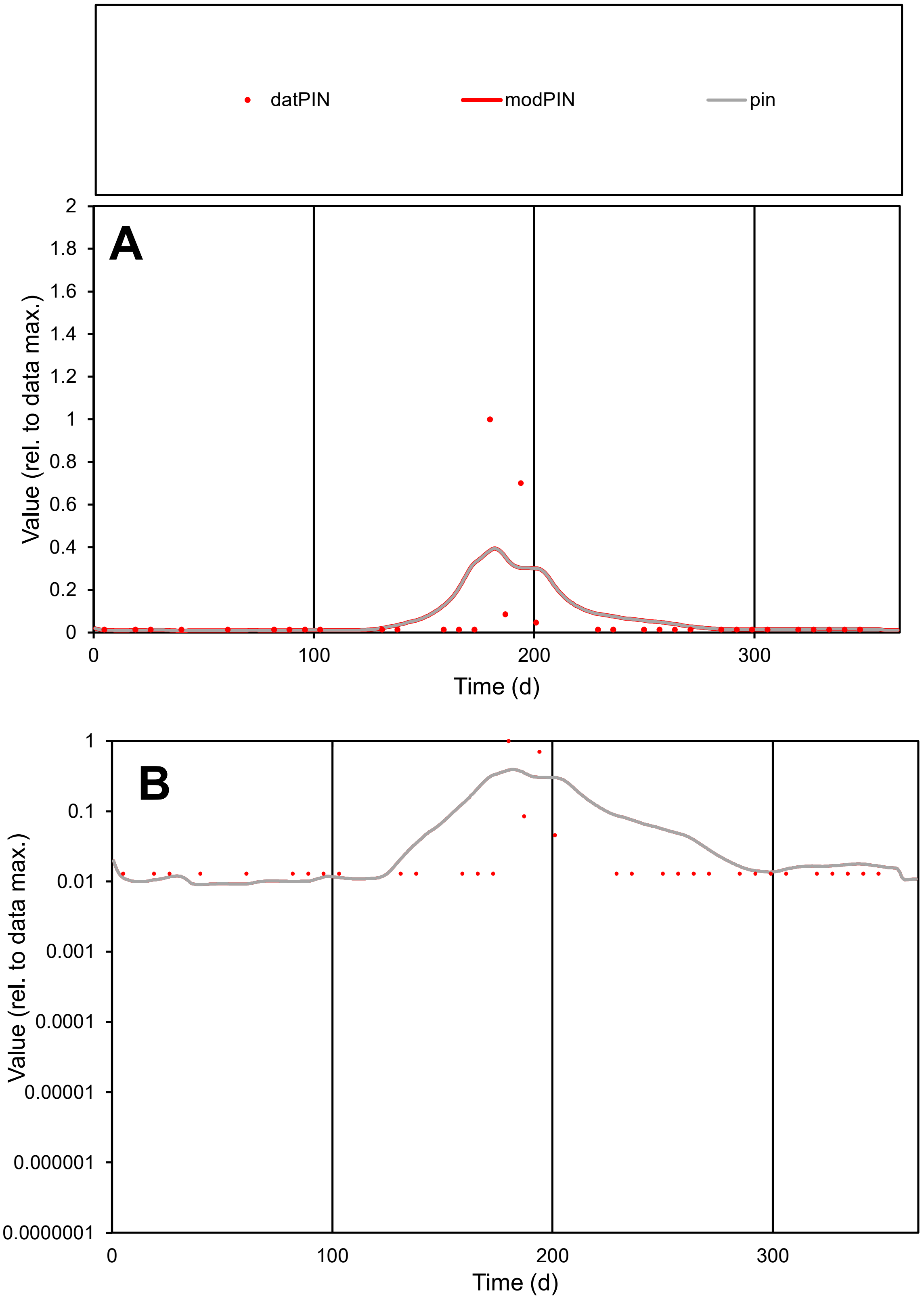


**C1**

Variables
PIN Provide jtd or Dst
CalibOnly?
2 1 to reproduce of
oNormalize?
2 1=no, 2=yes

ofxn calc*
2.34469
ofxn mod
2.37034

sRun
RunA1
zout
1

Ave
Data
2.1E-05
Model
2.4E-05

data/vMax
0.00031

ofxng
263.459

Min d/m
1.423

ave/min d
5.18288
ave/min m
8.64379

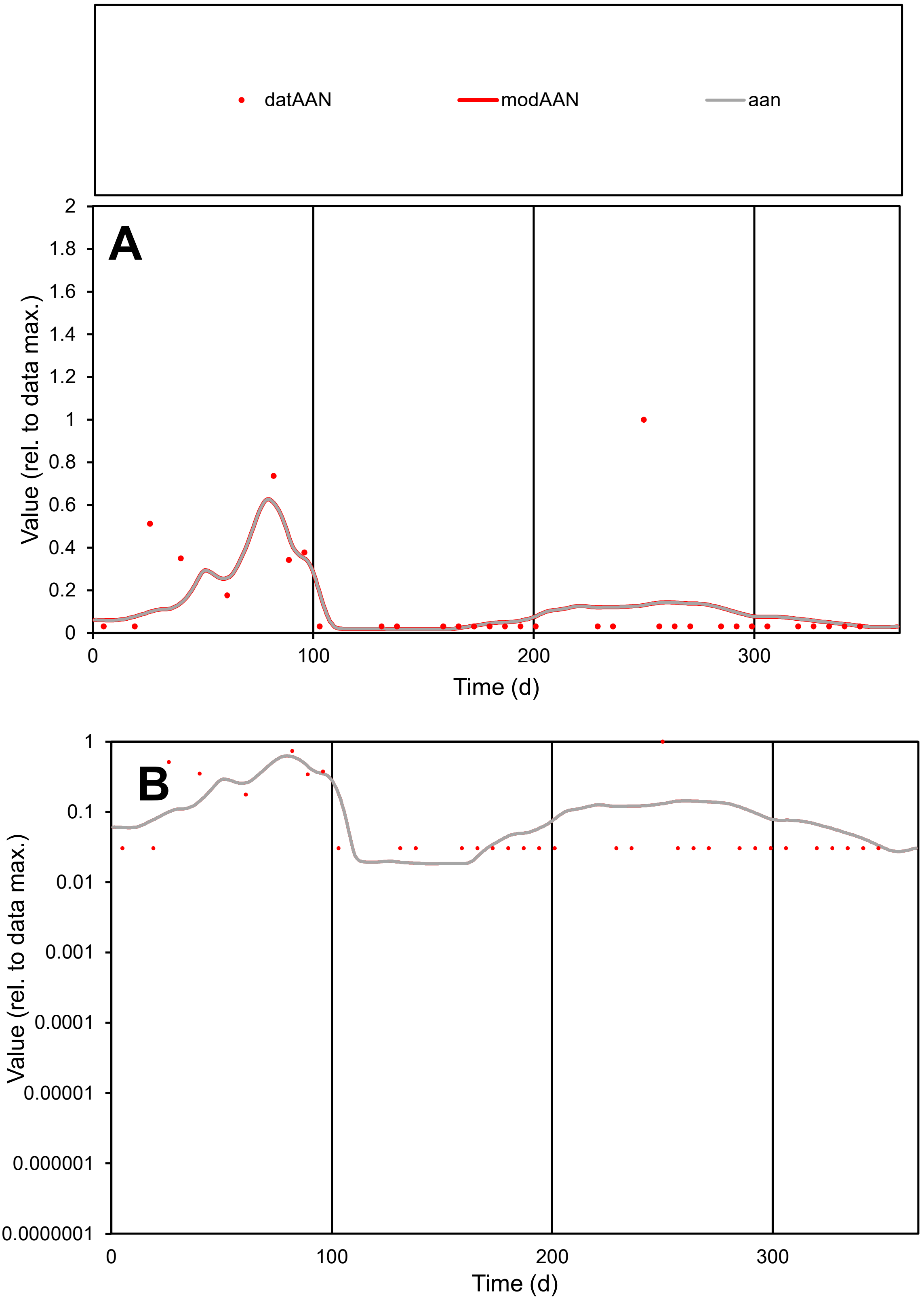


**C1**

Variables
AAN Provide jtd or Dst
CalibOnly?
2 1 to reproduce of
oNormalize?
2 1=no, 2=yes

ofxn calc*
1.39135
ofxn mod
1.40631

sRun
RunA1
zout
1

Ave
Data
2.4E-05
Model
2.2E-05

data/vMax
0.00018

ofxng
263.459

Min d/m
1.64107

ave/min d
4.27441
ave/min m
6.33898

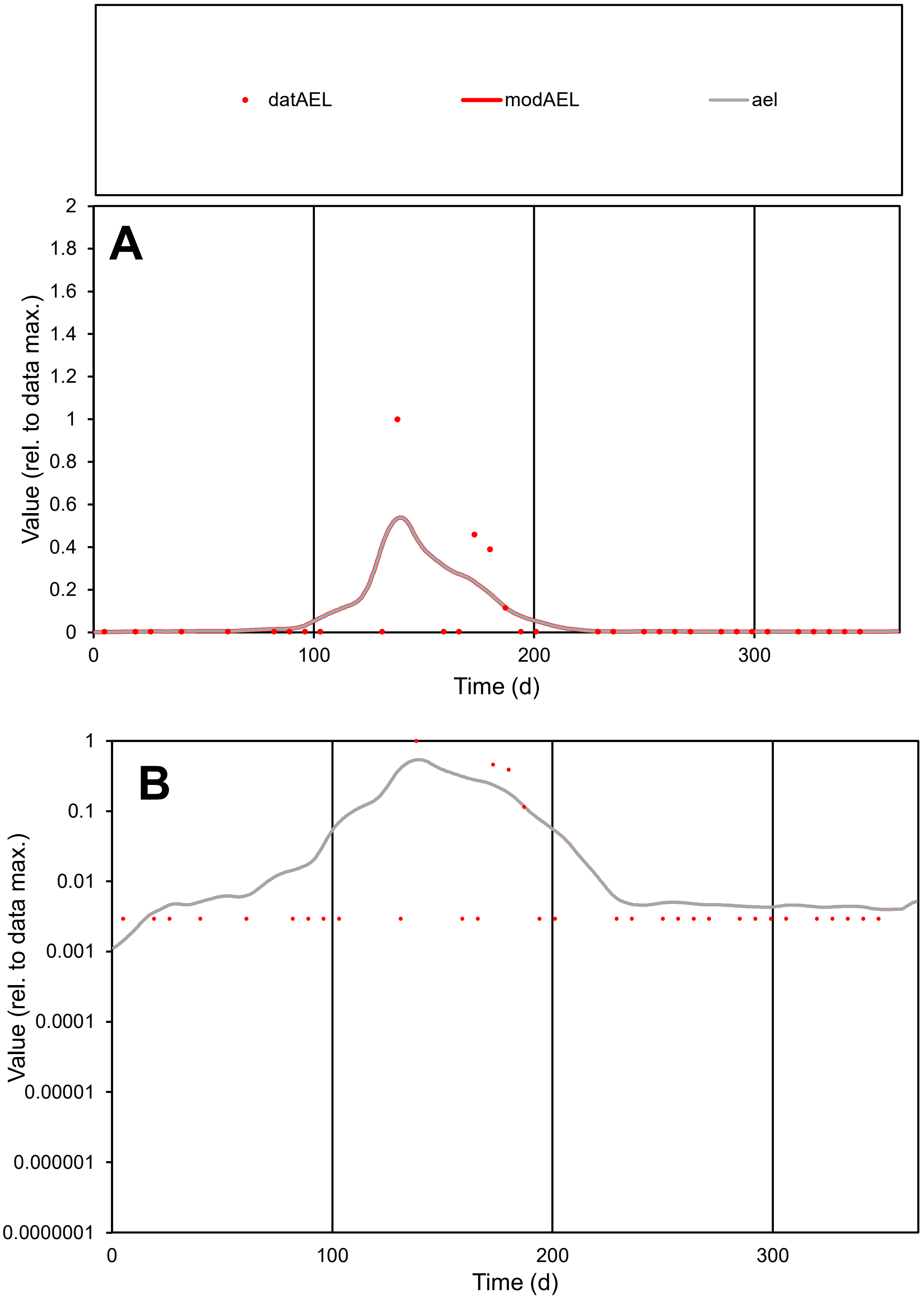


**C1**

Variables
AEL Provide jtd or Dst
CalibOnly?
2 1 to reproduce of
oNormalize?
2 1=no, 2=yes

ofxn calc*
2.11293
ofxn mod
2.26653

sRun
RunA1
zout
1

Ave
Data
1.5E-05
Model
1.6E-05

data/vMax
0.00024

ofxng
263.459

Min d/m
1.8657

ave/min d
21.1441
ave/min m
43.4787

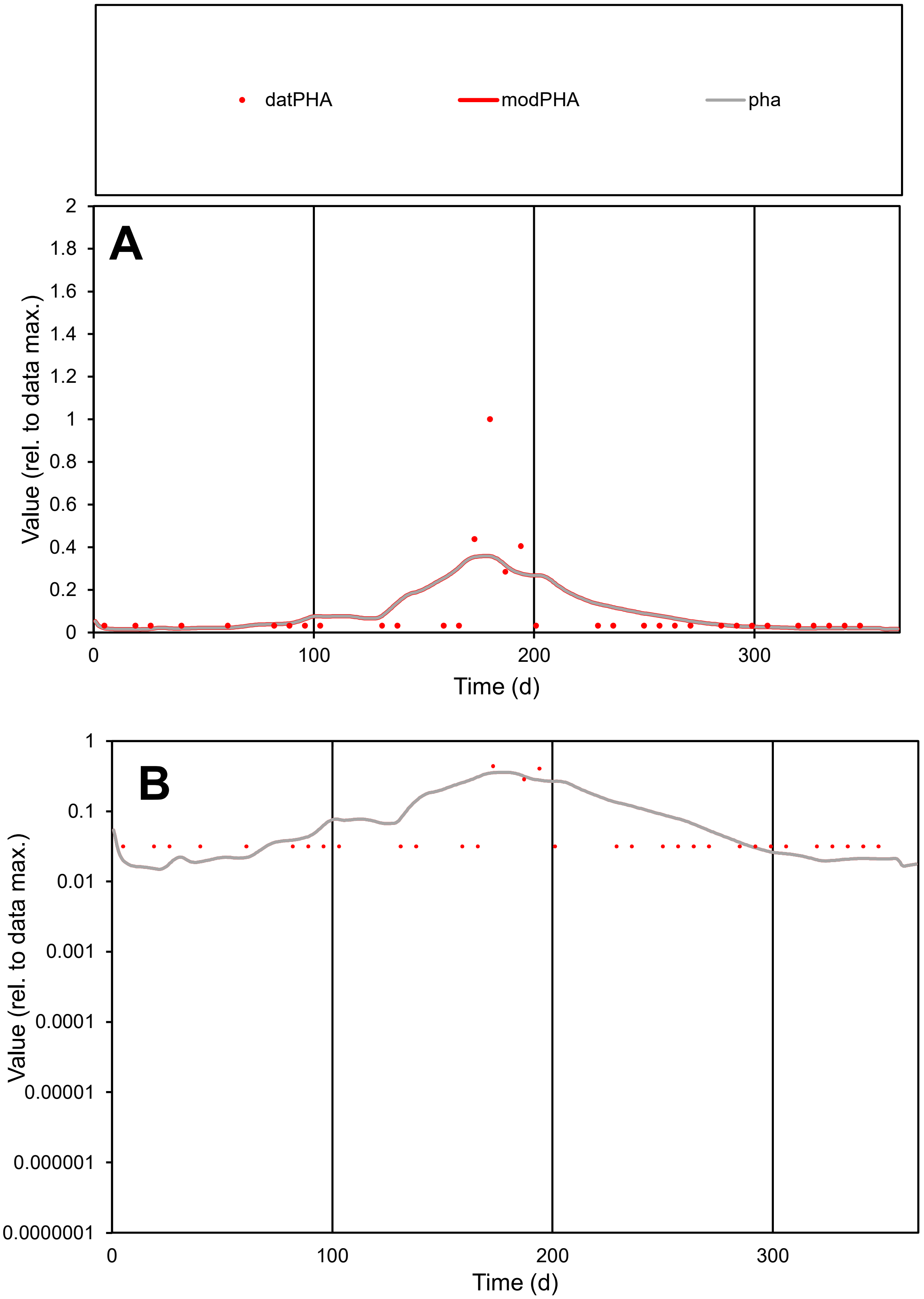


**C1**

Variables

PHA Provide jtd or Dst

CalibOnly?

2 1 to reproduce of

oNormalize?

2 1=no, 2=yes

ofxn calc*

1.53319

ofxn mod

1.55607

sRun

RunA1

zout

1

Ave

Data

3E-05

Model

3.4E-05

data/vMax

0.00033

ofxng

263.459

Min d/m

2.08743

ave/min d

2.91975

ave/min m

6.794

326_Calib_S_A

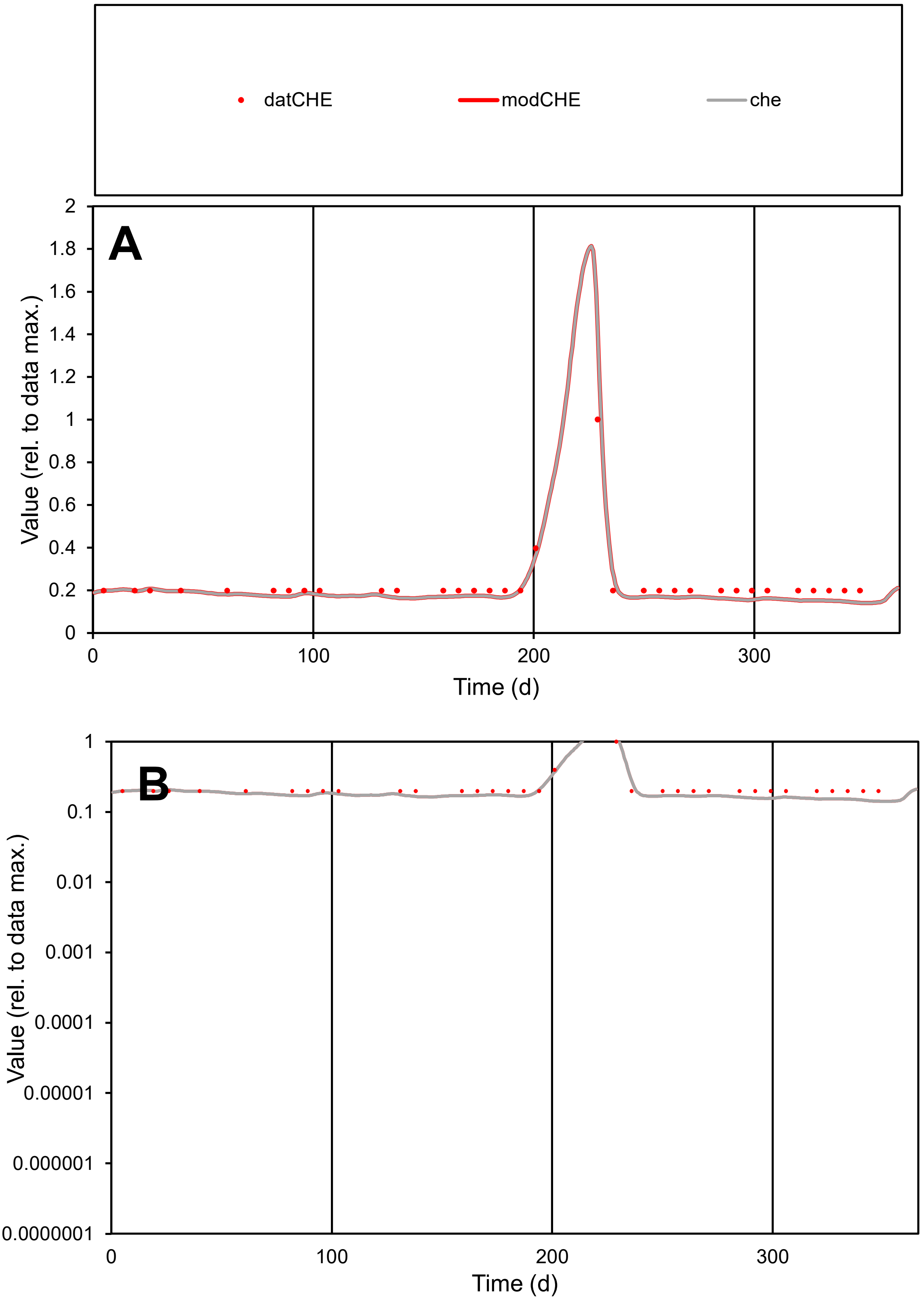


**C1**

Variables
CHE Provide jtd or Dst
CalibOnly?
2 1 to reproduce of
oNormalize?
2 1=no, 2=yes

ofxn calc*
0.13437
ofxn mod
0.1992

sRun
RunA1
zout
1

Ave
Data
8.1E-06
Model
7.3E-06

data/vMax
3.5E-05

ofxng
263.459

Min d/m
1.40681

ave/min d
1.15255
ave/min m
1.45649

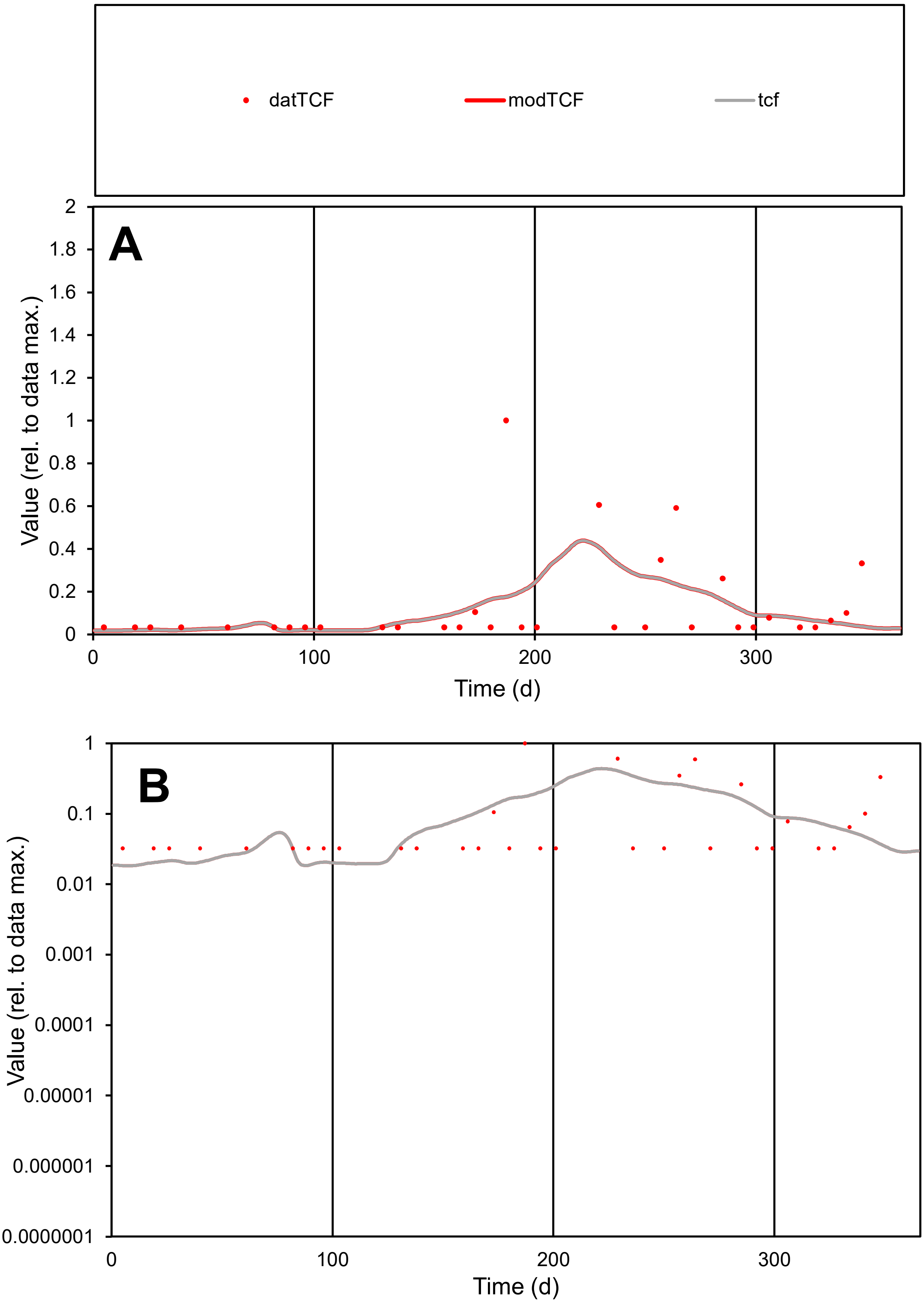


**C1**

Variables

| | |
|---|---|
| TCF | Provide jtd or Dst |
| CalibOnly? | |
| 2 | 1 to reproduce of |
| oNormalize? | |
| 2 | 1=no, 2=yes |

ofxn calc*
1.51669
ofxn mod
1.53286

sRun
RunA1
zout
1

Ave
Data
1.2E-05
Model
1.1E-05

data/vMax
9.6E-05

ofxng
263.459

Min d/m
1.76514

ave/min d
3.95464
ave/min m
6.3315

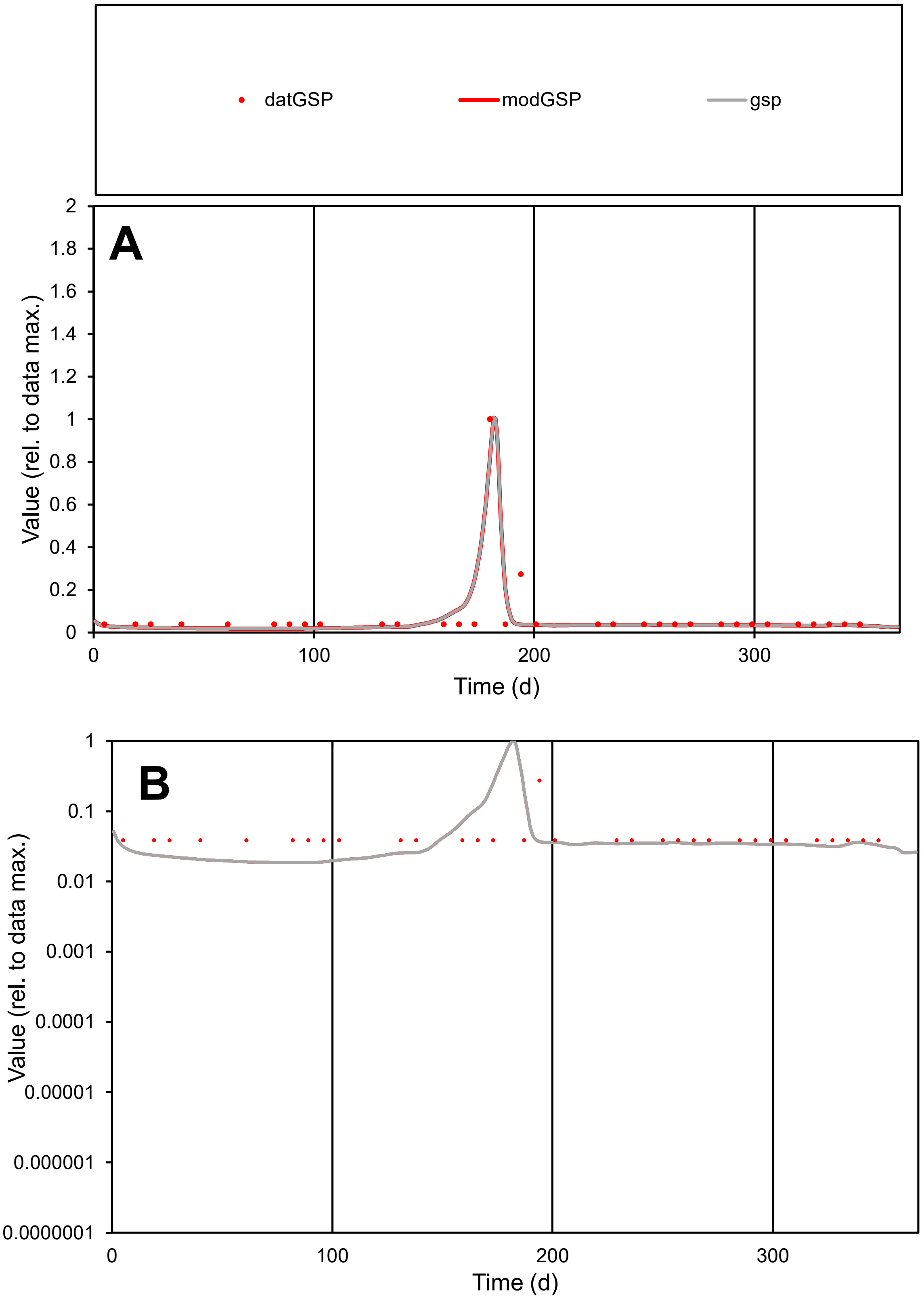


C1

Variables
GSP Provide jtd or Dst
CalibOnly?
2 1 to reproduce of
oNormalize?
2 1=no, 2=yes

ofxn calc*
0.72153
ofxn mod
0.79297

sRun
RunA1
zout
1

Ave
Data
1.7E-05
Model
1.6E-05

data/vMax
0.00023

ofxng
263.459

Min d/m
2.07352

ave/min d
1.94003
ave/min m
3.66244

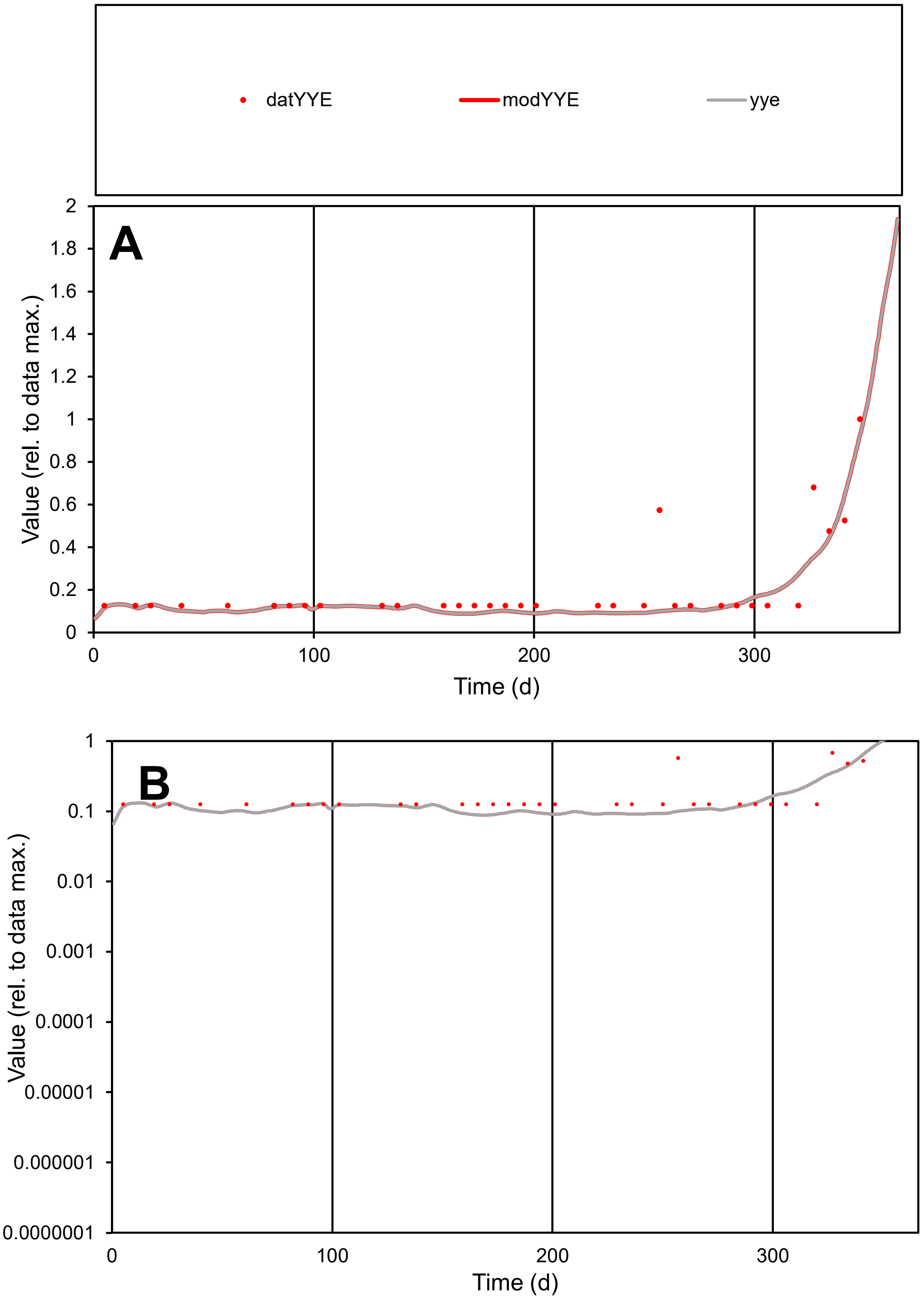


**C1**

Variables

YE Provide jtd or Dst

CalibOnly?

2 1 to reproduce of

oNormalize?

2 1=no, 2=yes

ofxn calc*

0.52255

ofxn mod

0.5457

sRun

RunA1

zout

1

Ave

Data

6.1E-06

Model

5.3E-06

data/vMax

3E-05

ofxng

263.459

Min d/m

1.427

ave/min d

1.63155

ave/min m

2.01387

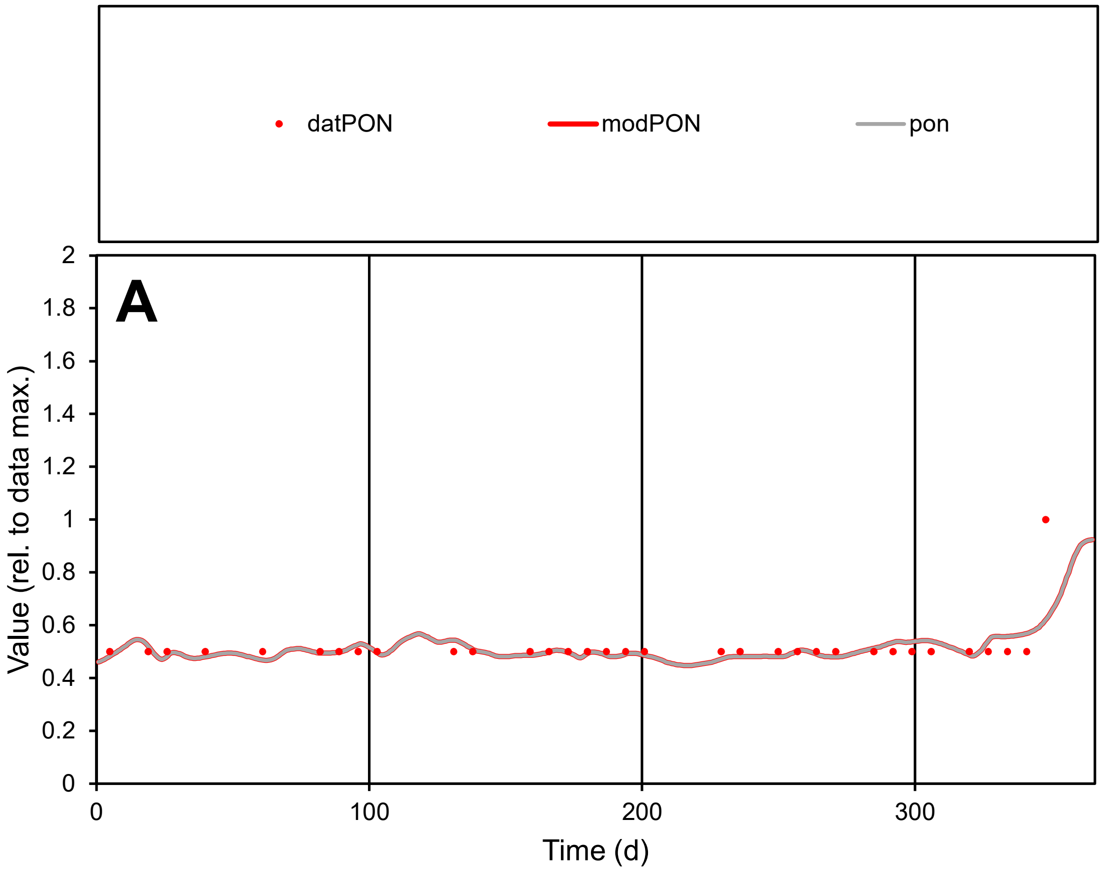


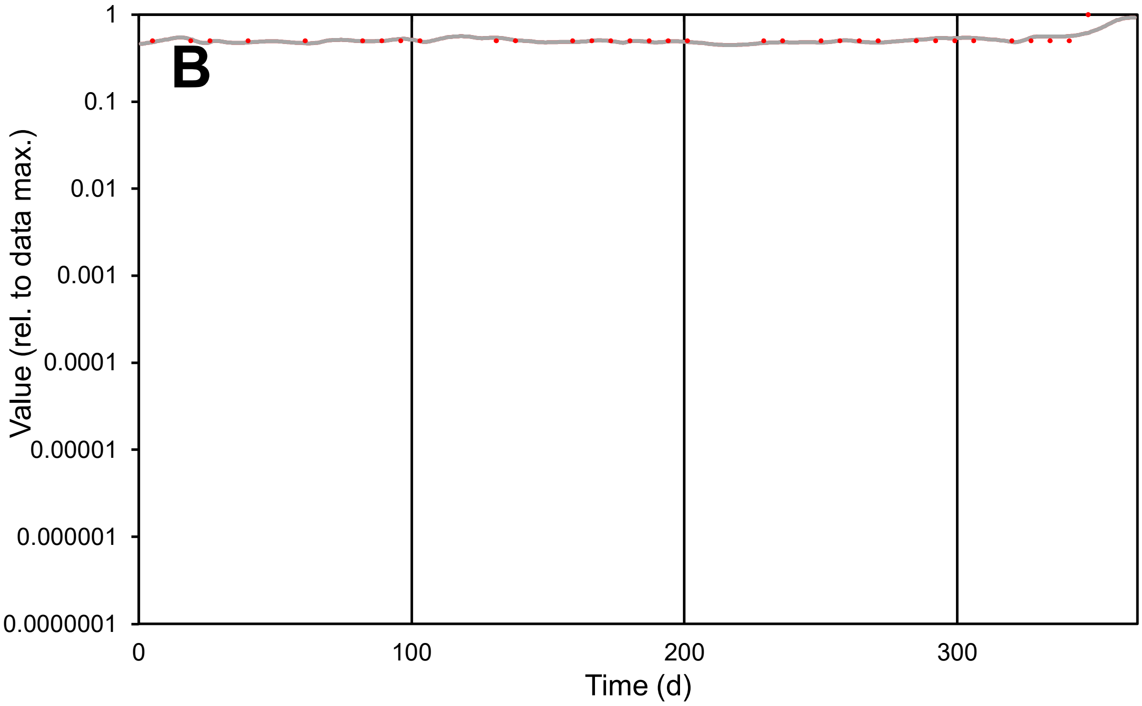


**C1**

Variables
PON Provide jtd or Dst
CalibOnly?
2 1 to reproduce of
oNormalize?
2 1=no, 2=yes

ofxn calc*
0.13352
ofxn mod
0.17383

sRun
RunA1
zout
1

Ave
Data
3E-06
Model
3E-06

data/vMax
5.9E-06

ofxng
263.459

Min d/m
1.07276

ave/min d
1.0303
ave/min m
1.09085

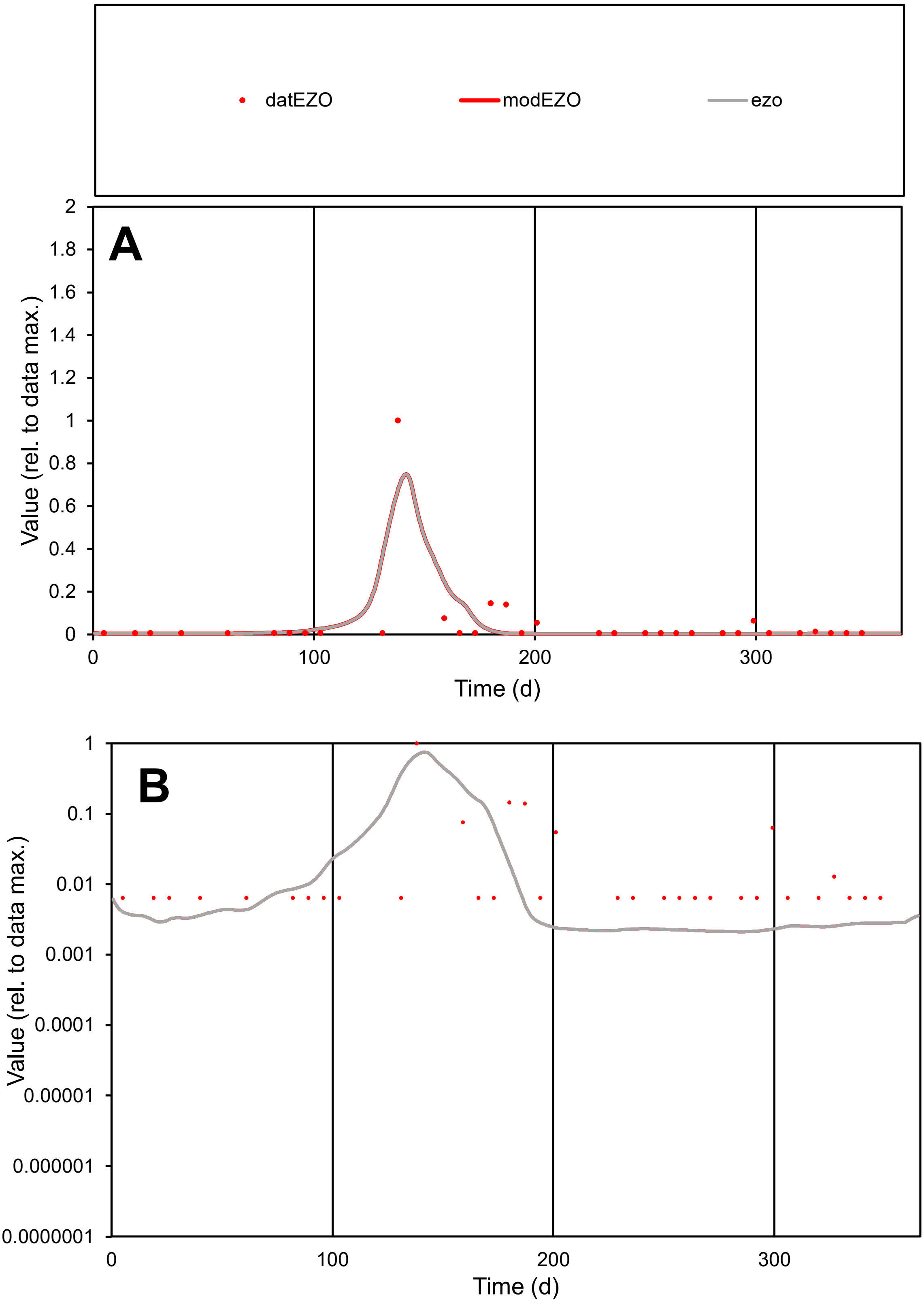


**C1**

Variables
EZO Provide jtd or Dst
CalibOnly?
2 1 to reproduce of
oNormalize?
2 1=no, 2=yes

ofxn calc*
1.5486
ofxn mod
1.816

sRun
RunA1
zout
1

Ave
Data
3.1E-05
Model
2.7E-05

data/vMax
0.00061

ofxng
263.459

Min d/m
3.04714

ave/min d
7.80666
ave/min m
21.0145

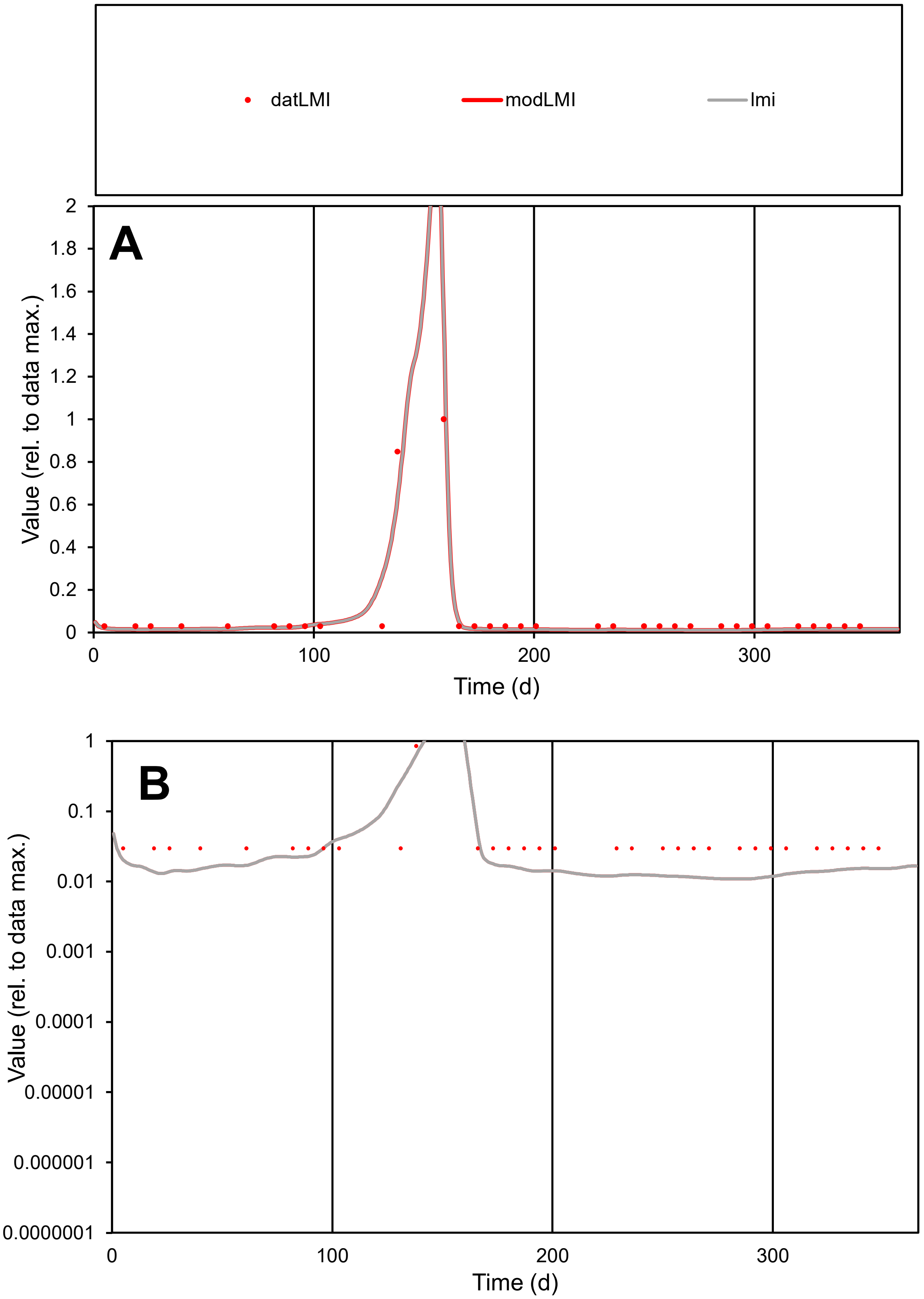


**C1**

Variables

LMI Provide jtd or Dst

CalibOnly?

2 1 to reproduce of

oNormalize?

2 1=no, 2=yes

ofxn calc*

0.48654

ofxn mod

0.56868

sRun

RunA1

zout

1

Ave

Data

8.5E-06

Model

7.7E-06

data/vMax

0.0001

ofxng

263.459

Min d/m

2.73909

ave/min d

2.82275

ave/min m

7.02444

326_Calib_S_A

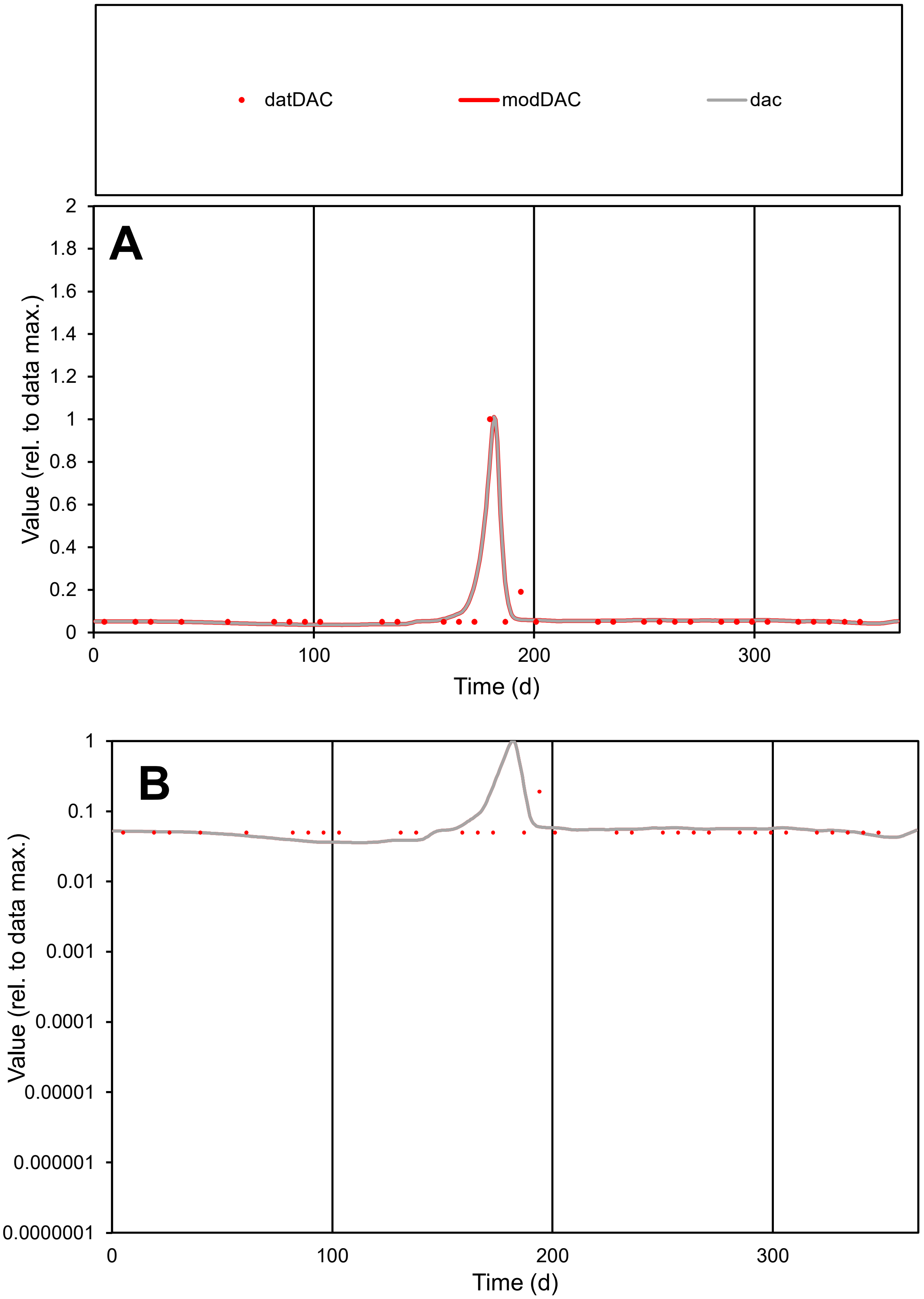


**C1**

Variables

DAC Provide jtd or Dst

CalibOnly?

2 1 to reproduce of

oNormalize?

2 1=no, 2=yes

ofxn calc*

0.46603

ofxn mod

0.52202

sRun

RunA1

zout

1

Ave

Data

1.3E-05

Model

1.3E-05

data/vMax

0.00015

ofxng

263.459

Min d/m

1.38398

ave/min d

1.66295

ave/min m

2.37849

326_Calib_S_A

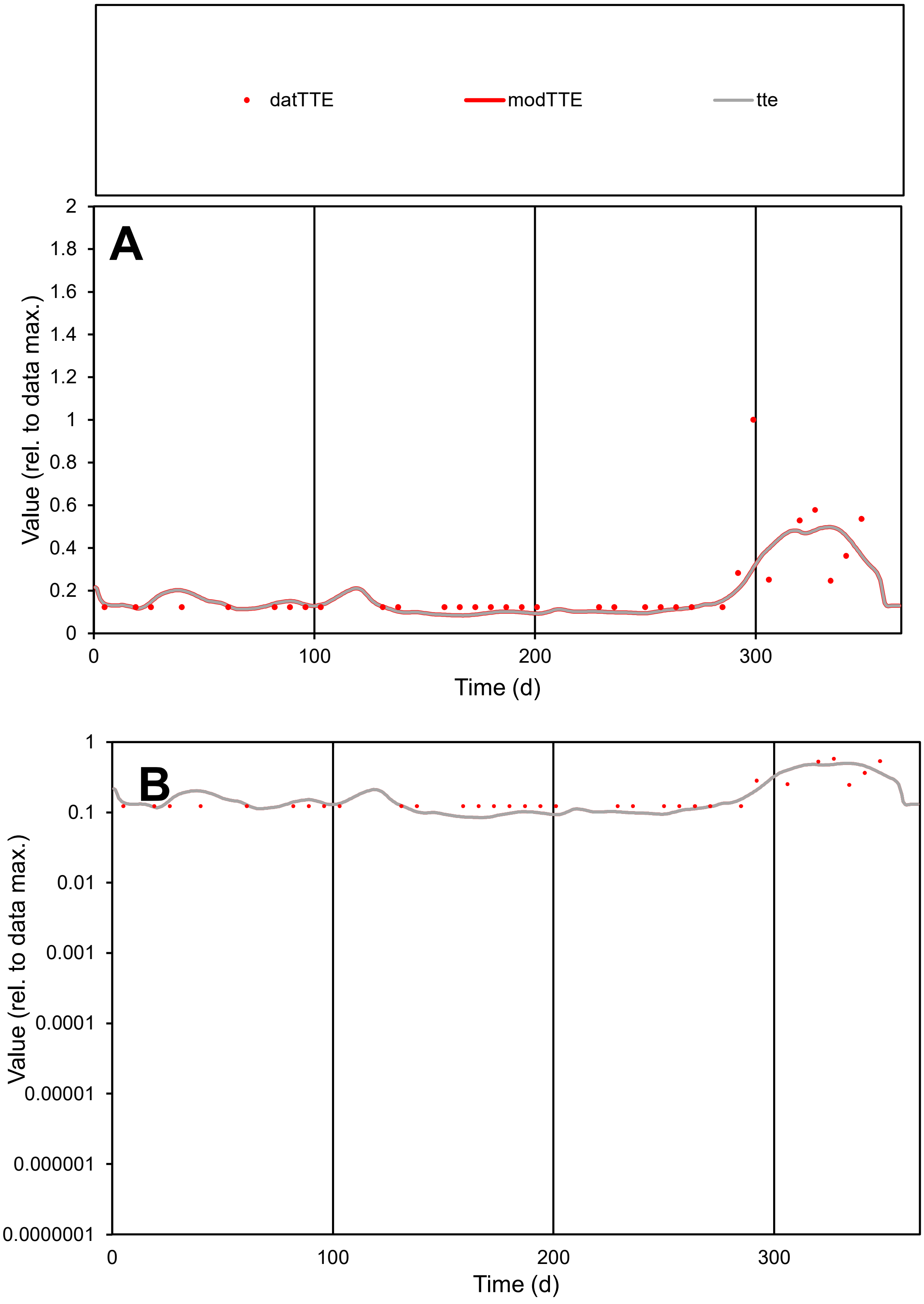


**C1**

Variables
TTE Provide jtd or Dst
CalibOnly?
2 1 to reproduce of
oNormalize?
2 1=no, 2=yes

ofxn calc*
0.65385
ofxn mod
0.66749

sRun
RunA1
zout
1

Ave
Data
6.4E-06
Model
5.7E-06

data/vMax
3.1E-05

ofxng
263.459

Min d/m
1.45945

ave/min d
1.69168
ave/min m
2.21507

326_Calib_S_A

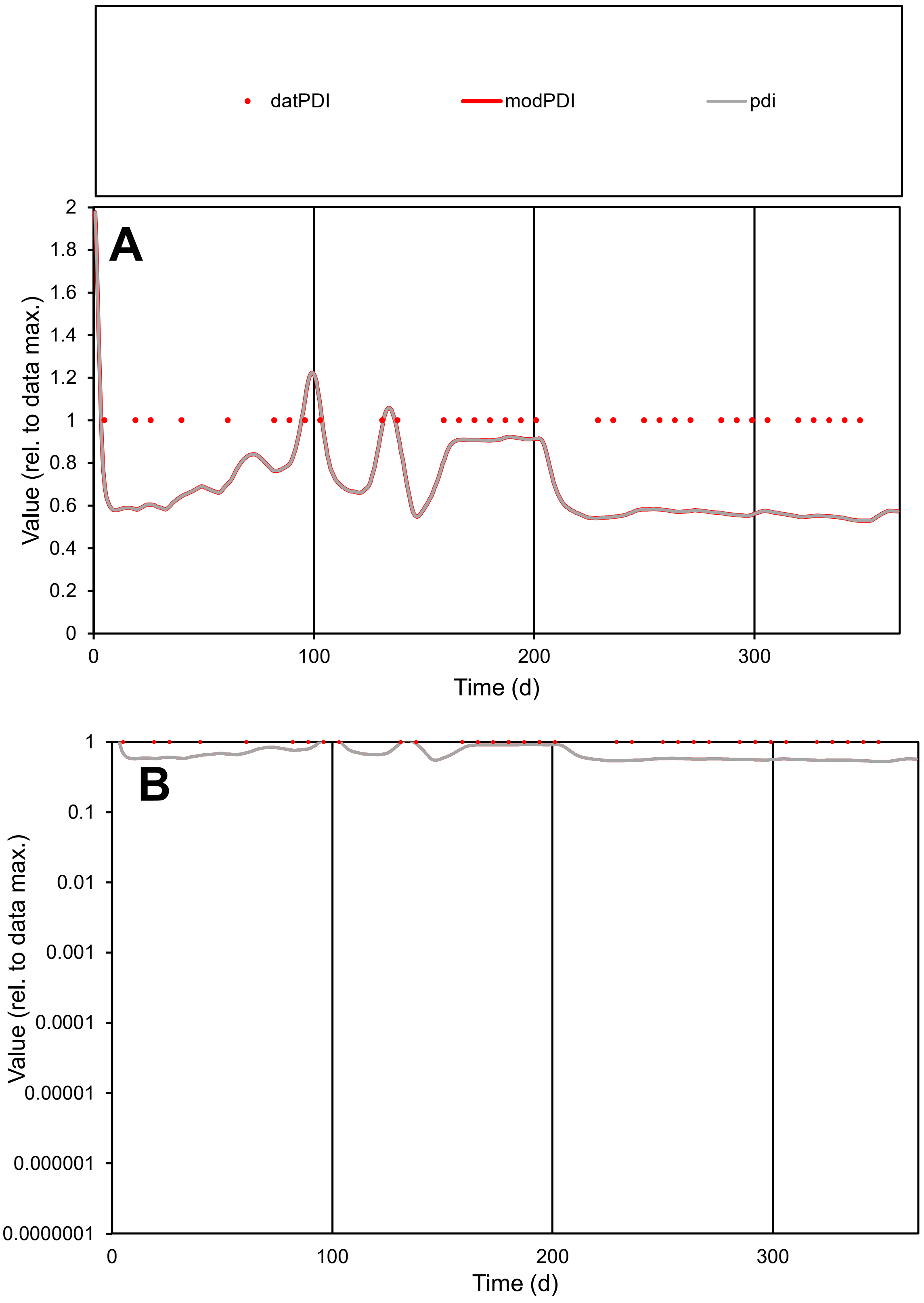


**C1**

Variables

PDI Provide jtd or Dst

CalibOnly?

2 1 to reproduce of

oNormalize?

2 1=no, 2=yes

ofxn calc*

0.33833

ofxn mod

0.35297

sRun

RunA1

zout

1

Ave

Data

4.4E-06

Model

3.1E-06

data/vMax

4.4E-06

ofxng

263.459

Min d/m

1.8867

ave/min d

1

ave/min m

1.3472

326_Calib_S_A

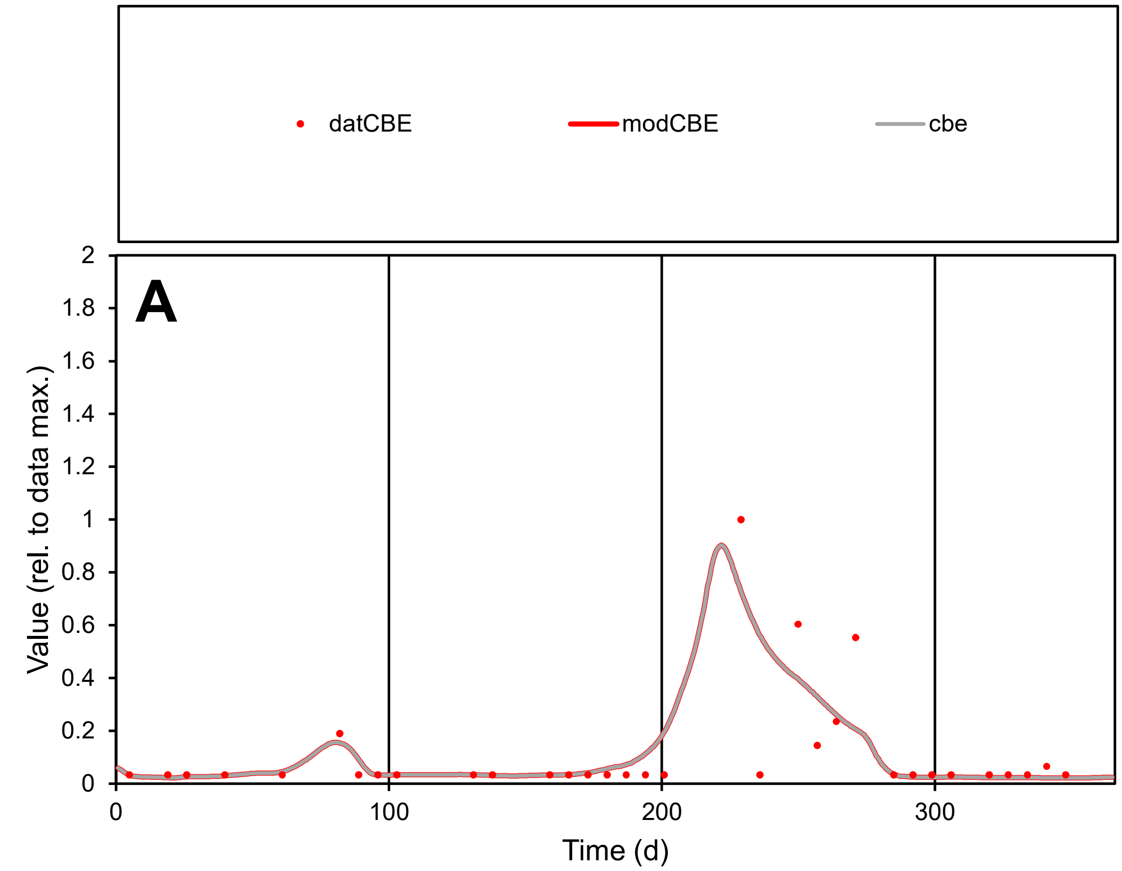


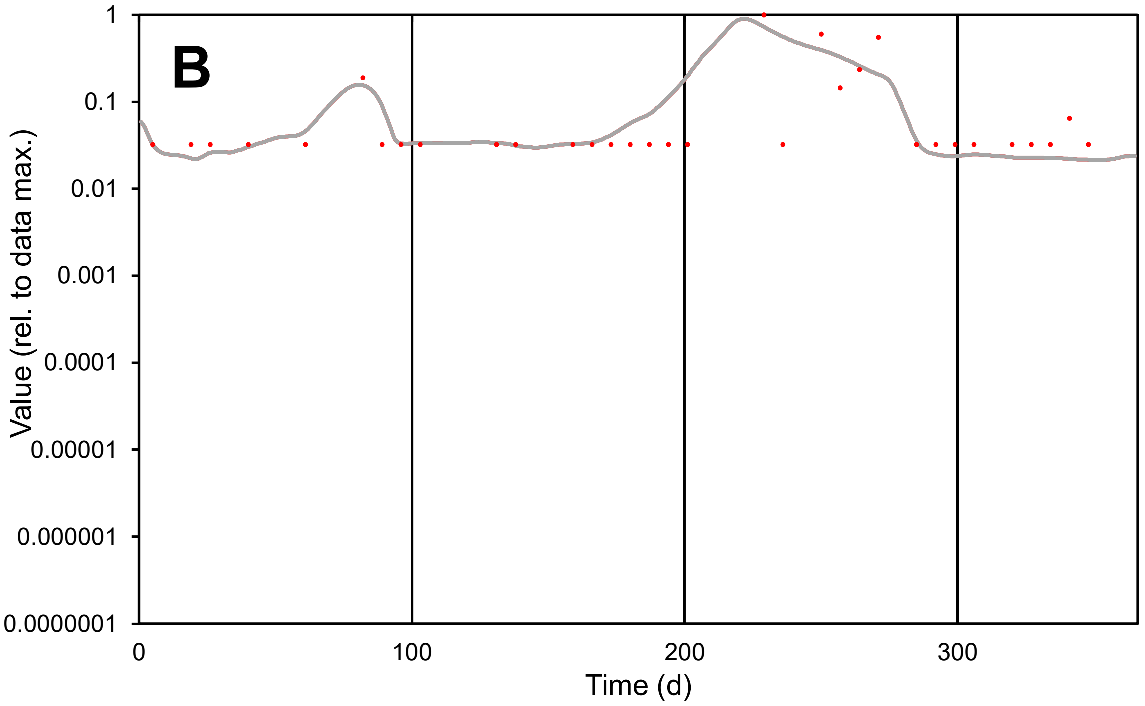


**C1**

Variables
CBE Provide jtd or Dst
CalibOnly?
2 1 to reproduce of
oNormalize?
2 1=no, 2=yes

ofxn calc*
1.18355
ofxn mod
1.24042

sRun
RunA1
zout
1

Ave
Data
5.7E-06
Model
5.6E-06

data/vMax
5.1E-05

ofxng
263.459

Min d/m
1.51499

ave/min d
3.38787
ave/min m
5.06961

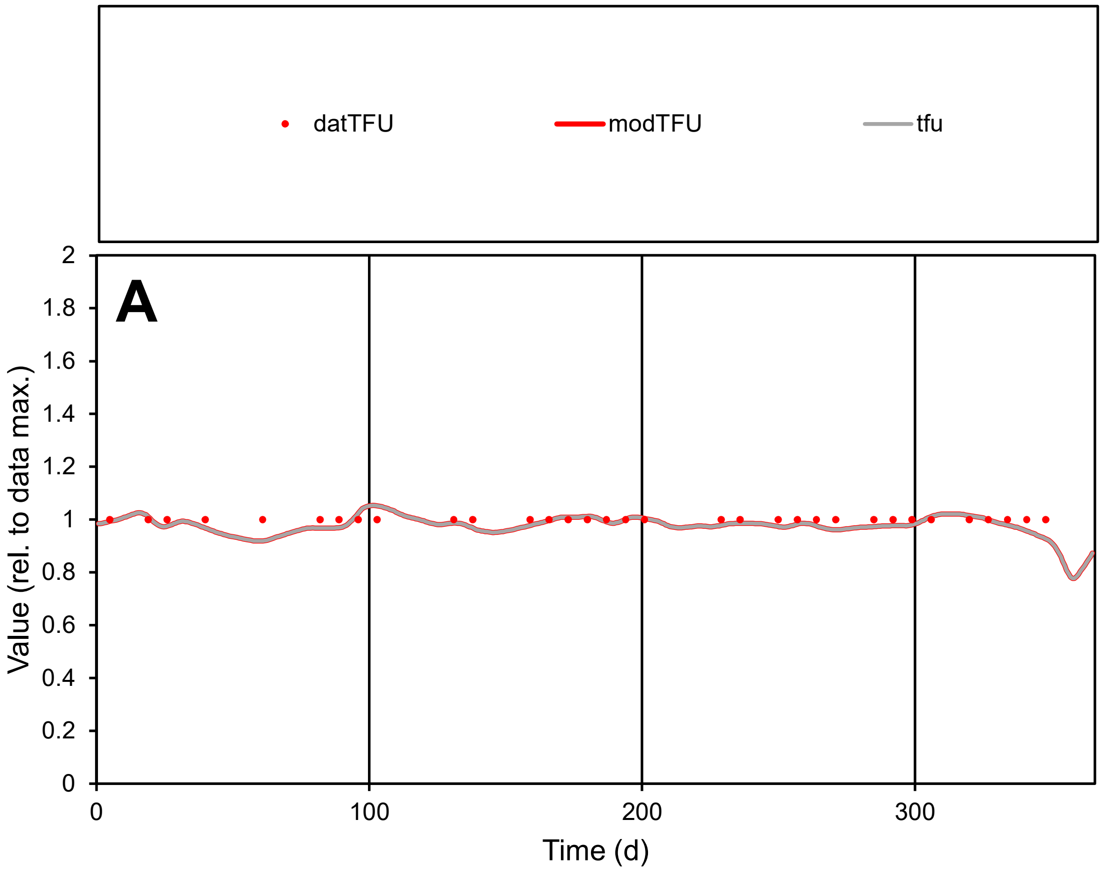


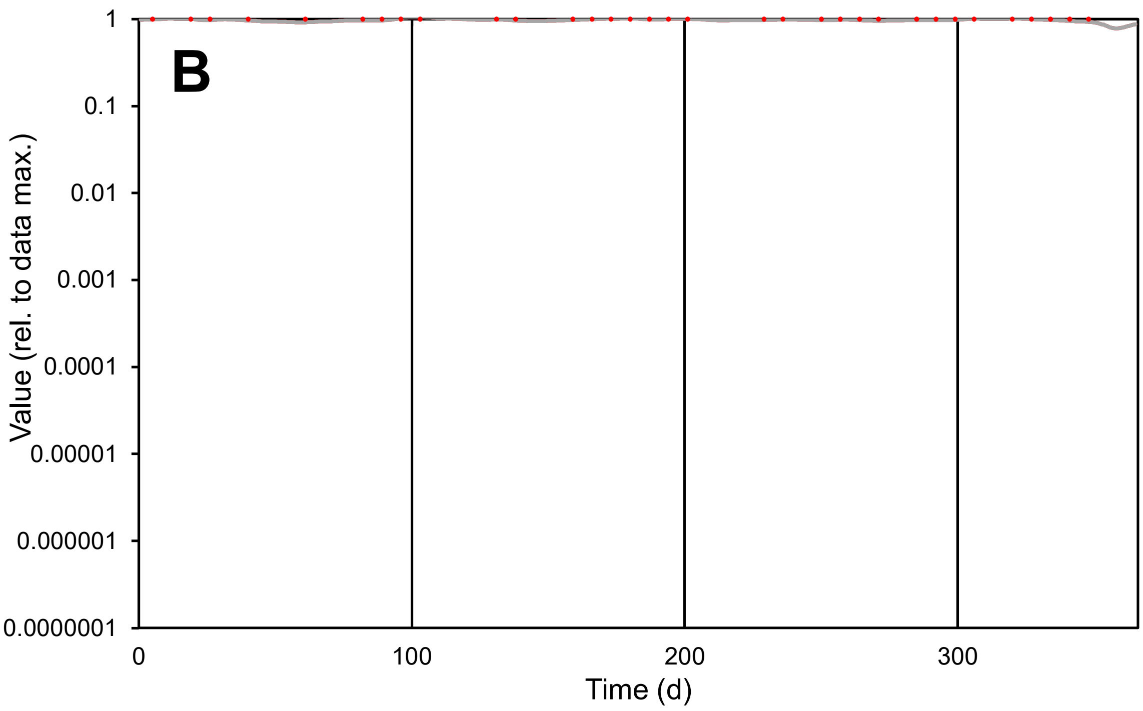


**C1**

Variables

TFU Provide jtd or Dst

CalibOnly?

2 1 to reproduce of

oNormalize?

2 1=no, 2=yes

ofxn calc*

0.03072

ofxn mod

0.11634

sRun

RunA1

zout

1

Ave

Data

1.3E-05

Model

1.3E-05

data/vMax

1.3E-05

ofxng

263.459

Min d/m

1.08603

ave/min d

1

ave/min m

1.06974

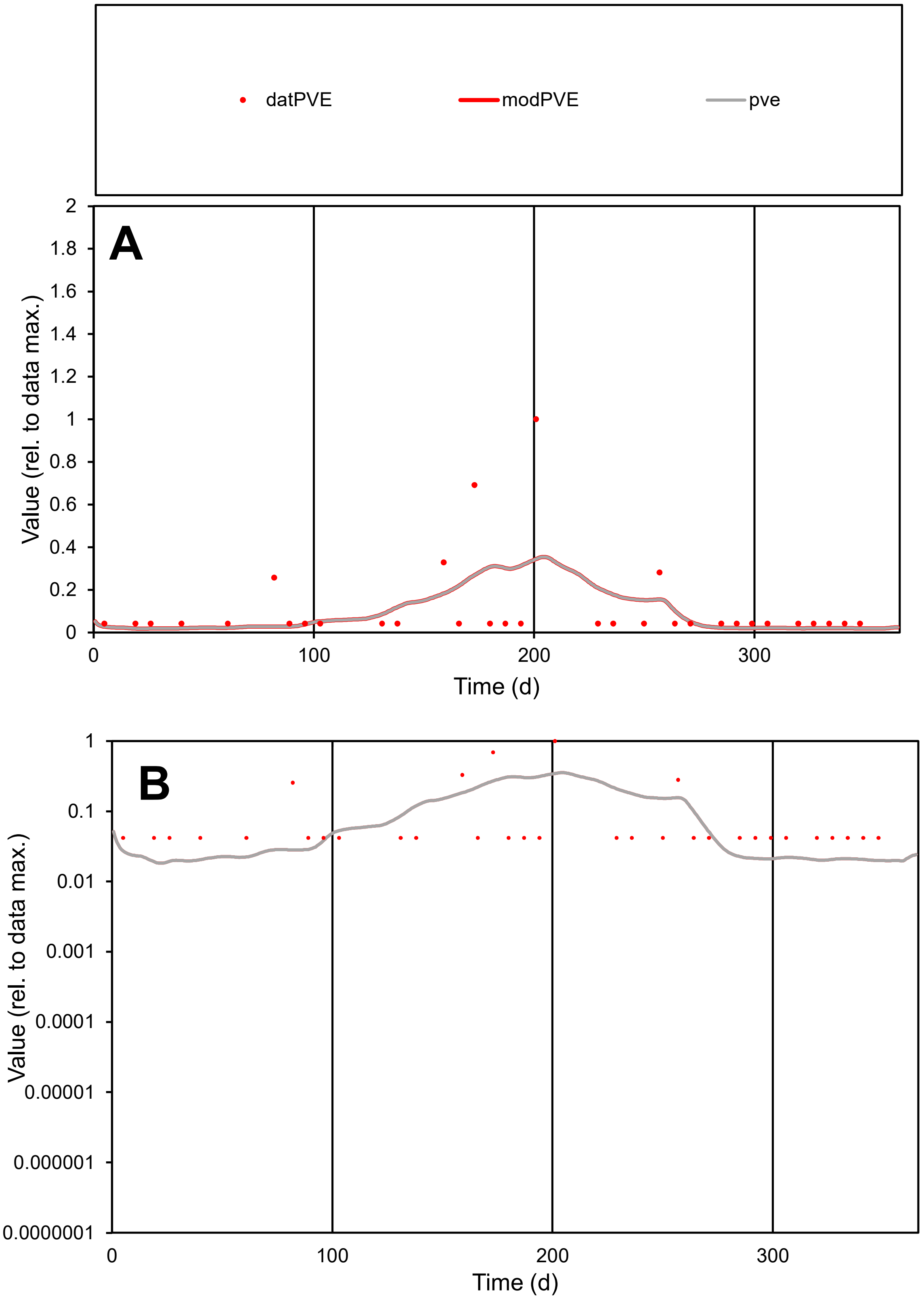


**C1**

Variables
PVE Provide jtd or Dst
CalibOnly?
2 1 to reproduce of
oNormalize?
2 1=no, 2=yes

ofxn calc*
1.54088
ofxn mod
1.55126

sRun
RunA1
zout
1

Ave
Data
5.7E-06
Model
5.2E-06

data/vMax
5E-05

ofxng
263.459

Min d/m
2.21775

ave/min d
2.70751
ave/min m
5.54221

326_Calib_S_A

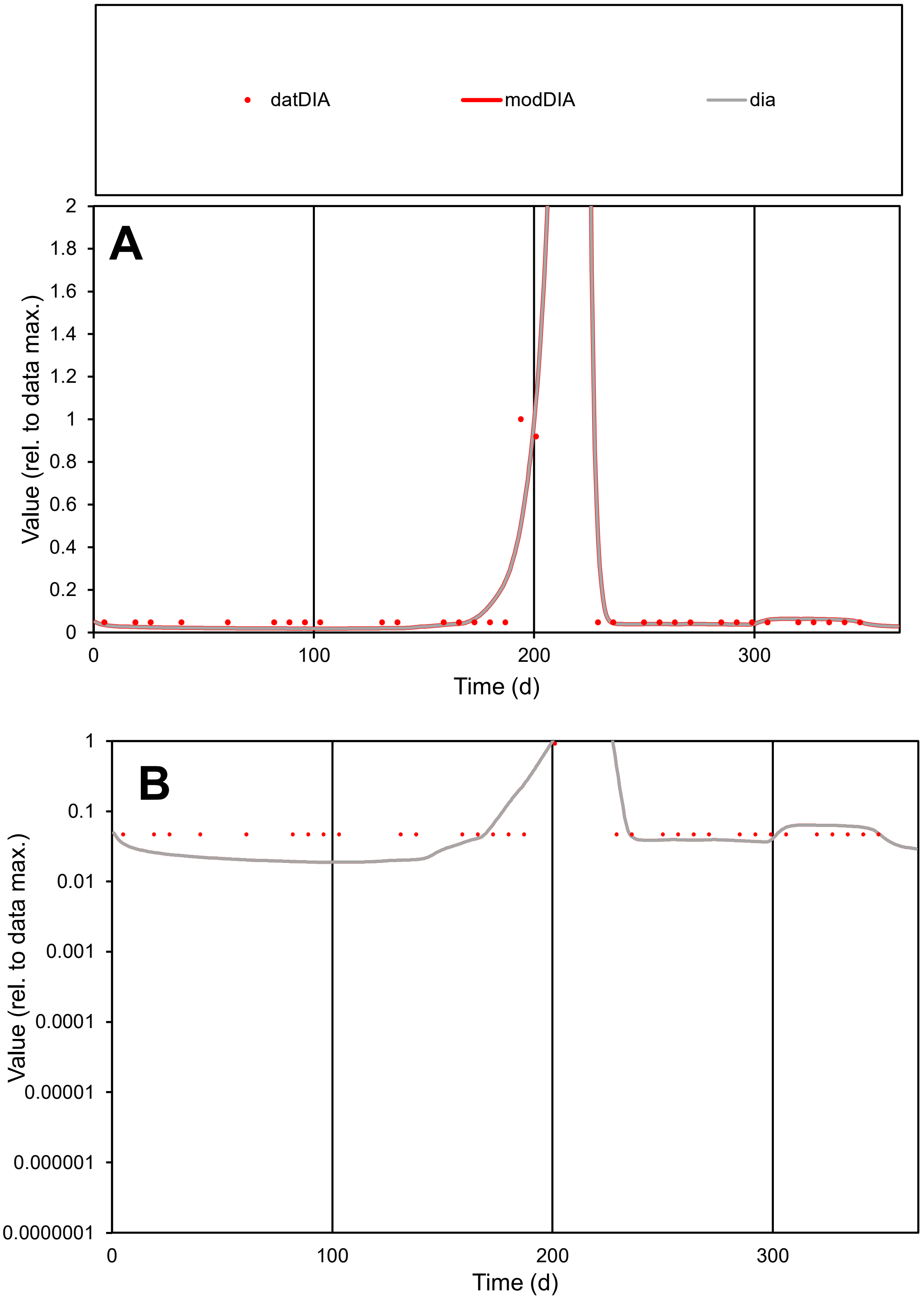


**C1**

Variables
DIA Provide jtd or Dst
CalibOnly?
2 1 to reproduce of
oNormalize?
2 1=no, 2=yes

ofxn calc*
0.88717
ofxn mod
0.97116

sRun
RunA1
zout
1

Ave
Data
1.4E-05
Model
1.3E-05

data/vMax
0.00013

ofxng
263.459

Min d/m
2.48088

ave/min d
2.18227
ave/min m
5.36667

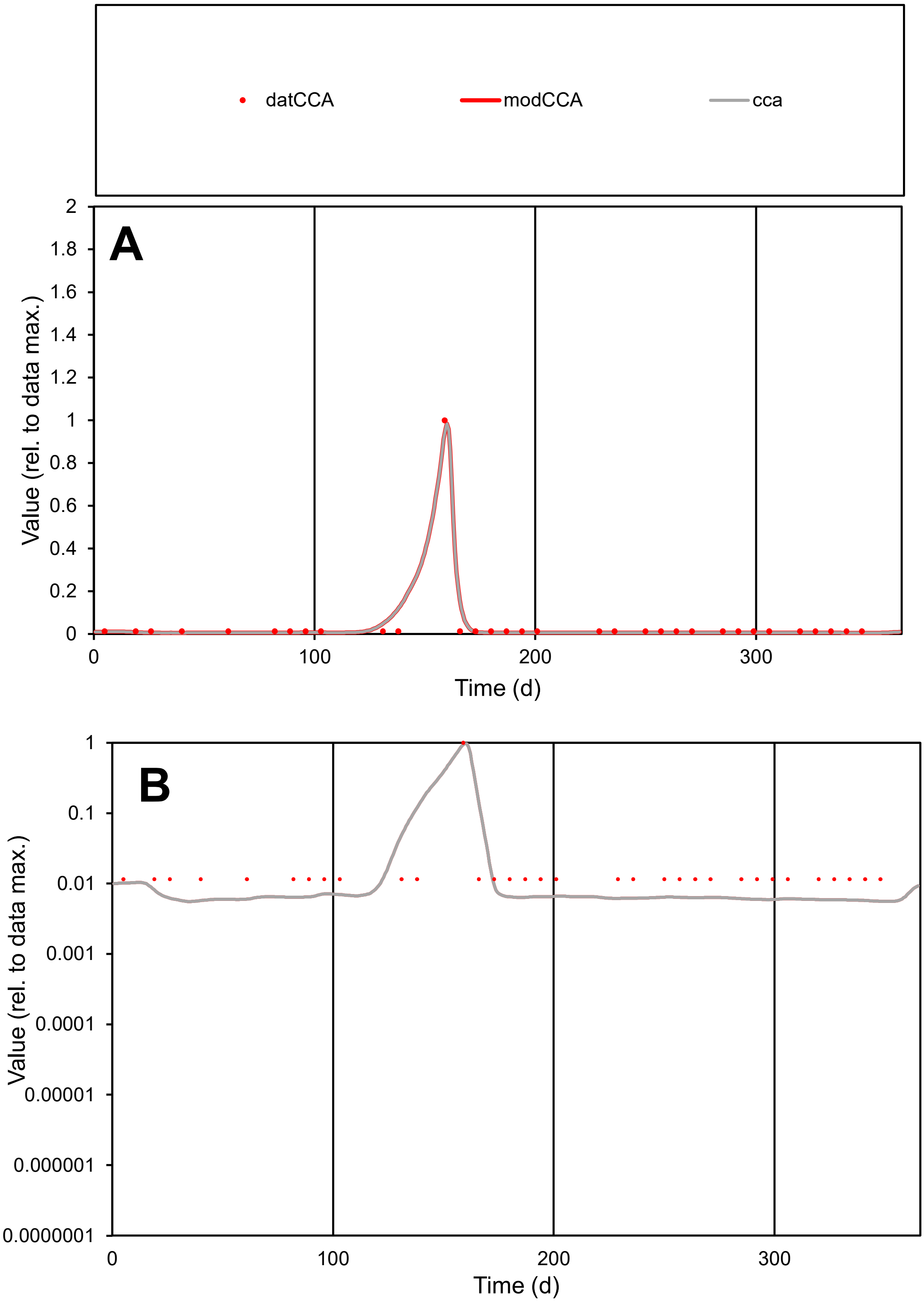


**C1**

Variables
CCA Provide jtd or Dst
CalibOnly?
2 1 to reproduce of
oNormalize?
2 1=no, 2=yes

ofxn calc*
0.40693
ofxn mod
0.55332

sRun
RunA1
zout
1

Ave
Data
6.6E-06
Model
6.5E-06

data/vMax
0.00016

ofxng
263.459

Min d/m
2.05232

ave/min d
3.60546
ave/min m
7.28636

326_Calib_S_A

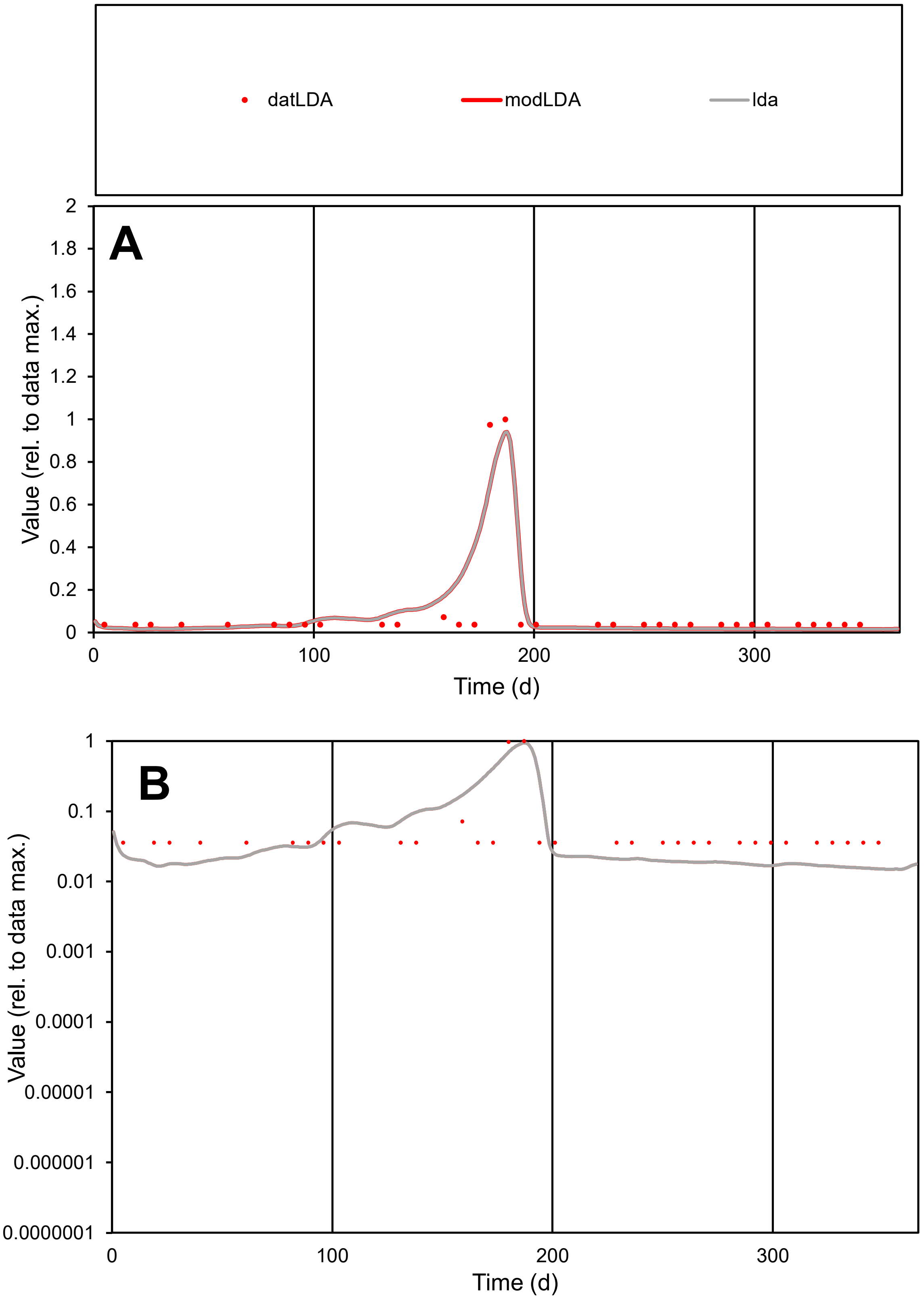


**C1**

Variables
LDA Provide jtd or Dst
CalibOnly?
2 1 to reproduce of
oNormalize?
2 1=no, 2=yes

ofxn calc*
0.92202
ofxn mod
0.99715

sRun
RunA1
zout
1

Ave
Data
1.8E-05
Model
1.9E-05

data/vMax
0.00019

ofxng
263.459

Min d/m
2.38393

ave/min d
2.63853
ave/min m
6.6581

326_Calib_S_A

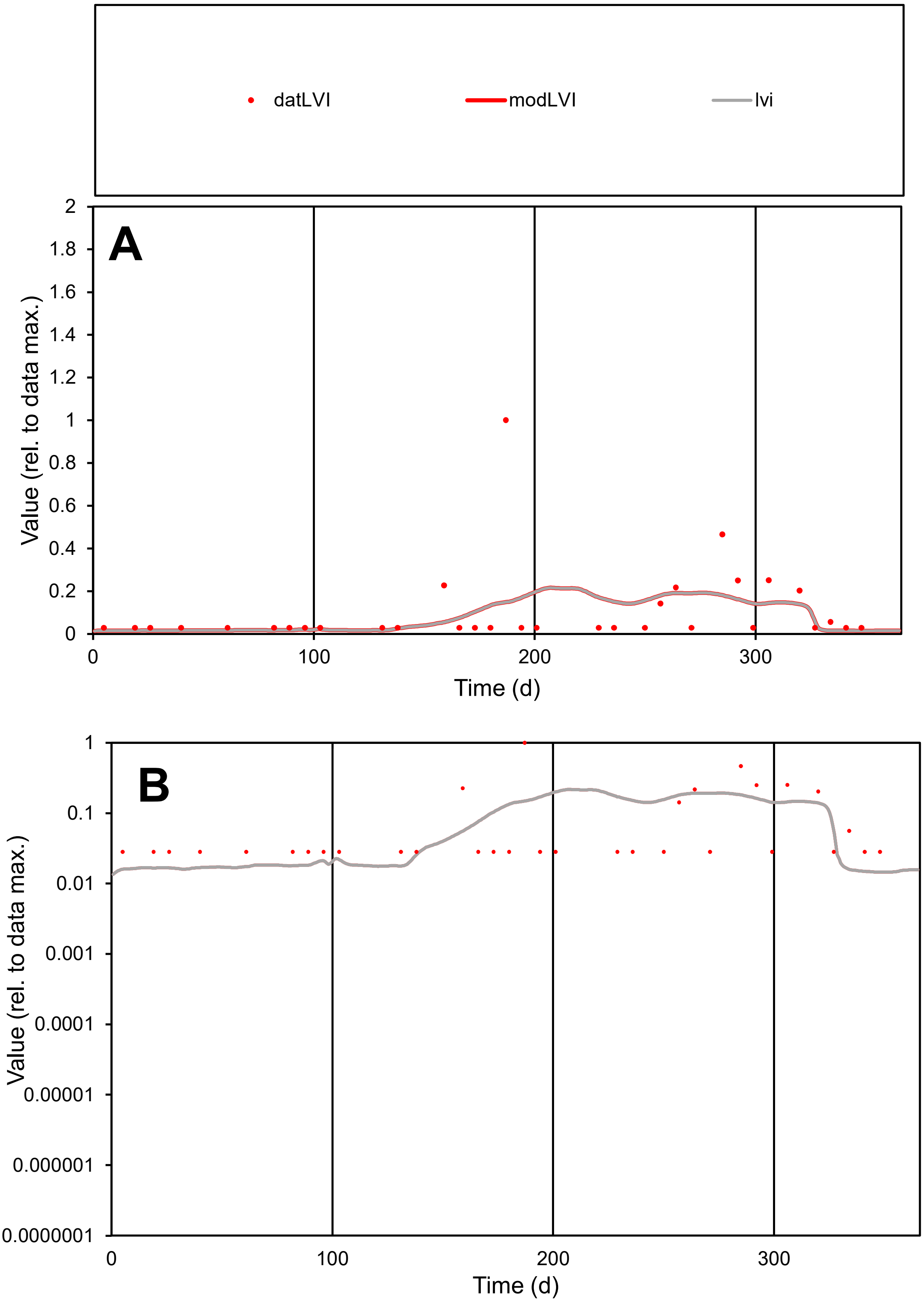


**C1**

Variables
LVI Provide jtd or Dst
CalibOnly?
2 1 to reproduce of
oNormalize?
2 1=no, 2=yes

ofxn calc*
1.66483
ofxn mod
1.67442

sRun
RunA1
zout
1

Ave
Data
5.3E-06
Model
4.6E-06

data/vMax
5.1E-05

ofxng
263.459

Min d/m
1.93285

ave/min d
3.76005
ave/min m
6.22202

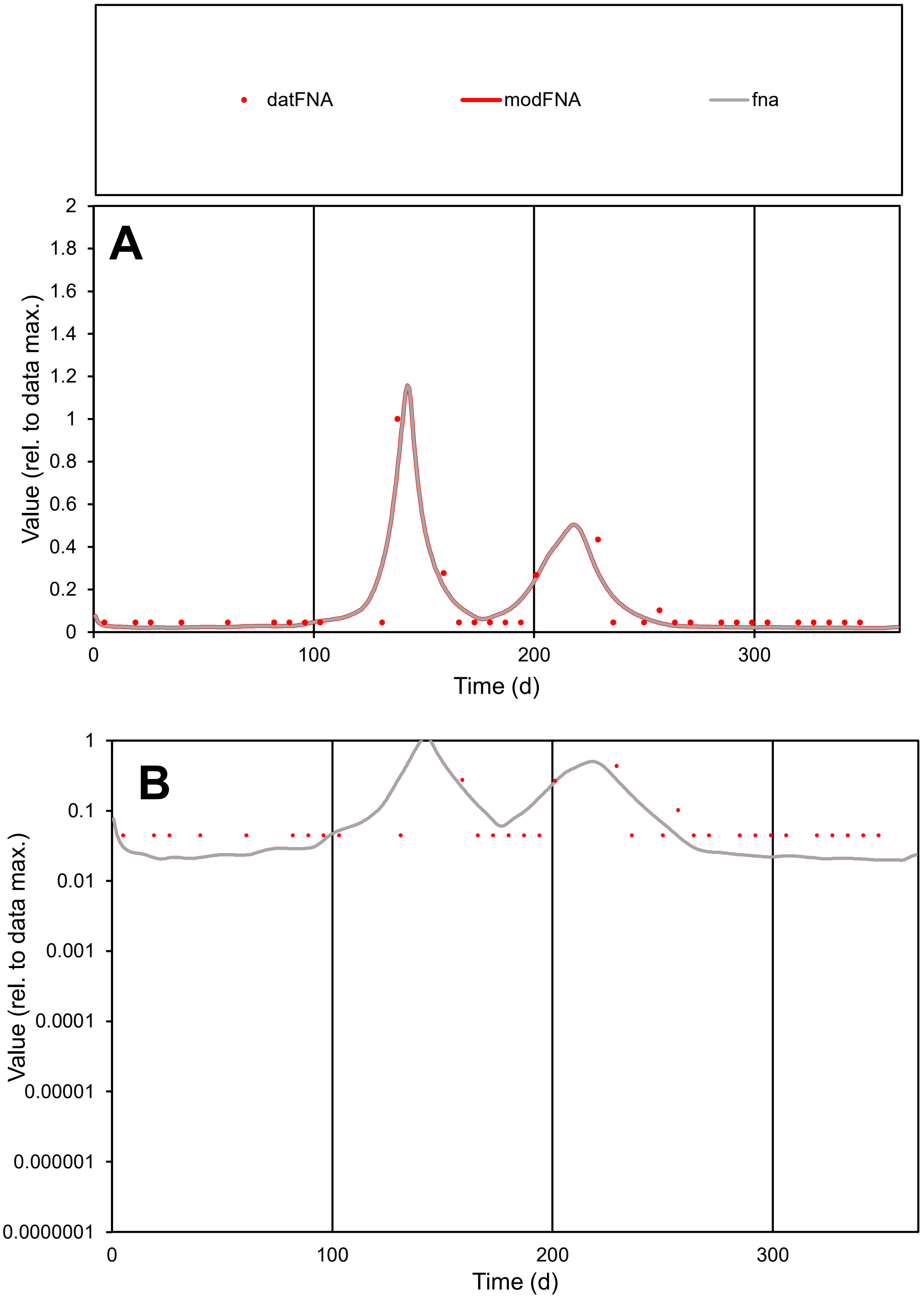


**C1**

Variables
FNA Provide jtd or Dst
CalibOnly?
2 1 to reproduce of
oNormalize?
2 1=no, 2=yes

ofxn calc*
0.6348
ofxn mod
0.71489

sRun
RunA1
zout
1

Ave
Data
7E-06
Model
6.8E-06

data/vMax
7E-05

ofxng
263.459

Min d/m
2.24317

ave/min d
2.26026
ave/min m
4.9104

326_Calib_S_A

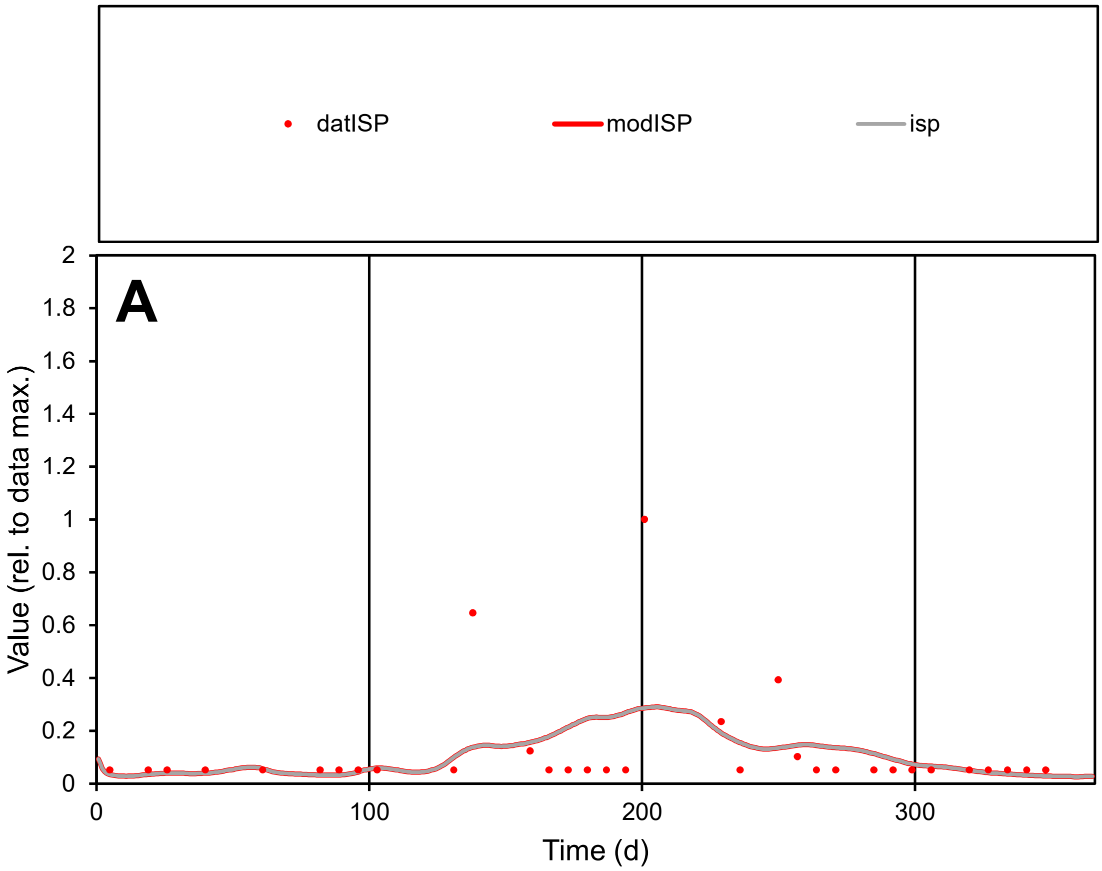


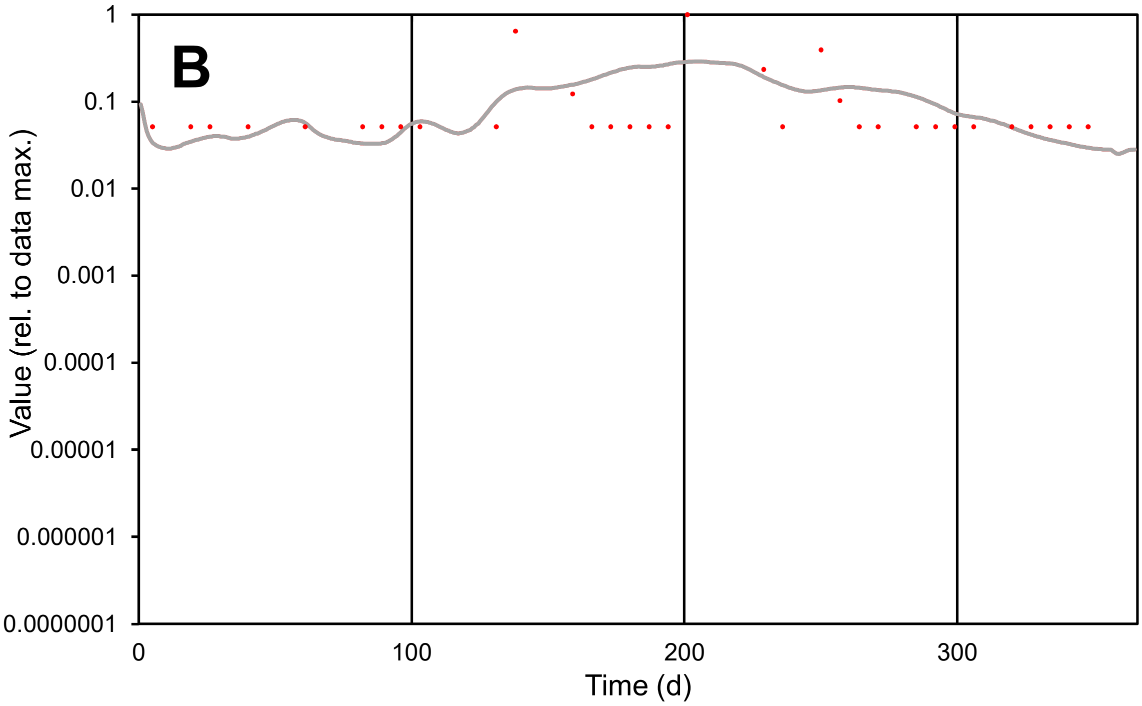


**C1**

Variables
ISP Provide jtd or Dst
CalibOnly?
2 1 to reproduce of
oNormalize?
2 1=no, 2=yes

ofxn calc*
1.50508
ofxn mod
1.51538

sRun
RunA1
zout
1

Ave
Data
1.2E-05
Model
1.1E-05

data/vMax
9.8E-05

ofxng
263.459

Min d/m
1.76232

ave/min d
2.29198
ave/min m
3.79449

326_Calib_S_A

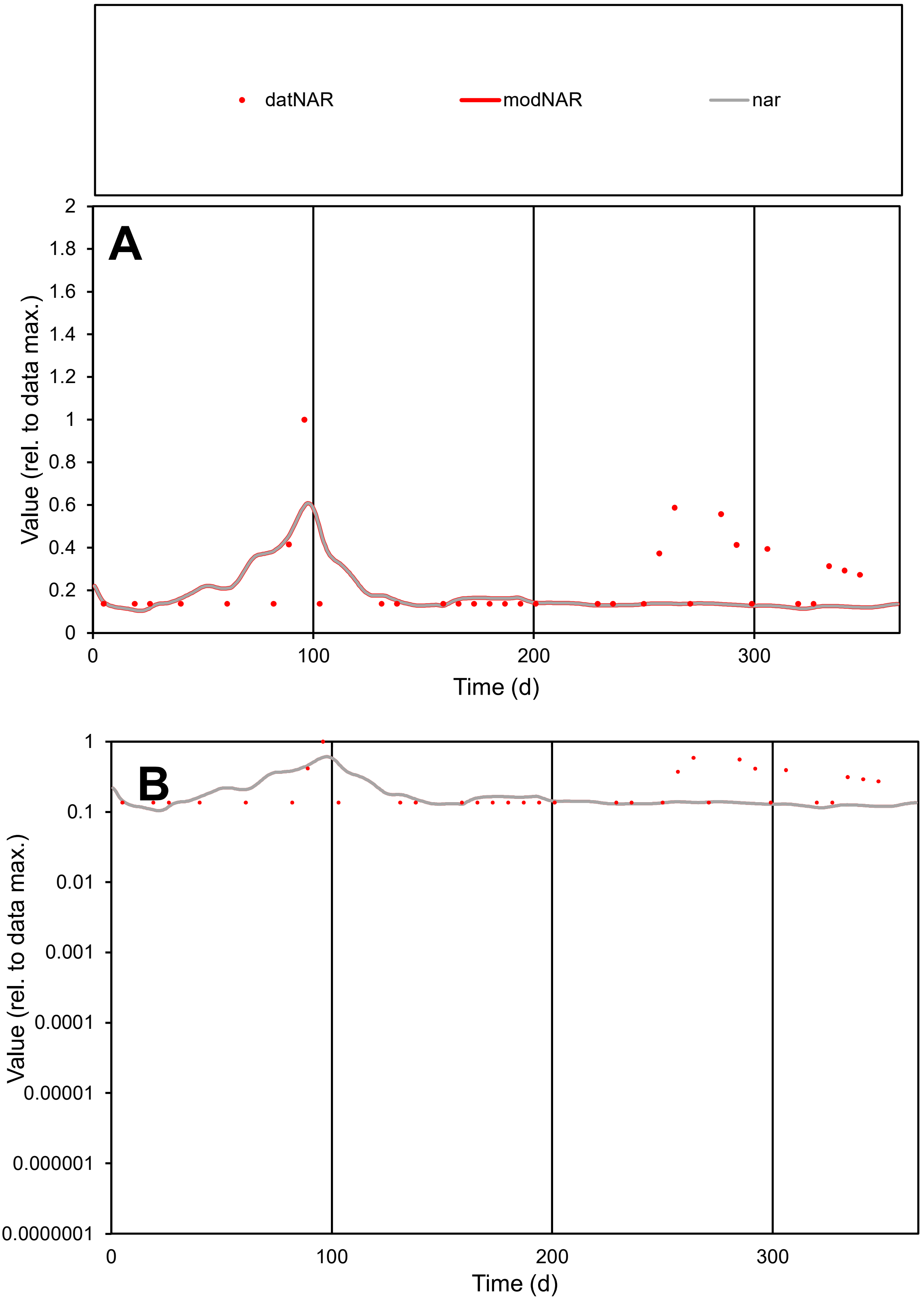


**C1**

| | |
|---|---|
| Variables | |
| NAR | Provide jtd or Dst |
| CalibOnly? | |
| 2 | 1 to reproduce of |
| oNormalize? | |
| 2 | 1=no, 2=yes |
| ofxn calc* | |
| 0.73321 | |
| ofxn mod | |
| 0.74736 | |
| sRun | |
| RunA1 | |
| zout | |
| 1 | |
| Ave Data | |
| 4.9E-06 | |
| Model | |
| 3.7E-06 | |
| data/vMax | |
| 2.1E-05 | |
| ofxng | |
| 263.459 | |
| Min d/m | |
| 1.30005 | |
| ave/min d | |
| 1.72364 | |
| ave/min m | |
| 1.72518 | |

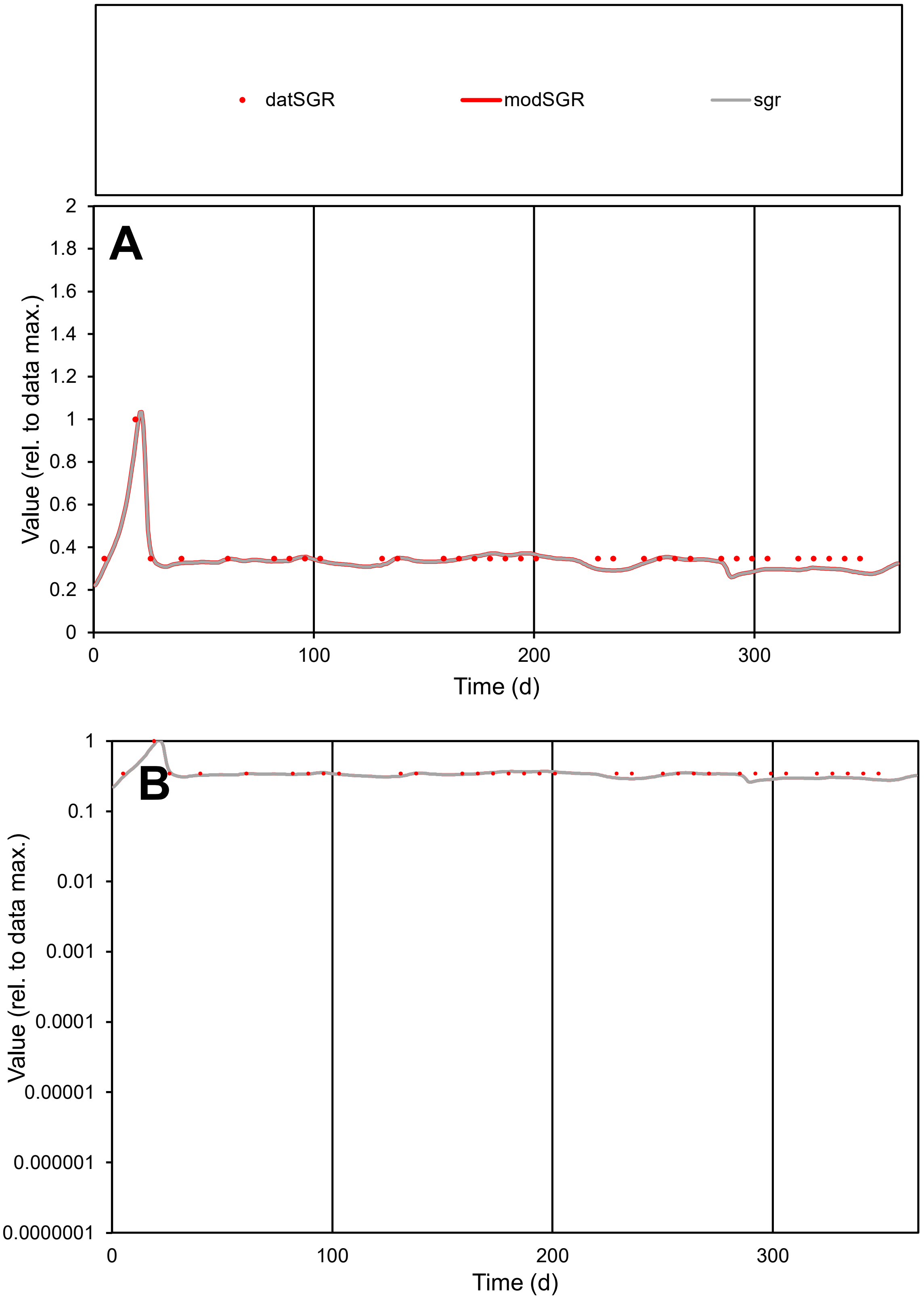


**C1**

Variables
SGR Provide jtd or Dst
CalibOnly?
2 1 to reproduce of
oNormalize?
2 1=no, 2=yes

ofxn calc*
0.09099
ofxn mod
0.14942

sRun
RunA1
zout
1

Ave
Data
4.8E-06
Model
4.5E-06

data/vMax
1.3E-05

ofxng
263.459

Min d/m
1.27173

ave/min d
1.0571
ave/min m
1.27592

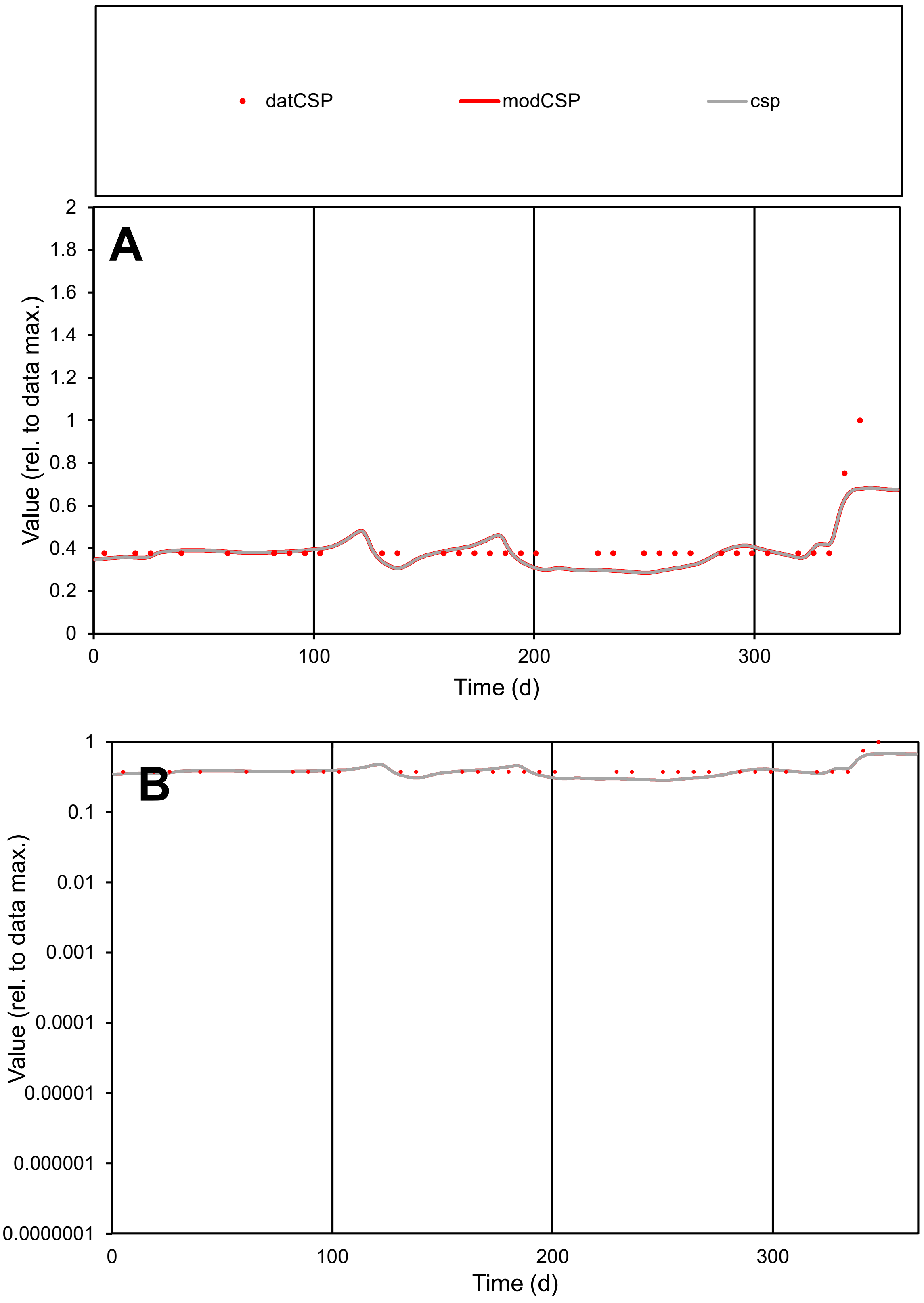


**C1**

Variables
CSP Provide jtd or Dst
CalibOnly?
2 1 to reproduce of
oNormalize?
2 1=no, 2=yes

ofxn calc*
0.18212
ofxn mod
0.21239

sRun
RunA1
zout
1

Ave
Data
2.6E-06
Model
2.4E-06

data/vMax
6.4E-06

ofxng
263.459

Min d/m
1.31588

ave/min d
1.0807
ave/min m
1.34556

326_Calib_S_A

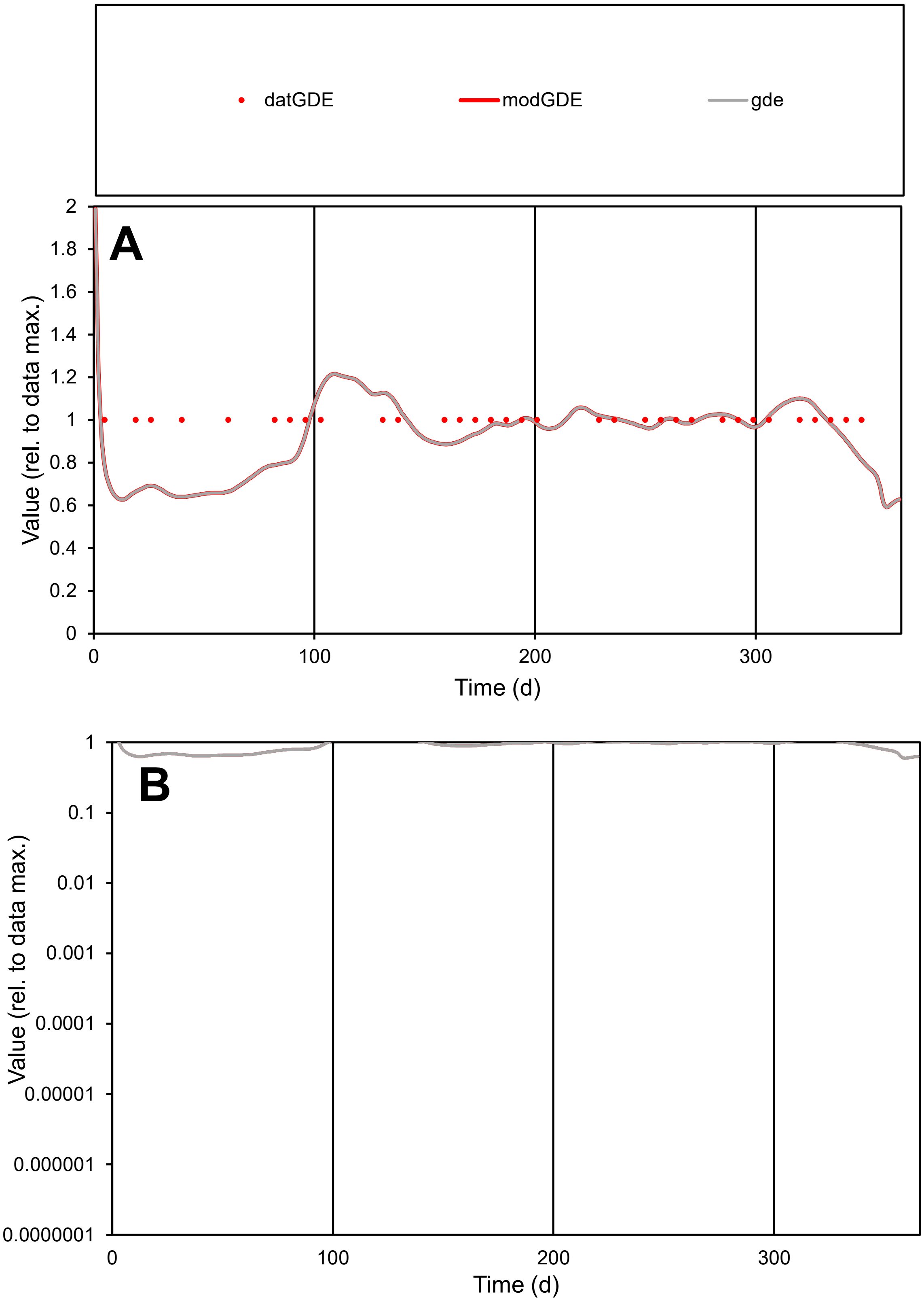


**C1**

Variables

GDE Provide jtd or Dst

CalibOnly?

2 1 to reproduce of

oNormalize?

2 1=no, 2=yes

ofxn calc*

0.14736

ofxn mod

0.18539

sRun

RunA1

zout

1

Ave

Data

2.7E-05

Model

2.6E-05

data/vMax

2.7E-05

ofxng

263.459

Min d/m

1.56048

ave/min d

1

ave/min m

1.45849

326_Calib_S_A

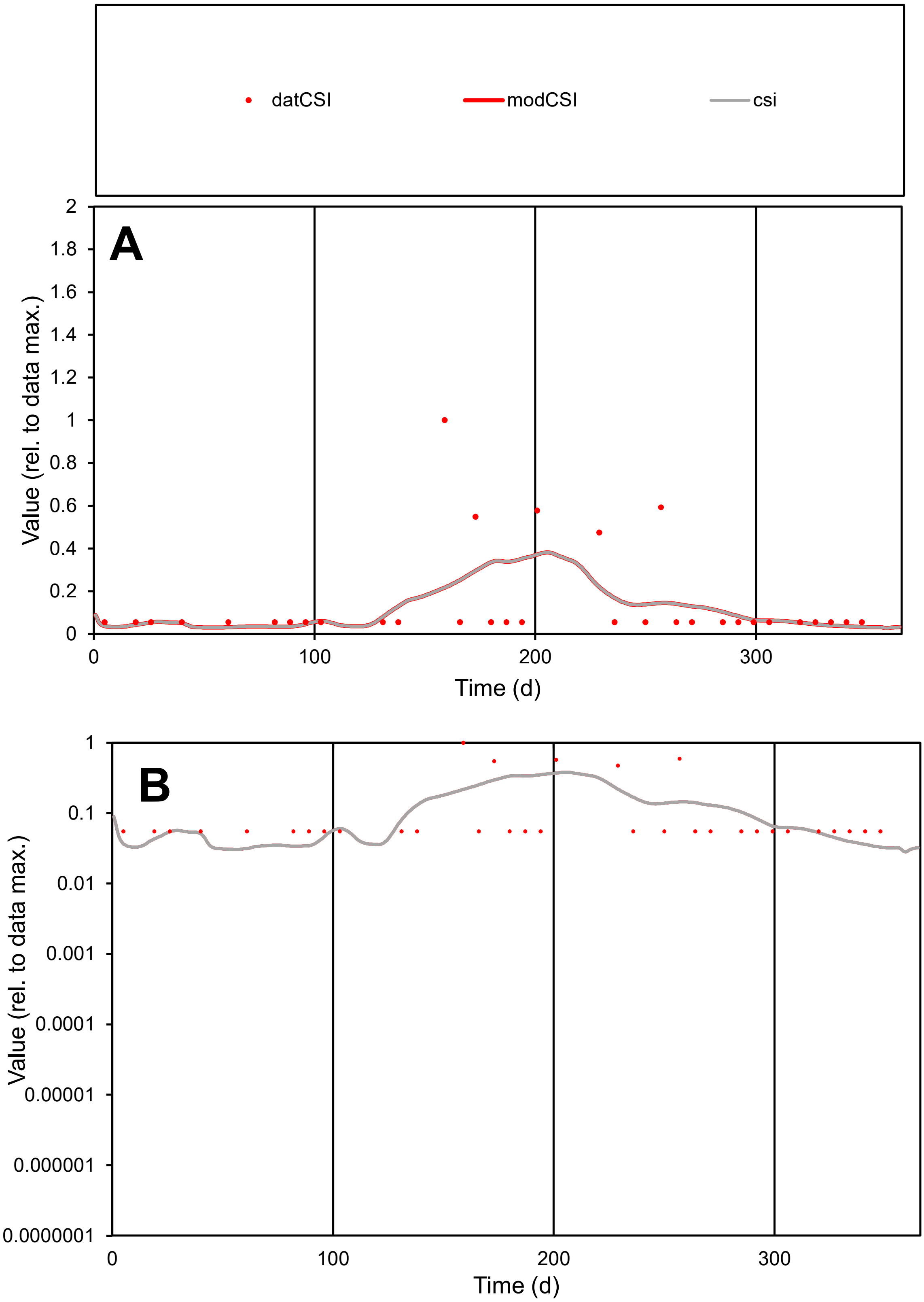


**C1**

Variables

CSI Provide jtd or Dst

CalibOnly?

2 1 to reproduce of

oNormalize?

2 1=no, 2=yes

ofxn calc*

1.38253

ofxn mod

1.39292

sRun

RunA1

zout

1

Ave

Data

6E-06

Model

5.4E-06

data/vMax

4.2E-05

ofxng

263.459

Min d/m

1.72131

ave/min d

2.59753

ave/min m

4.00029

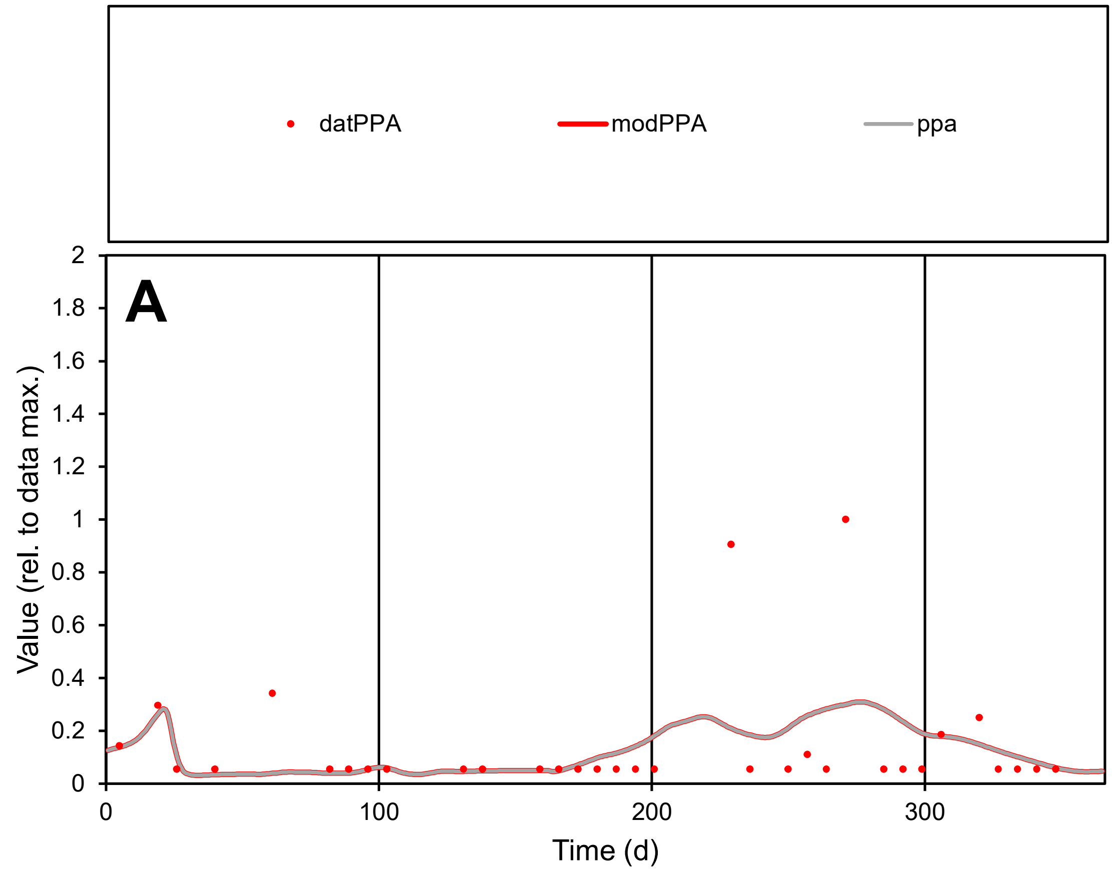


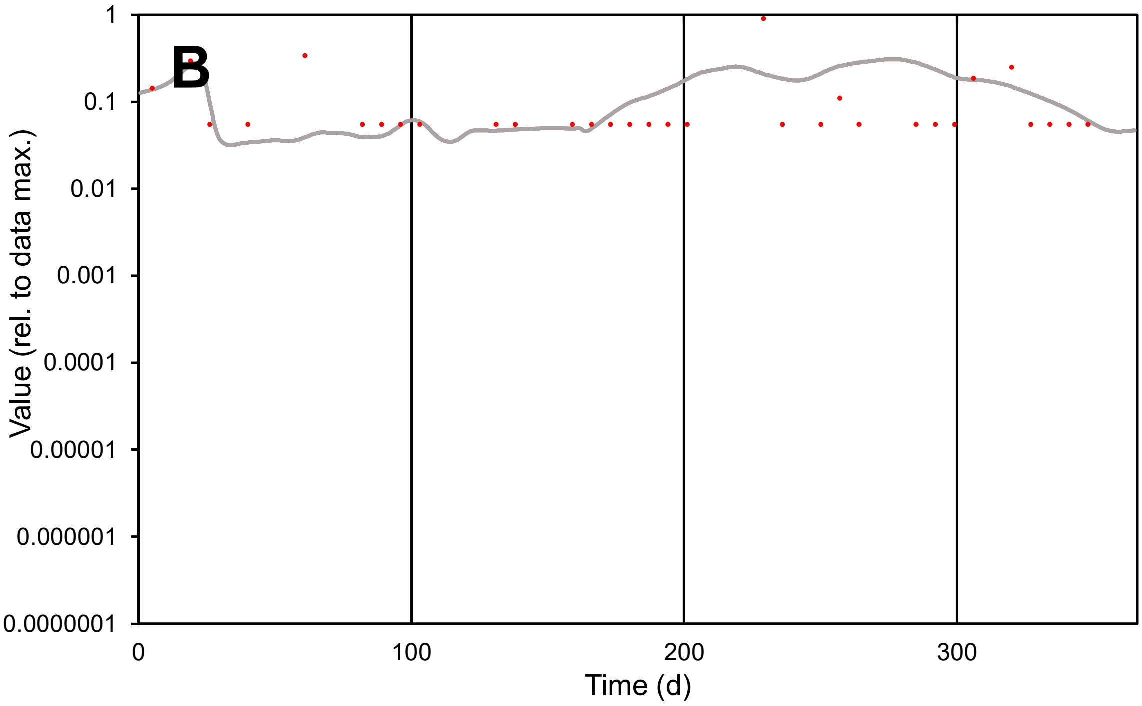


**C1**

Variables
PPA Provide jtd or Dst
CalibOnly?
2 1 to reproduce of
oNormalize?
2 1=no, 2=yes

ofxn calc*
1.43489
ofxn mod
1.44732

sRun
RunA1
zout
1

Ave
Data
8.1E-06
Model
7.7E-06

data/vMax
5.8E-05

ofxng
263.459

Min d/m
1.6055

ave/min d
2.53064
ave/min m
3.84979

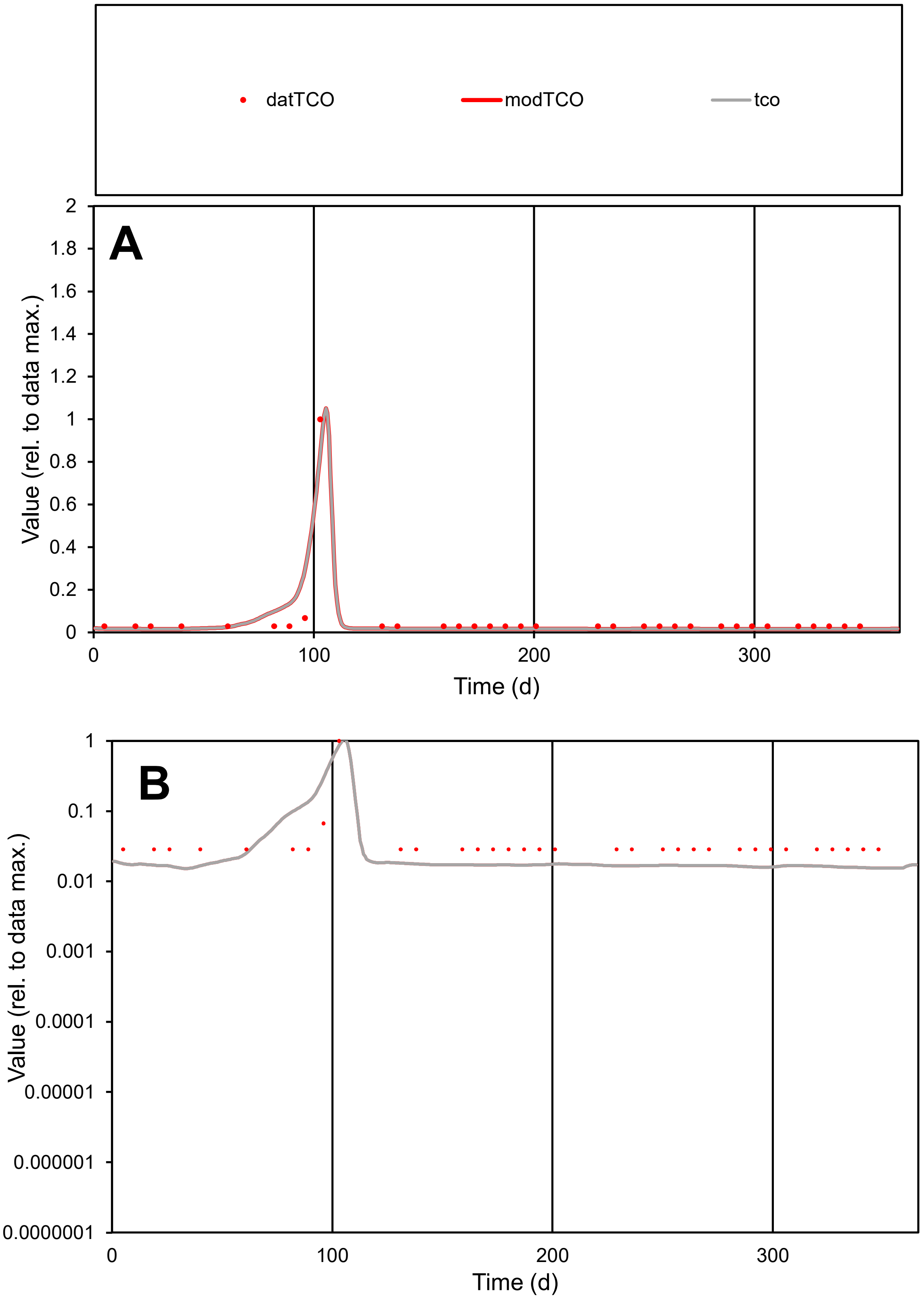


**C1**

Variables
TCO Provide jtd or Dst
CalibOnly?
2 1 to reproduce of
oNormalize?
2 1=no, 2=yes

ofxn calc*
0.71555
ofxn mod
0.7894

sRun
RunA1
zout
1

Ave
Data
1.8E-05
Model
1.8E-05

data/vMax
0.0003

ofxng
263.459

Min d/m
1.86484

ave/min d
2.06514
ave/min m
3.81896

326_Calib_S_A

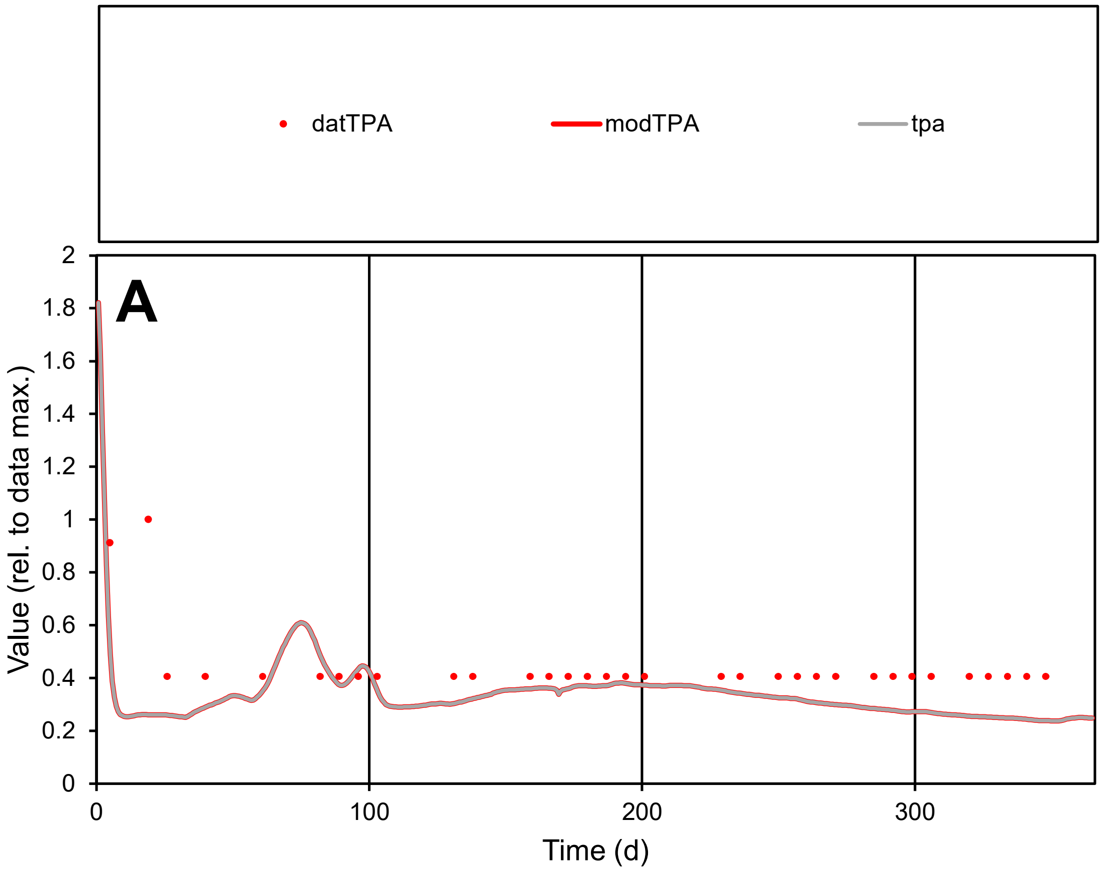


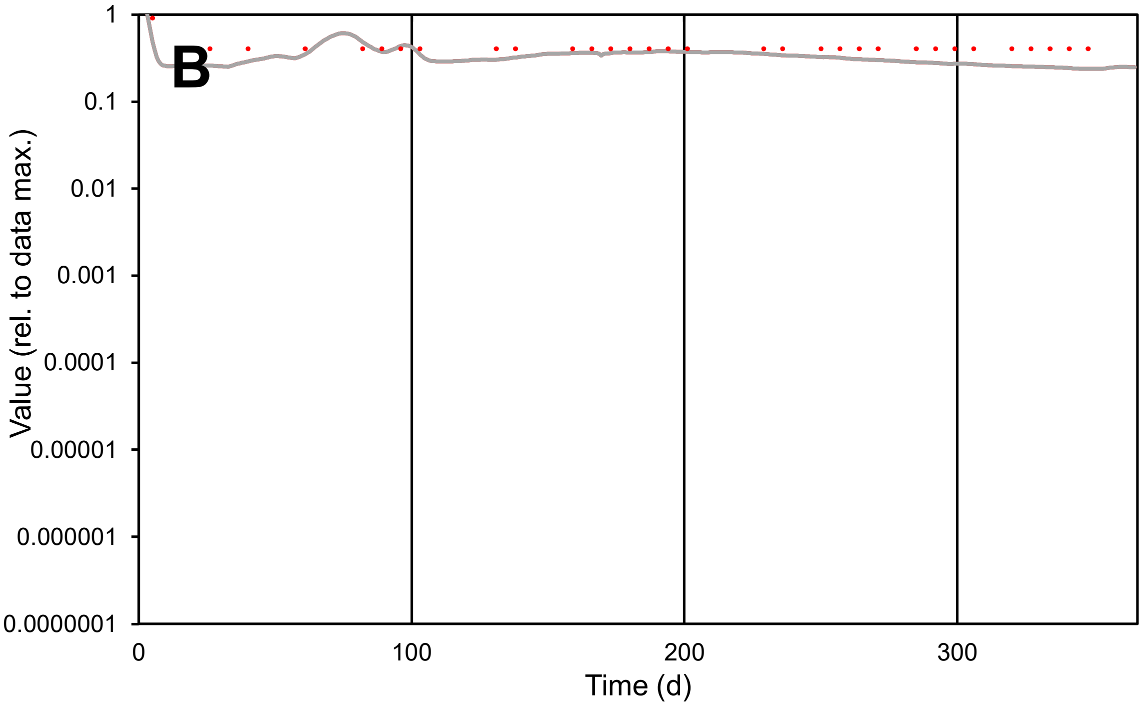


**C1**

Variables
TPA Provide jtd or Dst
CalibOnly?
2 1 to reproduce of
oNormalize?
2 1=no, 2=yes

ofxn calc*
0.38302
ofxn mod
0.41549

sRun
RunA1
zout
1

Ave
Data
5.1E-06
Model
3.9E-06

data/vMax
1.2E-05

ofxng
263.459

Min d/m
1.69928

ave/min d
1.08231
ave/min m
1.38531

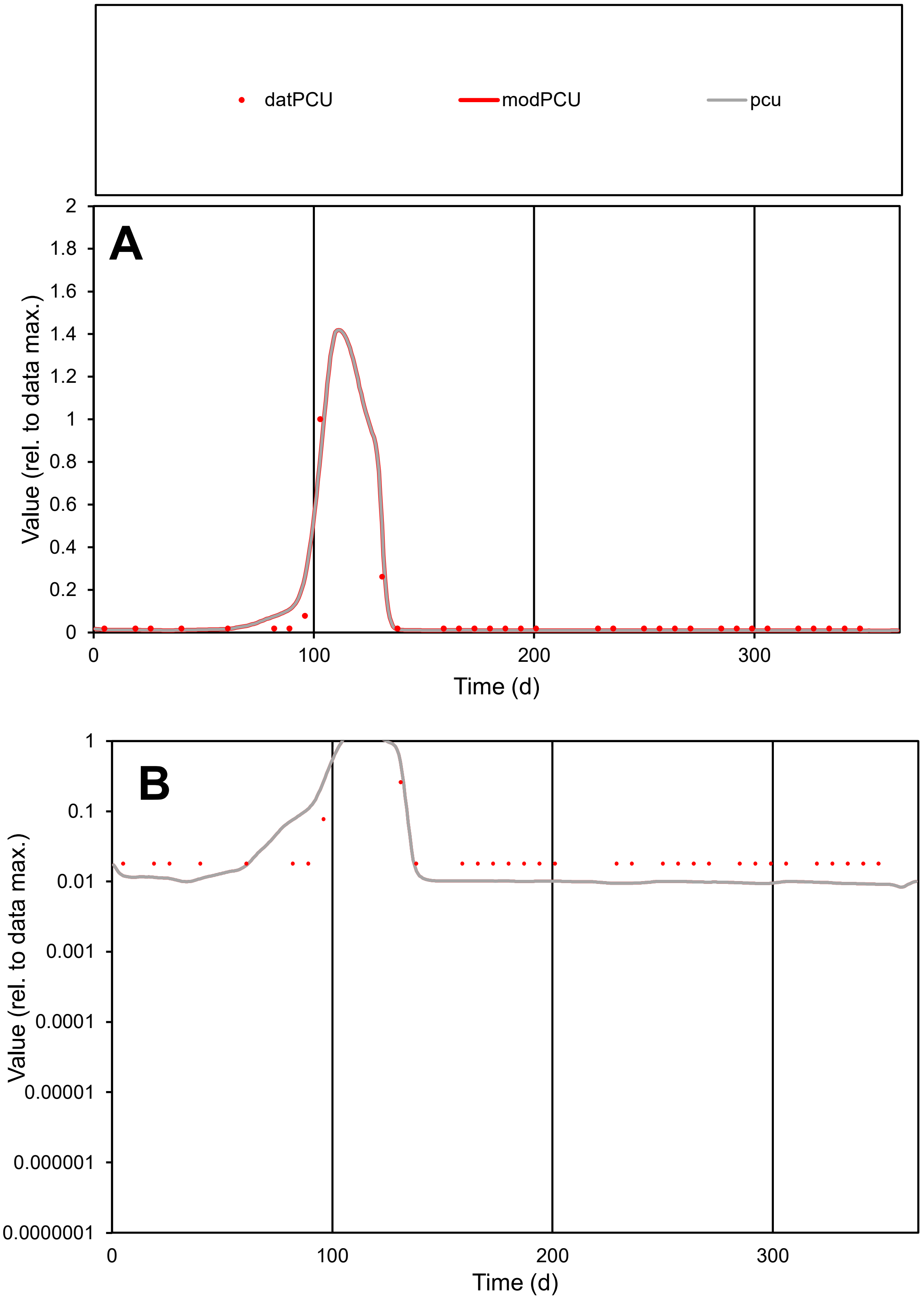


**C1**

Variables
PCU Provide jtd or Dst
CalibOnly?
2 1 to reproduce of
oNormalize?
2 1=no, 2=yes

ofxn calc*
0.61002
ofxn mod
0.69786

sRun
RunA1
zout
1

Ave
Data
2E-05
Model
2E-05

data/vMax
0.00034

ofxng
263.459

Min d/m
1.99036

ave/min d
3.15109
ave/min m
6.28007

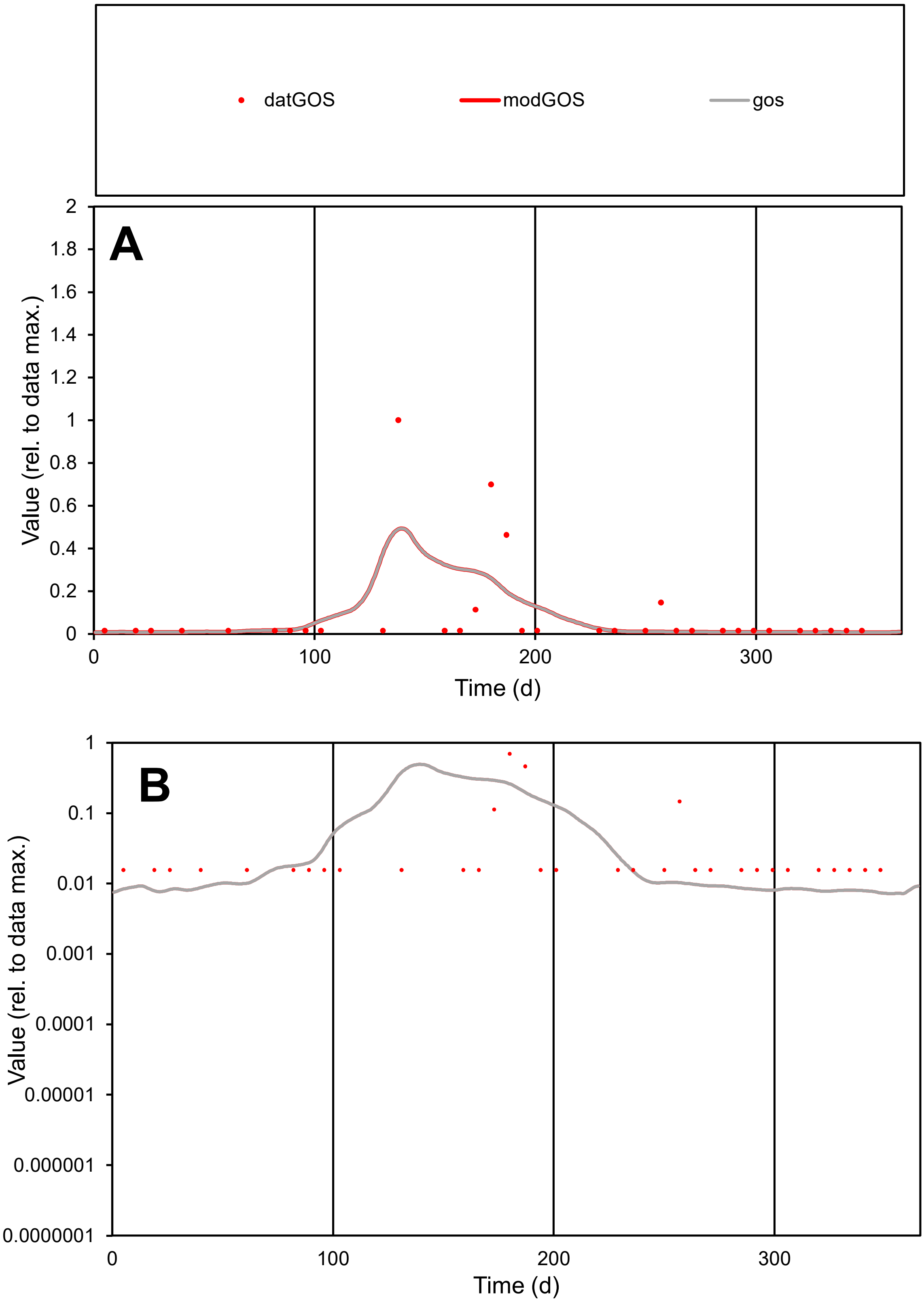


| | |
|---|---|
| **C1** | |
| Variables | |
| GOS | Provide jtd or Dst |
| CalibOnly? | |
| 2 | 1 to reproduce of |
| oNormalize? | |
| 2 | 1=no, 2=yes |
| ofxn calc* | |
| 1.90686 | |
| ofxn mod | |
| 1.9358 | |
| sRun | |
| RunA1 | |
| zout | |
| 1 | |
| Ave | |
| Data | |
| 1E-05 | |
| Model | |
| 9.8E-06 | |
| data/vMax | |
| 0.00012 | |
| ofxng | |
| 263.459 | |
| Min d/m | |
| 2.13252 | |
| ave/min d | |
| 5.54696 | |
| ave/min m | |
| 11.5738 | |

326_Calib_S_A

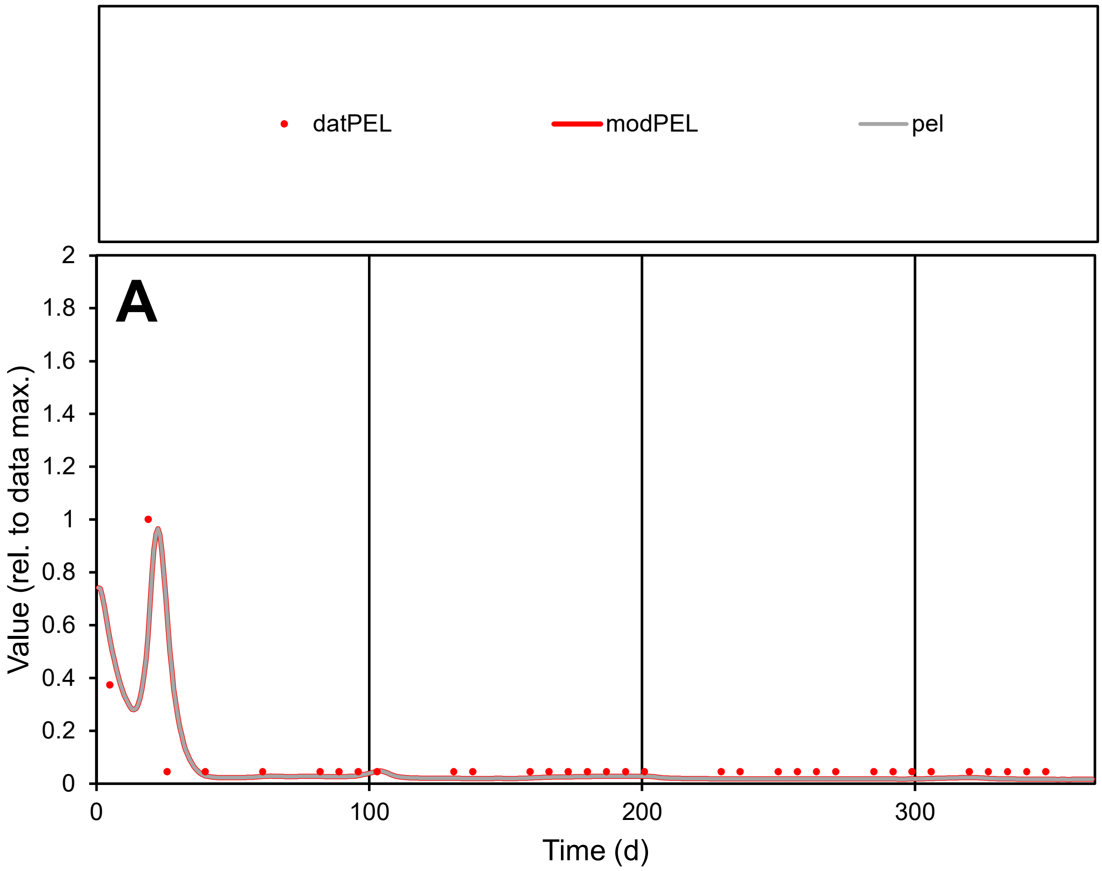


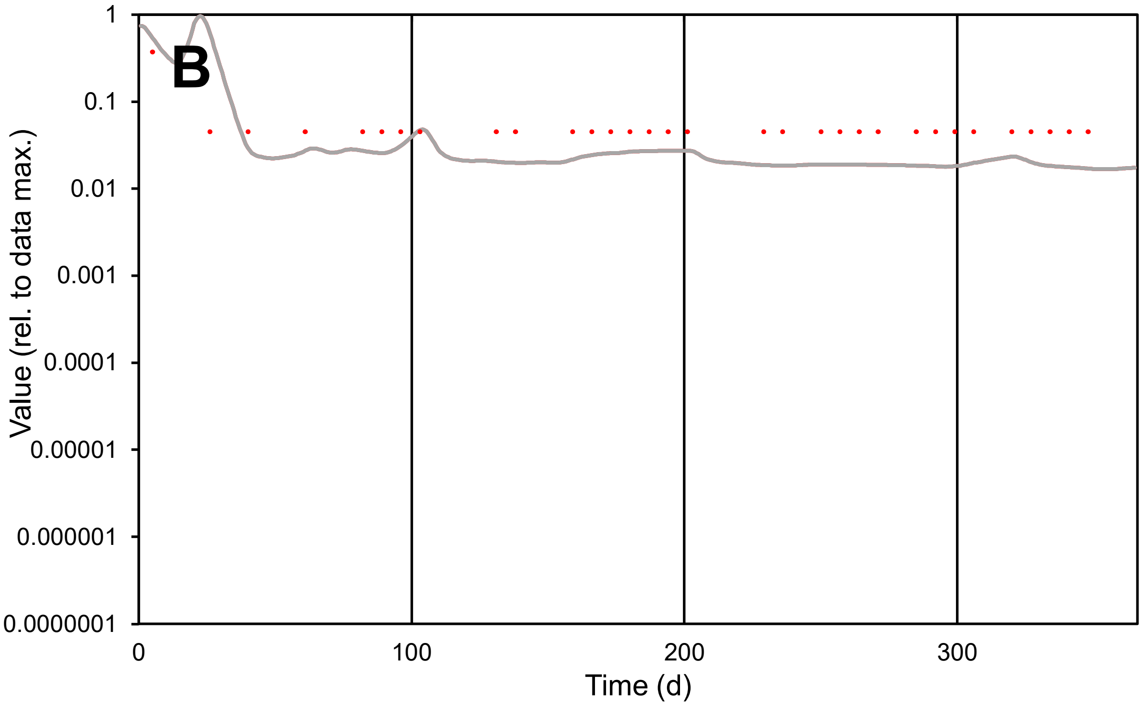


**C1**

Variables
PEL Provide jtd or Dst
CalibOnly?
2 1 to reproduce of
oNormalize?
2 1=no, 2=yes

ofxn calc*
0.76175
ofxn mod
0.92143

sRun
RunA1
zout
1

Ave
Data
6.1E-06
Model
5E-06

data/vMax
7.3E-05

ofxng
263.459

Min d/m
2.64677

ave/min d
1.8651
ave/min m
4.01618

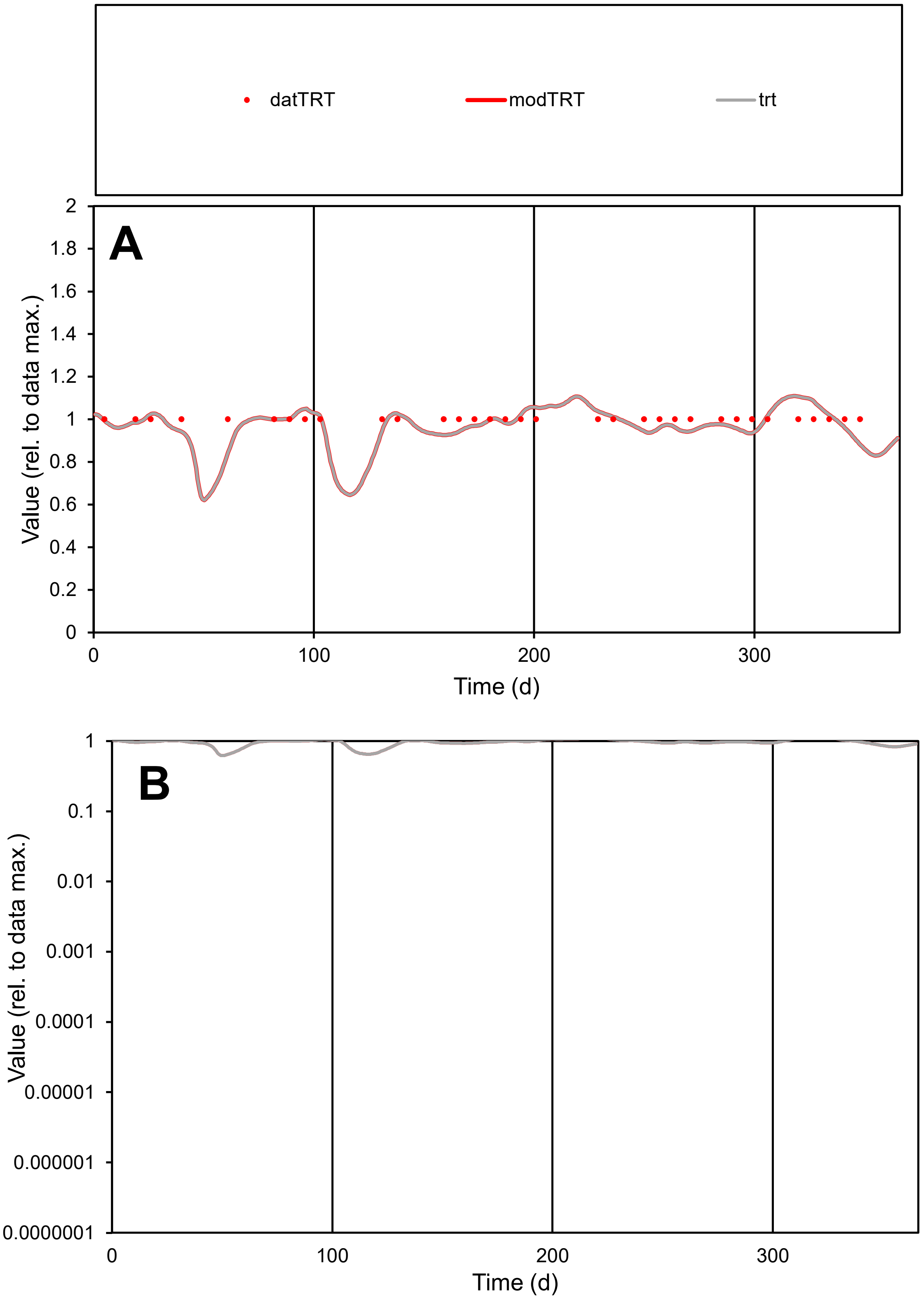


**C1**

Variables
TRT Provide jtd or Dst
CalibOnly?
2 1 to reproduce of
oNormalize?
2 1=no, 2=yes

ofxn calc*
0.05173
ofxn mod
0.12367

sRun
RunA1
zout
1

Ave
Data
2.3E-05
Model
2.3E-05

data/vMax
2.3E-05

ofxng
263.459

Min d/m
1.14817

ave/min d
1
ave/min m
1.13235

326_Calib_S_A

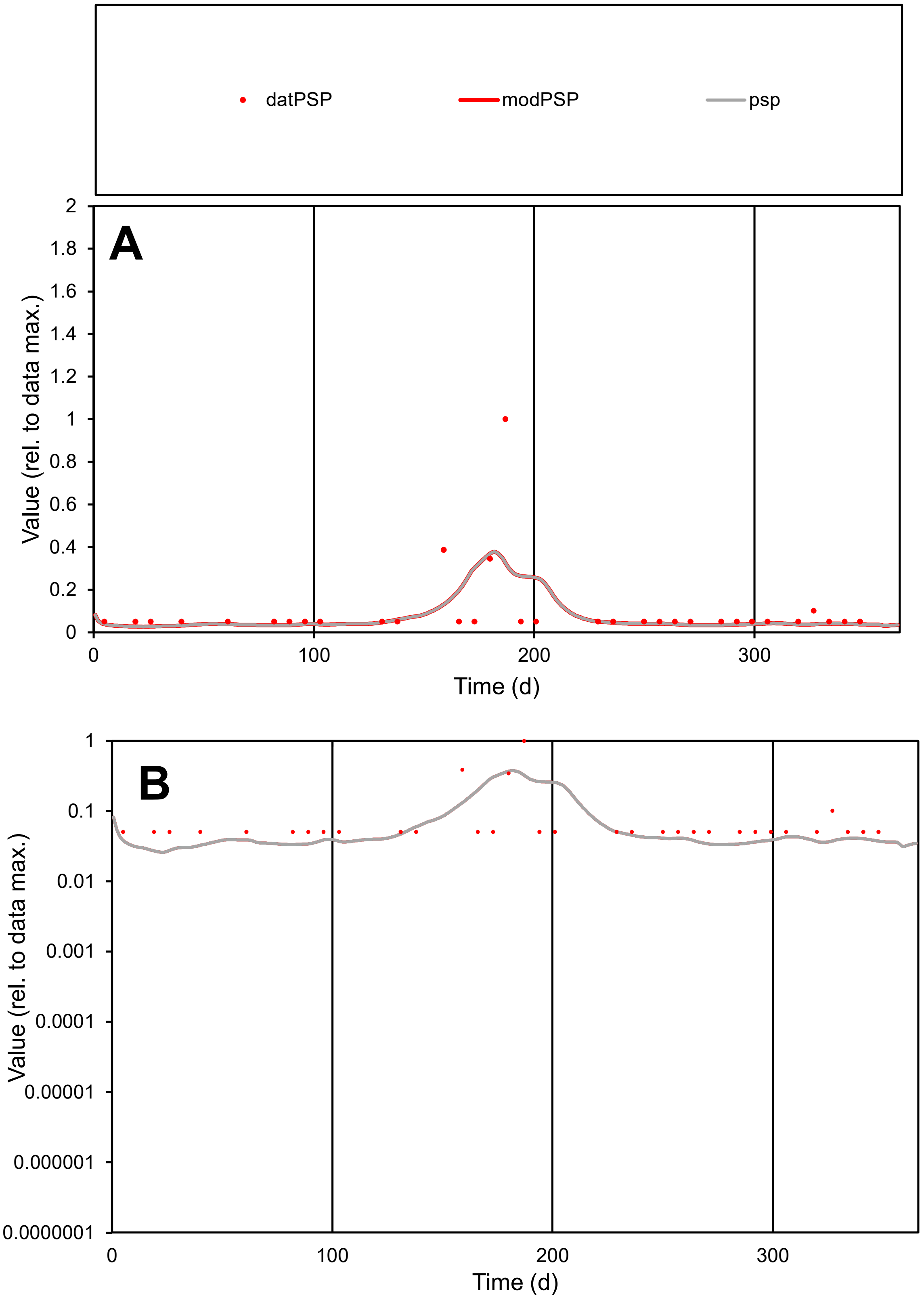


**C1**

Variables

| | |
|---|---|
| PSP | Provide jtd or Dst |
| CalibOnly? | |
| 2 | 1 to reproduce of |
| oNormalize? | |
| 2 | 1=no, 2=yes |
| ofxn calc* | |
| 1.46498 | |
| ofxn mod | |
| 1.48427 | |
| sRun | |
| RunA1 | |
| zout | |
| 1 | |
| Ave | |
| Data | |
| 8.5E-06 | |
| Model | |
| 7.4E-06 | |
| data/vMax | |
| 8.5E-05 | |
| ofxng | |
| 263.459 | |
| Min d/m | |
| 1.88274 | |
| ave/min d | |
| 1.9759 | |
| ave/min m | |
| 3.21809 | |

326_Calib_S_A

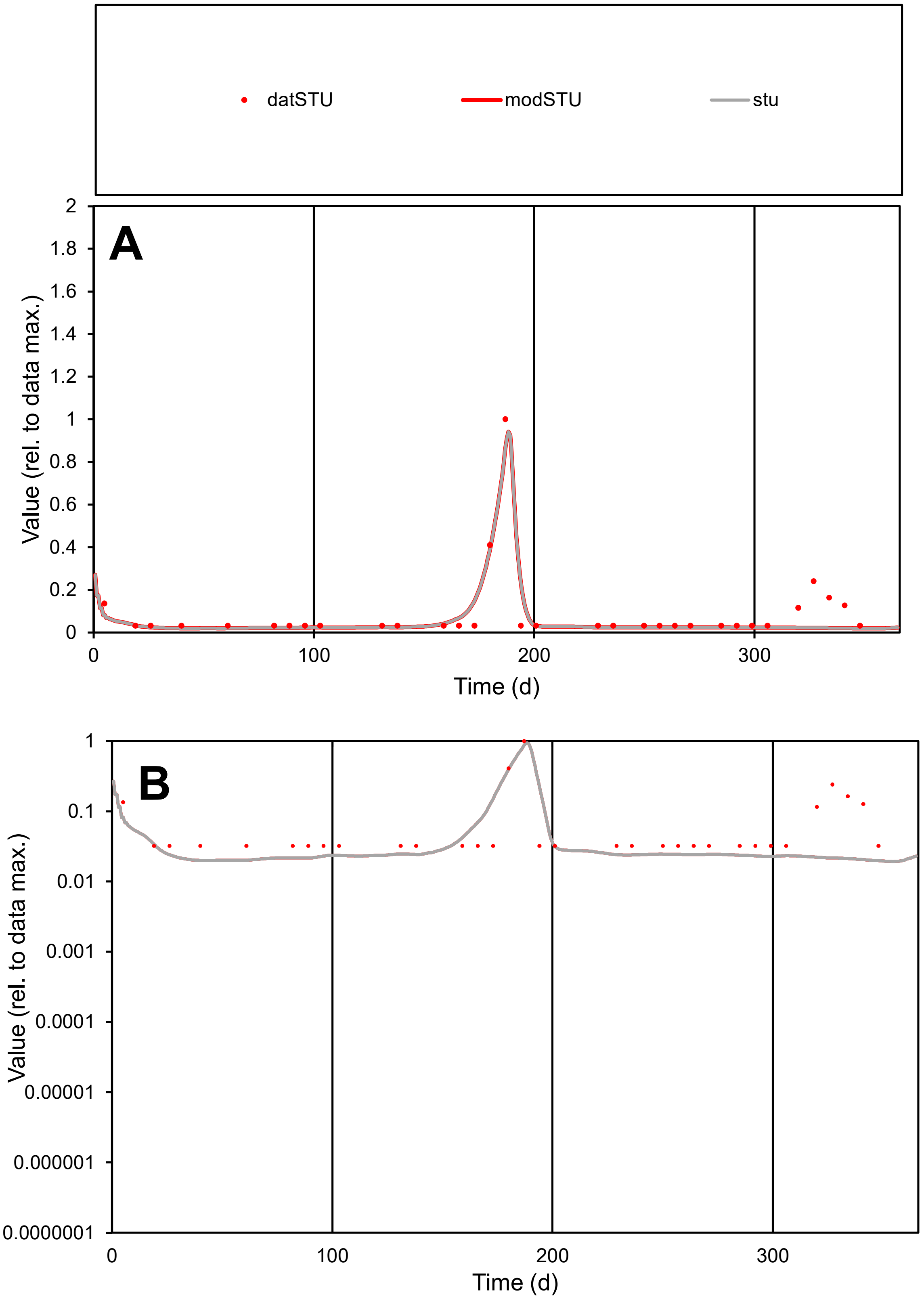


**C1**

Variables
STU Provide jtd or Dst
CalibOnly?
2 1 to reproduce of
oNormalize?
2 1=no, 2=yes

ofxn calc*
0.64422
ofxn mod
0.70216

sRun
RunA1
zout
1

Ave
Data
7.1E-06
Model
5.7E-06

data/vMax
7.7E-05

ofxng
263.459

Min d/m
1.64121

ave/min d
2.84852
ave/min m
3.76326

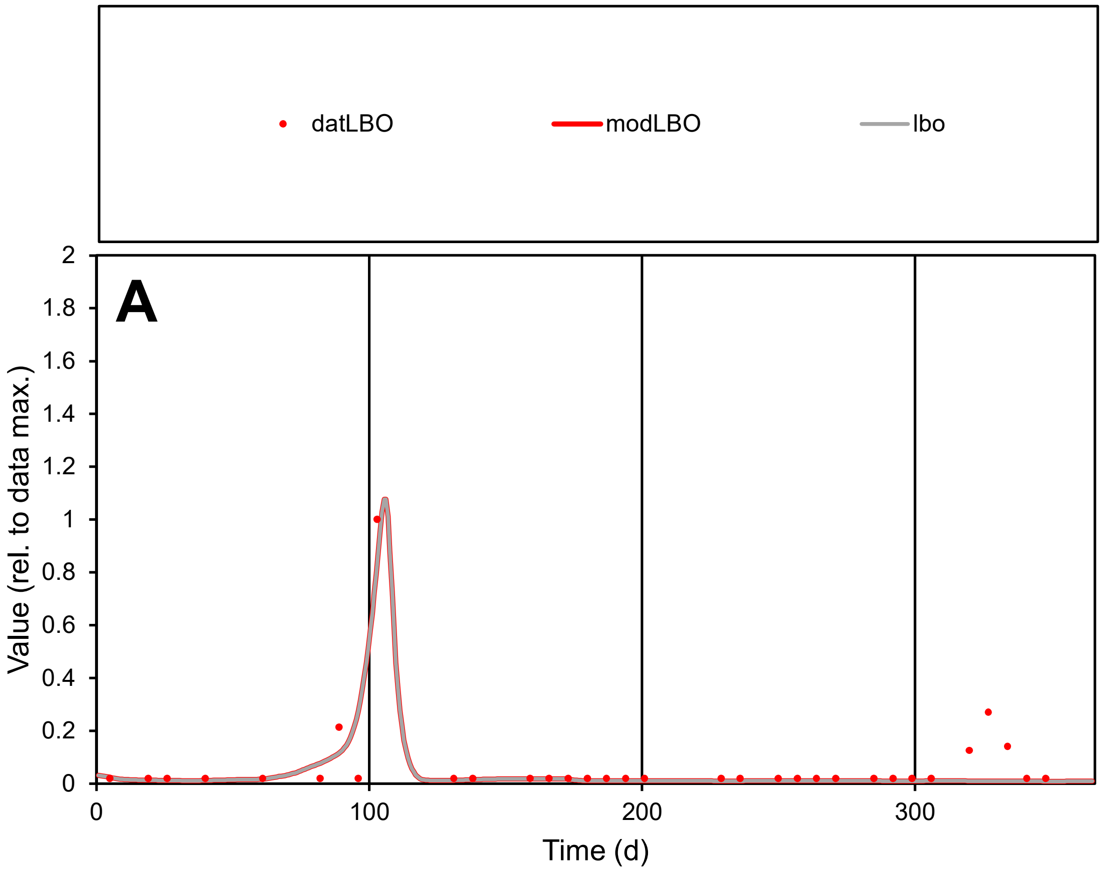


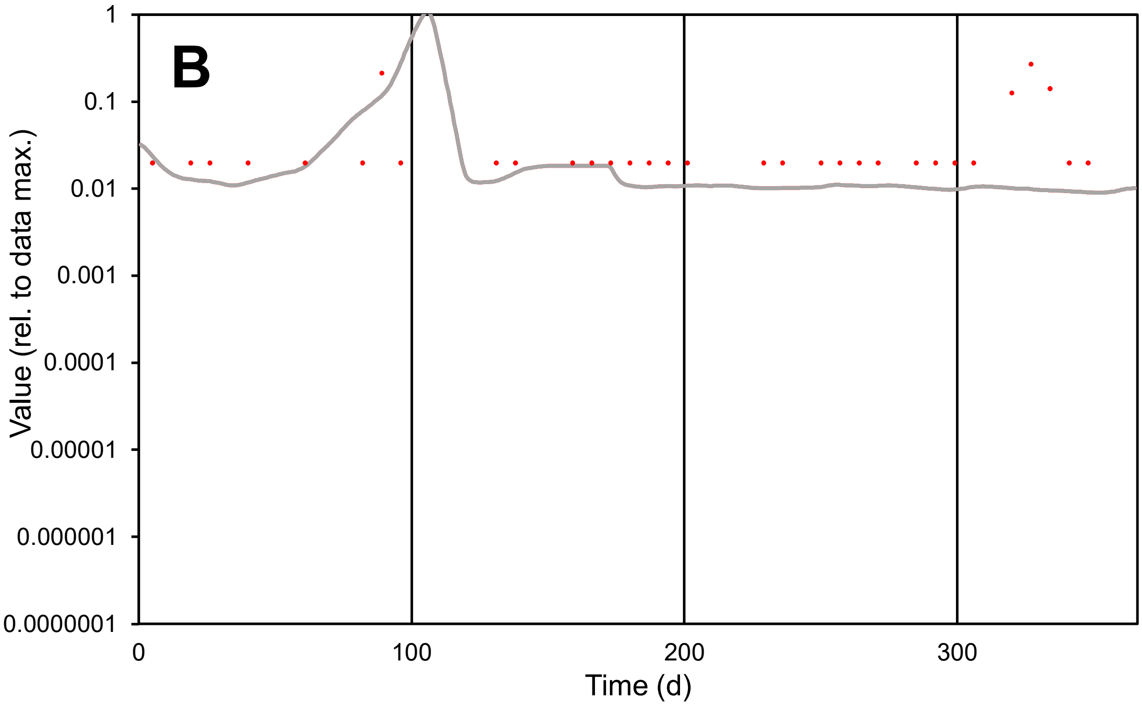


**C1**

Variables

LBO Provide jtd or Dst

CalibOnly?

2 1 to reproduce of

oNormalize?

2 1=no, 2=yes

ofxn calc*

0.92359

ofxn mod

1.02081

sRun

RunA1

zout

1

Ave

Data

4.9E-06

Model

3.6E-06

data/vMax

7E-05

ofxng

263.459

Min d/m

2.18391

ave/min d

3.53925

ave/min m

5.67503

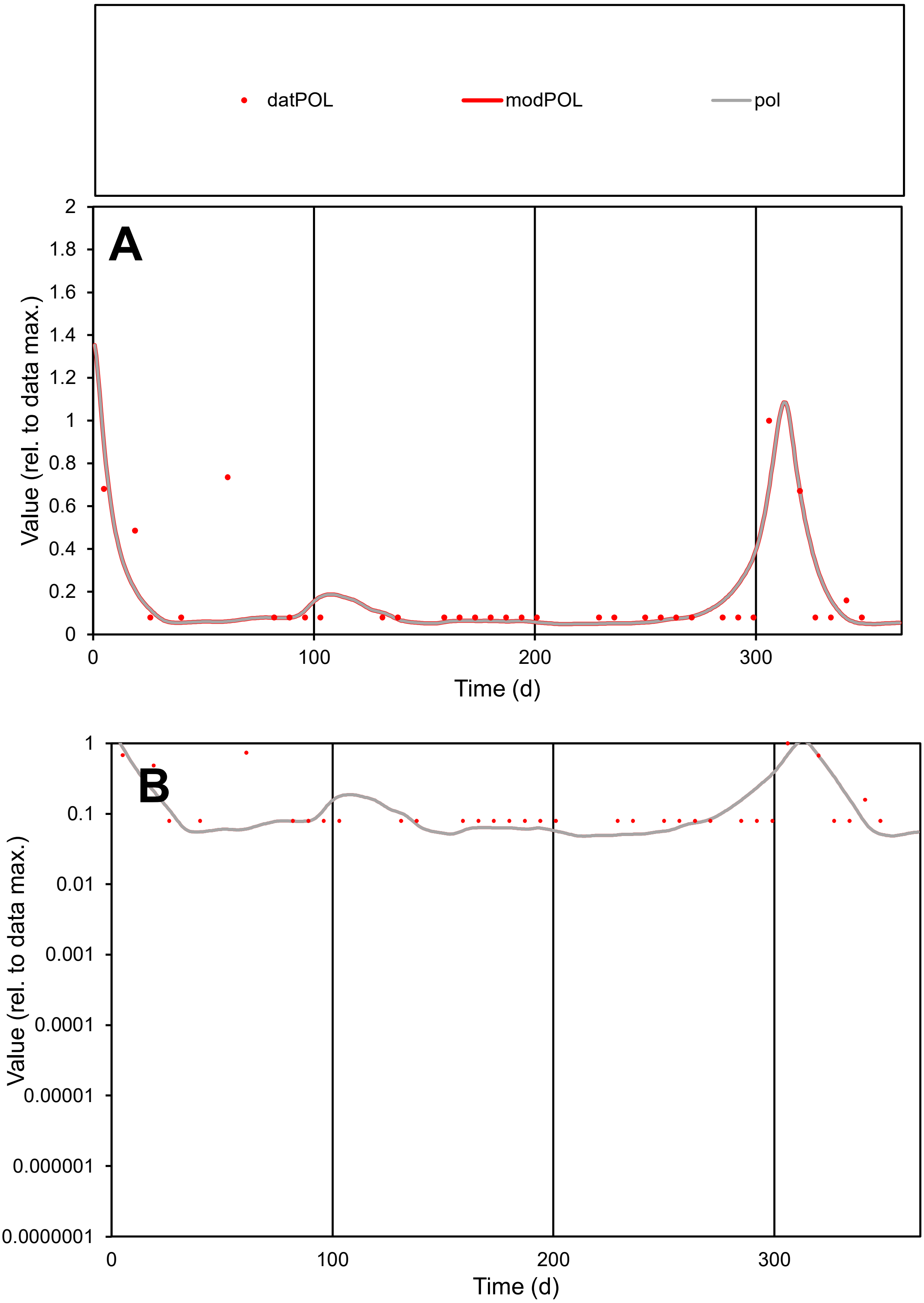


**C1**

Variables
POL Provide jtd or Dst
CalibOnly?
2 1 to reproduce of
oNormalize?
2 1=no, 2=yes

ofxn calc*
0.84626
ofxn mod
0.87768

sRun
RunA1
zout
1

Ave
Data
3.6E-06
Model
3.2E-06

data/vMax
2E-05

ofxng
263.459

Min d/m
1.57087

ave/min d
2.24358
ave/min m
3.13991

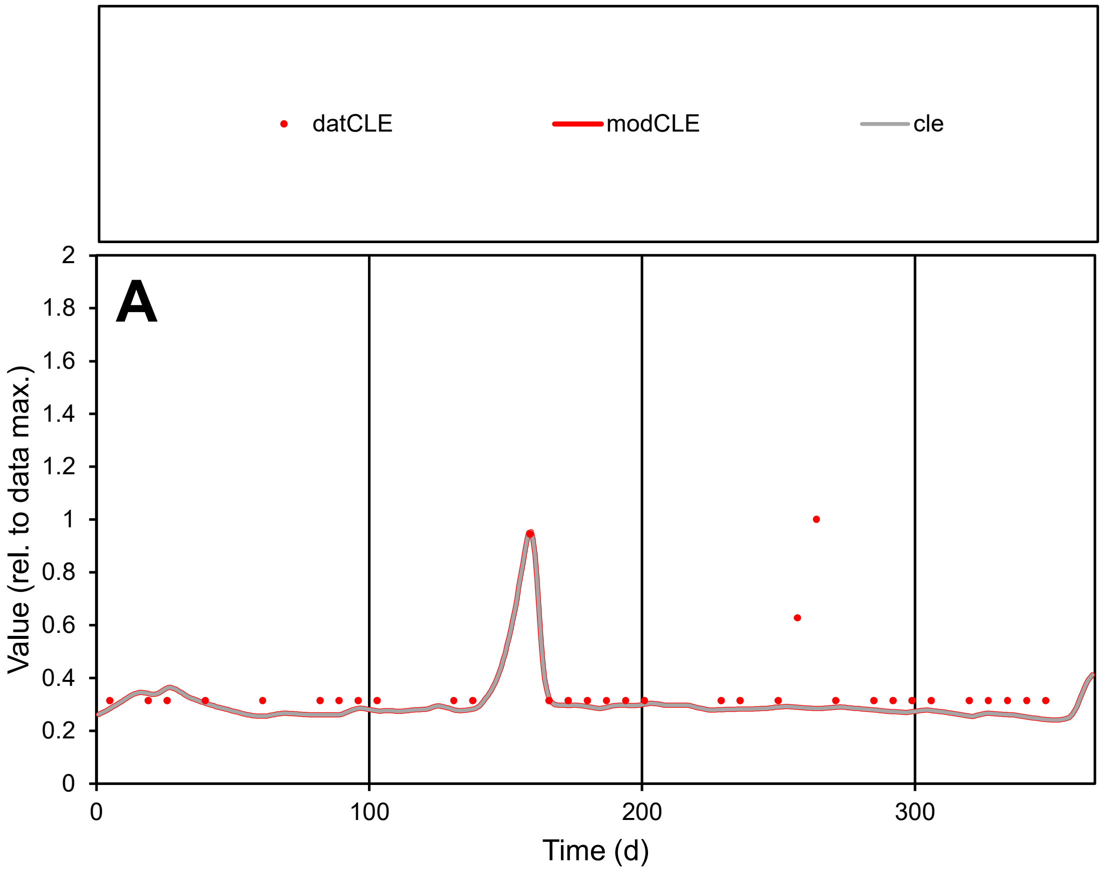


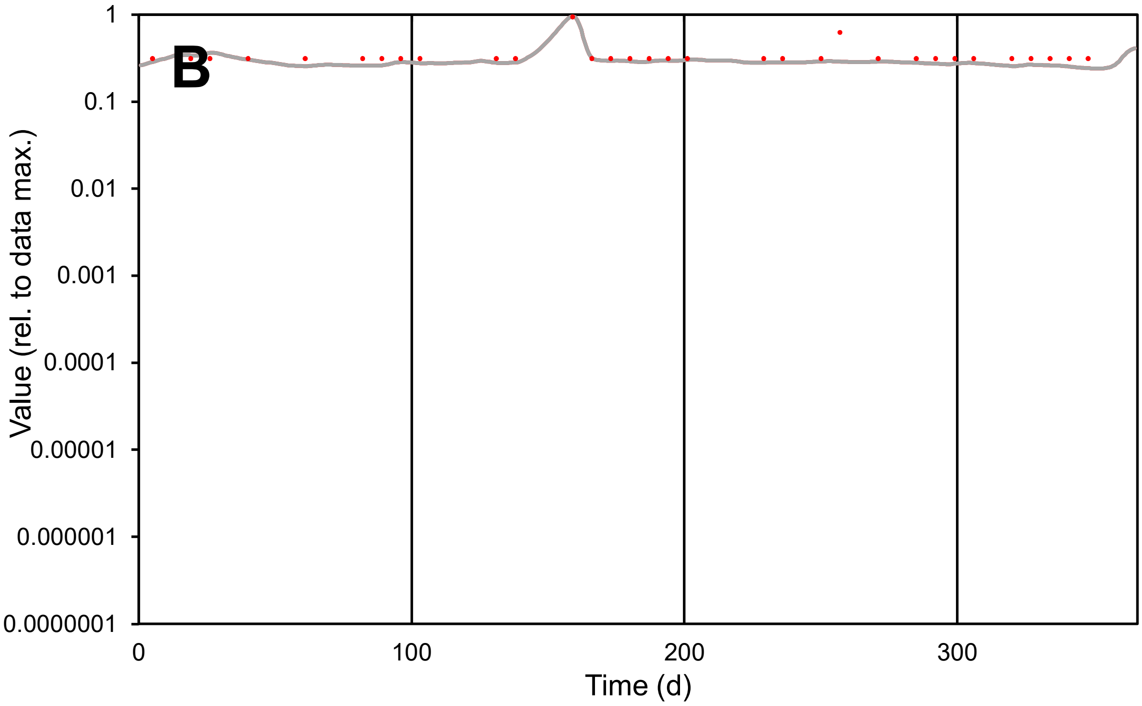


**C1**

Variables
CLE Provide jtd or Dst
CalibOnly?
2 1 to reproduce of
oNormalize?
2 1=no, 2=yes

ofxn calc*
0.39277
ofxn mod
0.40668

sRun
RunA1
zout
1

Ave
Data
5.4E-06
Model
4.5E-06

data/vMax
1.5E-05

ofxng
263.459

Min d/m
1.29473

ave/min d
1.15754
ave/min m
1.25131

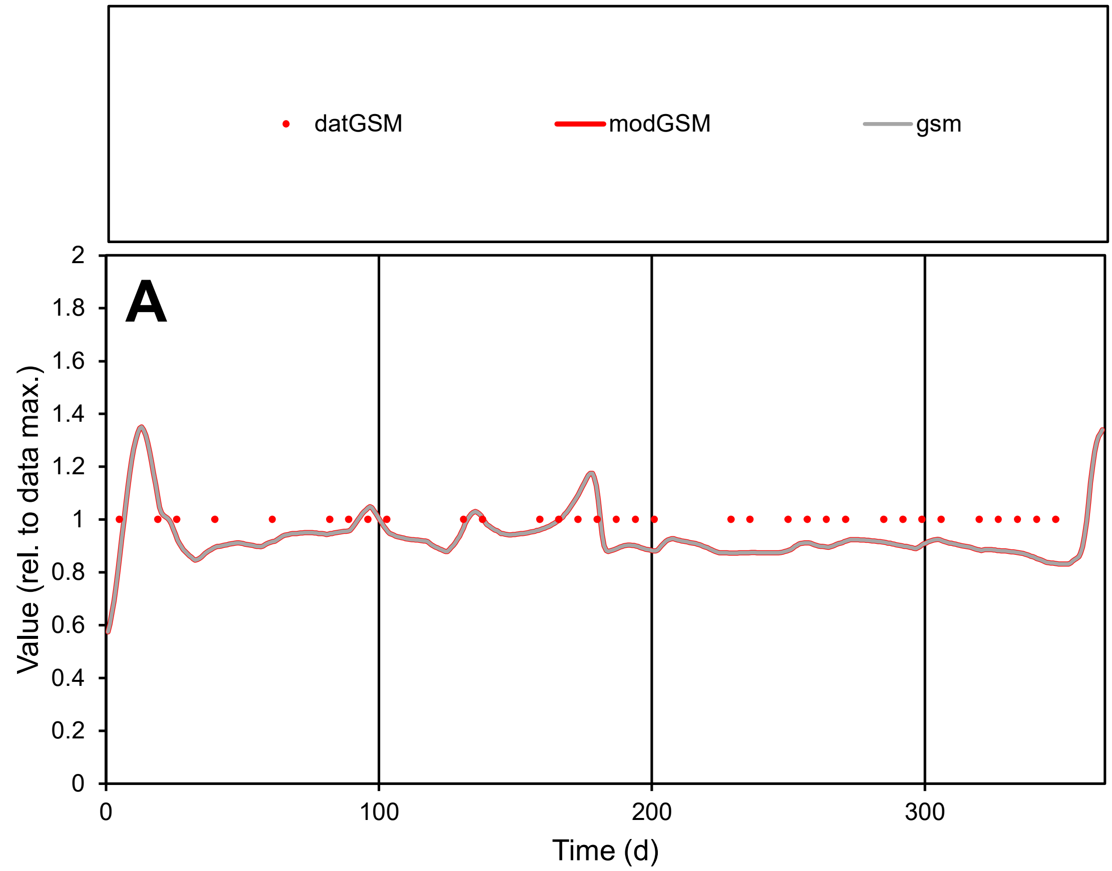


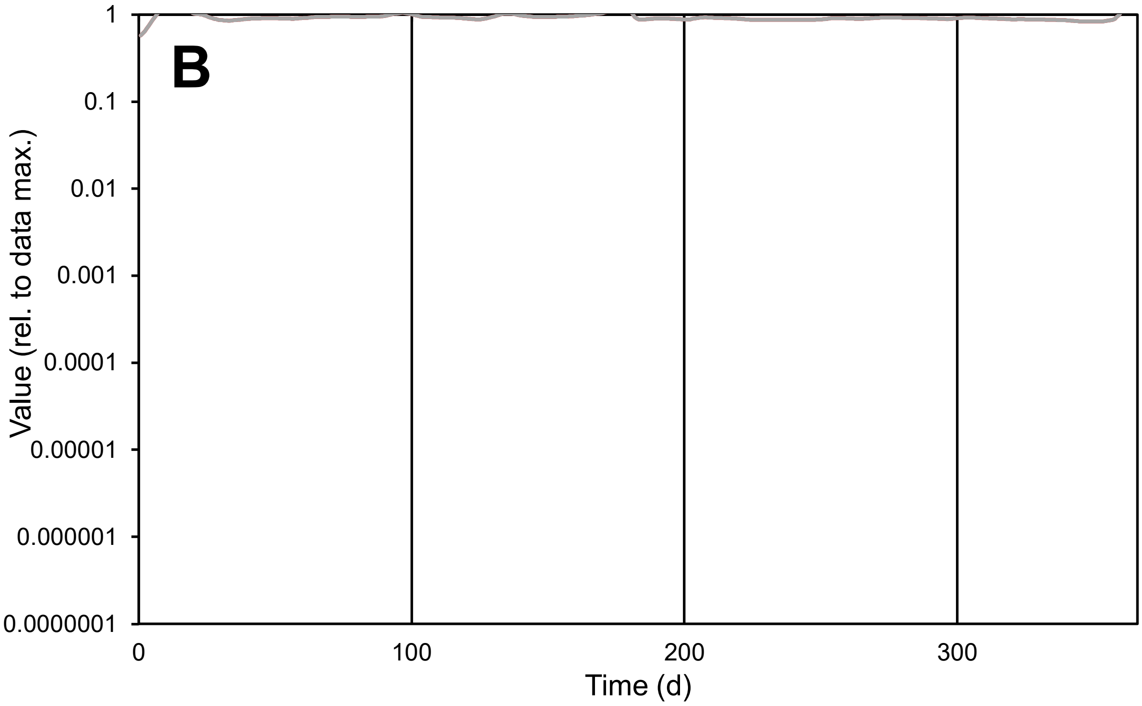


**C1**

Variables
GSM Provide jtd or Dst
CalibOnly?
2 1 to reproduce of
oNormalize?
2 1=no, 2=yes

ofxn calc*
0.09299
ofxn mod
0.14346

sRun
RunA1
zout
1

Ave
Data
1.6E-06
Model
1.5E-06

data/vMax
1.6E-06

ofxng
263.459

Min d/m
1.20167

ave/min d
1
ave/min m
1.11911

326_Calib_S_A

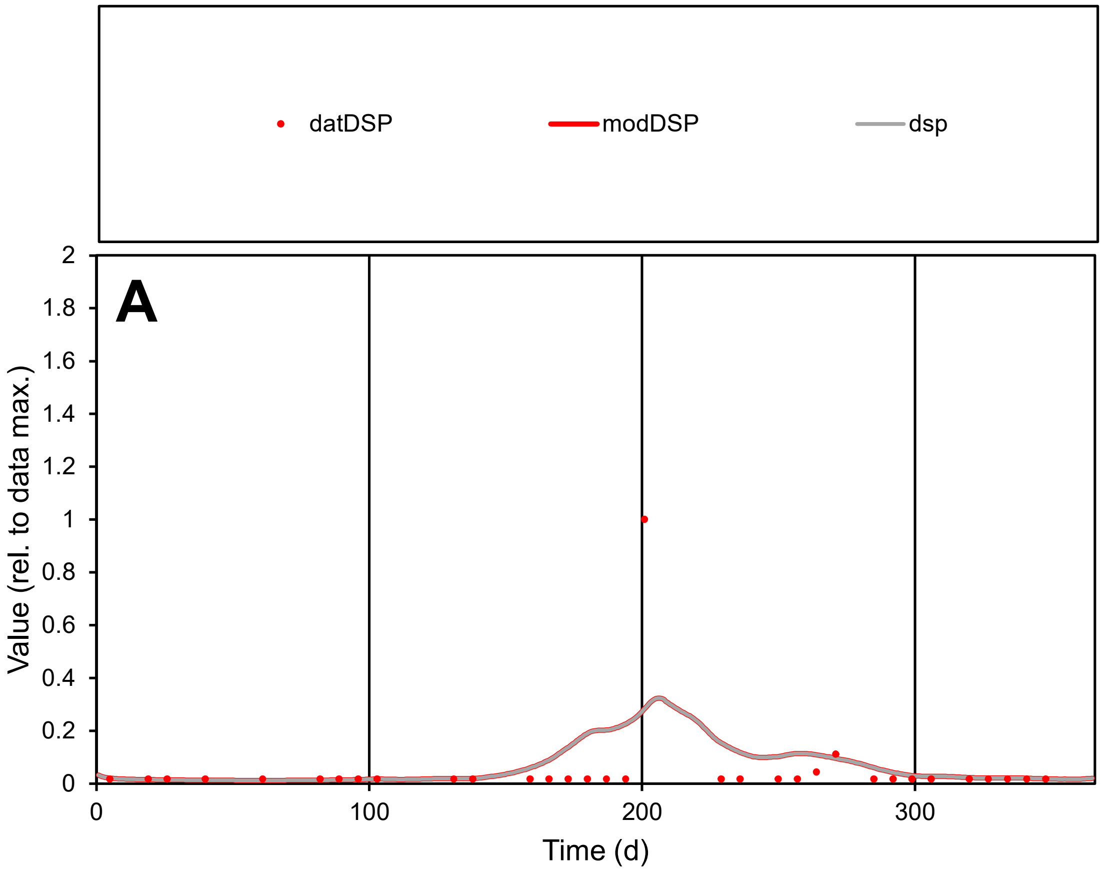


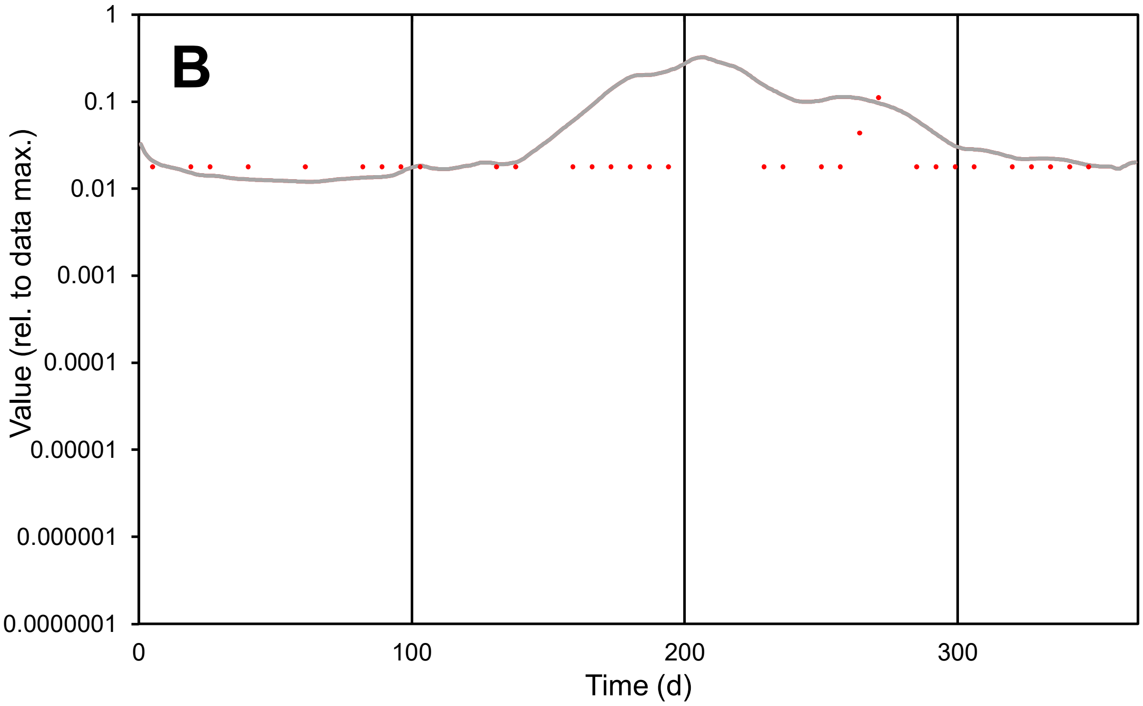


**C1**

Variables
DSP Provide jtd or Dst
CalibOnly?
2 1 to reproduce of
oNormalize?
2 1=no, 2=yes

ofxn calc*
2.77783
ofxn mod
2.79751

sRun
RunA1
zout
1

Ave
Data
1.4E-05
Model
2E-05

data/vMax
0.00028

ofxng
263.459

Min d/m
1.48384

ave/min d
2.86713
ave/min m
5.80938

326_Calib_S_A

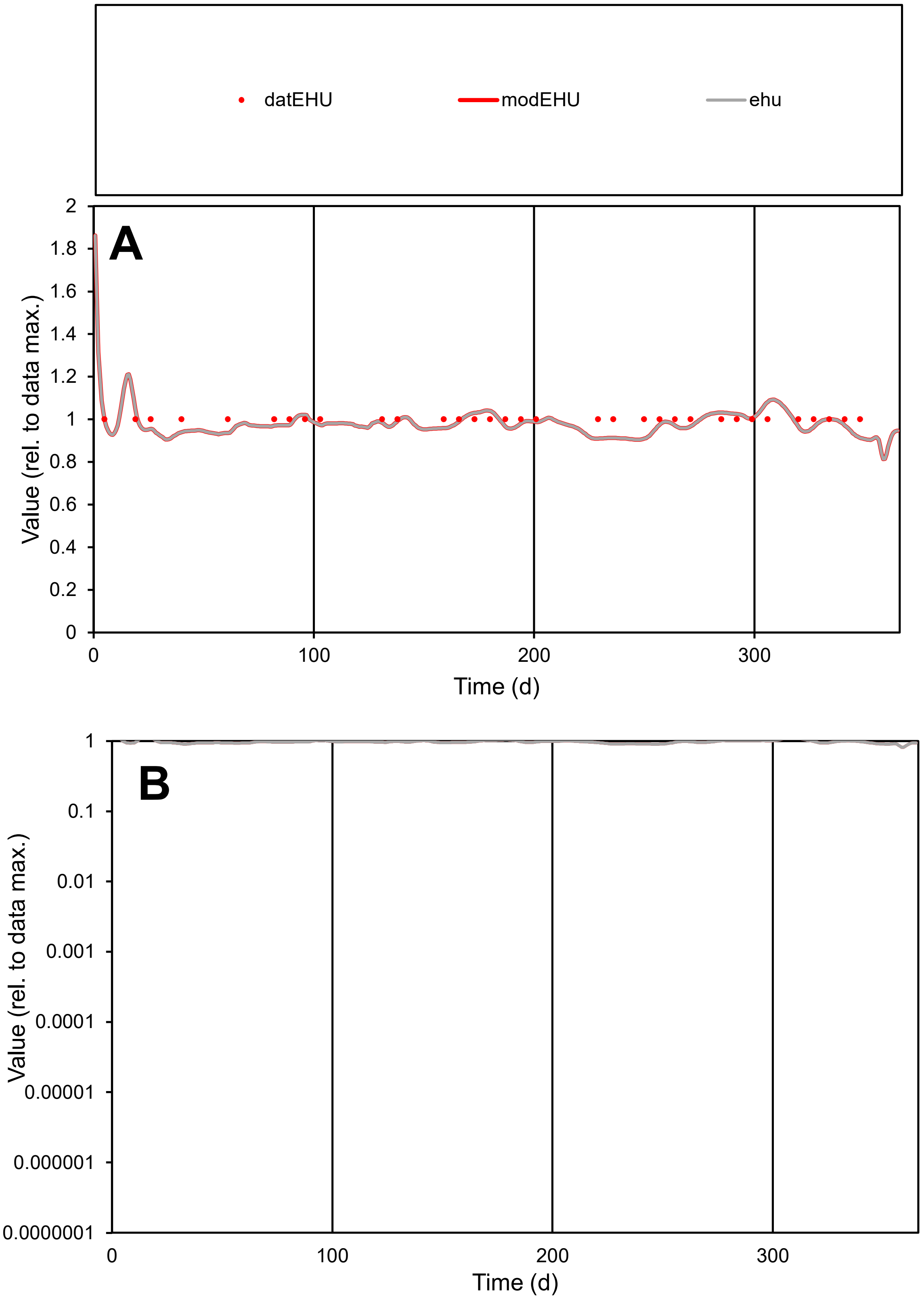


**C1**

Variables
EHU Provide jtd or Dst
CalibOnly?
2 1 to reproduce of
oNormalize?
2 1=no, 2=yes

ofxn calc*
0.04547
ofxn mod
0.12098

sRun
RunA1
zout
1

Ave
Data
5.8E-06
Model
5.6E-06

data/vMax
5.8E-06

ofxng
263.459

Min d/m
1.09912

ave/min d
1
ave/min m
1.07476

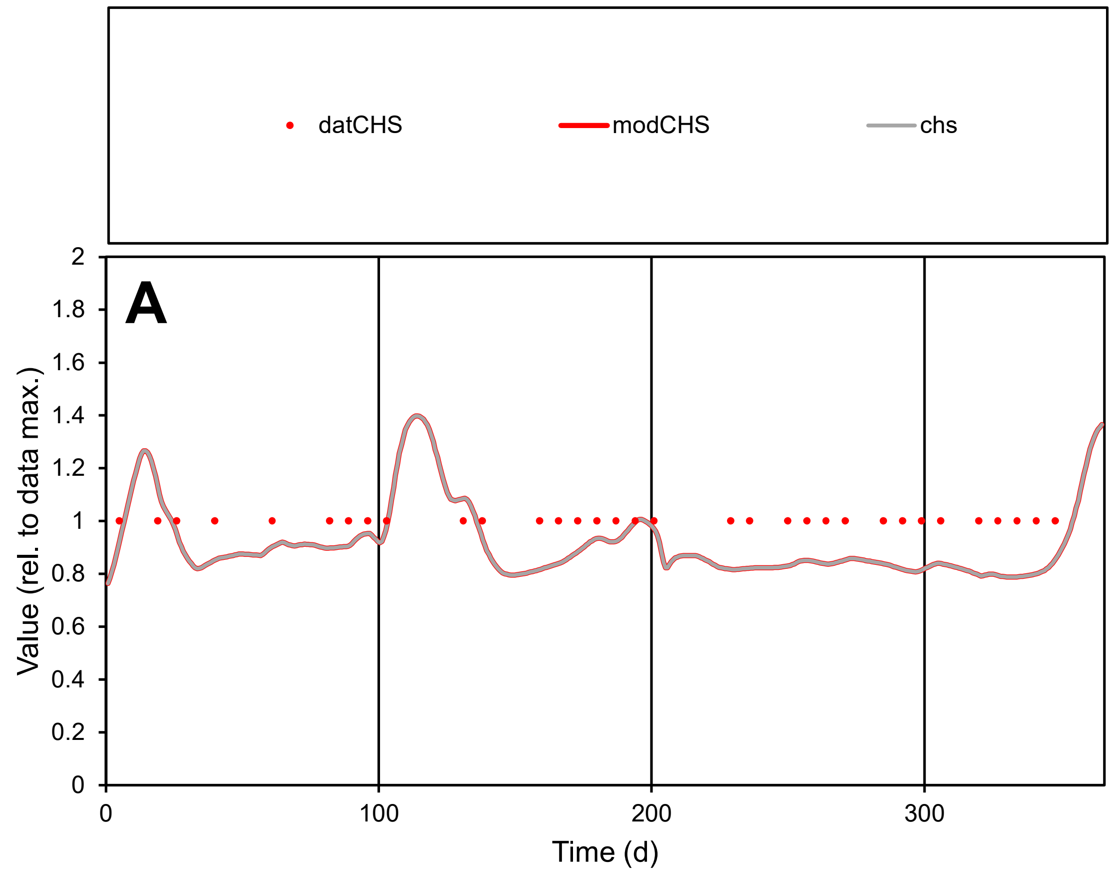


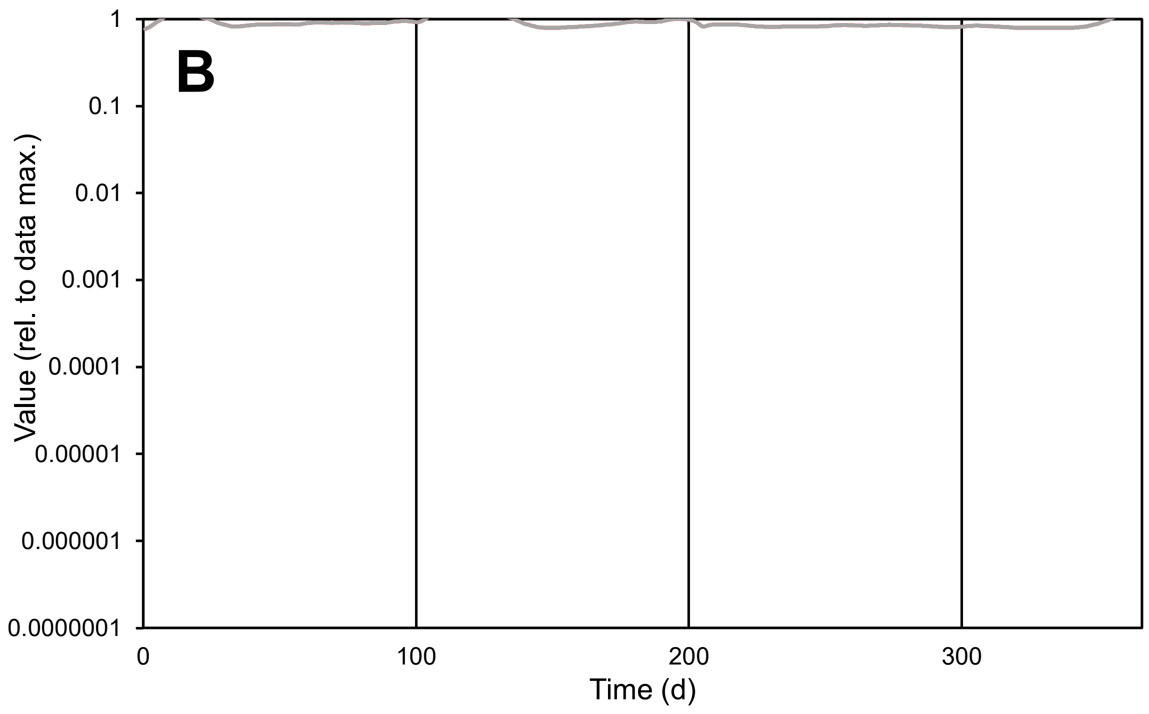


**C1**

Variables
CHS Provide jtd or Dst
CalibOnly?
2 1 to reproduce of
oNormalize?
2 1=no, 2=yes

ofxn calc*
0.13822
ofxn mod
0.17503

sRun
RunA1
zout
1

Ave
Data
2.1E-05
Model
1.9E-05

data/vMax
2.1E-05

ofxng
263.459

Min d/m
1.26493

ave/min d
1
ave/min m
1.12152

326_Calib_S_A

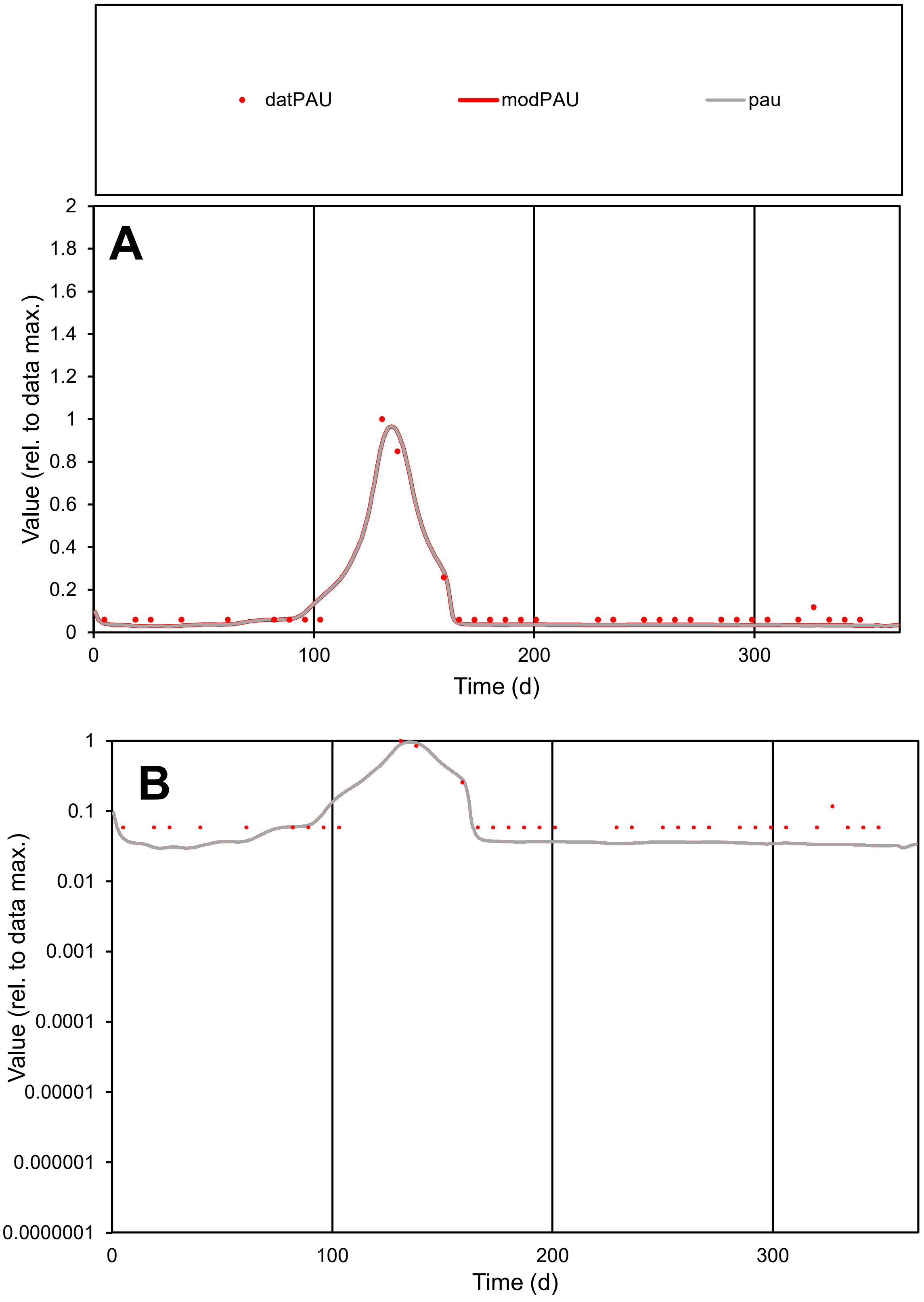


**C1**

Variables
PAU Provide jtd or Dst
CalibOnly?
2 1 to reproduce of
oNormalize?
2 1=no, 2=yes

ofxn calc*
0.29484
ofxn mod
0.38672

sRun
RunA1
zout
1

Ave
Data
8E-06
Model
6.9E-06

data/vMax
6.7E-05

ofxng
263.459

Min d/m
1.95198

ave/min d
2.02533
ave/min m
3.40575

326_Calib_S_A

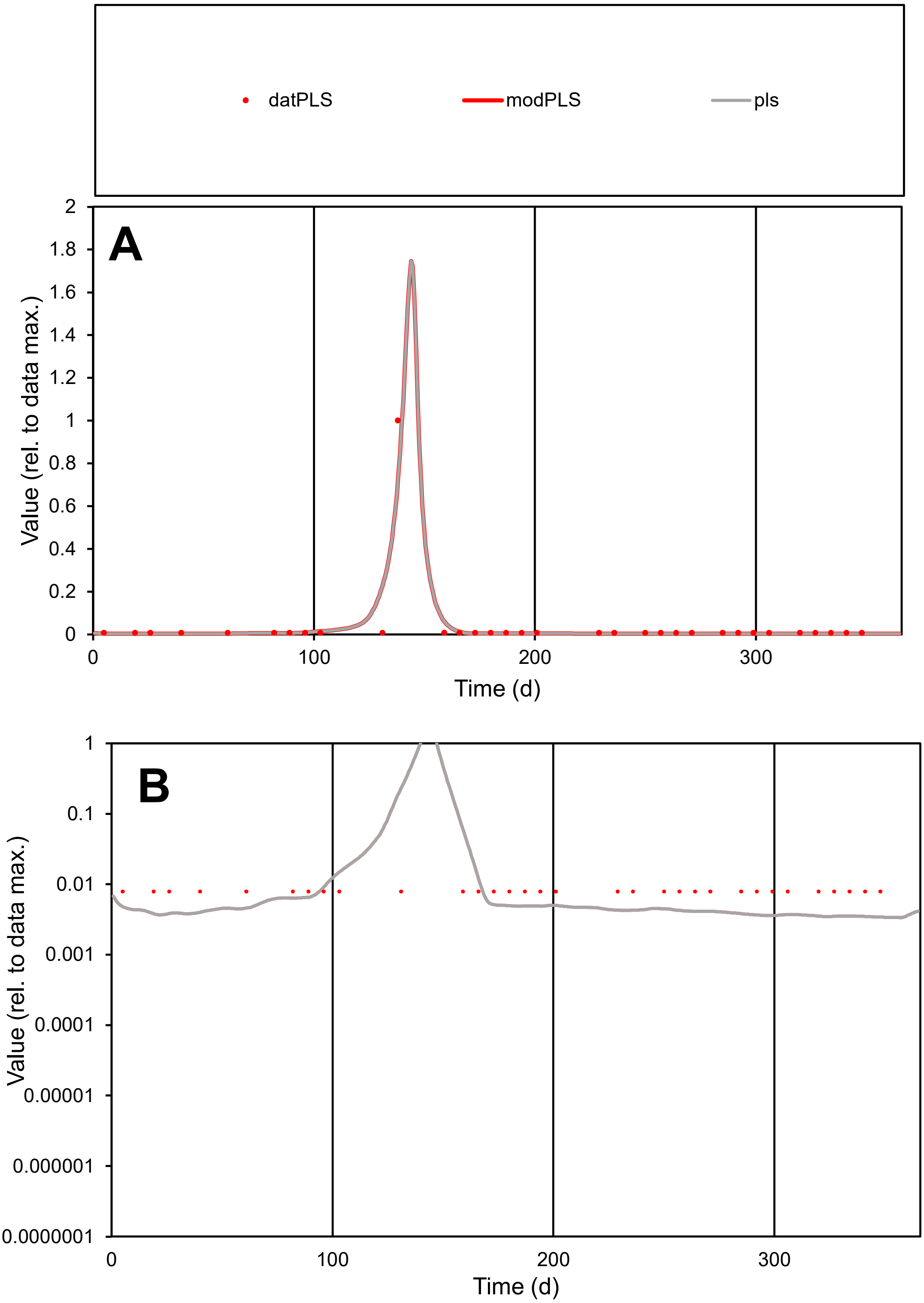


**C1**

Variables

PLS Provide jtd or Dst

CalibOnly?

2 1 to reproduce of

oNormalize?

2 1=no, 2=yes

ofxn calc*

0.65433

ofxn mod

1.08523

sRun

RunA1

zout

1

Ave

Data

1.9E-05

Model

1.9E-05

data/vMax

0.00049

ofxng

263.459

Min d/m

2.30829

ave/min d

4.81813

ave/min m

11.4589

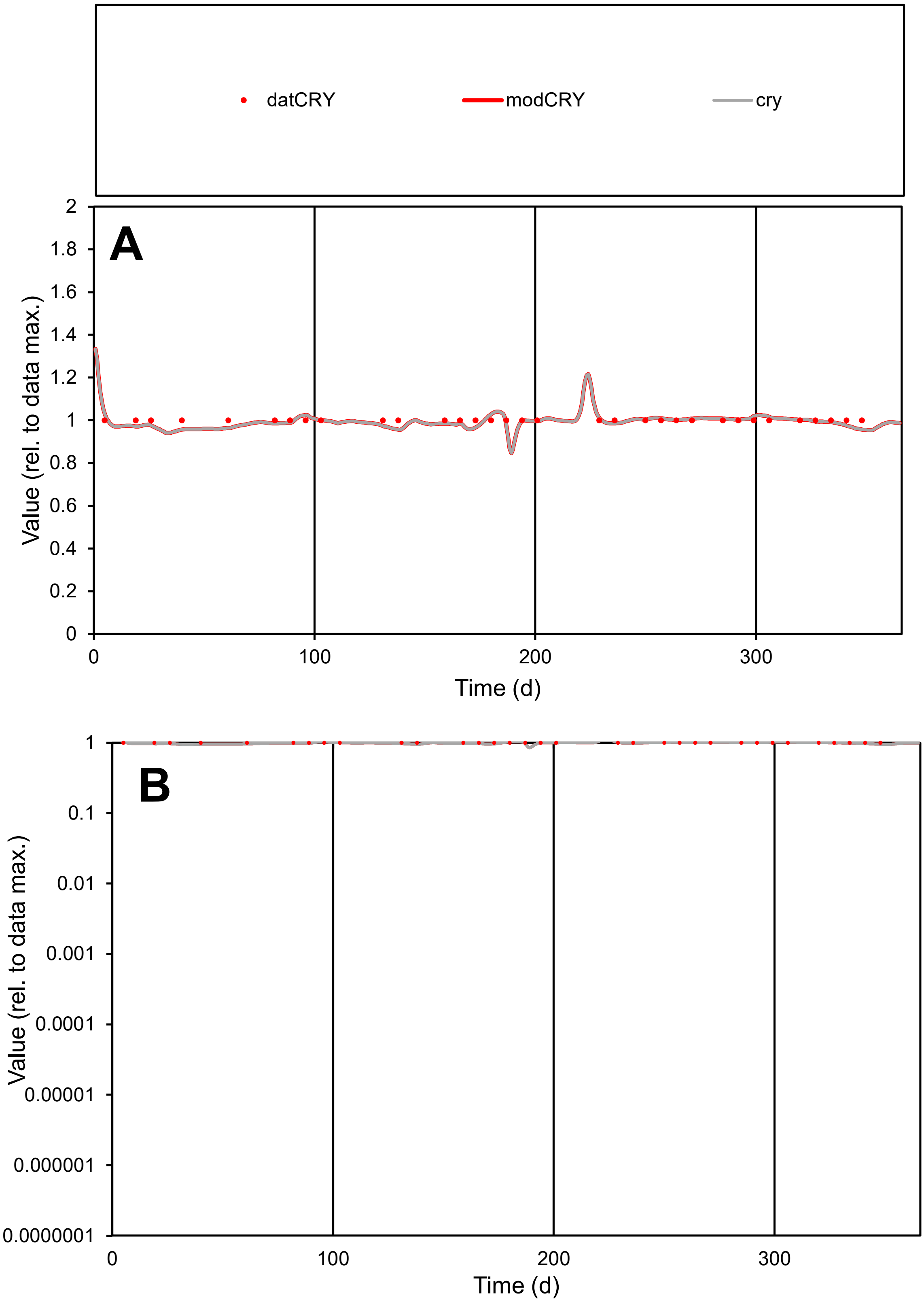


C1

Variables
CRY Provide jtd or Dst
CalibOnly?
2 1 to reproduce of
oNormalize?
2 1=no, 2=yes

ofxn calc*
0.02471
ofxn mod
0.11627

sRun
RunA1
zout
1

Ave
Data
3E-06
Model
3E-06

data/vMax
3E-06

ofxng
263.459

Min d/m
1.08032

ave/min d
1
ave/min m
1.06886

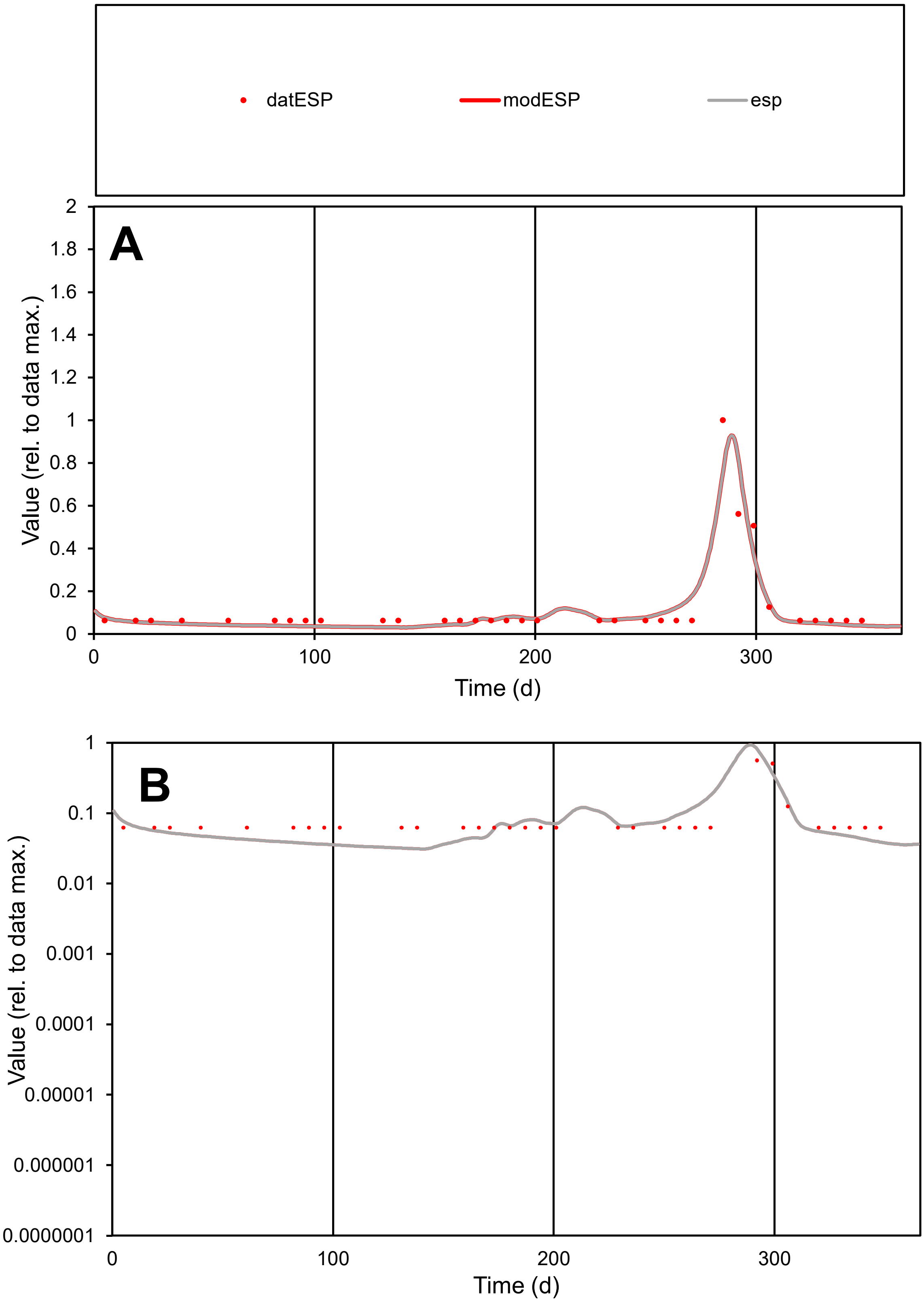


**C1**

Variables
ESP Provide jtd or Dst
CalibOnly?
2 1 to reproduce of
oNormalize?
2 1=no, 2=yes

ofxn calc*
0.4705
ofxn mod
0.51499

sRun
RunA1
zout
1

Ave
Data
1.6E-05
Model
1.5E-05

data/vMax
0.00013

ofxng
263.459

Min d/m
2.00853

ave/min d
1.94223
ave/min m
3.72635

326_Calib_S_A

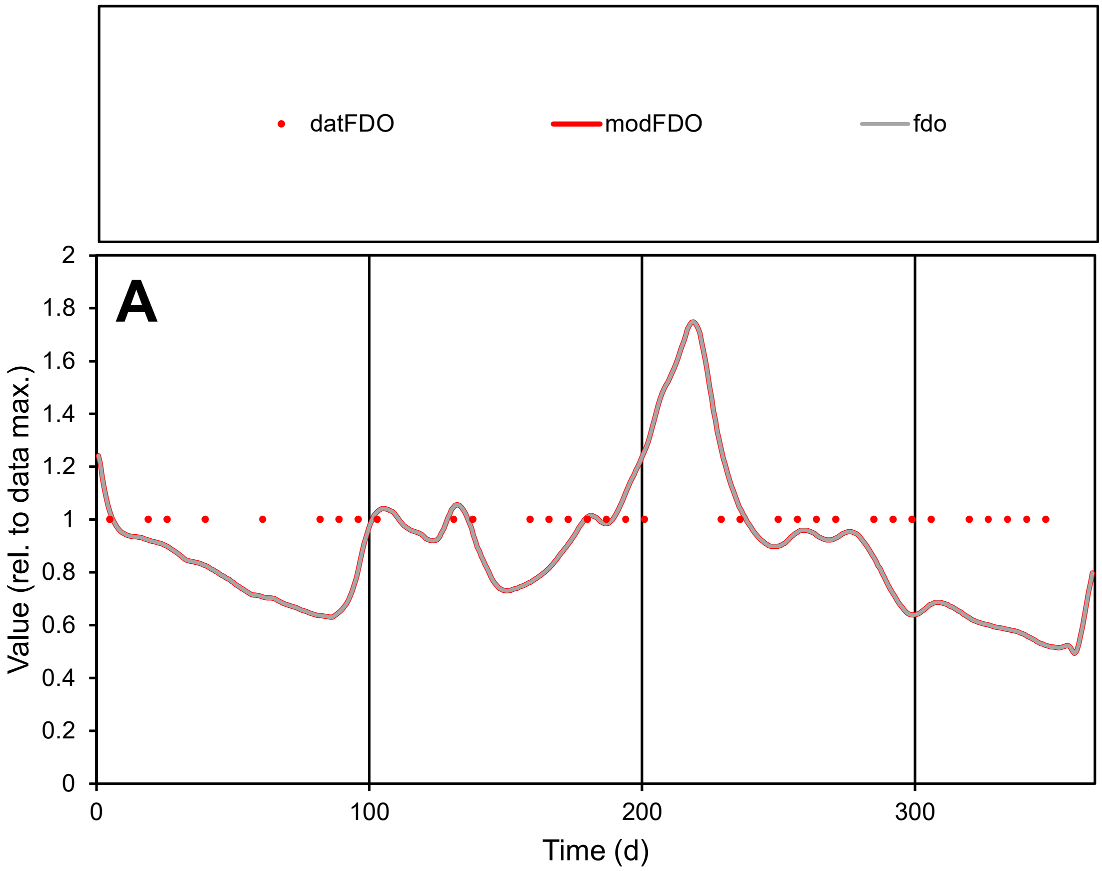


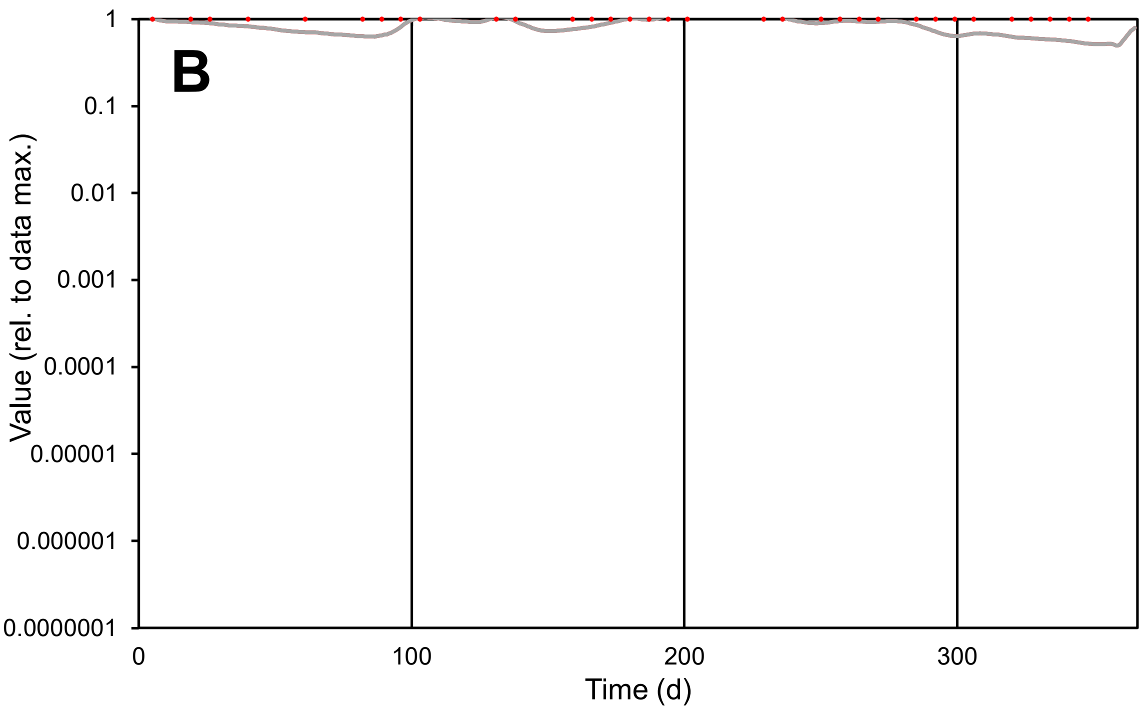


**C1**

Variables
FDO Provide jtd or Dst
CalibOnly?
2 1 to reproduce of
oNormalize?
2 1=no, 2=yes

ofxn calc*
0.24201
ofxn mod
0.26446

sRun
RunA1
zout
1

Ave
Data
1E-07
Model
8.5E-08

data/vMax
1E-07

ofxng
263.459

Min d/m
1.92188

ave/min d
1
ave/min m
1.63881

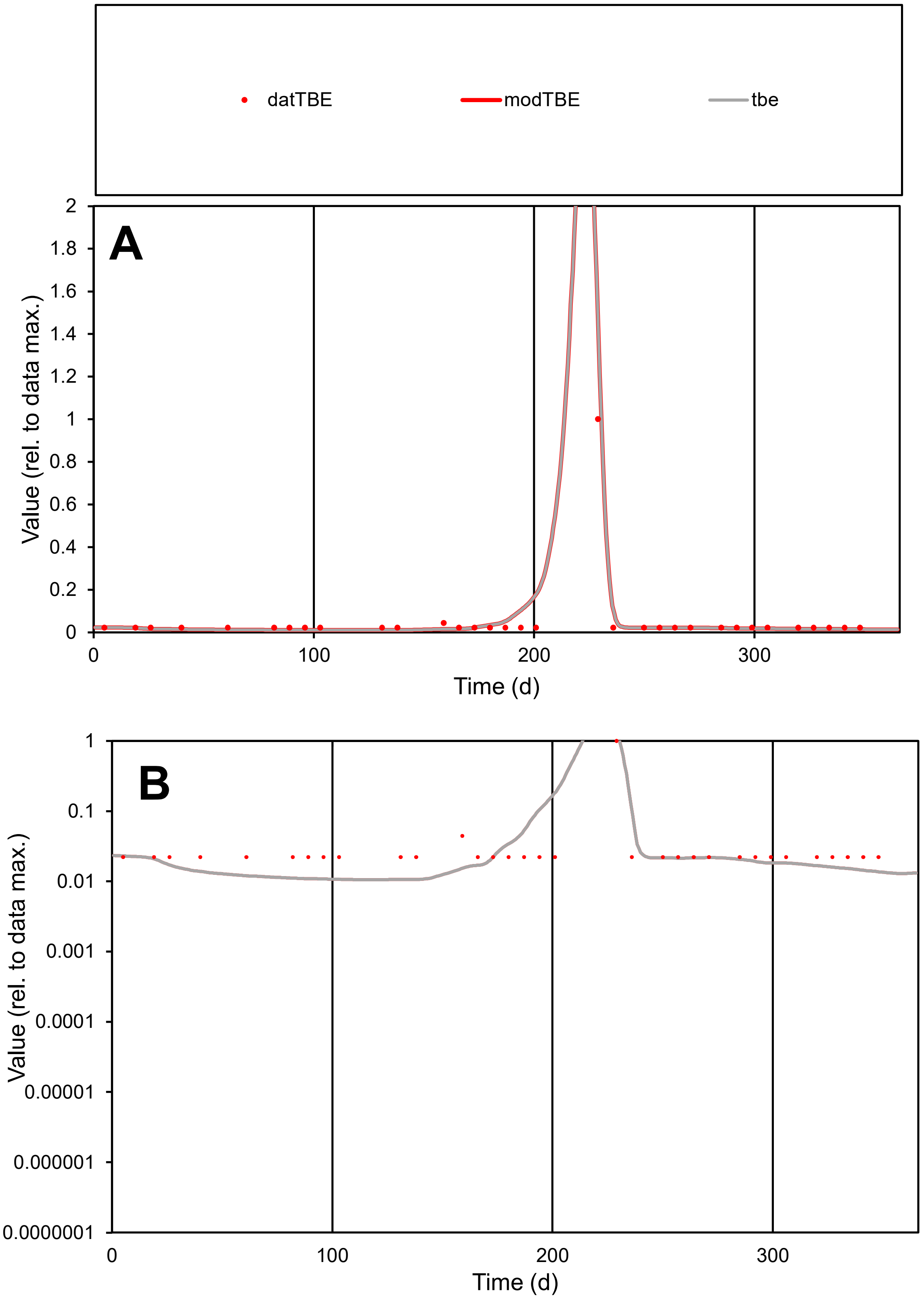


**C1**

Variables
TBE Provide jtd or Dst
CalibOnly?
2 1 to reproduce of
oNormalize?
2 1=no, 2=yes

ofxn calc*
0.548
ofxn mod
0.79958

sRun
RunA1
zout
1

Ave
Data
1.6E-05
Model
1.7E-05

data/vMax
0.0003

ofxng
263.459

Min d/m
2.08333

ave/min d
2.36874
ave/min m
5.17036

326_Calib_S_A

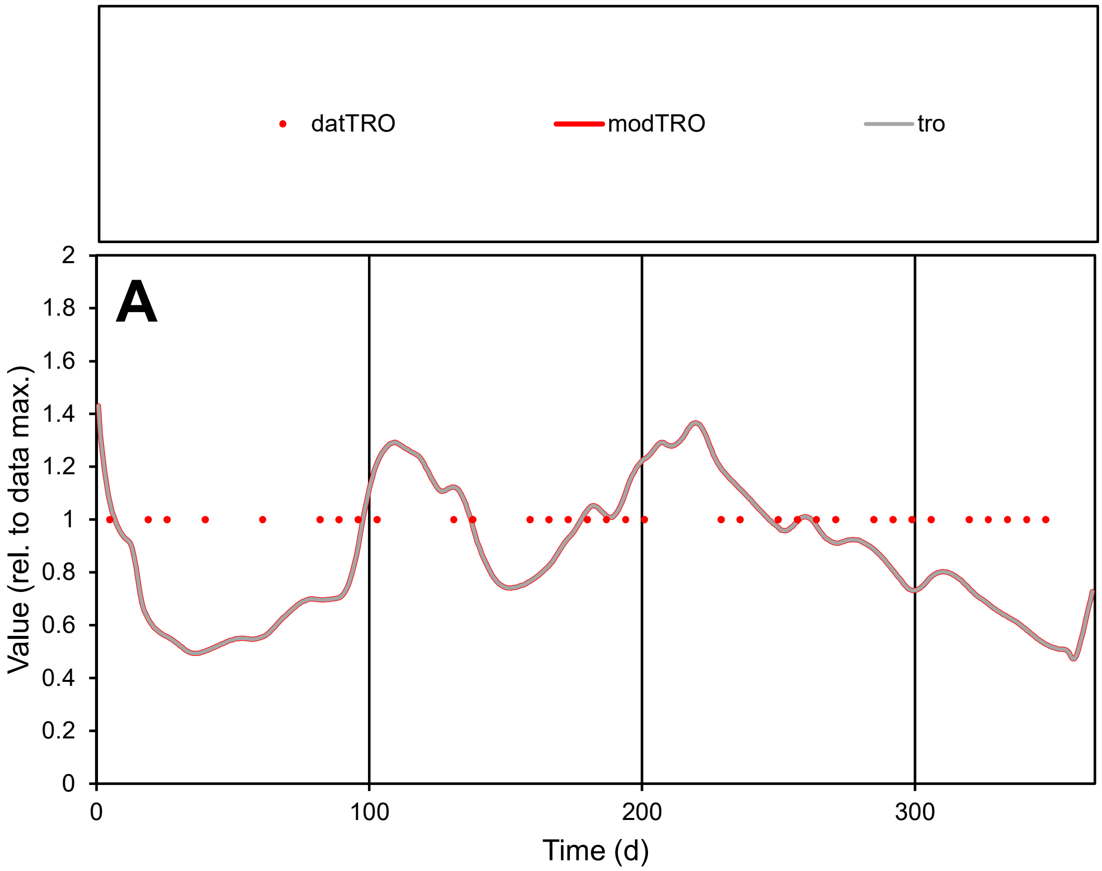


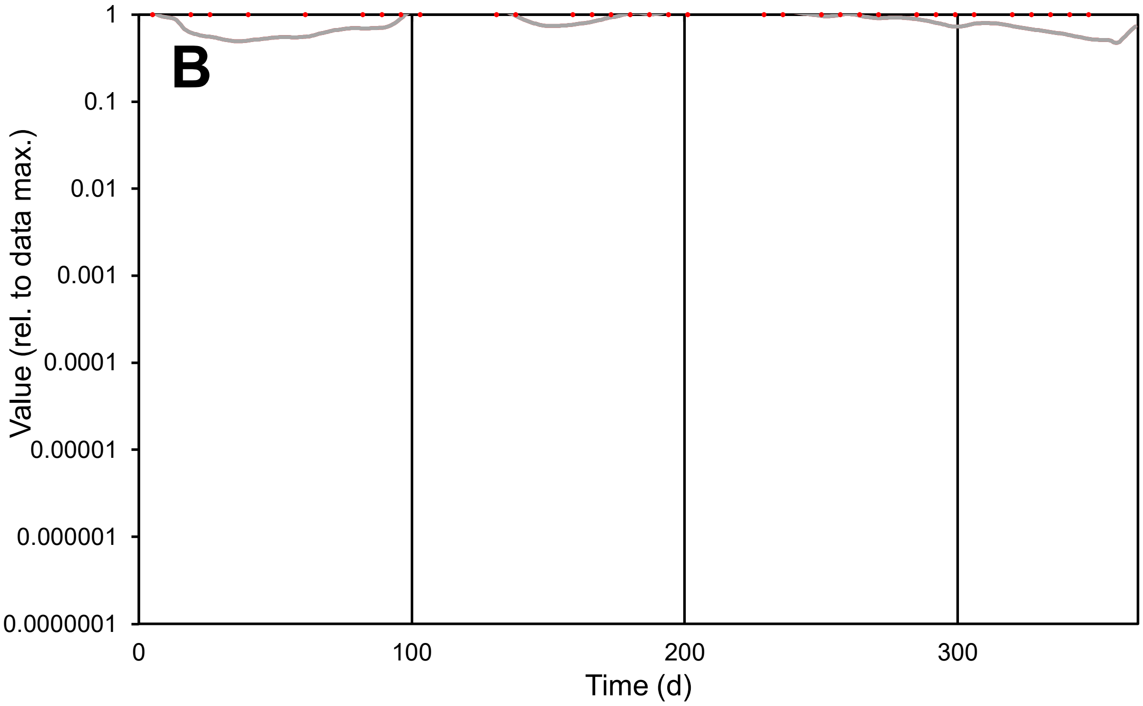


**C1**

Variables

| | |
|---|---|
| TRO | Provide jtd or Dst |
| CalibOnly? | |
| 2 | 1 to reproduce of |
| oNormalize? | |
| 2 | 1=no, 2=yes |

ofxn calc*
0.25604
ofxn mod
0.27734

sRun
RunA1
zout
1

Ave
Data
5E-06
Model
4.3E-06

data/vMax
5E-06

ofxng
263.459

Min d/m
1.96637

ave/min d
1
ave/min m
1.68841

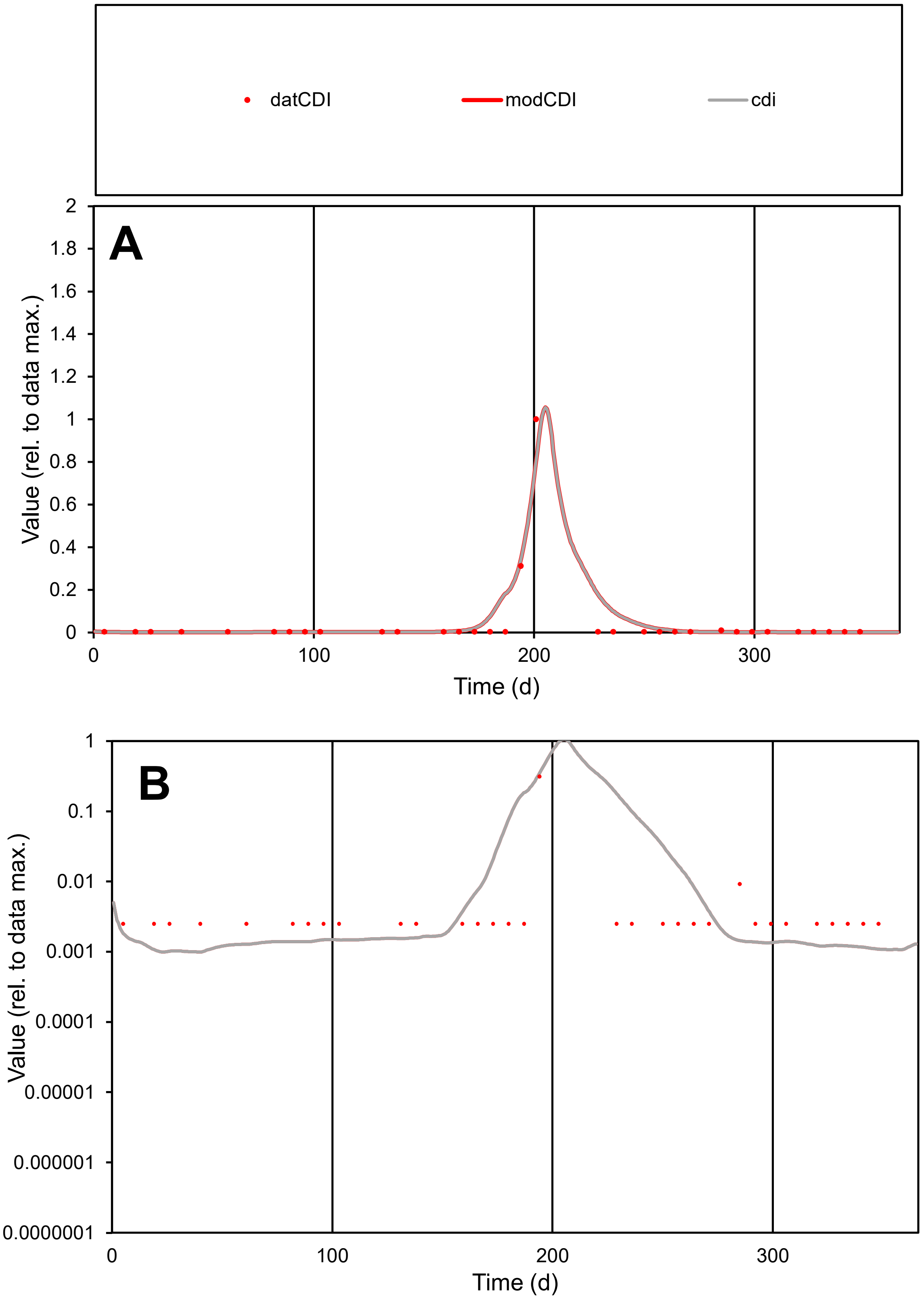


**C1**

Variables
CDI Provide jtd or Dst
CalibOnly?
2 1 to reproduce of
oNormalize?
2 1=no, 2=yes

ofxn calc*
1.02155
ofxn mod
1.20988

sRun
RunA1
zout
1

Ave
Data
2.9E-06
Model
3.8E-06

data/vMax
6.9E-05

ofxng
263.459

Min d/m
2.46408

ave/min d
16.9549
ave/min m
53.8661

326_Calib_S_A

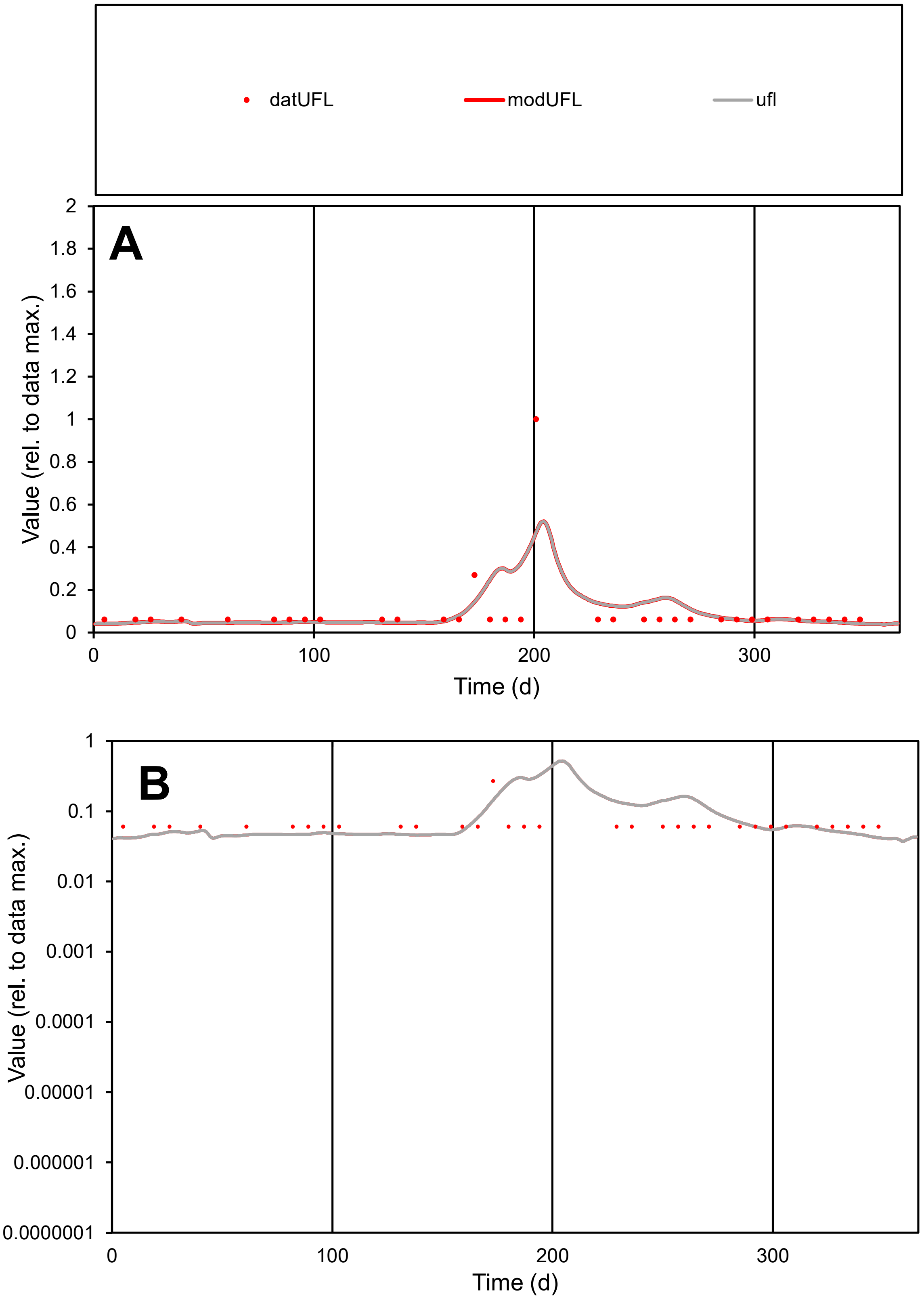


**C1**

Variables
UFL Provide jtd or Dst
CalibOnly?
2 1 to reproduce of
oNormalize?
2 1=no, 2=yes

ofxn calc*
1.23926
ofxn mod
1.25437

sRun
RunA1
zout
1

Ave
Data
4.8E-06
Model
5.3E-06

data/vMax
5.1E-05

ofxng
263.459

Min d/m
1.45671

ave/min d
1.57524
ave/min m
2.54234

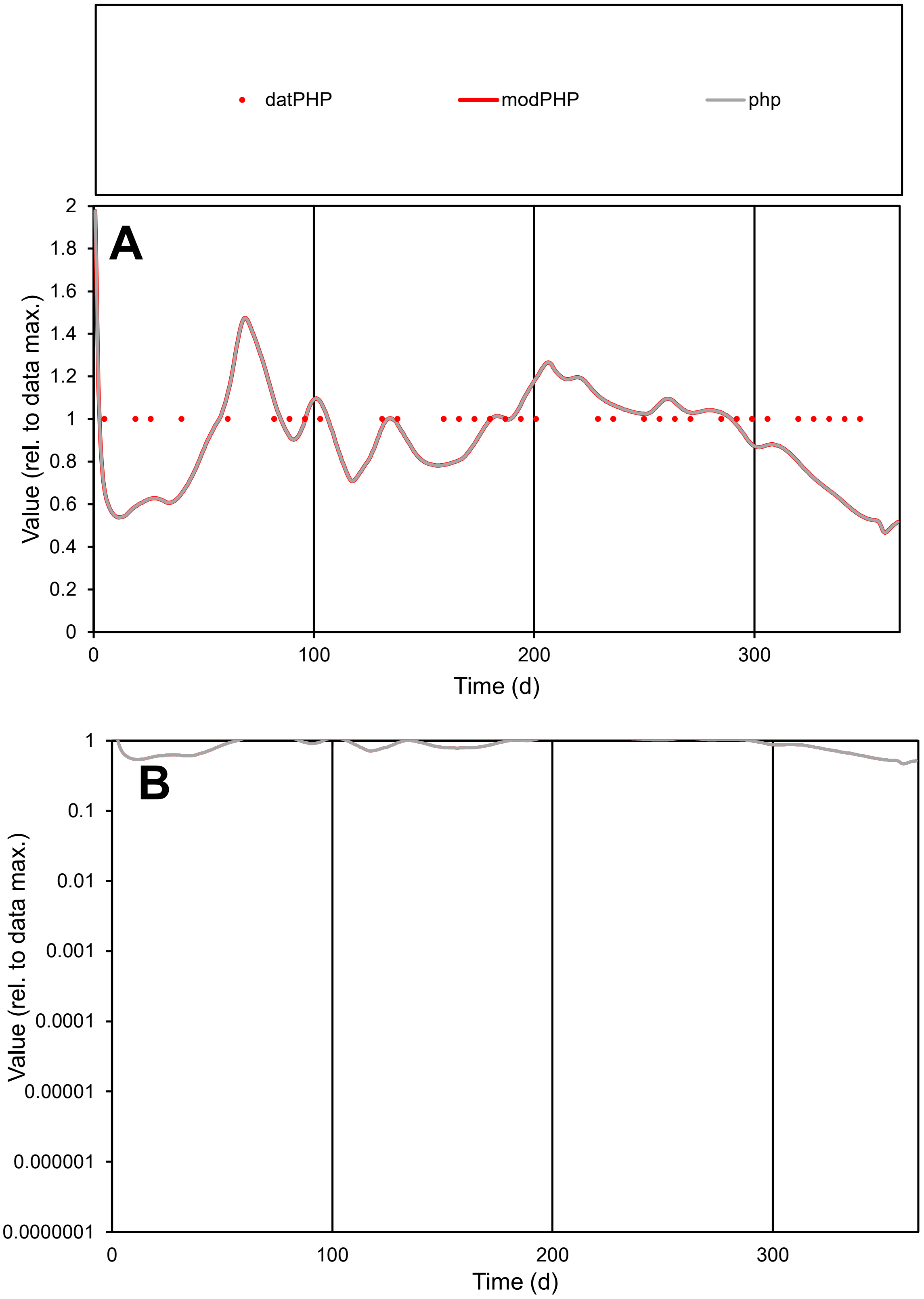


**C1**

Variables
PHP Provide jtd or Dst
CalibOnly?
2 1 to reproduce of
oNormalize?
2 1=no, 2=yes

ofxn calc*
0.20639
ofxn mod
0.2349

sRun
RunA1
zout
1

Ave
Data
6.9E-06
Model
6.3E-06

data/vMax
6.9E-06

ofxng
263.459

Min d/m
1.84111

ave/min d
1
ave/min m
1.6717

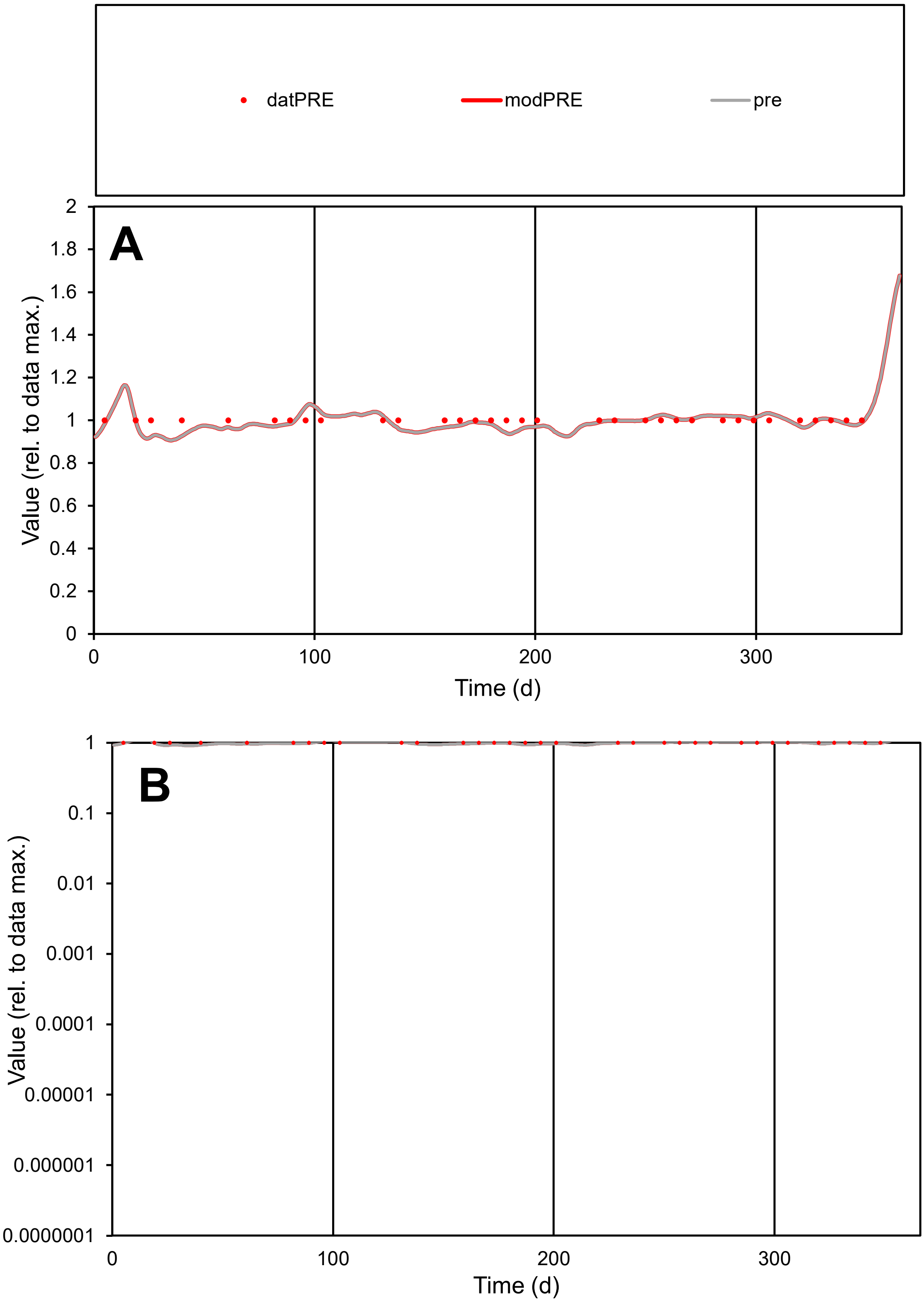


**C1**

Variables
PRE Provide jtd or Dst
CalibOnly?
2 1 to reproduce of
oNormalize?
2 1=no, 2=yes

ofxn calc*
0.03251
ofxn mod
0.11711

sRun
RunA1
zout
1

Ave
Data
1.3E-05
Model
1.3E-05

data/vMax
1.3E-05

ofxng
263.459

Min d/m
1.0744

ave/min d
1
ave/min m
1.06424

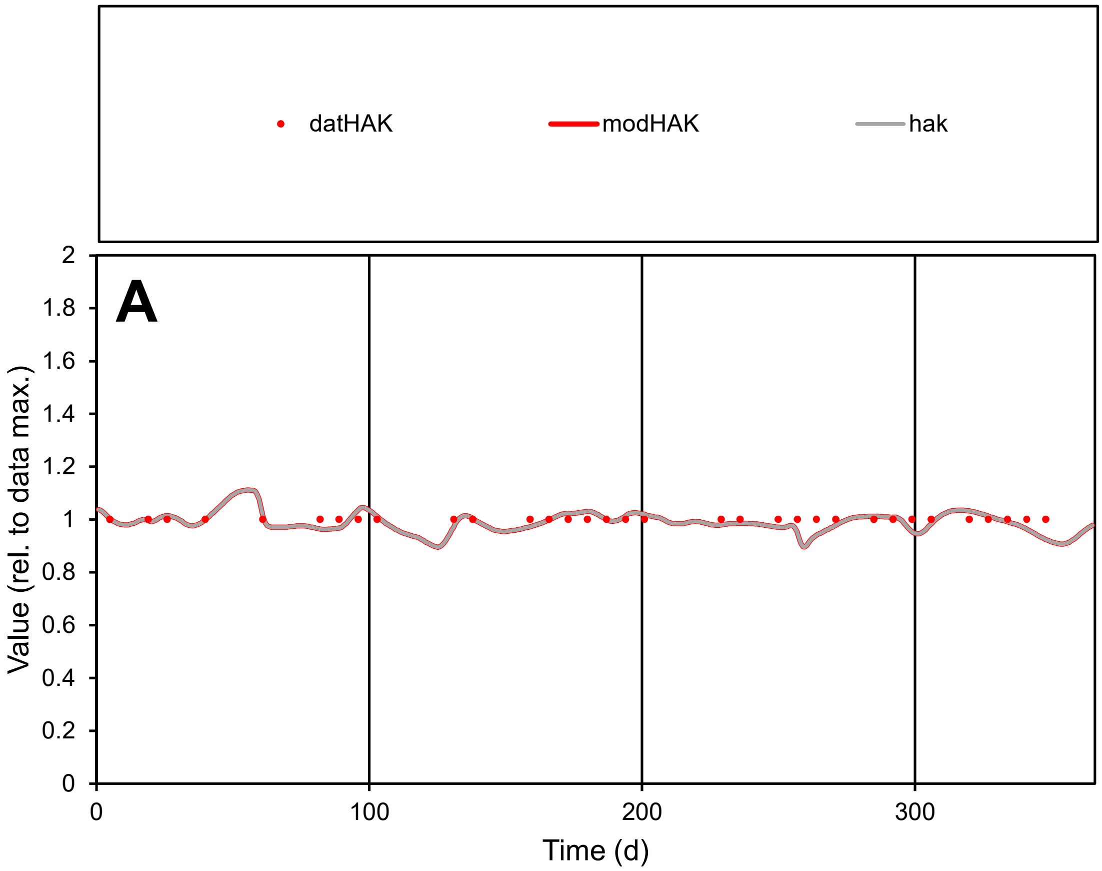


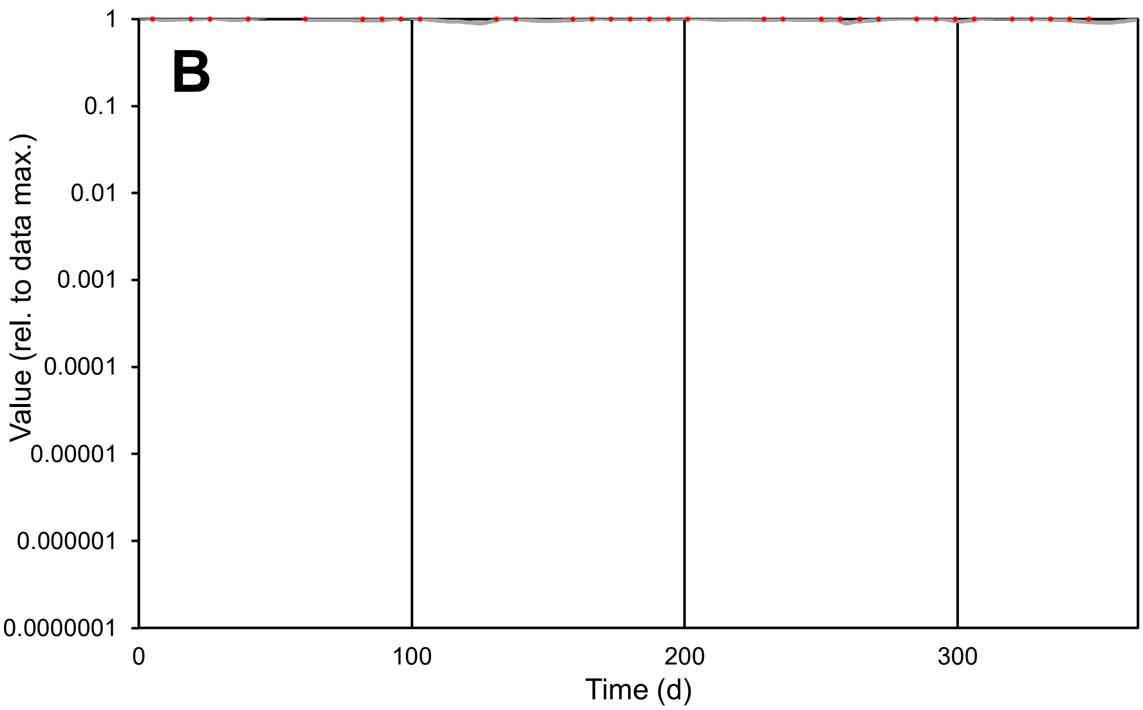


**C1**

Variables
HAK Provide jtd or Dst
CalibOnly?
2 1 to reproduce of
oNormalize?
2 1=no, 2=yes

ofxn calc*
0.03175
ofxn mod
0.11716

sRun
RunA1
zout
1

Ave
Data
5.1E-06
Model
5E-06

data/vMax
5.1E-06

ofxng
263.459

Min d/m
1.09912

ave/min d
1
ave/min m
1.08809

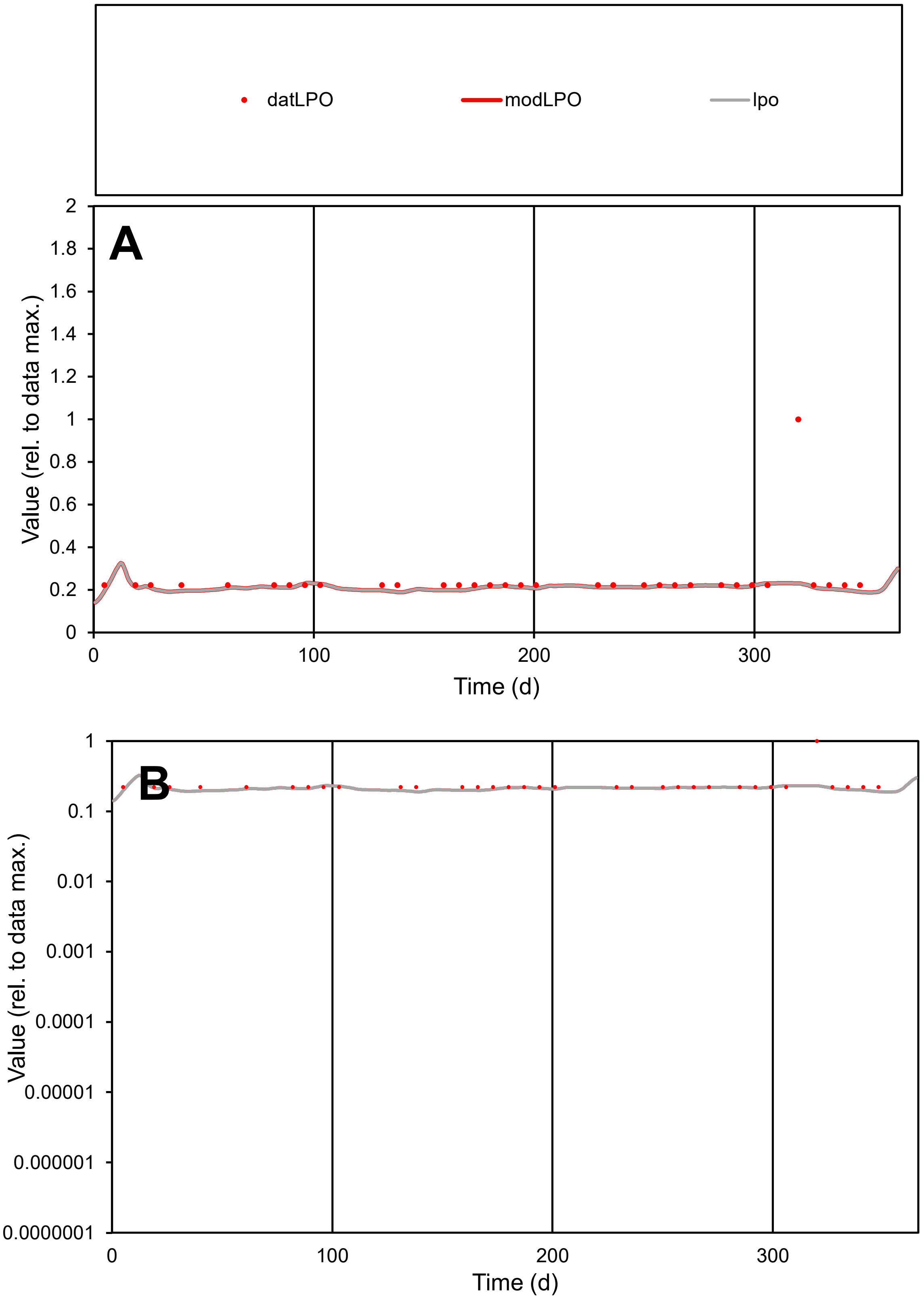


**C1**

| | |
|---|---|
| Variables | |
| LPO | Provide jtd or Dst |
| CalibOnly? | |
| 2 | 1 to reproduce of |
| oNormalize? | |
| 2 | 1=no, 2=yes |
| ofxn calc* | |
| 0.55134 | |
| ofxn mod | |
| 0.56151 | |
| sRun | |
| RunA1 | |
| zout | |
| 1 | |
| Ave | |
| Data | |
| 6.5E-06 | |
| Model | |
| 5.6E-06 | |
| data/vMax | |
| 2.6E-05 | |
| ofxng | |
| 263.459 | |
| Min d/m | |
| 1.1698 | |
| ave/min d | |
| 1.10657 | |
| ave/min m | |
| 1.11674 | |

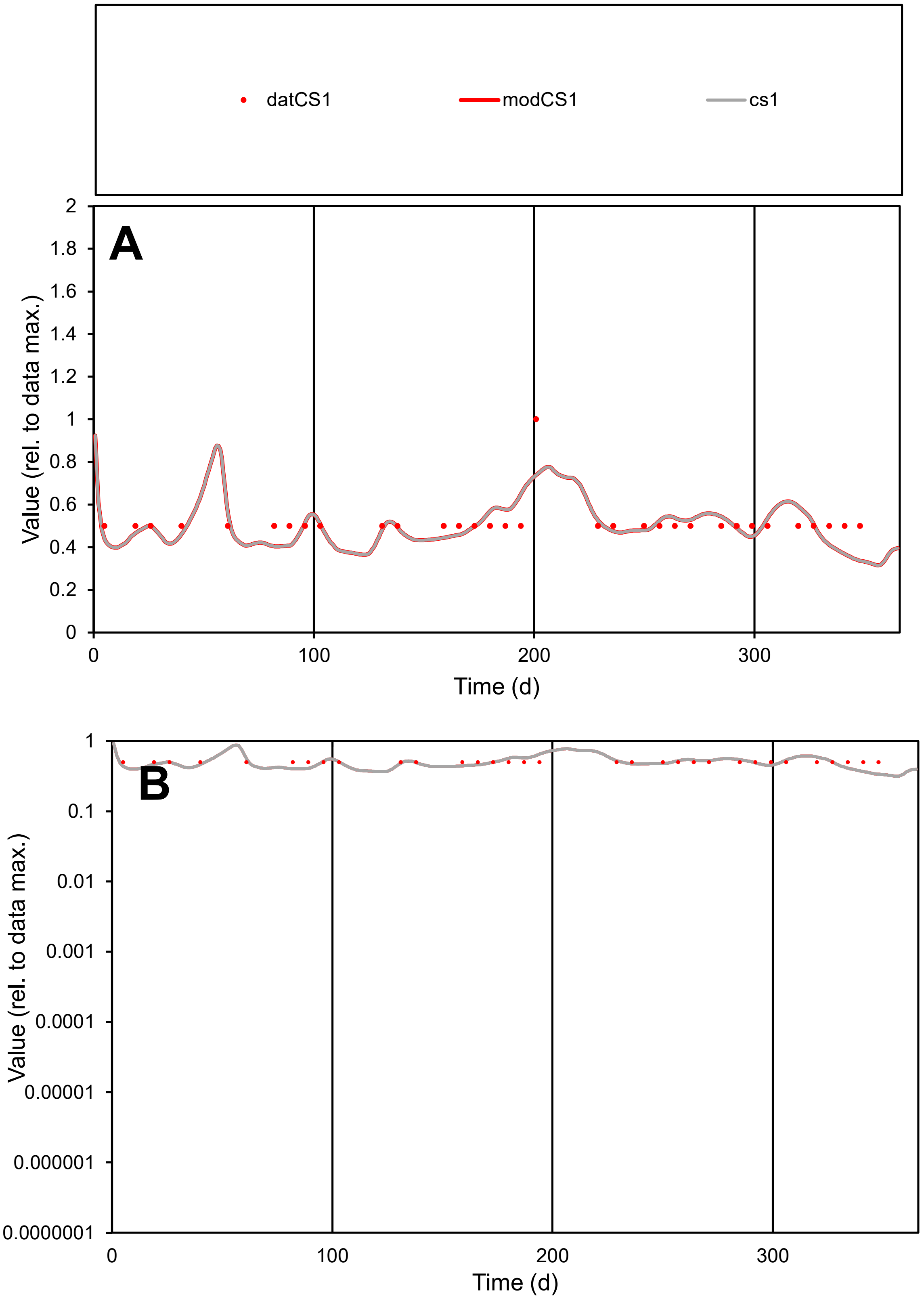


**C1**

Variables
CS1 Provide jtd or Dst
CalibOnly?
2 1 to reproduce of
oNormalize?
2 1=no, 2=yes

ofxn calc*
0.15363
ofxn mod
0.19241

sRun
RunA1
zout
1

Ave
Data
1.8E-05
Model
1.7E-05

data/vMax
3.5E-05

ofxng
263.459

Min d/m
1.49616

ave/min d
1.0303
ave/min m
1.48674

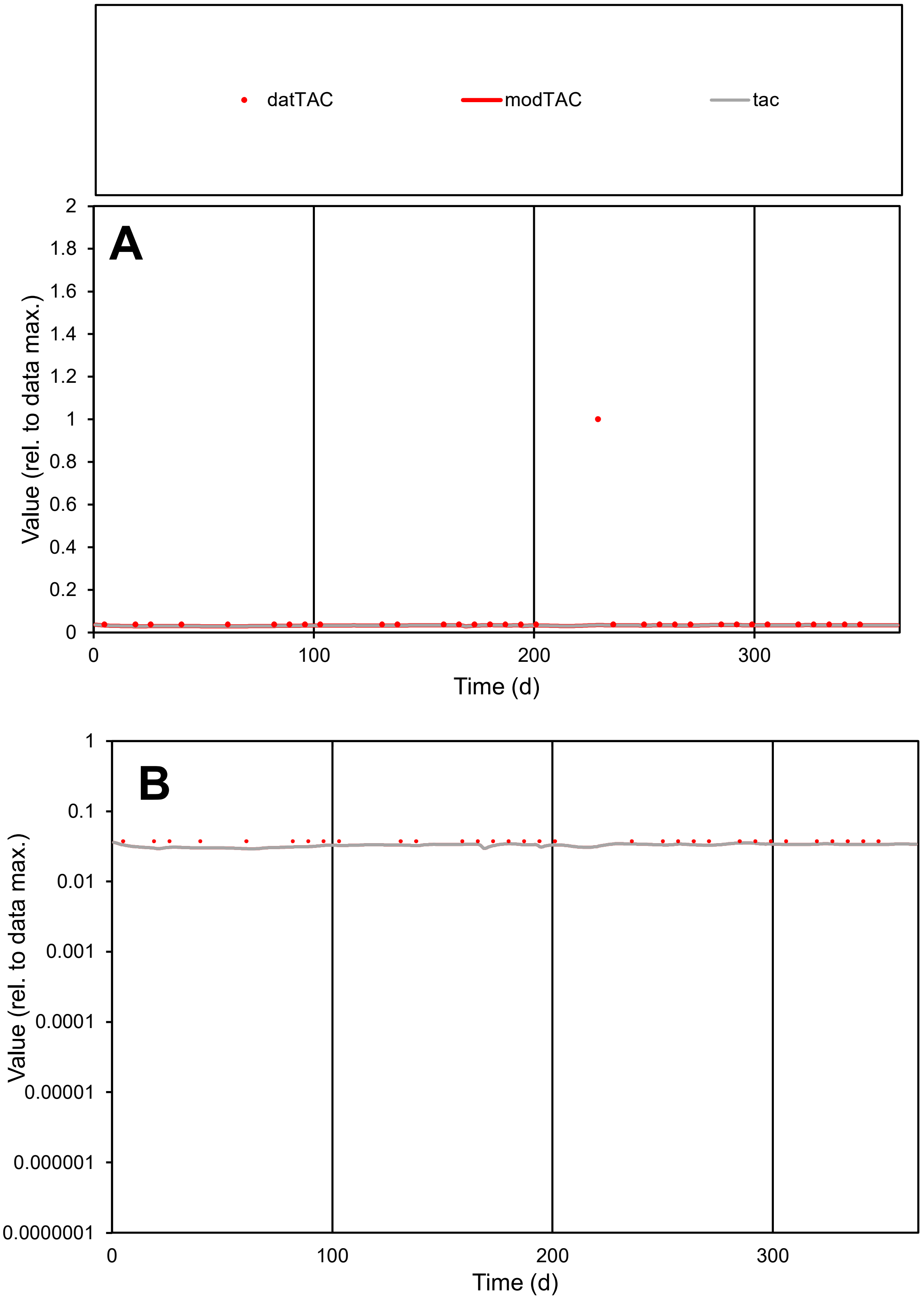


**C1**

Variables
TAC Provide jtd or Dst
CalibOnly?
2 1 to reproduce of
oNormalize?
2 1=no, 2=yes

ofxn calc*
2.51861
ofxn mod
2.5244

sRun
RunA1
zout
1

Ave
Data
2.3E-06
Model
1.1E-06

data/vMax
3.4E-05

ofxng
263.459

Min d/m
1.28472

ave/min d
1.77583
ave/min m
1.12626

326_Calib_S_A

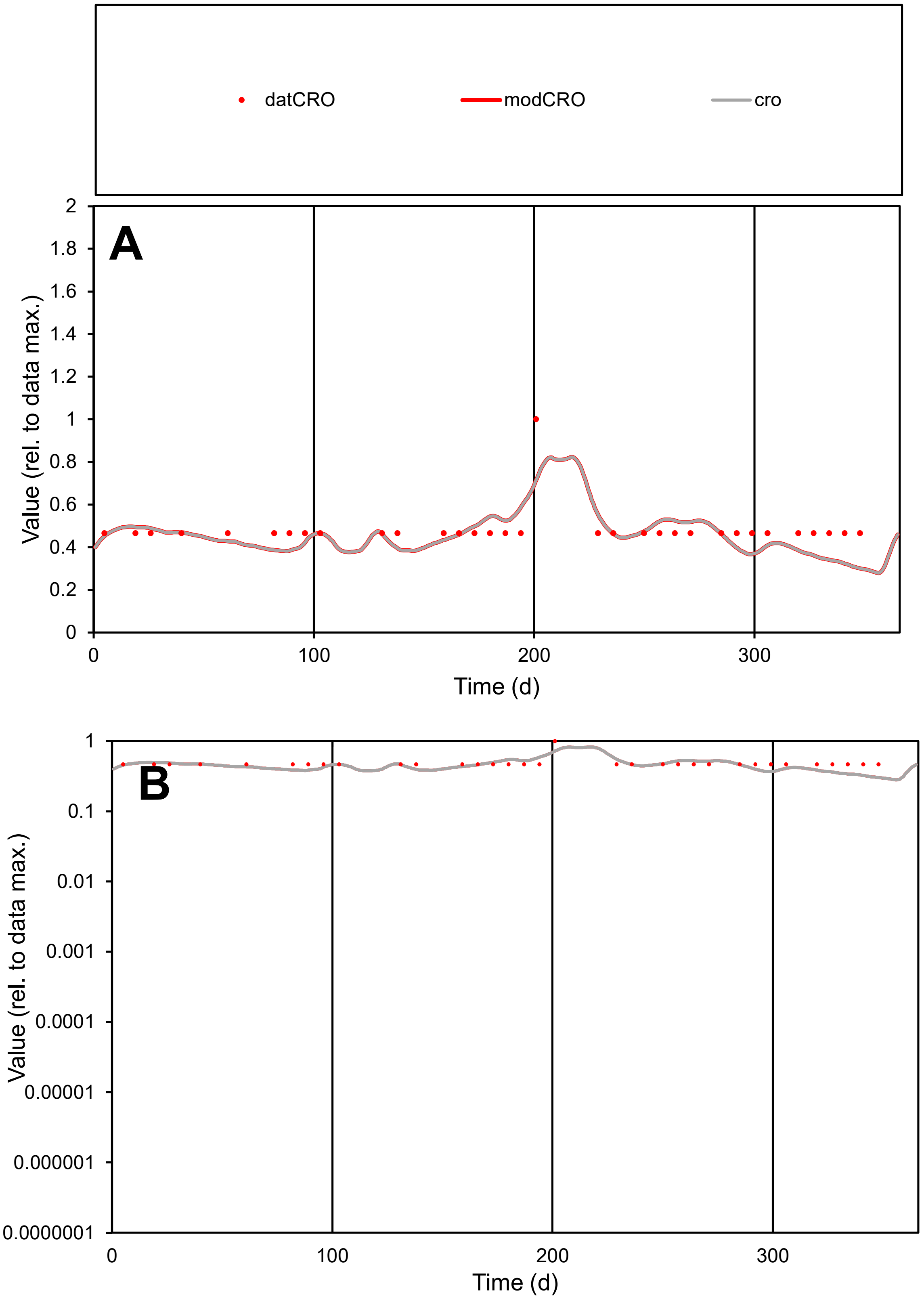


**C1**

Variables
CRO Provide jtd or Dst
CalibOnly?
2 1 to reproduce of
oNormalize?
2 1=no, 2=yes

ofxn calc*
0.1741
ofxn mod
0.20879

sRun
RunA1
zout
1

Ave
Data
1.1E-05
Model
1E-05

data/vMax
2.2E-05

ofxng
263.459

Min d/m
1.55931

ave/min d
1.0348
ave/min m
1.52565

326_Calib_S_A

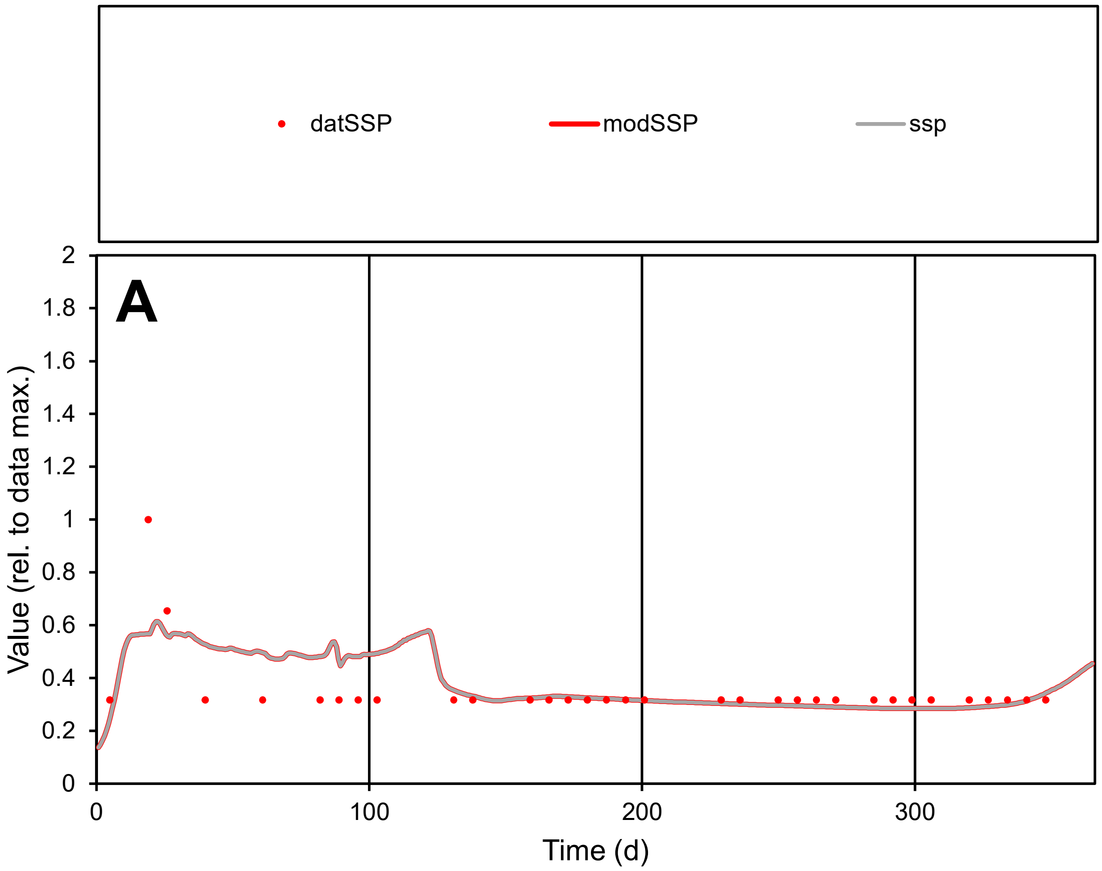


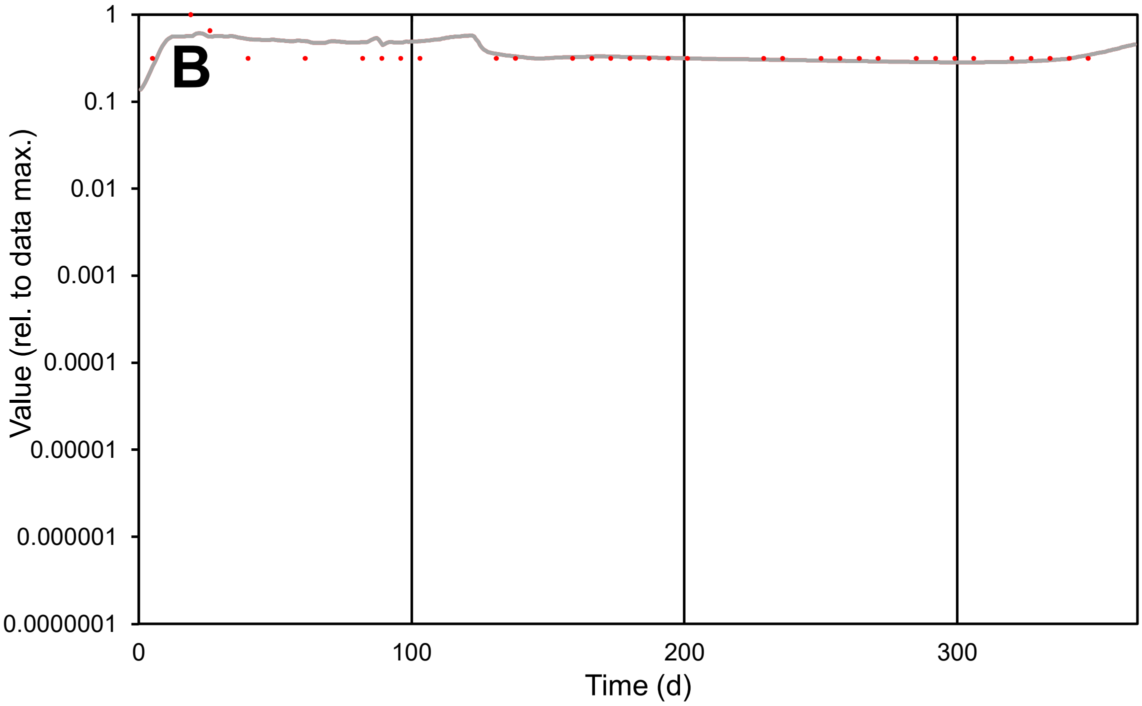


**C1**

Variables
SSP Provide jtd or Dst
CalibOnly?
2 1 to reproduce of
oNormalize?
2 1=no, 2=yes

ofxn calc*
0.30591
ofxn mod
0.32976

sRun
RunA1
zout
1

Ave
Data
1.5E-05
Model
1.5E-05

data/vMax
4.2E-05

ofxng
263.459

Min d/m
1.1127

ave/min d
1.09803
ave/min m
1.2595

326_Calib_S_A

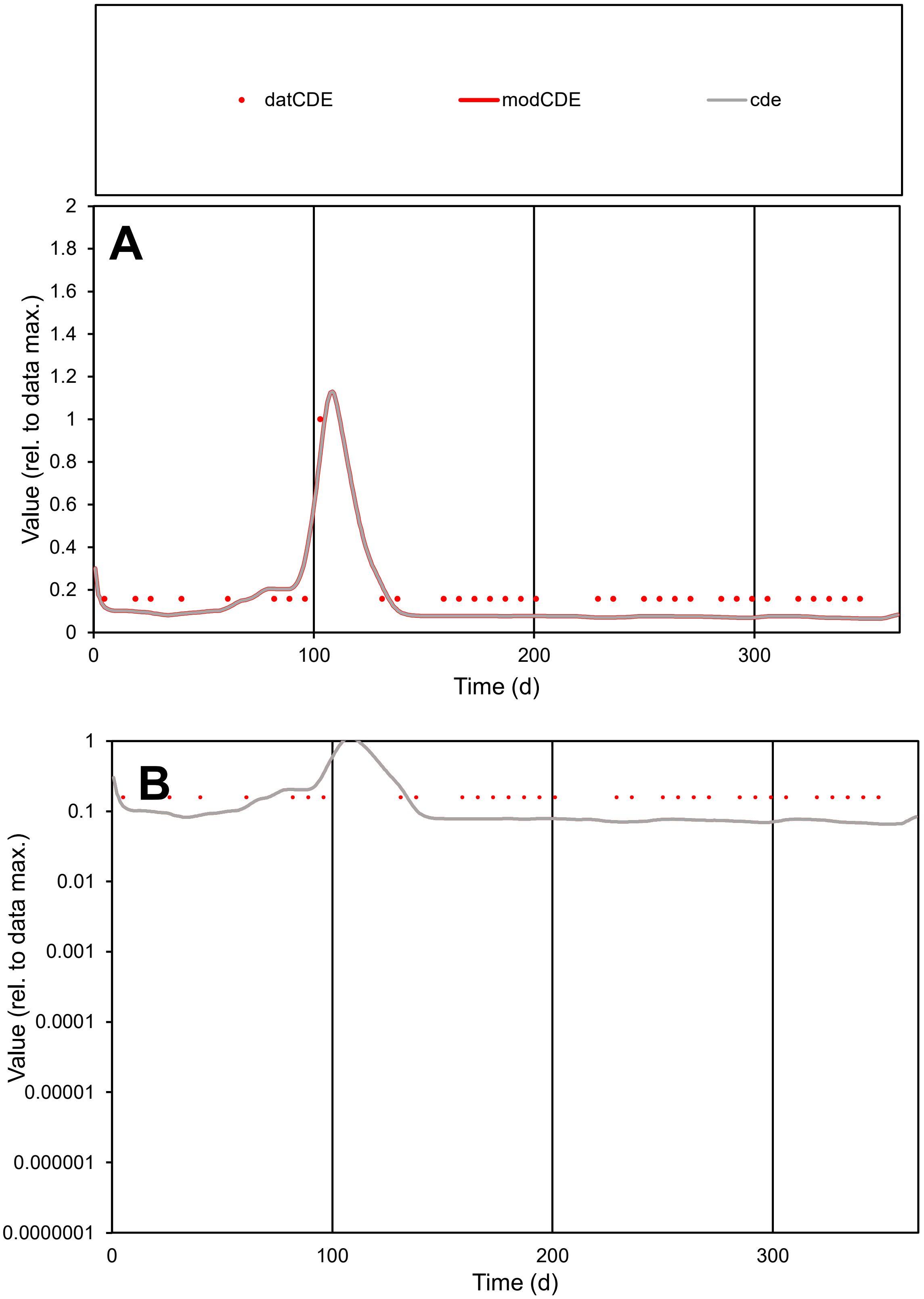


**C1**

Variables

| | |
|---|---|
| CDE | Provide jtd or Dst |
| CalibOnly? | |
| 2 | 1 to reproduce of |
| oNormalize? | |
| 2 | 1=no, 2=yes |

ofxn calc*
0.44455
ofxn mod
0.45873

sRun
RunA1
zout
1

Ave
Data
1.2E-05
Model
7.9E-06

data/vMax
6.3E-05

ofxng
263.459

Min d/m
2.38923

ave/min d
1.16146
ave/min m
1.88973

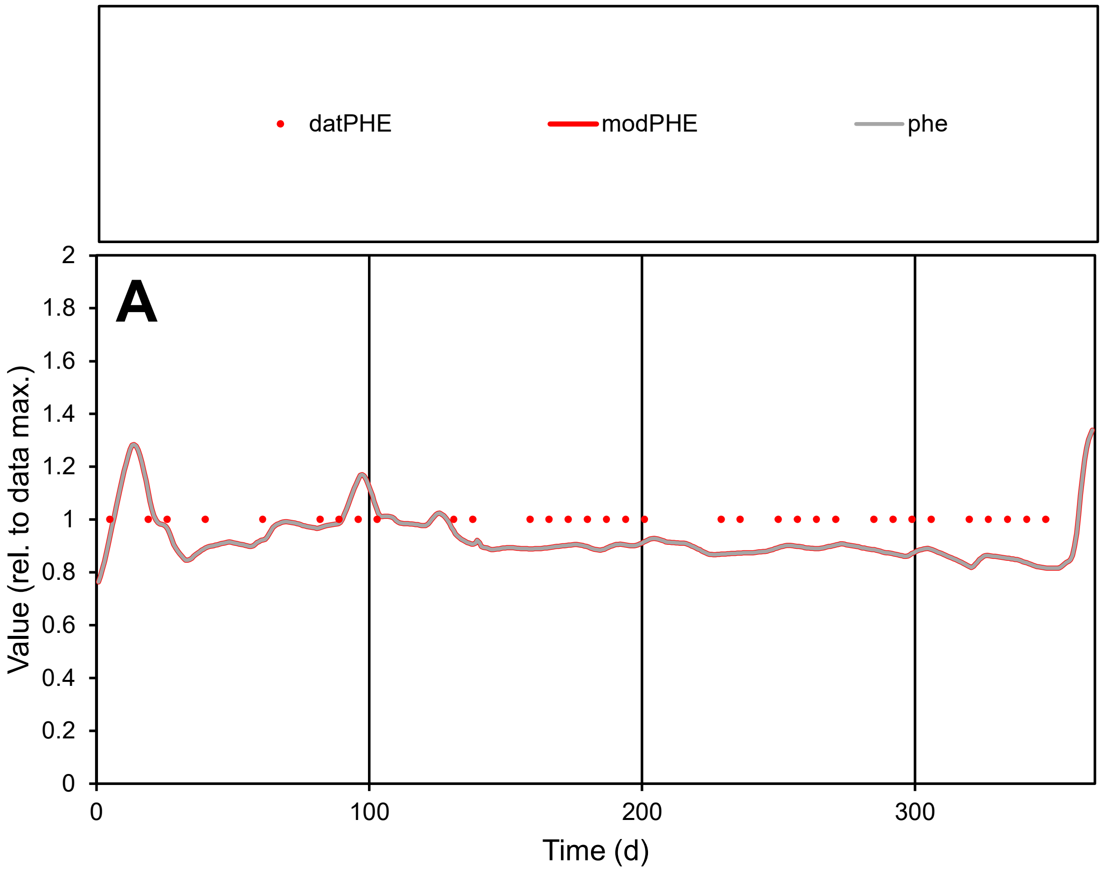


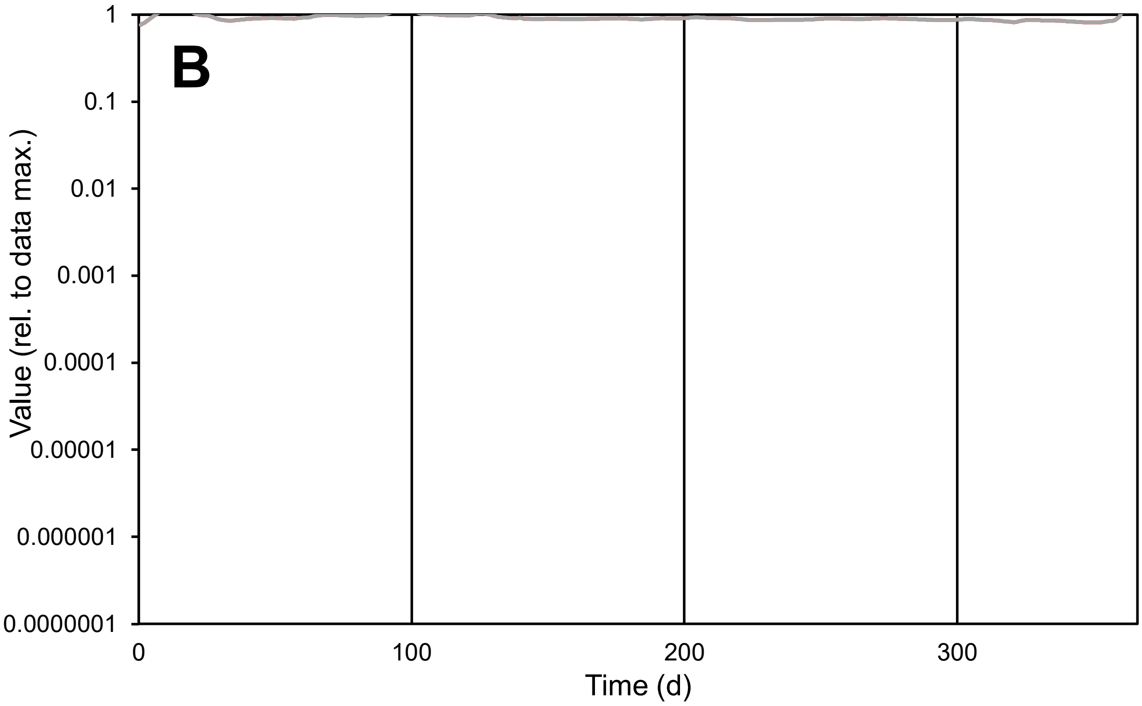


**C1**

Variables
PHE Provide jtd or Dst
CalibOnly?
2 1 to reproduce of
oNormalize?
2 1=no, 2=yes

ofxn calc*
0.11044
ofxn mod
0.15482

sRun
RunA1
zout
1

Ave
Data
4E-06
Model
3.6E-06

data/vMax
4E-06

ofxng
263.459

Min d/m
1.22552

ave/min d
1
ave/min m
1.11995

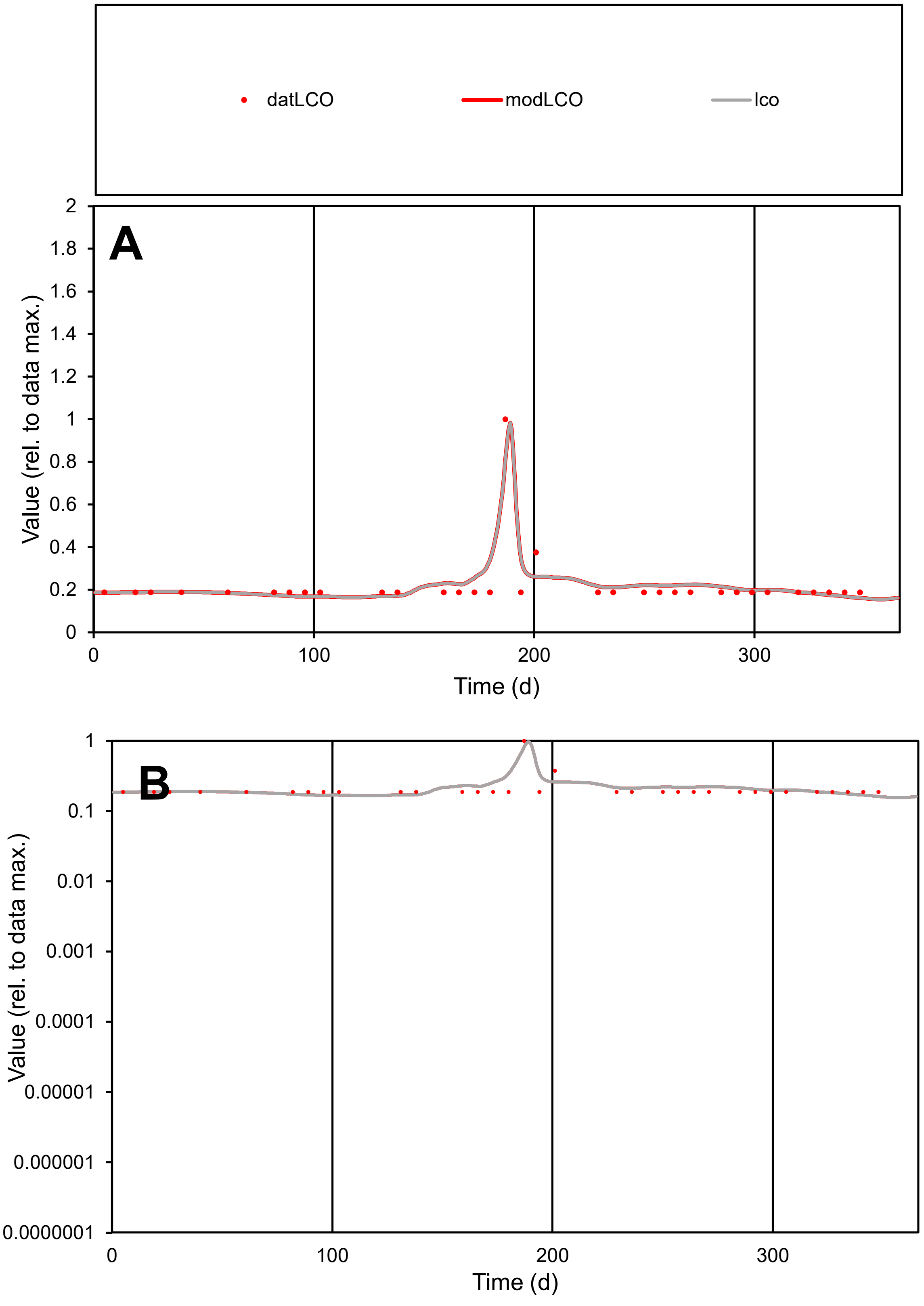


C1

Variables
LCO Provide jtd or Dst
CalibOnly?
2 1 to reproduce of
oNormalize?
2 1=no, 2=yes

ofxn calc*
0.21711
ofxn mod
0.26128

sRun
RunA1
zout
1

Ave
Data
8.6E-06
Model
9E-06

data/vMax
4E-05

ofxng
263.459

Min d/m
1.16774

ave/min d
1.16167
ave/min m
1.42251

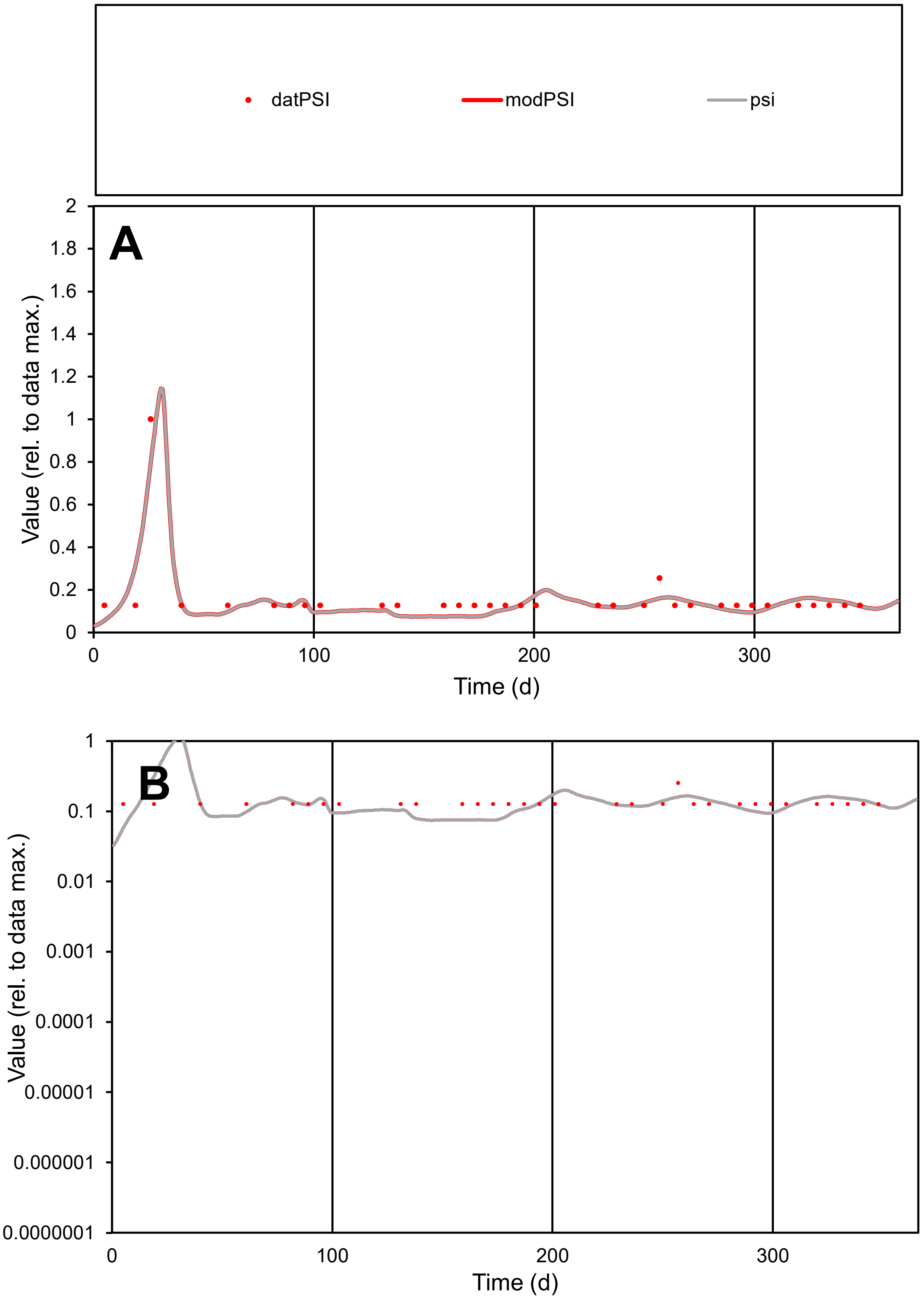


**C1**

Variables
PSI Provide jtd or Dst
CalibOnly?
2 1 to reproduce of
oNormalize?
2 1=no, 2=yes

ofxn calc*
0.27784
ofxn mod
0.3517

sRun
RunA1
zout
1

Ave
Data
4.4E-06
Model
4.2E-06

data/vMax
2.8E-05

ofxng
263.459

Min d/m
1.83162

ave/min d
1.23836
ave/min m
2.14927

326_Calib_S_A

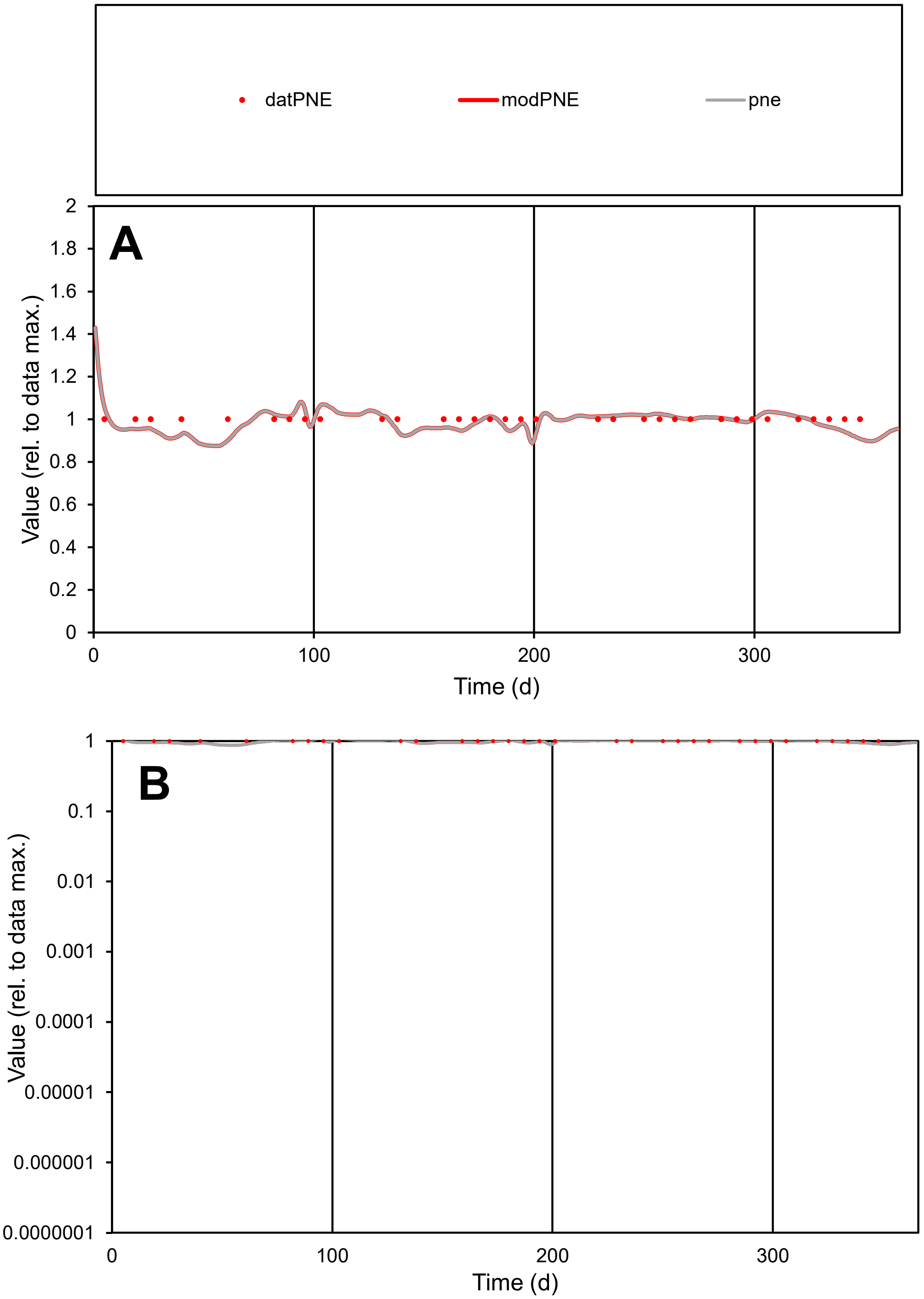


**C1**

Variables
PNE Provide jtd or Dst
CalibOnly?
2 1 to reproduce of
oNormalize?
2 1=no, 2=yes

ofxn calc*
0.0403
ofxn mod
0.11956

sRun
RunA1
zout
1

Ave
Data
1.5E-05
Model
1.4E-05

data/vMax
1.5E-05

ofxng
263.459

Min d/m
1.10502

ave/min d
1
ave/min m
1.08927

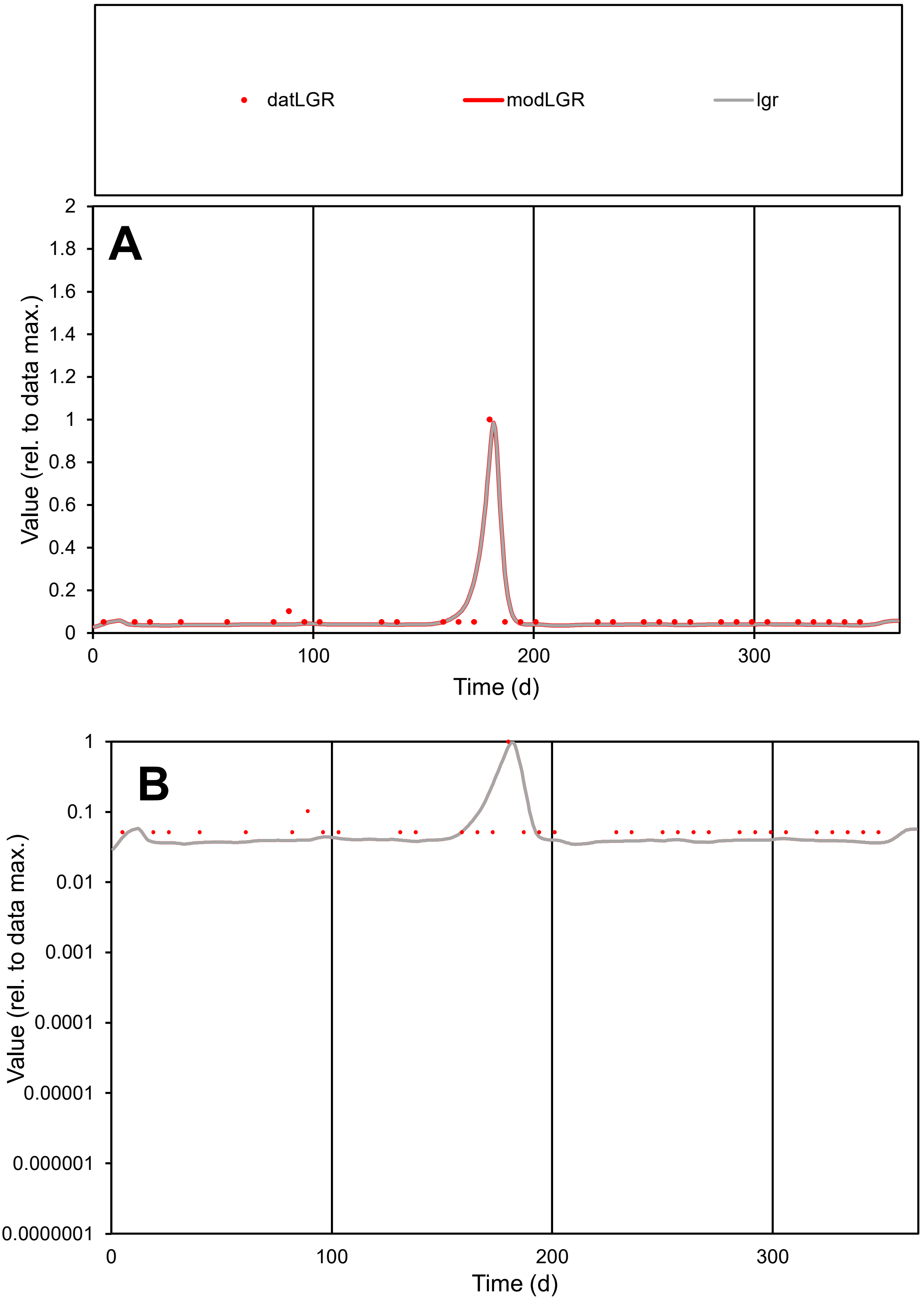


**C1**

Variables

LGR Provide jtd or Dst

CalibOnly?

2 1 to reproduce of

oNormalize?

2 1=no, 2=yes

ofxn calc*

0.46495

ofxn mod

0.52696

sRun

RunA1

zout

1

Ave

Data

8.2E-06

Model

7.8E-06

data/vMax

0.0001

ofxng

263.459

Min d/m

1.41309

ave/min d

1.59054

ave/min m

2.12911

326_Calib_S_A

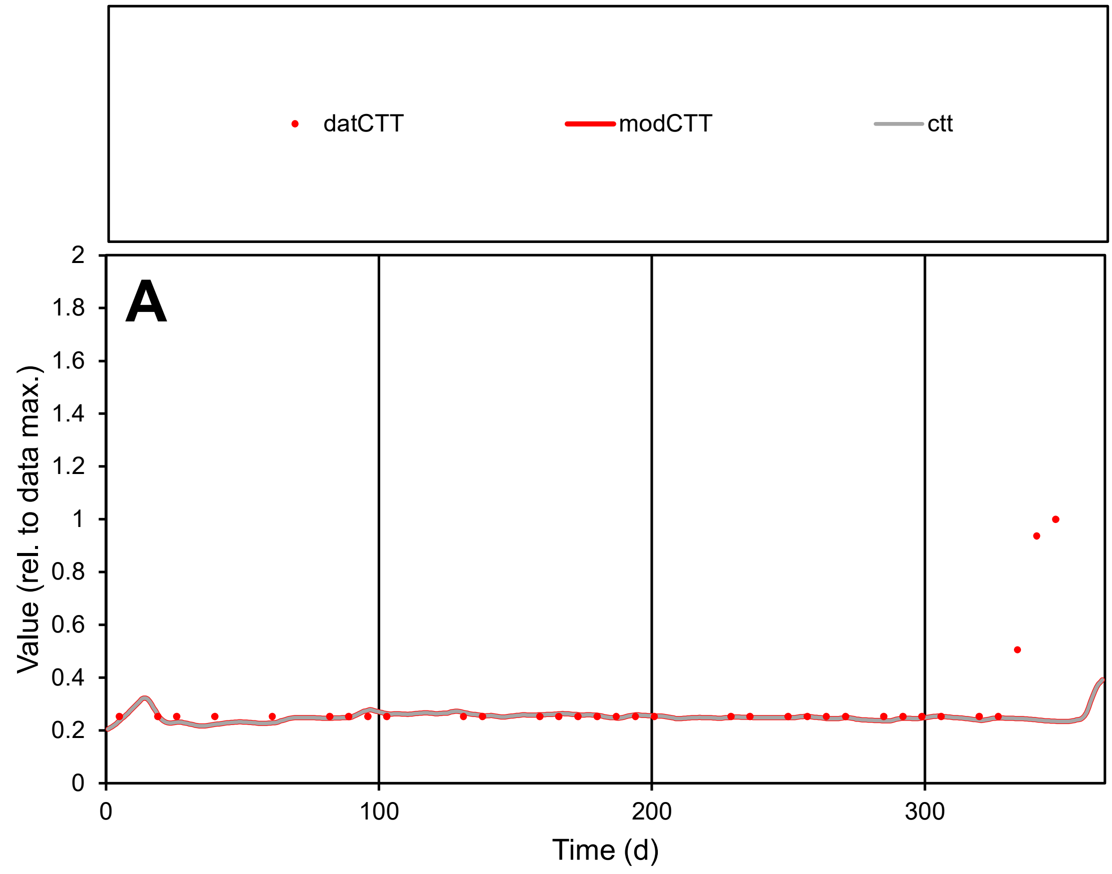


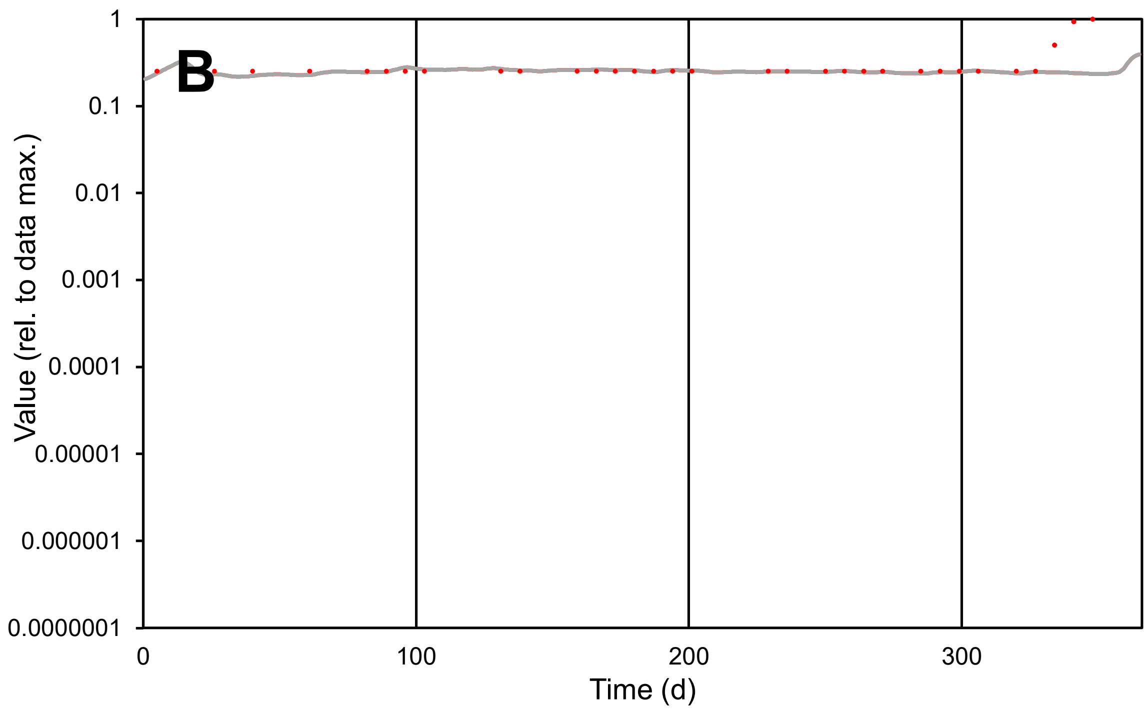


**C1**

Variables

CTT Provide jtd or Dst

CalibOnly?

2 1 to reproduce of

oNormalize?

2 1=no, 2=yes

ofxn calc*

0.61331

ofxn mod

0.62211

sRun

RunA1

zout

1

Ave

Data

1.8E-06

Model

1.5E-06

data/vMax

5.9E-06

ofxng

263.459

Min d/m

1.1269

ave/min d

1.20184

ave/min m

1.10979

326_Calib_S_A

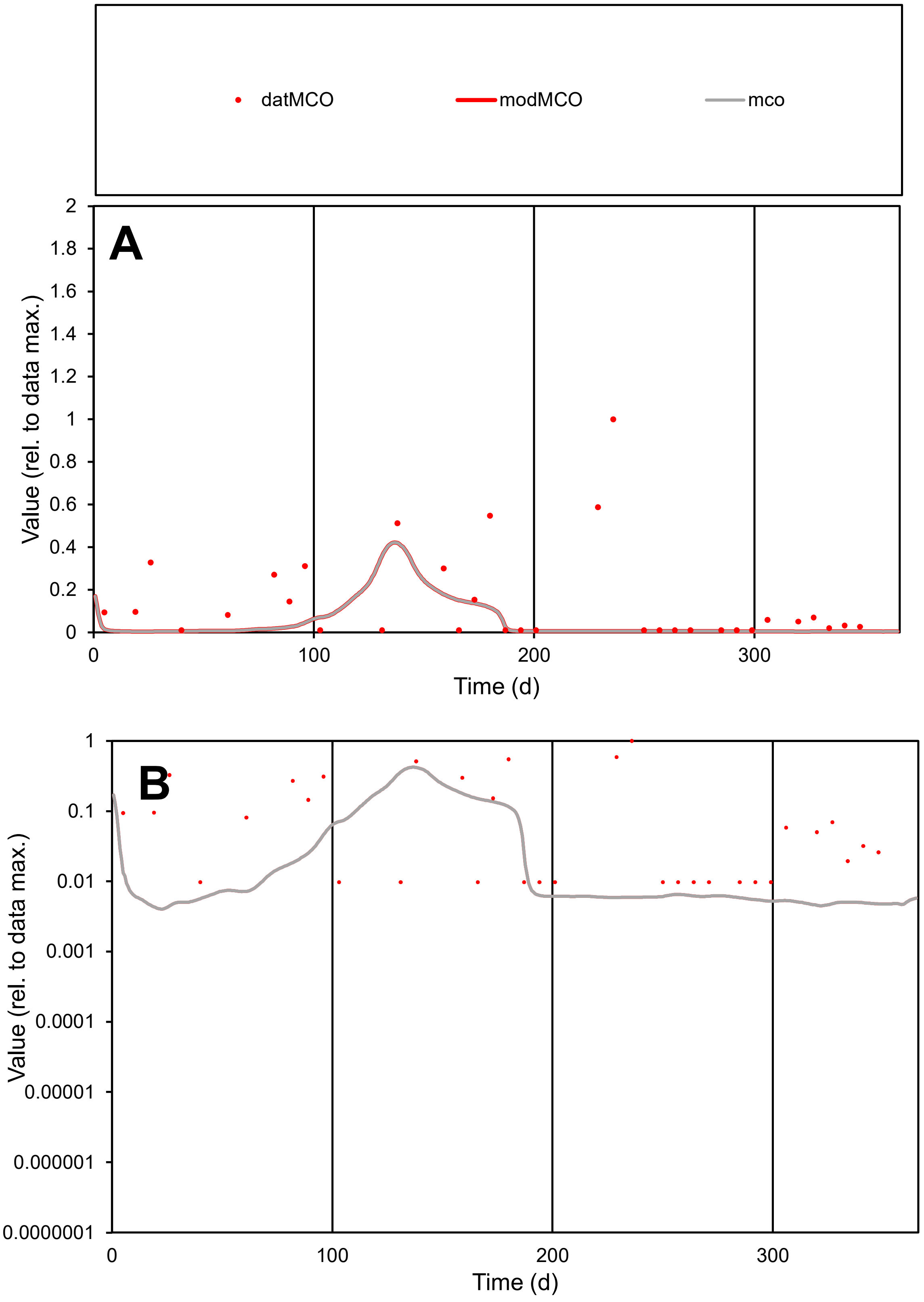


**C1**

Variables
MCO Provide jtd or Dst
CalibOnly?
2 1 to reproduce of
oNormalize?
2 1=no, 2=yes

ofxn calc*
1.66469
ofxn mod
1.67552

sRun
RunA1
zout
1

Ave
Data
9.2E-07
Model
3.3E-07

data/vMax
6.3E-06

ofxng
263.459

Min d/m
2.34201
ave/min d
15.0153
ave/min m
12.4603

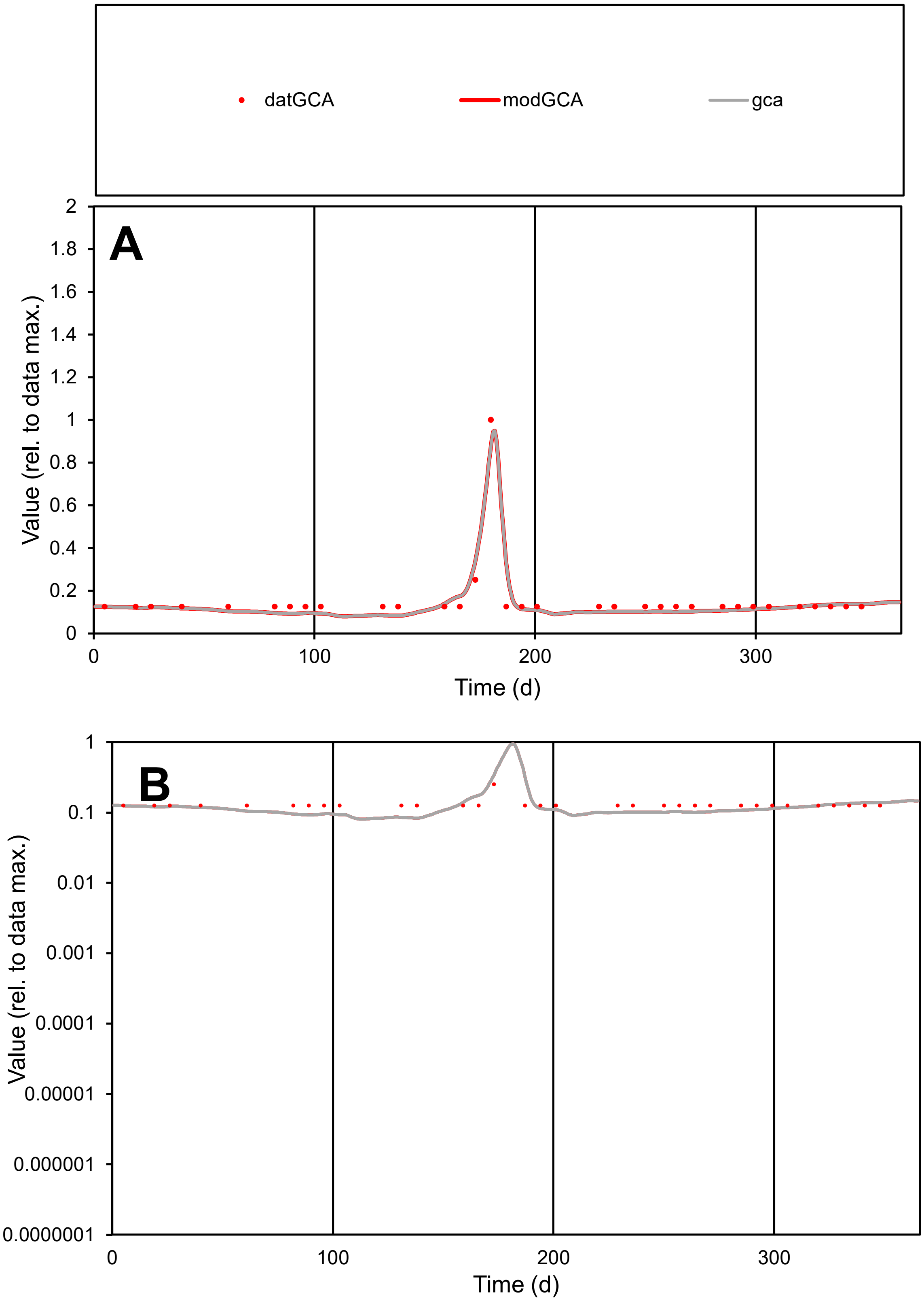


**C1**

Variables
GCA Provide jtd or Dst
CalibOnly?
2 1 to reproduce of
oNormalize?
2 1=no, 2=yes

ofxn calc*
0.20138
ofxn mod
0.25754

sRun
RunA1
zout
1

Ave
Data
1E-05
Model
9.9E-06

data/vMax
6.7E-05

ofxng
263.459

Min d/m
1.5017

ave/min d
1.2408
ave/min m
1.76937

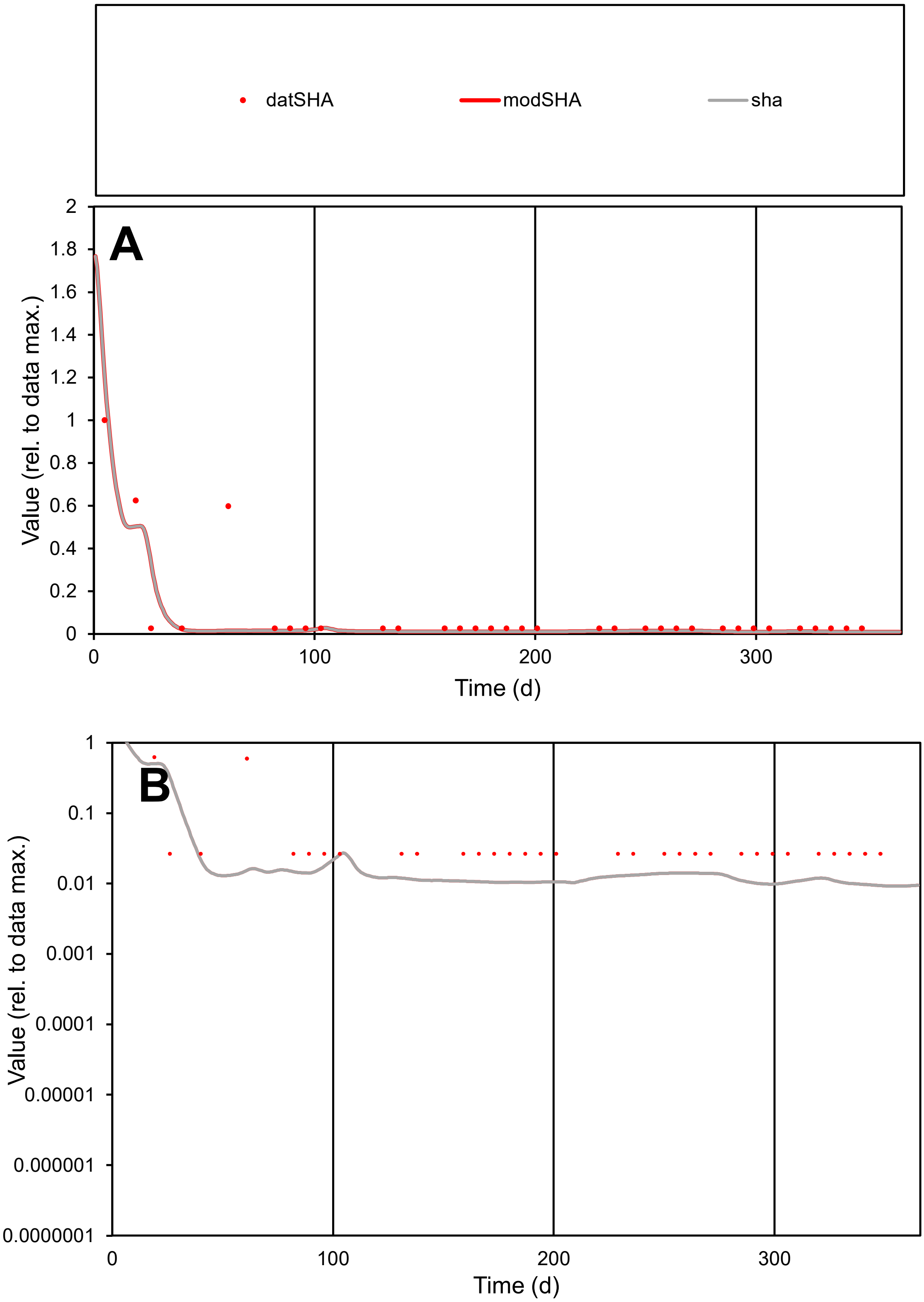


**C1**

Variables
SHA Provide jtd or Dst
CalibOnly?
2 1 to reproduce of
oNormalize?
2 1=no, 2=yes

ofxn calc*
1.17647
ofxn mod
1.23373

sRun
RunA1
zout
1

Ave
Data
2.4E-06
Model
1.7E-06

data/vMax
2.6E-05

ofxng
263.459

Min d/m
2.83266

ave/min d
3.45665
ave/min m
6.70763

326_Calib_S_A

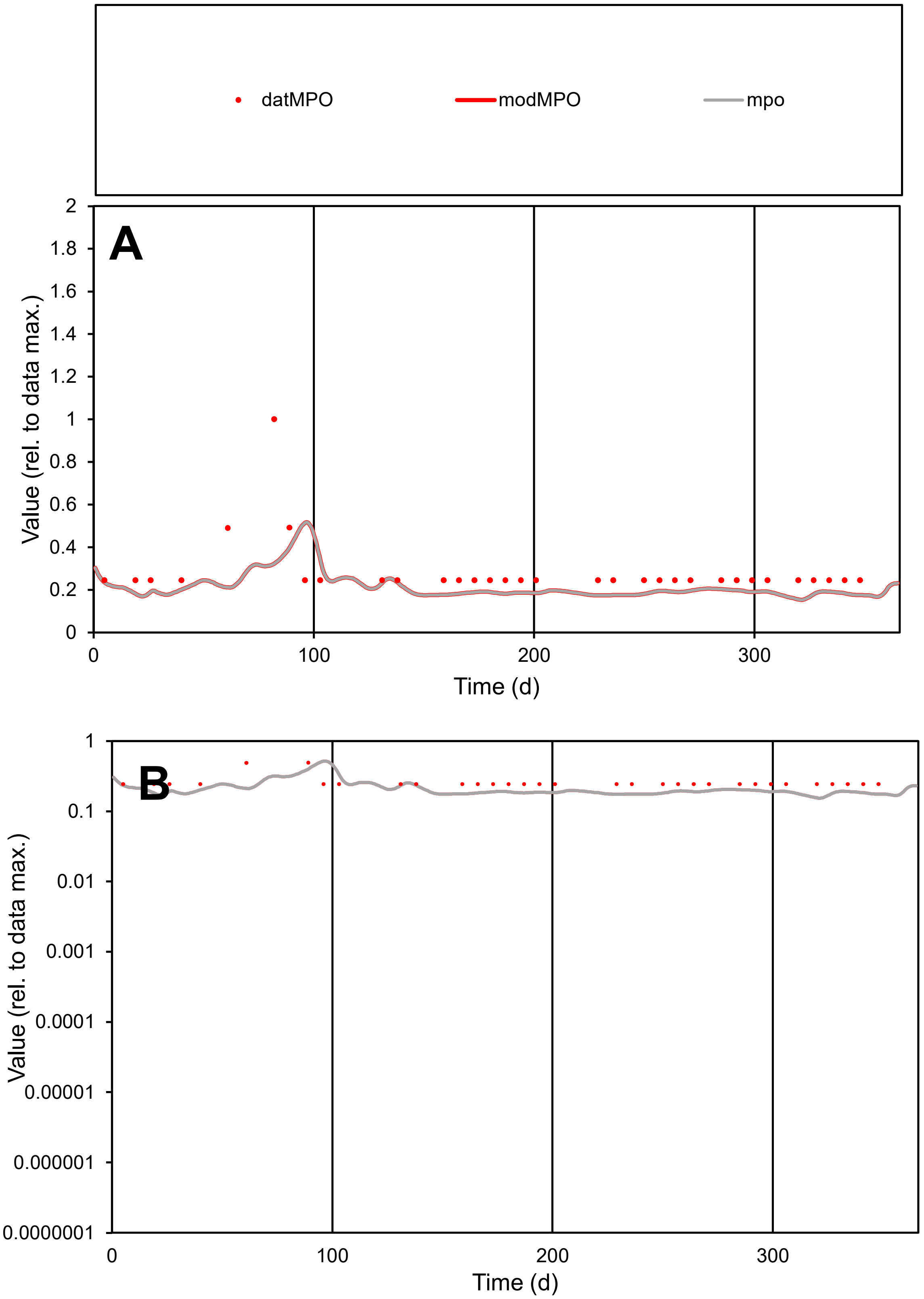


**C1**

Variables
MPO Provide jtd or Dst
CalibOnly?
2 1 to reproduce of
oNormalize?
2 1=no, 2=yes

ofxn calc*
0.51136
ofxn mod
0.52249

sRun
RunA1
zout
1

Ave
Data
1.4E-05
Model
1.1E-05

data/vMax
5.1E-05

ofxng
263.459

Min d/m
1.59389

ave/min d
1.15459
ave/min m
1.40877

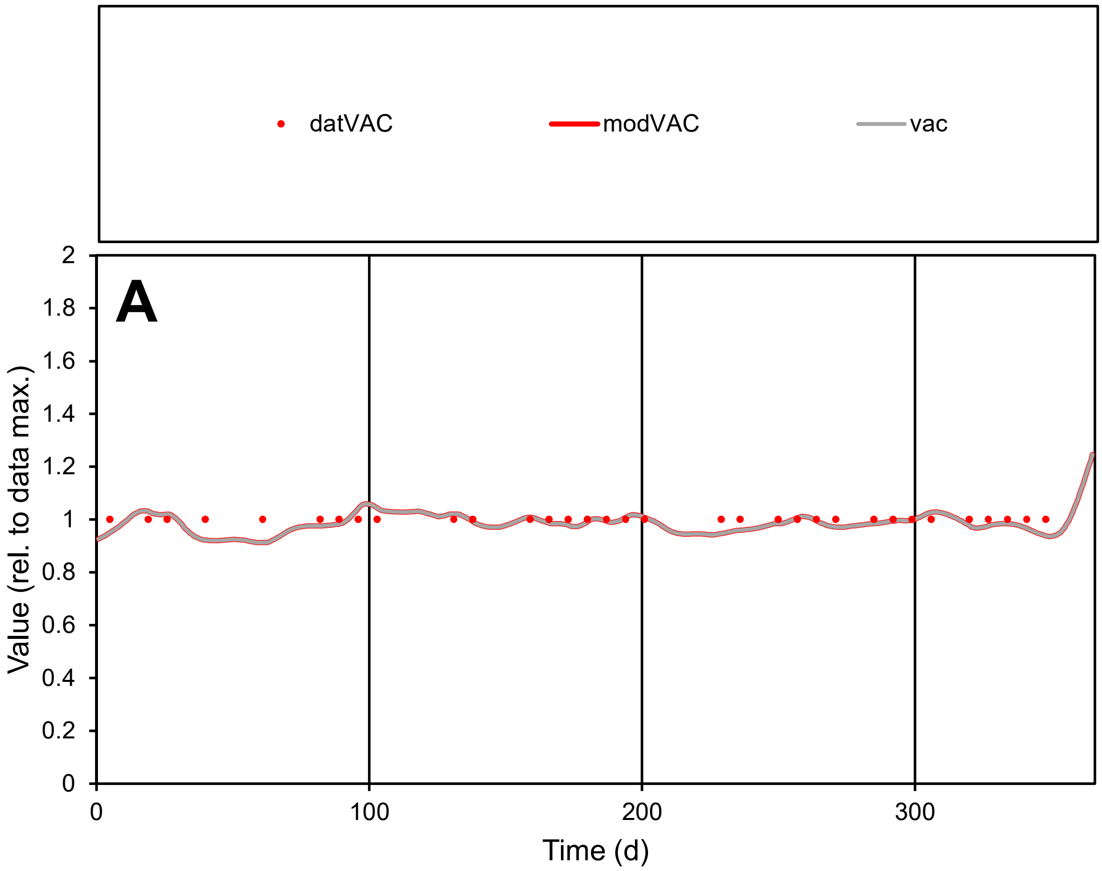


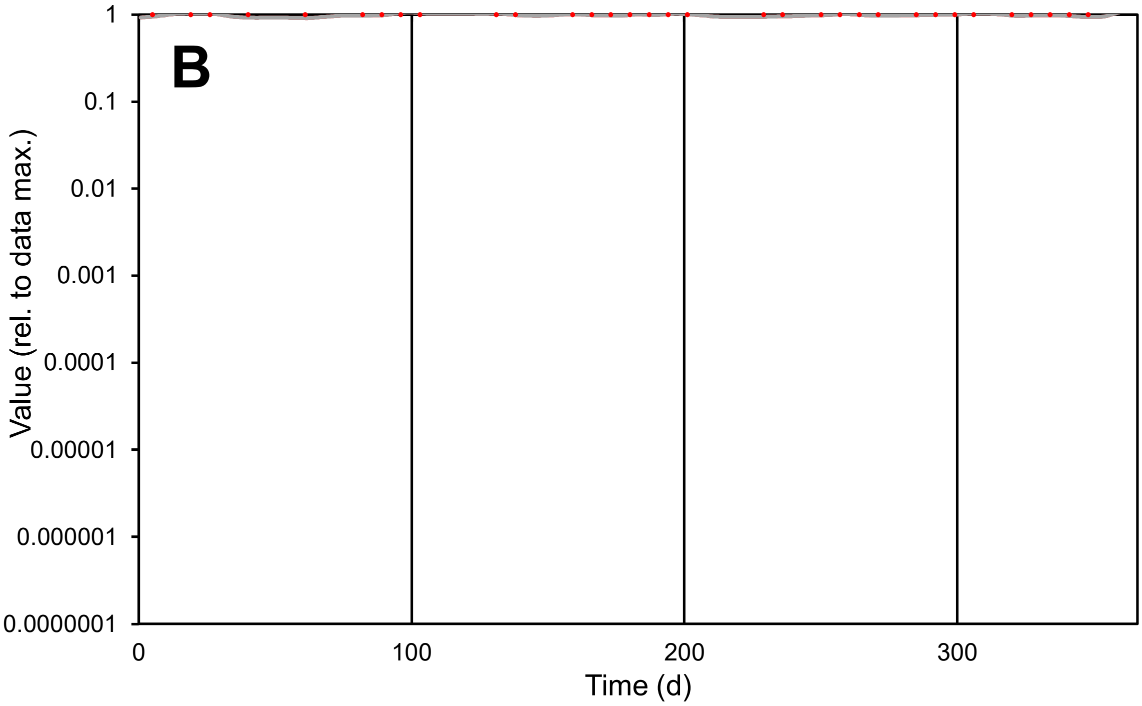


**C1**

Variables
VAC Provide jtd or Dst
CalibOnly?
2 1 to reproduce of
oNormalize?
2 1=no, 2=yes

ofxn calc*
0.03341
ofxn mod
0.11736

sRun
RunA1
zout
1

Ave
Data
9.5E-06
Model
9.4E-06

data/vMax
9.5E-06

ofxng
263.459

Min d/m
1.09447

ave/min d
1
ave/min m
1.08066

326_Calib_S_A

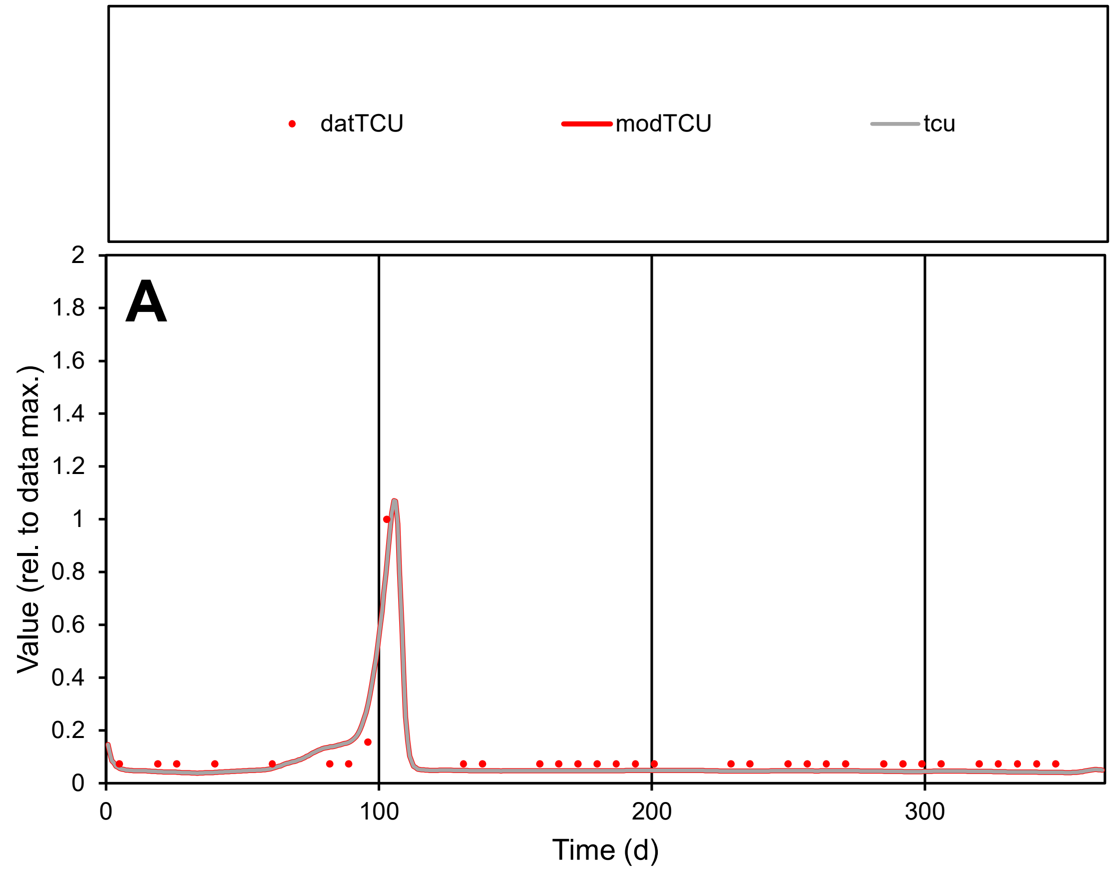


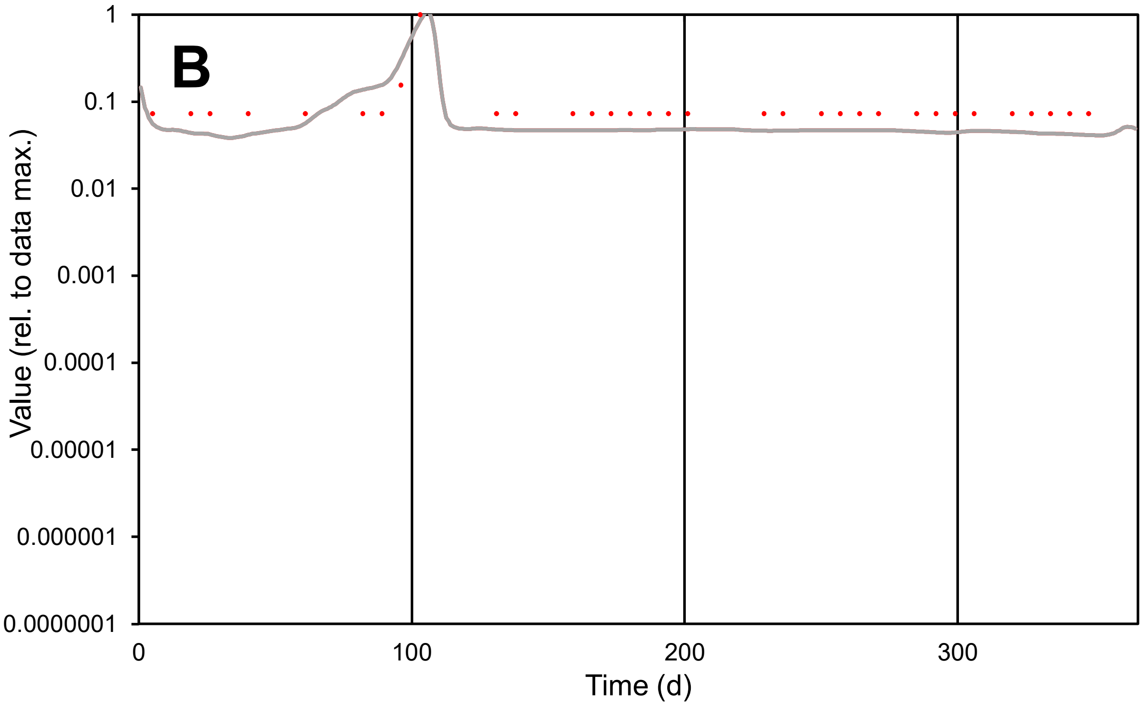


**C1**

Variables
TCU Provide jtd or Dst
CalibOnly?
2 1 to reproduce of
oNormalize?
2 1=no, 2=yes

ofxn calc*
0.395
ofxn mod
0.42306

sRun
RunA1
zout
1

Ave
Data
1.4E-05
Model
1.2E-05

data/vMax
0.00013

ofxng
263.459

Min d/m
1.77077

ave/min d
1.41902
ave/min m
2.1243

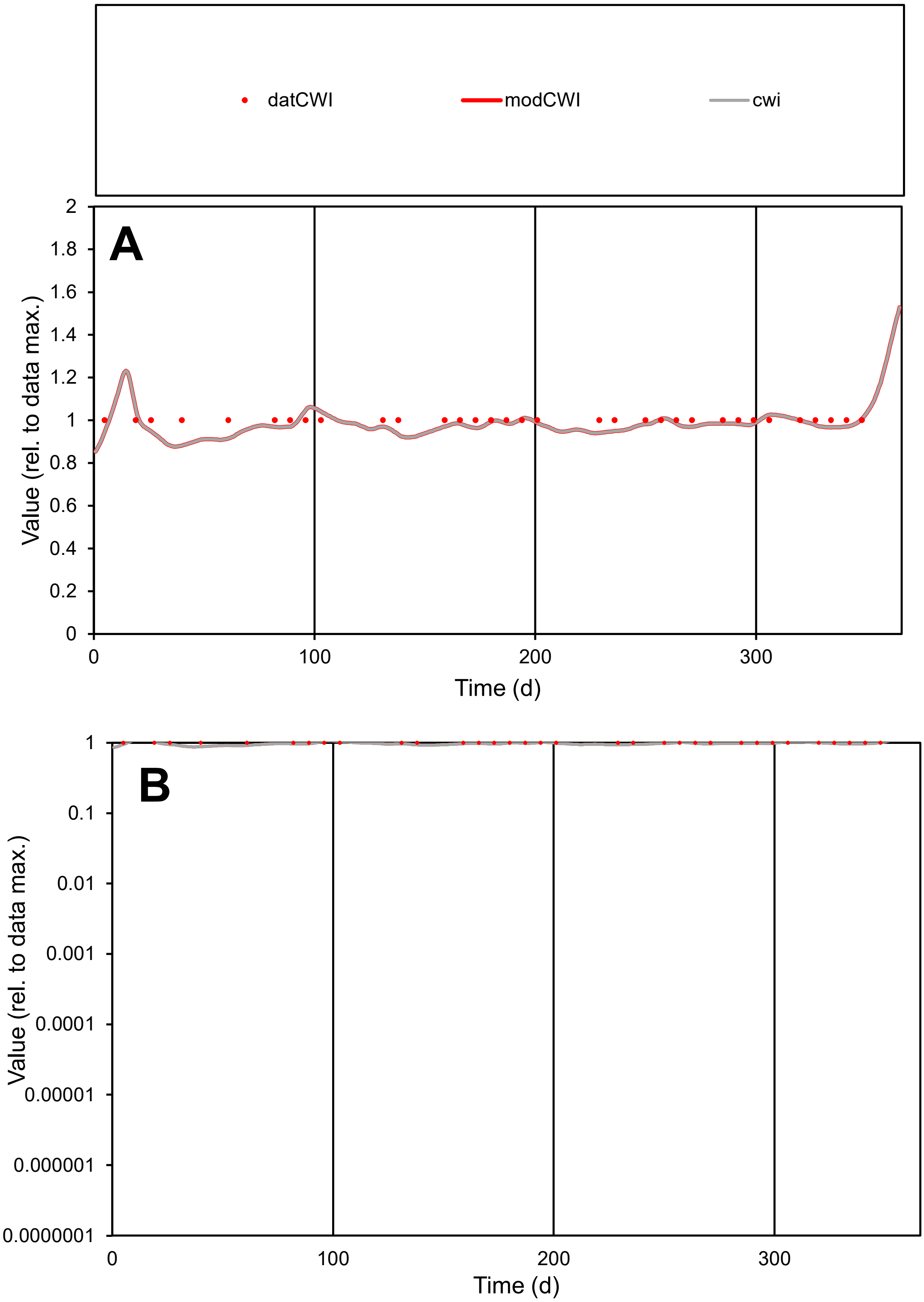


**C1**

Variables

CWI Provide jtd or Dst

CalibOnly?

2 1 to reproduce of

oNormalize?

2 1=no, 2=yes

ofxn calc*

0.03968

ofxn mod

0.11863

sRun

RunA1

zout

1

Ave

Data

1.3E-05

Model

1.3E-05

data/vMax

1.3E-05

ofxng

263.459

Min d/m

1.13106

ave/min d

1

ave/min m

1.10749

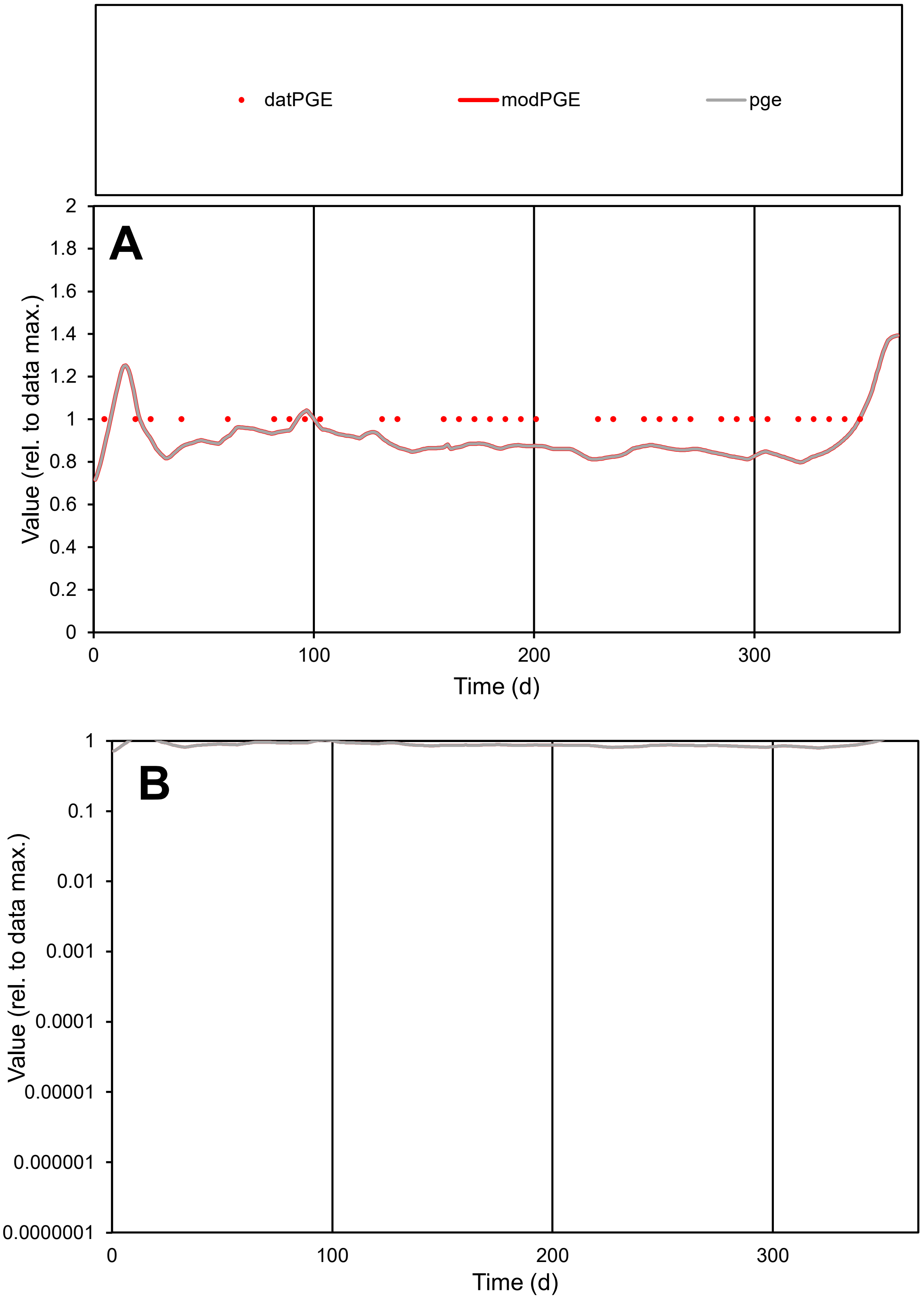


**C1**

Variables

PGE Provide jtd or Dst

CalibOnly?

2 1 to reproduce of

oNormalize?

2 1=no, 2=yes

ofxn calc*

0.12681

ofxn mod

0.16581

sRun

RunA1

zout

1

Ave

Data

5.9E-06

Model

5.2E-06

data/vMax

5.9E-06

ofxng

263.459

Min d/m

1.25266

ave/min d

1

ave/min m

1.11262

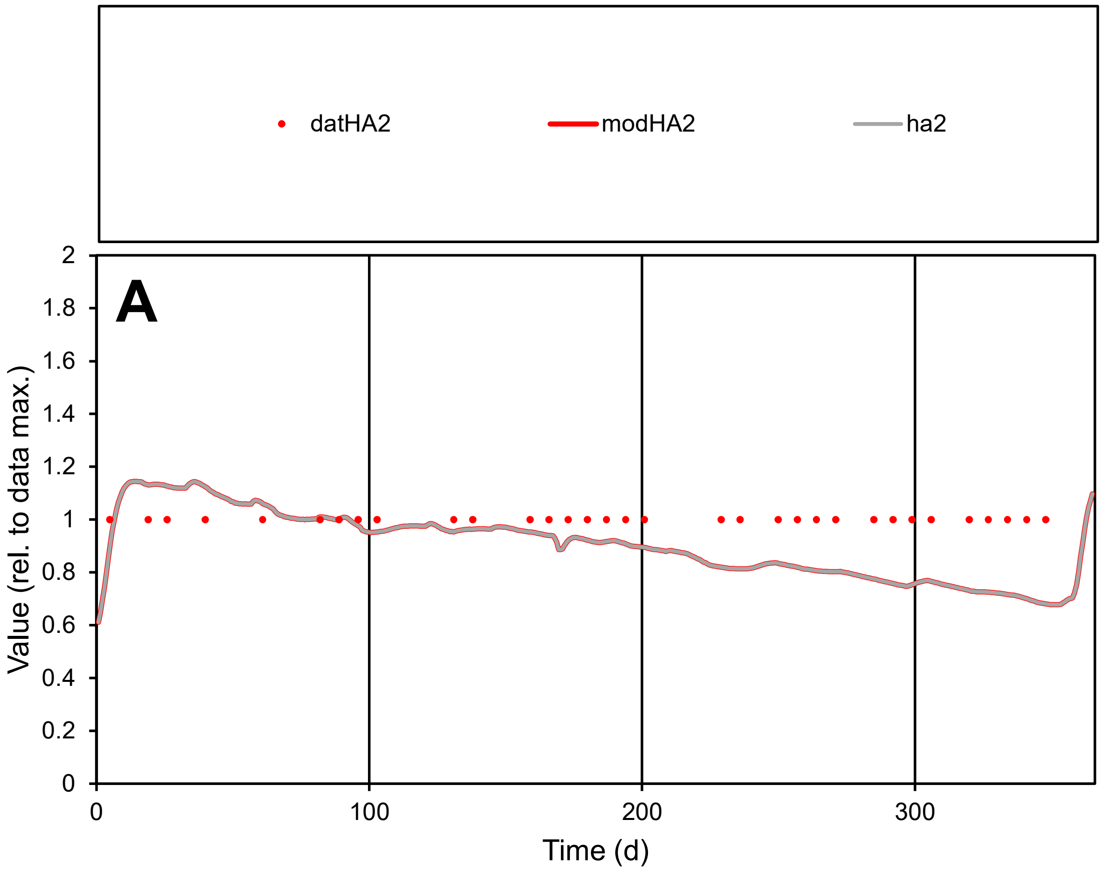


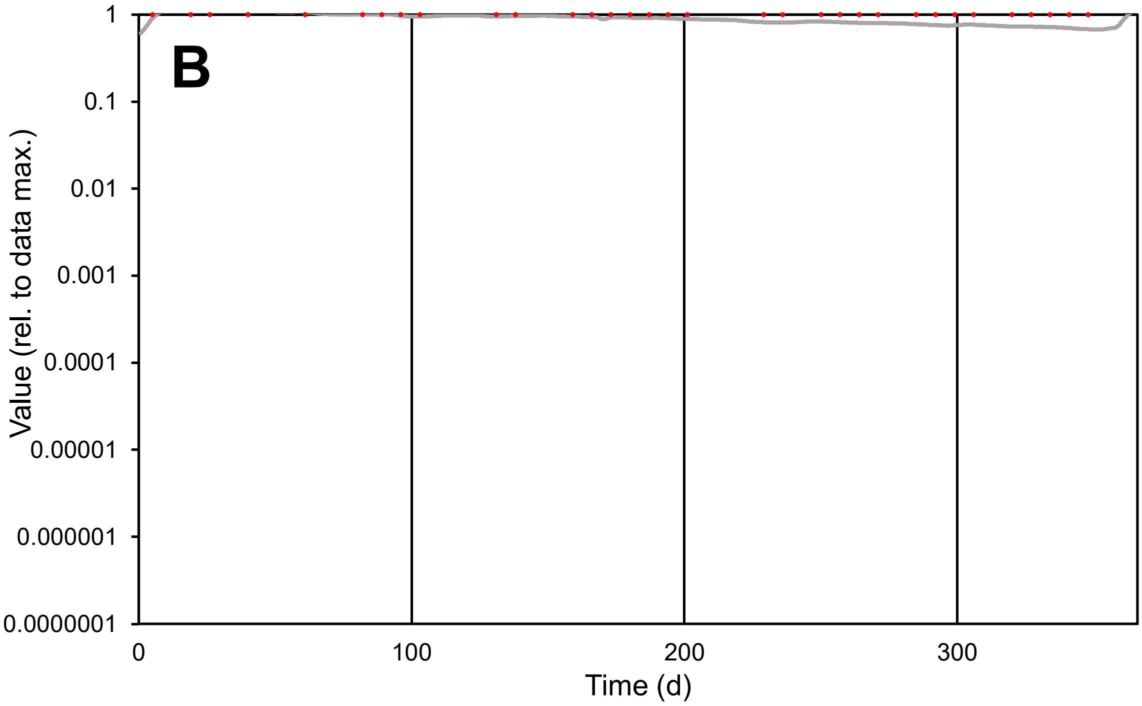


**C1**

| | |
|---|---|
| Variables | |
| HA2 | Provide jtd or Dst |
| CalibOnly? | |
| 2 | 1 to reproduce of |
| oNormalize? | |
| 2 | 1=no, 2=yes |
| ofxn calc* | |
| 0.17086 | |
| ofxn mod | |
| 0.20195 | |
| sRun | |
| RunA1 | |
| zout | |
| 1 | |
| Ave | |
| Data | |
| 2.6E-06 | |
| Model | |
| 2.3E-06 | |
| data/vMax | |
| 2.6E-06 | |
| ofxng | |
| 263.459 | |
| Min d/m | |
| 1.47187 | |
| ave/min d | |
| 1 | |
| ave/min m | |
| 1.30274 | |

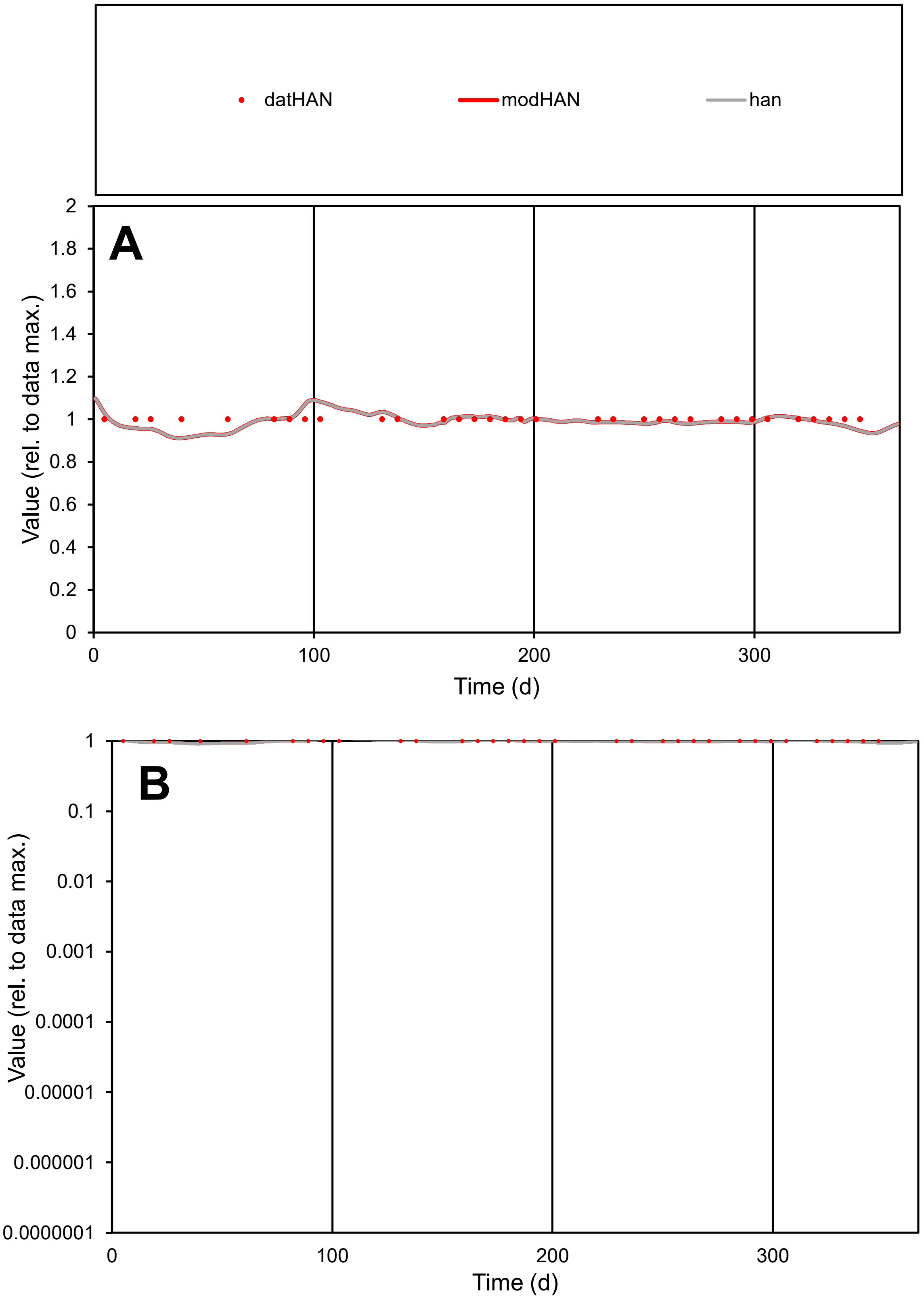


| | |
|---|---|
| **C1** | |
| Variables | |
| HAN | Provide jtd or Dst |
| CalibOnly? | |
| 2 | 1 to reproduce of |
| oNormalize? | |
| 2 | 1=no, 2=yes |
| ofxn calc* | |
| 0.03428 | |
| ofxn mod | |
| 0.11775 | |
| sRun | |
| RunA1 | |
| zout | |
| 1 | |
| Ave | |
| Data | |
| 5.1E-06 | |
| Model | |
| 5.1E-06 | |
| data/vMax | |
| 5.1E-06 | |
| ofxng | |
| 263.459 | |
| Min d/m | |
| 1.09511 | |
| ave/min d | |
| 1 | |
| ave/min m | |
| 1.08713 | |

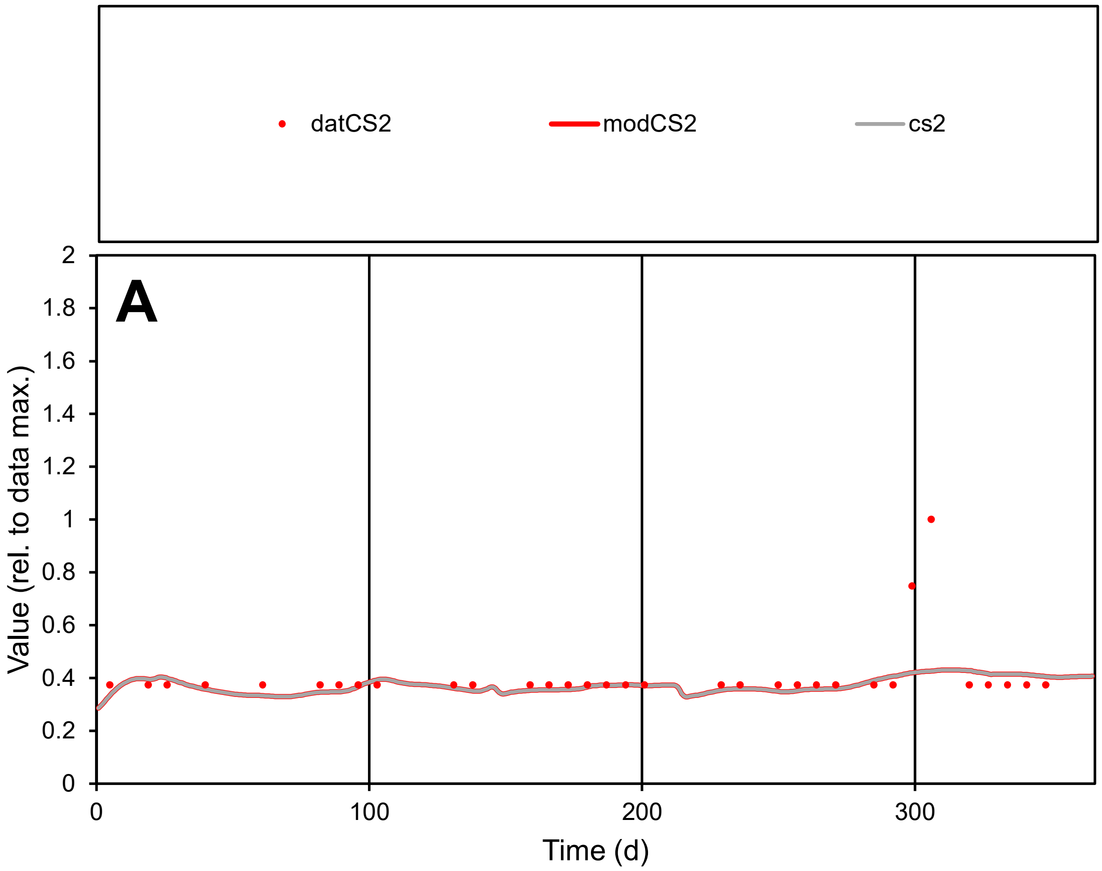


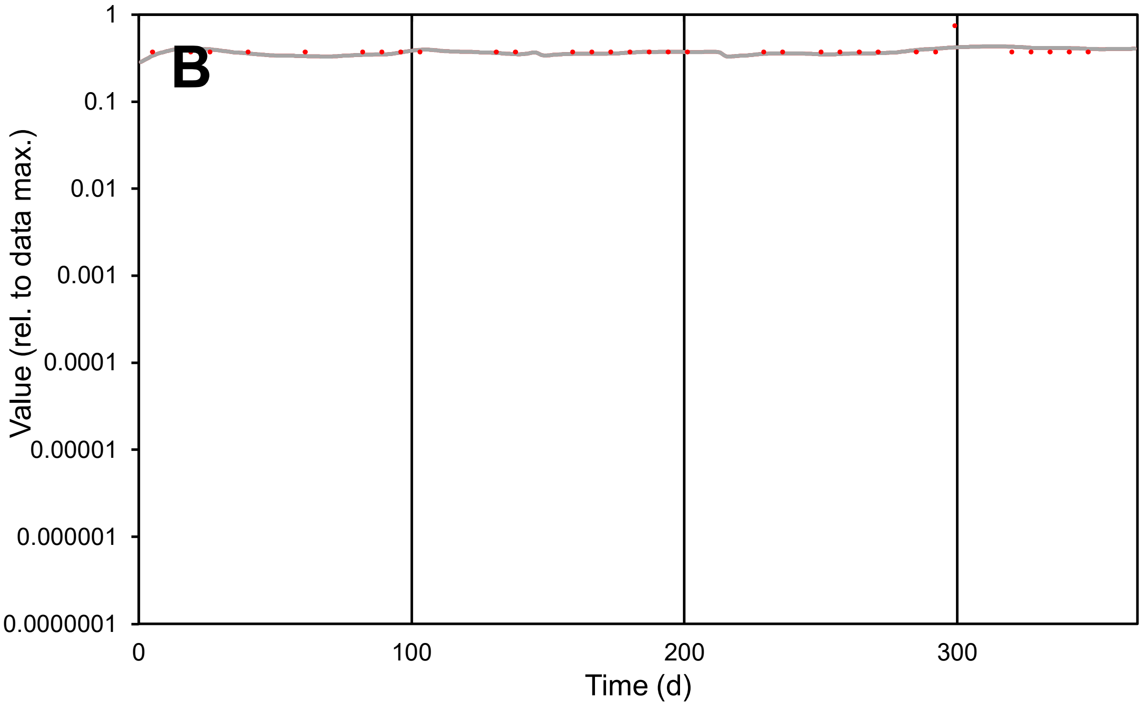


**C1**

Variables
CS2 Provide jtd or Dst
CalibOnly?
2 1 to reproduce of
oNormalize?
2 1=no, 2=yes

ofxn calc*
0.28972
ofxn mod
0.31069

sRun
RunA1
zout
1

Ave
Data
8.2E-06
Model
7.6E-06

data/vMax
2E-05

ofxng
263.459

Min d/m
1.12651

ave/min d
1.08107
ave/min m
1.13362

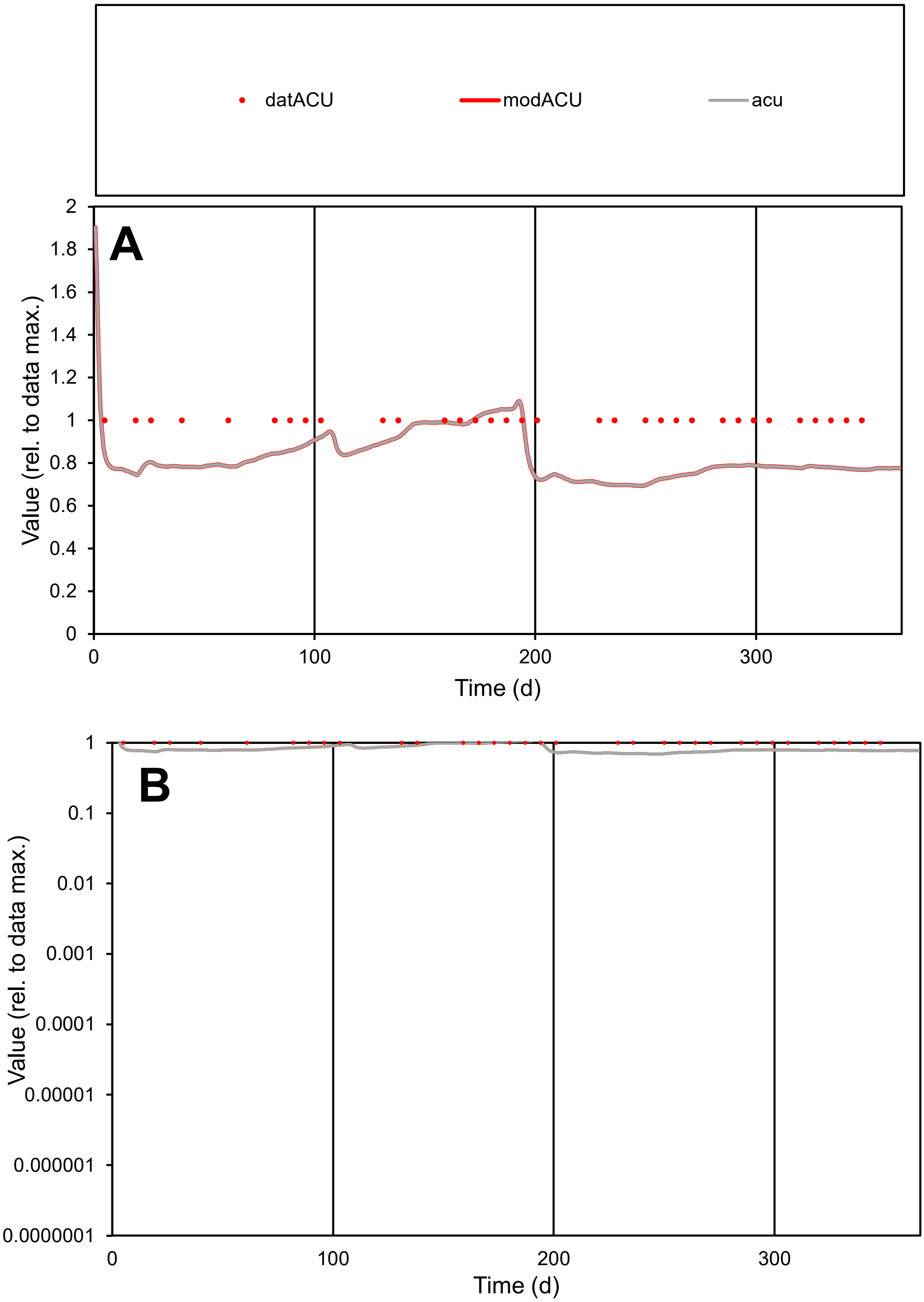


**C1**

Variables
ACU Provide jtd or Dst
CalibOnly?
2 1 to reproduce of
oNormalize?
2 1=no, 2=yes

ofxn calc*
0.19918
ofxn mod
0.22424

sRun
RunA1
zout
1

Ave
Data
1.1E-05
Model
9.2E-06

data/vMax
1.1E-05

ofxng
263.459

Min d/m
1.43392

ave/min d
1
ave/min m
1.18772

326_Calib_S_A

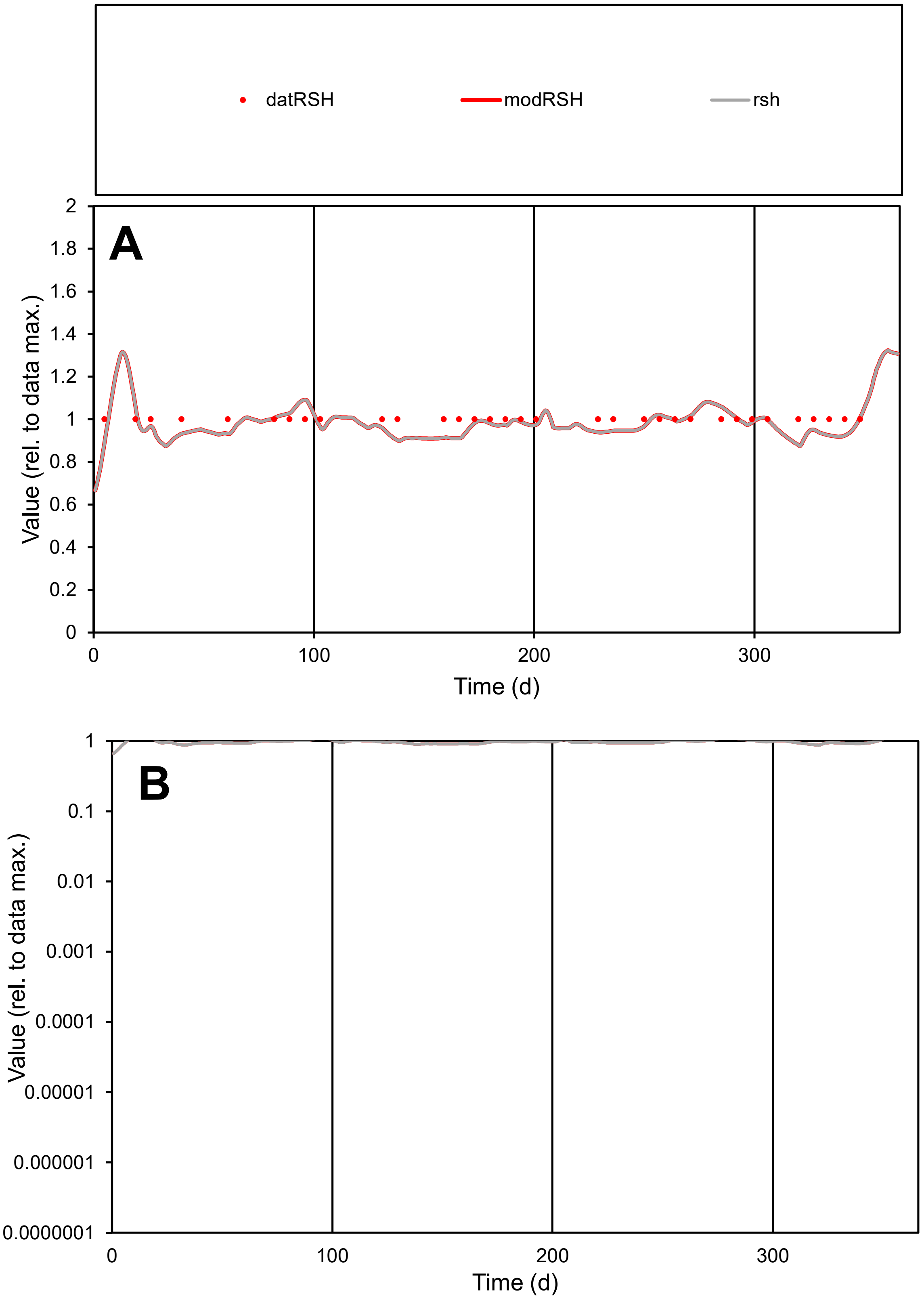


**C1**

Variables
RSH Provide jtd or Dst
CalibOnly?
2 1 to reproduce of
oNormalize?
2 1=no, 2=yes

ofxn calc*
0.0518
ofxn mod
0.1232

sRun
RunA1
zout
1

Ave
Data
1.8E-05
Model
1.7E-05

data/vMax
1.8E-05

ofxng
263.459

Min d/m
1.13038

ave/min d
1
ave/min m
1.10056

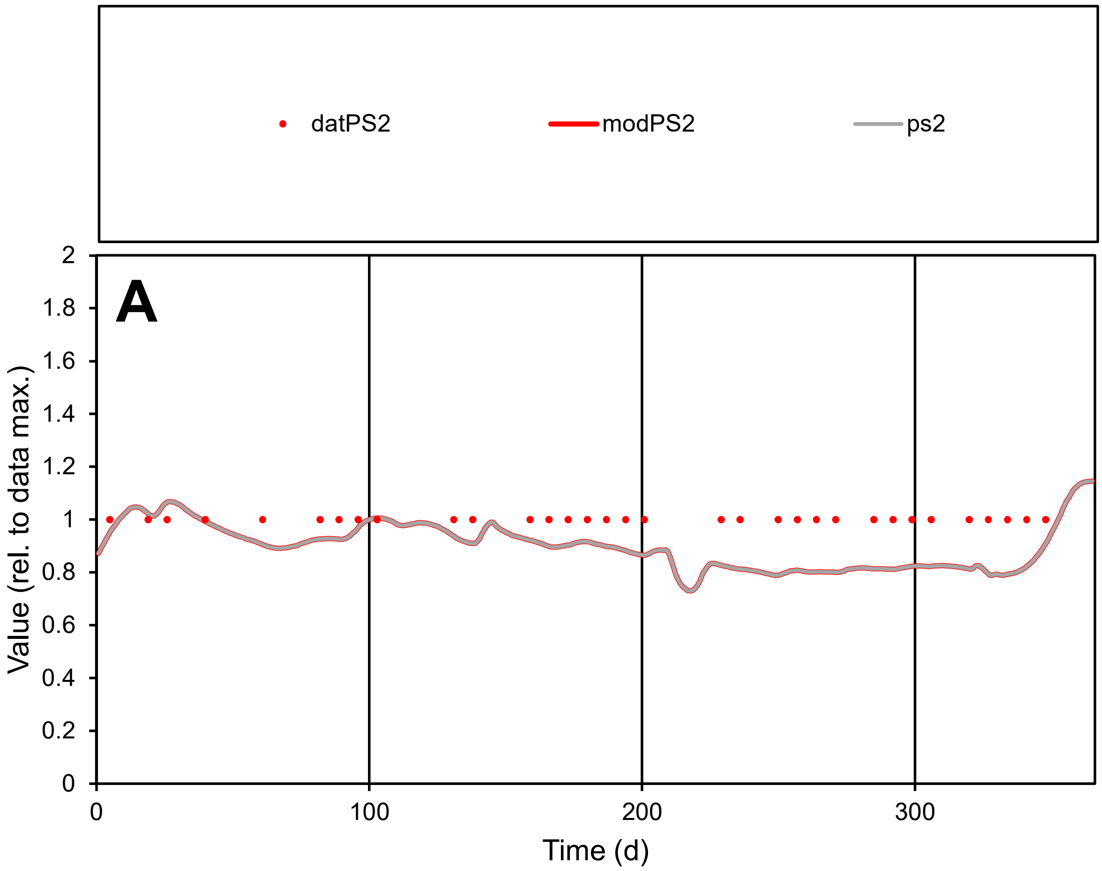


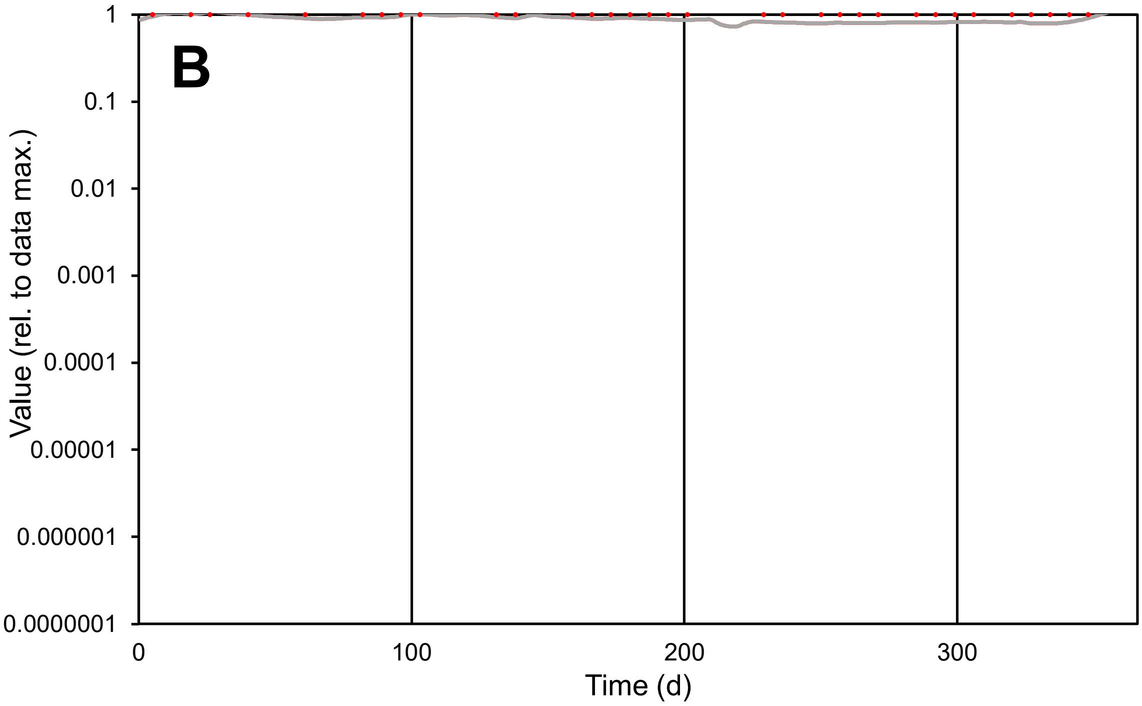


**C1**

Variables
PS2 Provide jtd or Dst
CalibOnly?
2 1 to reproduce of
oNormalize?
2 1=no, 2=yes

ofxn calc*
0.13745
ofxn mod
0.17397

sRun
RunA1
zout
1

Ave
Data
1.9E-05
Model
1.7E-05

data/vMax
1.9E-05

ofxng
263.459

Min d/m
1.26831

ave/min d
1
ave/min m
1.12194

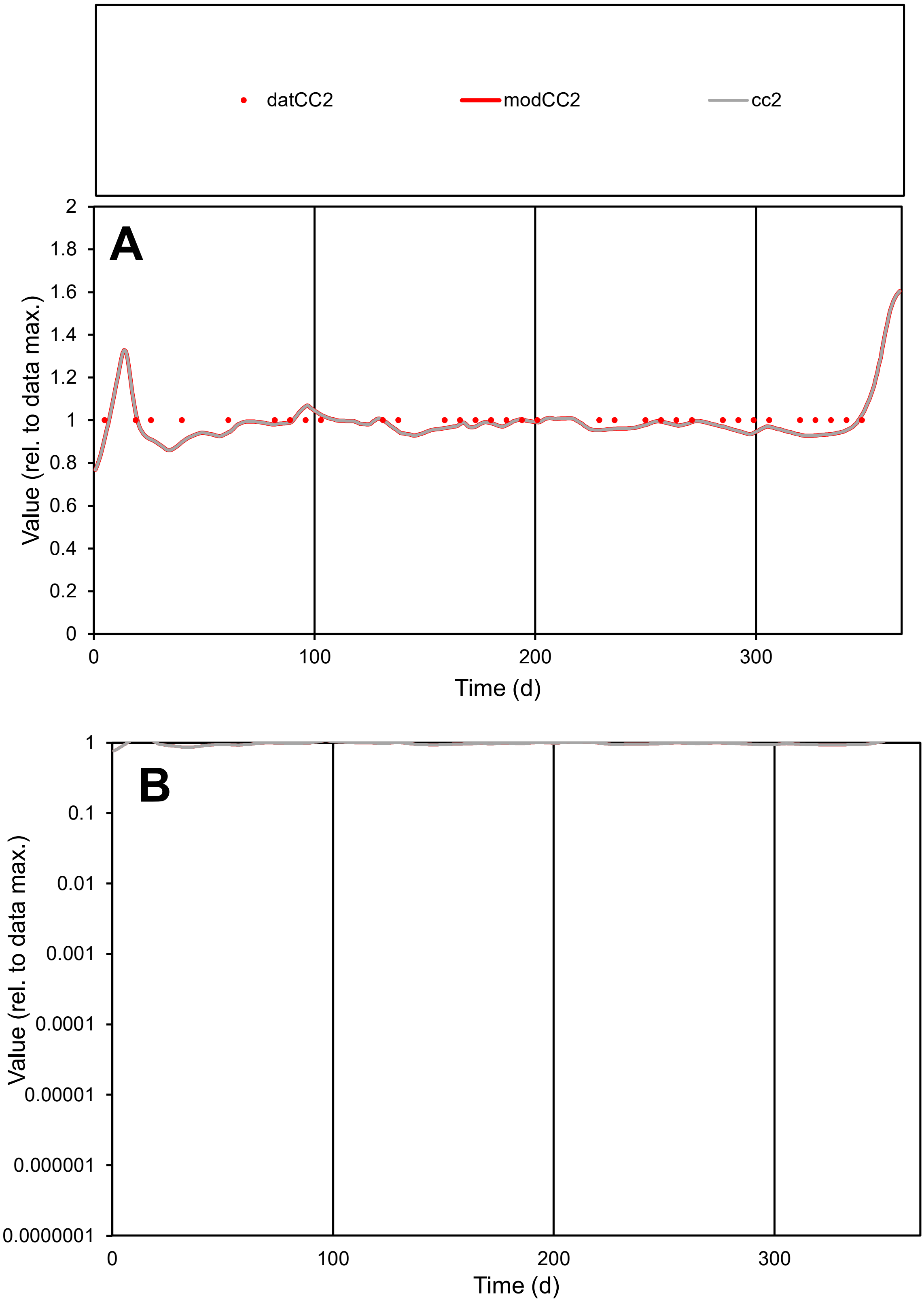


**C1**

Variables

CC2 Provide jtd or Dst

CalibOnly?

2 1 to reproduce of

oNormalize?

2 1=no, 2=yes

ofxn calc*

0.04357

ofxn mod

0.11971

sRun

RunA1

zout

1

Ave

Data

5.9E-06

Model

5.7E-06

data/vMax

5.9E-06

ofxng

263.459

Min d/m

1.10964

ave/min d

1

ave/min m

1.07982

326_Calib_S_A

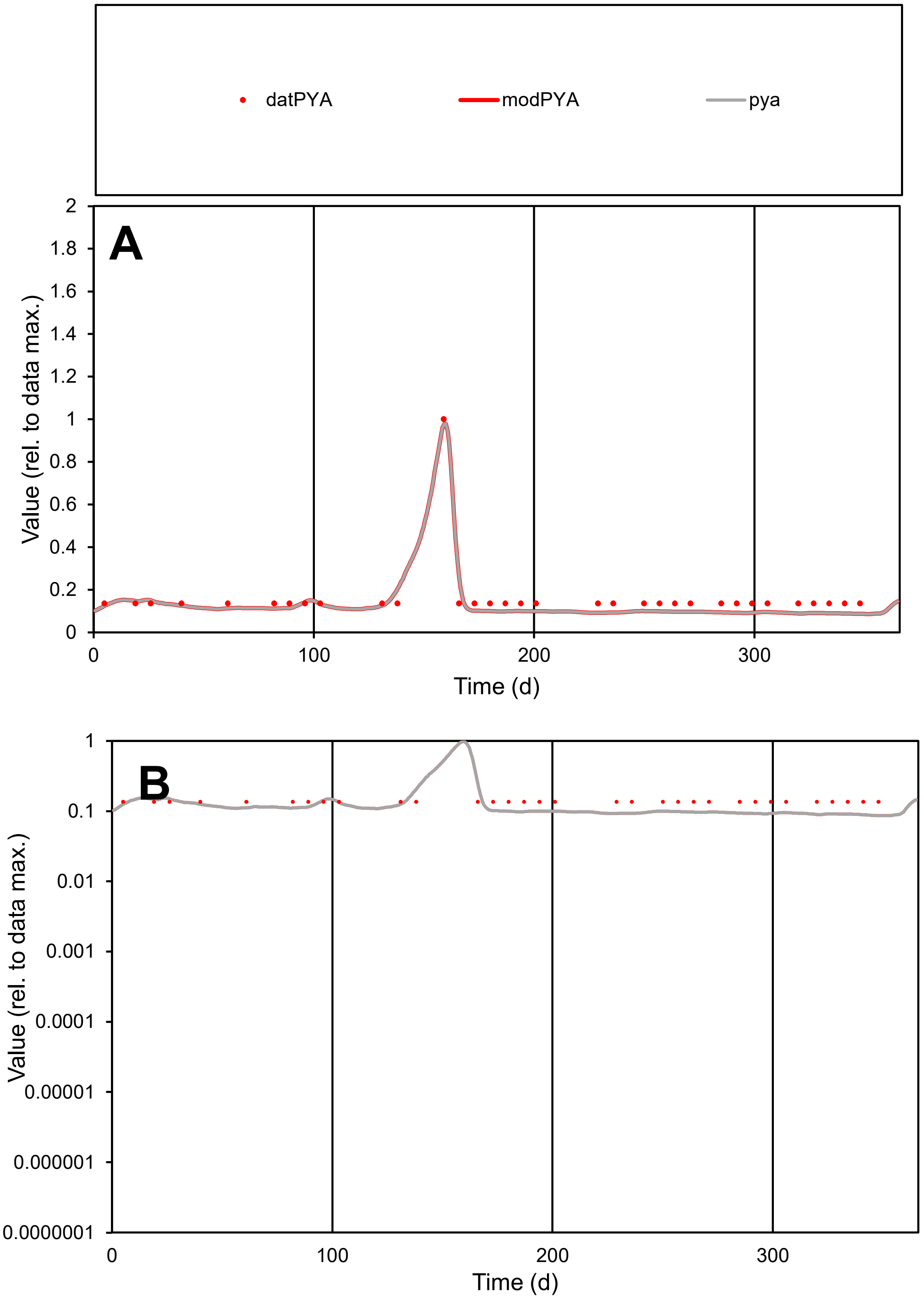


**C1**

Variables
PYA Provide jtd or Dst
CalibOnly?
2 1 to reproduce of
oNormalize?
2 1=no, 2=yes

ofxn calc*
0.22261
ofxn mod
0.24752

sRun
RunA1
zout
1

Ave
Data
2.2E-06
Model
1.8E-06

data/vMax
1.3E-05

ofxng
263.459

Min d/m
1.56883

ave/min d
1.19296
ave/min m
1.5936

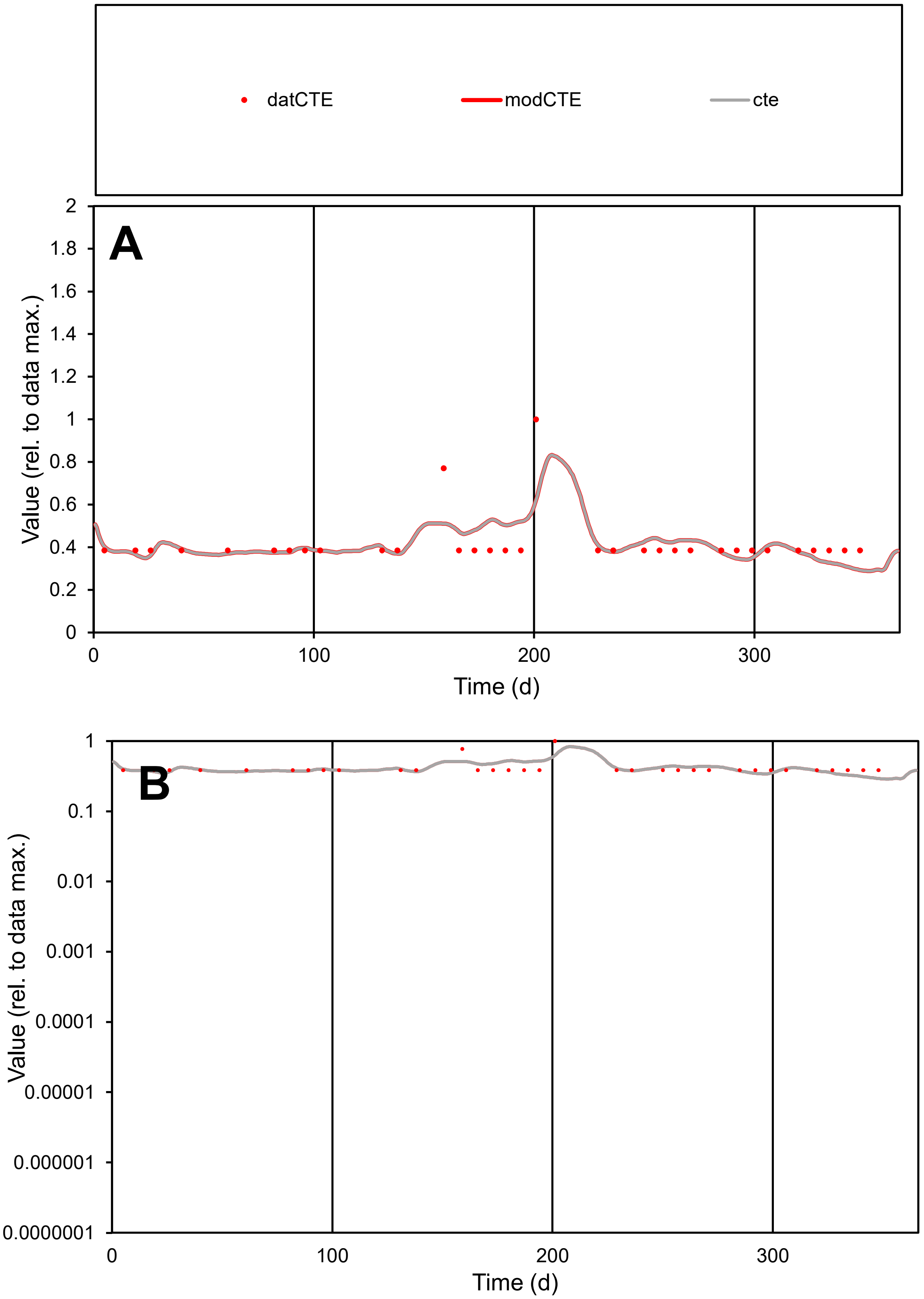


**C1**

Variables
CTE Provide jtd or Dst
CalibOnly?
2 1 to reproduce of
oNormalize?
2 1=no, 2=yes

ofxn calc*
0.22168
ofxn mod
0.24874

sRun
RunA1
zout
1

Ave
Data
6.6E-06
Model
6.5E-06

data/vMax
1.6E-05

ofxng
263.459

Min d/m
1.3214

ave/min d
1.07874
ave/min m
1.40581

326_Calib_S_A

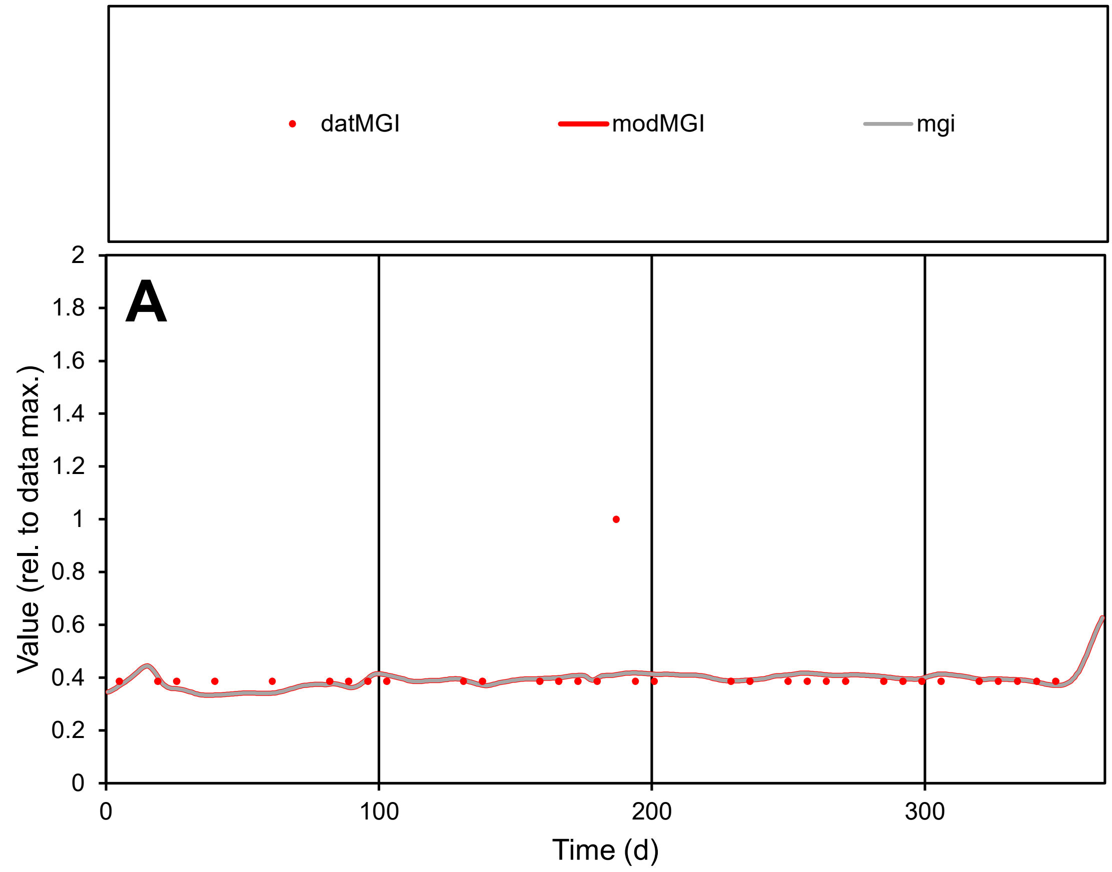


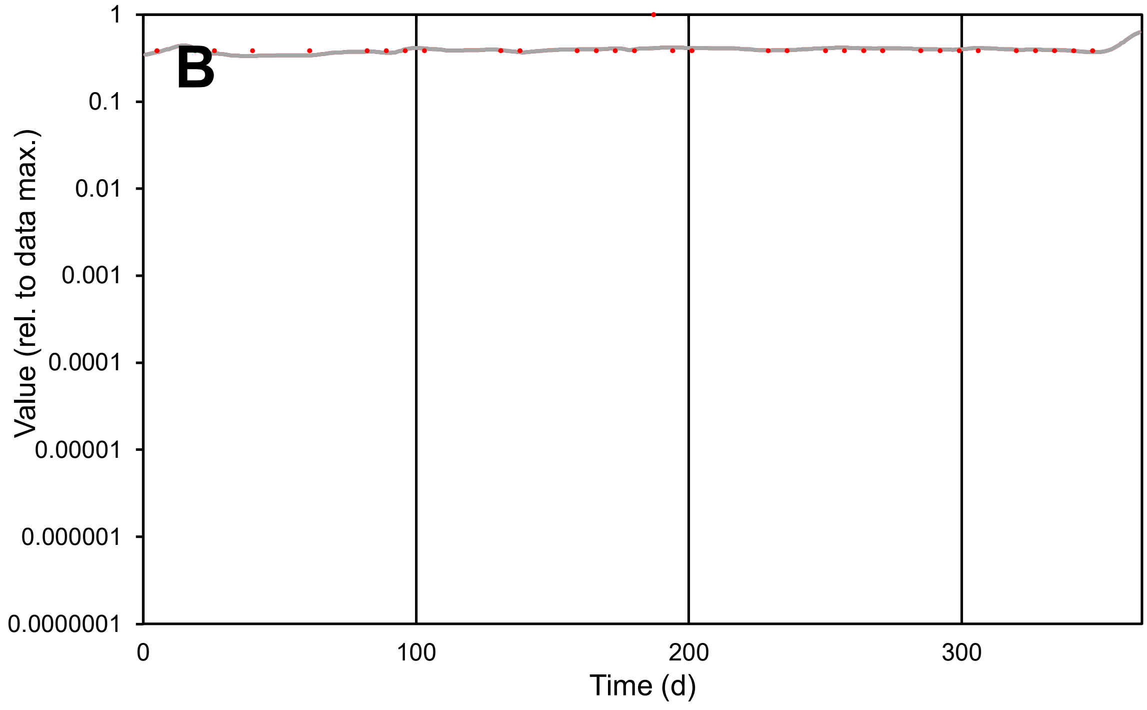


**C1**

Variables
MGI Provide jtd or Dst
CalibOnly?
2 1 to reproduce of
oNormalize?
2 1=no, 2=yes

ofxn calc*
0.25772
ofxn mod
0.28147

sRun
RunA1
zout
1

Ave
Data
1.1E-05
Model
1E-05

data/vMax
2.7E-05

ofxng
263.459

Min d/m
1.15488

ave/min d
1.04824
ave/min m
1.17309

326_Calib_S_A

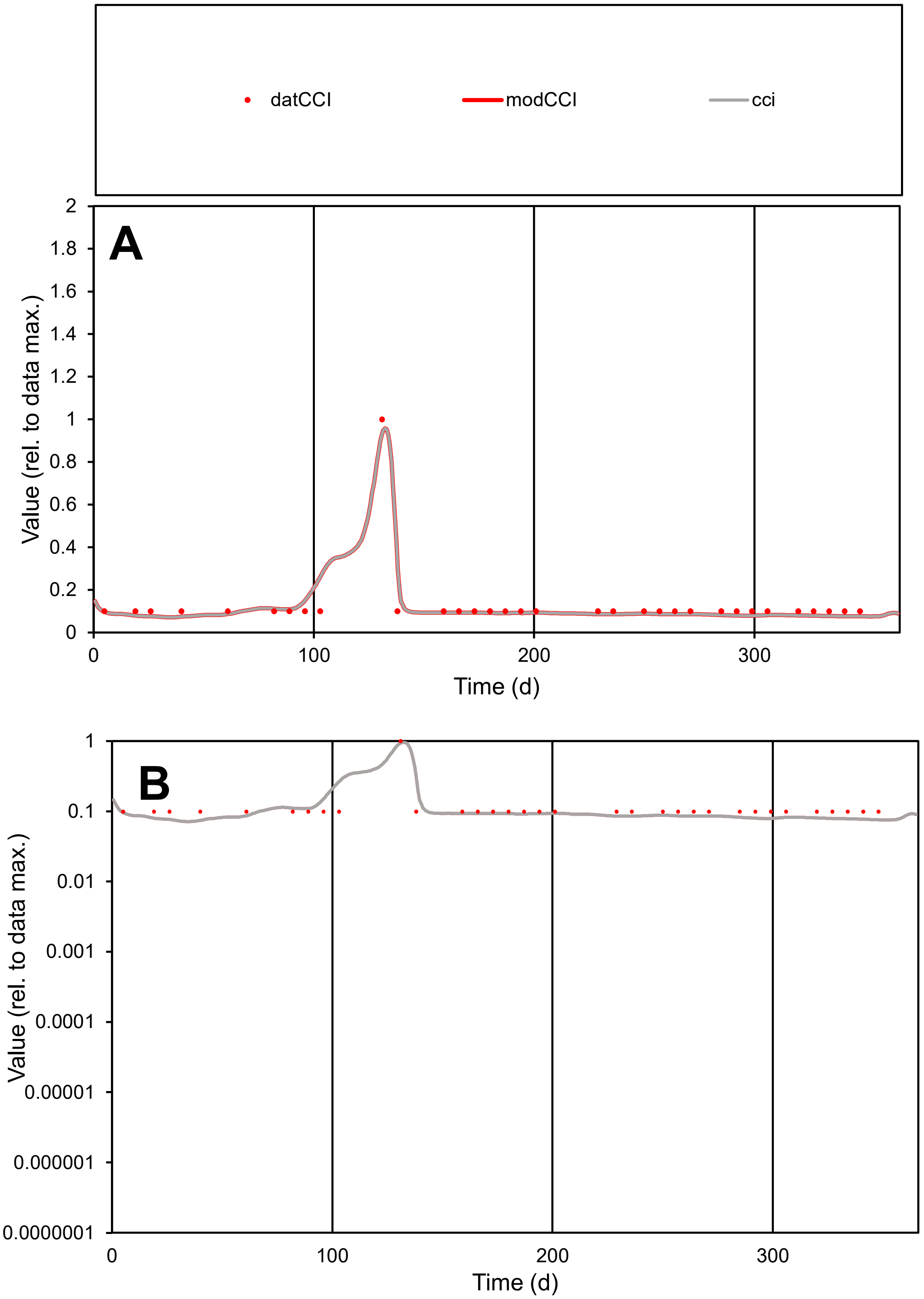


**C1**

Variables
CCI Provide jtd or Dst
CalibOnly?
2 1 to reproduce of
oNormalize?
2 1=no, 2=yes

ofxn calc*
0.28218
ofxn mod
0.34345

sRun
RunA1
zout
1

Ave
Data
1.5E-05
Model
1.4E-05

data/vMax
0.00012

ofxng
263.459

Min d/m
1.30886

ave/min d
1.27642
ave/min m
1.61794

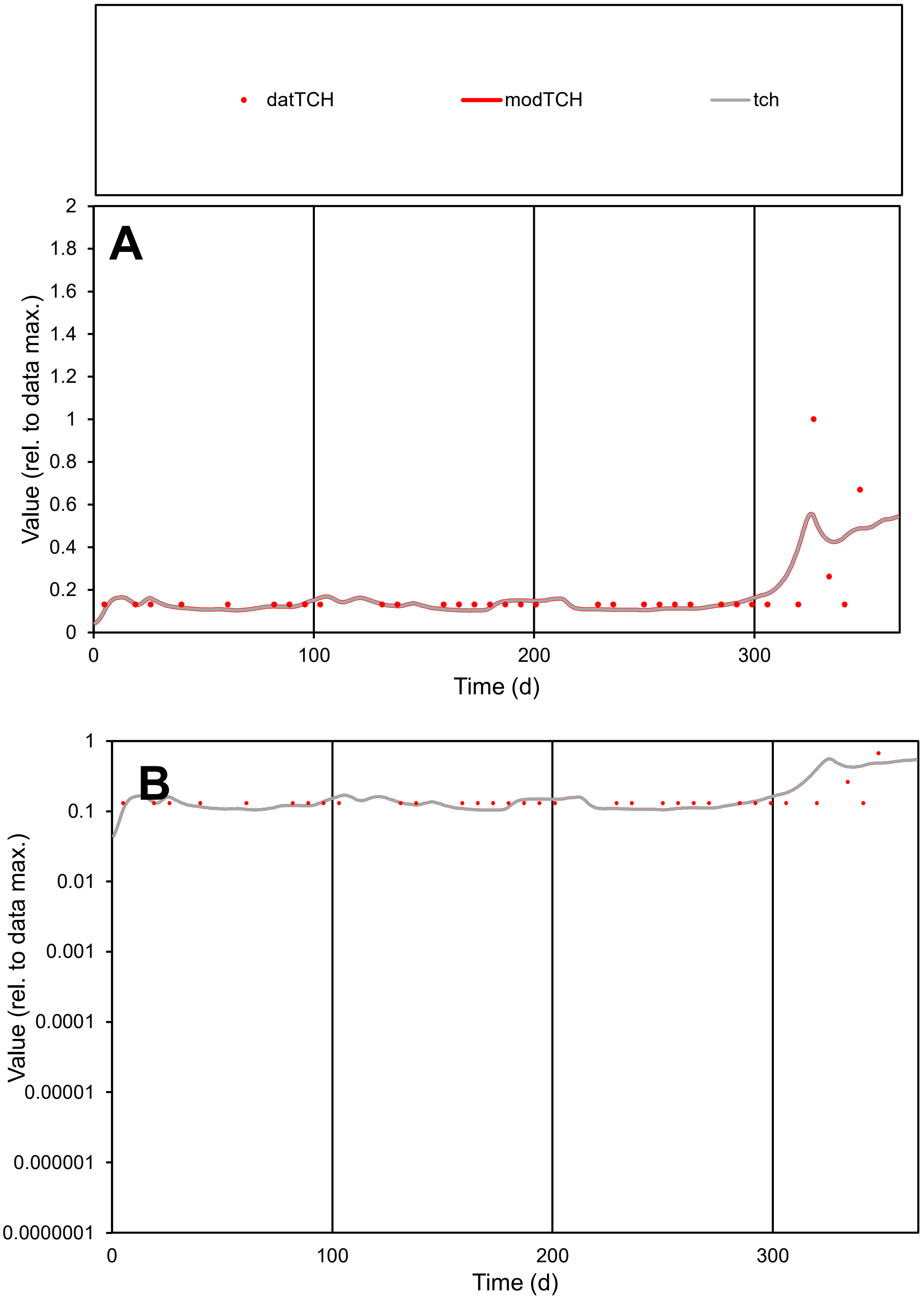


**C1**

Variables
TCH Provide jtd or Dst
CalibOnly?
2 1 to reproduce of
oNormalize?
2 1=no, 2=yes

ofxn calc*
0.67712
ofxn mod
0.69168

sRun
RunA1
zout
1

Ave
Data
2.3E-06
Model
2.3E-06

data/vMax
1.3E-05

ofxng
263.459

Min d/m
1.2571

ave/min d
1.35548
ave/min m
1.70823

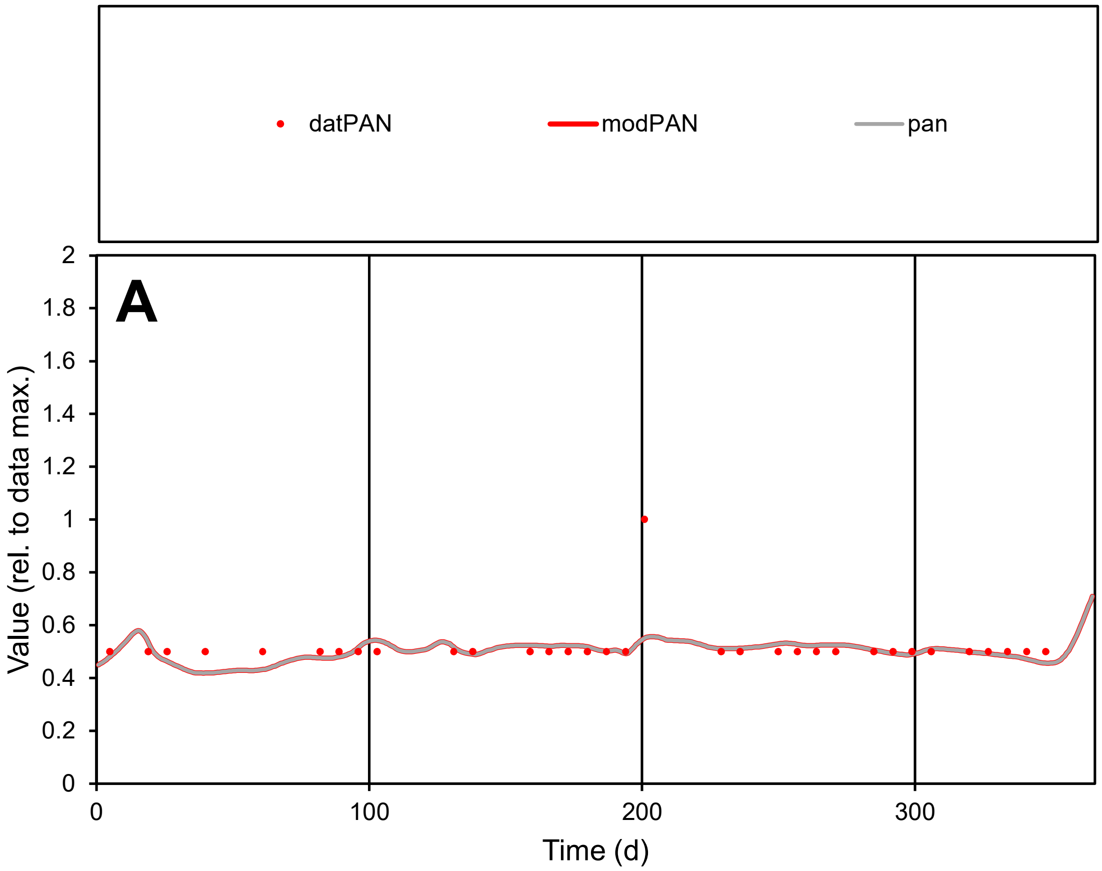


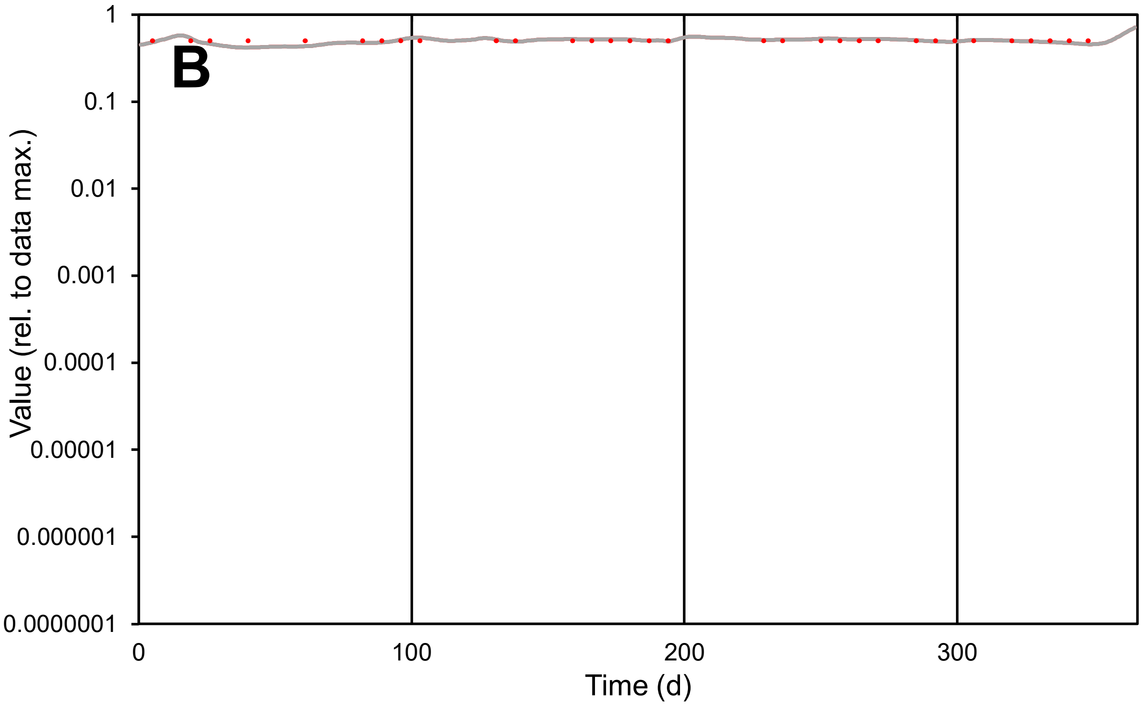


**C1**

| | |
|---|---|
| Variables | |
| PAN | Provide jtd or Dst |
| CalibOnly? | |
| 2 | 1 to reproduce of |
| oNormalize? | |
| 2 | 1=no, 2=yes |
| ofxn calc* | |
| 0.15925 | |
| ofxn mod | |
| 0.19438 | |
| sRun | |
| RunA1 | |
| zout | |
| 1 | |
| Ave | |
| Data | |
| 9.4E-06 | |
| Model | |
| 9.1E-06 | |
| data/vMax | |
| 1.8E-05 | |
| ofxng | |
| 263.459 | |
| Min d/m | |
| 1.19294 | |
| ave/min d | |
| 1.0303 | |
| ave/min m | |
| 1.19317 | |

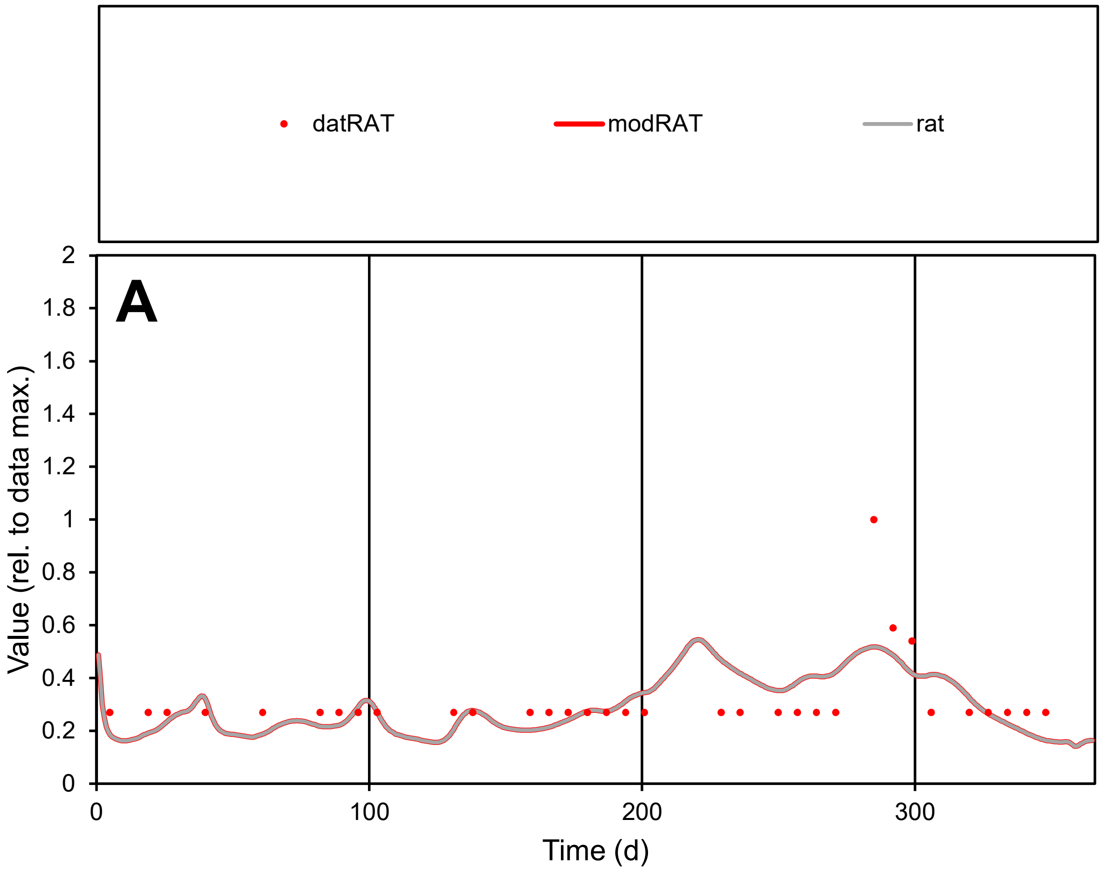


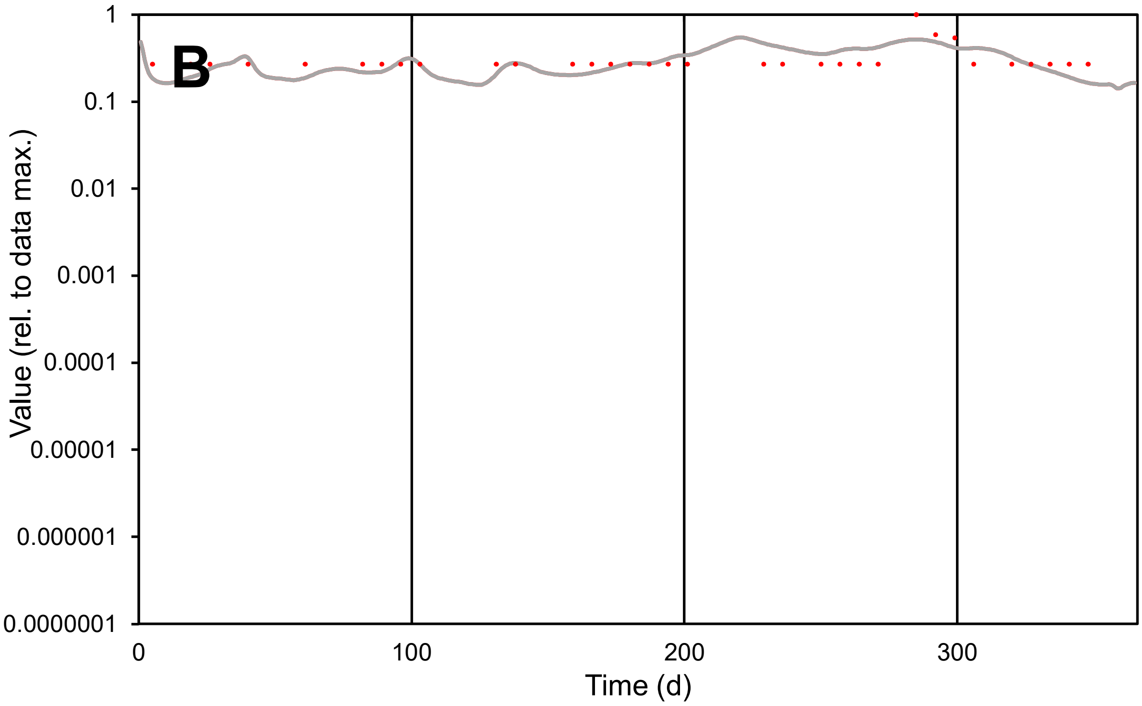


**C1**

Variables

RAT Provide jtd or Dst

CalibOnly?

2 1 to reproduce of

oNormalize?

2 1=no, 2=yes

ofxn calc*

0.3856

ofxn mod

0.40505

sRun

RunA1

zout

1

Ave

Data

1.3E-05

Model

1.3E-05

data/vMax

4.2E-05

ofxng

263.459

Min d/m

1.66325

ave/min d

1.14824

ave/min m

1.851

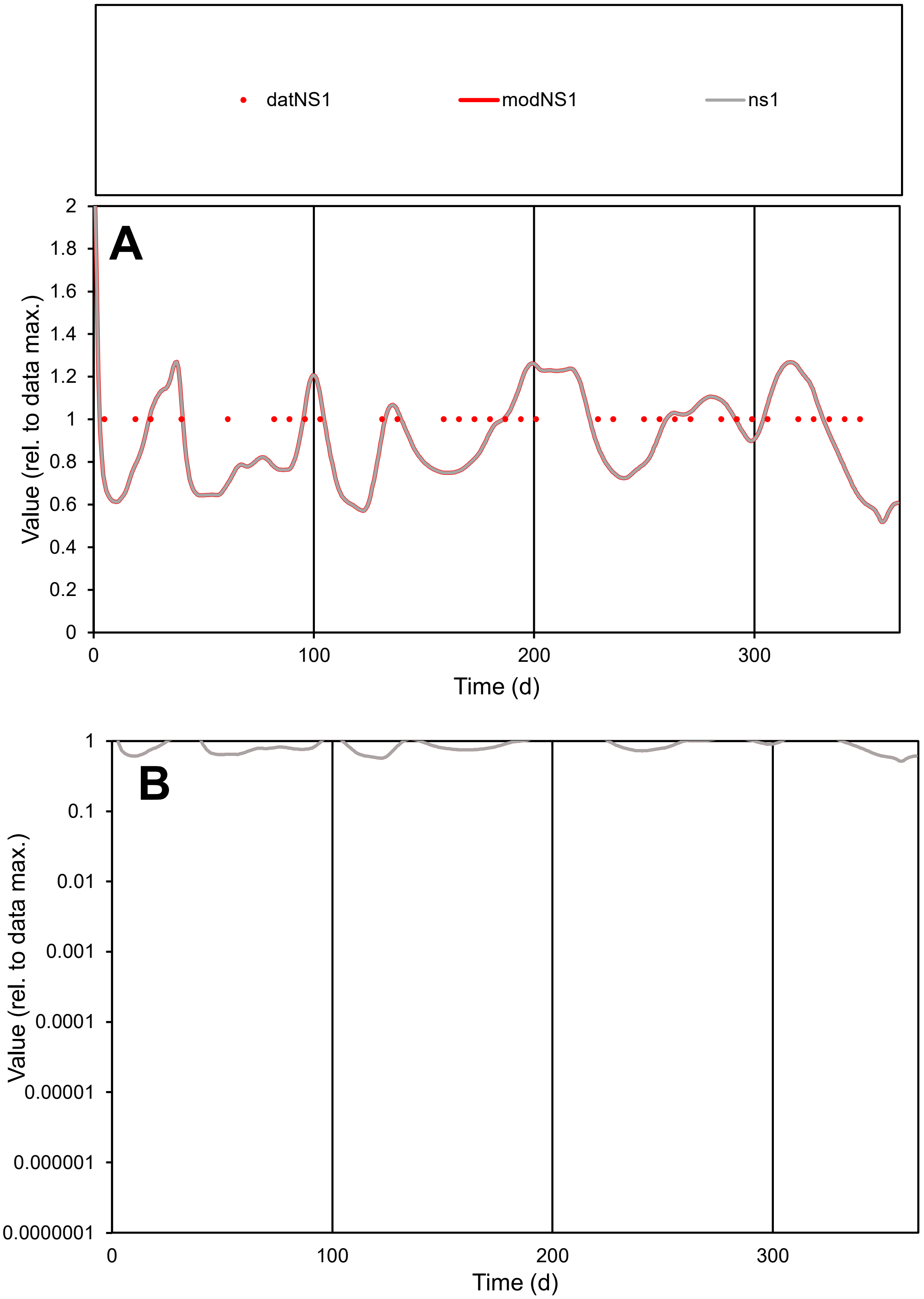


**C1**

Variables

NS1 Provide jtd or Dst

CalibOnly?

2 1 to reproduce of

oNormalize?

2 1=no, 2=yes

ofxn calc*

0.17669

ofxn mod

0.21002

sRun

RunA1

zout

1

Ave

Data

2.7E-05

Model

2.6E-05

data/vMax

2.7E-05

ofxng

263.459

Min d/m

1.61624

ave/min d

1

ave/min m

1.51383

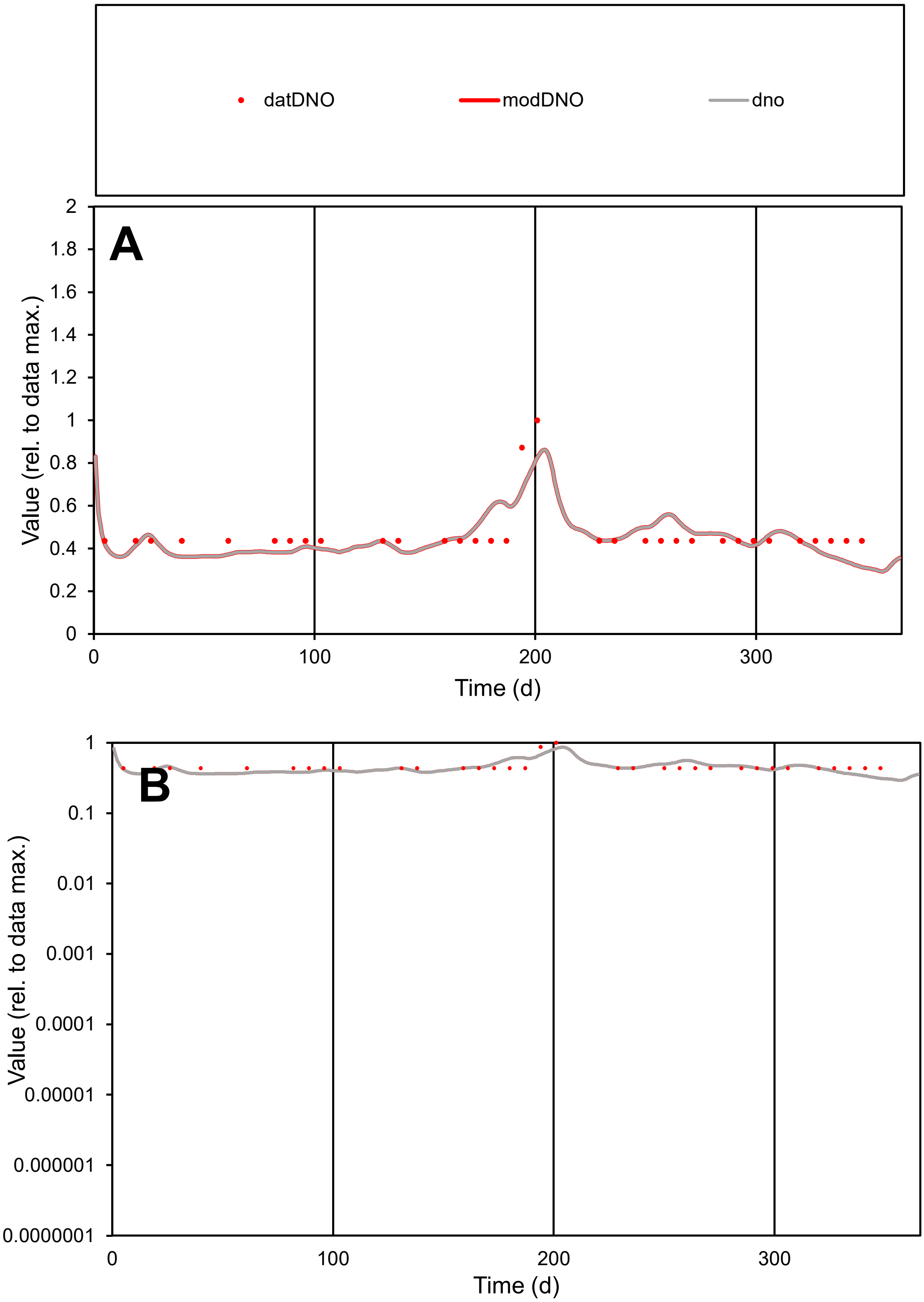


**C1**

Variables

DNO Provide jtd or Dst

CalibOnly?

2 1 to reproduce of

oNormalize?

2 1=no, 2=yes

ofxn calc*

0.16559

ofxn mod

0.20277

sRun

RunA1

zout

1

Ave

Data

1.9E-05

Model

1.9E-05

data/vMax

4.2E-05

ofxng

263.459

Min d/m

1.40201

ave/min d

1.0696

ave/min m

1.47119

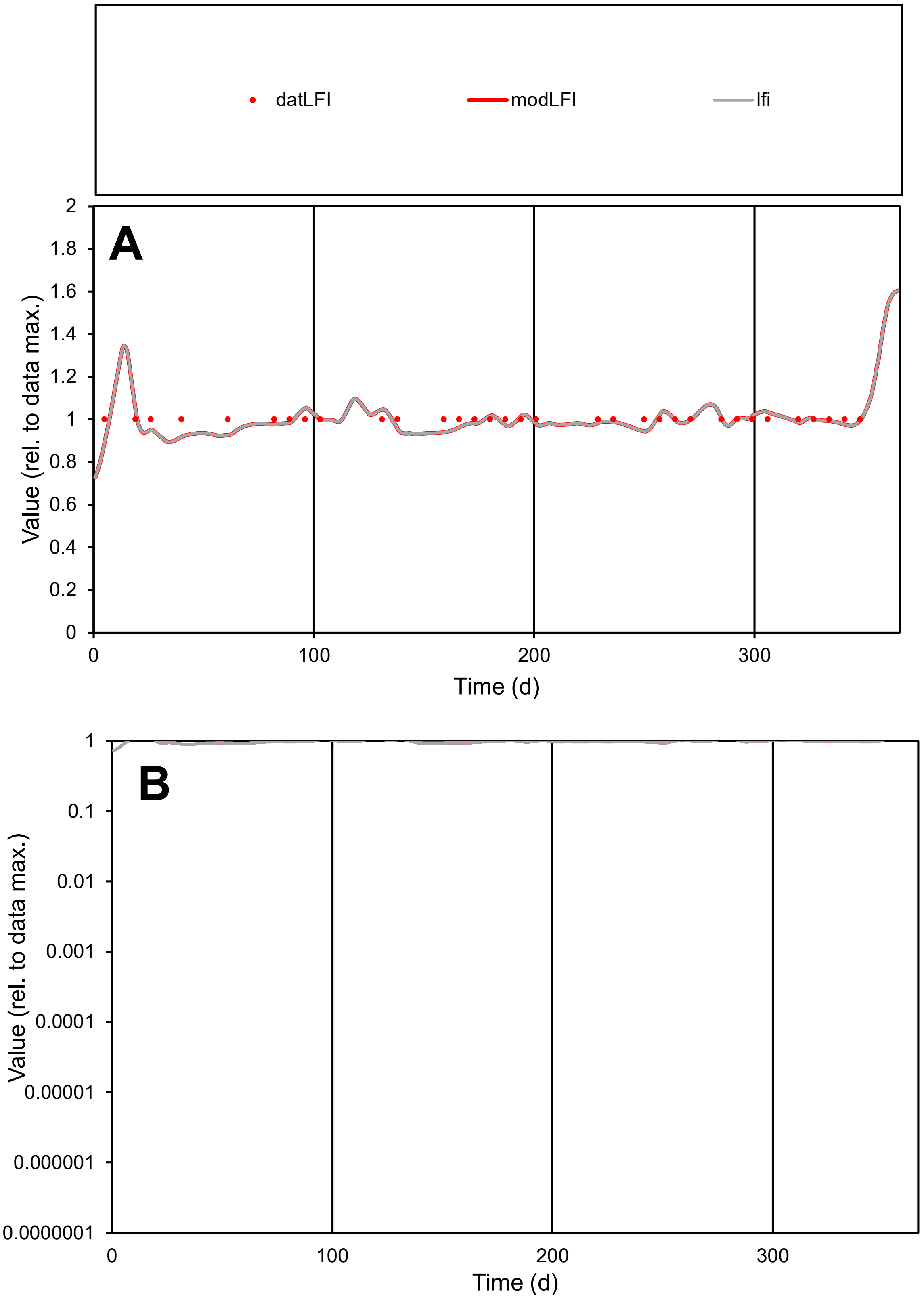


**C1**

Variables
LFI Provide jtd or Dst
CalibOnly?
2 1 to reproduce of
oNormalize?
2 1=no, 2=yes

ofxn calc*
0.03484
ofxn mod
0.1176

sRun
RunA1
zout
1

Ave
Data
8.3E-06
Model
8.2E-06

data/vMax
8.3E-06

ofxng
263.459

Min d/m
1.08558

ave/min d
1
ave/min m
1.07093

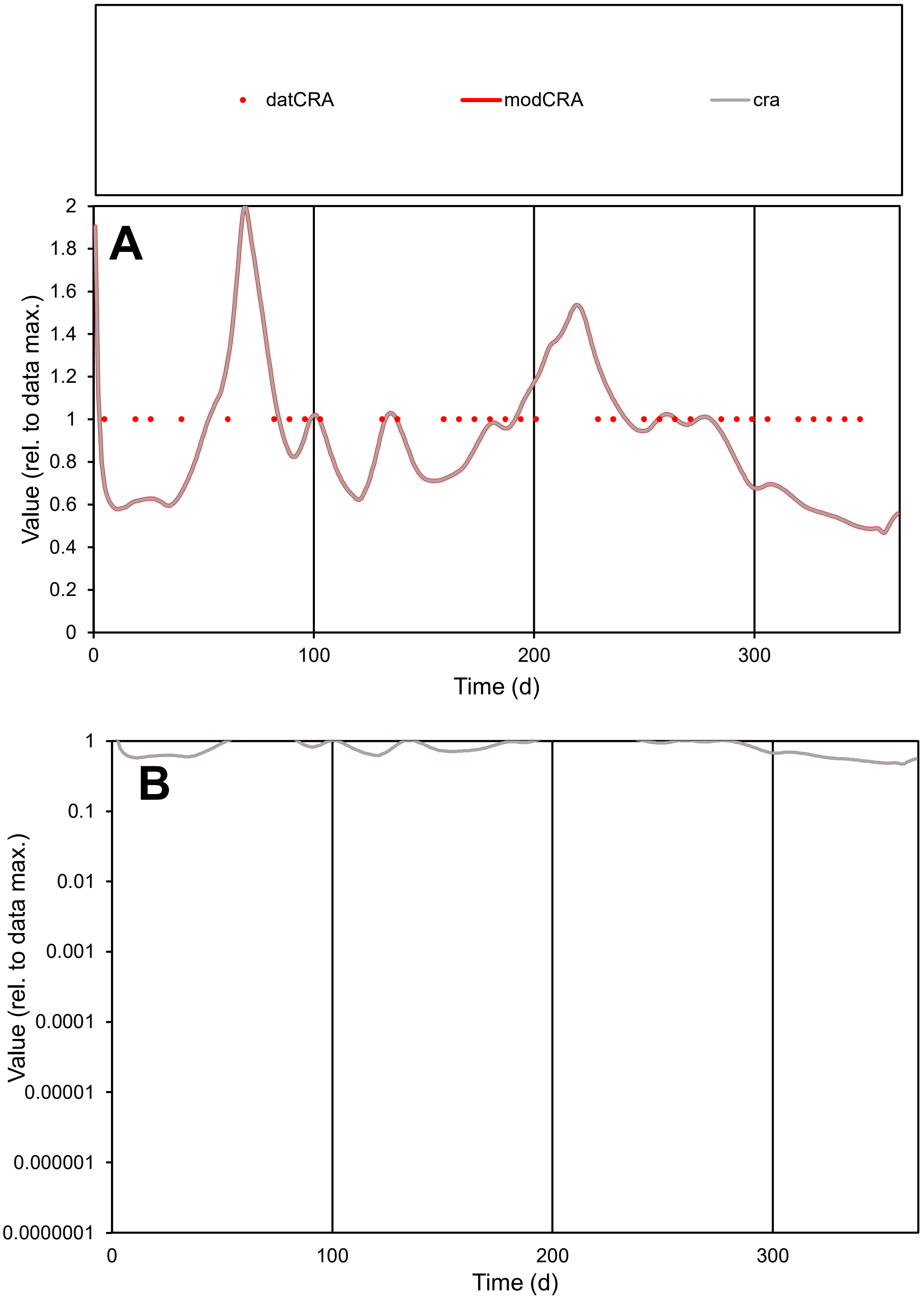


**C1**

Variables
CRA Provide jtd or Dst
CalibOnly?
2 1 to reproduce of
oNormalize?
2 1=no, 2=yes

ofxn calc*
0.25506
ofxn mod
0.28046

sRun
RunA1
zout
1

Ave
Data
4.9E-06
Model
4.2E-06

data/vMax
4.9E-06

ofxng
263.459

Min d/m
2.0343

ave/min d
1
ave/min m
1.74474

326_Calib_S_A

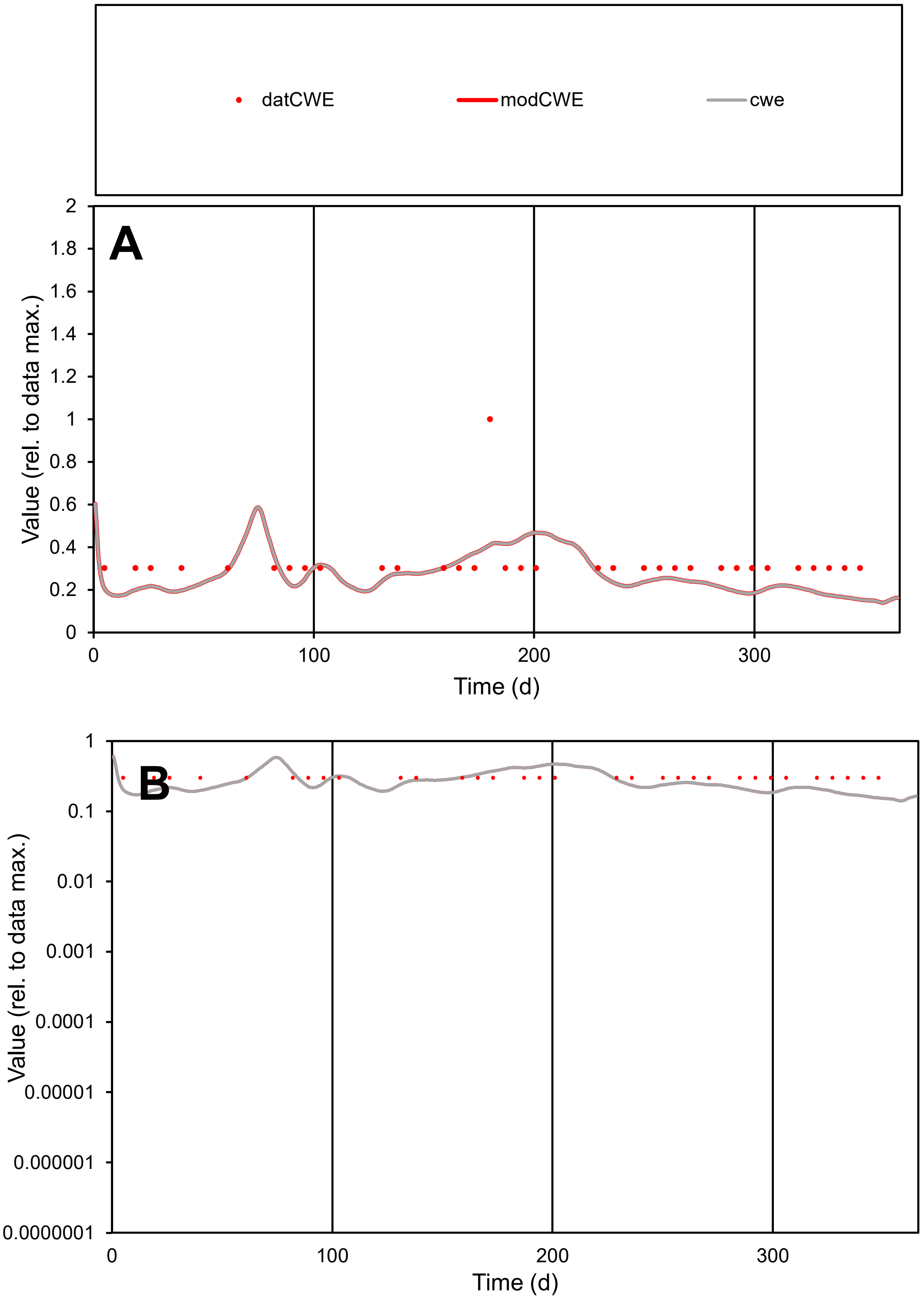


**C1**

Variables
CWE Provide jtd or Dst
CalibOnly?
2 1 to reproduce of
oNormalize?
2 1=no, 2=yes

ofxn calc*
0.41638
ofxn mod
0.43071

sRun
RunA1
zout
1

Ave
Data
2.7E-05
Model
2.2E-05

data/vMax
8.5E-05

ofxng
263.459

Min d/m
1.96456

ave/min d
1.07015
ave/min m
1.724

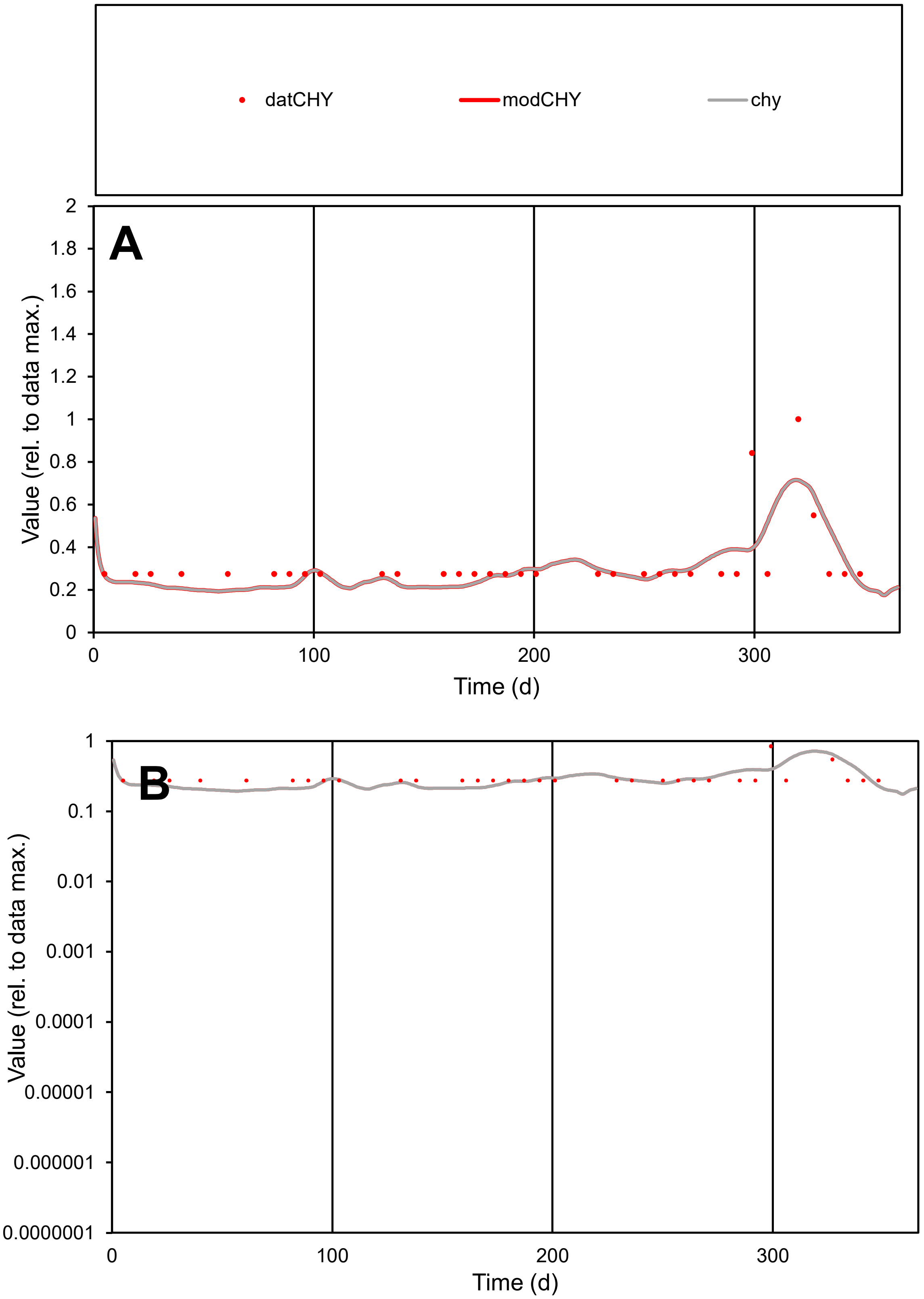


**C1**

Variables
CHY Provide jtd or Dst
CalibOnly?
2 1 to reproduce of
oNormalize?
2 1=no, 2=yes

ofxn calc*
0.36148
ofxn mod
0.3809

sRun
RunA1
zout
1

Ave
Data
7.9E-06
Model
7.5E-06

data/vMax
2.4E-05

ofxng
263.459

Min d/m
1.38913

ave/min d
1.17298
ave/min m
1.54672

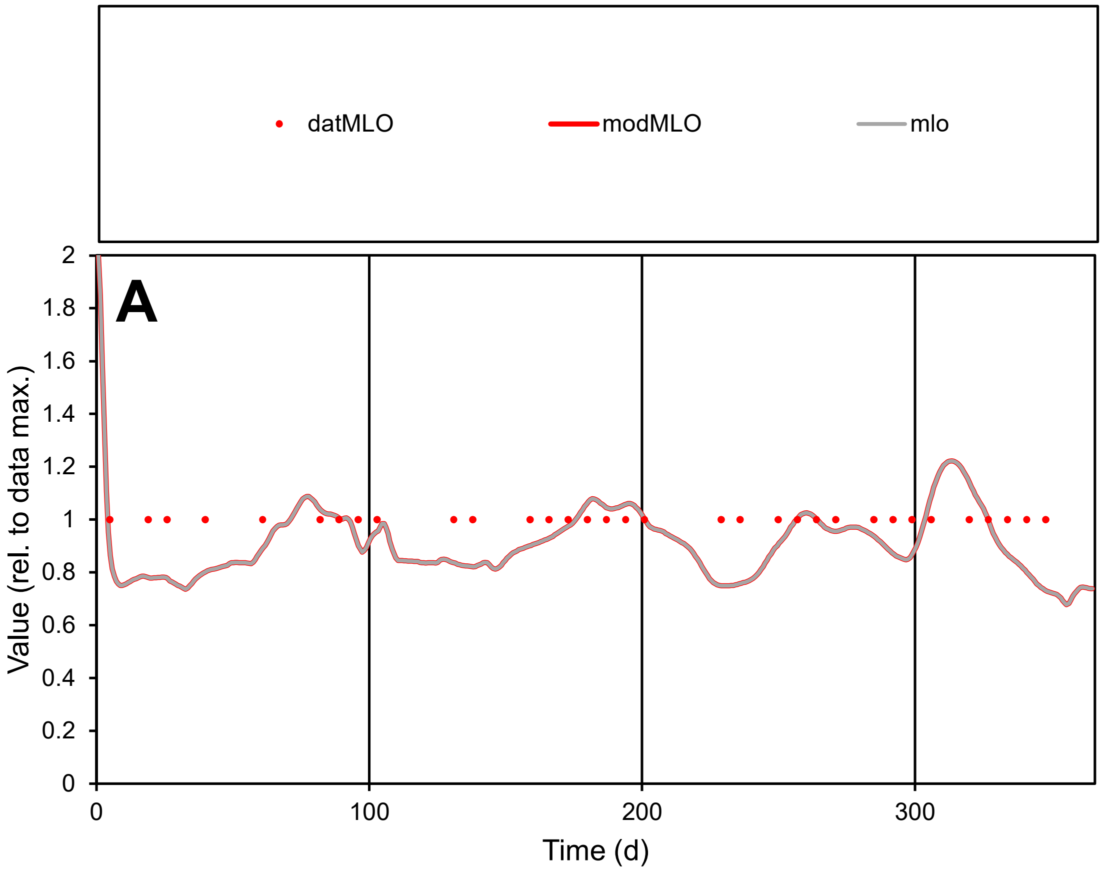


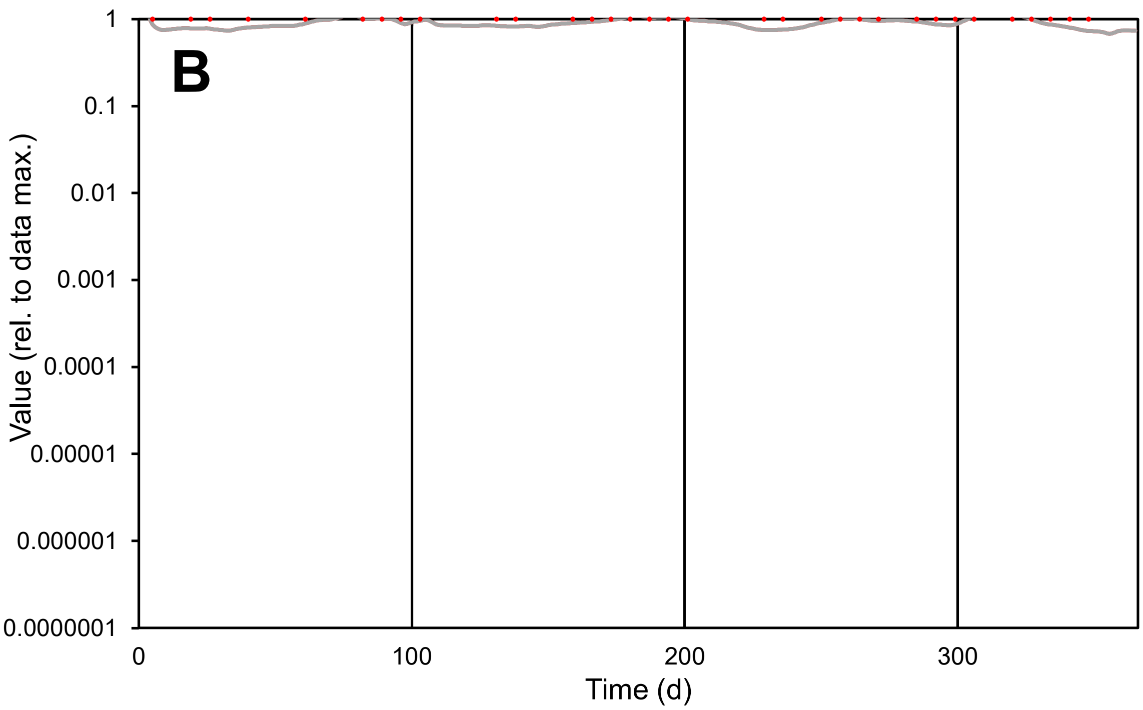


**C1**

Variables
MLO Provide jtd or Dst
CalibOnly?
2 1 to reproduce of
oNormalize?
2 1=no, 2=yes

ofxn calc*
0.13886
ofxn mod
0.17631

sRun
RunA1
zout
1

Ave
Data
1E-05
Model
9.6E-06

data/vMax
1E-05

ofxng
263.459

Min d/m
1.37763

ave/min d
1
ave/min m
1.26242

326_Calib_S_A

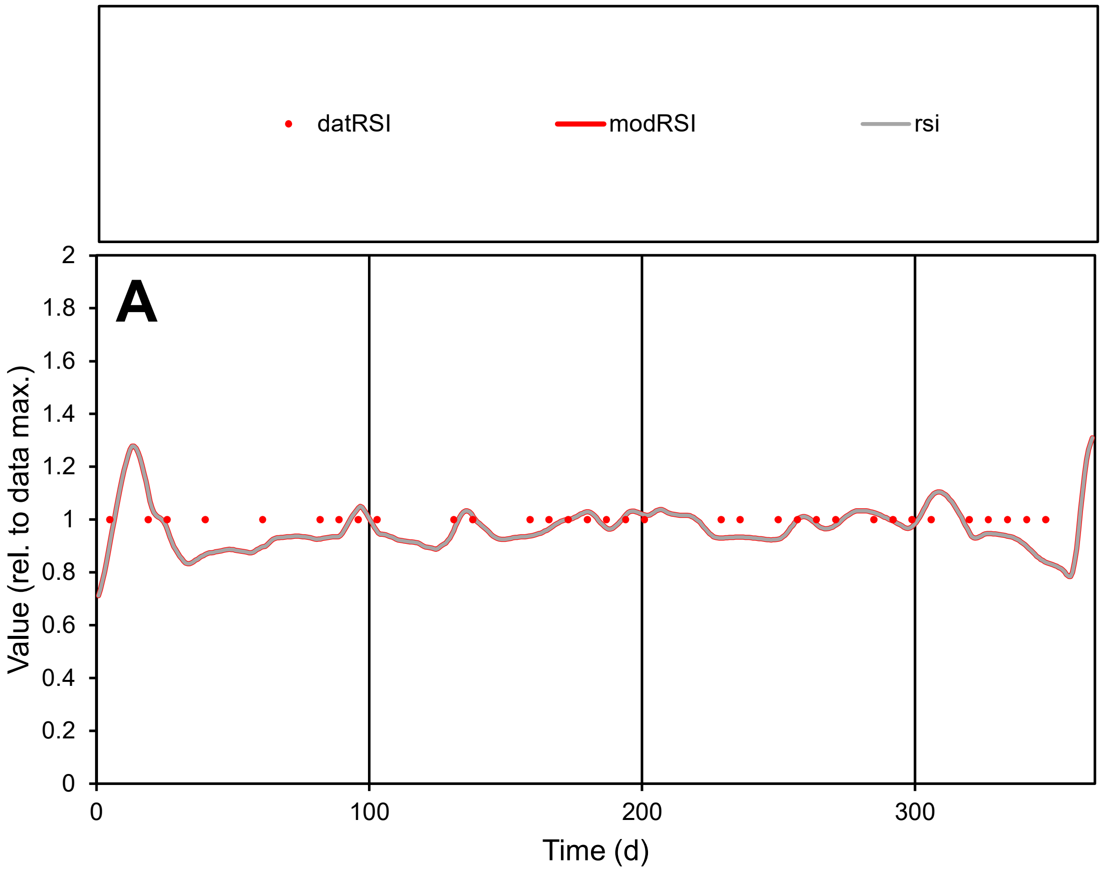


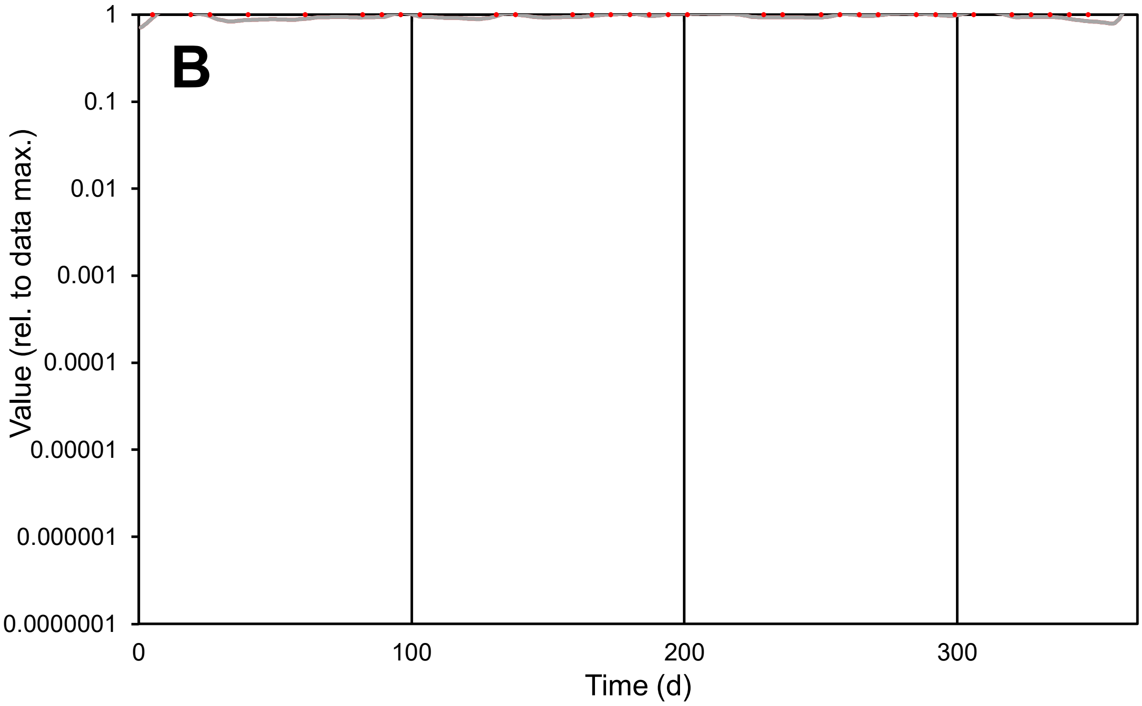


**C1**

Variables
RSI Provide jtd or Dst
CalibOnly?
2 1 to reproduce of
oNormalize?
2 1=no, 2=yes

ofxn calc*
0.06376
ofxn mod
0.12851

sRun
RunA1
zout
1

Ave
Data
1E-05
Model
1E-05

data/vMax
1E-05

ofxng
263.459

Min d/m
1.19907

ave/min d
1
ave/min m
1.16075

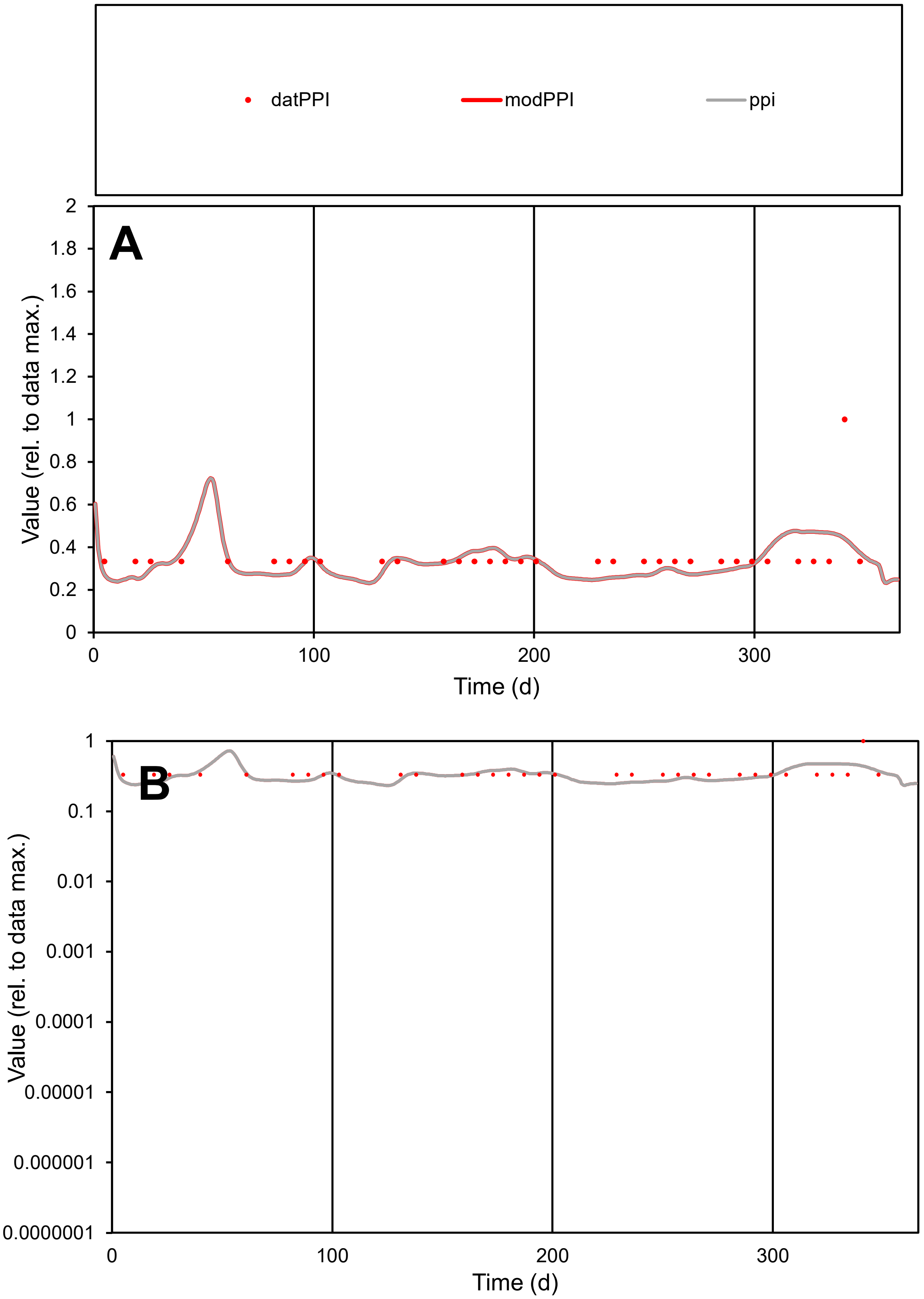


**C1**

Variables
PPI Provide jtd or Dst
CalibOnly?
2 1 to reproduce of
oNormalize?
2 1=no, 2=yes

ofxn calc*
0.33082
ofxn mod
0.34924

sRun
RunA1
zout
1

Ave
Data
1.5E-06
Model
1.5E-06

data/vMax
4.3E-06

ofxng
263.459

Min d/m
1.32418

ave/min d
1.06071
ave/min m
1.33164

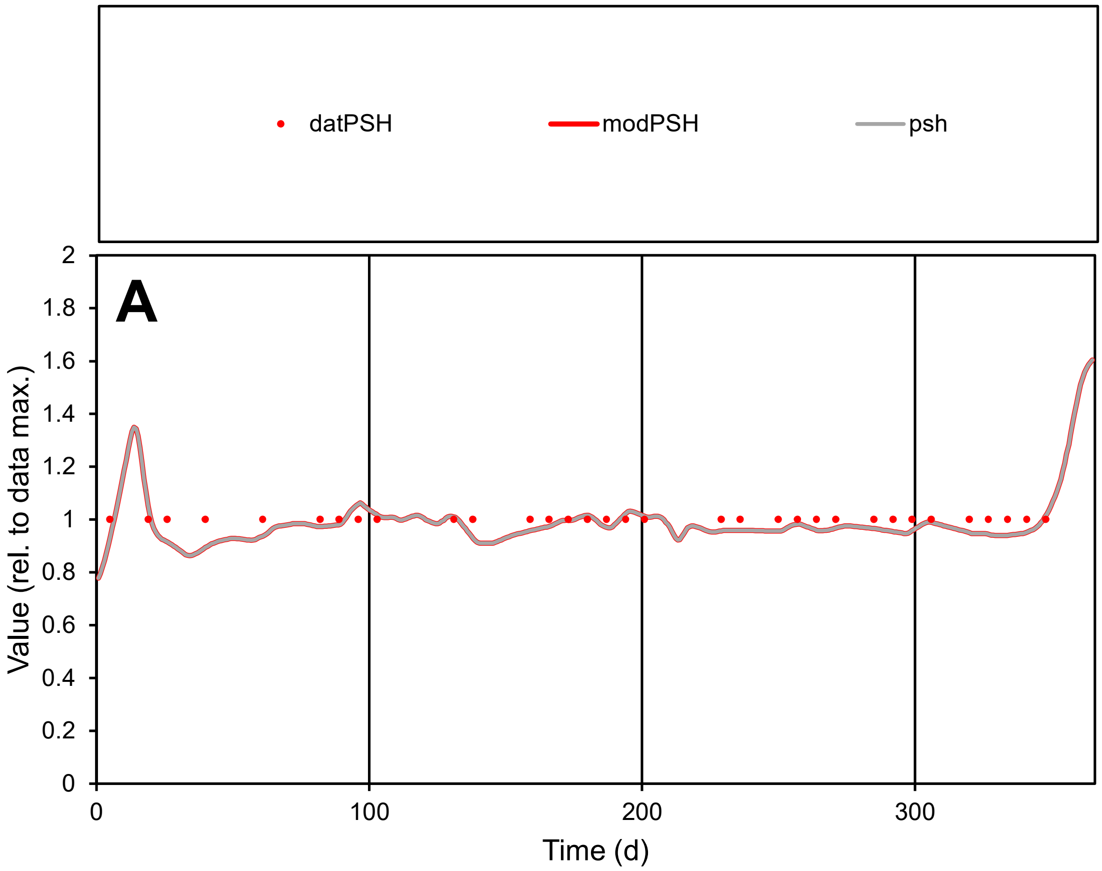


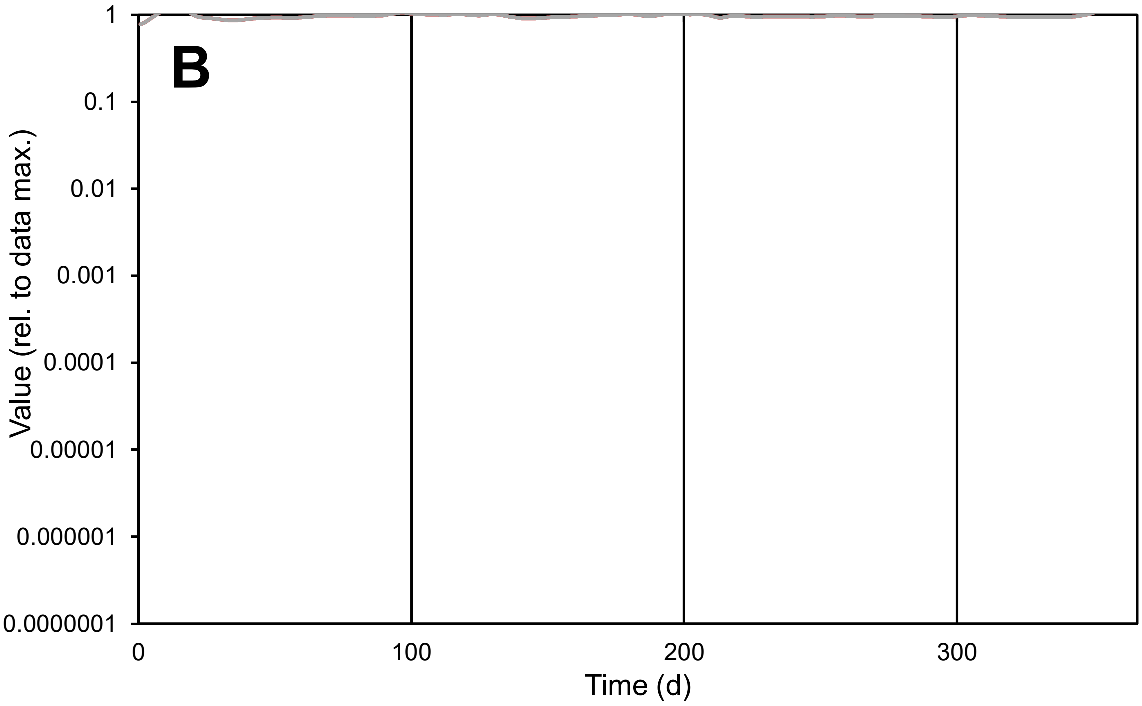


**C1**

Variables
PSH Provide jtd or Dst
CalibOnly?
2 1 to reproduce of
oNormalize?
2 1=no, 2=yes

ofxn calc*
0.04385
ofxn mod
0.11983

sRun
RunA1
zout
1

Ave
Data
1.3E-05
Model
1.3E-05

data/vMax
1.3E-05

ofxng
263.459

Min d/m
1.11207

ave/min d
1
ave/min m
1.08284

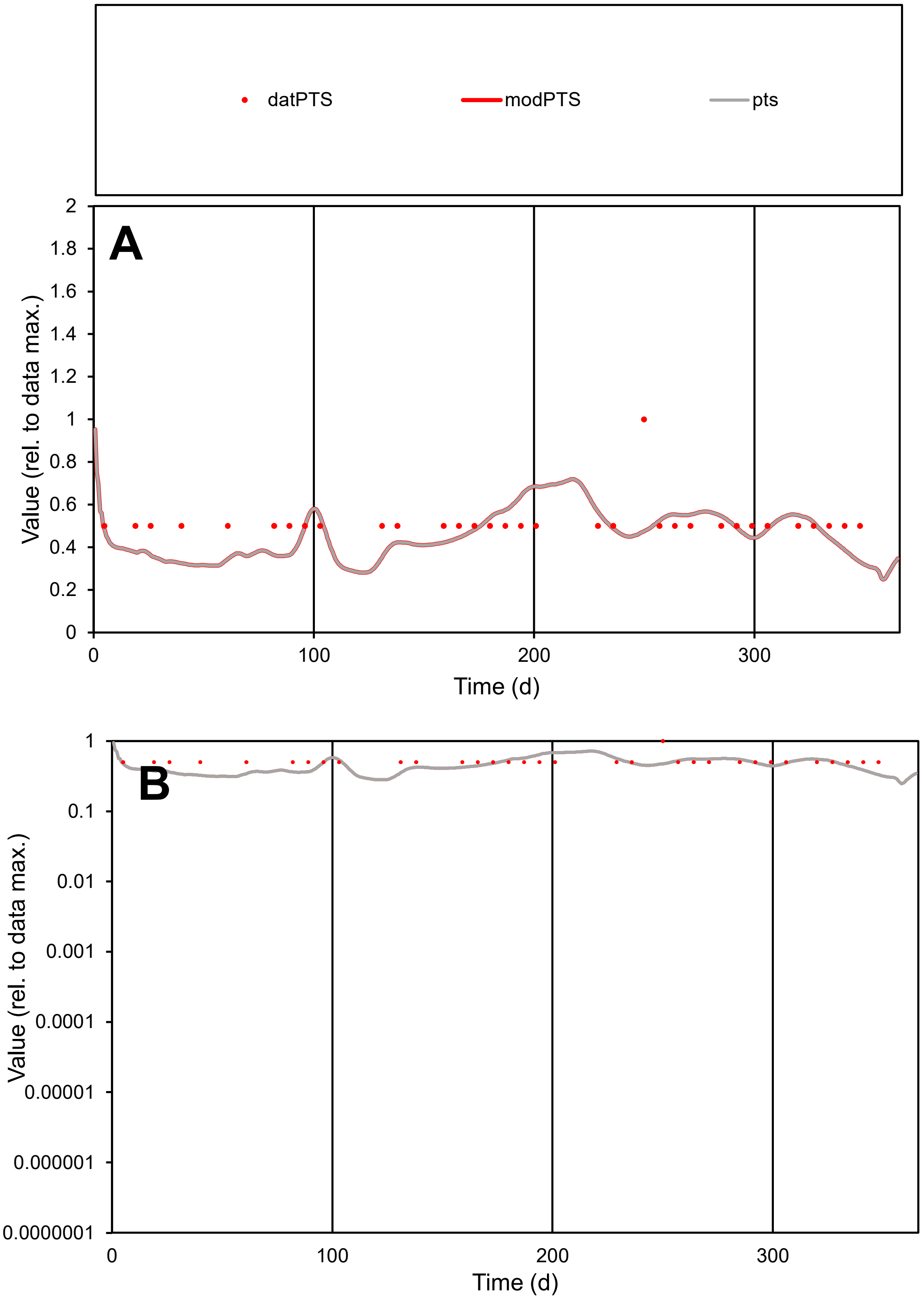


**C1**

Variables

PTS Provide jtd or Dst

CalibOnly?

2 1 to reproduce of

oNormalize?

2 1=no, 2=yes

ofxn calc*

0.25331

ofxn mod

0.27645

sRun

RunA1

zout

1

Ave

Data

1E-05

Model

9.1E-06

data/vMax

1.9E-05

ofxng

263.459

Min d/m

1.54409

ave/min d

1.03032

ave/min m

1.45833

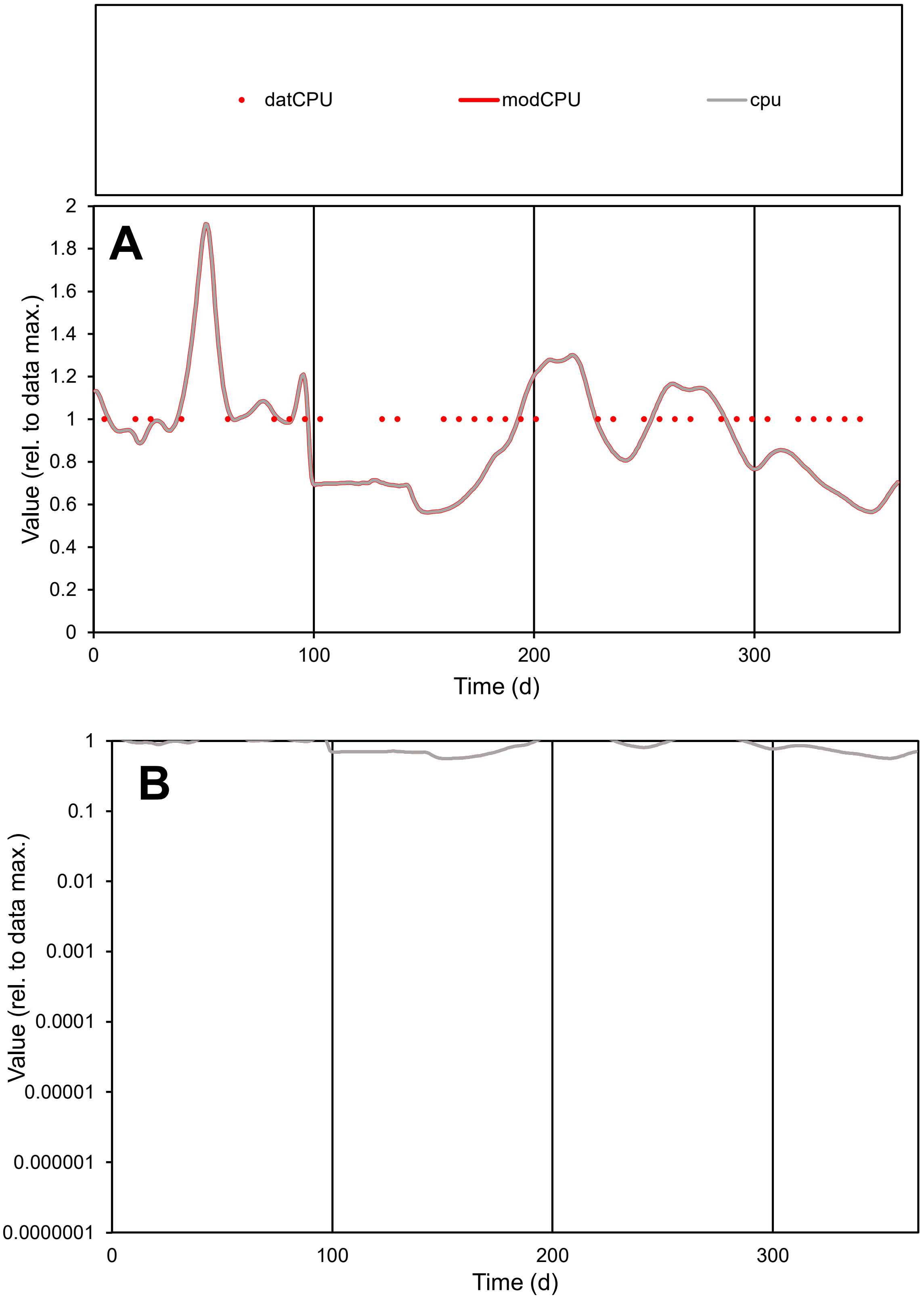


**C1**

Variables

CPU Provide jtd or Dst

CalibOnly?

2 1 to reproduce of

oNormalize?

2 1=no, 2=yes

ofxn calc*

0.21868

ofxn mod

0.2446

sRun

RunA1

zout

1

Ave

Data

4.4E-07

Model

3.9E-07

data/vMax

4.4E-07

ofxng

263.459

Min d/m

1.73821

ave/min d

1

ave/min m

1.52893

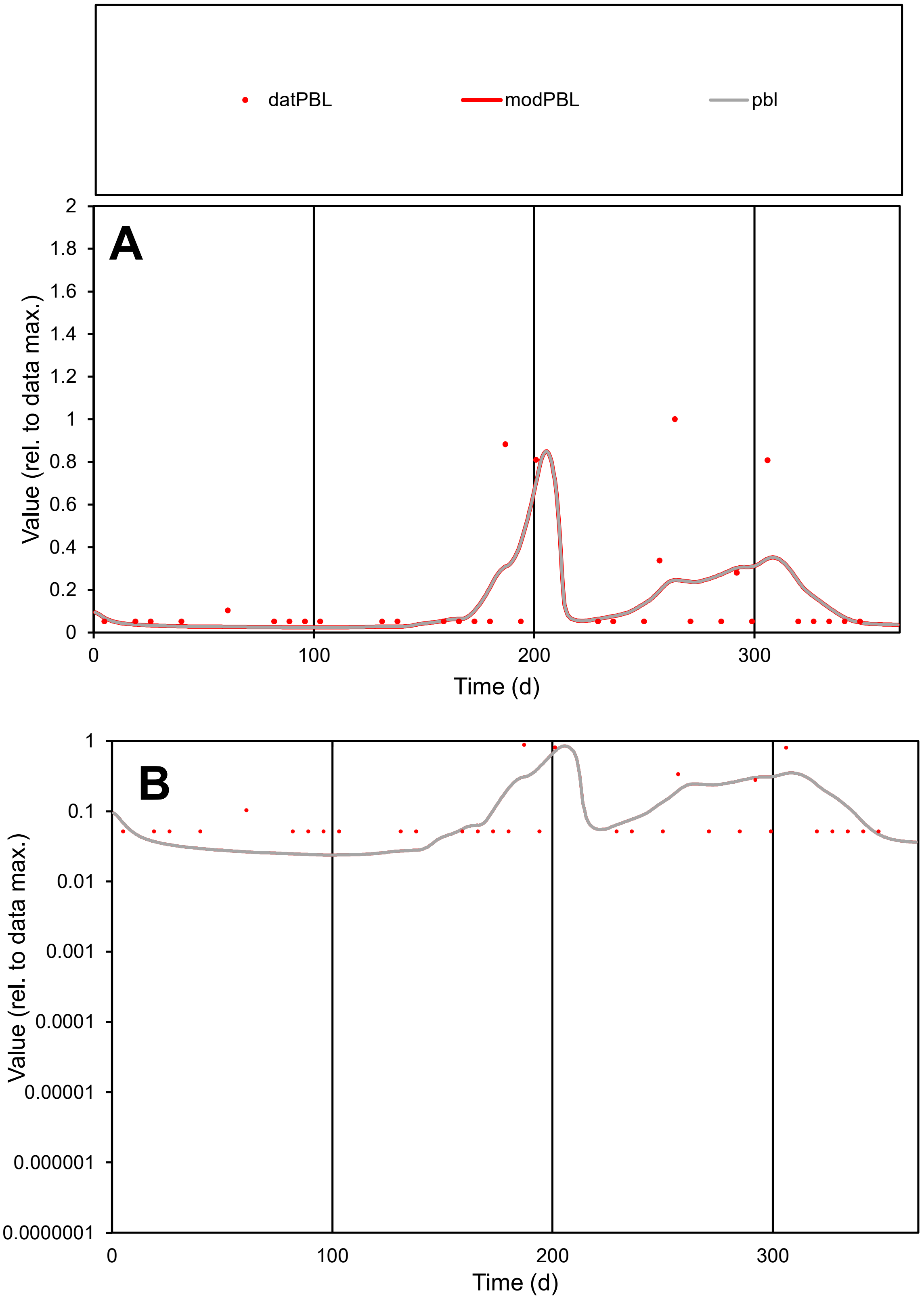


**C1**

Variables
PBL Provide jtd or Dst
CalibOnly?
2 1 to reproduce of
oNormalize?
2 1=no, 2=yes

ofxn calc*
1.25308
ofxn mod
1.2746

sRun
RunA1
zout
1

Ave
Data
1.4E-07
Model
1.3E-07

data/vMax
8.2E-07

ofxng
263.459

Min d/m
2.17476

ave/min d
3.24895
ave/min m
6.40074

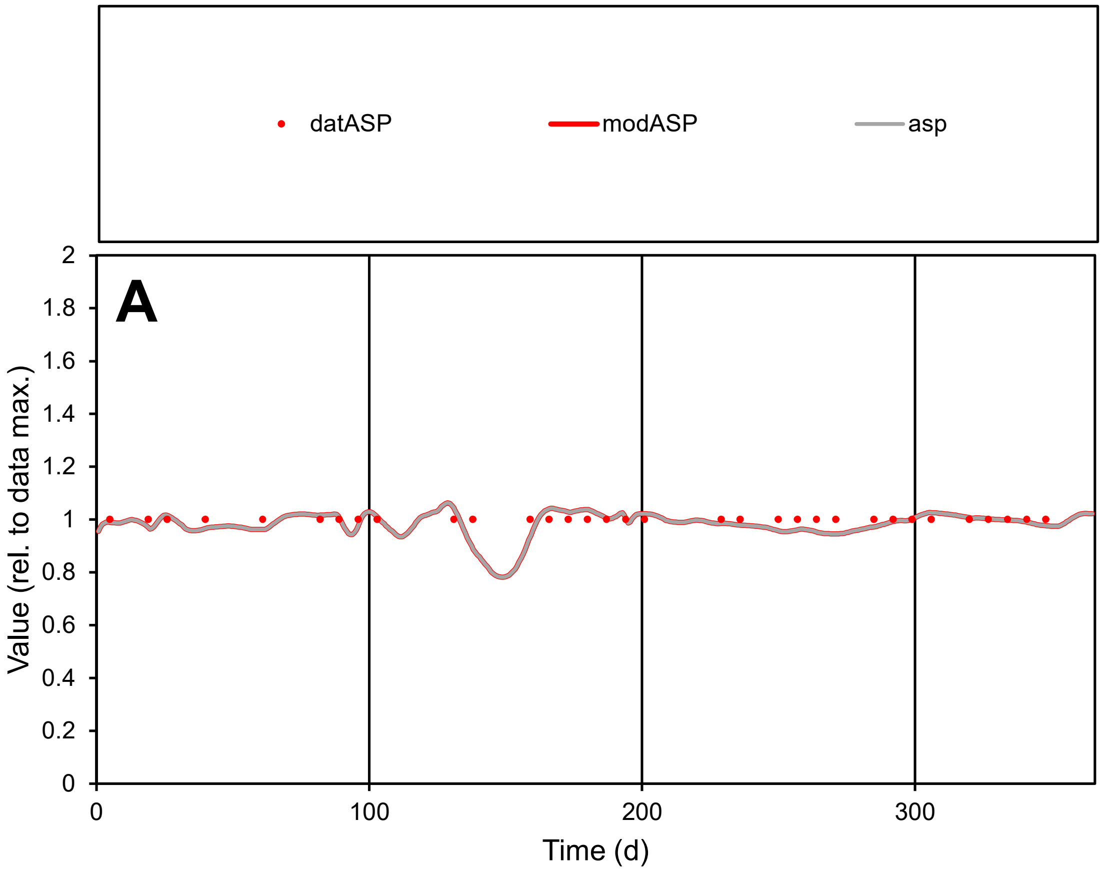


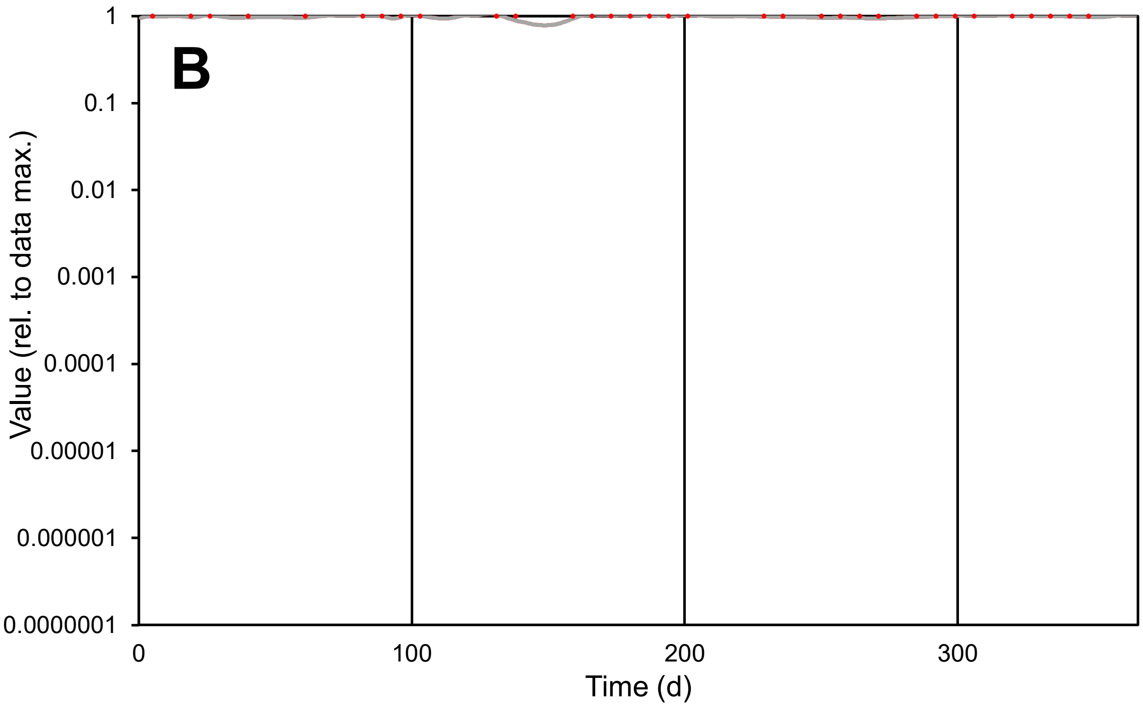


**C1**

Variables
ASP Provide jtd or Dst
CalibOnly?
2 1 to reproduce of
oNormalize?
2 1=no, 2=yes

ofxn calc*
0.03465
ofxn mod
0.11773

sRun
RunA1
zout
1

Ave
Data
5.8E-06
Model
5.7E-06

data/vMax
5.8E-06

ofxng
263.459

Min d/m
1.15383

ave/min d
1
ave/min m
1.14071

326_Calib_S_A

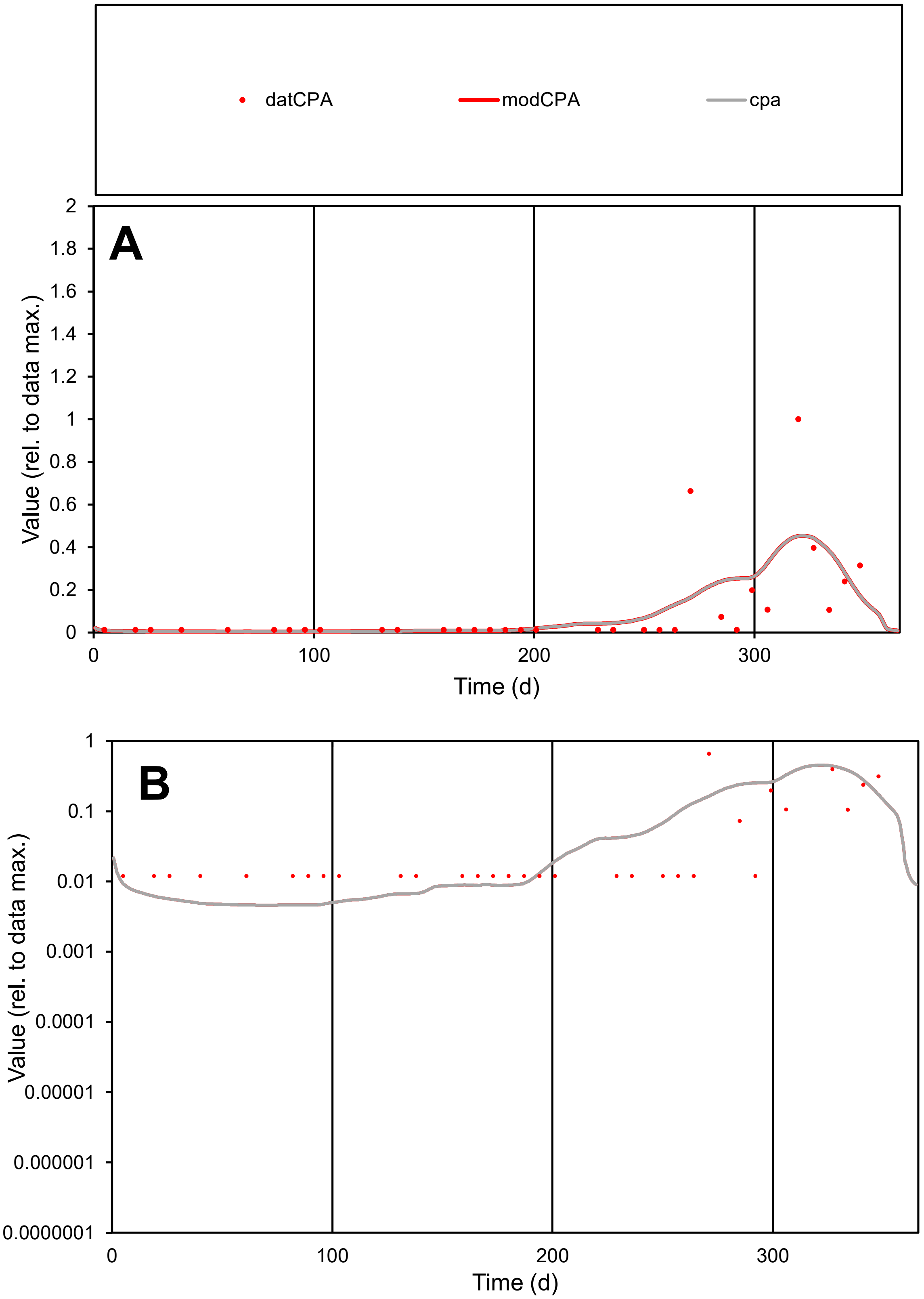


**C1**

Variables

| | |
|---|---|
| CPA | Provide jtd or Dst |
| CalibOnly? | |
| 2 | 1 to reproduce of |
| oNormalize? | |
| 2 | 1=no, 2=yes |

ofxn calc*
1.51566
ofxn mod
1.53401

sRun
RunA1
zout
1

Ave
Data
5.7E-07
Model
5.8E-07

data/vMax
5.5E-06

ofxng
263.459

Min d/m
2.60471

ave/min d
8.53224
ave/min m
22.7593

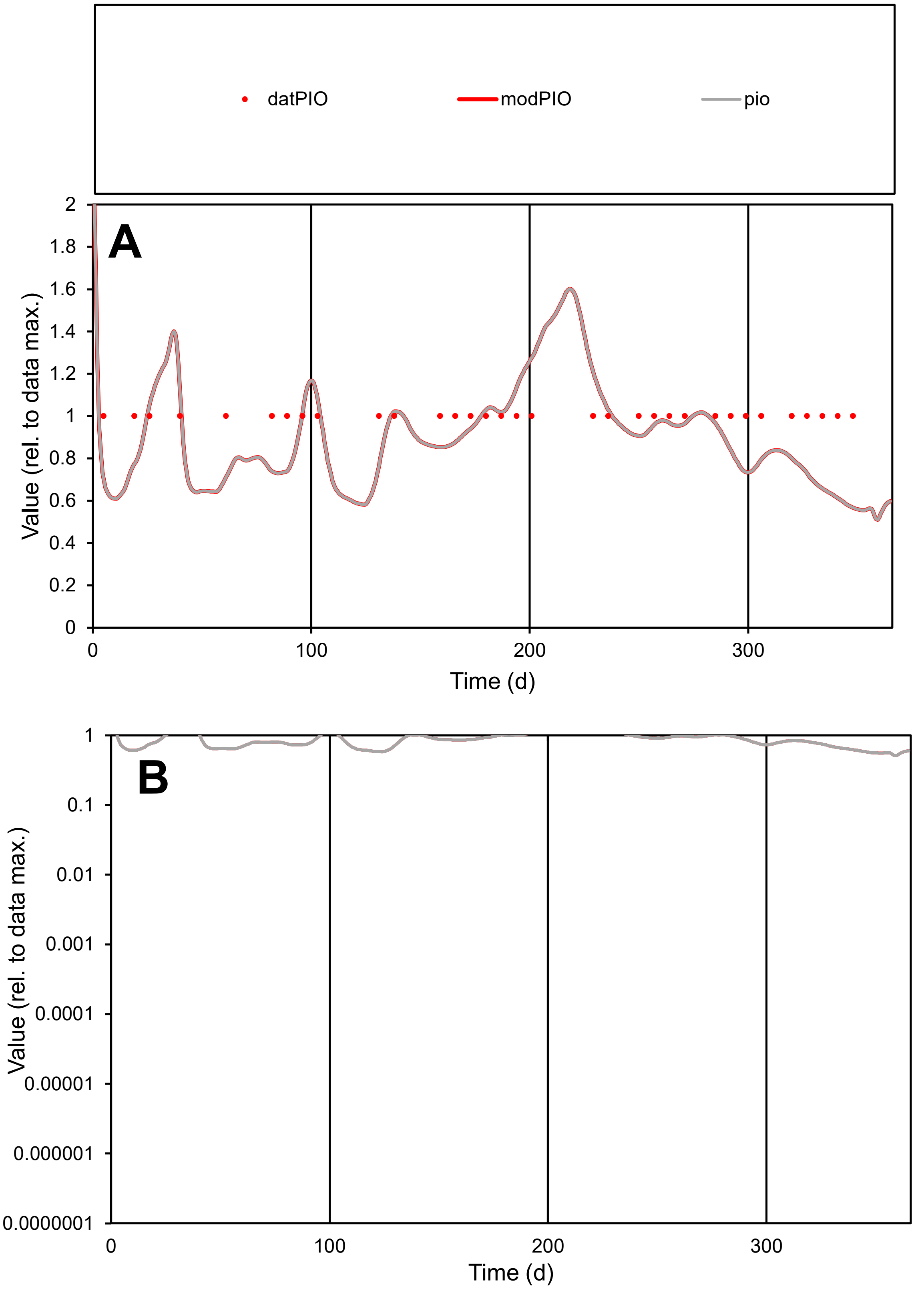


**C1**

Variables
PIO Provide jtd or Dst
CalibOnly?
2 1 to reproduce of
oNormalize?
2 1=no, 2=yes

ofxn calc*
0.19992
ofxn mod
0.22905

sRun
RunA1
zout
1

Ave
Data
1.5E-07
Model
1.3E-07

data/vMax
1.5E-07

ofxng
263.459

Min d/m
1.77658

ave/min d
1
ave/min m
1.59674

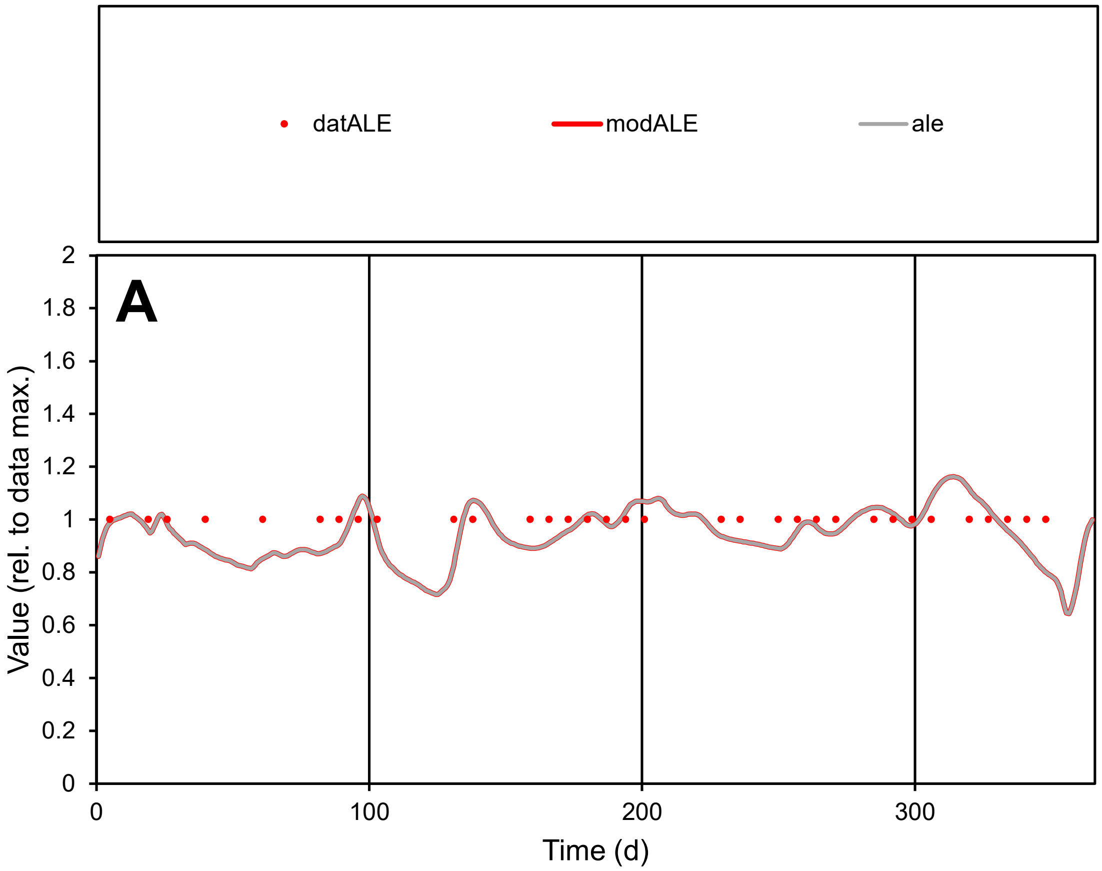


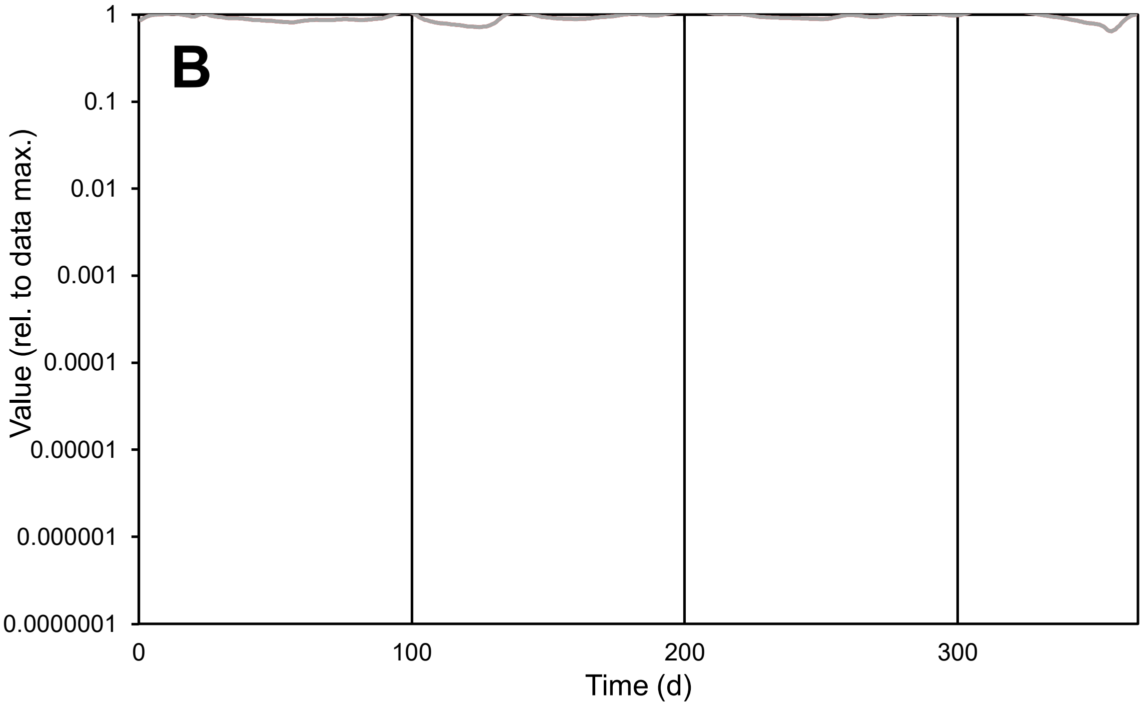


**C1**

Variables
ALE Provide jtd or Dst
CalibOnly?
2 1 to reproduce of
oNormalize?
2 1=no, 2=yes

ofxn calc*
0.08443
ofxn mod
0.13974

sRun
RunA1
zout
1

Ave
Data
1.1E-05
Model
1.1E-05

data/vMax
1.1E-05

ofxng
263.459

Min d/m
1.26198

ave/min d
1
ave/min m
1.21733

326_Calib_S_A

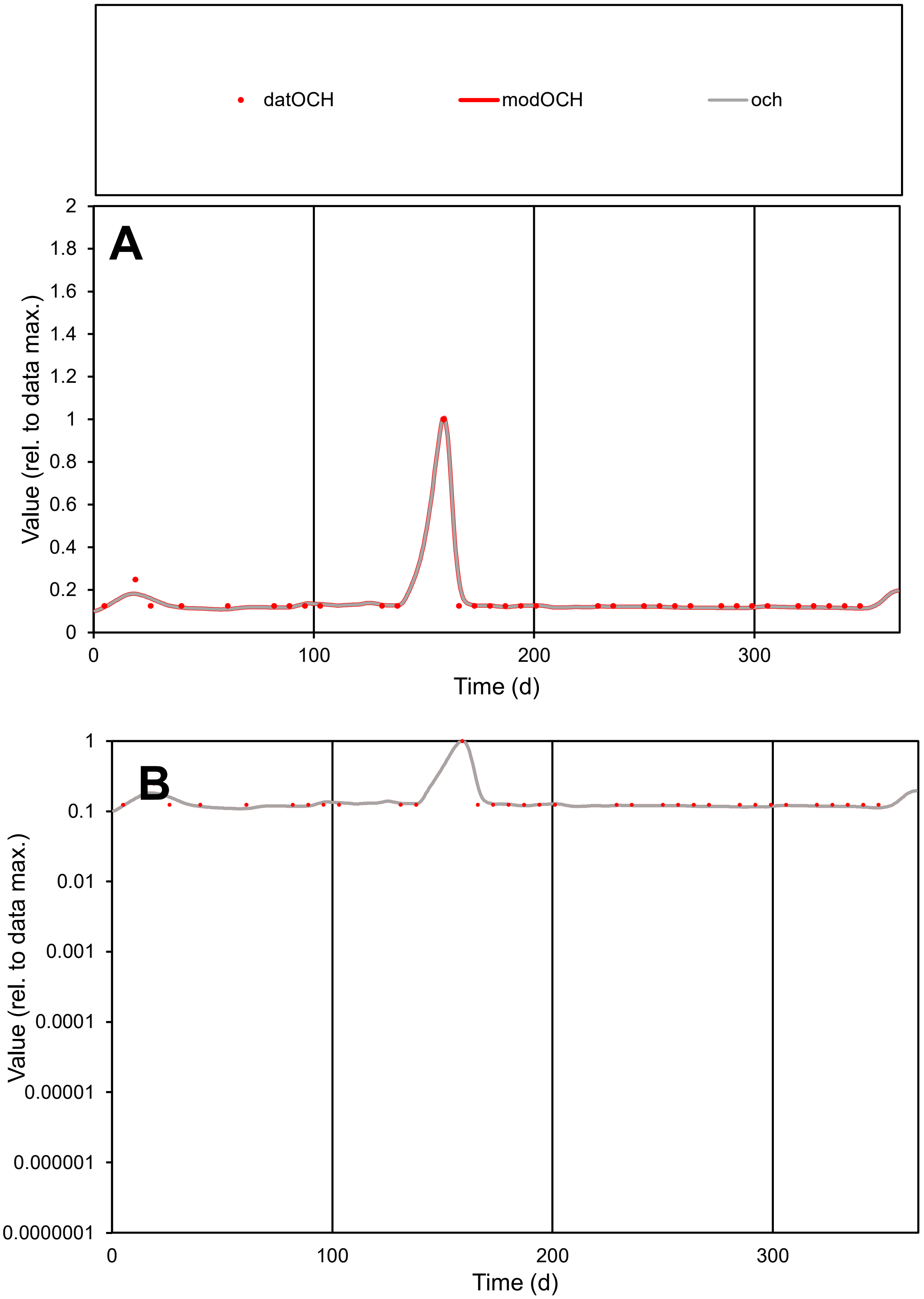


**C1**

Variables
OCH Provide jtd or Dst
CalibOnly?
2 1 to reproduce of
oNormalize?
2 1=no, 2=yes

ofxn calc*
0.11207
ofxn mod
0.15726

sRun
RunA1
zout
1

Ave
Data
5.5E-06
Model
5.4E-06

data/vMax
3.6E-05

ofxng
263.459

Min d/m
1.12369

ave/min d
1.24466
ave/min m
1.37567

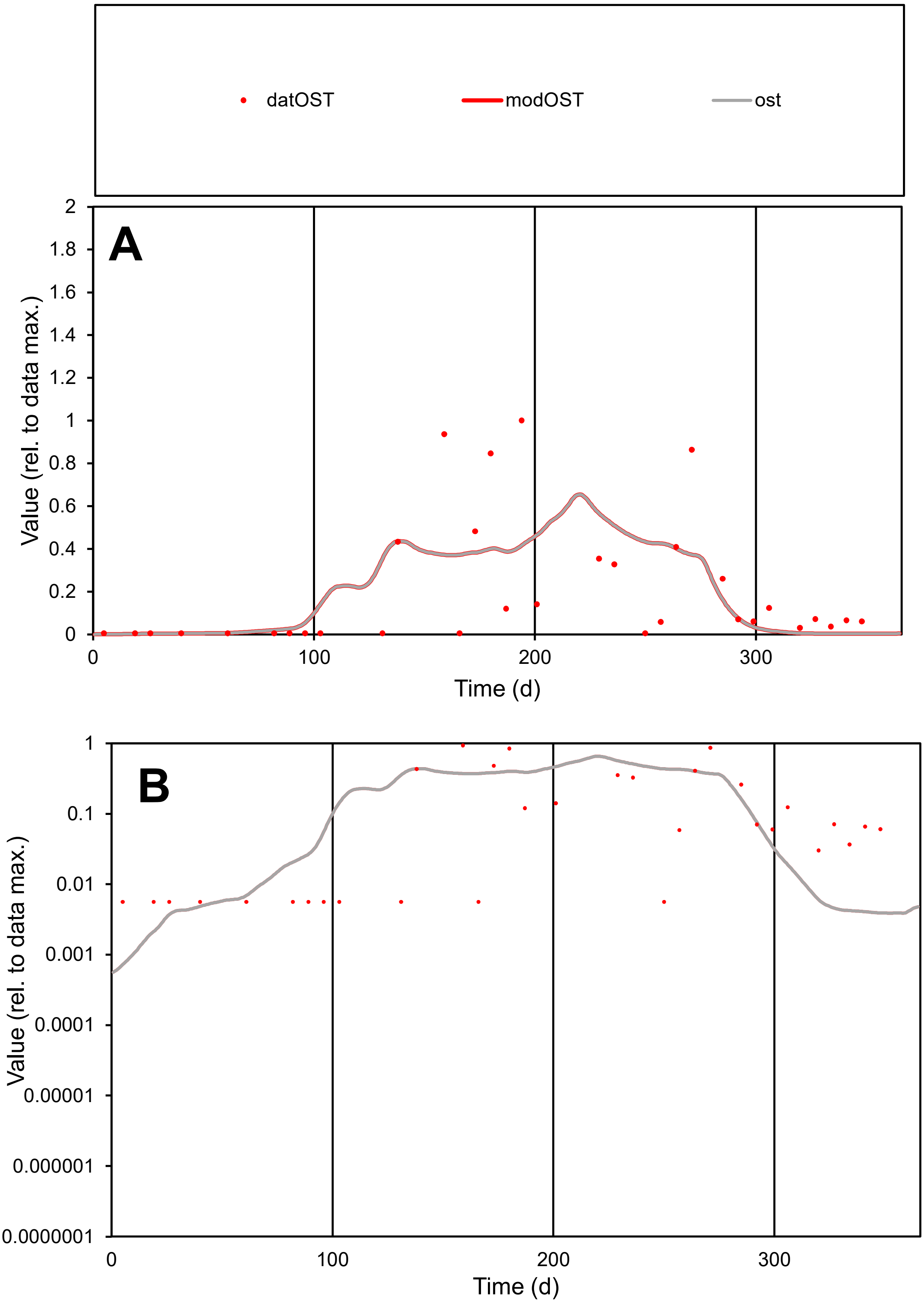


**C1**

Variables

OST Provide jtd or Dst

CalibOnly?

2 1 to reproduce of

oNormalize?

2 1=no, 2=yes

ofxn calc*

1.17735

ofxn mod

1.19519

sRun

RunA1

zout

1

Ave

Data

7.9E-05

Model

7.9E-05

data/vMax

0.00038

ofxng

263.459

Min d/m

7.038

ave/min d

36.6456

ave/min m

257.168

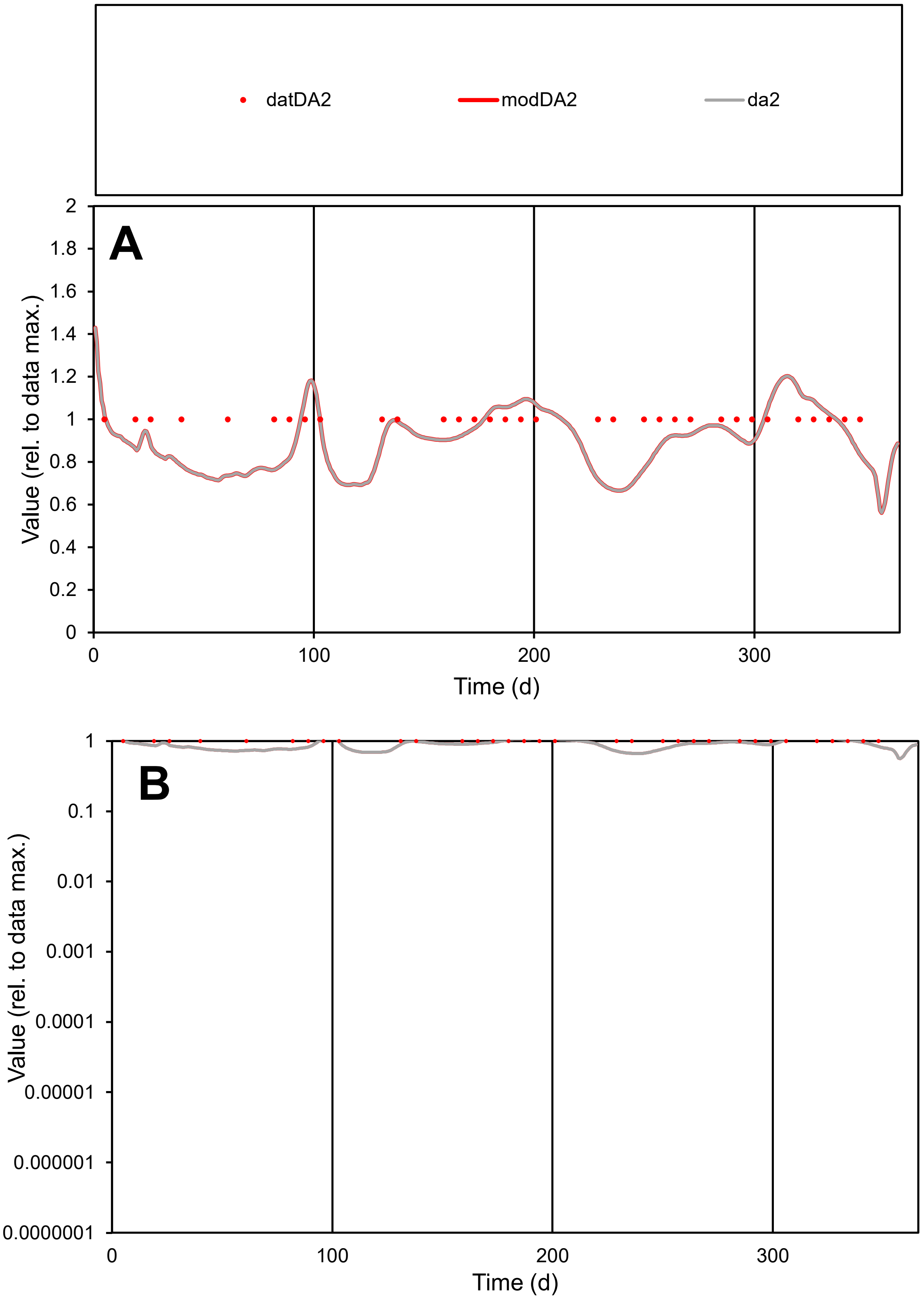


**C1**

Variables
DA2 Provide jtd or Dst
CalibOnly?
2 1 to reproduce of
oNormalize?
2 1=no, 2=yes

ofxn calc*
0.14138
ofxn mod
0.17851

sRun
RunA1
zout
1

Ave
Data
3.1E-06
Model
2.9E-06

data/vMax
3.1E-06

ofxng
263.459

Min d/m
1.49856

ave/min d
1
ave/min m
1.39004

326_Calib_S_A

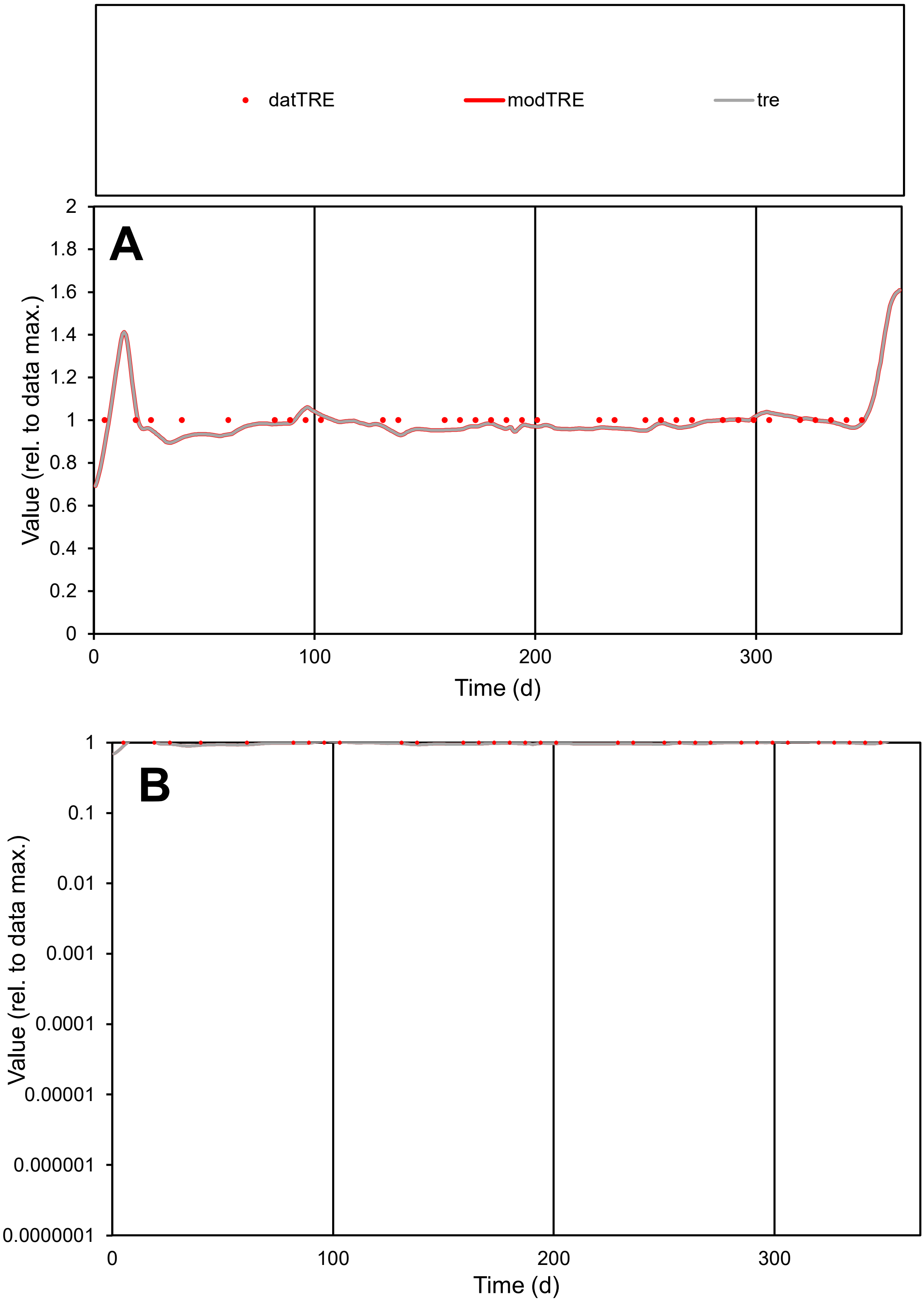


**C1**

Variables
TRE Provide jtd or Dst
CalibOnly?
2 1 to reproduce of
oNormalize?
2 1=no, 2=yes

ofxn calc*
0.03488
ofxn mod
0.11707

sRun
RunA1
zout
1

Ave
Data
6.9E-06
Model
6.8E-06

data/vMax
6.9E-06

ofxng
263.459

Min d/m
1.08562

ave/min d
1
ave/min m
1.06394

326_Calib_S_A

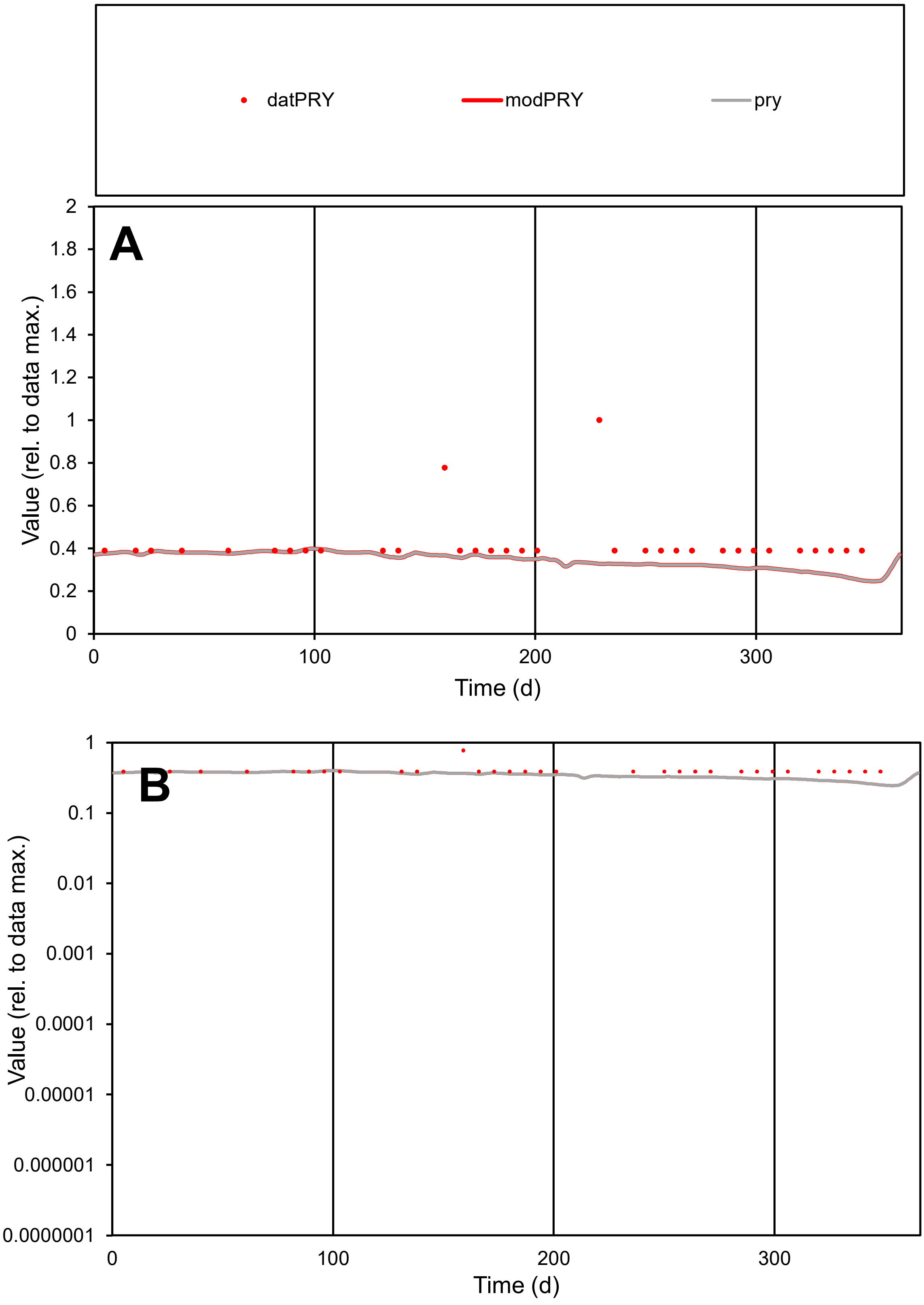


**C1**

Variables
PRY Provide jtd or Dst
CalibOnly?
2 1 to reproduce of
oNormalize?
2 1=no, 2=yes

ofxn calc*
0.35876
ofxn mod
0.37337

sRun
RunA1
zout
1

Ave
Data
9.5E-06
Model
7.7E-06

data/vMax
2.3E-05

ofxng
263.459

Min d/m
1.56207

ave/min d
1.07792
ave/min m
1.36693

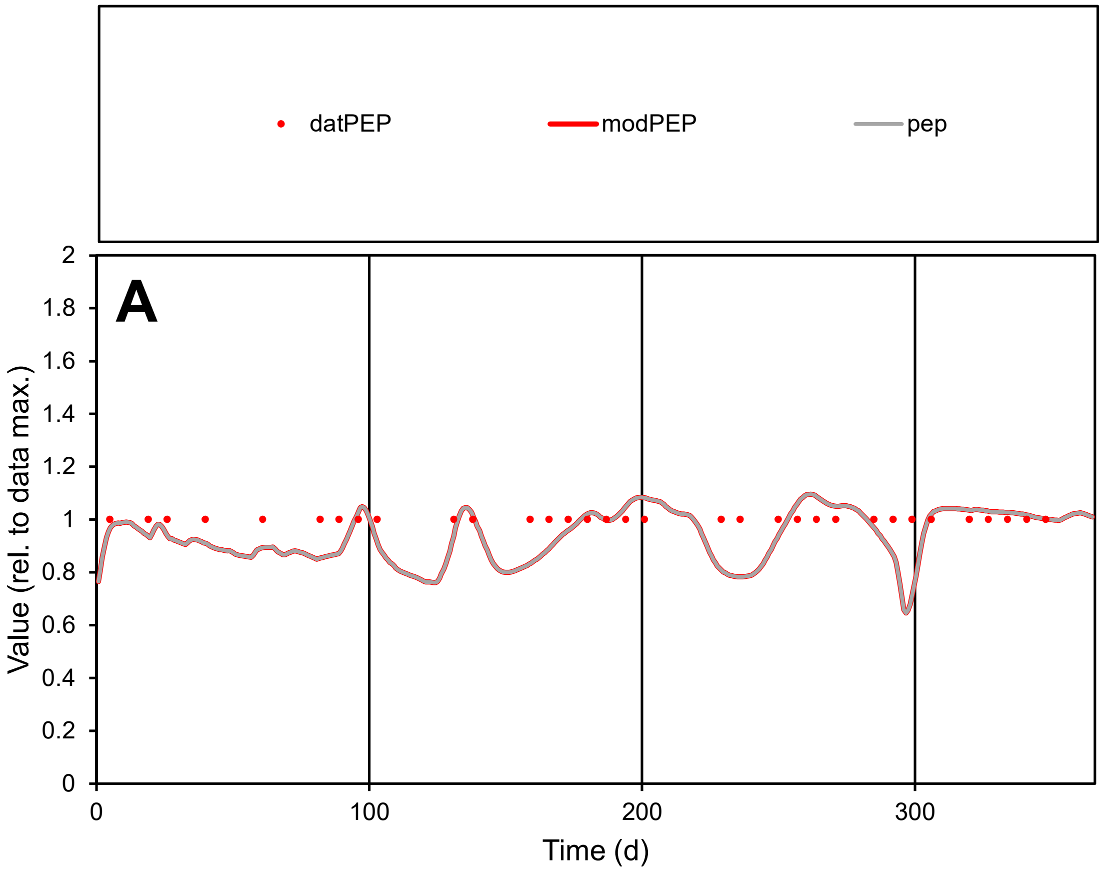


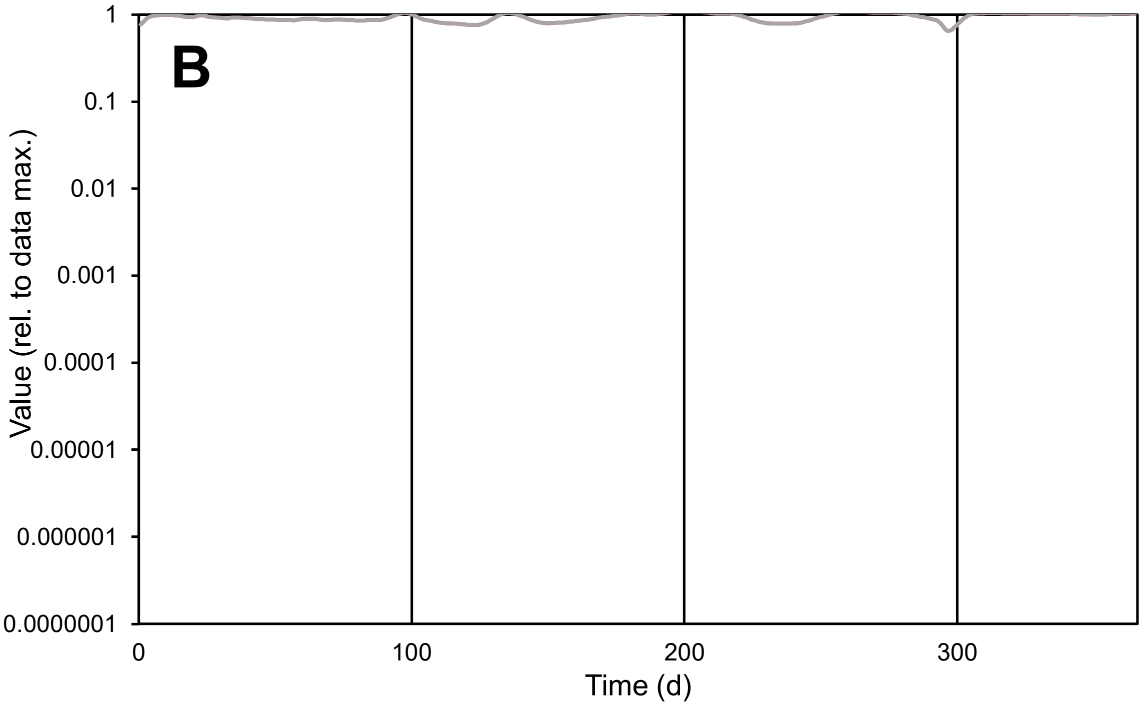


**C1**

Variables
PEP Provide jtd or Dst
CalibOnly?
2 1 to reproduce of
oNormalize?
2 1=no, 2=yes

ofxn calc*
0.09649
ofxn mod
0.14734

sRun
RunA1
zout
1

Ave
Data
4.9E-06
Model
4.7E-06

data/vMax
4.9E-06

ofxng
263.459

Min d/m
1.27725

ave/min d
1
ave/min m
1.2257

326_Calib_S_A

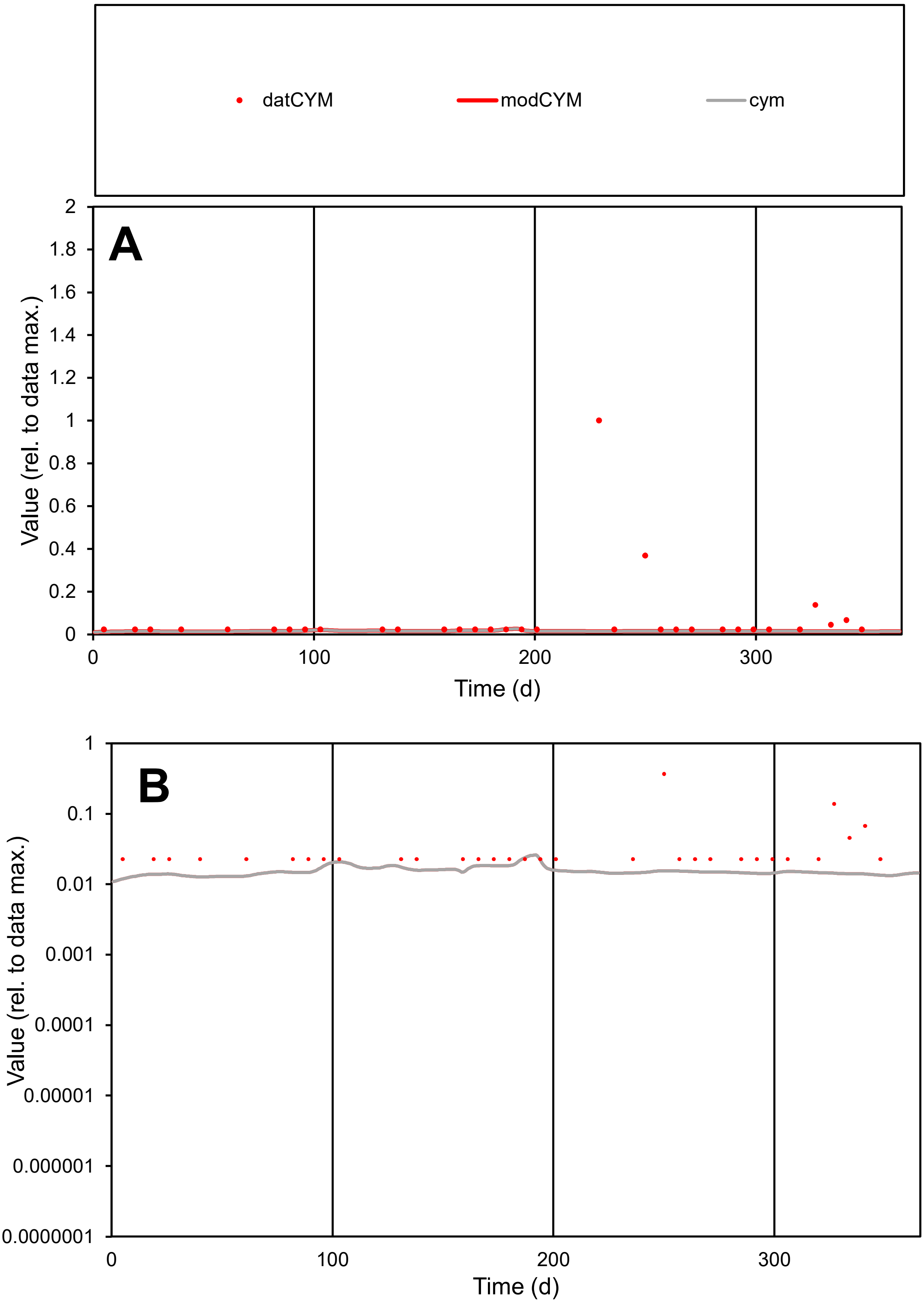


**C1**

Variables
CYM Provide jtd or Dst
CalibOnly?
2 1 to reproduce of
oNormalize?
2 1=no, 2=yes

ofxn calc*
2.69352
ofxn mod
2.6994

sRun
RunA1
zout
1

Ave
Data
4.1E-06
Model
9.4E-07

data/vMax
6E-05

ofxng
263.459

Min d/m
1.85195

ave/min d
3.01196
ave/min m
1.28584

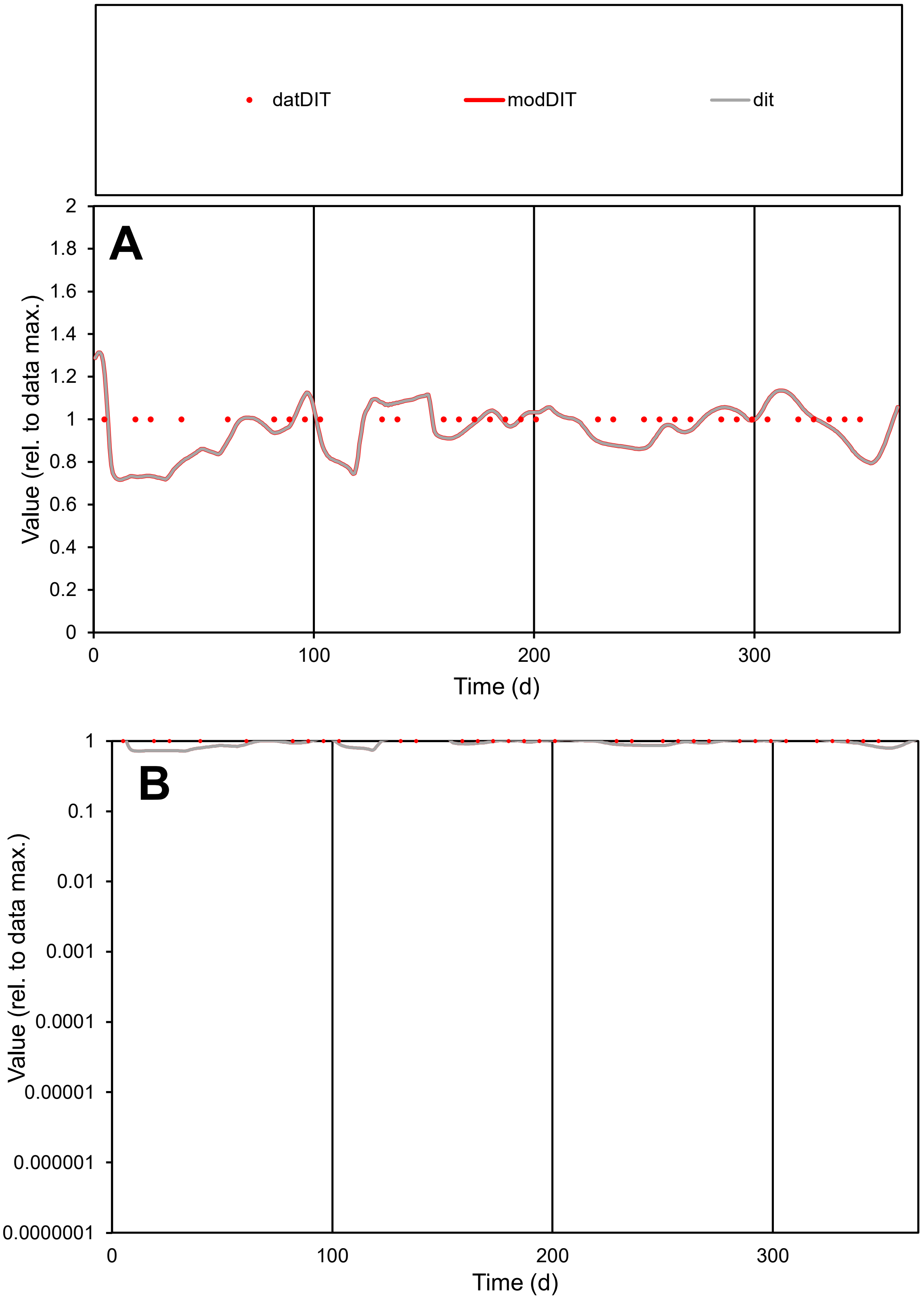


**C1**

Variables
DIT Provide jtd or Dst
CalibOnly?
2 1 to reproduce of
oNormalize?
2 1=no, 2=yes

ofxn calc*
0.10557
ofxn mod
0.15338

sRun
RunA1
zout
1

Ave
Data
5.1E-06
Model
4.9E-06

data/vMax
5.1E-06

ofxng
263.459

Min d/m
1.36867

ave/min d
1
ave/min m
1.31063

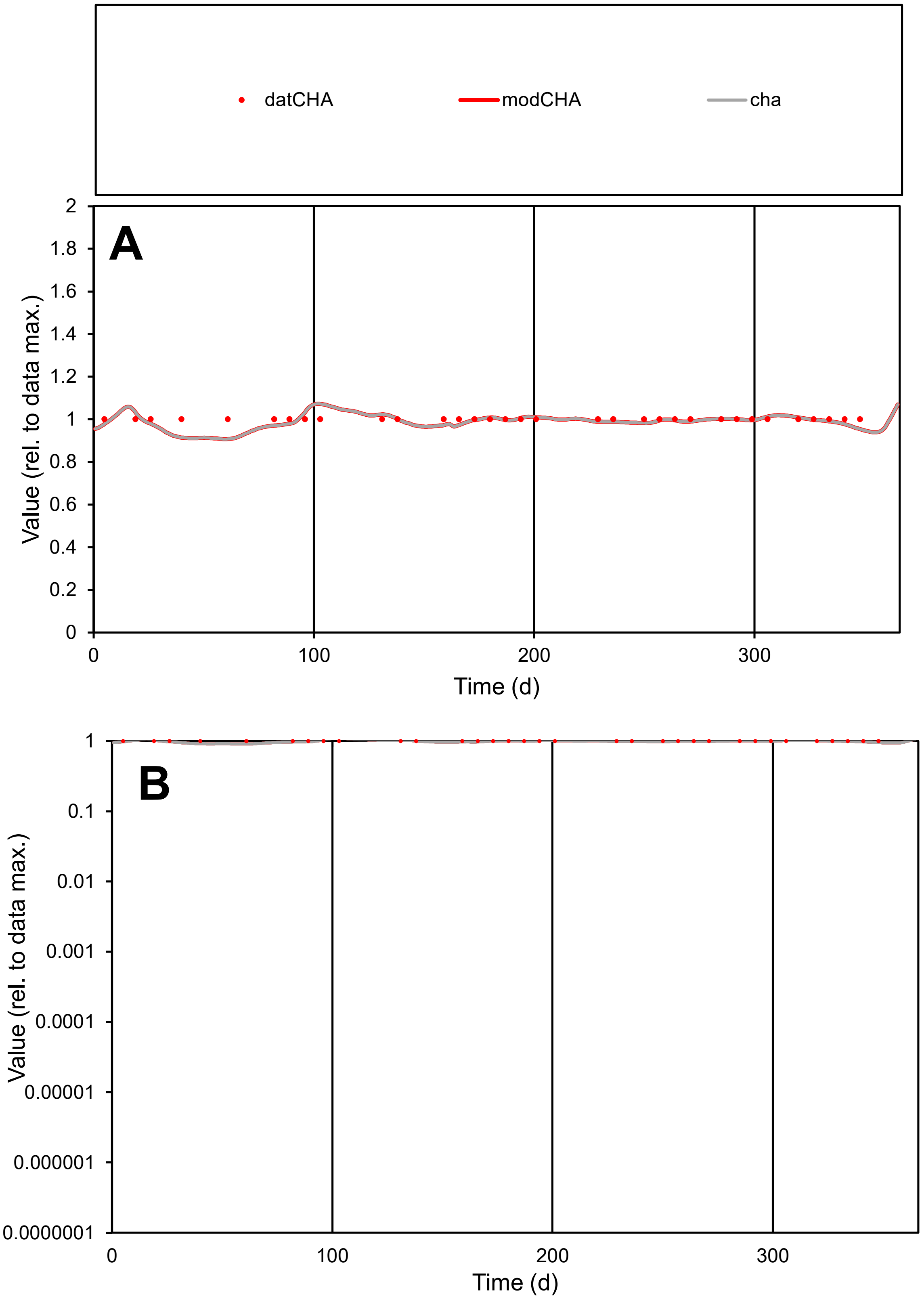


C1

Variables
CHA Provide jtd or Dst
CalibOnly?
2 1 to reproduce of
oNormalize?
2 1=no, 2=yes

ofxn calc*
0.03109
ofxn mod
0.11689

sRun
RunA1
zout
1

Ave
Data
8.7E-08
Model
8.6E-08

data/vMax
8.7E-08

ofxng
263.459

Min d/m
1.10058

ave/min d
1
ave/min m
1.09211

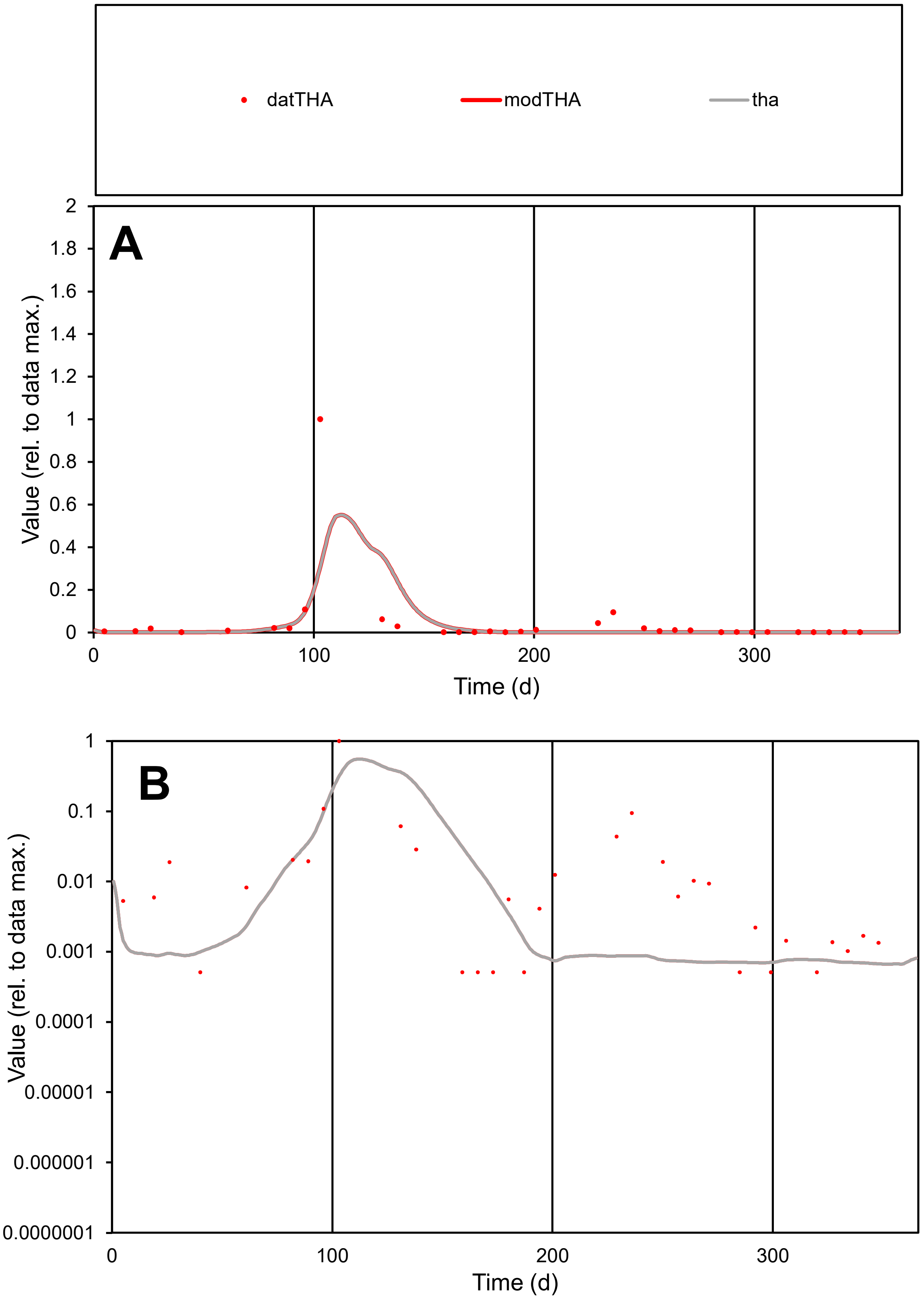


**C1**

Variables
THA Provide jtd or Dst
CalibOnly?
2 1 to reproduce of
oNormalize?
2 1=no, 2=yes

ofxn calc*
2.59667
ofxn mod
2.80703

sRun
RunA1
zout
1

Ave
Data
0.00024
Model
0.00019

data/vMax
0.00536

ofxng
263.459

Min d/m
0.76466

ave/min d
88.3419
ave/min m
54.1061

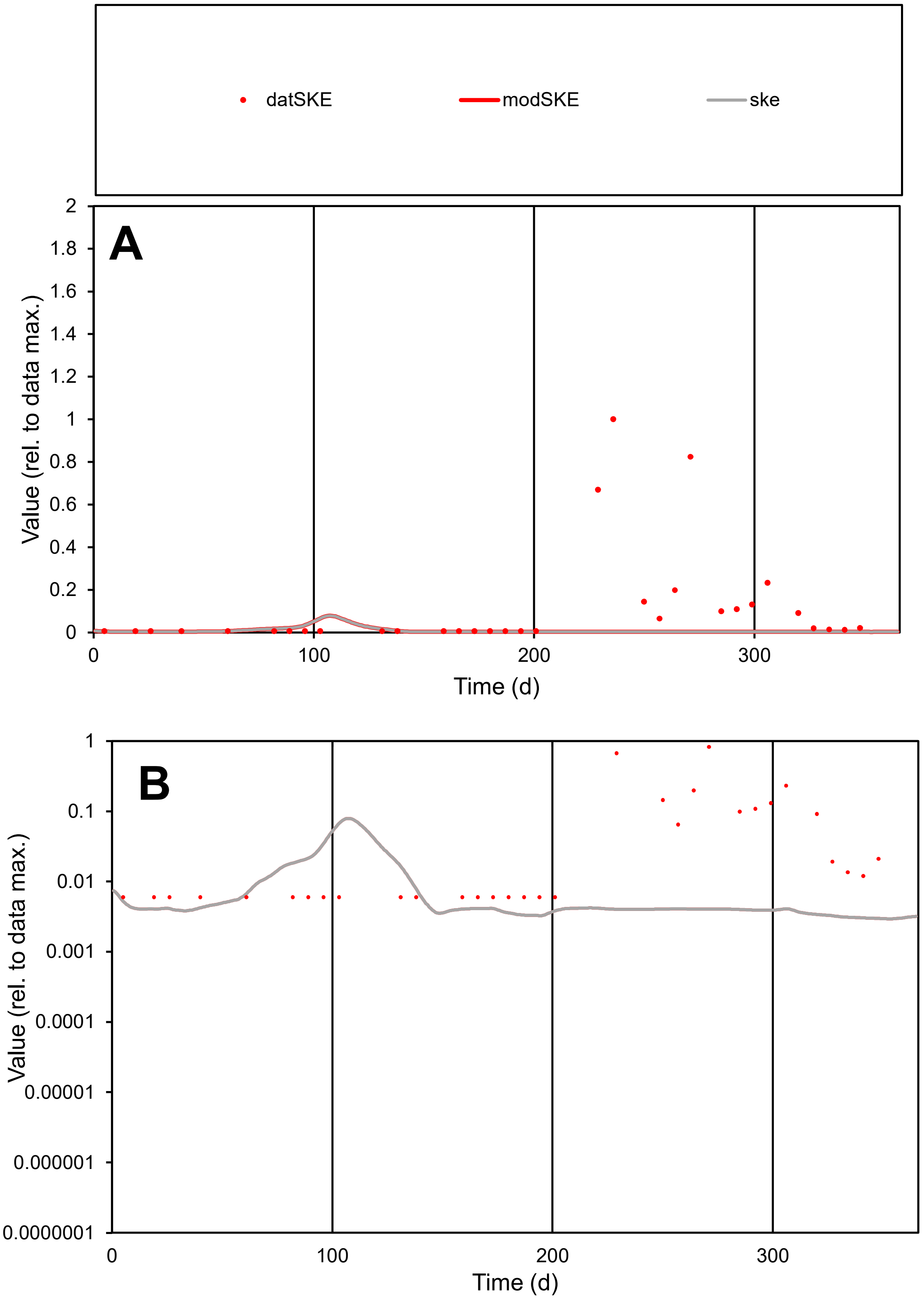


**C1**

Variables
SKE Provide jtd or Dst
CalibOnly?
2 1 to reproduce of
oNormalize?
2 1=no, 2=yes

ofxn calc*
2.3156
ofxn mod
2.32036

sRun
RunA1
zout
1

Ave
Data
4.4E-05
Model
3E-06

data/vMax
0.00039

ofxng
263.459

Min d/m
2.01489

ave/min d
18.9708
ave/min m
2.61508

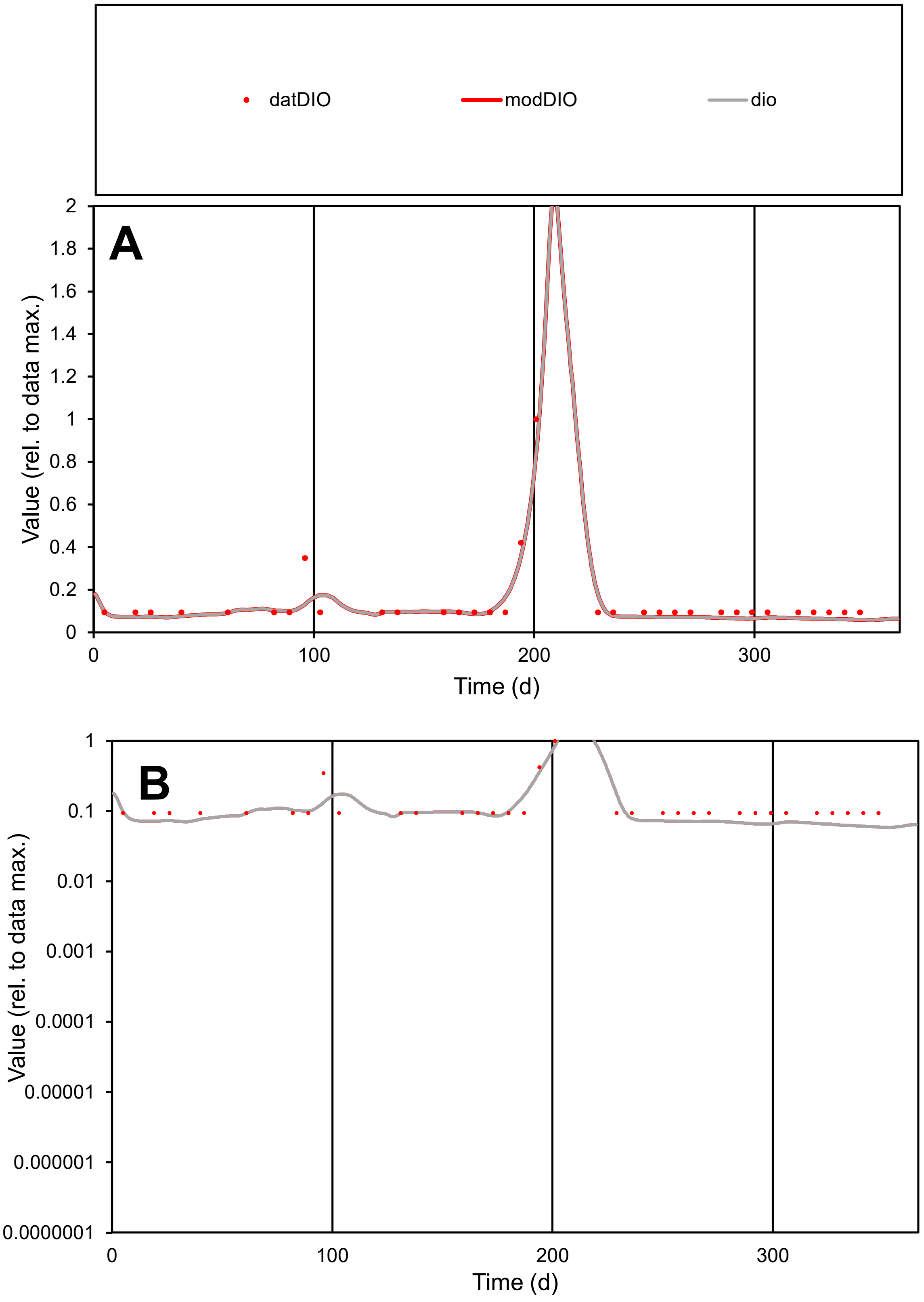


**C1**

Variables
DIO Provide jtd or Dst
CalibOnly?
2 1 to reproduce of
oNormalize?
2 1=no, 2=yes

ofxn calc*
0.32823
ofxn mod
0.36512

sRun
RunA1
zout
1

Ave
Data
6.9E-06
Model
6.3E-06

data/vMax
5E-05

ofxng
263.459

Min d/m
1.59406

ave/min d
1.4792
ave/min m
2.14483

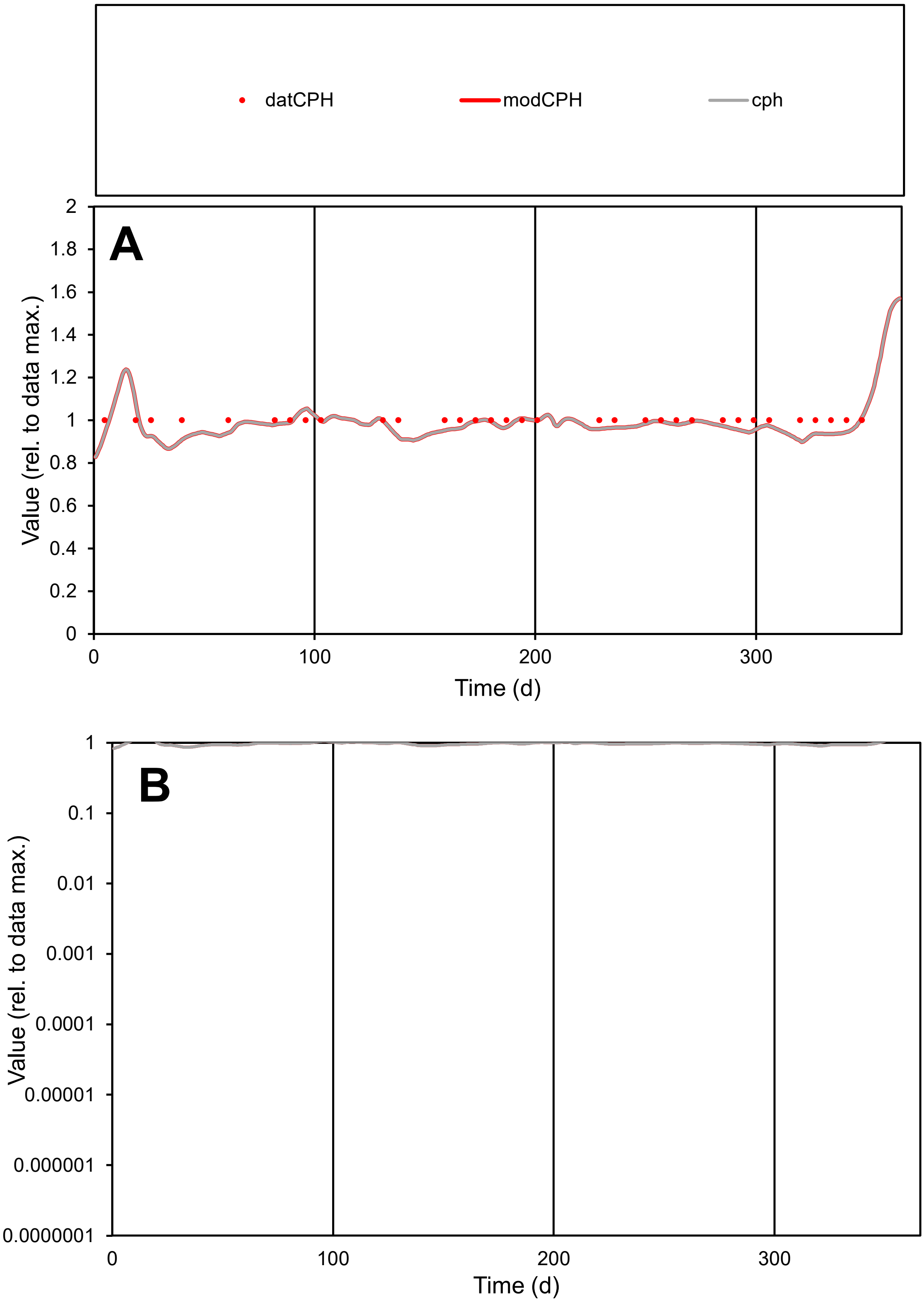


**C1**

Variables
CPH Provide jtd or Dst
CalibOnly?
2 1 to reproduce of
oNormalize?
2 1=no, 2=yes

ofxn calc*
0.04318
ofxn mod
0.11973

sRun
RunA1
zout
1

Ave
Data
9.6E-06
Model
9.3E-06

data/vMax
9.6E-06

ofxng
263.459

Min d/m
1.10954

ave/min d
1
ave/min m
1.07895

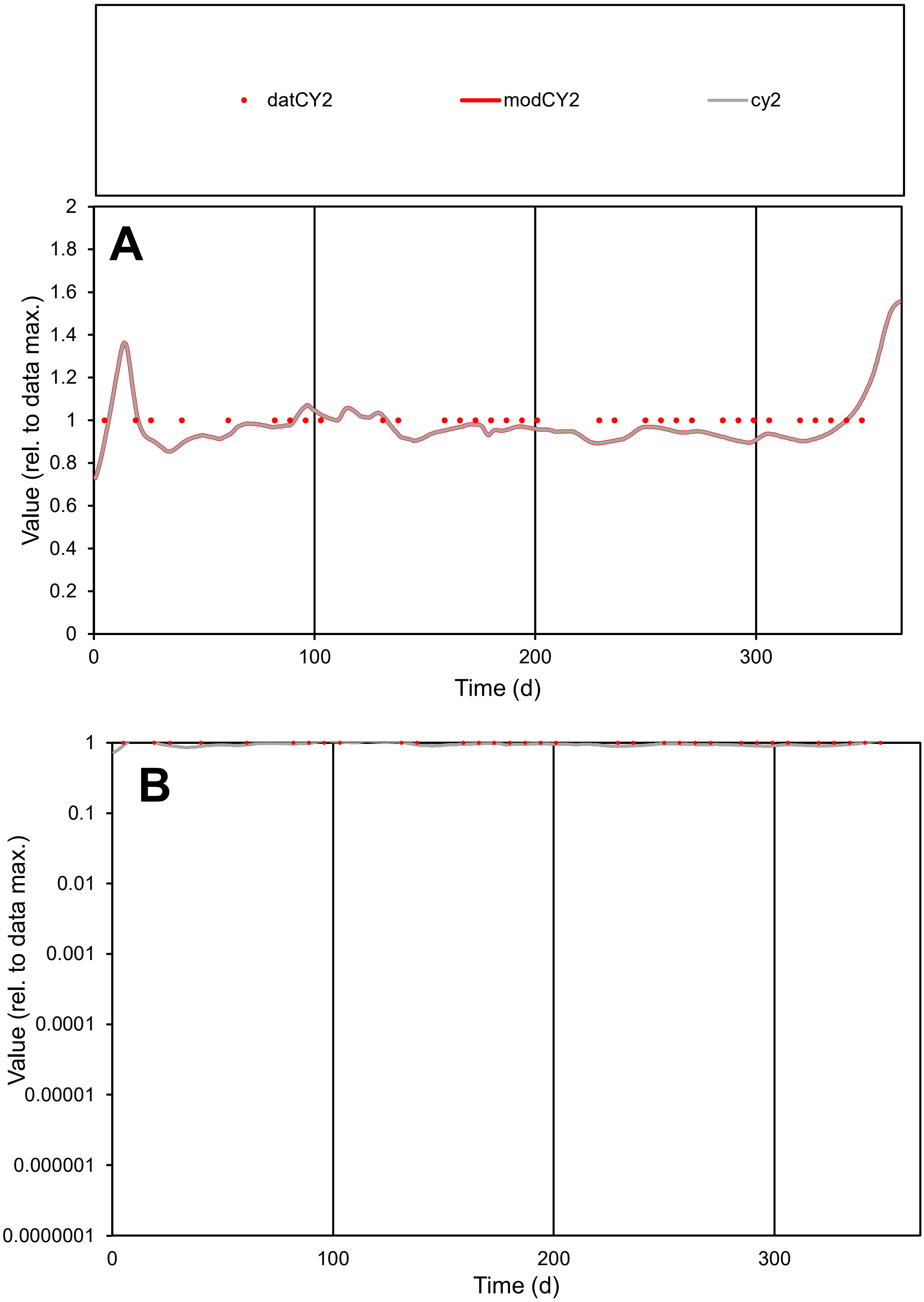


**C1**

Variables
CY2 Provide jtd or Dst
CalibOnly?
2 1 to reproduce of
oNormalize?
2 1=no, 2=yes

ofxn calc*
0.06437
ofxn mod
0.12788

sRun
RunA1
zout
1

Ave
Data
1.1E-05
Model
1.1E-05

data/vMax
1.1E-05

ofxng
263.459

Min d/m
1.11814

ave/min d
1
ave/min m
1.07179

326_Calib_S_A

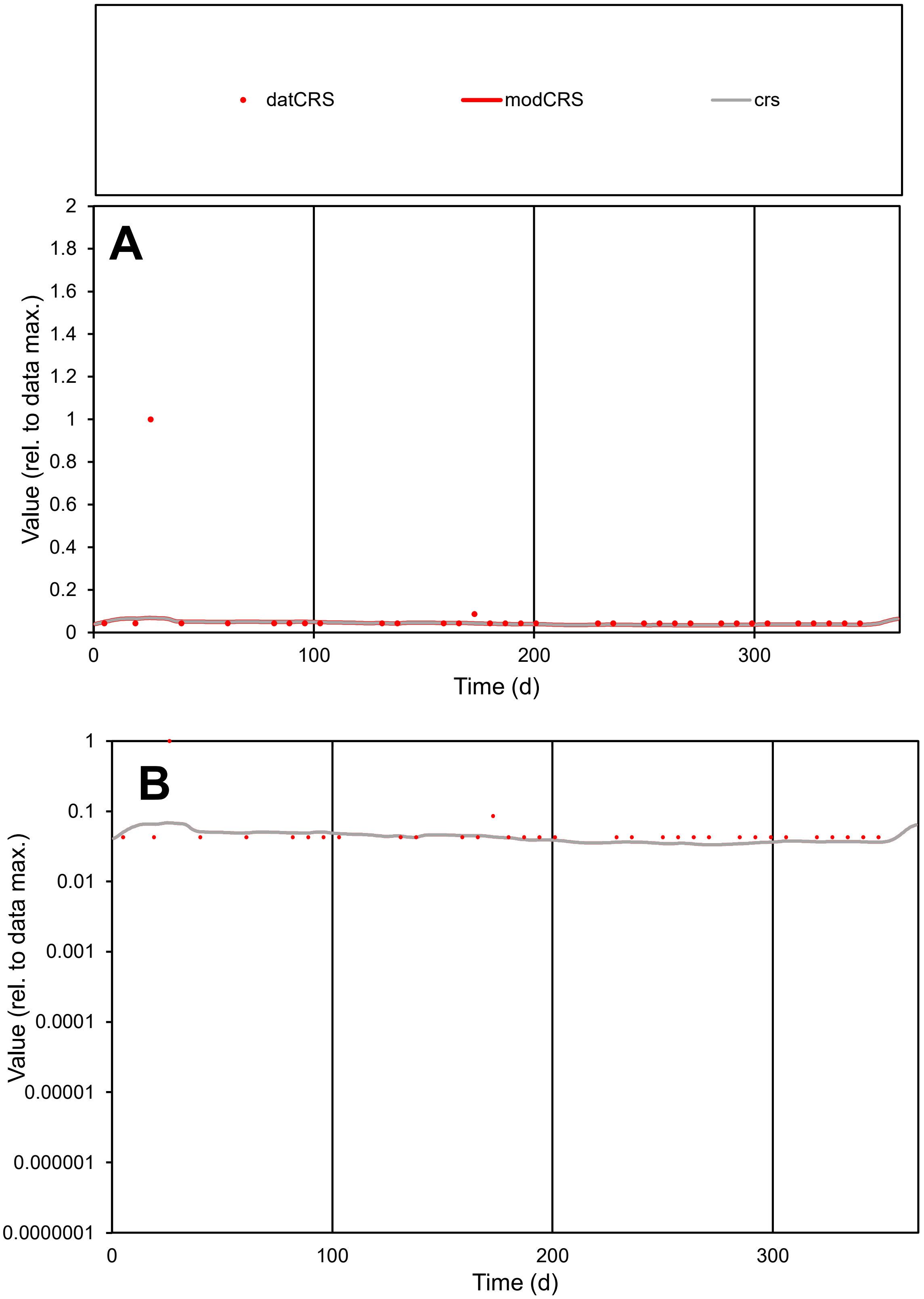


**C1**

Variables
CRS Provide jtd or Dst
CalibOnly?
2 1 to reproduce of
oNormalize?
2 1=no, 2=yes

ofxn calc*
2.22084
ofxn mod
2.22916

sRun
RunA1
zout
1

Ave
Data
9.2E-06
Model
5.4E-06

data/vMax
0.00013

ofxng
263.459

Min d/m
1.28443

ave/min d
1.70632
ave/min m
1.271

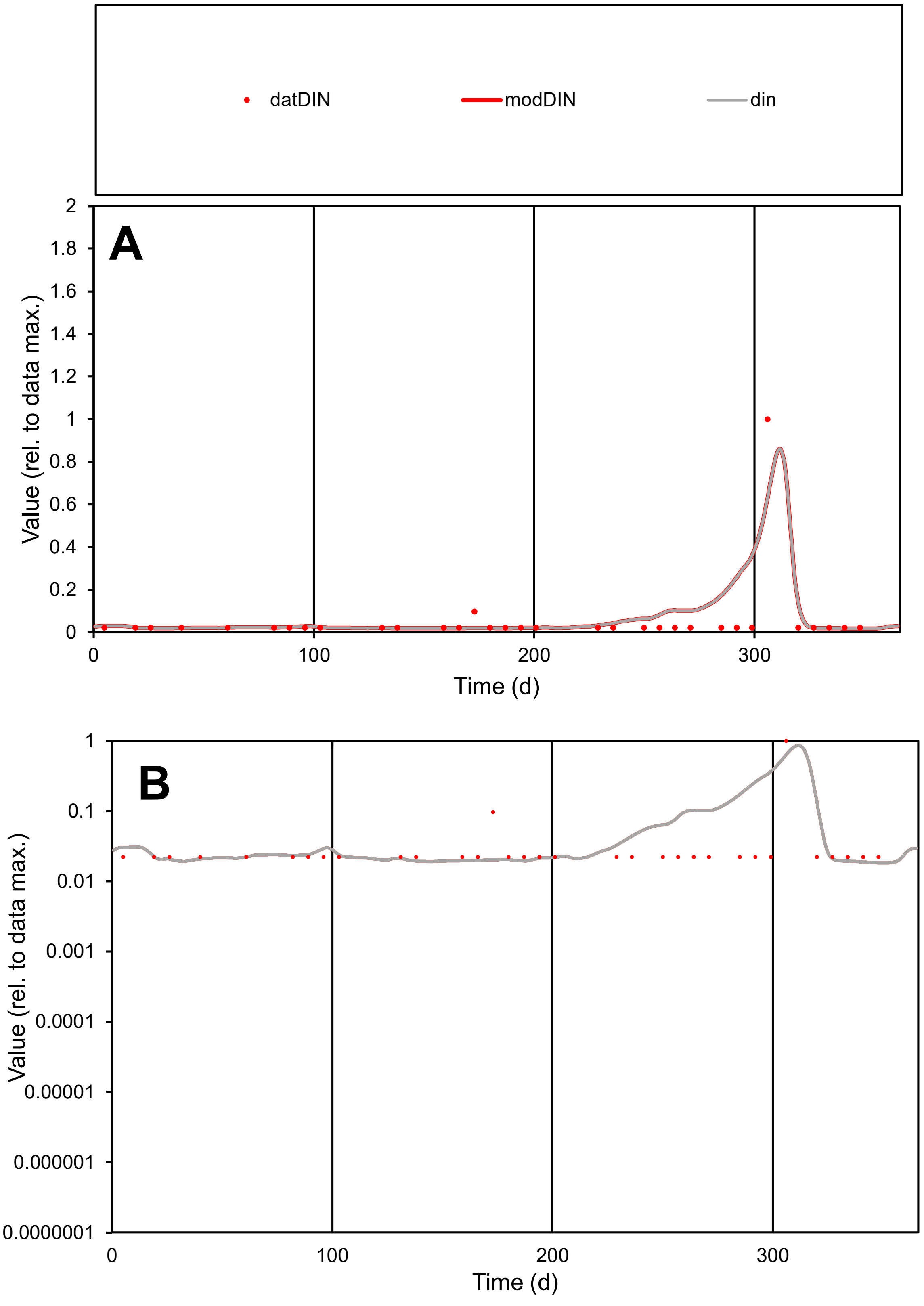


**C1**

Variables
DIN Provide jtd or Dst
CalibOnly?
2 1 to reproduce of
oNormalize?
2 1=no, 2=yes

ofxn calc*
1.71094
ofxn mod
1.80182

sRun
RunA1
zout
1

Ave
Data
7E-06
Model
9.6E-06

data/vMax
0.00013

ofxng
263.459

Min d/m
1.20922
ave/min d
2.43332
ave/min m
4.03847

326_Calib_S_A

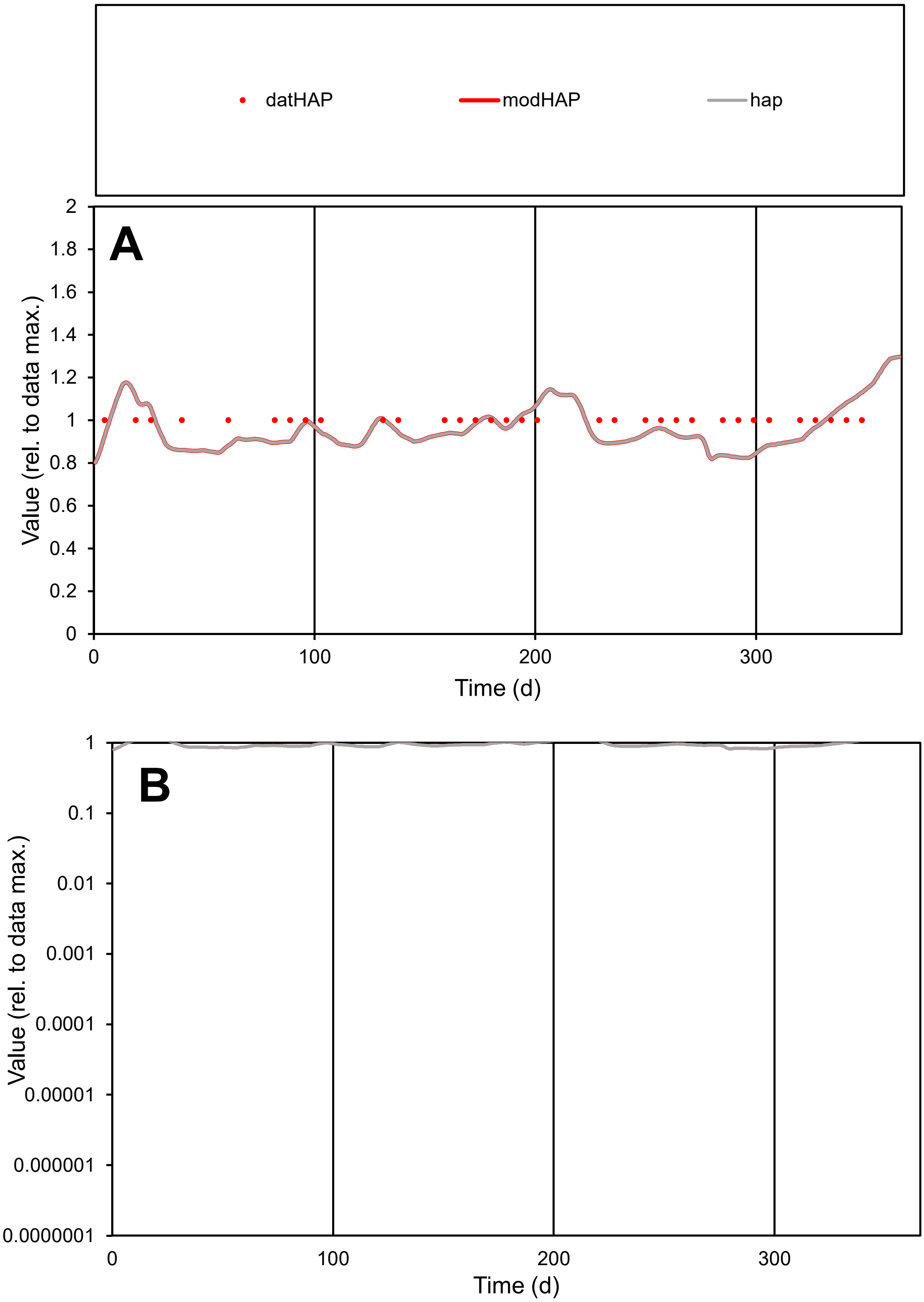


**C1**

Variables

| | |
|---|---|
| HAP | Provide jtd or Dst |
| CalibOnly? | |
| 2 | 1 to reproduce of |
| oNormalize? | |
| 2 | 1=no, 2=yes |

ofxn calc*
0.08754
ofxn mod
0.14114

sRun
RunA1
zout
1

Ave
Data
1.1E-05
Model
1.1E-05

data/vMax
1.1E-05

ofxng
263.459

Min d/m
1.21131

ave/min d
1
ave/min m
1.15859

326_Calib_S_A

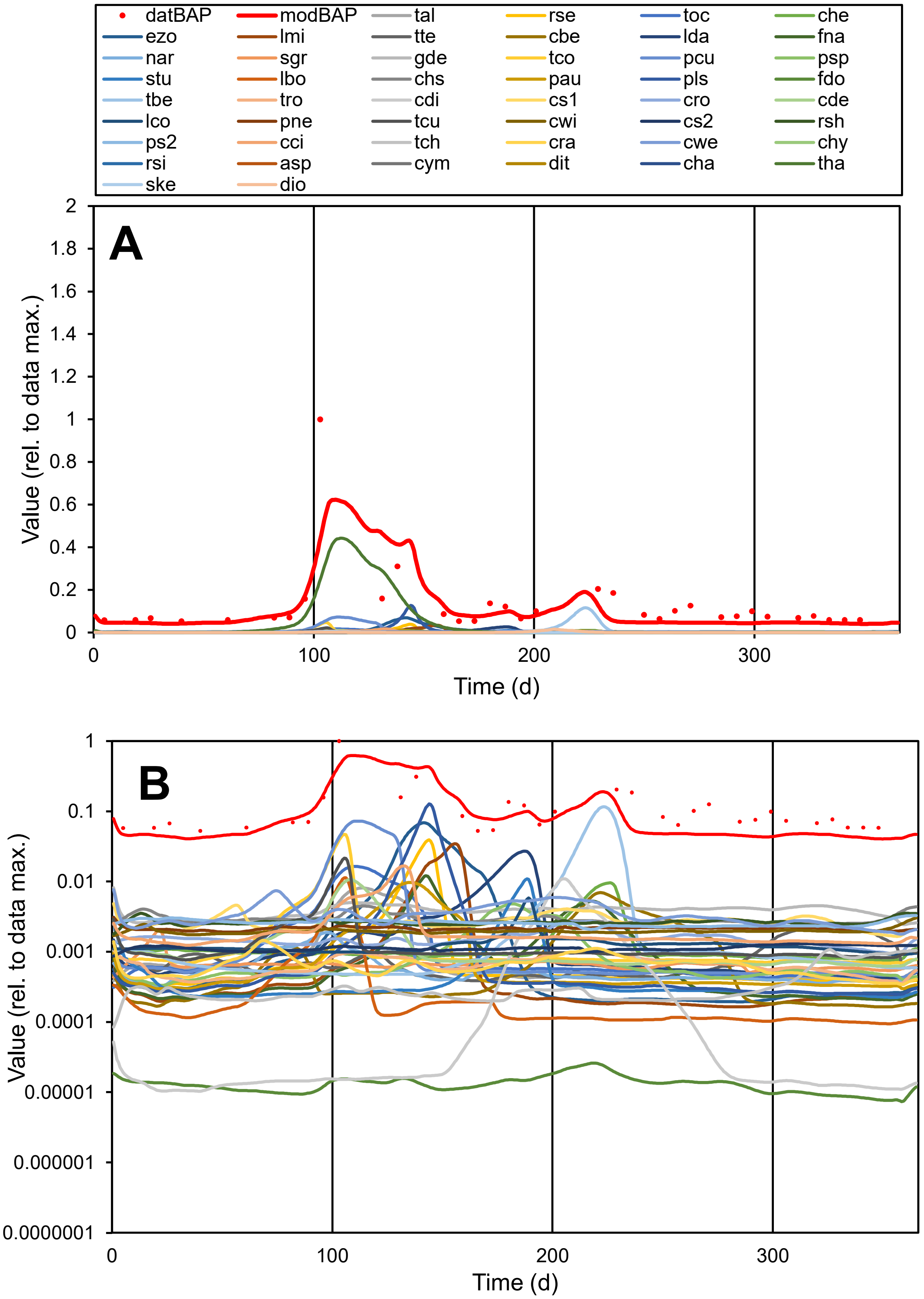


C1

Variables

BAP Provide jtd or Dst

CalibOnly?

2 1 to reproduce of

oNormalize?

2 1=no, 2=yes

ofxn calc*

0.89719

ofxn mod

0.90825

sRun

RunA1

zout

1

Ave

Data

0.00083

Model

0.00067

data/vMax

0.00669

ofxng

263.459

Min d/m

1.29764

ave/min d

2.34955

ave/min m

2.48411

326_Calib_S_A

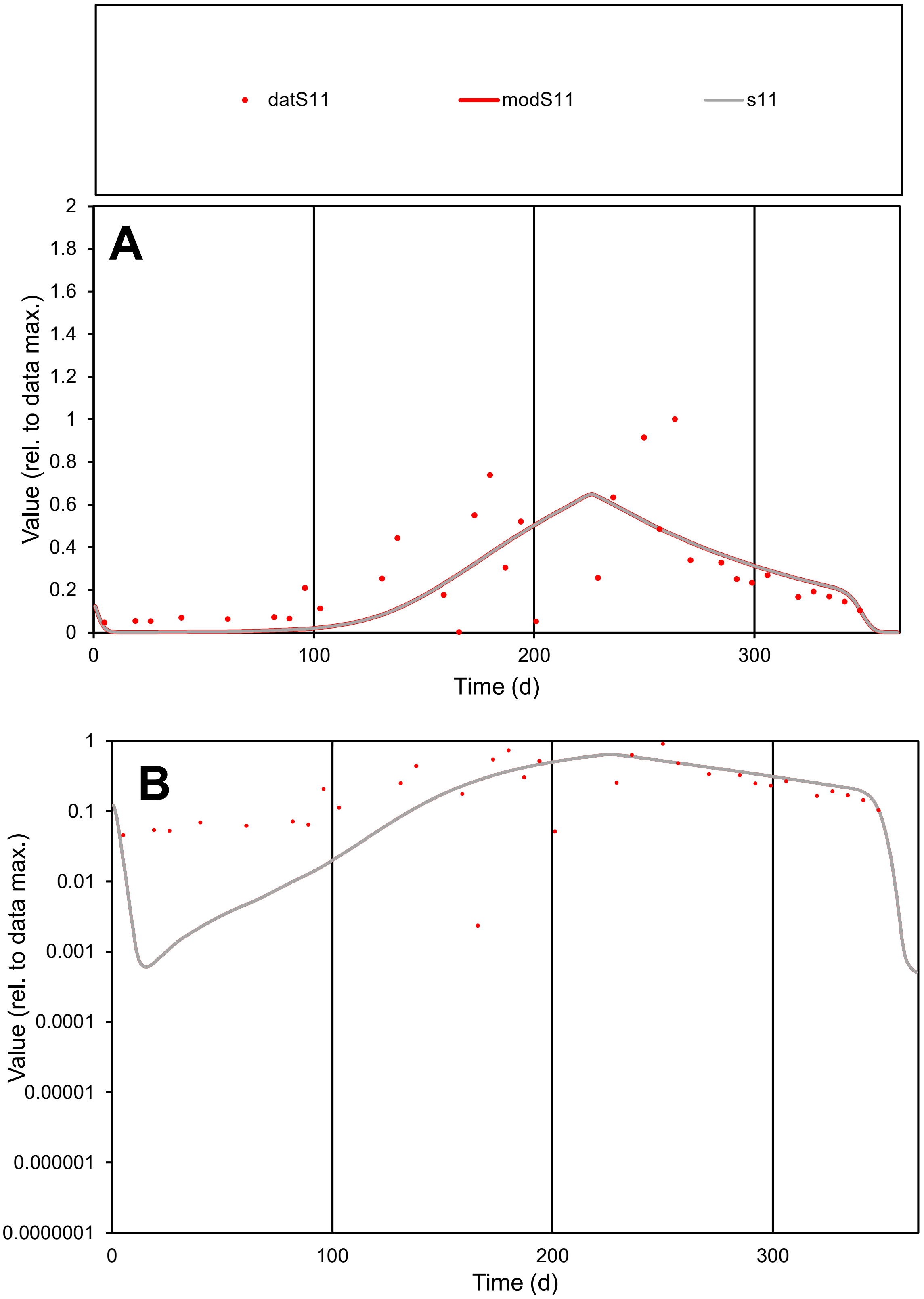


**C1**

Variables

| | |
|---|---|
| S11 | Provide jtd or Dst |
| CalibOnly? | |
| 2 | 1 to reproduce of |
| oNormalize? | |
| 2 | 1=no, 2=yes |

ofxn calc*
0.7093
ofxn mod
0.72576

sRun
RunA1
zout
1

Ave
Data
0.0821
Model
0.07354

data/vMax
0.2927

ofxng
263.459

Min d/m
3.10921

ave/min d
119.156
ave/min m
331.876

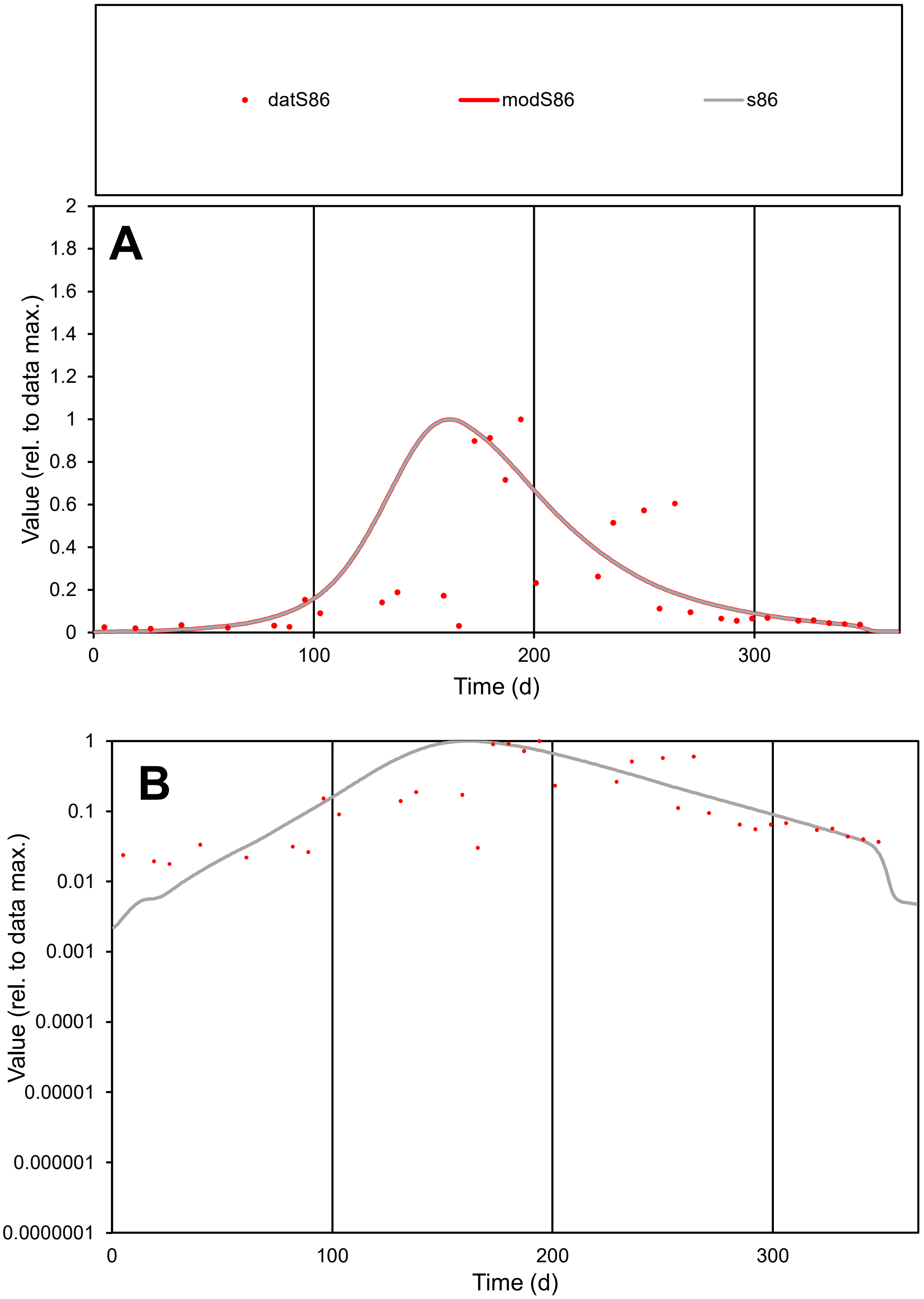


**C1**

Variables
S86 Provide jtd or Dst
CalibOnly?
2 1 to reproduce of
oNormalize?
2 1=no, 2=yes

ofxn calc*
1.29034
ofxn mod
1.30849

sRun
RunA1
zout
1

Ave
Data
0.04325
Model
0.05851

data/vMax
0.1942

ofxng
263.459

Min d/m
5.13289

ave/min d
12.5098
ave/min m
86.8713

326_Calib_S_A

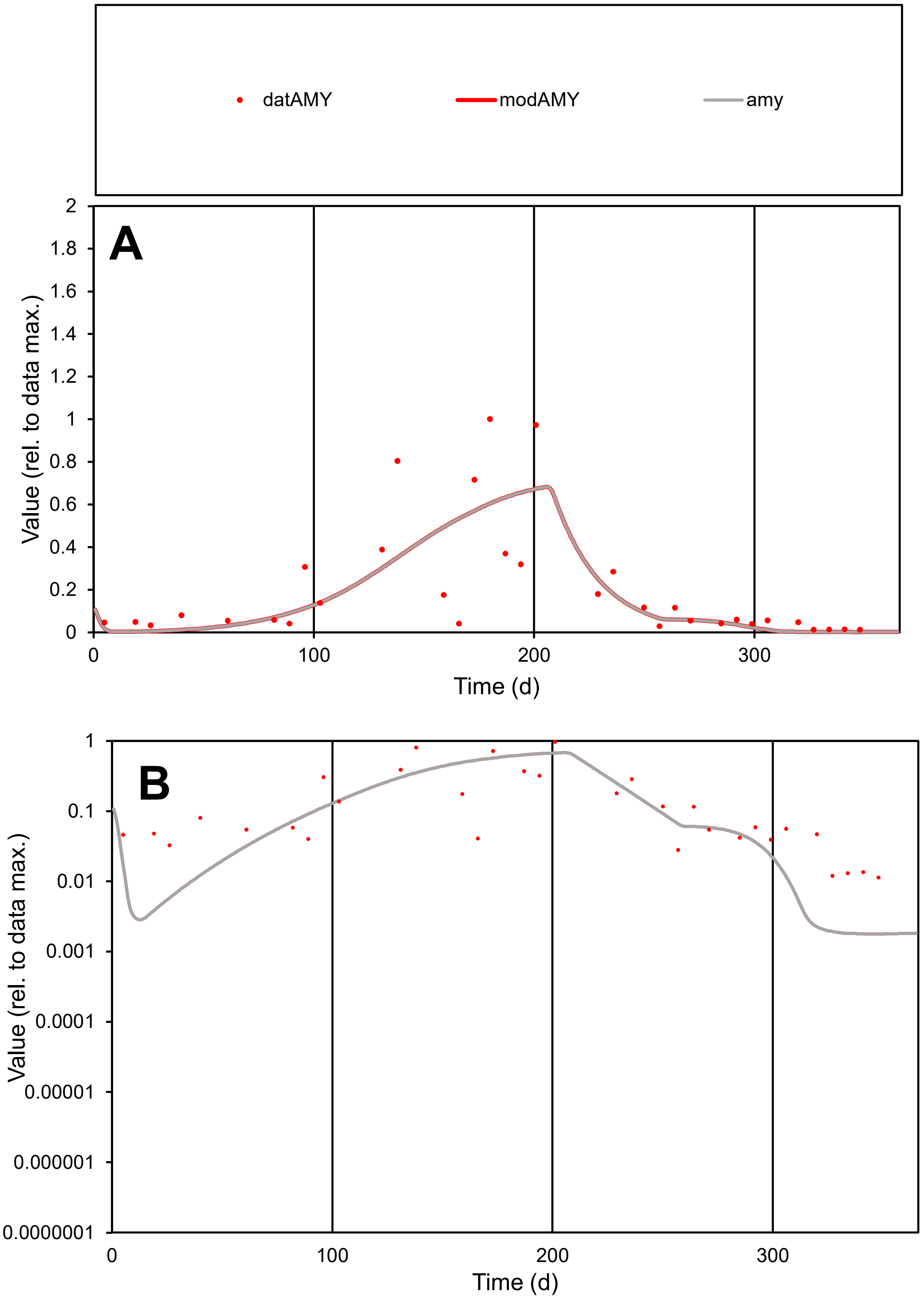


**C1**

Variables
AMY Provide jtd or Dst
CalibOnly?
2 1 to reproduce of
oNormalize?
2 1=no, 2=yes

ofxn calc*
0.89244
ofxn mod
0.90937

sRun
RunA1
zout
1

Ave
Data
0.0278
Model
0.02593

data/vMax
0.1377

ofxng
263.459

Min d/m
6.38628

ave/min d
17.7734
ave/min m
105.871

326_Calib_S_A

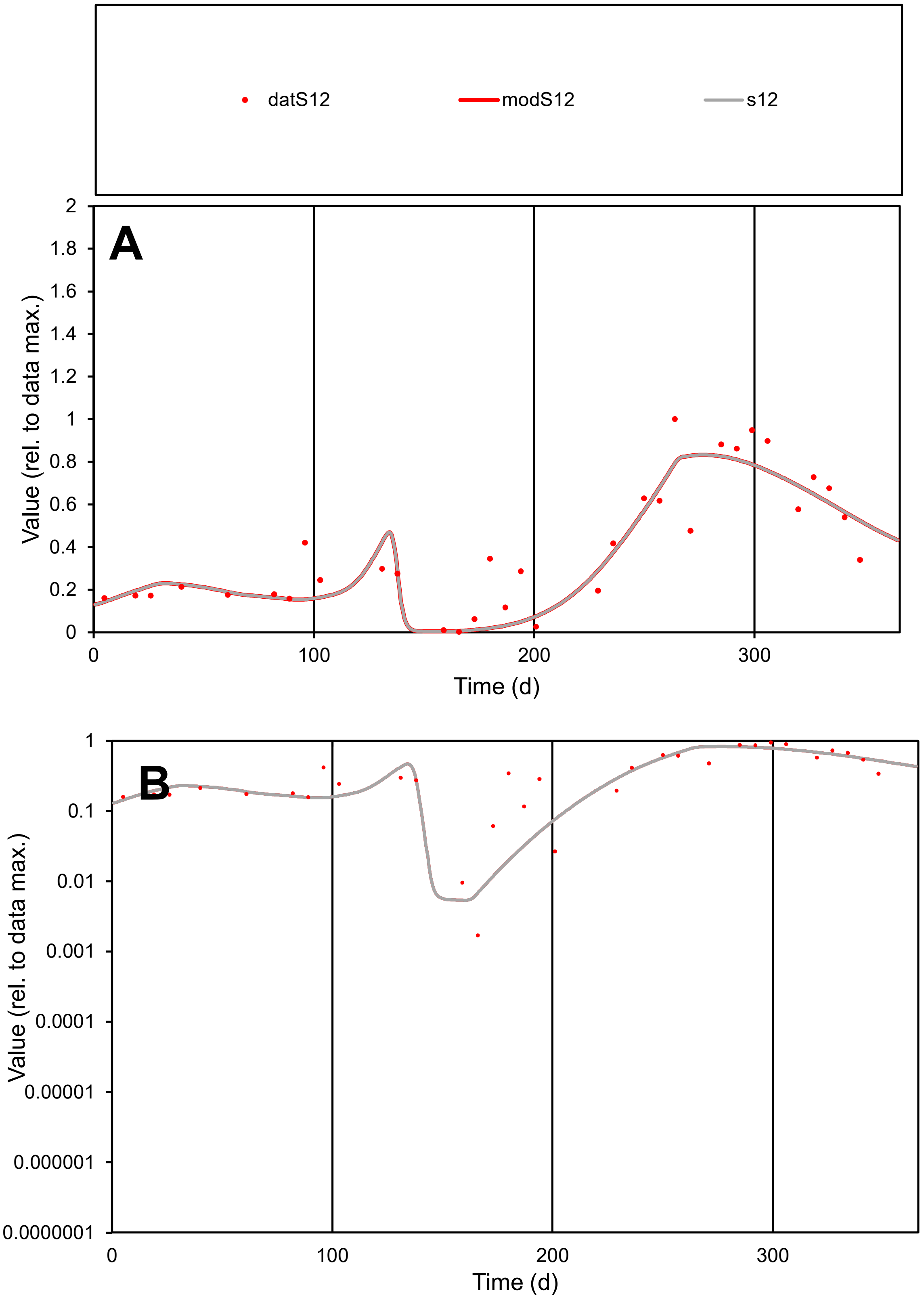


**C1**

Variables

S12 Provide jtd or Dst

CalibOnly?

2 1 to reproduce of

oNormalize?

2 1=no, 2=yes

ofxn calc*

0.32739

ofxn mod

0.35805

sRun

RunA1

zout

1

Ave

Data

0.0323

Model

0.03039

data/vMax

0.08138

ofxng

263.459

Min d/m

0.31732

ave/min d

232.897

ave/min m

69.5213

326_Calib_S_A

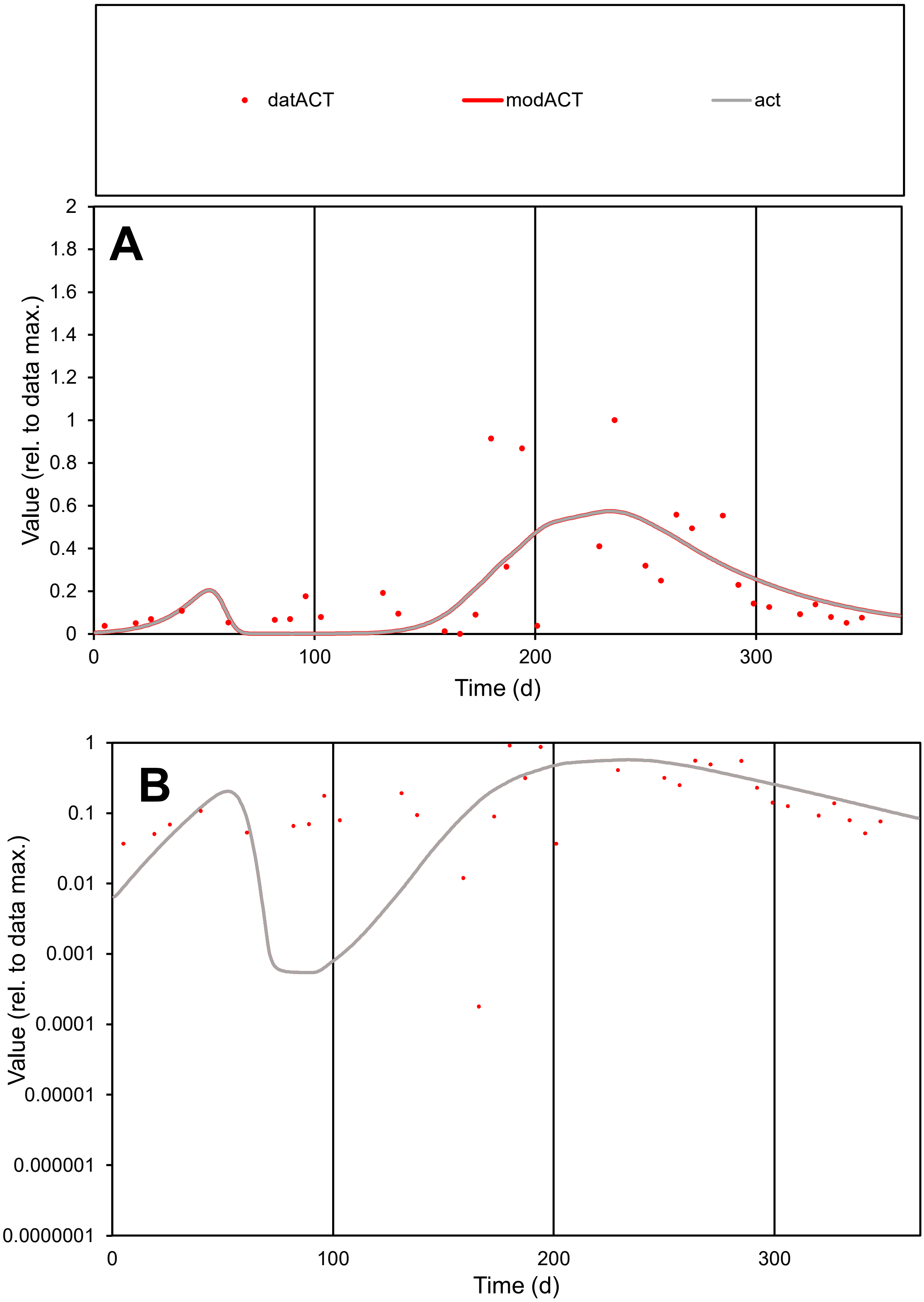


**C1**

Variables
ACT Provide jtd or Dst
CalibOnly?
2 1 to reproduce of
oNormalize?
2 1=no, 2=yes

ofxn calc*
0.85445
ofxn mod
0.86952

sRun
RunA1
zout
1

Ave
Data
0.03376
Model
0.03092

data/vMax
0.1438

ofxng
263.459

Min d/m
0.32697

ave/min d
1314.59
ave/min m
393.628

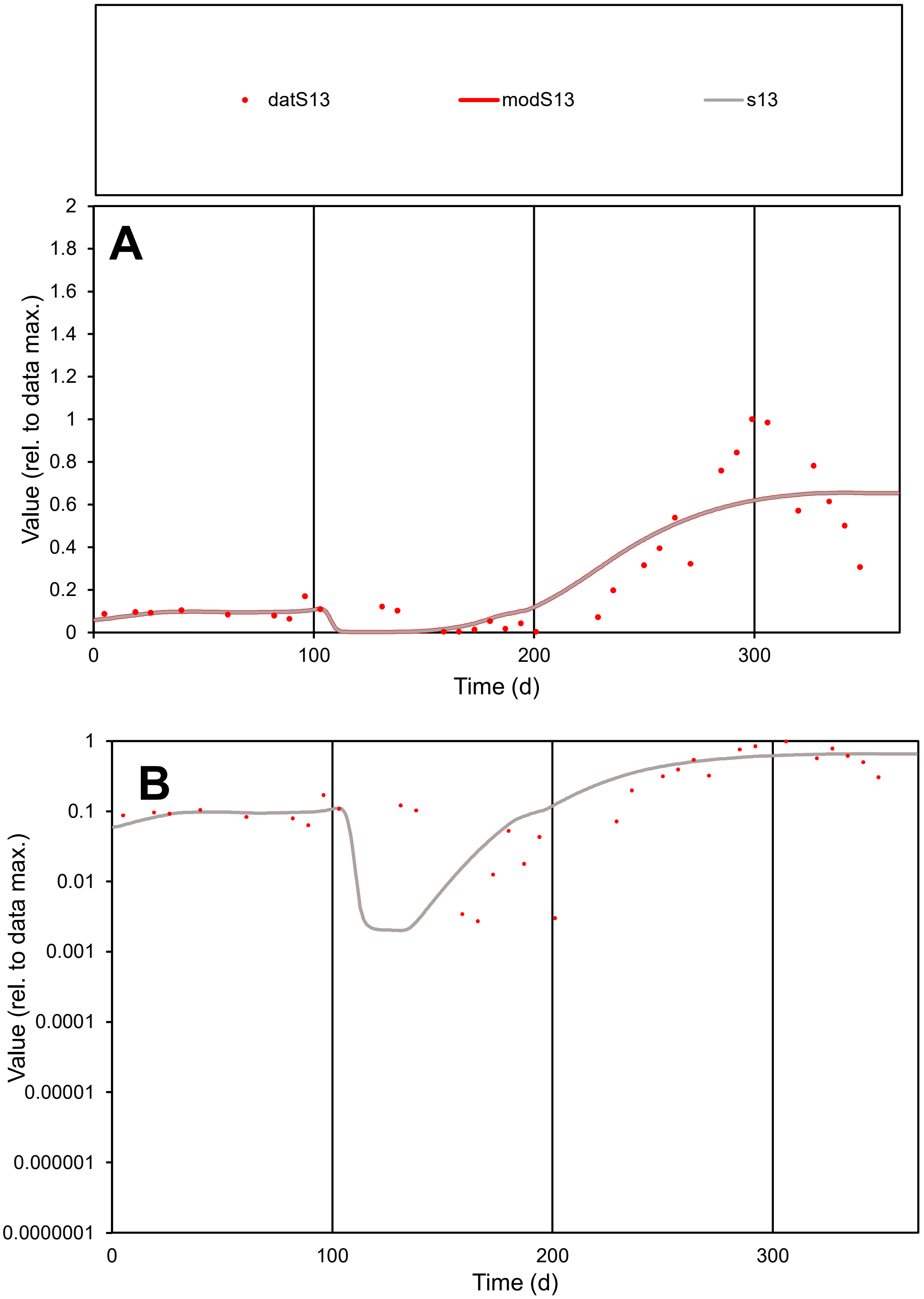


**C1**

Variables
S13 Provide jtd or Dst
CalibOnly?
2 1 to reproduce of
oNormalize?
2 1=no, 2=yes

ofxn calc*
0.51966
ofxn mod
0.54221

sRun
RunA1
zout
1

Ave
Data
0.02911
Model
0.02976

data/vMax
0.1017

ofxng
263.459

Min d/m
1.34208

ave/min d
105.402
ave/min m
144.621

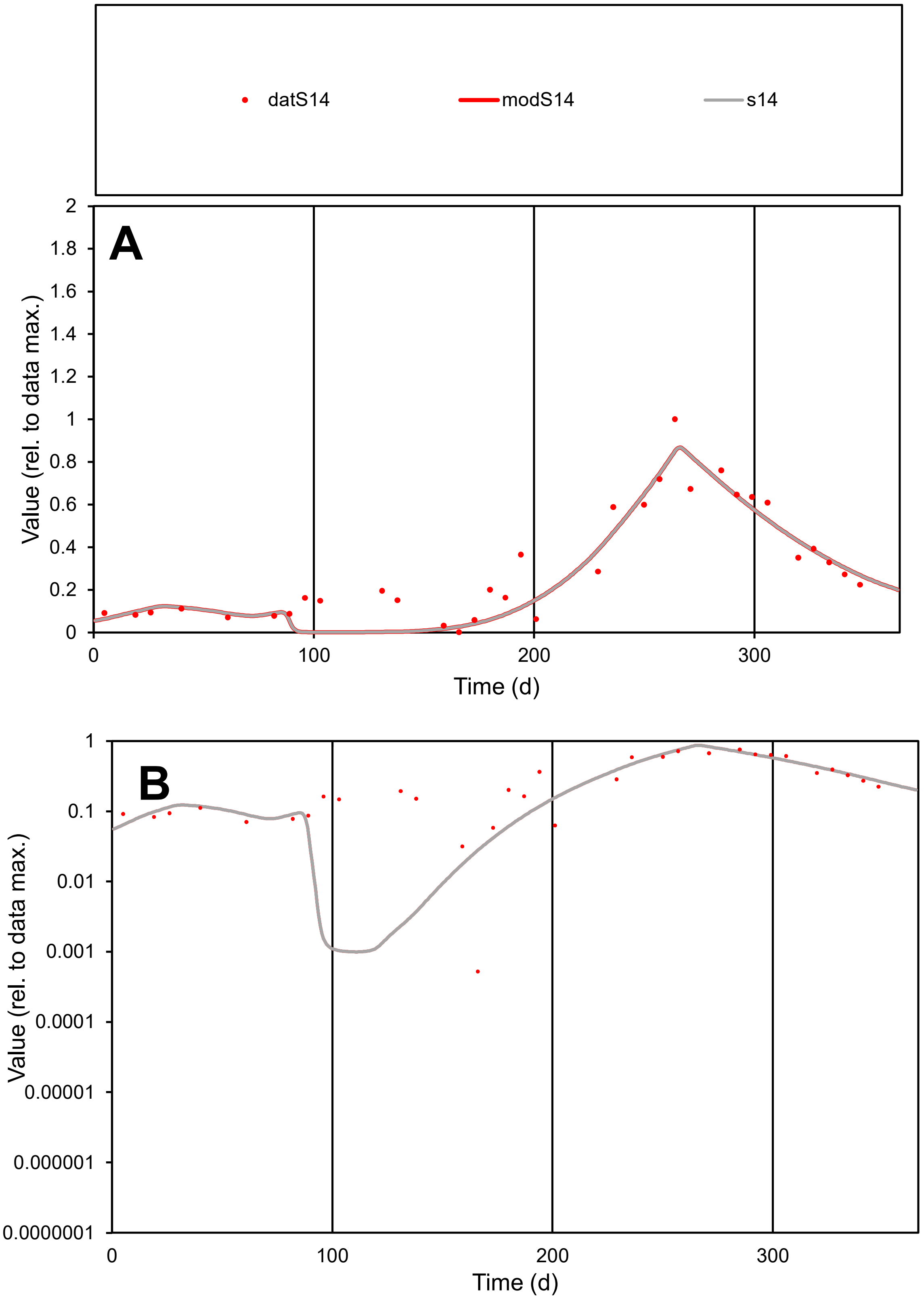


**C1**

Variables
S14 Provide jtd or Dst
CalibOnly?
2 1 to reproduce of
oNormalize?
2 1=no, 2=yes

ofxn calc*
0.30563
ofxn mod
0.34371

sRun
RunA1
zout
1

Ave
Data
0.02919
Model
0.02651

data/vMax
0.09409

ofxng
263.459

Min d/m
0.50998

ave/min d
591.651
ave/min m
274.069

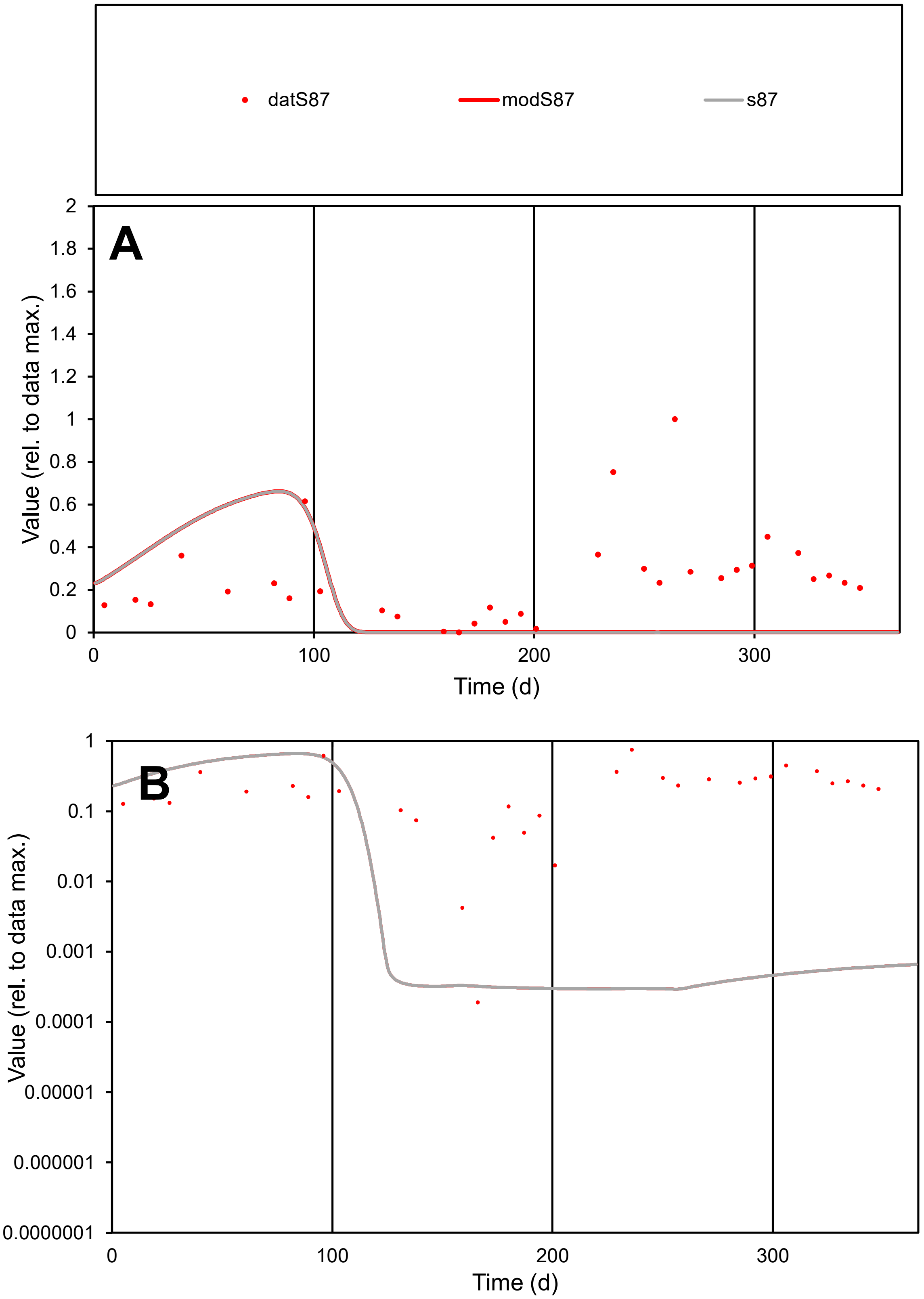


**C1**

Variables
S87 Provide jtd or Dst
CalibOnly?
2 1 to reproduce of
oNormalize?
2 1=no, 2=yes

ofxn calc*
1.31339
ofxn mod
1.32125

sRun
RunA1
zout
1

Ave
Data
0.023
Model
0.012

data/vMax
0.09219

ofxng
263.459

Min d/m
0.64307

ave/min d
1311.83
ave/min m
440.127

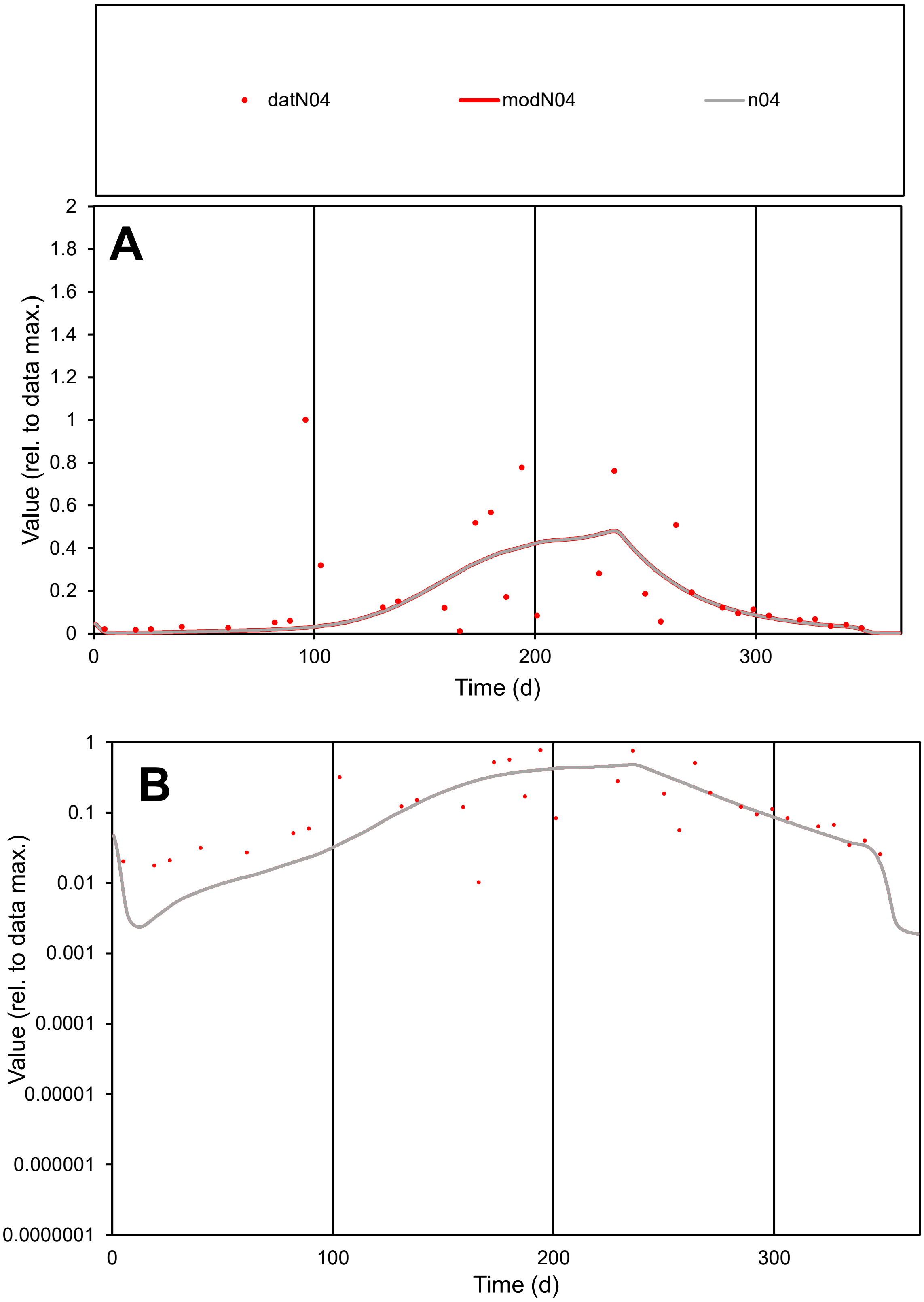


**C1**

Variables

N04 Provide jtd or Dst

CalibOnly?

2 1 to reproduce of

oNormalize?

2 1=no, 2=yes

ofxn calc*

1.13813

ofxn mod

1.14738

sRun

RunA1

zout

1

Ave

Data

0.01365

Model

0.01098

data/vMax

0.06724

ofxng

263.459

Min d/m

3.00218

ave/min d

19.8055

ave/min m

47.832

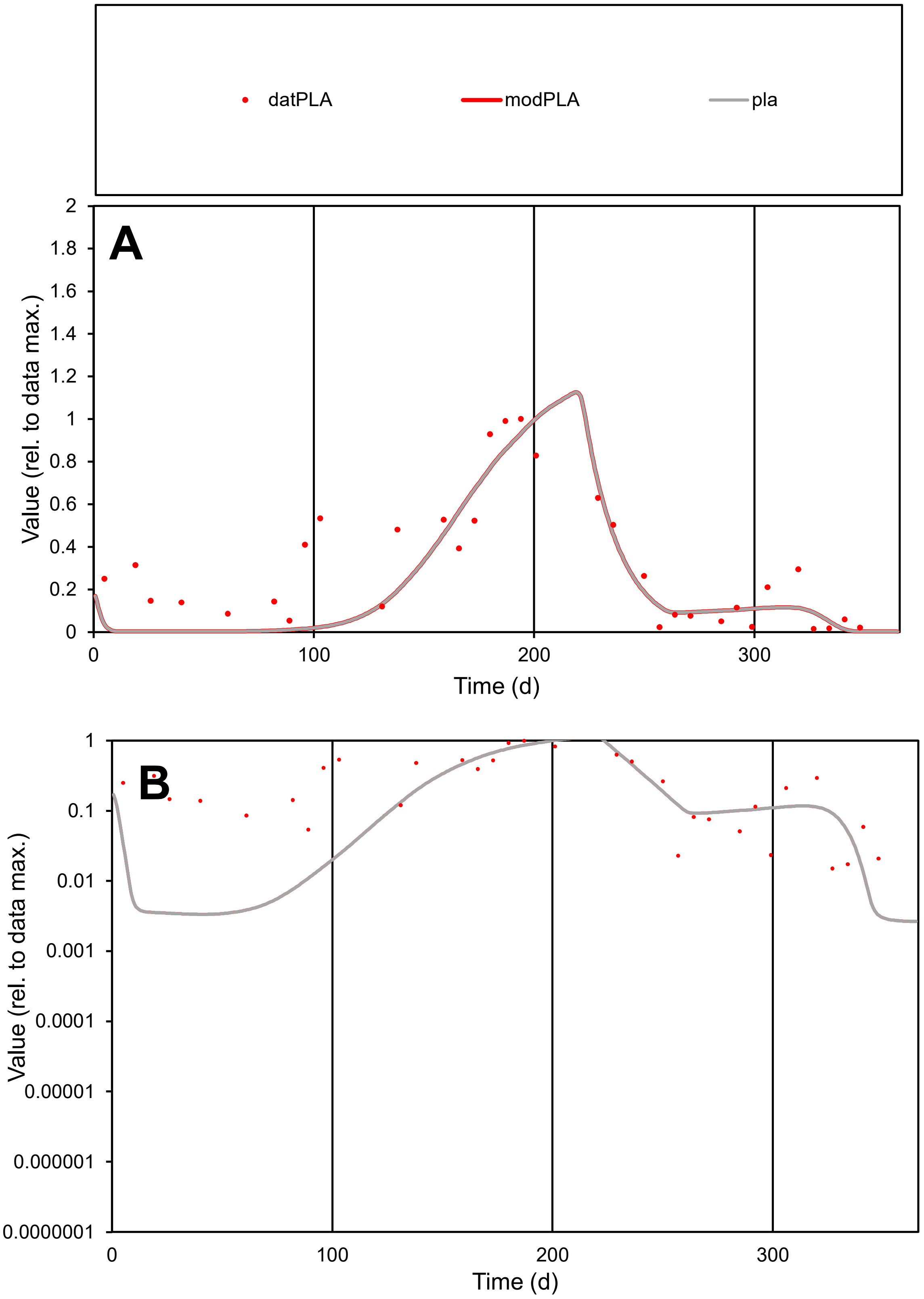


**C1**

Variables
PLA Provide jtd or Dst
CalibOnly?
2 1 to reproduce of
oNormalize?
2 1=no, 2=yes

ofxn calc*
0.52771
ofxn mod
0.55505

sRun
RunA1
zout
1

Ave
Data
0.01075
Model
0.00855

data/vMax
0.03462

ofxng
263.459

Min d/m
4.85634

ave/min d
20.6509
ave/min m
79.727

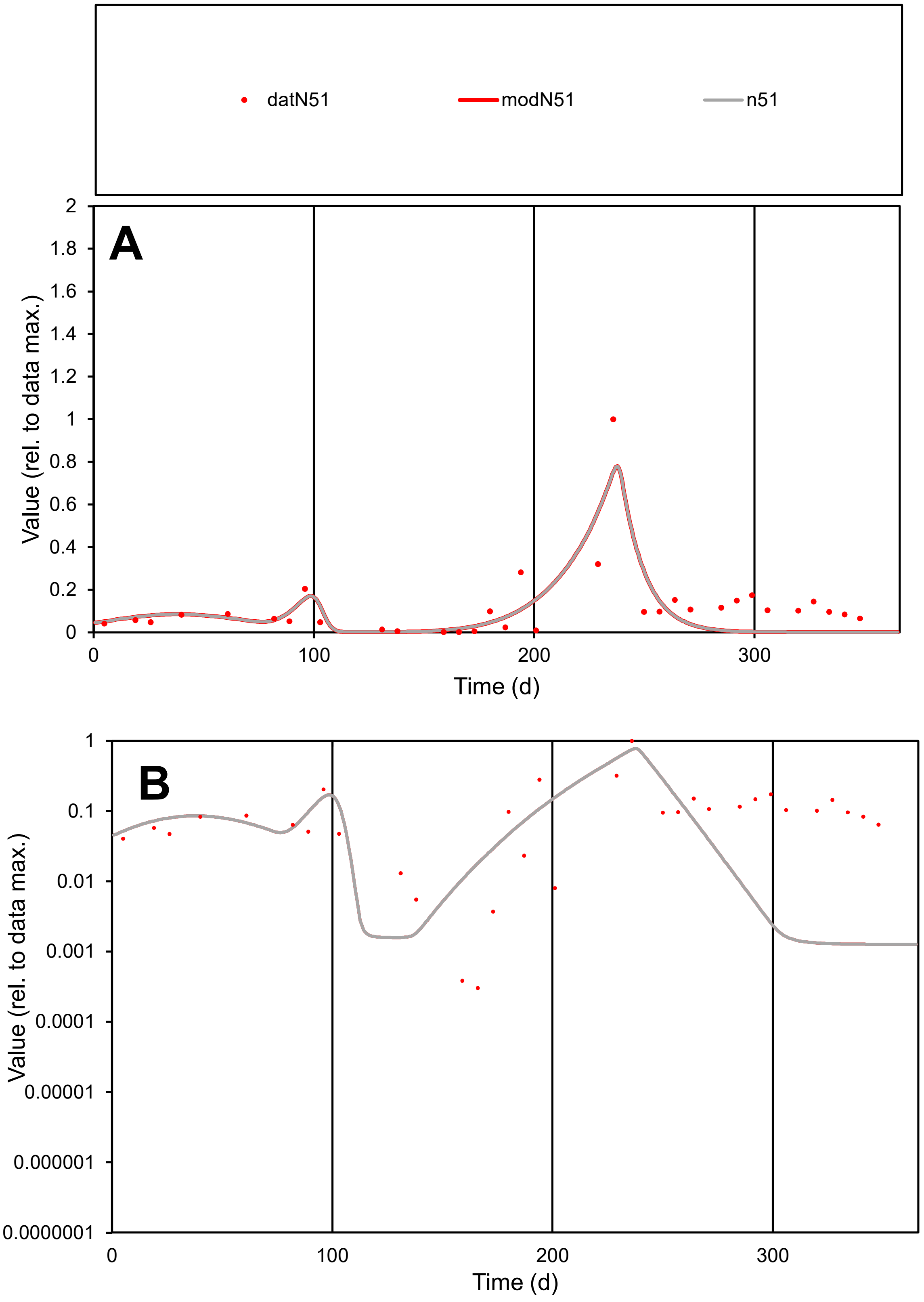


**C1**

Variables
N51 Provide jtd or Dst
CalibOnly?
2 1 to reproduce of
oNormalize?
2 1=no, 2=yes

ofxn calc*
0.83577
ofxn mod
0.86657

sRun
RunA1
zout
1

Ave
Data
0.01206
Model
0.00938

data/vMax
0.1016

ofxng
263.459

Min d/m
0.23897

ave/min d
393.262
ave/min m
73.1485

326_Calib_S_A

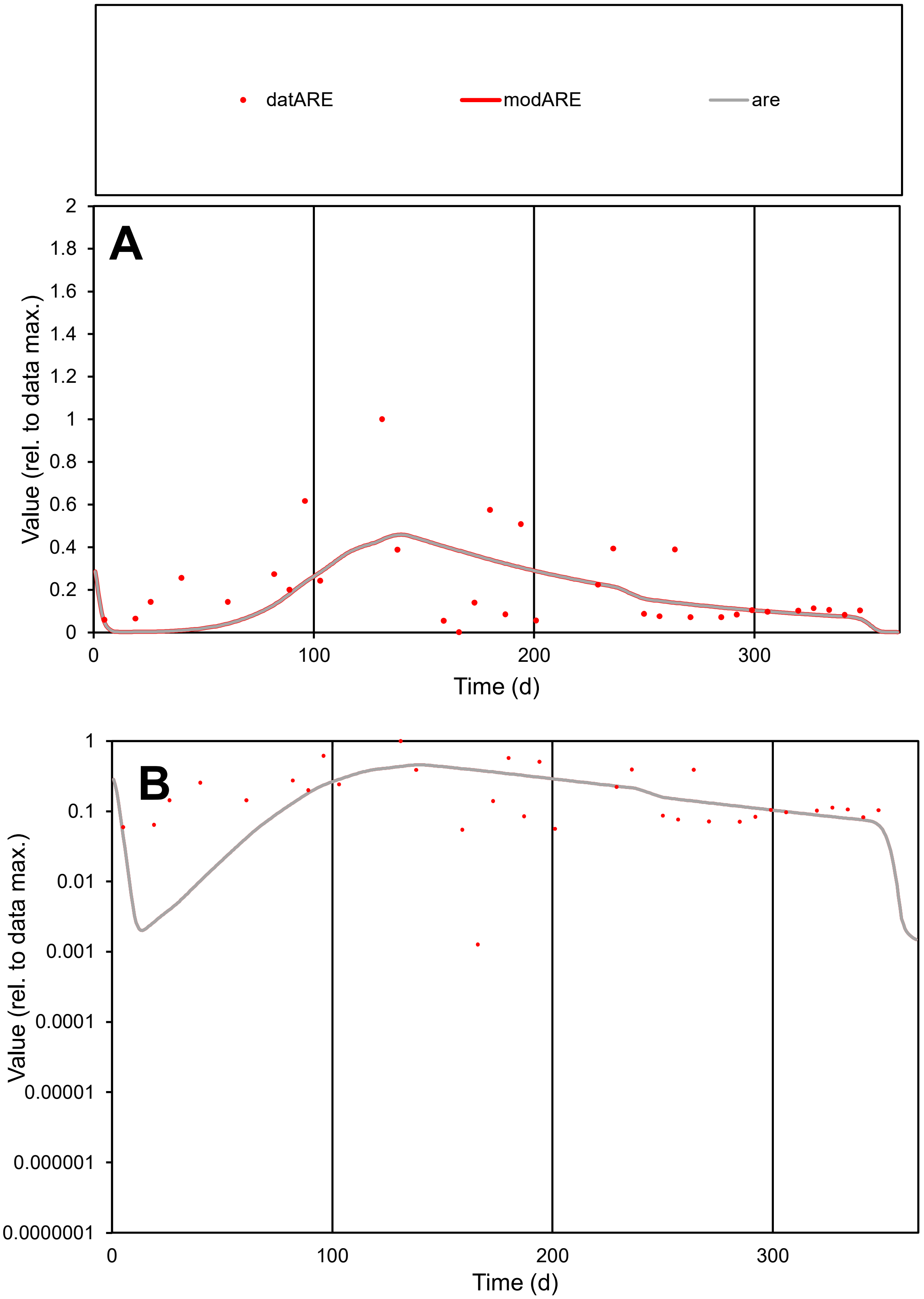


**C1**

Variables

ARE Provide jtd or Dst

CalibOnly?

2 1 to reproduce of

oNormalize?

2 1=no, 2=yes

ofxn calc*

0.91016

ofxn mod

0.91983

sRun

RunA1

zout

1

Ave

Data

0.00885

Model

0.00779

data/vMax

0.04227

ofxng

263.459

Min d/m

0.44322

ave/min d

165.056

ave/min m

64.4159

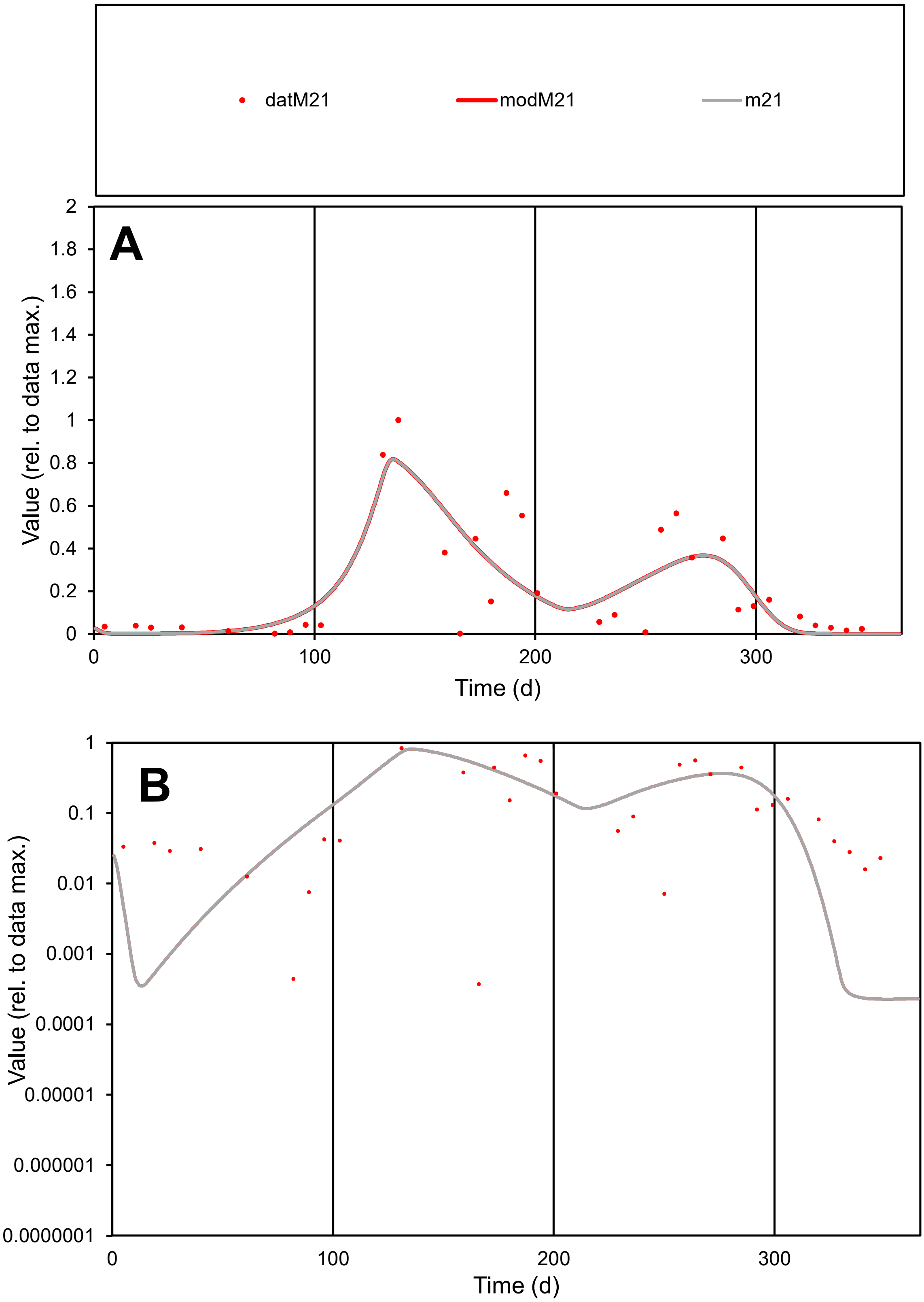


**C1**

Variables
M21 Provide jtd or Dst
CalibOnly?
2 1 to reproduce of
oNormalize?
2 1=no, 2=yes

ofxn calc*
0.74727
ofxn mod
0.77445

sRun
RunA1
zout
1

Ave
Data
0.01127
Model
0.01099

data/vMax
0.05272

ofxng
263.459

Min d/m
1.64387

ave/min d
571.686
ave/min m
916.876

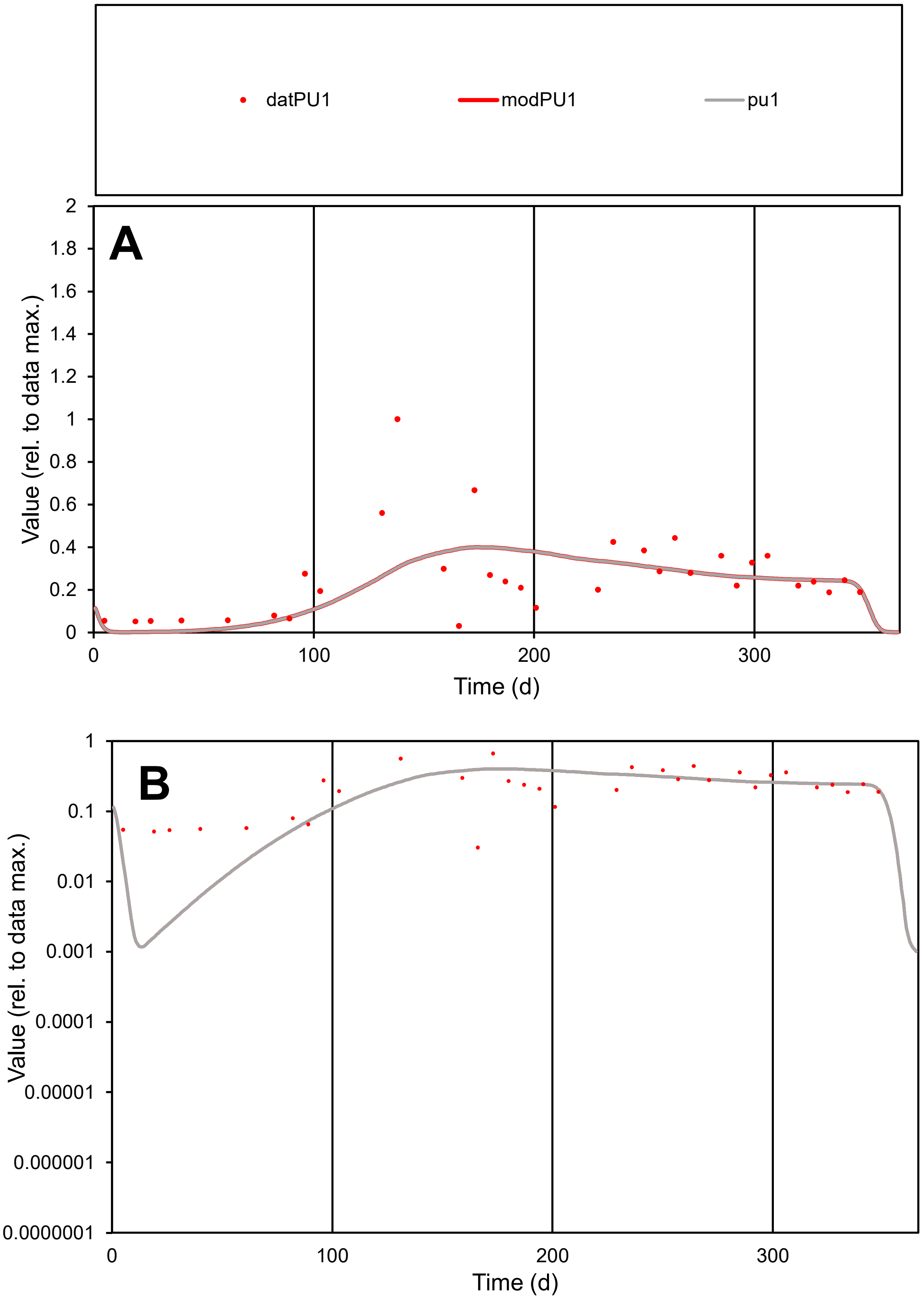


**C1**

Variables
PU1 Provide jtd or Dst
CalibOnly?
2 1 to reproduce of
oNormalize?
2 1=no, 2=yes

ofxn calc*
0.67707
ofxn mod
0.68925

sRun
RunA1
zout
1

Ave
Data
0.01067
Model
0.00963

data/vMax
0.04076

ofxng
263.459

Min d/m
17.2318

ave/min d
8.60662
ave/min m
133.798

326_Calib_S_A

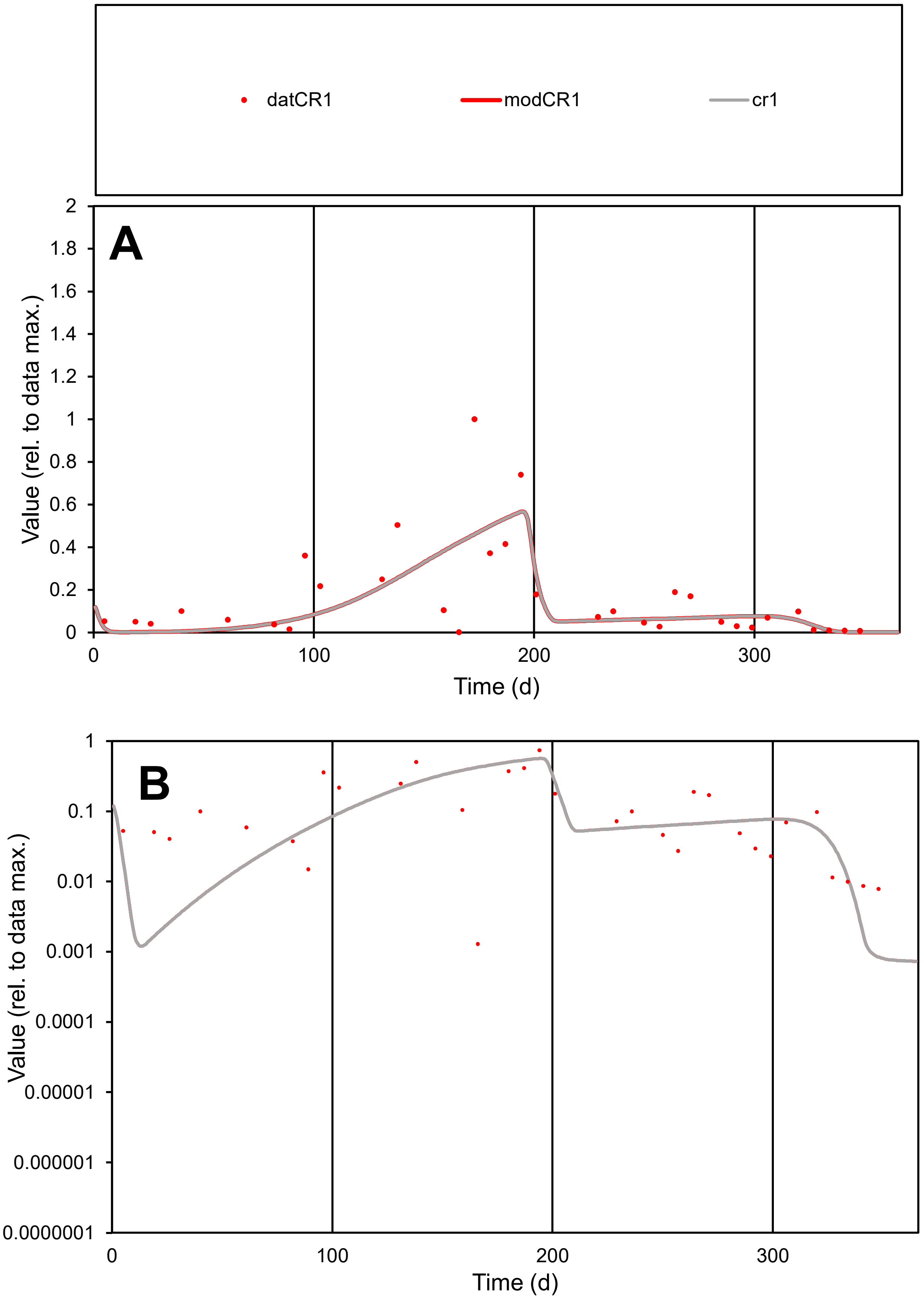


**C1**

Variables
CR1 Provide jtd or Dst
CalibOnly?
2 1 to reproduce of
oNormalize?
2 1=no, 2=yes

ofxn calc*
0.95651
ofxn mod
0.97538

sRun
RunA1
zout
1

Ave
Data
0.00824
Model
0.00719

data/vMax
0.05033

ofxng
263.459

Min d/m
1.57226

ave/min d
127.366
ave/min m
174.691

326_Calib_S_A

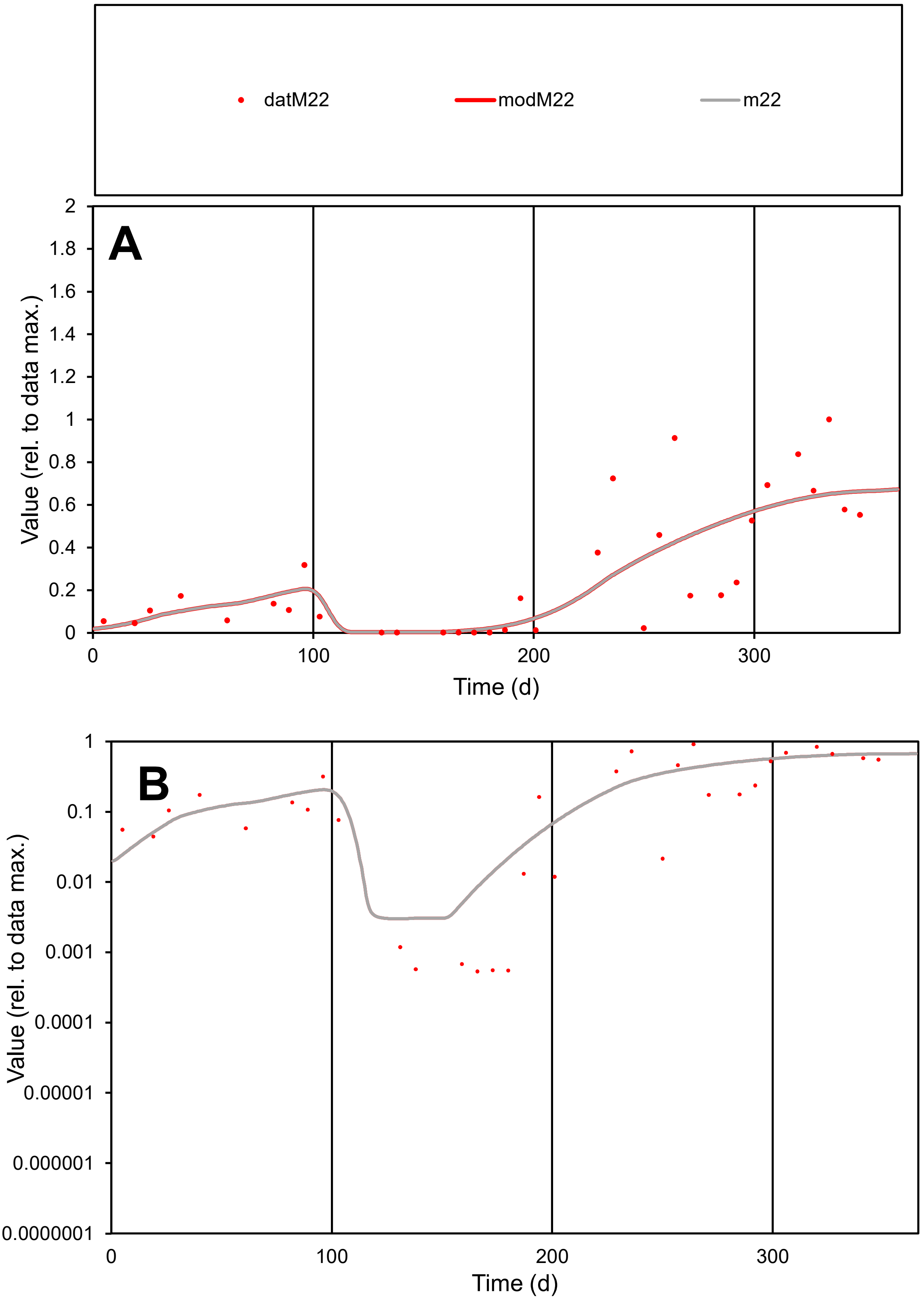


**C1**

Variables

M22 Provide jtd or Dst

CalibOnly?

2 1 to reproduce of

oNormalize?

2 1=no, 2=yes

ofxn calc*

0.65837

ofxn mod

0.67568

sRun

RunA1

zout

1

Ave

Data

0.01113

Model

0.01084

data/vMax

0.03993

ofxng

263.459

Min d/m

0.17745

ave/min d

521.699

ave/min m

90.1602

326_Calib_S_A

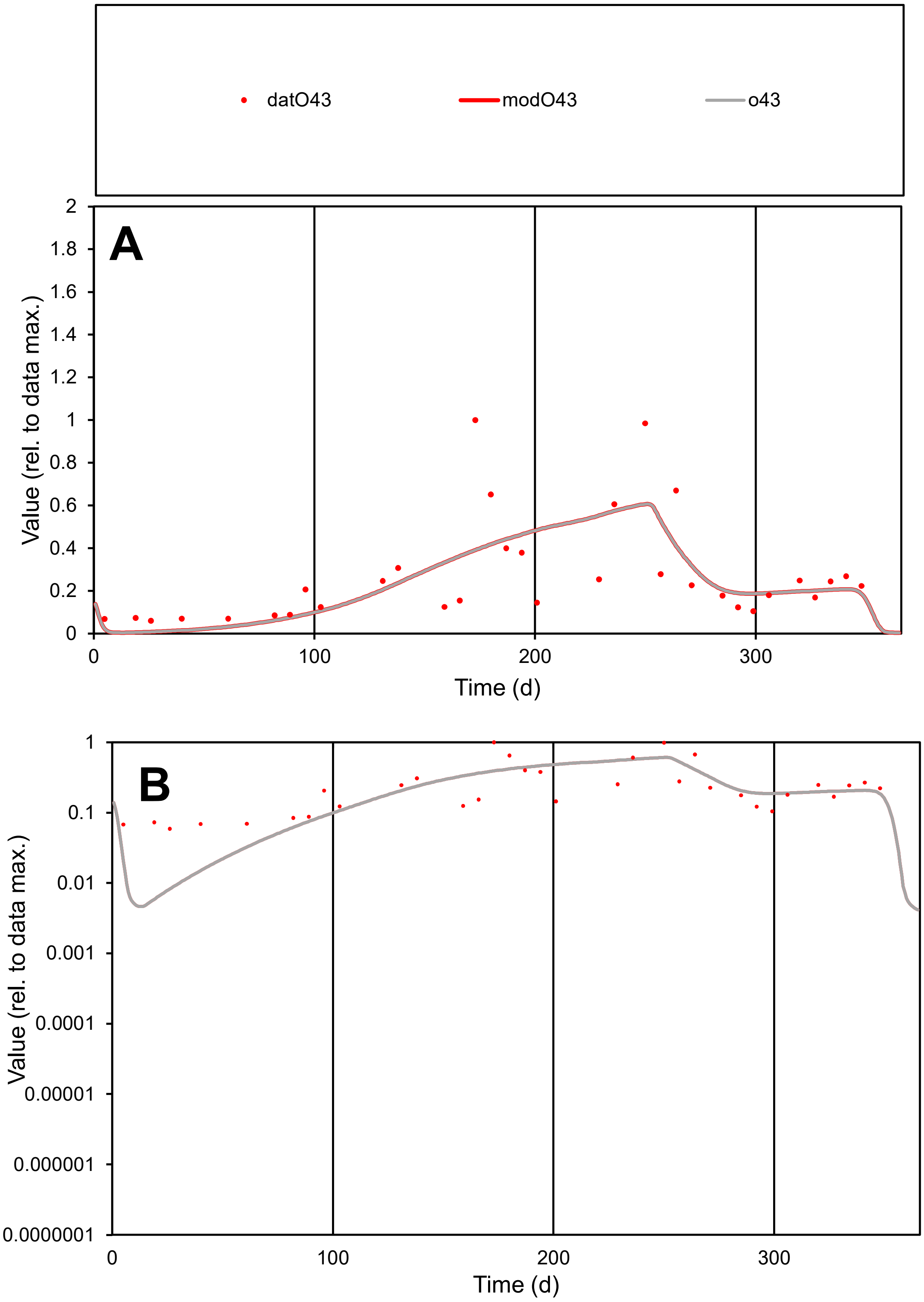


**C1**

Variables
O43 Provide jtd or Dst
CalibOnly?
2 1 to reproduce of
oNormalize?
2 1=no, 2=yes

ofxn calc*
0.65572
ofxn mod
0.67182

sRun
RunA1
zout
1

Ave
Data
0.01103
Model
0.01048

data/vMax
0.04049

ofxng
263.459

Min d/m
9.39442

ave/min d
4.61806
ave/min m
41.2057

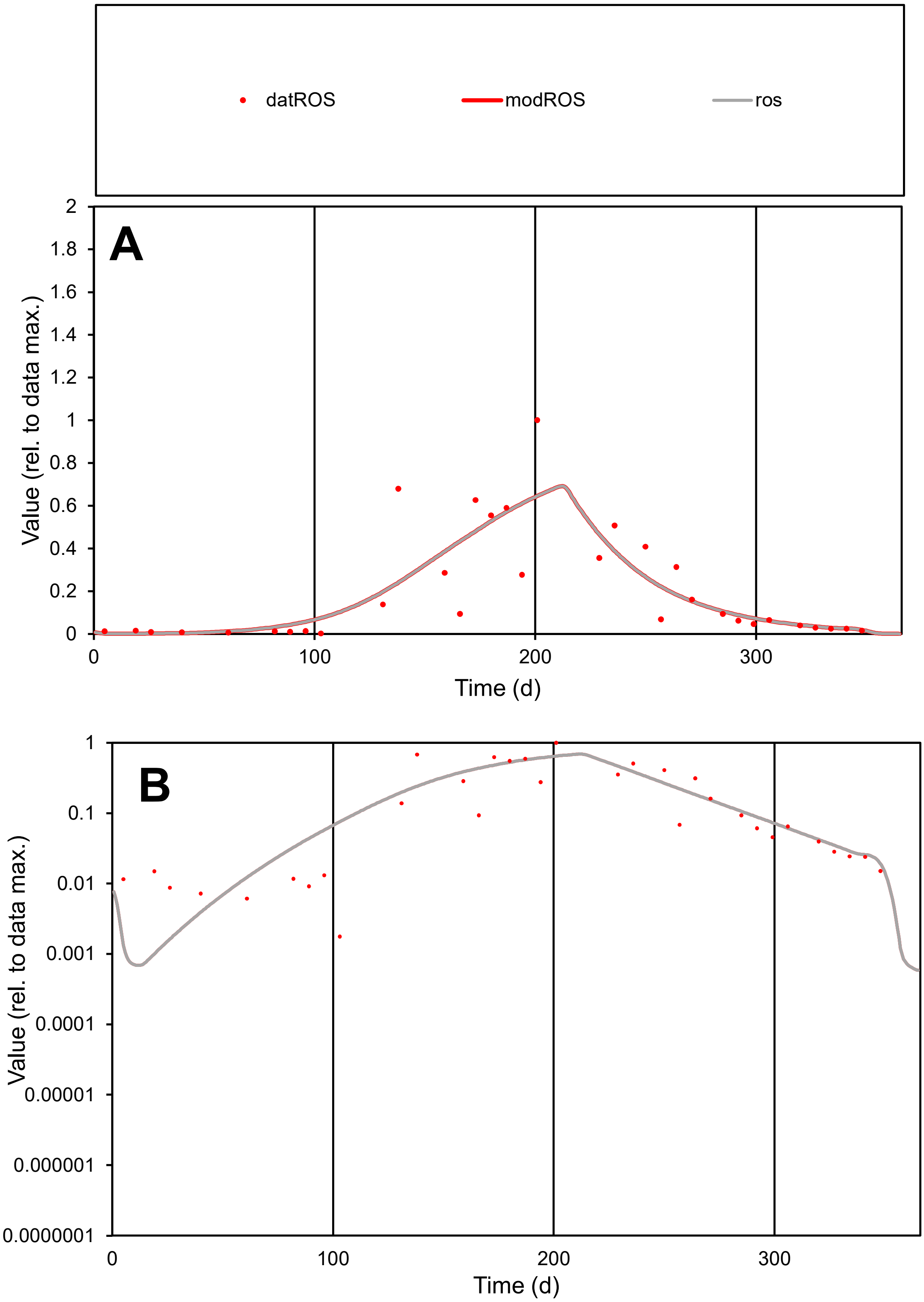


**C1**

Variables
ROS Provide jtd or Dst
CalibOnly?
2 1 to reproduce of
oNormalize?
2 1=no, 2=yes

ofxn calc*
0.71702
ofxn mod
0.74061

sRun
RunA1
zout
1

Ave
Data
0.0115
Model
0.0114

data/vMax
0.05804

ofxng
263.459

Min d/m
1.6015

ave/min d
112.28
ave/min m
178.334

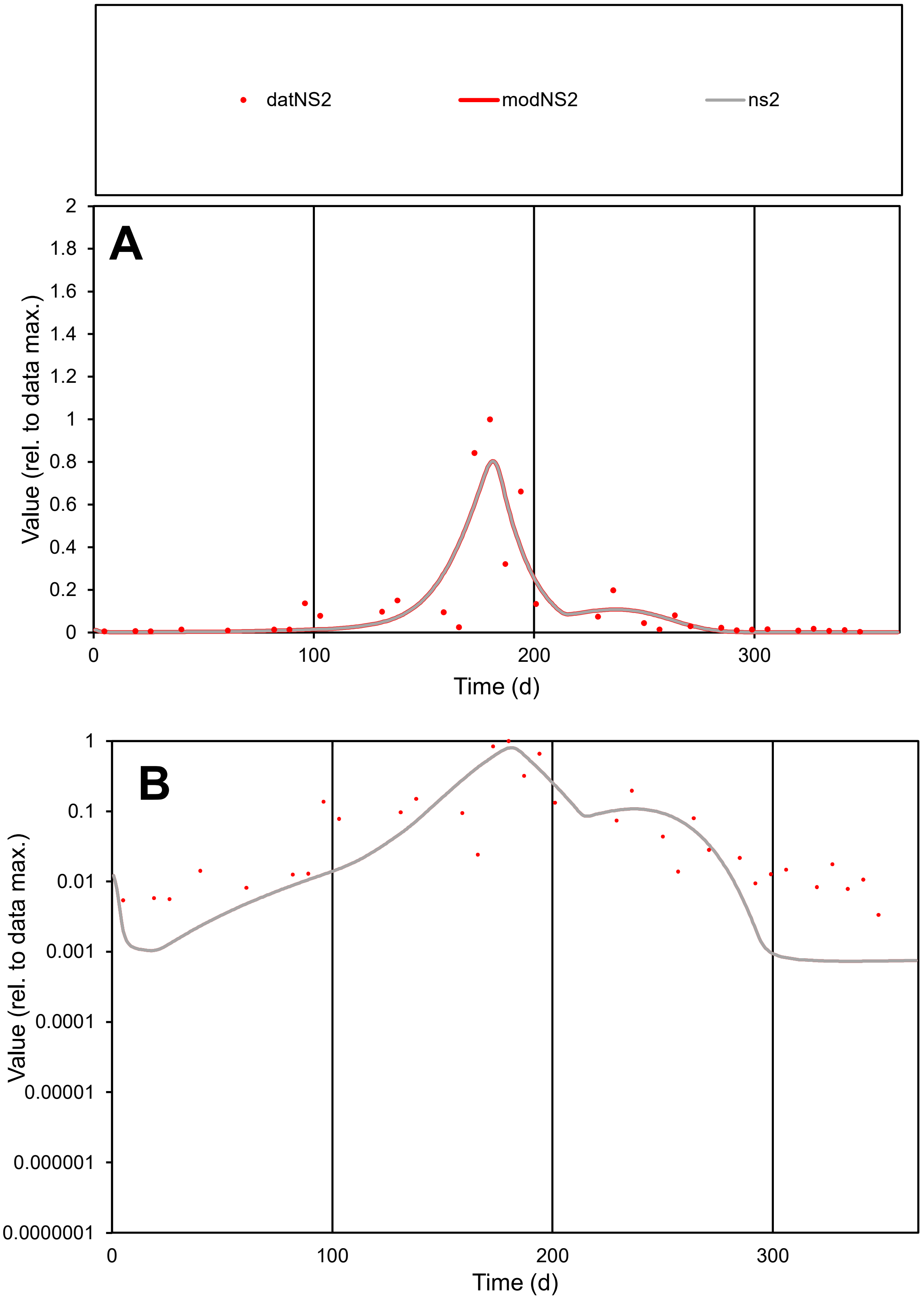


**C1**

Variables
NS2 Provide jtd or Dst
CalibOnly?
2 1 to reproduce of
oNormalize?
2 1=no, 2=yes

ofxn calc*
0.92146
ofxn mod
0.96594

sRun
RunA1
zout
1

Ave
Data
0.00666
Model
0.0064

data/vMax
0.05303

ofxng
263.459

Min d/m
4.57466

ave/min d
37.3924
ave/min m
164.551

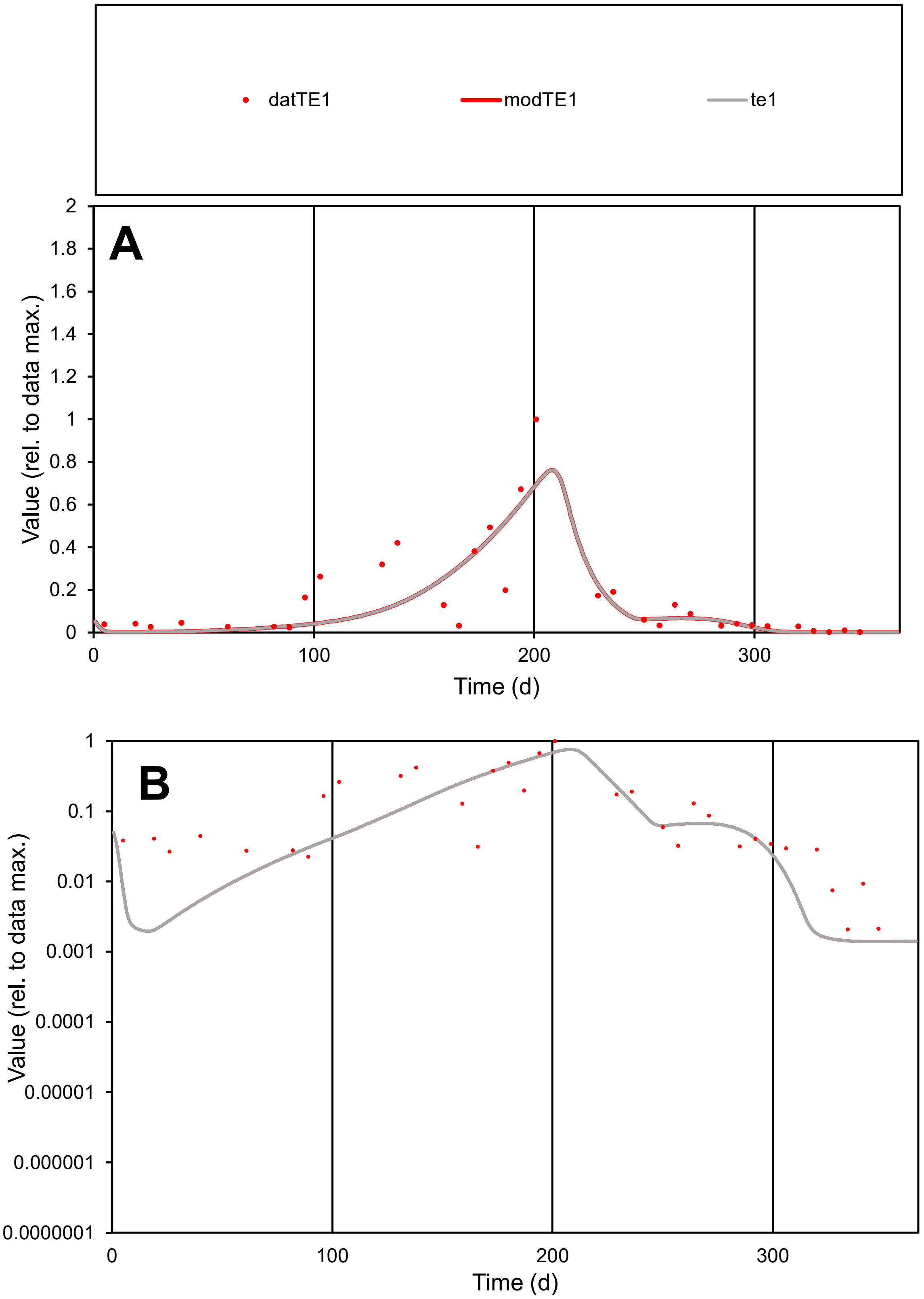


**C1**

Variables
TE1 Provide jtd or Dst
CalibOnly?
2 1 to reproduce of
oNormalize?
2 1=no, 2=yes

ofxn calc*
0.78204
ofxn mod
0.80528

sRun
RunA1
zout
1

Ave
Data
0.00411
Model
0.00355

data/vMax
0.0263

ofxng
263.459

Min d/m
1.48585

ave/min d
75.2468
ave/min m
96.6644

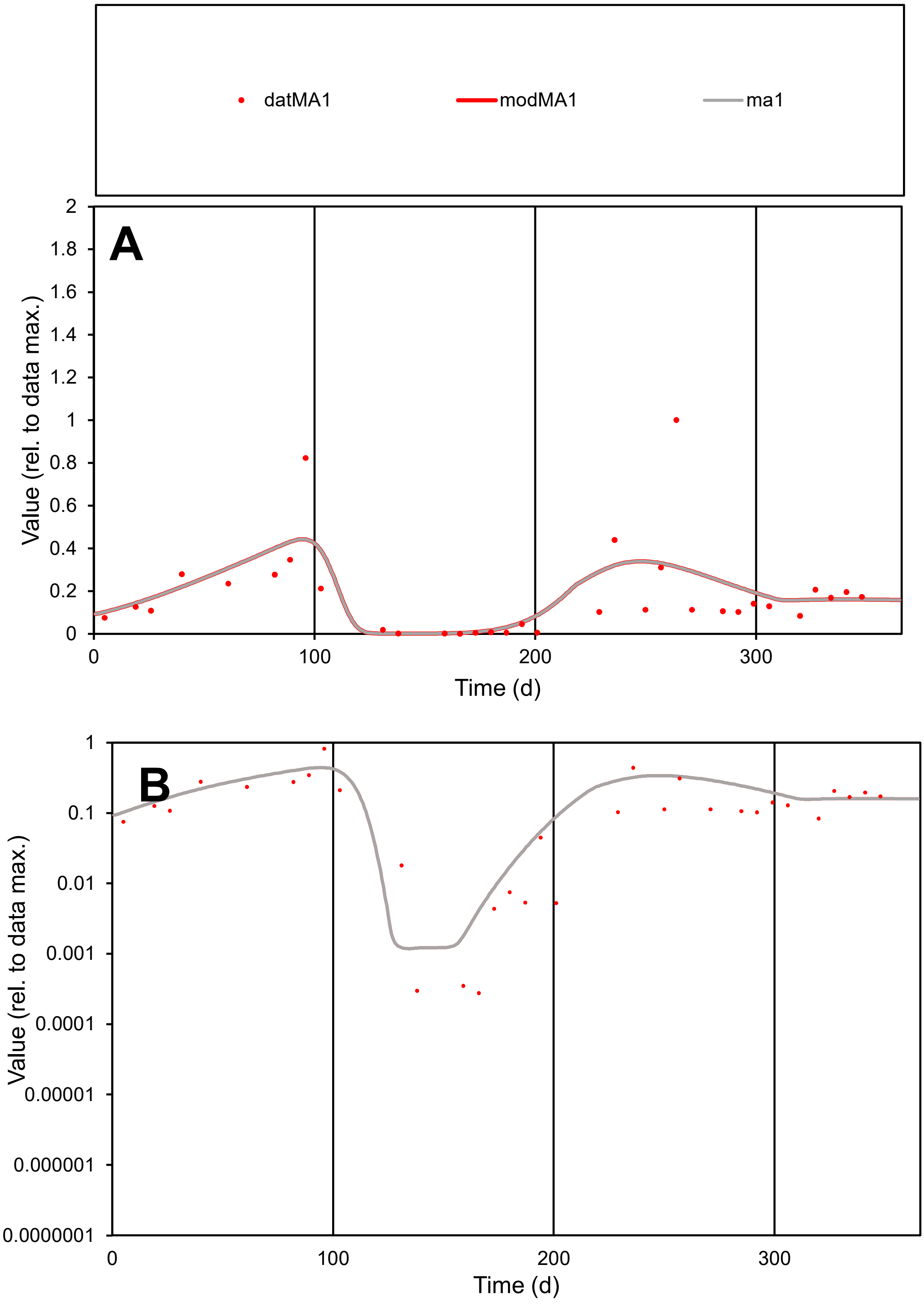


**C1**

Variables
MA1 Provide jtd or Dst
CalibOnly?
2 1 to reproduce of
oNormalize?
2 1=no, 2=yes

ofxn calc*
0.89915
ofxn mod
0.91395

sRun
RunA1
zout
1

Ave
Data
0.00931
Model
0.00987

data/vMax
0.05166

ofxng
263.459

Min d/m
0.23093

ave/min d
651.461
ave/min m
159.538

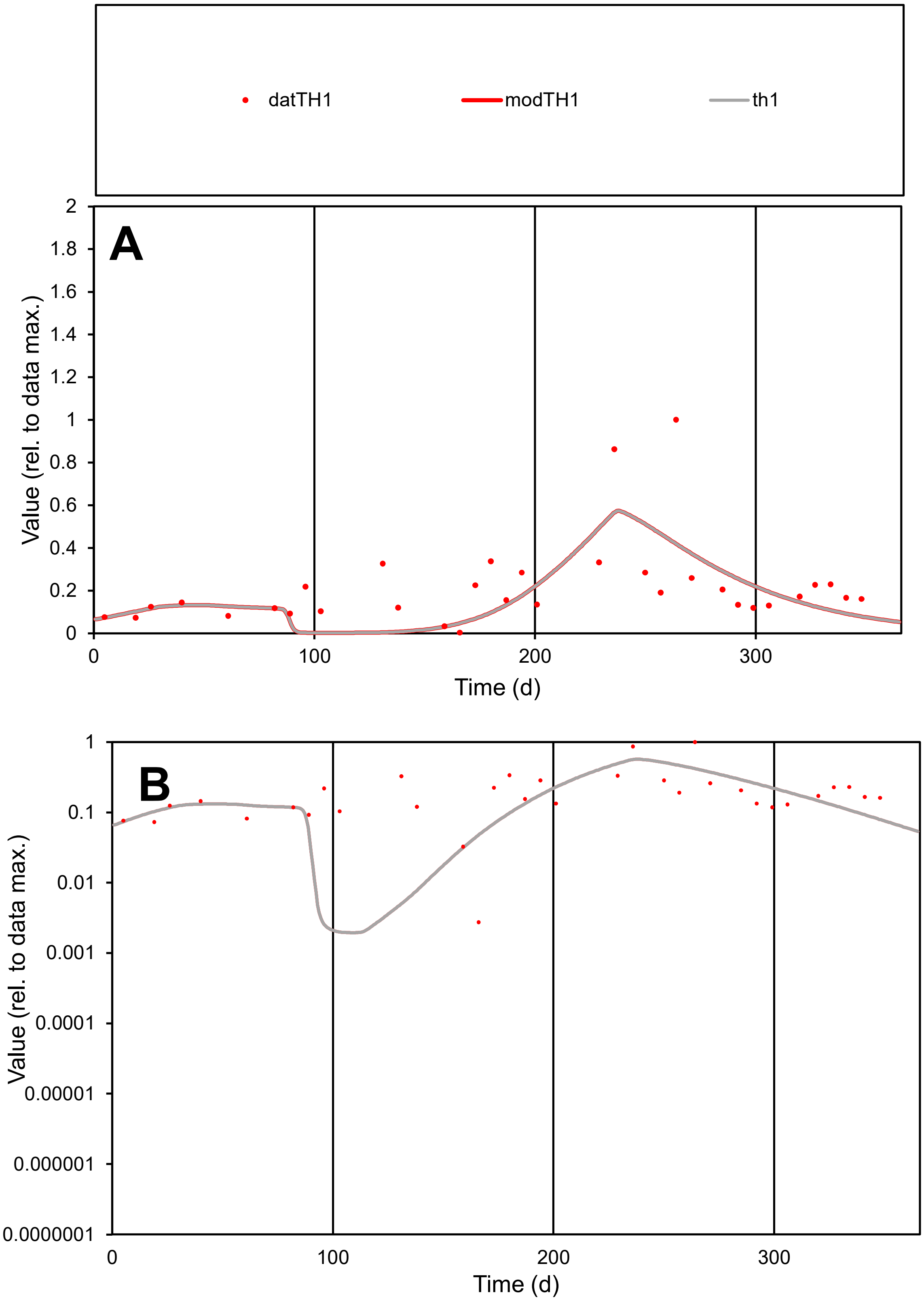


**C1**

Variables
TH1 Provide jtd or Dst
CalibOnly?
2 1 to reproduce of
oNormalize?
2 1=no, 2=yes

ofxn calc*
0.78033
ofxn mod
0.79343

sRun
RunA1
zout
1

Ave
Data
0.00723
Model
0.00598

data/vMax
0.03352

ofxng
263.459

Min d/m
1.38911

ave/min d
78.5384
ave/min m
90.2403

326_Calib_S_A

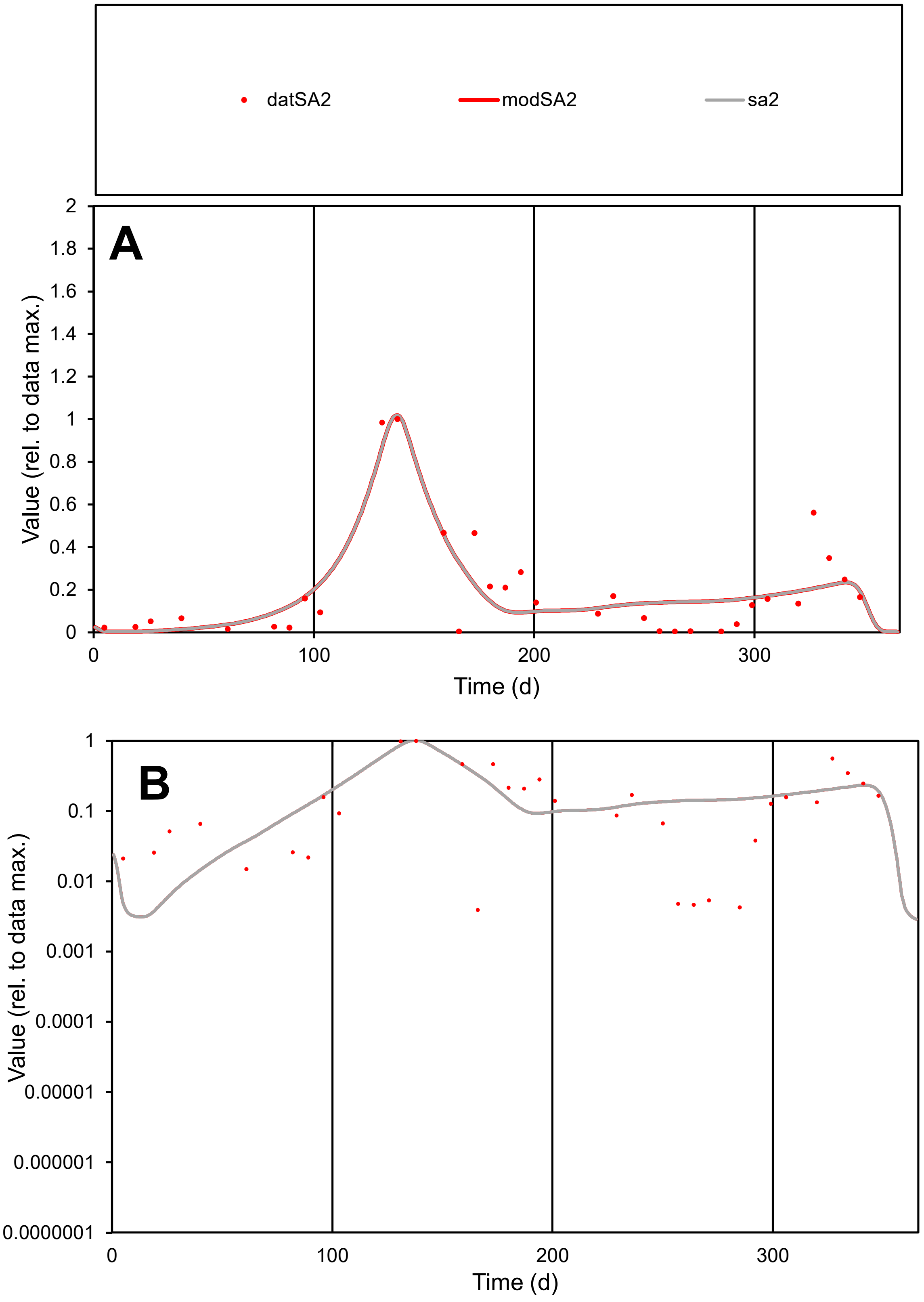


**C1**

Variables
SA2 Provide jtd or Dst
CalibOnly?
2 1 to reproduce of
oNormalize?
2 1=no, 2=yes

ofxn calc*
0.62197
ofxn mod
0.66026

sRun
RunA1
zout
1

Ave
Data
0.00496
Model
0.00505

data/vMax
0.0257

ofxng
263.459

Min d/m
0.94638

ave/min d
49.2954
ave/min m
47.4823

326_Calib_S_A

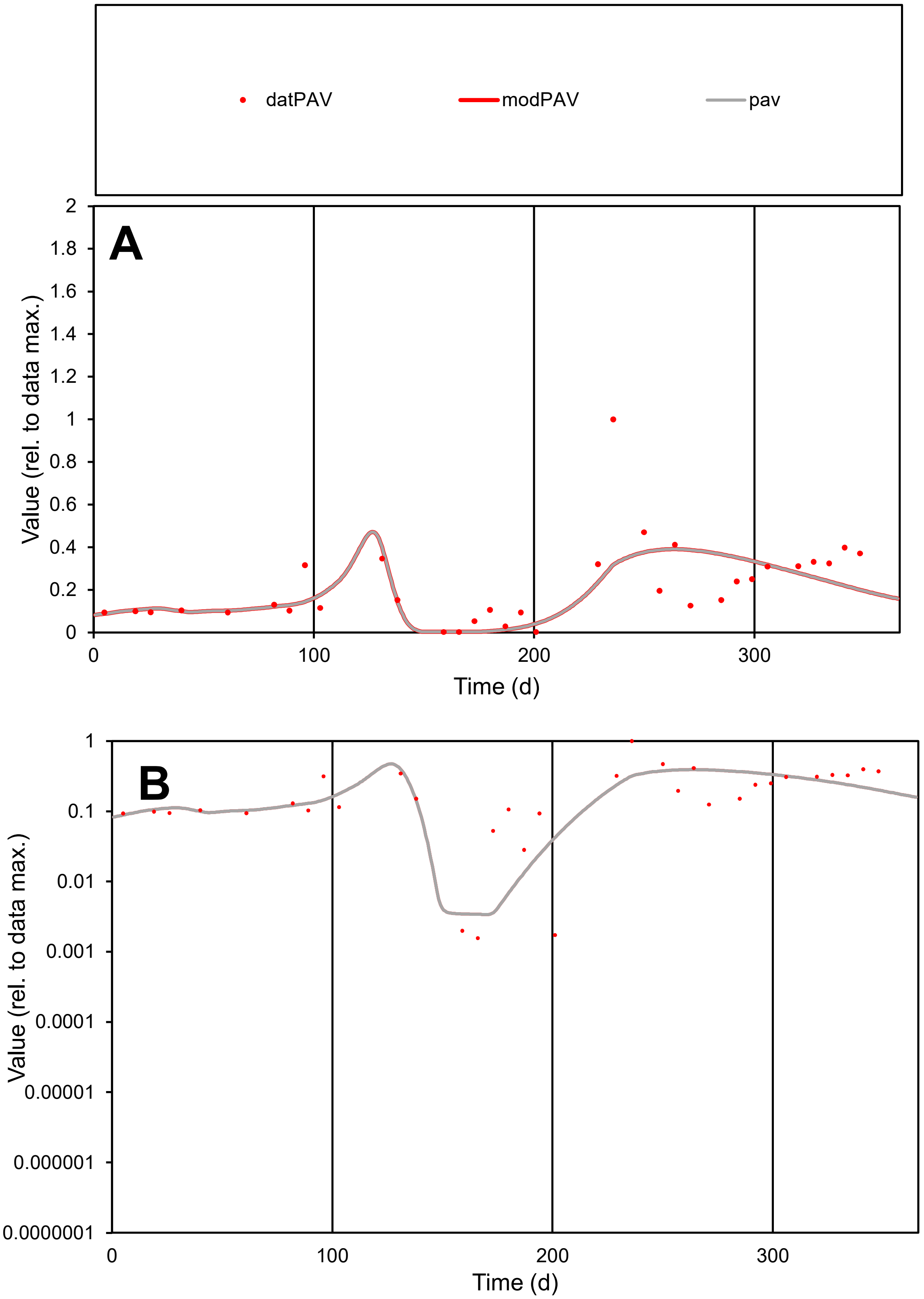


**C1**

Variables
PAV Provide jtd or Dst
CalibOnly?
2 1 to reproduce of
oNormalize?
2 1=no, 2=yes

ofxn calc*
0.70483
ofxn mod
0.71682

sRun
RunA1
zout
1

Ave
Data
0.00508
Model
0.00452

data/vMax
0.02349

ofxng
263.459

Min d/m
0.4598

ave/min d
137.901
ave/min m
56.4434

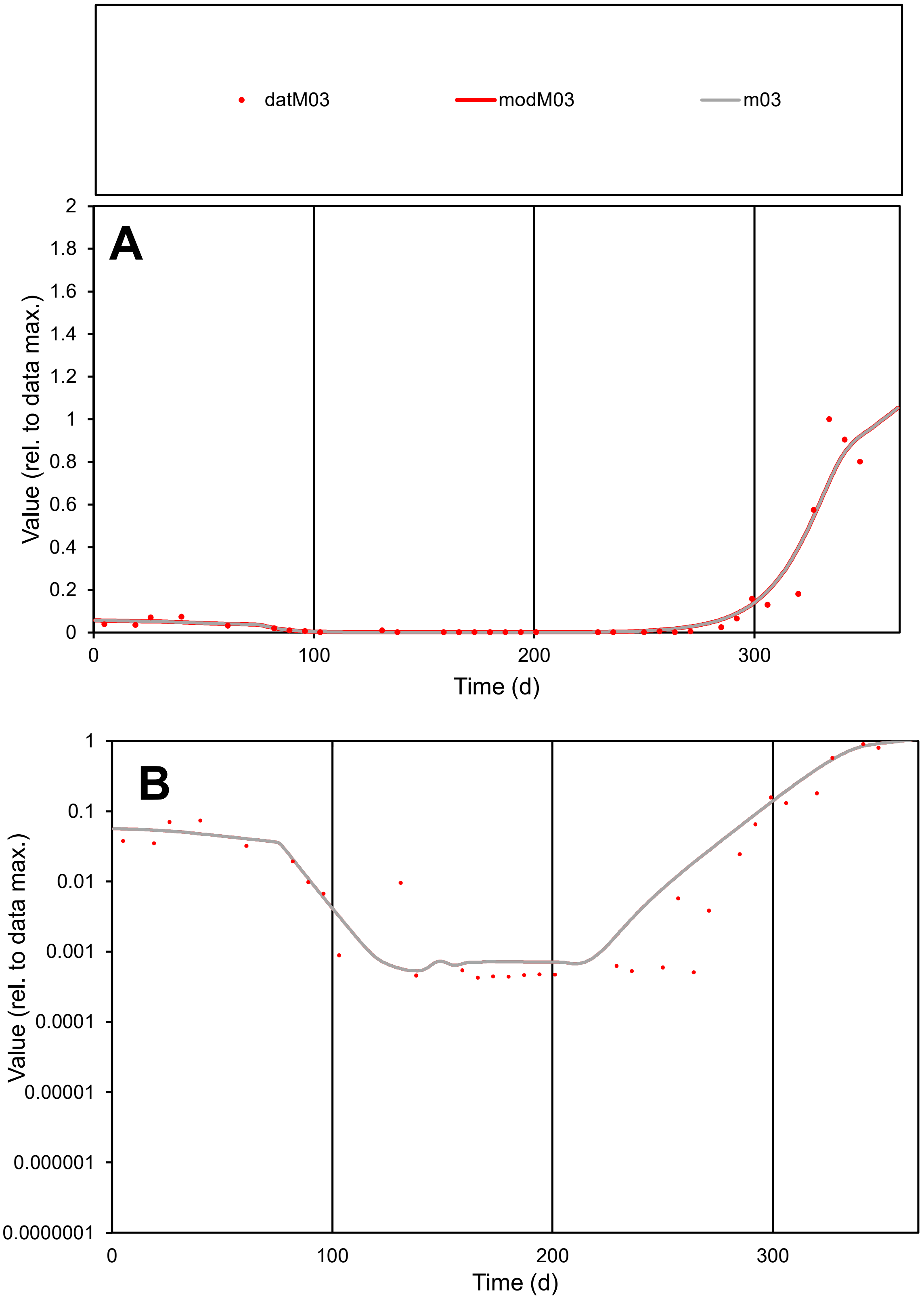


**C1**

Variables
M03 Provide jtd or Dst
CalibOnly?
2 1 to reproduce of
oNormalize?
2 1=no, 2=yes

ofxn calc*
0.52508
ofxn mod
0.57843

sRun
RunA1
zout
1

Ave
Data
0.0051
Model
0.00535

data/vMax
0.04056

ofxng
263.459

Min d/m
0.79652

ave/min d
293.461
ave/min m
245.007

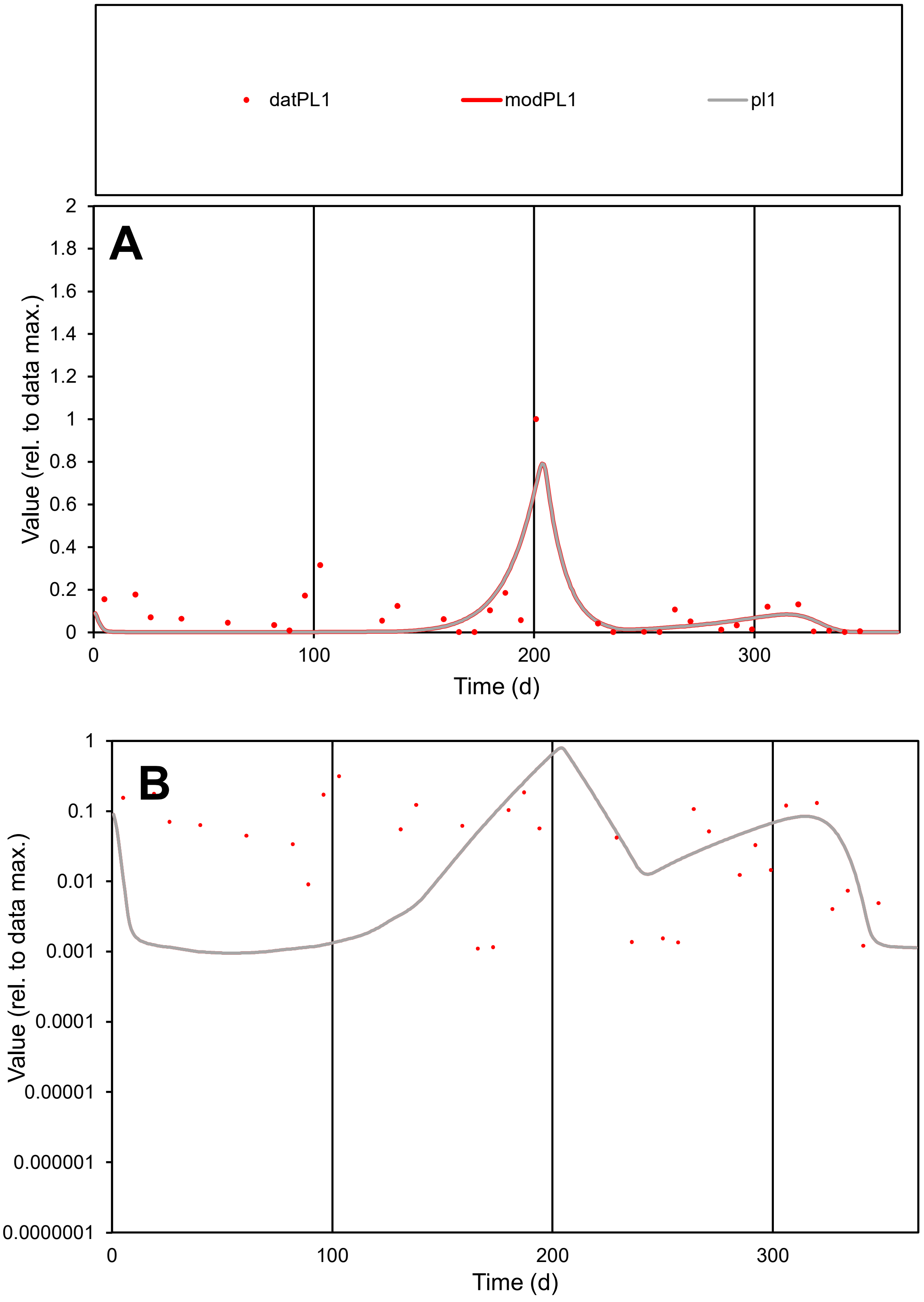


**C1**

Variables
PL1 Provide jtd or Dst
CalibOnly?
2 1 to reproduce of
oNormalize?
2 1=no, 2=yes

ofxn calc*
1.18376
ofxn mod
1.22826

sRun
RunA1
zout
1

Ave
Data
0.00297
Model
0.00222

data/vMax
0.03106

ofxng
263.459

Min d/m
1.15127

ave/min d
86.4604
ave/min m
74.2832

326_Calib_S_A

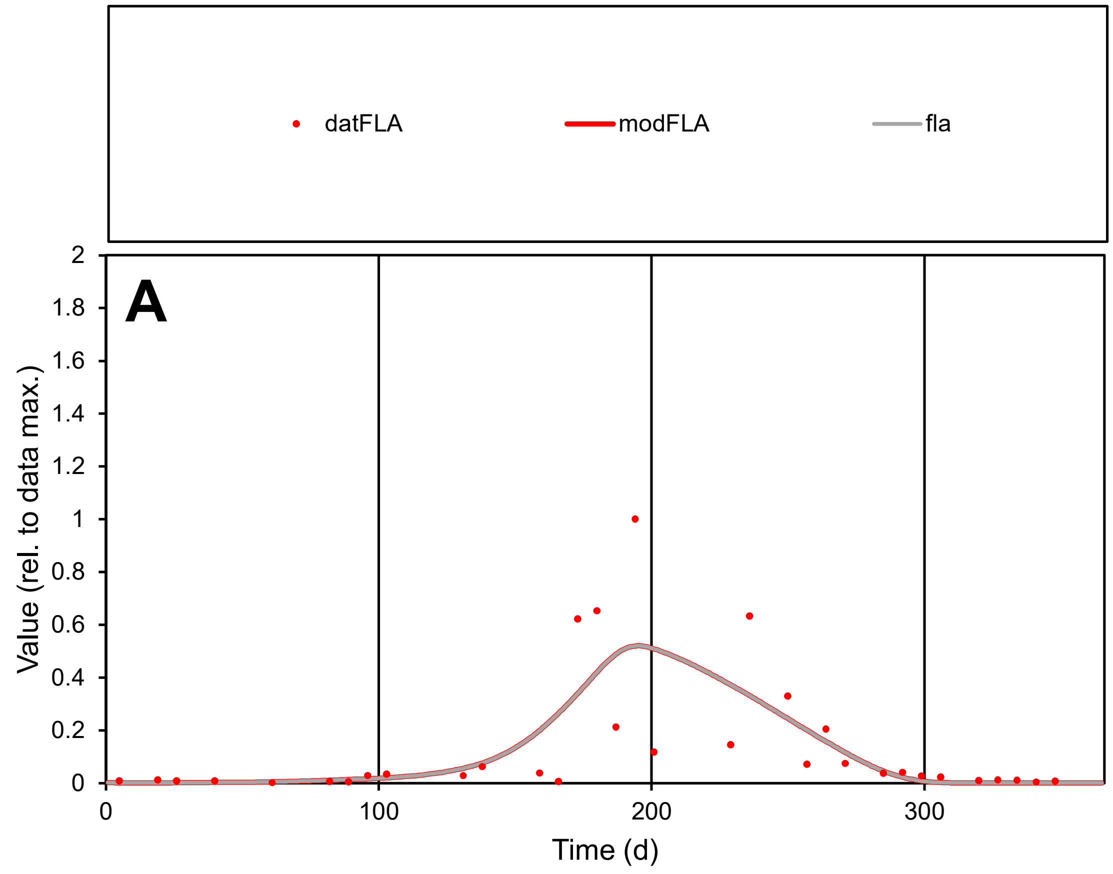


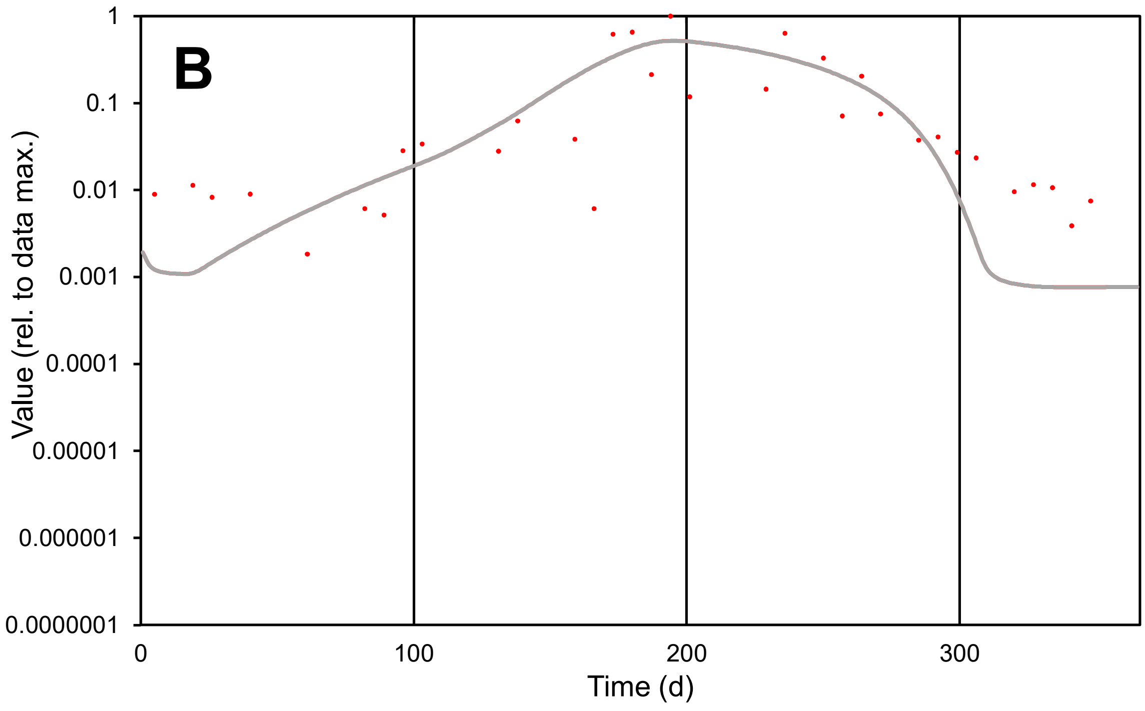


**C1**

Variables

FLA Provide jtd or Dst

CalibOnly?

2 1 to reproduce of

oNormalize?

2 1=no, 2=yes

ofxn calc*

1.17281

ofxn mod

1.1965

sRun

RunA1

zout

1

Ave

Data

0.00509

Model

0.00505

data/vMax

0.03746

ofxng

263.459

Min d/m

2.42254

ave/min d

73.9972

ave/min m

177.754

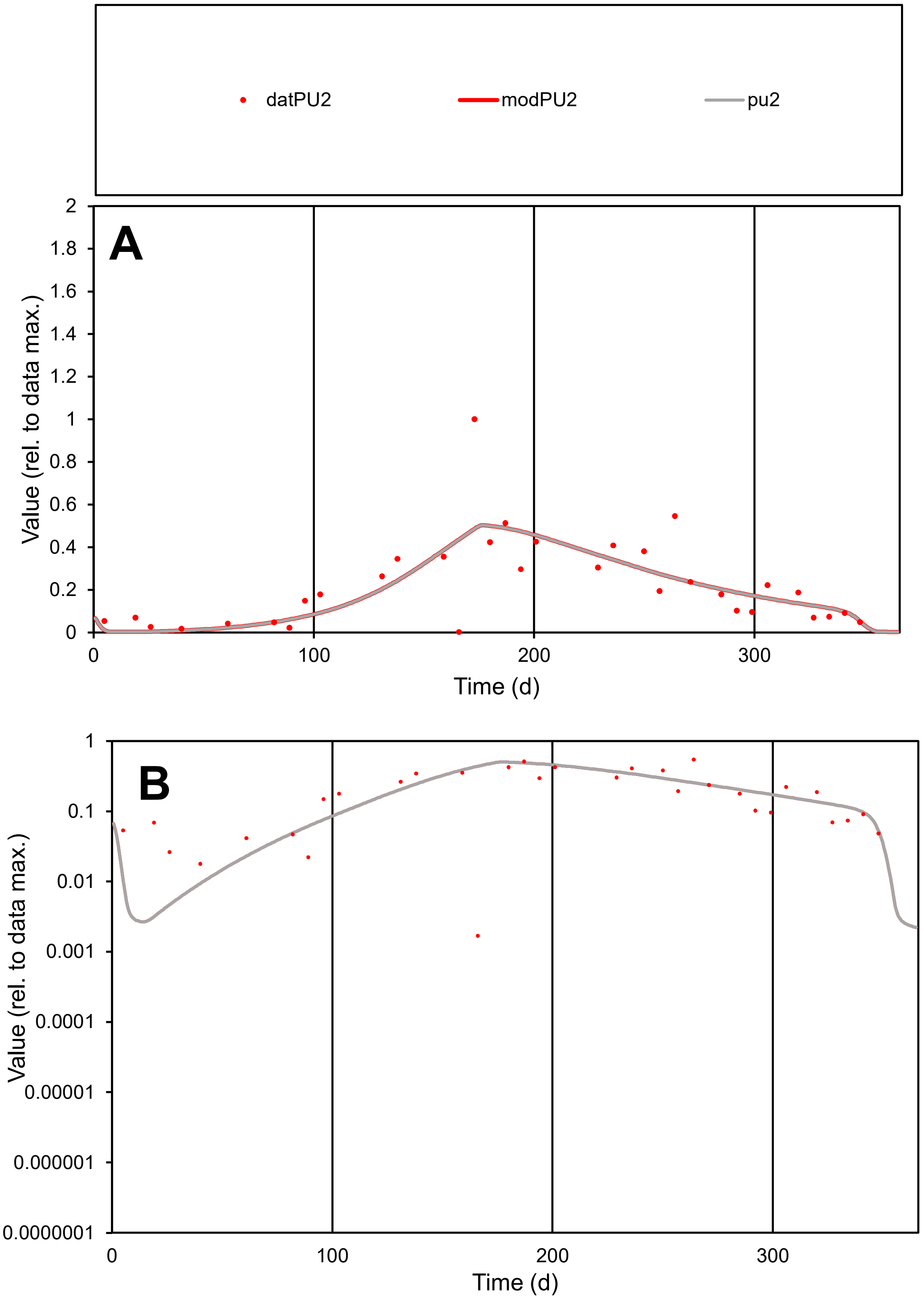


**C1**

Variables
PU2 Provide jtd or Dst
CalibOnly?
2 1 to reproduce of
oNormalize?
2 1=no, 2=yes

ofxn calc*
0.62689
ofxn mod
0.64417

sRun
RunA1
zout
1

Ave
Data
0.0072
Model
0.00689

data/vMax
0.03225

ofxng
263.459

Min d/m
0.49517

ave/min d
132.571
ave/min m
62.7721

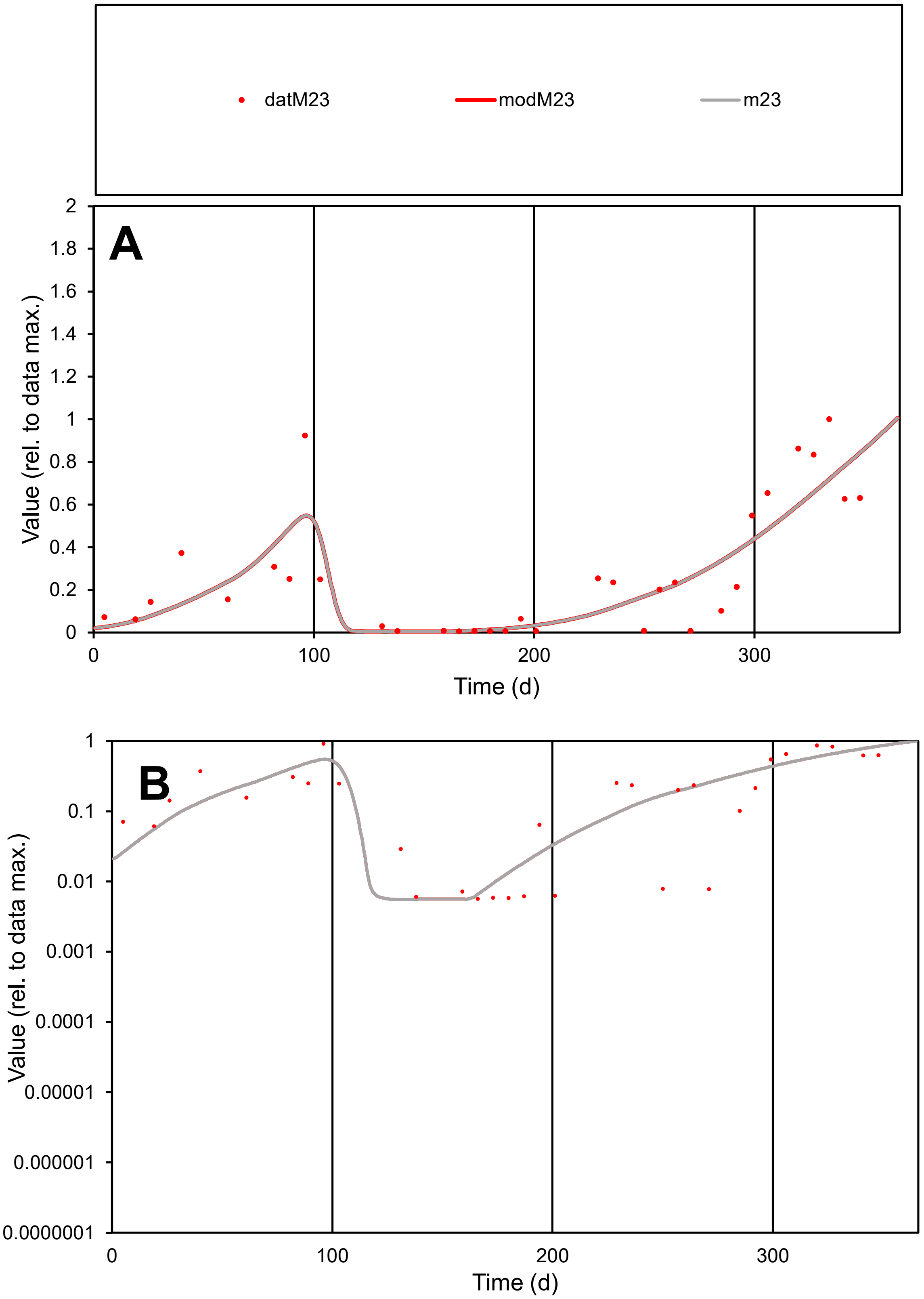


**C1**

Variables
M23 Provide jtd or Dst
CalibOnly?
2 1 to reproduce of
oNormalize?
2 1=no, 2=yes

ofxn calc*
0.56714
ofxn mod
0.58458

sRun
RunA1
zout
1

Ave
Data
0.00516
Model
0.0051

data/vMax
0.01876

ofxng
263.459

Min d/m
1.02111

ave/min d
48.5055
ave/min m
48.9056

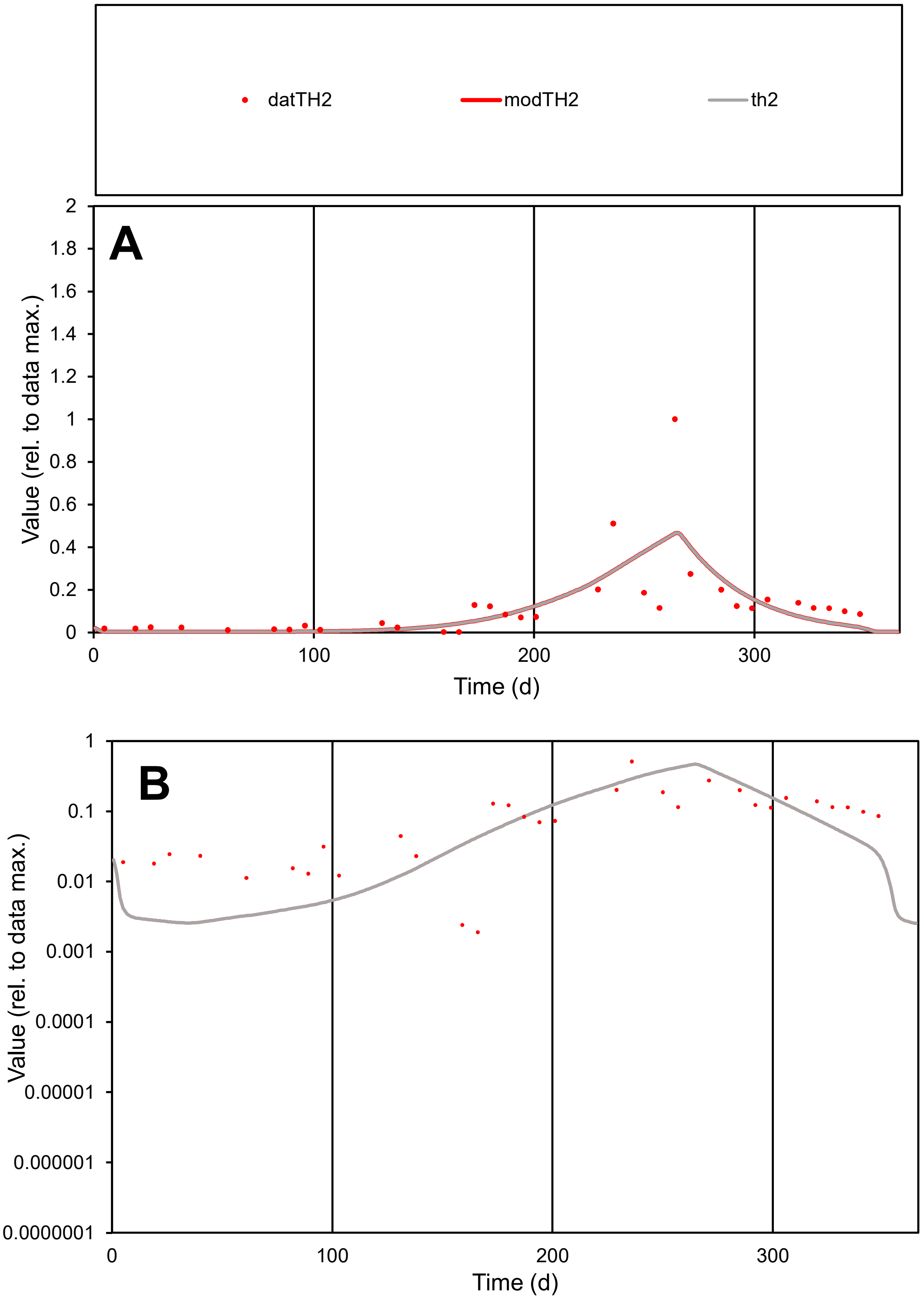


**C1**

Variables
TH2 Provide jtd or Dst
CalibOnly?
2 1 to reproduce of
oNormalize?
2 1=no, 2=yes

ofxn calc*
1.00409
ofxn mod
1.02619

sRun
RunA1
zout
1

Ave
Data
0.00695
Model
0.00627

data/vMax
0.0553

ofxng
263.459

Min d/m
0.71732

ave/min d
66.3343
ave/min m
42.8885

326_Calib_S_A

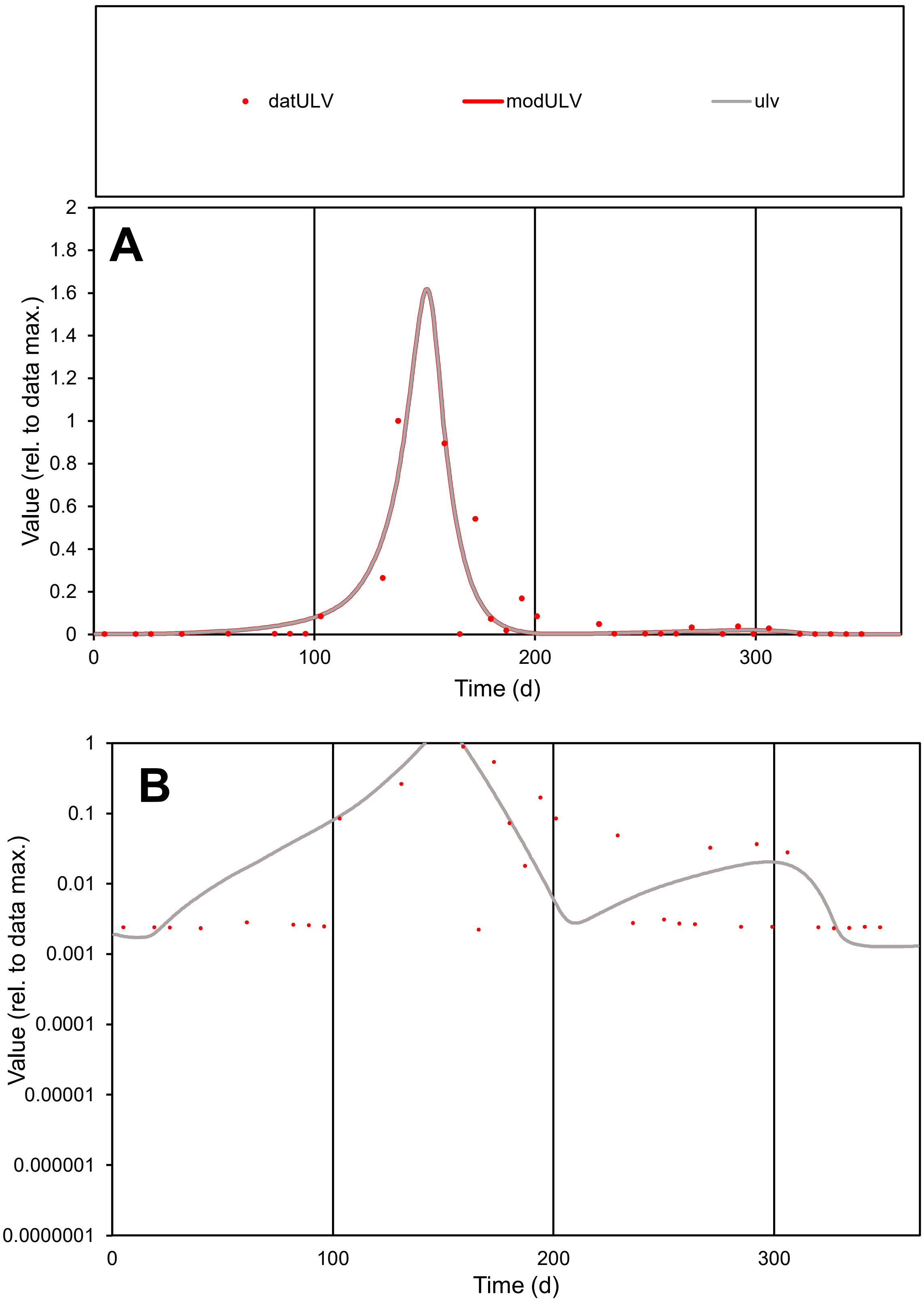


**C1**

Variables

ULV Provide jtd or Dst

CalibOnly?

2 1 to reproduce of

oNormalize?

2 1=no, 2=yes

ofxn calc*

0.9729

ofxn mod

1.04429

sRun

RunA1

zout

1

Ave

Data

0.00077

Model

0.00074

data/vMax

0.00763

ofxng

263.459

Min d/m

1.75128

ave/min d

45.1502

ave/min m

76.4763

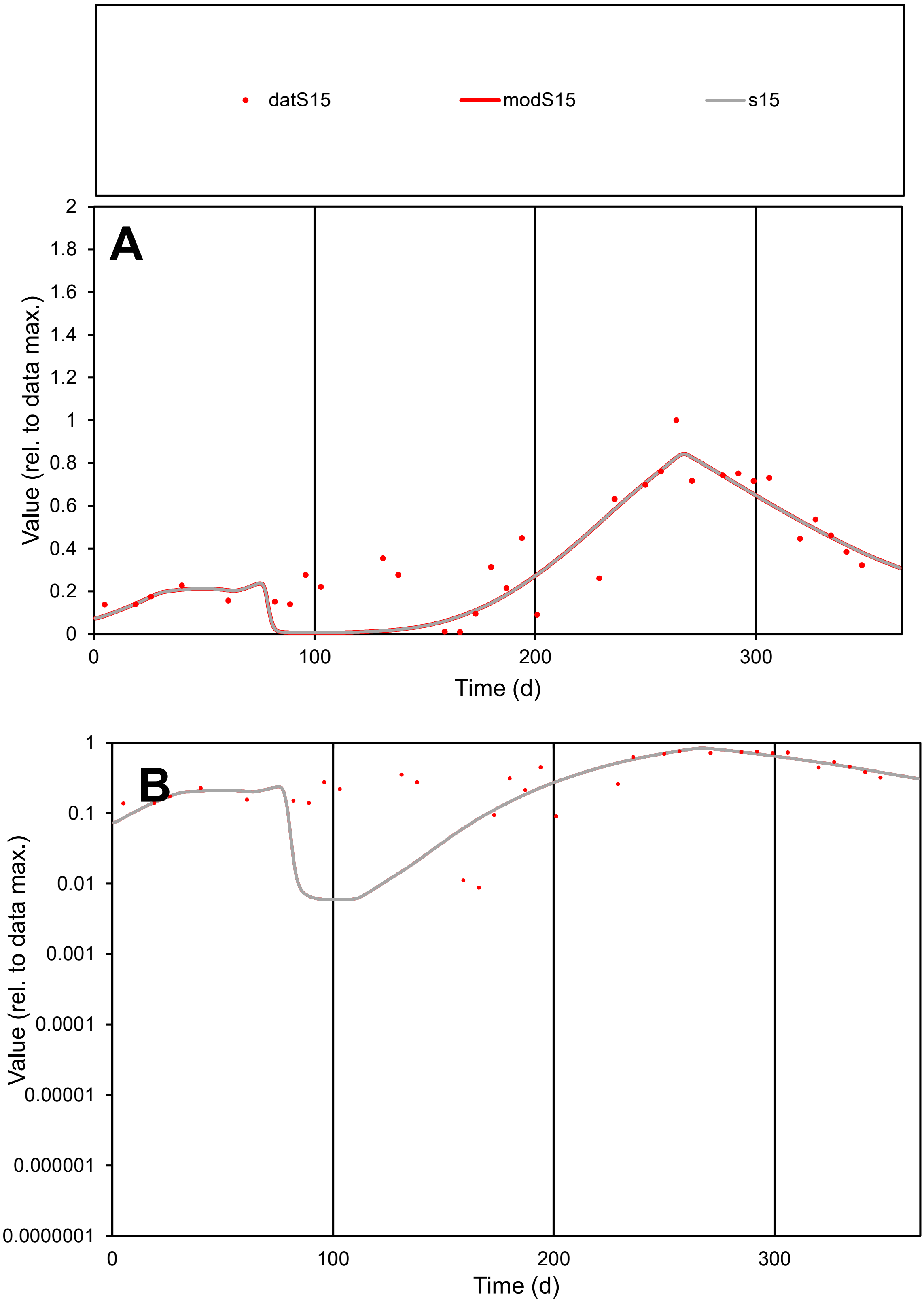


**C1**

Variables
S15 Provide jtd or Dst
CalibOnly?
2 1 to reproduce of
oNormalize?
2 1=no, 2=yes

ofxn calc*
0.34826
ofxn mod
0.37952

sRun
RunA1
zout
1

Ave
Data
0.00533
Model
0.00479

data/vMax
0.01396

ofxng
263.459

Min d/m
1.48211

ave/min d
43.3045
ave/min m
57.672

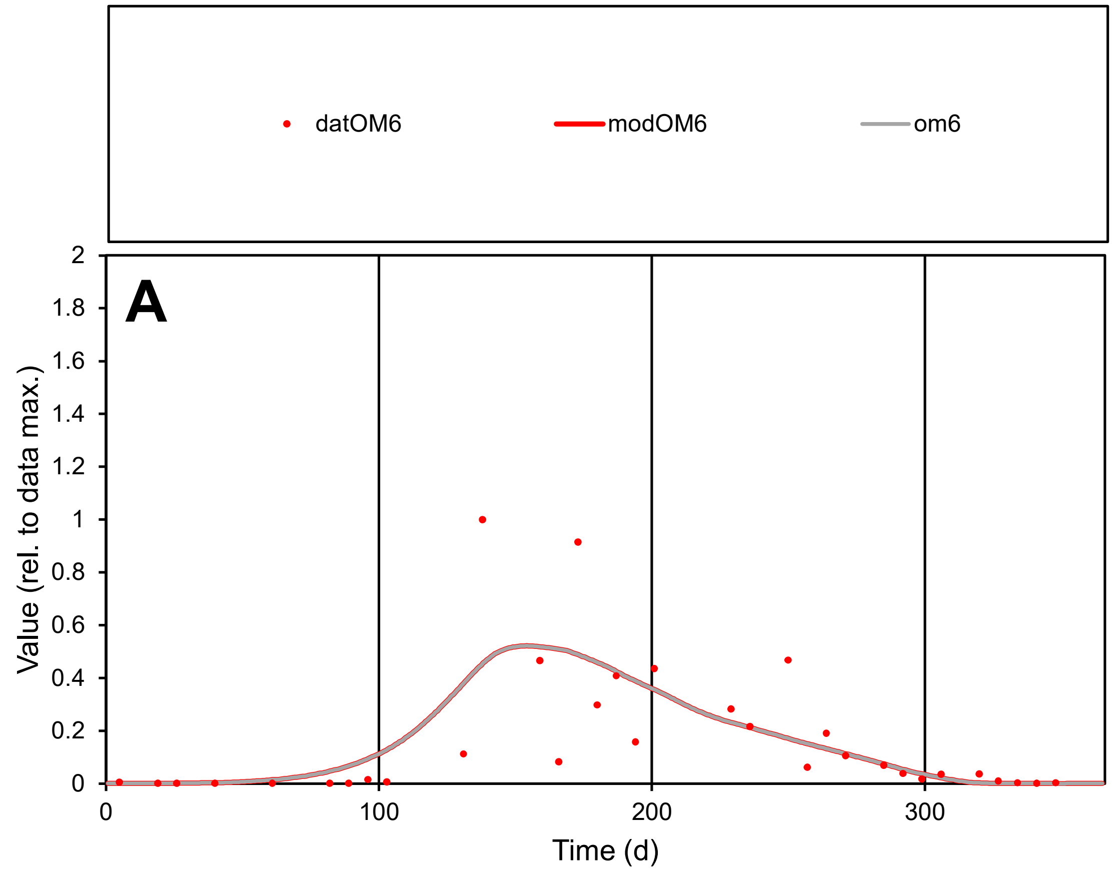


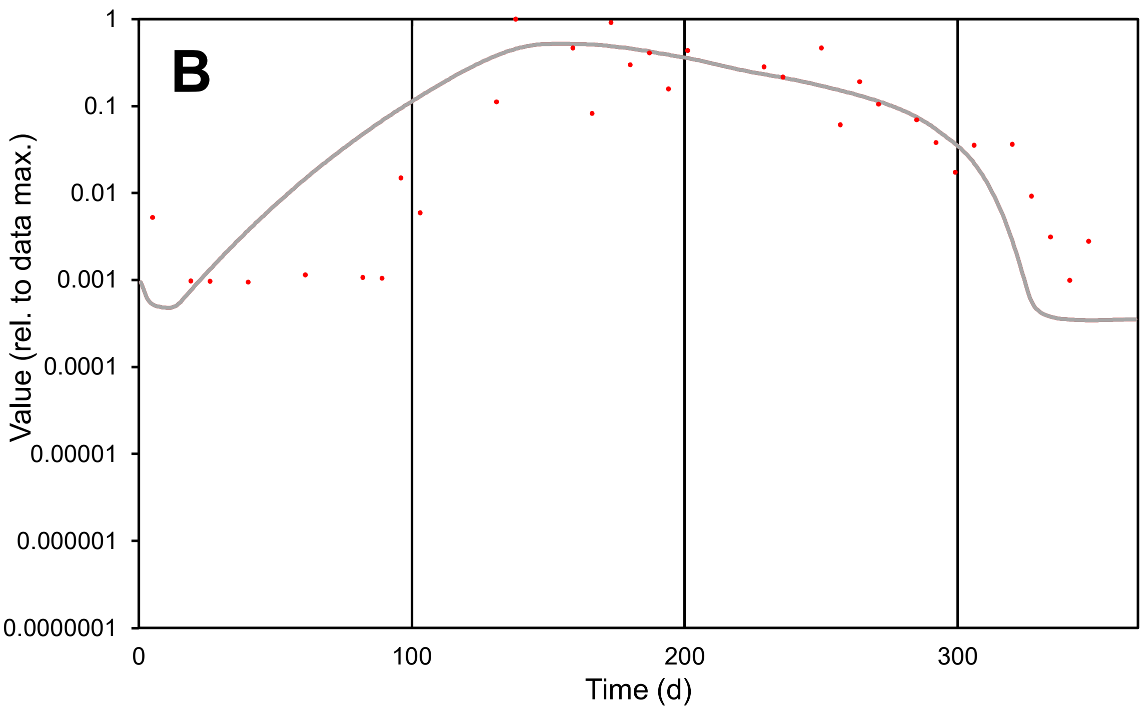


**C1**

Variables
OM6 Provide jtd or Dst
CalibOnly?
2 1 to reproduce of
oNormalize?
2 1=no, 2=yes

ofxn calc*
1.02356
ofxn mod
1.04235

sRun
RunA1
zout
1

Ave
Data
0.00451
Model
0.00455

data/vMax
0.02733

ofxng
263.459

Min d/m
2.75005

ave/min d
173.987
ave/min m
482.952

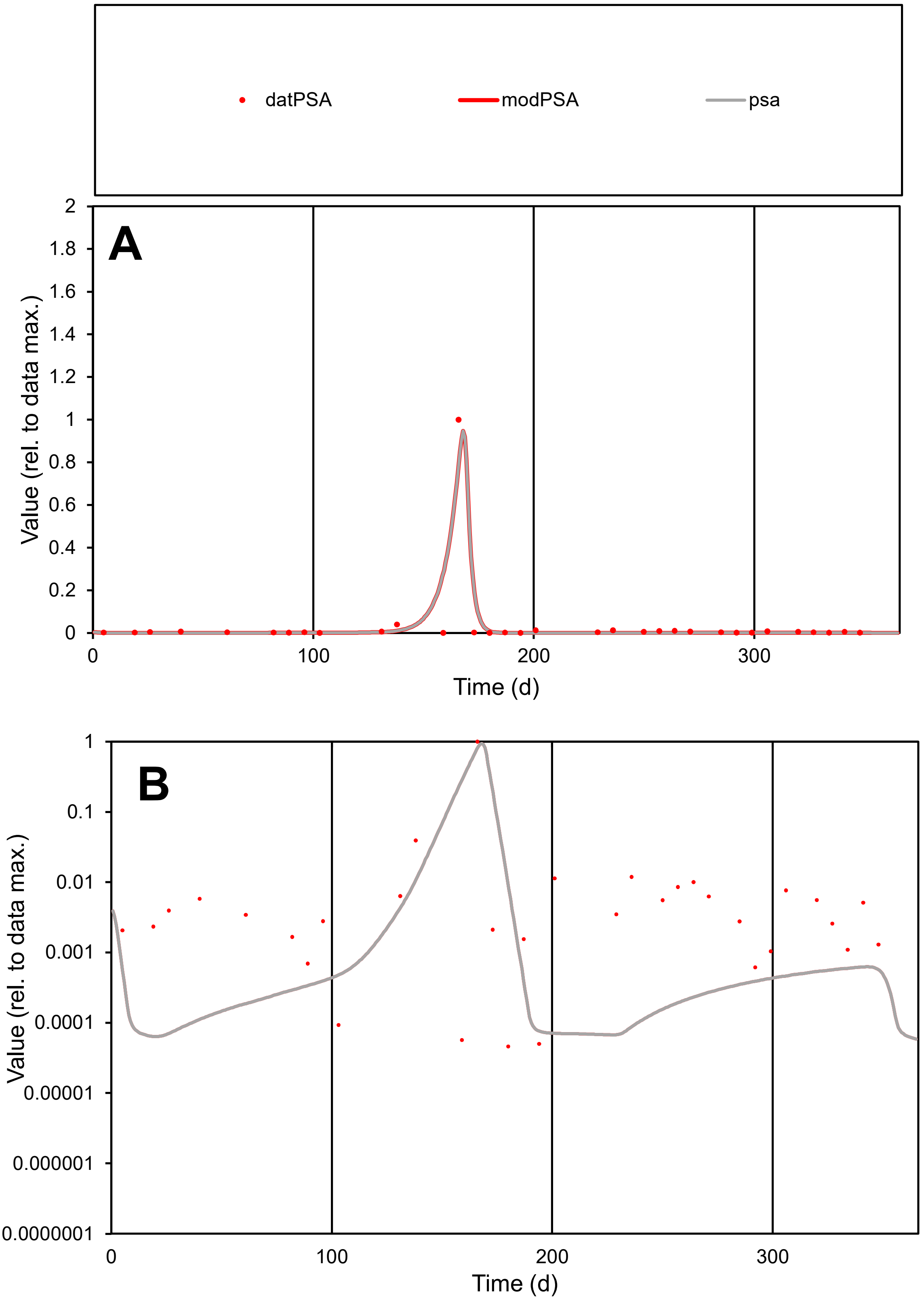


**C1**

Variables
PSA Provide jtd or Dst
CalibOnly?
2 1 to reproduce of
oNormalize?
2 1=no, 2=yes

ofxn calc*
1.16952
ofxn mod
1.42907

sRun
RunA1
zout
1

Ave
Data
0.00684
Model
0.00716

data/vMax
0.1951

ofxng
263.459

Min d/m
0.72623

ave/min d
757.784
ave/min m
575.811

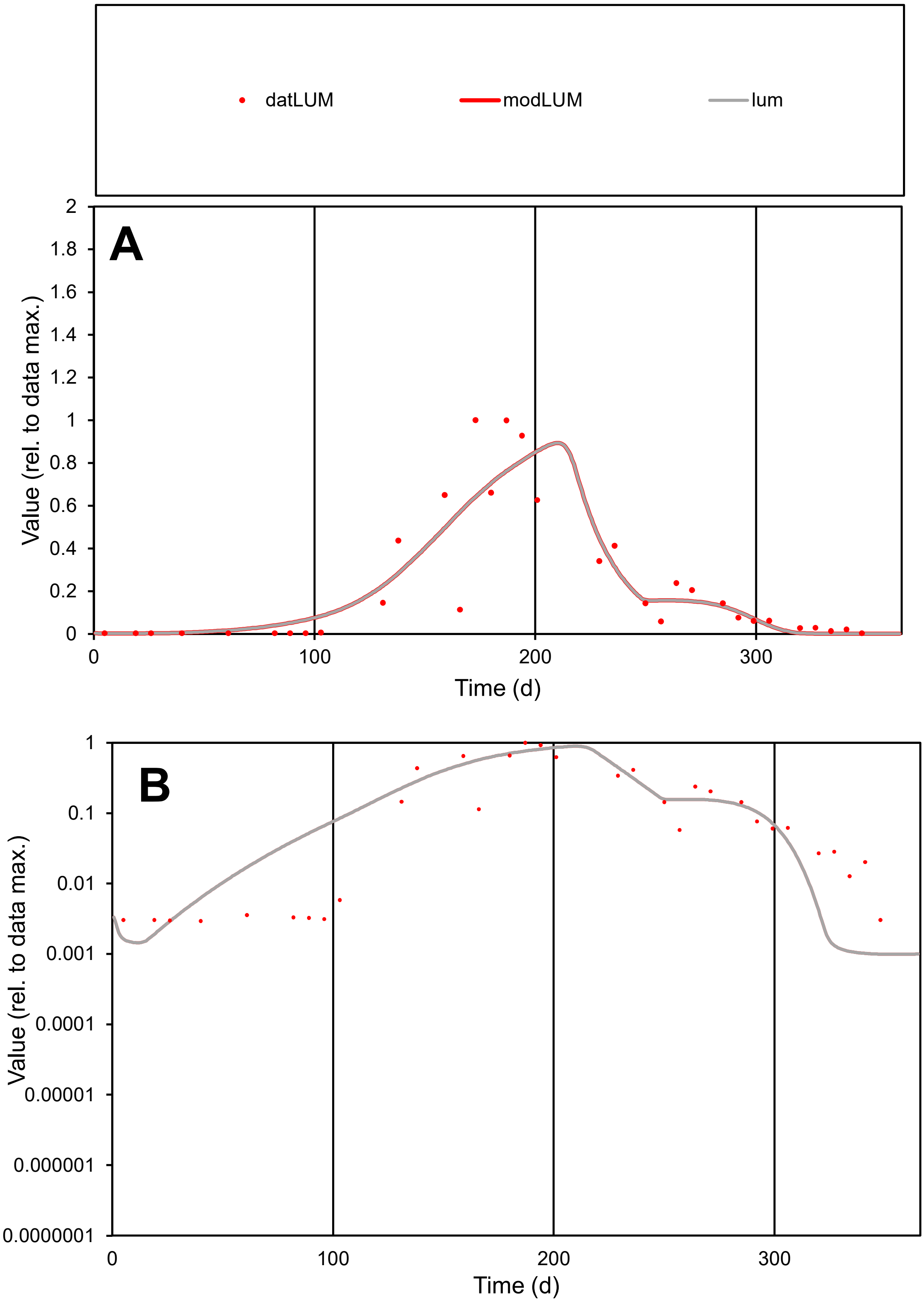


**C1**

Variables

LUM Provide jtd or Dst

CalibOnly?

2 1 to reproduce of

oNormalize?

2 1=no, 2=yes

ofxn calc*

0.57785

ofxn mod

0.60856

sRun

RunA1

zout

1

Ave

Data

0.00354

Model

0.00353

data/vMax

0.01575

ofxng

263.459

Min d/m

2.95156

ave/min d

76.4858

ave/min m

224.816

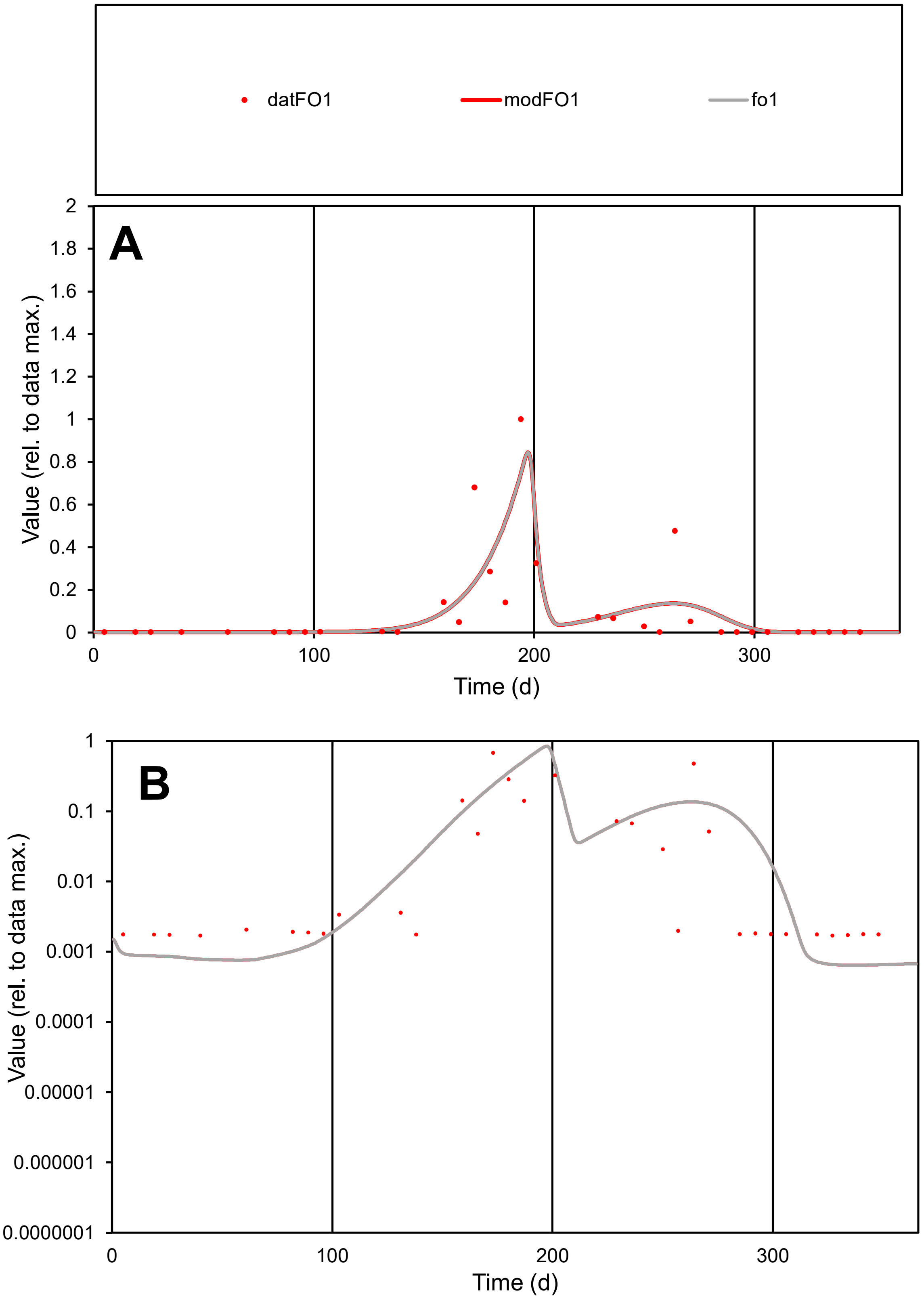


**C1**

Variables

FO1 Provide jtd or Dst

CalibOnly?

2 1 to reproduce of

oNormalize?

2 1=no, 2=yes

ofxn calc*

1.22656

ofxn mod

1.29225

sRun

RunA1

zout

1

Ave

Data

0.0036

Model

0.00359

data/vMax

0.03535

ofxng

263.459

Min d/m

2.63517

ave/min d

59.7386

ave/min m

157.039

326_Calib_S_A

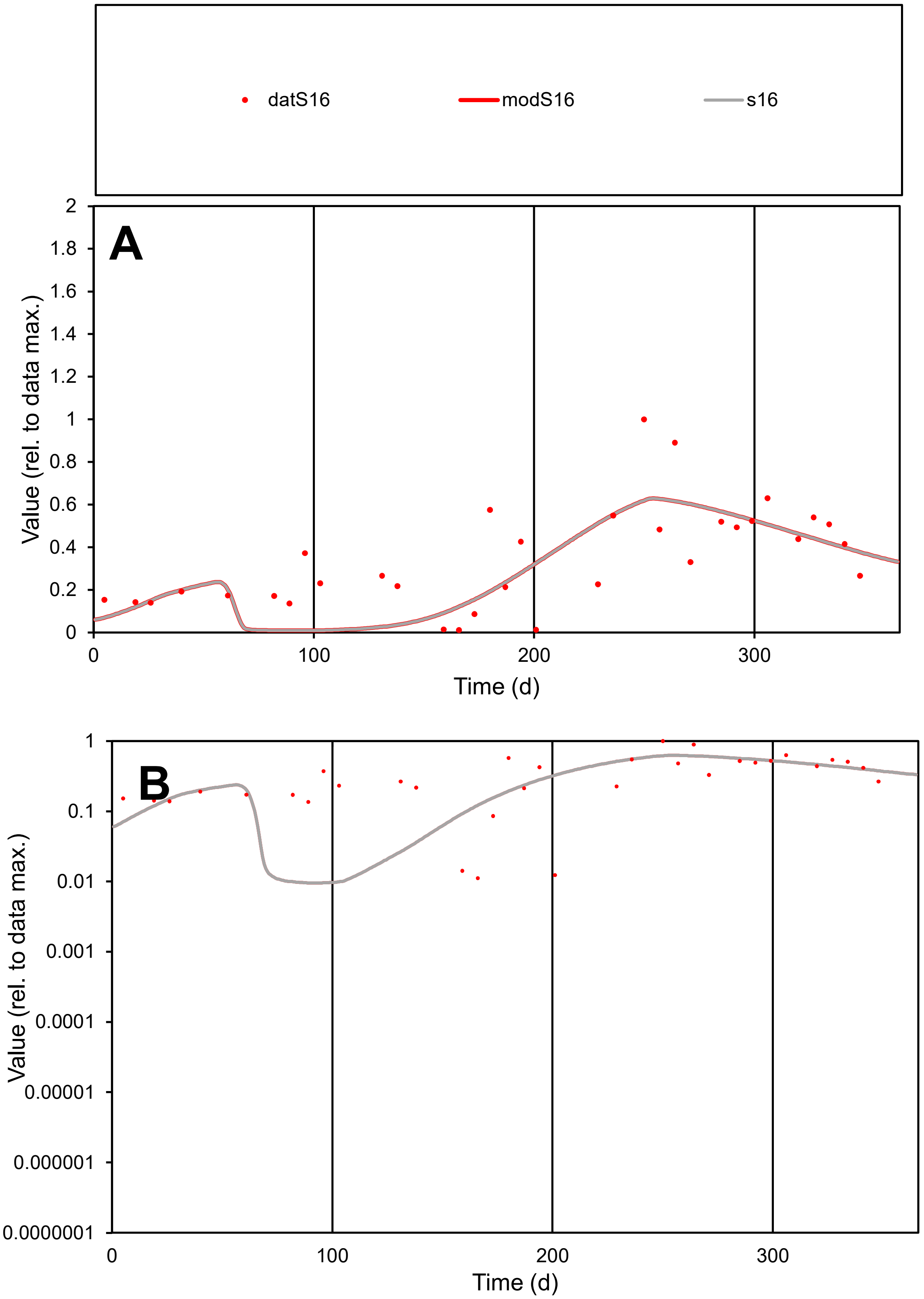


C1

Variables
S16 Provide jtd or Dst
CalibOnly?
2 1 to reproduce of
oNormalize?
2 1=no, 2=yes

ofxn calc*
0.51747
ofxn mod
0.53593

sRun
RunA1
zout
1

Ave
Data
0.00377
Model
0.00333

data/vMax
0.01099

ofxng
263.459

Min d/m
1.17479

ave/min d
30.6819
ave/min m
31.8419

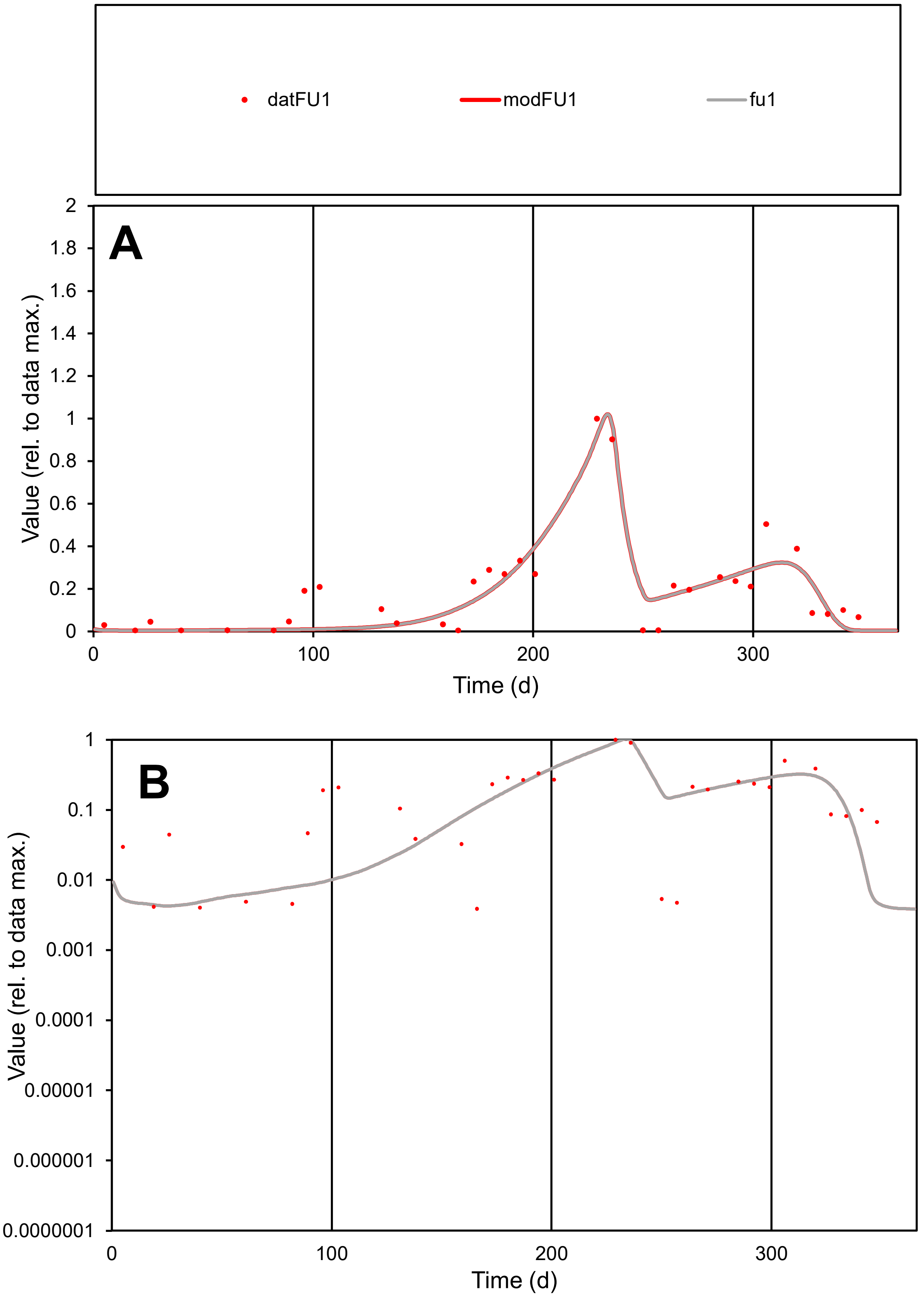


**C1**

Variables
FU1 Provide jtd or Dst
CalibOnly?
2 1 to reproduce of
oNormalize?
2 1=no, 2=yes

ofxn calc*
0.44814
ofxn mod
0.48391

sRun
RunA1
zout
1

Ave
Data
0.00229
Model
0.00213

data/vMax
0.01189

ofxng
263.459

Min d/m
0.90143

ave/min d
49.7227
ave/min m
41.713

326_Calib_S_A

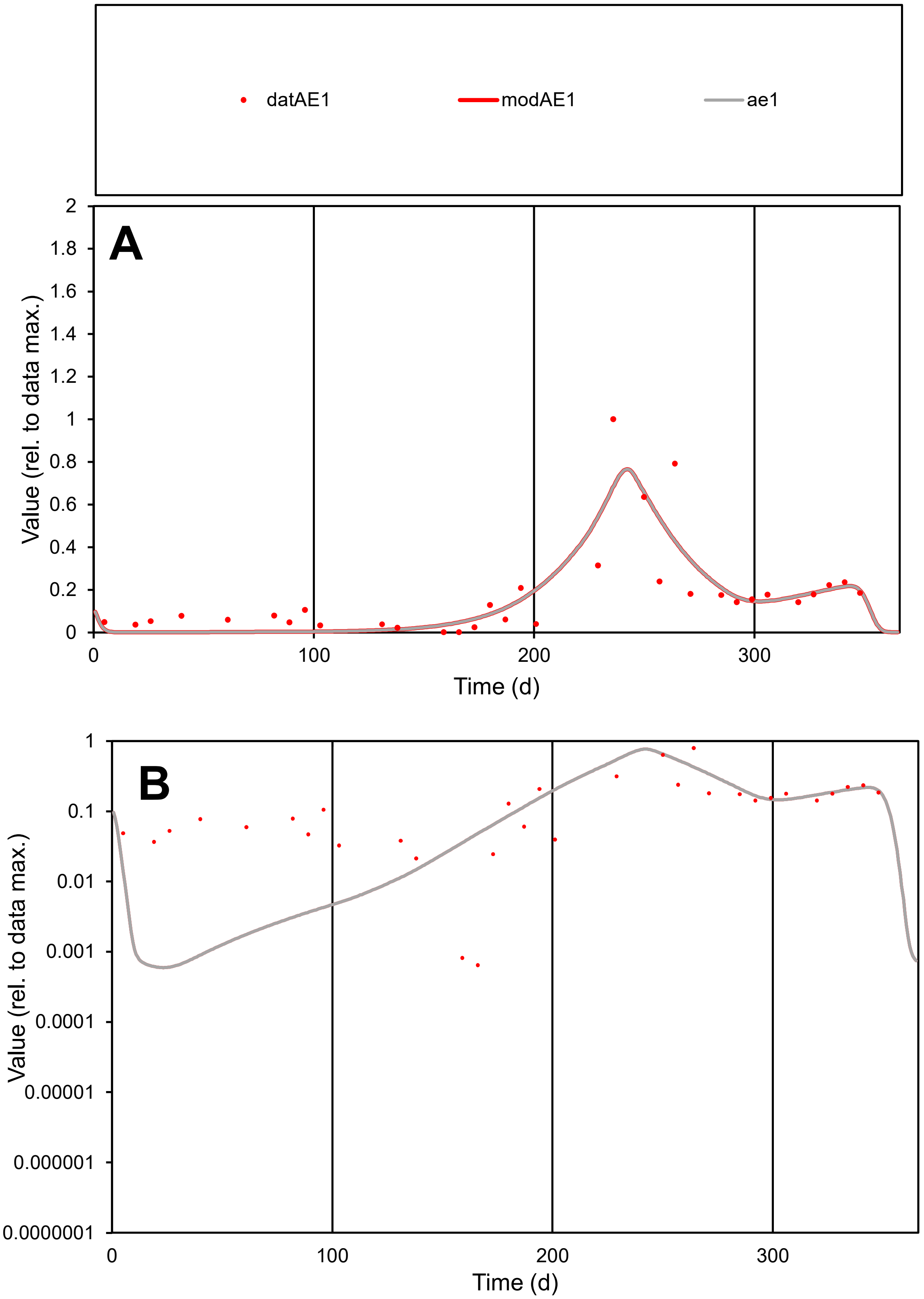


**C1**

Variables
AE1 Provide jtd or Dst
CalibOnly?
2 1 to reproduce of
oNormalize?
2 1=no, 2=yes

ofxn calc*
0.66092
ofxn mod
0.68858

sRun
RunA1
zout
1

Ave
Data
0.00379
Model
0.00362

data/vMax
0.02146

ofxng
263.459

Min d/m
1.06231

ave/min d
274.778
ave/min m
278.224

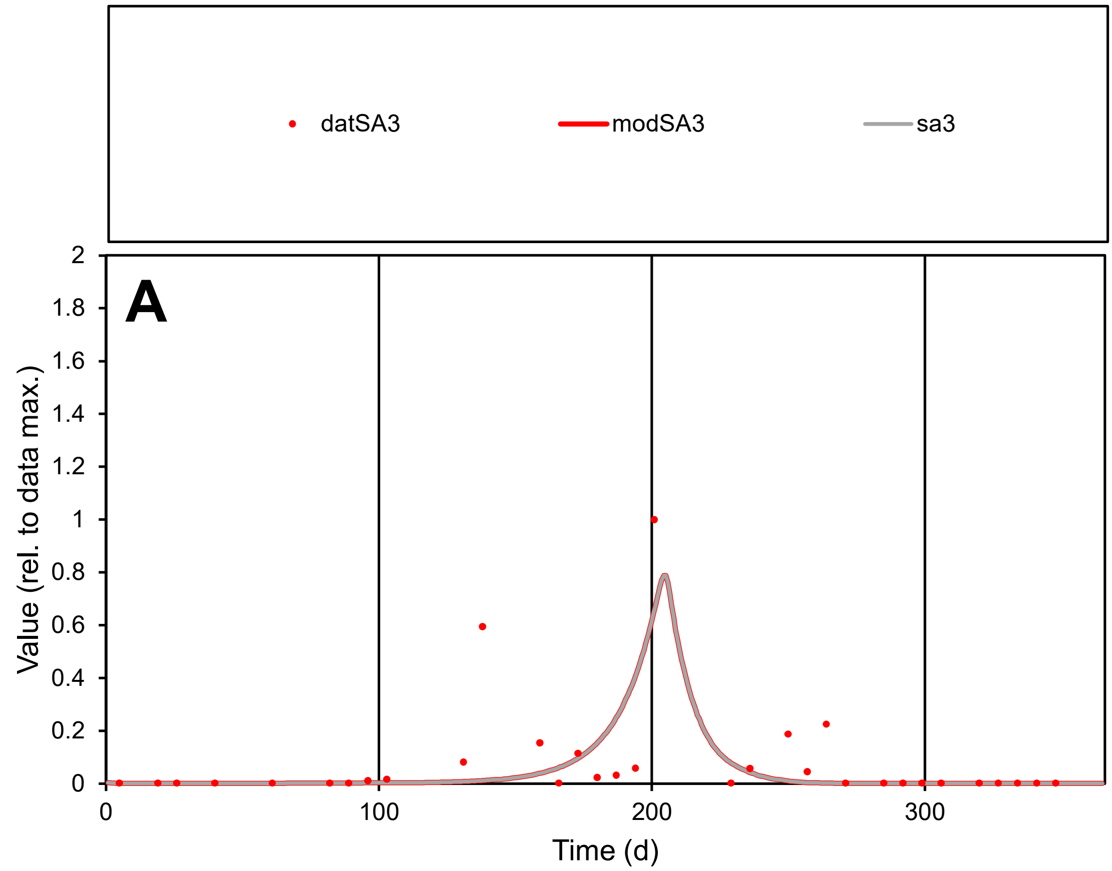


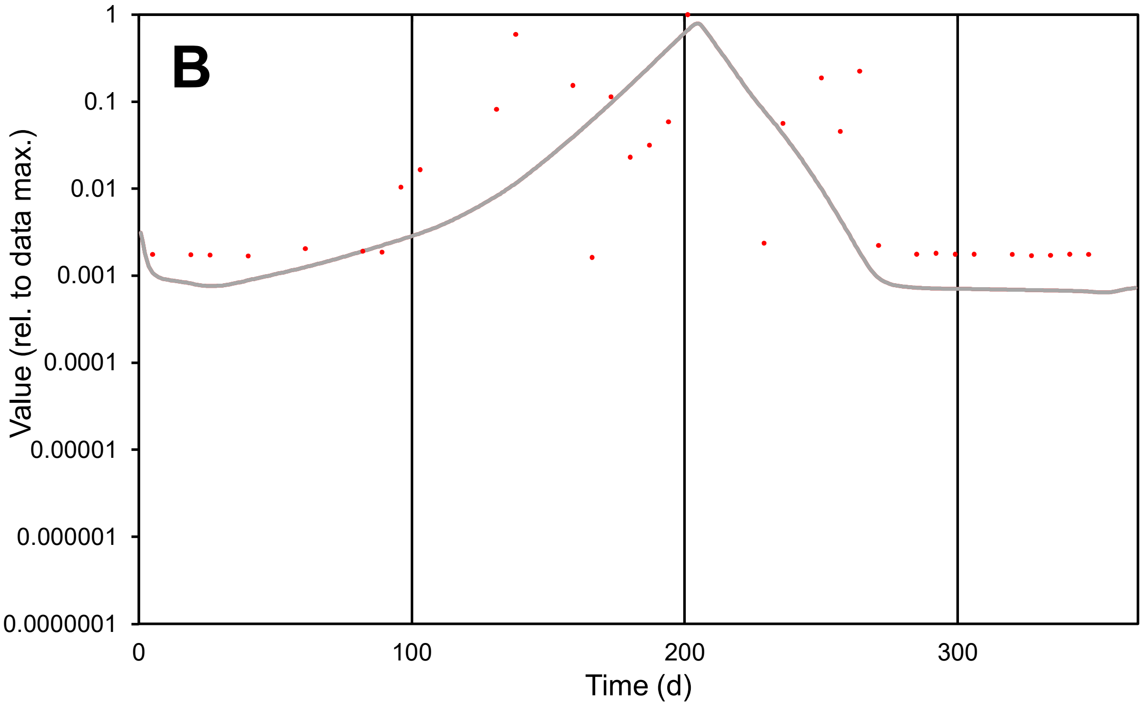


**C1**

Variables
SA3 Provide jtd or Dst
CalibOnly?
2 1 to reproduce of
oNormalize?
2 1=no, 2=yes

ofxn calc*
1.82321
ofxn mod
1.86147

sRun
RunA1
zout
1

Ave
Data
0.00479
Model
0.00338

data/vMax
0.06005

ofxng
263.459

Min d/m
2.46915

ave/min d
49.2467
ave/min m
85.7361

326_Calib_S_A

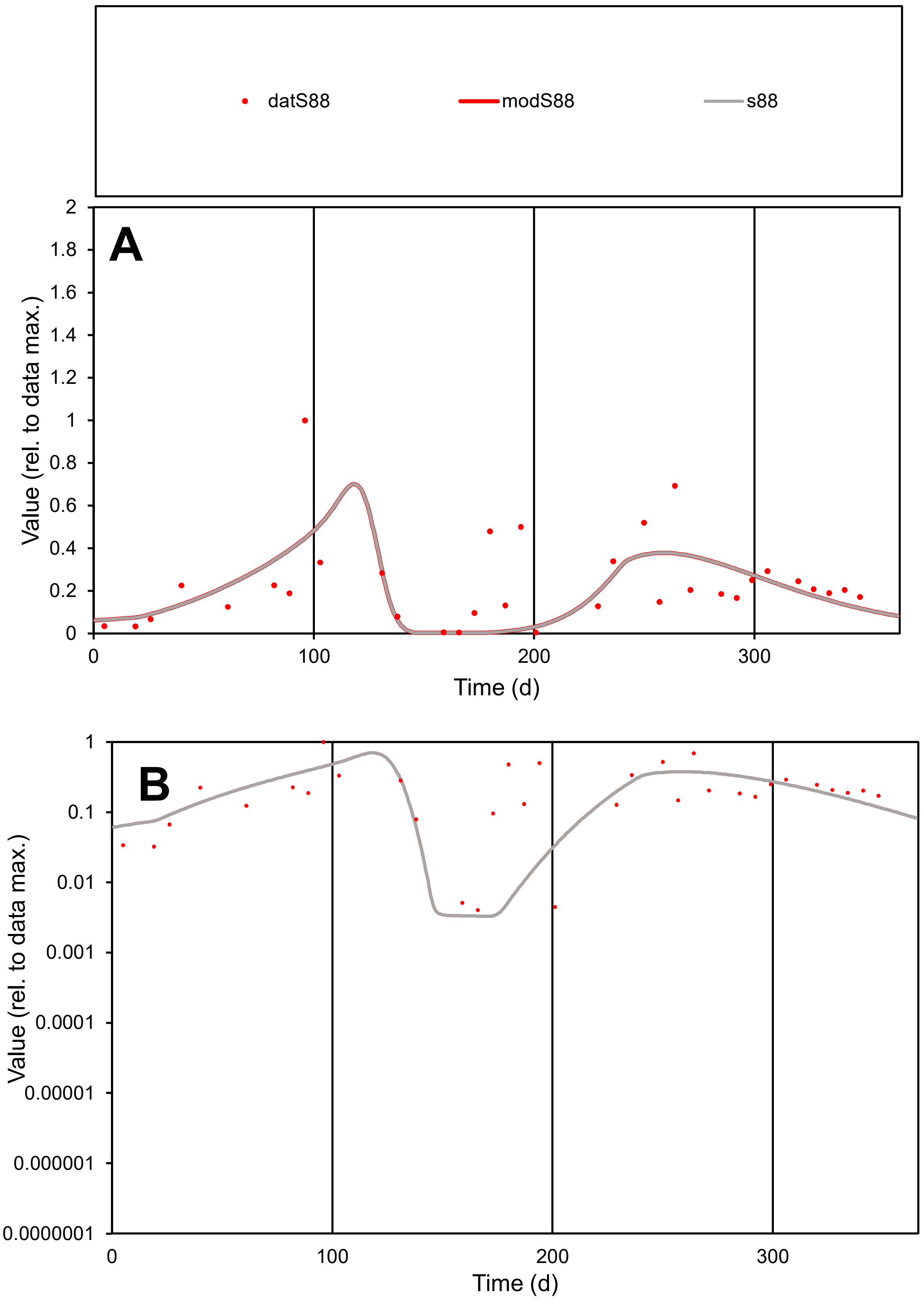


**C1**

Variables

| | |
|---|---|
| S88 | Provide jtd or Dst |

CalibOnly?

| | |
|---|---|
| 2 | 1 to reproduce of |

oNormalize?

| | |
|---|---|
| 2 | 1=no, 2=yes |

ofxn calc*
0.79419
ofxn mod
0.80558

sRun
RunA1
zout
1

Ave
Data
0.00331
Model
0.00283

data/vMax
0.01411

ofxng
263.459

Min d/m
1.22011

ave/min d
57.9764
ave/min m
60.432

326_Calib_S_A

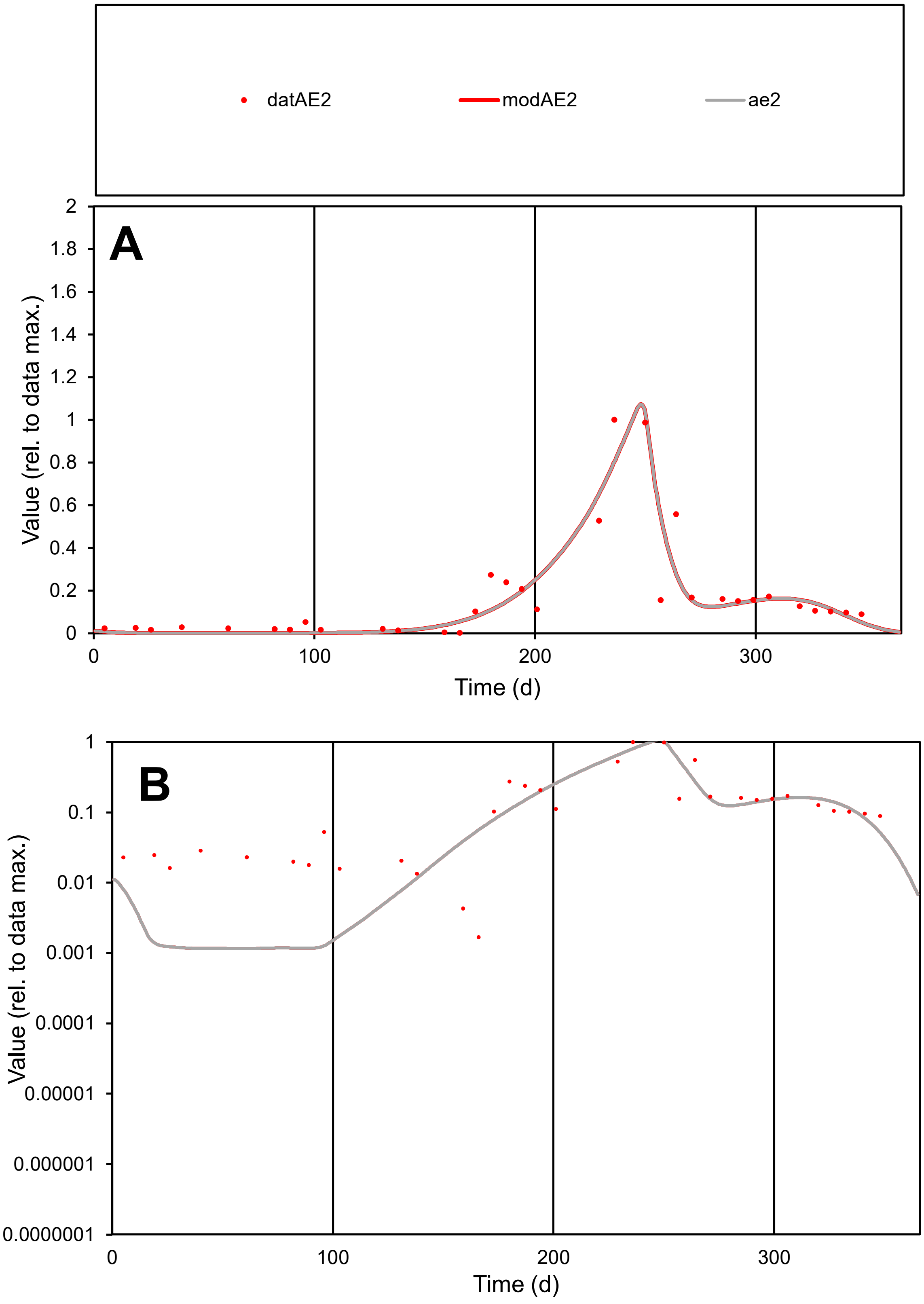


**C1**

Variables

AE2 Provide jtd or Dst

CalibOnly?

2 1 to reproduce of

oNormalize?

2 1=no, 2=yes

ofxn calc*

0.53876

ofxn mod

0.5947

sRun

RunA1

zout

1

Ave

Data

0.00593

Model

0.00551

data/vMax

0.03404

ofxng

263.459

Min d/m

1.45946

ave/min d

103.648

ave/min m

140.611

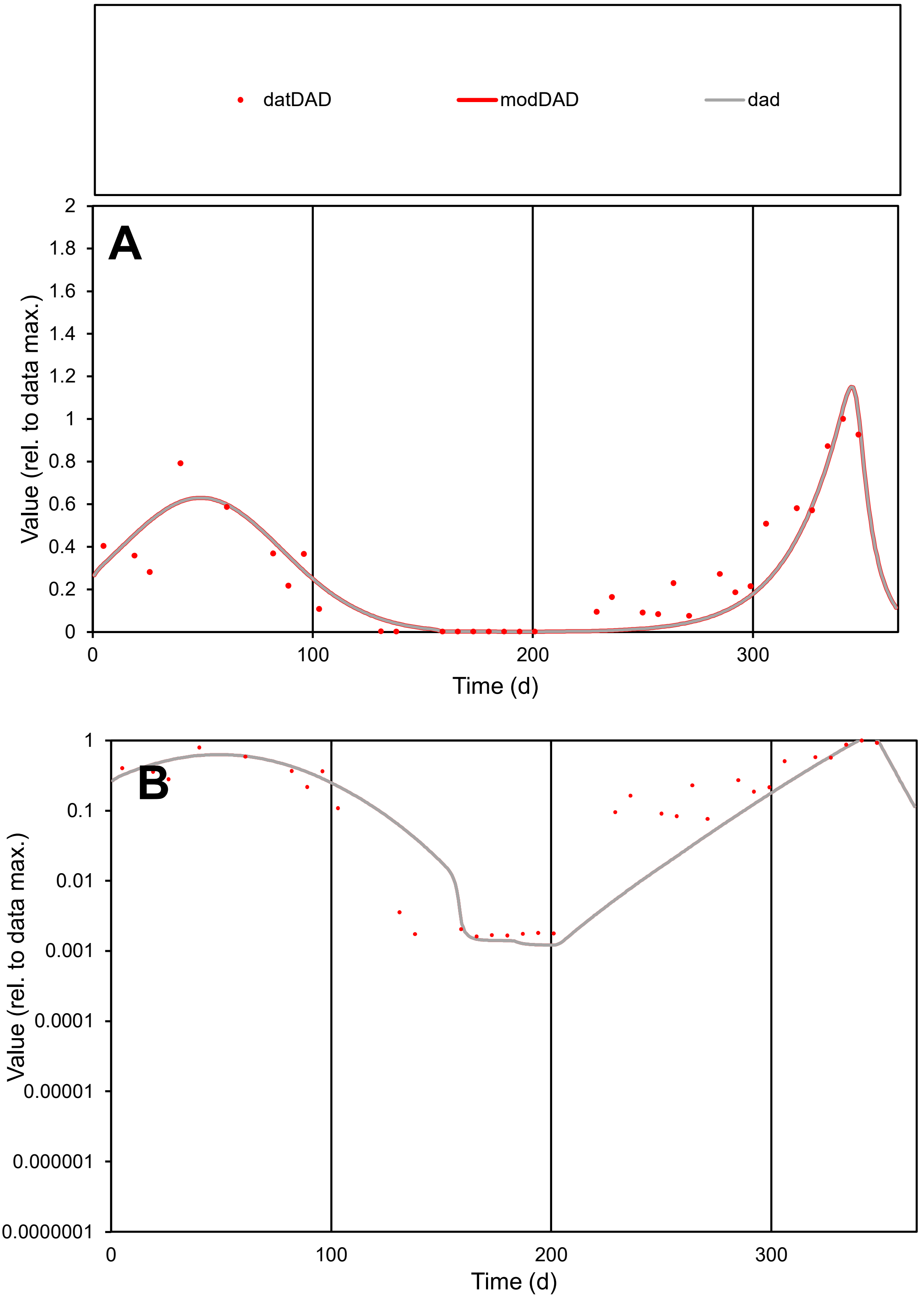


**C1**

Variables
DAD Provide jtd or Dst
CalibOnly?
2 1 to reproduce of
oNormalize?
2 1=no, 2=yes

ofxn calc*
0.36533
ofxn mod
0.39811

sRun
RunA1
zout
1

Ave
Data
0.00269
Model
0.00247

data/vMax
0.00946

ofxng
263.459

Min d/m
1.32239

ave/min d
175.606
ave/min m
213.537

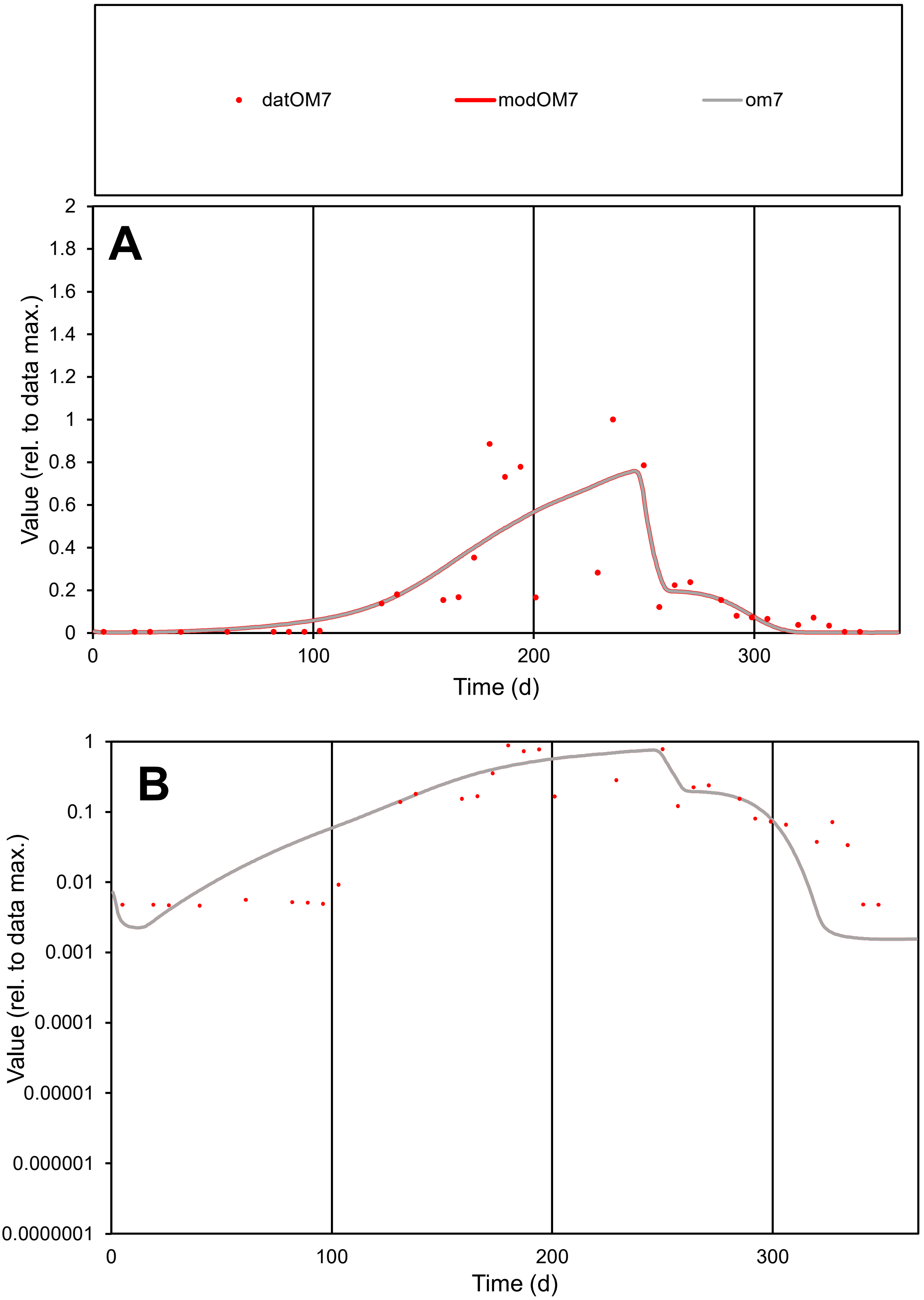


**C1**

Variables
OM7 Provide jtd or Dst
CalibOnly?
2 1 to reproduce of
oNormalize?
2 1=no, 2=yes

ofxn calc*
0.77604
ofxn mod
0.80176

sRun
RunA1
zout
1

Ave
Data
0.00429
Model
0.00429

data/vMax
0.02086

ofxng
263.459

Min d/m
2.99072

ave/min d
44.3389
ave/min m
132.638

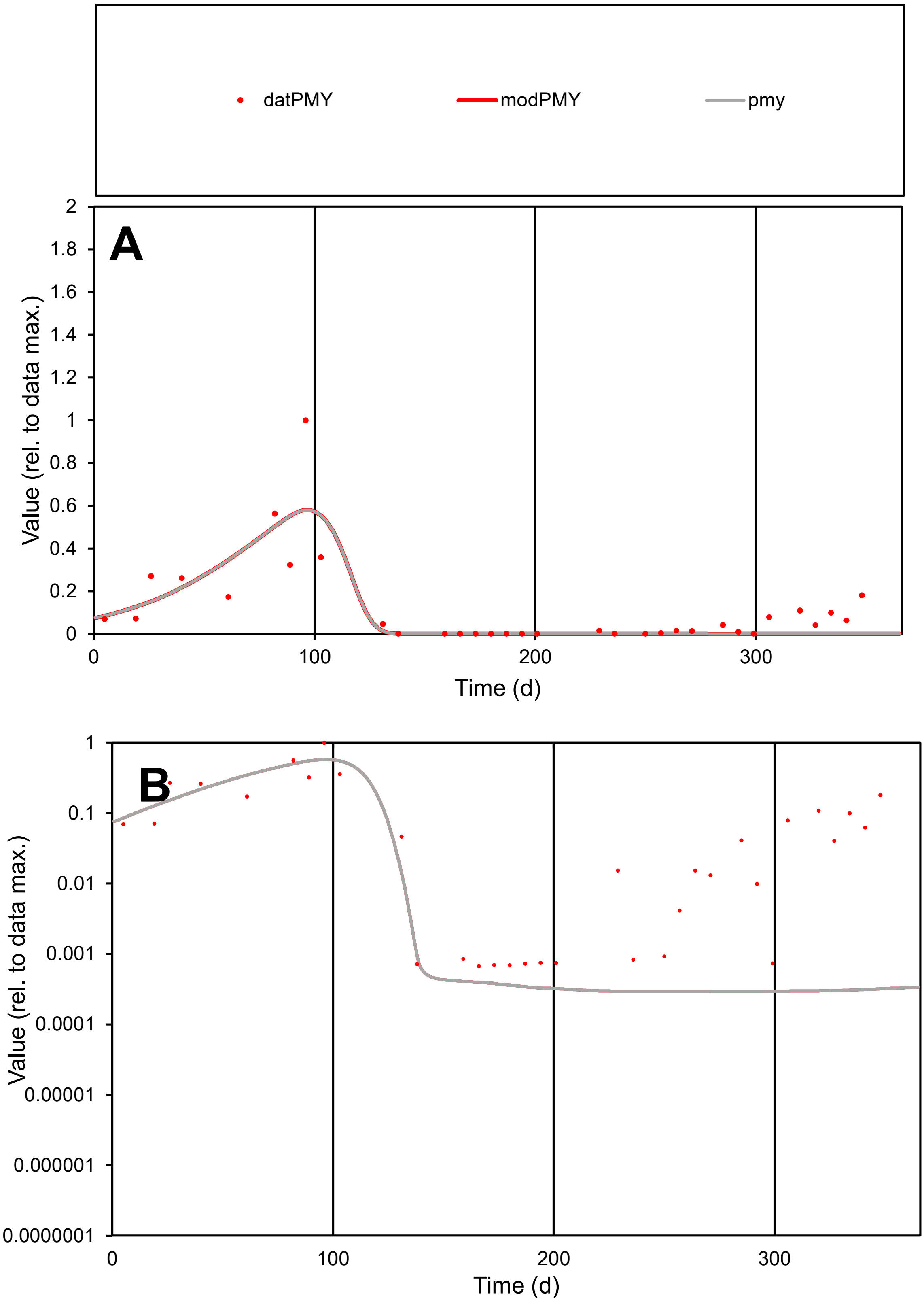


**C1**

Variables

PMY Provide jtd or Dst

CalibOnly?

2 1 to reproduce of

oNormalize?

2 1=no, 2=yes

ofxn calc*

0.93631

ofxn mod

0.95806

sRun

RunA1

zout

1

Ave

Data

0.00164

Model

0.00137

data/vMax

0.01422

ofxng

263.459

Min d/m

2.28997

ave/min d

172.469

ave/min m

329.001

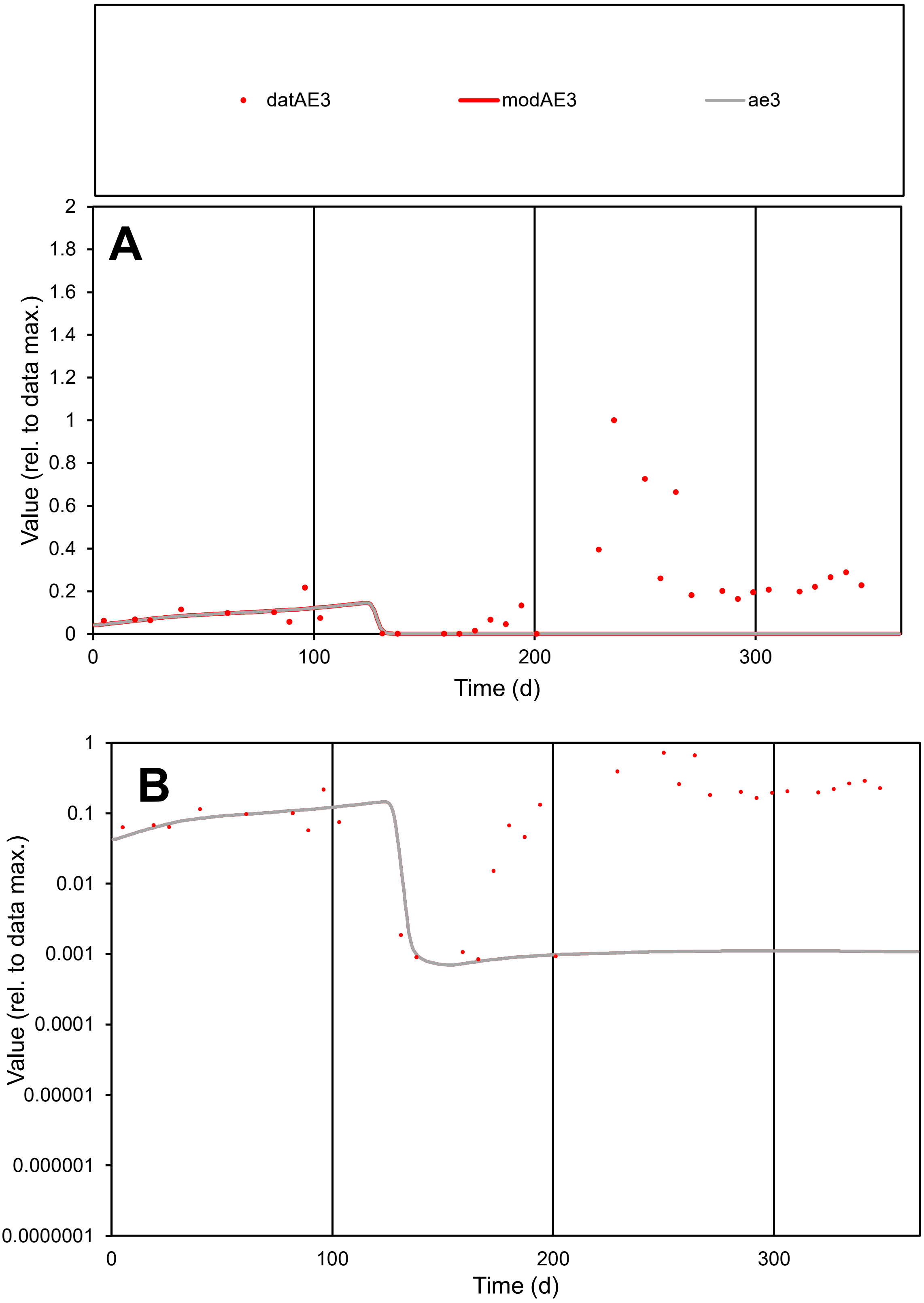


**C1**

Variables

| | |
|---|---|
| AE3 | Provide jtd or Dst |
| CalibOnly? | |
| 2 | 1 to reproduce of |
| oNormalize? | |
| 2 | 1=no, 2=yes |
| ofxn calc* | |
| 1.49246 | |
| ofxn mod | |
| 1.49521 | |
| sRun | |
| RunA1 | |
| zout | |
| 1 | |
| Ave Data | |
| 0.00378 | |
| Model | |
| 0.00052 | |
| data/vMax | |
| 0.01973 | |
| ofxng | |
| 263.459 | |
| Min d/m | |
| 1.14178 | |
| ave/min d | |
| 226.639 | |
| ave/min m | |
| 35.4391 | |

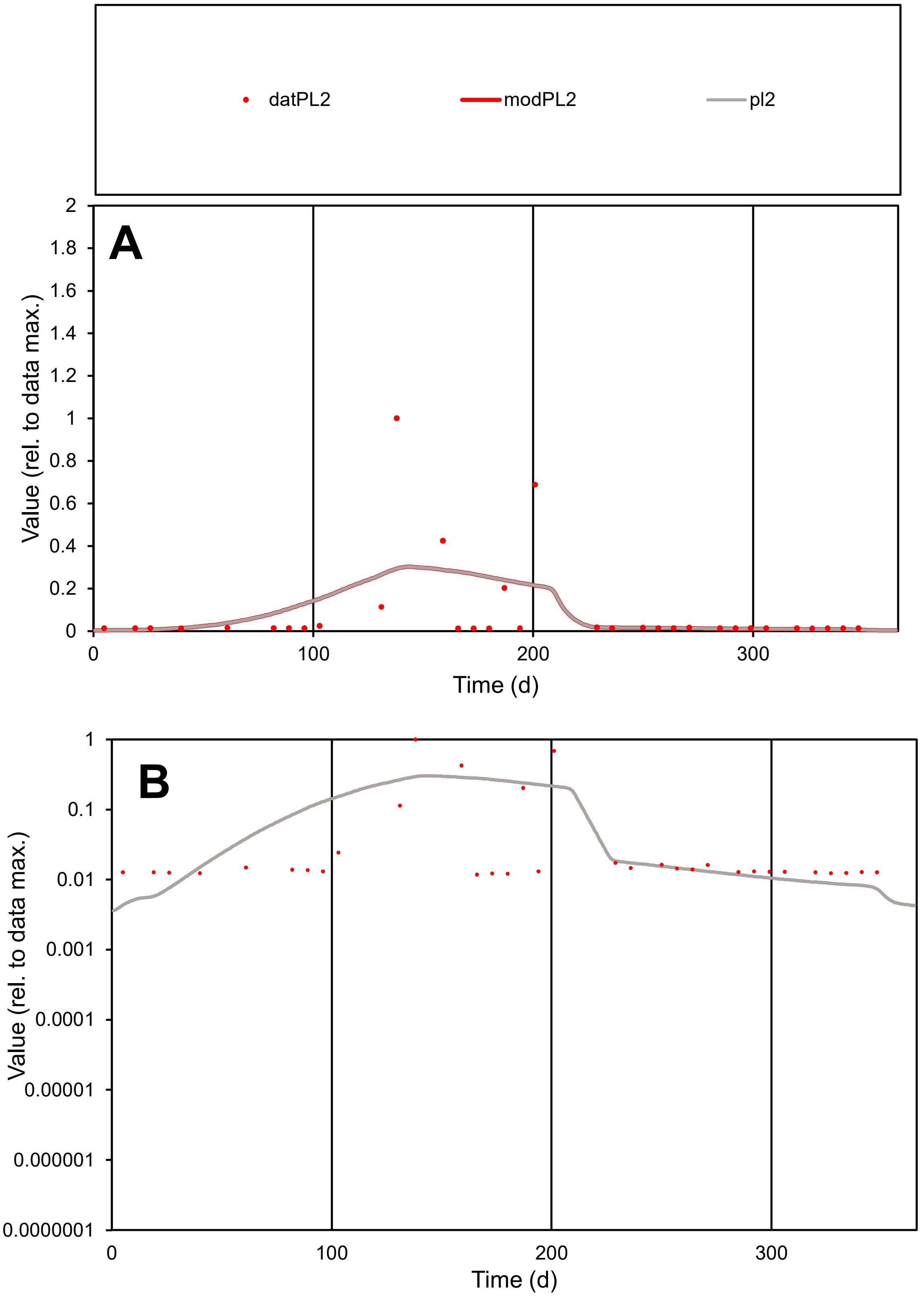


**C1**

Variables
PL2 Provide jtd or Dst
CalibOnly?
2 1 to reproduce of
oNormalize?
2 1=no, 2=yes

ofxn calc*
2.08329
ofxn mod
2.09664

sRun
RunA1
zout
1

Ave
Data
0.00117
Model
0.00126

data/vMax
0.01369

ofxng
263.459

Min d/m
2.5591

ave/min d
7.24428
ave/min m
20.0662

326_Calib_S_A

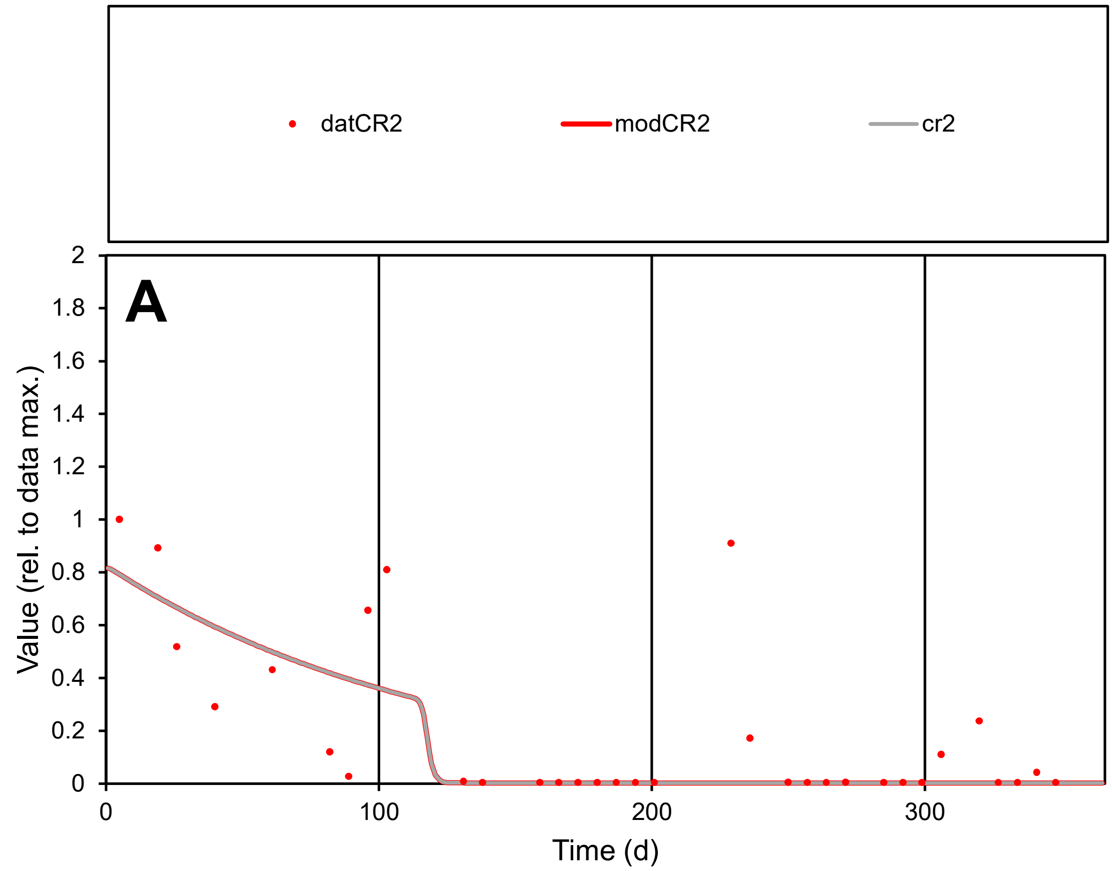


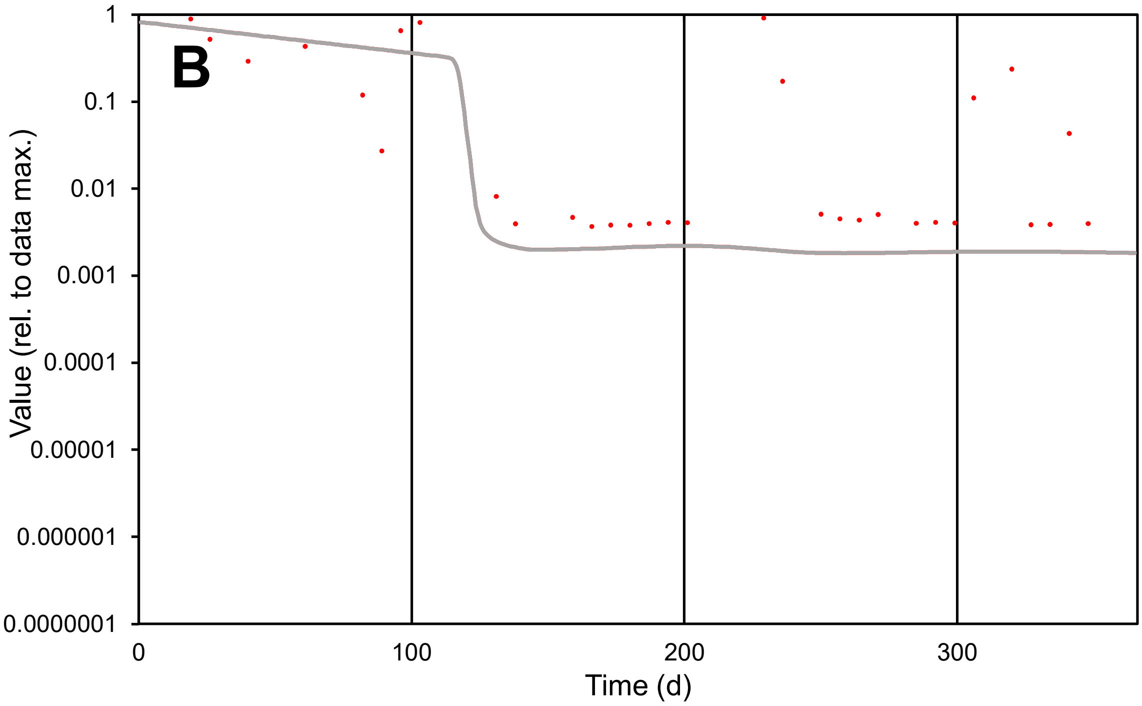


**C1**

Variables
CR2 Provide jtd or Dst
CalibOnly?
2 1 to reproduce of
oNormalize?
2 1=no, 2=yes

ofxn calc*
1.16429
ofxn mod
1.18034

sRun
RunA1
zout
1

Ave
Data
0.00205
Model
0.00156

data/vMax
0.01075

ofxng
263.459

Min d/m
2.01479

ave/min d
51.9507
ave/min m
79.5404

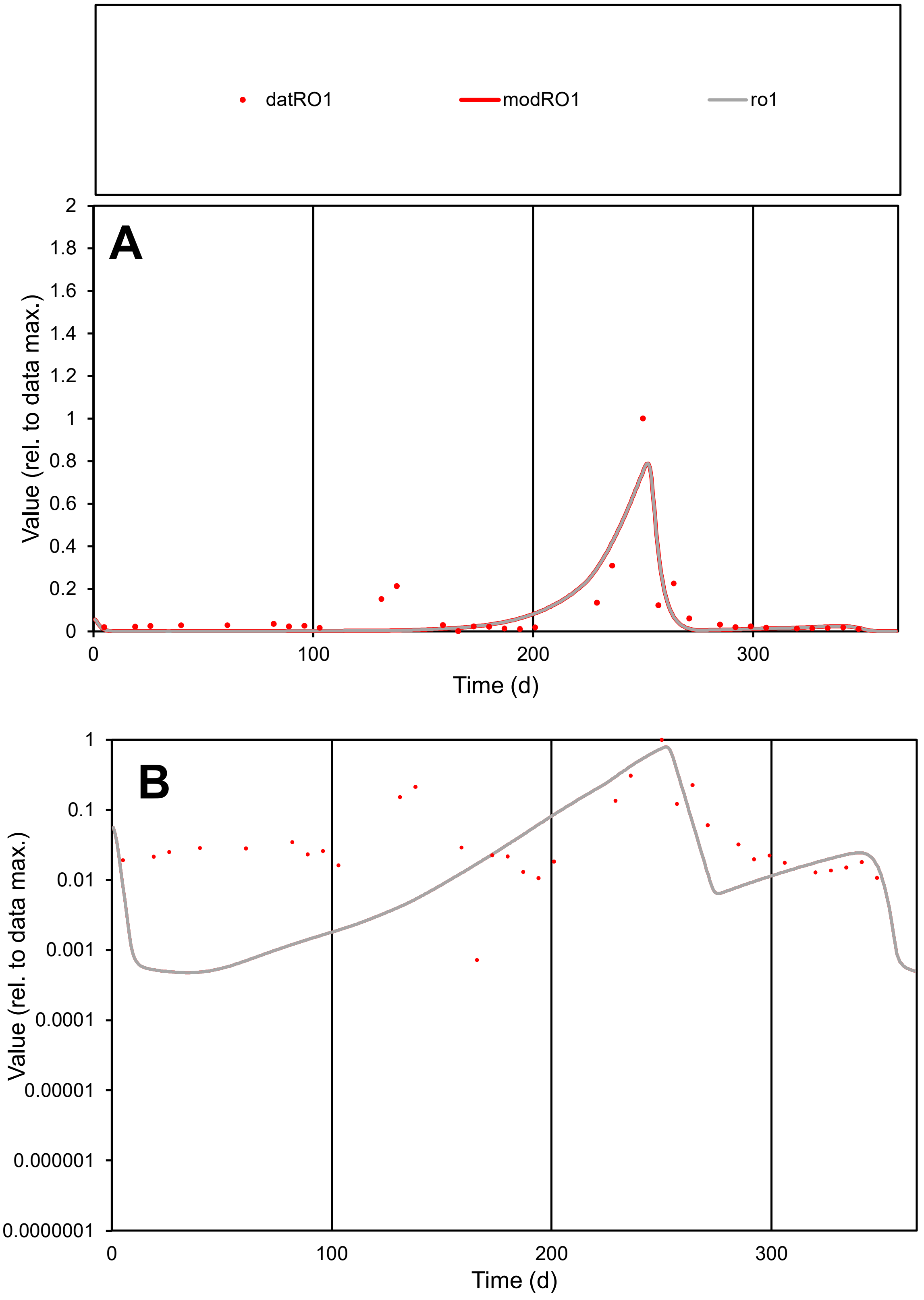


**C1**

Variables

RO1 Provide jtd or Dst

CalibOnly?

2 1 to reproduce of

oNormalize?

2 1=no, 2=yes

ofxn calc*

0.93296

ofxn mod

1.02102

sRun

RunA1

zout

1

Ave

Data

0.00399

Model

0.00331

data/vMax

0.04848

ofxng

263.459

Min d/m

1.48475

ave/min d

113.904

ave/min m

140.358

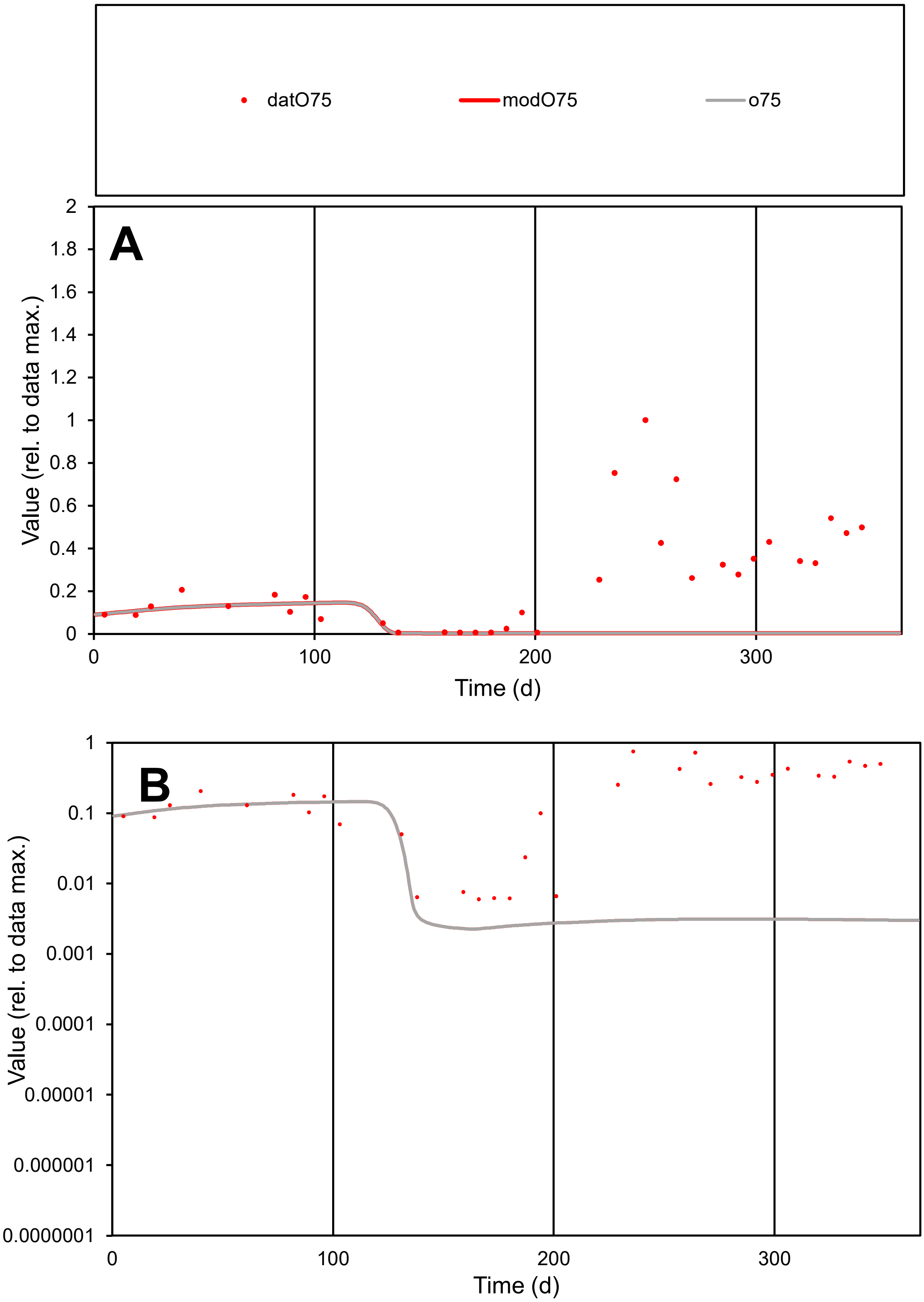


**C1**

Variables
O75 Provide jtd or Dst
CalibOnly?
2 1 to reproduce of
oNormalize?
2 1=no, 2=yes

ofxn calc*
1.35029
ofxn mod
1.3532

sRun
RunA1
zout
1

Ave
Data
0.00295
Model
0.00044

data/vMax
0.01163

ofxng
263.459

Min d/m
2.62798

ave/min d
42.4317
ave/min m
16.6998

326_Calib_S_A

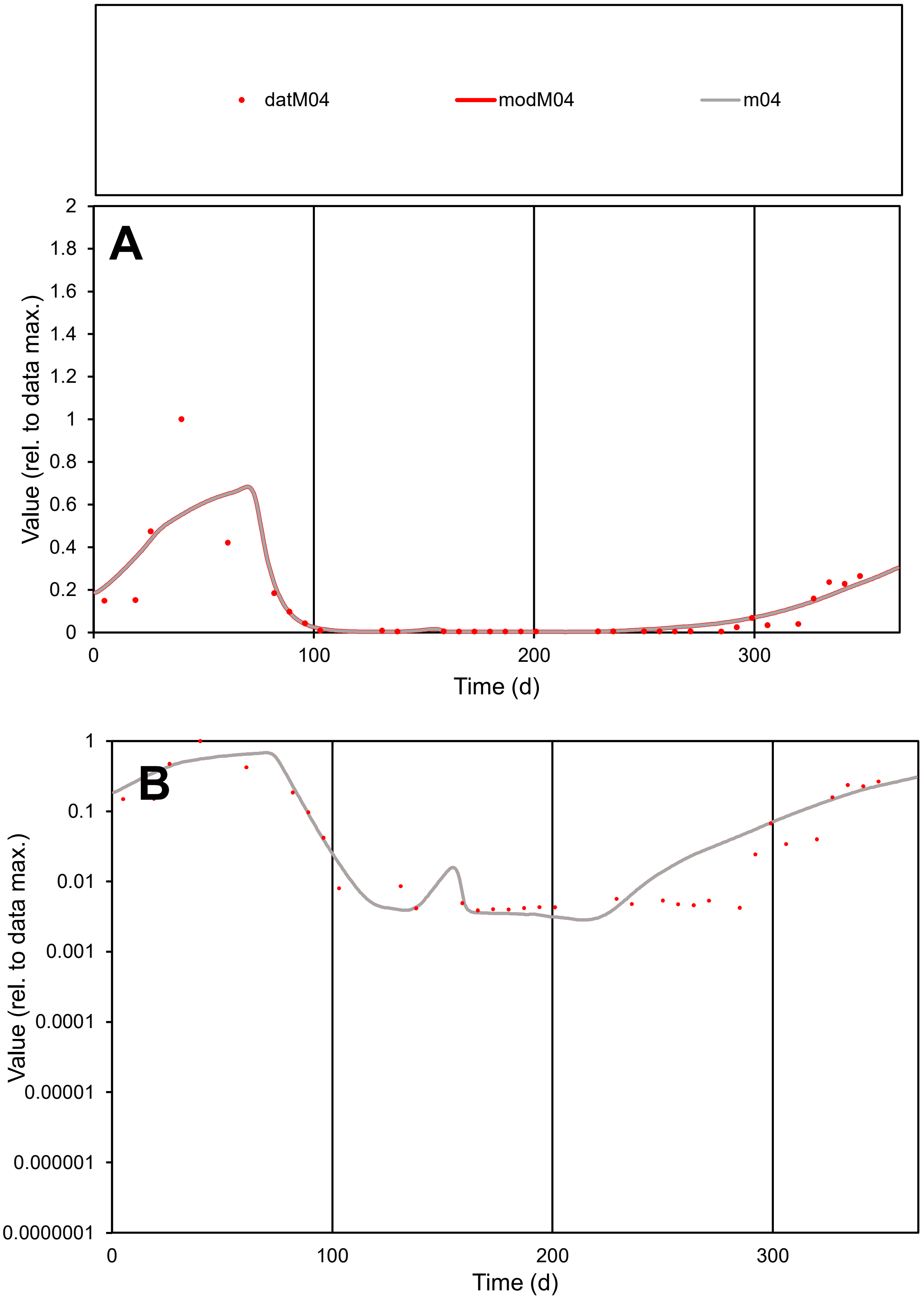


**C1**

Variables
M04 Provide jtd or Dst
CalibOnly?
2 1 to reproduce of
oNormalize?
2 1=no, 2=yes

ofxn calc*
0.87803
ofxn mod
0.91669

sRun
RunA1
zout
1

Ave
Data
0.001
Model
0.00105

data/vMax
0.00902

ofxng
263.459

Min d/m
1.24679

ave/min d
28.6225
ave/min m
37.5434

326_Calib_S_A

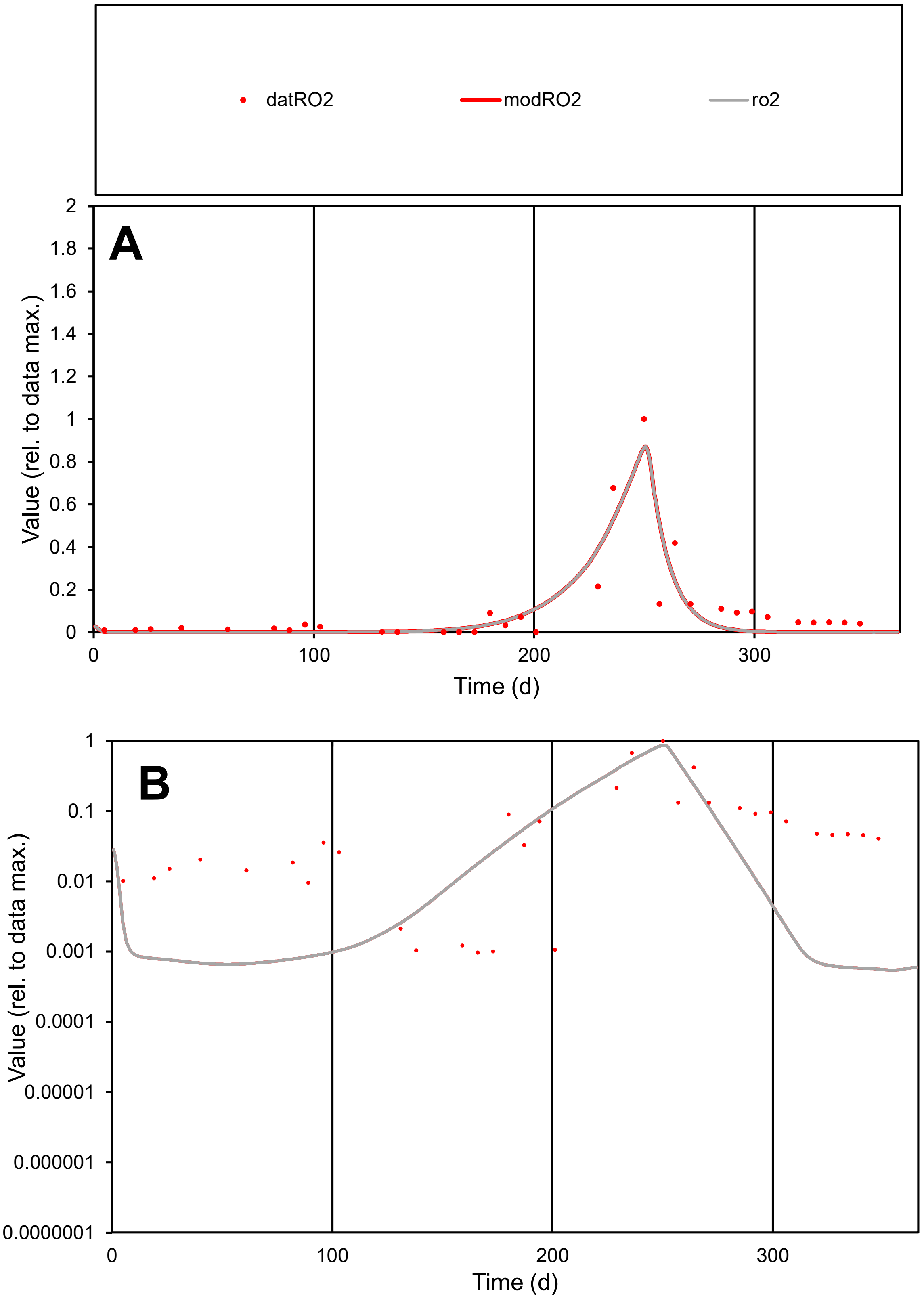


**C1**

Variables
RO2 Provide jtd or Dst
CalibOnly?
2 1 to reproduce of
oNormalize?
2 1=no, 2=yes

ofxn calc*
0.77187
ofxn mod
0.85318

sRun
RunA1
zout
1

Ave
Data
0.00423
Model
0.00349

data/vMax
0.03943

ofxng
263.459

Min d/m
1.72711

ave/min d
110.958
ave/min m
158.098

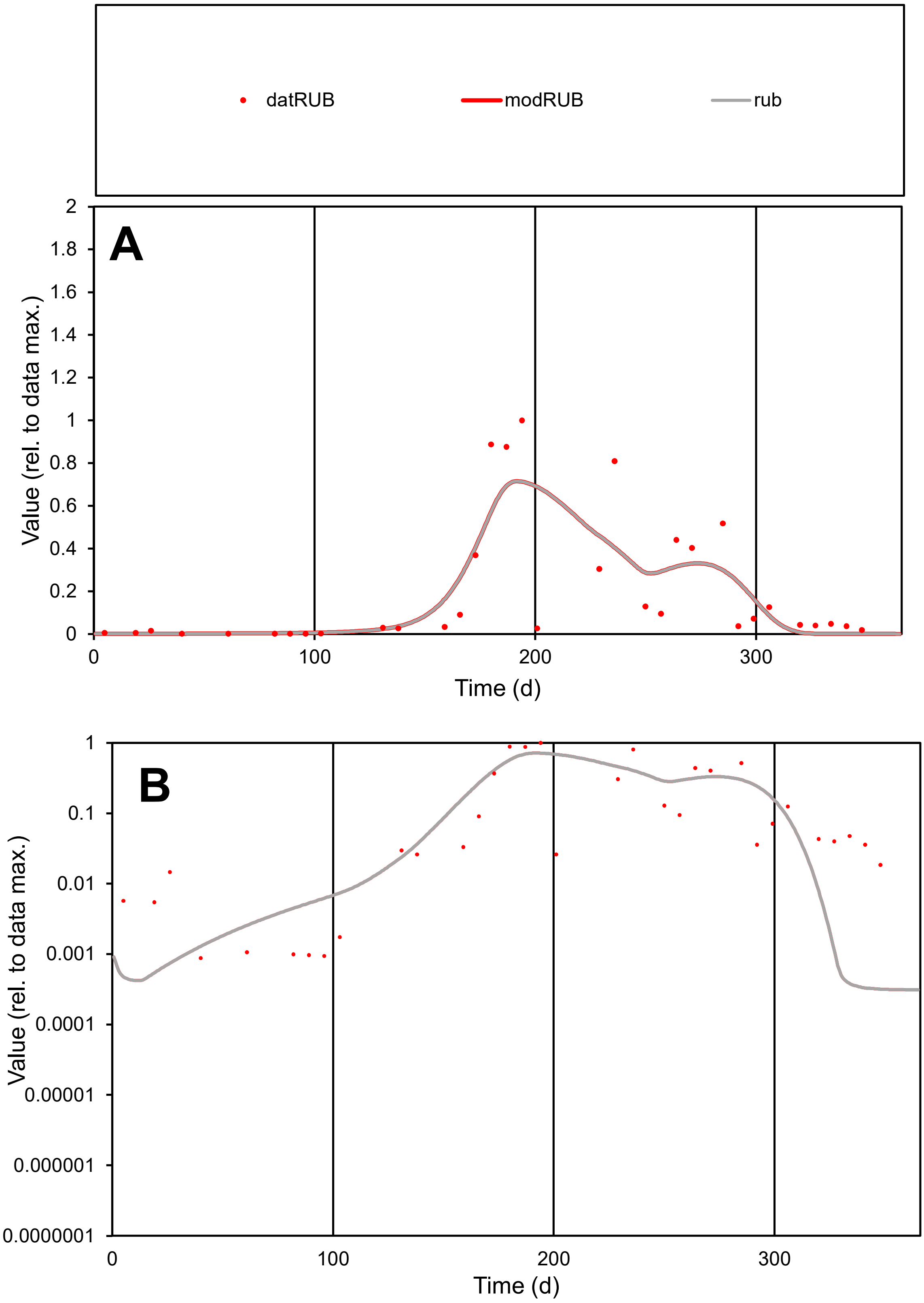


**C1**

Variables
RUB Provide jtd or Dst
CalibOnly?
2 1 to reproduce of
oNormalize?
2 1=no, 2=yes

ofxn calc*
0.90766
ofxn mod
0.93452

sRun
RunA1
zout
1

Ave
Data
0.0023
Model
0.0023

data/vMax
0.01173

ofxng
263.459

Min d/m
2.78092

ave/min d
223.27
ave/min m
618.893

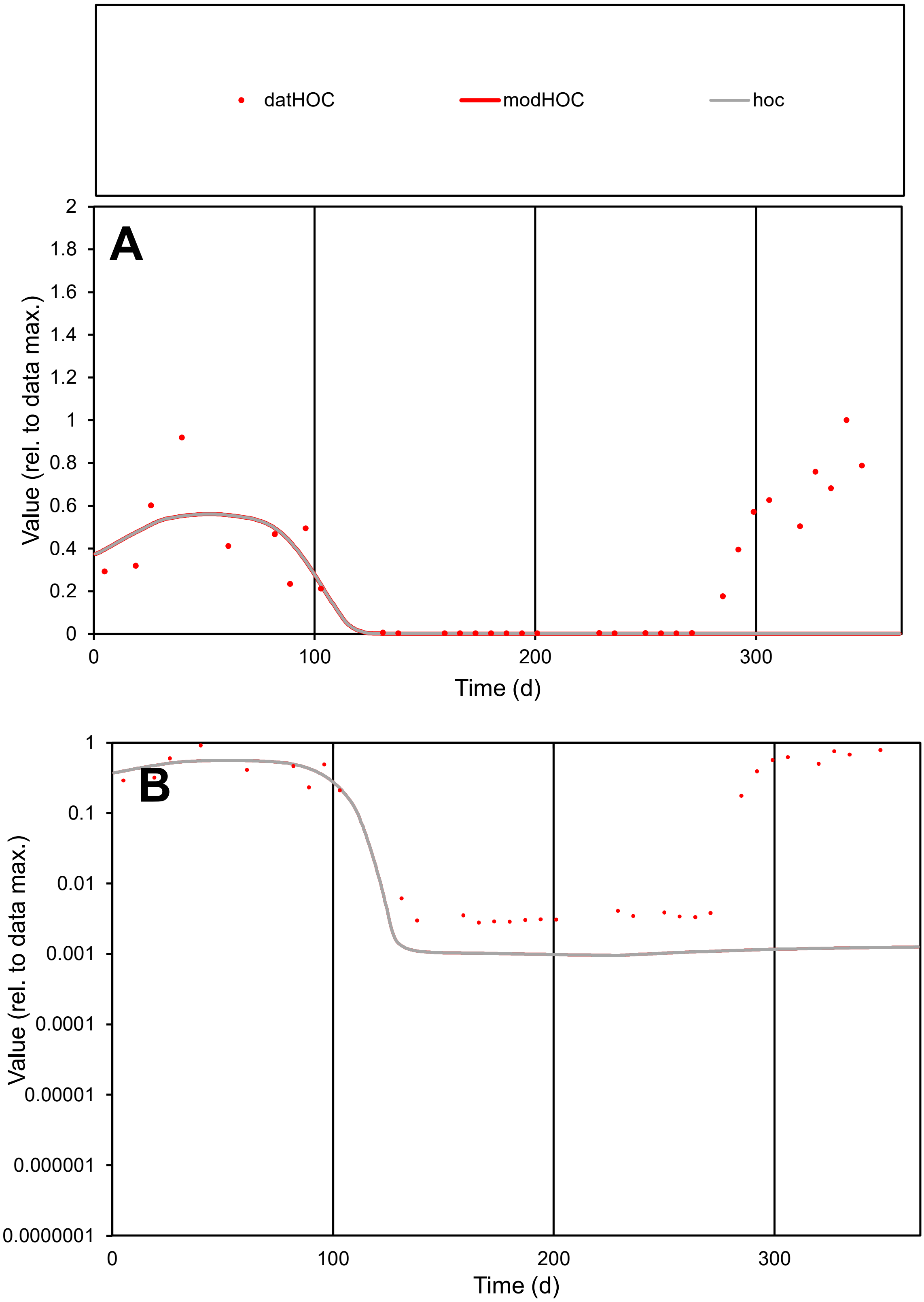


C1

Variables
HOC Provide jtd or Dst
CalibOnly?
2 1 to reproduce of
oNormalize?
2 1=no, 2=yes

ofxn calc*
1.22002
ofxn mod
1.22634

sRun
RunA1
zout
1

Ave
Data
0.00129
Model
0.00054

data/vMax
0.00447

ofxng
263.459

Min d/m
2.92174

ave/min d
103.182
ave/min m
125.94

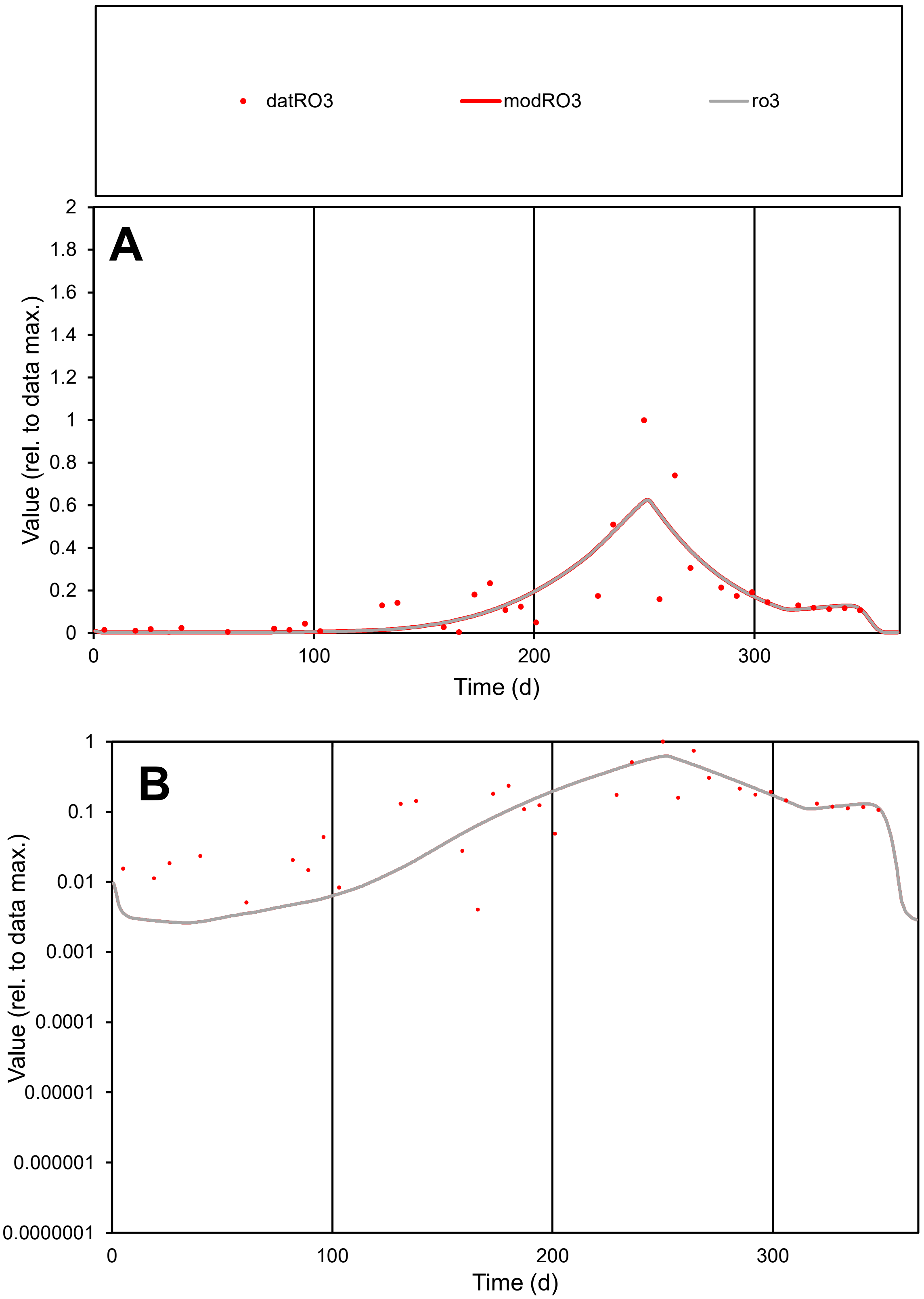


**C1**

Variables

RO3 Provide jtd or Dst

CalibOnly?

2 1 to reproduce of

oNormalize?

2 1=no, 2=yes

ofxn calc*

0.76831

ofxn mod

0.79496

sRun

RunA1

zout

1

Ave

Data

0.00384

Model

0.00368

data/vMax

0.02365

ofxng

263.459

Min d/m

1.51263

ave/min d

40.3002

ave/min m

58.4116

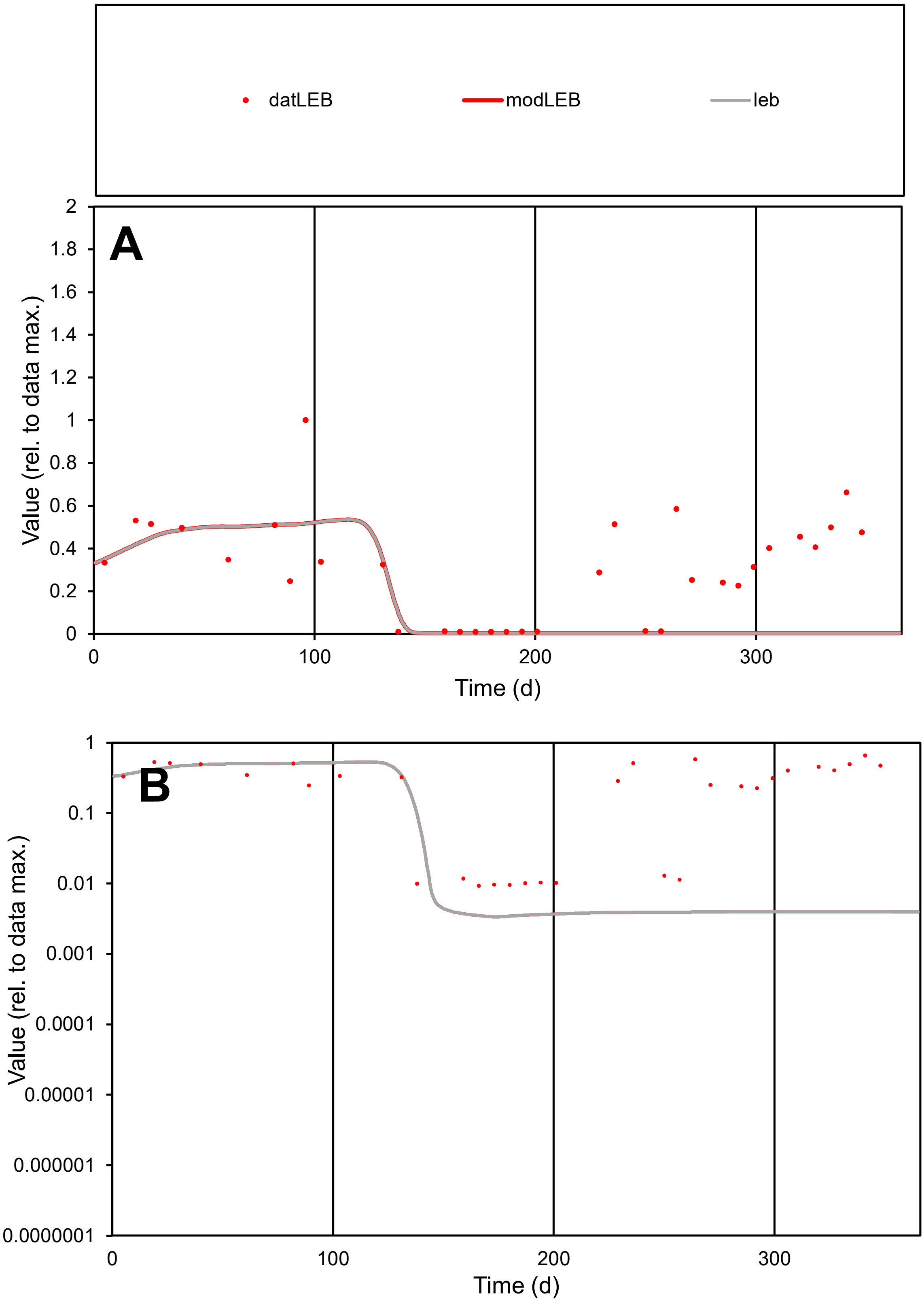


**C1**

Variables

LEB Provide jtd or Dst

CalibOnly?

2 1 to reproduce of

oNormalize?

2 1=no, 2=yes

ofxn calc*

0.94528

ofxn mod

0.9518

sRun

RunA1

zout

1

Ave

Data

0.00227

Model

0.00107

data/vMax

0.00744

ofxng

263.459

Min d/m

2.75968

ave/min d

32.783

ave/min m

42.5361

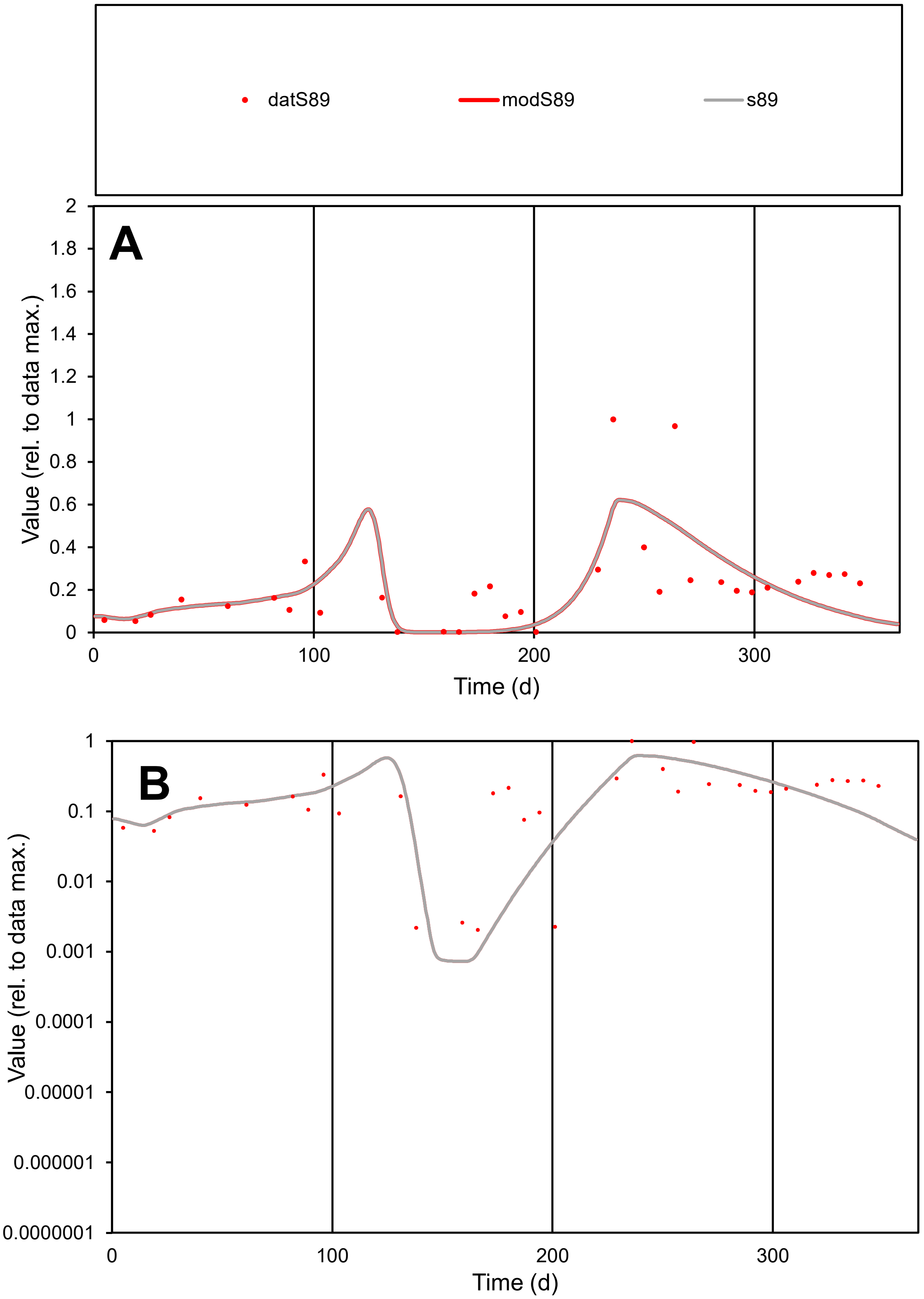


**C1**

Variables
S89 Provide jtd or Dst
CalibOnly?
2 1 to reproduce of
oNormalize?
2 1=no, 2=yes

ofxn calc*
0.74386
ofxn mod
0.76149

sRun
RunA1
zout
1

Ave
Data
0.00376
Model
0.00335

data/vMax
0.01741

ofxng
263.459

Min d/m
2.81039

ave/min d
105.14
ave/min m
263.397

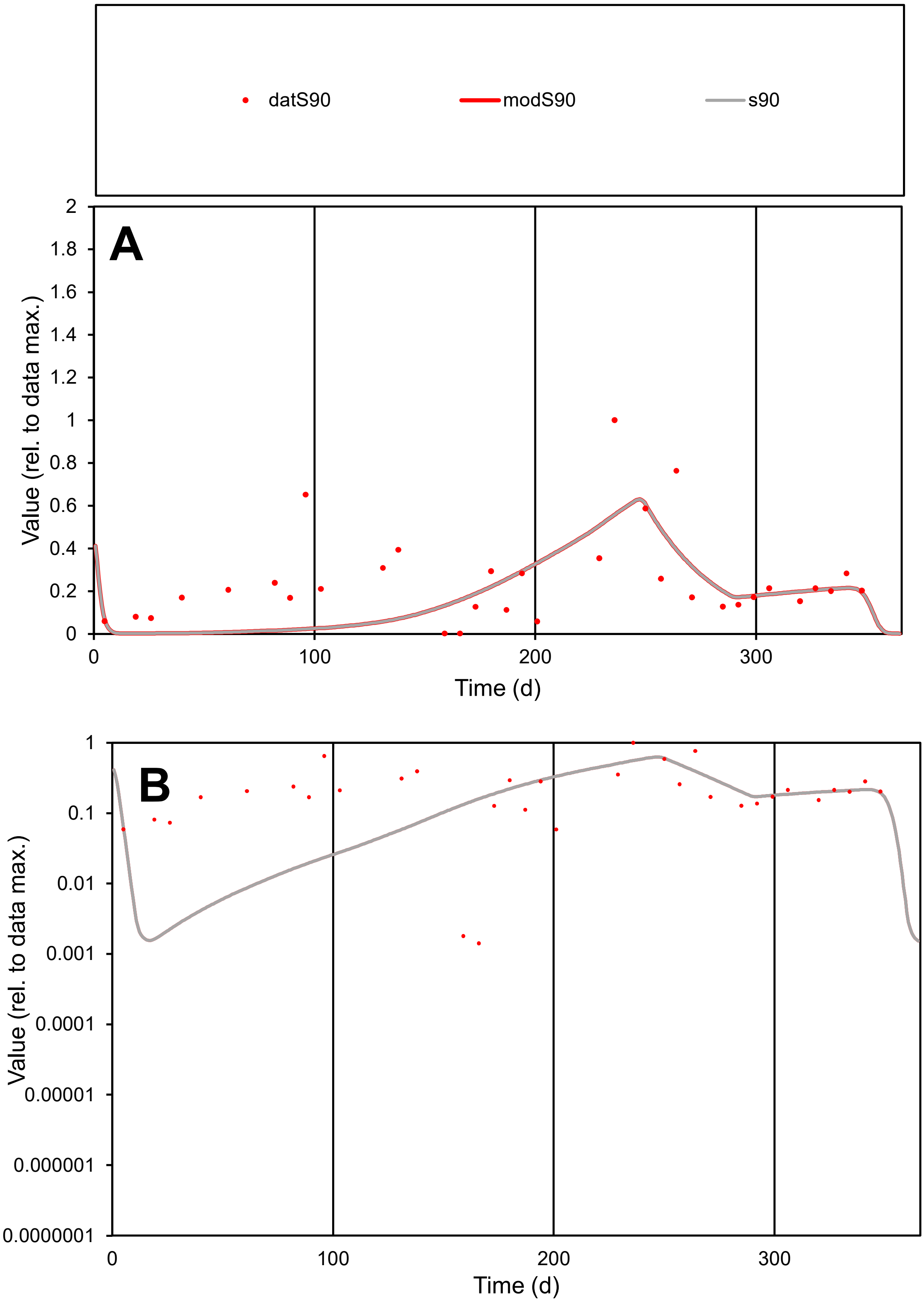


**C1**

Variables

S90 Provide jtd or Dst

CalibOnly?

2 1 to reproduce of

oNormalize?

2 1=no, 2=yes

ofxn calc*

0.79808

ofxn mod

0.80898

sRun

RunA1

zout

1

Ave

Data

0.0035

Model

0.00274

data/vMax

0.01395

ofxng

263.459

Min d/m

0.82914

ave/min d

177.153

ave/min m

115.023

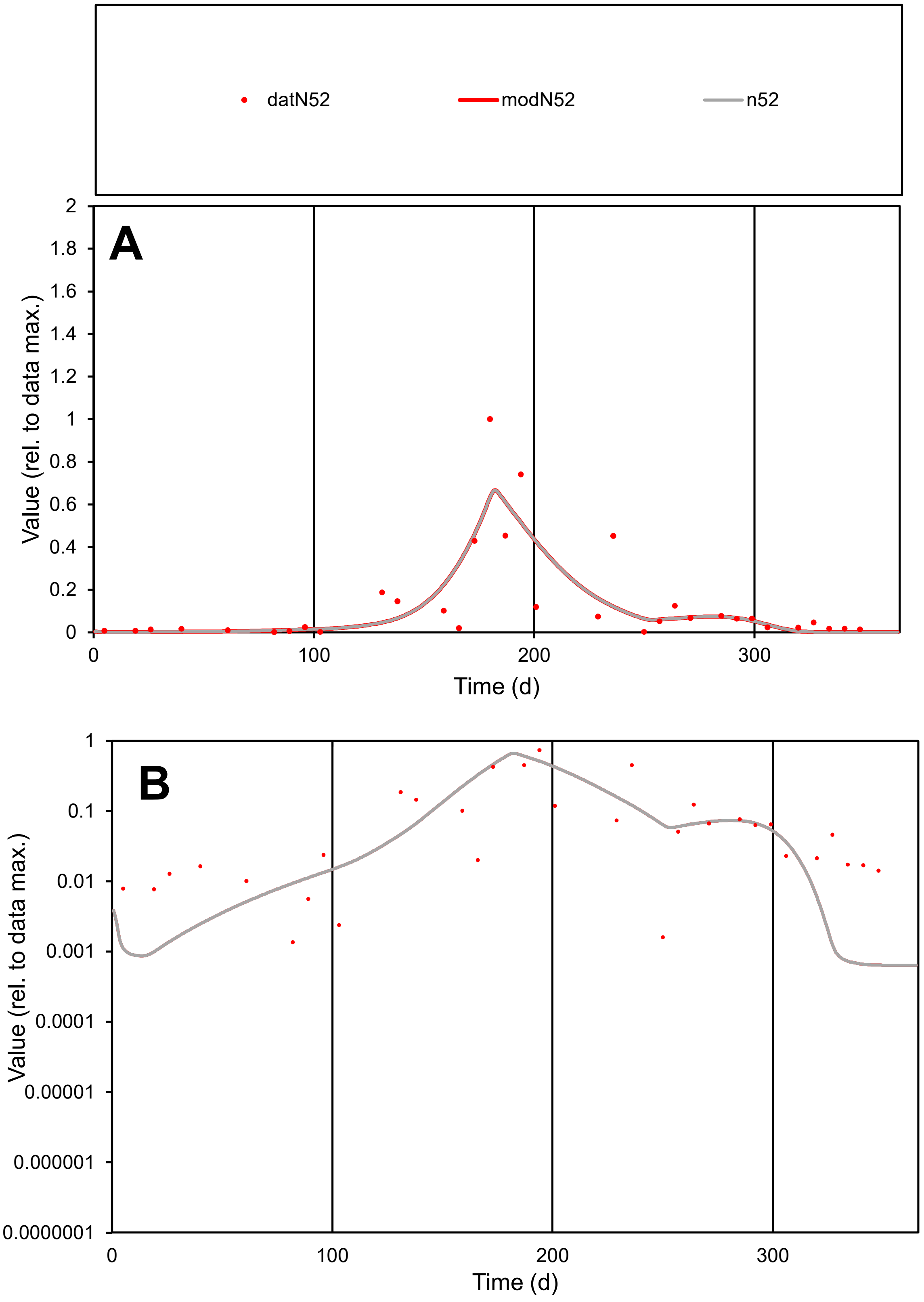


**C1**

Variables
N52 Provide jtd or Dst
CalibOnly?
2 1 to reproduce of
oNormalize?
2 1=no, 2=yes

ofxn calc*
0.95158
ofxn mod
0.98743

sRun
RunA1
zout
1

Ave
Data
0.00274
Model
0.00259

data/vMax
0.02059

ofxng
263.459

Min d/m
2.1306

ave/min d
97.7776
ave/min m
196.838

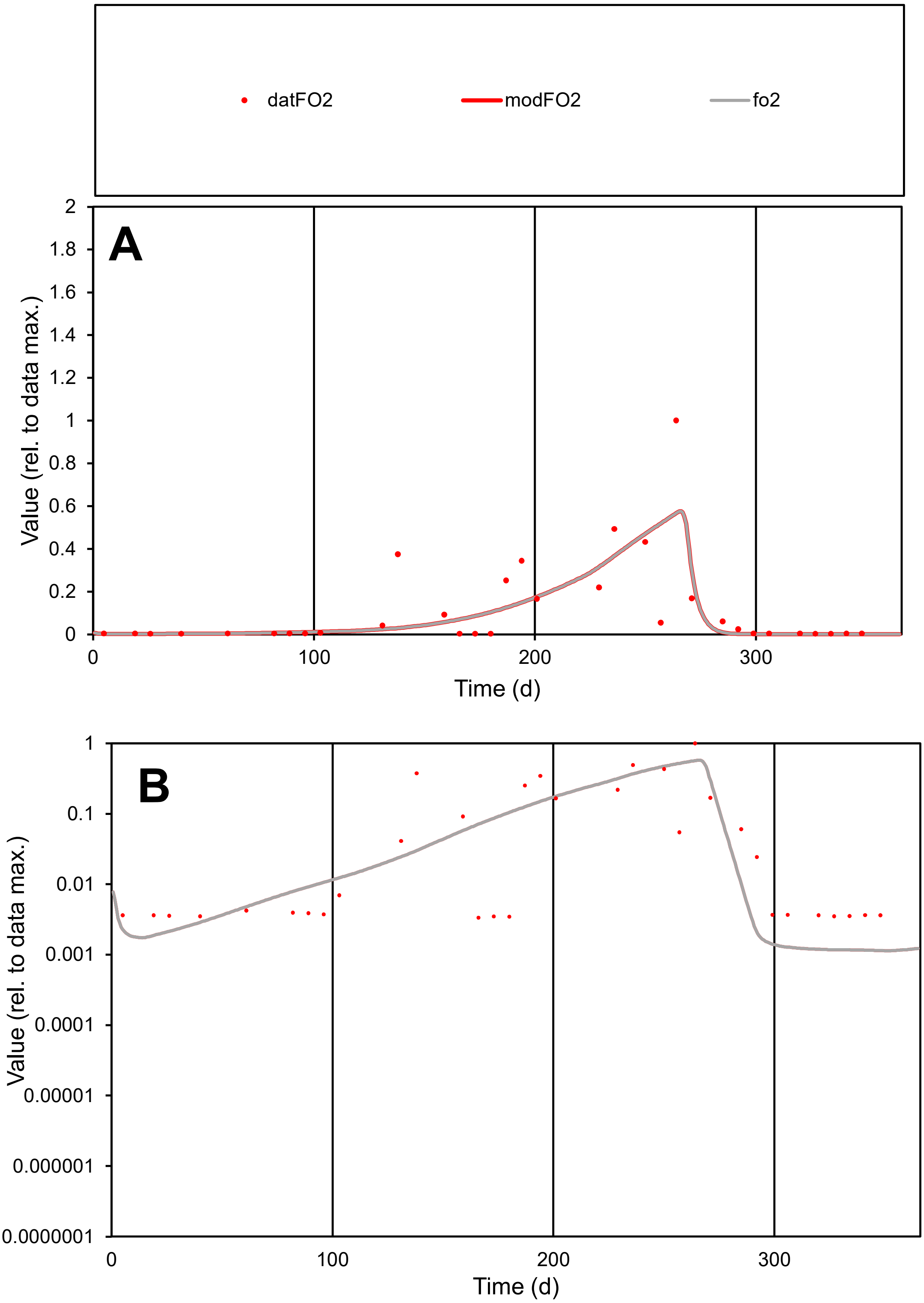


**C1**

Variables

FO2 Provide jtd or Dst

CalibOnly?

2 1 to reproduce of

oNormalize?

2 1=no, 2=yes

ofxn calc*

1.19417

ofxn mod

1.2197

sRun

RunA1

zout

1

Ave

Data

0.00228

Model

0.00204

data/vMax

0.01983

ofxng

263.459

Min d/m

2.94032

ave/min d

34.2963

ave/min m

90.3863

326_Calib_S_A

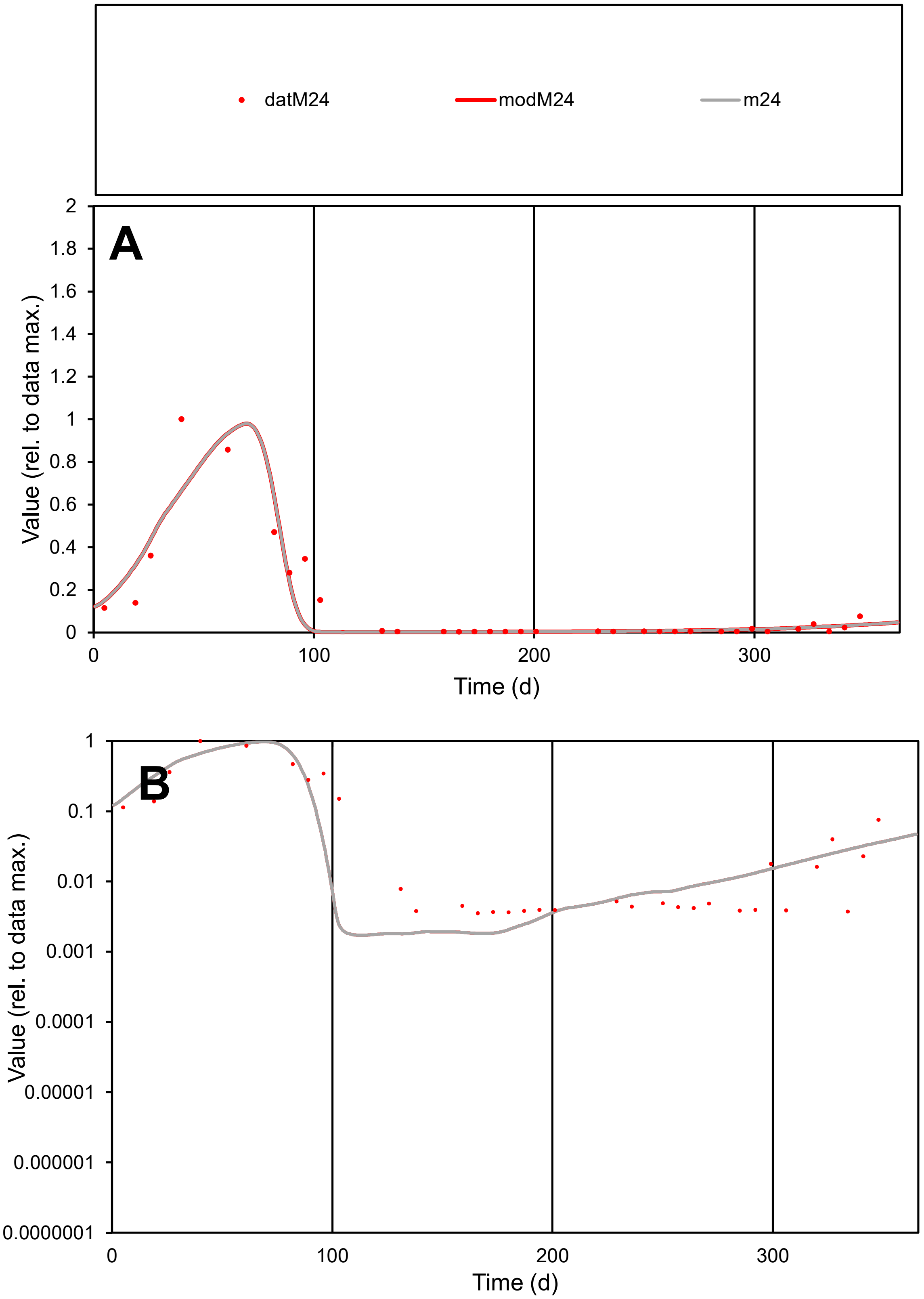


**C1**

Variables

M24 Provide jtd or Dst

CalibOnly?

2 1 to reproduce of

oNormalize?

2 1=no, 2=yes

ofxn calc*

0.76469

ofxn mod

0.82496

sRun

RunA1

zout

1

Ave

Data

0.00244

Model

0.00222

data/vMax

0.02023

ofxng

263.459

Min d/m

1.96782

ave/min d

34.0383

ave/min m

61.0427

326_Calib_S_A

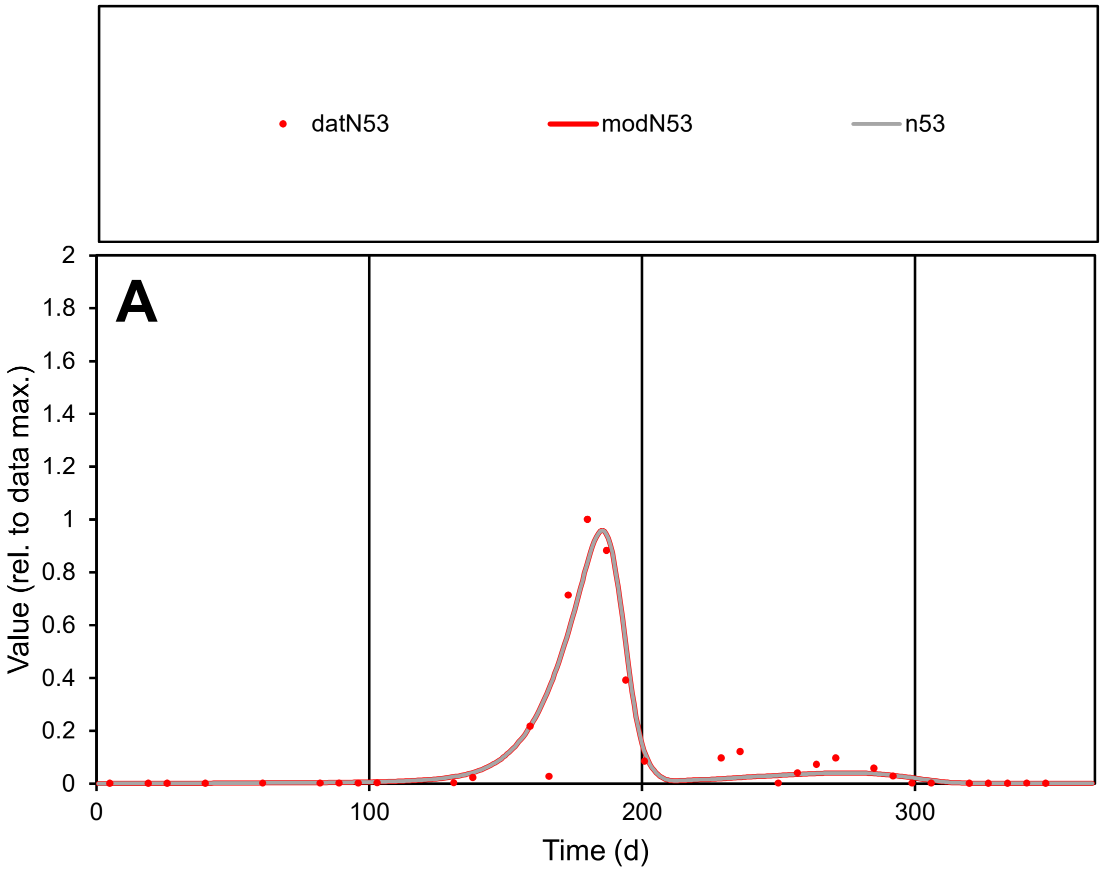


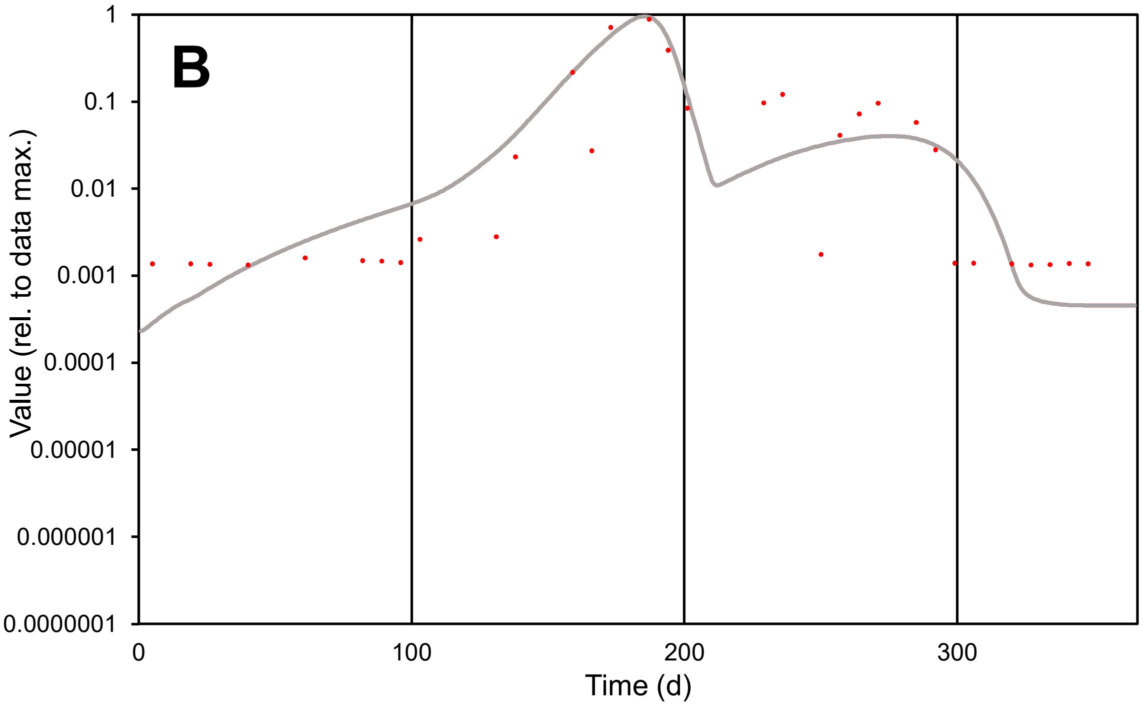


**C1**

Variables
N53 Provide jtd or Dst
CalibOnly?
2 1 to reproduce of
oNormalize?
2 1=no, 2=yes

ofxn calc*
0.56128
ofxn mod
0.63518

sRun
RunA1
zout
1

Ave
Data
0.00336
Model
0.00334

data/vMax
0.02856

ofxng
263.459

Min d/m
4.28377

ave/min d
88.7108
ave/min m
377.46

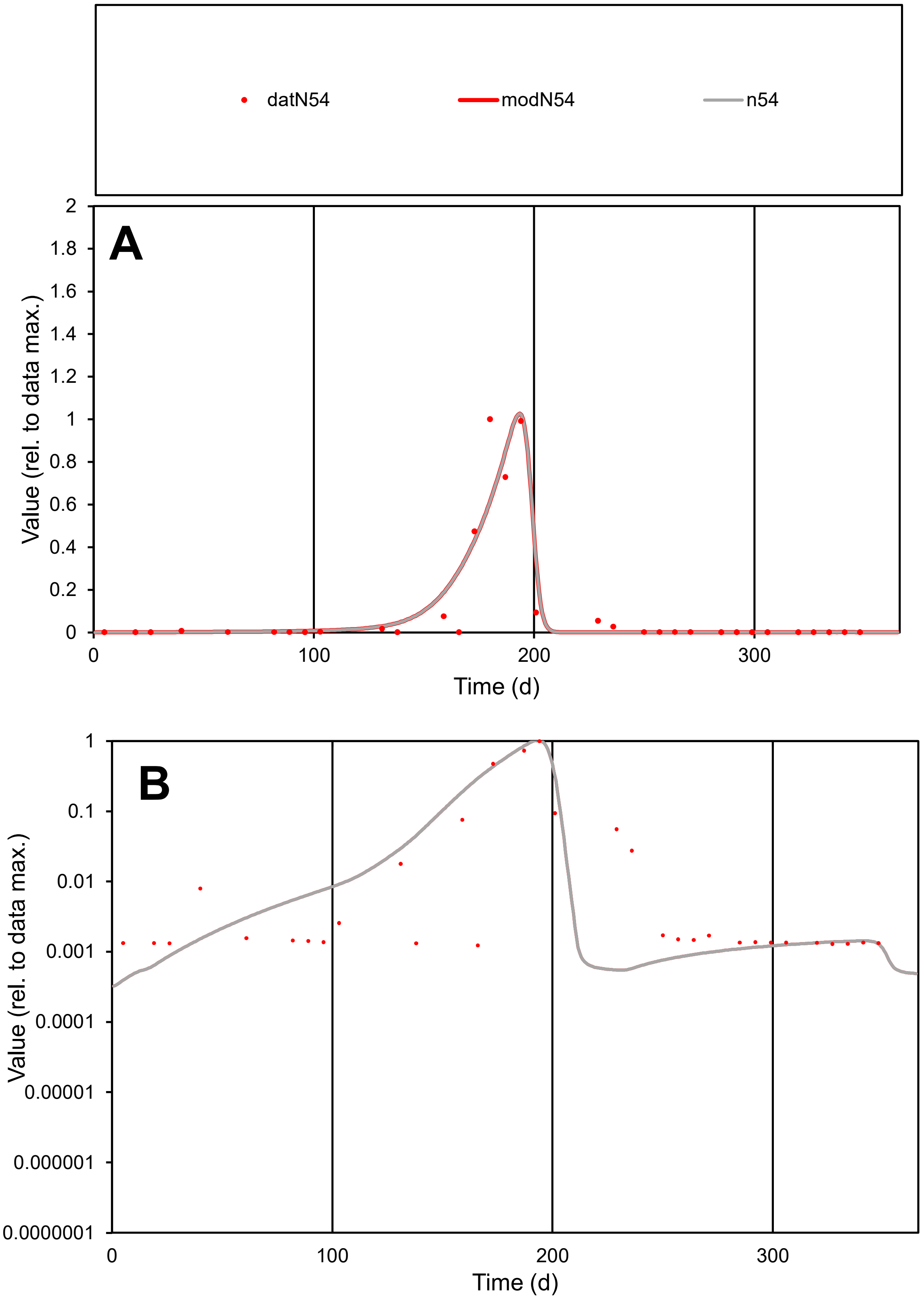


**C1**

Variables
N54 Provide jtd or Dst
CalibOnly?
2 1 to reproduce of
oNormalize?
2 1=no, 2=yes

ofxn calc*
0.80368
ofxn mod
0.89423

sRun
RunA1
zout
1

Ave
Data
0.00185
Model
0.00192

data/vMax
0.01738

ofxng
263.459

Min d/m
2.91446

ave/min d
86.1382
ave/min m
260.794

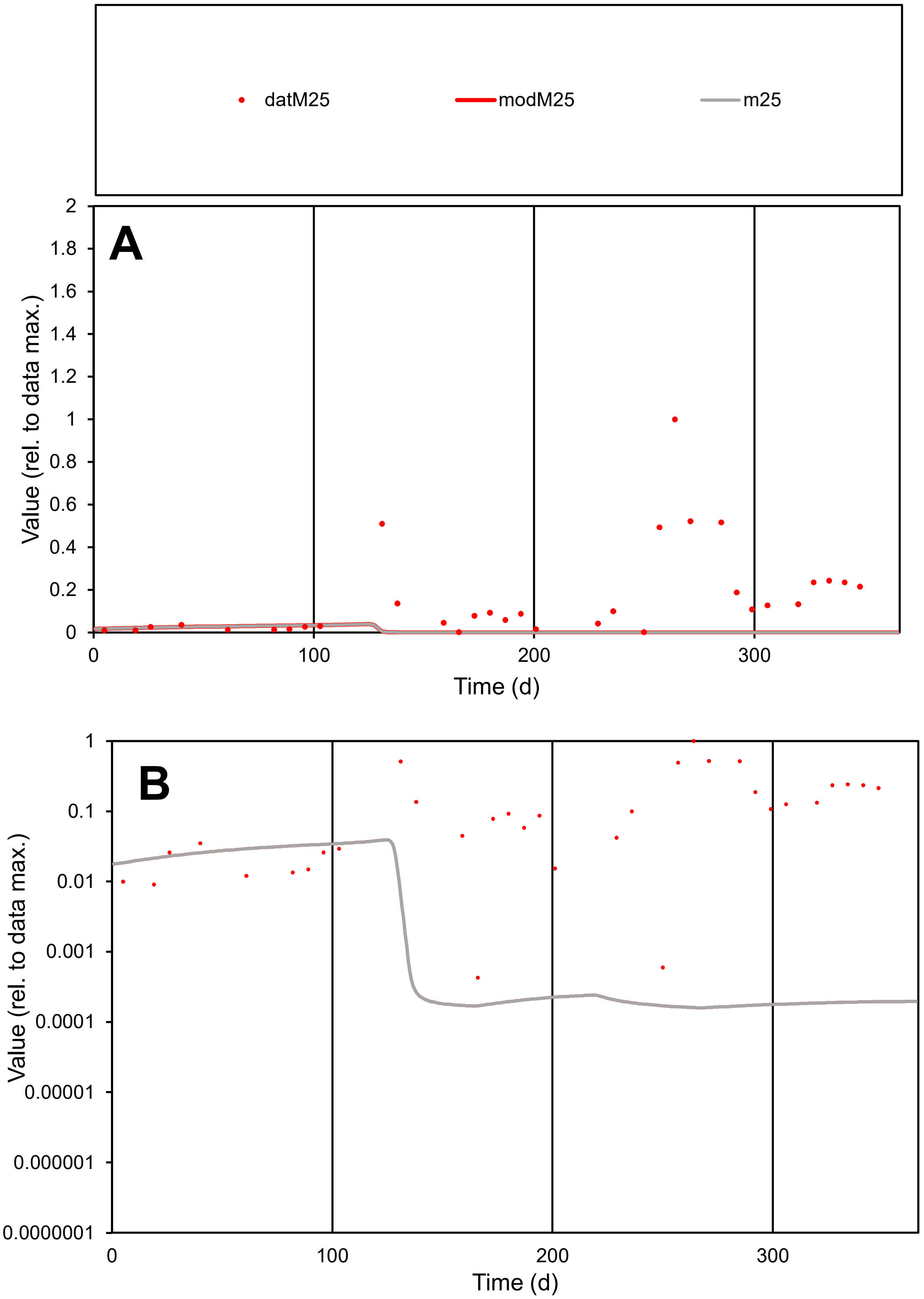


**C1**

Variables
M25 Provide jtd or Dst
CalibOnly?
2 1 to reproduce of
oNormalize?
2 1=no, 2=yes

ofxn calc*
1.65992
ofxn mod
1.66364

sRun
RunA1
zout
1

Ave
Data
0.00359
Model
0.00018

data/vMax
0.02217

ofxng
263.459

Min d/m
2.6866

ave/min d
377.411
ave/min m
49.7337

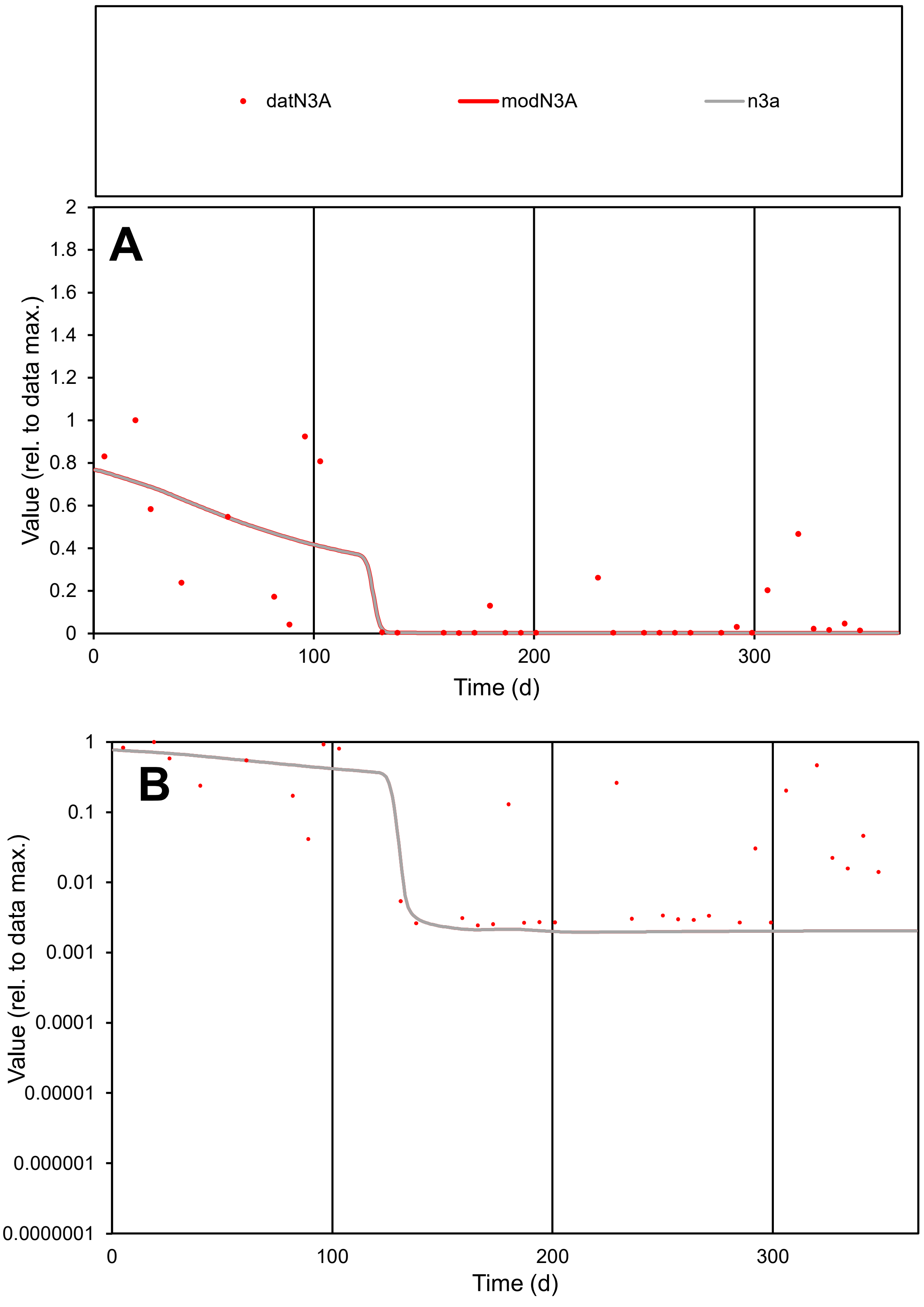


**C1**

Variables
N3A Provide jtd or Dst
CalibOnly?
2 1 to reproduce of
oNormalize?
2 1=no, 2=yes

ofxn calc*
1.00831
ofxn mod
1.02678

sRun
RunA1
zout
1

Ave
Data
0.0014
Model
0.00112

data/vMax
0.00725

ofxng
263.459

Min d/m
1.24073

ave/min d
78.9915
ave/min m
78.3452

326_Calib_S_A

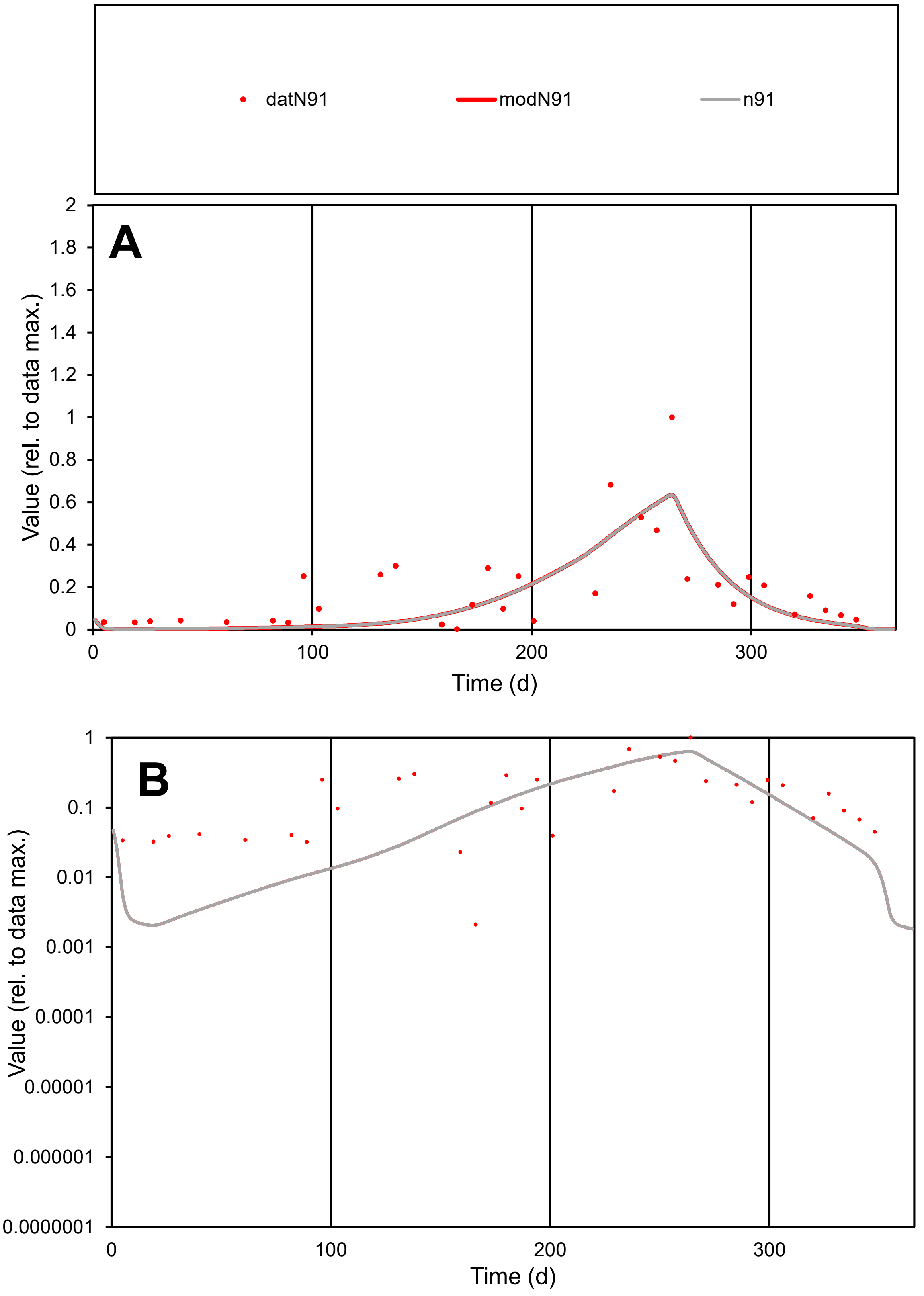


**C1**

Variables
N91 Provide jtd or Dst
CalibOnly?
2 1 to reproduce of
oNormalize?
2 1=no, 2=yes

ofxn calc*
0.72106
ofxn mod
0.74365

sRun
RunA1
zout
1

Ave
Data
0.00281
Model
0.00231

data/vMax
0.01478

ofxng
263.459

Min d/m
1.02457

ave/min d
89.8021
ave/min m
75.532

326_Calib_S_A

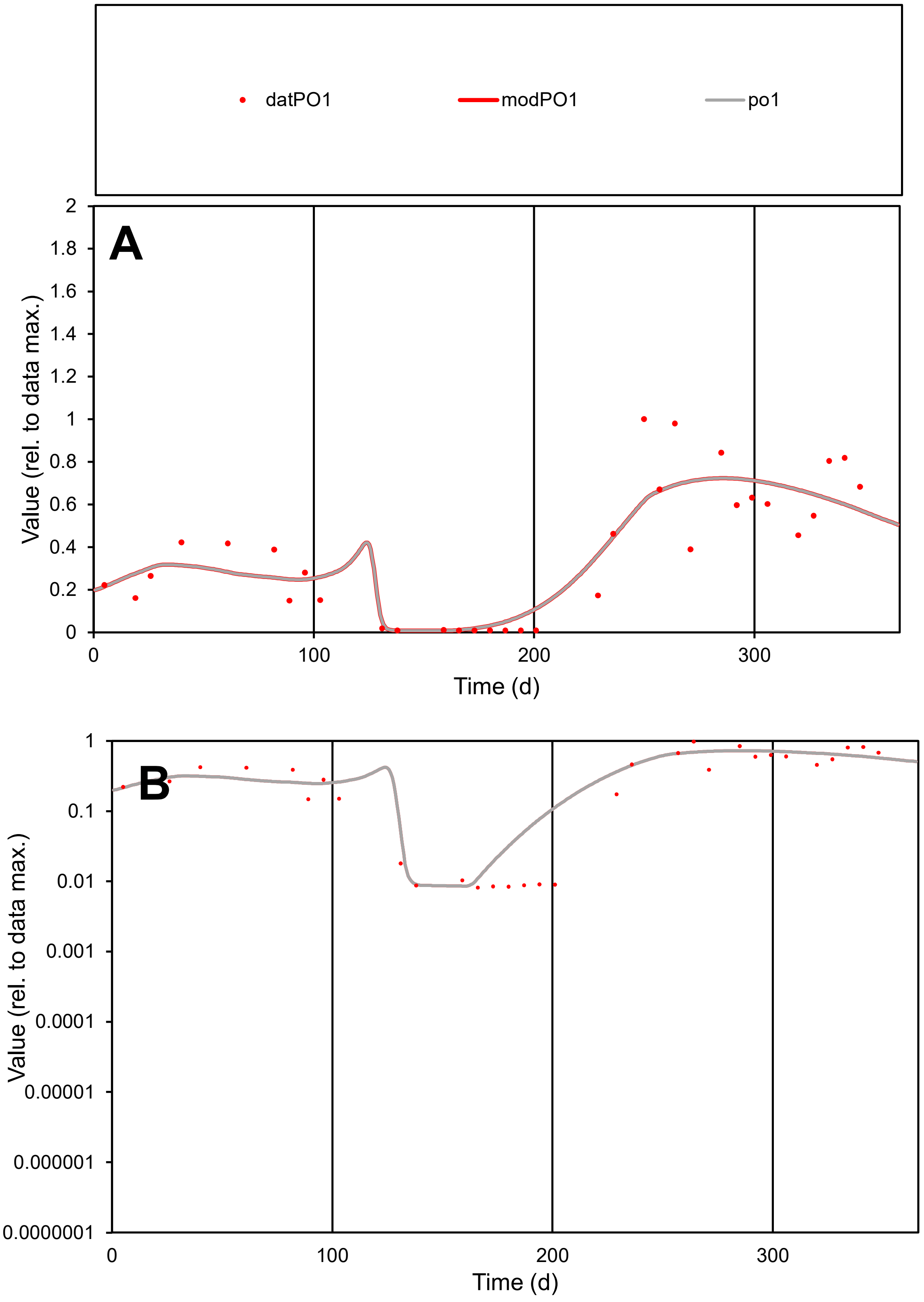


**C1**

Variables

PO1 Provide jtd or Dst

CalibOnly?

2 1 to reproduce of

oNormalize?

2 1=no, 2=yes

ofxn calc*

0.38135

ofxn mod

0.40954

sRun

RunA1

zout

1

Ave

Data

0.00205

Model

0.00206

data/vMax

0.00555

ofxng

263.459

Min d/m

0.94774

ave/min d

45.406

ave/min m

43.2

326_Calib_S_A

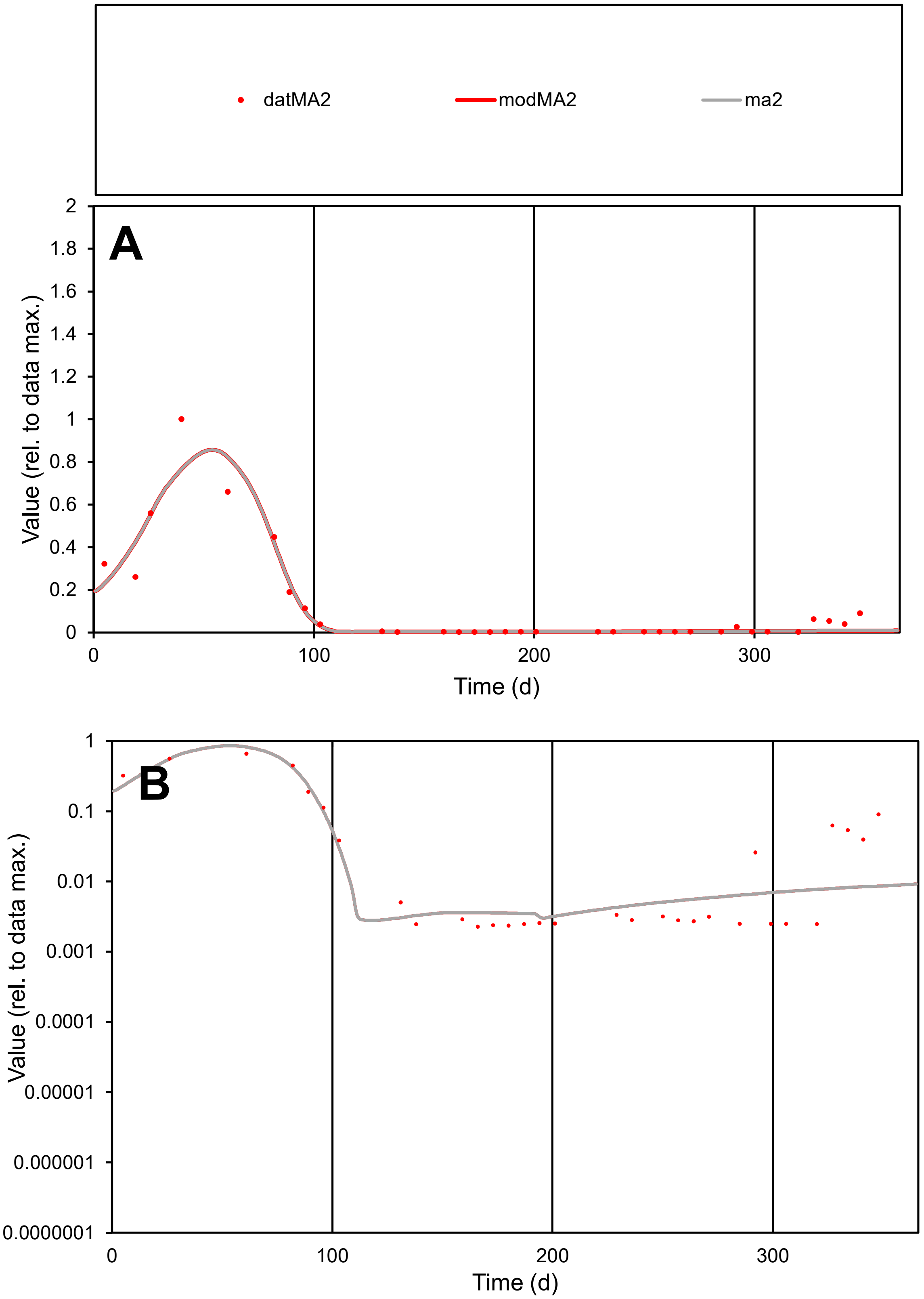


**C1**

Variables
MA2 Provide jtd or Dst
CalibOnly?
2 1 to reproduce of
oNormalize?
2 1=no, 2=yes

ofxn calc*
0.51906
ofxn mod
0.59663

sRun
RunA1
zout
1

Ave
Data
0.00092
Model
0.00086

data/vMax
0.00772

ofxng
263.459

Min d/m
0.75772

ave/min d
51.8078
ave/min m
37.0843

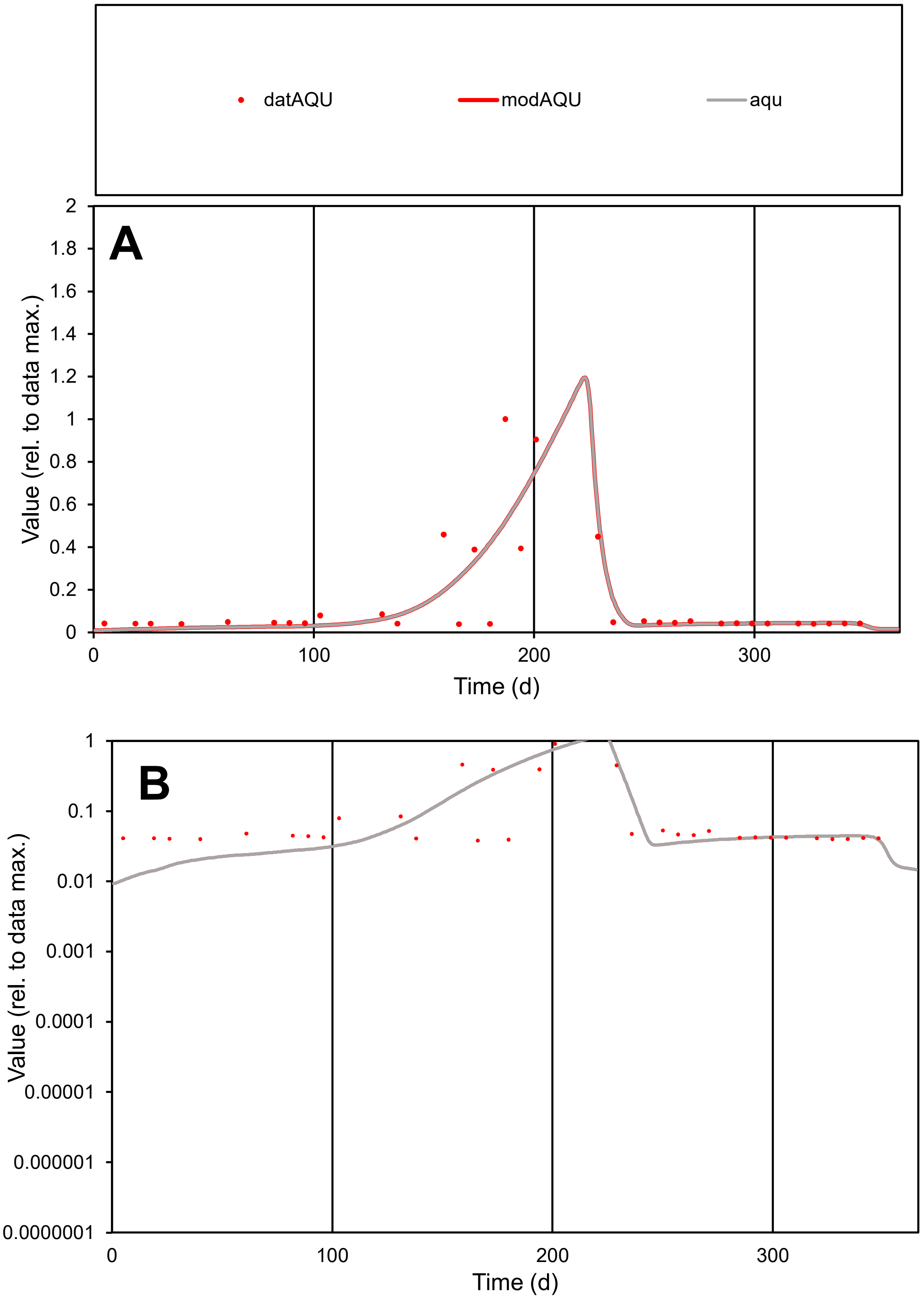


**C1**

Variables

AQU Provide jtd or Dst

CalibOnly?

2 1 to reproduce of

oNormalize?

2 1=no, 2=yes

ofxn calc*

0.88578

ofxn mod

0.91967

sRun

RunA1

zout

1

Ave

Data

0.0005

Model

0.00047

data/vMax

0.00339

ofxng

263.459

Min d/m

3.50748

ave/min d

3.84844

ave/min m

12.8479

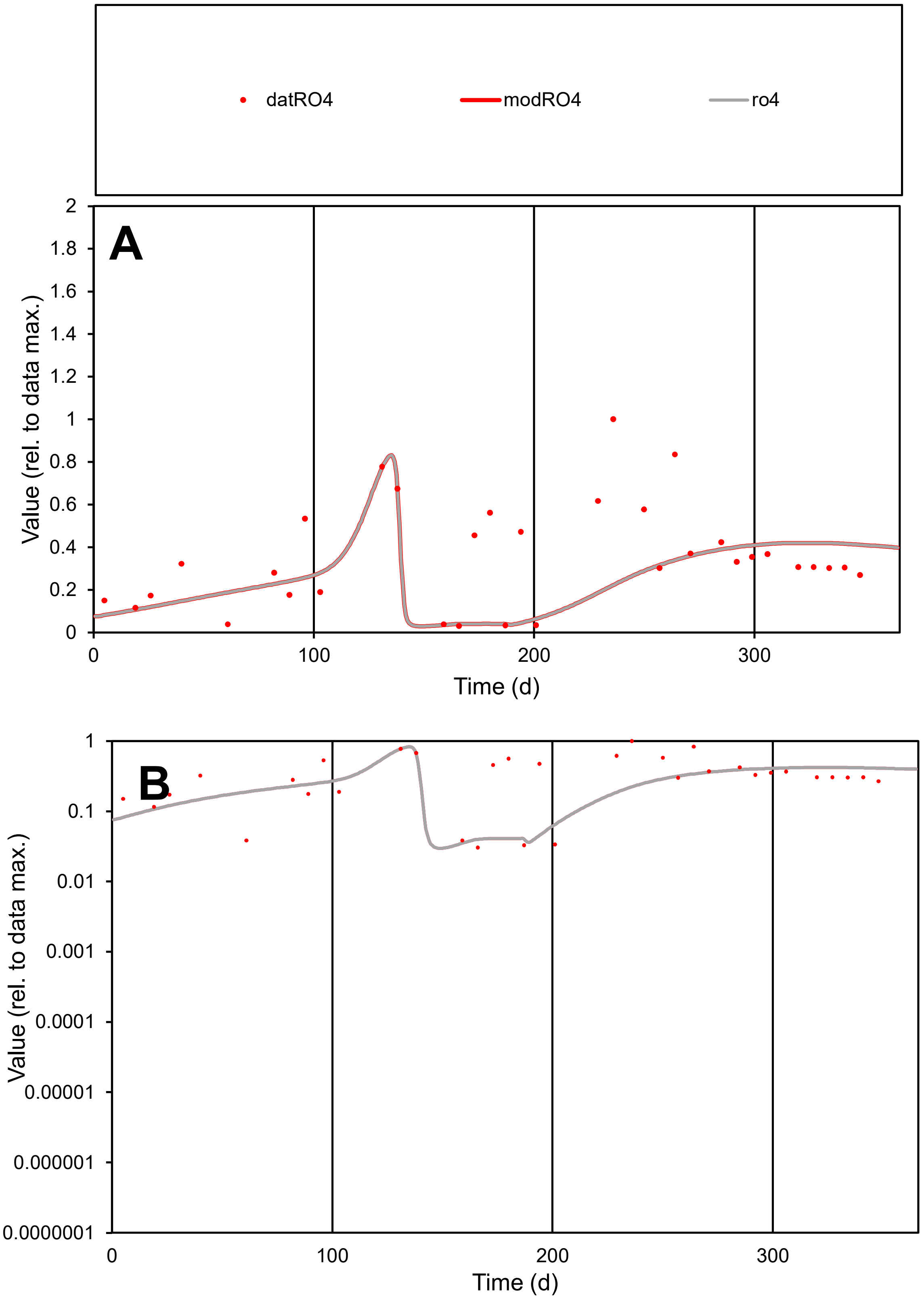


**C1**

Variables
RO4 Provide jtd or Dst
CalibOnly?
2 1 to reproduce of
oNormalize?
2 1=no, 2=yes

ofxn calc*
0.68484
ofxn mod
0.69753

sRun
RunA1
zout
1

Ave
Data
0.00252
Model
0.00191

data/vMax
0.00709

ofxng
263.459

Min d/m
0.84531

ave/min d
11.7018
ave/min m
7.49713

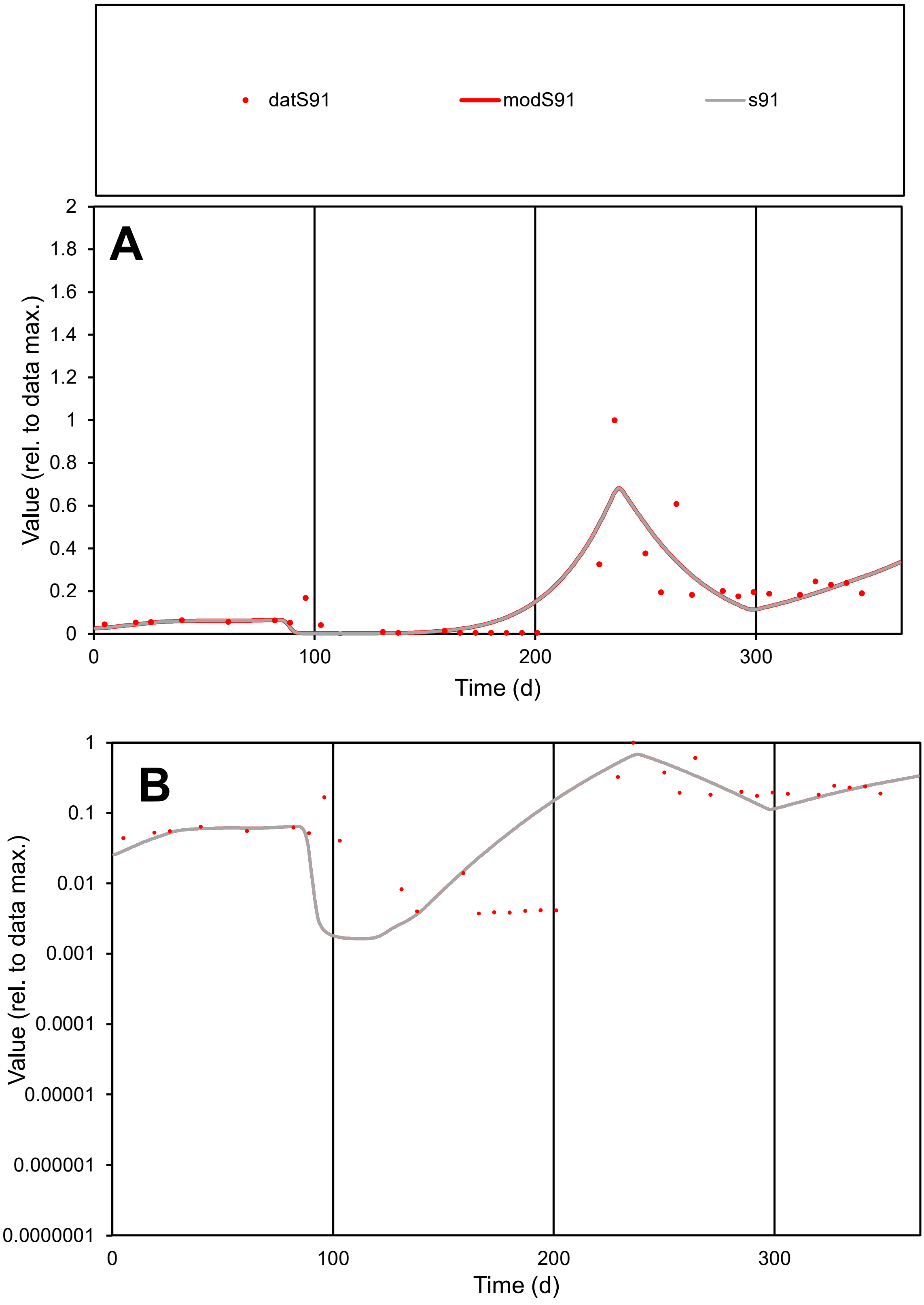


**C1**

Variables
S91 Provide jtd or Dst
CalibOnly?
2 1 to reproduce of
oNormalize?
2 1=no, 2=yes

ofxn calc*
0.67507
ofxn mod
0.70505

sRun
RunA1
zout
1

Ave
Data
0.00272
Model
0.00271

data/vMax
0.01735

ofxng
263.459

Min d/m
2.20197

ave/min d
41.9075
ave/min m
91.8236

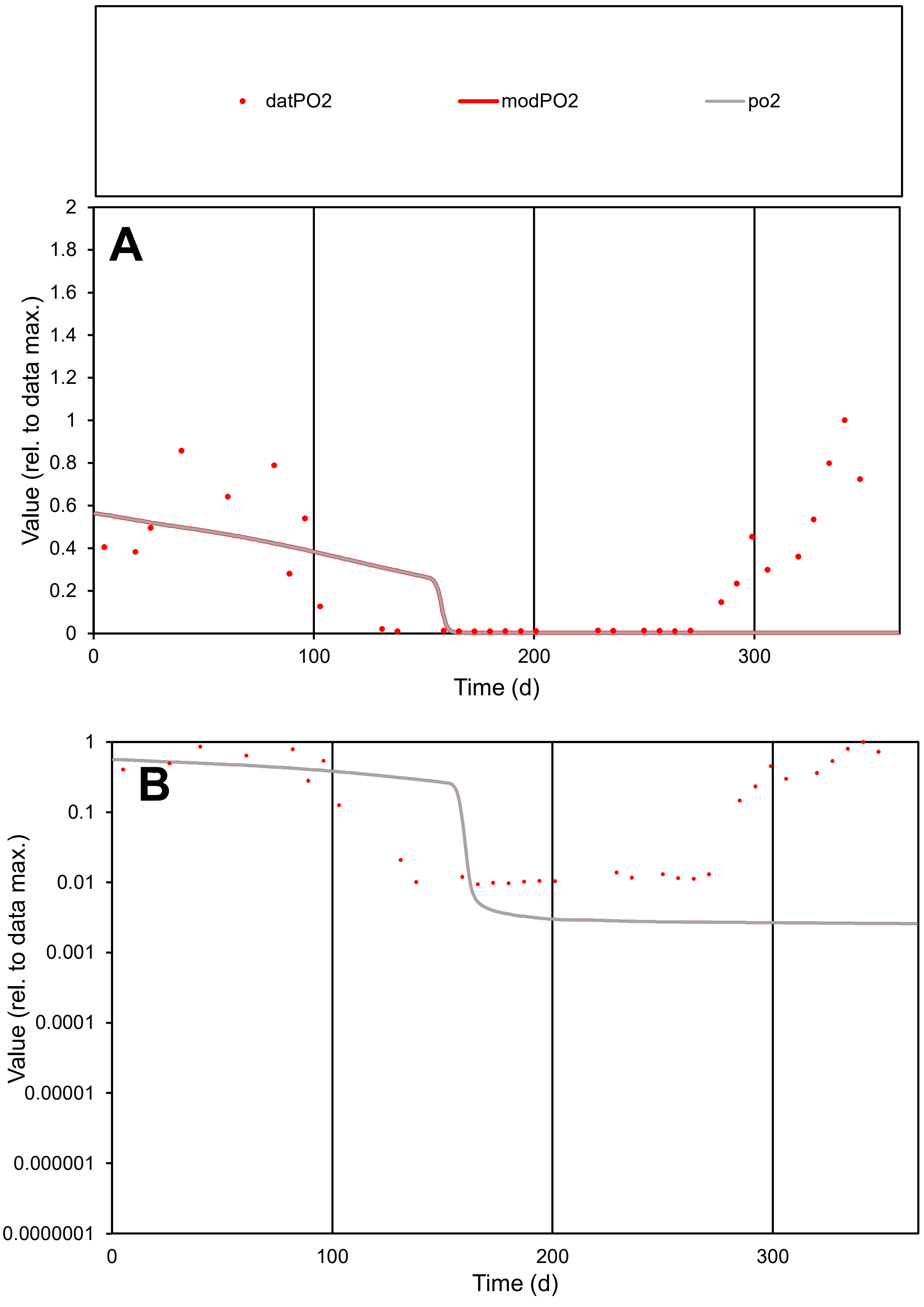


**C1**

Variables

PO2 Provide jtd or Dst

CalibOnly?

2 1 to reproduce of

oNormalize?

2 1=no, 2=yes

ofxn calc*

1.16718

ofxn mod

1.17441

sRun

RunA1

zout

1

Ave

Data

0.00233

Model

0.00122

data/vMax

0.00833

ofxng

263.459

Min d/m

3.61868

ave/min d

29.651

ave/min m

56.0405

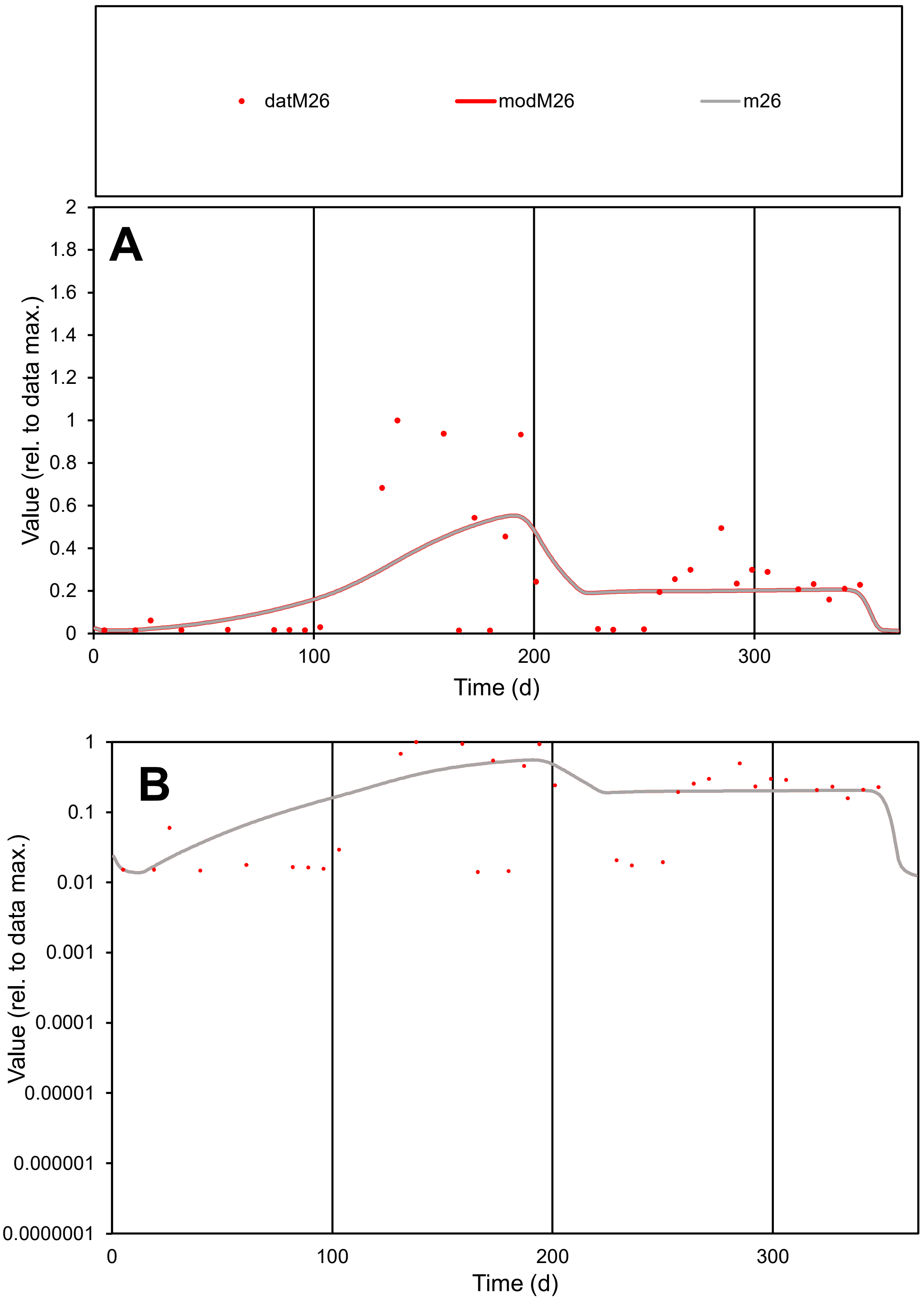


**C1**

Variables

M26 Provide jtd or Dst

CalibOnly?

2 1 to reproduce of

oNormalize?

2 1=no, 2=yes

ofxn calc*

0.93454

ofxn mod

0.9473

sRun

RunA1

zout

1

Ave

Data

0.00143

Model

0.00137

data/vMax

0.00575

ofxng

263.459

Min d/m

0.96496

ave/min d

17.6005

ave/min m

16.3749

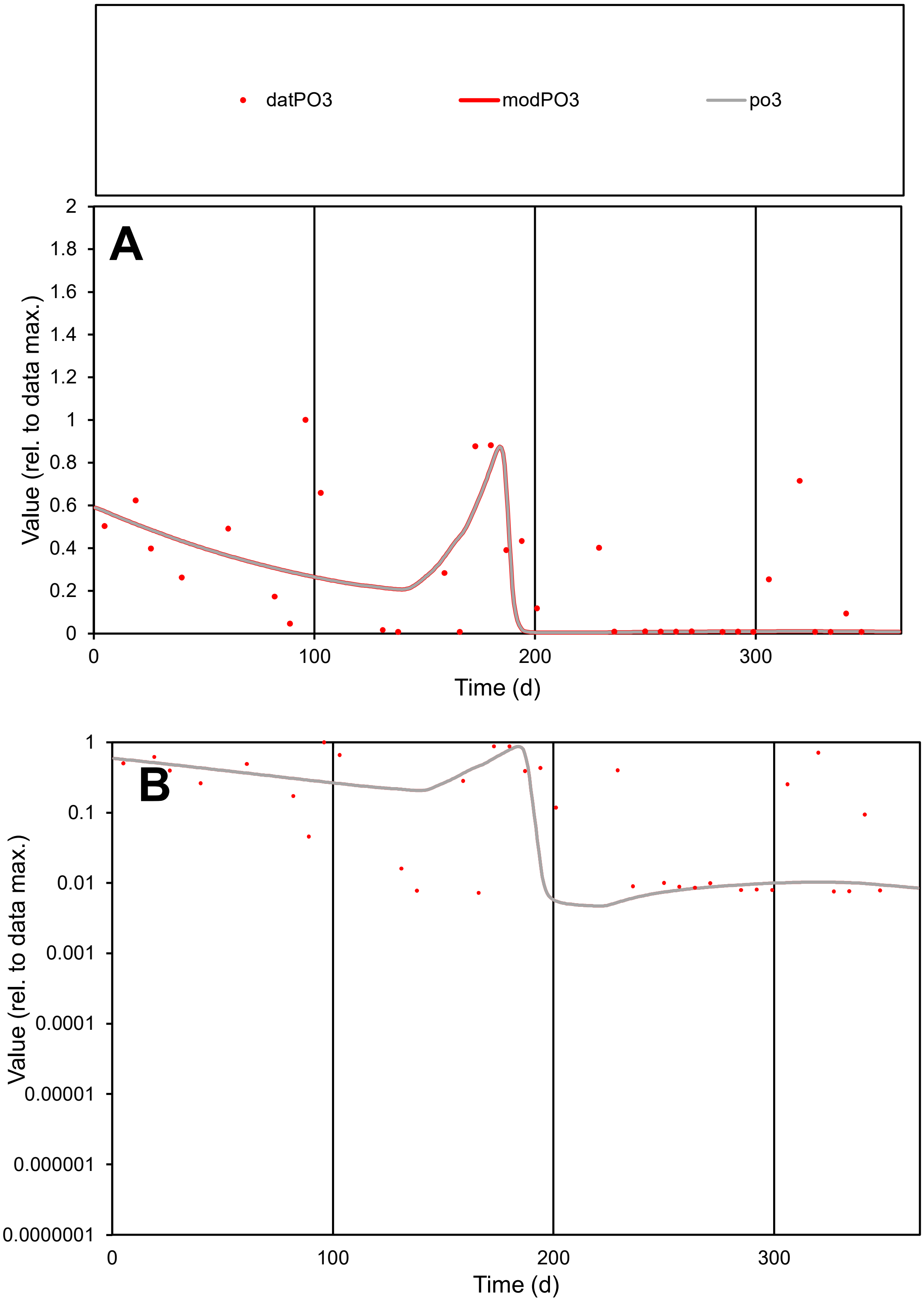


**C1**

Variables

PO3 Provide jtd or Dst

CalibOnly?

2 1 to reproduce of

oNormalize?

2 1=no, 2=yes

ofxn calc*

0.91456

ofxn mod

0.96268

sRun

RunA1

zout

1

Ave

Data

0.00168

Model

0.00135

data/vMax

0.00635

ofxng

263.459

Min d/m

1.34394

ave/min d

36.454

ave/min m

39.547

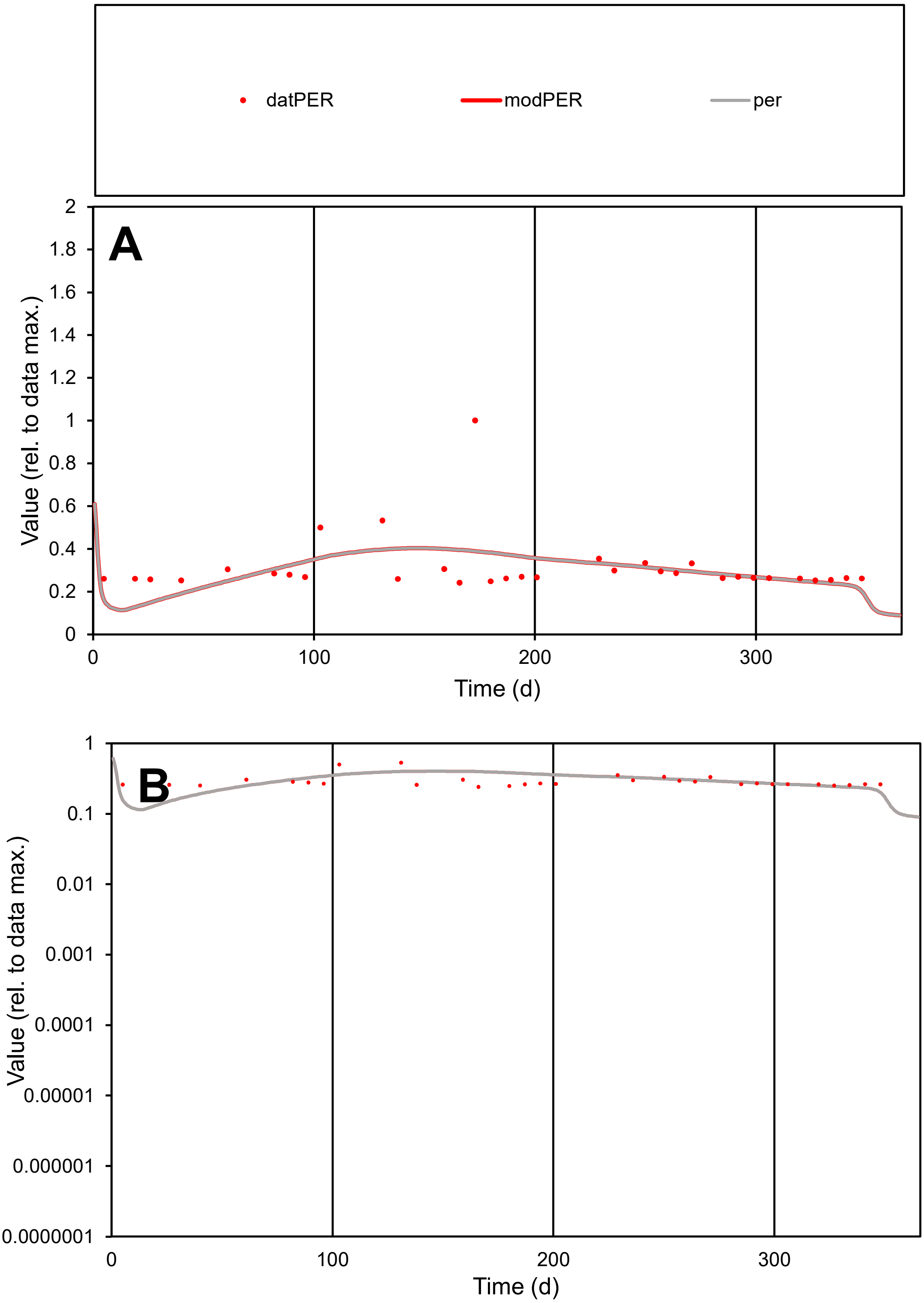


**C1**

Variables
PER Provide jtd or Dst
CalibOnly?
2 1 to reproduce of
oNormalize?
2 1=no, 2=yes

ofxn calc*
0.42274
ofxn mod
0.4385

sRun
RunA1
zout
1

Ave
Data
2.3E-05
Model
2.2E-05

data/vMax
7.5E-05

ofxng
263.459

Min d/m
1.8223

ave/min d
1.29545
ave/min m
2.24464

326_Calib_S_A

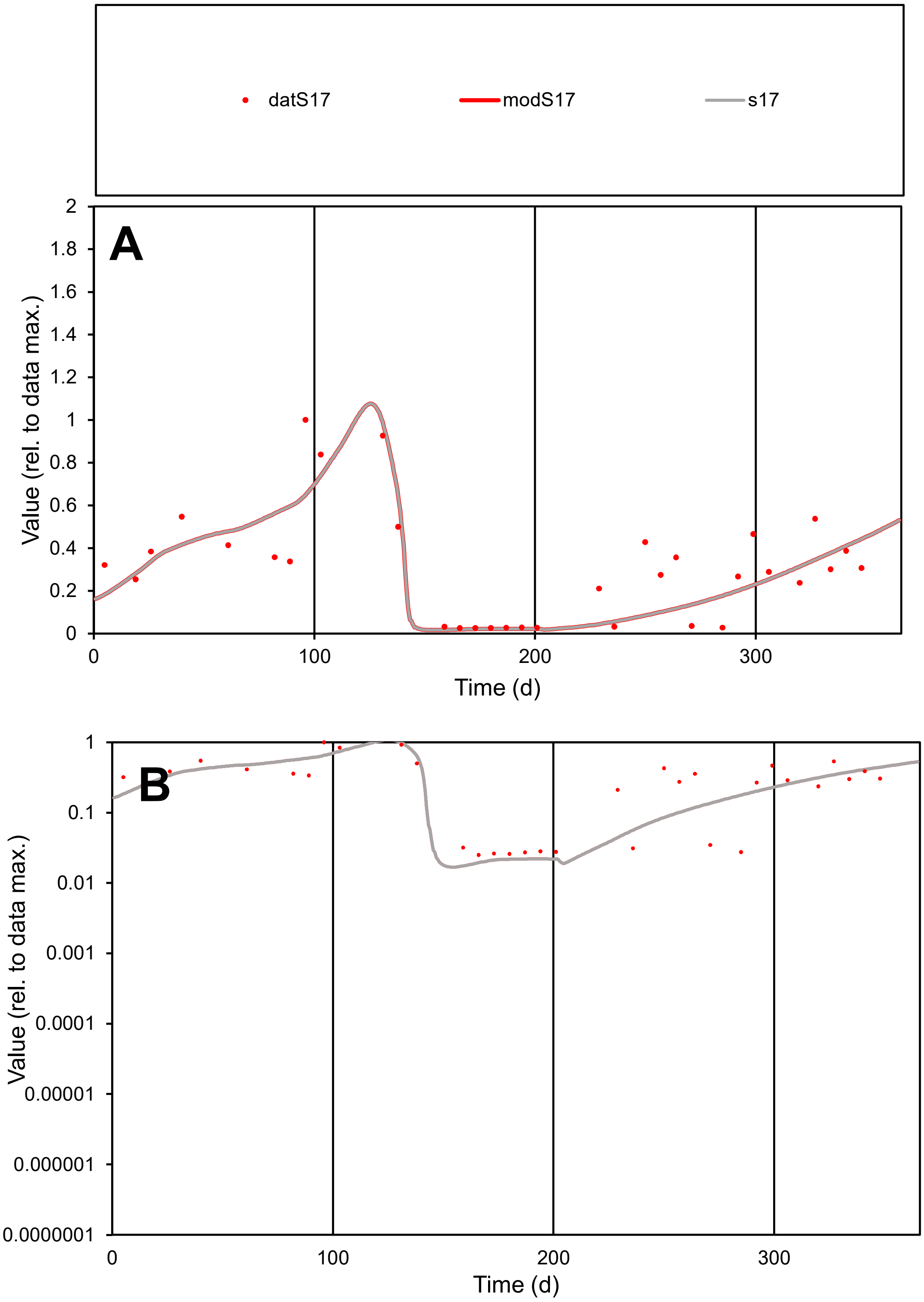


**C1**

Variables
S17 Provide jtd or Dst
CalibOnly?
2 1 to reproduce of
oNormalize?
2 1=no, 2=yes

ofxn calc*
0.45707
ofxn mod
0.4763

sRun
RunA1
zout
1

Ave
Data
0.00132
Model
0.0012

data/vMax
0.00428

ofxng
263.459

Min d/m
1.42234

ave/min d
12.3095
ave/min m
15.8857

326_Calib_S_A

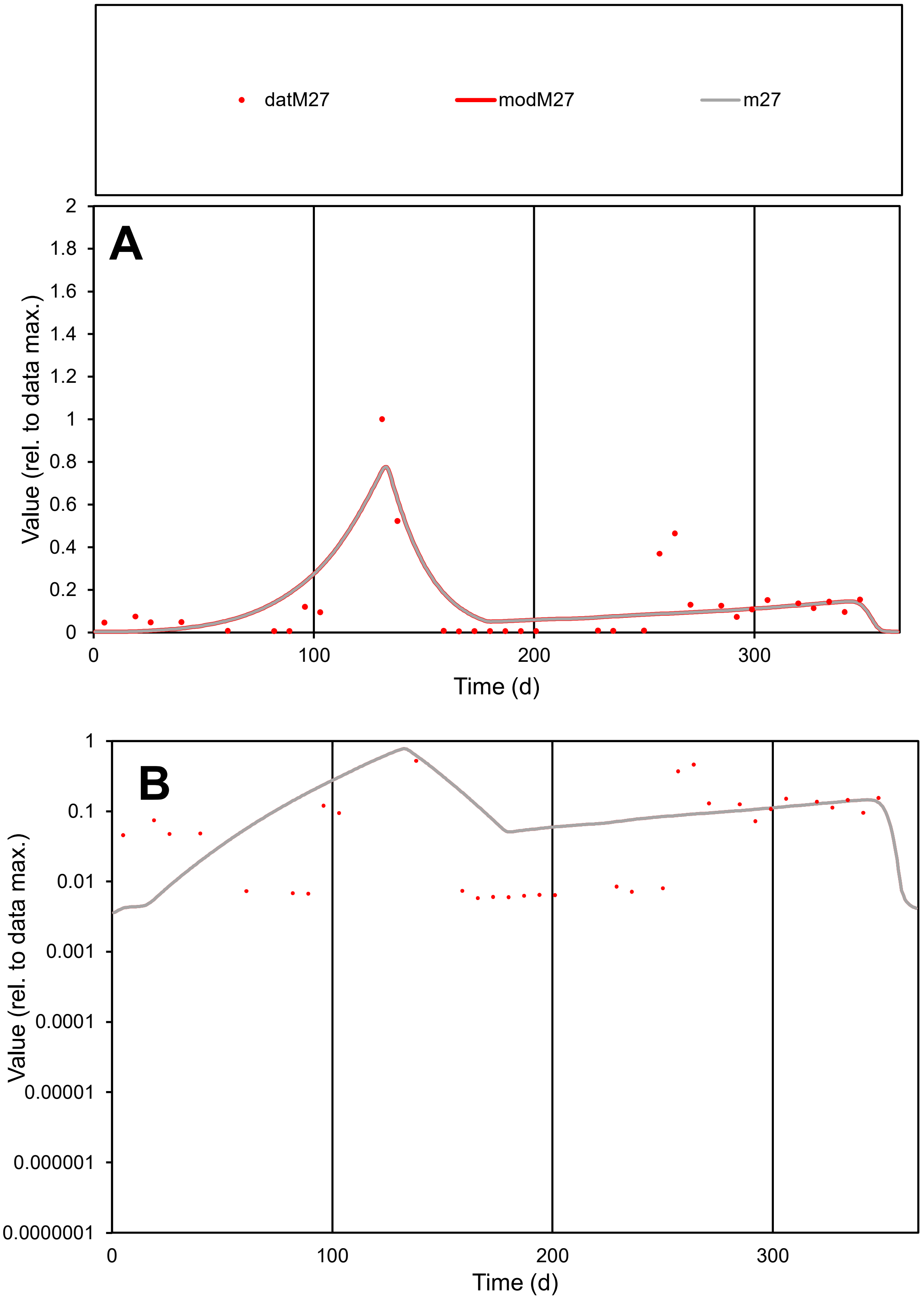


**C1**

Variables
M27 Provide jtd or Dst
CalibOnly?
2 1 to reproduce of
oNormalize?
2 1=no, 2=yes

ofxn calc*
0.94377
ofxn mod
0.97632

sRun
RunA1
zout
1

Ave
Data
0.00138
Model
0.00151

data/vMax
0.01111

ofxng
263.459

Min d/m
1.34999

ave/min d
21.4996
ave/min m
31.6239

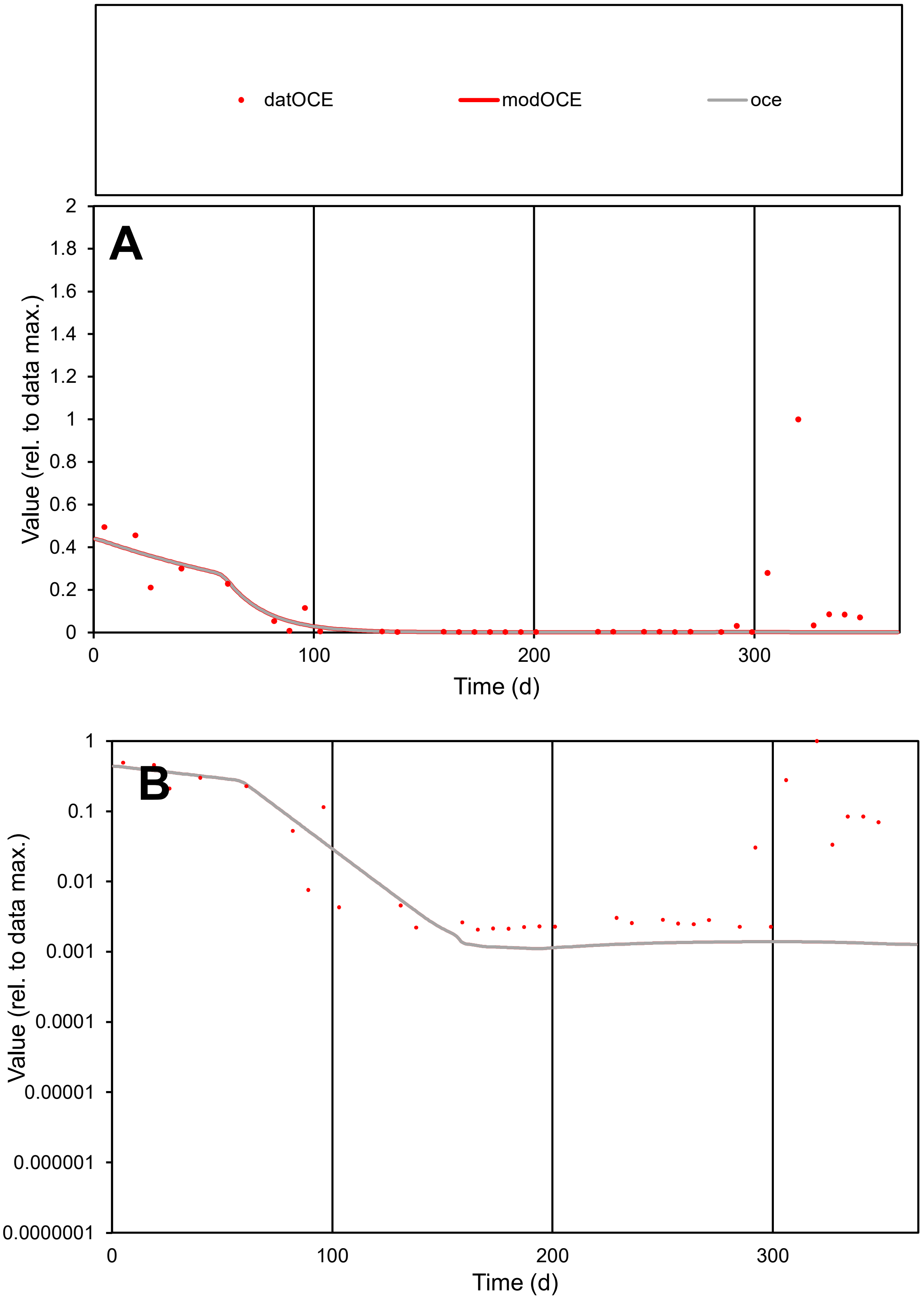


**C1**

Variables
OCE Provide jtd or Dst
CalibOnly?
2 1 to reproduce of
oNormalize?
2 1=no, 2=yes

ofxn calc*
1.75351
ofxn mod
1.76388

sRun
RunA1
zout
1

Ave
Data
0.00188
Model
0.00103

data/vMax
0.01779

ofxng
263.459

Min d/m
1.85693

ave/min d
51.2312
ave/min m
51.8892

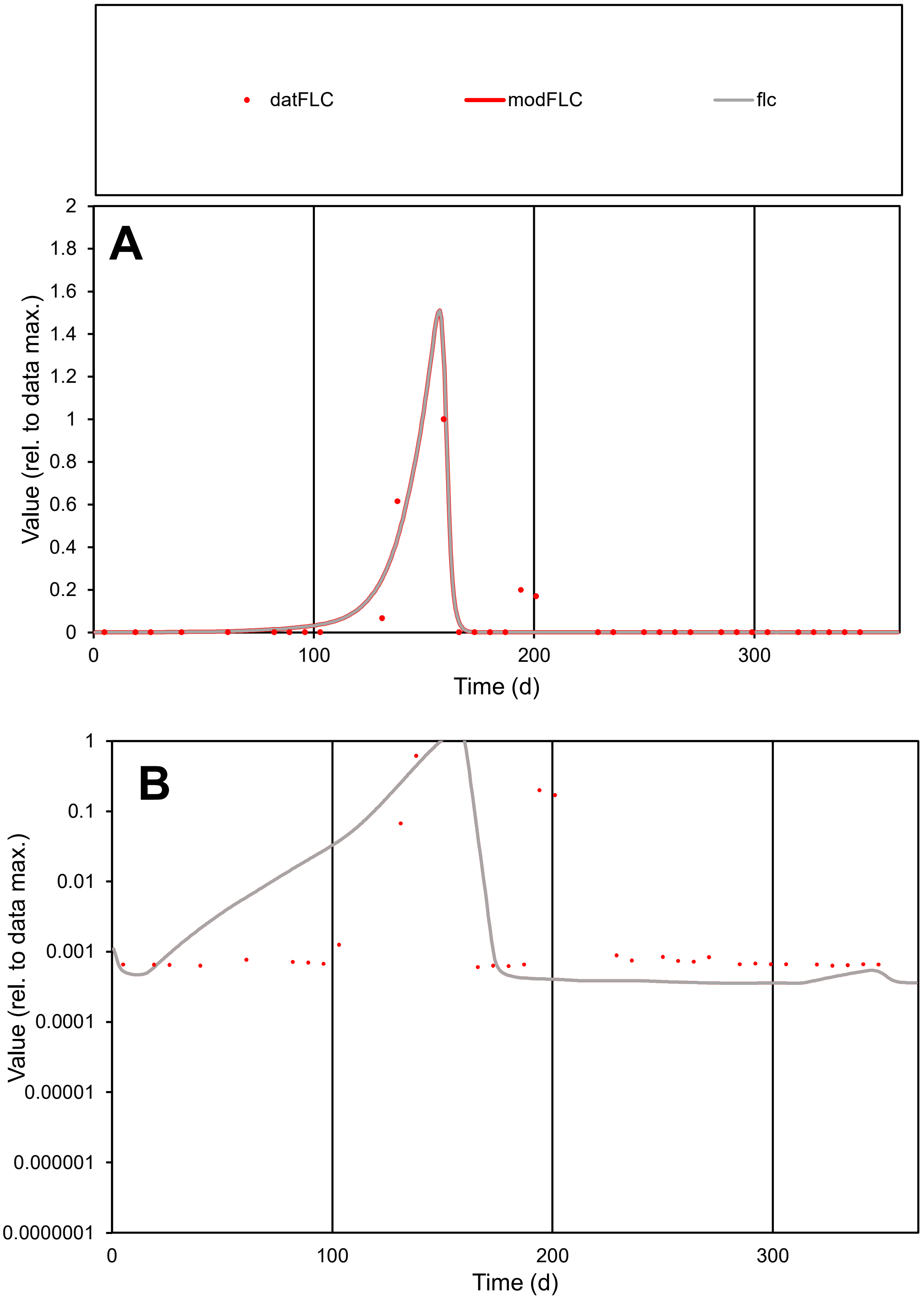


**C1**

Variables
FLC Provide jtd or Dst
CalibOnly?
2 1 to reproduce of
oNormalize?
2 1=no, 2=yes

ofxn calc*
0.92961
ofxn mod
0.99508

sRun
RunA1
zout
1

Ave
Data
0.00123
Model
0.00111

data/vMax
0.0196

ofxng
263.459

Min d/m
1.69973

ave/min d
103.339
ave/min m
158.389

326_Calib_S_A

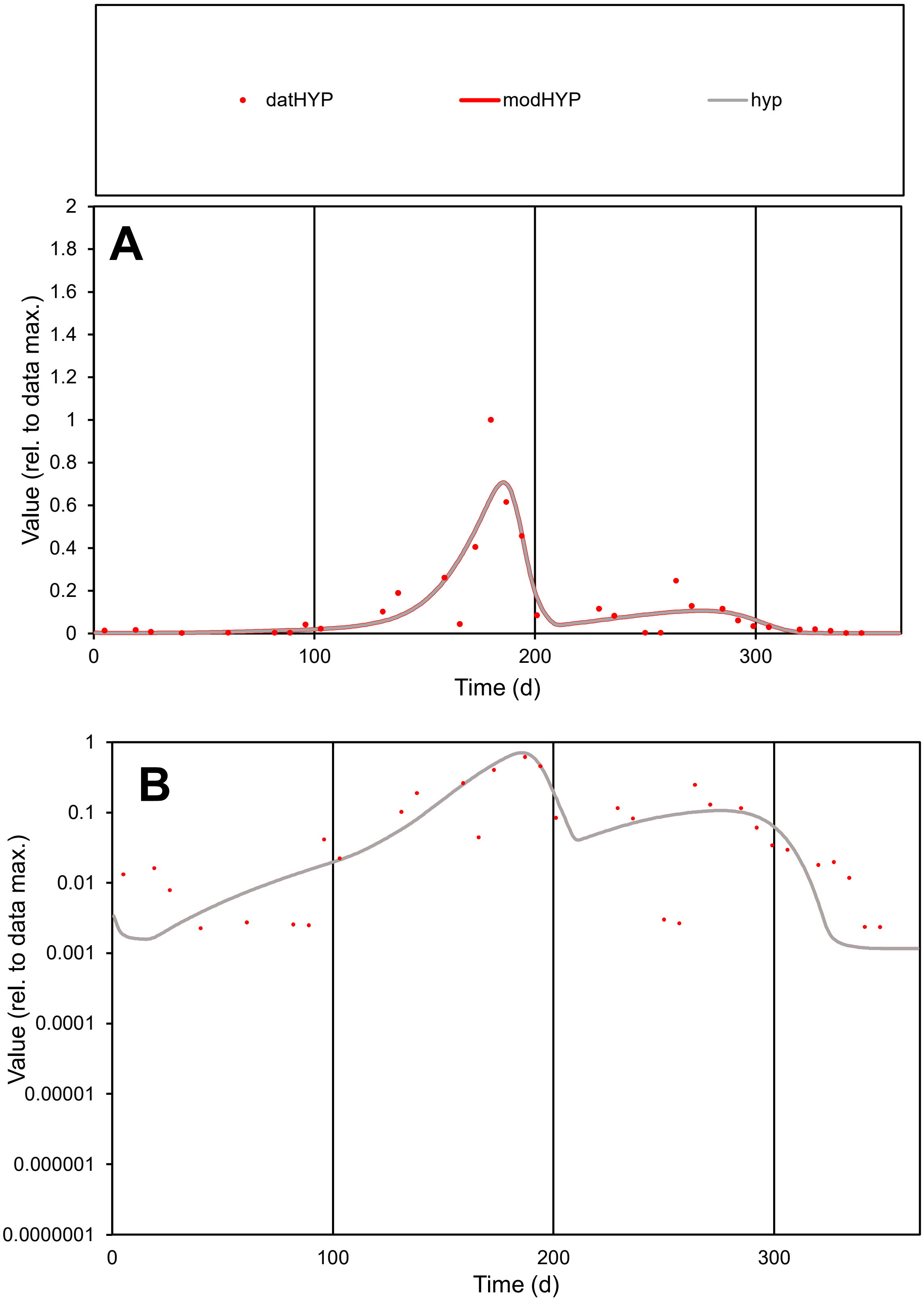


**C1**

| | |
|---|---|
| Variables | |
| HYP | Provide jtd or Dst |
| CalibOnly? | |
| 2 | 1 to reproduce of |
| oNormalize? | |
| 2 | 1=no, 2=yes |
| ofxn calc* | |
| 0.73387 | |
| ofxn mod | |
| 0.779 | |
| sRun | |
| RunA1 | |
| zout | |
| 1 | |
| Ave | |
| Data | |
| 0.00091 | |
| Model | |
| 0.0009 | |
| data/vMax | |
| 0.00725 | |
| ofxng | |
| 263.459 | |
| Min d/m | |
| 1.94747 | |
| ave/min d | |
| 55.2951 | |
| ave/min m | |
| 106.749 | |

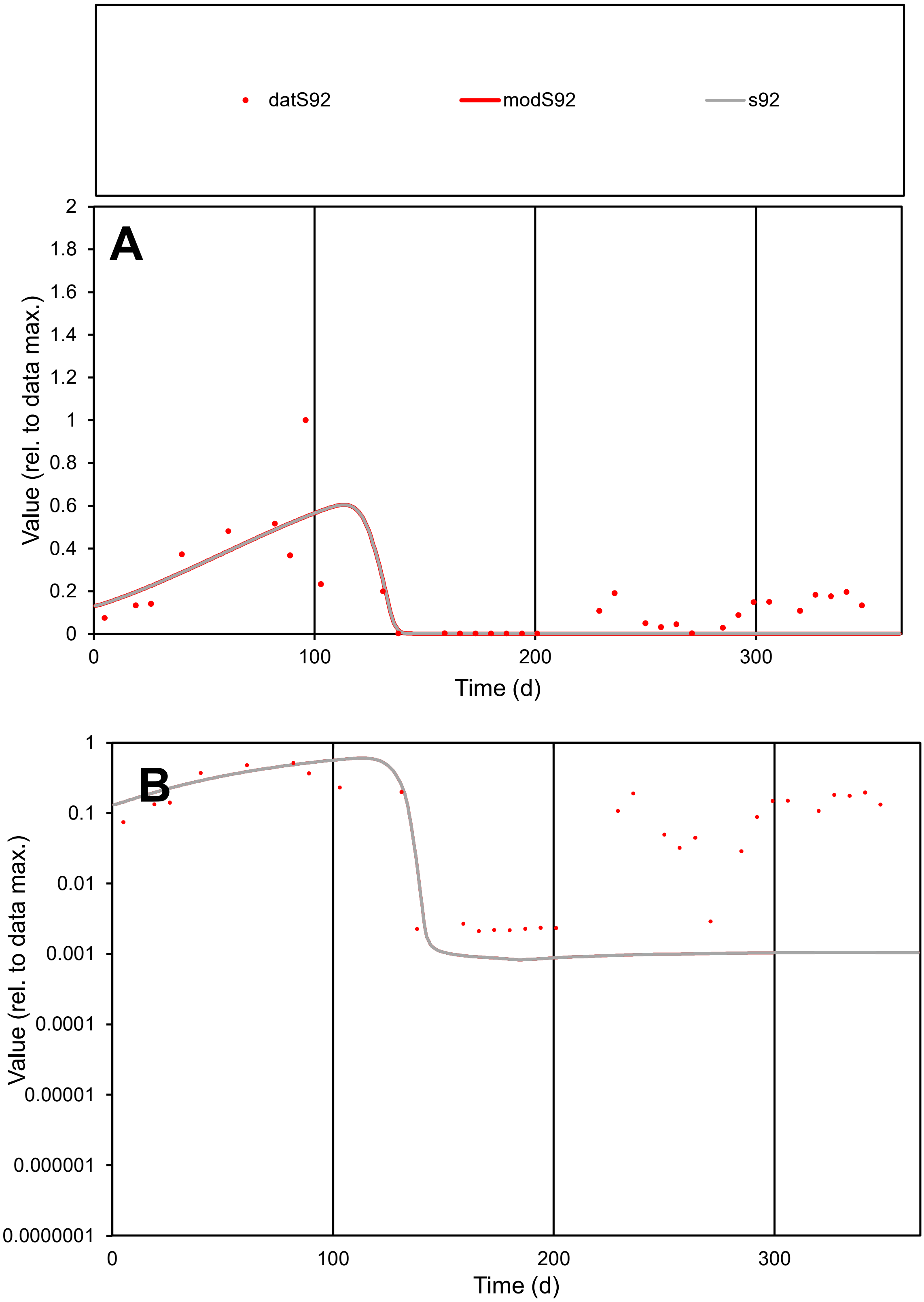


**C1**

Variables
S92 Provide jtd or Dst
CalibOnly?
2 1 to reproduce of
oNormalize?
2 1=no, 2=yes

ofxn calc*
0.86727
ofxn mod
0.8815

sRun
RunA1
zout
1

Ave
Data
0.00171
Model
0.0012

data/vMax
0.01092

ofxng
263.459

Min d/m
2.52109

ave/min d
74.4252
ave/min m
131.796

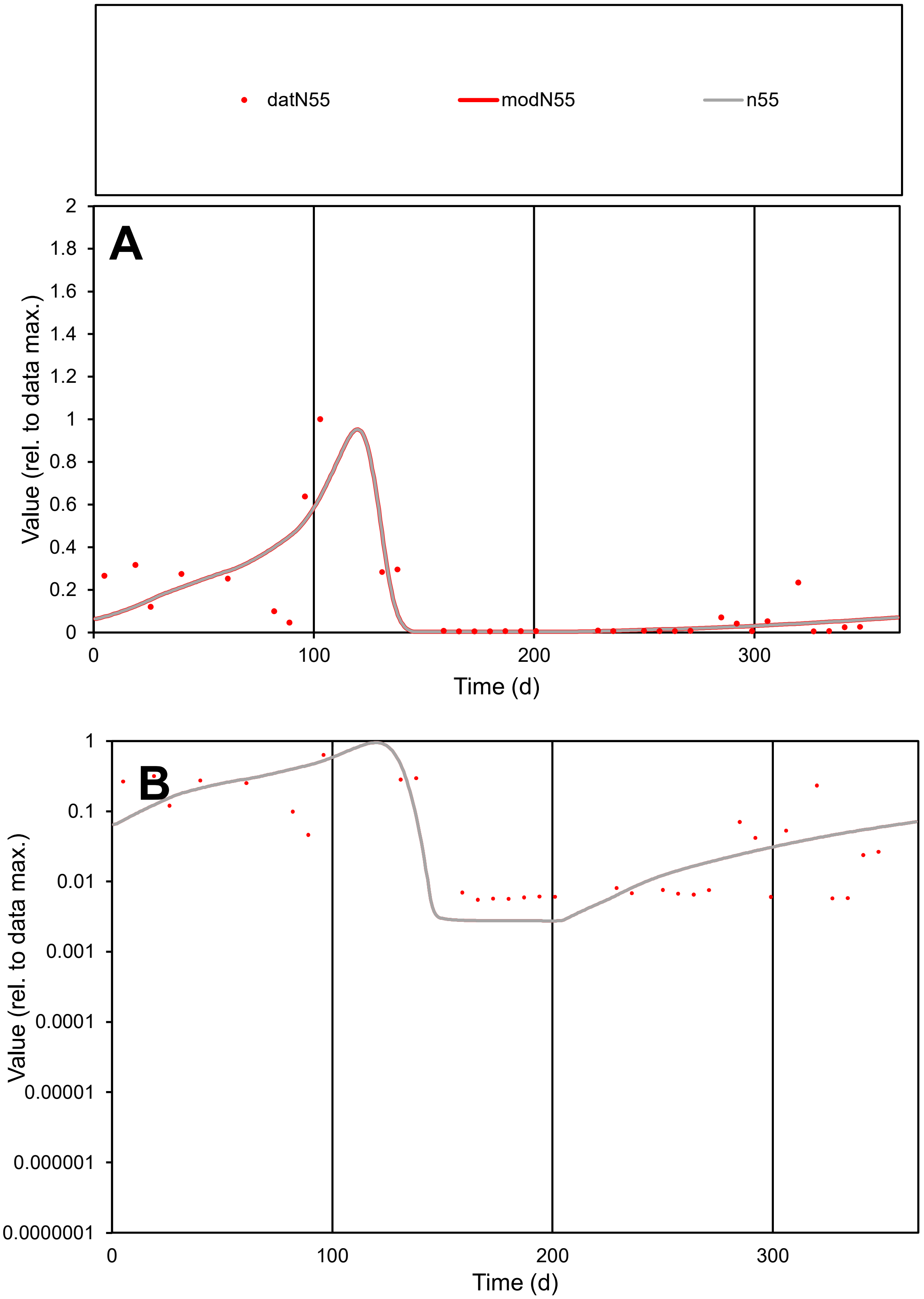


**C1**

Variables
N55 Provide jtd or Dst
CalibOnly?
2 1 to reproduce of
oNormalize?
2 1=no, 2=yes

ofxn calc*
1.01152
ofxn mod
1.05154

sRun
RunA1
zout
1

Ave
Data
0.00084
Model
0.0008

data/vMax
0.00671

ofxng
263.459

Min d/m
2.00054
ave/min d
22.8523
ave/min m
43.3747

326_Calib_S_A

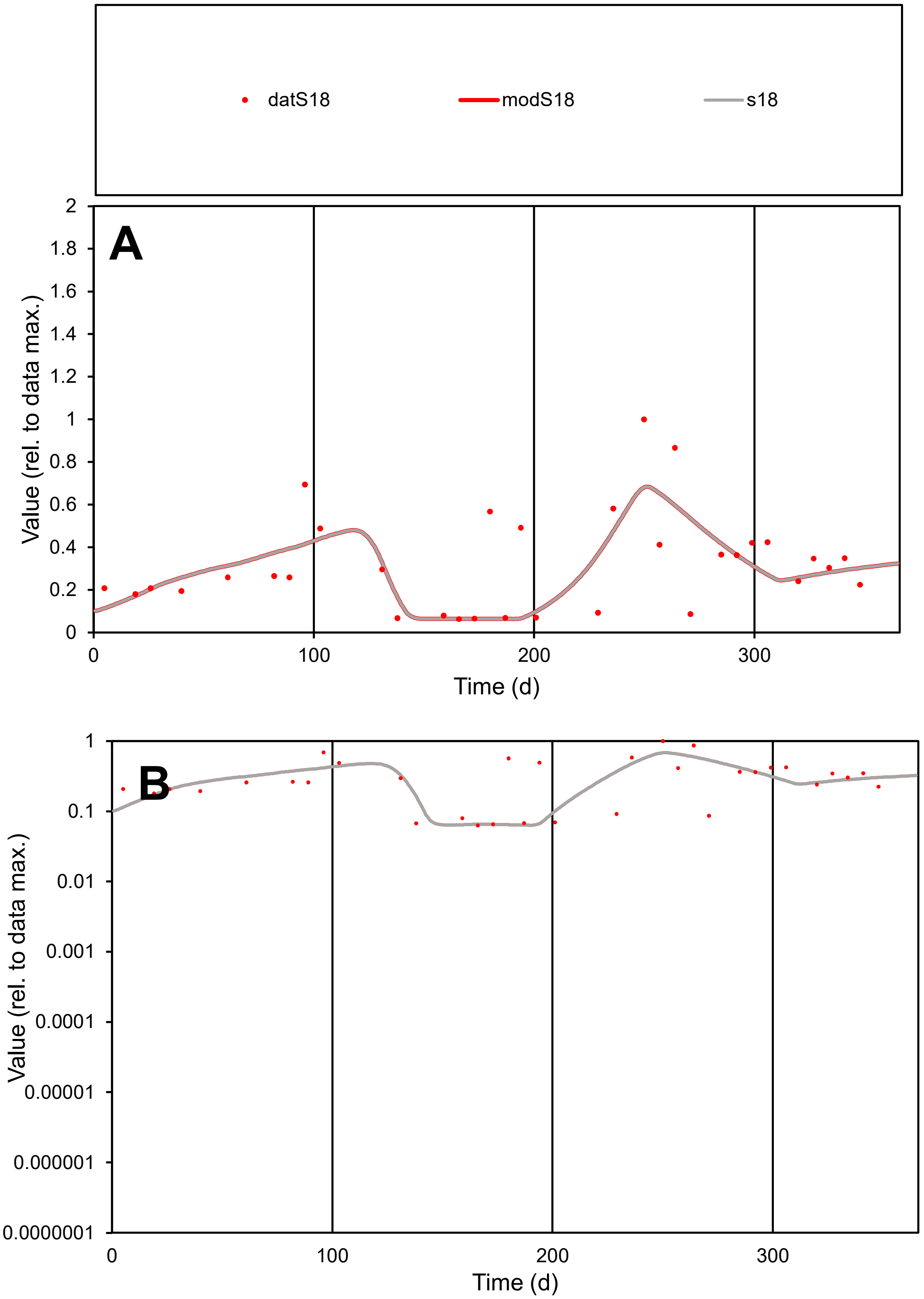


**C1**

Variables
S18 Provide jtd or Dst
CalibOnly?
2 1 to reproduce of
oNormalize?
2 1=no, 2=yes

ofxn calc*
0.5732
ofxn mod
0.58826

sRun
RunA1
zout
1

Ave
Data
0.00135
Model
0.00124

data/vMax
0.0042

ofxng
263.459

Min d/m
0.98874

ave/min d
5.11514
ave/min m
4.63437

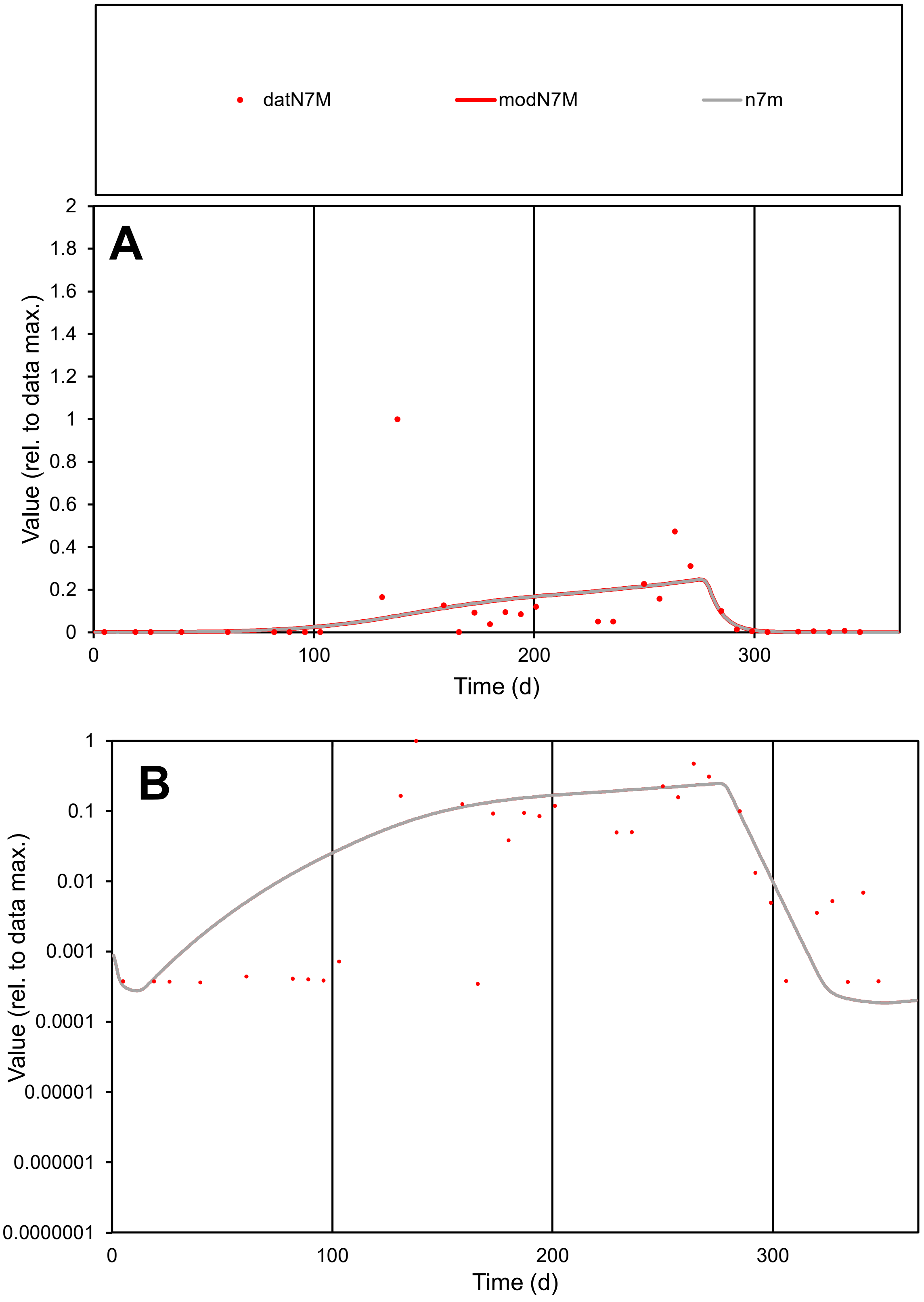


**C1**

Variables

N7M Provide jtd or Dst

CalibOnly?

2 1 to reproduce of

oNormalize?

2 1=no, 2=yes

ofxn calc*

1.84433

ofxn mod

1.85664

sRun

RunA1

zout

1

Ave

Data

0.00259

Model

0.00224

data/vMax

0.02733

ofxng

263.459

Min d/m

1.8737

ave/min d

272.06

ave/min m

440.241

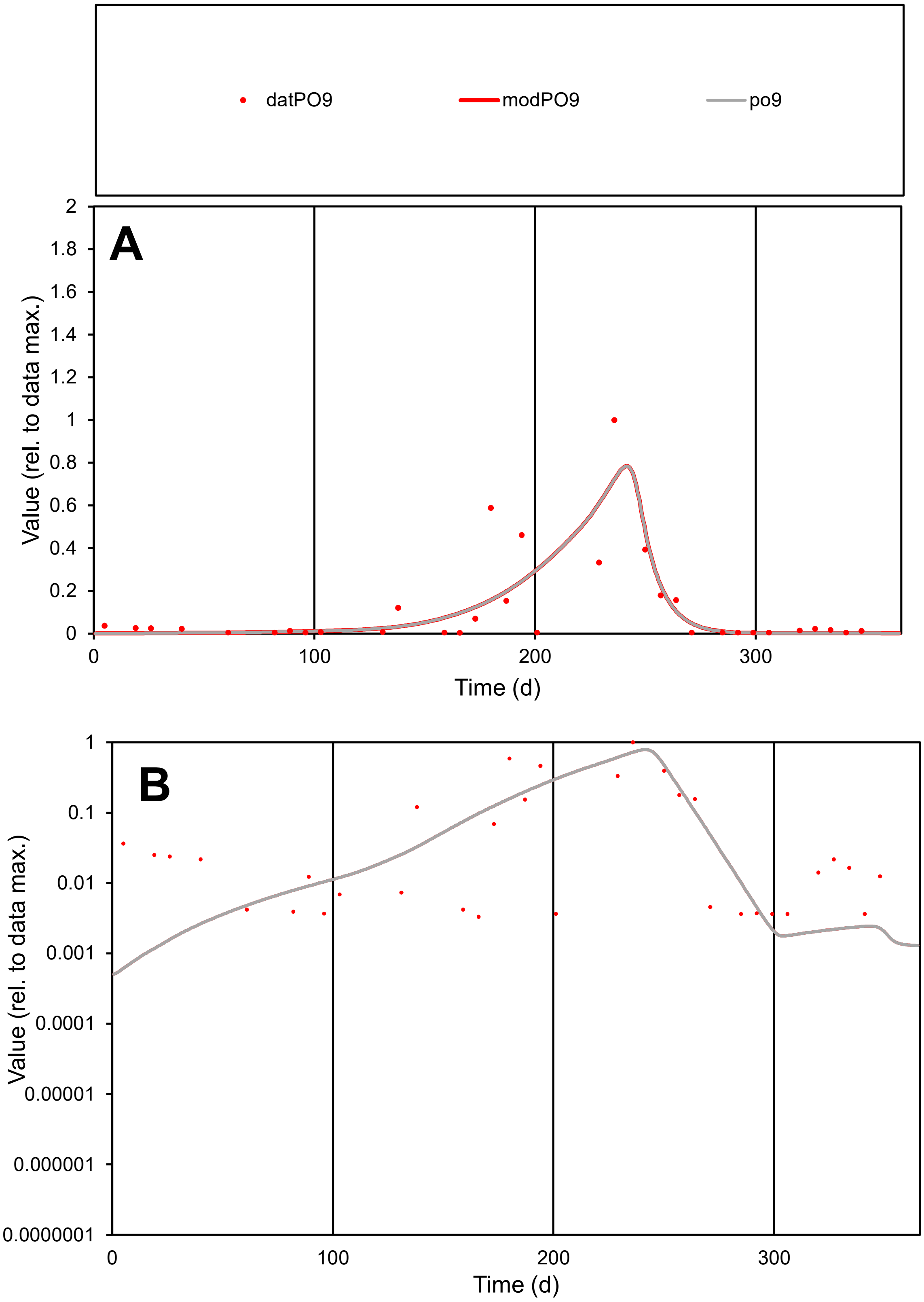


**C1**

Variables

PO9 Provide jtd or Dst

CalibOnly?

2 1 to reproduce of

oNormalize?

2 1=no, 2=yes

ofxn calc*

1.09233

ofxn mod

1.12117

sRun

RunA1

zout

1

Ave

Data

0.00195

Model

0.00182

data/vMax

0.01741

ofxng

263.459

Min d/m

5.01913

ave/min d

33.7671

ave/min m

158.043

326_Calib_S_A

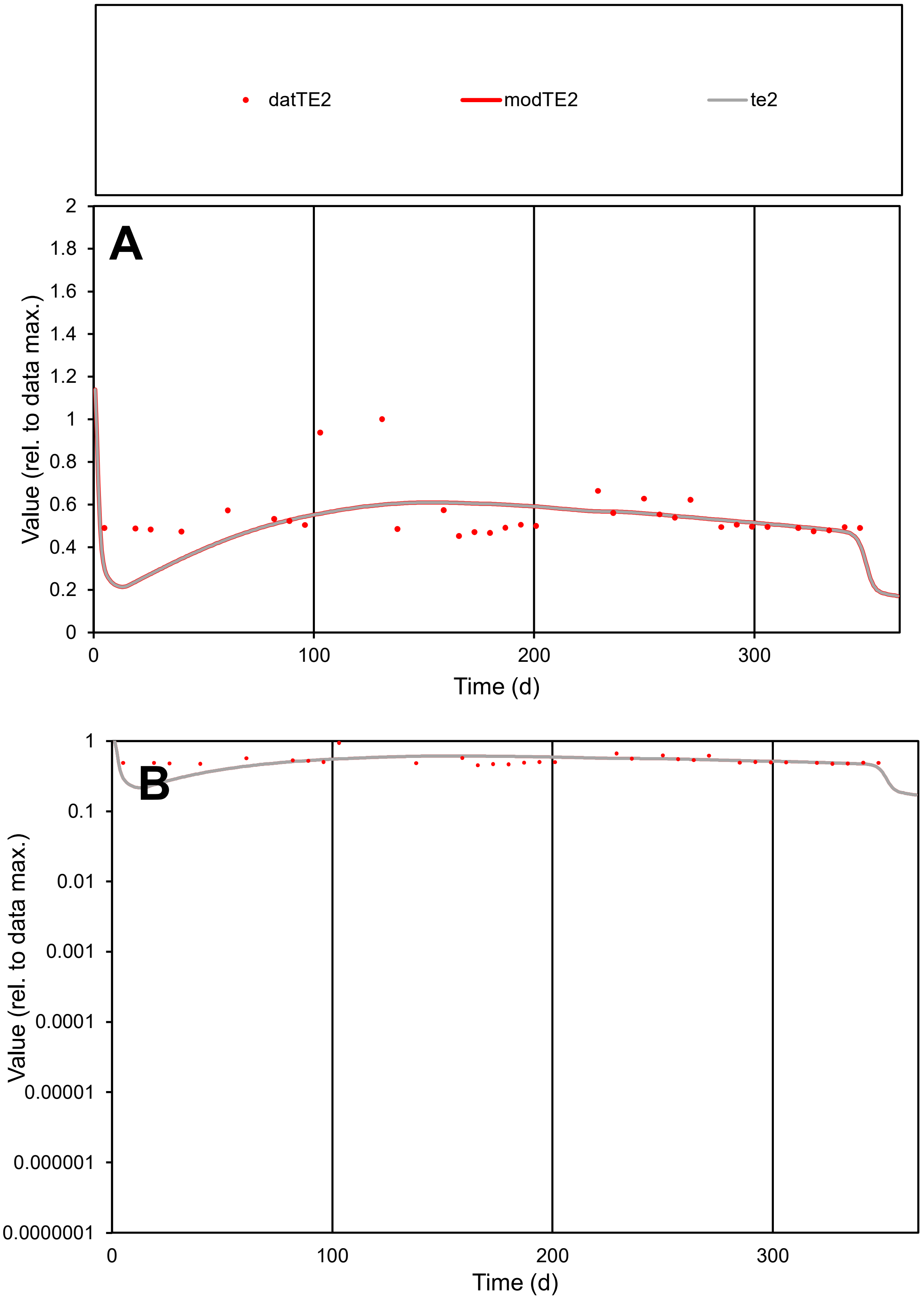


**C1**

Variables
TE2 Provide jtd or Dst
CalibOnly?
2 1 to reproduce of
oNormalize?
2 1=no, 2=yes

ofxn calc*
0.25047
ofxn mod
0.2746

sRun
RunA1
zout
1

Ave
Data
0.0001
Model
9.6E-05

data/vMax
0.00019

ofxng
263.459

Min d/m
1.84461

ave/min d
1.20135
ave/min m
2.08935

326_Calib_S_A

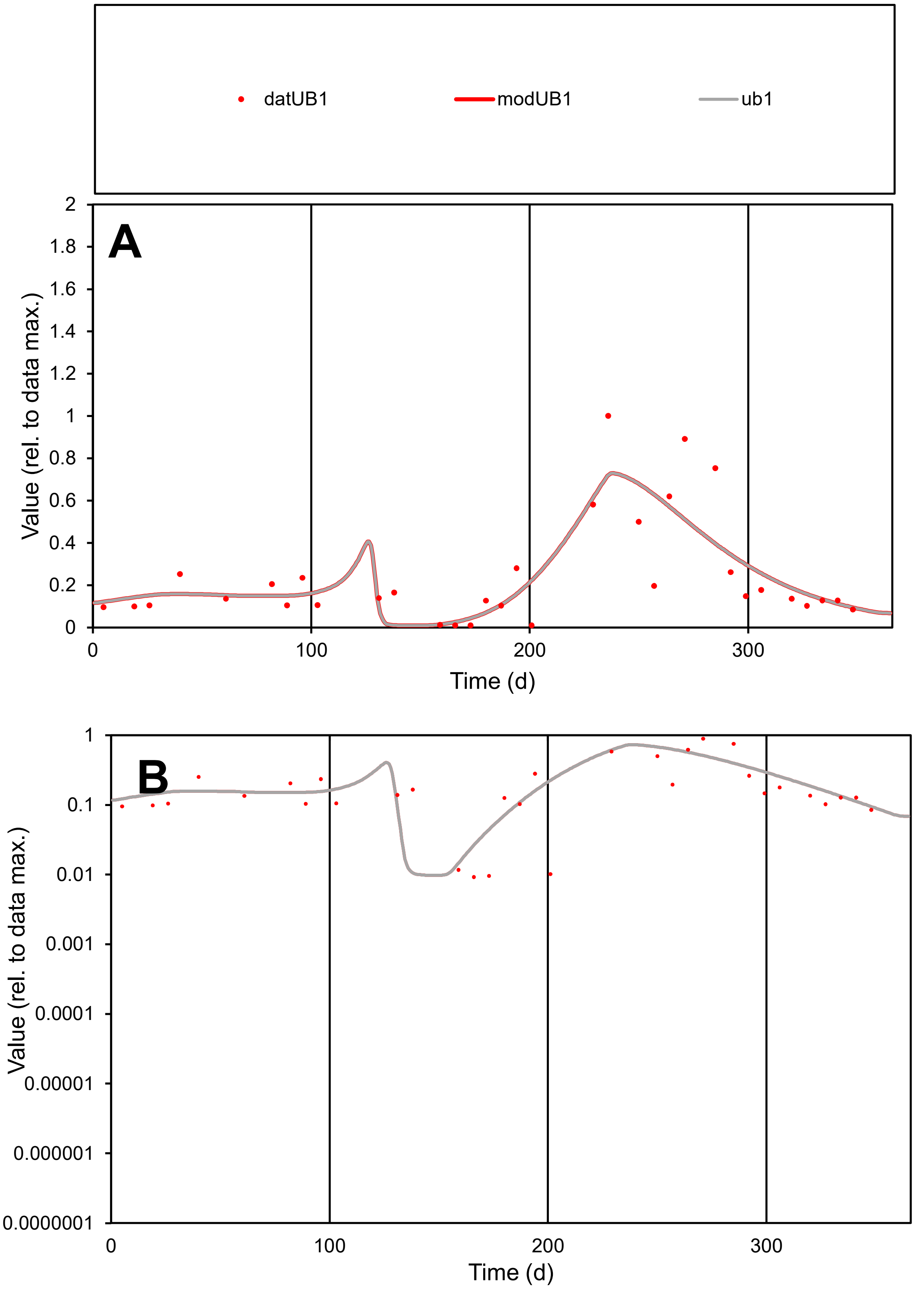


**C1**

Variables
UB1 Provide jtd or Dst
CalibOnly?
2 1 to reproduce of
oNormalize?
2 1=no, 2=yes

ofxn calc*
0.61706
ofxn mod
0.64352

sRun
RunA1
zout
1

Ave
Data
0.00147
Model
0.00146

data/vMax
0.00614

ofxng
263.459

Min d/m
0.90484

ave/min d
26.0102
ave/min m
23.3618

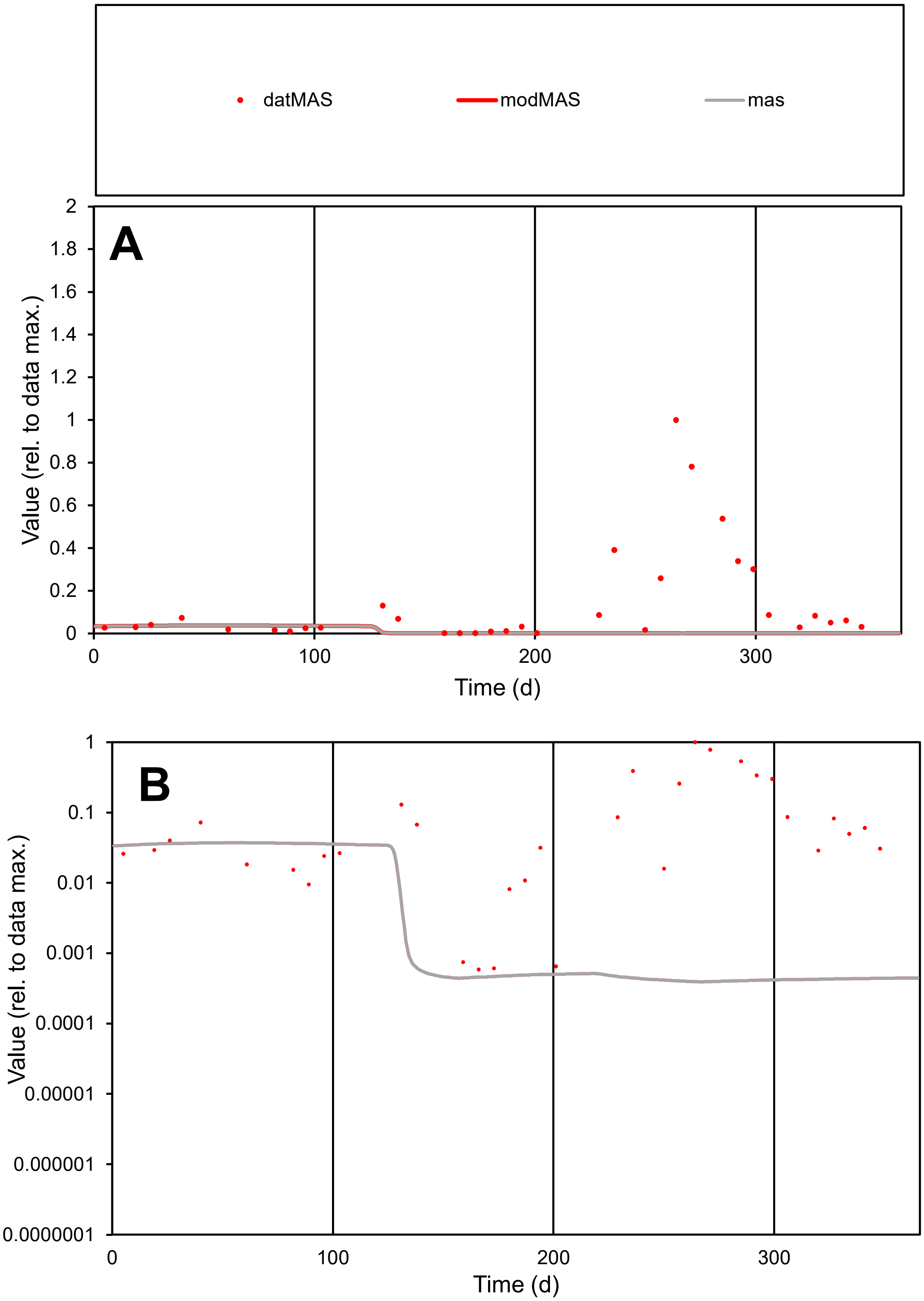


**C1**

Variables
MAS Provide jtd or Dst
CalibOnly?
2 1 to reproduce of
oNormalize?
2 1=no, 2=yes

ofxn calc*
1.9411
ofxn mod
1.94604

sRun
RunA1
zout
1

Ave
Data
0.00281
Model
0.00021

data/vMax
0.02032

ofxng
263.459

Min d/m
1.48708

ave/min d
235.674
ave/min m
26.4084

326_Calib_S_A

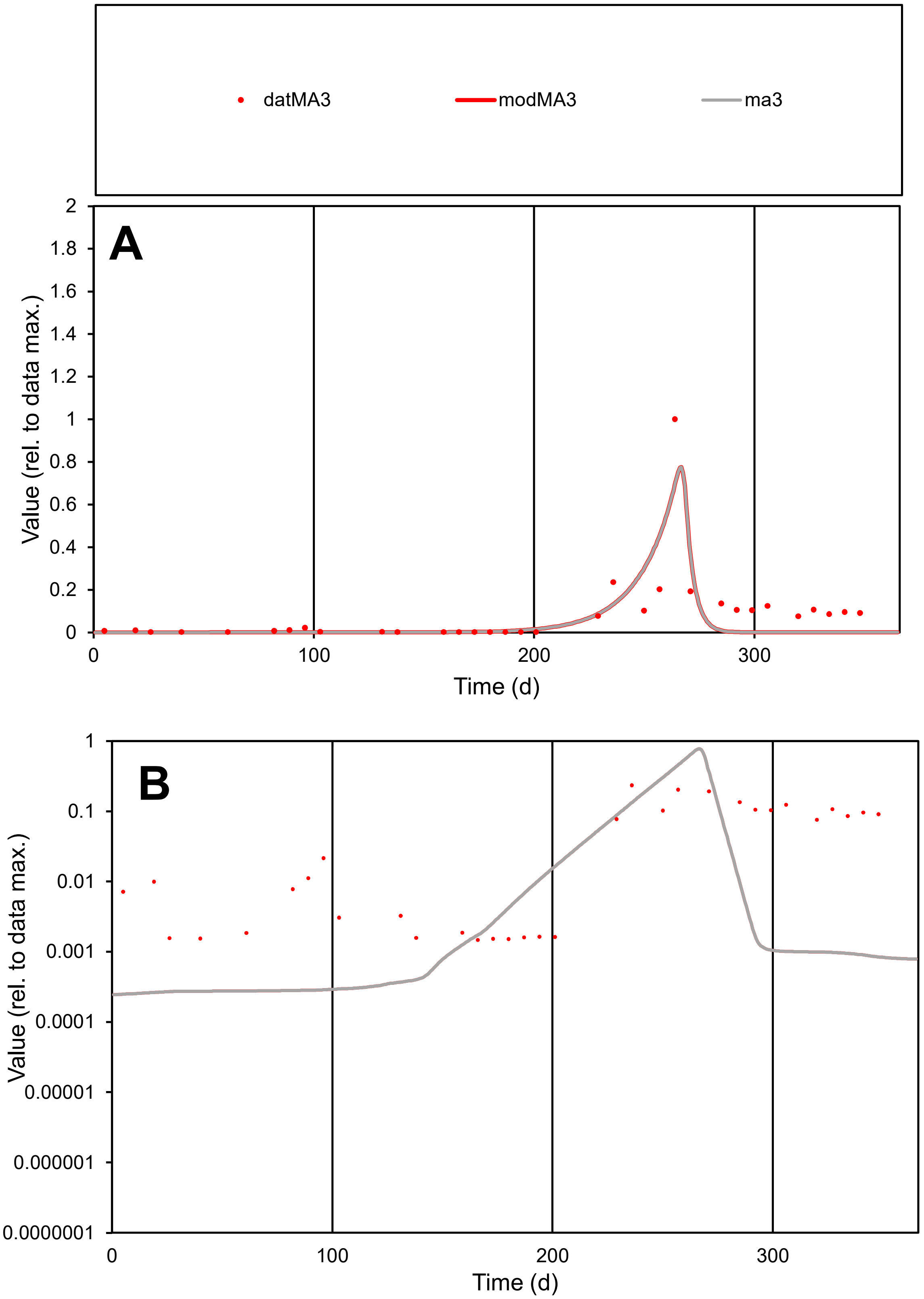


**C1**

Variables
MA3 Provide jtd or Dst
CalibOnly?
2 1 to reproduce of
oNormalize?
2 1=no, 2=yes

ofxn calc*
1.07667
ofxn mod
1.12053

sRun
RunA1
zout
1

Ave
Data
0.00356
Model
0.00261

data/vMax
0.04172

ofxng
263.459

Min d/m
5.87452

ave/min d
58.0396
ave/min m
249.549

326_Calib_S_A

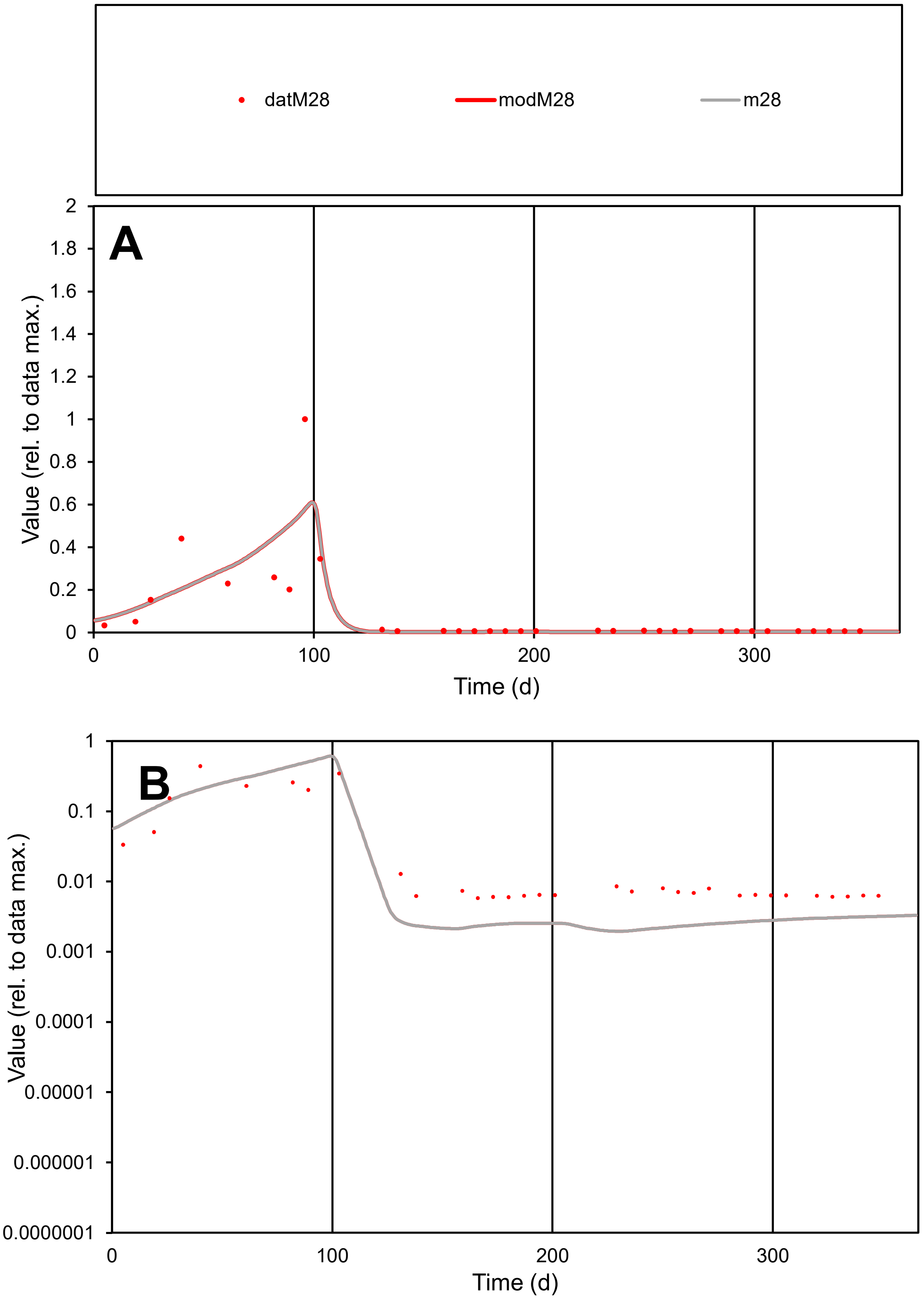


**C1**

Variables
M28 Provide jtd or Dst
CalibOnly?
2 1 to reproduce of
oNormalize?
2 1=no, 2=yes

ofxn calc*
1.20049
ofxn mod
1.22819

sRun
RunA1
zout
1

Ave
Data
0.0008
Model
0.00079

data/vMax
0.00921

ofxng
263.459

Min d/m
2.98099

ave/min d
15.0456
ave/min m
43.9636

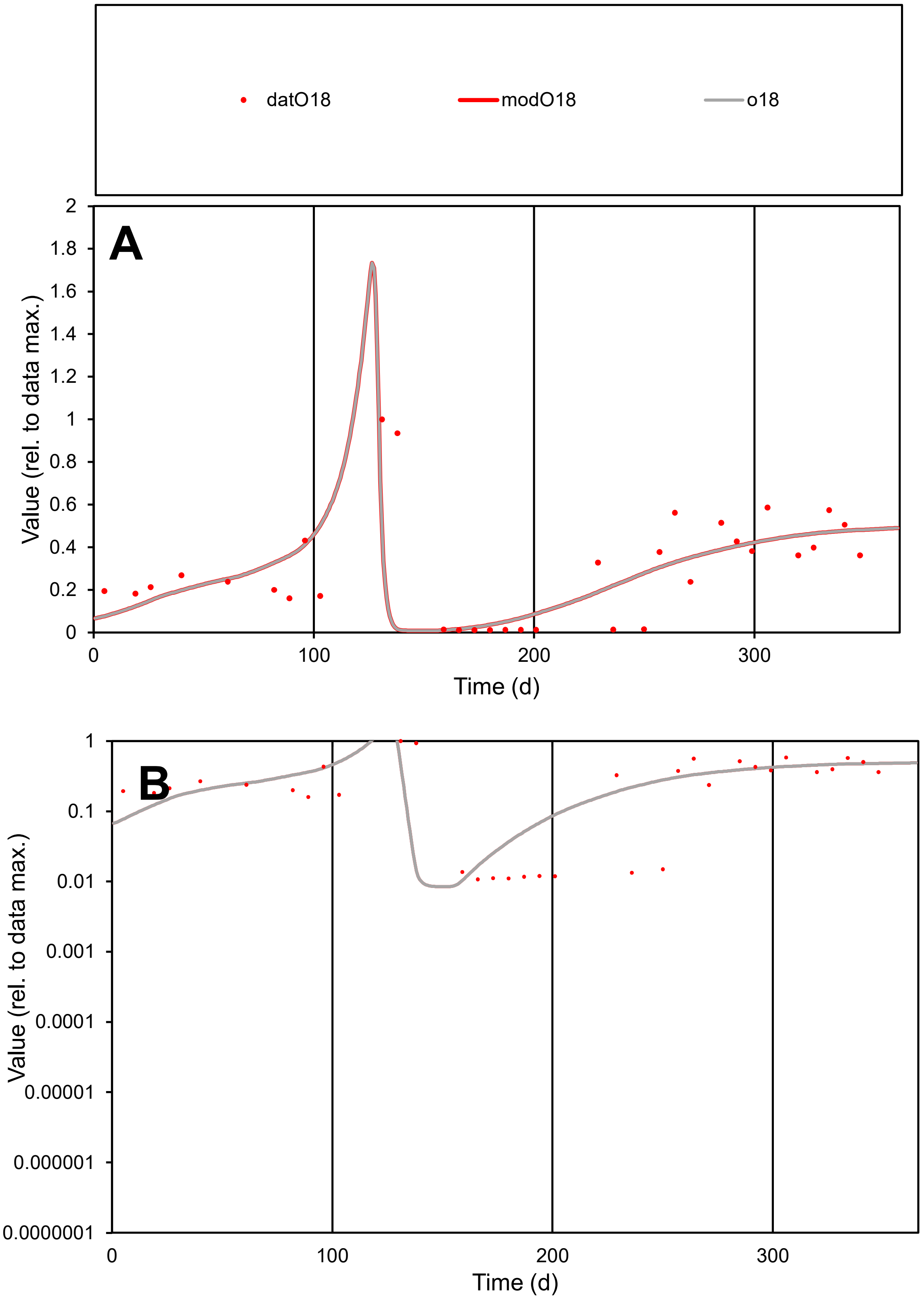


**C1**

Variables

| | |
|---|---|
| O18 | Provide jtd or Dst |
| CalibOnly? | |
| 2 | 1 to reproduce of |
| oNormalize? | |
| 2 | 1=no, 2=yes |

ofxn calc*
0.70705
ofxn mod
0.74766

sRun
RunA1
zout
1

Ave
Data
0.00139
Model
0.00132

data/vMax
0.00474

ofxng
263.459

Min d/m
1.00631

ave/min d
27.3344
ave/min m
26.0565

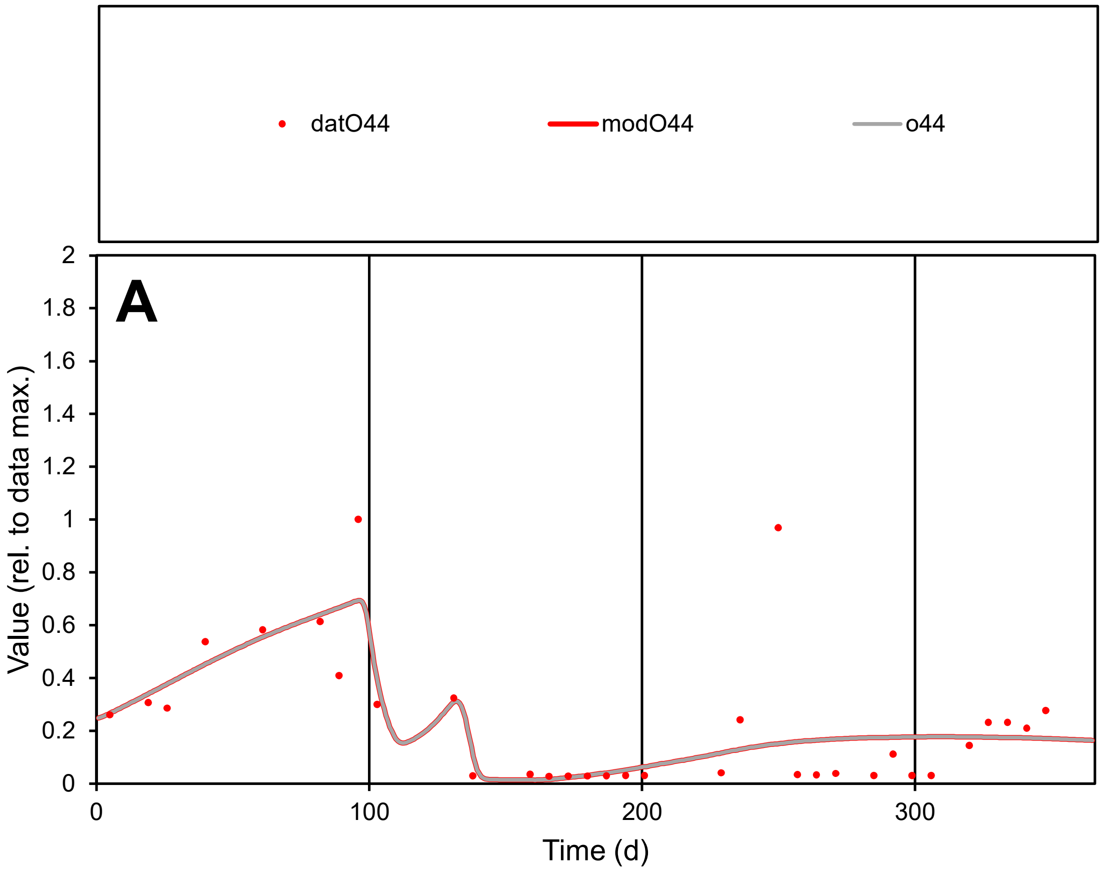


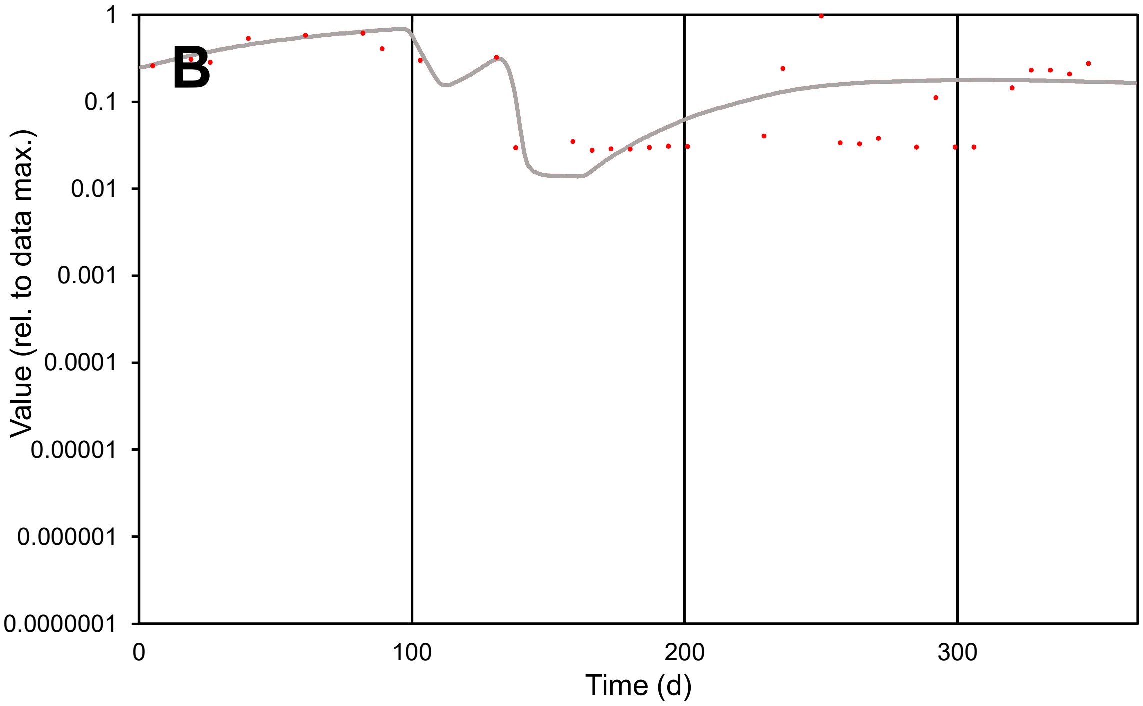


**C1**

Variables
O44 Provide jtd or Dst
CalibOnly?
2 1 to reproduce of
oNormalize?
2 1=no, 2=yes

ofxn calc*
0.76878
ofxn mod
0.7847

sRun
RunA1
zout
1

Ave
Data
0.0009
Model
0.00089

data/vMax
0.00395

ofxng
263.459

Min d/m
1.99562

ave/min d
8.22537
ave/min m
16.3096

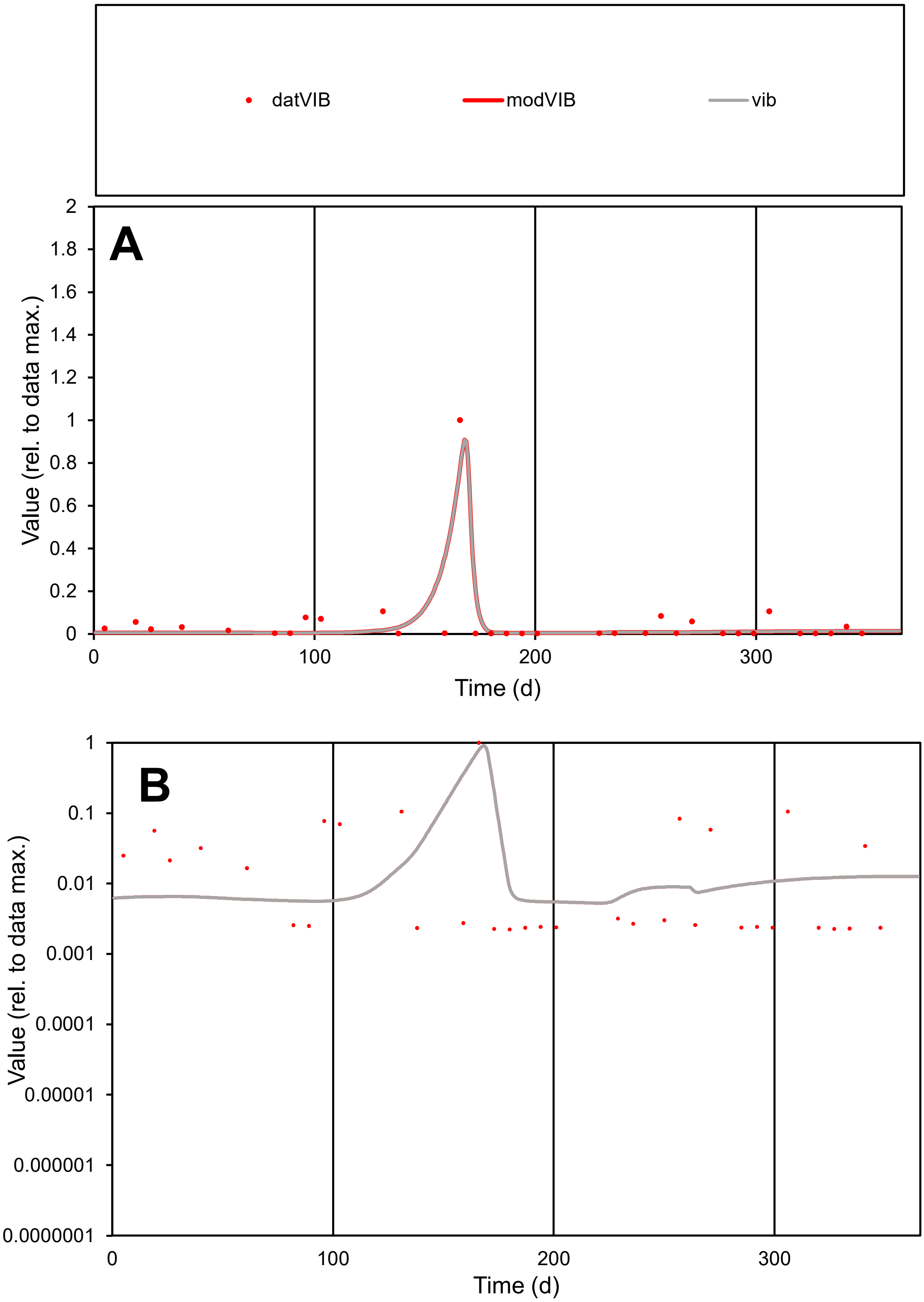


**C1**

Variables

VIB Provide jtd or Dst

CalibOnly?

2 1 to reproduce of

oNormalize?

2 1=no, 2=yes

ofxn calc*

1.23969

ofxn mod

1.38219

sRun

RunA1

zout

1

Ave

Data

0.00043

Model

0.00037

data/vMax

0.00808

ofxng

263.459

Min d/m

0.40943

ave/min d

23.5358

ave/min m

8.37112

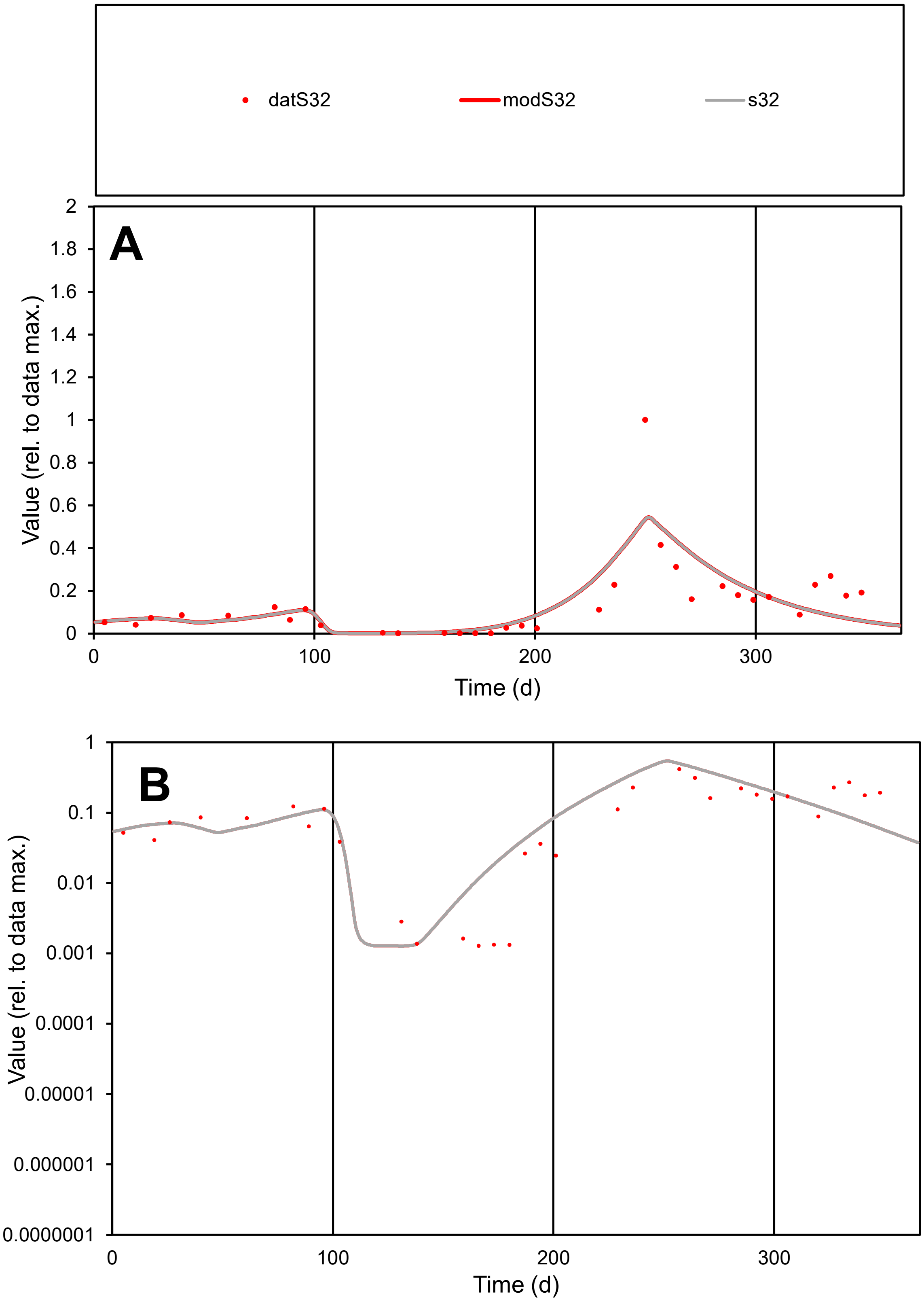


**C1**

Variables
S32 Provide jtd or Dst
CalibOnly?
2 1 to reproduce of
oNormalize?
2 1=no, 2=yes

ofxn calc*
0.78267
ofxn mod
0.80969

sRun
RunA1
zout
1

Ave
Data
0.00118
Model
0.00118

data/vMax
0.00835

ofxng
263.459

Min d/m
1.00471

ave/min d
111.004
ave/min m
111.064

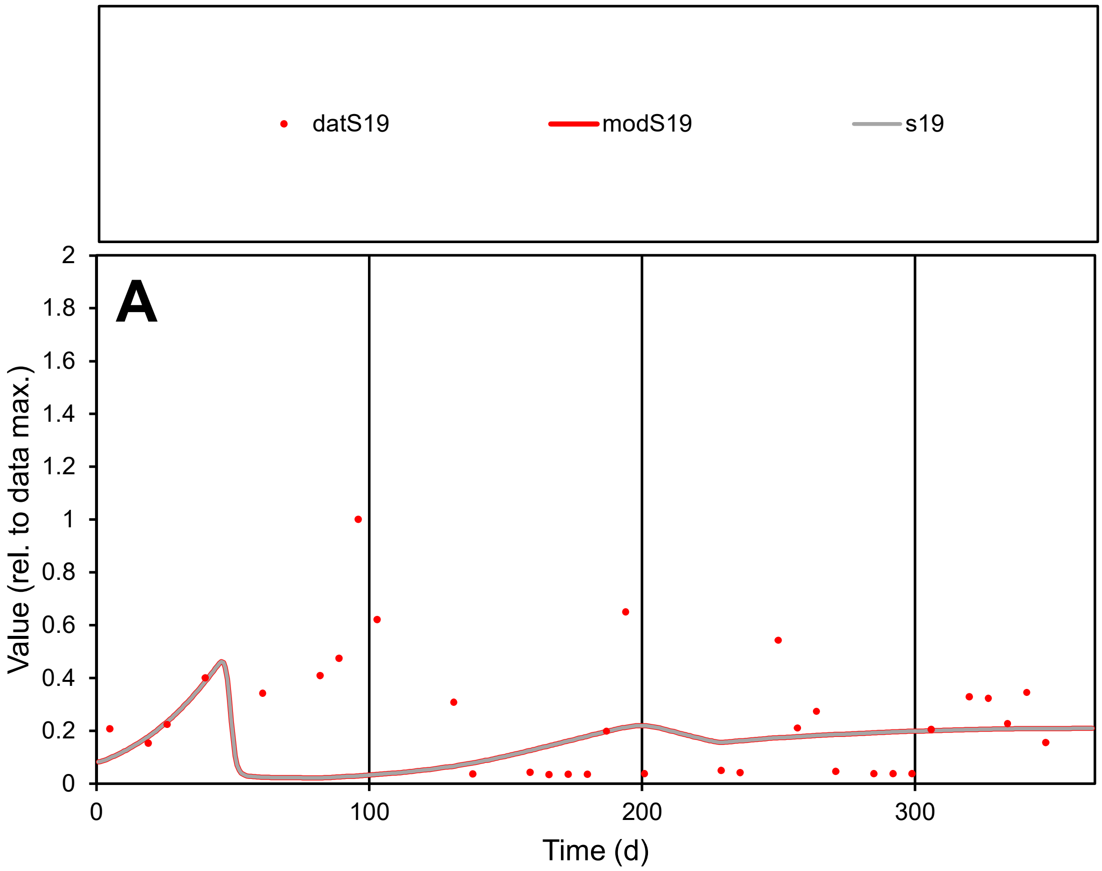


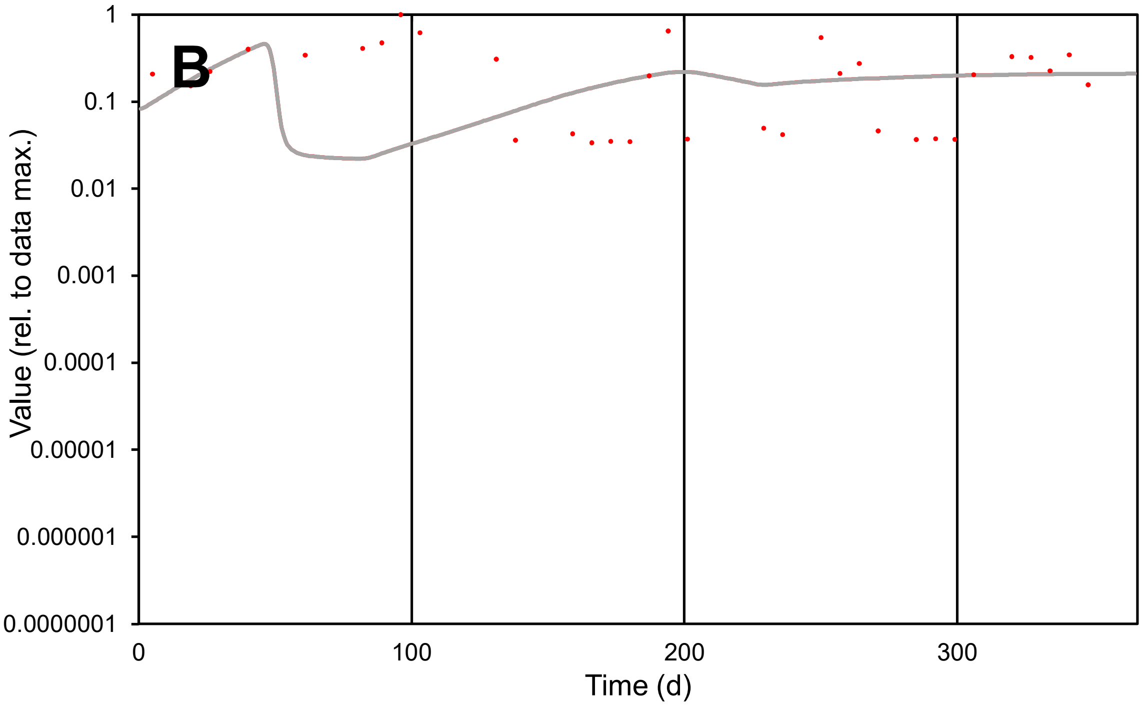


**C1**

Variables
S19 Provide jtd or Dst
CalibOnly?
2 1 to reproduce of
oNormalize?
2 1=no, 2=yes

ofxn calc*
1.10343
ofxn mod
1.10917

sRun
RunA1
zout
1

Ave
Data
0.00034
Model
0.00022

data/vMax
0.00139

ofxng
263.459

Min d/m
1.49856

ave/min d
7.27084
ave/min m
7.21787

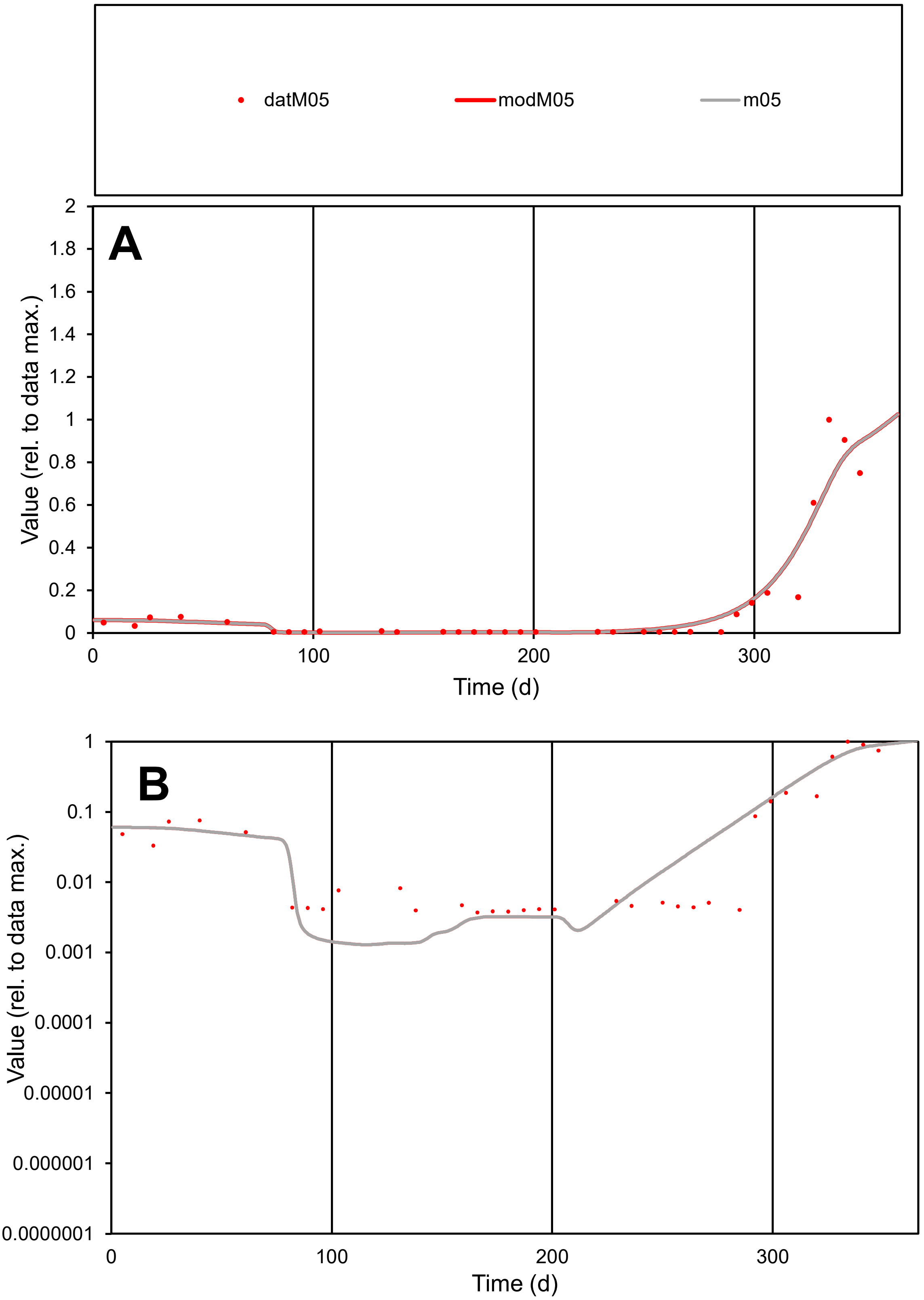


**C1**

Variables
M05 Provide jtd or Dst
CalibOnly?
2 1 to reproduce of
oNormalize?
2 1=no, 2=yes

ofxn calc*
0.57084
ofxn mod
0.61704

sRun
RunA1
zout
1

Ave
Data
0.00147
Model
0.00157

data/vMax
0.01153

ofxng
263.459

Min d/m
2.74005

ave/min d
34.5469
ave/min m
100.475

326_Calib_S_A

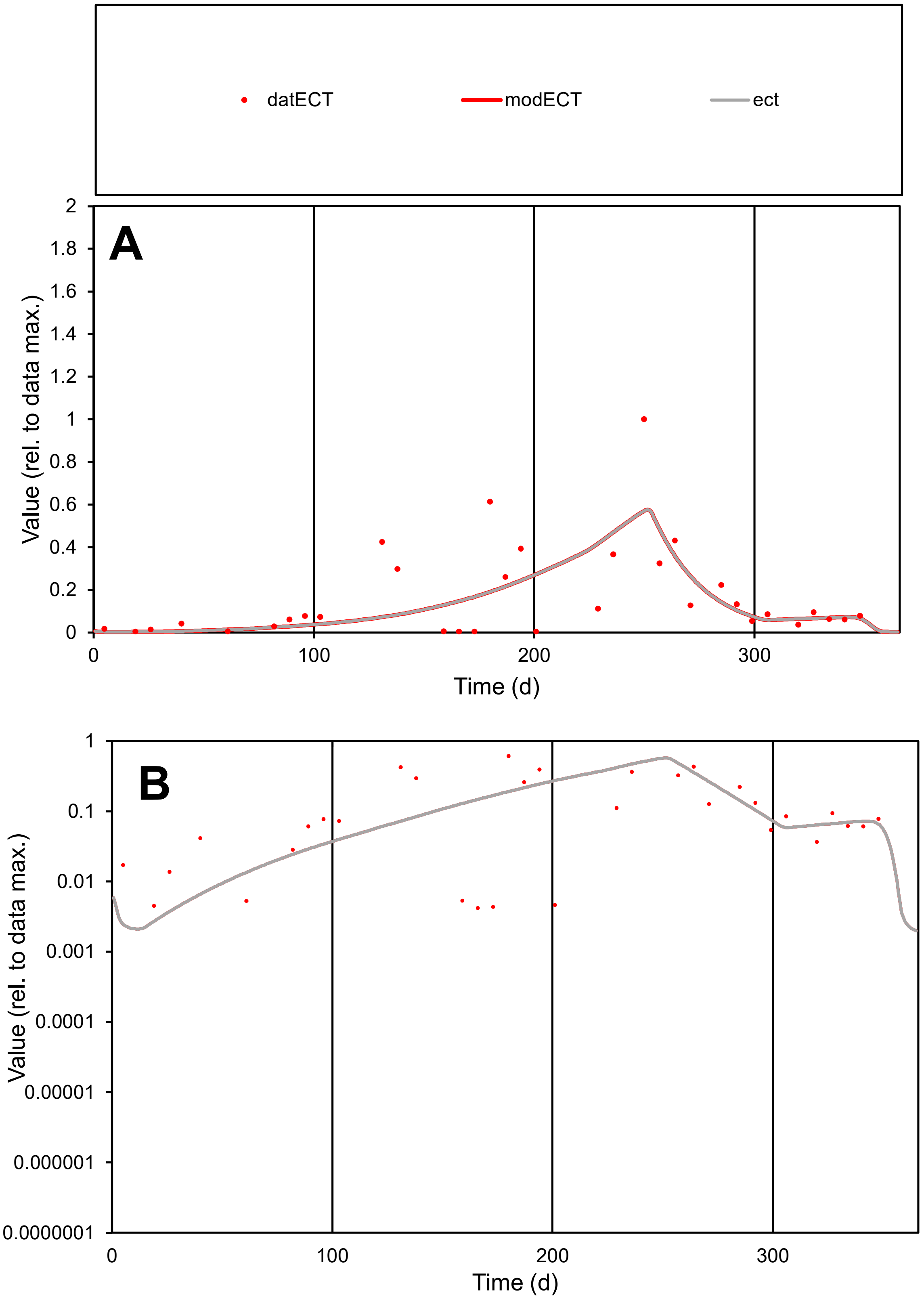


**C1**

Variables
ECT Provide jtd or Dst
CalibOnly?
2 1 to reproduce of
oNormalize?
2 1=no, 2=yes

ofxn calc*
0.95635
ofxn mod
0.97446

sRun
RunA1
zout
1

Ave
Data
0.00112
Model
0.001

data/vMax
0.00668

ofxng
263.459

Min d/m
1.70377

ave/min d
39.835
ave/min m
60.8629

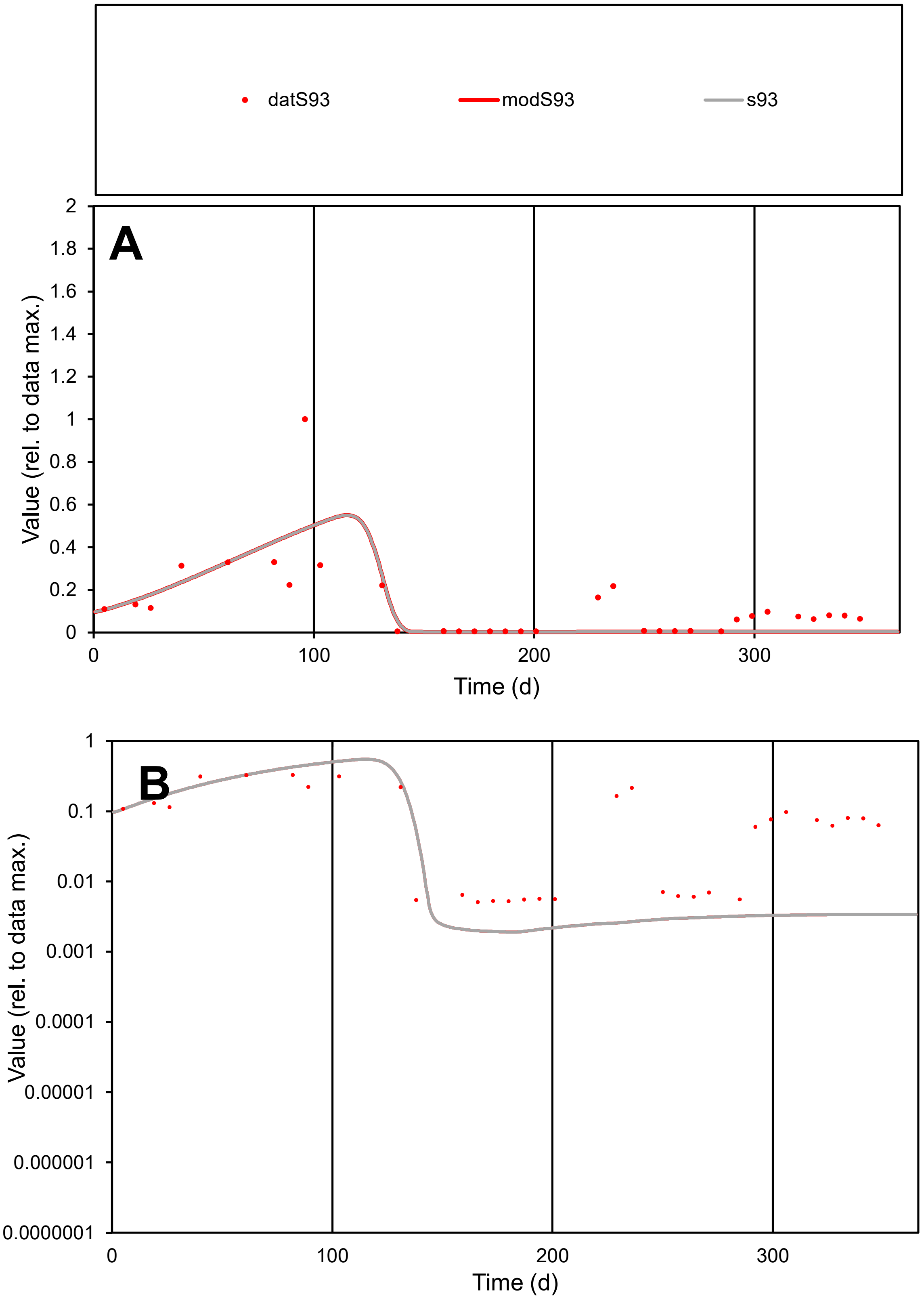


**C1**

Variables
S93 Provide jtd or Dst
CalibOnly?
2 1 to reproduce of
oNormalize?
2 1=no, 2=yes

ofxn calc*
0.97329
ofxn mod
0.98749

sRun
RunA1
zout
1

Ave
Data
0.00159
Model
0.00124

data/vMax
0.01266

ofxng
263.459

Min d/m
2.67621

ave/min d
24.6059
ave/min m
51.5182

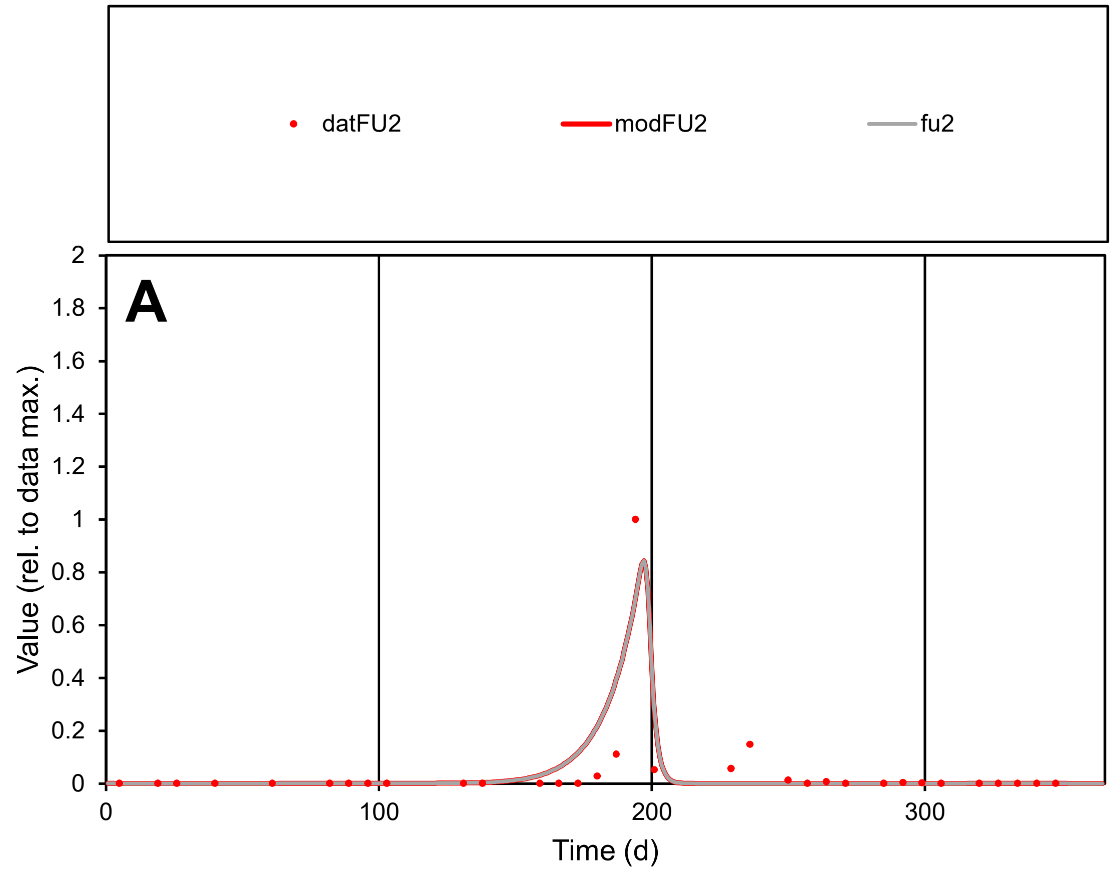


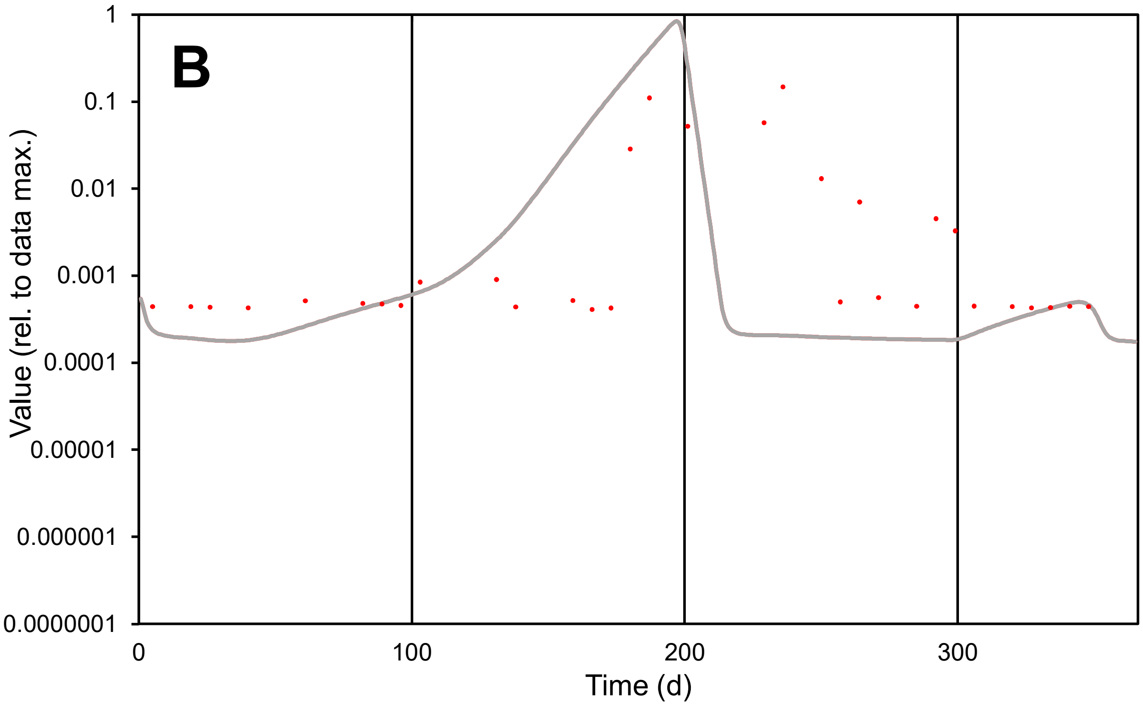


**C1**

Variables
FU2 Provide jtd or Dst
CalibOnly?
2 1 to reproduce of
oNormalize?
2 1=no, 2=yes

ofxn calc*
1.64977
ofxn mod
1.88799

sRun
RunA1
zout
1

Ave
Data
0.00269
Model
0.00301

data/vMax
0.0619

ofxng
263.459

Min d/m
2.25939

ave/min d
106.656
ave/min m
269.353

326_Calib_S_A

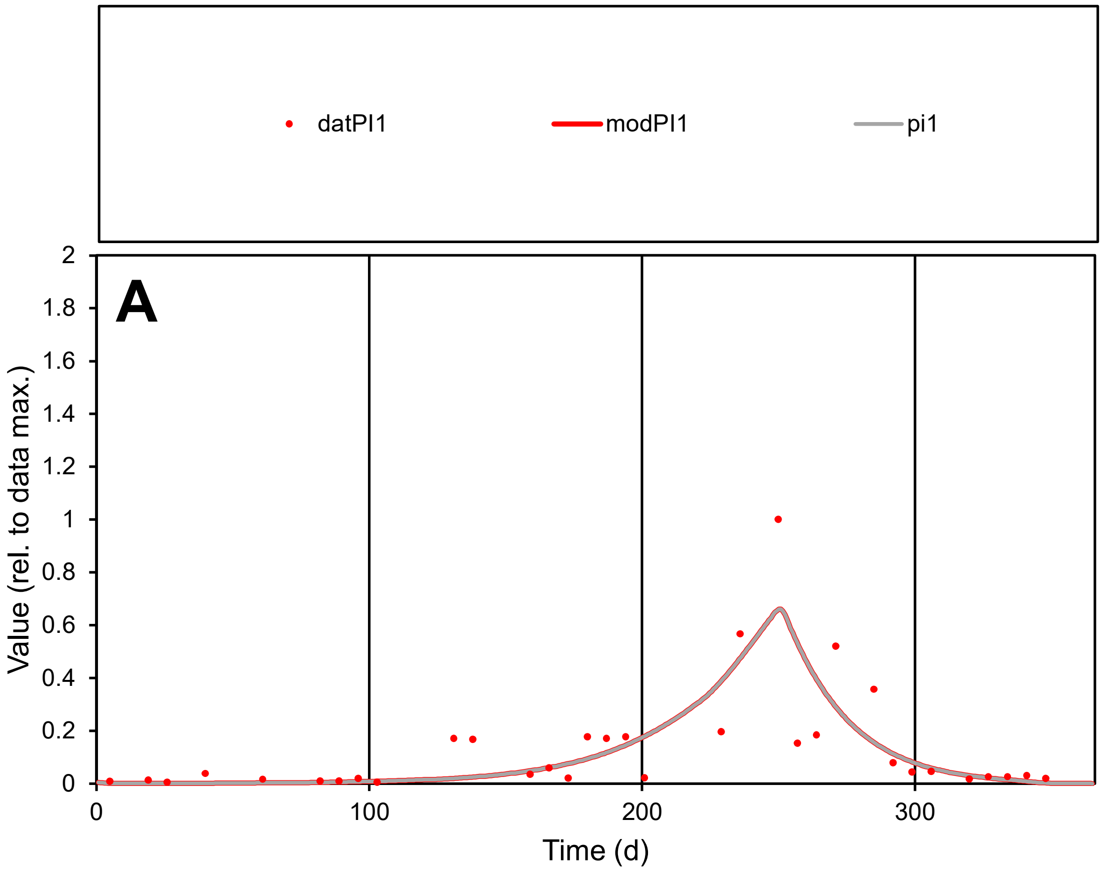


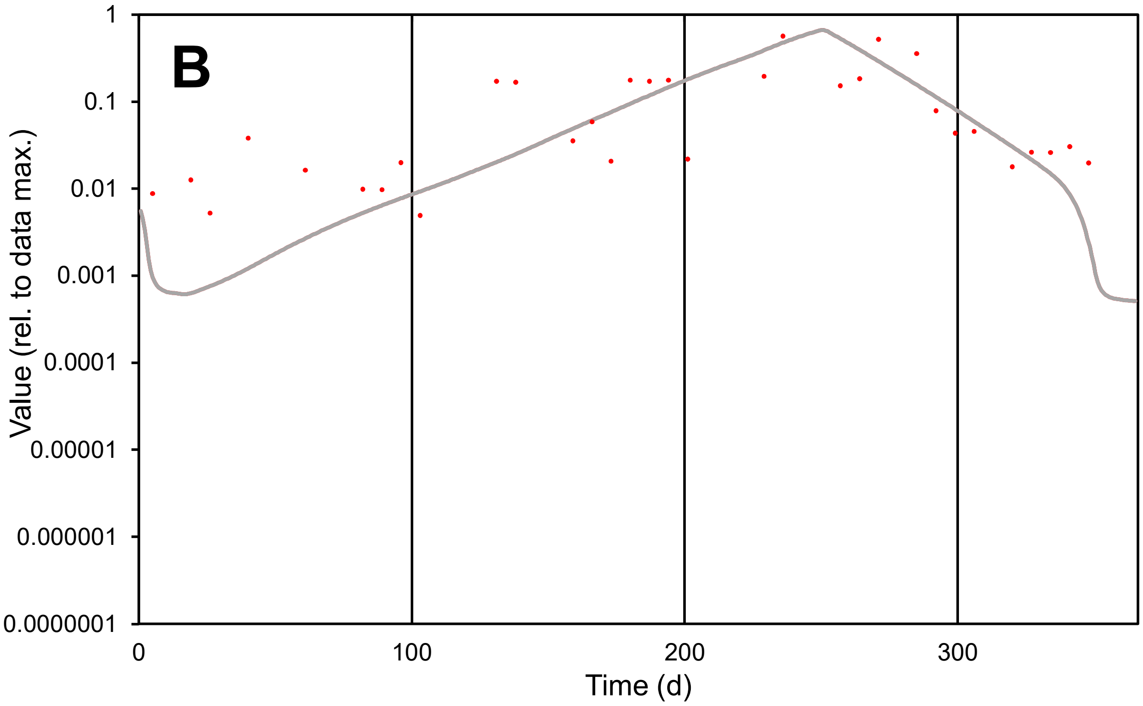


**C1**

Variables
PI1 Provide jtd or Dst
CalibOnly?
2 1 to reproduce of
oNormalize?
2 1=no, 2=yes

ofxn calc*
0.92711
ofxn mod
0.95914

sRun
RunA1
zout
1

Ave
Data
0.00145
Model
0.00132

data/vMax
0.01093

ofxng
263.459

Min d/m
7.60079

ave/min d
26.9795
ave/min m
186.662

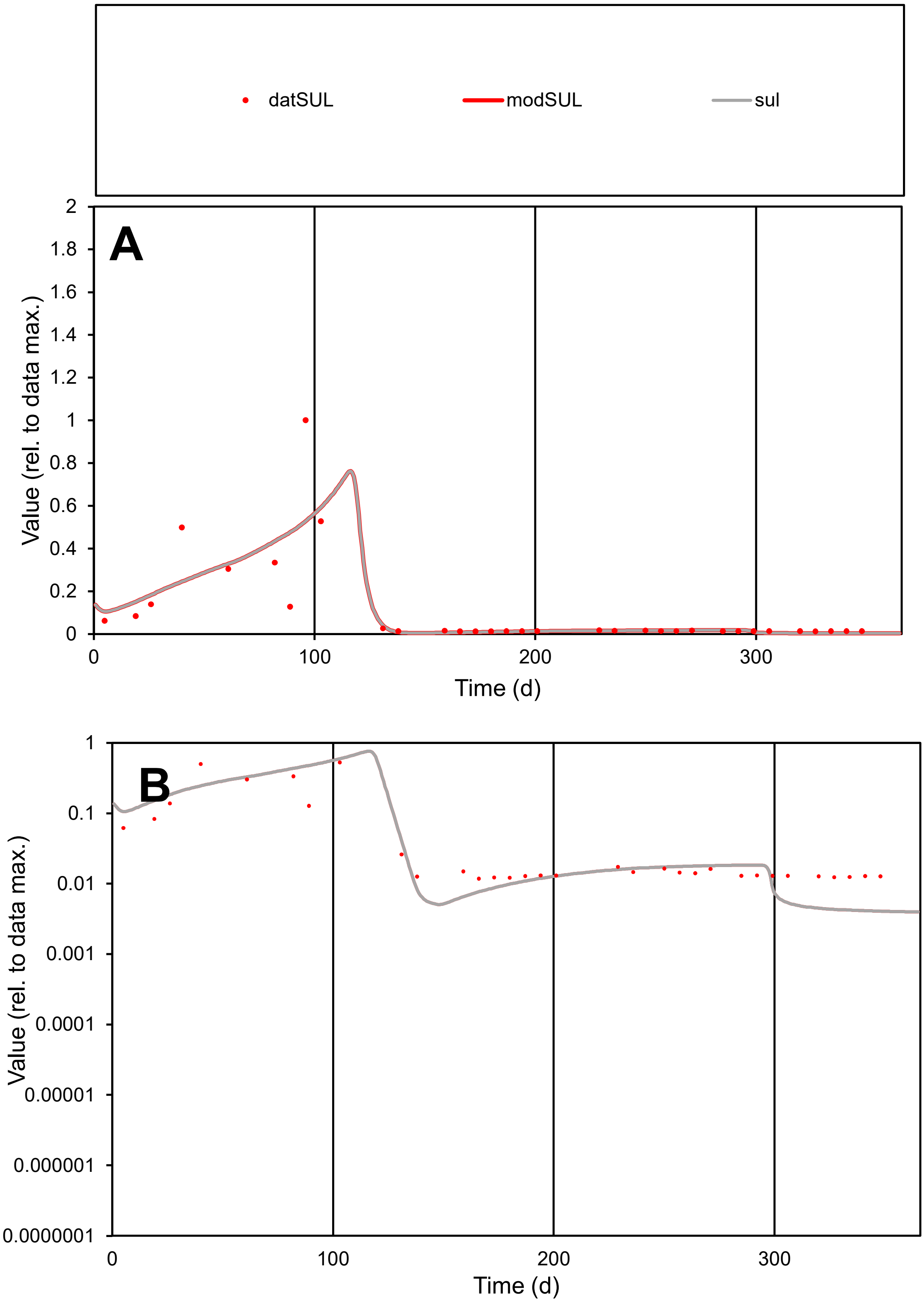


**C1**

Variables
SUL Provide jtd or Dst
CalibOnly?
2 1 to reproduce of
oNormalize?
2 1=no, 2=yes

ofxn calc*
1.09977
ofxn mod
1.11911

sRun
RunA1
zout
1

Ave
Data
0.00069
Model
0.00068

data/vMax
0.00065

ofxng
263.459

Min d/m
2.9066

ave/min d
8.76283
ave/min m
25.3036

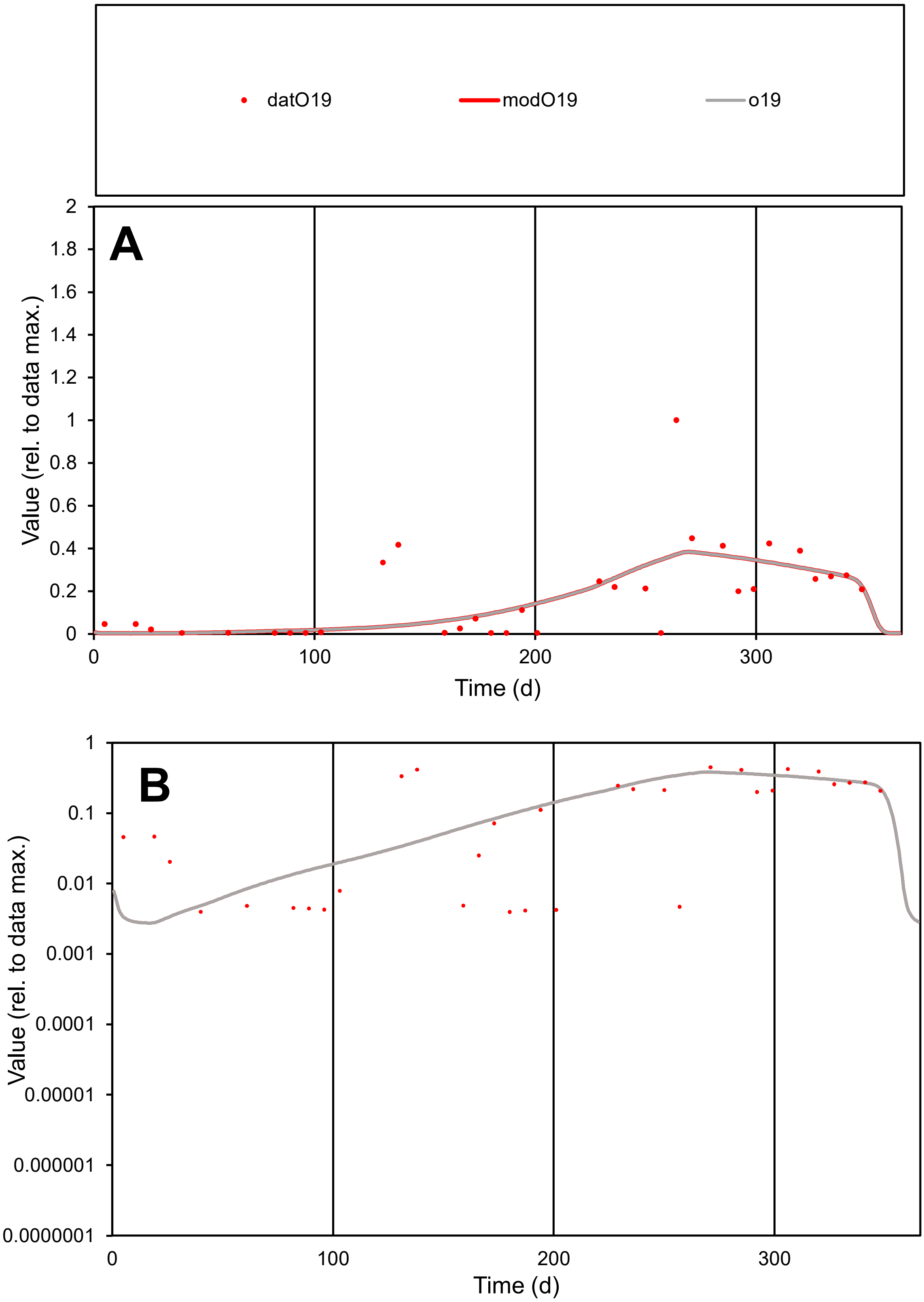


**C1**

Variables
O19 Provide jtd or Dst
CalibOnly?
2 1 to reproduce of
oNormalize?
2 1=no, 2=yes

ofxn calc*
0.90922
ofxn mod
0.92482

sRun
RunA1
zout
1

Ave
Data
0.00131
Model
0.00124

data/vMax
0.00733

ofxng
263.459

Min d/m
1.37464

ave/min d
45.3979
ave/min m
58.9906

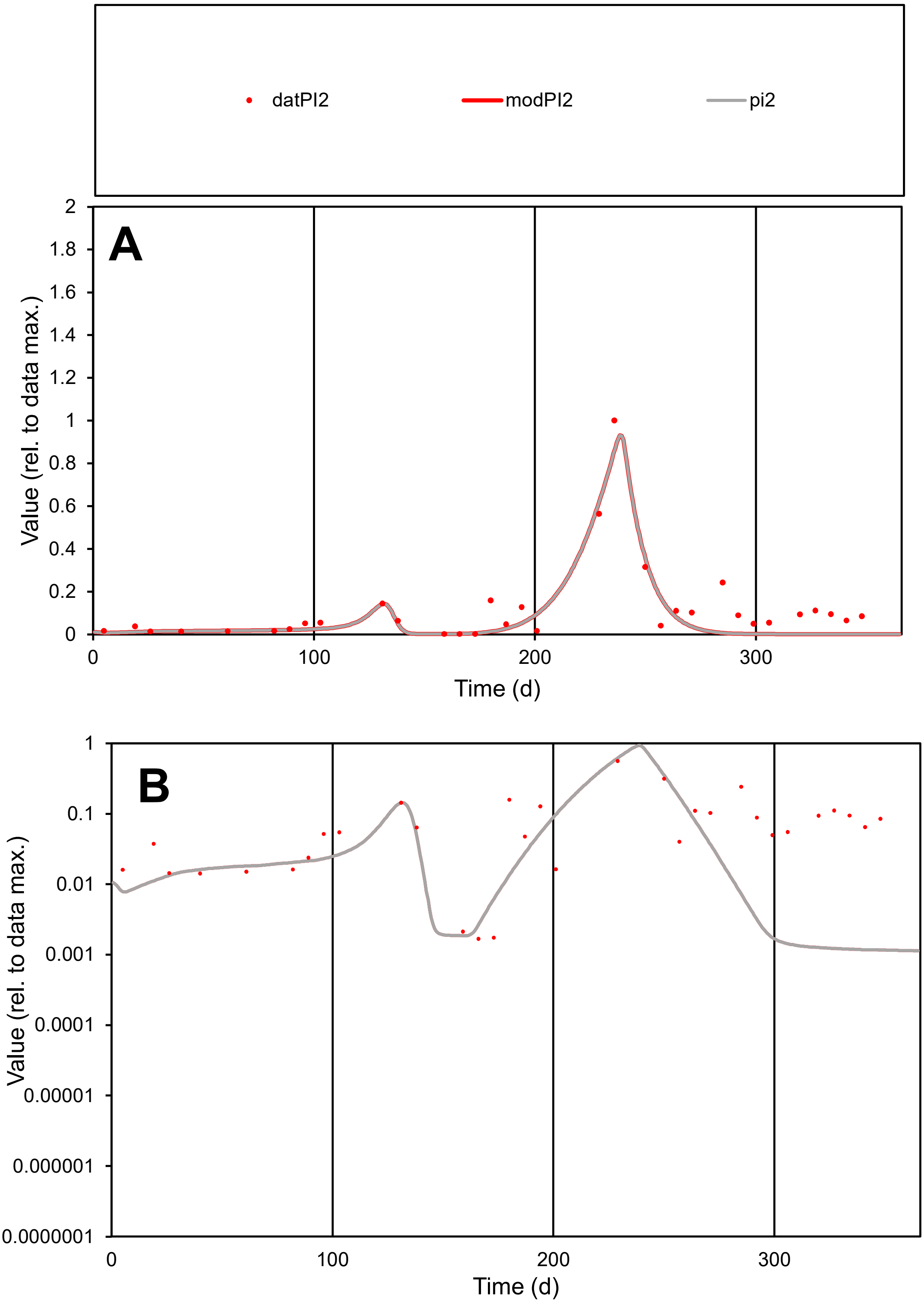


C1

Variables
PI2 Provide jtd or Dst
CalibOnly?
2 1 to reproduce of
oNormalize?
2 1=no, 2=yes

ofxn calc*
0.63171
ofxn mod
0.6637

sRun
RunA1
zout
1

Ave
Data
0.00194
Model
0.00136

data/vMax
0.01675

ofxng
263.459

Min d/m
1.43959

ave/min d
69.3452
ave/min m
69.8166

326_Calib_S_A

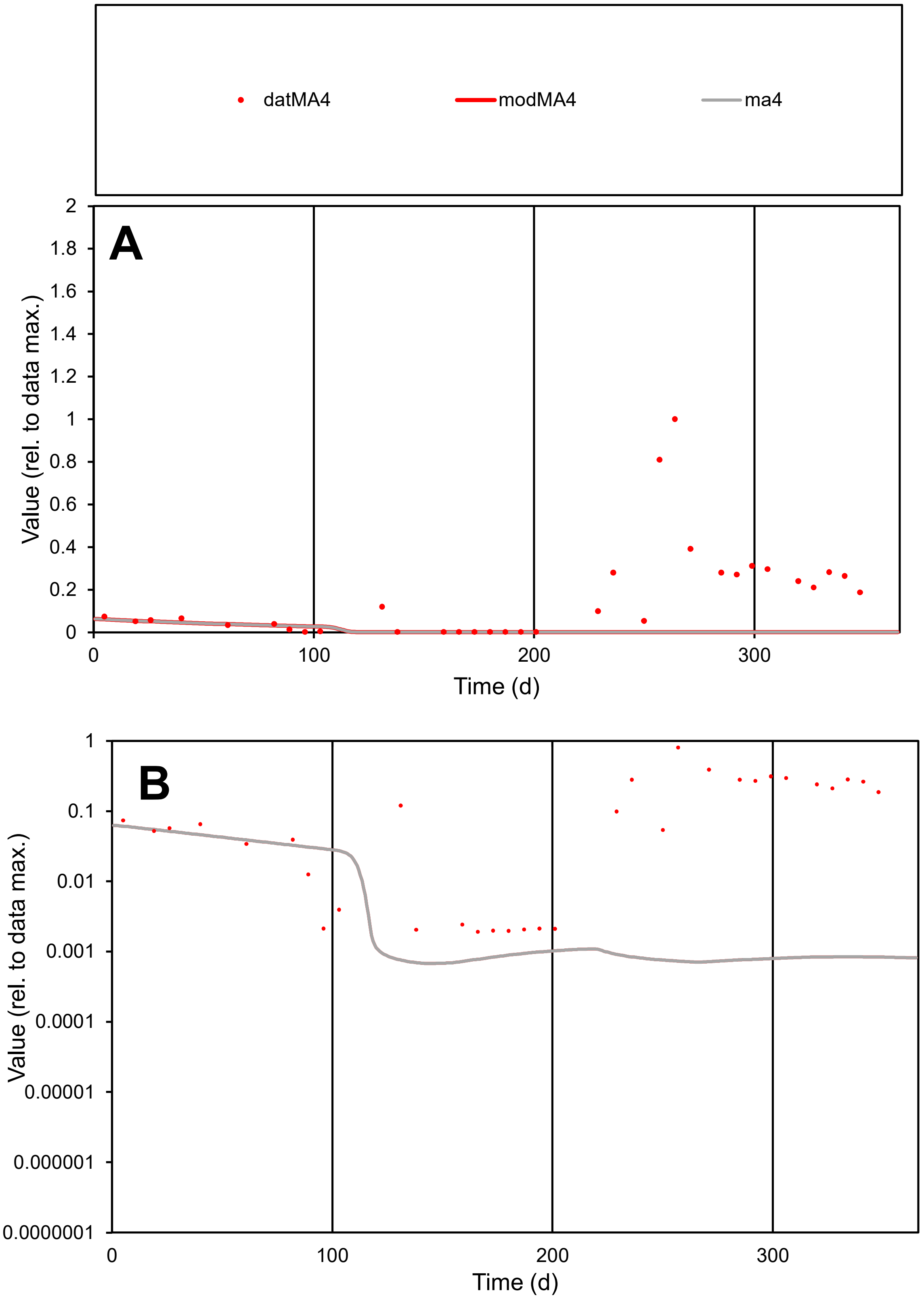


**C1**

Variables
MA4 Provide jtd or Dst
CalibOnly?
2 1 to reproduce of
oNormalize?
2 1=no, 2=yes

ofxn calc*
1.67553
ofxn mod
1.67881

sRun
RunA1
zout
1

Ave
Data
0.00165
Model
0.00012

data/vMax
0.01

ofxng
263.459

Min d/m
2.78053

ave/min d
86.5851
ave/min m
17.0832

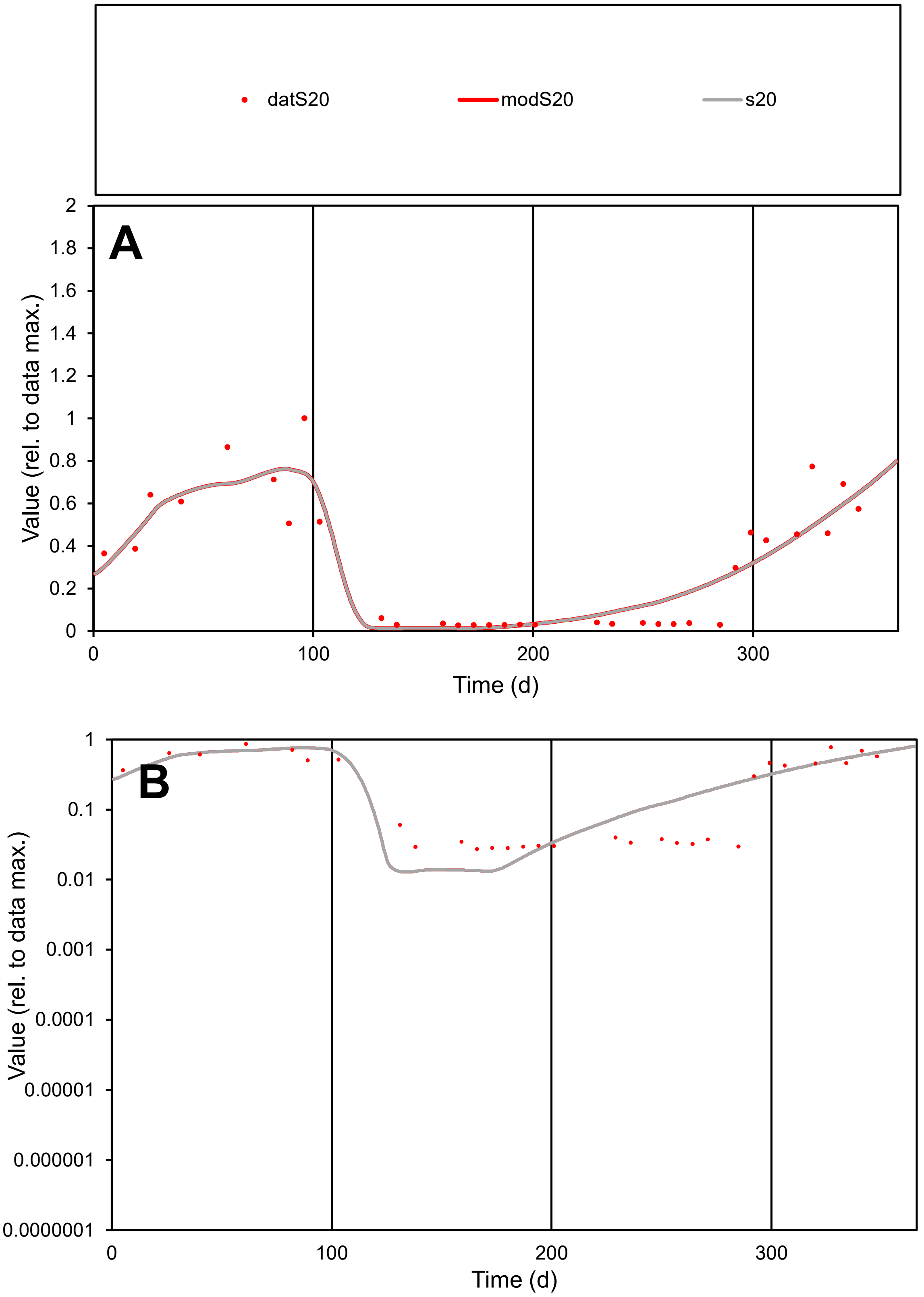


**C1**

Variables

S20 Provide jtd or Dst

CalibOnly?

2 1 to reproduce of

oNormalize?

2 1=no, 2=yes

ofxn calc*

0.36302

ofxn mod

0.39198

sRun

RunA1

zout

1

Ave

Data

0.00088

Model

0.0009

data/vMax

0.00283

ofxng

263.459

Min d/m

2.11647

ave/min d

11.4265

ave/min m

24.6599

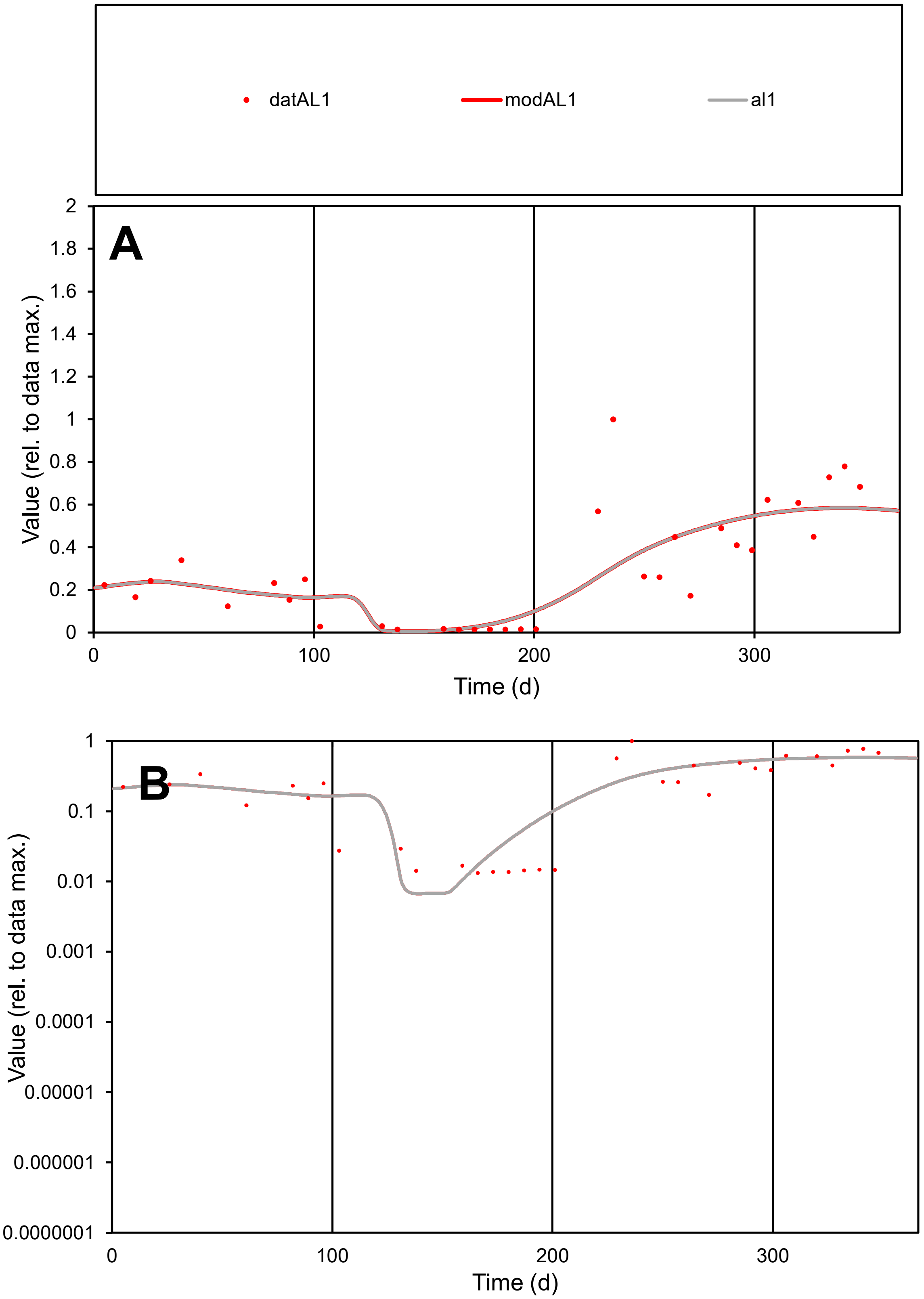


**C1**

Variables

AL1 Provide jtd or Dst

CalibOnly?

2 1 to reproduce of

oNormalize?

2 1=no, 2=yes

ofxn calc*

0.55767

ofxn mod

0.57414

sRun

RunA1

zout

1

Ave

Data

0.00097

Model

0.00095

data/vMax

0.00328

ofxng

263.459

Min d/m

1.98447

ave/min d

22.3182

ave/min m

43.2825

326_Calib_S_A

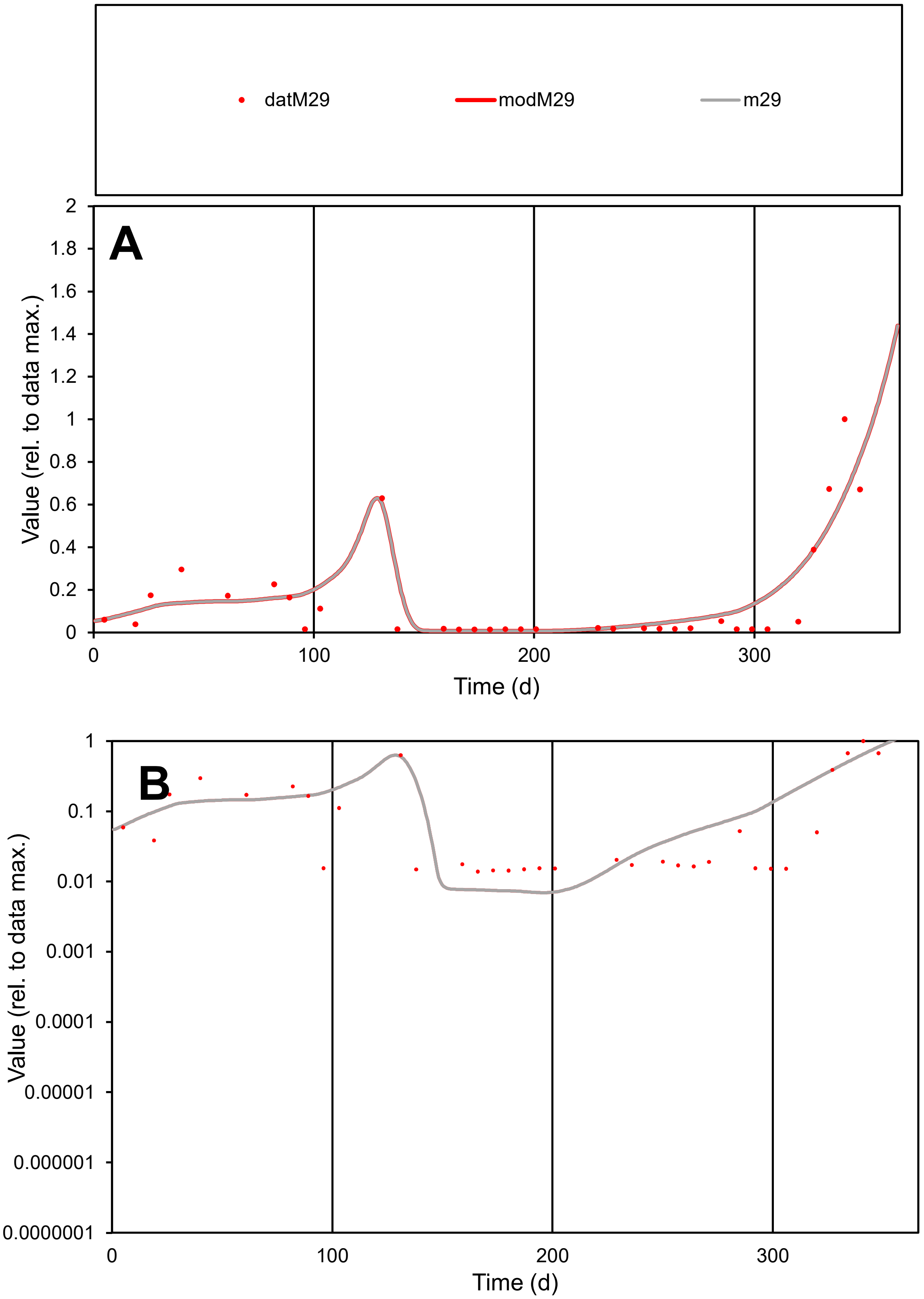


**C1**

Variables
M29 Provide jtd or Dst
CalibOnly?
2 1 to reproduce of
oNormalize?
2 1=no, 2=yes

ofxn calc*
0.69012
ofxn mod
0.72713

sRun
RunA1
zout
1

Ave
Data
0.00059
Model
0.00065

data/vMax
0.00389

ofxng
263.459

Min d/m
2.00223

ave/min d
10.9247
ave/min m
24.2652

326_Calib_S_A

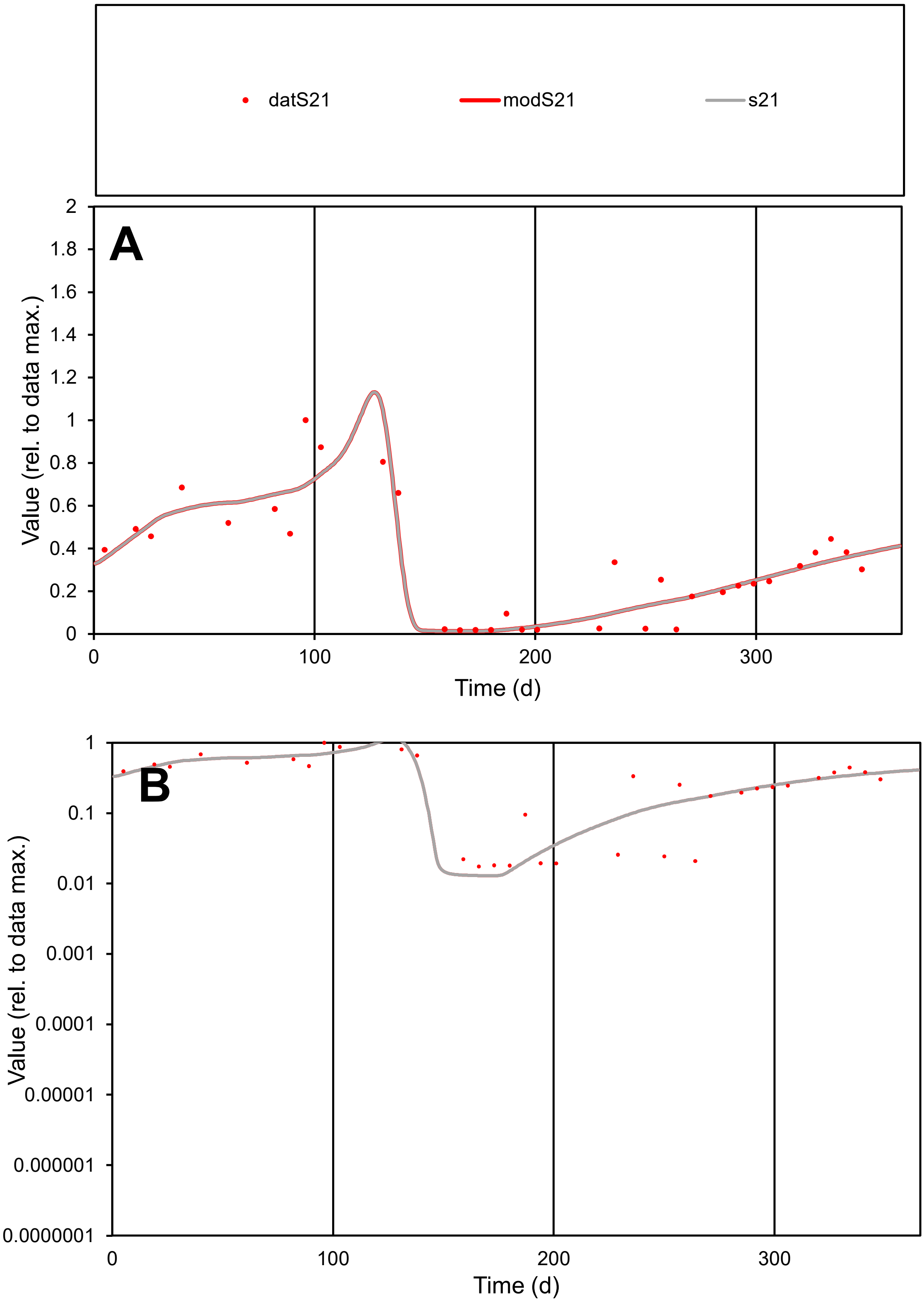


**C1**

Variables
S21 Provide jtd or Dst
CalibOnly?
2 1 to reproduce of
oNormalize?
2 1=no, 2=yes

ofxn calc*
0.32414
ofxn mod
0.35714

sRun
RunA1
zout
1

Ave
Data
0.00075
Model
0.00073

data/vMax
0.00232

ofxng
263.459

Min d/m
1.34485

ave/min d
18.5575
ave/min m
24.3151

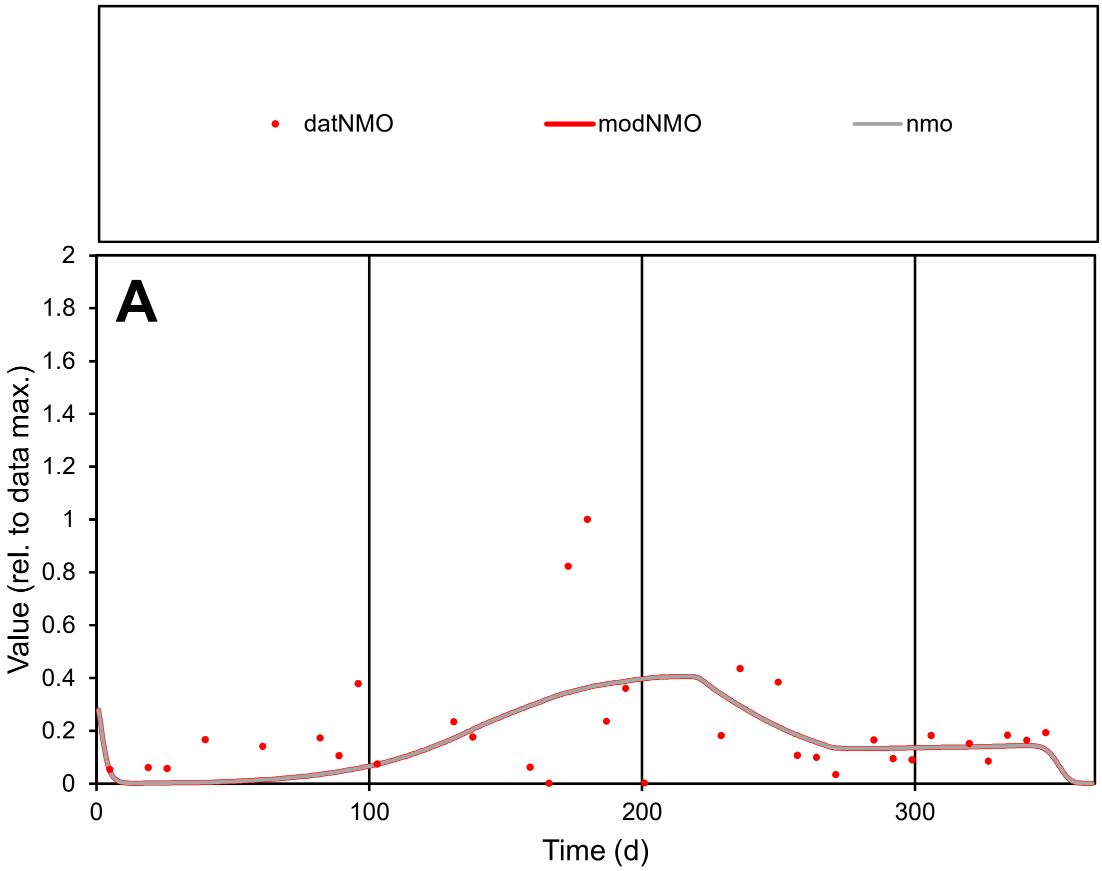


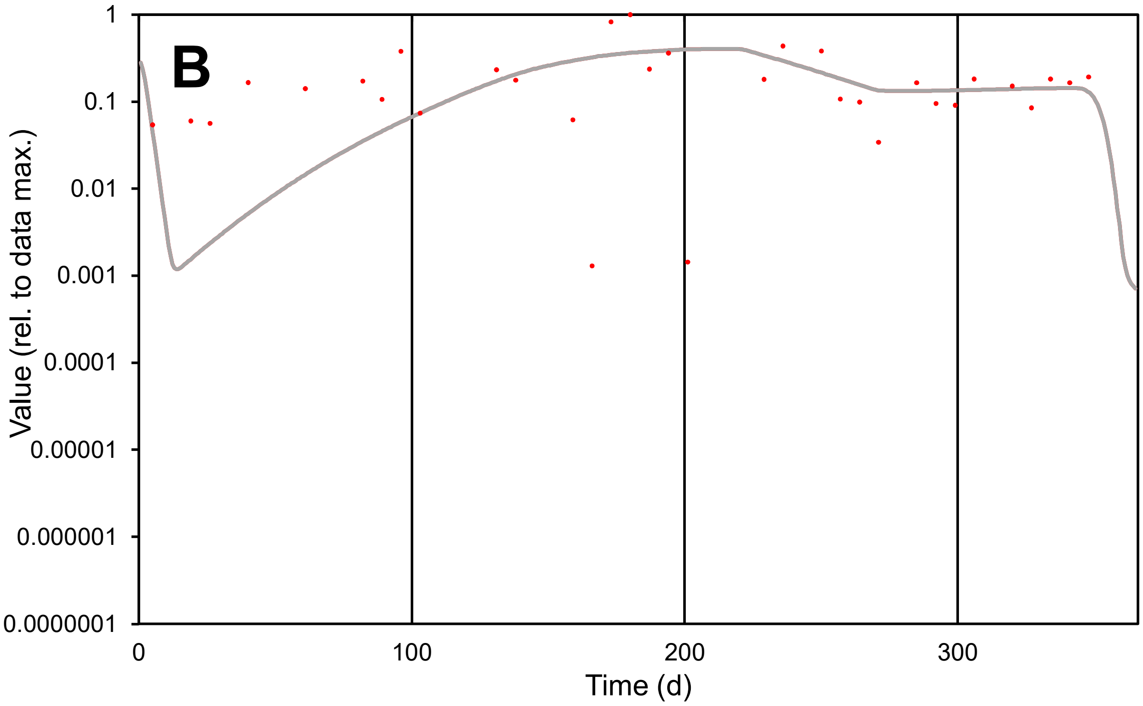


**C1**

Variables
NMO Provide jtd or Dst
CalibOnly?
2 1 to reproduce of
oNormalize?
2 1=no, 2=yes

ofxn calc*
0.96368
ofxn mod
0.97366

sRun
RunA1
zout
1

Ave
Data
0.0011
Model
0.00095

data/vMax
0.00548

ofxng
263.459

Min d/m
0.77481

ave/min d
154.661
ave/min m
103.007

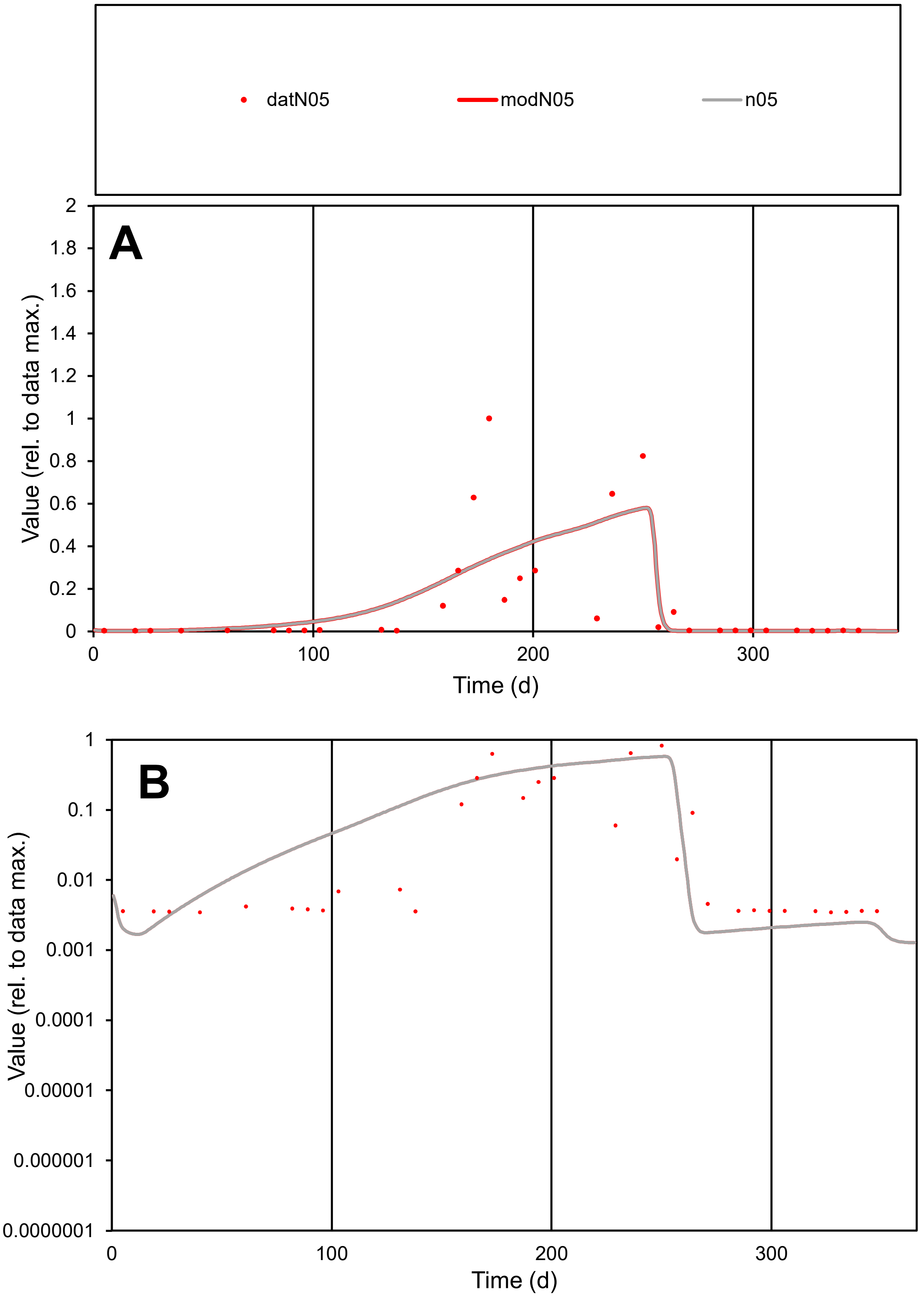


**C1**

Variables
N05 Provide jtd or Dst
CalibOnly?
2 1 to reproduce of
oNormalize?
2 1=no, 2=yes

ofxn calc*
1.26473
ofxn mod
1.285

sRun
RunA1
zout
1

Ave
Data
0.0016
Model
0.00163

data/vMax
0.0119

ofxng
263.459

Min d/m
1.93841

ave/min d
38.8619
ave/min m
76.5494

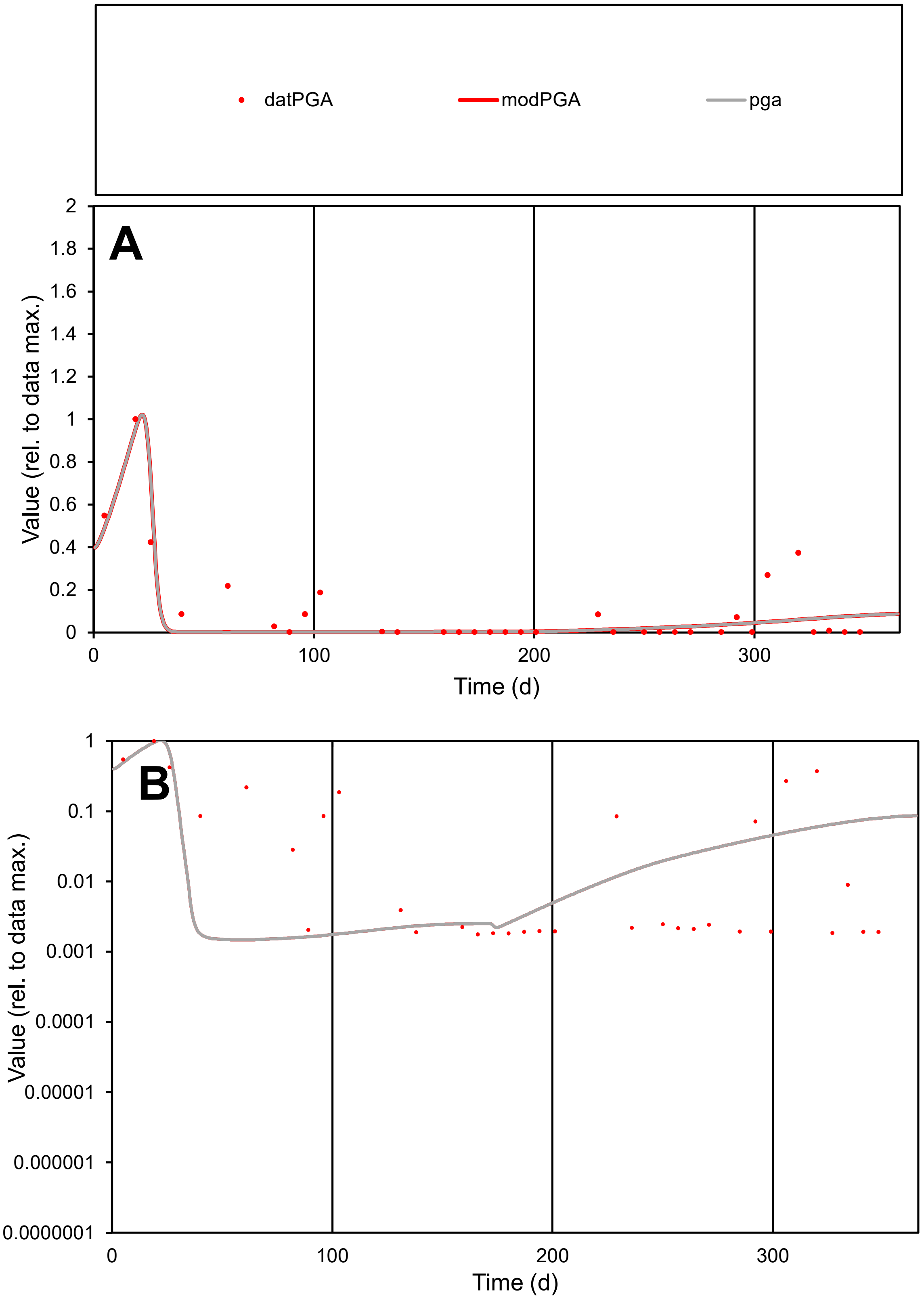


**C1**

Variables
PGA Provide jtd or Dst
CalibOnly?
2 1 to reproduce of
oNormalize?
2 1=no, 2=yes

ofxn calc*
0.87542
ofxn mod
0.919

sRun
RunA1
zout
1

Ave
Data
0.00099
Model
0.00076

data/vMax
0.00951

ofxng
263.459

Min d/m
1.20114

ave/min d
58.681
ave/min m
54.0694

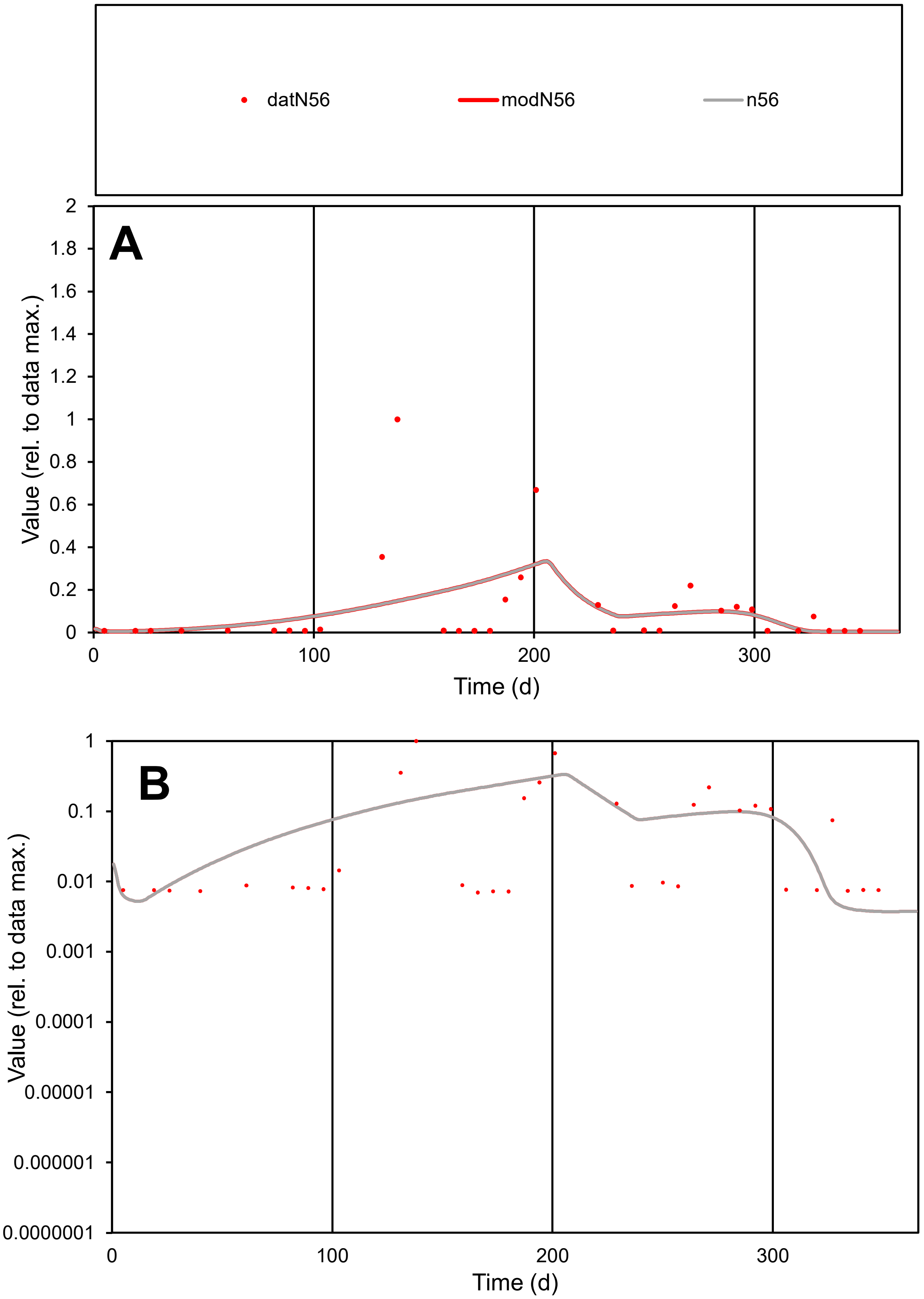


**C1**

Variables
N56 Provide jtd or Dst
CalibOnly?
2 1 to reproduce of
oNormalize?
2 1=no, 2=yes

ofxn calc*
1.76549
ofxn mod
1.77805

sRun
RunA1
zout
1

Ave
Data
0.00051
Model
0.00049

data/vMax
0.00484

ofxng
263.459

Min d/m
1.86504

ave/min d
15.1293
ave/min m
26.8923

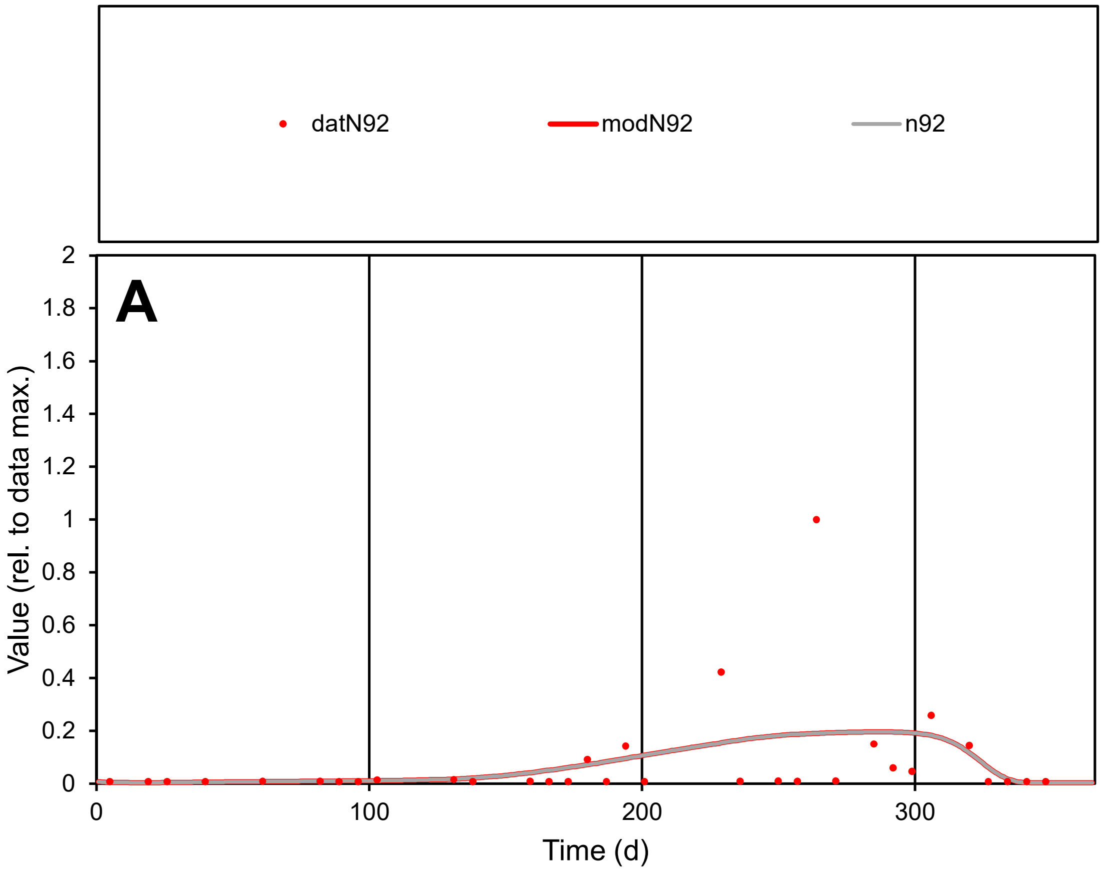


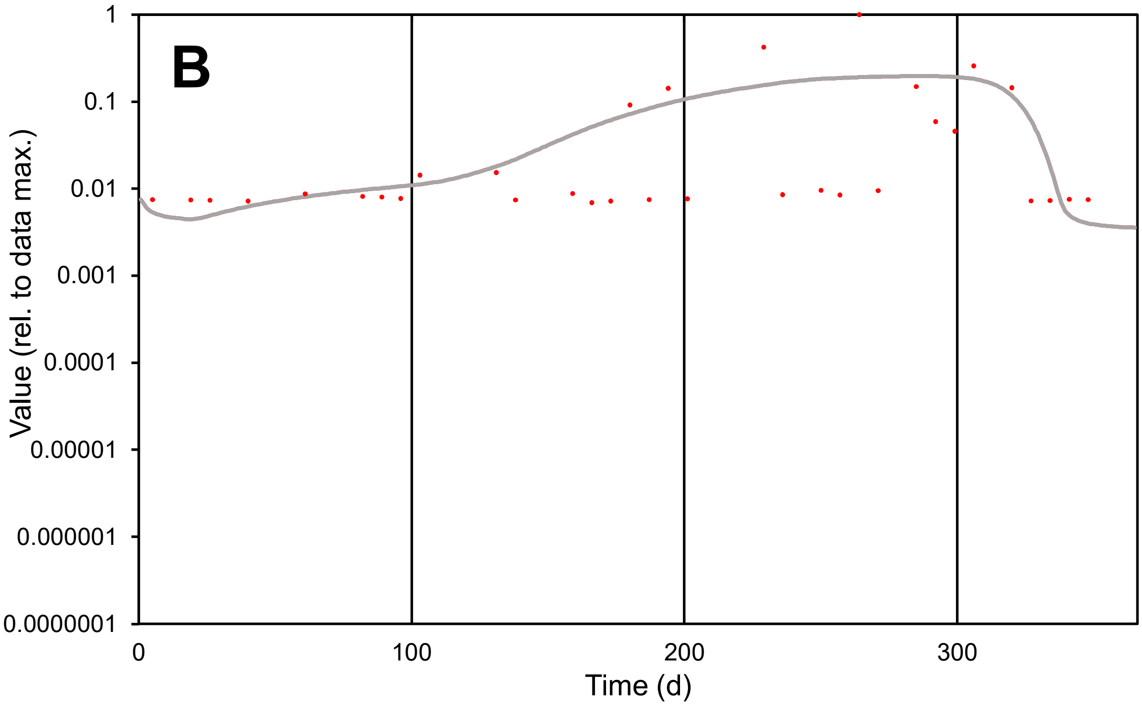


**C1**

Variables
N92 | Provide jtd or Dst
CalibOnly?
2 | 1 to reproduce of
oNormalize?
2 | 1=no, 2=yes

ofxn calc*
2.19171
ofxn mod
2.20564

sRun
RunA1
zout
1

Ave
Data
0.00041
Model
0.00043

data/vMax
0.00543

ofxng
263.459

Min d/m
1.75585

ave/min d
11.0326
ave/min m
20.3417

326_Calib_S_A

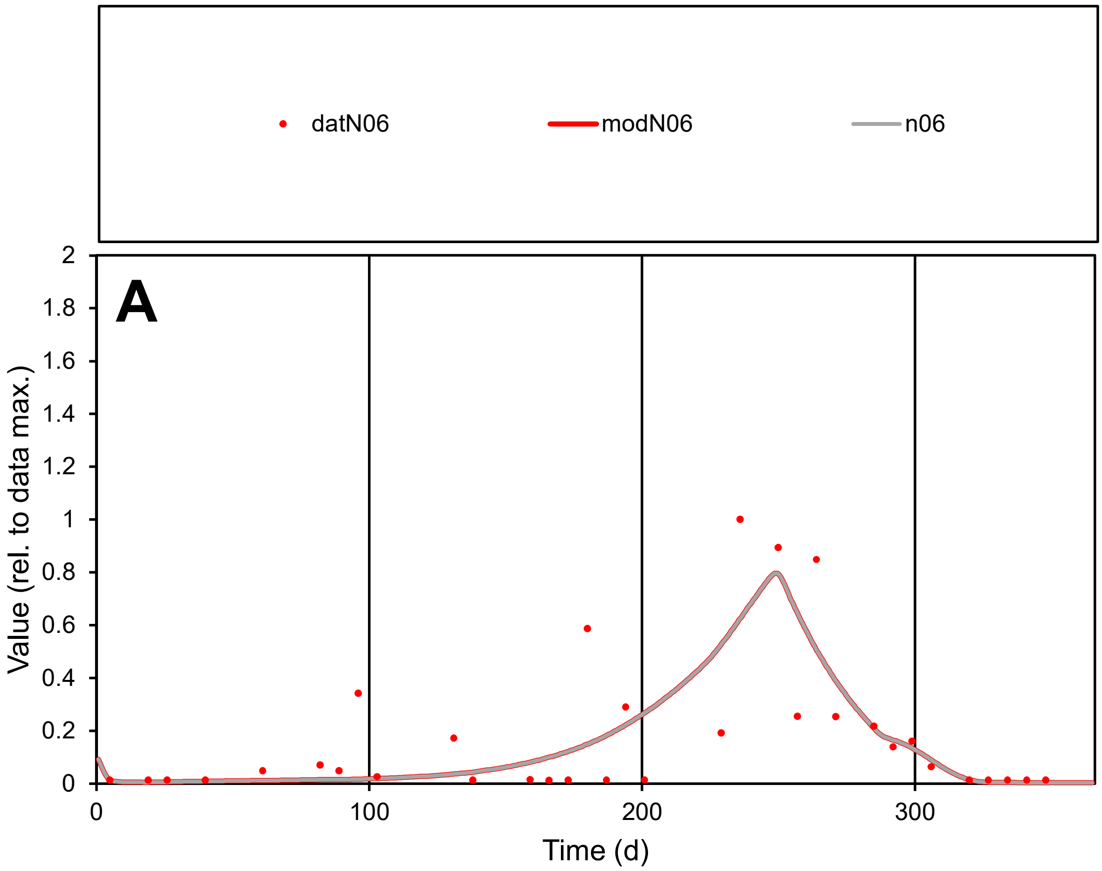


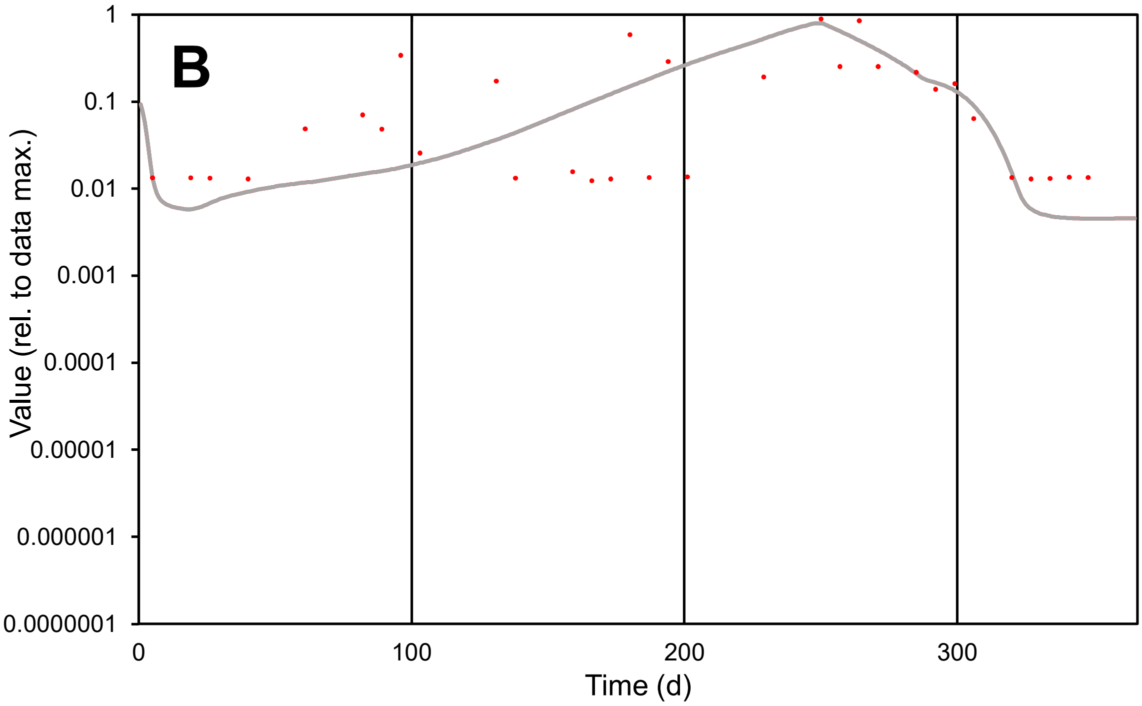


**C1**

Variables

N06 Provide jtd or Dst

CalibOnly?

2 1 to reproduce of

oNormalize?

2 1=no, 2=yes

ofxn calc*

0.97438

ofxn mod

0.99744

sRun

RunA1

zout

1

Ave

Data

0.00091

Model

0.00084

data/vMax

0.00519

ofxng

263.459

Min d/m

2.74199

ave/min d

14.2177

ave/min m

35.9123

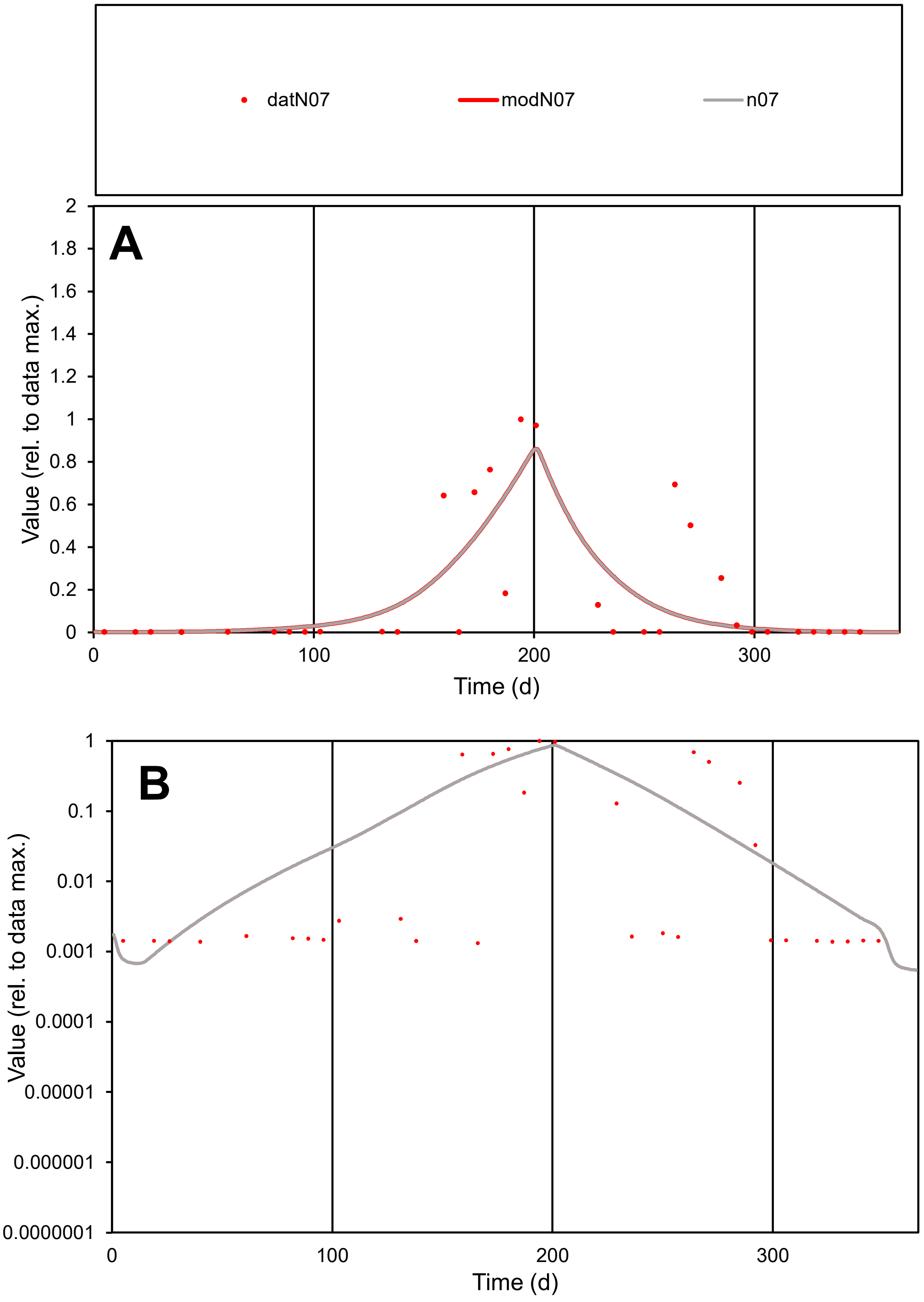


**C1**

Variables

| | |
|---|---|
| N07 | Provide jtd or Dst |
| CalibOnly? | |
| 2 | 1 to reproduce of |
| oNormalize? | |
| 2 | 1=no, 2=yes |

ofxn calc*
1.14582
ofxn mod
1.17076

sRun
RunA1
zout
1

Ave
Data
0.00227
Model
0.00207

data/vMax
0.01276

ofxng
263.459

Min d/m
1.70084

ave/min d
134.639
ave/min m
208.715

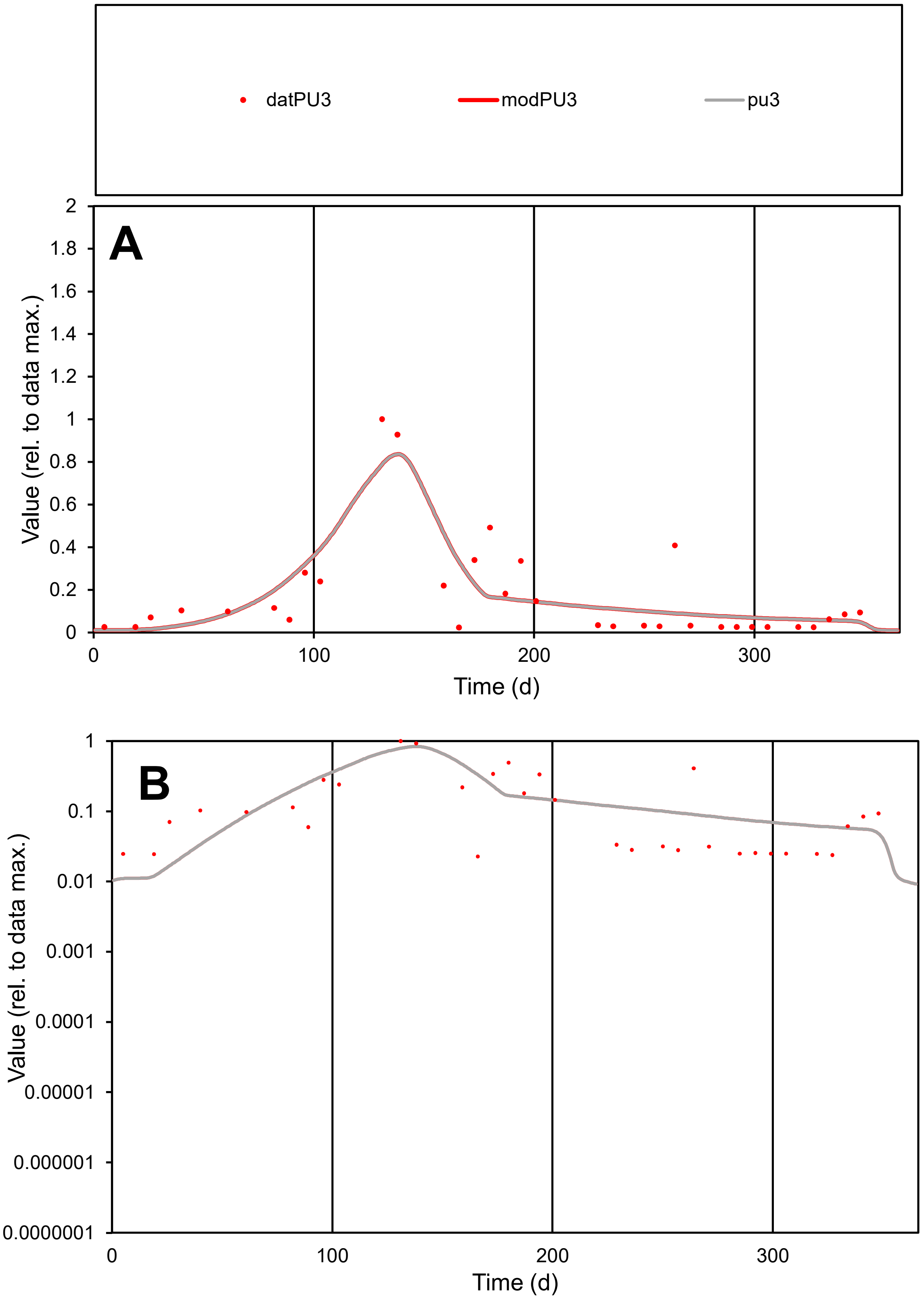


**C1**

Variables
PU3 Provide jtd or Dst
CalibOnly?
2 1 to reproduce of
oNormalize?
2 1=no, 2=yes

ofxn calc*
0.75227
ofxn mod
0.78775

sRun
RunA1
zout
1

Ave
Data
0.00111
Model
0.00113

data/vMax
0.00653

ofxng
263.459

Min d/m
2.04934

ave/min d
7.48422
ave/min m
15.6102

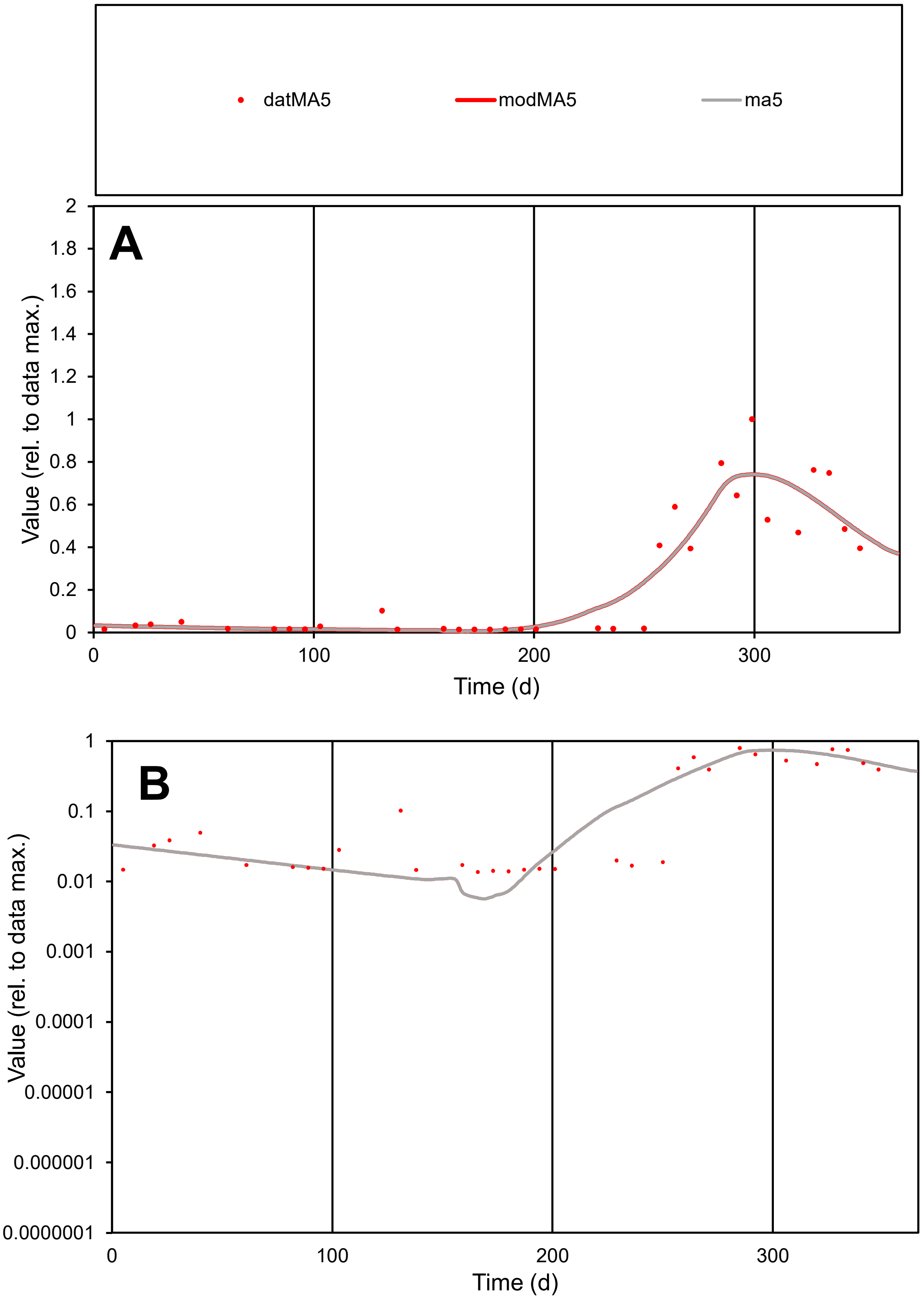


**C1**

Variables
MA5 Provide jtd or Dst
CalibOnly?
2 1 to reproduce of
oNormalize?
2 1=no, 2=yes

ofxn calc*
0.44703
ofxn mod
0.48077

sRun
RunA1
zout
1

Ave
Data
0.0016
Model
0.00159

data/vMax
0.00683

ofxng
263.459

Min d/m
2.37302

ave/min d
17.1901
ave/min m
40.6595

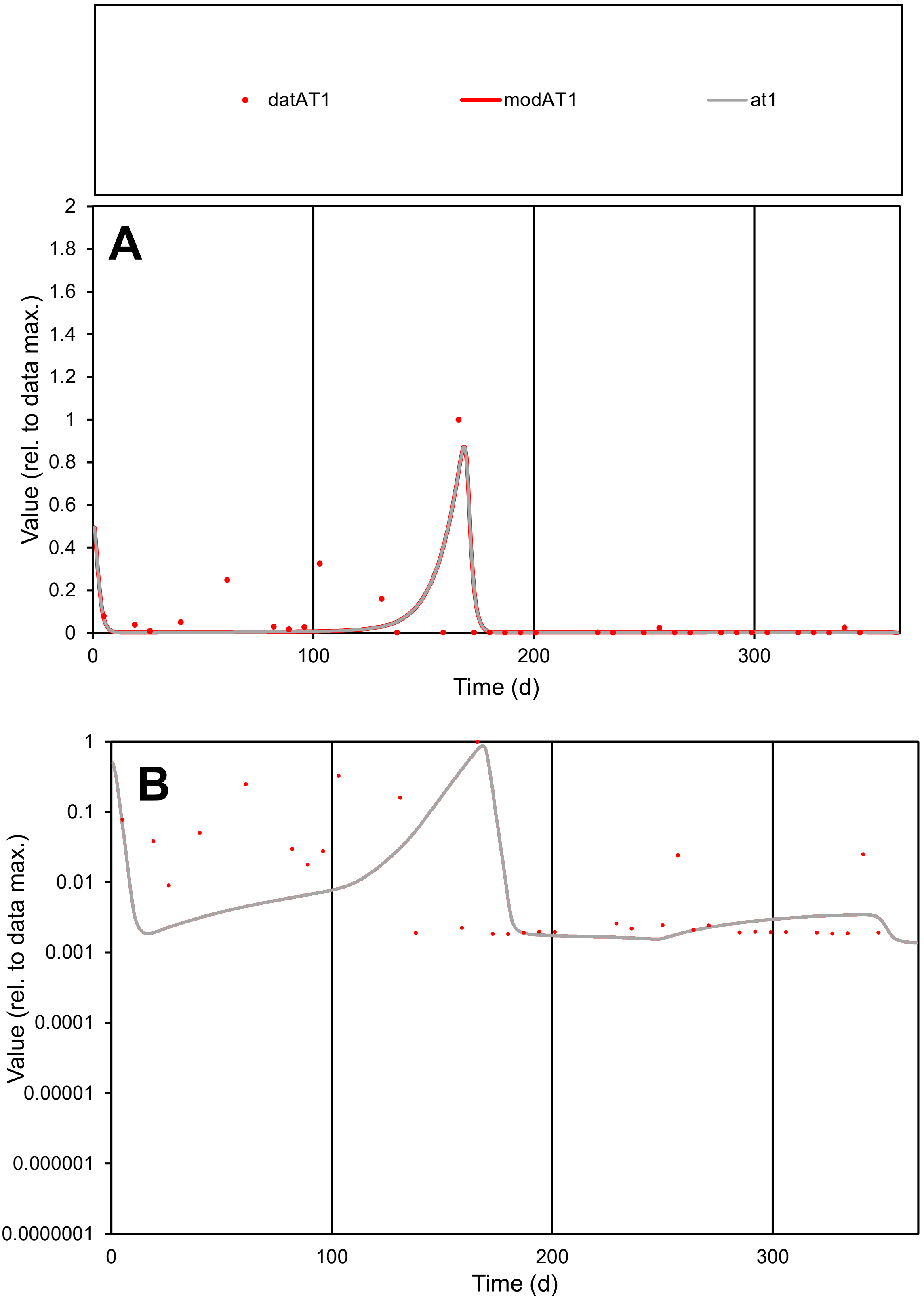


**C1**

Variables
AT1 Provide jtd or Dst
CalibOnly?
2 1 to reproduce of
oNormalize?
2 1=no, 2=yes

ofxn calc*
1.57907
ofxn mod
1.67477

sRun
RunA1
zout
1

Ave
Data
0.00168
Model
0.00119

data/vMax
0.02678

ofxng
263.459

Min d/m
1.13607

ave/min d
34.5026
ave/min m
27.7987

326_Calib_S_A

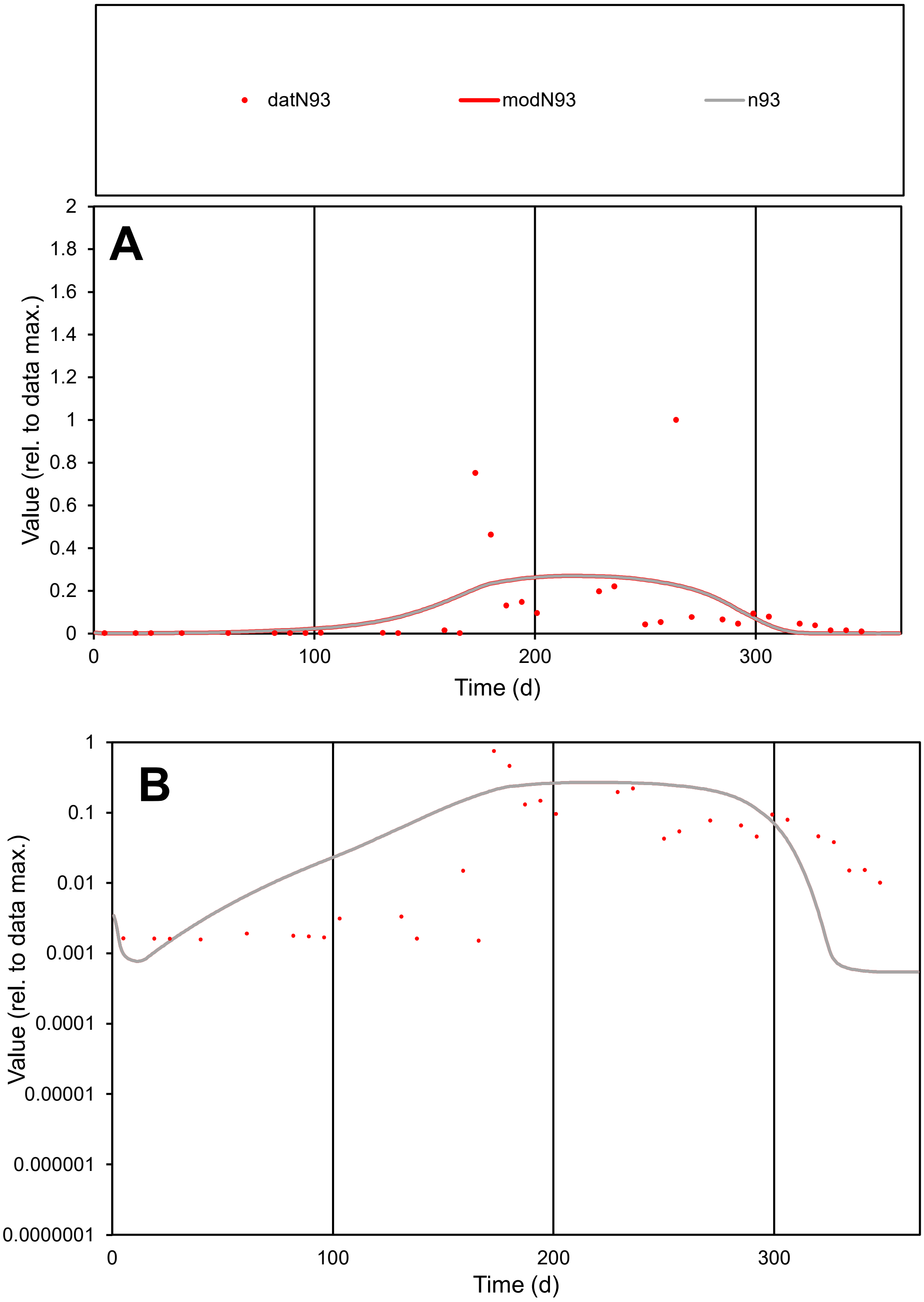


**C1**

Variables
N93 Provide jtd or Dst
CalibOnly?
2 1 to reproduce of
oNormalize?
2 1=no, 2=yes

ofxn calc*
1.70996
ofxn mod
1.72439

sRun
RunA1
zout
1

Ave
Data
0.00125
Model
0.00124

data/vMax
0.01135

ofxng
263.459

Min d/m
2.77221

ave/min d
72.7232
ave/min m
199.622

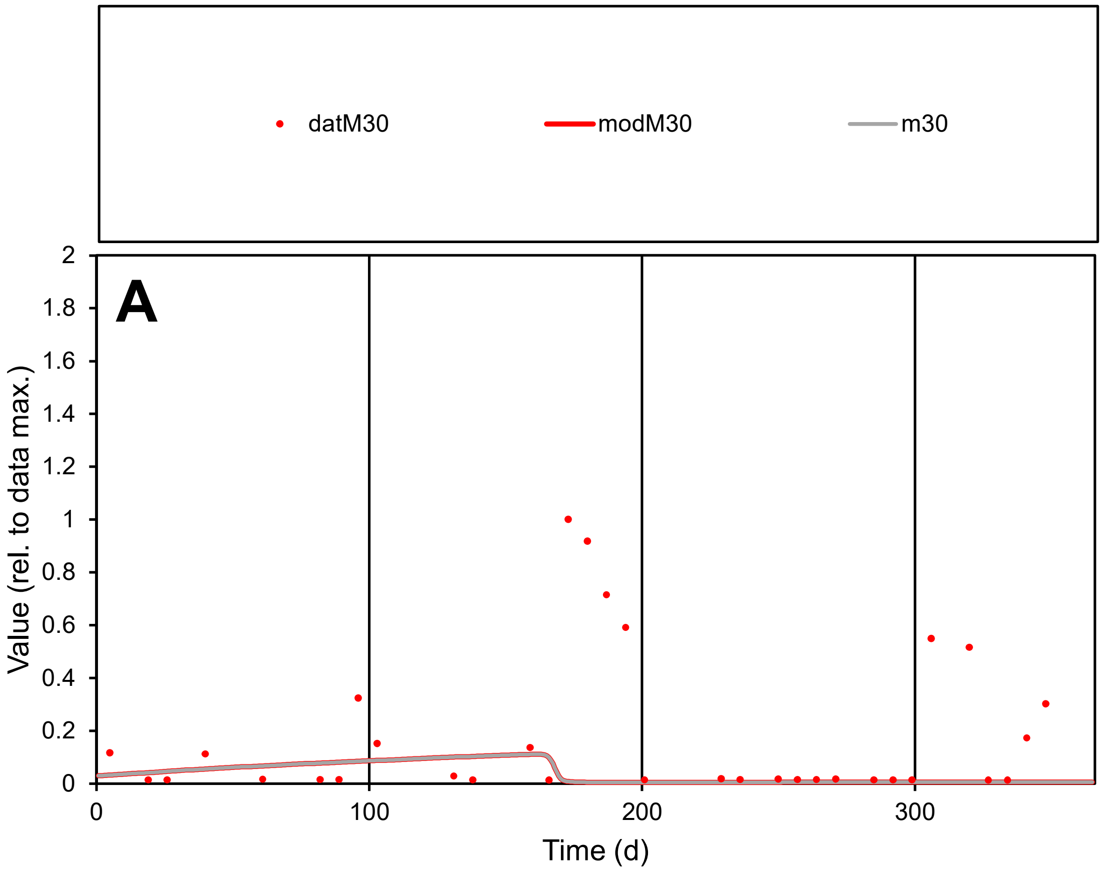


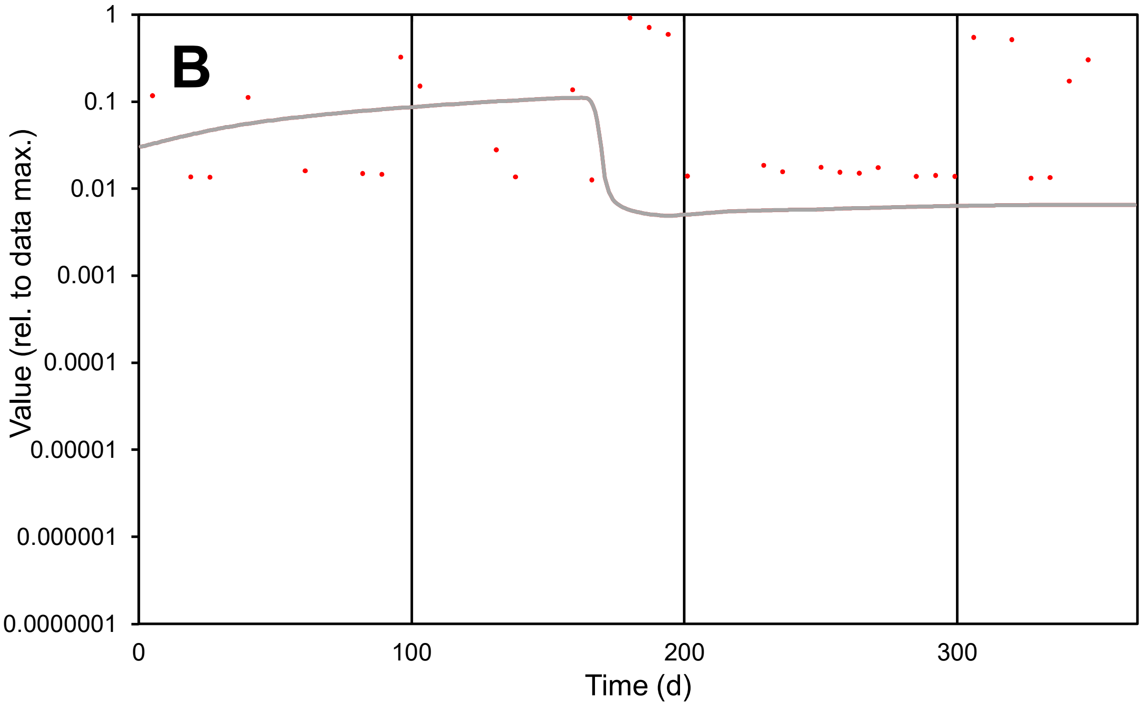


**C1**

Variables
M30 Provide jtd or Dst
CalibOnly?
2 1 to reproduce of
oNormalize?
2 1=no, 2=yes

ofxn calc*
1.80004
ofxn mod
1.80351

sRun
RunA1
zout
1

Ave
Data
0.00057
Model
0.0001

data/vMax
0.0032

ofxng
263.459

Min d/m
2.59155

ave/min d
14.1489
ave/min m
6.48983

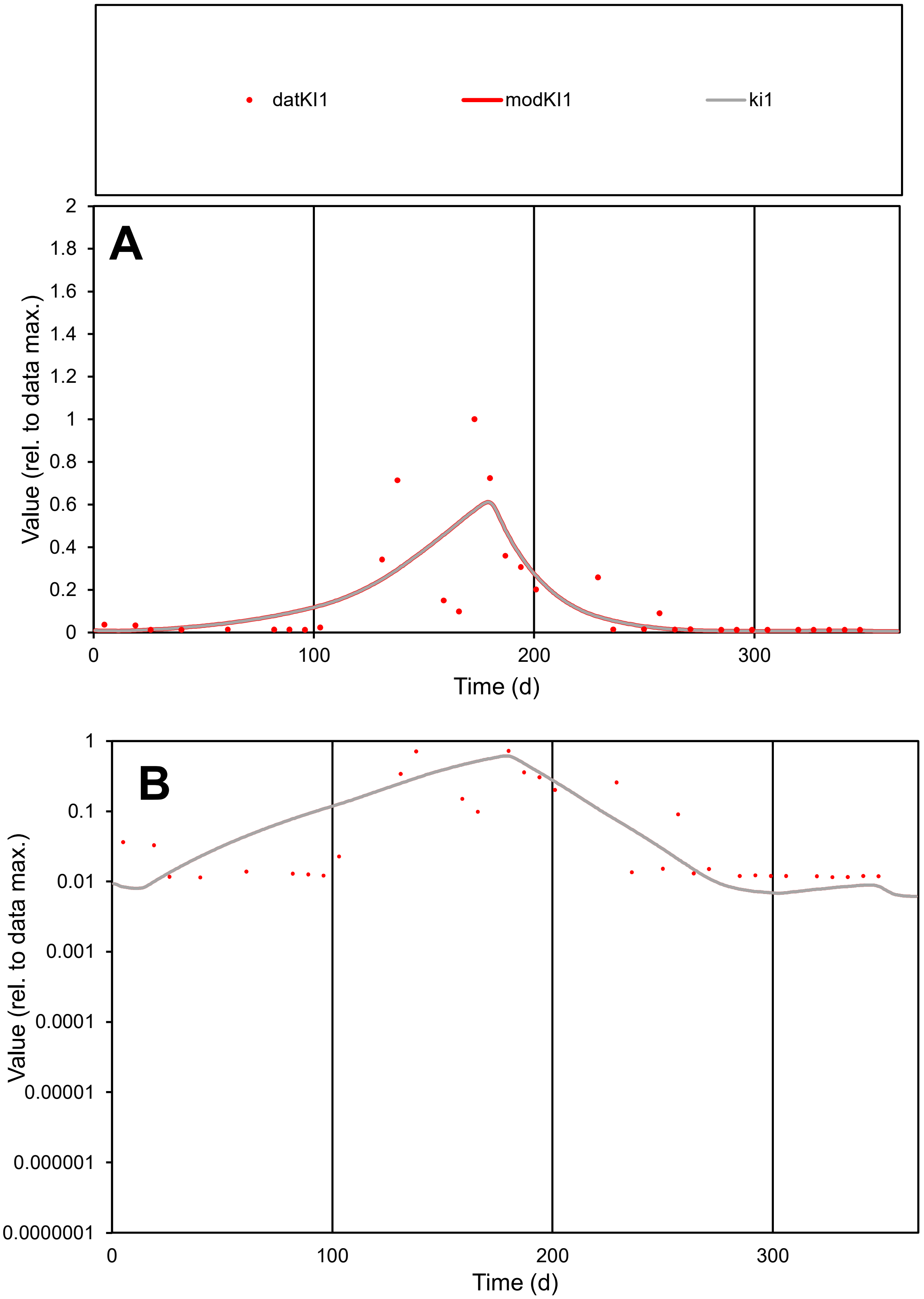


**C1**

Variables
KI1 Provide jtd or Dst
CalibOnly?
2 1 to reproduce of
oNormalize?
2 1=no, 2=yes

ofxn calc*
1.06493
ofxn mod
1.08842

sRun
RunA1
zout
1

Ave
Data
0.00058
Model
0.00058

data/vMax
0.00421

ofxng
263.459

Min d/m
1.67221

ave/min d
12.0879
ave/min m
20.1187

326_Calib_S_A

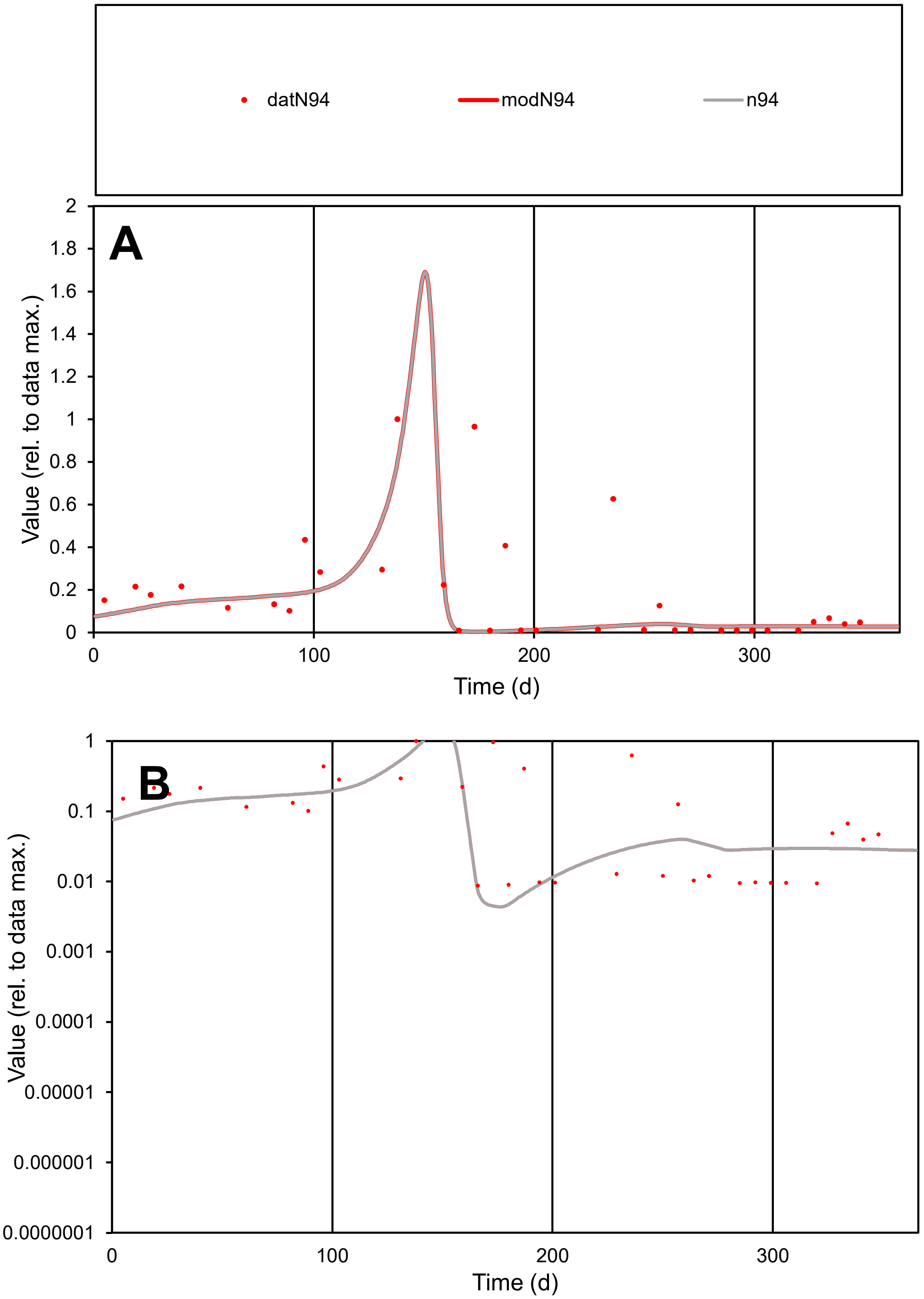


**C1**

Variables

N94 Provide jtd or Dst

CalibOnly?

2 1 to reproduce of

oNormalize?

2 1=no, 2=yes

ofxn calc*

1.26235

ofxn mod

1.27389

sRun

RunA1

zout

1

Ave

Data

0.0009

Model

0.00055

data/vMax

0.00514

ofxng

263.459

Min d/m

1.97837

ave/min d

20.1625

ave/min m

24.3764

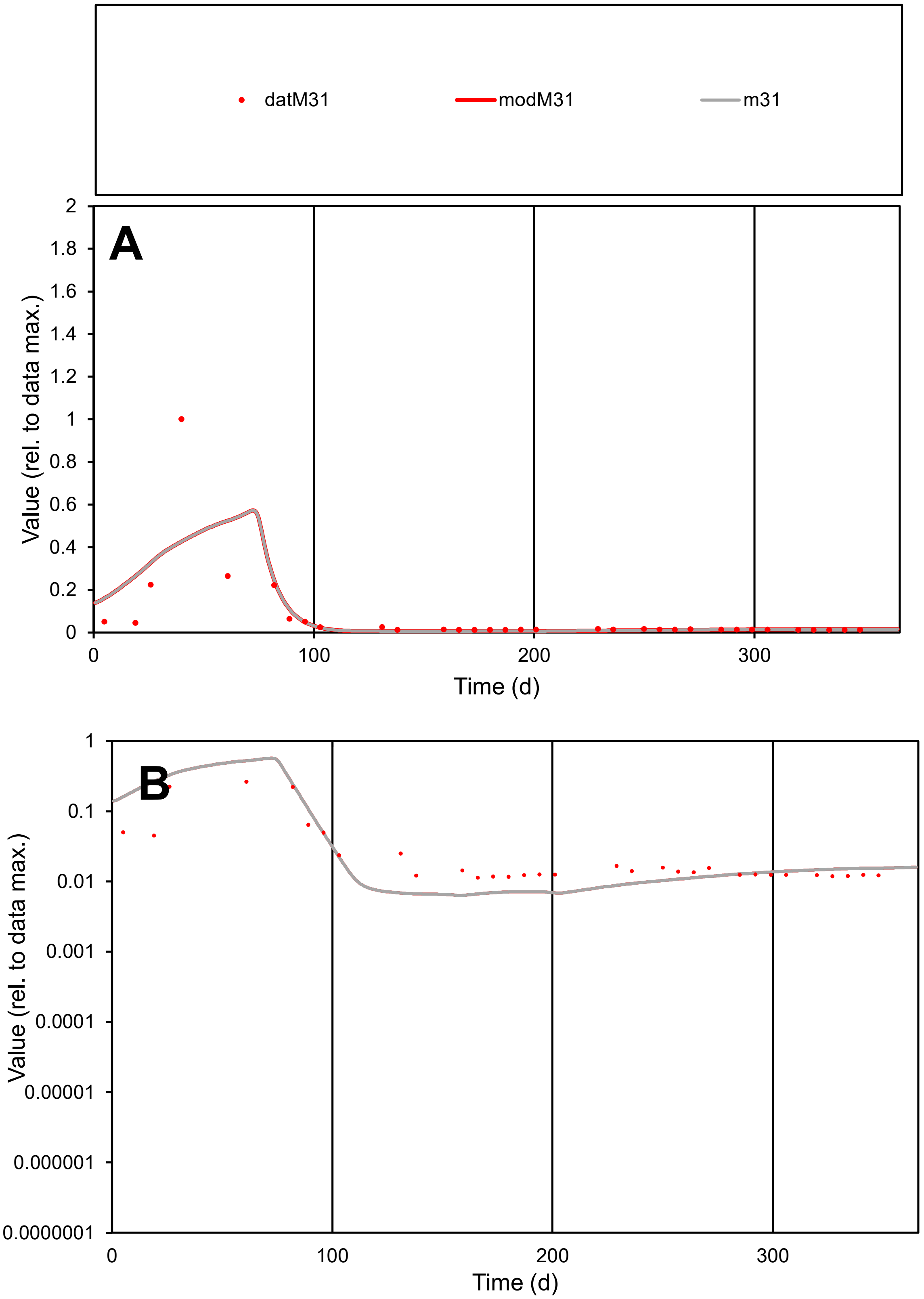


**C1**

Variables
M31 Provide jtd or Dst
CalibOnly?
2 1 to reproduce of
oNormalize?
2 1=no, 2=yes

ofxn calc*
1.72966
ofxn mod
1.76397

sRun
RunA1
zout
1

Ave
Data
0.00049
Model
0.0005

data/vMax
0.00708

ofxng
263.459

Min d/m
1.78473

ave/min d
6.04886
ave/min m
11.1465

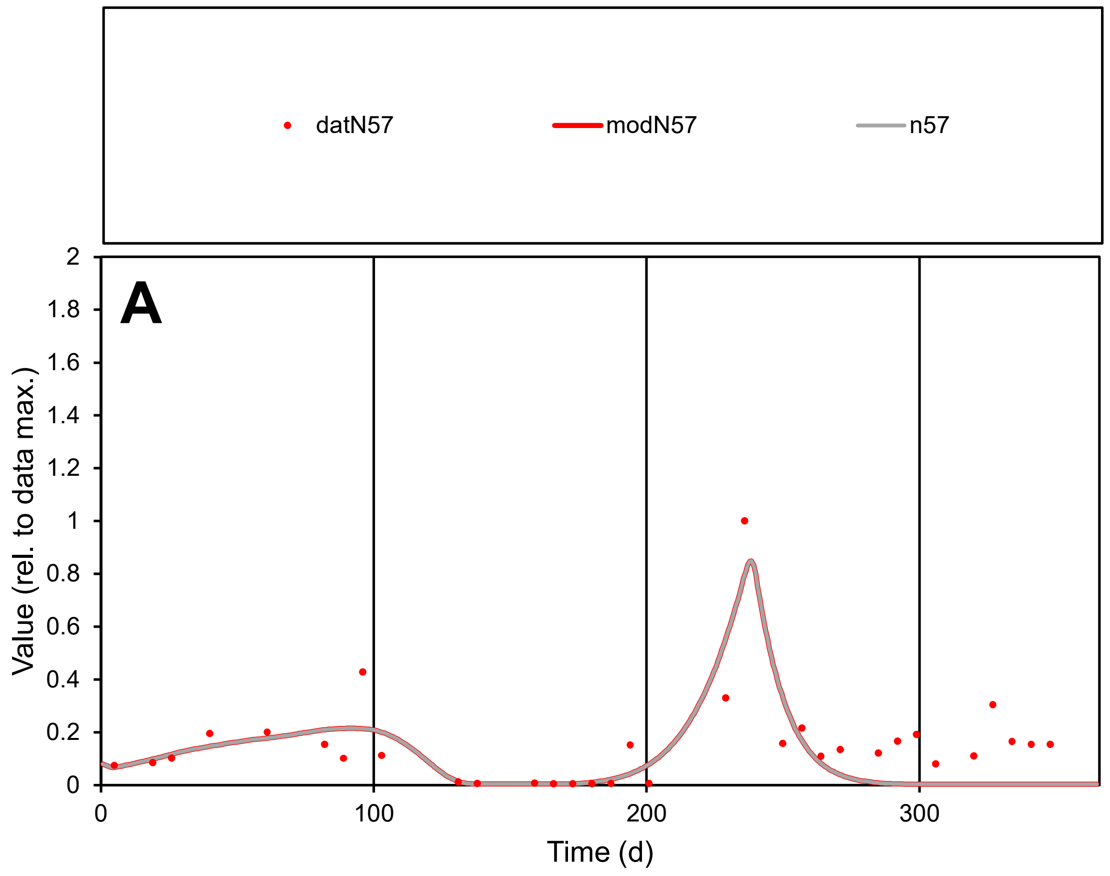


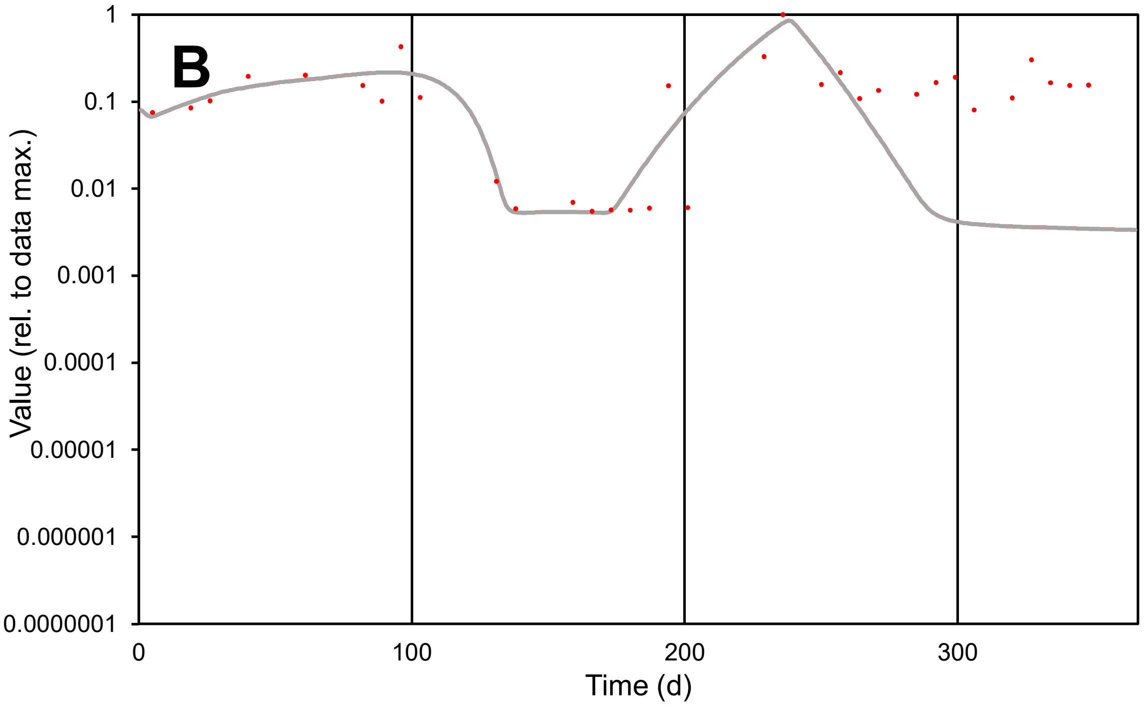


**C1**

Variables
N57 Provide jtd or Dst
CalibOnly?
2 1 to reproduce of
oNormalize?
2 1=no, 2=yes

ofxn calc*
0.76129
ofxn mod
0.78025

sRun
RunA1
zout
1

Ave
Data
0.001
Model
0.00072

data/vMax
0.0065

ofxng
263.459

Min d/m
1.5918

ave/min d
27.8652
ave/min m
31.8688

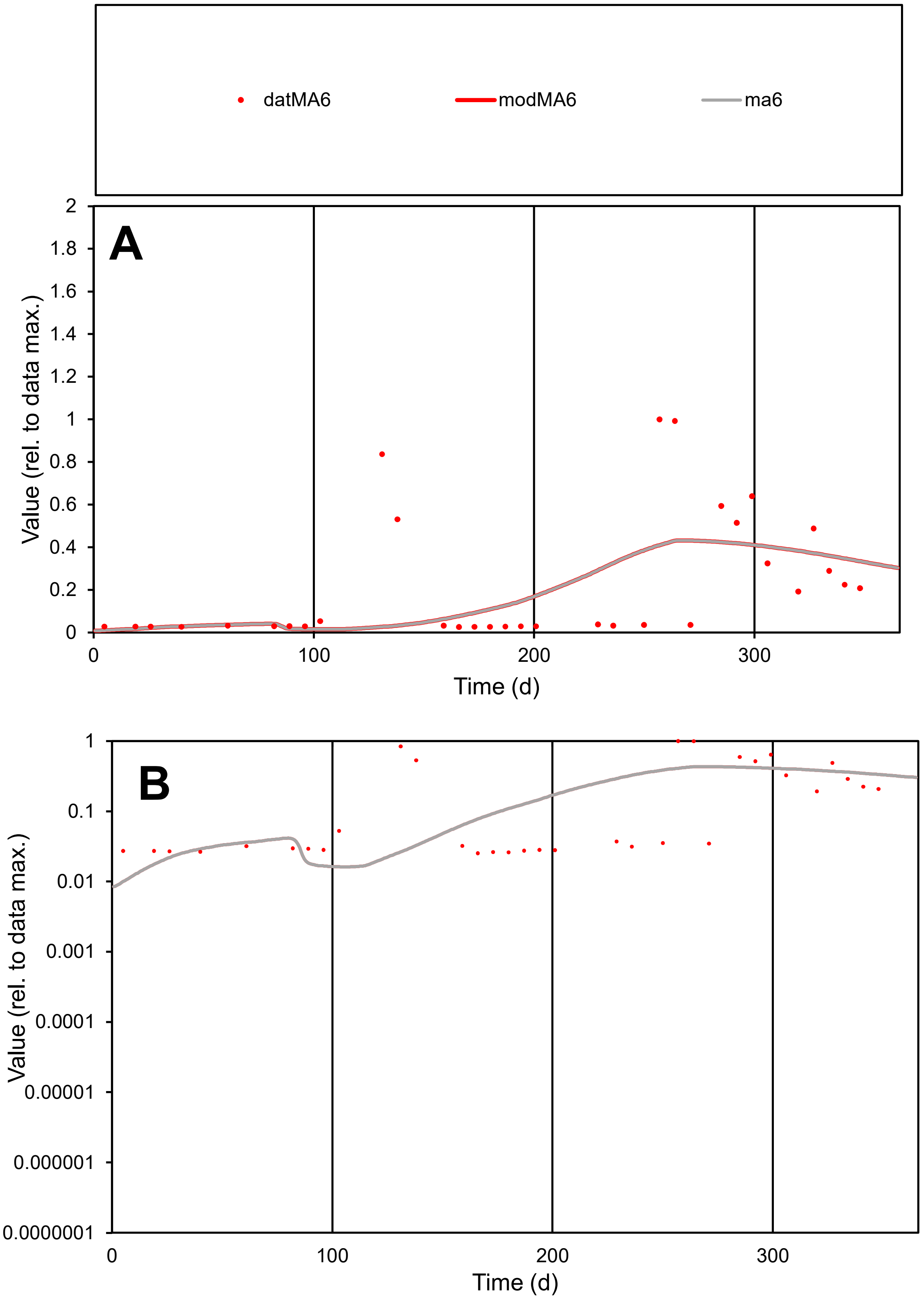


**C1**

Variables
MA6 Provide jtd or Dst
CalibOnly?
2 1 to reproduce of
oNormalize?
2 1=no, 2=yes

ofxn calc*
1.15457
ofxn mod
1.16506

sRun
RunA1
zout
1

Ave
Data
0.00104
Model
0.00095

data/vMax
0.0046

ofxng
263.459

Min d/m
2.37842

ave/min d
8.91232
ave/min m
19.3218

326_Calib_S_A

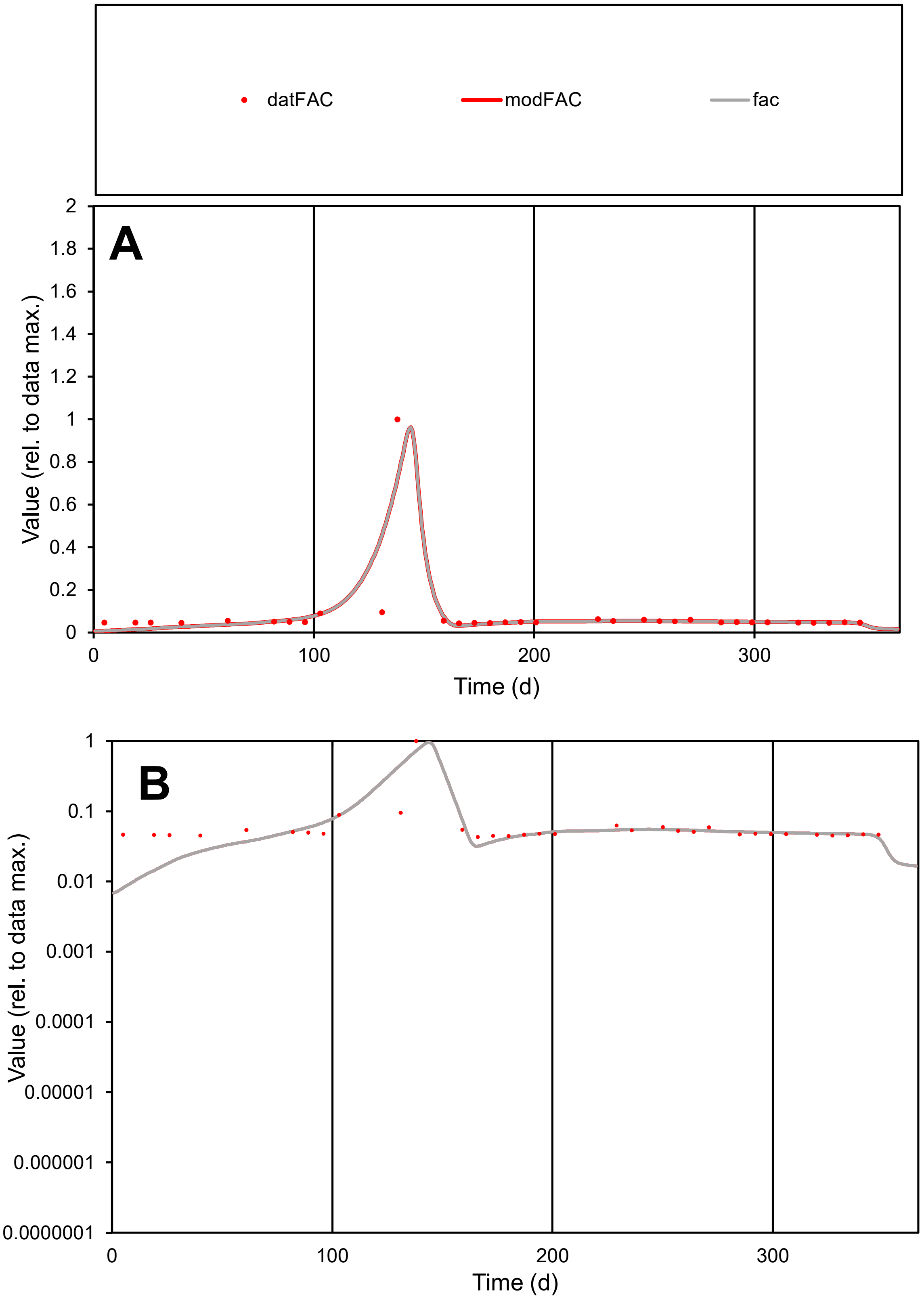


**C1**

Variables
FAC Provide jtd or Dst
CalibOnly?
2 1 to reproduce of
oNormalize?
2 1=no, 2=yes

ofxn calc*
0.86657
ofxn mod
0.96048

sRun
RunA1
zout
1

Ave
Data
0.00016
Model
0.00016

data/vMax
0.00199

ofxng
263.459

Min d/m
4.95183

ave/min d
1.87383
ave/min m
9.40115

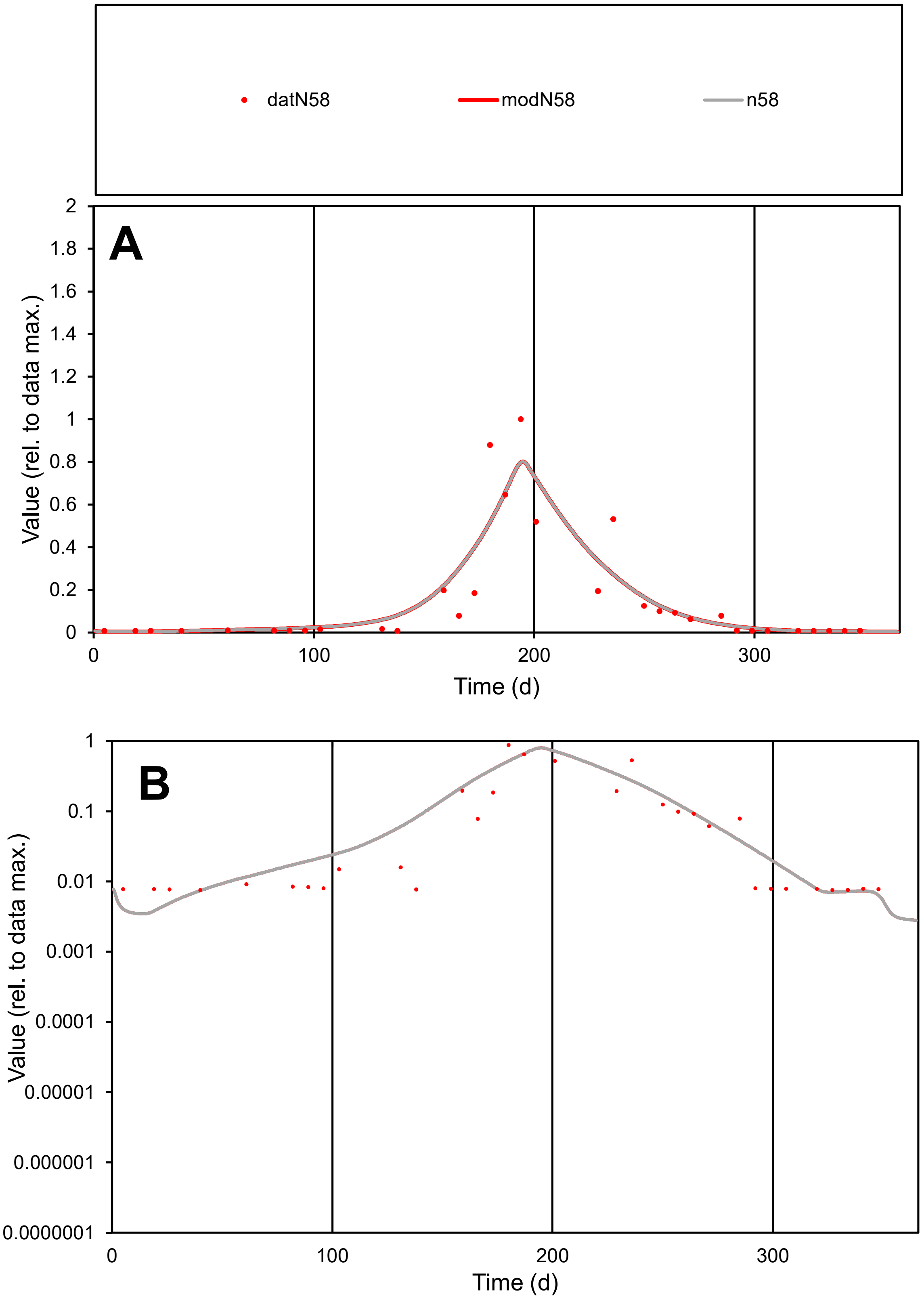


C1

Variables
N58 Provide jtd or Dst
CalibOnly?
2 1 to reproduce of
oNormalize?
2 1=no, 2=yes

ofxn calc*
0.74216
ofxn mod
0.77903

sRun
RunA1
zout
1

Ave
Data
0.00165
Model
0.00173

data/vMax
0.01121

ofxng
263.459

Min d/m
1.88765

ave/min d
19.4992
ave/min m
38.5609

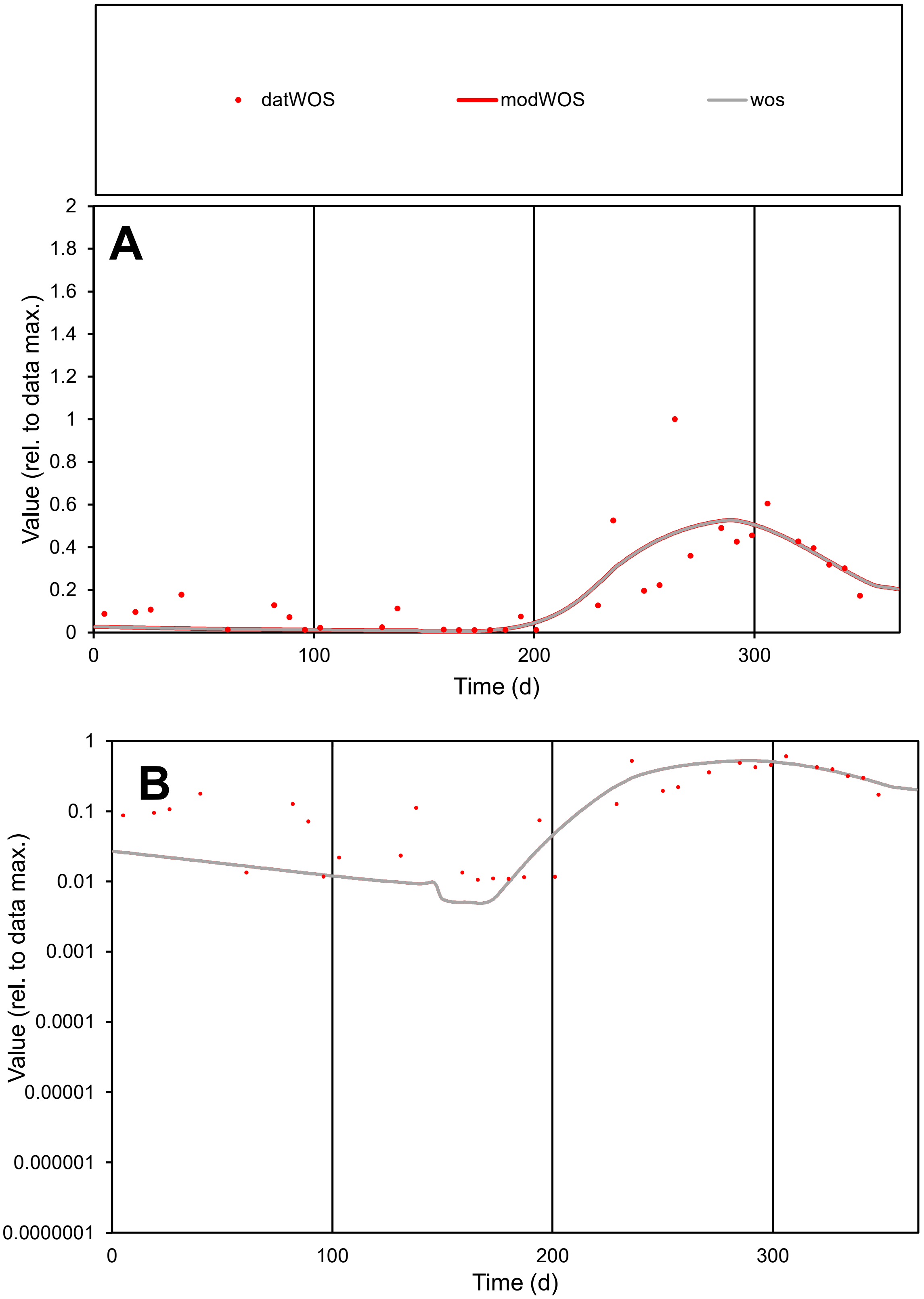


C1

Variables
WOS Provide jtd or Dst
CalibOnly?
2 1 to reproduce of
oNormalize?
2 1=no, 2=yes

ofxn calc*
0.60876
ofxn mod
0.63257

sRun
RunA1
zout
1

Ave
Data
0.00135
Model
0.00122

data/vMax
0.00636

ofxng
263.459

Min d/m
2.17634

ave/min d
19.9936
ave/min m
39.1758

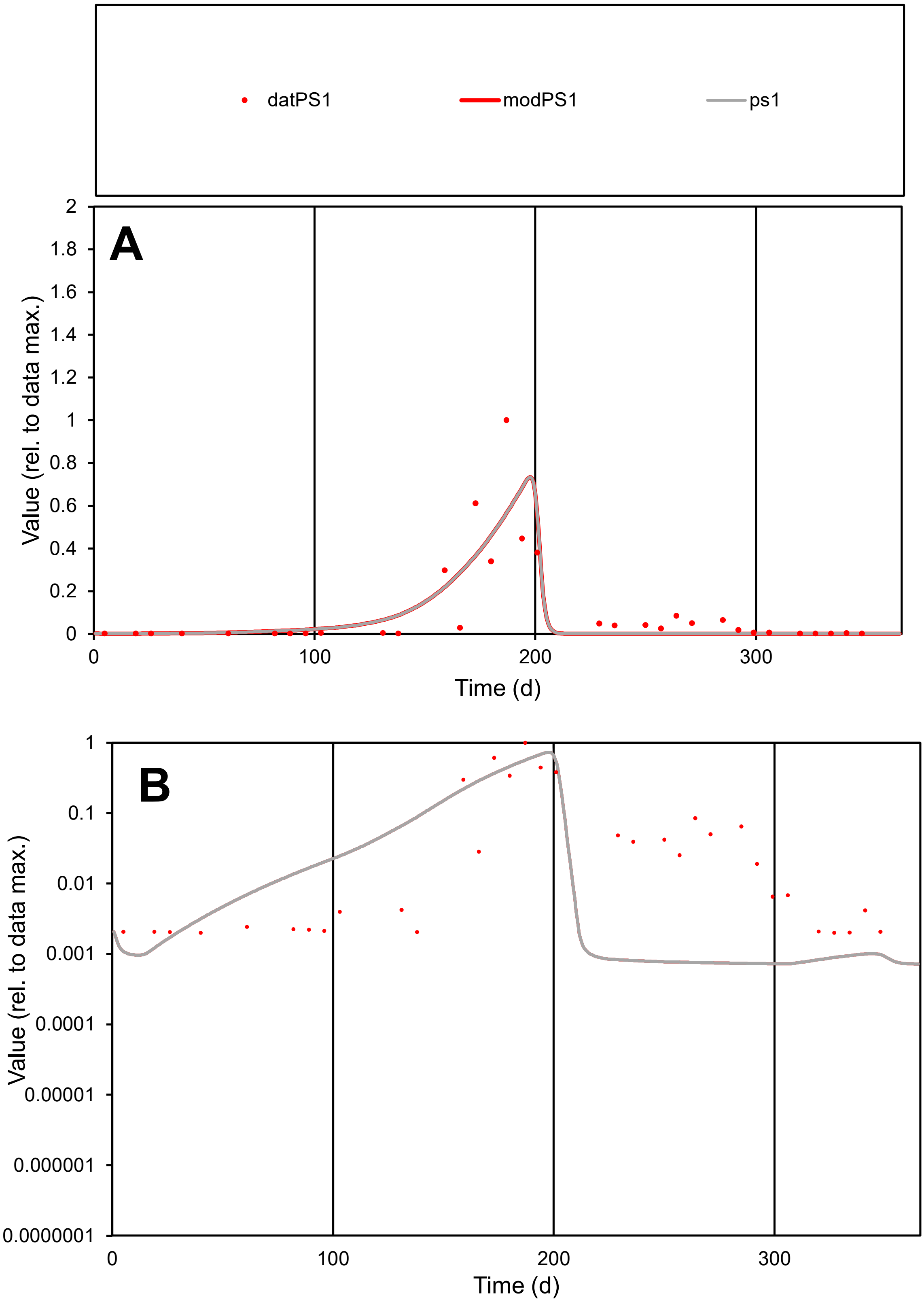


**C1**

Variables

PS1 Provide jtd or Dst

CalibOnly?

2 1 to reproduce of

oNormalize?

2 1=no, 2=yes

ofxn calc*

1.02276

ofxn mod

1.07025

sRun

RunA1

zout

1

Ave

Data

0.0026

Model

0.00241

data/vMax

0.02431

ofxng

263.459

Min d/m

2.75212

ave/min d

53.4941

ave/min m

136.383

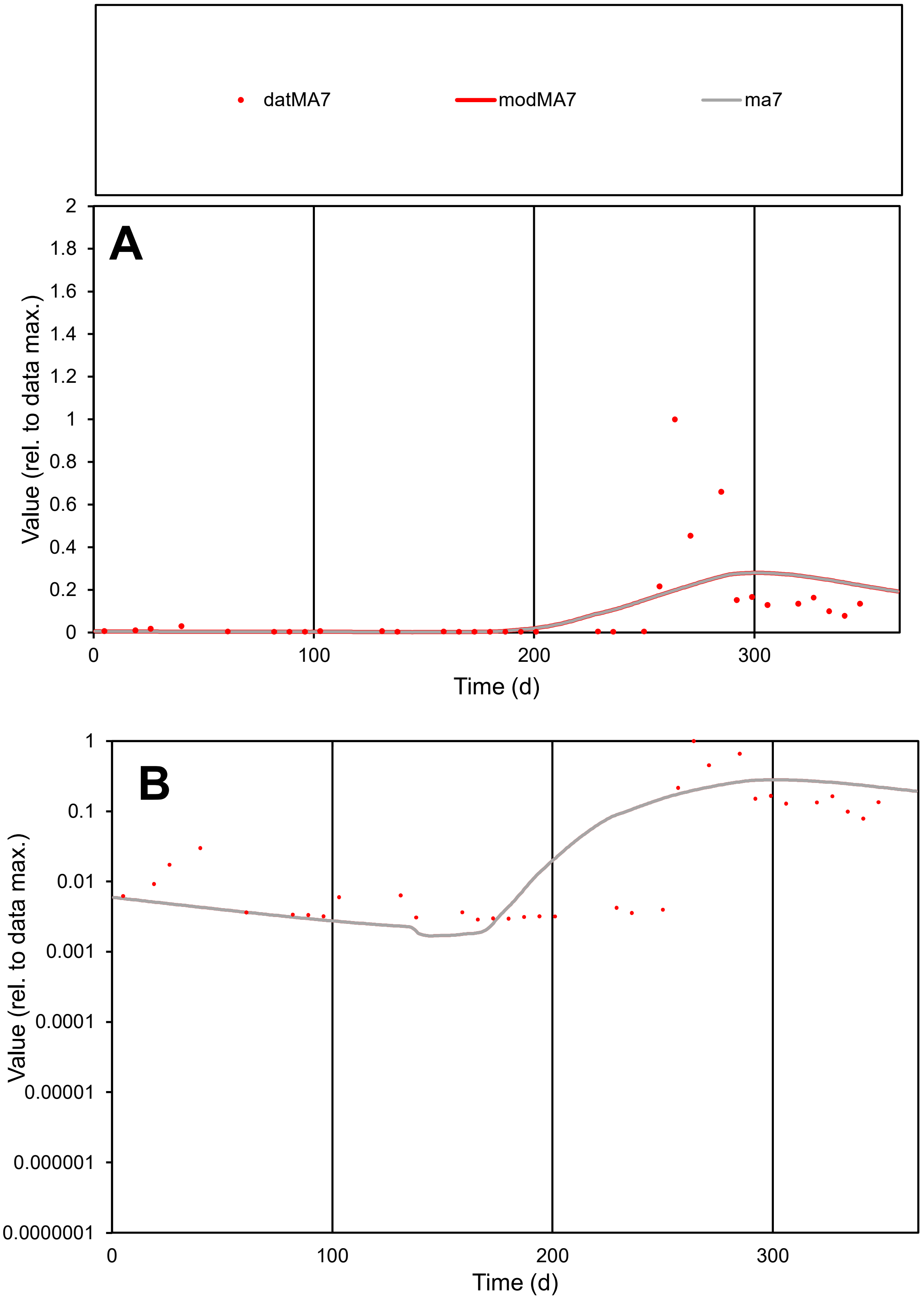


**C1**

Variables
MA7 Provide jtd or Dst
CalibOnly?
2 1 to reproduce of
oNormalize?
2 1=no, 2=yes

ofxn calc*
1.64522
ofxn mod
1.65961

sRun
RunA1
zout
1

Ave
Data
0.00049
Model
0.00047

data/vMax
0.00462

ofxng
263.459

Min d/m
1.61091

ave/min d
36.974
ave/min m
57.1447

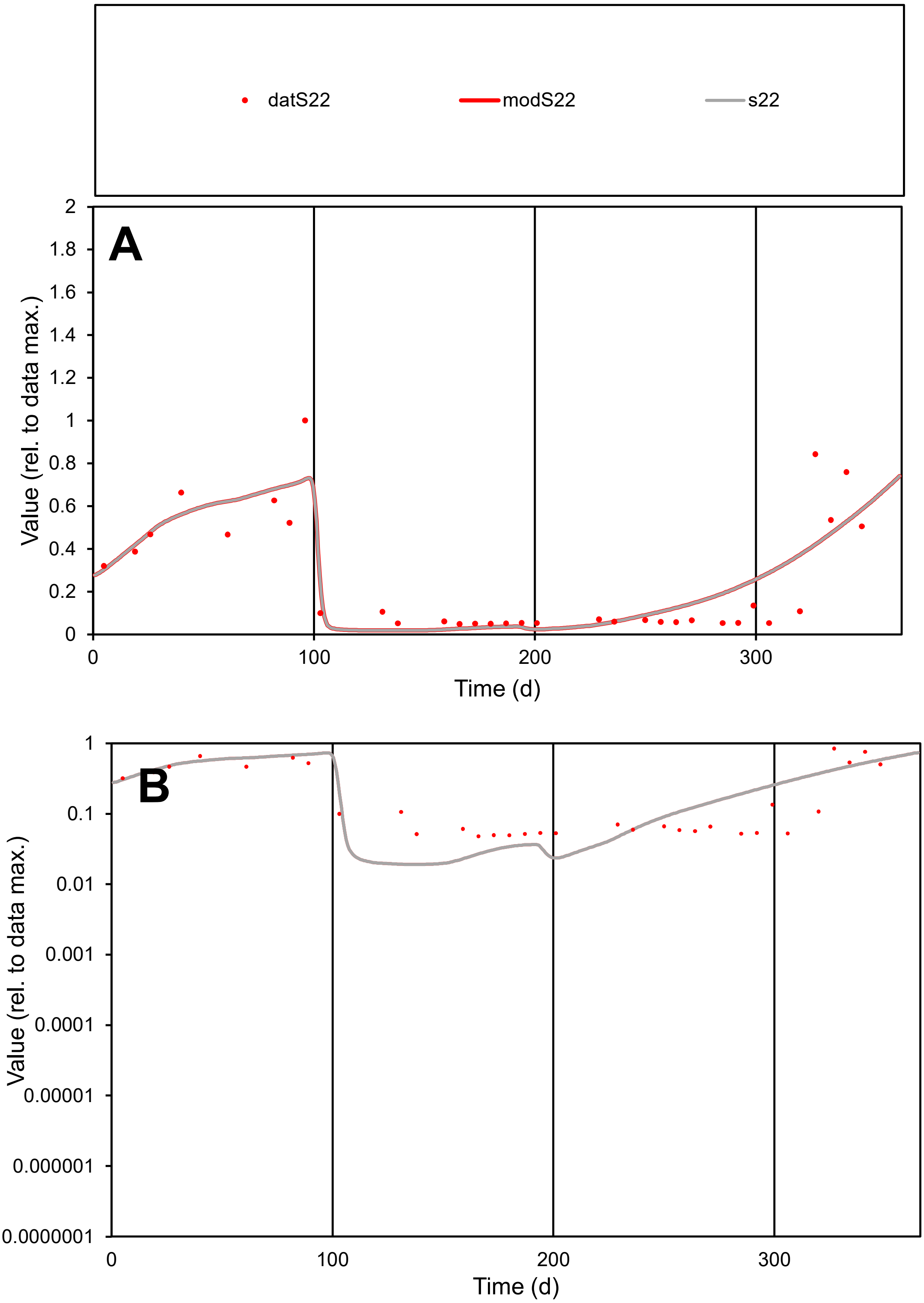


**C1**

Variables
S22 Provide jtd or Dst
CalibOnly?
2 1 to reproduce of
oNormalize?
2 1=no, 2=yes

ofxn calc*
0.52075
ofxn mod
0.54504

sRun
RunA1
zout
1

Ave
Data
0.00078
Model
0.00081

data/vMax
0.00302

ofxng
263.459

Min d/m
2.51039

ave/min d
5.36723
ave/min m
14.0907

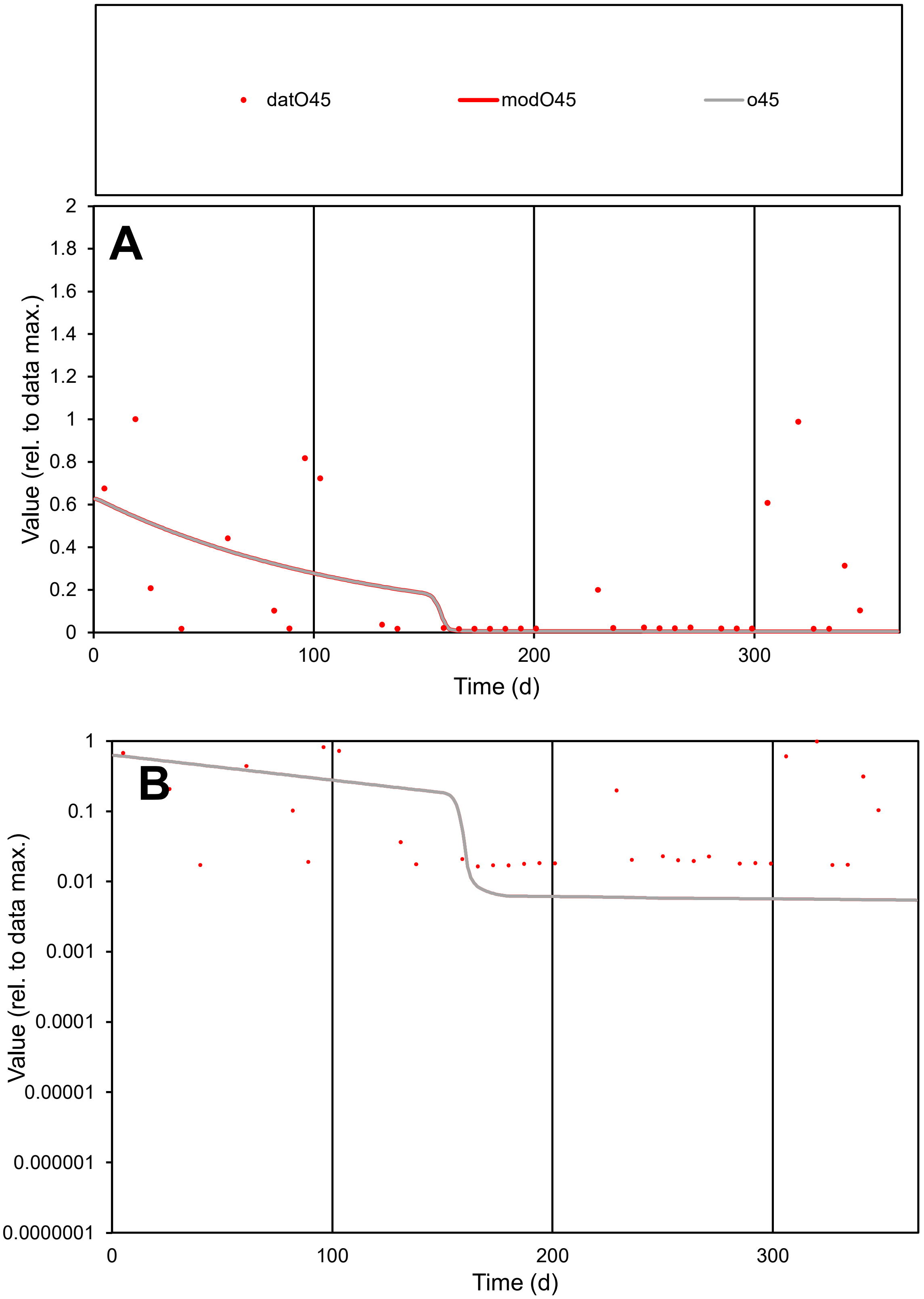


**C1**

Variables
O45 Provide jtd or Dst
CalibOnly?
2 1 to reproduce of
oNormalize?
2 1=no, 2=yes

ofxn calc*
1.42061
ofxn mod
1.42904

sRun
RunA1
zout
1

Ave
Data
0.00069
Model
0.00044

data/vMax
0.00343

ofxng
263.459

Min d/m
2.99629

ave/min d
12.135
ave/min m
23.2721

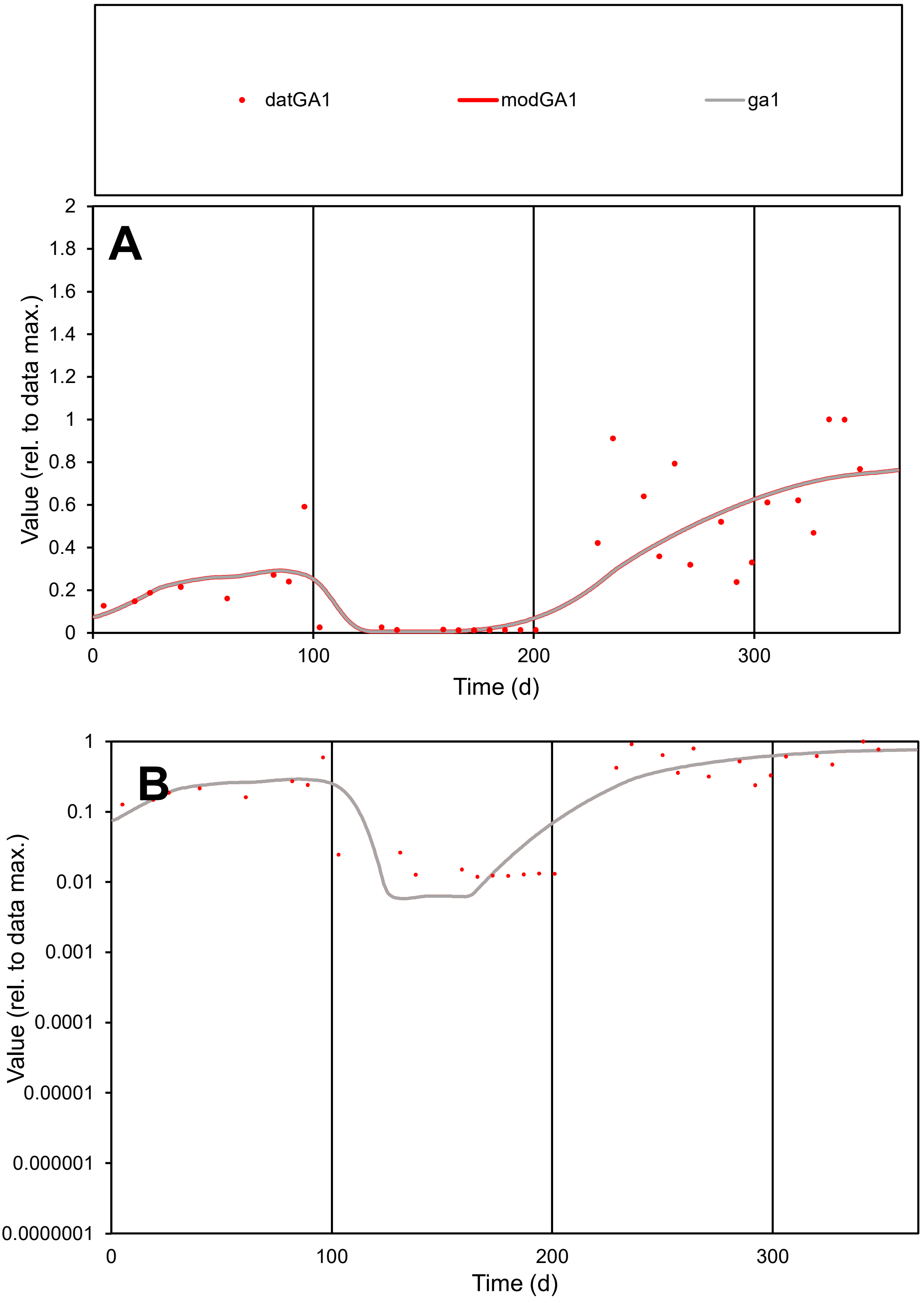


C1

Variables
GA1 Provide jtd or Dst
CalibOnly?
2 1 to reproduce of
oNormalize?
2 1=no, 2=yes

ofxn calc*
0.56761
ofxn mod
0.58537

sRun
RunA1
zout
1

Ave
Data
0.00099
Model
0.00095

data/vMax
0.00295

ofxng
263.459

Min d/m
2.04684

ave/min d
28.3335
ave/min m
55.469

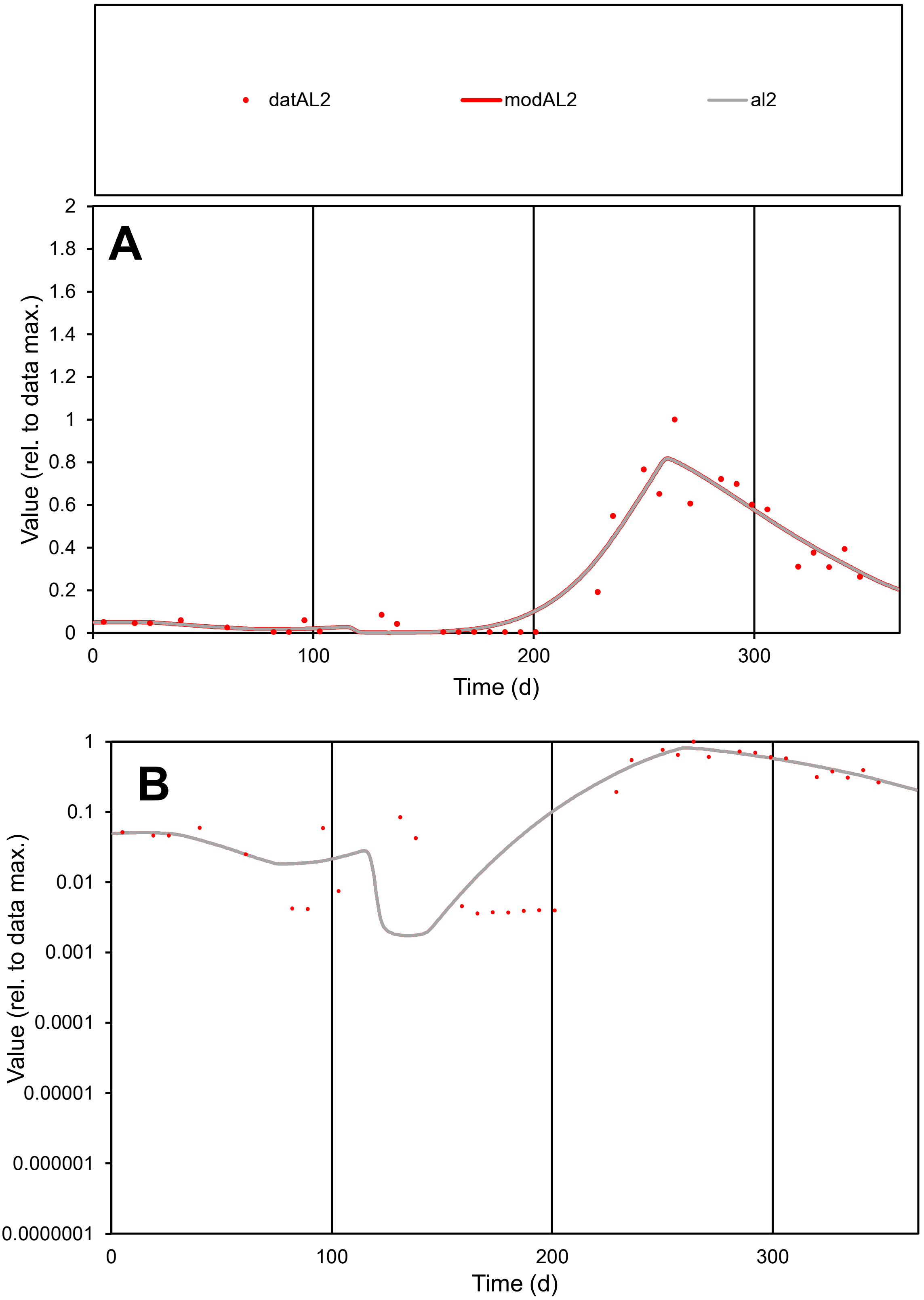


**C1**

Variables

AL2 Provide jtd or Dst

CalibOnly?

2 1 to reproduce of

oNormalize?

2 1=no, 2=yes

ofxn calc*

0.29136

ofxn mod

0.34702

sRun

RunA1

zout

1

Ave

Data

0.00153

Model

0.00156

data/vMax

0.00598

ofxng

263.459

Min d/m

2.06334

ave/min d

71.3936

ave/min m

149.836

326_Calib_S_A

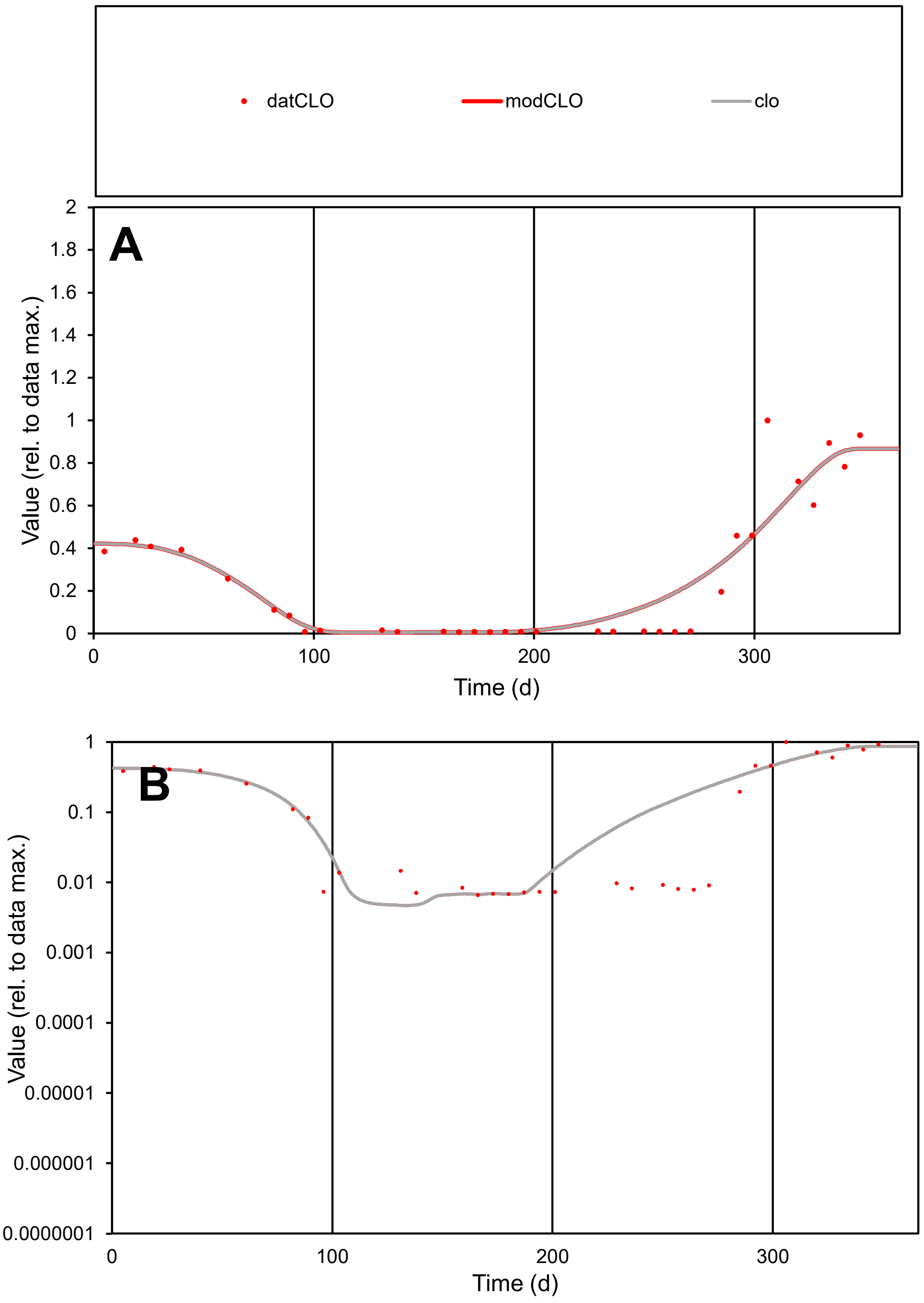


**C1**

Variables
CLO Provide jtd or Dst
CalibOnly?
2 1 to reproduce of
oNormalize?
2 1=no, 2=yes

ofxn calc*
0.44062
ofxn mod
0.47315

sRun
RunA1
zout
1

Ave
Data
0.00049
Model
0.00052

data/vMax
0.00197

ofxng
263.459

Min d/m
1.40606

ave/min d
37.772
ave/min m
56.4799

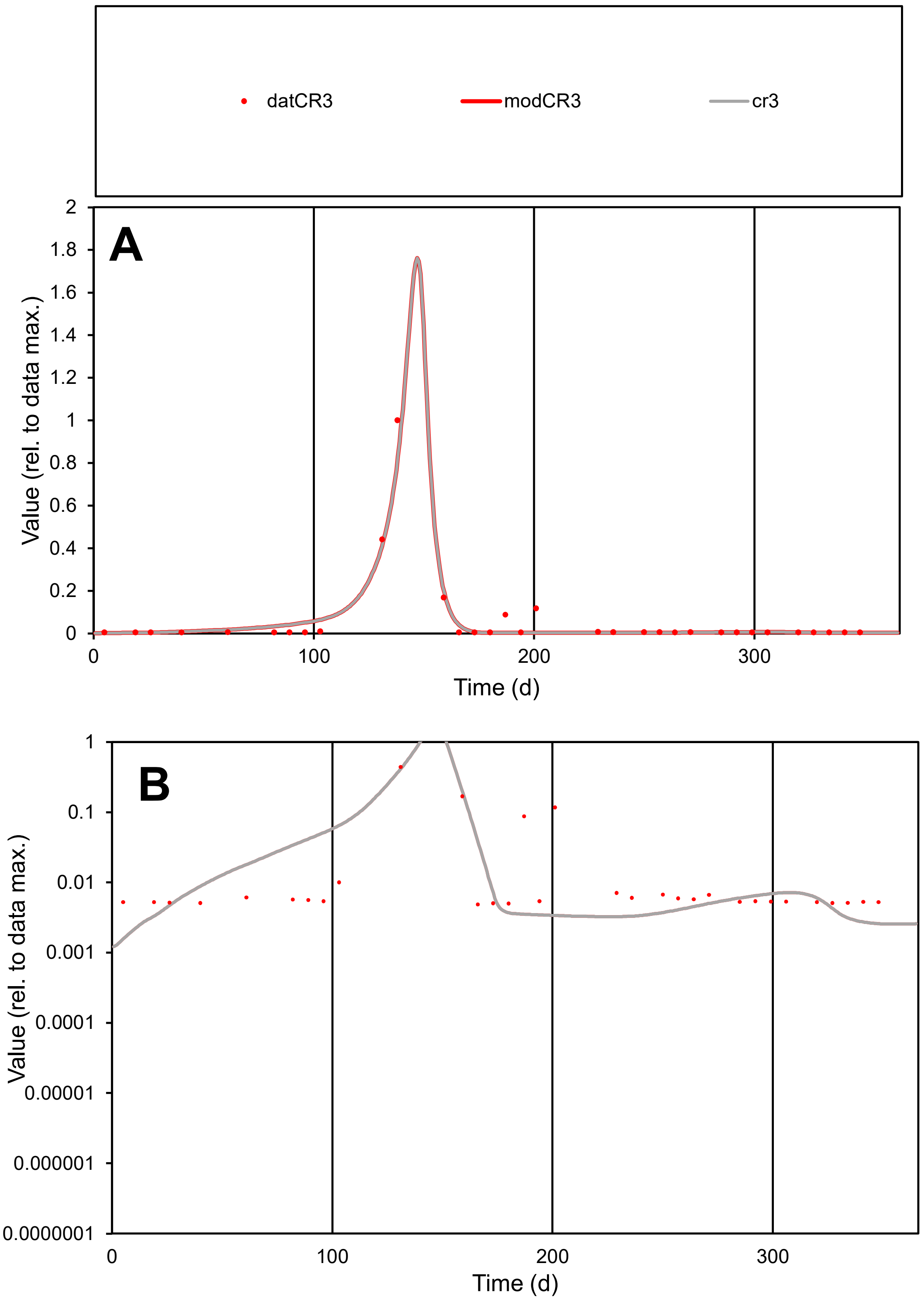


**C1**

Variables
CR3 Provide jtd or Dst
CalibOnly?
2 1 to reproduce of
oNormalize?
2 1=no, 2=yes

ofxn calc*
0.49767
ofxn mod
0.63264

sRun
RunA1
zout
1

Ave
Data
0.00043
Model
0.00042

data/vMax
0.00723

ofxng
263.459

Min d/m
2.81672

ave/min d
12.3353
ave/min m
34.0377

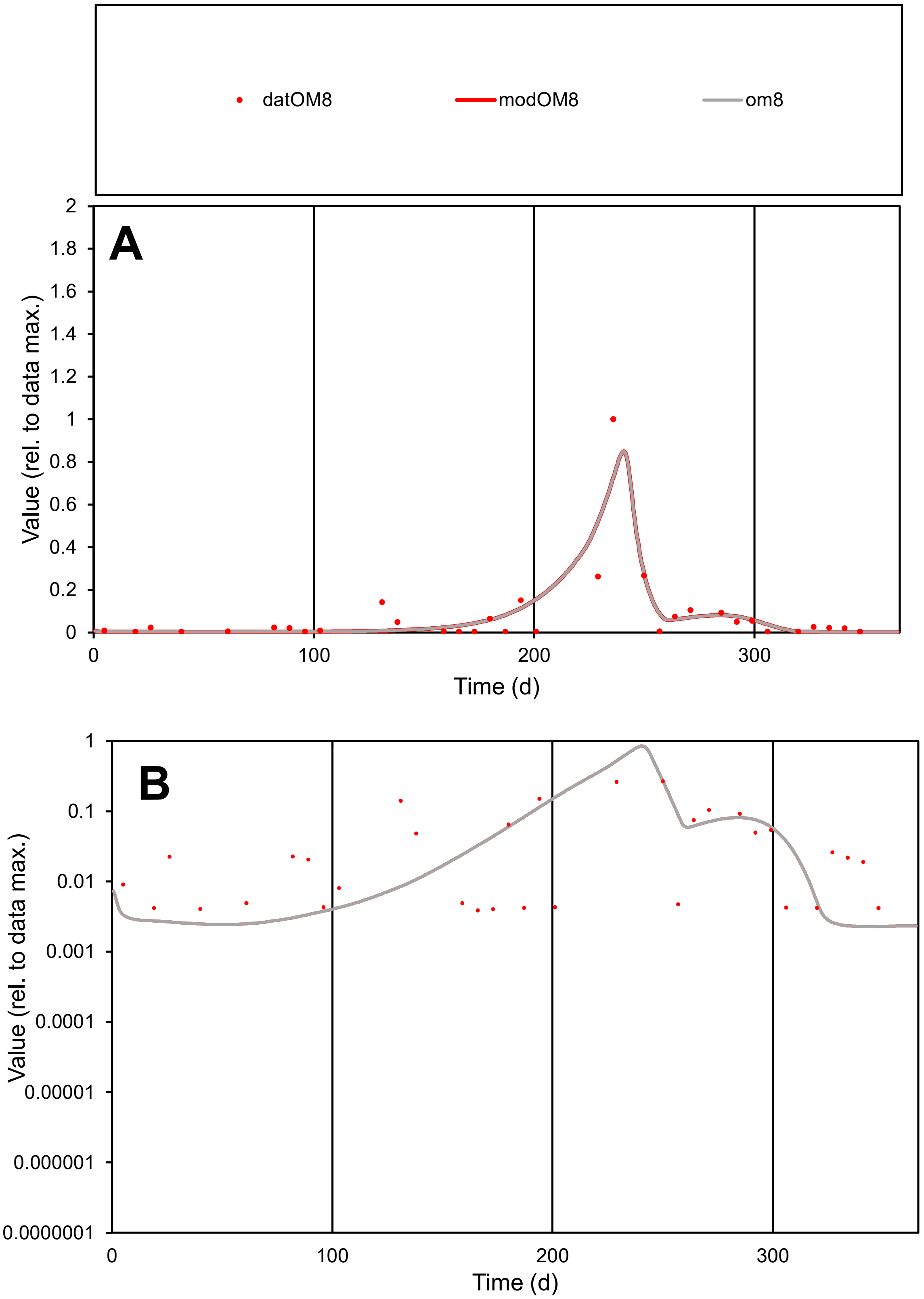


**C1**

Variables
OM8 Provide jtd or Dst
CalibOnly?
2 1 to reproduce of
oNormalize?
2 1=no, 2=yes

ofxn calc*
0.9738
ofxn mod
1.02205

sRun
RunA1
zout
1

Ave
Data
0.00105
Model
0.00107

data/vMax
0.01371

ofxng
263.459

Min d/m
1.70624
ave/min d
19.6175
ave/min m
34.2192

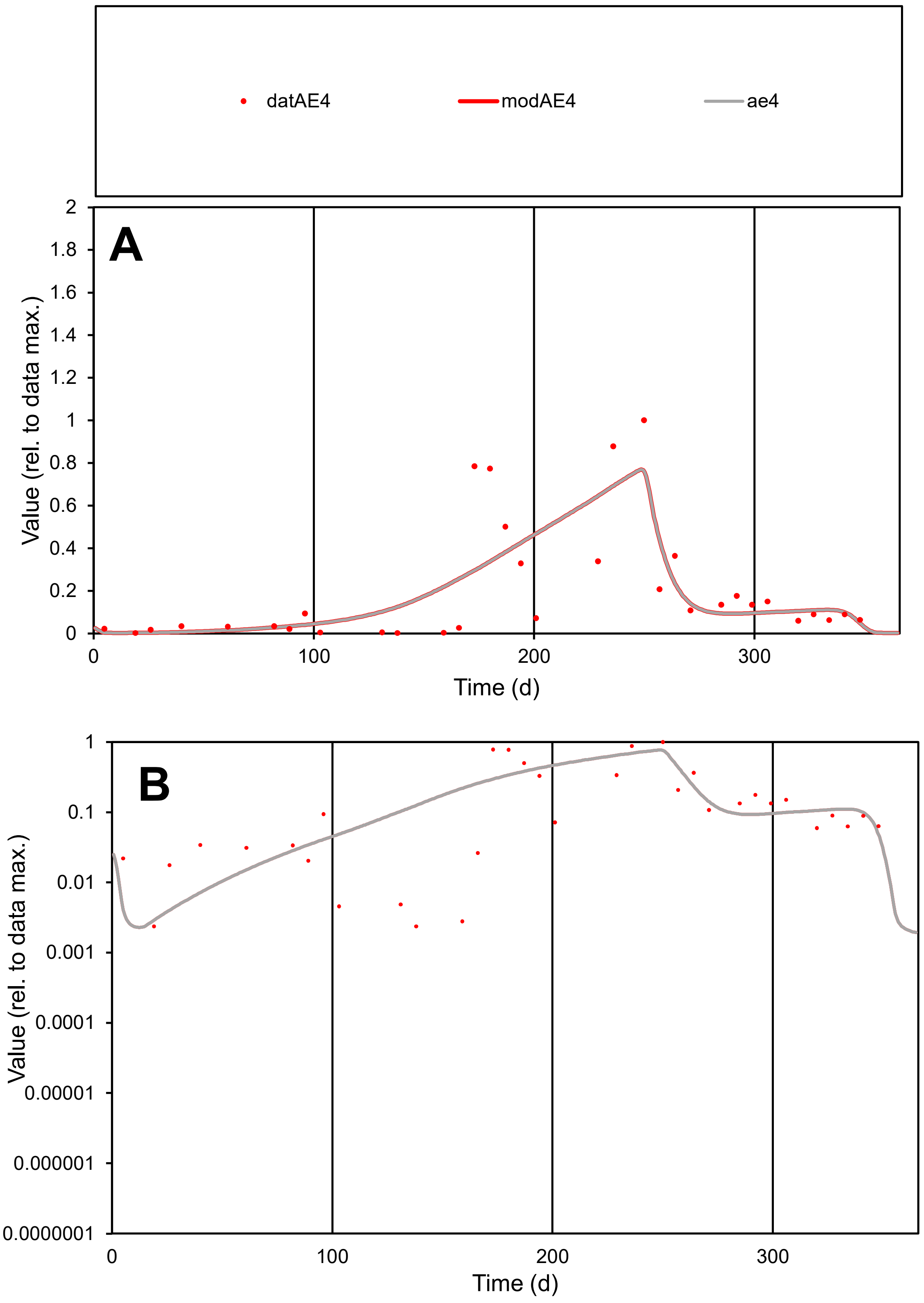


**C1**

Variables

AE4 Provide jtd or Dst

CalibOnly?

2 1 to reproduce of

oNormalize?

2 1=no, 2=yes

ofxn calc*

0.86625

ofxn mod

0.89122

sRun

RunA1

zout

1

Ave

Data

0.00105

Model

0.00104

data/vMax

0.00523

ofxng

263.459

Min d/m

0.79448

ave/min d

84.5892

ave/min m

66.7494

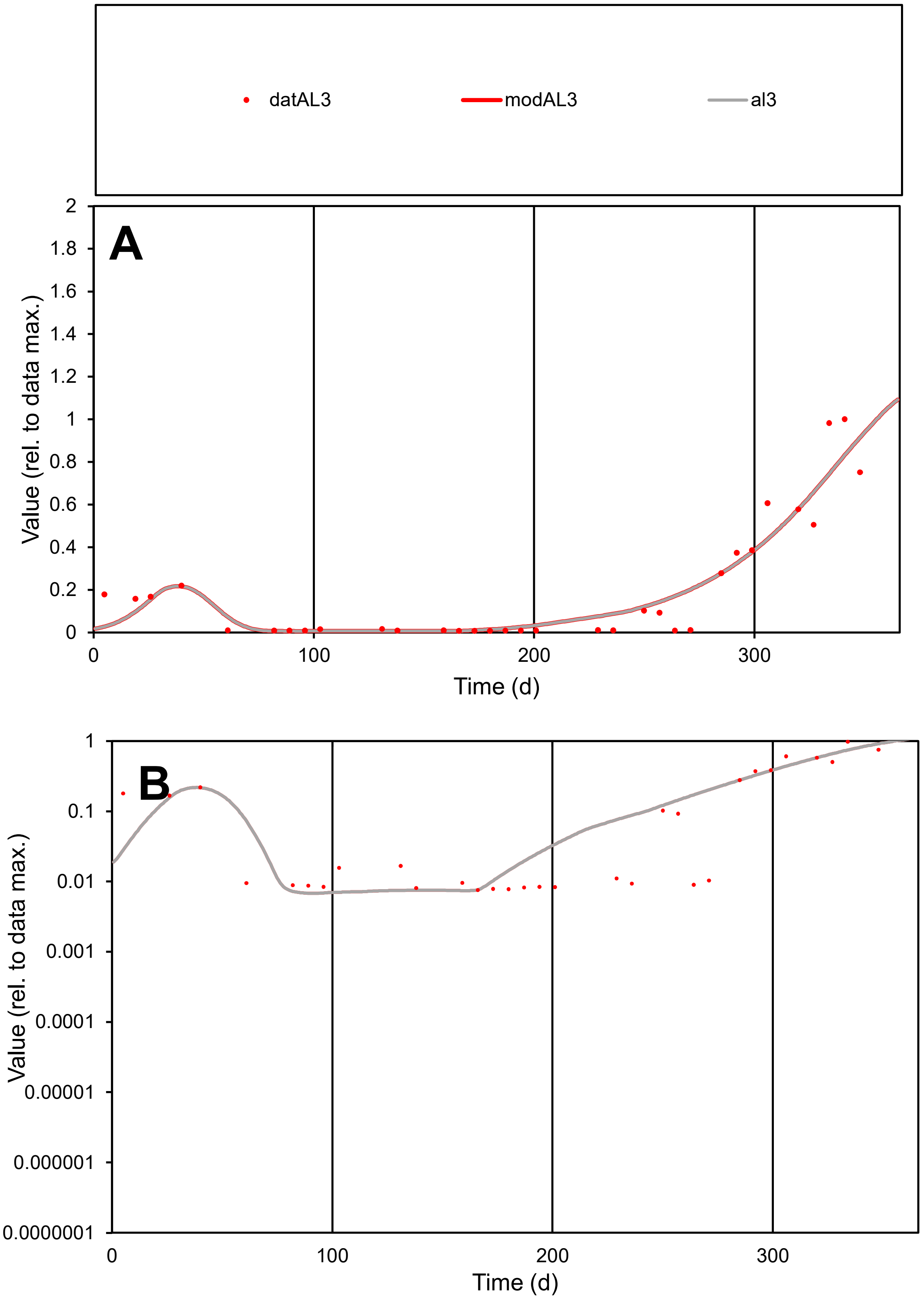


C1

Variables
AL3 Provide jtd or Dst
CalibOnly?
2 1 to reproduce of
oNormalize?
2 1=no, 2=yes

ofxn calc*
0.44599
ofxn mod
0.48086

sRun
RunA1
zout
1

Ave
Data
0.00063
Model
0.00065

data/vMax
0.00316

ofxng
263.459

Min d/m
1.11345

ave/min d
26.3344
ave/min m
30.4937

326_Calib_S_A

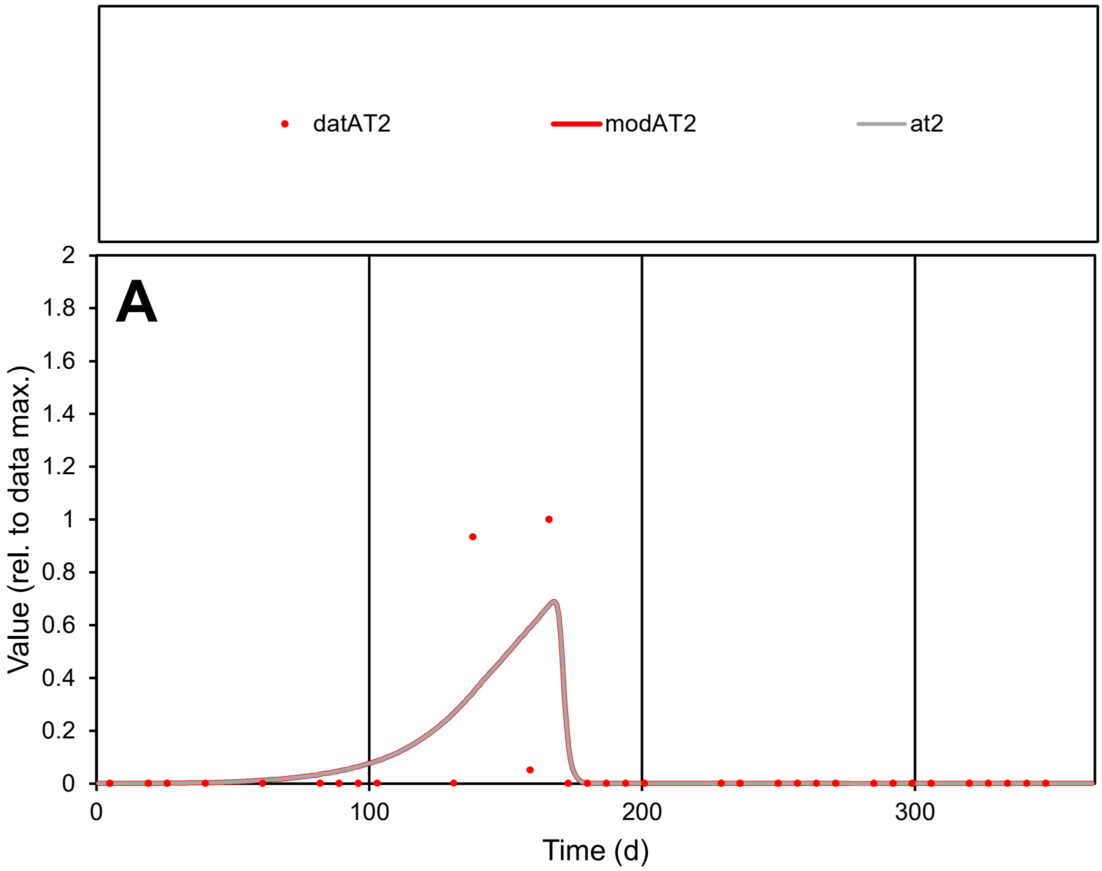


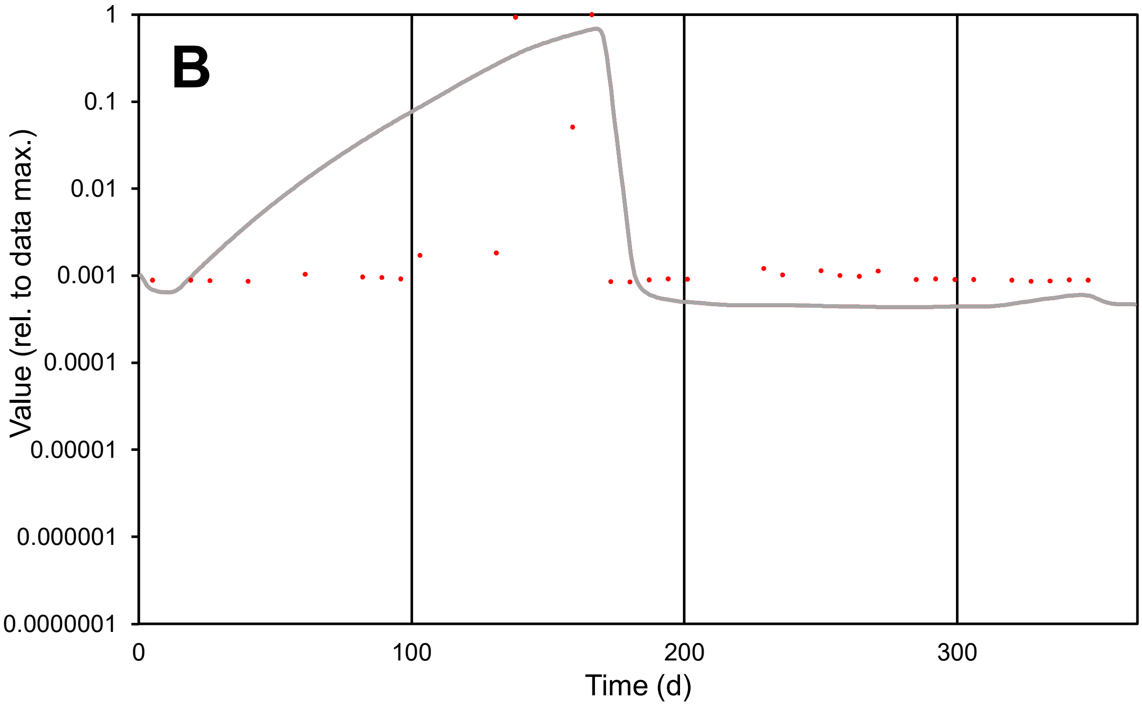


**C1**

Variables
AT2 Provide jtd or Dst
CalibOnly?
2 1 to reproduce of
oNormalize?
2 1=no, 2=yes

ofxn calc*
2.54857
ofxn mod
2.58908

sRun
RunA1
zout
1

Ave
Data
0.00583
Model
0.00637

data/vMax
0.0954

ofxng
263.459

Min d/m
1.9481

ave/min d
71.8511
ave/min m
153.073

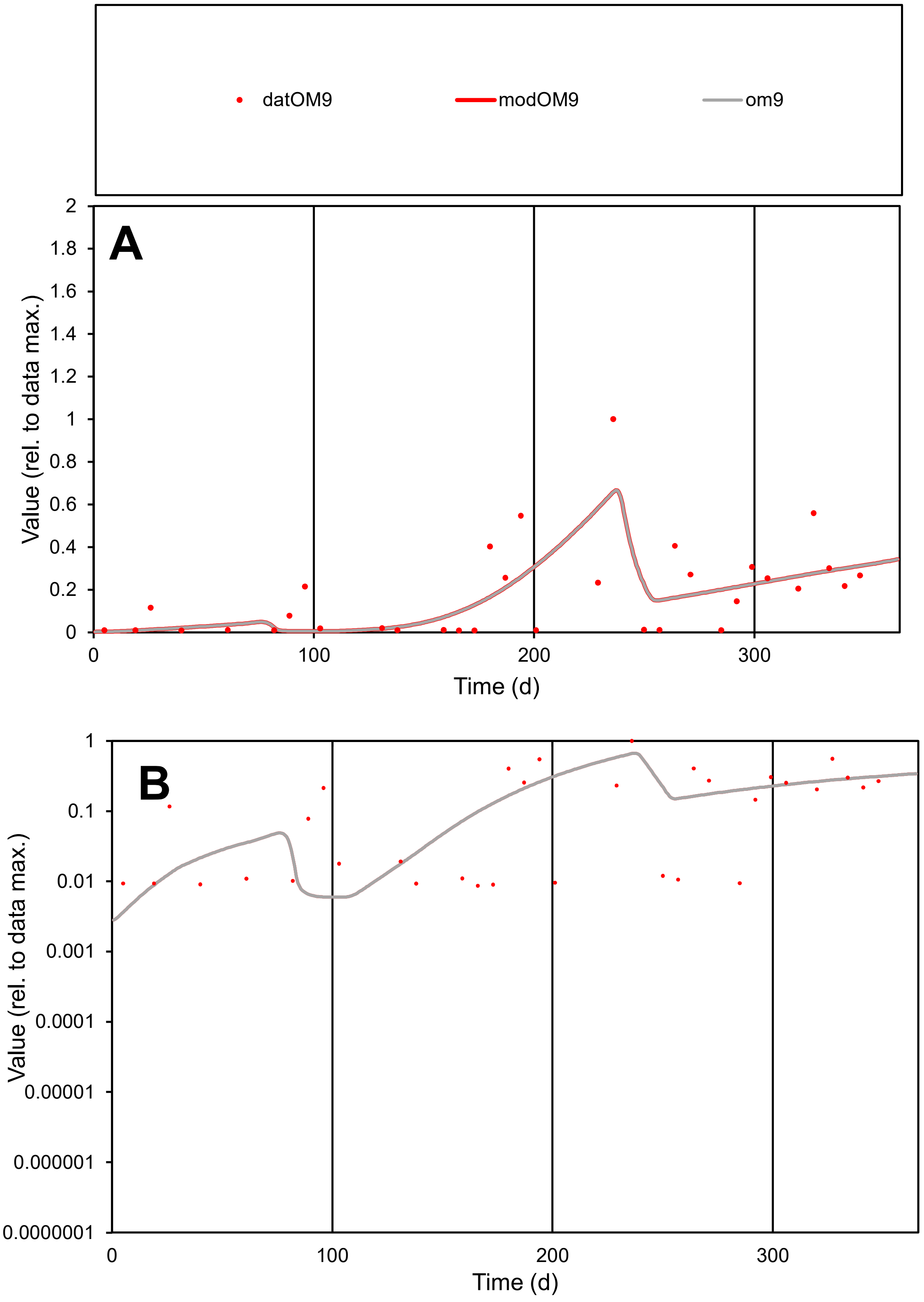


**C1**

Variables
OM9 Provide jtd or Dst
CalibOnly?
2 1 to reproduce of
oNormalize?
2 1=no, 2=yes

ofxn calc*
0.86354
ofxn mod
0.88017

sRun
RunA1
zout
1

Ave
Data
0.00089
Model
0.00085

data/vMax
0.00495

ofxng
263.459

Min d/m
2.14429

ave/min d
20.8219
ave/min m
42.8339

326_Calib_S_A

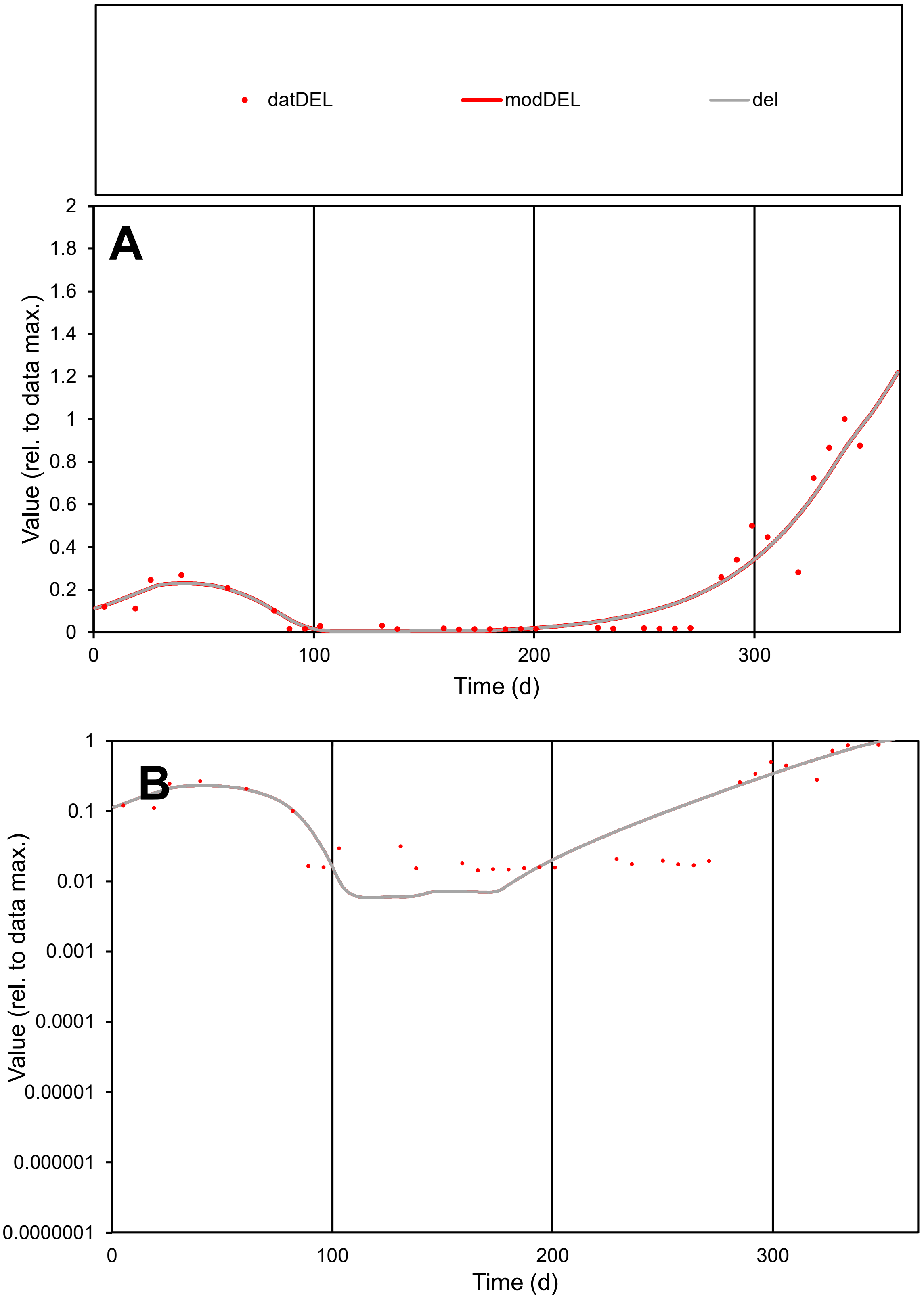


**C1**

Variables
DEL Provide jtd or Dst
CalibOnly?
2 1 to reproduce of
oNormalize?
2 1=no, 2=yes

ofxn calc*
0.38572
ofxn mod
0.42897

sRun
RunA1
zout
1

Ave
Data
0.00087
Model
0.00091

data/vMax
0.0043

ofxng
263.459

Min d/m
2.37007

ave/min d
14.1759
ave/min m
35.1017

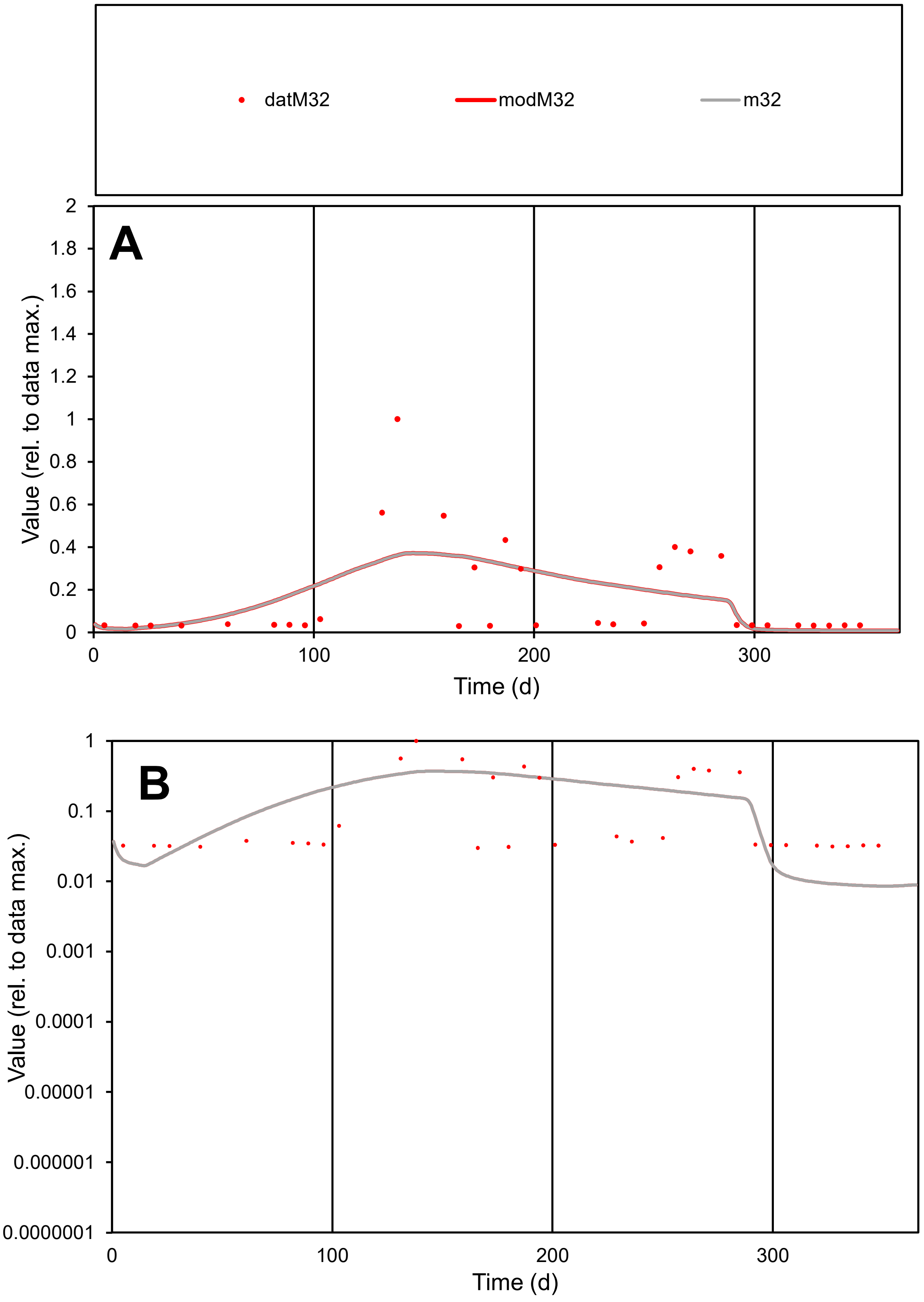


**C1**

Variables
M32 Provide jtd or Dst
CalibOnly?
2 1 to reproduce of
oNormalize?
2 1=no, 2=yes

ofxn calc*
1.11489
ofxn mod
1.1288

sRun
RunA1
zout
1

Ave
Data
0.00113
Model
0.00114

data/vMax
0.00692

ofxng
263.459

Min d/m
3.48544

ave/min d
5.46533
ave/min m
19.1129

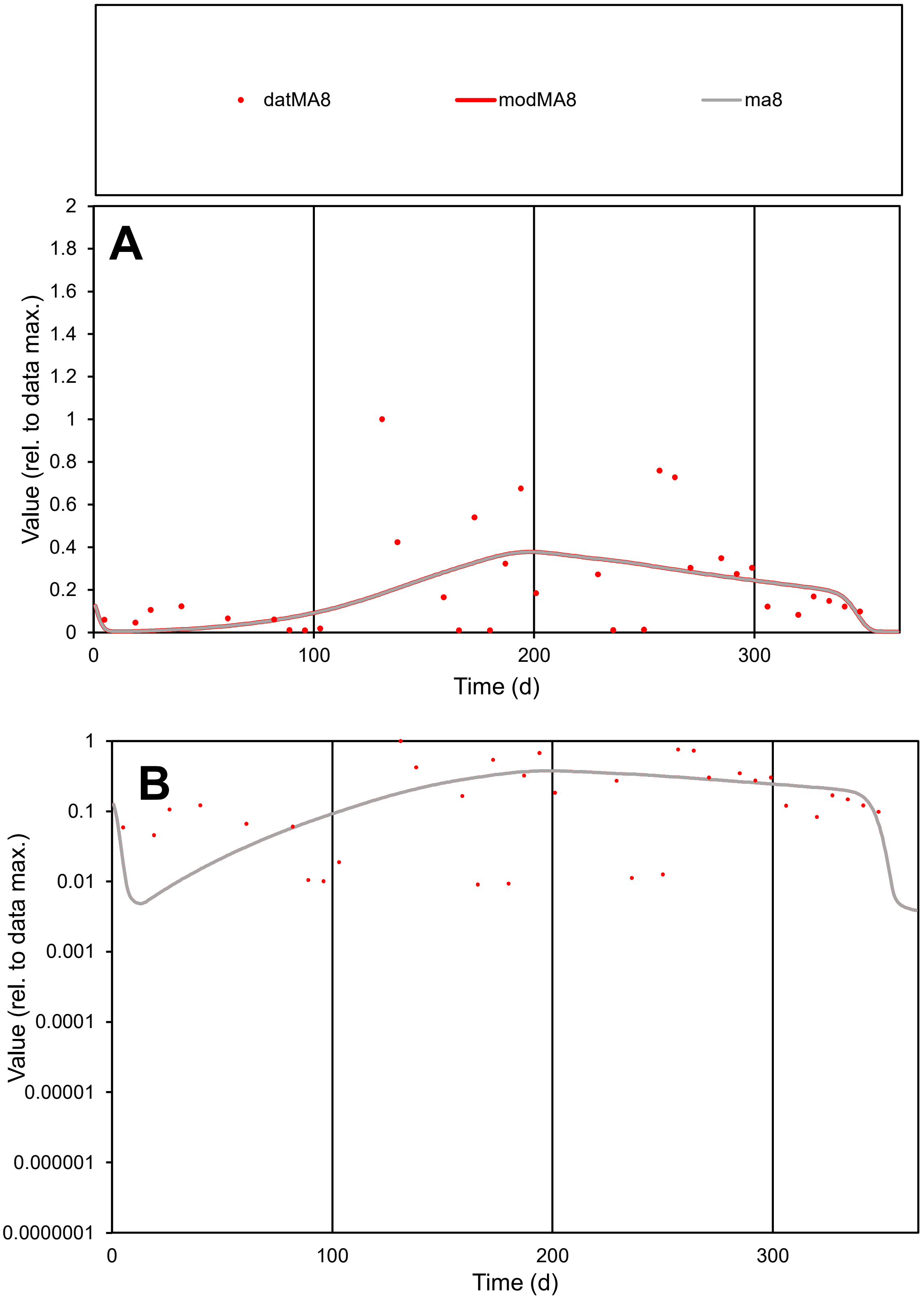


C1

Variables
MA8 Provide jtd or Dst
CalibOnly?
2 1 to reproduce of
oNormalize?
2 1=no, 2=yes

ofxn calc*
1.01064
ofxn mod
1.02151

sRun
RunA1
zout
1

Ave
Data
0.00149
Model
0.00136

data/vMax
0.00649

ofxng
263.459

Min d/m
1.39815

ave/min d
25.2122
ave/min m
32.1472

326_Calib_S_A

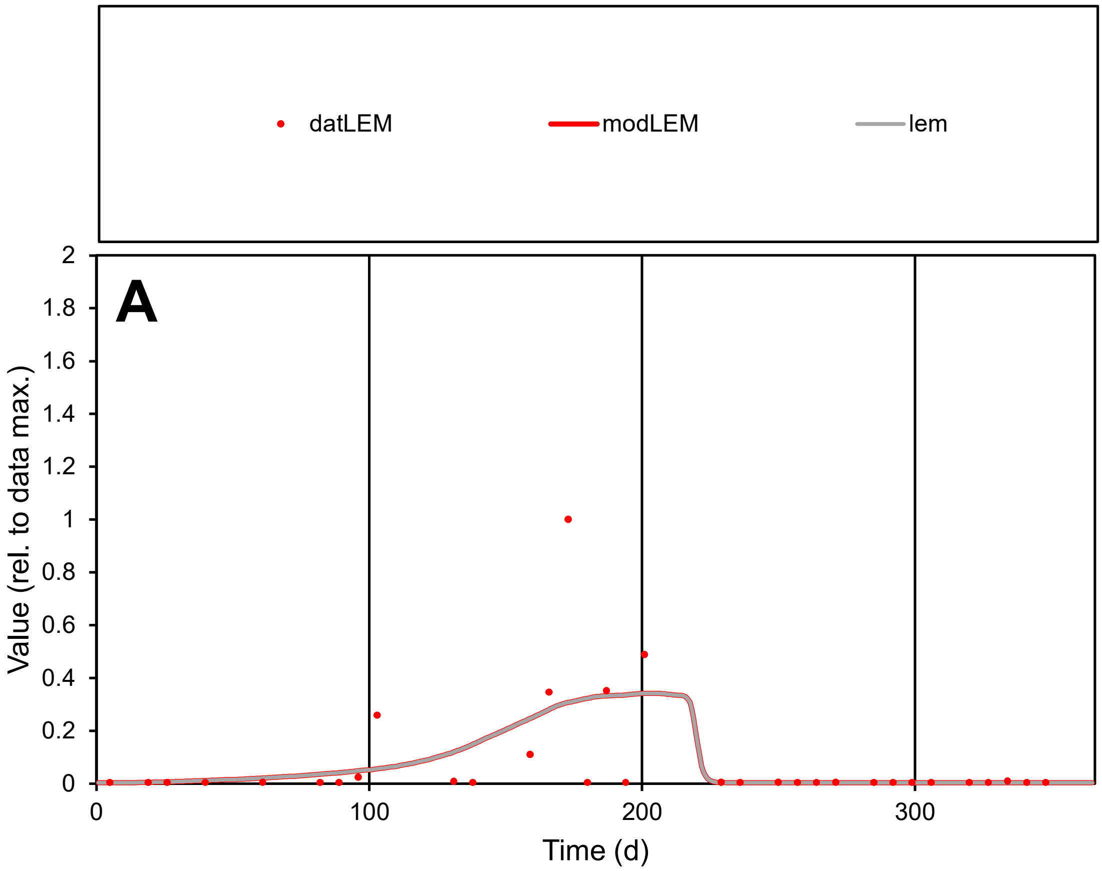


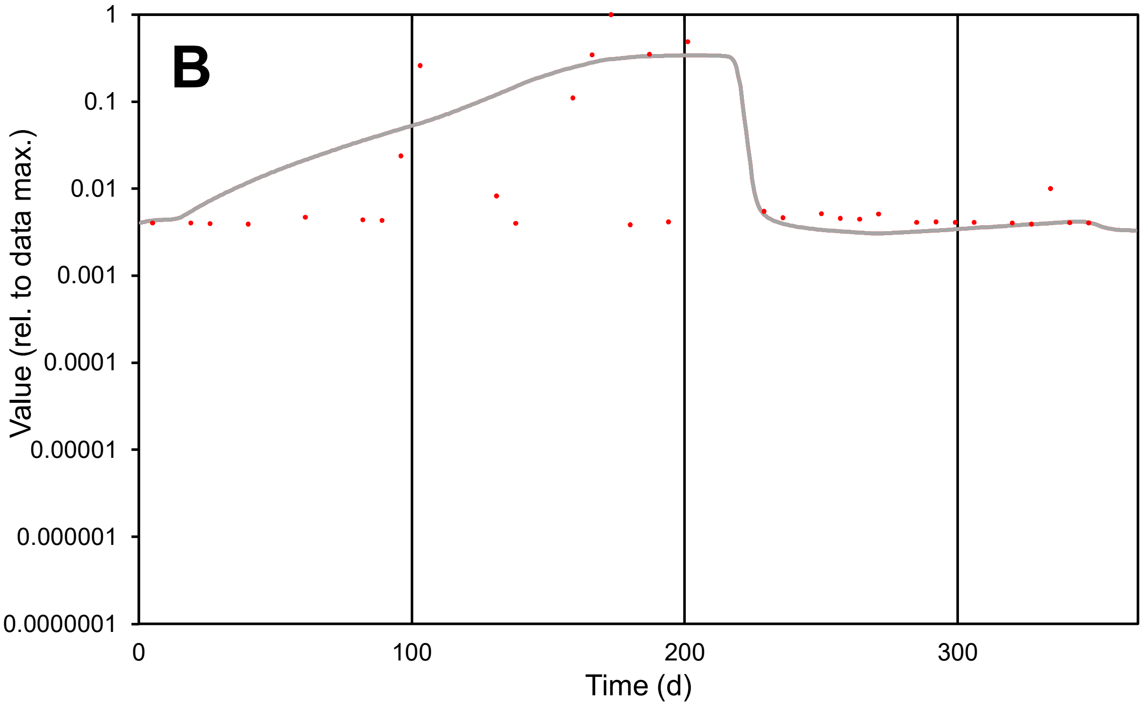


**C1**

Variables
LEM Provide jtd or Dst
CalibOnly?
2 1 to reproduce of
oNormalize?
2 1=no, 2=yes

ofxn calc*
1.91352
ofxn mod
1.93065

sRun
RunA1
zout
1

Ave
Data
0.00047
Model
0.00047

data/vMax
0.00571

ofxng
263.459

Min d/m
1.25128

ave/min d
21.2513
ave/min m
26.9952

326_Calib_S_A

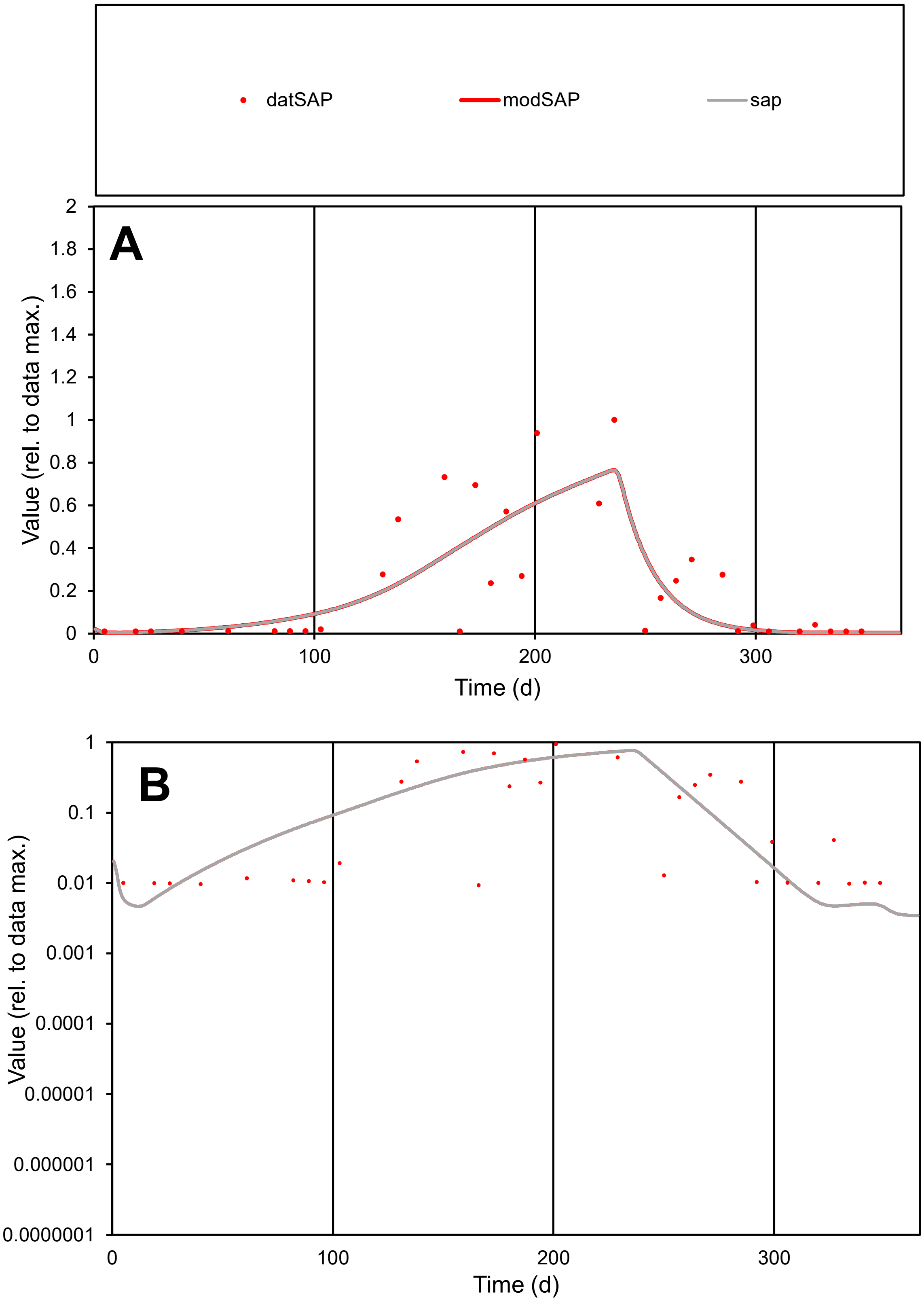


**C1**

Variables
SAP Provide jtd or Dst
CalibOnly?
2 1 to reproduce of
oNormalize?
2 1=no, 2=yes

ofxn calc*
0.81199
ofxn mod
0.83226

sRun
RunA1
zout
1

Ave
Data
0.00076
Model
0.00072

data/vMax
0.00352

ofxng
263.459

Min d/m
1.98536

ave/min d
23.4591
ave/min m
43.6696

326_Calib_S_A

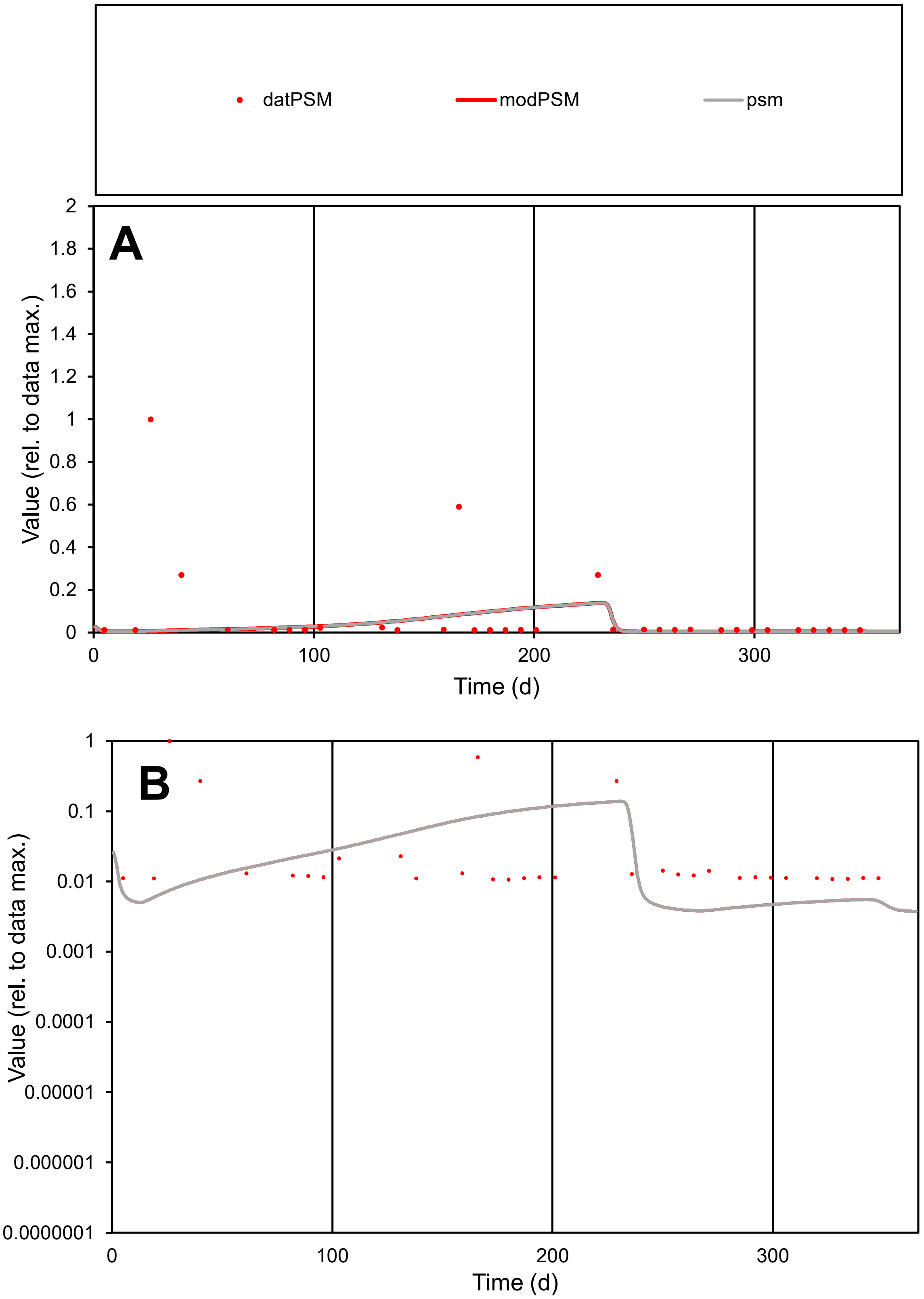


**C1**

| | |
|---|---|
| Variables | |
| PSM | Provide jtd or Dst |
| CalibOnly? | |
| 2 | 1 to reproduce of |
| oNormalize? | |
| 2 | 1=no, 2=yes |
| ofxn calc* | |
| 2.70609 | |
| ofxn mod | |
| 2.71436 | |
| sRun | |
| RunA1 | |
| zout | |
| 1 | |
| Ave | |
| Data | |
| 0.0002 | |
| Model | |
| 9.3E-05 | |
| data/vMax | |
| 0.00265 | |
| ofxng | |
| 263.459 | |
| Min d/m | |
| 2.77657 | |
| ave/min d | |
| 7.09234 | |
| ave/min m | |
| 9.1474 | |

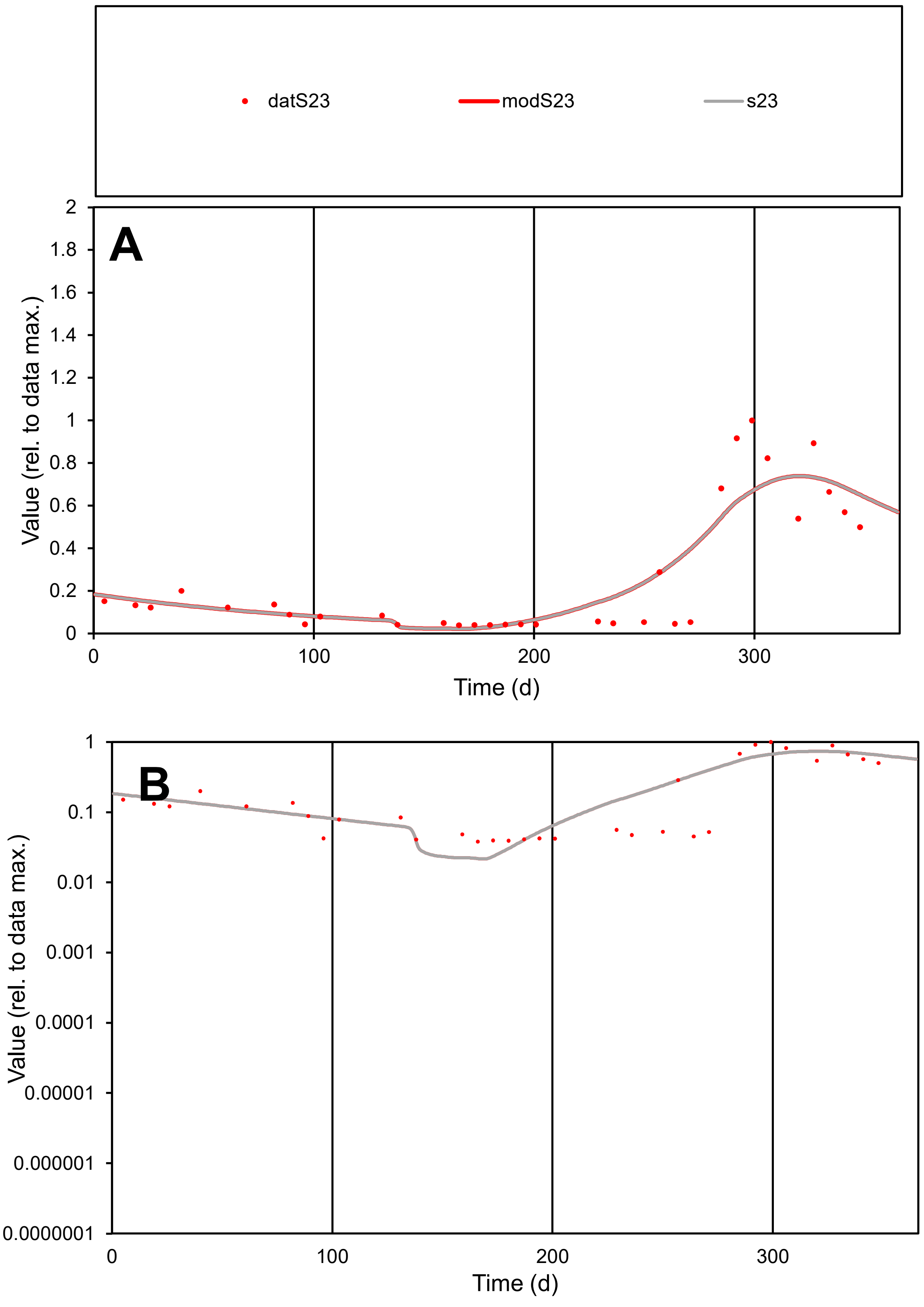


**C1**

Variables
S23 Provide jtd or Dst
CalibOnly?
2 1 to reproduce of
oNormalize?
2 1=no, 2=yes

ofxn calc*
0.51527
ofxn mod
0.53825

sRun
RunA1
zout
1

Ave
Data
0.00109
Model
0.00115

data/vMax
0.00418

ofxng
263.459

Min d/m
1.75497

ave/min d
6.85885
ave/min m
12.7333

326_Calib_S_A

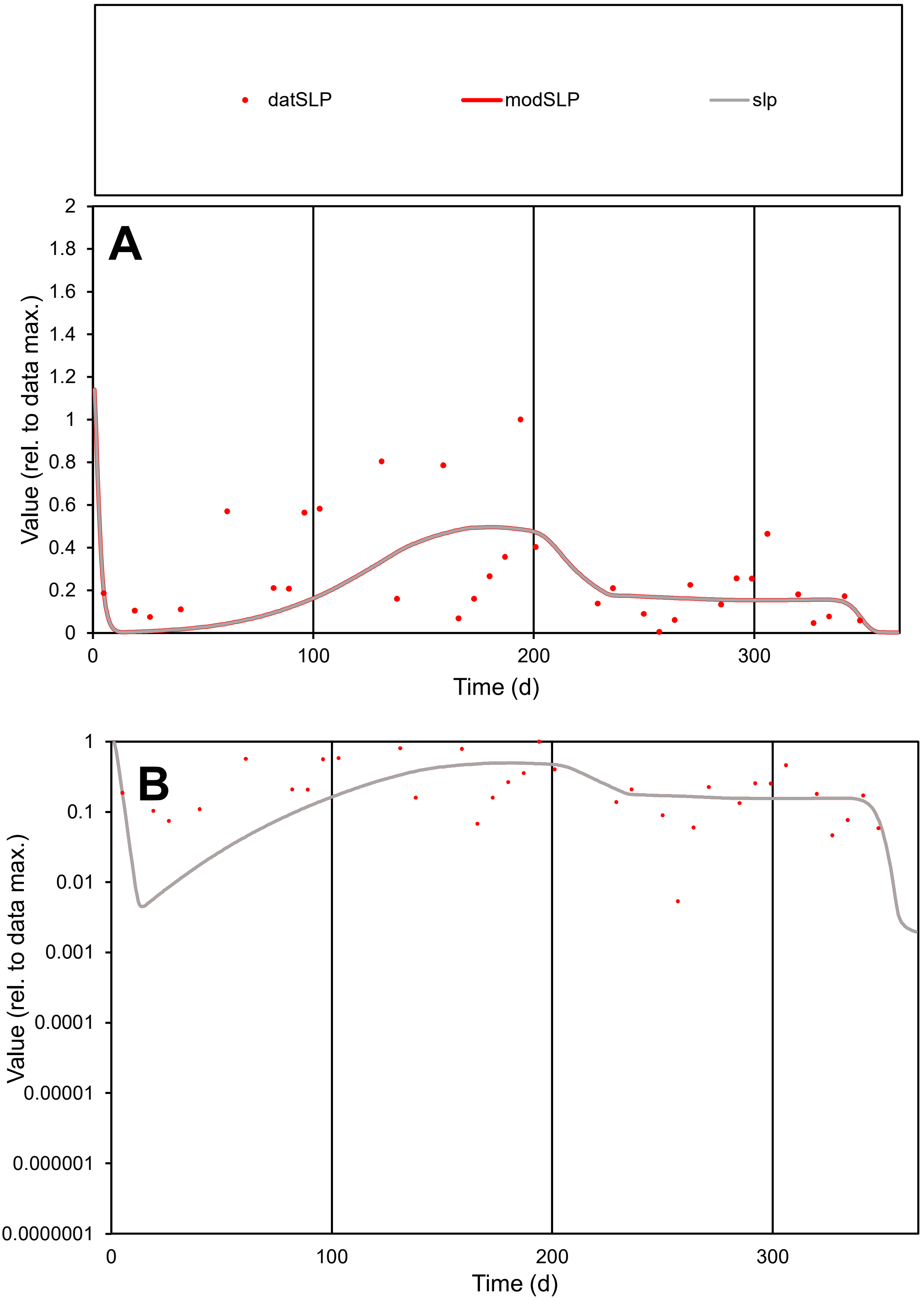


**C1**

| | |
|---|---|
| Variables | |
| SLP | Provide jtd or Dst |
| CalibOnly? | |
| 2 | 1 to reproduce of |
| oNormalize? | |
| 2 | 1=no, 2=yes |
| ofxn calc* | |
| 0.86441 | |
| ofxn mod | |
| 0.8739 | |
| sRun | |
| RunA1 | |
| zout | |
| 1 | |
| Ave Data | |
| 0.00147 | |
| Model | |
| 0.00119 | |
| data/vMax | |
| 0.00541 | |
| ofxng | |
| 263.459 | |
| Min d/m | |
| 0.86408 | |
| ave/min d | |
| 50.7992 | |
| ave/min m | |
| 35.459 | |

326_Calib_S_A

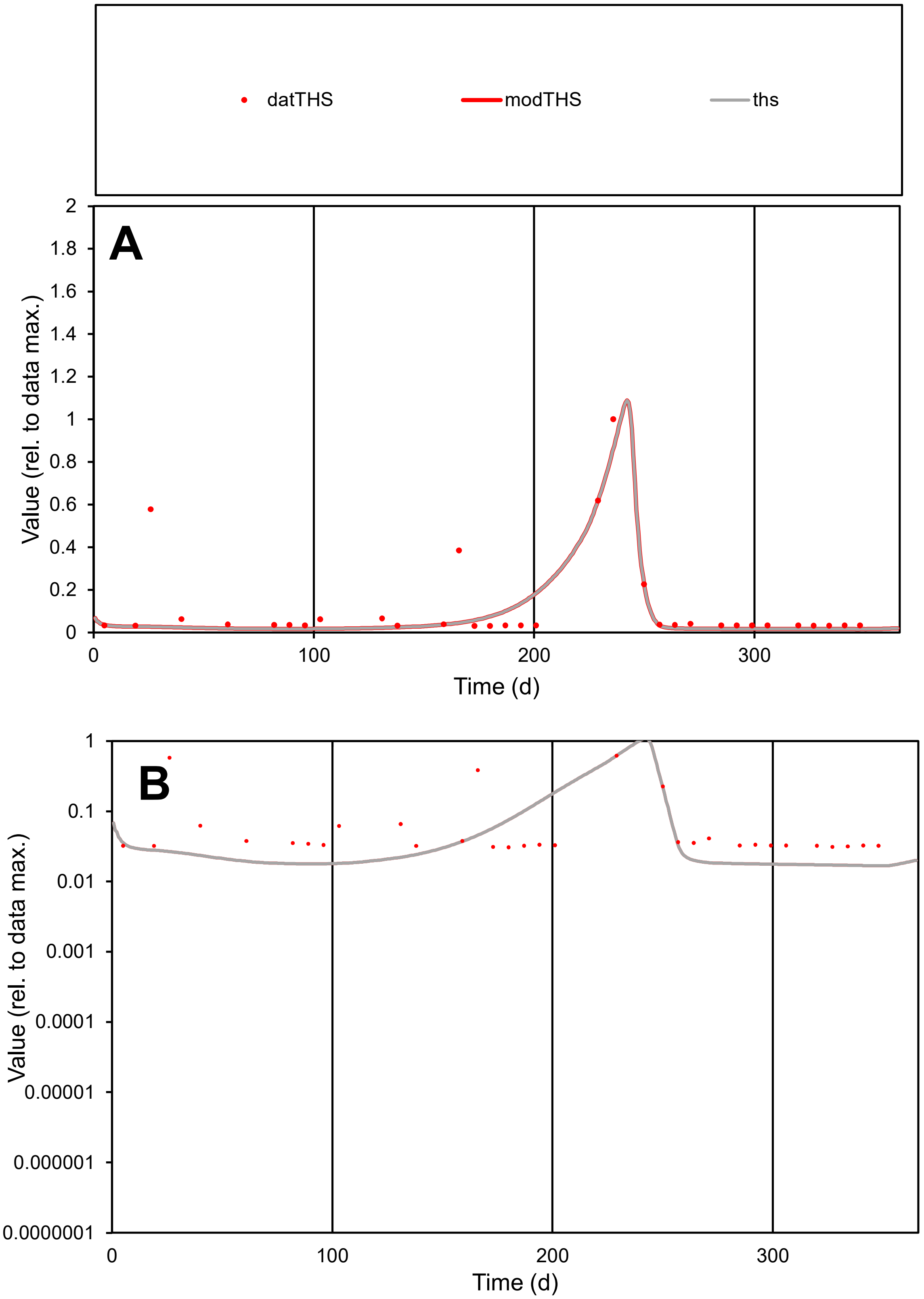


**C1**

Variables
THS Provide jtd or Dst
CalibOnly?
2 1 to reproduce of
oNormalize?
2 1=no, 2=yes

ofxn calc*
1.03936
ofxn mod
1.05686

sRun
RunA1
zout
1

Ave
Data
0.00011
Model
8.4E-05

data/vMax
0.00095

ofxng
263.459

Min d/m
1.83021

ave/min d
3.78275
ave/min m
5.24161

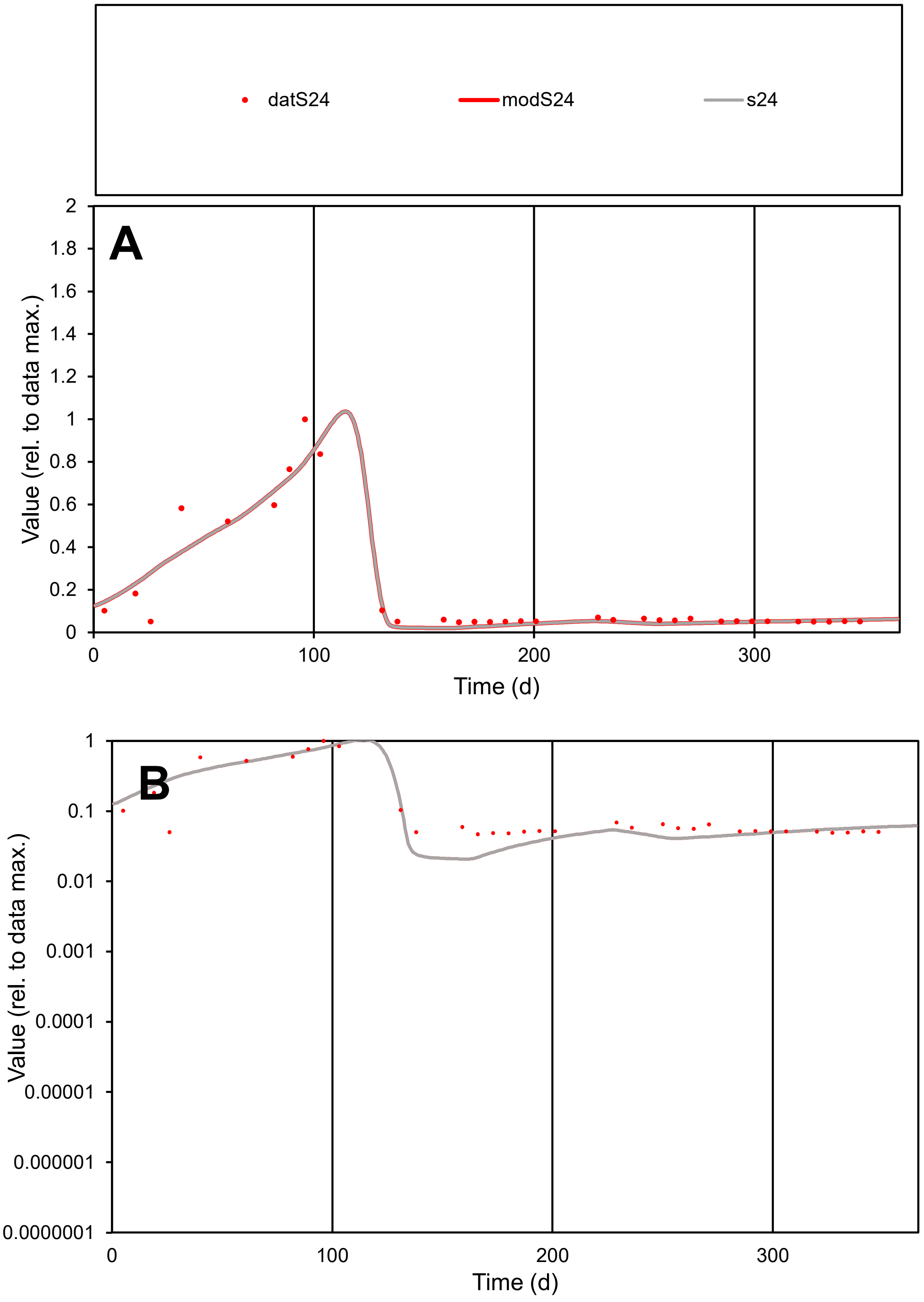


**C1**

Variables
S24 Provide jtd or Dst
CalibOnly?
2 1 to reproduce of
oNormalize?
2 1=no, 2=yes

ofxn calc*
0.37139
ofxn mod
0.40904

sRun
RunA1
zout
1

Ave
Data
0.00022
Model
0.00022

data/vMax
0.00123

ofxng
263.459

Min d/m
2.26169

ave/min d
3.85761
ave/min m
8.47787

326_Calib_S_A

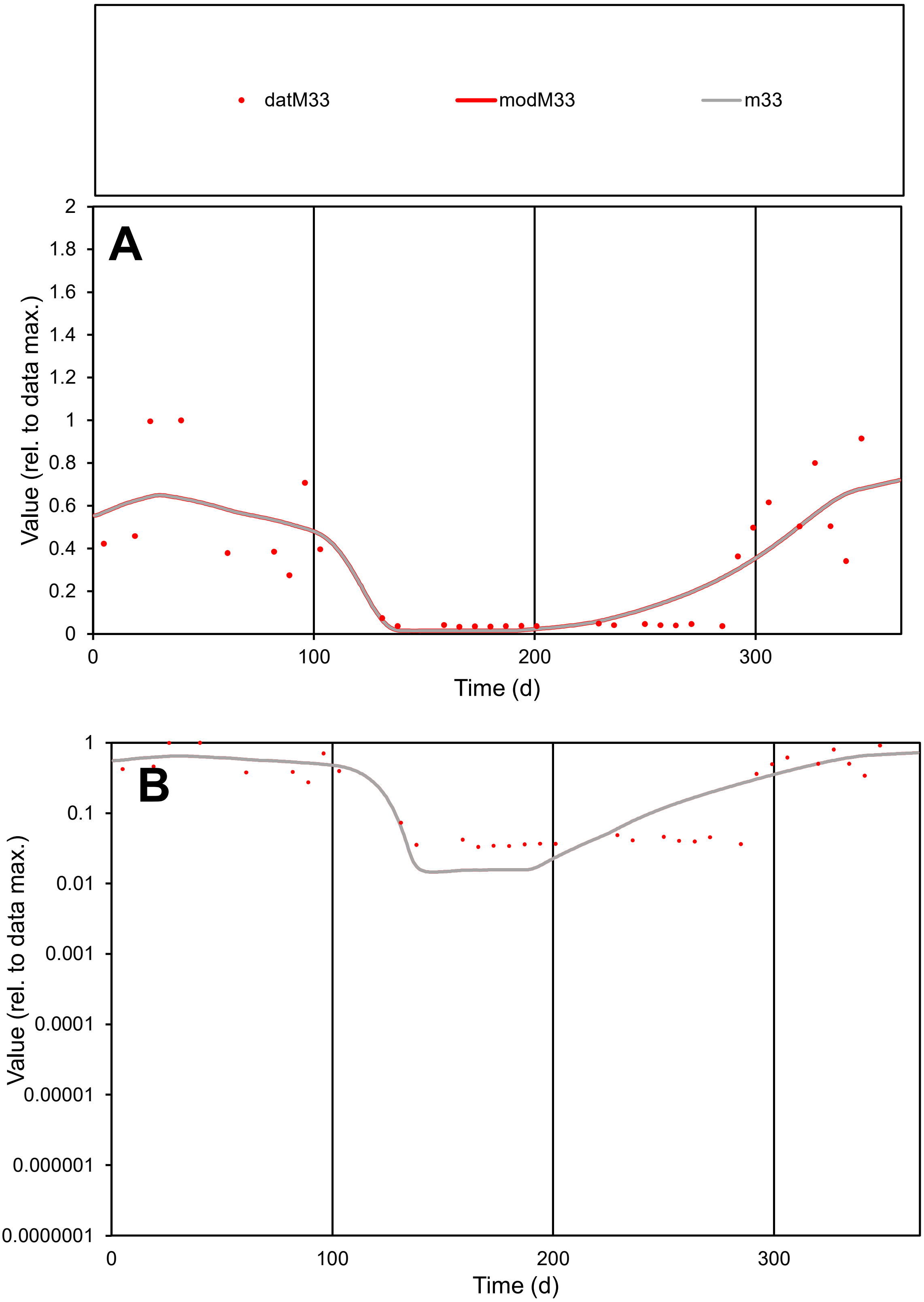


**C1**

Variables
M33 Provide jtd or Dst
CalibOnly?
2 1 to reproduce of
oNormalize?
2 1=no, 2=yes

ofxn calc*
0.52731
ofxn mod
0.54815

sRun
RunA1
zout
1

Ave
Data
0.00063
Model
0.00064

data/vMax
0.00204

ofxng
263.459

Min d/m
2.14707

ave/min d
9.33374
ave/min m
20.4284

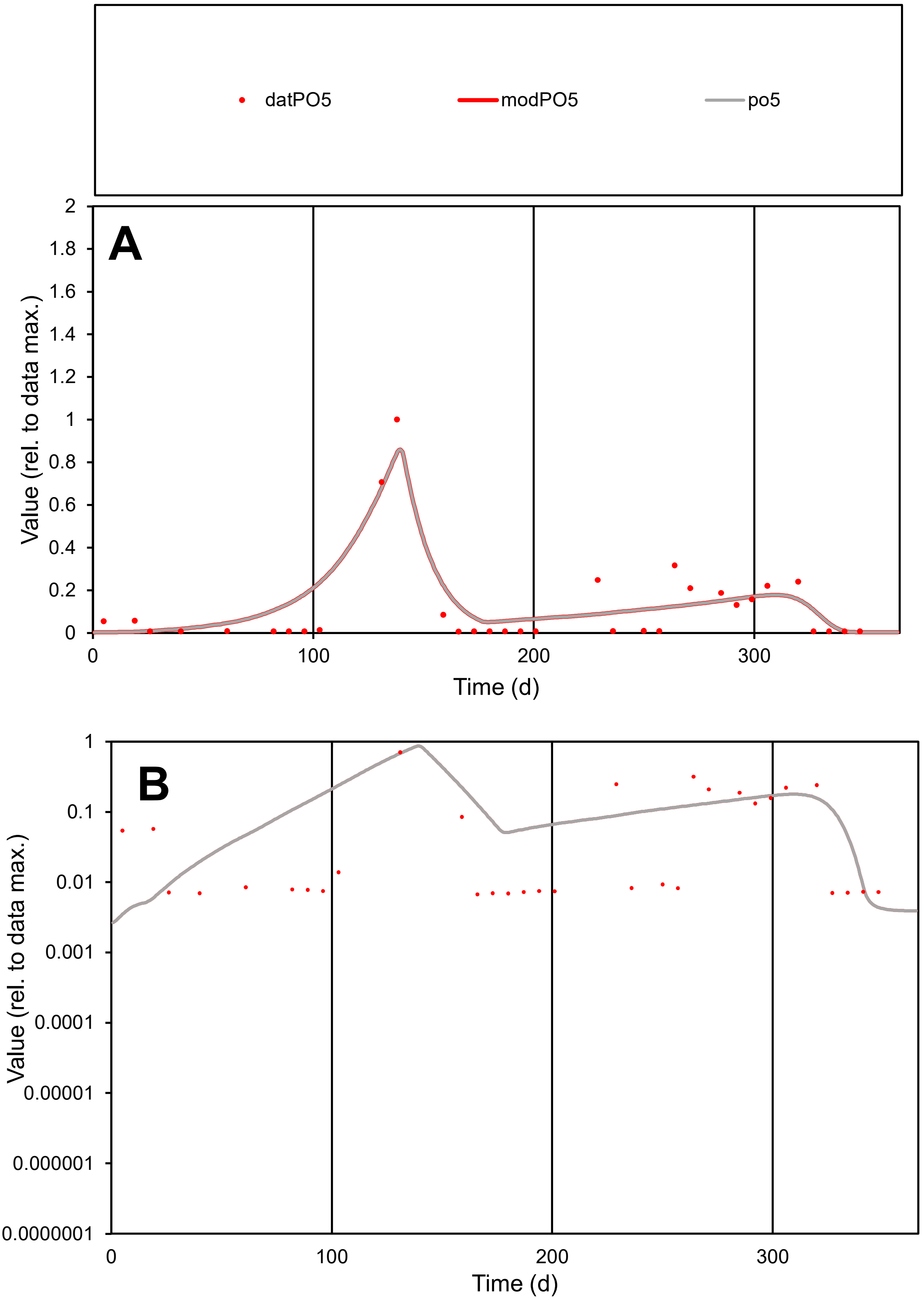


**C1**

Variables
PO5 Provide jtd or Dst
CalibOnly?
2 1 to reproduce of
oNormalize?
2 1=no, 2=yes

ofxn calc*
0.80294
ofxn mod
0.85751

sRun
RunA1
zout
1

Ave
Data
0.00079
Model
0.00094

data/vMax
0.00687

ofxng
263.459

Min d/m
1.75019

ave/min d
17.0577
ave/min m
35.5884

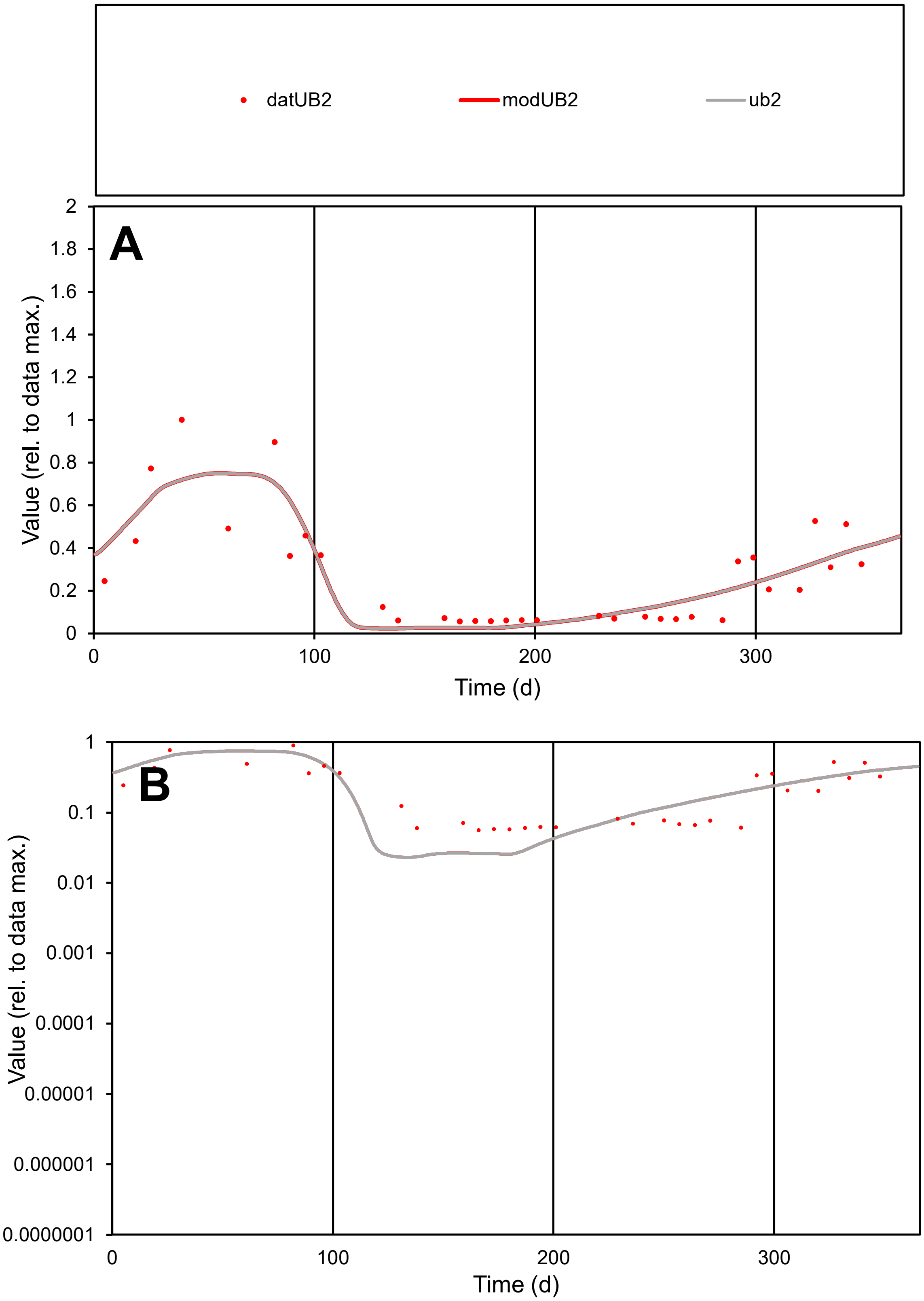


**C1**

Variables

UB2 Provide jtd or Dst

CalibOnly?

2 1 to reproduce of

oNormalize?

2 1=no, 2=yes

ofxn calc*

0.44004

ofxn mod

0.4709

sRun

RunA1

zout

1

Ave

Data

0.0005

Model

0.00049

data/vMax

0.00184

ofxng

263.459

Min d/m

2.42931

ave/min d

4.82753

ave/min m

11.6577

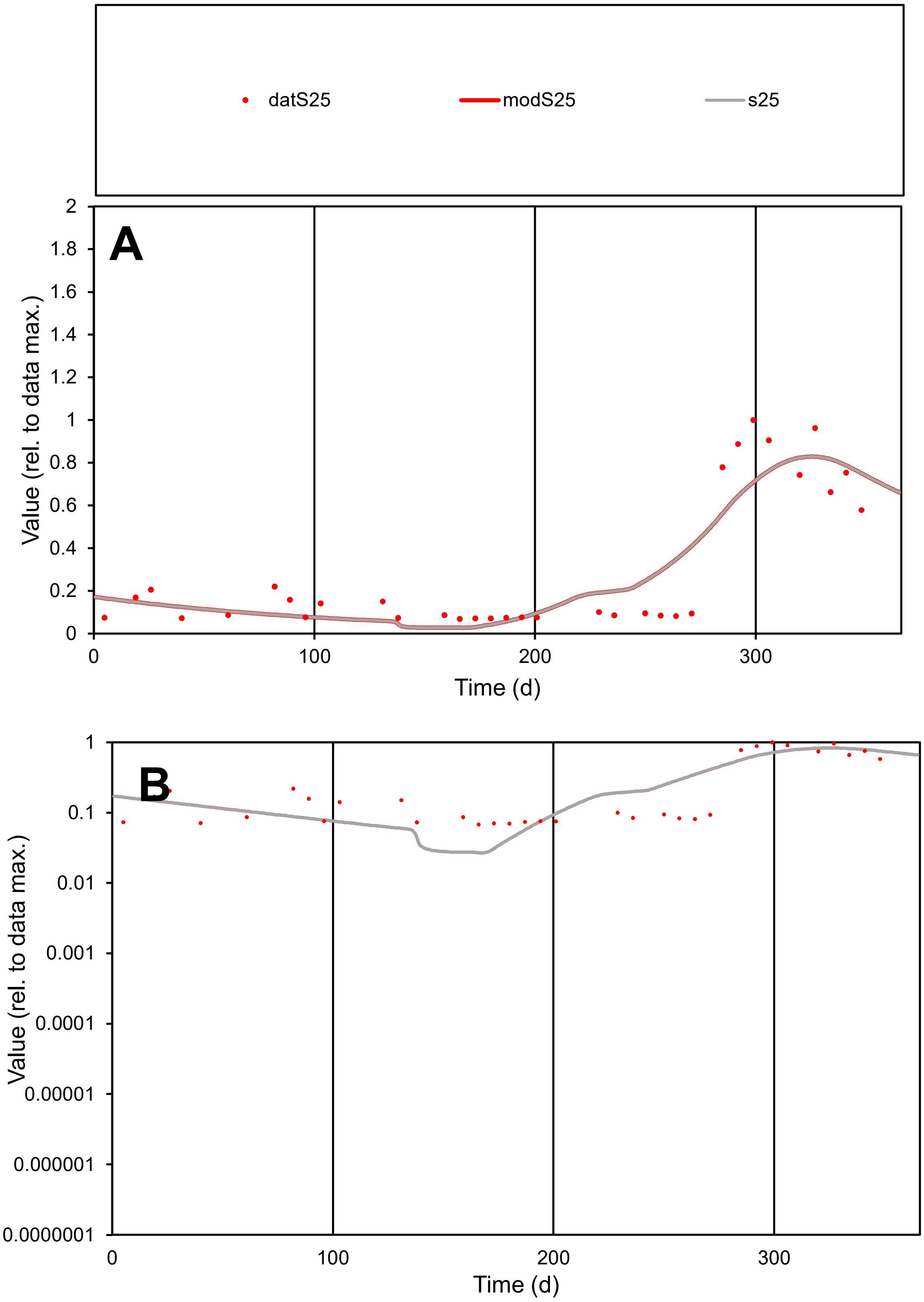


**C1**

Variables
S25 Provide jtd or Dst
CalibOnly?
2 1 to reproduce of
oNormalize?
2 1=no, 2=yes

ofxn calc*
0.45432
ofxn mod
0.47859

sRun
RunA1
zout
1

Ave
Data
0.00073
Model
0.00074

data/vMax
0.00246

ofxng
263.459

Min d/m
2.54291

ave/min d
4.34528
ave/min m
11.2213

326_Calib_S_A

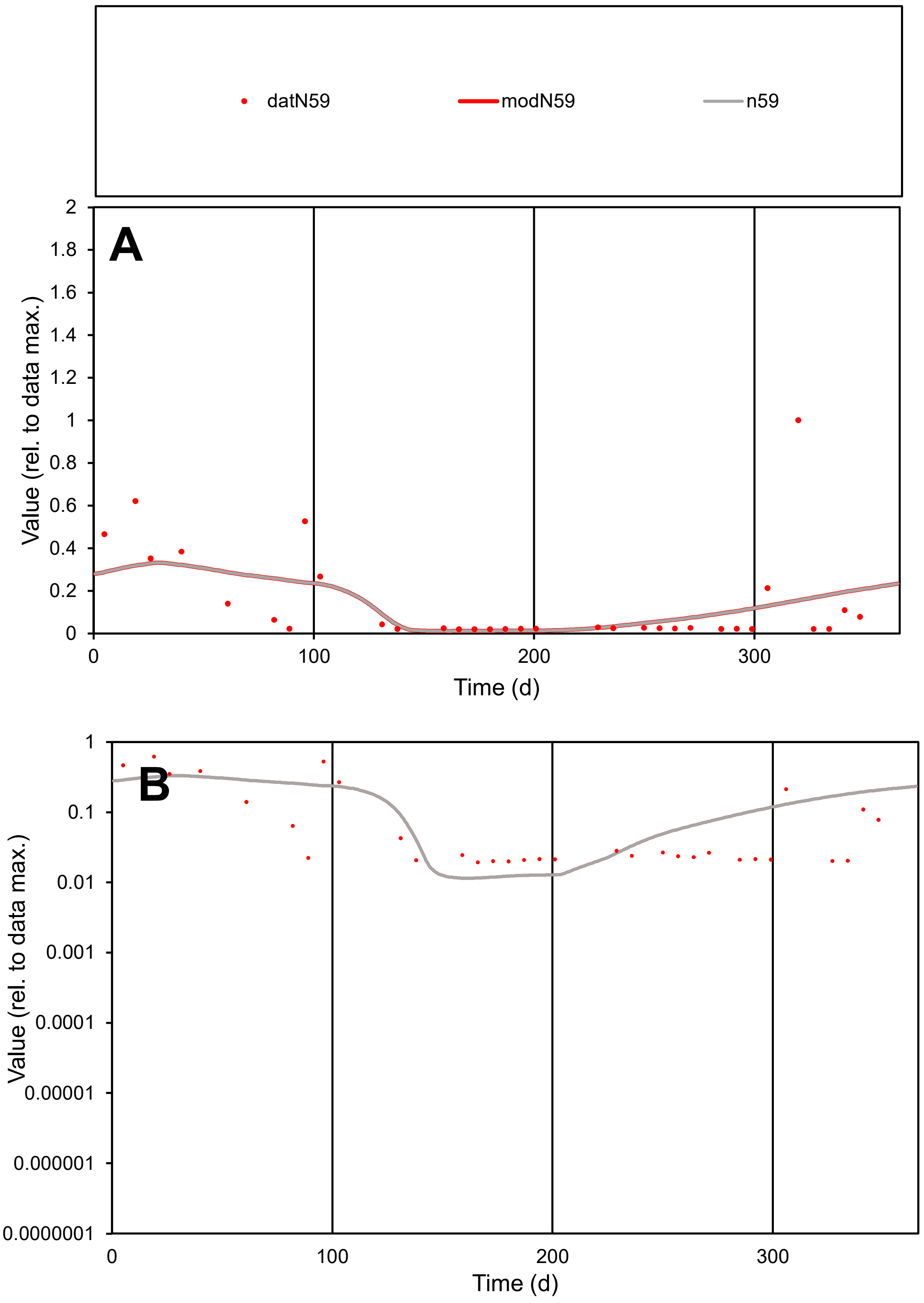


**C1**

Variables

N59 Provide jtd or Dst

CalibOnly?

2 1 to reproduce of

oNormalize?

2 1=no, 2=yes

ofxn calc*

1.30025

ofxn mod

1.31104

sRun

RunA1

zout

1

Ave

Data

0.0003

Model

0.00028

data/vMax

0.00212

ofxng

263.459

Min d/m

1.67857

ave/min d

7.40115

ave/min m

11.5996

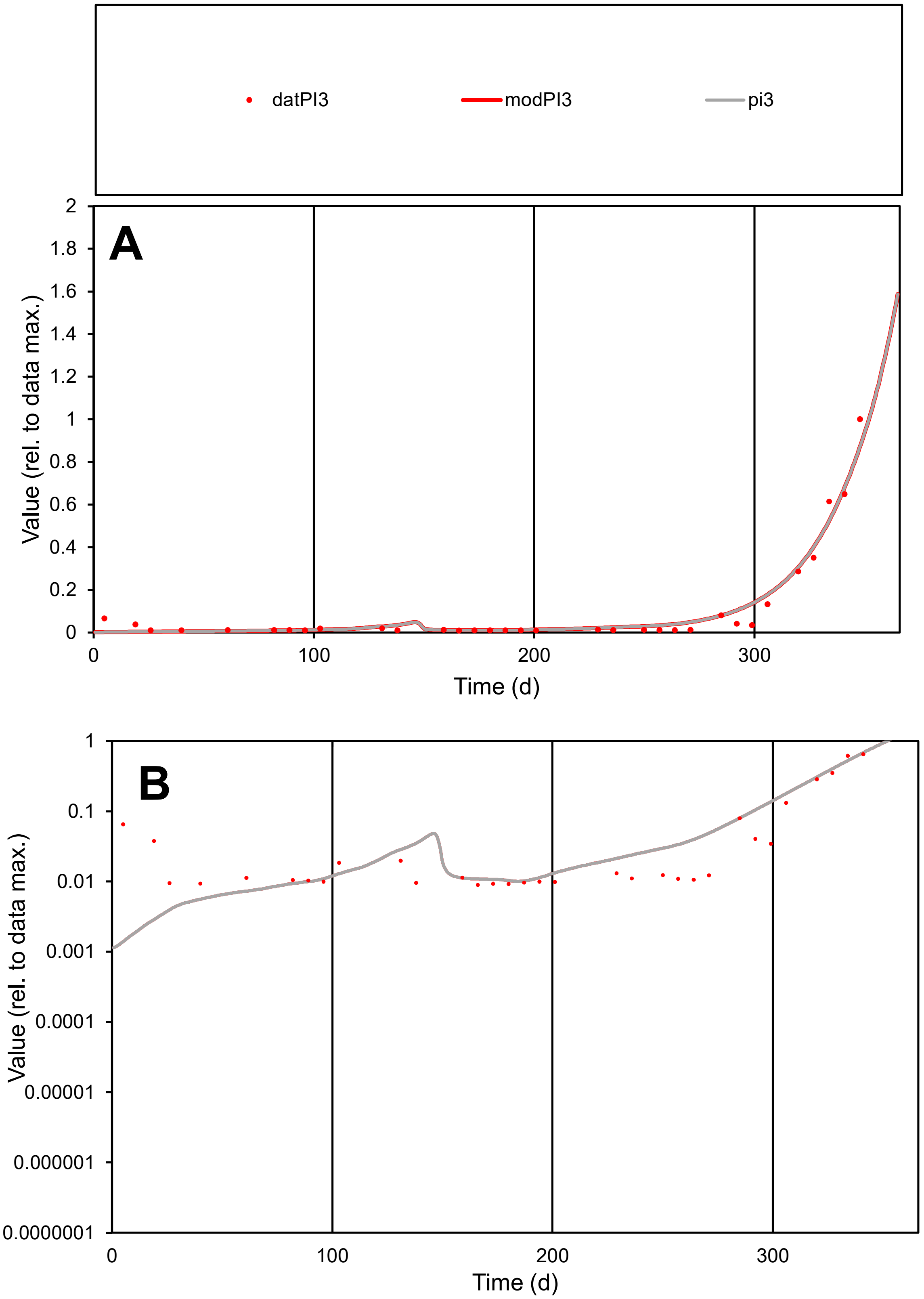


**C1**

Variables
PI3 Provide jtd or Dst
CalibOnly?
2 1 to reproduce of
oNormalize?
2 1=no, 2=yes

ofxn calc*
0.36774
ofxn mod
0.42575

sRun
RunA1
zout
1

Ave
Data
0.00085
Model
0.00092

data/vMax
0.00789

ofxng
263.459

Min d/m
5.8774

ave/min d
12.0039
ave/min m
76.5853

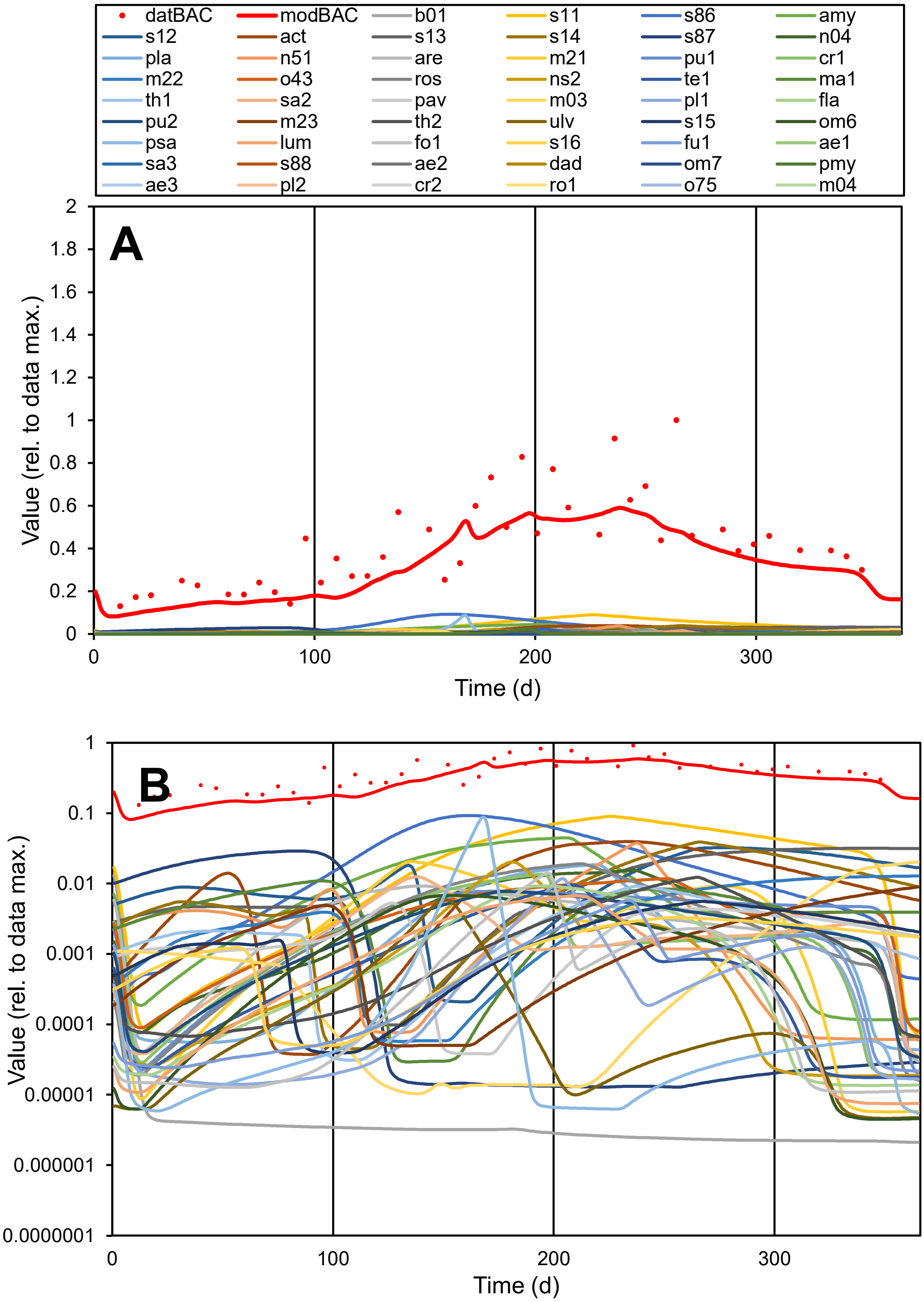


C1

Variables

BAC Provide jtd or Dst

CalibOnly?

2 1 to reproduce of

oNormalize?

2 1=no, 2=yes

ofxn calc*

0.36631

ofxn mod

0.38595

sRun

RunA1

zout

1

Ave

Data

0.88962

Model

0.70494

data/vMax

2.102

ofxng

263.459

Min d/m

1.49891

ave/min d

3.2468

ave/min m

3.85635

327_Calib_S_A

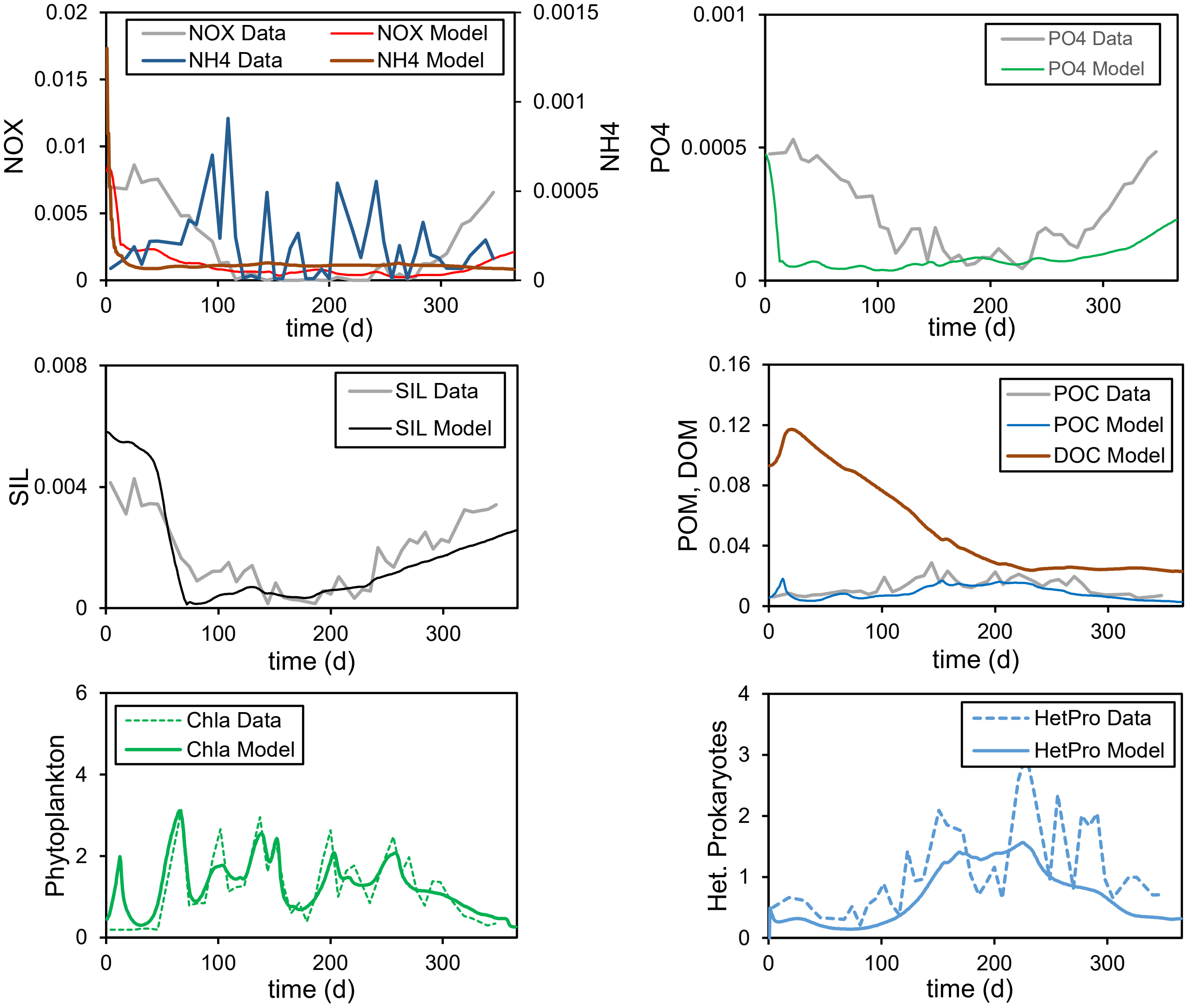

NOX Data
NOX Model
NH4 Data
NH4 Model
NOX
NH4
time (d)
PO4 Data
PO4 Model
PO4
time (d)
SIL Data
SIL Model
SIL
time (d)
POC Data
POC Model
DOC Model
POM, DOM
time (d)
Chla Data
Chla Model
Phytoplankton
time (d)
HetPro Data
HetPro Model
Het. Prokaryotes
time (d)

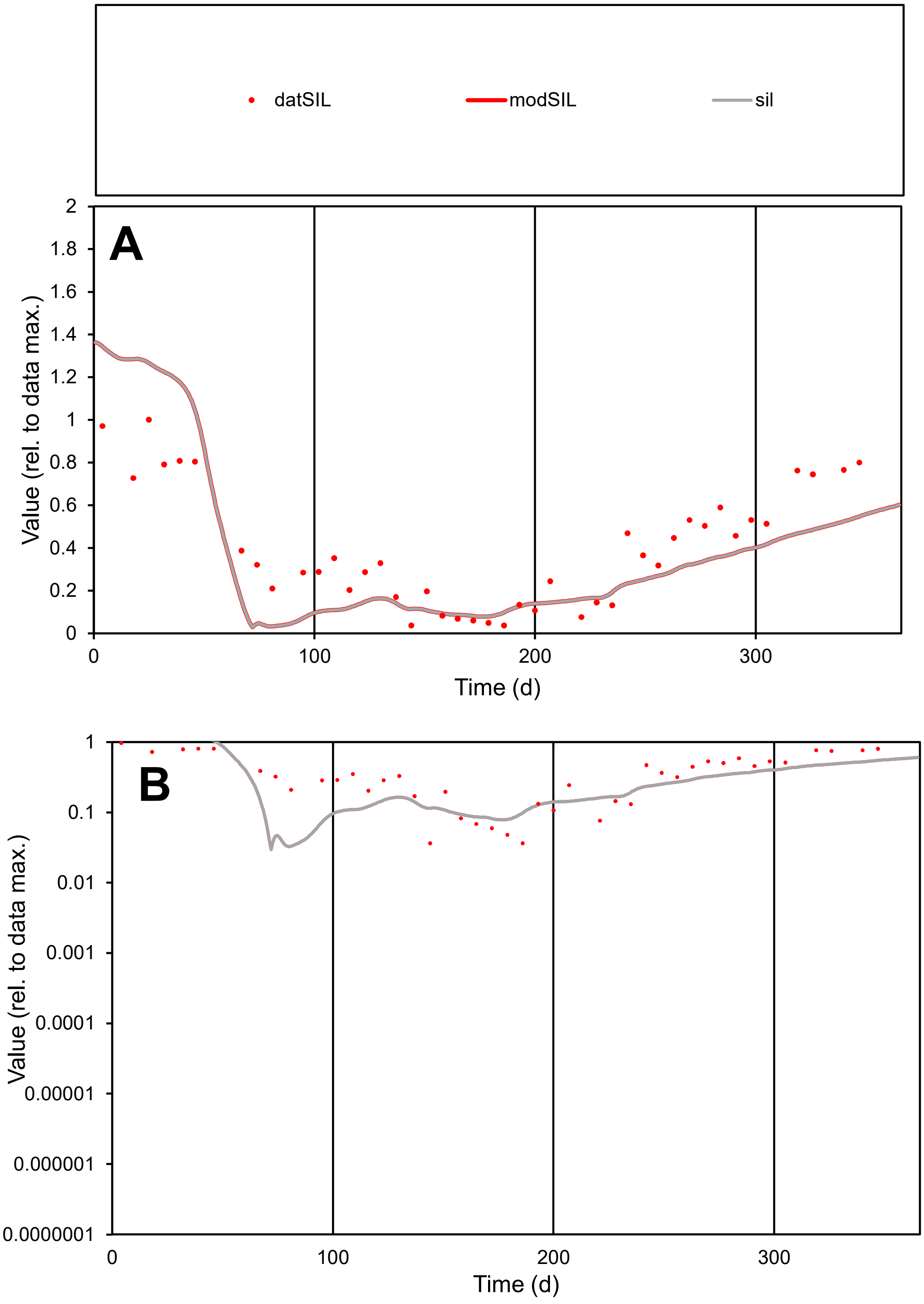


**C1**

Variables
SIL Provide jtd or Dst
CalibOnly?
2 1 to reproduce of
oNormalize?
2 1=no, 2=yes

ofxn calc*
1.51835
ofxn mod
1.5861

sRun
RunA1
zout
1

Ave
Data
0.0017
Model
0.00152

data/vMax
0.00427

ofxng
334.07

Min d/m
1.0697

ave/min d
10.9364
ave/min m
10.4946

327_Calib_S_A

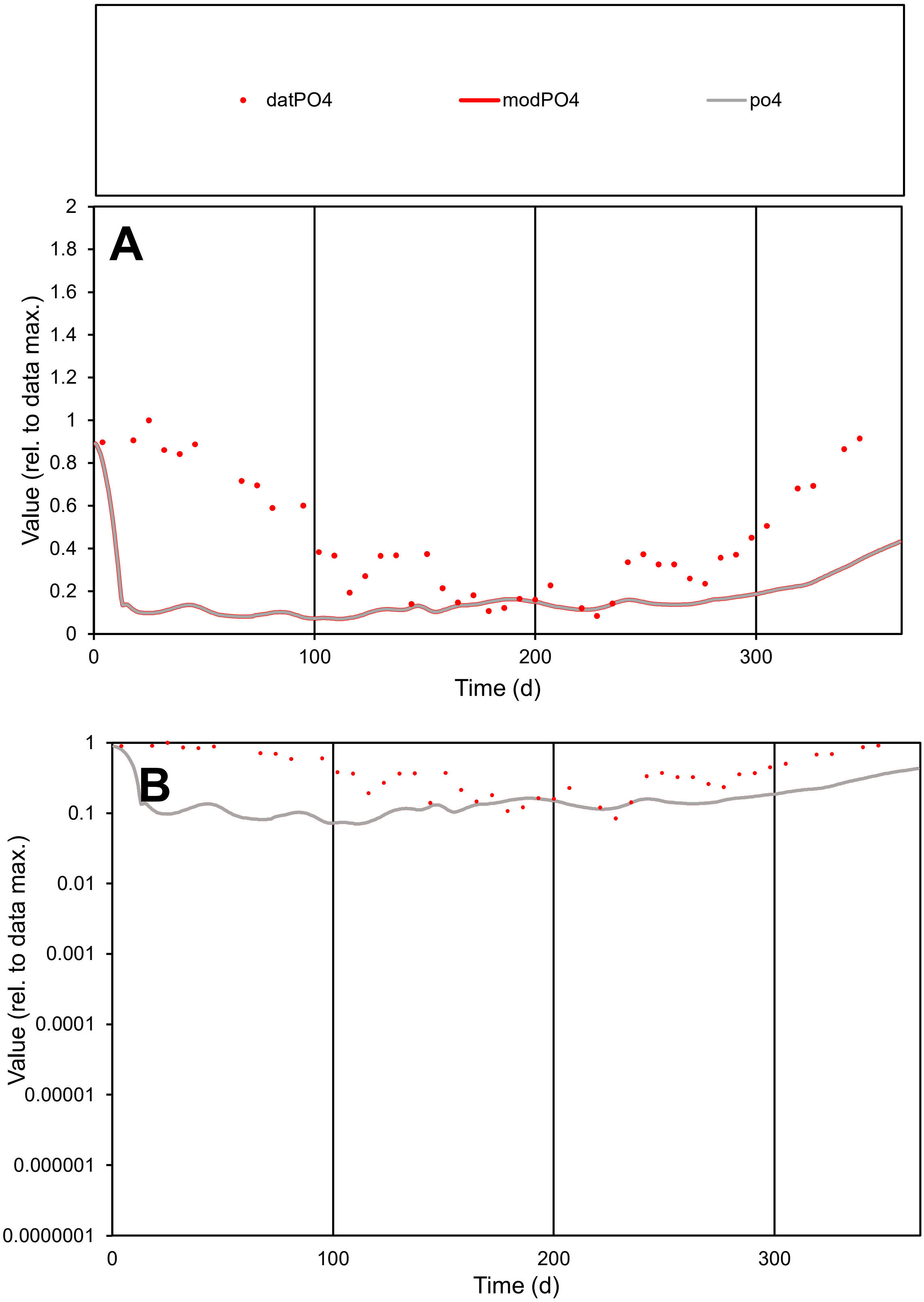


**C1**

Variables

PO4 Provide jtd or Dst

CalibOnly?

2 1 to reproduce of

oNormalize?

2 1=no, 2=yes

ofxn calc*

2.61757

ofxn mod

2.63116

sRun

RunA1

zout

1

Ave

Data

0.00023

Model

8.3E-05

data/vMax

0.00053

ofxng

334.07

Min d/m

1.19855

ave/min d

5.20235

ave/min m

2.23894

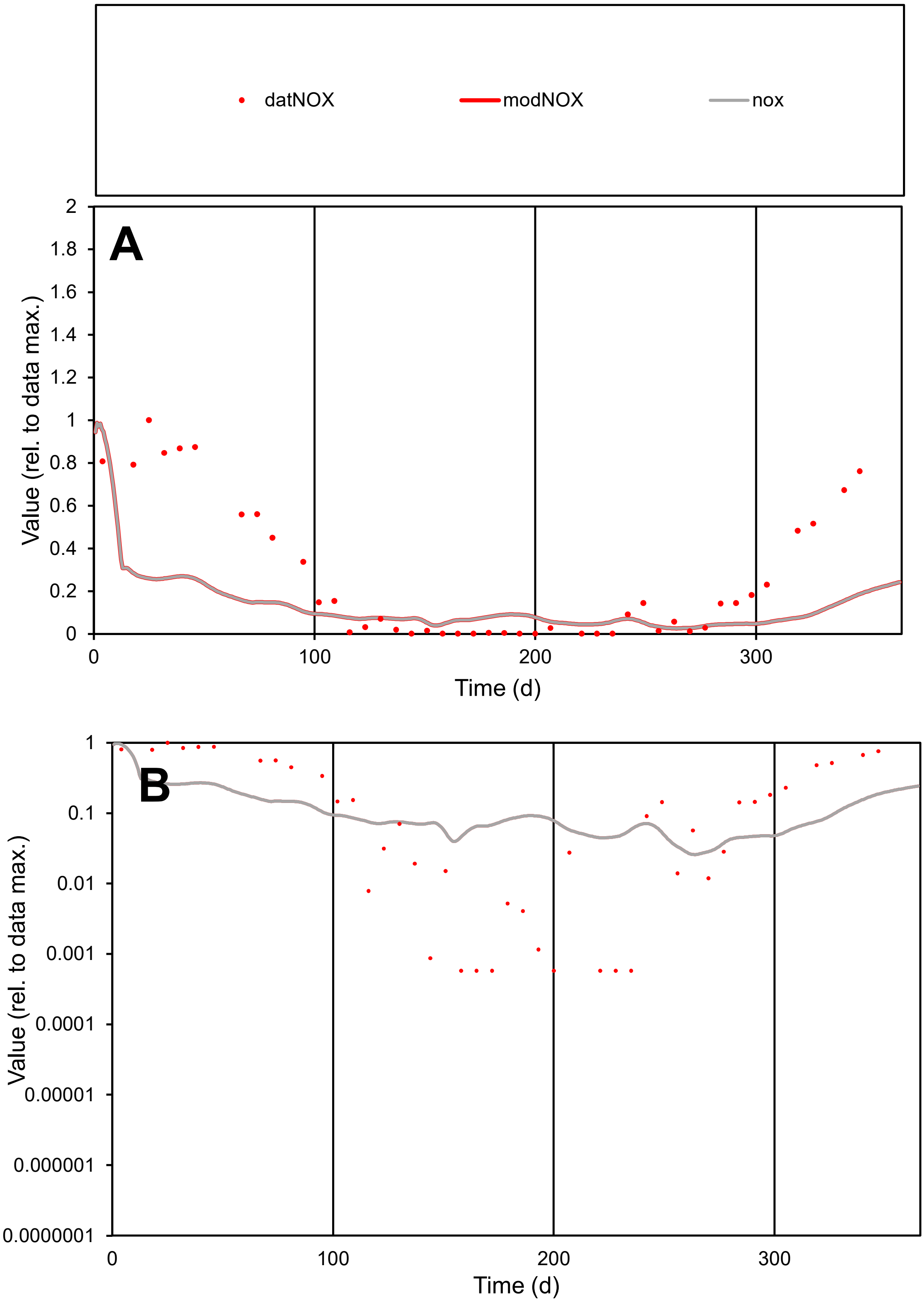


**C1**

Variables
NOX Provide jtd or Dst
CalibOnly?
2 1 to reproduce of
oNormalize?
2 1=no, 2=yes

ofxn calc*
3.30585
ofxn mod
3.33135

sRun
RunA1
zout
1

Ave
Data
0.00222
Model
0.00101

data/vMax
0.00862

ofxng
334.07

Min d/m
0.02229

ave/min d
443.44
ave/min m
4.51528

327_Calib_S_A

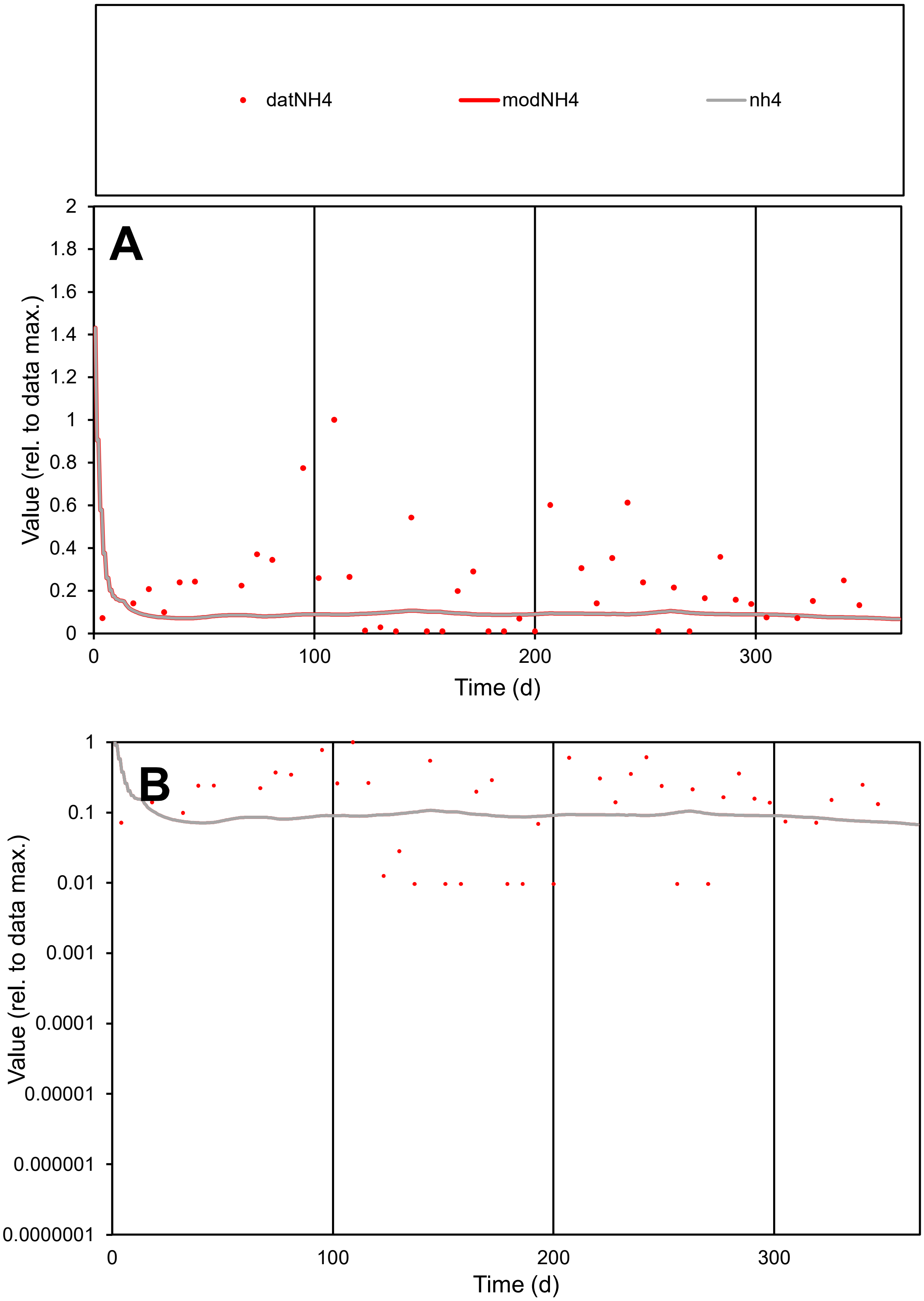


**C1**

Variables
NH4 Provide jtd or Dst
CalibOnly?
2 1 to reproduce of
oNormalize?
2 1=no, 2=yes

ofxn calc*
3.45237
ofxn mod
3.48088

sRun
RunA1
zout
1

Ave
Data
0.0002
Model
8.4E-05

data/vMax
0.00091

ofxng
334.07

Min d/m
0.13585

ave/min d
22.7078
ave/min m
1.30294

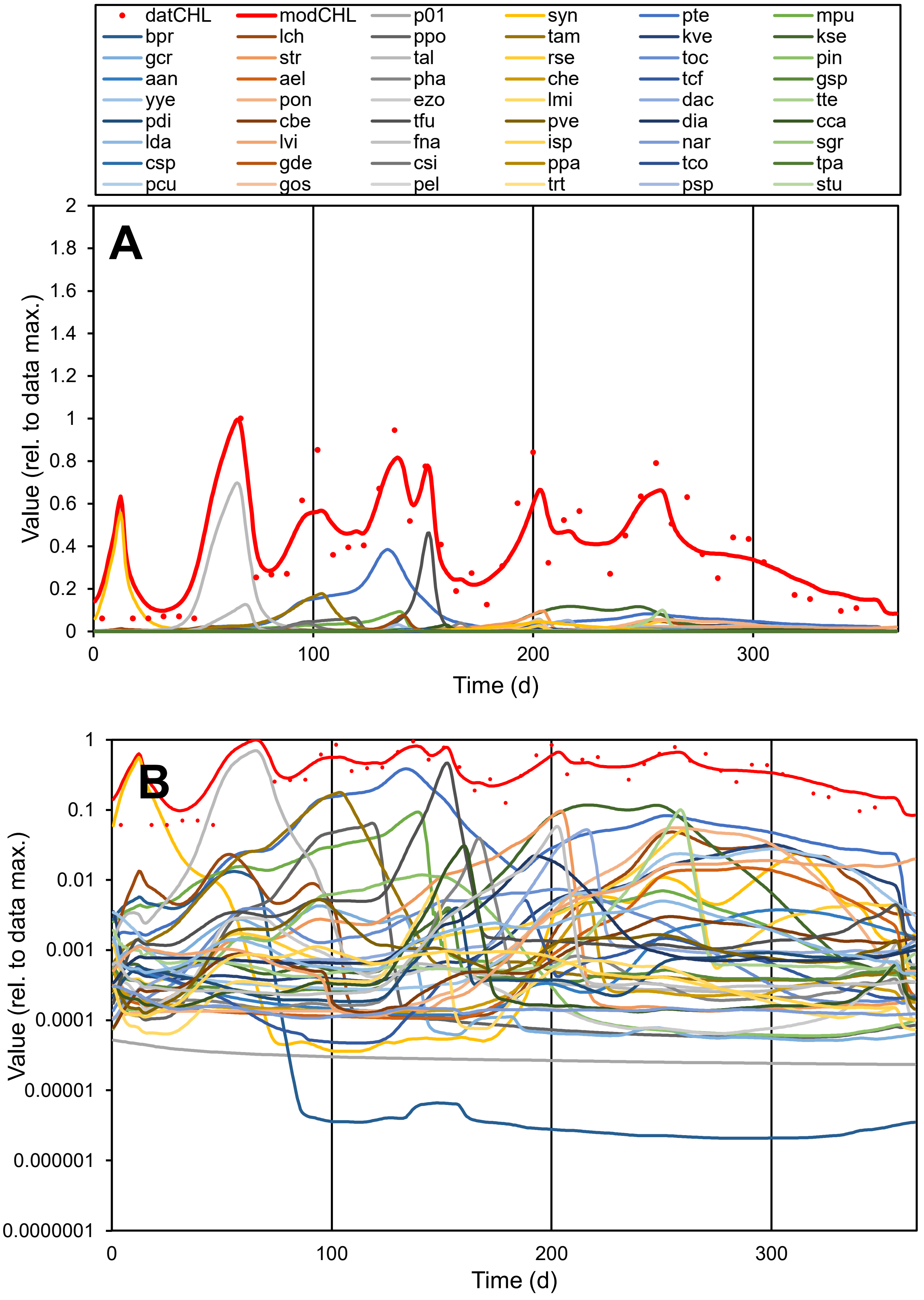


**C1**

Variables

CHL Provide jtd or Dst

CalibOnly?

2 1 to reproduce of

oNormalize?

2 1=no, 2=yes

ofxn calc*

2.71147

ofxn mod

3.10075

sRun

RunA1

zout

1

Ave

Data

1.24336

Model

1.25597

data/vMax

3.128

ofxng

334.07

Min d/m

0.6197

ave/min d

6.54402

ave/min m

4.09645

327_Calib_S_A

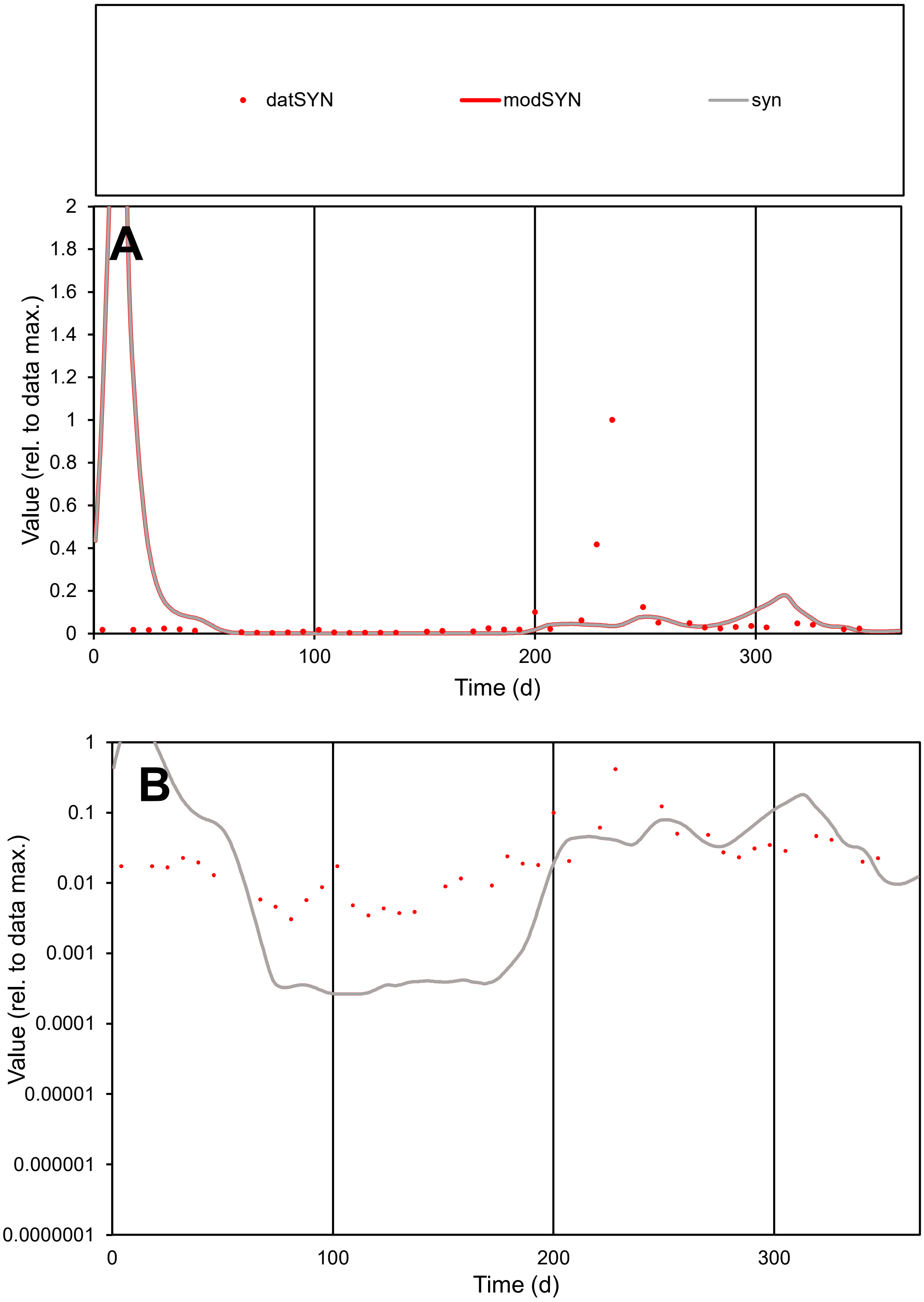


**C1**

Variables
SYN Provide jtd or Dst
CalibOnly?
2 1 to reproduce of
oNormalize?
2 1=no, 2=yes

ofxn calc*
3.59894
ofxn mod
4.27556

sRun
RunA1
zout
1

Ave
Data
15.004
Model
16.8841

data/vMax
254.6

ofxng
334.07

Min d/m
11.6142

ave/min d
19.2556
ave/min m
251.663

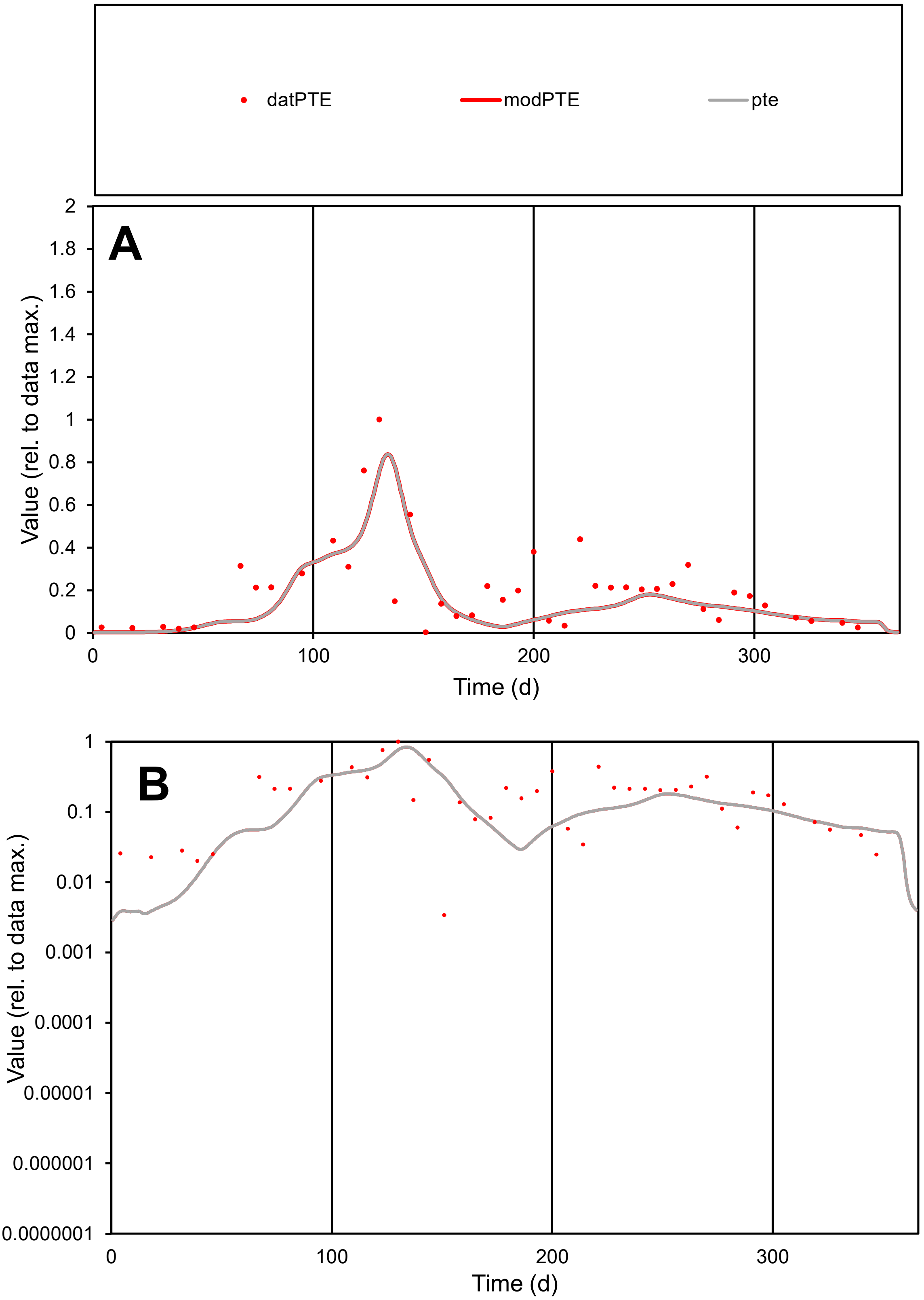


**C1**

Variables
PTE Provide jtd or Dst
CalibOnly?
2 1 to reproduce of
oNormalize?
2 1=no, 2=yes

ofxn calc*
0.71408
ofxn mod
0.74686

sRun
RunA1
zout
1

Ave
Data
0.0005
Model
0.00039

data/vMax
0.00245

ofxng
334.07

Min d/m
0.87539

ave/min d
60.0116
ave/min m
41.0433

327_Calib_S_A

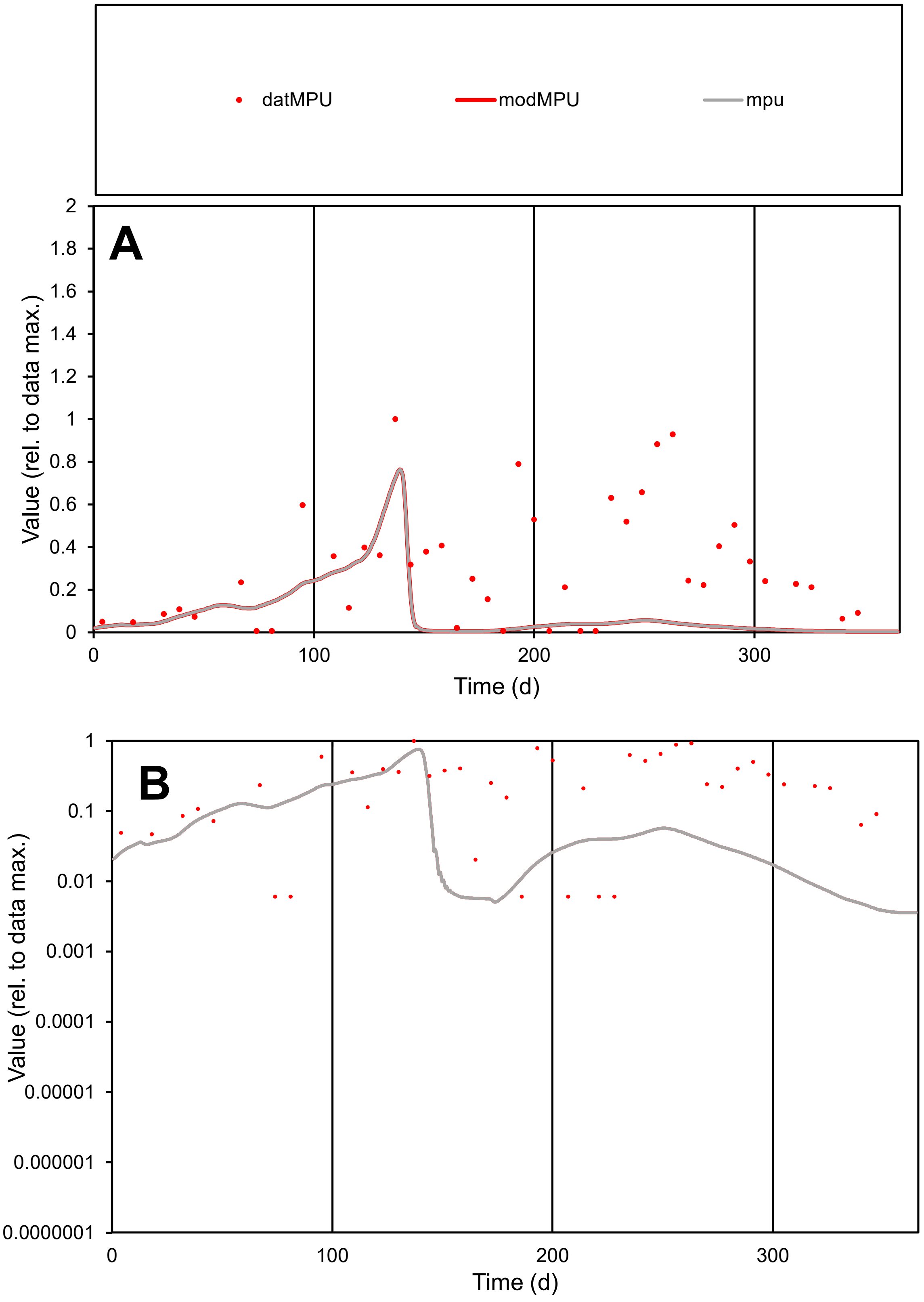


**C1**

Variables

MPU Provide jtd or Dst

CalibOnly?

2 1 to reproduce of

oNormalize?

2 1=no, 2=yes

ofxn calc*

1.10504

ofxn mod

1.11198

sRun

RunA1

zout

1

Ave

Data

0.0002

Model

6.5E-05

data/vMax

0.00067

ofxng

334.07

Min d/m

1.54014

ave/min d

49.6911

ave/min m

24.5795

327_Calib_S_A

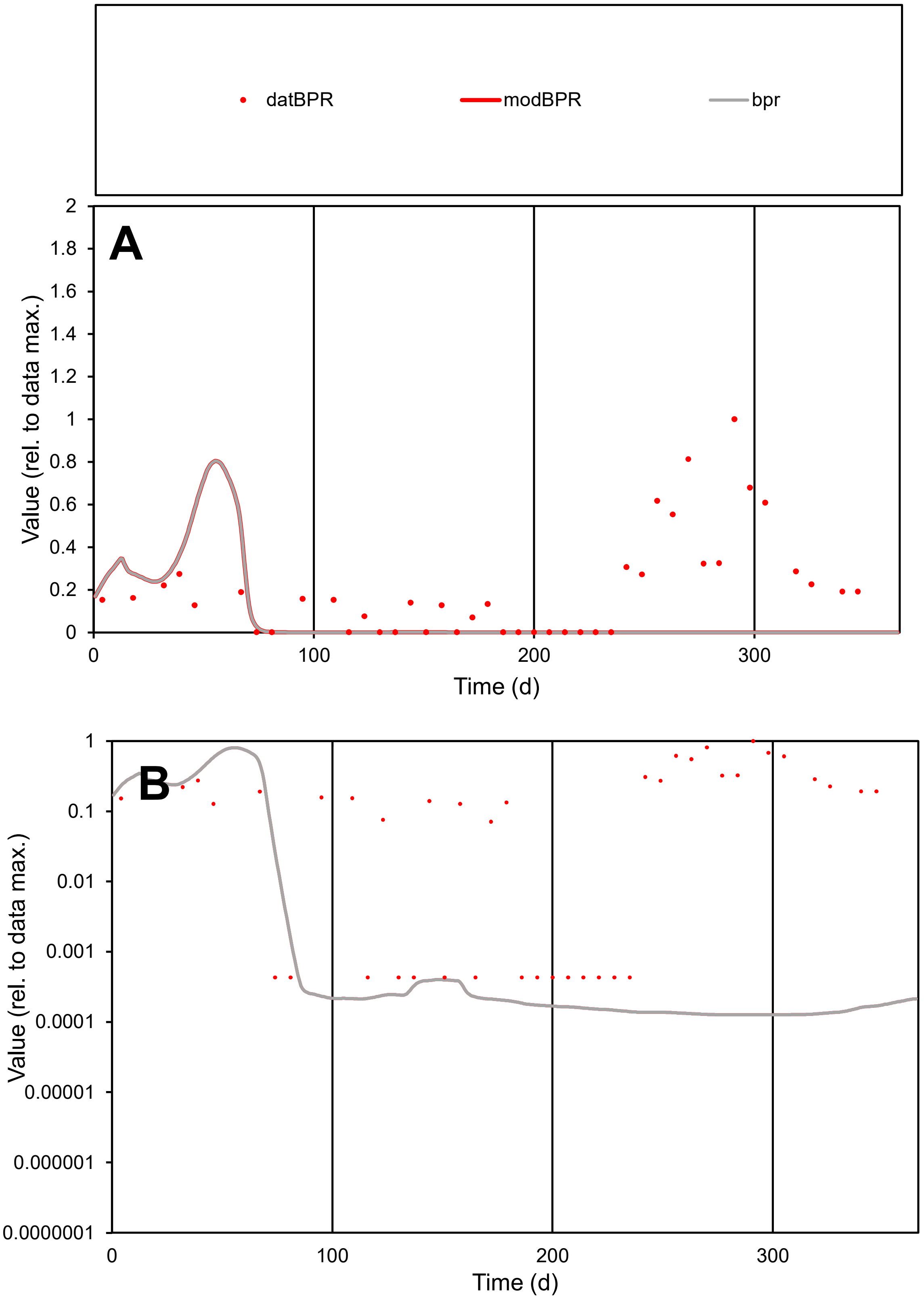


**C1**

Variables
BPR Provide jtd or Dst
CalibOnly?
2 1 to reproduce of
oNormalize?
2 1=no, 2=yes

ofxn calc*
1.56247
ofxn mod
1.57596

sRun
RunA1
zout
1

Ave
Data
8.6E-05
Model
2E-05

data/vMax
0.00043

ofxng
334.07

Min d/m
3.41945

ave/min d
462.23
ave/min m
370.047

327_Calib_S_A

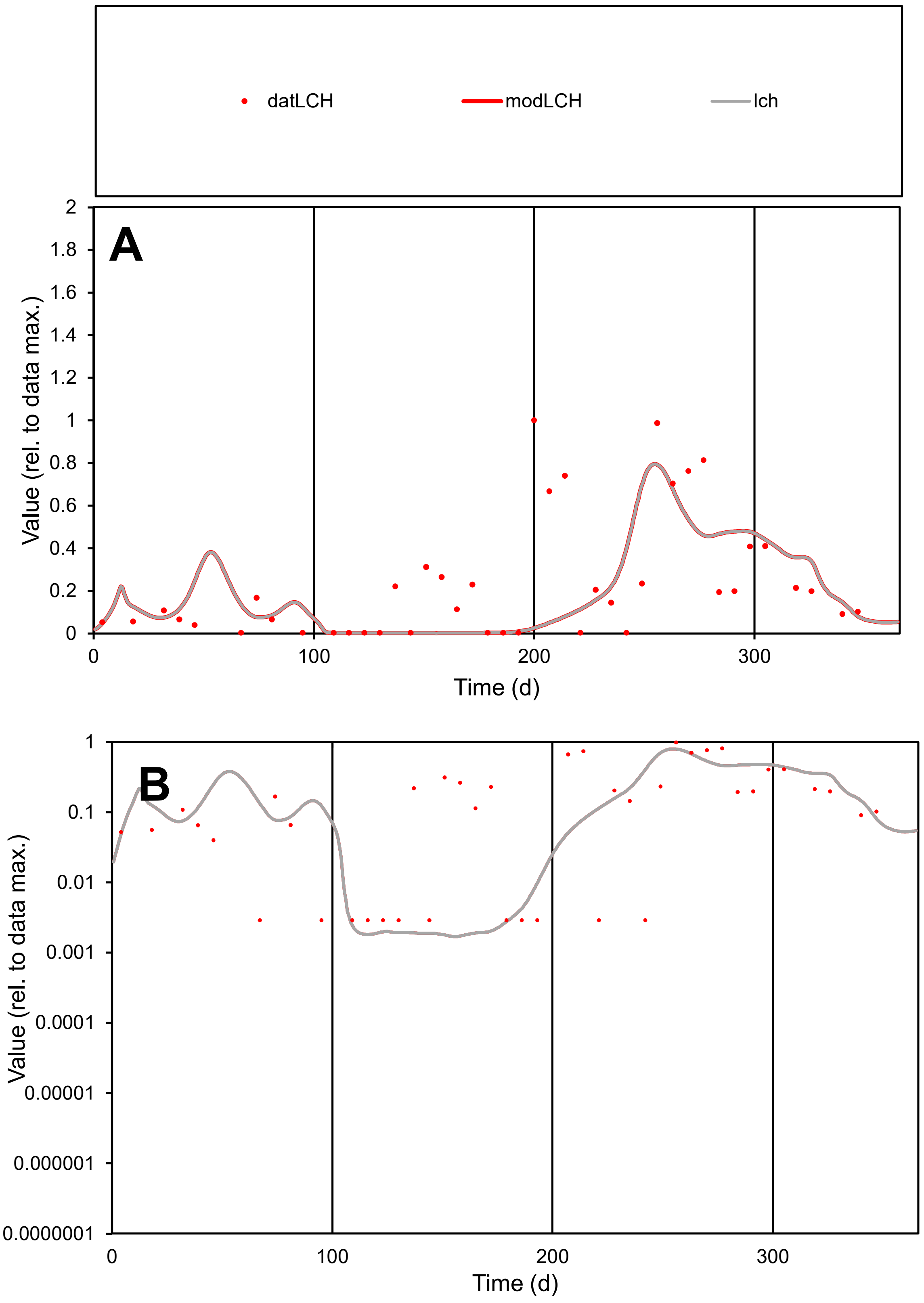


**C1**

Variables

LCH Provide jtd or Dst

CalibOnly?

2 1 to reproduce of

oNormalize?

2 1=no, 2=yes

ofxn calc*

1.1178

ofxn mod

1.14226

sRun

RunA1

zout

1

Ave

Data

0.0002

Model

0.00016

data/vMax

0.00085

ofxng

334.07

Min d/m

1.64936

ave/min d

80.6734

ave/min m

106.151

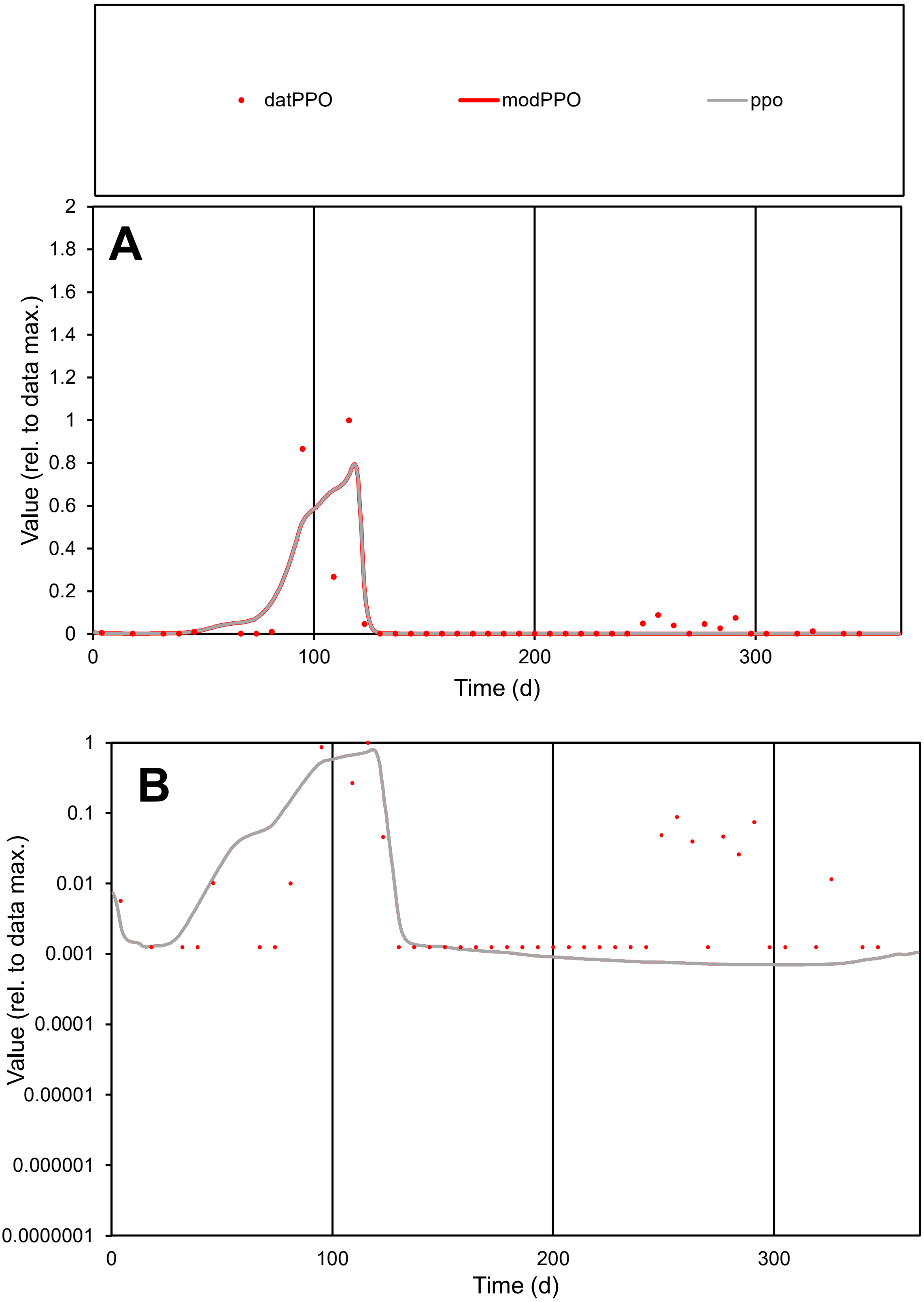


**C1**

Variables
PPO Provide jtd or Dst
CalibOnly?
2 1 to reproduce of
oNormalize?
2 1=no, 2=yes

ofxn calc*
1.53194
ofxn mod
1.55951

sRun
RunA1
zout
1

Ave
Data
9.4E-05
Model
8.6E-05

data/vMax
0.00153

ofxng
334.07

Min d/m
1.78119

ave/min d
49.1302
ave/min m
80.4163

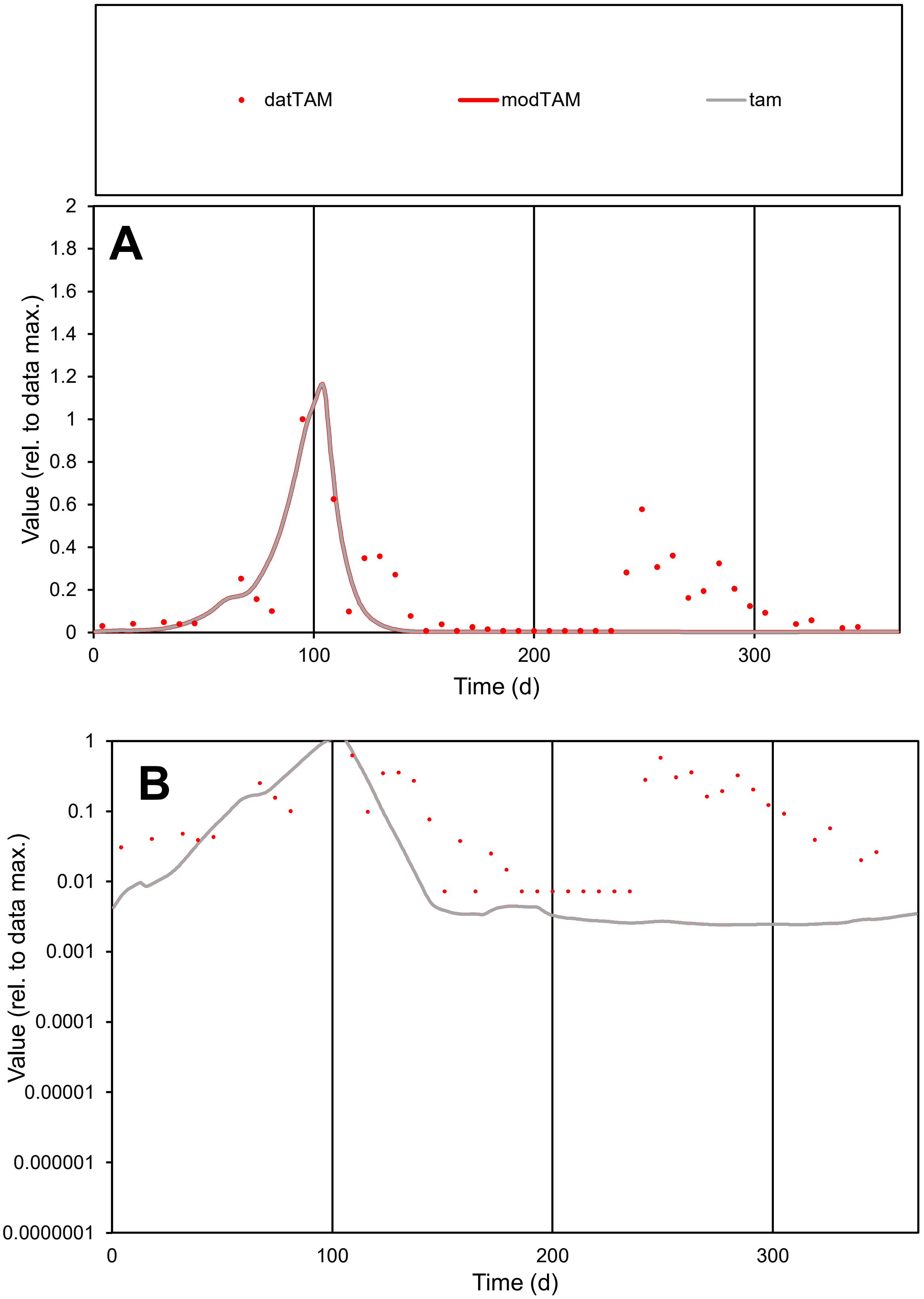


**C1**

Variables
TAM Provide jtd or Dst
CalibOnly?
2 1 to reproduce of
oNormalize?
2 1=no, 2=yes

ofxn calc*
1.10588
ofxn mod
1.12498

sRun
RunA1
zout
1

Ave
Data
0.00012
Model
5.4E-05

data/vMax
0.0008

ofxng
334.07

Min d/m
2.99013
ave/min d
21.0829
ave/min m
28.0988

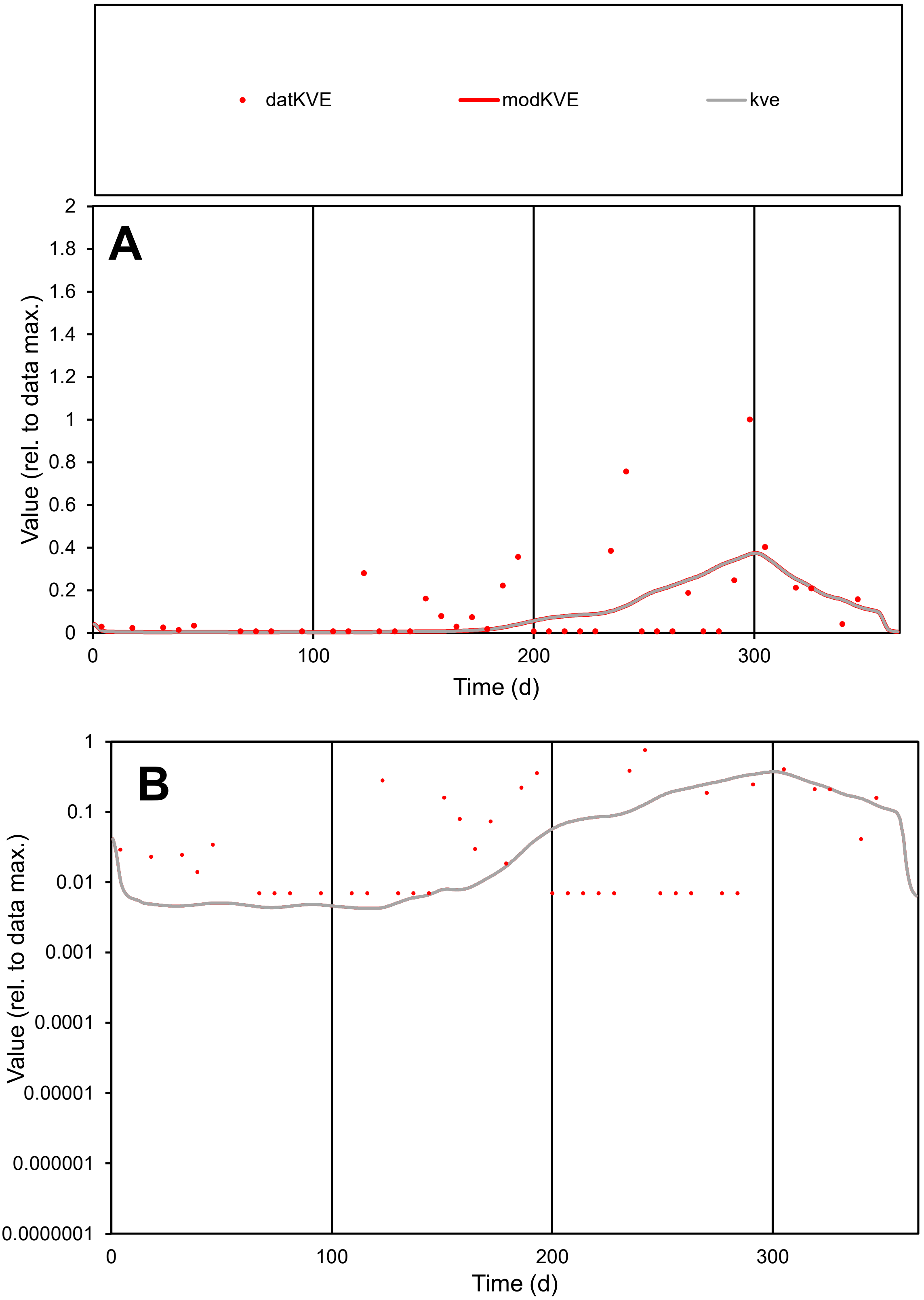


**C1**

Variables
KVE Provide jtd or Dst
CalibOnly?
2 1 to reproduce of
oNormalize?
2 1=no, 2=yes

ofxn calc*
1.5301
ofxn mod
1.54217

sRun
RunA1
zout
1

Ave
Data
5.4E-05
Model
4.4E-05

data/vMax
0.00045

ofxng
334.07

Min d/m
1.64443

ave/min d
17.3302
ave/min m
22.9165

327_Calib_S_A

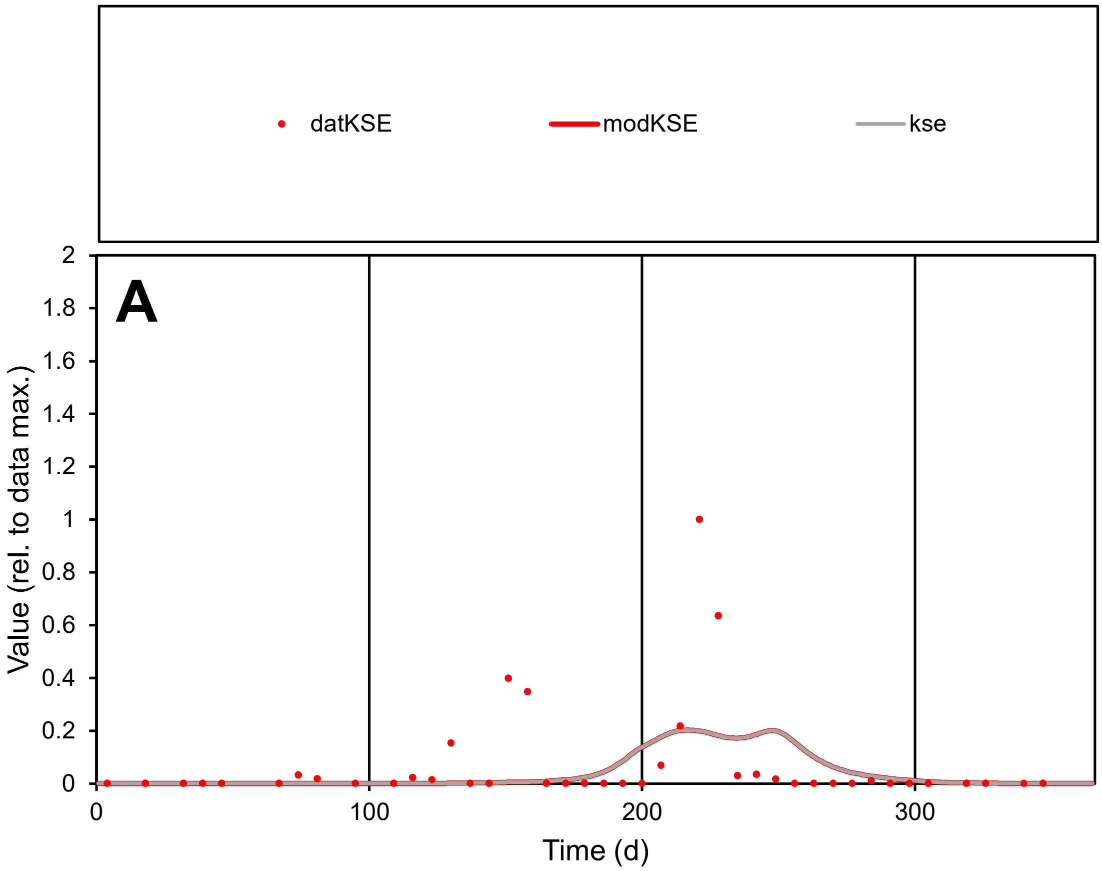


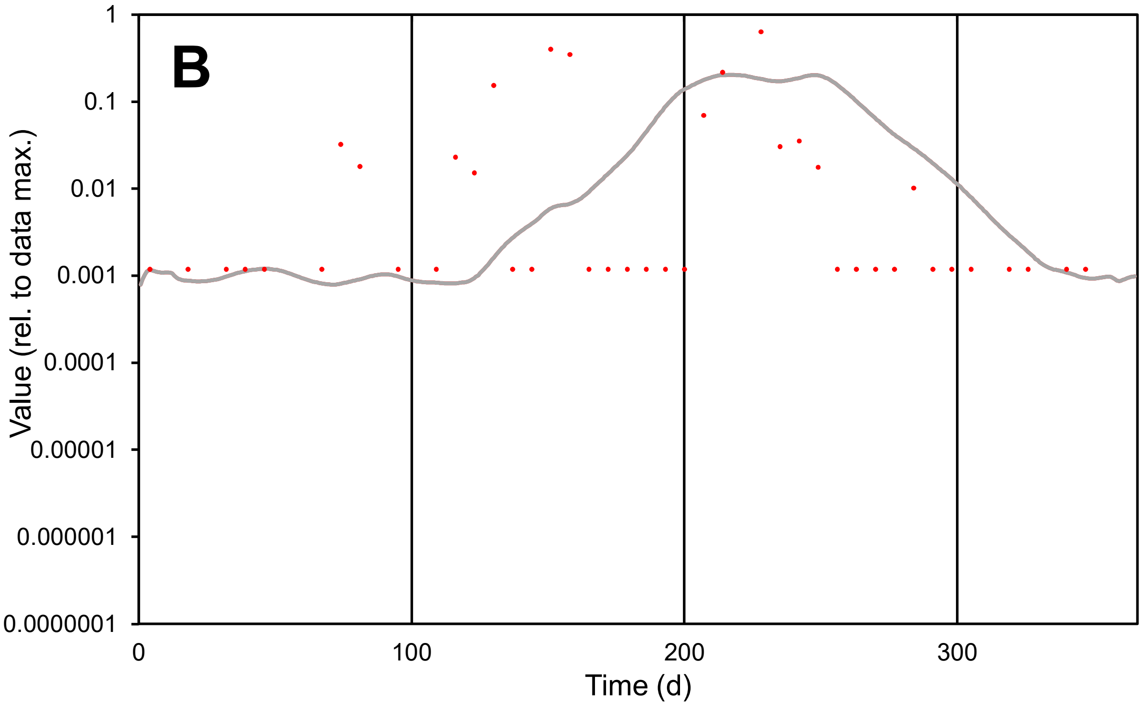


**C1**

Variables
KSE Provide jtd or Dst
CalibOnly?
2 1 to reproduce of
oNormalize?
2 1=no, 2=yes

ofxn calc*
2.41947
ofxn mod
2.4368

sRun
RunA1
zout
1

Ave
Data
0.00023
Model
0.00015

data/vMax
0.00316

ofxng
334.07

Min d/m
1.47501

ave/min d
61.062
ave/min m
59.2194

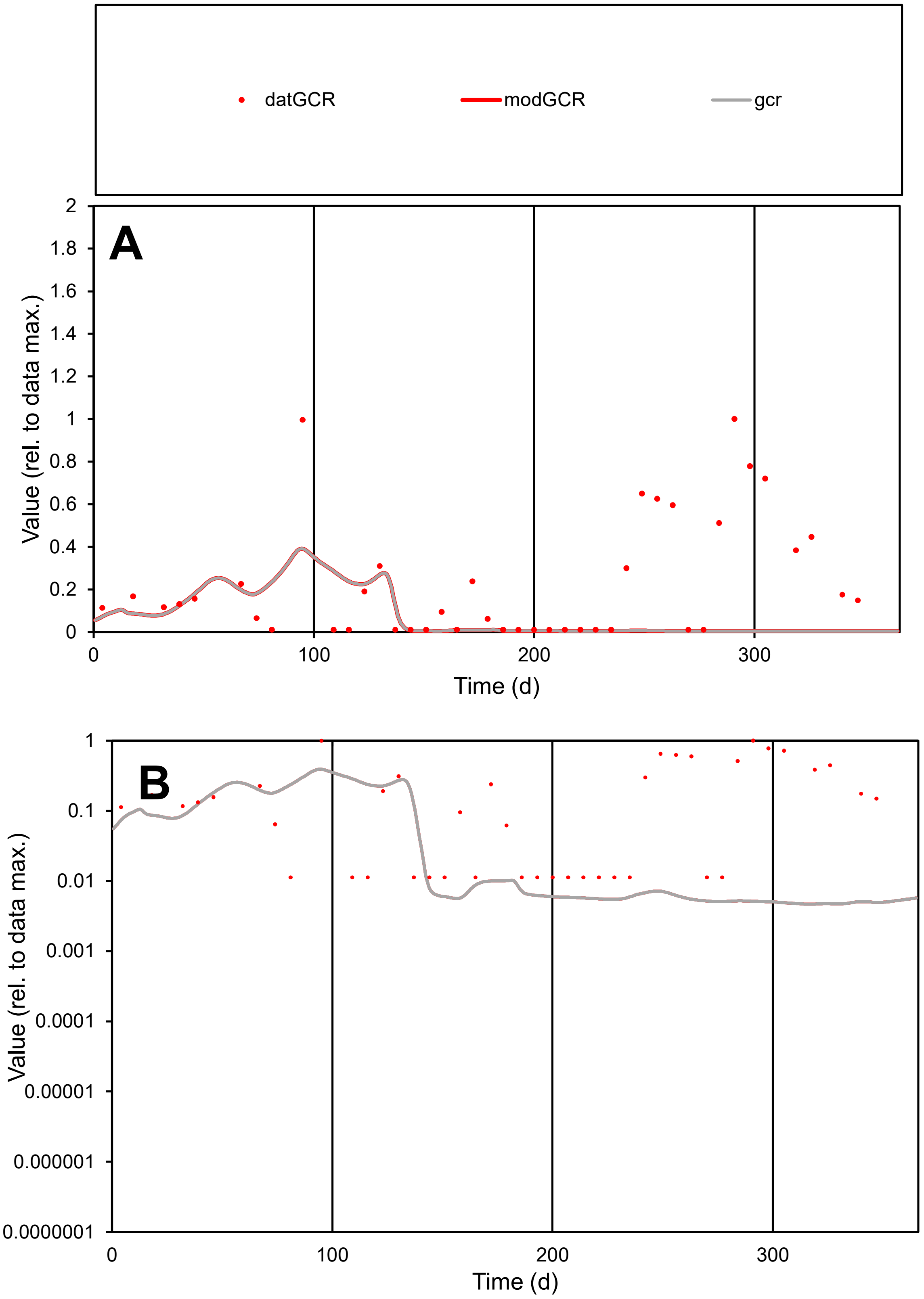


**C1**

Variables
GCR Provide jtd or Dst
CalibOnly?
2 1 to reproduce of
oNormalize?
2 1=no, 2=yes

ofxn calc*
1.48023
ofxn mod
1.48446

sRun
RunA1
zout
1

Ave
Data
6.2E-05
Model
1.9E-05

data/vMax
0.00028

ofxng
334.07

Min d/m
2.37965

ave/min d
19.9094
ave/min m
14.1215

327_Calib_S_A

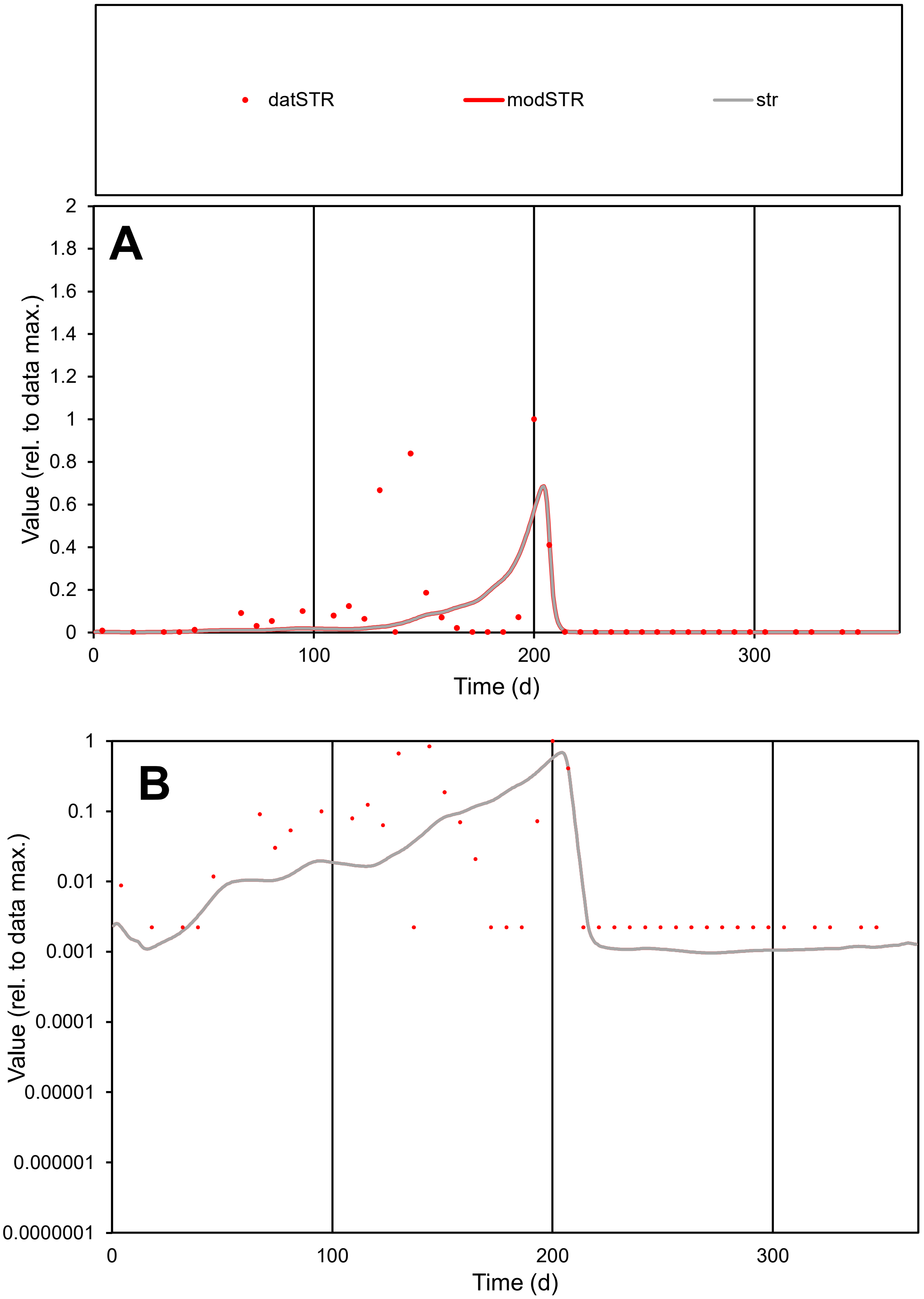


**C1**

Variables
STR Provide jtd or Dst
CalibOnly?
2 1 to reproduce of
oNormalize?
2 1=no, 2=yes

ofxn calc*
1.97506
ofxn mod
2.00339

sRun
RunA1
zout
1

Ave
Data
6.8E-05
Model
4.4E-05

data/vMax
0.00073

ofxng
334.07

Min d/m
2.31754

ave/min d
41.5849
ave/min m
62.1646

327_Calib_S_A

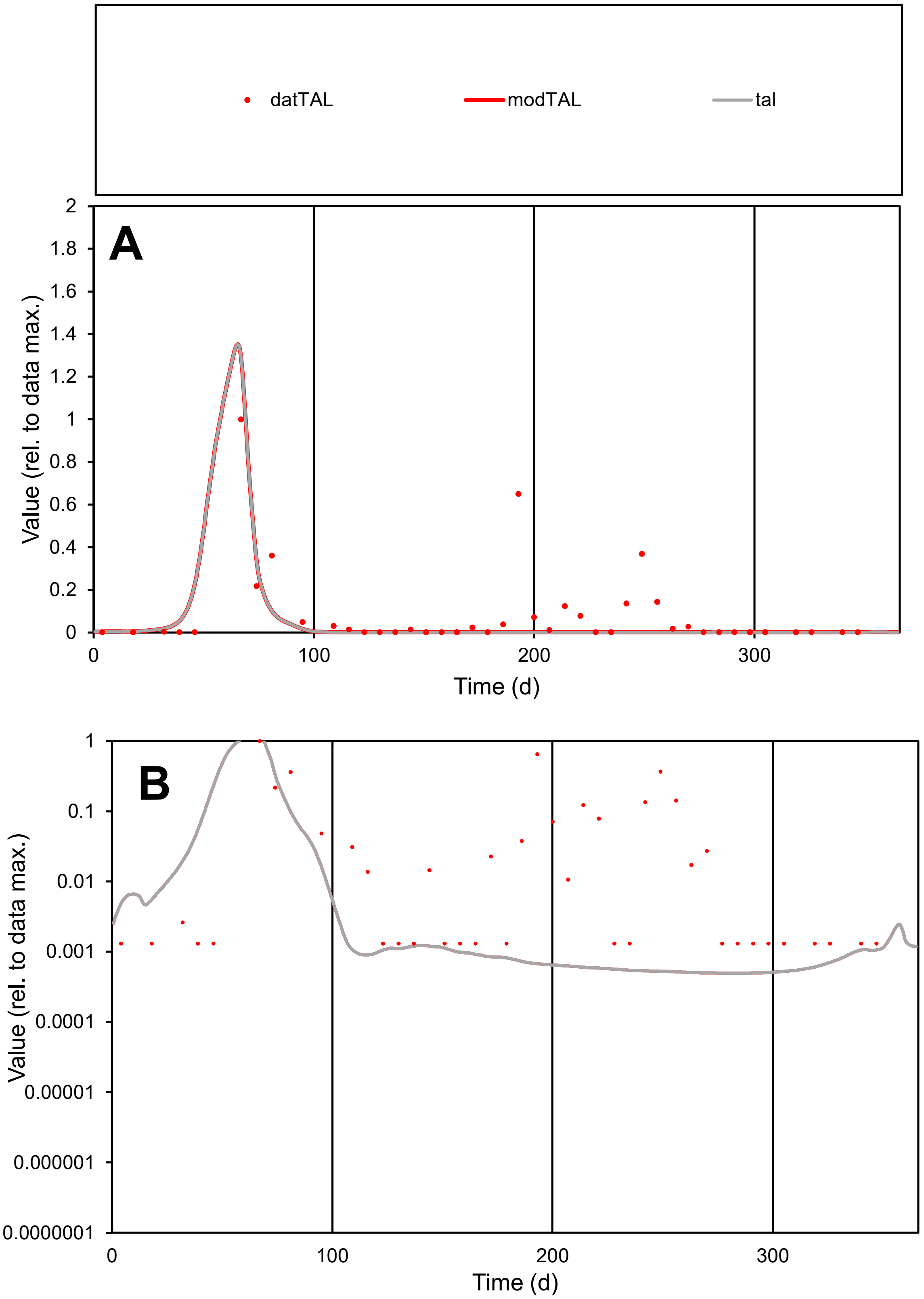


**C1**

Variables
TAL Provide jtd or Dst
CalibOnly?
2 1 to reproduce of
oNormalize?
2 1=no, 2=yes

ofxn calc*
1.63588
ofxn mod
1.67522

sRun
RunA1
zout
1

Ave
Data
0.00026
Model
0.00013

data/vMax
0.00317

ofxng
334.07

Min d/m
2.65301

ave/min d
61.9445
ave/min m
82.5974

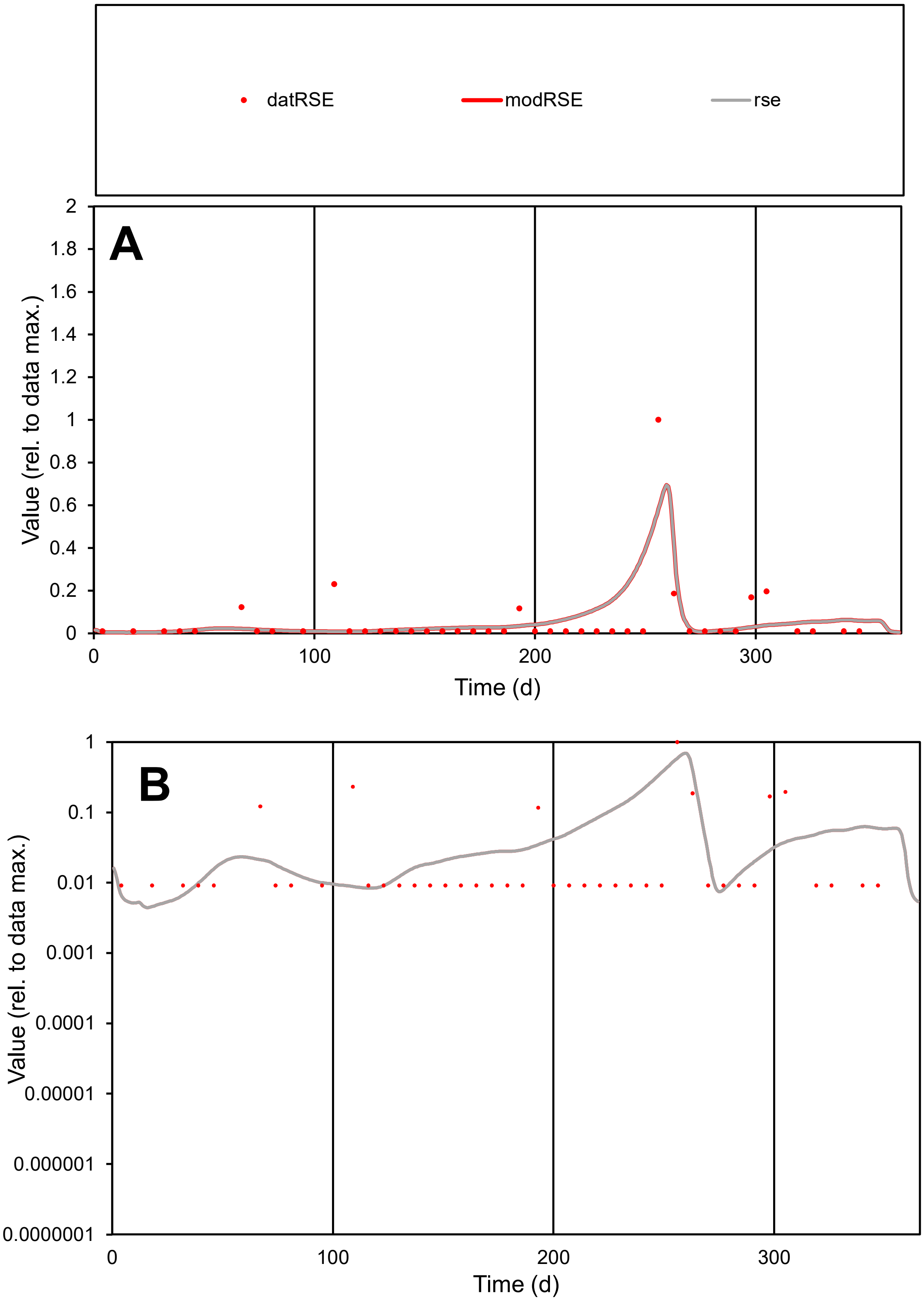


**C1**

Variables
RSE Provide jtd or Dst
CalibOnly?
2 1 to reproduce of
oNormalize?
2 1=no, 2=yes

ofxn calc*
1.84905
ofxn mod
1.94502

sRun
RunA1
zout
1

Ave
Data
2.8E-05
Model
3.2E-05

data/vMax
0.00051

ofxng
334.07

Min d/m
1.92051

ave/min d
6.10161
ave/min m
13.1746

327_Calib_S_A

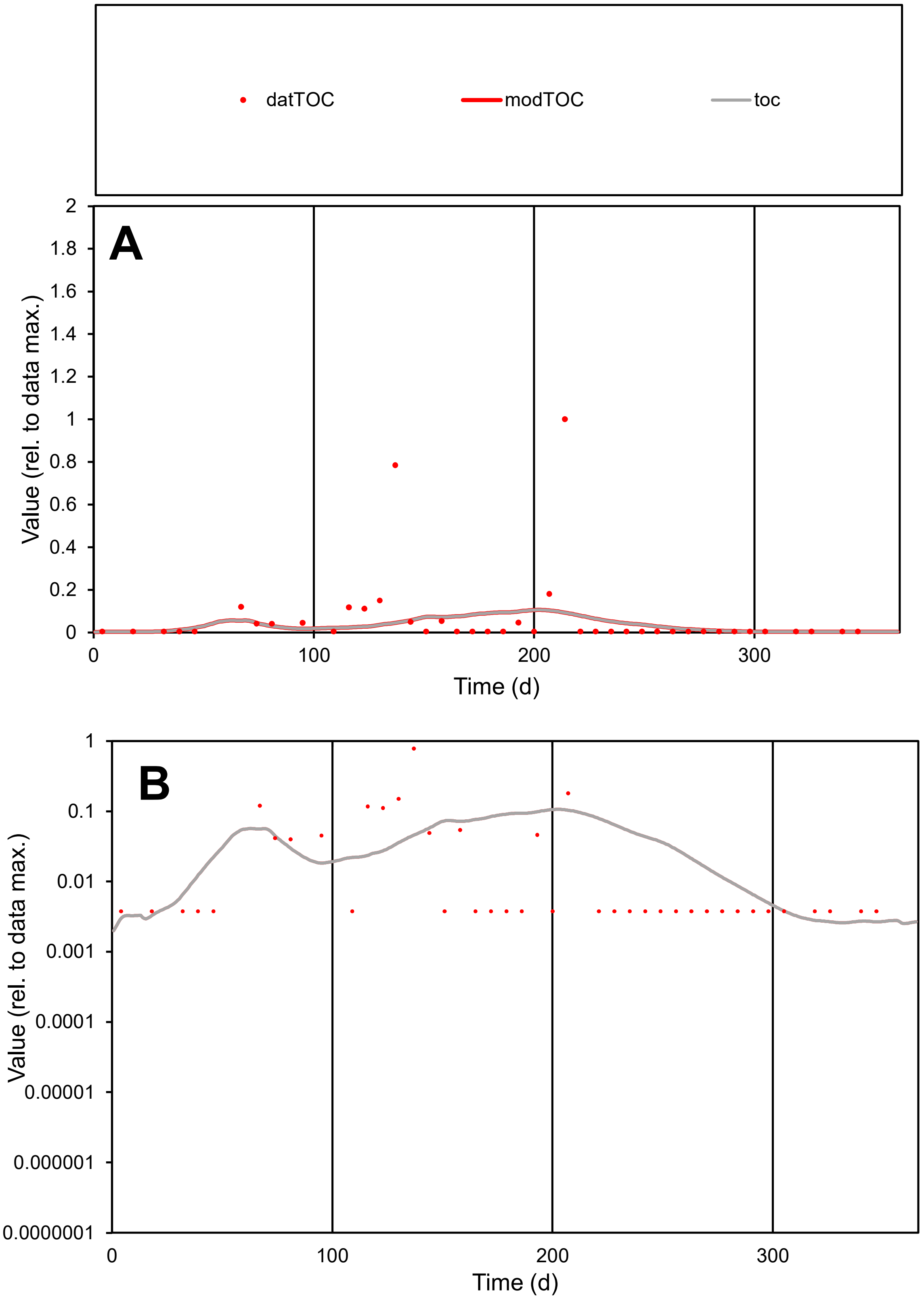


**C1**

Variables
TOC Provide jtd or Dst
CalibOnly?
2 1 to reproduce of
oNormalize?
2 1=no, 2=yes

ofxn calc*
2.75768
ofxn mod
2.76365

sRun
RunA1
zout
1

Ave
Data
7.8E-05
Model
4.6E-05

data/vMax
0.00115

ofxng
334.07

Min d/m
1.42405
ave/min d
17.9982
ave/min m
15.0116

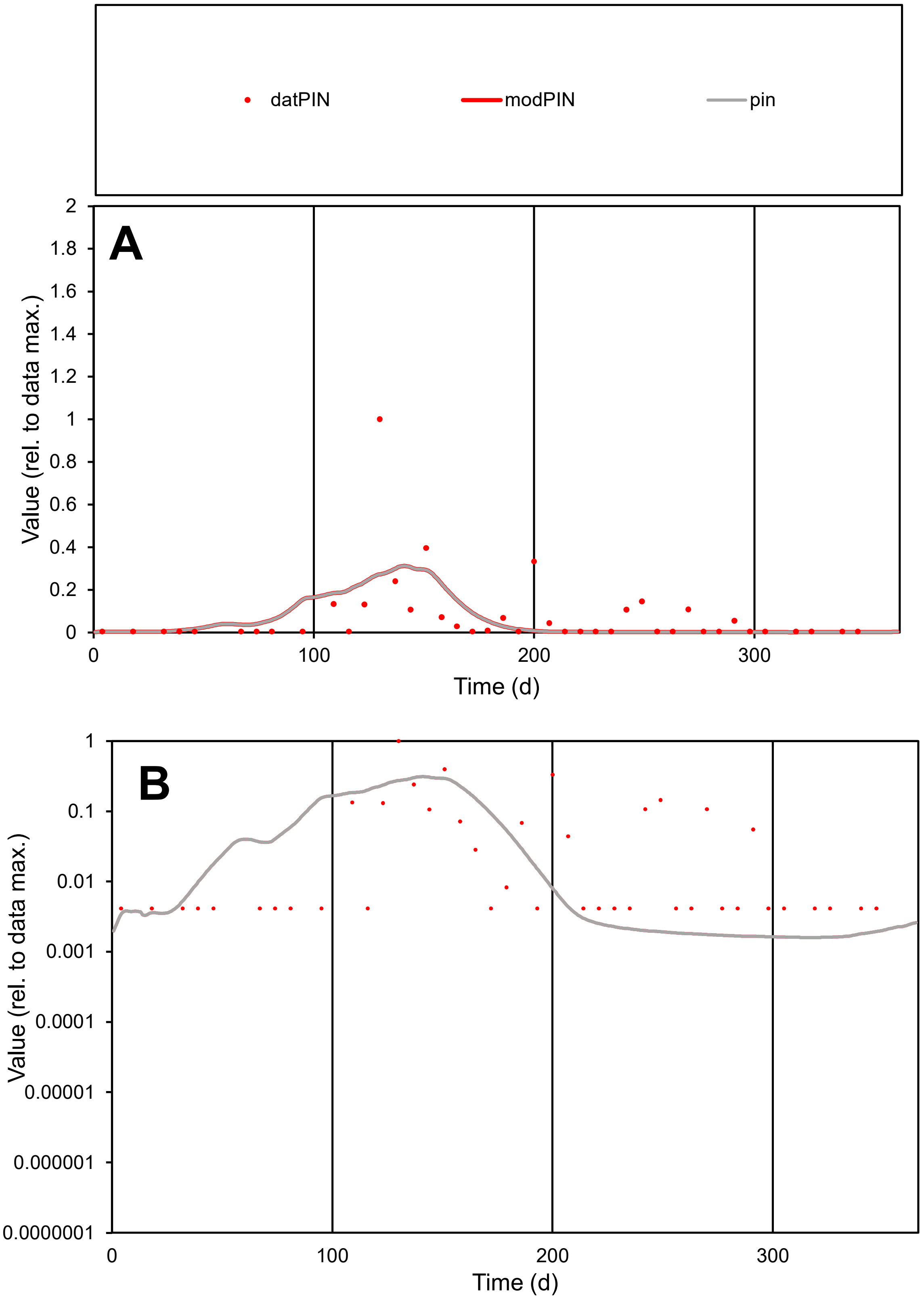


**C1**

Variables
PIN Provide jtd or Dst
CalibOnly?
2 1 to reproduce of
oNormalize?
2 1=no, 2=yes

ofxn calc*
1.93313
ofxn mod
1.95356

sRun
RunA1
zout
1

Ave
Data
7.1E-05
Model
6.1E-05

data/vMax
0.00097

ofxng
334.07

Min d/m
2.5871

ave/min d
17.7519
ave/min m
39.6483

327_Calib_S_A

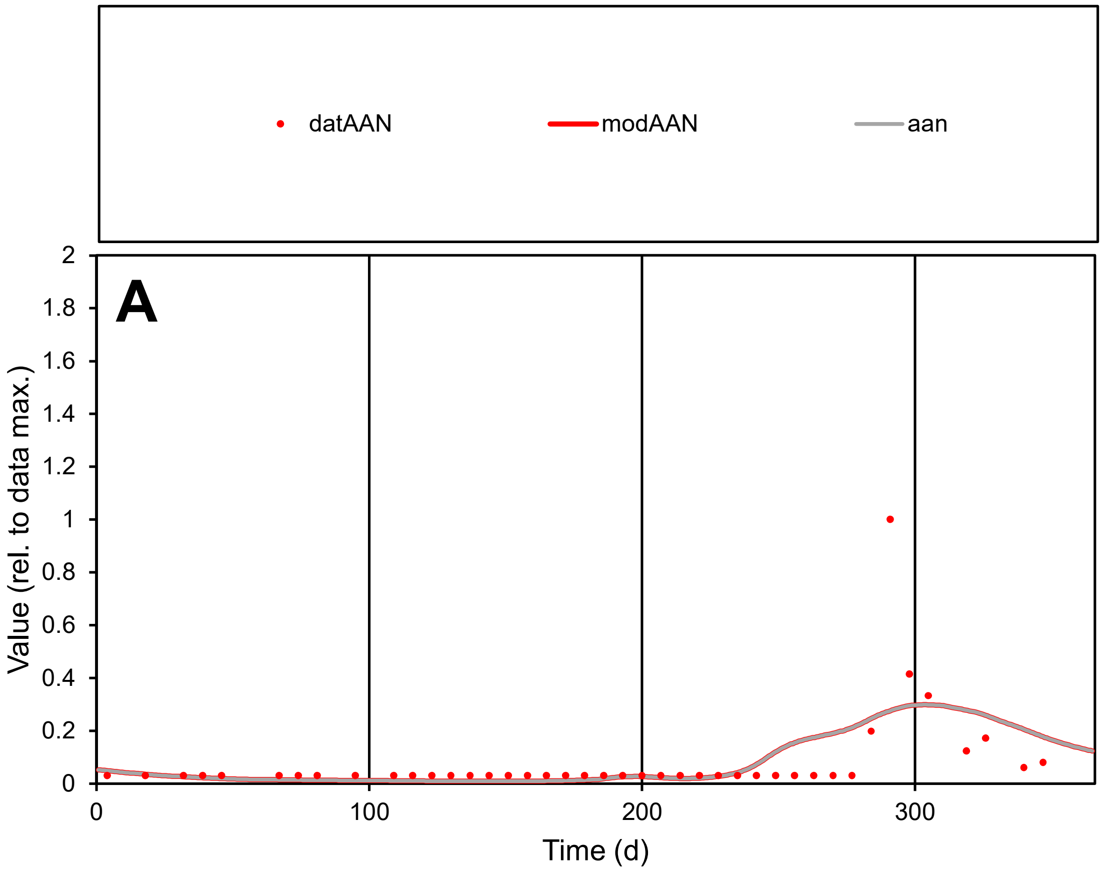


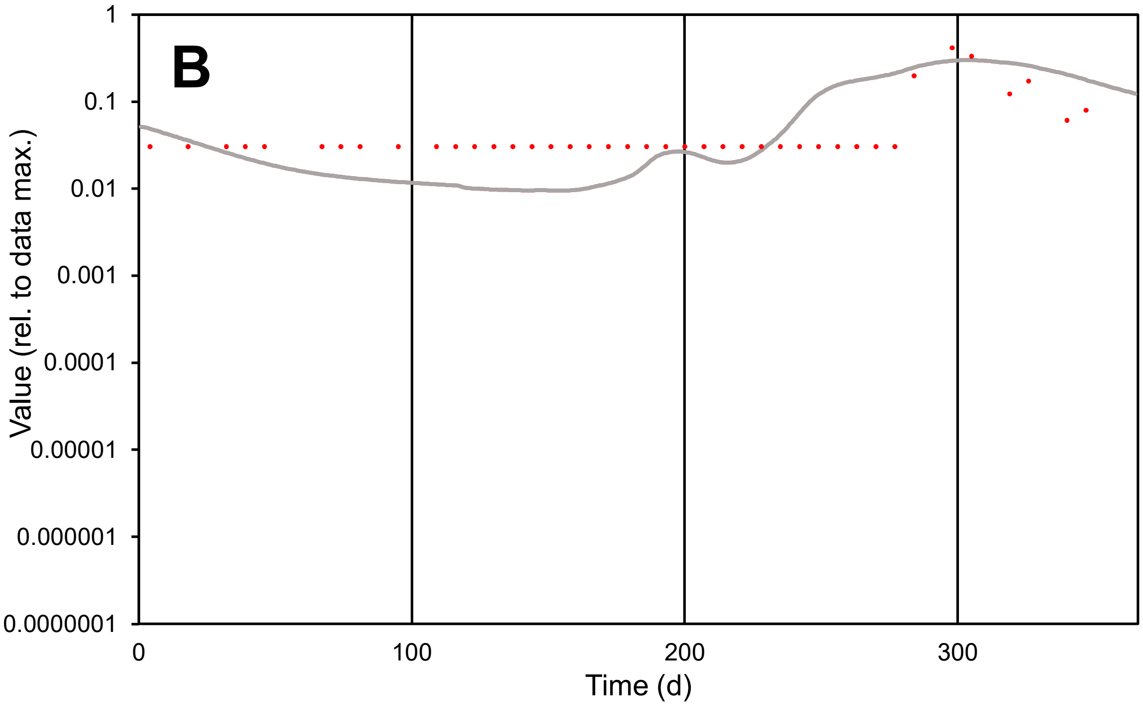


**C1**

Variables
AAN Provide jtd or Dst
CalibOnly?
2 1 to reproduce of
oNormalize?
2 1=no, 2=yes

ofxn calc*
1.5938
ofxn mod
1.6076

sRun
RunA1
zout
1

Ave
Data
1.5E-05
Model
1.5E-05

data/vMax
0.00018

ofxng
334.07

Min d/m
3.19473

ave/min d
2.67049
ave/min m
8.73129

327_Calib_S_A

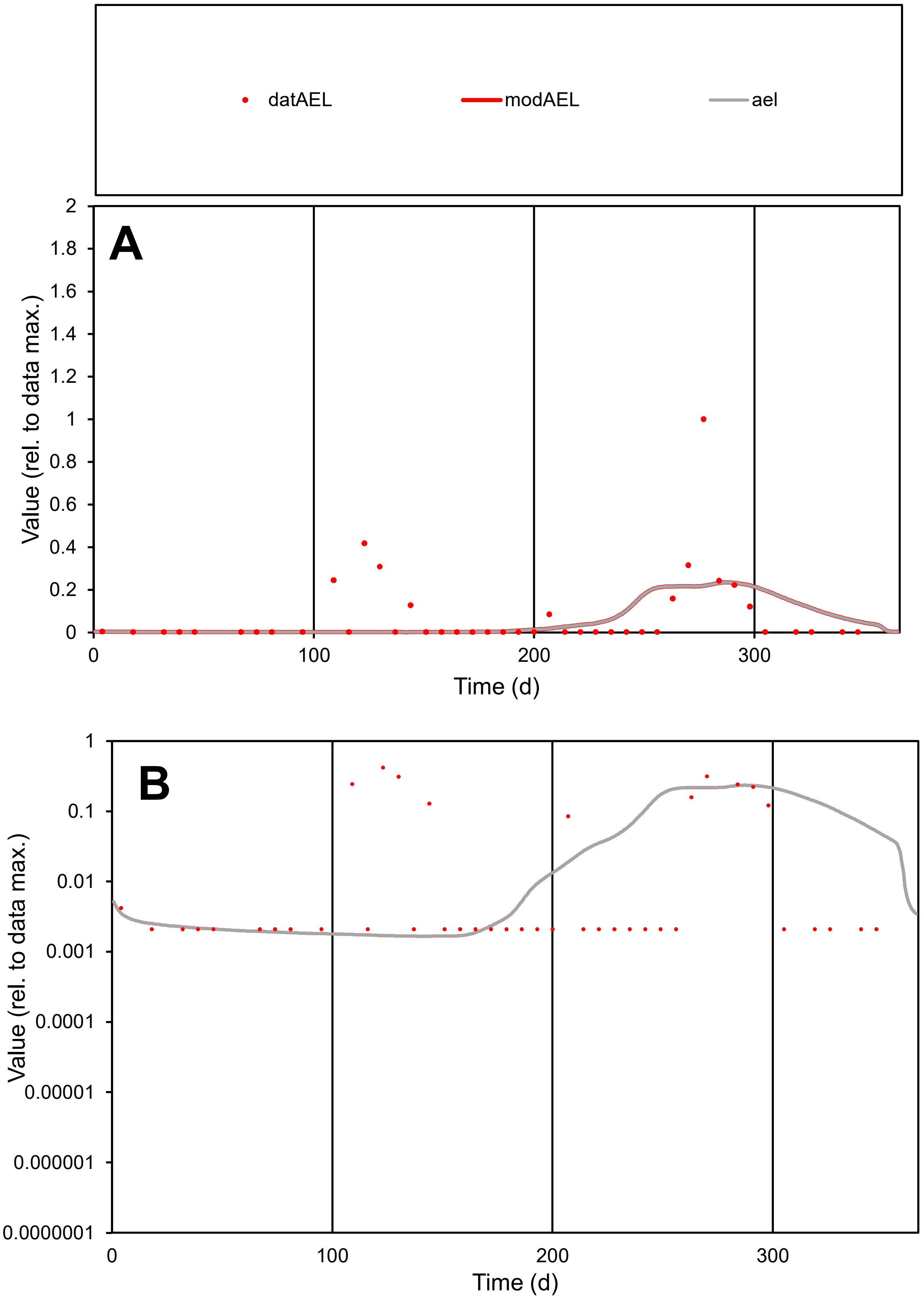


**C1**

Variables
AEL Provide jtd or Dst
CalibOnly?
2 1 to reproduce of
oNormalize?
2 1=no, 2=yes

ofxn calc*
2.07501
ofxn mod
2.09397

sRun
RunA1
zout
1

Ave
Data
2.6E-05
Model
2E-05

data/vMax
0.00033

ofxng
334.07

Min d/m
1.25162

ave/min d
37.6138
ave/min m
36.7362

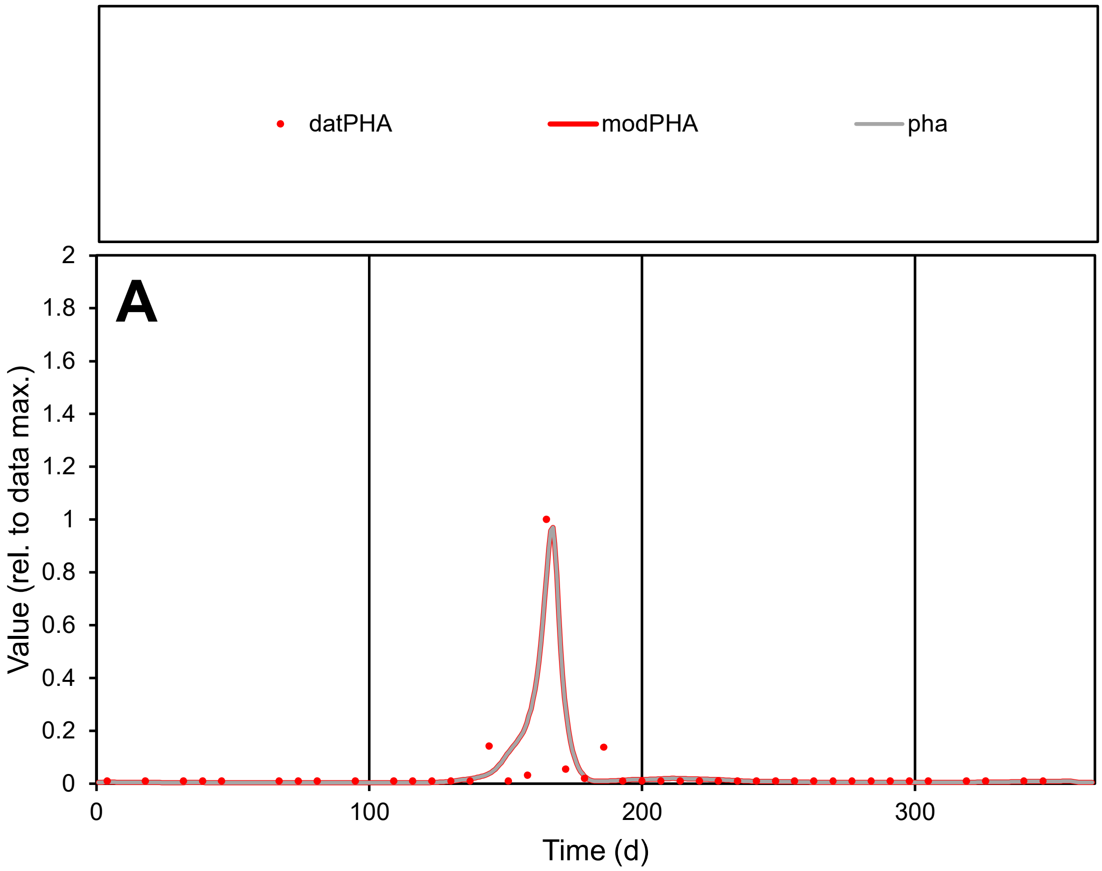


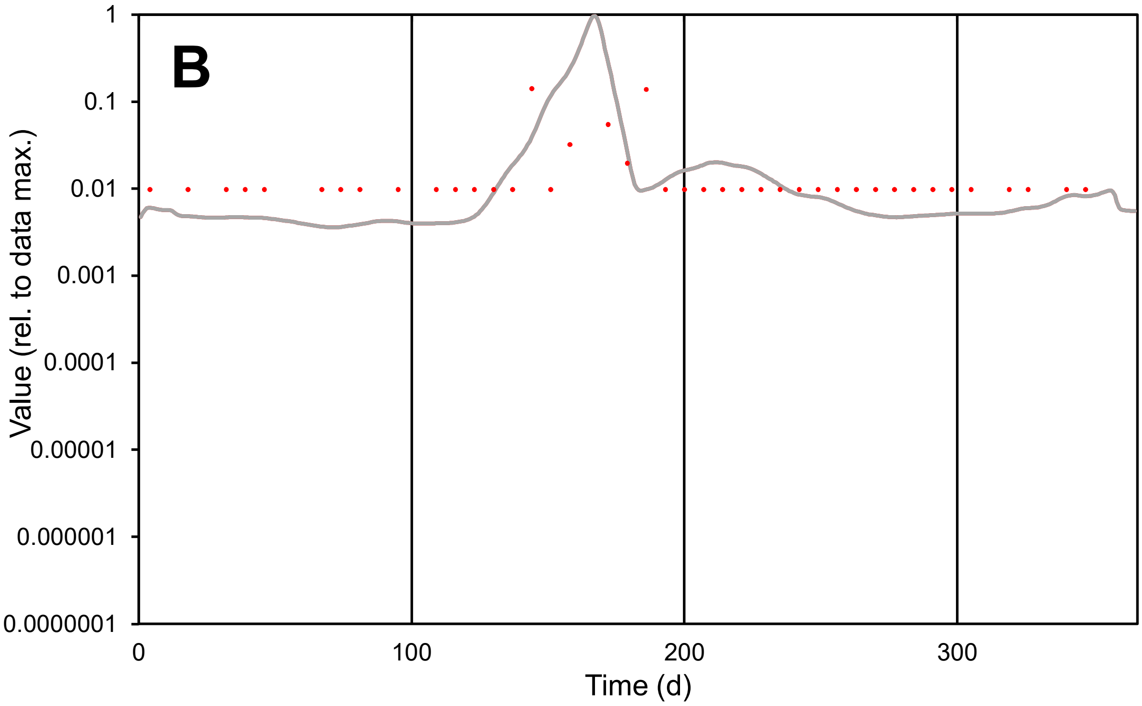


**C1**

Variables
PHA Provide jtd or Dst
CalibOnly?
2 1 to reproduce of
oNormalize?
2 1=no, 2=yes

ofxn calc*
0.98706
ofxn mod
1.15303

sRun
RunA1
zout
1

Ave
Data
4.4E-05
Model
4.4E-05

data/vMax
0.00106

ofxng
334.07

Min d/m
2.70109

ave/min d
4.21664
ave/min m
11.432

327_Calib_S_A

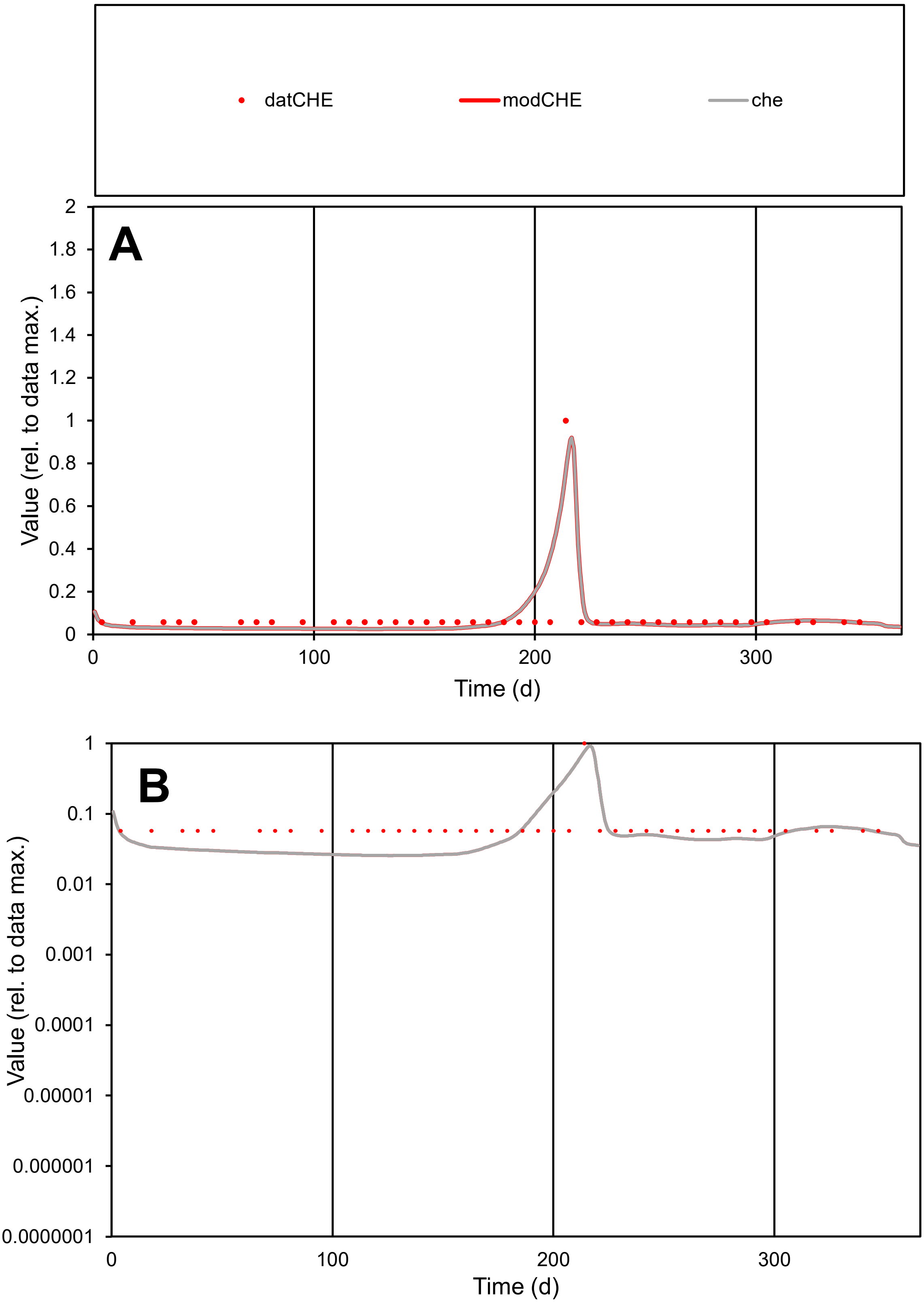


**C1**

Variables
CHE Provide jtd or Dst
CalibOnly?
2 1 to reproduce of
oNormalize?
2 1=no, 2=yes

ofxn calc*
0.67336
ofxn mod
0.79227

sRun
RunA1
zout
1

Ave
Data
9.8E-06
Model
8.9E-06

data/vMax
0.00012

ofxng
334.07

Min d/m
2.2464

ave/min d
1.39167
ave/min m
2.85504

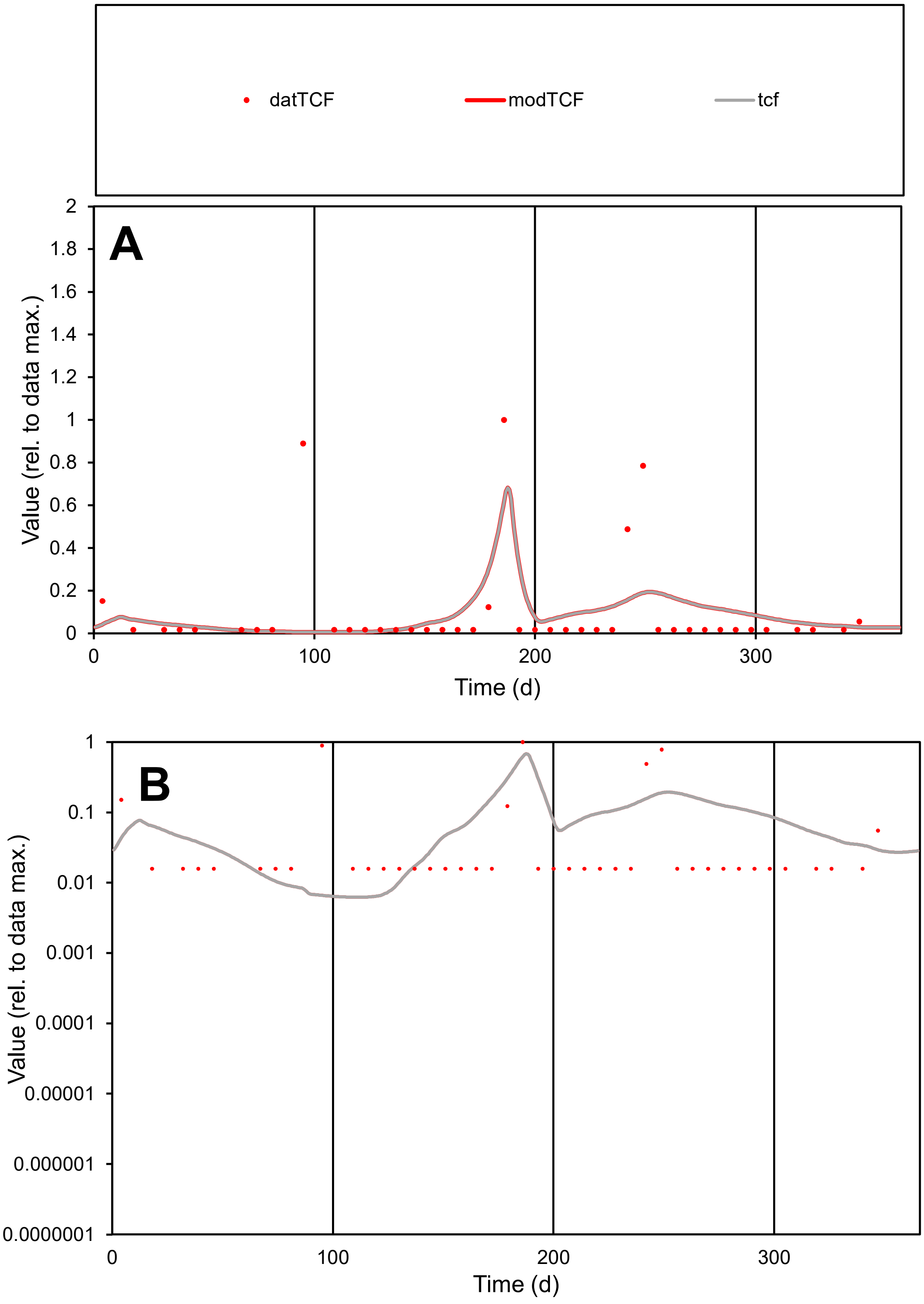


**C1**

Variables
TCF Provide jtd or Dst
CalibOnly?
2 1 to reproduce of
oNormalize?
2 1=no, 2=yes

ofxn calc*
1.99926
ofxn mod
2.03295

sRun
RunA1
zout
1

Ave
Data
1.9E-05
Model
1.8E-05

data/vMax
0.0002

ofxng
334.07

Min d/m
2.53577

ave/min d
6.09189
ave/min m
14.5856

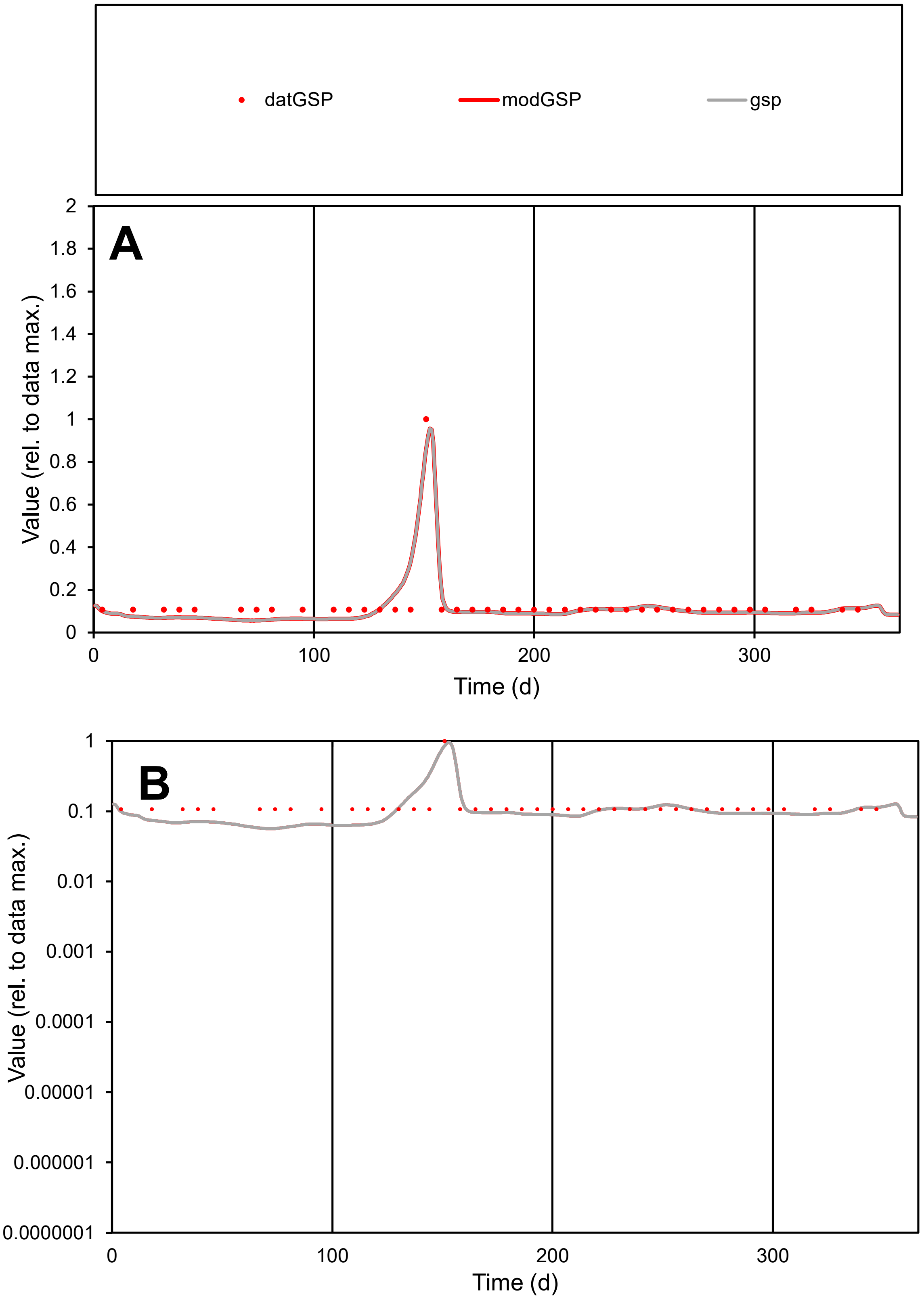


**C1**

Variables
GSP Provide jtd or Dst
CalibOnly?
2 1 to reproduce of
oNormalize?
2 1=no, 2=yes

ofxn calc*
0.32975
ofxn mod
0.37214

sRun
RunA1
zout
1

Ave
Data
1.1E-05
Model
1E-05

data/vMax
8.4E-05

ofxng
334.07

Min d/m
1.87398

ave/min d
1.1983
ave/min m
2.08083

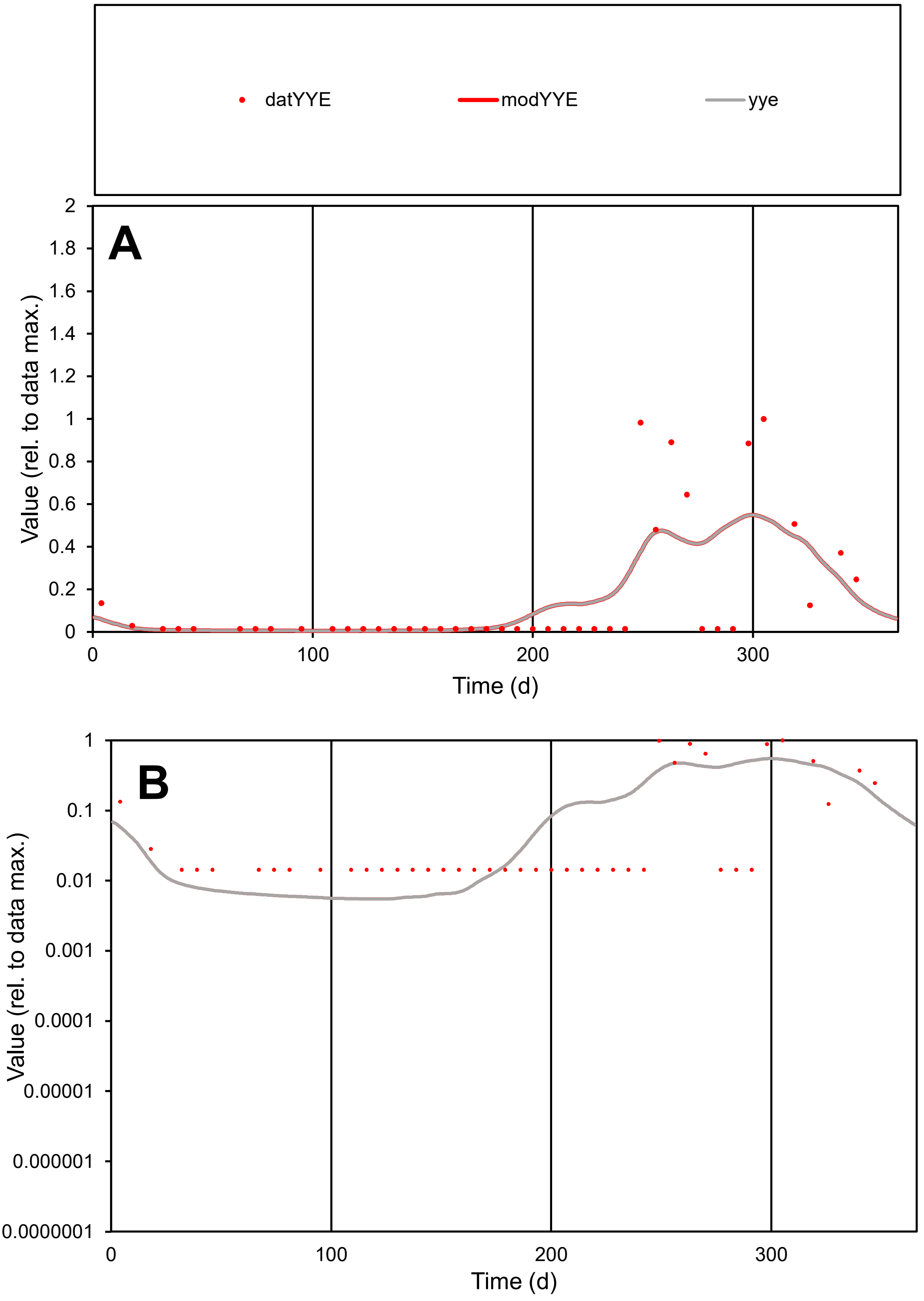


**C1**

Variables

YYE Provide jtd or Dst

CalibOnly?

2 1 to reproduce of

oNormalize?

2 1=no, 2=yes

ofxn calc*

1.28467

ofxn mod

1.30582

sRun

RunA1

zout

1

Ave

Data

4.2E-05

Model

4.2E-05

data/vMax

0.00026

ofxng

334.07

Min d/m

2.58981

ave/min d

11.2218

ave/min m

29.1679

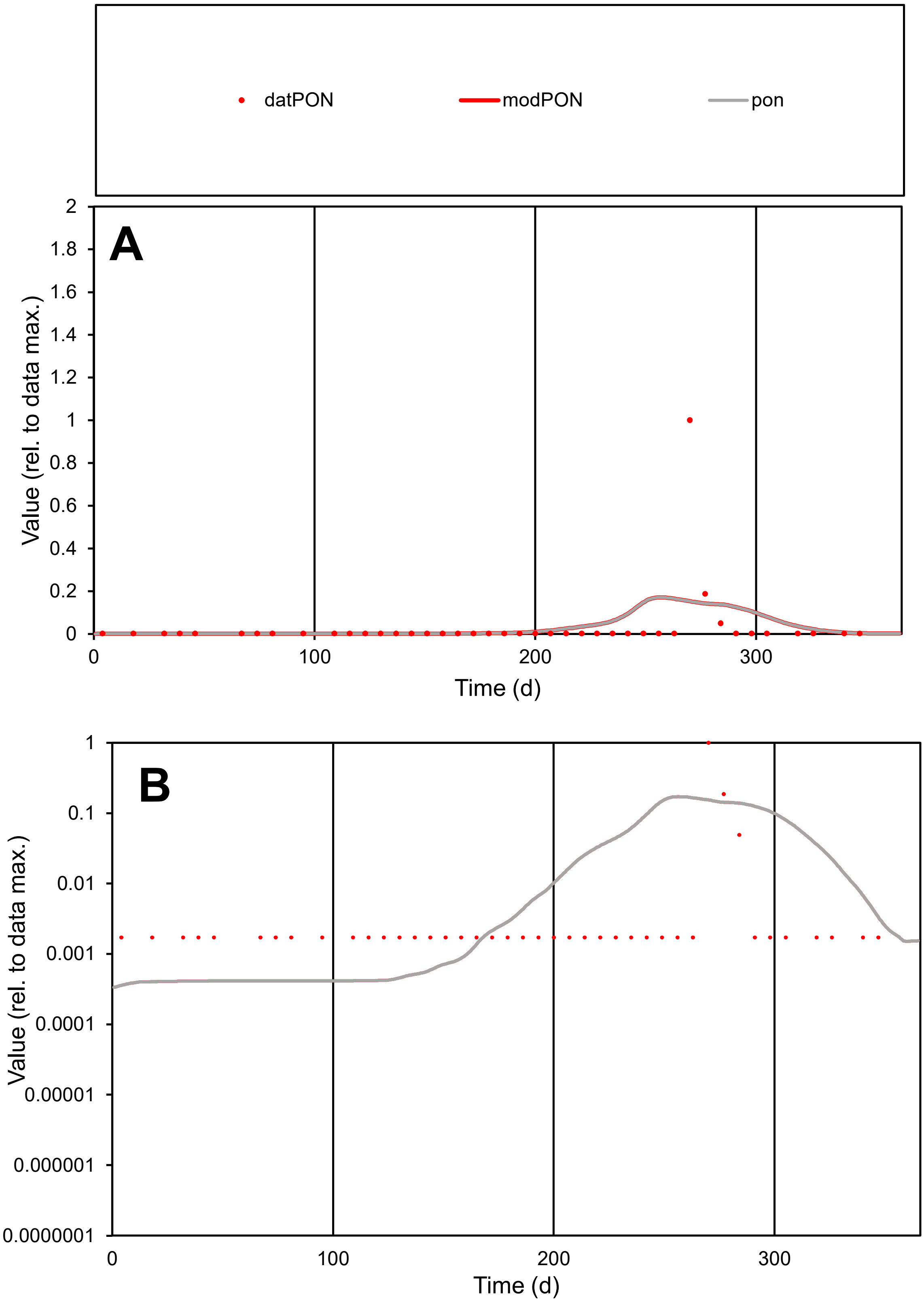


**C1**

Variables
PON Provide jtd or Dst
CalibOnly?
2 1 to reproduce of
oNormalize?
2 1=no, 2=yes

ofxn calc*
4.5772
ofxn mod
4.60397

sRun
RunA1
zout
1

Ave
Data
5.3E-05
Model
6.3E-05

data/vMax
0.00171

ofxng
334.07

Min d/m
4.71704

ave/min d
18.0494
ave/min m
101.203

327_Calib_S_A

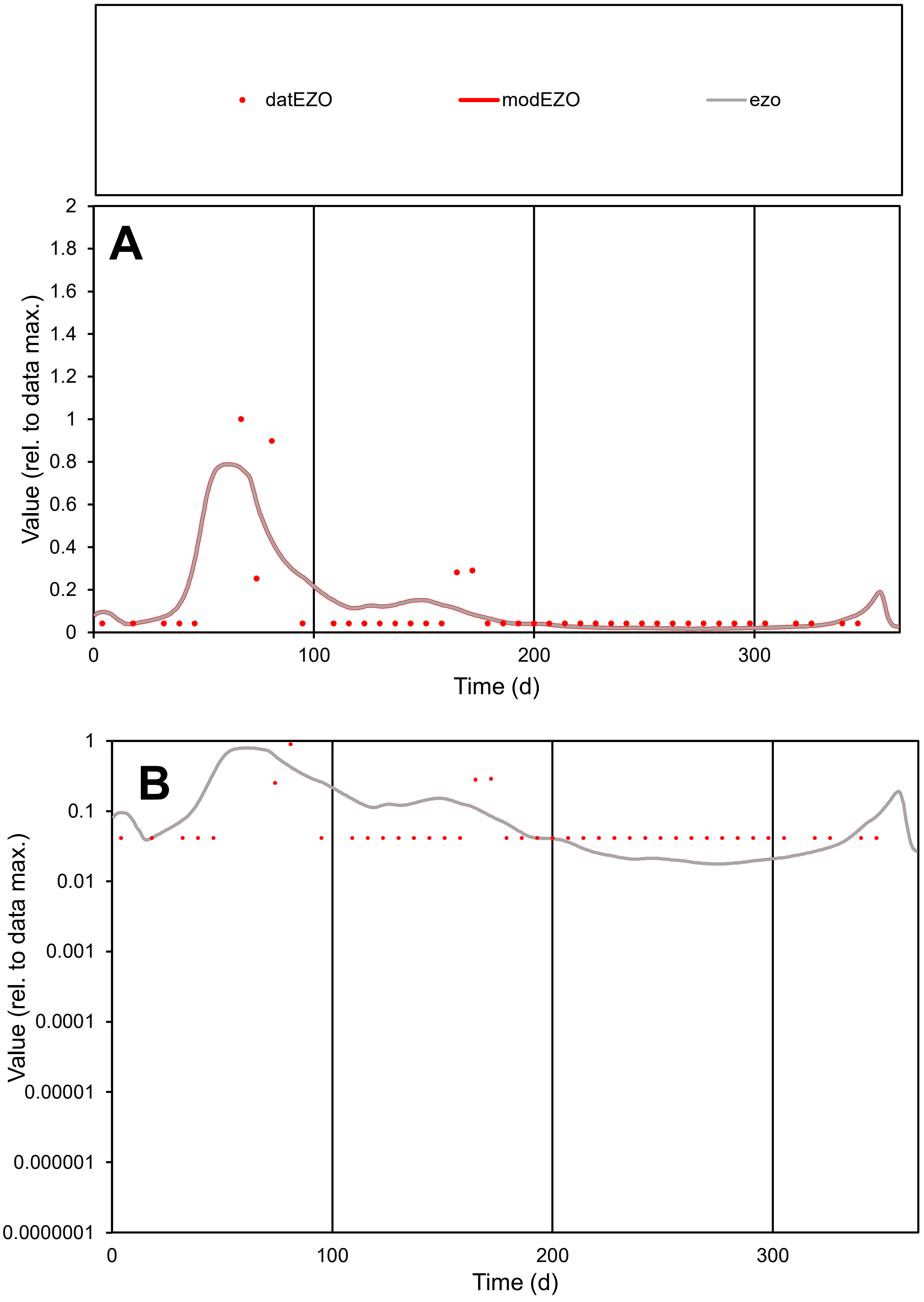


**C1**

Variables
EZO Provide jtd or Dst
CalibOnly?
2 1 to reproduce of
oNormalize?
2 1=no, 2=yes

ofxn calc*
1.22382
ofxn mod
1.27272

sRun
RunA1
zout
1

Ave
Data
9.6E-06
Model
1E-05

data/vMax
9.5E-05

ofxng
334.07

Min d/m
2.3359

ave/min d
2.44261
ave/min m
6.08487

327_Calib_S_A

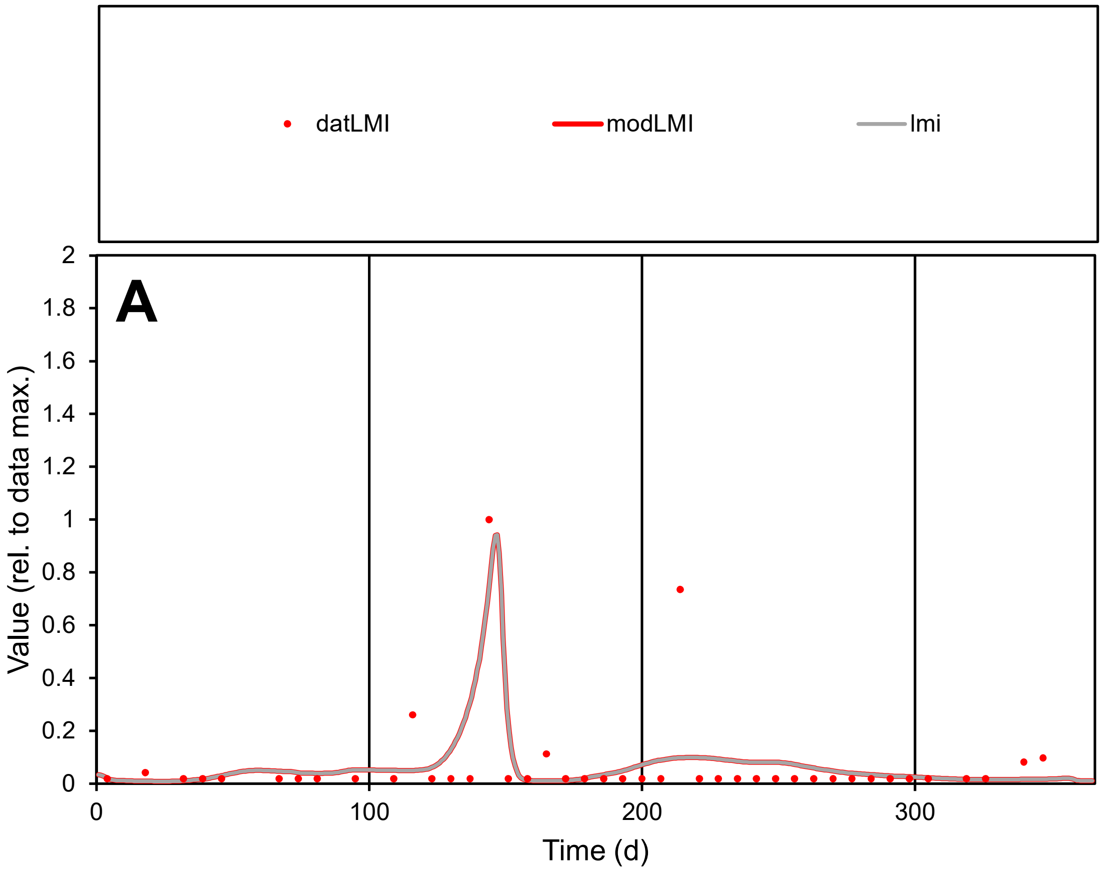


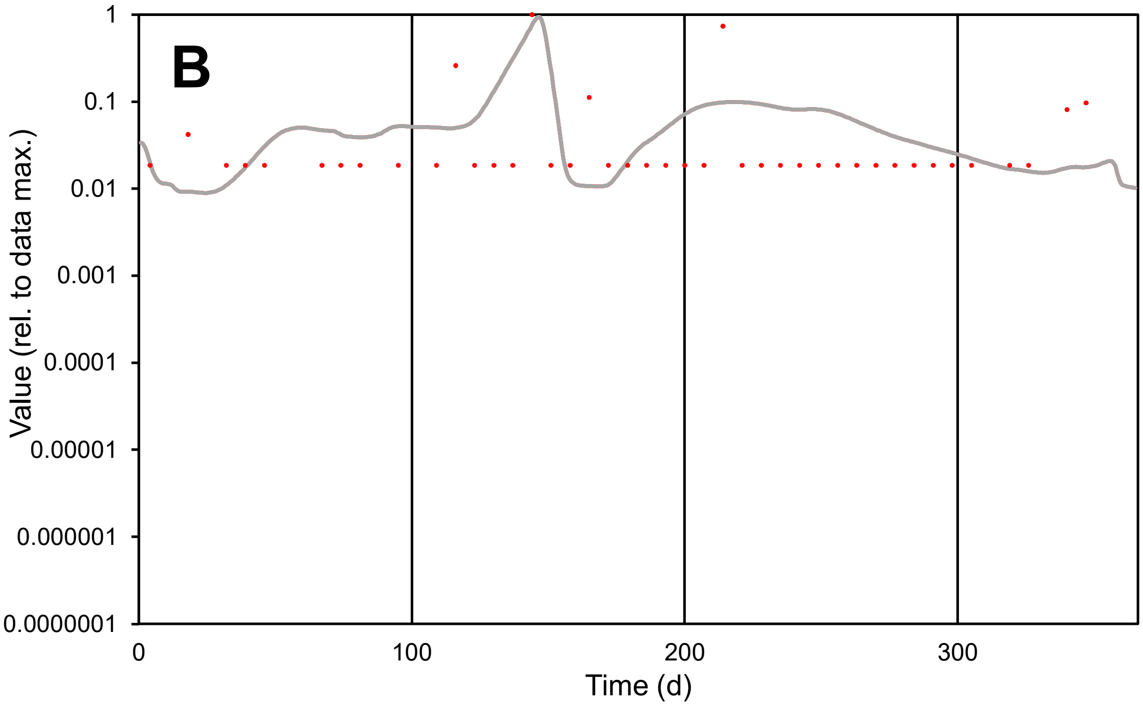


**C1**

Variables
LMI Provide jtd or Dst
CalibOnly?
2 1 to reproduce of
oNormalize?
2 1=no, 2=yes

ofxn calc*
1.67243
ofxn mod
1.72136

sRun
RunA1
zout
1

Ave
Data
1.1E-05
Model
1.1E-05

data/vMax
0.00016

ofxng
334.07

Min d/m
2.02759

ave/min d
3.8104
ave/min m
7.68719

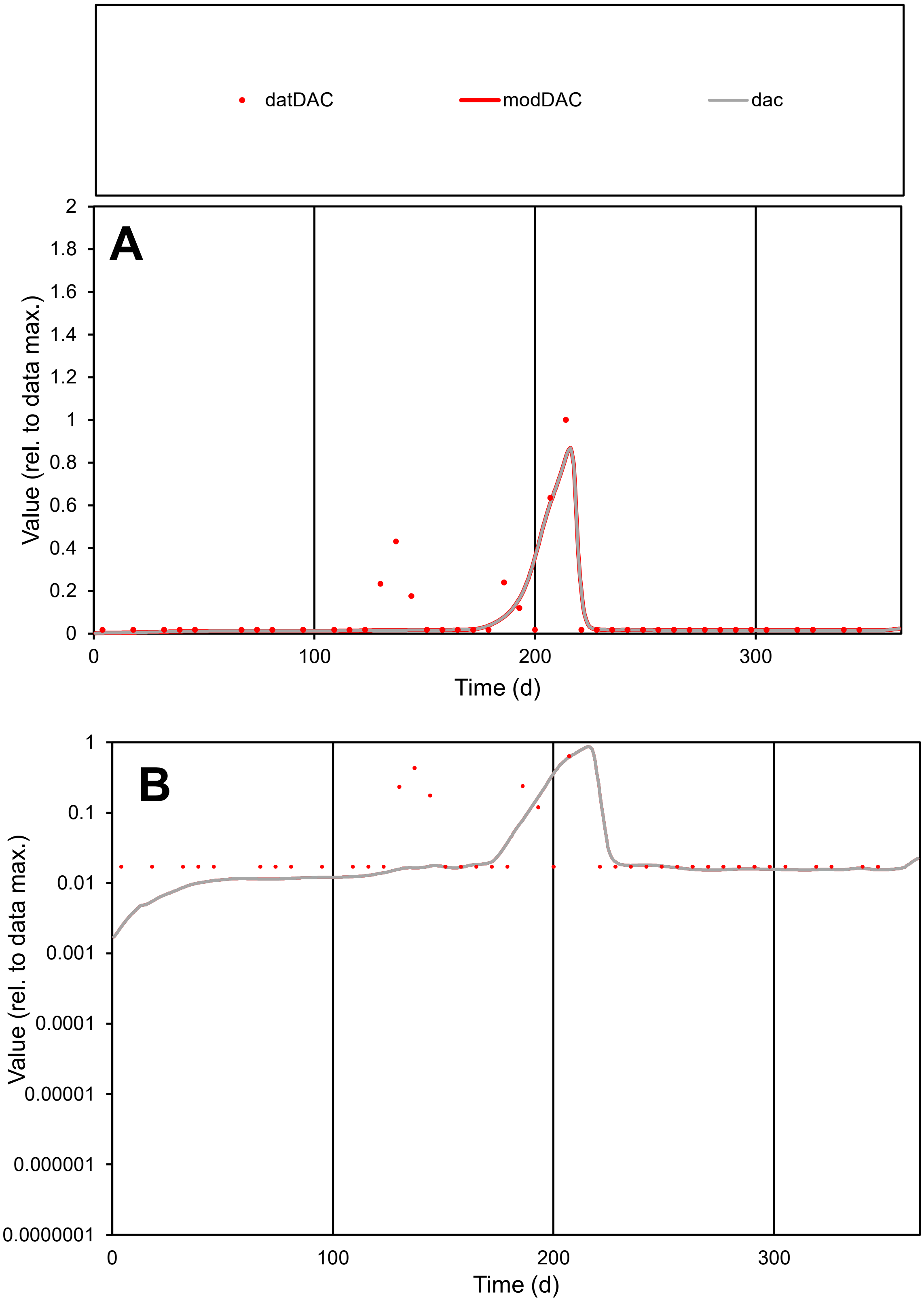


**C1**

Variables

DAC Provide jtd or Dst

CalibOnly?

2 1 to reproduce of

oNormalize?

2 1=no, 2=yes

ofxn calc*

1.15529

ofxn mod

1.22583

sRun

RunA1

zout

1

Ave

Data

3.7E-05

Model

2.9E-05

data/vMax

0.00045

ofxng

334.07

Min d/m

6.50295

ave/min d

4.77794

ave/min m

24.4306

327_Calib_S_A

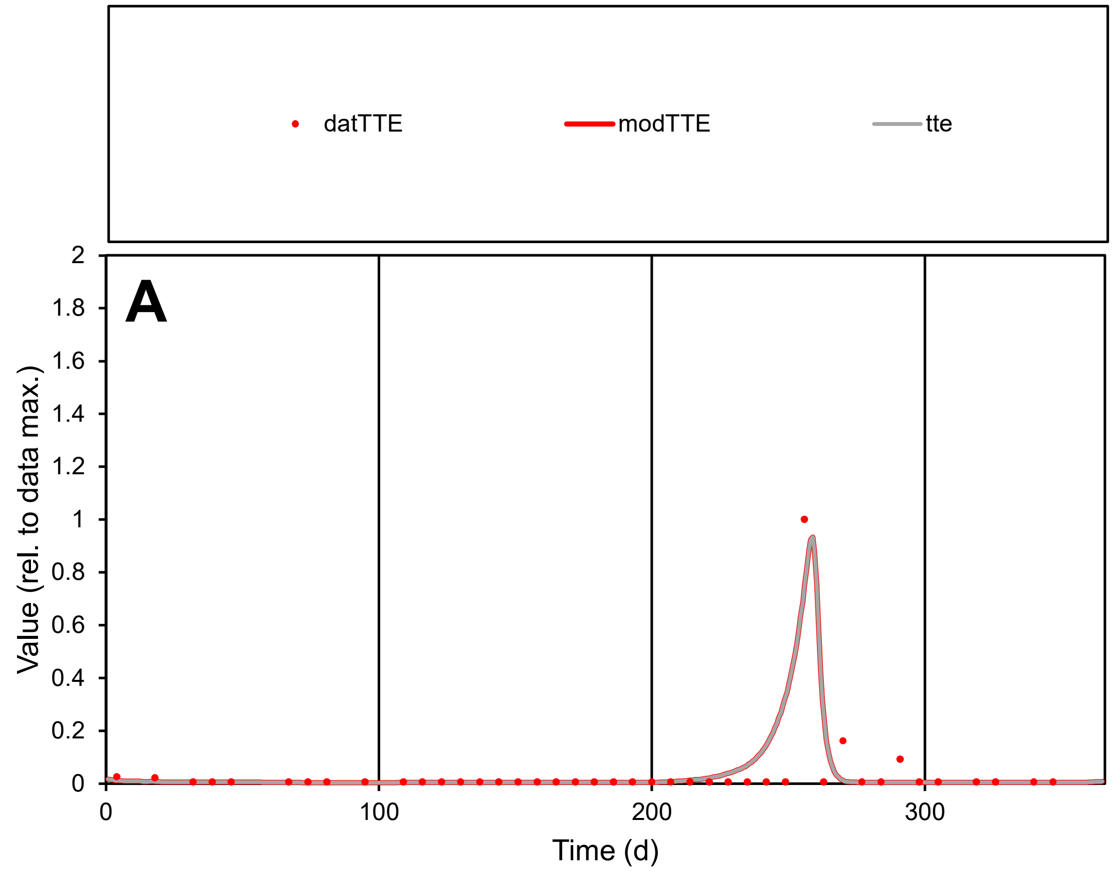


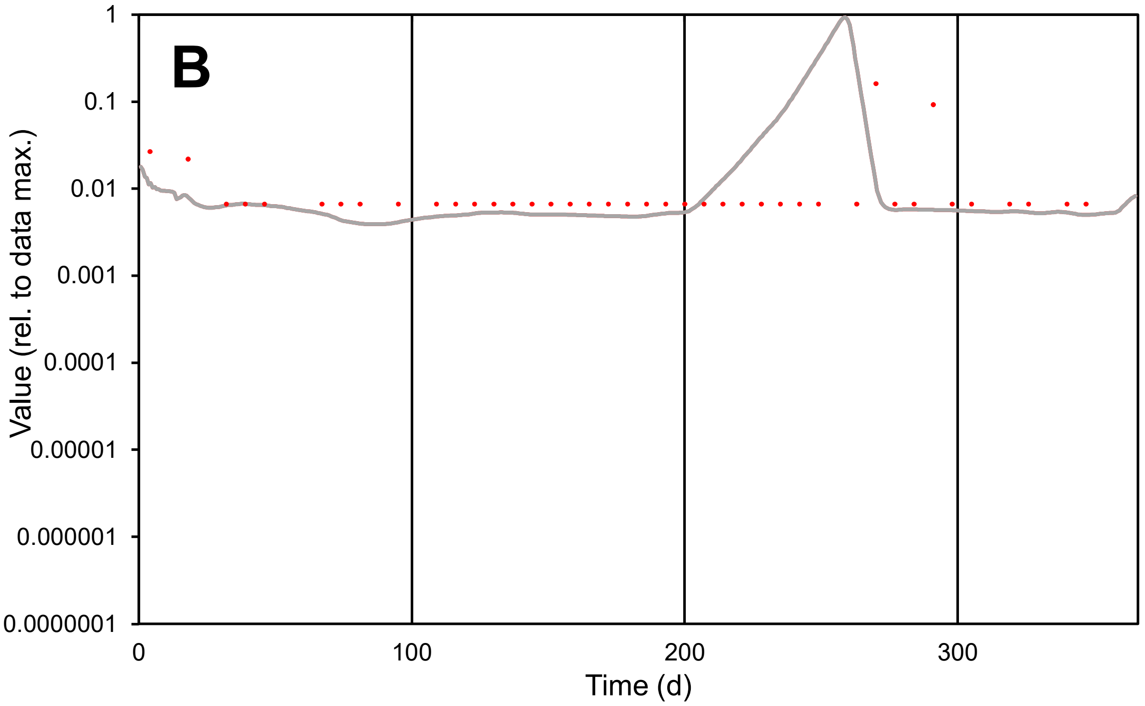


**C1**

Variables
TTE Provide jtd or Dst
CalibOnly?
2 1 to reproduce of
oNormalize?
2 1=no, 2=yes

ofxn calc*
1.53196
ofxn mod
1.80022

sRun
RunA1
zout
1

Ave
Data
2.1E-05
Model
2.2E-05

data/vMax
0.00056

ofxng
334.07

Min d/m
1.69034

ave/min d
5.52359
ave/min m
10.0998

327_Calib_S_A

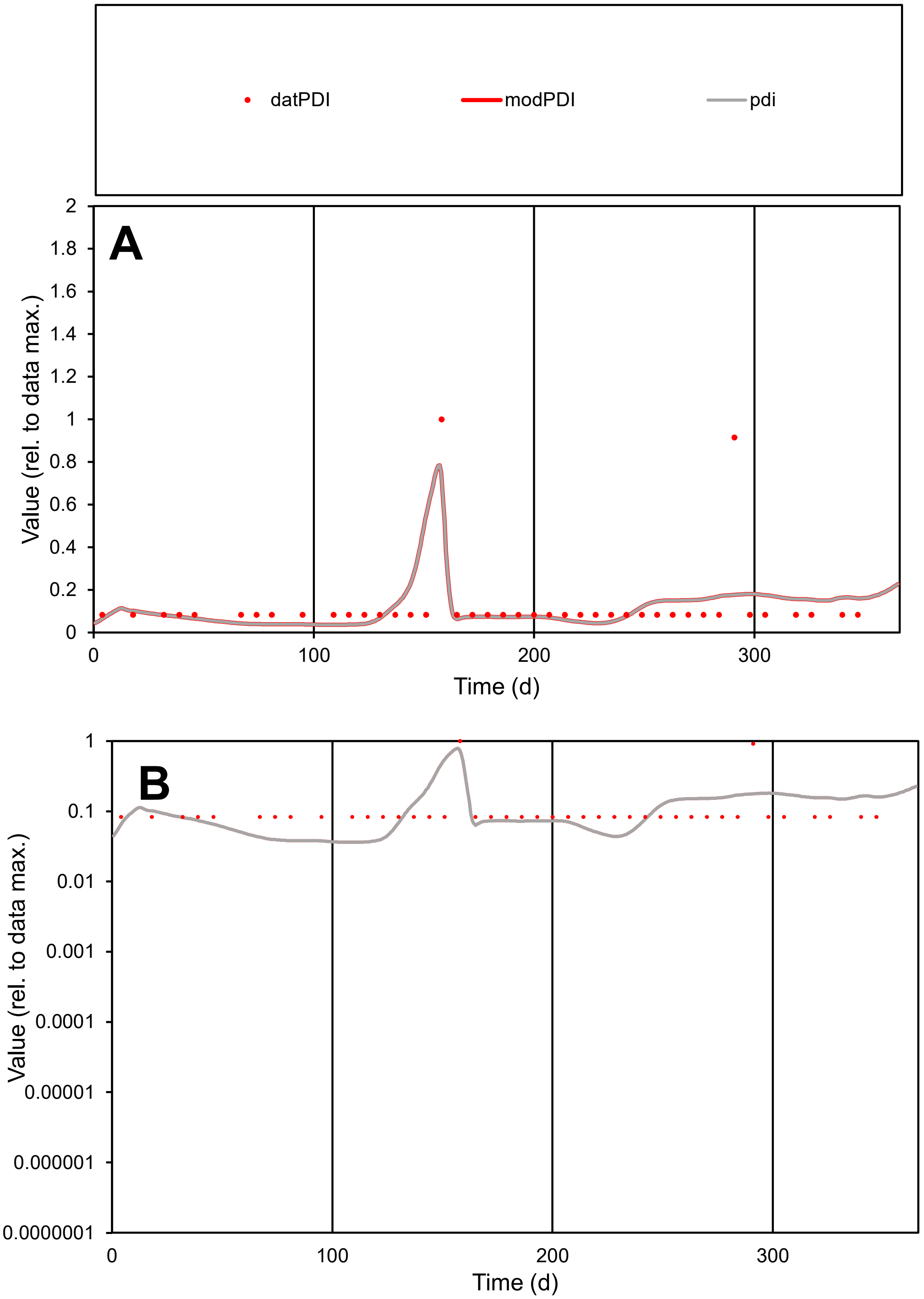


**C1**

Variables
PDI Provide jtd or Dst
CalibOnly?
2 1 to reproduce of
oNormalize?
2 1=no, 2=yes

ofxn calc*
1.17722
ofxn mod
1.24607

sRun
RunA1
zout
1

Ave
Data
6.6E-06
Model
6.4E-06

data/vMax
5.3E-05

ofxng
334.07

Min d/m
2.27301

ave/min d
1.50196
ave/min m
3.29787

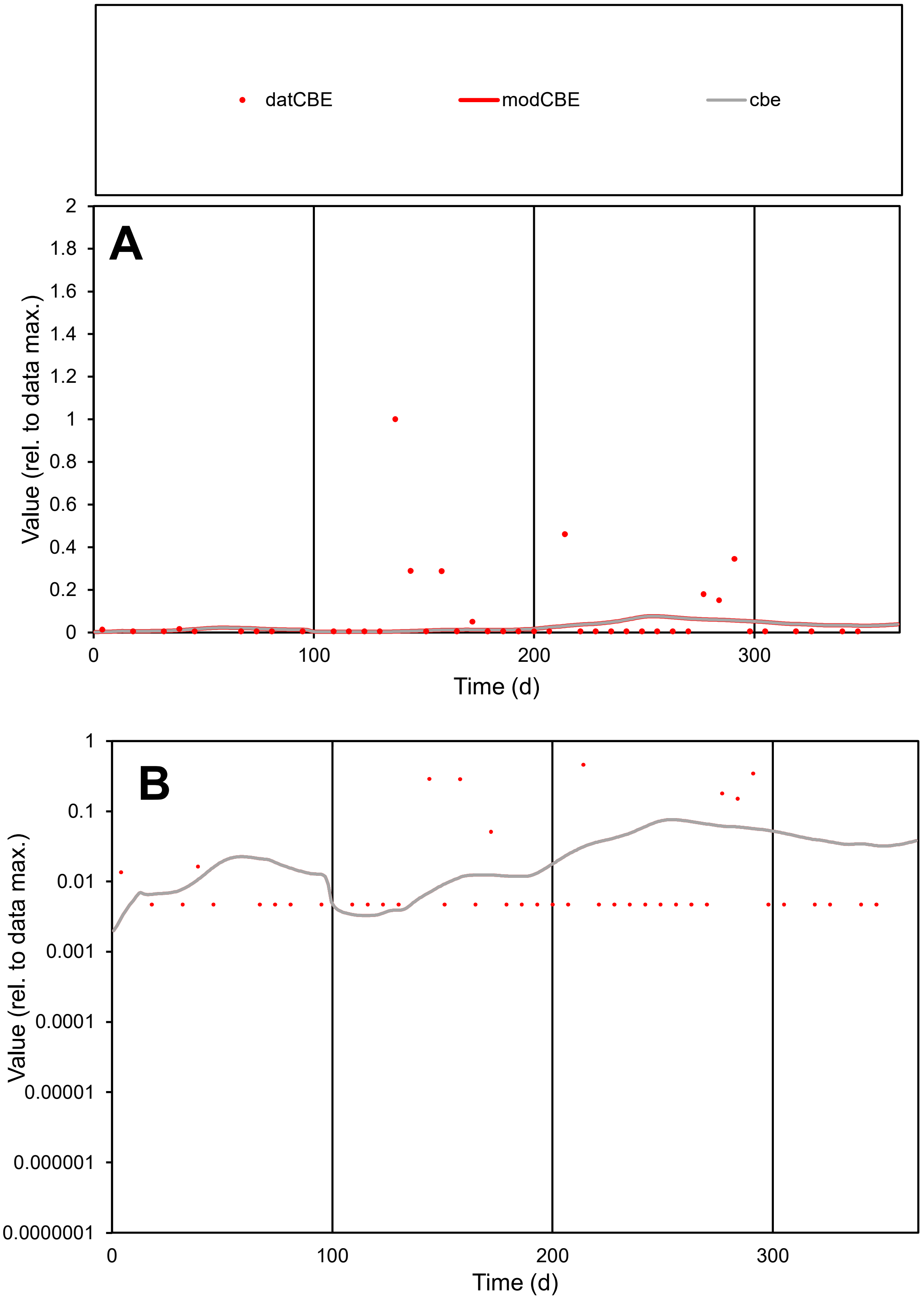


**C1**

Variables
CBE Provide jtd or Dst
CalibOnly?
2 1 to reproduce of
oNormalize?
2 1=no, 2=yes

ofxn calc*
2.66882
ofxn mod
2.67221

sRun
RunA1
zout
1

Ave
Data
2.5E-05
Model
1E-05

data/vMax
0.00035

ofxng
334.07

Min d/m
1.44166

ave/min d
14.8696
ave/min m
8.63629

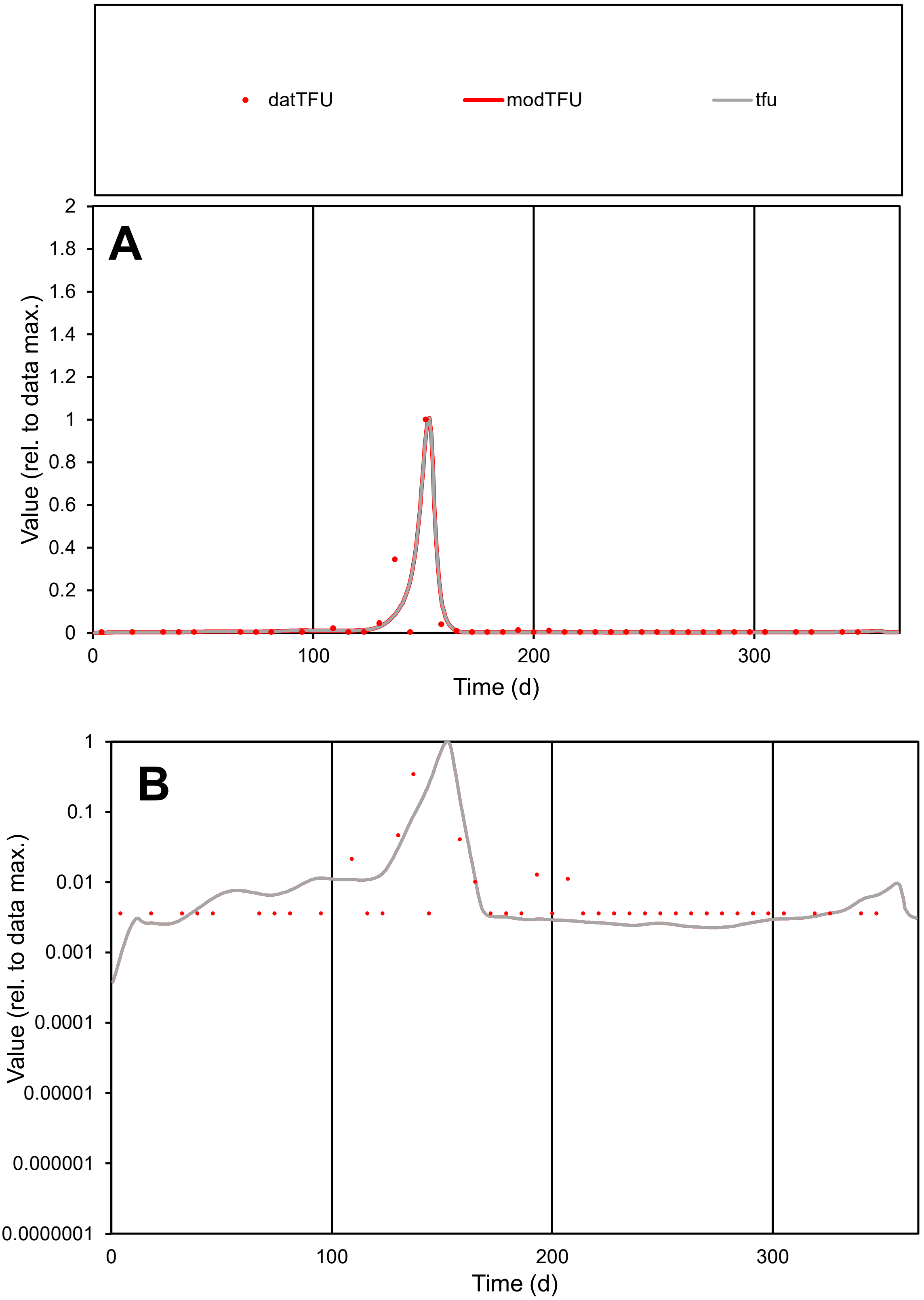


**C1**

Variables
TFU Provide jtd or Dst
CalibOnly?
2 1 to reproduce of
oNormalize?
2 1=no, 2=yes

ofxn calc*
1.08482
ofxn mod
1.3016

sRun
RunA1
zout
1

Ave
Data
0.00014
Model
0.00013

data/vMax
0.00355

ofxng
334.07

Min d/m
2.34074

ave/min d
10.6776
ave/min m
24.2733

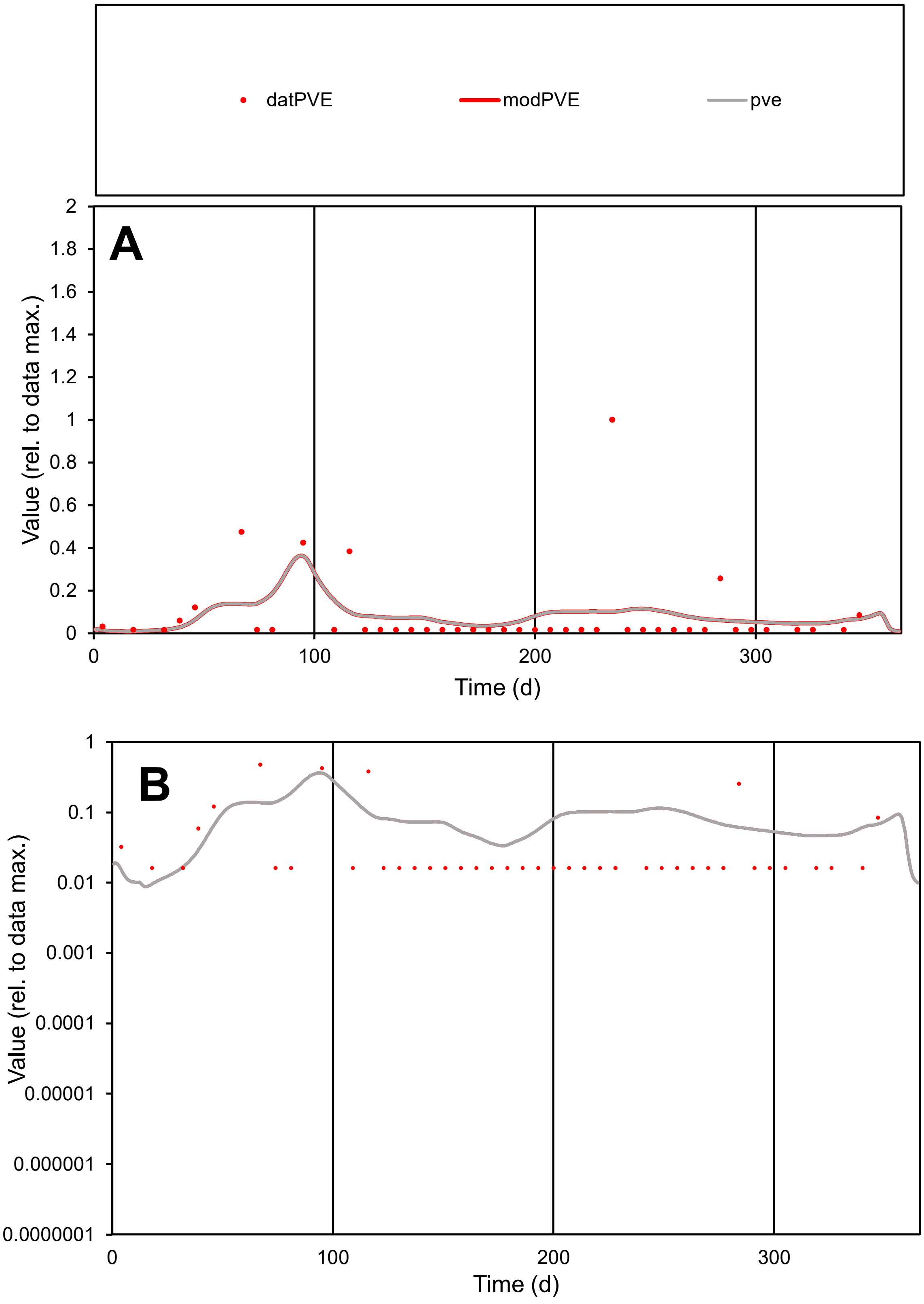


**C1**

Variables
PVE Provide jtd or Dst
CalibOnly?
2 1 to reproduce of
oNormalize?
2 1=no, 2=yes

ofxn calc*
2.11175
ofxn mod
2.12583

sRun
RunA1
zout
1

Ave
Data
1E-05
Model
1.1E-05

data/vMax
0.00013

ofxng
334.07

Min d/m
1.57293

ave/min d
4.97234
ave/min m
7.93636

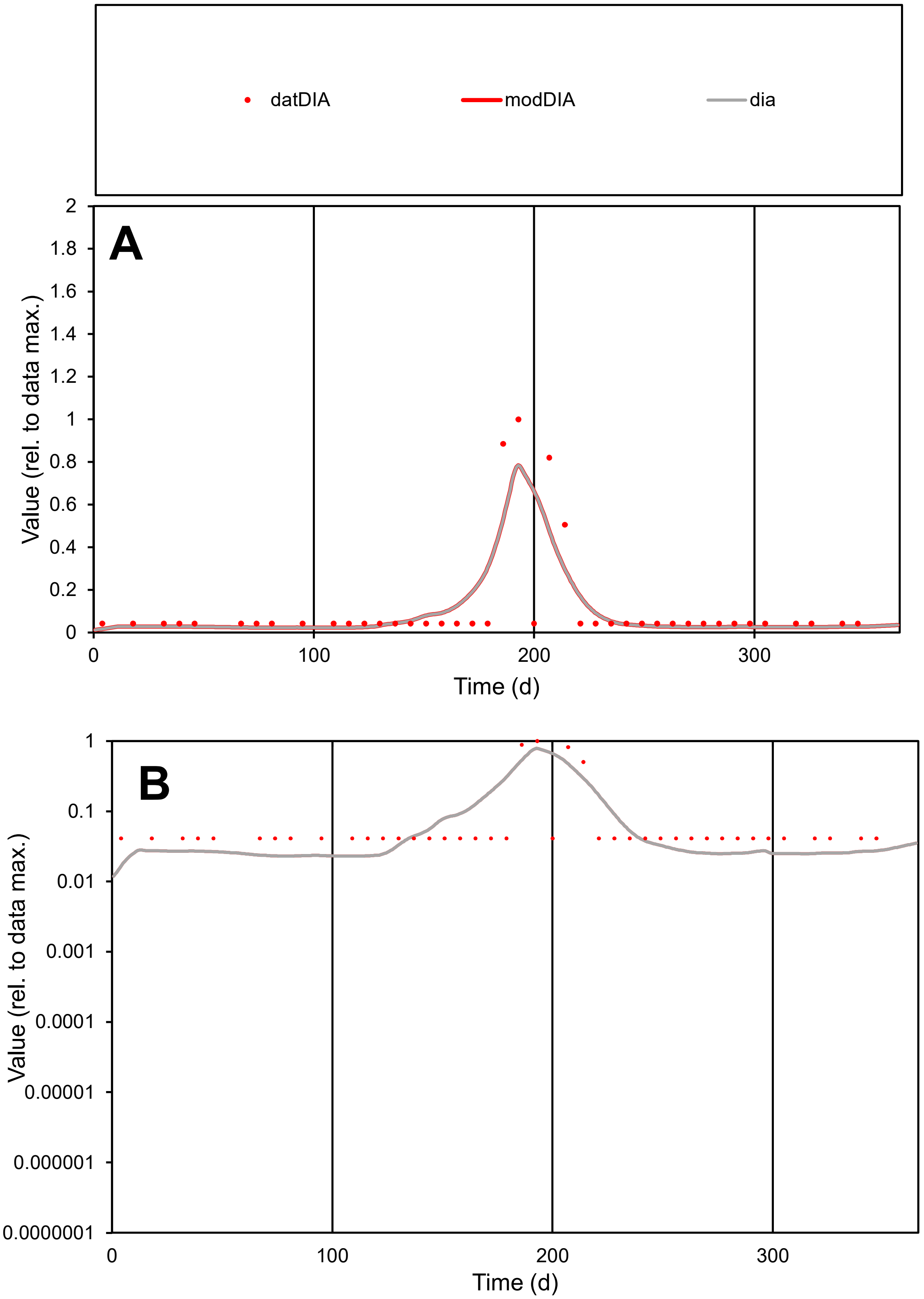


**C1**

Variables
DIA Provide jtd or Dst
CalibOnly?
2 1 to reproduce of
oNormalize?
2 1=no, 2=yes

ofxn calc*
1.17237
ofxn mod
1.2275

sRun
RunA1
zout
1

Ave
Data
1.7E-05
Model
1.7E-05

data/vMax
0.00015

ofxng
334.07

Min d/m
2.26849

ave/min d
2.7571
ave/min m
6.02542

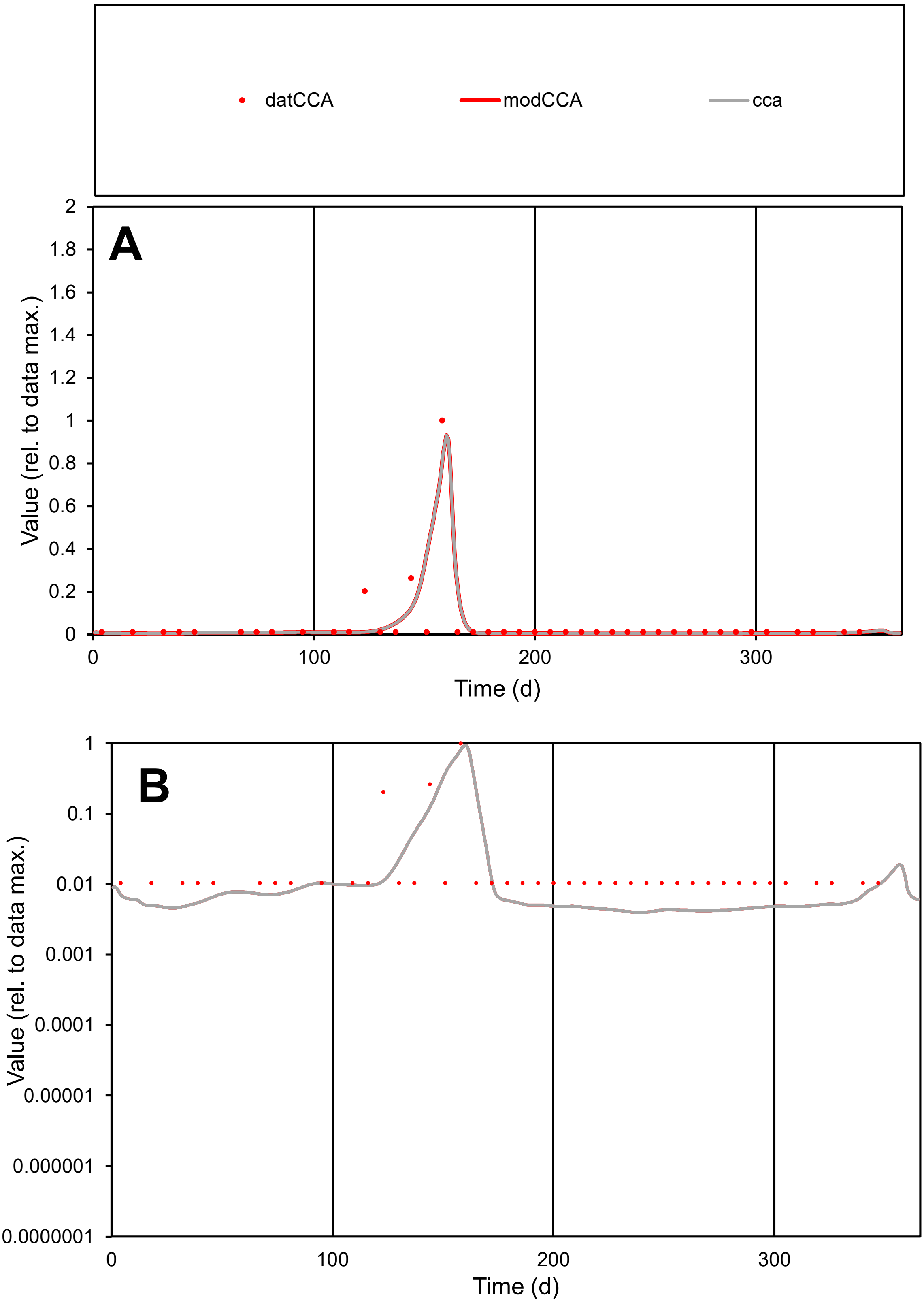


**C1**

Variables

CCA Provide jtd or Dst

CalibOnly?

2 1 to reproduce of

oNormalize?

2 1=no, 2=yes

ofxn calc*

1.29009

ofxn mod

1.45931

sRun

RunA1

zout

1

Ave

Data

7.8E-06

Model

7.1E-06

data/vMax

0.00018

ofxng

334.07

Min d/m

2.59034

ave/min d

4.28822

ave/min m

10.1098

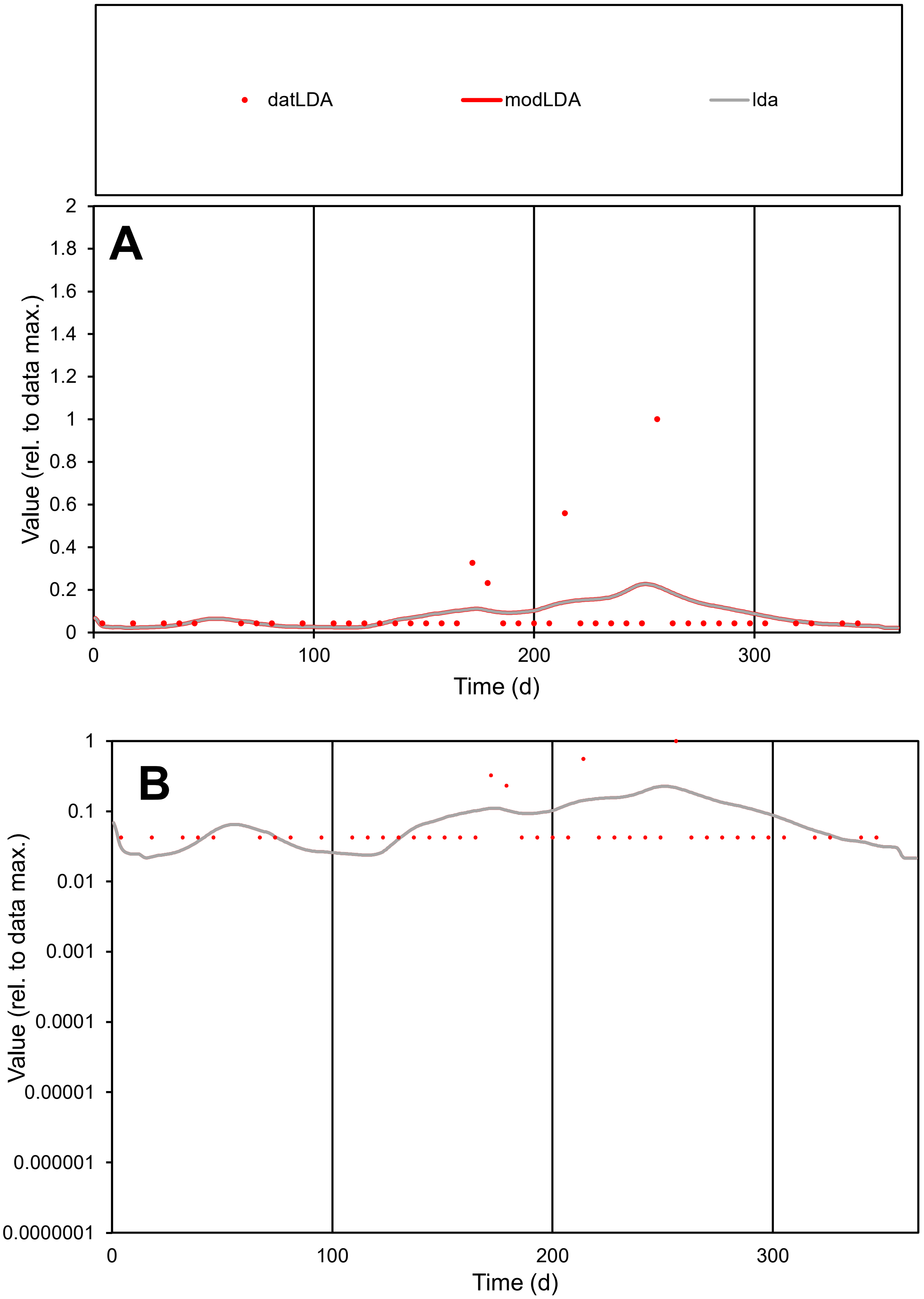


**C1**

Variables
LDA Provide jtd or Dst
CalibOnly?
2 1 to reproduce of
oNormalize?
2 1=no, 2=yes

ofxn calc*
1.76701
ofxn mod
1.77774

sRun
RunA1
zout
1

Ave
Data
1.4E-05
Model
1.5E-05

data/vMax
0.00016

ofxng
334.07

Min d/m
1.83901

ave/min d
2.09489
ave/min m
3.88584

327_Calib_S_A

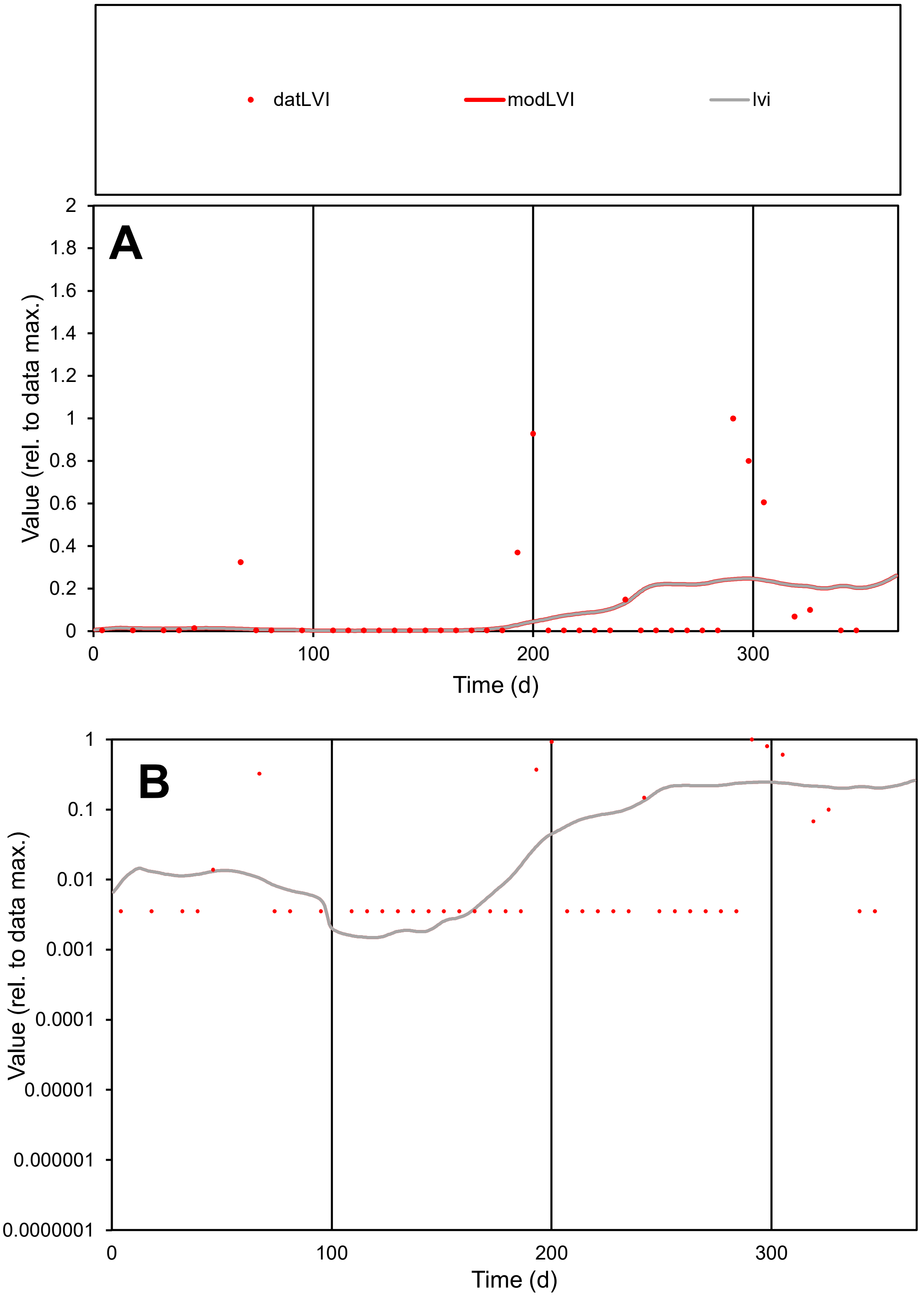


**C1**

Variables
LVI Provide jtd or Dst
CalibOnly?
2 1 to reproduce of
oNormalize?
2 1=no, 2=yes

ofxn calc*
2.25495
ofxn mod
2.26762

sRun
RunA1
zout
1

Ave
Data
4.3E-05
Model
3.5E-05

data/vMax
0.0004

ofxng
334.07

Min d/m
2.37753

ave/min d
30.0642
ave/min m
57.9973

327_Calib_S_A

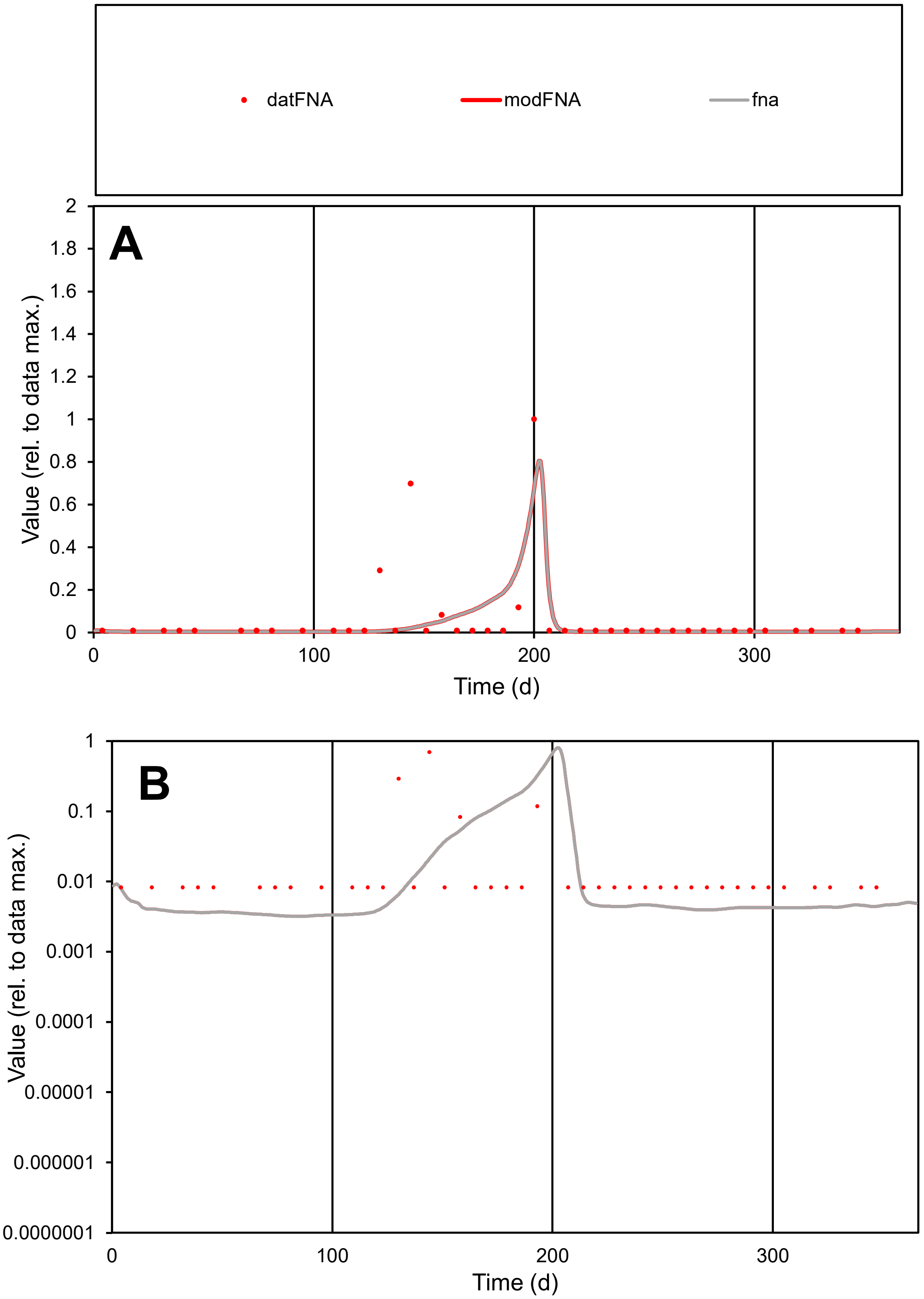


**C1**

Variables
FNA Provide jtd or Dst
CalibOnly?
2 1 to reproduce of
oNormalize?
2 1=no, 2=yes

ofxn calc*
2.15528
ofxn mod
2.20691

sRun
RunA1
zout
1

Ave
Data
2.2E-05
Model
1.7E-05

data/vMax
0.00038

ofxng
334.07

Min d/m
2.58541
ave/min d
7.19429
ave/min m
14.1402

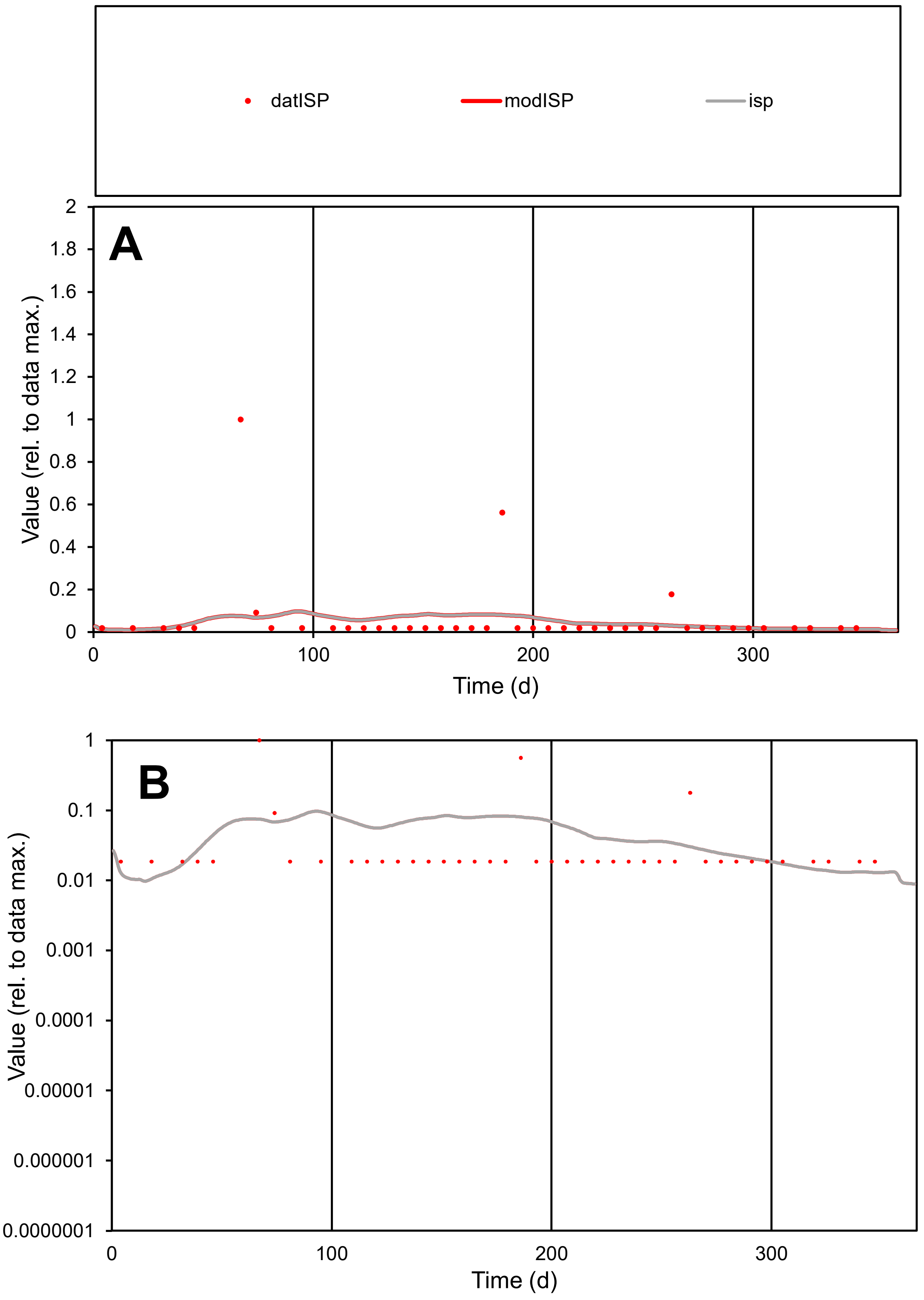


**C1**

Variables
ISP Provide jtd or Dst
CalibOnly?
2 1 to reproduce of
oNormalize?
2 1=no, 2=yes

ofxn calc*
2.75805
ofxn mod
2.76768

sRun
RunA1
zout
1

Ave
Data
1.6E-05
Model
1.3E-05

data/vMax
0.00027

ofxng
334.07

Min d/m
1.67397

ave/min d
3.24926
ave/min m
4.2661

327_Calib_S_A

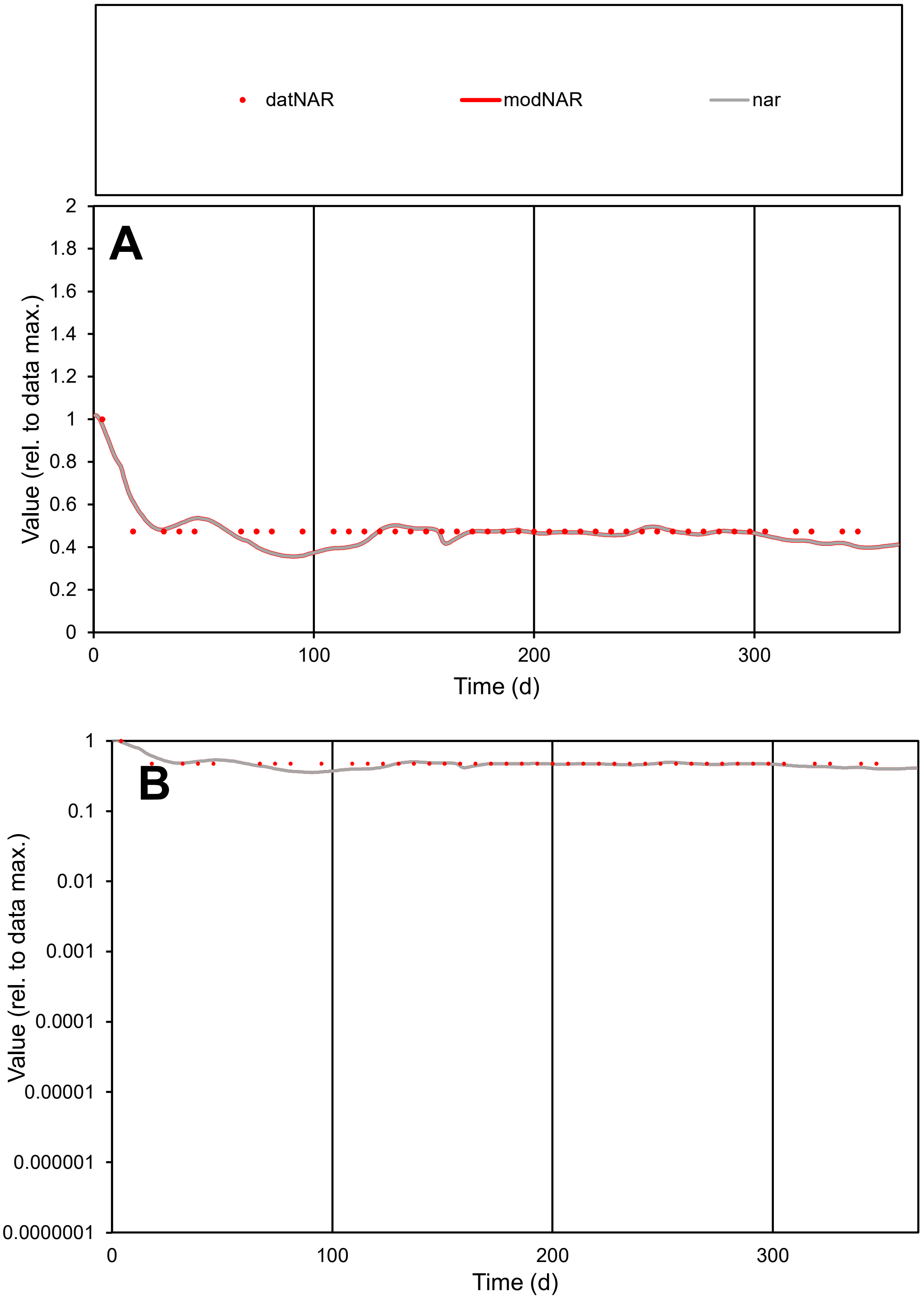


**C1**

Variables
NAR Provide jtd or Dst
CalibOnly?
2 1 to reproduce of
oNormalize?
2 1=no, 2=yes

ofxn calc*
0.0925
ofxn mod
0.14536

sRun
RunA1
zout
1

Ave
Data
2.9E-06
Model
2.8E-06

data/vMax
6E-06

ofxng
334.07

Min d/m
1.30304

ave/min d
1.02657
ave/min m
1.29752

327_Calib_S_A

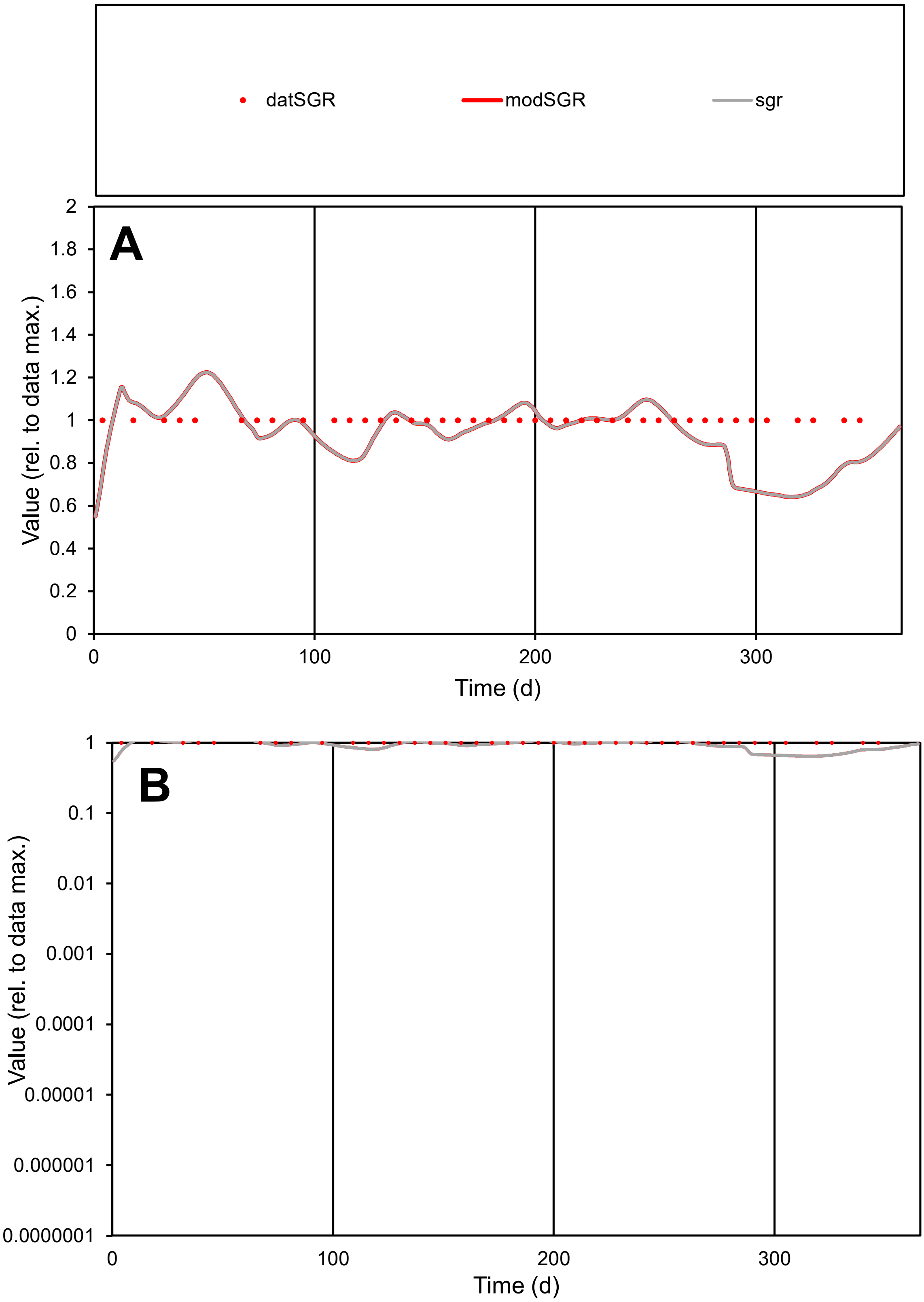


C1

Variables
SGR Provide jtd or Dst
CalibOnly?
2 1 to reproduce of
oNormalize?
2 1=no, 2=yes

ofxn calc*
0.14631
ofxn mod
0.18367

sRun
RunA1
zout
1

Ave
Data
4.5E-06
Model
4.2E-06

data/vMax
4.5E-06

ofxng
334.07

Min d/m
1.54937

ave/min d
1
ave/min m
1.45063

327_Calib_S_A

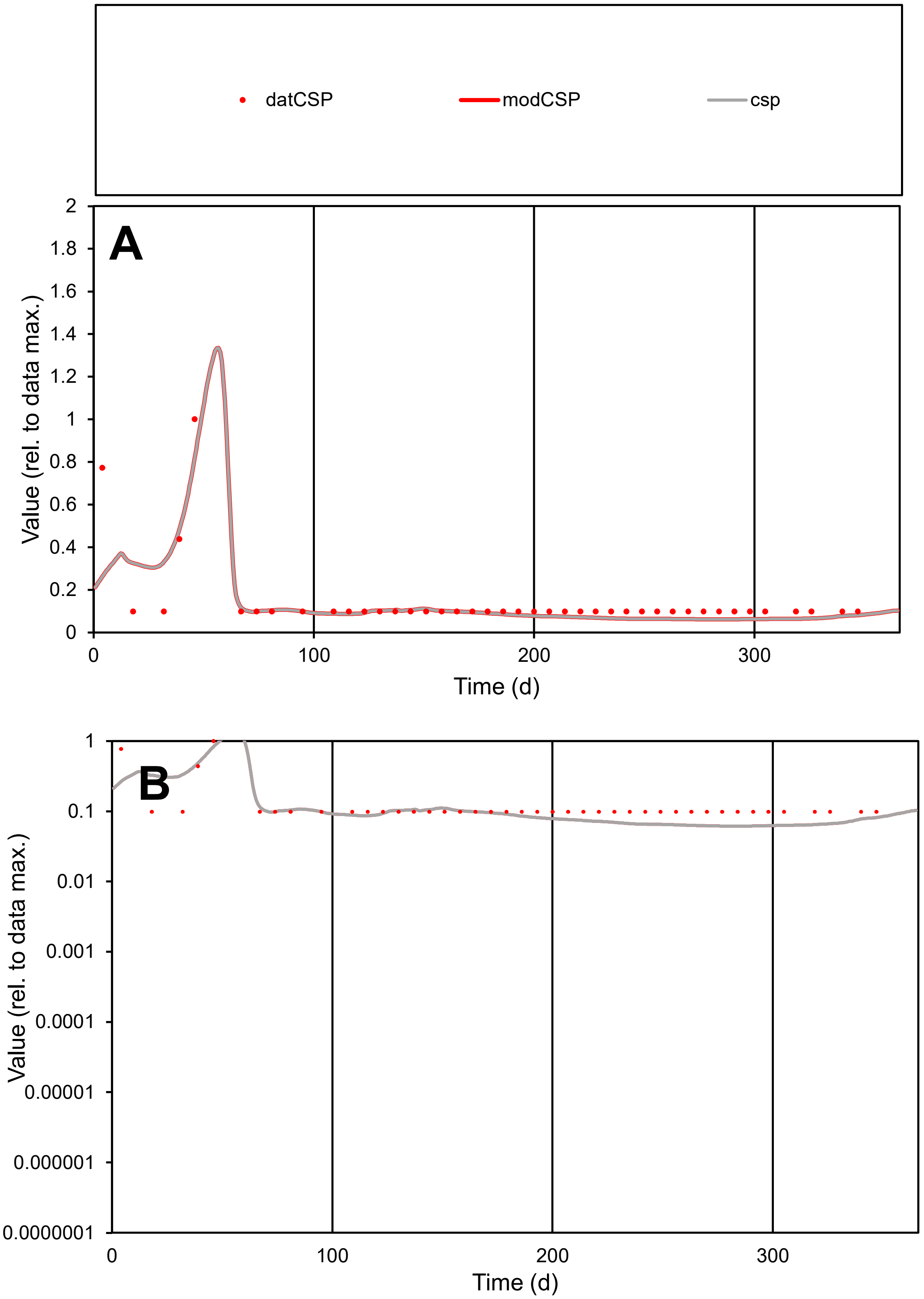


**C1**

Variables
CSP Provide jtd or Dst
CalibOnly?
2 1 to reproduce of
oNormalize?
2 1=no, 2=yes

ofxn calc*
0.66608
ofxn mod
0.68706

sRun
RunA1
zout
1

Ave
Data
3.5E-06
Model
3.1E-06

data/vMax
2.4E-05

ofxng
334.07

Min d/m
1.58816

ave/min d
1.46521
ave/min m
2.06248

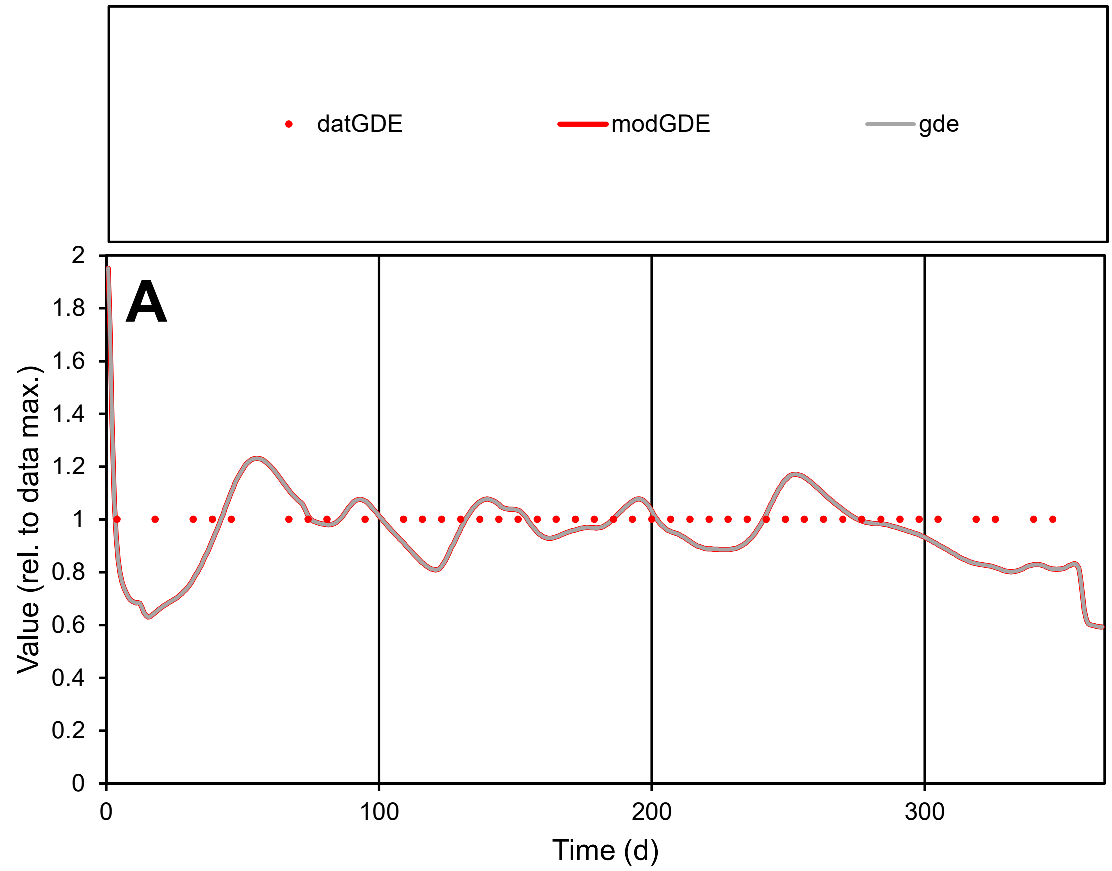


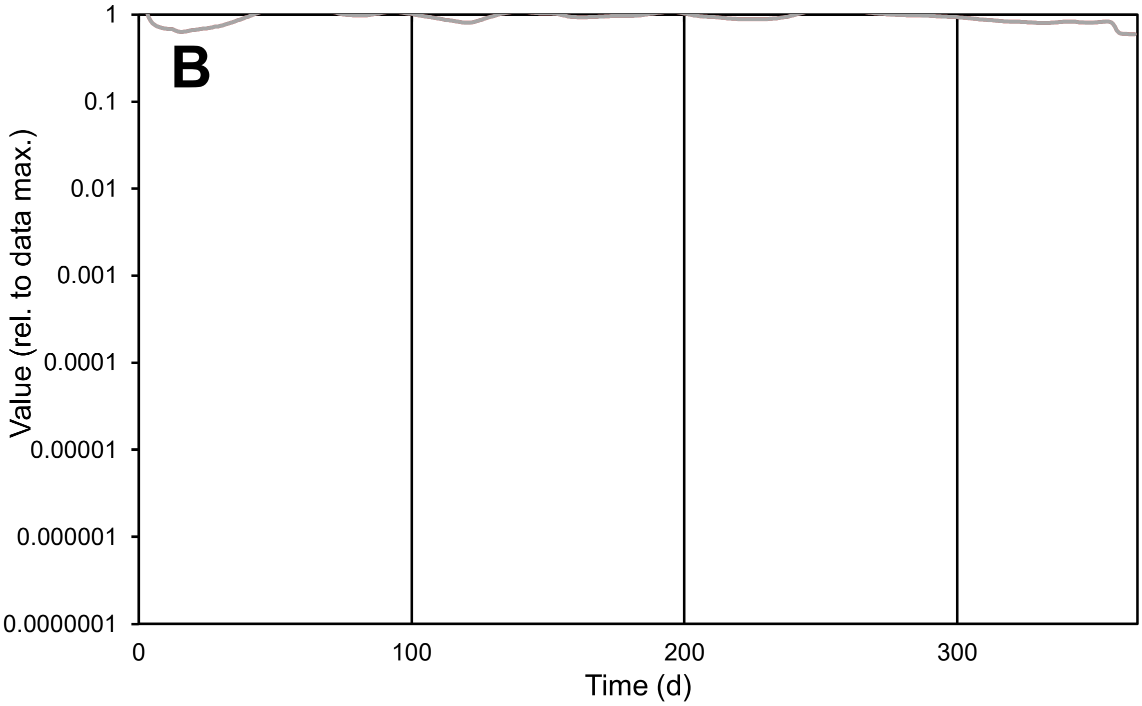


**C1**

Variables
GDE Provide jtd or Dst
CalibOnly?
2 1 to reproduce of
oNormalize?
2 1=no, 2=yes

ofxn calc*
0.11406
ofxn mod
0.16068

sRun
RunA1
zout
1

Ave
Data
2.7E-05
Model
2.6E-05

data/vMax
2.7E-05

ofxng
334.07

Min d/m
1.50989

ave/min d
1
ave/min m
1.44802

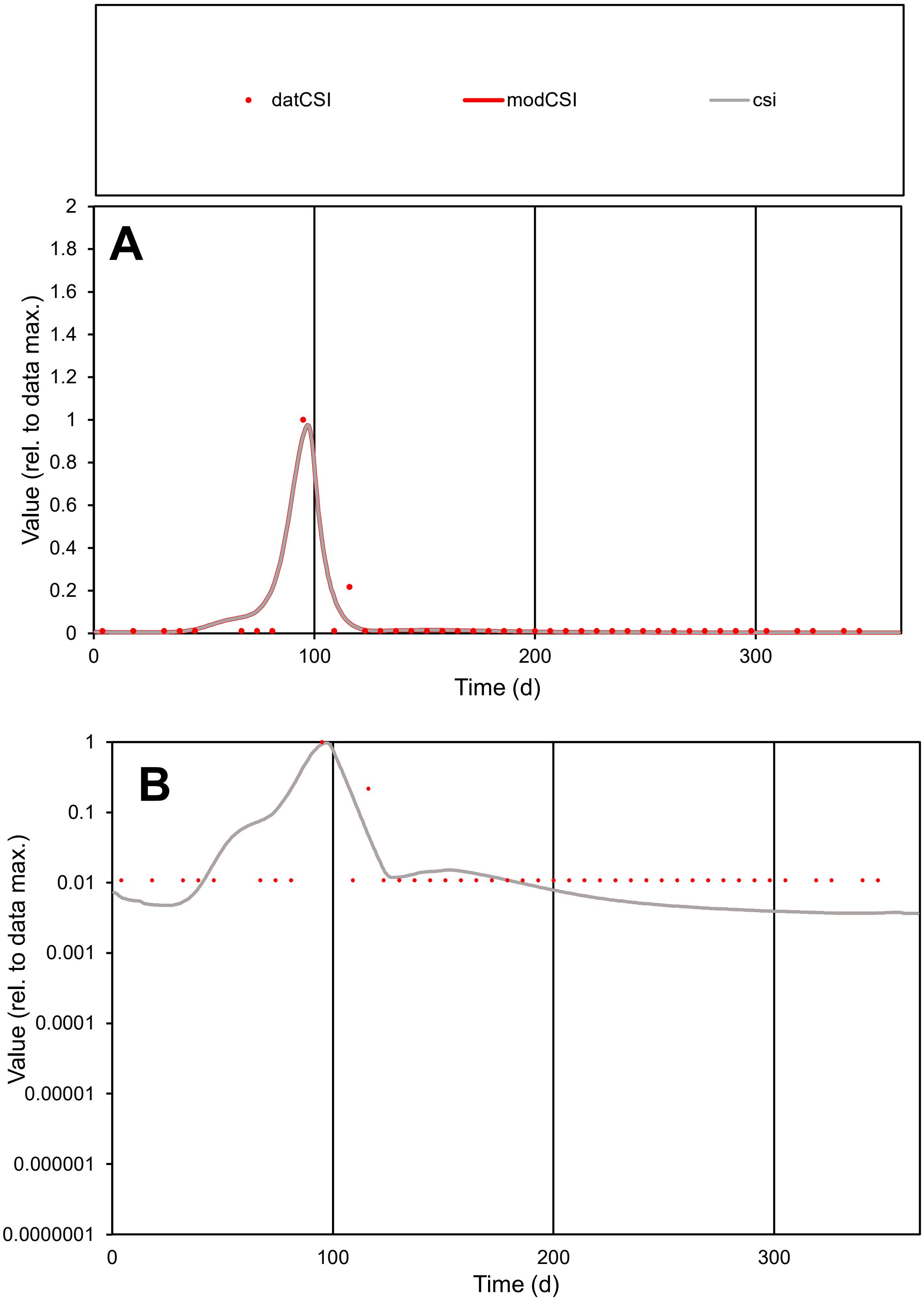


**C1**

Variables

CSI Provide jtd or Dst

CalibOnly?

2 1 to reproduce of

oNormalize?

2 1=no, 2=yes

ofxn calc*

1.10236

ofxn mod

1.20908

sRun

RunA1

zout

1

Ave

Data

8.4E-06

Model

8.8E-06

data/vMax

0.00021

ofxng

334.07

Min d/m

2.93557

ave/min d

3.62327

ave/min m

11.1869

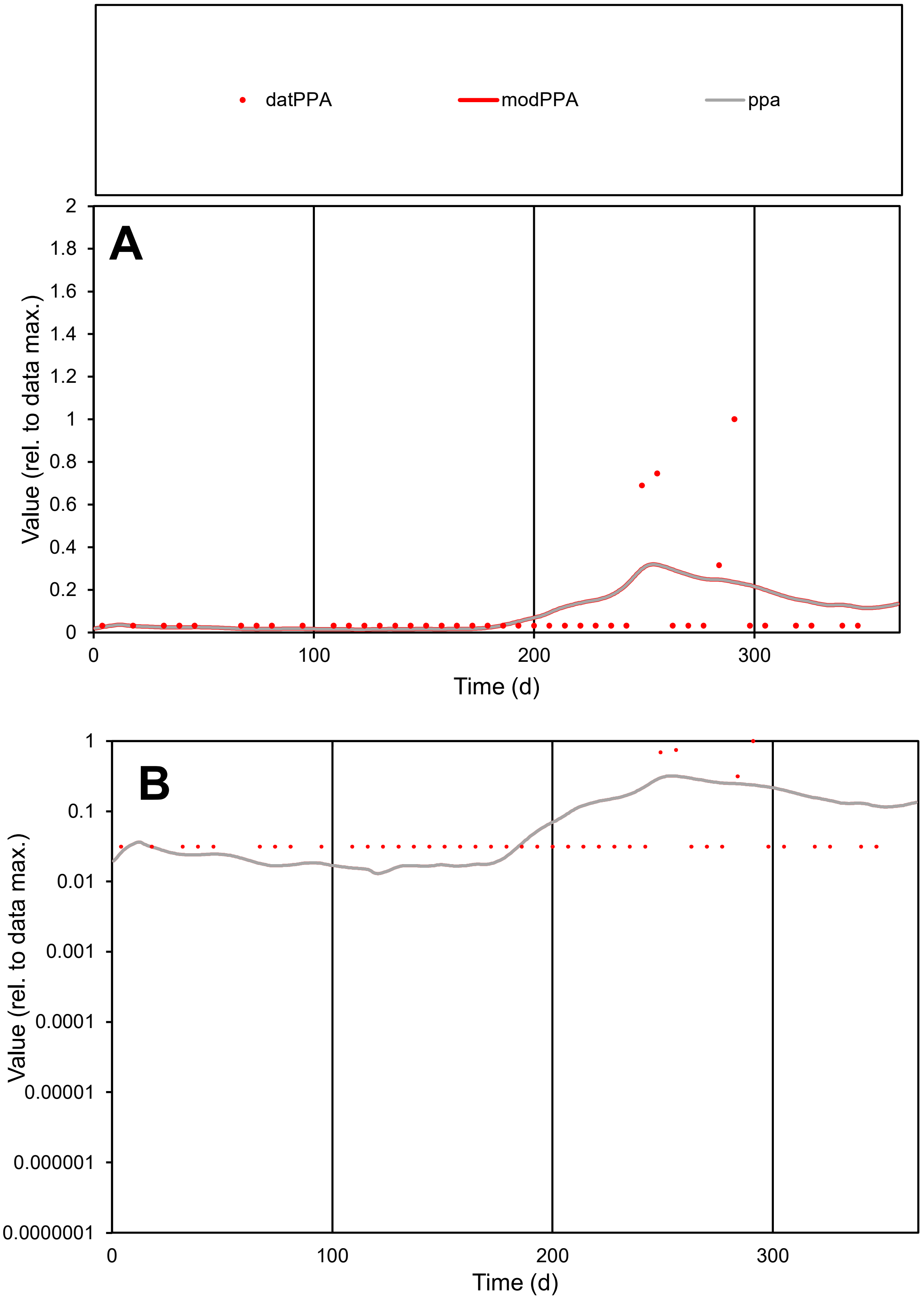


**C1**

Variables
PPA Provide jtd or Dst
CalibOnly?
2 1 to reproduce of
oNormalize?
2 1=no, 2=yes

ofxn calc*
1.87494
ofxn mod
1.88787

sRun
RunA1
zout
1

Ave
Data
9.6E-06
Model
1.1E-05

data/vMax
0.0001

ofxng
334.07

Min d/m
2.27247

ave/min d
2.99297
ave/min m
7.46086

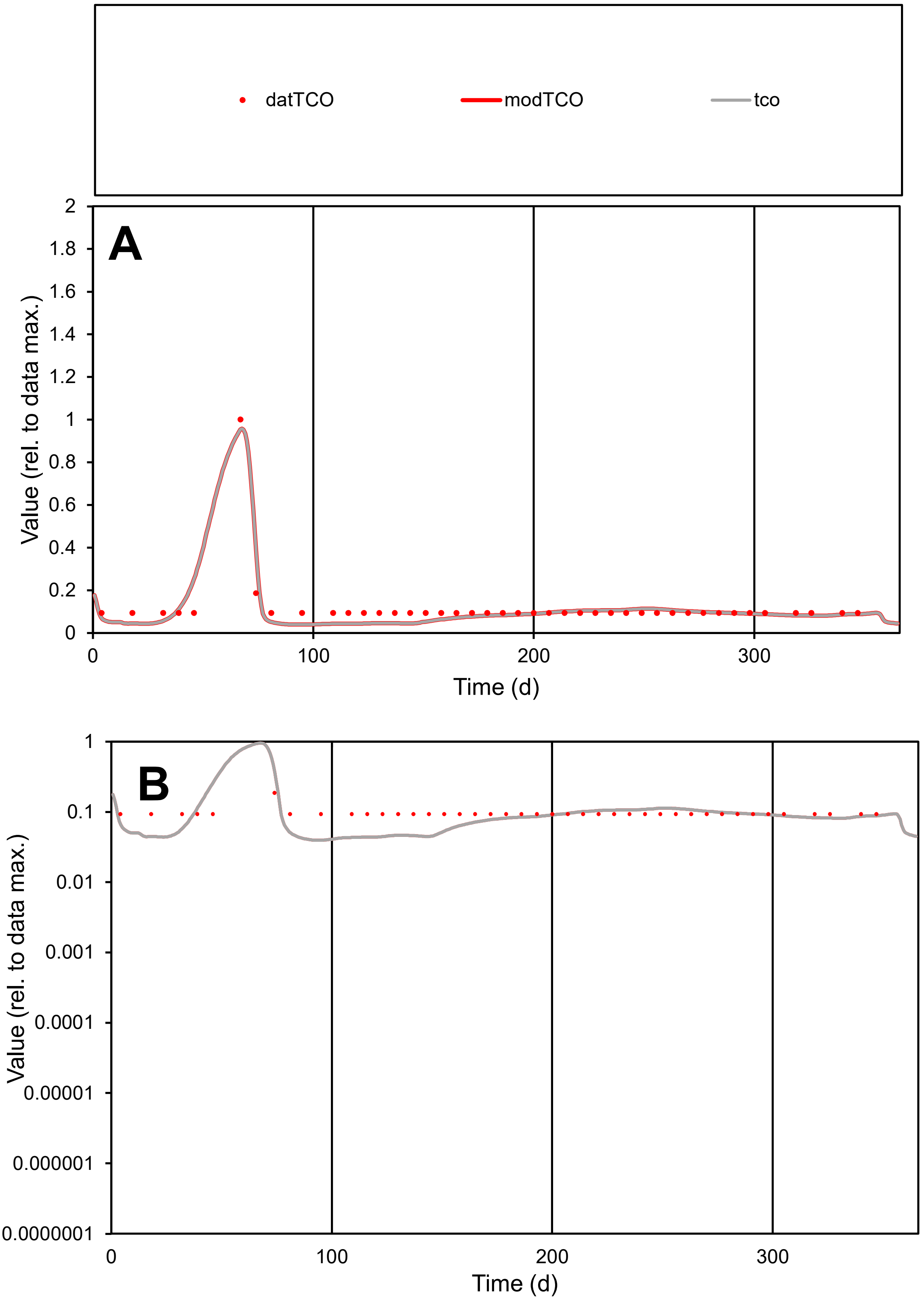


C1

Variables
TCO Provide jtd or Dst
CalibOnly?
2 1 to reproduce of
oNormalize?
2 1=no, 2=yes

ofxn calc*
0.29422
ofxn mod
0.37318

sRun
RunA1
zout
1

Ave
Data
1.1E-05
Model
9.9E-06

data/vMax
9.2E-05

ofxng
334.07

Min d/m
2.34576

ave/min d
1.25514
ave/min m
2.69227

327_Calib_S_A

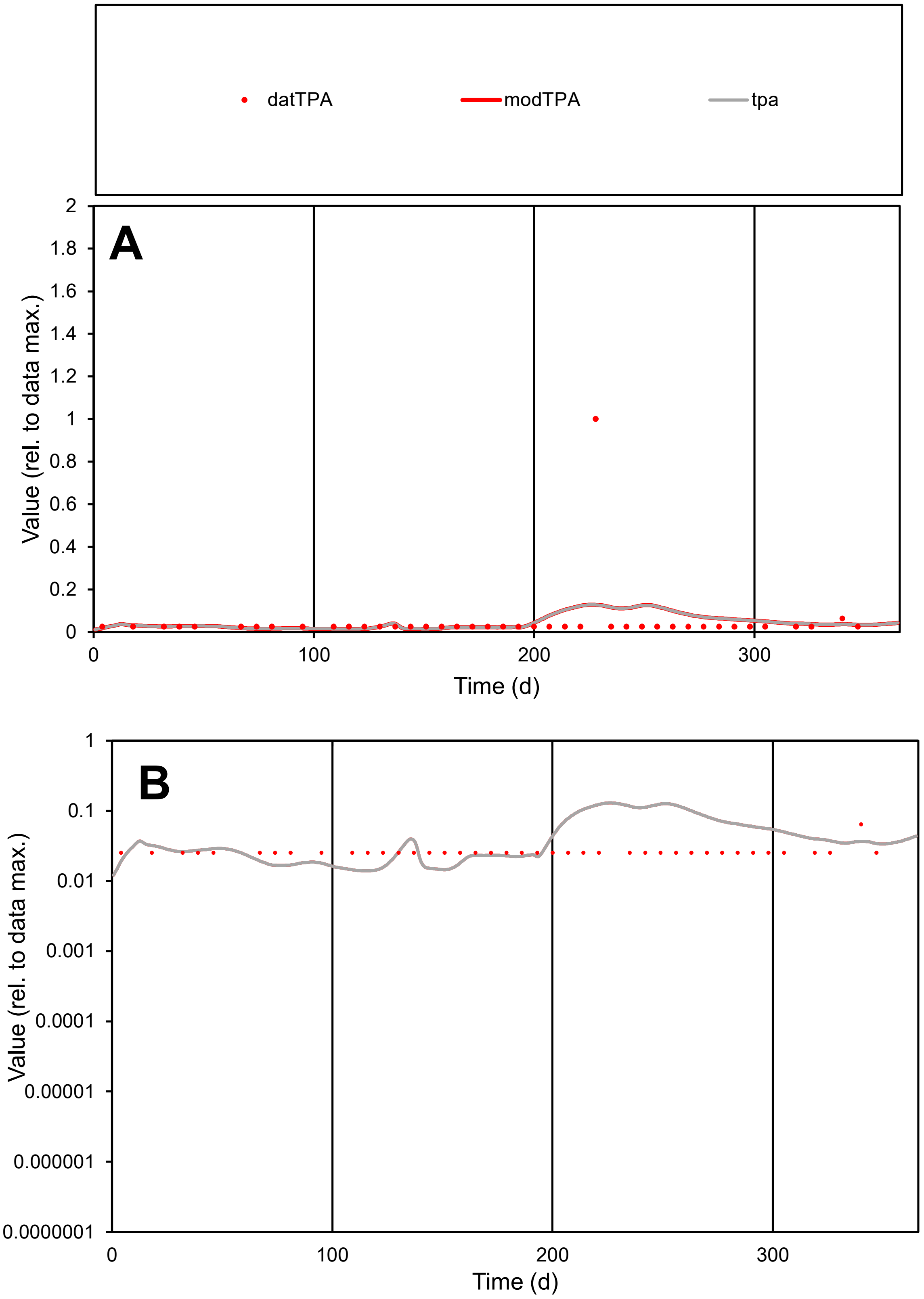


**C1**

Variables

TPA Provide jtd or Dst

CalibOnly?

2 1 to reproduce of

oNormalize?

2 1=no, 2=yes

ofxn calc*

2.84805

ofxn mod

2.85437

sRun

RunA1

zout

1

Ave

Data

9.2E-06

Model

9.1E-06

data/vMax

0.00019

ofxng

334.07

Min d/m

1.80719

ave/min d

1.95587

ave/min m

3.4837

327_Calib_S_A

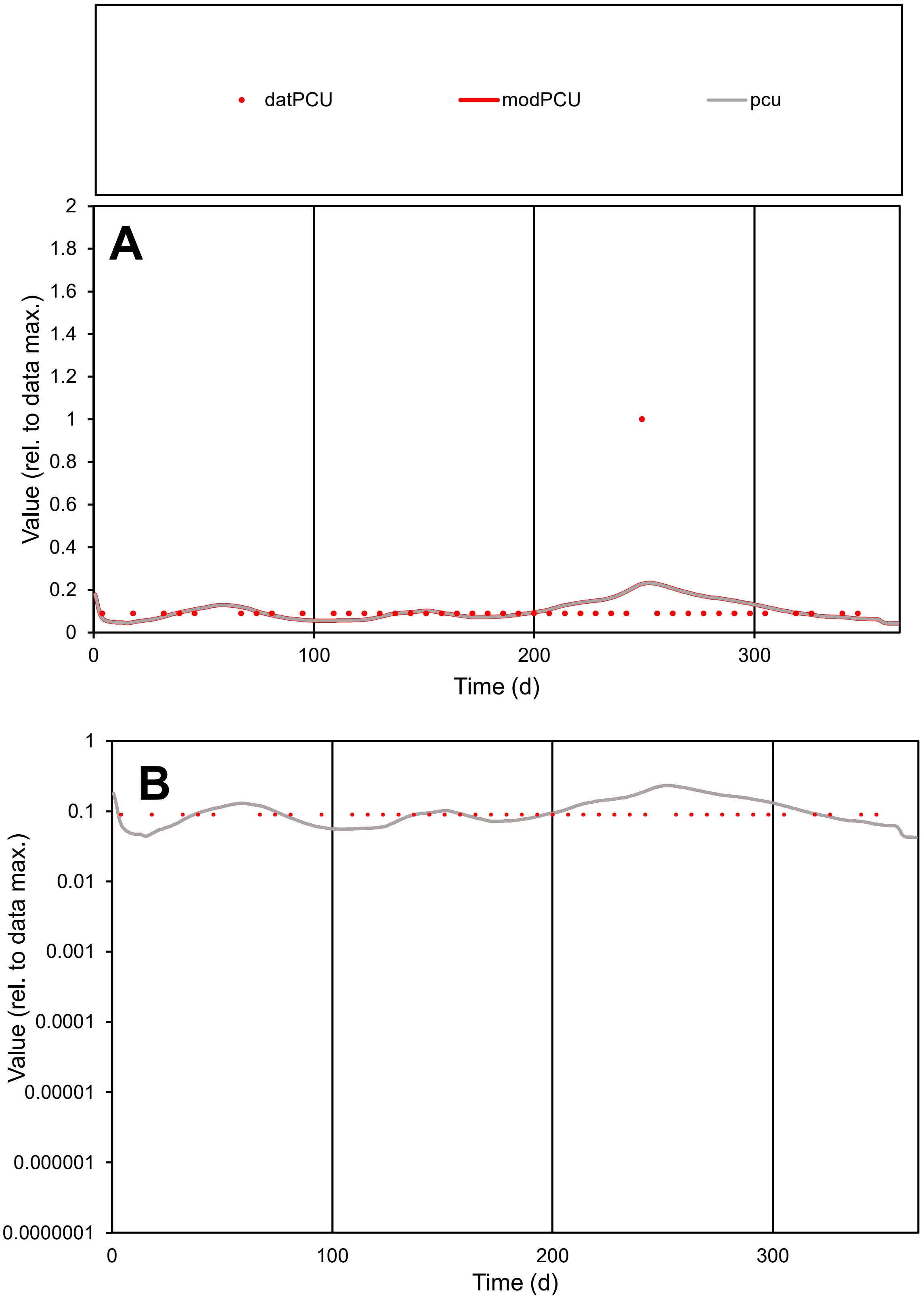


**C1**

Variables
PCU Provide jtd or Dst
CalibOnly?
2 1 to reproduce of
oNormalize?
2 1=no, 2=yes

ofxn calc*
1.14716
ofxn mod
1.15586

sRun
RunA1
zout
1

Ave
Data
7.7E-06
Model
7.6E-06

data/vMax
6.9E-05

ofxng
334.07

Min d/m
1.74972

ave/min d
1.24231
ave/min m
2.15426

327_Calib_S_A

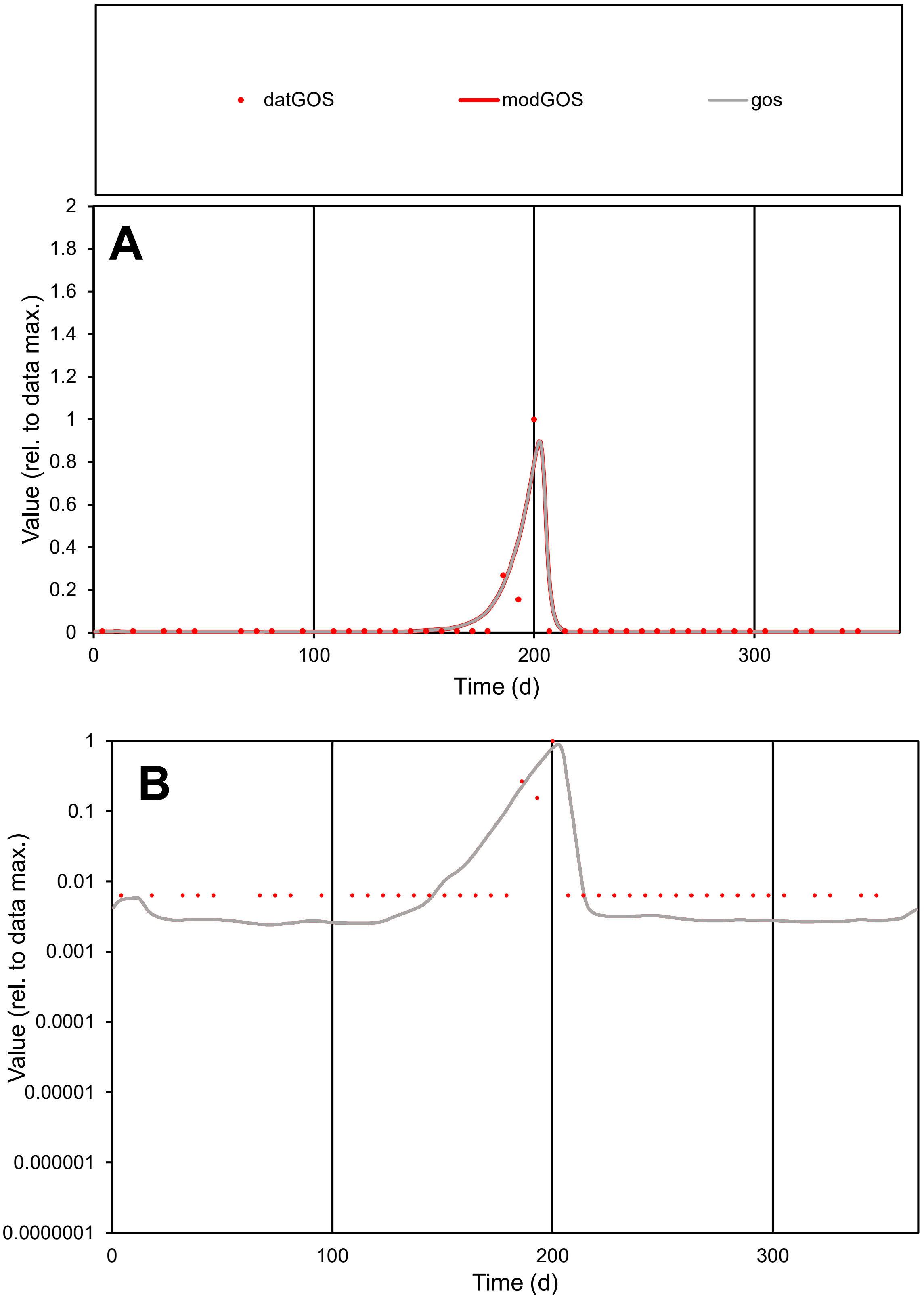


**C1**

Variables

GOS Provide jtd or Dst

CalibOnly?

2 1 to reproduce of

oNormalize?

2 1=no, 2=yes

ofxn calc*

1.2646

ofxn mod

1.49257

sRun

RunA1

zout

1

Ave

Data

1.1E-05

Model

1.3E-05

data/vMax

0.00028

ofxng

334.07

Min d/m

2.60945

ave/min d

6.24327

ave/min m

18.7724

327_Calib_S_A

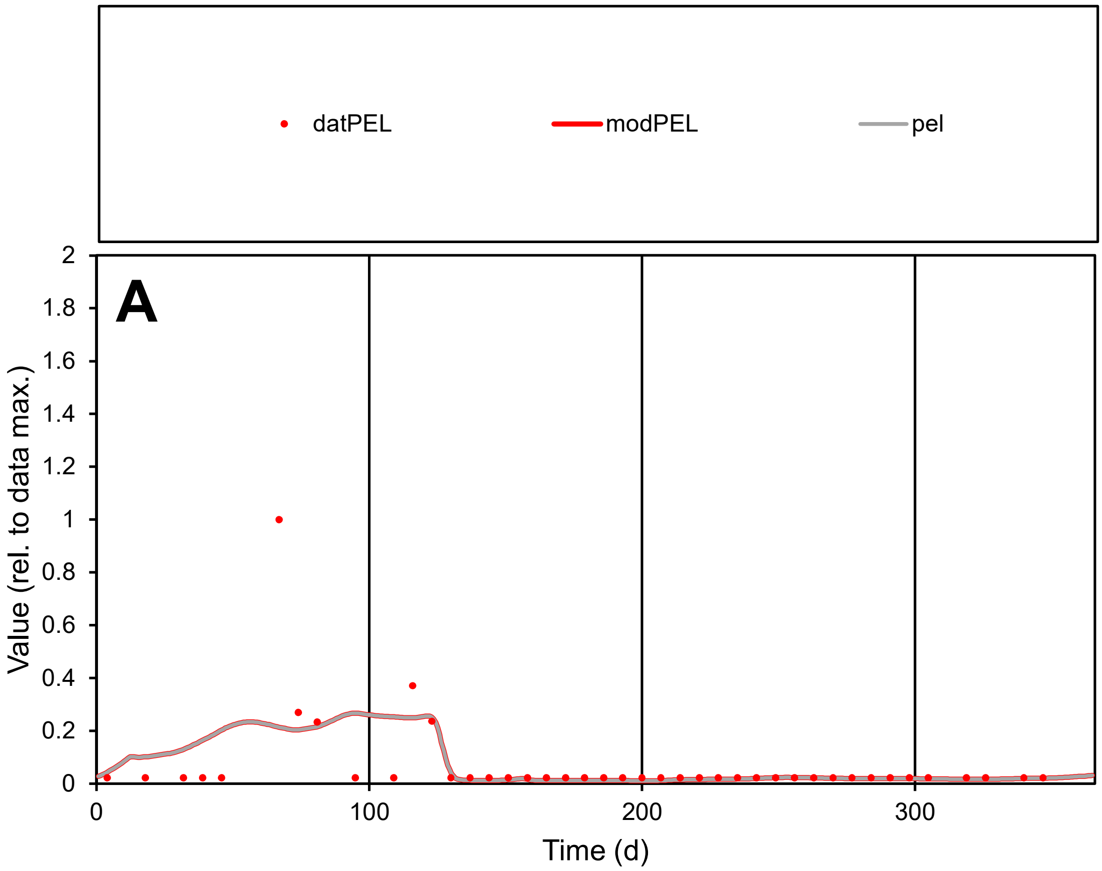


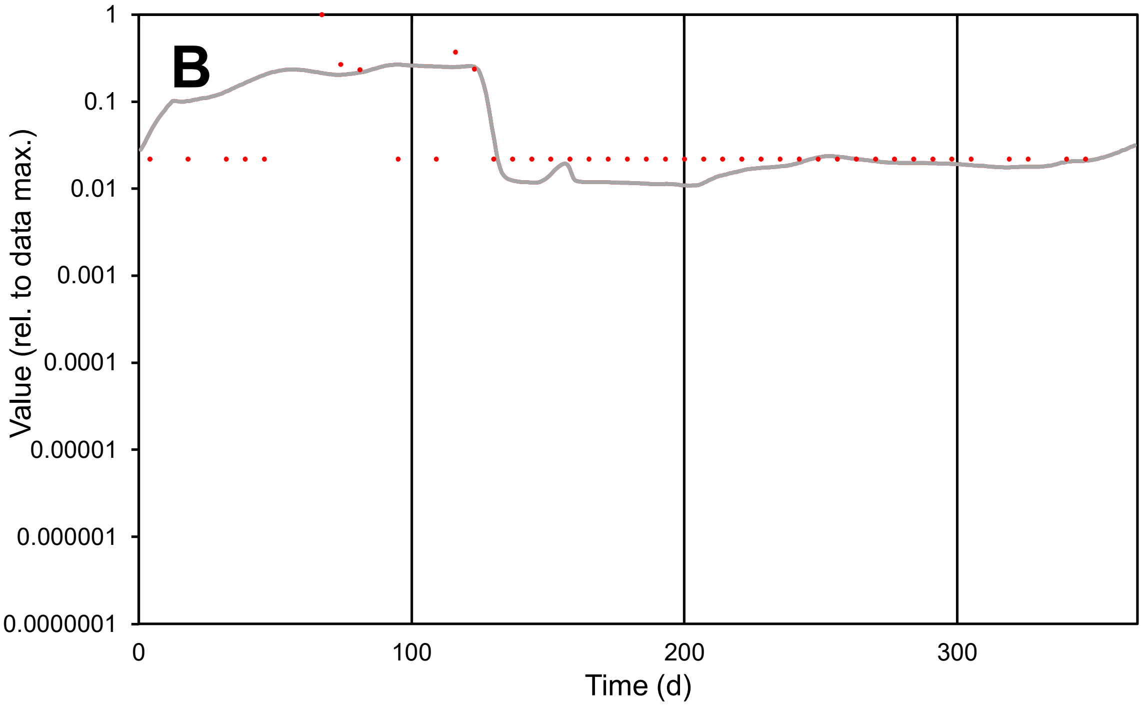


**C1**

Variables
PEL Provide jtd or Dst
CalibOnly?
2 1 to reproduce of
oNormalize?
2 1=no, 2=yes

ofxn calc*
2.02511
ofxn mod
2.03708

sRun
RunA1
zout
1

Ave
Data
1E-05
Model
1E-05

data/vMax
0.00015

ofxng
334.07

Min d/m
2.01597

ave/min d
3.17681
ave/min m
6.14312

327_Calib_S_A

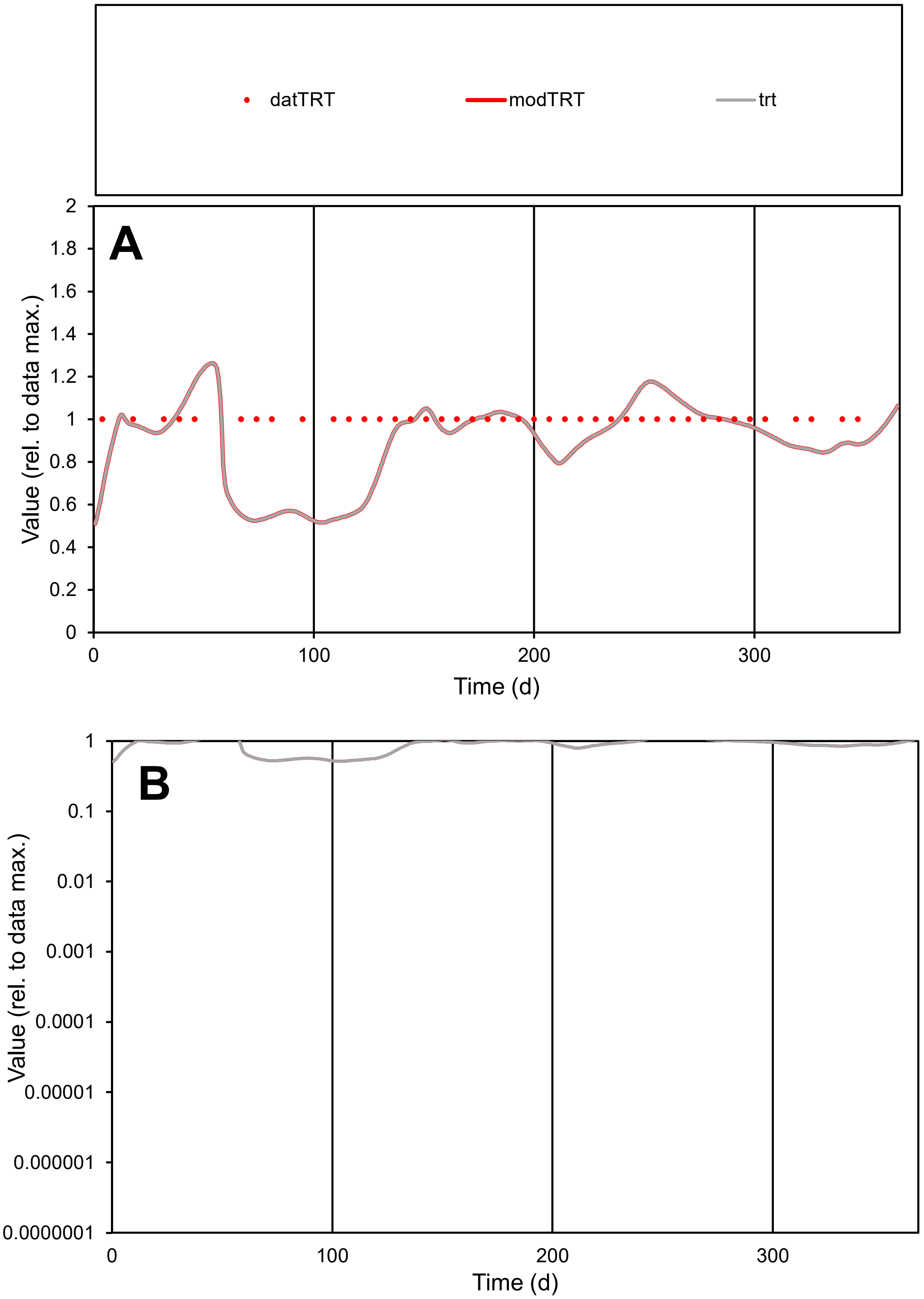


**C1**

Variables

TRT Provide jtd or Dsı

CalibOnly?

2 1 to reproduce of

oNormalize?

2 1=no, 2=yes

ofxn calc*

0.20647

ofxn mod

0.23378

sRun

RunA1

zout

1

Ave

Data

2.3E-05

Model

2.1E-05

data/vMax

2.3E-05

ofxng

334.07

Min d/m

1.88989

ave/min d

1

ave/min m

1.70557

327_Calib_S_A

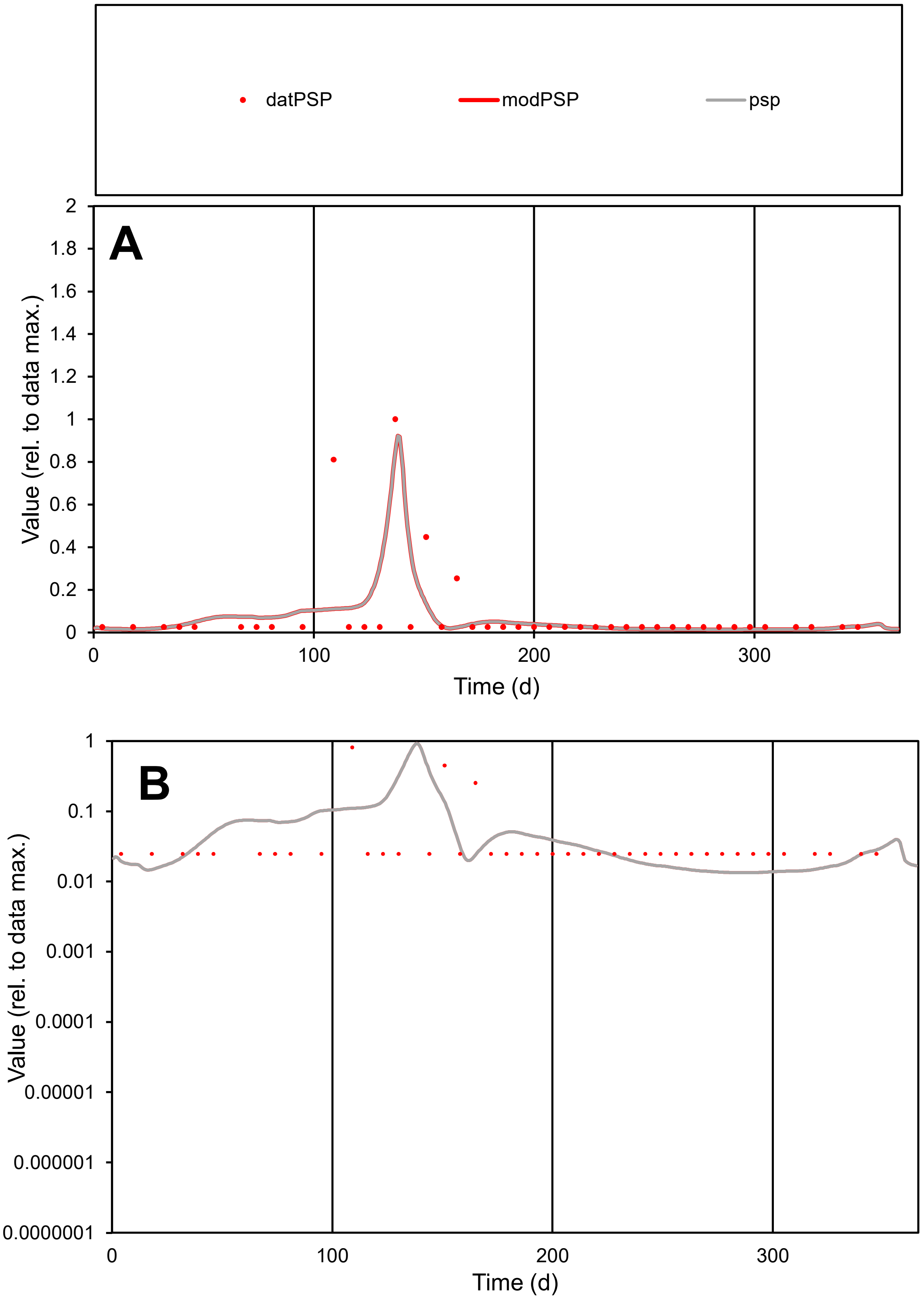


| | |
|---|---|
| **C1** | |
| Variables | |
| PSP | Provide jtd or Dst |
| CalibOnly? | |
| 2 | 1 to reproduce of |
| oNormalize? | |
| 2 | 1=no, 2=yes |
| ofxn calc* | |
| 1.6436 | |
| ofxn mod | |
| 1.70471 | |
| sRun | |
| RunA1 | |
| zout | |
| 1 | |
| Ave | |
| Data | |
| 1.4E-05 | |
| Model | |
| 1.2E-05 | |
| data/vMax | |
| 0.00018 | |
| ofxng | |
| 334.07 | |
| Min d/m | |
| 1.84336 | |
| ave/min d | |
| 3.3296 | |
| ave/min m | |
| 5.26512 | |

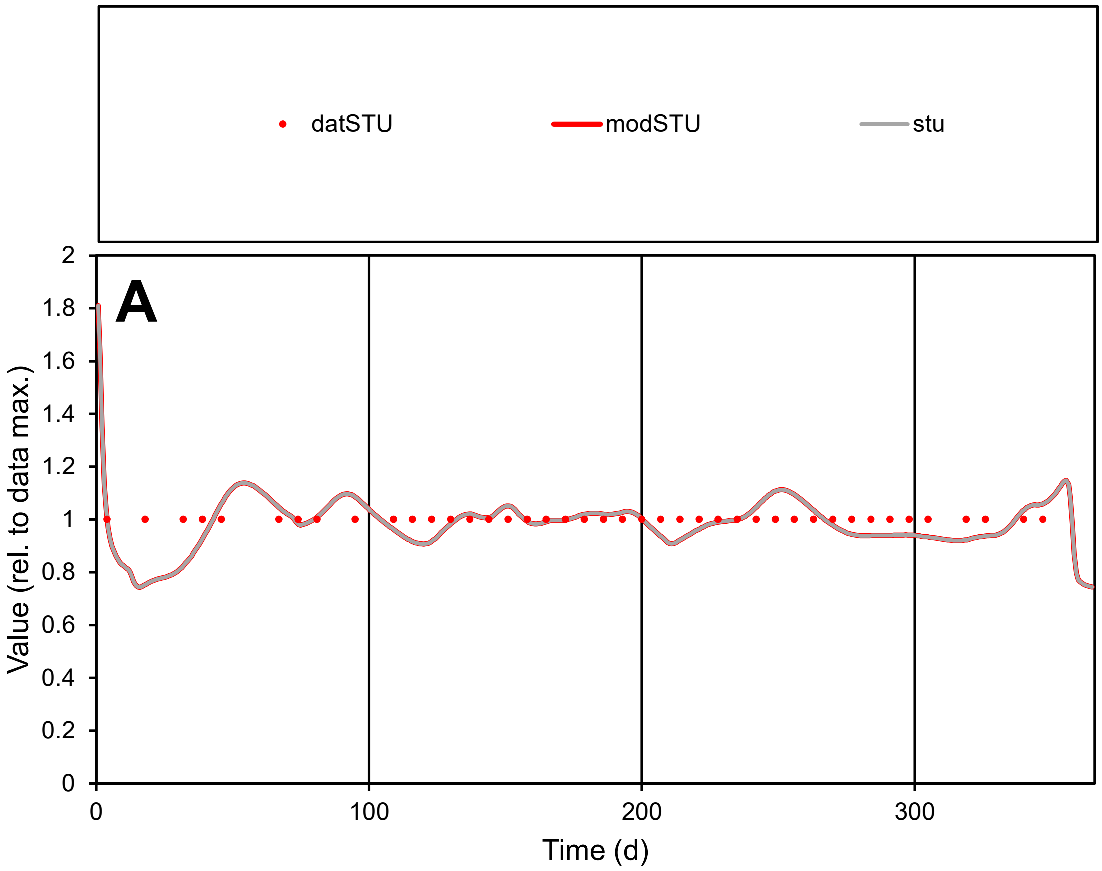


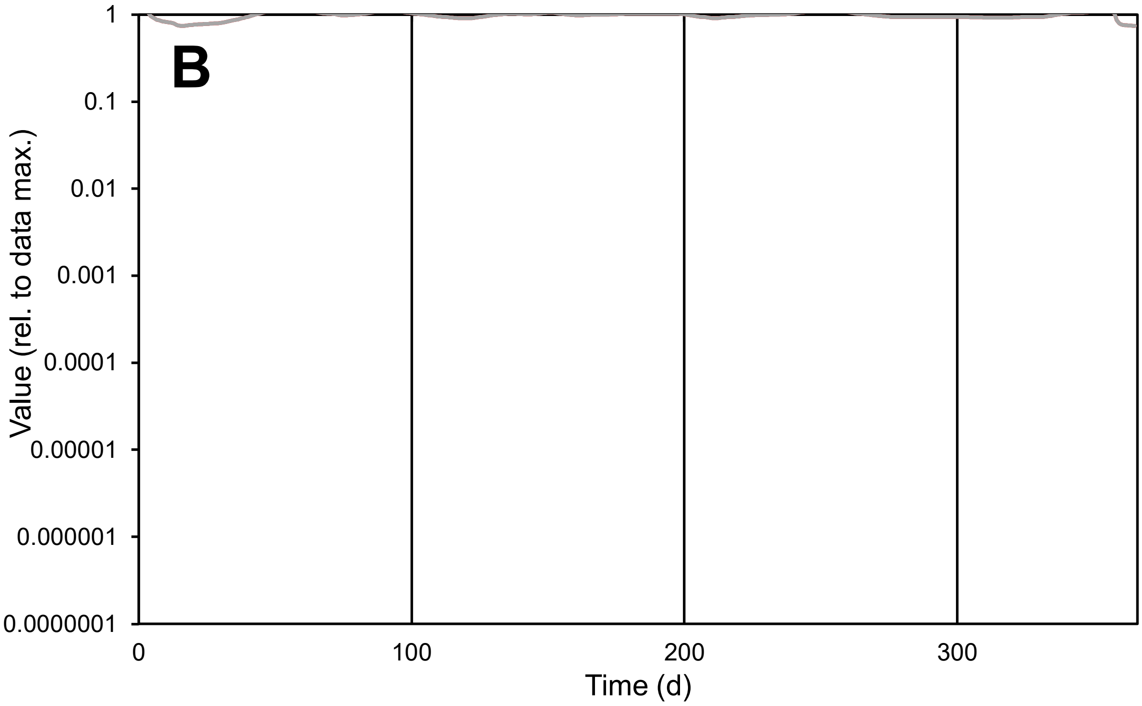


**C1**

Variables
STU Provide jtd or Dst
CalibOnly?
2 1 to reproduce of
oNormalize?
2 1=no, 2=yes

ofxn calc*
0.06833
ofxn mod
0.13148

sRun
RunA1
zout
1

Ave
Data
2.5E-06
Model
2.5E-06

data/vMax
2.5E-06

ofxng
334.07

Min d/m
1.31142

ave/min d
1
ave/min m
1.29102

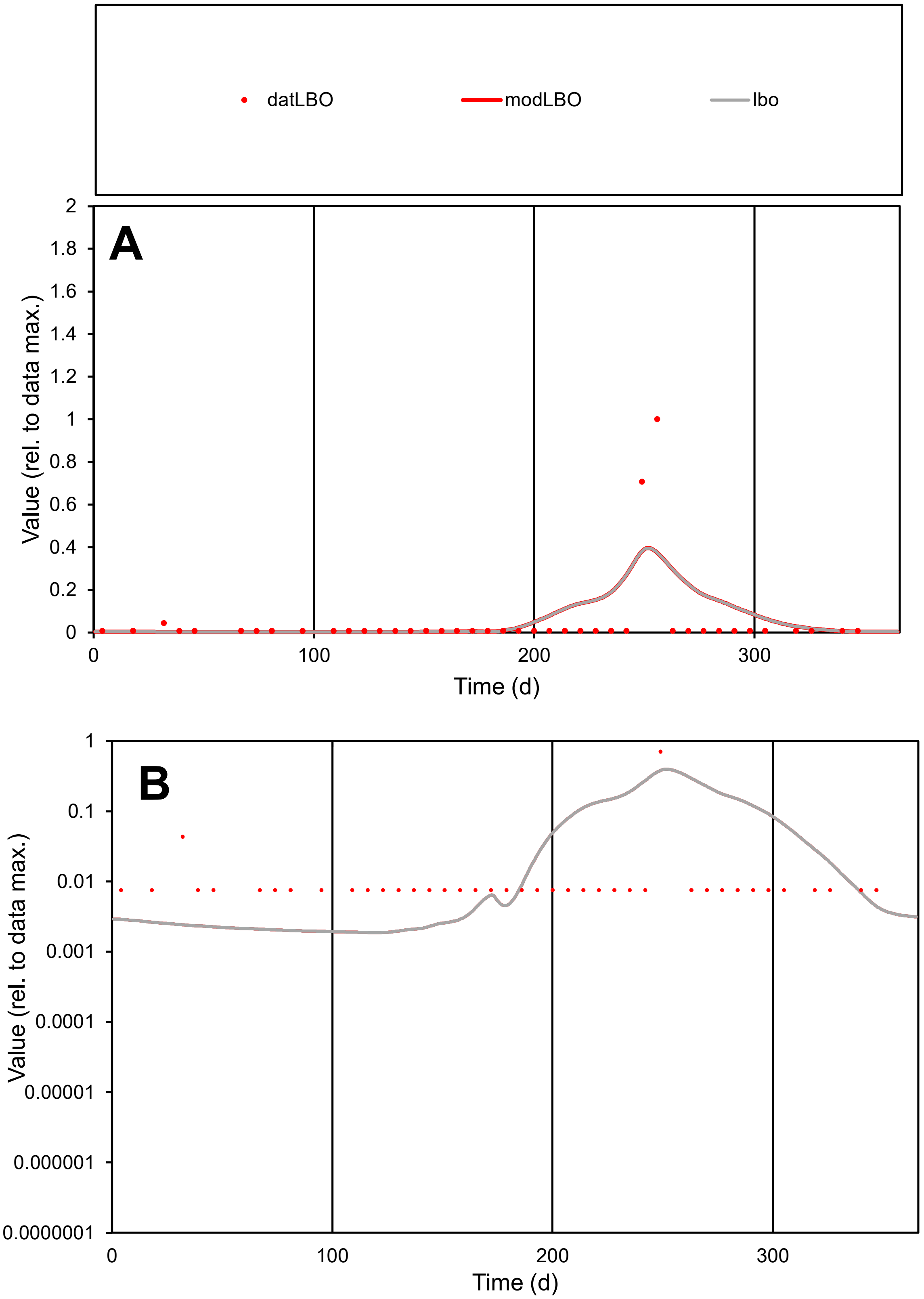


**C1**

Variables
LBO Provide jtd or Dst
CalibOnly?
2 1 to reproduce of
oNormalize?
2 1=no, 2=yes

ofxn calc*
2.87634
ofxn mod
2.93127

sRun
RunA1
zout
1

Ave
Data
8.9E-06
Model
1.3E-05

data/vMax
0.00018

ofxng
334.07

Min d/m
4.0308

ave/min d
6.43631
ave/min m
36.6842

327_Calib_S_A

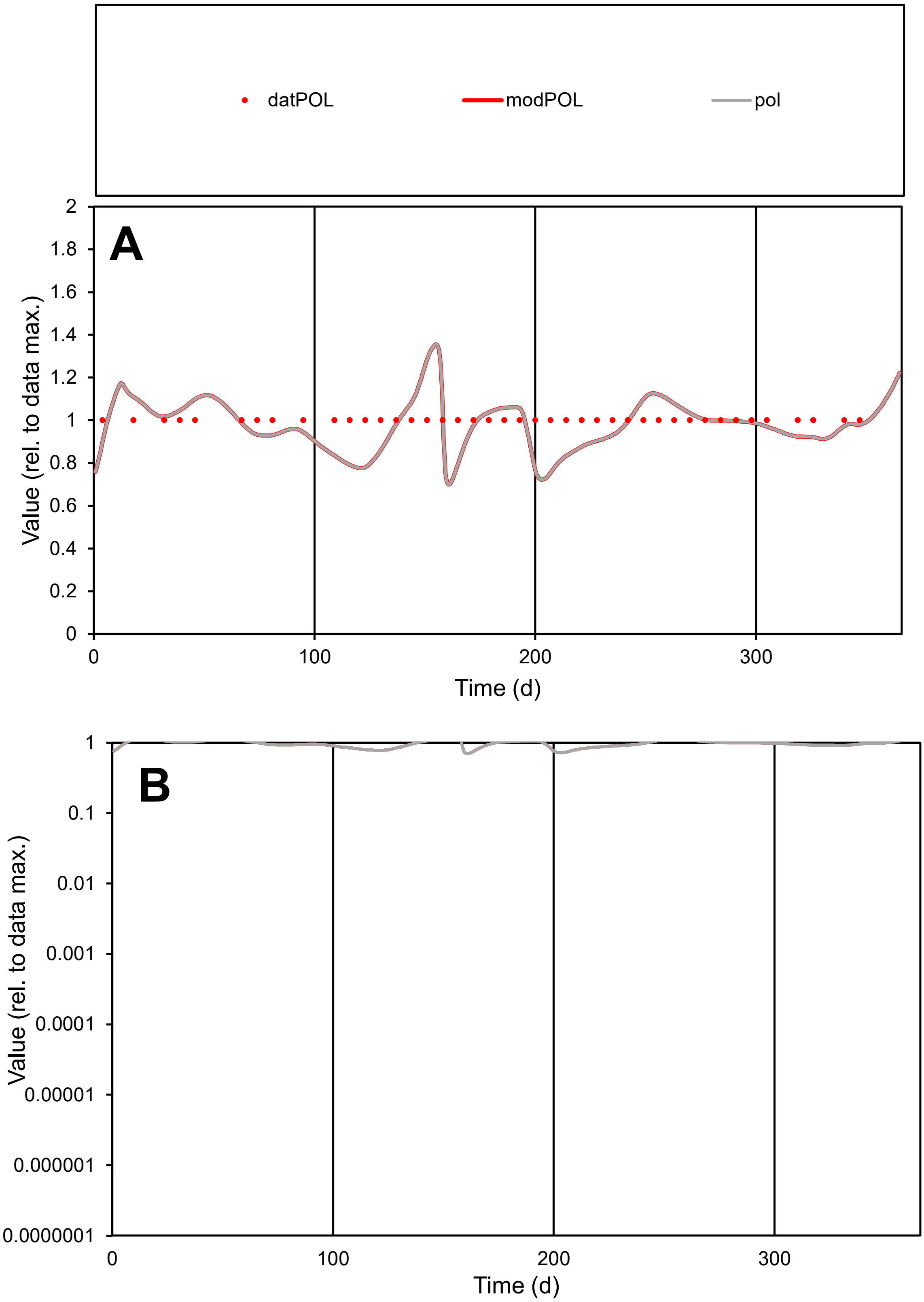


**C1**

Variables
POL Provide jtd or Dst
CalibOnly?
2 1 to reproduce of
oNormalize?
2 1=no, 2=yes

ofxn calc*
0.11278
ofxn mod
0.16017

sRun
RunA1
zout
1

Ave
Data
1.6E-06
Model
1.6E-06

data/vMax
1.6E-06

ofxng
334.07

Min d/m
1.3423

ave/min d
1
ave/min m
1.3041

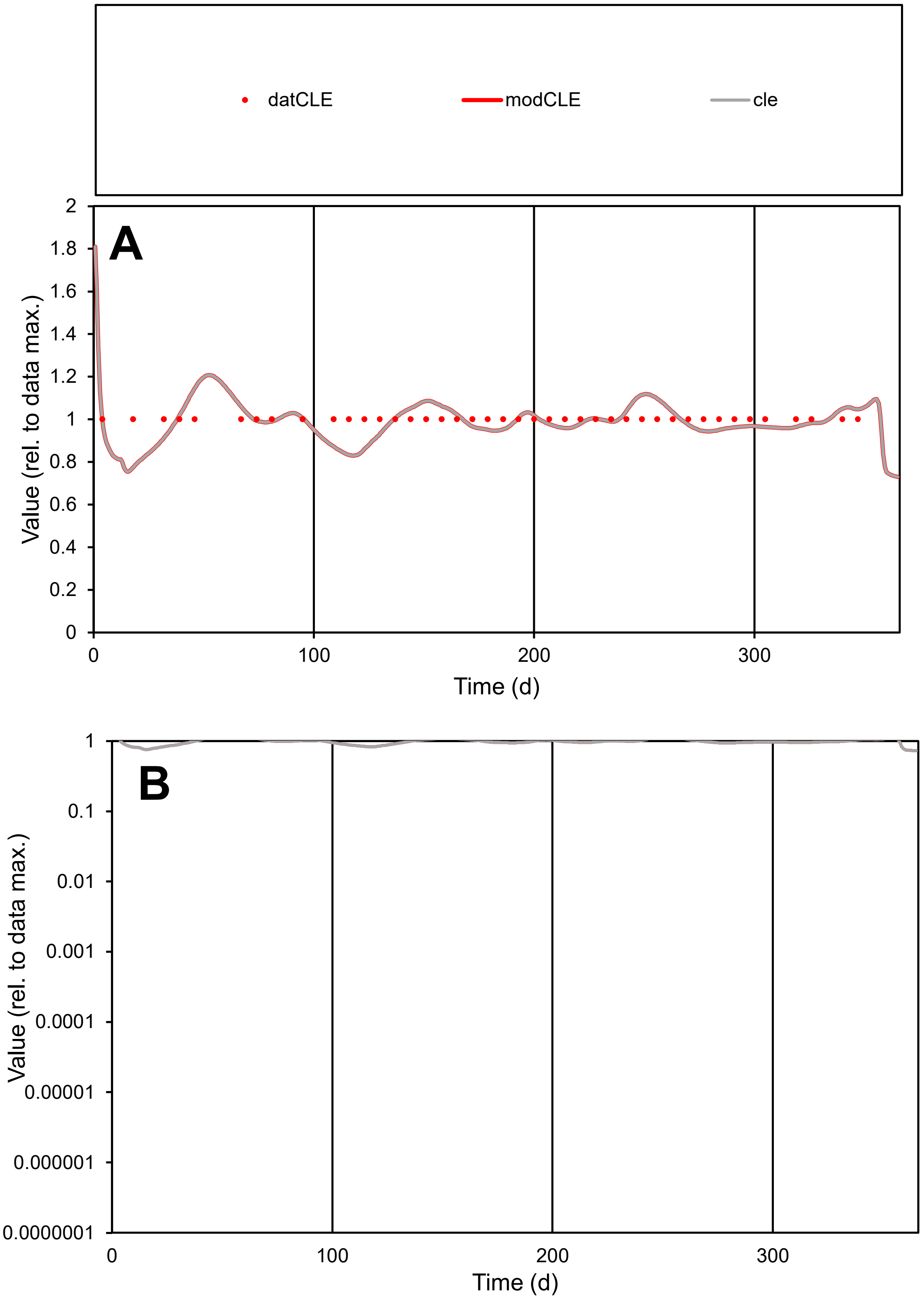


**C1**

Variables
CLE Provide jtd or Dst
CalibOnly?
2 1 to reproduce of
oNormalize?
2 1=no, 2=yes

ofxn calc*
0.07256
ofxn mod
0.13393

sRun
RunA1
zout
1

Ave
Data
4.7E-06
Model
4.6E-06

data/vMax
4.7E-06

ofxng
334.07

Min d/m
1.26135

ave/min d
1
ave/min m
1.24765

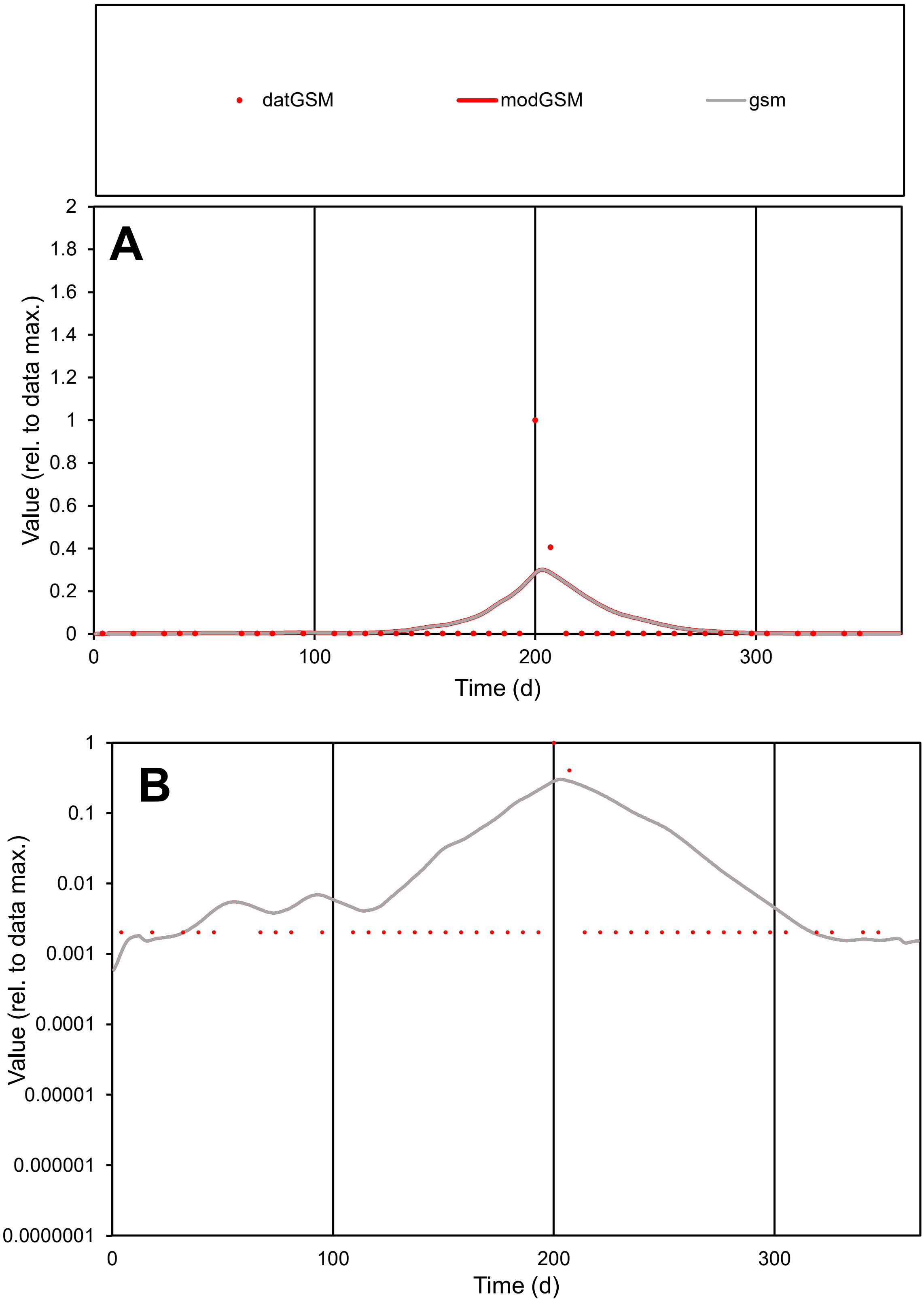


C1

| | |
|---|---|
| Variables | |
| GSM | Provide jtd or Dst |
| CalibOnly? | |
| 2 | 1 to reproduce of |
| oNormalize? | |
| 2 | 1=no, 2=yes |
| ofxn calc* | |
| 3.67926 | |
| ofxn mod | |
| 3.74514 | |
| sRun | |
| RunA1 | |
| zout | |
| 1 | |
| Ave | |
| Data | |
| 2.8E-05 | |
| Model | |
| 4.1E-05 | |
| data/vMax | |
| 0.00078 | |
| ofxng | |
| 334.07 | |
| Min d/m | |
| 1.63458 | |
| ave/min d | |
| 17.4592 | |
| ave/min m | |
| 42.7189 | |

327_Calib_S_A

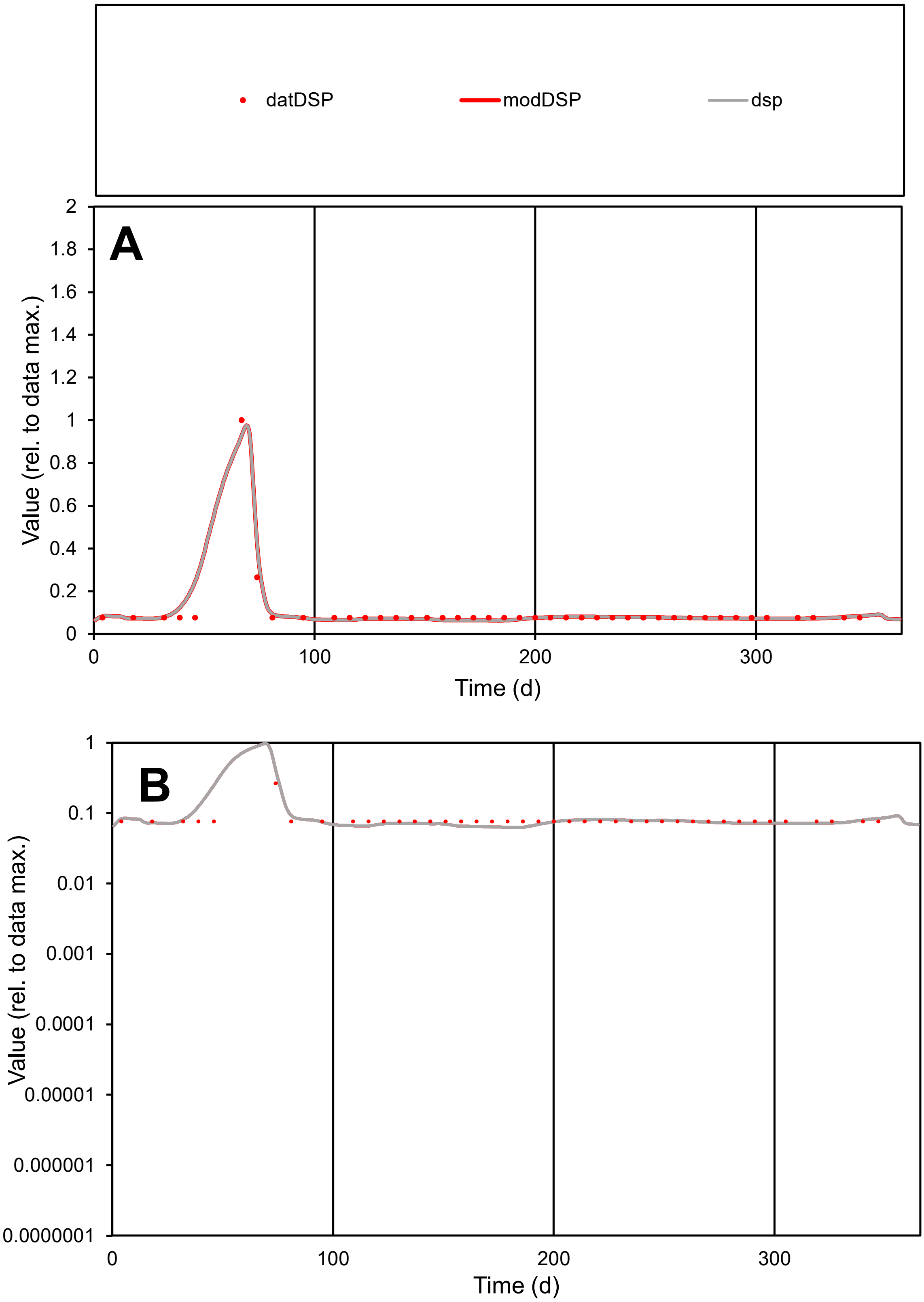


**C1**

Variables

DSP Provide jtd or Dst

CalibOnly?

2 1 to reproduce of

oNormalize?

2 1=no, 2=yes

ofxn calc*

0.25063

ofxn mod

0.35577

sRun

RunA1

zout

1

Ave

Data

6.7E-06

Model

6.9E-06

data/vMax

6.6E-05

ofxng

334.07

Min d/m

1.2104

ave/min d

1.34751

ave/min m

1.6603

327_Calib_S_A

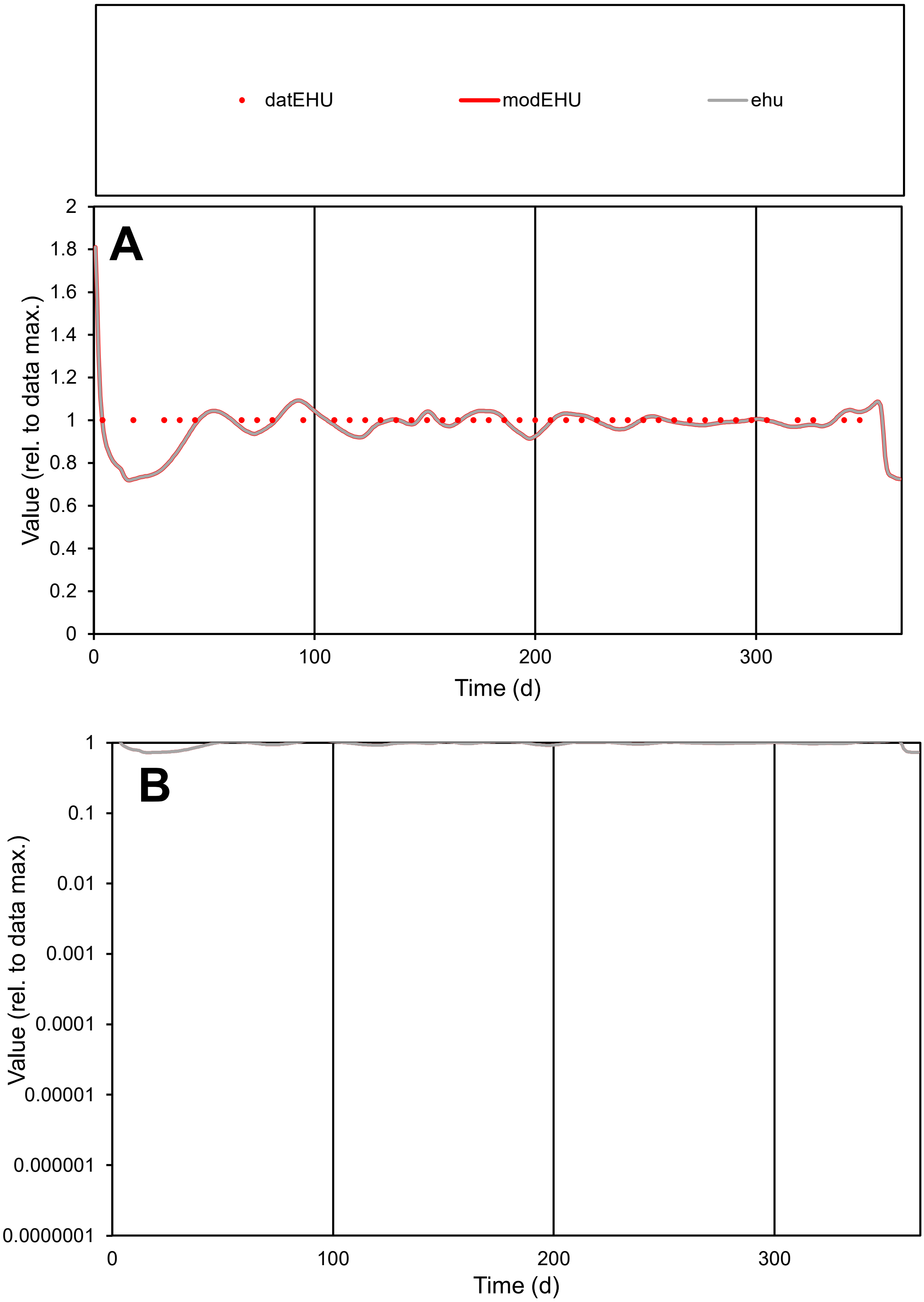


**C1**

Variables
EHU Provide jtd or Dst
CalibOnly?
2 1 to reproduce of
oNormalize?
2 1=no, 2=yes

ofxn calc*
0.06551
ofxn mod
0.12938

sRun
RunA1
zout
1

Ave
Data
5.8E-06
Model
5.6E-06

data/vMax
5.8E-06

ofxng
334.07

Min d/m
1.37089

ave/min d
1
ave/min m
1.33996

327_Calib_S_A

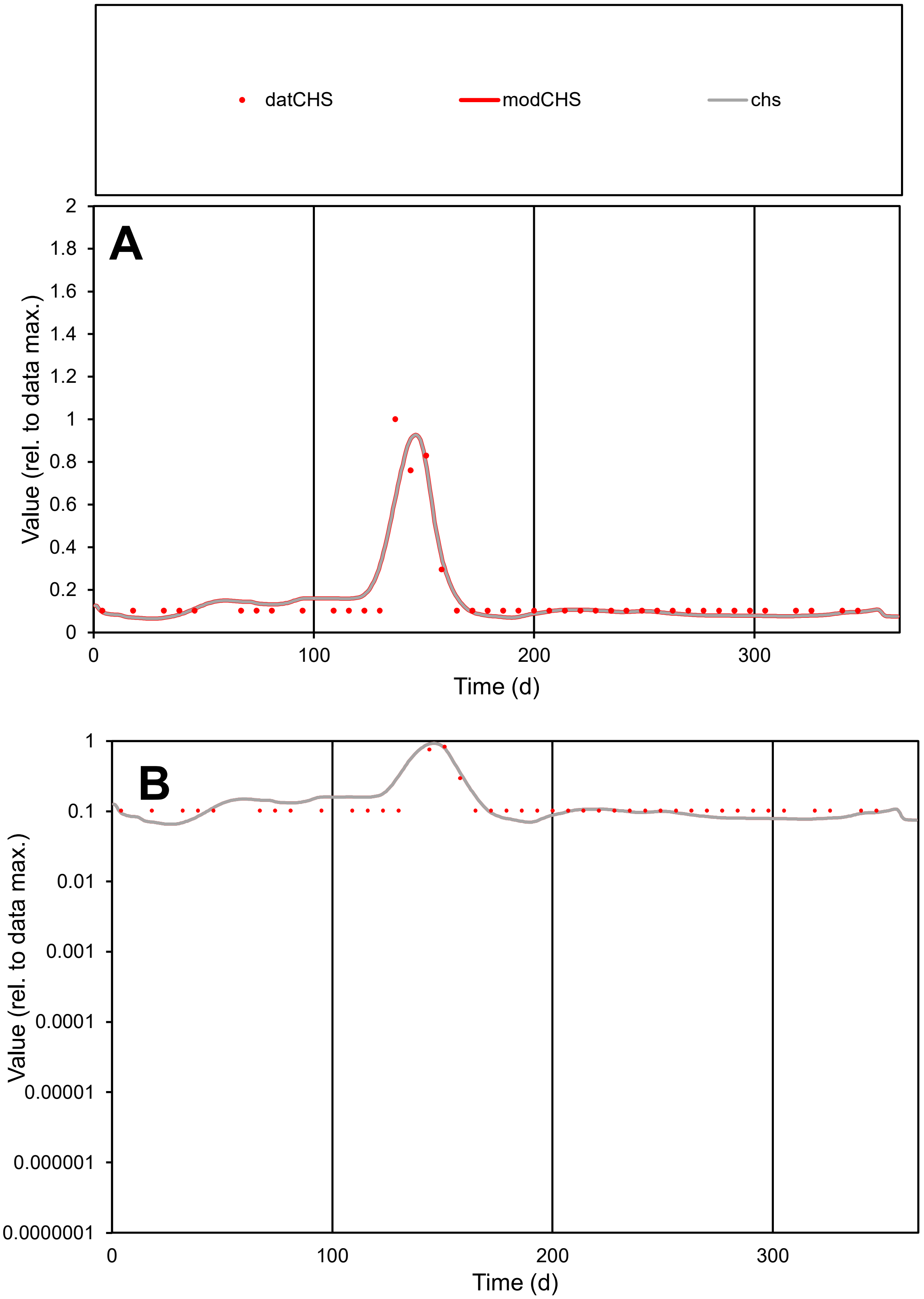


C1

Variables

CHS Provide jtd or Dst

CalibOnly?

2 1 to reproduce of

oNormalize?

2 1=no, 2=yes

ofxn calc*

0.43488

ofxn mod

0.46475

sRun

RunA1

zout

1

Ave

Data

3.4E-05

Model

3.4E-05

data/vMax

0.00021

ofxng

334.07

Min d/m

1.4701

ave/min d

1.57954

ave/min m

2.31283

327_Calib_S_A

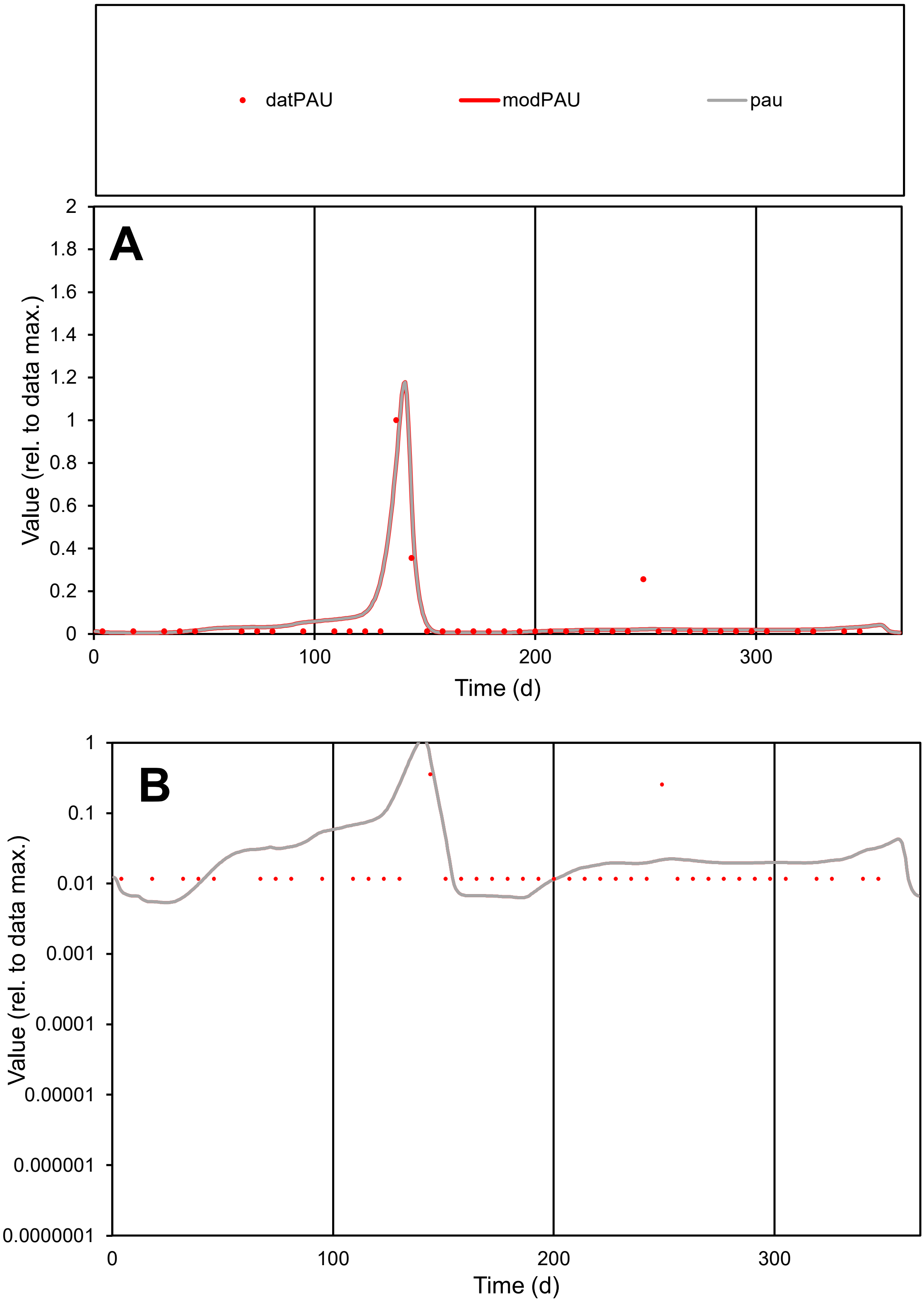


**C1**

Variables

| | |
|---|---|
| PAU | Provide jtd or Dst |
| CalibOnly? | |
| 2 | 1 to reproduce of |
| oNormalize? | |
| 2 | 1=no, 2=yes |

ofxn calc*
1.02213
ofxn mod
1.12244

sRun
RunA1
zout
1

Ave
Data
1.7E-05
Model
1.9E-05

data/vMax
0.00034

ofxng
334.07

Min d/m
2.1279

ave/min d
4.20563
ave/min m
10.3668

327_Calib_S_A

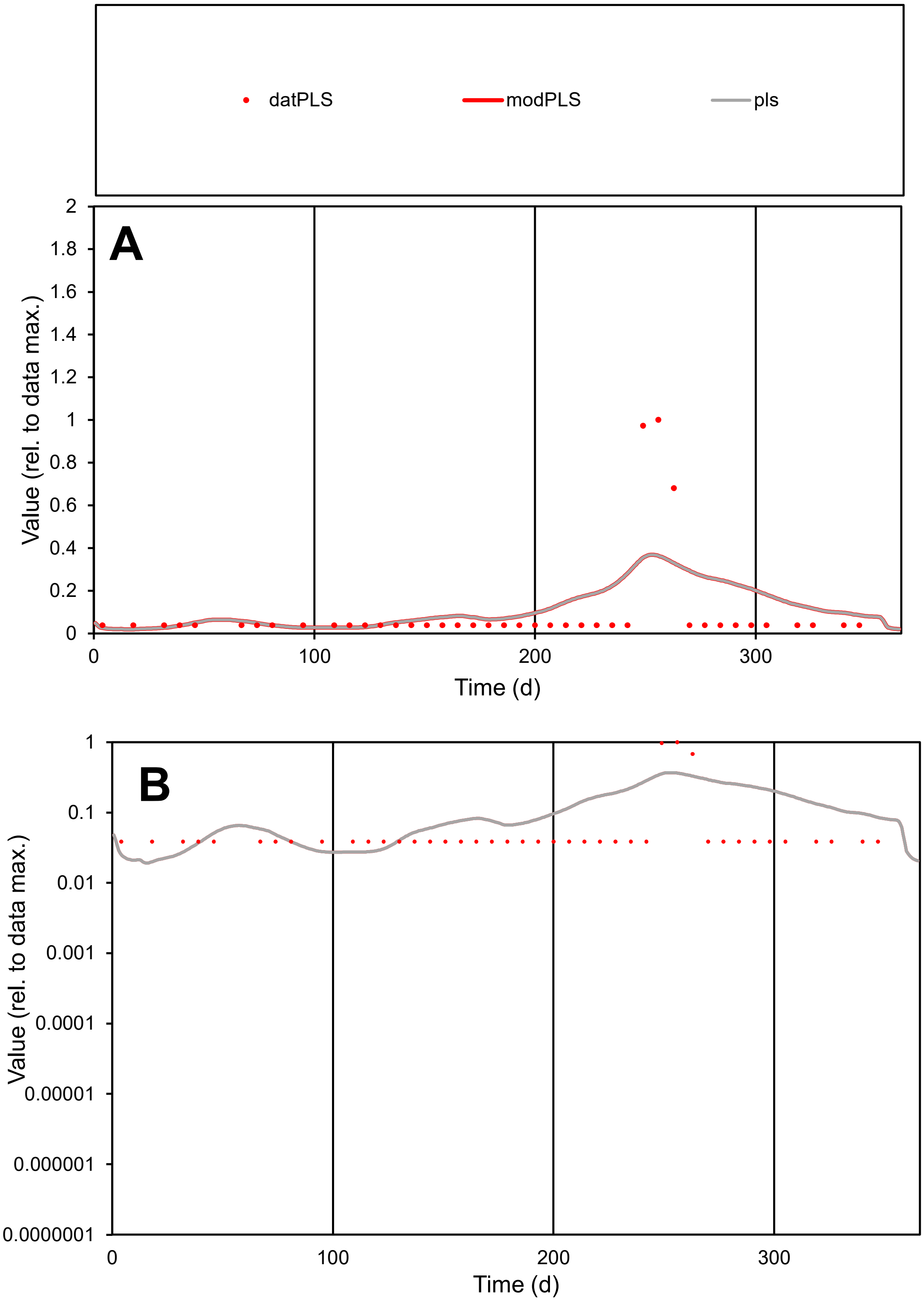


**C1**

Variables

PLS Provide jtd or Dst

CalibOnly?

2 1 to reproduce of

oNormalize?

2 1=no, 2=yes

ofxn calc*

1.79649

ofxn mod

1.81588

sRun

RunA1

zout

1

Ave

Data

9.8E-06

Model

1.2E-05

data/vMax

9.9E-05

ofxng

334.07

Min d/m

1.86185

ave/min d

2.55798

ave/min m

5.97681

327_Calib_S_A

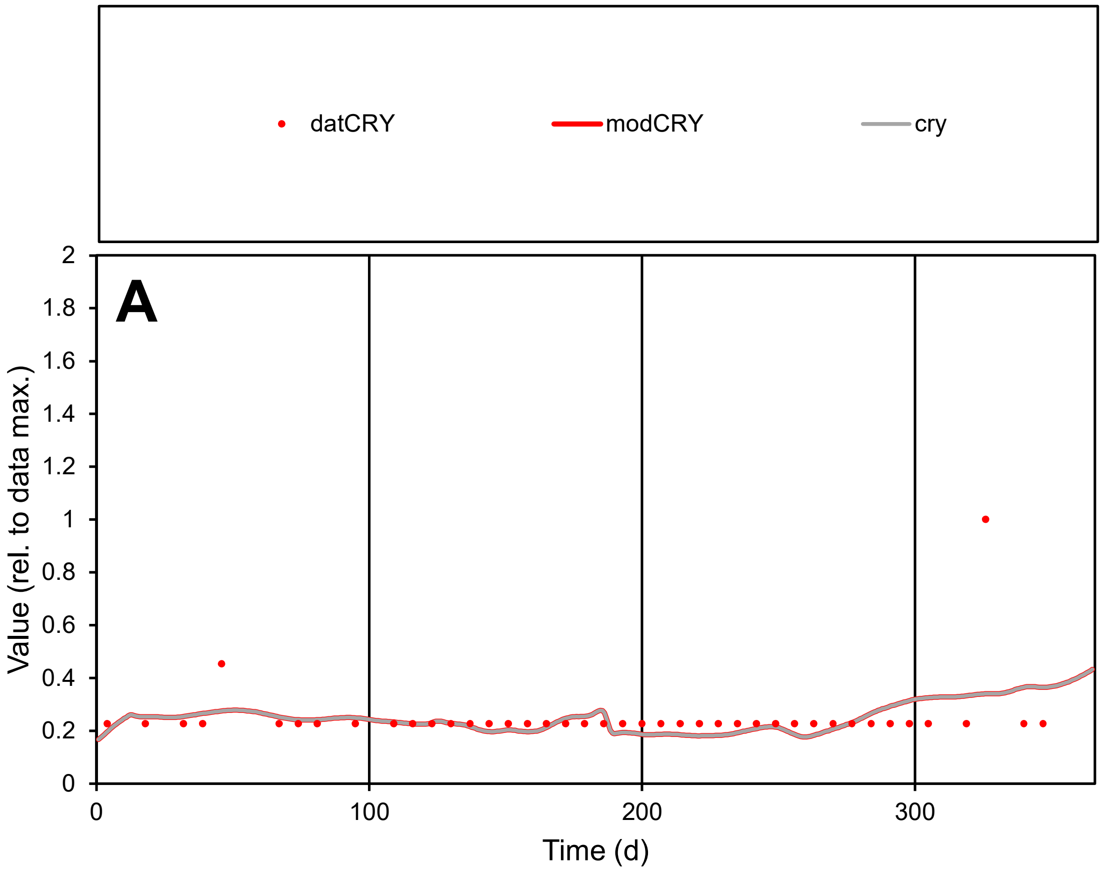


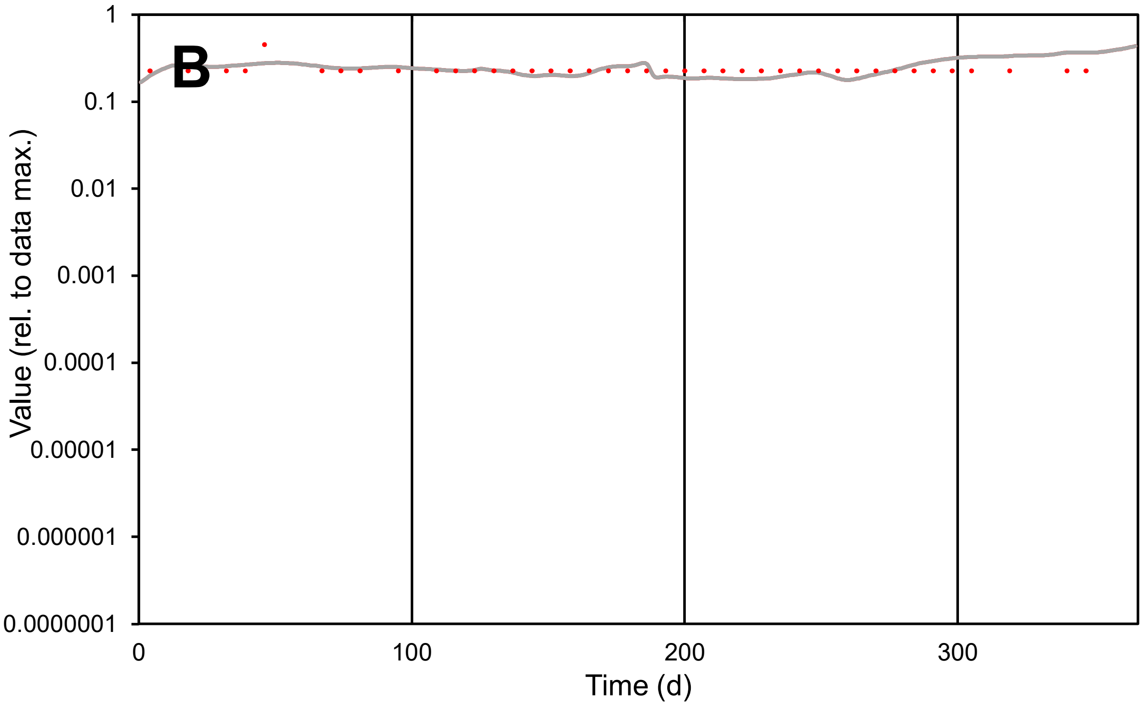


**C1**

Variables
CRY Provide jtd or Dst
CalibOnly?
2 1 to reproduce of
oNormalize?
2 1=no, 2=yes

ofxn calc*
0.46299
ofxn mod
0.47679

sRun
RunA1
zout
1

Ave
Data
3.3E-06
Model
3.2E-06

data/vMax
1.3E-05

ofxng
334.07

Min d/m
1.24728

ave/min d
1.10493
ave/min m
1.32265

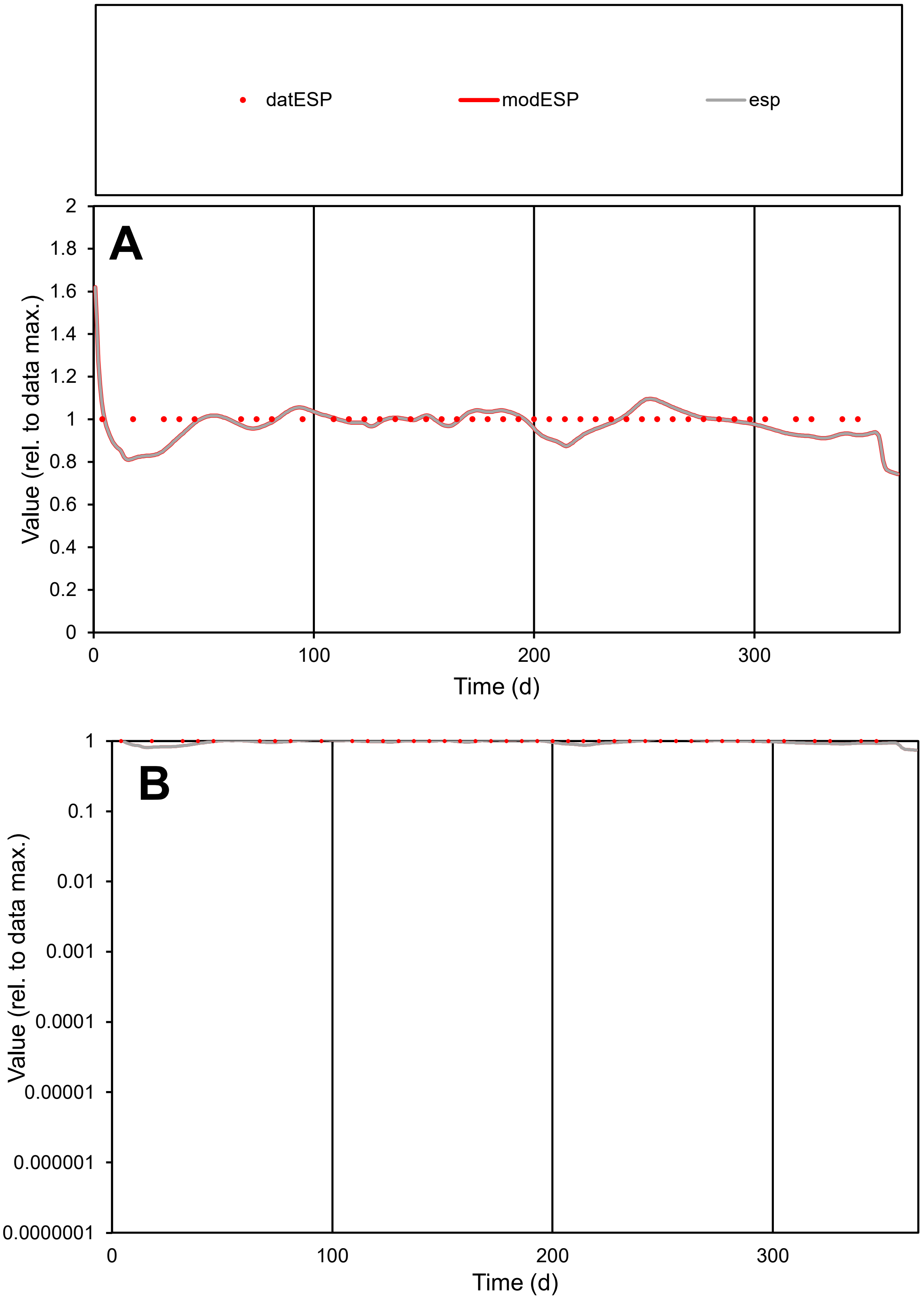


**C1**

Variables
ESP Provide jtd or Dst
CalibOnly?
2 1 to reproduce of
oNormalize?
2 1=no, 2=yes

ofxn calc*
0.05884
ofxn mod
0.12651

sRun
RunA1
zout
1

Ave
Data
8E-06
Model
7.9E-06

data/vMax
8E-06

ofxng
334.07

Min d/m
1.21653

ave/min d
1
ave/min m
1.19297

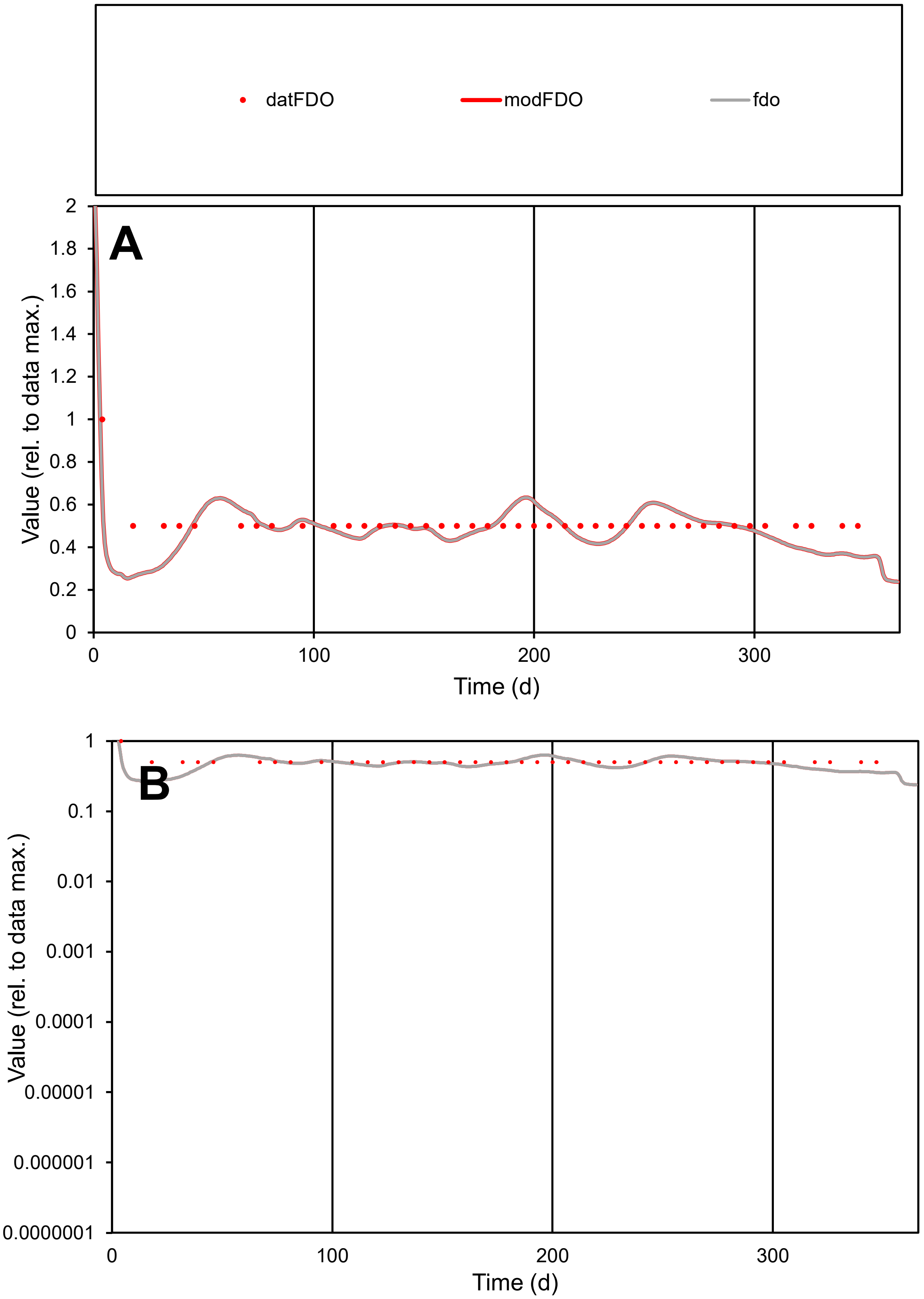


**C1**

Variables

FDO Provide jtd or Dst

CalibOnly?

2 1 to reproduce of

oNormalize?

2 1=no, 2=yes

ofxn calc*

0.17149

ofxn mod

0.22659

sRun

RunA1

zout

1

Ave

Data

1E-07

Model

9.7E-08

data/vMax

2E-07

ofxng

334.07

Min d/m

1.856

ave/min d

1.02382

ave/min m

1.81103

327_Calib_S_A

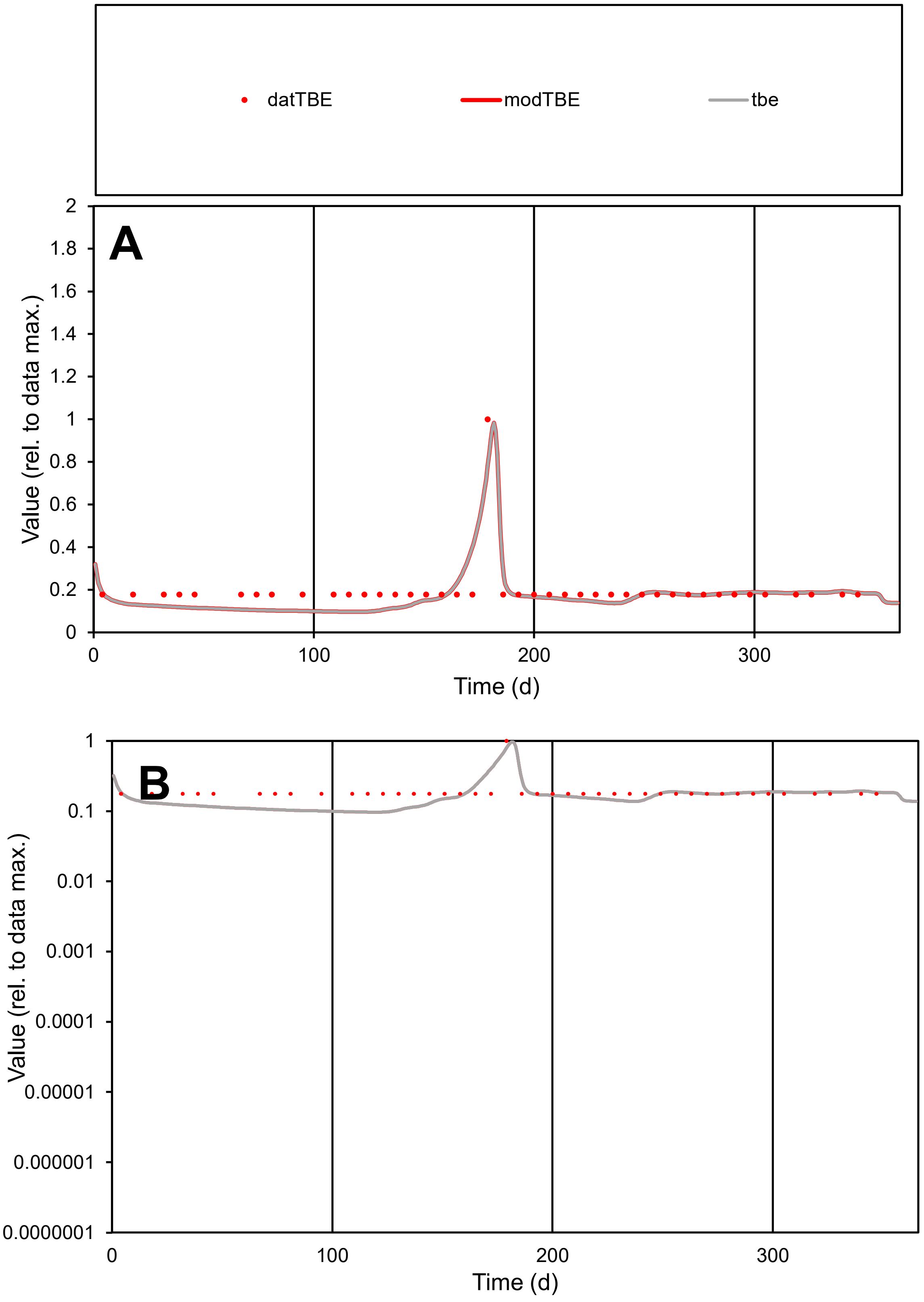


**C1**

Variables
TBE Provide jtd or Dst
CalibOnly?
2 1 to reproduce of
oNormalize?
2 1=no, 2=yes

ofxn calc*
0.2929
ofxn mod
0.32417

sRun
RunA1
zout
1

Ave
Data
7.4E-06
Model
6.7E-06

data/vMax
3.8E-05

ofxng
334.07

Min d/m
1.82927

ave/min d
1.11081
ave/min m
1.83147

327_Calib_S_A

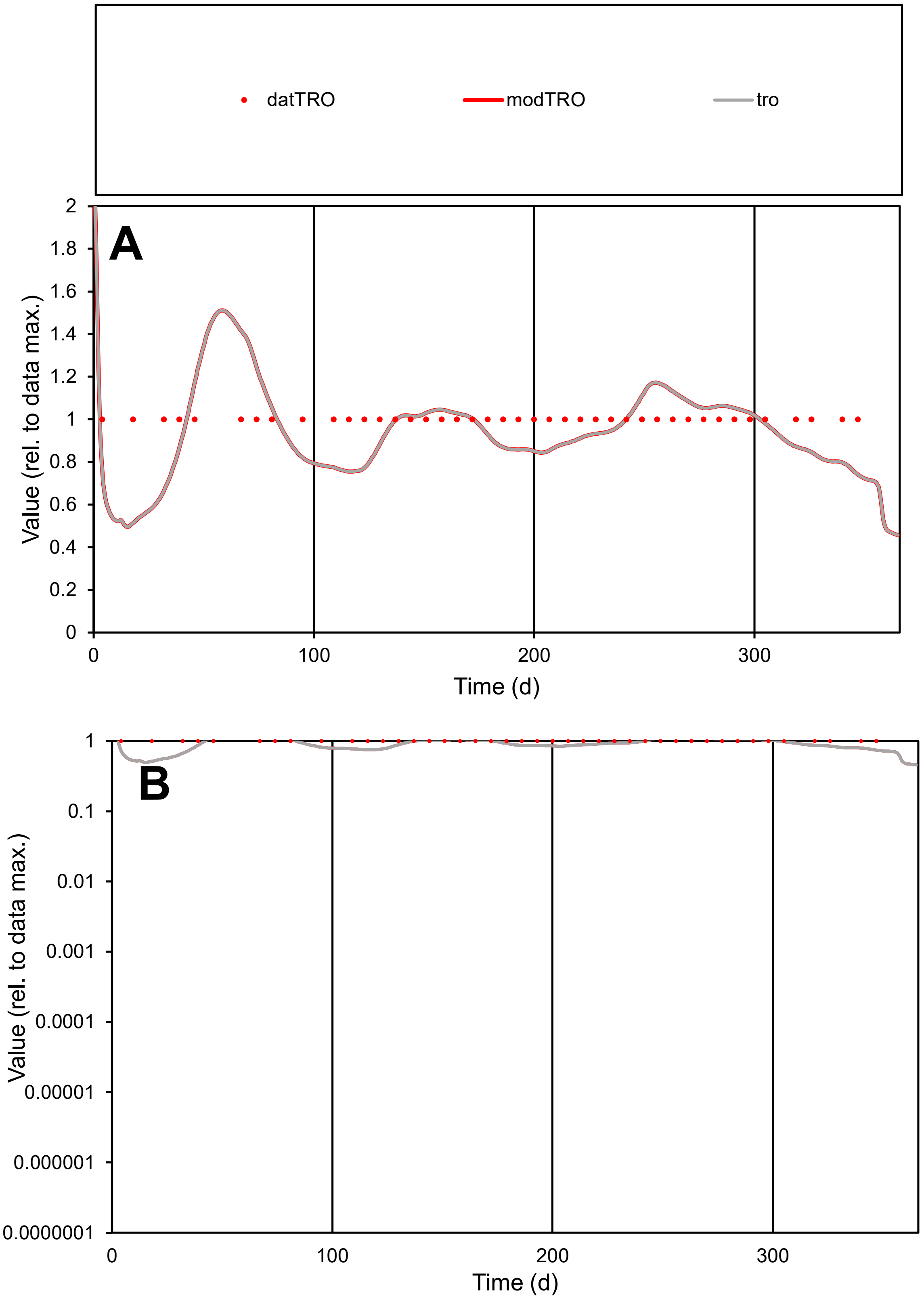


**C1**

Variables
TRO Provide jtd or Dst
CalibOnly?
2 1 to reproduce of
oNormalize?
2 1=no, 2=yes

ofxn calc*
0.16979
ofxn mod
0.20363

sRun
RunA1
zout
1

Ave
Data
5E-06
Model
4.8E-06

data/vMax
5E-06

ofxng
334.07

Min d/m
1.8881

ave/min d
1
ave/min m
1.78996

327_Calib_S_A

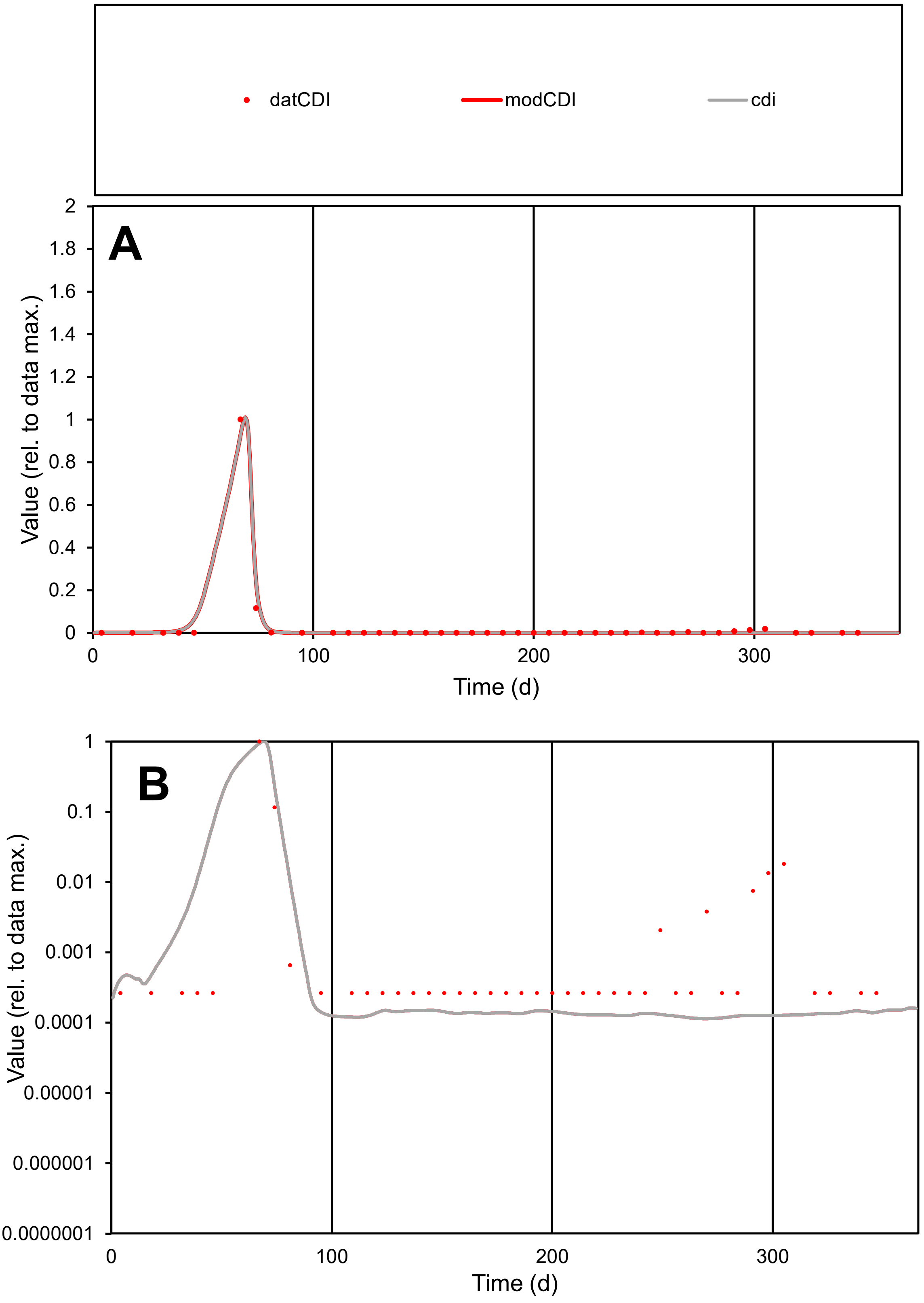


**C1**

Variables
CDI Provide jtd or Dst
CalibOnly?
2 1 to reproduce of
oNormalize?
2 1=no, 2=yes

ofxn calc*
0.27648
ofxn mod
0.77399

sRun
RunA1
zout
1

Ave
Data
1.8E-05
Model
1.8E-05

data/vMax
0.00066

ofxng
334.07

Min d/m
2.30749

ave/min d
106.087
ave/min m
244.182

327_Calib_S_A

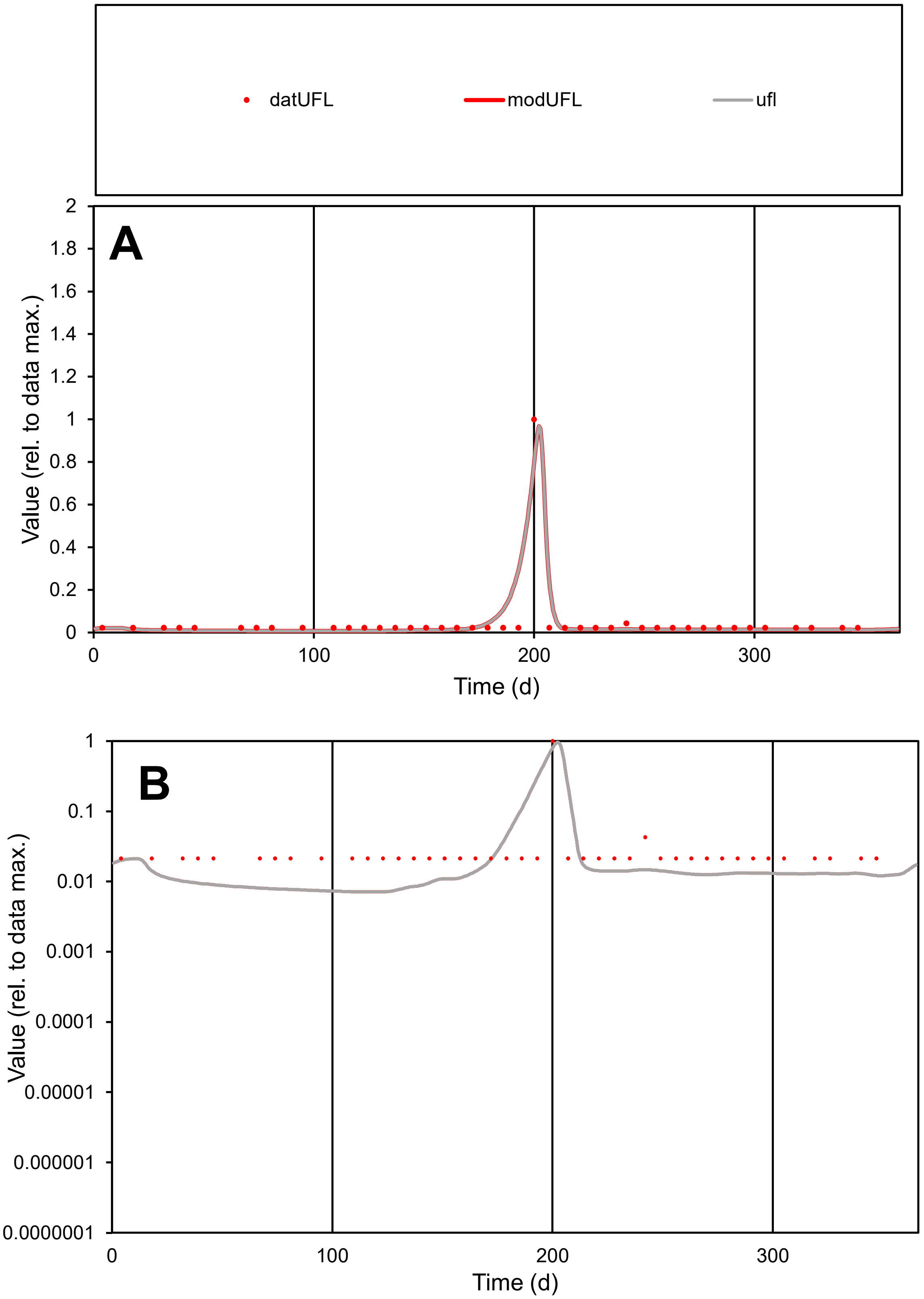


**C1**

Variables
UFL Provide jtd or Dst
CalibOnly?
2 1 to reproduce of
oNormalize?
2 1=no, 2=yes

ofxn calc*
0.85917
ofxn mod
1.10545

sRun
RunA1
zout
1

Ave
Data
6.5E-06
Model
6.2E-06

data/vMax
0.00014

ofxng
334.07

Min d/m
3.00786

ave/min d
2.1135
ave/min m
6.08839

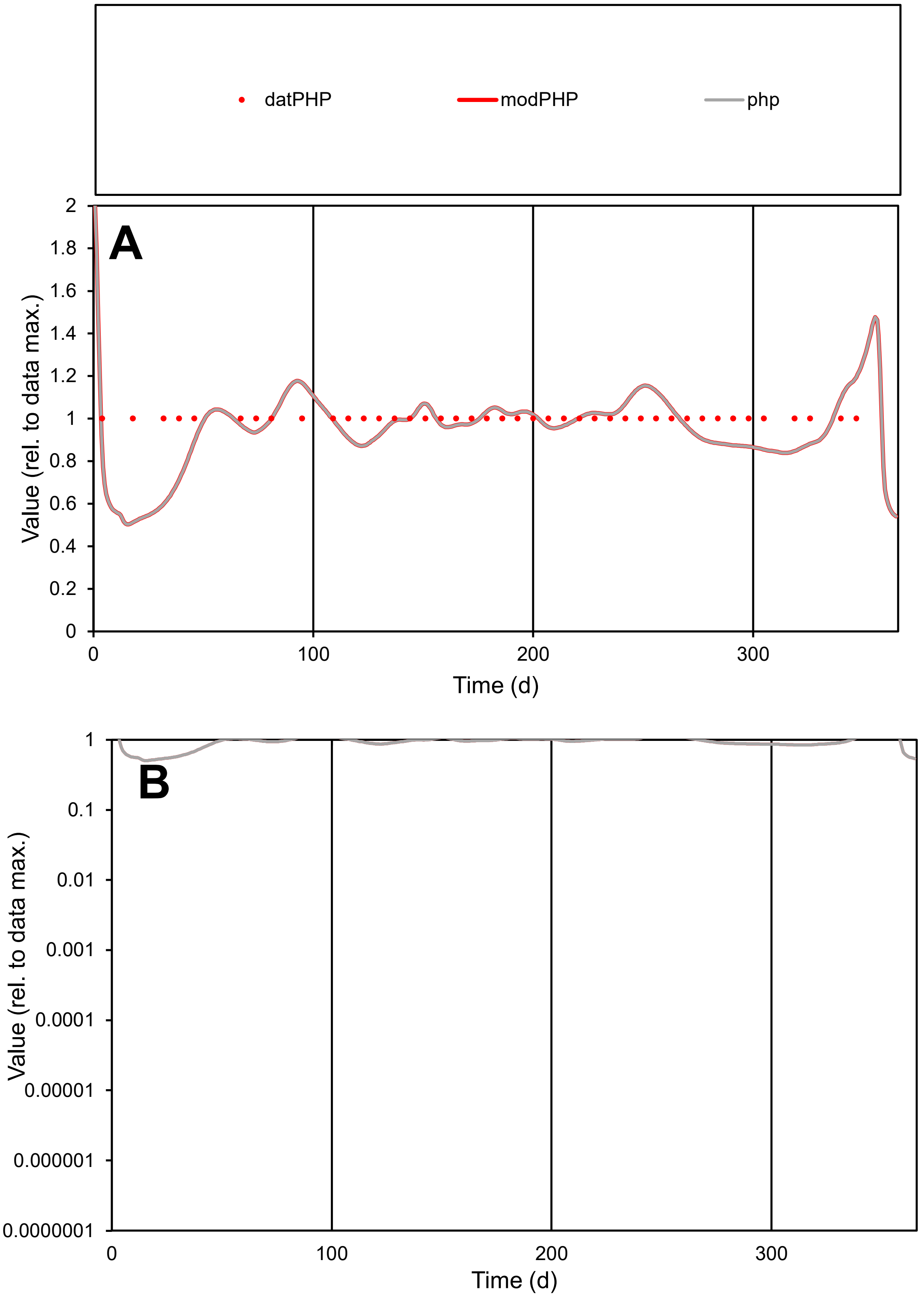


**C1**

Variables
PHP Provide jtd or Dst
CalibOnly?
2 1 to reproduce of
oNormalize?
2 1=no, 2=yes

ofxn calc*
0.13782
ofxn mod
0.17773

sRun
RunA1
zout
1

Ave
Data
6.9E-06
Model
6.7E-06

data/vMax
6.9E-06

ofxng
334.07

Min d/m
1.91687

ave/min d
1
ave/min m
1.8373

327_Calib_S_A

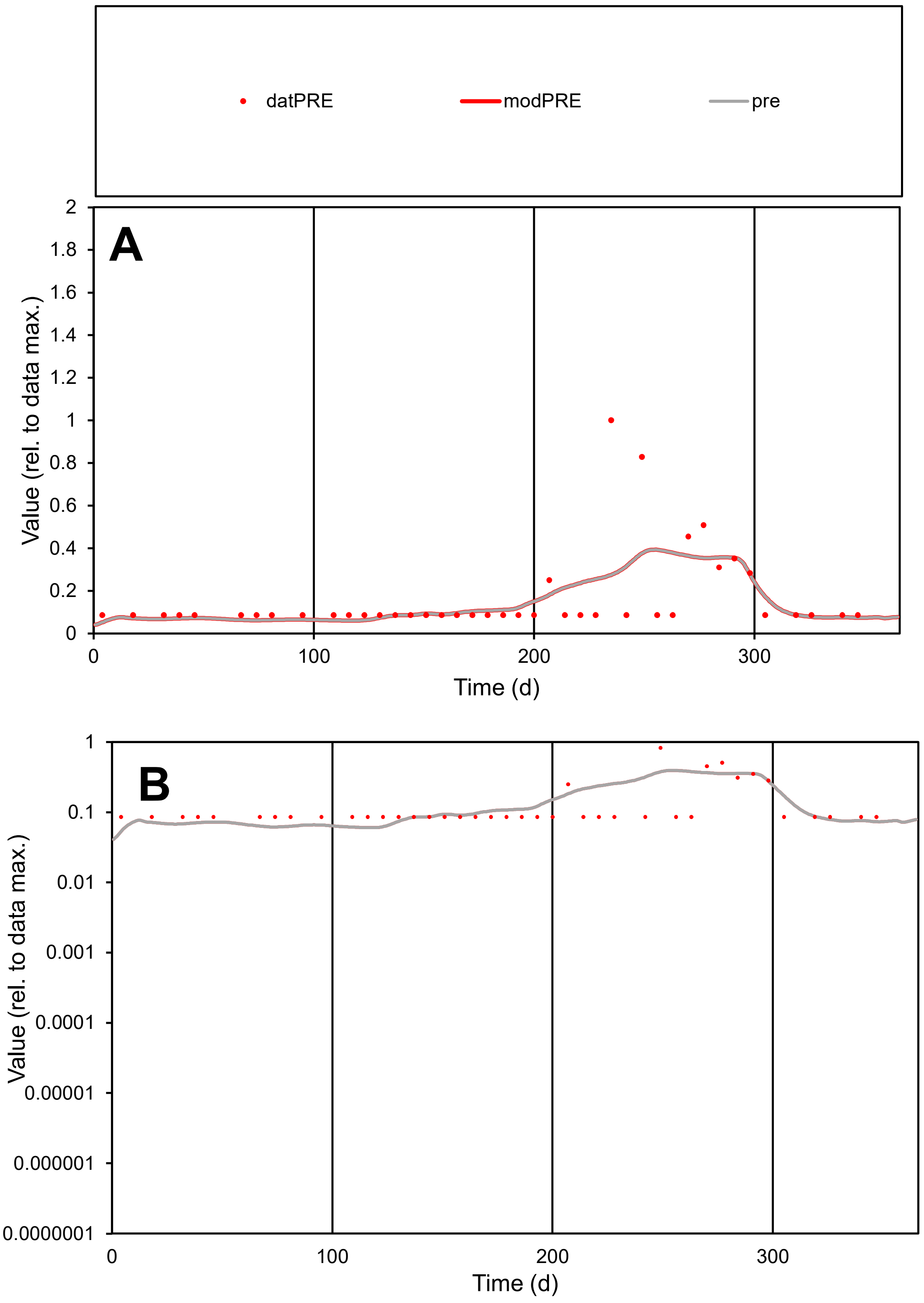


**C1**

Variables

PRE Provide jtd or Dst

CalibOnly?

2 1 to reproduce of

oNormalize?

2 1=no, 2=yes

ofxn calc*

0.9758

ofxn mod

0.98896

sRun

RunA1

zout

1

Ave

Data

2.6E-05

Model

2.5E-05

data/vMax

0.00016

ofxng

334.07

Min d/m

1.48891

ave/min d

1.92175

ave/min m

2.78817

327_Calib_S_A

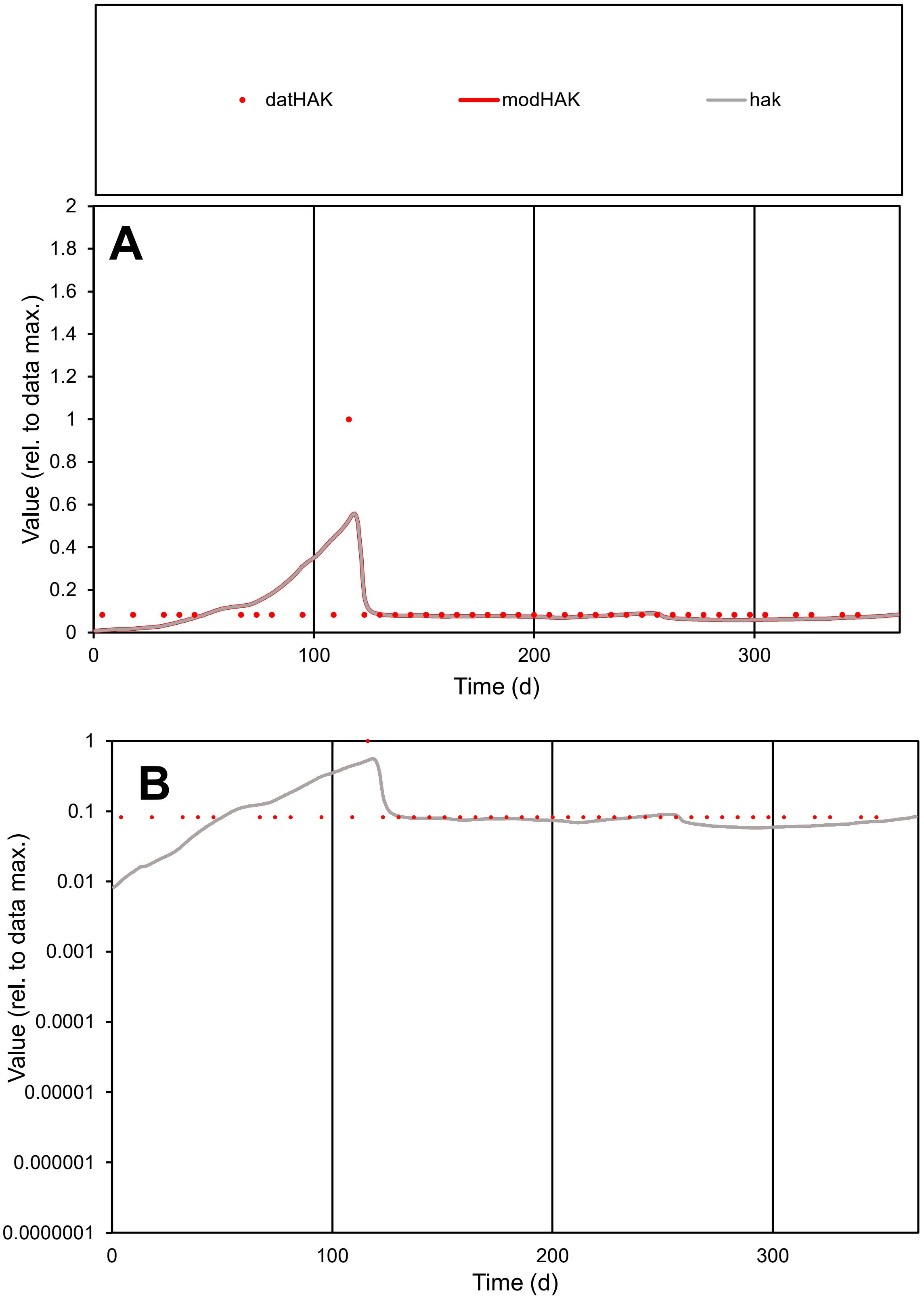


**C1**

Variables
HAK Provide jtd or Dst
CalibOnly?
2 1 to reproduce of
oNormalize?
2 1=no, 2=yes

ofxn calc*
0.96947
ofxn mod
0.98399

sRun
RunA1
zout
1

Ave
Data
6.5E-06
Model
6.3E-06

data/vMax
6.2E-05

ofxng
334.07

Min d/m
7.60095

ave/min d
1.26596
ave/min m
9.37001

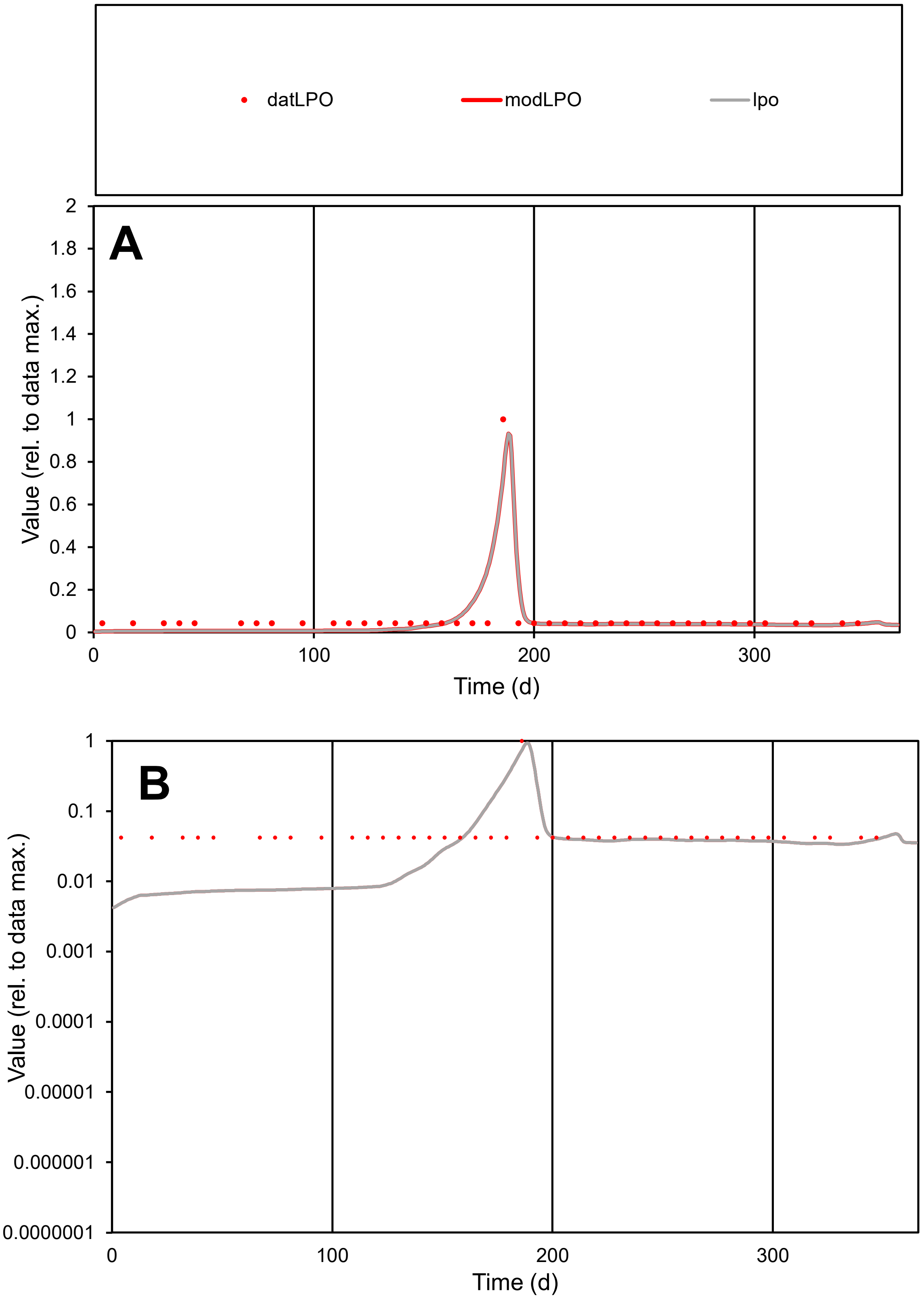


**C1**

Variables
LPO Provide jtd or Dst
CalibOnly?
2 1 to reproduce of
oNormalize?
2 1=no, 2=yes

ofxn calc*
0.72445
ofxn mod
0.83755

sRun
RunA1
zout
1

Ave
Data
9E-06
Model
8E-06

data/vMax
0.00014

ofxng
334.07

Min d/m
8.35021

ave/min d
1.54012
ave/min m
11.3991

327_Calib_S_A

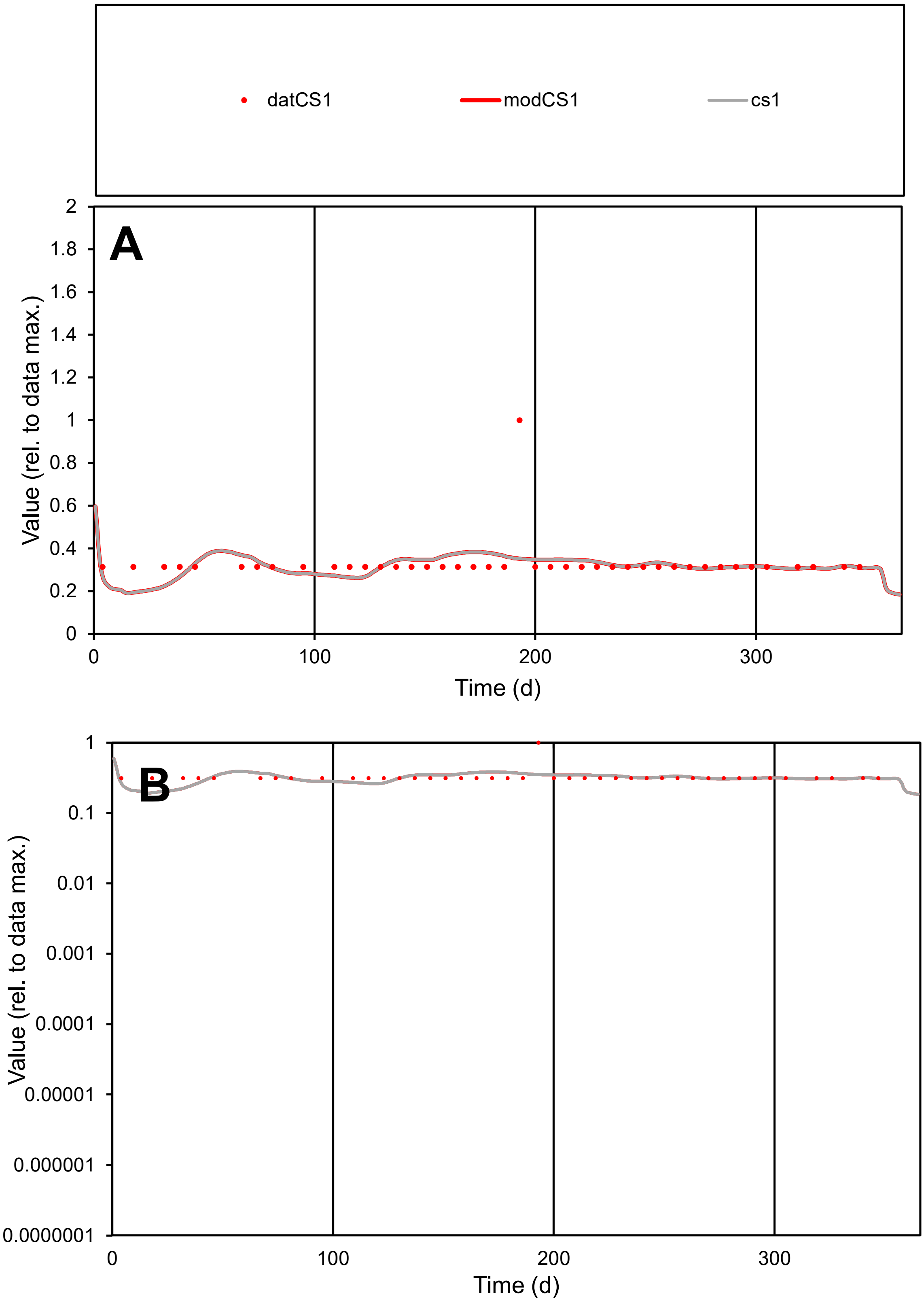


**C1**

Variables
CS1 Provide jtd or Dst
CalibOnly?
2 1 to reproduce of
oNormalize?
2 1=no, 2=yes

ofxn calc*
0.32553
ofxn mod
0.3448

sRun
RunA1
zout
1

Ave
Data
1.8E-05
Model
1.8E-05

data/vMax
5.6E-05

ofxng
334.07

Min d/m
1.59418

ave/min d
1.05214
ave/min m
1.62488

327_Calib_S_A

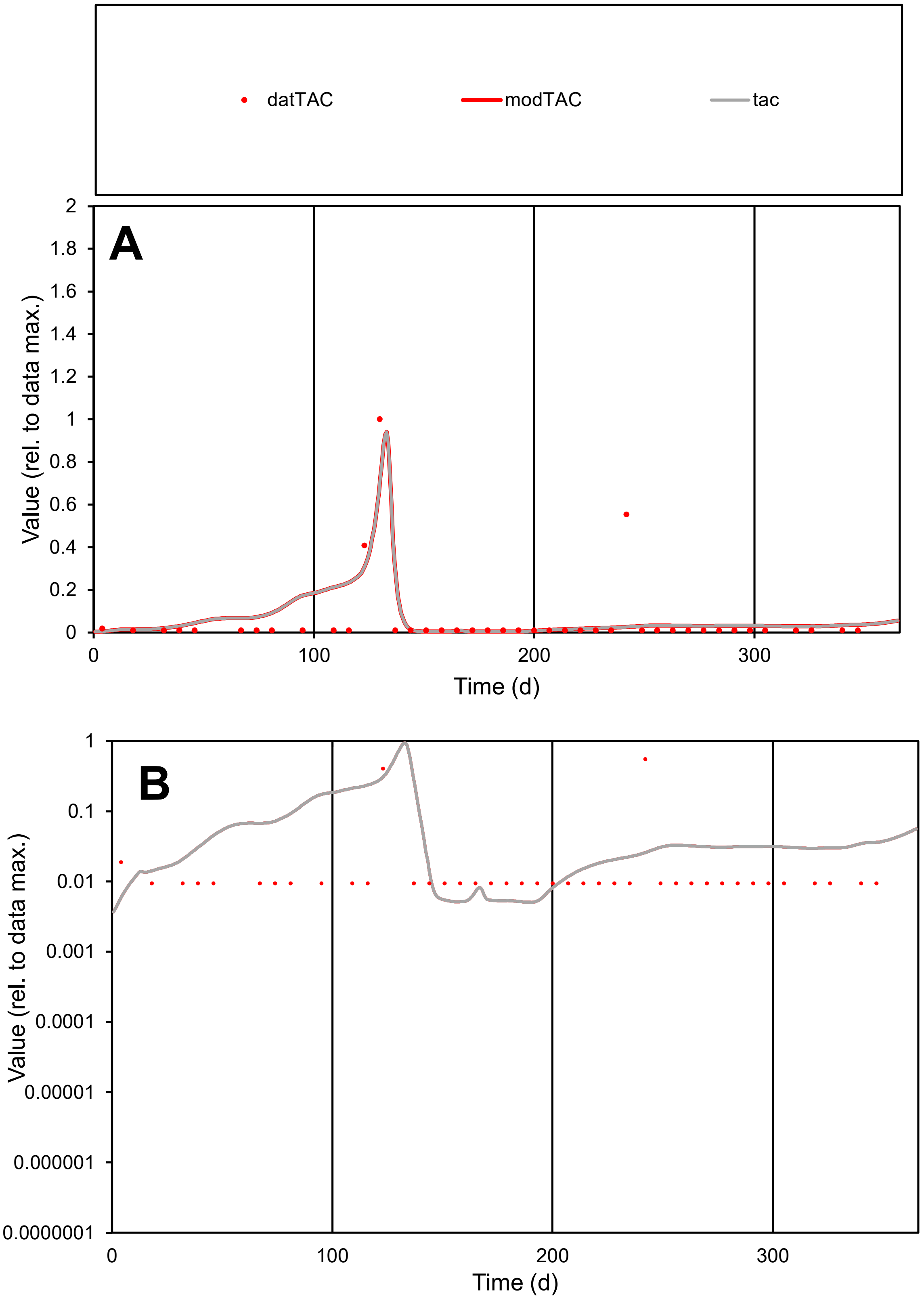


**C1**

Variables
TAC Provide jtd or Dst
CalibOnly?
2 1 to reproduce of
oNormalize?
2 1=no, 2=yes

ofxn calc*
1.85474
ofxn mod
1.90059

sRun
RunA1
zout
1

Ave
Data
7.7E-06
Model
9.4E-06

data/vMax
0.00014

ofxng
334.07

Min d/m
1.84394
ave/min d
5.90734
ave/min m
13.3608

327_Calib_S_A

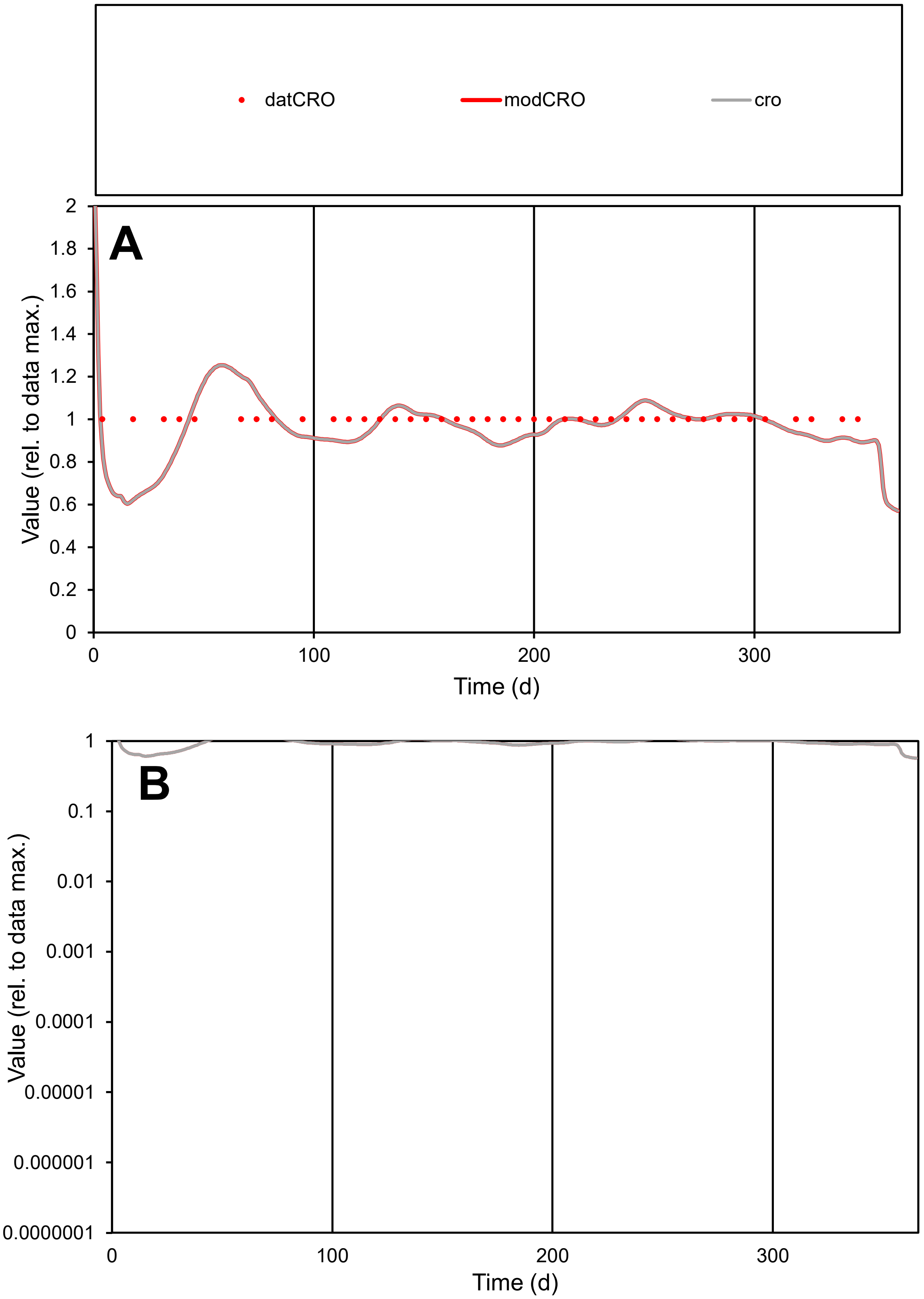


**C1**

Variables
CRO Provide jtd or Dst
CalibOnly?
2 1 to reproduce of
oNormalize?
2 1=no, 2=yes

ofxn calc*
0.09907
ofxn mod
0.15058

sRun
RunA1
zout
1

Ave
Data
1E-05
Model
9.9E-06

data/vMax
1E-05

ofxng
334.07

Min d/m
1.57272

ave/min d
1
ave/min m
1.52593

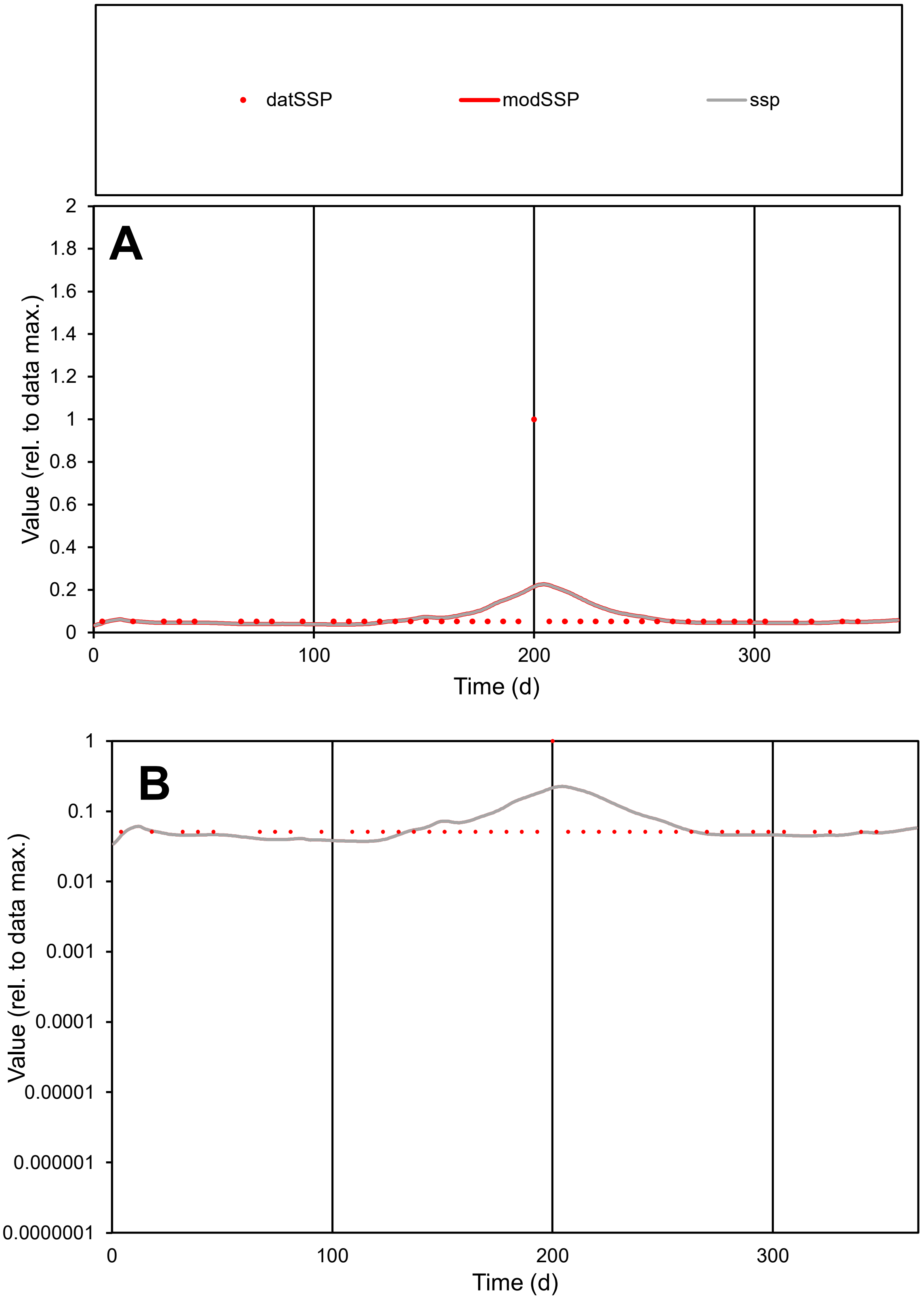


**C1**

Variables
SSP Provide jtd or Dst
CalibOnly?
2 1 to reproduce of
oNormalize?
2 1=no, 2=yes

ofxn calc*
1.76453
ofxn mod
1.77858

sRun
RunA1
zout
1

Ave
Data
1.9E-05
Model
2E-05

data/vMax
0.00026

ofxng
334.07

Min d/m
1.36125

ave/min d
1.44266
ave/min m
2.01944

327_Calib_S_A

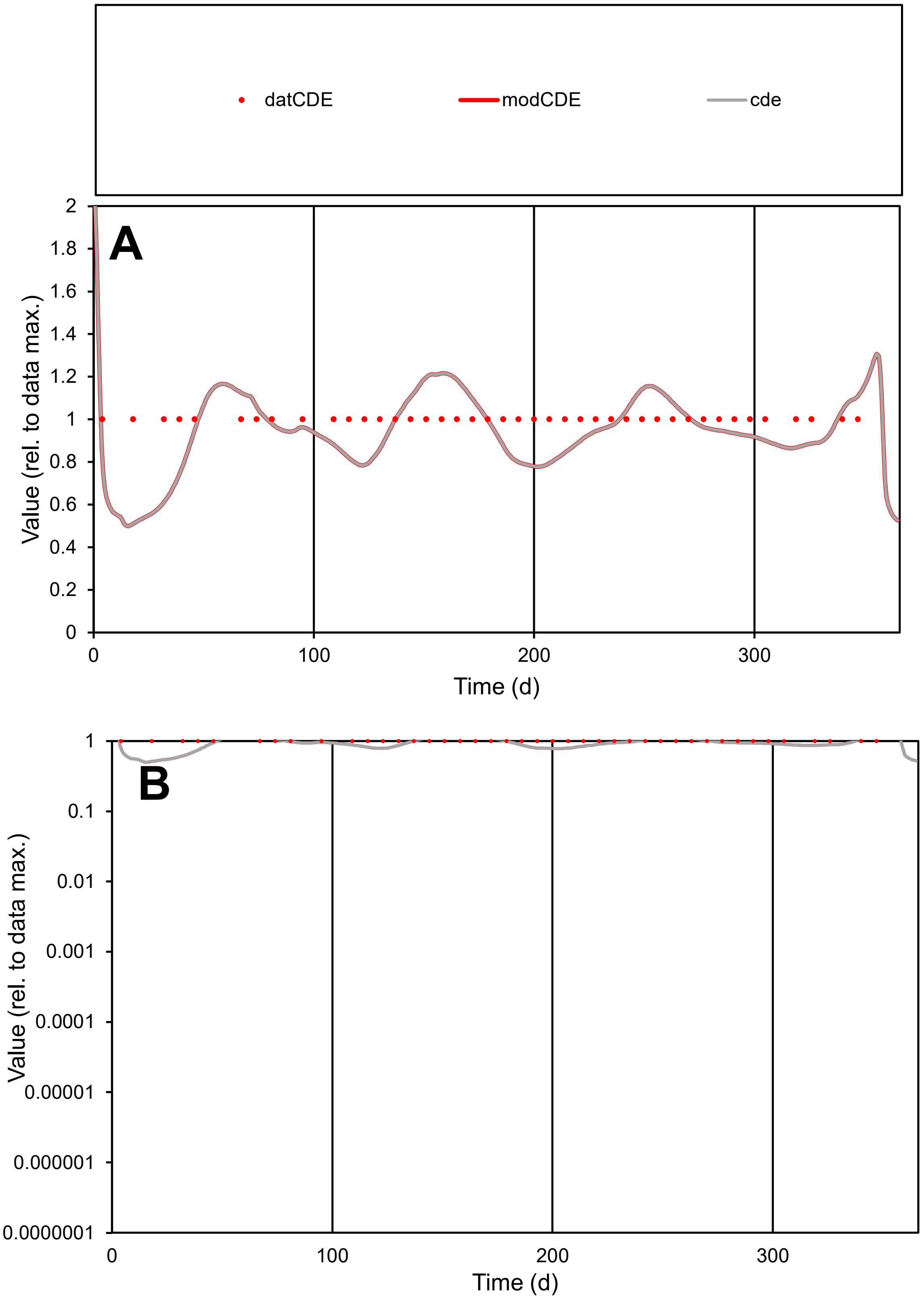


**C1**

Variables
CDE Provide jtd or Dst
CalibOnly?
2 1 to reproduce of
oNormalize?
2 1=no, 2=yes

ofxn calc*
0.15807
ofxn mod
0.19339

sRun
RunA1
zout
1

Ave
Data
1E-05
Model
9.5E-06

data/vMax
1E-05

ofxng
334.07

Min d/m
1.91984

ave/min d
1
ave/min m
1.82161

327_Calib_S_A

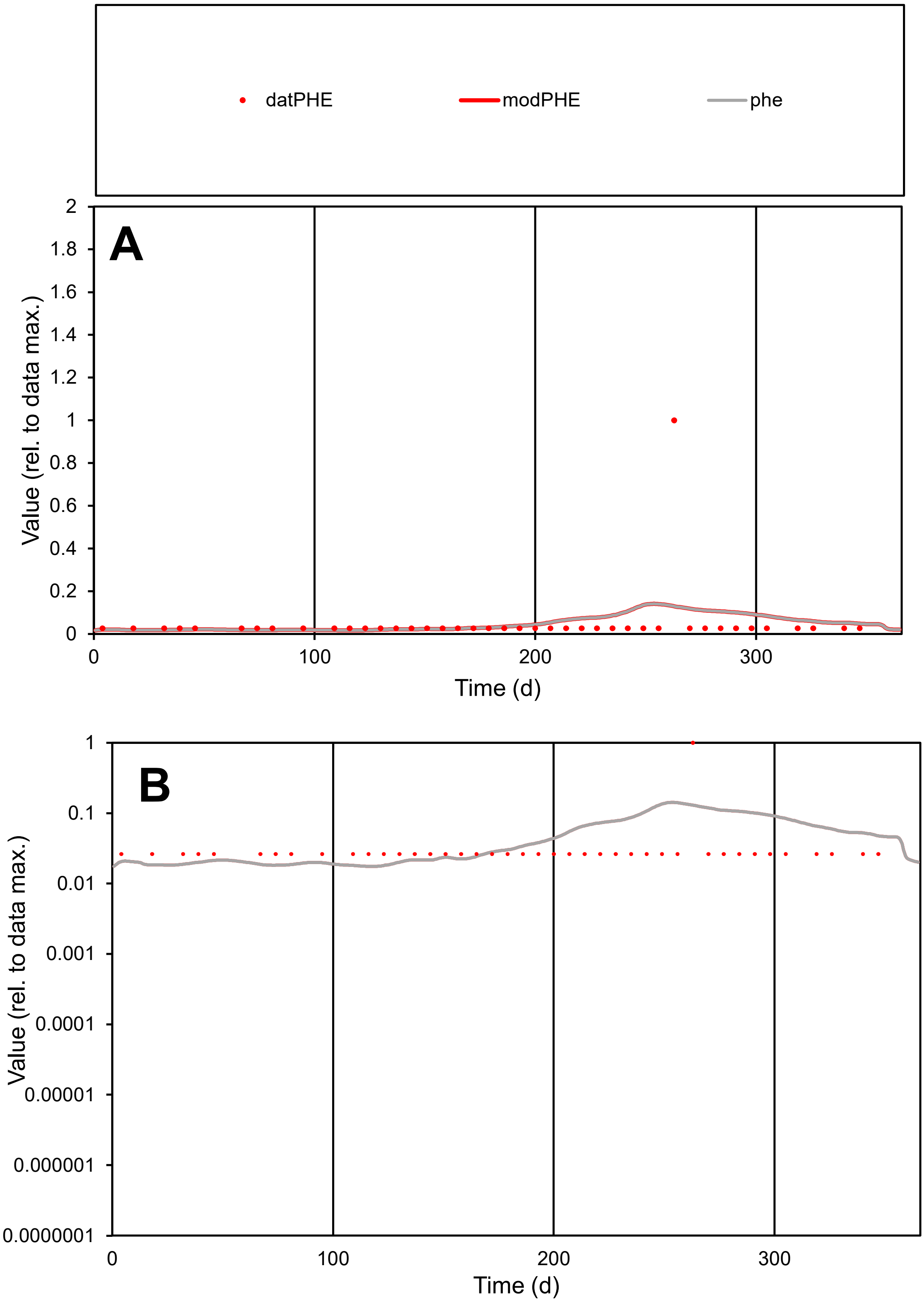


C1

Variables

PHE Provide jtd or Dst

CalibOnly?

2 1 to reproduce of

oNormalize?

2 1=no, 2=yes

ofxn calc*

2.86569

ofxn mod

2.88152

sRun

RunA1

zout

1

Ave

Data

7.5E-06

Model

8.1E-06

data/vMax

0.00015

ofxng

334.07

Min d/m

1.49924

ave/min d

1.88248

ave/min m

3.04244

327_Calib_S_A

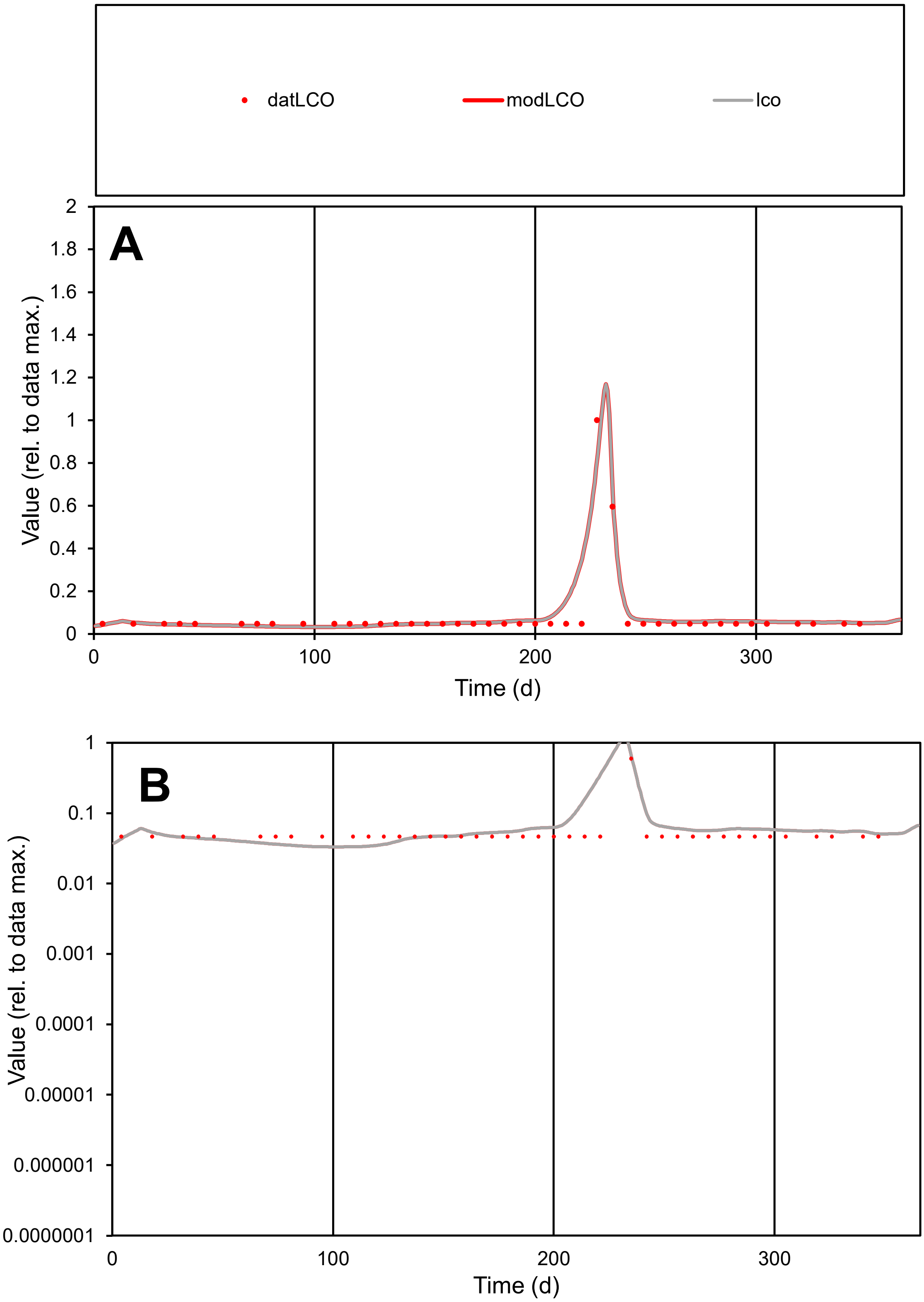


**C1**

Variables
LCO Provide jtd or Dst
CalibOnly?
2 1 to reproduce of
oNormalize?
2 1=no, 2=yes

ofxn calc*
0.54415
ofxn mod
0.66026

sRun
RunA1
zout
1

Ave
Data
1.3E-05
Model
1.5E-05

data/vMax
0.00016

ofxng
334.07

Min d/m
1.40604

ave/min d
1.7658
ave/min m
2.77852

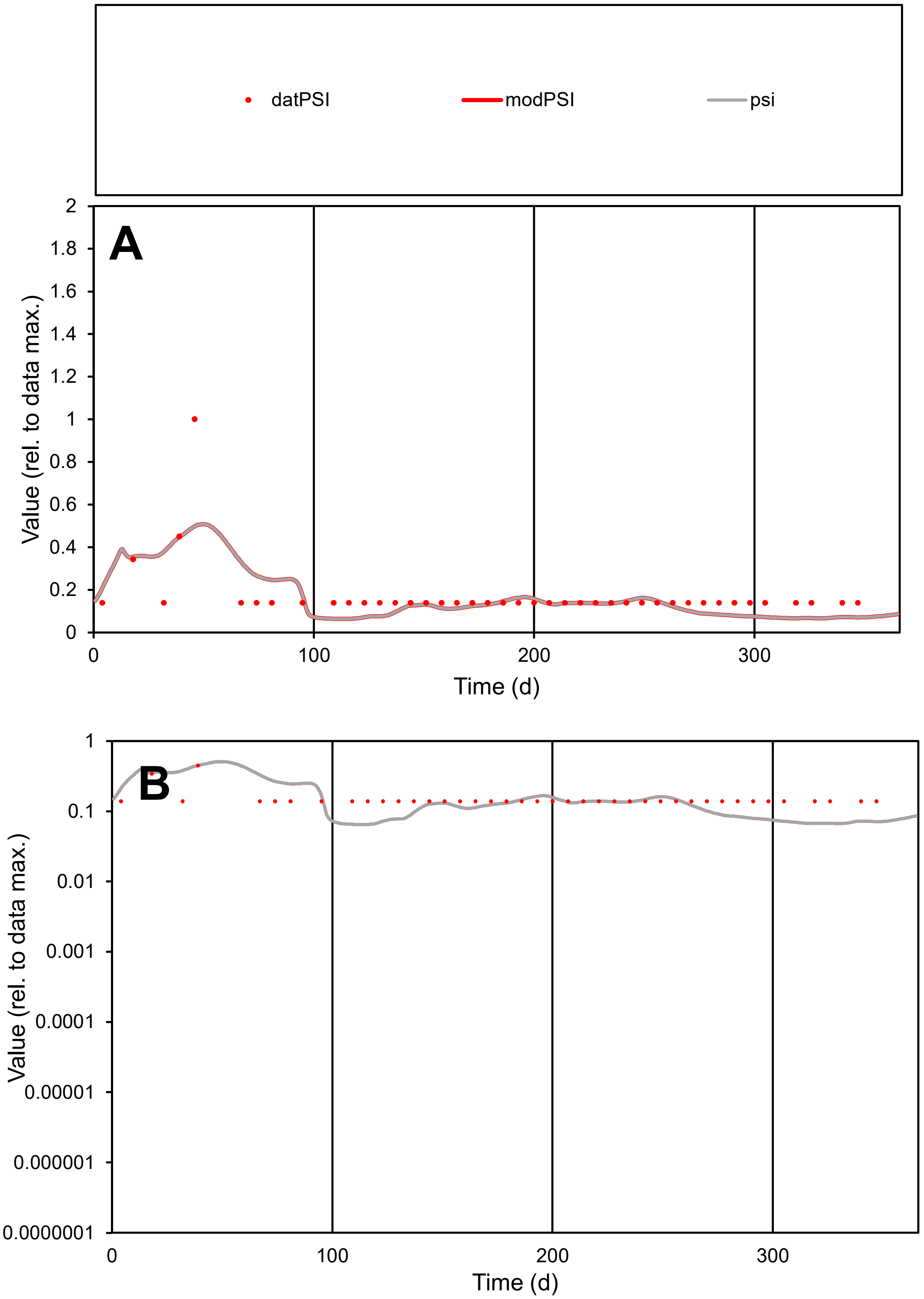


C1

Variables
PSI Provide jtd or Dst
CalibOnly?
2 1 to reproduce of
oNormalize?
2 1=no, 2=yes

ofxn calc*
0.59266
ofxn mod
0.60933

sRun
RunA1
zout
1

Ave
Data
4.4E-06
Model
4E-06

data/vMax
2.6E-05

ofxng
334.07

Min d/m
2.14019

ave/min d
1.2365
ave/min m
2.39329

327_Calib_S_A

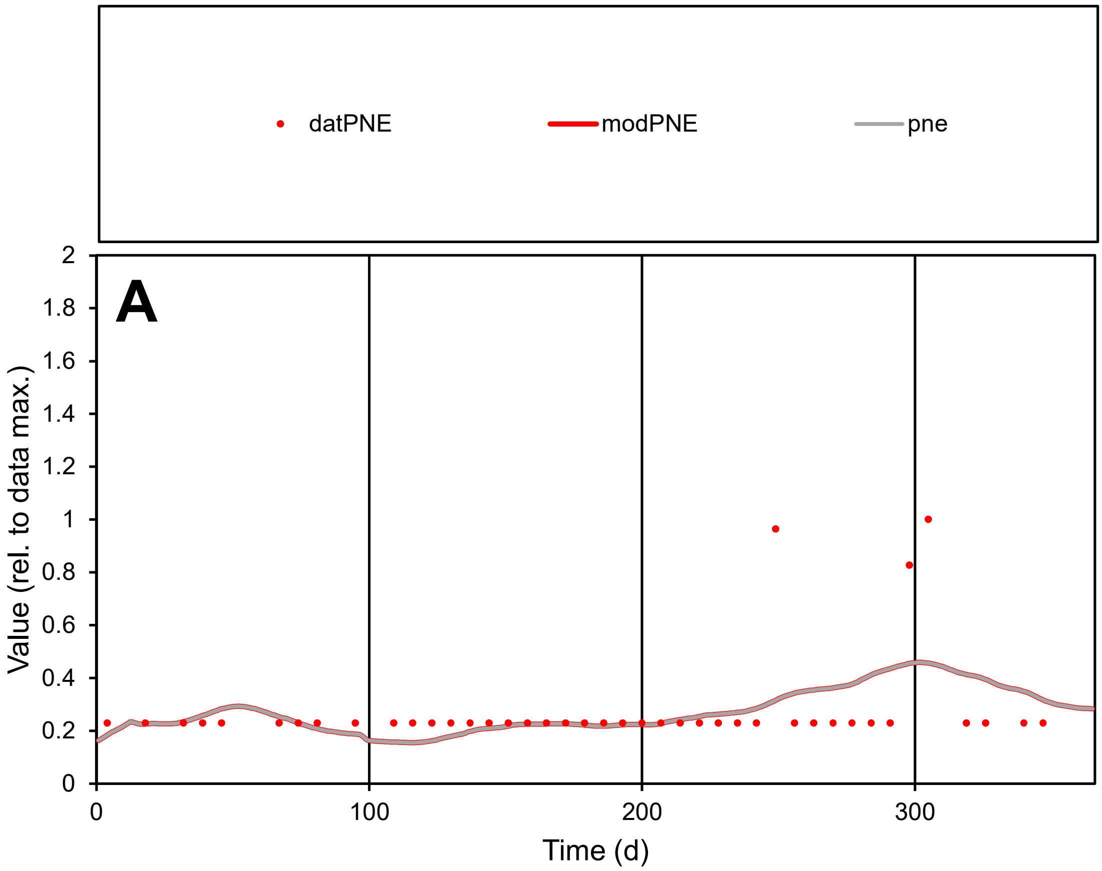


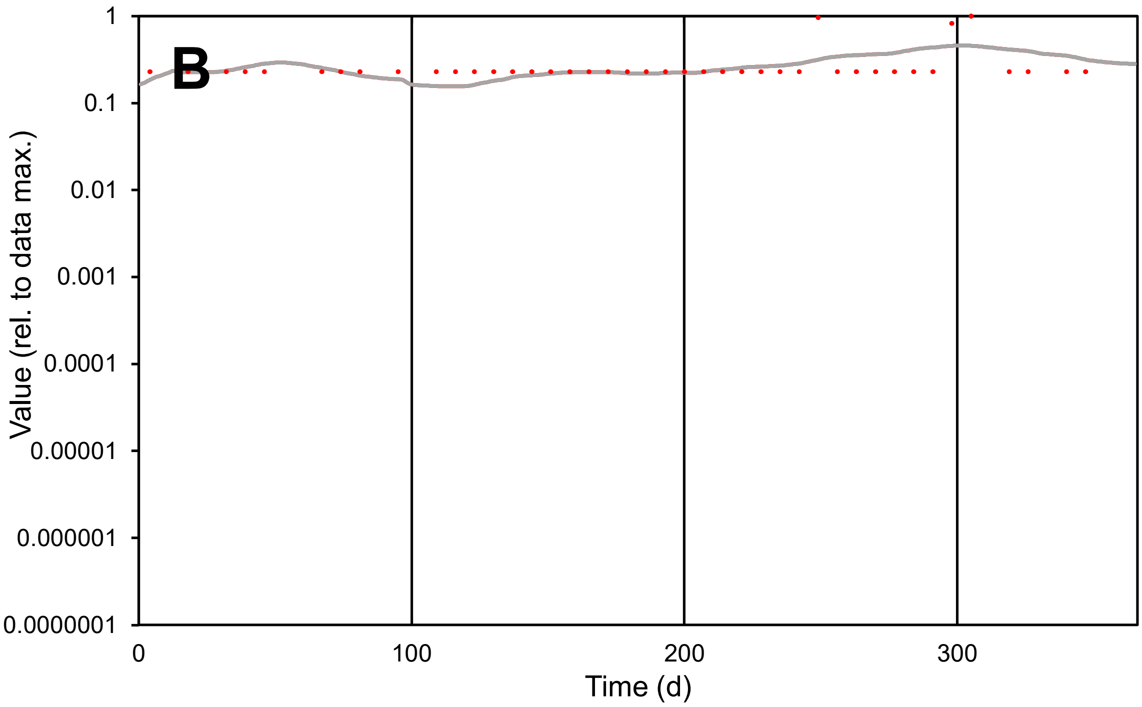


**C1**

Variables
PNE Provide jtd or Dst
CalibOnly?
2 1 to reproduce of
oNormalize?
2 1=no, 2=yes

ofxn calc*
0.5827
ofxn mod
0.59451

sRun
RunA1
zout
1

Ave
Data
1.8E-05
Model
1.7E-05

data/vMax
6.3E-05

ofxng
334.07

Min d/m
1.47112

ave/min d
1.21848
ave/min m
1.7555

327_Calib_S_A

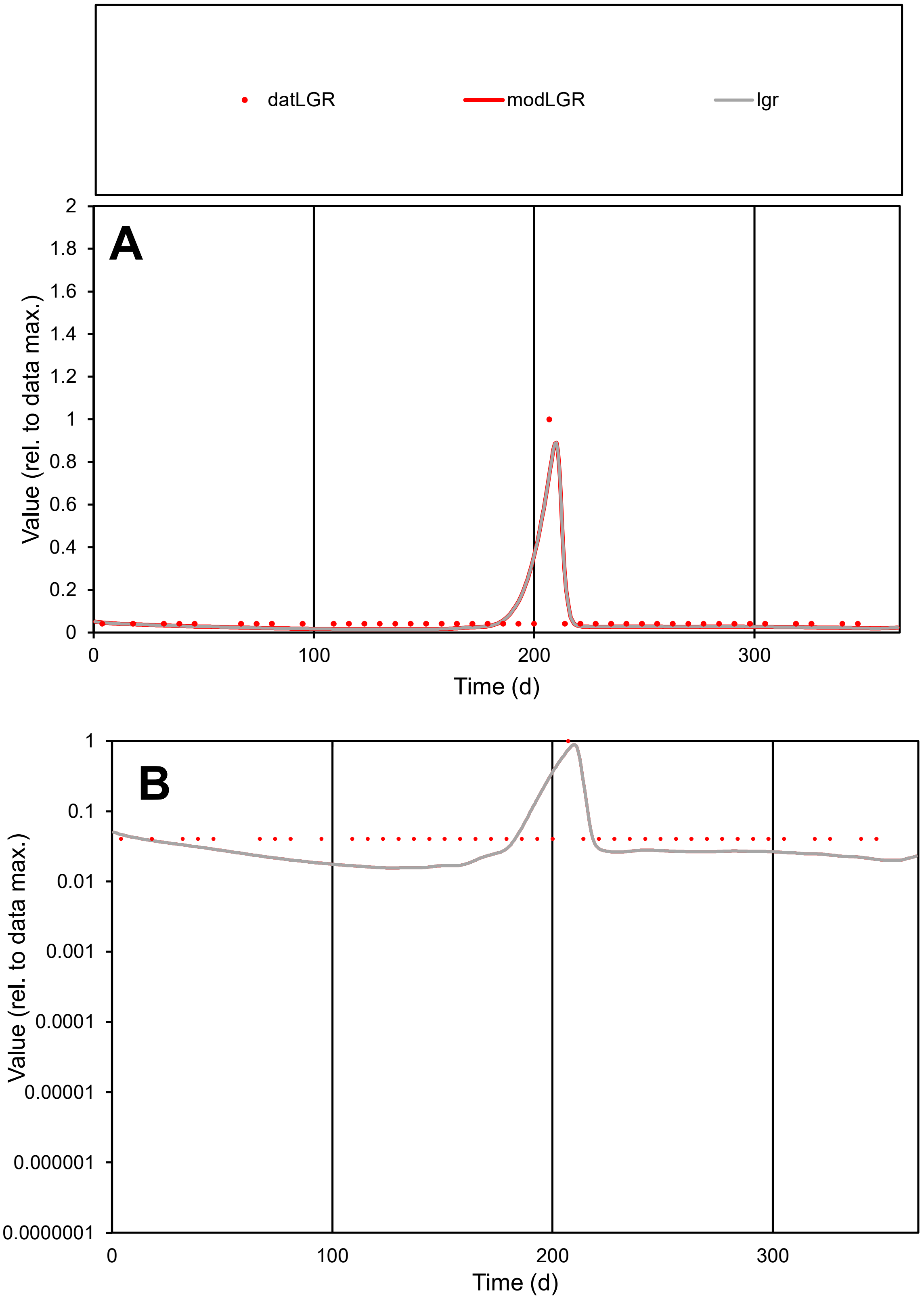


**C1**

Variables
LGR Provide jtd or Dst
CalibOnly?
2 1 to reproduce of
oNormalize?
2 1=no, 2=yes

ofxn calc*
0.85489
ofxn mod
0.98818

sRun
RunA1
zout
1

Ave
Data
8.1E-06
Model
7.1E-06

data/vMax
0.00013

ofxng
334.07

Min d/m
2.57713

ave/min d
1.56566
ave/min m
3.55047

327_Calib_S_A

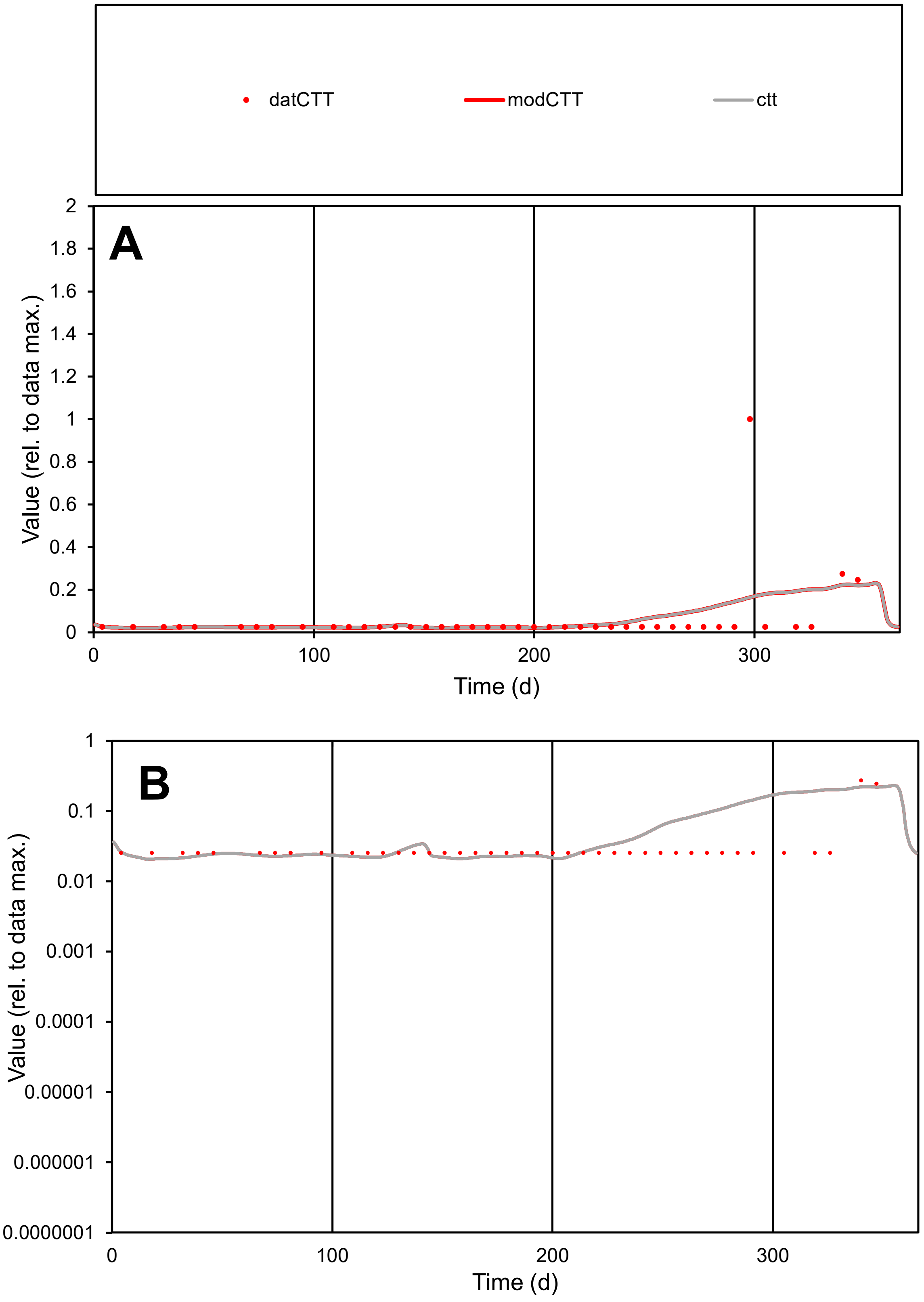


**C1**

Variables

CTT Provide jtd or Dst

CalibOnly?

2 1 to reproduce of

oNormalize?

2 1=no, 2=yes

ofxn calc*

2.33842

ofxn mod

2.35061

sRun

RunA1

zout

1

Ave

Data

3.5E-06

Model

3.6E-06

data/vMax

5.8E-05

ofxng

334.07

Min d/m

1.22158

ave/min d

2.34735

ave/min m

3.00167

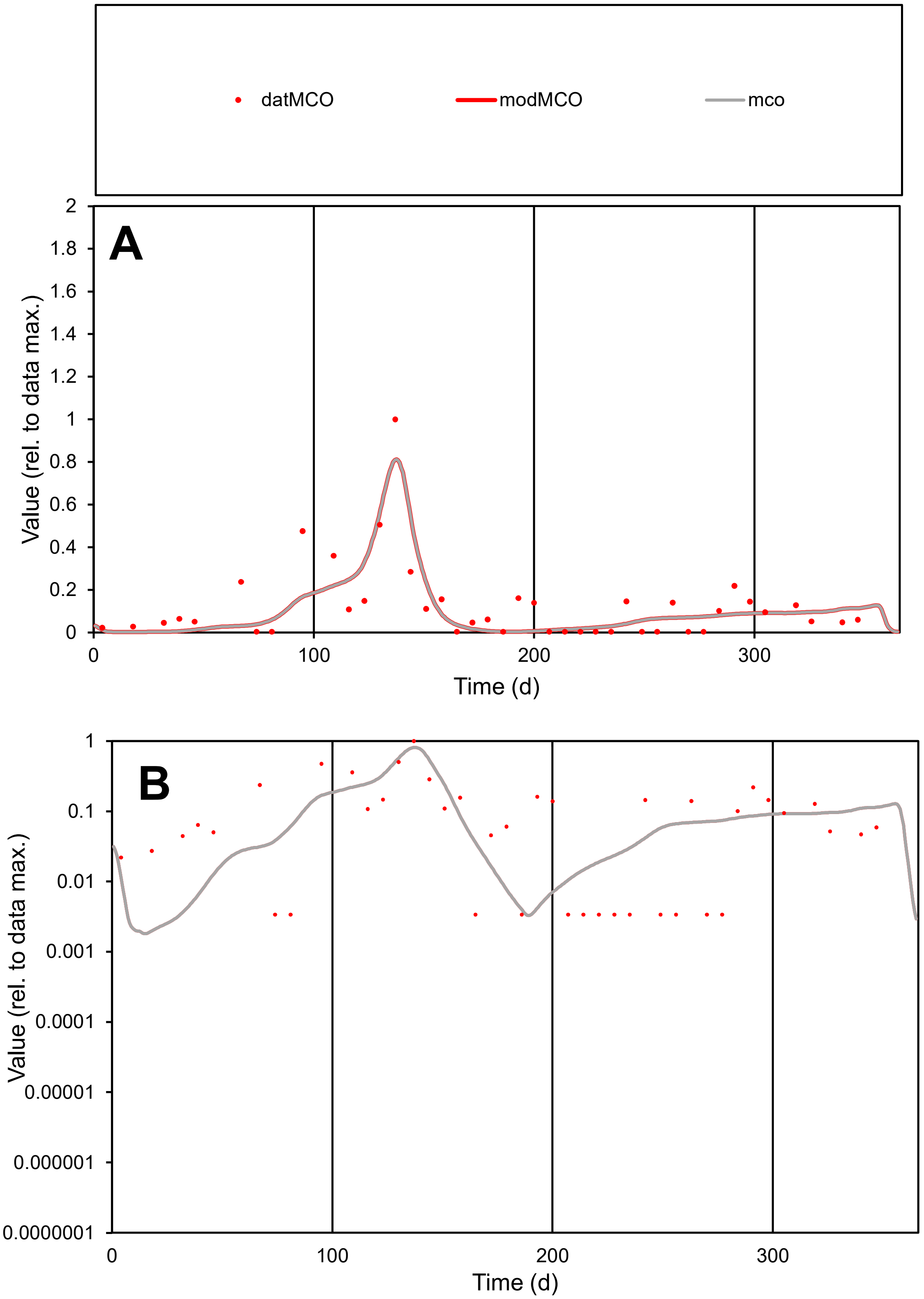


**C1**

Variables
MCO Provide jtd or Dst
CalibOnly?
2 1 to reproduce of
oNormalize?
2 1=no, 2=yes

ofxn calc*
0.80512
ofxn mod
0.83657

sRun
RunA1
zout
1

Ave
Data
2.2E-06
Model
2E-06

data/vMax
1.8E-05

ofxng
334.07

Min d/m
1.59786

ave/min d
36.4577
ave/min m
51.4766

327_Calib_S_A

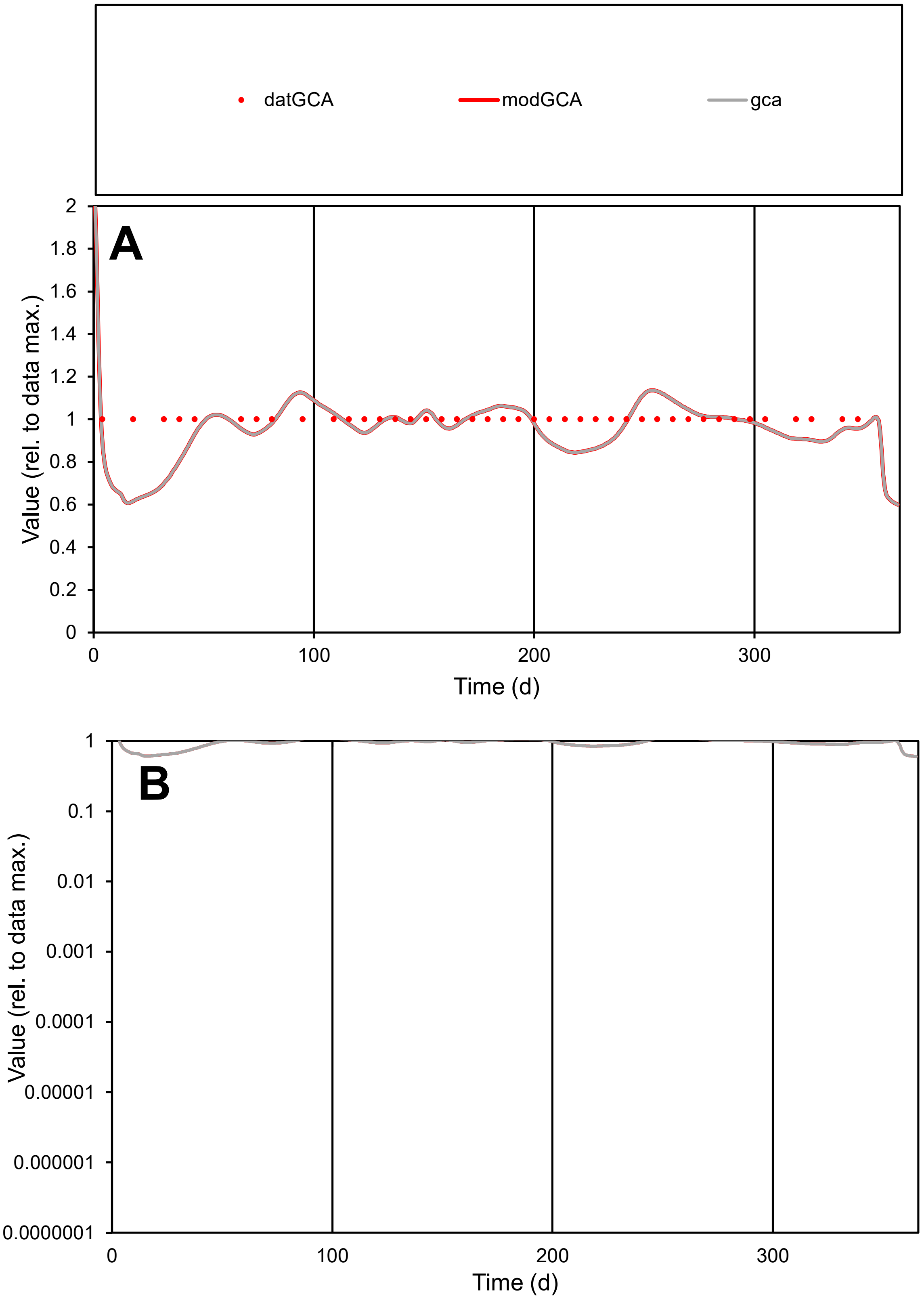


**C1**

Variables
GCA Provide jtd or Dst
CalibOnly?
2 1 to reproduce of
oNormalize?
2 1=no, 2=yes

ofxn calc*
0.10461
ofxn mod
0.15379

sRun
RunA1
zout
1

Ave
Data
8.4E-06
Model
8.1E-06

data/vMax
8.4E-06

ofxng
334.07

Min d/m
1.5995

ave/min d
1
ave/min m
1.54131

327_Calib_S_A

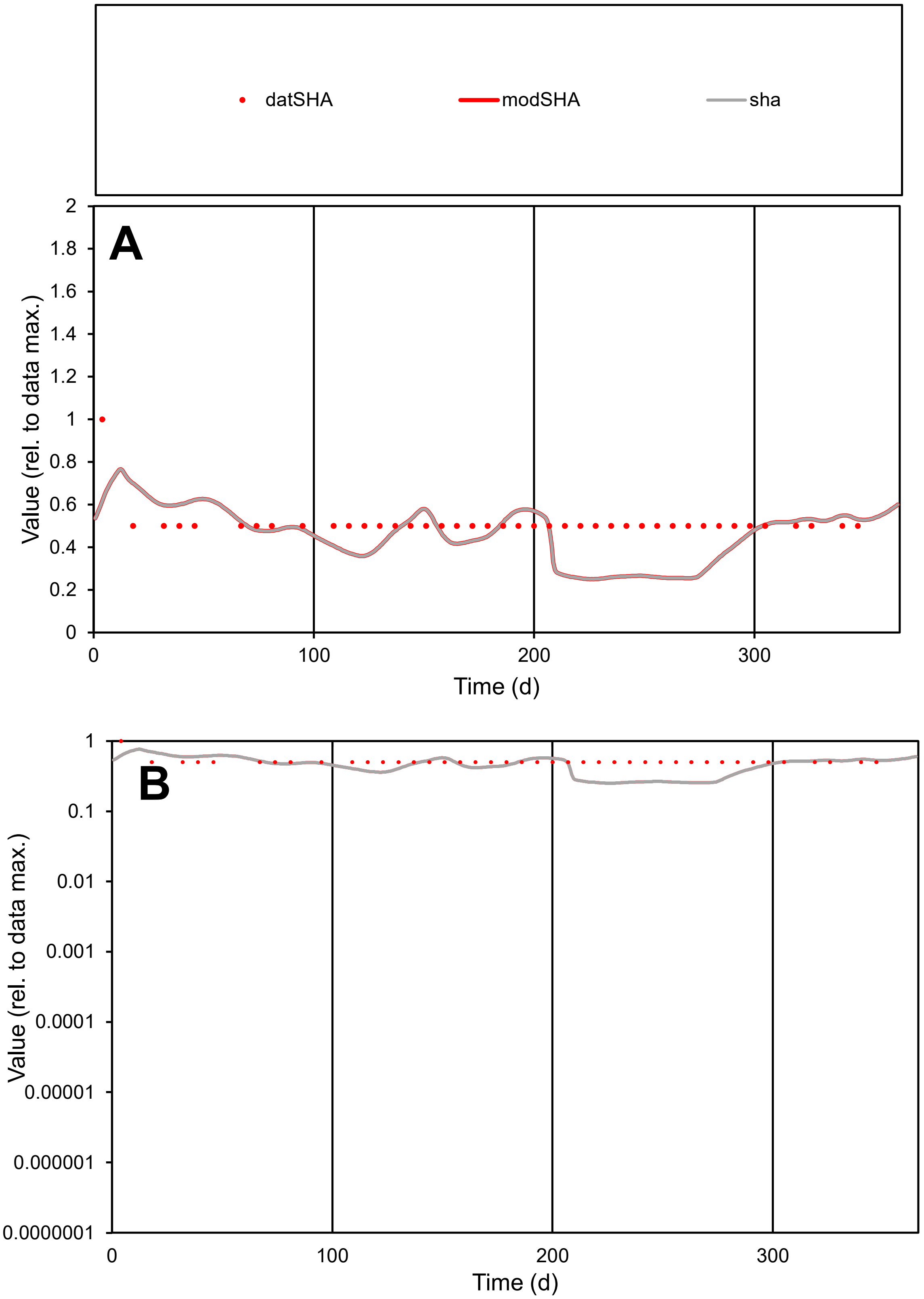


**C1**

Variables
SHA Provide jtd or Dst
CalibOnly?
2 1 to reproduce of
oNormalize?
2 1=no, 2=yes

ofxn calc*
0.28255
ofxn mod
0.30225

sRun
RunA1
zout
1

Ave
Data
7.1E-07
Model
6.2E-07

data/vMax
1.4E-06

ofxng
334.07

Min d/m
1.9852

ave/min d
1.02382
ave/min m
1.76747

327_Calib_S_A

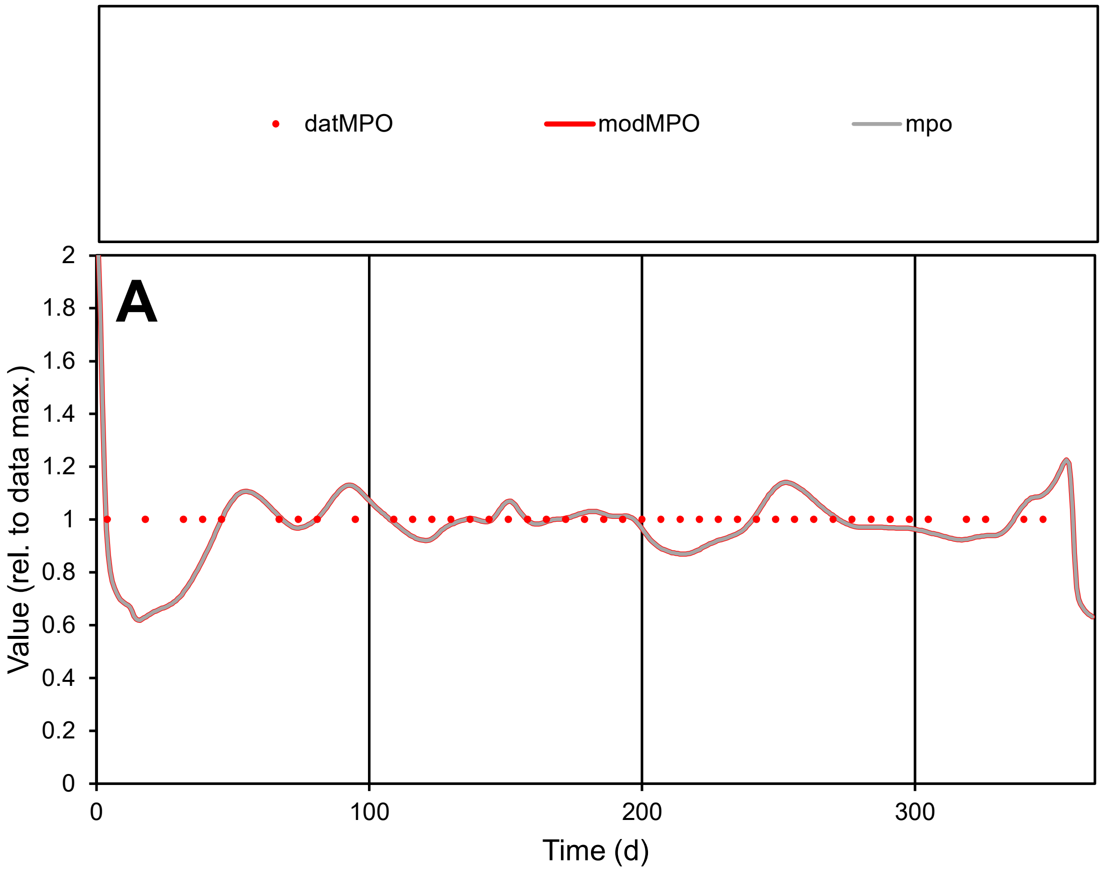


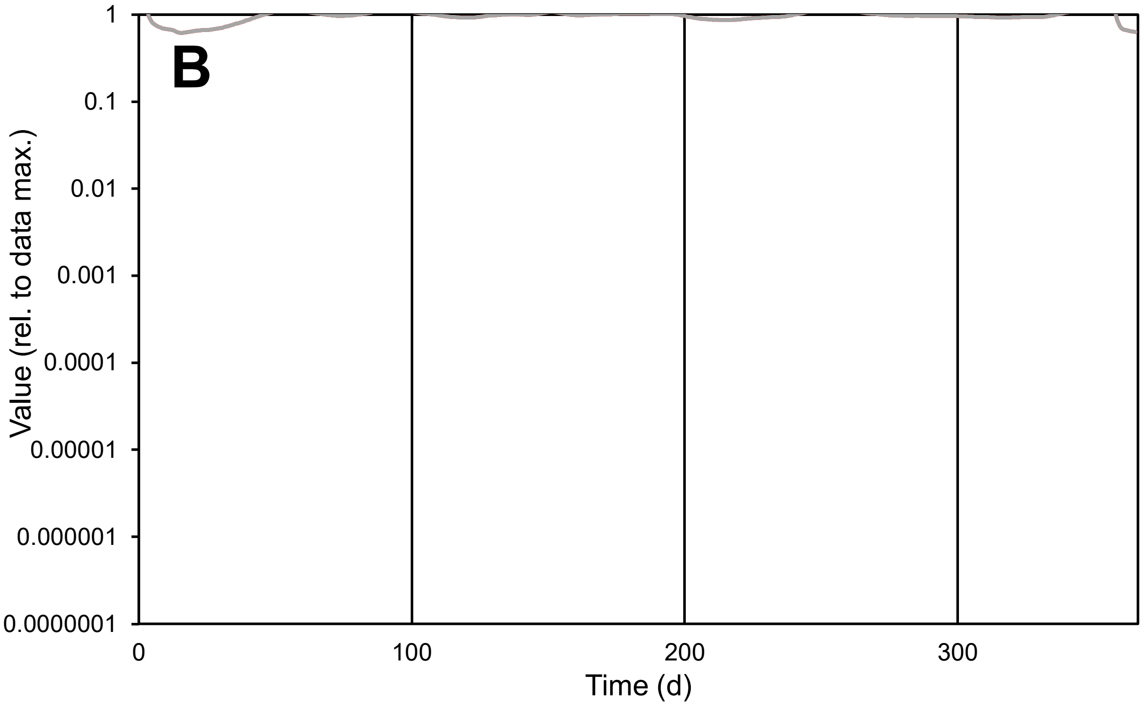


**C1**

Variables
MPO Provide jtd or Dst
CalibOnly?
2 1 to reproduce of
oNormalize?
2 1=no, 2=yes

ofxn calc*
0.09469
ofxn mod
0.14769

sRun
RunA1
zout
1

Ave
Data
1.2E-05
Model
1.2E-05

data/vMax
1.2E-05

ofxng
334.07

Min d/m
1.55728

ave/min d
1
ave/min m
1.51817

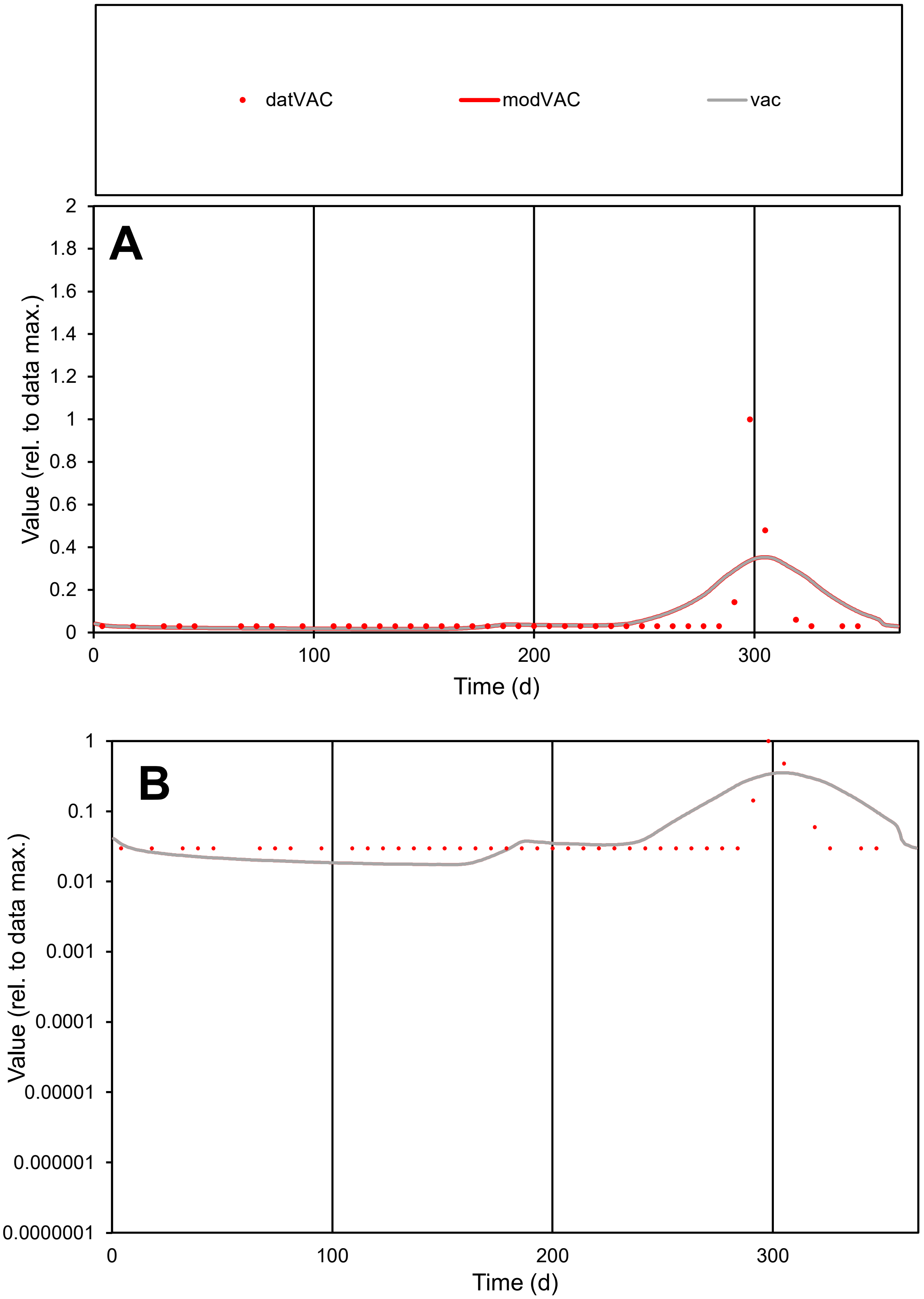


**C1**

Variables
VAC Provide jtd or Dst
CalibOnly?
2 1 to reproduce of
oNormalize?
2 1=no, 2=yes

ofxn calc*
1.86946
ofxn mod
1.89108

sRun
RunA1
zout
1

Ave
Data
2.1E-05
Model
2.5E-05

data/vMax
0.00032

ofxng
334.07

Min d/m
1.70526

ave/min d
2.25065
ave/min m
4.42485

327_Calib_S_A

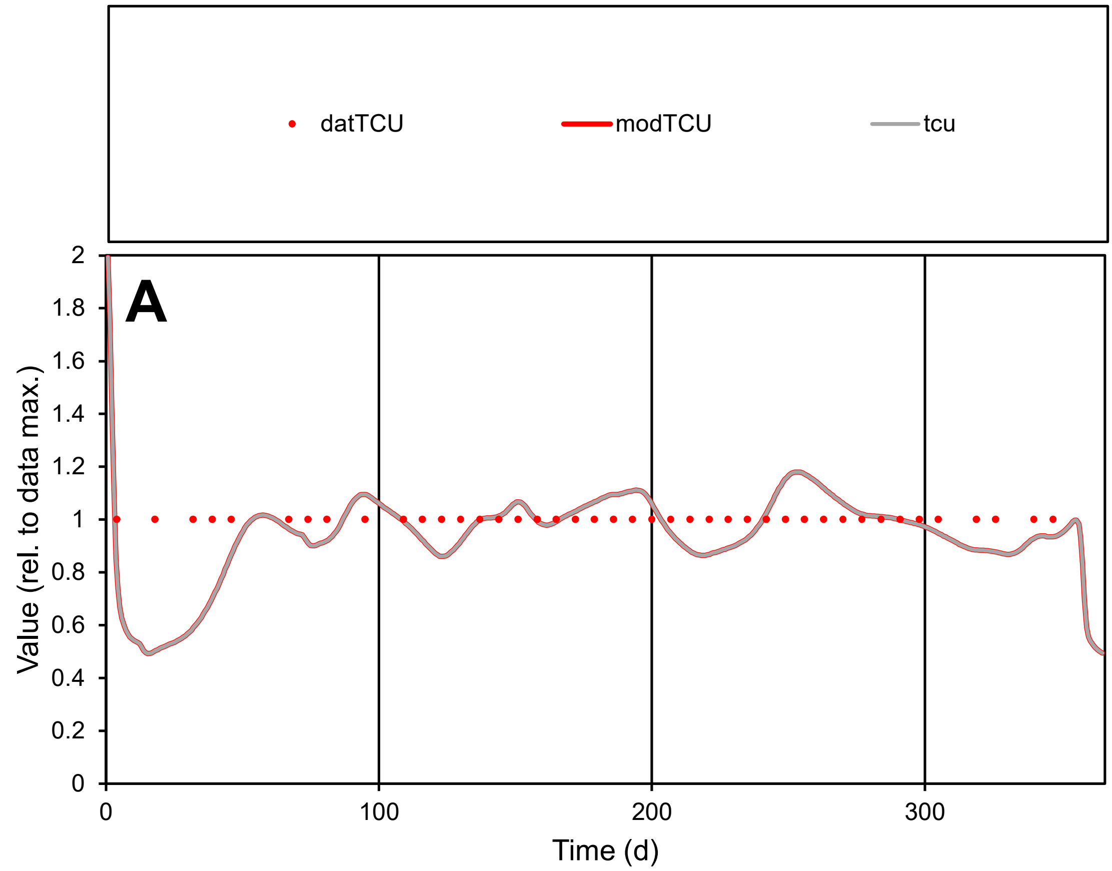


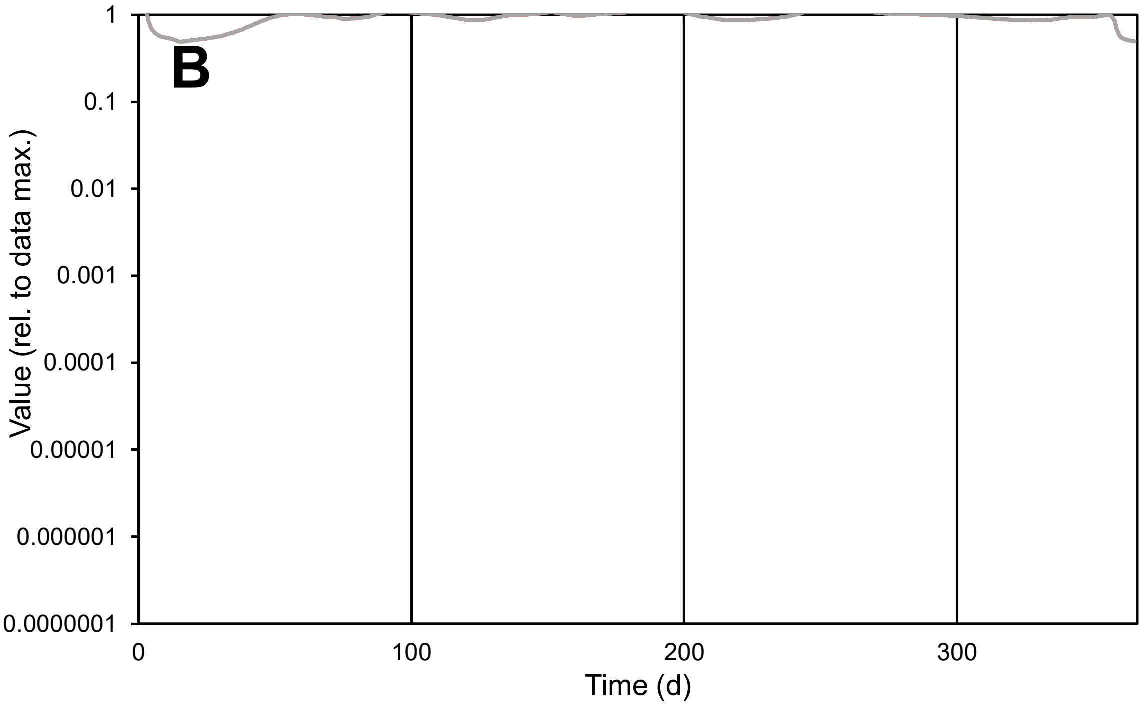


**C1**

Variables
TCU Provide jtd or Dst
CalibOnly?
2 1 to reproduce of
oNormalize?
2 1=no, 2=yes

ofxn calc*
0.13774
ofxn mod
0.17768

sRun
RunA1
zout
1

Ave
Data
9.8E-06
Model
9.3E-06

data/vMax
9.8E-06

ofxng
334.07

Min d/m
1.95515

ave/min d
1
ave/min m
1.86867

327_Calib_S_A

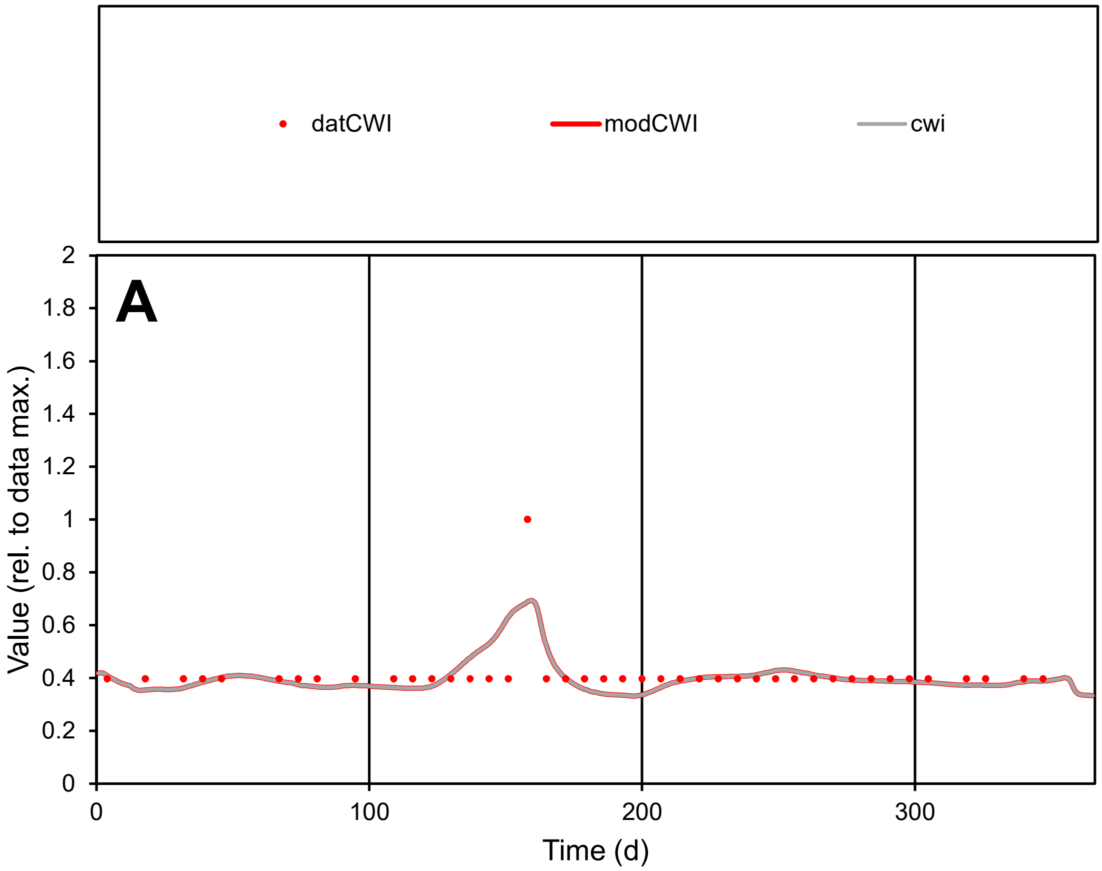


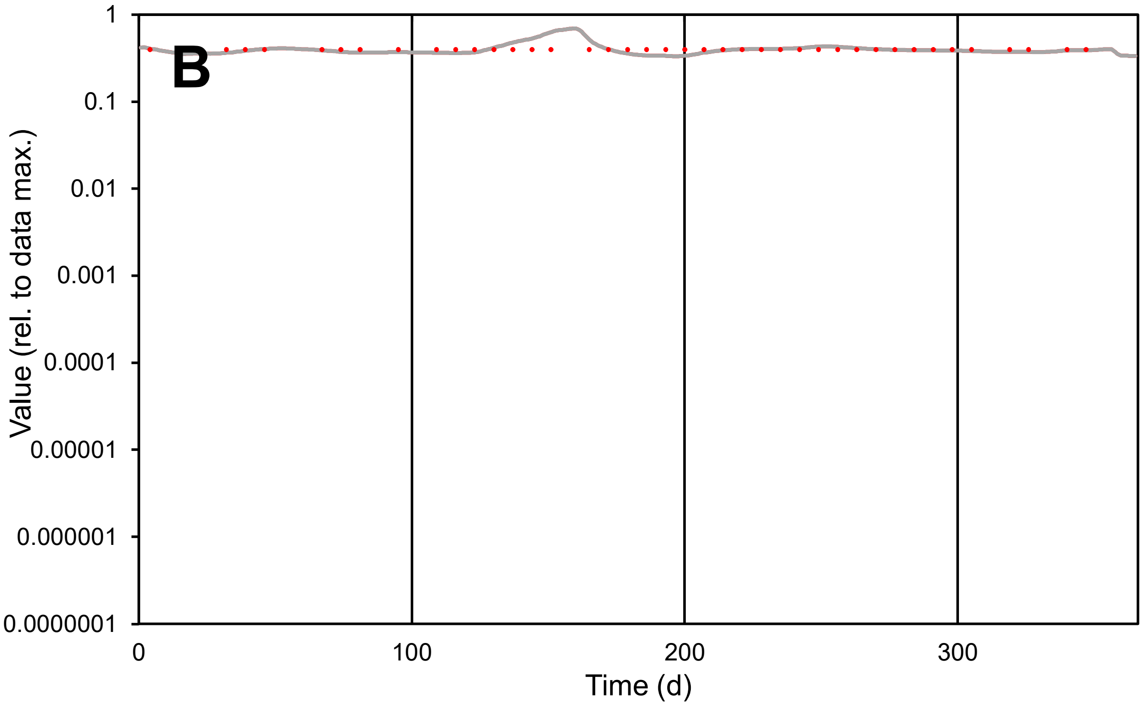


**C1**

Variables
CWI Provide jtd or Dst
CalibOnly?
2 1 to reproduce of
oNormalize?
2 1=no, 2=yes

ofxn calc*
0.17509
ofxn mod
0.20805

sRun
RunA1
zout
1

Ave
Data
1.4E-05
Model
1.4E-05

data/vMax
3.3E-05

ofxng
334.07

Min d/m
1.18873

ave/min d
1.0361
ave/min m
1.21467

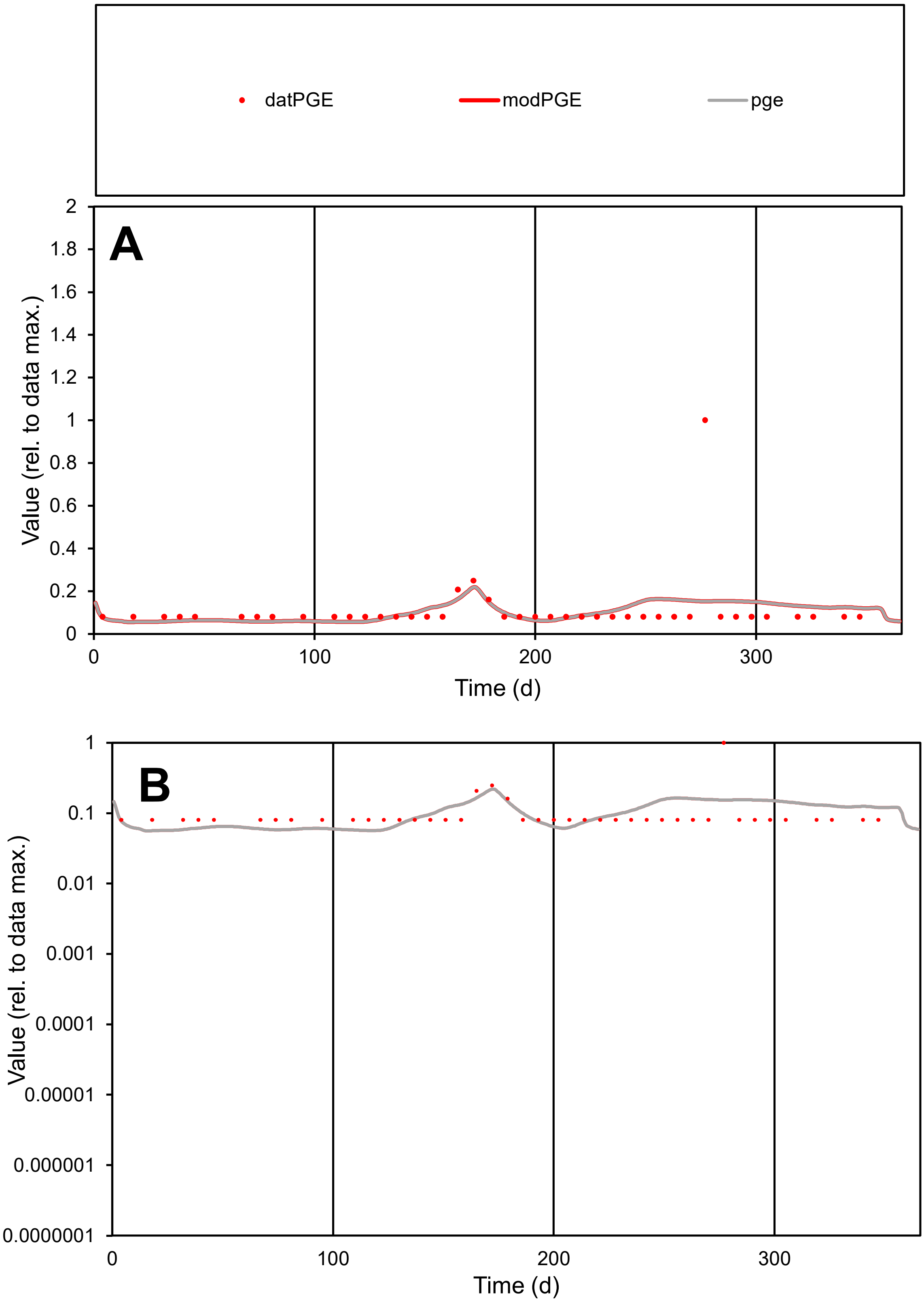


**C1**

Variables
PGE Provide jtd or Dst
CalibOnly?
2 1 to reproduce of
oNormalize?
2 1=no, 2=yes

ofxn calc*
1.22905
ofxn mod
1.23703

sRun
RunA1
zout
1

Ave
Data
8.1E-06
Model
7.8E-06

data/vMax
7.3E-05

ofxng
334.07

Min d/m
1.42047

ave/min d
1.38285
ave/min m
1.87291

327_Calib_S_A

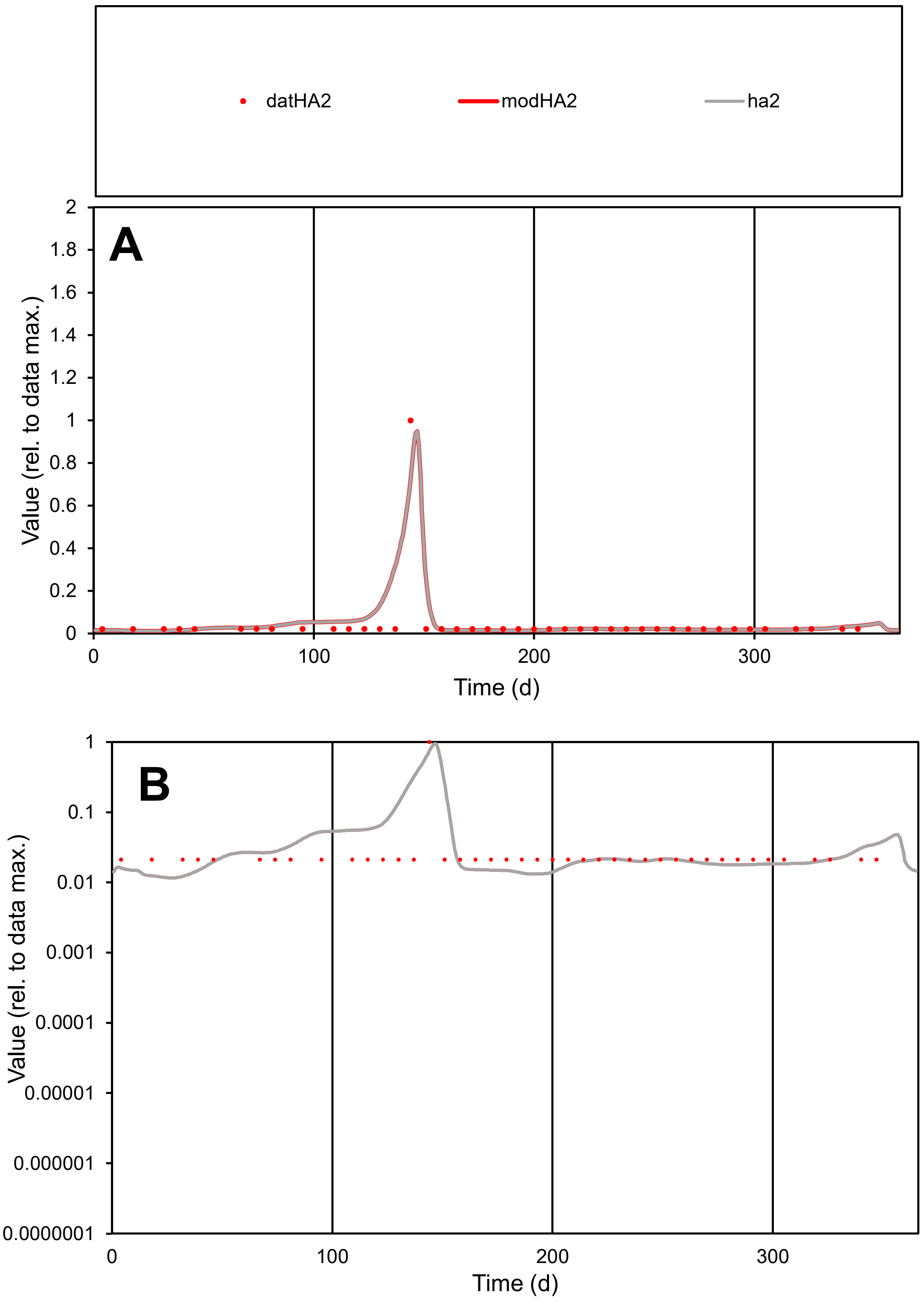


**C1**

Variables
HA2 Provide jtd or Dst
CalibOnly?
2 1 to reproduce of
oNormalize?
2 1=no, 2=yes

ofxn calc*
1.03274
ofxn mod
1.21851

sRun
RunA1
zout
1

Ave
Data
5.6E-06
Model
6.7E-06

data/vMax
0.00013

ofxng
334.07

Min d/m
1.7167

ave/min d
2.10268
ave/min m
4.32815

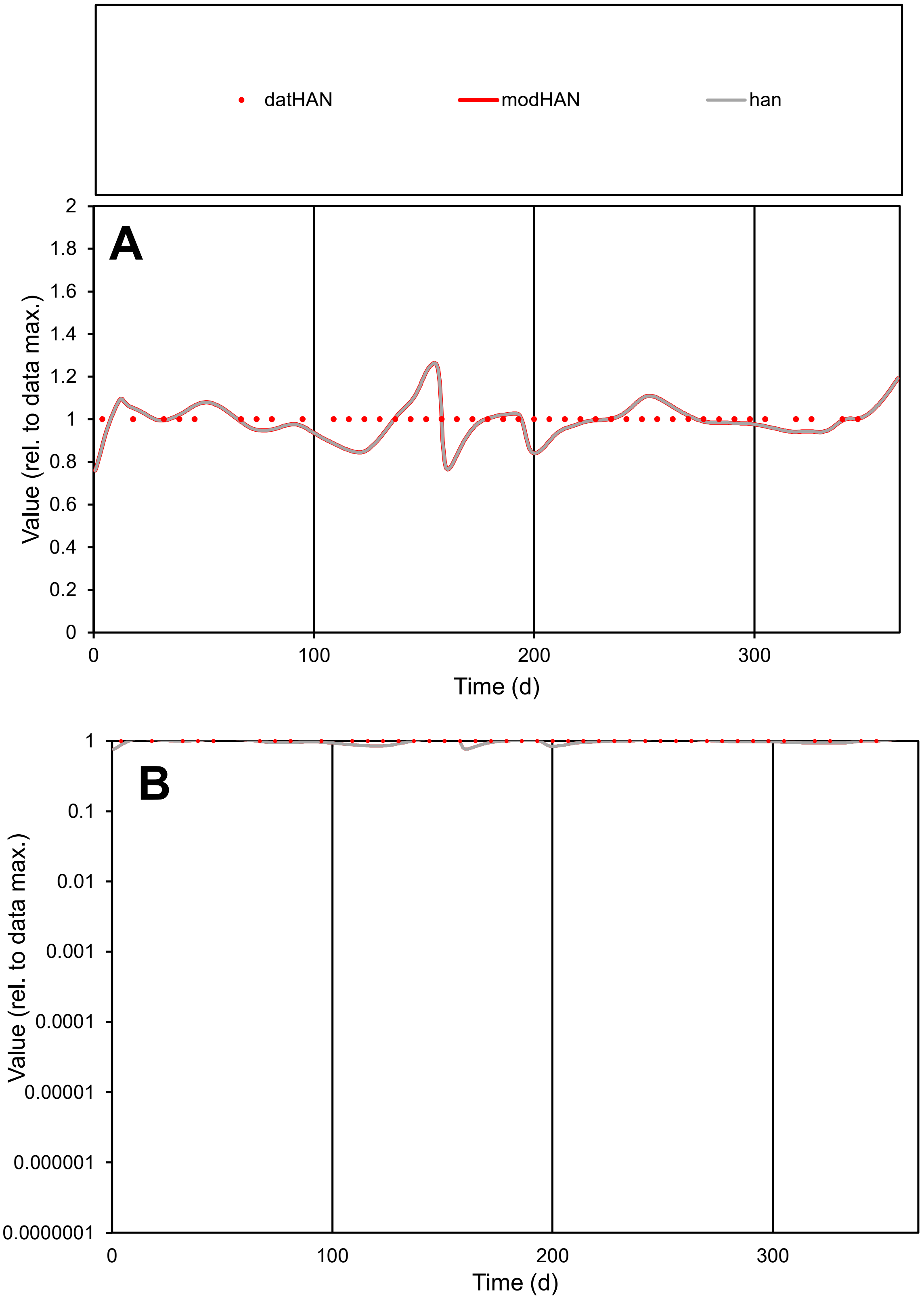


**C1**

Variables
HAN Provide jtd or Dst
CalibOnly?
2 1 to reproduce of
oNormalize?
2 1=no, 2=yes

ofxn calc*
0.07699
ofxn mod
0.13799

sRun
RunA1
zout
1

Ave
Data
5.1E-06
Model
5E-06

data/vMax
5.1E-06

ofxng
334.07

Min d/m
1.18886

ave/min d
1
ave/min m
1.17103

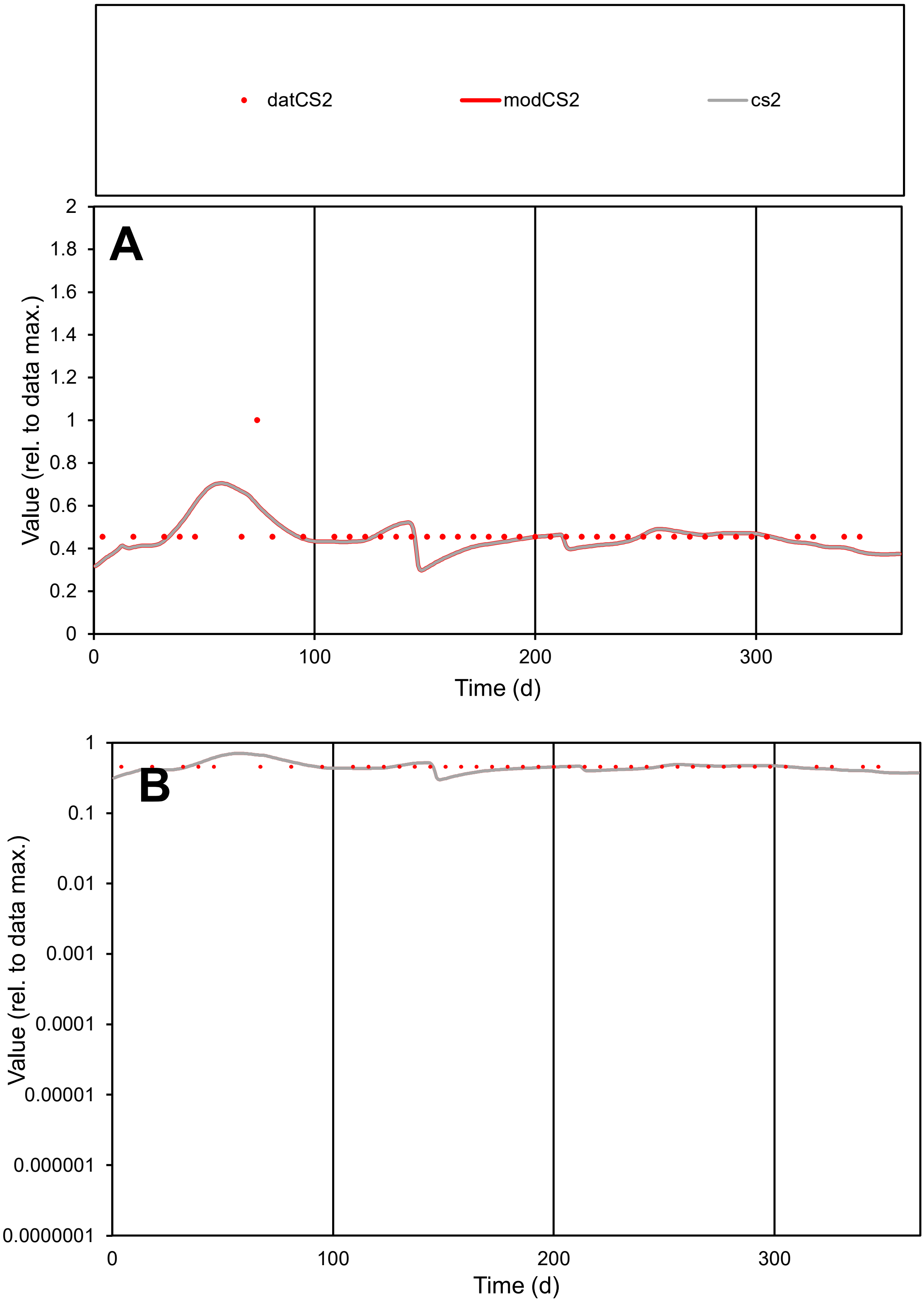


**C1**

Variables
CS2 Provide jtd or Dst
CalibOnly?
2 1 to reproduce of
oNormalize?
2 1=no, 2=yes

ofxn calc*
0.18982
ofxn mod
0.22152

sRun
RunA1
zout
1

Ave
Data
7.8E-06
Model
7.5E-06

data/vMax
1.7E-05

ofxng
334.07

Min d/m
1.43659

ave/min d
1.02853
ave/min m
1.42839

327_Calib_S_A

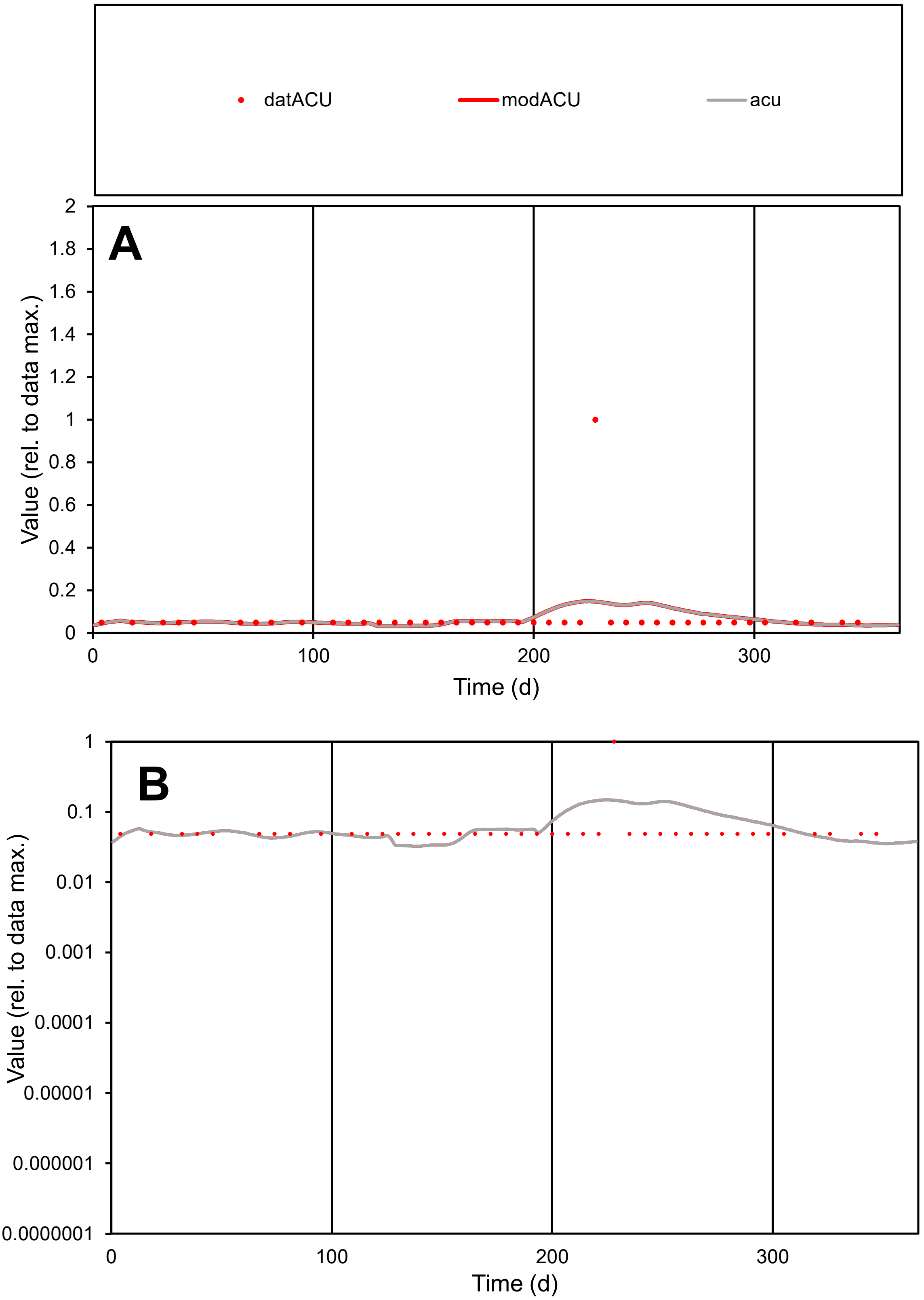


**C1**

Variables
ACU Provide jtd or Dst
CalibOnly?
2 1 to reproduce of
oNormalize?
2 1=no, 2=yes

ofxn calc*
1.92425
ofxn mod
1.93018

sRun
RunA1
zout
1

Ave
Data
1.6E-05
Model
1.6E-05

data/vMax
0.00023

ofxng
334.07

Min d/m
1.50014

ave/min d
1.46481
ave/min m
2.12846

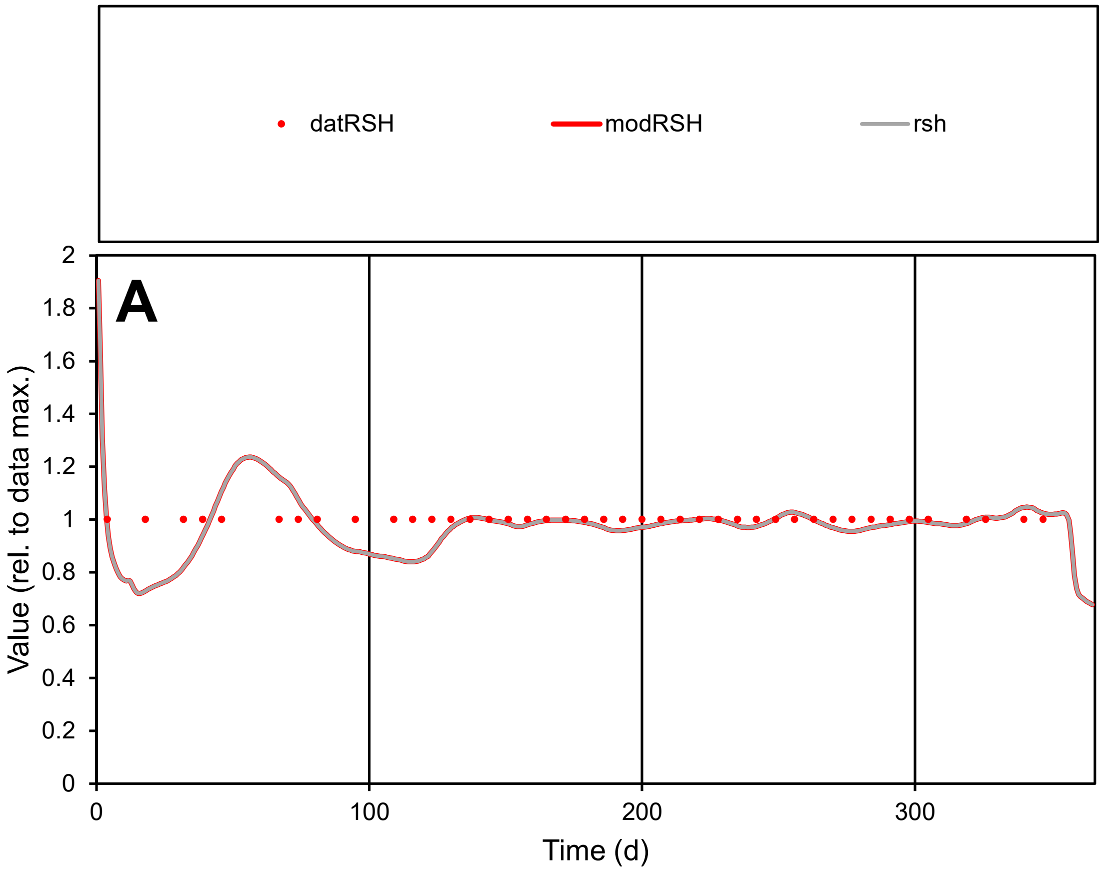


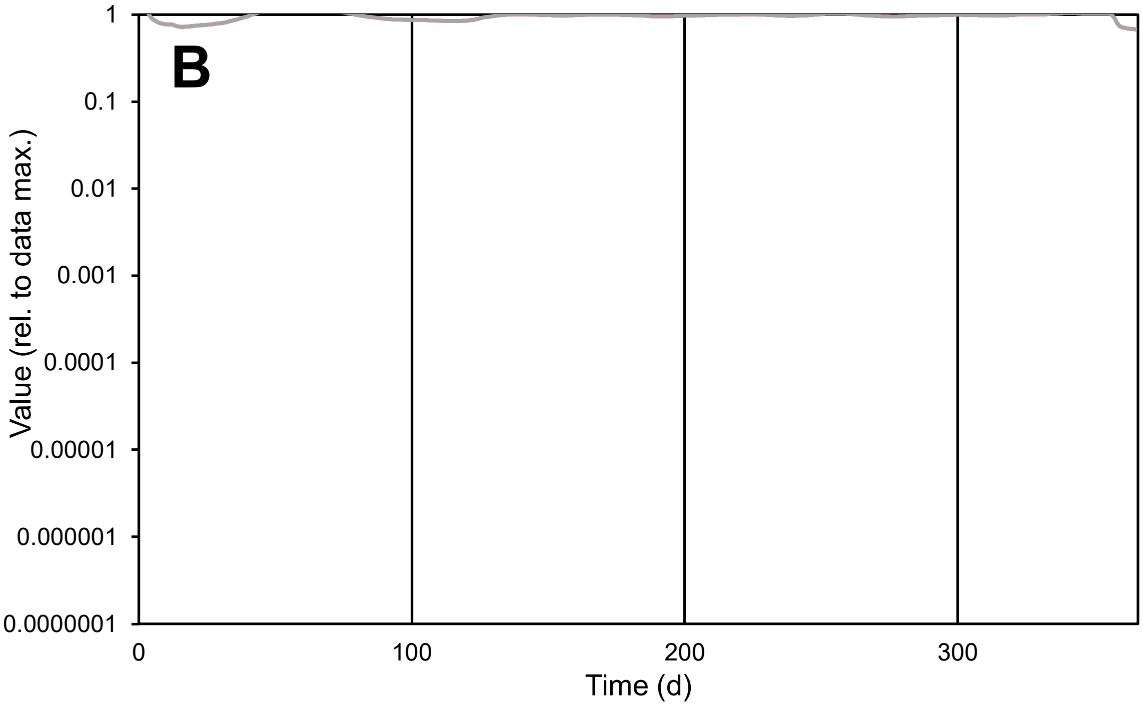


**C1**

Variables
RSH Provide jtd or Dst
CalibOnly?
2 1 to reproduce of
oNormalize?
2 1=no, 2=yes

ofxn calc*
0.07544
ofxn mod
0.13507

sRun
RunA1
zout
1

Ave
Data
1.8E-05
Model
1.7E-05

data/vMax
1.8E-05

ofxng
334.07

Min d/m
1.35201

ave/min d
1
ave/min m
1.31751

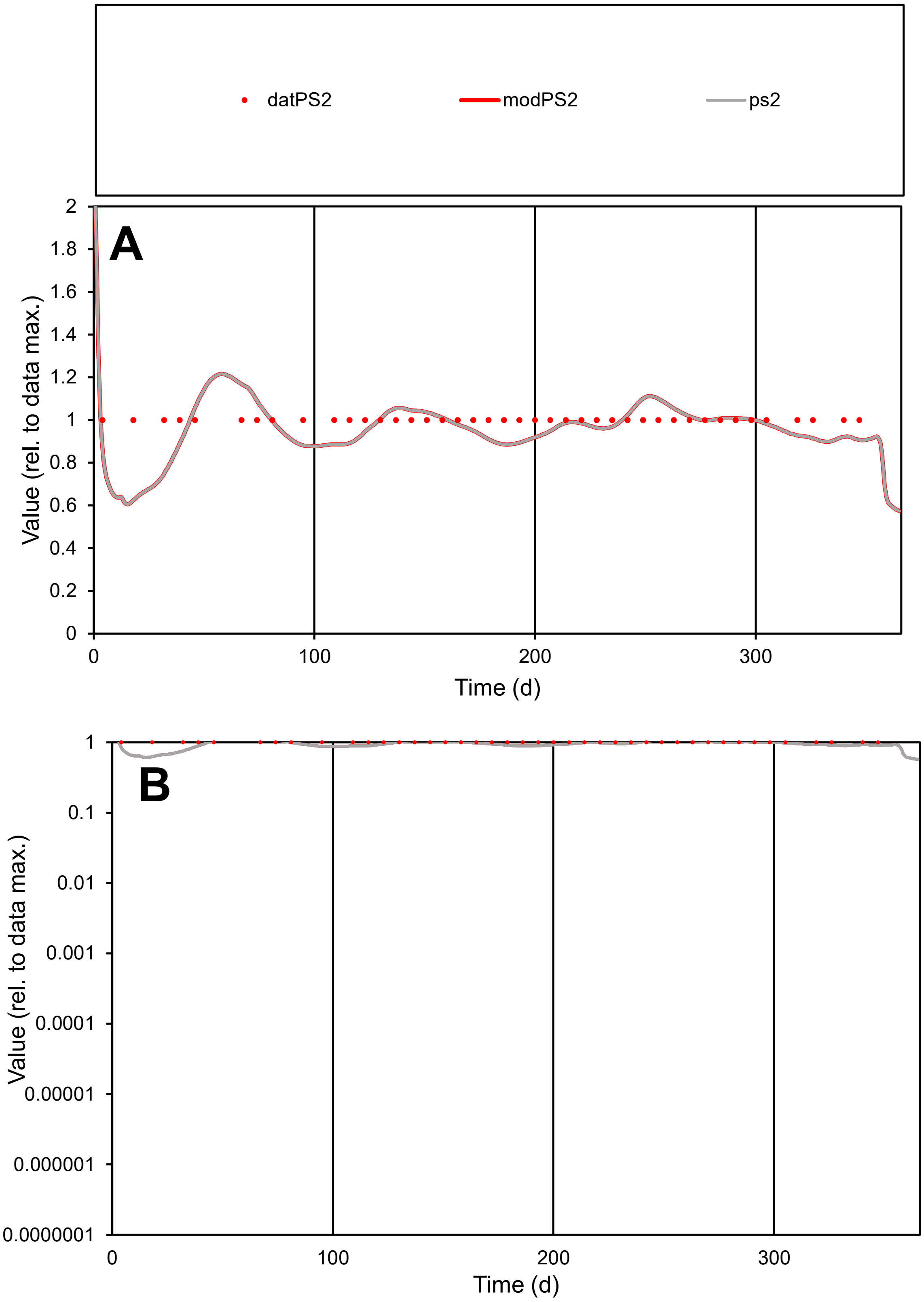


**C1**

Variables
PS2 Provide jtd or Dst
CalibOnly?
2 1 to reproduce of
oNormalize?
2 1=no, 2=yes

ofxn calc*
0.09808
ofxn mod
0.14979

sRun
RunA1
zout
1

Ave
Data
1.9E-05
Model
1.8E-05

data/vMax
1.9E-05

ofxng
334.07

Min d/m
1.56148

ave/min d
1
ave/min m
1.50909

327_Calib_S_A

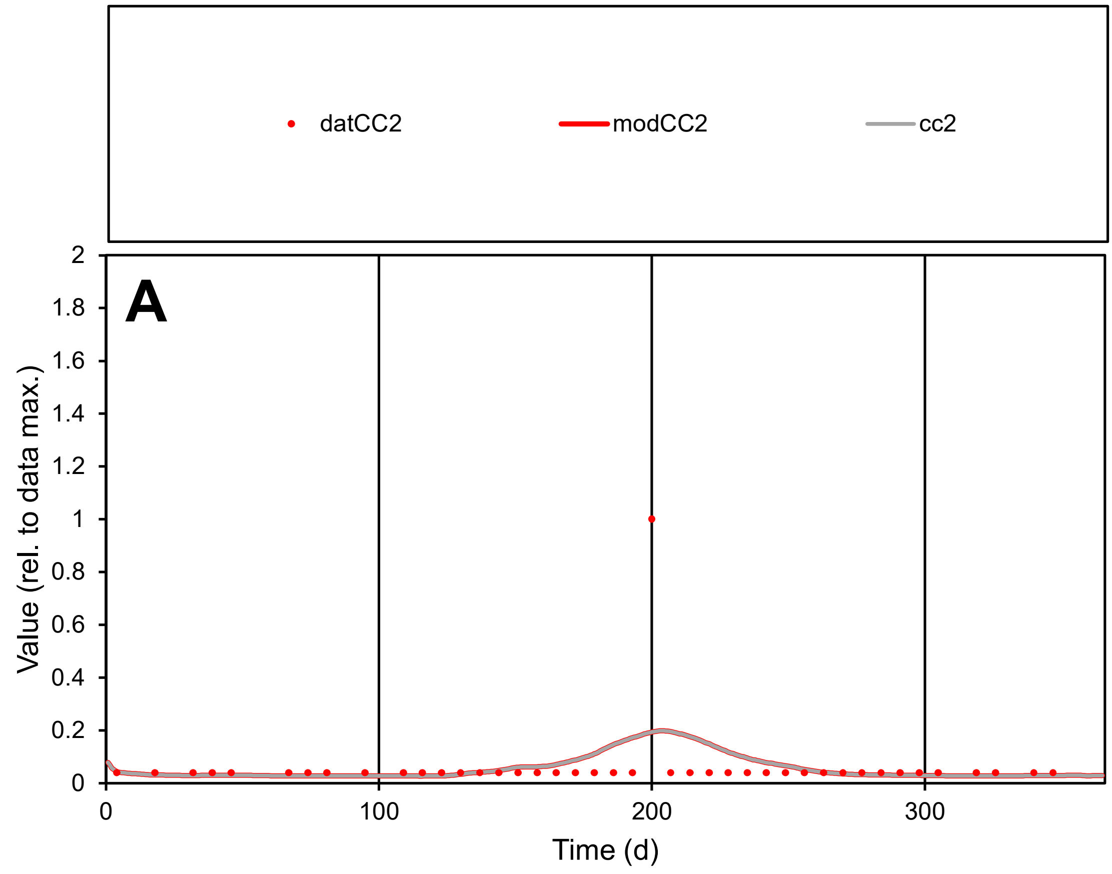


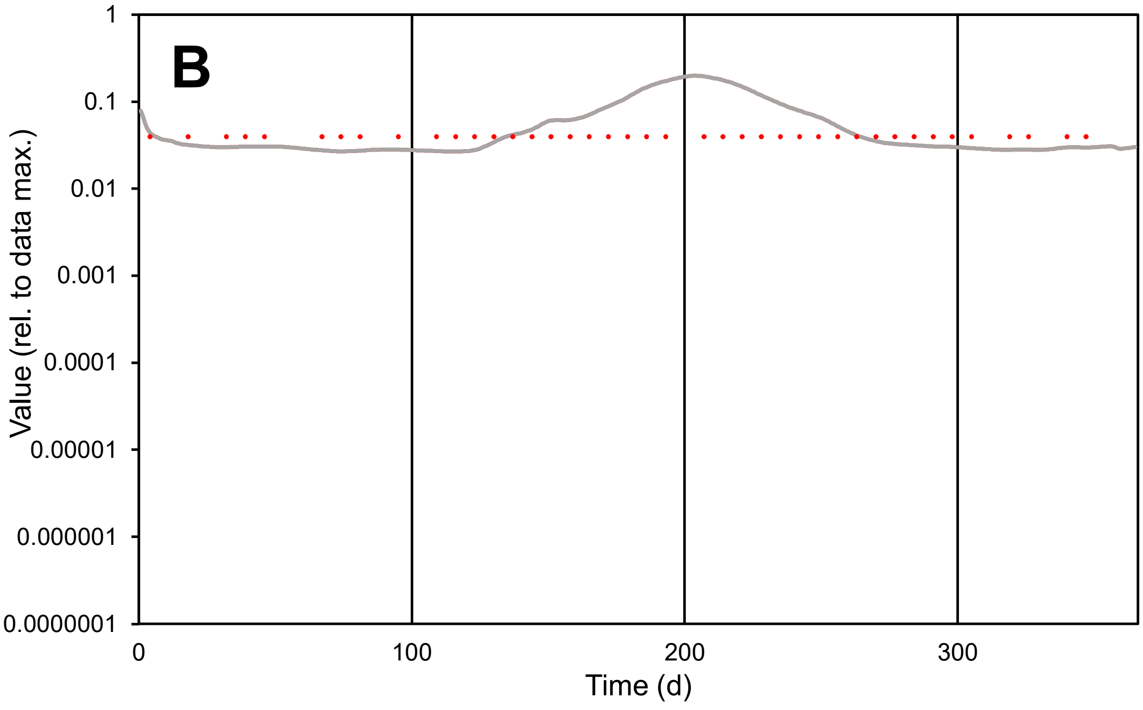


C1

Variables
CC2 Provide jtd or Dst
CalibOnly?
2 1 to reproduce of
oNormalize?
2 1=no, 2=yes

ofxn calc*
2.14234
ofxn mod
2.15714

sRun
RunA1
zout
1

Ave
Data
9.3E-06
Model
9.4E-06

data/vMax
0.00015

ofxng
334.07

Min d/m
1.47265

ave/min d
1.57934
ave/min m
2.34898

327_Calib_S_A

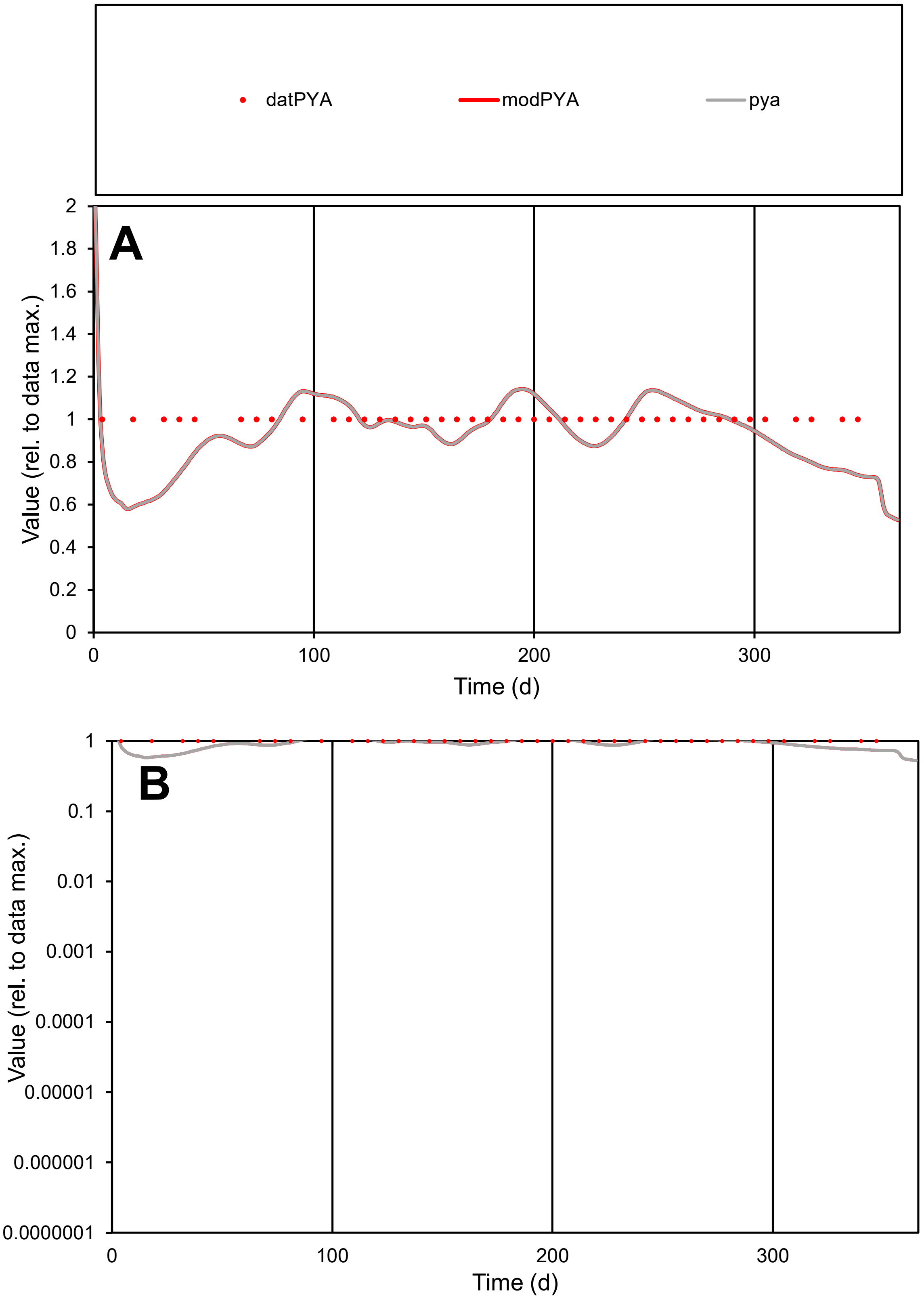


**C1**

Variables

PYA Provide jtd or Dst

CalibOnly?

2 1 to reproduce of

oNormalize?

2 1=no, 2=yes

ofxn calc*

0.13766

ofxn mod

0.17772

sRun

RunA1

zout

1

Ave

Data

1.8E-06

Model

1.7E-06

data/vMax

1.8E-06

ofxng

334.07

Min d/m

1.67623

ave/min d

1

ave/min m

1.58973

327_Calib_S_A

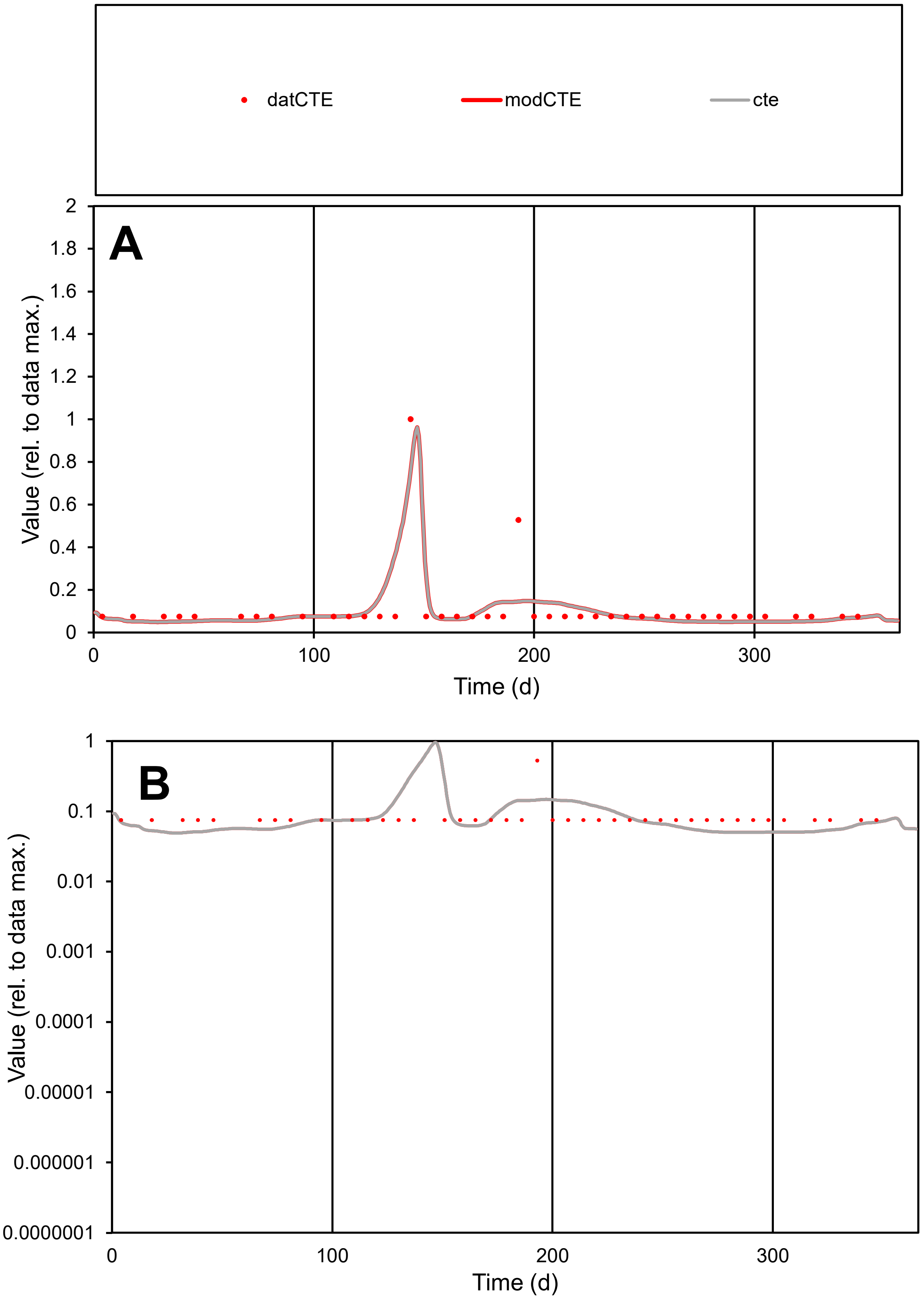


**C1**

Variables
CTE Provide jtd or Dst
CalibOnly?
2 1 to reproduce of
oNormalize?
2 1=no, 2=yes

ofxn calc*
0.73165
ofxn mod
0.7749

sRun
RunA1
zout
1

Ave
Data
8.8E-06
Model
8.5E-06

data/vMax
8.1E-05

ofxng
334.07

Min d/m
1.50998

ave/min d
1.43576
ave/min m
2.09389

327_Calib_S_A

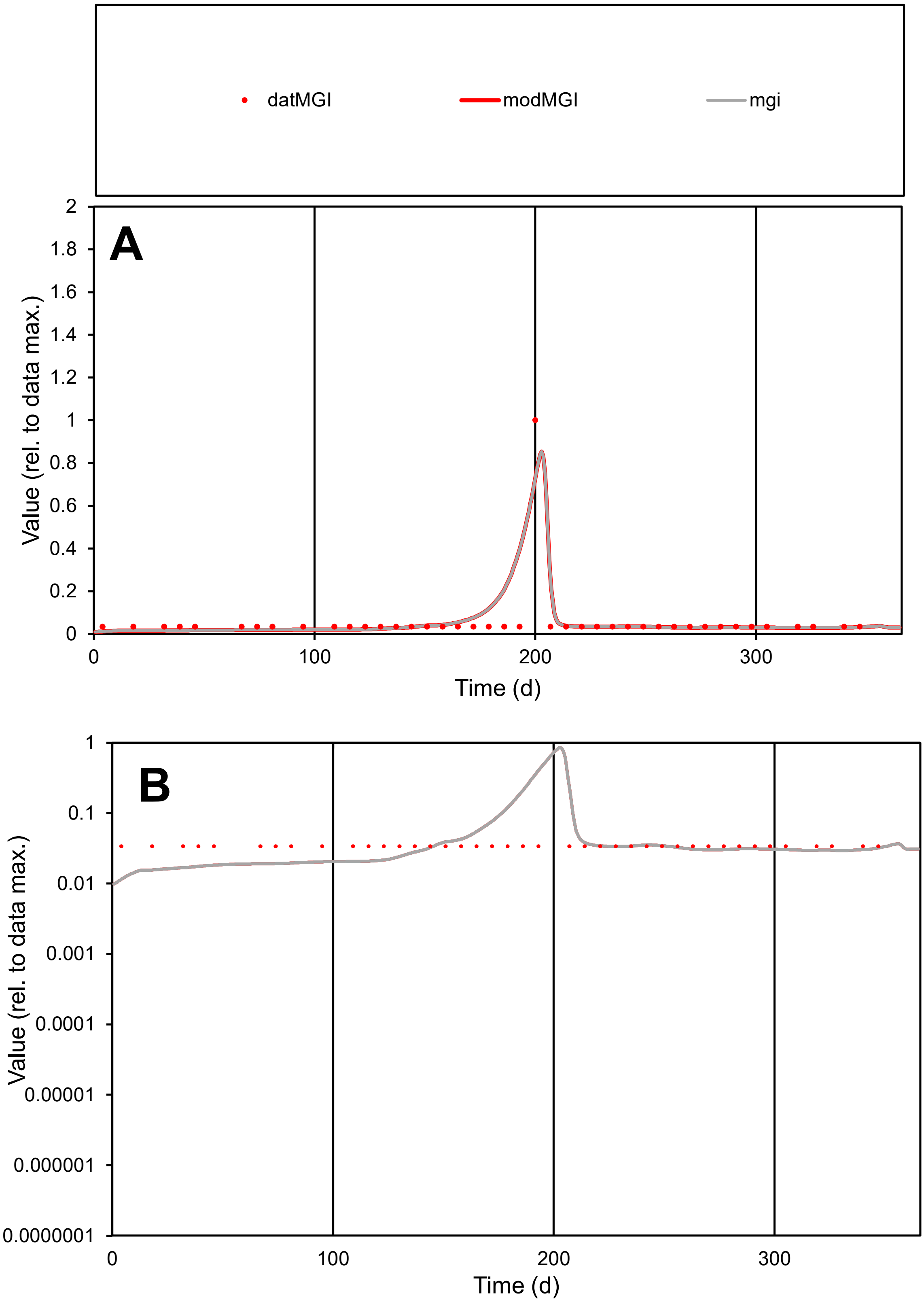


**C1**

Variables
MGI Provide jtd or Dst
CalibOnly?
2 1 to reproduce of
oNormalize?
2 1=no, 2=yes

ofxn calc*
1.12634
ofxn mod
1.30164

sRun
RunA1
zout
1

Ave
Data
1.7E-05
Model
1.9E-05

data/vMax
0.0003

ofxng
334.07

Min d/m
2.82662
ave/min d
1.6764
ave/min m
5.2156

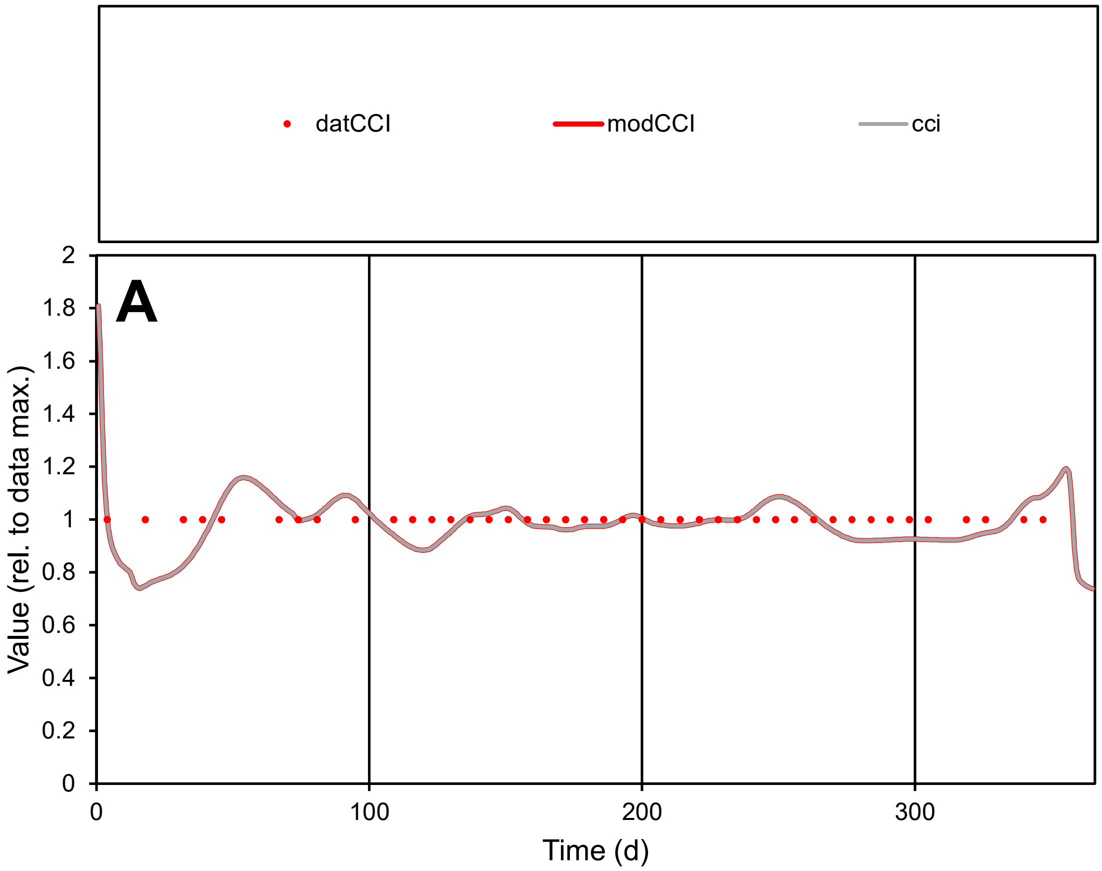


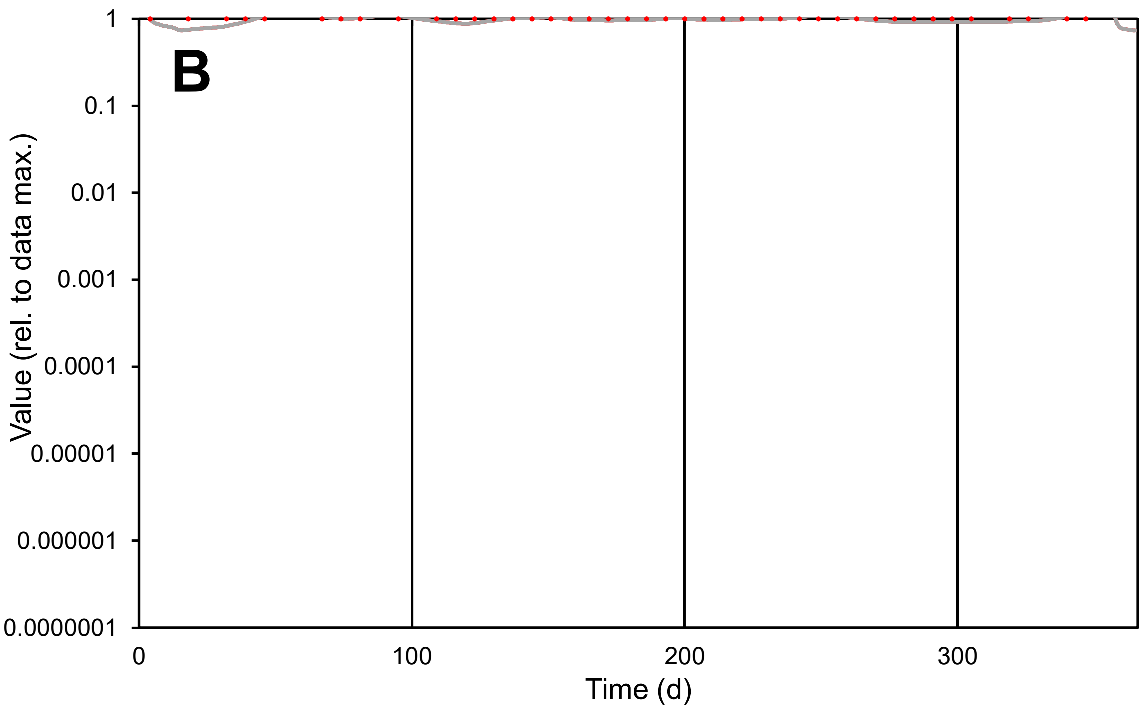


**C1**

Variables
CCI Provide jtd or Dst
CalibOnly?
2 1 to reproduce of
oNormalize?
2 1=no, 2=yes

ofxn calc*
0.07071
ofxn mod
0.13247

sRun
RunA1
zout
1

Ave
Data
1.1E-05
Model
1.1E-05

data/vMax
1.1E-05

ofxng
334.07

Min d/m
1.31636

ave/min d
1
ave/min m
1.28973

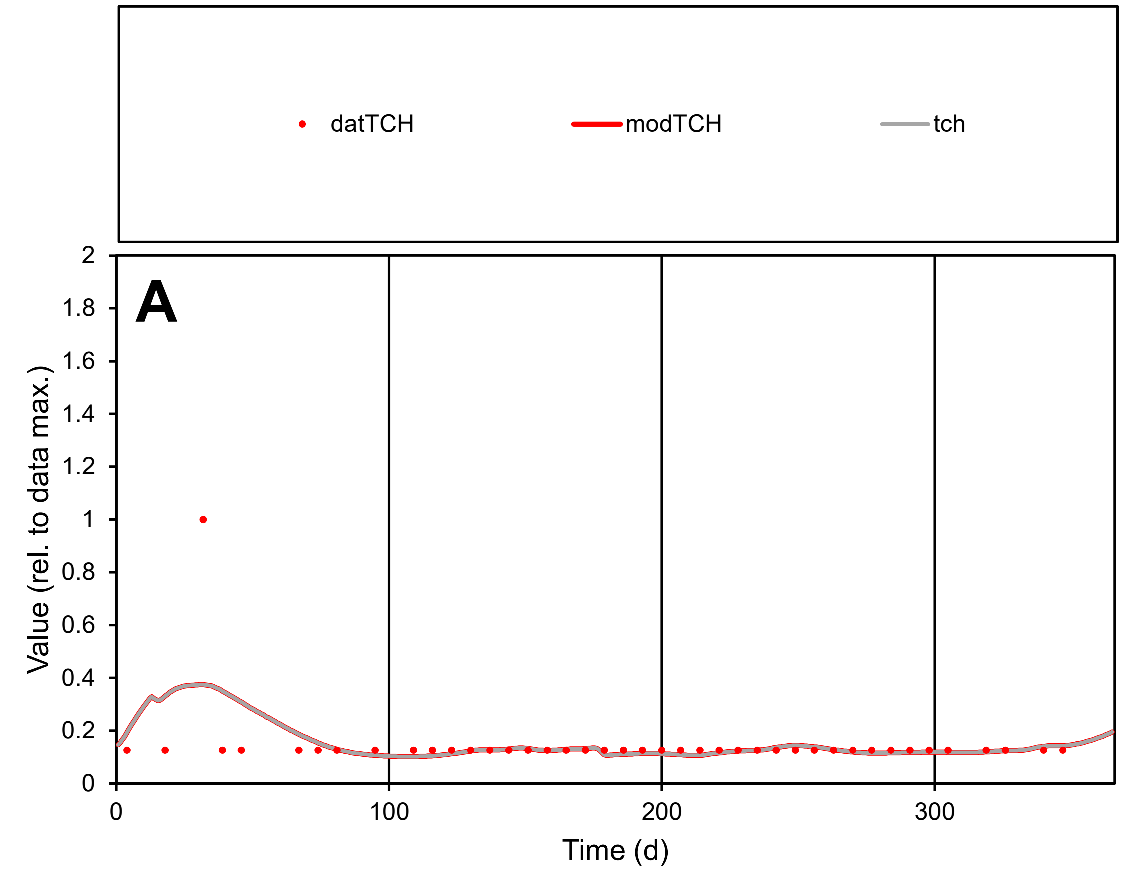


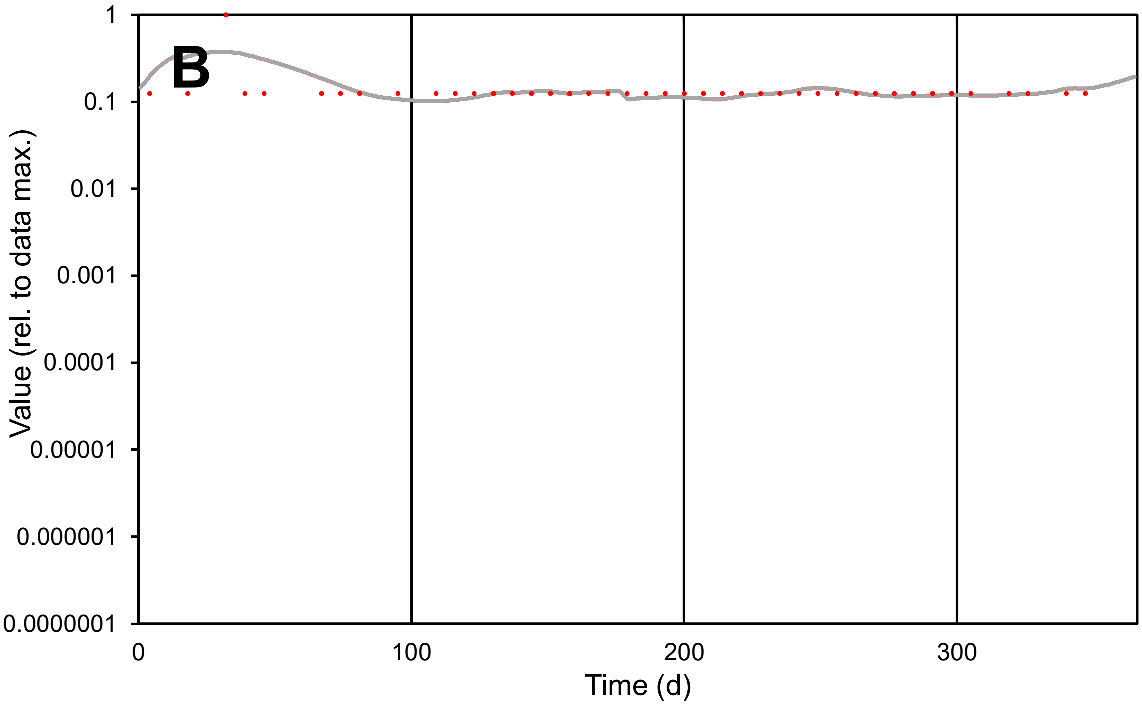


**C1**

Variables
TCH Provide jtd or Dst
CalibOnly?
2 1 to reproduce of
oNormalize?
2 1=no, 2=yes

ofxn calc*
0.76557
ofxn mod
0.7807

sRun
RunA1
zout
1

Ave
Data
2E-06
Model
2E-06

data/vMax
1.3E-05

ofxng
334.07

Min d/m
1.21884

ave/min d
1.1663
ave/min m
1.42345

327_Calib_S_A

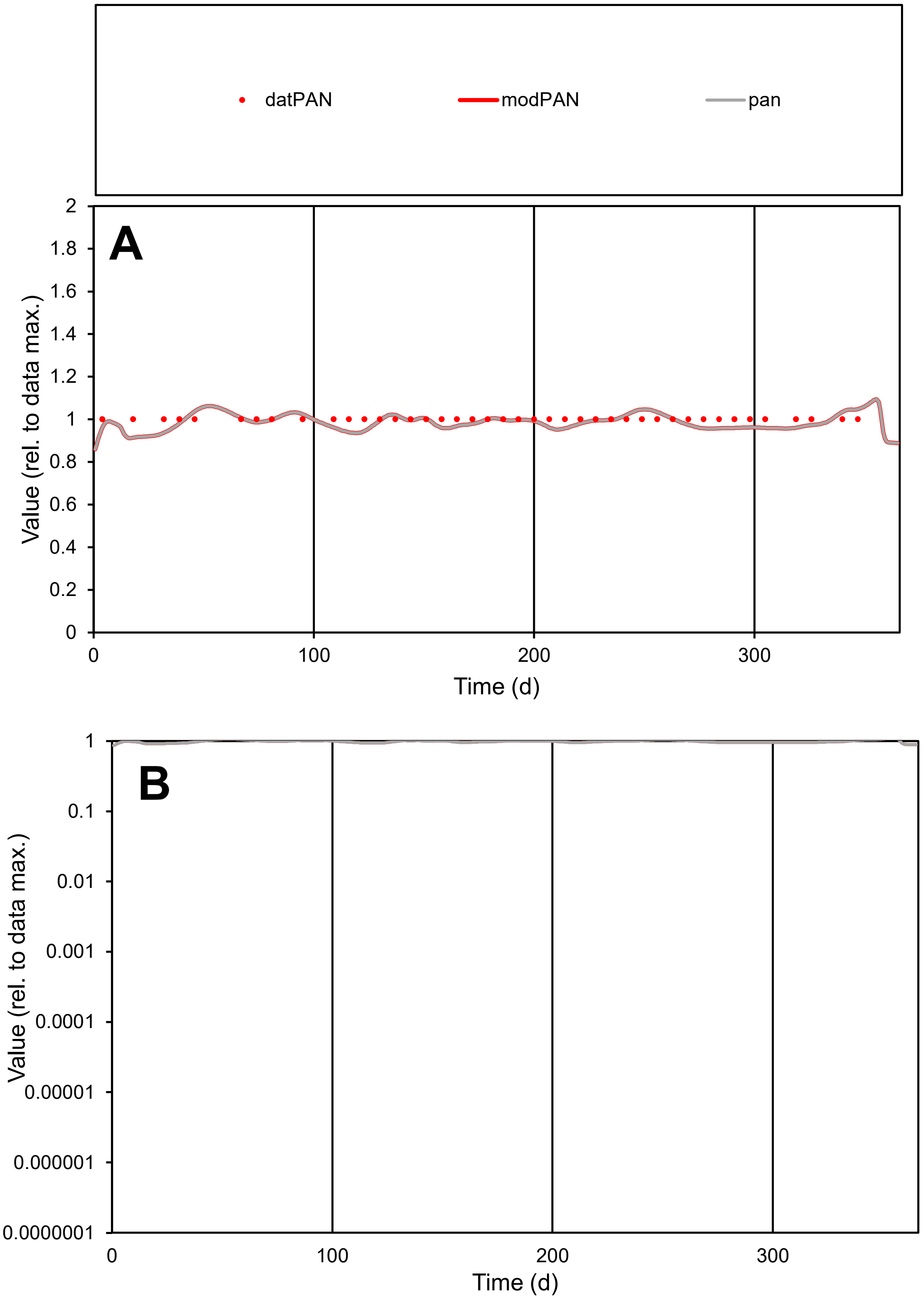


**C1**

Variables
PAN Provide jtd or Dst
CalibOnly?
2 1 to reproduce of
oNormalize?
2 1=no, 2=yes

ofxn calc*
0.03396
ofxn mod
0.11748

sRun
RunA1
zout
1

Ave
Data
9.1E-06
Model
9E-06

data/vMax
9.1E-06

ofxng
334.07

Min d/m
1.09117

ave/min d
1
ave/min m
1.07686

327_Calib_S_A

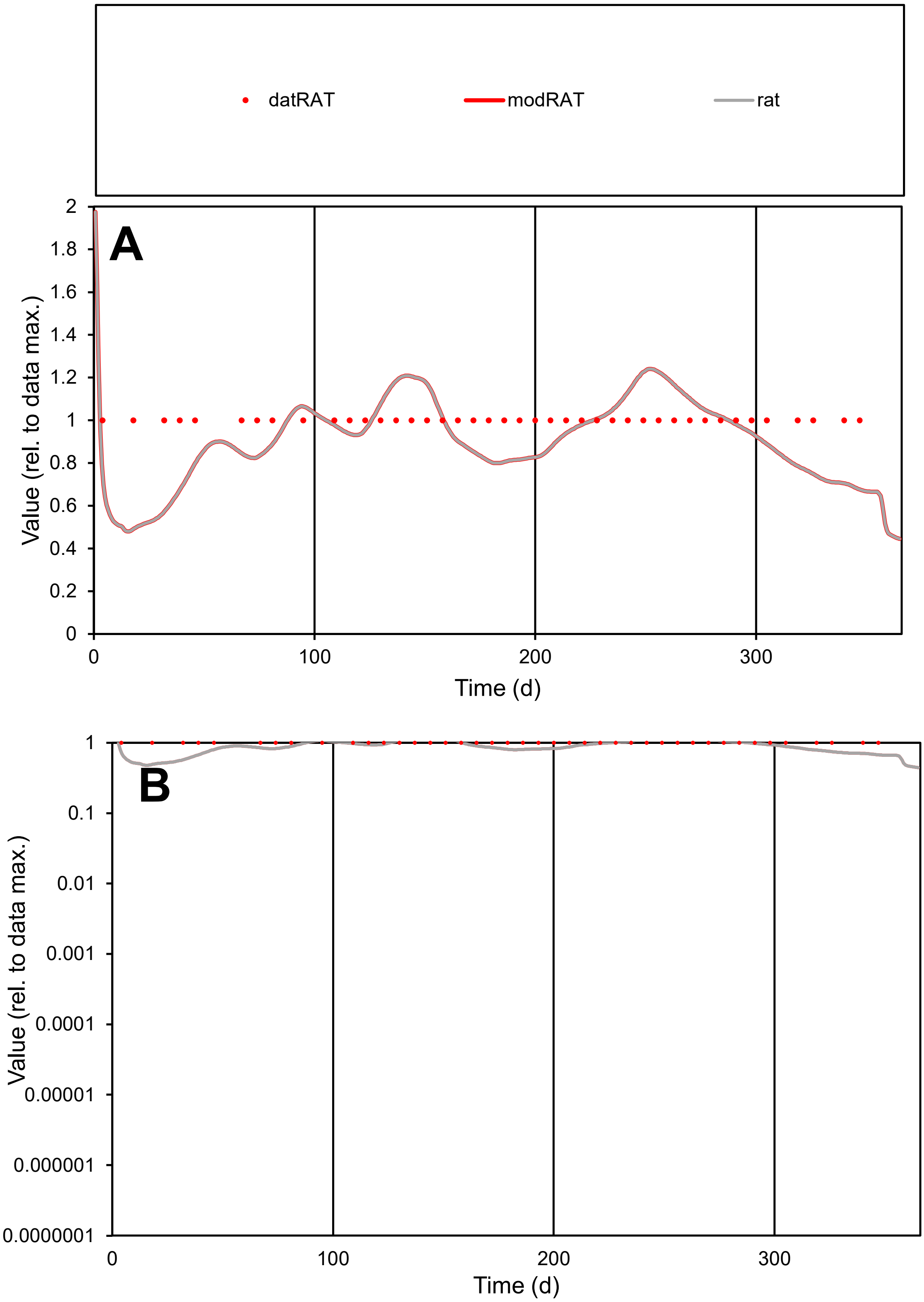


**C1**

Variables
RAT Provide jtd or Dst
CalibOnly?
2 1 to reproduce of
oNormalize?
2 1=no, 2=yes

ofxn calc*
0.18877
ofxn mod
0.21878

sRun
RunA1
zout
1

Ave
Data
1.1E-05
Model
1.1E-05

data/vMax
1.1E-05

ofxng
334.07

Min d/m
1.99439

ave/min d
1
ave/min m
1.85001

327_Calib_S_A

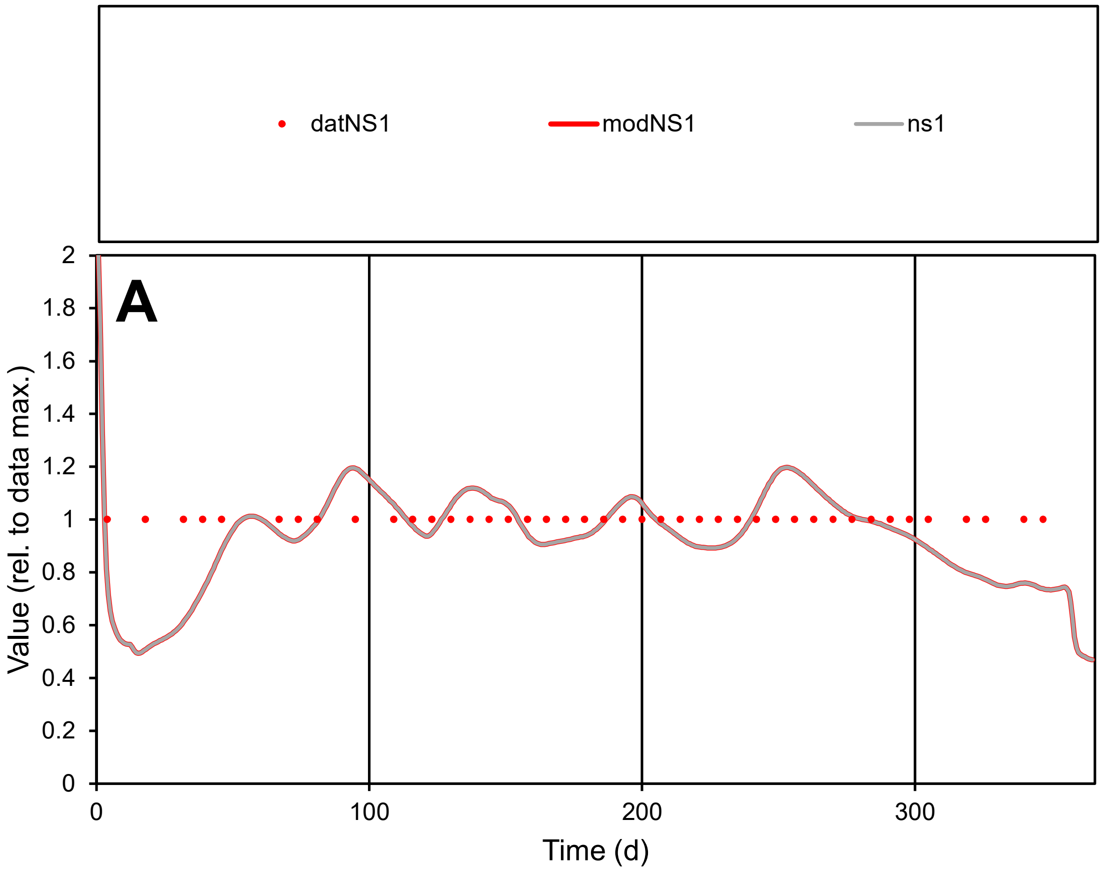


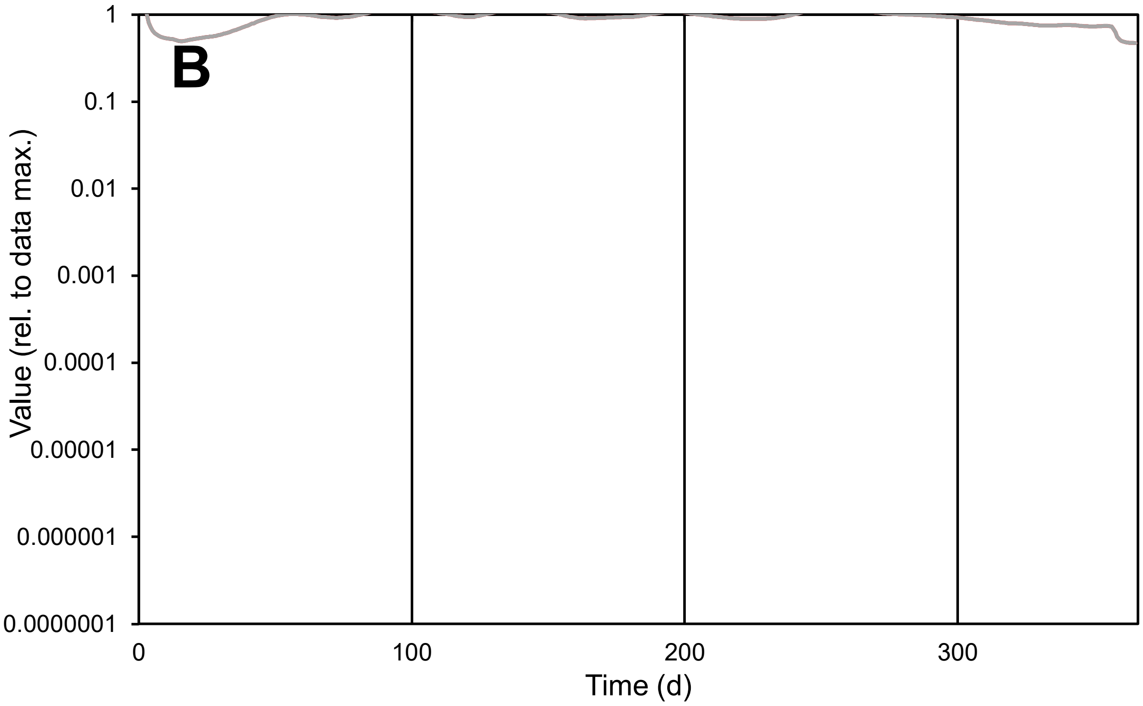


**C1**

Variables
NS1 Provide jtd or Dst
CalibOnly?
2 1 to reproduce of
oNormalize?
2 1=no, 2=yes

ofxn calc*
0.15286
ofxn mod
0.18953

sRun
RunA1
zout
1

Ave
Data
2.7E-05
Model
2.6E-05

data/vMax
2.7E-05

ofxng
334.07

Min d/m
1.92162

ave/min d
1

ave/min m
1.82174

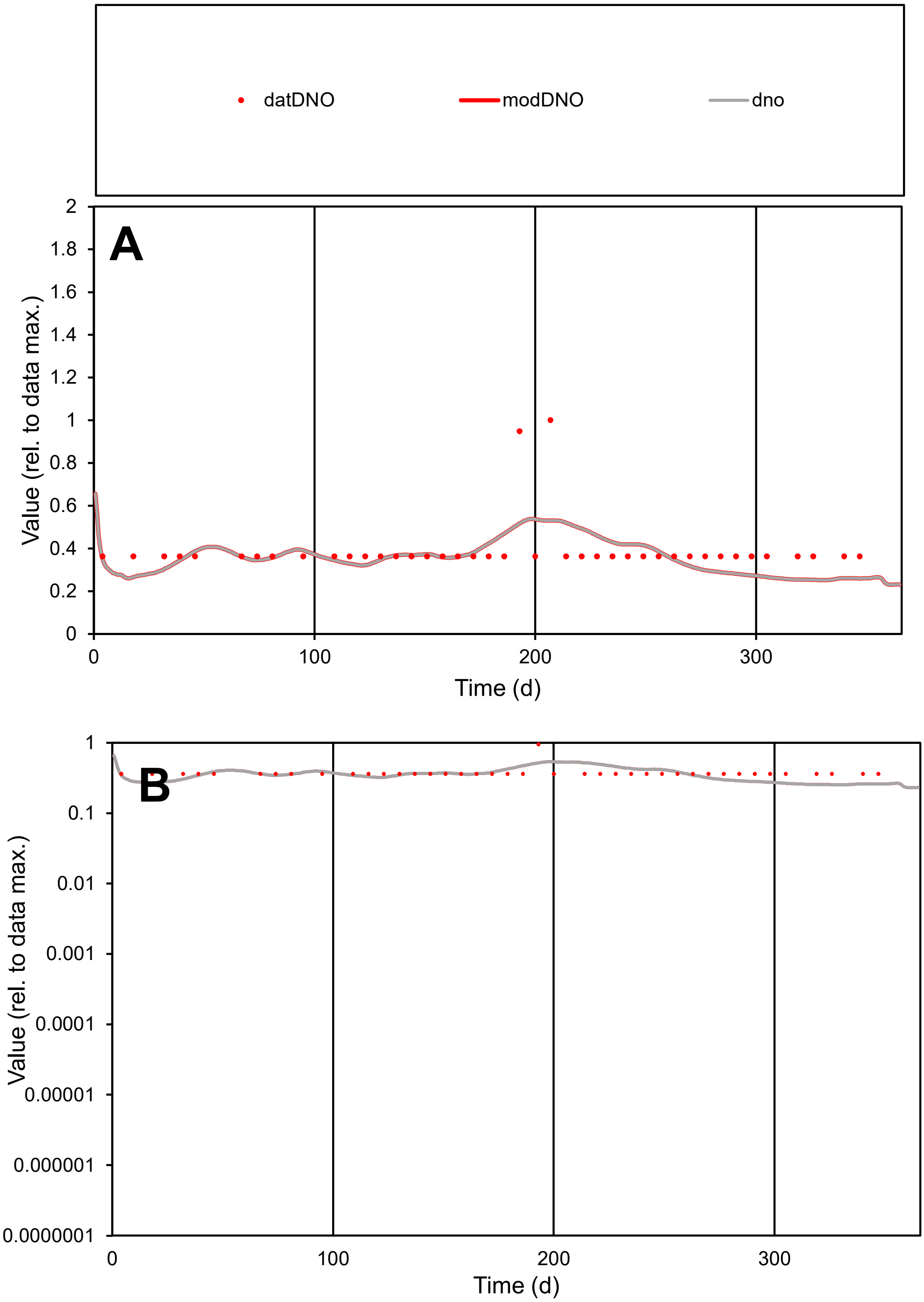


**C1**

Variables
DNO Provide jtd or Dst
CalibOnly?
2 1 to reproduce of
oNormalize?
2 1=no, 2=yes

ofxn calc*
0.30667
ofxn mod
0.32813

sRun
RunA1
zout
1

Ave
Data
2E-05
Model
1.8E-05

data/vMax
5E-05

ofxng
334.07

Min d/m
1.43083

ave/min d
1.08047
ave/min m
1.44425

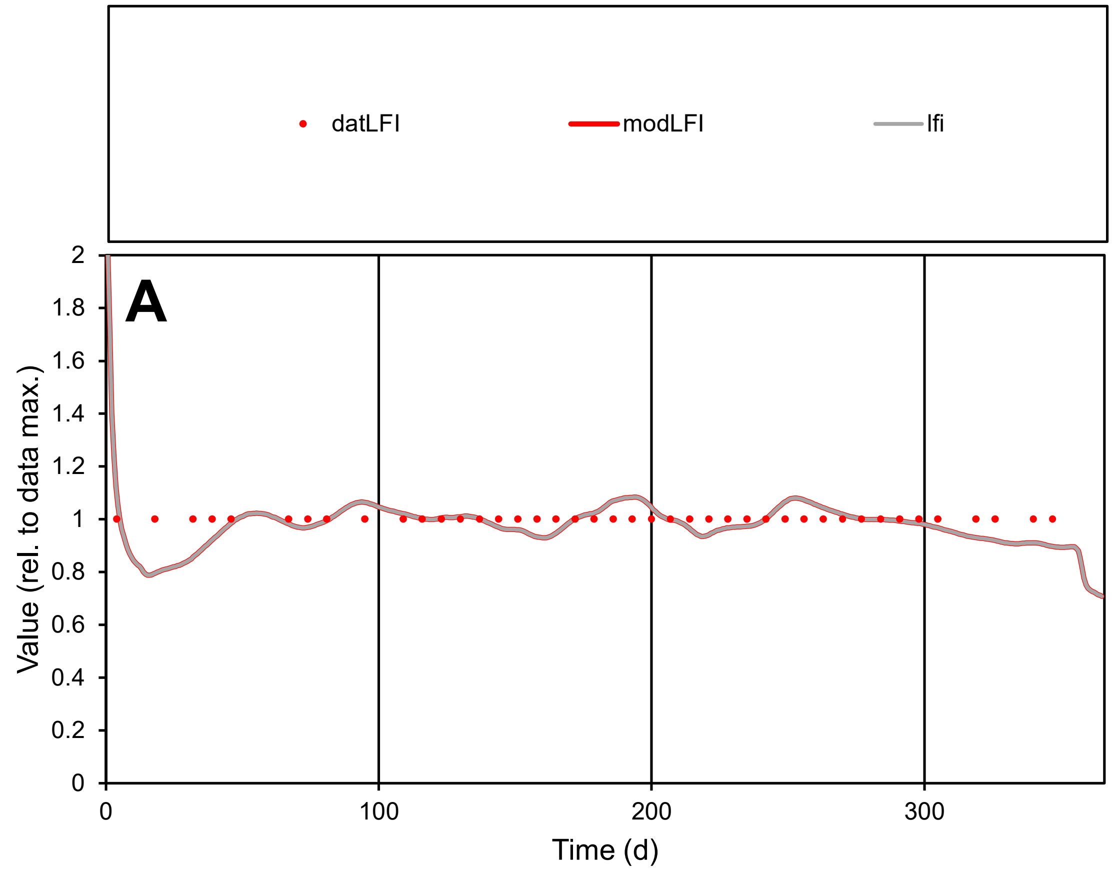


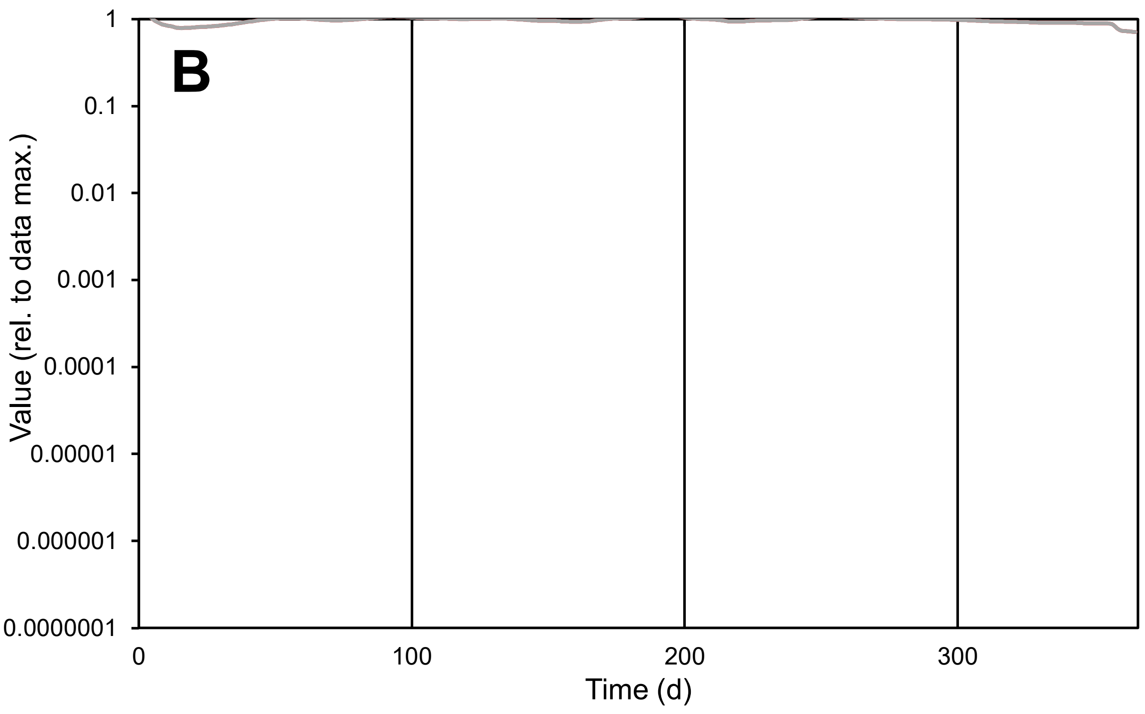


**C1**

Variables
LFI Provide jtd or Dst
CalibOnly?
2 1 to reproduce of
oNormalize?
2 1=no, 2=yes

ofxn calc*
0.0583
ofxn mod
0.12664

sRun
RunA1
zout
1

Ave
Data
8.3E-06
Model
8.2E-06

data/vMax
8.3E-06

ofxng
334.07

Min d/m
1.24517

ave/min d
1
ave/min m
1.22543

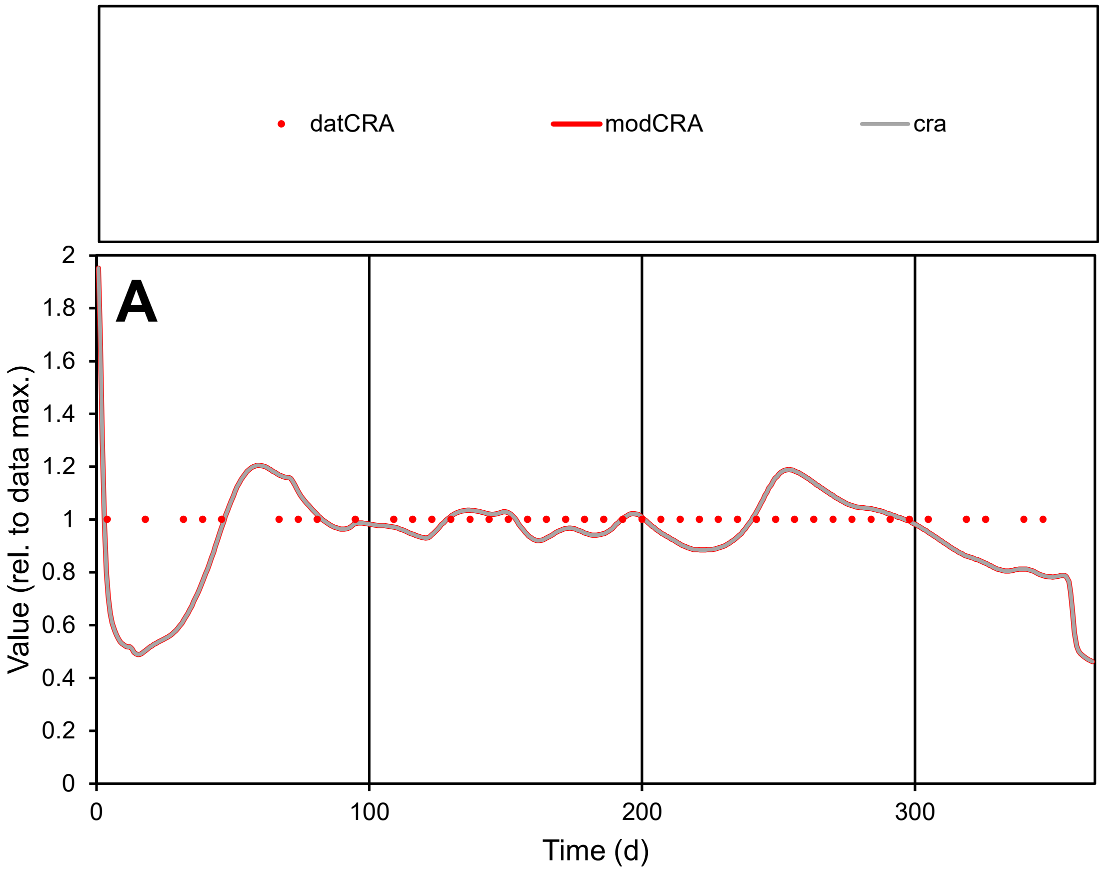


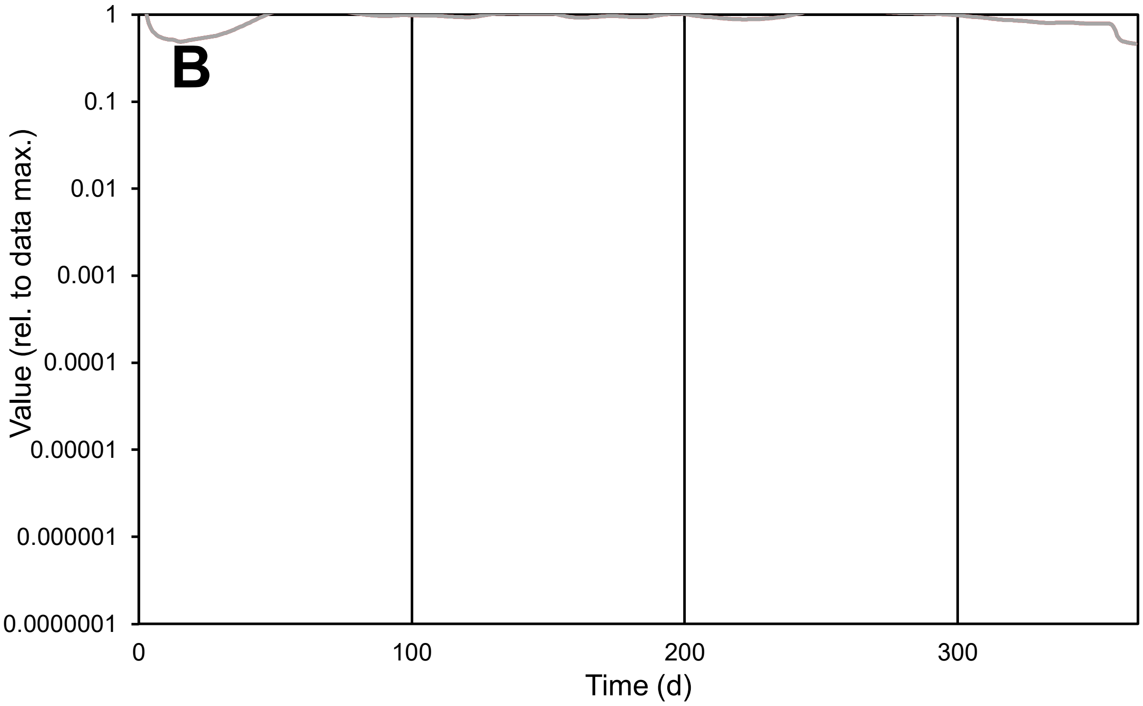


**C1**

Variables
CRA Provide jtd or Dst
CalibOnly?
2 1 to reproduce of
oNormalize?
2 1=no, 2=yes

ofxn calc*
0.13875
ofxn mod
0.17874

sRun
RunA1
zout
1

Ave
Data
4.9E-06
Model
4.6E-06

data/vMax
4.9E-06

ofxng
334.07

Min d/m
1.93477

ave/min d
1
ave/min m
1.84908

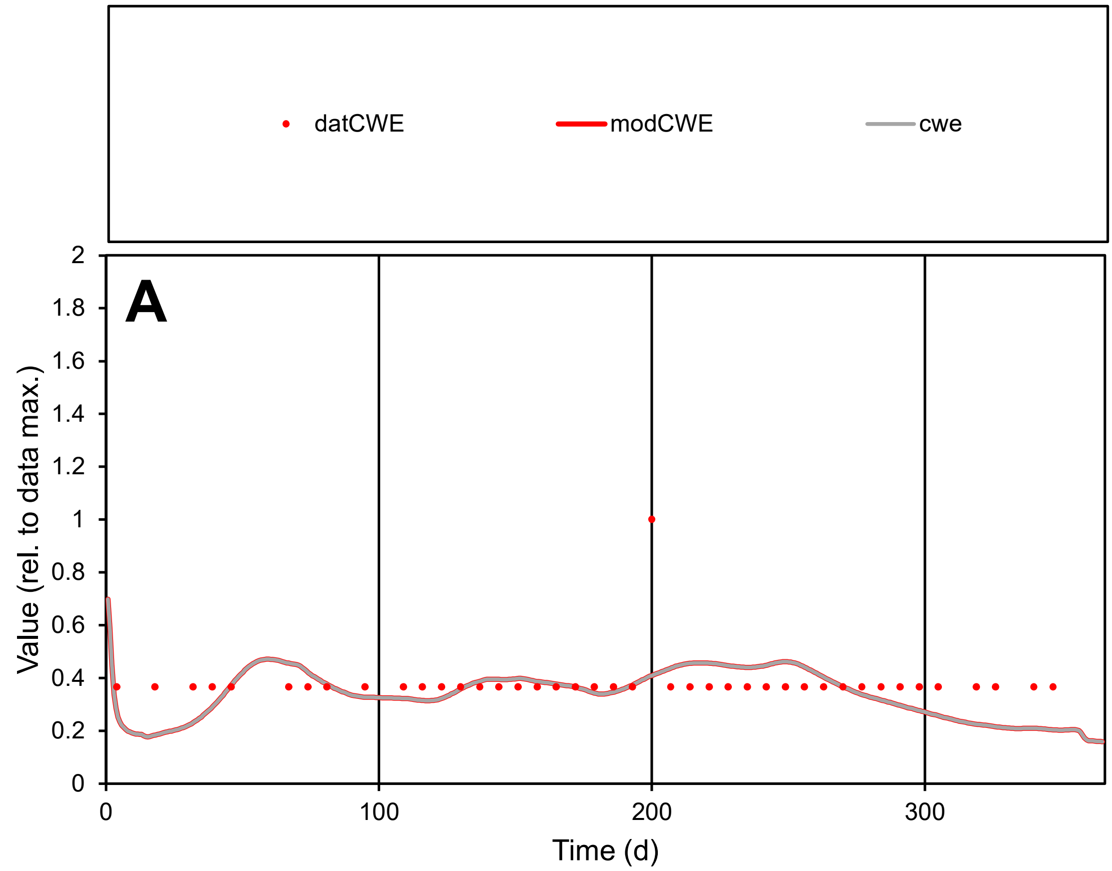


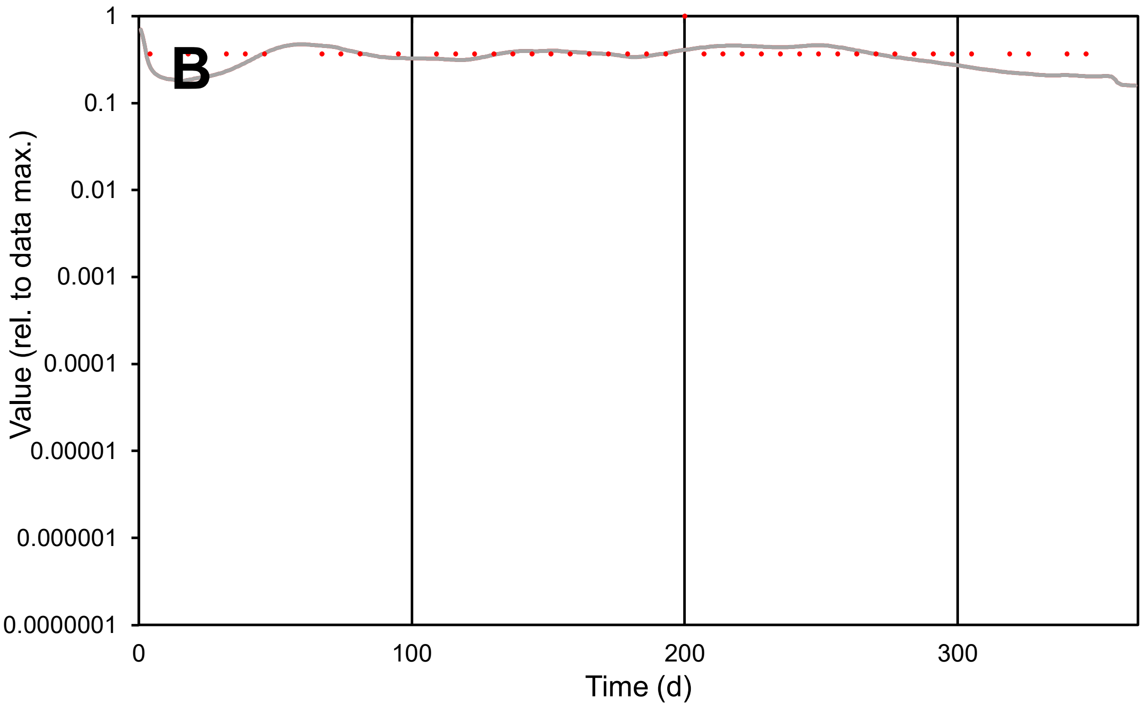


**C1**

Variables
CWE Provide jtd or Dst
CalibOnly?
2 1 to reproduce of
oNormalize?
2 1=no, 2=yes

ofxn calc*
0.31539
ofxn mod
0.33593

sRun
RunA1
zout
1

Ave
Data
2.7E-05
Model
2.4E-05

data/vMax
7E-05

ofxng
334.07

Min d/m
1.94064

ave/min d
1.04116
ave/min m
1.85685

327_Calib_S_A

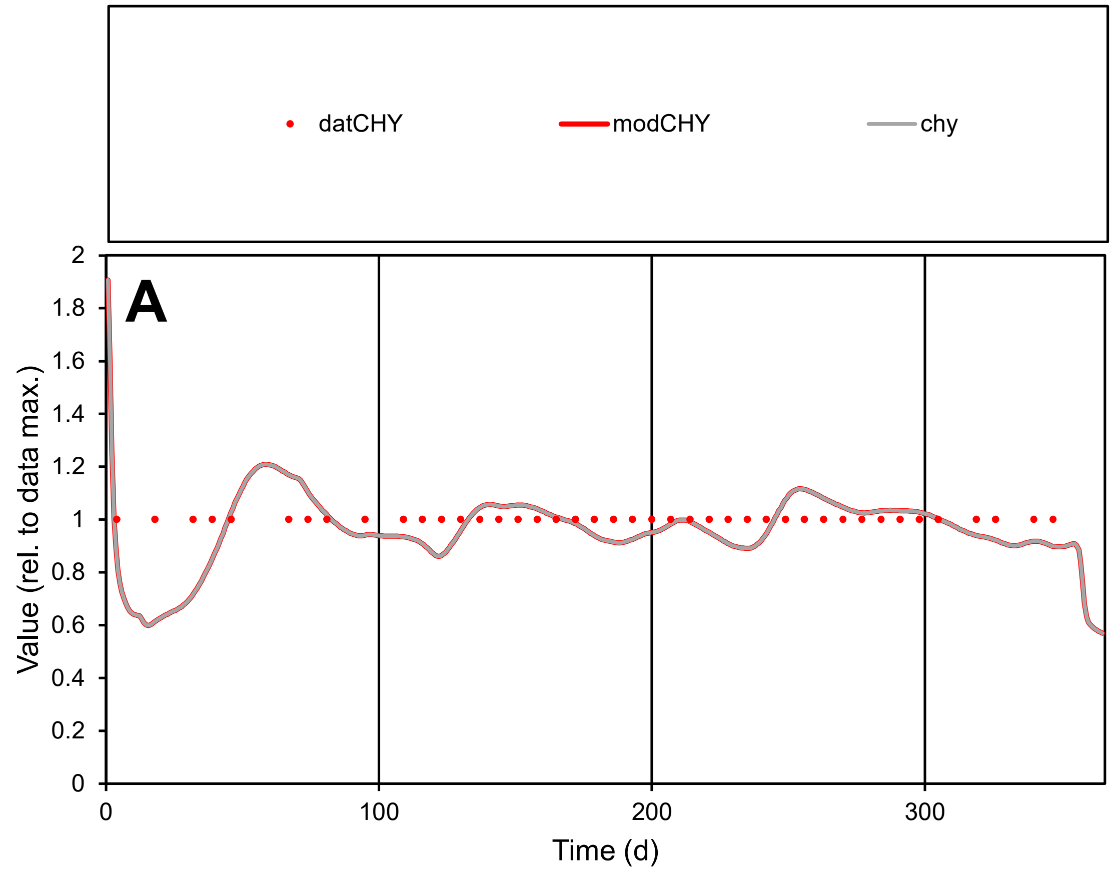


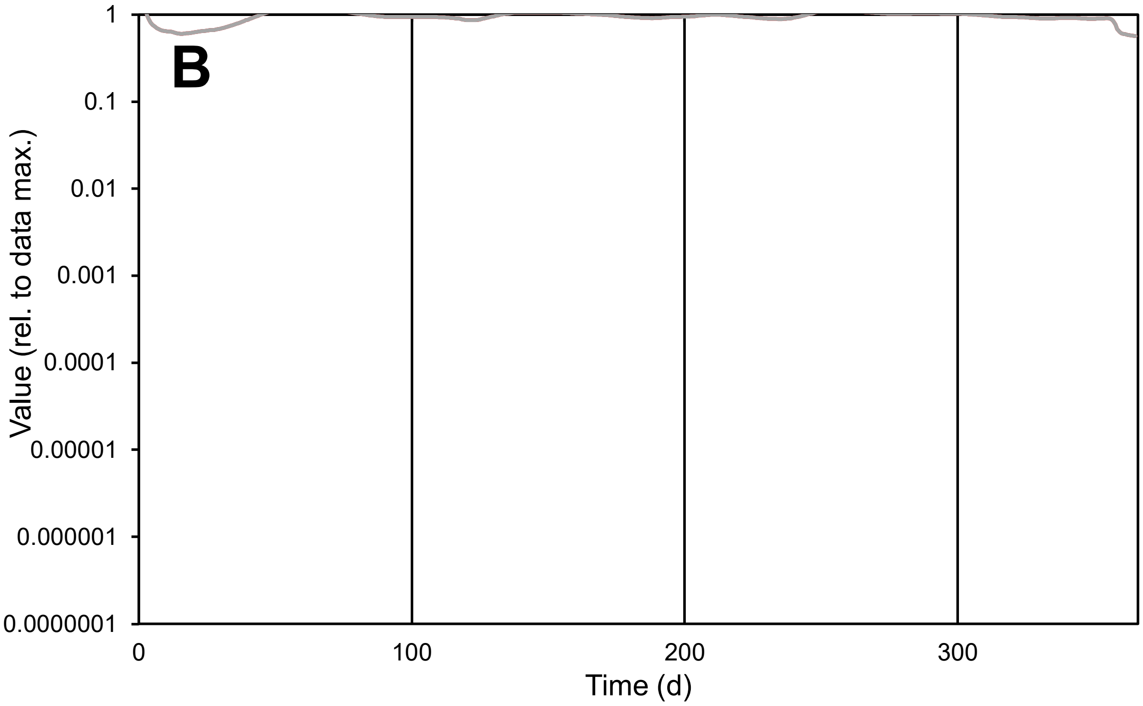


**C1**

Variables

CHY Provide jtd or Dst

CalibOnly?

2 1 to reproduce of

oNormalize?

2 1=no, 2=yes

ofxn calc*

0.10234

ofxn mod

0.15262

sRun

RunA1

zout

1

Ave

Data

6.7E-06

Model

6.5E-06

data/vMax

6.7E-06

ofxng

334.07

Min d/m

1.59909

ave/min d

1

ave/min m

1.54997

327_Calib_S_A

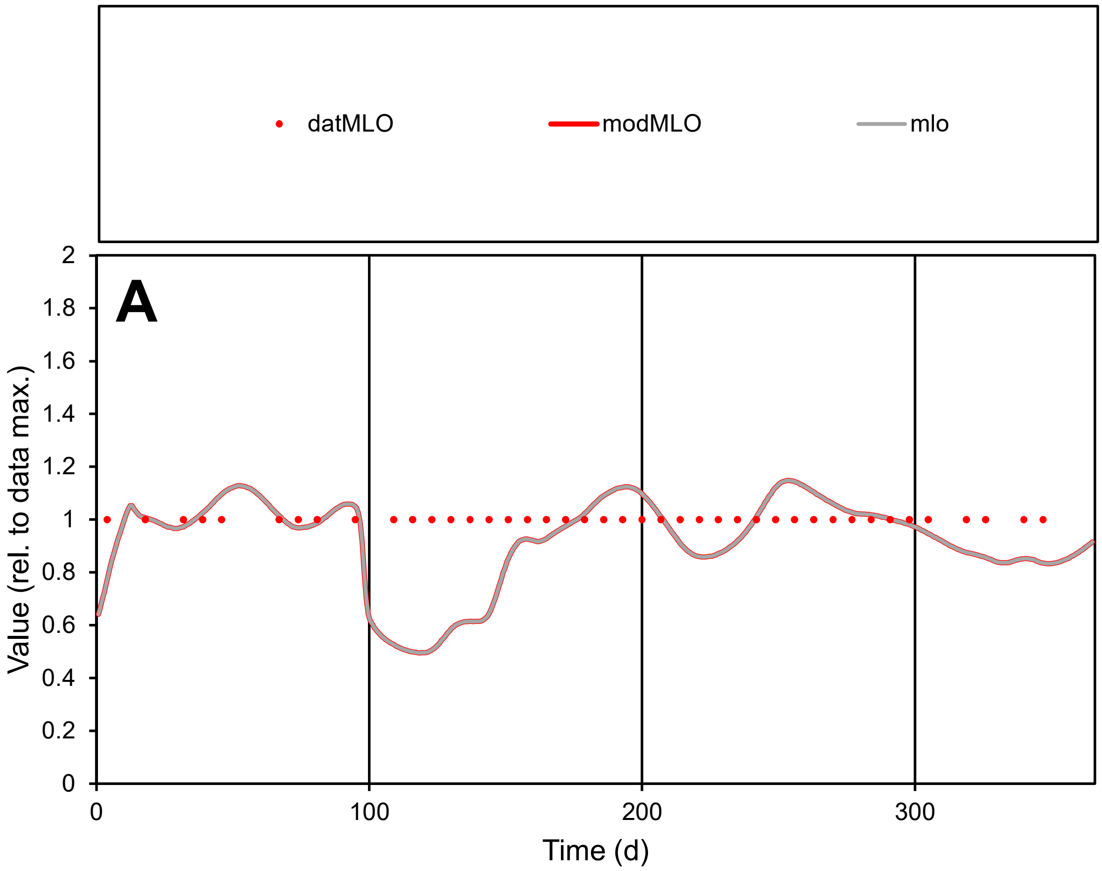


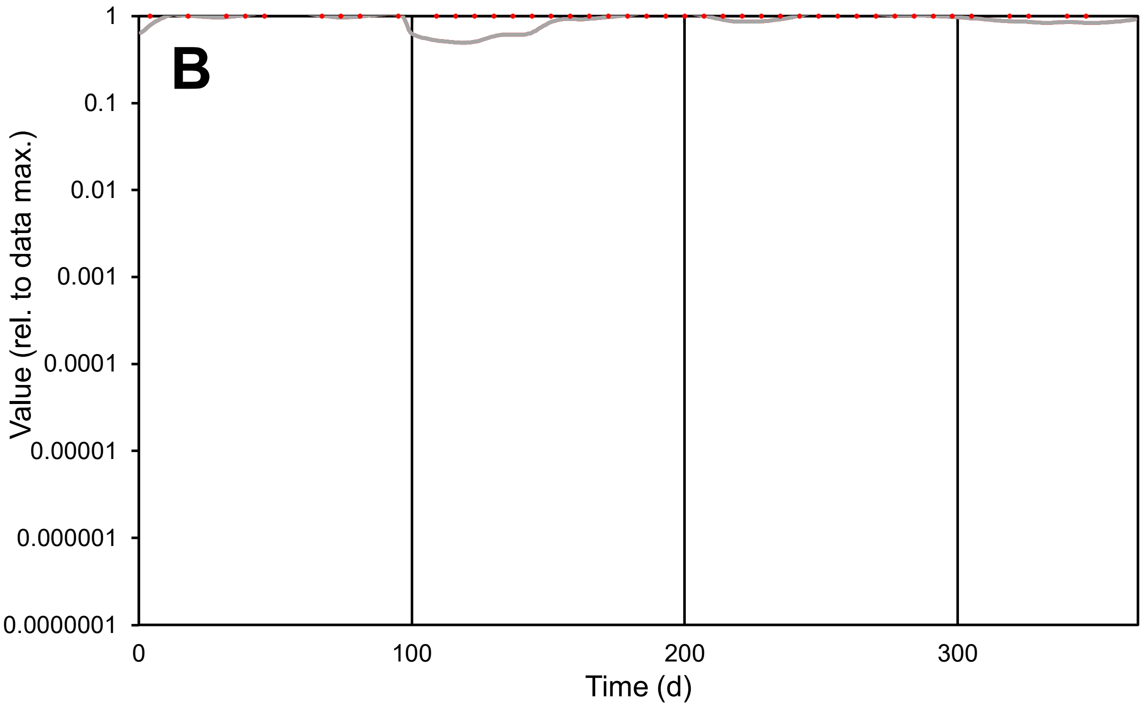


**C1**

Variables

MLO Provide jtd or Dst

CalibOnly?

2 1 to reproduce of

oNormalize?

2 1=no, 2=yes

ofxn calc*

0.1847

ofxn mod

0.2153

sRun

RunA1

zout

1

Ave

Data

1E-05

Model

9.6E-06

data/vMax

1E-05

ofxng

334.07

Min d/m

2.01235

ave/min d

1

ave/min m

1.85929

327_Calib_S_A

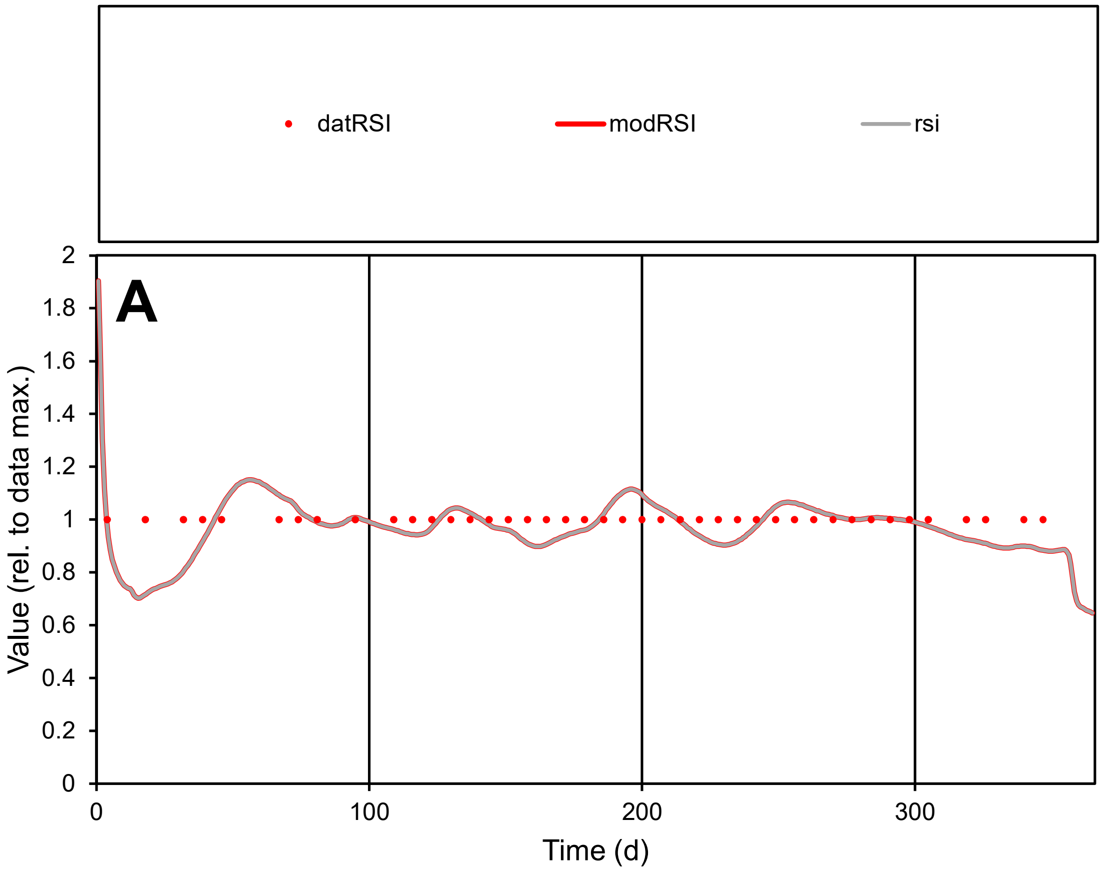


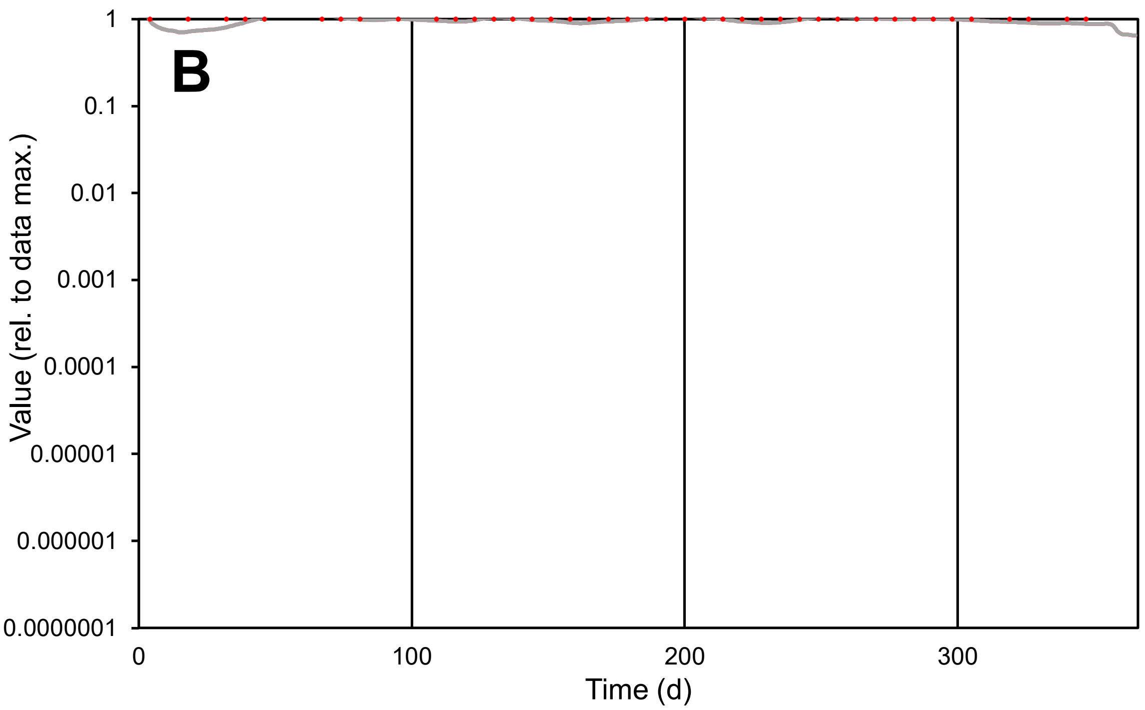


**C1**

Variables
RSI Provide jtd or Dst
CalibOnly?
2 1 to reproduce of
oNormalize?
2 1=no, 2=yes

ofxn calc*
0.07801
ofxn mod
0.13668

sRun
RunA1
zout
1

Ave
Data
1E-05
Model
1E-05

data/vMax
1E-05

ofxng
334.07

Min d/m
1.37239

ave/min d
1
ave/min m
1.33682

327_Calib_S_A

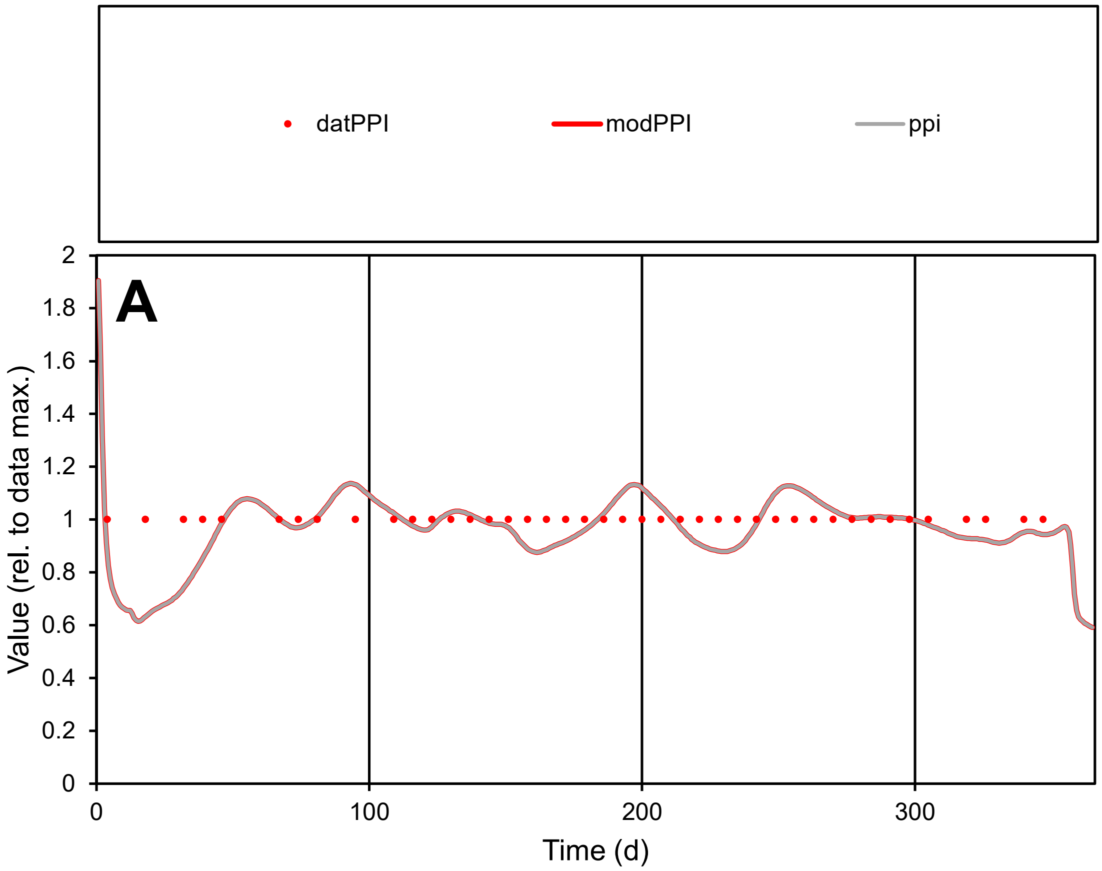


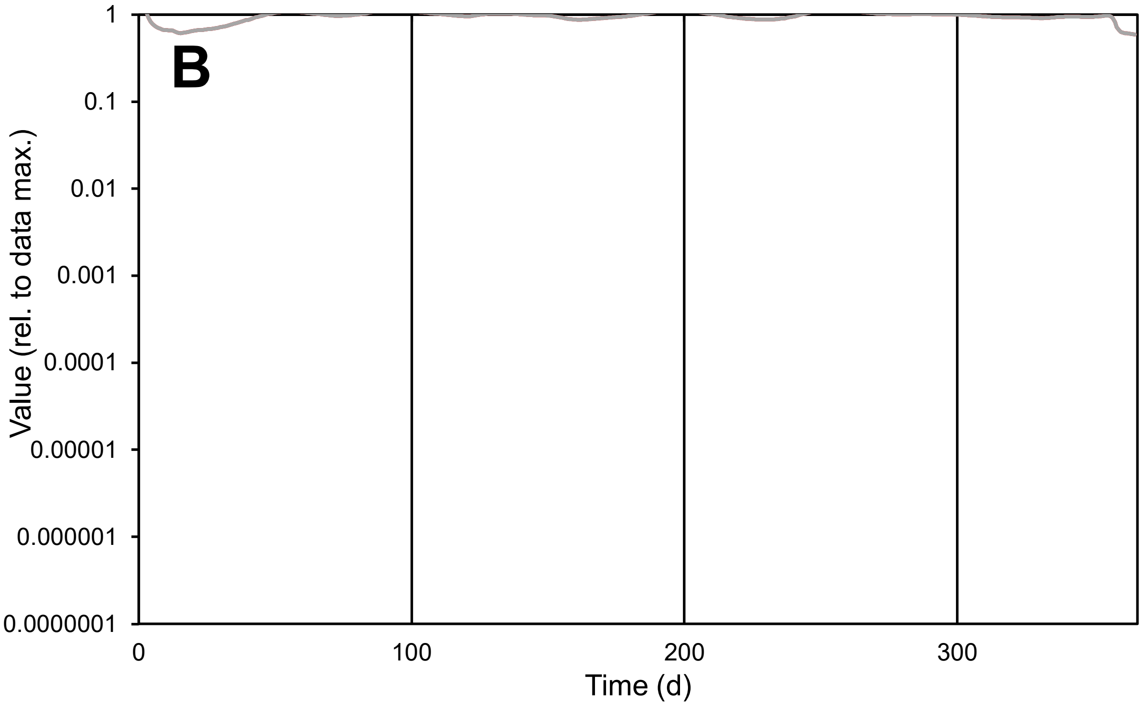


**C1**

Variables
PPI Provide jtd or Dst
CalibOnly?
2 1 to reproduce of
oNormalize?
2 1=no, 2=yes

ofxn calc*
0.09707
ofxn mod
0.14952

sRun
RunA1
zout
1

Ave
Data
1.4E-06
Model
1.4E-06

data/vMax
1.4E-06

ofxng
334.07

Min d/m
1.54735

ave/min d
1
ave/min m
1.50703

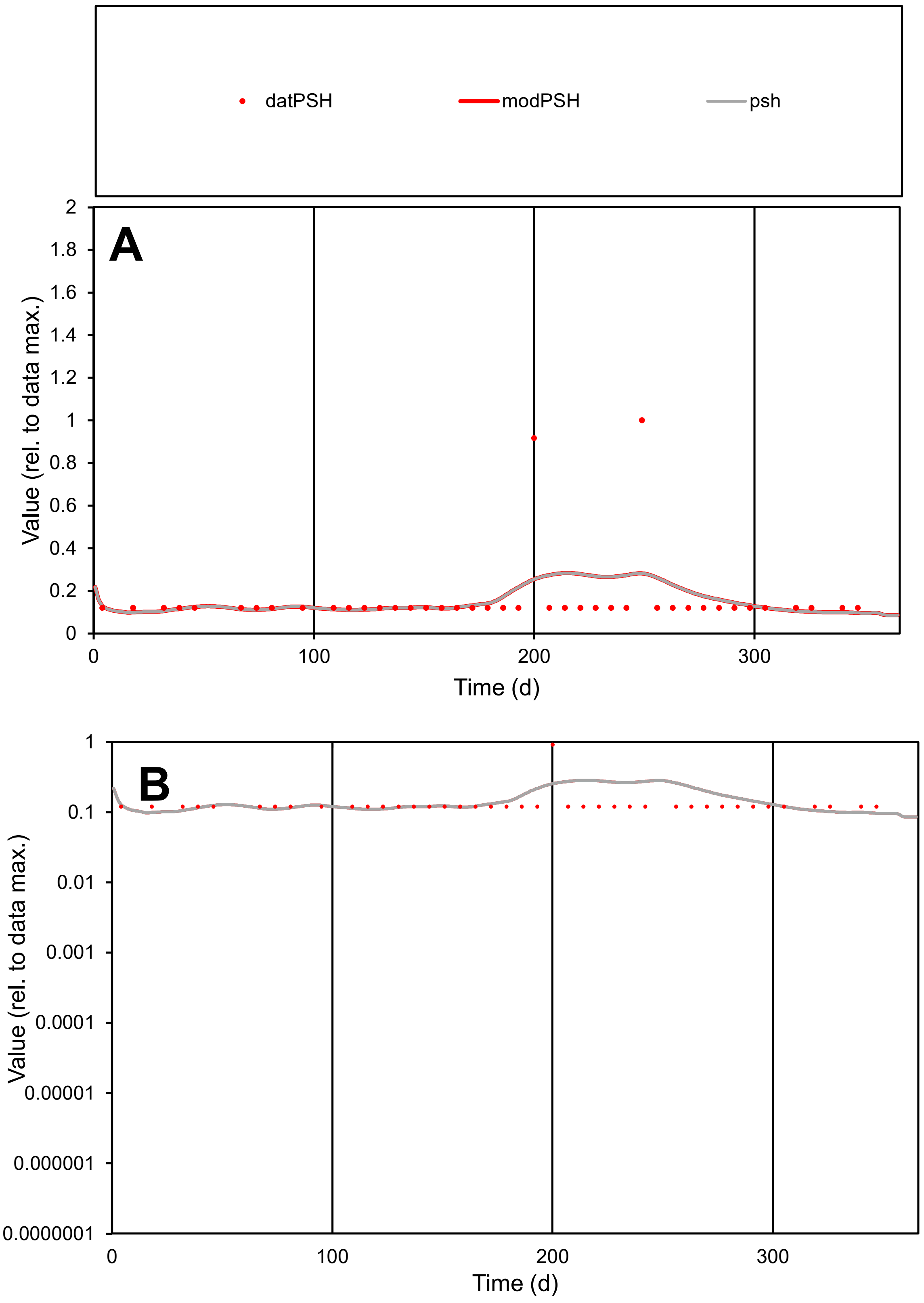


**C1**

Variables
PSH Provide jtd or Dst
CalibOnly?
2 1 to reproduce of
oNormalize?
2 1=no, 2=yes

ofxn calc*
1.03083
ofxn mod
1.04182

sRun
RunA1
zout
1

Ave
Data
1.7E-05
Model
1.7E-05

data/vMax
0.00011

ofxng
334.07

Min d/m
1.25

ave/min d
1.33171
ave/min m
1.66796

327_Calib_S_A

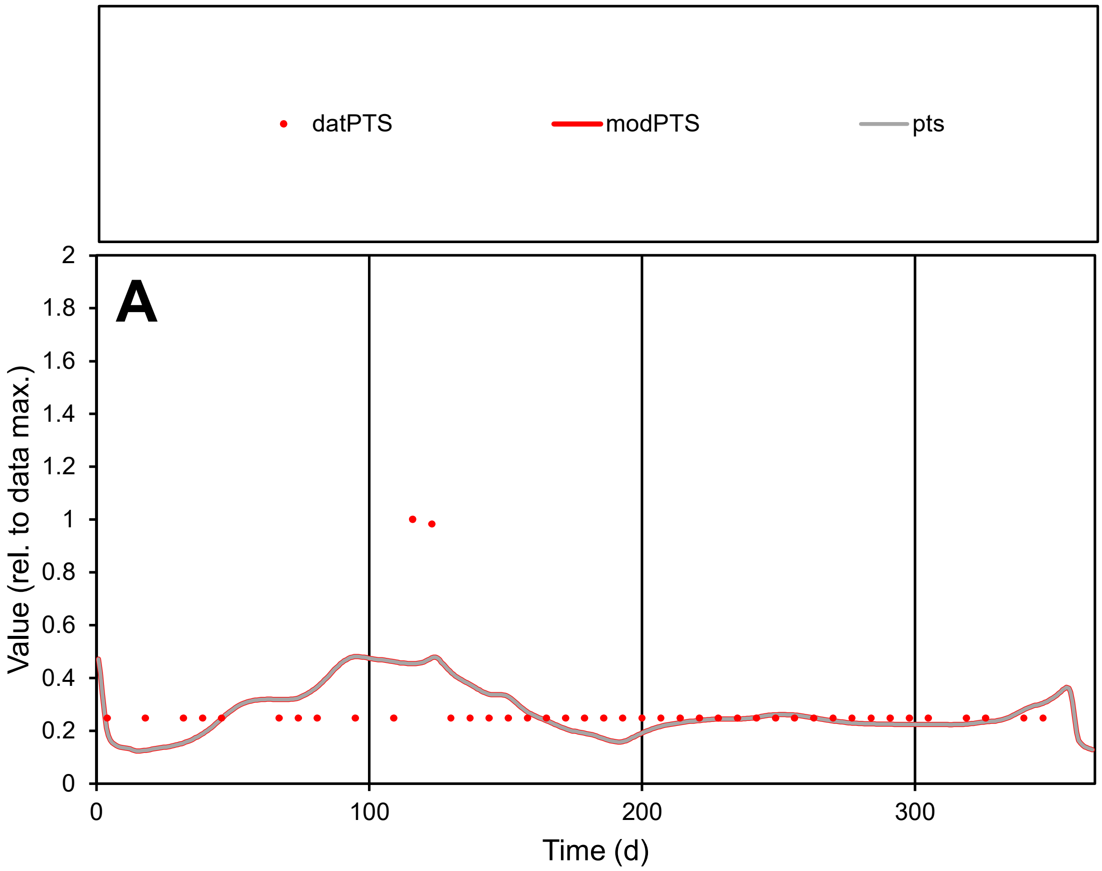


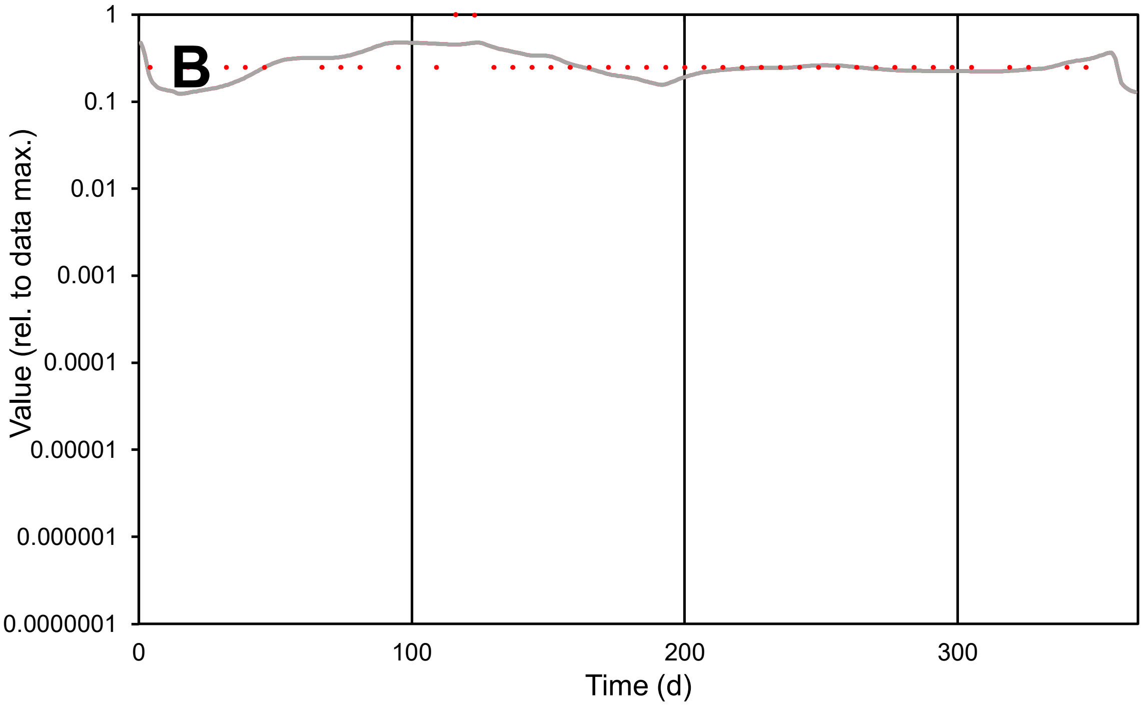


**C1**

Variables
PTS Provide jtd or Dst
CalibOnly?
2 1 to reproduce of
oNormalize?
2 1=no, 2=yes

ofxn calc*
0.48542
ofxn mod
0.4988

sRun
RunA1
zout
1

Ave
Data
1.1E-05
Model
1.1E-05

data/vMax
3.9E-05

ofxng
334.07

Min d/m
1.91667

ave/min d
1.14285
ave/min m
2.08327

327_Calib_S_A

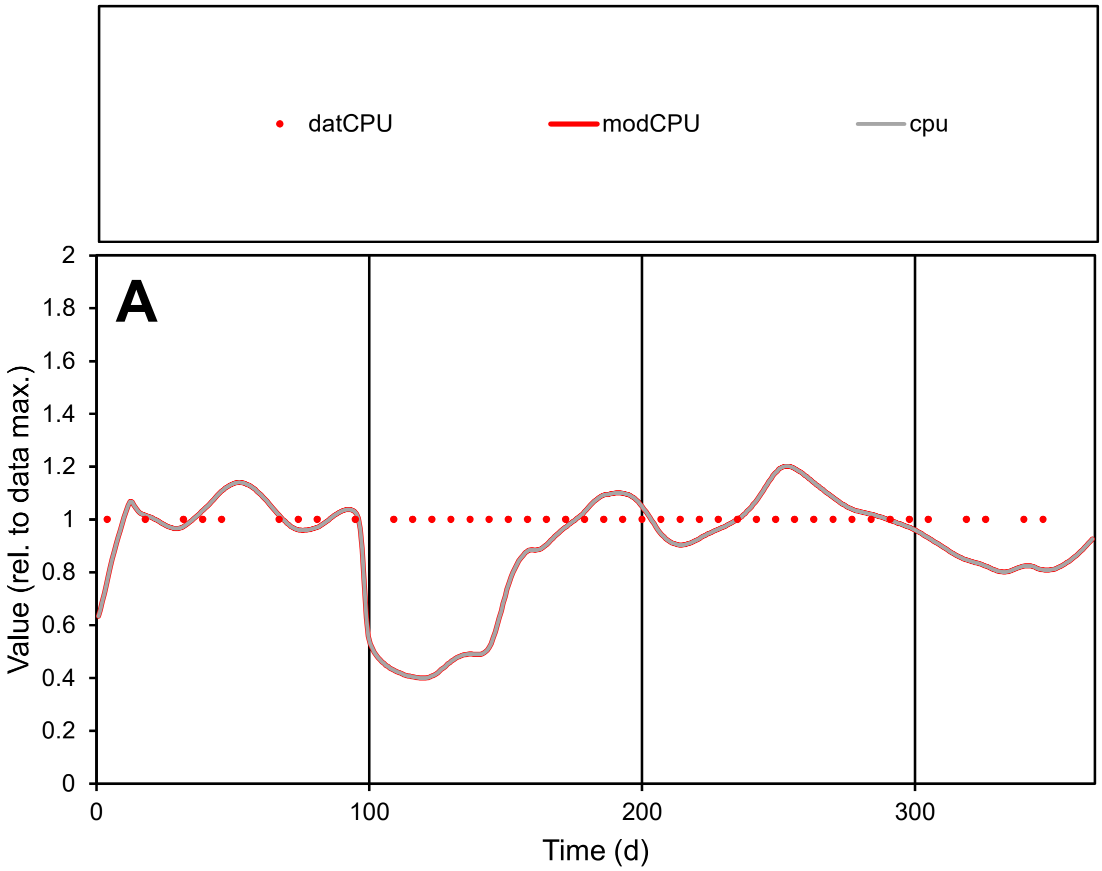


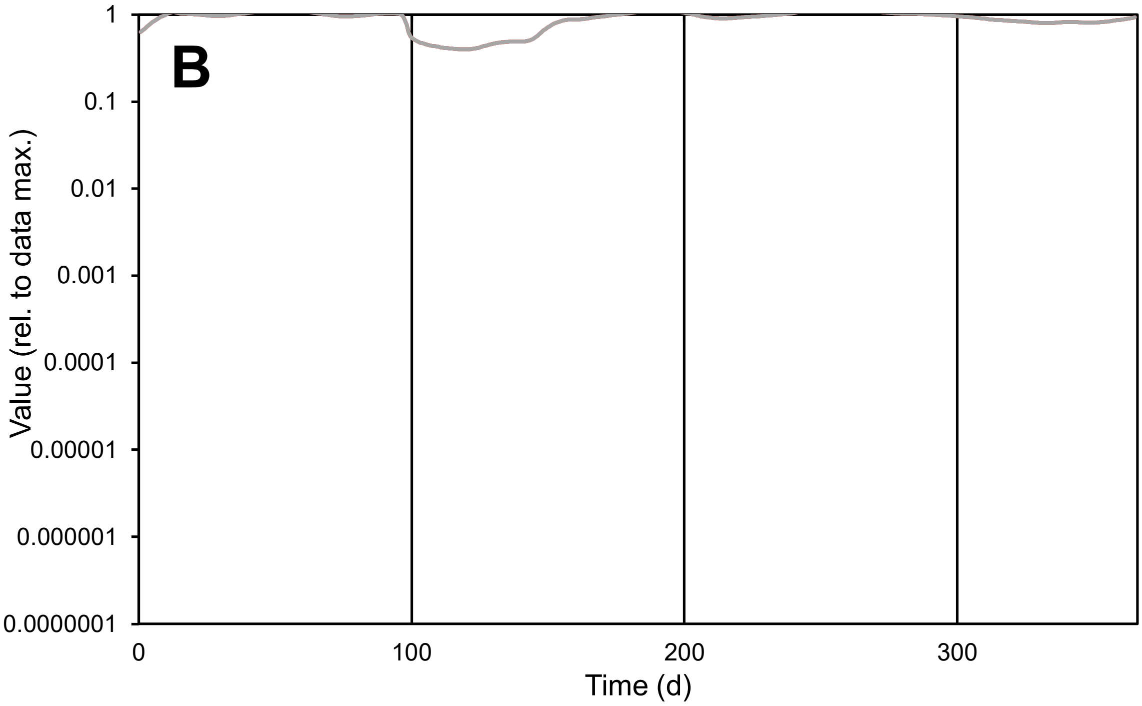


**C1**

Variables

CPU Provide jtd or Dst

CalibOnly?

2 1 to reproduce of

oNormalize?

2 1=no, 2=yes

ofxn calc*

0.22728

ofxn mod

0.25269

sRun

RunA1

zout

1

Ave

Data

4.4E-07

Model

4E-07

data/vMax

4.4E-07

ofxng

334.07

Min d/m

2.48567

ave/min d

1

ave/min m

2.25913

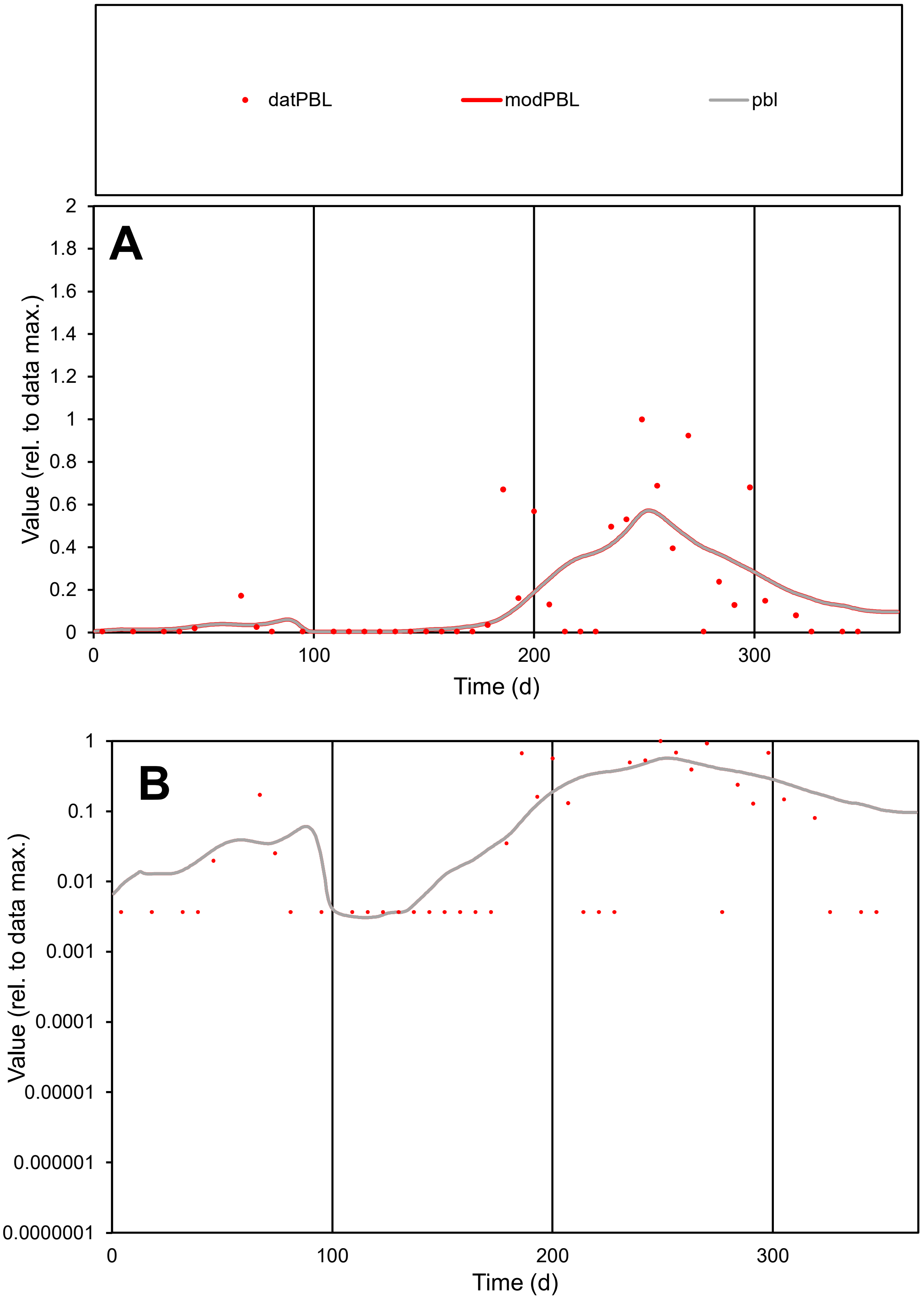


**C1**

Variables
PBL Provide jtd or Dst
CalibOnly?
2 1 to reproduce of
oNormalize?
2 1=no, 2=yes

ofxn calc*
1.19597
ofxn mod
1.21581

sRun
RunA1
zout
1

Ave
Data
2E-06
Model
2E-06

data/vMax
1.2E-05

ofxng
334.07

Min d/m
1.19383

ave/min d
46.3317
ave/min m
55.5847

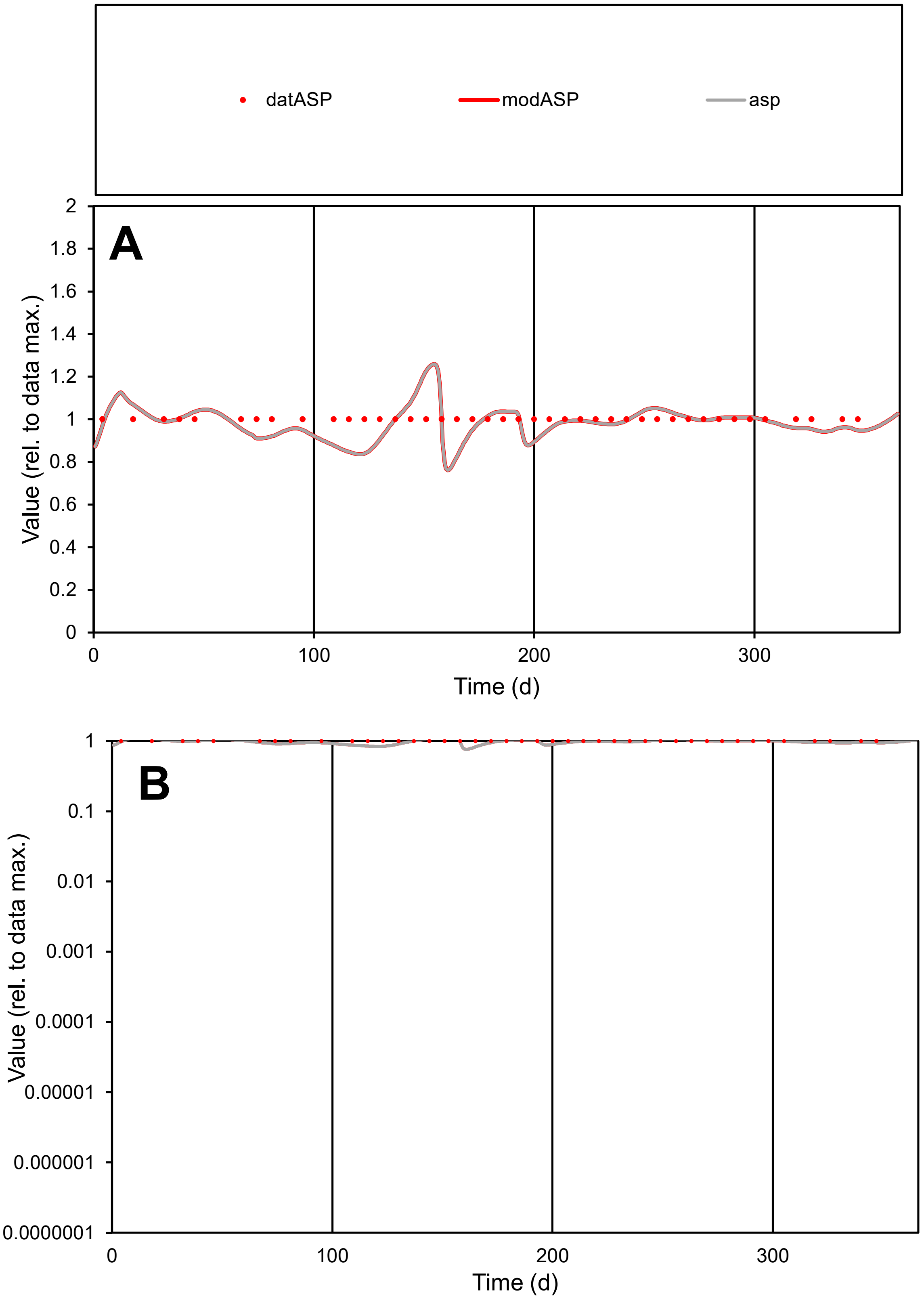


**C1**

Variables
ASP Provide jtd or Dst
CalibOnly?
2 1 to reproduce of
oNormalize?
2 1=no, 2=yes

ofxn calc*
0.07353
ofxn mod
0.13588

sRun
RunA1
zout
1

Ave
Data
5.8E-06
Model
5.7E-06

data/vMax
5.8E-06

ofxng
334.07

Min d/m
1.18569

ave/min d
1
ave/min m
1.1626

327_Calib_S_A

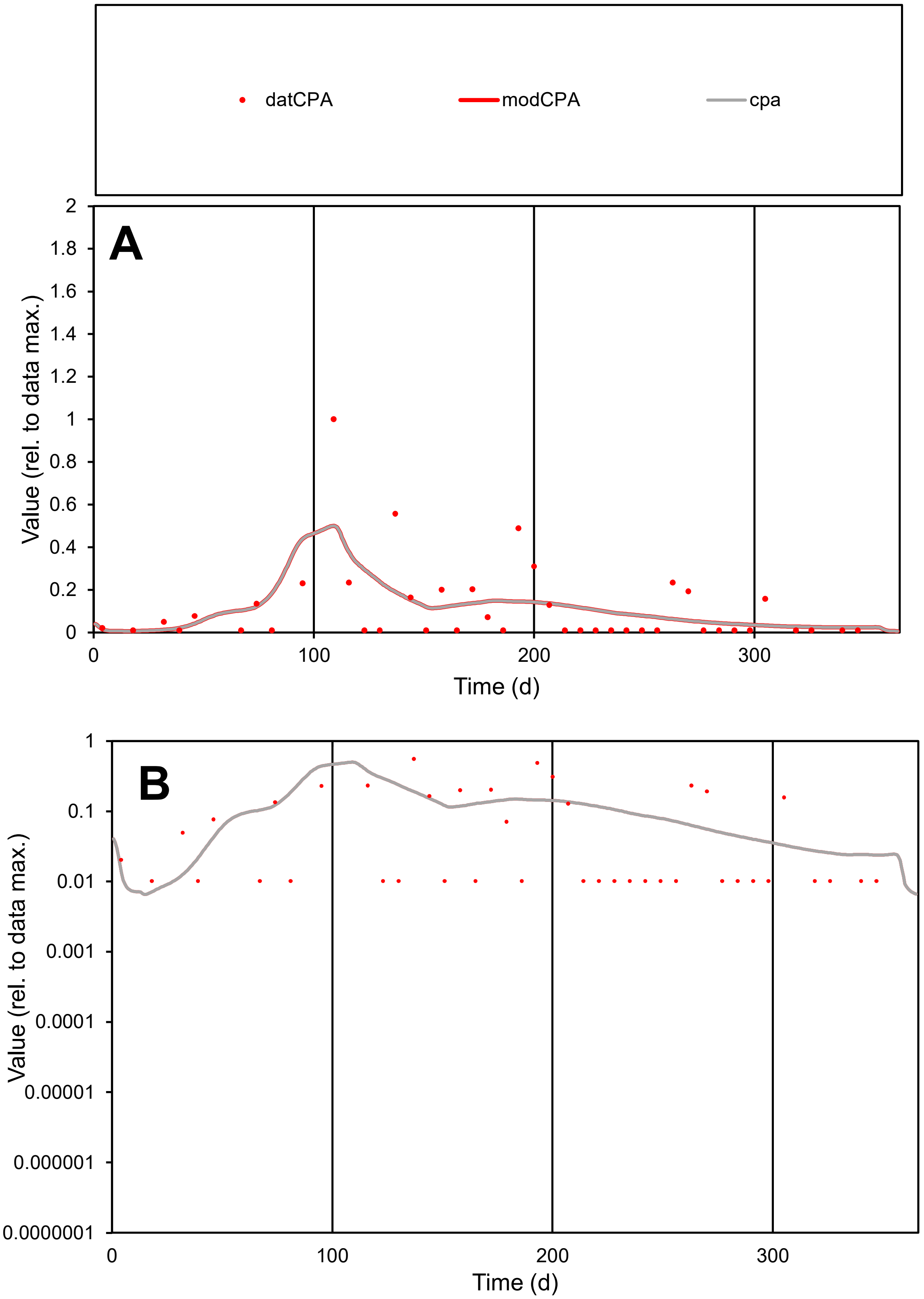


**C1**

Variables
CPA Provide jtd or Dst
CalibOnly?
2 1 to reproduce of
oNormalize?
2 1=no, 2=yes

ofxn calc*
1.34071
ofxn mod
1.356

sRun
RunA1
zout
1

Ave
Data
7.3E-07
Model
7.7E-07

data/vMax
6.5E-06

ofxng
334.07

Min d/m
1.3489

ave/min d
10.9585
ave/min m
15.5925

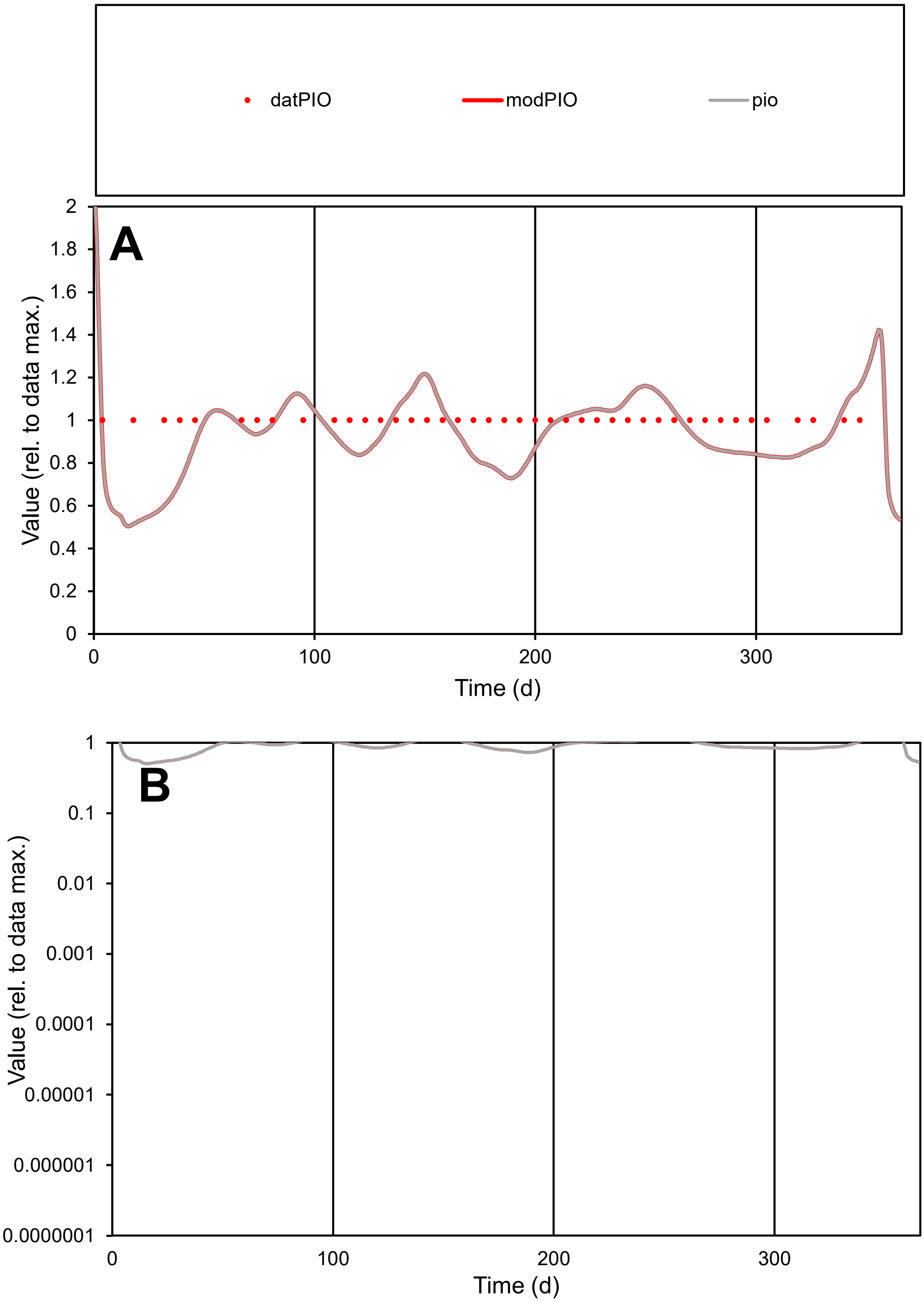


**C1**

Variables

PIO Provide jtd or Dst

CalibOnly?

2 1 to reproduce of

oNormalize?

2 1=no, 2=yes

ofxn calc*

0.16246

ofxn mod

0.19678

sRun

RunA1

zout

1

Ave

Data

1.5E-07

Model

1.4E-07

data/vMax

1.5E-07

ofxng

334.07

Min d/m

1.90802

ave/min d

1

ave/min m

1.78631

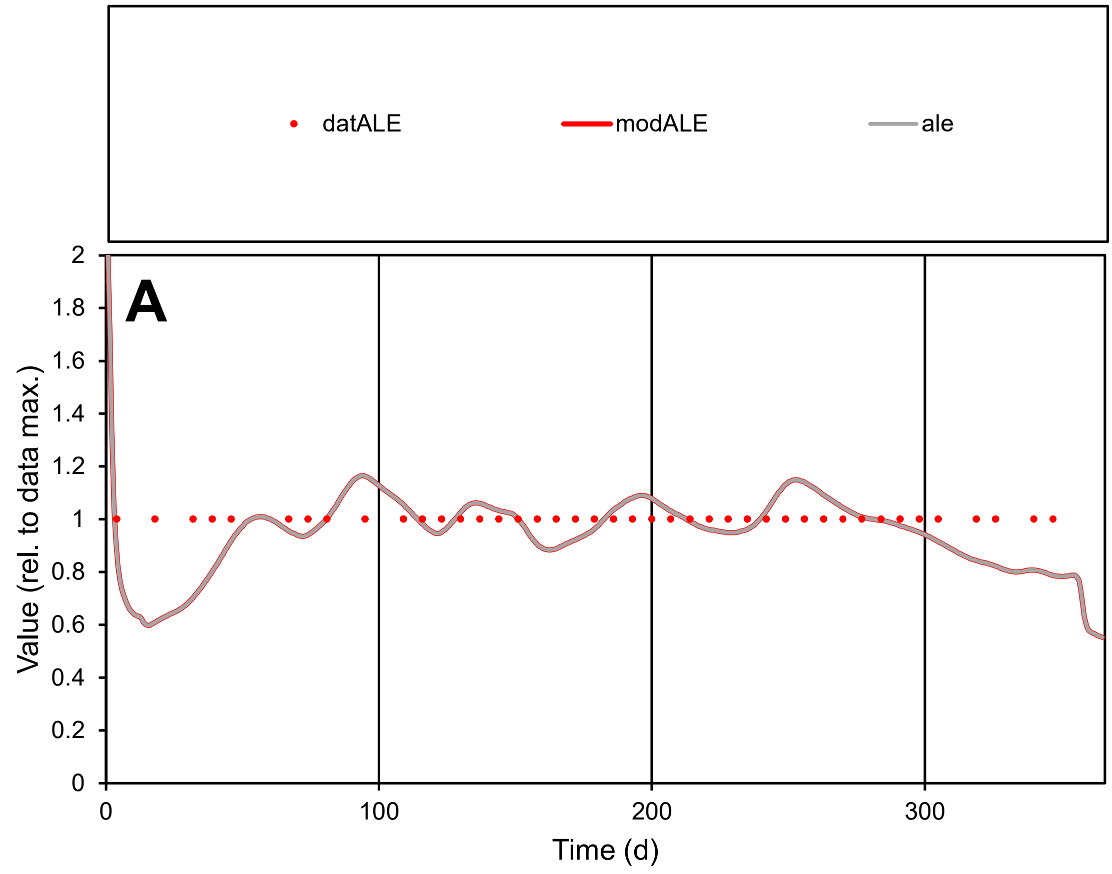


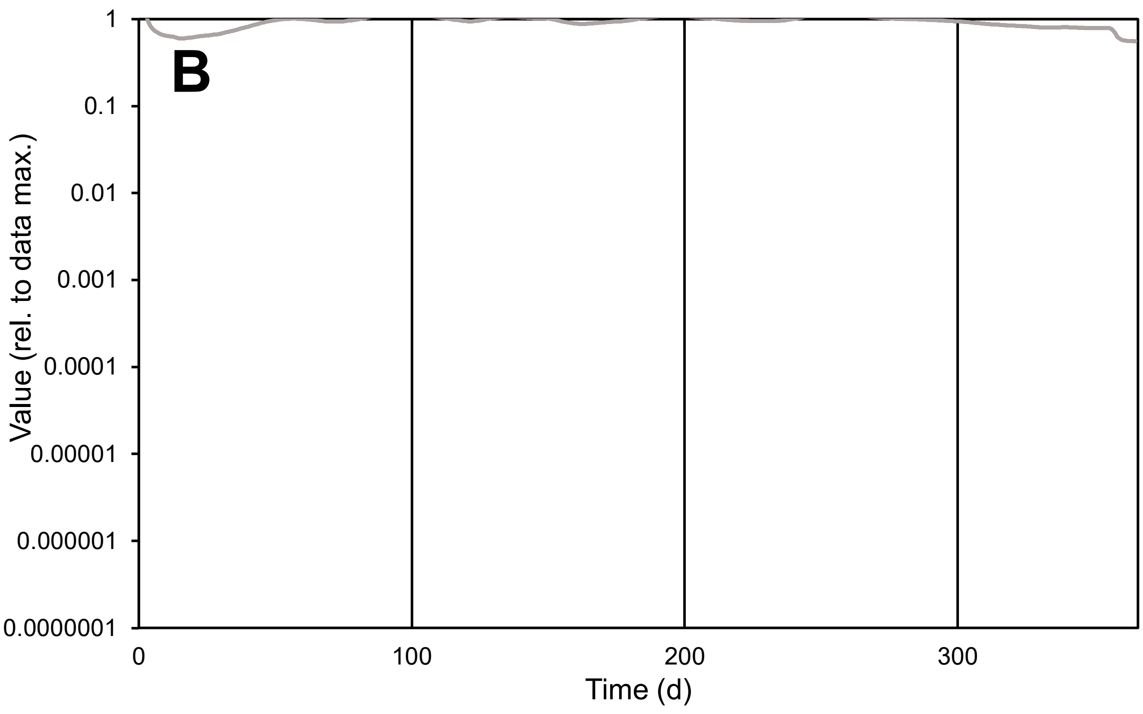


**C1**

Variables
ALE Provide jtd or Dst
CalibOnly?
2 1 to reproduce of
oNormalize?
2 1=no, 2=yes

ofxn calc*
0.11741
ofxn mod
0.1631

sRun
RunA1
zout
1

Ave
Data
1.1E-05
Model
1.1E-05

data/vMax
1.1E-05

ofxng
334.07

Min d/m
1.61248

ave/min d
1
ave/min m
1.5516

327_Calib_S_A

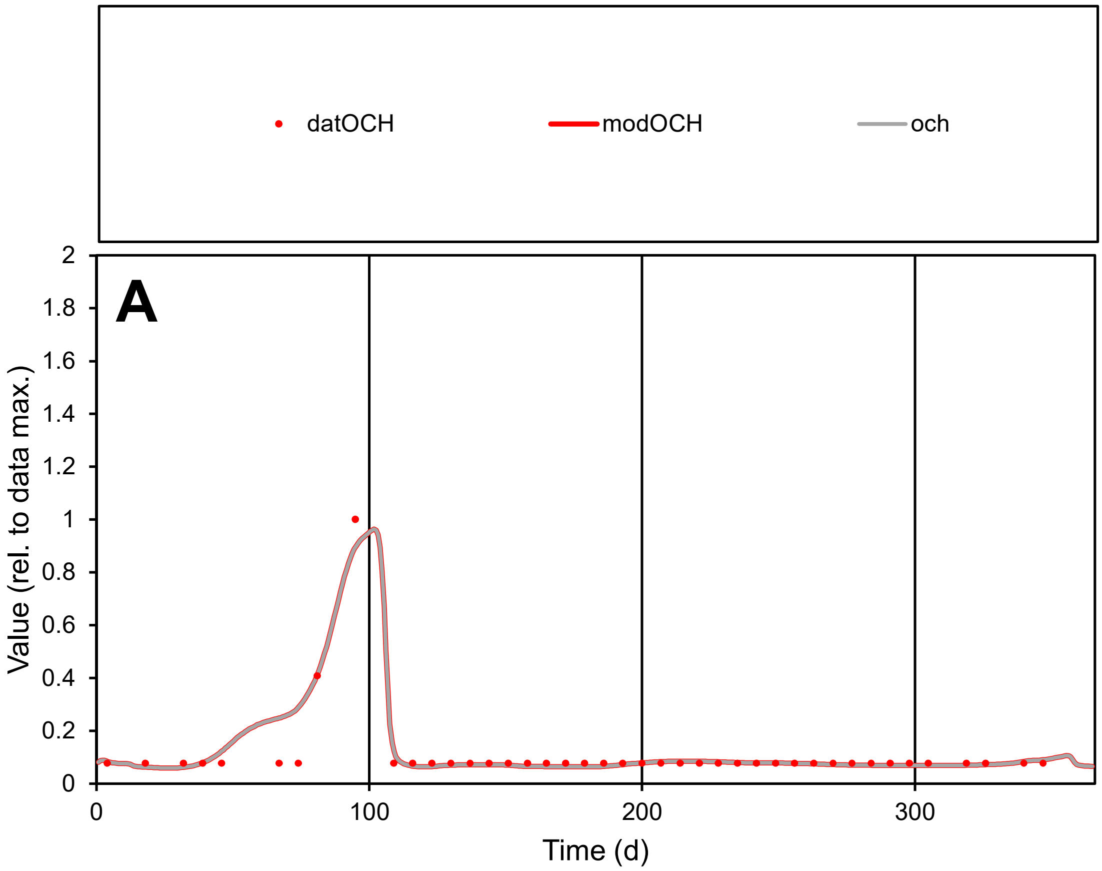


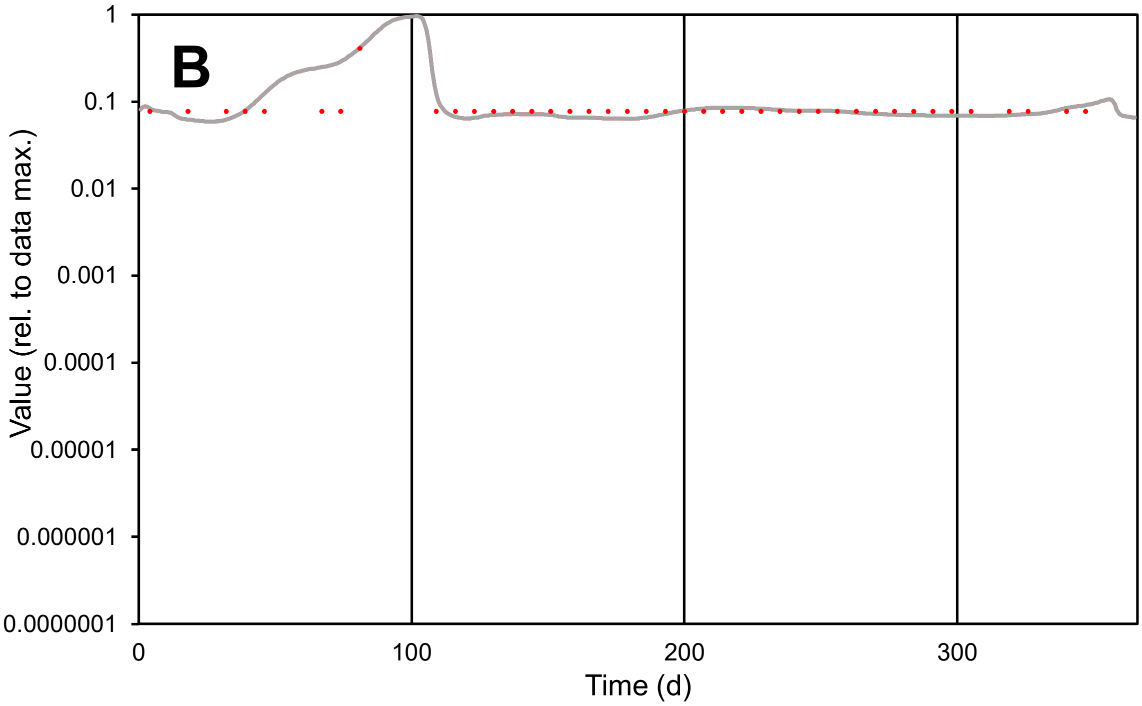


**C1**

Variables
OCH Provide jtd or Dst
CalibOnly?
2 1 to reproduce of
oNormalize?
2 1=no, 2=yes

ofxn calc*
0.43204
ofxn mod
0.45258

sRun
RunA1
zout
1

Ave
Data
6.1E-06
Model
6.5E-06

data/vMax
5.7E-05

ofxng
334.07

Min d/m
1.25629

ave/min d
1.38606
ave/min m
1.84568

327_Calib_S_A

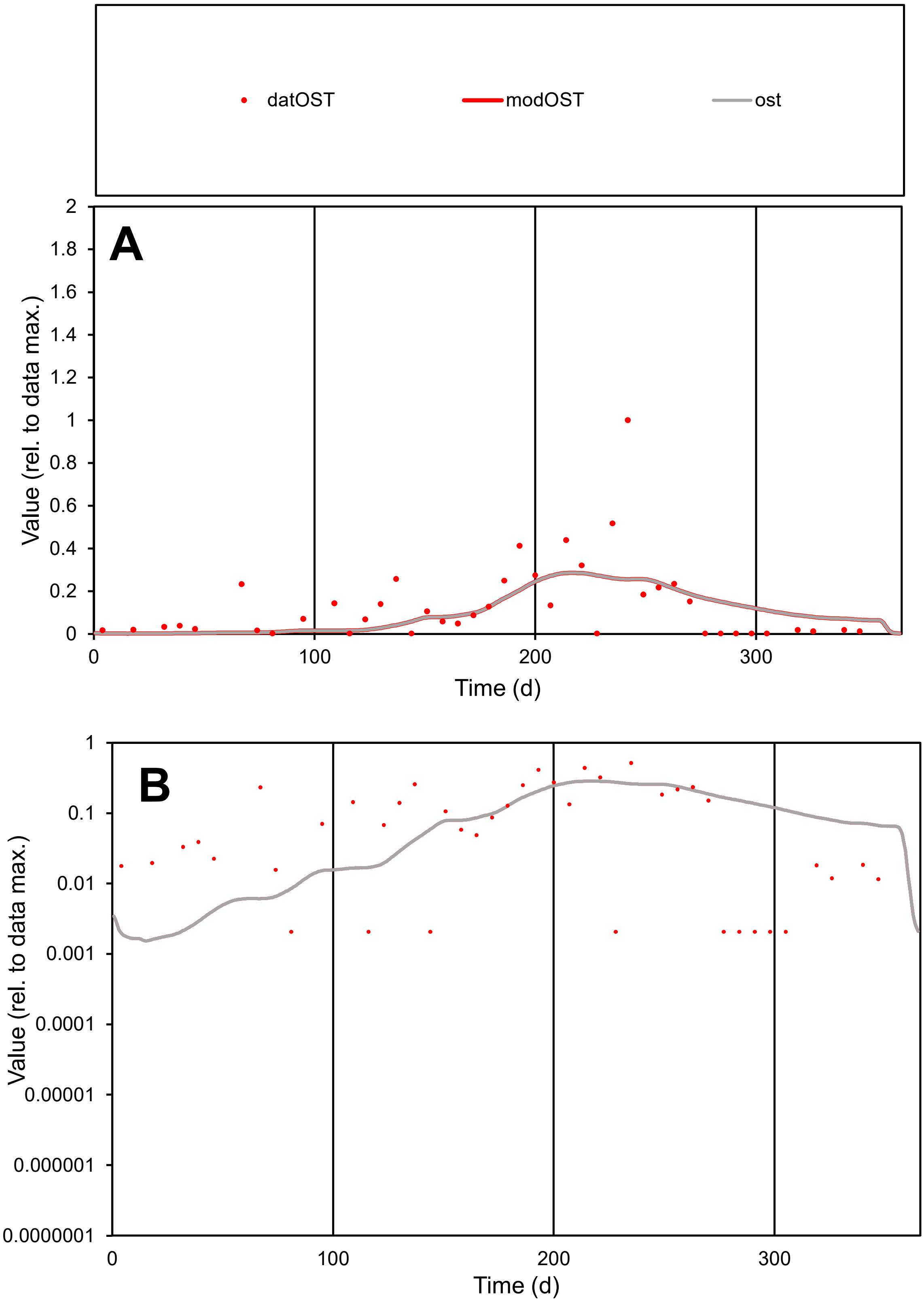


**C1**

Variables

OST Provide jtd or Dst

CalibOnly?

2 1 to reproduce of

oNormalize?

2 1=no, 2=yes

ofxn calc*

1.15613

ofxn mod

1.1695

sRun

RunA1

zout

1

Ave

Data

0.00014

Model

0.00012

data/vMax

0.00105

ofxng

334.07

Min d/m

1.25188

ave/min d

65.6878

ave/min m

69.9127

327_Calib_S_A

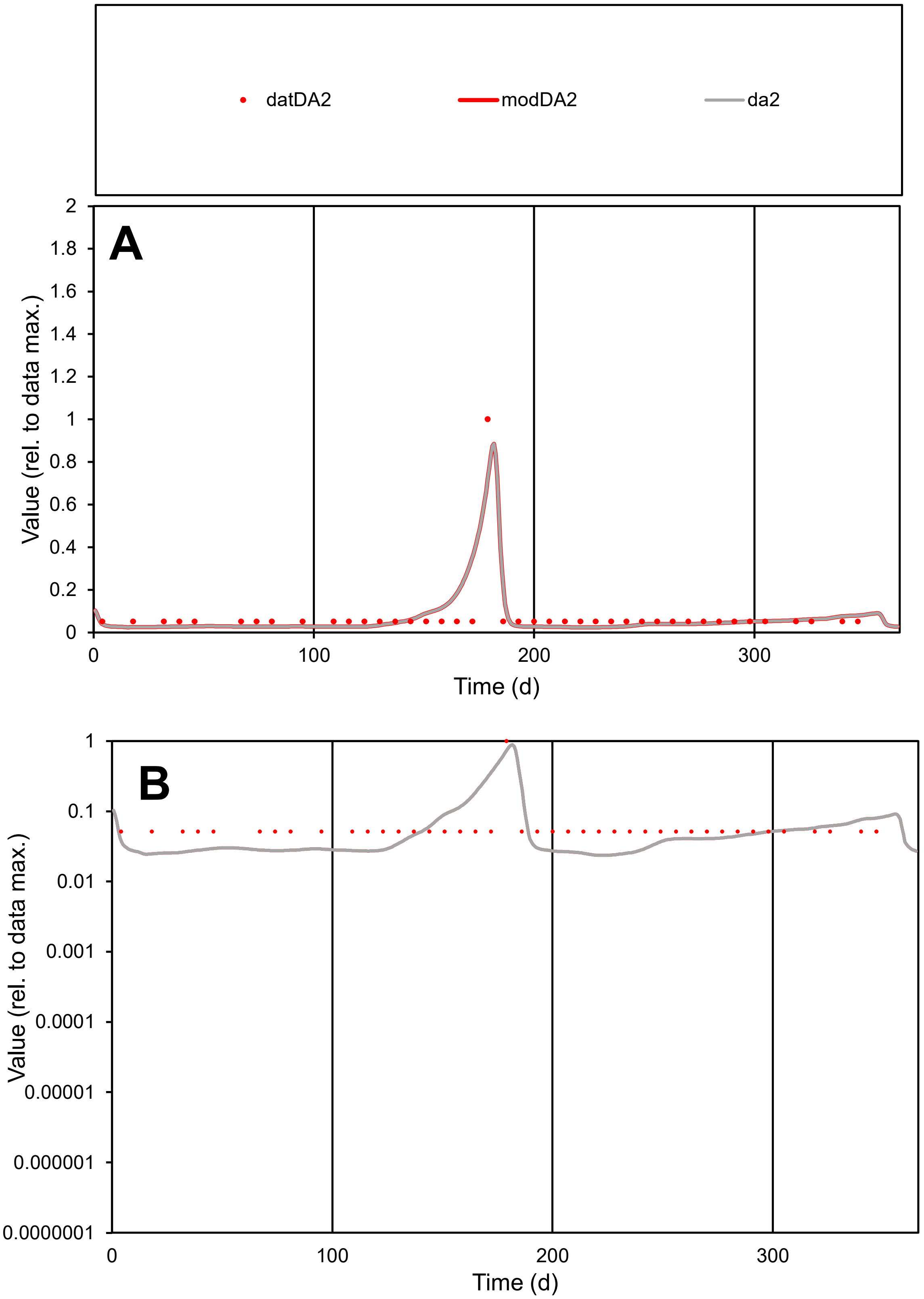


**C1**

Variables
DA2 Provide jtd or Dst
CalibOnly?
2 1 to reproduce of
oNormalize?
2 1=no, 2=yes

ofxn calc*
0.74034
ofxn mod
0.86452

sRun
RunA1
zout
1

Ave
Data
4.5E-06
Model
4.3E-06

data/vMax
6E-05

ofxng
334.07

Min d/m
2.17877

ave/min d
1.43765
ave/min m
3.00515

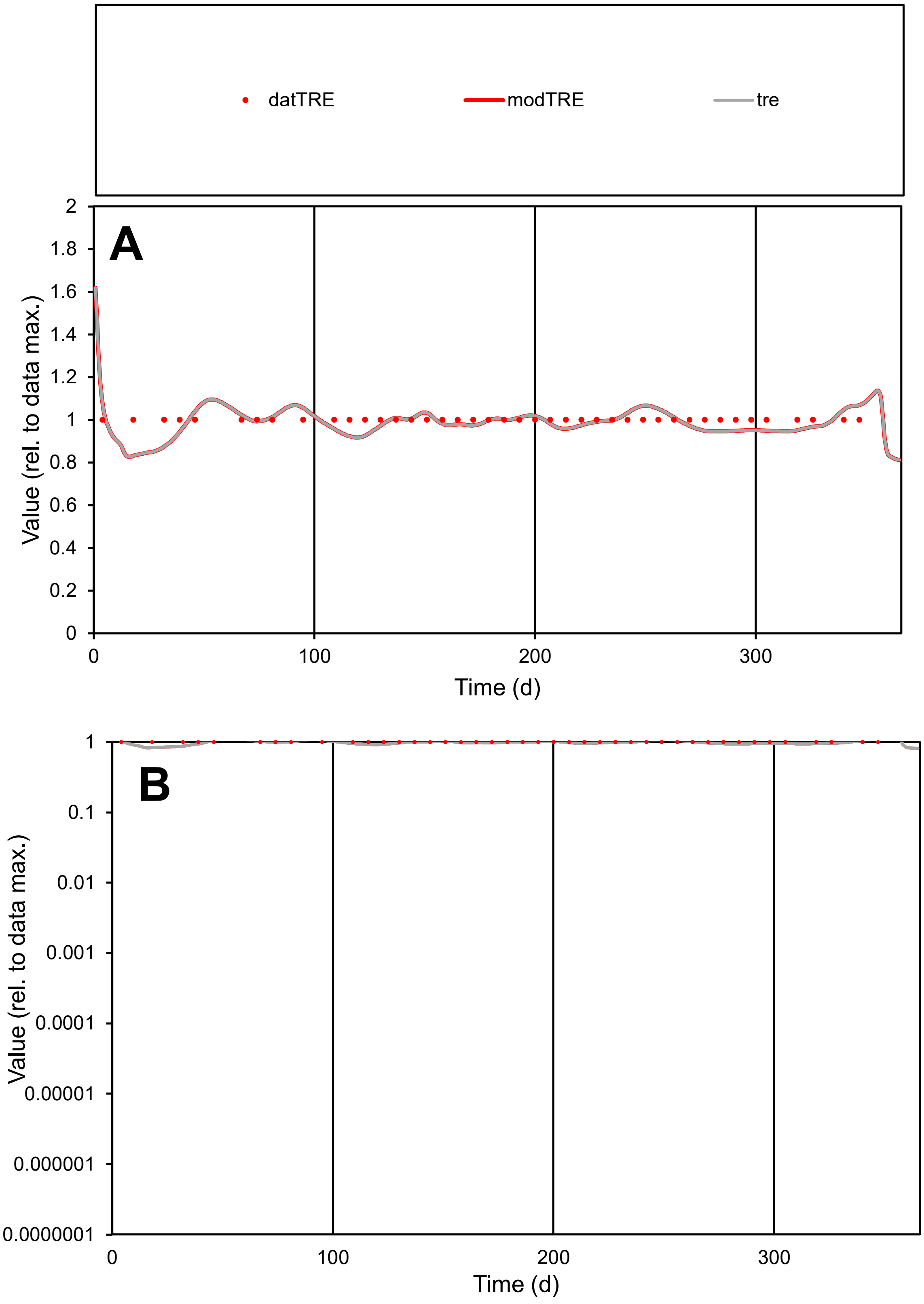


**C1**

Variables
TRE Provide jtd or Dst
CalibOnly?
2 1 to reproduce of
oNormalize?
2 1=no, 2=yes

ofxn calc*
0.04946
ofxn mod
0.12278

sRun
RunA1
zout
1

Ave
Data
6.9E-06
Model
6.8E-06

data/vMax
6.9E-06

ofxng
334.07

Min d/m
1.19506

ave/min d
1
ave/min m
1.18013

327_Calib_S_A

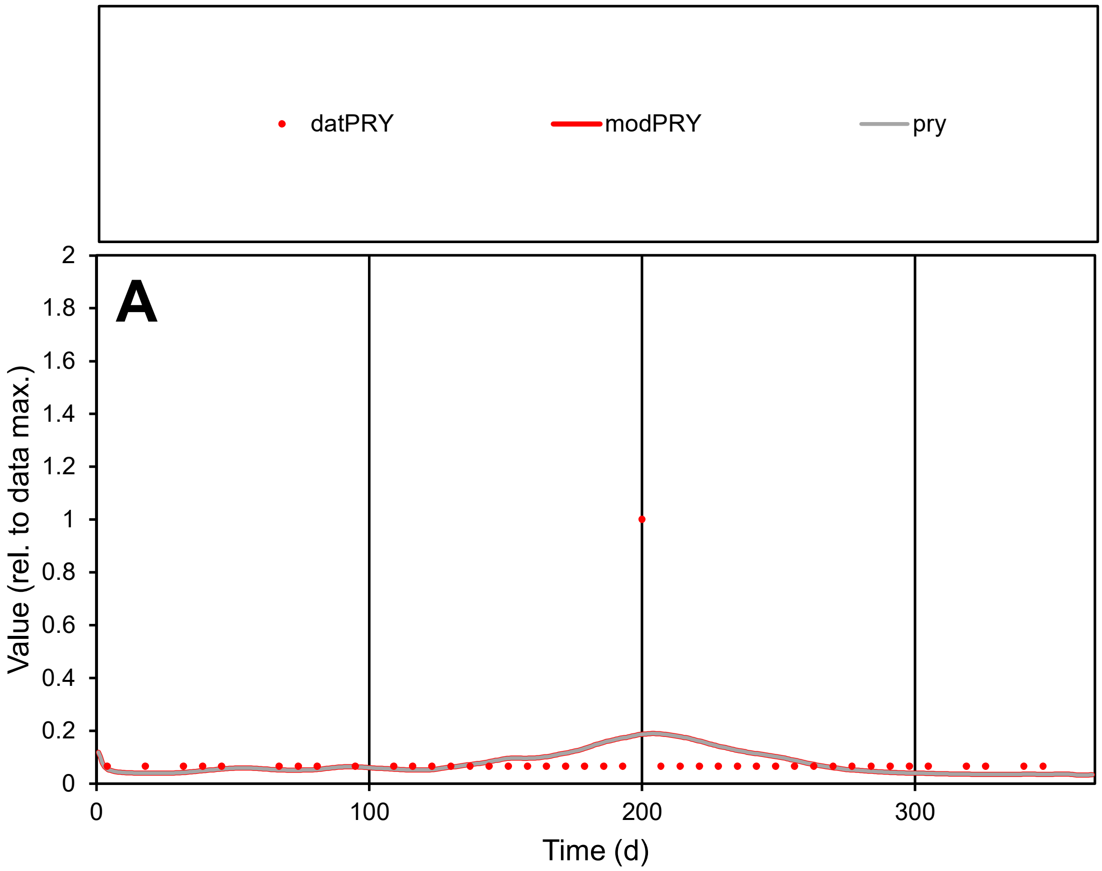


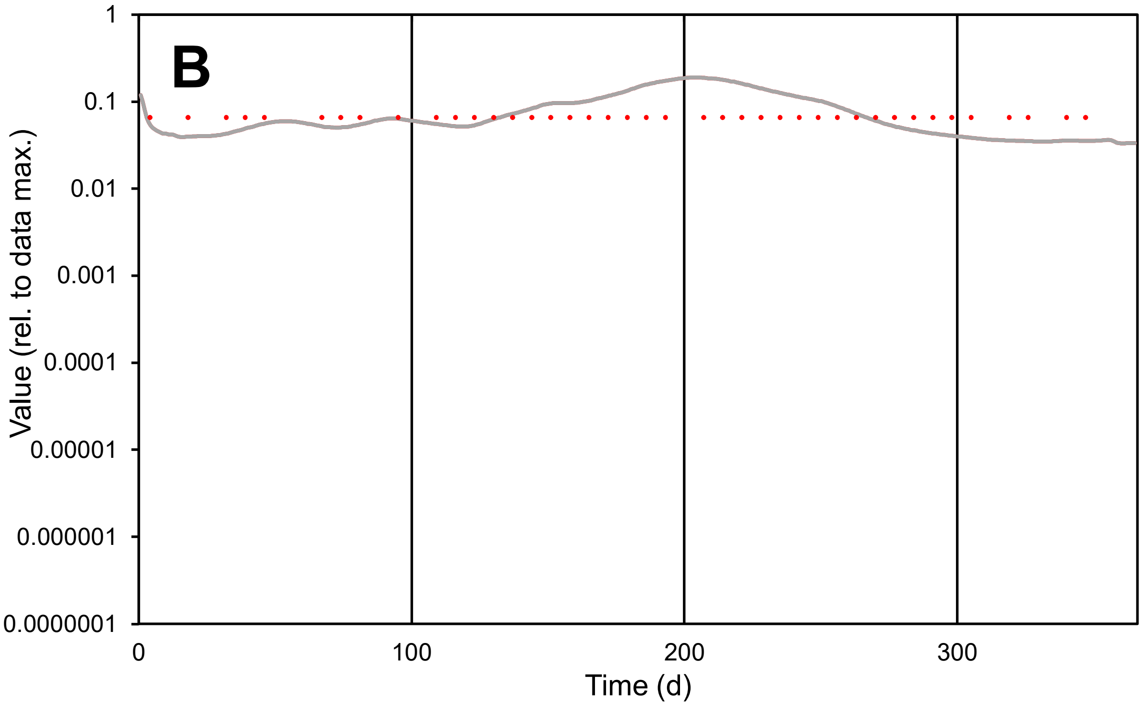


**C1**

Variables
PRY Provide jtd or Dst
CalibOnly?
2 1 to reproduce of
oNormalize?
2 1=no, 2=yes

ofxn calc*
1.51305
ofxn mod
1.52484

sRun
RunA1
zout
1

Ave
Data
1.2E-05
Model
1.1E-05

data/vMax
0.00013

ofxng
334.07

Min d/m
1.87989

ave/min d
1.33783
ave/min m
2.37303

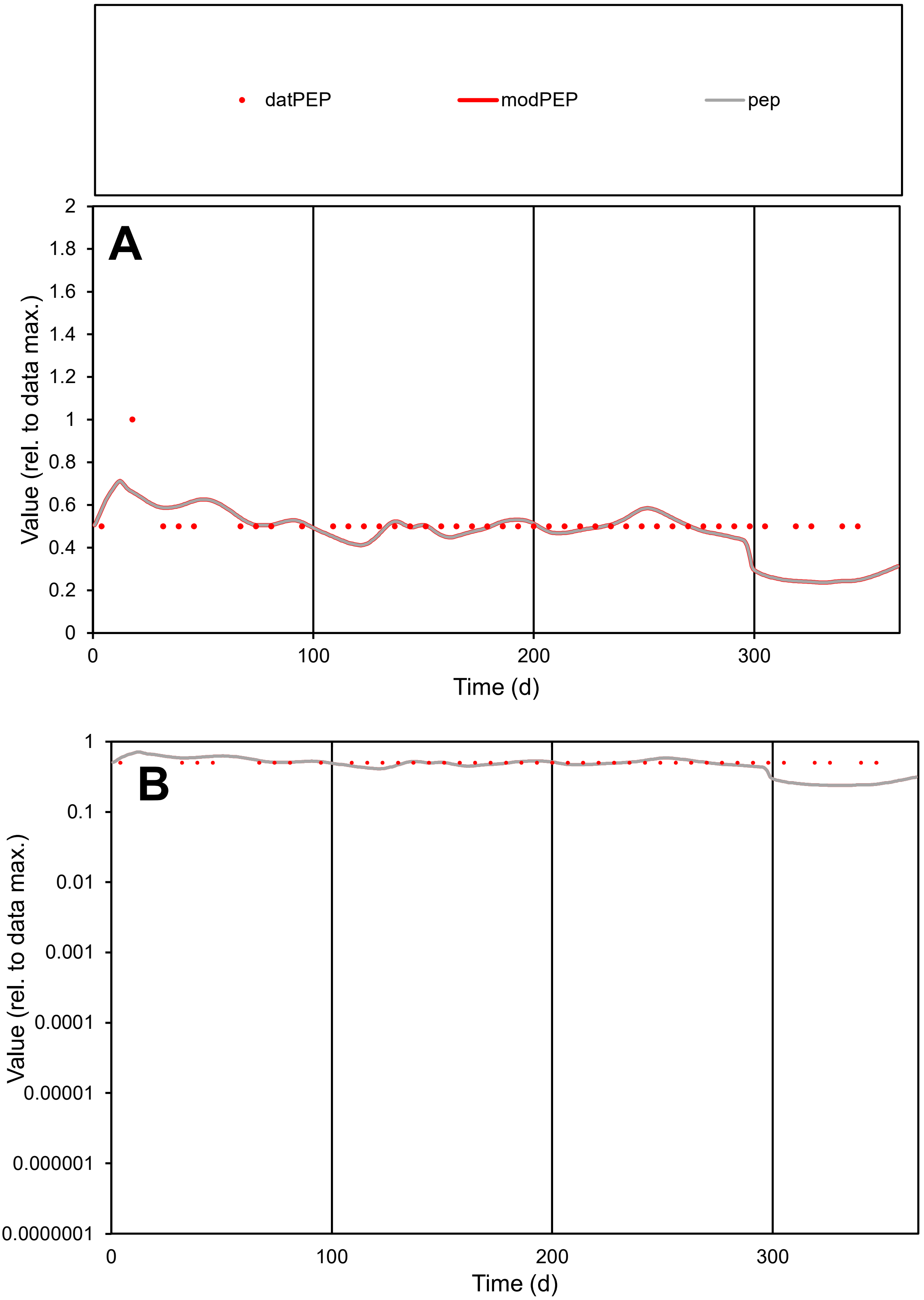


**C1**

Variables
PEP Provide jtd or Dst
CalibOnly?
2 1 to reproduce of
oNormalize?
2 1=no, 2=yes

ofxn calc*
0.22445
ofxn mod
0.25259

sRun
RunA1
zout
1

Ave
Data
5.1E-06
Model
4.7E-06

data/vMax
9.9E-06

ofxng
334.07

Min d/m
2.09334

ave/min d
1.02381
ave/min m
1.99303

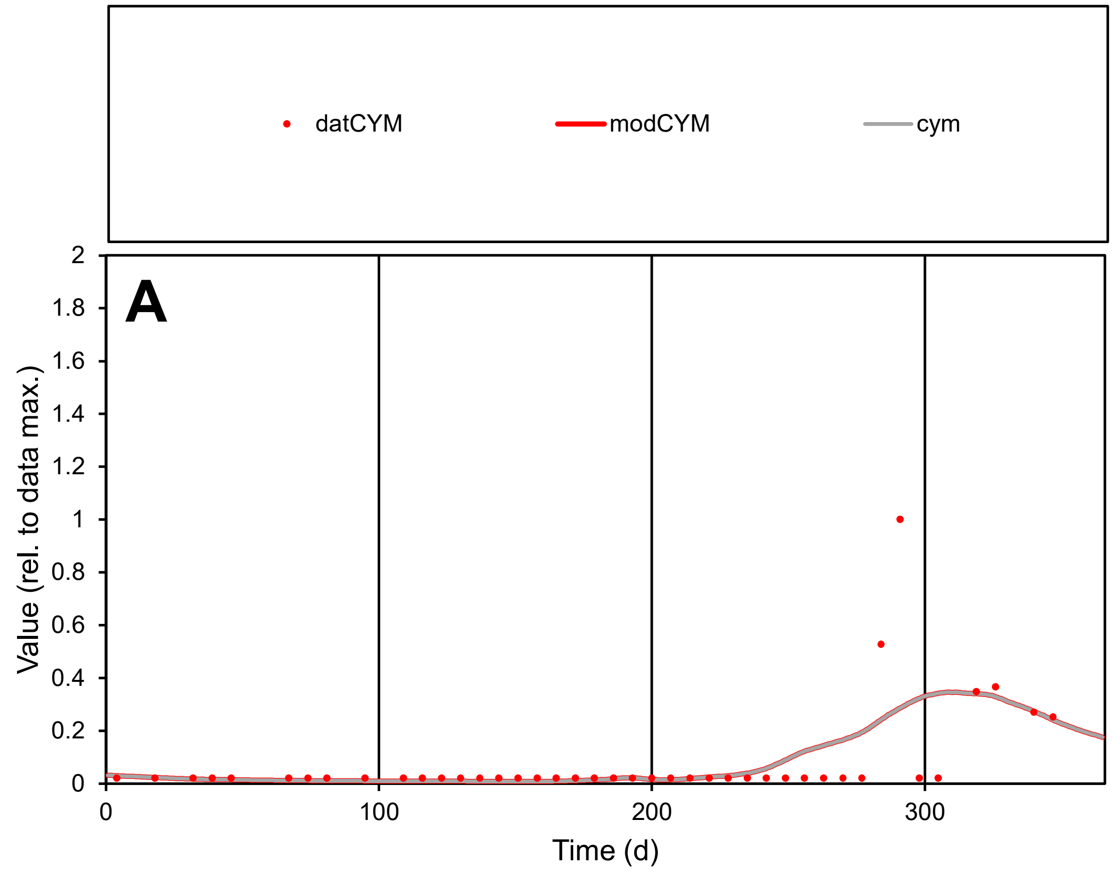


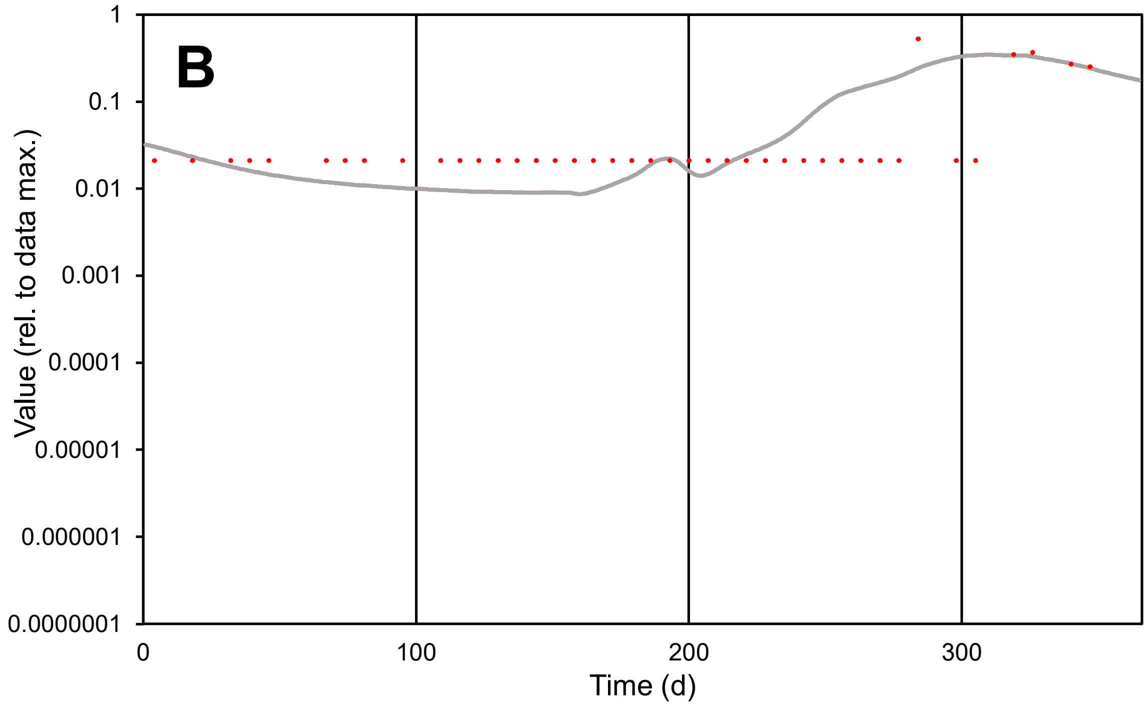


**C1**

Variables
CYM Provide jtd or Dst
CalibOnly?
2 1 to reproduce of
oNormalize?
2 1=no, 2=yes

ofxn calc*
1.70712
ofxn mod
1.72001

sRun
RunA1
zout
1

Ave
Data
5.4E-06
Model
5.6E-06

data/vMax
6.5E-05

ofxng
334.07

Min d/m
2.40842

ave/min d
3.98999
ave/min m
9.85587

327_Calib_S_A

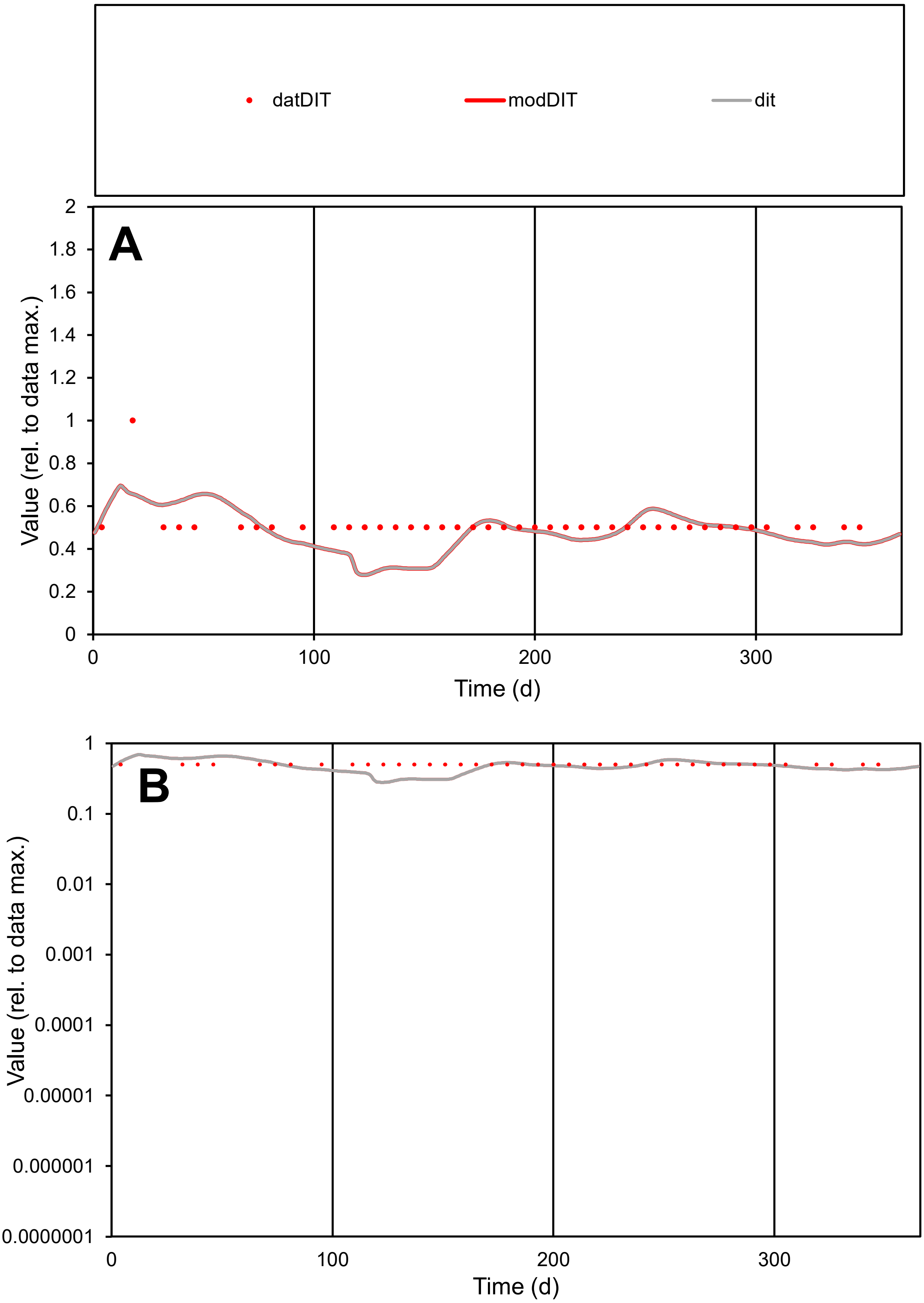


**C1**

Variables
DIT Provide jtd or Dst
CalibOnly?
2 1 to reproduce of
oNormalize?
2 1=no, 2=yes

ofxn calc*
0.21055
ofxn mod
0.23923

sRun
RunA1
zout
1

Ave
Data
5.2E-06
Model
4.8E-06

data/vMax
1E-05

ofxng
334.07

Min d/m
1.78492

ave/min d
1.02379
ave/min m
1.6921

327_Calib_S_A

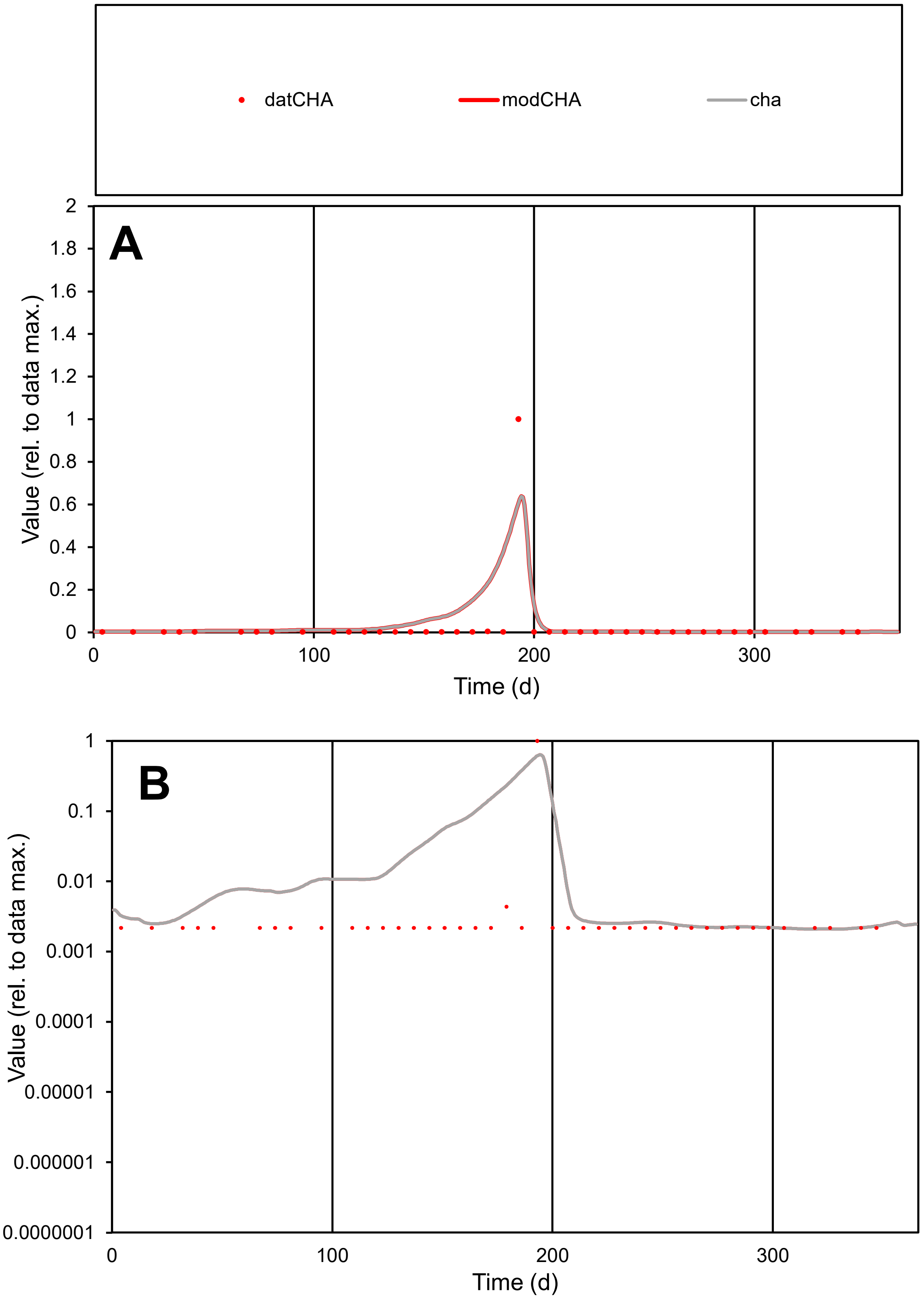


**C1**

Variables

CHA Provide jtd or Dst

CalibOnly?

2 1 to reproduce of

oNormalize?

2 1=no, 2=yes

ofxn calc*

3.41742

ofxn mod

3.613

sRun

RunA1

zout

1

Ave

Data

1E-06

Model

1.7E-06

data/vMax

4E-05

ofxng

334.07

Min d/m

1.04064

ave/min d

11.9295

ave/min m

20.4439

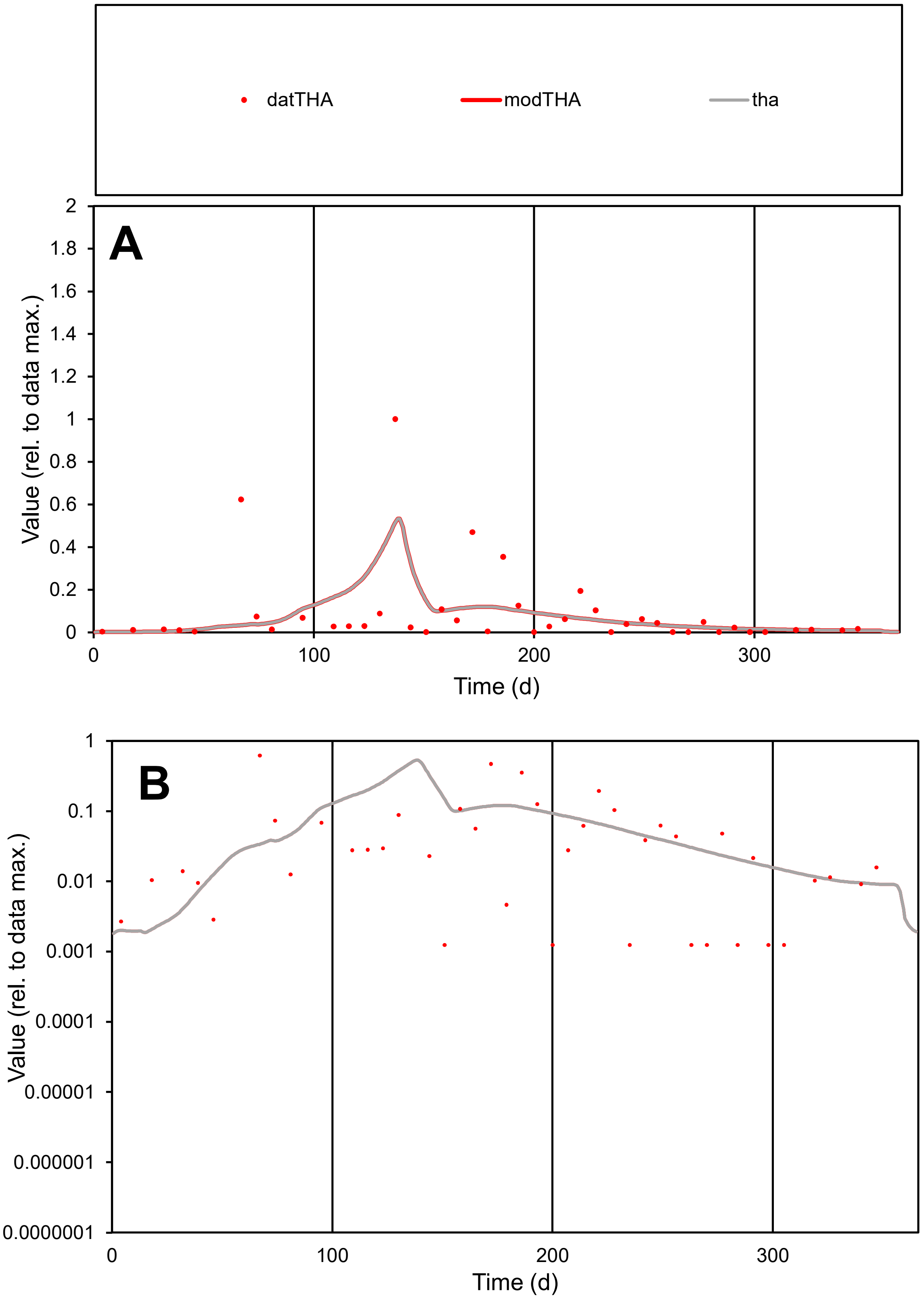


**C1**

Variables
THA Provide jtd or Dst
CalibOnly?
2 1 to reproduce of
oNormalize?
2 1=no, 2=yes

ofxn calc*
1.73824
ofxn mod
1.76795

sRun
RunA1
zout
1

Ave
Data
0.0002
Model
0.00019

data/vMax
0.00222

ofxng
334.07

Min d/m
0.61676

ave/min d
72.8737
ave/min m
41.6441

327_Calib_S_A

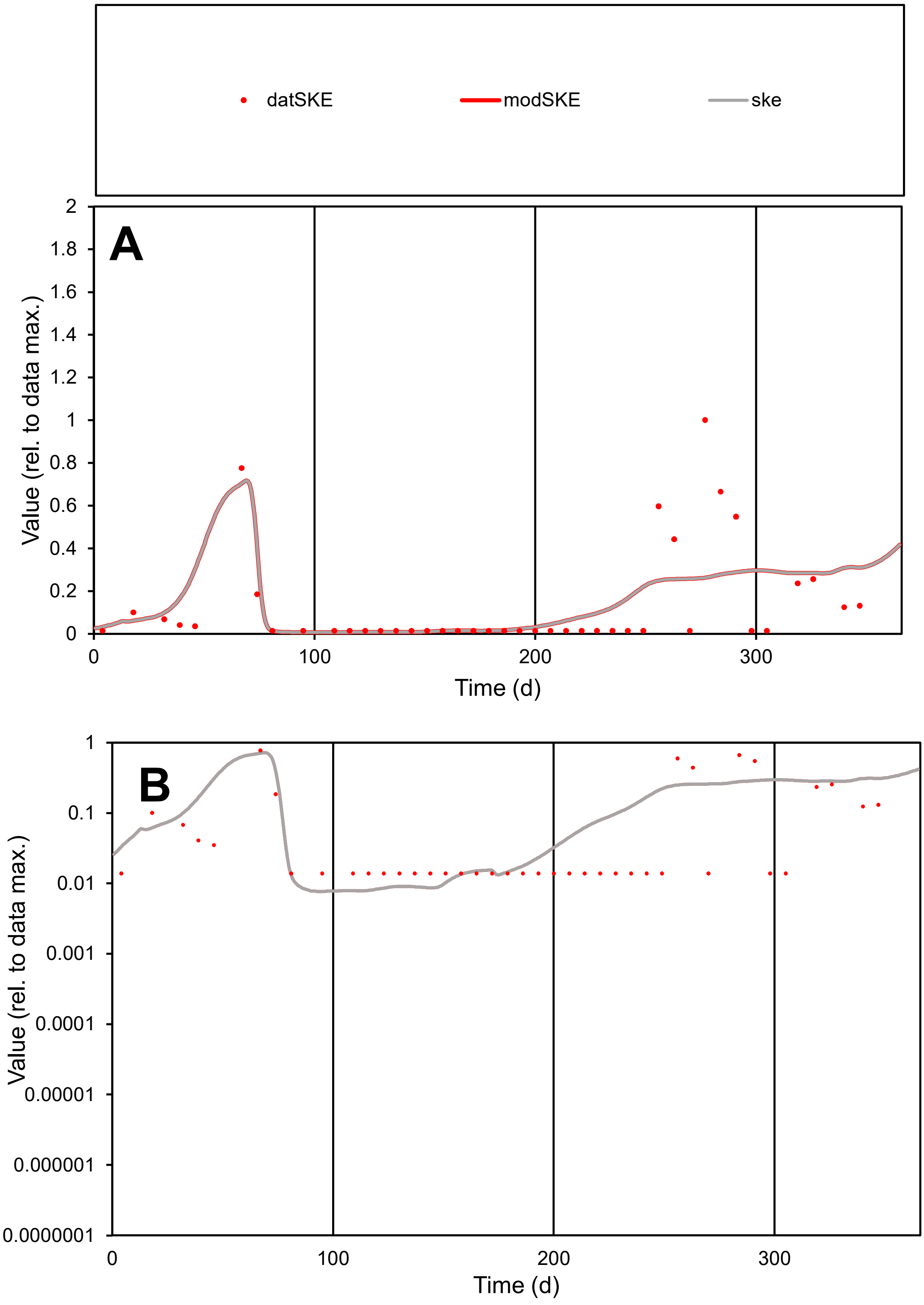


**C1**

Variables

SKE Provide jtd or Dst

CalibOnly?

2 1 to reproduce of

oNormalize?

2 1=no, 2=yes

ofxn calc*

1.3617

ofxn mod

1.38382

sRun

RunA1

zout

1

Ave

Data

2.2E-05

Model

2.4E-05

data/vMax

0.00017

ofxng

334.07

Min d/m

1.80754

ave/min d

9.57992

ave/min m

18.491

327_Calib_S_A

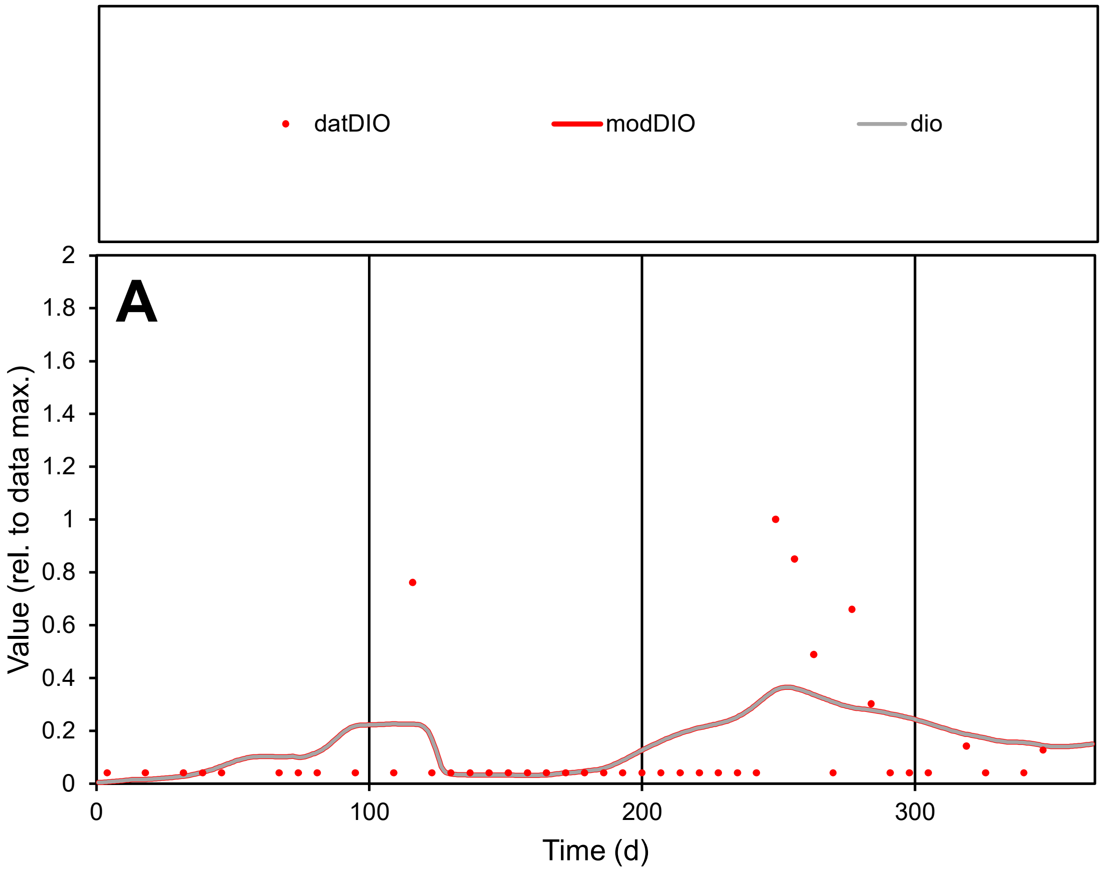


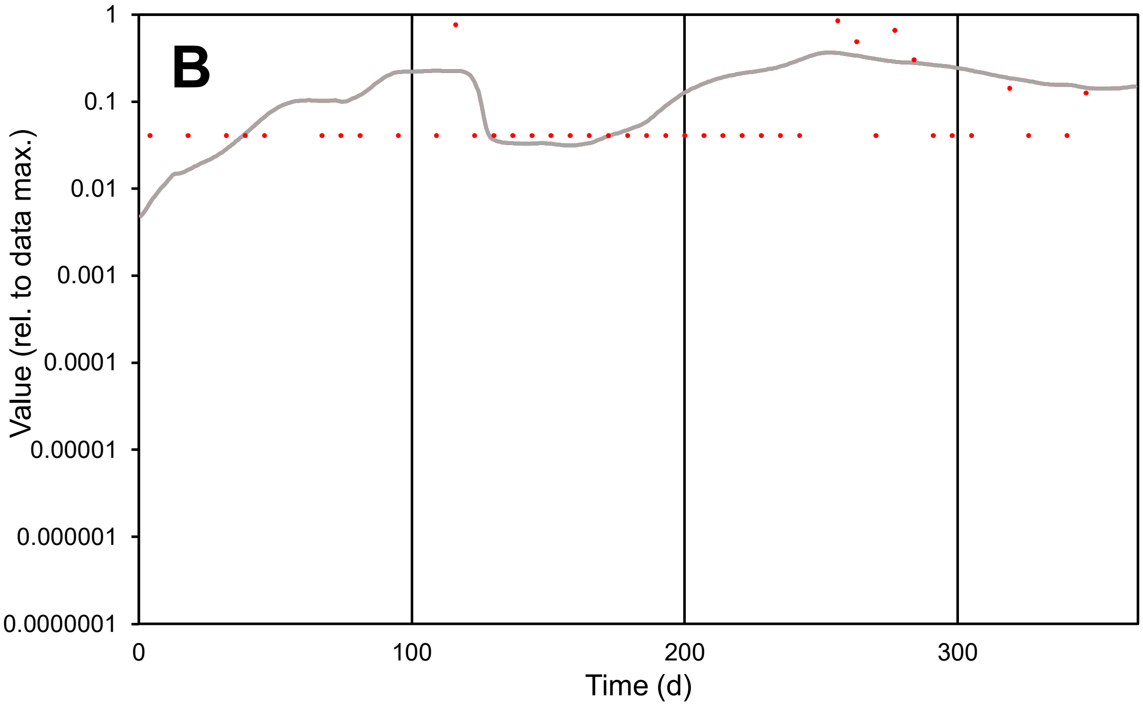


**C1**

Variables
DIO Provide jtd or Dst
CalibOnly?
2 1 to reproduce of
oNormalize?
2 1=no, 2=yes

ofxn calc*
1.45019
ofxn mod
1.46478

sRun
RunA1
zout
1

Ave
Data
1.6E-05
Model
1.8E-05

data/vMax
0.00011

ofxng
334.07

Min d/m
5.37841

ave/min d
3.32775
ave/min m
20.1813

327_Calib_S_A

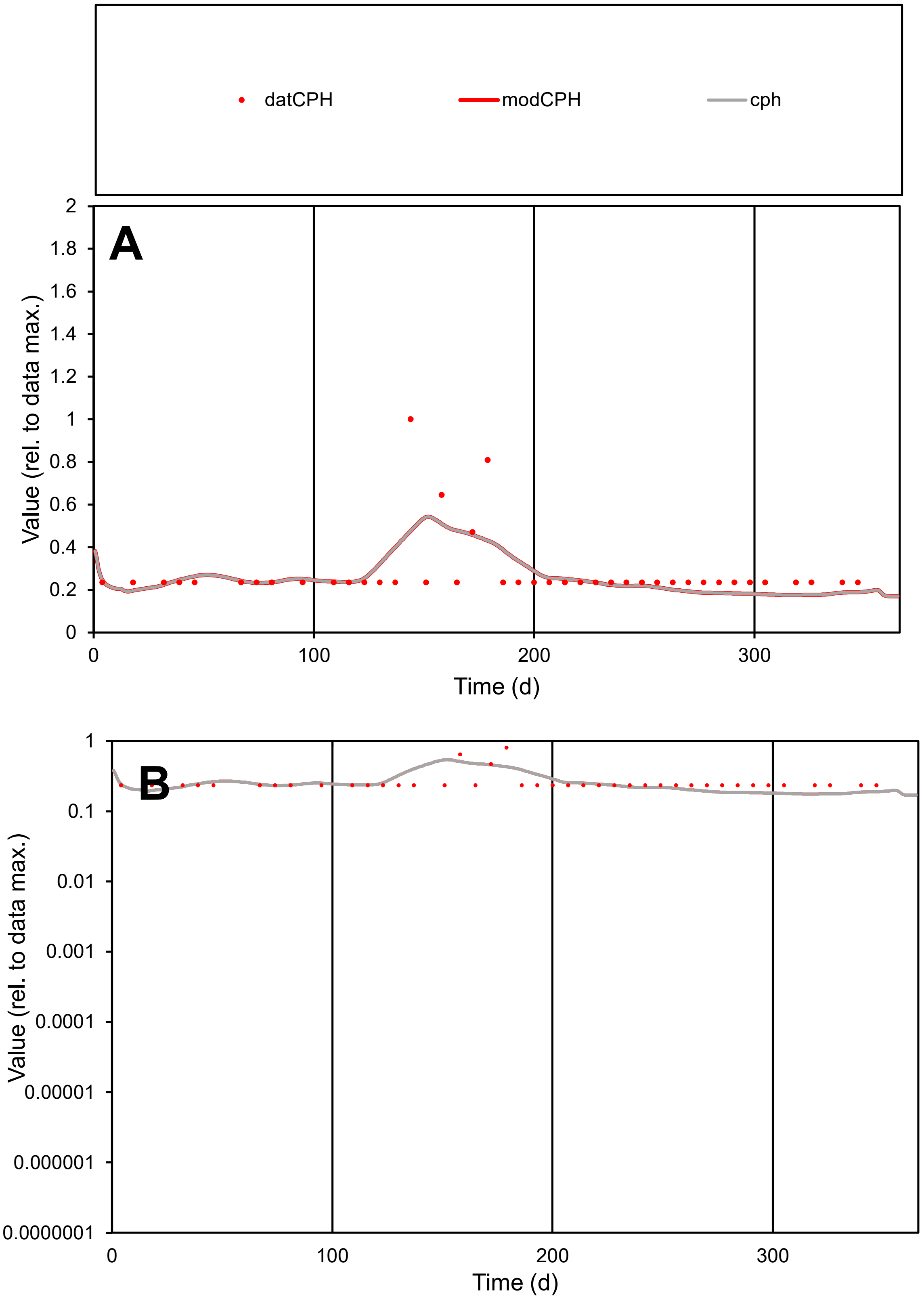


**C1**

Variables

| CPH | Provide jtd or Dst |
|---|---|

CalibOnly?

| 2 | 1 to reproduce of |
|---|---|

oNormalize?

| 2 | 1=no, 2=yes |
|---|---|

ofxn calc*
0.45081
ofxn mod
0.46617

sRun
RunA1
zout
1

Ave
Data
1.2E-05
Model
1.1E-05

data/vMax
4.1E-05

ofxng
334.07

Min d/m
1.33435

ave/min d
1.20075
ave/min m
1.53604

327_Calib_S_A

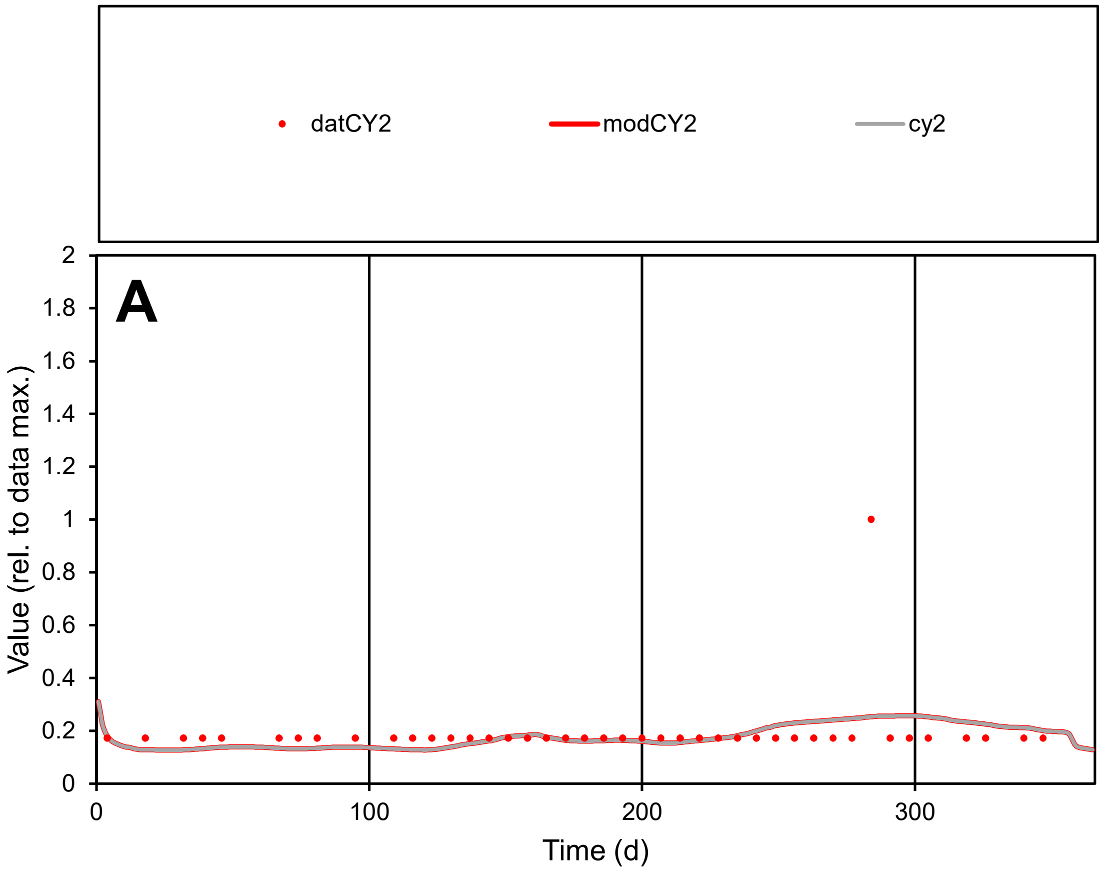


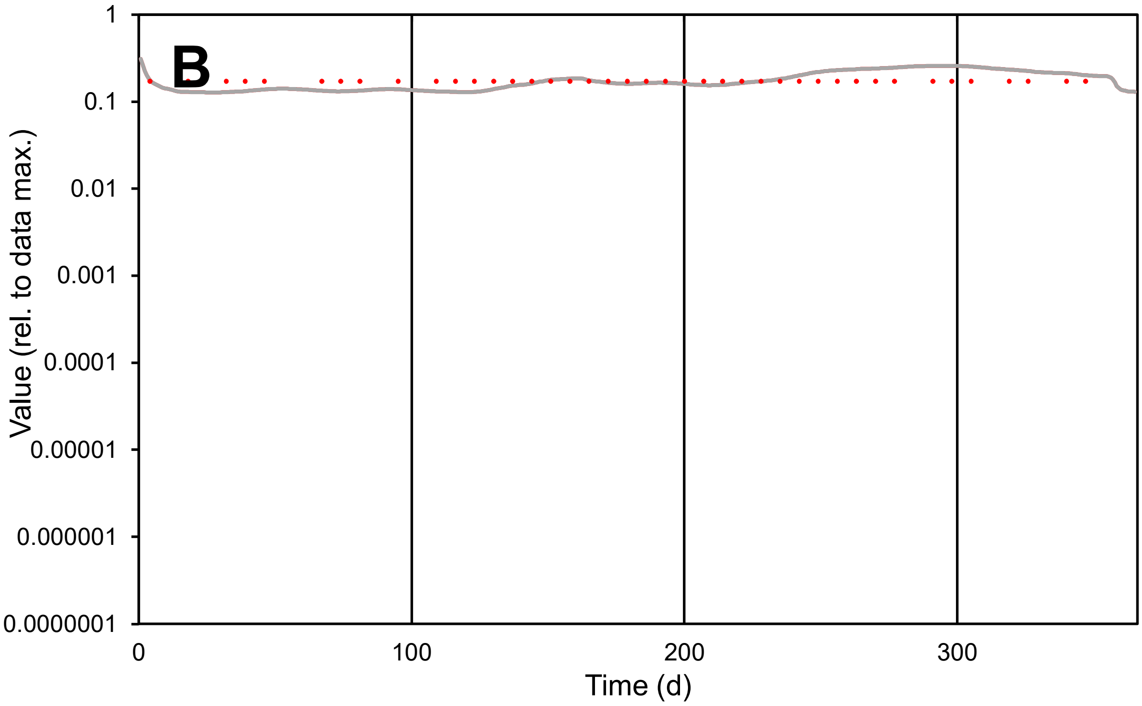


**C1**

Variables
CY2 Provide jtd or Dst
CalibOnly?
2 1 to reproduce of
oNormalize?
2 1=no, 2=yes

ofxn calc*
0.63903
ofxn mod
0.64868

sRun
RunA1
zout
1

Ave
Data
1.2E-05
Model
1.2E-05

data/vMax
6.5E-05

ofxng
334.07

Min d/m
1.33333

ave/min d
1.11503
ave/min m
1.39496

327_Calib_S_A

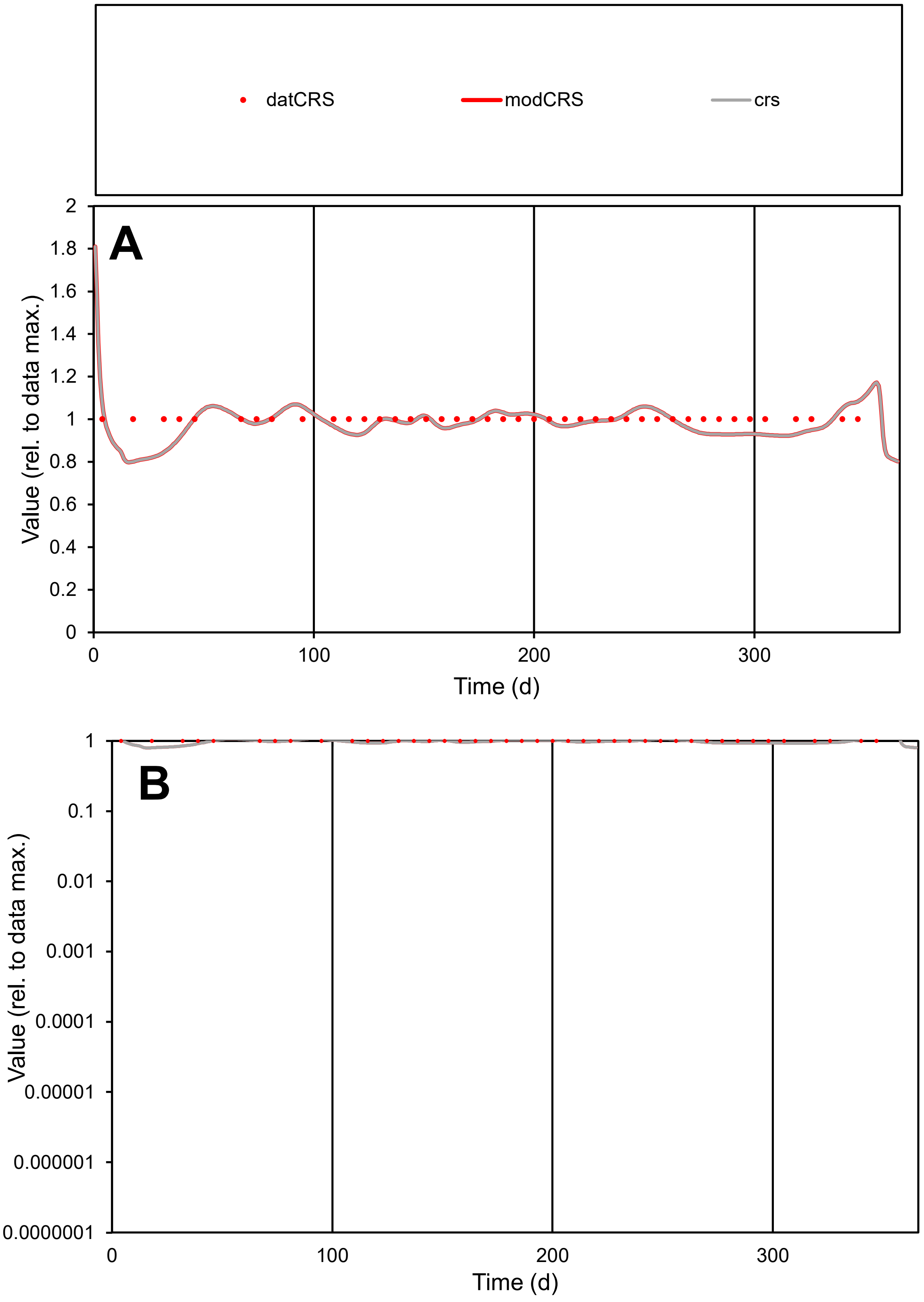


**C1**

Variables

CRS Provide jtd or Dst

CalibOnly?

2 1 to reproduce of

oNormalize?

2 1=no, 2=yes

ofxn calc*

0.05735

ofxn mod

0.12586

sRun

RunA1

zout

1

Ave

Data

5.4E-06

Model

5.3E-06

data/vMax

5.4E-06

ofxng

334.07

Min d/m

1.24054

ave/min d

1

ave/min m

1.21683

327_Calib_S_A

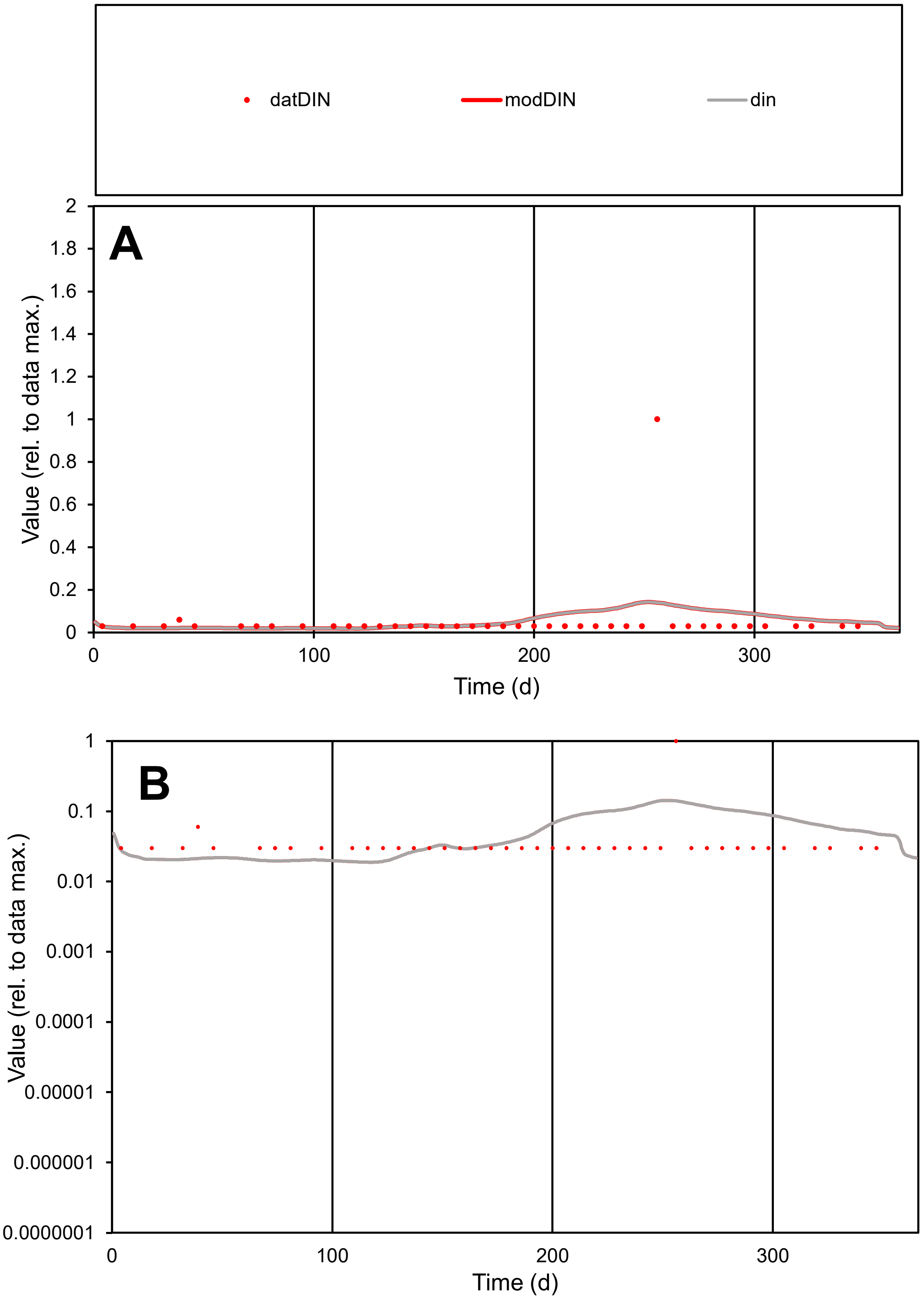


**C1**

Variables
DIN Provide jtd or Dst
CalibOnly?
2 1 to reproduce of
oNormalize?
2 1=no, 2=yes

ofxn calc*
2.61771
ofxn mod
2.63049

sRun
RunA1
zout
1

Ave
Data
5.2E-06
Model
5.7E-06

data/vMax
9.6E-05

ofxng
334.07

Min d/m
1.58549

ave/min d
1.79618
ave/min m
3.12914

327_Calib_S_A

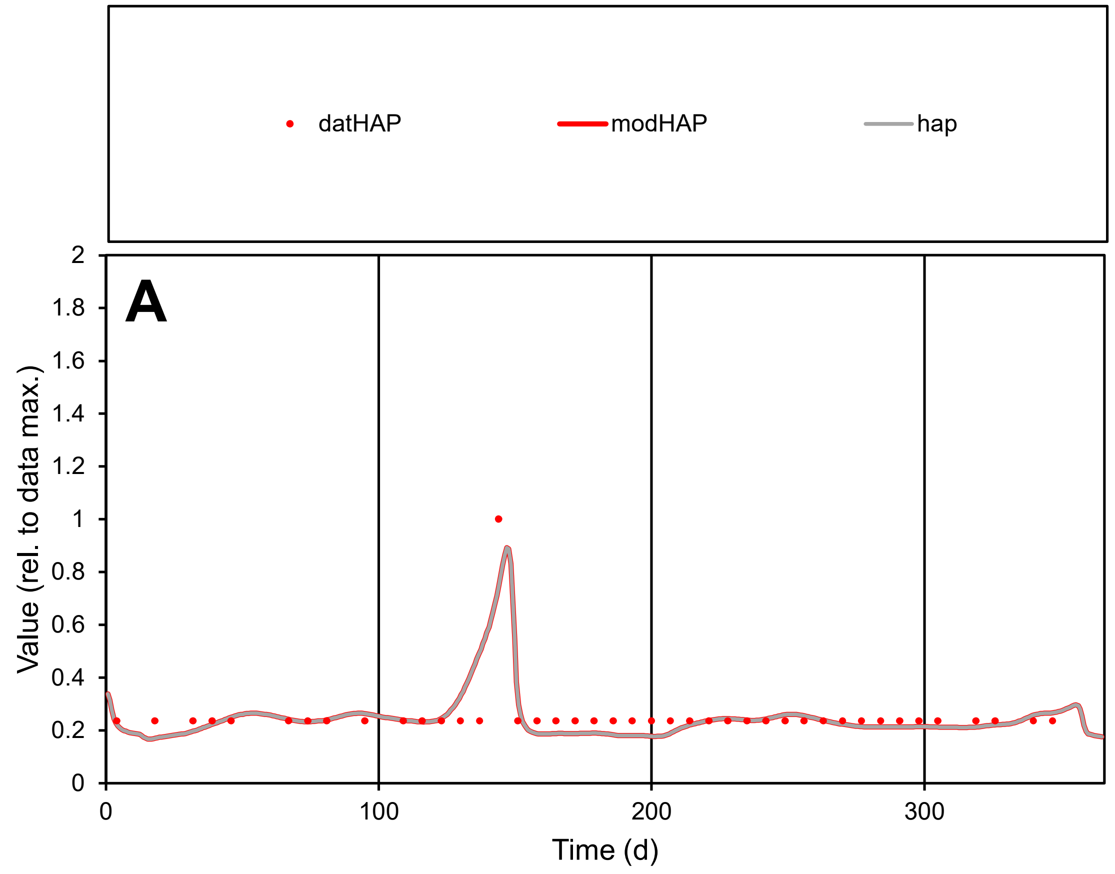


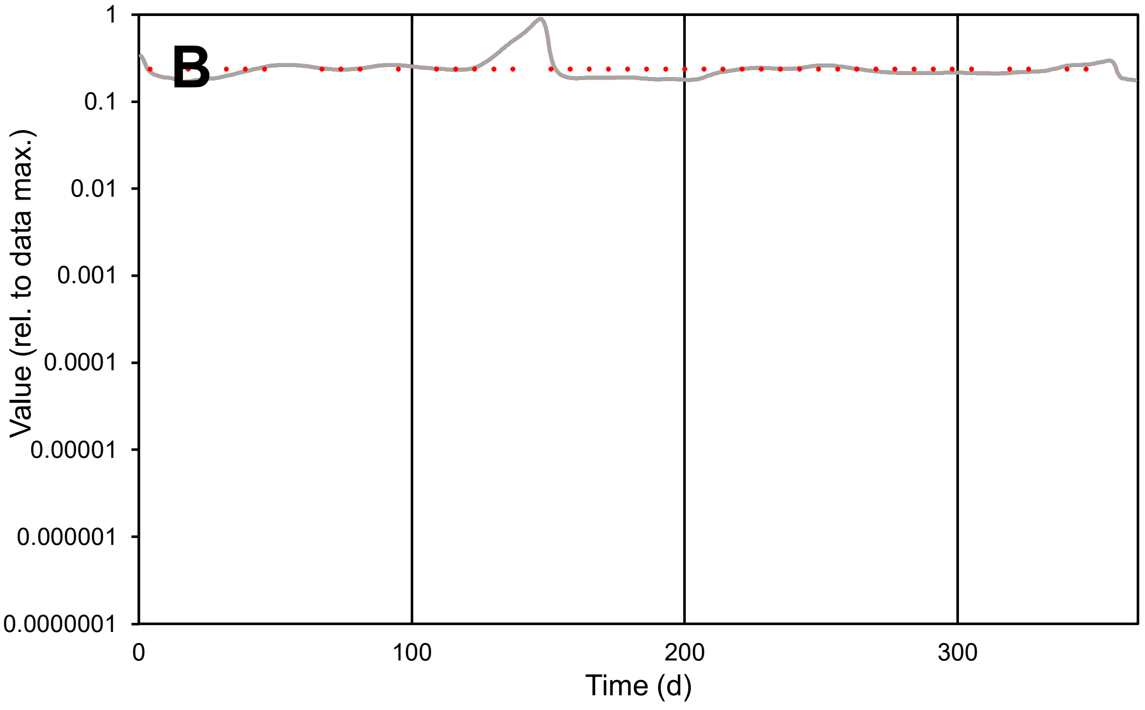


**C1**

Variables
HAP Provide jtd or Dst
CalibOnly?
2 1 to reproduce of
oNormalize?
2 1=no, 2=yes

ofxn calc*
0.23314
ofxn mod
0.26504

sRun
RunA1
zout
1

Ave
Data
1.2E-05
Model
1.2E-05

data/vMax
4.8E-05

ofxng
334.07

Min d/m
1.35944

ave/min d
1.07692
ave/min m
1.42344

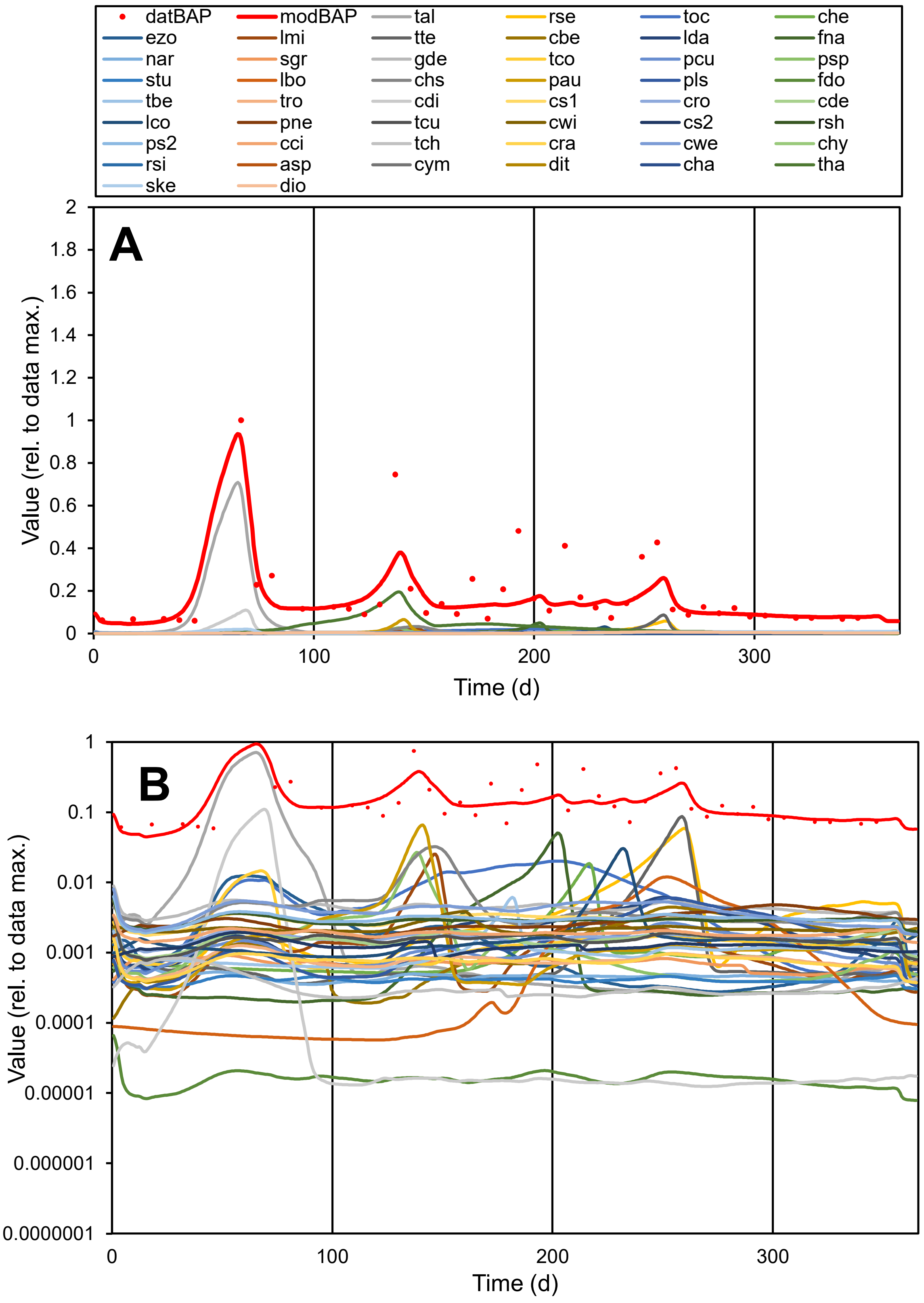


**C1**

Variables

BAP Provide jtd or Dst

CalibOnly?

2 1 to reproduce of

oNormalize?

2 1=no, 2=yes

ofxn calc*

0.59405

ofxn mod

0.61949

sRun

RunA1

zout

1

Ave

Data

0.0011

Model

0.00094

data/vMax

0.00605

ofxng

334.07

Min d/m

1.2758

ave/min d

3.07346

ave/min m

3.34659

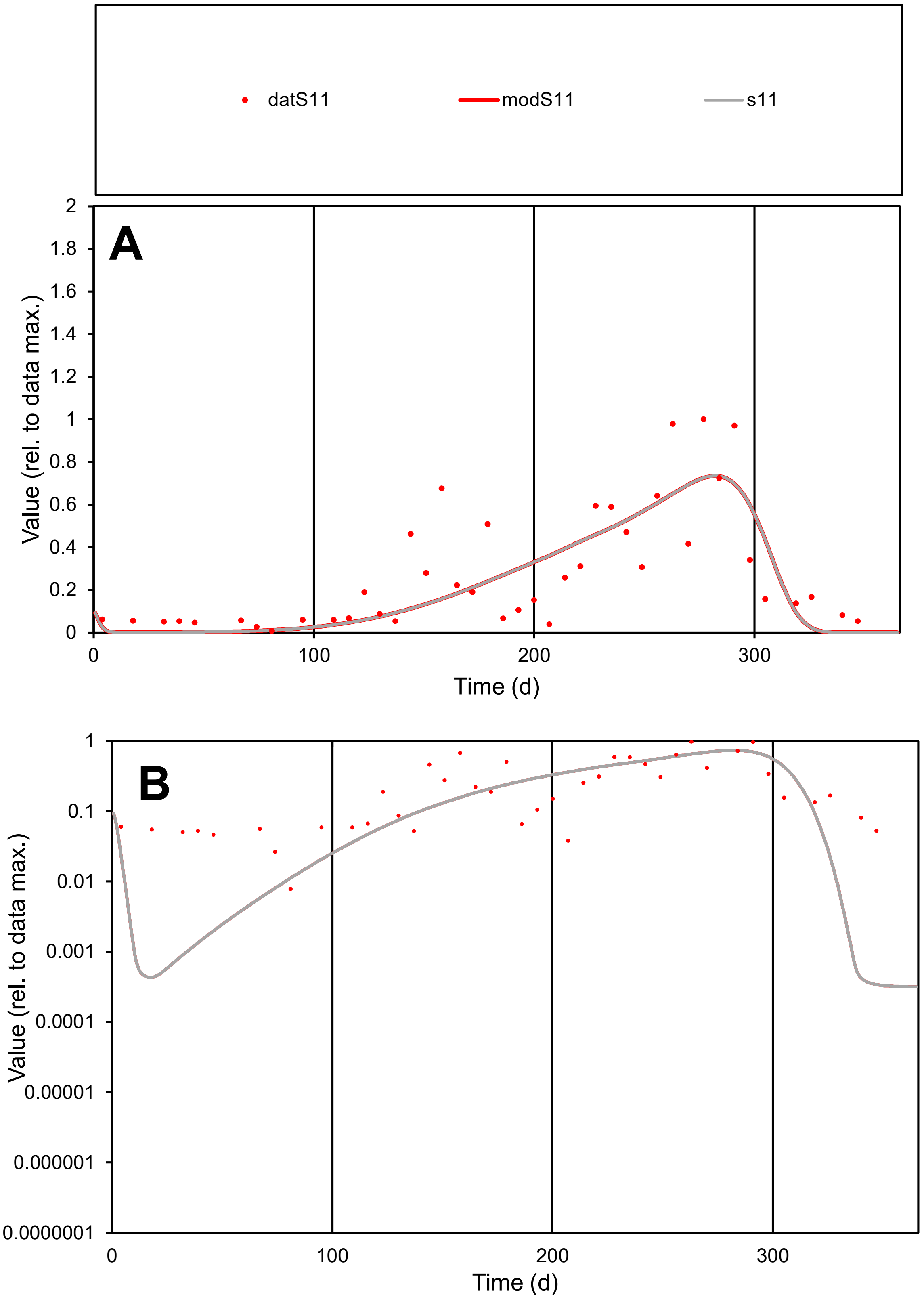


**C1**

Variables
S11 Provide jtd or Dst
CalibOnly?
2 1 to reproduce of
oNormalize?
2 1=no, 2=yes

ofxn calc*
0.63398
ofxn mod
0.65809

sRun
RunA1
zout
1

Ave
Data
0.10213
Model
0.09419

data/vMax
0.3649

ofxng
334.07

Min d/m
23.4069

ave/min d
35.6486
ave/min m
769.521

327_Calib_S_A

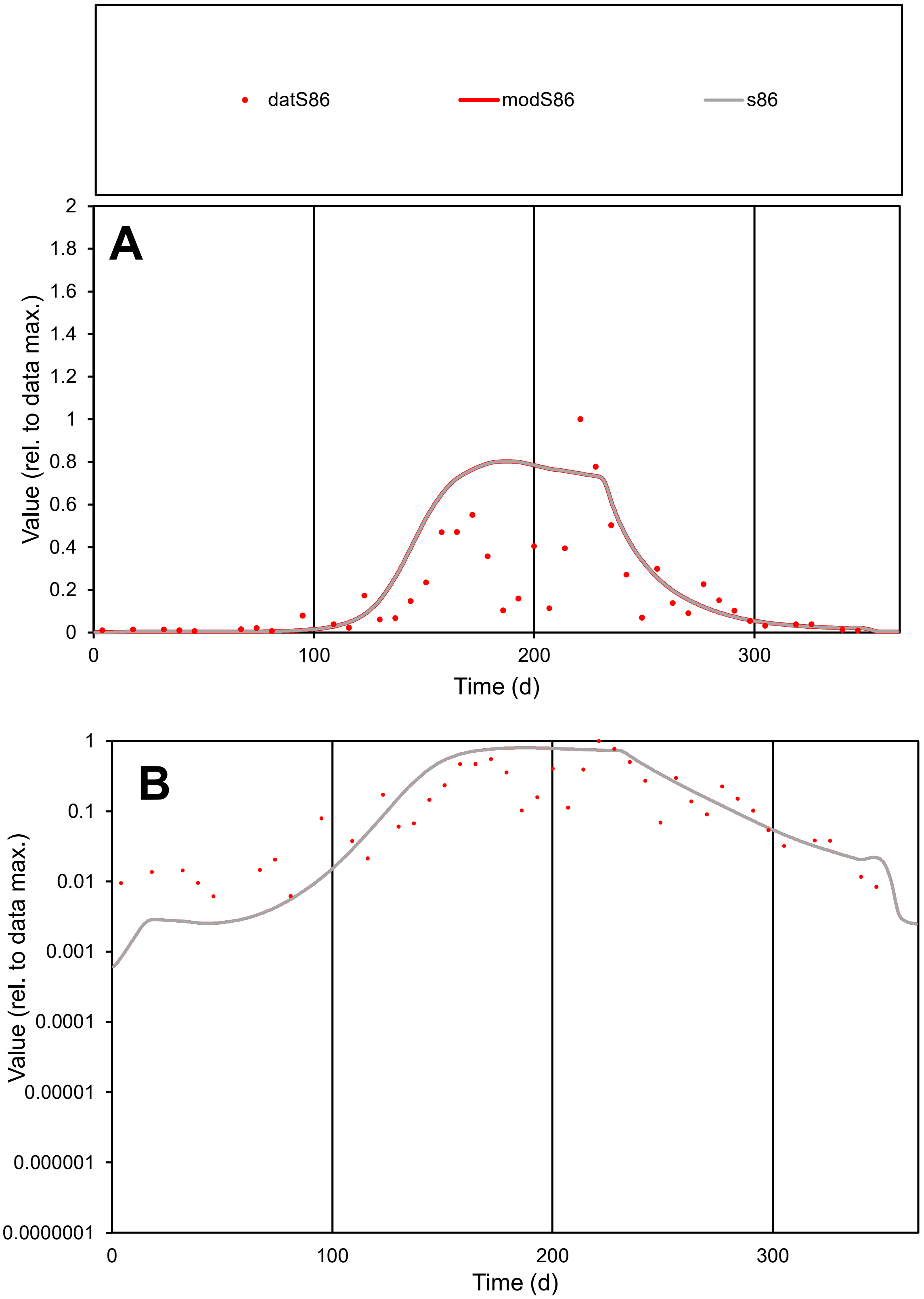


C1

Variables

S86 Provide jtd or Dst

CalibOnly?

2 1 to reproduce of

oNormalize?

2 1=no, 2=yes

ofxn calc*

1.27719

ofxn mod

1.30232

sRun

RunA1

zout

1

Ave

Data

0.07546

Model

0.11945

data/vMax

0.4092

ofxng

334.07

Min d/m

6.71558

ave/min d

29.9801

ave/min m

318.71

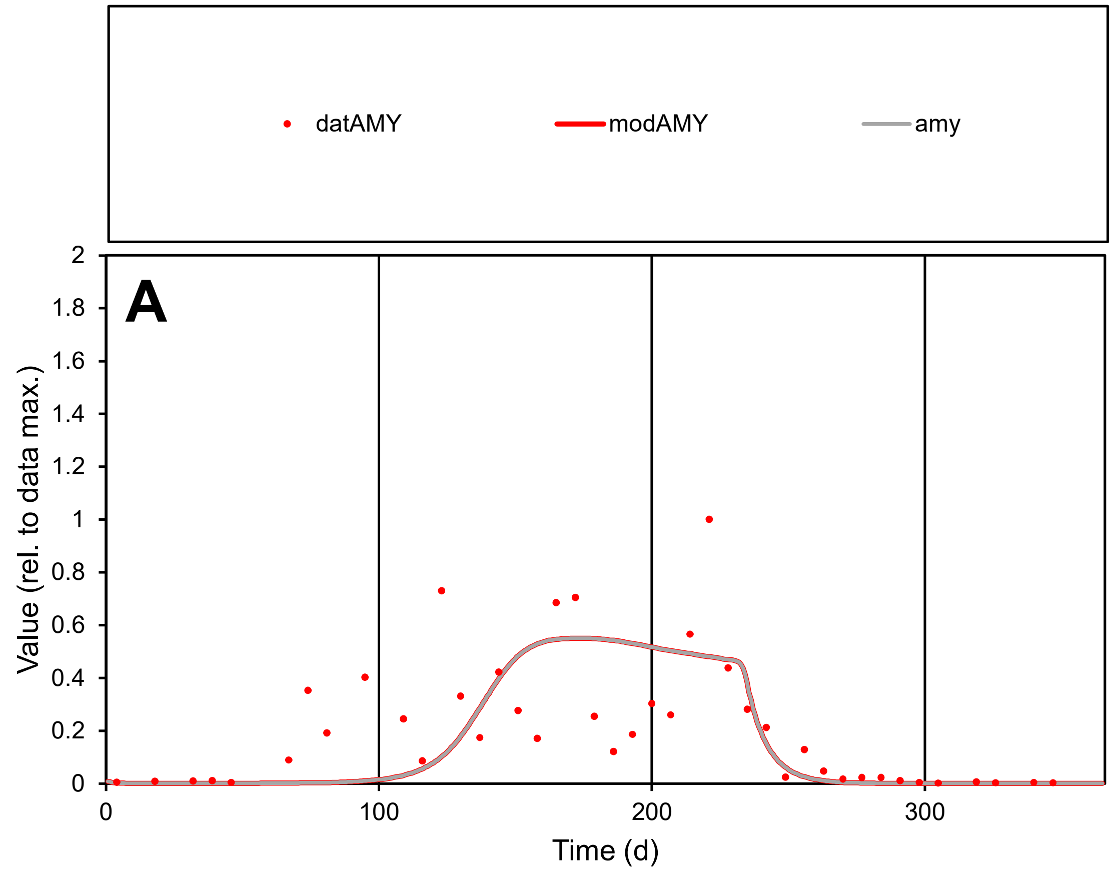


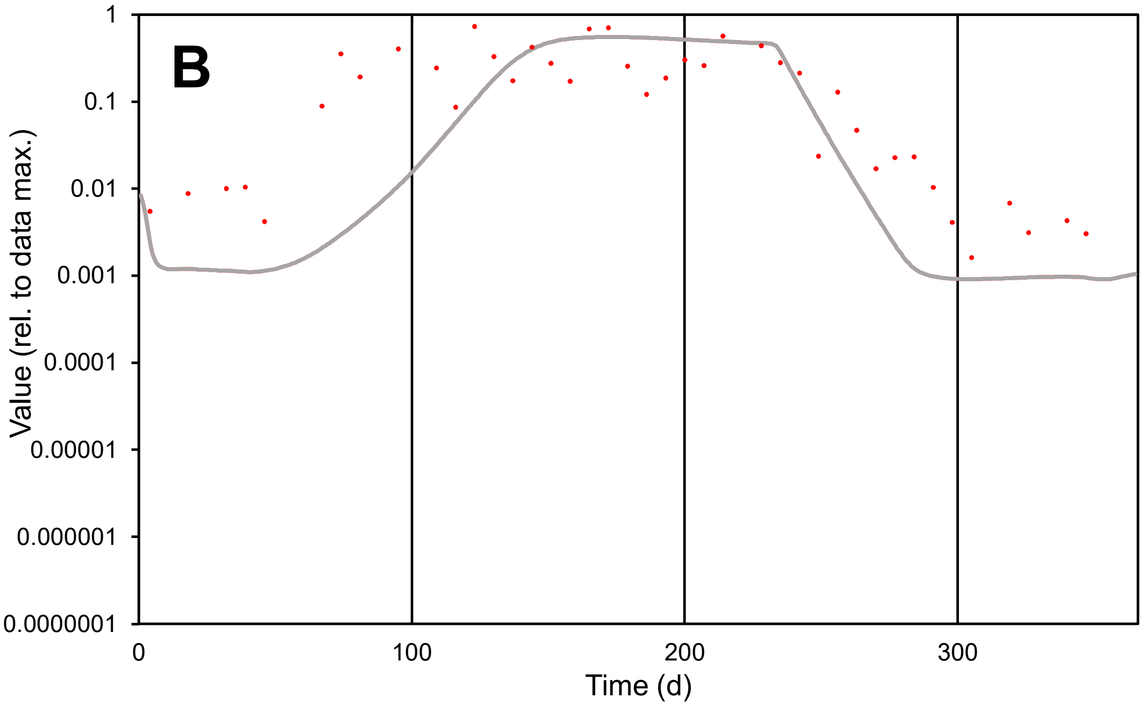


**C1**

Variables
AMY Provide jtd or Dst
CalibOnly?
2 1 to reproduce of
oNormalize?
2 1=no, 2=yes

ofxn calc*
0.97853
ofxn mod
0.99134

sRun
RunA1
zout
1

Ave
Data
0.06911
Model
0.06197

data/vMax
0.3291

ofxng
334.07

Min d/m
1.75895

ave/min d
130.266
ave/min m
205.483

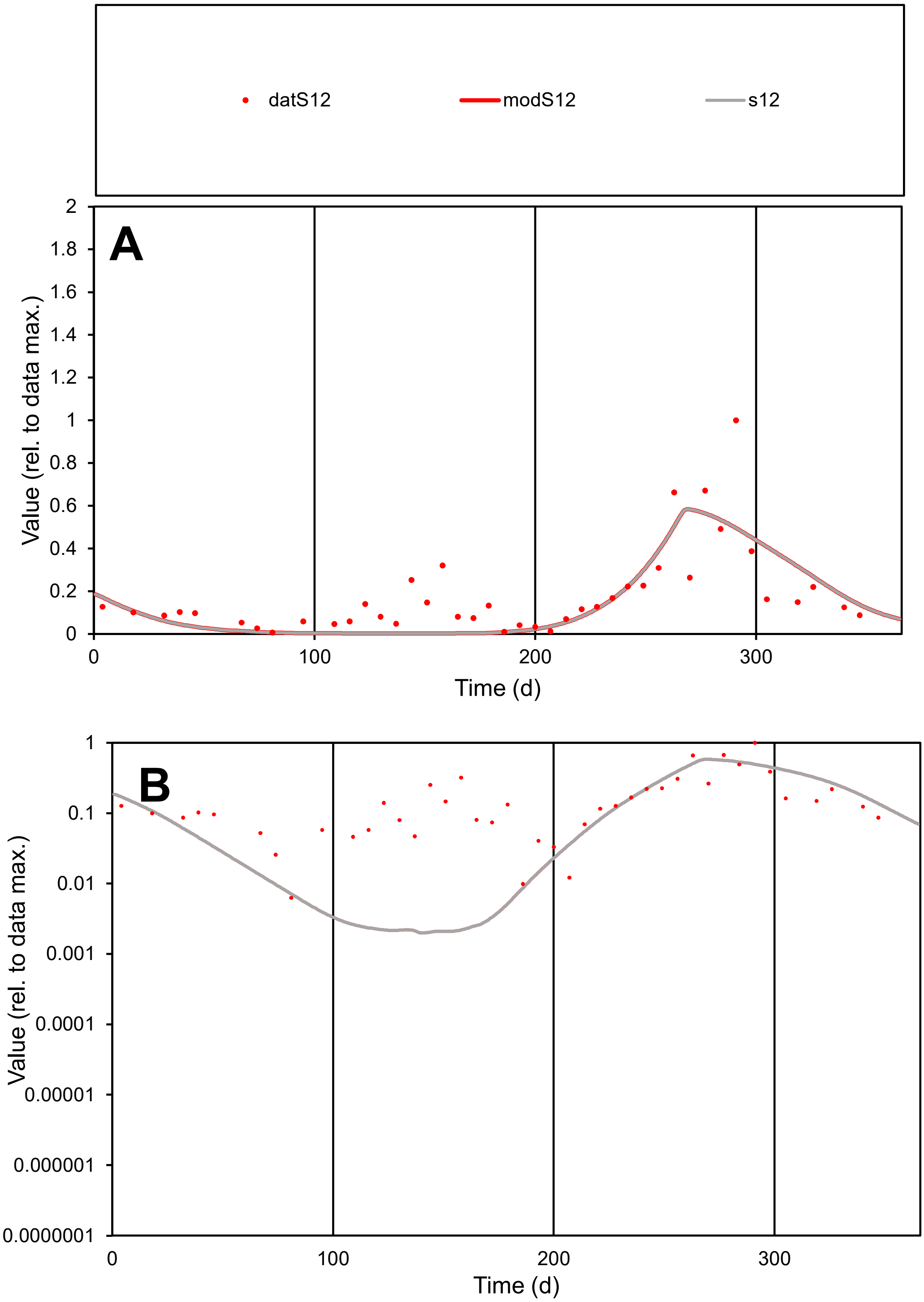


**C1**

Variables
S12 Provide jtd or Dst
CalibOnly?
2 1 to reproduce of
oNormalize?
2 1=no, 2=yes

ofxn calc*
0.74519
ofxn mod
0.76345

sRun
RunA1
zout
1

Ave
Data
0.0284
Model
0.0238

data/vMax
0.1575

ofxng
334.07

Min d/m
3.03606

ave/min d
28.8299
ave/min m
73.3417

327_Calib_S_A

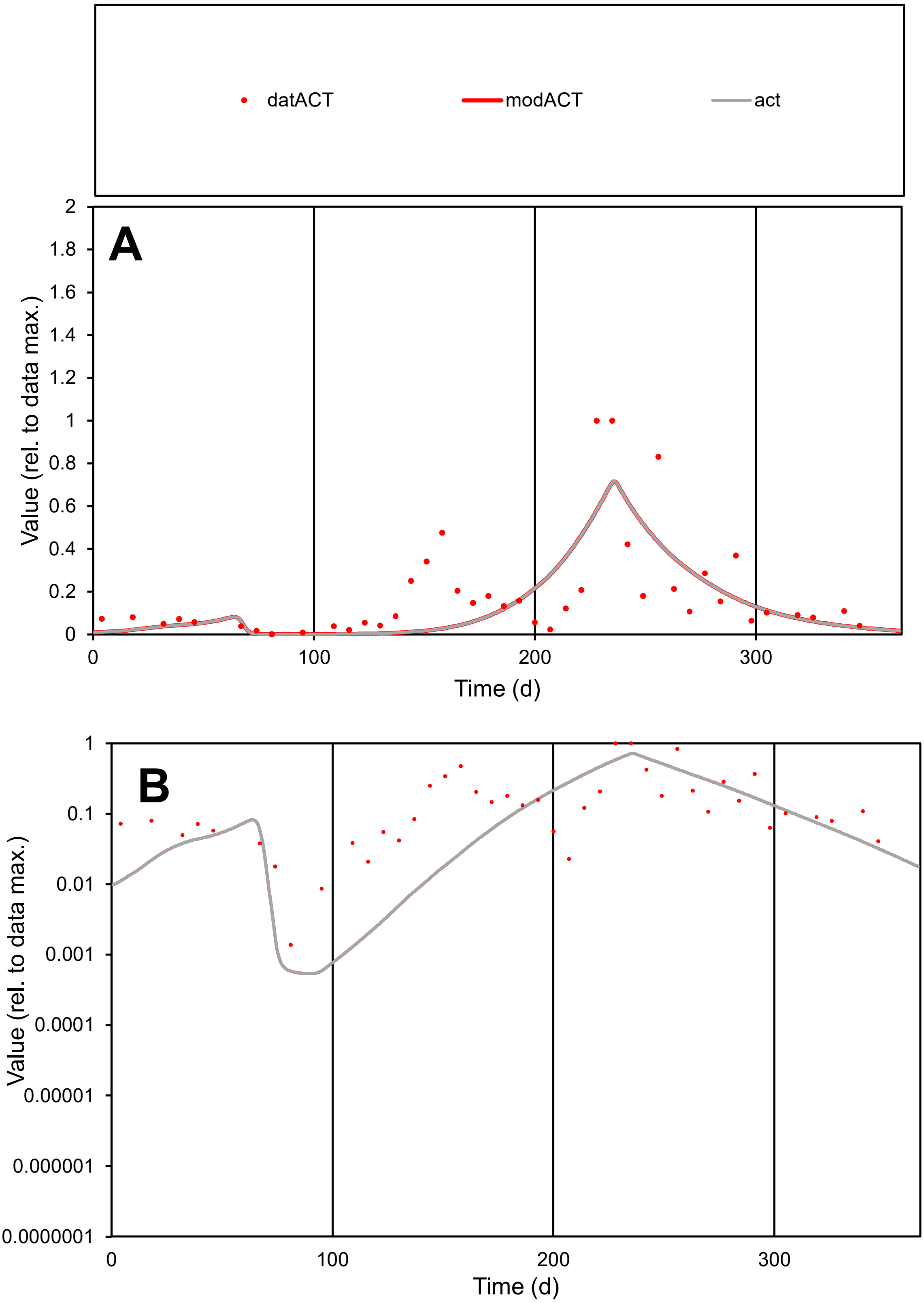


**C1**

Variables
ACT Provide jtd or Dst
CalibOnly?
2 1 to reproduce of
oNormalize?
2 1=no, 2=yes

ofxn calc*
0.92213
ofxn mod
0.93914

sRun
RunA1
zout
1

Ave
Data
0.02612
Model
0.02182

data/vMax
0.1376

ofxng
334.07

Min d/m
2.42123

ave/min d
137.599
ave/min m
278.373

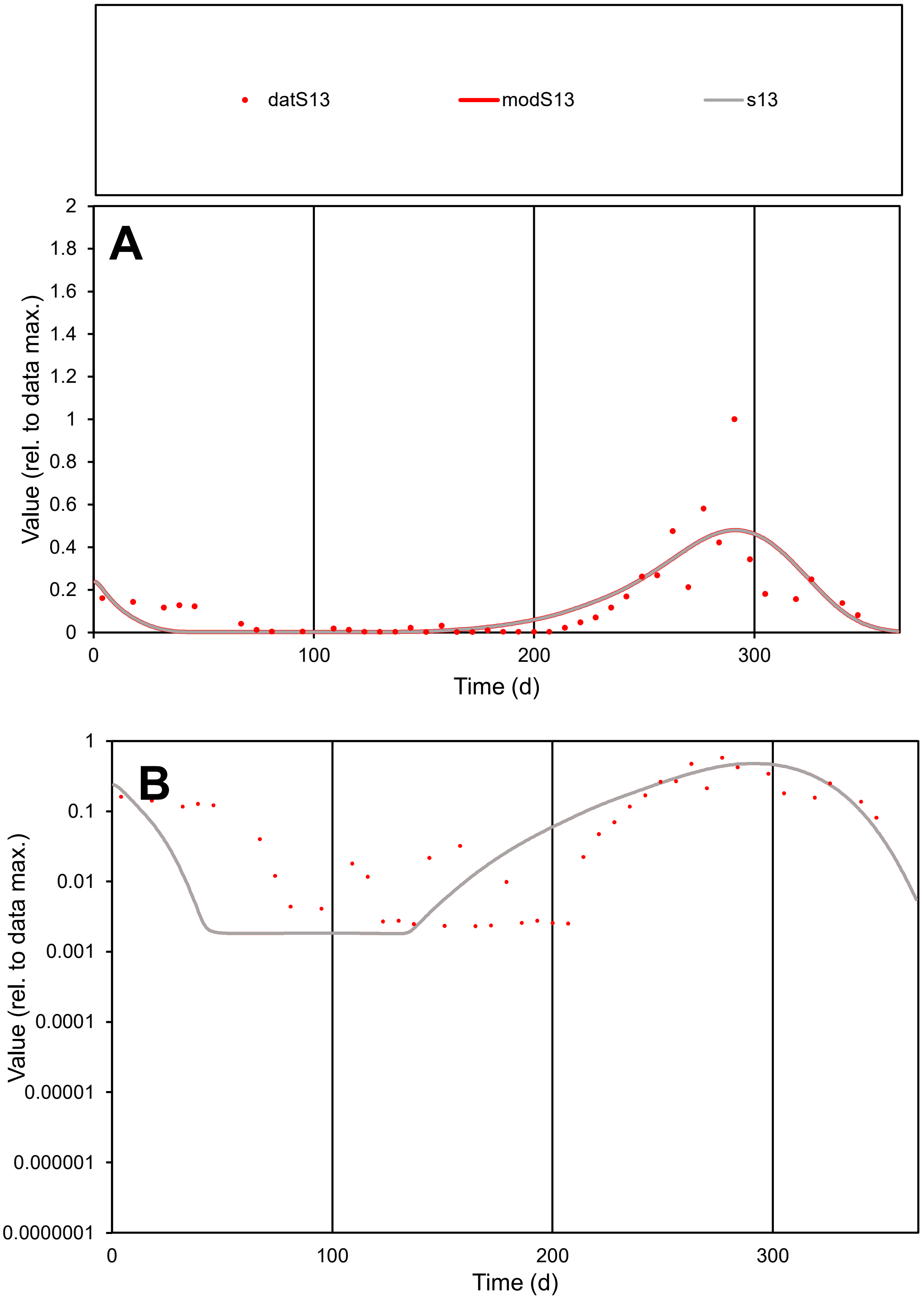


**C1**

Variables
S13 Provide jtd or Dst
CalibOnly?
2 1 to reproduce of
oNormalize?
2 1=no, 2=yes

ofxn calc*
0.83512
ofxn mod
0.85819

sRun
RunA1
zout
1

Ave
Data
0.01604
Model
0.01602

data/vMax
0.1194

ofxng
334.07

Min d/m
1.27852

ave/min d
57.9536
ave/min m
73.9938

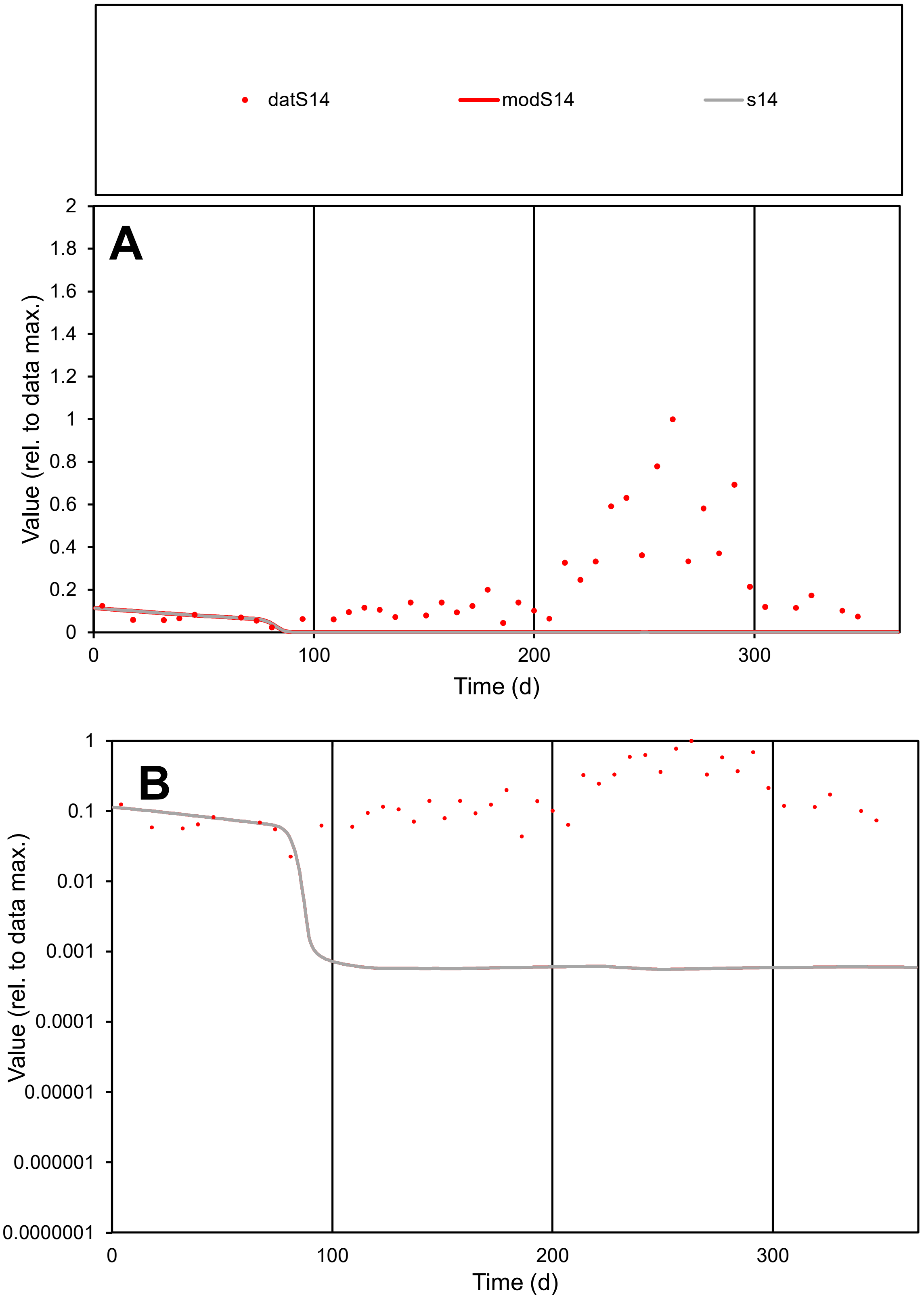


**C1**

Variables
S14 Provide jtd or Dst
CalibOnly?
2 1 to reproduce of
oNormalize?
2 1=no, 2=yes

ofxn calc*
1.43072
ofxn mod
1.43394

sRun
RunA1
zout
1

Ave
Data
0.02486
Model
0.00173

data/vMax
0.1134

ofxng
334.07

Min d/m
40.3279

ave/min d
9.71678
ave/min m
27.284

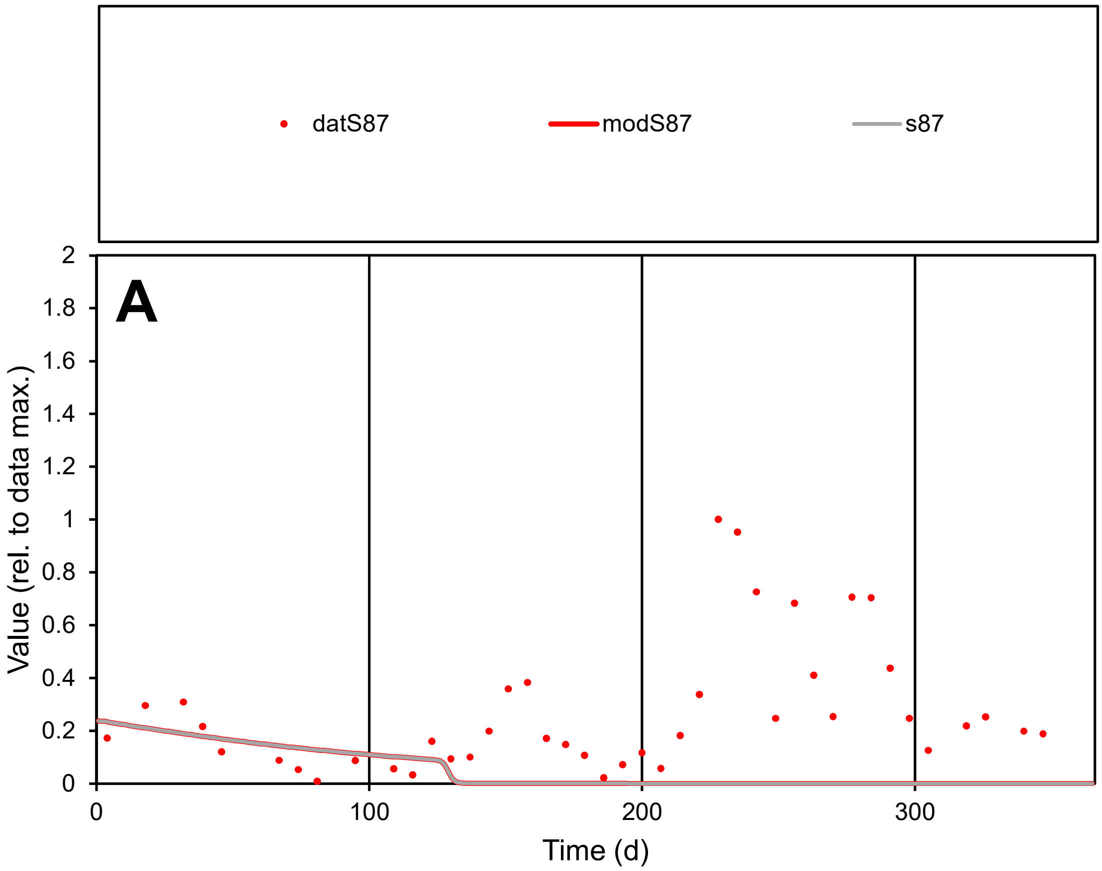


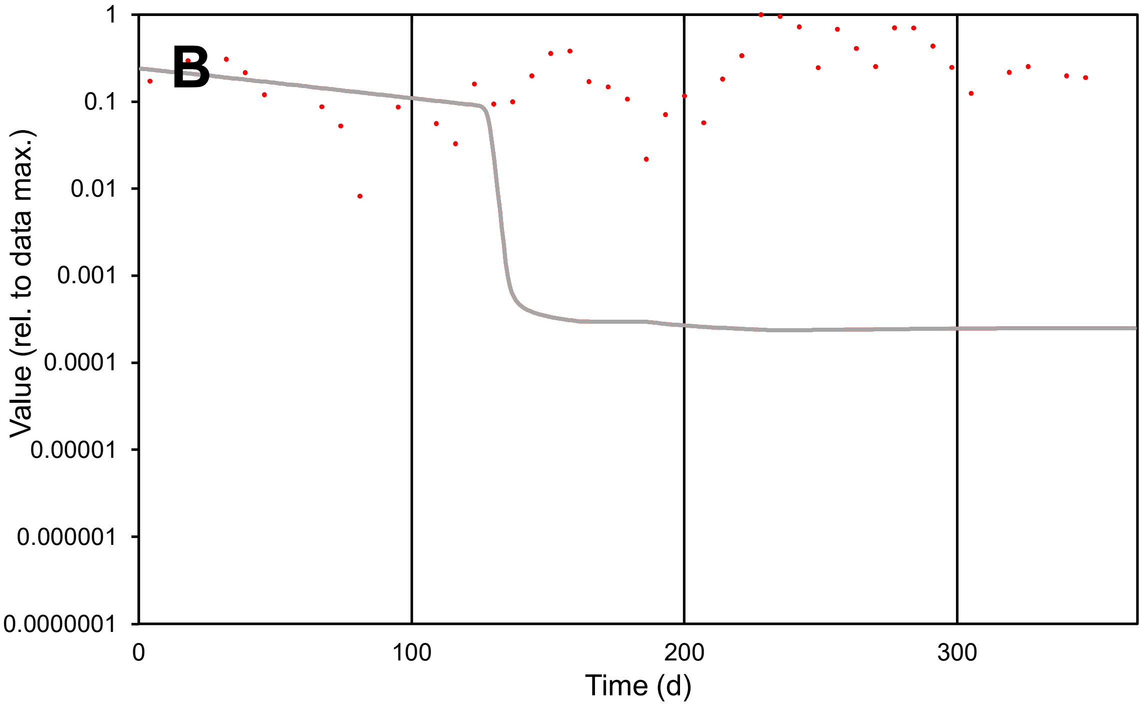


**C1**

Variables

S87 Provide jtd or Dst

CalibOnly?

2 1 to reproduce of

oNormalize?

2 1=no, 2=yes

ofxn calc*

1.31619

ofxn mod

1.3192

sRun

RunA1

zout

1

Ave

Data

0.02168

Model

0.00355

data/vMax

0.08068

ofxng

334.07

Min d/m

34.4903

ave/min d

32.8657

ave/min m

185.605

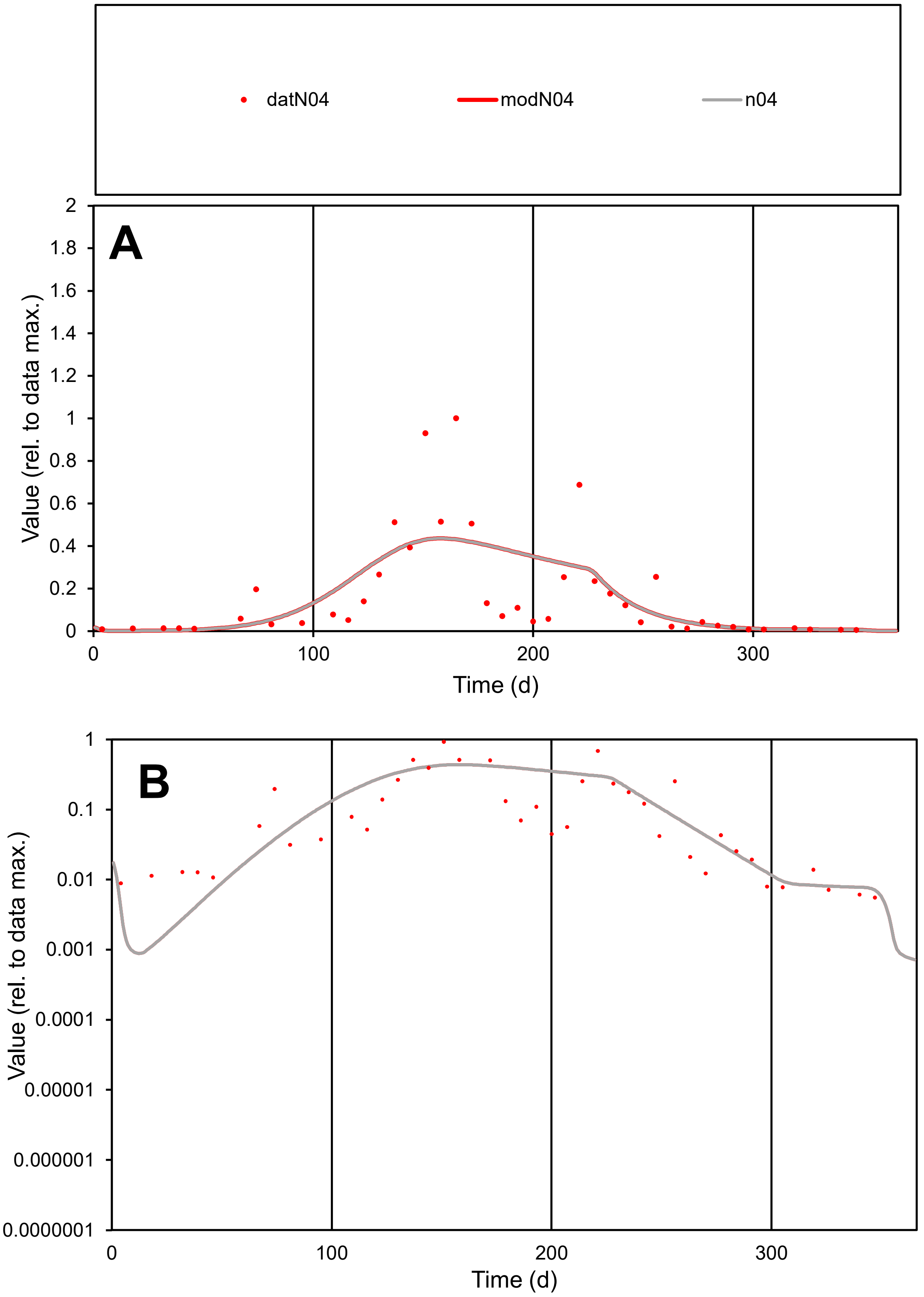


**C1**

Variables
N04 Provide jtd or Dst
CalibOnly?
2 1 to reproduce of
oNormalize?
2 1=no, 2=yes

ofxn calc*
1.041
ofxn mod
1.053

sRun
RunA1
zout
1

Ave
Data
0.02944
Model
0.03

data/vMax
0.1737

ofxng
334.07

Min d/m
4.54541

ave/min d
30.4825
ave/min m
141.185

327_Calib_S_A

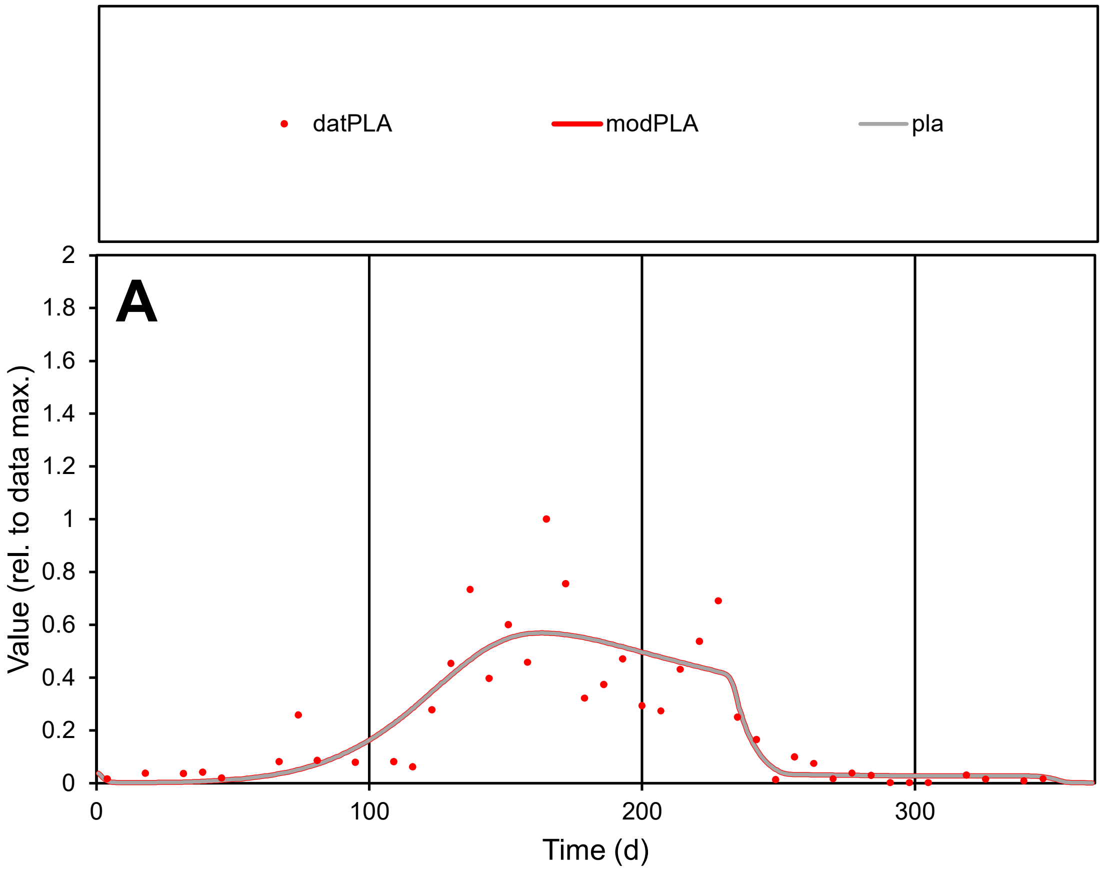


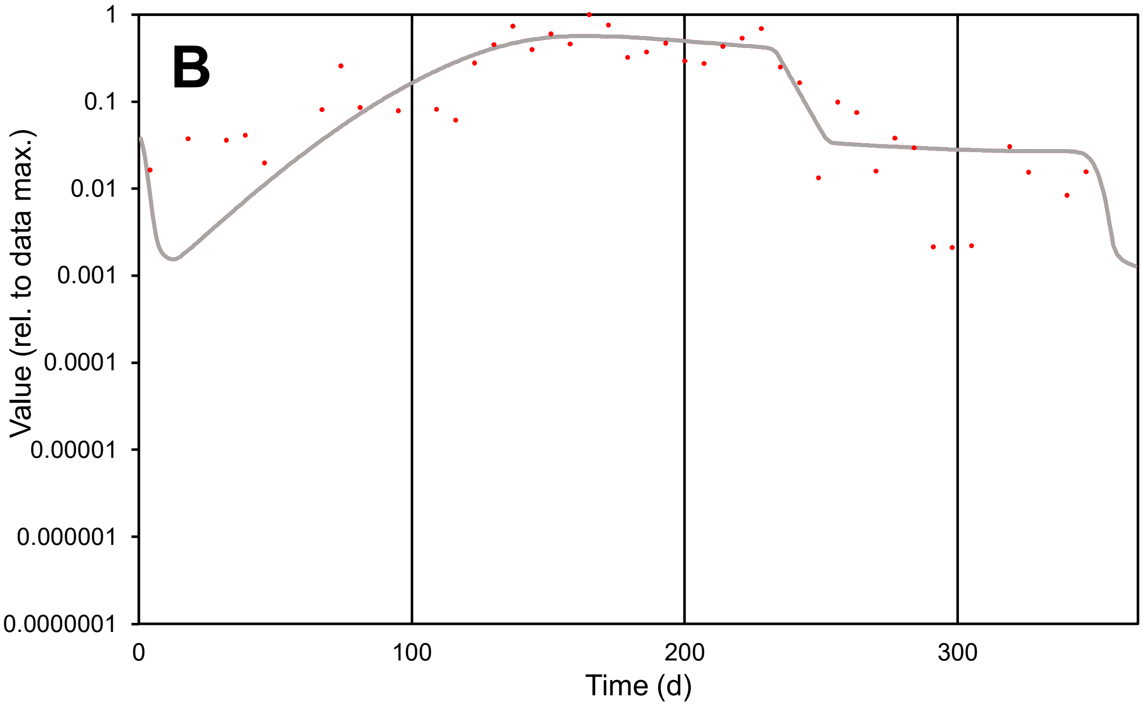


**C1**

Variables
PLA Provide jtd or Dst
CalibOnly?
2 1 to reproduce of
oNormalize?
2 1=no, 2=yes

ofxn calc*
0.56434
ofxn mod
0.58506

sRun
RunA1
zout
1

Ave
Data
0.01813
Model
0.01794

data/vMax
0.07909

ofxng
334.07

Min d/m
0.98128

ave/min d
108.095
ave/min m
104.965

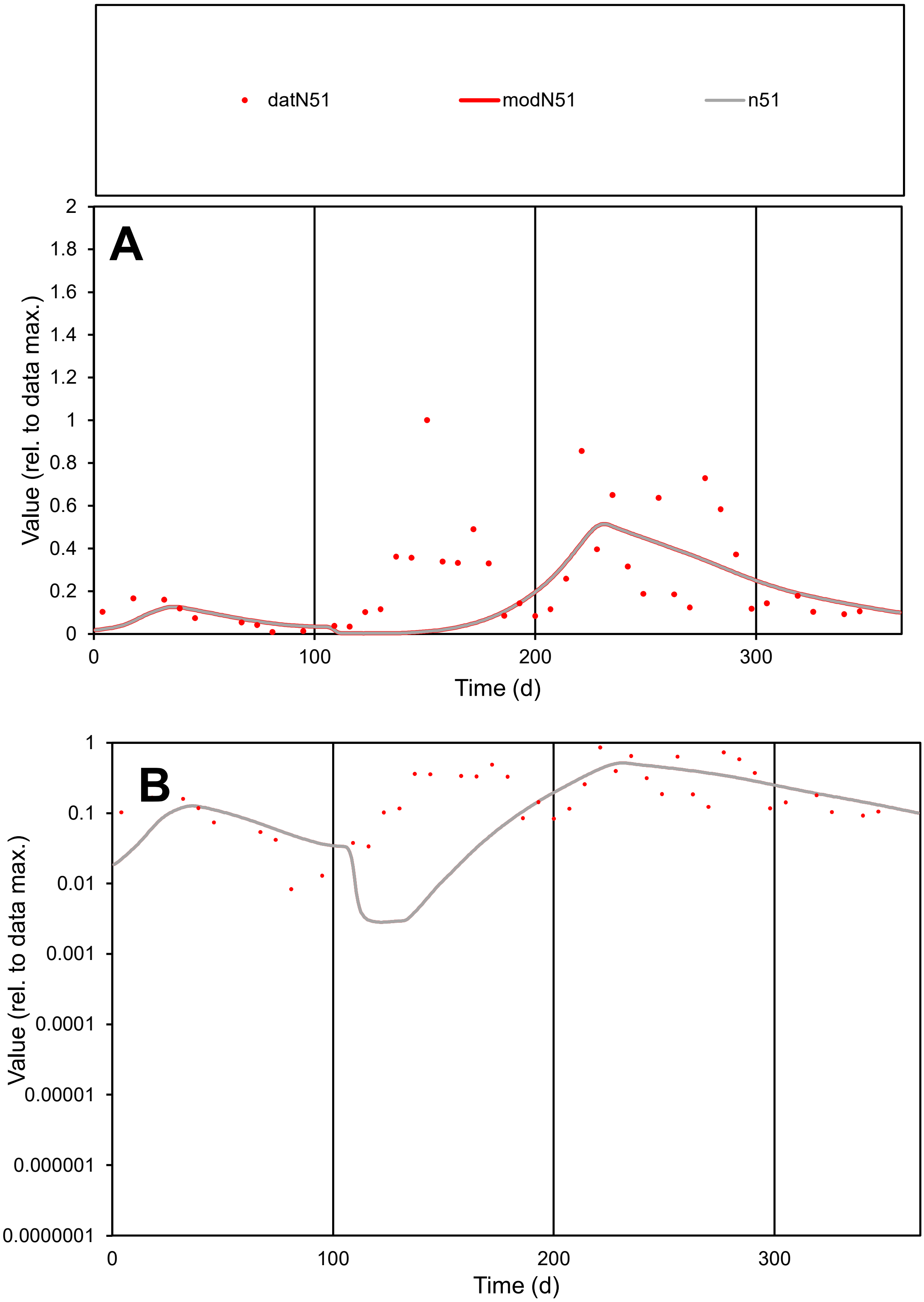


C1

Variables
N51 Provide jtd or Dst
CalibOnly?
2 1 to reproduce of
oNormalize?
2 1=no, 2=yes

ofxn calc*
0.95172
ofxn mod
0.95974

sRun
RunA1
zout
1

Ave
Data
0.01438
Model
0.01018

data/vMax
0.05647

ofxng
334.07

Min d/m
2.9247

ave/min d
30.5891
ave/min m
63.332

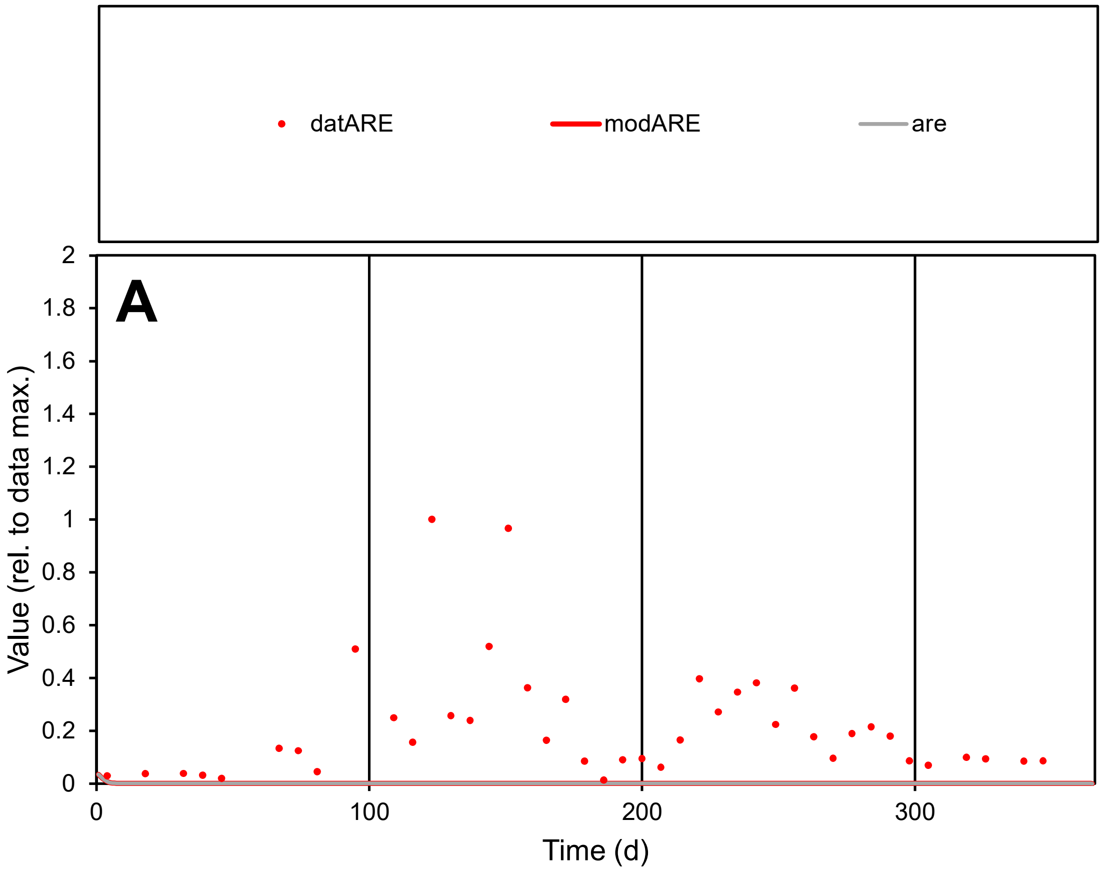


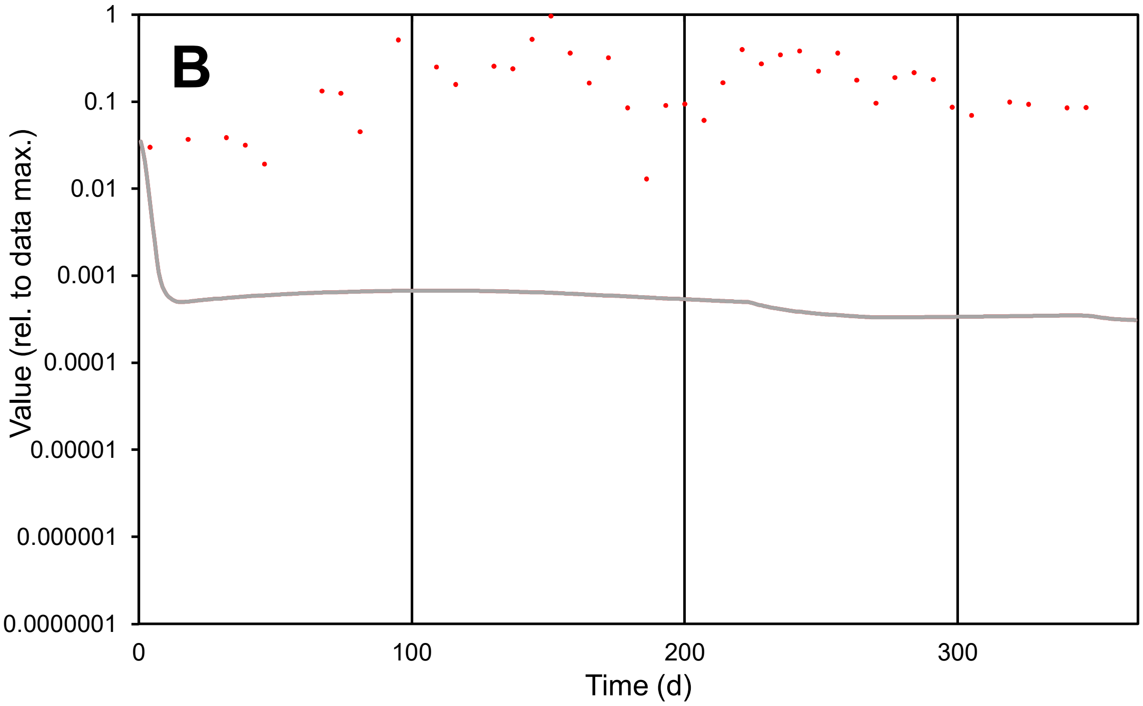


**C1**

Variables
ARE Provide jtd or Dst
CalibOnly?
2 1 to reproduce of
oNormalize?
2 1=no, 2=yes

ofxn calc*
1.41092
ofxn mod
1.41277

sRun
RunA1
zout
1

Ave
Data
0.02832
Model
0.00013

data/vMax
0.1312

ofxng
334.07

Min d/m
38.8953

ave/min d
16.6905
ave/min m
2.96176

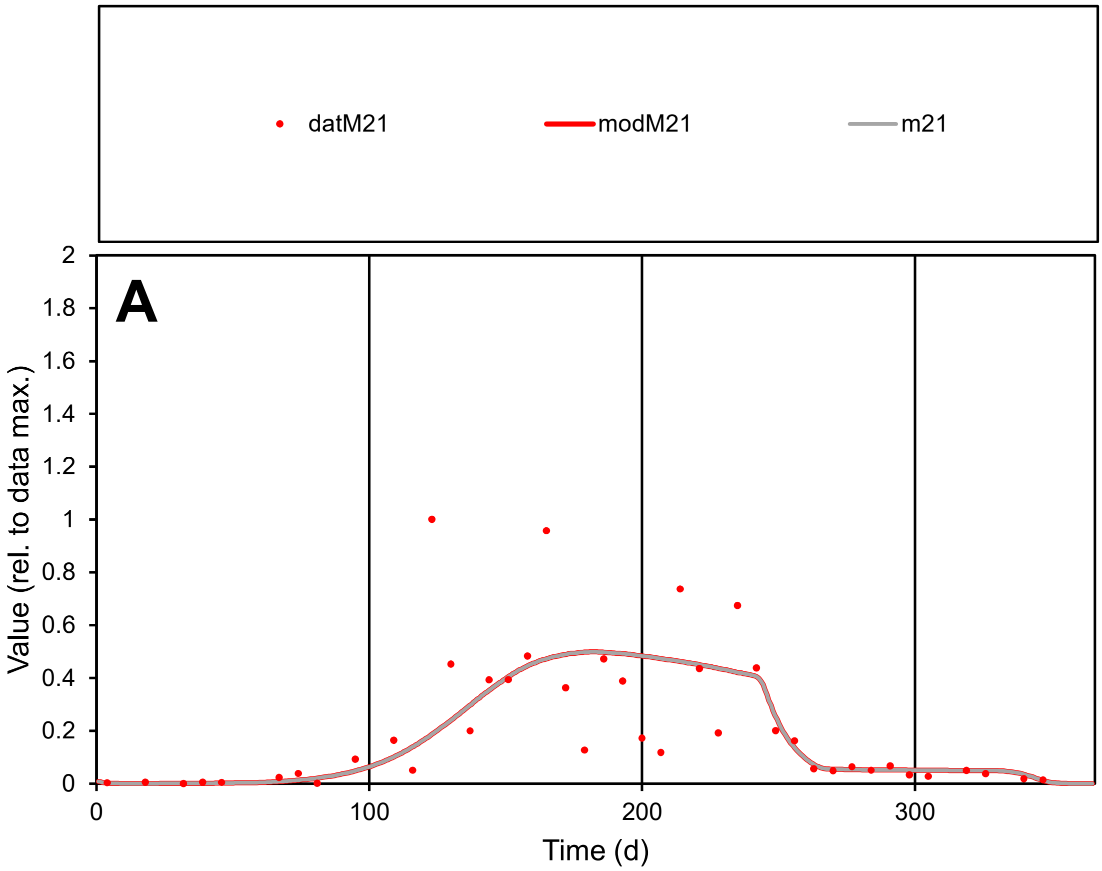


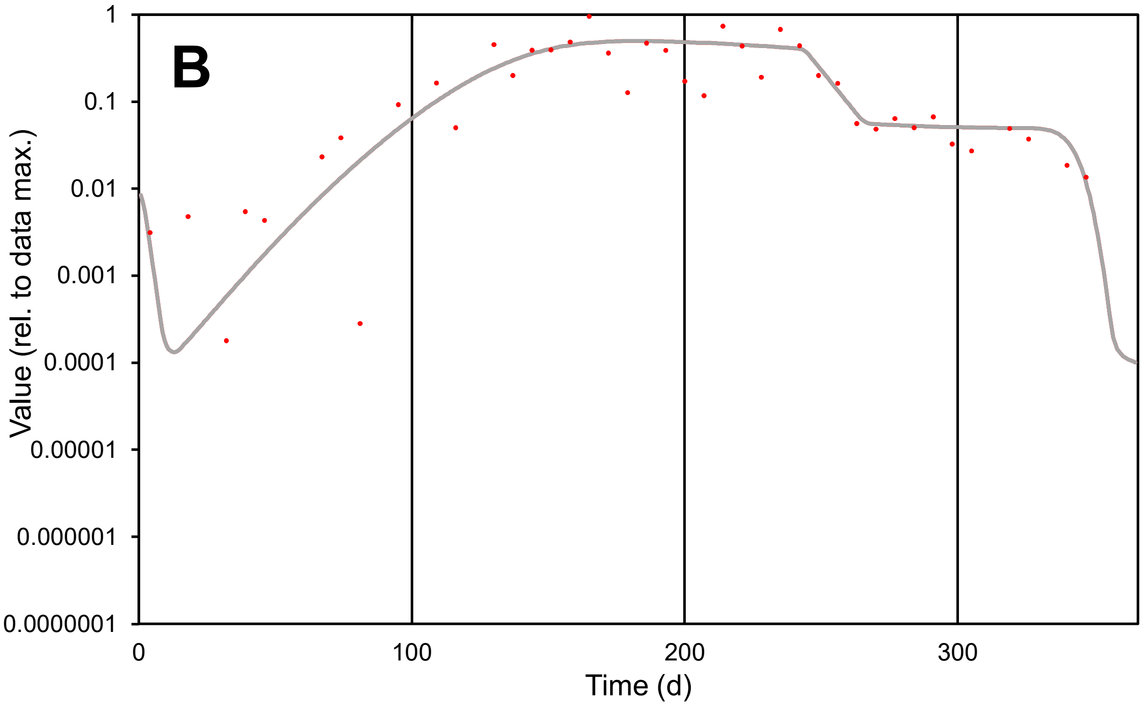


**C1**

Variables

M21 Provide jtd or Dst

CalibOnly?

2 1 to reproduce of

oNormalize?

2 1=no, 2=yes

ofxn calc*

0.87285

ofxn mod

0.887

sRun

RunA1

zout

1

Ave

Data

0.02906

Model

0.02769

data/vMax

0.1326

ofxng

334.07

Min d/m

0.86812

ave/min d

1219.5

ave/min m

1008.61

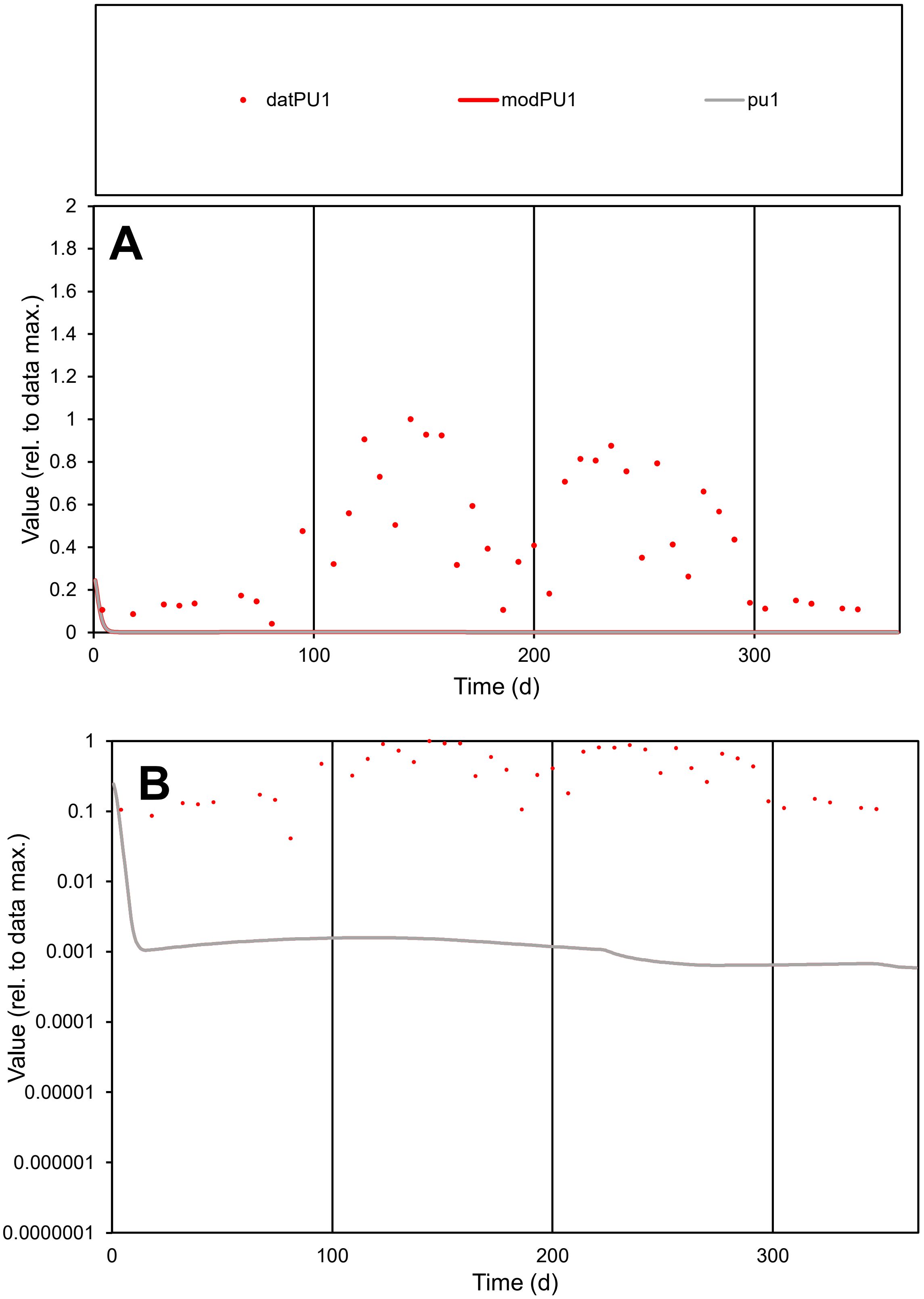


**C1**

Variables
PU1 Provide jtd or Dst
CalibOnly?
2 1 to reproduce of
oNormalize?
2 1=no, 2=yes

ofxn calc*
1.21479
ofxn mod
1.21665

sRun
RunA1
zout
1

Ave
Data
0.01692
Model
0.00011

data/vMax
0.0399

ofxng
334.07

Min d/m
64.1066

ave/min d
10.344
ave/min m
4.12559

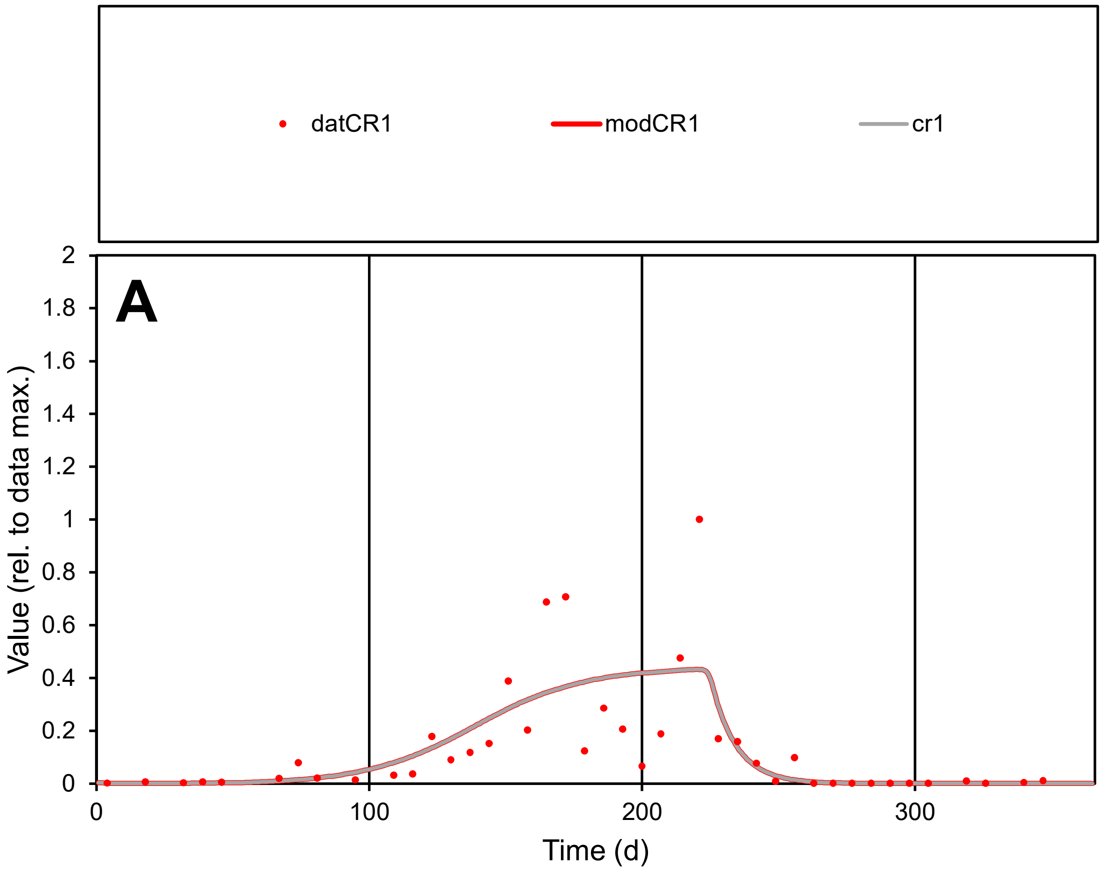


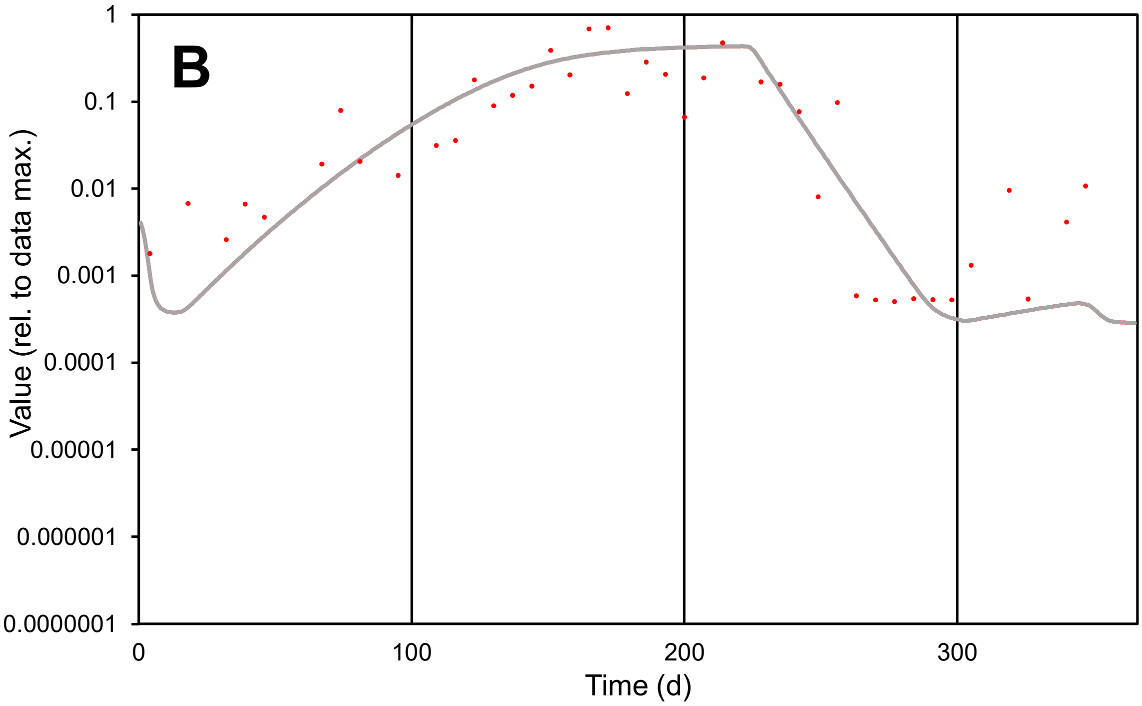


**C1**

Variables
CR1 Provide jtd or Dst
CalibOnly?
2 1 to reproduce of
oNormalize?
2 1=no, 2=yes

ofxn calc*
1.11687
ofxn mod
1.13355

sRun
RunA1
zout
1

Ave
Data
0.01799
Model
0.01858

data/vMax
0.1342

ofxng
334.07

Min d/m
1.61932

ave/min d
265.707
ave/min m
444.386

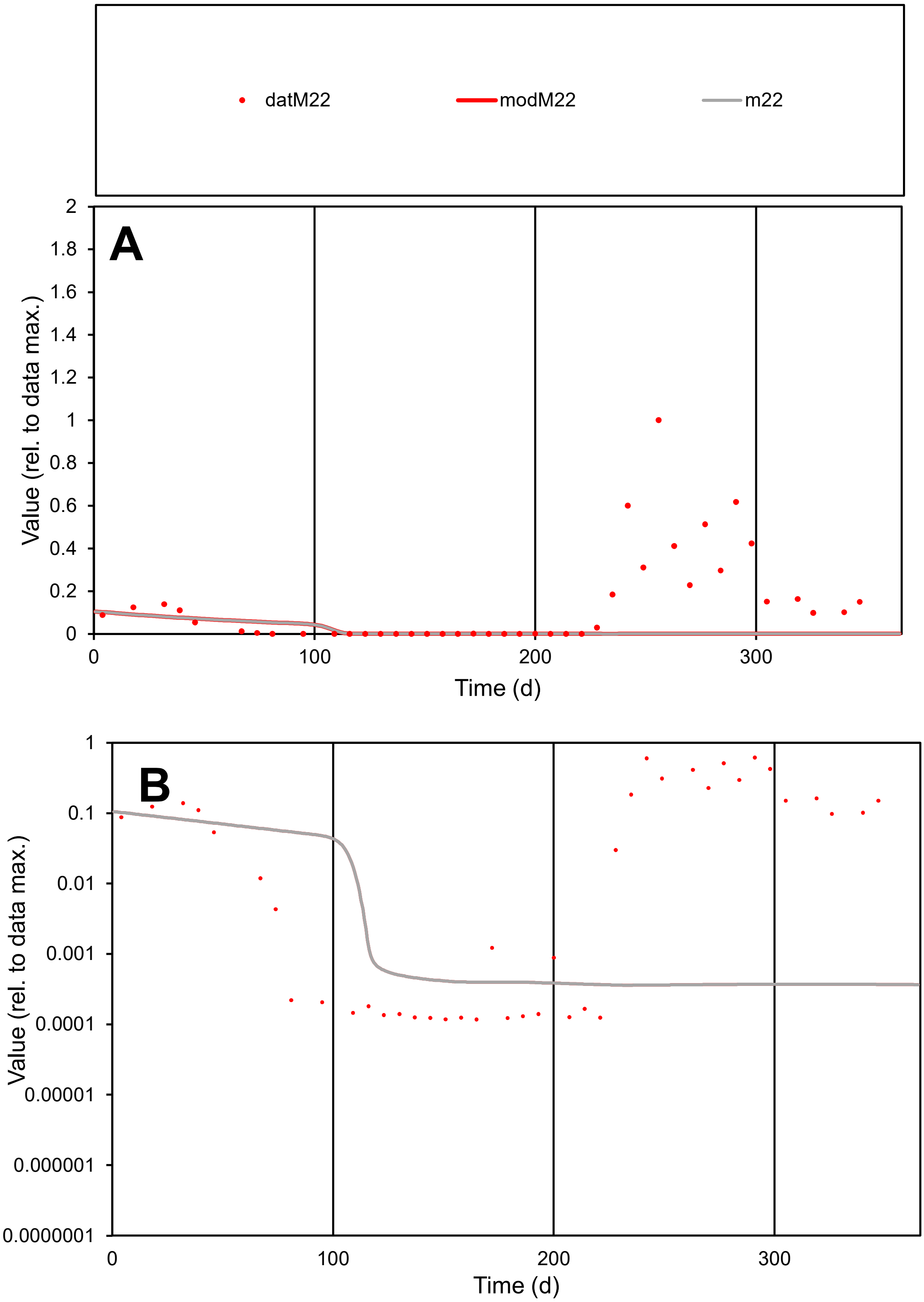


**C1**

Variables
M22 Provide jtd or Dst
CalibOnly?
2 1 to reproduce of
oNormalize?
2 1=no, 2=yes

ofxn calc*
1.84192
ofxn mod
1.84627

sRun
RunA1
zout
1

Ave
Data
0.02531
Model
0.00286

data/vMax
0.1829

ofxng
334.07

Min d/m
0.3232

ave/min d
1184.29
ave/min m
43.2599

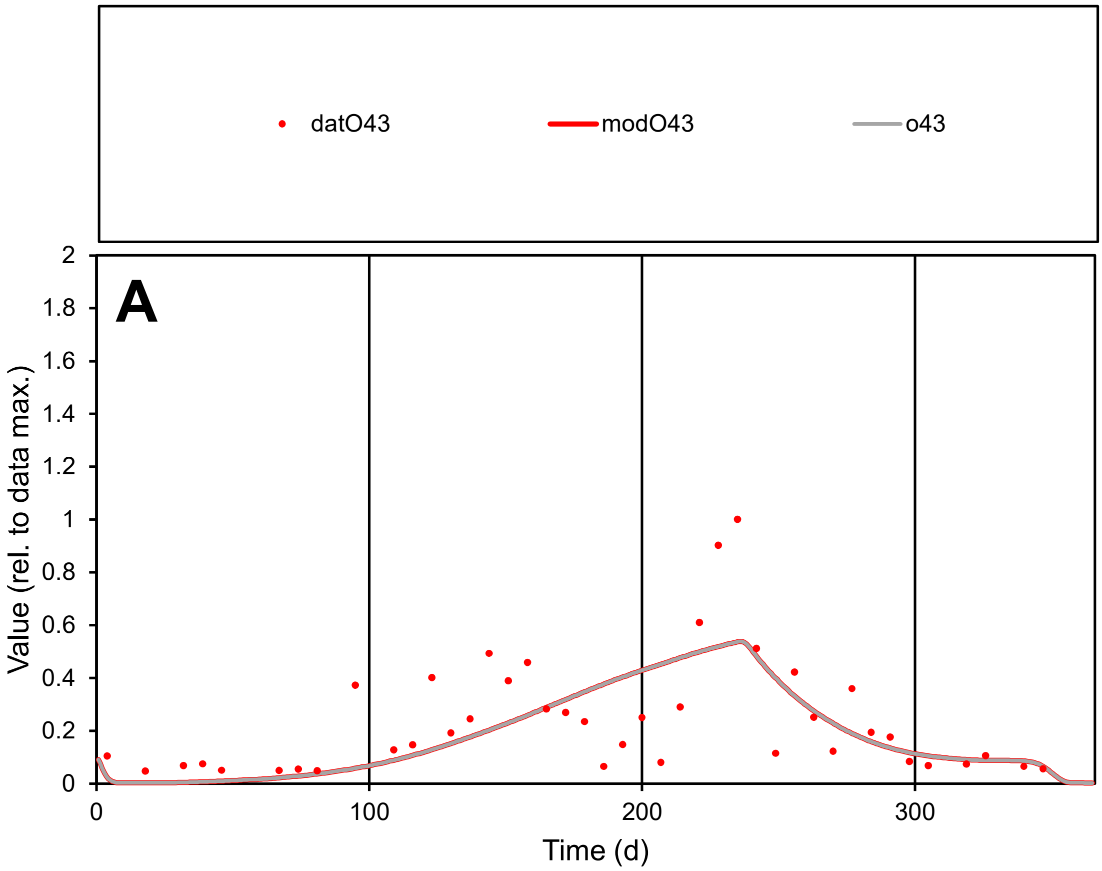


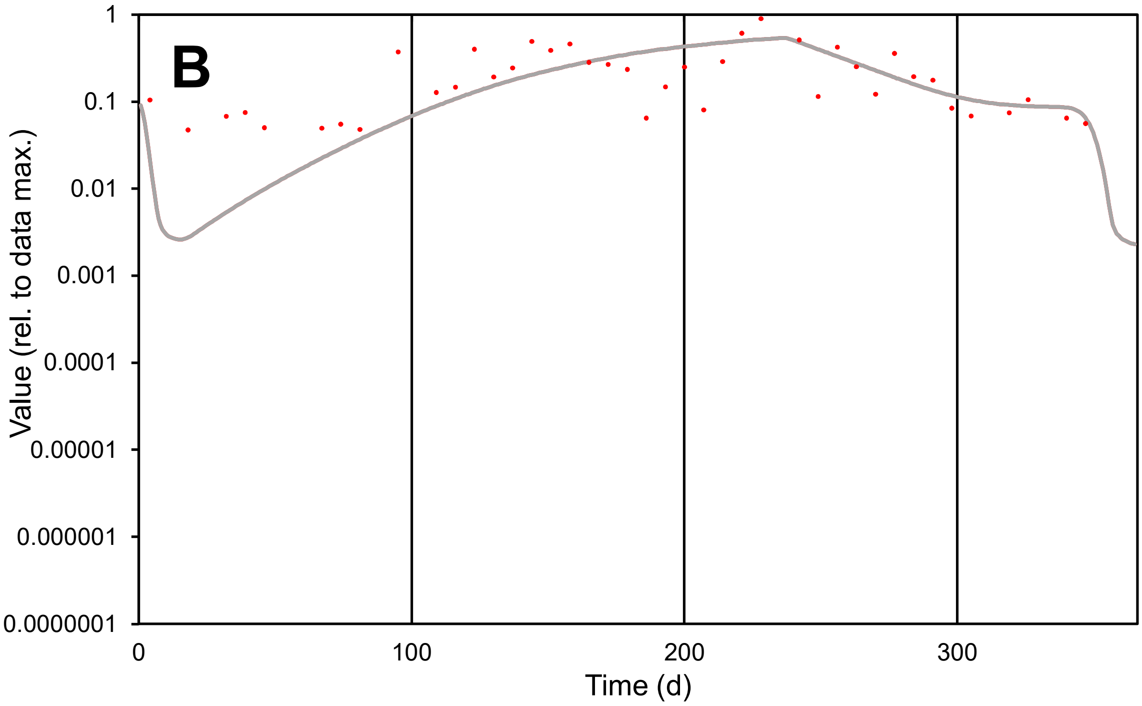


**C1**

Variables
O43 Provide jtd or Dst
CalibOnly?
2 1 to reproduce of
oNormalize?
2 1=no, 2=yes

ofxn calc*
0.71437
ofxn mod
0.72883

sRun
RunA1
zout
1

Ave
Data
0.01631
Model
0.01454

data/vMax
0.0681

ofxng
334.07

Min d/m
16.046

ave/min d
5.08741
ave/min m
72.7964

327_Calib_S_A

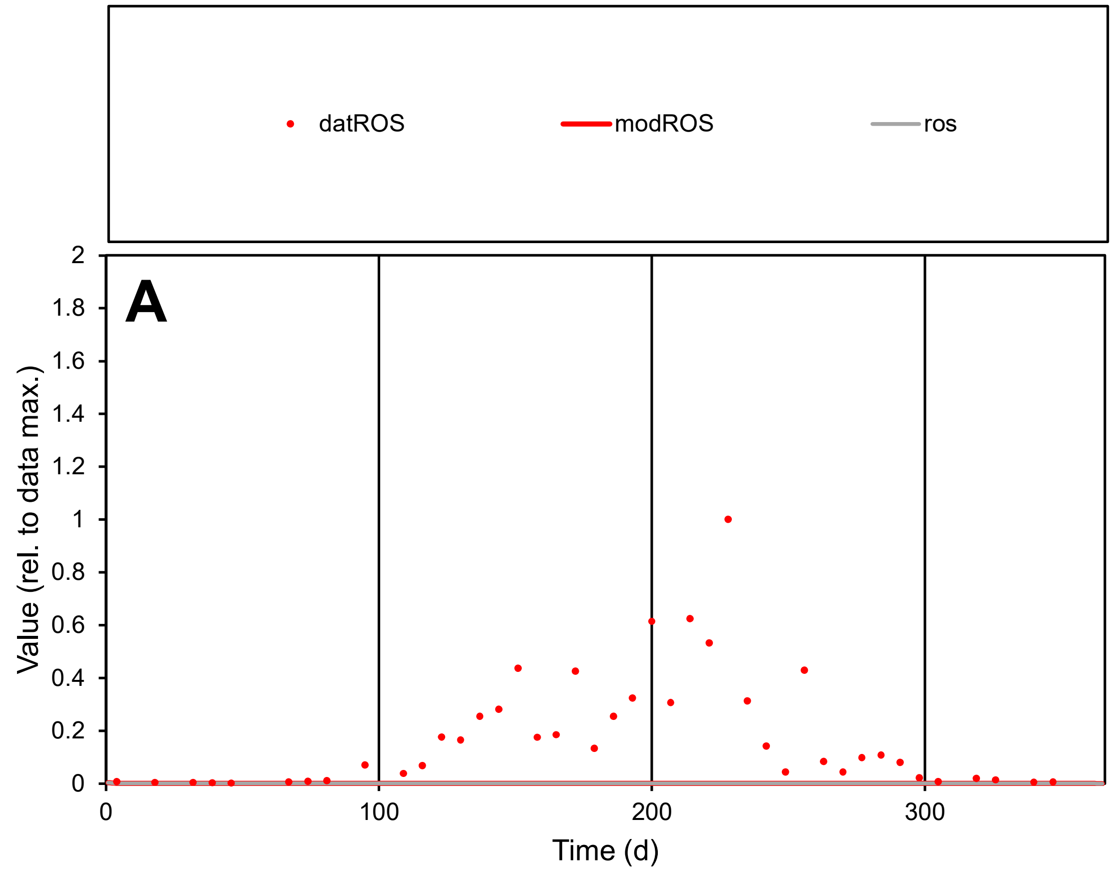


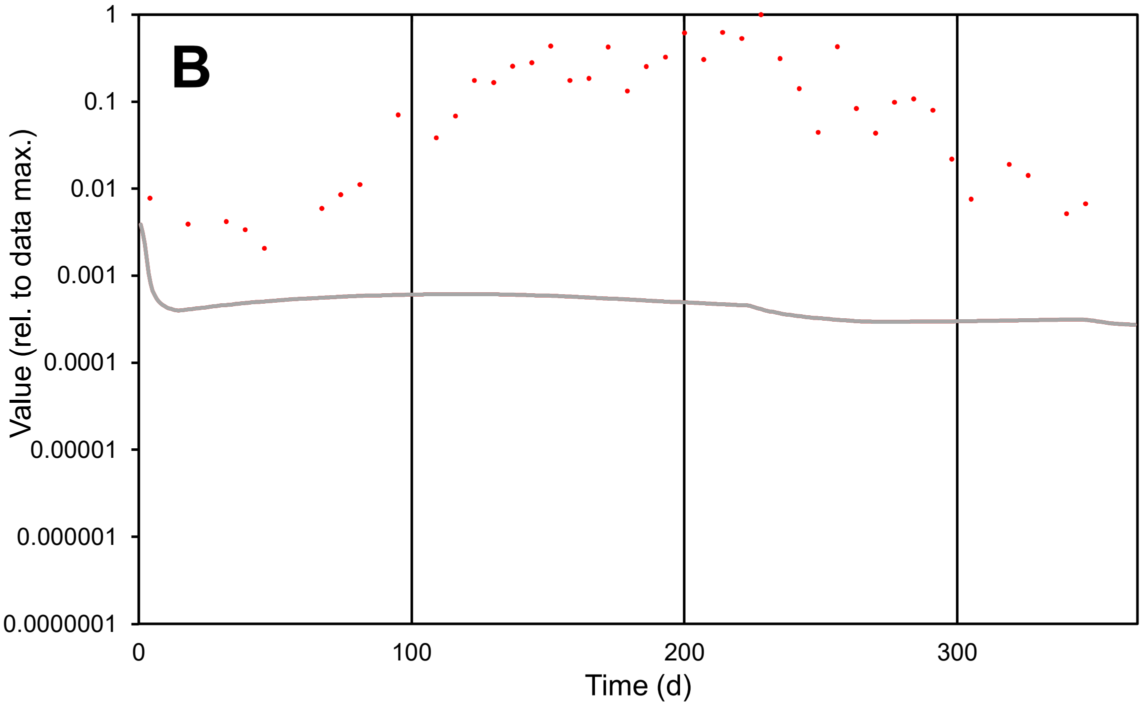


**C1**

Variables
ROS Provide jtd or Dst
CalibOnly?
2 1 to reproduce of
oNormalize?
2 1=no, 2=yes

ofxn calc*
1.56782
ofxn mod
1.57001

sRun
RunA1
zout
1

Ave
Data
0.01489
Model
4.3E-05

data/vMax
0.08307

ofxng
334.07

Min d/m
7.02084

ave/min d
86.669
ave/min m
1.76282

327_Calib_S_A

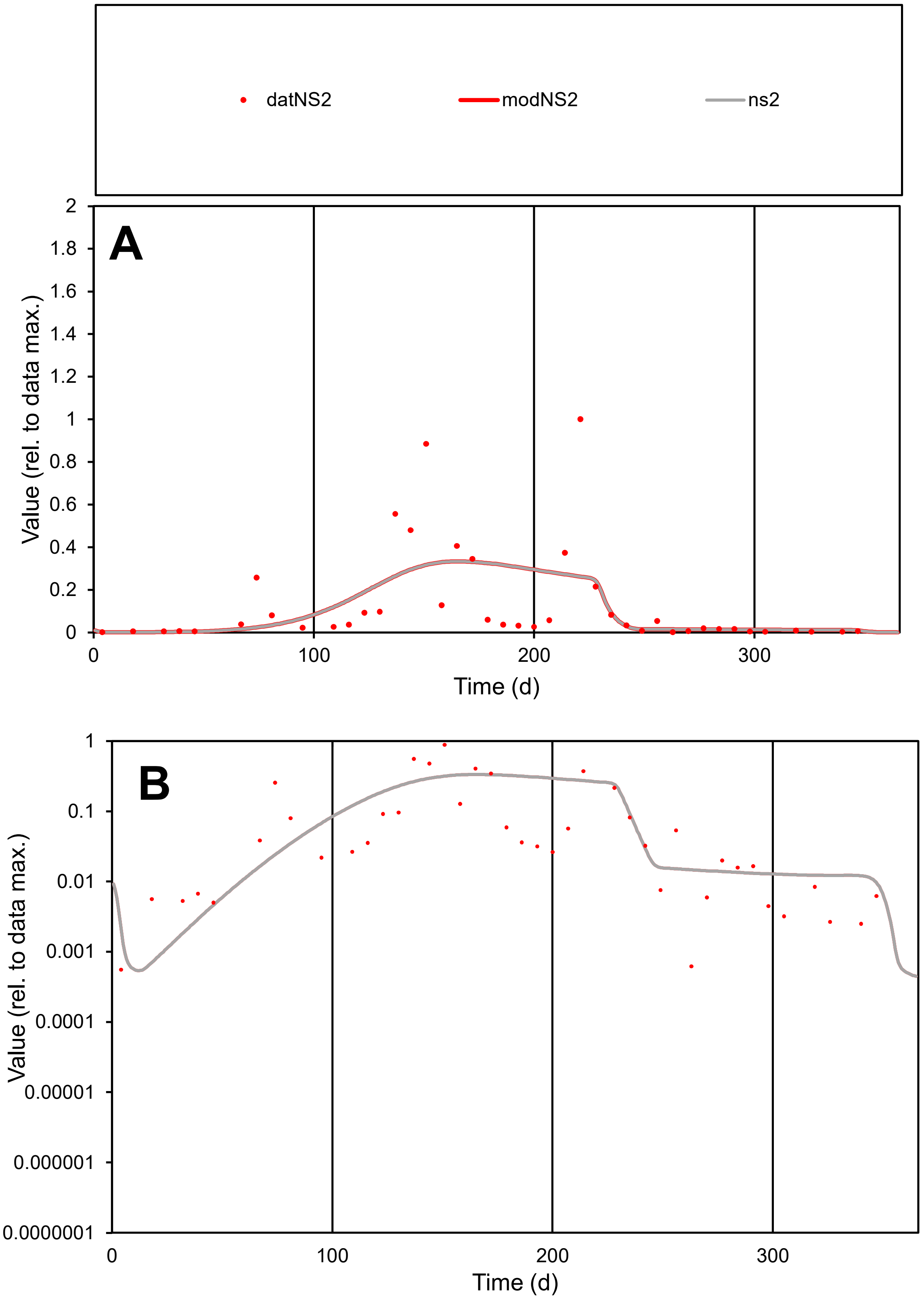


**C1**

Variables
NS2 Provide jtd or Dst
CalibOnly?
2 1 to reproduce of
oNormalize?
2 1=no, 2=yes

ofxn calc*
1.43169
ofxn mod
1.44094

sRun
RunA1
zout
1

Ave
Data
0.01457
Model
0.01395

data/vMax
0.1107

ofxng
334.07

Min d/m
0.71394

ave/min d
235.933
ave/min m
161.243

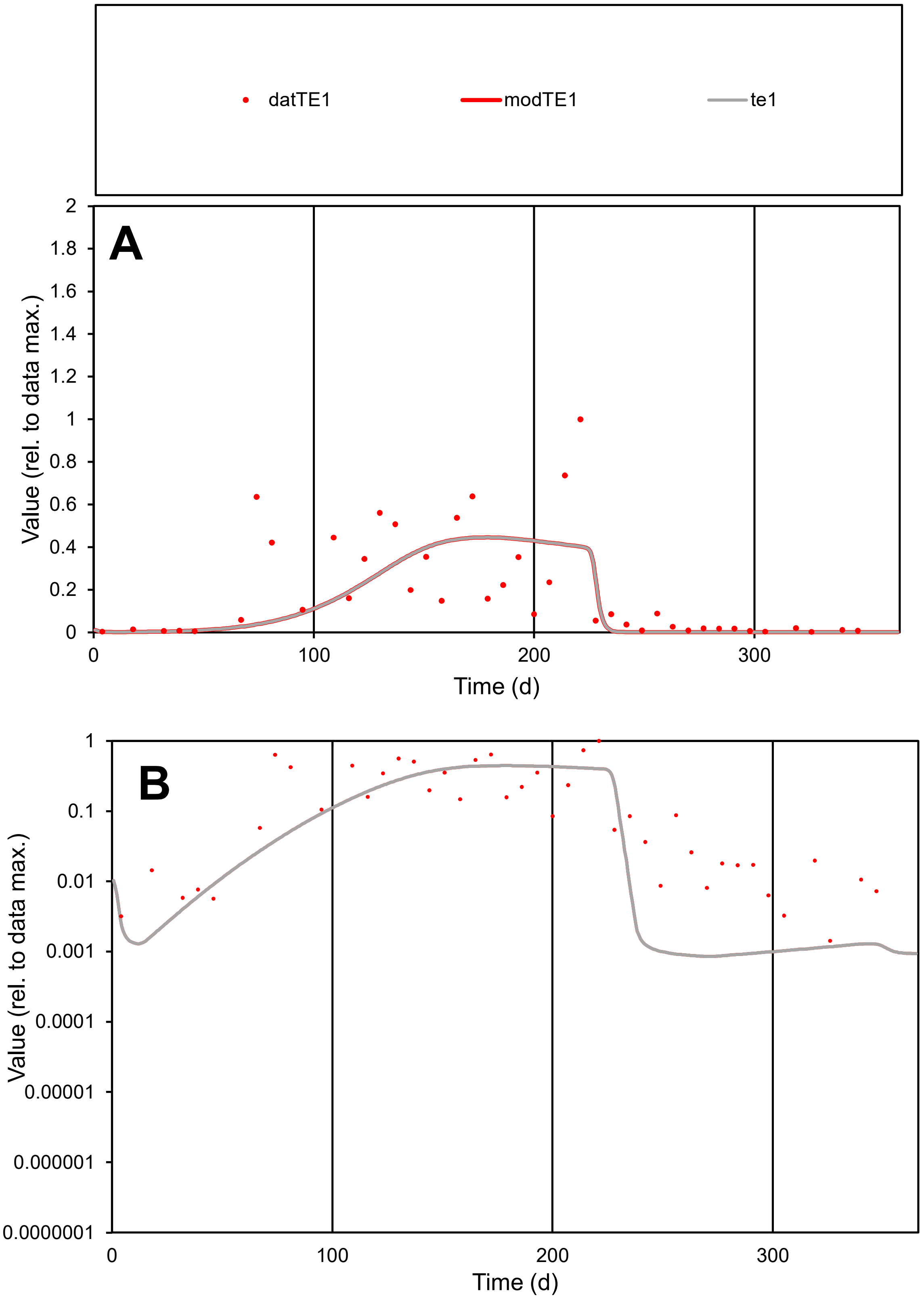


**C1**

Variables

TE1 Provide jtd or Dst

CalibOnly?

2 1 to reproduce of

oNormalize?

2 1=no, 2=yes

ofxn calc*

0.98657

ofxn mod

0.99972

sRun

RunA1

zout

1

Ave

Data

0.00808

Model

0.00647

data/vMax

0.04069

ofxng

334.07

Min d/m

1.66839

ave/min d

139.397

ave/min m

186.339

327_Calib_S_A

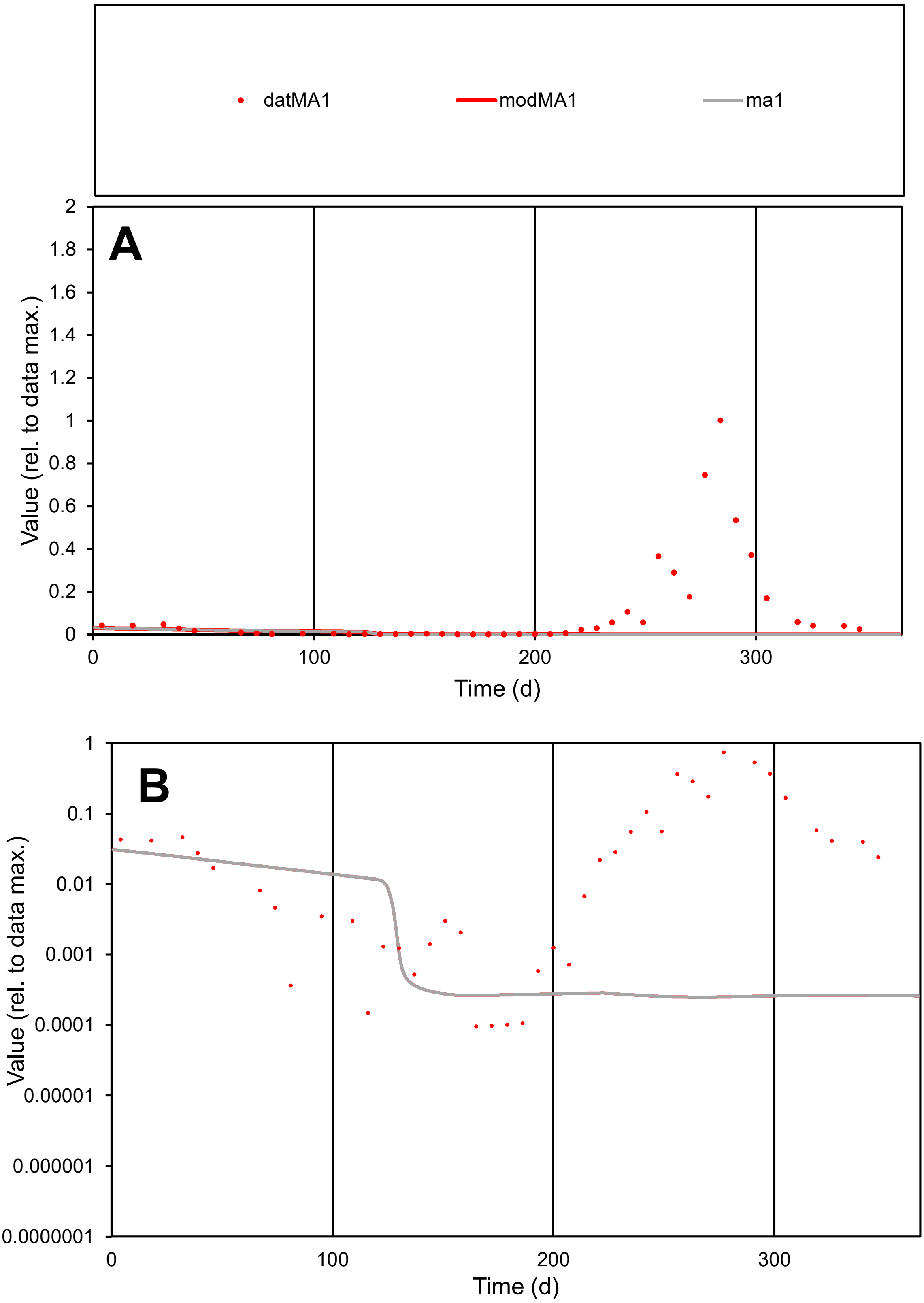


**C1**

Variables
MA1 Provide jtd or Dst
CalibOnly?
2 1 to reproduce of
oNormalize?
2 1=no, 2=yes

ofxn calc*
2.27891
ofxn mod
2.28228

sRun
RunA1
zout
1

Ave
Data
0.01525
Model
0.00082

data/vMax
0.1492

ofxng
334.07

Min d/m
0.38724

ave/min d
1065.19
ave/min m
22.1004

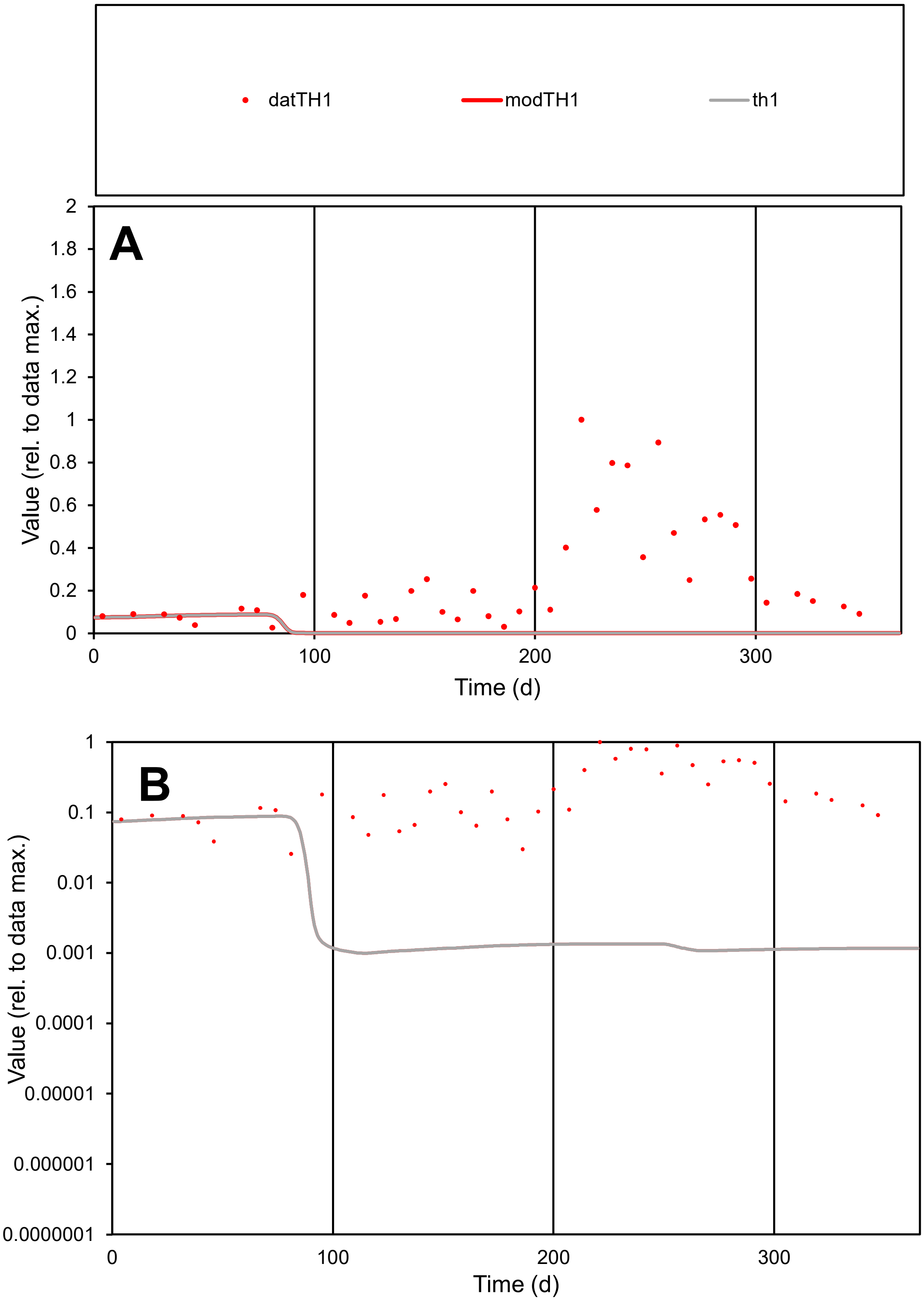


**C1**

Variables

TH1 Provide jtd or Dst

CalibOnly?

2 1 to reproduce of

oNormalize?

2 1=no, 2=yes

ofxn calc*

1.39542

ofxn mod

1.39825

sRun

RunA1

zout

1

Ave

Data

0.01344

Model

0.00087

data/vMax

0.05295

ofxng

334.07

Min d/m

25.6242

ave/min d

9.84516

ave/min m

16.3162

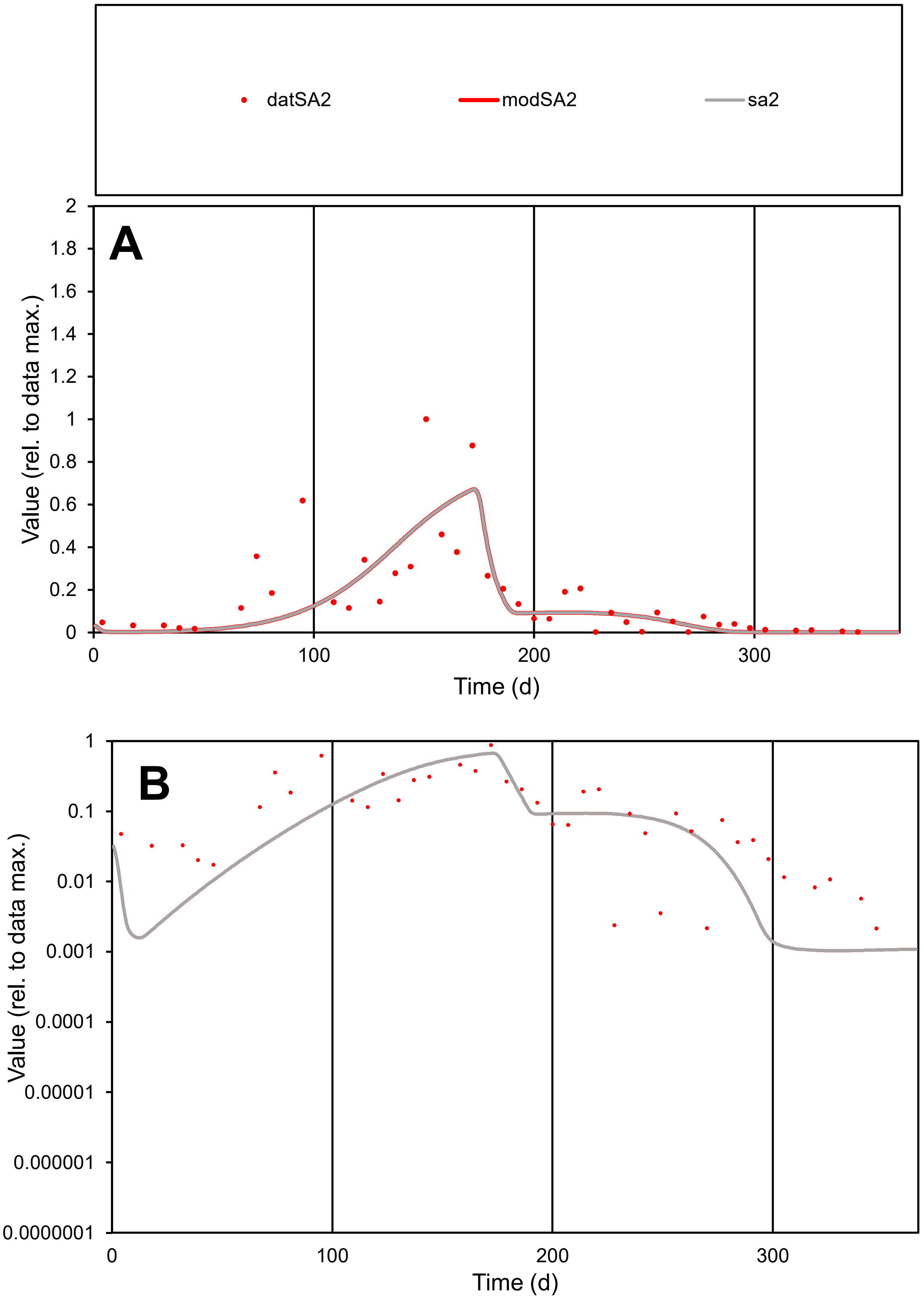


**C1**

| | |
|---|---|
| Variables | |
| SA2 | Provide jtd or Dst |
| CalibOnly? | |
| 2 | 1 to reproduce of |
| oNormalize? | |
| 2 | 1=no, 2=yes |
| ofxn calc* | |
| 0.8581 | |
| ofxn mod | |
| 0.87038 | |
| sRun | |
| RunA1 | |
| zout | |
| 1 | |
| Ave | |
| Data | |
| 0.00863 | |
| Model | |
| 0.00747 | |
| data/vMax | |
| 0.05106 | |
| ofxng | |
| 334.07 | |
| Min d/m | |
| 2.07601 | |
| ave/min d | |
| 78.6261 | |
| ave/min m | |
| 141.194 | |

327_Calib_S_A

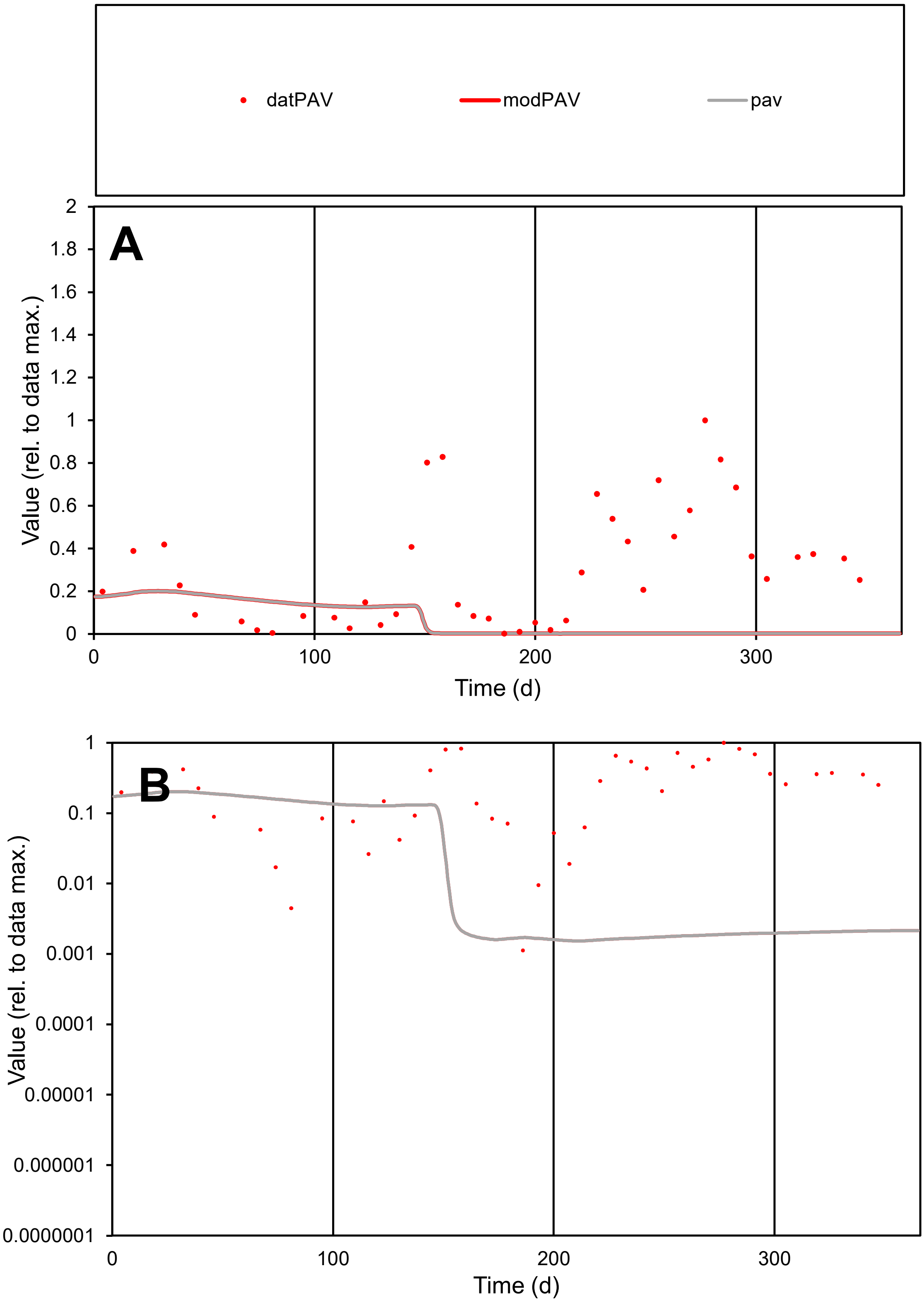


**C1**

Variables
PAV Provide jtd or Dst
CalibOnly?
2 1 to reproduce of
oNormalize?
2 1=no, 2=yes

ofxn calc*
1.2968
ofxn mod
1.29994

sRun
RunA1
zout
1

Ave
Data
0.01103
Model
0.00209

data/vMax
0.03656

ofxng
334.07

Min d/m
0.73248

ave/min d
267.859
ave/min m
37.238

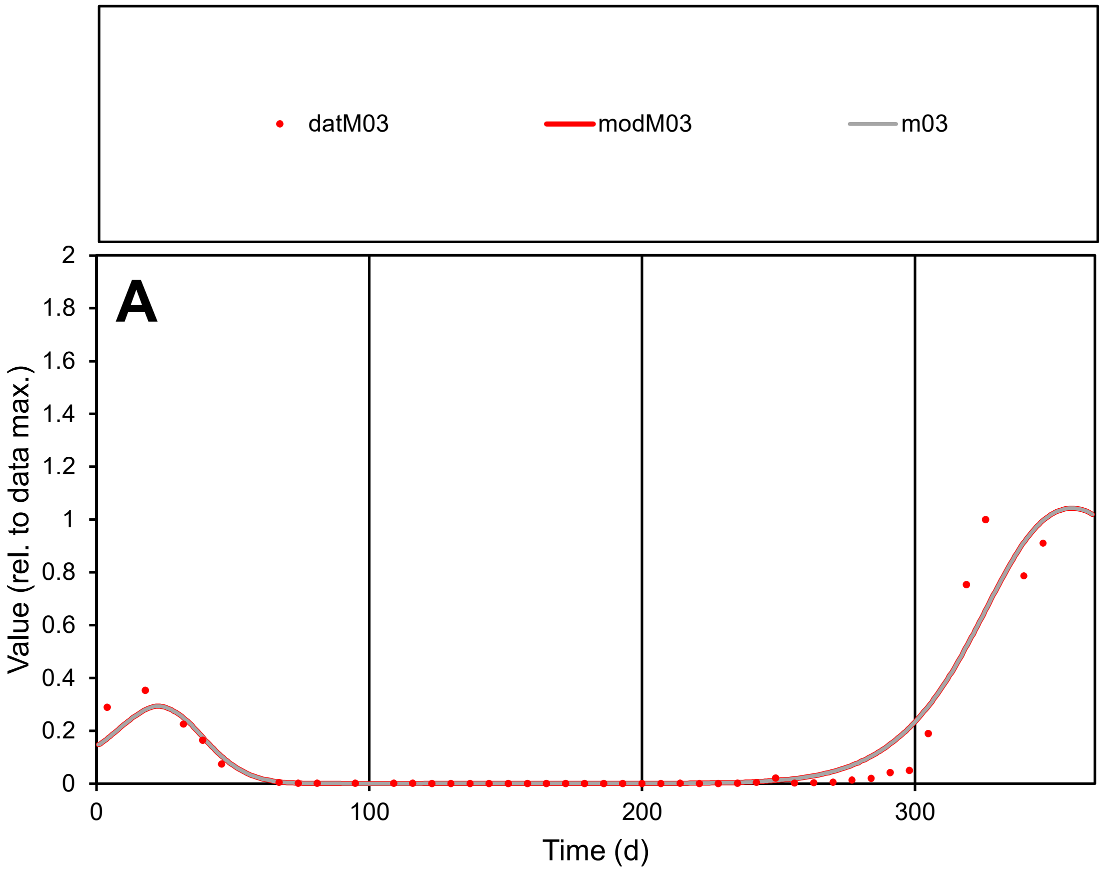


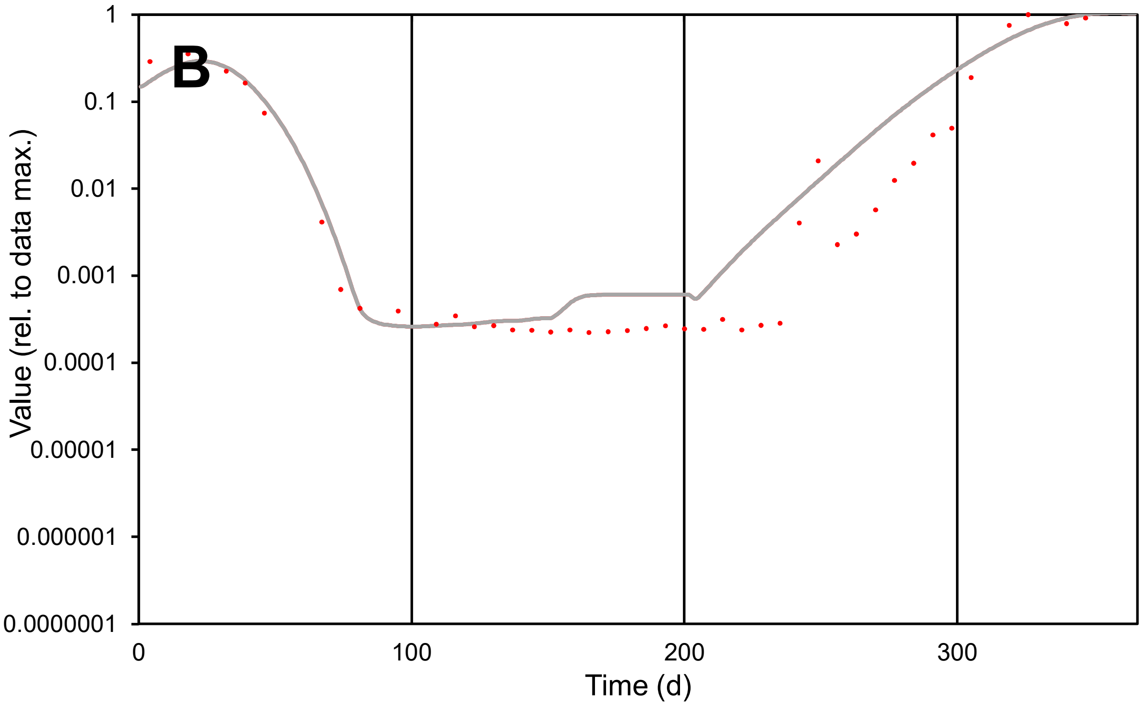


**C1**

Variables
M03 Provide jtd or Dst
CalibOnly?
2 1 to reproduce of
oNormalize?
2 1=no, 2=yes

ofxn calc*
0.66485
ofxn mod
0.71136

sRun
RunA1
zout
1

Ave
Data
0.00915
Model
0.00946

data/vMax
0.07822

ofxng
334.07

Min d/m
0.84769

ave/min d
525.319
ave/min m
460.226

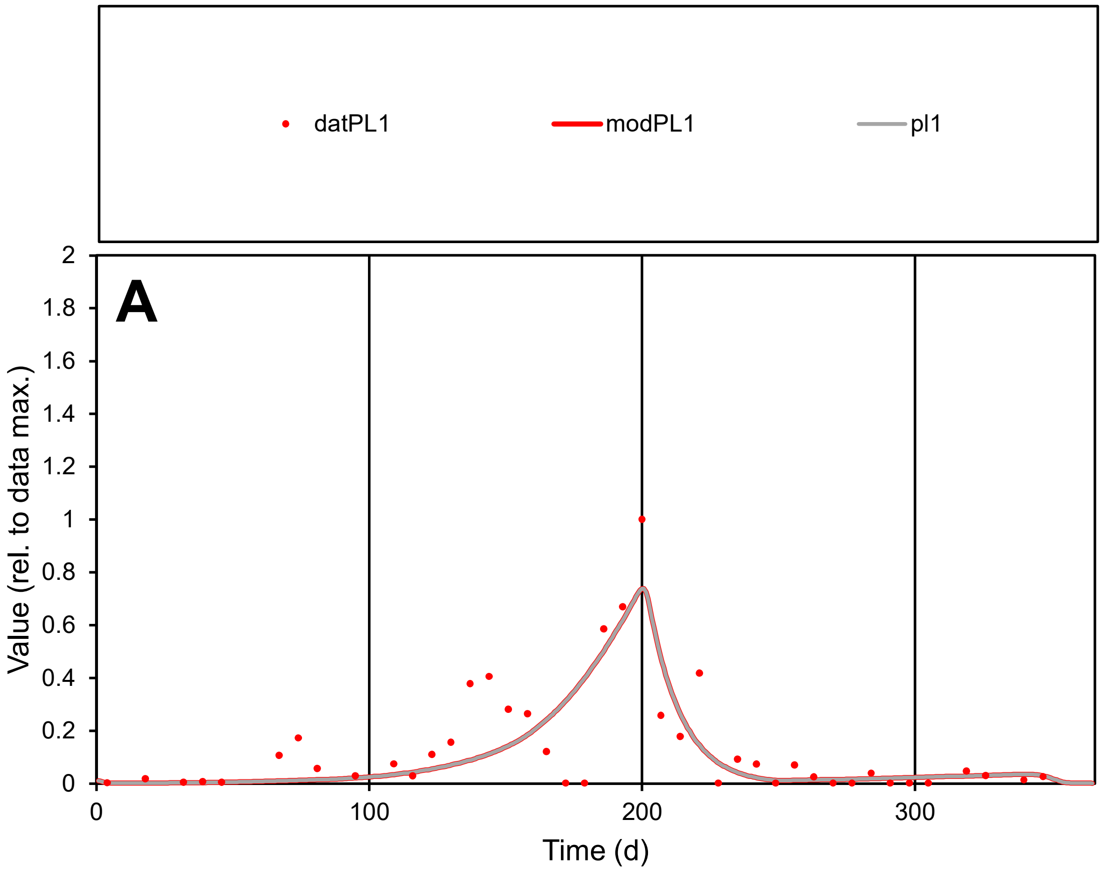


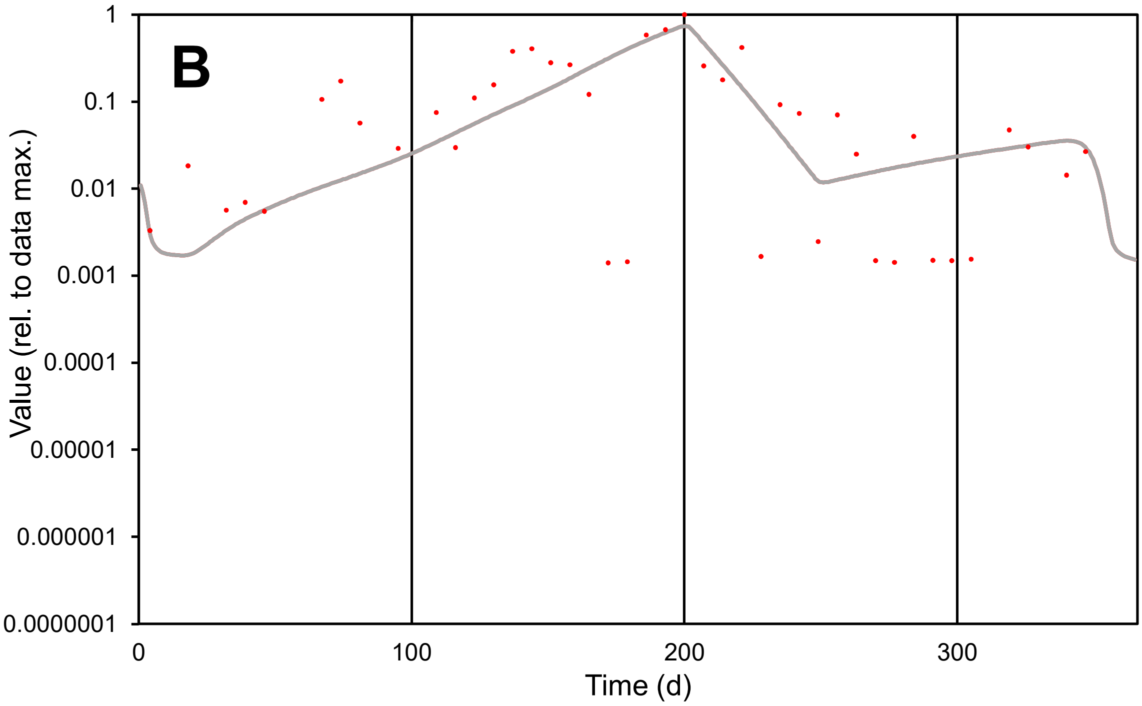


**C1**

Variables
PL1 Provide jtd or Dst
CalibOnly?
2 1 to reproduce of
oNormalize?
2 1=no, 2=yes

ofxn calc*
0.92964
ofxn mod
0.96622

sRun
RunA1
zout
1

Ave
Data
0.00345
Model
0.00294

data/vMax
0.02515

ofxng
334.07

Min d/m
0.78004

ave/min d
98.2041
ave/min m
65.1212

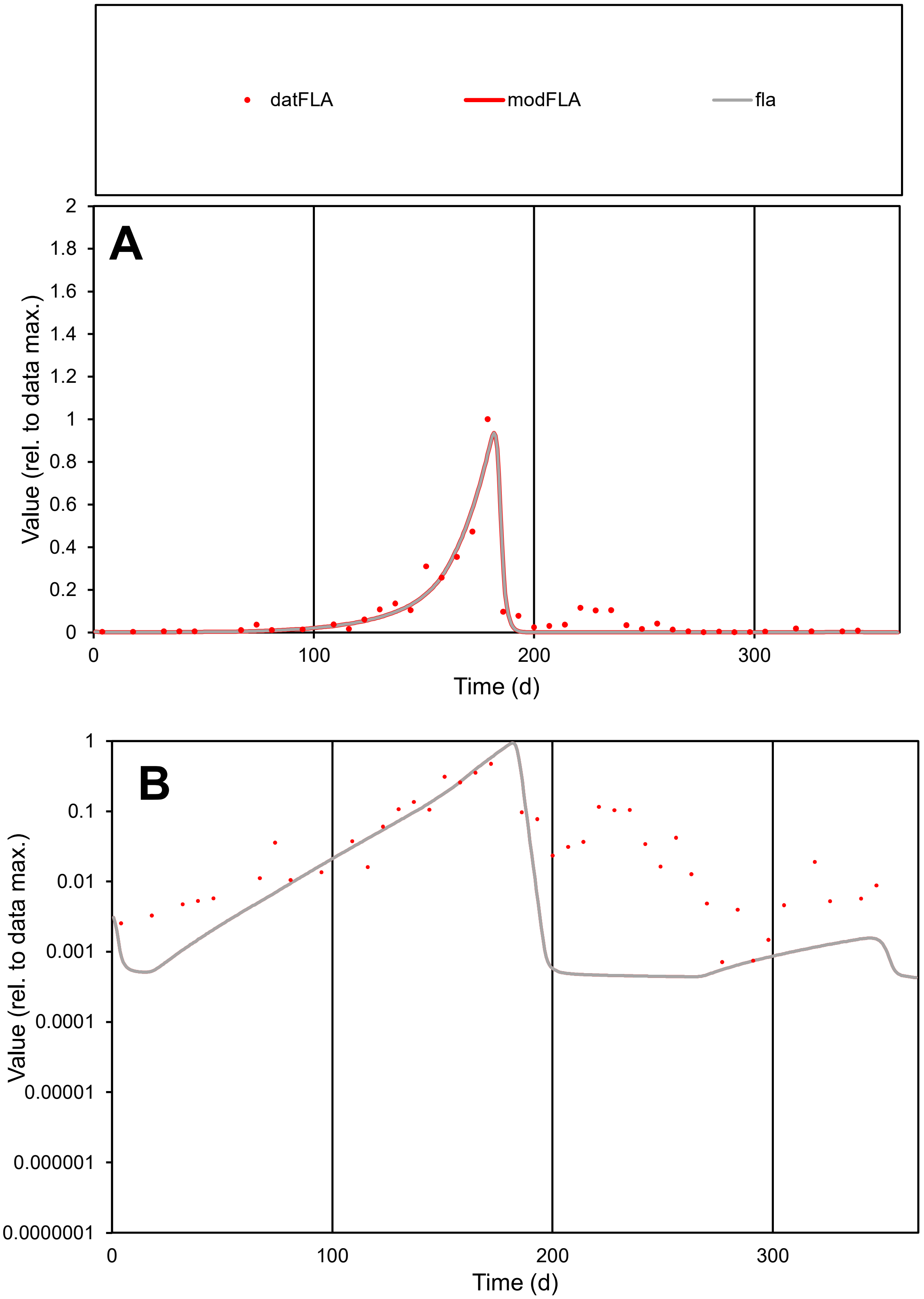


**C1**

Variables

FLA Provide jtd or Dst

CalibOnly?

2 1 to reproduce of

oNormalize?

2 1=no, 2=yes

ofxn calc*

0.55237

ofxn mod

0.61002

sRun

RunA1

zout

1

Ave

Data

0.00706

Model

0.00592

data/vMax

0.08017

ofxng

334.07

Min d/m

1.61172

ave/min d

124.045

ave/min m

167.78

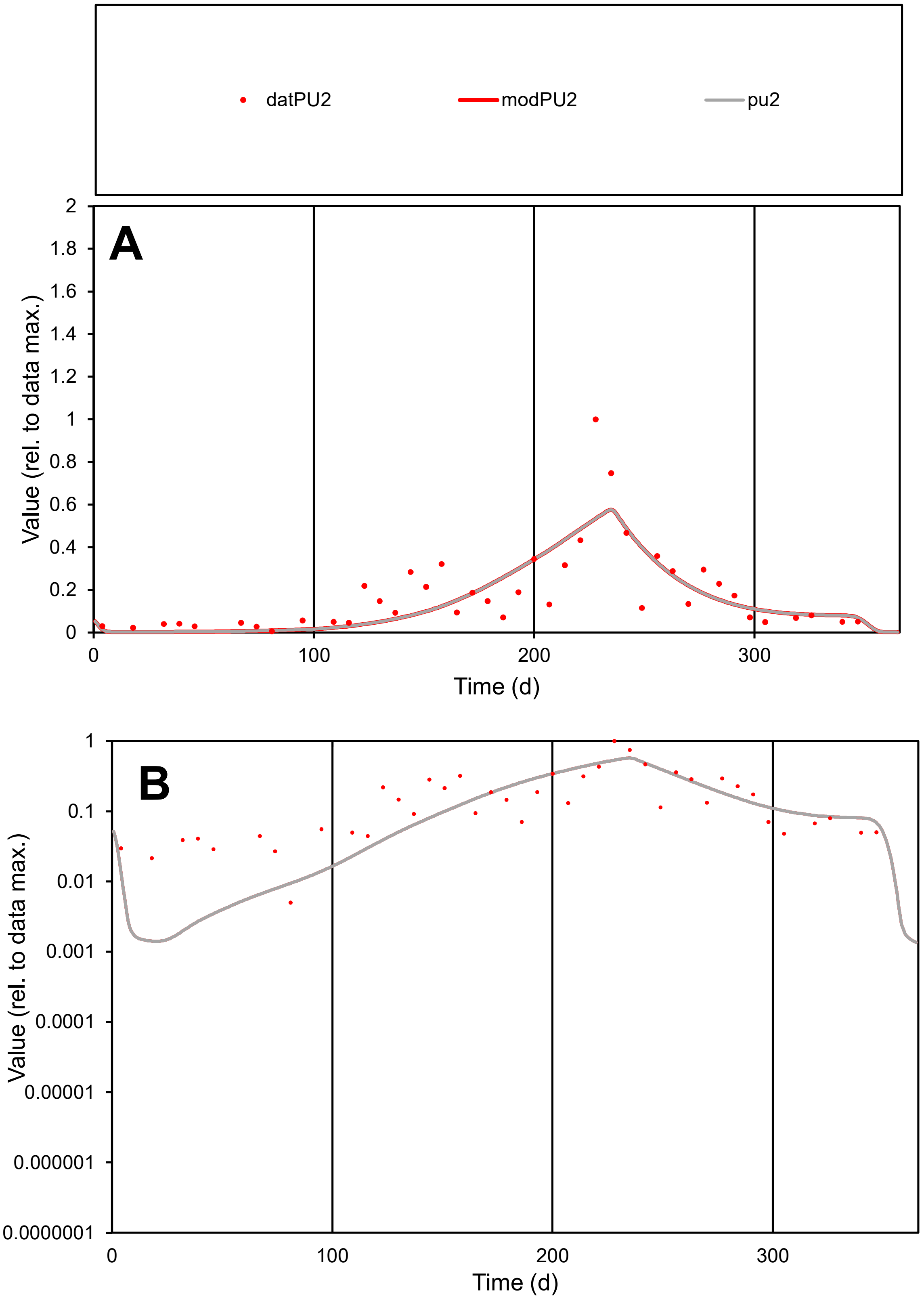


**C1**

Variables
PU2 Provide jtd or Dst
CalibOnly?
2 1 to reproduce of
oNormalize?
2 1=no, 2=yes

ofxn calc*
0.6688
ofxn mod
0.68832

sRun
RunA1
zout
1

Ave
Data
0.0106
Model
0.00977

data/vMax
0.05754

ofxng
334.07

Min d/m
3.56553

ave/min d
36.6453
ave/min m
120.478

327_Calib_S_A

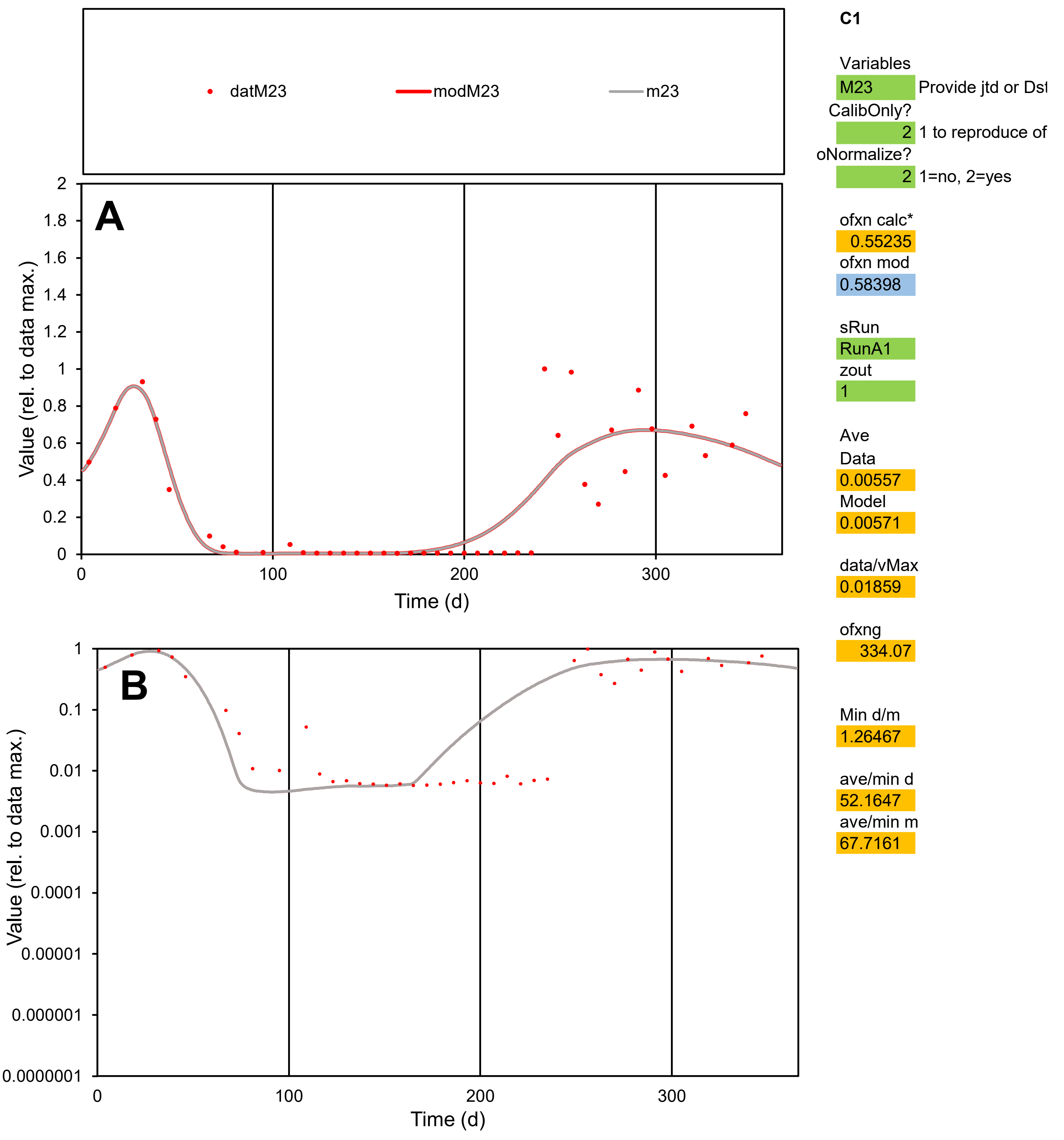

datM23
modM23
m23
A
Value (rel. to data max.)
Time (d)
B
Value (rel. to data max.)
Time (d)
C1
Variables
M23
Provide jtd or Dst
CalibOnly?
2
1 to reproduce of
oNormalize?
2
1=no, 2=yes
ofxn calc*
0.55235
ofxn mod
0.58398
sRun
RunA1
zout
1
Ave
Data
0.00557
Model
0.00571
data/vMax
0.01859
ofxng
334.07
Min d/m
1.26467
ave/min d
52.1647
ave/min m
67.7161

327_Calib_S_A

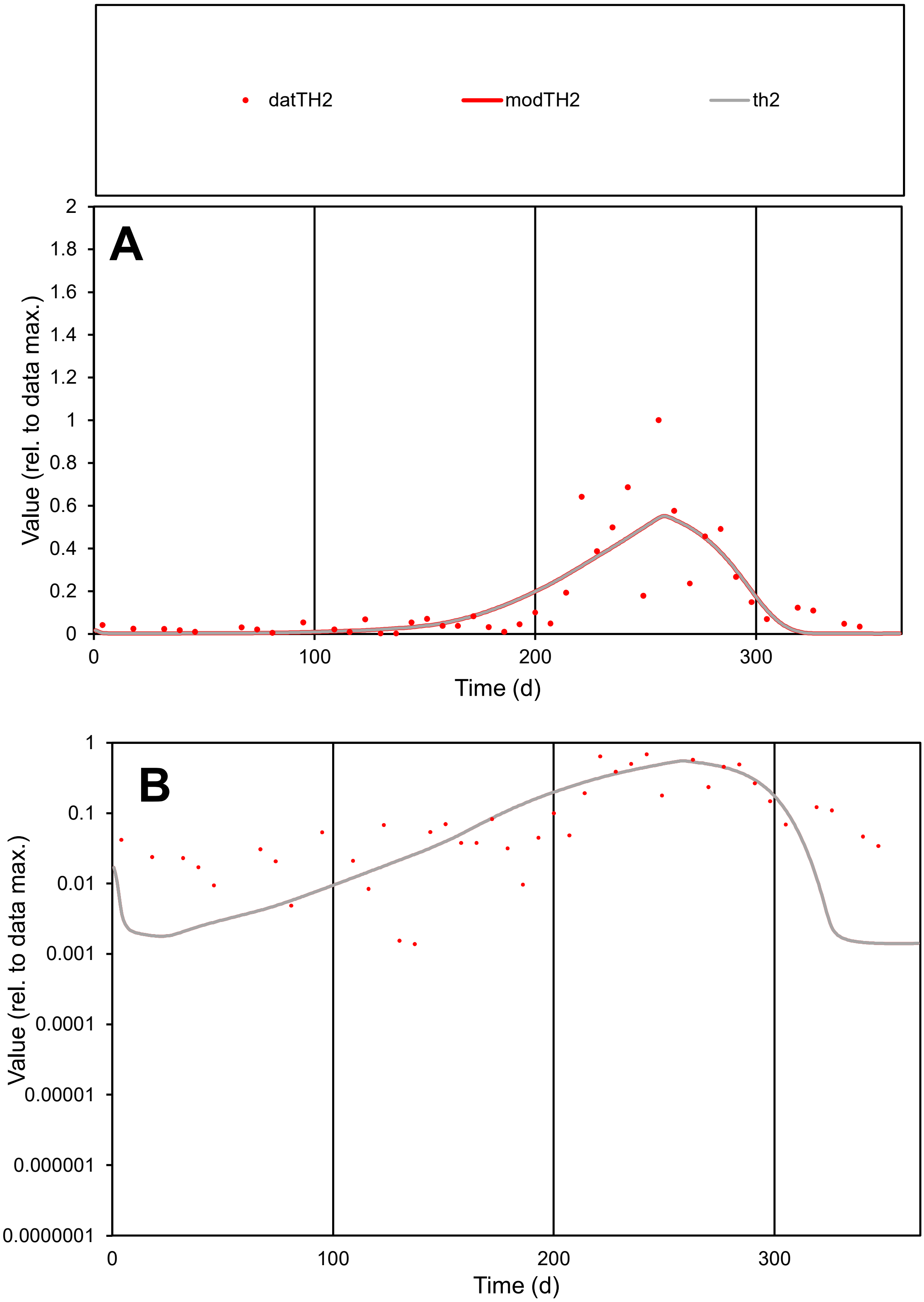


**C1**

Variables

TH2 Provide jtd or Dst

CalibOnly?

2 1 to reproduce of

oNormalize?

2 1=no, 2=yes

ofxn calc*

0.76252

ofxn mod

0.78963

sRun

RunA1

zout

1

Ave

Data

0.01355

Model

0.01277

data/vMax

0.08154

ofxng

334.07

Min d/m

0.97489

ave/min d

120.377

ave/min m

110.568

327_Calib_S_A

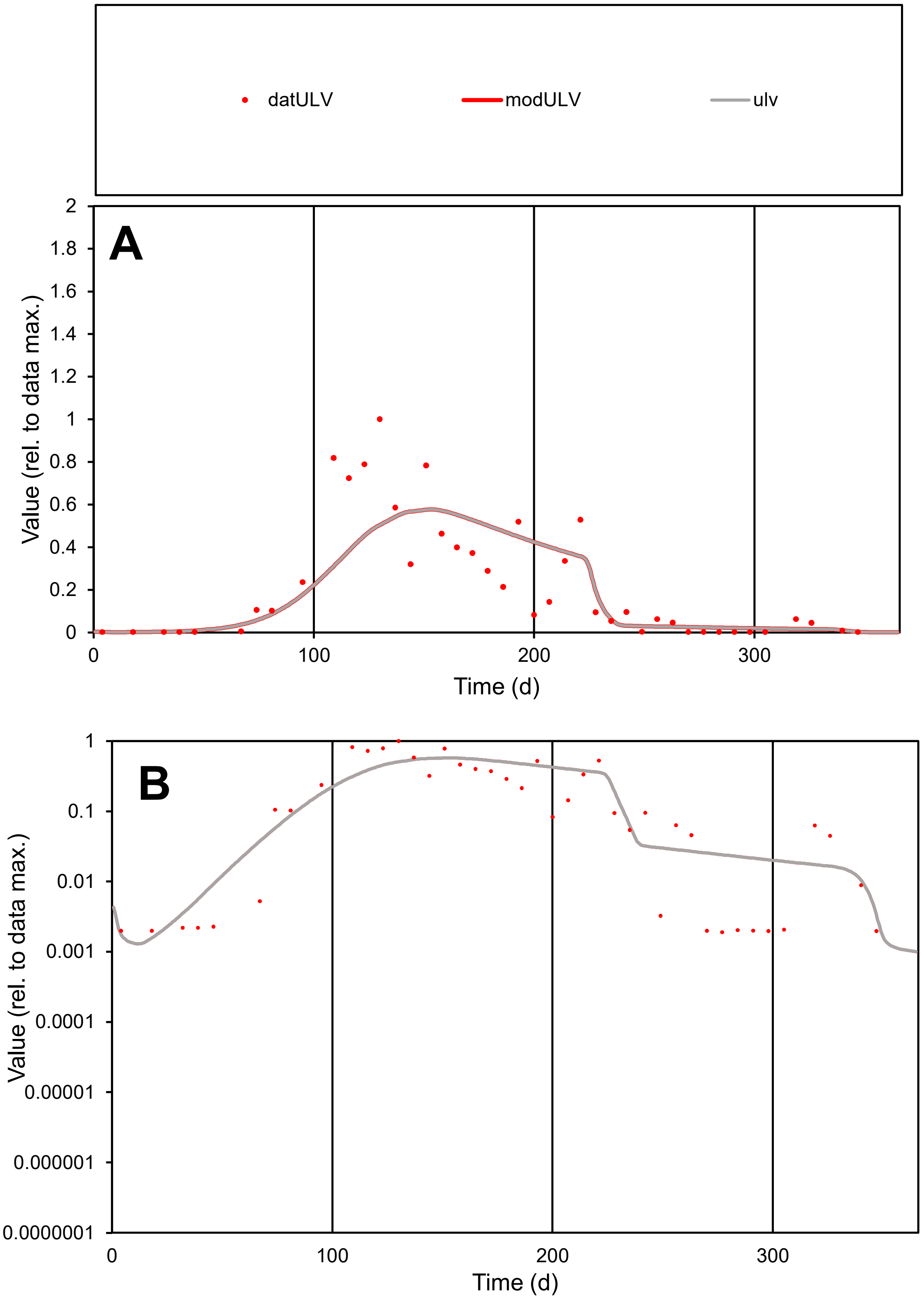


**C1**

Variables

ULV Provide jtd or Dst

CalibOnly?

2 1 to reproduce of

oNormalize?

2 1=no, 2=yes

ofxn calc*

0.76066

ofxn mod

0.77707

sRun

RunA1

zout

1

Ave

Data

0.00208

Model

0.00198

data/vMax

0.0094

ofxng

334.07

Min d/m

1.11584

ave/min d

116.989

ave/min m

124.234

327_Calib_S_A

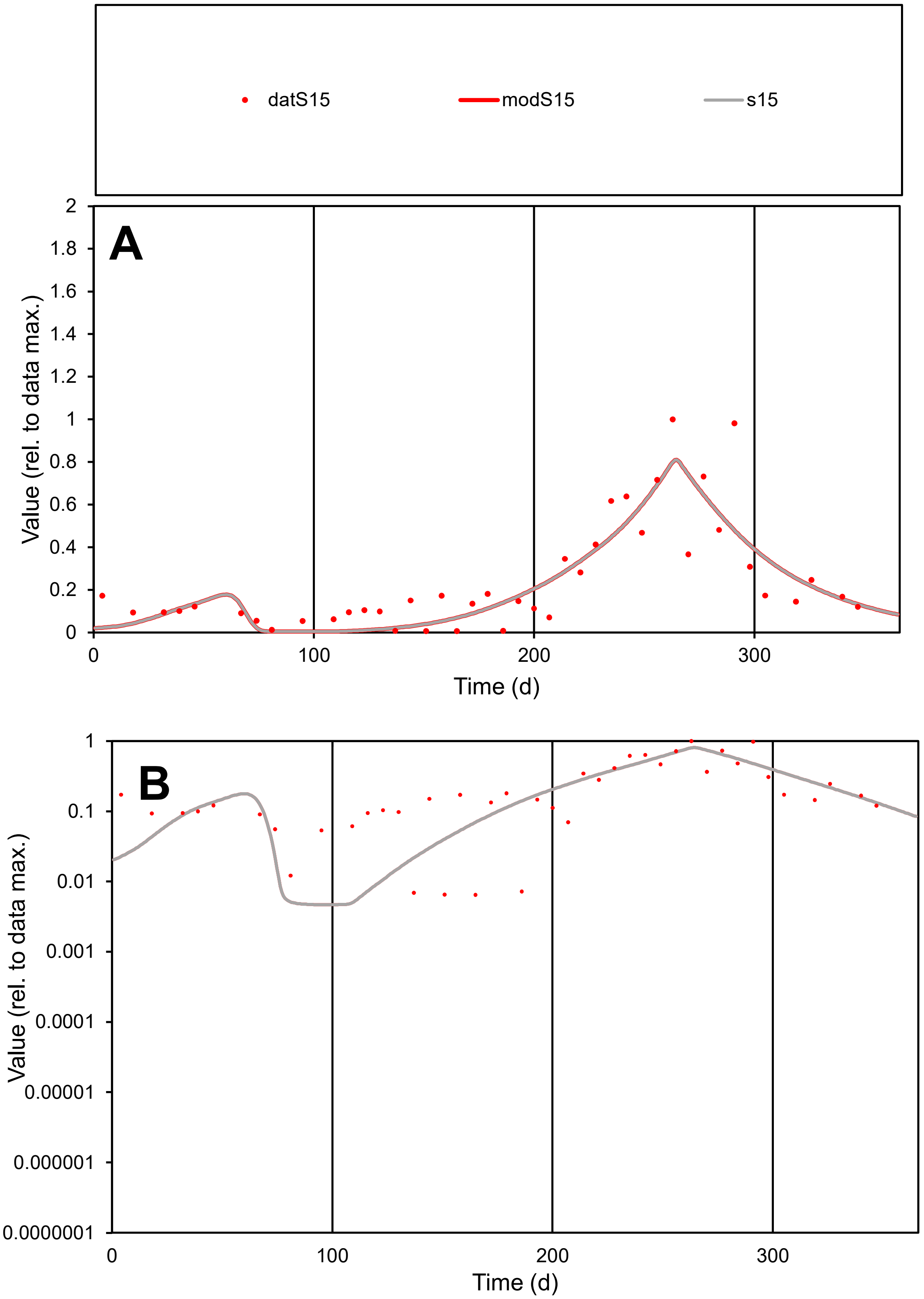


**C1**

Variables

S15 Provide jtd or Dst

CalibOnly?

2 1 to reproduce of

oNormalize?

2 1=no, 2=yes

ofxn calc*

0.52476

ofxn mod

0.55496

sRun

RunA1

zout

1

Ave

Data

0.00471

Model

0.00447

data/vMax

0.01914

ofxng

334.07

Min d/m

1.37046

ave/min d

38.2036

ave/min m

49.6864

327_Calib_S_A

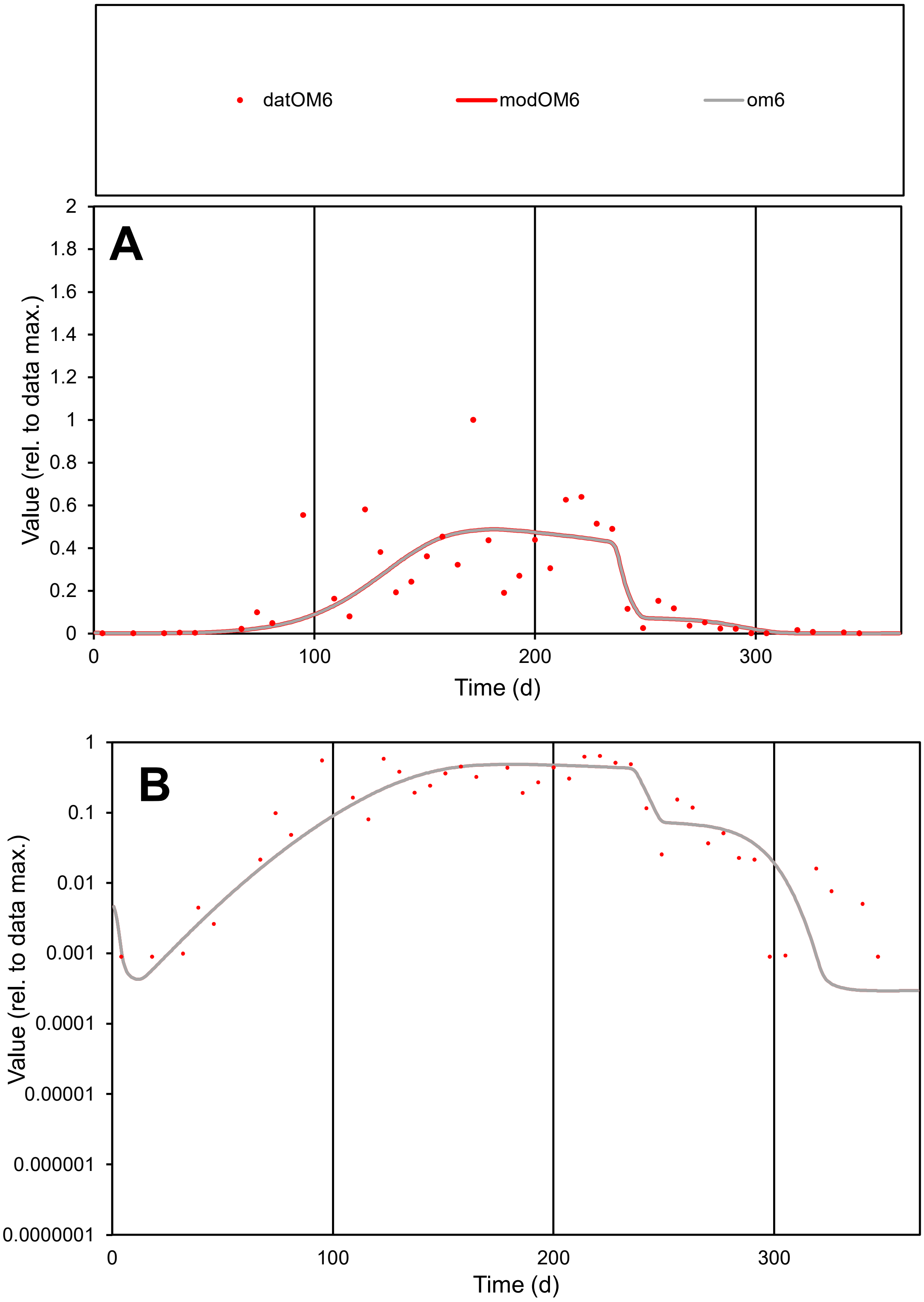


**C1**

Variables

OM6 Provide jtd or Dst

CalibOnly?

2 1 to reproduce of

oNormalize?

2 1=no, 2=yes

ofxn calc*

0.70736

ofxn mod

0.72327

sRun

RunA1

zout

1

Ave

Data

0.00647

Model

0.00596

data/vMax

0.0302

ofxng

334.07

Min d/m

3.05104

ave/min d

239.46

ave/min m

673.003

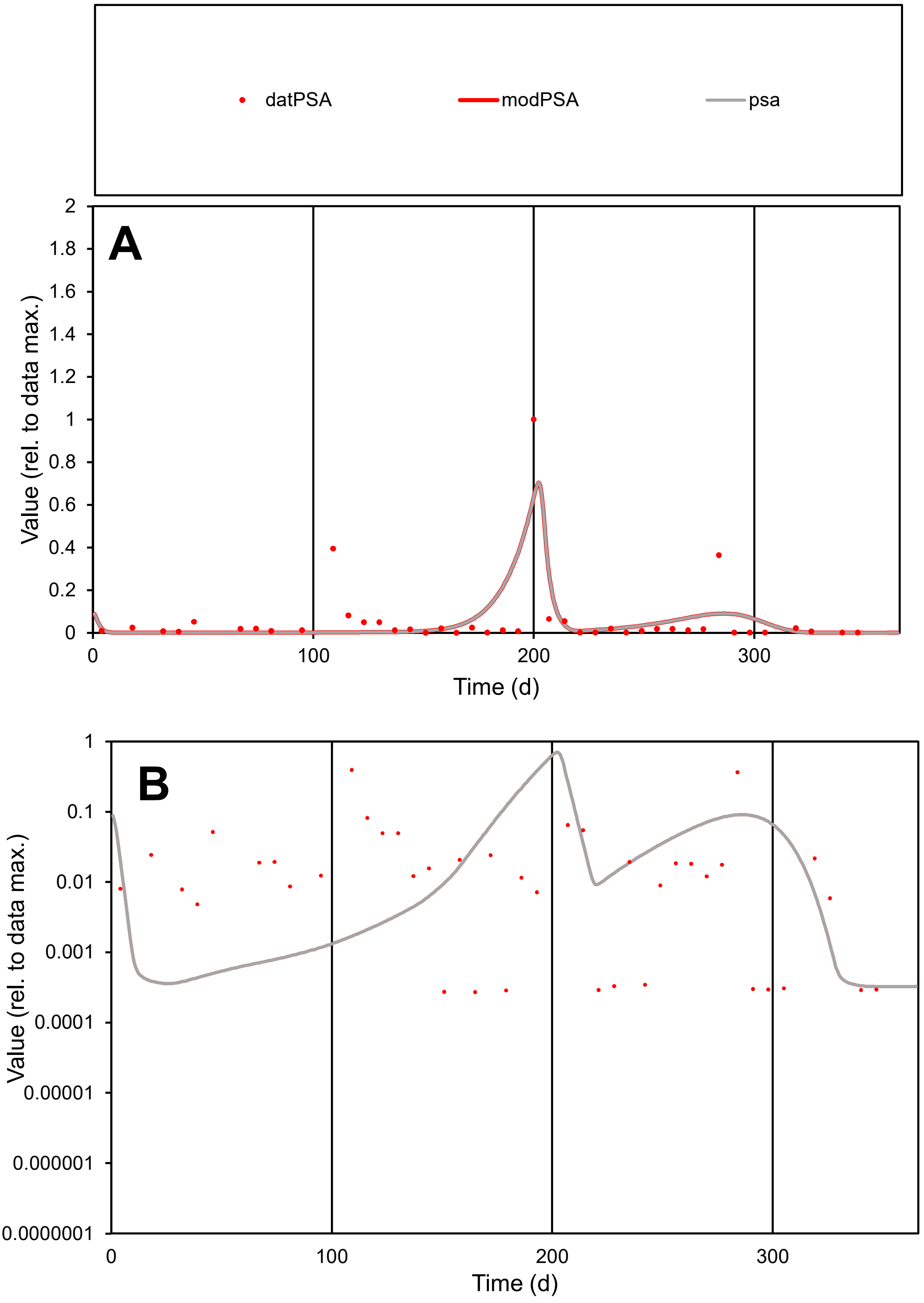


**C1**

Variables
PSA Provide jtd or Dst
CalibOnly?
2 1 to reproduce of
oNormalize?
2 1=no, 2=yes

ofxn calc*
1.92368
ofxn mod
2.02883

sRun
RunA1
zout
1

Ave
Data
0.00187
Model
0.00177

data/vMax
0.03227

ofxng
334.07

Min d/m
0.83981

ave/min d
212.795
ave/min m
169.418

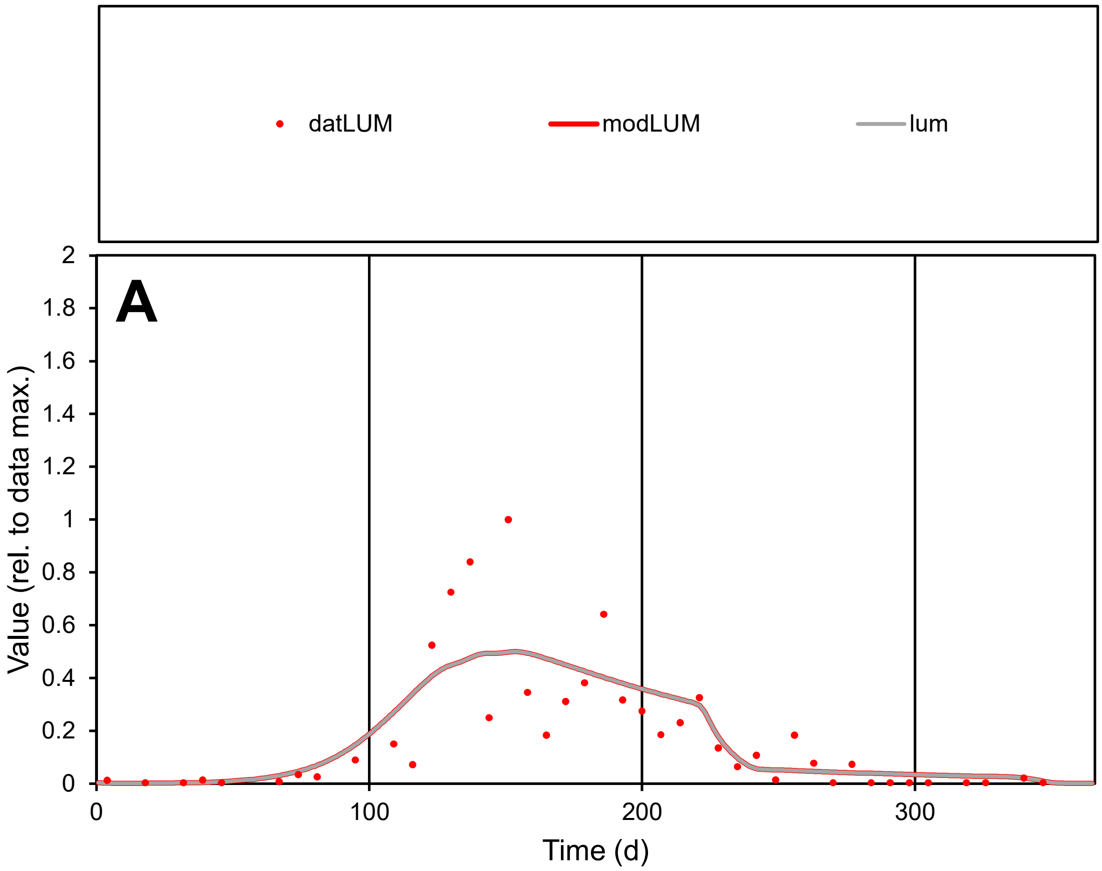


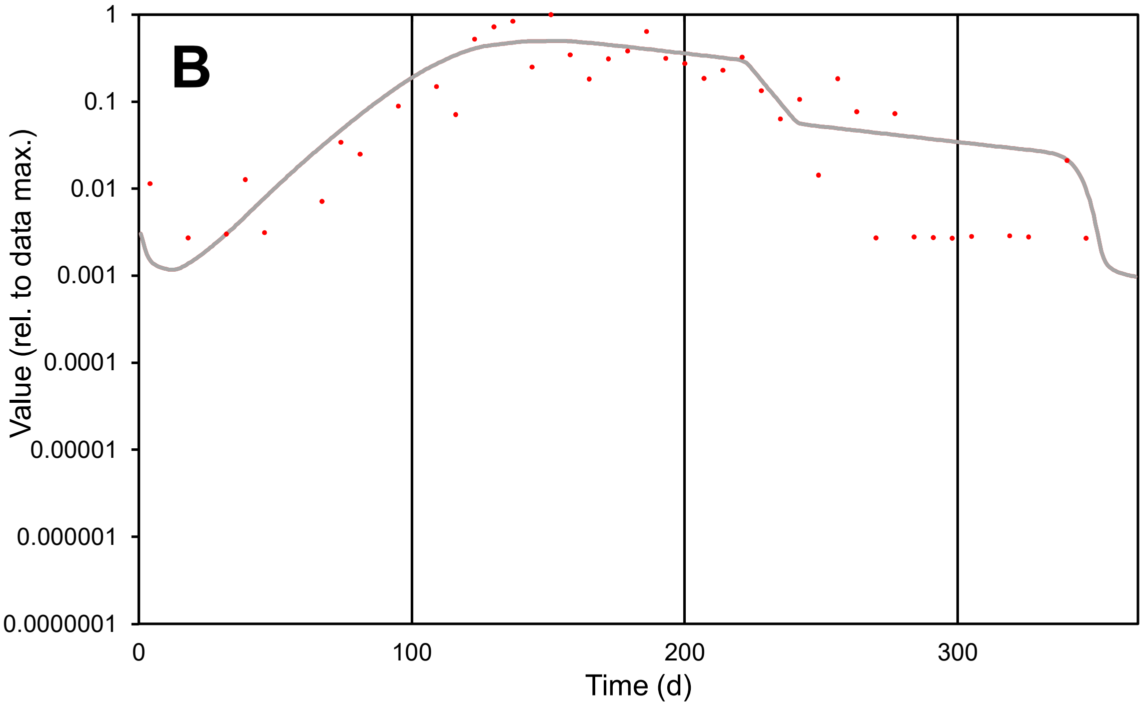


**C1**

Variables
LUM Provide jtd or Dst
CalibOnly?
2 1 to reproduce of
oNormalize?
2 1=no, 2=yes

ofxn calc*
0.79456
ofxn mod
0.81213

sRun
RunA1
zout
1

Ave
Data
0.00324
Model
0.00337

data/vMax
0.01784

ofxng
334.07

Min d/m
1.83378

ave/min d
67.0998
ave/min m
127.796

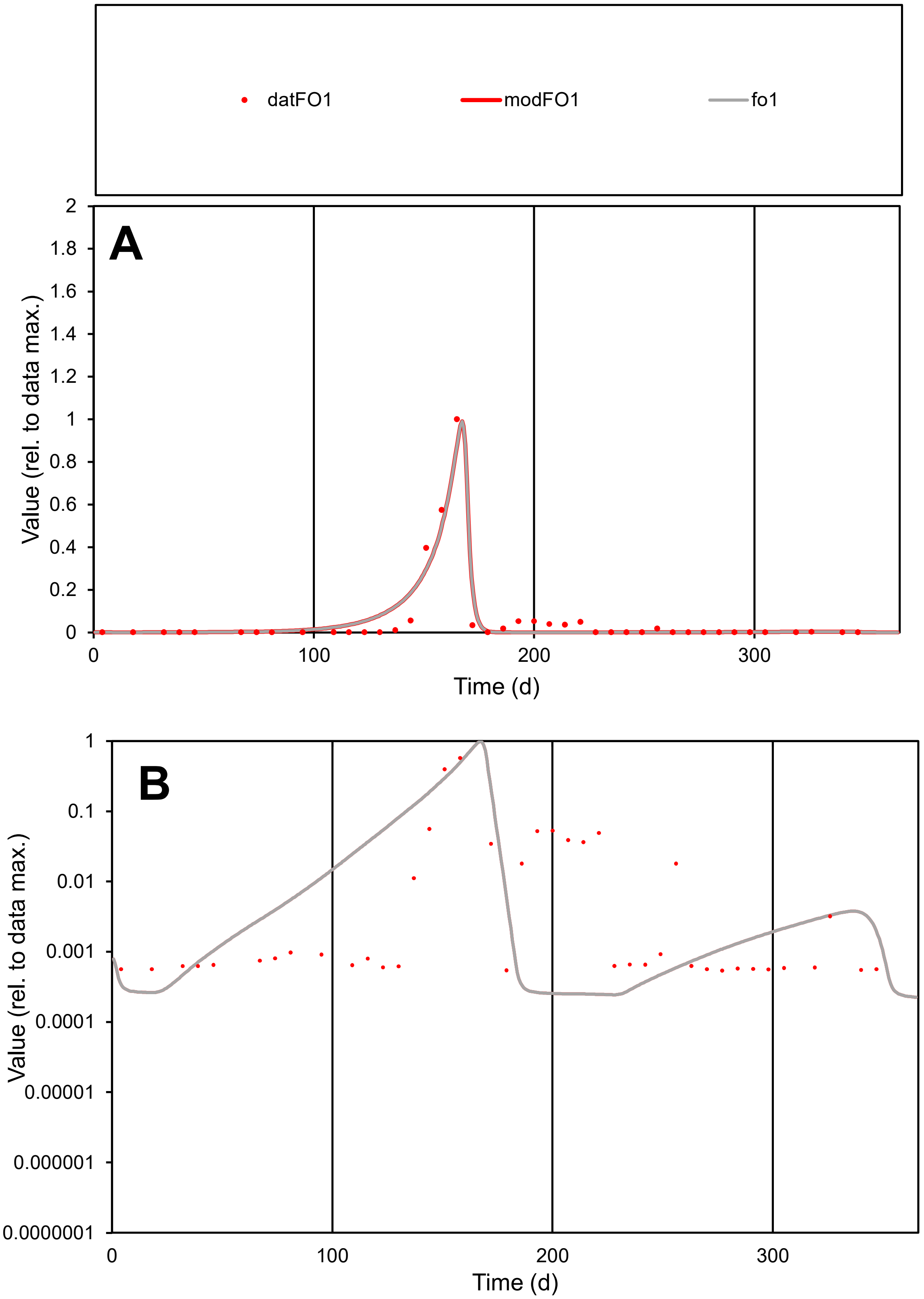


**C1**

Variables
FO1 Provide jtd or Dst
CalibOnly?
2 1 to reproduce of
oNormalize?
2 1=no, 2=yes

ofxn calc*
0.65479
ofxn mod
0.77075

sRun
RunA1
zout
1

Ave
Data
0.00628
Model
0.00646

data/vMax
0.1117

ofxng
334.07

Min d/m
2.20608

ave/min d
104.167
ave/min m
236.598

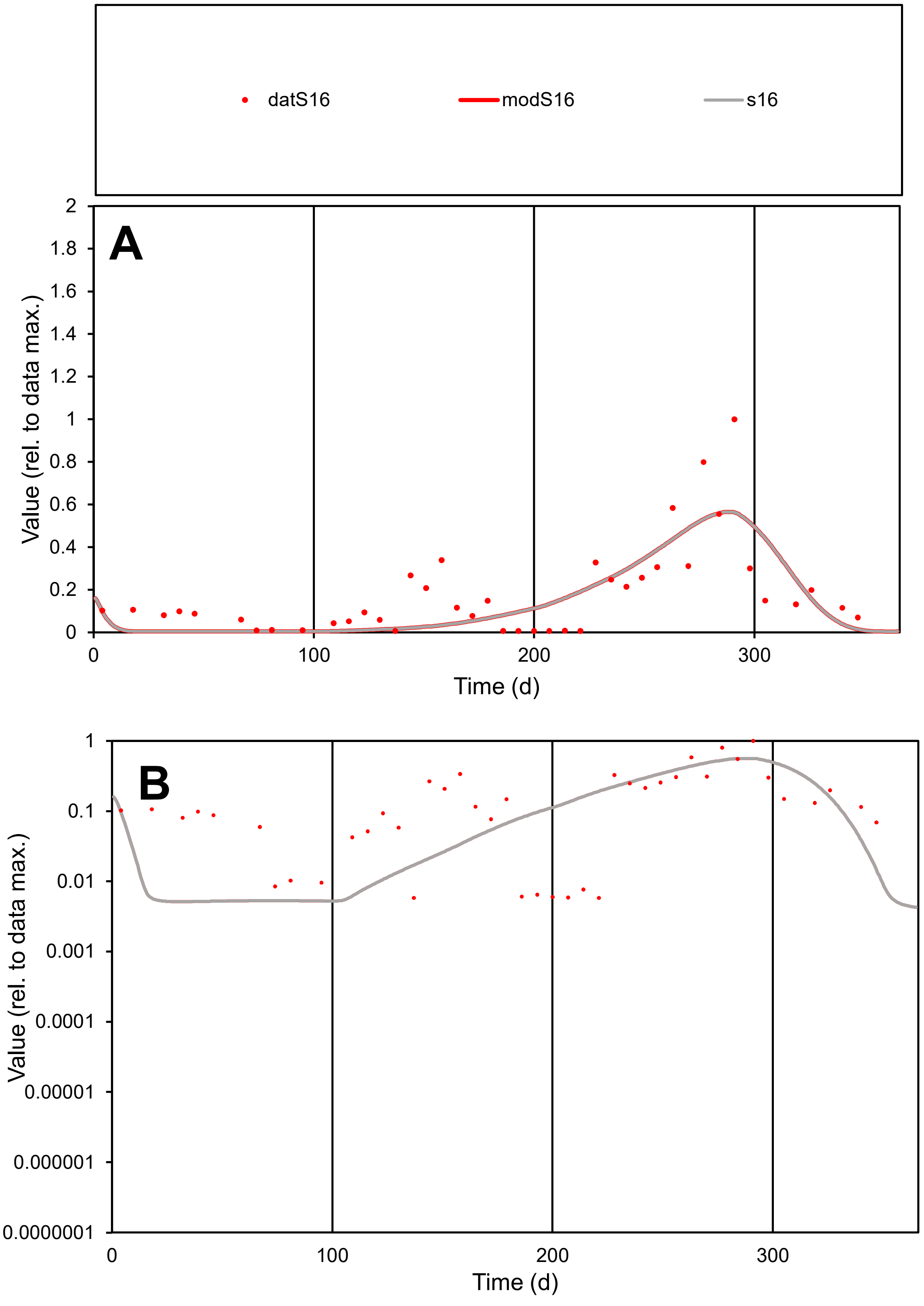


**C1**

Variables
S16 Provide jtd or Dst
CalibOnly?
2 1 to reproduce of
oNormalize?
2 1=no, 2=yes

ofxn calc*
0.7796
ofxn mod
0.79911

sRun
RunA1
zout
1

Ave
Data
0.00409
Model
0.00364

data/vMax
0.02272

ofxng
334.07

Min d/m
1.12543

ave/min d
31.0254
ave/min m
31.0335

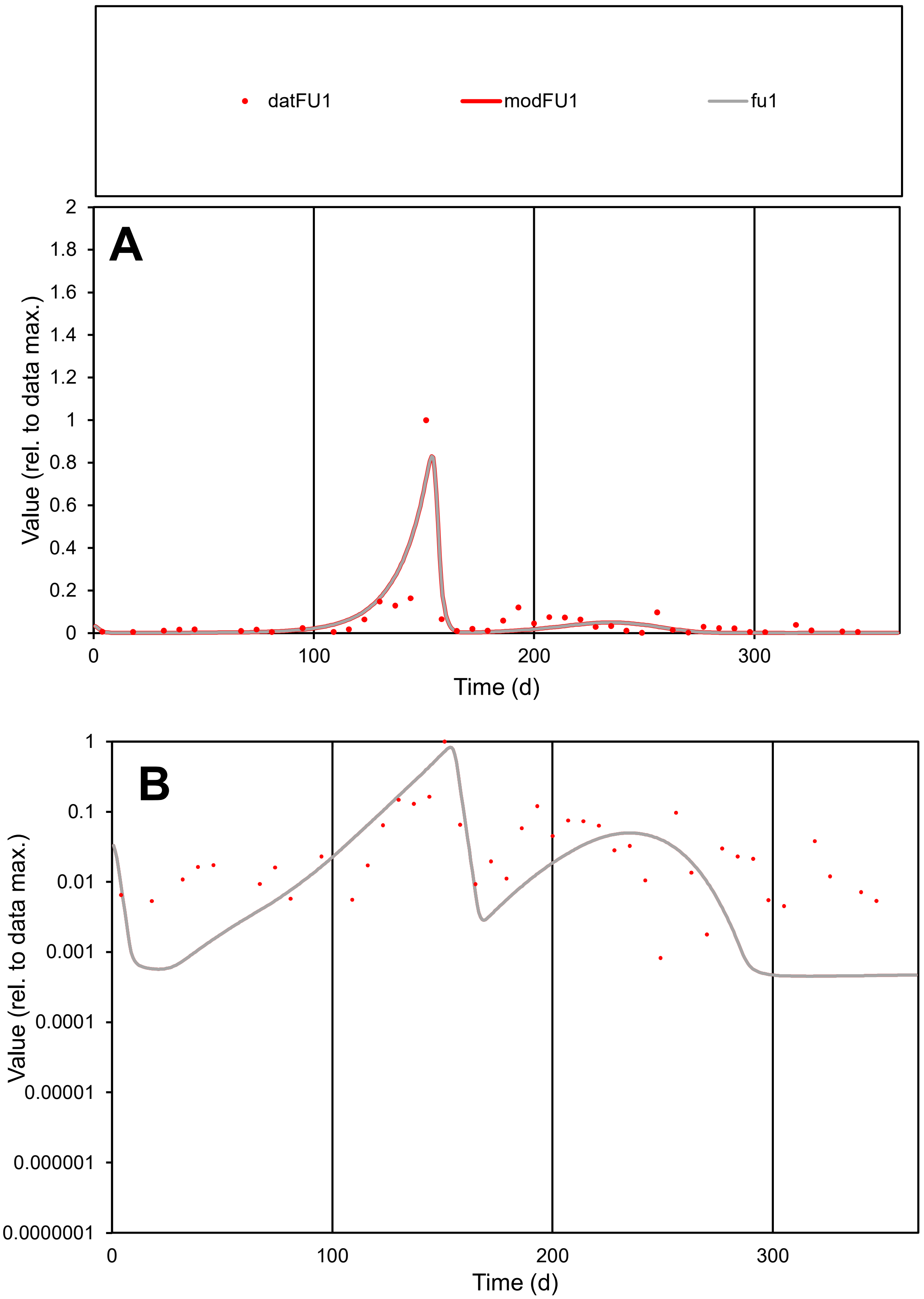


**C1**

Variables
FU1 Provide jtd or Dst
CalibOnly?
2 1 to reproduce of
oNormalize?
2 1=no, 2=yes

ofxn calc*
1.10753
ofxn mod
1.14631

sRun
RunA1
zout
1

Ave
Data
0.00602
Model
0.00578

data/vMax
0.1007

ofxng
334.07

Min d/m
1.82064

ave/min d
72.5596
ave/min m
126.791

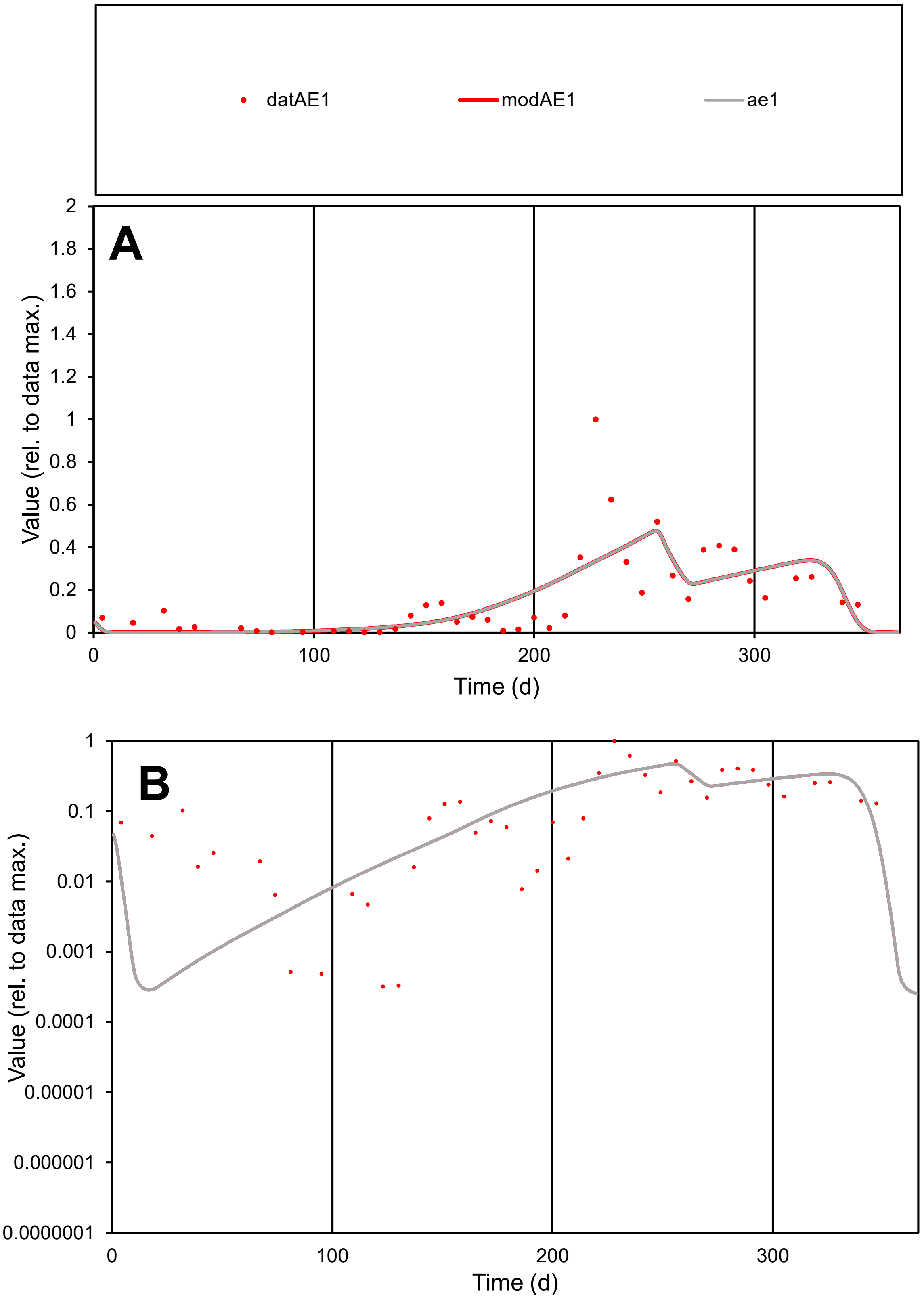


**C1**

Variables
AE1 Provide jtd or Dst
CalibOnly?
2 1 to reproduce of
oNormalize?
2 1=no, 2=yes

ofxn calc*
0.8727
ofxn mod
0.89006

sRun
RunA1
zout
1

Ave
Data
0.0082
Model
0.00807

data/vMax
0.05036

ofxng
334.07

Min d/m
1.05851

ave/min d
509.277
ave/min m
530.854

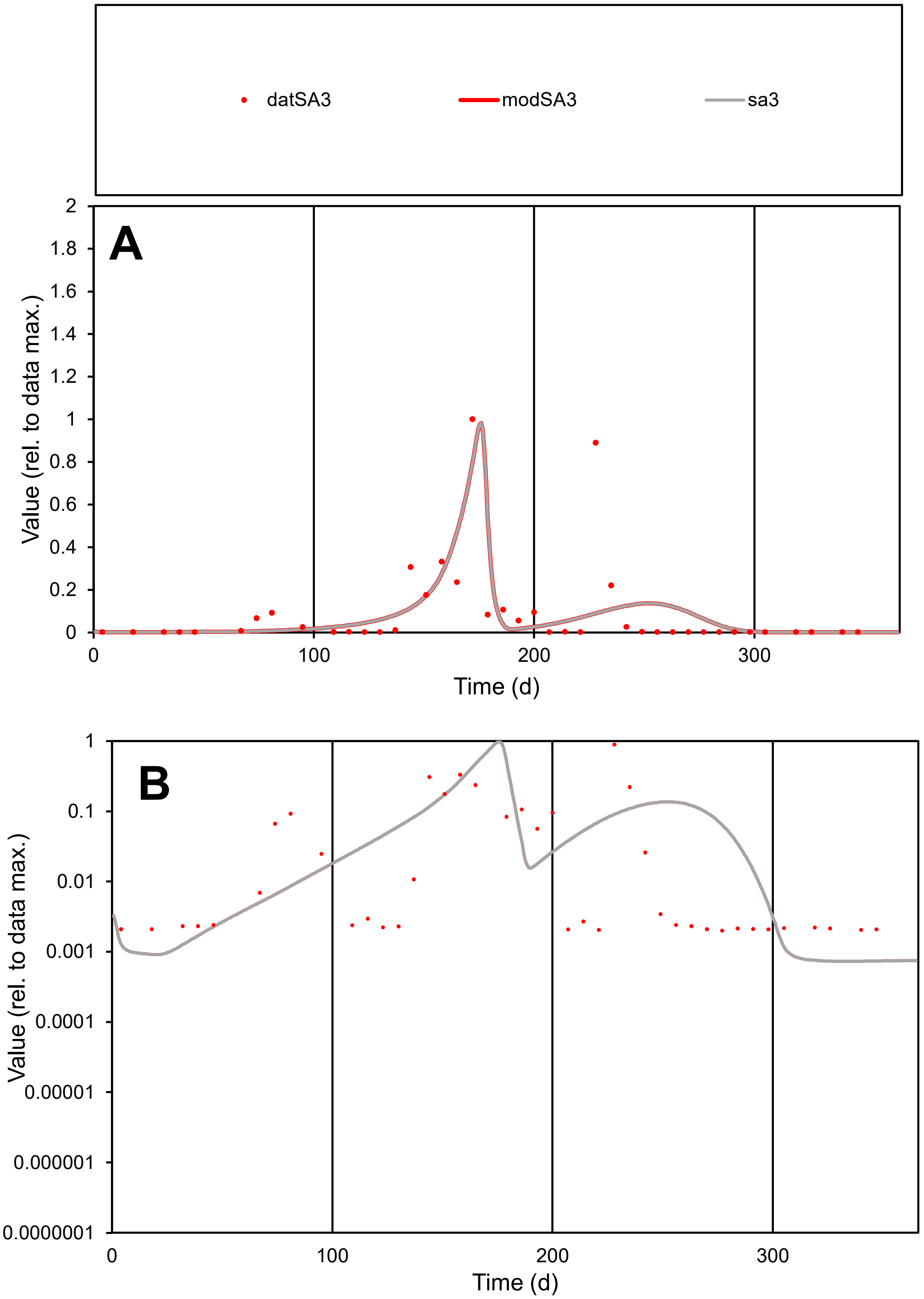


**C1**

Variables
SA3 Provide jtd or Dst
CalibOnly?
2 1 to reproduce of
oNormalize?
2 1=no, 2=yes

ofxn calc*
1.61209
ofxn mod
1.63708

sRun
RunA1
zout
1

Ave
Data
0.00459
Model
0.00438

data/vMax
0.05091

ofxng
334.07

Min d/m
2.7263

ave/min d
45.09
ave/min m
117.194

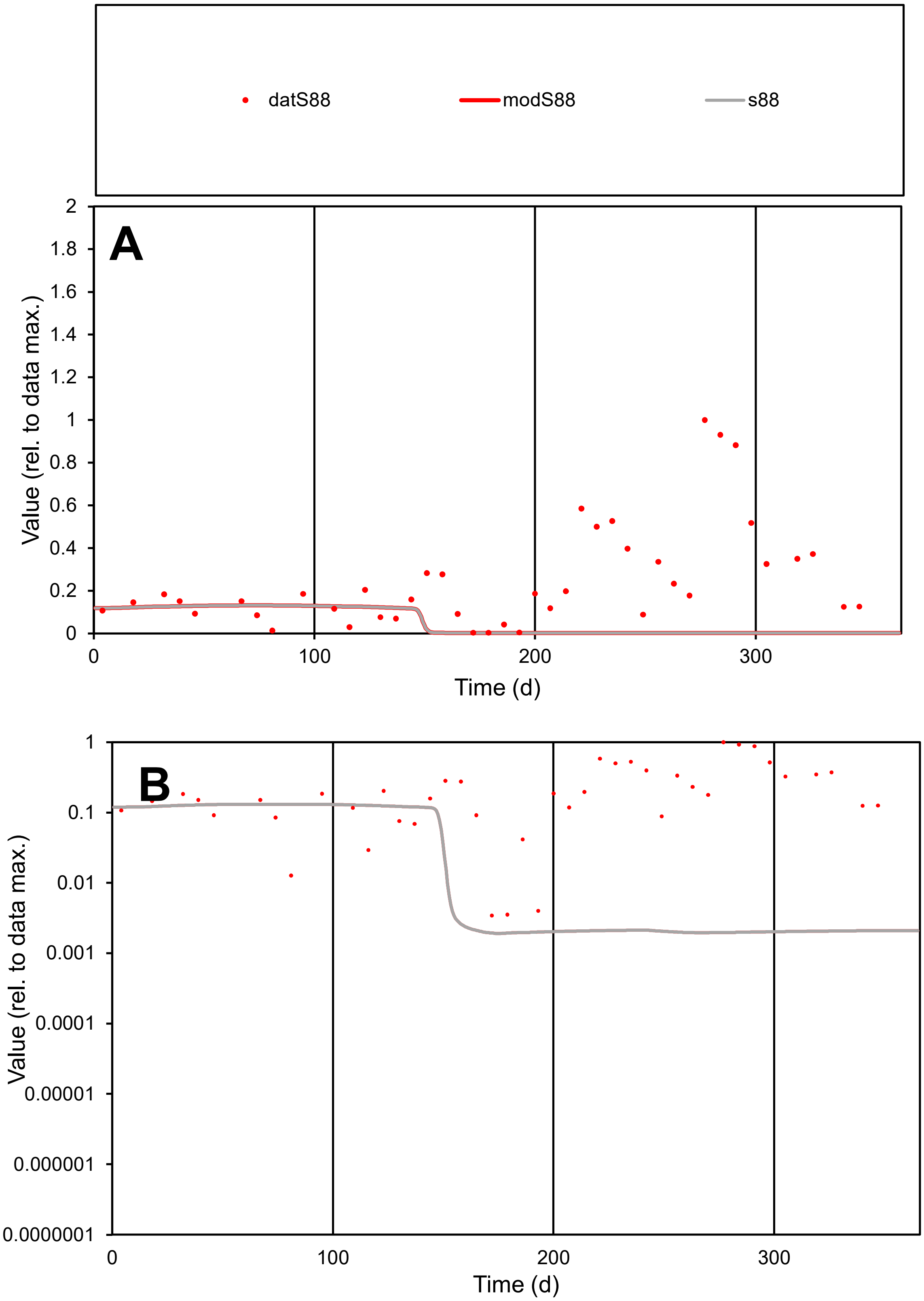


**C1**

Variables

S88 Provide jtd or Dst

CalibOnly?

2 1 to reproduce of

oNormalize?

2 1=no, 2=yes

ofxn calc*

1.35718

ofxn mod

1.36183

sRun

RunA1

zout

1

Ave

Data

0.00423

Model

0.00081

data/vMax

0.01702

ofxng

334.07

Min d/m

1.7902

ave/min d

72.3612

ave/min m

24.7843

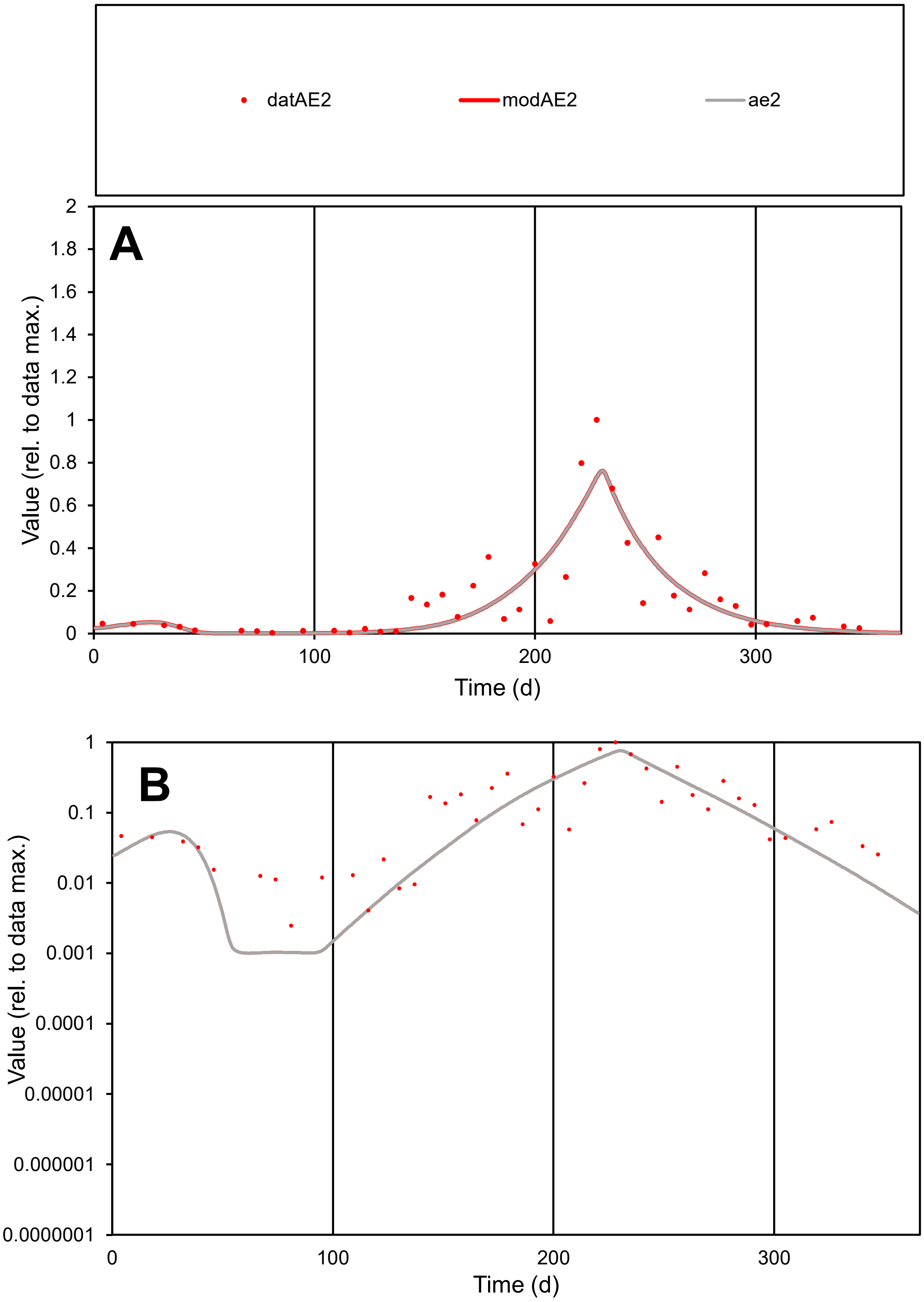


**C1**

Variables
AE2 Provide jtd or Dst
CalibOnly?
2 1 to reproduce of
oNormalize?
2 1=no, 2=yes

ofxn calc*
0.67356
ofxn mod
0.70117

sRun
RunA1
zout
1

Ave
Data
0.00714
Model
0.00656

data/vMax
0.04362

ofxng
334.07

Min d/m
2.43171

ave/min d
65.7685
ave/min m
146.976

327_Calib_S_A

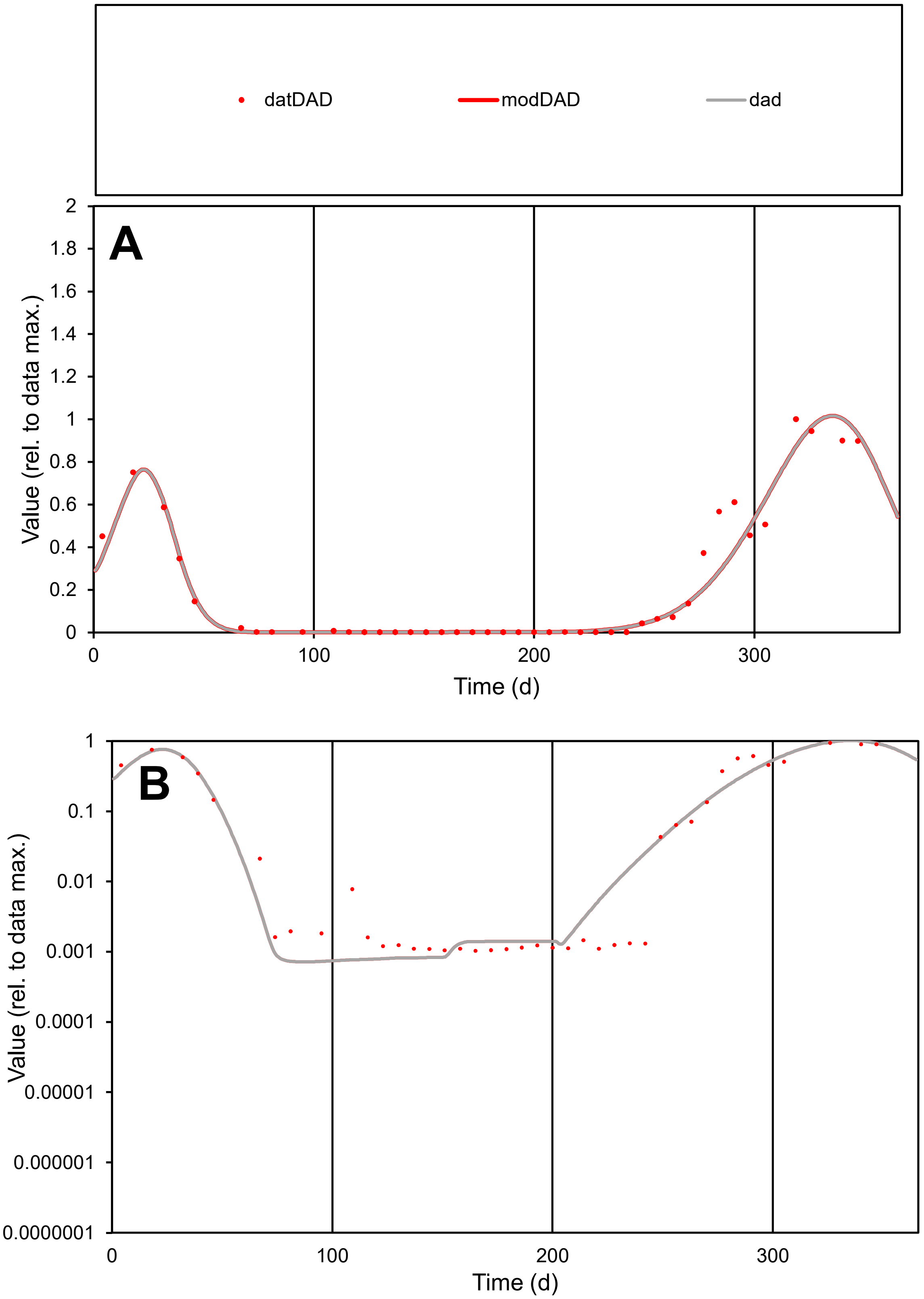


**C1**

Variables
DAD Provide jtd or Dst
CalibOnly?
2 1 to reproduce of
oNormalize?
2 1=no, 2=yes

ofxn calc*
0.31929
ofxn mod
0.36836

sRun
RunA1
zout
1

Ave
Data
0.00315
Model
0.00299

data/vMax
0.01484

ofxng
334.07

Min d/m
1.4234

ave/min d
205.29
ave/min m
277.796

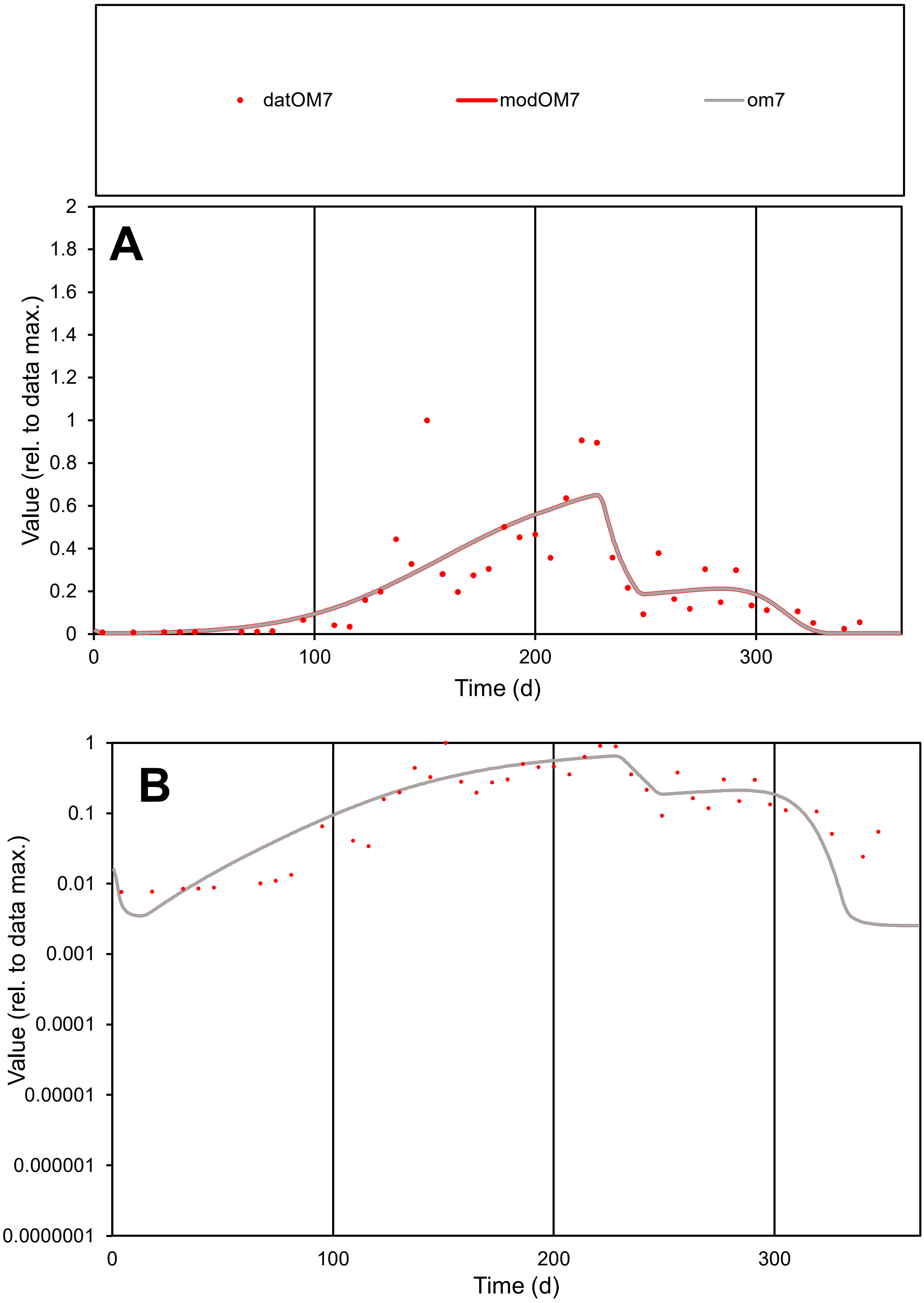


**C1**

Variables
OM7 Provide jtd or Dst
CalibOnly?
2 1 to reproduce of
oNormalize?
2 1=no, 2=yes

ofxn calc*
0.61187
ofxn mod
0.62974

sRun
RunA1
zout
1

Ave
Data
0.00319
Model
0.00316

data/vMax
0.01318

ofxng
334.07

Min d/m
2.93639

ave/min d
31.5425
ave/min m
91.6445

327_Calib_S_A

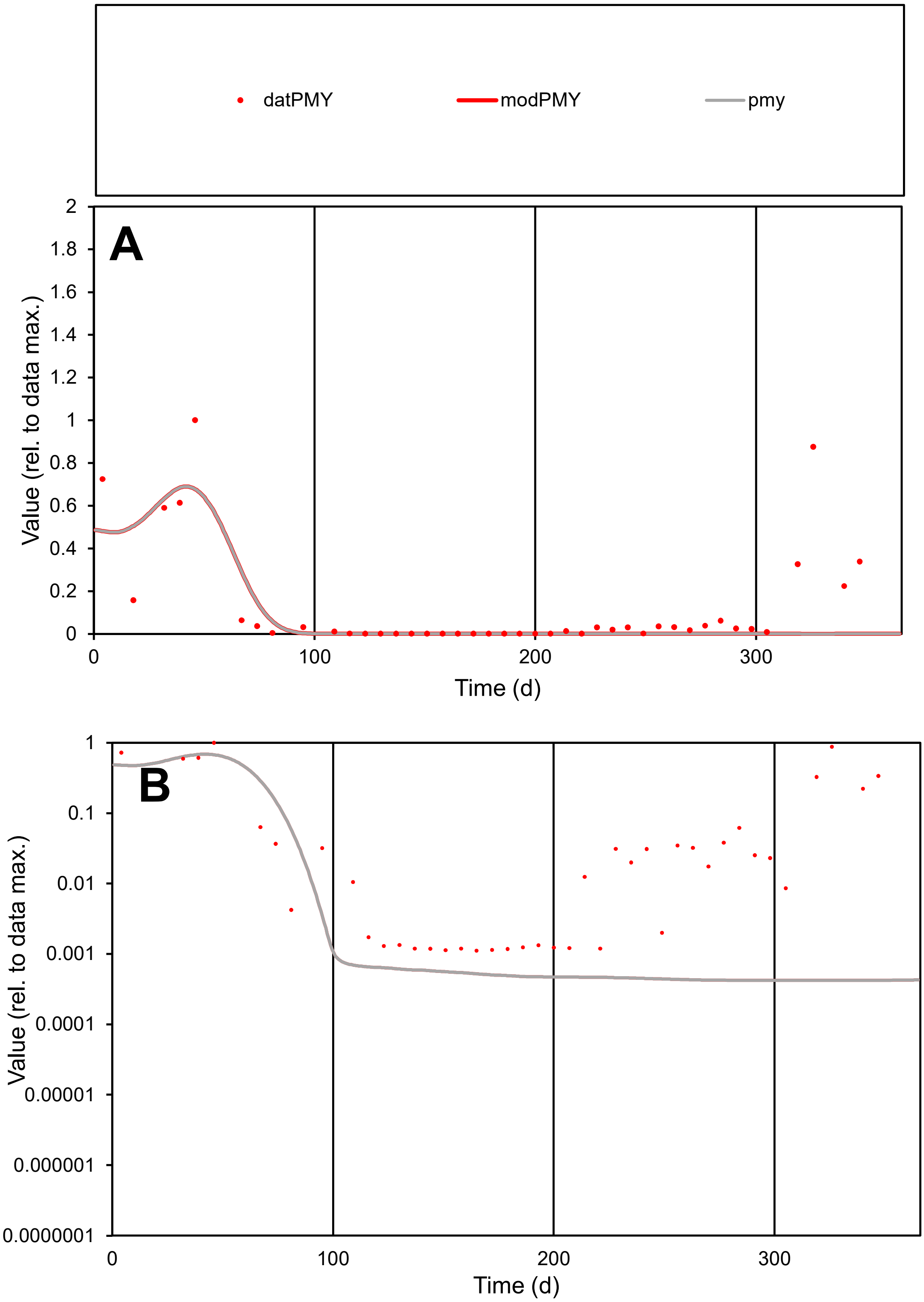


**C1**

Variables

PMY Provide jtd or Dst

CalibOnly?

2 1 to reproduce of

oNormalize?

2 1=no, 2=yes

ofxn calc*

1.42688

ofxn mod

1.44752

sRun

RunA1

zout

1

Ave

Data

0.00109

Model

0.00069

data/vMax

0.00856

ofxng

334.07

Min d/m

2.65628

ave/min d

114.228

ave/min m

192.79

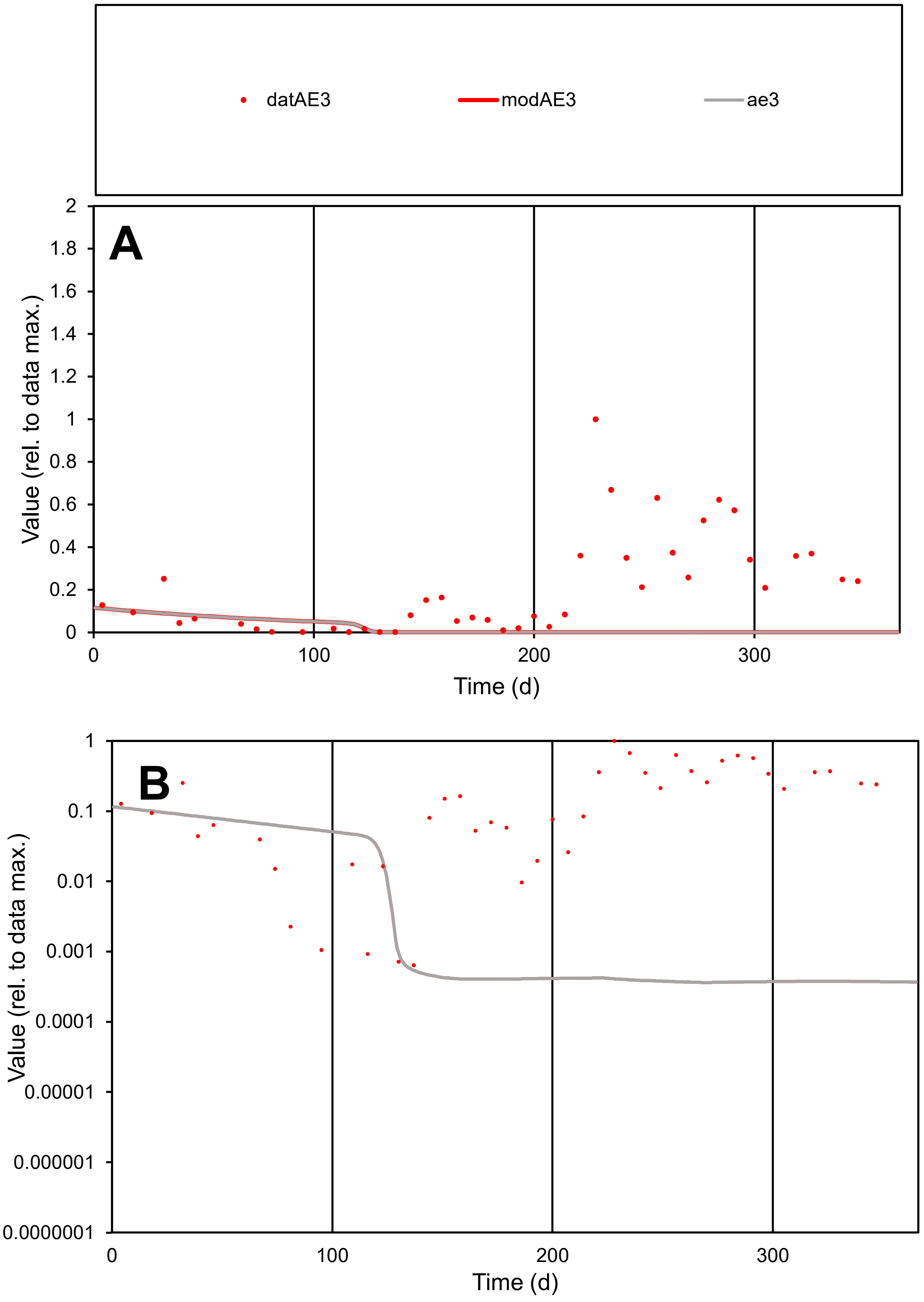


**C1**

Variables
AE3 Provide jtd or Dst
CalibOnly?
2 1 to reproduce of
oNormalize?
2 1=no, 2=yes

ofxn calc*
1.47184
ofxn mod
1.47423

sRun
RunA1
zout
1

Ave
Data
0.00585
Model
0.00054

data/vMax
0.02793

ofxng
334.07

Min d/m
1.76106

ave/min d
326.704
ave/min m
53

327_Calib_S_A

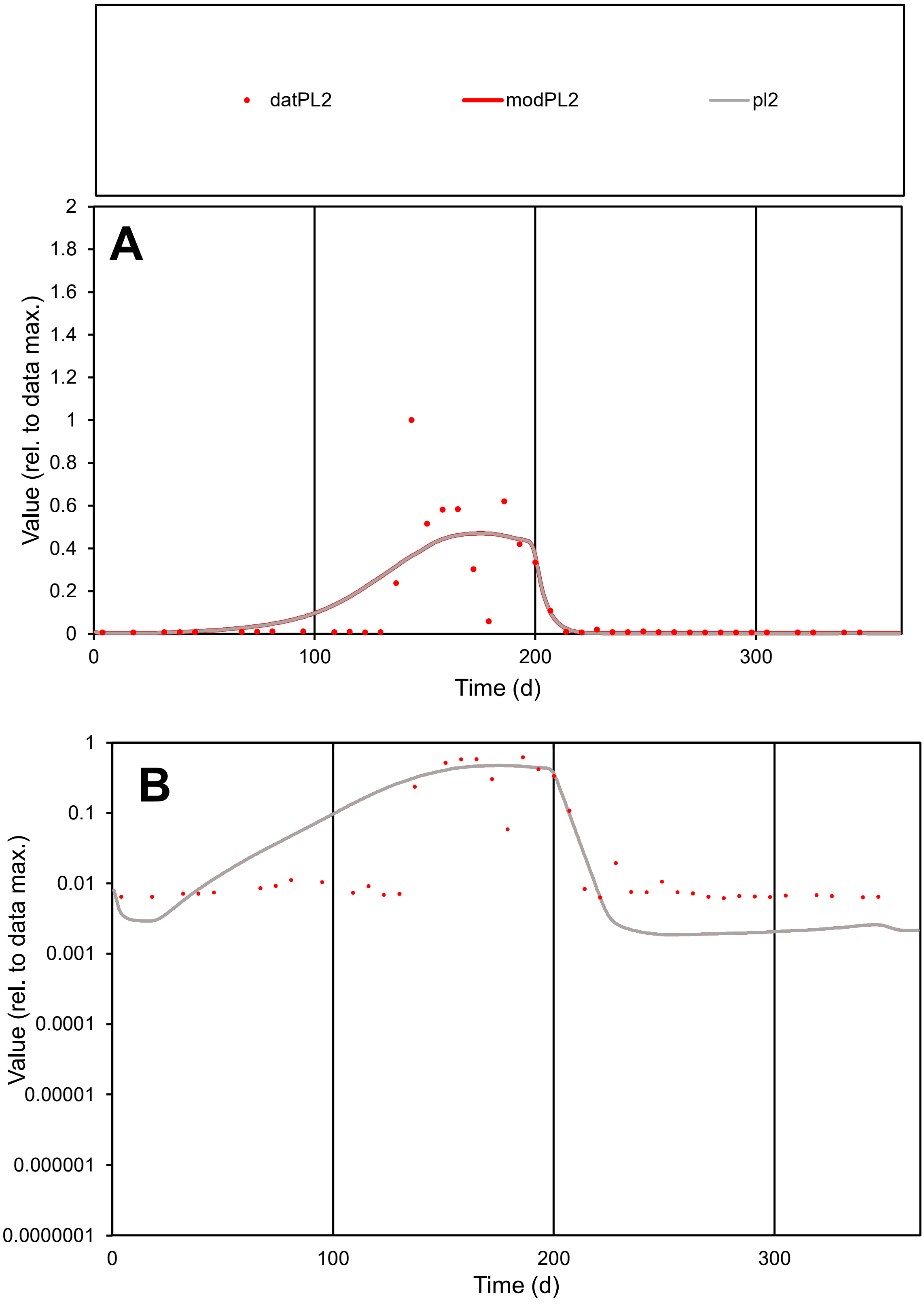


**C1**

Variables
PL2 Provide jtd or Dst
CalibOnly?
2 1 to reproduce of
oNormalize?
2 1=no, 2=yes

ofxn calc*
1.18079
ofxn mod
1.2018

sRun
RunA1
zout
1

Ave
Data
0.00326
Model
0.00349

data/vMax
0.02733

ofxng
334.07

Min d/m
3.31174

ave/min d
19.2973
ave/min m
68.4506

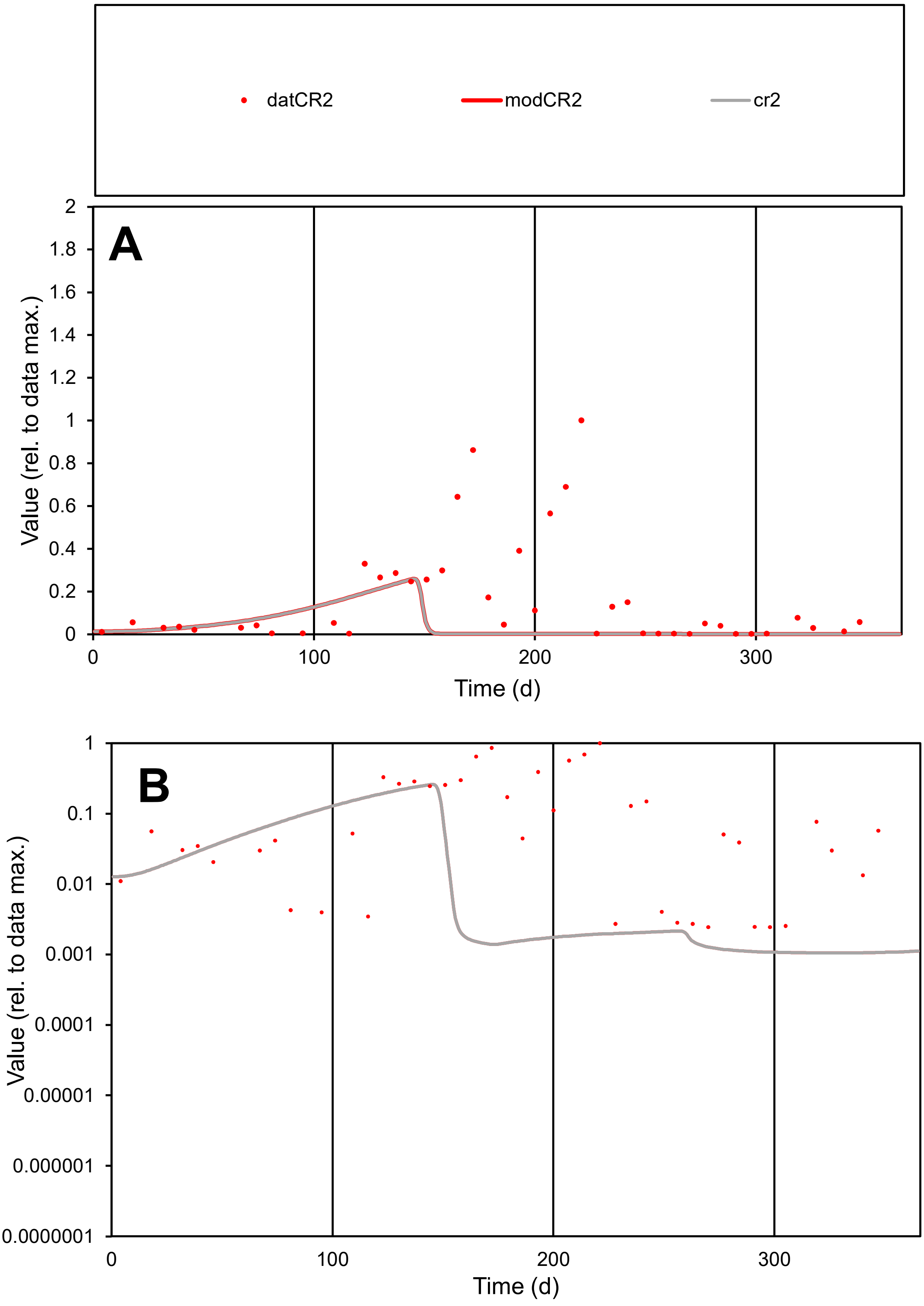


| | |
|---|---|
| **C1** | |
| Variables | |
| CR2 | Provide jtd or Dst |
| CalibOnly? | |
| 2 | 1 to reproduce of |
| oNormalize? | |
| 2 | 1=no, 2=yes |
| ofxn calc* | |
| 1.69508 | |
| ofxn mod | |
| 1.70247 | |
| sRun | |
| RunA1 | |
| zout | |
| 1 | |
| Ave | |
| Data | |
| 0.00295 | |
| Model | |
| 0.00079 | |
| data/vMax | |
| 0.01765 | |
| ofxng | |
| 334.07 | |
| Min d/m | |
| 2.3247 | |
| ave/min d | |
| 68.3841 | |
| ave/min m | |
| 42.7066 | |

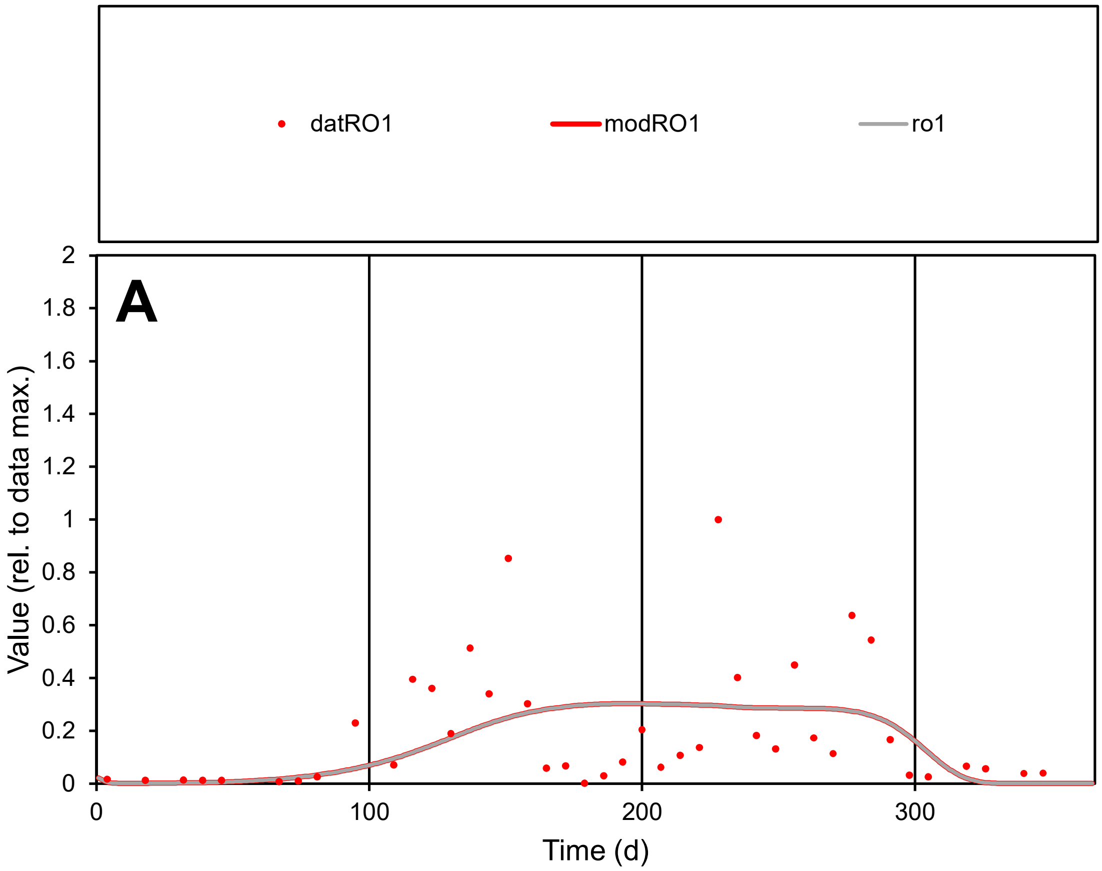


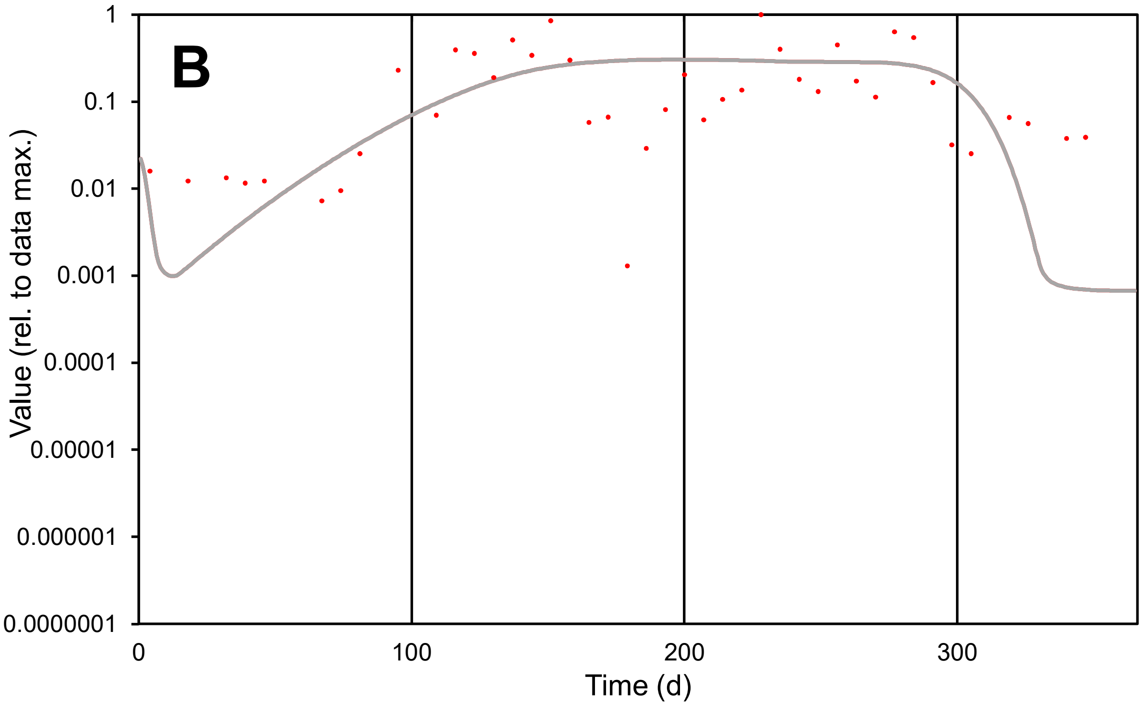


**C1**

Variables
RO1 Provide jtd or Dst
CalibOnly?
2 1 to reproduce of
oNormalize?
2 1=no, 2=yes

ofxn calc*
1.10173
ofxn mod
1.10943

sRun
RunA1
zout
1

Ave
Data
0.00554
Model
0.00503

data/vMax
0.02855

ofxng
334.07

Min d/m
1.88878

ave/min d
149.694
ave/min m
256.857

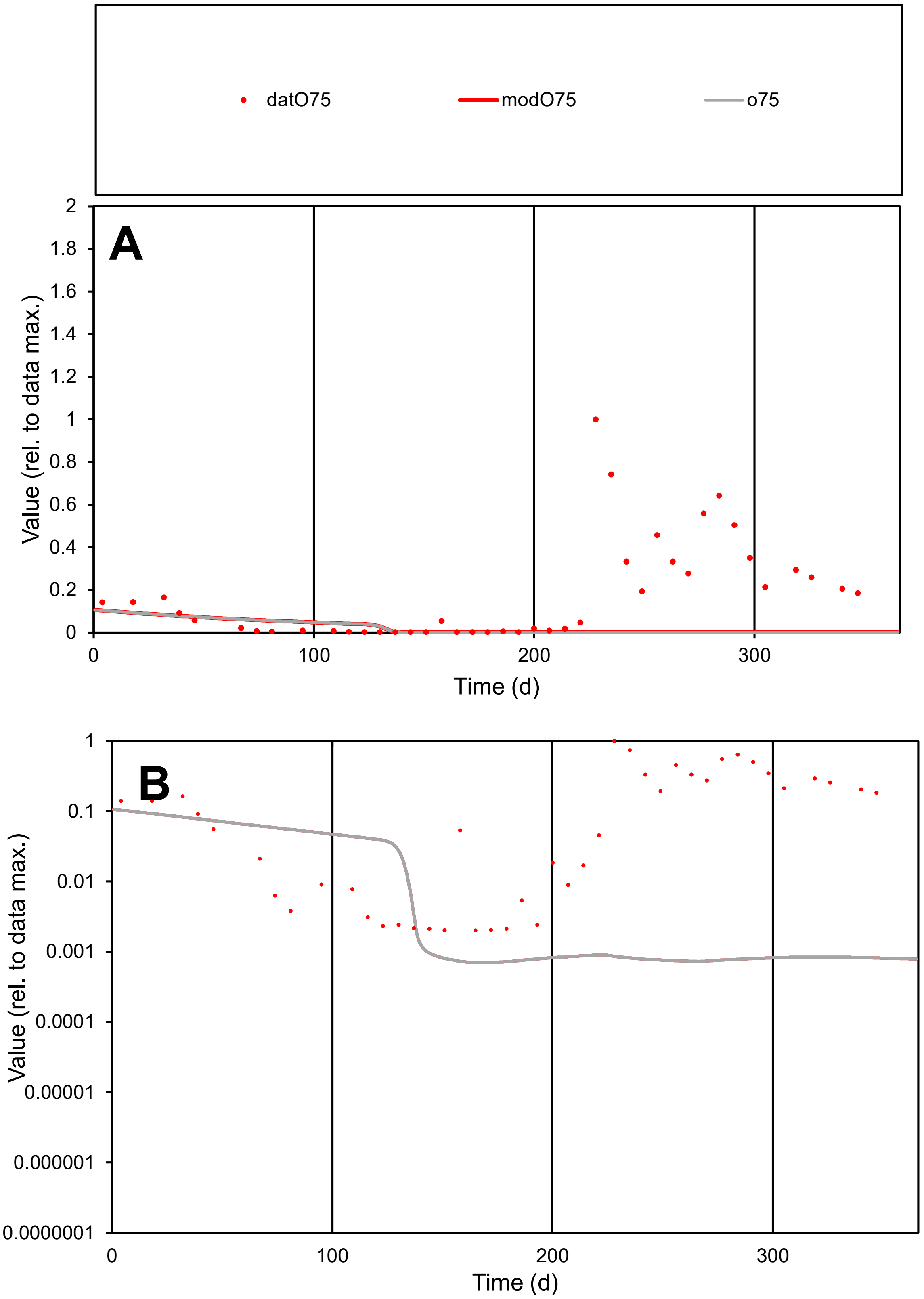


**C1**

Variables

O75 Provide jtd or Dst

CalibOnly?

2 1 to reproduce of

oNormalize?

2 1=no, 2=yes

ofxn calc*

1.64564

ofxn mod

1.64825

sRun

RunA1

zout

1

Ave

Data

0.00605

Model

0.00067

data/vMax

0.03458

ofxng

334.07

Min d/m

2.88884

ave/min d

86.8731

ave/min m

27.6932

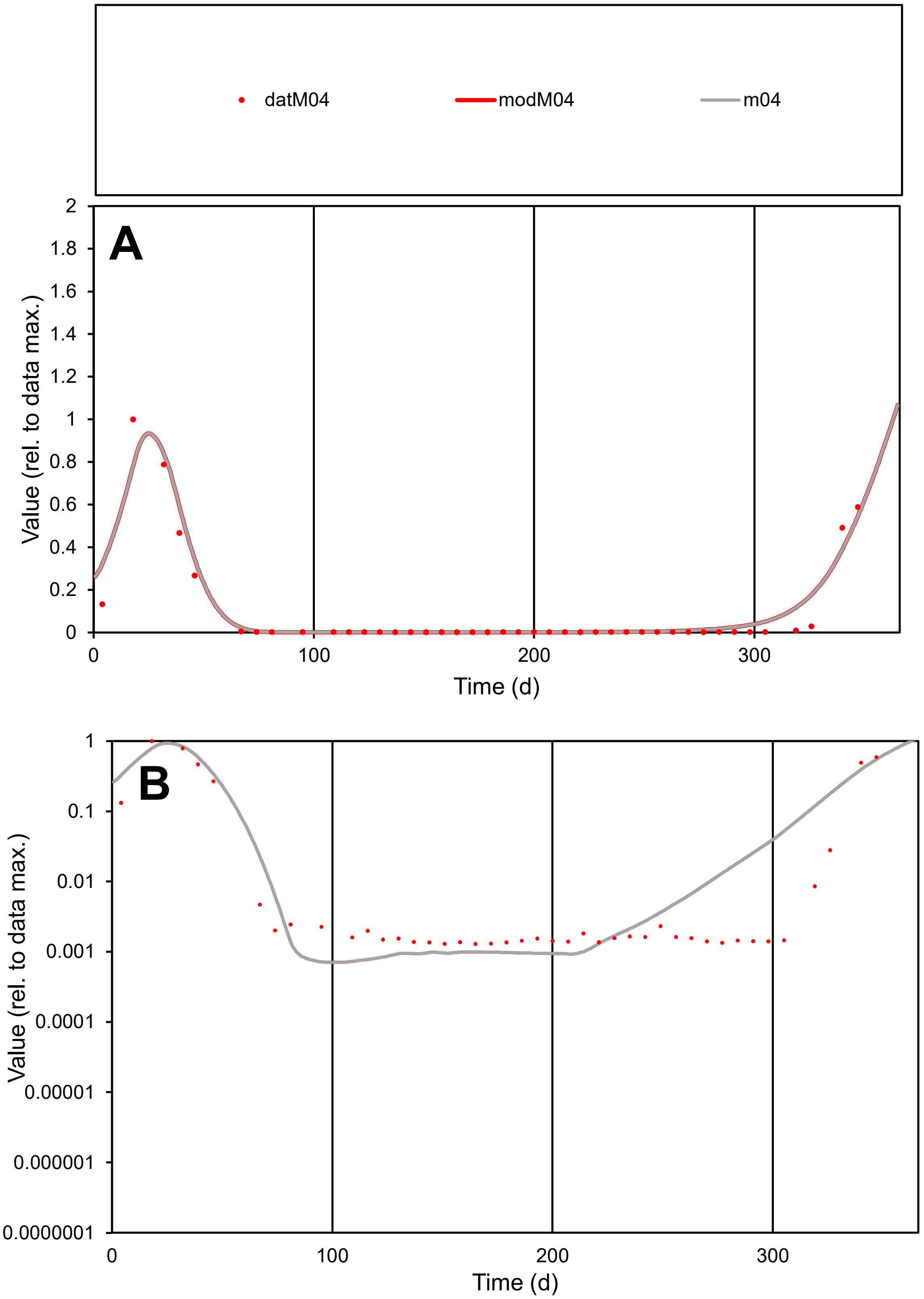


**C1**

Variables
M04 Provide jtd or Dst
CalibOnly?
2 1 to reproduce of
oNormalize?
2 1=no, 2=yes

ofxn calc*
0.56139
ofxn mod
0.68022

sRun
RunA1
zout
1

Ave
Data
0.00248
Model
0.00278

data/vMax
0.02722

ofxng
334.07

Min d/m
1.80103

ave/min d
70.7506
ave/min m
143.054

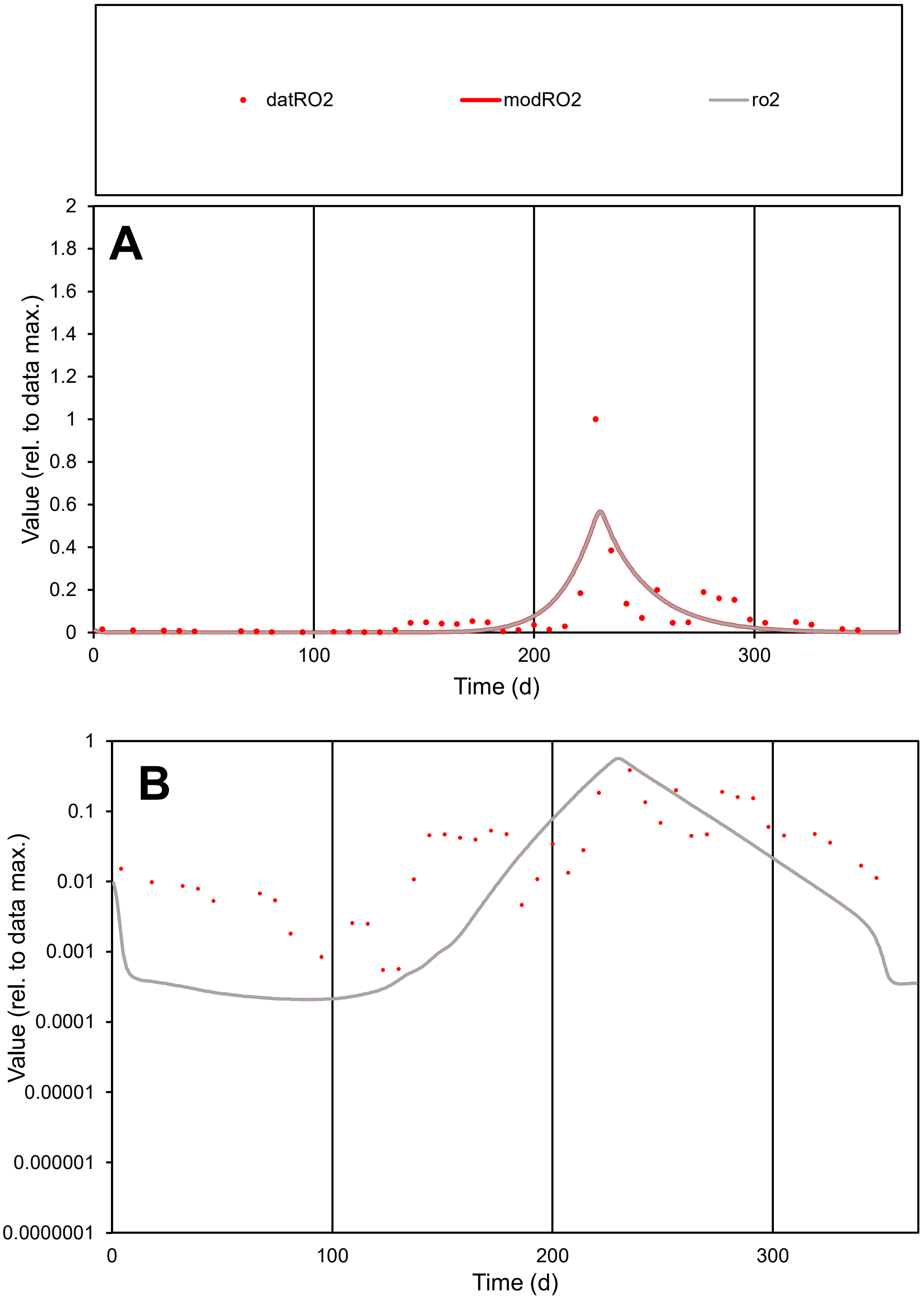


**C1**

Variables
RO2 Provide jtd or Dst
CalibOnly?
2 1 to reproduce of
oNormalize?
2 1=no, 2=yes

ofxn calc*
1.24528
ofxn mod
1.28722

sRun
RunA1
zout
1

Ave
Data
0.00616
Model
0.0056

data/vMax
0.08014

ofxng
334.07

Min d/m
2.63777

ave/min d
138.568
ave/min m
332.414

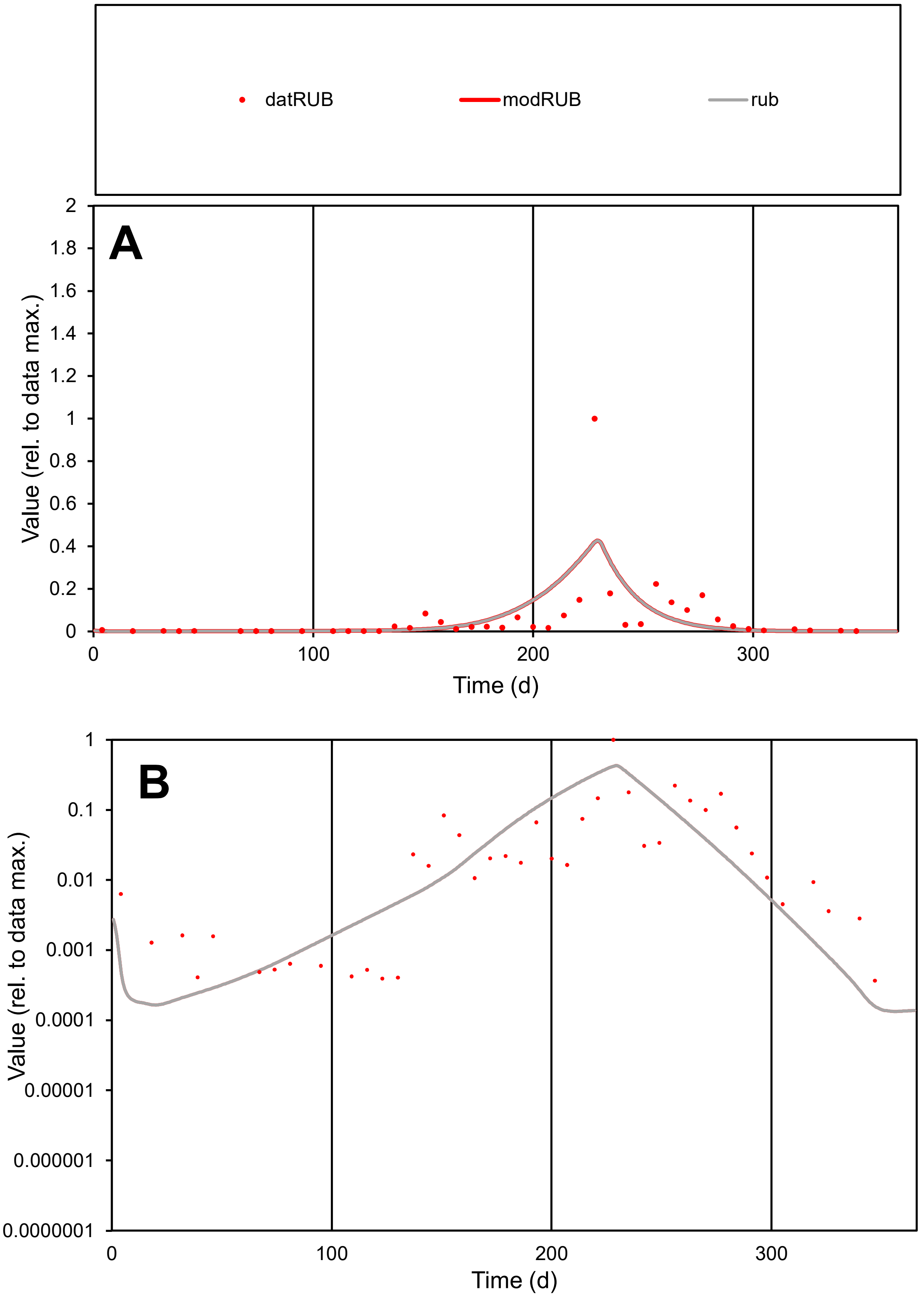


**C1**

Variables

RUB Provide jtd or Dst

CalibOnly?

2 1 to reproduce of

oNormalize?

2 1=no, 2=yes

ofxn calc*

1.8966

ofxn mod

1.92383

sRun

RunA1

zout

1

Ave

Data

0.00178

Model

0.00181

data/vMax

0.02926

ofxng

334.07

Min d/m

2.39024

ave/min d

165.534

ave/min m

400.598

327_Calib_S_A

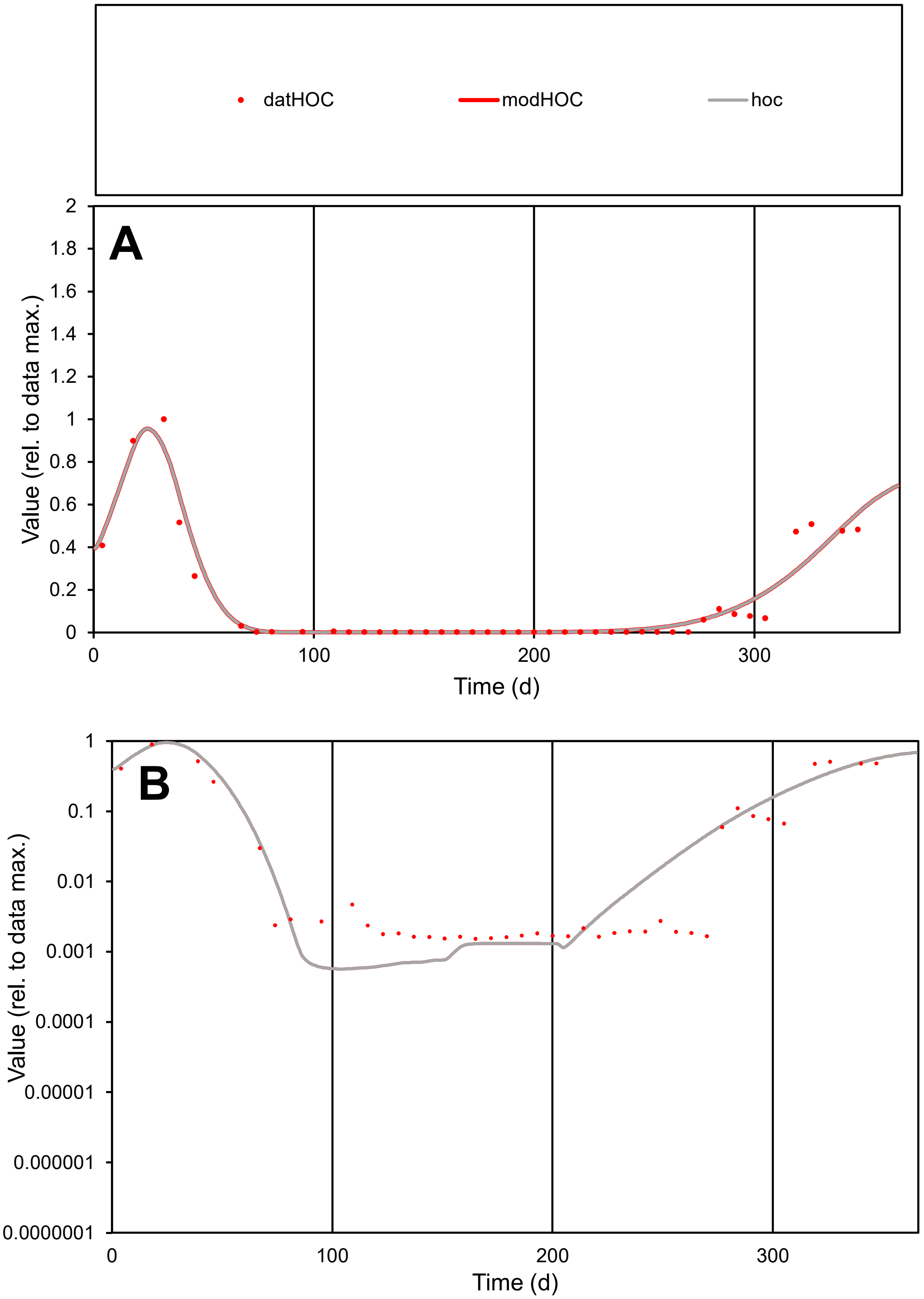


**C1**

Variables
HOC Provide jtd or Dst
CalibOnly?
2 1 to reproduce of
oNormalize?
2 1=no, 2=yes

ofxn calc*
0.41861
ofxn mod
0.49961

sRun
RunA1
zout
1

Ave
Data
0.00107
Model
0.00111

data/vMax
0.00817

ofxng
334.07

Min d/m
2.63948

ave/min d
85.8817
ave/min m
235.43

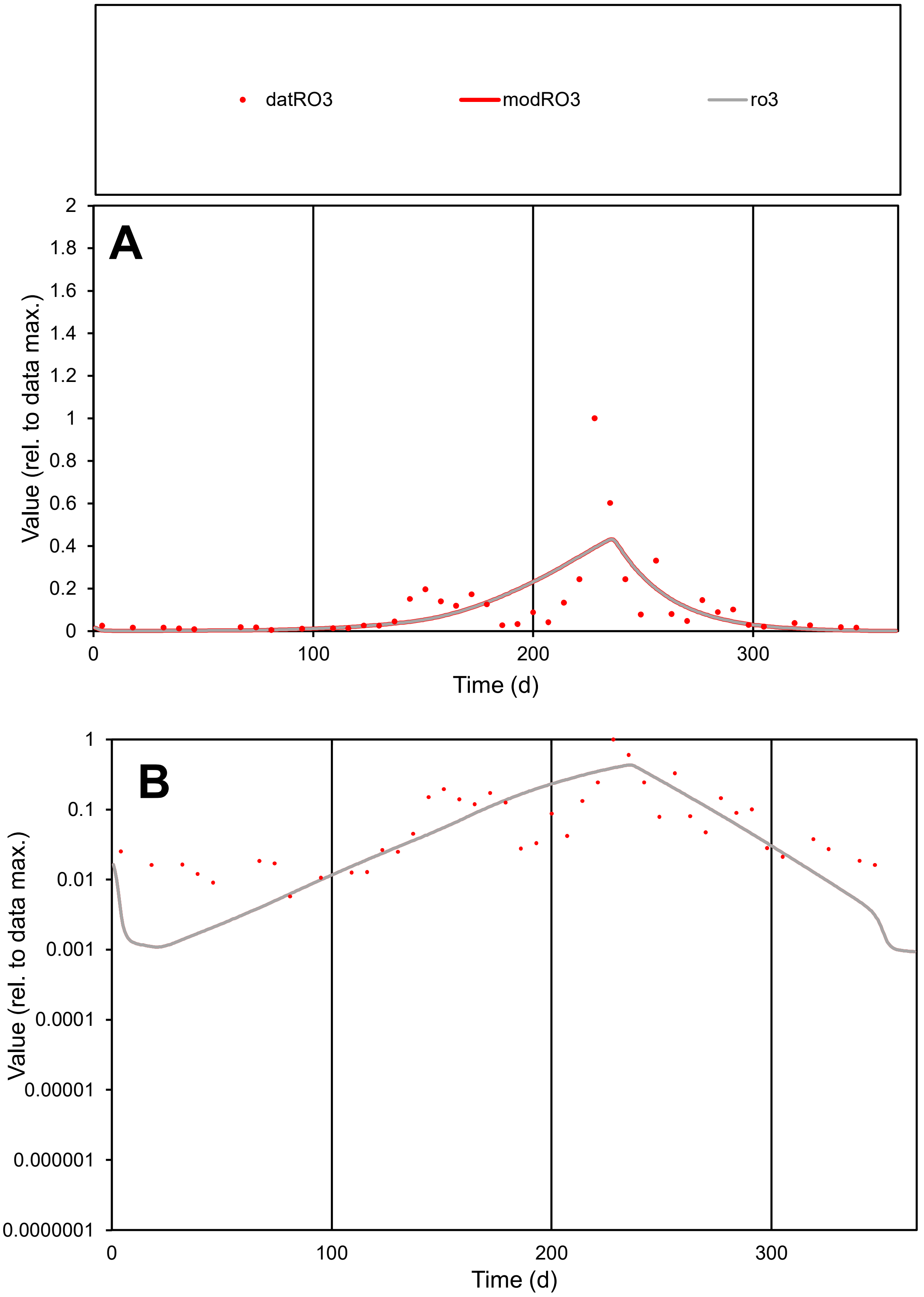


**C1**

Variables
RO3 Provide jtd or Dst
CalibOnly?
2 1 to reproduce of
oNormalize?
2 1=no, 2=yes

ofxn calc*
1.15649
ofxn mod
1.17189

sRun
RunA1
zout
1

Ave
Data
0.00685
Model
0.00651

data/vMax
0.06268

ofxng
334.07

Min d/m
5.26346

ave/min d
18.9486
ave/min m
94.7119

327_Calib_S_A

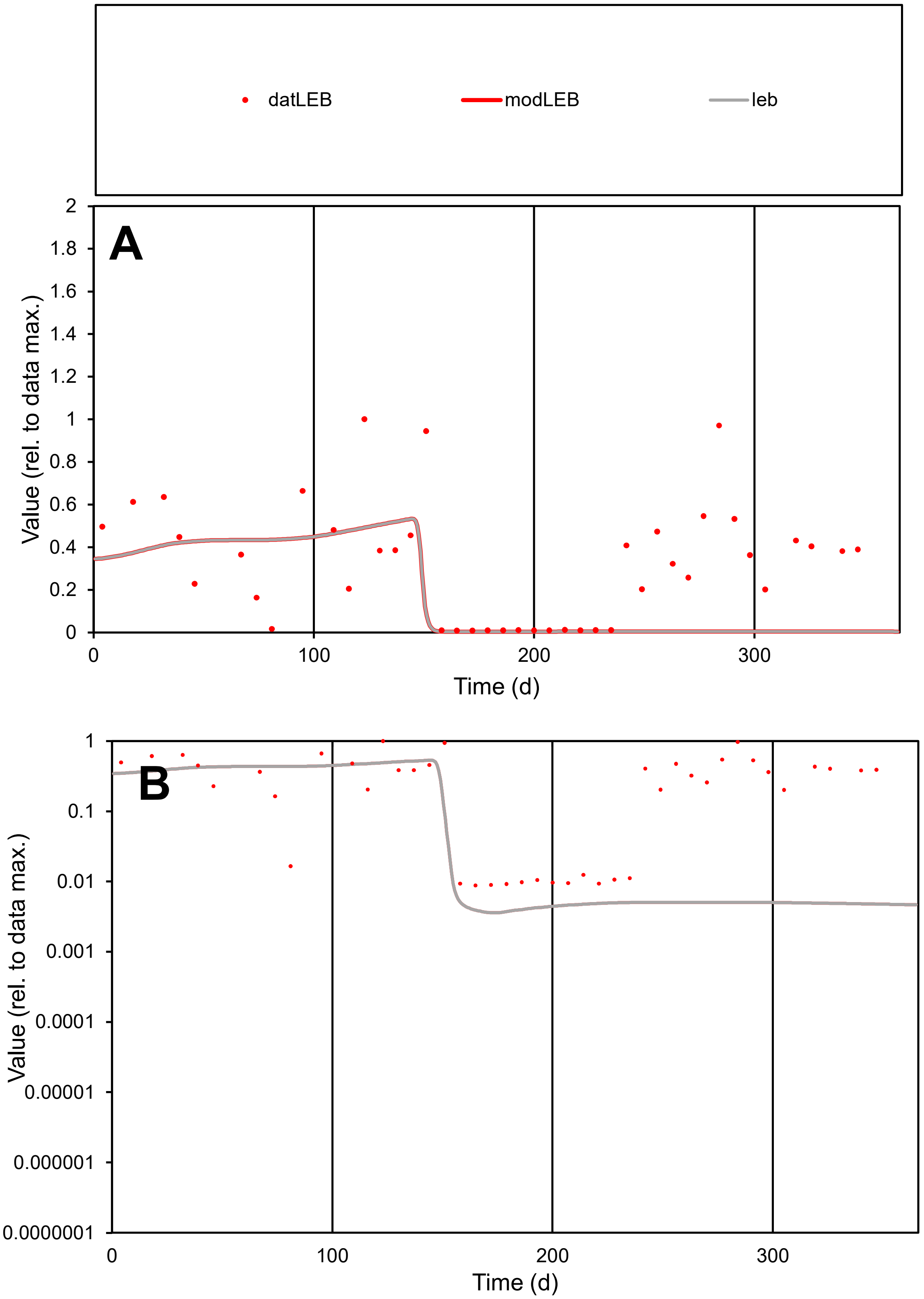


**C1**

Variables
LEB Provide jtd or Dst
CalibOnly?
2 1 to reproduce of
oNormalize?
2 1=no, 2=yes

ofxn calc*
0.98043
ofxn mod
1.00706

sRun
RunA1
zout
1

Ave
Data
0.00254
Model
0.00135

data/vMax
0.0079

ofxng
334.07

Min d/m
2.44339

ave/min d
36.6306
ave/min m
47.4924

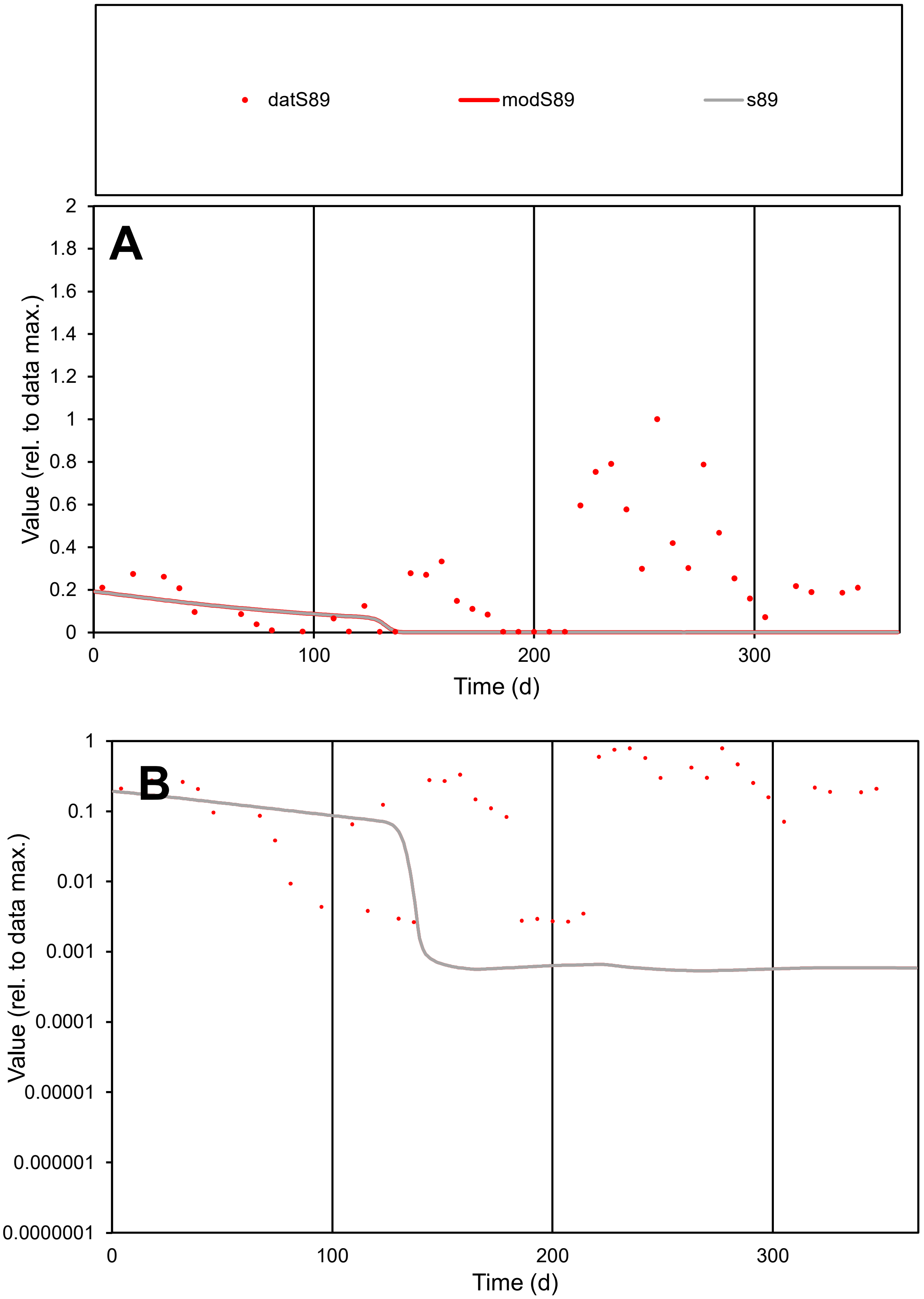


**C1**

Variables
S89 Provide jtd or Dst
CalibOnly?
2 1 to reproduce of
oNormalize?
2 1=no, 2=yes

ofxn calc*
1.41624
ofxn mod
1.41947

sRun
RunA1
zout
1

Ave
Data
0.00341
Model
0.0005

data/vMax
0.01448

ofxng
334.07

Min d/m
4.9593

ave/min d
88.922
ave/min m
64.8197

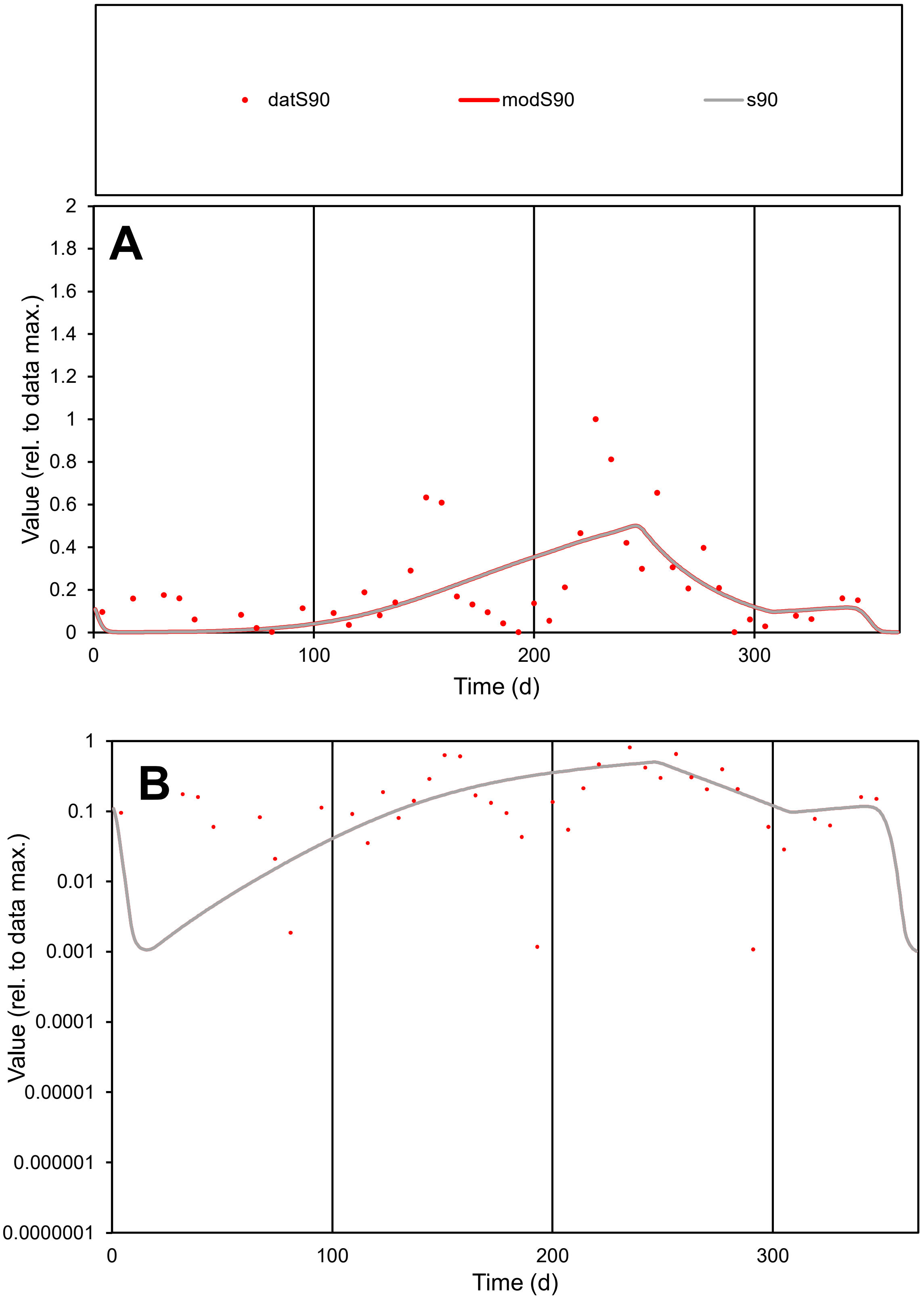


**C1**

Variables
S90 Provide jtd or Dst
CalibOnly?
2 1 to reproduce of
oNormalize?
2 1=no, 2=yes

ofxn calc*
0.87822
ofxn mod
0.89092

sRun
RunA1
zout
1

Ave
Data
0.00435
Model
0.00393

data/vMax
0.02012

ofxng
334.07

Min d/m
0.92015

ave/min d
199.854
ave/min m
165.95

327_Calib_S_A

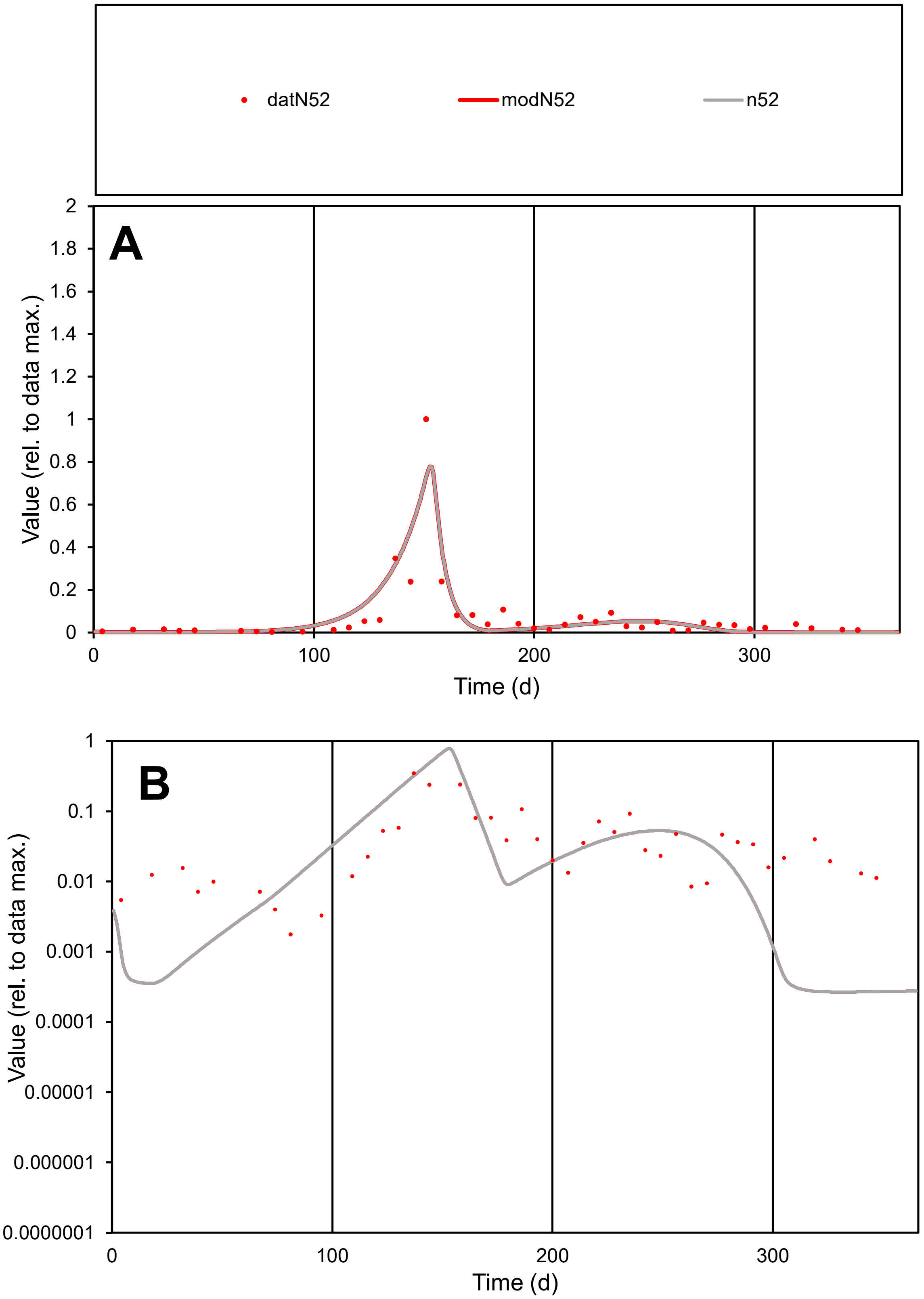


**C1**

Variables
N52 Provide jtd or Dst
CalibOnly?
2 1 to reproduce of
oNormalize?
2 1=no, 2=yes

ofxn calc*
0.9247
ofxn mod
0.95859

sRun
RunA1
zout
1

Ave
Data
0.00366
Model
0.00373

data/vMax
0.05091

ofxng
334.07

Min d/m
6.64632

ave/min d
40.5458
ave/min m
274.53

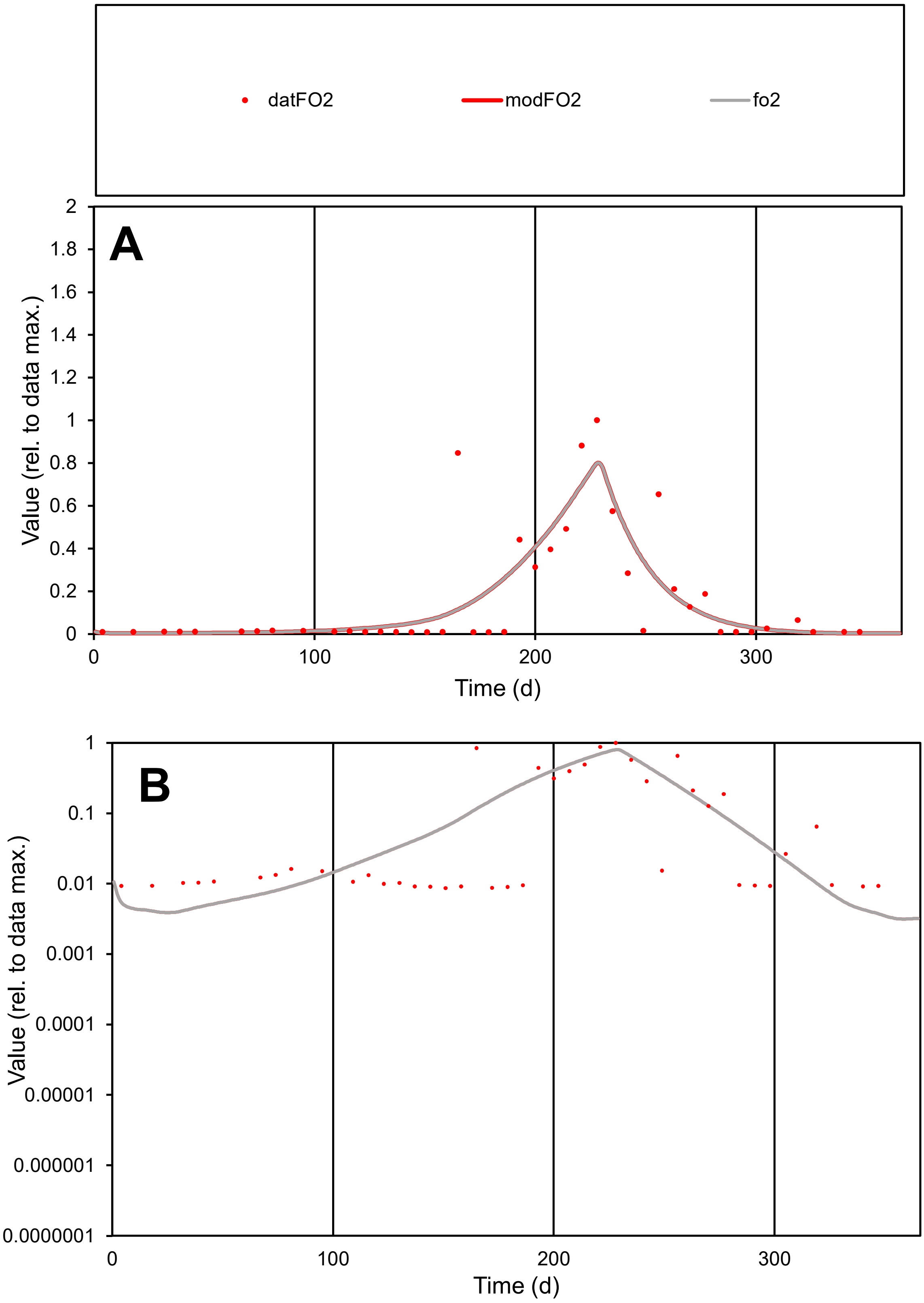


**C1**

Variables
FO2 Provide jtd or Dst
CalibOnly?
2 1 to reproduce of
oNormalize?
2 1=no, 2=yes

ofxn calc*
0.98307
ofxn mod
1.01214

sRun
RunA1
zout
1

Ave
Data
0.00126
Model
0.00124

data/vMax
0.0078

ofxng
334.07

Min d/m
2.34179

ave/min d
18.7182
ave/min m
43.0272

327_Calib_S_A

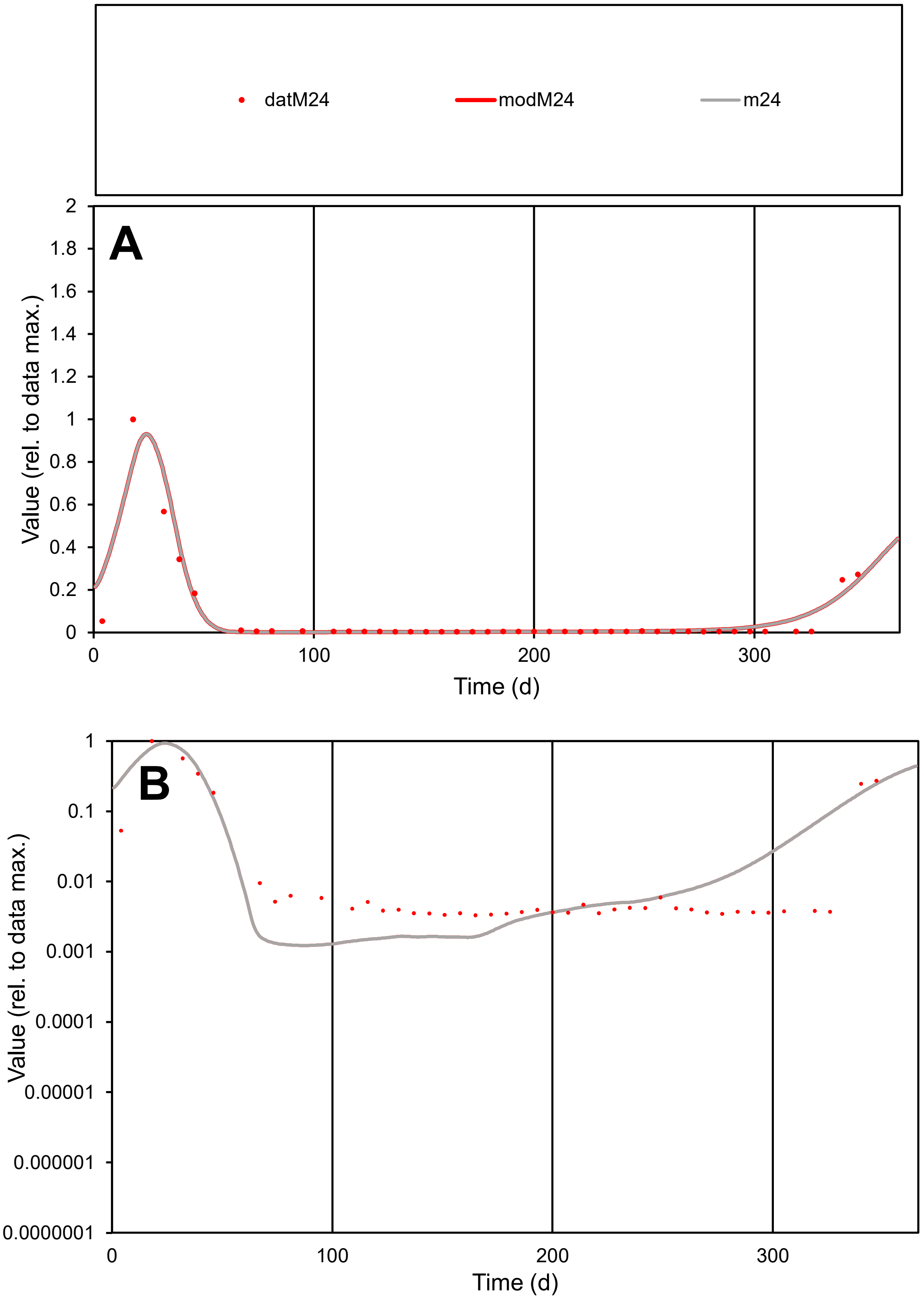


**C1**

Variables
M24 Provide jtd or Dst
CalibOnly?
2 1 to reproduce of
oNormalize?
2 1=no, 2=yes

ofxn calc*
0.67372
ofxn mod
0.84884

sRun
RunA1
zout
1

Ave
Data
0.00145
Model
0.00157

data/vMax
0.02167

ofxng
334.07

Min d/m
2.66803

ave/min d
20.2427
ave/min m
58.4772

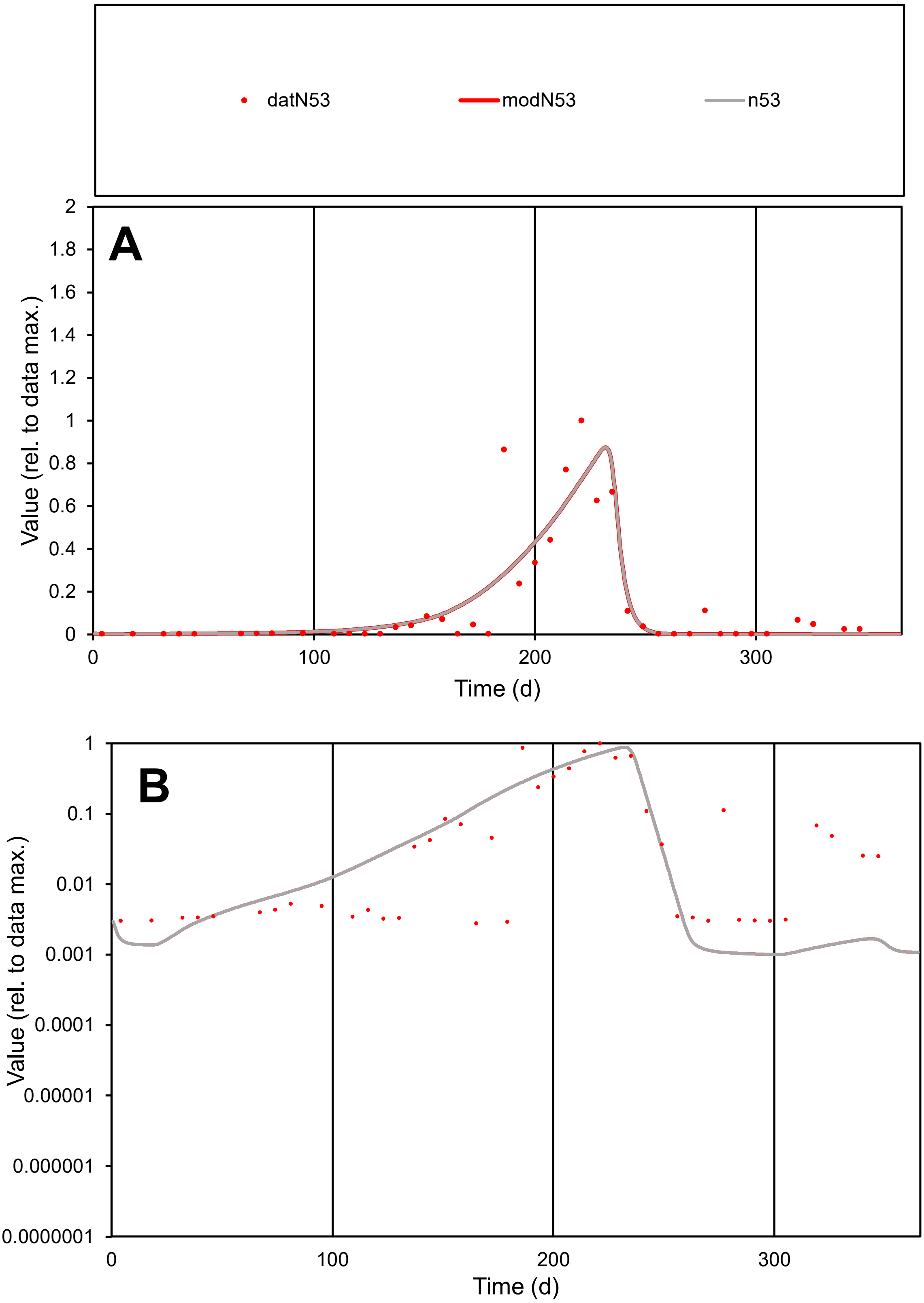


**C1**

Variables

N53 Provide jtd or Dst

CalibOnly?

2 1 to reproduce of

oNormalize?

2 1=no, 2=yes

ofxn calc*

0.86432

ofxn mod

0.90375

sRun

RunA1

zout

1

Ave

Data

0.00177

Model

0.00174

data/vMax

0.01301

ofxng

334.07

Min d/m

2.76104

ave/min d

48.9054

ave/min m

132.168

327_Calib_S_A

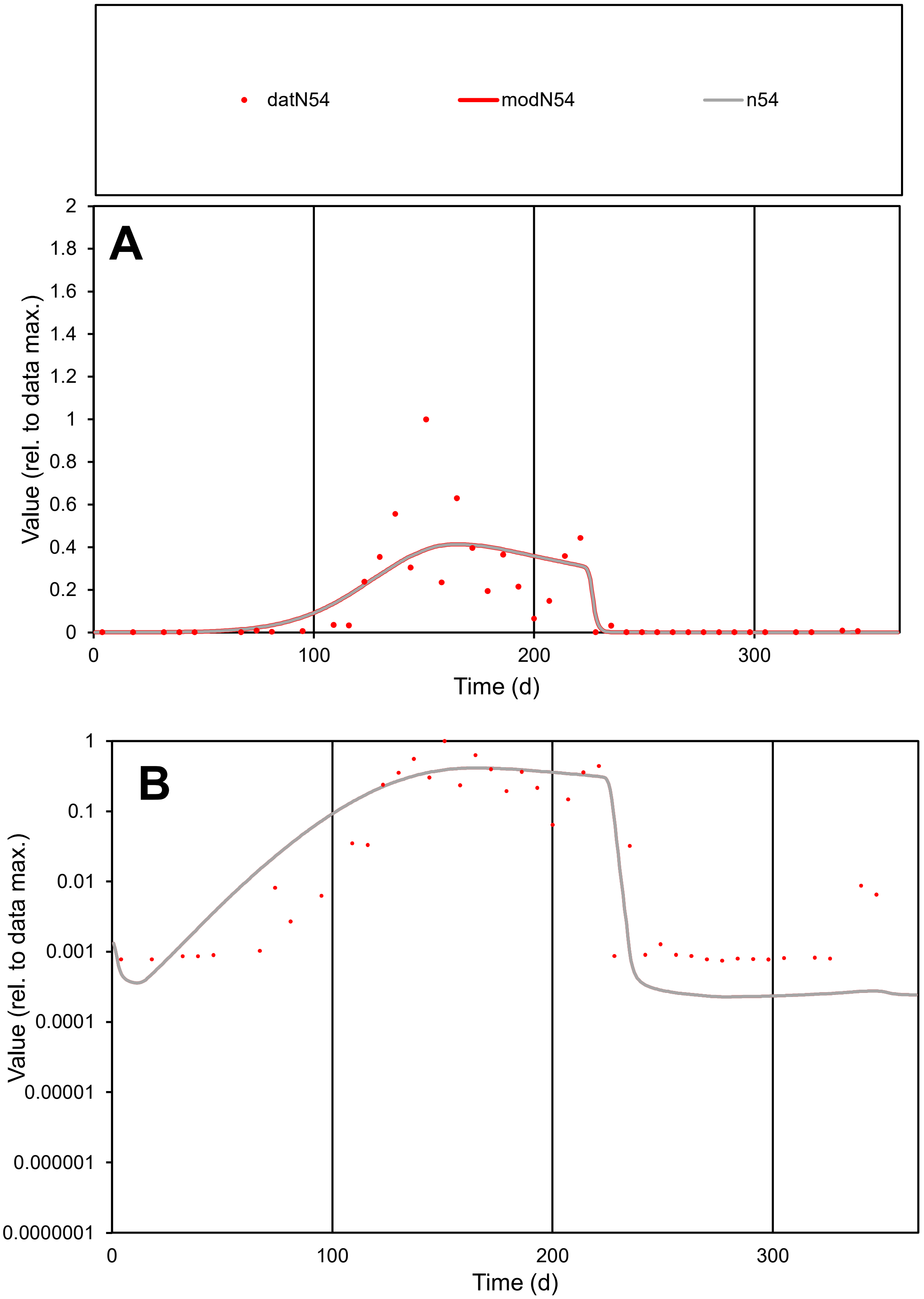


**C1**

Variables

N54 Provide jtd or Dst

CalibOnly?

2 1 to reproduce of

oNormalize?

2 1=no, 2=yes

ofxn calc*

0.99611

ofxn mod

1.01185

sRun

RunA1

zout

1

Ave

Data

0.00405

Model

0.00416

data/vMax

0.03007

ofxng

334.07

Min d/m

3.28114

ave/min d

180.445

ave/min m

608.384

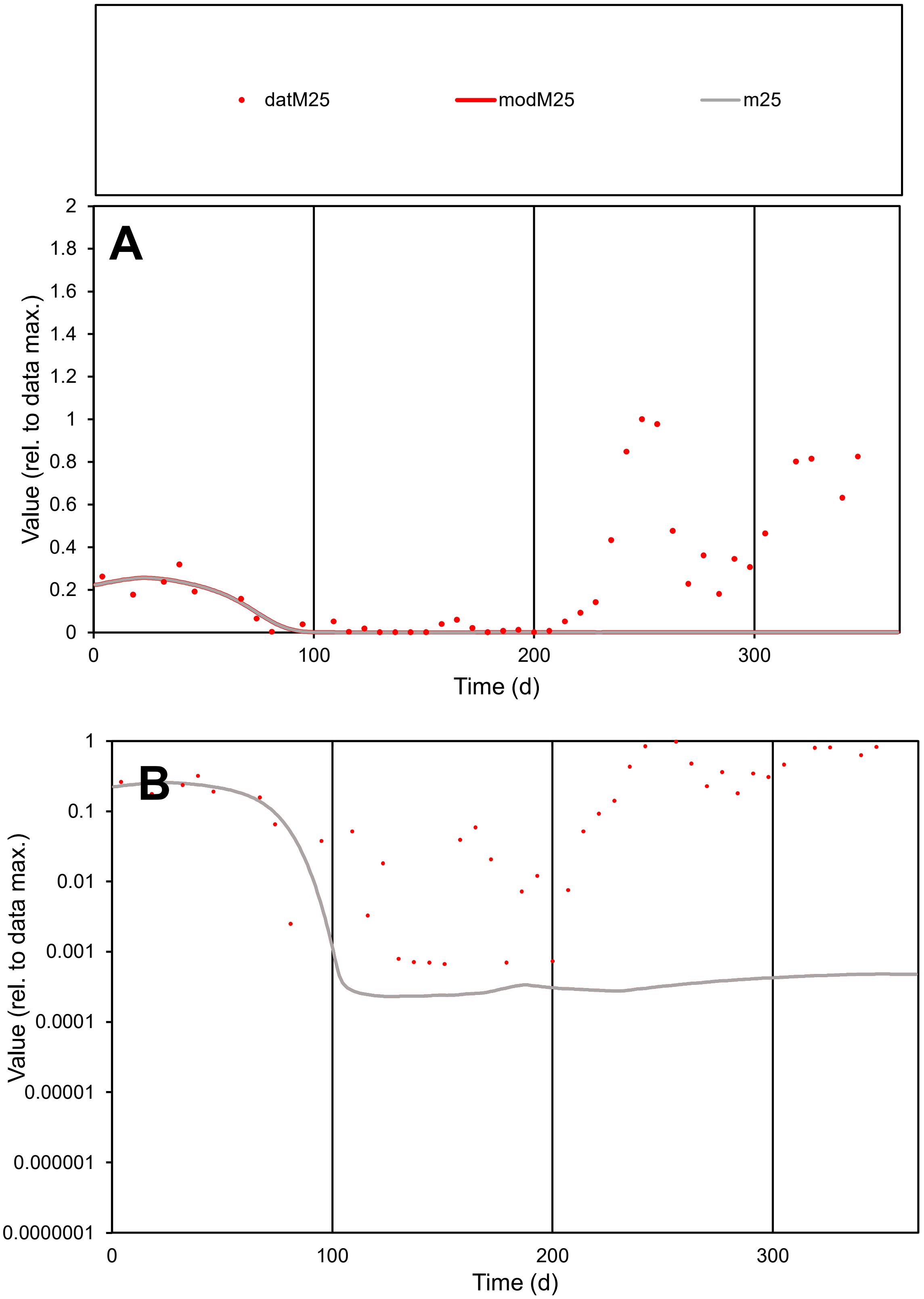


**C1**

Variables
M25 Provide jtd or Dst
CalibOnly?
2 1 to reproduce of
oNormalize?
2 1=no, 2=yes

ofxn calc*
1.51126
ofxn mod
1.5145

sRun
RunA1
zout
1

Ave
Data
0.00365
Model
0.0005

data/vMax
0.01439

ofxng
334.07

Min d/m
2.88686

ave/min d
378.446
ave/min m
149.659

327_Calib_S_A

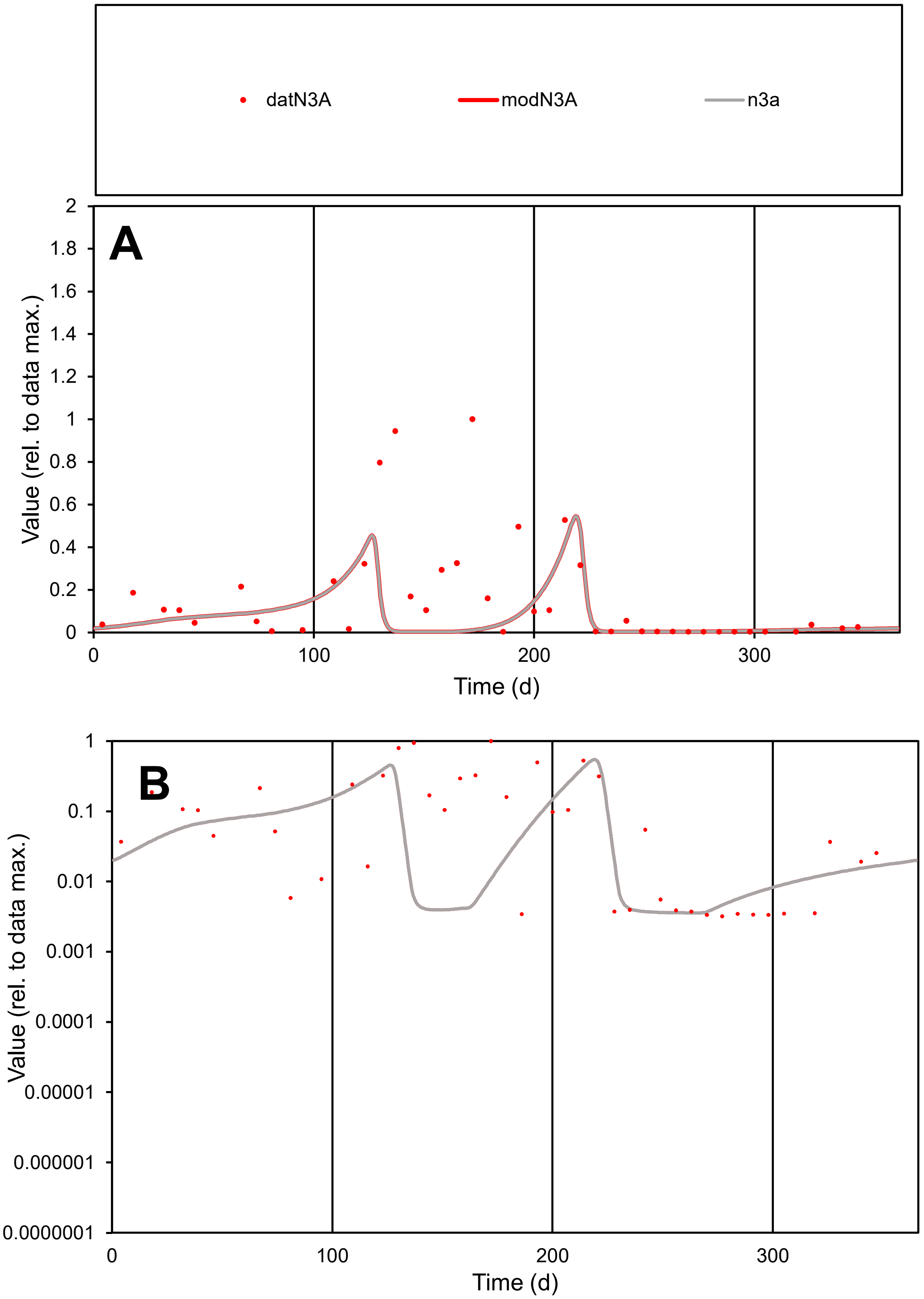


**C1**

Variables
N3A Provide jtd or Dst
CalibOnly?
2 1 to reproduce of
oNormalize?
2 1=no, 2=yes

ofxn calc*
1.52414
ofxn mod
1.57916

sRun
RunA1
zout
1

Ave
Data
0.00094
Model
0.00048

data/vMax
0.00577

ofxng
334.07

Min d/m
0.90709

ave/min d
50.8023
ave/min m
23.6441

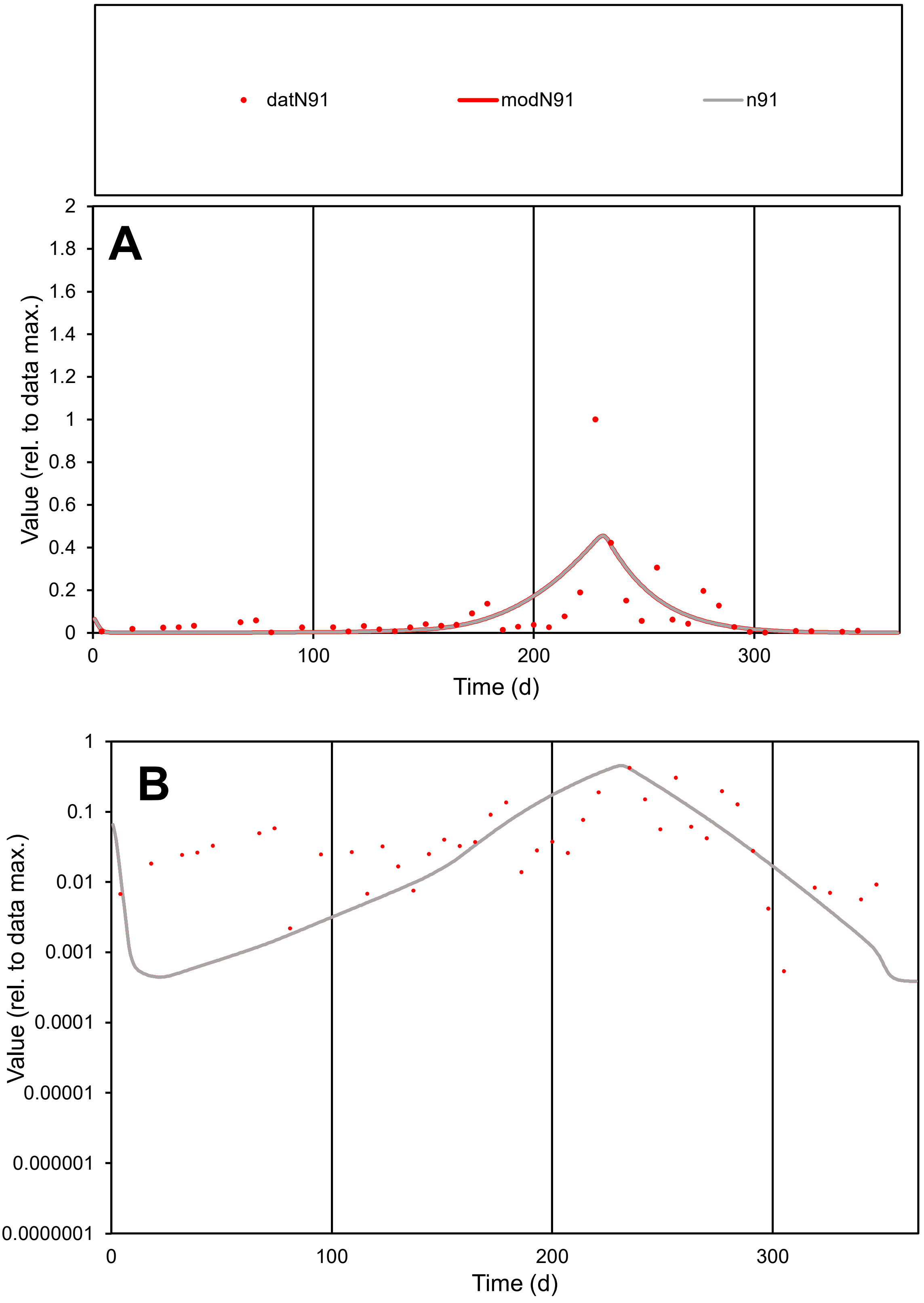


**C1**

Variables
N91 Provide jtd or Dst
CalibOnly?
2 1 to reproduce of
oNormalize?
2 1=no, 2=yes

ofxn calc*
1.39972
ofxn mod
1.4236

sRun
RunA1
zout
1

Ave
Data
0.00548
Model
0.00526

data/vMax
0.06587

ofxng
334.07

Min d/m
1.19858

ave/min d
154.099
ave/min m
177.183

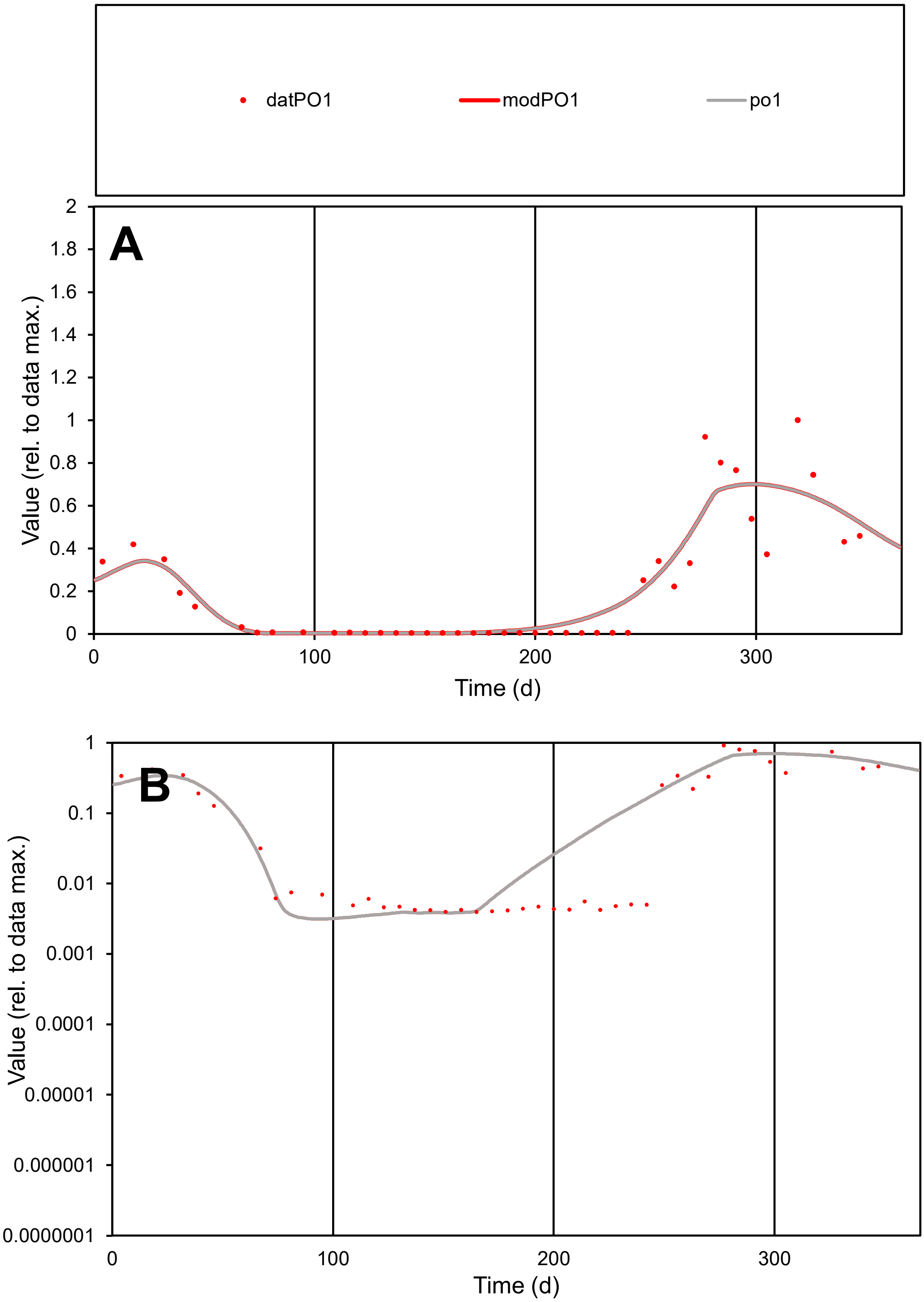


**C1**

Variables
PO1 Provide jtd or Dst
CalibOnly?
2 1 to reproduce of
oNormalize?
2 1=no, 2=yes

ofxn calc*
0.52481
ofxn mod
0.55605

sRun
RunA1
zout
1

Ave
Data
0.00239
Model
0.00249

data/vMax
0.01148

ofxng
334.07

Min d/m
1.2559

ave/min d
52.8421
ave/min m
69.091

327_Calib_S_A

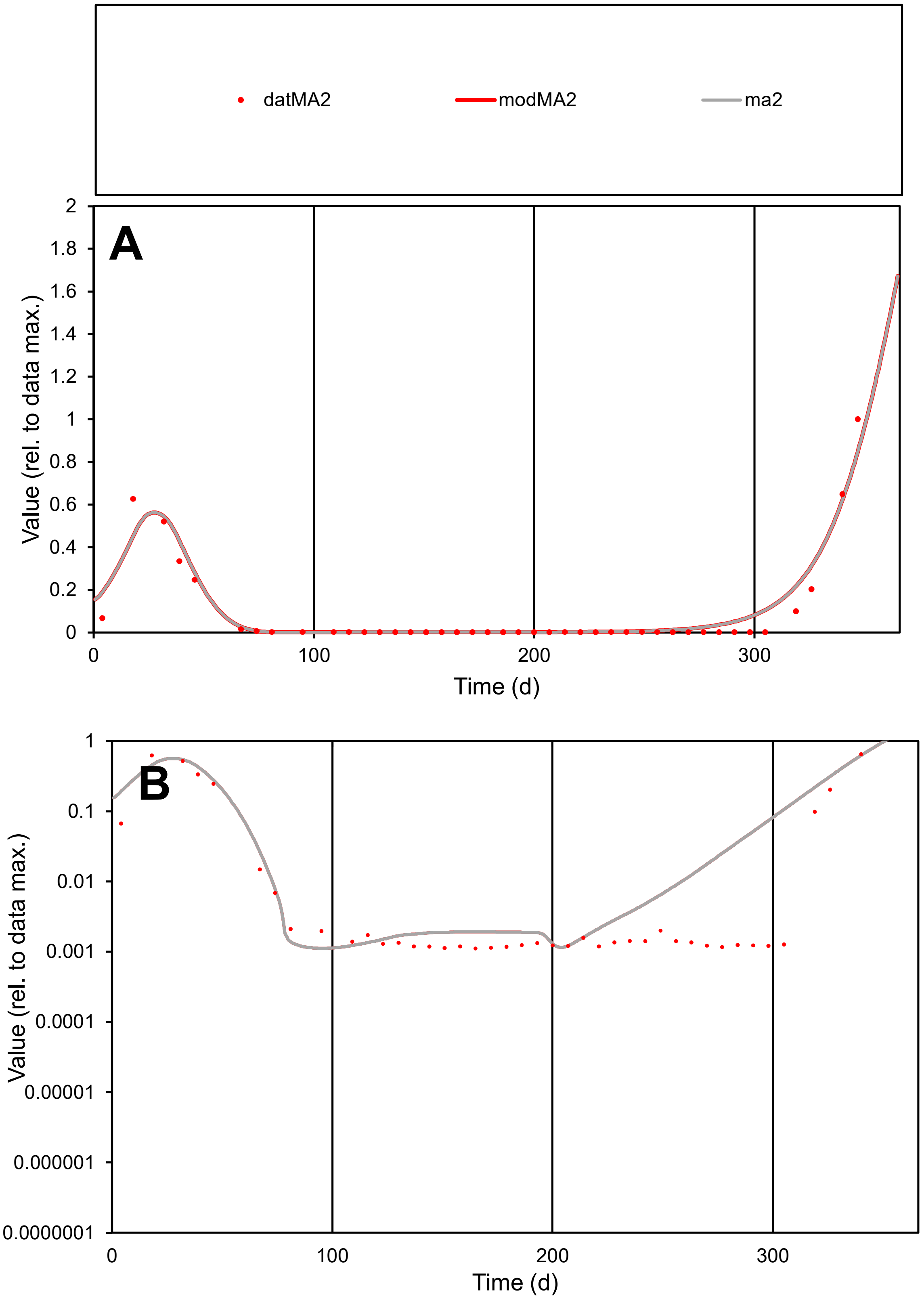


**C1**

Variables
MA2 Provide jtd or Dst
CalibOnly?
2 1 to reproduce of
oNormalize?
2 1=no, 2=yes

ofxn calc*
0.53154
ofxn mod
0.62505

sRun
RunA1
zout
1

Ave
Data
0.00144
Model
0.00164

data/vMax
0.01586

ofxng
334.07

Min d/m
0.99775

ave/min d
81.1841
ave/min m
92.5872

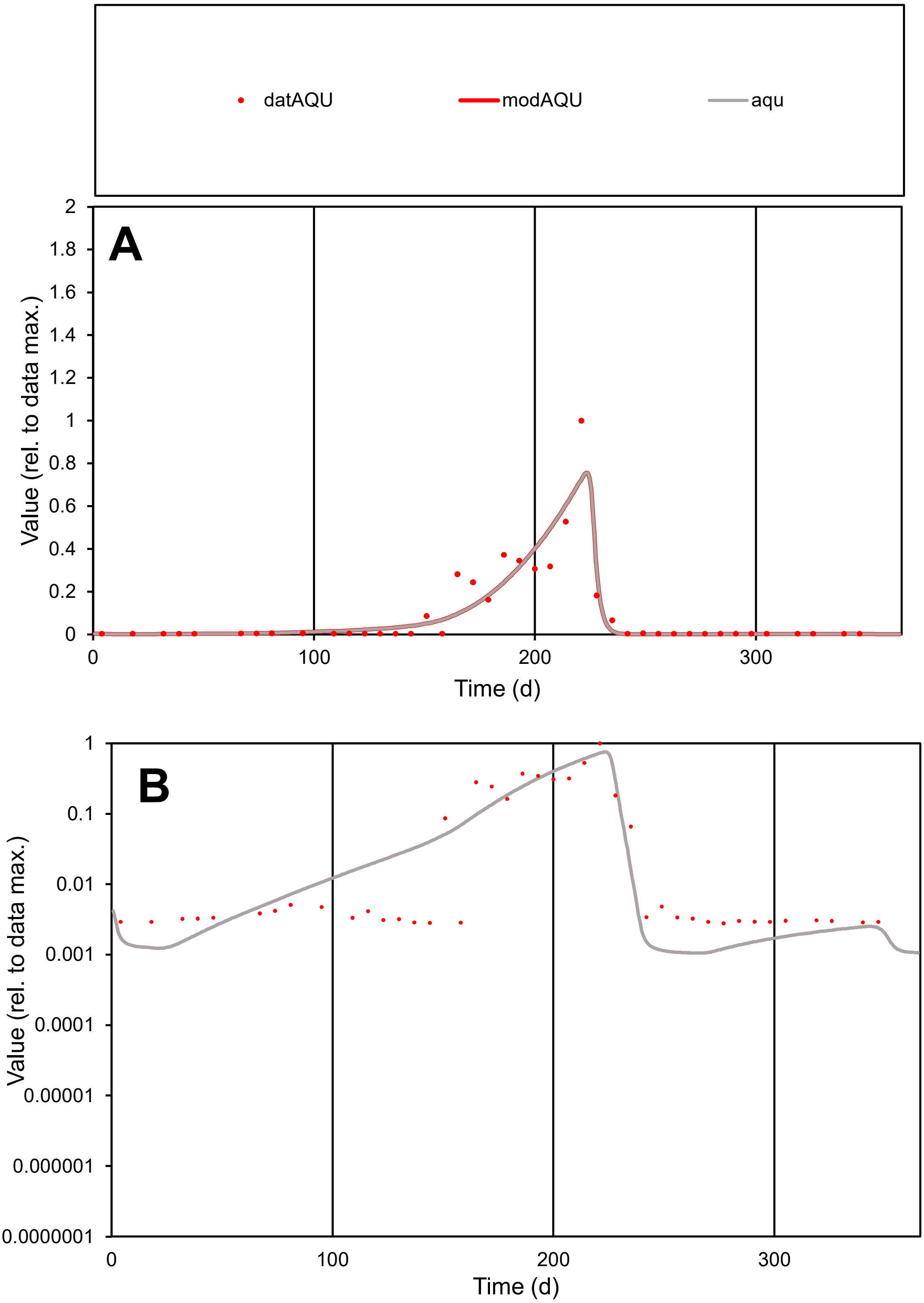


**C1**

Variables
AQU Provide jtd or Dst
CalibOnly?
2 1 to reproduce of
oNormalize?
2 1=no, 2=yes

ofxn calc*
0.69313
ofxn mod
0.74669

sRun
RunA1
zout
1

Ave
Data
0.00458
Model
0.00446

data/vMax
0.04823

ofxng
334.07

Min d/m
2.66653

ave/min d
33.9598
ave/min m
88.0776

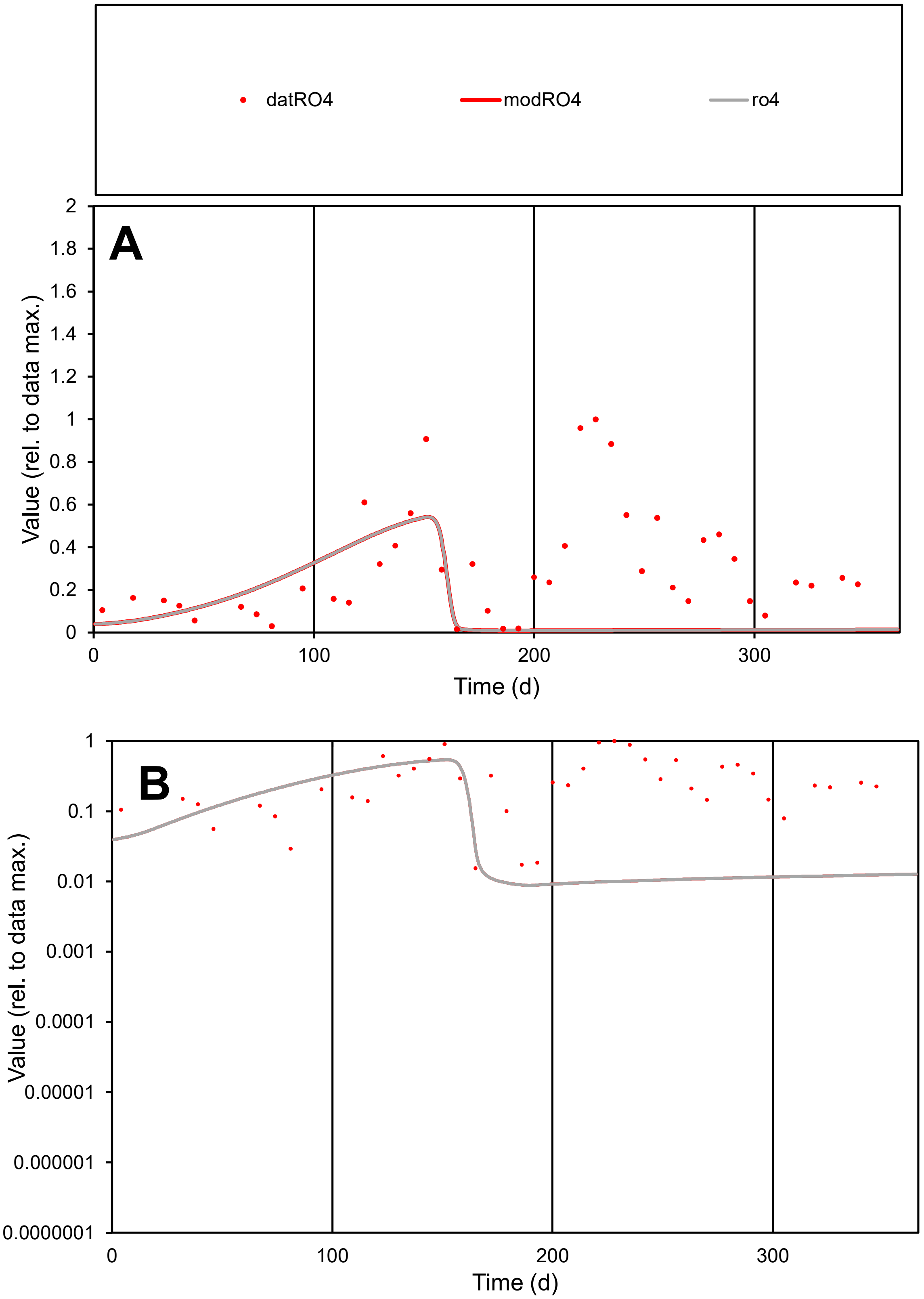


**C1**

Variables
RO4 Provide jtd or Dst
CalibOnly?
2 1 to reproduce of
oNormalize?
2 1=no, 2=yes

ofxn calc*
1.11635
ofxn mod
1.12055

sRun
RunA1
zout
1

Ave
Data
0.00424
Model
0.00174

data/vMax
0.01392

ofxng
334.07

Min d/m
1.75223

ave/min d
19.6386
ave/min m
14.127

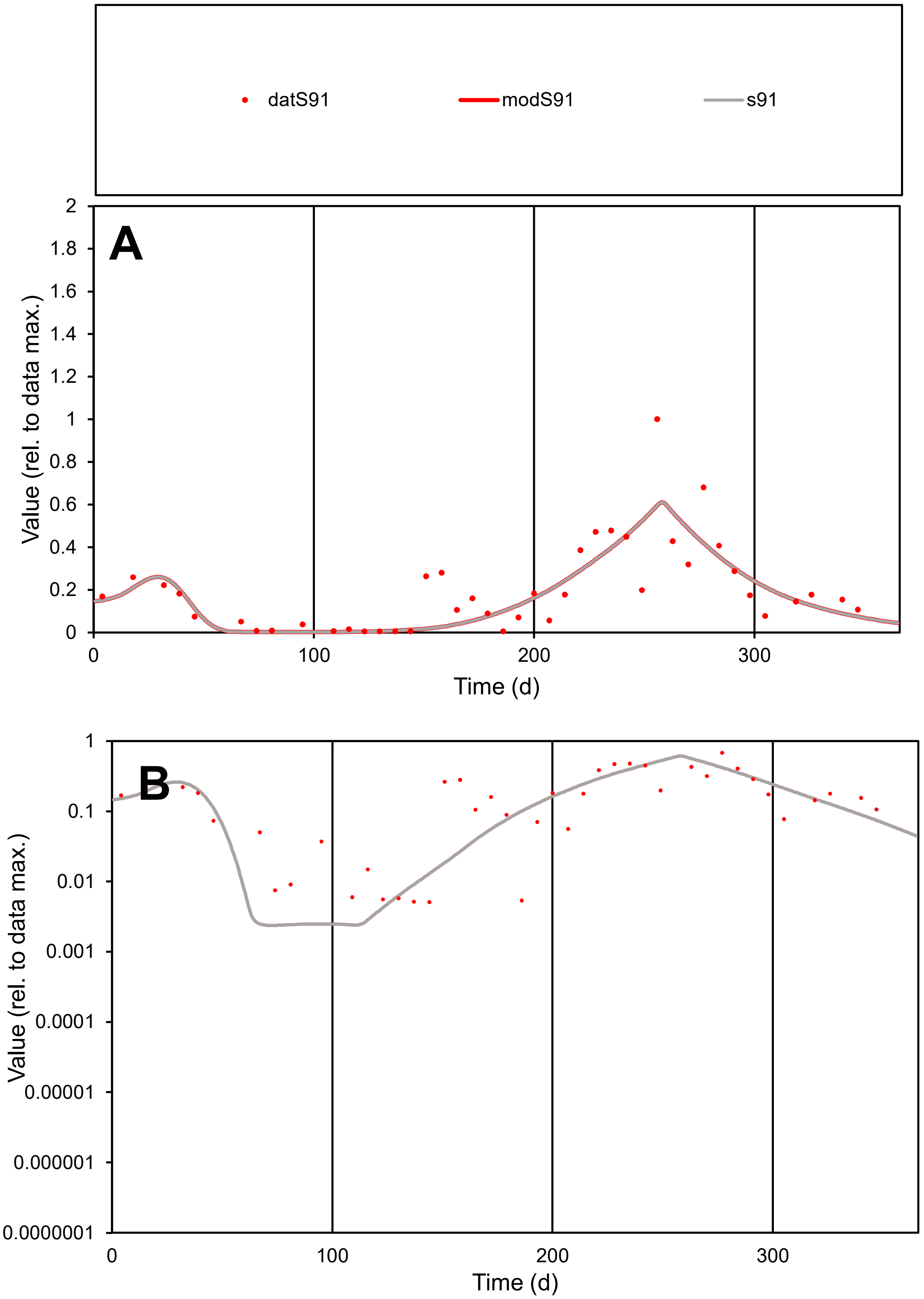


**C1**

Variables

S91 Provide jtd or Dst

CalibOnly?

2 1 to reproduce of

oNormalize?

2 1=no, 2=yes

ofxn calc*

0.60885

ofxn mod

0.63291

sRun

RunA1

zout

1

Ave

Data

0.0027

Model

0.00246

data/vMax

0.01353

ofxng

334.07

Min d/m

2.13199

ave/min d

39.1232

ave/min m

76.1248

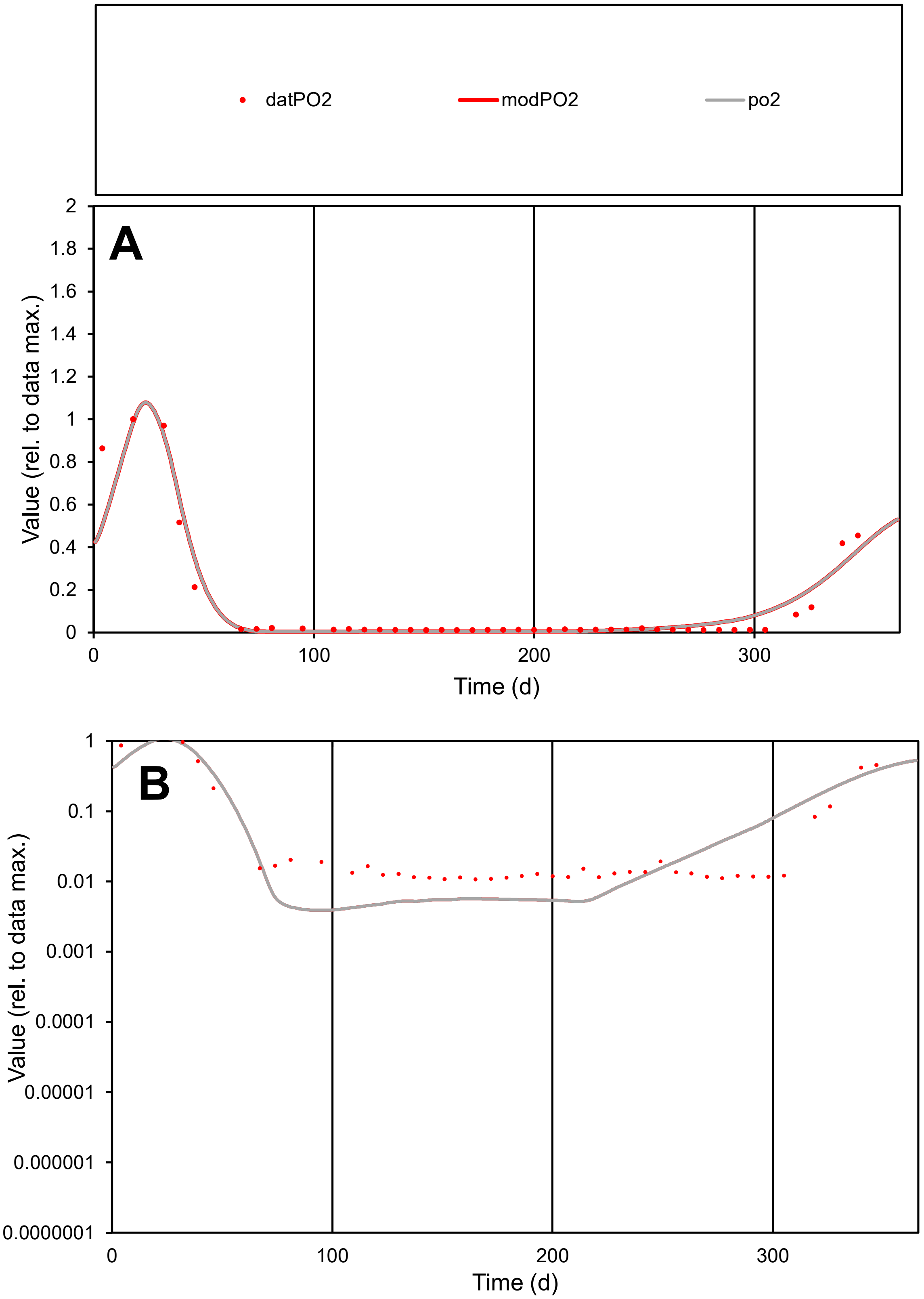


**C1**

| | |
|---|---|
| Variables | |
| PO2 | Provide jtd or Dst |
| CalibOnly? | |
| 2 | 1 to reproduce of |
| oNormalize? | |
| 2 | 1=no, 2=yes |
| ofxn calc* | |
| 0.51743 | |
| ofxn mod | |
| 0.59973 | |
| sRun | |
| RunA1 | |
| zout | |
| 1 | |
| Ave | |
| Data | |
| 0.00089 | |
| Model | |
| 0.00087 | |
| data/vMax | |
| 0.00735 | |
| ofxng | |
| 334.07 | |
| Min d/m | |
| 2.74956 | |
| ave/min d | |
| 11.259 | |
| ave/min m | |
| 30.3477 | |

327_Calib_S_A

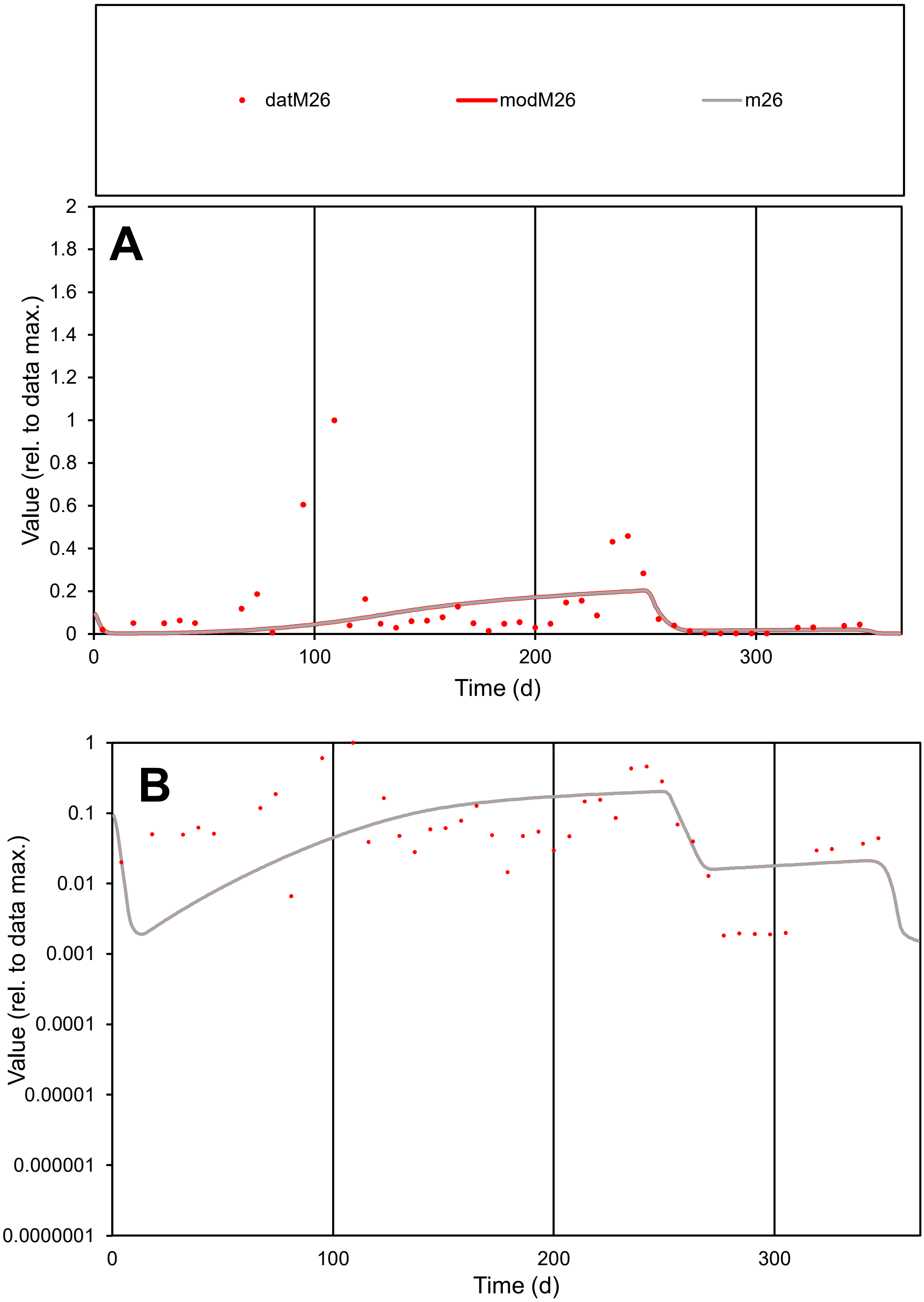


**C1**

Variables
M26 Provide jtd or Dst
CalibOnly?
2 1 to reproduce of
oNormalize?
2 1=no, 2=yes

ofxn calc*
1.65127
ofxn mod
1.65813

sRun
RunA1
zout
1

Ave
Data
0.00534
Model
0.00386

data/vMax
0.04647

ofxng
334.07

Min d/m
0.74634

ave/min d
63.0322
ave/min m
33.9797

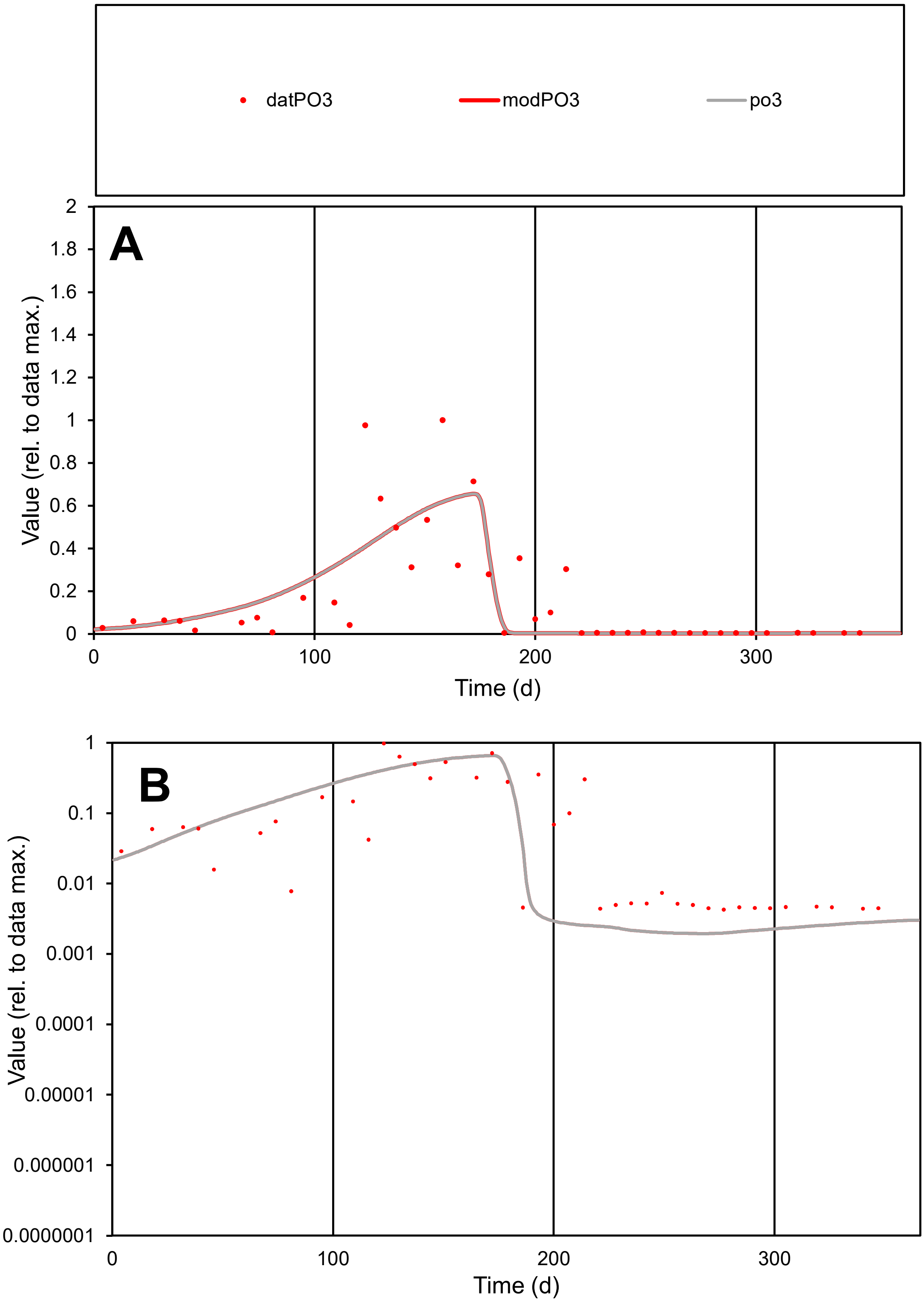


**C1**

| | |
|---|---|
| Variables | |
| PO3 | Provide jtd or Dst |
| CalibOnly? | |
| 2 | 1 to reproduce of |
| oNormalize? | |
| 2 | 1=no, 2=yes |
| ofxn calc* | |
| 0.96876 | |
| ofxn mod | |
| 0.98504 | |
| sRun | |
| RunA1 | |
| zout | |
| 1 | |
| Ave | |
| Data | |
| 0.00185 | |
| Model | |
| 0.00173 | |
| data/vMax | |
| 0.01127 | |
| ofxng | |
| 334.07 | |
| Min d/m | |
| 2.2011 | |
| ave/min d | |
| 38.4542 | |
| ave/min m | |
| 79.1829 | |

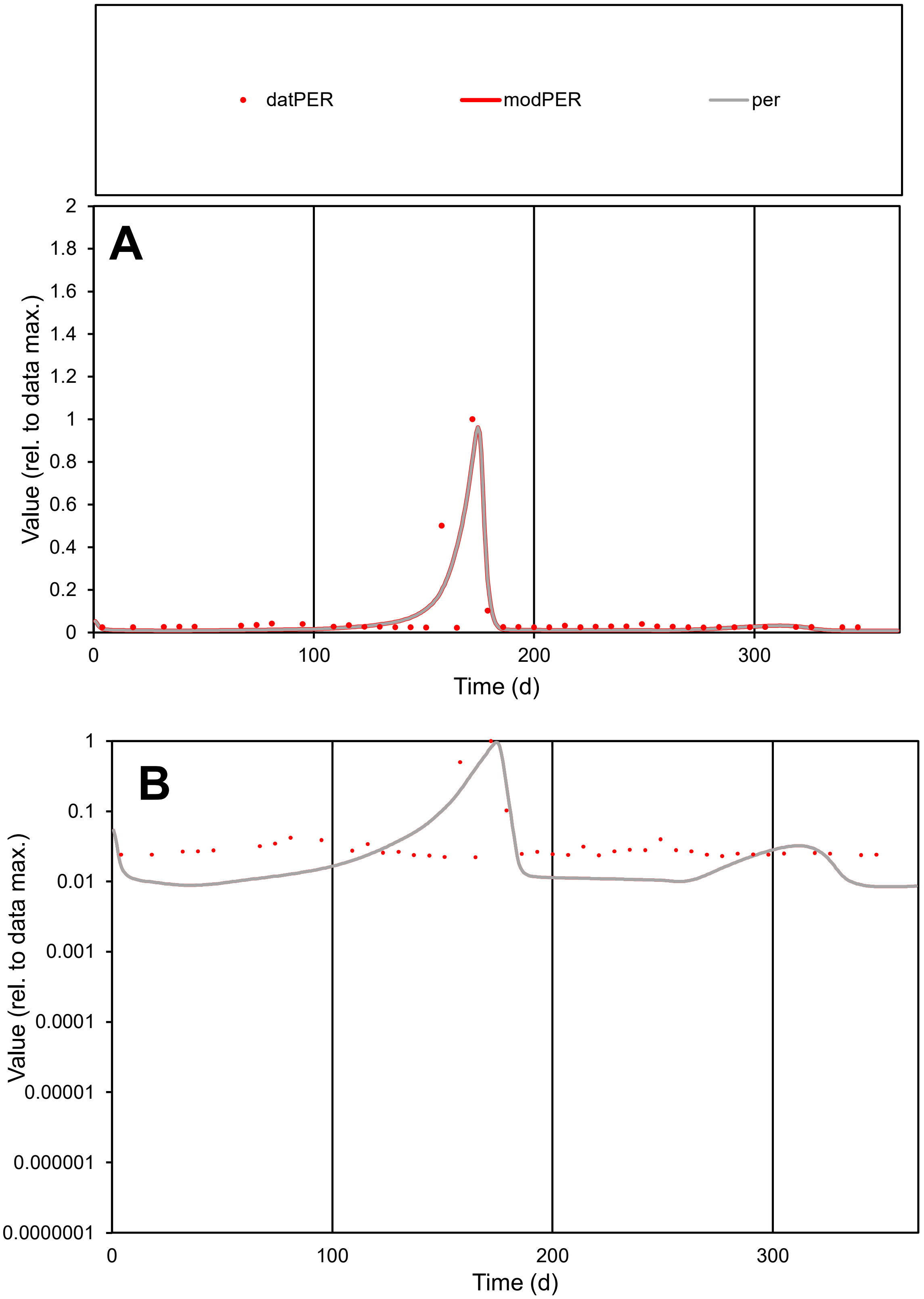


**C1**

Variables
PER Provide jtd or Dst
CalibOnly?
2 1 to reproduce of
oNormalize?
2 1=no, 2=yes

ofxn calc*
1.12855
ofxn mod
1.21628

sRun
RunA1
zout
1

Ave
Data
5.2E-05
Model
4.6E-05

data/vMax
0.00082

ofxng
334.07

Min d/m
2.62245

ave/min d
2.86232
ave/min m
6.67494

327_Calib_S_A

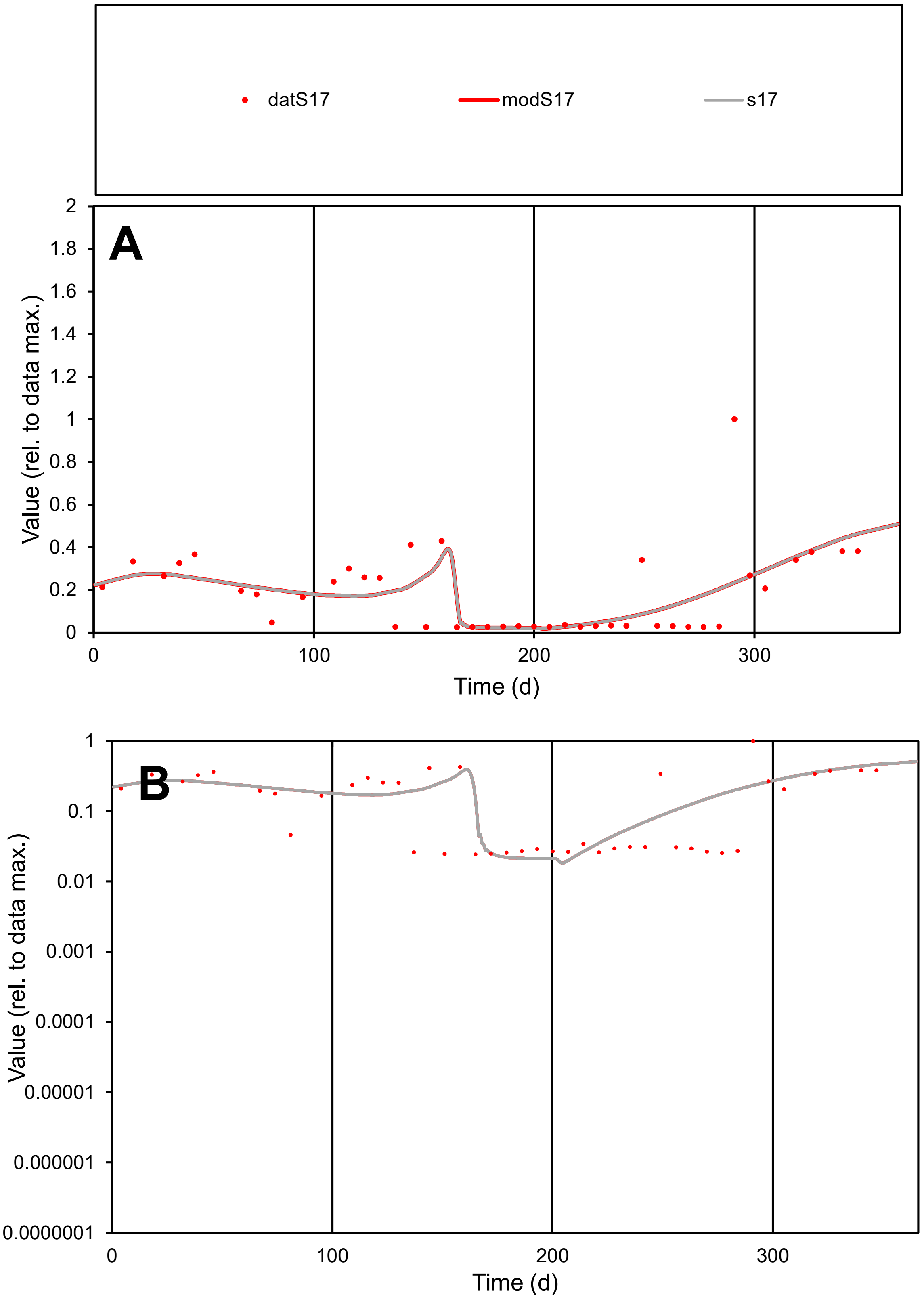


**C1**

Variables

S17 Provide jtd or Dst

CalibOnly?

2 1 to reproduce of

oNormalize?

2 1=no, 2=yes

ofxn calc*

0.81221

ofxn mod

0.8247

sRun

RunA1

zout

1

Ave

Data

0.00082

Model

0.0008

data/vMax

0.00443

ofxng

334.07

Min d/m

1.15872

ave/min d

7.61331

ave/min m

8.63274

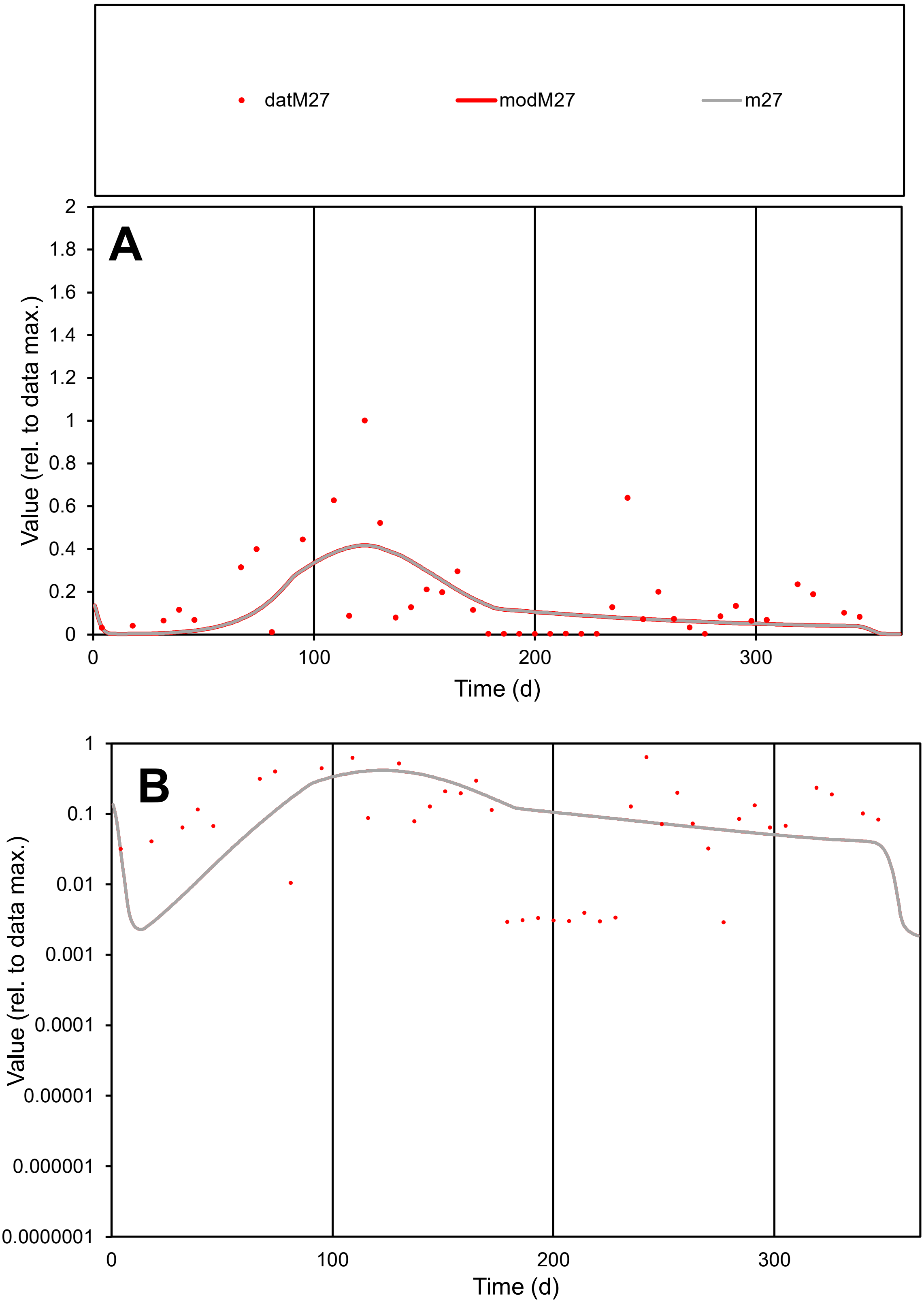


**C1**

Variables
M27 Provide jtd or Dst
CalibOnly?
2 1 to reproduce of
oNormalize?
2 1=no, 2=yes

ofxn calc*
1.09416
ofxn mod
1.10517

sRun
RunA1
zout
1

Ave
Data
0.00379
Model
0.00311

data/vMax
0.02315

ofxng
334.07

Min d/m
0.94163

ave/min d
56.3442
ave/min m
43.4848

327_Calib_S_A

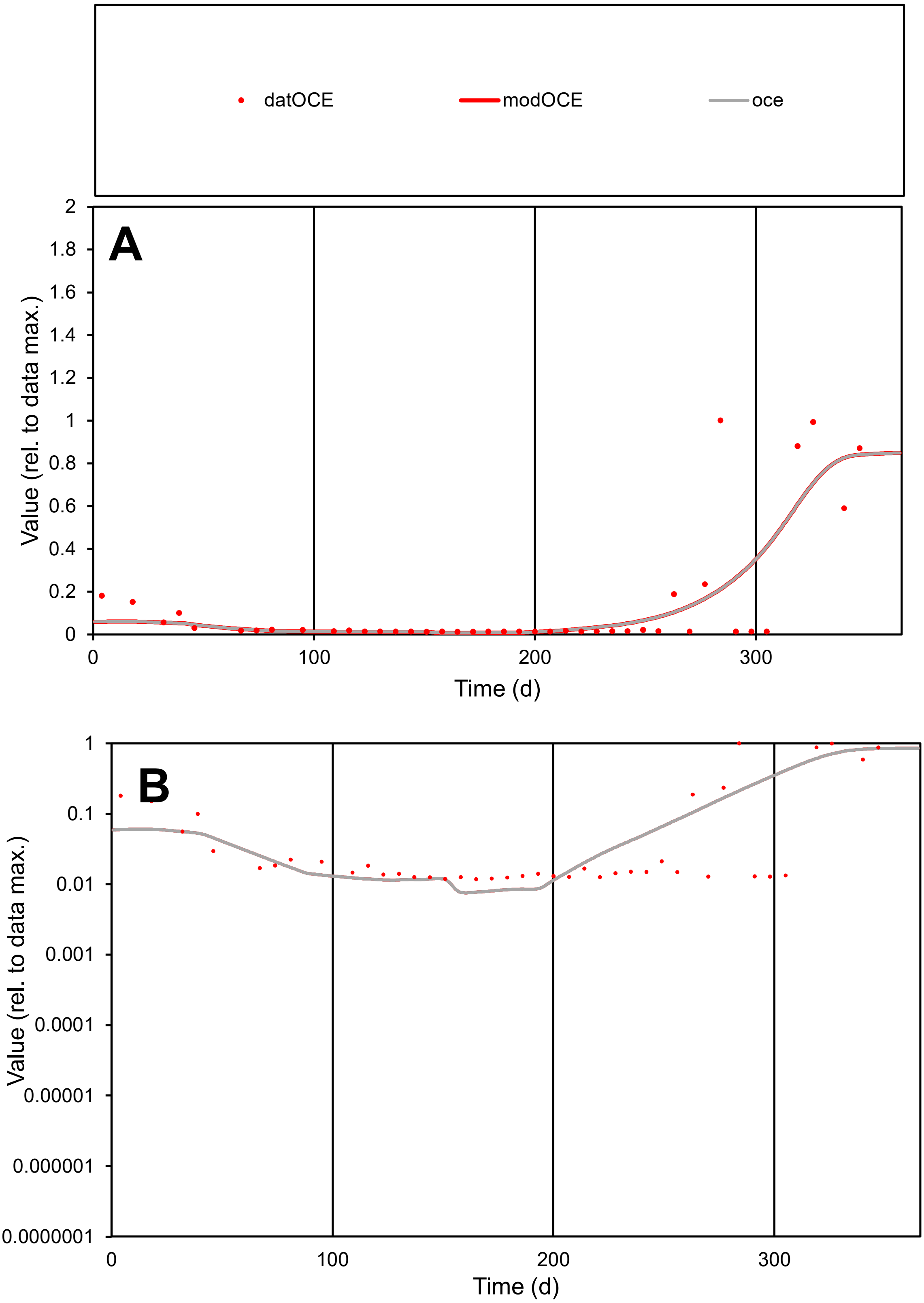


**C1**

Variables
OCE Provide jtd or Dst
CalibOnly?
2 1 to reproduce of
oNormalize?
2 1=no, 2=yes

ofxn calc*
1.22928
ofxn mod
1.25459

sRun
RunA1
zout
1

Ave
Data
0.00042
Model
0.00041

data/vMax
0.00312

ofxng
334.07

Min d/m
1.56064

ave/min d
11.5448
ave/min m
17.2737

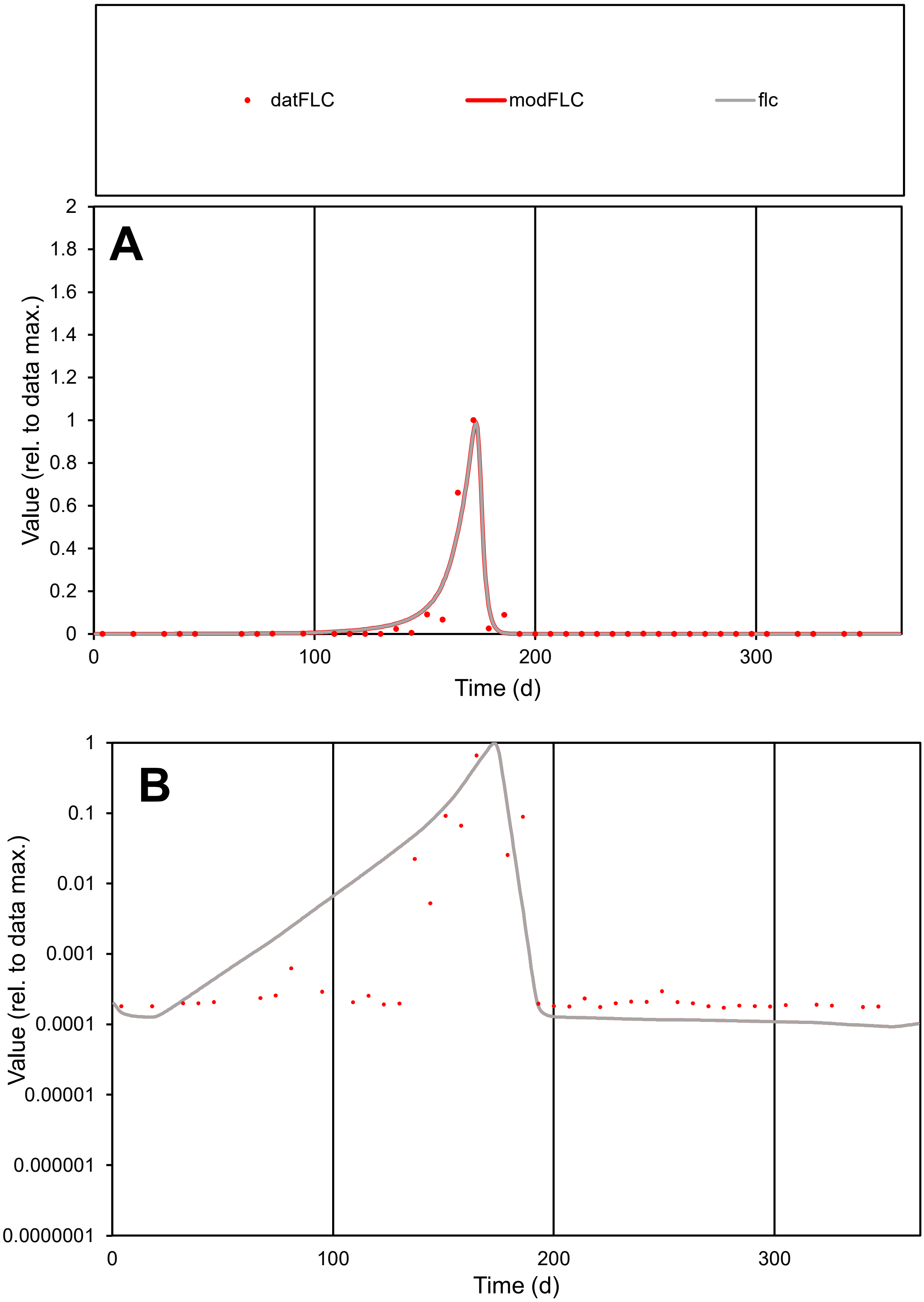


**C1**

Variables

FLC Provide jtd or Dst

CalibOnly?

2 1 to reproduce of

oNormalize?

2 1=no, 2=yes

ofxn calc*

0.71606

ofxn mod

0.8448

sRun

RunA1

zout

1

Ave

Data

0.00339

Model

0.00371

data/vMax

0.07236

ofxng

334.07

Min d/m

1.83047

ave/min d

272.041

ave/min m

545.369

327_Calib_S_A

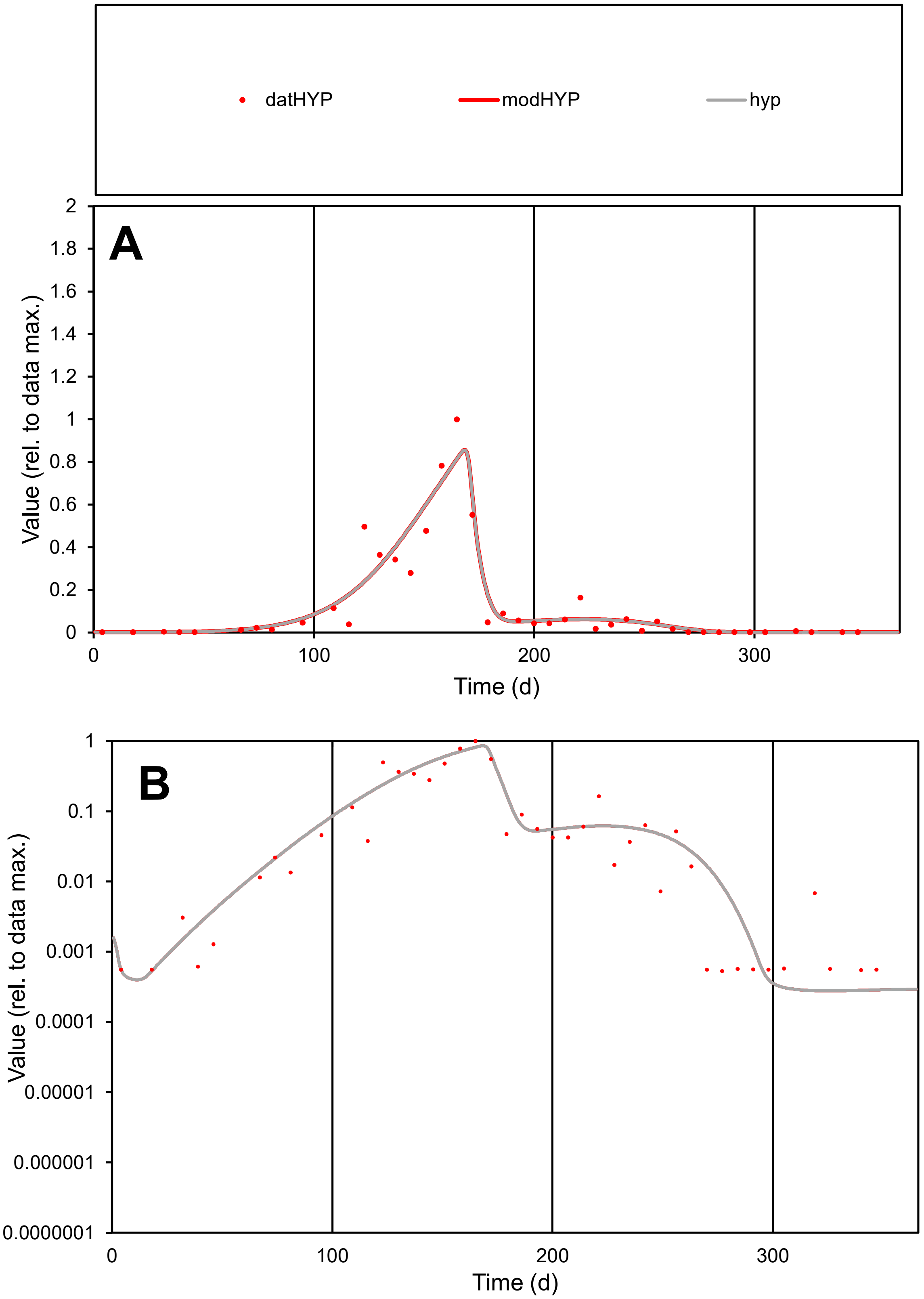


**C1**

Variables
HYP Provide jtd or Dst
CalibOnly?
2 1 to reproduce of
oNormalize?
2 1=no, 2=yes

ofxn calc*
0.56268
ofxn mod
0.61092

sRun
RunA1
zout
1

Ave
Data
0.00387
Model
0.00402

data/vMax
0.03095

ofxng
334.07

Min d/m
1.90868

ave/min d
234.709
ave/min m
465.884

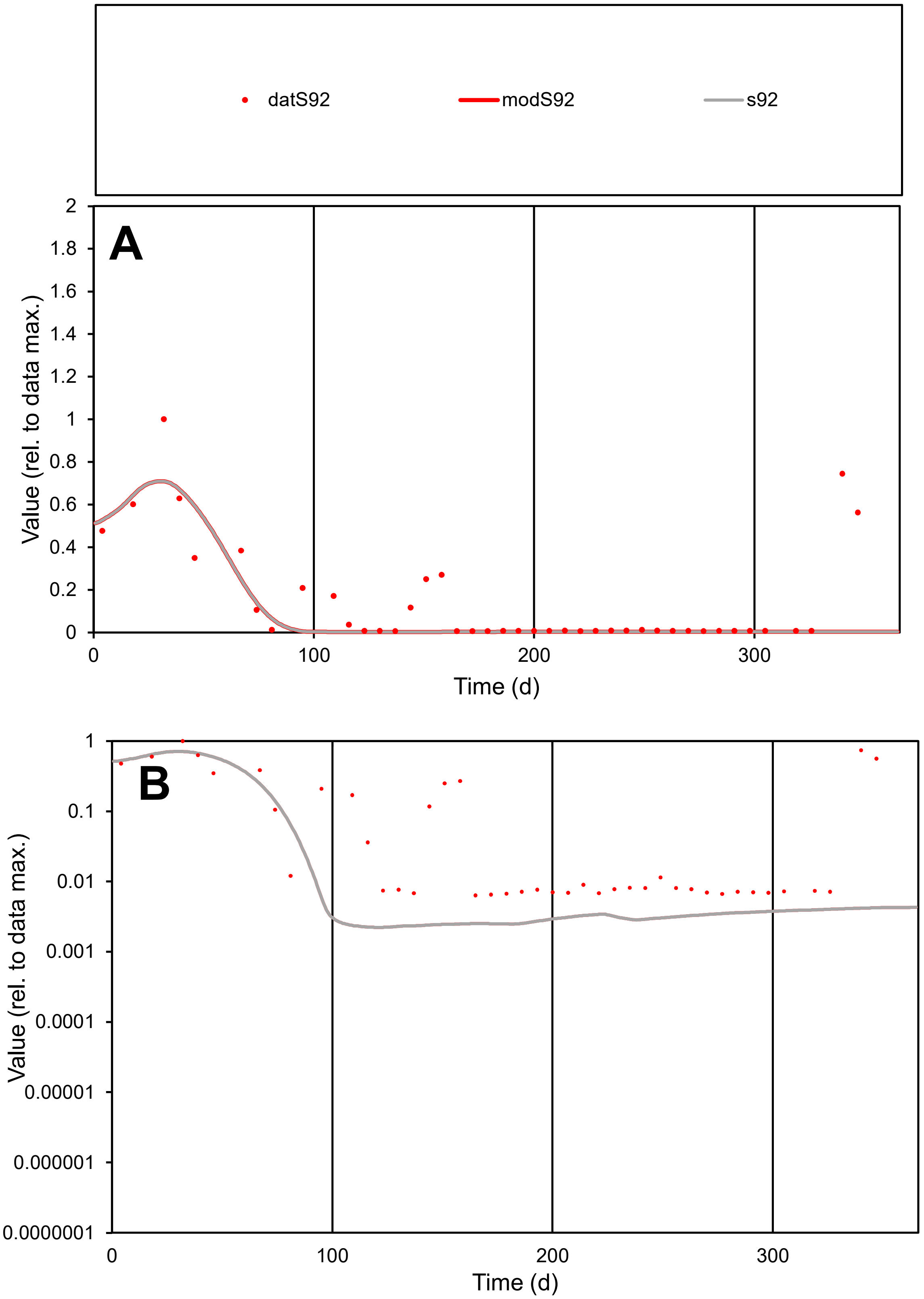


**C1**

Variables
S92 Provide jtd or Dst
CalibOnly?
2 1 to reproduce of
oNormalize?
2 1=no, 2=yes

ofxn calc*
1.18275
ofxn mod
1.20267

sRun
RunA1
zout
1

Ave
Data
0.00053
Model
0.00031

data/vMax
0.00361

ofxng
334.07

Min d/m
2.8575

ave/min d
22.7769
ave/min m
38.813

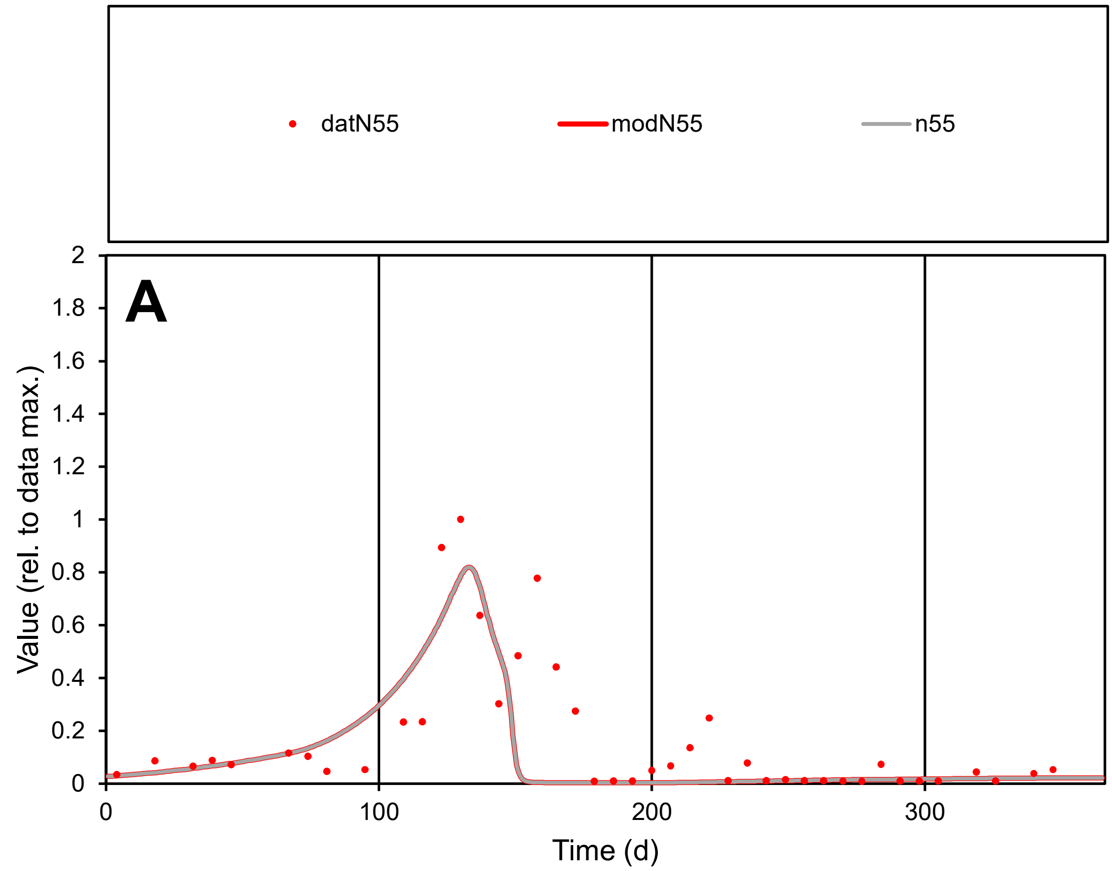


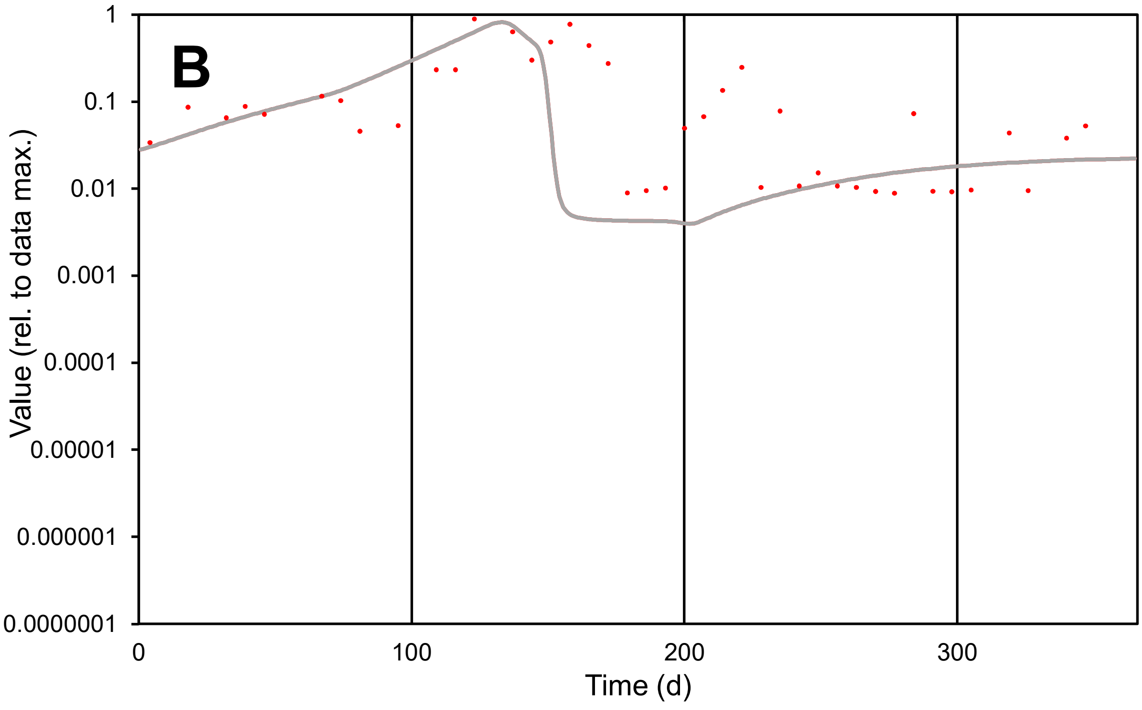


**C1**

Variables
N55 Provide jtd or Dst
CalibOnly?
2 1 to reproduce of
oNormalize?
2 1=no, 2=yes

ofxn calc*
1.06663
ofxn mod
1.11632

sRun
RunA1
zout
1

Ave
Data
0.00071
Model
0.00052

data/vMax
0.00435

ofxng
334.07

Min d/m
2.23006

ave/min d
18.426
ave/min m
30.1969

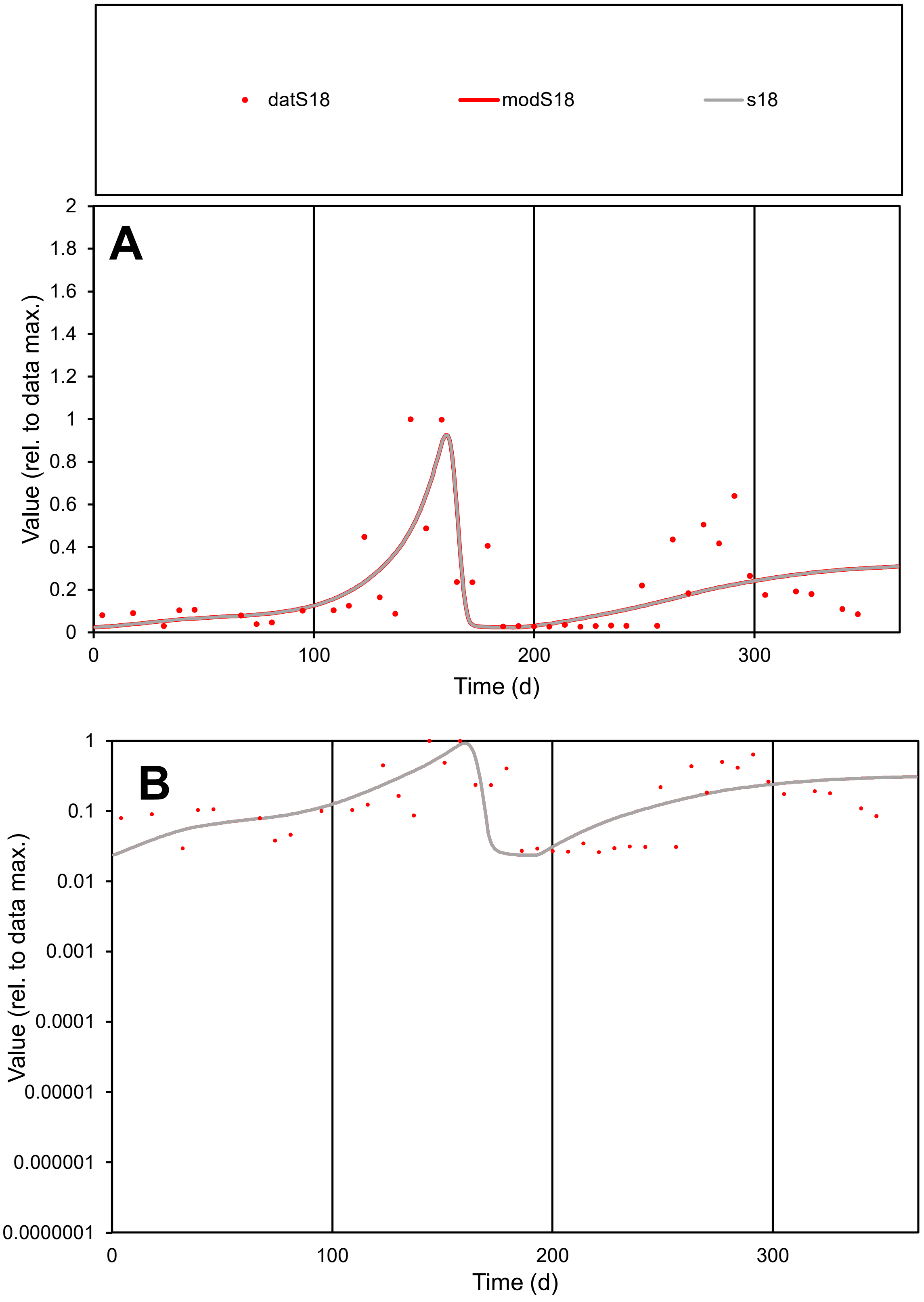


**C1**

Variables
S18 Provide jtd or Dst
CalibOnly?
2 1 to reproduce of
oNormalize?
2 1=no, 2=yes

ofxn calc*
0.79491
ofxn mod
0.82099

sRun
RunA1
zout
1

Ave
Data
0.00222
Model
0.00206

data/vMax
0.01079

ofxng
334.07

Min d/m
1.10958

ave/min d
7.87563
ave/min m
8.07162

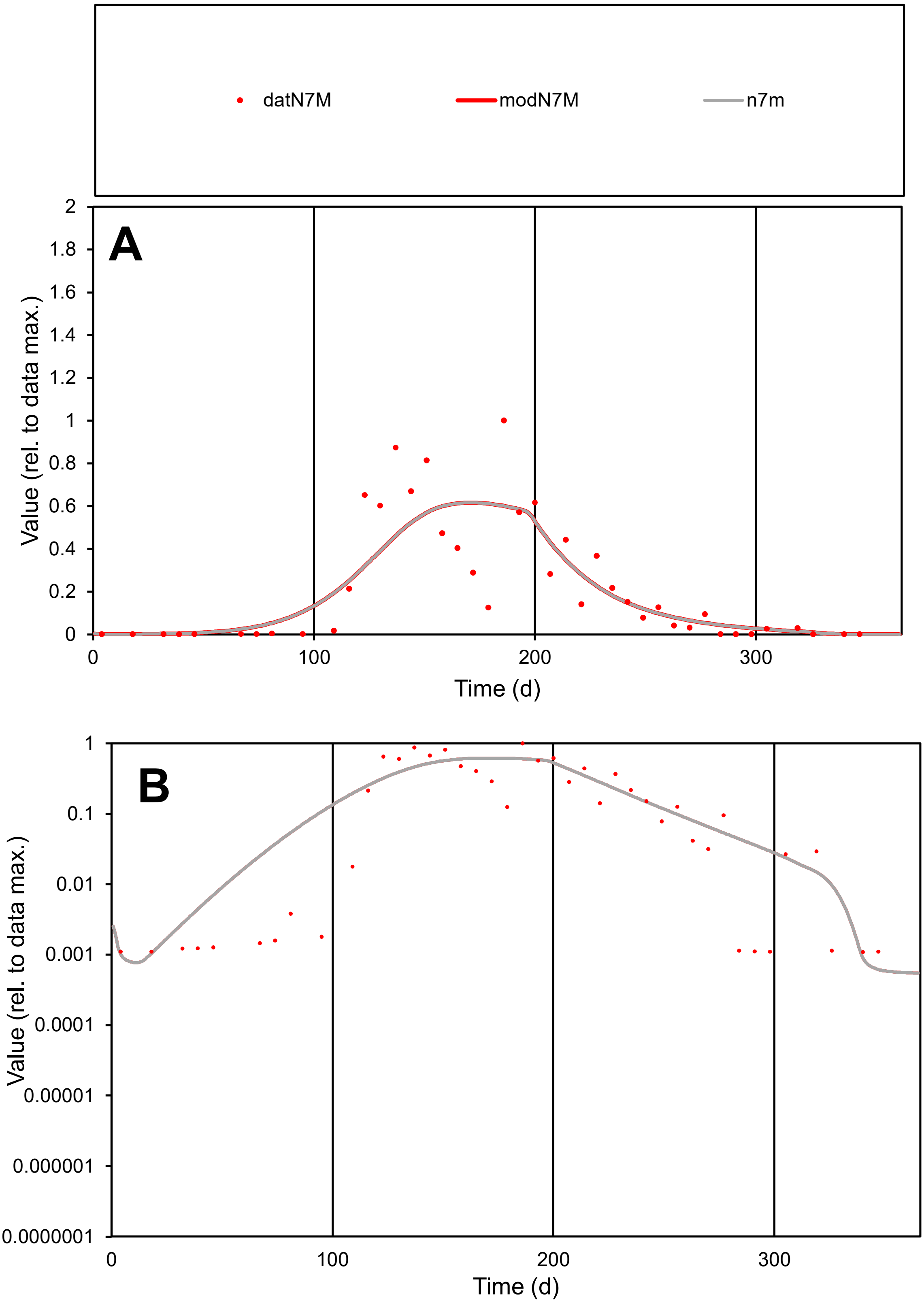


**C1**

Variables
N7M Provide jtd or Dst
CalibOnly?
2 1 to reproduce of
oNormalize?
2 1=no, 2=yes

ofxn calc*
0.71706
ofxn mod
0.73604

sRun
RunA1
zout
1

Ave
Data
0.0021
Model
0.00207

data/vMax
0.00941

ofxng
334.07

Min d/m
1.80409

ave/min d
205.132
ave/min m
365.369

327_Calib_S_A

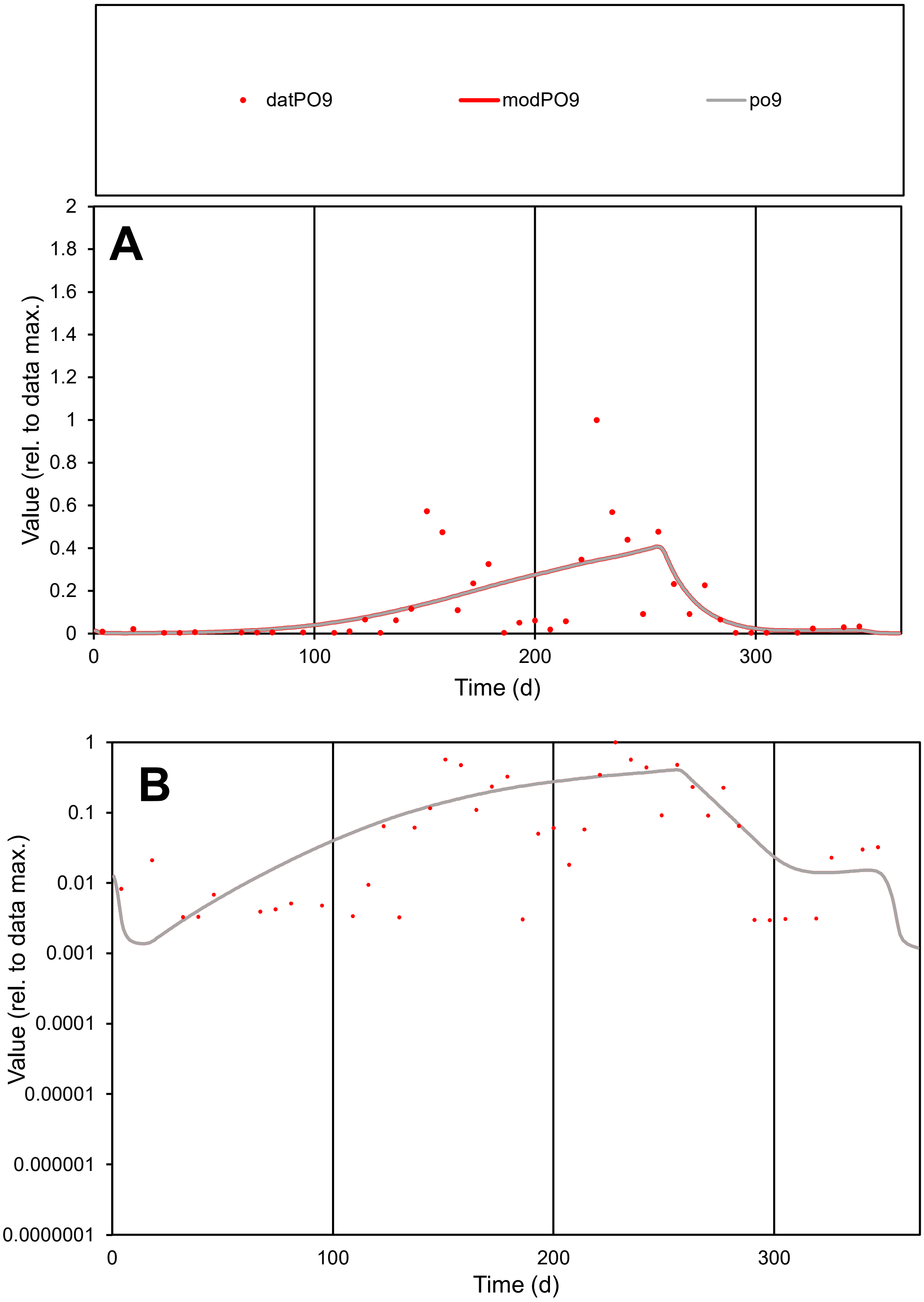


**C1**

Variables
PO9 Provide jtd or Dst
CalibOnly?
2 1 to reproduce of
oNormalize?
2 1=no, 2=yes

ofxn calc*
1.20948
ofxn mod
1.22316

sRun
RunA1
zout
1

Ave
Data
0.00297
Model
0.00297

data/vMax
0.02128

ofxng
334.07

Min d/m
1.85643

ave/min d
47.1241
ave/min m
87.4716

327_Calib_S_A

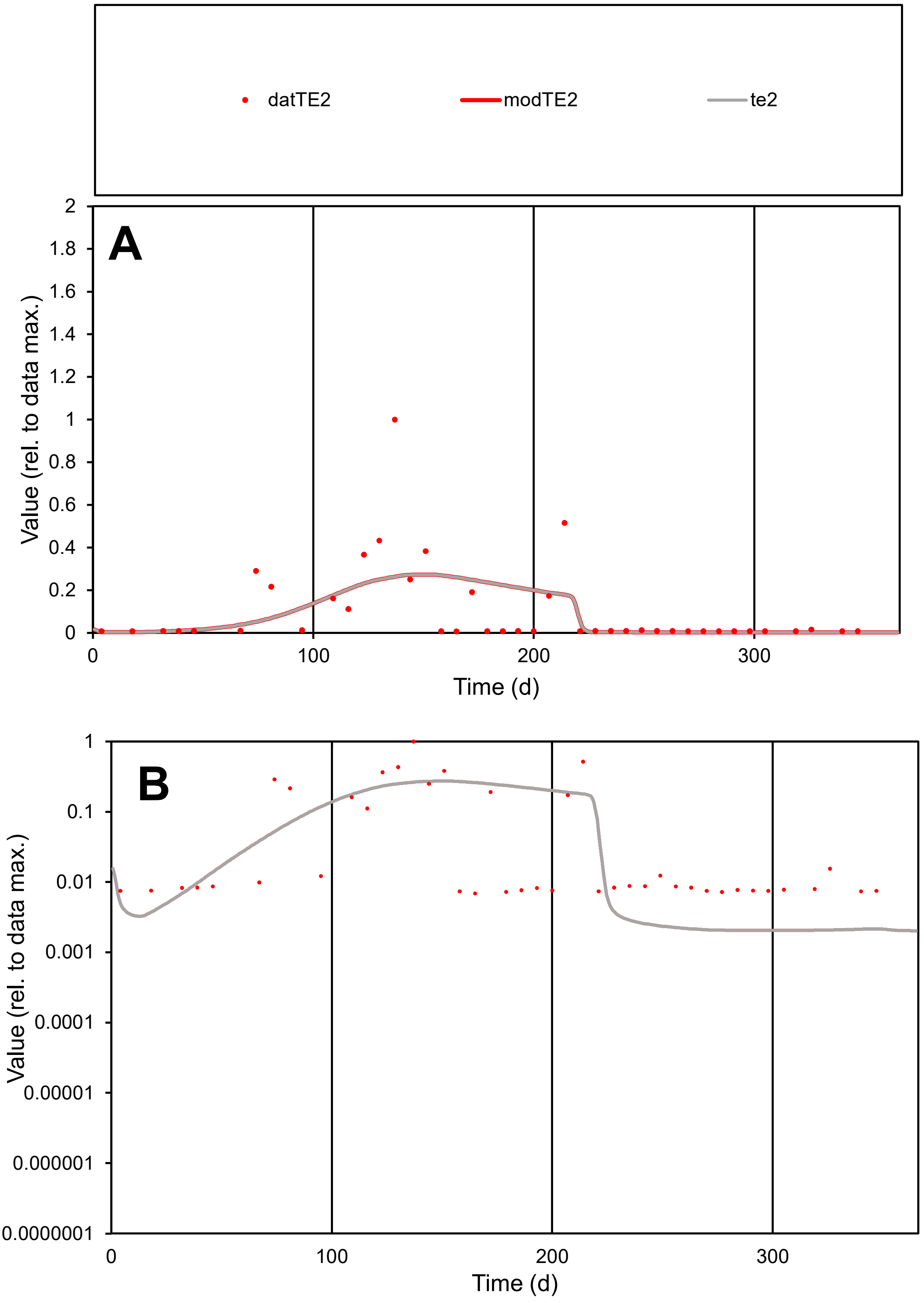


**C1**

Variables

TE2 Provide jtd or Dst

CalibOnly?

2 1 to reproduce of

oNormalize?

2 1=no, 2=yes

ofxn calc*

1.58025

ofxn mod

1.5882

sRun

RunA1

zout

1

Ave

Data

0.00127

Model

0.00119

data/vMax

0.01232

ofxng

334.07

Min d/m

3.34386

ave/min d

15.0383

ave/min m

47.1029

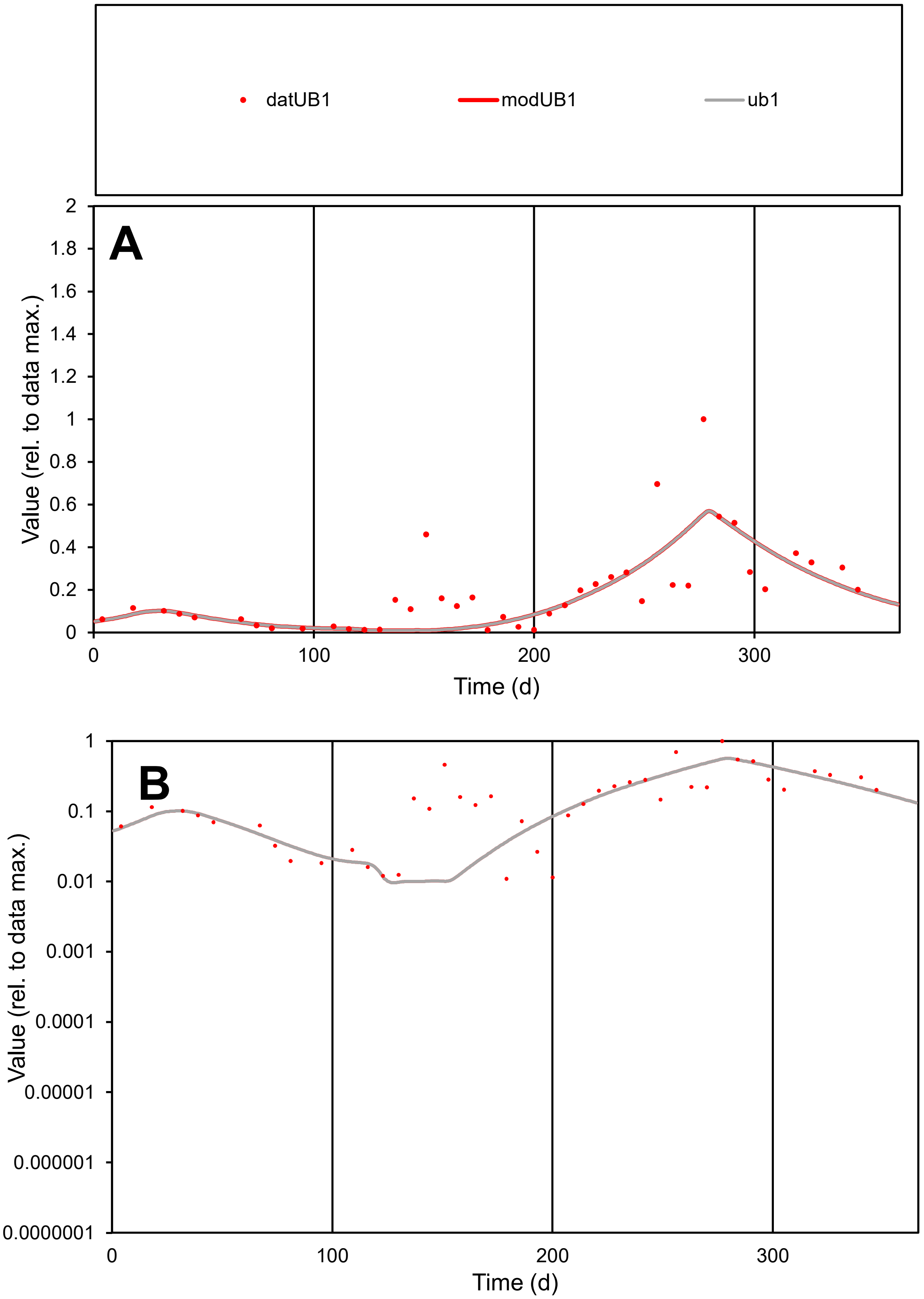


**C1**

Variables
UB1 Provide jtd or Dst
CalibOnly?
2 1 to reproduce of
oNormalize?
2 1=no, 2=yes

ofxn calc*
0.71321
ofxn mod
0.73116

sRun
RunA1
zout
1

Ave
Data
0.00106
Model
0.00092

data/vMax
0.00546

ofxng
334.07

Min d/m
1.10744

ave/min d
17.7151
ave/min m
17.066

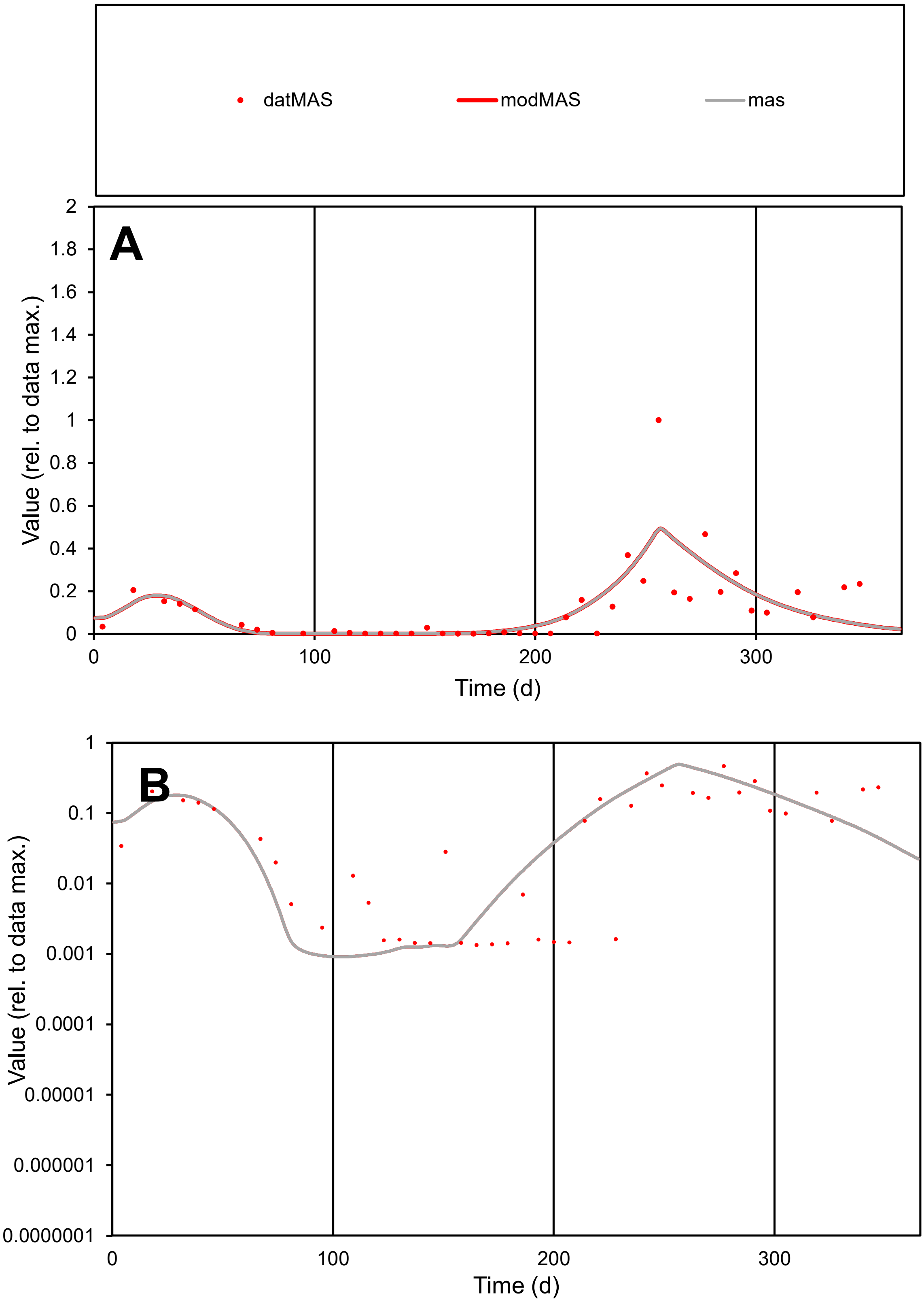


C1

Variables
MAS Provide jtd or Dst
CalibOnly?
2 1 to reproduce of
oNormalize?
2 1=no, 2=yes

ofxn calc*
0.94757
ofxn mod
0.97595

sRun
RunA1
zout
1

Ave
Data
0.00106
Model
0.00104

data/vMax
0.0089

ofxng
334.07

Min d/m
1.43562

ave/min d
88.95
ave/min m
124.867

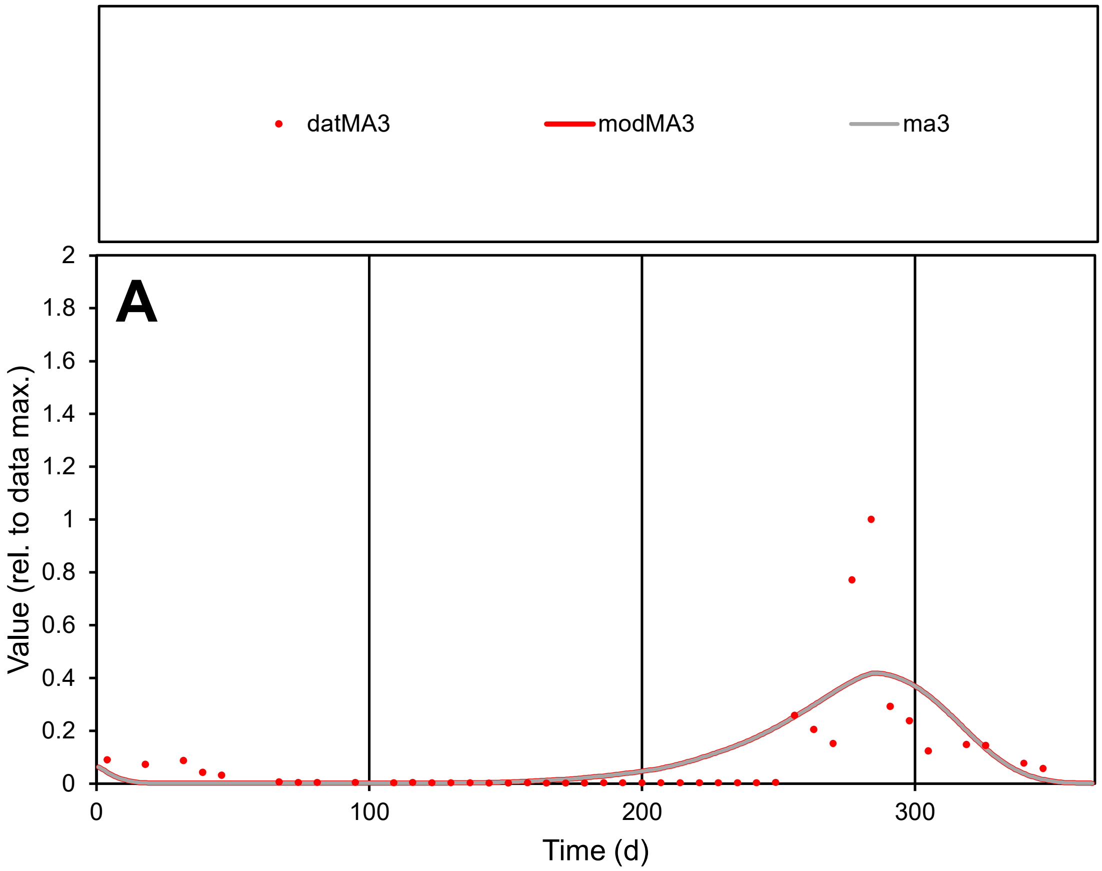


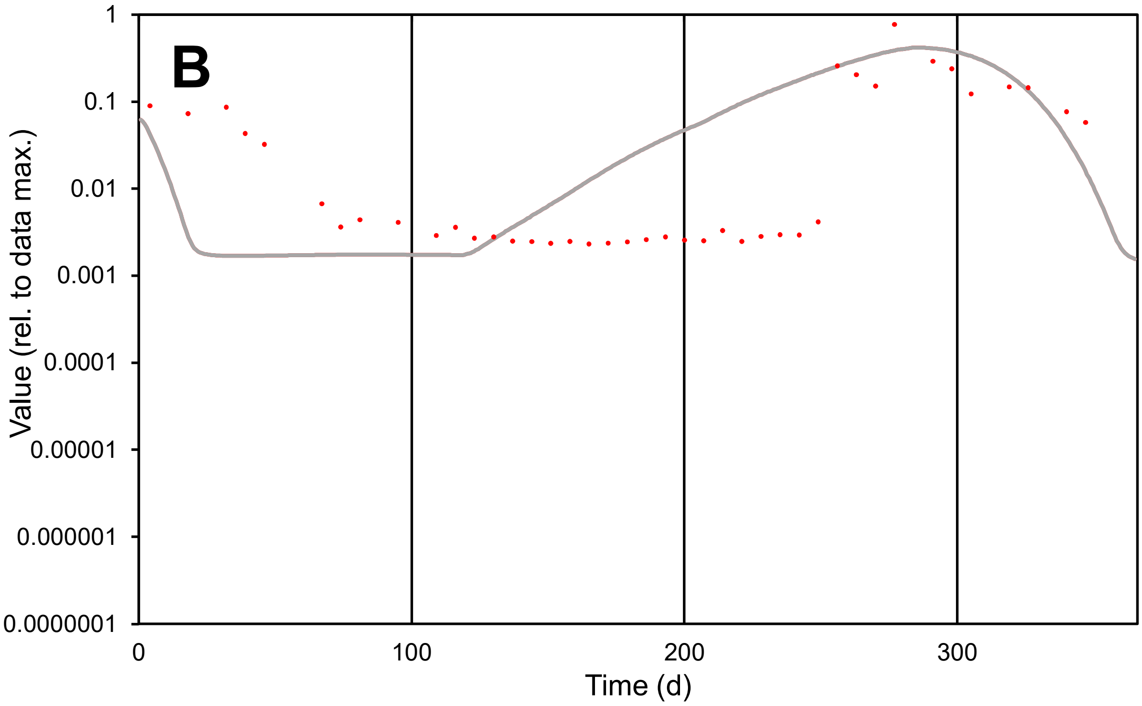


**C1**

Variables
MA3 Provide jtd or Dst
CalibOnly?
2 1 to reproduce of
oNormalize?
2 1=no, 2=yes

ofxn calc*
1.45255
ofxn mod
1.4734

sRun
RunA1
zout
1

Ave
Data
0.00243
Model
0.00272

data/vMax
0.02641

ofxng
334.07

Min d/m
1.37073

ave/min d
39.5704
ave/min m
60.648

327_Calib_S_A

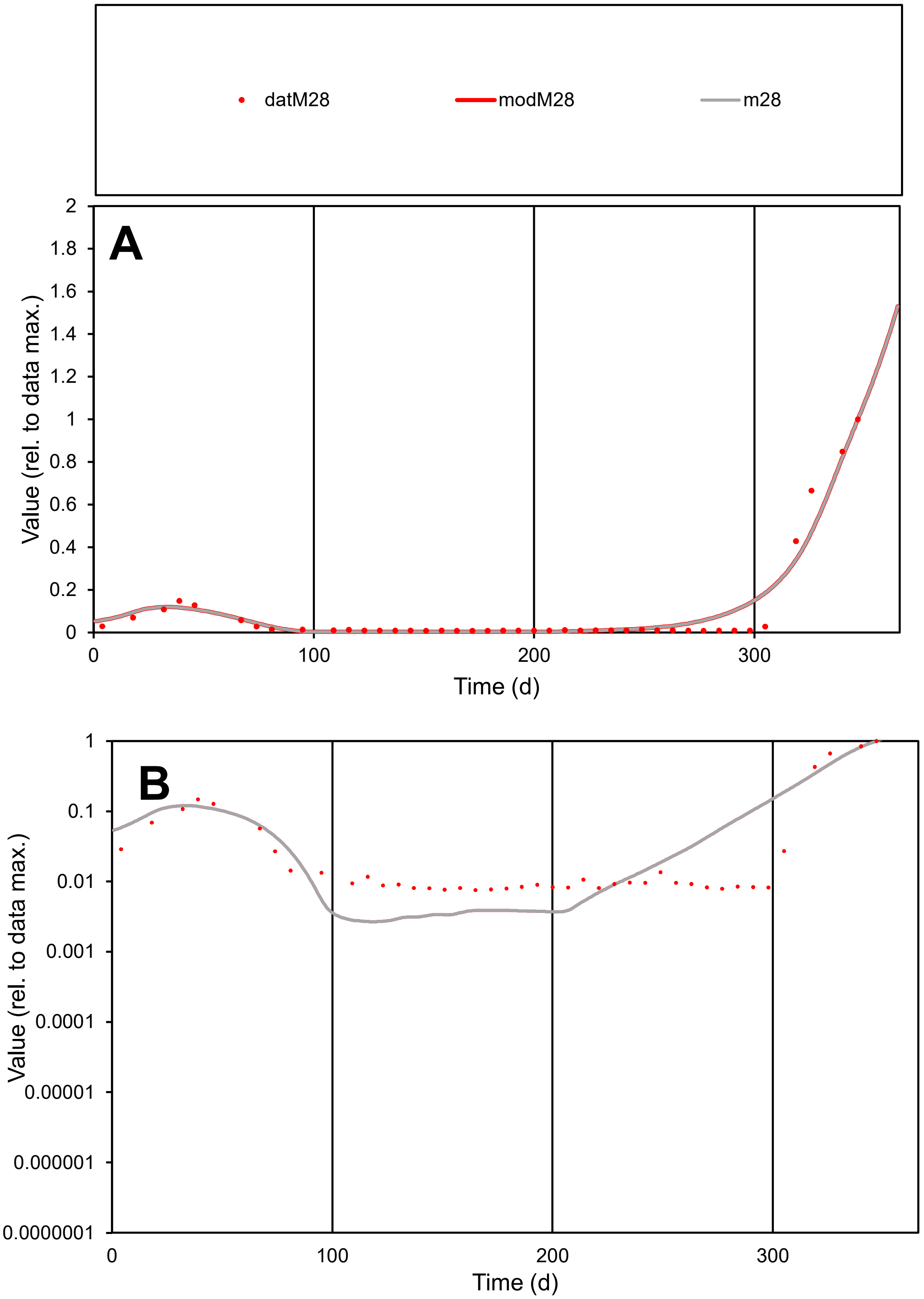


**C1**

Variables

M28 Provide jtd or Dst

CalibOnly?

2 1 to reproduce of

oNormalize?

2 1=no, 2=yes

ofxn calc*

0.52359

ofxn mod

0.59776

sRun

RunA1

zout

1

Ave

Data

0.00064

Model

0.0007

data/vMax

0.00707

ofxng

334.07

Min d/m

2.8156

ave/min d

12.0023

ave/min m

36.7304

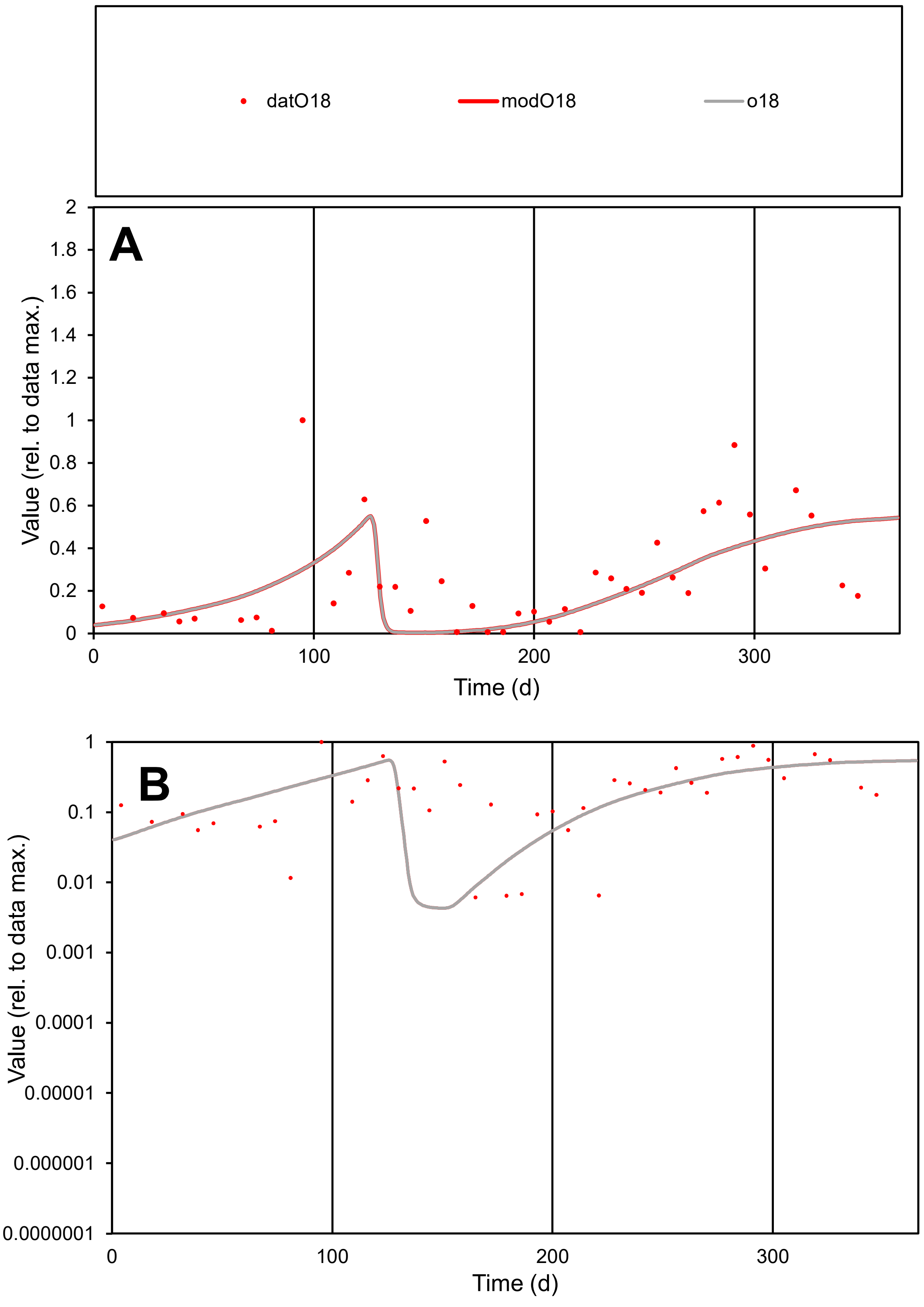


**C1**

Variables
O18 Provide jtd or Dst
CalibOnly?
2 1 to reproduce of
oNormalize?
2 1=no, 2=yes

ofxn calc*
0.79792
ofxn mod
0.80806

sRun
RunA1
zout
1

Ave
Data
0.00216
Model
0.00181

data/vMax
0.00837

ofxng
334.07

Min d/m
1.40467

ave/min d
42.2304
ave/min m
49.7871

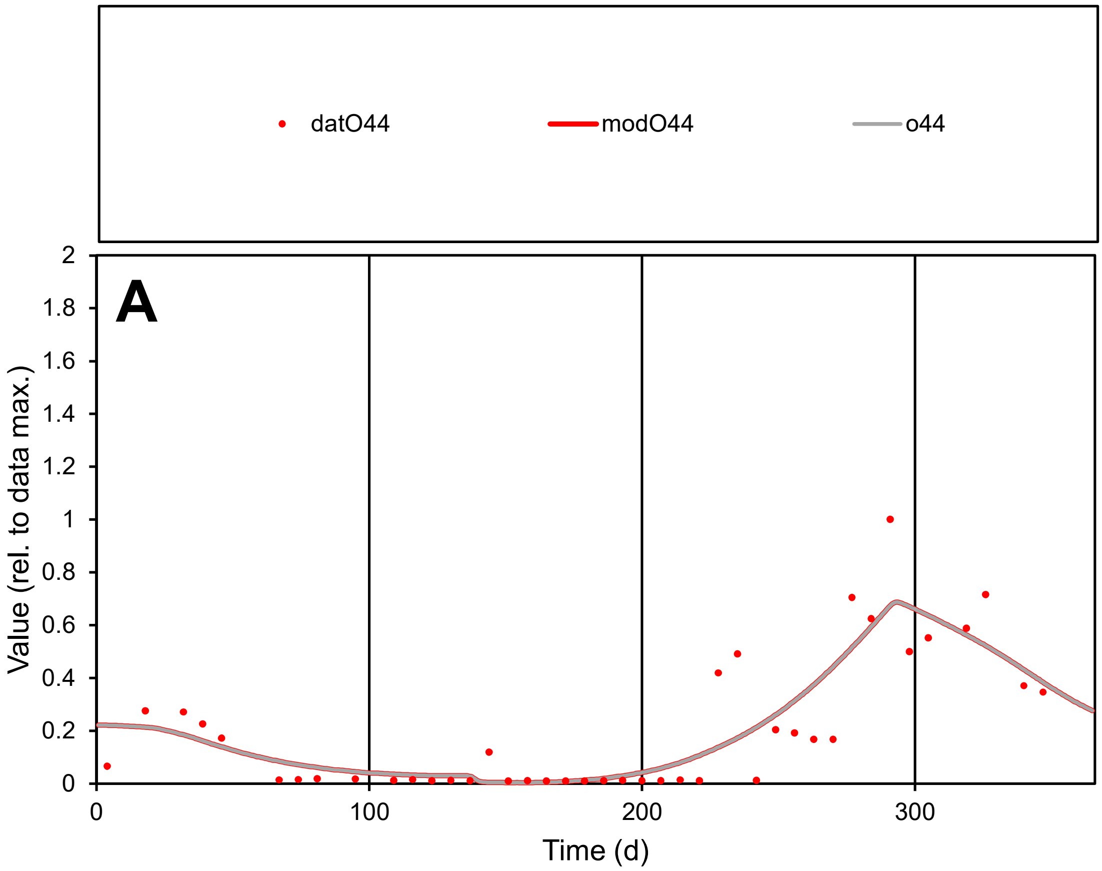


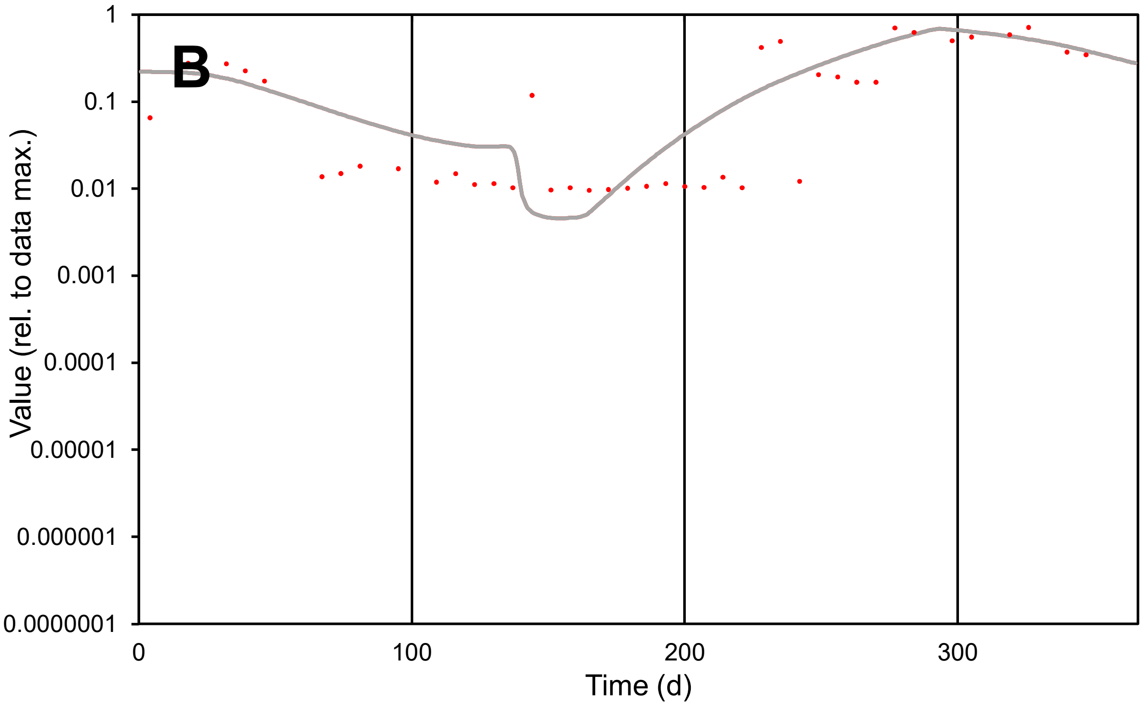


**C1**

Variables
O44 Provide jtd or Dst
CalibOnly?
2 1 to reproduce of
oNormalize?
2 1=no, 2=yes

ofxn calc*
0.61885
ofxn mod
0.64204

sRun
RunA1
zout
1

Ave
Data
0.00229
Model
0.00235

data/vMax
0.01144

ofxng
334.07

Min d/m
2.09249

ave/min d
20.947
ave/min m
44.8991

327_Calib_S_A

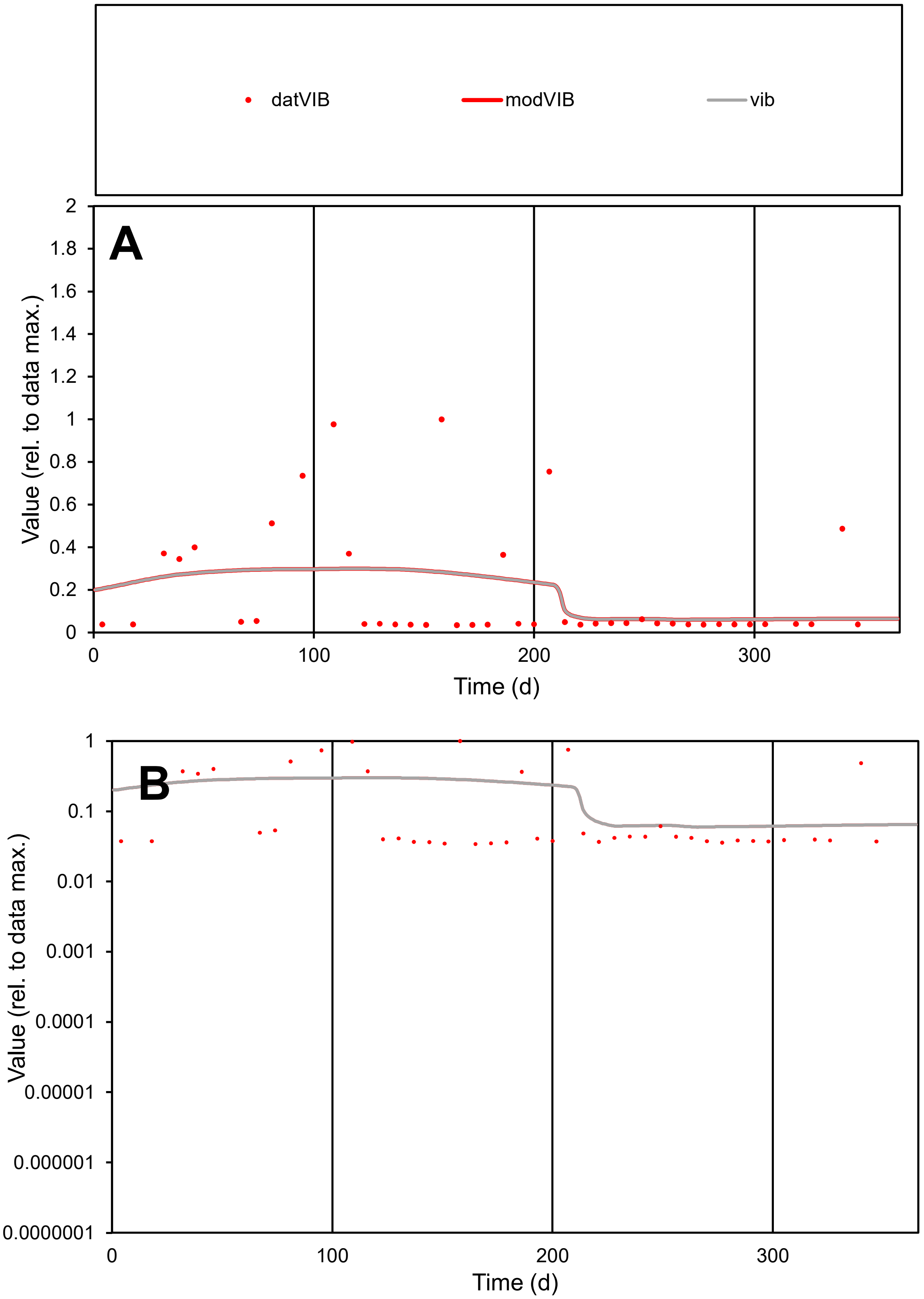


**C1**

Variables

VIB Provide jtd or Dst

CalibOnly?

2 1 to reproduce of

oNormalize?

2 1=no, 2=yes

ofxn calc*

1.35149

ofxn mod

1.36013

sRun

RunA1

zout

1

Ave

Data

9.2E-05

Model

9.4E-05

data/vMax

0.00051

ofxng

334.07

Min d/m

0.57536

ave/min d

5.25466

ave/min m

3.07835

327_Calib_S_A

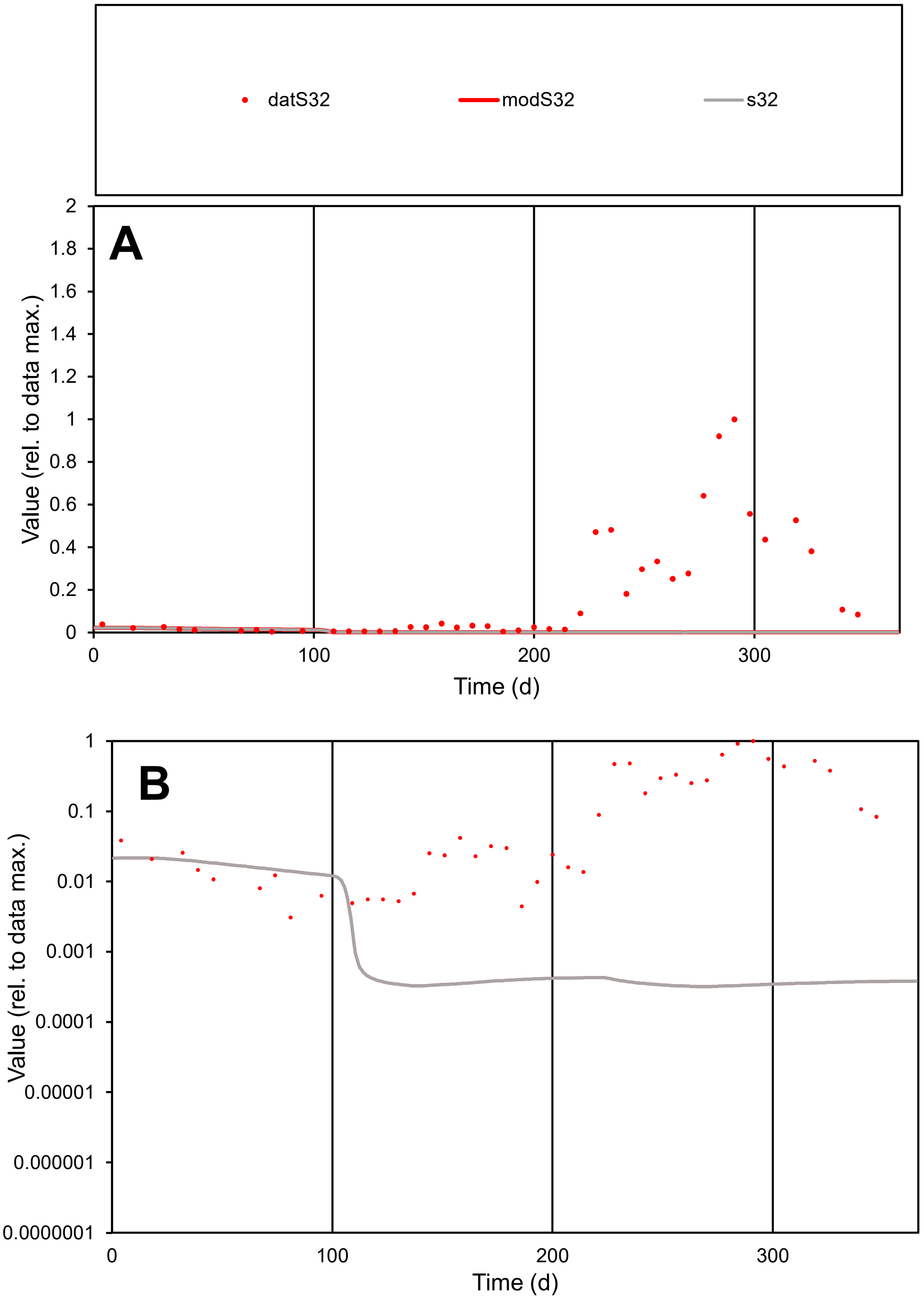


**C1**

Variables

S32 Provide jtd or Dst

CalibOnly?

2 1 to reproduce of

oNormalize?

2 1=no, 2=yes

ofxn calc*

1.74877

ofxn mod

1.75123

sRun

RunA1

zout

1

Ave

Data

0.00414

Model

9.5E-05

data/vMax

0.02337

ofxng

334.07

Min d/m

9.70083

ave/min d

57.225

ave/min m

12.8108

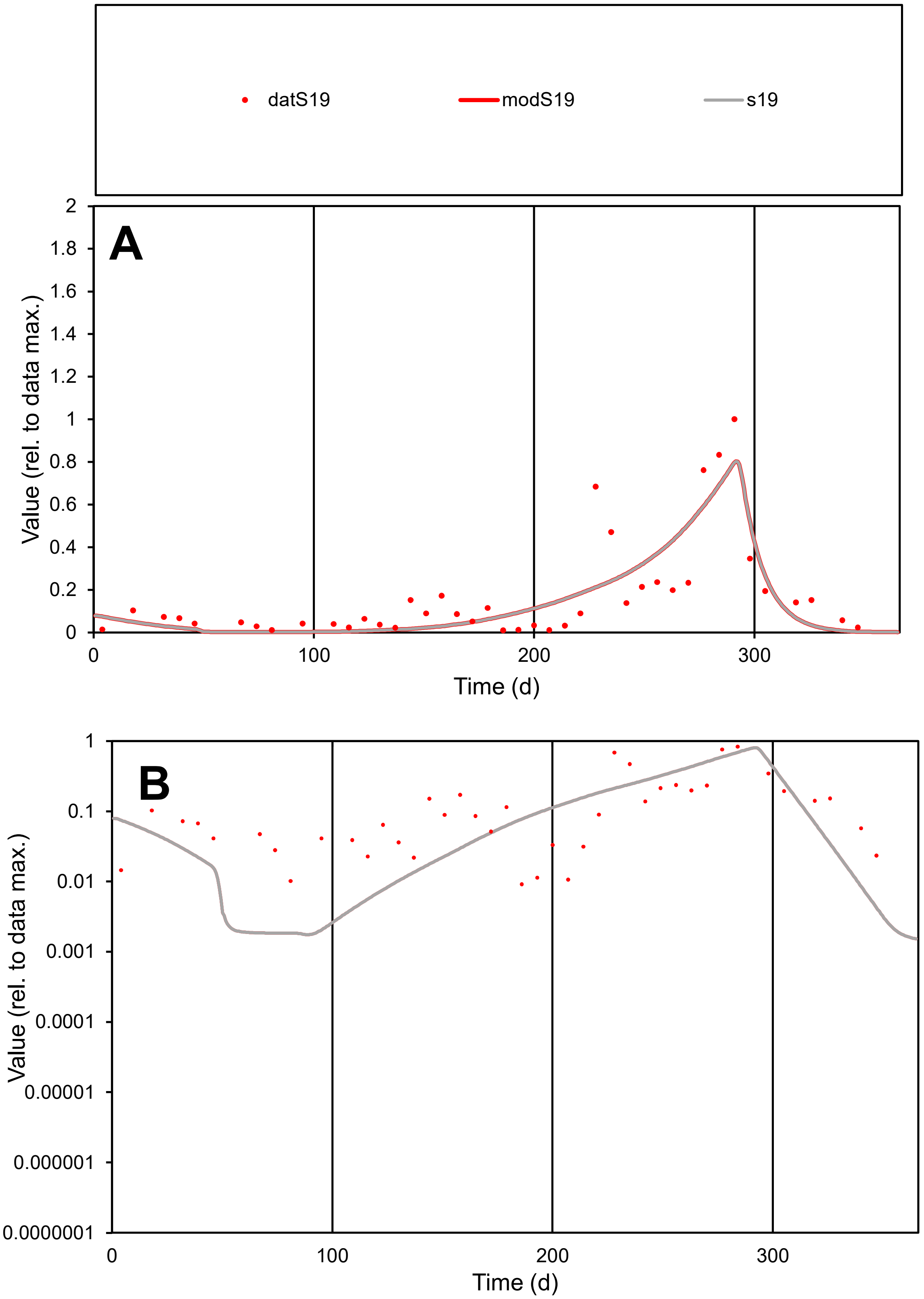


**C1**

Variables
S19 Provide jtd or Dst
CalibOnly?
2 1 to reproduce of
oNormalize?
2 1=no, 2=yes

ofxn calc*
0.75475
ofxn mod
0.77932

sRun
RunA1
zout
1

Ave
Data
0.00346
Model
0.00317

data/vMax
0.02034

ofxng
334.07

Min d/m
4.96401

ave/min d
18.5697
ave/min m
84.4442

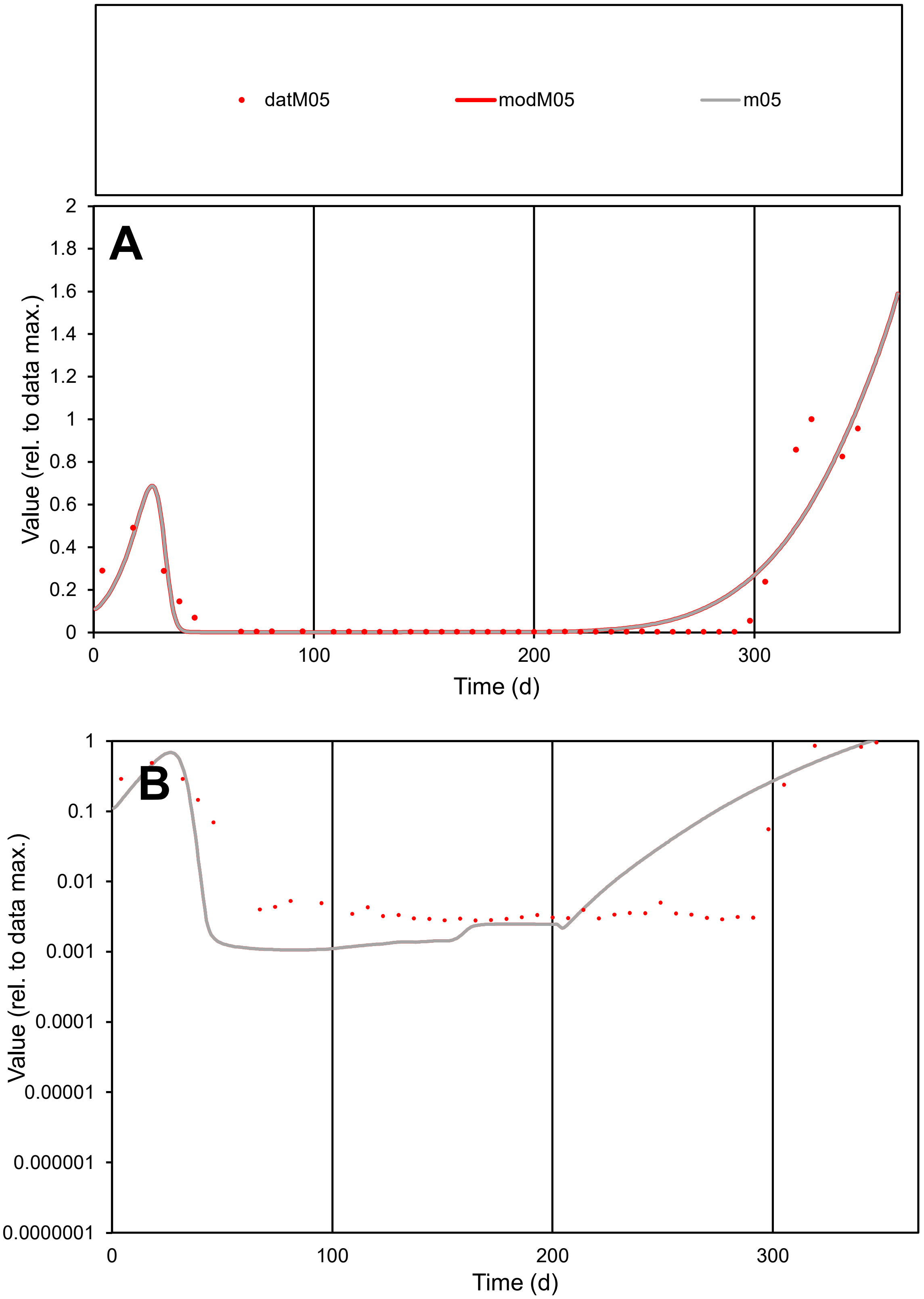


C1

Variables
M05 Provide jtd or Dst
CalibOnly?
2 1 to reproduce of
oNormalize?
2 1=no, 2=yes

ofxn calc*
0.80022
ofxn mod
0.84524

sRun
RunA1
zout
1

Ave
Data
0.00195
Model
0.00203

data/vMax
0.01536

ofxng
334.07

Min d/m
2.63362

ave/min d
45.5252
ave/min m
125.048

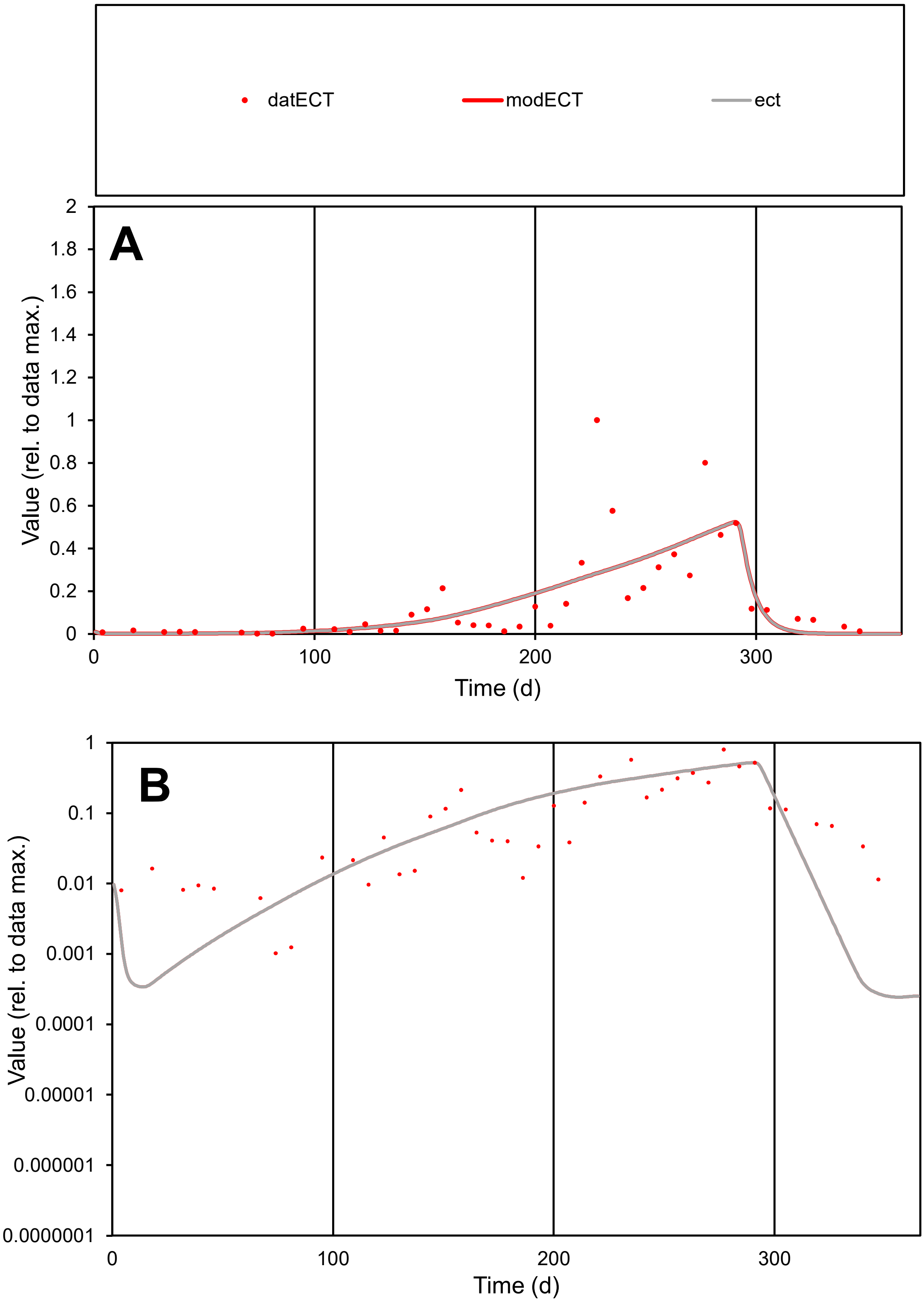


**C1**

Variables
ECT Provide jtd or Dst
CalibOnly?
2 1 to reproduce of
oNormalize?
2 1=no, 2=yes

ofxn calc*
0.94566
ofxn mod
0.96268

sRun
RunA1
zout
1

Ave
Data
0.00666
Model
0.00637

data/vMax
0.04282

ofxng
334.07

Min d/m
3.86596

ave/min d
152.017
ave/min m
561.74

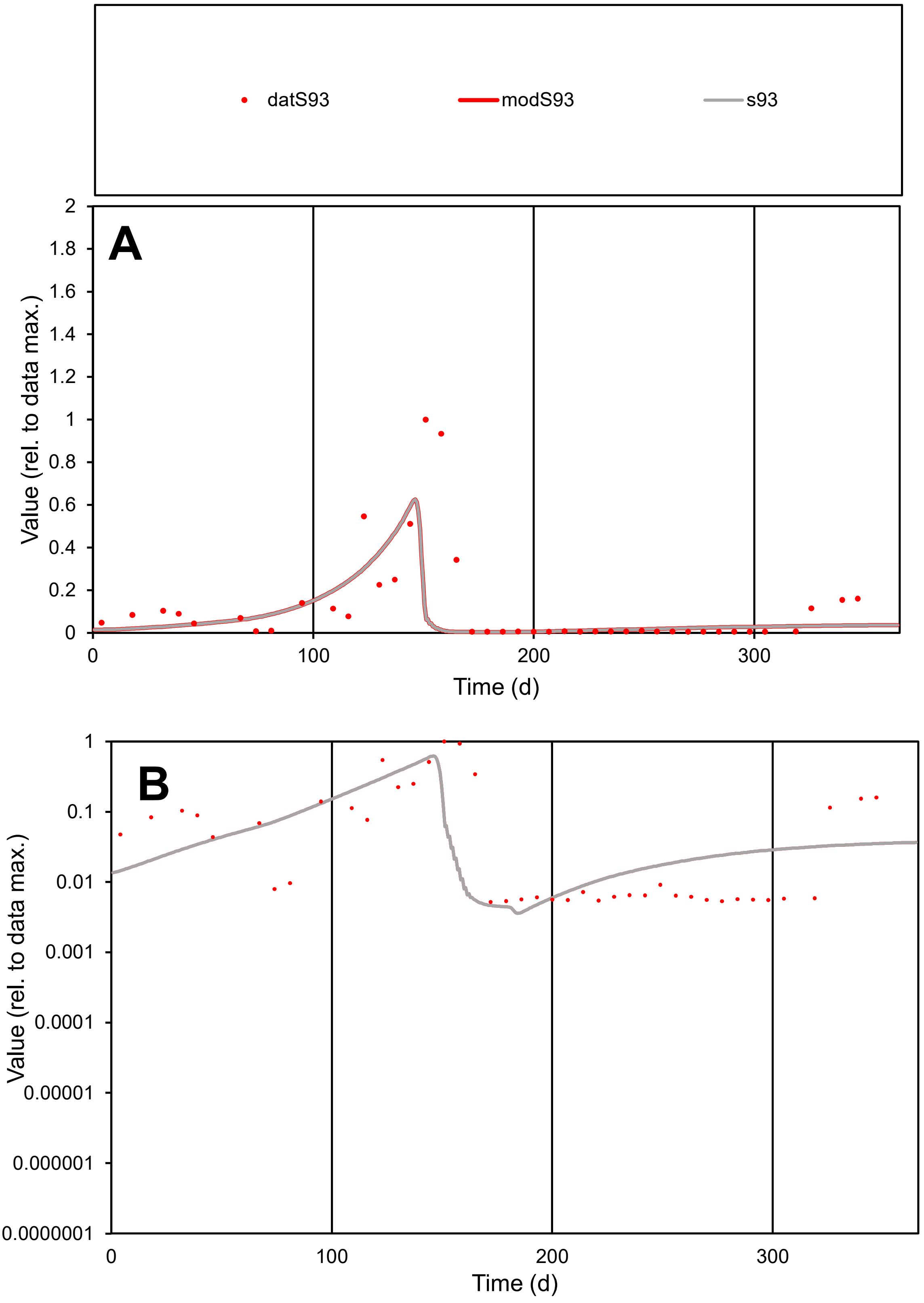


**C1**

Variables
S93 Provide jtd or Dst
CalibOnly?
2 1 to reproduce of
oNormalize?
2 1=no, 2=yes

ofxn calc*
1.52811
ofxn mod
1.69964

sRun
RunA1
zout
1

Ave
Data
0.00155
Model
0.00111

data/vMax
0.01268

ofxng
334.07

Min d/m
1.3281

ave/min d
23.557
ave/min m
22.3984

327_Calib_S_A

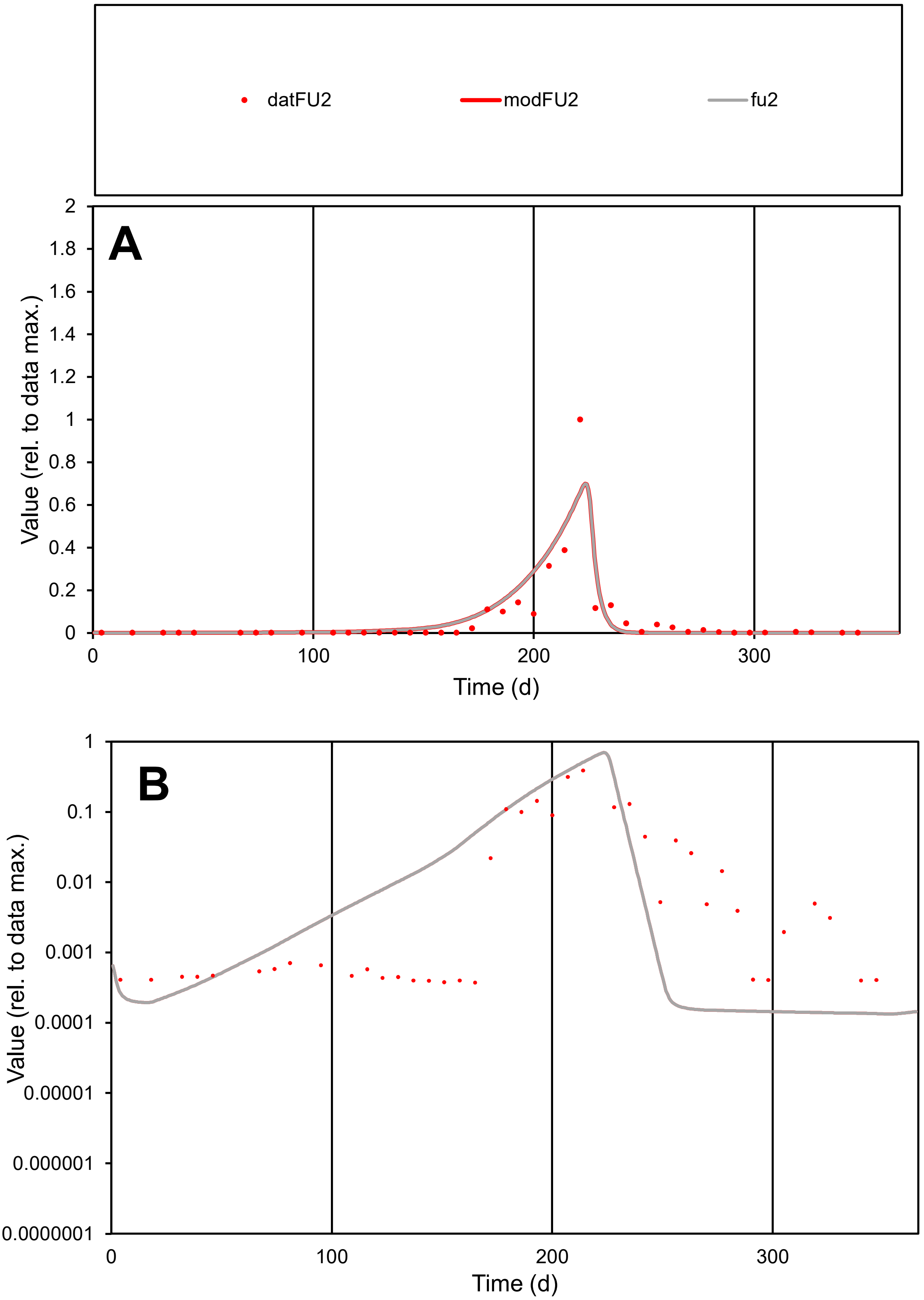


**C1**

Variables
FU2 Provide jtd or Dst
CalibOnly?
2 1 to reproduce of
oNormalize?
2 1=no, 2=yes

ofxn calc*
1.1113
ofxn mod
1.19013

sRun
RunA1
zout
1

Ave
Data
0.00415
Model
0.00461

data/vMax
0.06778

ofxng
334.07

Min d/m
2.78101

ave/min d
163.935
ave/min m
506.217

327_Calib_S_A

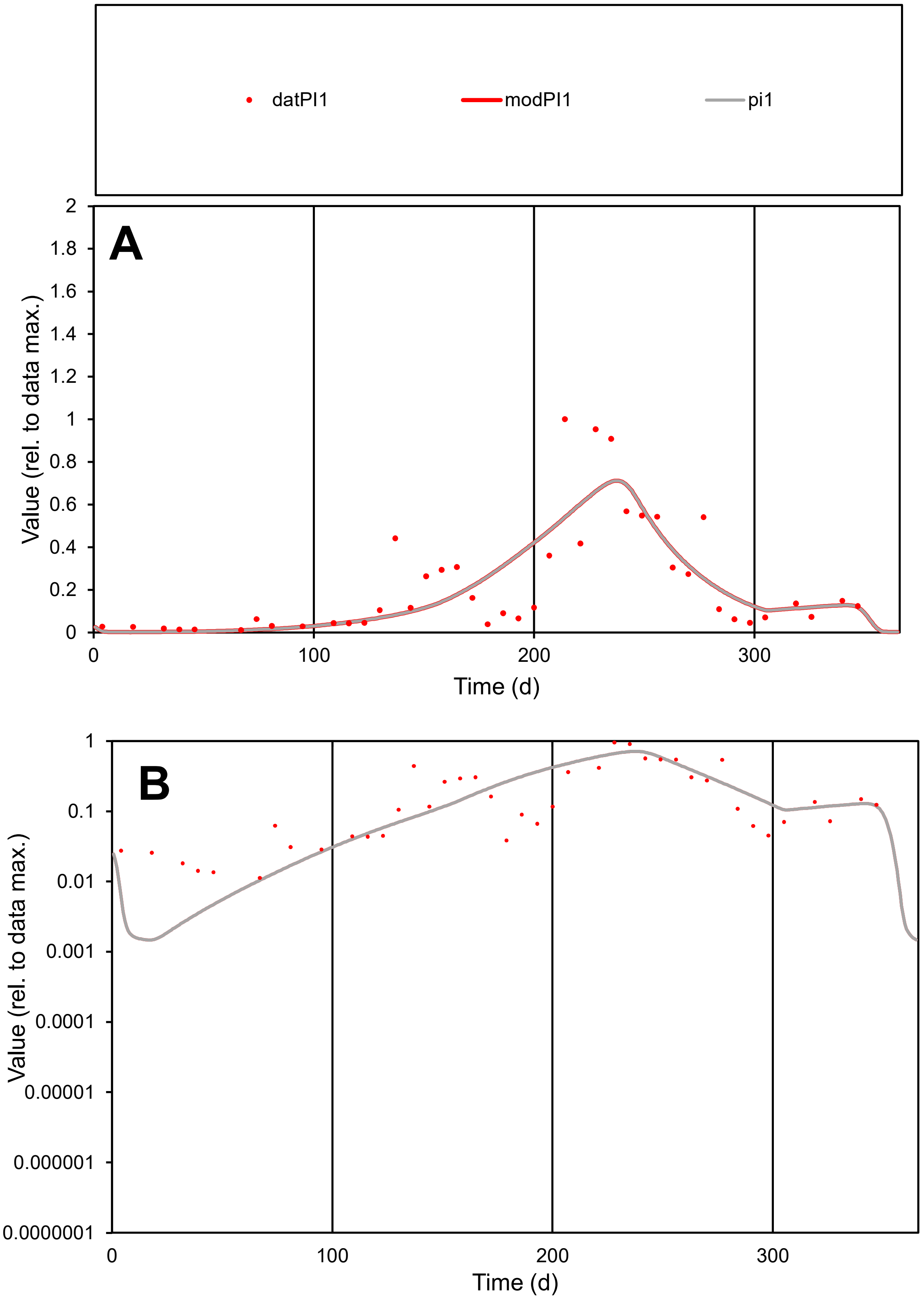


**C1**

Variables

PI1 Provide jtd or Dst

CalibOnly?

2 1 to reproduce of

oNormalize?

2 1=no, 2=yes

ofxn calc*

0.67517

ofxn mod

0.698

sRun

RunA1

zout

1

Ave

Data

0.00099

Model

0.00097

data/vMax

0.00437

ofxng

334.07

Min d/m

7.42413

ave/min d

20.2107

ave/min m

146.054

327_Calib_S_A

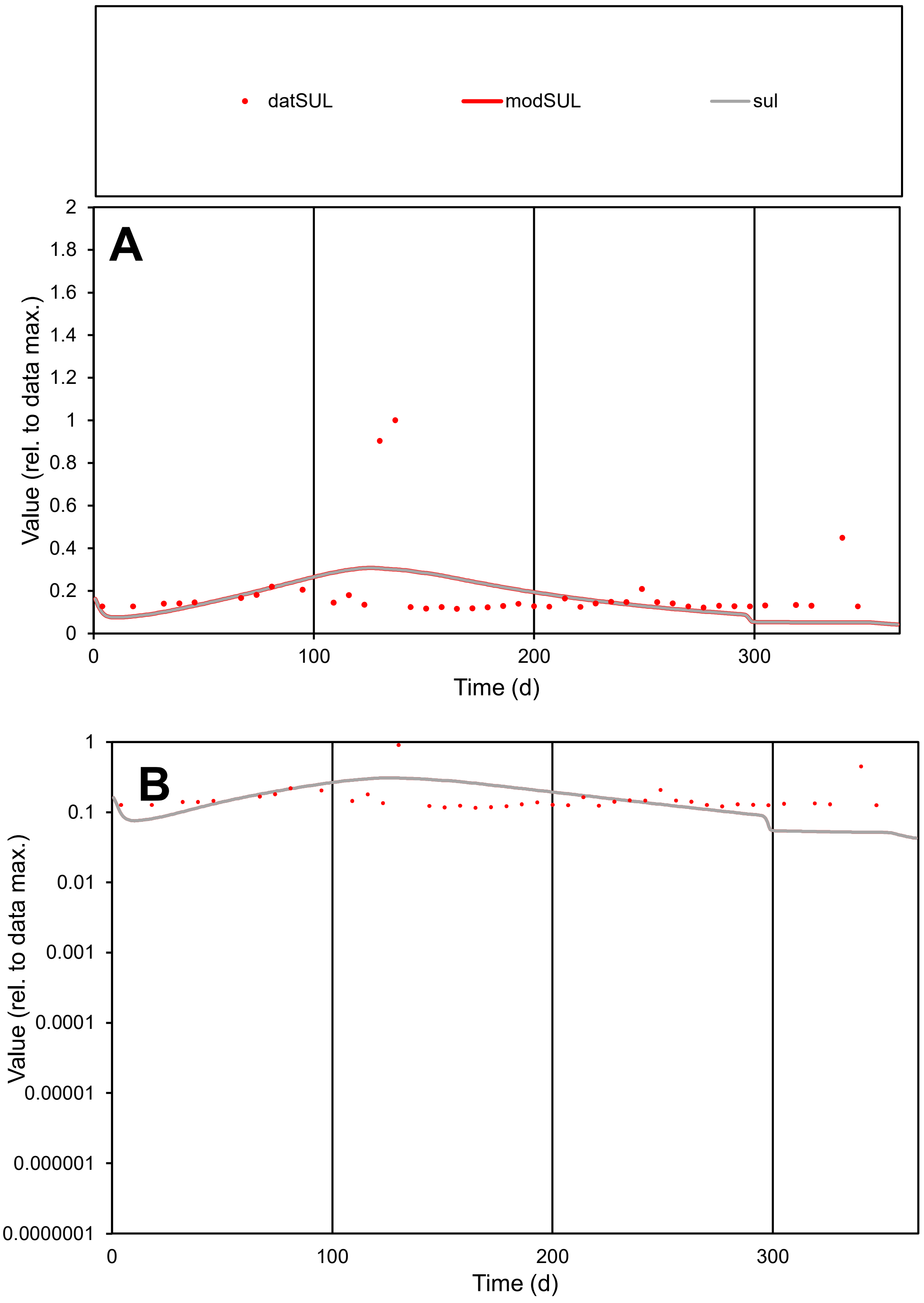


**C1**

Variables
SUL Provide jtd or Dst
CalibOnly?
2 1 to reproduce of
oNormalize?
2 1=no, 2=yes

ofxn calc*
0.92453
ofxn mod
0.93241

sRun
RunA1
zout
1

Ave
Data
0.00013
Model
0.00011

data/vMax
0.00068

ofxng
334.07

Min d/m
2.22985

ave/min d
1.6183
ave/min m
3.25538

327_Calib_S_A

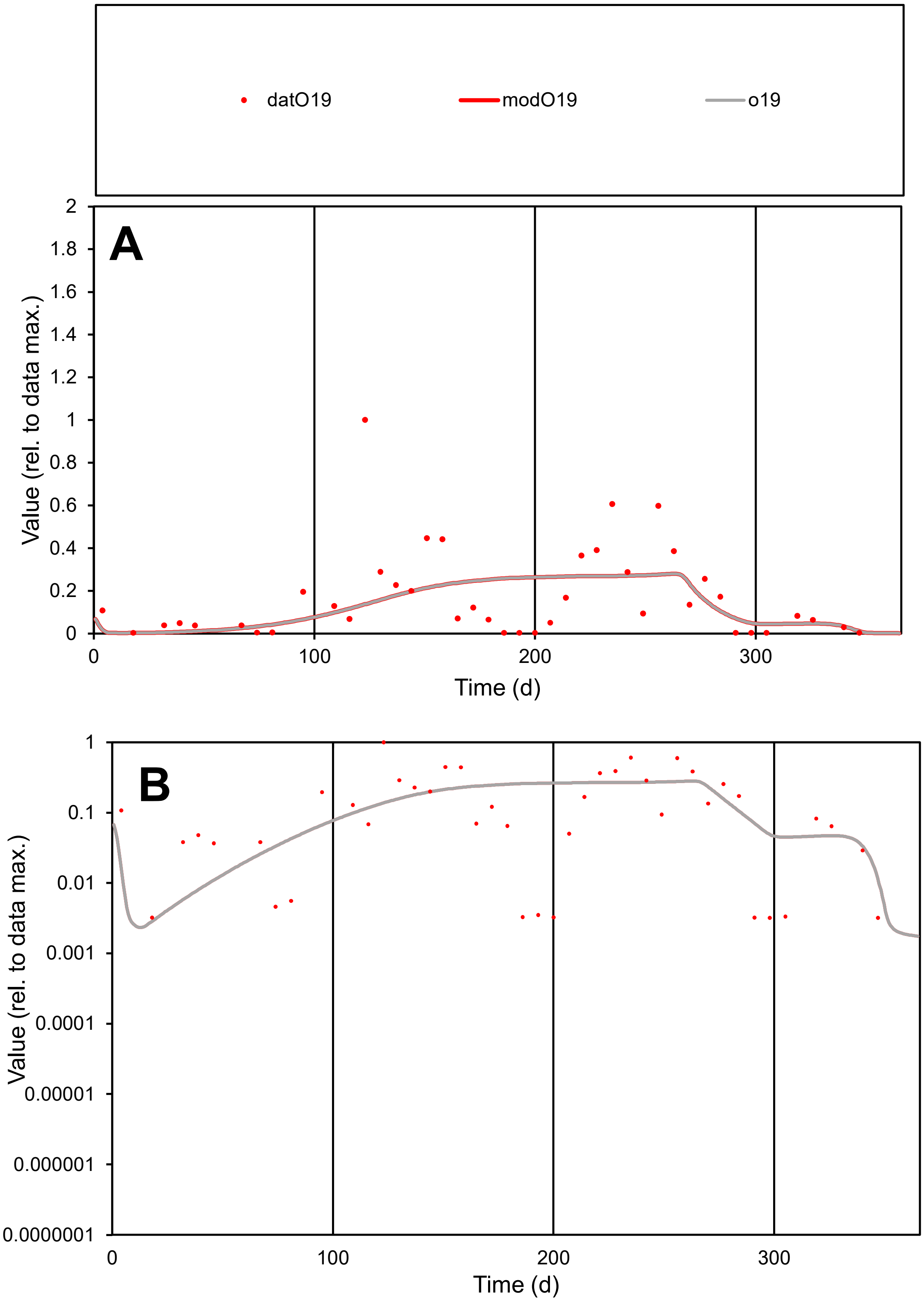


**C1**

Variables
O19 Provide jtd or Dst
CalibOnly?
2 1 to reproduce of
oNormalize?
2 1=no, 2=yes

ofxn calc*
1.1077
ofxn mod
1.11934

sRun
RunA1
zout
1

Ave
Data
0.00164
Model
0.00144

data/vMax
0.00954

ofxng
334.07

Min d/m
1.03142

ave/min d
53.844
ave/min m
48.6362

327_Calib_S_A

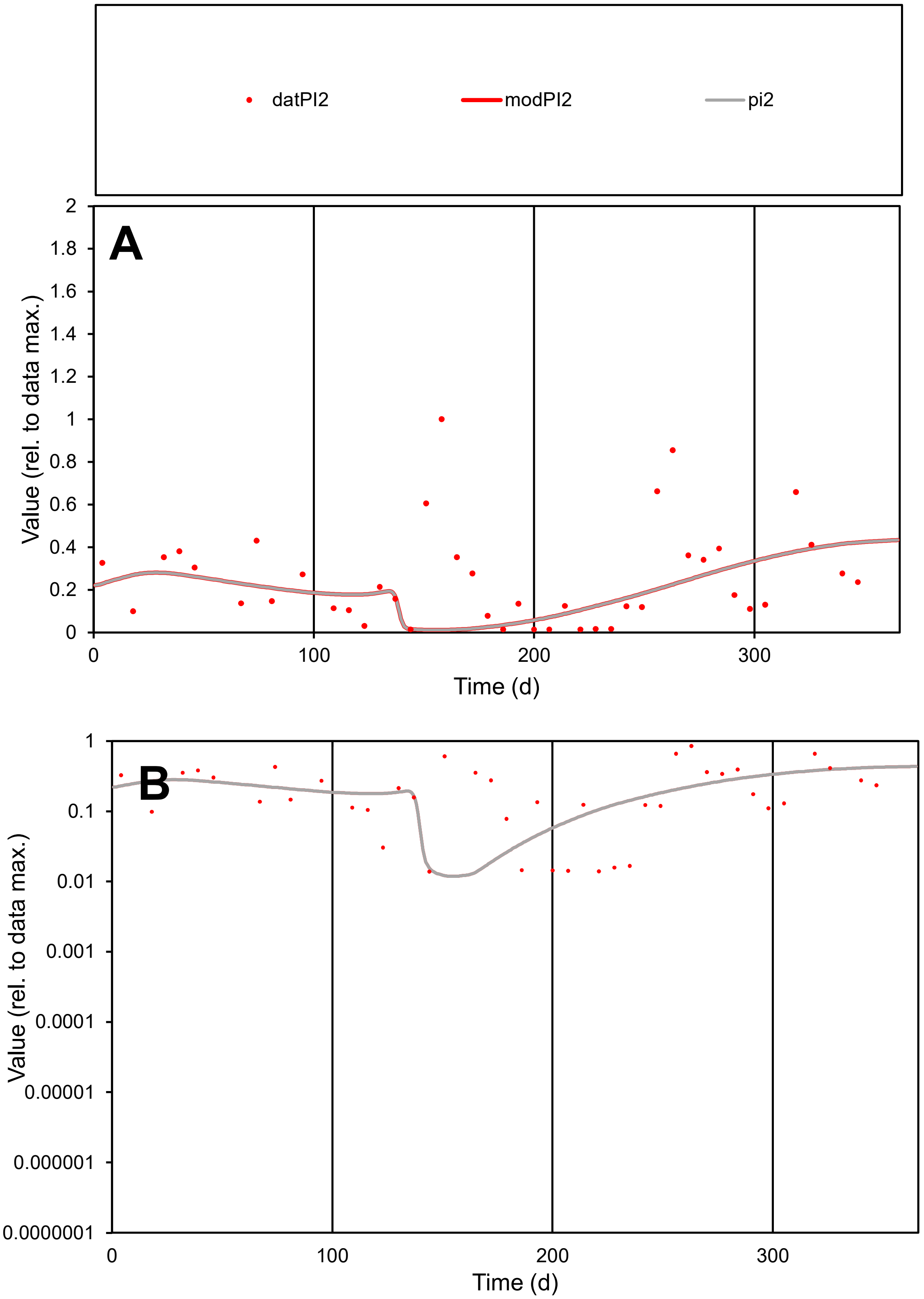


**C1**

Variables
PI2 Provide jtd or Dst
CalibOnly?
2 1 to reproduce of
oNormalize?
2 1=no, 2=yes

ofxn calc*
0.98483
ofxn mod
0.99253

sRun
RunA1
zout
1

Ave
Data
0.00054
Model
0.00041

data/vMax
0.00215

ofxng
334.07

Min d/m
1.15432

ave/min d
18.236
ave/min m
15.9863

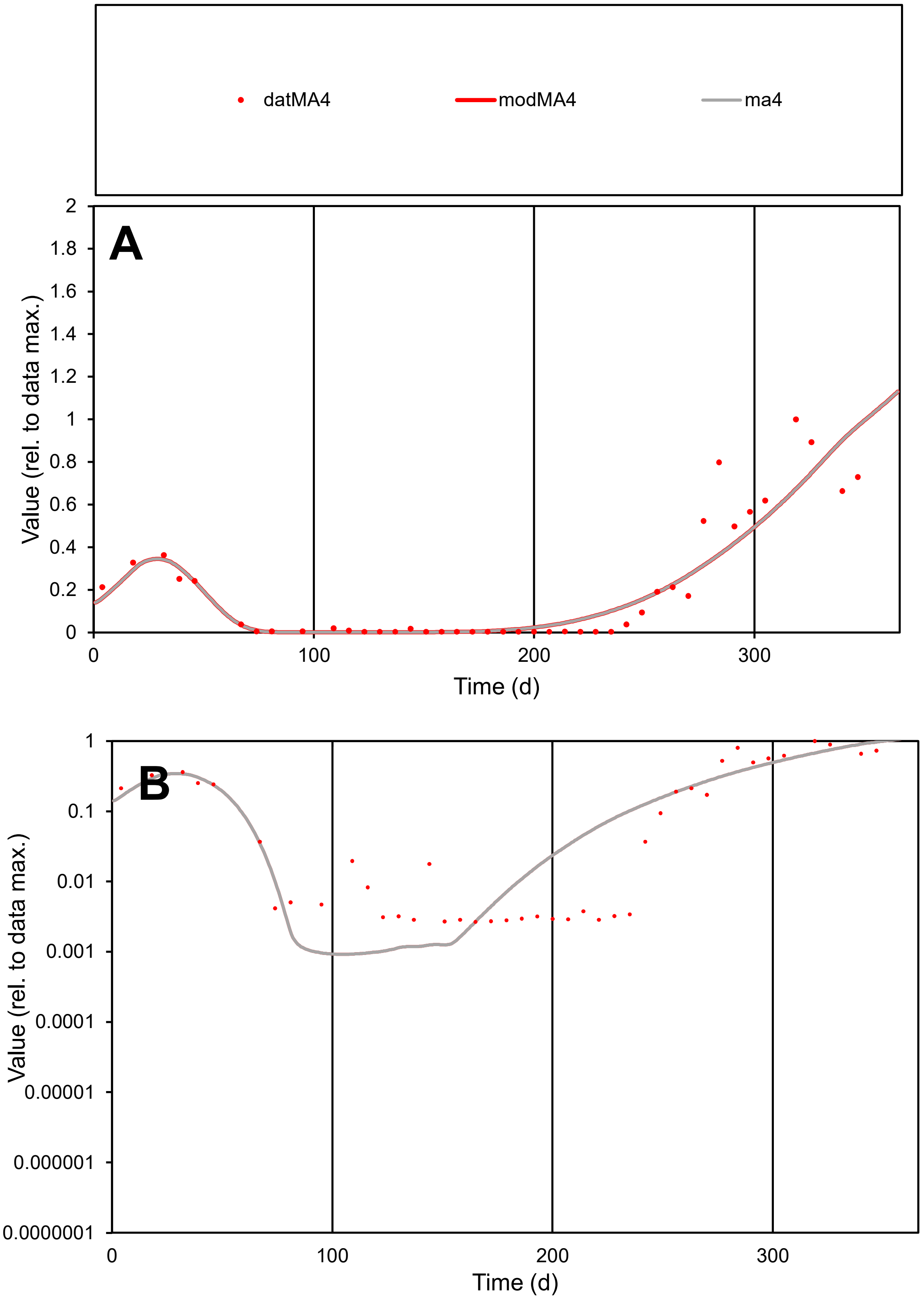


C1

Variables
MA4 Provide jtd or Dst
CalibOnly?
2 1 to reproduce of
oNormalize?
2 1=no, 2=yes

ofxn calc*
0.55048
ofxn mod
0.57555

sRun
RunA1
zout
1

Ave
Data
0.00146
Model
0.00141

data/vMax
0.00717

ofxng
334.07

Min d/m
2.85629

ave/min d
76.1403
ave/min m
210.74

327_Calib_S_A

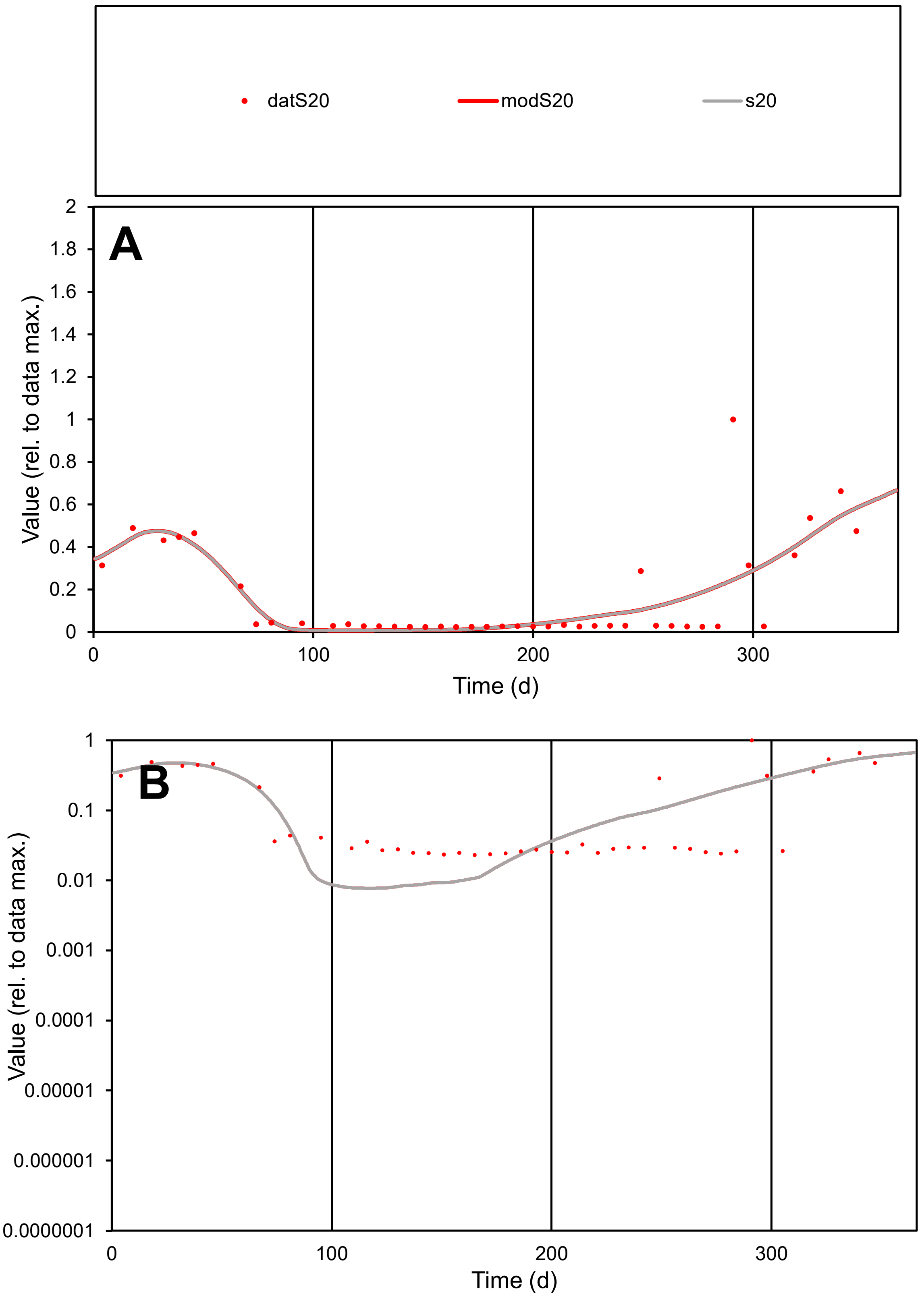


**C1**

Variables
S20 Provide jtd or Dst
CalibOnly?
2 1 to reproduce of
oNormalize?
2 1=no, 2=yes

ofxn calc*
0.87558
ofxn mod
0.89477

sRun
RunA1
zout
1

Ave
Data
0.00054
Model
0.00055

data/vMax
0.00335

ofxng
334.07

Min d/m
3.00622

ave/min d
7.00642
ave/min m
21.3749

327_Calib_S_A

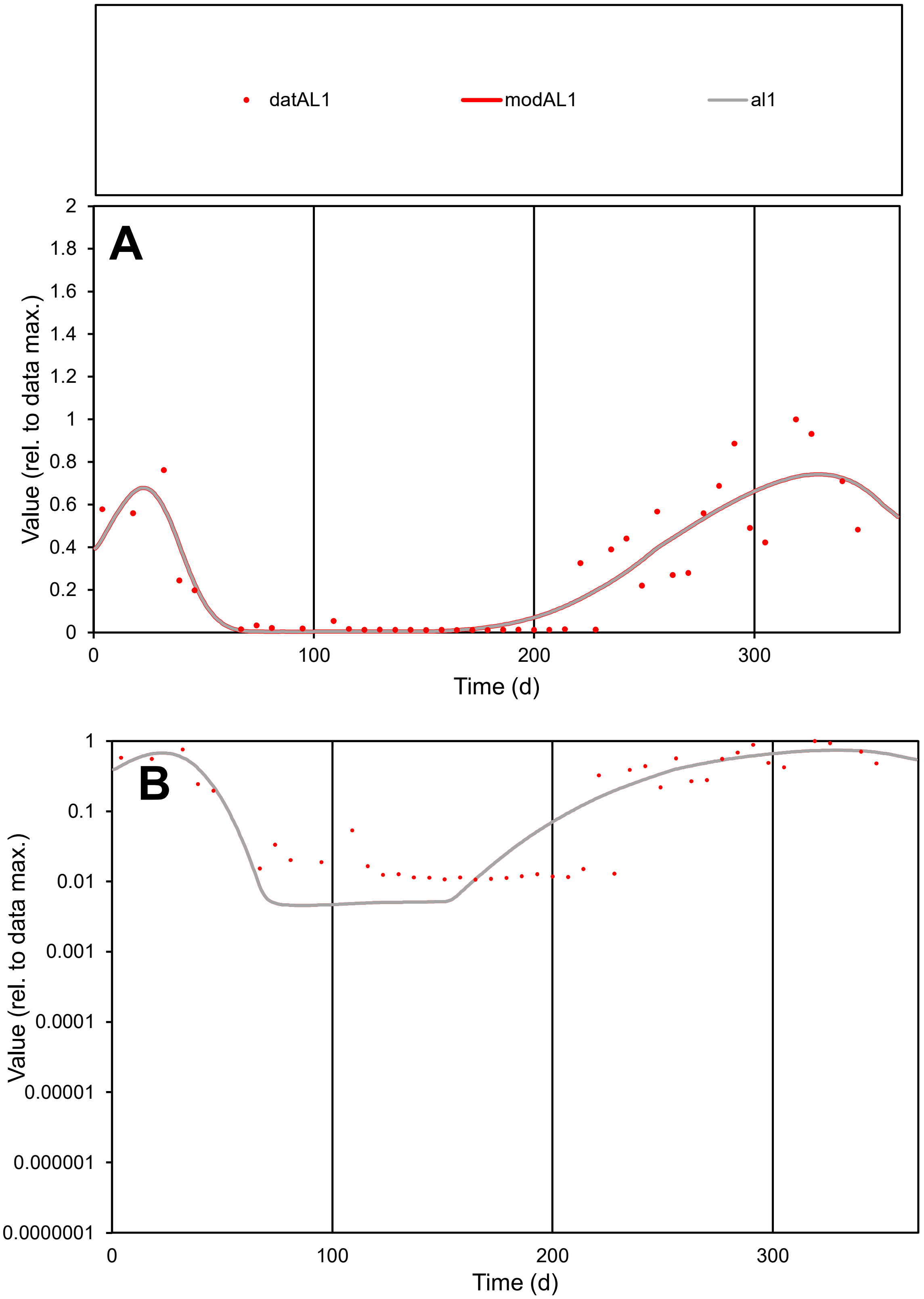


**C1**

Variables

AL1 Provide jtd or Dst

CalibOnly?

2 1 to reproduce of

oNormalize?

2 1=no, 2=yes

ofxn calc*

0.462

ofxn mod

0.4901

sRun

RunA1

zout

1

Ave

Data

0.0011

Model

0.0011

data/vMax

0.00408

ofxng

334.07

Min d/m

2.33405

ave/min d

25.2894

ave/min m

59.062

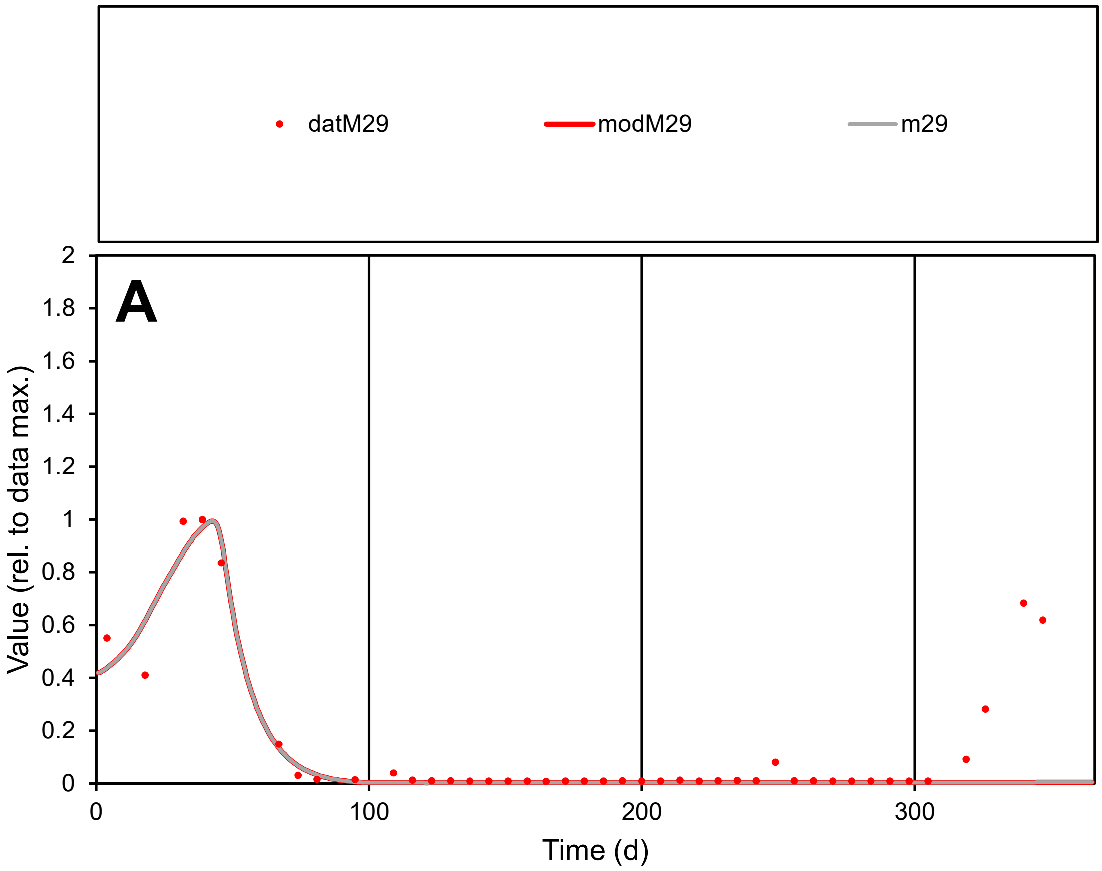


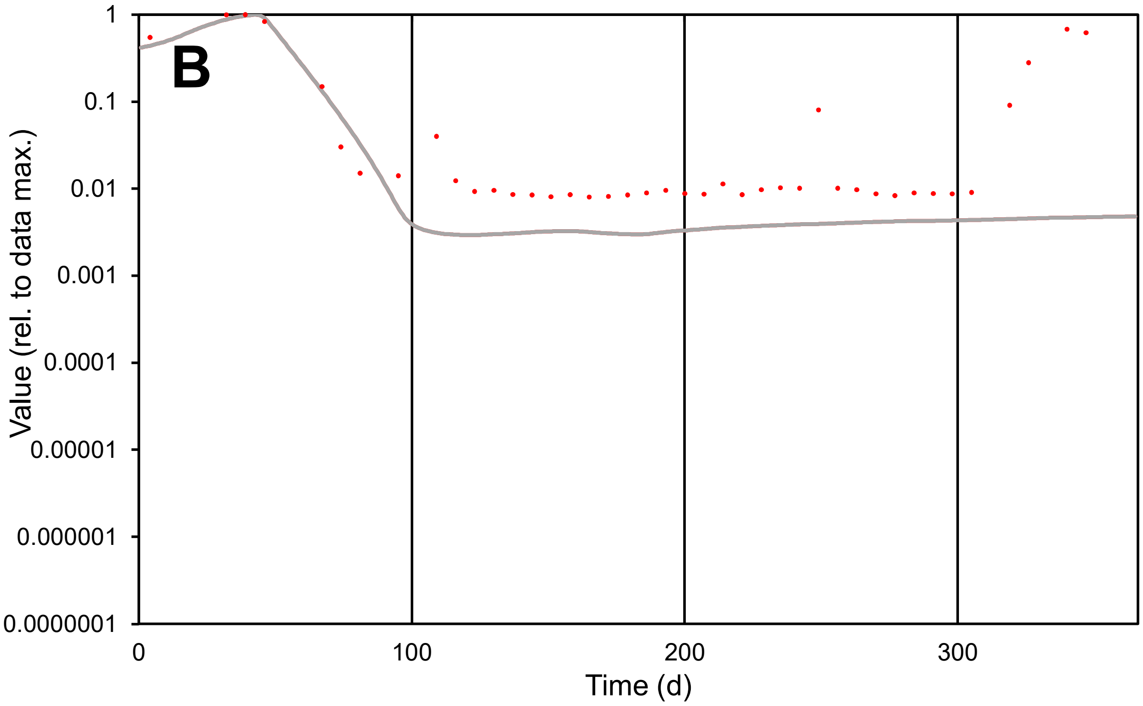


**C1**

Variables
M29 Provide jtd or Dst
CalibOnly?
2 1 to reproduce of
oNormalize?
2 1=no, 2=yes

ofxn calc*
1.07036
ofxn mod
1.10576

sRun
RunA1
zout
1

Ave
Data
0.00097
Model
0.00066

data/vMax
0.00675

ofxng
334.07

Min d/m
2.72264

ave/min d
17.9719
ave/min m
33.4075

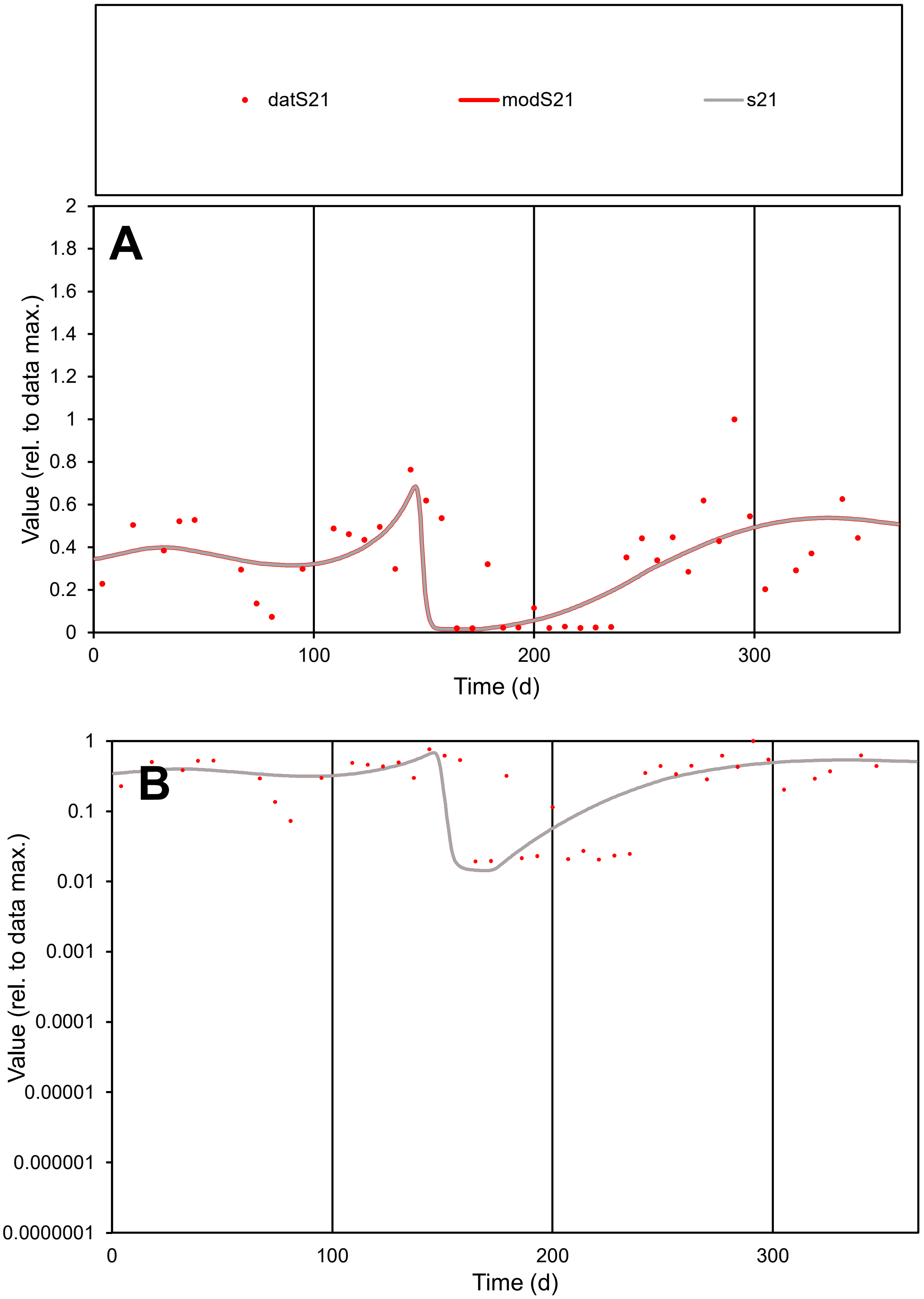


**C1**

Variables
S21 Provide jtd or Dst
CalibOnly?
2 1 to reproduce of
oNormalize?
2 1=no, 2=yes

ofxn calc*
0.52712
ofxn mod
0.55974

sRun
RunA1
zout
1

Ave
Data
0.00071
Model
0.00067

data/vMax
0.00211

ofxng
334.07

Min d/m
1.33421

ave/min d
17.3982
ave/min m
22.0052

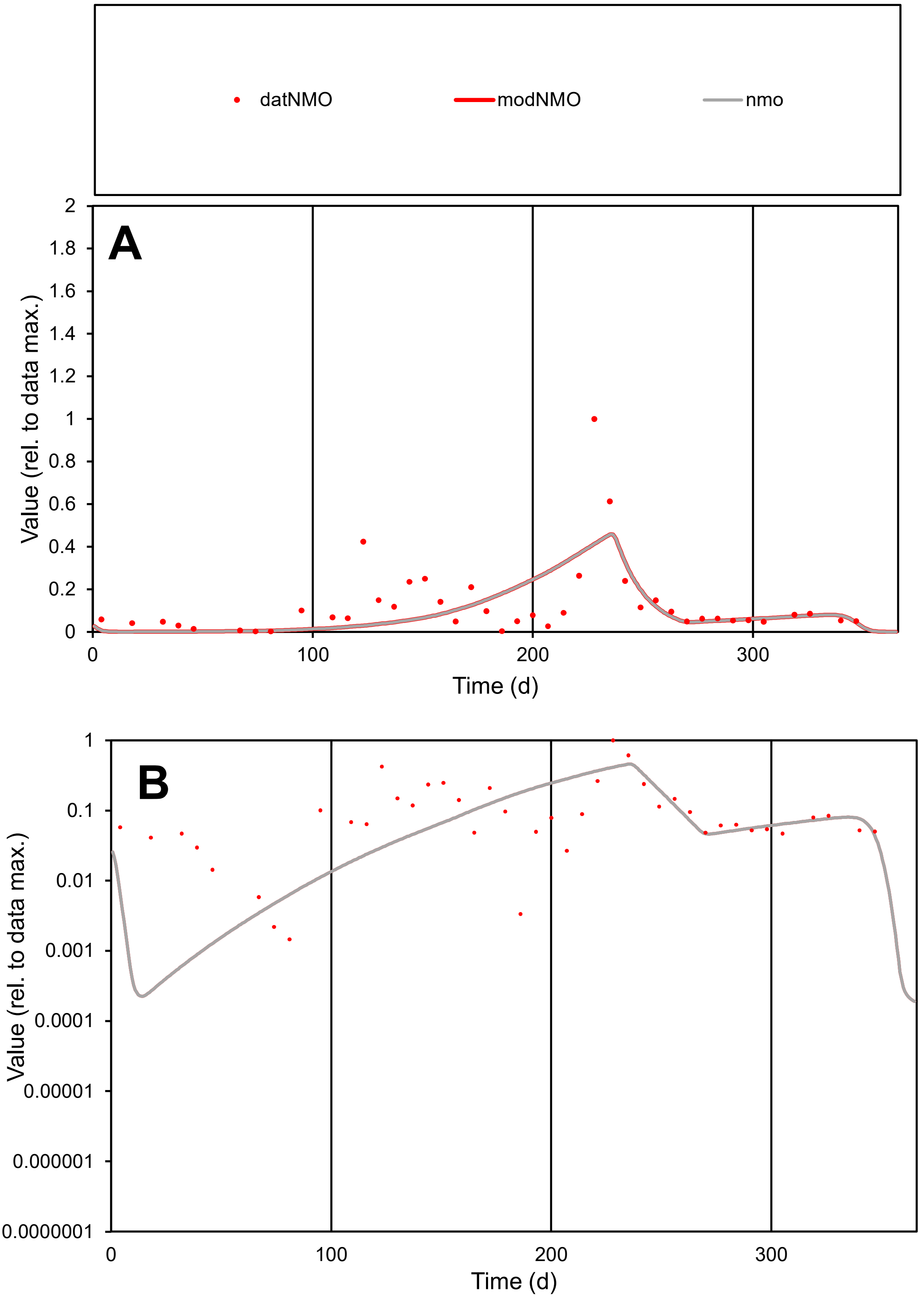


**C1**

Variables
NMO Provide jtd or Dst
CalibOnly?
2 1 to reproduce of
oNormalize?
2 1=no, 2=yes

ofxn calc*
1.11165
ofxn mod
1.12371

sRun
RunA1
zout
1

Ave
Data
0.00239
Model
0.00202

data/vMax
0.01853

ofxng
334.07

Min d/m
4.98896

ave/min d
88.0708
ave/min m
371.348

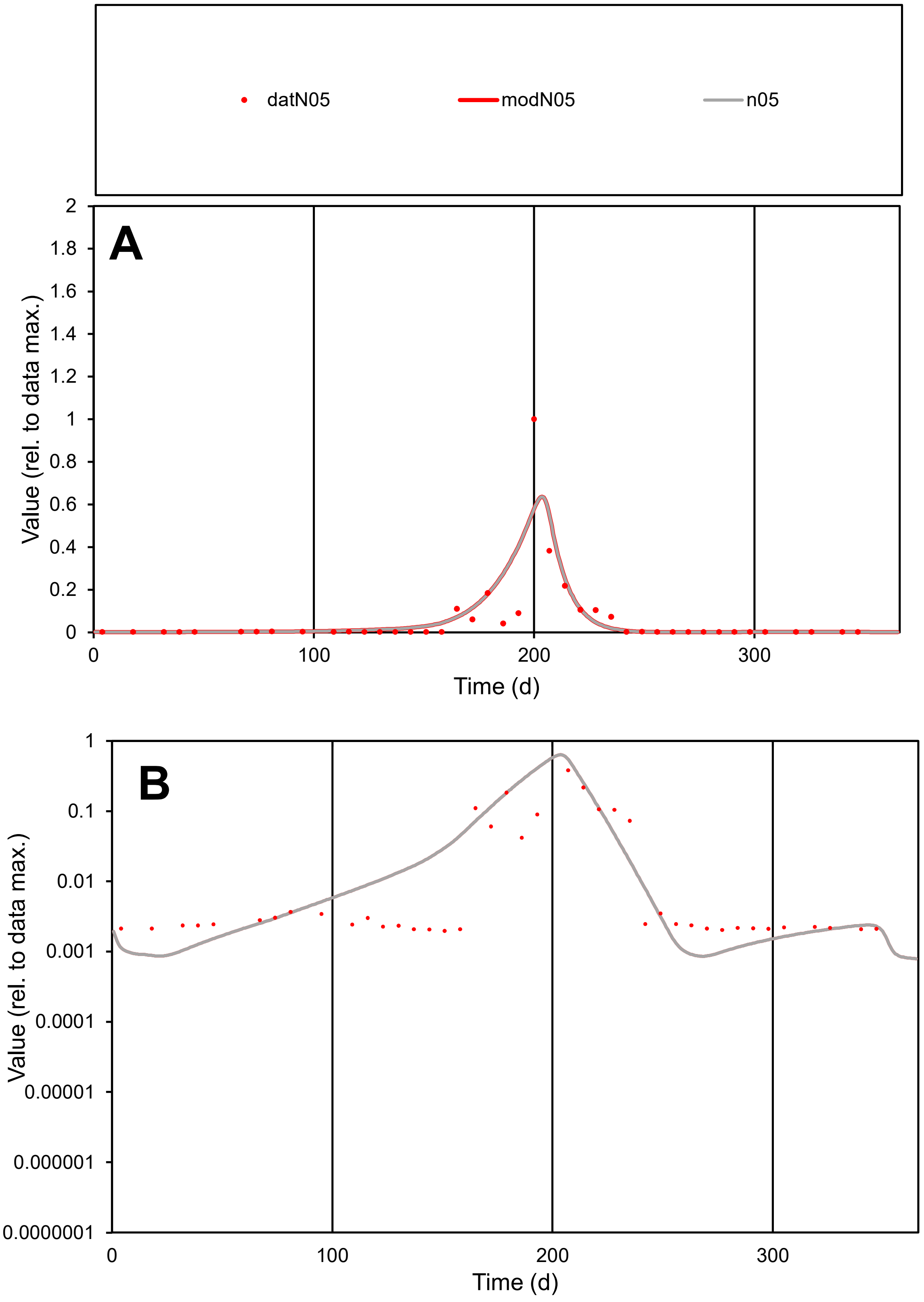


**C1**

Variables
N05 Provide jtd or Dst
CalibOnly?
2 1 to reproduce of
oNormalize?
2 1=no, 2=yes

ofxn calc*
1.52102
ofxn mod
1.58484

sRun
RunA1
zout
1

Ave
Data
0.00118
Model
0.00132

data/vMax
0.02031

ofxng
334.07

Min d/m
2.2592

ave/min d
29.6118
ave/min m
74.6163

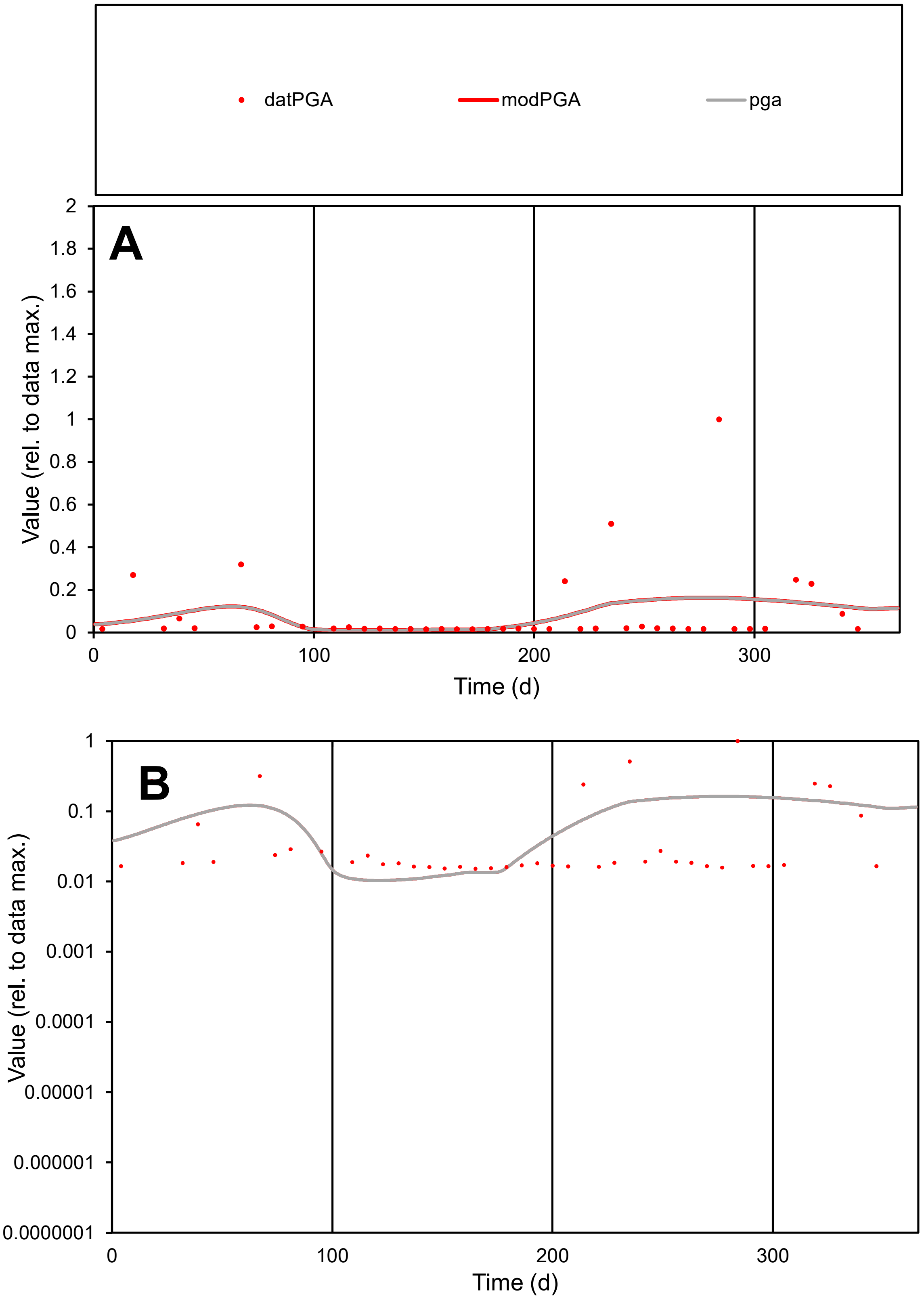


C1

Variables
PGA Provide jtd or Dst
CalibOnly?
2 1 to reproduce of
oNormalize?
2 1=no, 2=yes

ofxn calc*
1.98008
ofxn mod
1.98701

sRun
RunA1
zout
1

Ave
Data
9.5E-05
Model
9.3E-05

data/vMax
0.00111

ofxng
334.07

Min d/m
1.46401

ave/min d
5.60445
ave/min m
8.03316

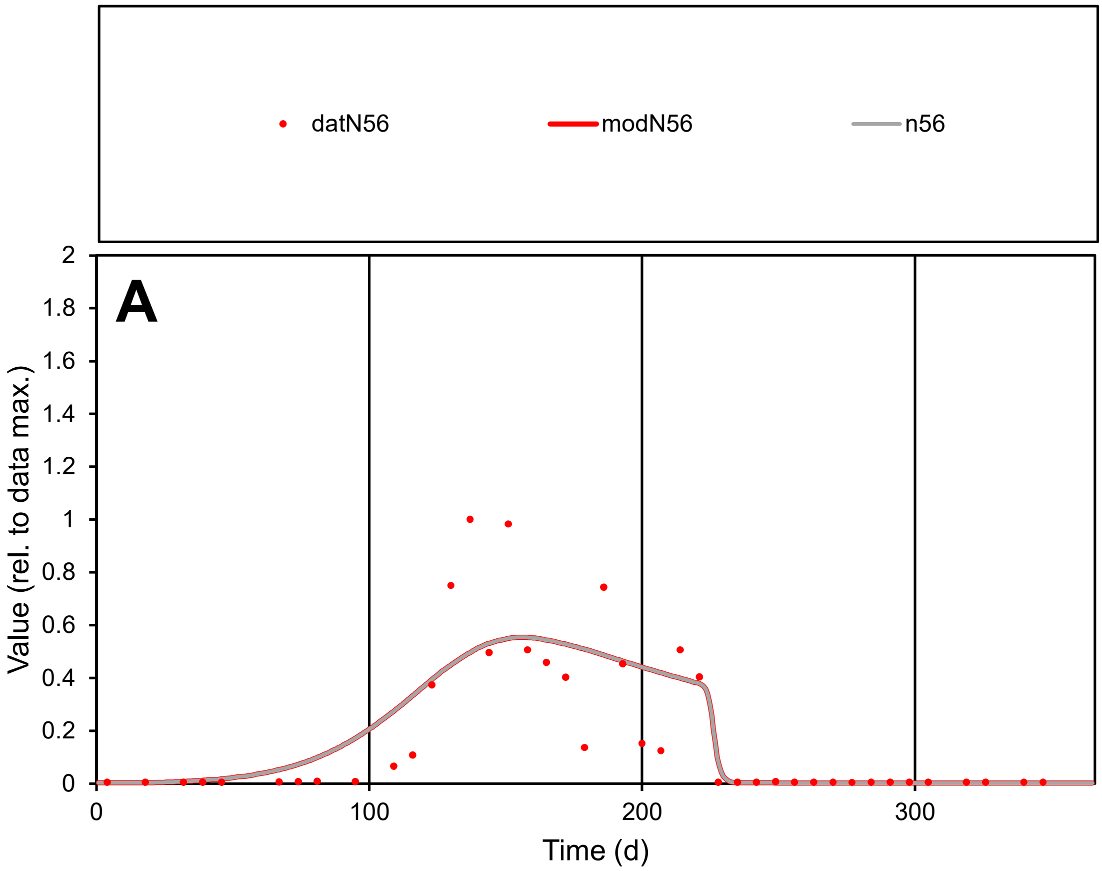


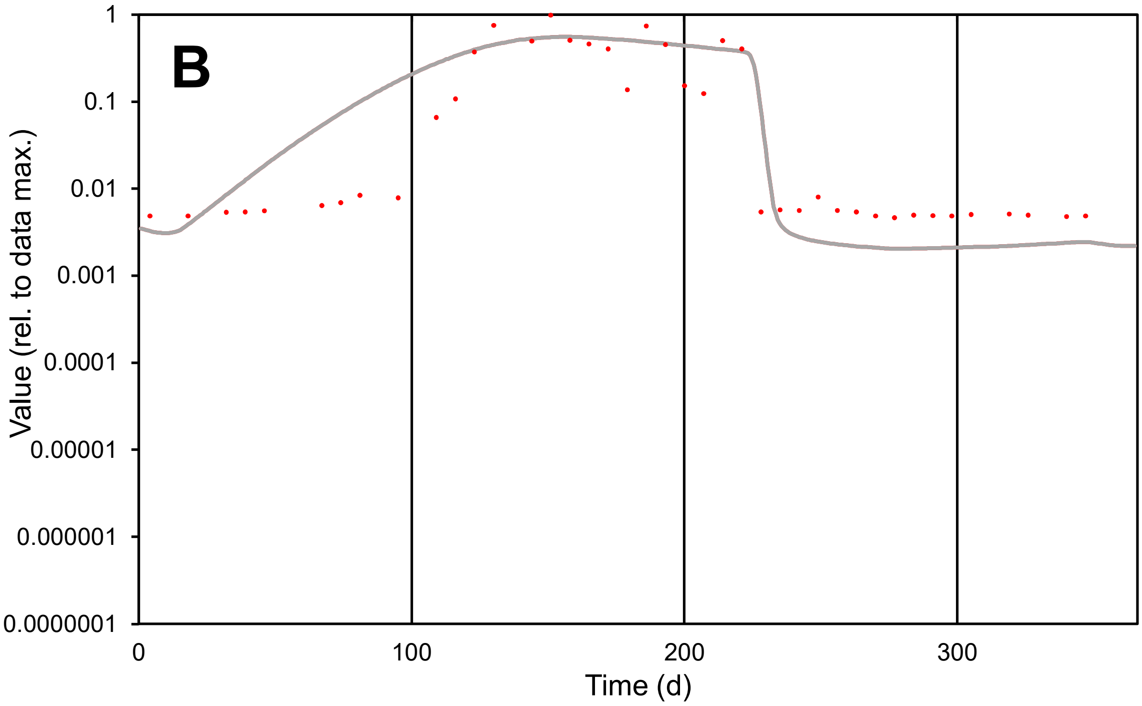


**C1**

| | |
|---|---|
| Variables | |
| N56 | Provide jtd or Dst |
| CalibOnly? | |
| 2 | 1 to reproduce of |
| oNormalize? | |
| 2 | 1=no, 2=yes |
| ofxn calc* | |
| 0.85692 | |
| ofxn mod | |
| 0.87585 | |
| sRun | |
| RunA1 | |
| zout | |
| 1 | |
| Ave | |
| Data | |
| 0.00141 | |
| Model | |
| 0.00149 | |
| data/vMax | |
| 0.00759 | |
| ofxng | |
| 334.07 | |
| Min d/m | |
| 2.27613 | |
| ave/min d | |
| 39.991 | |
| ave/min m | |
| 96.2321 | |

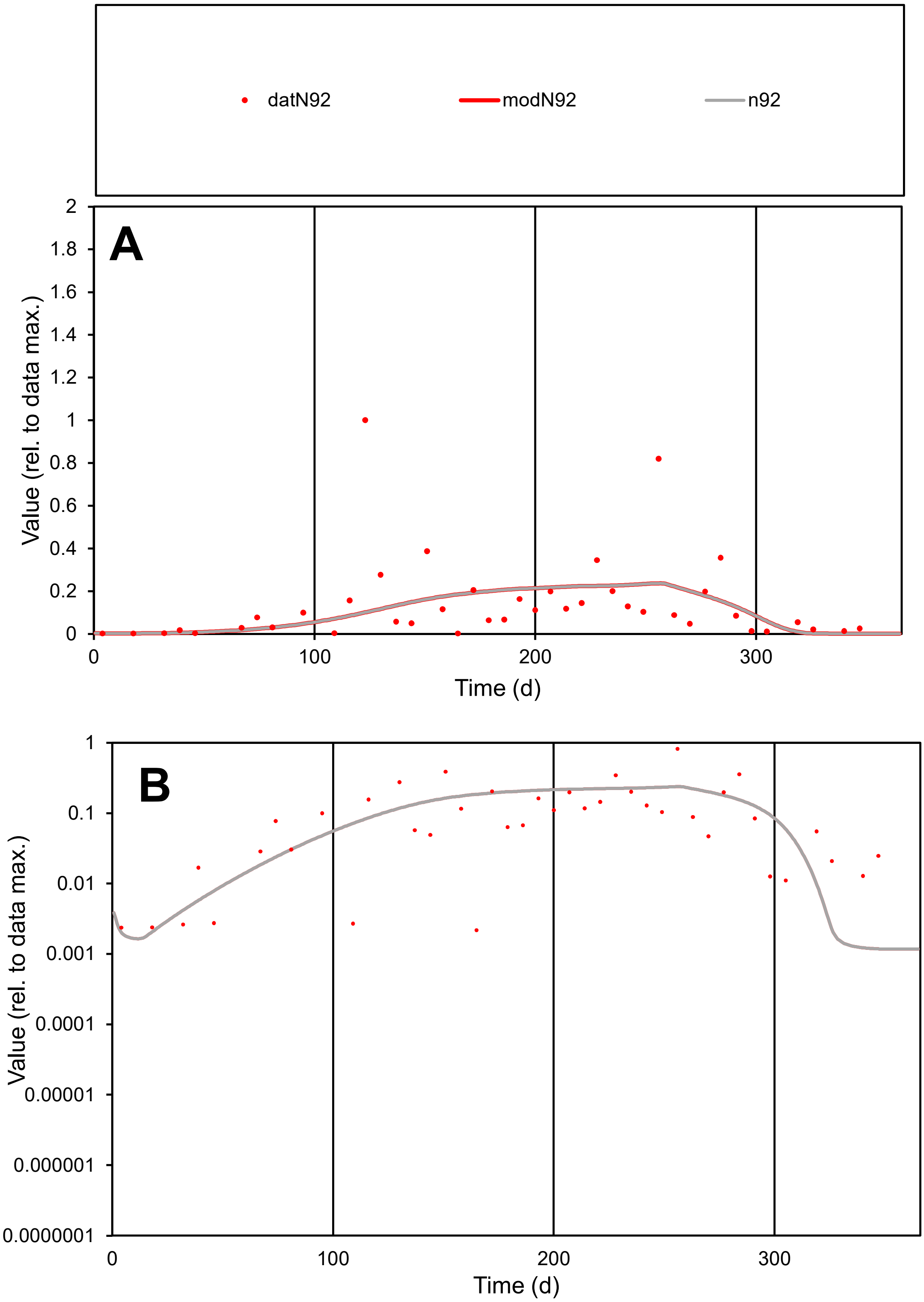


**C1**

Variables
N92 Provide jtd or Dst
CalibOnly?
2 1 to reproduce of
oNormalize?
2 1=no, 2=yes

ofxn calc*
1.3423
ofxn mod
1.35184

sRun
RunA1
zout
1

Ave
Data
0.00242
Model
0.00213

data/vMax
0.01724

ofxng
334.07

Min d/m
1.84363

ave/min d
64.2618
ave/min m
104.635

327_Calib_S_A

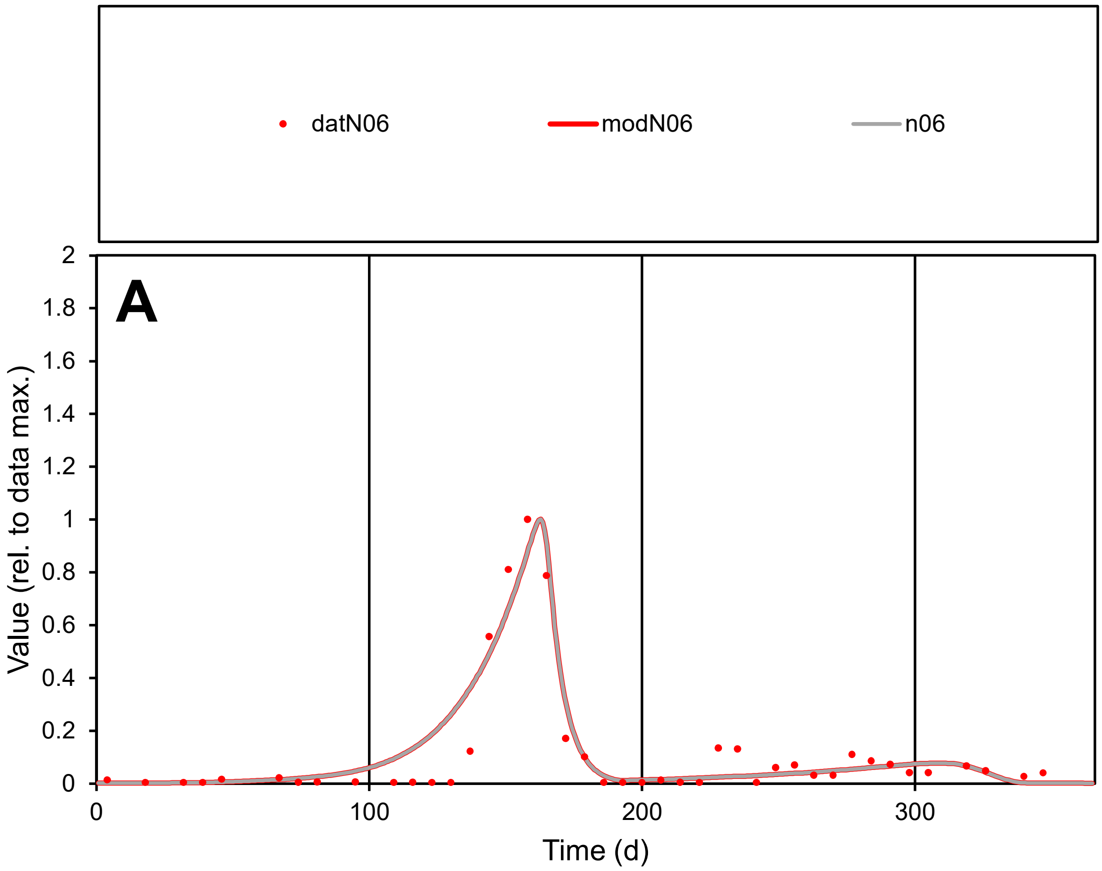


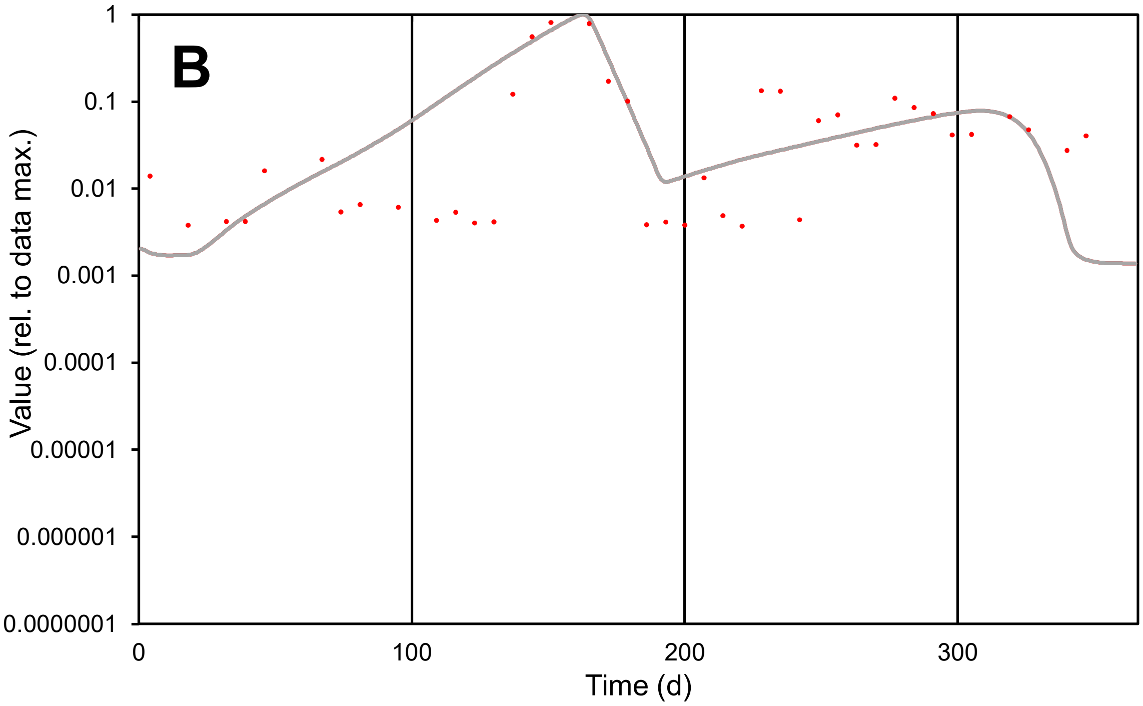


**C1**

Variables
N06 Provide jtd or Dst
CalibOnly?
2 1 to reproduce of
oNormalize?
2 1=no, 2=yes

ofxn calc*
0.68779
ofxn mod
0.73602

sRun
RunA1
zout
1

Ave
Data
0.00207
Model
0.0023

data/vMax
0.01855

ofxng
334.07

Min d/m
2.49982

ave/min d
30.0402
ave/min m
83.3829

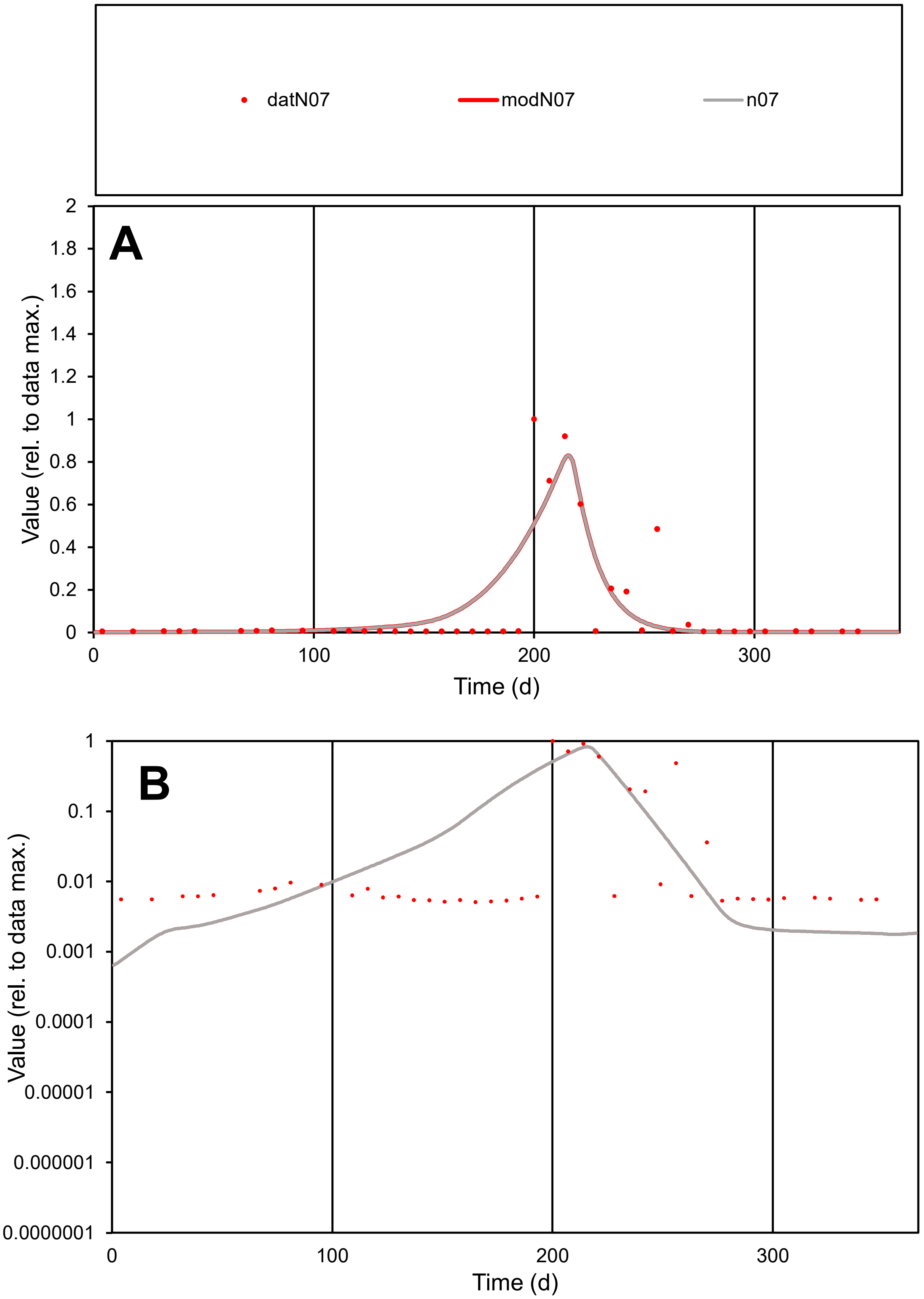


**C1**

Variables
N07 Provide jtd or Dst
CalibOnly?
2 1 to reproduce of
oNormalize?
2 1=no, 2=yes

ofxn calc*
1.33726
ofxn mod
1.39161

sRun
RunA1
zout
1

Ave
Data
0.00034
Model
0.00037

data/vMax
0.00331

ofxng
334.07

Min d/m
6.47239

ave/min d
20.3383
ave/min m
140.595

327_Calib_S_A

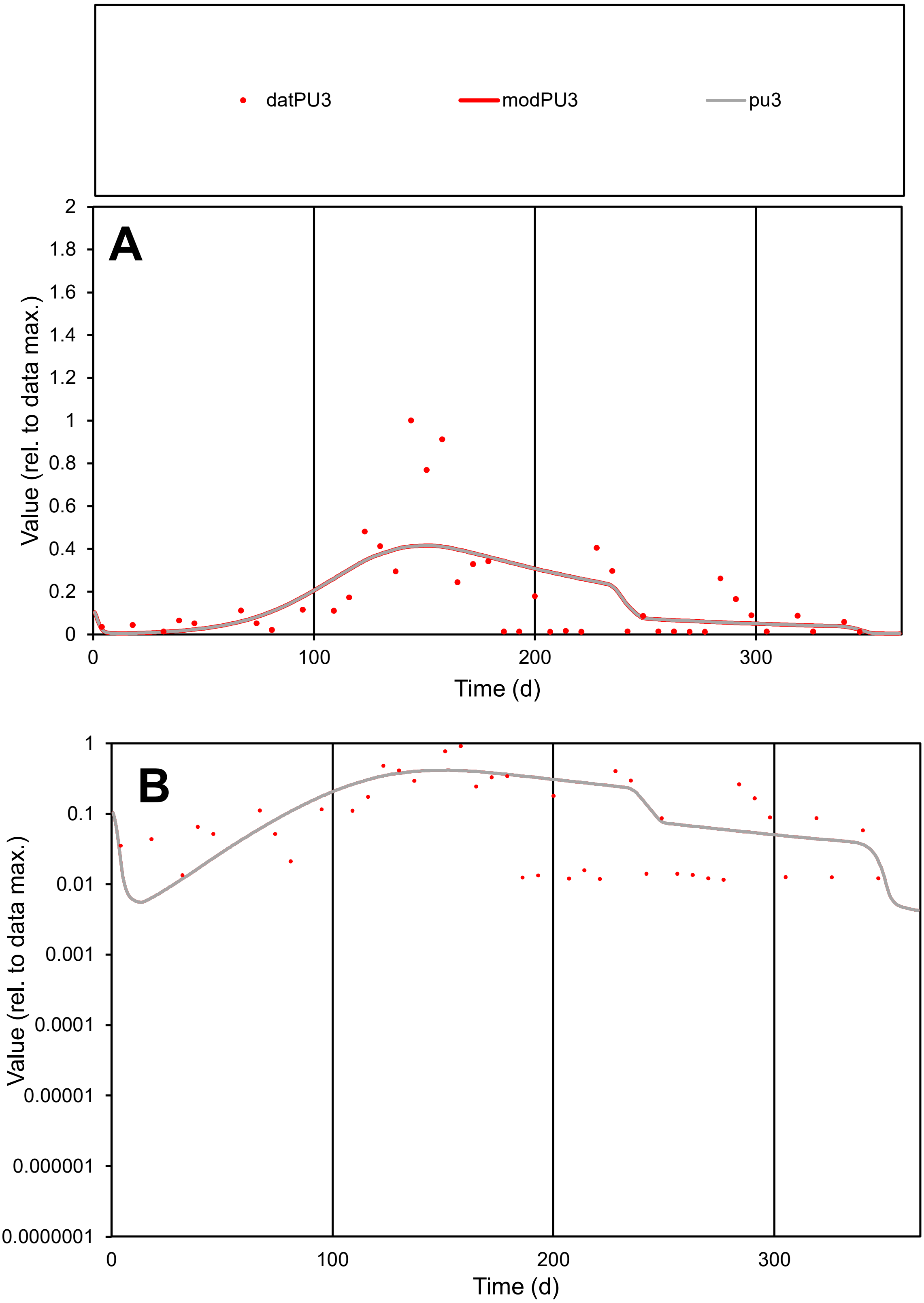


**C1**

Variables
PU3 Provide jtd or Dst
CalibOnly?
2 1 to reproduce of
oNormalize?
2 1=no, 2=yes

ofxn calc*
1.02924
ofxn mod
1.04024

sRun
RunA1
zout
1

Ave
Data
0.00235
Model
0.00243

data/vMax
0.01337

ofxng
334.07

Min d/m
1.70689

ave/min d
15.0812
ave/min m
26.6981

327_Calib_S_A

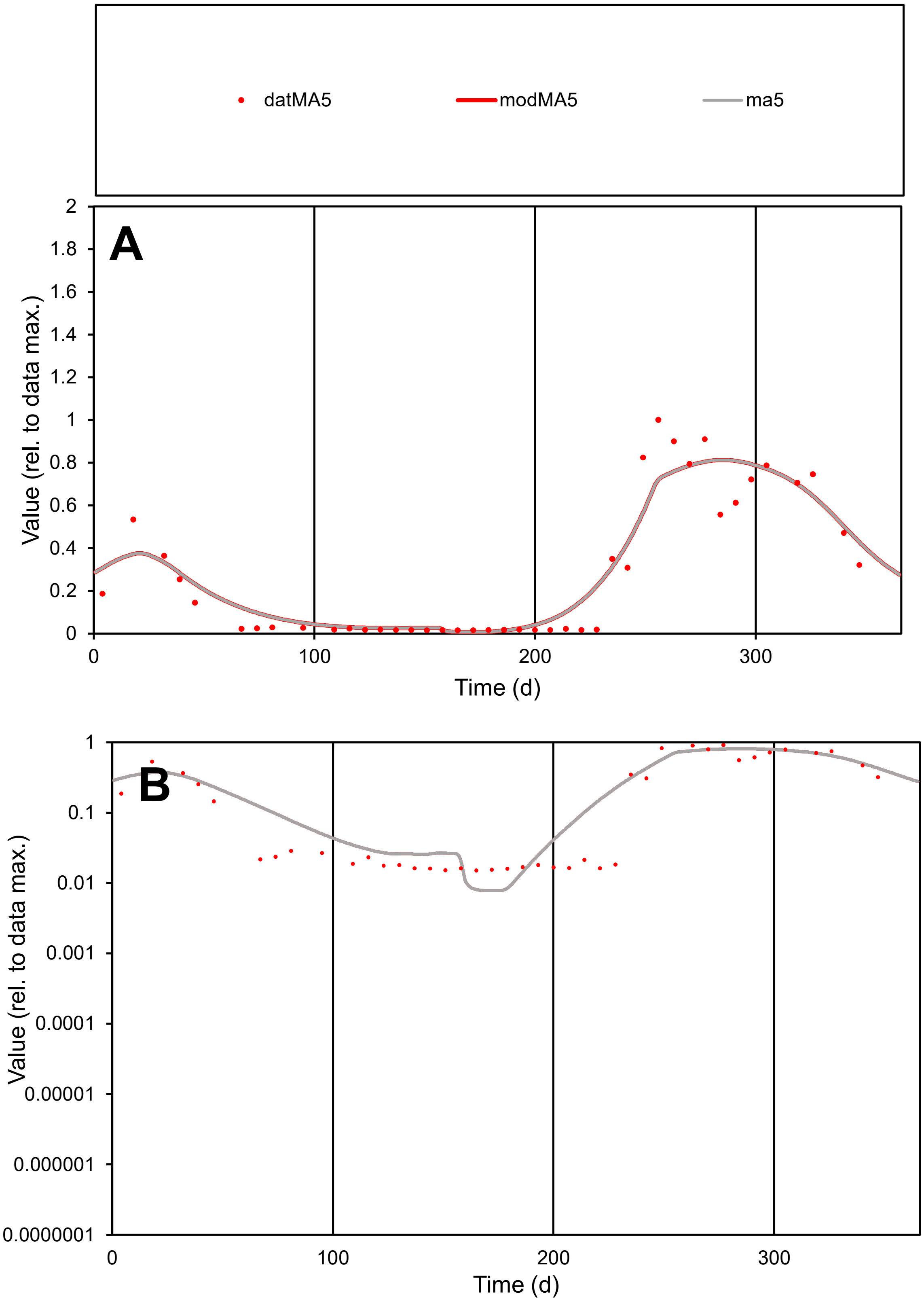


**C1**

Variables
MA5 Provide jtd or Dst
CalibOnly?
2 1 to reproduce of
oNormalize?
2 1=no, 2=yes

ofxn calc*
0.35669
ofxn mod
0.40043

sRun
RunA1
zout
1

Ave
Data
0.00175
Model
0.00184

data/vMax
0.00617

ofxng
334.07

Min d/m
1.94301

ave/min d
18.792
ave/min m
38.495

327_Calib_S_A

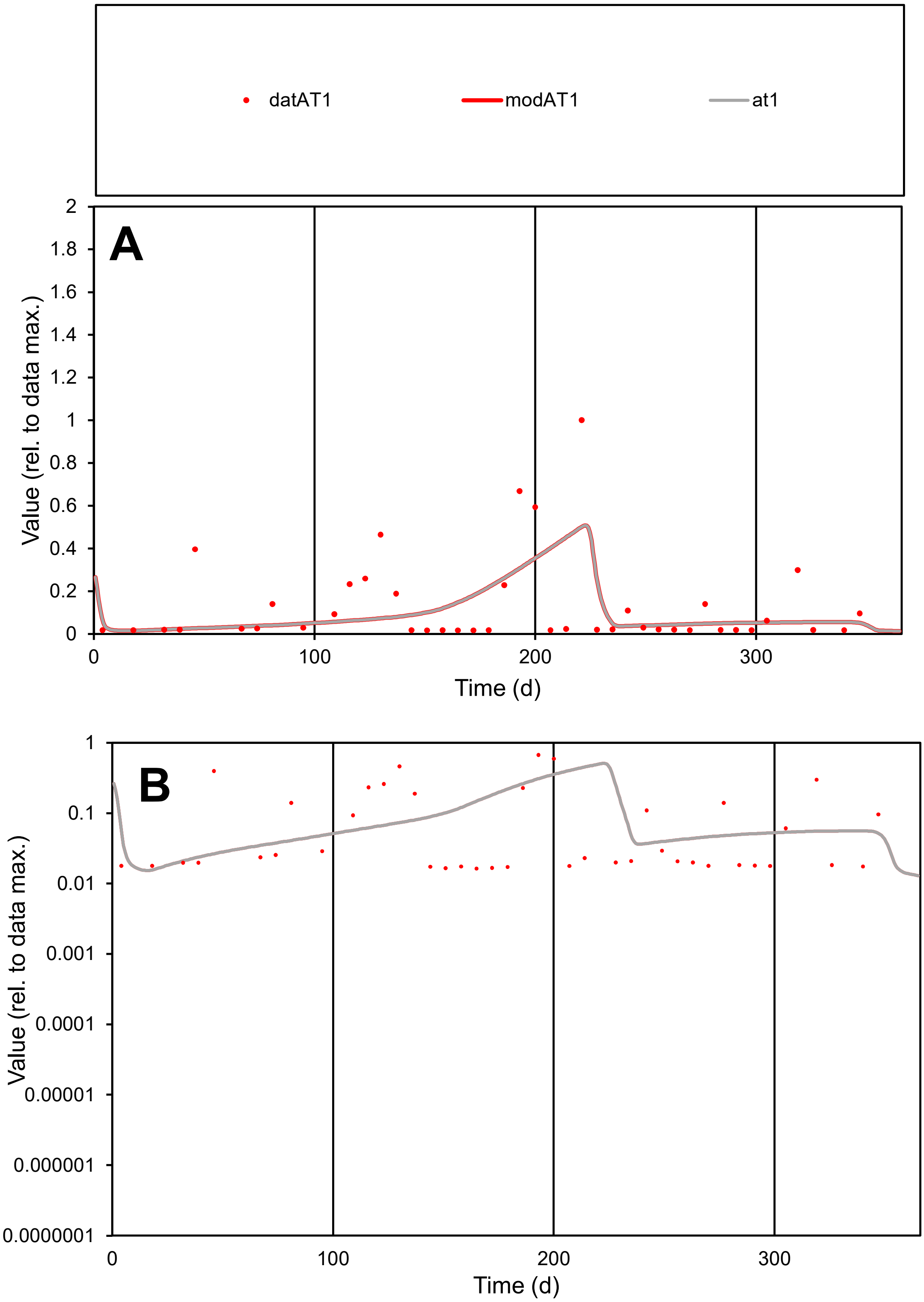


**C1**

Variables
AT1 Provide jtd or Dst
CalibOnly?
2 1 to reproduce of
oNormalize?
2 1=no, 2=yes

ofxn calc*
1.3694
ofxn mod
1.38376

sRun
RunA1
zout
1

Ave
Data
0.00038
Model
0.00032

data/vMax
0.0029

ofxng
334.07

Min d/m
1.01608

ave/min d
7.98814
ave/min m
6.9174

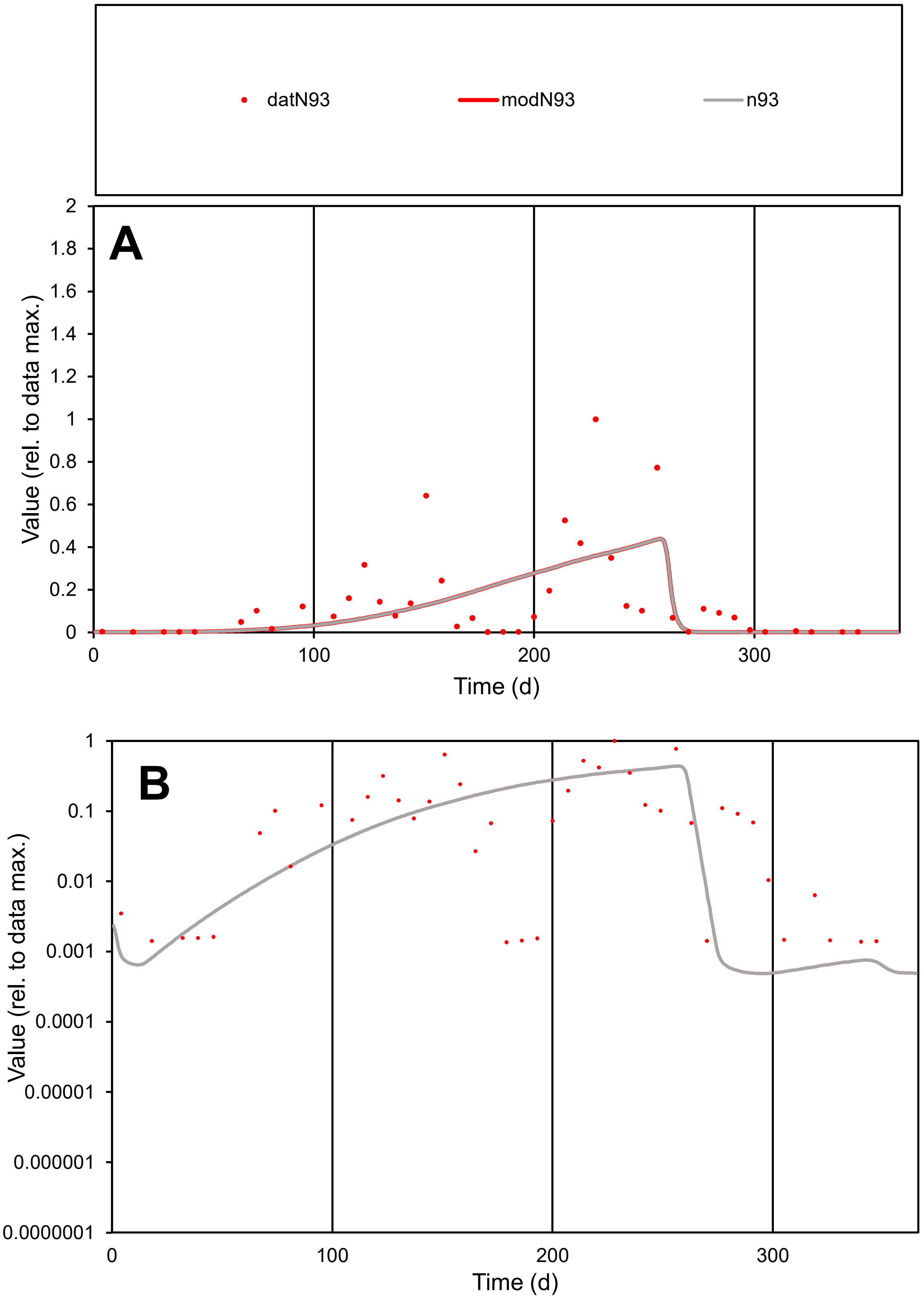


**C1**

Variables

N93 Provide jtd or Dst

CalibOnly?

2 1 to reproduce of

oNormalize?

2 1=no, 2=yes

ofxn calc*

1.24699

ofxn mod

1.25938

sRun

RunA1

zout

1

Ave

Data

0.00193

Model

0.00165

data/vMax

0.0133

ofxng

334.07

Min d/m

2.79009

ave/min d

106.684

ave/min m

254.402

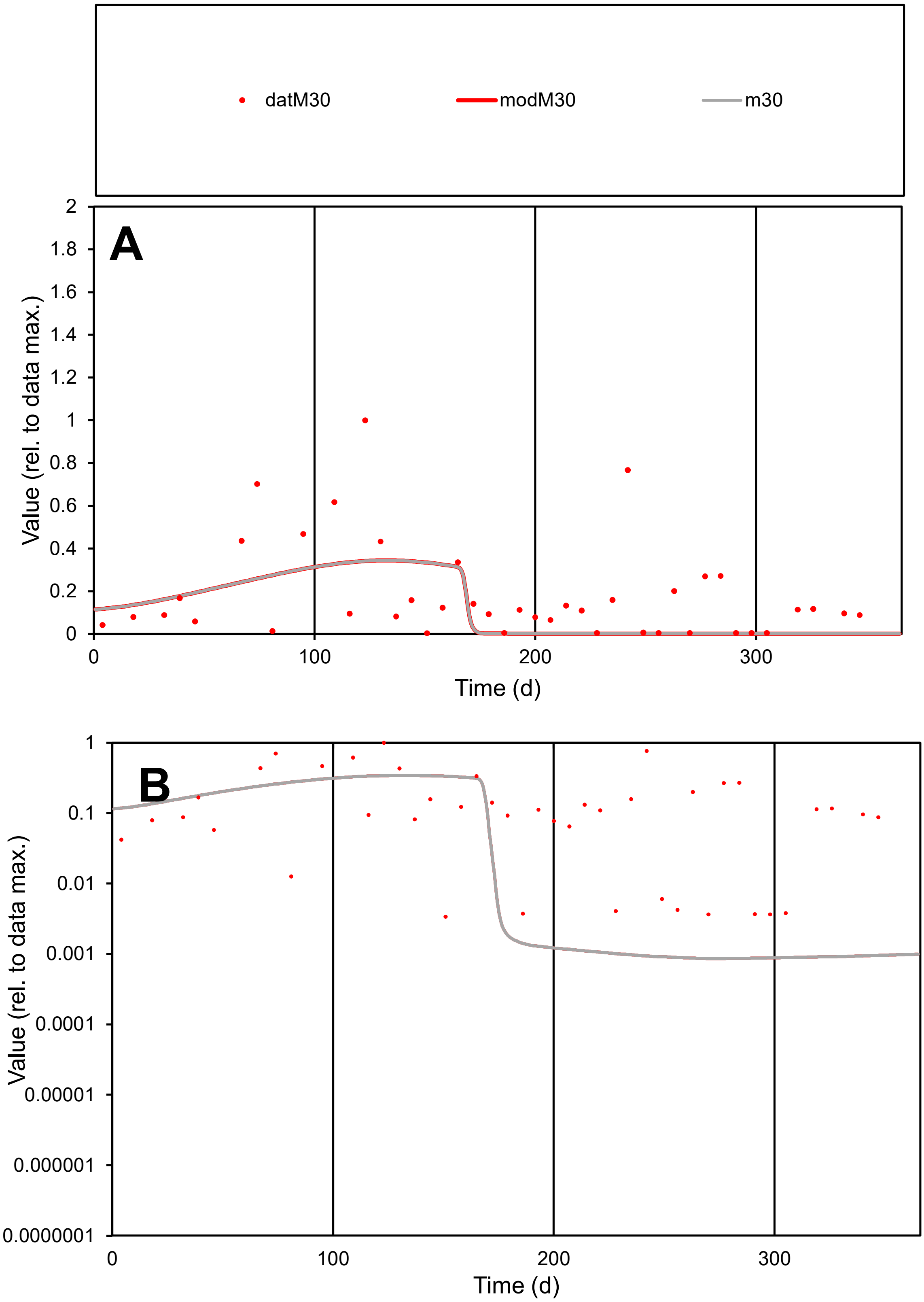


**C1**

Variables
M30 Provide jtd or Dst
CalibOnly?
2 1 to reproduce of
oNormalize?
2 1=no, 2=yes

ofxn calc*
1.21153
ofxn mod
1.22107

sRun
RunA1
zout
1

Ave
Data
0.00223
Model
0.00144

data/vMax
0.01212

ofxng
334.07

Min d/m
3.91874

ave/min d
54.4747
ave/min m
137.437

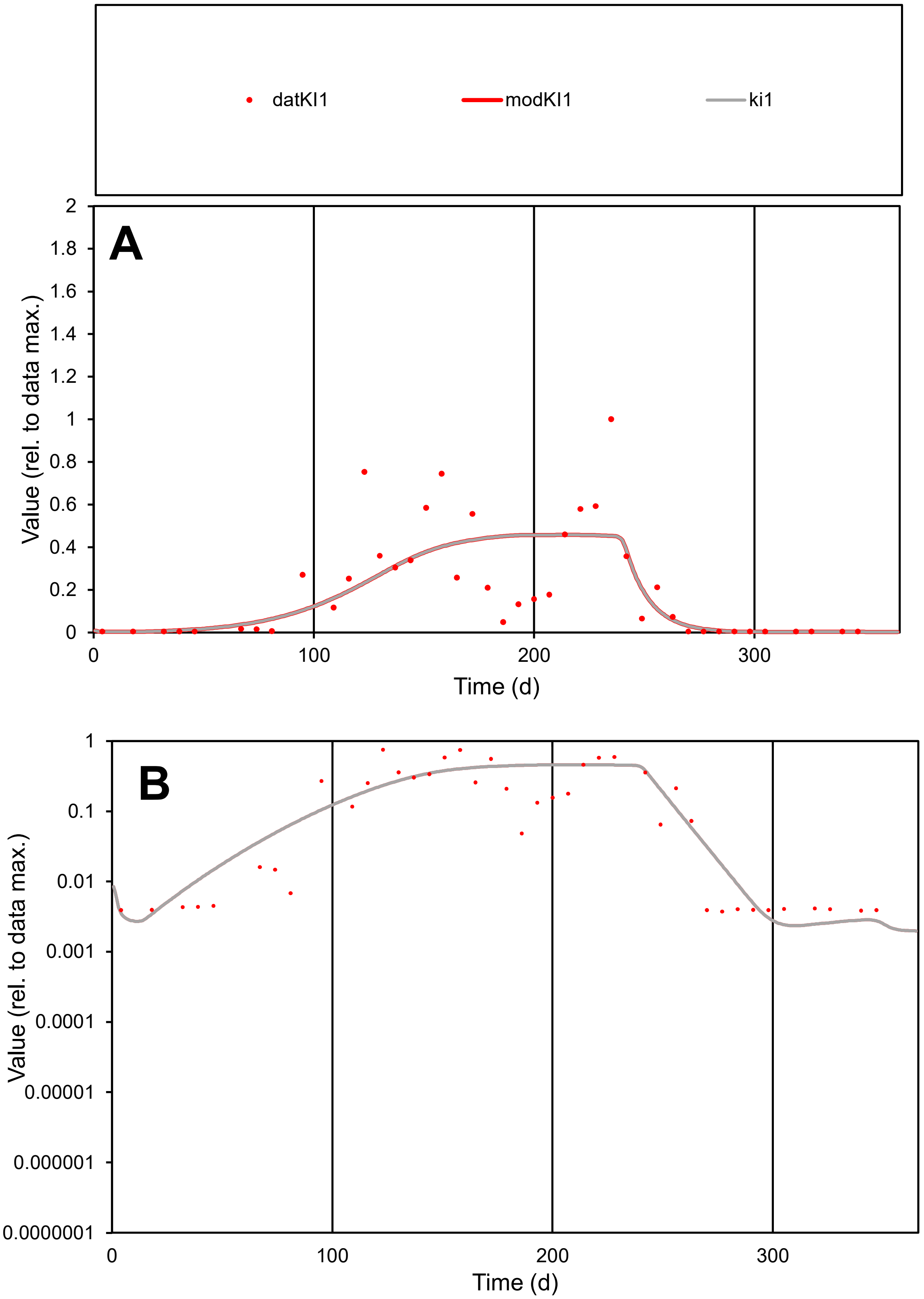


**C1**

Variables
KI1 Provide jtd or Dst
CalibOnly?
2 1 to reproduce of
oNormalize?
2 1=no, 2=yes

ofxn calc*
0.86701
ofxn mod
0.88182

sRun
RunA1
zout
1

Ave
Data
0.00266
Model
0.00255

data/vMax
0.01287

ofxng
334.07

Min d/m
1.56481

ave/min d
55.1833
ave/min m
82.6656

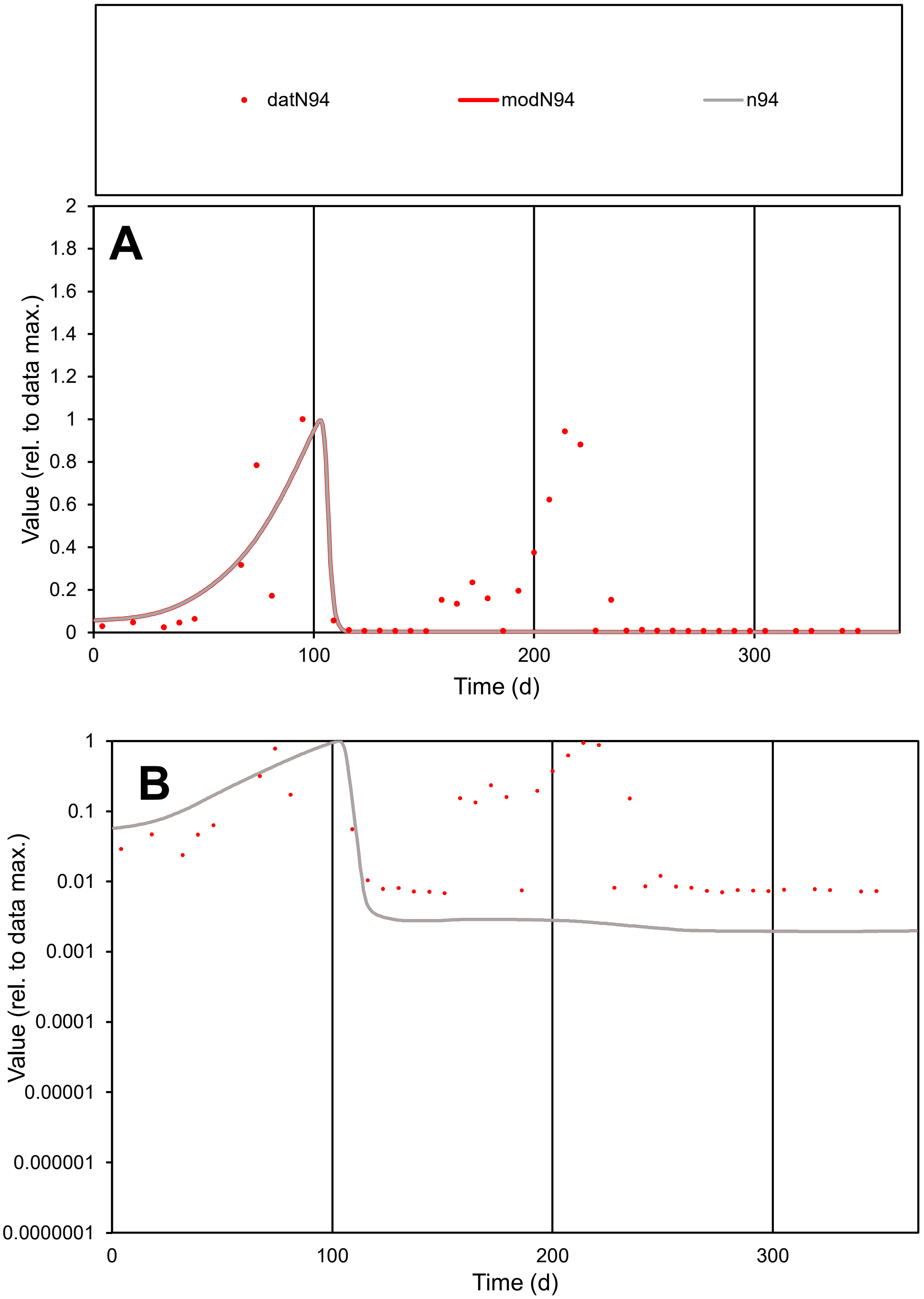


**C1**

Variables
N94 Provide jtd or Dst
CalibOnly?
2 1 to reproduce of
oNormalize?
2 1=no, 2=yes

ofxn calc*
1.61053
ofxn mod
1.6185

sRun
RunA1
zout
1

Ave
Data
0.00104
Model
0.00047

data/vMax
0.00667

ofxng
334.07

Min d/m
3.51783

ave/min d
22.9854
ave/min m
36.1394

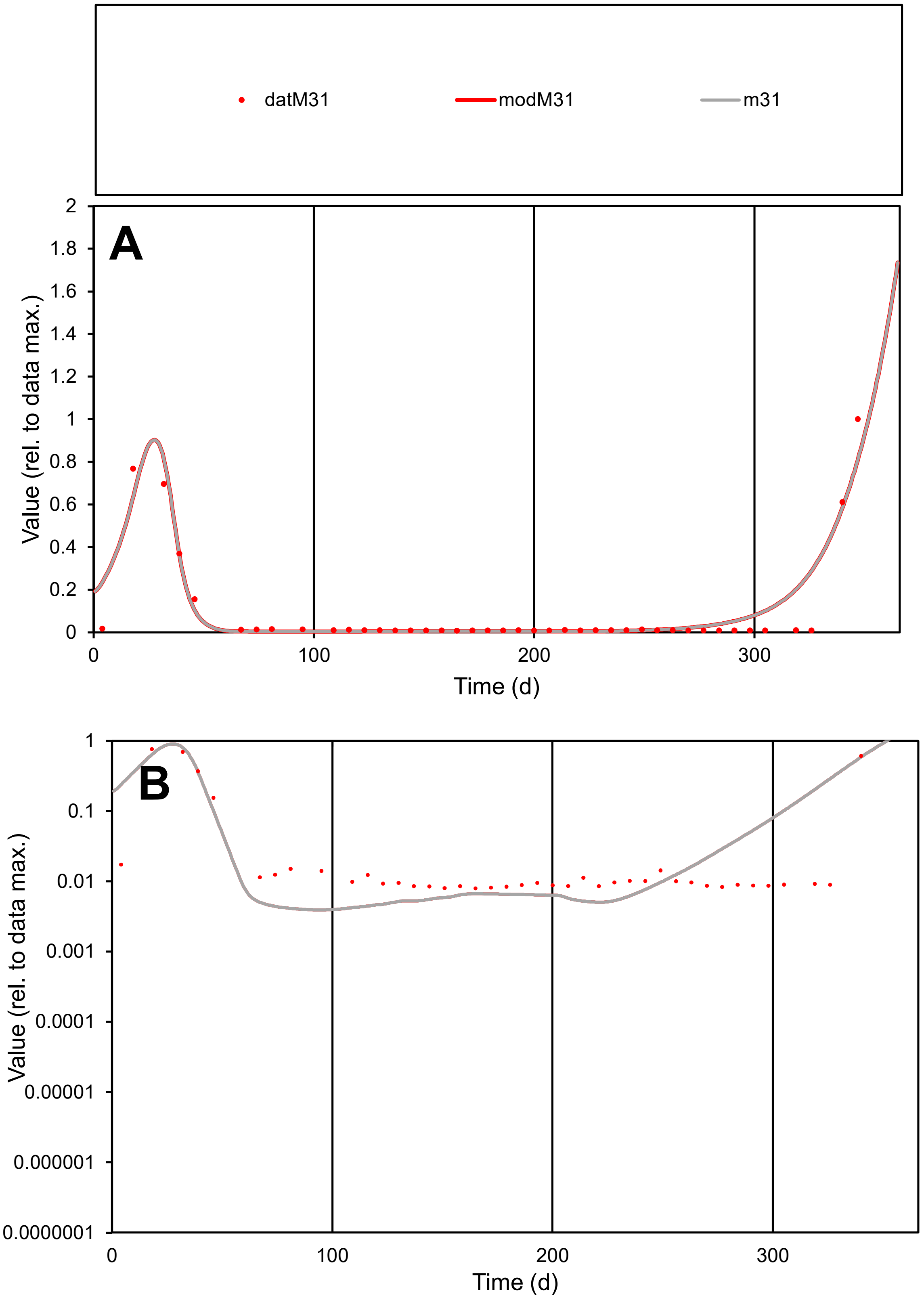


**C1**

Variables
M31 Provide jtd or Dst
CalibOnly?
2 1 to reproduce of
oNormalize?
2 1=no, 2=yes

ofxn calc*
0.71135
ofxn mod
0.82117

sRun
RunA1
zout
1

Ave
Data
0.00095
Model
0.00109

data/vMax
0.01013

ofxng
334.07

Min d/m
2.01955

ave/min d
11.8463
ave/min m
27.3571

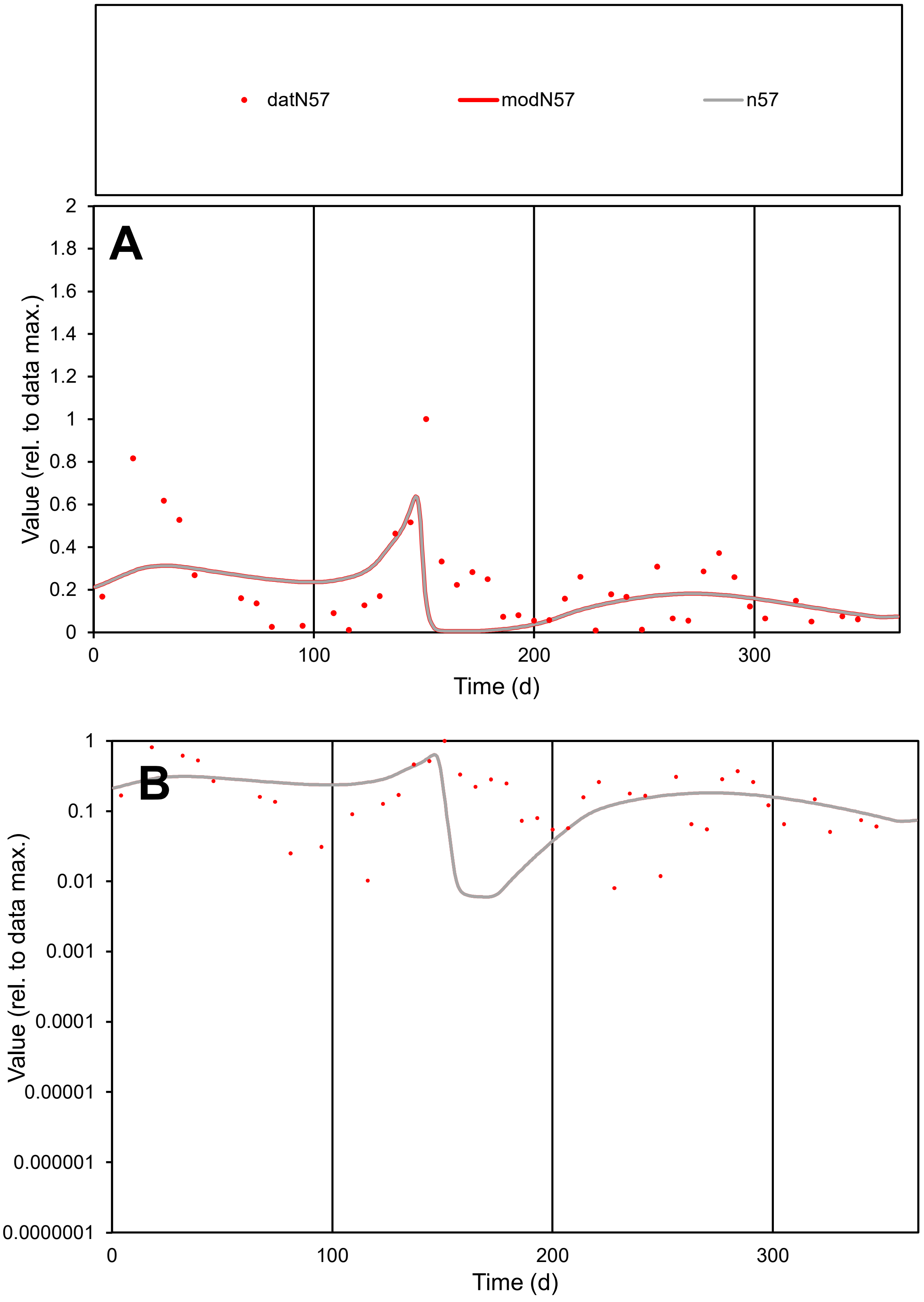


**C1**

Variables
N57 Provide jtd or Dst
CalibOnly?
2 1 to reproduce of
oNormalize?
2 1=no, 2=yes

ofxn calc*
0.87621
ofxn mod
0.94606

sRun
RunA1
zout
1

Ave
Data
0.00117
Model
0.00101

data/vMax
0.00541

ofxng
334.07

Min d/m
1.32601

ave/min d
27.0217
ave/min m
30.7248

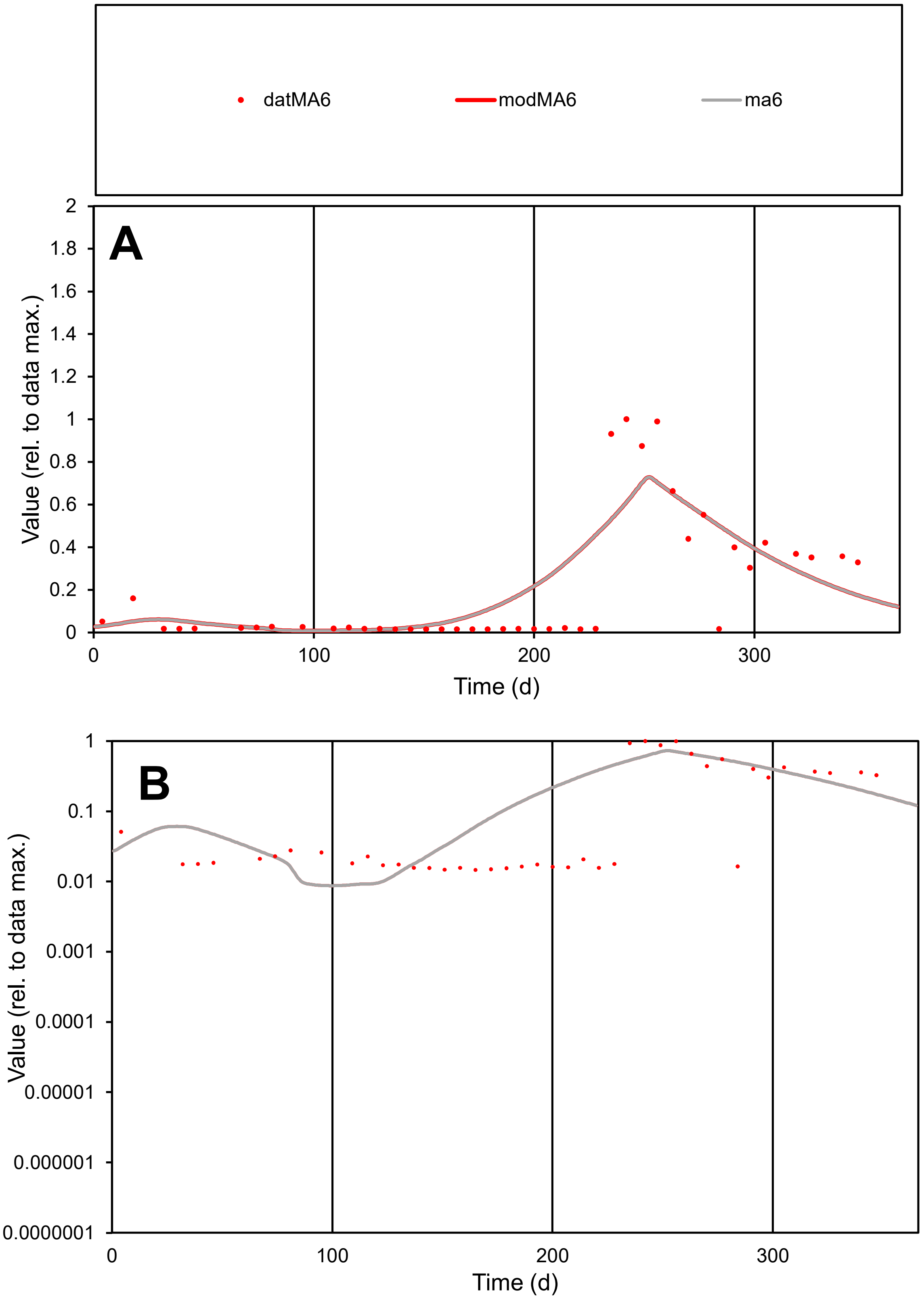


**C1**

Variables
MA6 Provide jtd or Dst
CalibOnly?
2 1 to reproduce of
oNormalize?
2 1=no, 2=yes

ofxn calc*
0.86173
ofxn mod
0.88999

sRun
RunA1
zout
1

Ave
Data
0.00164
Model
0.00183

data/vMax
0.00798

ofxng
334.07

Min d/m
1.66786

ave/min d
14.0873
ave/min m
26.1853

327_Calib_S_A

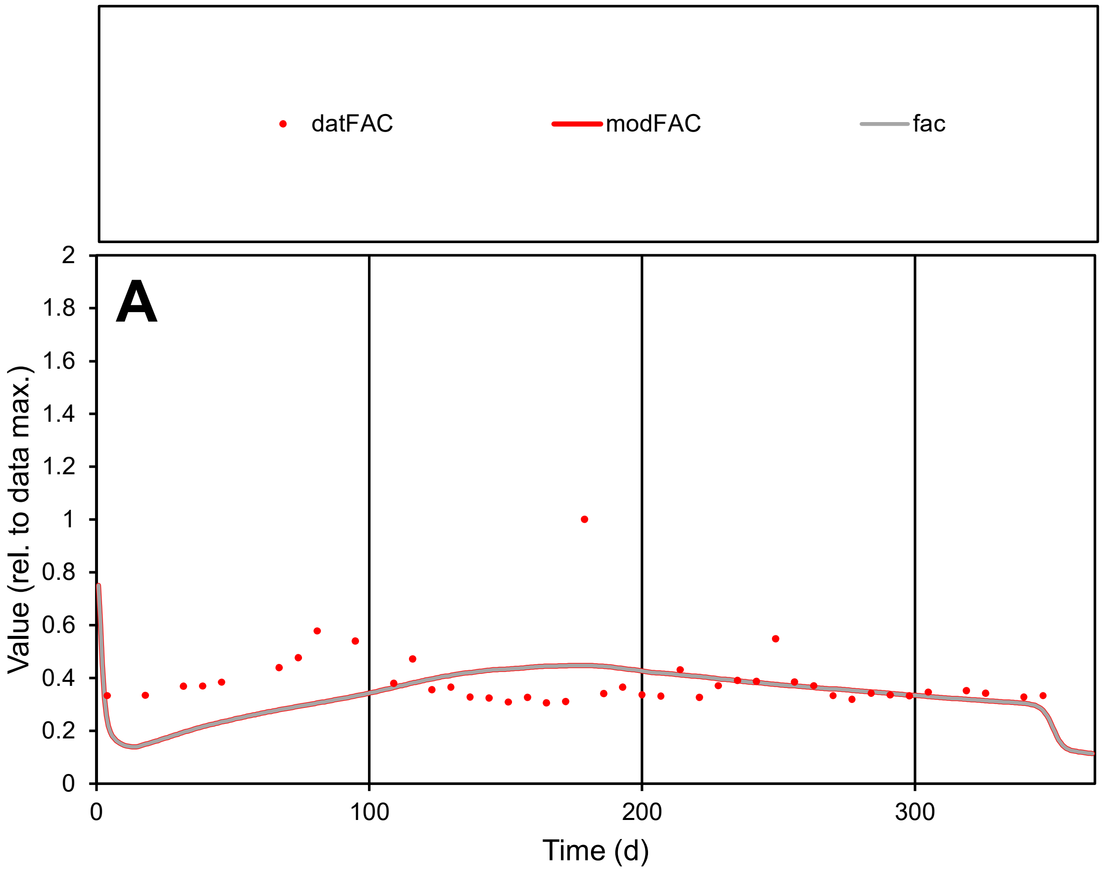


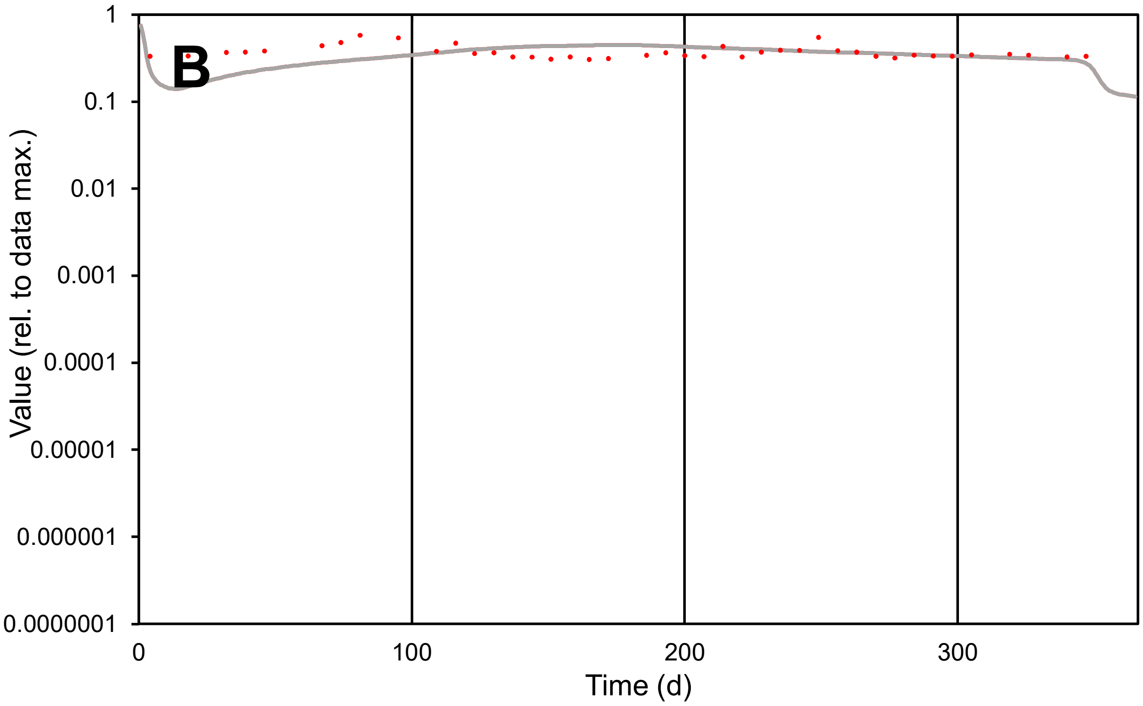


**C1**

Variables
FAC Provide jtd or Dst
CalibOnly?
2 1 to reproduce of
oNormalize?
2 1=no, 2=yes

ofxn calc*
0.35054
ofxn mod
0.36802

sRun
RunA1
zout
1

Ave
Data
0.00011
Model
1E-04

data/vMax
0.00028

ofxng
334.07

Min d/m
1.97939

ave/min d
1.26736
ave/min m
2.31097

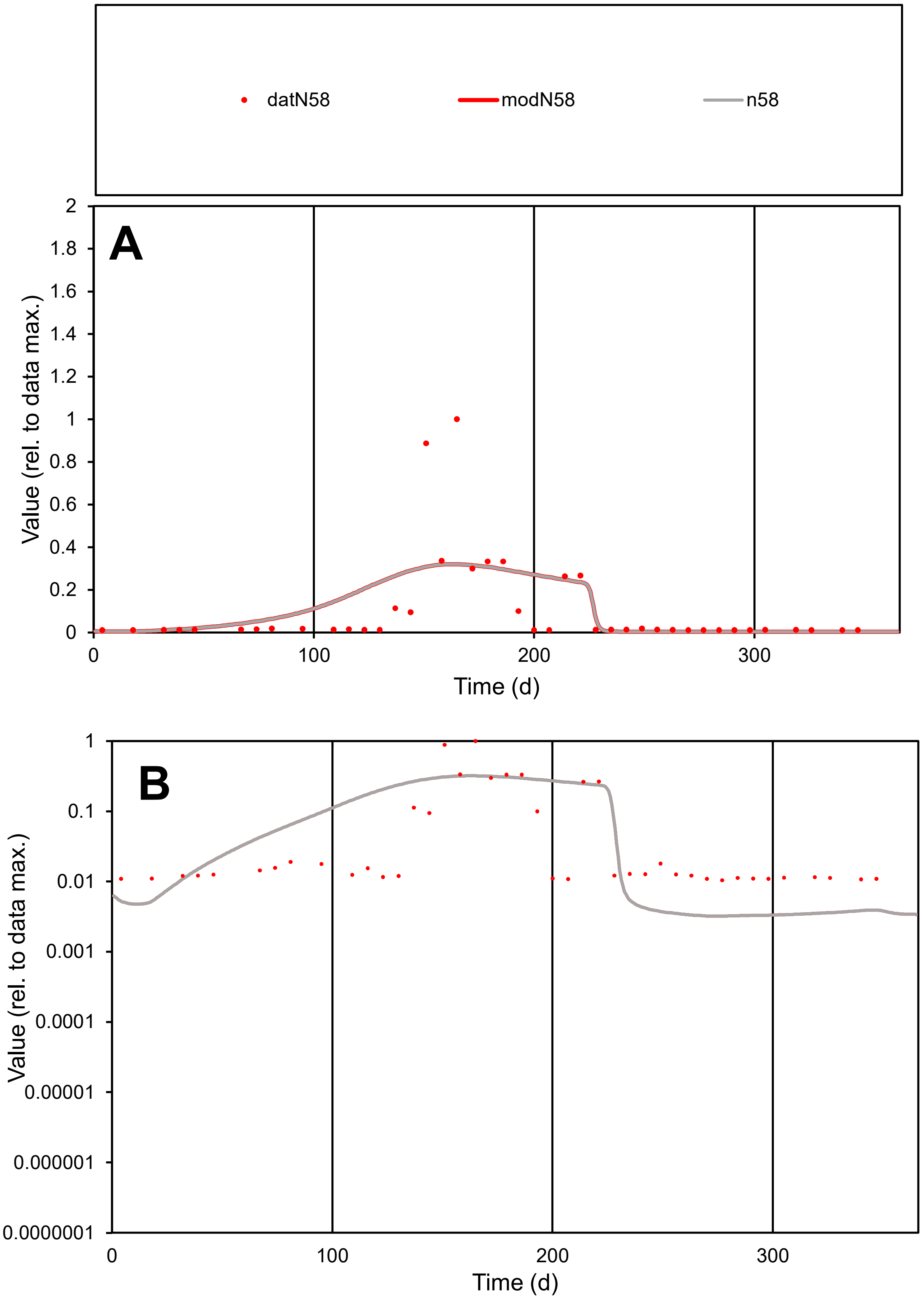


C1

Variables
N58 Provide jtd or Dst
CalibOnly?
2 1 to reproduce of
oNormalize?
2 1=no, 2=yes

ofxn calc*
1.58715
ofxn mod
1.59924

sRun
RunA1
zout
1

Ave
Data
0.00085
Model
0.00094

data/vMax
0.00808

ofxng
334.07

Min d/m
3.25568

ave/min d
10.0504
ave/min m
36.0655

327_Calib_S_A

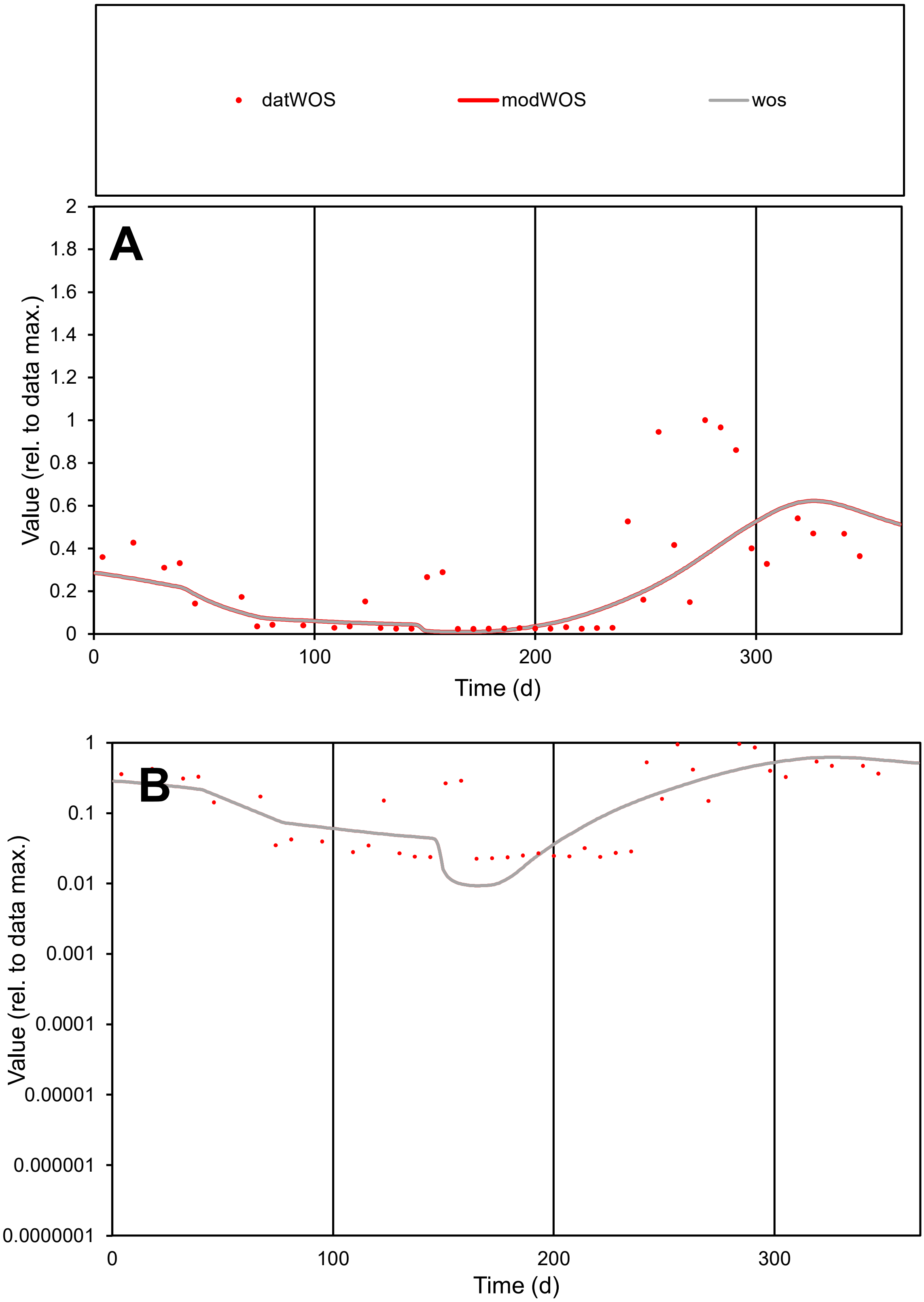


**C1**

Variables
WOS Provide jtd or Dst
CalibOnly?
2 1 to reproduce of
oNormalize?
2 1=no, 2=yes

ofxn calc*
0.8418
ofxn mod
0.85374

sRun
RunA1
zout
1

Ave
Data
0.00076
Model
0.0006

data/vMax
0.00301

ofxng
334.07

Min d/m
2.40064

ave/min d
11.2155
ave/min m
21.2933

327_Calib_S_A

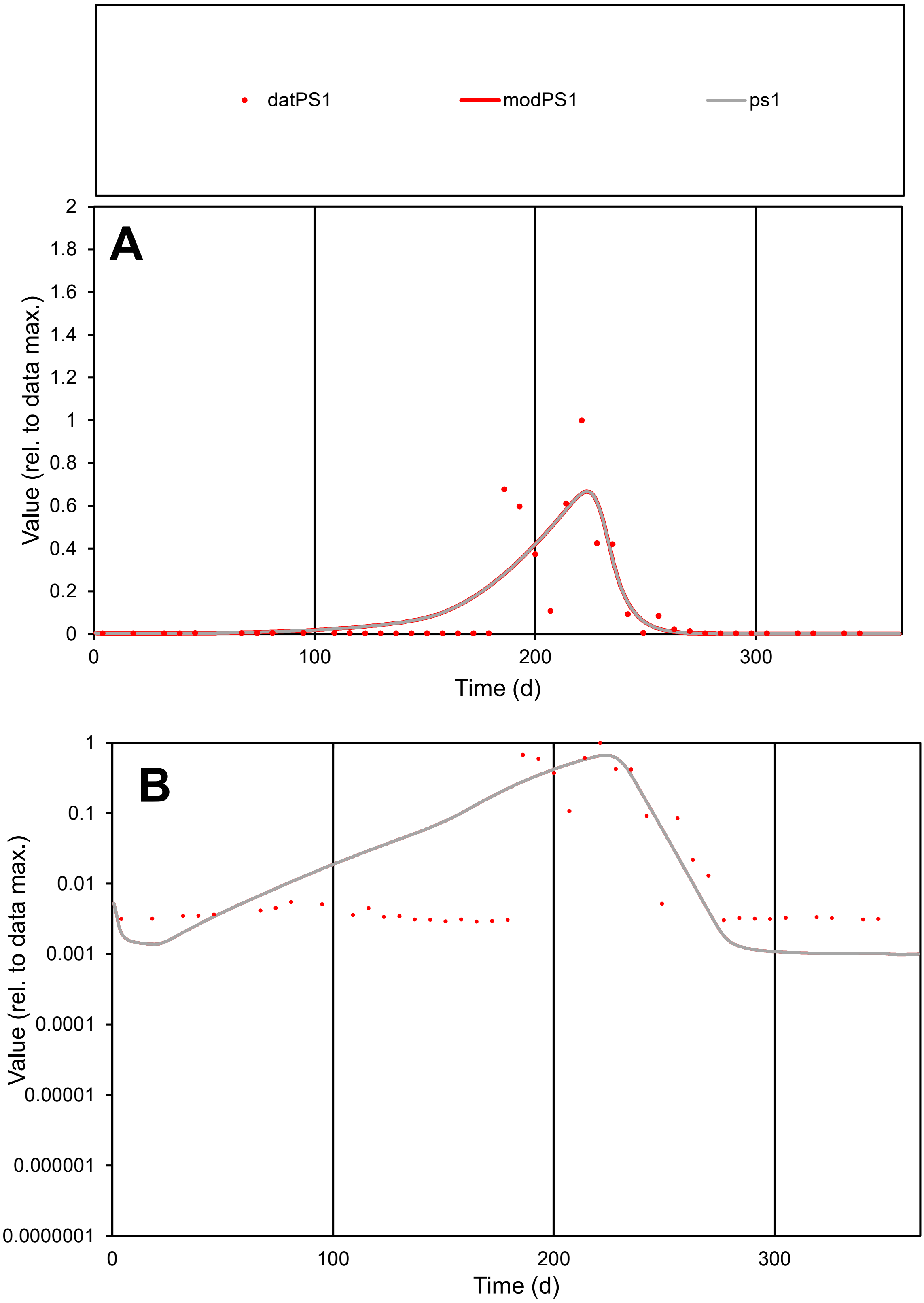


**C1**

Variables

PS1 Provide jtd or Dst

CalibOnly?

2 1 to reproduce of

oNormalize?

2 1=no, 2=yes

ofxn calc*

1.13336

ofxn mod

1.1697

sRun

RunA1

zout

1

Ave

Data

0.00173

Model

0.0019

data/vMax

0.01608

ofxng

334.07

Min d/m

2.8432

ave/min d

37.2022

ave/min m

116.029

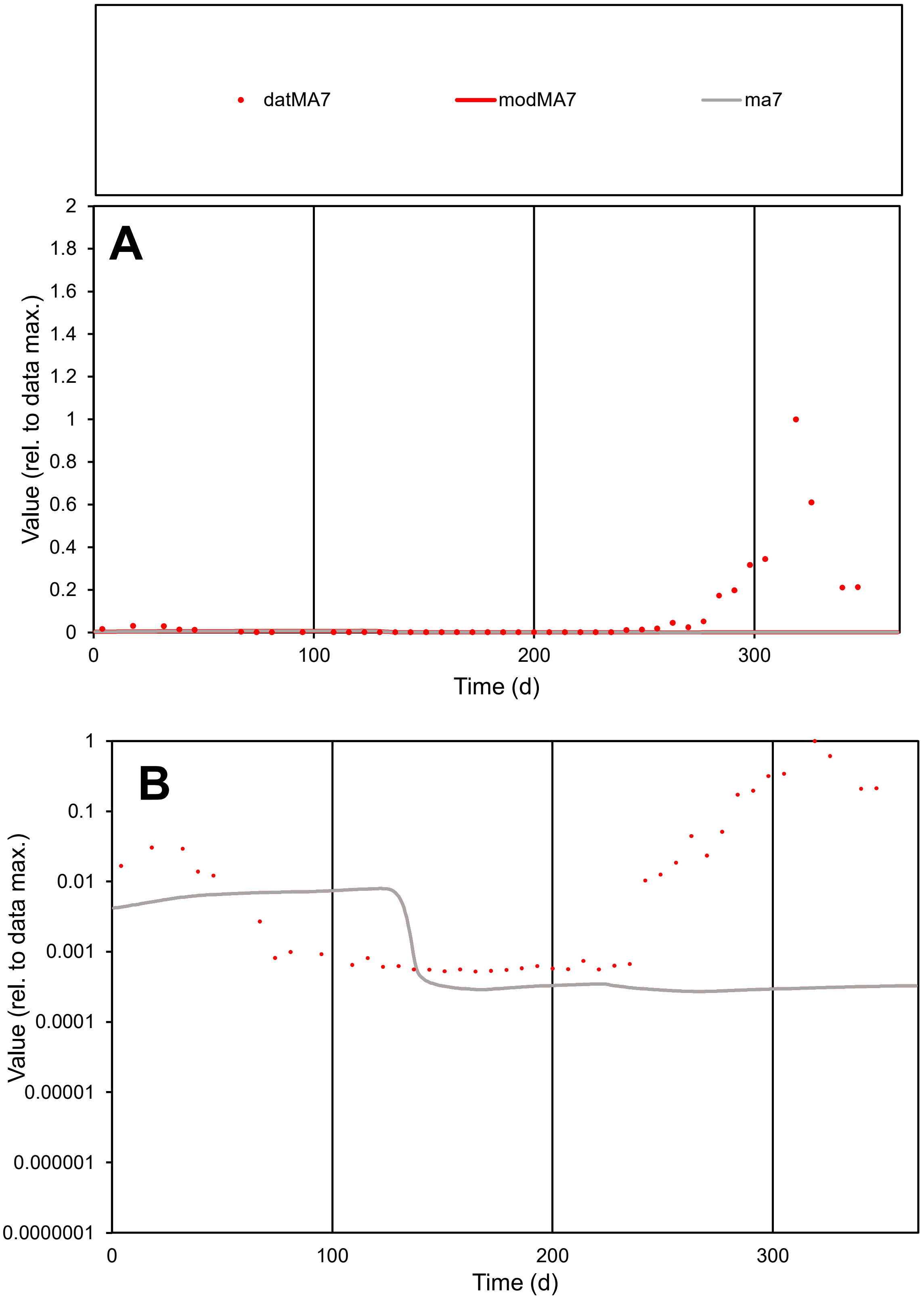


**C1**

Variables
MA7 Provide jtd or Dst
CalibOnly?
2 1 to reproduce of
oNormalize?
2 1=no, 2=yes

ofxn calc*
2.56763
ofxn mod
2.56948

sRun
RunA1
zout
1

Ave
Data
0.00203
Model
5.7E-05

data/vMax
0.02546

ofxng
334.07

Min d/m
1.92759

ave/min d
152.235
ave/min m
8.18941

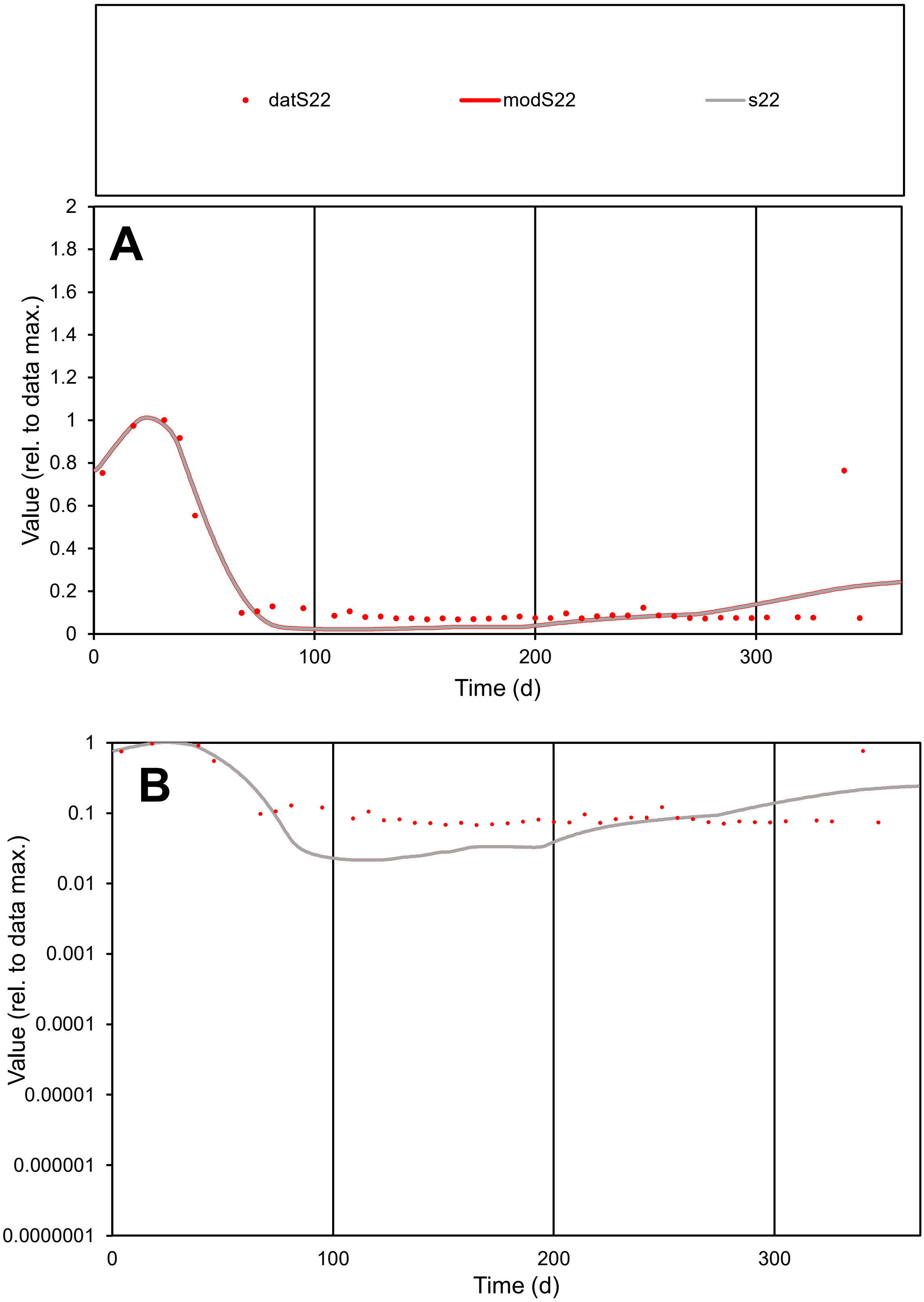


**C1**

Variables
S22 Provide jtd or Dst
CalibOnly?
2 1 to reproduce of
oNormalize?
2 1=no, 2=yes

ofxn calc*
0.54162
ofxn mod
0.5842

sRun
RunA1
zout
1

Ave
Data
0.0004
Model
0.00036

data/vMax
0.00214

ofxng
334.07

Min d/m
3.15309

ave/min d
2.78875
ave/min m
7.86885

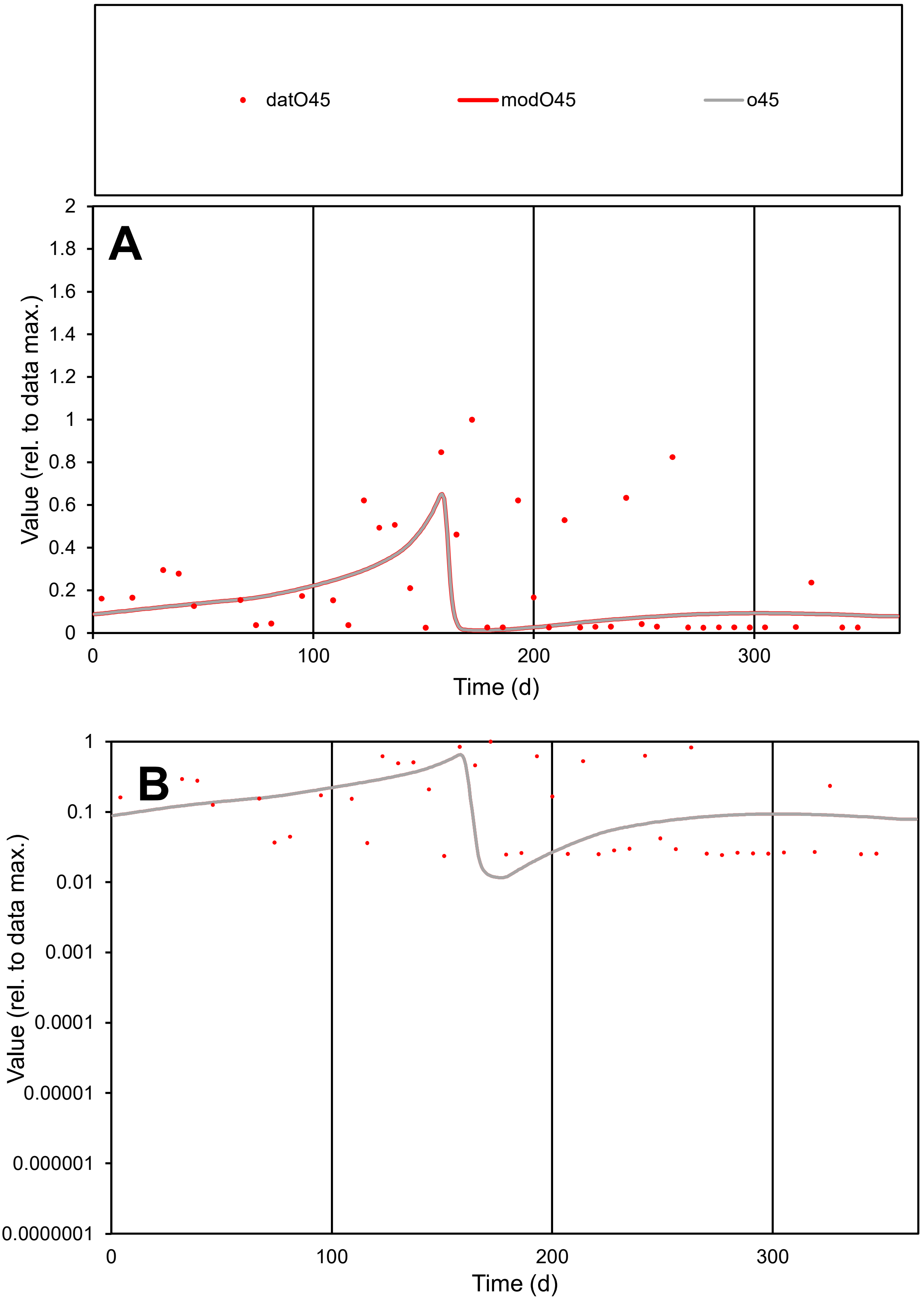


**C1**

Variables
O45 Provide jtd or Dst
CalibOnly?
2 1 to reproduce of
oNormalize?
2 1=no, 2=yes

ofxn calc*
1.2558
ofxn mod
1.27923

sRun
RunA1
zout
1

Ave
Data
0.00053
Model
0.00036

data/vMax
0.00242

ofxng
334.07

Min d/m
1.96631

ave/min d
9.32538
ave/min m
12.2414

327_Calib_S_A

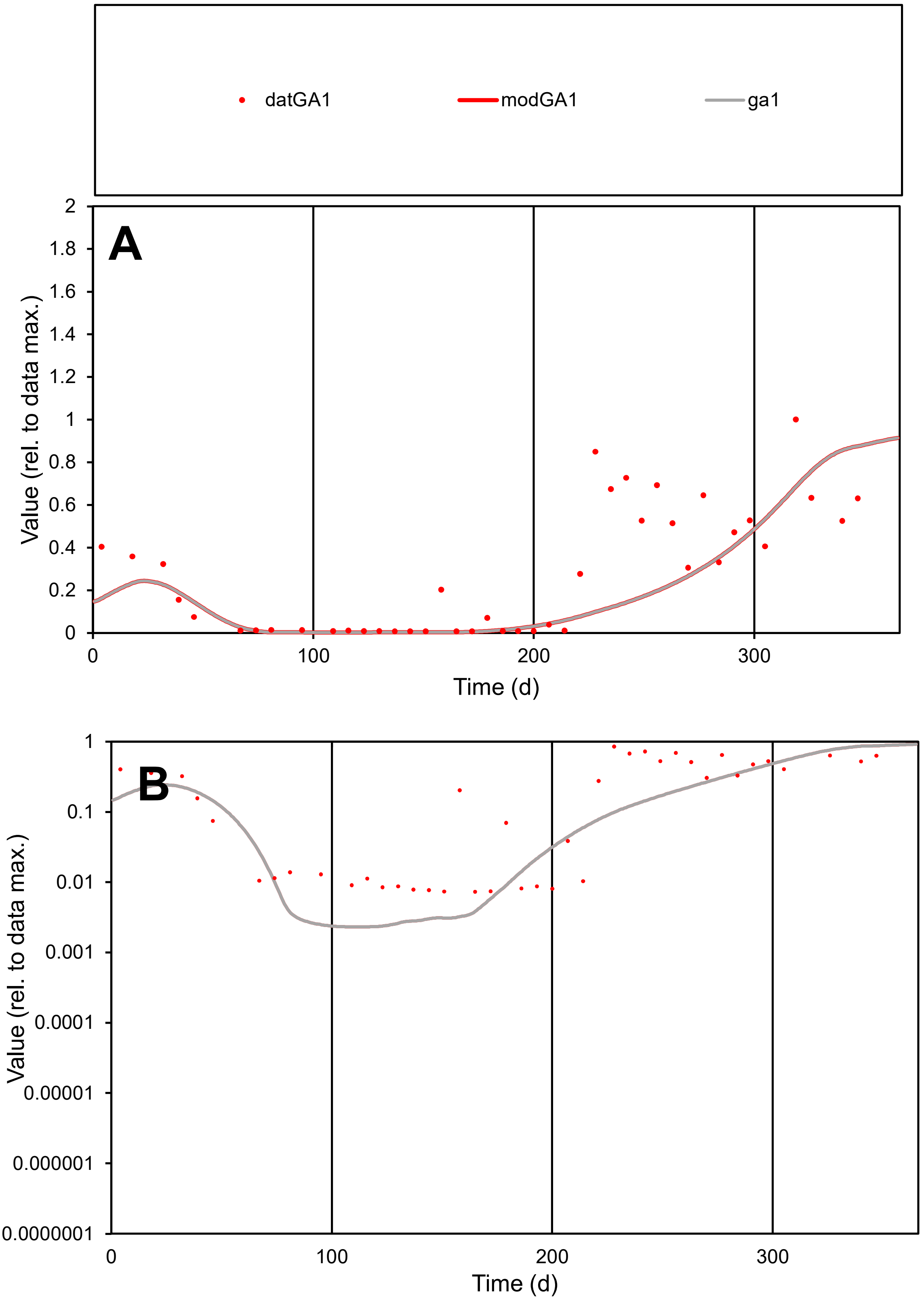


C1

Variables
GA1 Provide jtd or Dst
CalibOnly?
2 1 to reproduce of
oNormalize?
2 1=no, 2=yes

ofxn calc*
0.83805
ofxn mod
0.84817

sRun
RunA1
zout
1

Ave
Data
0.00132
Model
0.0009

data/vMax
0.0048

ofxng
334.07

Min d/m
3.17301

ave/min d
37.5751
ave/min m
81.5595

327_Calib_S_A

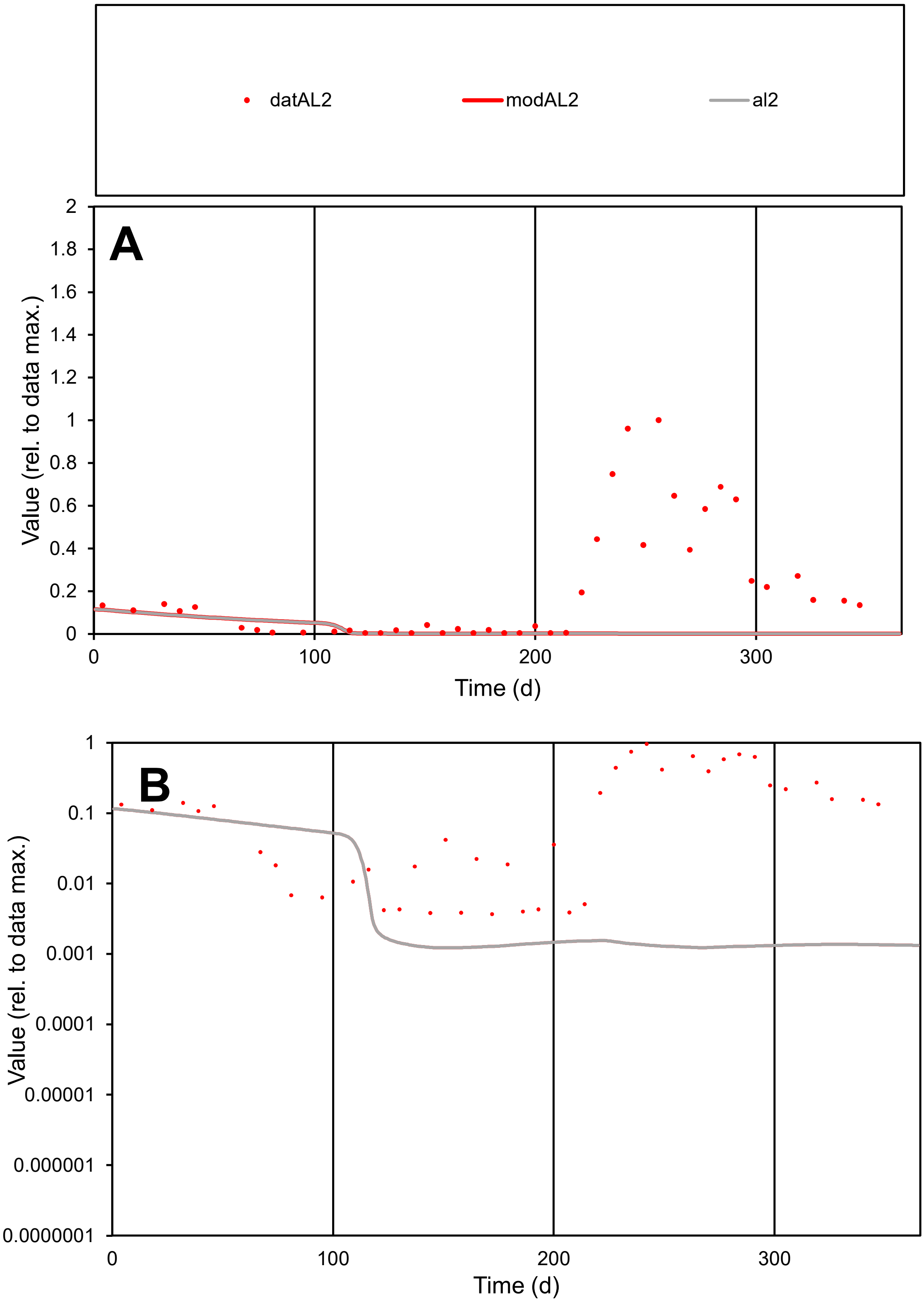


**C1**

Variables
AL2 Provide jtd or Dst
CalibOnly?
2 1 to reproduce of
oNormalize?
2 1=no, 2=yes

ofxn calc*
1.64047
ofxn mod
1.64399

sRun
RunA1
zout
1

Ave
Data
0.00125
Model
0.00011

data/vMax
0.00599

ofxng
334.07

Min d/m
2.99224

ave/min d
56.8828
ave/min m
15.5017

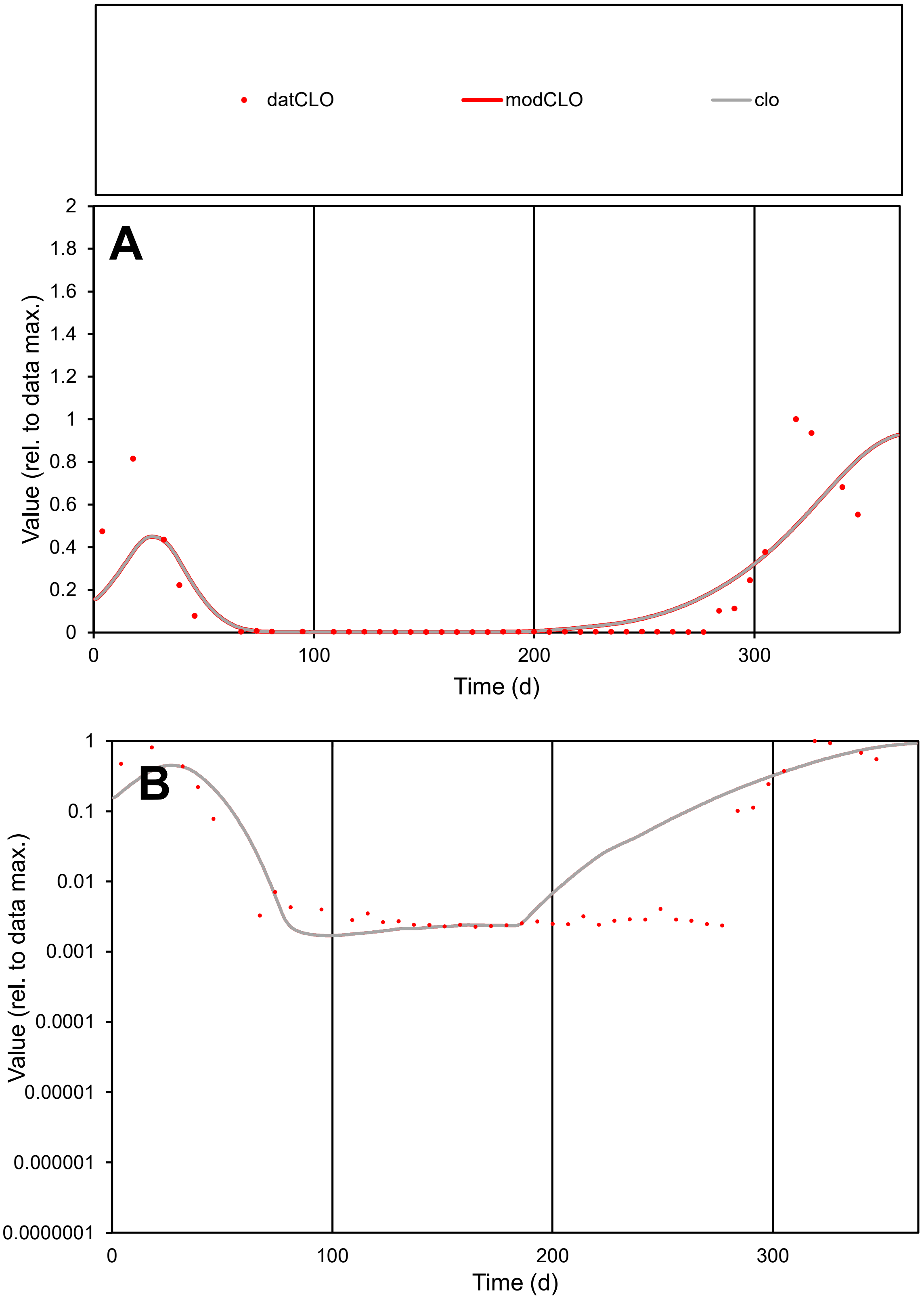


**C1**

Variables
CLO Provide jtd or Dst
CalibOnly?
2 1 to reproduce of
oNormalize?
2 1=no, 2=yes

ofxn calc*
0.95071
ofxn mod
0.9751

sRun
RunA1
zout
1

Ave
Data
0.00084
Model
0.00084

data/vMax
0.00575

ofxng
334.07

Min d/m
1.33306

ave/min d
64.1122
ave/min m
85.5609

327_Calib_S_A

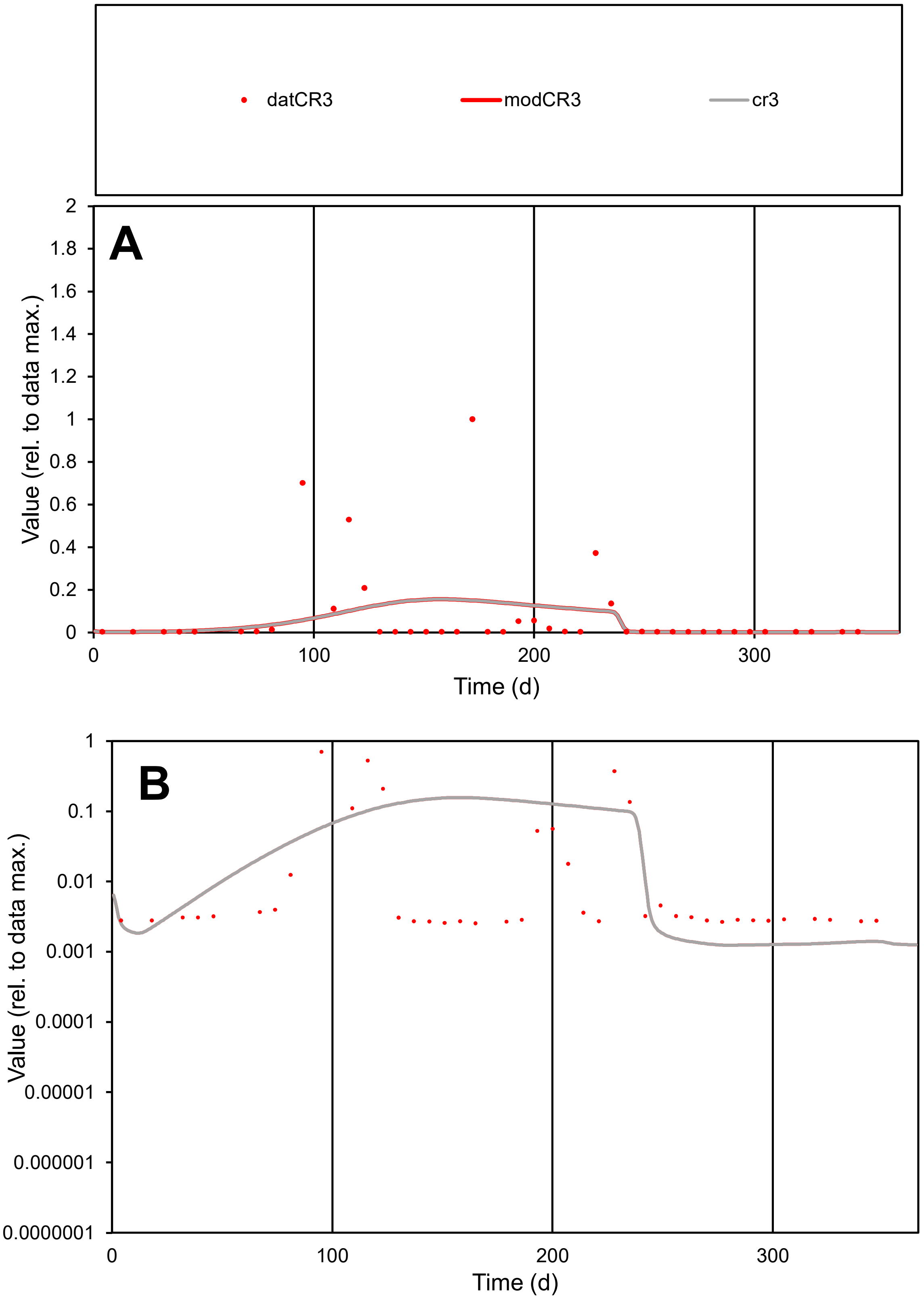


**C1**

Variables
CR3 Provide jtd or Dst
CalibOnly?
2 1 to reproduce of
oNormalize?
2 1=no, 2=yes

ofxn calc*
2.49793
ofxn mod
2.50224

sRun
RunA1
zout
1

Ave
Data
0.00108
Model
0.00086

data/vMax
0.01378

ofxng
334.07

Min d/m
2.05442

ave/min d
30.7546
ave/min m
50.3592

327_Calib_S_A

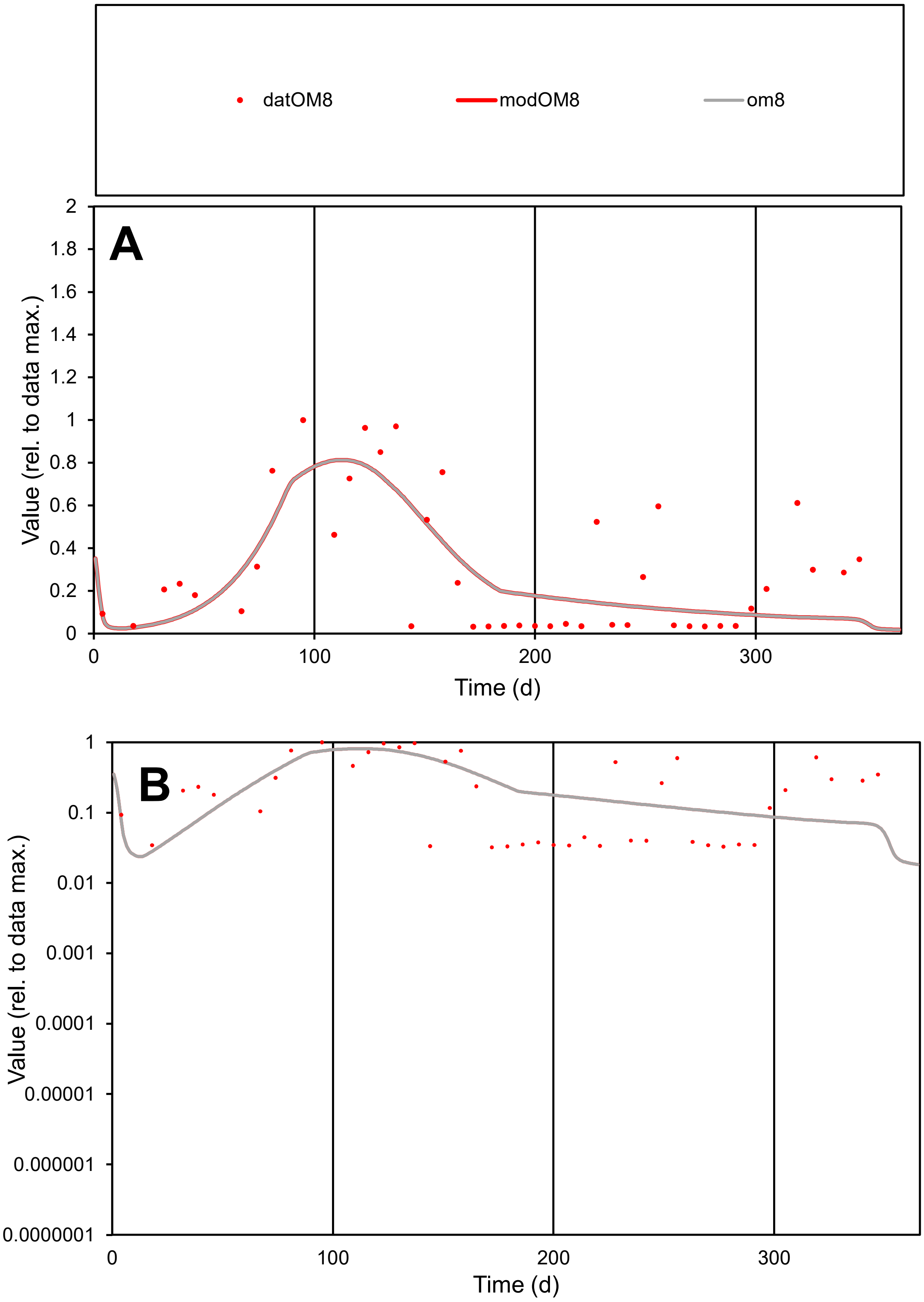


**C1**

Variables
OM8 Provide jtd or Dst
CalibOnly?
2 1 to reproduce of
oNormalize?
2 1=no, 2=yes

ofxn calc*
0.75027
ofxn mod
0.76258

sRun
RunA1
zout
1

Ave
Data
0.00049
Model
0.00045

data/vMax
0.0017

ofxng
334.07

Min d/m
1.0613

ave/min d
9.0621
ave/min m
8.73771

327_Calib_S_A

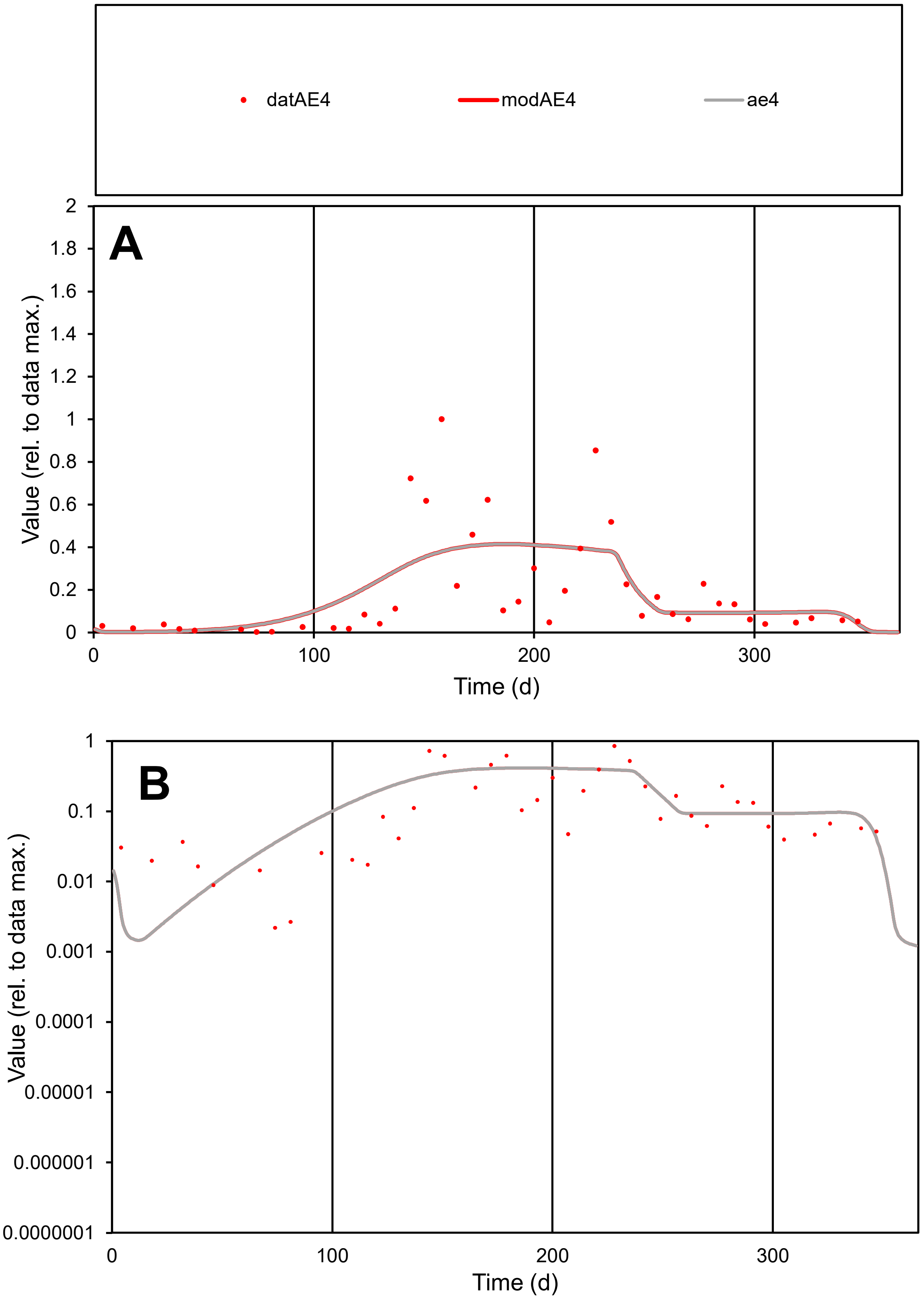


**C1**

Variables
AE4 Provide jtd or Dst
CalibOnly?
2 1 to reproduce of
oNormalize?
2 1=no, 2=yes

ofxn calc*
0.96691
ofxn mod
0.97857

sRun
RunA1
zout
1

Ave
Data
0.00157
Model
0.00161

data/vMax
0.0082

ofxng
334.07

Min d/m
1.07198

ave/min d
87.3634
ave/min m
95.8359

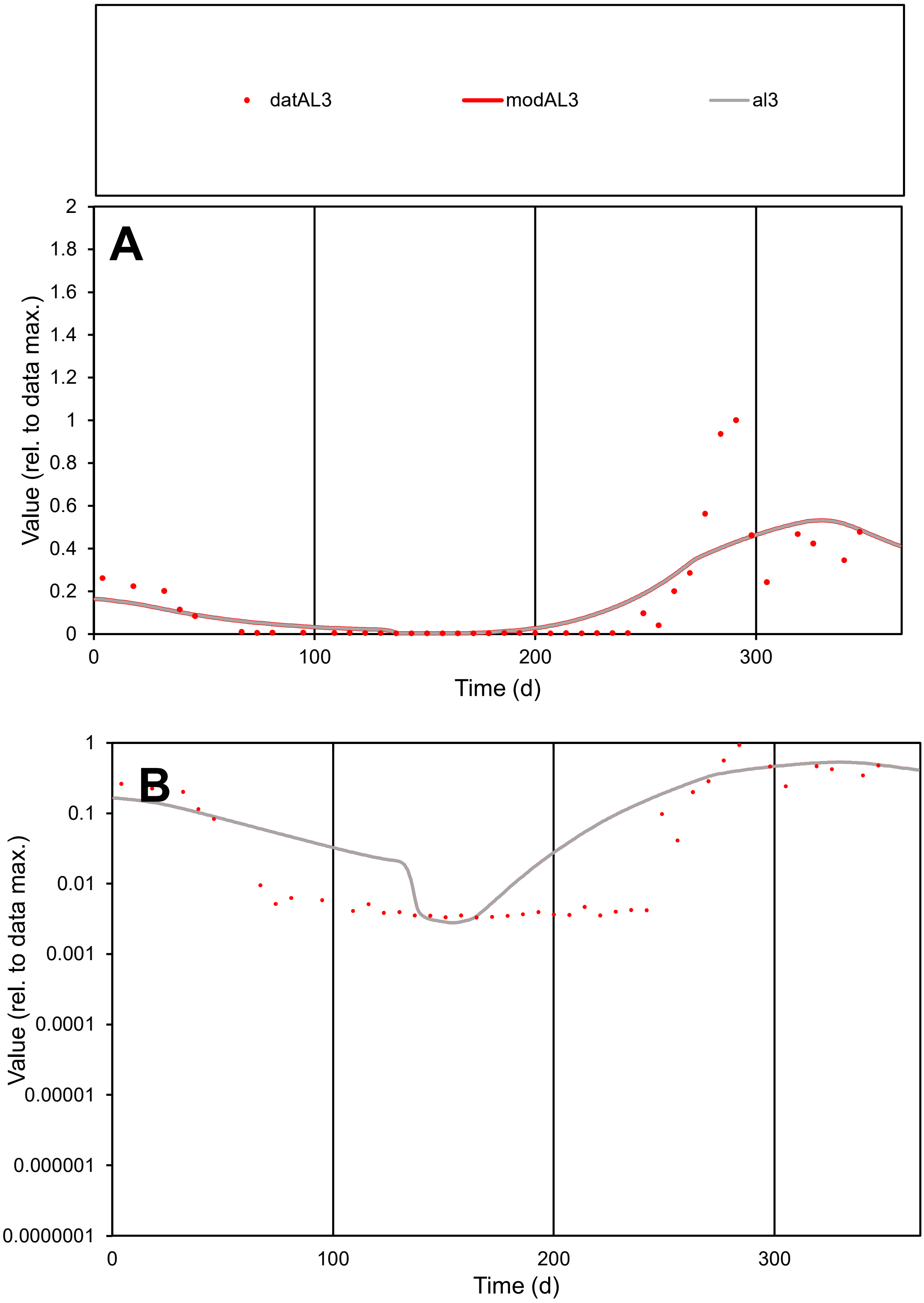


**C1**

Variables
AL3 Provide jtd or Dst
CalibOnly?
2 1 to reproduce of
oNormalize?
2 1=no, 2=yes

ofxn calc*
0.92788
ofxn mod
0.94556

sRun
RunA1
zout
1

Ave
Data
0.00112
Model
0.00117

data/vMax
0.00723

ofxng
334.07

Min d/m
1.17329

ave/min d
47.0399
ave/min m
57.1936

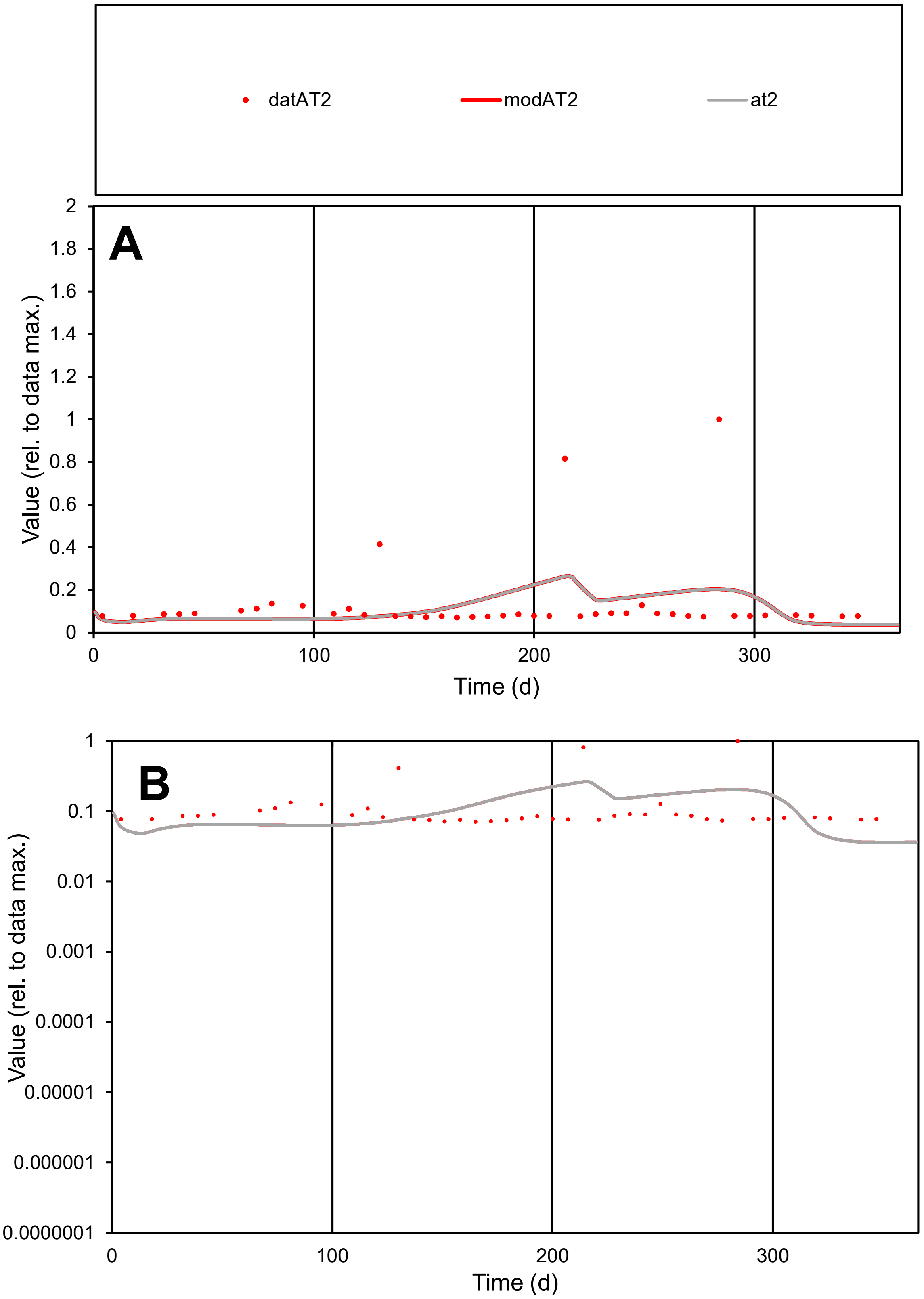


**C1**

Variables
AT2 Provide jtd or Dst
CalibOnly?
2 1 to reproduce of
oNormalize?
2 1=no, 2=yes

ofxn calc*
1.31127
ofxn mod
1.31848

sRun
RunA1
zout
1

Ave
Data
0.00015
Model
0.00014

data/vMax
0.00111

ofxng
334.07

Min d/m
1.96829

ave/min d
1.87387
ave/min m
3.48183

327_Calib_S_A

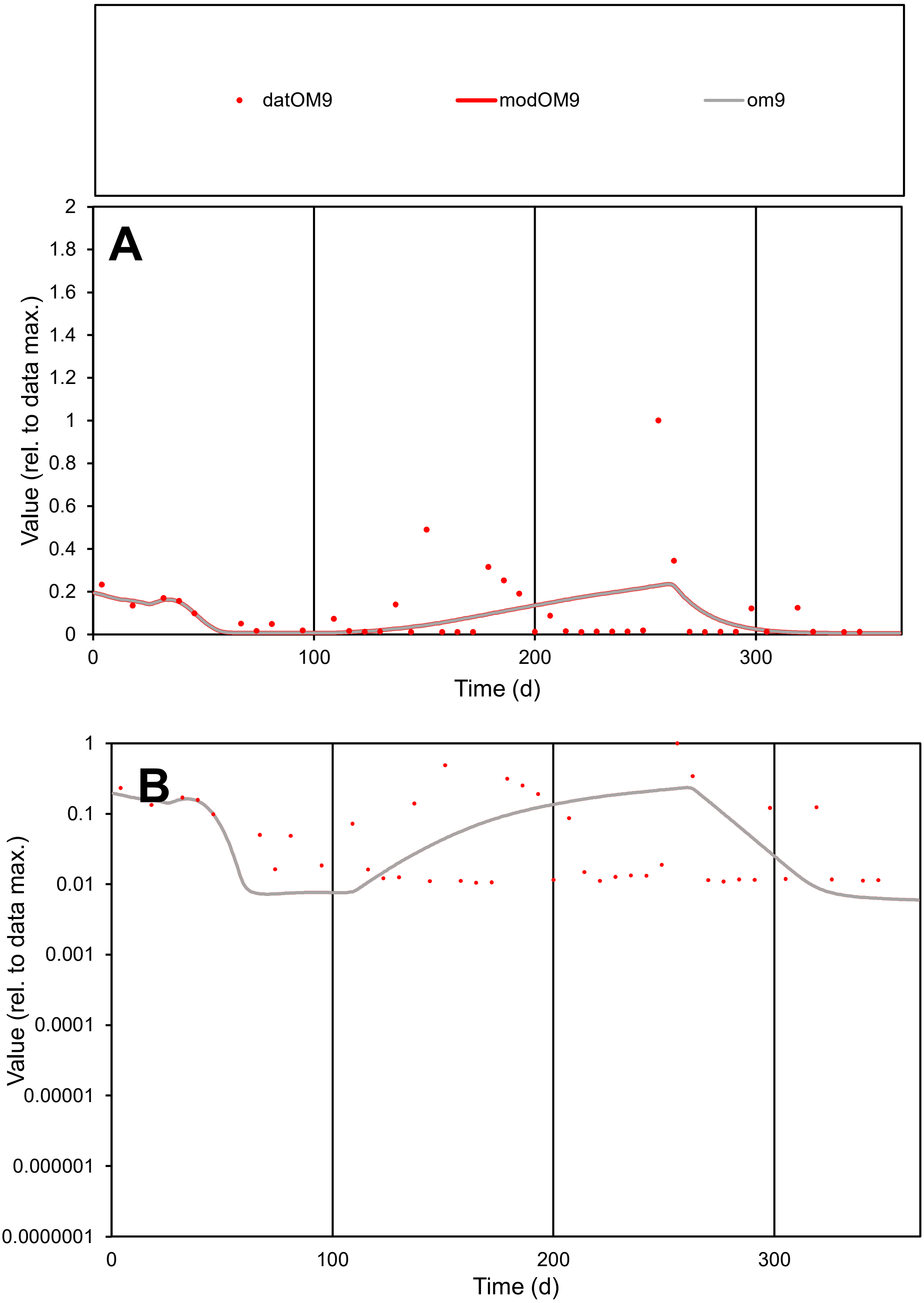


**C1**

Variables
OM9 Provide jtd or Dst
CalibOnly?
2 1 to reproduce of
oNormalize?
2 1=no, 2=yes

ofxn calc*
1.61251
ofxn mod
1.62682

sRun
RunA1
zout
1

Ave
Data
0.00042
Model
0.00036

data/vMax
0.0041

ofxng
334.07

Min d/m
1.67082

ave/min d
9.85089
ave/min m
14.2128

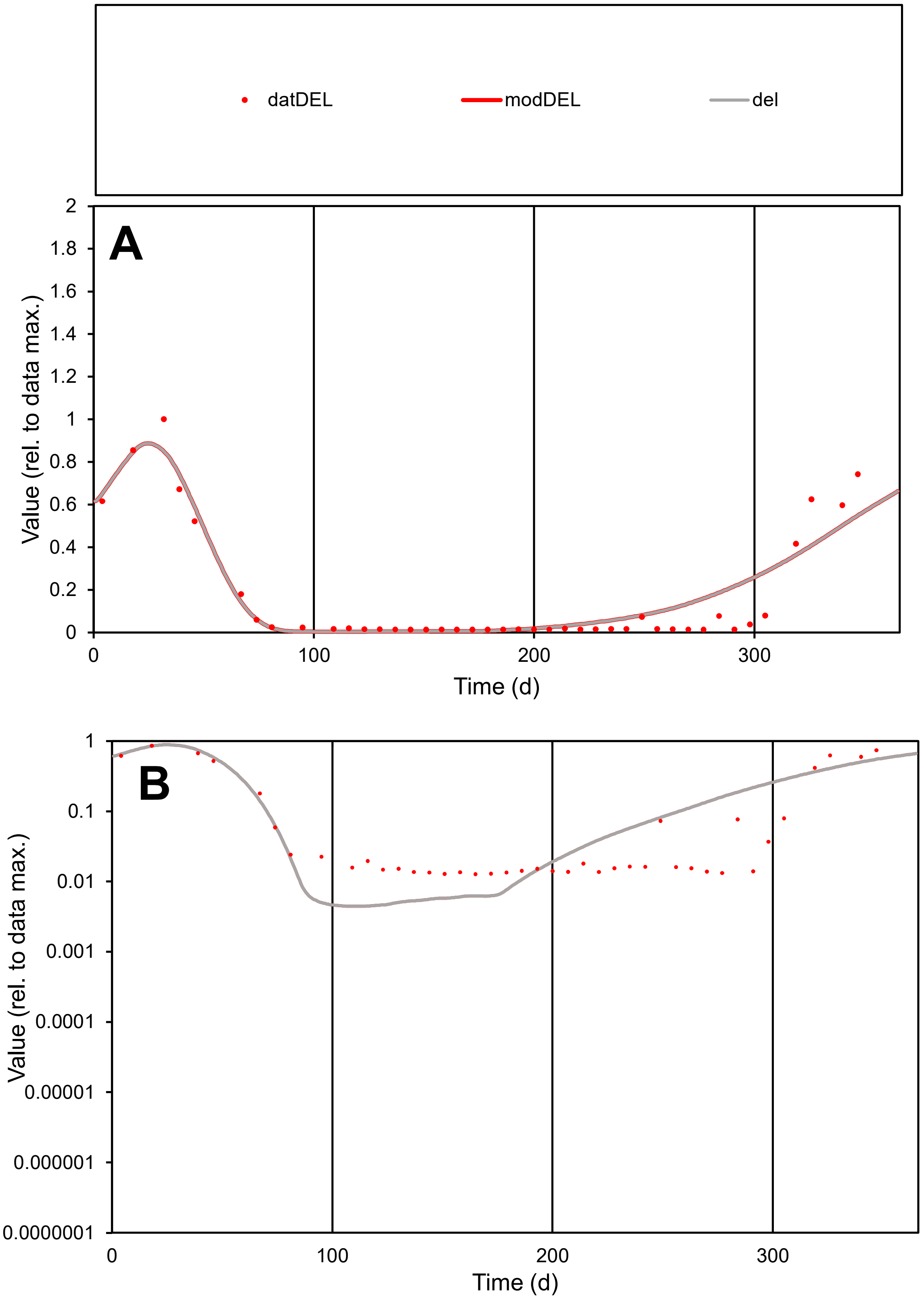


**C1**

Variables
DEL Provide jtd or Dst
CalibOnly?
2 1 to reproduce of
oNormalize?
2 1=no, 2=yes

ofxn calc*
0.53014
ofxn mod
0.58215

sRun
RunA1
zout
1

Ave
Data
0.0008
Model
0.00088

data/vMax
0.00483

ofxng
334.07

Min d/m
2.86829

ave/min d
13.0338
ave/min m
40.9471

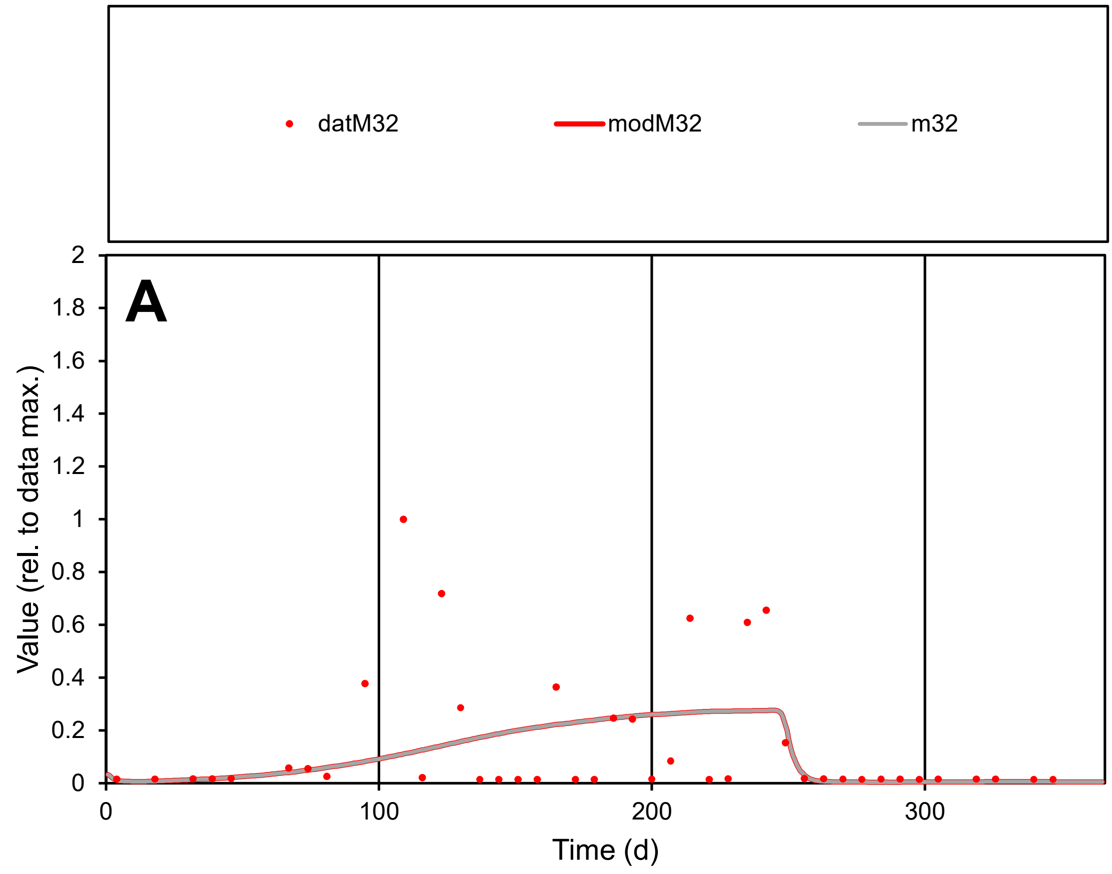


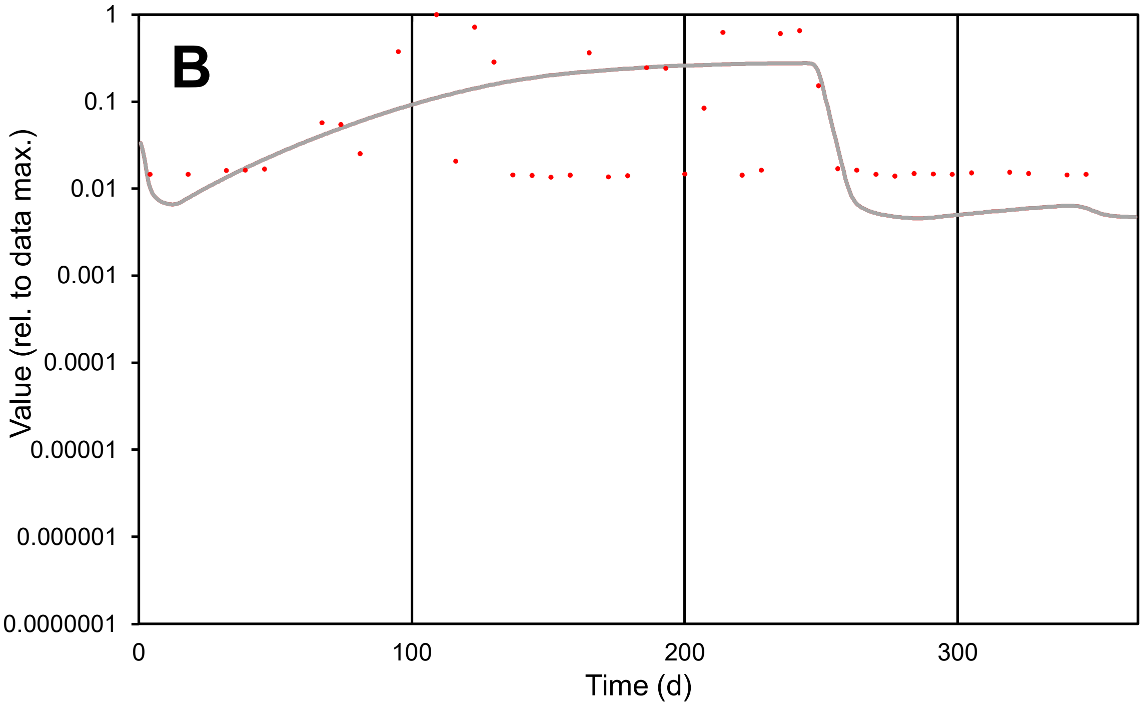


**C1**

| | |
|---|---|
| Variables | |
| M32 | Provide jtd or Dst |
| CalibOnly? | |
| 2 | 1 to reproduce of |
| oNormalize? | |
| 2 | 1=no, 2=yes |
| ofxn calc* | |
| 1.58404 | |
| ofxn mod | |
| 1.59391 | |
| sRun | |
| RunA1 | |
| zout | |
| 1 | |
| Ave | |
| Data | |
| 0.00217 | |
| Model | |
| 0.00183 | |
| data/vMax | |
| 0.01547 | |
| ofxng | |
| 334.07 | |
| Min d/m | |
| 2.98151 | |
| ave/min d | |
| 10.3765 | |
| ave/min m | |
| 26.0413 | |

327_Calib_S_A

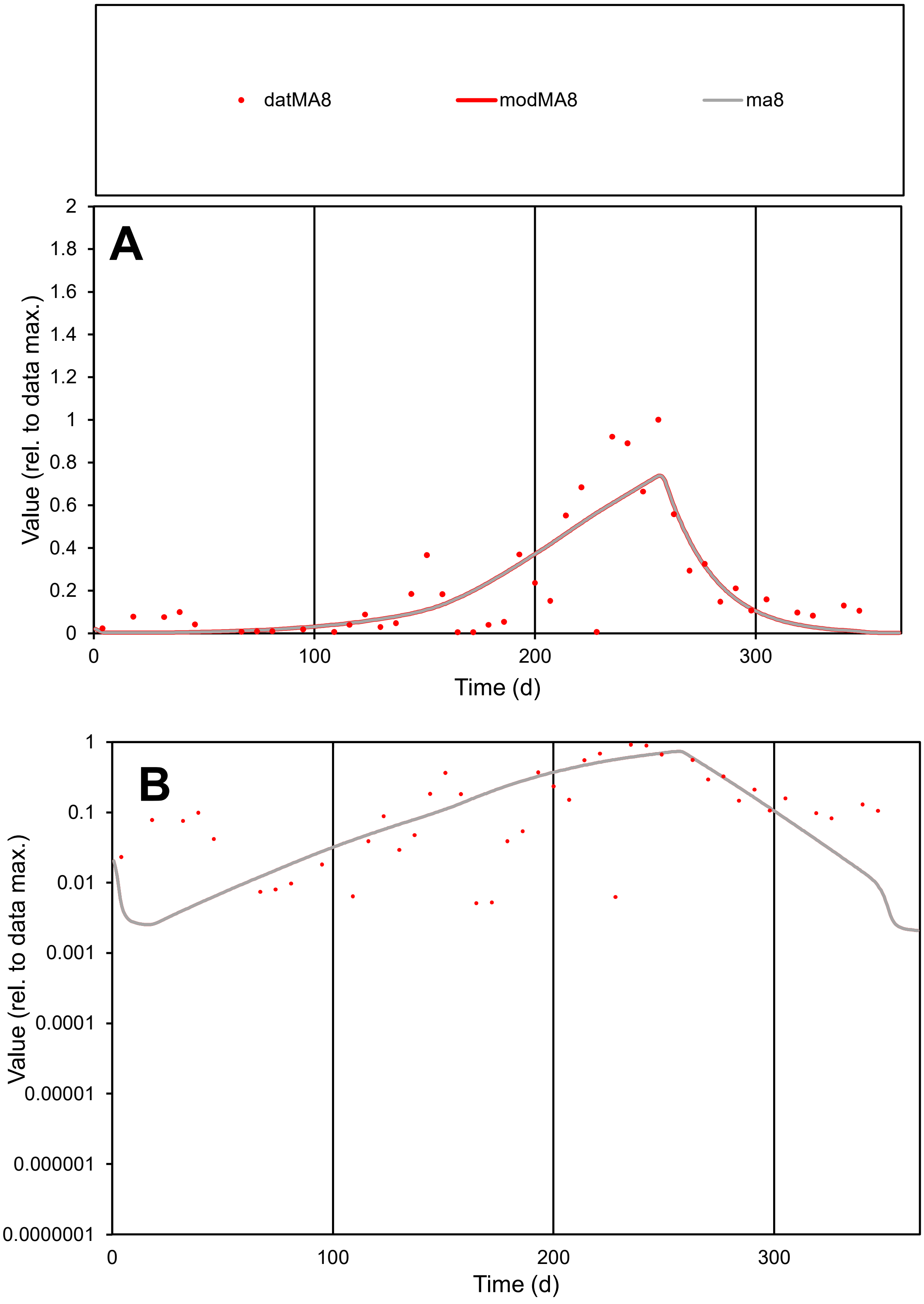


**C1**

Variables
MA8 Provide jtd or Dst
CalibOnly?
2 1 to reproduce of
oNormalize?
2 1=no, 2=yes

ofxn calc*
0.70009
ofxn mod
0.72686

sRun
RunA1
zout
1

Ave
Data
0.00249
Model
0.00246

data/vMax
0.01151

ofxng
334.07

Min d/m
1.94291

ave/min d
42.0931
ave/min m
80.7091

327_Calib_S_A

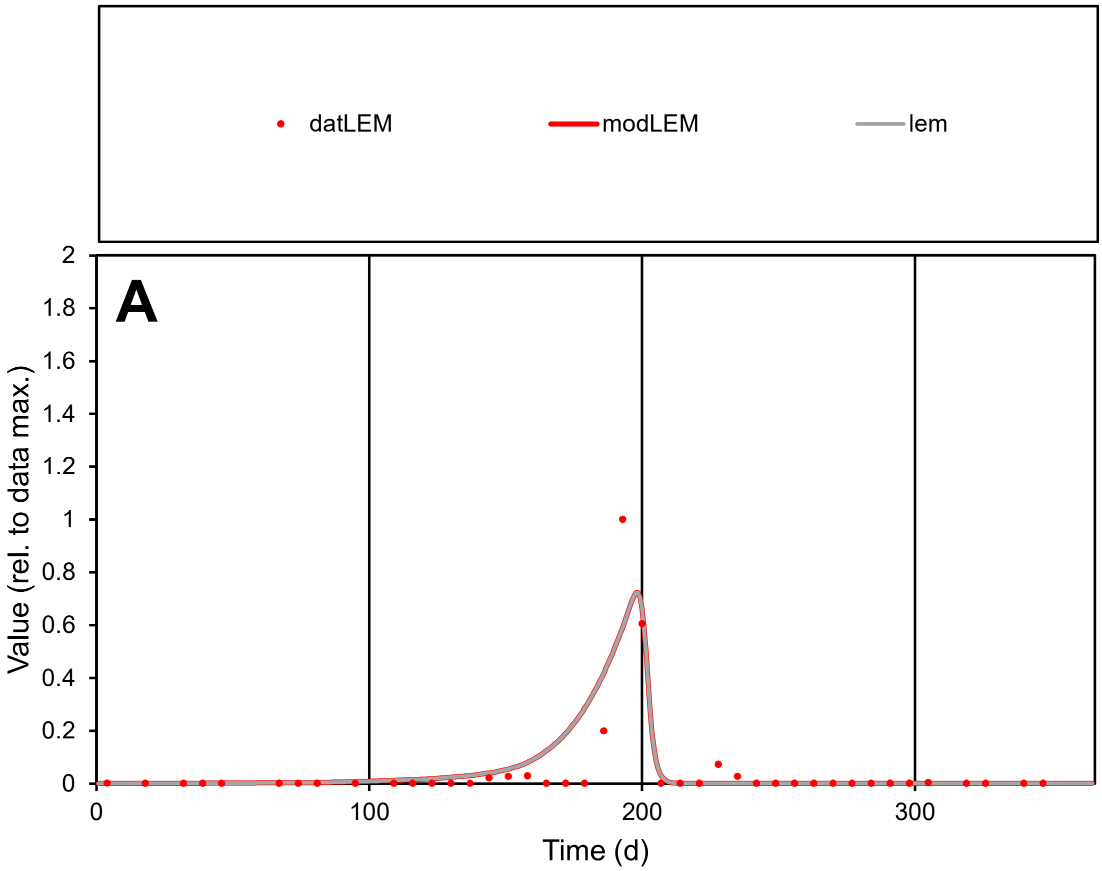


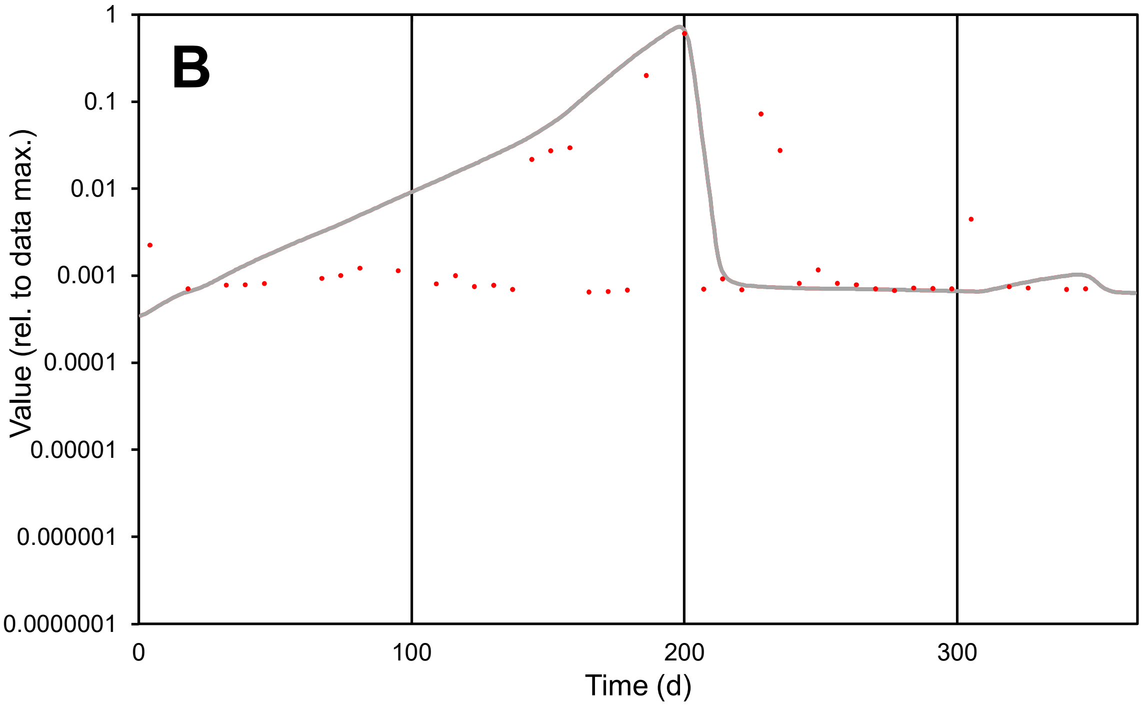


**C1**

Variables

LEM Provide jtd or Dst

CalibOnly?

2 1 to reproduce of

oNormalize?

2 1=no, 2=yes

ofxn calc*

1.80915

ofxn mod

1.897

sRun

RunA1

zout

1

Ave

Data

0.00159

Model

0.002

data/vMax

0.03304

ofxng

334.07

Min d/m

1.58401

ave/min d

74.0732

ave/min m

147.873

327_Calib_S_A

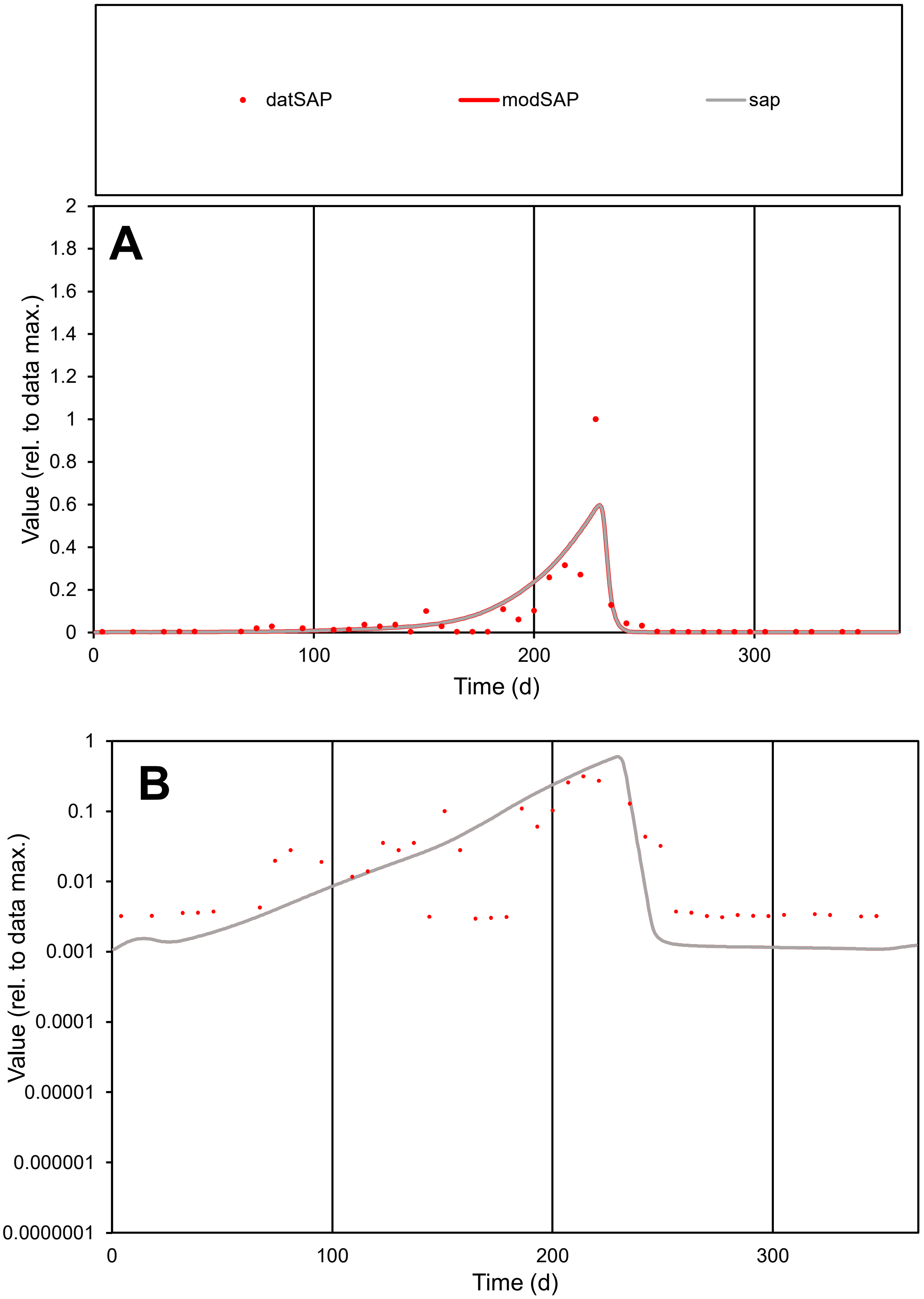


**C1**

Variables

SAP Provide jtd or Dst

CalibOnly?

2 1 to reproduce of

oNormalize?

2 1=no, 2=yes

ofxn calc*

1.25966

ofxn mod

1.30331

sRun

RunA1

zout

1

Ave

Data

0.00071

Model

0.00078

data/vMax

0.01099

ofxng

334.07

Min d/m

2.71893

ave/min d

21.7918

ave/min m

64.8259

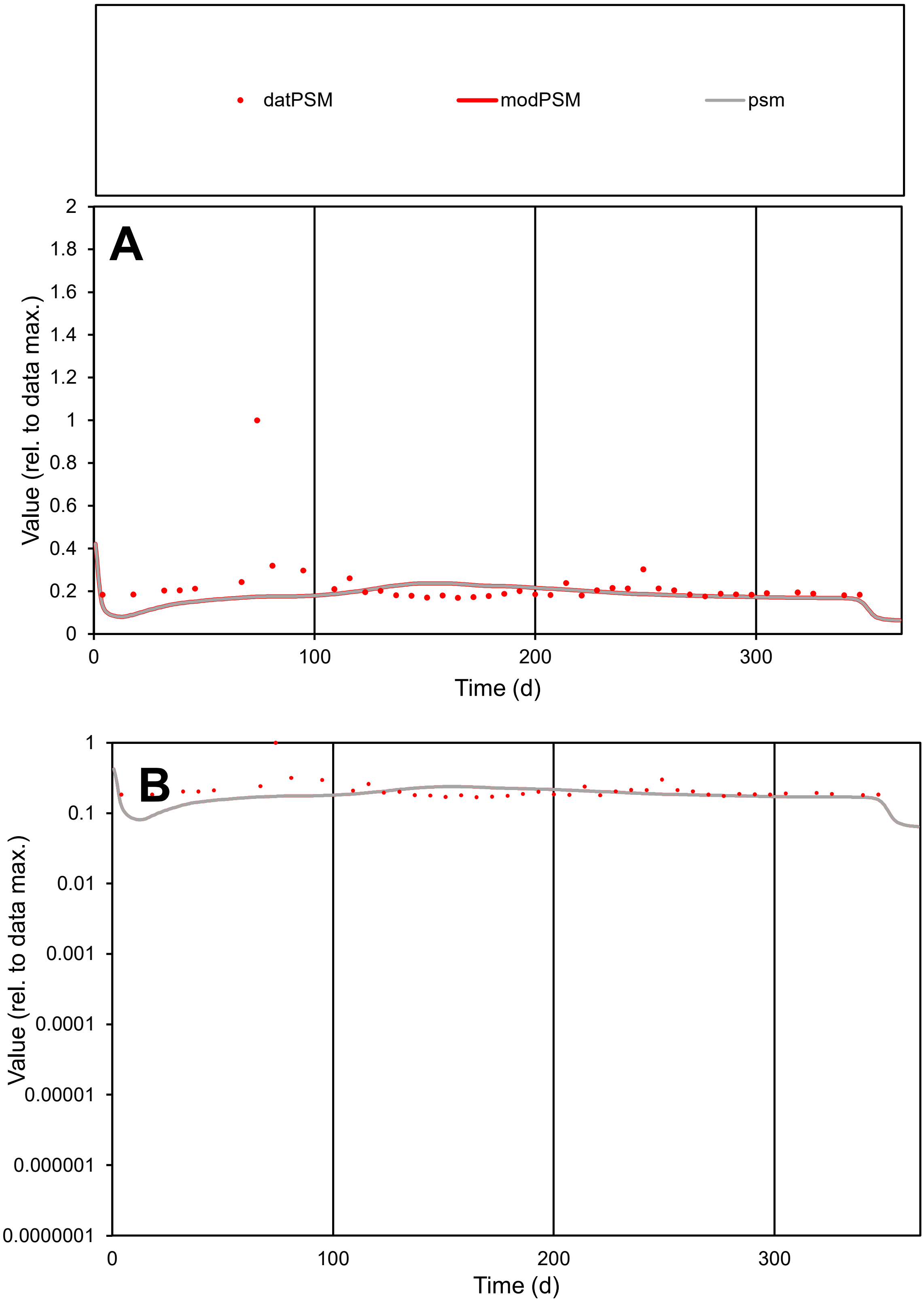


**C1**

Variables

PSM Provide jtd or Dst

CalibOnly?

2 1 to reproduce of

oNormalize?

2 1=no, 2=yes

ofxn calc*

0.62089

ofxn mod

0.63108

sRun

RunA1

zout

1

Ave

Data

3.6E-05

Model

3.1E-05

data/vMax

0.00016

ofxng

334.07

Min d/m

1.74539

ave/min d

1.319

ave/min m

1.95414

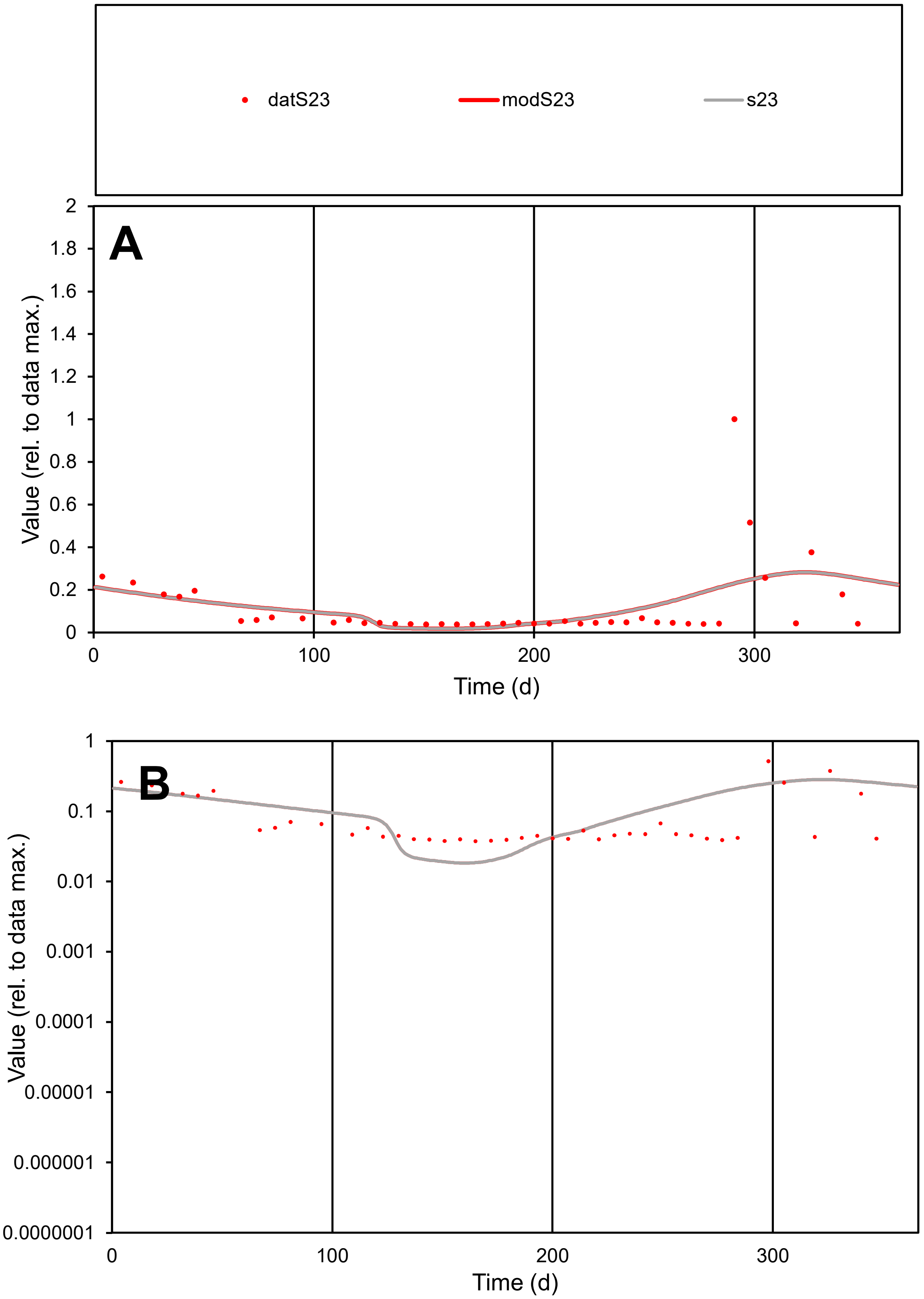


**C1**

Variables
S23 Provide jtd or Dst
CalibOnly?
2 1 to reproduce of
oNormalize?
2 1=no, 2=yes

ofxn calc*
1.26452
ofxn mod
1.27619

sRun
RunA1
zout
1

Ave
Data
0.00049
Model
0.00051

data/vMax
0.00426

ofxng
334.07

Min d/m
2.03213

ave/min d
3.0852
ave/min m
6.54729

327_Calib_S_A

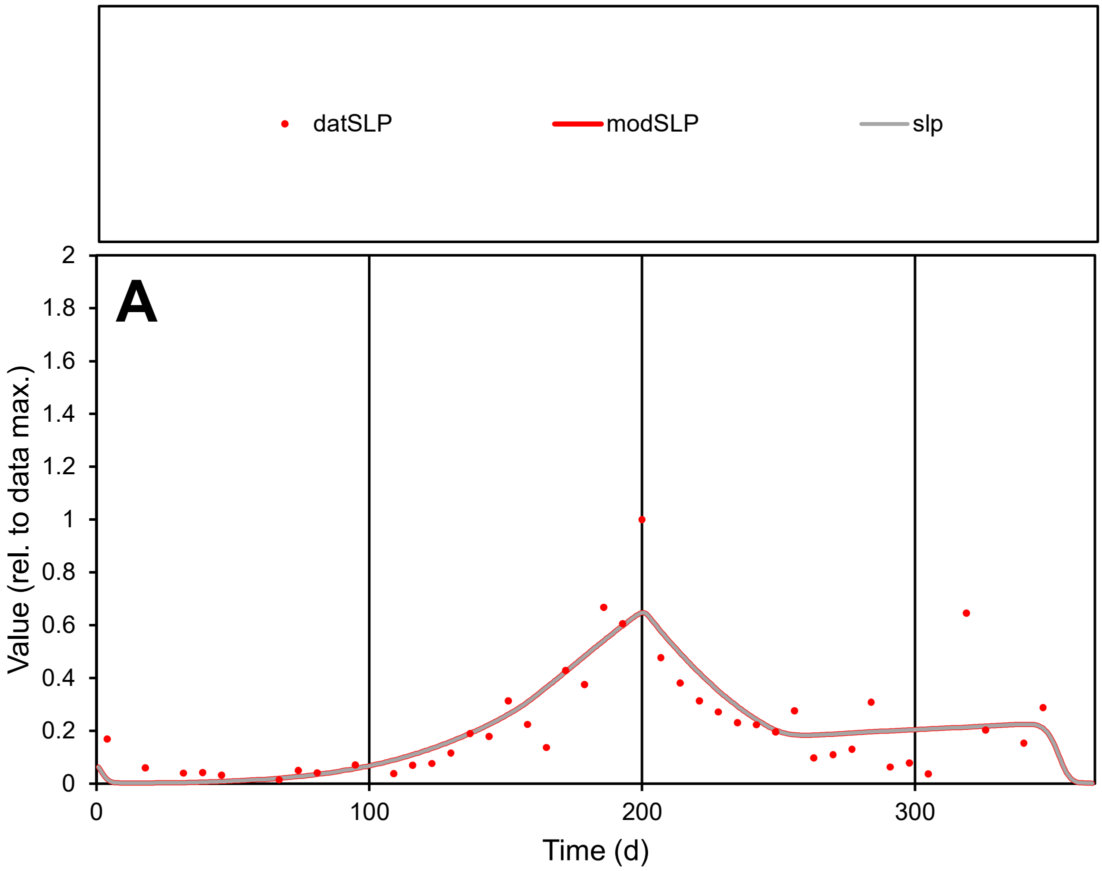


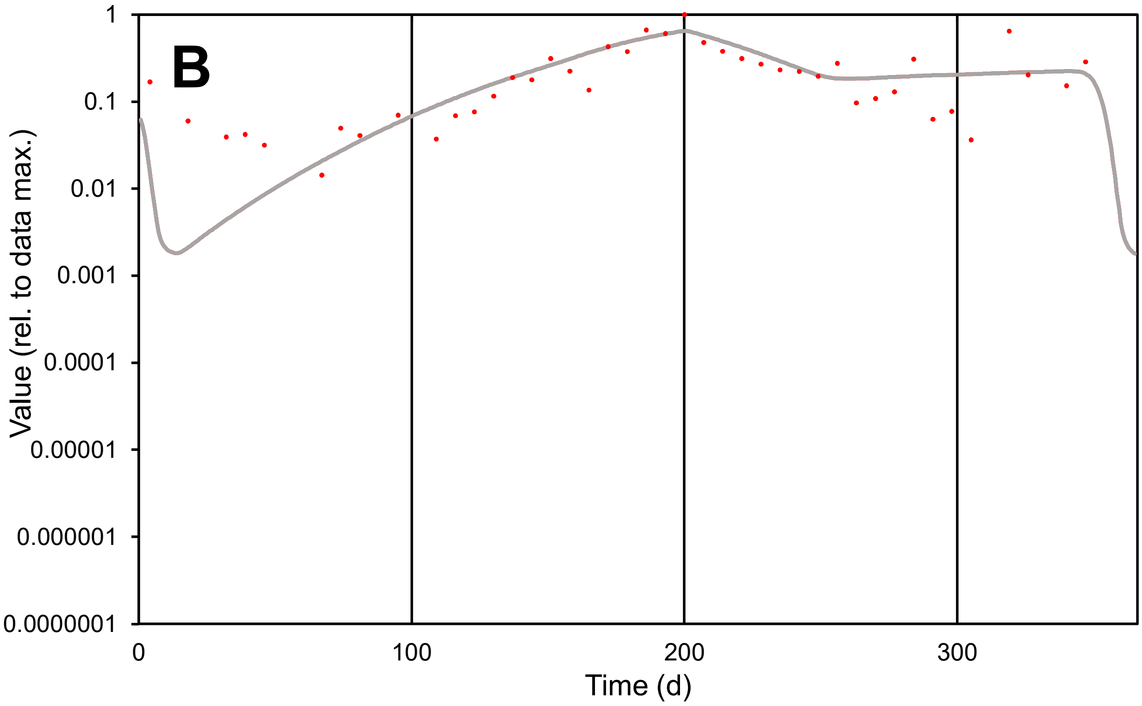


**C1**

Variables
SLP Provide jtd or Dst
CalibOnly?
2 1 to reproduce of
oNormalize?
2 1=no, 2=yes

ofxn calc*
0.52753
ofxn mod
0.55355

sRun
RunA1
zout
1

Ave
Data
0.00154
Model
0.0016

data/vMax
0.00688

ofxng
334.07

Min d/m
6.23921

ave/min d
15.6637
ave/min m
101.322

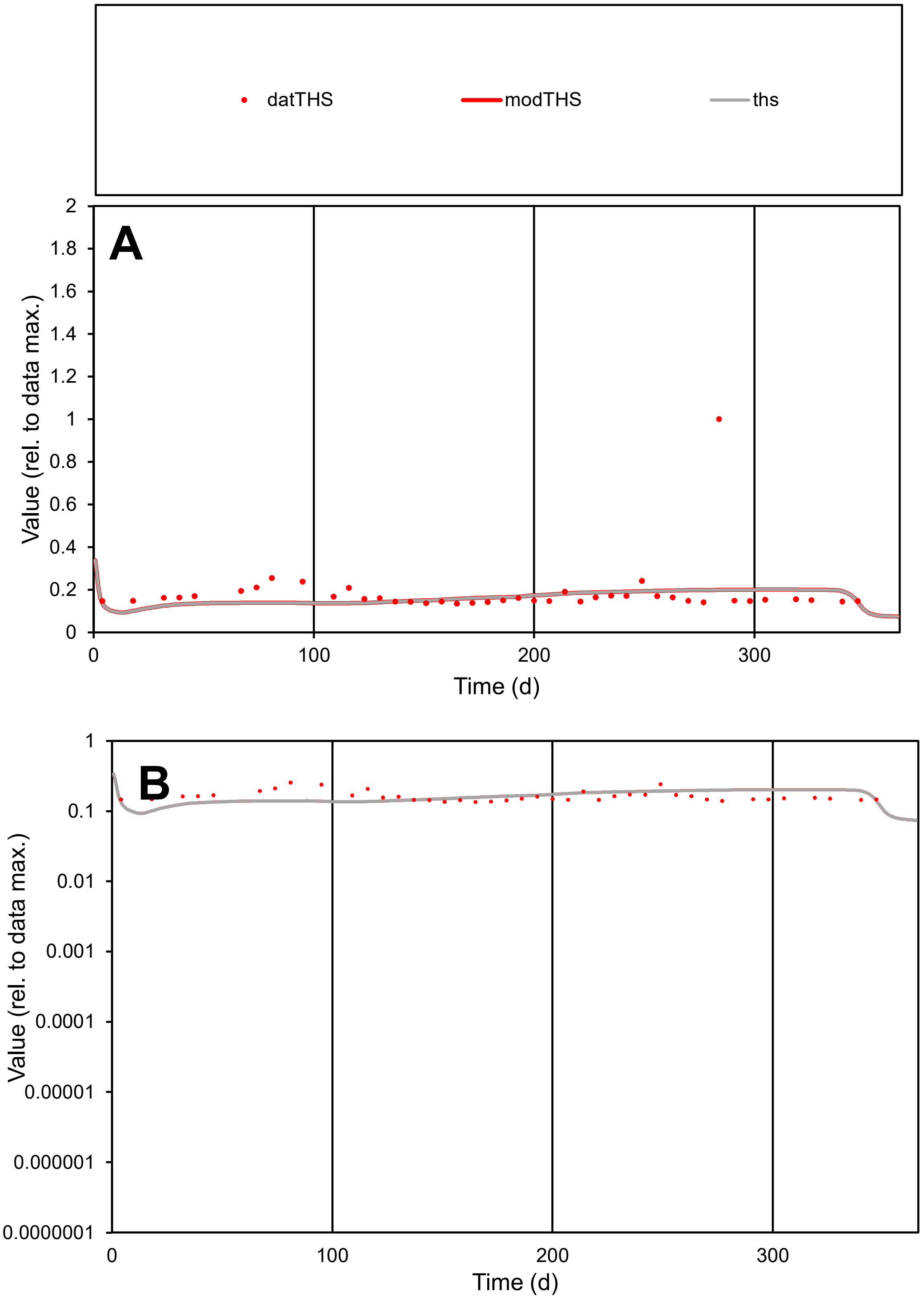


**C1**

Variables
THS Provide jtd or Dst
CalibOnly?
2 1 to reproduce of
oNormalize?
2 1=no, 2=yes

ofxn calc*
0.71023
ofxn mod
0.7188

sRun
RunA1
zout
1

Ave
Data
3.9E-05
Model
3.5E-05

data/vMax
0.00021

ofxng
334.07

Min d/m
1.29063

ave/min d
1.36443
ave/min m
1.58876

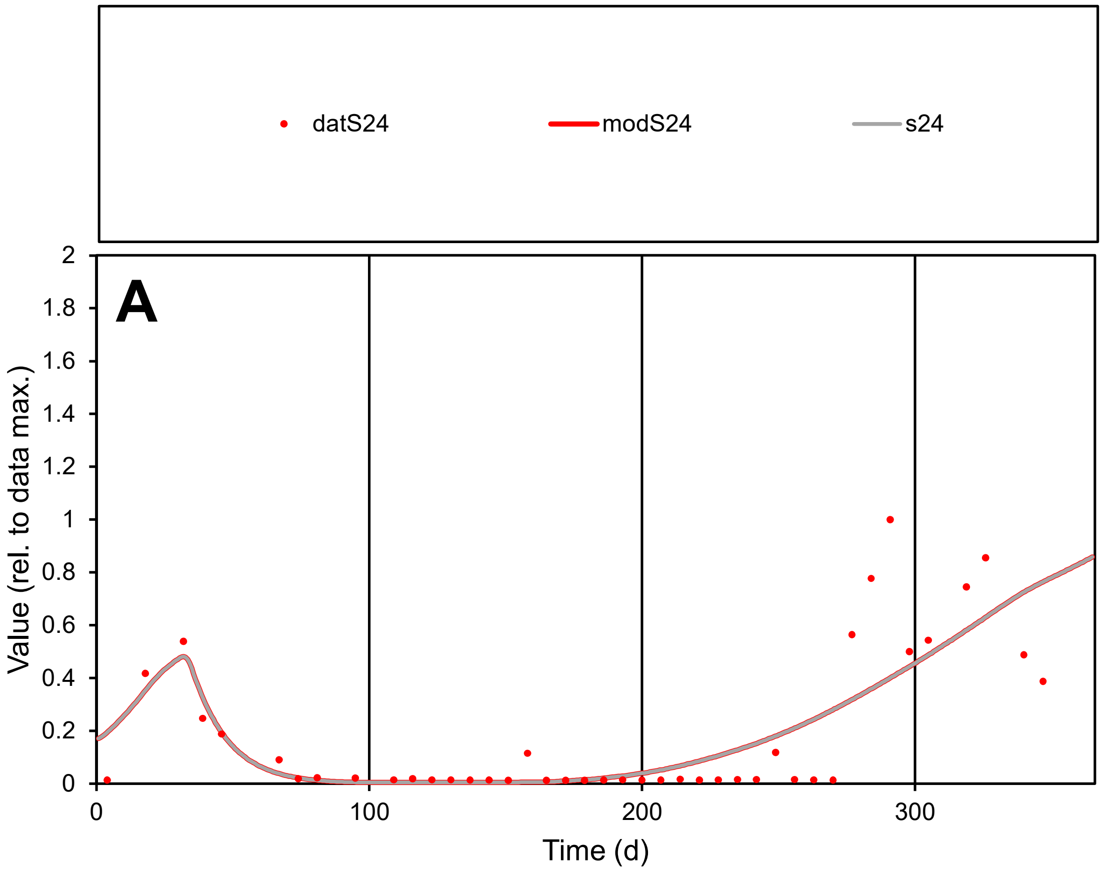


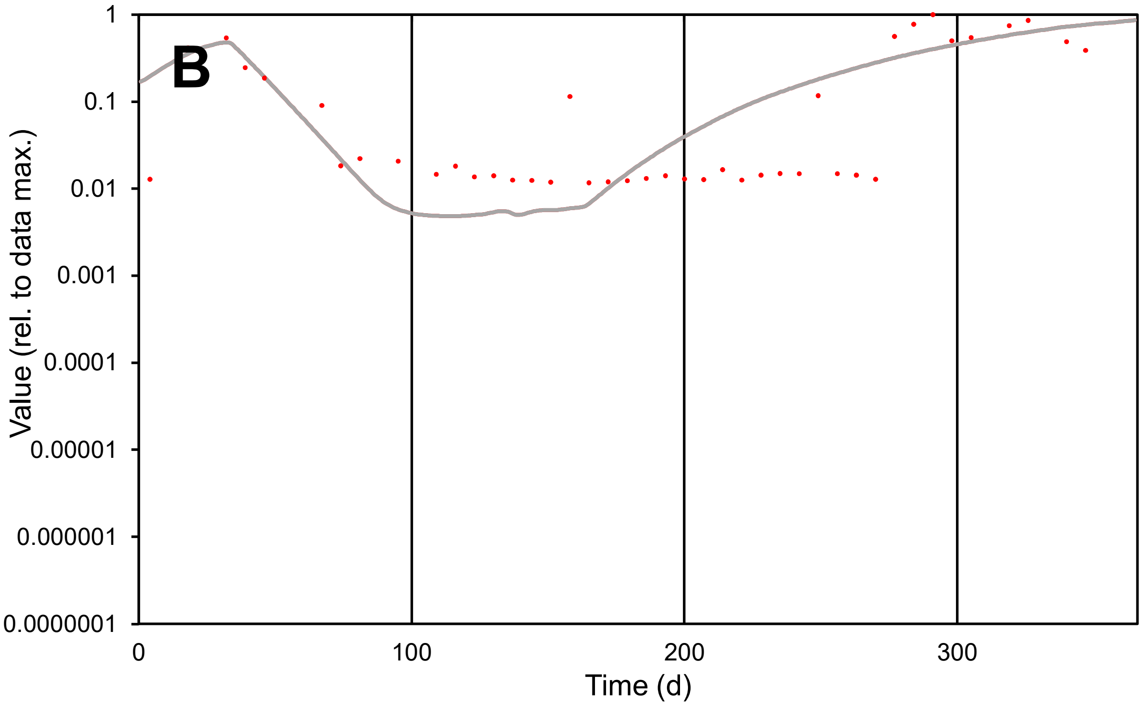


**C1**

Variables
S24 Provide jtd or Dst
CalibOnly?
2 1 to reproduce of
oNormalize?
2 1=no, 2=yes

ofxn calc*
0.86859
ofxn mod
0.88636

sRun
RunA1
zout
1

Ave
Data
0.00093
Model
0.00094

data/vMax
0.00491

ofxng
334.07

Min d/m
2.42107

ave/min d
16.1064
ave/min m
39.5247

327_Calib_S_A

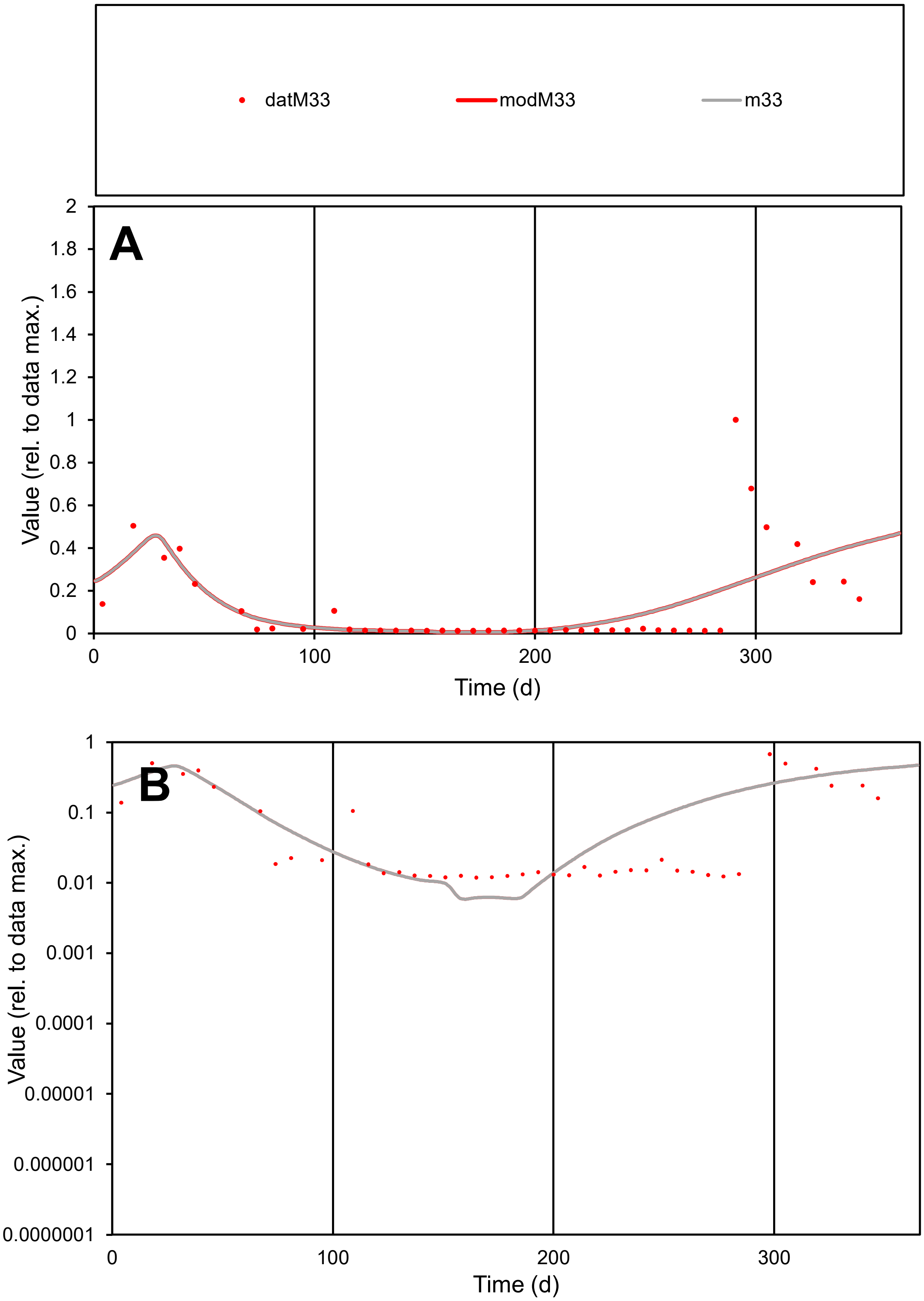


**C1**

Variables
M33 Provide jtd or Dst
CalibOnly?
2 1 to reproduce of
oNormalize?
2 1=no, 2=yes

ofxn calc*
1.23479
ofxn mod
1.25127

sRun
RunA1
zout
1

Ave
Data
0.00075
Model
0.00075

data/vMax
0.00572

ofxng
334.07

Min d/m
2.02869

ave/min d
10.9953
ave/min m
22.271

327_Calib_S_A

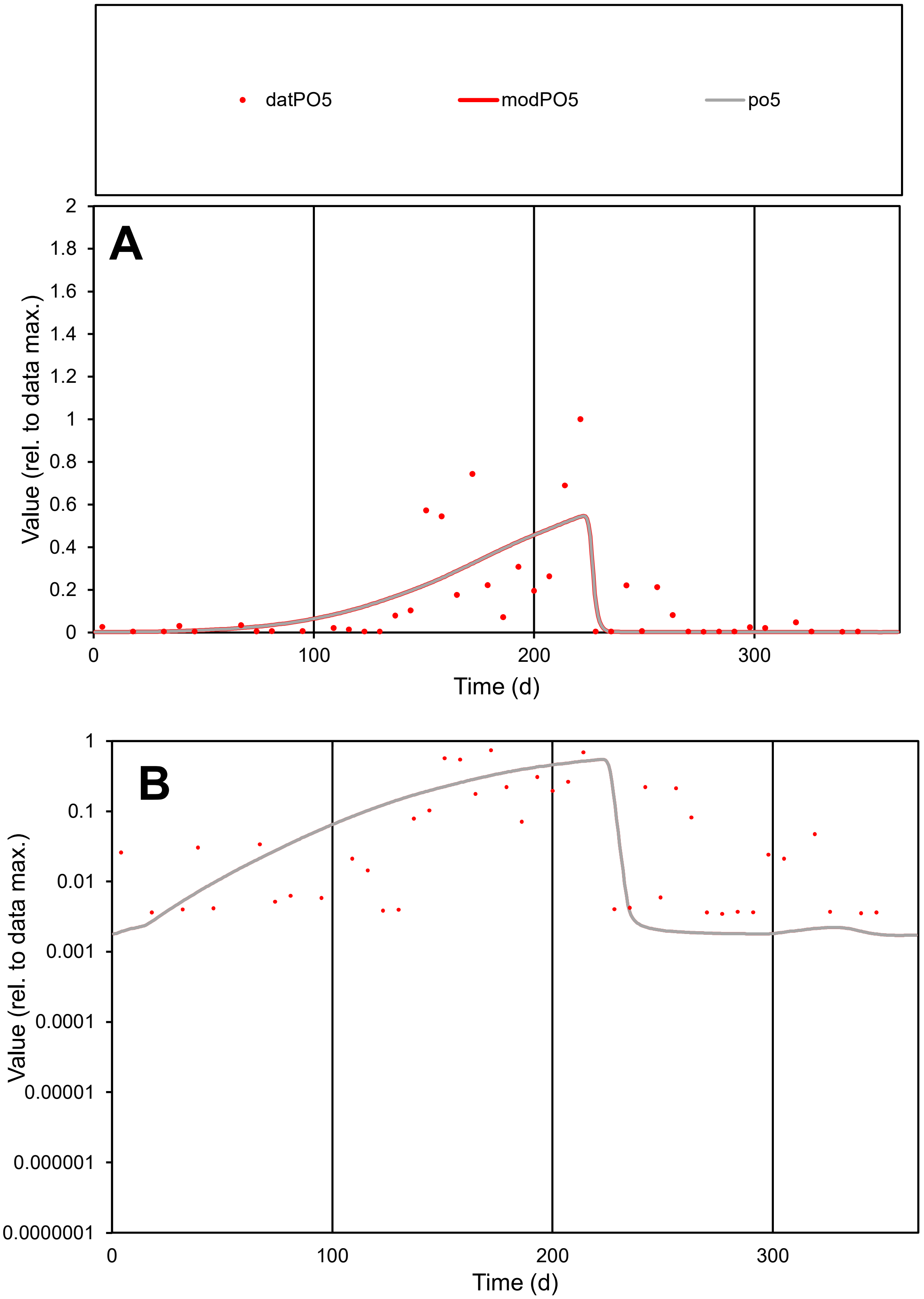


**C1**

Variables
PO5 Provide jtd or Dst
CalibOnly?
2 1 to reproduce of
oNormalize?
2 1=no, 2=yes

ofxn calc*
1.15054
ofxn mod
1.16827

sRun
RunA1
zout
1

Ave
Data
0.00191
Model
0.00179

data/vMax
0.01392

ofxng
334.07

Min d/m
1.96411

ave/min d
39.7572
ave/min m
72.8672

327_Calib_S_A

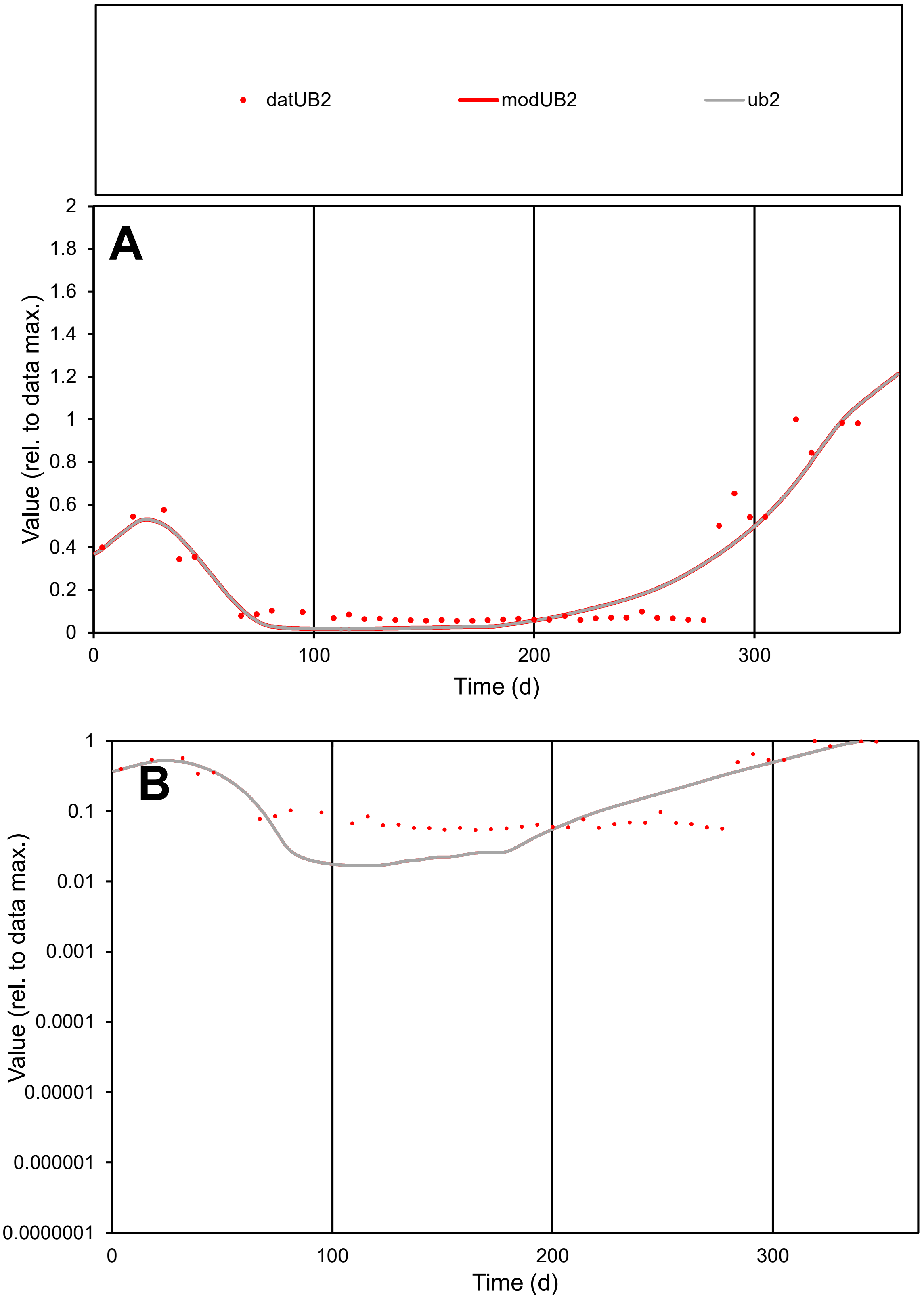


**C1**

Variables
UB2 Provide jtd or Dst
CalibOnly?
2 1 to reproduce of
oNormalize?
2 1=no, 2=yes

ofxn calc*
0.39943
ofxn mod
0.43107

sRun
RunA1
zout
1

Ave
Data
0.00046
Model
0.00046

data/vMax
0.0019

ofxng
334.07

Min d/m
3.22309

ave/min d
4.48403
ave/min m
14.3671

327_Calib_S_A

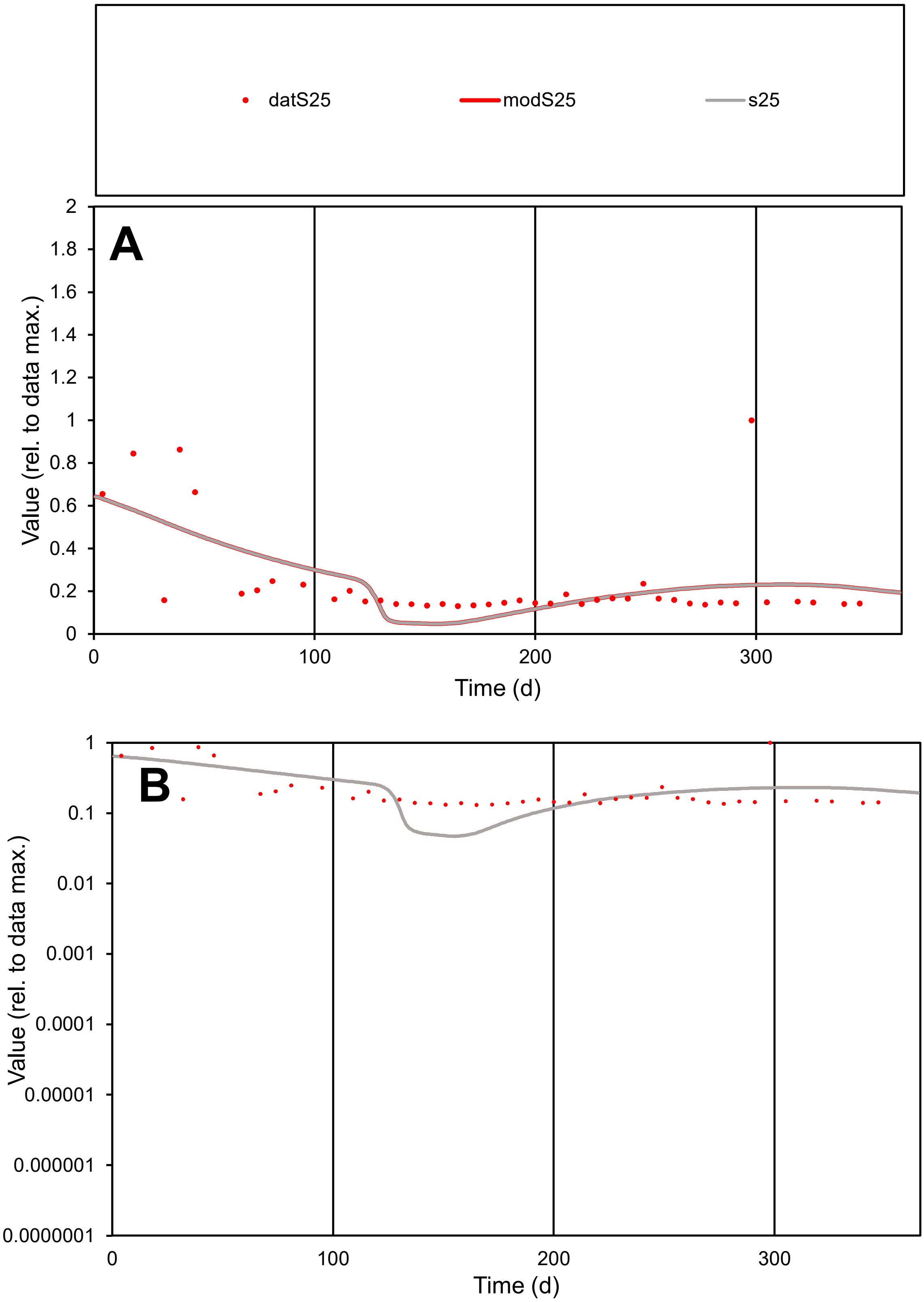


**C1**

Variables
S25 Provide jtd or Dst
CalibOnly?
2 1 to reproduce of
oNormalize?
2 1=no, 2=yes

ofxn calc*
0.71461
ofxn mod
0.72999

sRun
RunA1
zout
1

Ave
Data
0.0003
Model
0.00029

data/vMax
0.00129

ofxng
334.07

Min d/m
2.74273

ave/min d
1.81154
ave/min m
4.74421

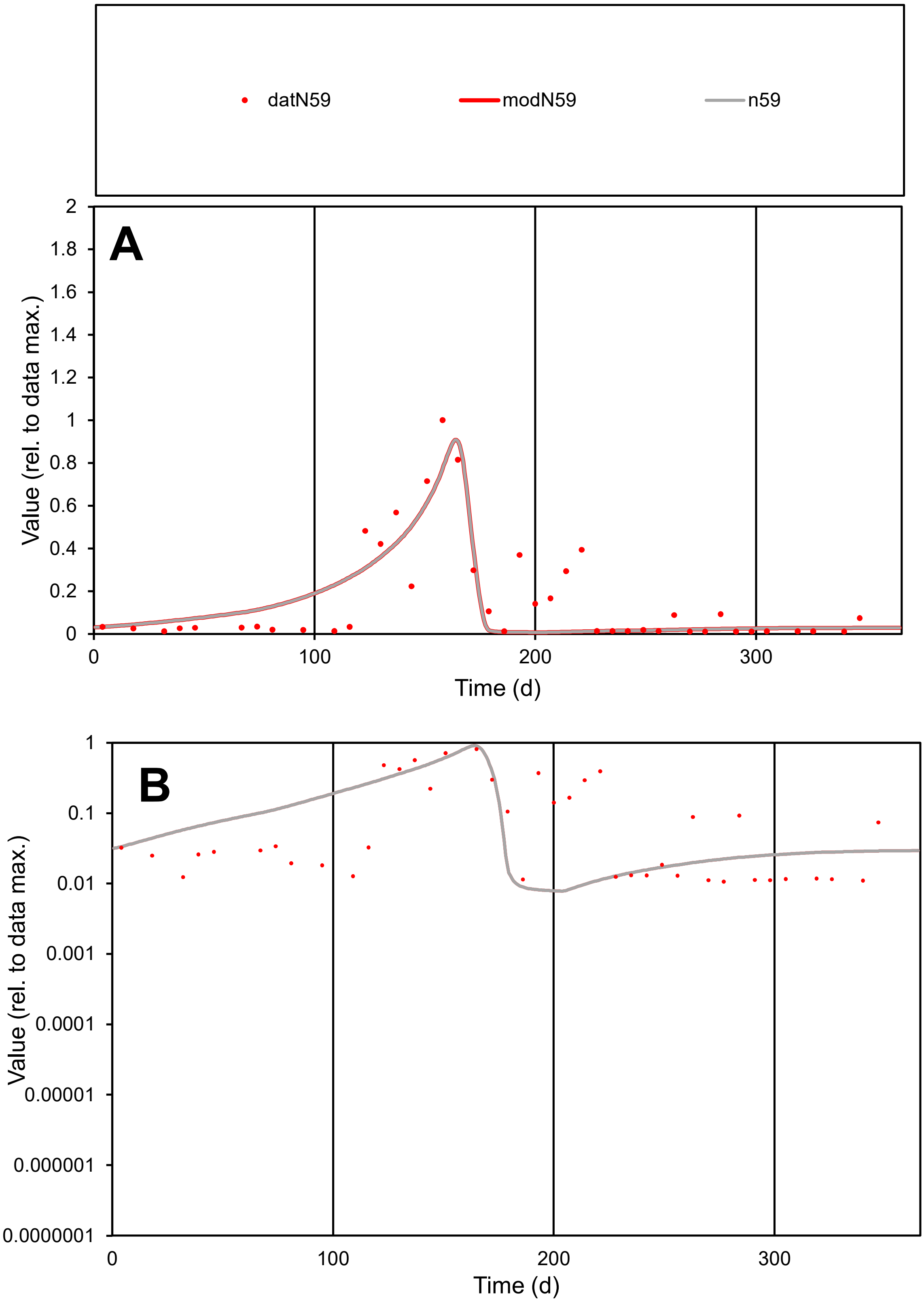


**C1**

Variables

N59 Provide jtd or Dst

CalibOnly?

2 1 to reproduce of

oNormalize?

2 1=no, 2=yes

ofxn calc*

0.83287

ofxn mod

0.85735

sRun

RunA1

zout

1

Ave

Data

0.00064

Model

0.00057

data/vMax

0.004

ofxng

334.07

Min d/m

1.3568

ave/min d

14.8766

ave/min m

18.2225

327_Calib_S_A

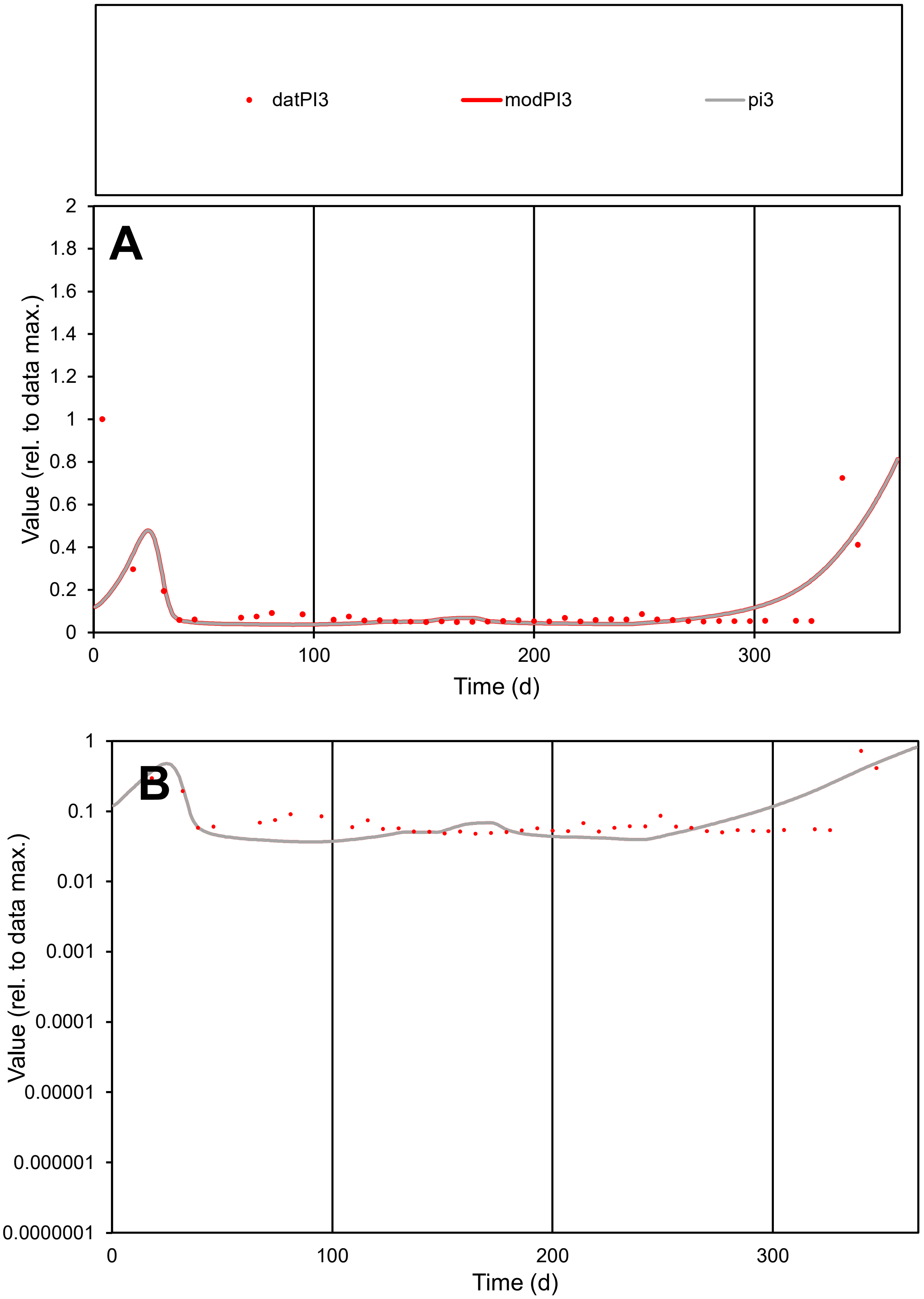


**C1**

Variables
PI3 Provide jtd or Dst
CalibOnly?
2 1 to reproduce of
oNormalize?
2 1=no, 2=yes

ofxn calc*
1.28785
ofxn mod
1.29913

sRun
RunA1
zout
1

Ave
Data
0.00017
Model
0.00014

data/vMax
0.00147

ofxng
334.07

Min d/m
1.30007

ave/min d
2.38492
ave/min m
2.59131

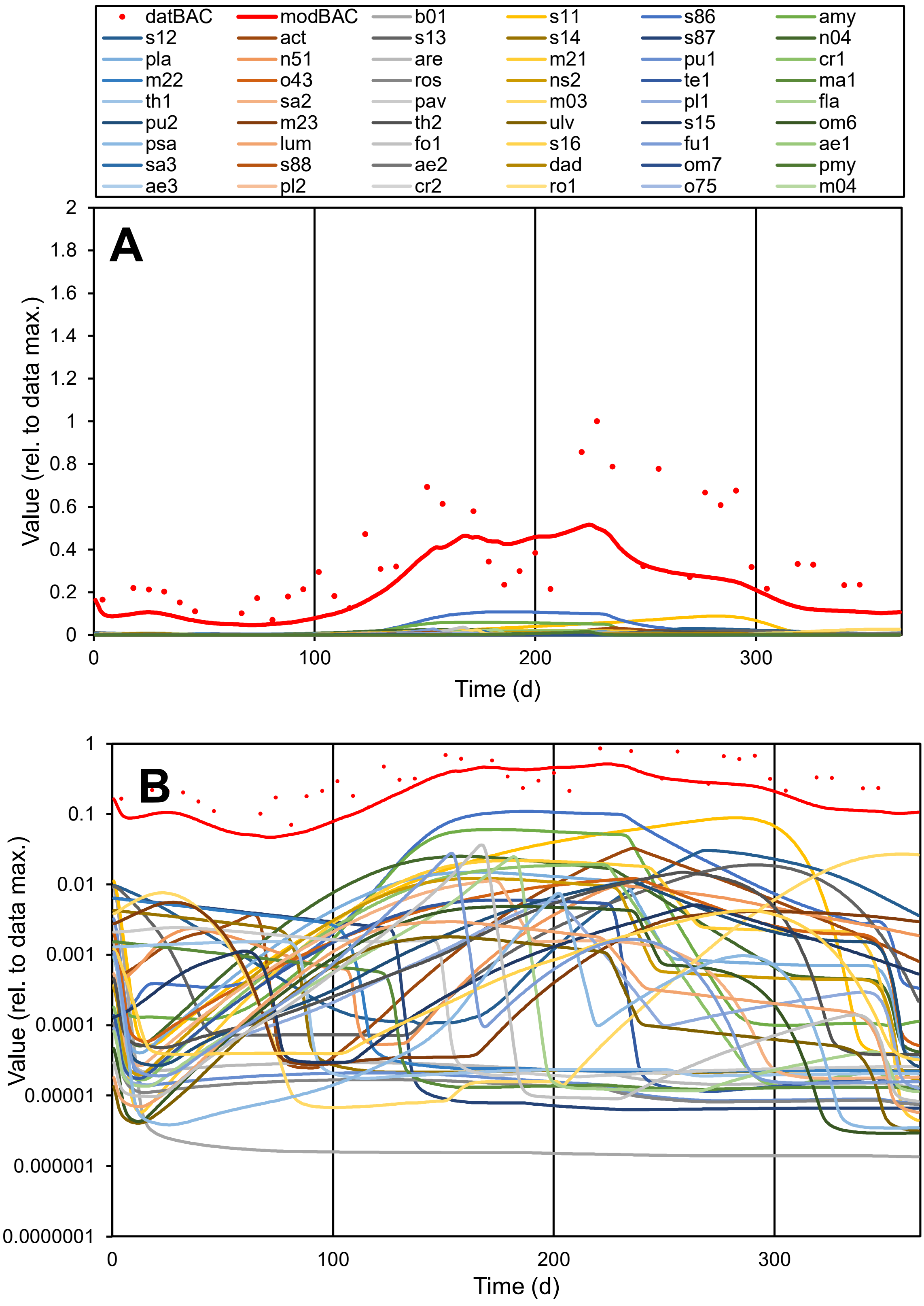


C1

Variables

BAC Provide jtd or Dst

CalibOnly?

2 1 to reproduce of

oNormalize?

2 1=no, 2=yes

ofxn calc*

0.59319

ofxn mod

0.6028

sRun

RunA1

zout

1

Ave

Data

1.09306

Model

0.69903

data/vMax

3.019

ofxng

334.07

Min d/m

1.46552

ave/min d

5.14379

ave/min m

4.82086

328_Calib_S_A

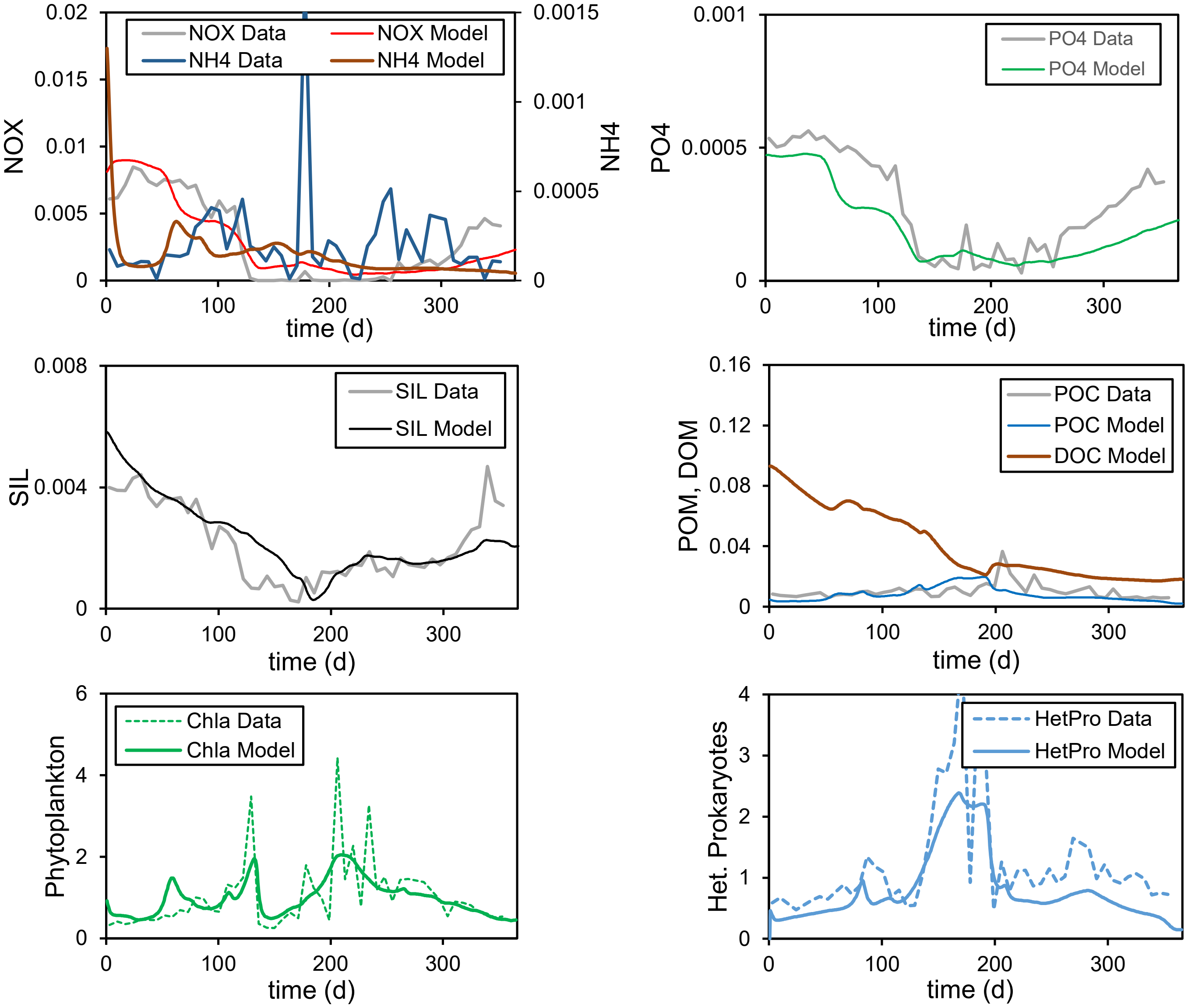
NOX Data
NOX Model
NH4 Data
NH4 Model
NOX
NH4
time (d)
PO4 Data
PO4 Model
PO4
time (d)
SIL Data
SIL Model
SIL
time (d)
POC Data
POC Model
DOC Model
POM, DOM
time (d)
Chla Data
Chla Model
Phytoplankton
time (d)
HetPro Data
HetPro Model
Het. Prokaryotes
time (d)

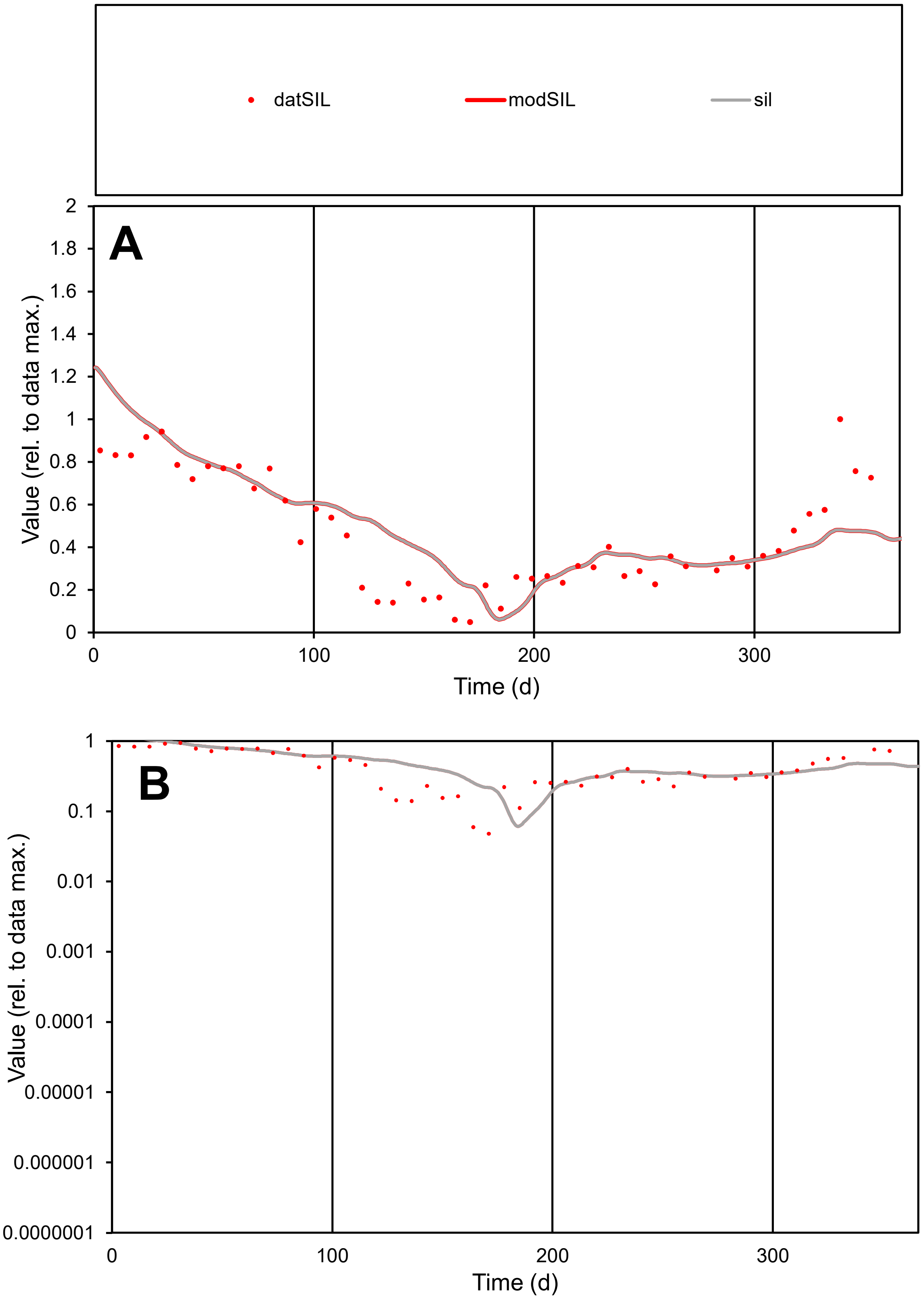


**C1**

Variables
SIL Provide jtd or Dst
CalibOnly?
2 1 to reproduce of
oNormalize?
2 1=no, 2=yes

ofxn calc*
1.06863
ofxn mod
1.14607

sRun
RunA1
zout
1

Ave
Data
0.00216
Model
0.00232

data/vMax
0.00469

ofxng
369.894

Min d/m
0.72464

ave/min d
9.58178
ave/min m
7.47688

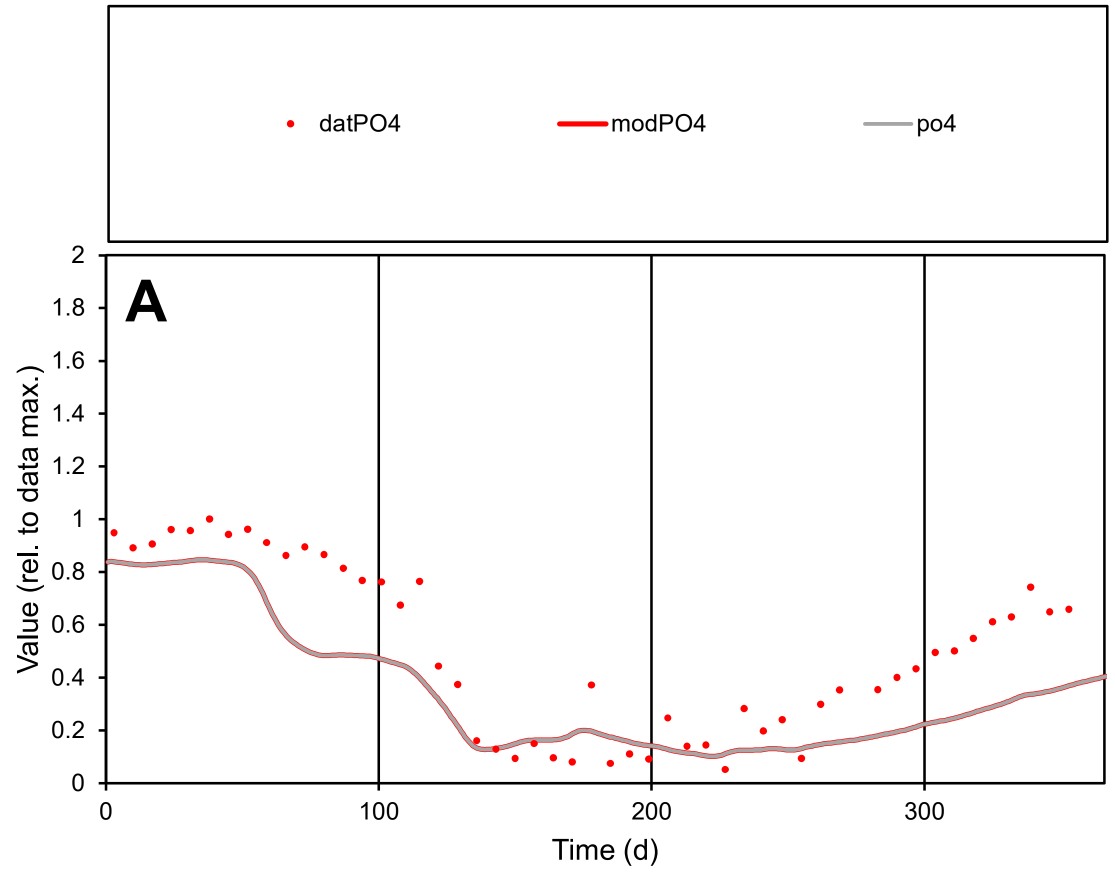


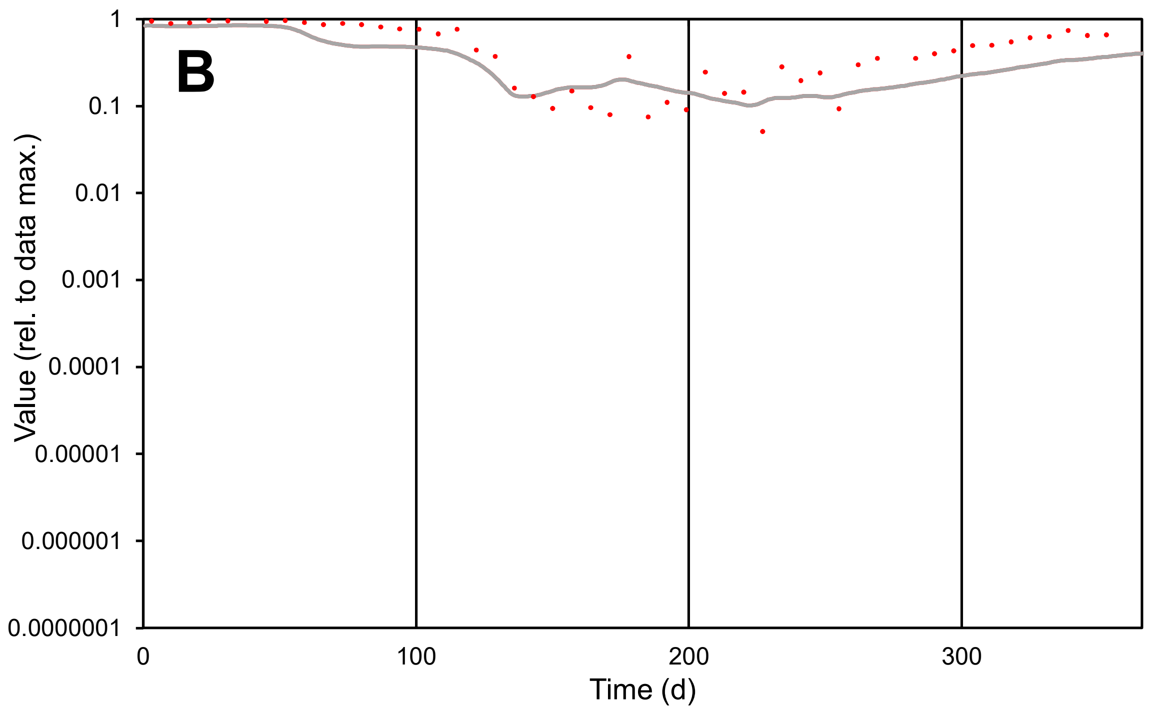


**C1**

Variables
PO4 Provide jtd or Dst
CalibOnly?
2 1 to reproduce of
oNormalize?
2 1=no, 2=yes

ofxn calc*
1.23068
ofxn mod
1.28719

sRun
RunA1
zout
1

Ave
Data
0.00028
Model
0.0002

data/vMax
0.00056

ofxng
369.894

Min d/m
0.49853

ave/min d
9.84803
ave/min m
3.4299

328_Calib_S_A

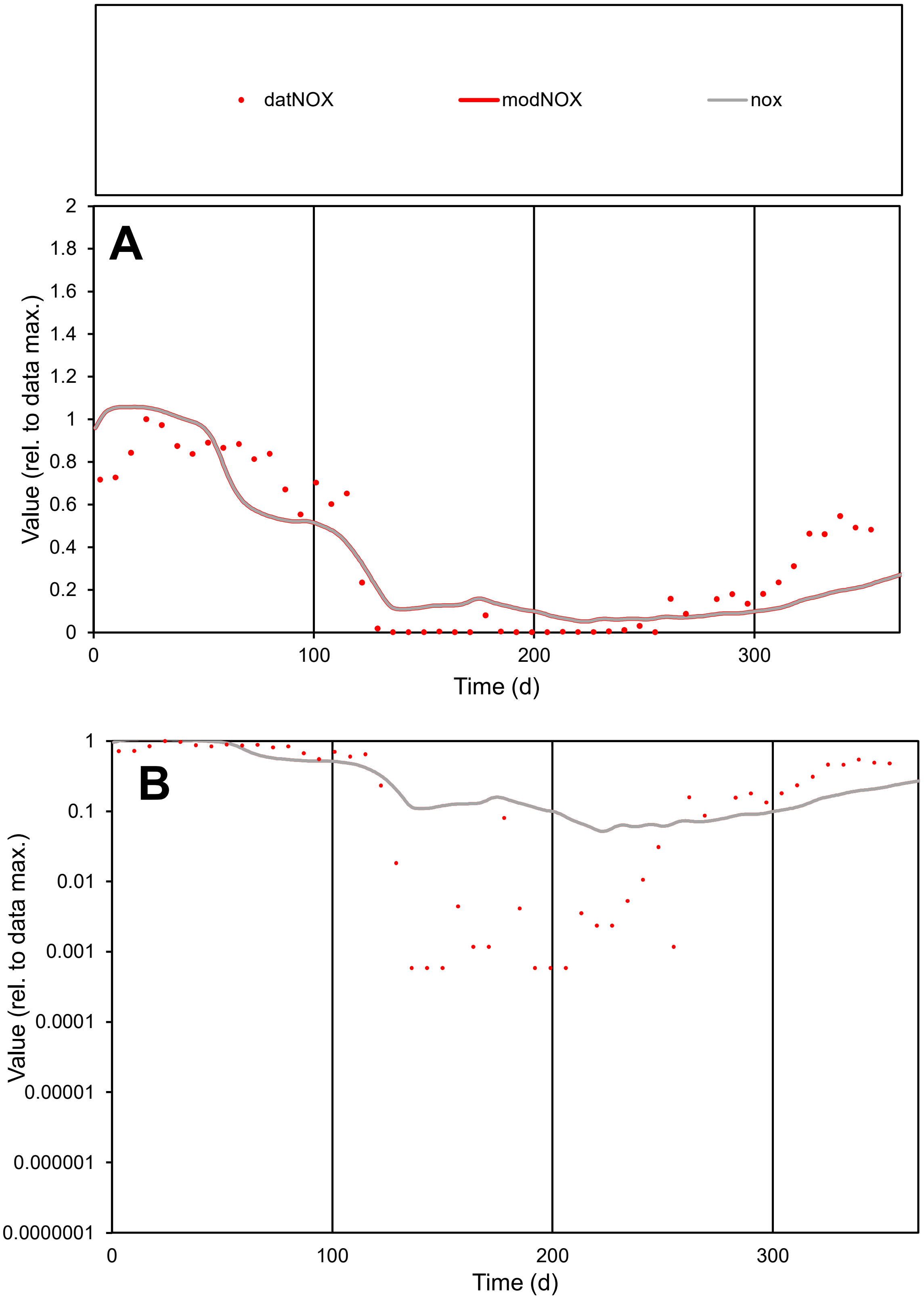


**C1**

Variables

NOX Provide jtd or Dst

CalibOnly?

2 1 to reproduce of

oNormalize?

2 1=no, 2=yes

ofxn calc*

1.38353

ofxn mod

1.4705

sRun

RunA1

zout

1

Ave

Data

0.00301

Model

0.00288

data/vMax

0.00848

ofxng

369.894

Min d/m

0.01119

ave/min d

601.322

ave/min m

6.44786

328_Calib_S_A

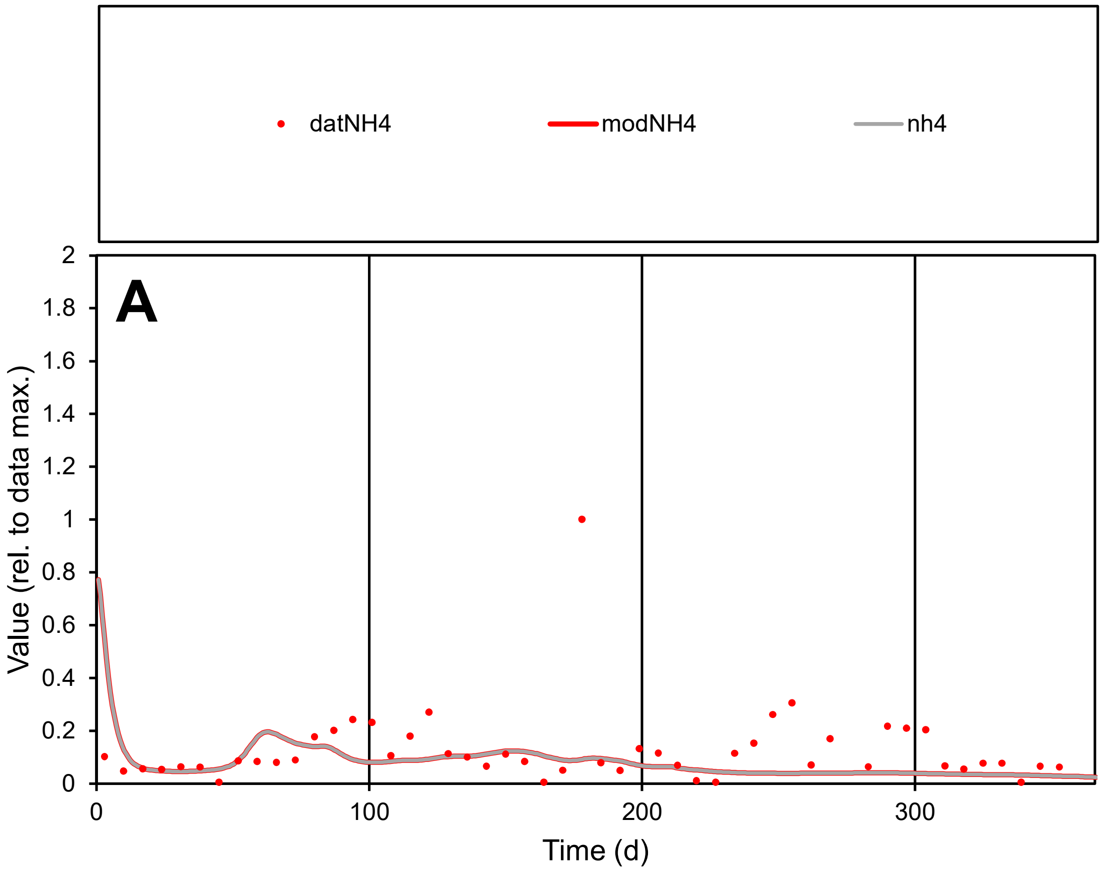


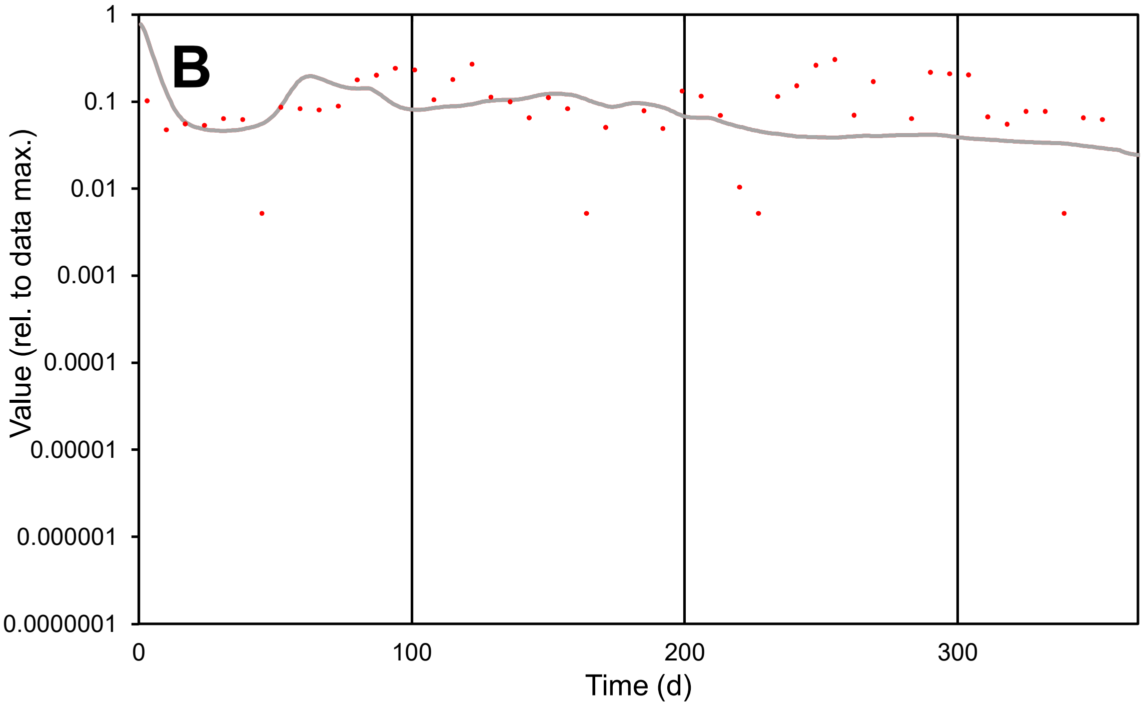


**C1**

Variables

NH4 Provide jtd or Dst

CalibOnly?

2 1 to reproduce of

oNormalize?

2 1=no, 2=yes

ofxn calc*

3.74893

ofxn mod

3.83291

sRun

RunA1

zout

1

Ave

Data

0.00021

Model

0.00013

data/vMax

0.00168

ofxng

369.894

Min d/m

0.17989

ave/min d

24.2421

ave/min m

2.67309

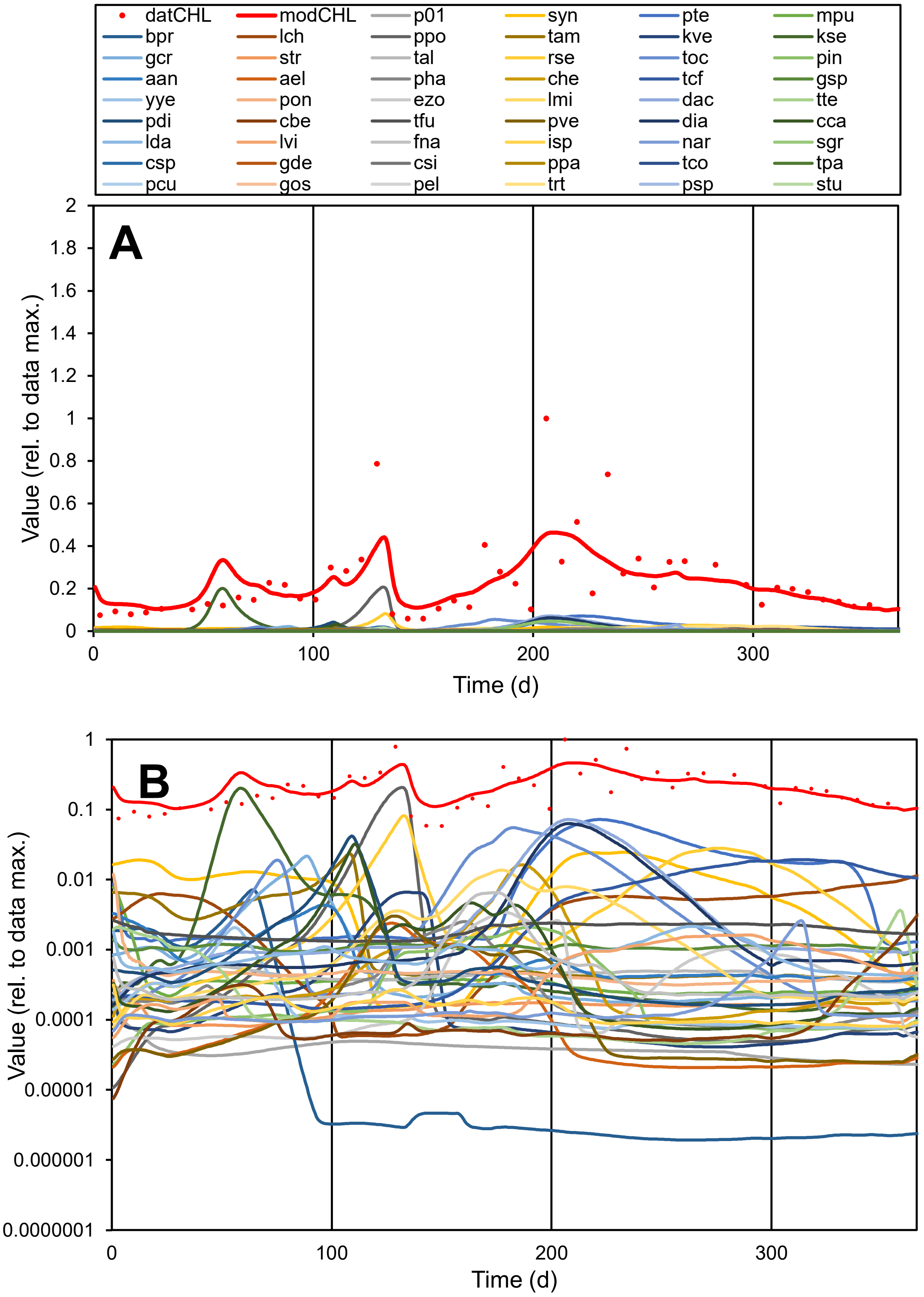


**C1**

Variables

CHL Provide jtd or Dst

CalibOnly?

2 1 to reproduce of

oNormalize?

2 1=no, 2=yes

ofxn calc*

5.94195

ofxn mod

6.11282

sRun

RunA1

zout

1

Ave

Data

1.02276

Model

0.9878

data/vMax

4.427

ofxng

369.894

Min d/m

0.55135

ave/min d

3.96879

ave/min m

2.1134

328_Calib_S_A

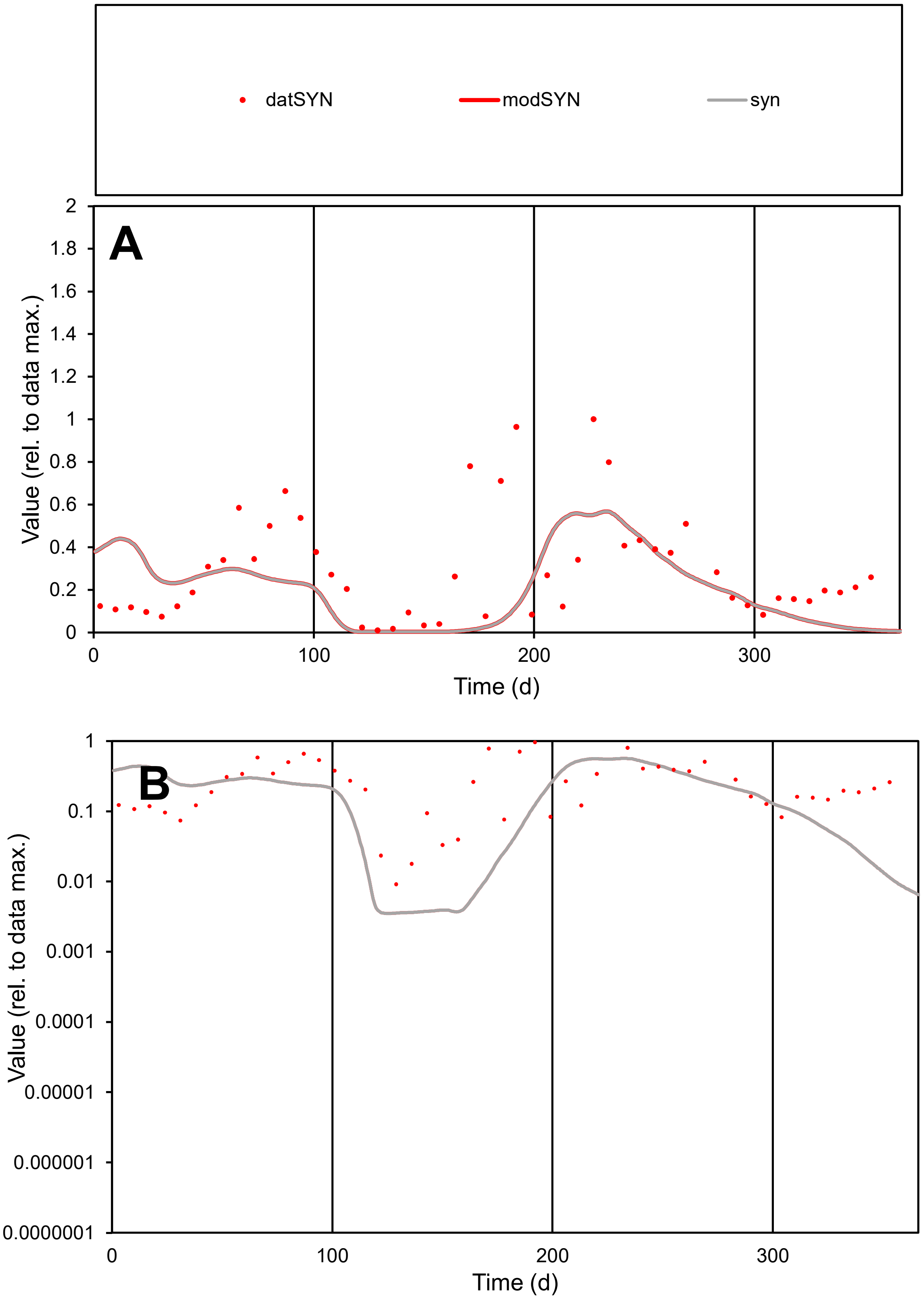


**C1**

Variables
SYN Provide jtd or Dst
CalibOnly?
2 1 to reproduce of
oNormalize?
2 1=no, 2=yes

ofxn calc*
0.89167
ofxn mod
0.90257

sRun
RunA1
zout
1

Ave
Data
6.69811
Model
4.77344

data/vMax
22.84

ofxng
369.894

Min d/m
2.60085

ave/min d
32.0637
ave/min m
59.4303

328_Calib_S_A

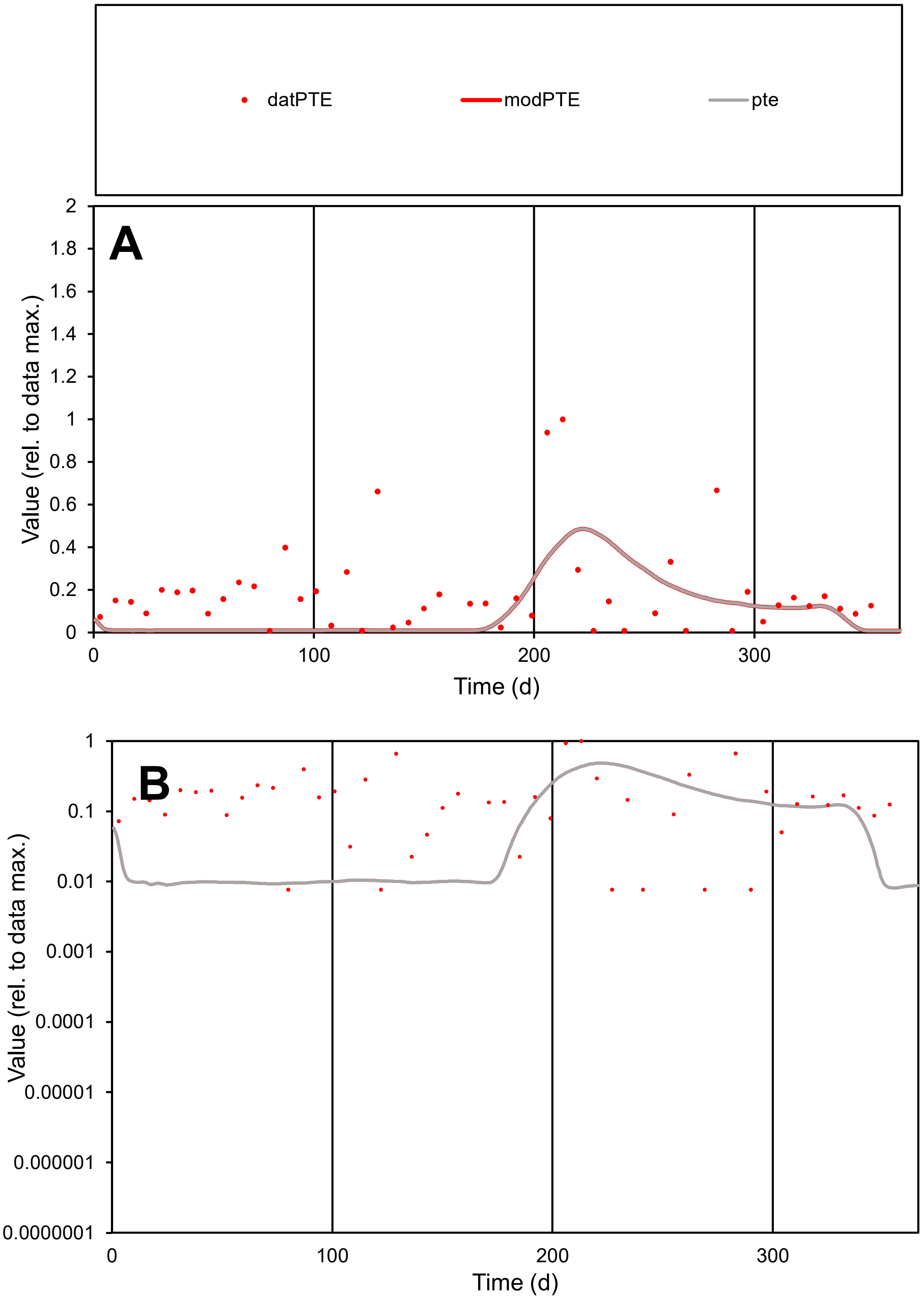


**C1**

Variables
PTE Provide jtd or Dst
CalibOnly?
2 1 to reproduce of
oNormalize?
2 1=no, 2=yes

ofxn calc*
1.22065
ofxn mod
1.22895

sRun
RunA1
zout
1

Ave
Data
0.0002
Model
0.00011

data/vMax
0.00109

ofxng
369.894

Min d/m
0.94795

ave/min d
24.4552
ave/min m
12.9709

328_Calib_S_A

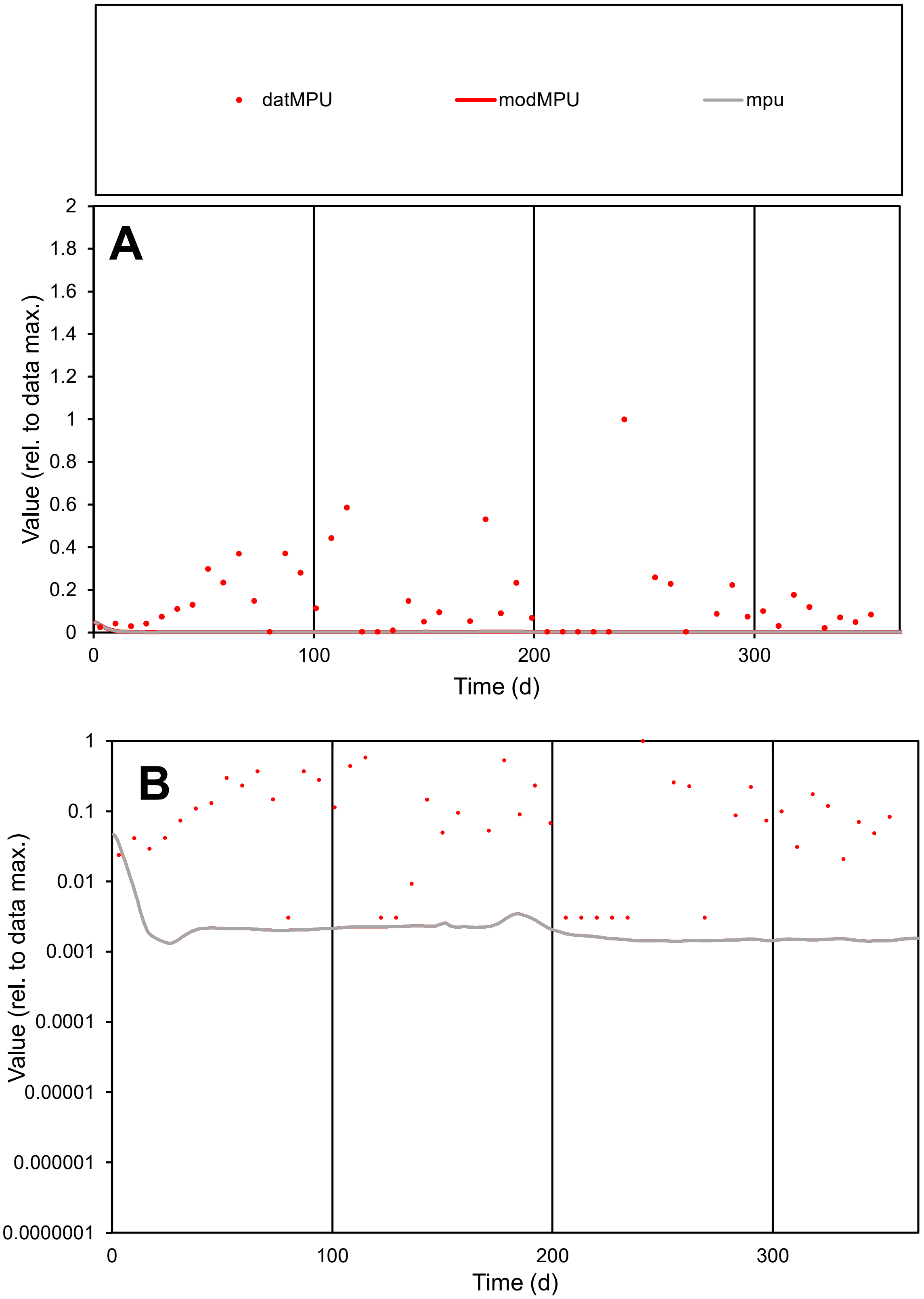


C1

Variables

MPU Provide jtd or Dst

CalibOnly?

2 1 to reproduce of

oNormalize?

2 1=no, 2=yes

ofxn calc*

1.60477

ofxn mod

1.60717

sRun

RunA1

zout

1

Ave

Data

0.0002

Model

3.4E-06

data/vMax

0.00133

ofxng

369.894

Min d/m

2.28396

ave/min d

48.311

ave/min m

1.89546

328_Calib_S_A

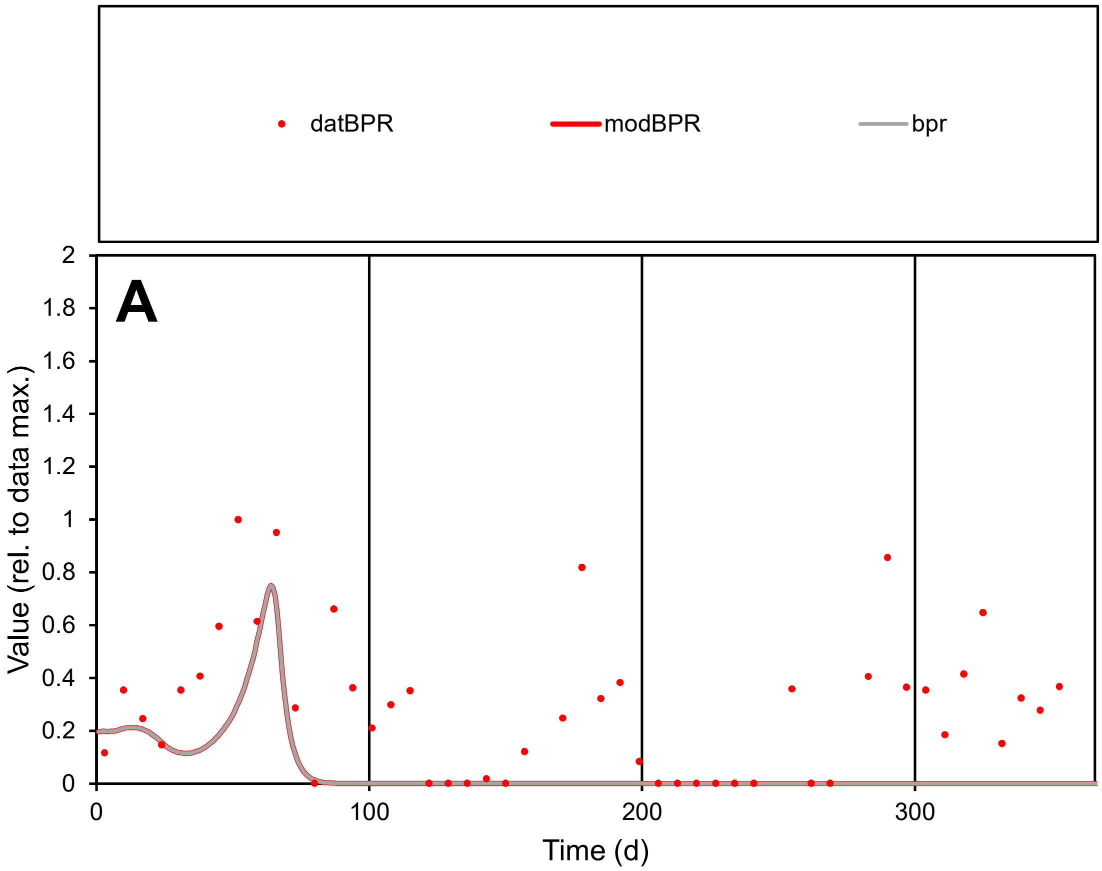


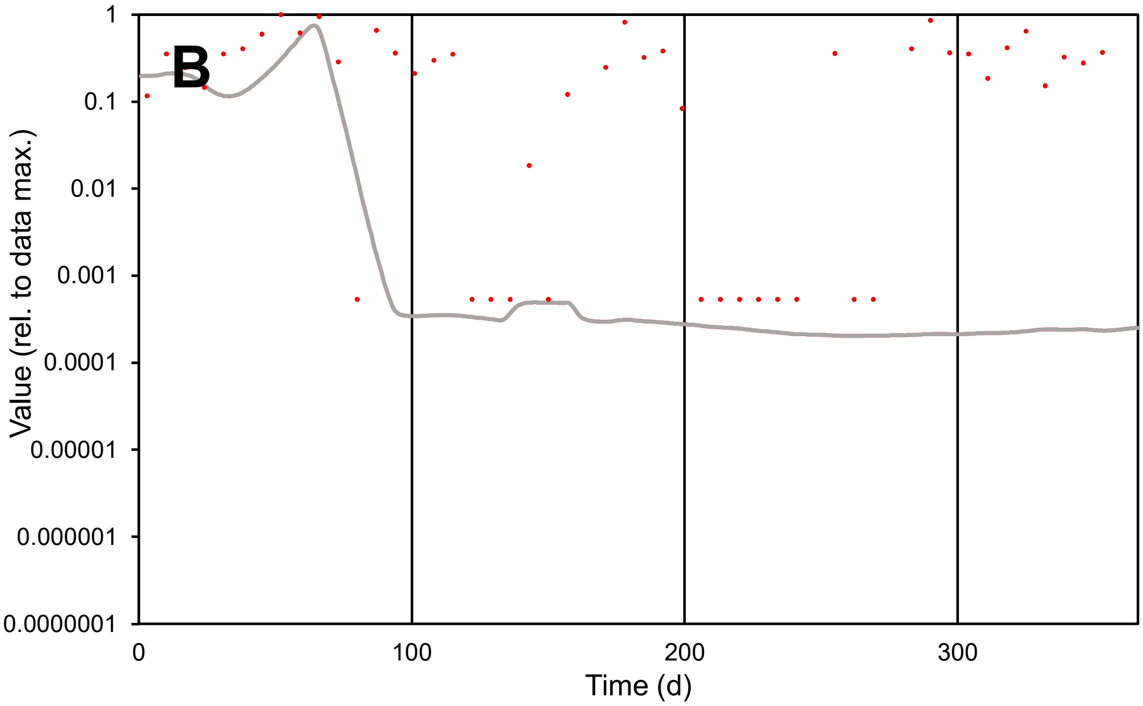


**C1**

Variables
BPR Provide jtd or Dst
CalibOnly?
2 1 to reproduce of
oNormalize?
2 1=no, 2=yes

ofxn calc*
1.12696
ofxn mod
1.13442

sRun
RunA1
zout
1

Ave
Data
9.8E-05
Model
2.1E-05

data/vMax
0.00035

ofxng
369.894

Min d/m
2.62465

ave/min d
531.292
ave/min m
302.64

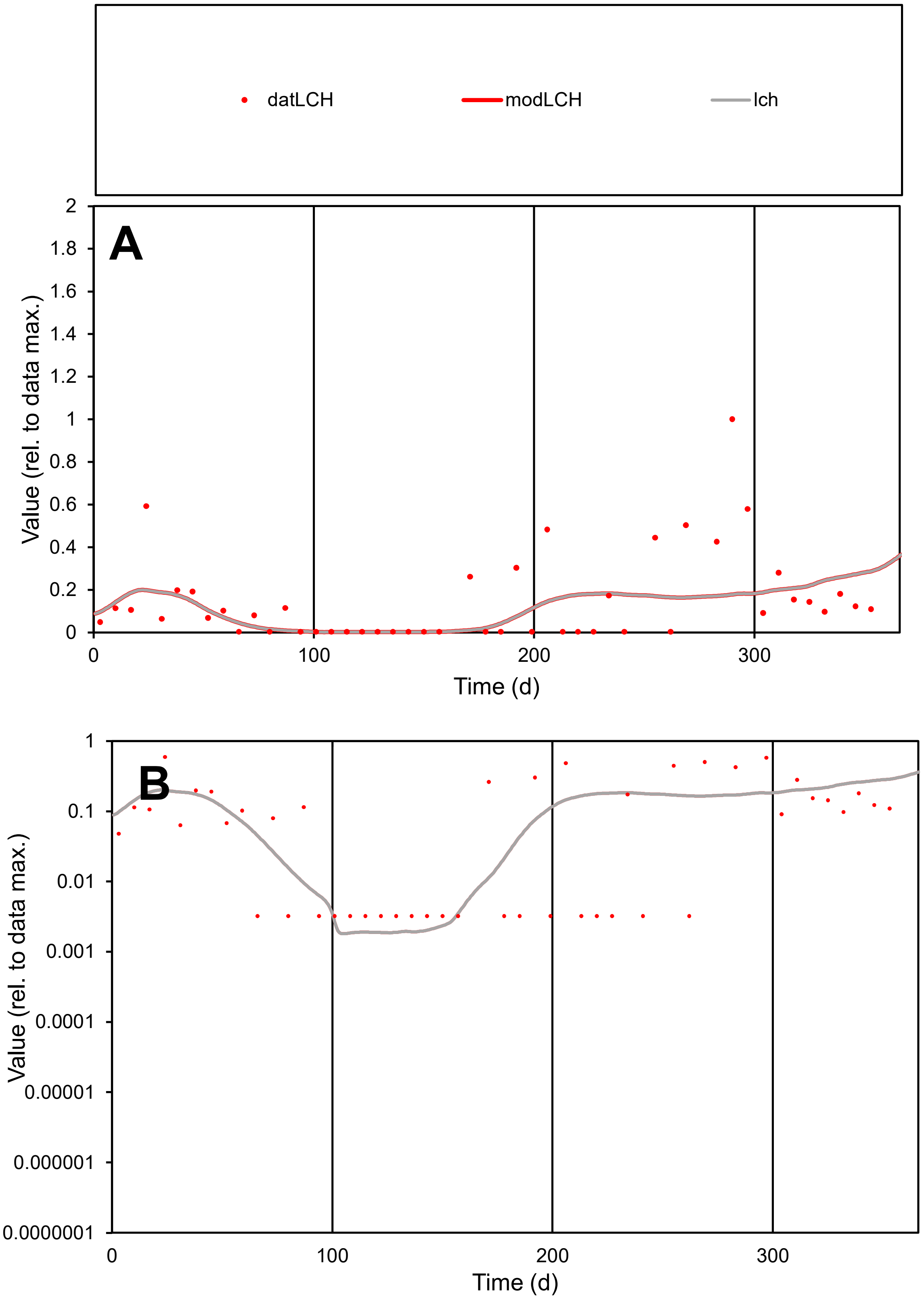


C1

Variables
LCH Provide jtd or Dst
CalibOnly?
2 1 to reproduce of
oNormalize?
2 1=no, 2=yes

ofxn calc*
1.30187
ofxn mod
1.31206

sRun
RunA1
zout
1

Ave
Data
0.00011
Model
8.9E-05

data/vMax
0.00076

ofxng
369.894

Min d/m
1.72496

ave/min d
45.7322
ave/min m
62.6031

328_Calib_S_A

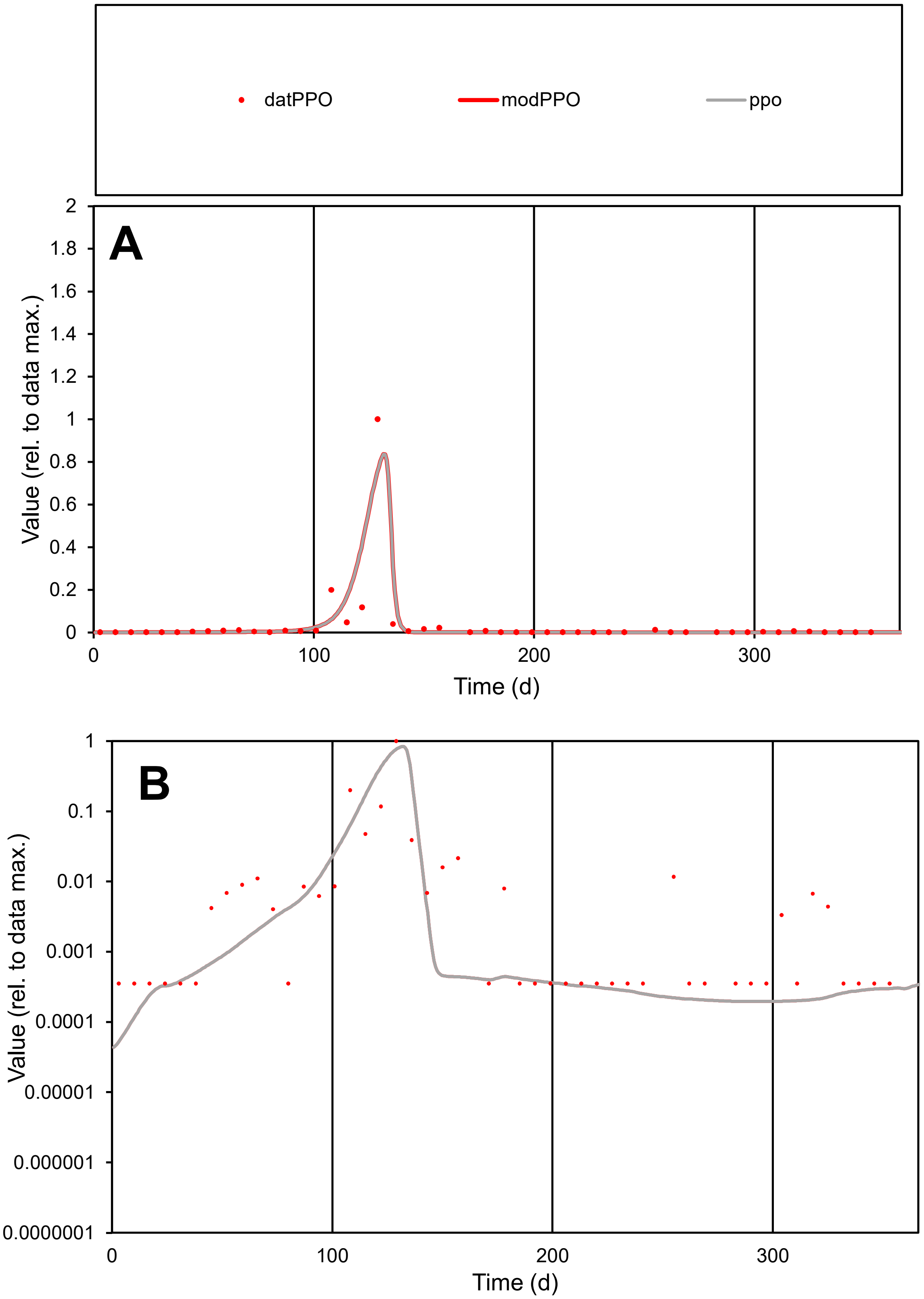


**C1**

Variables

| | |
|---|---|
| PPO | Provide jtd or Dst |
| CalibOnly? | |
| 2 | 1 to reproduce of |
| oNormalize? | |
| 2 | 1=no, 2=yes |

ofxn calc*
1.6882
ofxn mod
1.883

sRun
RunA1
zout
1

Ave
Data
0.00018
Model
0.00018

data/vMax
0.00542

ofxng
369.894

Min d/m
5.86091

ave/min d
91.5531
ave/min m
549.703

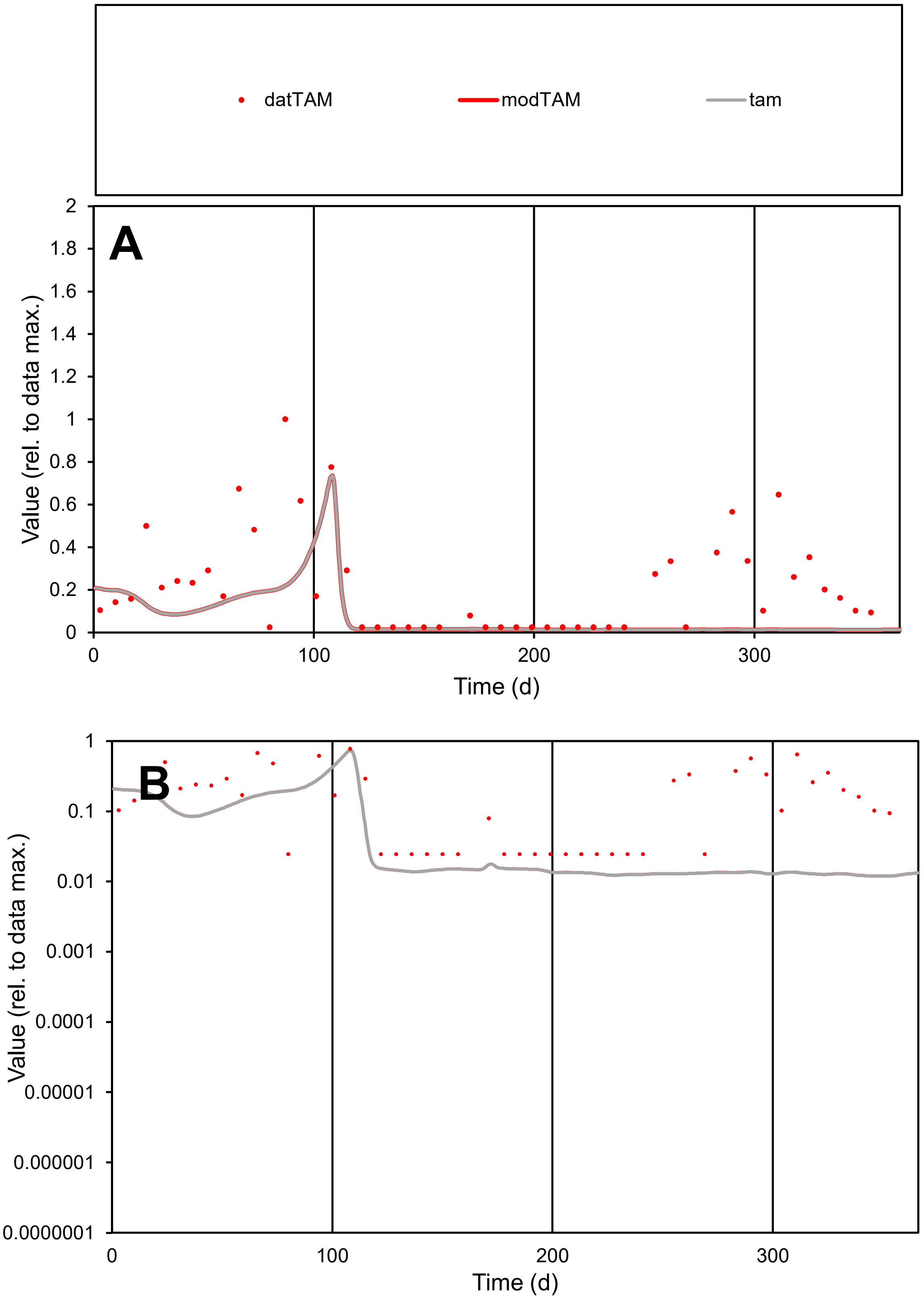


**C1**

Variables
TAM Provide jtd or Dst
CalibOnly?
2 1 to reproduce of
oNormalize?
2 1=no, 2=yes

ofxn calc*
1.10908
ofxn mod
1.12173

sRun
RunA1
zout
1

Ave
Data
5.1E-05
Model
2E-05

data/vMax
0.00023

ofxng
369.894

Min d/m
2.04292

ave/min d
8.81852
ave/min m
7.25314

328_Calib_S_A

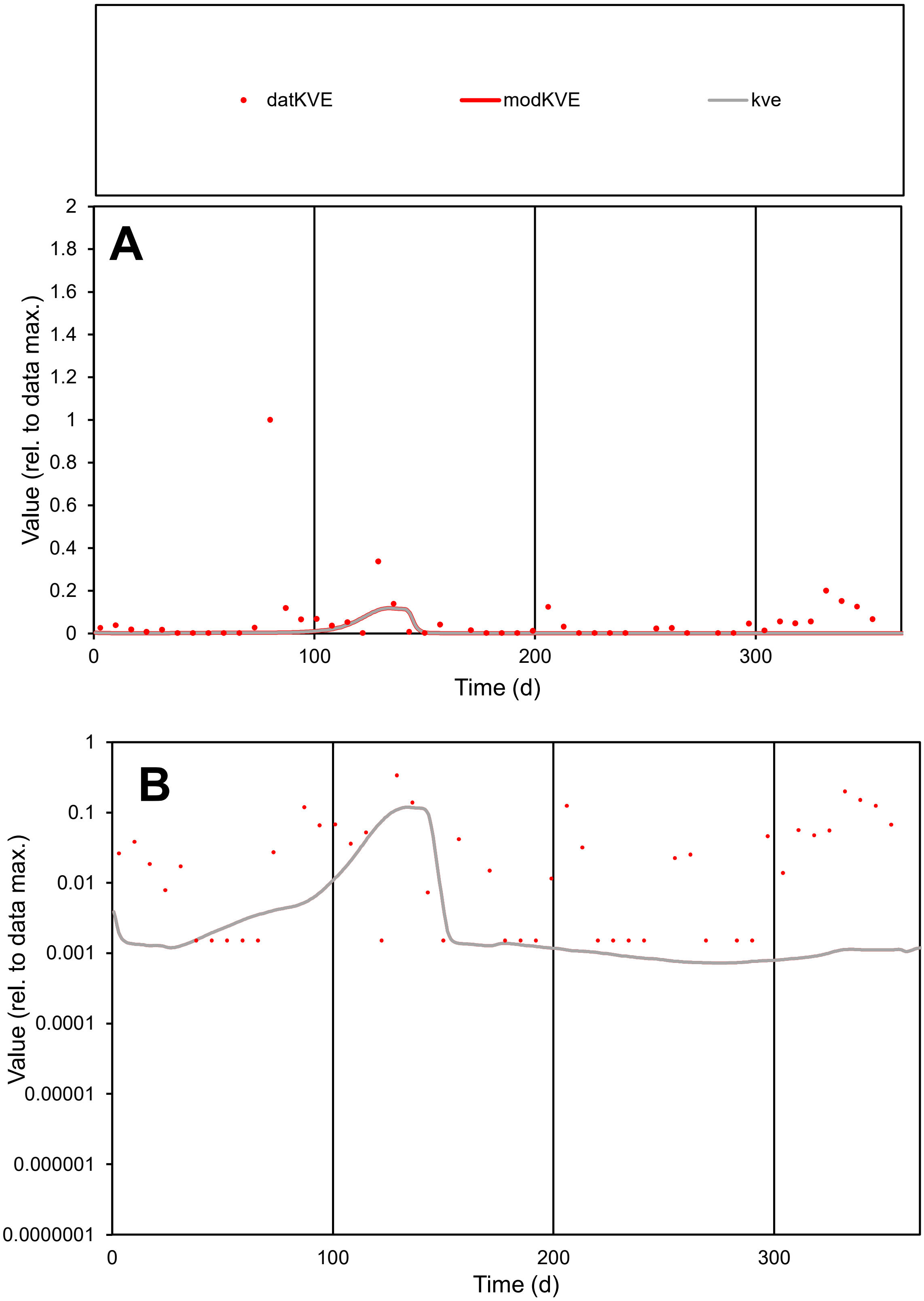


**C1**

Variables

KVE Provide jtd or Dst

CalibOnly?

2 1 to reproduce of

oNormalize?

2 1=no, 2=yes

ofxn calc*

2.49062

ofxn mod

2.49616

sRun

RunA1

zout

1

Ave

Data

0.00013

Model

2E-05

data/vMax

0.00206

ofxng

369.894

Min d/m

2.07902

ave/min d

41.4851

ave/min m

13.5695

328_Calib_S_A

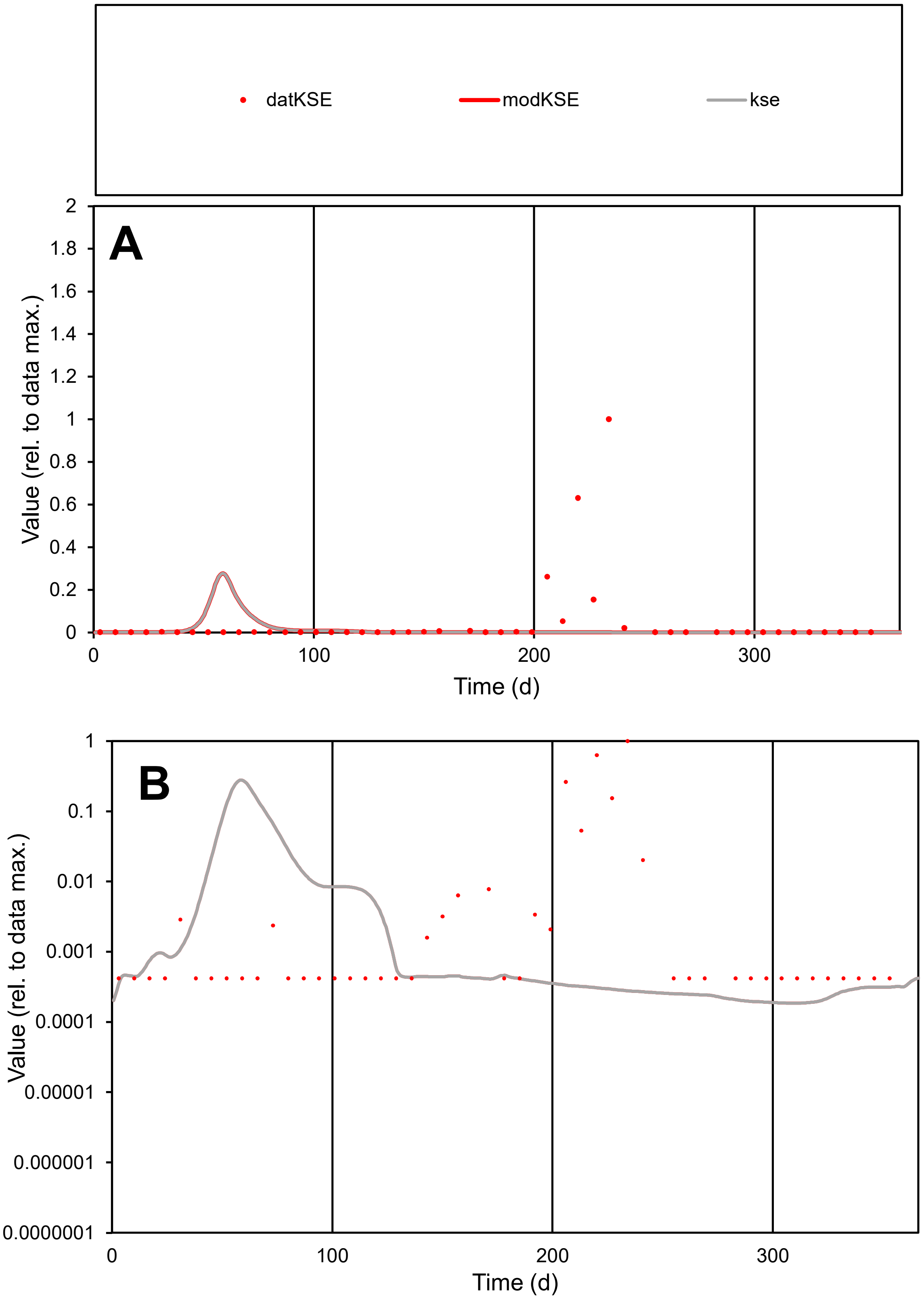


**C1**

Variables
KSE Provide jtd or Dst
CalibOnly?
2 1 to reproduce of
oNormalize?
2 1=no, 2=yes

ofxn calc*
3.99874
ofxn mod
4.0269

sRun
RunA1
zout
1

Ave
Data
0.0004
Model
1E-04

data/vMax
0.00893

ofxng
369.894

Min d/m
2.25783

ave/min d
107.325
ave/min m
60.0561

328_Calib_S_A

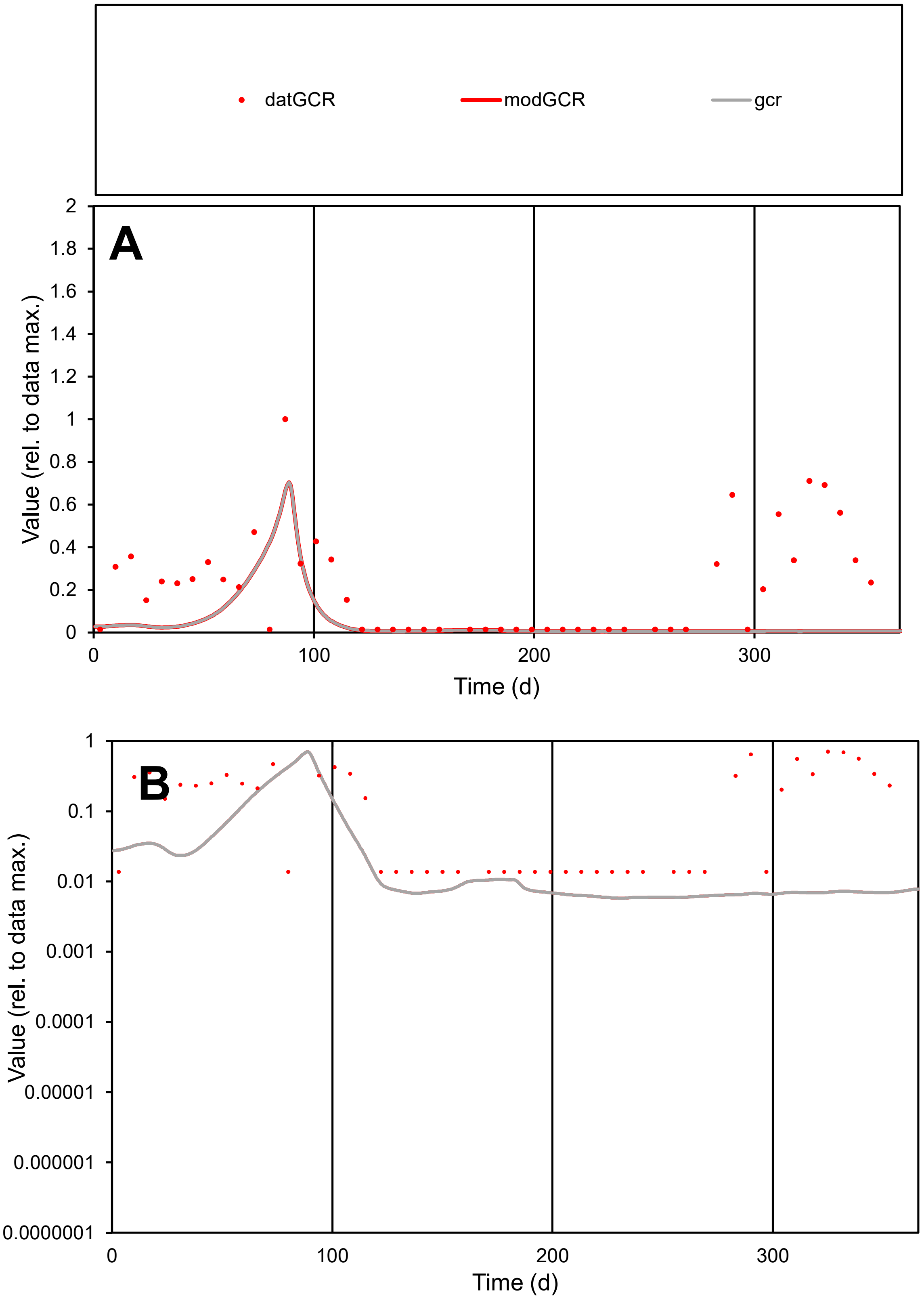


**C1**

Variables
GCR Provide jtd or Dst
CalibOnly?
2 1 to reproduce of
oNormalize?
2 1=no, 2=yes

ofxn calc*
1.25542
ofxn mod
1.26149

sRun
RunA1
zout
1

Ave
Data
4.7E-05
Model
1.3E-05

data/vMax
0.00023

ofxng
369.894

Min d/m
2.3635

ave/min d
15.0859
ave/min m
9.94452

328_Calib_S_A

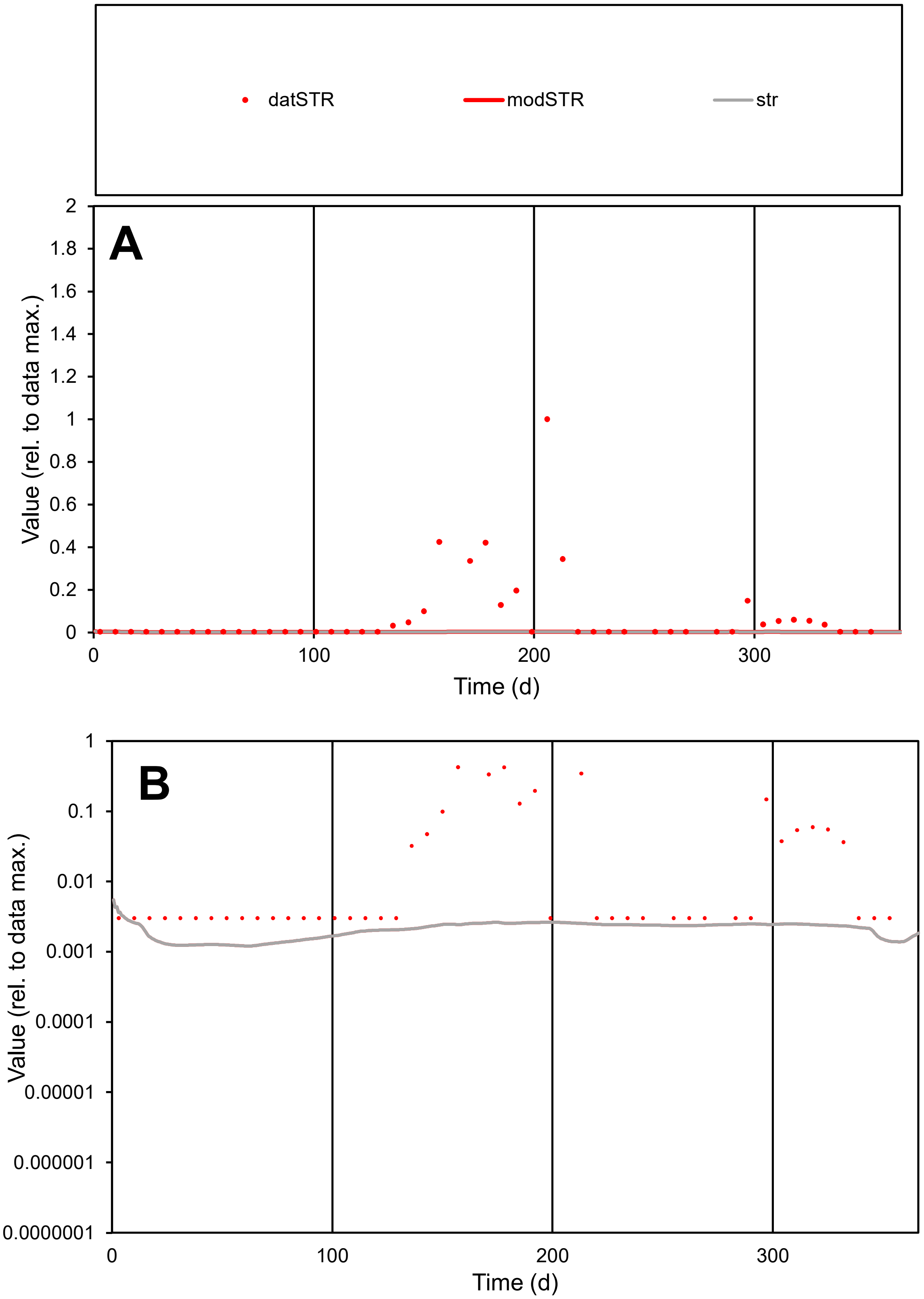


**C1**

Variables
STR Provide jtd or Dst
CalibOnly?
2 1 to reproduce of
oNormalize?
2 1=no, 2=yes

ofxn calc*
2.54904
ofxn mod
2.55301

sRun
RunA1
zout
1

Ave
Data
3.9E-05
Model
1.1E-06

data/vMax
0.00054

ofxng
369.894

Min d/m
2.49502

ave/min d
24.2247
ave/min m
1.73449

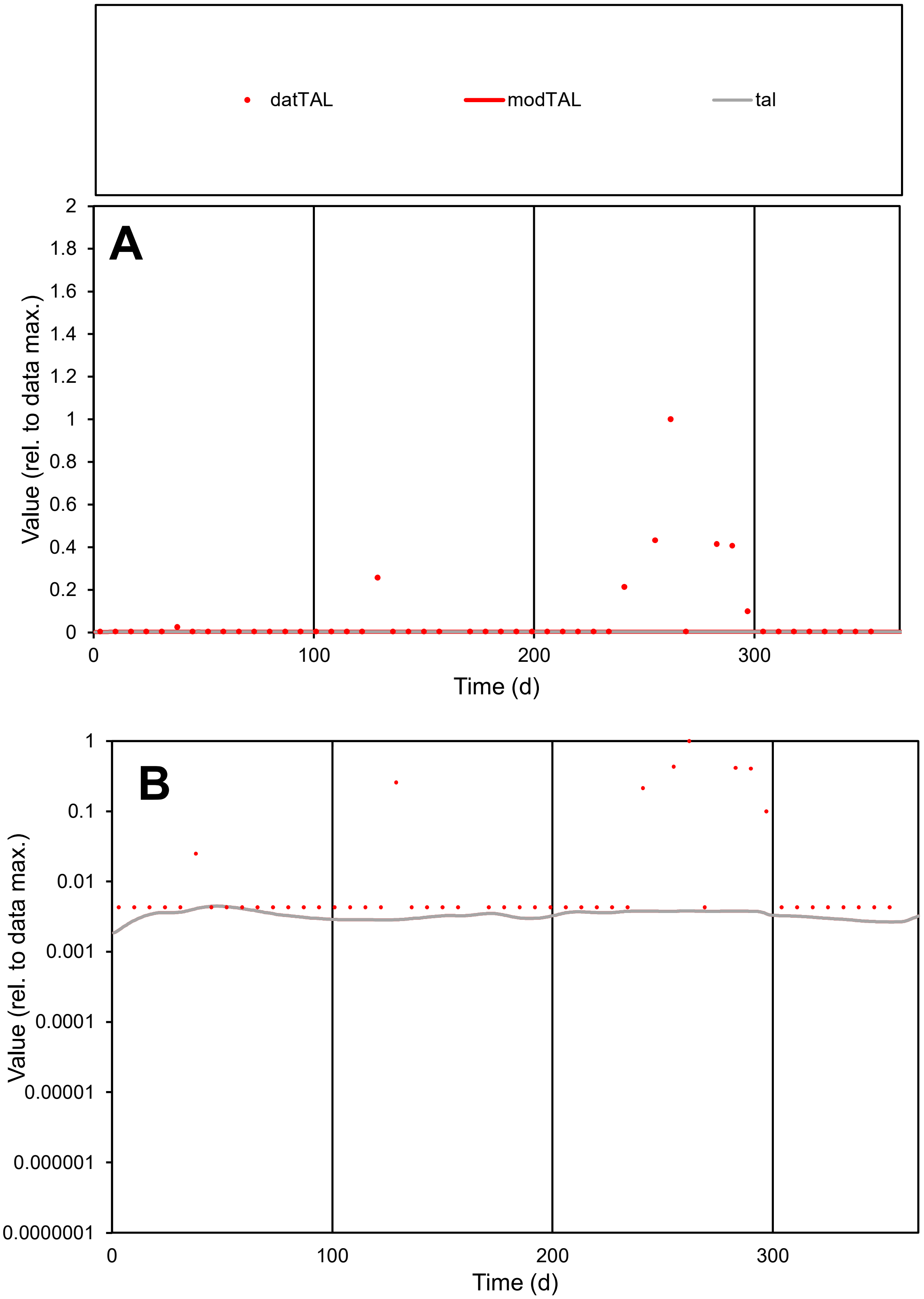


**C1**

Variables
TAL Provide jtd or Dst
CalibOnly?
2 1 to reproduce of
oNormalize?
2 1=no, 2=yes

ofxn calc*
2.92455
ofxn mod
2.92948

sRun
RunA1
zout
1

Ave
Data
6.1E-05
Model
3.2E-06

data/vMax
0.00097

ofxng
369.894

Min d/m
1.97804

ave/min d
14.6651
ave/min m
1.5387

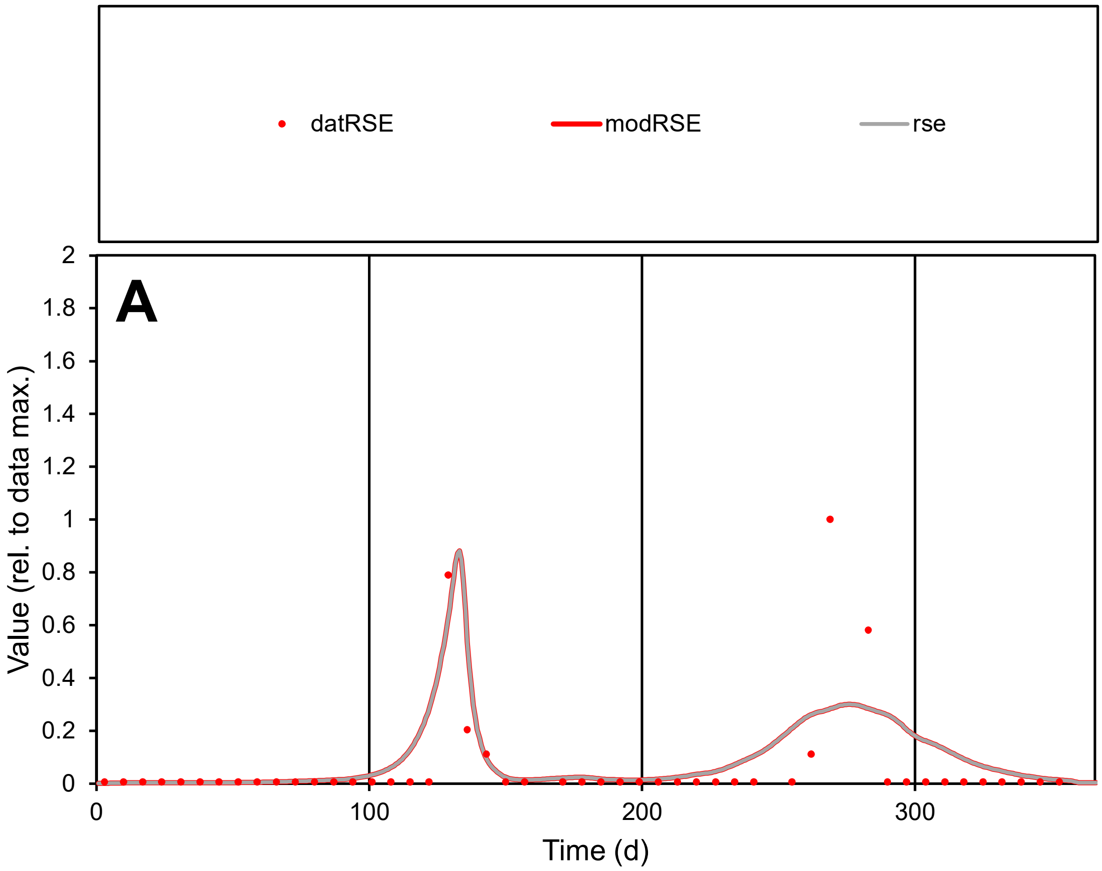


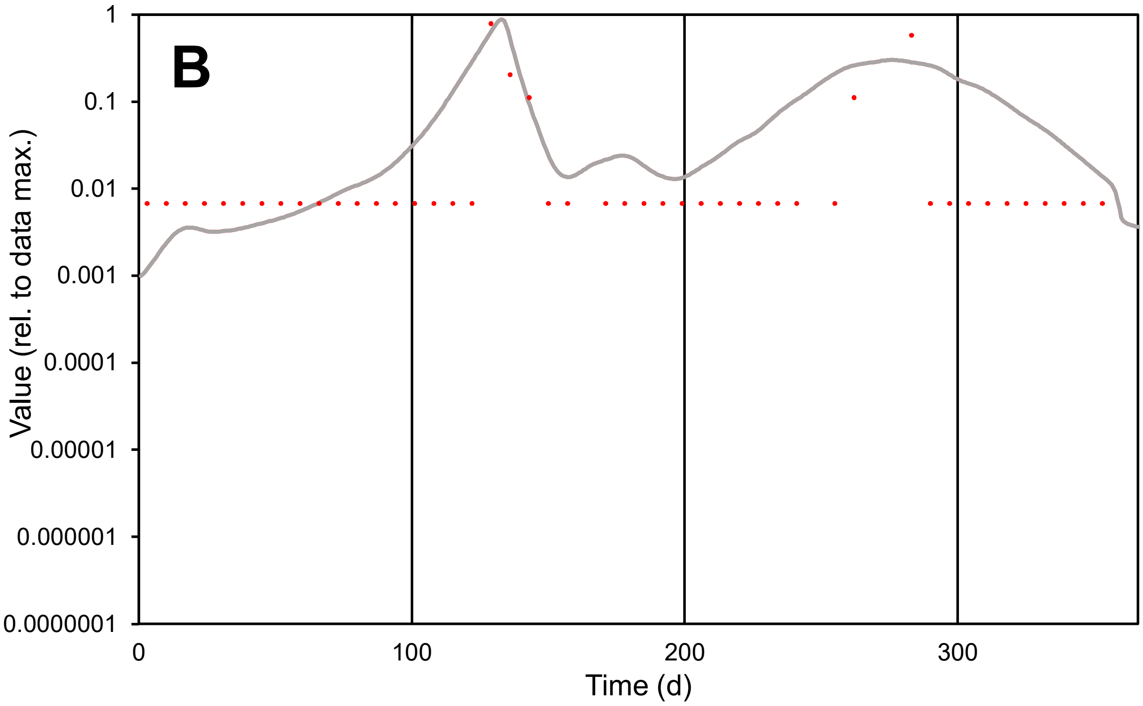


**C1**

Variables
RSE Provide jtd or Dst
CalibOnly?
2 1 to reproduce of
oNormalize?
2 1=no, 2=yes

ofxn calc*
2.13715
ofxn mod
2.18751

sRun
RunA1
zout
1

Ave
Data
4.4E-05
Model
5.9E-05

data/vMax
0.00069

ofxng
369.894

Min d/m
5.02858

ave/min d
9.49722
ave/min m
63.5259

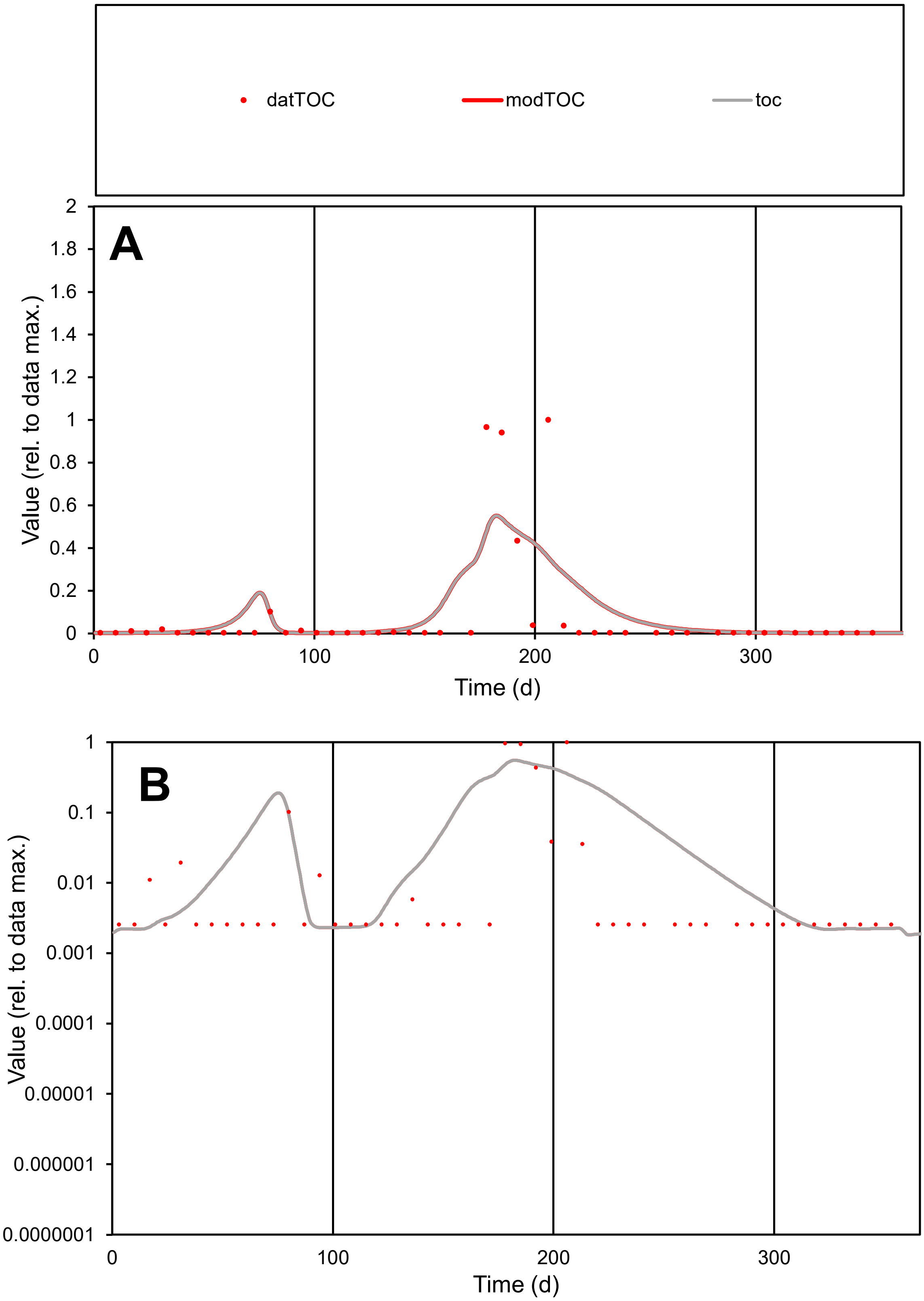


**C1**

Variables
TOC Provide jtd or Dst
CalibOnly?
2 1 to reproduce of
oNormalize?
2 1=no, 2=yes

ofxn calc*
2.0642
ofxn mod
2.11862

sRun
RunA1
zout
1

Ave
Data
0.00013
Model
0.00014

data/vMax
0.00169

ofxng
369.894

Min d/m
1.17159

ave/min d
29.7564
ave/min m
38.6301

328_Calib_S_A

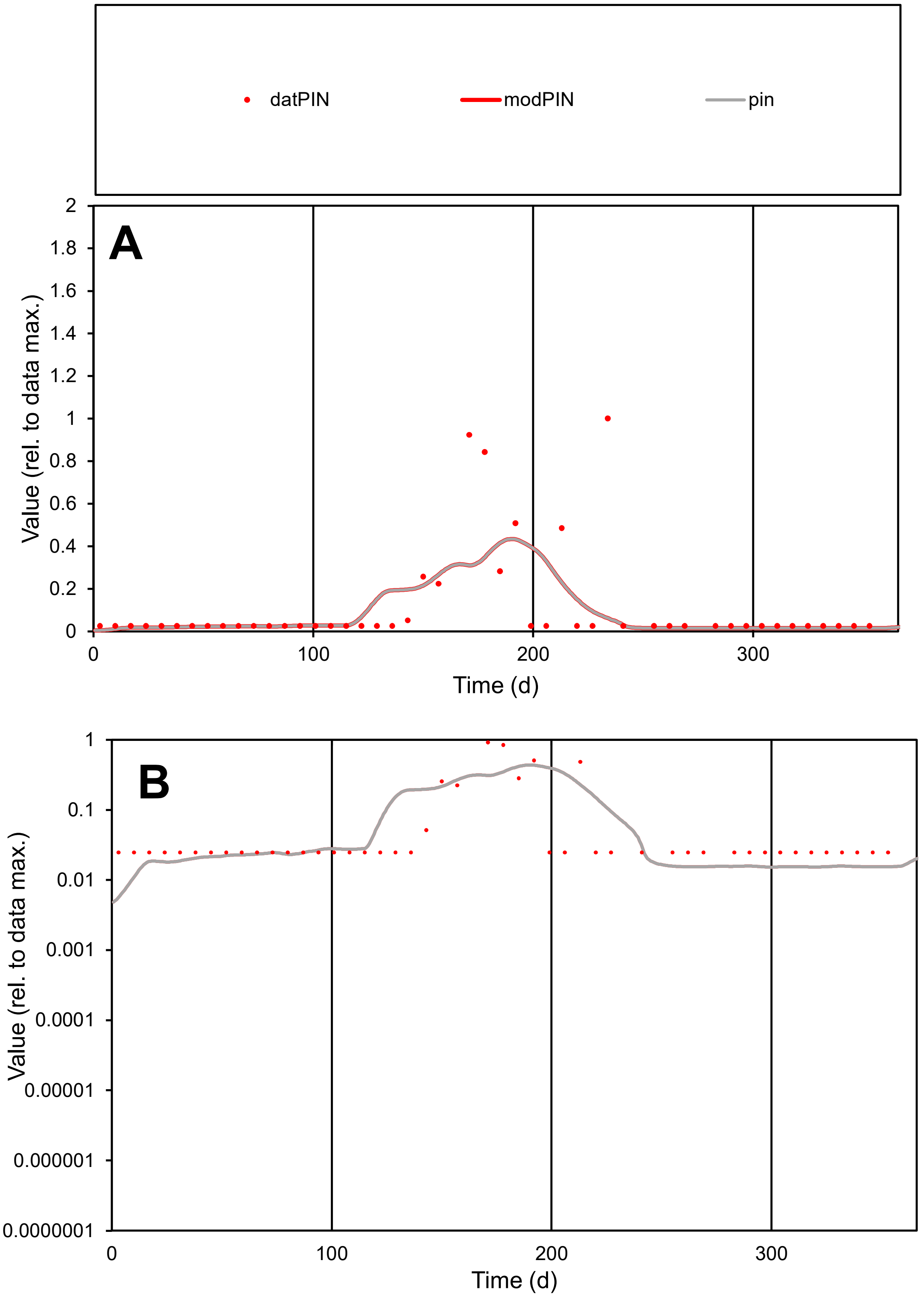


**C1**

Variables
PIN Provide jtd or Dst
CalibOnly?
2 1 to reproduce of
oNormalize?
2 1=no, 2=yes

ofxn calc*
1.70451
ofxn mod
1.72704

sRun
RunA1
zout
1

Ave
Data
1.9E-05
Model
1.5E-05

data/vMax
0.00016

ofxng
369.894

Min d/m
3.92752

ave/min d
4.65188
ave/min m
14.5758

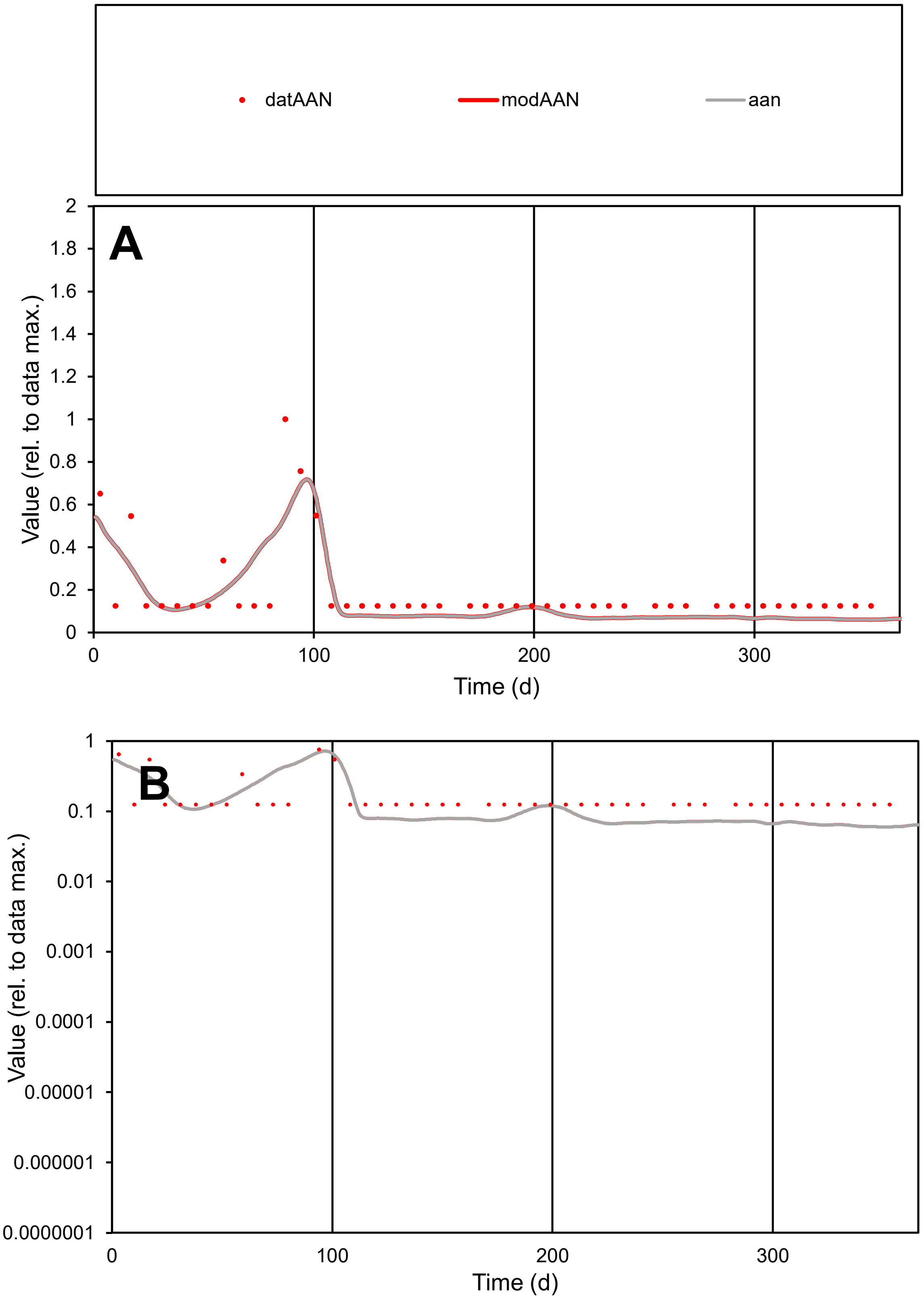


**C1**

Variables
AAN Provide jtd or Dst
CalibOnly?
2 1 to reproduce of
oNormalize?
2 1=no, 2=yes

ofxn calc*
0.60454
ofxn mod
0.62323

sRun
RunA1
zout
1

Ave
Data
8.5E-06
Model
7.1E-06

data/vMax
4.5E-05

ofxng
369.894

Min d/m
2.08057

ave/min d
1.51613
ave/min m
2.62975

328_Calib_S_A

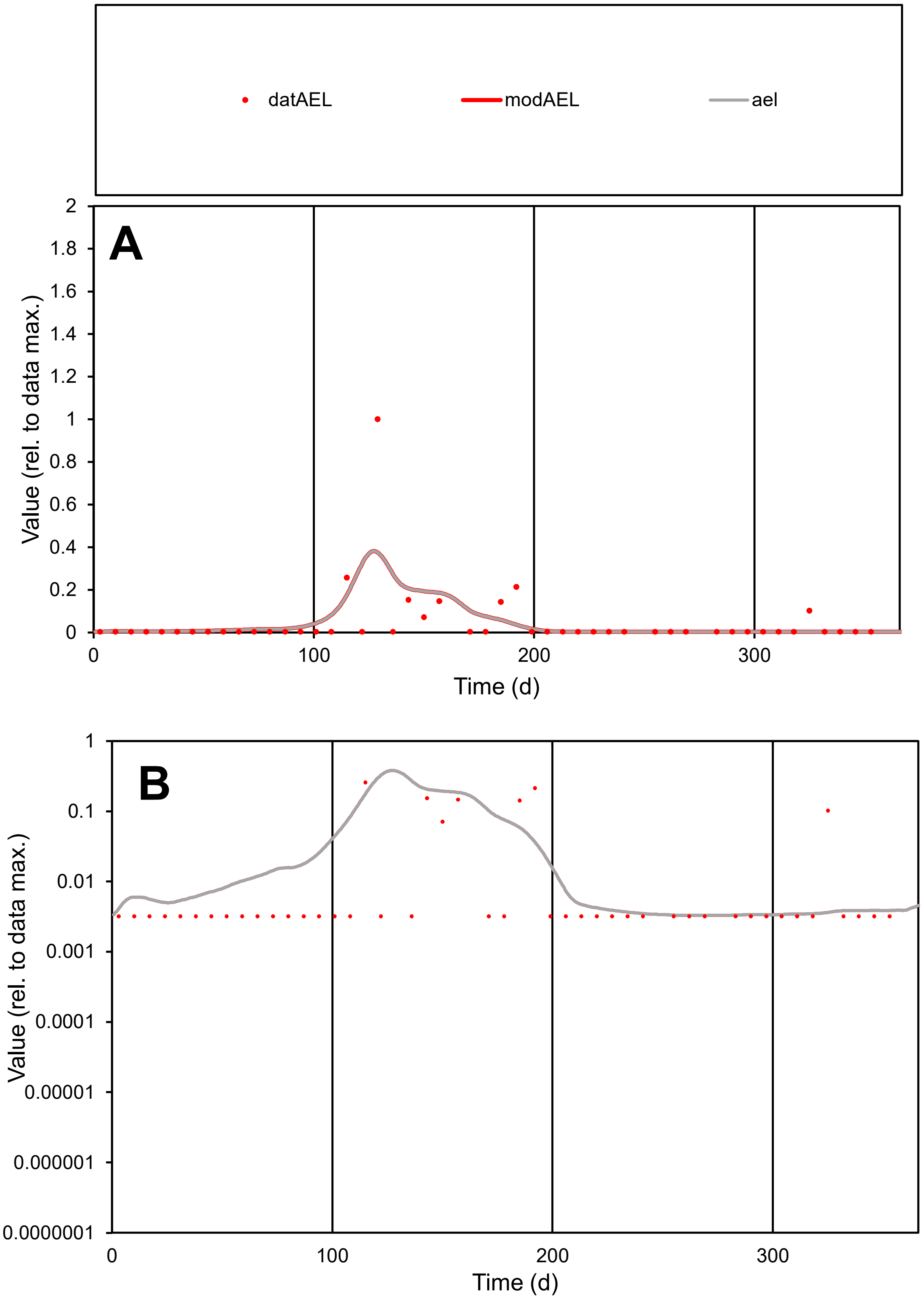


**C1**

Variables

AEL Provide jtd or Dst

CalibOnly?

2 1 to reproduce of

oNormalize?

2 1=no, 2=yes

ofxn calc*

2.48175

ofxn mod

2.53843

sRun

RunA1

zout

1

Ave

Data

1E-05

Model

1E-05

data/vMax

0.00022

ofxng

369.894

Min d/m

0.97348

ave/min d

14.3884

ave/min m

14.3456

328_Calib_S_A

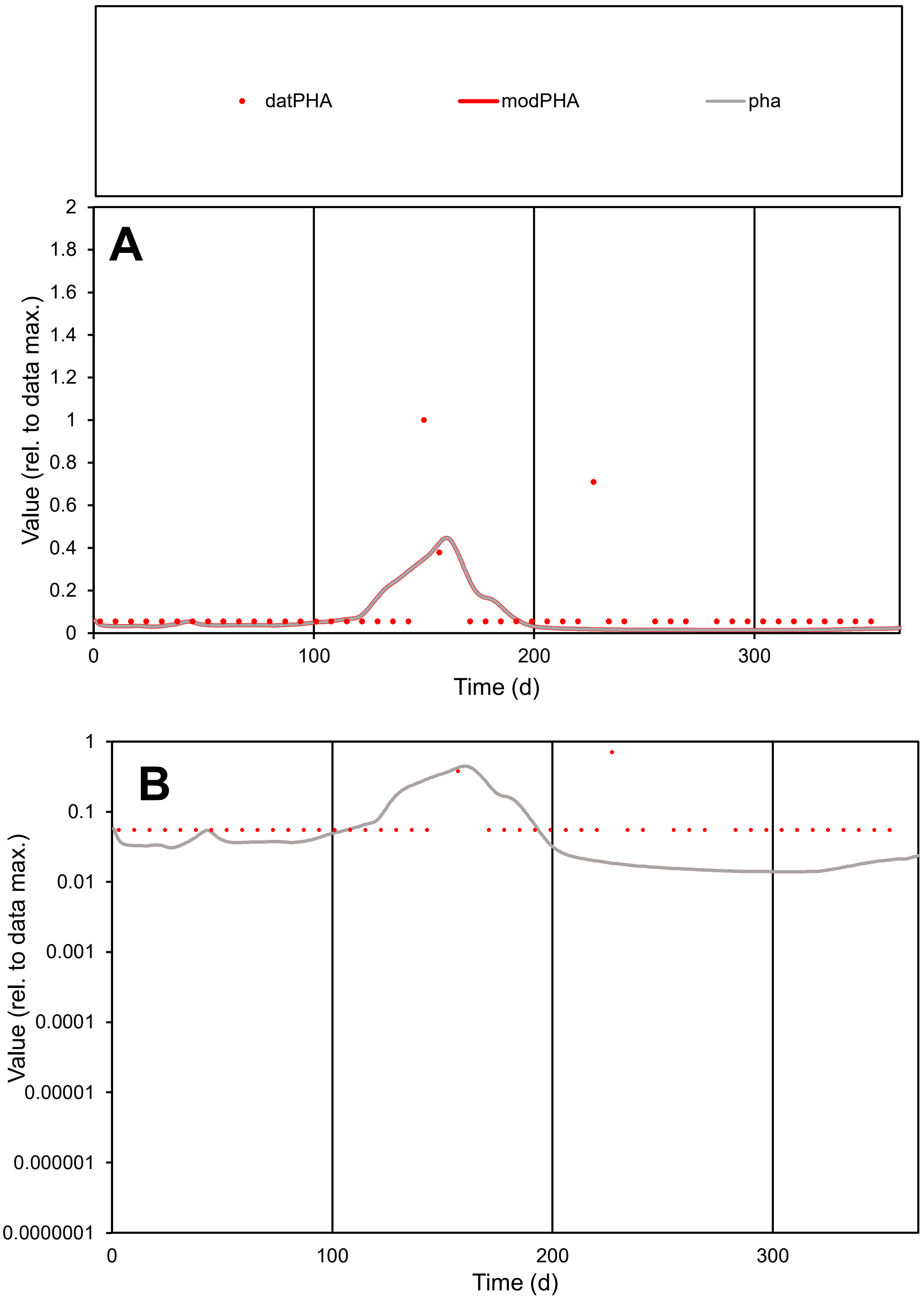


**C1**

Variables
PHA Provide jtd or Dst
CalibOnly?
2 1 to reproduce of
oNormalize?
2 1=no, 2=yes

ofxn calc*
1.56596
ofxn mod
1.58025

sRun
RunA1
zout
1

Ave
Data
1.8E-05
Model
1.3E-05

data/vMax
0.00019

ofxng
369.894

Min d/m
3.95817

ave/min d
1.72709
ave/min m
4.77327

328_Calib_S_A

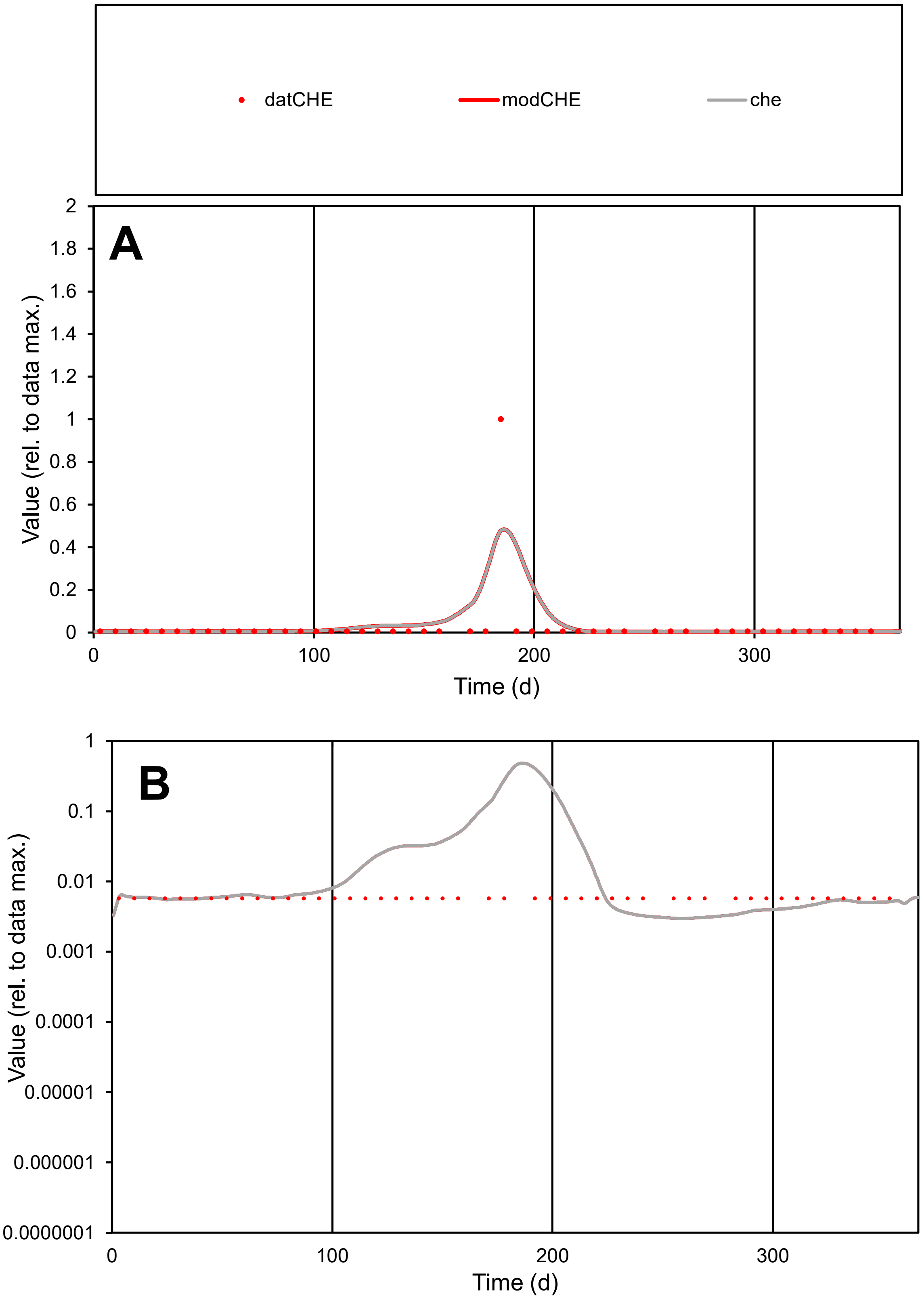


**C1**

Variables
CHE Provide jtd or Dst
CalibOnly?
2 1 to reproduce of
oNormalize?
2 1=no, 2=yes

ofxn calc*
3.66936
ofxn mod
3.8707

sRun
RunA1
zout
1

Ave
Data
3.2E-05
Model
4.6E-05

data/vMax
0.00122

ofxng
369.894

Min d/m
1.94137

ave/min d
4.59087
ave/min m
12.6069

328_Calib_S_A

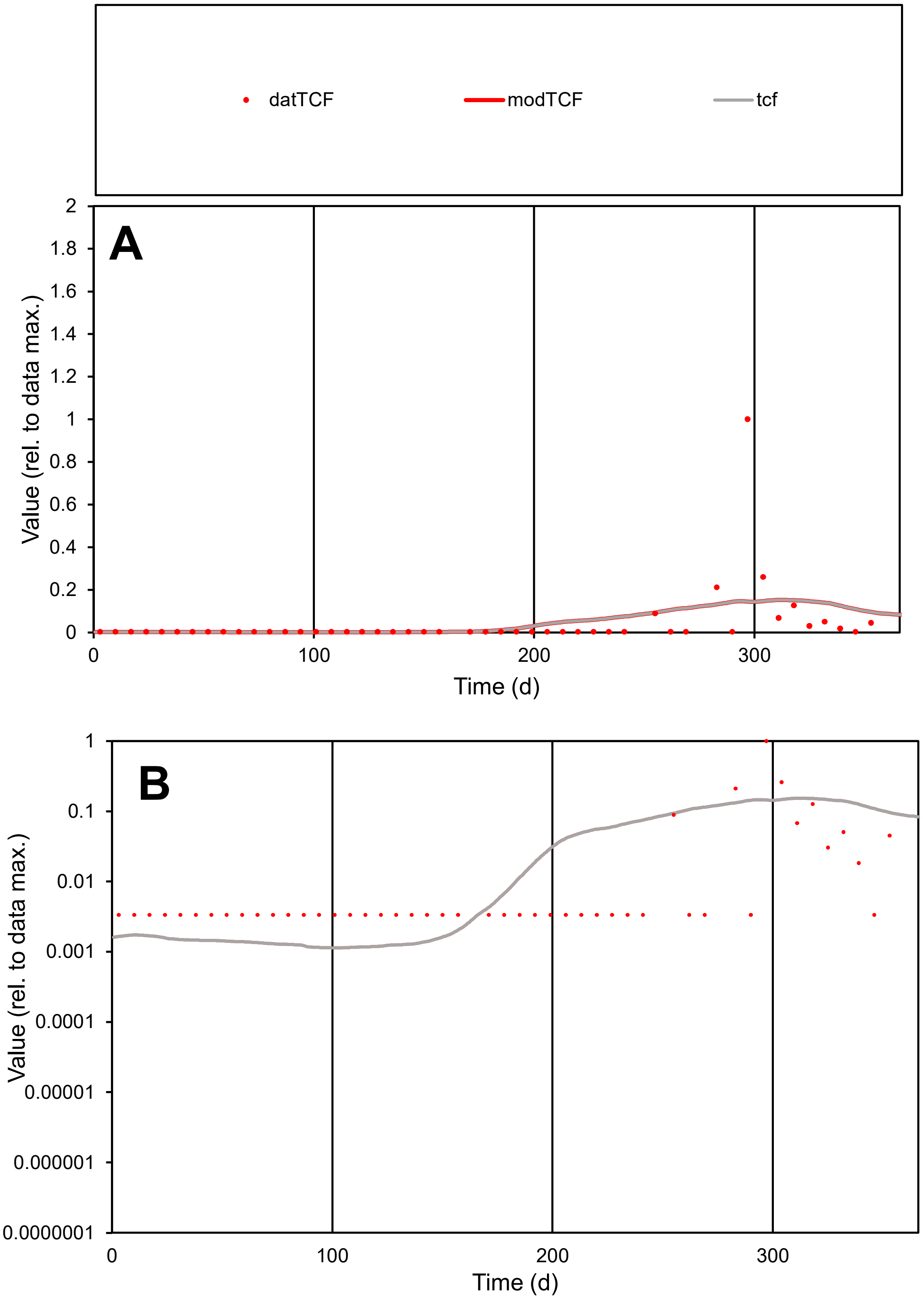


**C1**

Variables
TCF Provide jtd or Dst
CalibOnly?
2 1 to reproduce of
oNormalize?
2 1=no, 2=yes

ofxn calc*
3.18151
ofxn mod
3.18971

sRun
RunA1
zout
1

Ave
Data
3.9E-05
Model
4.4E-05

data/vMax
0.00093

ofxng
369.894

Min d/m
2.93691

ave/min d
12.6247
ave/min m
41.1734

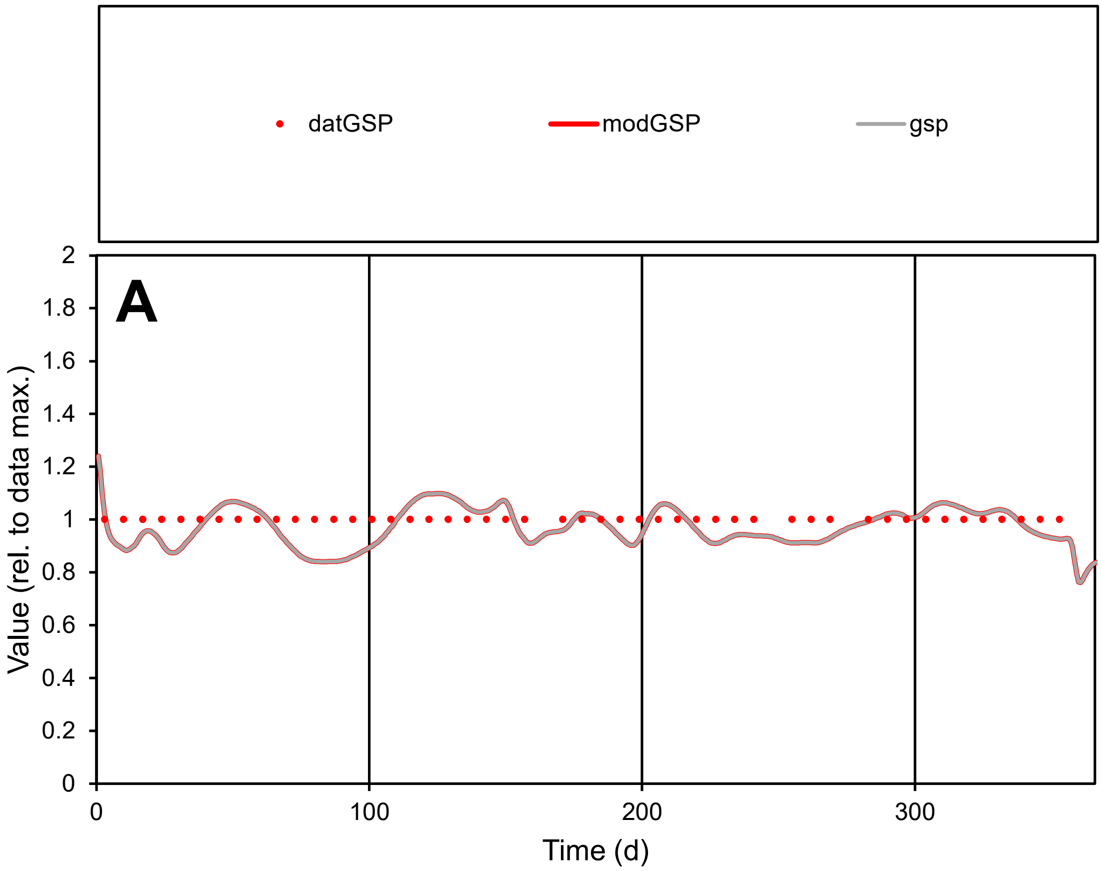


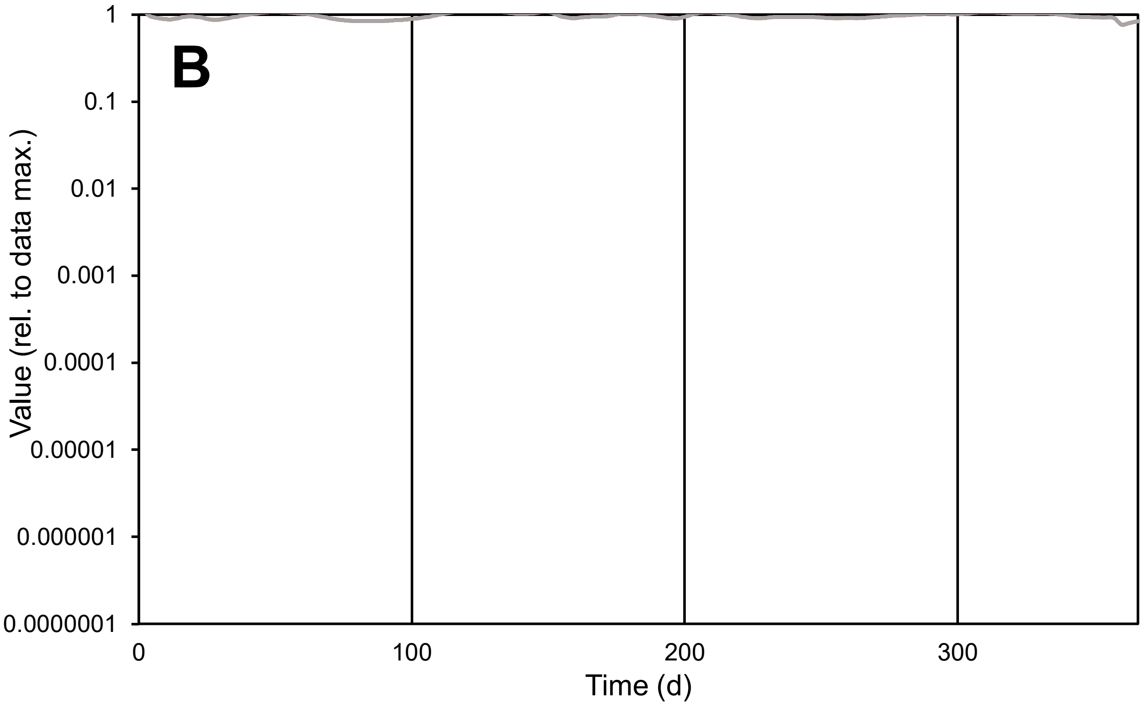


**C1**

Variables
GSP Provide jtd or Dst
CalibOnly?
2 1 to reproduce of
oNormalize?
2 1=no, 2=yes

ofxn calc*
0.07129
ofxn mod
0.13221

sRun
RunA1
zout
1

Ave
Data
9E-06
Model
8.8E-06

data/vMax
9E-06

ofxng
369.894

Min d/m
1.18882

ave/min d
1
ave/min m
1.15974

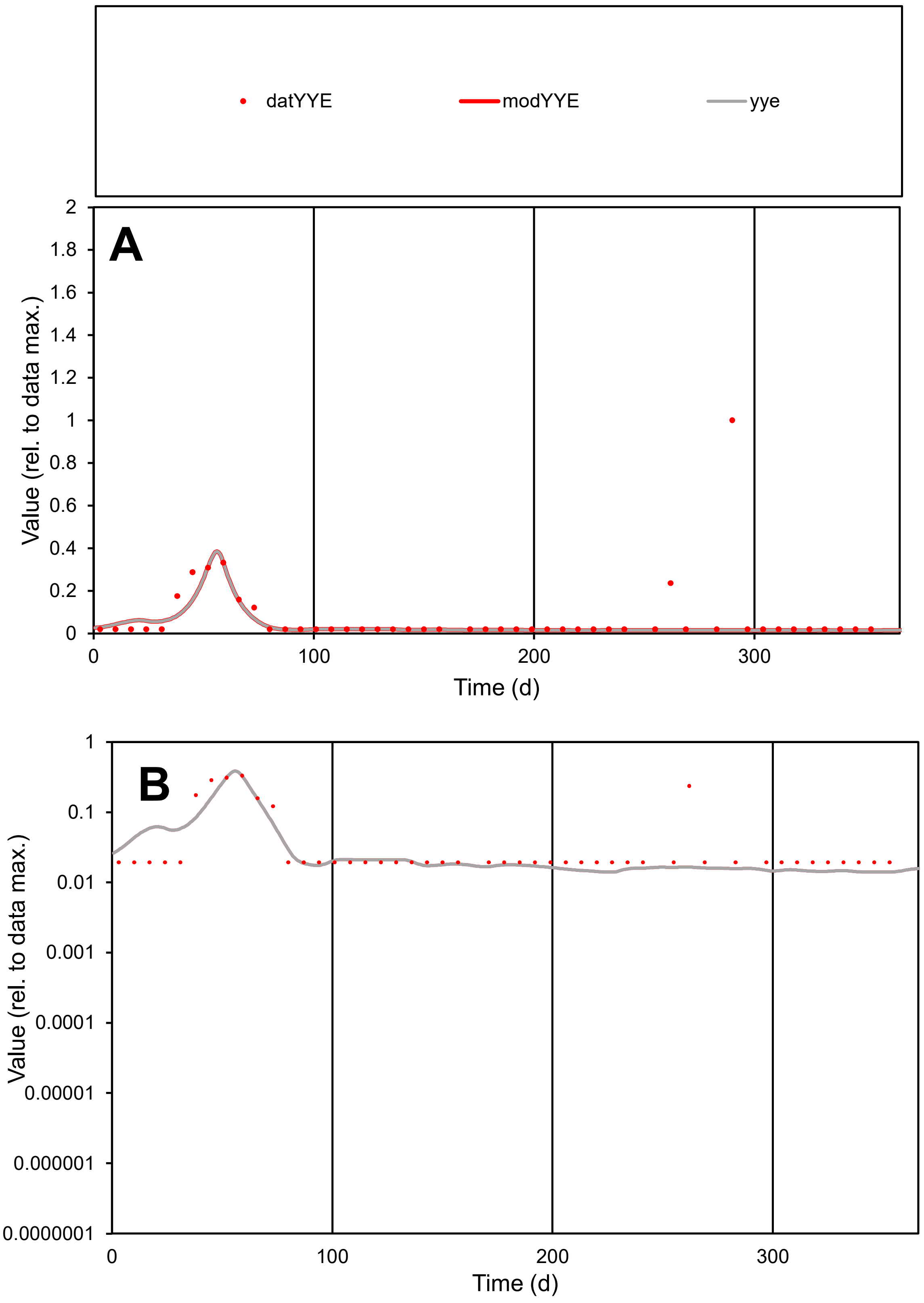


**C1**

Variables
YYE Provide jtd or Dst
CalibOnly?
2 1 to reproduce of
oNormalize?
2 1=no, 2=yes

ofxn calc*
2.08939
ofxn mod
2.09987

sRun
RunA1
zout
1

Ave
Data
1.4E-05
Model
8.2E-06

data/vMax
0.0002

ofxng
369.894

Min d/m
1.37286

ave/min d
3.66823
ave/min m
3.00579

328_Calib_S_A

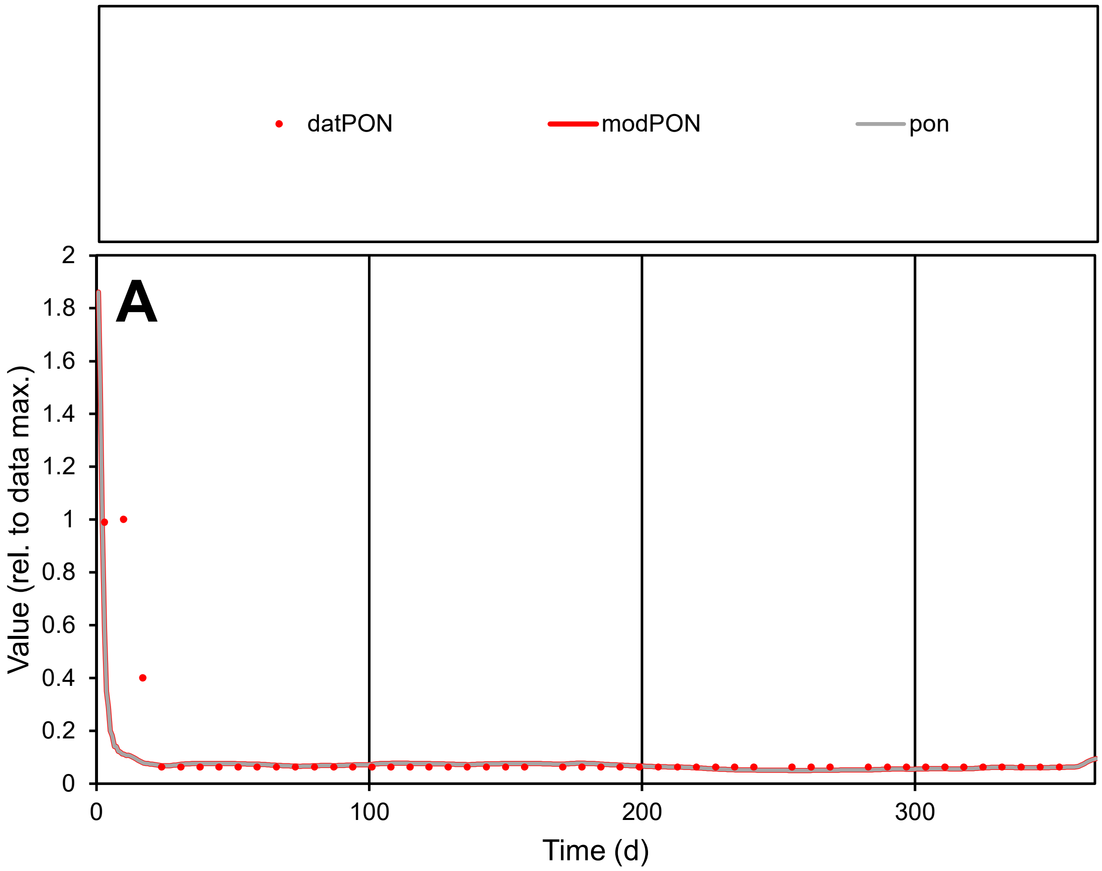


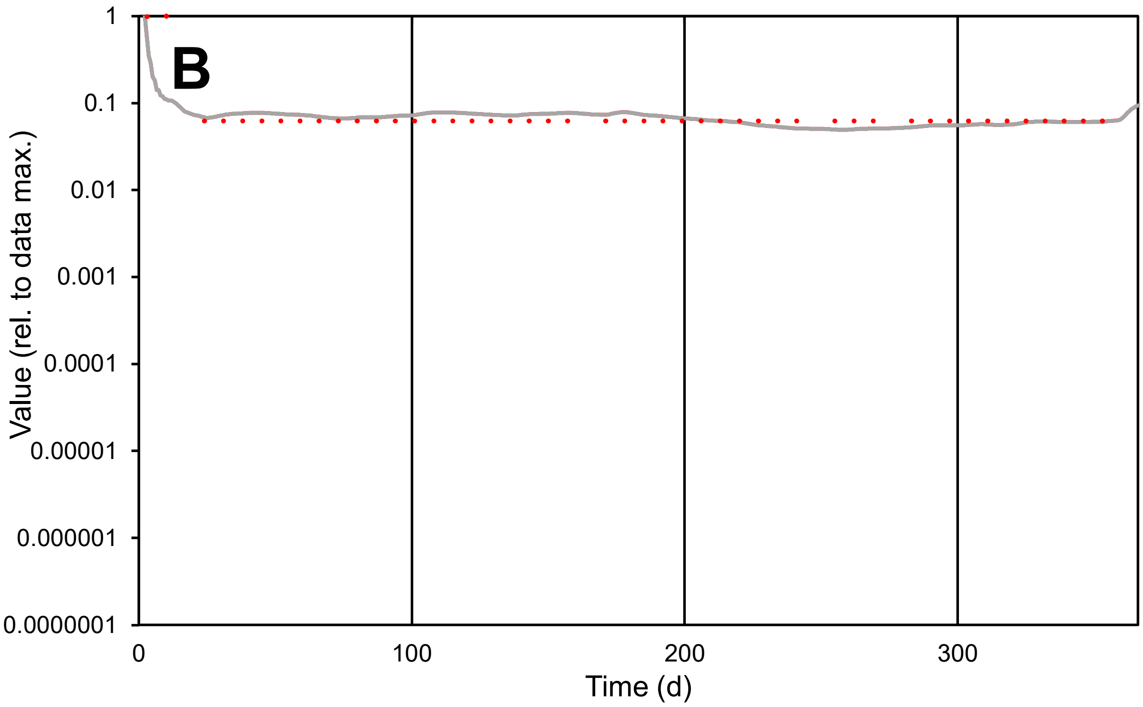


**C1**

Variables
PON Provide jtd or Dst
CalibOnly?
2 1 to reproduce of
oNormalize?
2 1=no, 2=yes

ofxn calc*
1.26858
ofxn mod
1.34049

sRun
RunA1
zout
1

Ave
Data
5.1E-06
Model
4E-06

data/vMax
4.7E-05

ofxng
369.894

Min d/m
1.26302

ave/min d
1.7364
ave/min m
1.7432

328_Calib_S_A

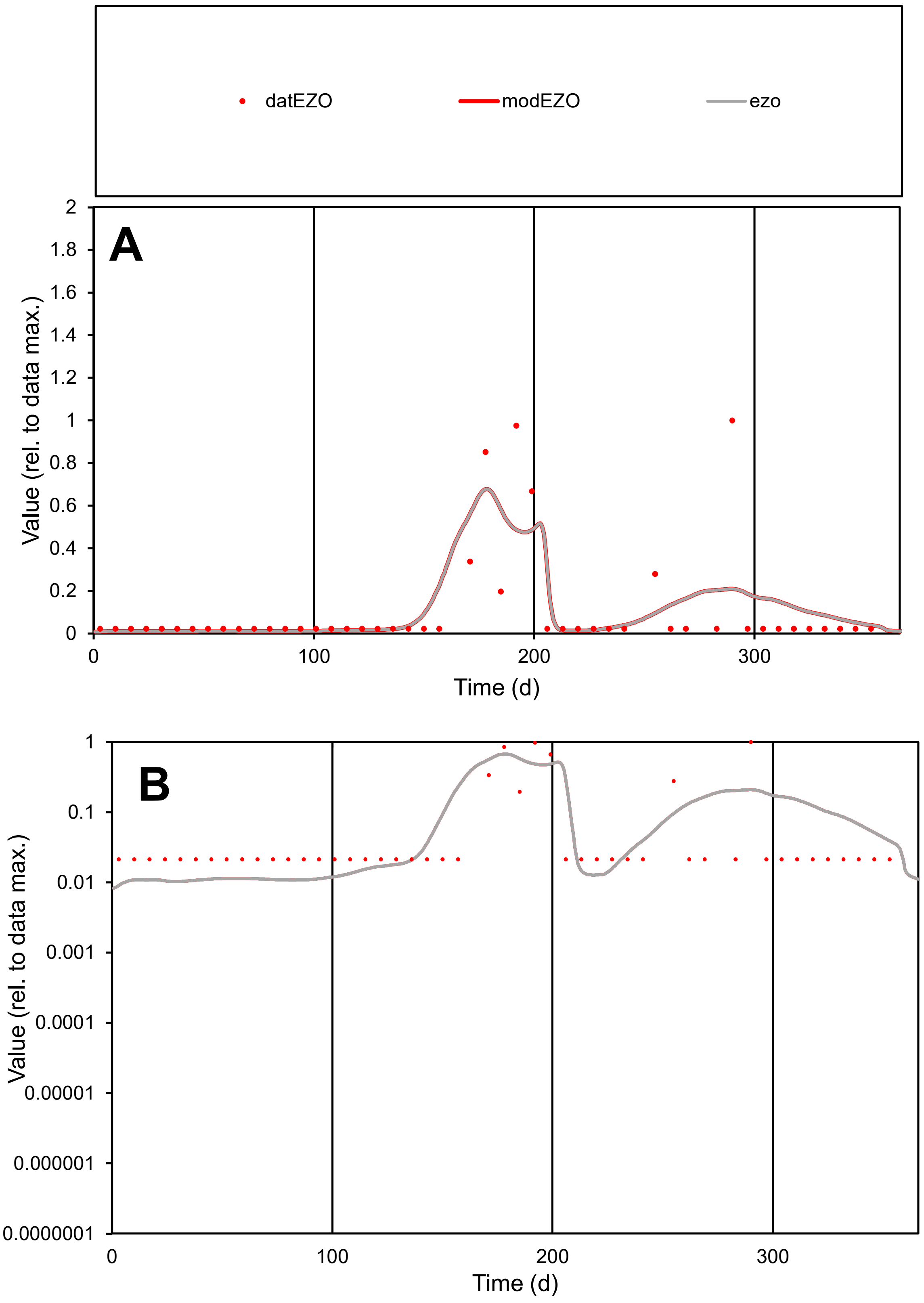


**C1**

Variables
EZO Provide jtd or Dst
CalibOnly?
2 1 to reproduce of
oNormalize?
2 1=no, 2=yes

ofxn calc*
1.5448
ofxn mod
1.58006

sRun
RunA1
zout
1

Ave
Data
2E-05
Model
2E-05

data/vMax
0.00018

ofxng
369.894

Min d/m
2.22768

ave/min d
5.05808
ave/min m
11.495

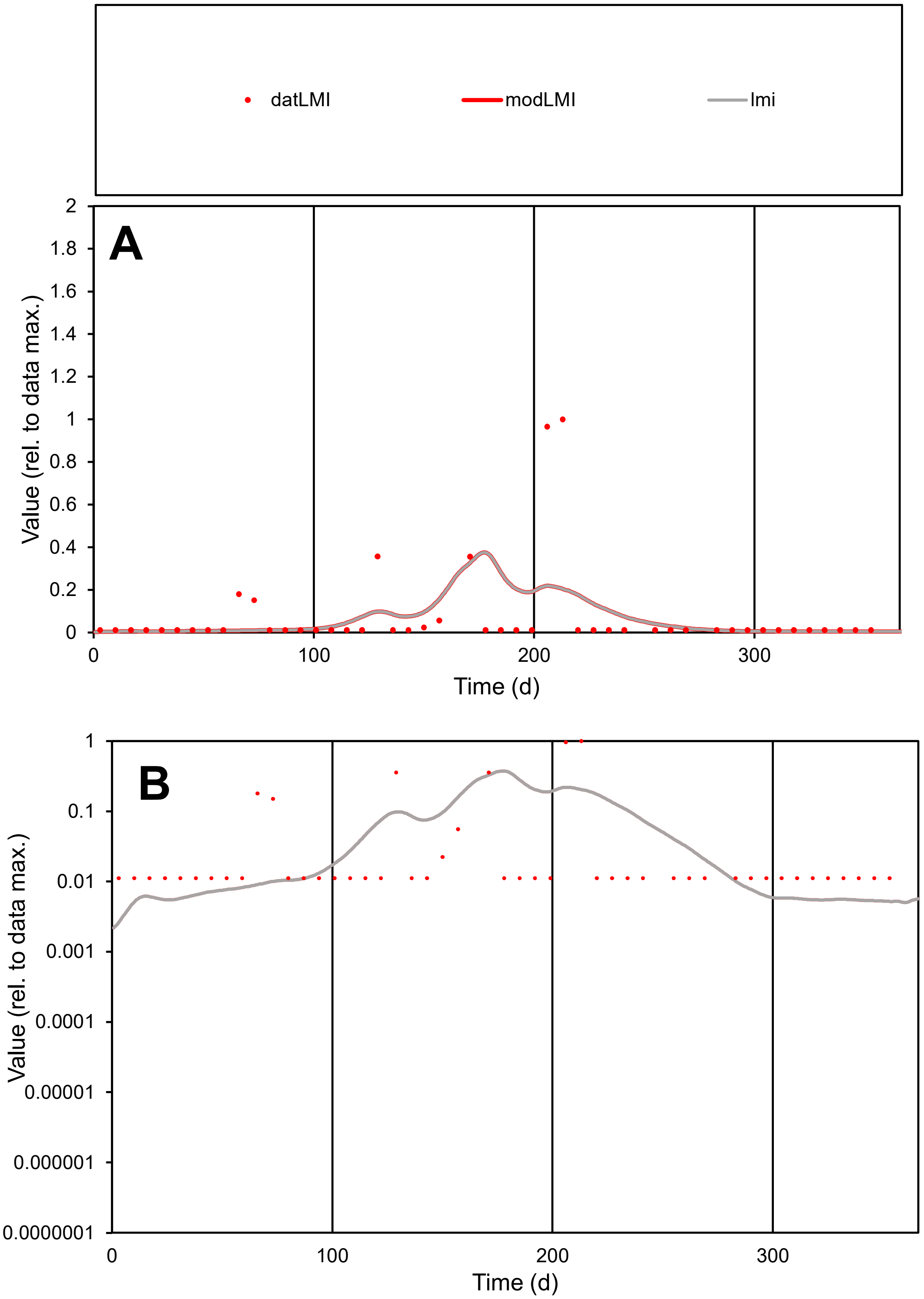


**C1**

Variables

| | |
|---|---|
| LMI | Provide jtd or Dst |
| CalibOnly? | |
| 2 | 1 to reproduce of |
| oNormalize? | |
| 2 | 1=no, 2=yes |

ofxn calc*
2.50792
ofxn mod
2.52432

sRun
RunA1
zout
1

Ave
Data
2E-05
Model
1.8E-05

data/vMax
0.00027

ofxng
369.894

Min d/m
3.75826

ave/min d
6.56534
ave/min m
21.8705

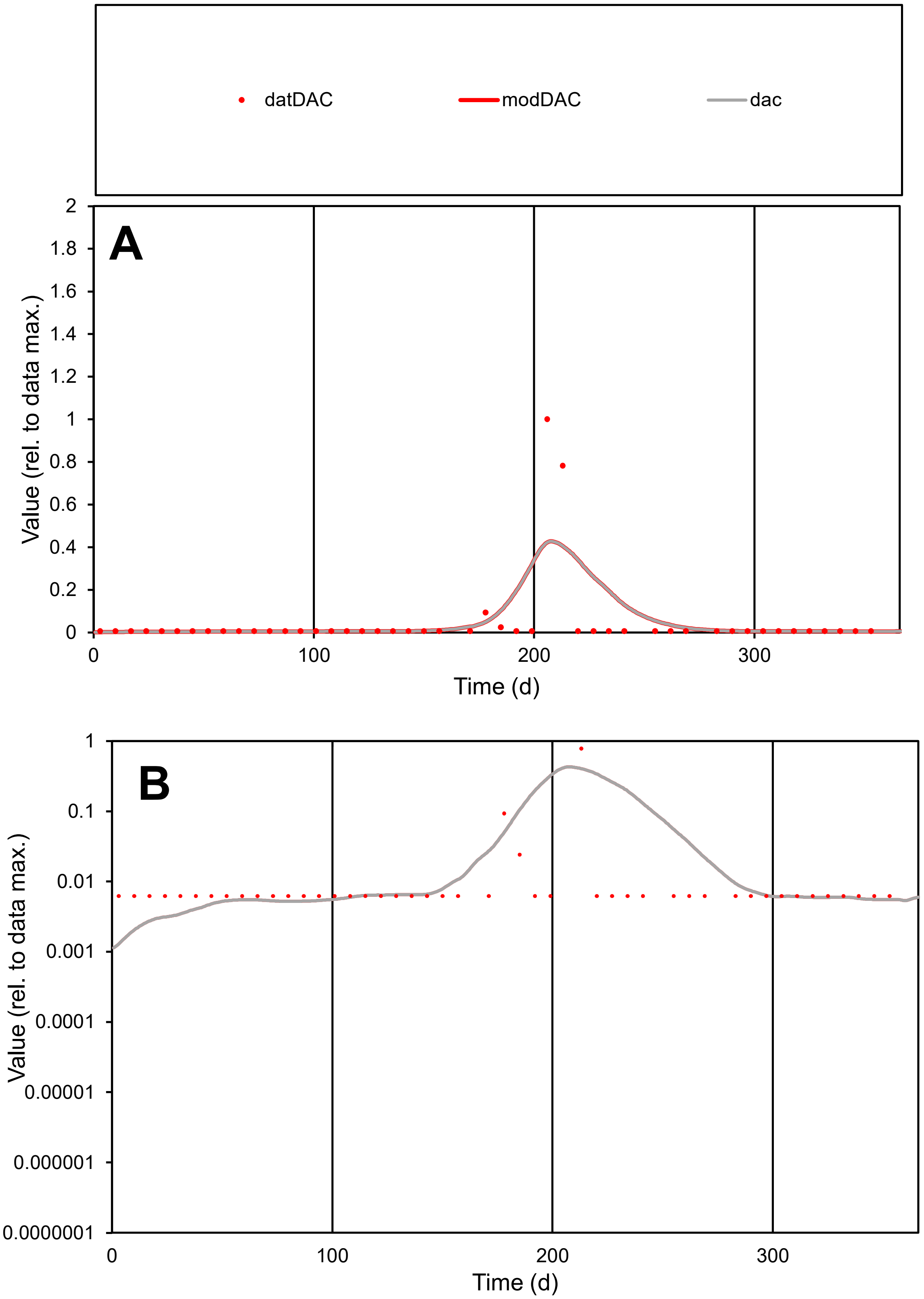


**C1**

Variables
DAC Provide jtd or Dst
CalibOnly?
2 1 to reproduce of
oNormalize?
2 1=no, 2=yes

ofxn calc*
2.81929
ofxn mod
2.90015

sRun
RunA1
zout
1

Ave
Data
5.6E-05
Model
6.7E-05

data/vMax
0.00124

ofxng
369.894

Min d/m
4.38339

ave/min d
7.2787
ave/min m
37.968

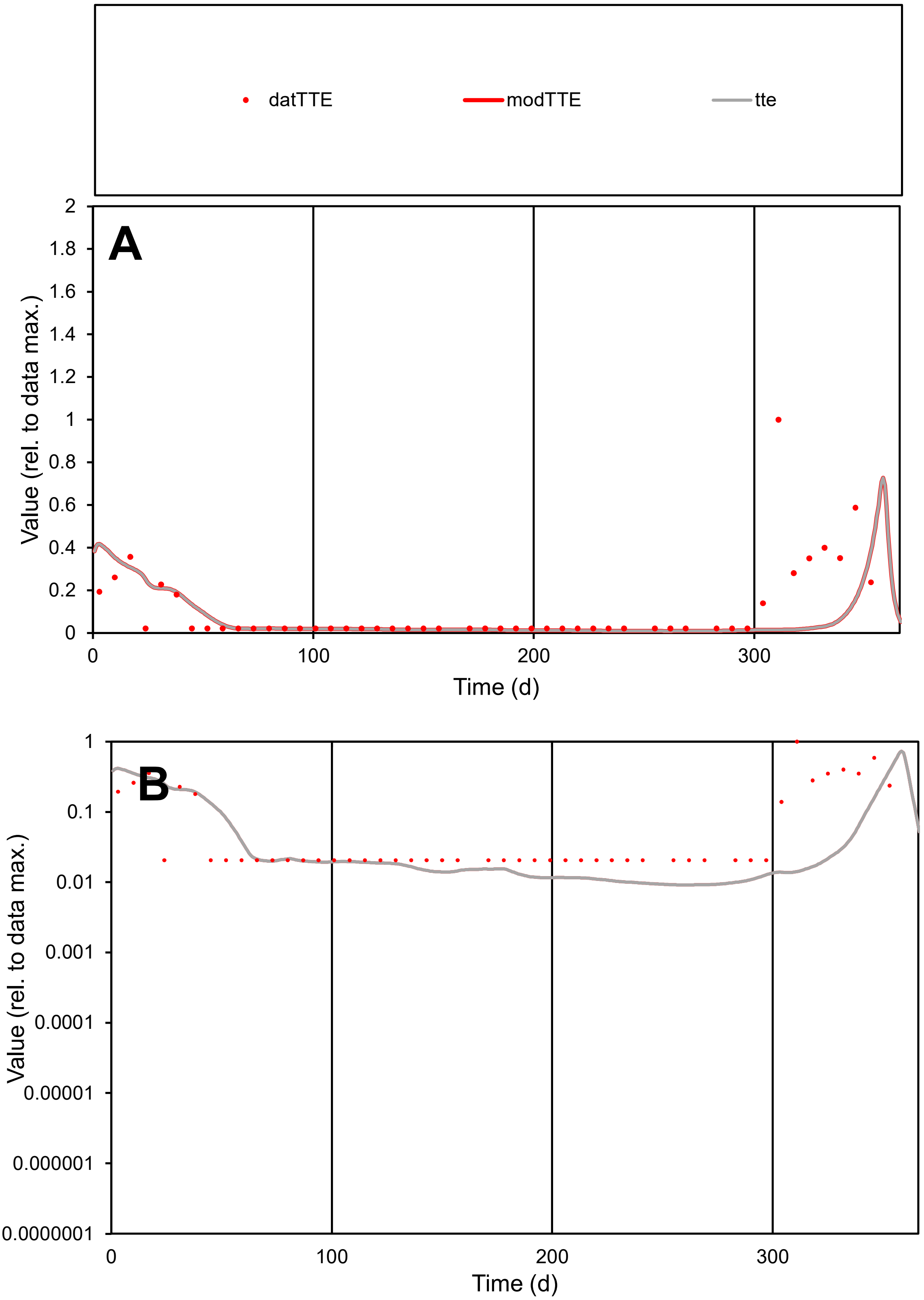


**C1**

Variables
TTE Provide jtd or Dst
CalibOnly?
2 1 to reproduce of
oNormalize?
2 1=no, 2=yes

ofxn calc*
1.69393
ofxn mod
1.70376

sRun
RunA1
zout
1

Ave
Data
2E-05
Model
1.2E-05

data/vMax
0.00018

ofxng
369.894

Min d/m
2.25886

ave/min d
5.34103
ave/min m
7.10294

328_Calib_S_A

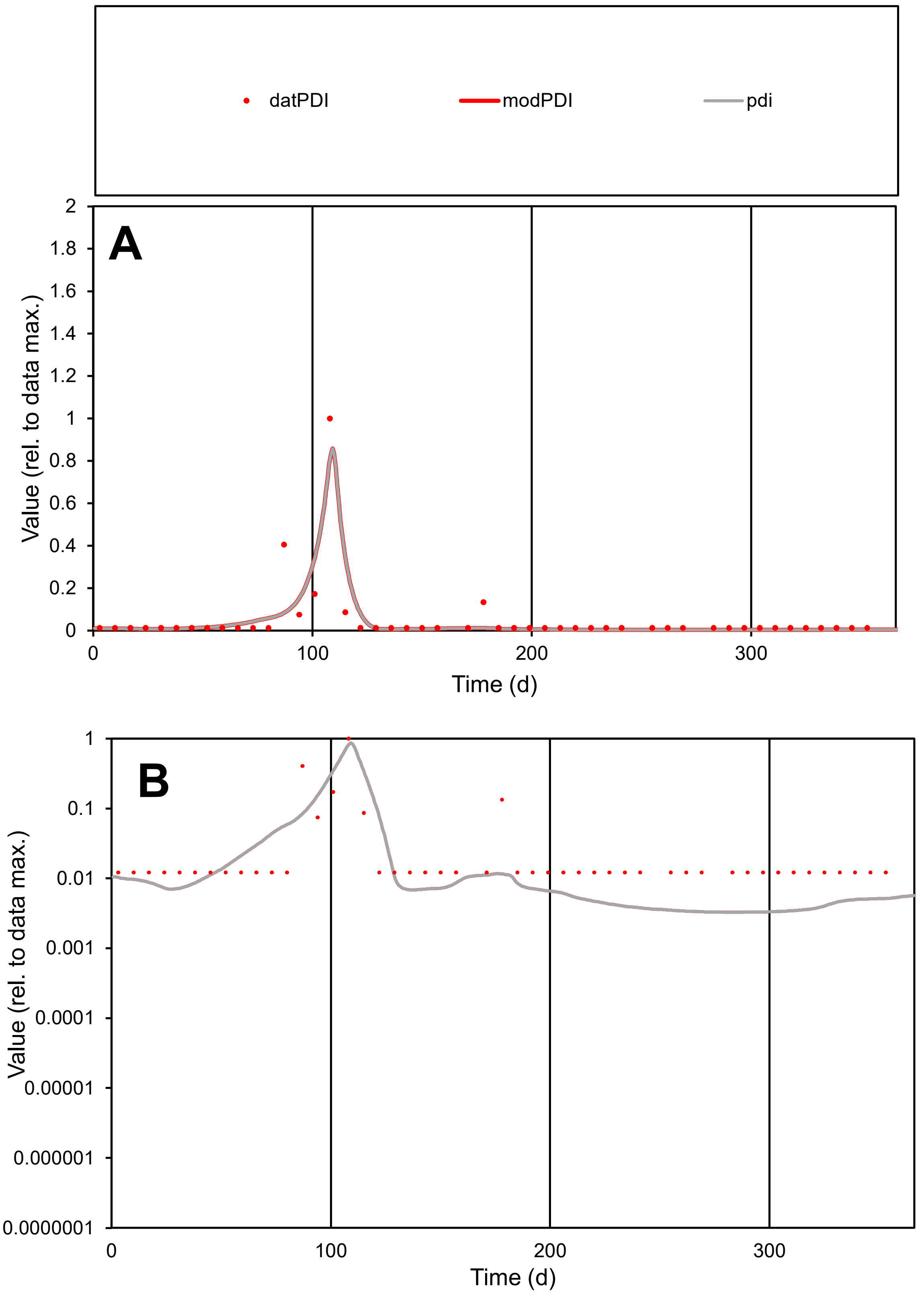


**C1**

Variables
PDI Provide jtd or Dst
CalibOnly?
2 1 to reproduce of
oNormalize?
2 1=no, 2=yes

ofxn calc*
1.28603
ofxn mod
1.34439

sRun
RunA1
zout
1

Ave
Data
1.8E-05
Model
1.6E-05

data/vMax
0.00036

ofxng
369.894

Min d/m
3.73492

ave/min d
4.07512
ave/min m
13.4449

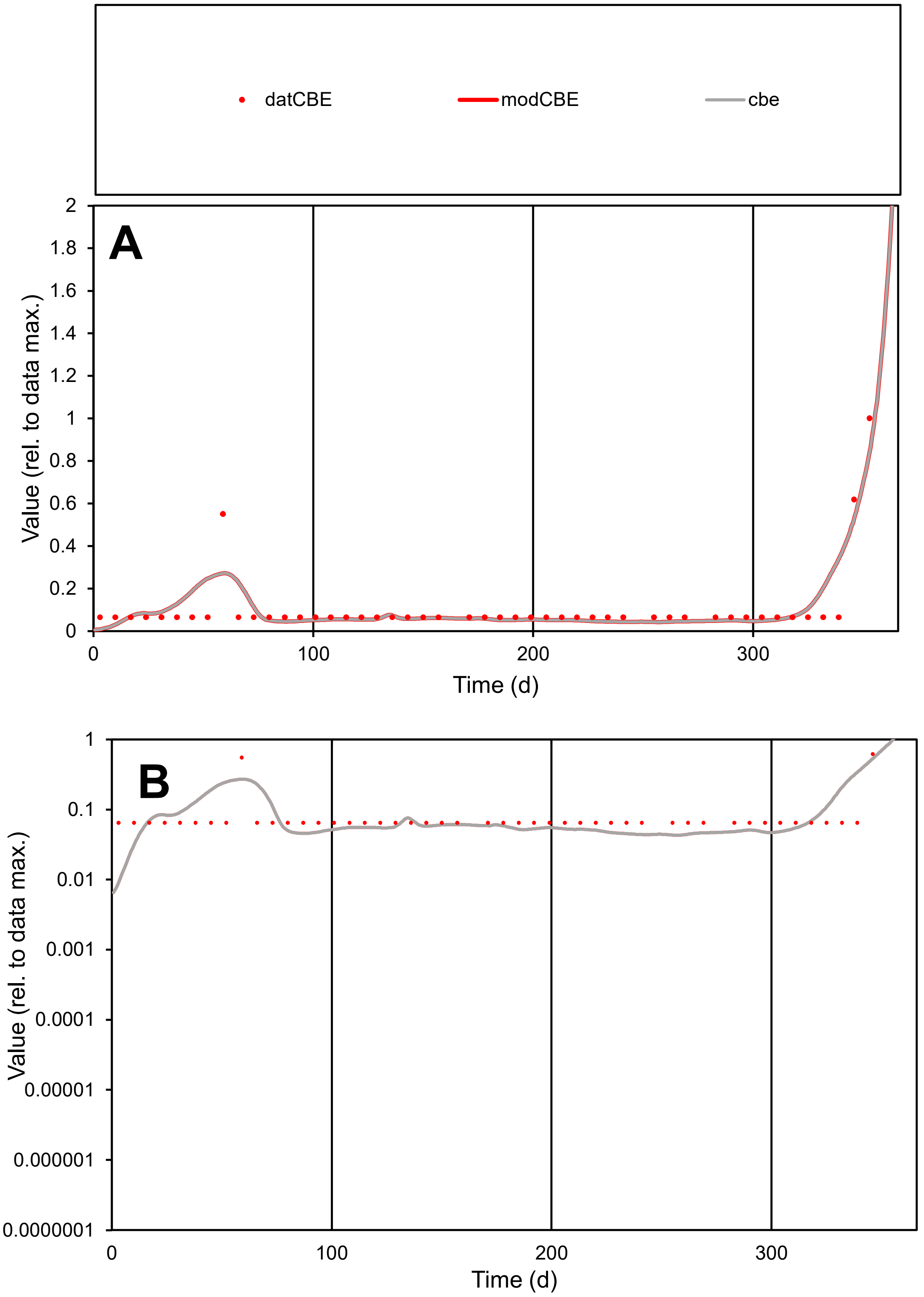


**C1**

Variables
CBE Provide jtd or Dst
CalibOnly?
2 1 to reproduce of
oNormalize?
2 1=no, 2=yes

ofxn calc*
0.69205
ofxn mod
0.73667

sRun
RunA1
zout
1

Ave
Data
2.7E-06
Model
2.8E-06

data/vMax
2.6E-05

ofxng
369.894

Min d/m
5.63514

ave/min d
1.63532
ave/min m
9.5599

328_Calib_S_A

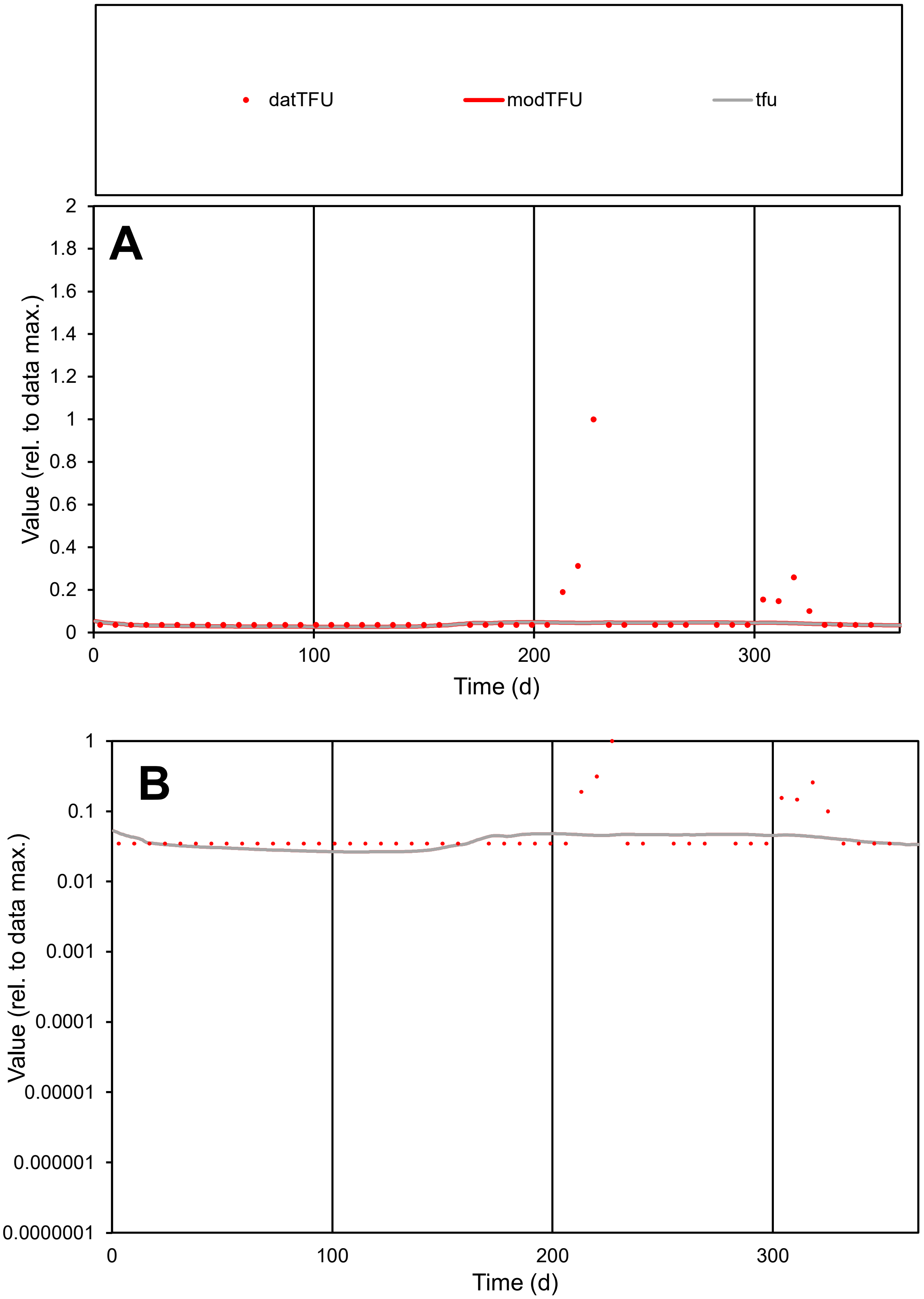


**C1**

Variables
TFU Provide jtd or Dst
CalibOnly?
2 1 to reproduce of
oNormalize?
2 1=no, 2=yes

ofxn calc*
2.00743
ofxn mod
2.01318

sRun
RunA1
zout
1

Ave
Data
2.7E-05
Model
1.4E-05

data/vMax
0.00037

ofxng
369.894

Min d/m
1.3085

ave/min d
2.15423
ave/min m
1.42379

328_Calib_S_A

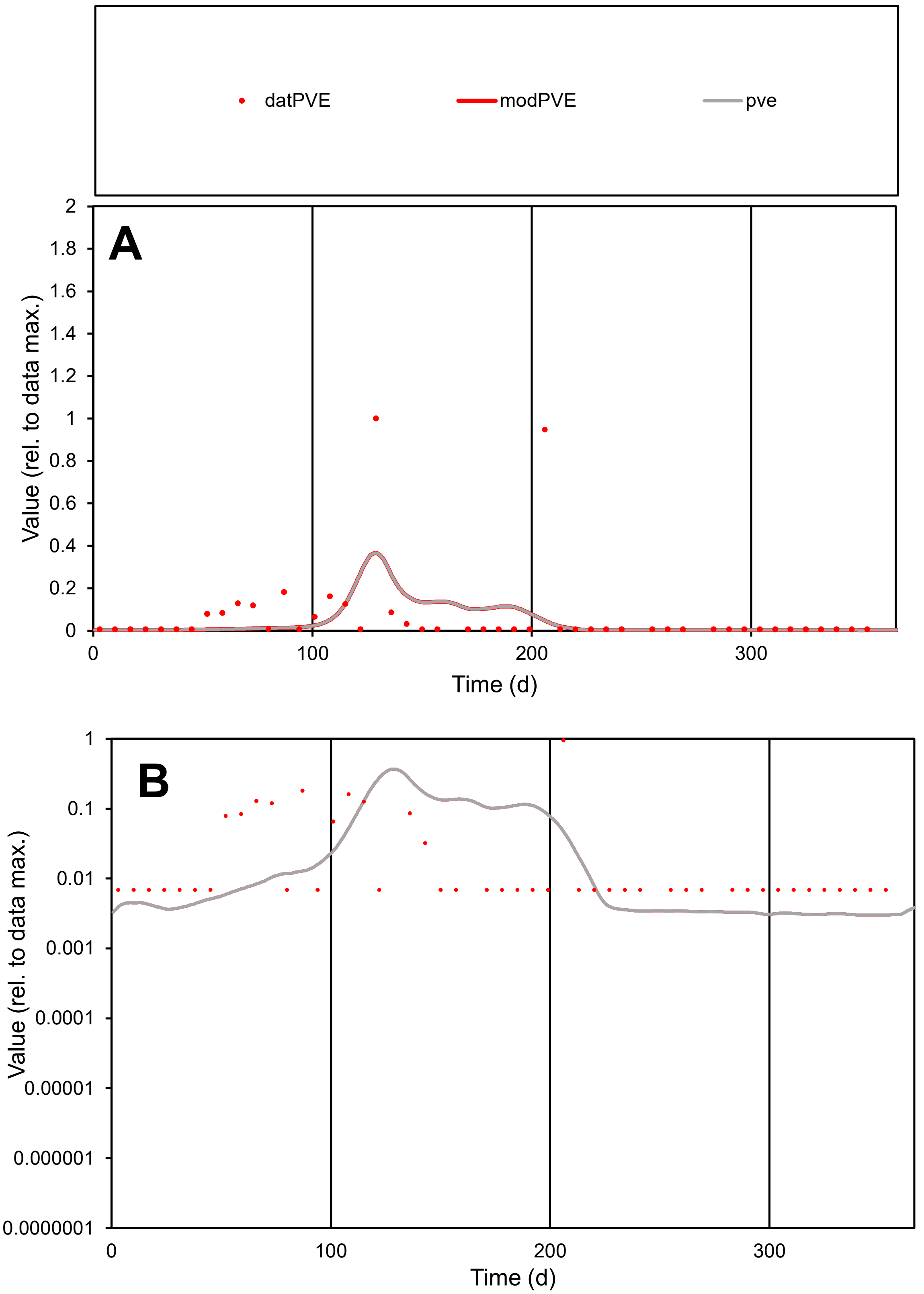


**C1**

Variables
PVE Provide jtd or Dst
CalibOnly?
2 1 to reproduce of
oNormalize?
2 1=no, 2=yes

ofxn calc*
2.57077
ofxn mod
2.59788

sRun
RunA1
zout
1

Ave
Data
2.1E-05
Model
1.4E-05

data/vMax
0.0003

ofxng
369.894

Min d/m
2.28559
ave/min d
9.86218
ave/min m
15.2401

328_Calib_S_A

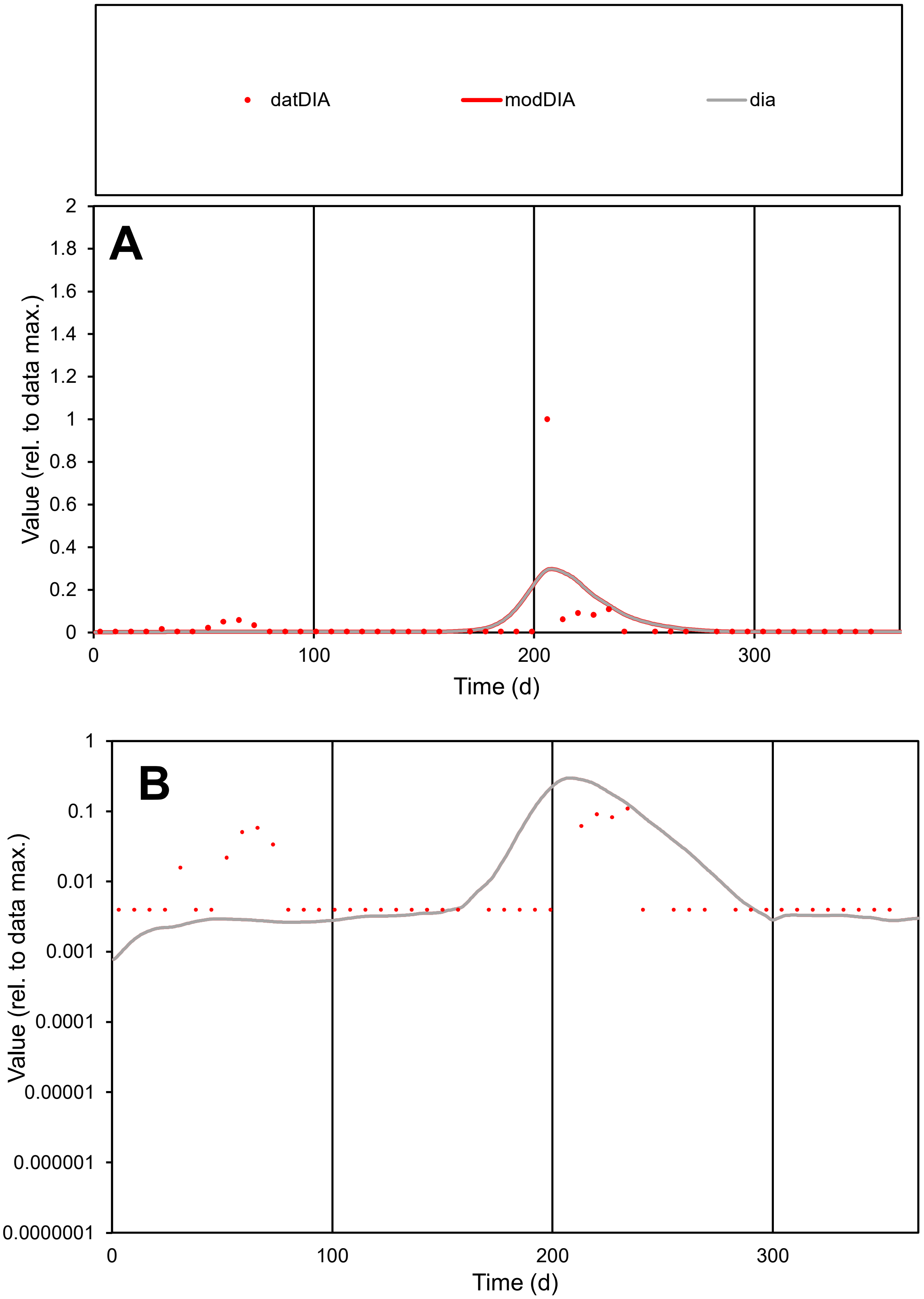


**C1**

Variables
DIA Provide jtd or Dst
CalibOnly?
2 1 to reproduce of
oNormalize?
2 1=no, 2=yes

ofxn calc*
3.25456
ofxn mod
3.2998

sRun
RunA1
zout
1

Ave
Data
5.5E-05
Model
5.5E-05

data/vMax
0.00157

ofxng
369.894

Min d/m
3.98146
ave/min d
8.77474
ave/min m
34.9811

328_Calib_S_A

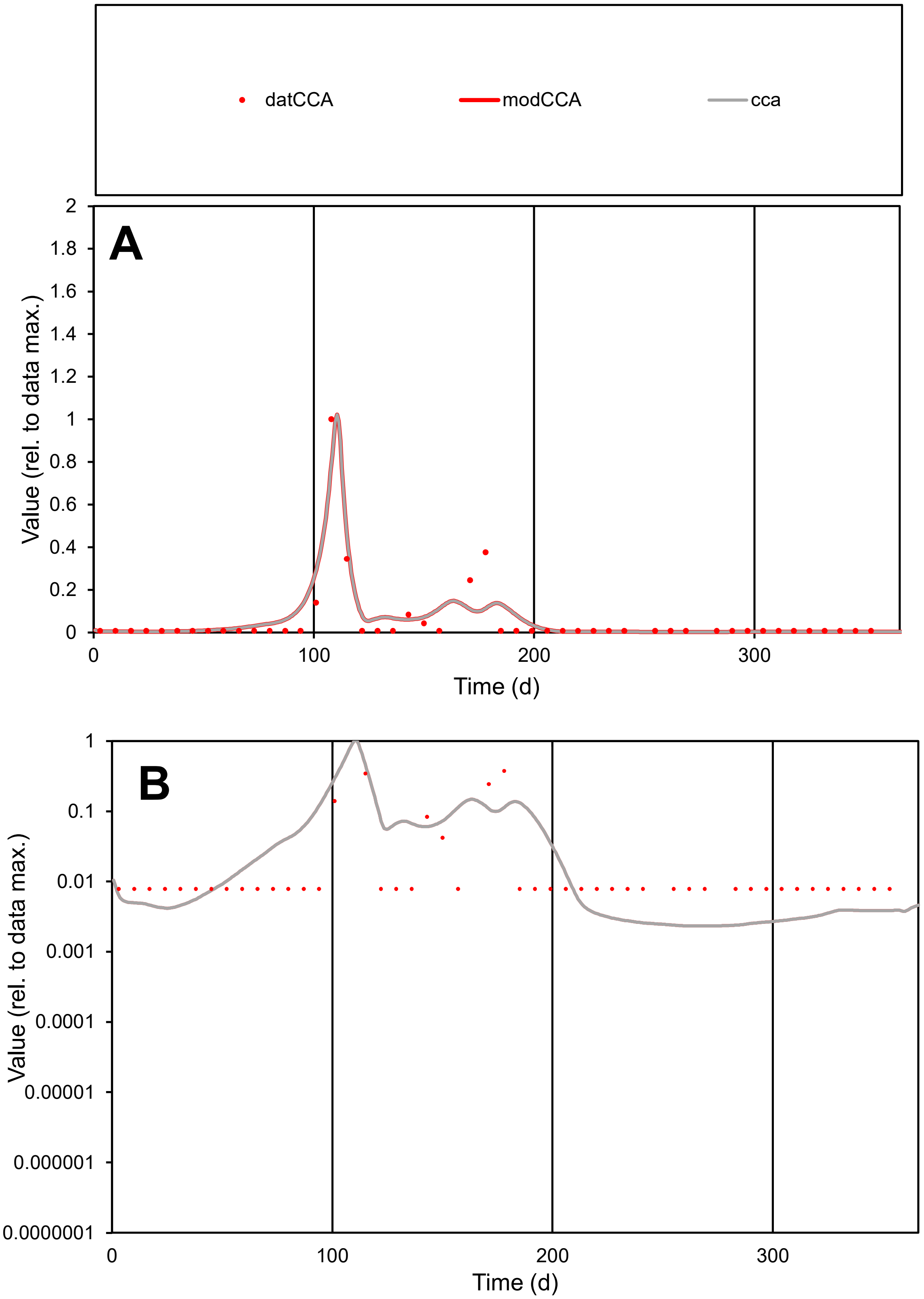


**C1**

Variables
CCA Provide jtd or Dst
CalibOnly?
2 1 to reproduce of
oNormalize?
2 1=no, 2=yes

ofxn calc*
1.08762
ofxn mod
1.14574

sRun
RunA1
zout
1

Ave
Data
1.2E-05
Model
1.4E-05

data/vMax
0.00023

ofxng
369.894

Min d/m
3.34615

ave/min d
6.79641
ave/min m
24.8313

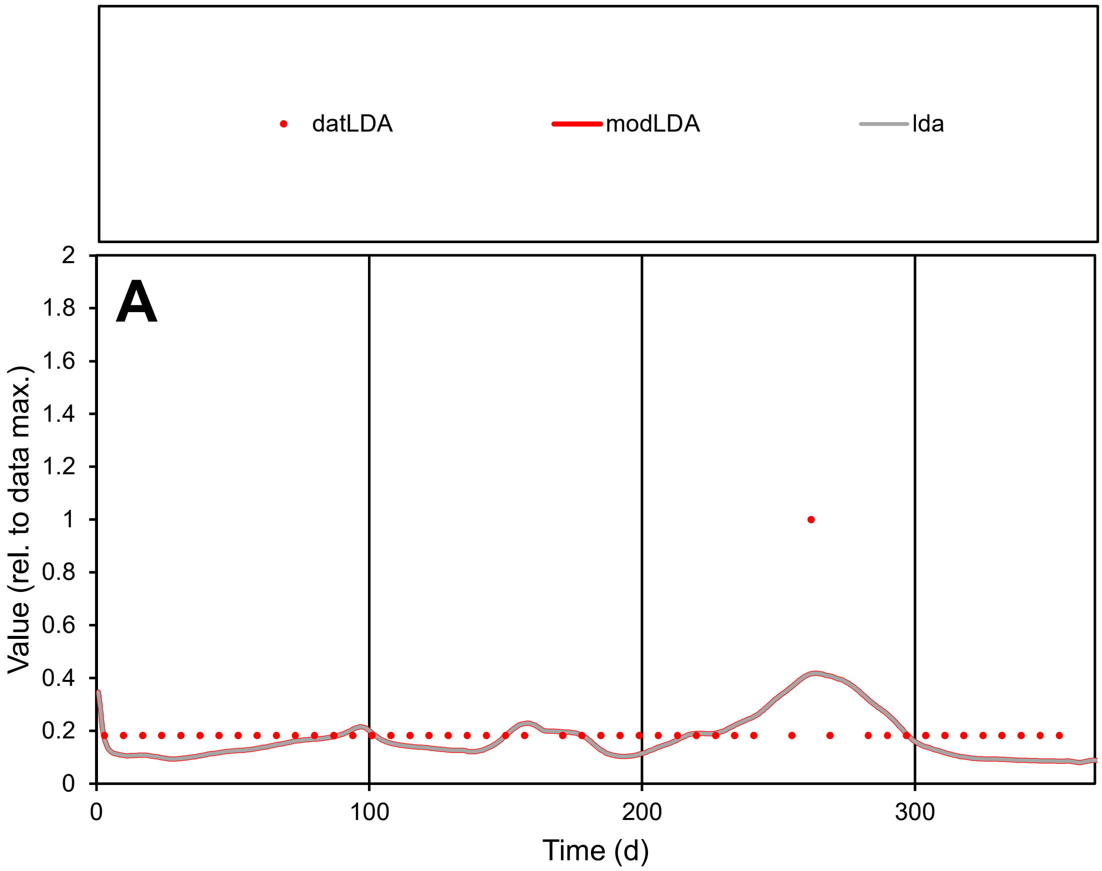


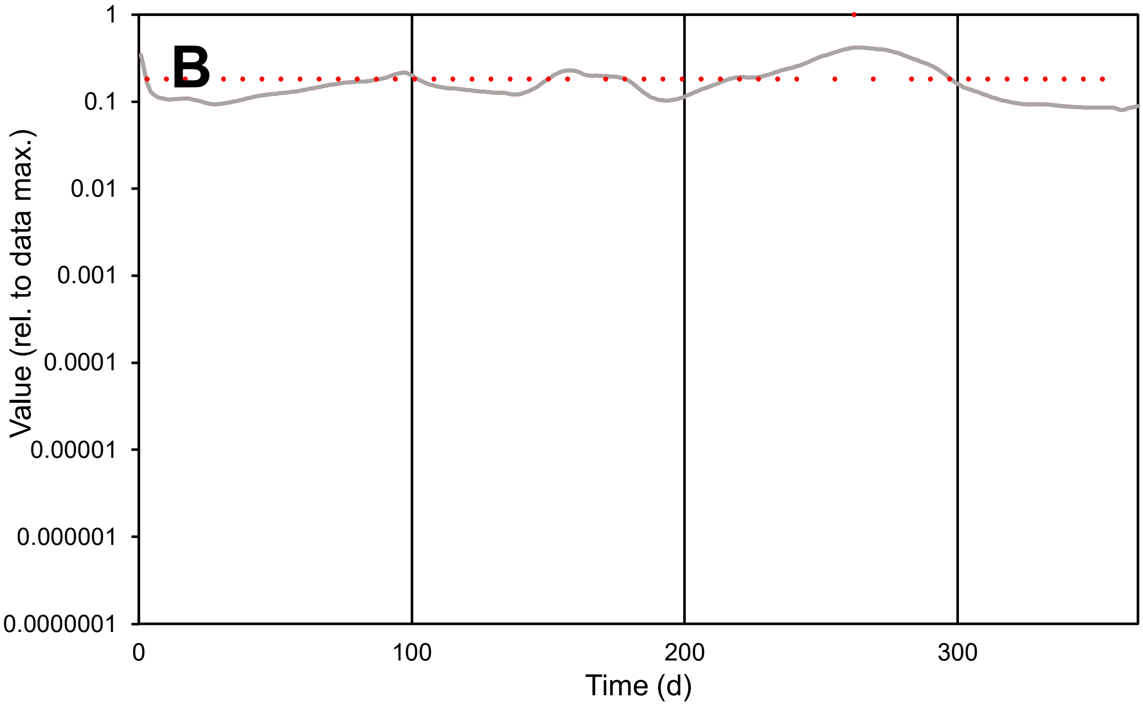


**C1**

Variables
LDA Provide jtd or Dst
CalibOnly?
2 1 to reproduce of
oNormalize?
2 1=no, 2=yes

ofxn calc*
0.55332
ofxn mod
0.5656

sRun
RunA1
zout
1

Ave
Data
7.6E-06
Model
6.3E-06

data/vMax
3.8E-05

ofxng
369.894

Min d/m
2.14228

ave/min d
1.09345
ave/min m
1.94431

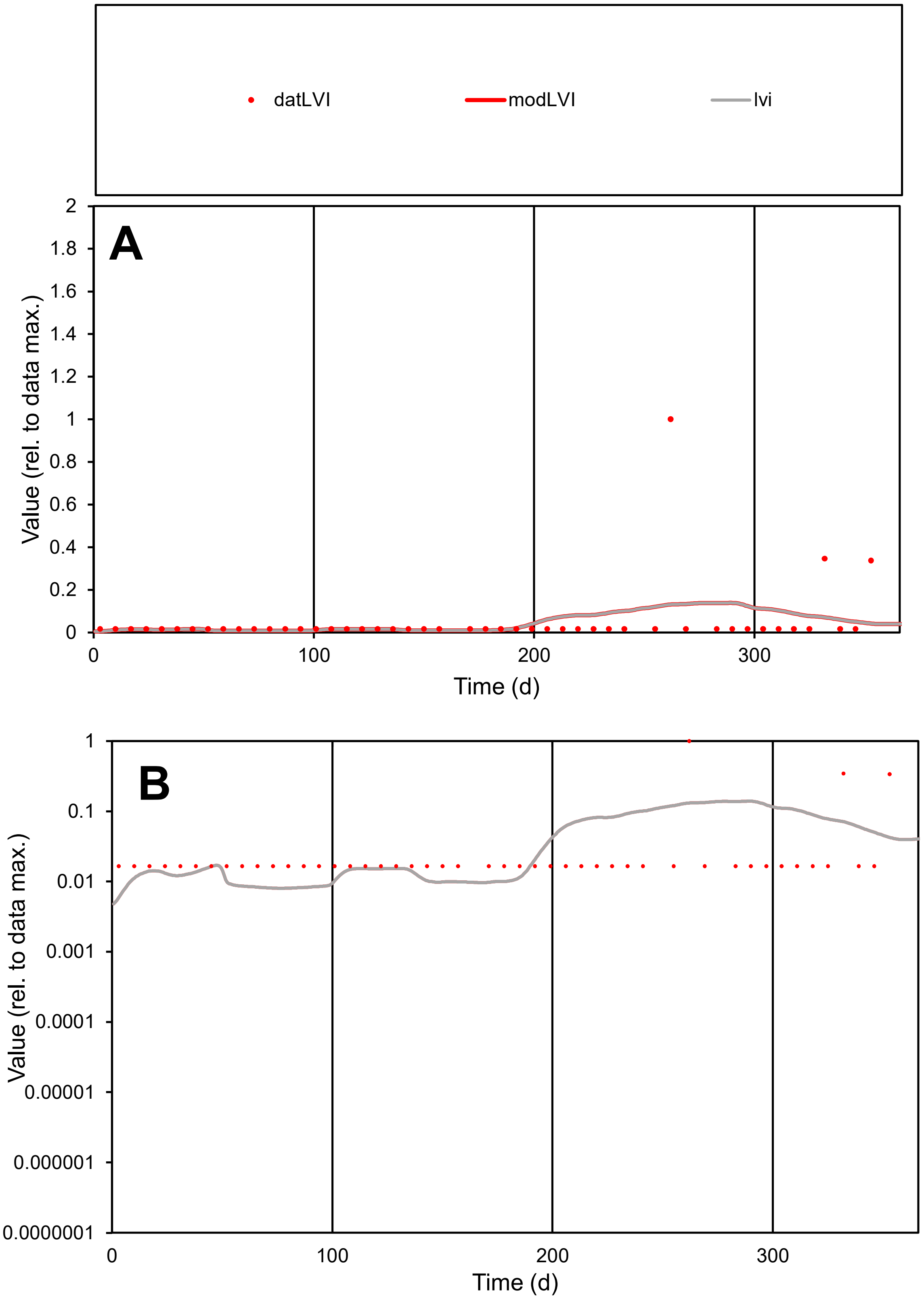


**C1**

Variables
LVI Provide jtd or Dst
CalibOnly?
2 1 to reproduce of
oNormalize?
2 1=no, 2=yes

ofxn calc*
2.90698
ofxn mod
2.91378

sRun
RunA1
zout
1

Ave
Data
4.3E-06
Model
4E-06

data/vMax
8.6E-05

ofxng
369.894

Min d/m
2.42538

ave/min d
3.05805
ave/min m
6.81646

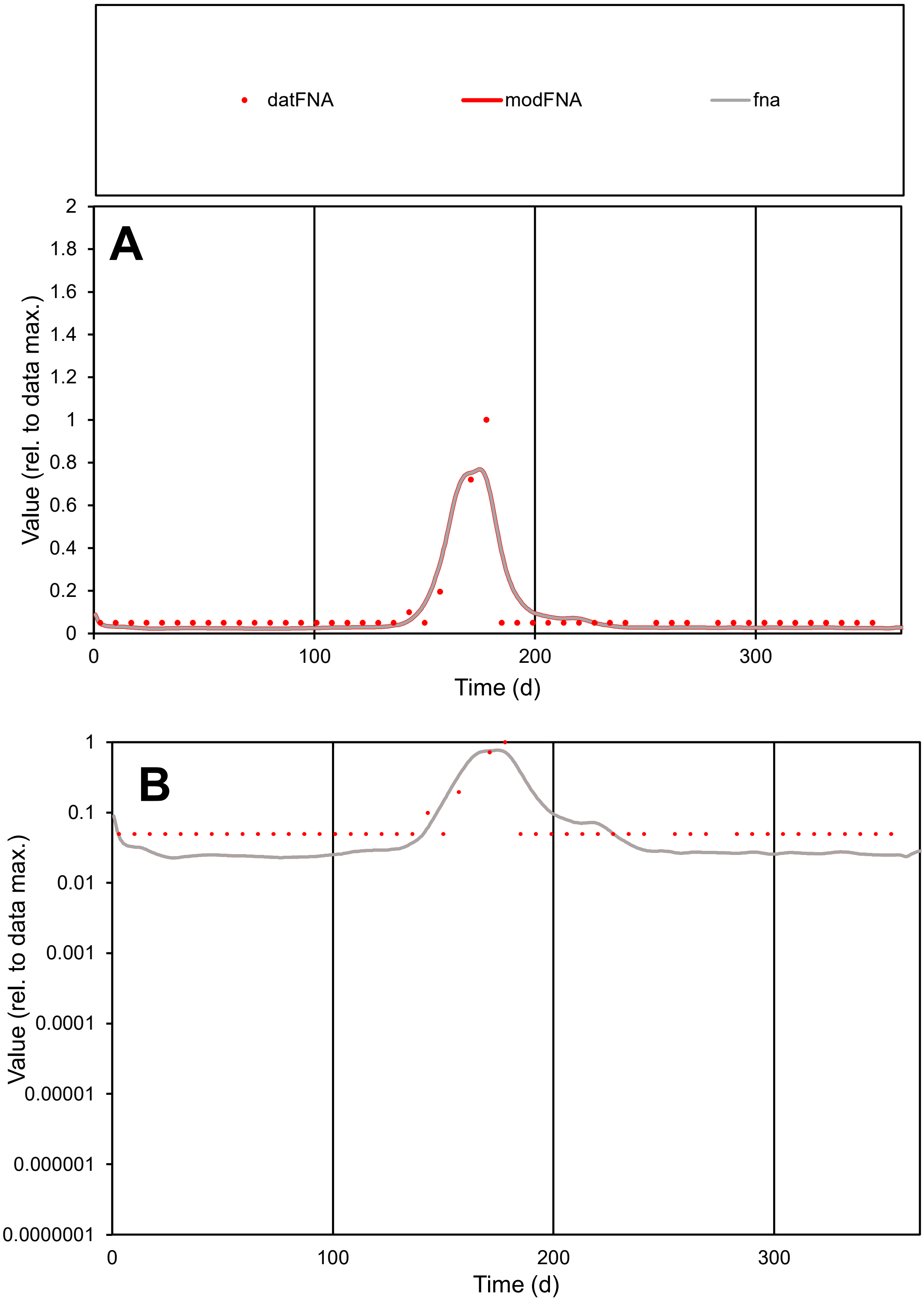


**C1**

Variables
FNA Provide jtd or Dst
CalibOnly?
2 1 to reproduce of
oNormalize?
2 1=no, 2=yes

ofxn calc*
0.71669
ofxn mod
0.82819

sRun
RunA1
zout
1

Ave
Data
5.5E-06
Model
4.9E-06

data/vMax
6.3E-05

ofxng
369.894

Min d/m
2.15331

ave/min d
1.76654
ave/min m
3.40637

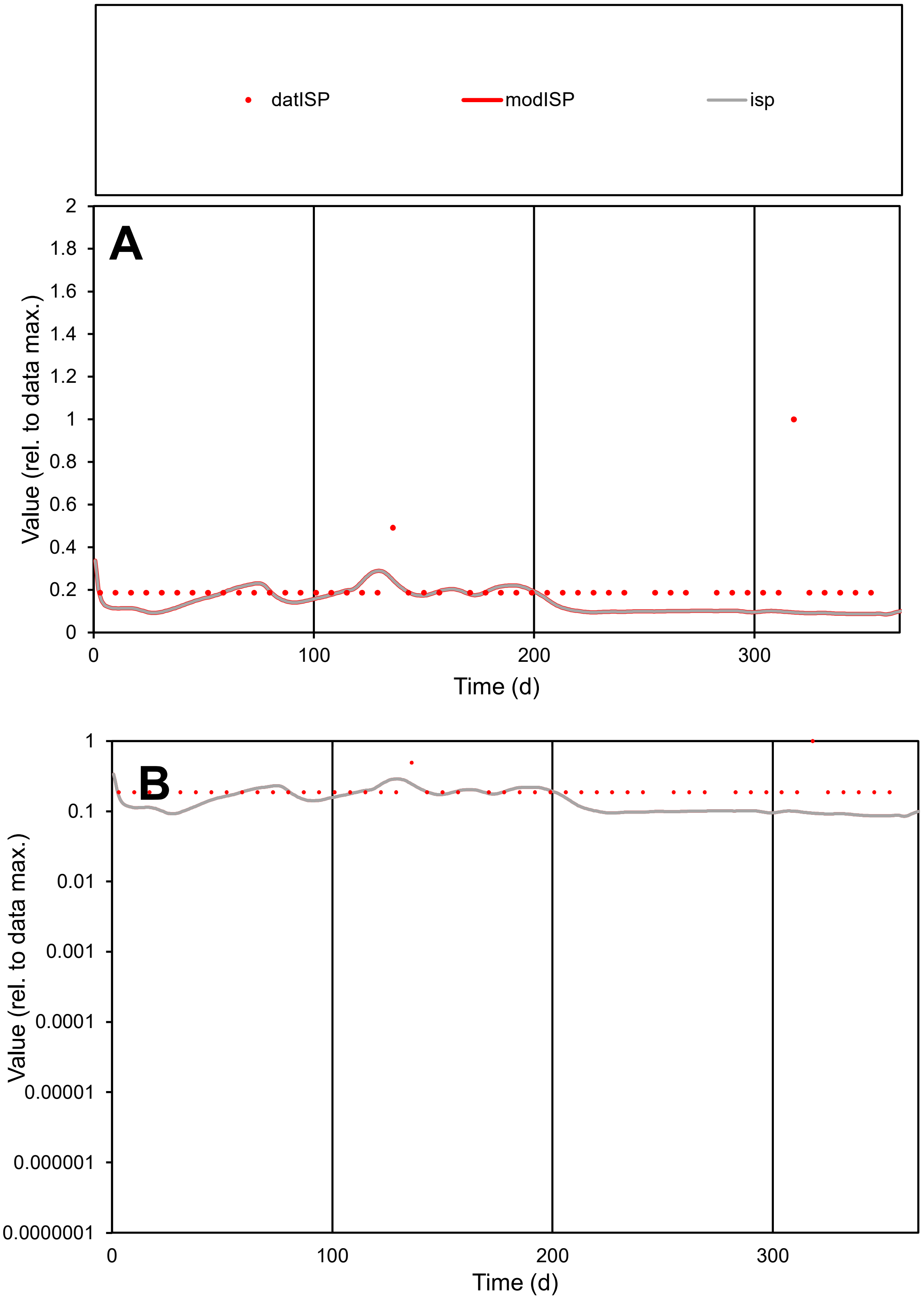


**C1**

Variables
ISP Provide jtd or Dst
CalibOnly?
2 1 to reproduce of
oNormalize?
2 1=no, 2=yes

ofxn calc*
0.71798
ofxn mod
0.72765

sRun
RunA1
zout
1

Ave
Data
5.7E-06
Model
3.9E-06

data/vMax
2.7E-05

ofxng
369.894

Min d/m
2.14827

ave/min d
1.12518
ave/min m
1.67842

328_Calib_S_A

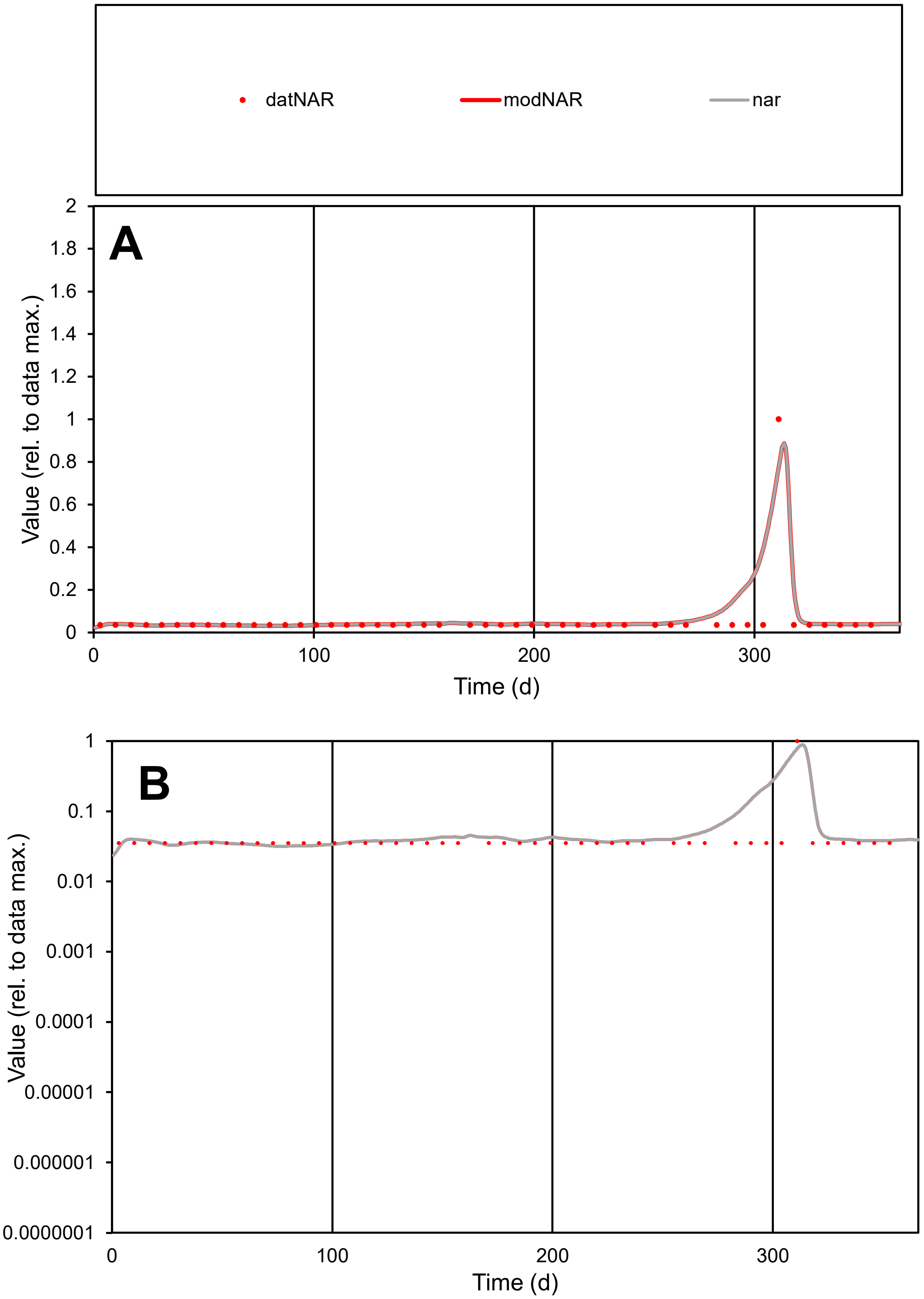


**C1**

Variables
NAR Provide jtd or Dst
CalibOnly?
2 1 to reproduce of
oNormalize?
2 1=no, 2=yes

ofxn calc*
1.12778
ofxn mod
1.19005

sRun
RunA1
zout
1

Ave
Data
4.4E-06
Model
5.6E-06

data/vMax
8E-05

ofxng
369.894

Min d/m
1.10697

ave/min d
1.57002
ave/min m
2.17693

328_Calib_S_A

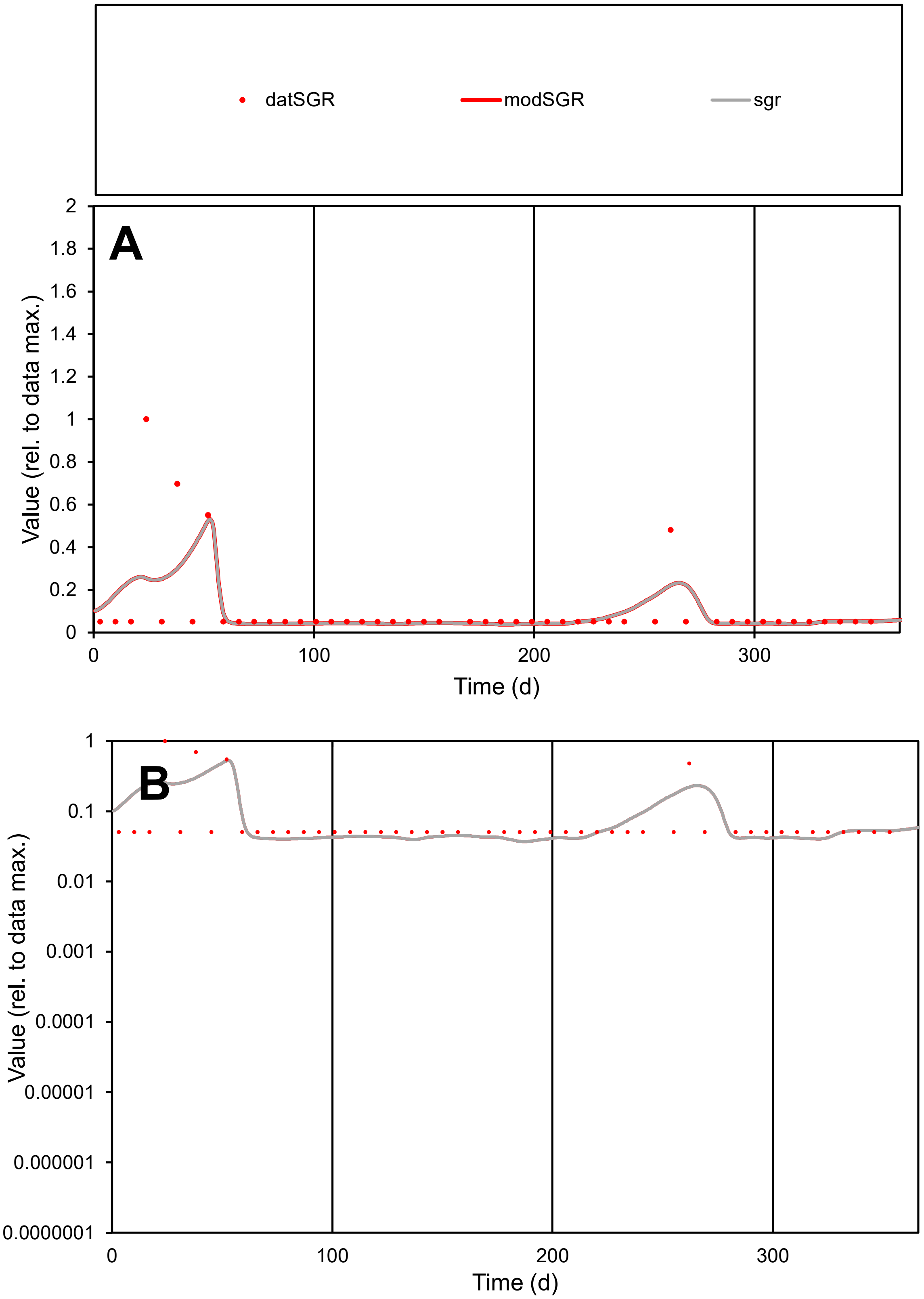


**C1**

Variables
SGR Provide jtd or Dst
CalibOnly?
2 1 to reproduce of
oNormalize?
2 1=no, 2=yes

ofxn calc*
1.42728
ofxn mod
1.44097

sRun
RunA1
zout
1

Ave
Data
9.3E-06
Model
8.7E-06

data/vMax
9E-05

ofxng
369.894

Min d/m
1.36145

ave/min d
2.0418
ave/min m
2.60109

328_Calib_S_A

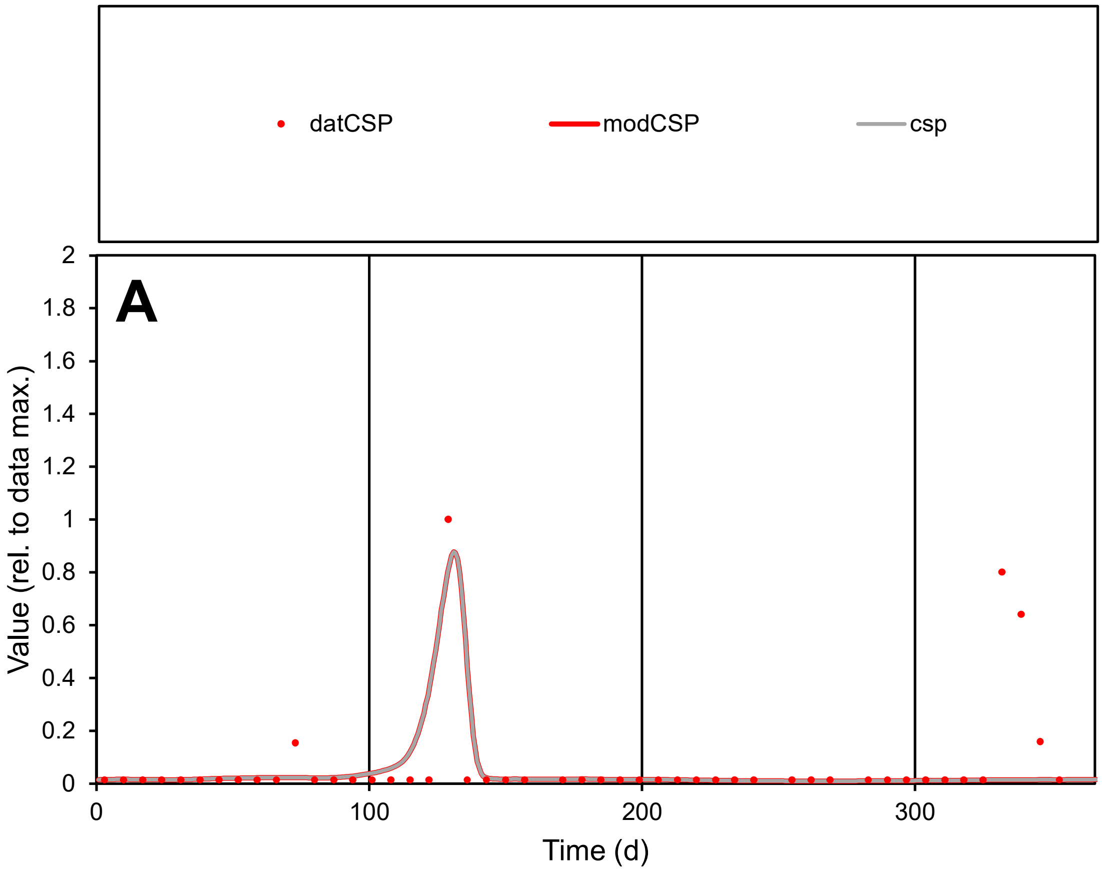


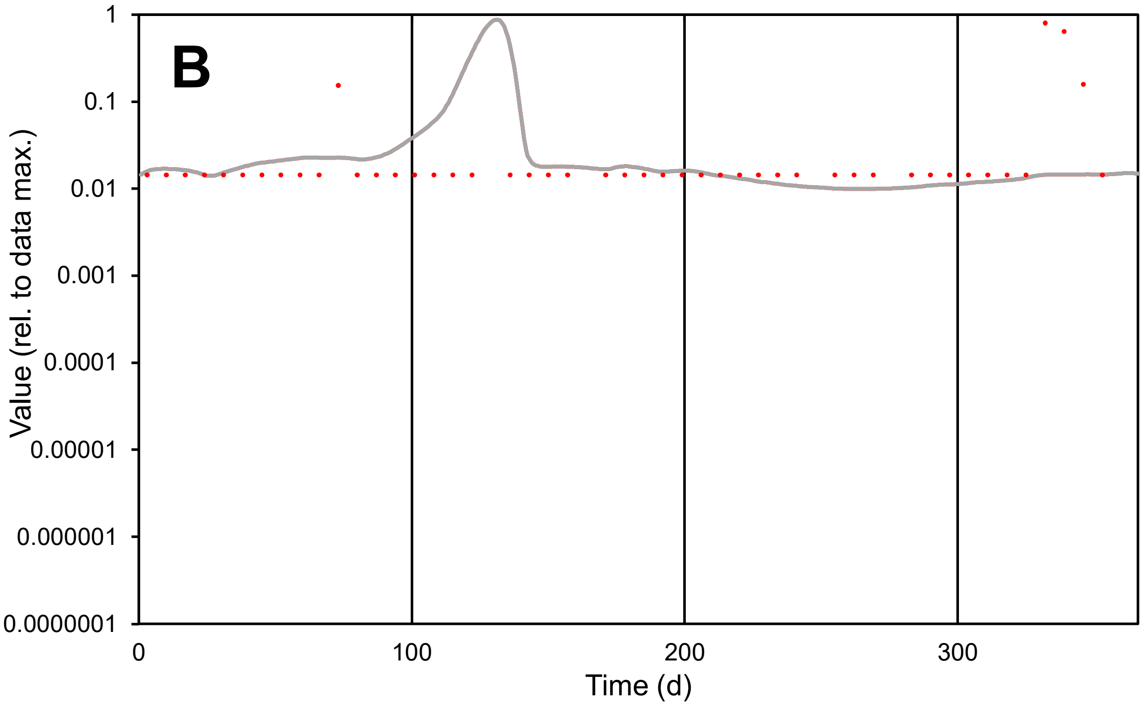


**C1**

| | |
|---|---|
| Variables | |
| CSP | Provide jtd or Dst |
| CalibOnly? | |
| 2 | 1 to reproduce of |
| oNormalize? | |
| 2 | 1=no, 2=yes |
| ofxn calc* | |
| 2.19929 | |
| ofxn mod | |
| 2.25987 | |
| sRun | |
| RunA1 | |
| zout | |
| 1 | |
| Ave | |
| Data | |
| 1.2E-05 | |
| Model | |
| 7.2E-06 | |
| data/vMax | |
| 0.00017 | |
| ofxng | |
| 369.894 | |
| Min d/m | |
| 1.45994 | |
| ave/min d | |
| 4.8711 | |
| ave/min m | |
| 4.40449 | |

328_Calib_S_A

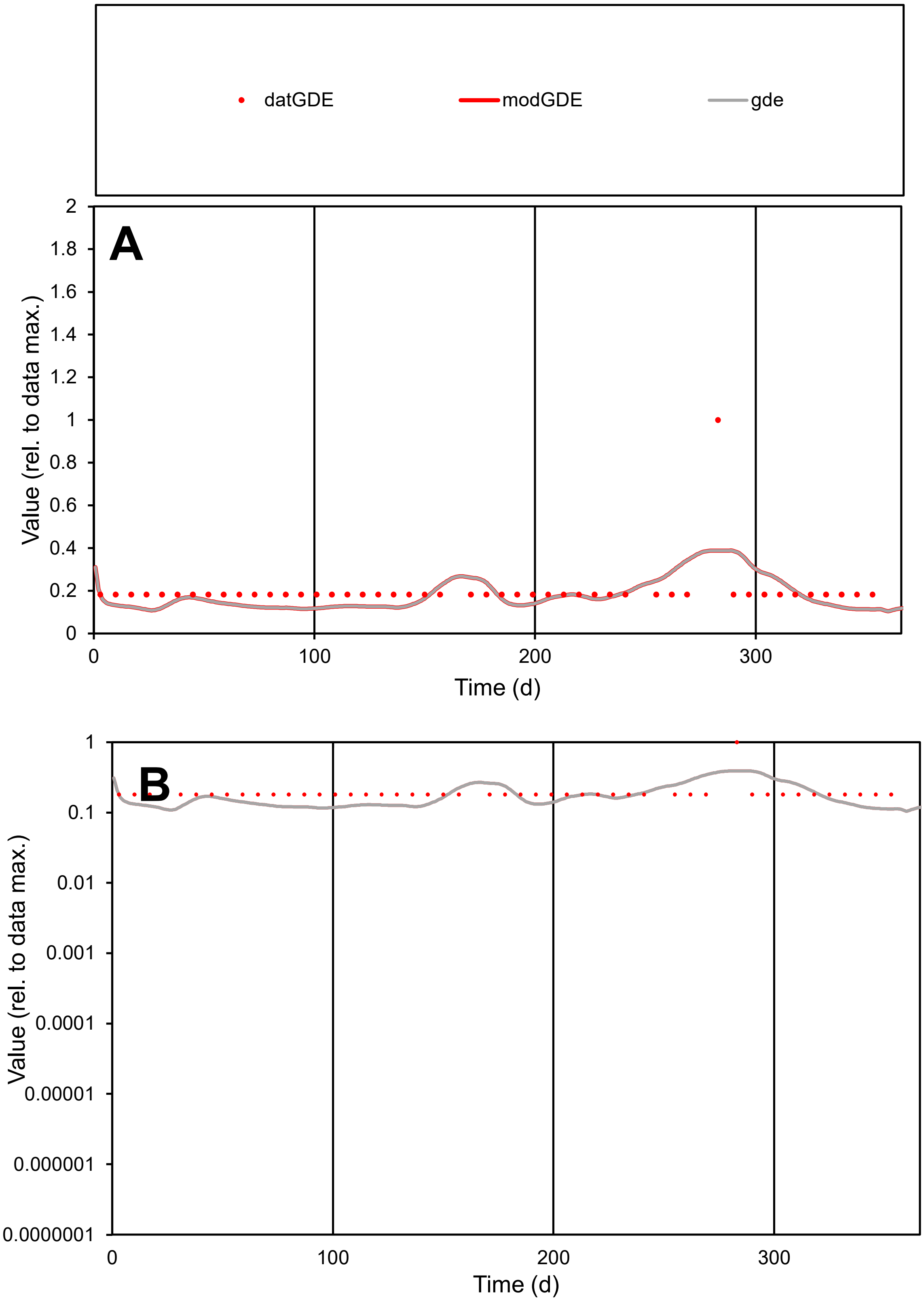


**C1**

Variables
GDE Provide jtd or Dst
CalibOnly?
2 1 to reproduce of
oNormalize?
2 1=no, 2=yes

ofxn calc*
0.55337
ofxn mod
0.5681

sRun
RunA1
zout
1

Ave
Data
3E-05
Model
2.7E-05

data/vMax
0.00015

ofxng
369.894

Min d/m
1.64946

ave/min d
1.0941
ave/min m
1.60314

328_Calib_S_A

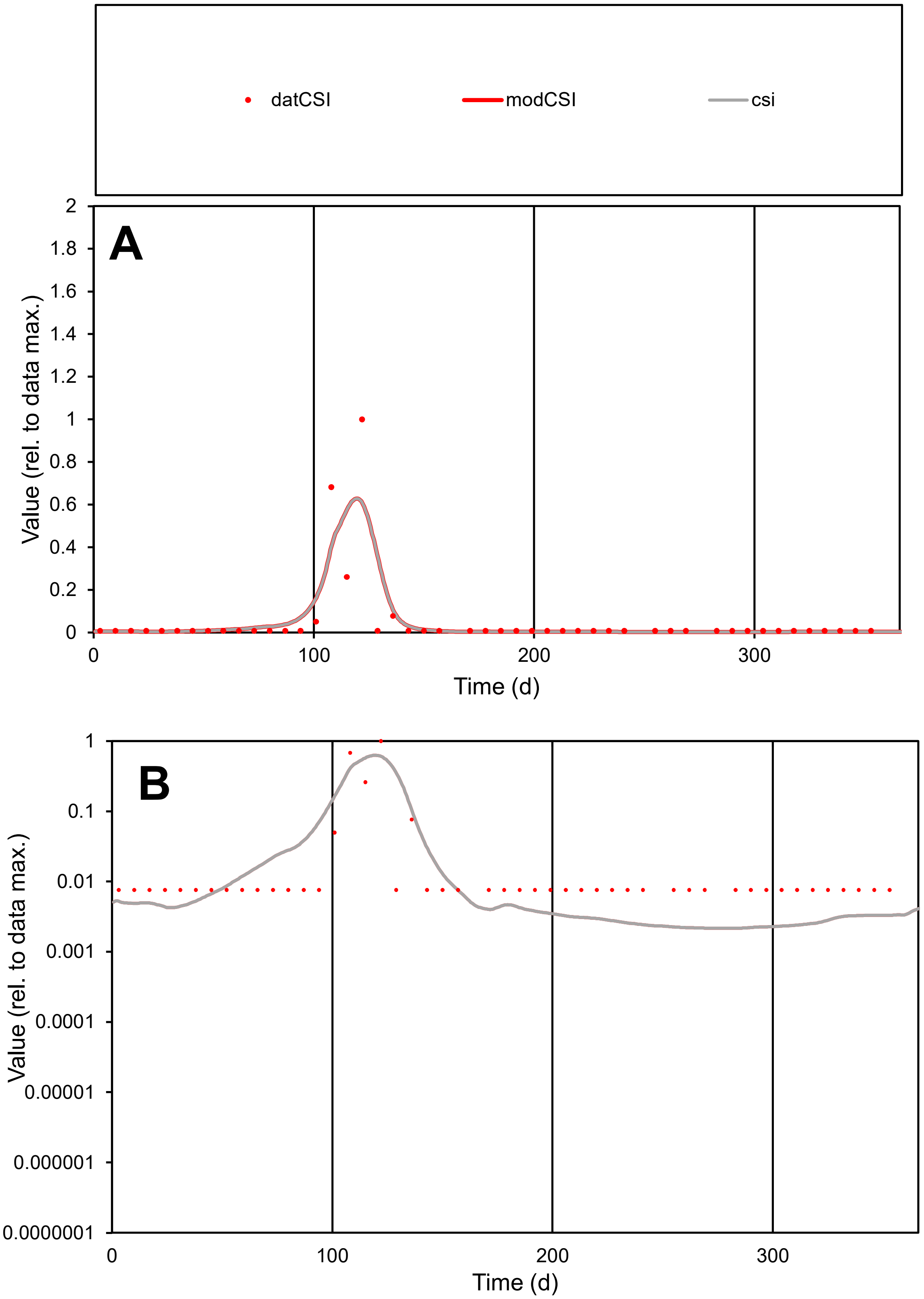


**C1**

Variables
CSI Provide jtd or Dst
CalibOnly?
2 1 to reproduce of
oNormalize?
2 1=no, 2=yes

ofxn calc*
1.77867
ofxn mod
1.91006

sRun
RunA1
zout
1

Ave
Data
1.5E-05
Model
1.5E-05

data/vMax
0.0003

ofxng
369.894

Min d/m
3.50797

ave/min d
6.55285
ave/min m
23.2168

328_Calib_S_A

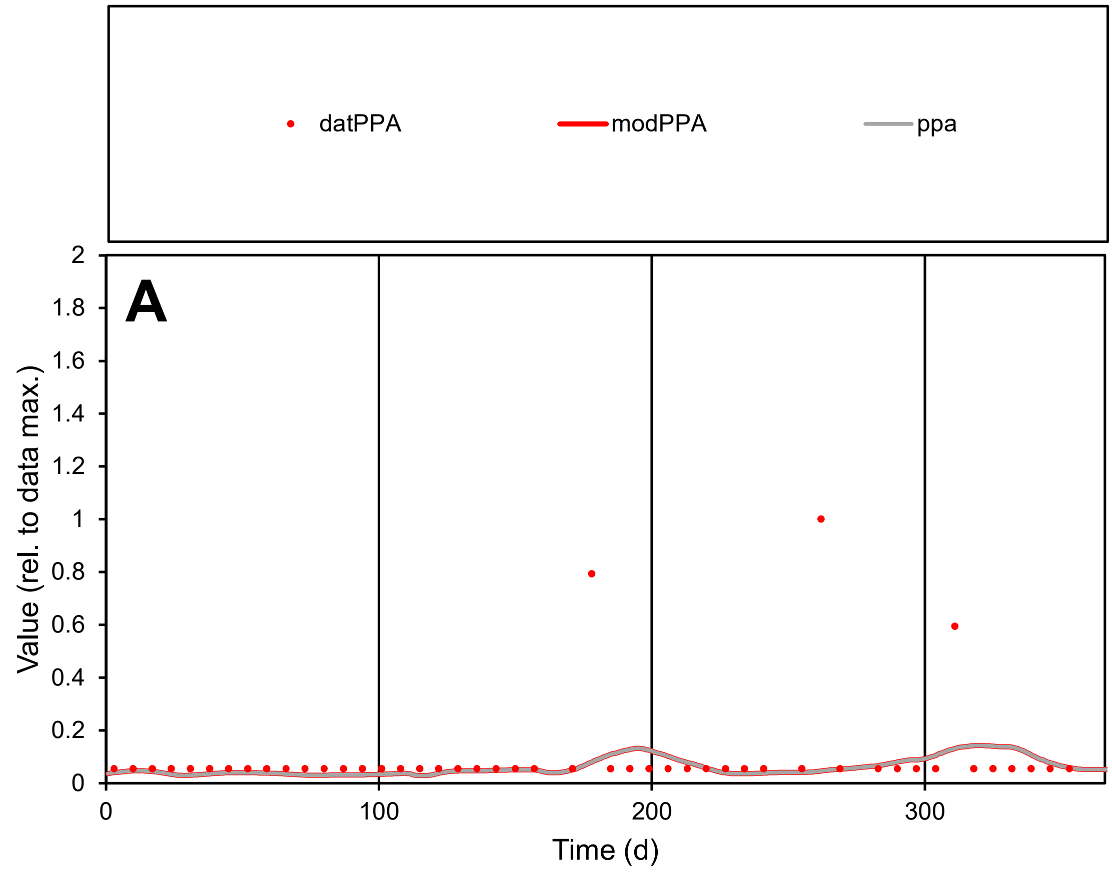


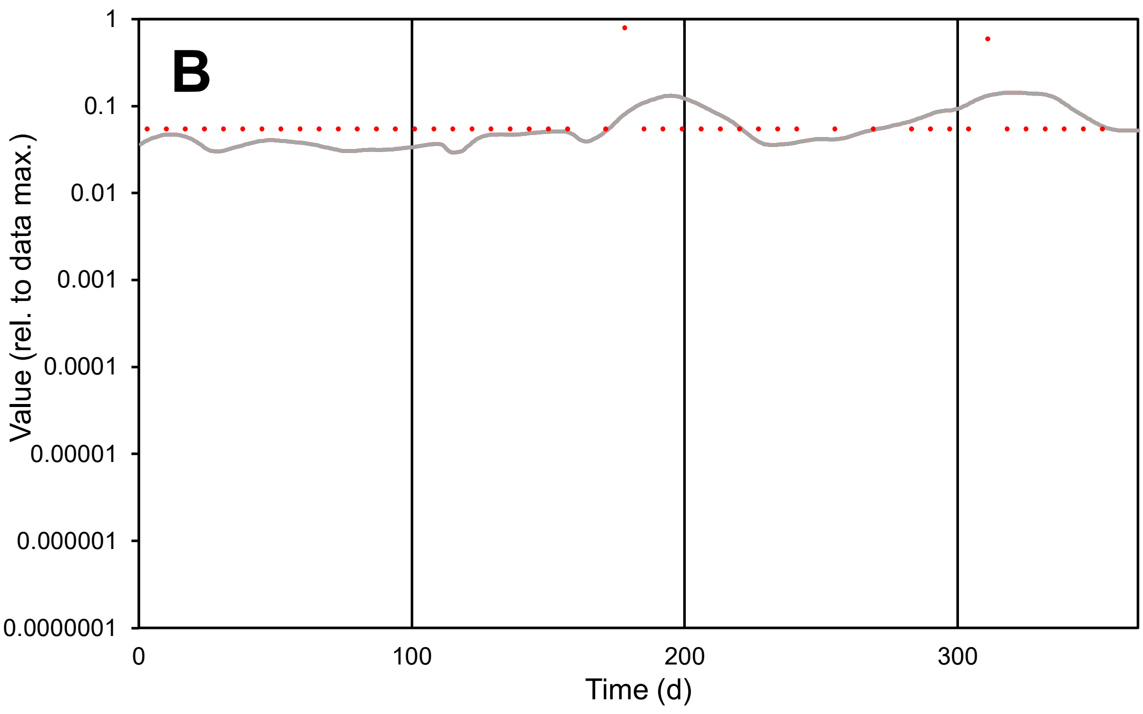


**C1**

Variables
PPA Provide jtd or Dst
CalibOnly?
2 1 to reproduce of
oNormalize?
2 1=no, 2=yes

ofxn calc*
1.84893
ofxn mod
1.85395

sRun
RunA1
zout
1

Ave
Data
5.9E-06
Model
3.7E-06

data/vMax
5.9E-05

ofxng
369.894

Min d/m
1.85393

ave/min d
1.85025
ave/min m
2.12931

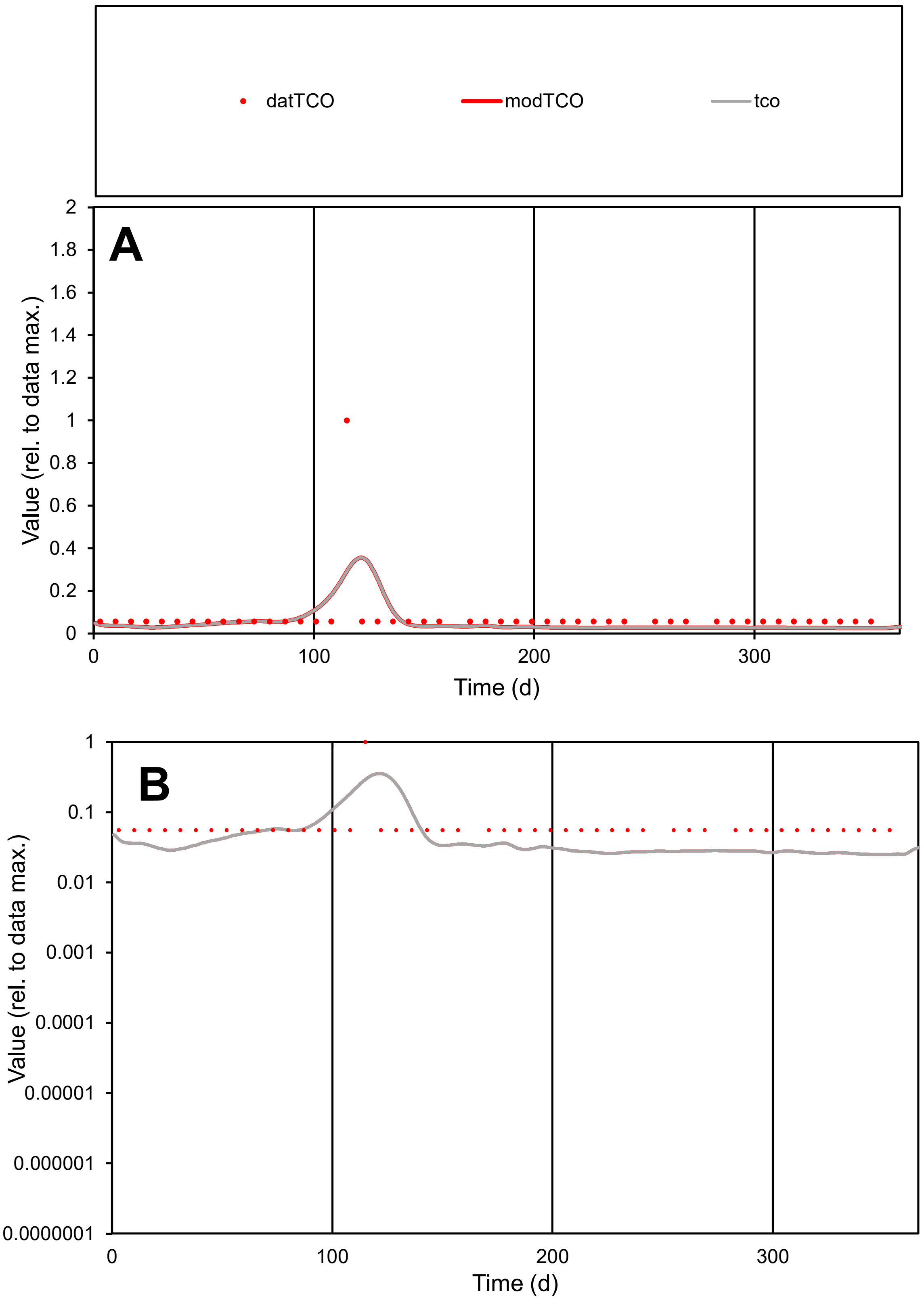


**C1**

Variables
TCO Provide jtd or Dst
CalibOnly?
2 1 to reproduce of
oNormalize?
2 1=no, 2=yes

ofxn calc*
1.50735
ofxn mod
1.5229

sRun
RunA1
zout
1

Ave
Data
1.2E-05
Model
8.7E-06

data/vMax
0.00015

ofxng
369.894

Min d/m
2.22759

ave/min d
1.35317
ave/min m
2.2421

328_Calib_S_A

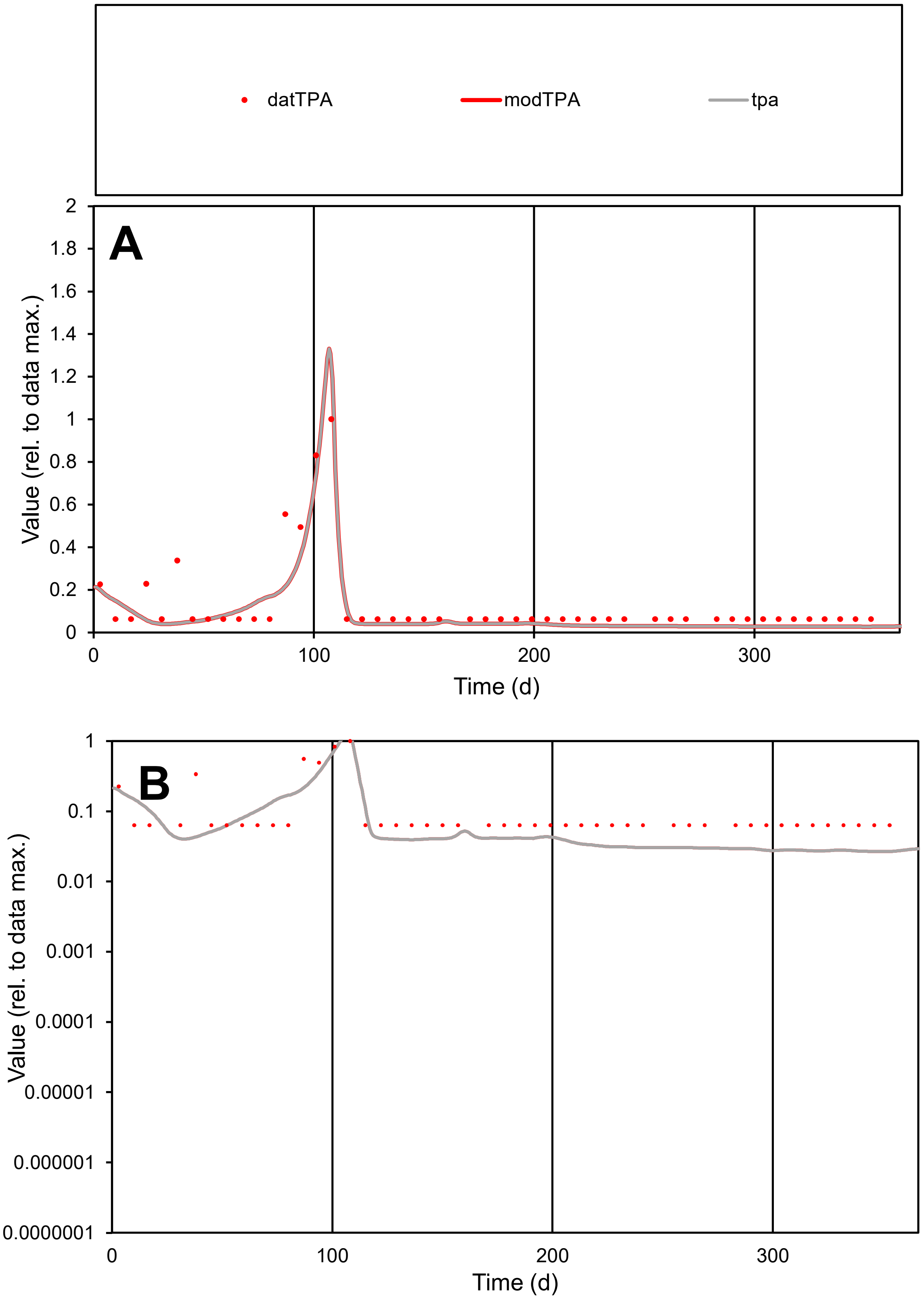


**C1**

Variables

TPA Provide jtd or Dst

CalibOnly?

2 1 to reproduce of

oNormalize?

2 1=no, 2=yes

ofxn calc*

0.59361

ofxn mod

0.6163

sRun

RunA1

zout

1

Ave

Data

9.8E-06

Model

7.5E-06

data/vMax

7.5E-05

ofxng

369.894

Min d/m

2.35494

ave/min d

2.06545

ave/min m

3.73382

328_Calib_S_A

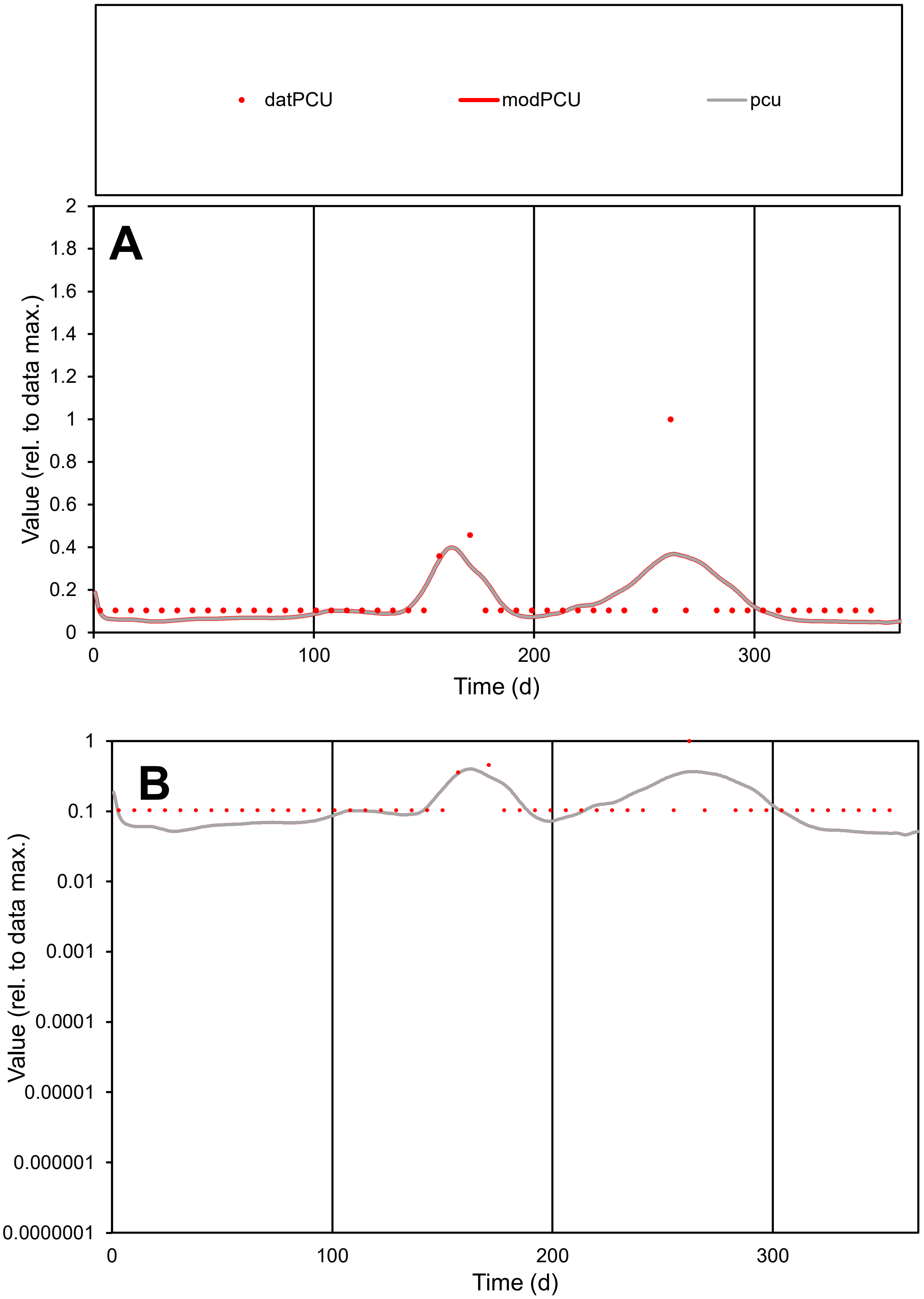


**C1**

Variables
PCU Provide jtd or Dst
CalibOnly?
2 1 to reproduce of
oNormalize?
2 1=no, 2=yes

ofxn calc*
0.85997
ofxn mod
0.8734

sRun
RunA1
zout
1

Ave
Data
8.1E-06
Model
7.4E-06

data/vMax
6E-05

ofxng
369.894

Min d/m
2.12999

ave/min d
1.3017
ave/min m
2.53551

328_Calib_S_A

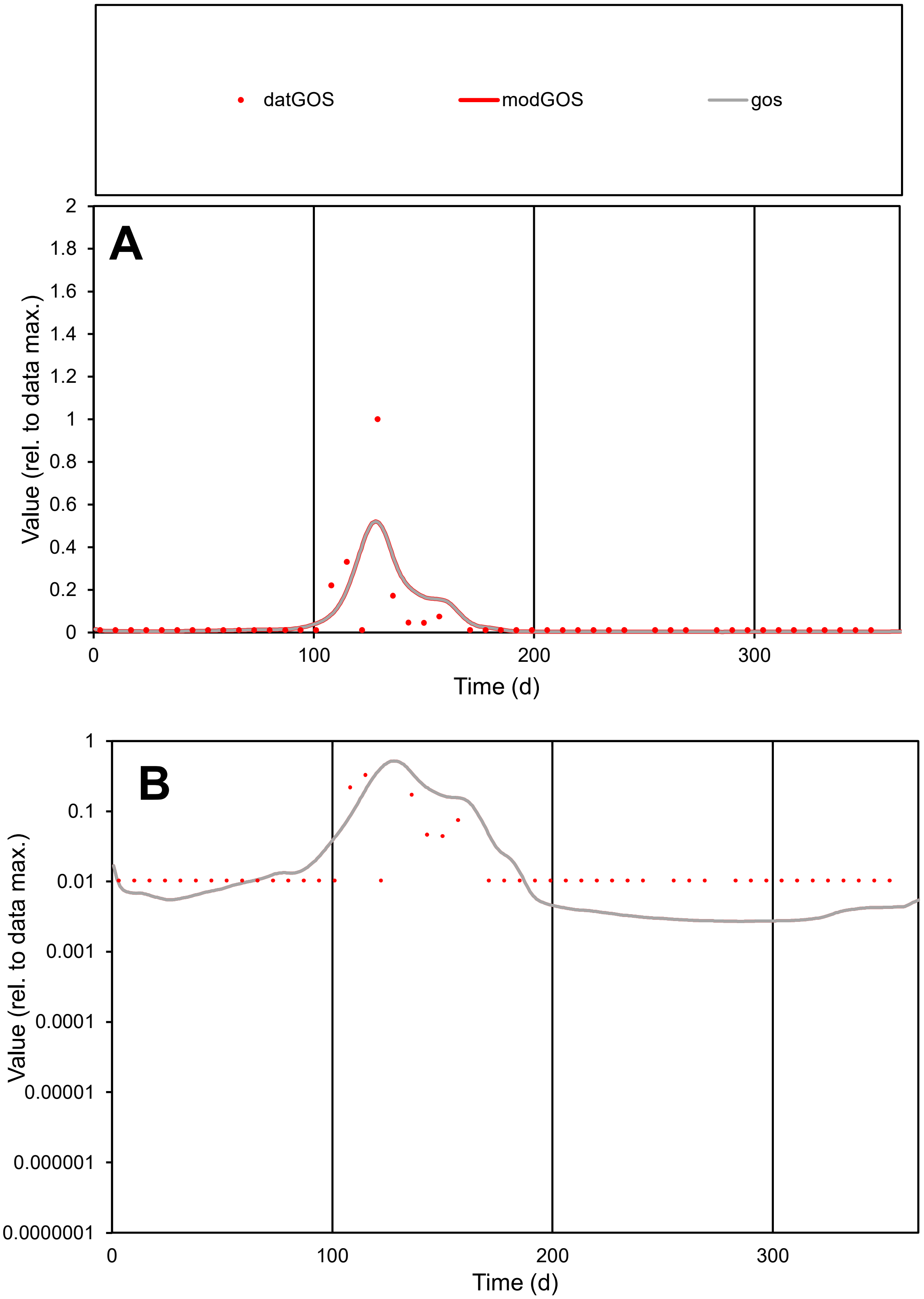


C1

Variables
GOS Provide jtd or Dst
CalibOnly?
2 1 to reproduce of
oNormalize?
2 1=no, 2=yes

ofxn calc*
1.99044
ofxn mod
2.08928

sRun
RunA1
zout
1

Ave
Data
8.4E-06
Model
8.6E-06

data/vMax
0.00017

ofxng
369.894

Min d/m
3.83898

ave/min d
4.64688
ave/min m
18.1451

328_Calib_S_A

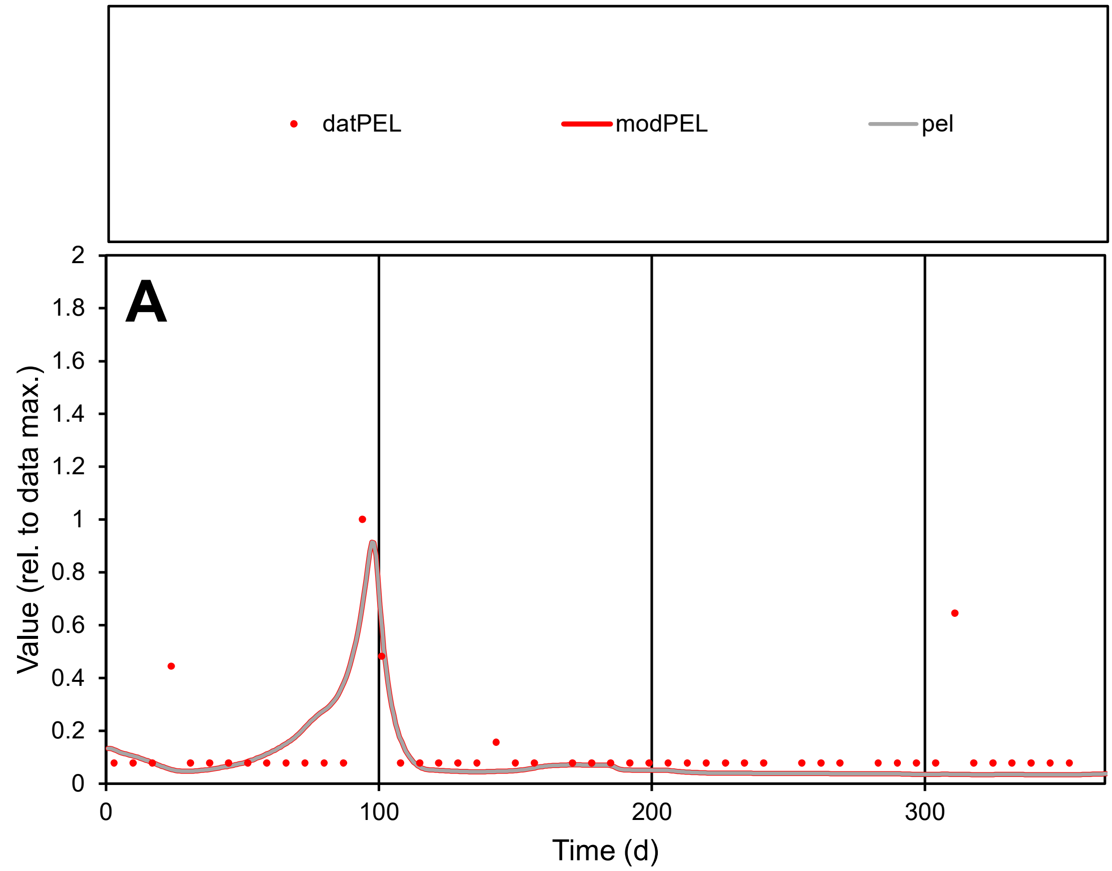


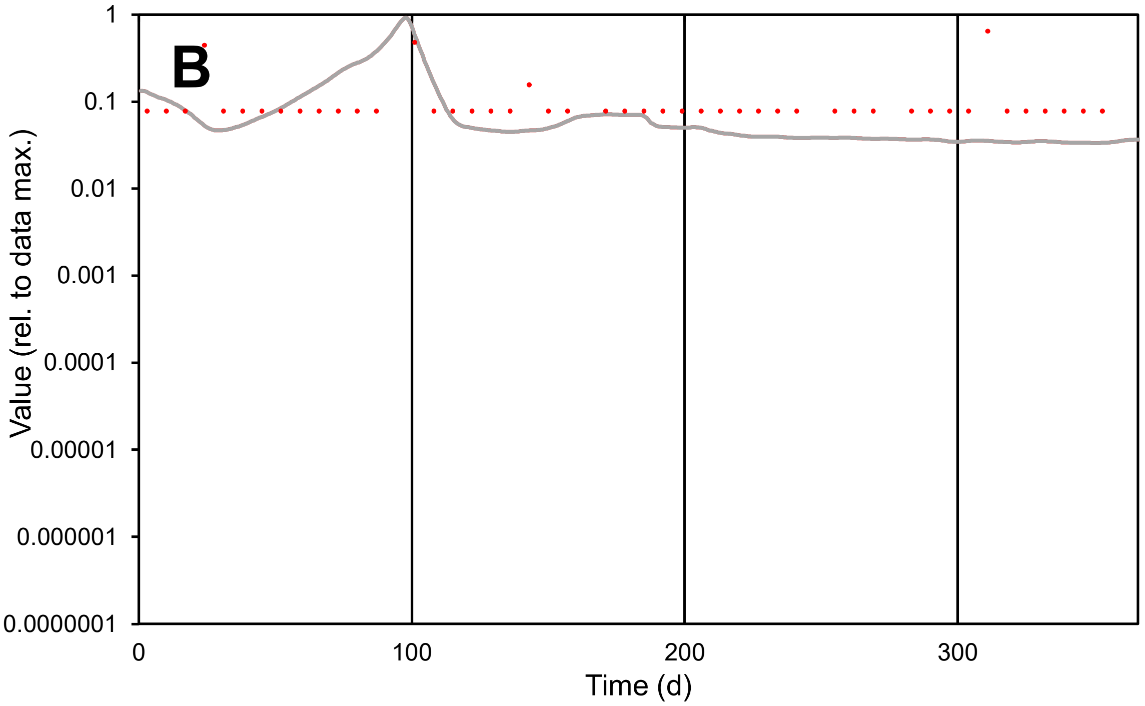


**C1**

Variables
PEL Provide jtd or Dst
CalibOnly?
2 1 to reproduce of
oNormalize?
2 1=no, 2=yes

ofxn calc*
1.01044
ofxn mod
1.03177

sRun
RunA1
zout
1

Ave
Data
5.3E-06
Model
3.9E-06

data/vMax
4.2E-05

ofxng
369.894

Min d/m
2.3178

ave/min d
1.62445
ave/min m
2.77629

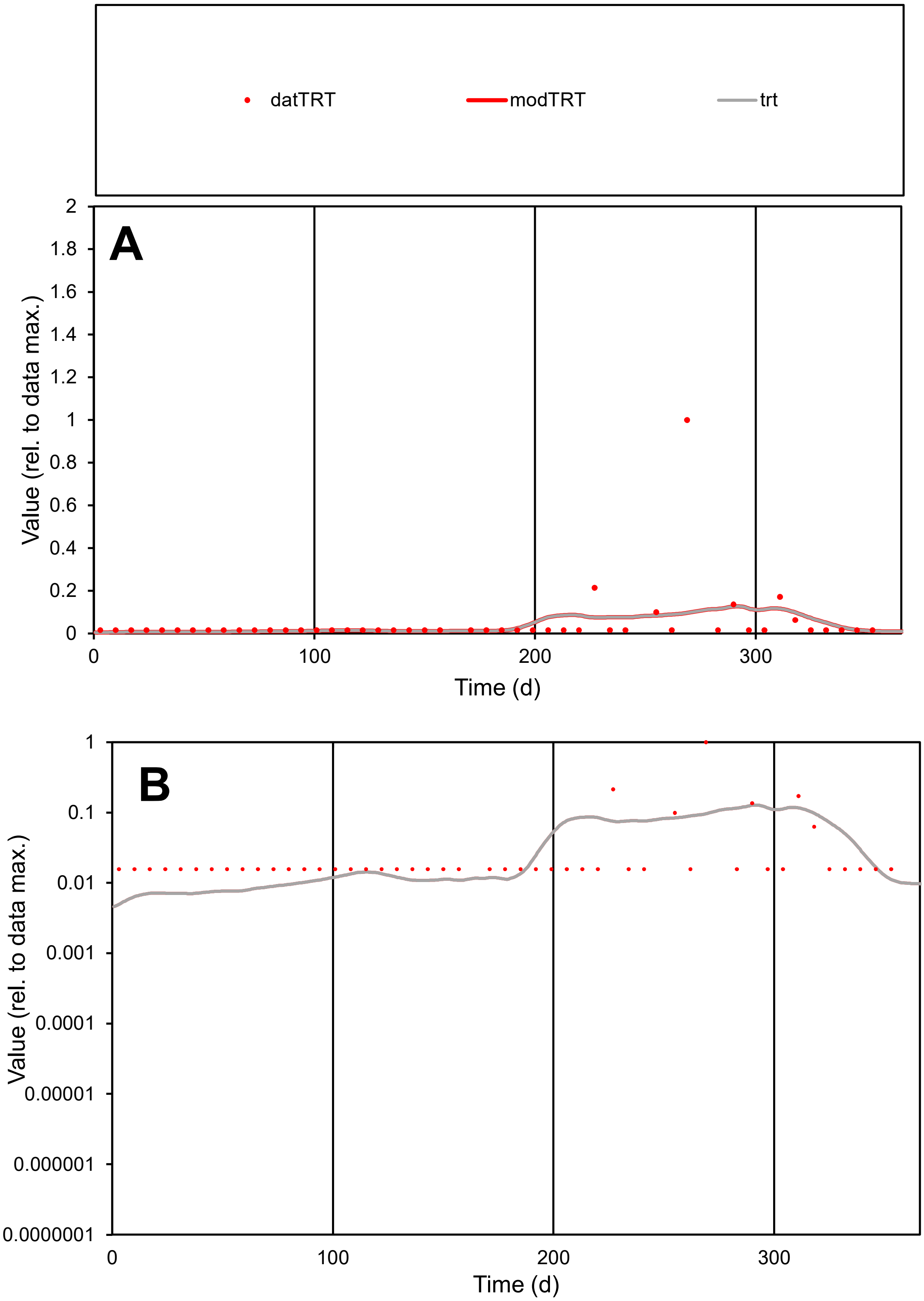


**C1**

Variables
TRT Provide jtd or Dst
CalibOnly?
2 1 to reproduce of
oNormalize?
2 1=no, 2=yes

ofxn calc*
2.80449
ofxn mod
2.81421

sRun
RunA1
zout
1

Ave
Data
7.2E-05
Model
5.9E-05

data/vMax
0.00148

ofxng
369.894

Min d/m
2.99354
ave/min d
3.11152
ave/min m
7.65396

328_Calib_S_A

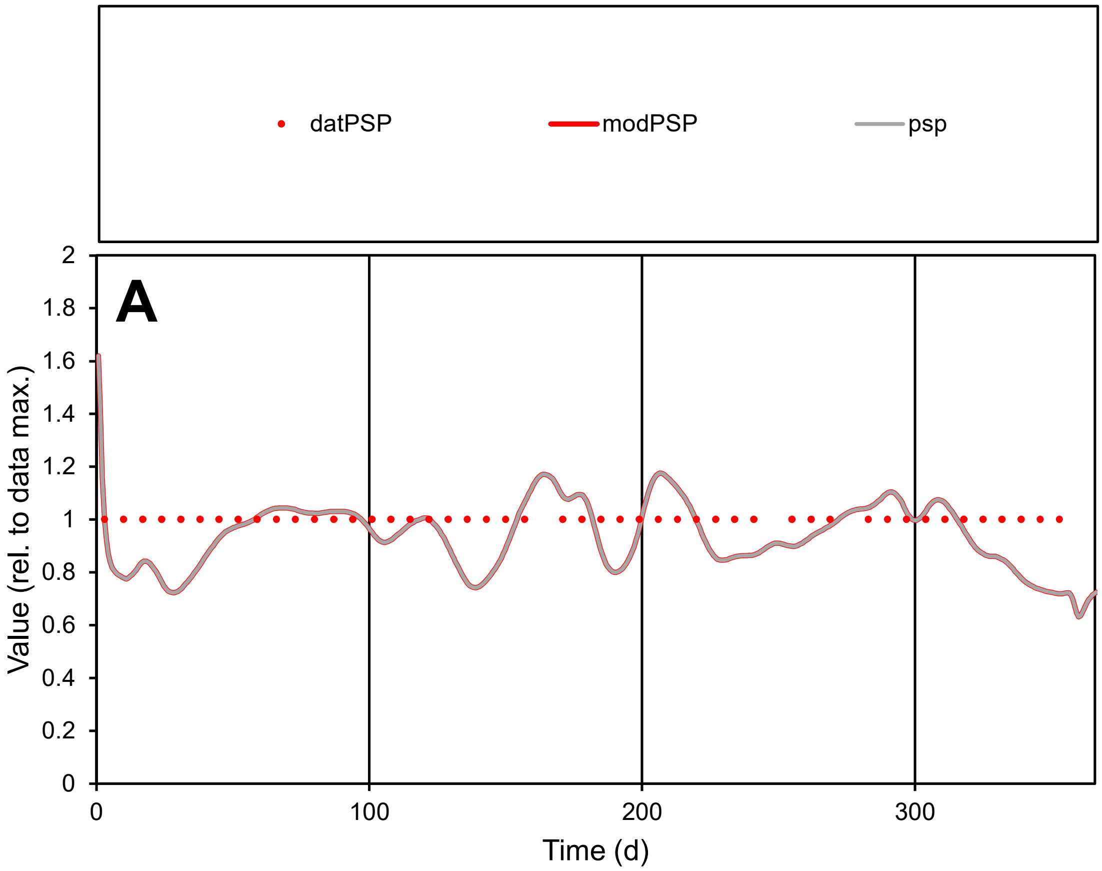


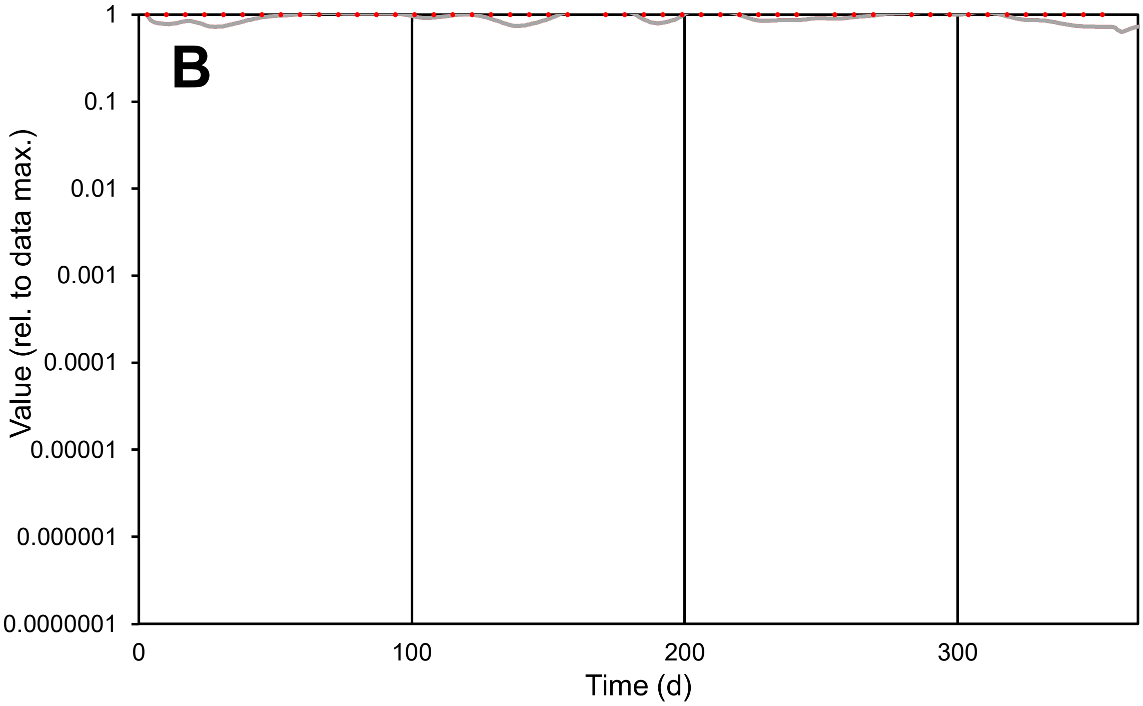


**C1**

Variables

PSP Provide jtd or Dst

CalibOnly?

2 1 to reproduce of

oNormalize?

2 1=no, 2=yes

ofxn calc*

0.1324

ofxn mod

0.17176

sRun

RunA1

zout

1

Ave

Data

4.3E-06

Model

4E-06

data/vMax

4.3E-06

ofxng

369.894

Min d/m

1.38964

ave/min d

1

ave/min m

1.29976

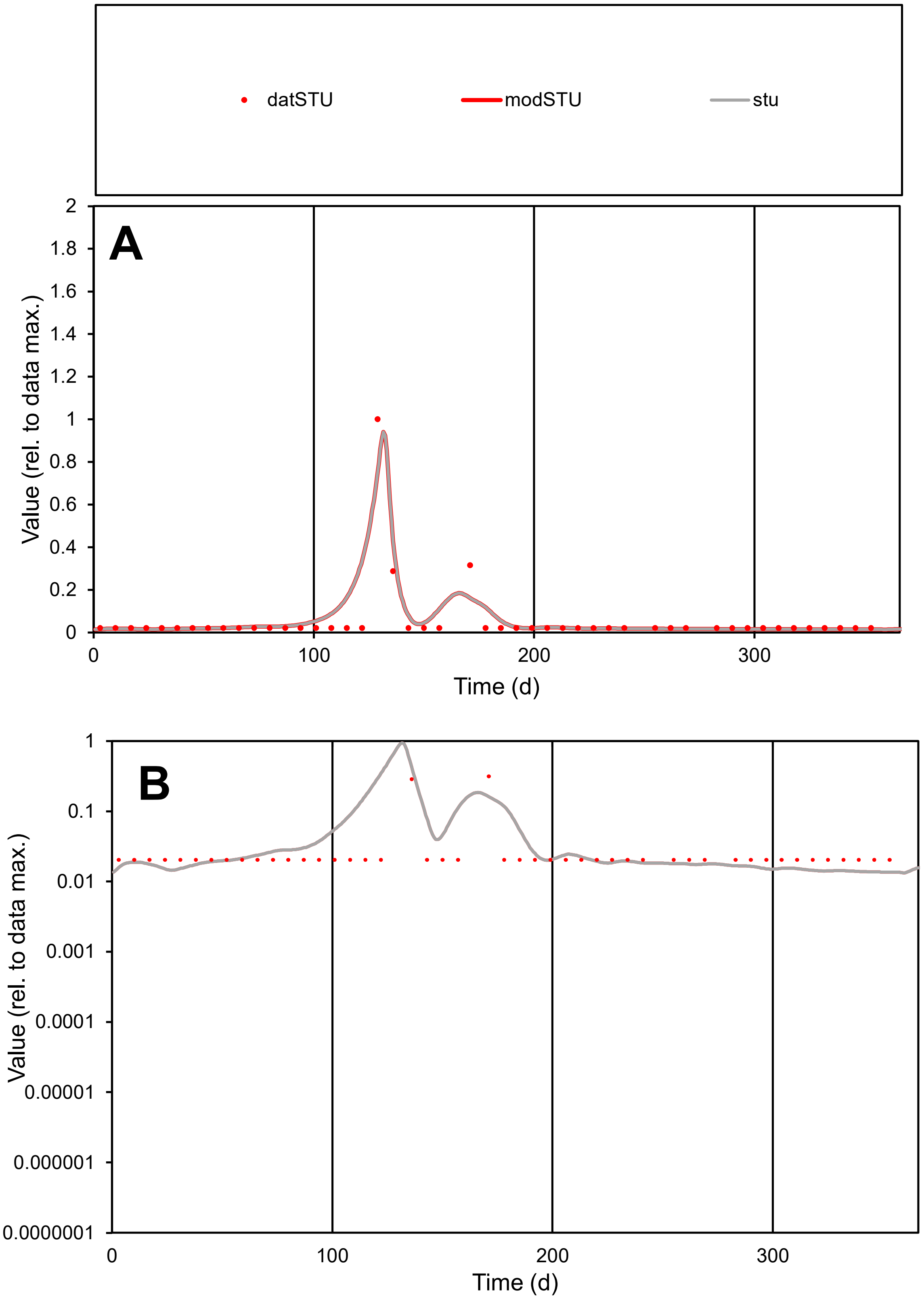


**C1**

Variables
STU Provide jtd or Dst
CalibOnly?
2 1 to reproduce of
oNormalize?
2 1=no, 2=yes

ofxn calc*
1.05787
ofxn mod
1.18627

sRun
RunA1
zout
1

Ave
Data
6.4E-06
Model
7.4E-06

data/vMax
0.00012

ofxng
369.894

Min d/m
1.5009

ave/min d
2.57751
ave/min m
4.46508

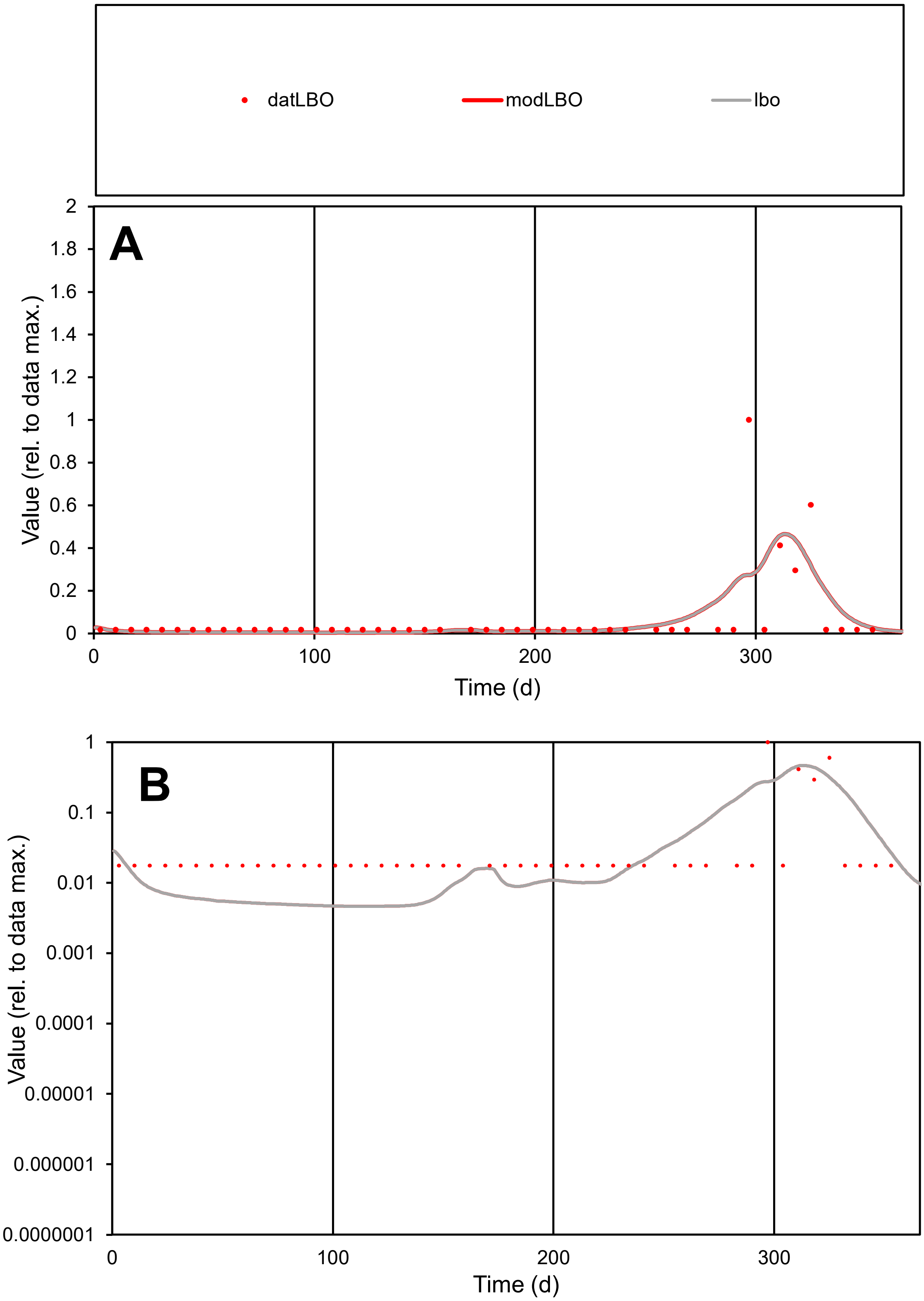


**C1**

Variables
LBO Provide jtd or Dst
CalibOnly?
2 1 to reproduce of
oNormalize?
2 1=no, 2=yes

ofxn calc*
2.02783
ofxn mod
2.06845

sRun
RunA1
zout
1

Ave
Data
5.1E-06
Model
4.8E-06

data/vMax
7.9E-05

ofxng
369.894

Min d/m
3.7721

ave/min d
3.65174
ave/min m
13.1872

328_Calib_S_A

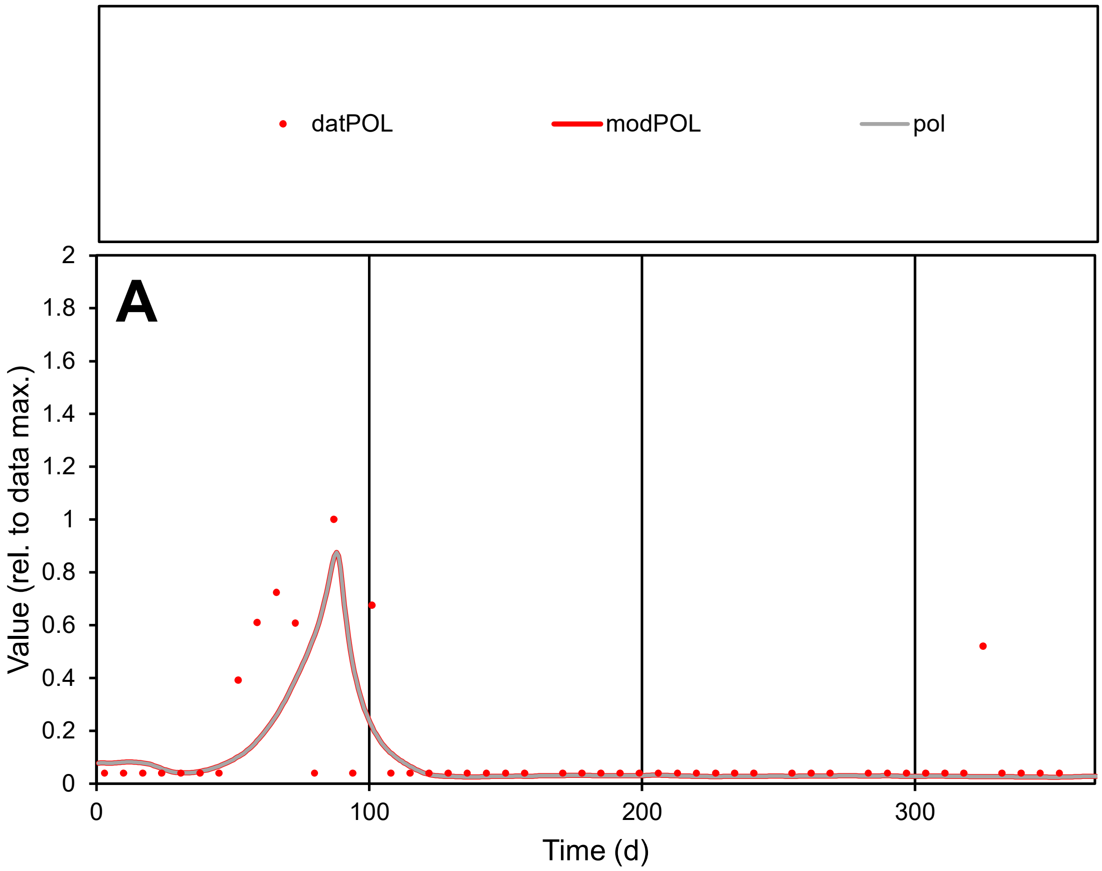


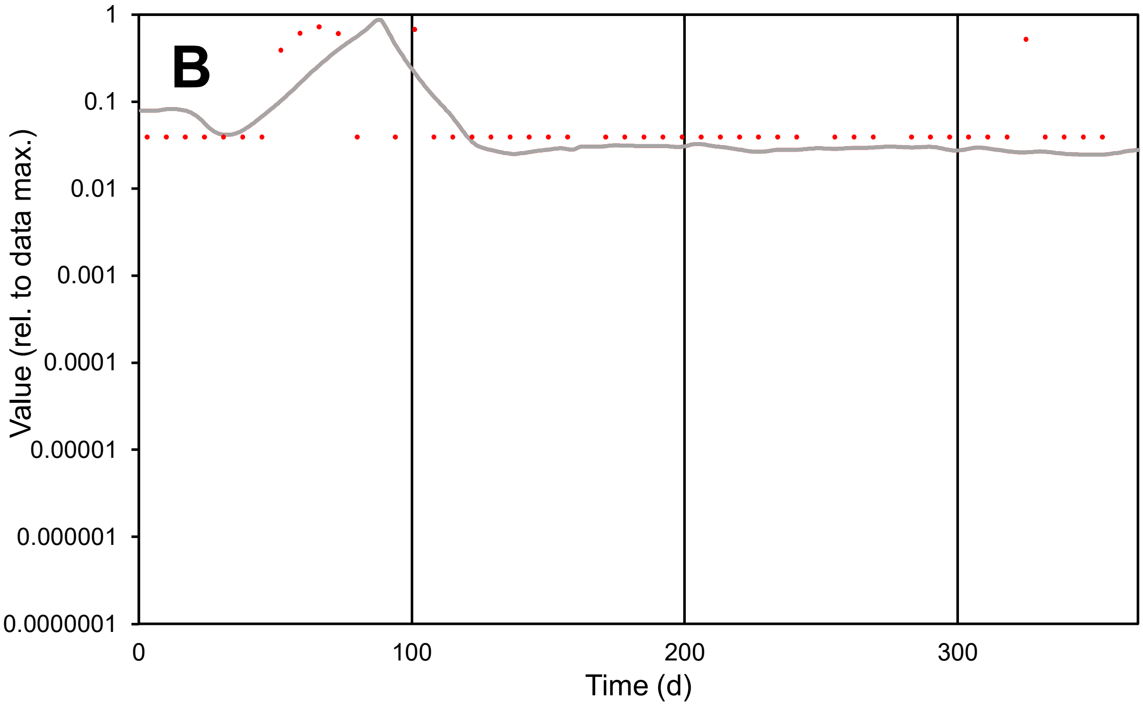


**C1**

Variables
POL Provide jtd or Dst
CalibOnly?
2 1 to reproduce of
oNormalize?
2 1=no, 2=yes

ofxn calc*
1.28876
ofxn mod
1.32446

sRun
RunA1
zout
1

Ave
Data
5.2E-06
Model
3.8E-06

data/vMax
4.1E-05

ofxng
369.894

Min d/m
1.58964

ave/min d
3.26136
ave/min m
3.73616

328_Calib_S_A

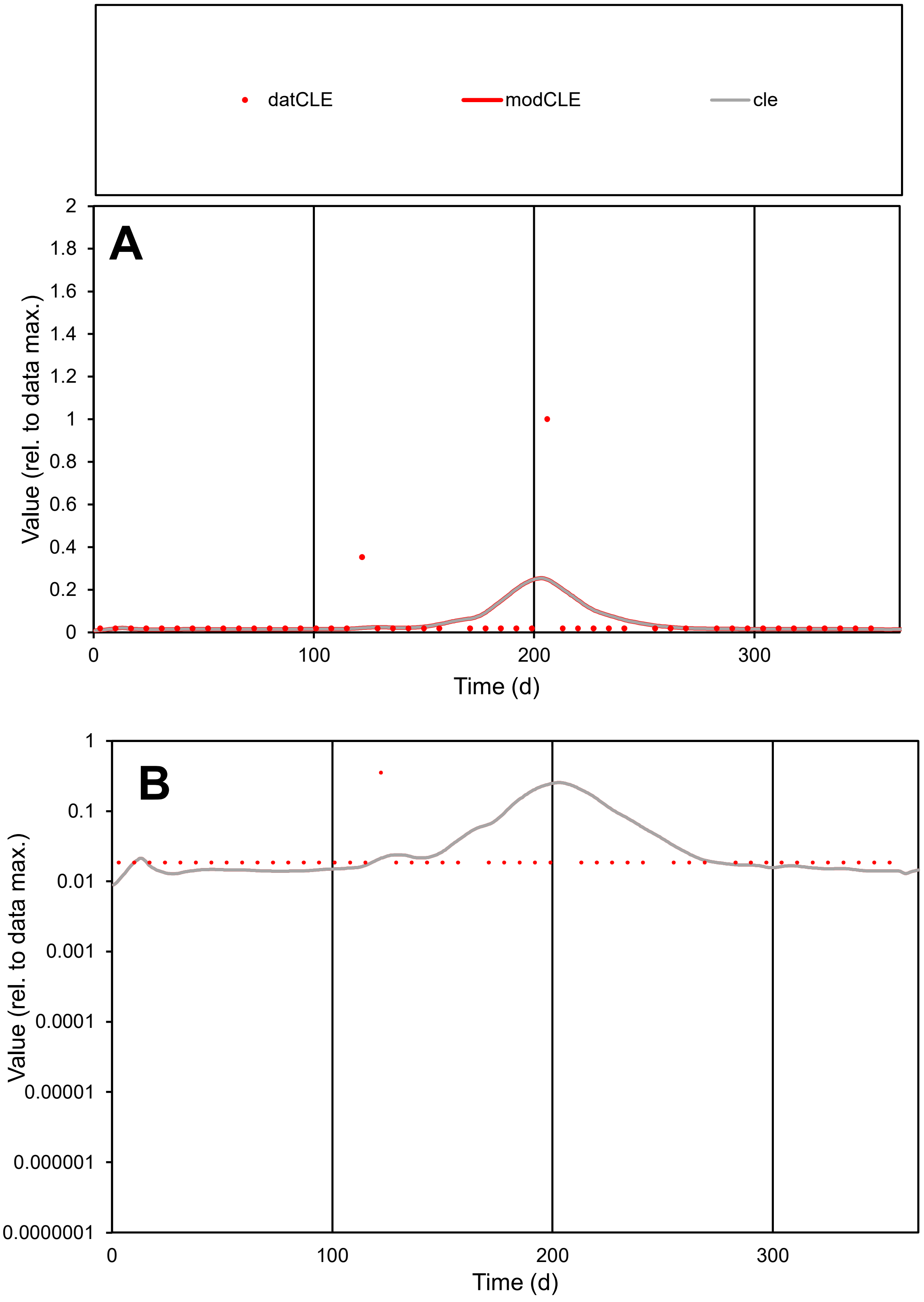


**C1**

Variables

CLE Provide jtd or Dst

CalibOnly?

2 1 to reproduce of

oNormalize?

2 1=no, 2=yes

ofxn calc*

2.88355

ofxn mod

2.90881

sRun

RunA1

zout

1

Ave

Data

1.2E-05

Model

1.2E-05

data/vMax

0.00025

ofxng

369.894

Min d/m

1.60767

ave/min d

2.4754

ave/min m

3.9392

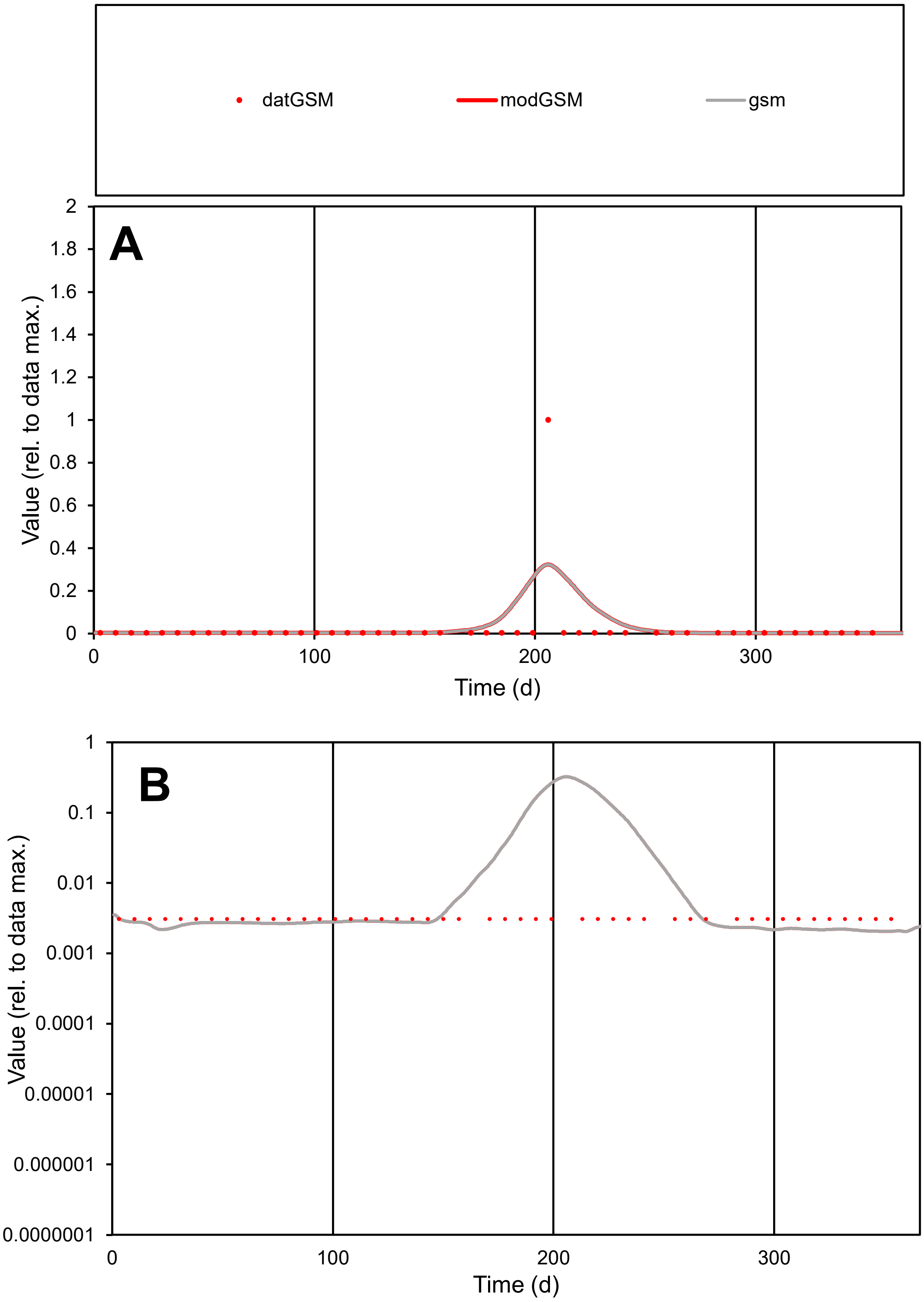


**C1**

Variables
GSM Provide jtd or Dst
CalibOnly?
2 1 to reproduce of
oNormalize?
2 1=no, 2=yes

ofxn calc*
4.82436
ofxn mod
4.93742

sRun
RunA1
zout
1

Ave
Data
1.2E-05
Model
1.6E-05

data/vMax
0.00051

ofxng
369.894

Min d/m
1.49385

ave/min d
7.73582
ave/min m
15.1842

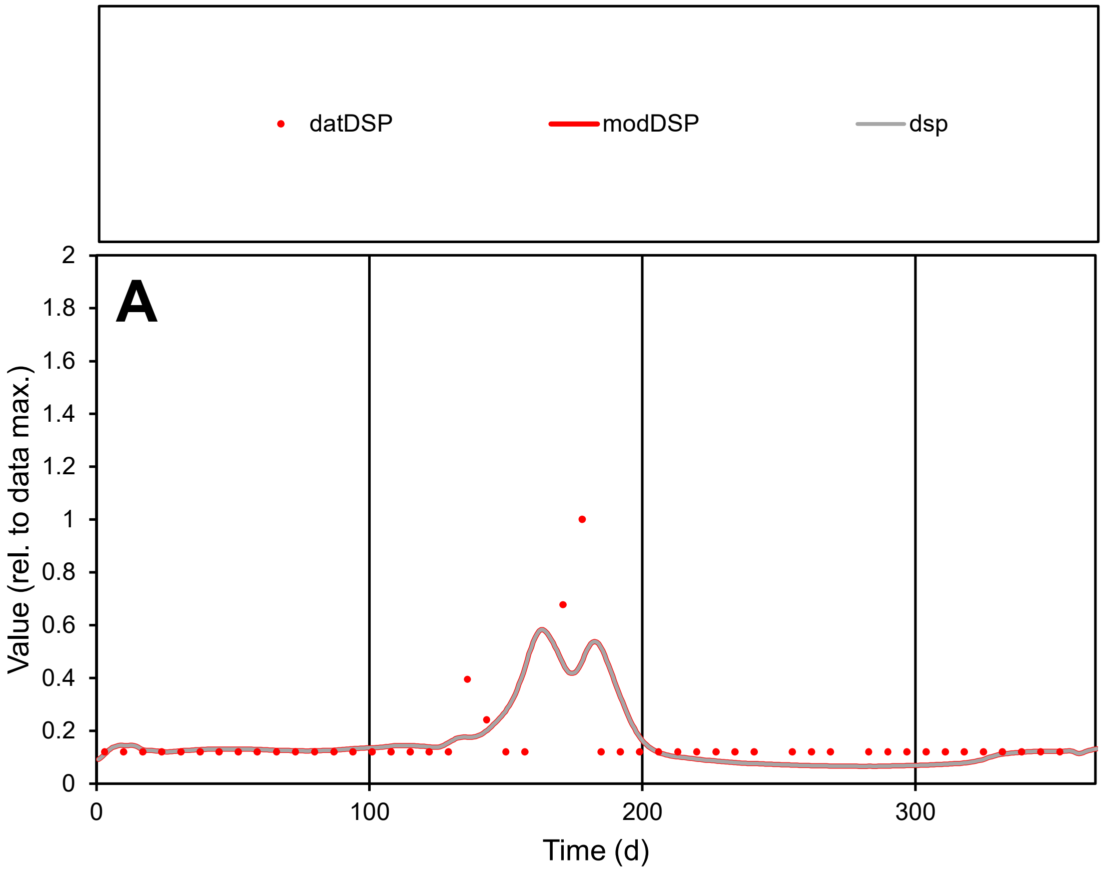


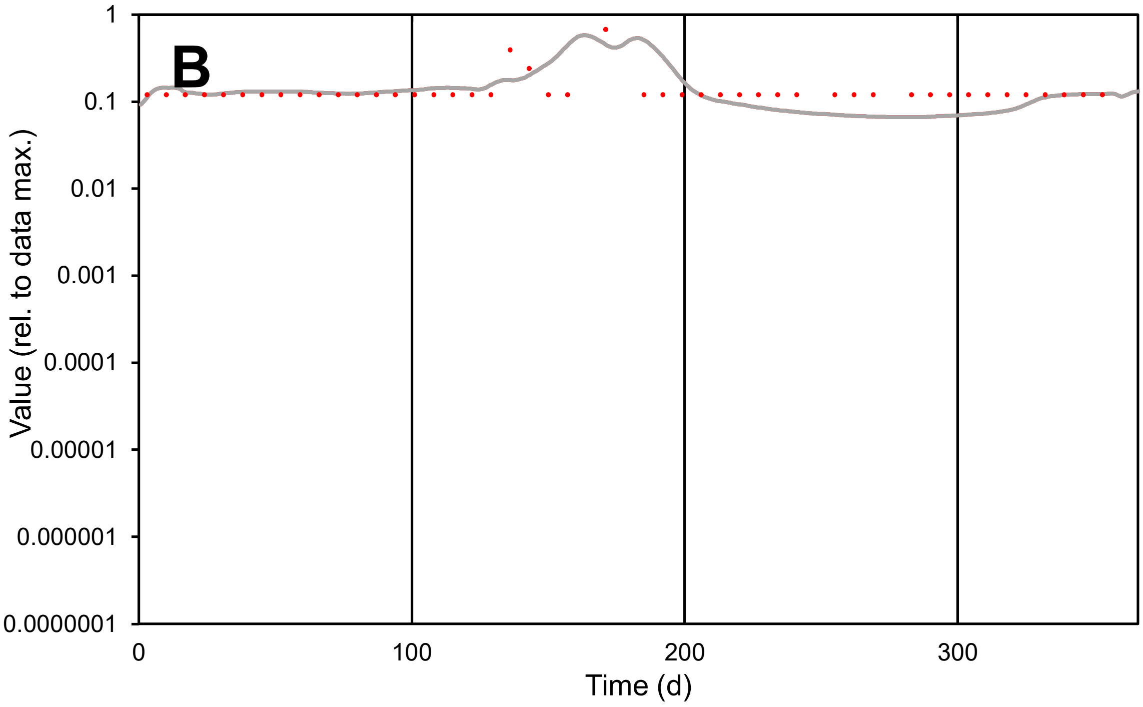


**C1**

Variables
DSP Provide jtd or Dst
CalibOnly?
2 1 to reproduce of
oNormalize?
2 1=no, 2=yes

ofxn calc*
0.72666
ofxn mod
0.76354

sRun
RunA1
zout
1

Ave
Data
6.6E-06
Model
6.2E-06

data/vMax
4.2E-05

ofxng
369.894

Min d/m
1.81472

ave/min d
1.3166
ave/min m
2.2603

328_Calib_S_A

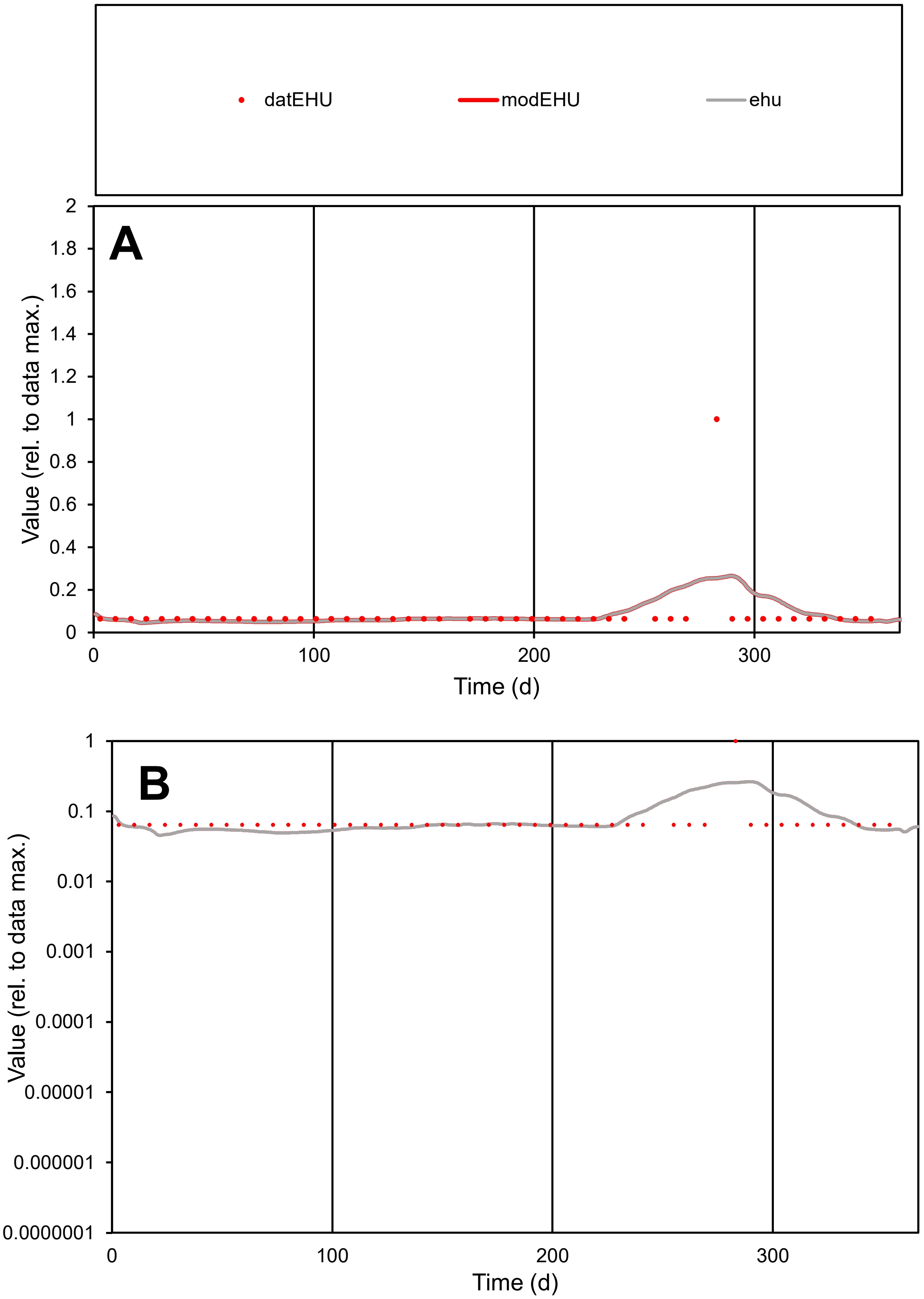


**C1**

Variables
EHU Provide jtd or Dst
CalibOnly?
2 1 to reproduce of
oNormalize?
2 1=no, 2=yes

ofxn calc*
1.42577
ofxn mod
1.43767

sRun
RunA1
zout
1

Ave
Data
7.5E-06
Model
7.6E-06

data/vMax
9E-05

ofxng
369.894

Min d/m
1.36504

ave/min d
1.30349
ave/min m
1.80464

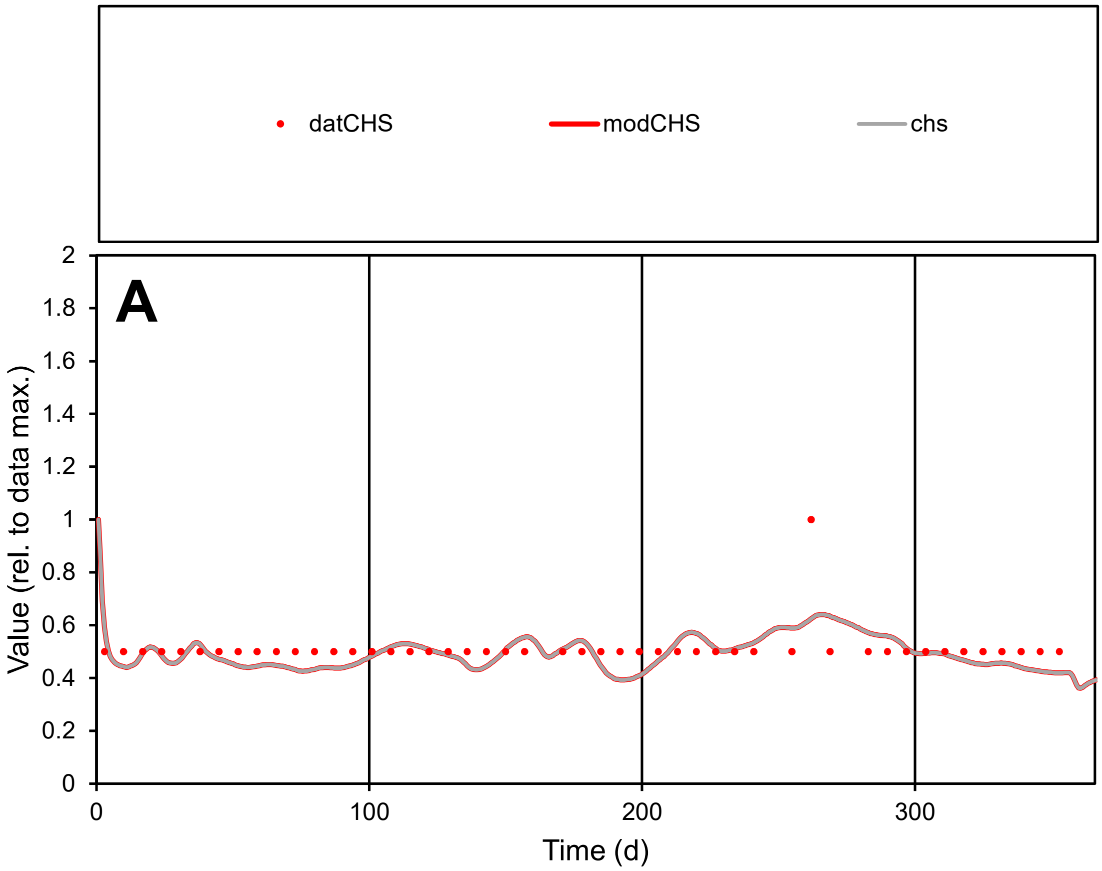


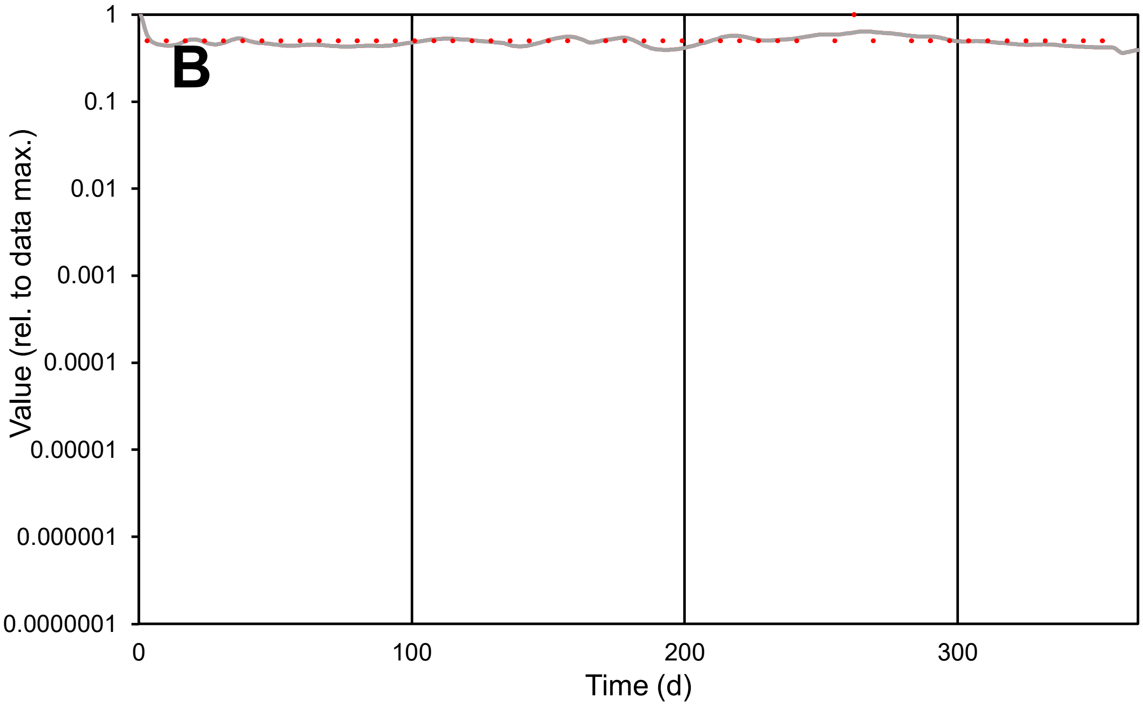


**C1**

Variables
CHS Provide jtd or Dst
CalibOnly?
2 1 to reproduce of
oNormalize?
2 1=no, 2=yes

ofxn calc*
0.14504
ofxn mod
0.18294

sRun
RunA1
zout
1

Ave
Data
2.2E-05
Model
2.1E-05

data/vMax
4.3E-05

ofxng
369.894

Min d/m
1.27246

ave/min d
1.02084
ave/min m
1.24772

328_Calib_S_A

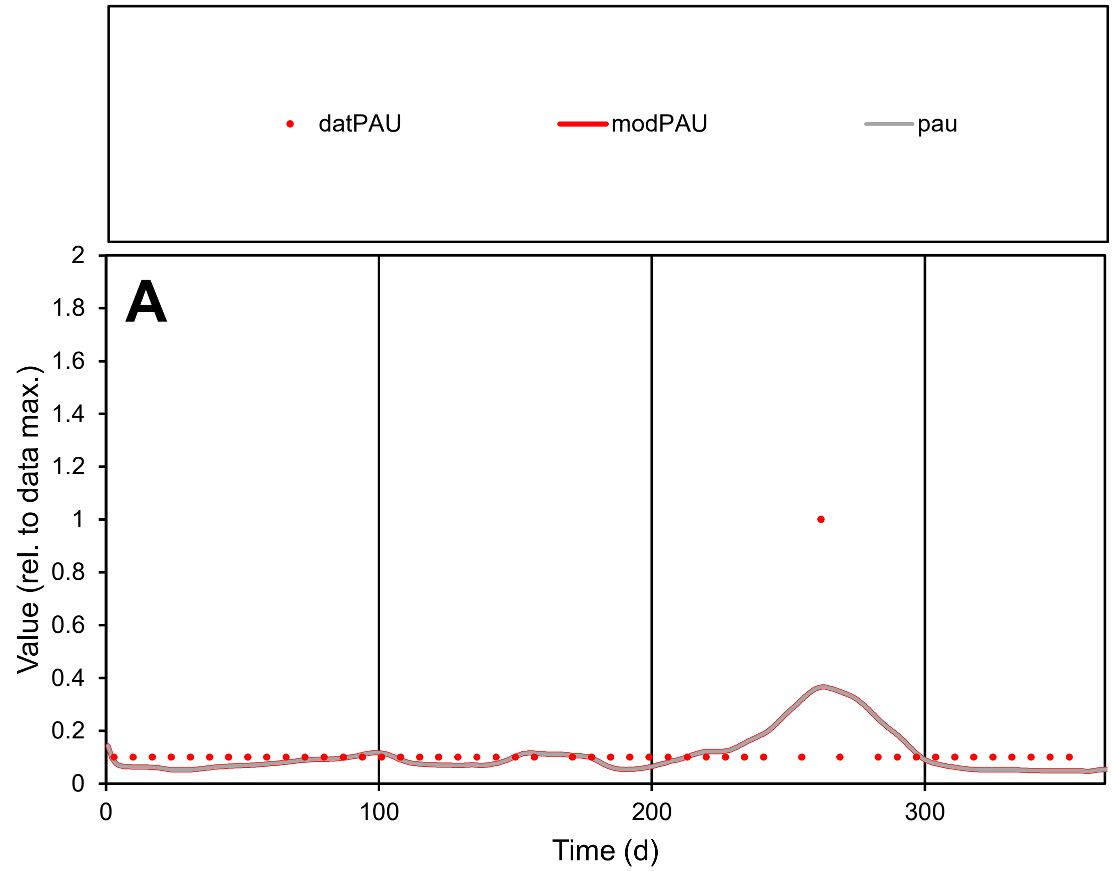


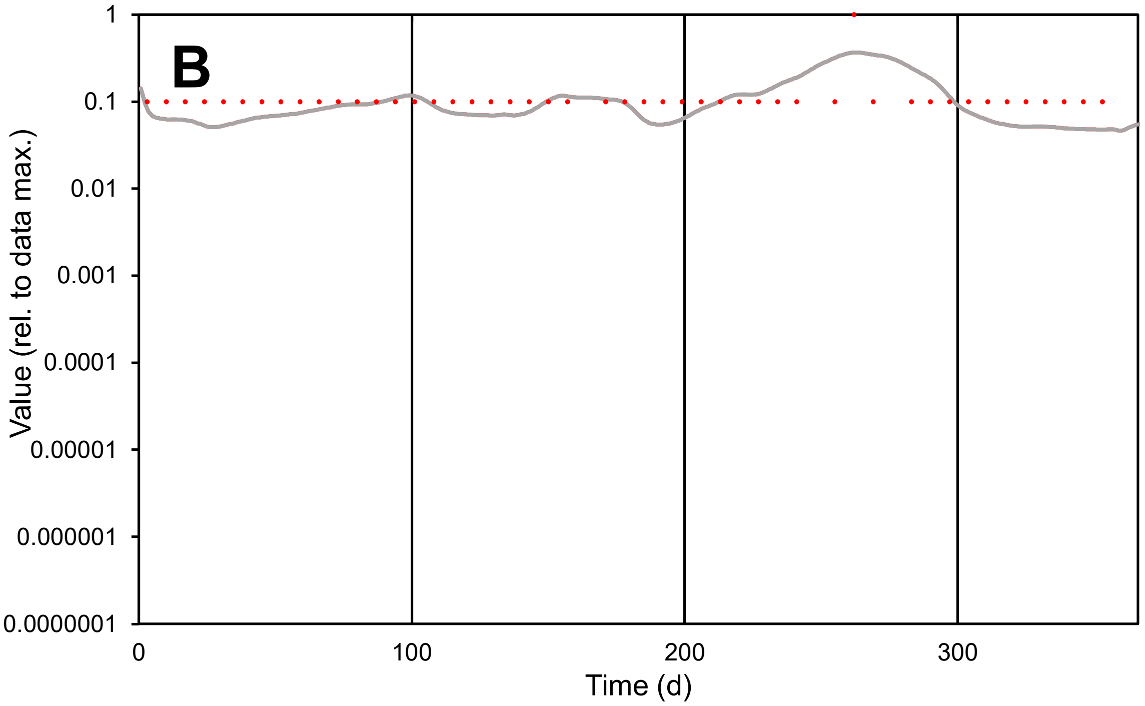


**C1**

Variables
PAU Provide jtd or Dst
CalibOnly?
2 1 to reproduce of
oNormalize?
2 1=no, 2=yes

ofxn calc*
0.92806
ofxn mod
0.93844

sRun
RunA1
zout
1

Ave
Data
4.7E-06
Model
4.1E-06

data/vMax
3.9E-05

ofxng
369.894

Min d/m
2.08514

ave/min d
1.18739
ave/min m
2.15778

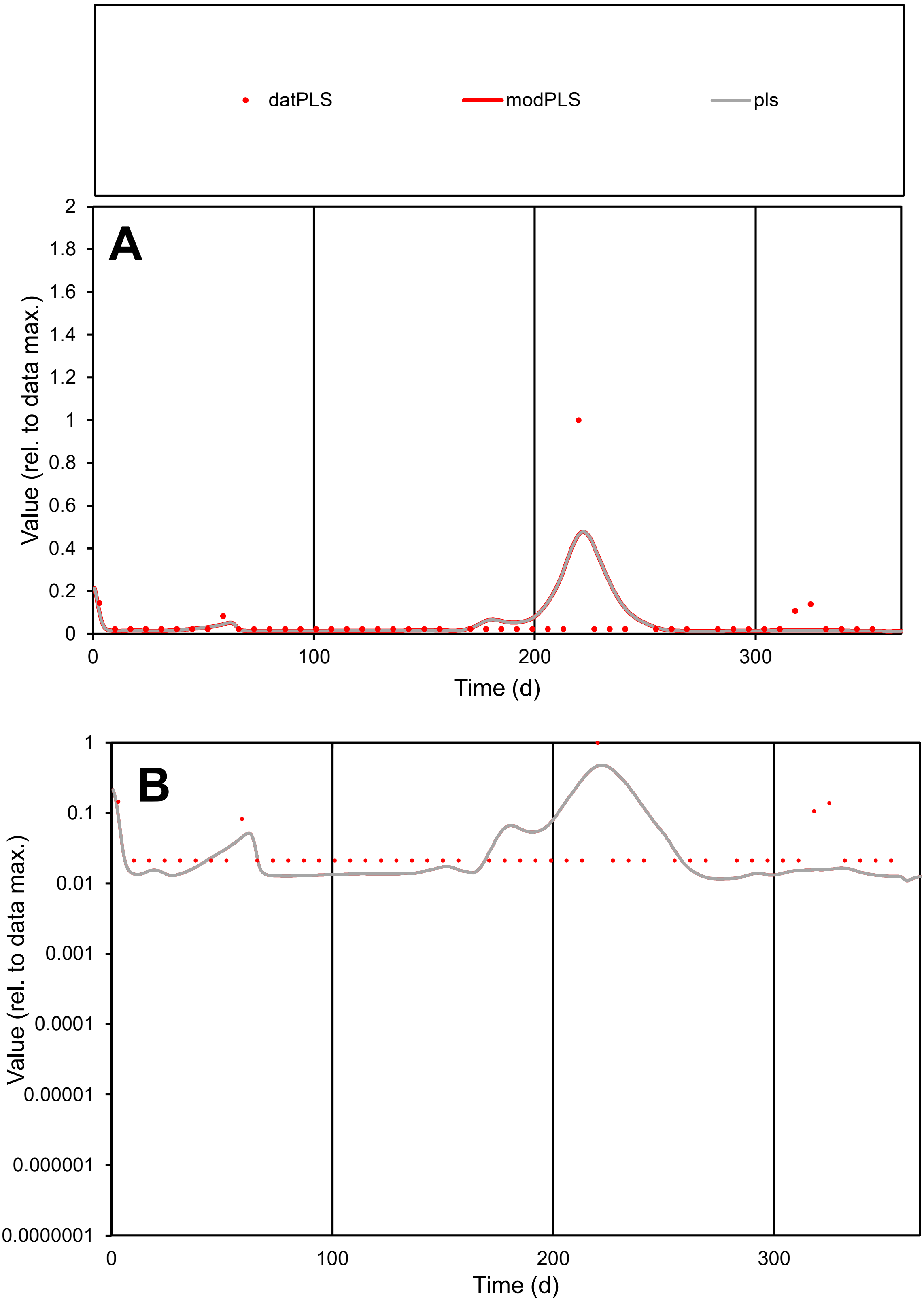


**C1**

Variables
PLS Provide jtd or Dst
CalibOnly?
2 1 to reproduce of
oNormalize?
2 1=no, 2=yes

ofxn calc*
2.05544
ofxn mod
2.15317

sRun
RunA1
zout
1

Ave
Data
9E-06
Model
9.3E-06

data/vMax
0.00018

ofxng
369.894

Min d/m
1.78984

ave/min d
2.33817
ave/min m
4.3507

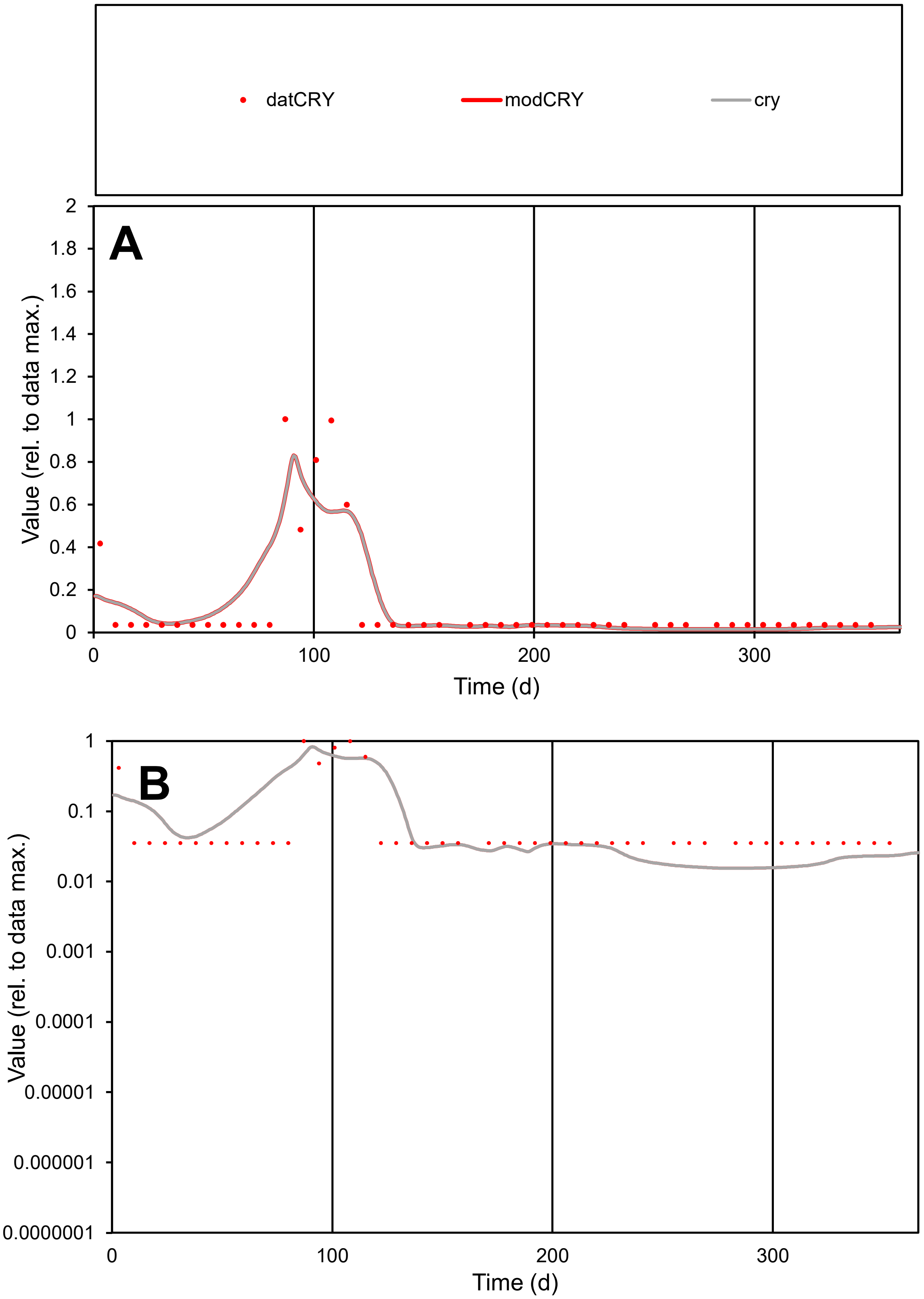


**C1**

Variables
CRY Provide jtd or Dst
CalibOnly?
2 1 to reproduce of
oNormalize?
2 1=no, 2=yes

ofxn calc*
1.04692
ofxn mod
1.09796

sRun
RunA1
zout
1

Ave
Data
1E-05
Model
1E-05

data/vMax
8.5E-05

ofxng
369.894

Min d/m
2.29516

ave/min d
3.41498
ave/min m
8.00796

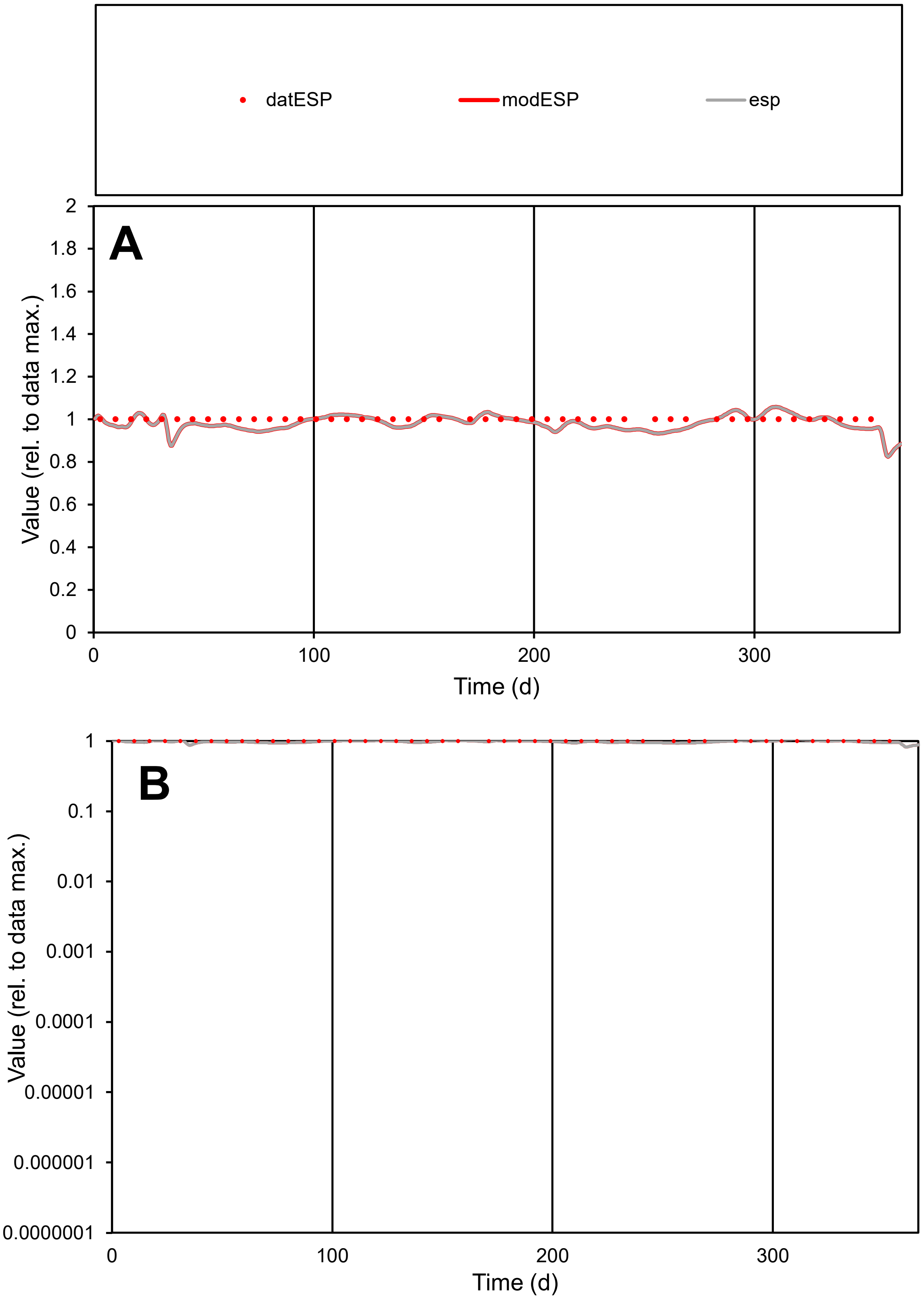


**C1**

Variables
ESP Provide jtd or Dst
CalibOnly?
2 1 to reproduce of
oNormalize?
2 1=no, 2=yes

ofxn calc*
0.0319
ofxn mod
0.11671

sRun
RunA1
zout
1

Ave
Data
8E-06
Model
7.9E-06

data/vMax
8E-06

ofxng
369.894

Min d/m
1.07118

ave/min d
1
ave/min m
1.05637

328_Calib_S_A

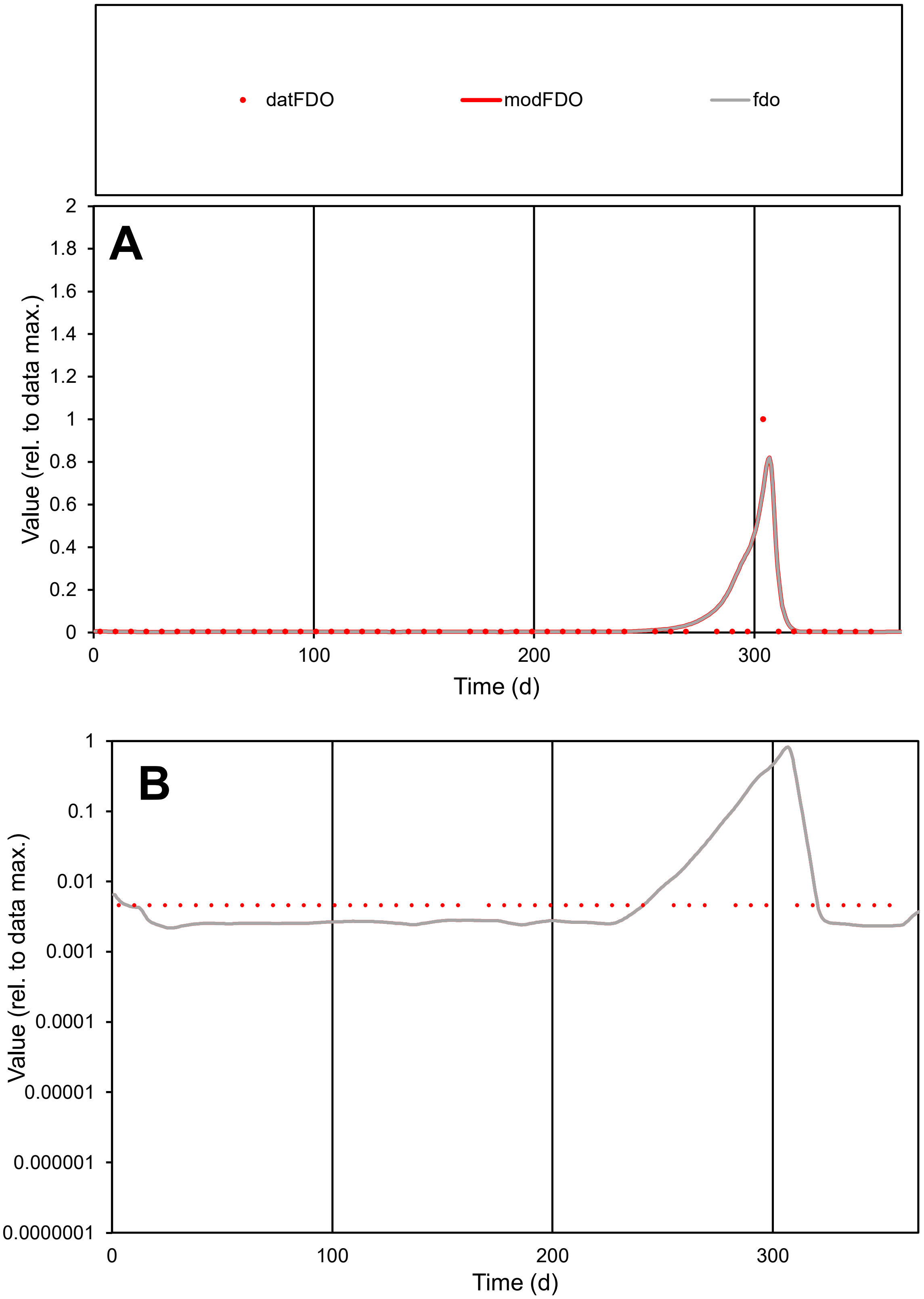


**C1**

Variables
FDO Provide jtd or Dst
CalibOnly?
2 1 to reproduce of
oNormalize?
2 1=no, 2=yes

ofxn calc*
2.72681
ofxn mod
2.91462

sRun
RunA1
zout
1

Ave
Data
5.5E-07
Model
7.6E-07

data/vMax
2.2E-05

ofxng
369.894

Min d/m
2.07519

ave/min d
5.51888
ave/min m
15.9153

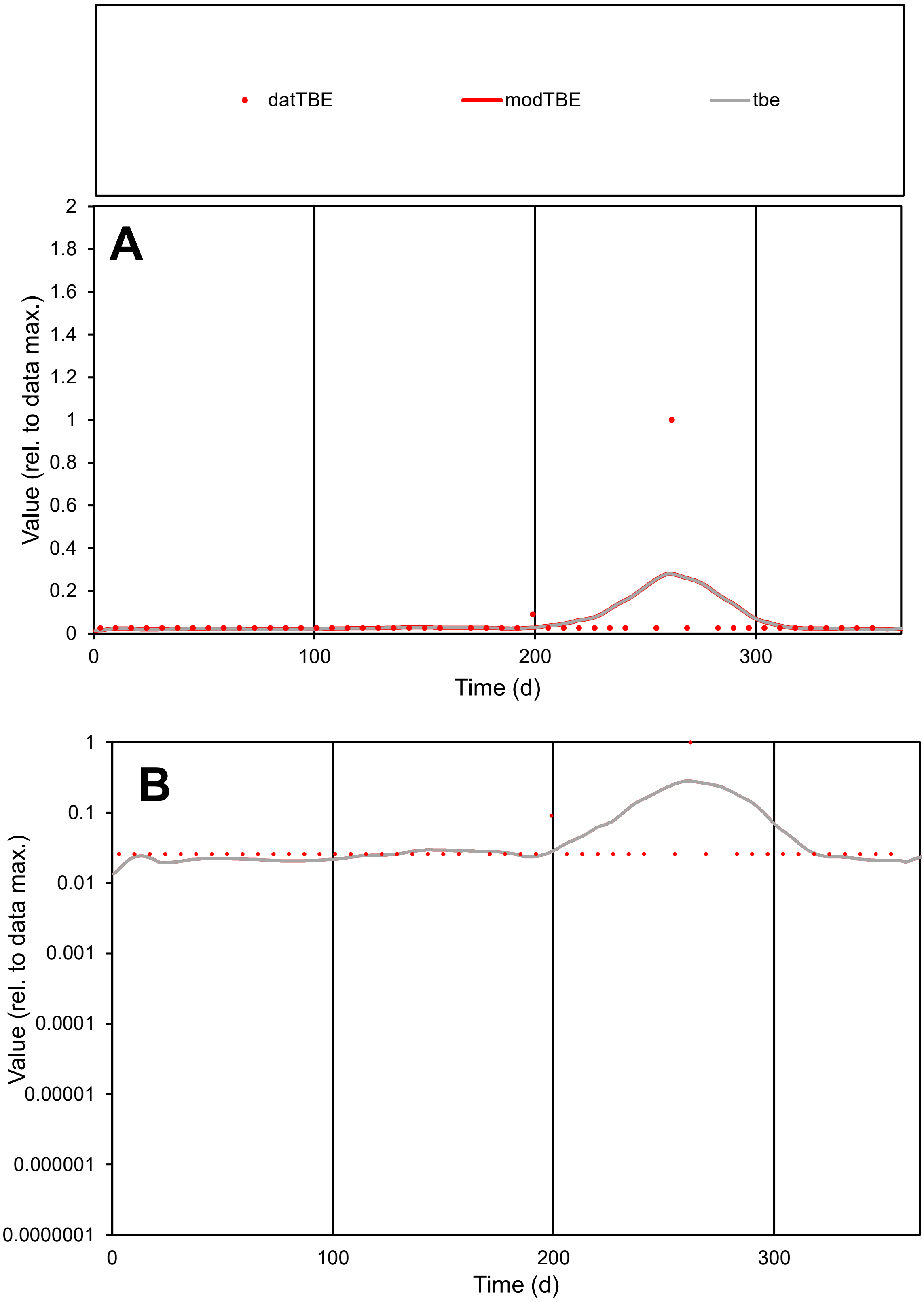


**C1**

Variables
TBE Provide jtd or Dst
CalibOnly?
2 1 to reproduce of
oNormalize?
2 1=no, 2=yes

ofxn calc*
2.54393
ofxn mod
2.56066

sRun
RunA1
zout
1

Ave
Data
1.2E-05
Model
1.4E-05

data/vMax
0.00026

ofxng
369.894

Min d/m
1.47318
ave/min d
1.84341
ave/min m
3.03934

328_Calib_S_A

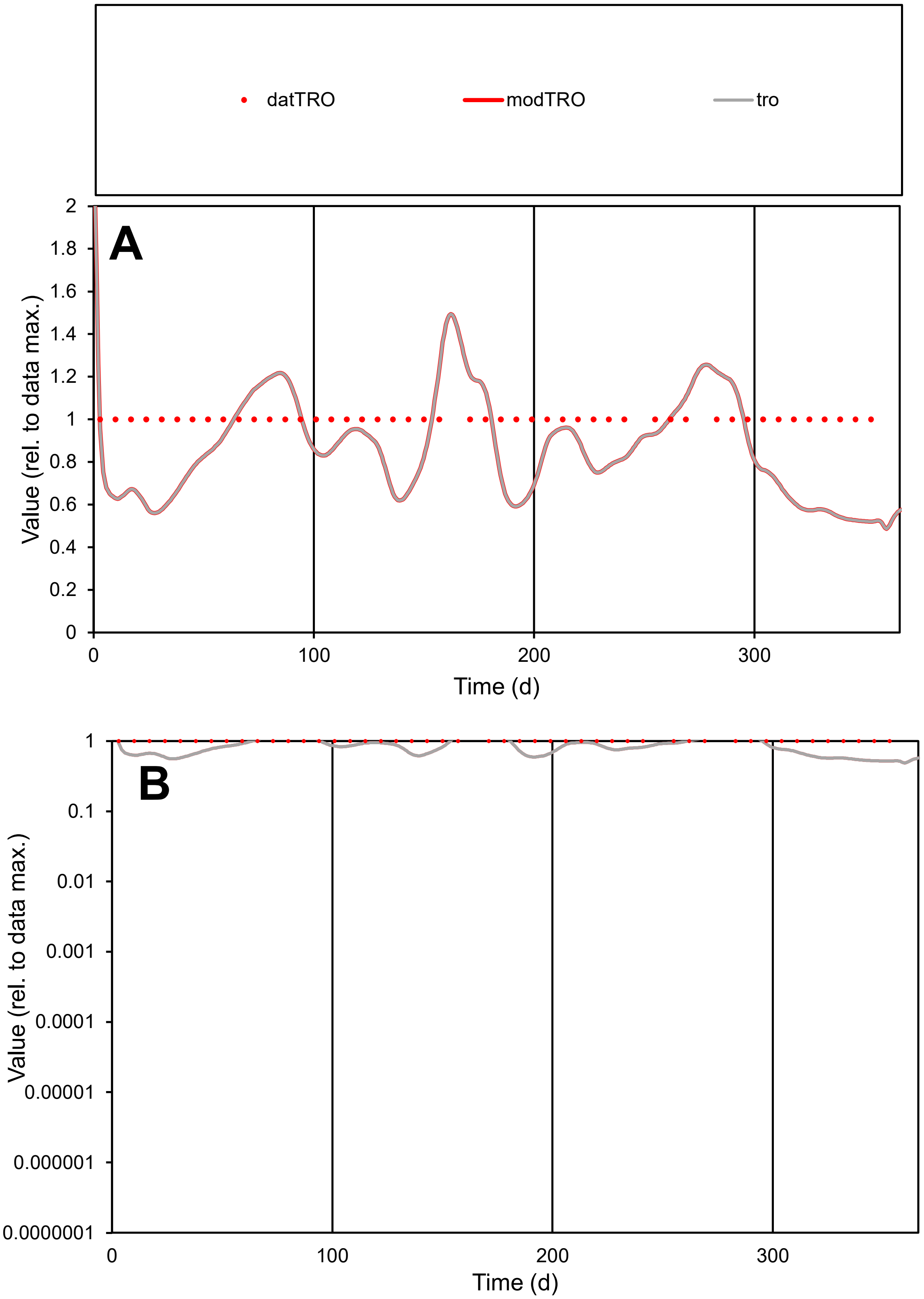


**C1**

Variables

TRO Provide jtd or Dst

CalibOnly?

2 1 to reproduce of

oNormalize?

2 1=no, 2=yes

ofxn calc*

0.26115

ofxn mod

0.28348

sRun

RunA1

zout

1

Ave

Data

5E-06

Model

4.2E-06

data/vMax

5E-06

ofxng

369.894

Min d/m

1.91762

ave/min d

1

ave/min m

1.61891

328_Calib_S_A

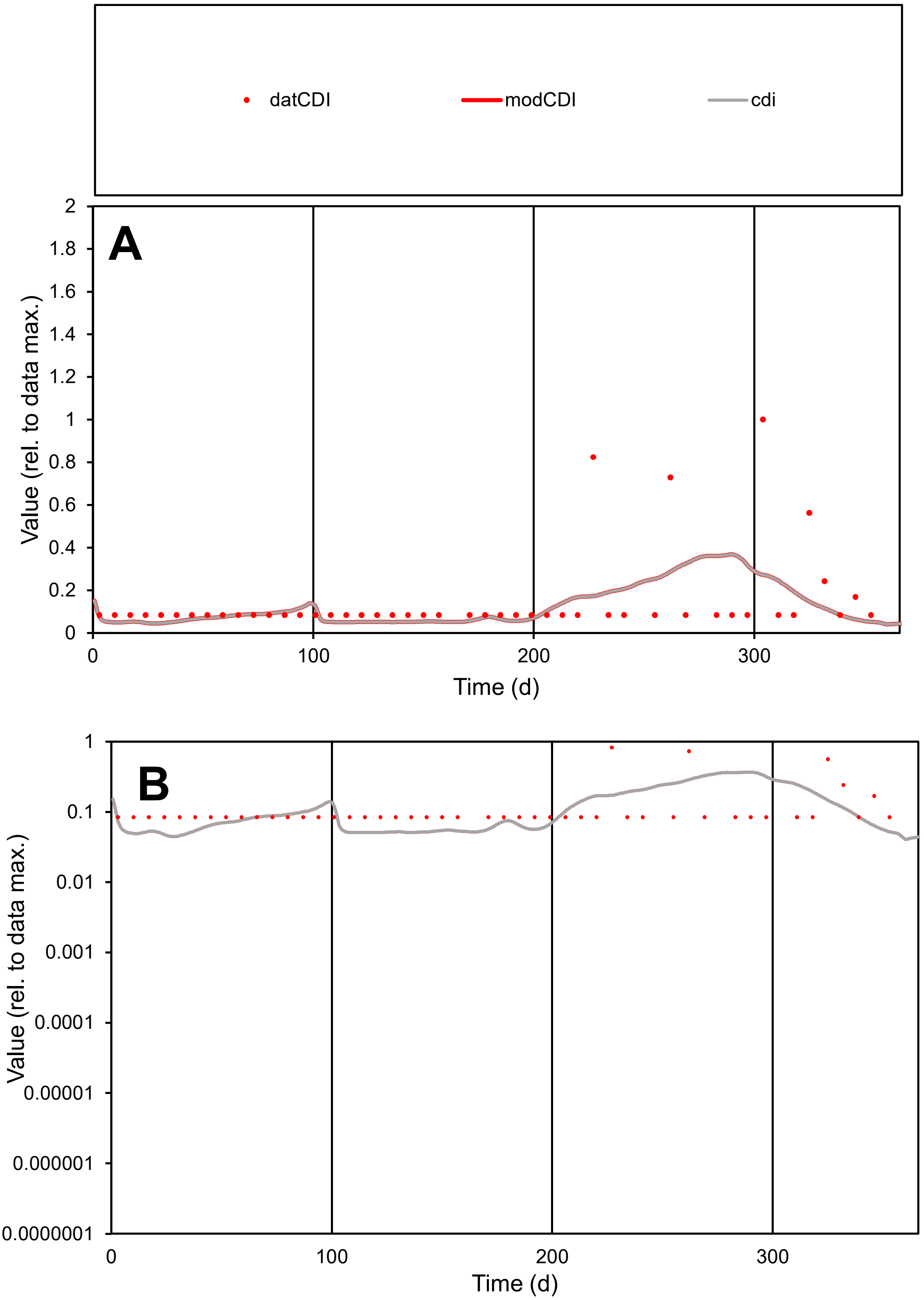


**C1**

| | |
|---|---|
| Variables | |
| CDI | Provide jtd or Dst |
| CalibOnly? | |
| 2 | 1 to reproduce of |
| oNormalize? | |
| 2 | 1=no, 2=yes |
| ofxn calc* | |
| 1.29001 | |
| ofxn mod | |
| 1.30023 | |
| sRun | |
| RunA1 | |
| zout | |
| 1 | |
| Ave | |
| Data | |
| 3E-07 | |
| Model | |
| 2.5E-07 | |
| data/vMax | |
| 2.1E-06 | |
| ofxng | |
| 369.894 | |
| Min d/m | |
| 1.8036 | |
| ave/min d | |
| 1.74844 | |
| ave/min m | |
| 2.63561 | |

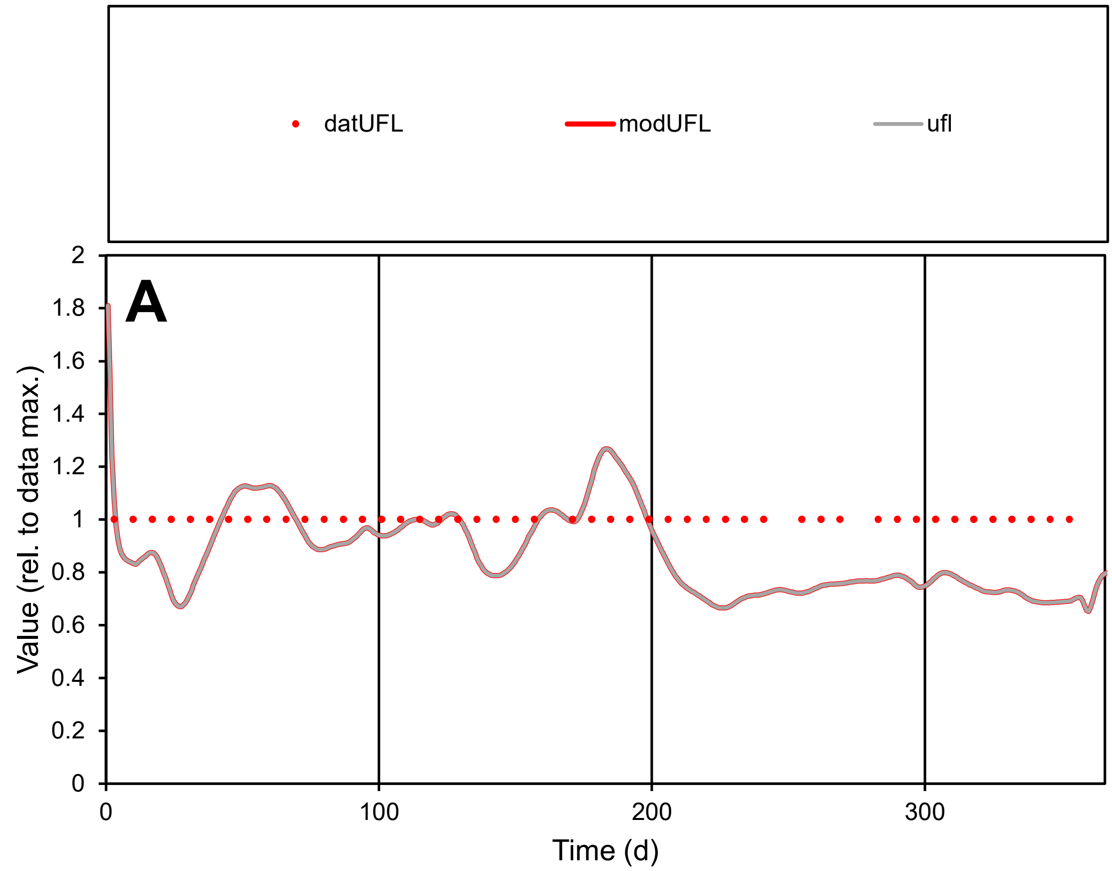


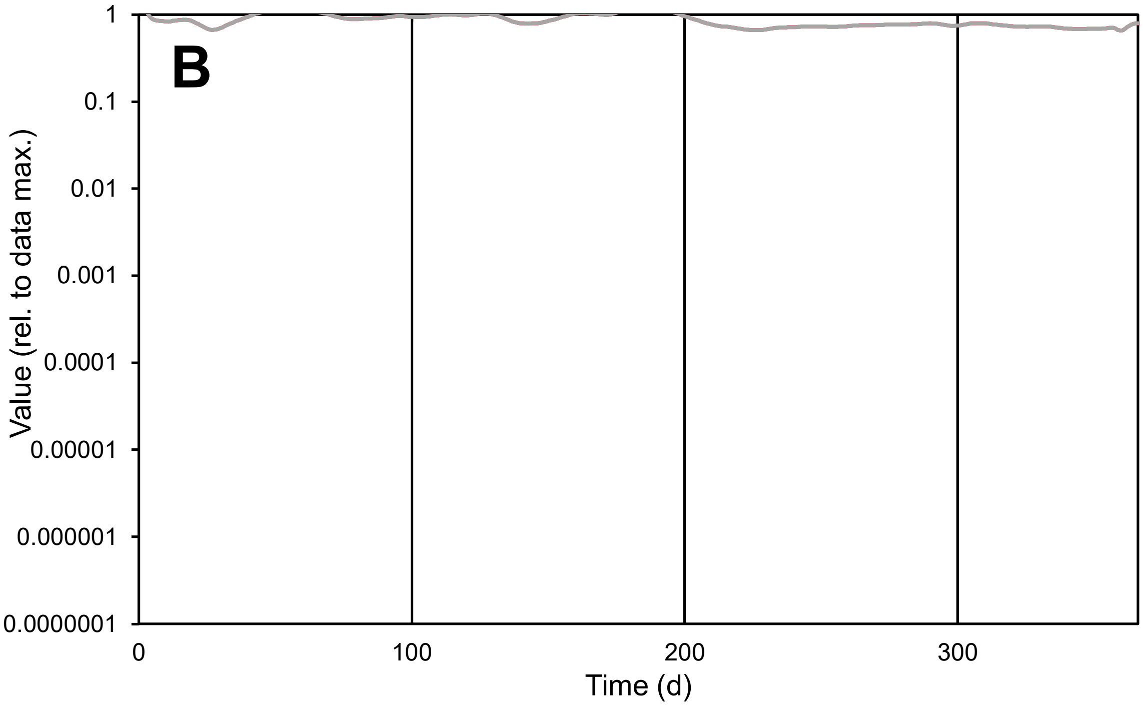


**C1**

Variables
UFL Provide jtd or Dst
CalibOnly?
2 1 to reproduce of
oNormalize?
2 1=no, 2=yes

ofxn calc*
0.19993
ofxn mod
0.22651

sRun
RunA1
zout
1

Ave
Data
3.1E-06
Model
2.7E-06

data/vMax
3.1E-06

ofxng
369.894

Min d/m
1.4922

ave/min d
1
ave/min m
1.29825

328_Calib_S_A

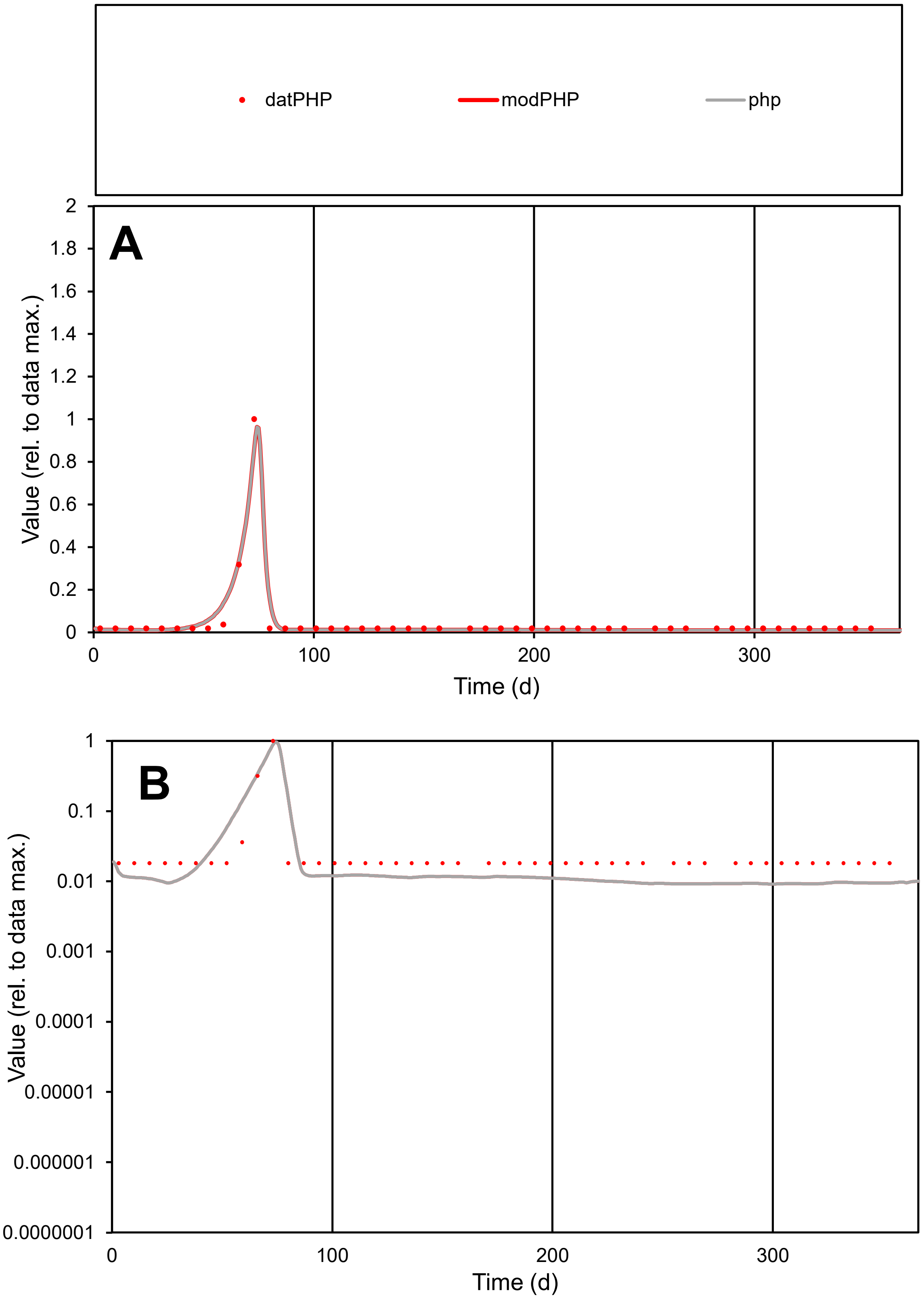


**C1**

Variables
PHP Provide jtd or Dst
CalibOnly?
2 1 to reproduce of
oNormalize?
2 1=no, 2=yes

ofxn calc*
0.39131
ofxn mod
0.53422

sRun
RunA1
zout
1

Ave
Data
1.7E-05
Model
1.6E-05

data/vMax
0.00038

ofxng
369.894

Min d/m
1.98088

ave/min d
2.49641
ave/min m
4.6186

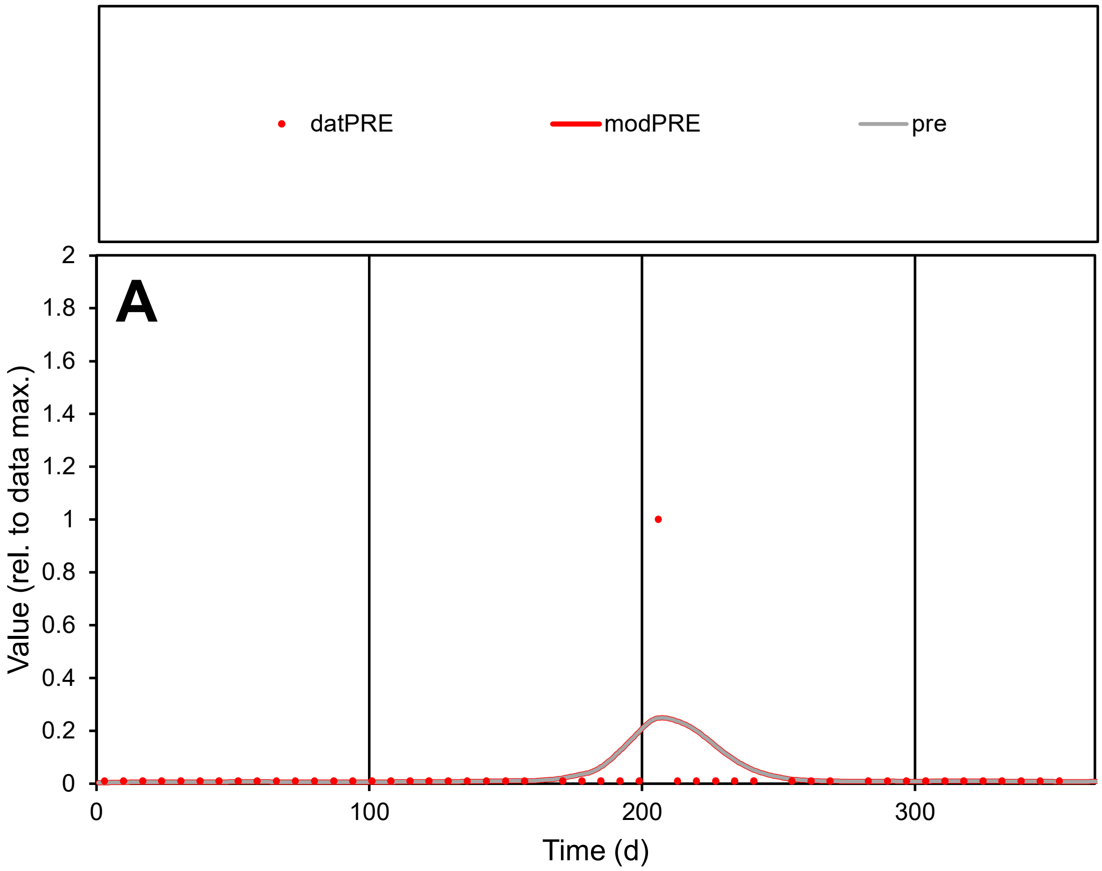


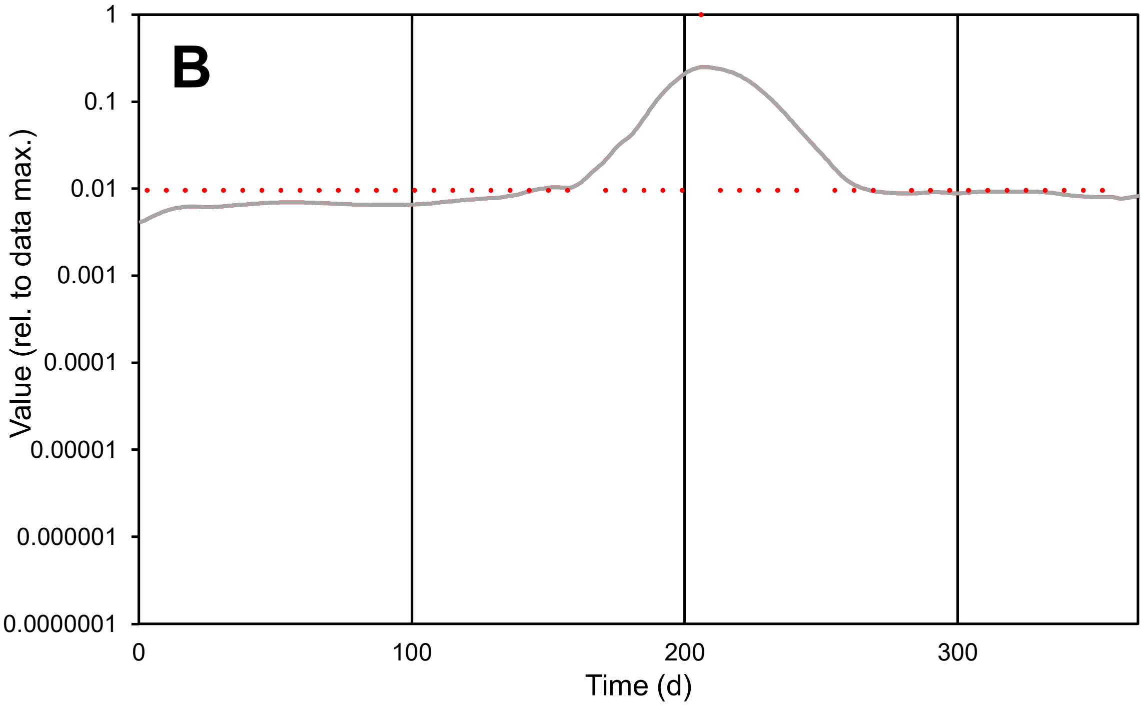


**C1**

Variables
PRE Provide jtd or Dst
CalibOnly?
2 1 to reproduce of
oNormalize?
2 1=no, 2=yes

ofxn calc*
3.99445
ofxn mod
4.04941

sRun
RunA1
zout
1

Ave
Data
4.2E-05
Model
4.7E-05

data/vMax
0.0014

ofxng
369.894

Min d/m
2.05258

ave/min d
3.14937
ave/min m
7.10754

328_Calib_S_A

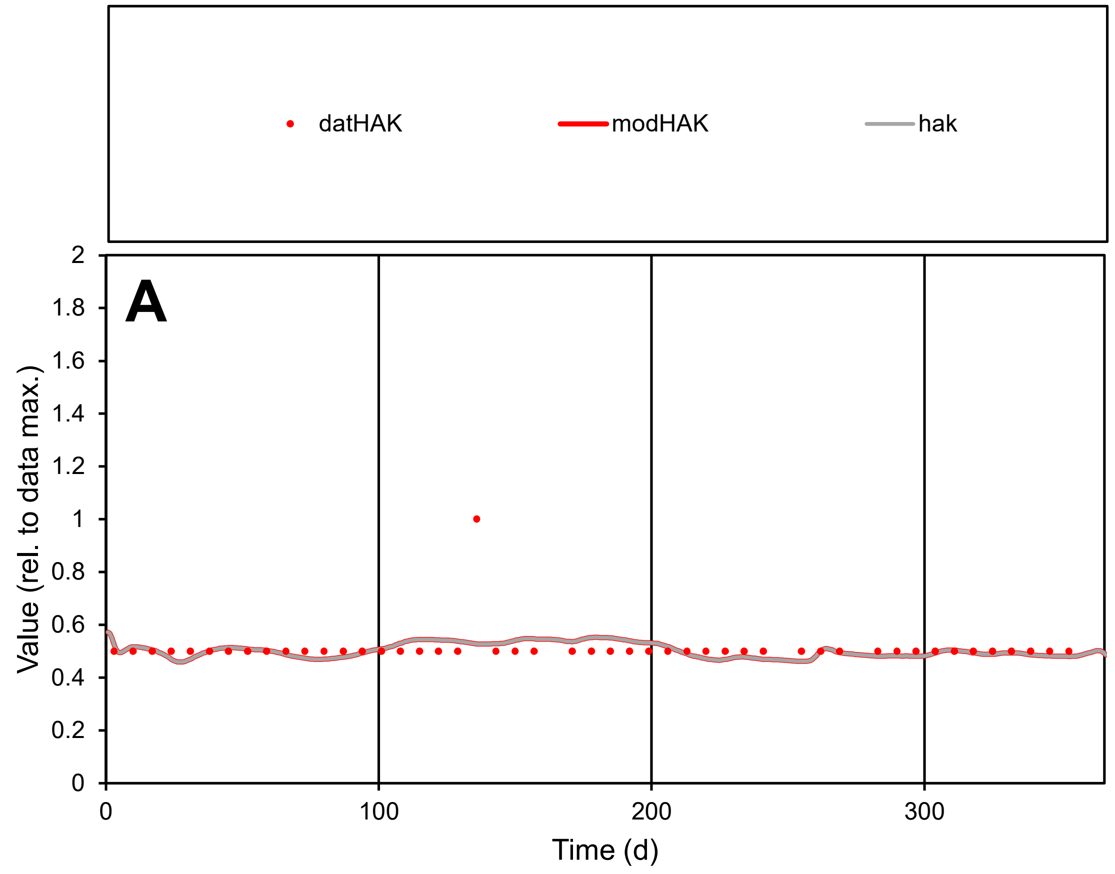


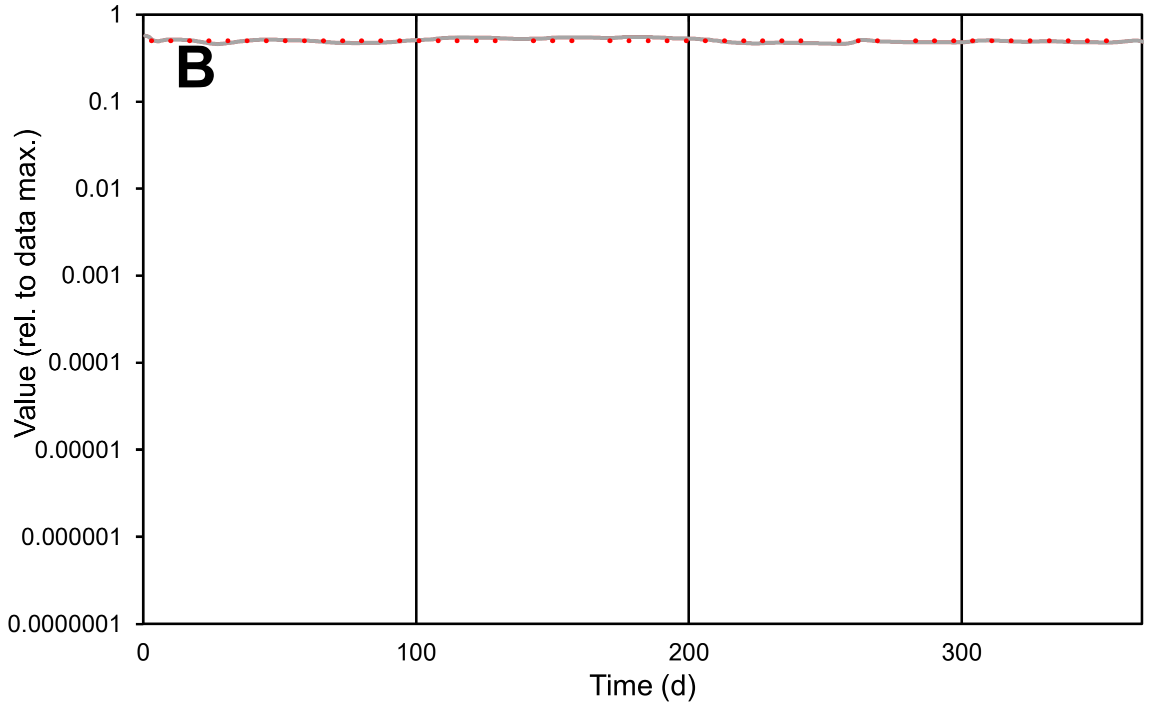


**C1**

Variables
HAK Provide jtd or Dst
CalibOnly?
2 1 to reproduce of
oNormalize?
2 1=no, 2=yes

ofxn calc*
0.14304
ofxn mod
0.18173

sRun
RunA1
zout
1

Ave
Data
5.2E-06
Model
5.1E-06

data/vMax
1E-05

ofxng
369.894

Min d/m
1.08141

ave/min d
1.02083
ave/min m
1.08711

328_Calib_S_A

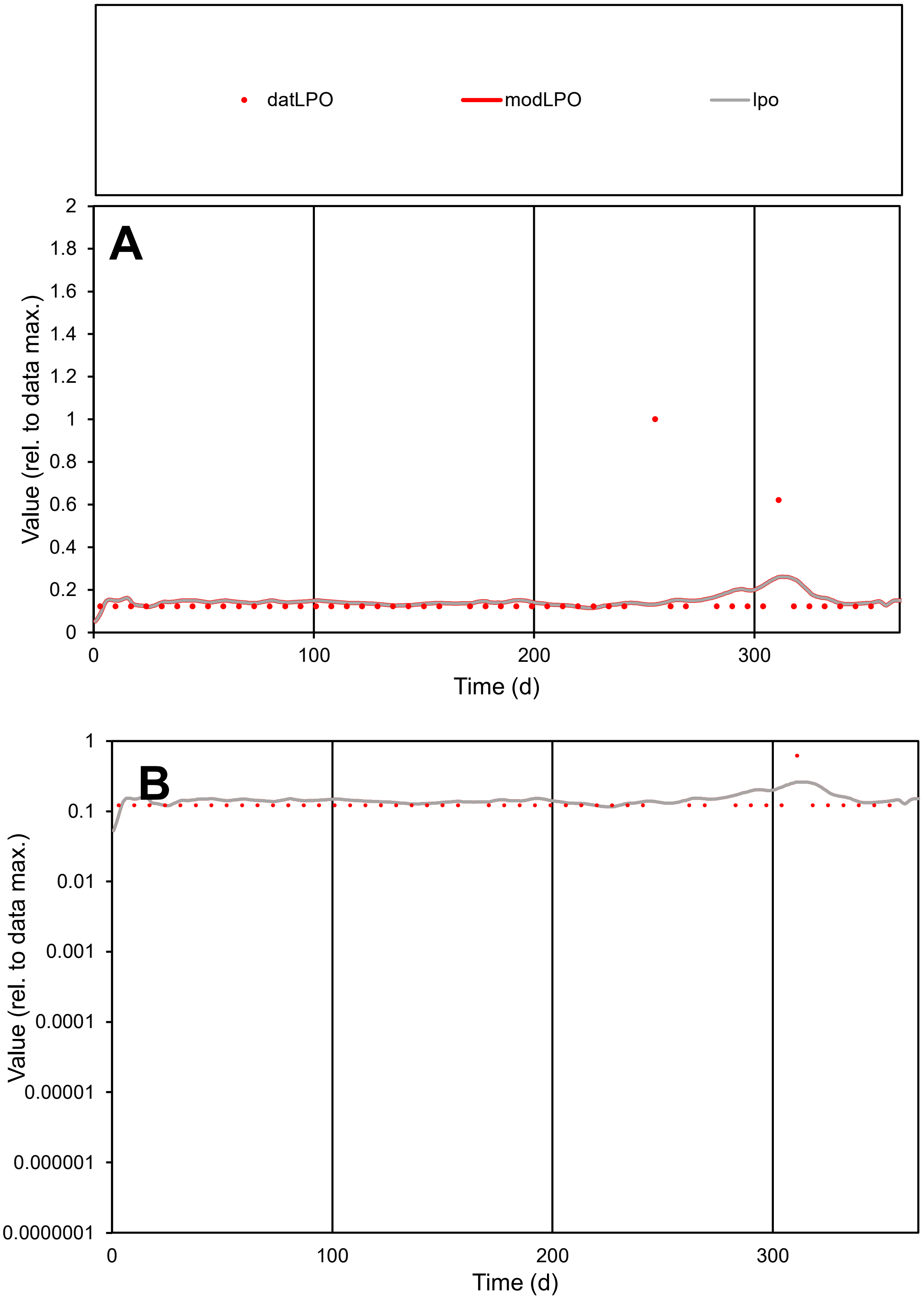


**C1**

Variables
LPO Provide jtd or Dst
CalibOnly?
2 1 to reproduce of
oNormalize?
2 1=no, 2=yes

ofxn calc*
0.92832
ofxn mod
0.93859

sRun
RunA1
zout
1

Ave
Data
7.2E-06
Model
7.2E-06

data/vMax
4.8E-05

ofxng
369.894

Min d/m
1.0383

ave/min d
1.23561
ave/min m
1.27113

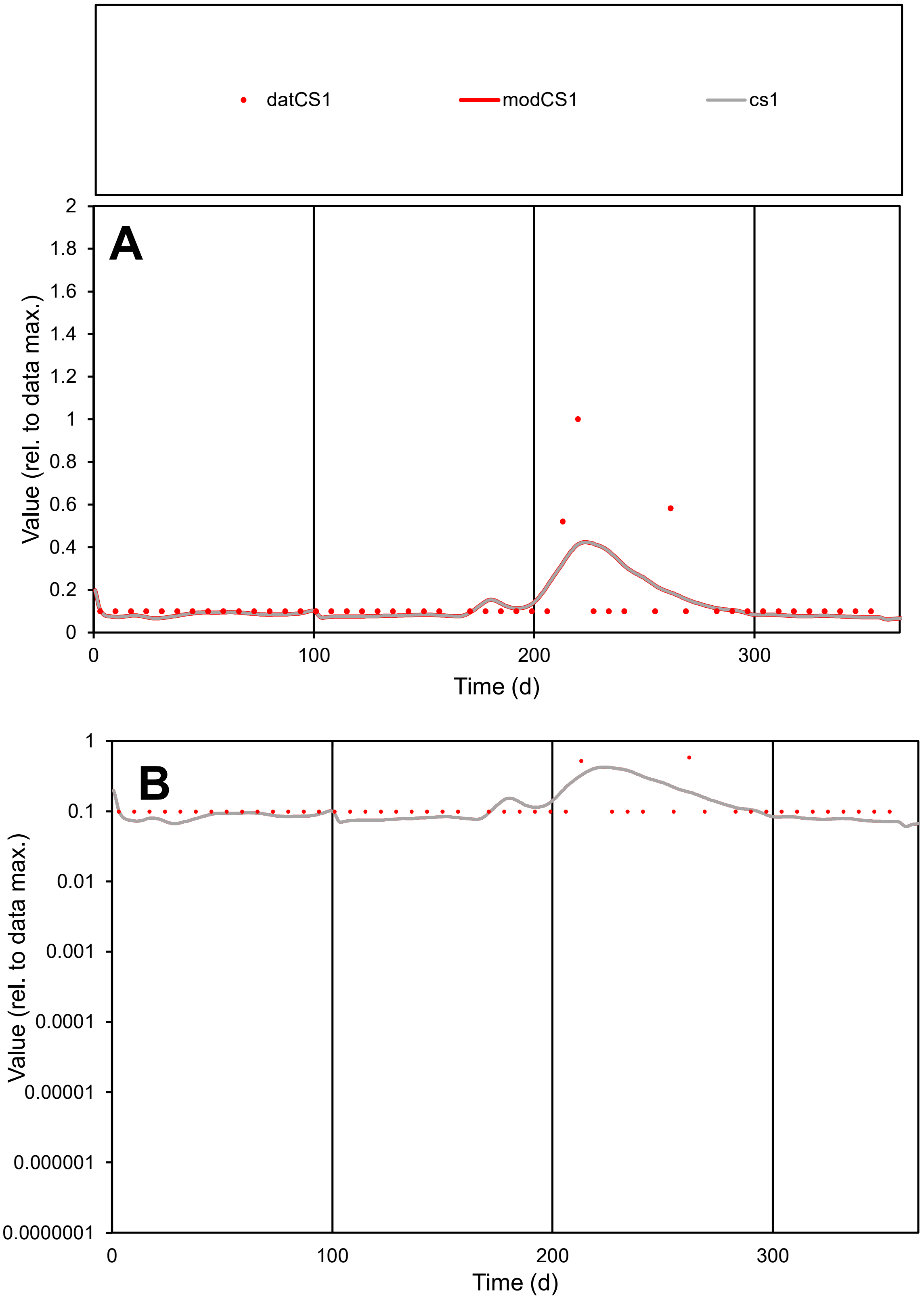


**C1**

Variables
CS1 Provide jtd or Dst
CalibOnly?
2 1 to reproduce of
oNormalize?
2 1=no, 2=yes

ofxn calc*
0.93745
ofxn mod
0.94755

sRun
RunA1
zout
1

Ave
Data
2.4E-05
Model
2.2E-05

data/vMax
0.00018

ofxng
369.894

Min d/m
1.4016

ave/min d
1.38108
ave/min m
1.79738

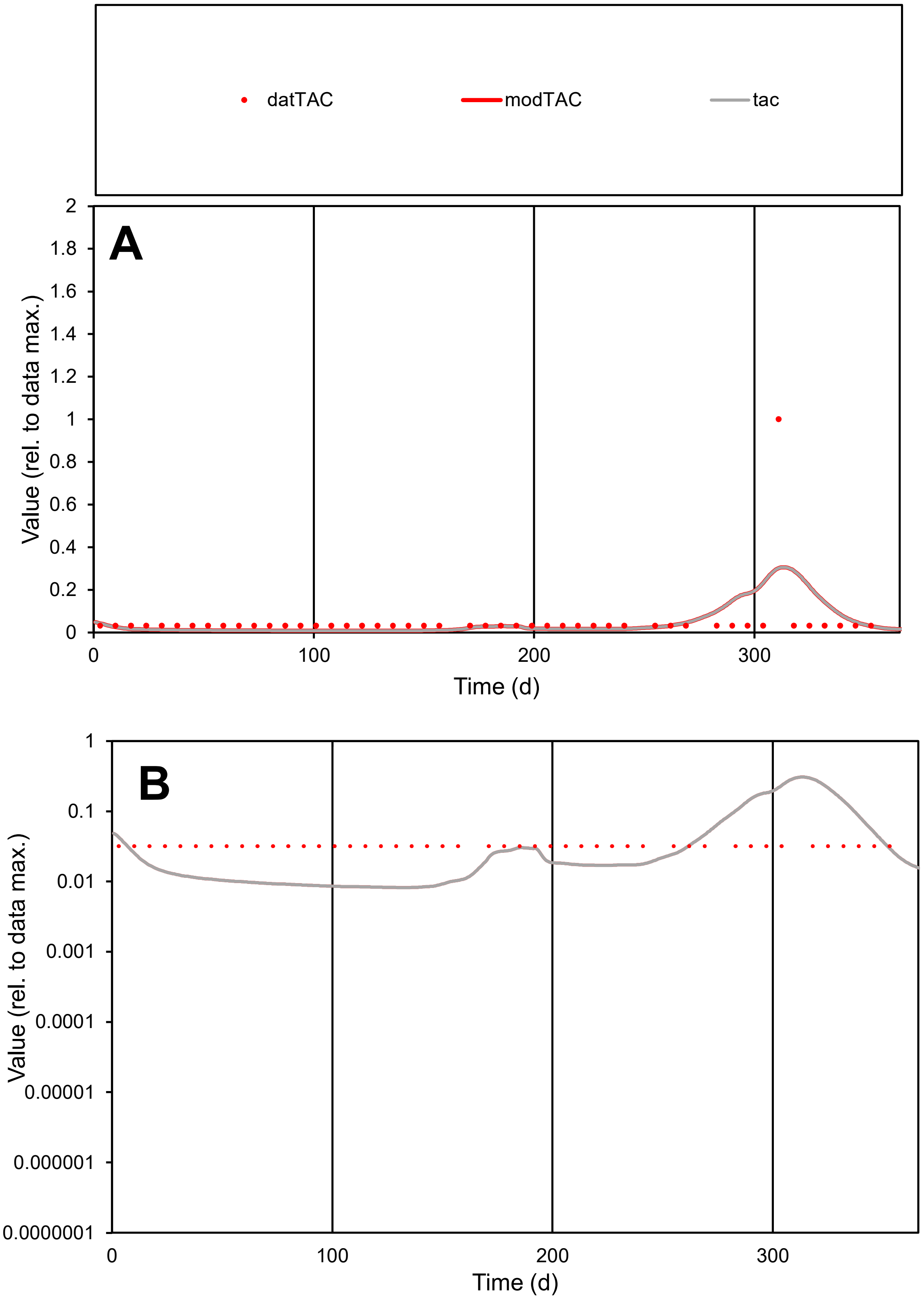


**C1**

Variables
TAC Provide jtd or Dst
CalibOnly?
2 1 to reproduce of
oNormalize?
2 1=no, 2=yes

ofxn calc*
2.26369
ofxn mod
2.28866

sRun
RunA1
zout
1

Ave
Data
2.1E-06
Model
2E-06

data/vMax
4.1E-05

ofxng
369.894

Min d/m
3.88655

ave/min d
1.63264
ave/min m
5.85609

328_Calib_S_A

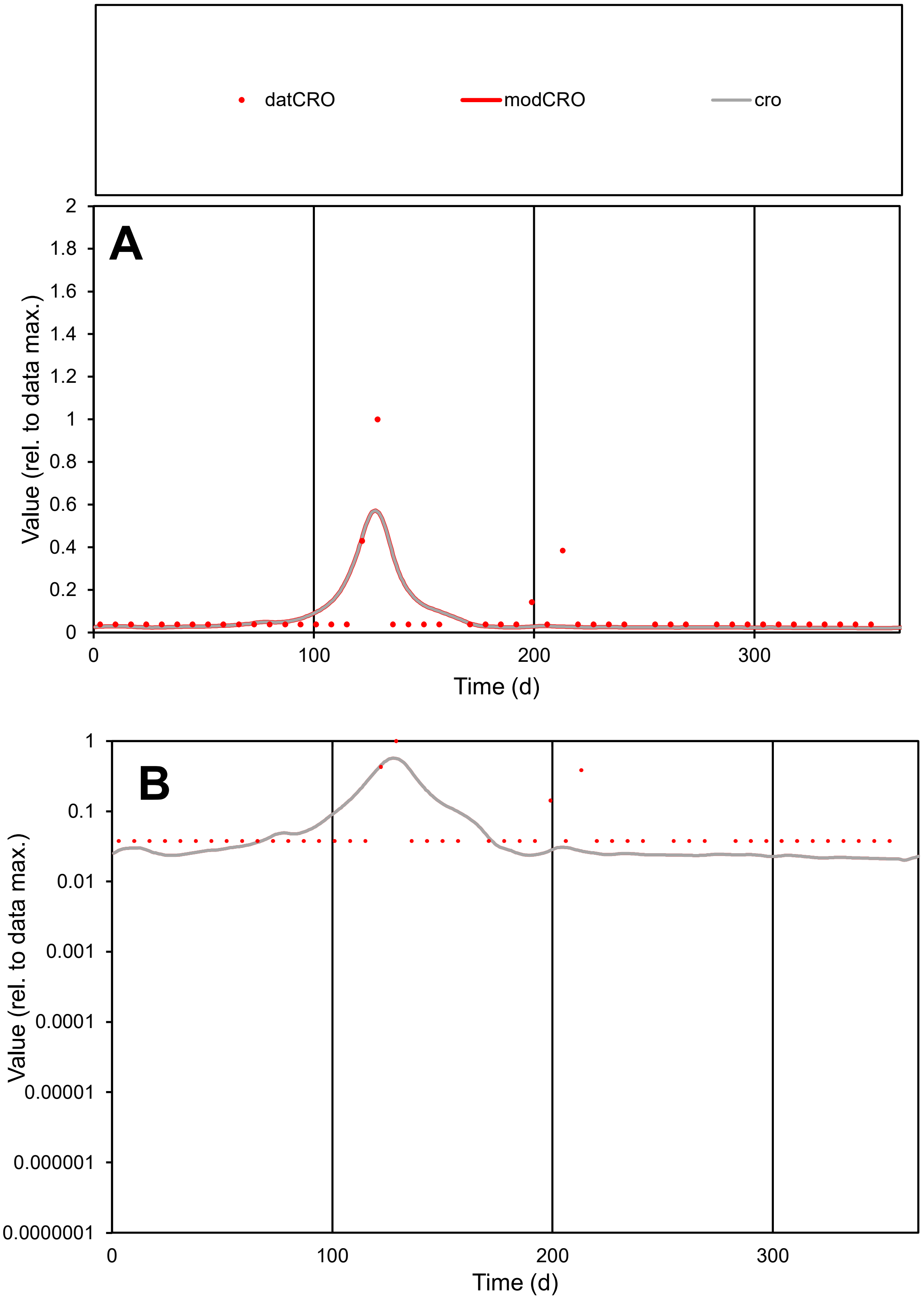


**C1**

Variables
CRO Provide jtd or Dst
CalibOnly?
2 1 to reproduce of
oNormalize?
2 1=no, 2=yes

ofxn calc*
1.33457
ofxn mod
1.3867

sRun
RunA1
zout
1

Ave
Data
2E-05
Model
1.9E-05

data/vMax
0.00027

ofxng
369.894

Min d/m
1.79744

ave/min d
1.99799
ave/min m
3.26882

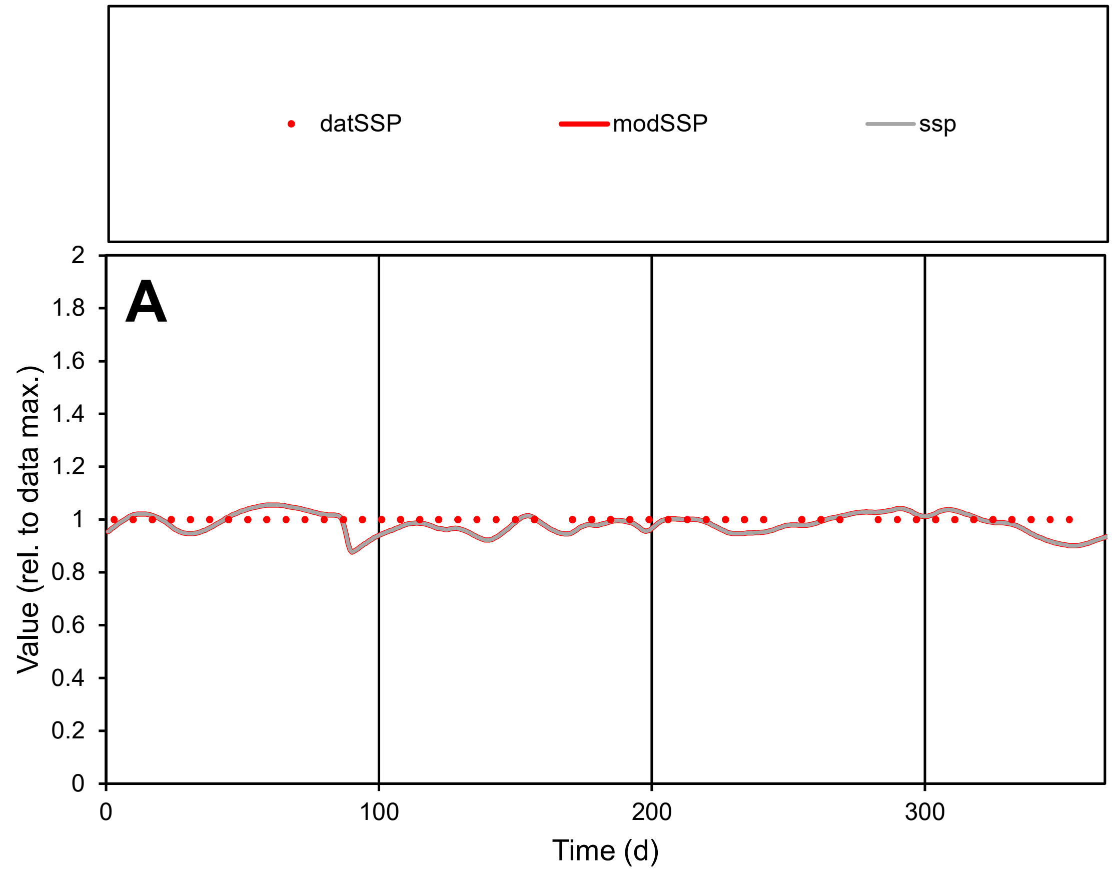


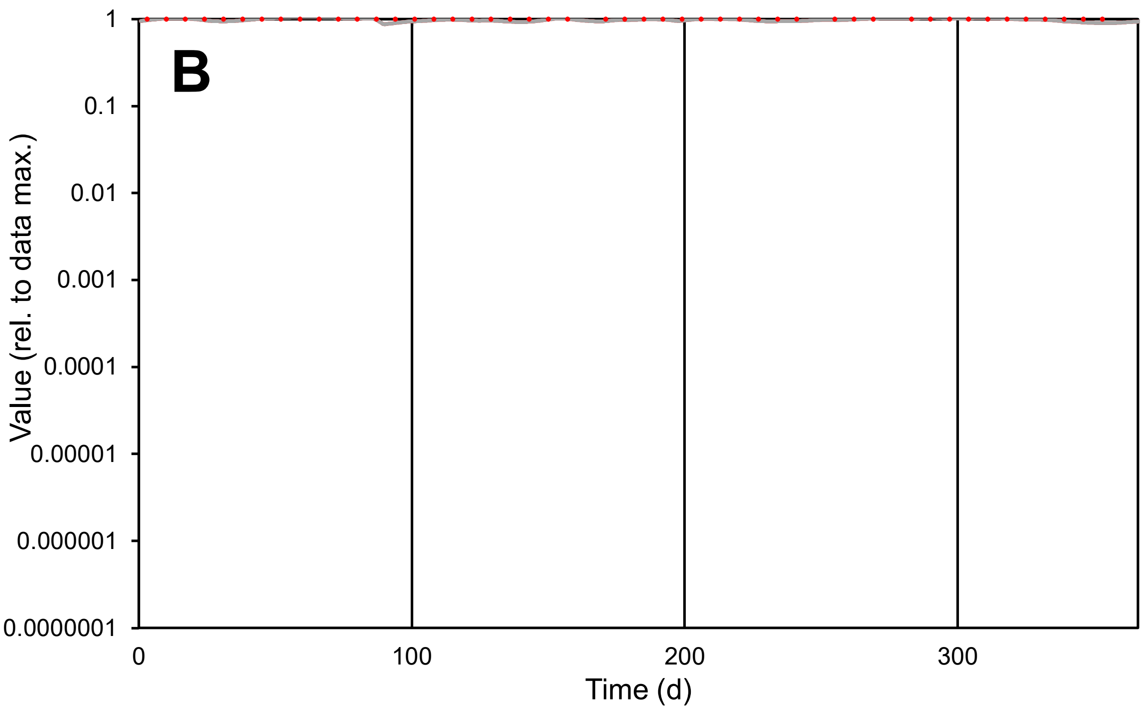


**C1**

Variables
SSP Provide jtd or Dst
CalibOnly?
2 1 to reproduce of
oNormalize?
2 1=no, 2=yes

ofxn calc*
0.04017
ofxn mod
0.11915

sRun
RunA1
zout
1

Ave
Data
1.3E-05
Model
1.3E-05

data/vMax
1.3E-05

ofxng
369.894

Min d/m
1.1099

ave/min d
1
ave/min m
1.09324

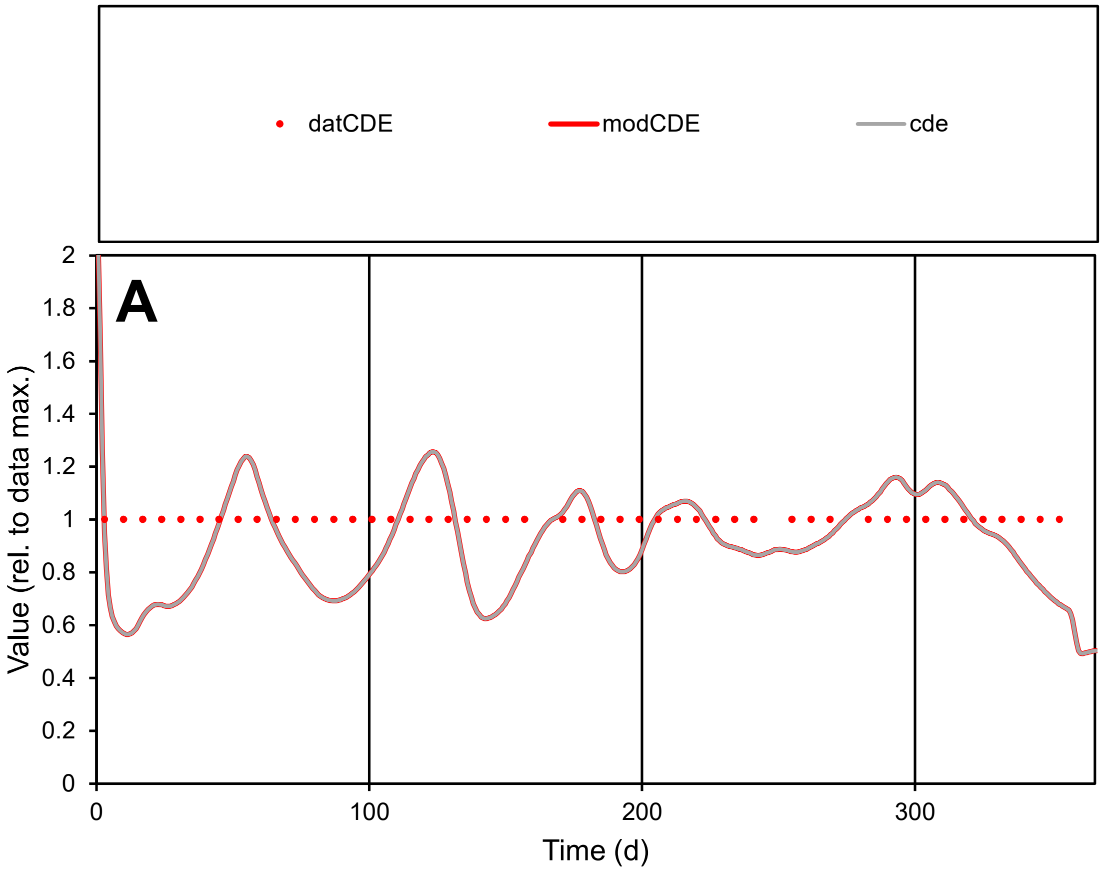


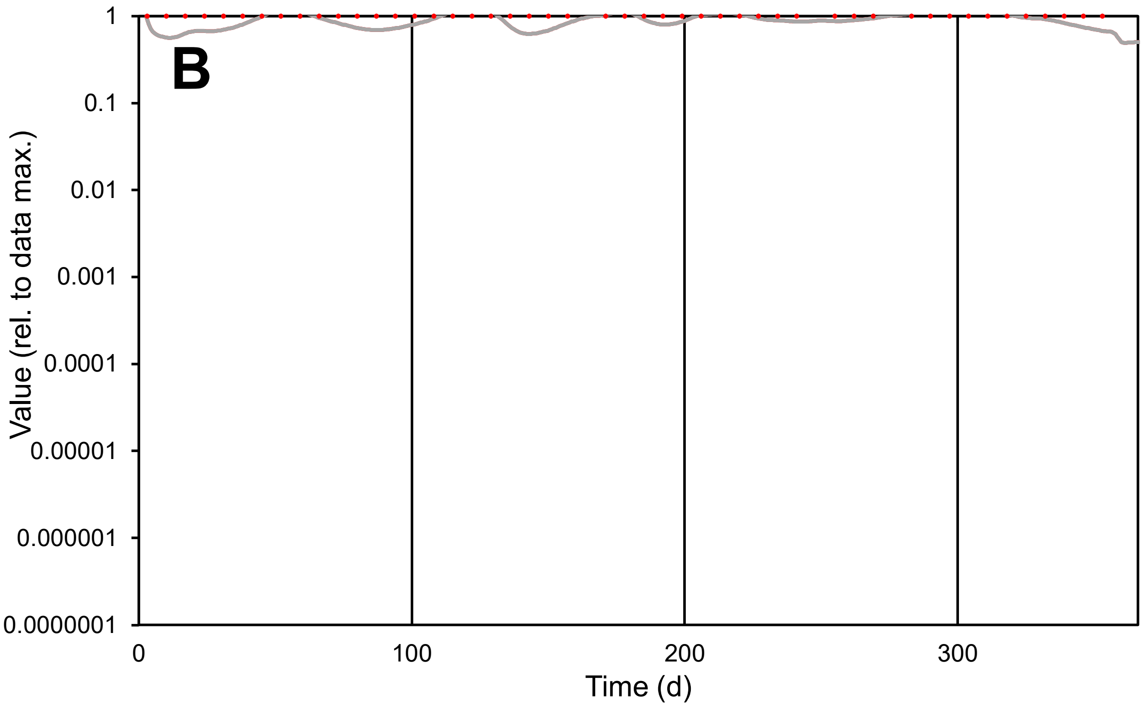


**C1**

Variables
CDE Provide jtd or Dst
CalibOnly?
2 1 to reproduce of
oNormalize?
2 1=no, 2=yes

ofxn calc*
0.19447
ofxn mod
0.22252

sRun
RunA1
zout
1

Ave
Data
1E-05
Model
9.1E-06

data/vMax
1E-05

ofxng
369.894

Min d/m
1.77043

ave/min d
1
ave/min m
1.6088

328_Calib_S_A

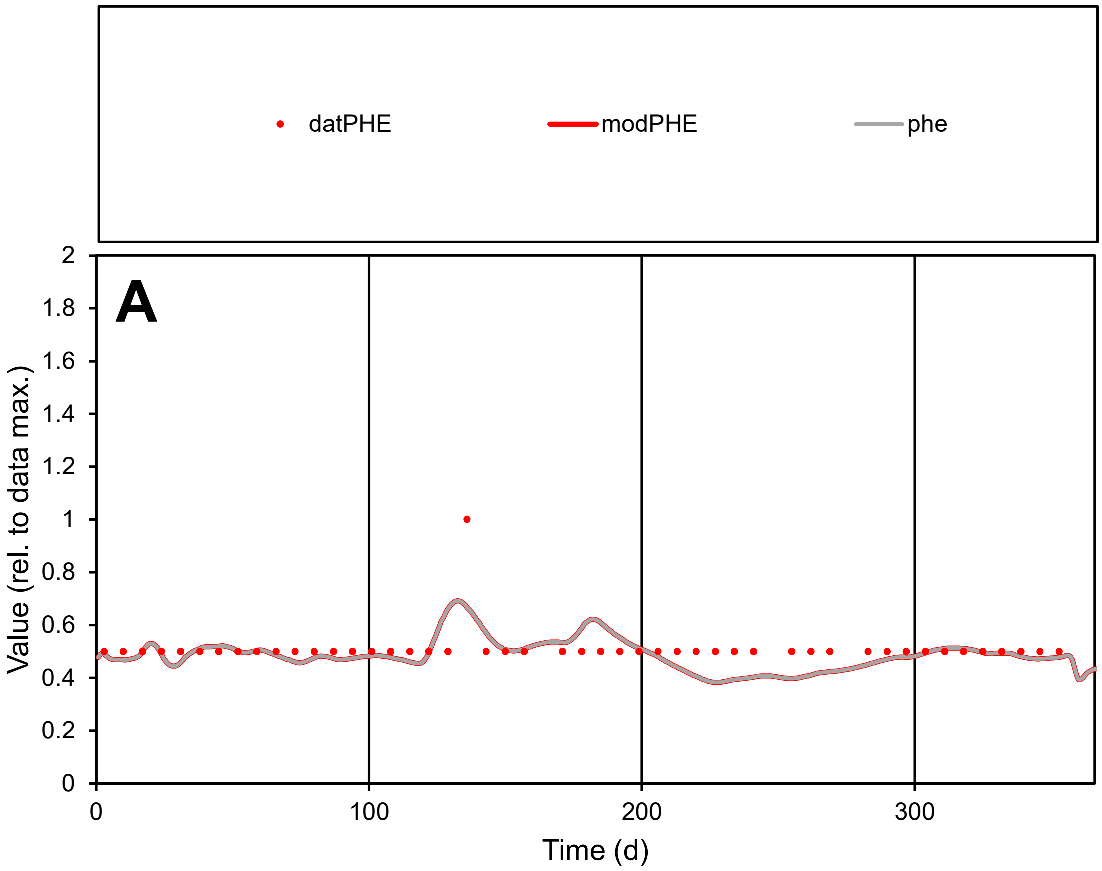


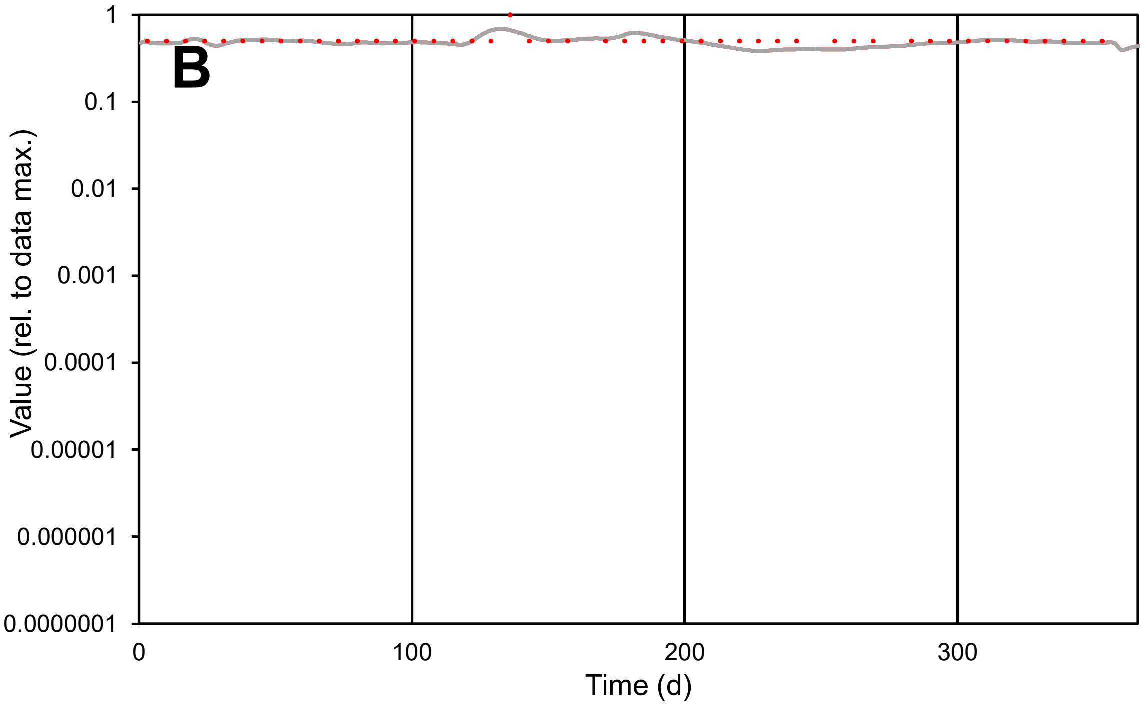


**C1**

Variables
PHE Provide jtd or Dst
CalibOnly?
2 1 to reproduce of
oNormalize?
2 1=no, 2=yes

ofxn calc*
0.14755
ofxn mod
0.18416

sRun
RunA1
zout
1

Ave
Data
4E-06
Model
3.9E-06

data/vMax
7.9E-06

ofxng
369.894

Min d/m
1.30365

ave/min d
1.02084
ave/min m
1.28004

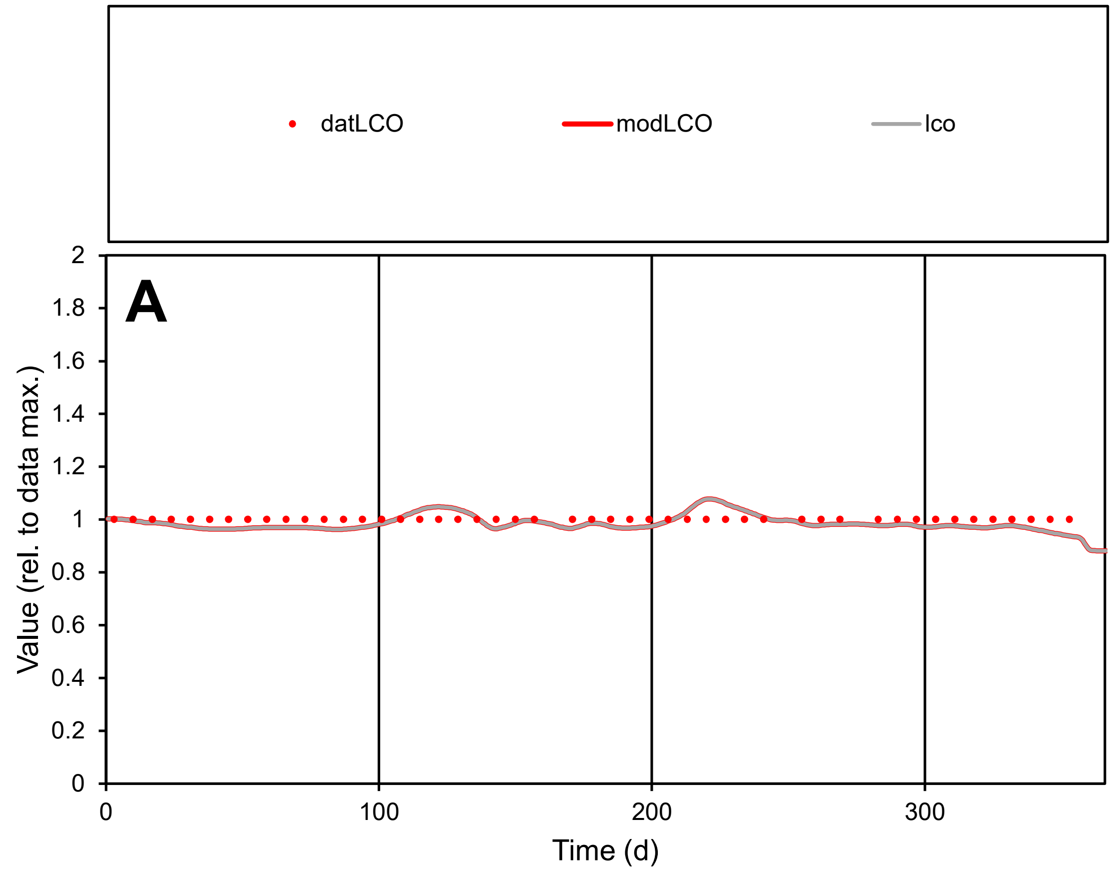


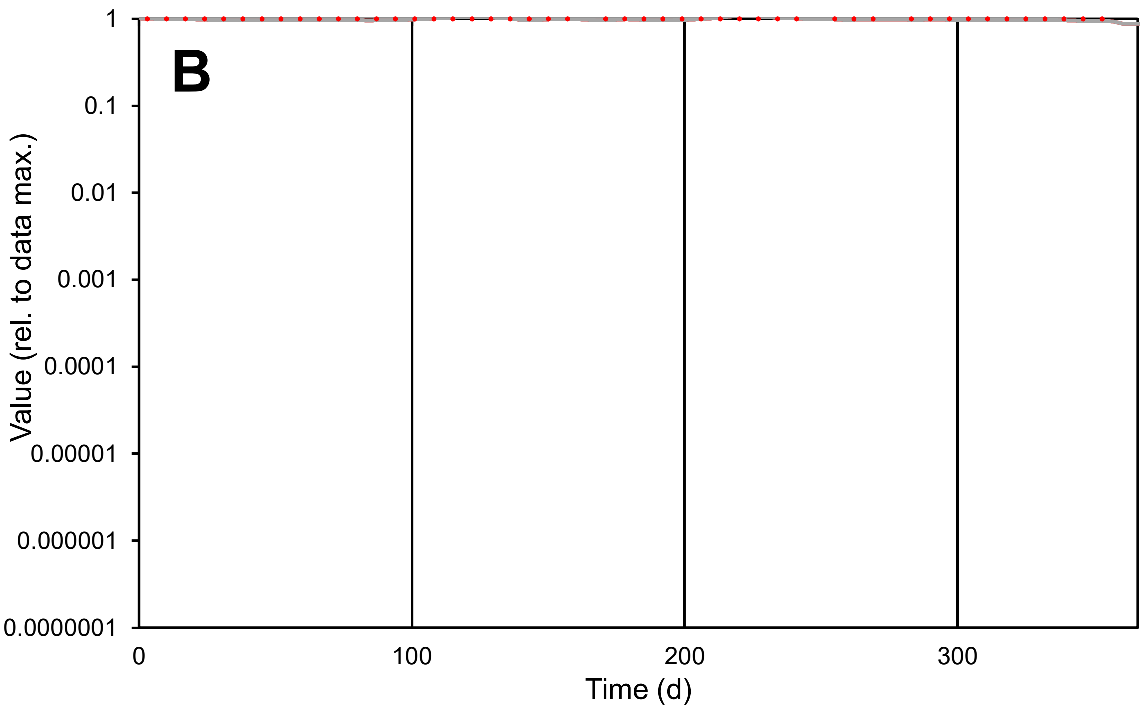


**C1**

Variables
LCO Provide jtd or Dst
CalibOnly?
2 1 to reproduce of
oNormalize?
2 1=no, 2=yes

ofxn calc*
0.03157
ofxn mod
0.11673

sRun
RunA1
zout
1

Ave
Data
7.4E-06
Model
7.3E-06

data/vMax
7.4E-06

ofxng
369.894

Min d/m
1.06791

ave/min d
1
ave/min m
1.05402

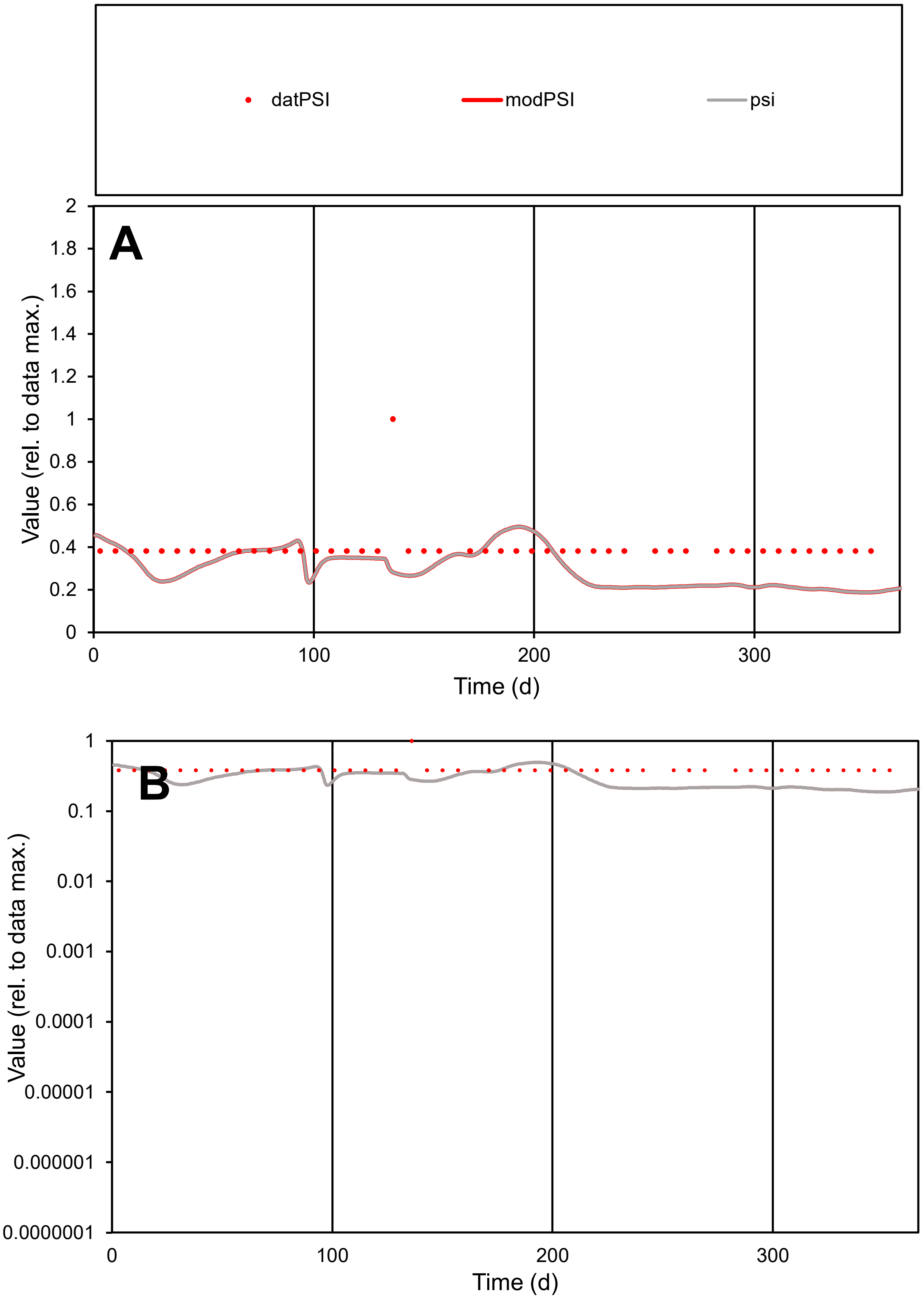


**C1**

Variables

PSI Provide jtd or Dst

CalibOnly?

2 1 to reproduce of

oNormalize?

2 1=no, 2=yes

ofxn calc*

0.3975

ofxn mod

0.41021

sRun

RunA1

zout

1

Ave

Data

3.7E-06

Model

2.8E-06

data/vMax

9.3E-06

ofxng

369.894

Min d/m

2.01873

ave/min d

1.03379

ave/min m

1.59907

328_Calib_S_A

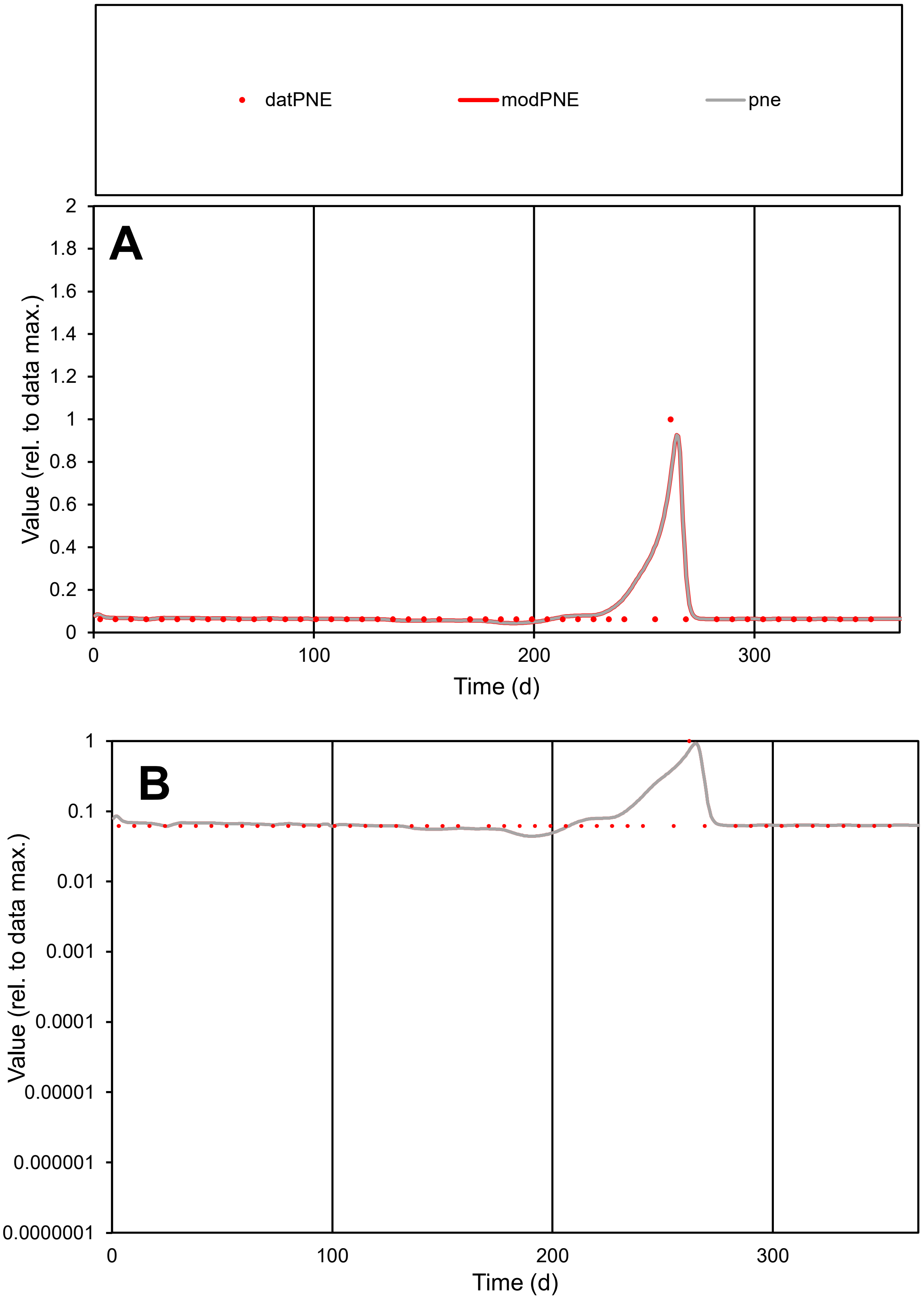


**C1**

Variables
PNE Provide jtd or Dst
CalibOnly?
2 1 to reproduce of
oNormalize?
2 1=no, 2=yes

ofxn calc*
0.70743
ofxn mod
0.75895

sRun
RunA1
zout
1

Ave
Data
1.9E-05
Model
2.1E-05

data/vMax
0.00023

ofxng
369.894

Min d/m
1.38418

ave/min d
1.31577
ave/min m
2.03023

328_Calib_S_A

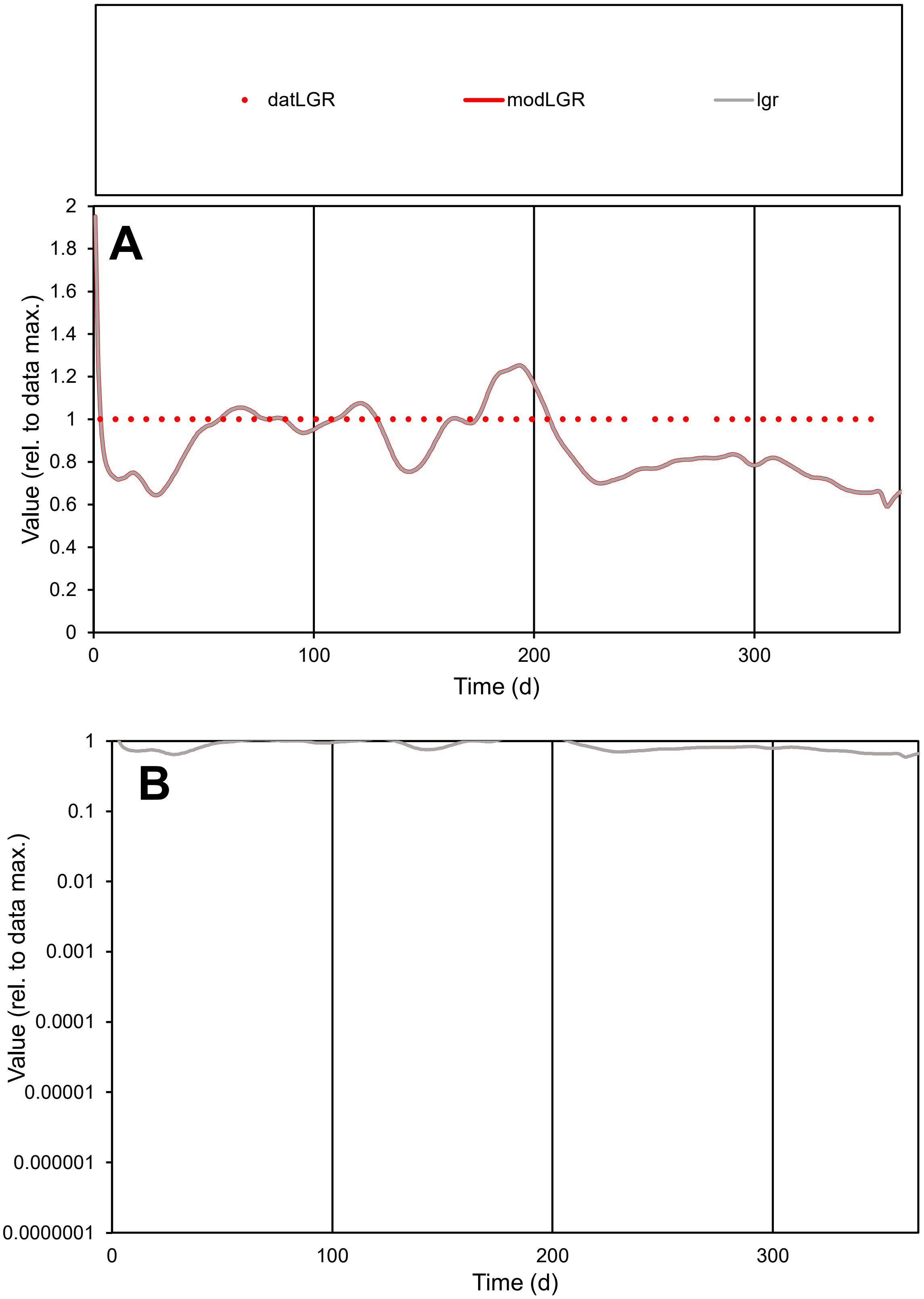


**C1**

Variables
LGR Provide jtd or Dst
CalibOnly?
2 1 to reproduce of
oNormalize?
2 1=no, 2=yes

ofxn calc*
0.19767
ofxn mod
0.22466

sRun
RunA1
zout
1

Ave
Data
5.2E-06
Model
4.5E-06

data/vMax
5.2E-06

ofxng
369.894

Min d/m
1.52137

ave/min d
1
ave/min m
1.3326

328_Calib_S_A

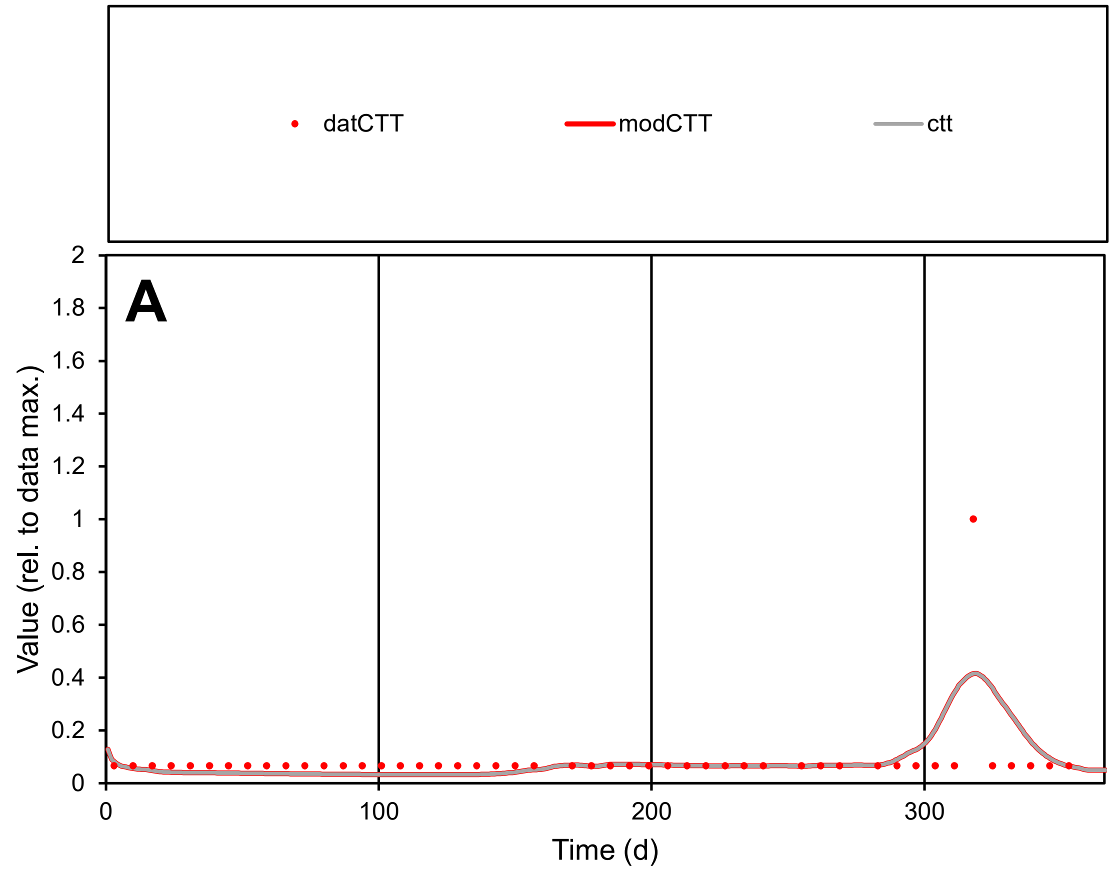


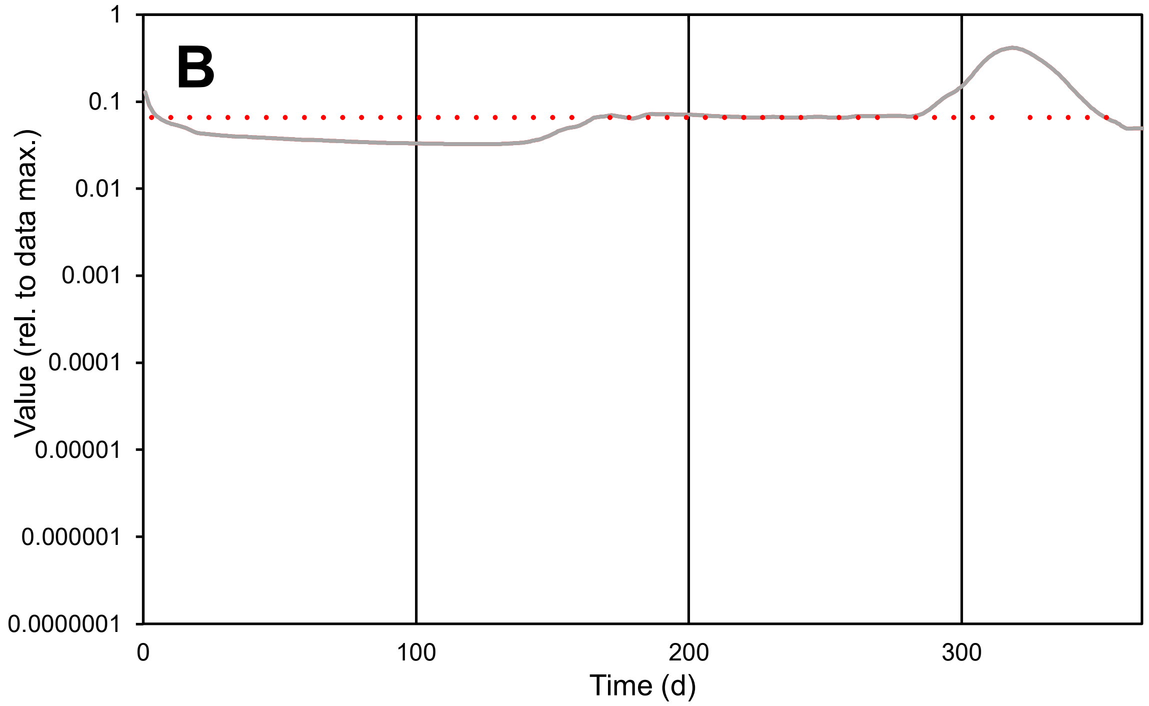


**C1**

Variables
CTT Provide jtd or Dst
CalibOnly?
2 1 to reproduce of
oNormalize?
2 1=no, 2=yes

ofxn calc*
1.27402
ofxn mod
1.29194

sRun
RunA1
zout
1

Ave
Data
1.9E-06
Model
1.9E-06

data/vMax
2.3E-05

ofxng
369.894

Min d/m
2.02568

ave/min d
1.29581
ave/min m
2.54042

328_Calib_S_A

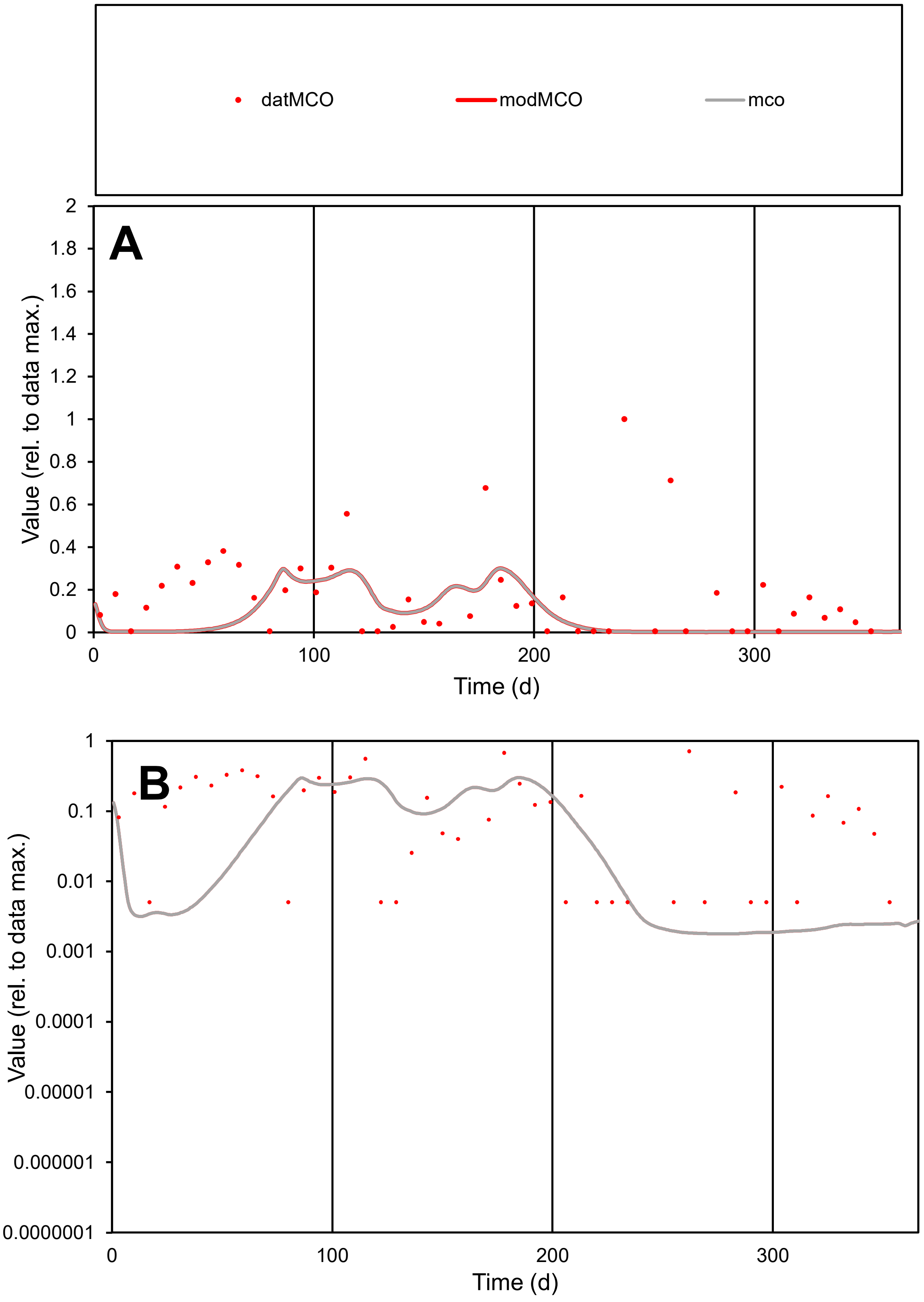


**C1**

Variables

MCO Provide jtd or Dst

CalibOnly?

2 1 to reproduce of

oNormalize?

2 1=no, 2=yes

ofxn calc*

1.34837

ofxn mod

1.36019

sRun

RunA1

zout

1

Ave

Data

2.1E-06

Model

1E-06

data/vMax

1.2E-05

ofxng

369.894

Min d/m

2.8102

ave/min d

33.9431

ave/min m

45.7761

328_Calib_S_A

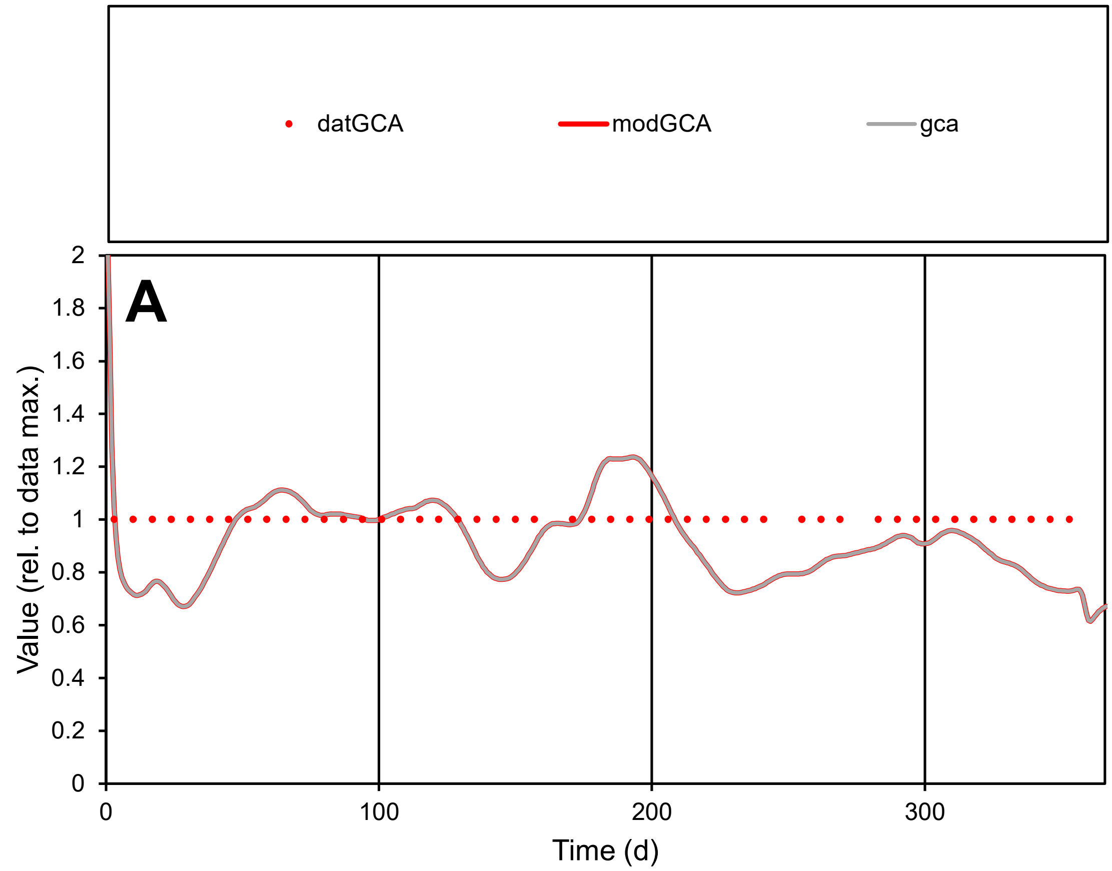


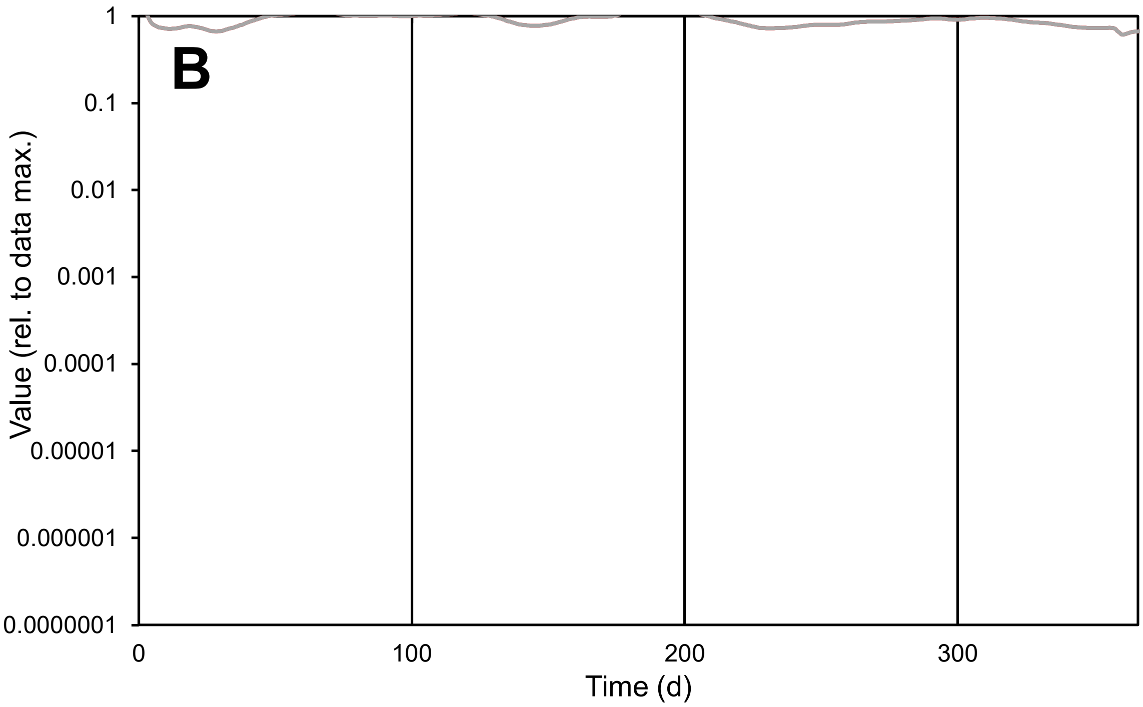


**C1**

Variables
GCA Provide jtd or Dst
CalibOnly?
2 1 to reproduce of
oNormalize?
2 1=no, 2=yes

ofxn calc*
0.16267
ofxn mod
0.1957

sRun
RunA1
zout
1

Ave
Data
8.4E-06
Model
7.7E-06

data/vMax
8.4E-06

ofxng
369.894

Min d/m
1.4326

ave/min d
1
ave/min m
1.31791

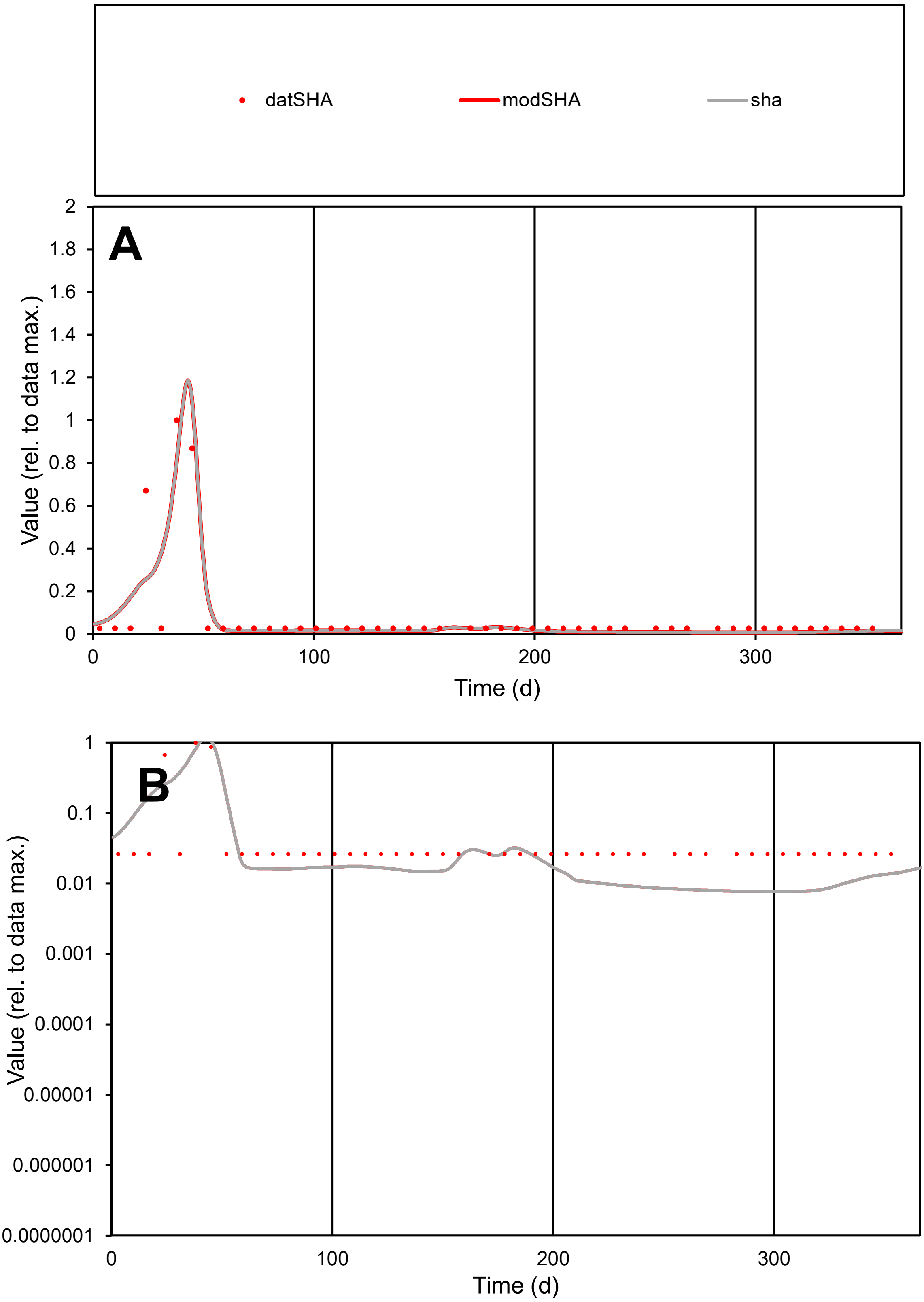


**C1**

Variables
SHA Provide jtd or Dst
CalibOnly?
2 1 to reproduce of
oNormalize?
2 1=no, 2=yes

ofxn calc*
0.99234
ofxn mod
1.10167

sRun
RunA1
zout
1

Ave
Data
2.1E-06
Model
1.8E-06

data/vMax
2.6E-05

ofxng
369.894

Min d/m
3.42703

ave/min d
2.94227
ave/min m
9.05464

328_Calib_S_A

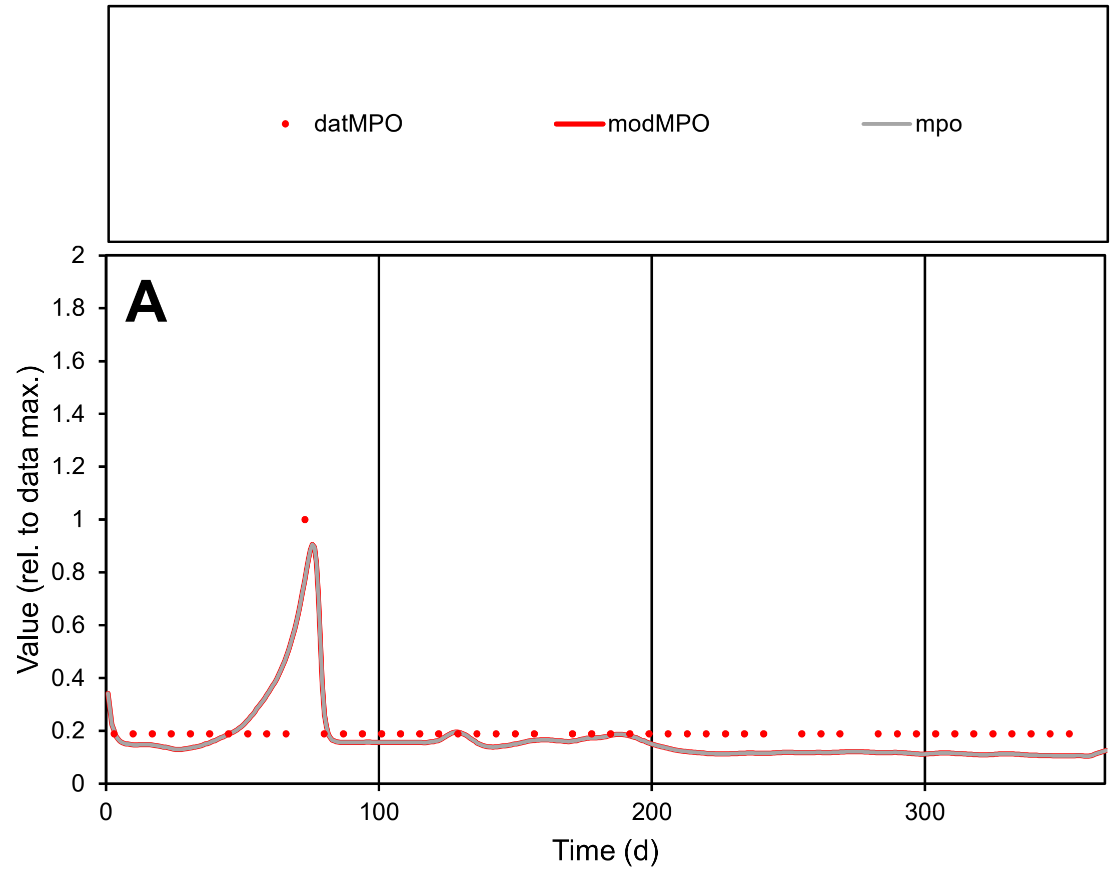


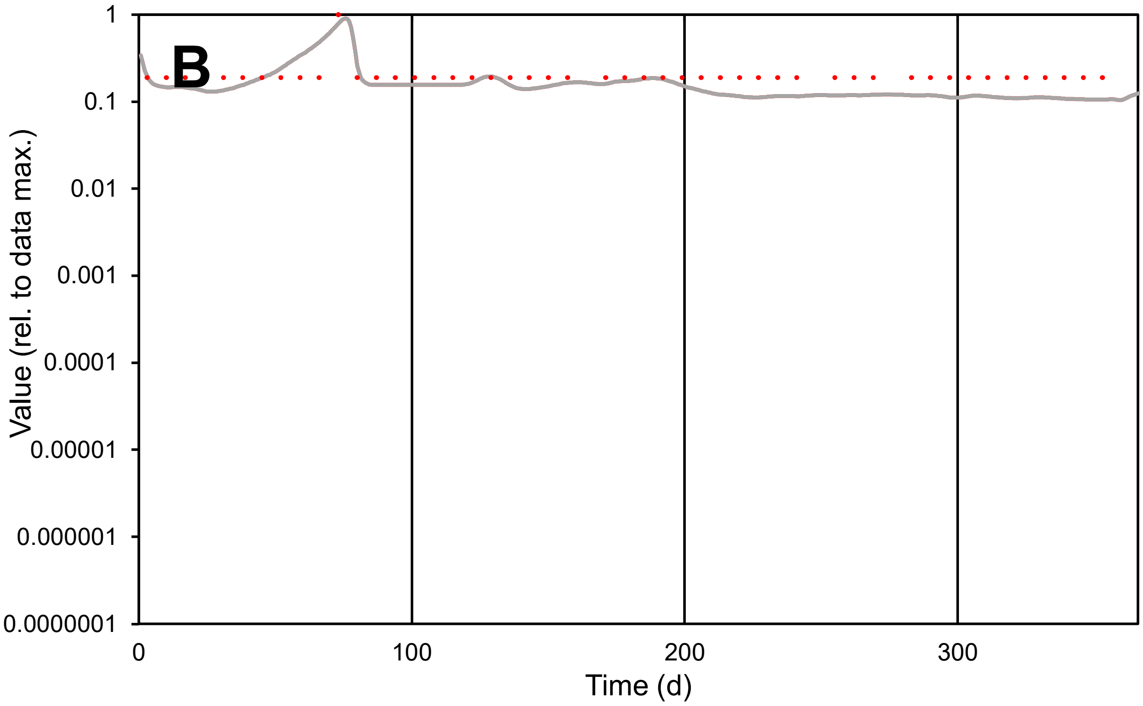


**C1**

Variables
MPO Provide jtd or Dst
CalibOnly?
2 1 to reproduce of
oNormalize?
2 1=no, 2=yes

ofxn calc*
0.36716
ofxn mod
0.38451

sRun
RunA1
zout
1

Ave
Data
1.4E-05
Model
1.1E-05

data/vMax
6.6E-05

ofxng
369.894

Min d/m
1.78253

ave/min d
1.08965
ave/min m
1.59823

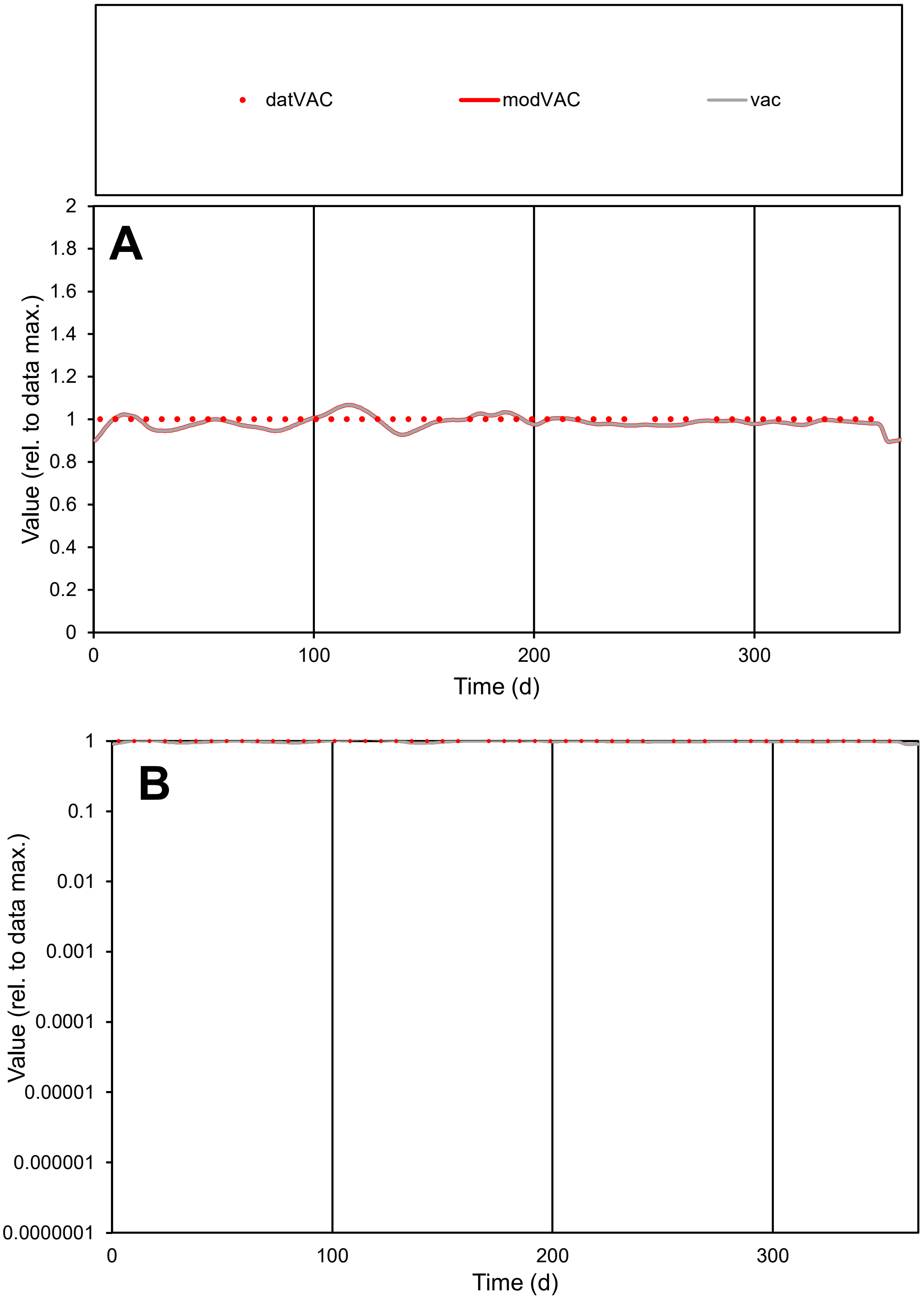


**C1**

Variables
VAC Provide jtd or Dst
CalibOnly?
2 1 to reproduce of
oNormalize?
2 1=no, 2=yes

ofxn calc*
0.02992
ofxn mod
0.11628

sRun
RunA1
zout
1

Ave
Data
9.5E-06
Model
9.4E-06

data/vMax
9.5E-06

ofxng
369.894

Min d/m
1.07066

ave/min d
1
ave/min m
1.05712

328_Calib_S_A

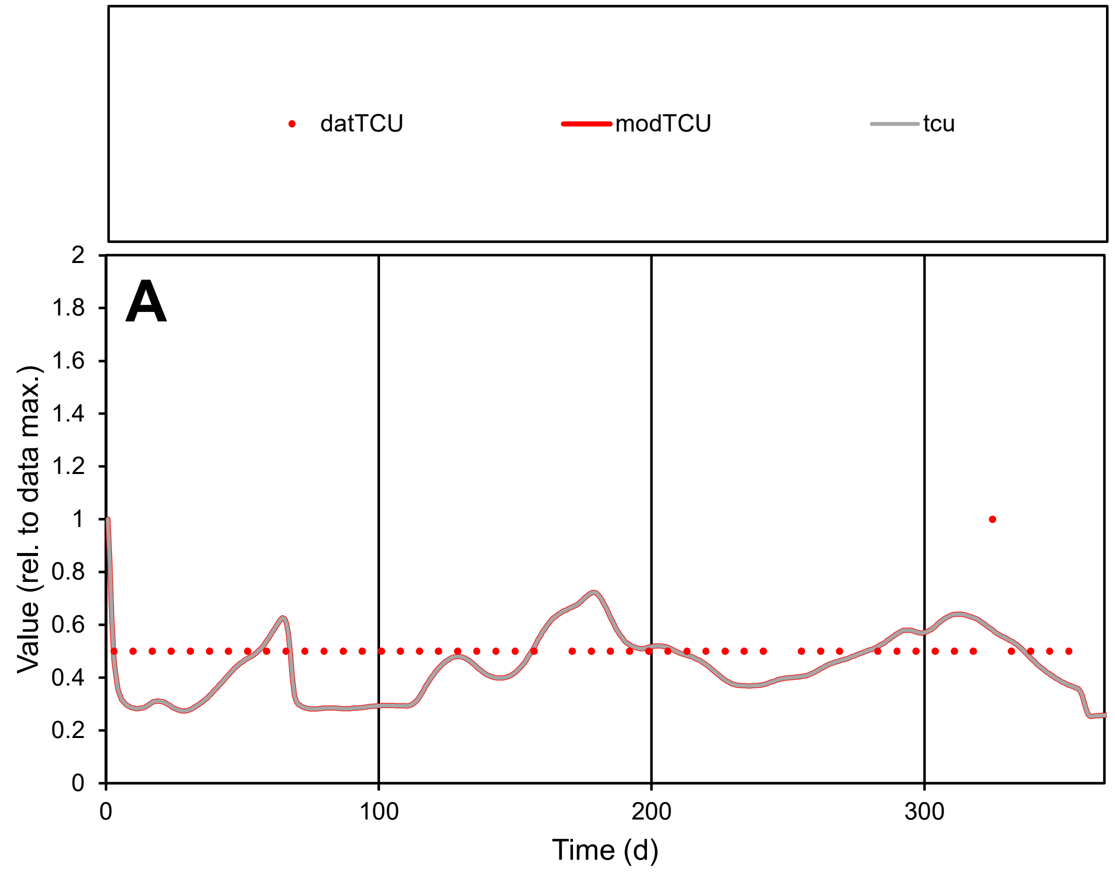


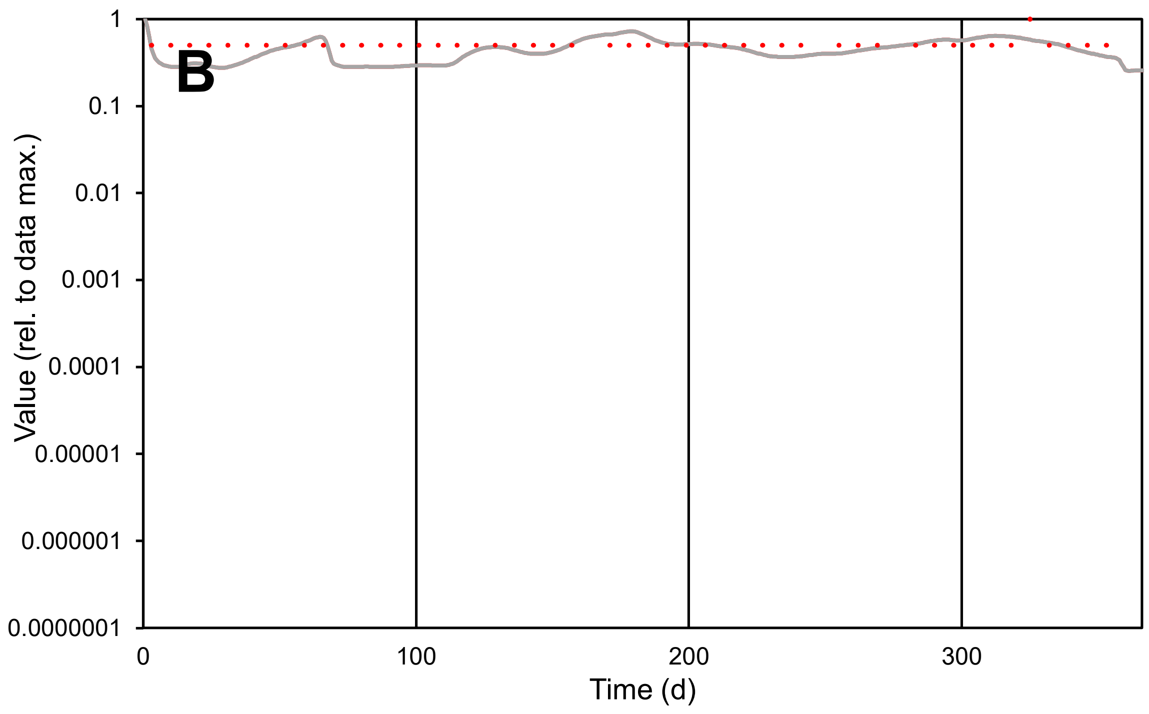


**C1**

Variables
TCU Provide jtd or Dst
CalibOnly?
2 1 to reproduce of
oNormalize?
2 1=no, 2=yes

ofxn calc*
0.27717
ofxn mod
0.29741

sRun
RunA1
zout
1

Ave
Data
1E-05
Model
8.7E-06

data/vMax
2E-05

ofxng
369.894

Min d/m
1.77463

ave/min d
1.02084
ave/min m
1.58783

328_Calib_S_A

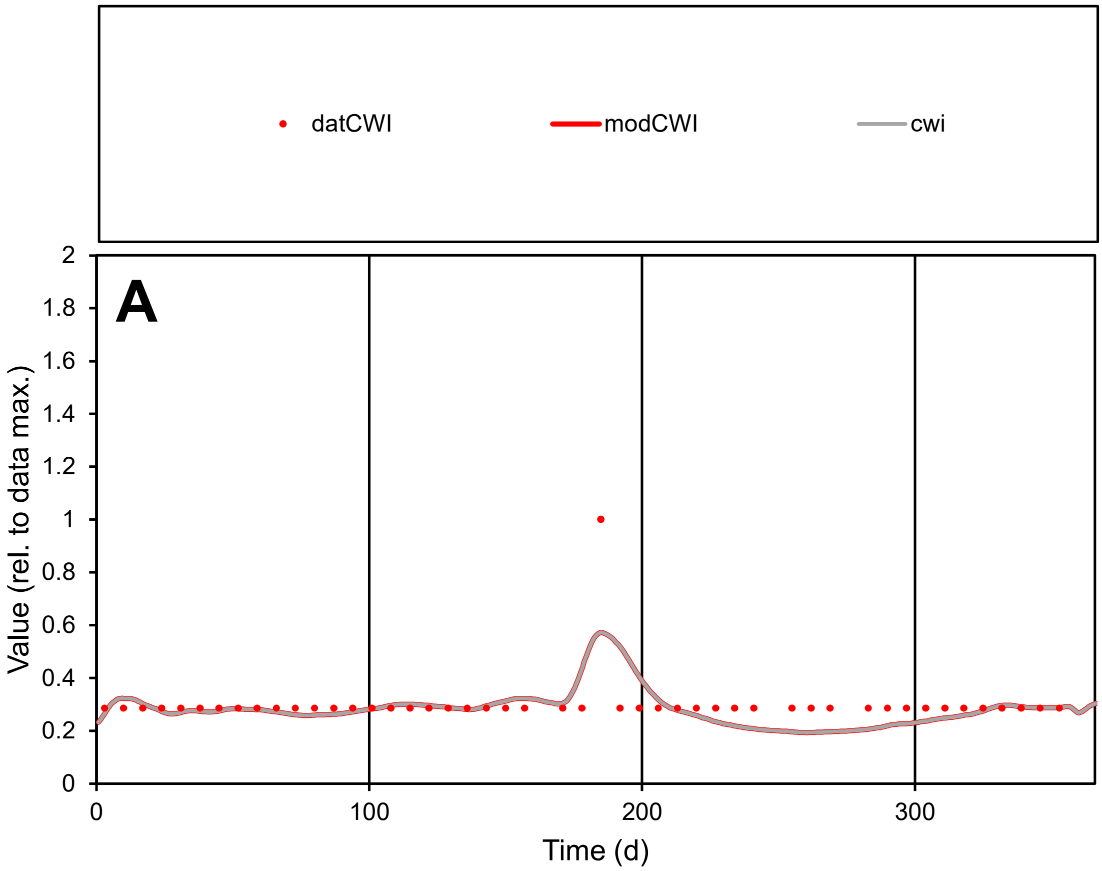


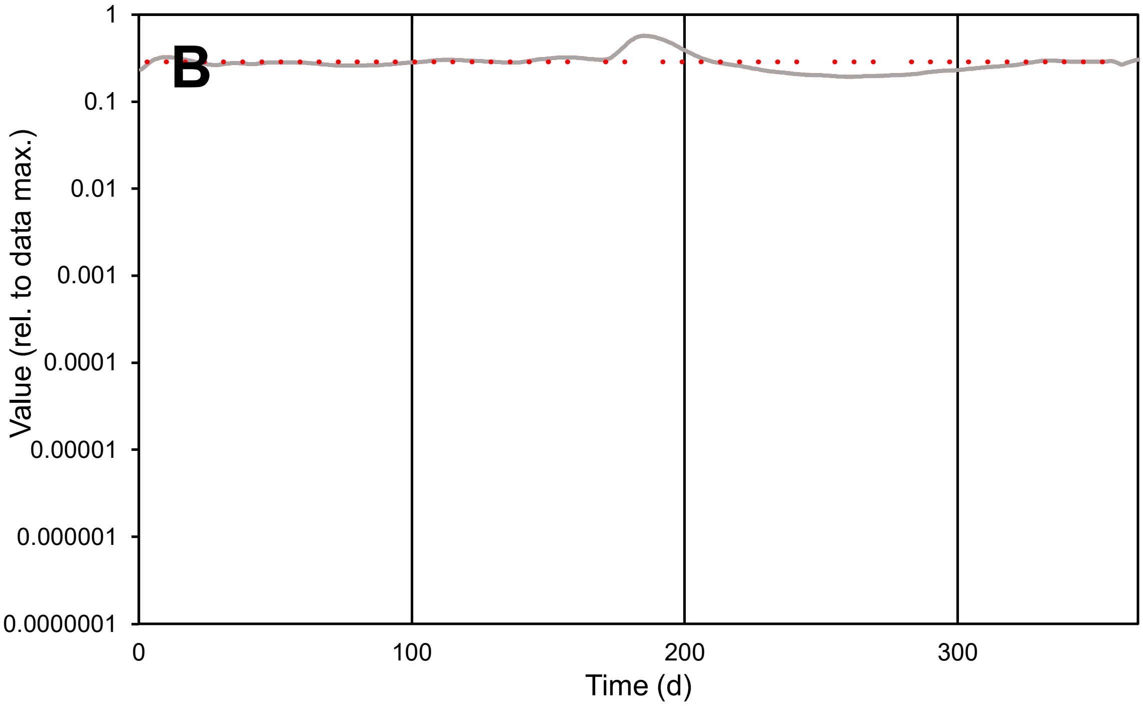


**C1**

Variables
CWI Provide jtd or Dst
CalibOnly?
2 1 to reproduce of
oNormalize?
2 1=no, 2=yes

ofxn calc*
0.27509
ofxn mod
0.29944

sRun
RunA1
zout
1

Ave
Data
1.4E-05
Model
1.3E-05

data/vMax
4.6E-05

ofxng
369.894

Min d/m
1.47143

ave/min d
1.052
ave/min m
1.46815

328_Calib_S_A

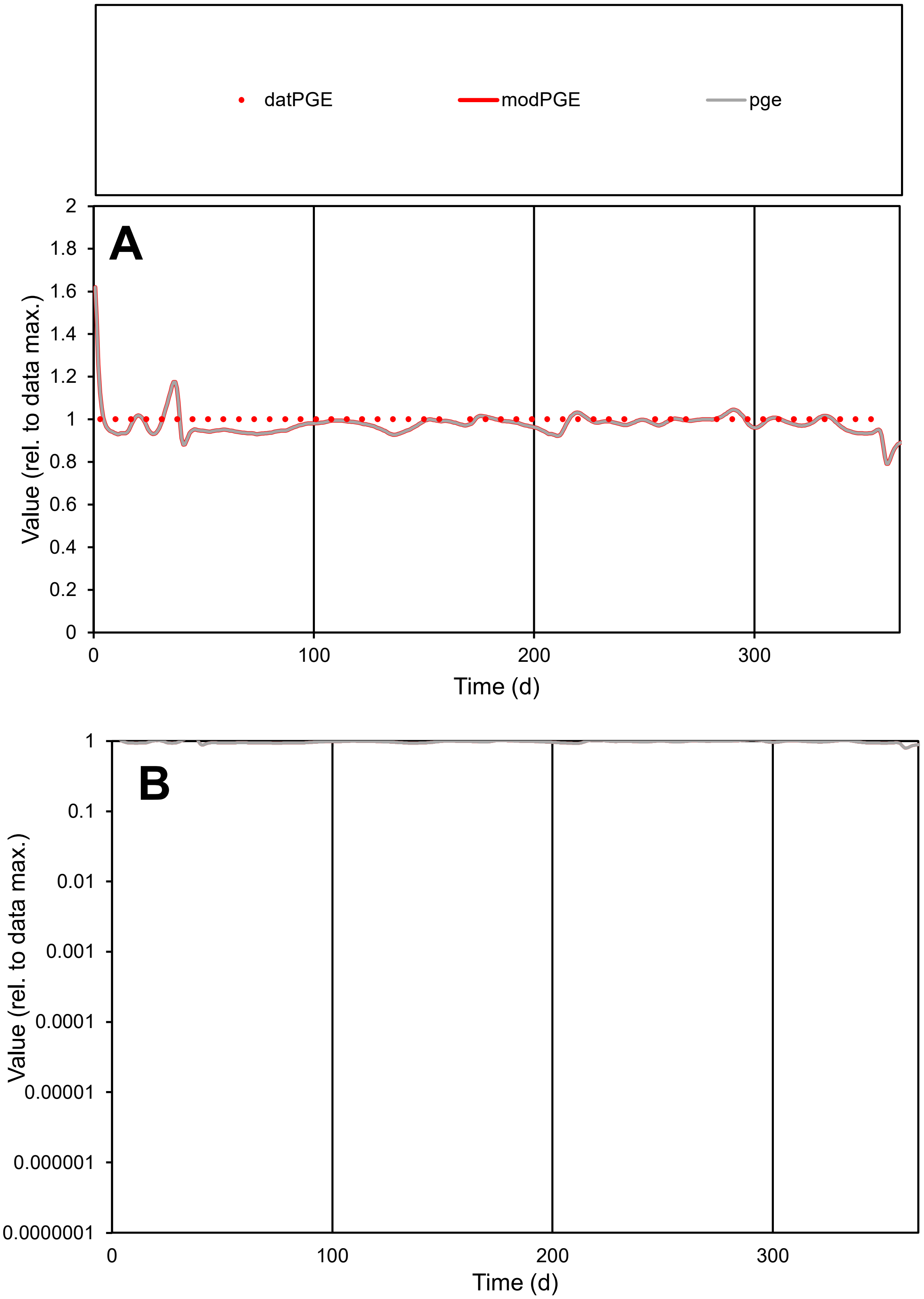


**C1**

Variables

| | |
|---|---|
| PGE | Provide jtd or Dst |
| CalibOnly? | |
| 2 | 1 to reproduce of |
| oNormalize? | |
| 2 | 1=no, 2=yes |

ofxn calc*
0.03725
ofxn mod
0.11793

sRun
RunA1
zout
1

Ave
Data
5.9E-06
Model
5.7E-06

data/vMax
5.9E-06

ofxng
369.894

Min d/m
1.07685

ave/min d
1
ave/min m
1.05178

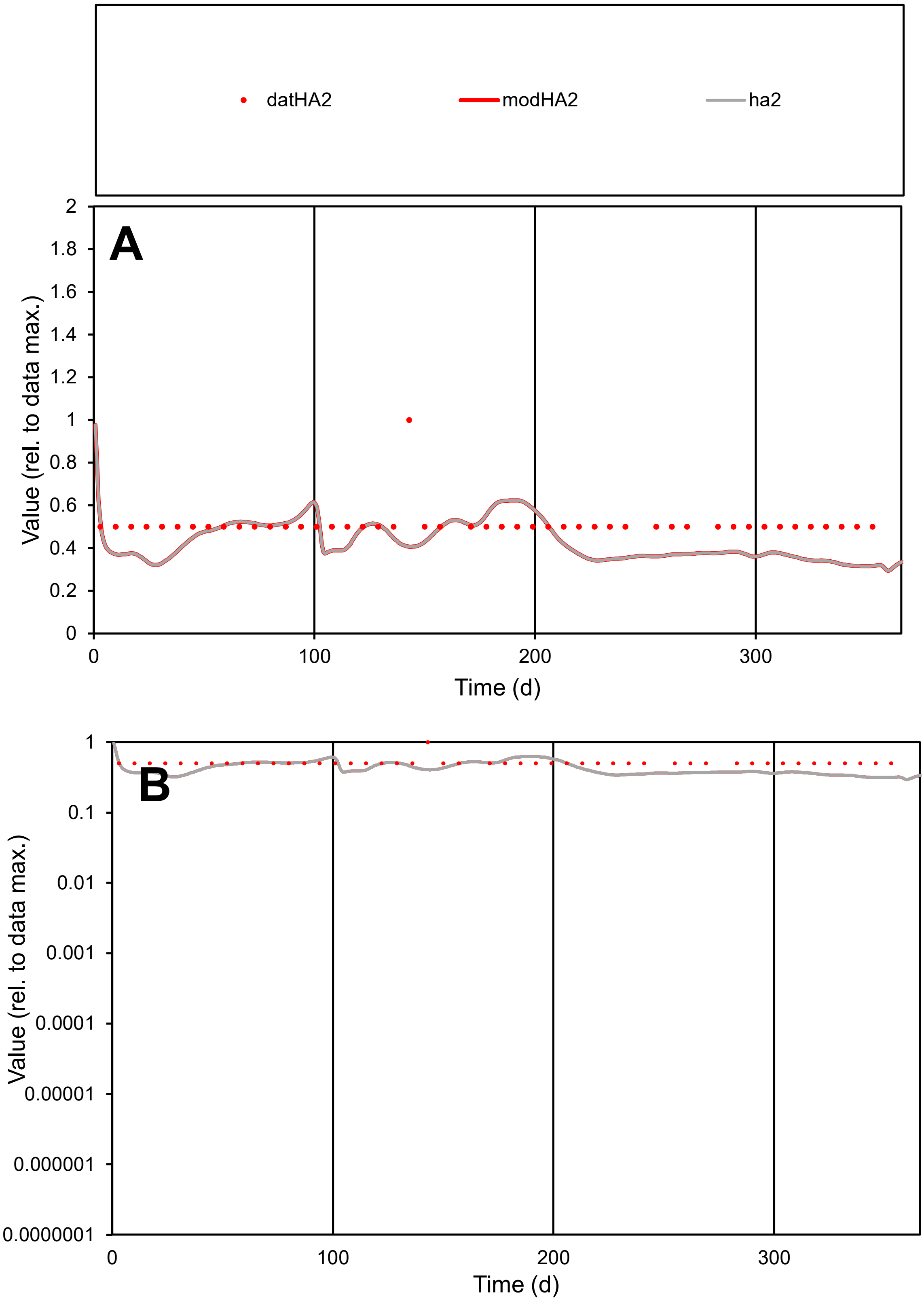


**C1**

Variables
HA2 Provide jtd or Dst
CalibOnly?
2 1 to reproduce of
oNormalize?
2 1=no, 2=yes

ofxn calc*
0.2718
ofxn mod
0.29101

sRun
RunA1
zout
1

Ave
Data
2.7E-06
Model
2.3E-06

data/vMax
5.3E-06

ofxng
369.894

Min d/m
1.58393

ave/min d
1.02083
ave/min m
1.36258

328_Calib_S_A

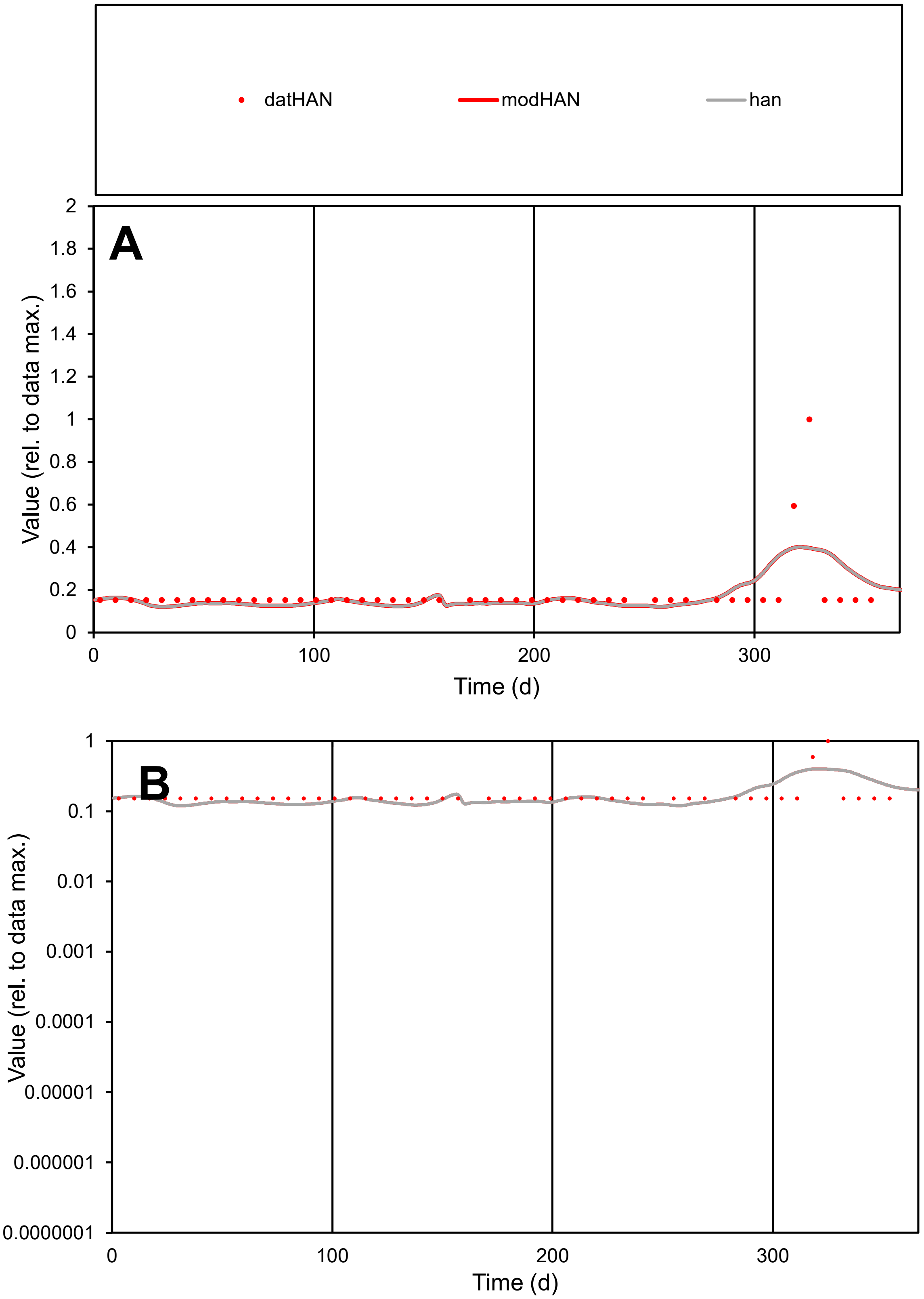


**C1**

Variables
HAN Provide jtd or Dst
CalibOnly?
2 1 to reproduce of
oNormalize?
2 1=no, 2=yes

ofxn calc*
0.61636
ofxn mod
0.62996

sRun
RunA1
zout
1

Ave
Data
6E-06
Model
5.8E-06

data/vMax
3.4E-05

ofxng
369.894

Min d/m
1.26957

ave/min d
1.17594
ave/min m
1.44566

328_Calib_S_A

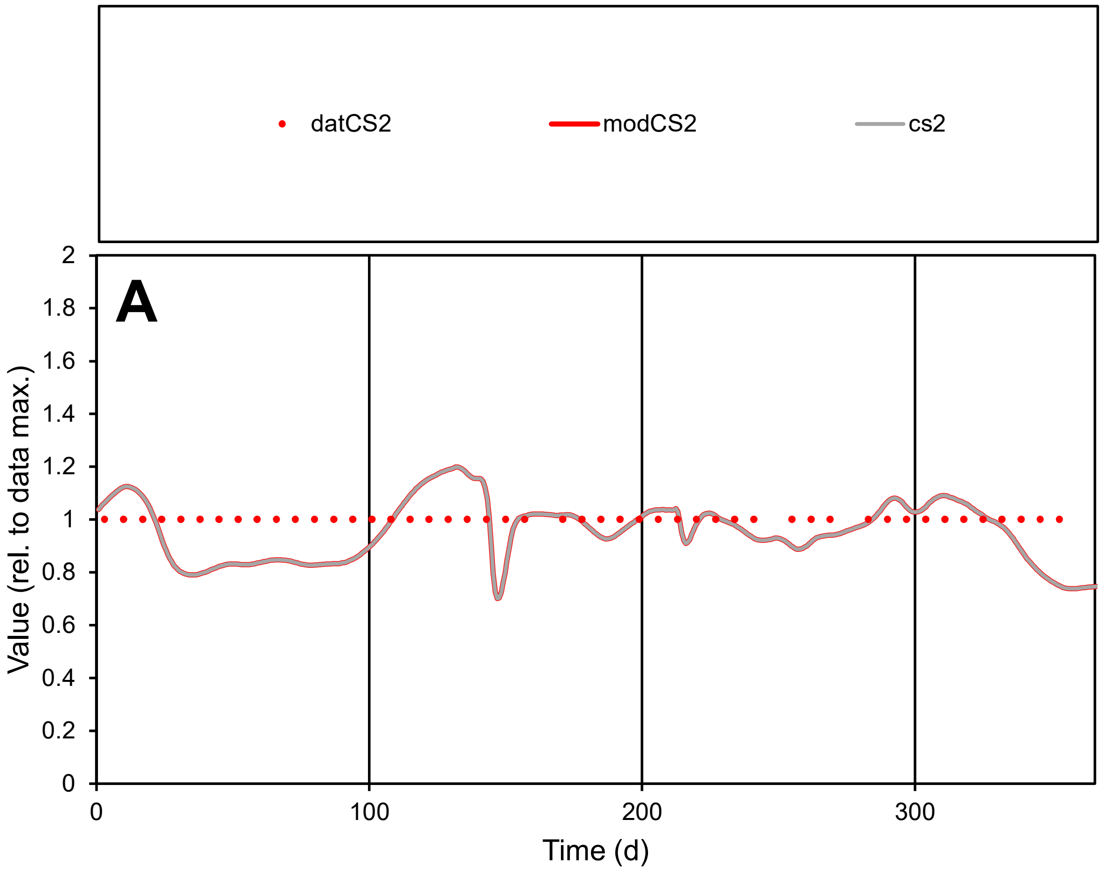


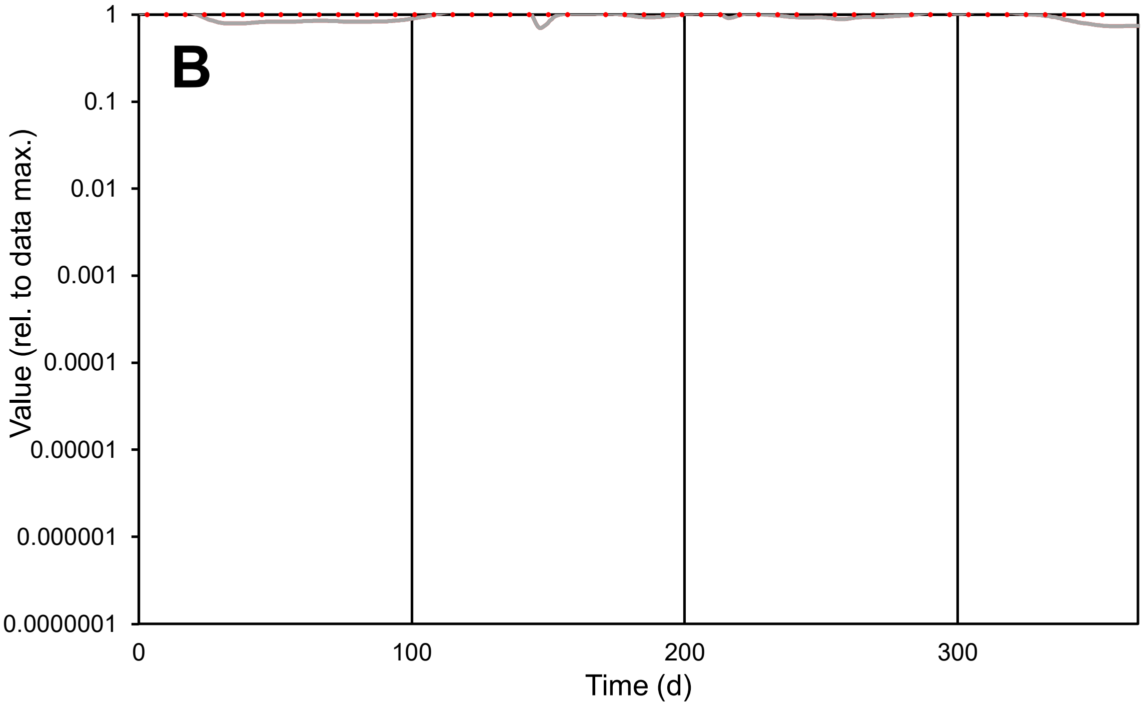


**C1**

Variables
CS2 Provide jtd or Dst
CalibOnly?
2 1 to reproduce of
oNormalize?
2 1=no, 2=yes

ofxn calc*
0.11601
ofxn mod
0.16054

sRun
RunA1
zout
1

Ave
Data
7.6E-06
Model
7.3E-06

data/vMax
7.6E-06

ofxng
369.894

Min d/m
1.34433

ave/min d
1
ave/min m
1.2928

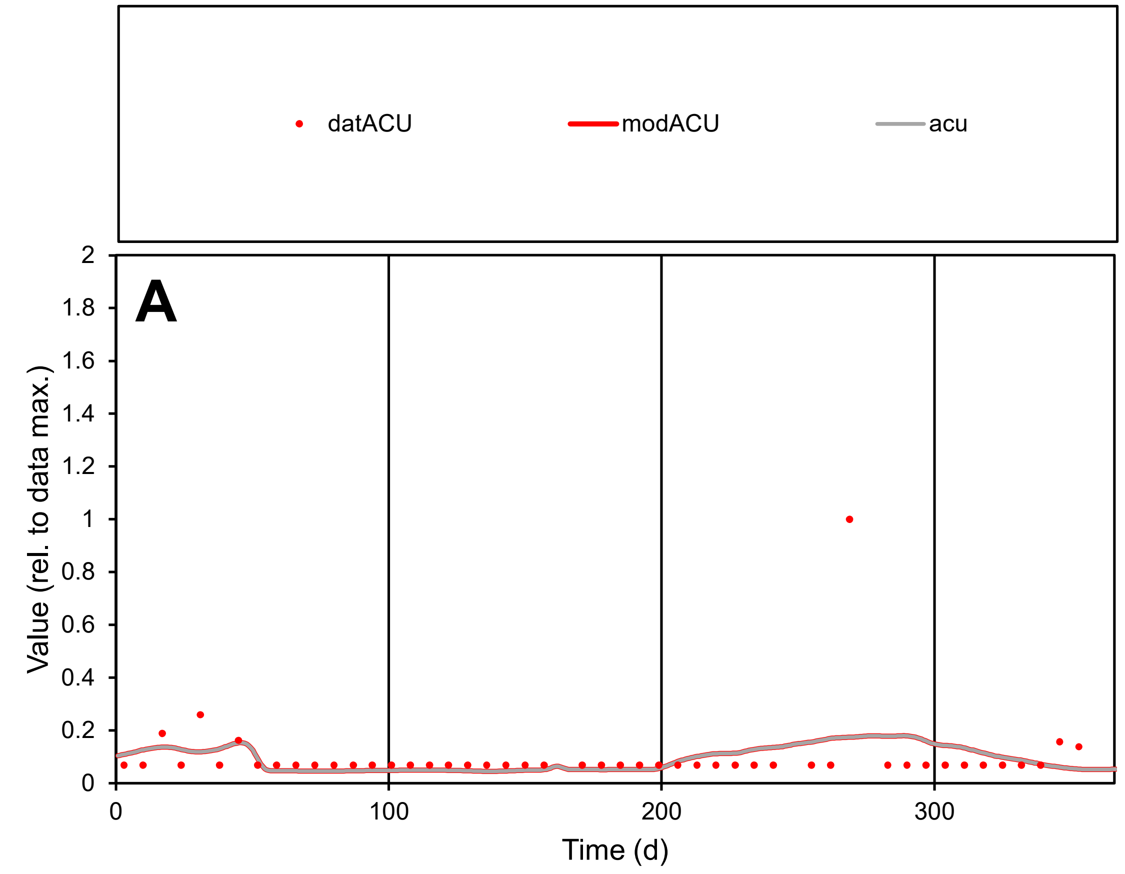


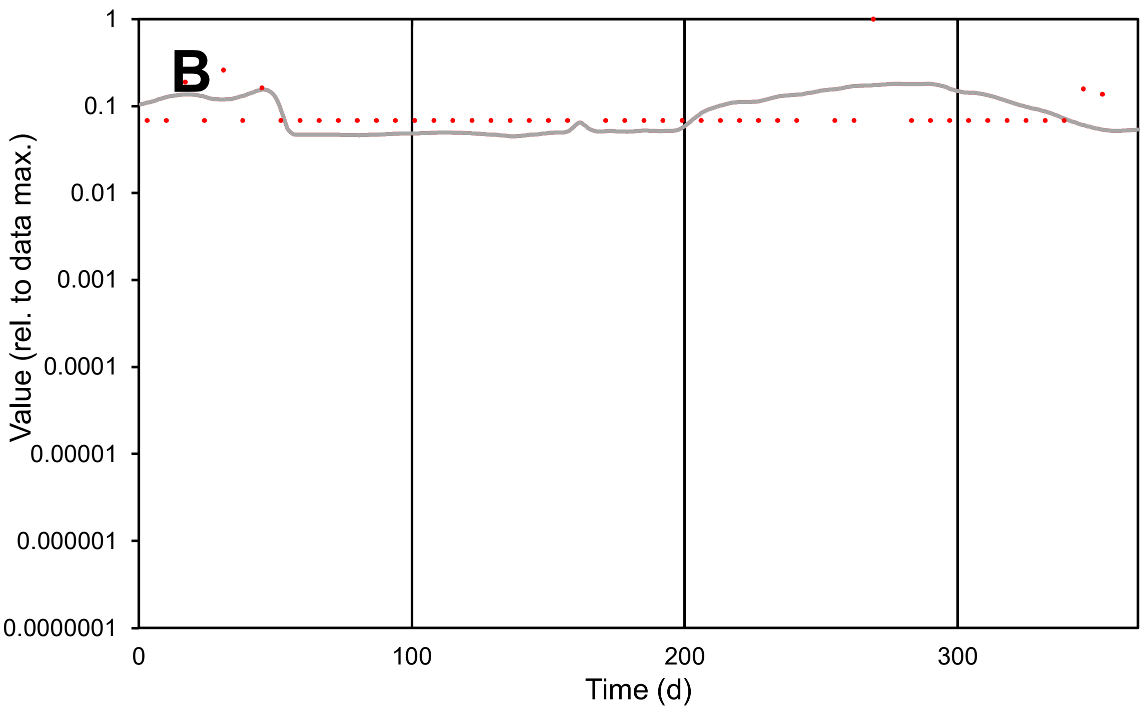


**C1**

Variables
ACU Provide jtd or Dst
CalibOnly?
2 1 to reproduce of
oNormalize?
2 1=no, 2=yes

ofxn calc*
1.30663
ofxn mod
1.31589

sRun
RunA1
zout
1

Ave
Data
1.6E-05
Model
1.5E-05

data/vMax
0.00016

ofxng
369.894

Min d/m
1.52505

ave/min d
1.45314
ave/min m
2.04036

328_Calib_S_A

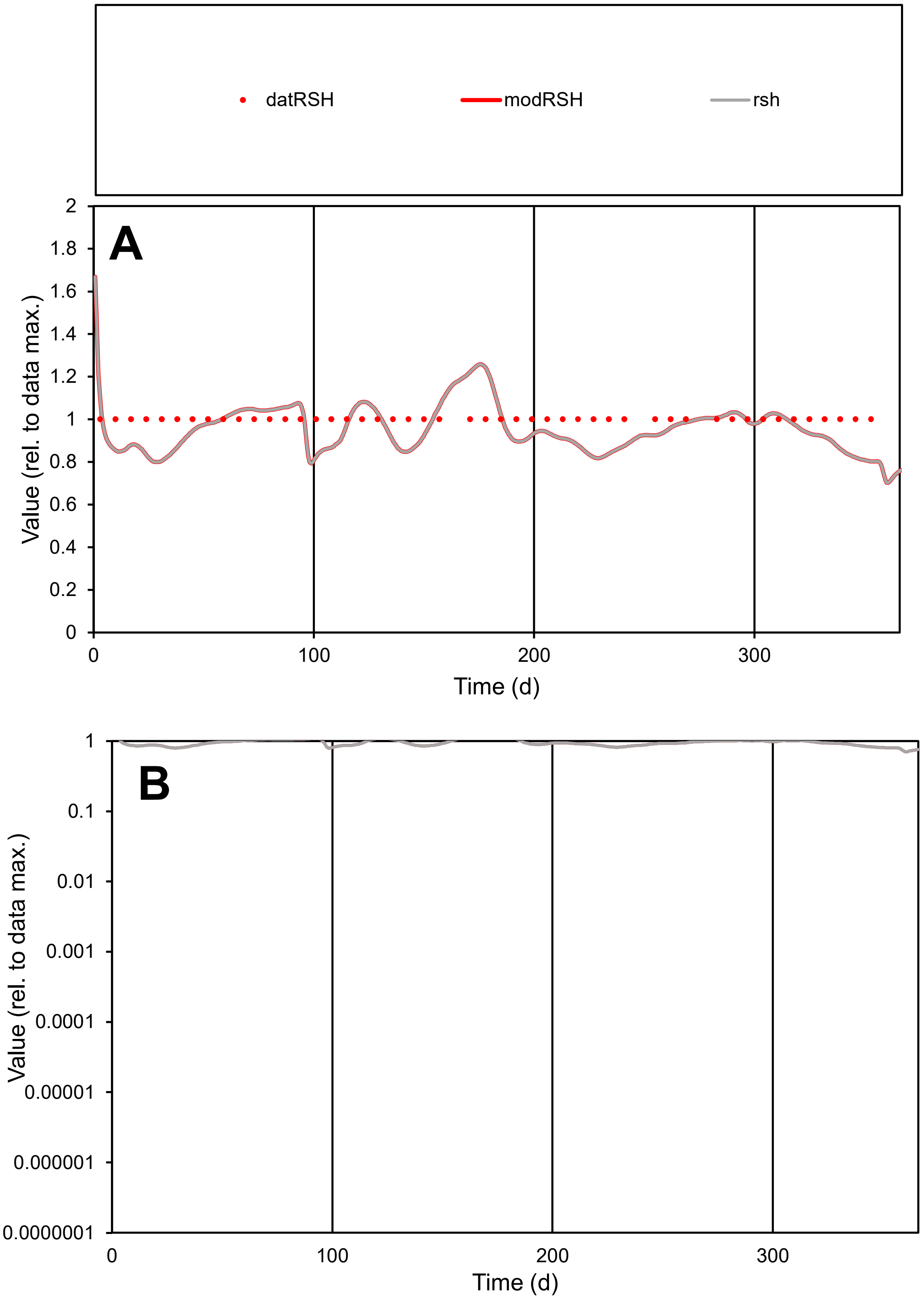


**C1**

Variables
RSH Provide jtd or Dst
CalibOnly?
2 1 to reproduce of
oNormalize?
2 1=no, 2=yes

ofxn calc*
0.10733
ofxn mod
0.15395

sRun
RunA1
zout
1

Ave
Data
1.8E-05
Model
1.7E-05

data/vMax
1.8E-05

ofxng
369.894

Min d/m
1.24547

ave/min d
1
ave/min m
1.18778

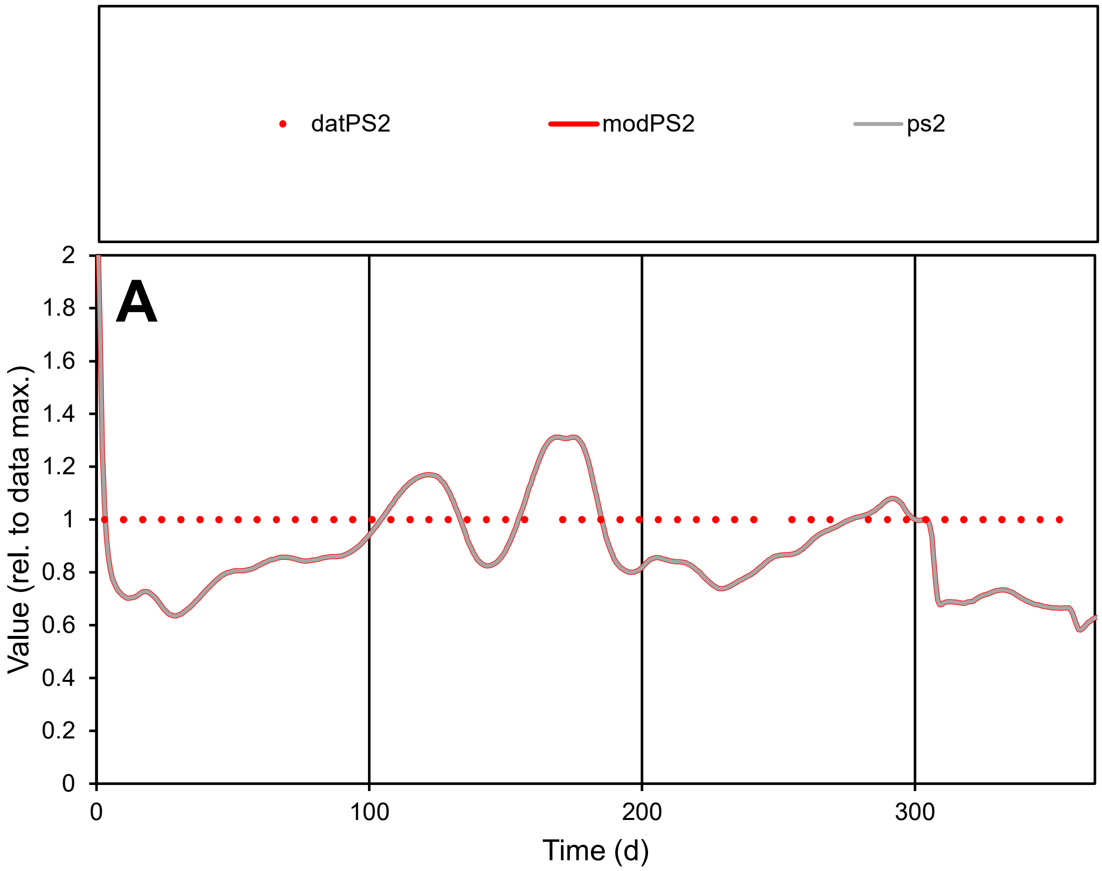


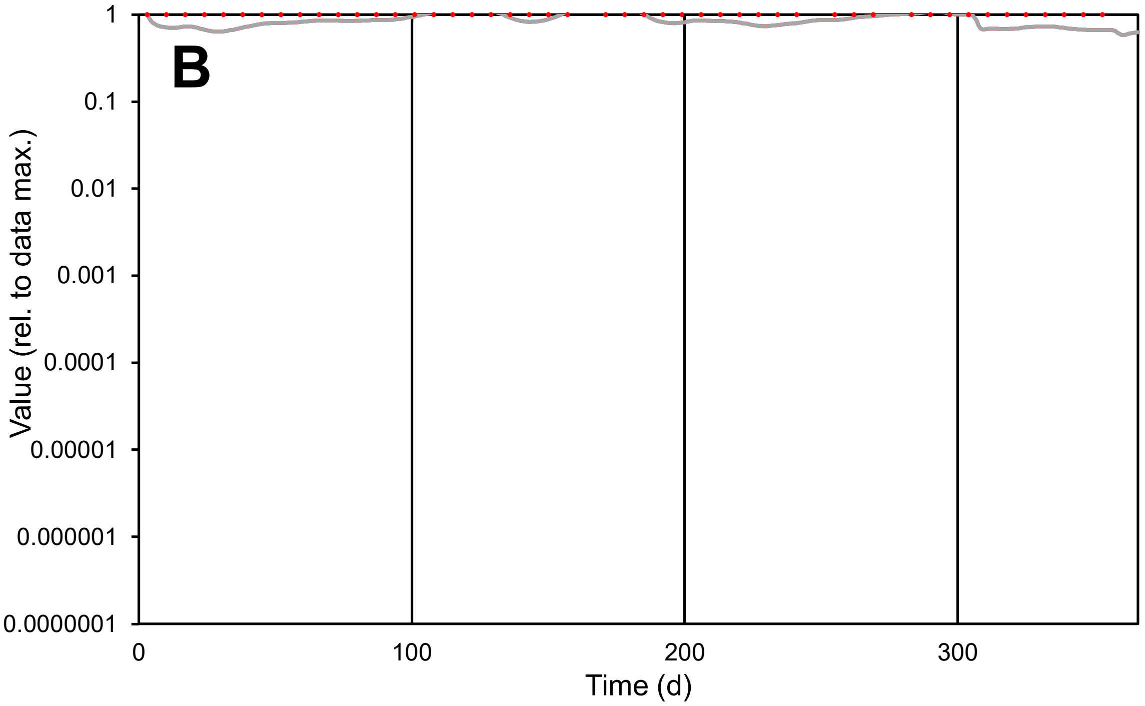


**C1**

Variables
PS2 Provide jtd or Dst
CalibOnly?
2 1 to reproduce of
oNormalize?
2 1=no, 2=yes

ofxn calc*
0.20258
ofxn mod
0.22875

sRun
RunA1
zout
1

Ave
Data
1.9E-05
Model
1.7E-05

data/vMax
1.9E-05

ofxng
369.894

Min d/m
1.52644

ave/min d
1
ave/min m
1.33599

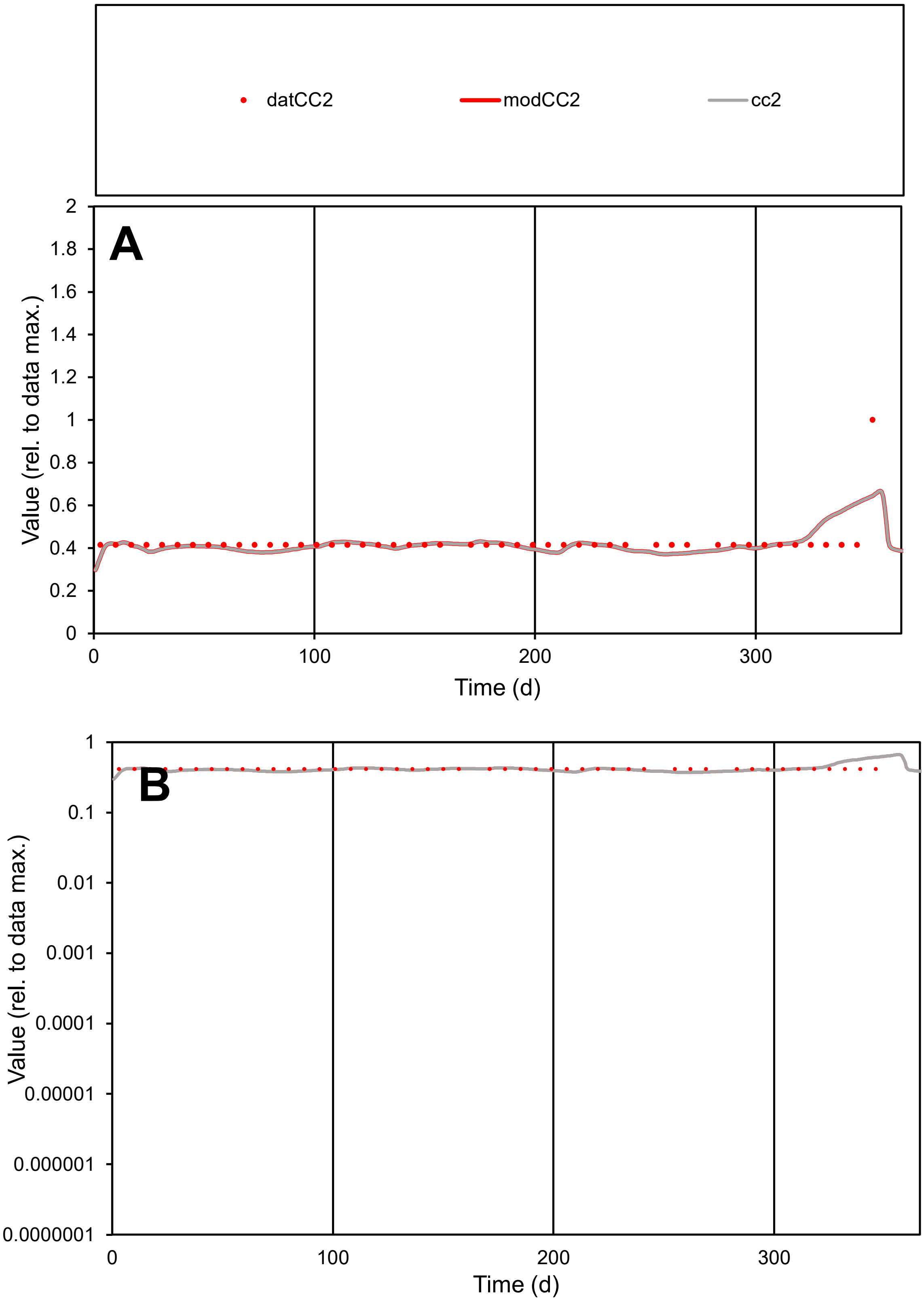


**C1**

Variables
CC2 Provide jtd or Dst
CalibOnly?
2 1 to reproduce of
oNormalize?
2 1=no, 2=yes

ofxn calc*
0.16113
ofxn mod
0.19598

sRun
RunA1
zout
1

Ave
Data
6E-06
Model
6E-06

data/vMax
1.4E-05

ofxng
369.894

Min d/m
1.1109

ave/min d
1.02946
ave/min m
1.13227

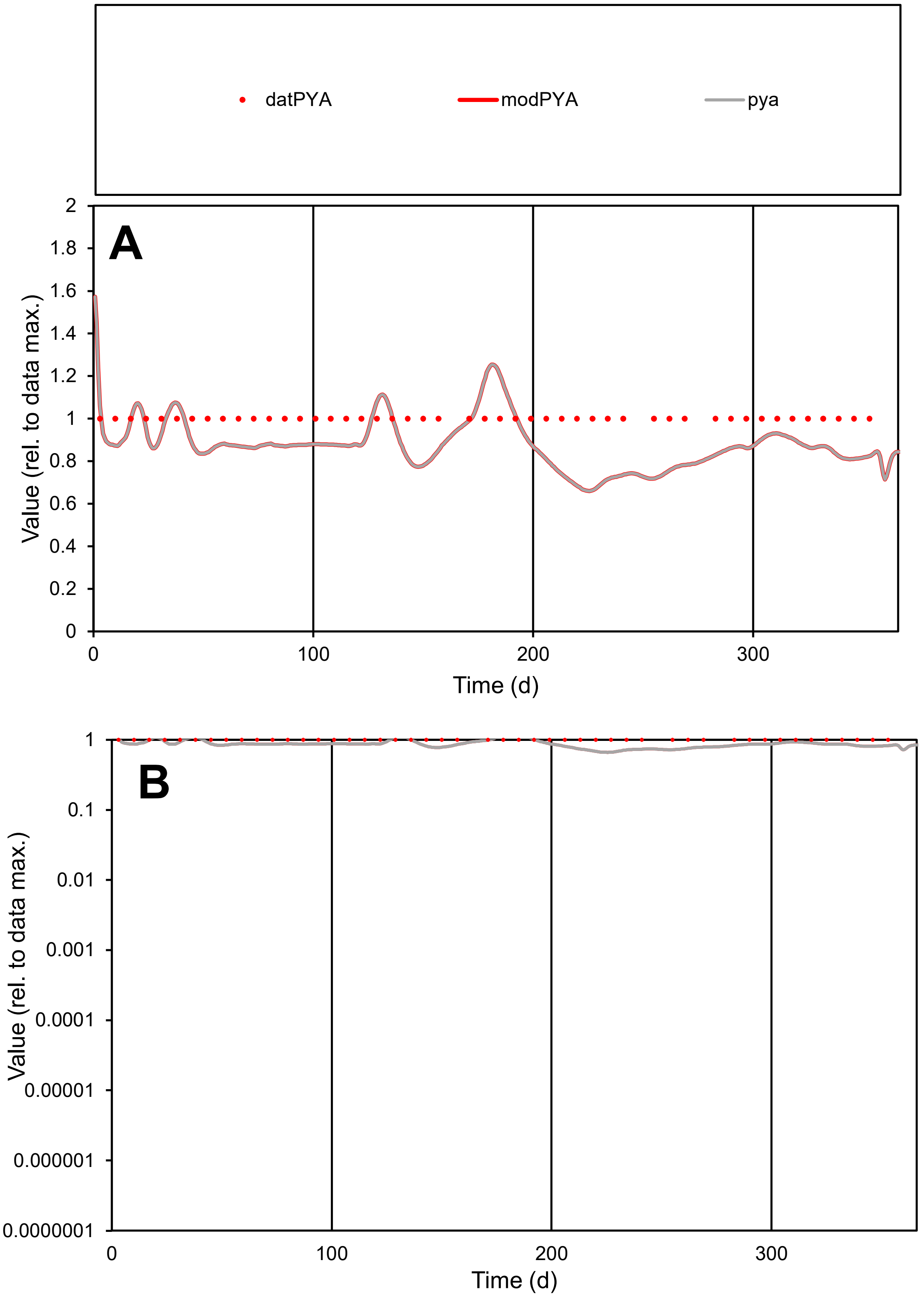


**C1**

Variables
PYA Provide jtd or Dst
CalibOnly?
2 1 to reproduce of
oNormalize?
2 1=no, 2=yes

ofxn calc*
0.16554
ofxn mod
0.19665

sRun
RunA1
zout
1

Ave
Data
1.8E-06
Model
1.6E-06

data/vMax
1.8E-06

ofxng
369.894

Min d/m
1.49136

ave/min d
1
ave/min m
1.31187

328_Calib_S_A

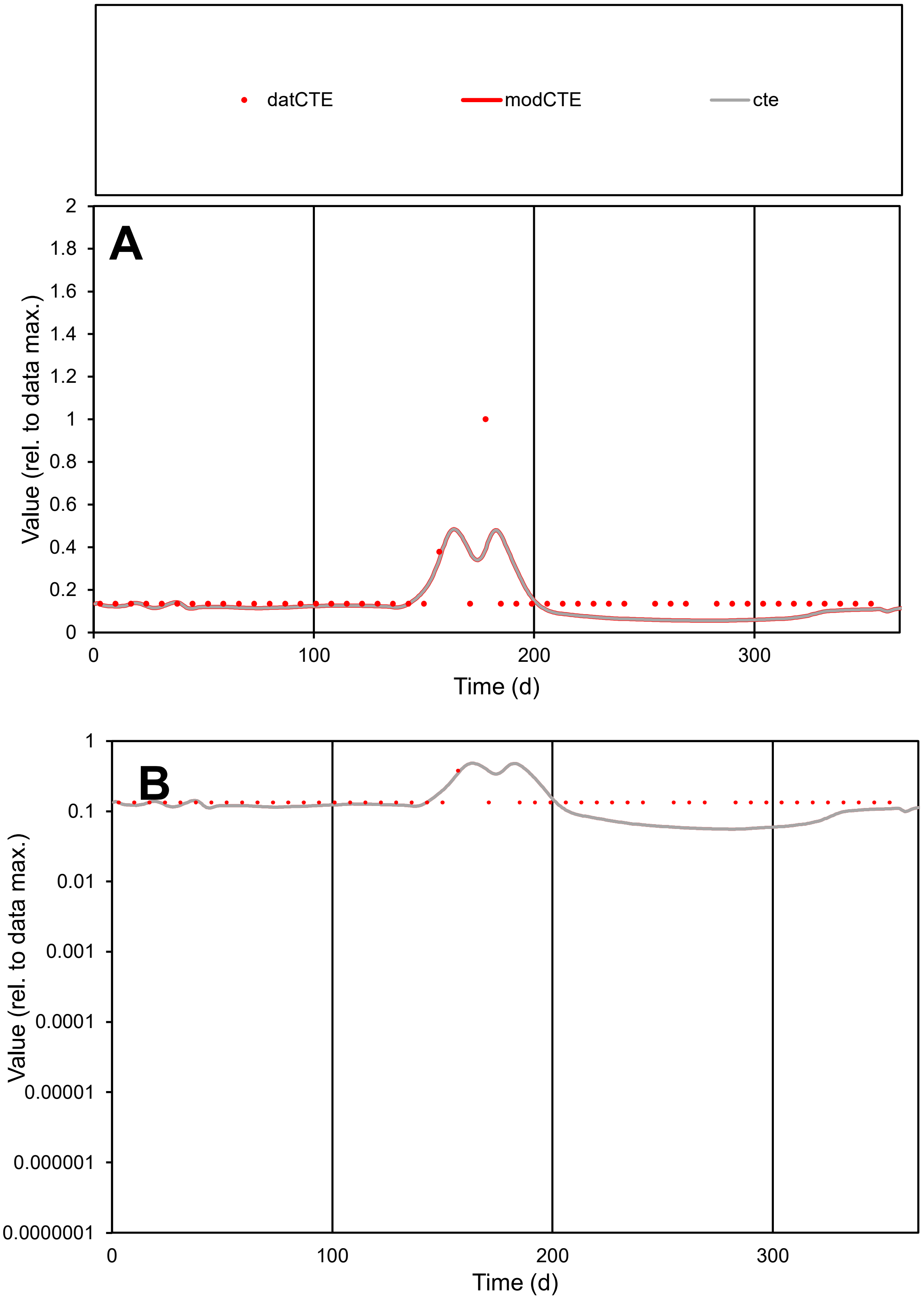


**C1**

Variables
CTE Provide jtd or Dst
CalibOnly?
2 1 to reproduce of
oNormalize?
2 1=no, 2=yes

ofxn calc*
0.66974
ofxn mod
0.69294

sRun
RunA1
zout
1

Ave
Data
7.2E-06
Model
6E-06

data/vMax
4.6E-05

ofxng
369.894

Min d/m
2.39578

ave/min d
1.17231
ave/min m
2.34521

328_Calib_S_A

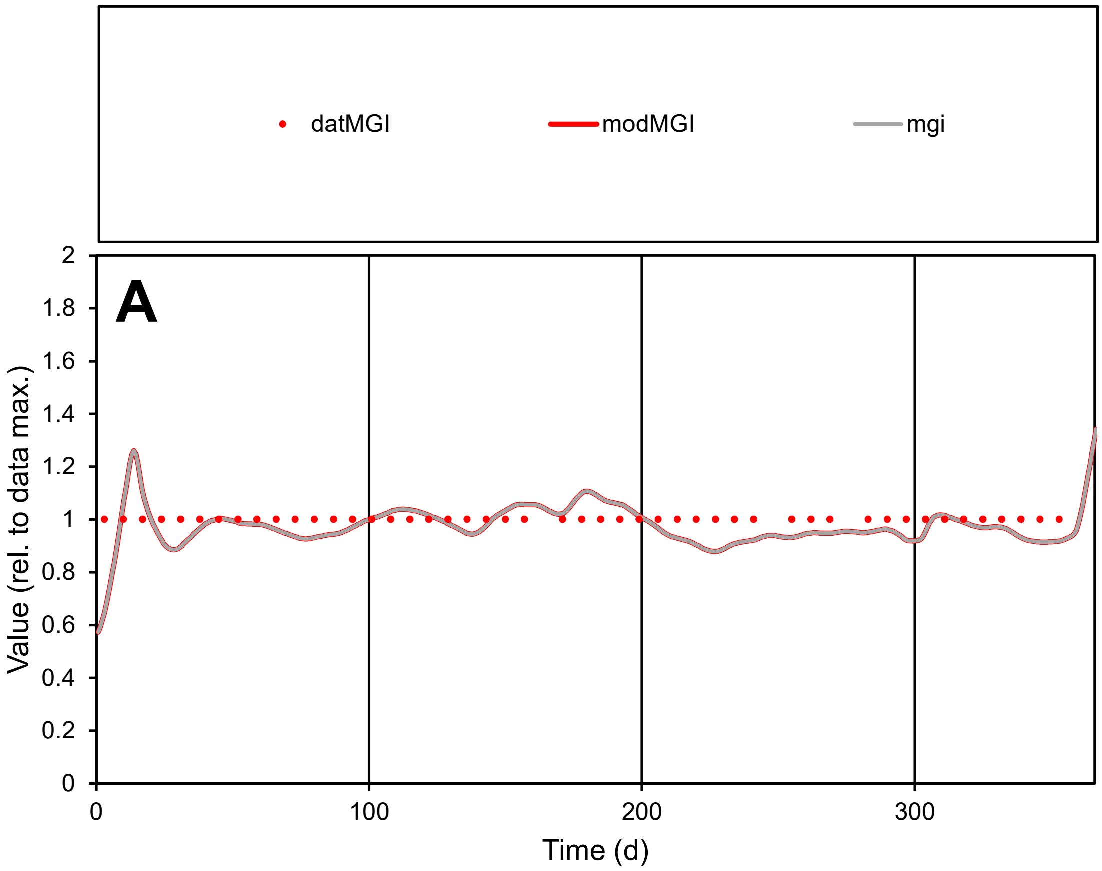


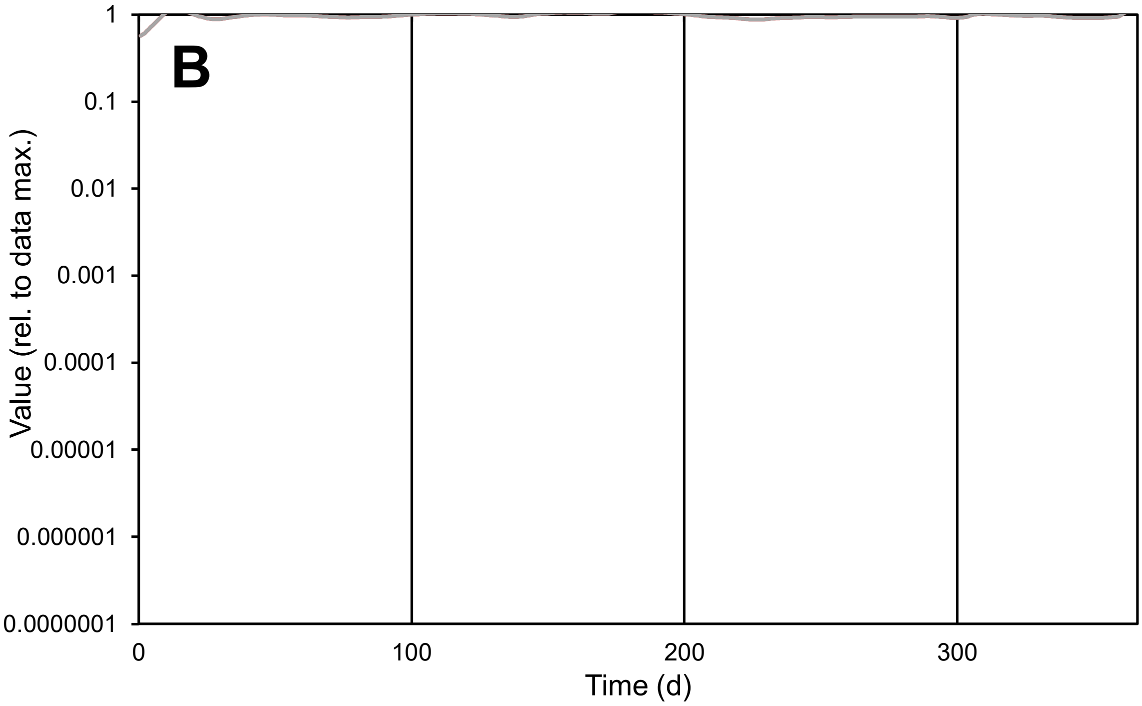


**C1**

Variables

| | |
|---|---|
| MGI | Provide jtd or Dst |
| CalibOnly? | |
| 2 | 1 to reproduce of |
| oNormalize? | |
| 2 | 1=no, 2=yes |

ofxn calc*
0.07412
ofxn mod
0.13368

sRun
RunA1
zout
1

Ave
Data
1E-05
Model
1E-05

data/vMax
1E-05

ofxng
369.894

Min d/m
1.40046

ave/min d
1
ave/min m
1.36147

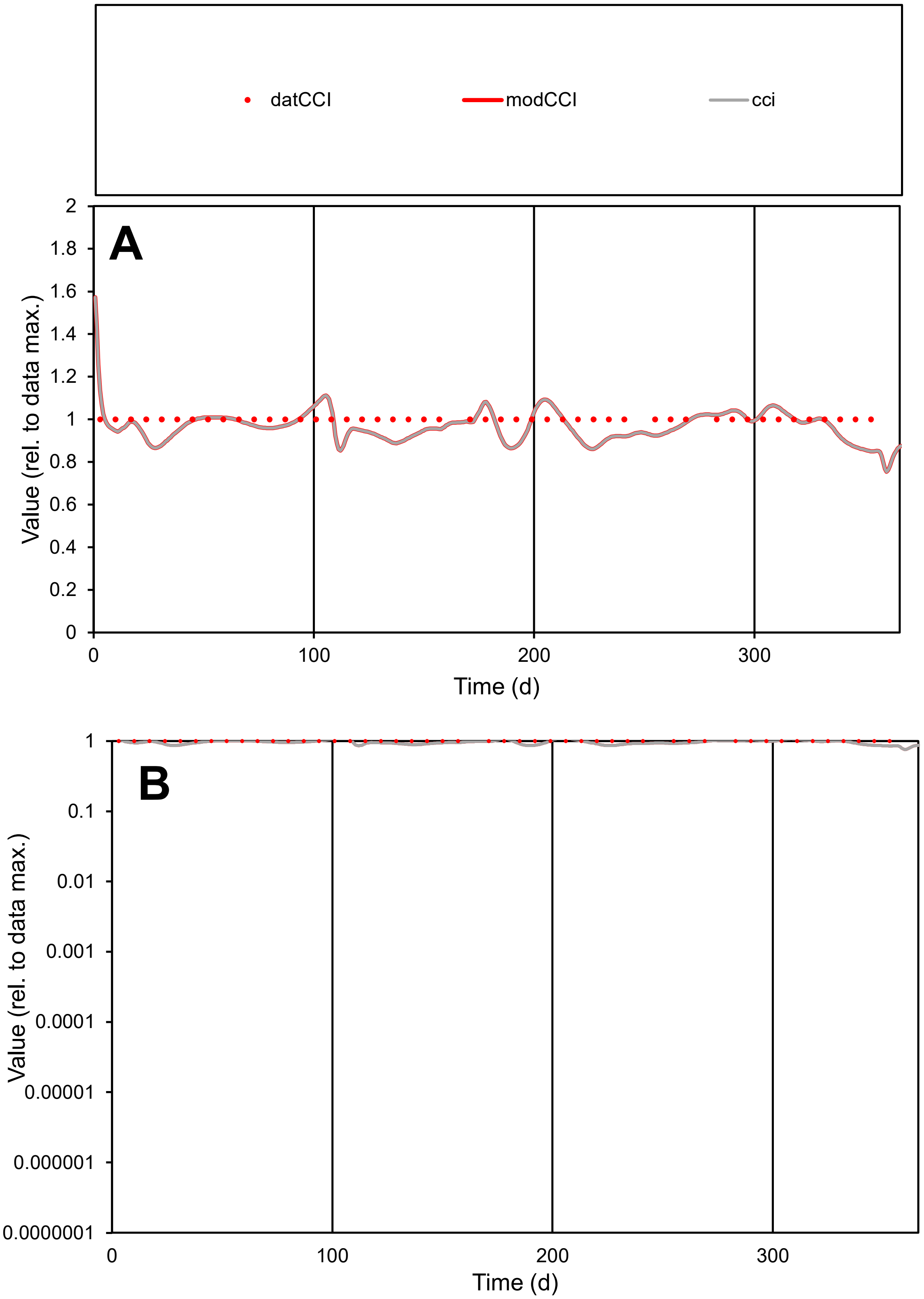


**C1**

Variables
CCI Provide jtd or Dst
CalibOnly?
2 1 to reproduce of
oNormalize?
2 1=no, 2=yes

ofxn calc*
0.06794
ofxn mod
0.13024

sRun
RunA1
zout
1

Ave
Data
1.1E-05
Model
1.1E-05

data/vMax
1.1E-05

ofxng
369.894

Min d/m
1.17563

ave/min d
1
ave/min m
1.13607

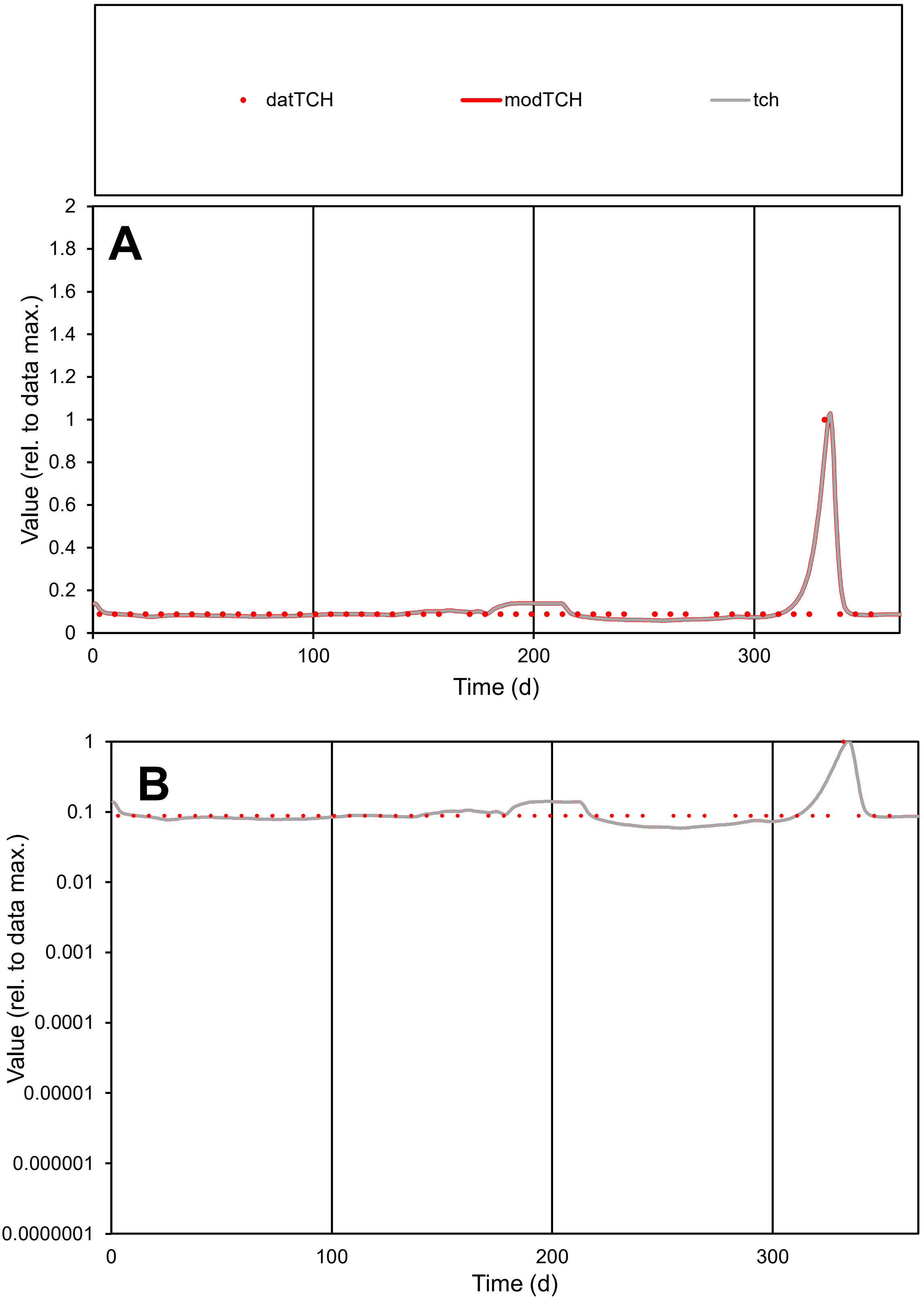


**C1**

Variables
TCH Provide jtd or Dst
CalibOnly?
2 1 to reproduce of
oNormalize?
2 1=no, 2=yes

ofxn calc*
0.31845
ofxn mod
0.37481

sRun
RunA1
zout
1

Ave
Data
2E-06
Model
2.1E-06

data/vMax
1.9E-05

ofxng
369.894

Min d/m
1.49511

ave/min d
1.21611
ave/min m
1.89122

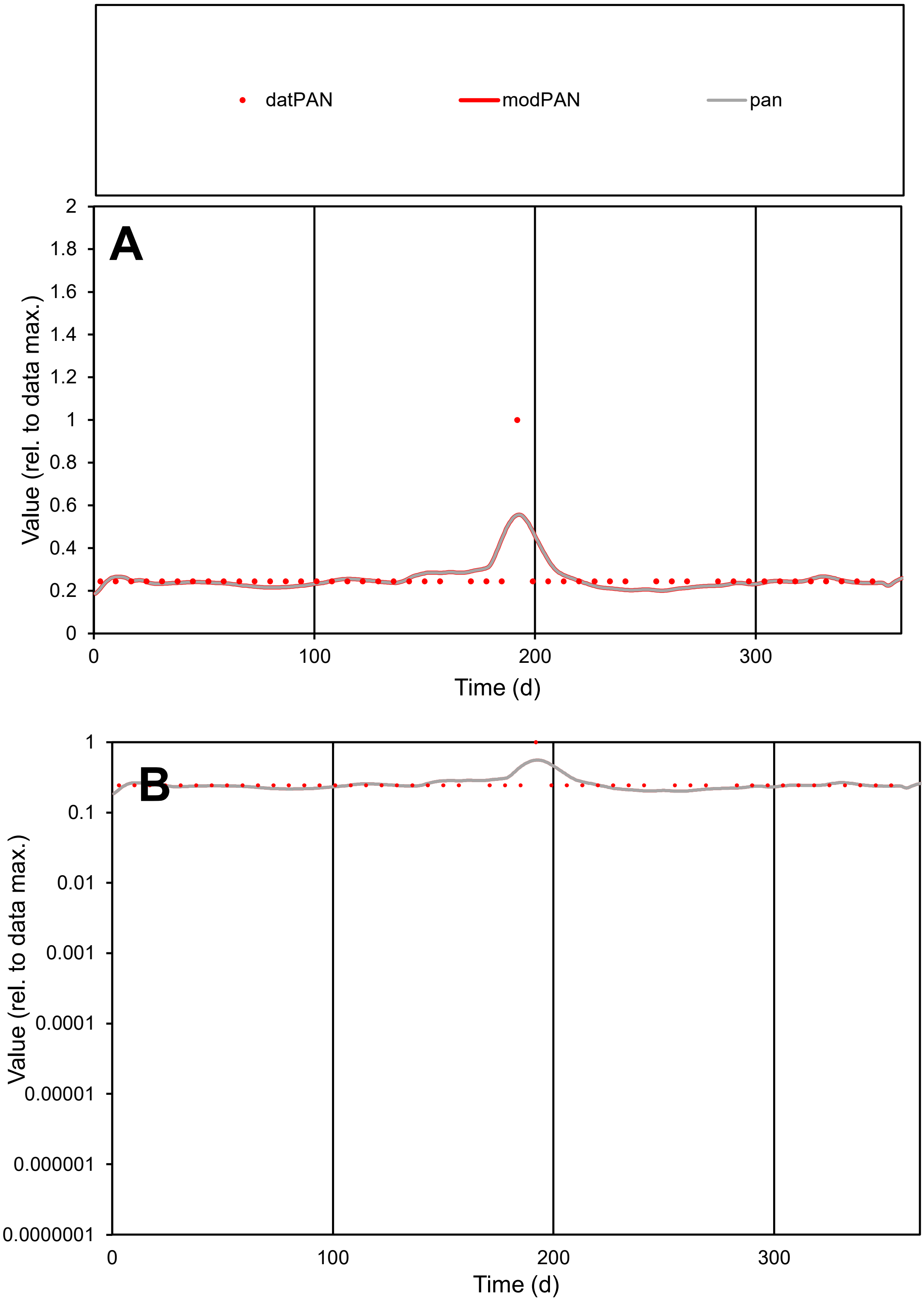


**C1**

Variables
PAN Provide jtd or Dst
CalibOnly?
2 1 to reproduce of
oNormalize?
2 1=no, 2=yes

ofxn calc*
0.31191
ofxn mod
0.33621

sRun
RunA1
zout
1

Ave
Data
9.7E-06
Model
9.7E-06

data/vMax
3.7E-05

ofxng
369.894

Min d/m
1.21132

ave/min d
1.06472
ave/min m
1.28409

328_Calib_S_A

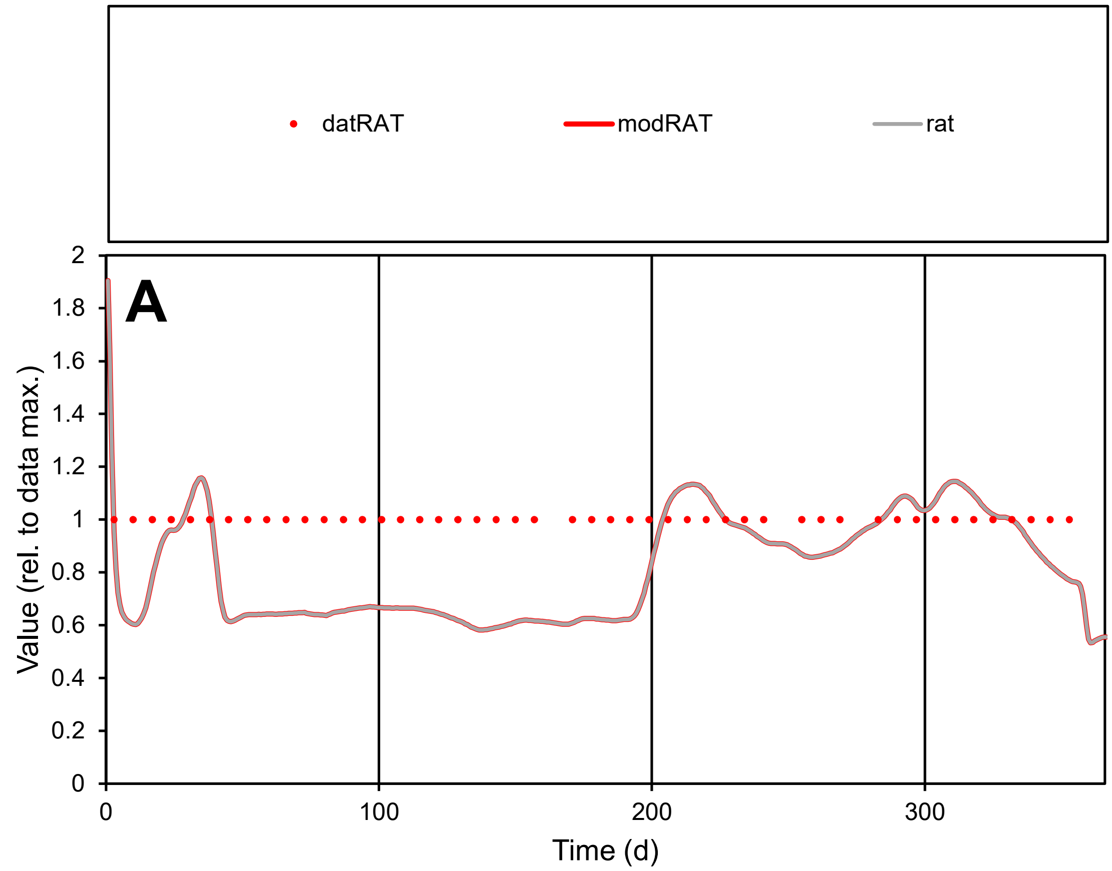


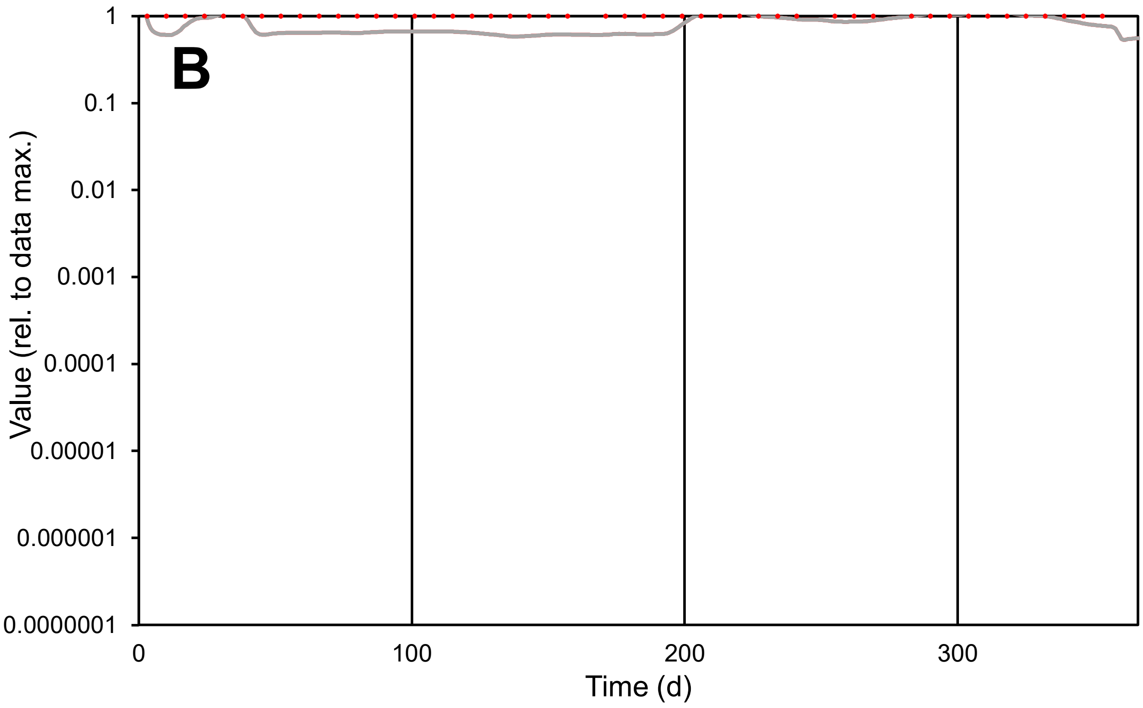


**C1**

Variables
RAT Provide jtd or Dst
CalibOnly?
2 1 to reproduce of
oNormalize?
2 1=no, 2=yes

ofxn calc*
0.26367
ofxn mod
0.28314

sRun
RunA1
zout
1

Ave
Data
1.1E-05
Model
9.3E-06

data/vMax
1.1E-05

ofxng
369.894

Min d/m
1.71726

ave/min d
1
ave/min m
1.40106

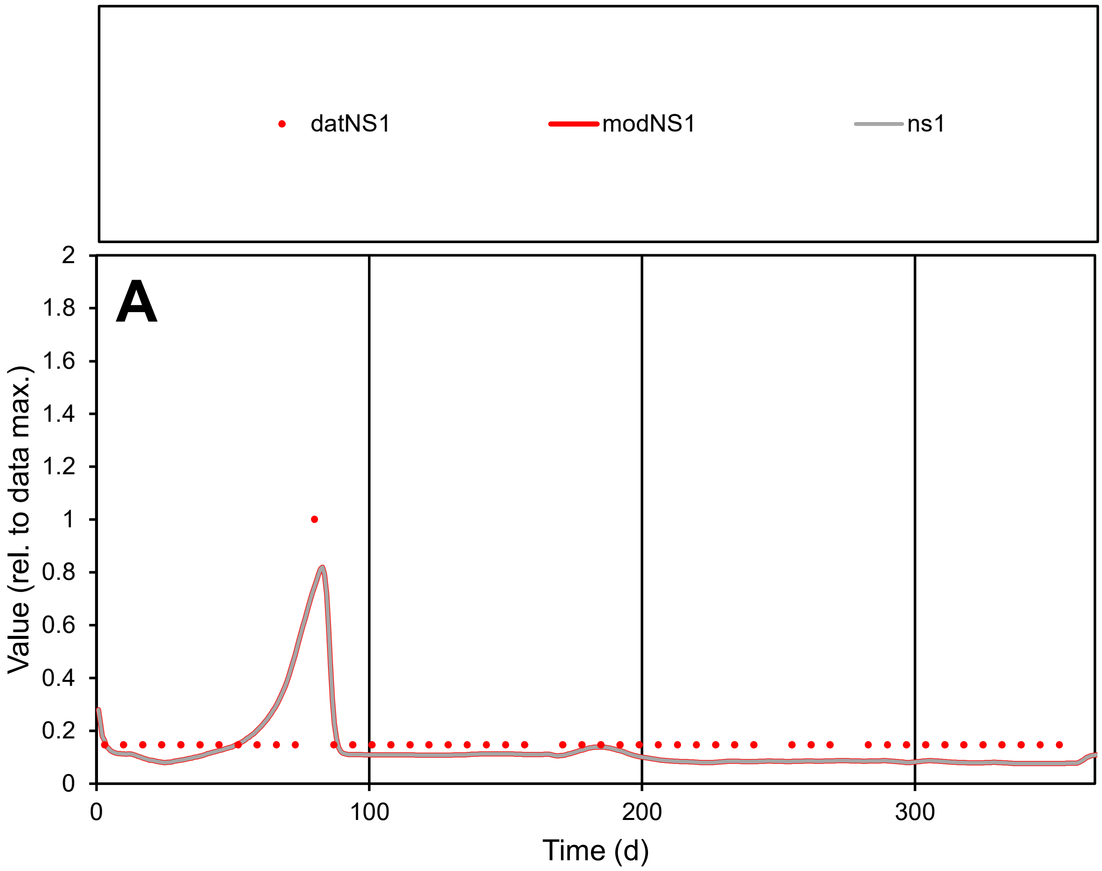


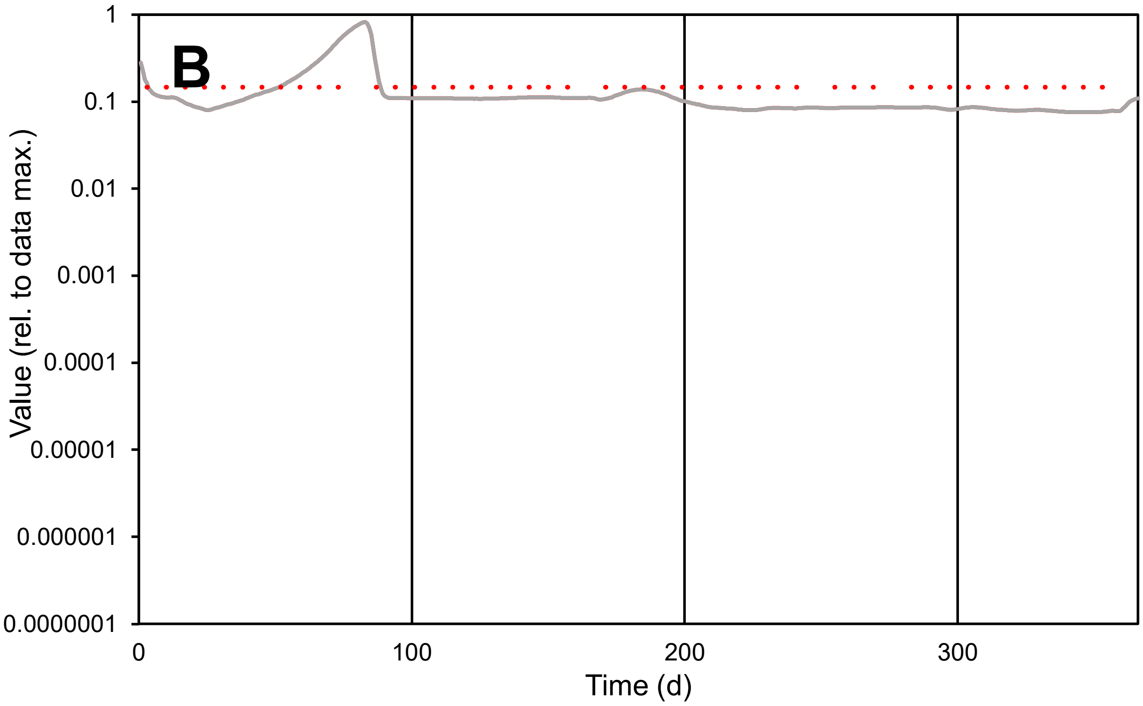


**C1**

Variables

NS1 Provide jtd or Dst

CalibOnly?

2 1 to reproduce of

oNormalize?

2 1=no, 2=yes

ofxn calc*

0.46553

ofxn mod

0.50599

sRun

RunA1

zout

1

Ave

Data

3.1E-05

Model

2.4E-05

data/vMax

0.00019

ofxng

369.894

Min d/m

1.93244

ave/min d

1.12096

ave/min m

1.6988

328_Calib_S_A

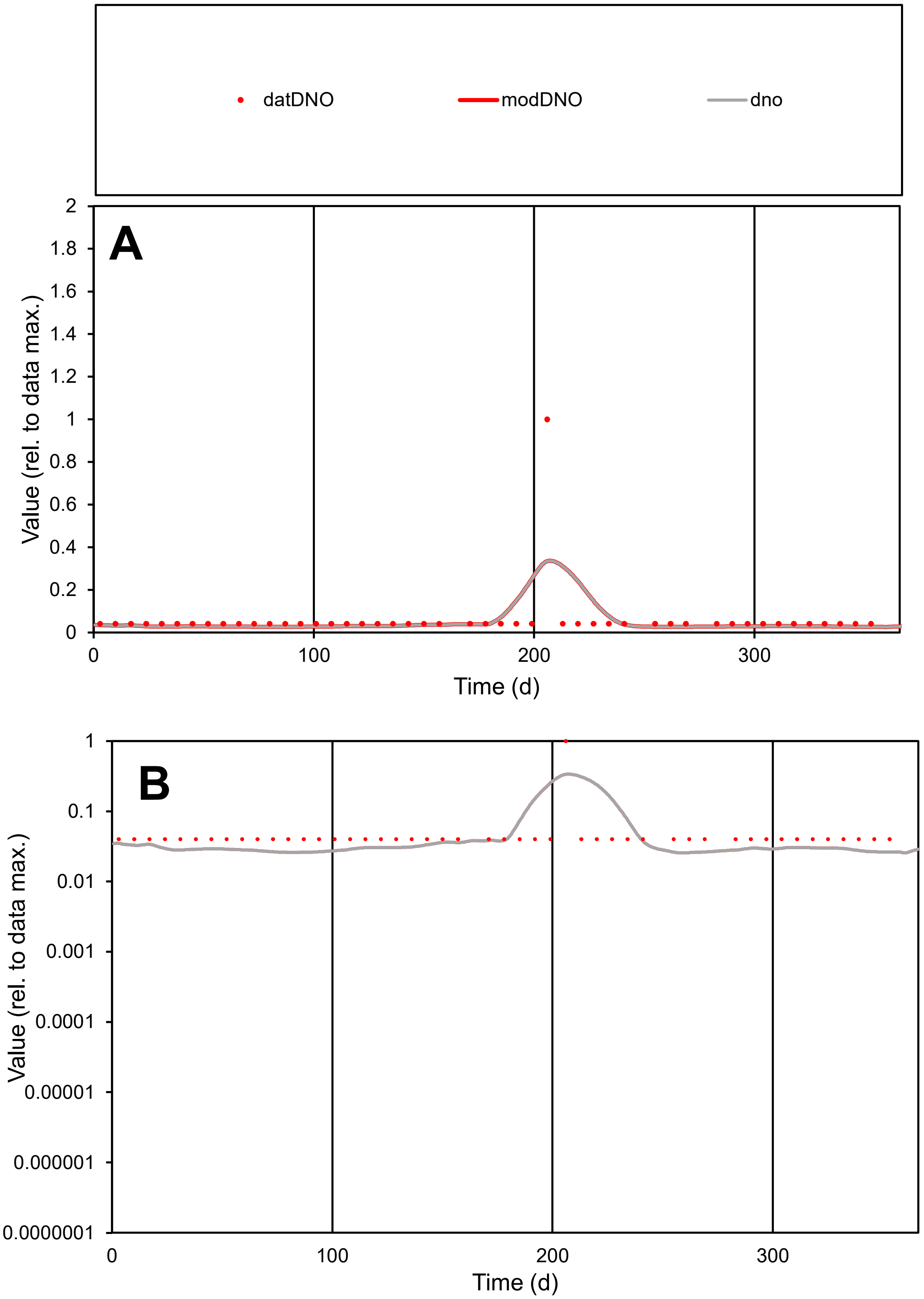


**C1**

Variables
DNO Provide jtd or Dst
CalibOnly?
2 1 to reproduce of
oNormalize?
2 1=no, 2=yes

ofxn calc*
1.85598
ofxn mod
1.88287

sRun
RunA1
zout
1

Ave
Data
2.7E-05
Model
2.5E-05

data/vMax
0.00045

ofxng
369.894

Min d/m
1.55498

ave/min d
1.49724
ave/min m
2.16849

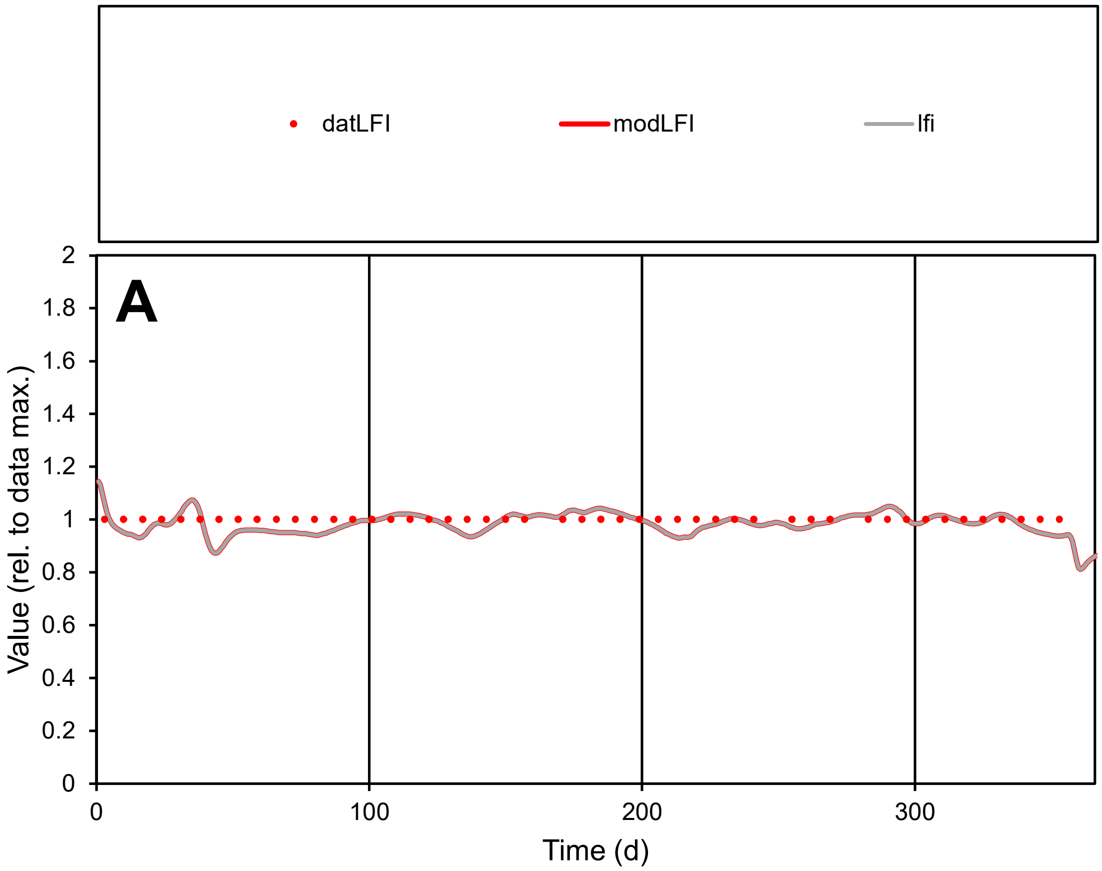


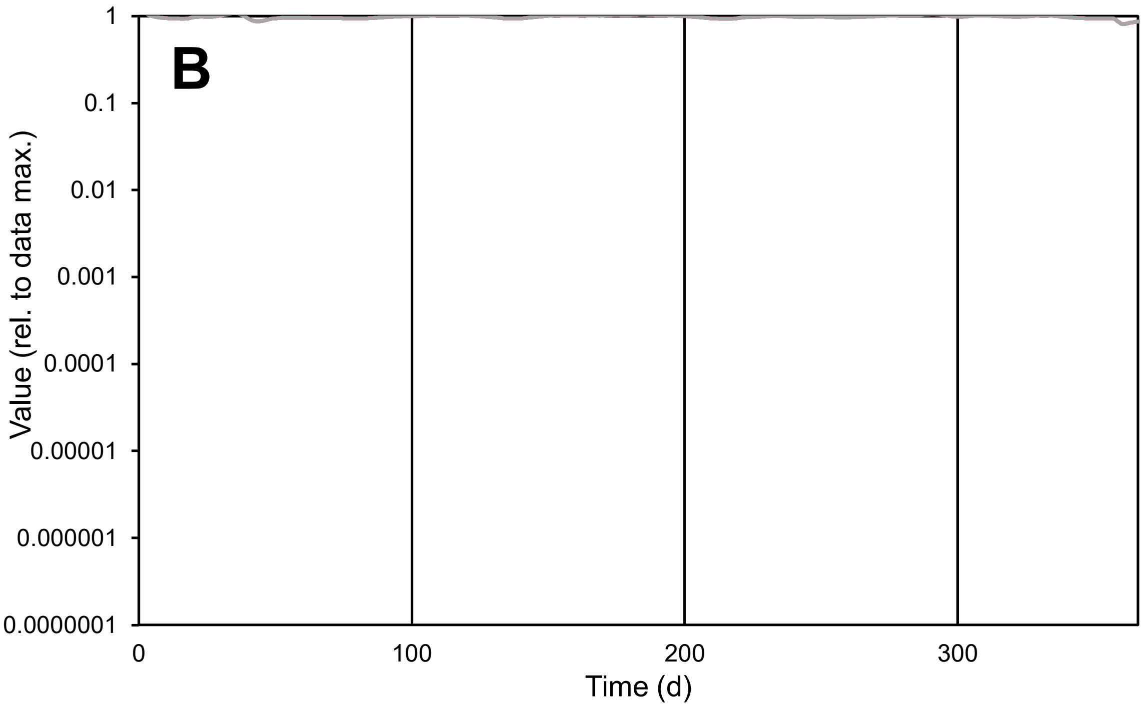


**C1**

Variables
LFI Provide jtd or Dst
CalibOnly?
2 1 to reproduce of
oNormalize?
2 1=no, 2=yes

ofxn calc*
0.03673
ofxn mod
0.11826

sRun
RunA1
zout
1

Ave
Data
8.3E-06
Model
8.2E-06

data/vMax
8.3E-06

ofxng
369.894

Min d/m
1.1092

ave/min d
1
ave/min m
1.09233

328_Calib_S_A

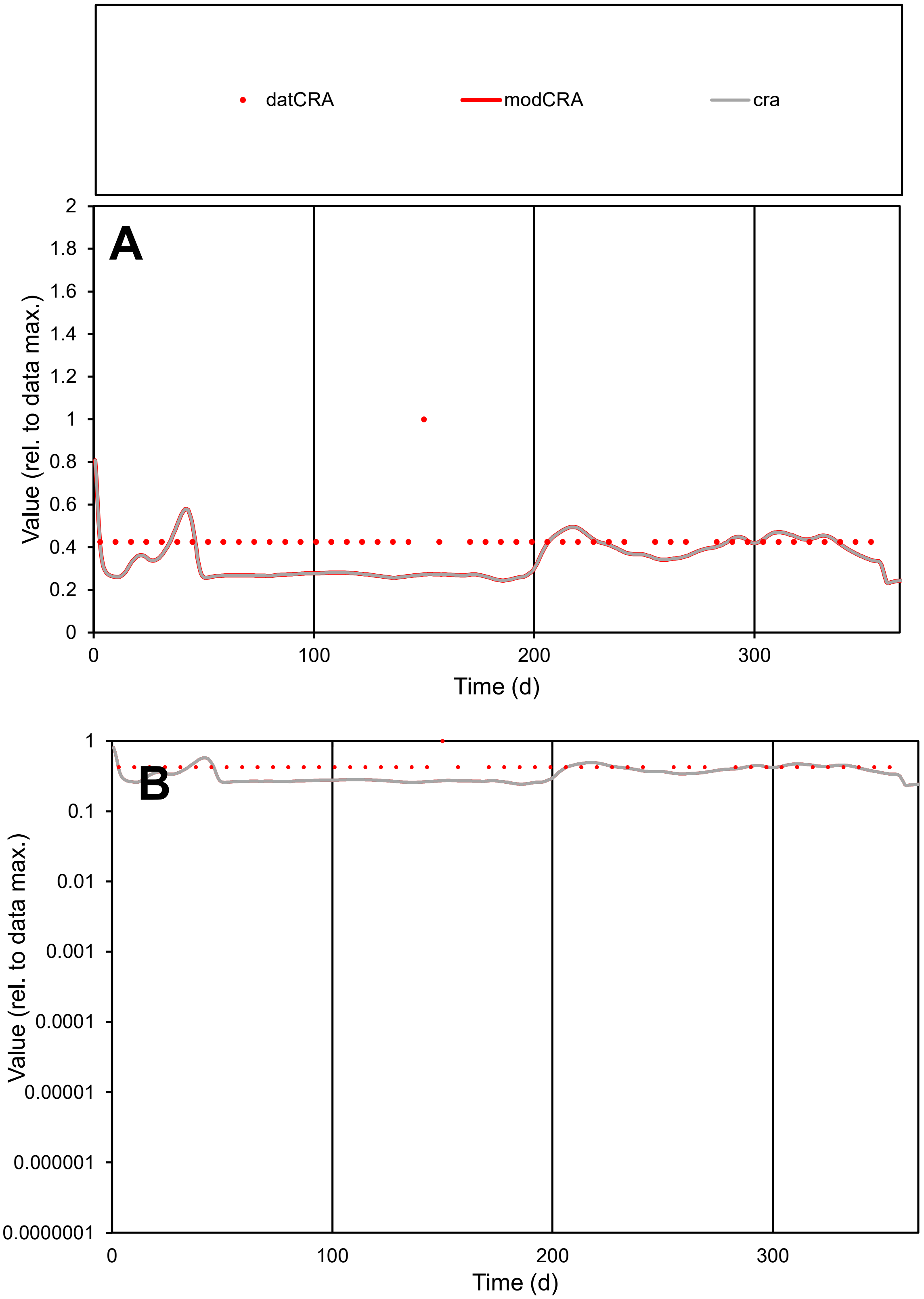


**C1**

Variables
CRA Provide jtd or Dst
CalibOnly?
2 1 to reproduce of
oNormalize?
2 1=no, 2=yes

ofxn calc*
0.349
ofxn mod
0.36349

sRun
RunA1
zout
1

Ave
Data
5E-06
Model
4E-06

data/vMax
1.1E-05

ofxng
369.894

Min d/m
1.74025

ave/min d
1.02829
ave/min m
1.41893

328_Calib_S_A

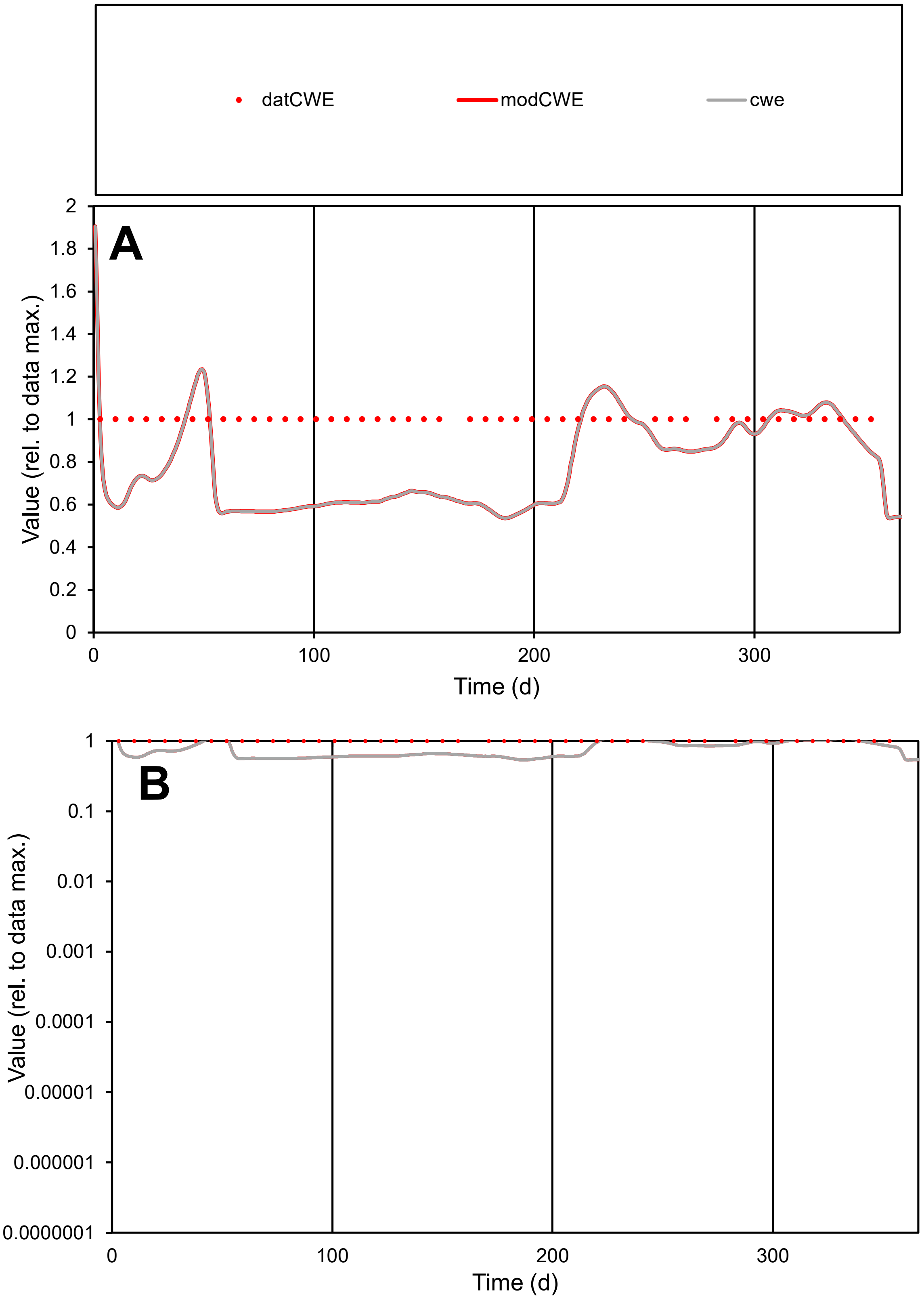


**C1**

Variables

CWE Provide jtd or Dst

CalibOnly?

2 1 to reproduce of

oNormalize?

2 1=no, 2=yes

ofxn calc*

0.29394

ofxn mod

0.31084

sRun

RunA1

zout

1

Ave

Data

2.6E-05

Model

2E-05

data/vMax

2.6E-05

ofxng

369.894

Min d/m

1.86267

ave/min d

1

ave/min m

1.45384

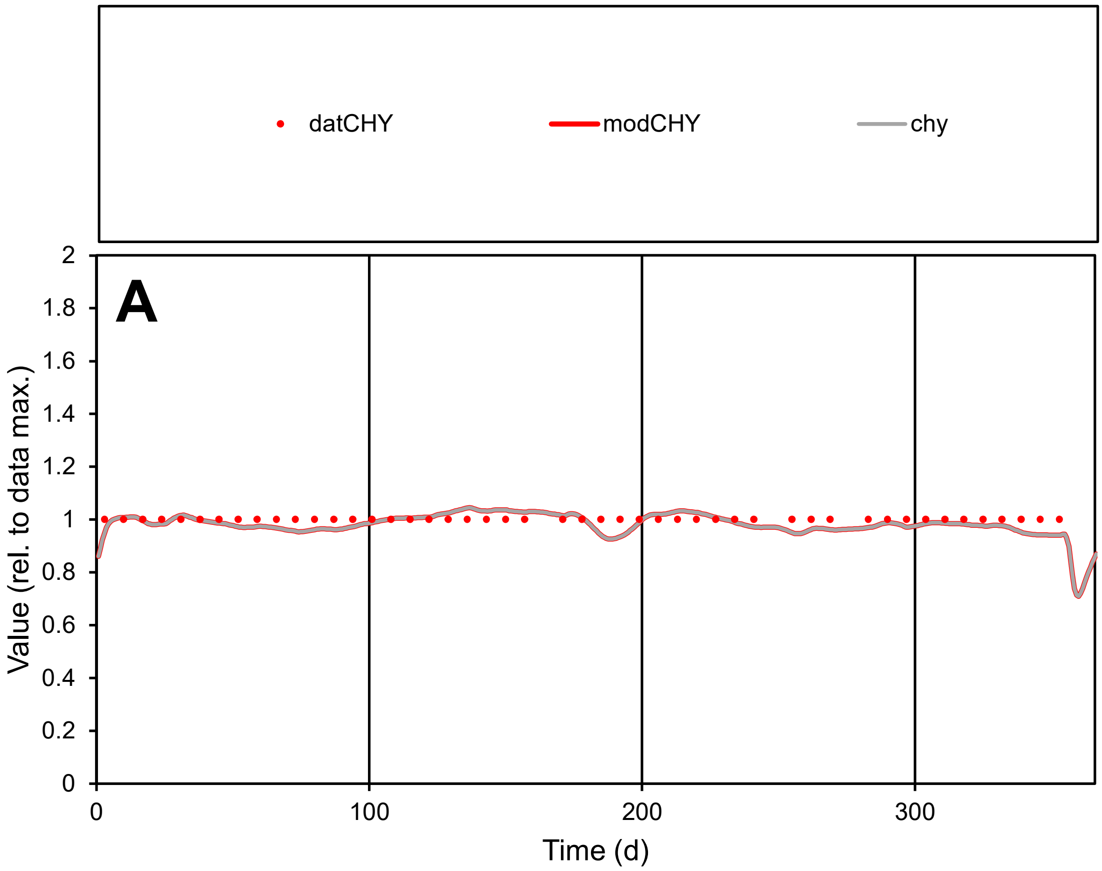


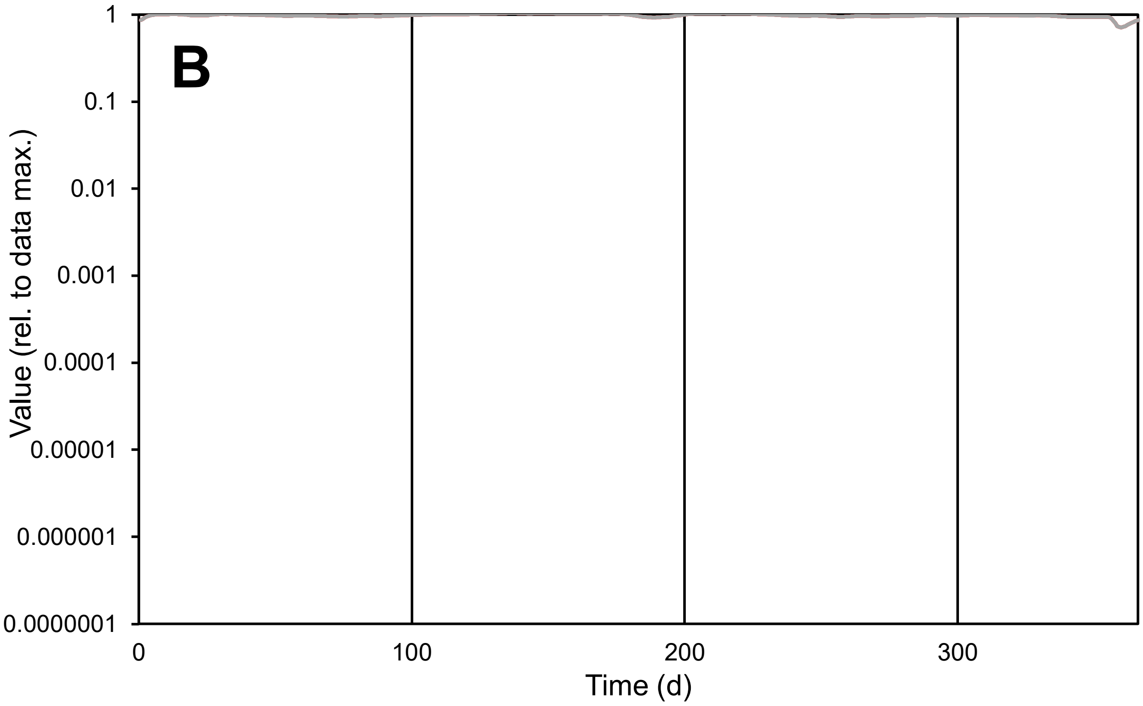


**C1**

Variables
CHY Provide jtd or Dst
CalibOnly?
2 1 to reproduce of
oNormalize?
2 1=no, 2=yes

ofxn calc*
0.03087
ofxn mod
0.11643

sRun
RunA1
zout
1

Ave
Data
6.7E-06
Model
6.6E-06

data/vMax
6.7E-06

ofxng
369.894

Min d/m
1.07255

ave/min d
1
ave/min m
1.05863

328_Calib_S_A

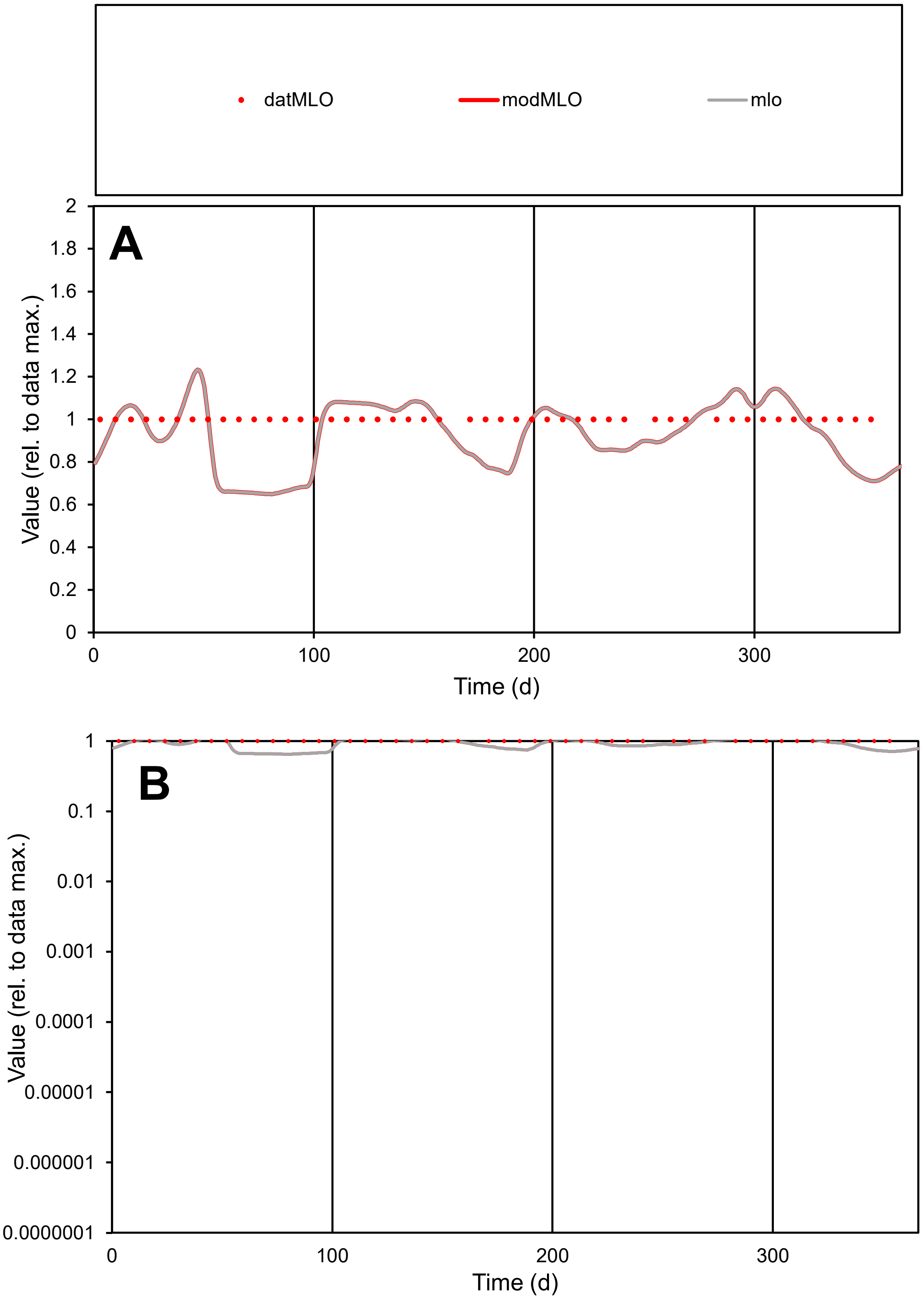


**C1**

Variables
MLO Provide jtd or Dst
CalibOnly?
2 1 to reproduce of
oNormalize?
2 1=no, 2=yes

ofxn calc*
0.16599
ofxn mod
0.19862

sRun
RunA1
zout
1

Ave
Data
1E-05
Model
9.7E-06

data/vMax
1E-05

ofxng
369.894

Min d/m
1.53948

ave/min d
1
ave/min m
1.43326

328_Calib_S_A

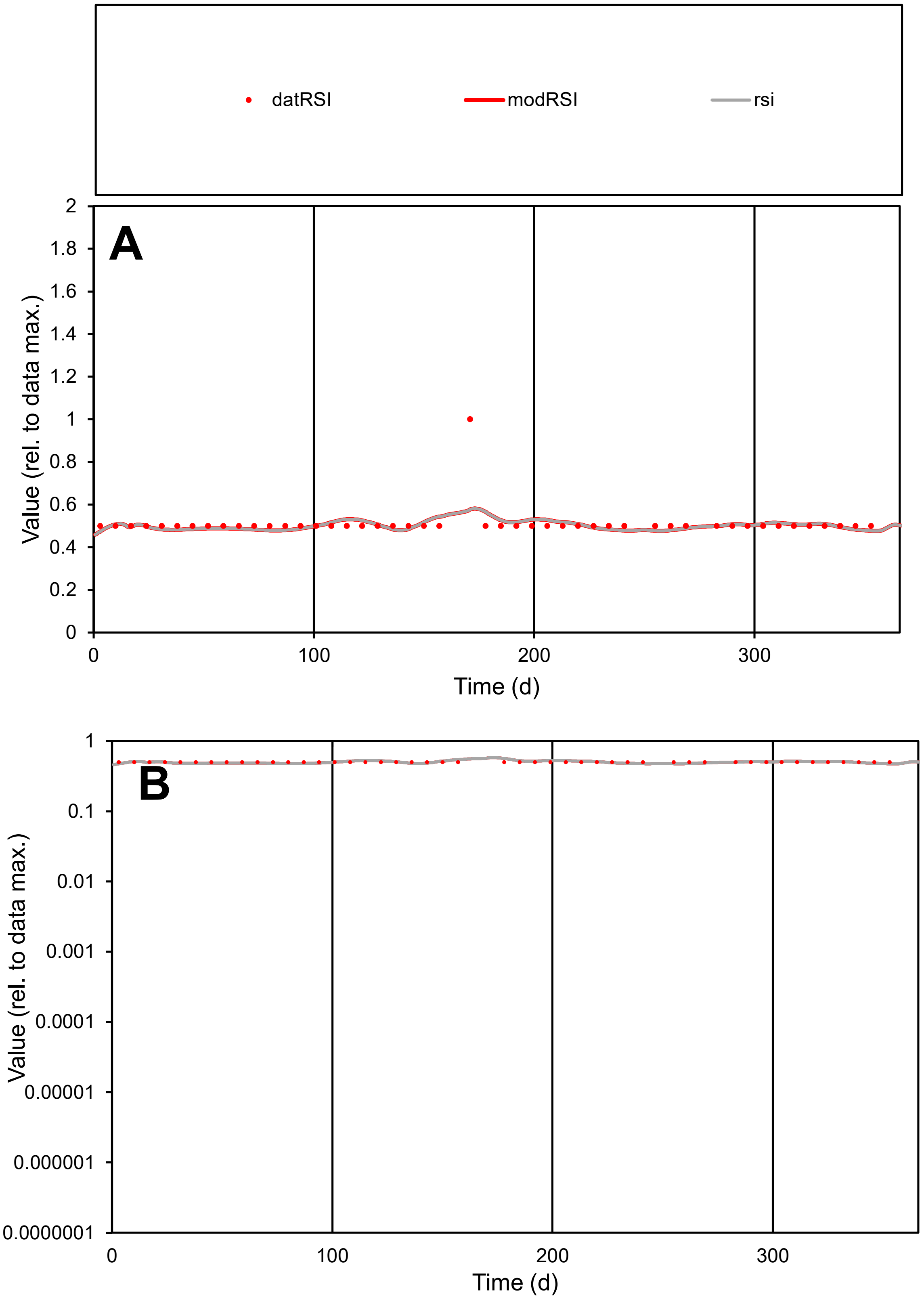


**C1**

Variables
RSI Provide jtd or Dst
CalibOnly?
2 1 to reproduce of
oNormalize?
2 1=no, 2=yes

ofxn calc*
0.12409
ofxn mod
0.16785

sRun
RunA1
zout
1

Ave
Data
1.1E-05
Model
1E-05

data/vMax
2.1E-05

ofxng
369.894

Min d/m
1.05044

ave/min d
1.02081
ave/min m
1.05522

328_Calib_S_A

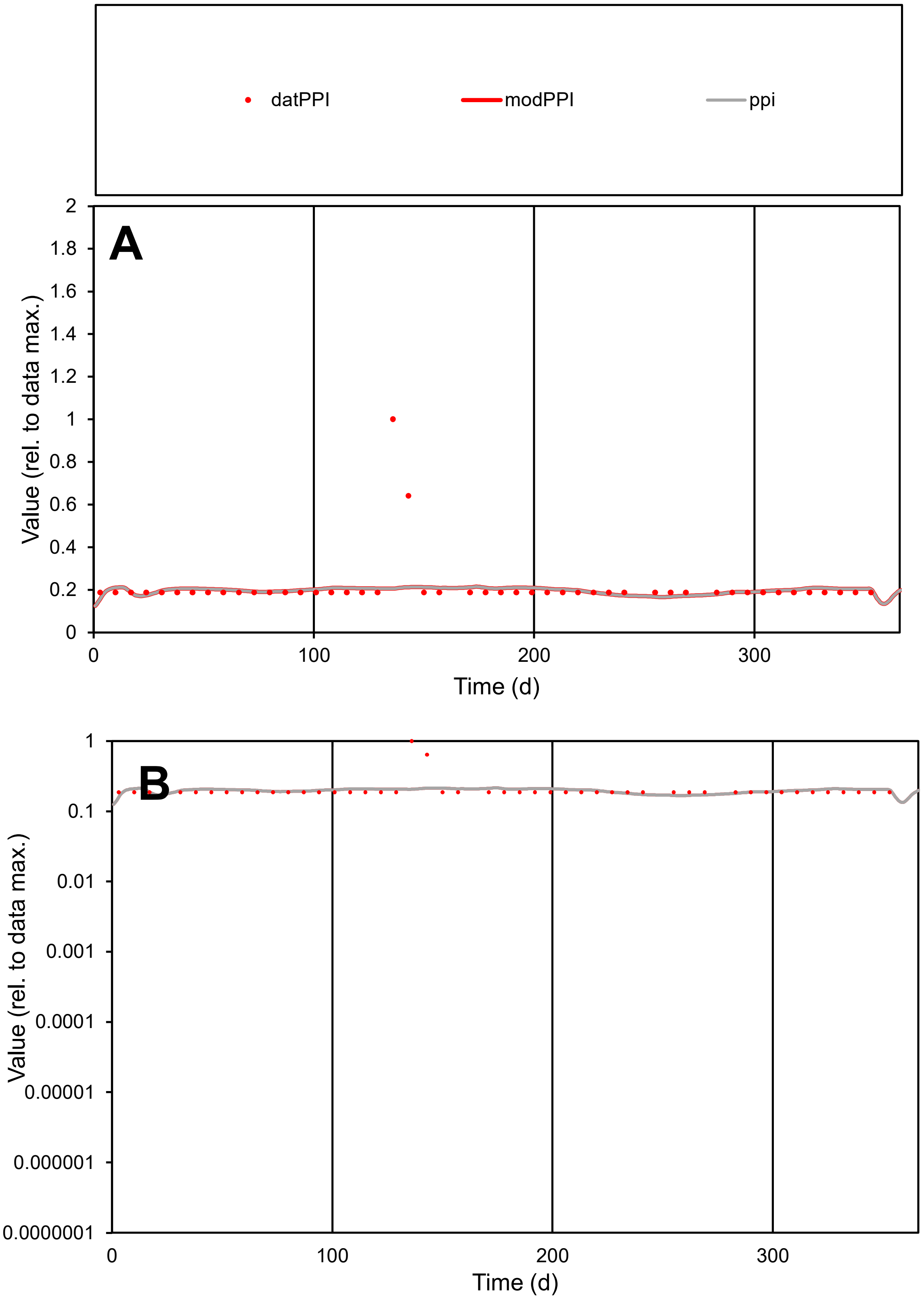


C1

Variables
PPI Provide jtd or Dst
CalibOnly?
2 1 to reproduce of
oNormalize?
2 1=no, 2=yes

ofxn calc*
0.6136
ofxn mod
0.62303

sRun
RunA1
zout
1

Ave
Data
1.7E-06
Model
1.5E-06

data/vMax
7.7E-06

ofxng
369.894

Min d/m
1.12267

ave/min d
1.14108
ave/min m
1.18465

328_Calib_S_A

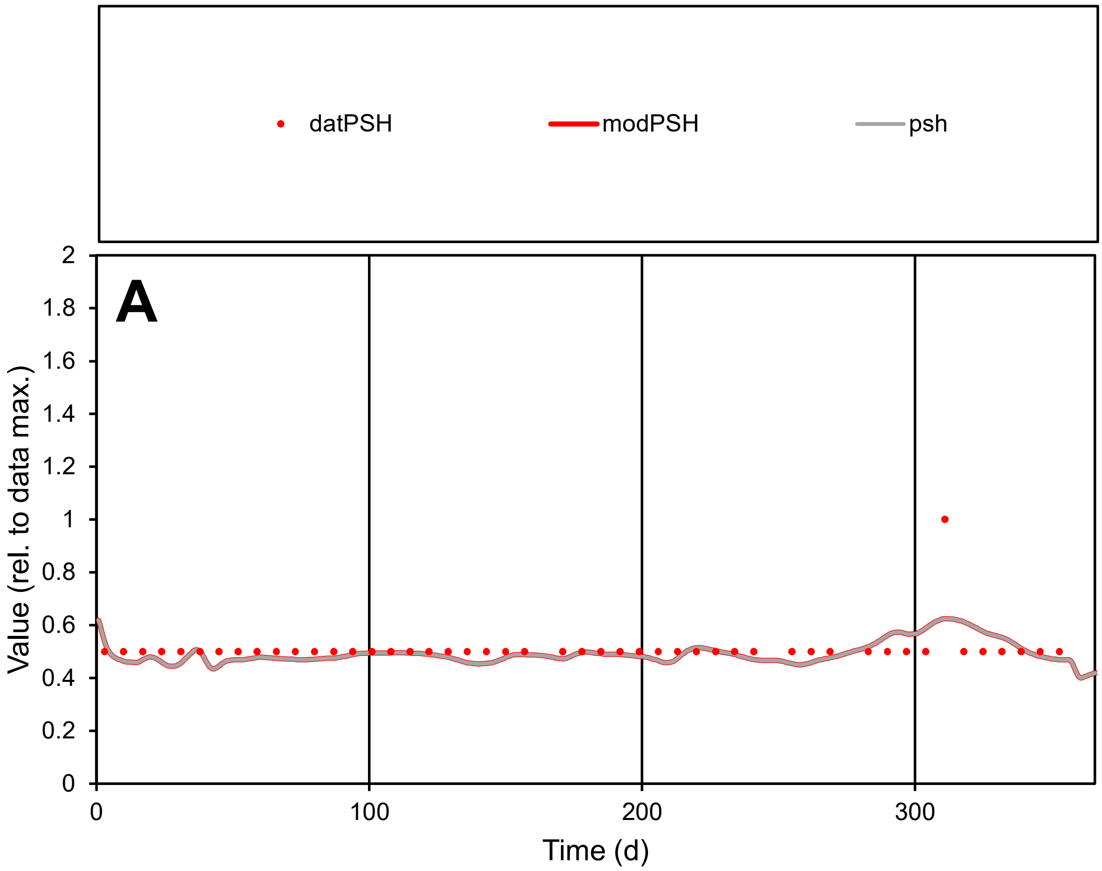


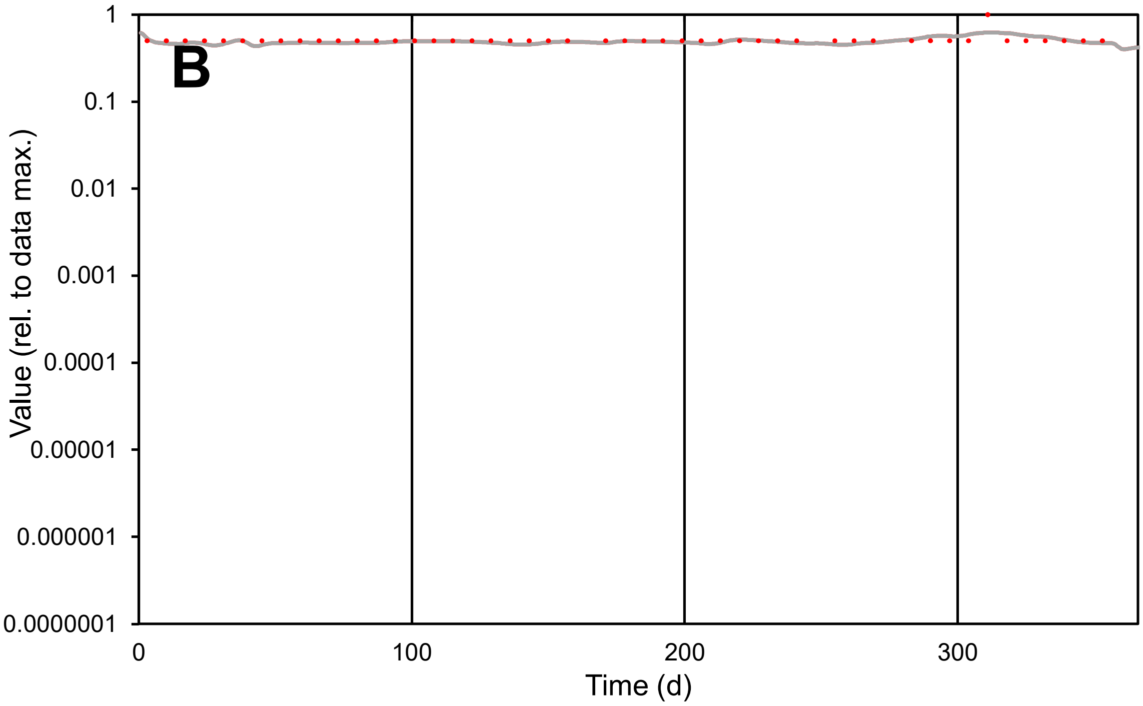


**C1**

Variables
PSH Provide jtd or Dst
CalibOnly?
2 1 to reproduce of
oNormalize?
2 1=no, 2=yes

ofxn calc*
0.12828
ofxn mod
0.16901

sRun
RunA1
zout
1

Ave
Data
1.3E-05
Model
1.3E-05

data/vMax
2.6E-05

ofxng
369.894

Min d/m
1.1092

ave/min d
1.02085
ave/min m
1.09913

328_Calib_S_A

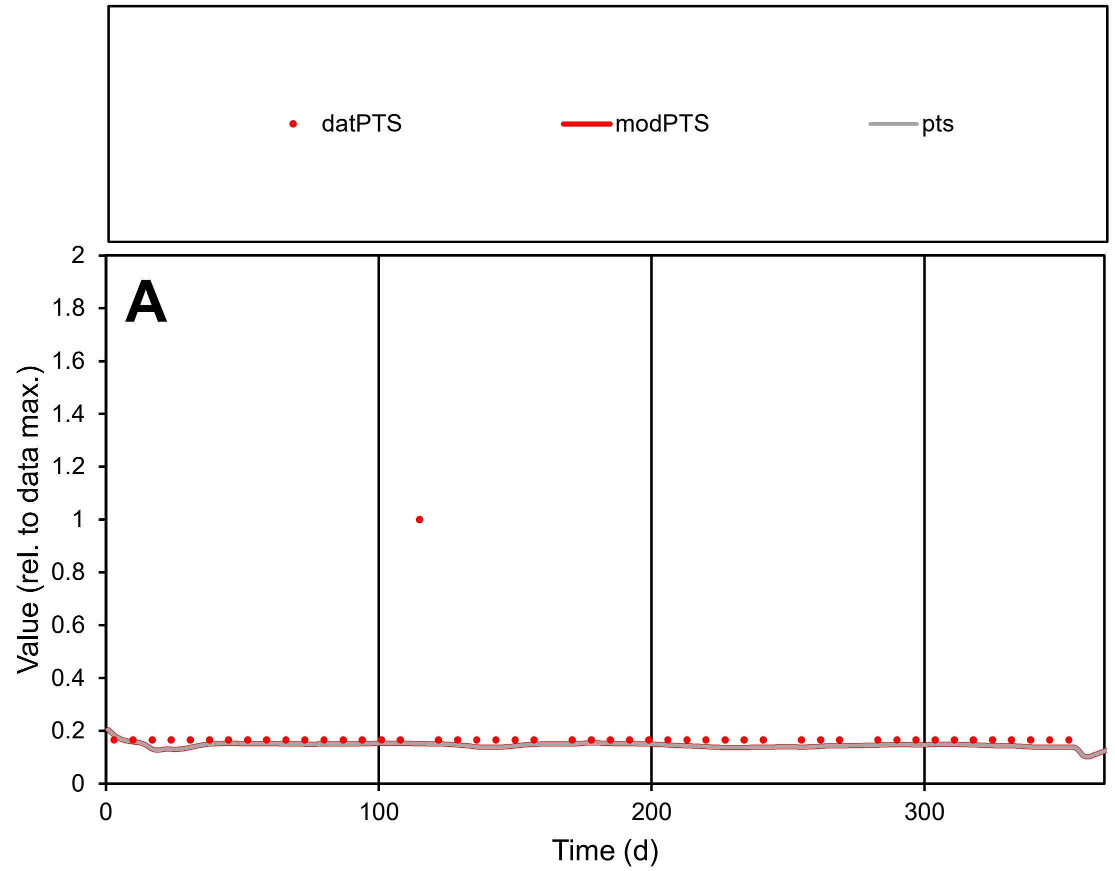


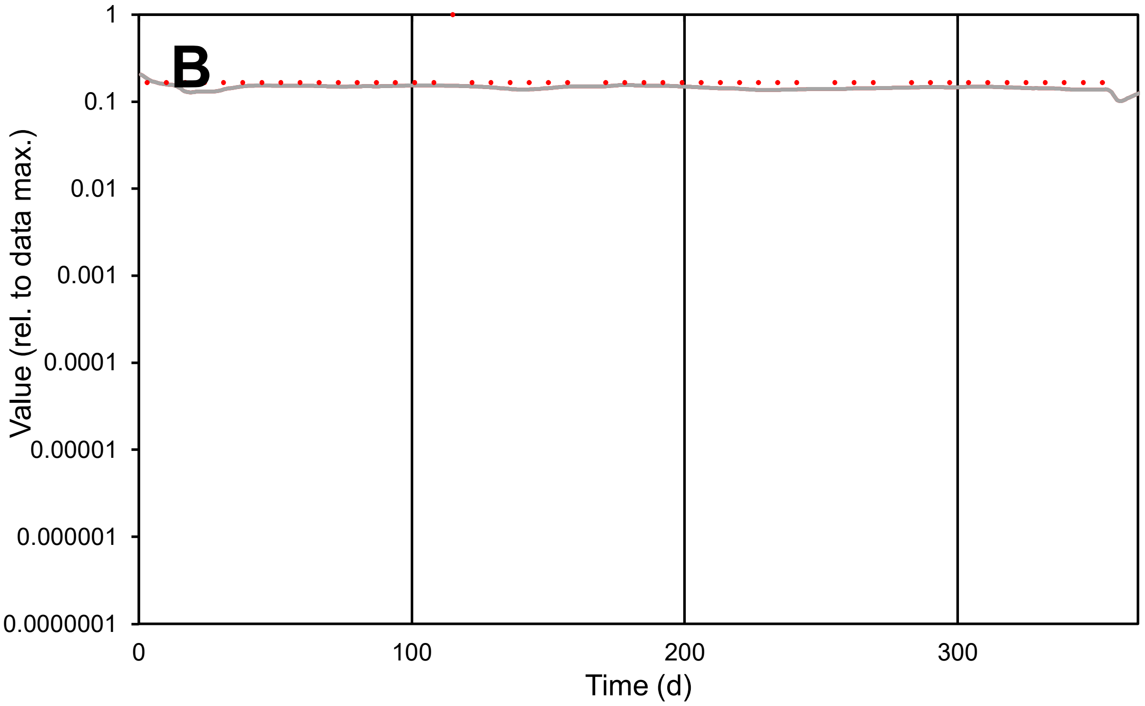


**C1**

Variables
PTS Provide jtd or Dst
CalibOnly?
2 1 to reproduce of
oNormalize?
2 1=no, 2=yes

ofxn calc*
0.67841
ofxn mod
0.68722

sRun
RunA1
zout
1

Ave
Data
1.1E-05
Model
8.5E-06

data/vMax
5.8E-05

ofxng
369.894

Min d/m
1.2966

ave/min d
1.1049
ave/min m
1.144

328_Calib_S_A

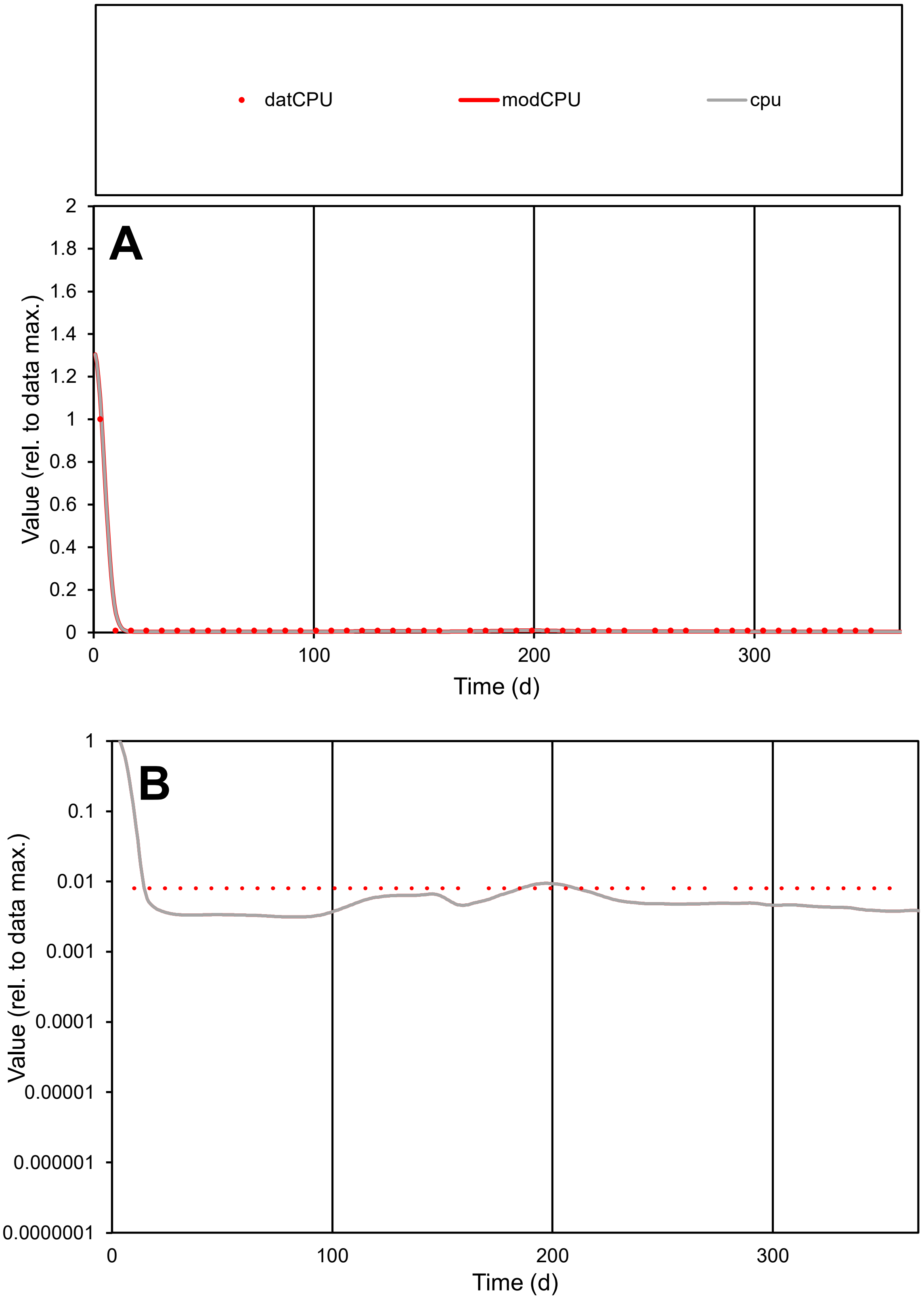


**C1**

Variables
CPU Provide jtd or Dst
CalibOnly?
2 1 to reproduce of
oNormalize?
2 1=no, 2=yes

ofxn calc*
0.1559
ofxn mod
0.70754

sRun
RunA1
zout
1

Ave
Data
1.6E-06
Model
1.4E-06

data/vMax
5.5E-05

ofxng
369.894

Min d/m
2.56645

ave/min d
3.56661
ave/min m
8.28019

328_Calib_S_A

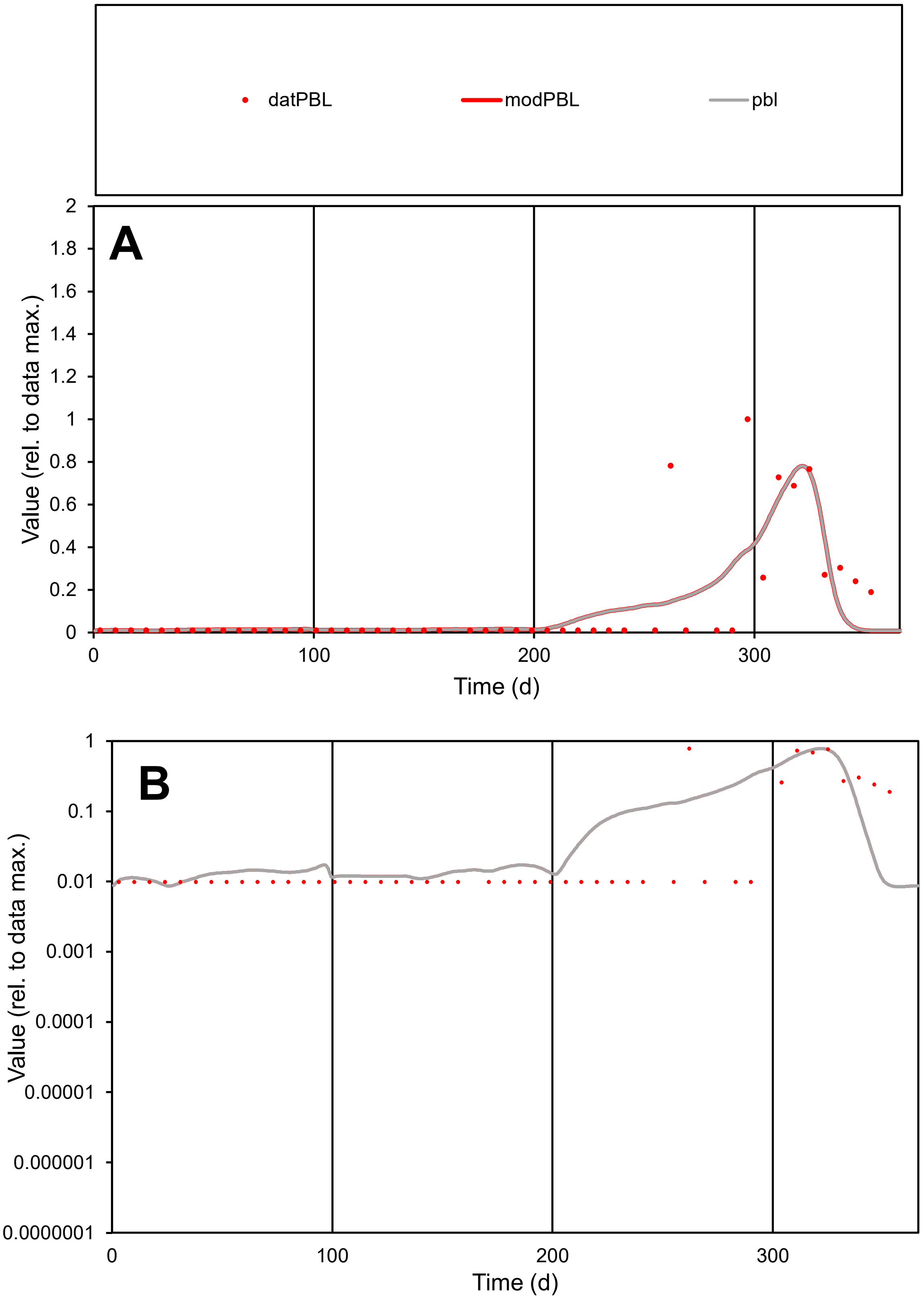


**C1**

Variables
PBL Provide jtd or Dst
CalibOnly?
2 1 to reproduce of
oNormalize?
2 1=no, 2=yes

ofxn calc*
1.33293
ofxn mod
1.36315

sRun
RunA1
zout
1

Ave
Data
5E-07
Model
4.8E-07

data/vMax
4.3E-06

ofxng
369.894

Min d/m
1.13827

ave/min d
11.8327
ave/min m
12.7528

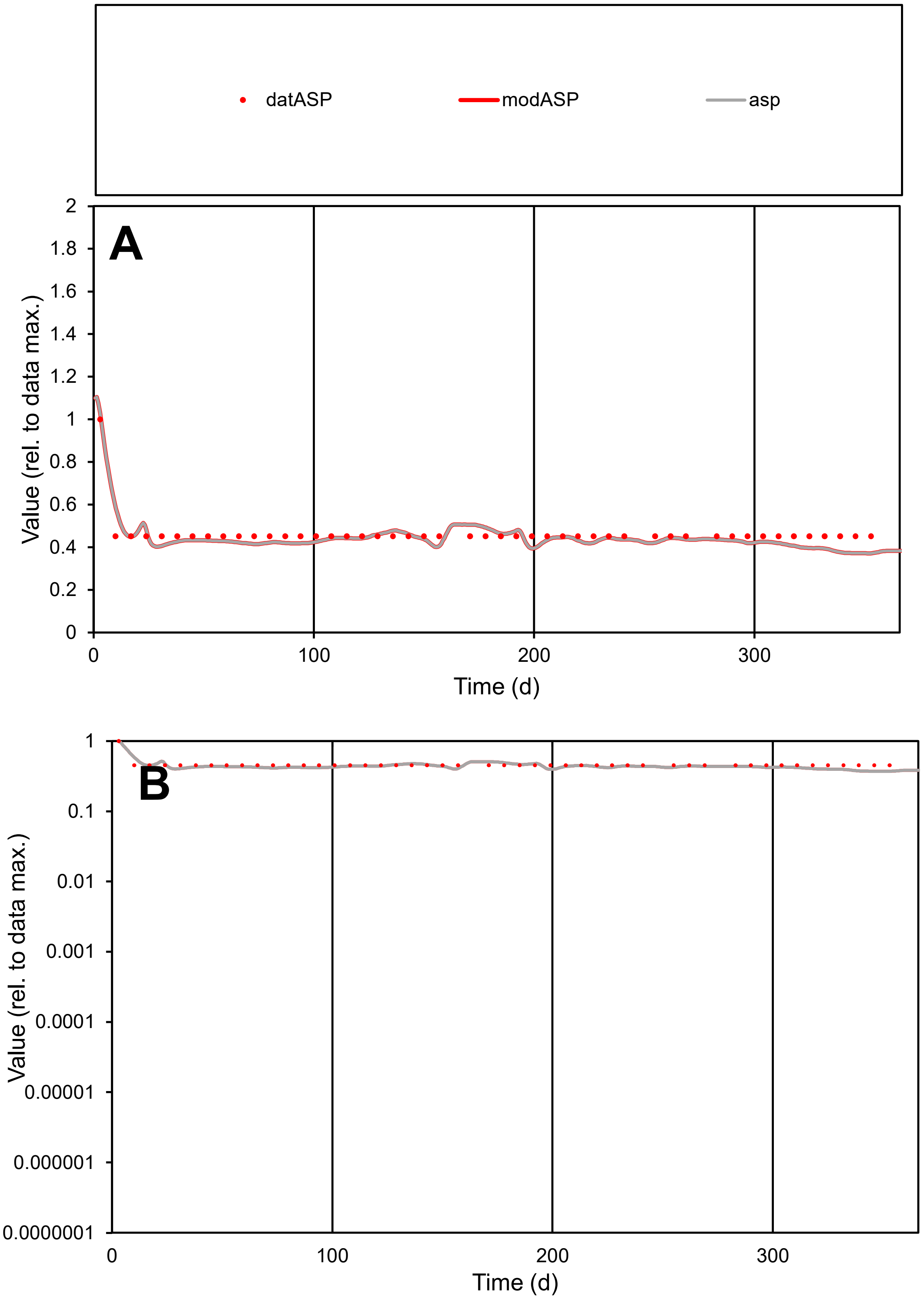


**C1**

Variables
ASP Provide jtd or Dst
CalibOnly?
2 1 to reproduce of
oNormalize?
2 1=no, 2=yes

ofxn calc*
0.07527
ofxn mod
0.13725

sRun
RunA1
zout
1

Ave
Data
5.9E-06
Model
5.7E-06

data/vMax
1.3E-05

ofxng
369.894

Min d/m
1.21094

ave/min d
1.02534
ave/min m
1.19432

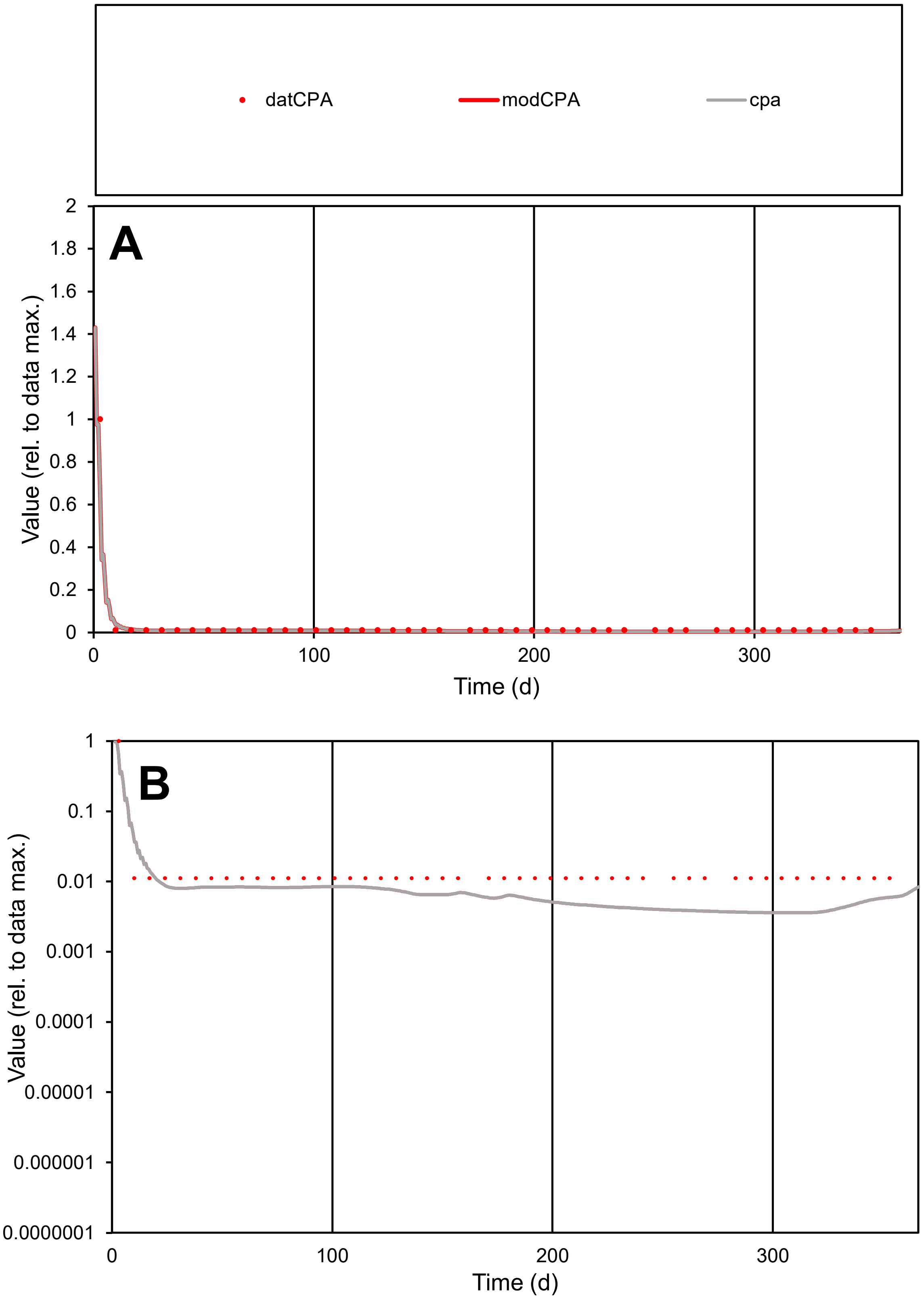


**C1**

Variables
CPA Provide jtd or Dst
CalibOnly?
2 1 to reproduce of
oNormalize?
2 1=no, 2=yes

ofxn calc*
0.20942
ofxn mod
1.42658

sRun
RunA1
zout
1

Ave
Data
1.9E-07
Model
1.6E-07

data/vMax
5.9E-06

ofxng
369.894

Min d/m
3.11831

ave/min d
2.83917
ave/min m
7.45752

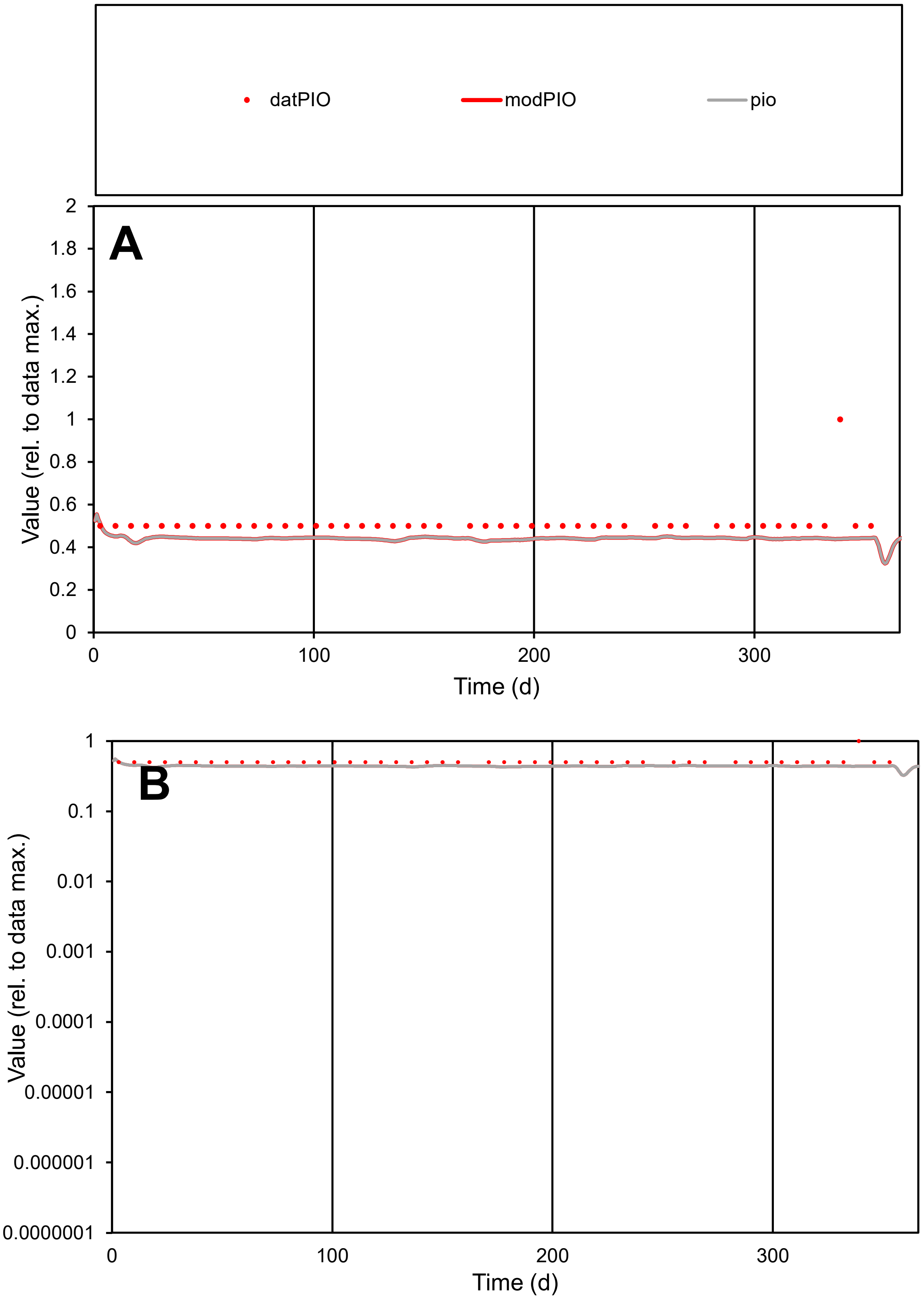


C1

Variables
PIO Provide jtd or Dst
CalibOnly?
2 1 to reproduce of
oNormalize?
2 1=no, 2=yes

ofxn calc*
0.19471
ofxn mod
0.22178

sRun
RunA1
zout
1

Ave
Data
1.5E-07
Model
1.3E-07

data/vMax
2.9E-07

ofxng
369.894

Min d/m
1.18627

ave/min d
1.02085
ave/min m
1.04958

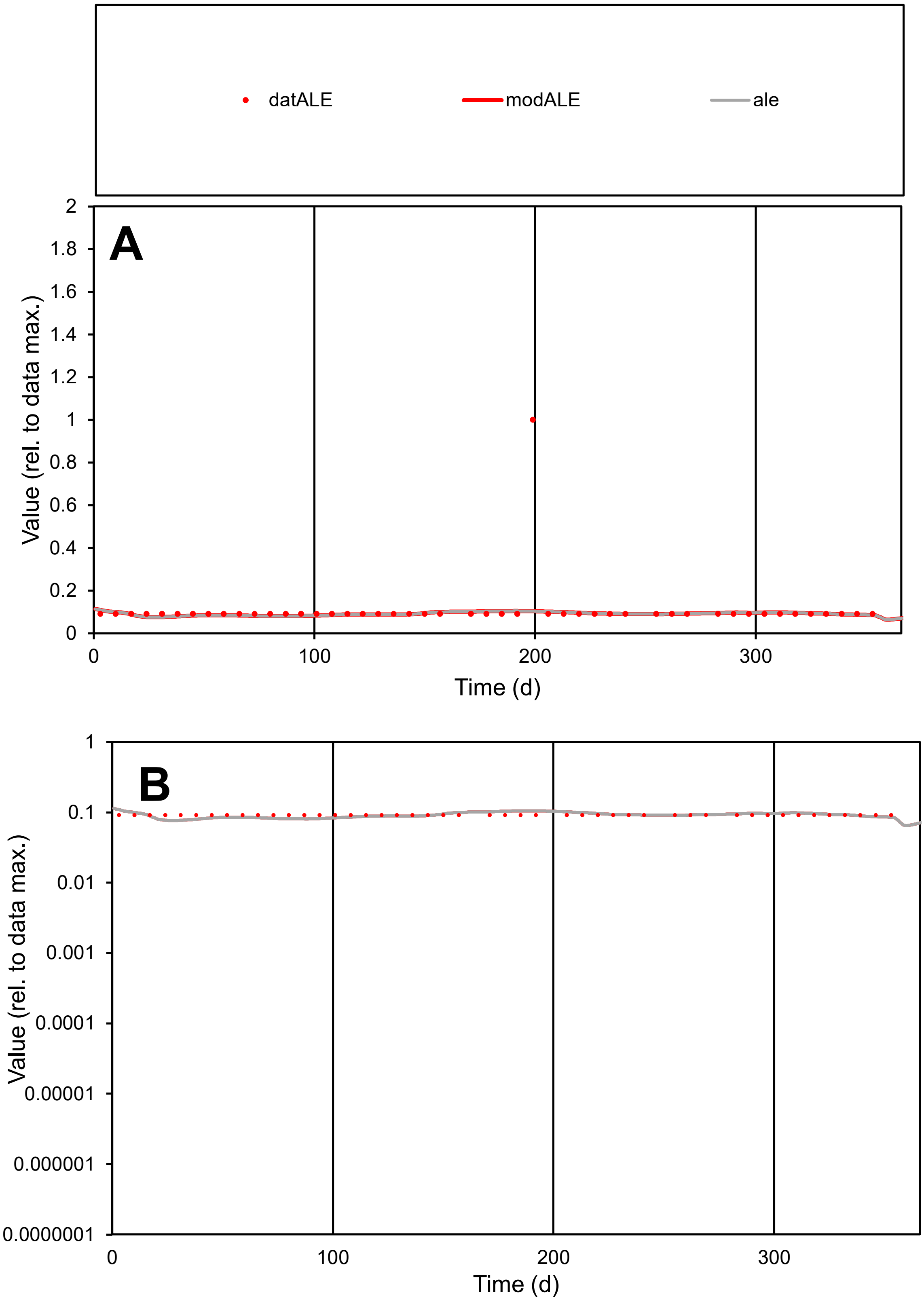


**C1**

Variables
ALE Provide jtd or Dst
CalibOnly?
2 1 to reproduce of
oNormalize?
2 1=no, 2=yes

ofxn calc*
1.17032
ofxn mod
1.17916

sRun
RunA1
zout
1

Ave
Data
1.3E-05
Model
1.1E-05

data/vMax
0.00012

ofxng
369.894

Min d/m
1.19271

ave/min d
1.20615
ave/min m
1.1904

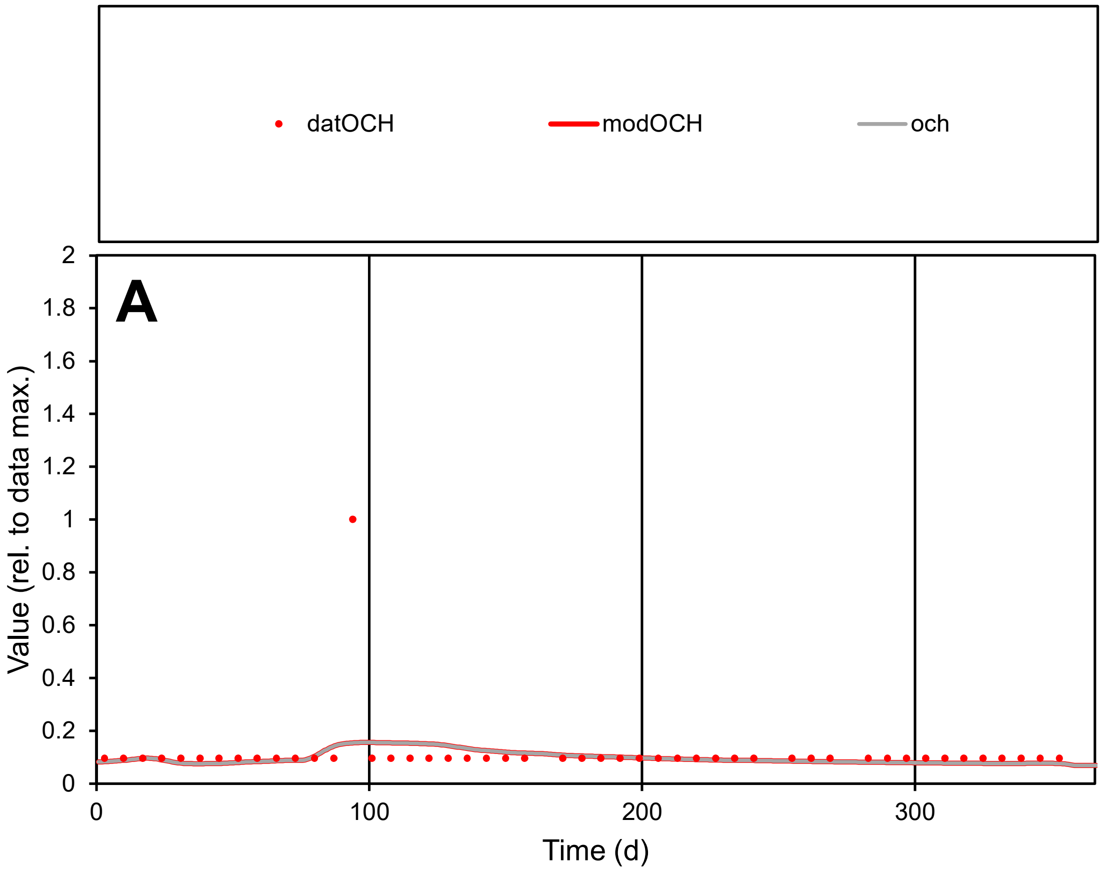


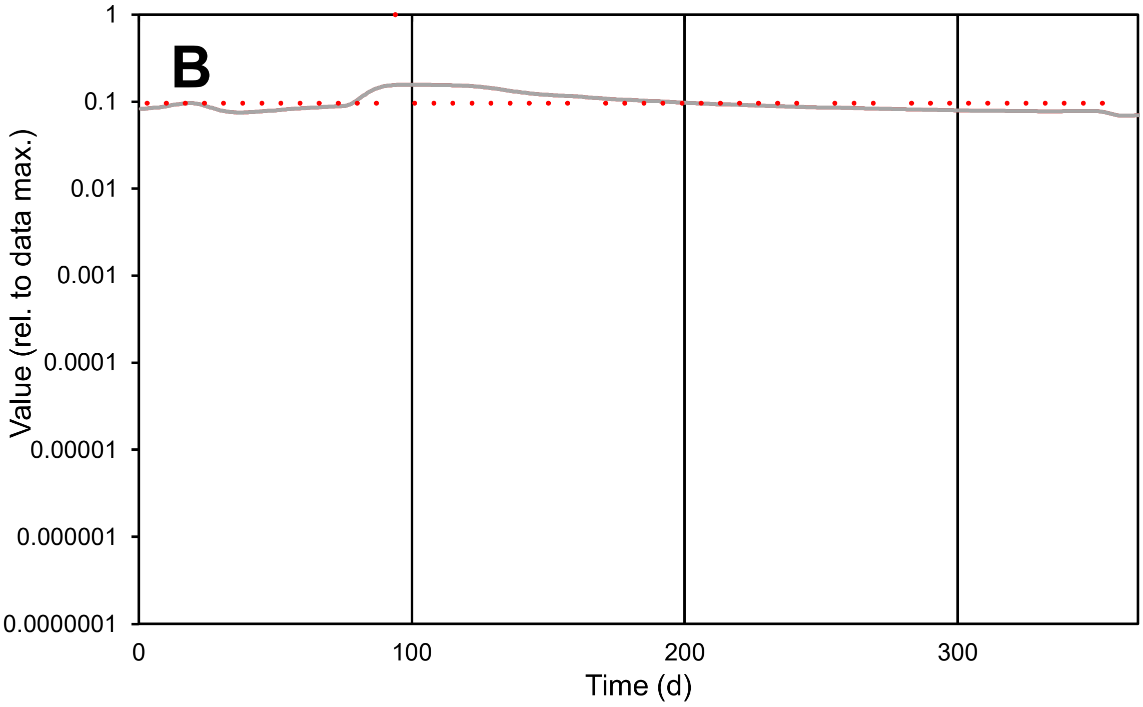


**C1**

Variables
OCH Provide jtd or Dst
CalibOnly?
2 1 to reproduce of
oNormalize?
2 1=no, 2=yes

ofxn calc*
1.08298
ofxn mod
1.08945

sRun
RunA1
zout
1

Ave
Data
5.3E-06
Model
4.5E-06

data/vMax
4.6E-05

ofxng
369.894

Min d/m
1.27486

ave/min d
1.1965
ave/min m
1.31843

328_Calib_S_A

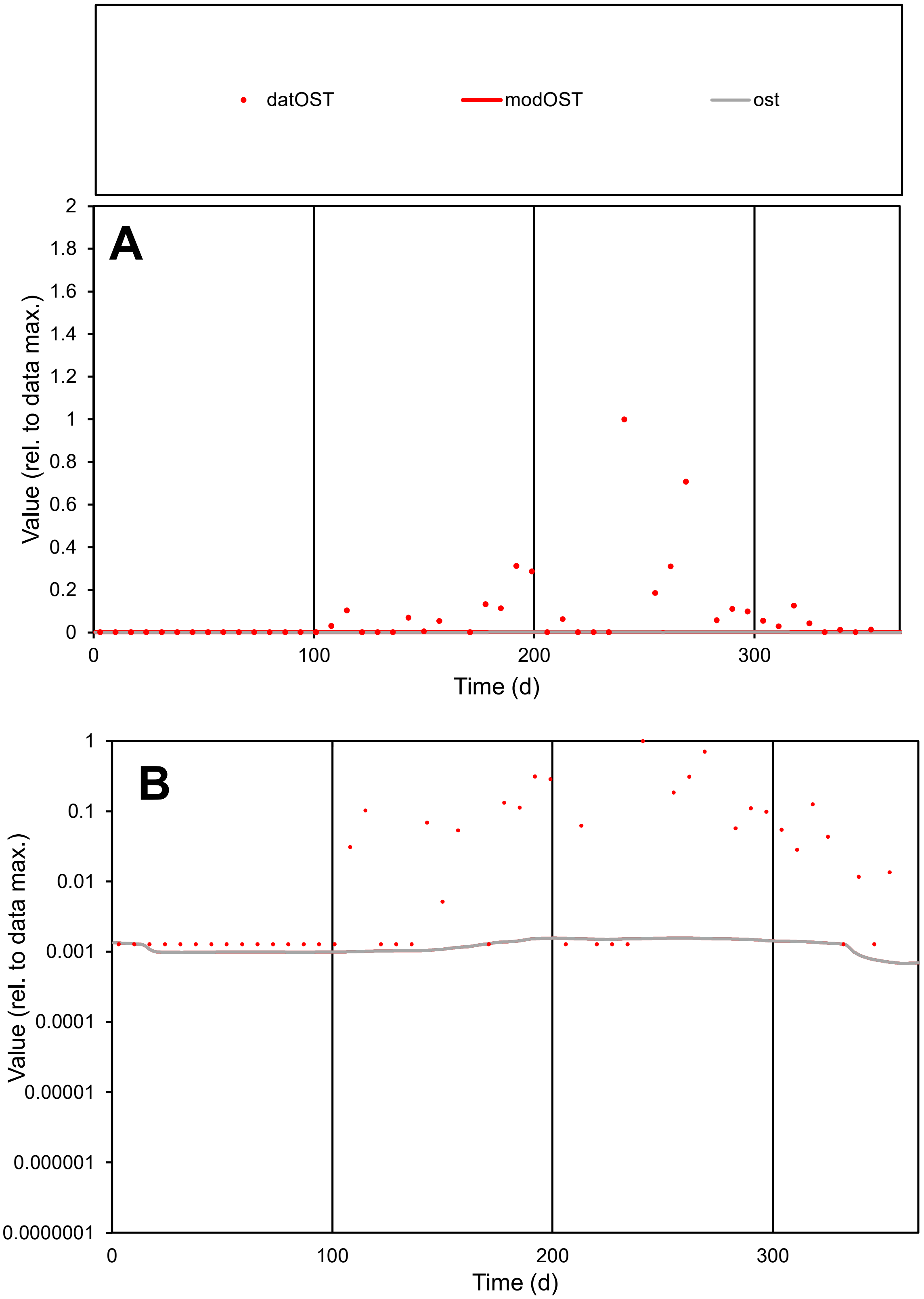


**C1**

Variables
OST Provide jtd or Dst
CalibOnly?
2 1 to reproduce of
oNormalize?
2 1=no, 2=yes

ofxn calc*
2.41962
ofxn mod
2.42328

sRun
RunA1
zout
1

Ave
Data
0.00014
Model
2.1E-06

data/vMax
0.00169

ofxng
369.894

Min d/m
1.81795
ave/min d
64.255
ave/min m
1.73264

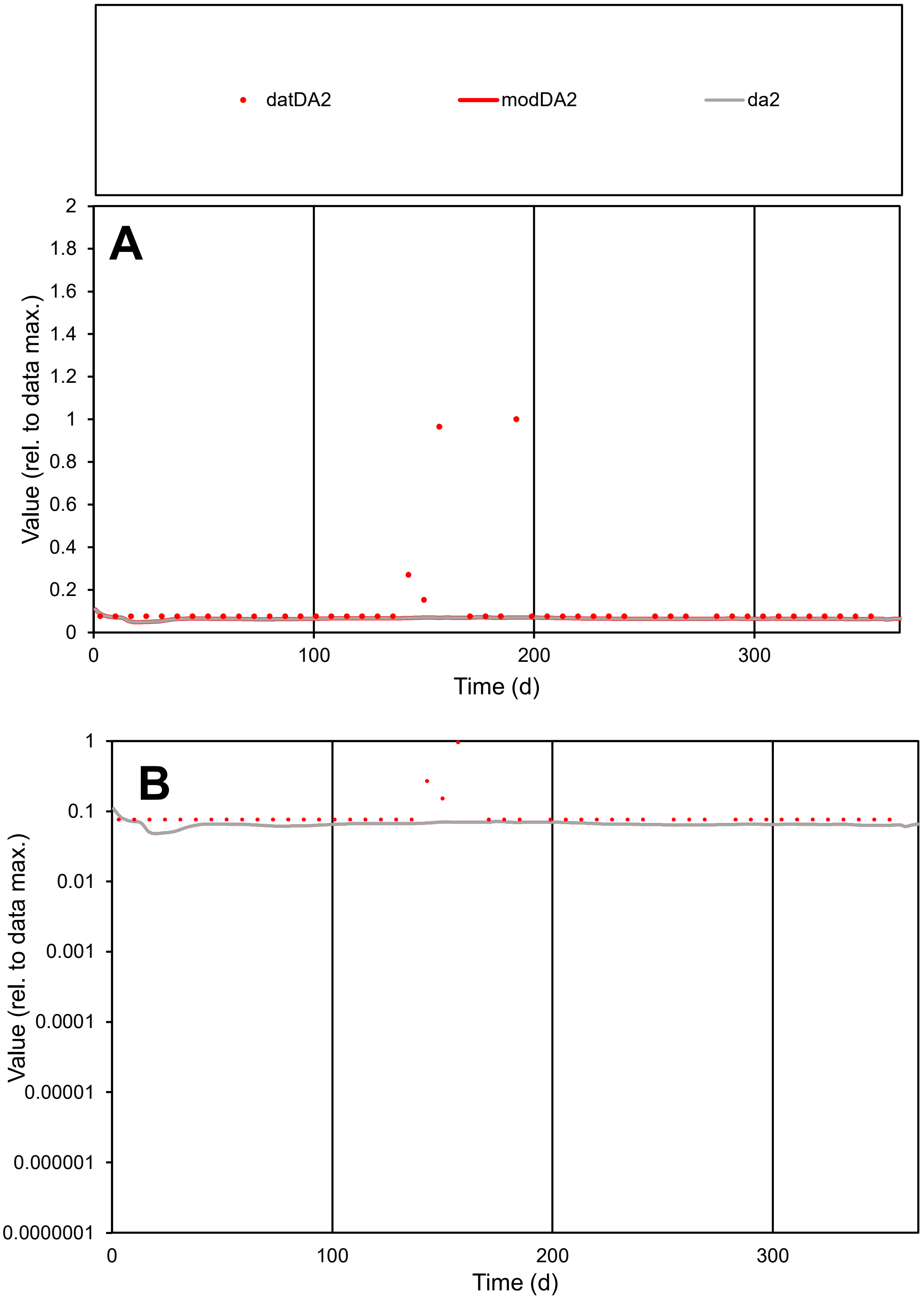


**C1**

Variables
DA2 Provide jtd or Dst
CalibOnly?
2 1 to reproduce of
oNormalize?
2 1=no, 2=yes

ofxn calc*
1.58112
ofxn mod
1.58691

sRun
RunA1
zout
1

Ave
Data
4.9E-06
Model
2.7E-06

data/vMax
4.1E-05

ofxng
369.894

Min d/m
1.56863

ave/min d
1.56804
ave/min m
1.34372

328_Calib_S_A

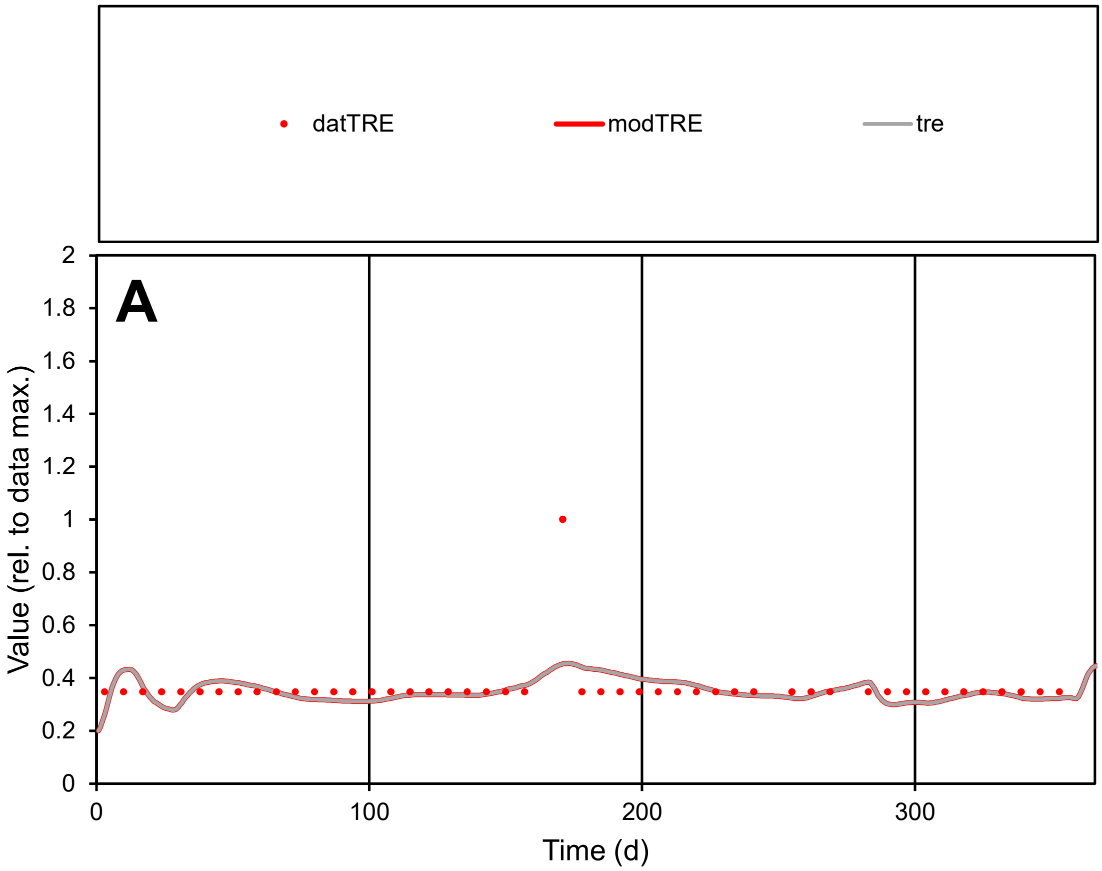


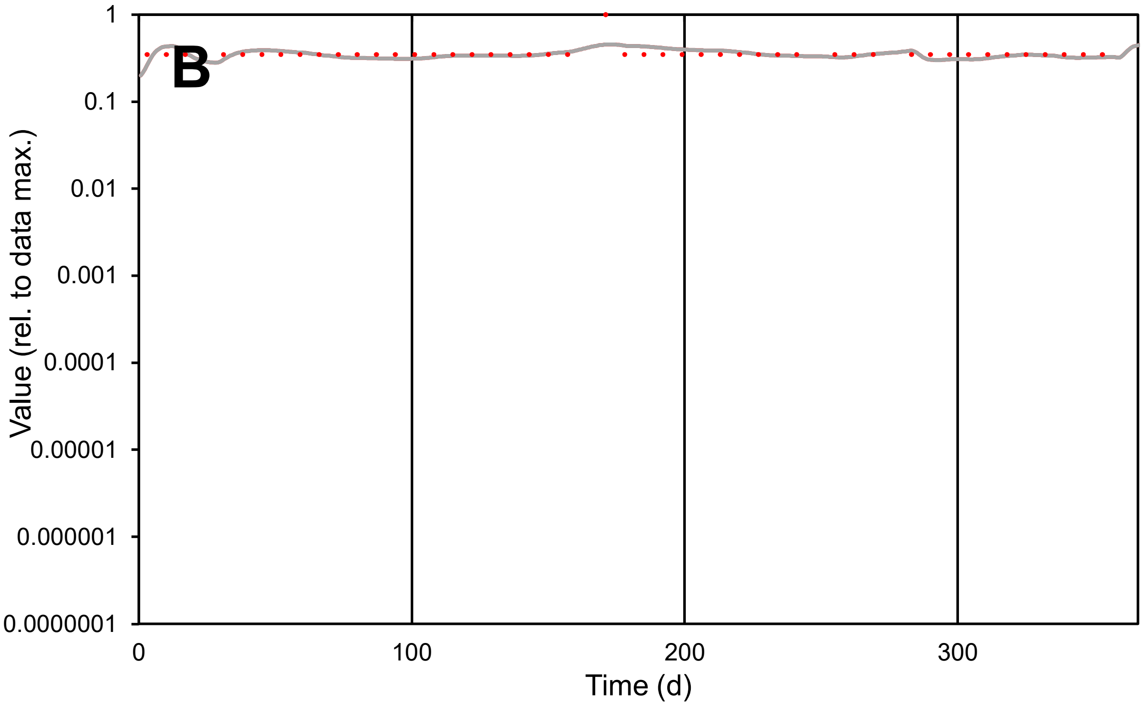


**C1**

Variables
TRE Provide jtd or Dst
CalibOnly?
2 1 to reproduce of
oNormalize?
2 1=no, 2=yes

ofxn calc*
0.23799
ofxn mod
0.26369

sRun
RunA1
zout
1

Ave
Data
7.2E-06
Model
7E-06

data/vMax
2E-05

ofxng
369.894

Min d/m
1.20484

ave/min d
1.03911
ave/min m
1.21205

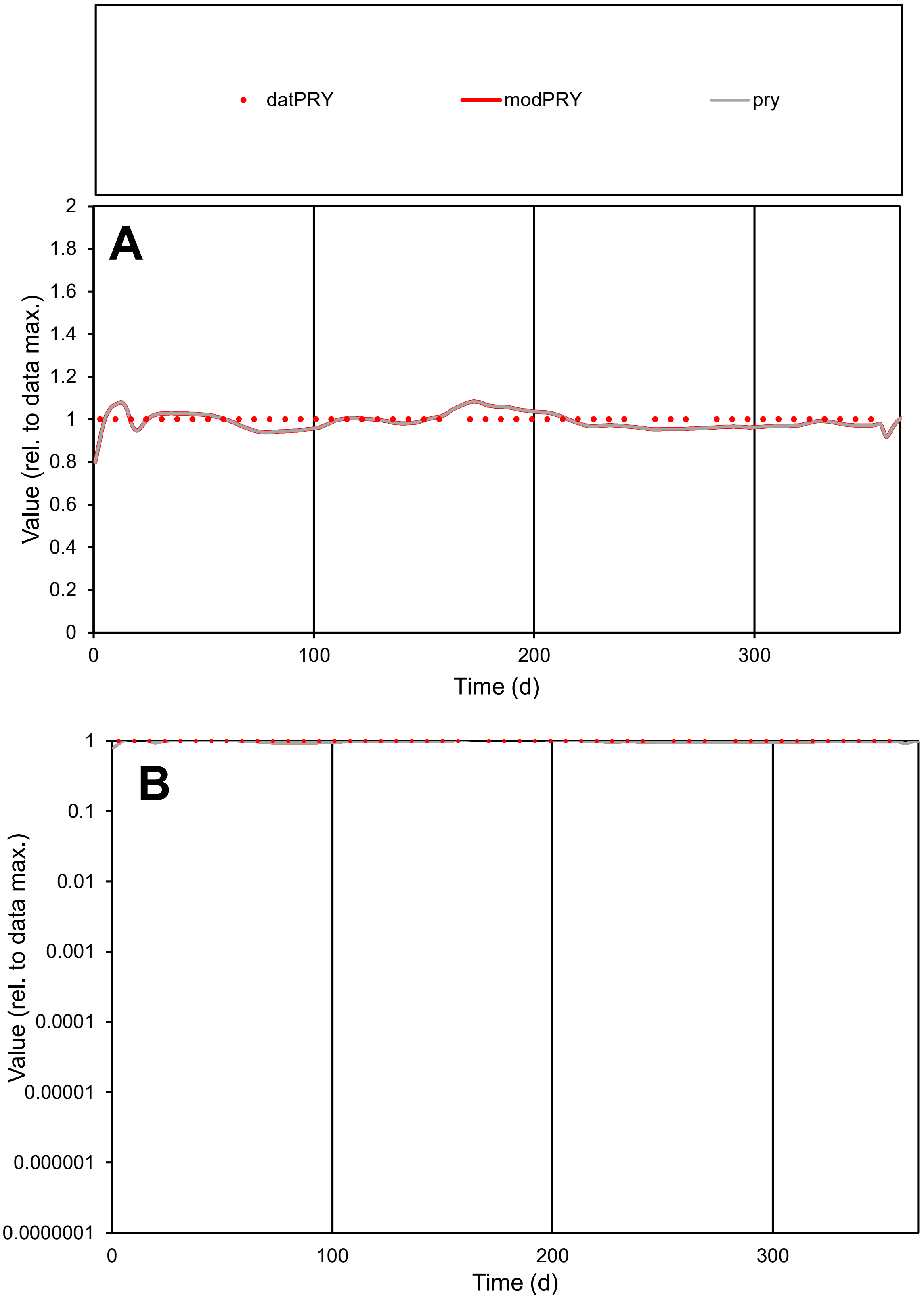


C1

Variables
PRY Provide jtd or Dst
CalibOnly?
2 1 to reproduce of
oNormalize?
2 1=no, 2=yes

ofxn calc*
0.03659
ofxn mod
0.11859

sRun
RunA1
zout
1

Ave
Data
8.8E-06
Model
8.7E-06

data/vMax
8.8E-06

ofxng
369.894

Min d/m
1.06296

ave/min d
1
ave/min m
1.05331

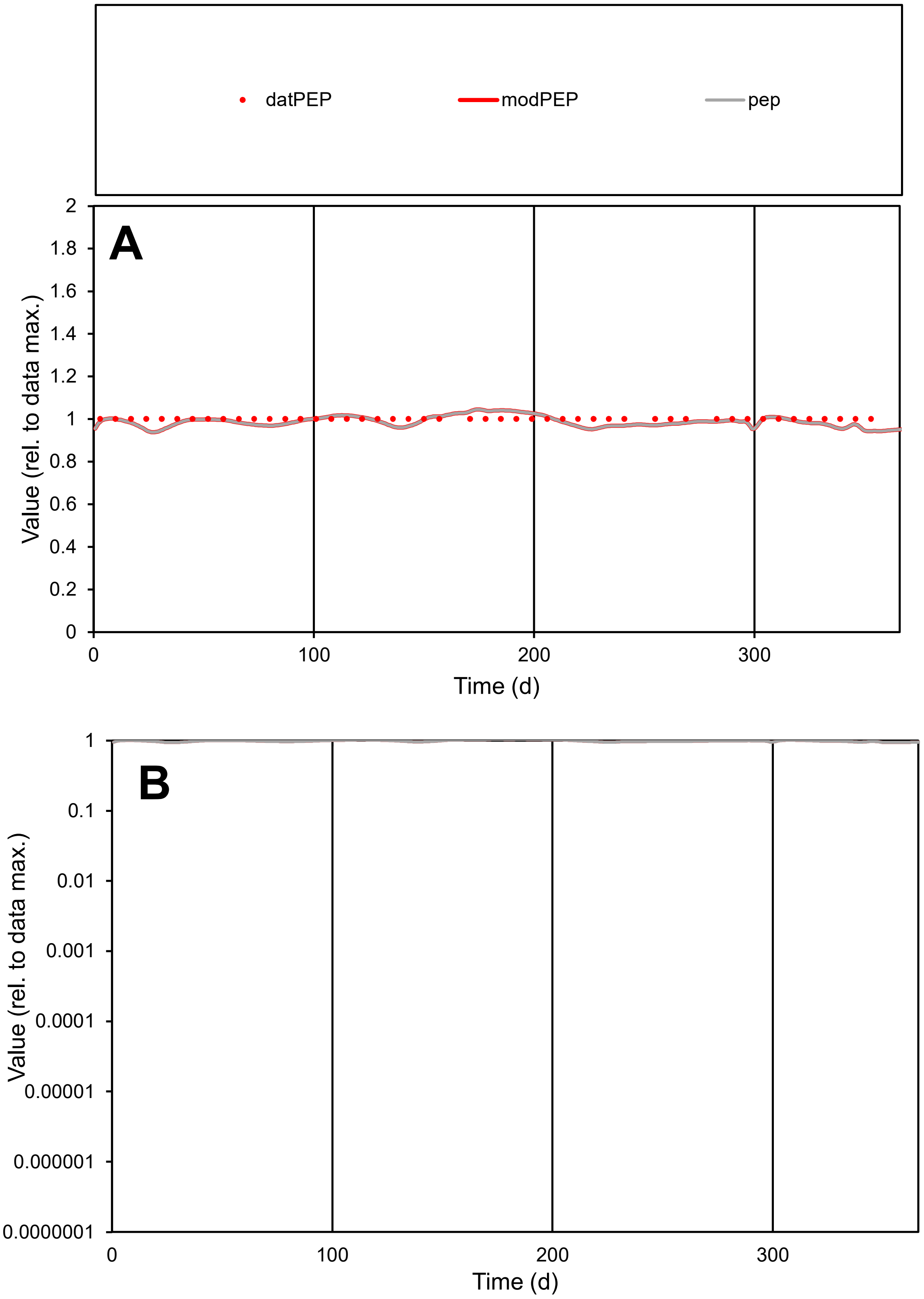


**C1**

Variables
PEP Provide jtd or Dst
CalibOnly?
2 1 to reproduce of
oNormalize?
2 1=no, 2=yes

ofxn calc*
0.02699
ofxn mod
0.11568

sRun
RunA1
zout
1

Ave
Data
4.9E-06
Model
4.9E-06

data/vMax
4.9E-06

ofxng
369.894

Min d/m
1.06062

ave/min d
1
ave/min m
1.04846

328_Calib_S_A

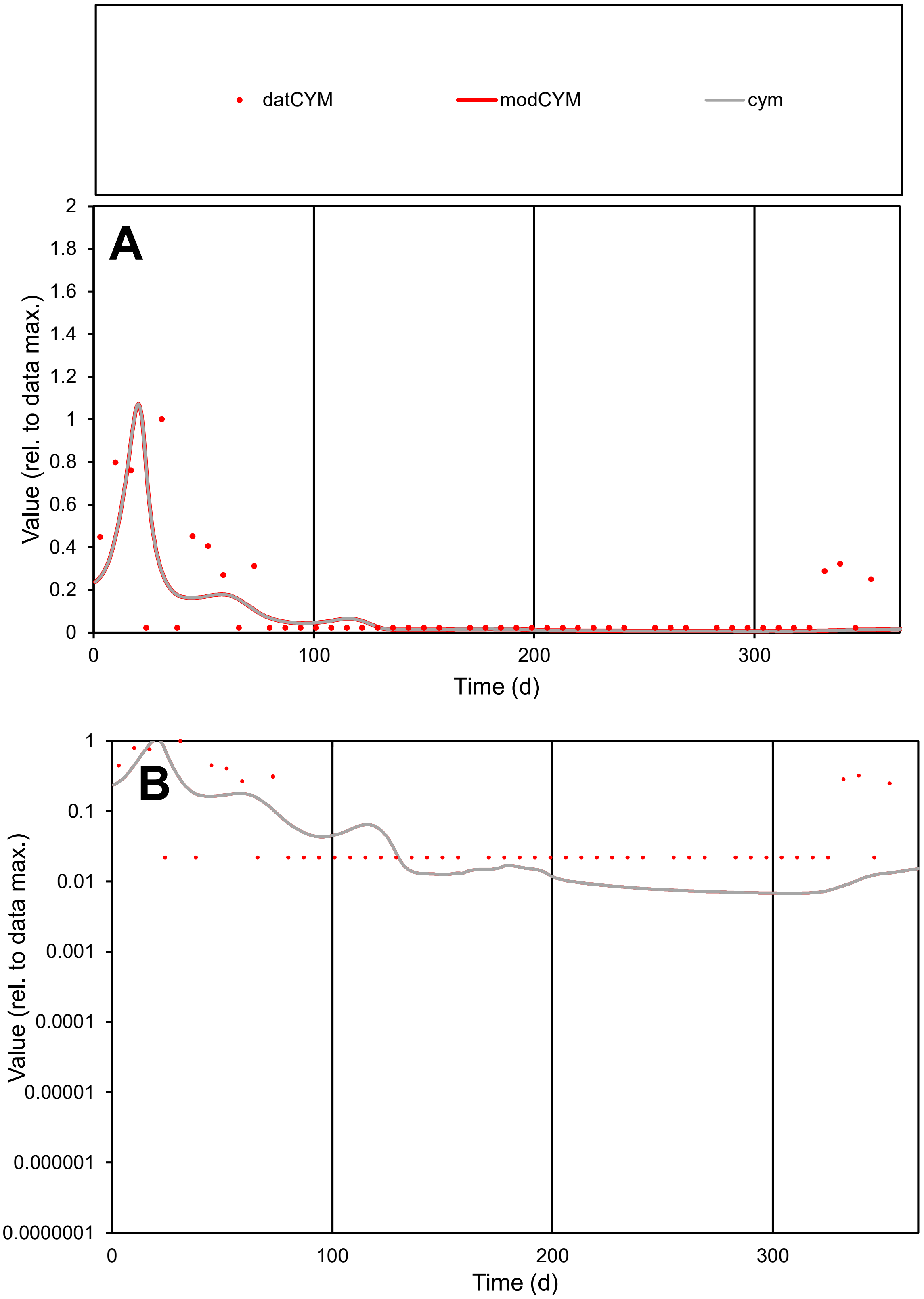


**C1**

Variables
CYM Provide jtd or Dst
CalibOnly?
2 1 to reproduce of
oNormalize?
2 1=no, 2=yes

ofxn calc*
1.28261
ofxn mod
1.33844

sRun
RunA1
zout
1

Ave
Data
7.9E-06
Model
5.2E-06

data/vMax
6.2E-05

ofxng
369.894

Min d/m
3.23508

ave/min d
5.78568
ave/min m
12.4349

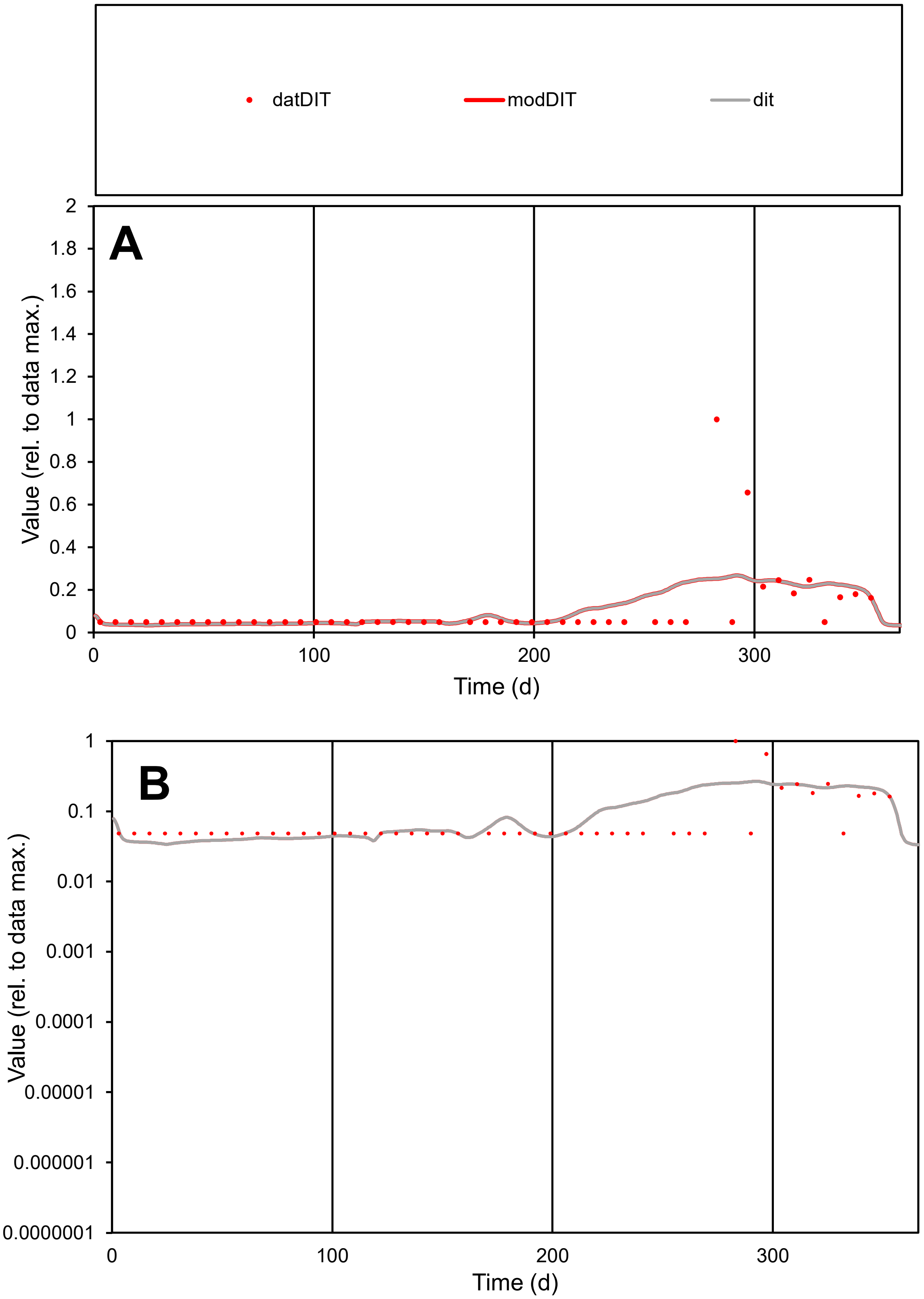


**C1**

Variables
DIT Provide jtd or Dst
CalibOnly?
2 1 to reproduce of
oNormalize?
2 1=no, 2=yes

ofxn calc*
1.34014
ofxn mod
1.35152

sRun
RunA1
zout
1

Ave
Data
1.1E-05
Model
1.1E-05

data/vMax
0.00011

ofxng
369.894

Min d/m
1.43354

ave/min d
2.12257
ave/min m
3.07489

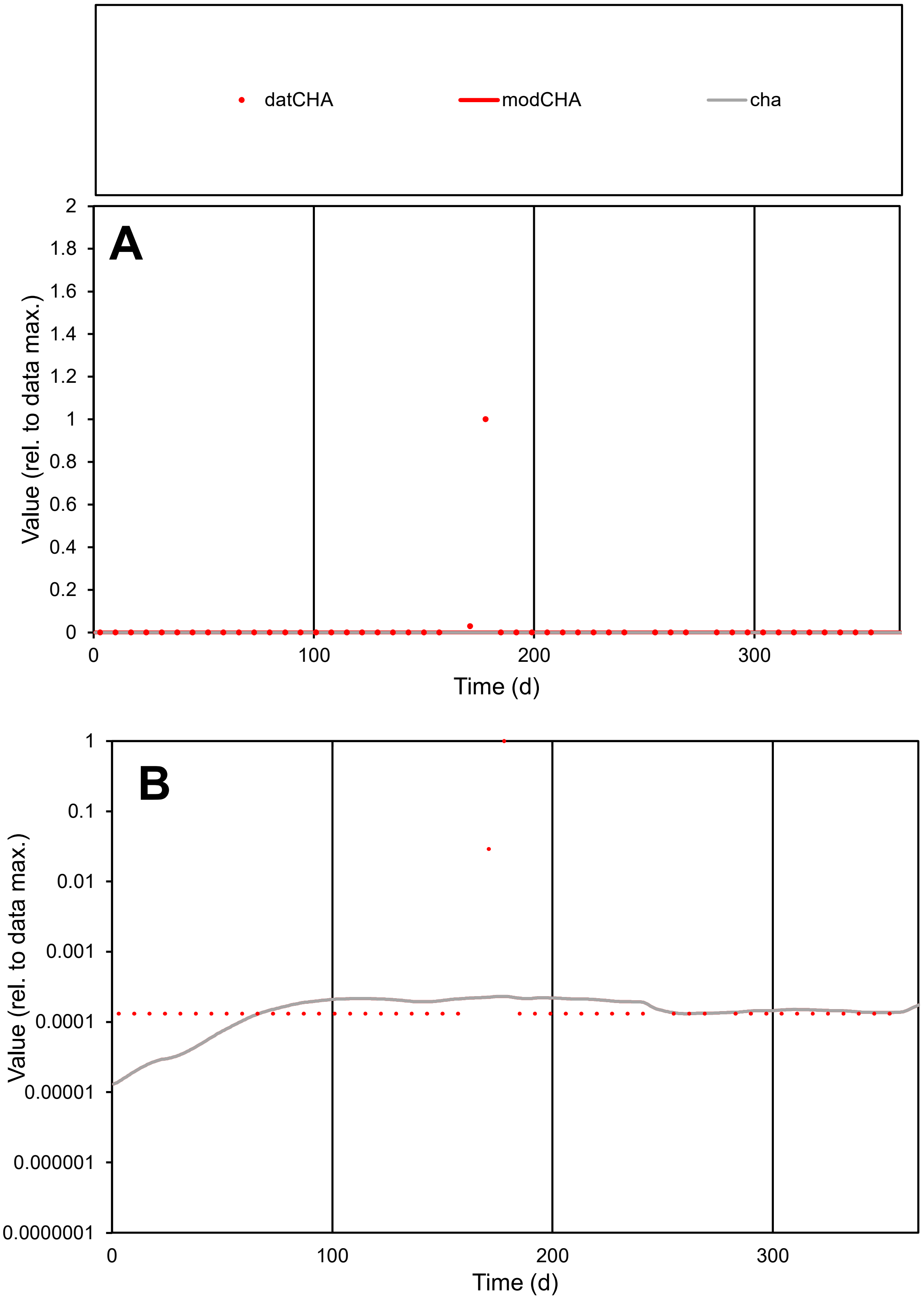


**C1**

Variables
CHA Provide jtd or Dst
CalibOnly?
2 1 to reproduce of
oNormalize?
2 1=no, 2=yes

ofxn calc*
6.69427
ofxn mod
6.70893

sRun
RunA1
zout
1

Ave
Data
1.4E-05
Model
1E-07

data/vMax
0.00066

ofxng
369.894

Min d/m
8.71513

ave/min d
164.301
ave/min m
10.315

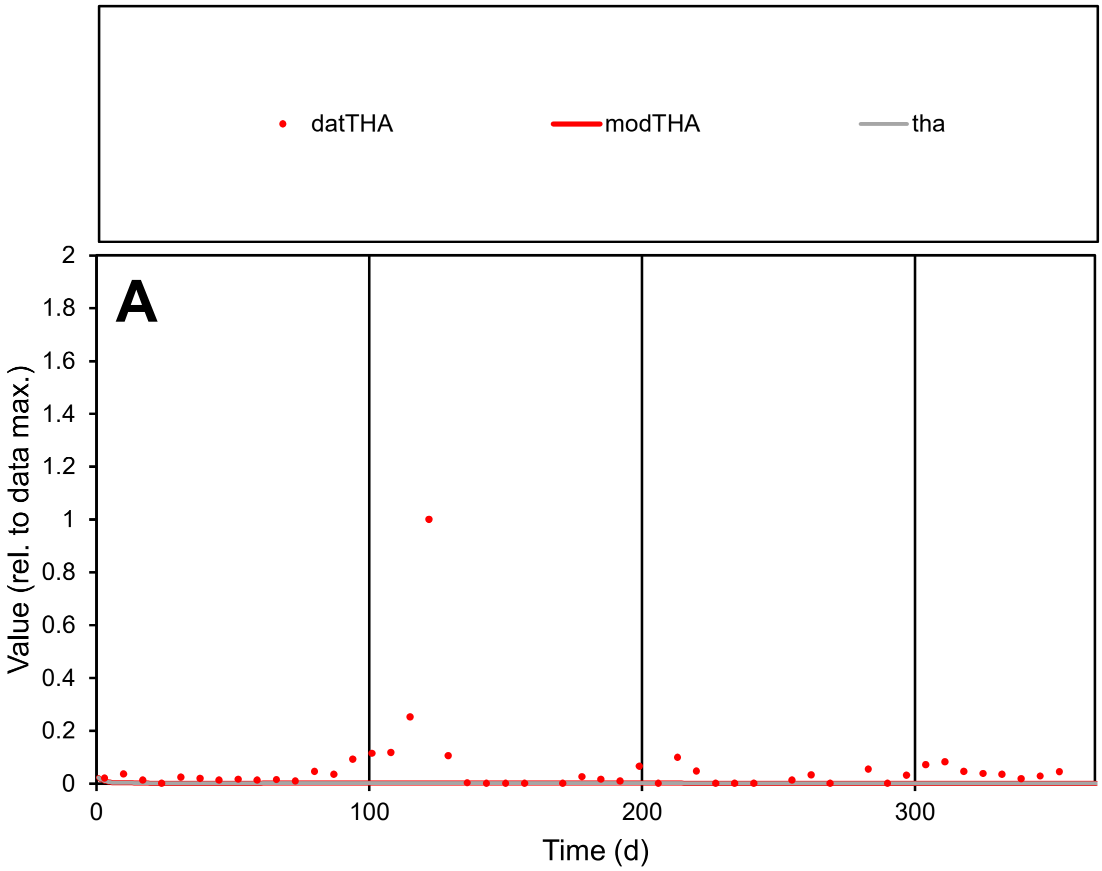


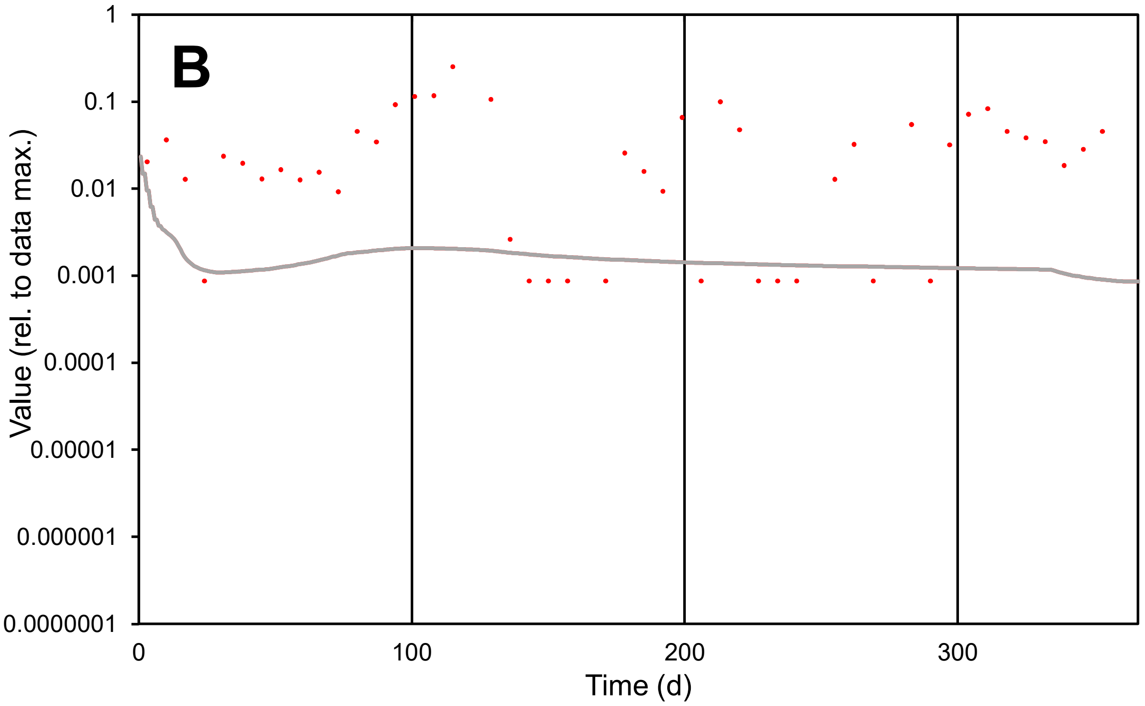


**C1**

Variables
THA Provide jtd or Dst
CalibOnly?
2 1 to reproduce of
oNormalize?
2 1=no, 2=yes

ofxn calc*
2.73521
ofxn mod
2.7413

sRun
RunA1
zout
1

Ave
Data
0.00018
Model
5.6E-06

data/vMax
0.00315

ofxng
369.894

Min d/m
0.97589

ave/min d
64.8519
ave/min m
1.97551

328_Calib_S_A

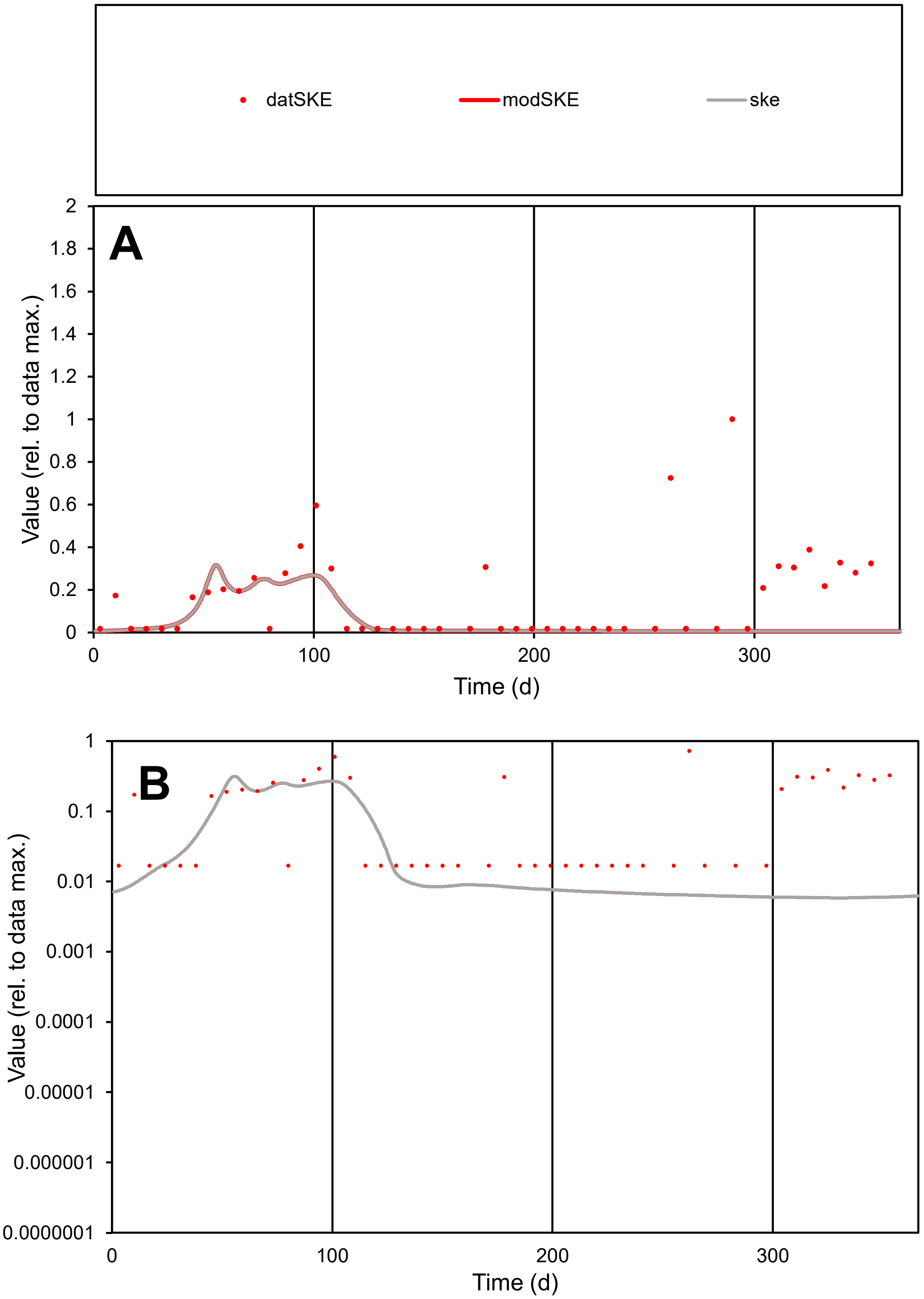


**C1**

Variables
SKE Provide jtd or Dst
CalibOnly?
2 1 to reproduce of
oNormalize?
2 1=no, 2=yes

ofxn calc*
1.4453
ofxn mod
1.45115

sRun
RunA1
zout
1

Ave
Data
2.2E-05
Model
7.6E-06

data/vMax
0.00014

ofxng
369.894

Min d/m
2.88853

ave/min d
9.38421
ave/min m
9.48207

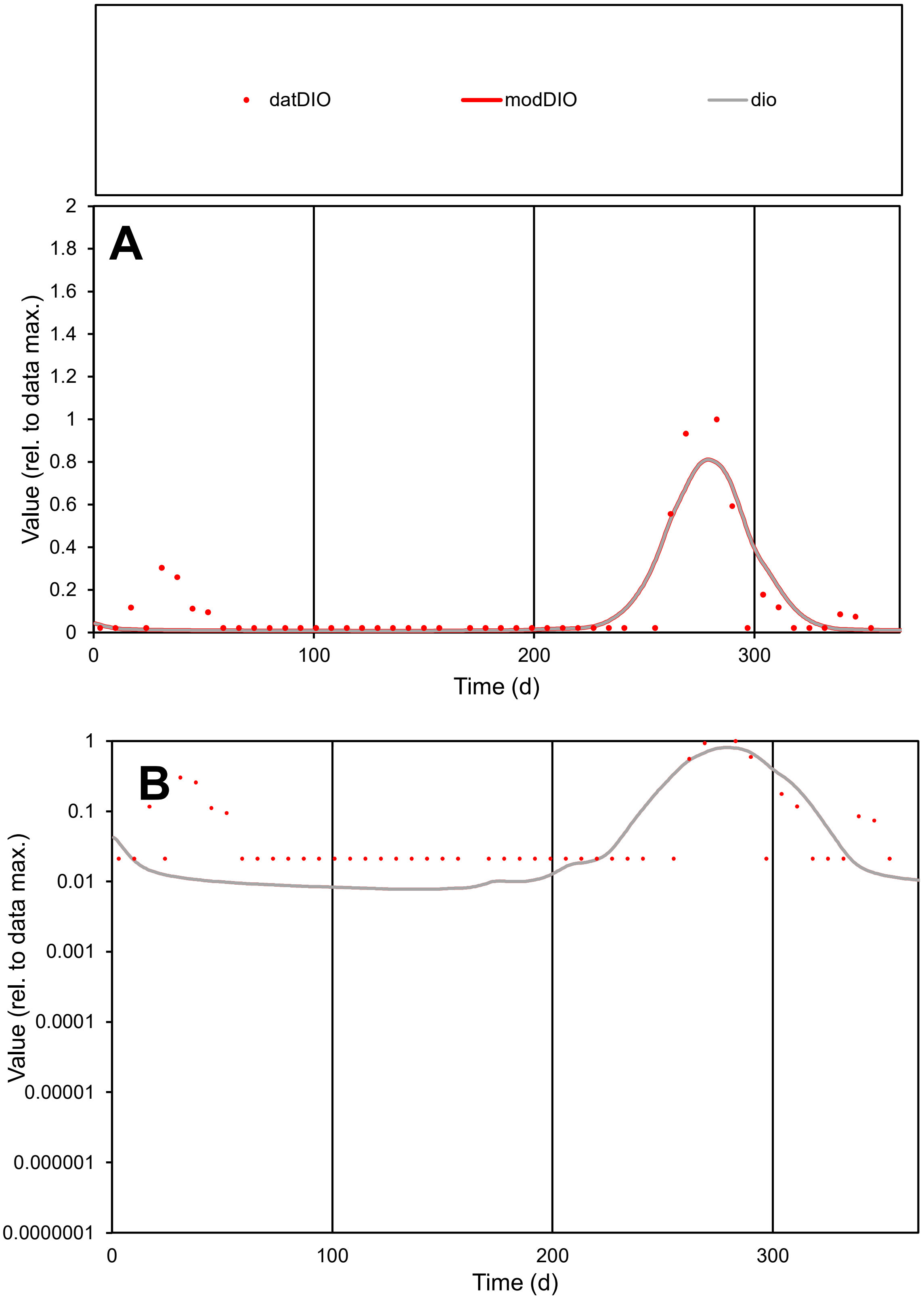


**C1**

Variables
DIO Provide jtd or Dst
CalibOnly?
2 1 to reproduce of
oNormalize?
2 1=no, 2=yes

ofxn calc*
0.98263
ofxn mod
1.04753

sRun
RunA1
zout
1

Ave
Data
2.4E-05
Model
2.1E-05

data/vMax
0.00022

ofxng
369.894

Min d/m
2.73361

ave/min d
5.08494
ave/min m
12.4445

328_Calib_S_A

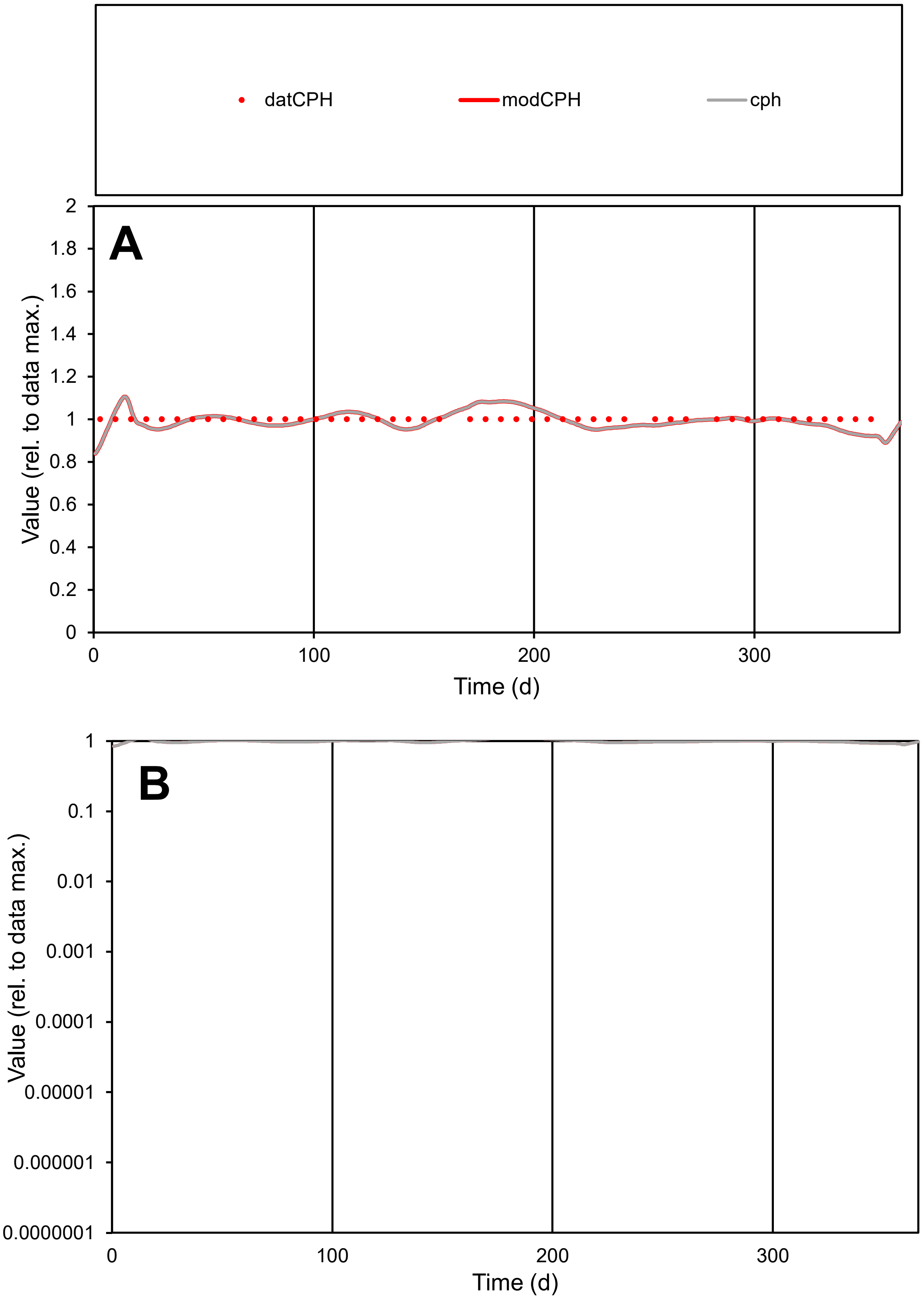


**C1**

Variables
CPH Provide jtd or Dst
CalibOnly?
2 1 to reproduce of
oNormalize?
2 1=no, 2=yes

ofxn calc*
0.04006
ofxn mod
0.11977

sRun
RunA1
zout
1

Ave
Data
9.6E-06
Model
9.5E-06

data/vMax
9.6E-06

ofxng
369.894

Min d/m
1.09936

ave/min d
1
ave/min m
1.0934

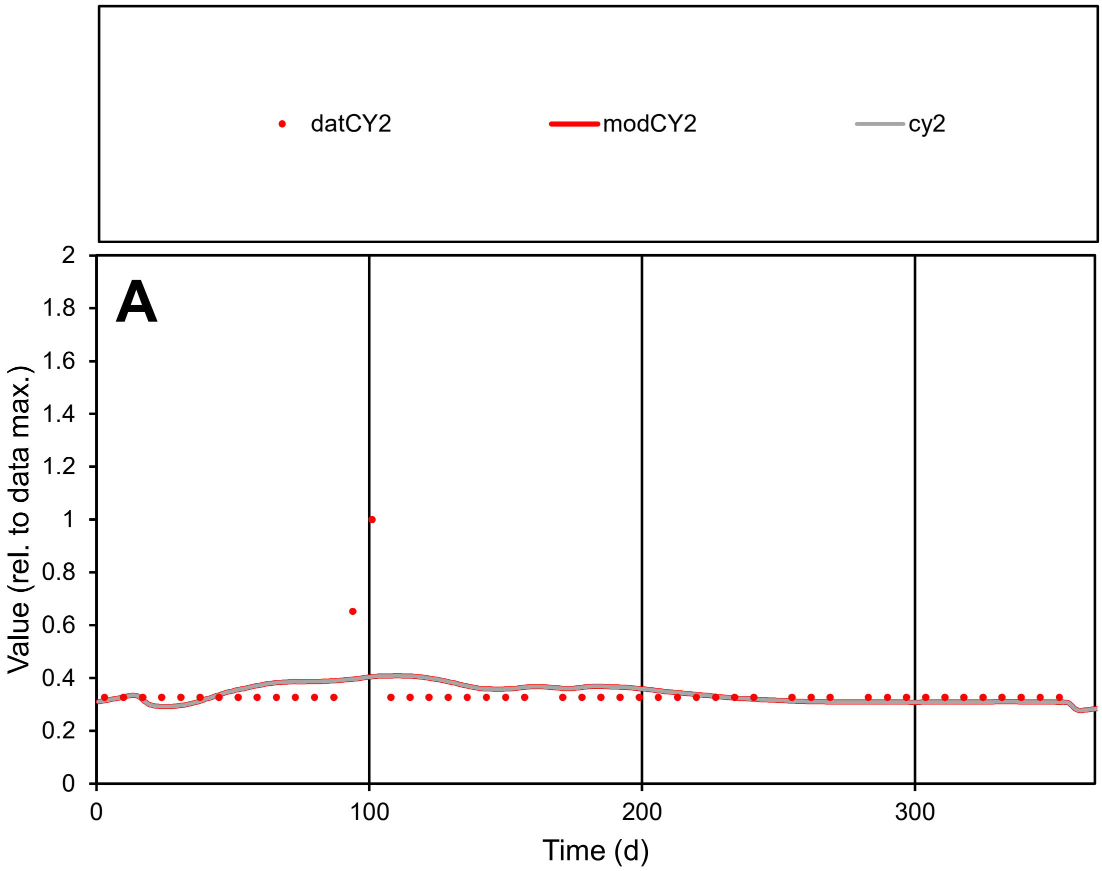


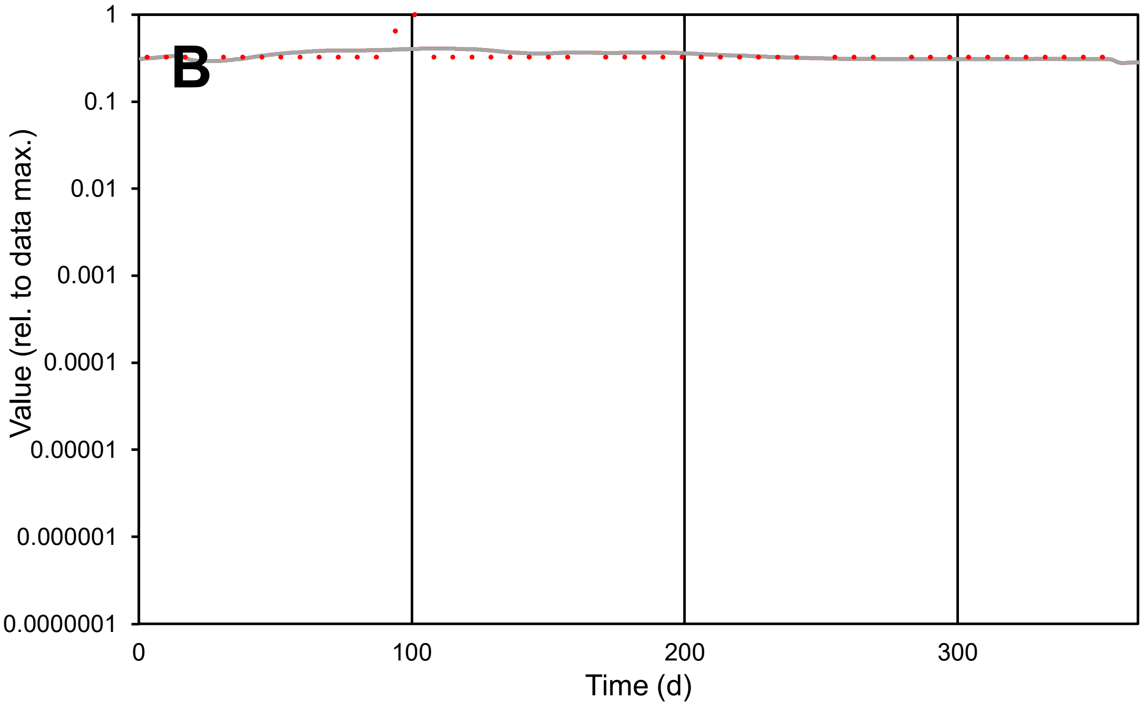


**C1**

Variables
CY2 Provide jtd or Dst
CalibOnly?
2 1 to reproduce of
oNormalize?
2 1=no, 2=yes

ofxn calc*
0.28754
ofxn mod
0.30933

sRun
RunA1
zout
1

Ave
Data
1.2E-05
Model
1.2E-05

data/vMax
3.4E-05

ofxng
369.894

Min d/m
1.11702

ave/min d
1.0639
ave/min m
1.17706

328_Calib_S_A

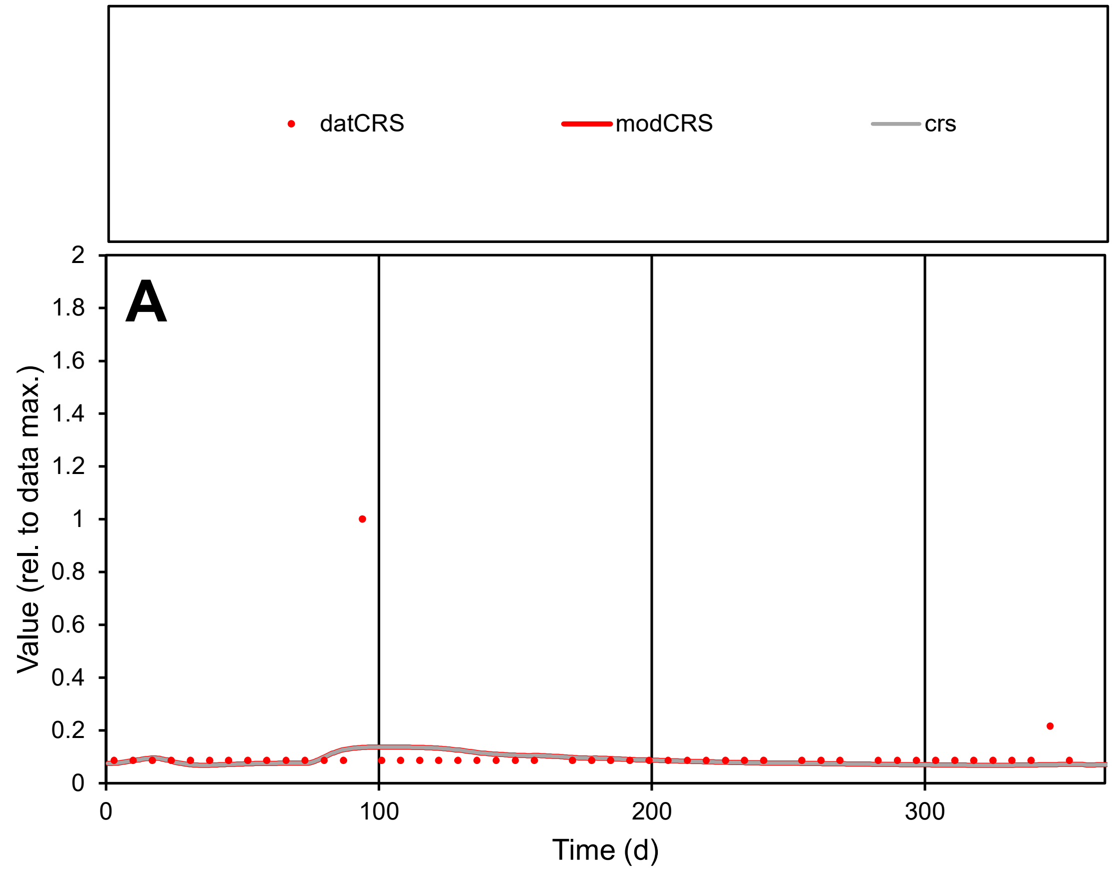


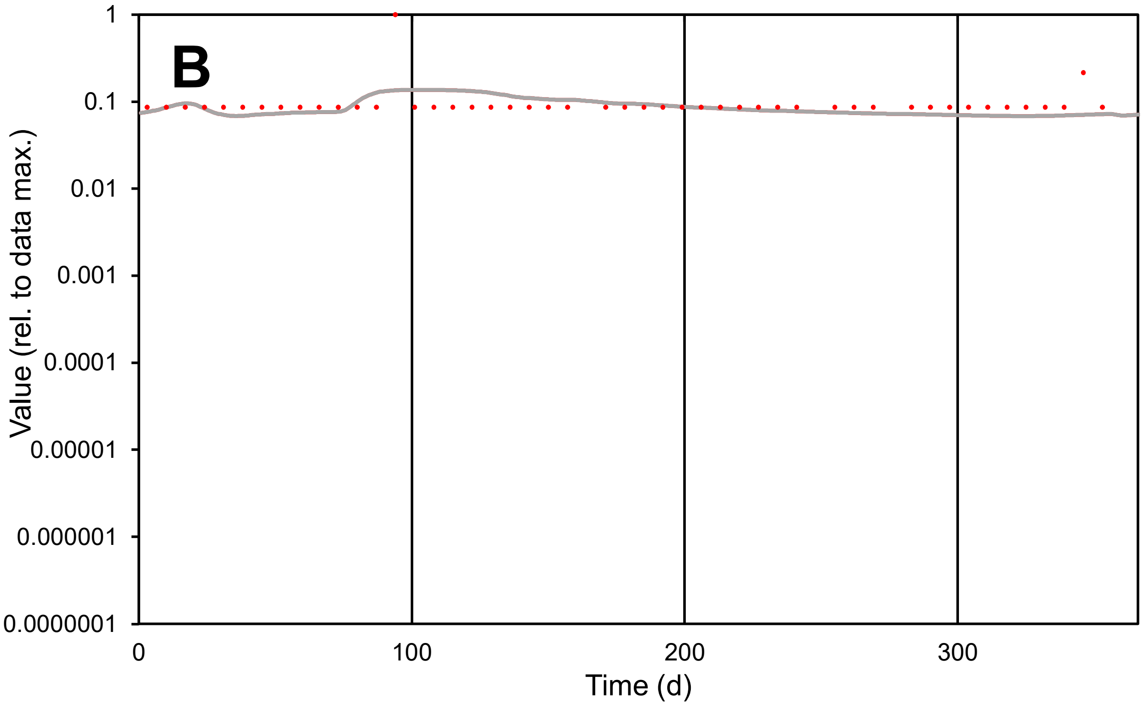


**C1**

Variables
CRS Provide jtd or Dst
CalibOnly?
2 1 to reproduce of
oNormalize?
2 1=no, 2=yes

ofxn calc*
1.18638
ofxn mod
1.1924

sRun
RunA1
zout
1

Ave
Data
6.8E-06
Model
5.6E-06

data/vMax
6.3E-05

ofxng
369.894

Min d/m
1.26284

ave/min d
1.25181
ave/min m
1.29898

328_Calib_S_A

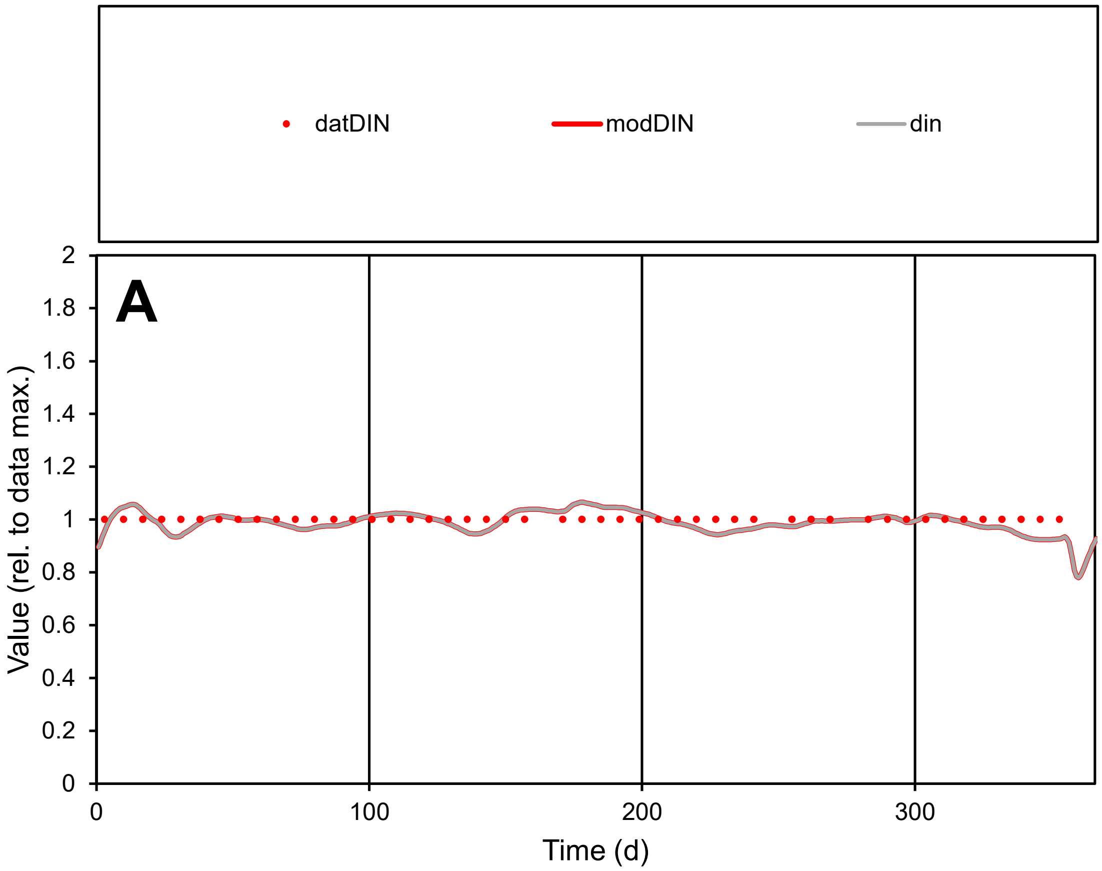


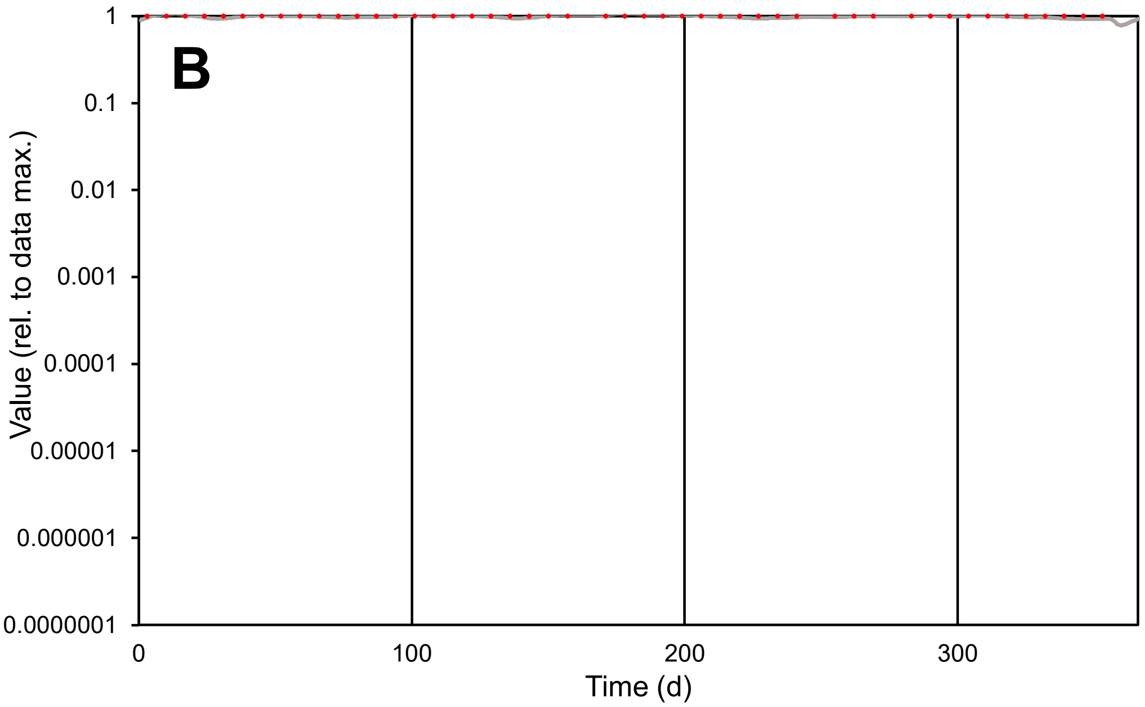


**C1**

Variables
DIN Provide jtd or Dst
CalibOnly?
2 1 to reproduce of
oNormalize?
2 1=no, 2=yes

ofxn calc*
0.03365
ofxn mod
0.11758

sRun
RunA1
zout
1

Ave
Data
2.9E-06
Model
2.9E-06

data/vMax
2.9E-06

ofxng
369.894

Min d/m
1.08218

ave/min d
1
ave/min m
1.07333

328_Calib_S_A

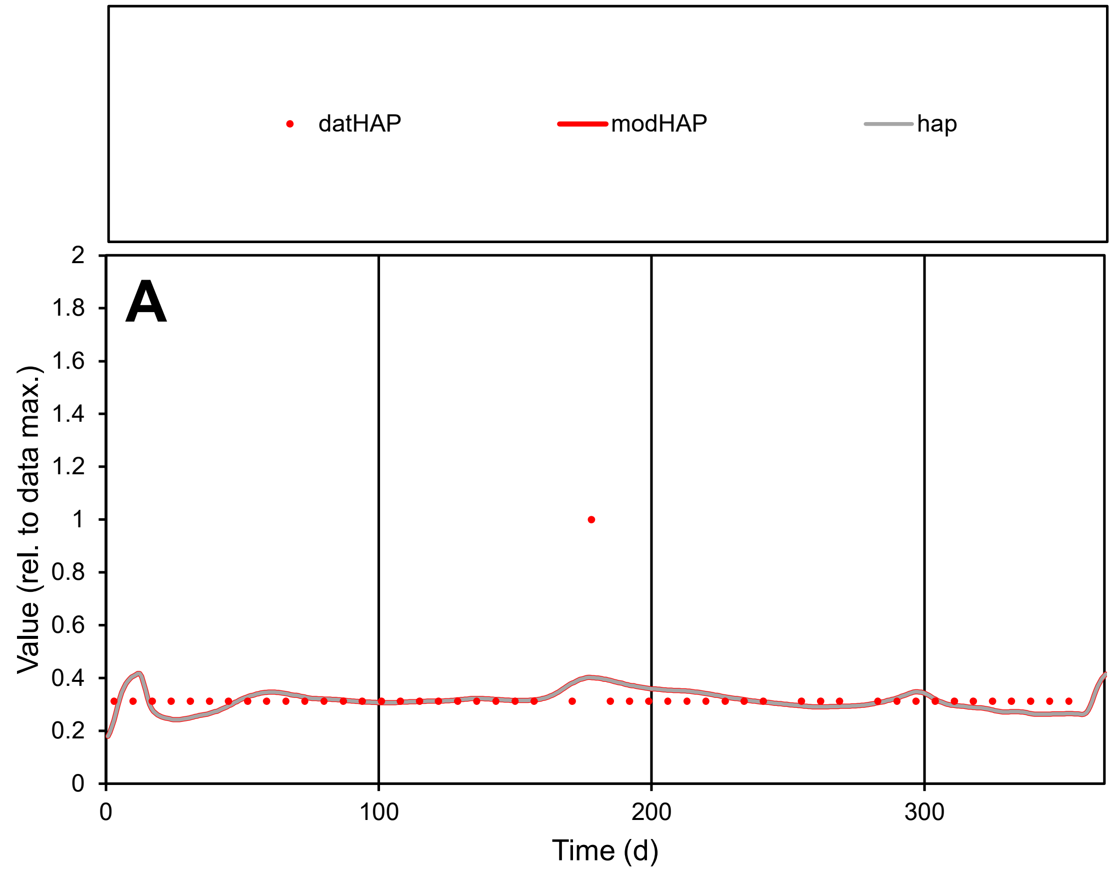


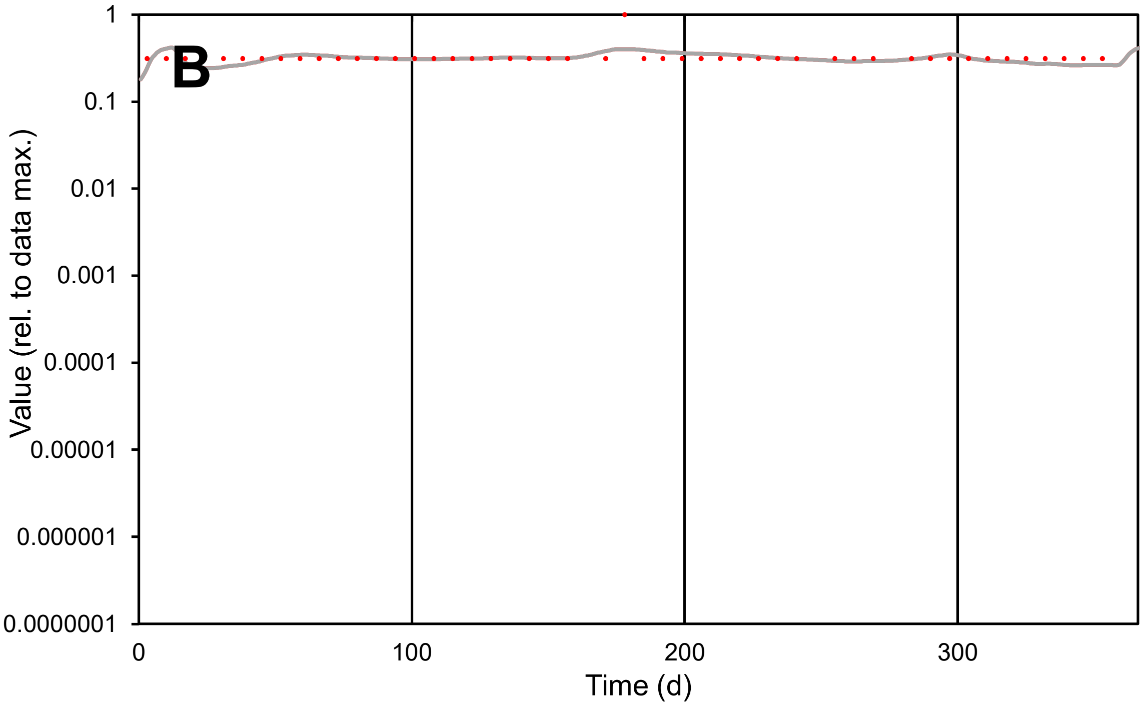


**C1**

Variables
HAP Provide jtd or Dst
CalibOnly?
2 1 to reproduce of
oNormalize?
2 1=no, 2=yes

ofxn calc*
0.28642
ofxn mod
0.3085

sRun
RunA1
zout
1

Ave
Data
1.2E-05
Model
1.1E-05

data/vMax
3.6E-05

ofxng
369.894

Min d/m
1.28641

ave/min d
1.04591
ave/min m
1.30236

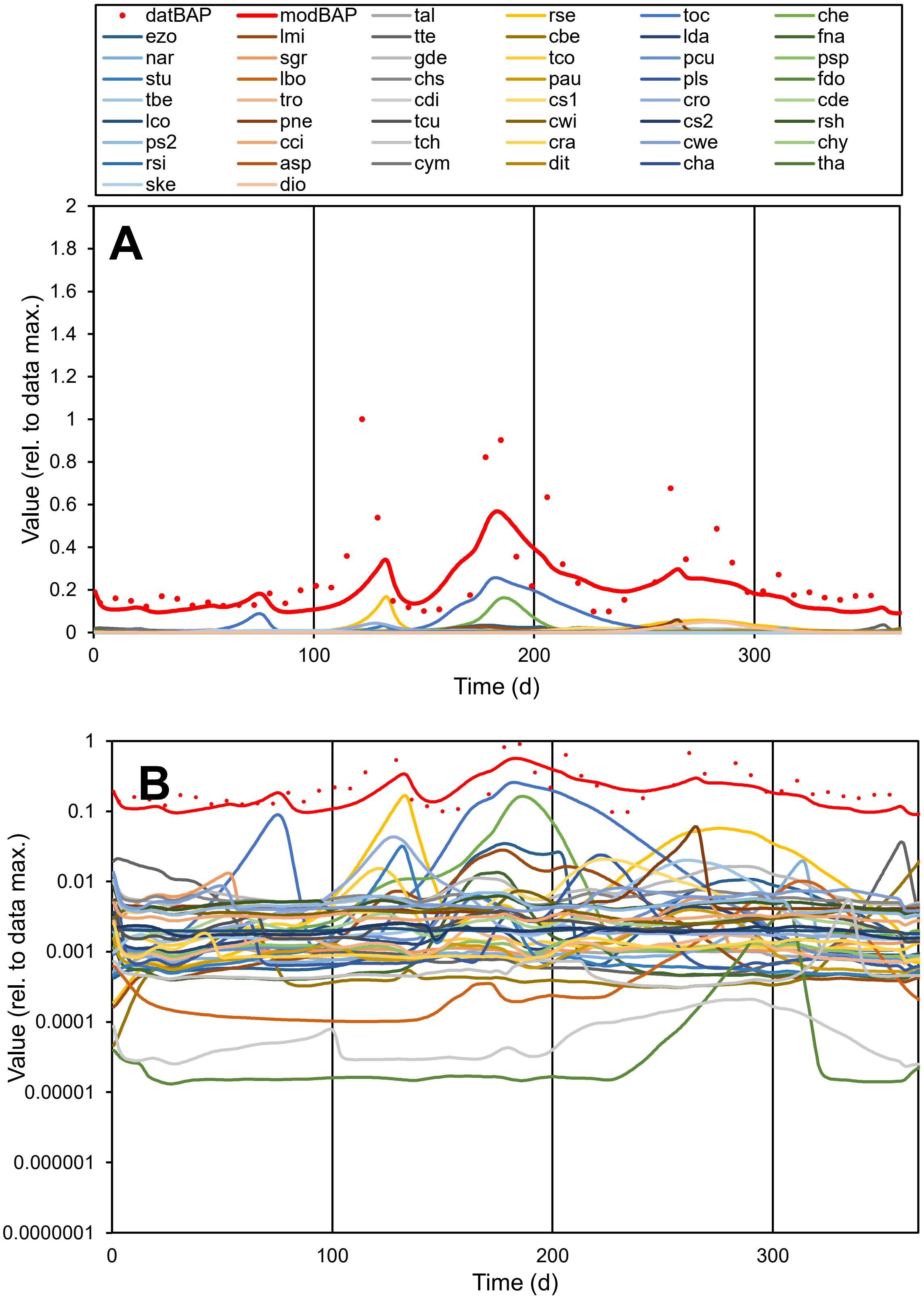


**C1**

Variables

BAP Provide jtd or Dst

CalibOnly?

2 1 to reproduce of

oNormalize?

2 1=no, 2=yes

ofxn calc*

0.65408

ofxn mod

0.66731

sRun

RunA1

zout

1

Ave

Data

0.00095

Model

0.00072

data/vMax

0.00361

ofxng

369.894

Min d/m

1.0121

ave/min d

2.69779

ave/min m

2.06267

328_Calib_S_A

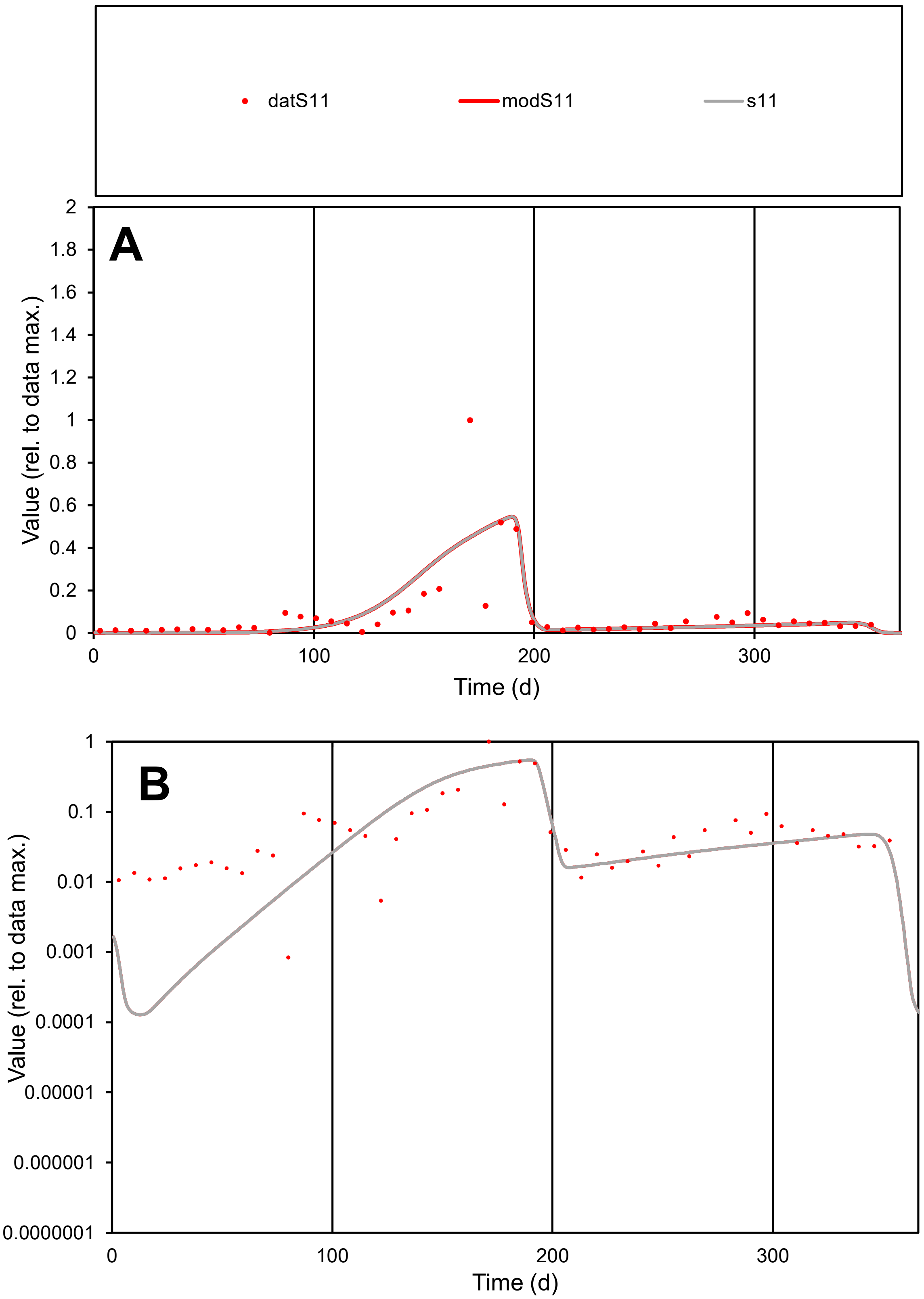

datS11
modS11
s11
A
Value (rel. to data max.)
Time (d)
B


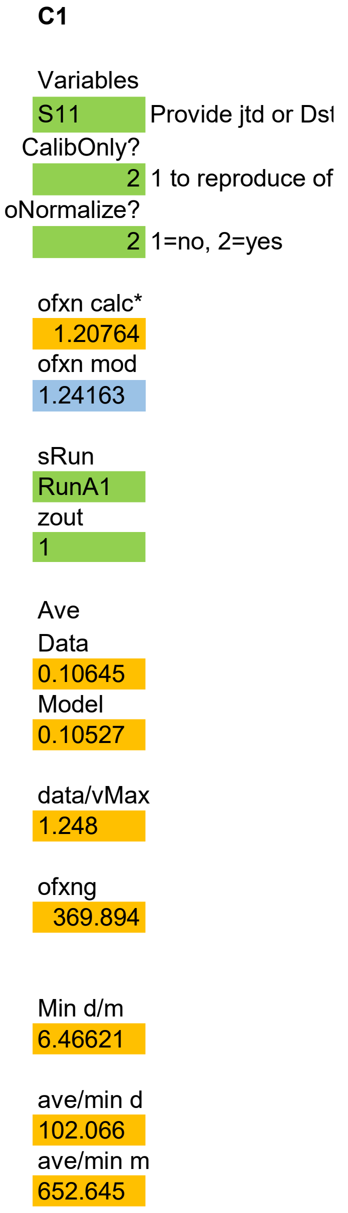

C1
Variables
S11
Provide jtd or Dst
CalibOnly?
2
1 to reproduce of
oNormalize?
2
1=no, 2=yes
ofxn calc*
1.20764
ofxn mod
1.24163
sRun
RunA1
zout
1
Ave
Data
0.10645
Model
0.10527
data/vMax
1.248
ofxng
369.894
Min d/m
6.46621
ave/min d
102.066
ave/min m
652.645

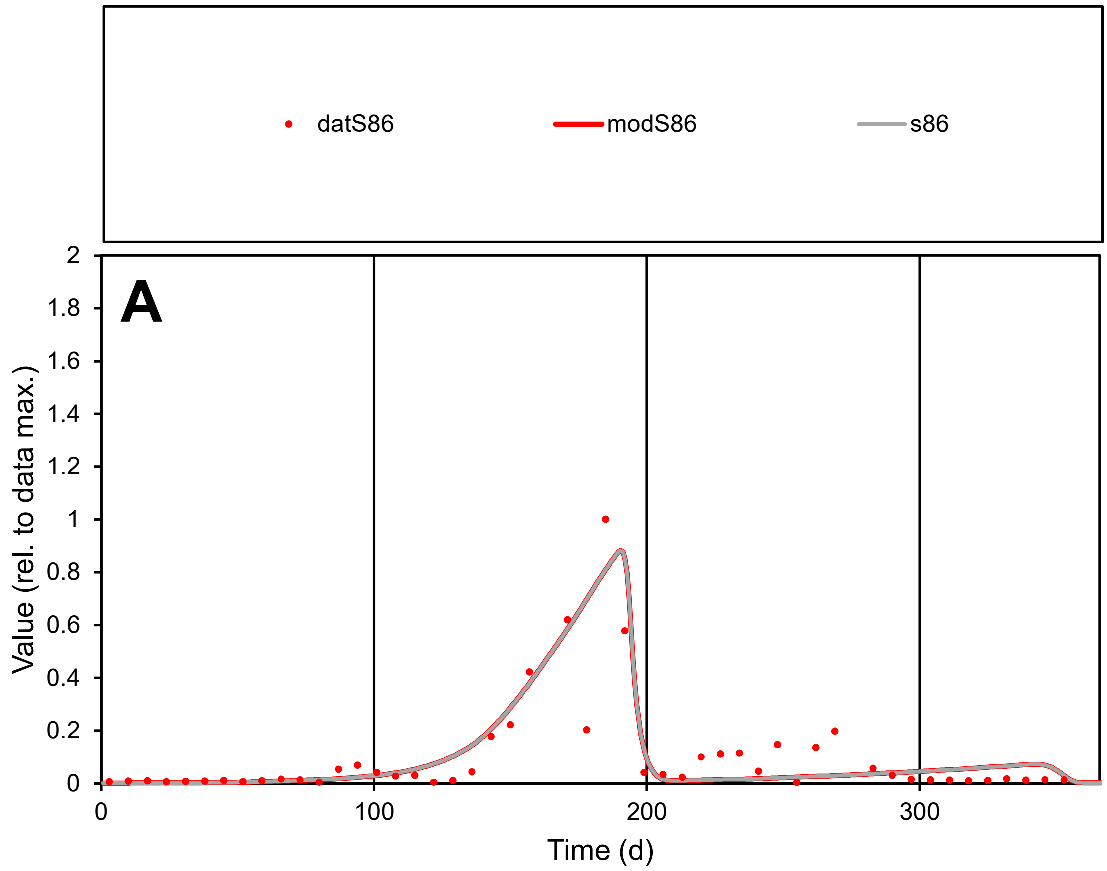


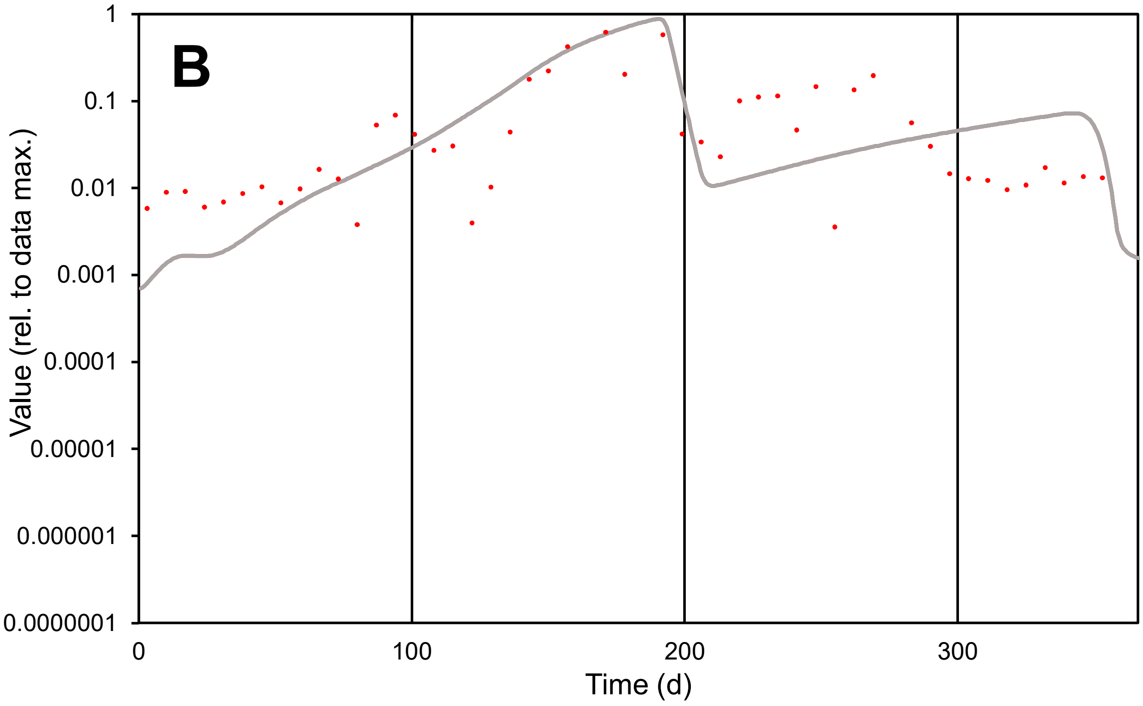


**C1**

Variables

S86 Provide jtd or Dst

CalibOnly?

2 1 to reproduce of

oNormalize?

2 1=no, 2=yes

ofxn calc*

0.96485

ofxn mod

1.01008

sRun

RunA1

zout

1

Ave

Data

0.06287

Model

0.06799

data/vMax

0.6468

ofxng

369.894

Min d/m

3.97829

ave/min d

27.23

ave/min m

117.15

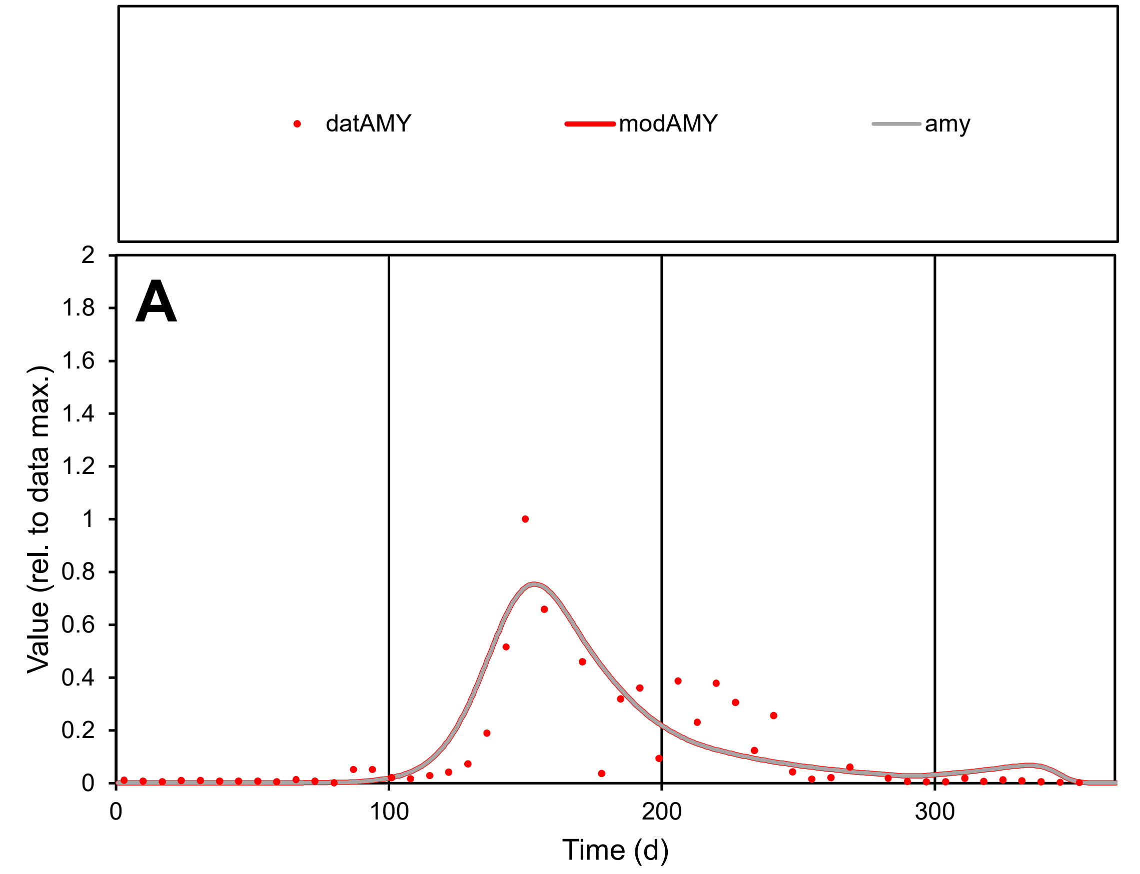


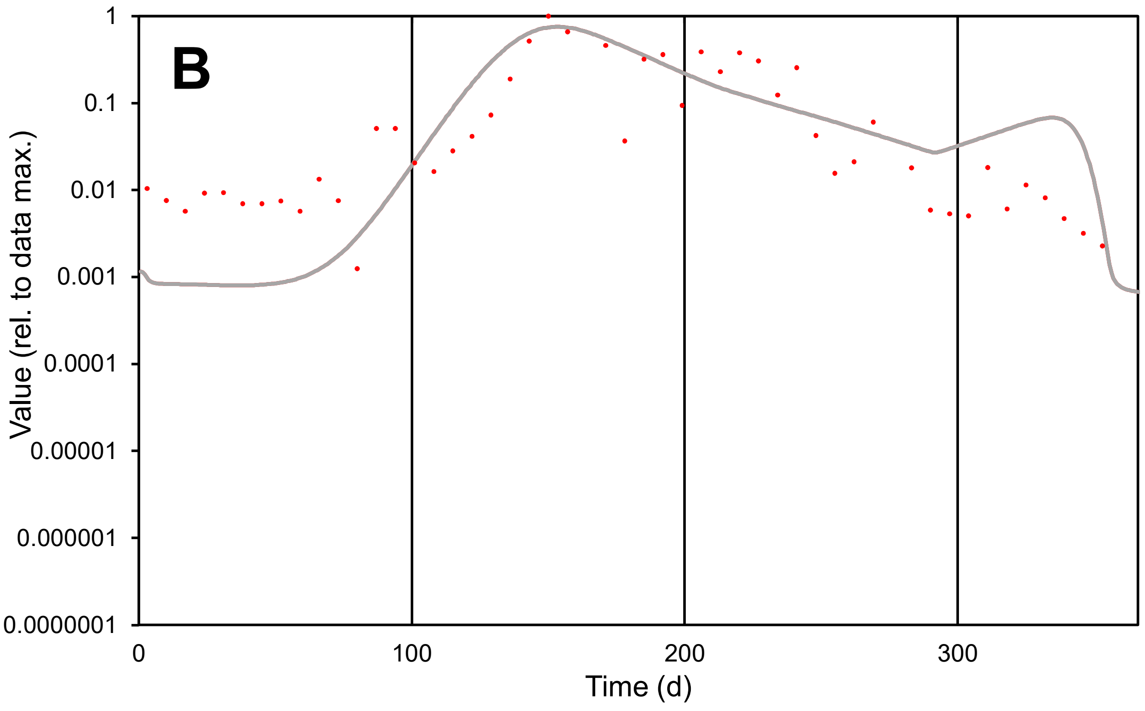


**C1**

Variables
AMY Provide jtd or Dst
CalibOnly?
2 1 to reproduce of
oNormalize?
2 1=no, 2=yes

ofxn calc*
0.91977
ofxn mod
0.95697

sRun
RunA1
zout
1

Ave
Data
0.05814
Model
0.06236

data/vMax
0.4817

ofxng
369.894

Min d/m
1.56491

ave/min d
96.2793
ave/min m
161.602

328_Calib_S_A

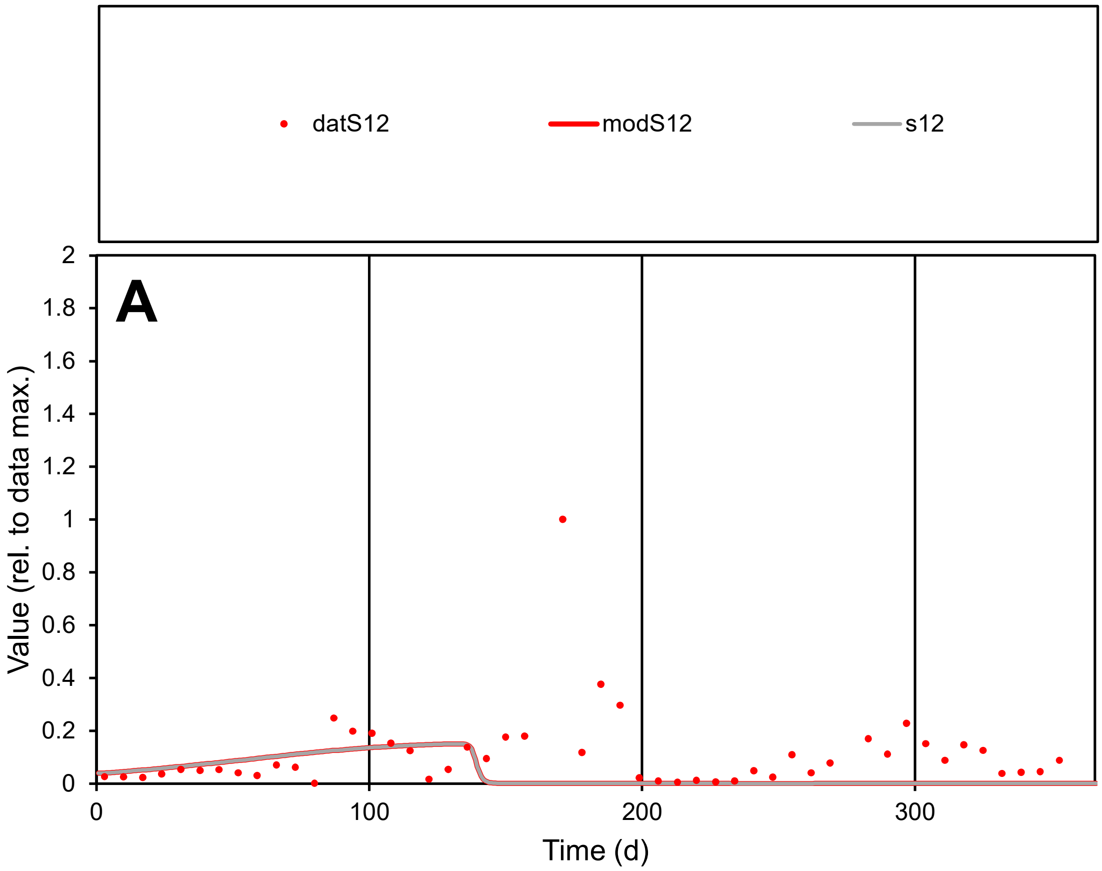


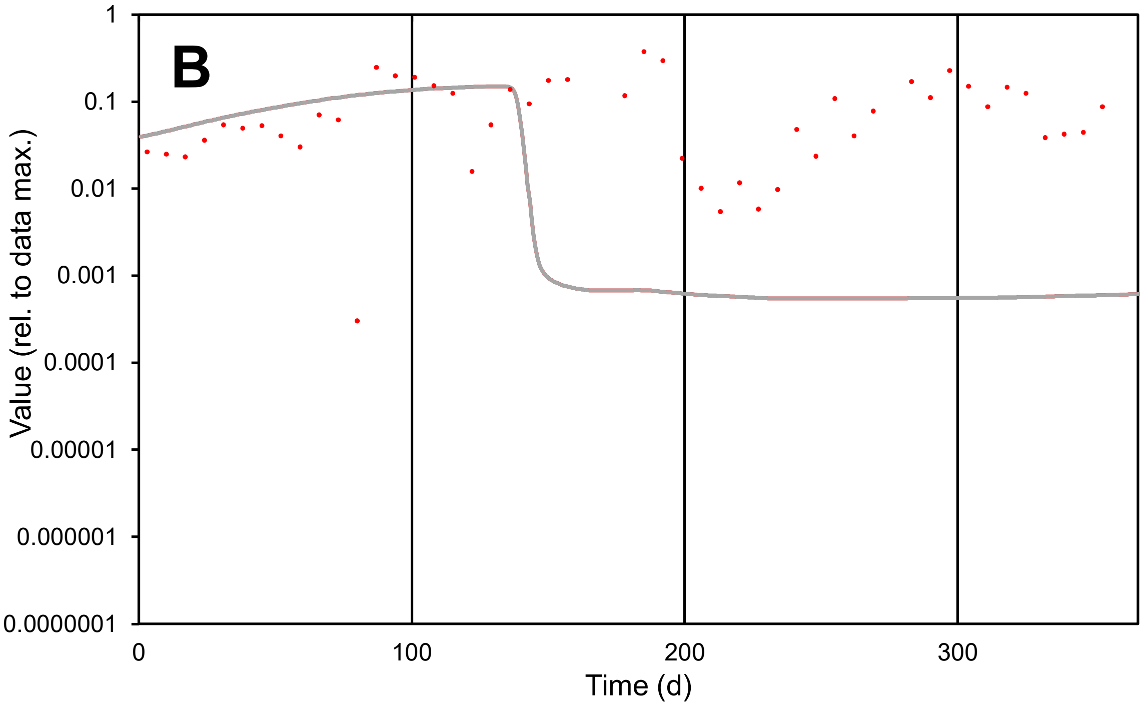


**C1**

Variables
S12 Provide jtd or Dst
CalibOnly?
2 1 to reproduce of
oNormalize?
2 1=no, 2=yes

ofxn calc*
1.61895
ofxn mod
1.62536

sRun
RunA1
zout
1

Ave
Data
0.05142
Model
0.02027

data/vMax
0.4641

ofxng
369.894

Min d/m
0.5519

ave/min d
364.908
ave/min m
79.3919

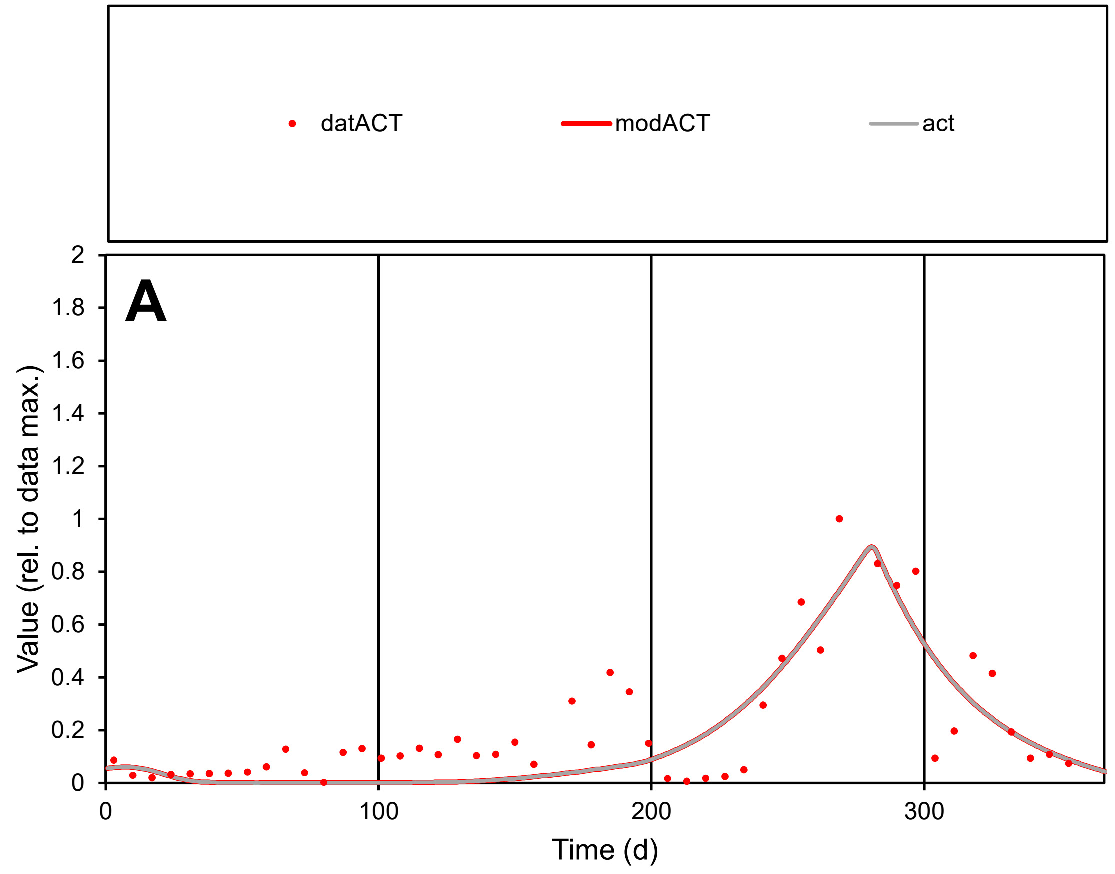


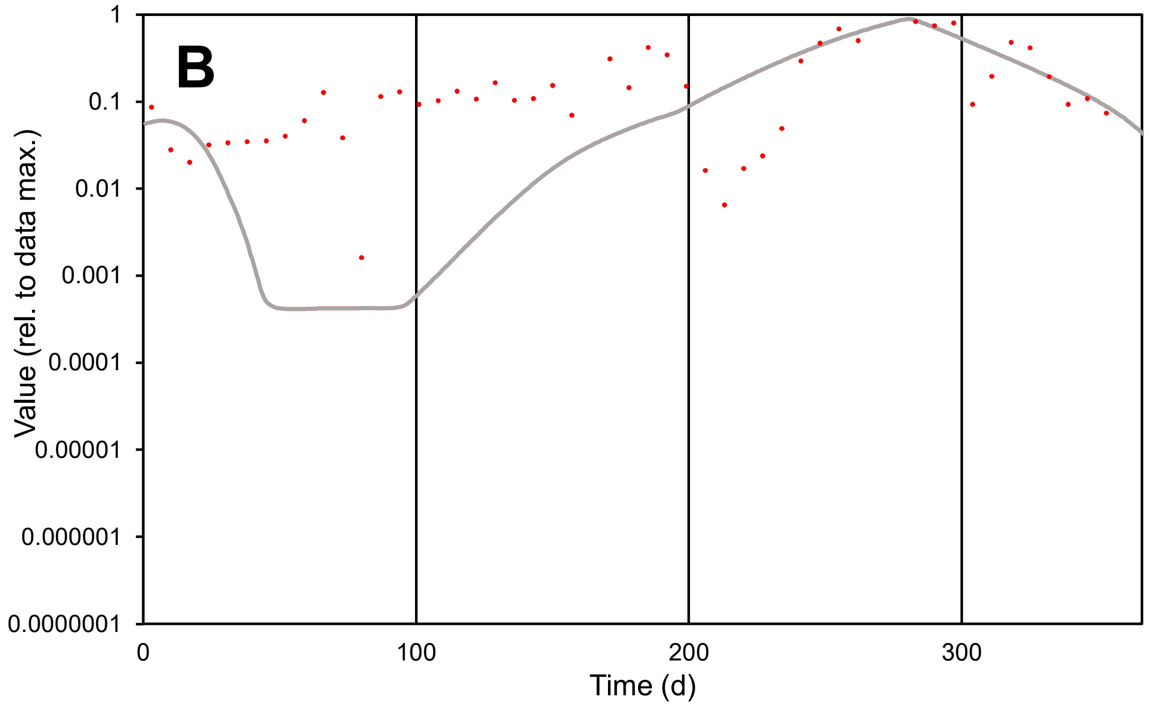


**C1**

Variables
ACT Provide jtd or Dst
CalibOnly?
2 1 to reproduce of
oNormalize?
2 1=no, 2=yes

ofxn calc*
0.68441
ofxn mod
0.70721

sRun
RunA1
zout
1

Ave
Data
0.03821
Model
0.03069

data/vMax
0.182

ofxng
369.894

Min d/m
3.89413

ave/min d
130.484
ave/min m
408.104

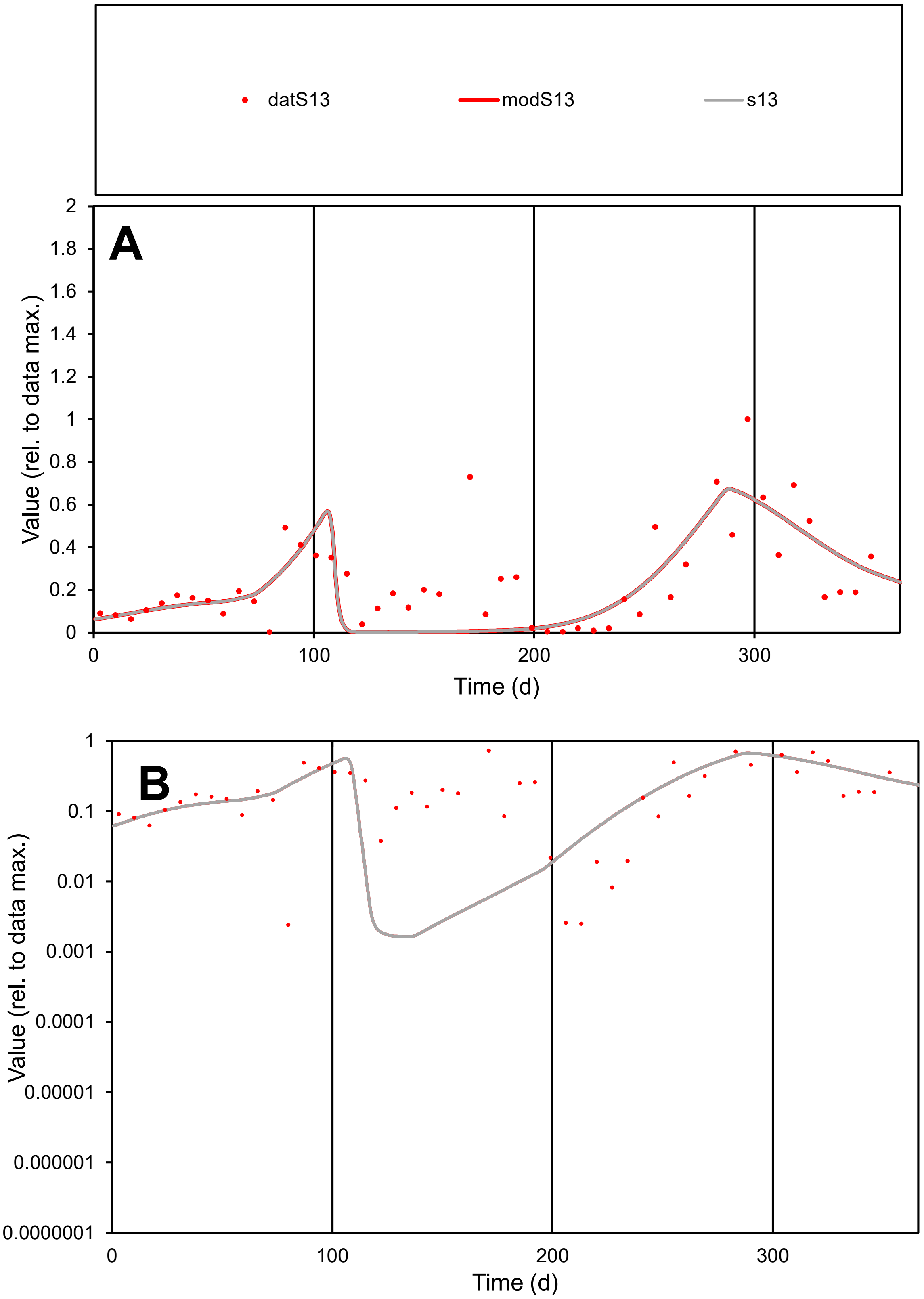


**C1**

Variables
S13 Provide jtd or Dst
CalibOnly?
2 1 to reproduce of
oNormalize?
2 1=no, 2=yes

ofxn calc*
0.69162
ofxn mod
0.71612

sRun
RunA1
zout
1

Ave
Data
0.02859
Model
0.02506

data/vMax
0.1167

ofxng
369.894

Min d/m
1.46122

ave/min d
101.837
ave/min m
130.476

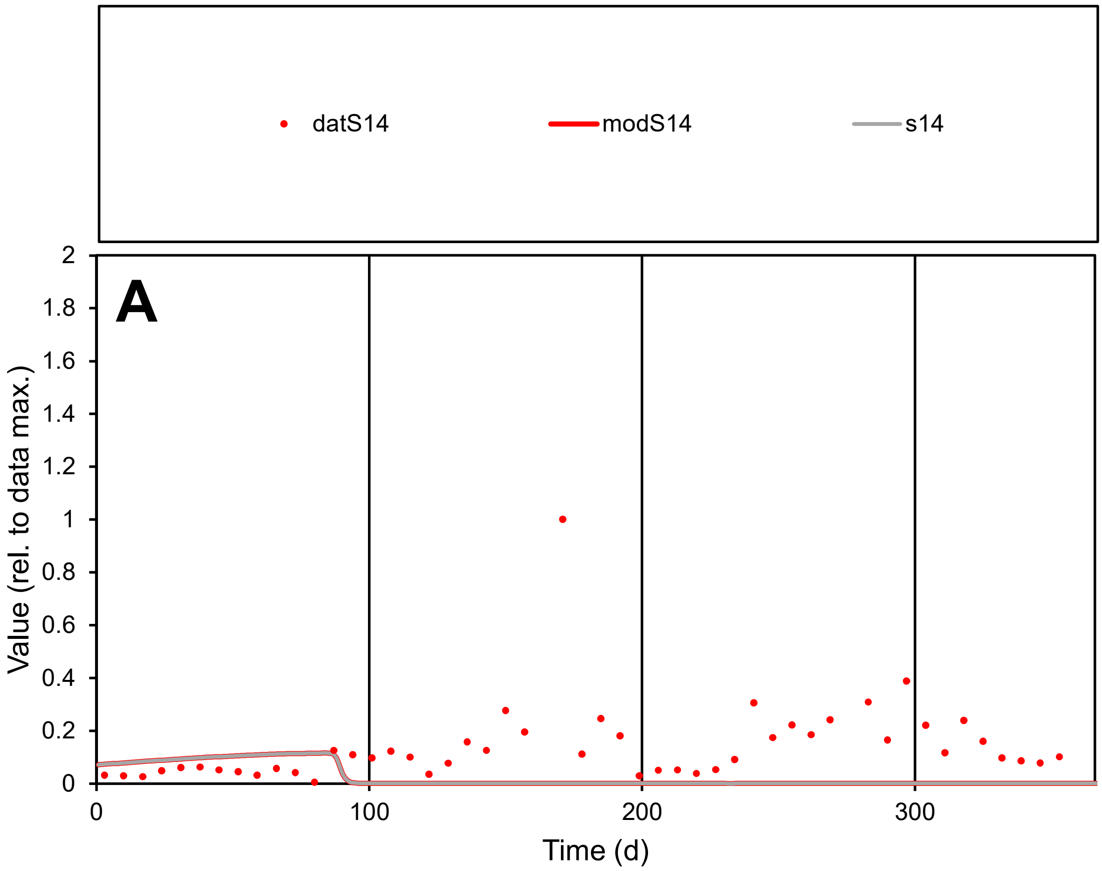


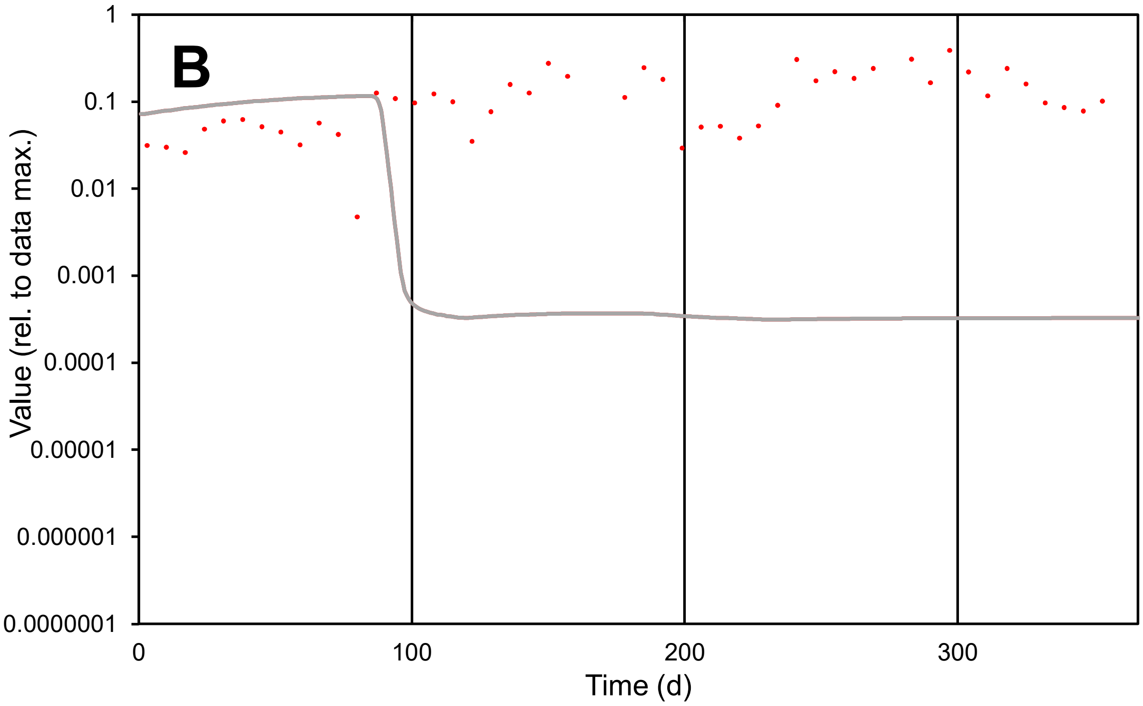


**C1**

Variables
S14 Provide jtd or Dst
CalibOnly?
2 1 to reproduce of
oNormalize?
2 1=no, 2=yes

ofxn calc*
1.47498
ofxn mod
1.4802

sRun
RunA1
zout
1

Ave
Data
0.03187
Model
0.00623

data/vMax
0.228

ofxng
369.894

Min d/m
15.1028

ave/min d
29.5059
ave/min m
87.1312

328_Calib_S_A

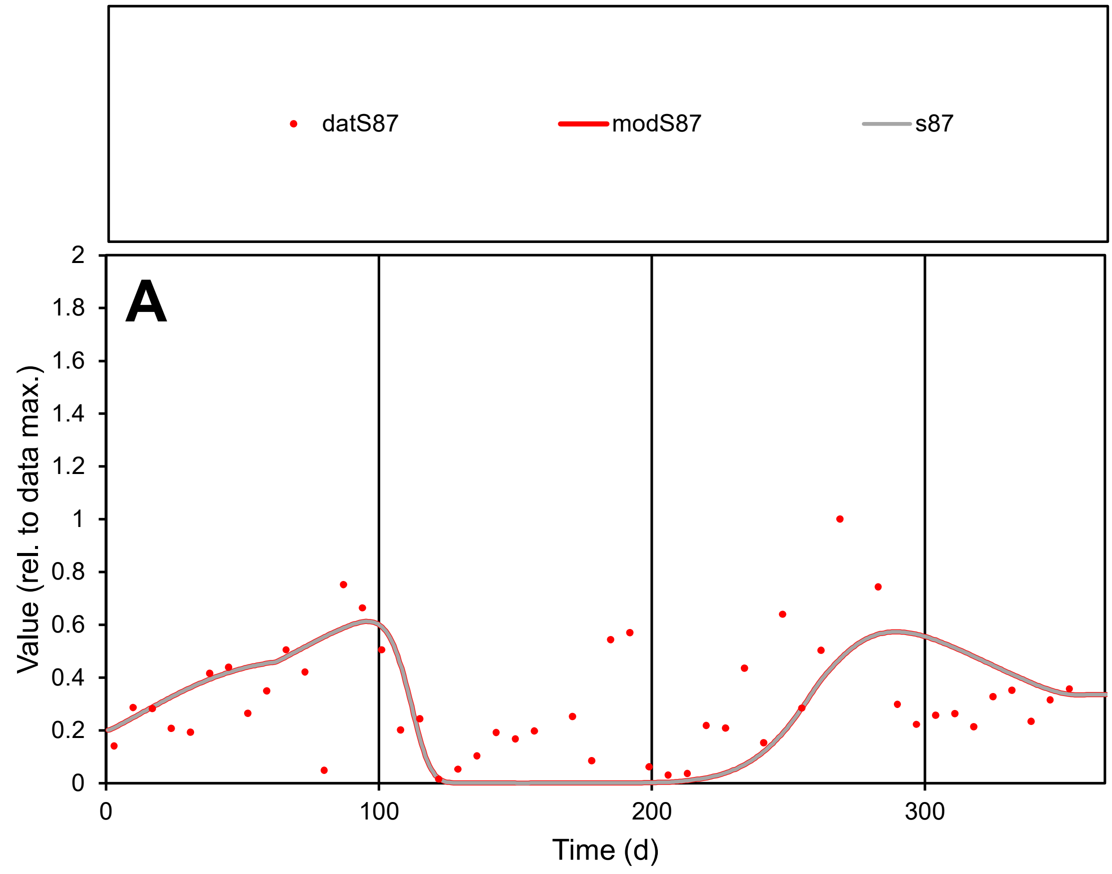


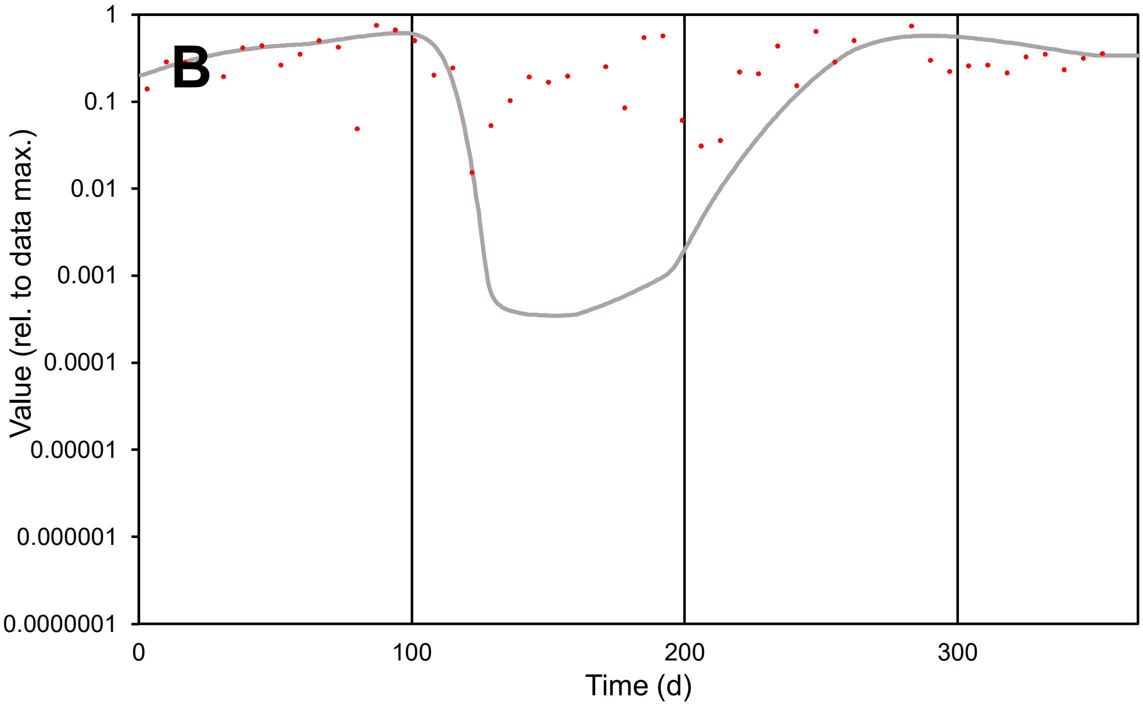


**C1**

Variables

S87 Provide jtd or Dst

CalibOnly?

2 1 to reproduce of

oNormalize?

2 1=no, 2=yes

ofxn calc*

0.70352

ofxn mod

0.71792

sRun

RunA1

zout

1

Ave

Data

0.02533

Model

0.02306

data/vMax

0.08137

ofxng

369.894

Min d/m

44.2872

ave/min d

20.2307

ave/min m

815.864

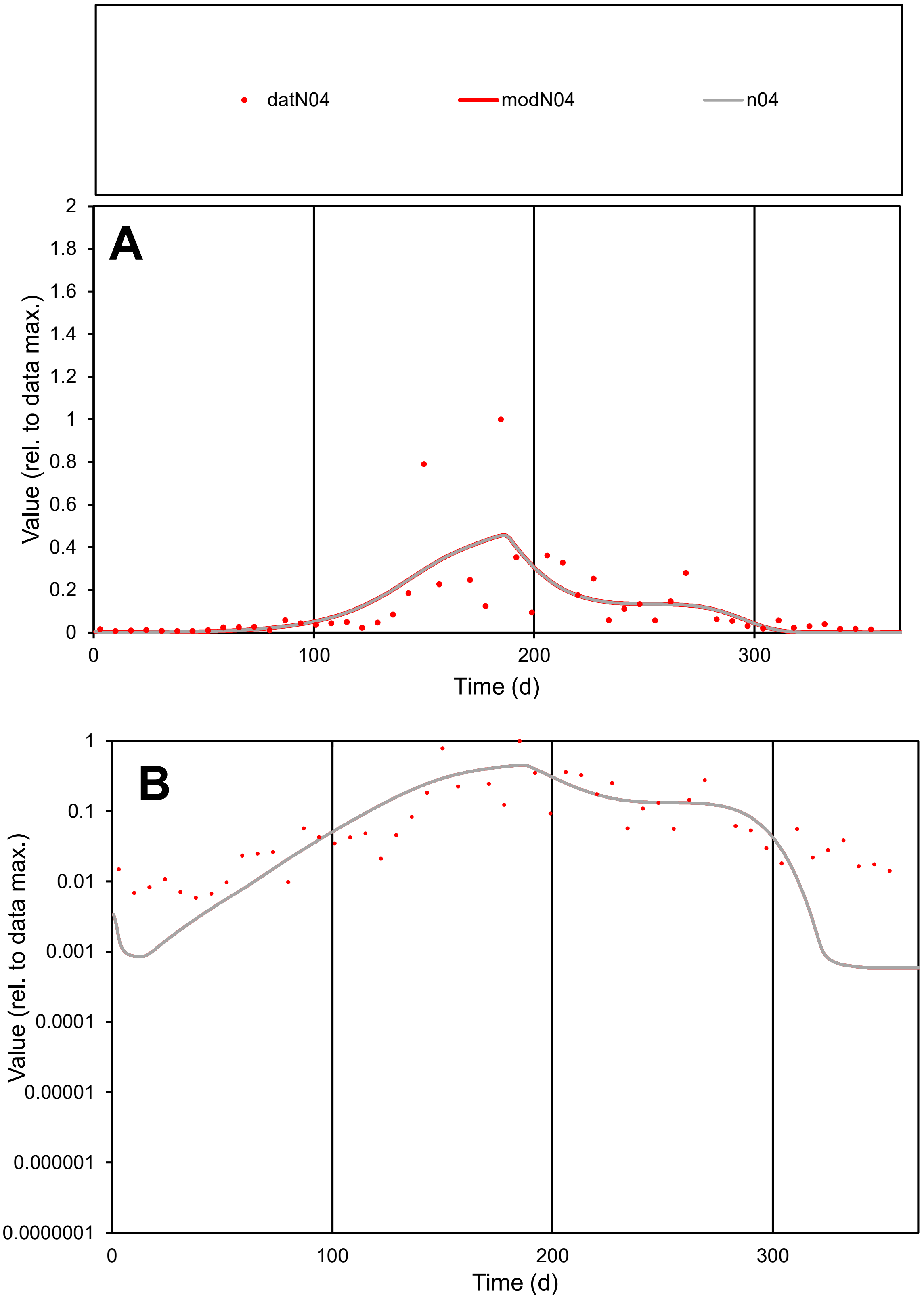


**C1**

Variables
N04 Provide jtd or Dst
CalibOnly?
2 1 to reproduce of
oNormalize?
2 1=no, 2=yes

ofxn calc*
1.11229
ofxn mod
1.1295

sRun
RunA1
zout
1

Ave
Data
0.02103
Model
0.02019

data/vMax
0.1777

ofxng
369.894

Min d/m
9.97137

ave/min d
20.1209
ave/min m
192.644

328_Calib_S_A

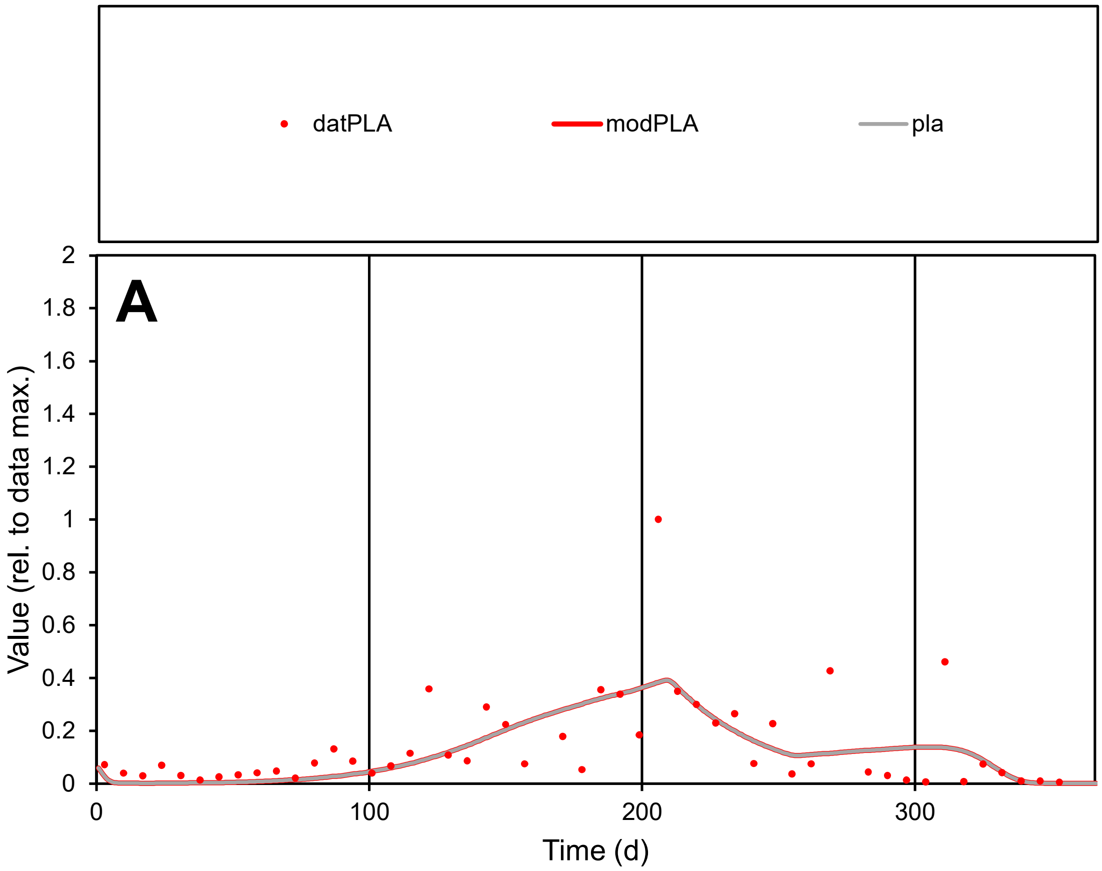


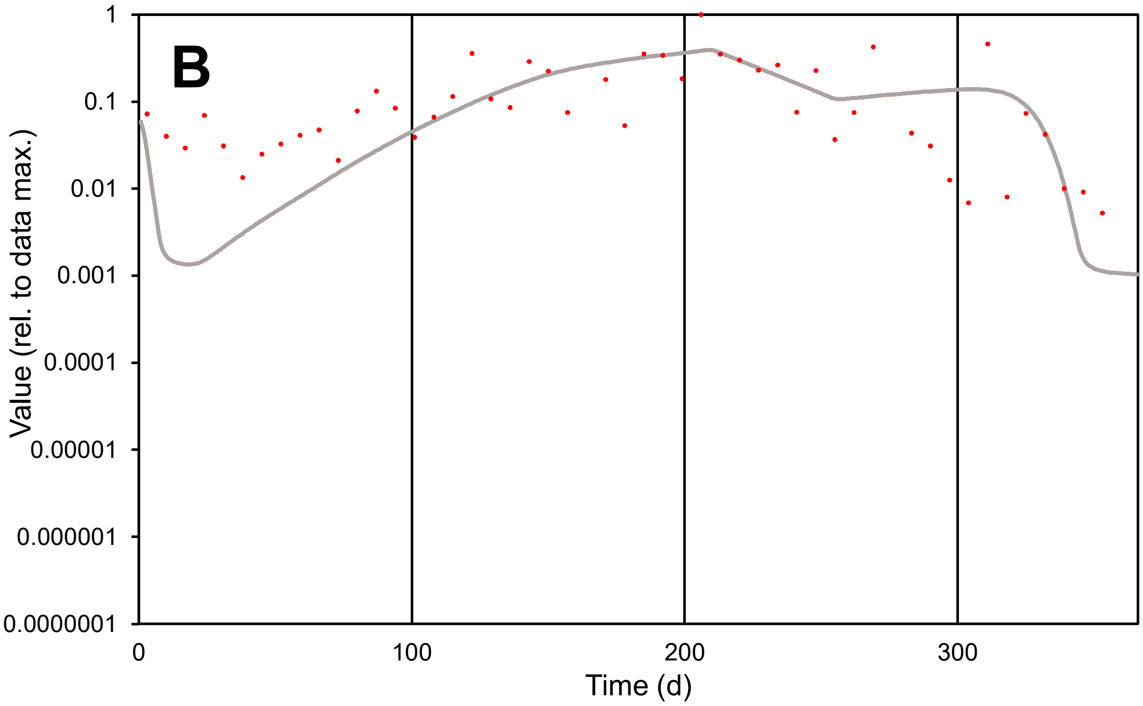


**C1**

Variables
PLA Provide jtd or Dst
CalibOnly?
2 1 to reproduce of
oNormalize?
2 1=no, 2=yes

ofxn calc*
0.96982
ofxn mod
0.9843

sRun
RunA1
zout
1

Ave
Data
0.01201
Model
0.01046

data/vMax
0.0856

ofxng
369.894

Min d/m
4.73375

ave/min d
26.6458
ave/min m
109.819

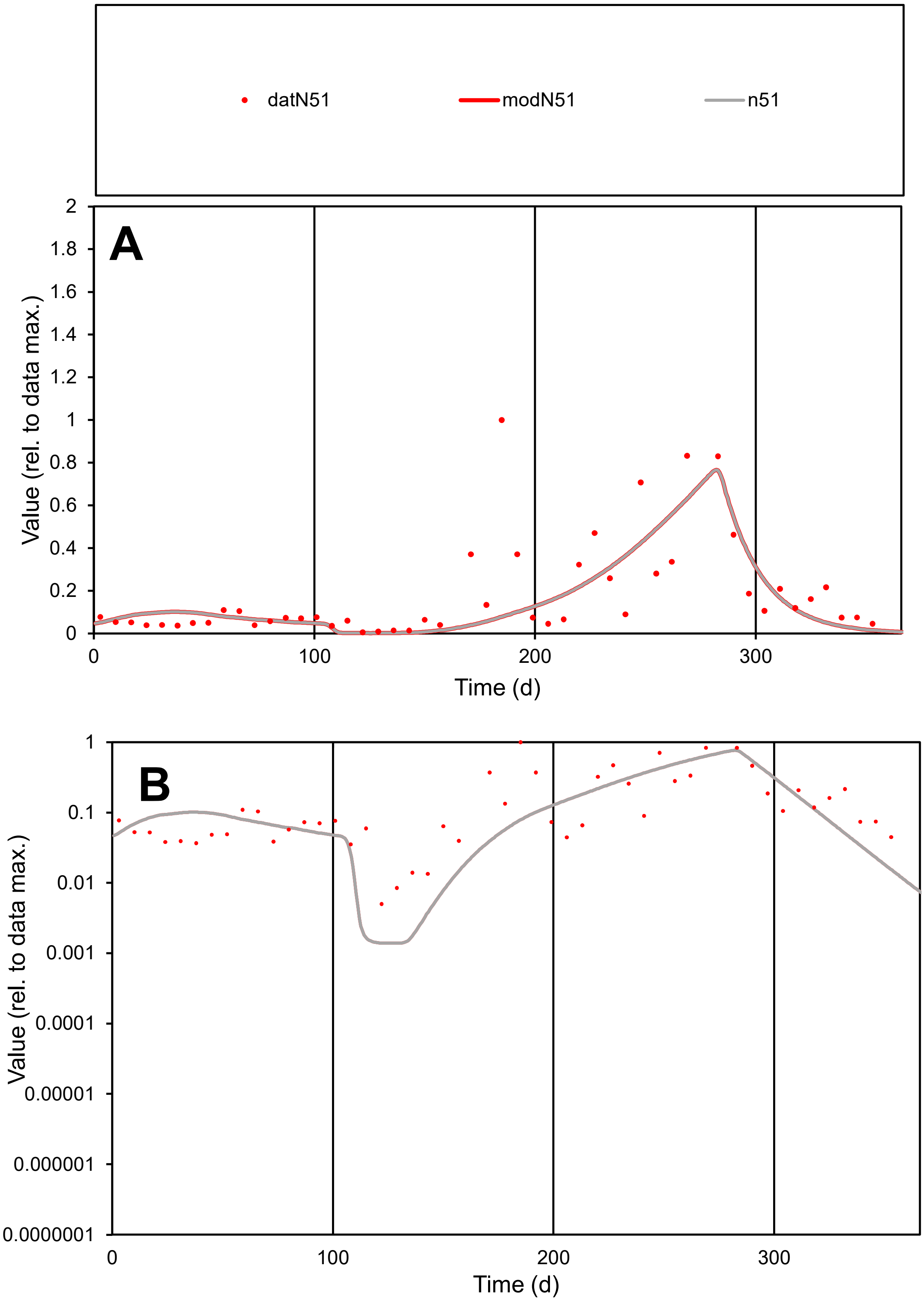


**C1**

Variables
N51 Provide jtd or Dst
CalibOnly?
2 1 to reproduce of
oNormalize?
2 1=no, 2=yes

ofxn calc*
0.95735
ofxn mod
0.97401

sRun
RunA1
zout
1

Ave
Data
0.02179
Model
0.01825

data/vMax
0.1187

ofxng
369.894

Min d/m
3.59612

ave/min d
36.7408
ave/min m
110.654

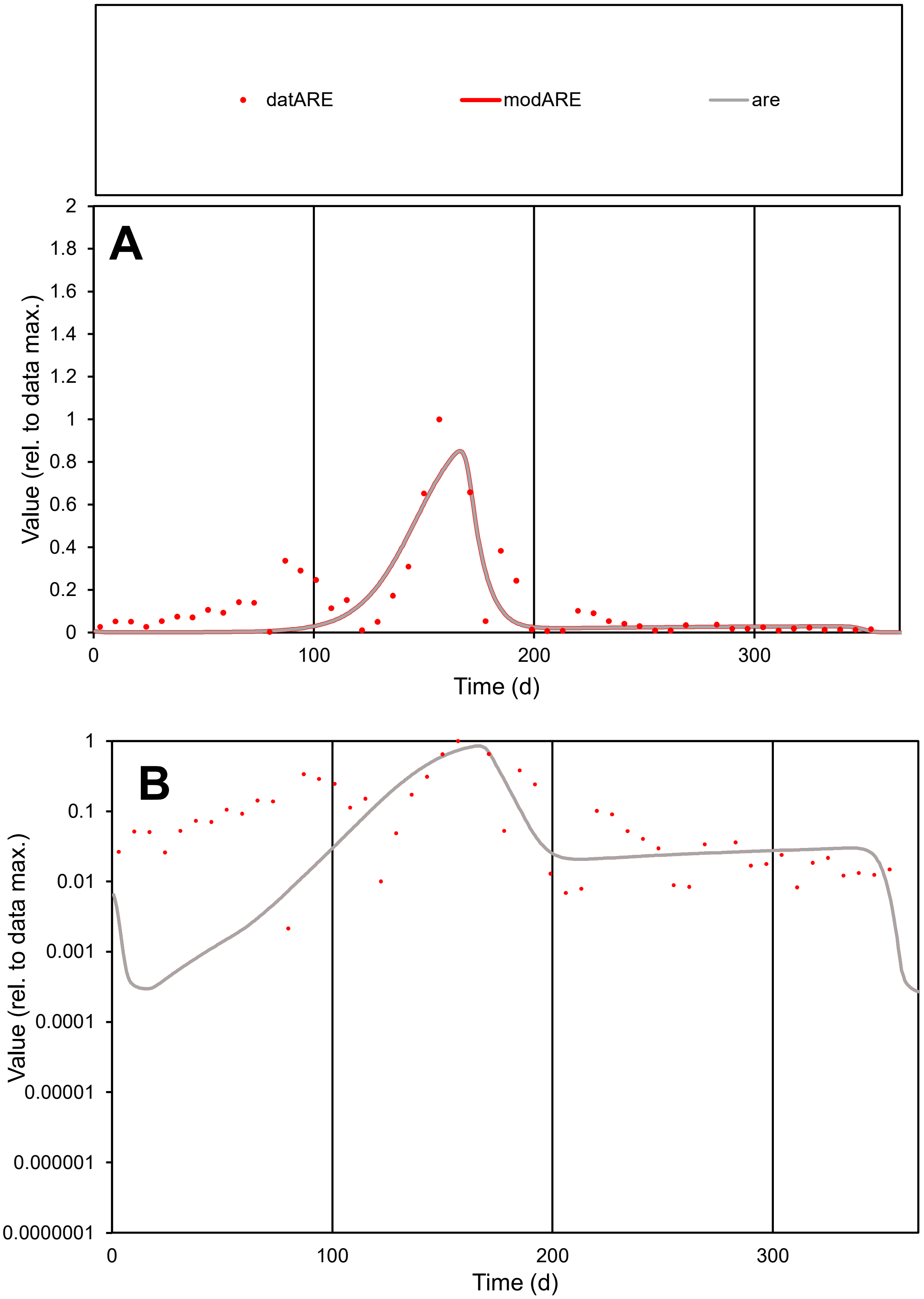


**C1**

Variables

ARE Provide jtd or Dst

CalibOnly?

2 1 to reproduce of

oNormalize?

2 1=no, 2=yes

ofxn calc*

0.90047

ofxn mod

0.93157

sRun

RunA1

zout

1

Ave

Data

0.02855

Model

0.01999

data/vMax

0.2299

ofxng

369.894

Min d/m

6.99759

ave/min d

57.7884

ave/min m

283.066

328_Calib_S_A

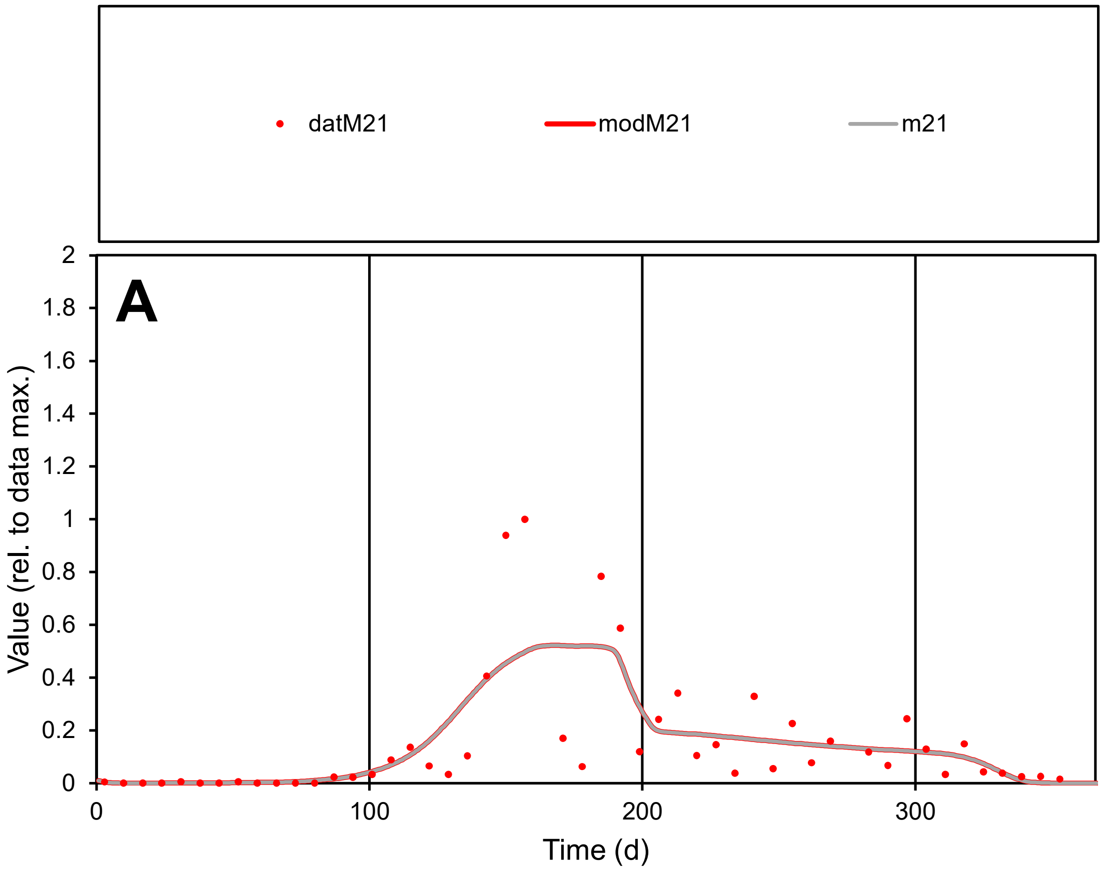


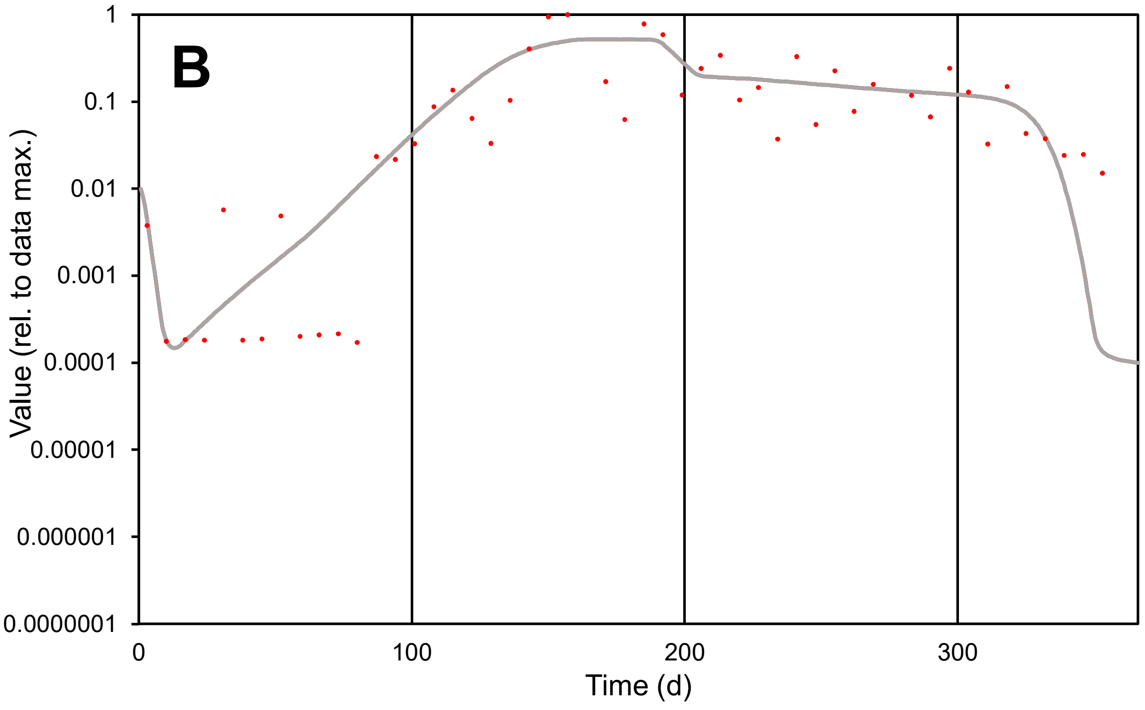


**C1**

Variables
M21 Provide jtd or Dst
CalibOnly?
2 1 to reproduce of
oNormalize?
2 1=no, 2=yes

ofxn calc*
1.03952
ofxn mod
1.05861

sRun
RunA1
zout
1

Ave
Data
0.01722
Model
0.0171

data/vMax
0.1175

ofxng
369.894

Min d/m
1.35246

ave/min d
859.701
ave/min m
1154.84

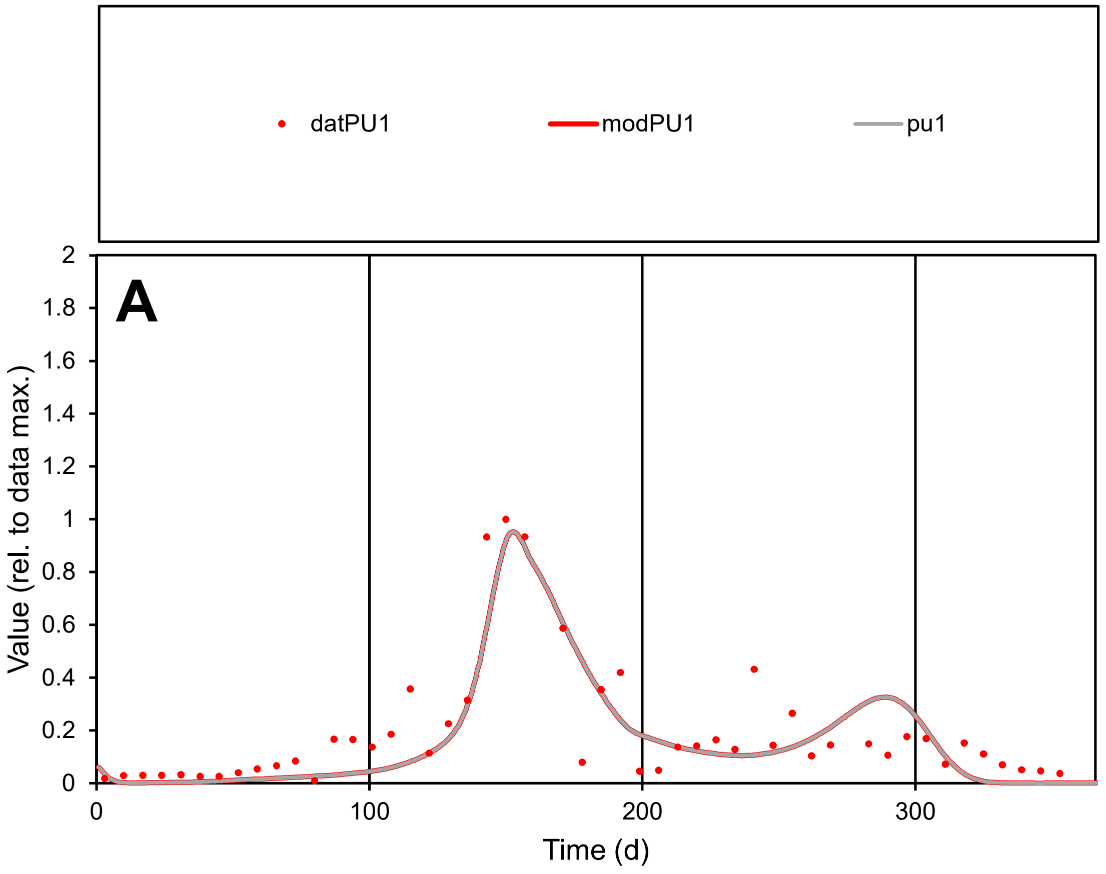


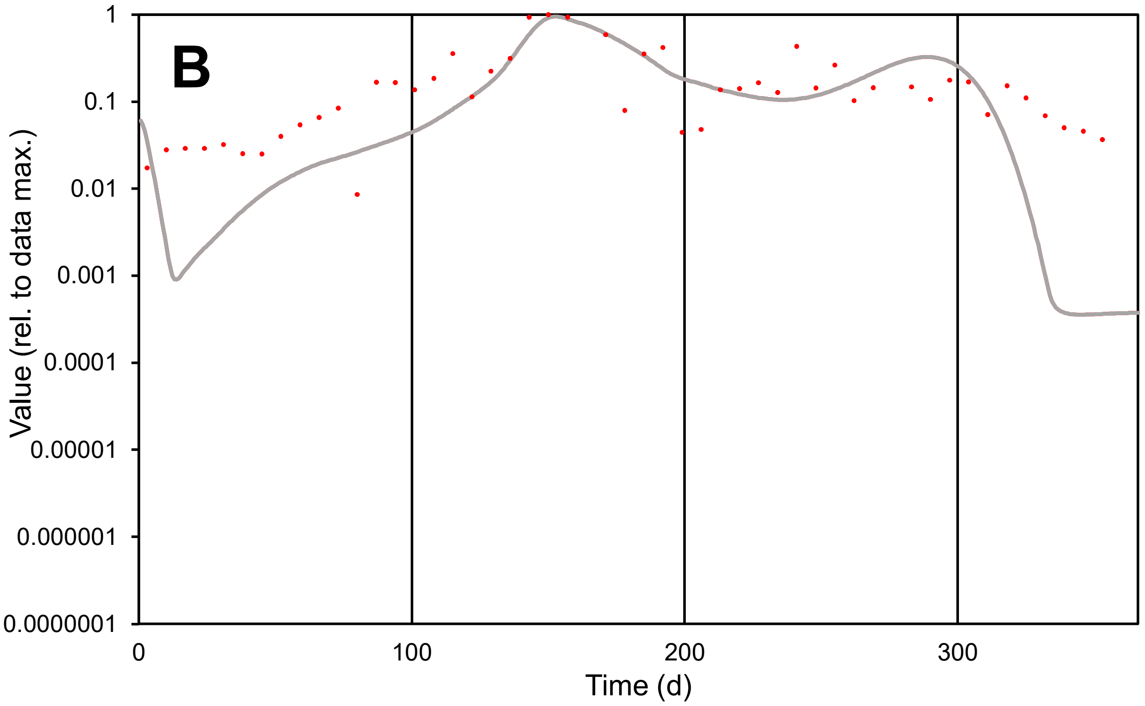


**C1**

Variables

PU1 Provide jtd or Dst

CalibOnly?

2 1 to reproduce of

oNormalize?

2 1=no, 2=yes

ofxn calc*

0.61933

ofxn mod

0.63996

sRun

RunA1

zout

1

Ave

Data

0.01932

Model

0.01635

data/vMax

0.1019

ofxng

369.894

Min d/m

24.0066

ave/min d

21.9948

ave/min m

446.951

328_Calib_S_A

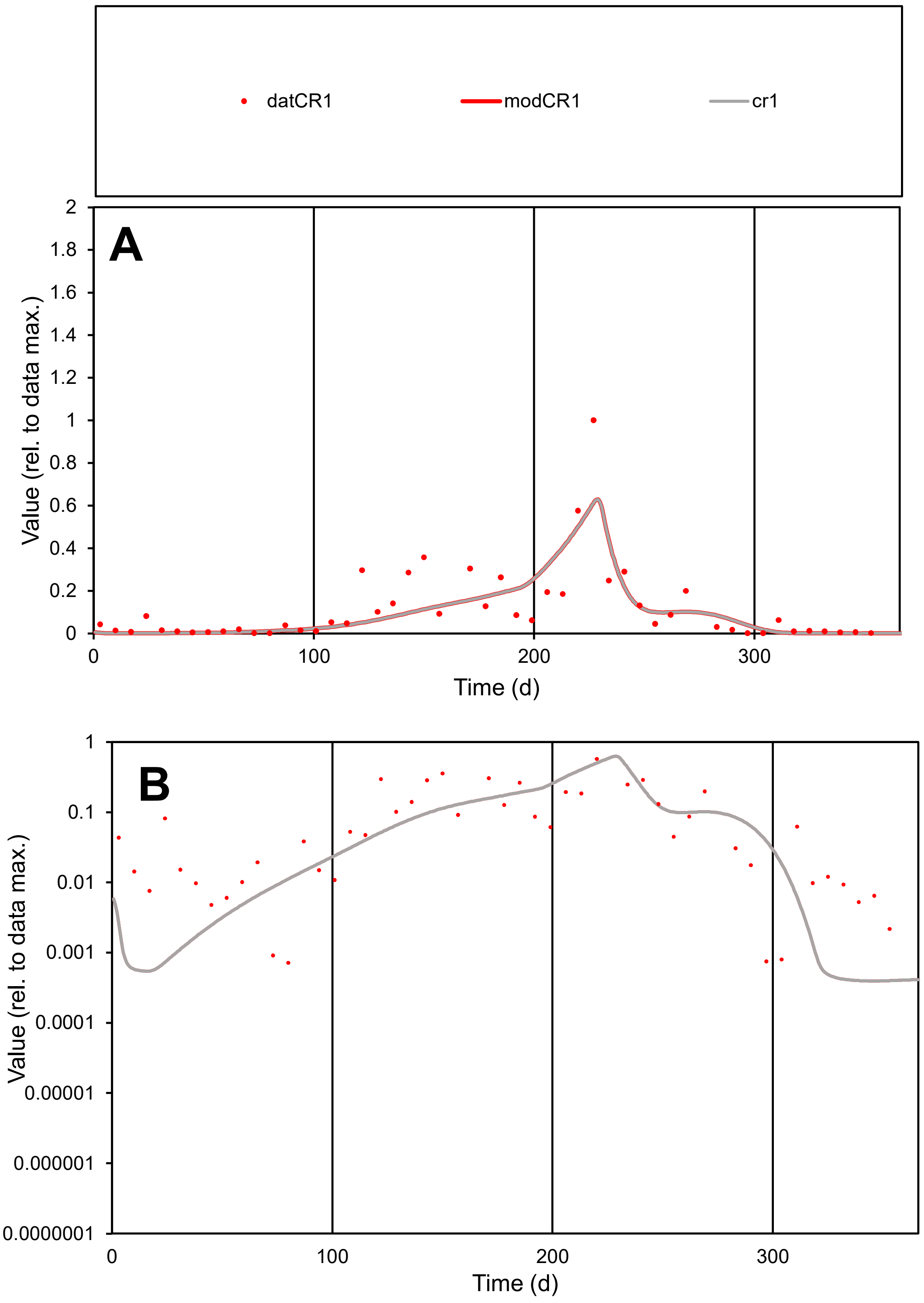


**C1**

Variables

CR1 Provide jtd or Dst

CalibOnly?

2 1 to reproduce of

oNormalize?

2 1=no, 2=yes

ofxn calc*

0.88029

ofxn mod

0.90562

sRun

RunA1

zout

1

Ave

Data

0.01047

Model

0.00896

data/vMax

0.09154

ofxng

369.894

Min d/m

1.81412

ave/min d

159.201

ave/min m

246.976

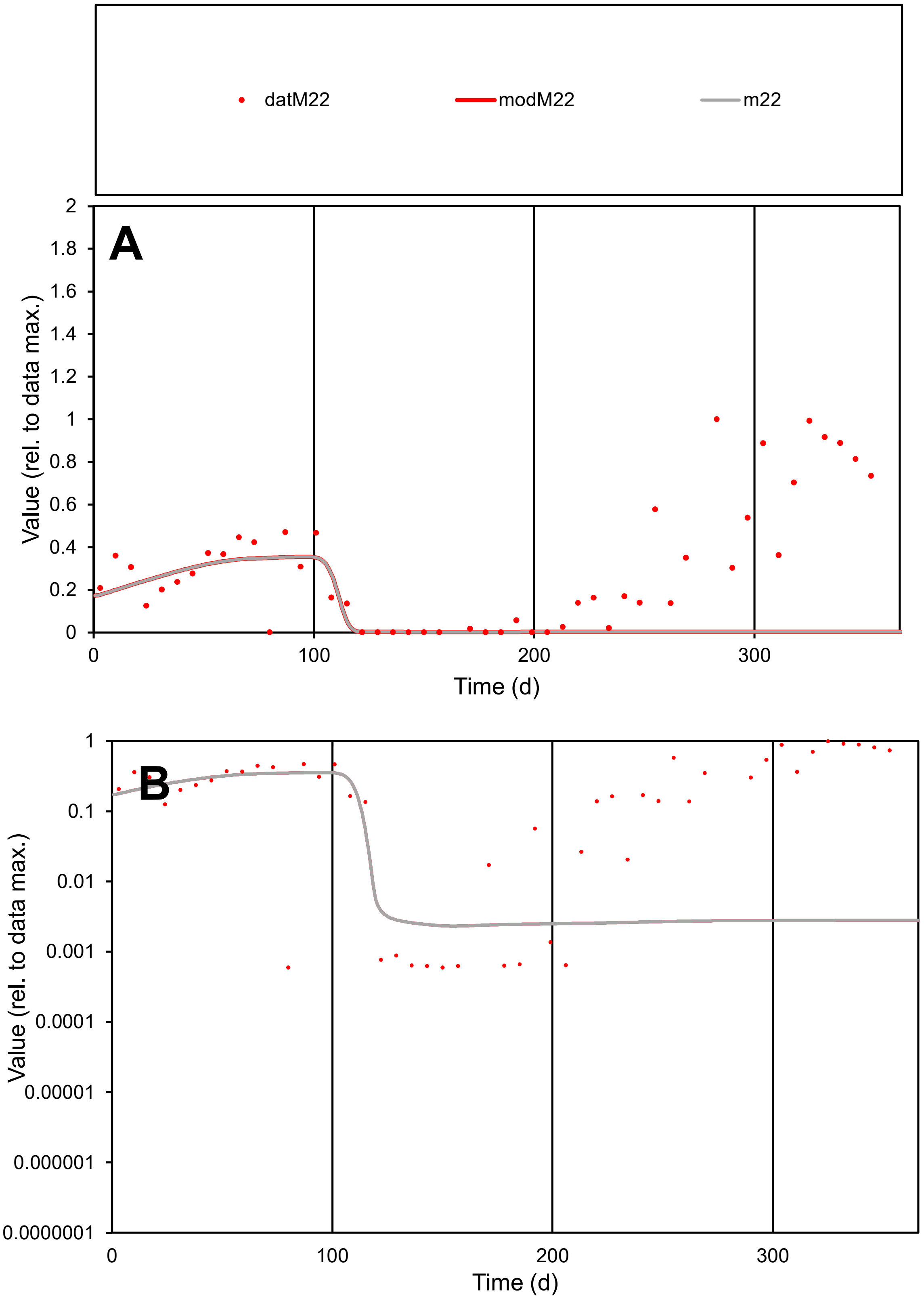


**C1**

Variables
M22 Provide jtd or Dst
CalibOnly?
2 1 to reproduce of
oNormalize?
2 1=no, 2=yes

ofxn calc*
1.28215
ofxn mod
1.2859

sRun
RunA1
zout
1

Ave
Data
0.01095
Model
0.00362

data/vMax
0.0362

ofxng
369.894

Min d/m
0.2556

ave/min d
507.991
ave/min m
42.8677

328_Calib_S_A

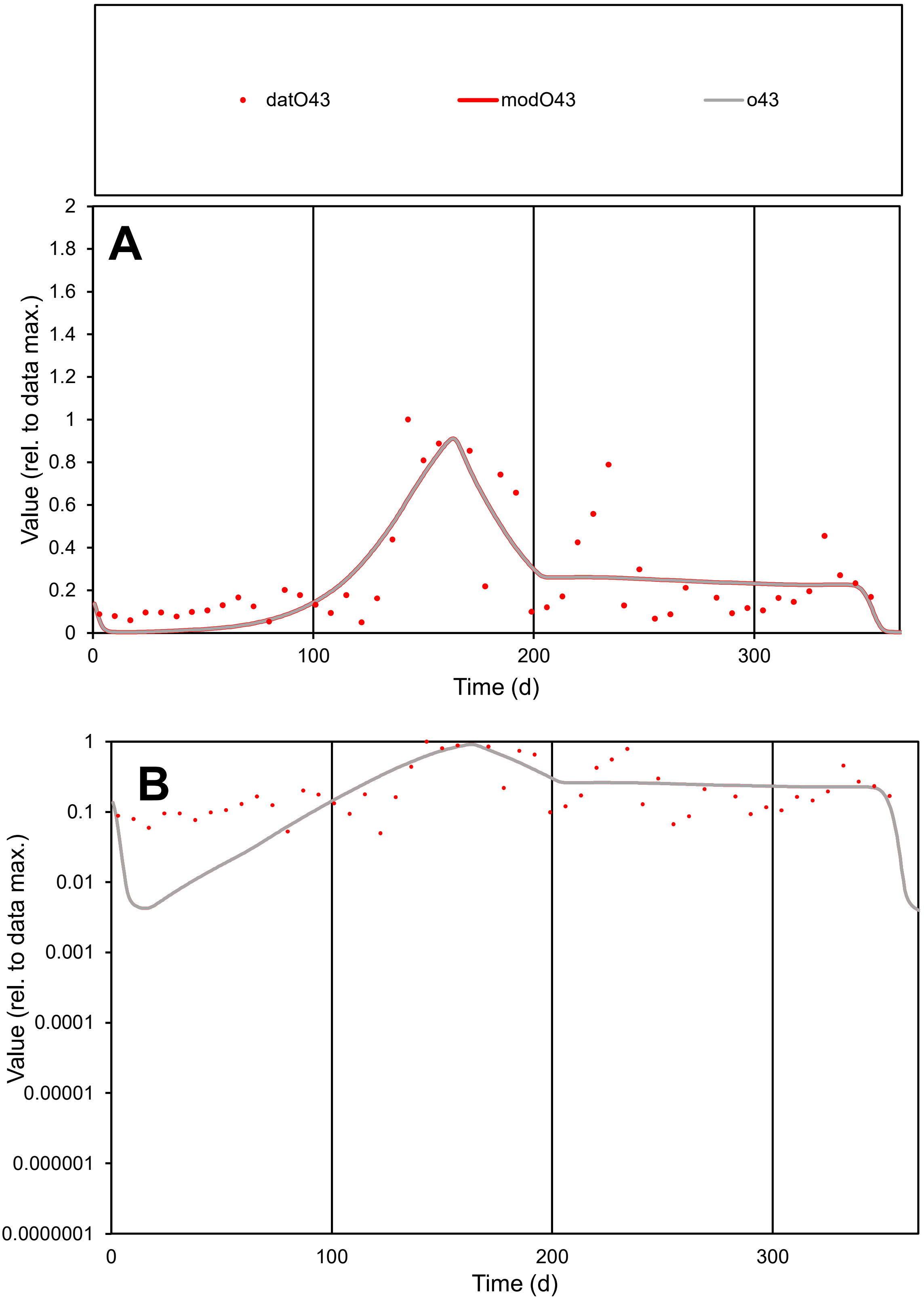


**C1**

Variables

O43 Provide jtd or Dst

CalibOnly?

2 1 to reproduce of

oNormalize?

2 1=no, 2=yes

ofxn calc*

0.63192

ofxn mod

0.64998

sRun

RunA1

zout

1

Ave

Data

0.01099

Model

0.01042

data/vMax

0.04196

ofxng

369.894

Min d/m

11.2265

ave/min d

5.26733

ave/min m

56.0428

328_Calib_S_A

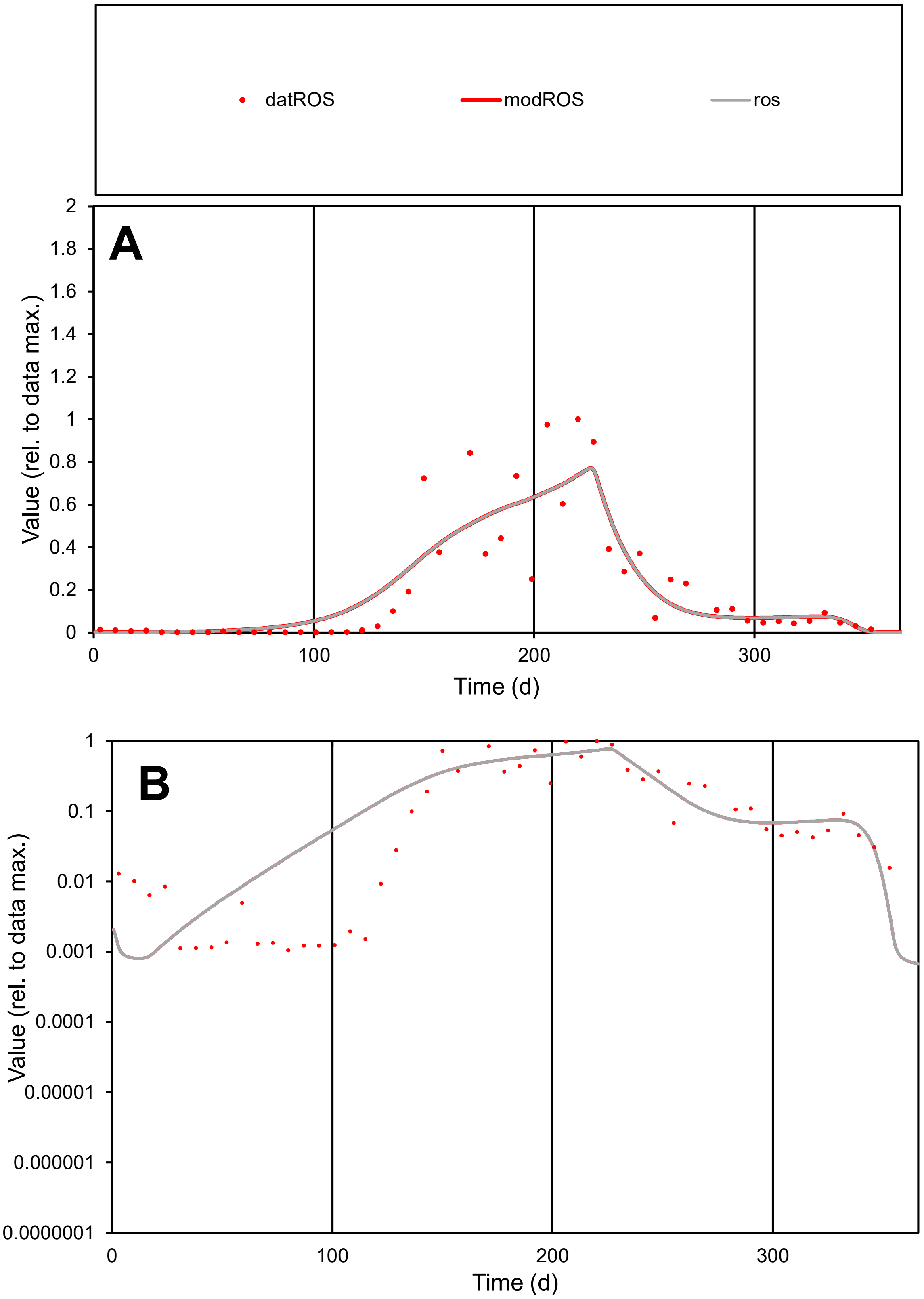


**C1**

Variables

| | |
|---|---|
| ROS | Provide jtd or Dst |
| CalibOnly? | |
| 2 | 1 to reproduce of |
| oNormalize? | |
| 2 | 1=no, 2=yes |

ofxn calc*
0.64043
ofxn mod
0.67404

sRun
RunA1
zout
1

Ave
Data
0.00952
Model
0.00968

data/vMax
0.04744

ofxng
369.894

Min d/m
1.31767

ave/min d
189.685
ave/min m
254.207

328_Calib_S_A

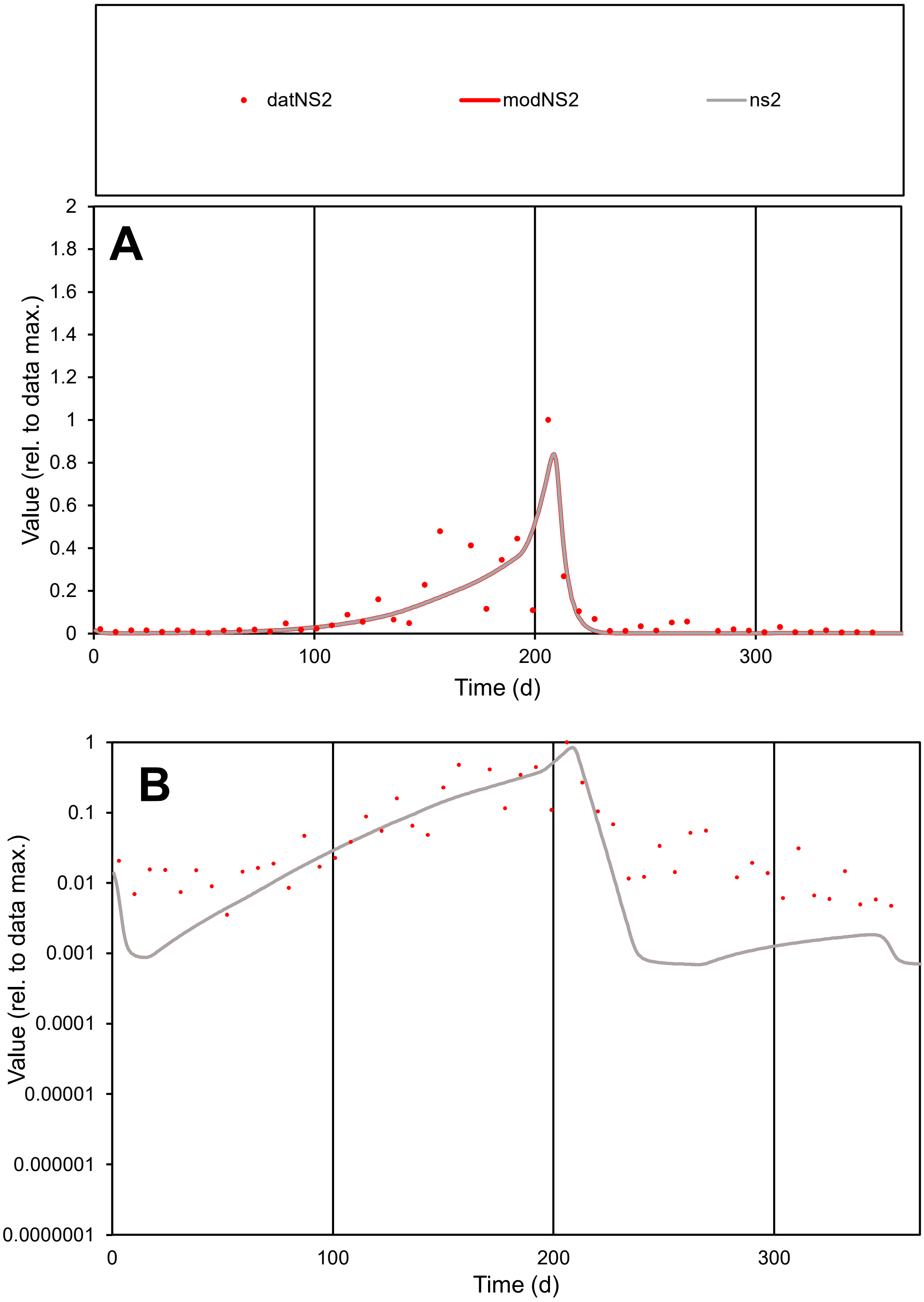


**C1**

Variables

NS2 Provide jtd or Dst

CalibOnly?

2 1 to reproduce of

oNormalize?

2 1=no, 2=yes

ofxn calc*

0.9306

ofxn mod

0.96117

sRun

RunA1

zout

1

Ave

Data

0.00619

Model

0.00515

data/vMax

0.06632

ofxng

369.894

Min d/m

5.09898

ave/min d

26.3952

ave/min m

111.957

328_Calib_S_A

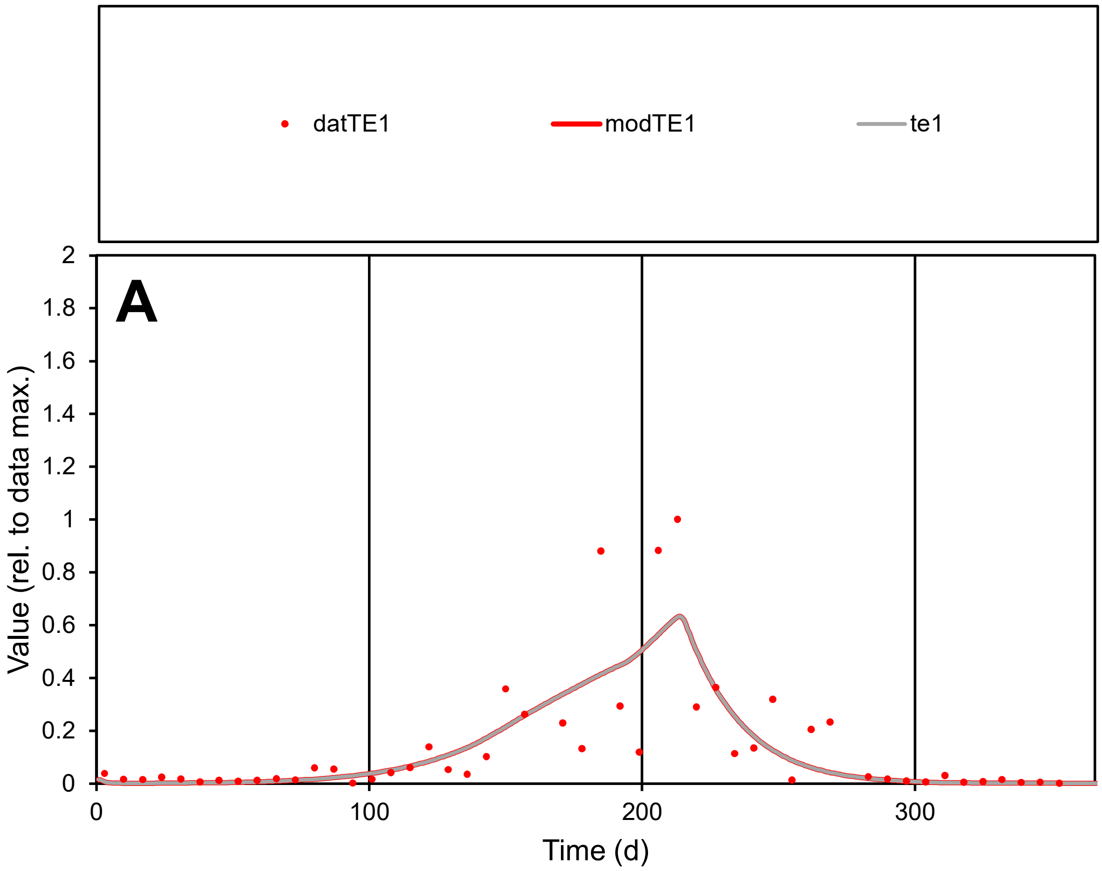


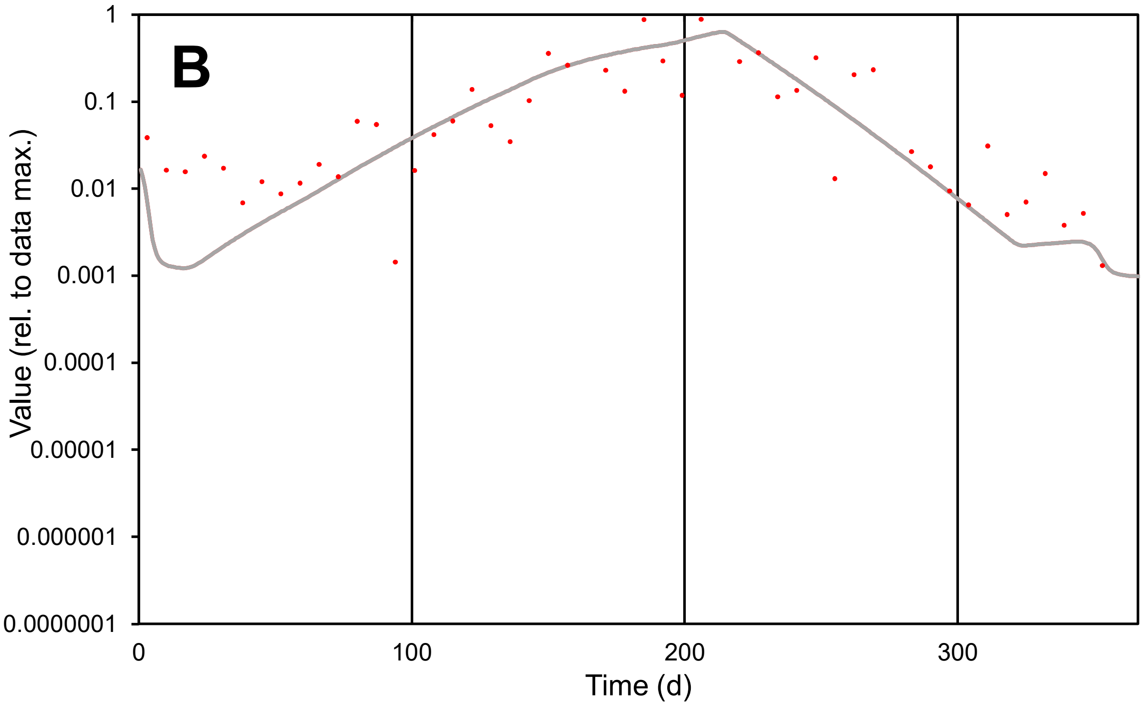


**C1**

Variables
TE1 Provide jtd or Dst
CalibOnly?
2 1 to reproduce of
oNormalize?
2 1=no, 2=yes

ofxn calc*
0.98242
ofxn mod
1.00457

sRun
RunA1
zout
1

Ave
Data
0.00578
Model
0.00534

data/vMax
0.04225

ofxng
369.894

Min d/m
1.0569
ave/min d
104.728
ave/min m
102.348

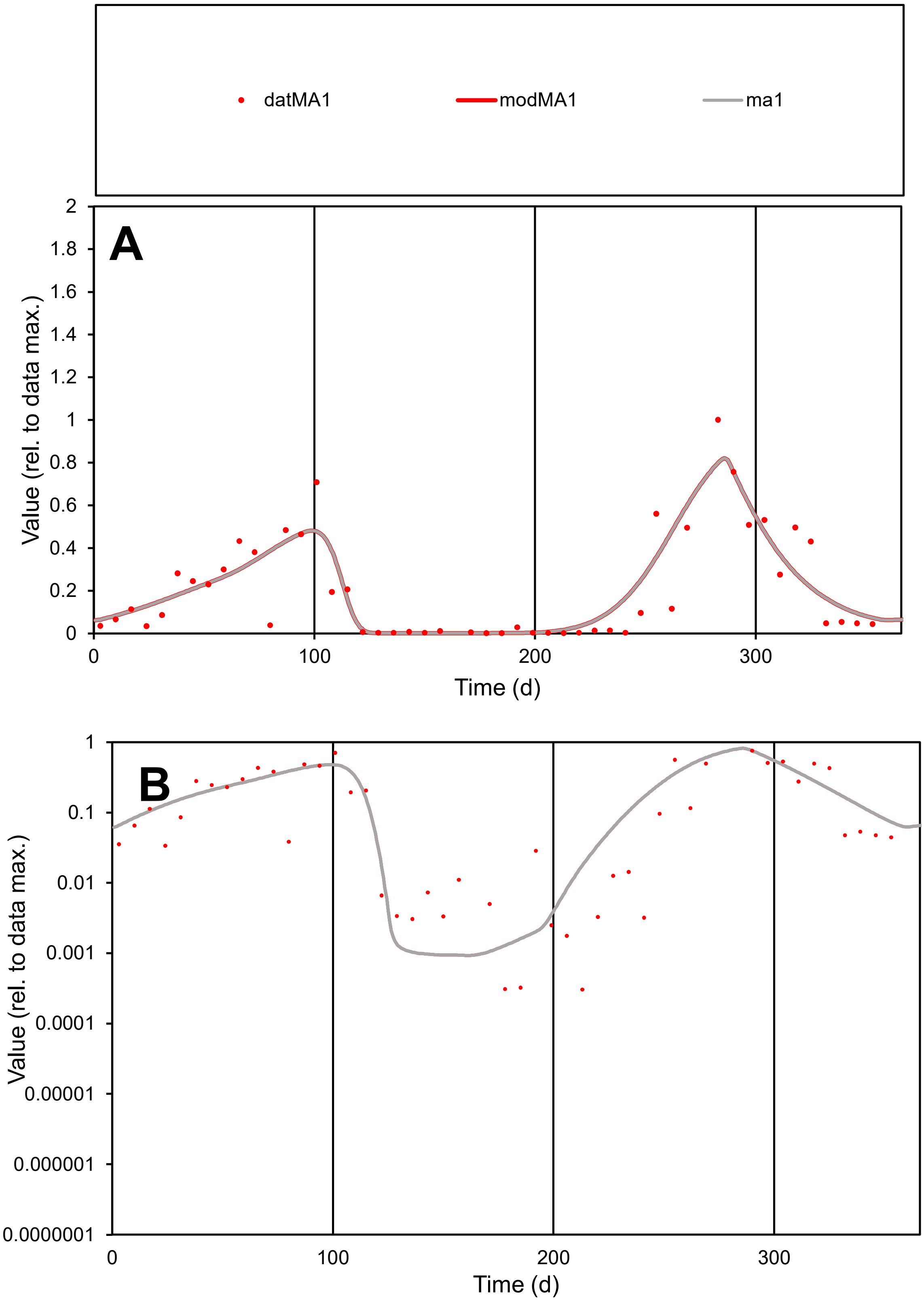


**C1**

Variables
MA1 Provide jtd or Dst
CalibOnly?
2 1 to reproduce of
oNormalize?
2 1=no, 2=yes

ofxn calc*
0.58193
ofxn mod
0.61296

sRun
RunA1
zout
1

Ave
Data
0.00999
Model
0.01051

data/vMax
0.04964

ofxng
369.894

Min d/m
0.32544

ave/min d
661.042
ave/min m
226.408

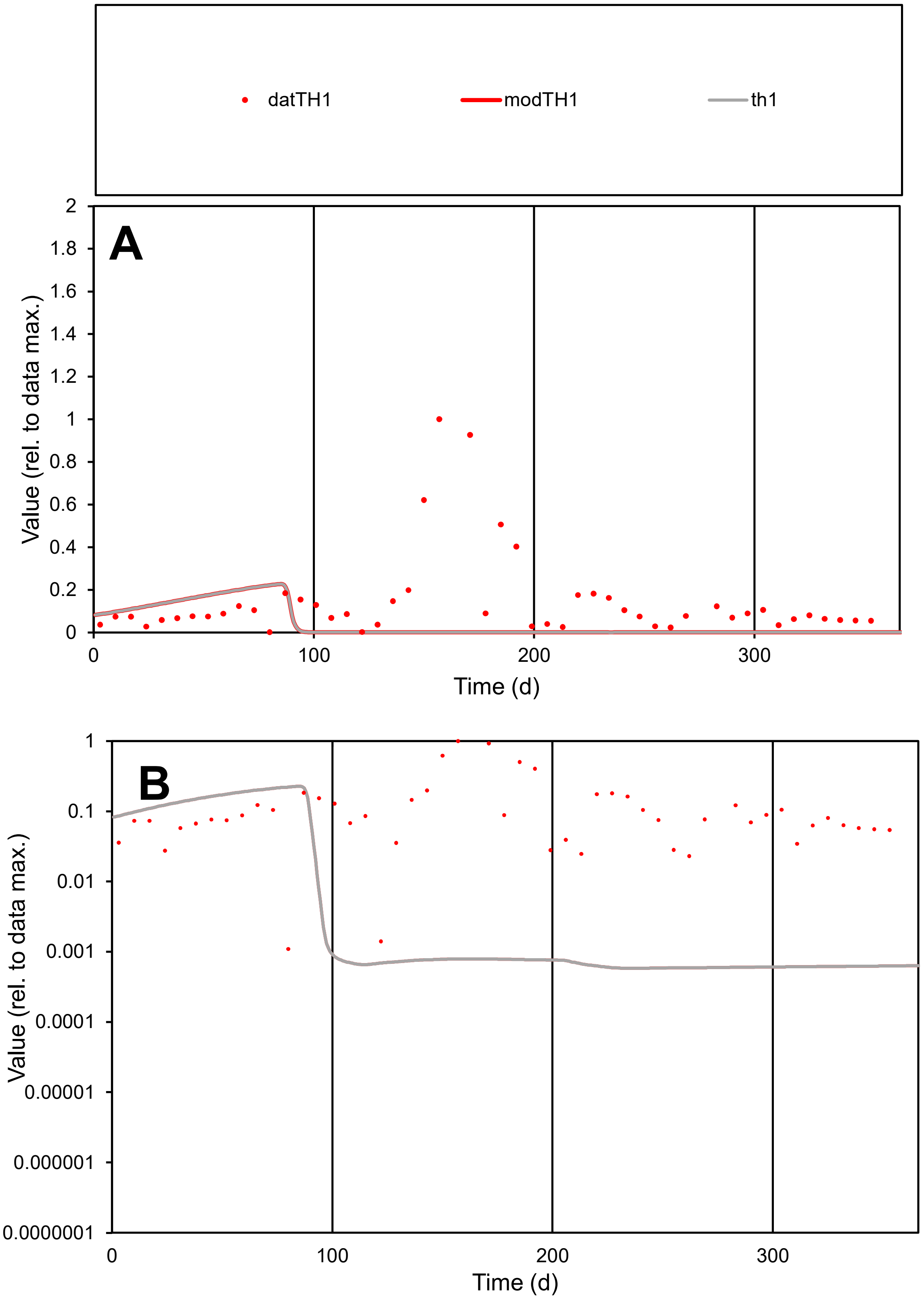


**C1**

Variables
TH1 Provide jtd or Dst
CalibOnly?
2 1 to reproduce of
oNormalize?
2 1=no, 2=yes

ofxn calc*
1.73433
ofxn mod
1.74169

sRun
RunA1
zout
1

Ave
Data
0.01231
Model
0.00379

data/vMax
0.08543

ofxng
369.894

Min d/m
1.89221

ave/min d
131.56
ave/min m
76.7052

328_Calib_S_A

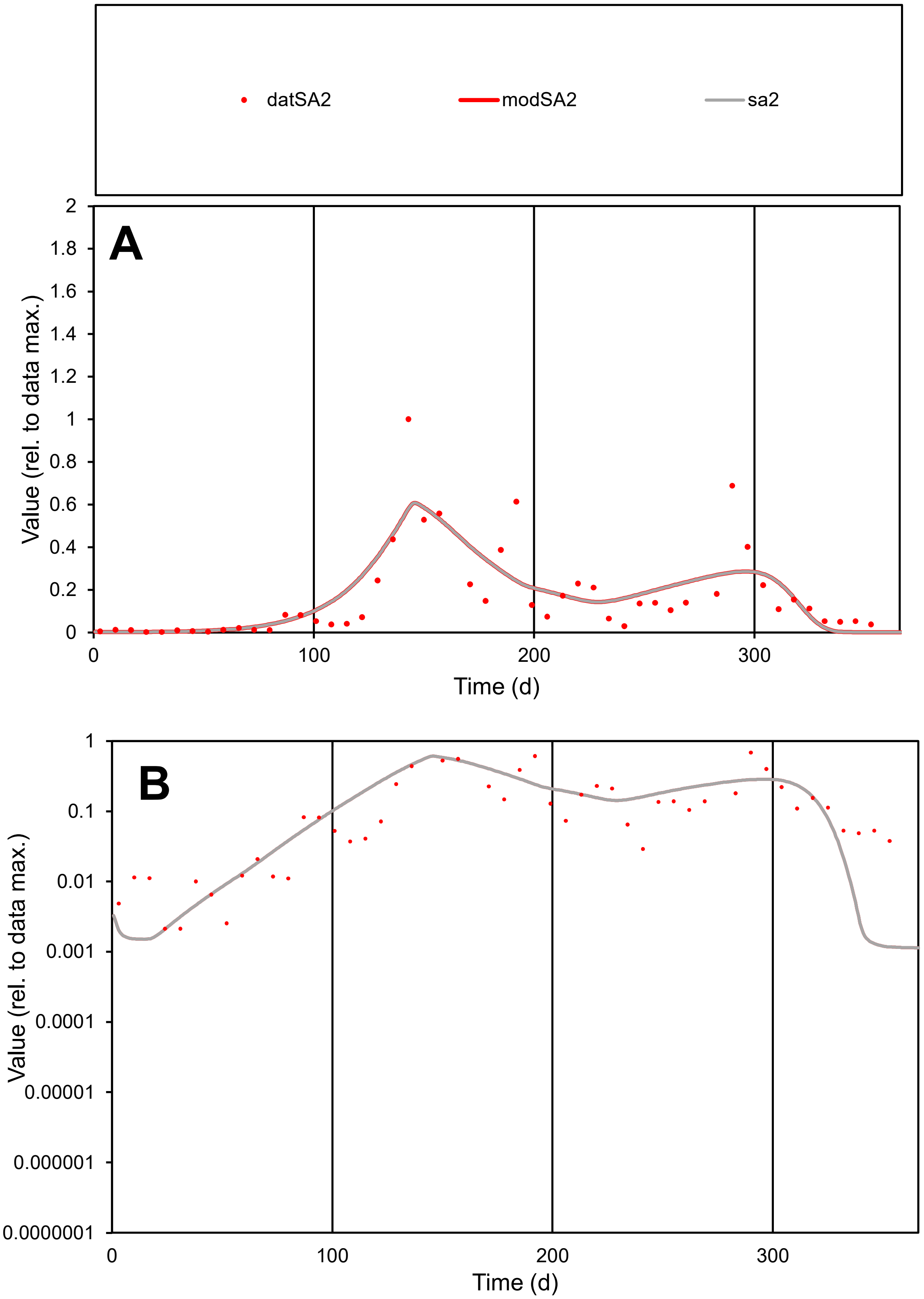


**C1**

Variables
SA2 Provide jtd or Dst
CalibOnly?
2 1 to reproduce of
oNormalize?
2 1=no, 2=yes

ofxn calc*
0.75905
ofxn mod
0.77702

sRun
RunA1
zout
1

Ave
Data
0.00845
Model
0.00874

data/vMax
0.05111

ofxng
369.894

Min d/m
1.82489

ave/min d
77.6683
ave/min m
146.599

328_Calib_S_A

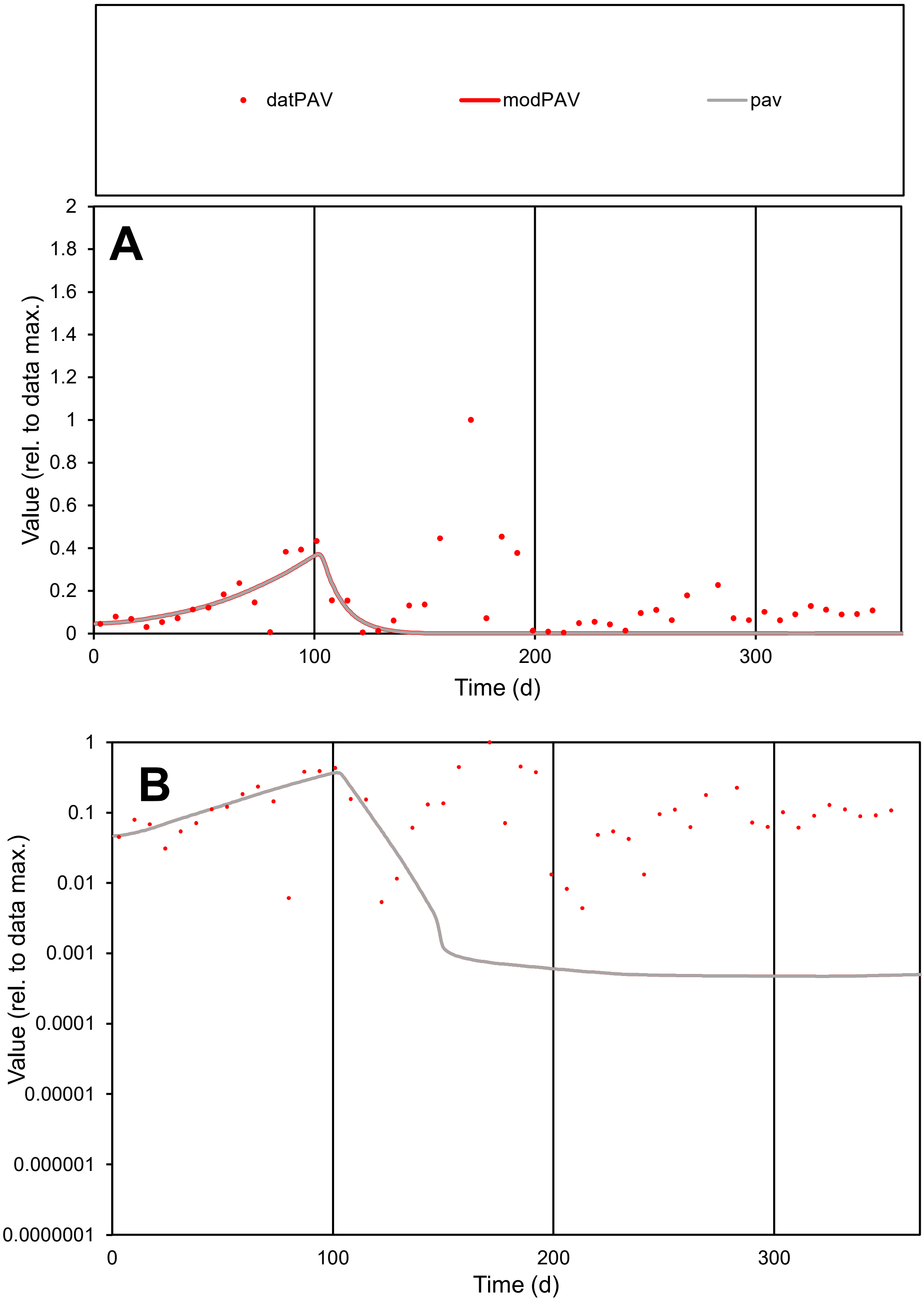


**C1**

Variables

PAV Provide jtd or Dst

CalibOnly?

2 1 to reproduce of

oNormalize?

2 1=no, 2=yes

ofxn calc*

1.34511

ofxn mod

1.35027

sRun

RunA1

zout

1

Ave

Data

0.01128

Model

0.00457

data/vMax

0.0775

ofxng

369.894

Min d/m

9.31316

ave/min d

33.0215

ave/min m

124.543

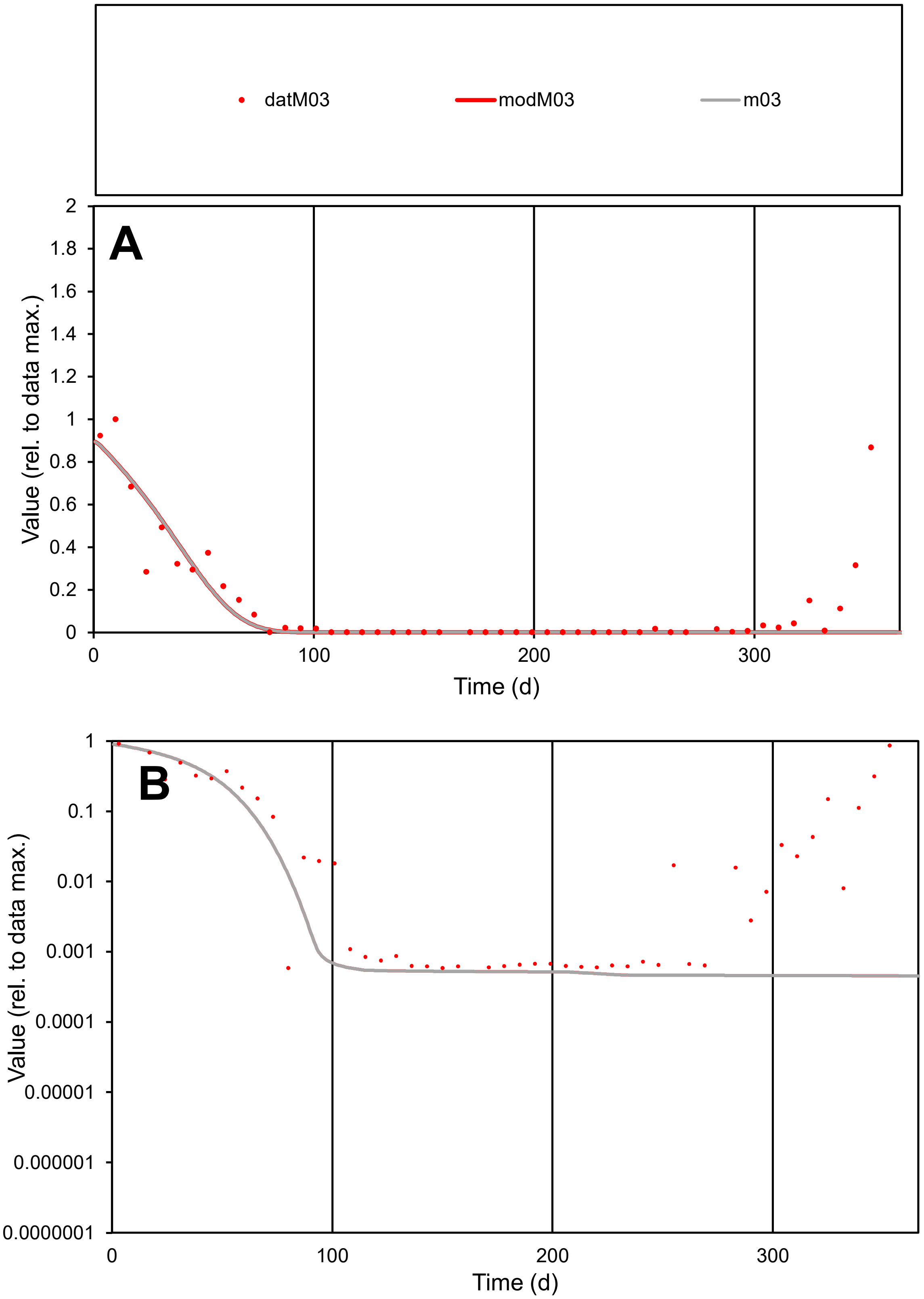


**C1**

Variables
M03 Provide jtd or Dst
CalibOnly?
2 1 to reproduce of
oNormalize?
2 1=no, 2=yes

ofxn calc*
1.12688
ofxn mod
1.154

sRun
RunA1
zout
1

Ave
Data
0.00398
Model
0.00284

data/vMax
0.02999

ofxng
369.894

Min d/m
1.29477

ave/min d
226.357
ave/min m
209.264

328_Calib_S_A

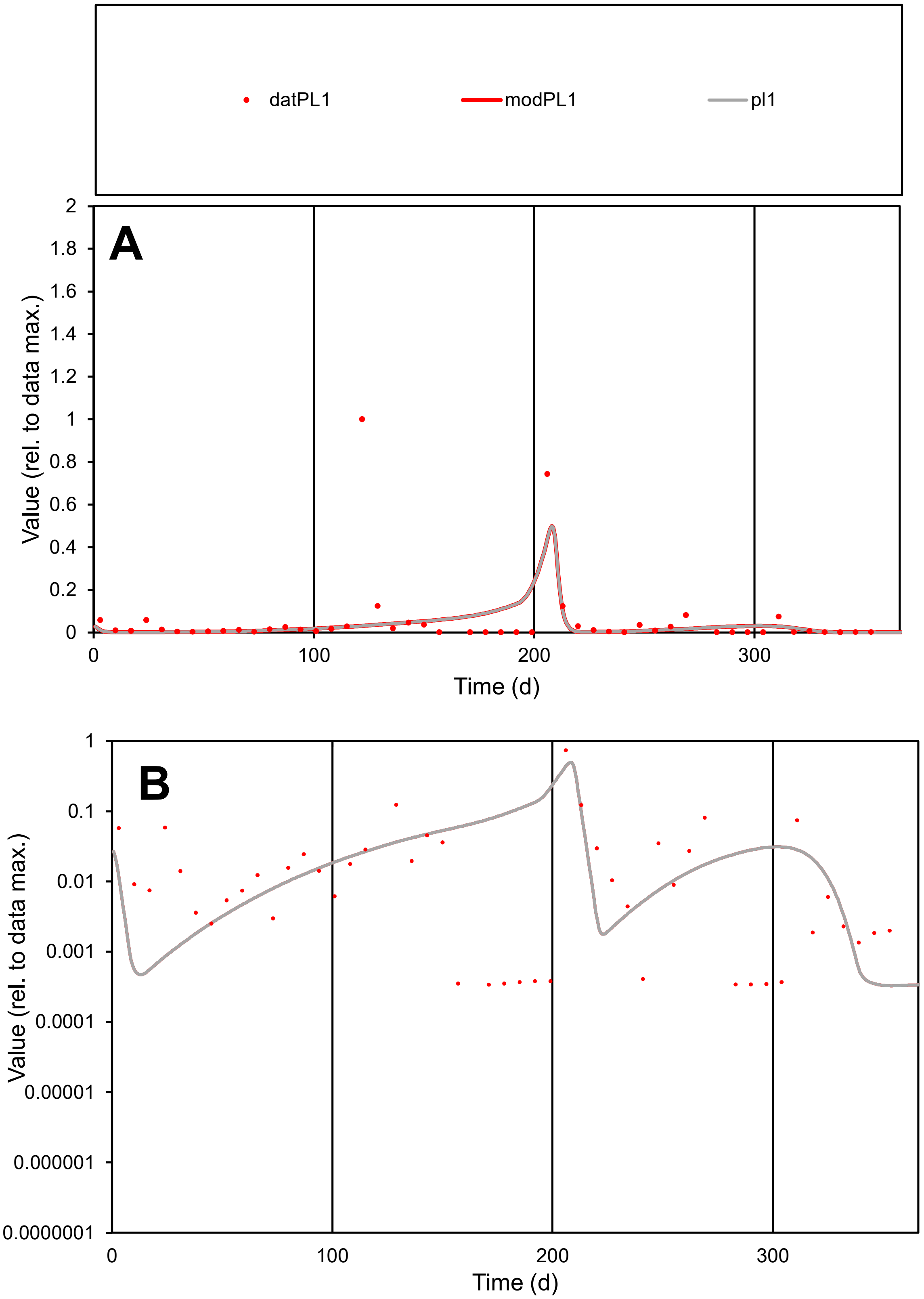


**C1**

Variables
PL1 Provide jtd or Dst
CalibOnly?
2 1 to reproduce of
oNormalize?
2 1=no, 2=yes

ofxn calc*
2.75819
ofxn mod
2.78286

sRun
RunA1
zout
1

Ave
Data
0.00571
Model
0.00397

data/vMax
0.1048

ofxng
369.894

Min d/m
1.03158

ave/min d
160.457
ave/min m
114.925

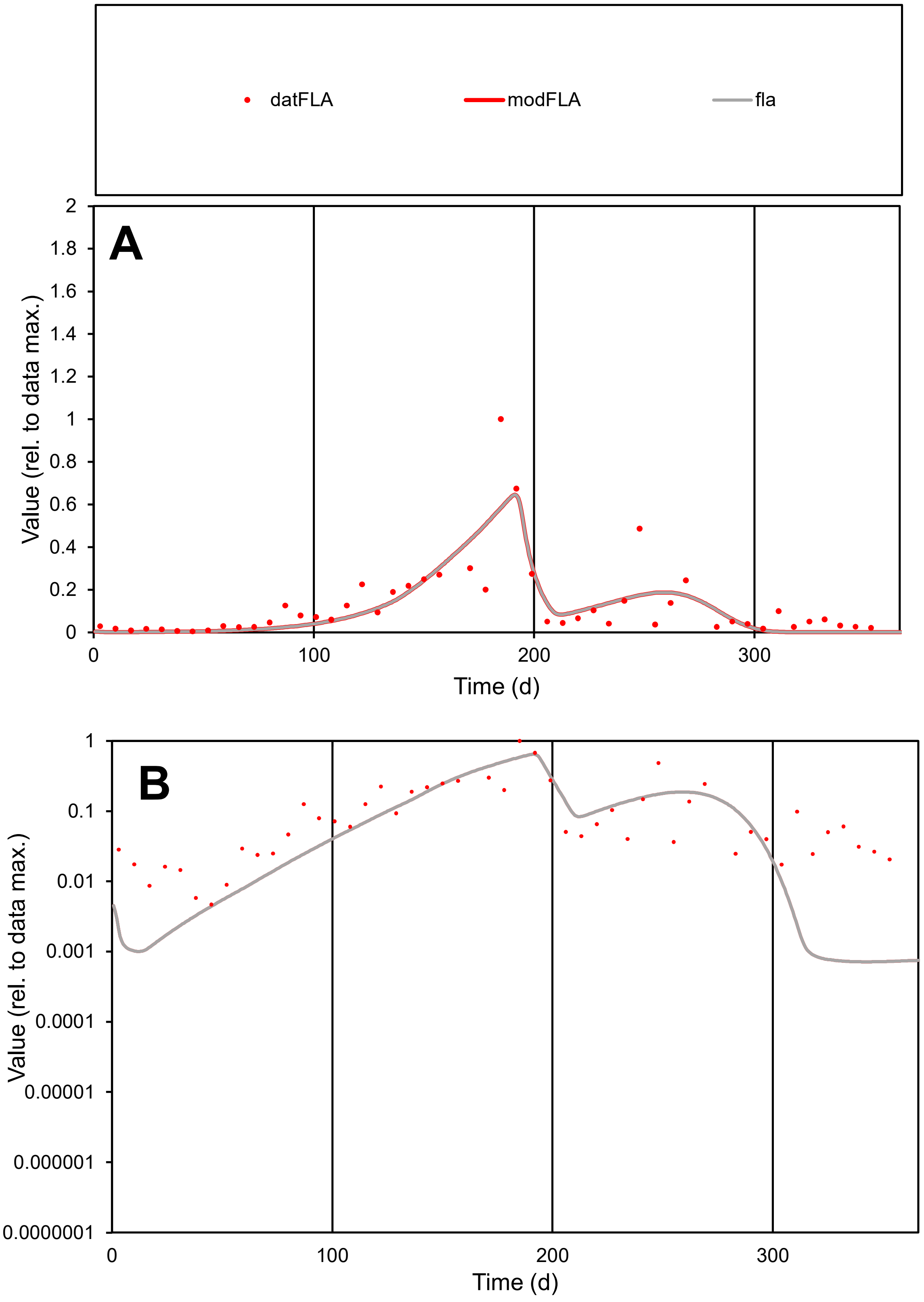


**C1**

Variables
FLA Provide jtd or Dst
CalibOnly?
2 1 to reproduce of
oNormalize?
2 1=no, 2=yes

ofxn calc*
0.78582
ofxn mod
0.81575

sRun
RunA1
zout
1

Ave
Data
0.00522
Model
0.00464

data/vMax
0.04138

ofxng
369.894

Min d/m
6.51178

ave/min d
27.0051
ave/min m
156.139

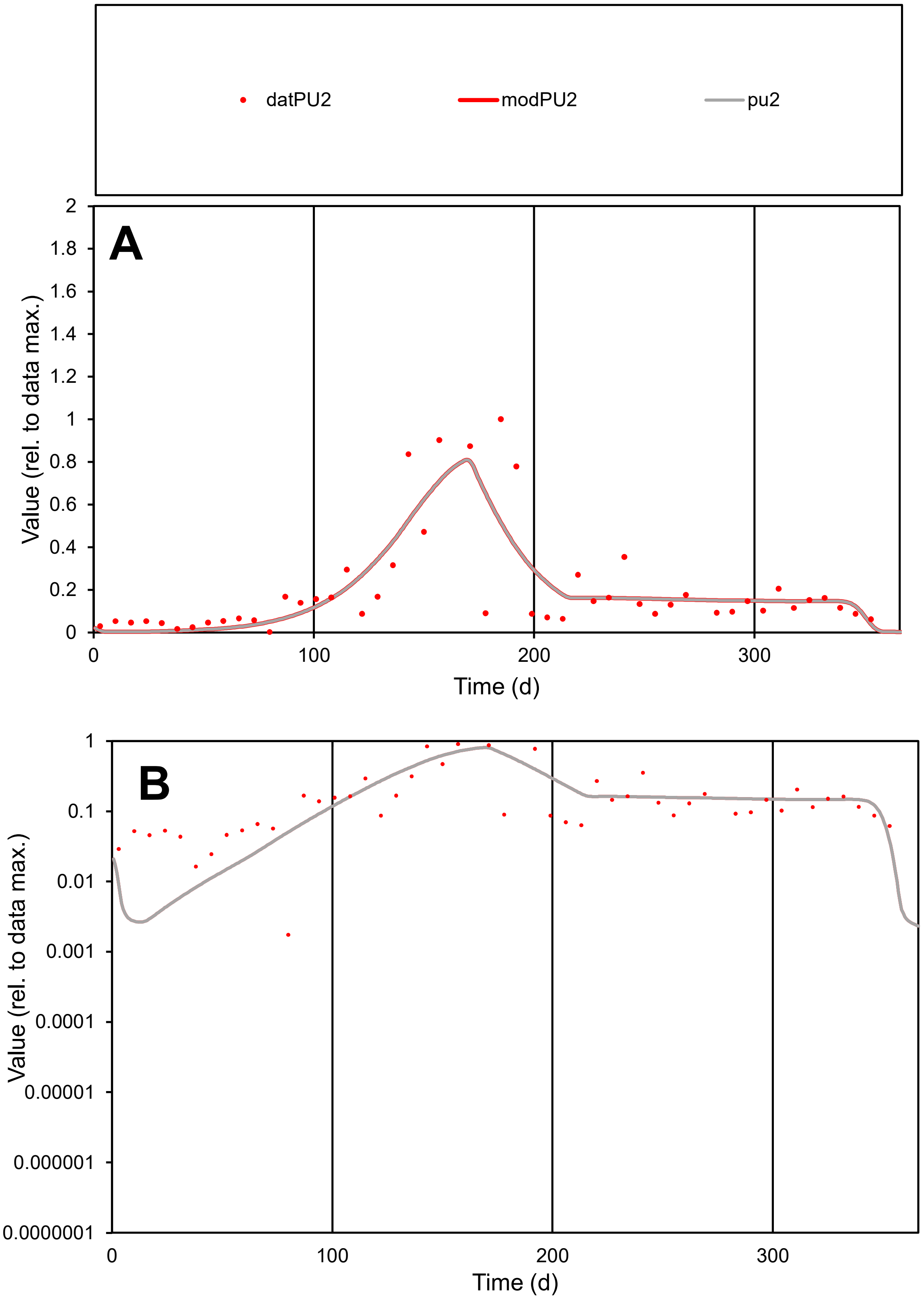


**C1**

Variables
PU2 Provide jtd or Dst
CalibOnly?
2 1 to reproduce of
oNormalize?
2 1=no, 2=yes

ofxn calc*
0.73513
ofxn mod
0.75538

sRun
RunA1
zout
1

Ave
Data
0.00641
Model
0.00619

data/vMax
0.0316

ofxng
369.894

Min d/m
0.658

ave/min d
116.208
ave/min m
73.8344

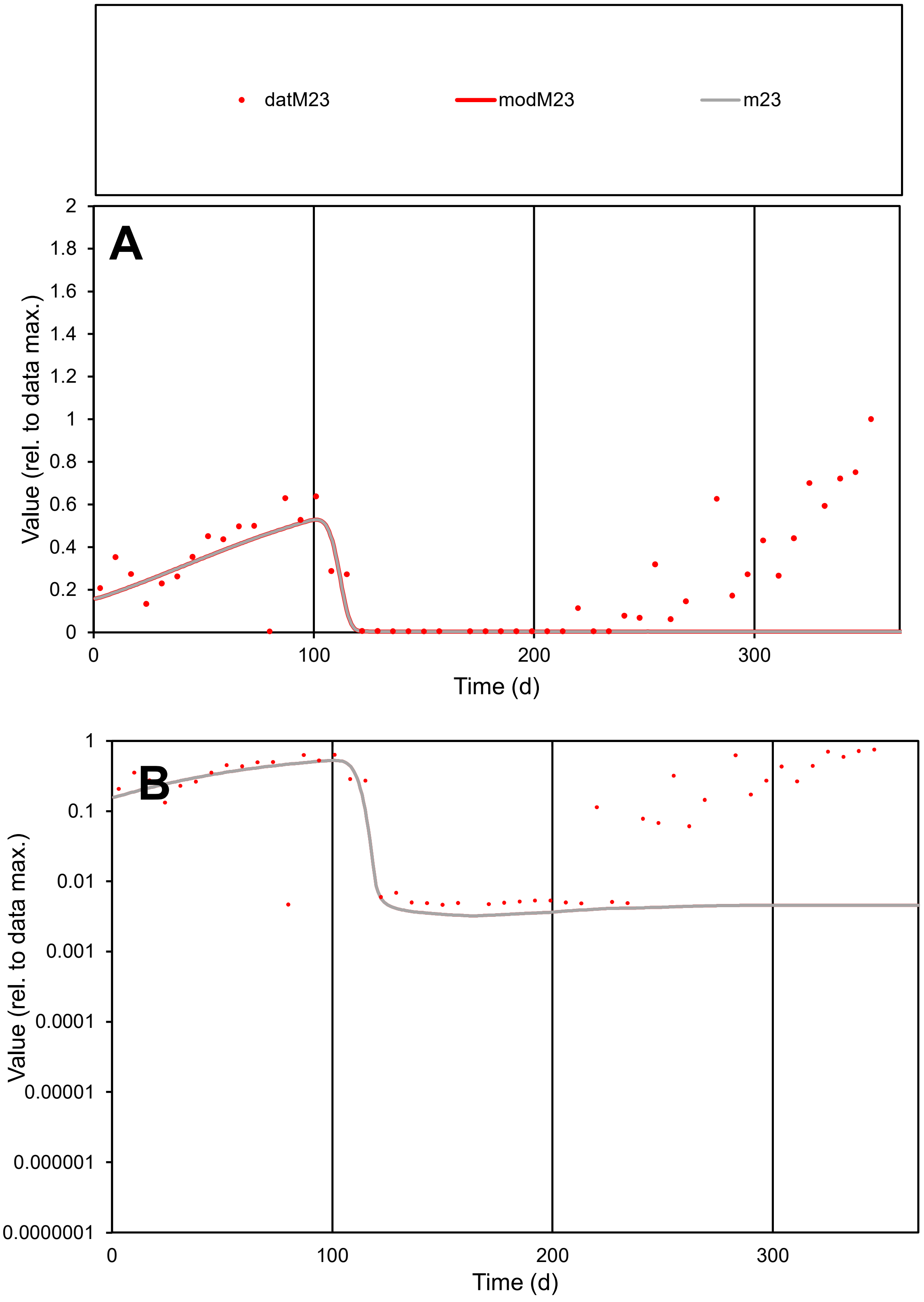


**C1**

Variables
M23 Provide jtd or Dst
CalibOnly?
2 1 to reproduce of
oNormalize?
2 1=no, 2=yes

ofxn calc*
1.12648
ofxn mod
1.13293

sRun
RunA1
zout
1

Ave
Data
0.00609
Model
0.00285

data/vMax
0.02315

ofxng
369.894

Min d/m
1.42084

ave/min d
56.5997
ave/min m
37.67

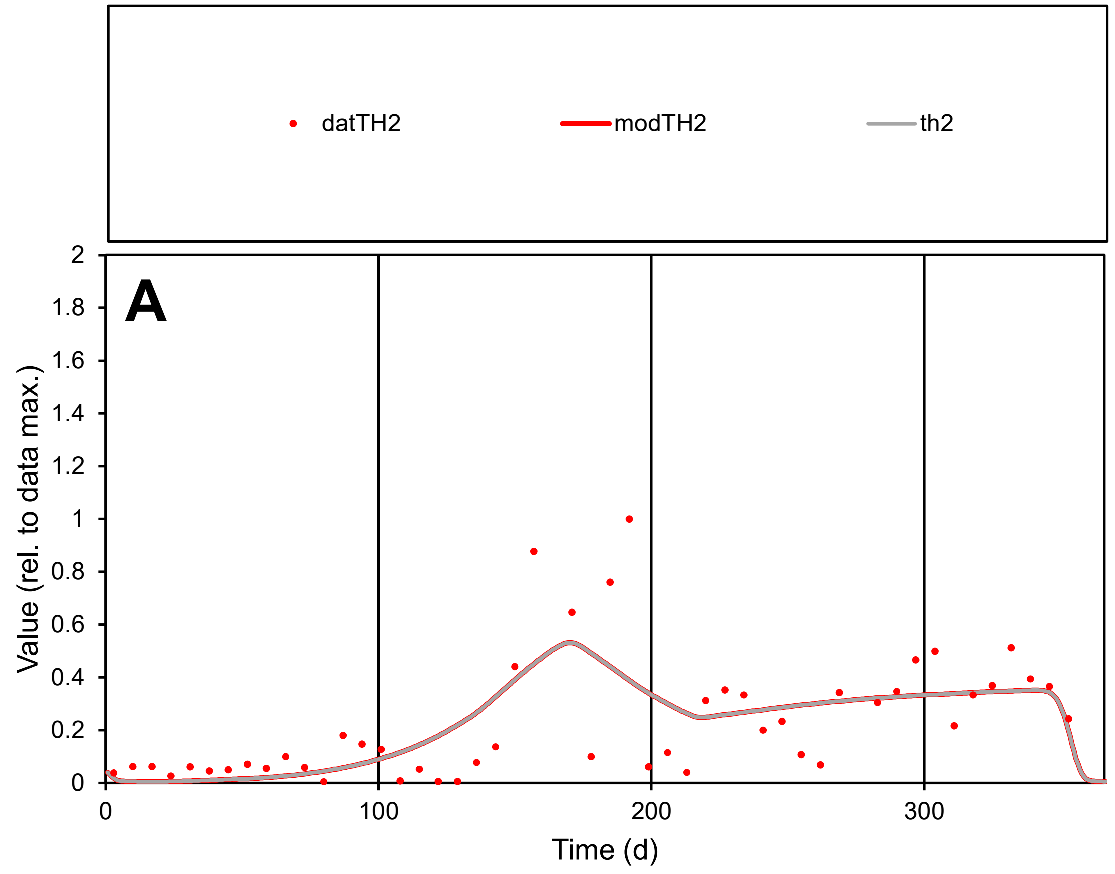


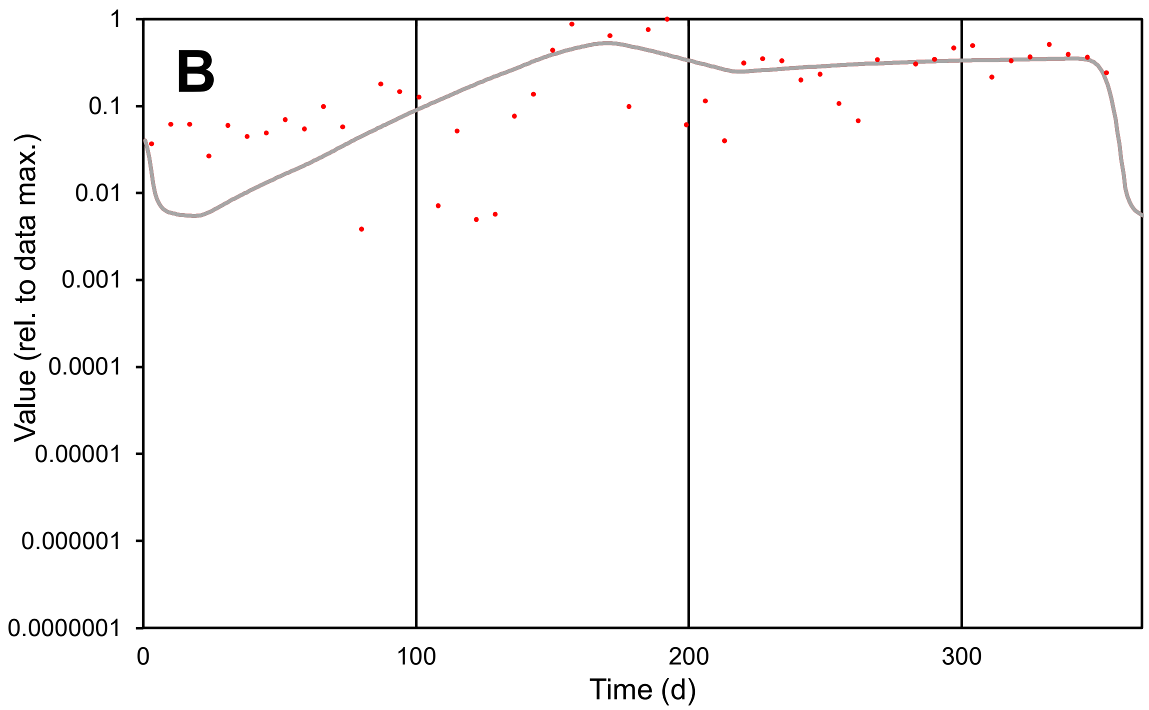


**C1**

Variables

TH2 Provide jtd or Dst

CalibOnly?

2 1 to reproduce of

oNormalize?

2 1=no, 2=yes

ofxn calc*

0.72558

ofxn mod

0.74215

sRun

RunA1

zout

1

Ave

Data

0.0064

Model

0.00623

data/vMax

0.02752

ofxng

369.894

Min d/m

0.71095

ave/min d

60.0677

ave/min m

41.5839

328_Calib_S_A

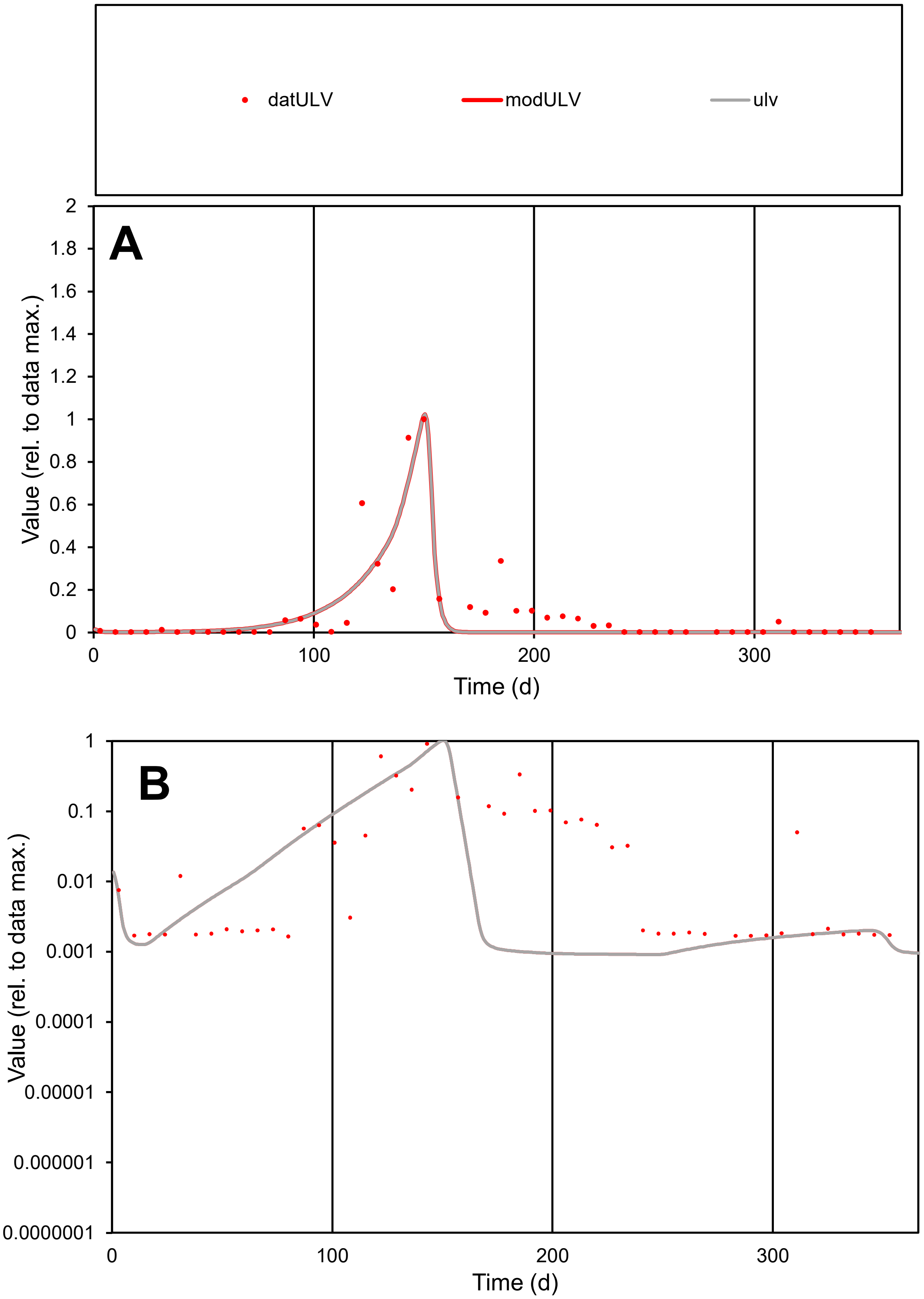


**C1**

Variables
ULV Provide jtd or Dst
CalibOnly?
2 1 to reproduce of
oNormalize?
2 1=no, 2=yes

ofxn calc*
1.00481
ofxn mod
1.0355

sRun
RunA1
zout
1

Ave
Data
0.00097
Model
0.00079

data/vMax
0.0105

ofxng
369.894

Min d/m
1.79882

ave/min d
56.2643
ave/min m
81.9563

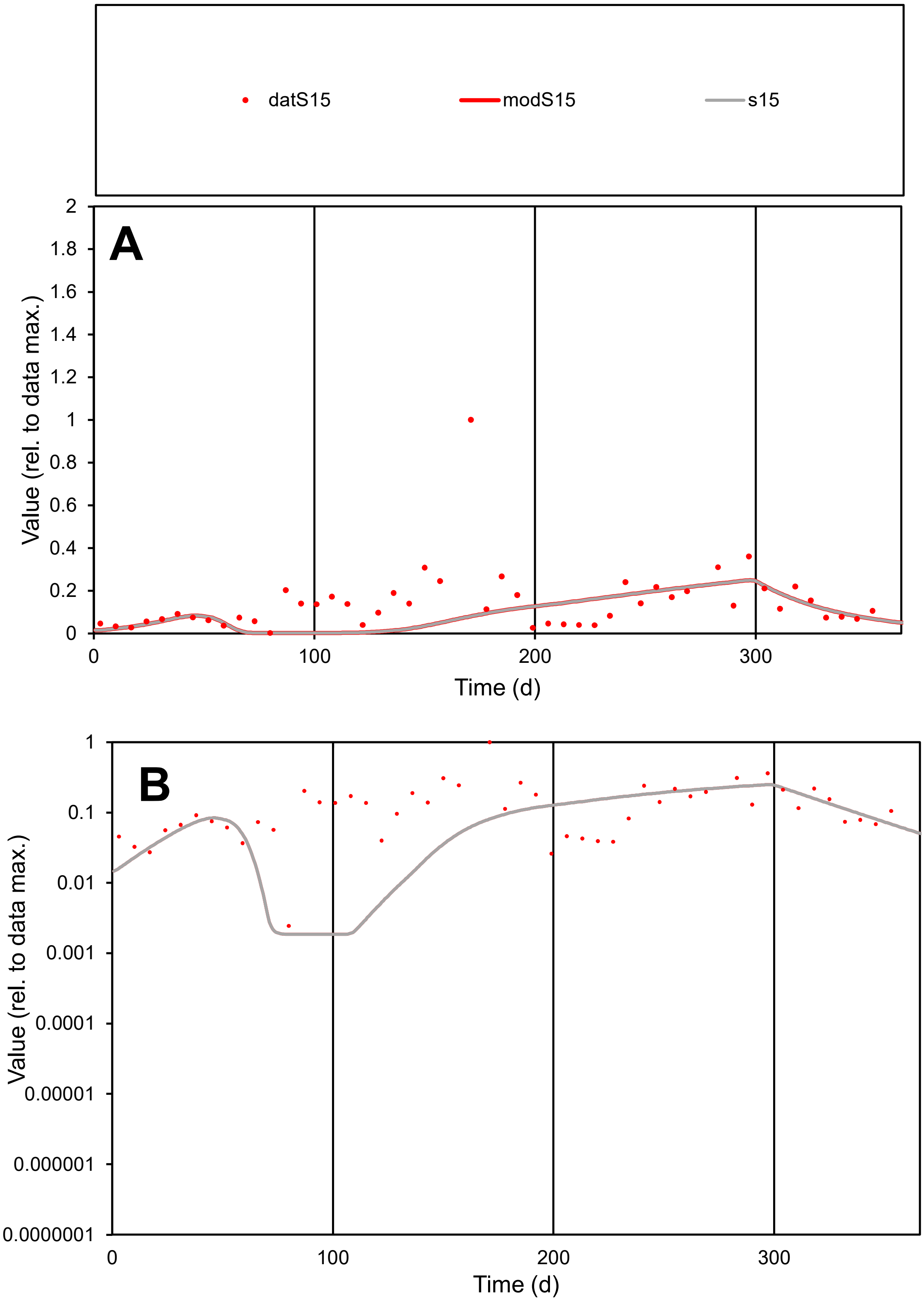


**C1**

Variables
S15 Provide jtd or Dst
CalibOnly?
2 1 to reproduce of
oNormalize?
2 1=no, 2=yes

ofxn calc*
1.12187
ofxn mod
1.12877

sRun
RunA1
zout
1

Ave
Data
0.00736
Model
0.00491

data/vMax
0.05113

ofxng
369.894

Min d/m
1.31372

ave/min d
58.9125
ave/min m
51.5915

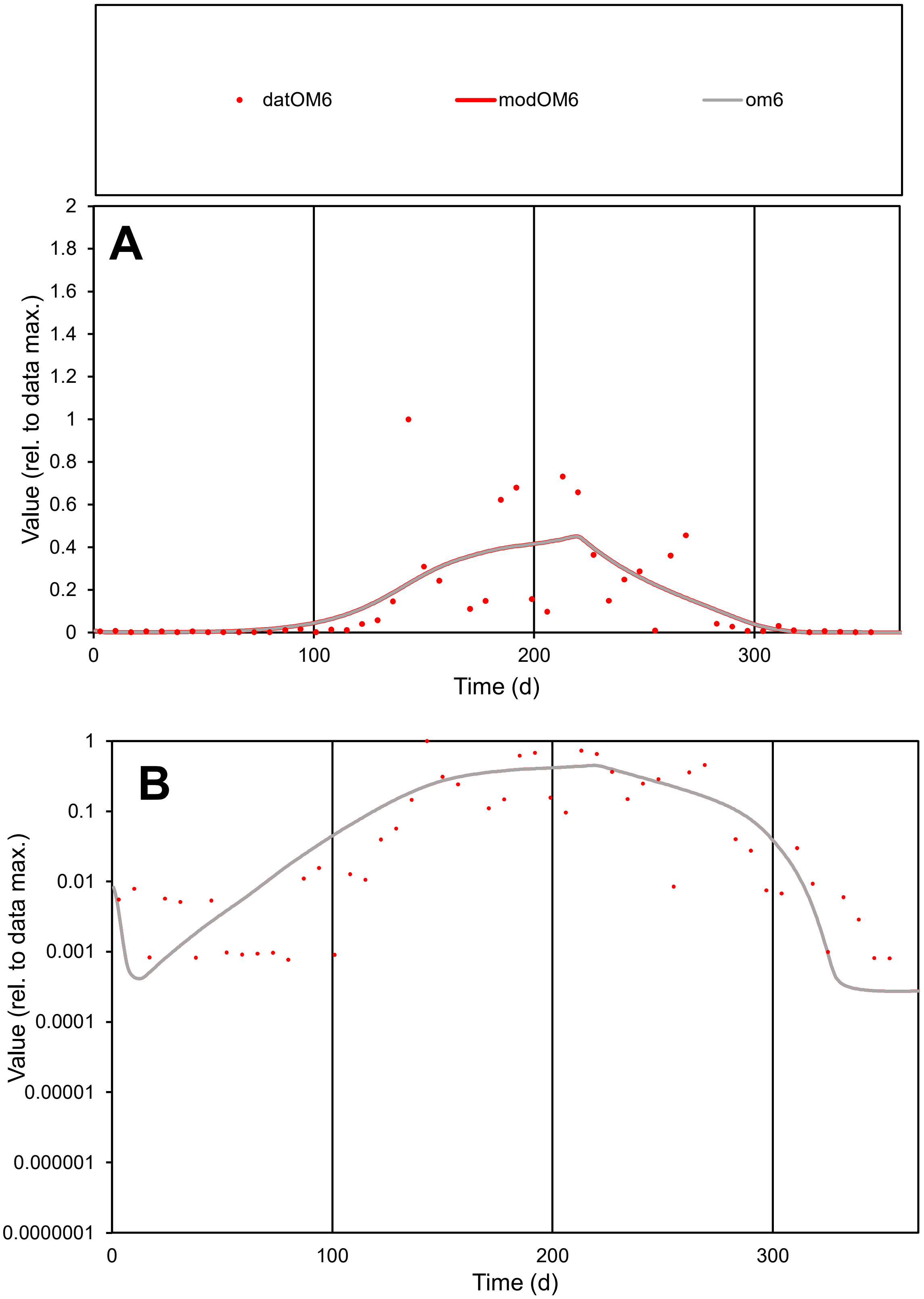


**C1**

Variables
OM6 Provide jtd or Dst
CalibOnly?
2 1 to reproduce of
oNormalize?
2 1=no, 2=yes

ofxn calc*
1.15697
ofxn mod
1.17257

sRun
RunA1
zout
1

Ave
Data
0.00473
Model
0.00465

data/vMax
0.03274

ofxng
369.894

Min d/m
2.79436
ave/min d
187.987
ave/min m
515.912

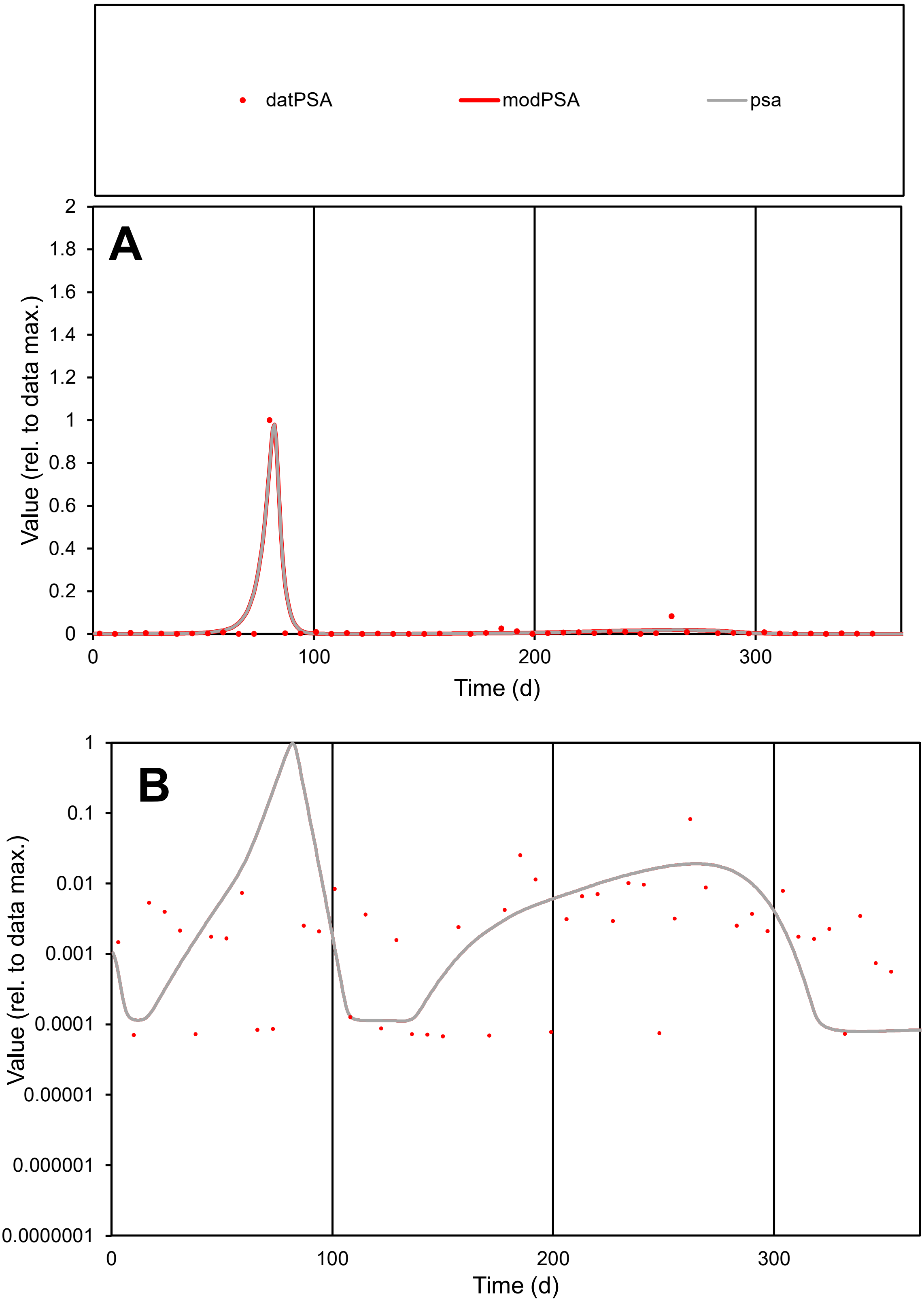


**C1**

Variables
PSA Provide jtd or Dst
CalibOnly?
2 1 to reproduce of
oNormalize?
2 1=no, 2=yes

ofxn calc*
1.02876
ofxn mod
1.31263

sRun
RunA1
zout
1

Ave
Data
0.00332
Model
0.00382

data/vMax
0.1304

ofxng
369.894

Min d/m
0.85971

ave/min d
374.781
ave/min m
371.29

328_Calib_S_A

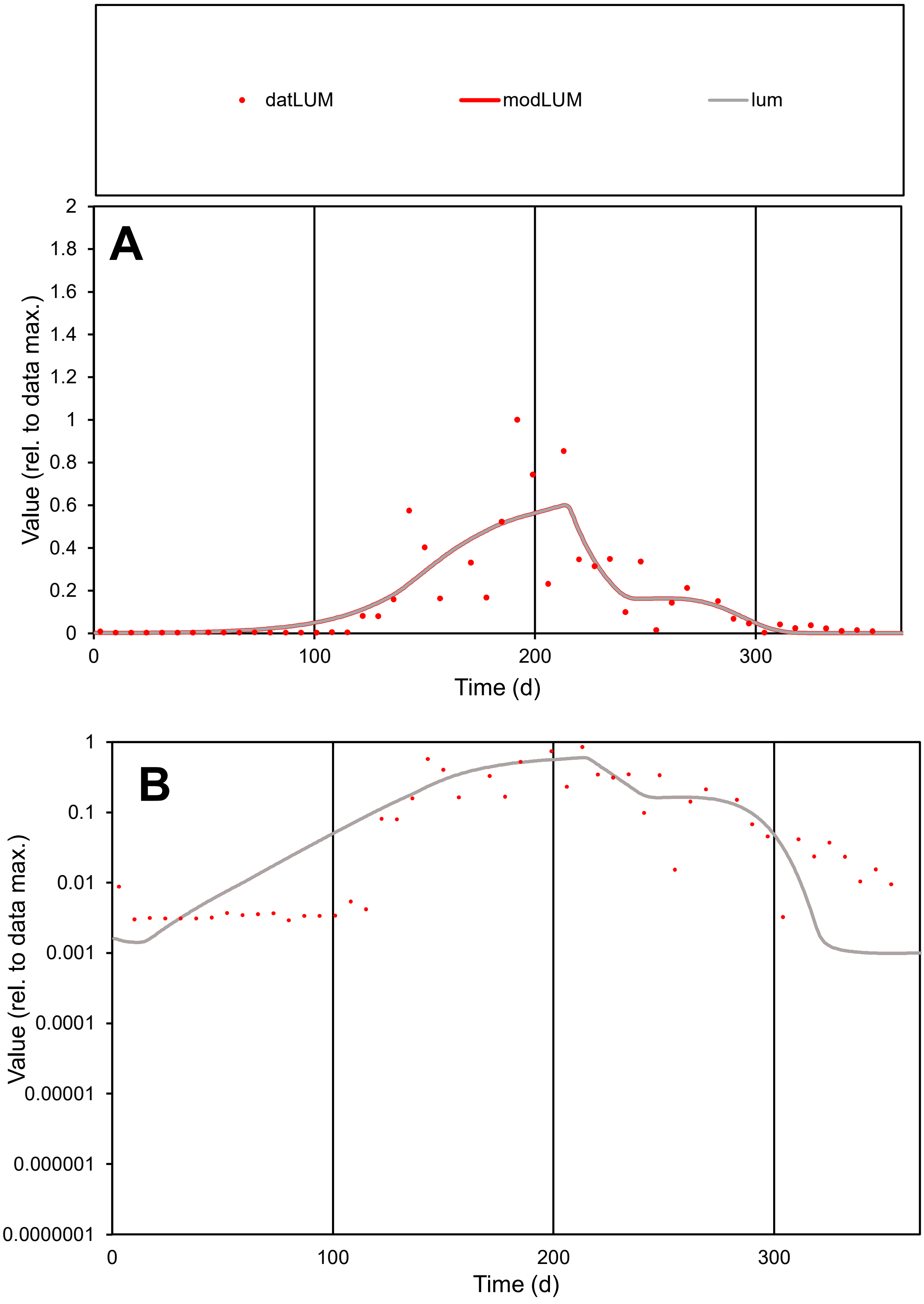


**C1**

Variables
LUM Provide jtd or Dst
CalibOnly?
2 1 to reproduce of
oNormalize?
2 1=no, 2=yes

ofxn calc*
0.8274
ofxn mod
0.85743

sRun
RunA1
zout
1

Ave
Data
0.0024
Model
0.00239

data/vMax
0.01543

ofxng
369.894

Min d/m
2.92653

ave/min d
53.2915
ave/min m
155.398

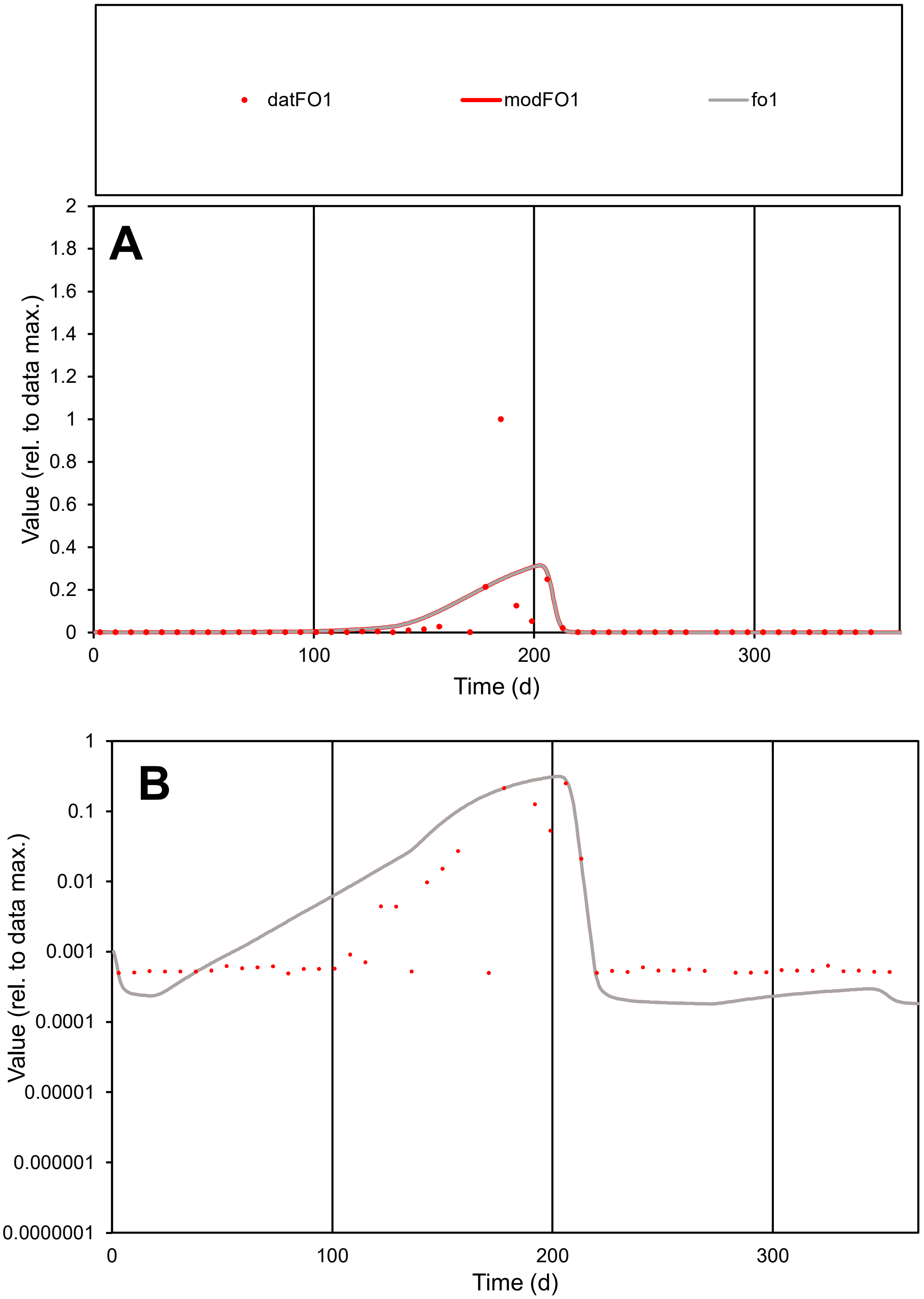


**C1**

Variables
FO1 Provide jtd or Dst
CalibOnly?
2 1 to reproduce of
oNormalize?
2 1=no, 2=yes

ofxn calc*
3.31892
ofxn mod
3.34544

sRun
RunA1
zout
1

Ave
Data
0.00424
Model
0.00442

data/vMax
0.1191

ofxng
369.894

Min d/m
2.70564

ave/min d
72.4533
ave/min m
204.086

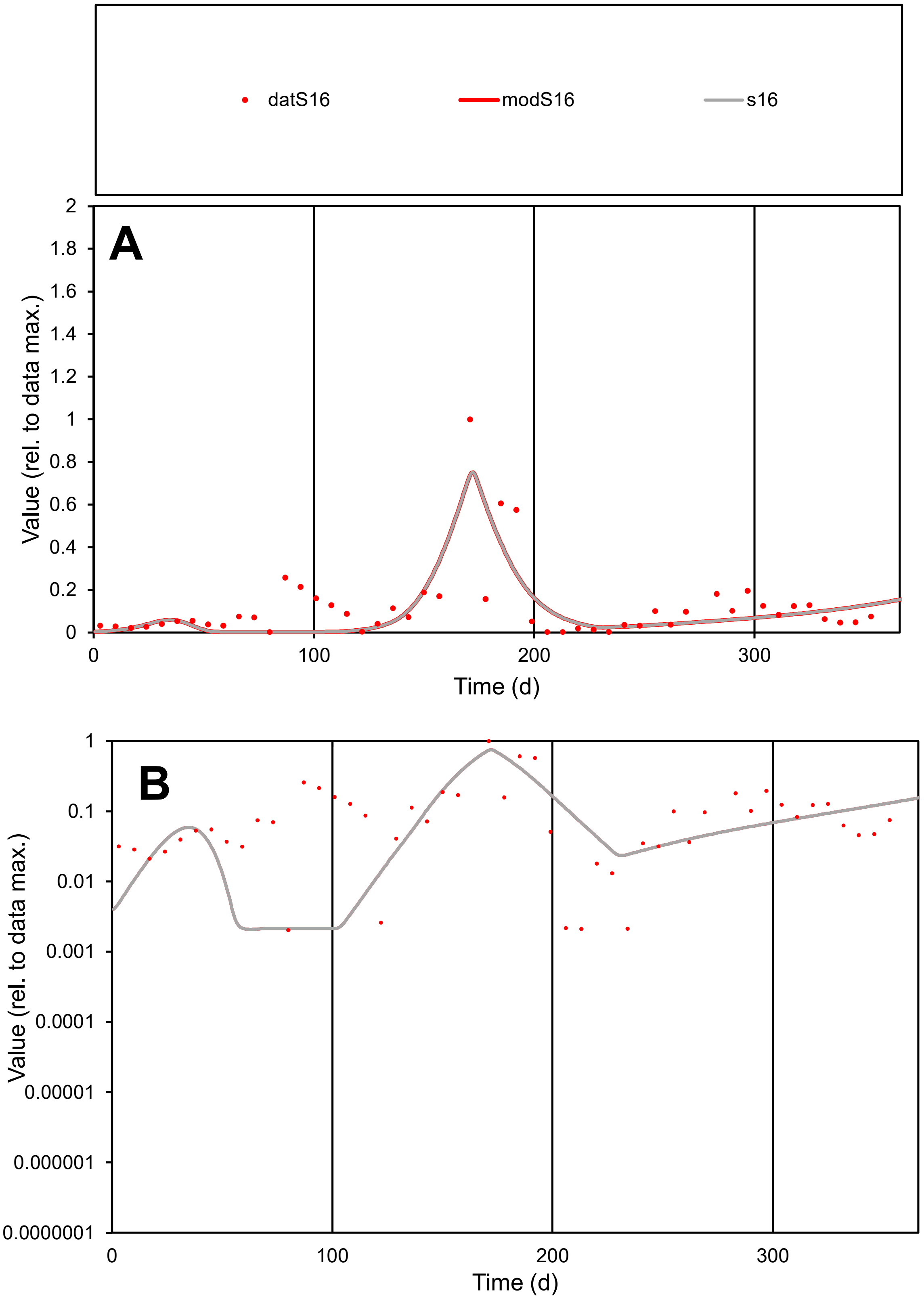


**C1**

Variables
S16 Provide jtd or Dst
CalibOnly?
2 1 to reproduce of
oNormalize?
2 1=no, 2=yes

ofxn calc*
0.9684
ofxn mod
0.99165

sRun
RunA1
zout
1

Ave
Data
0.00728
Model
0.00574

data/vMax
0.06158

ofxng
369.894

Min d/m
0.96228

ave/min d
58.2565
ave/min m
44.1507

328_Calib_S_A

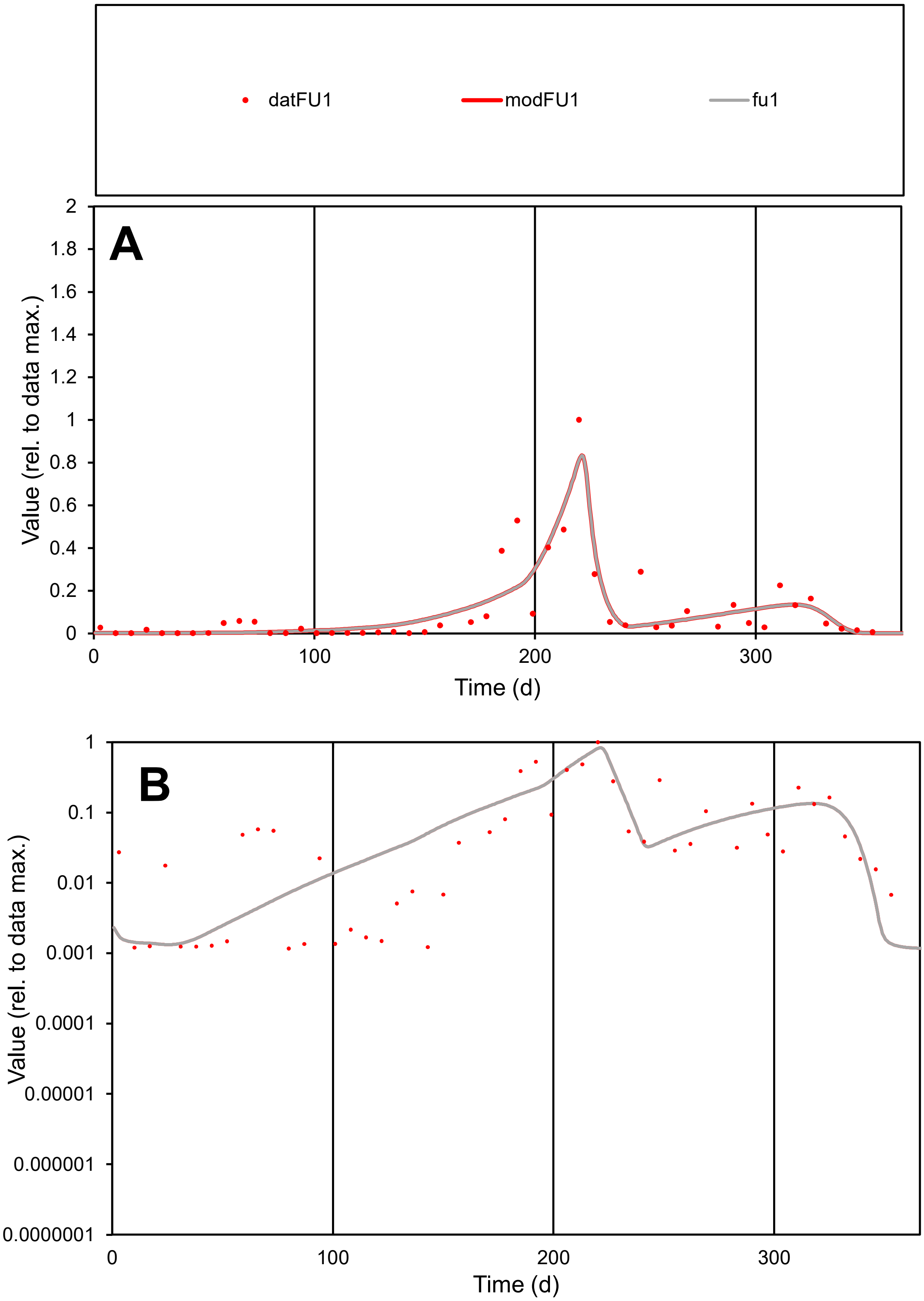


**C1**

Variables
FU1 Provide jtd or Dst
CalibOnly?
2 1 to reproduce of
oNormalize?
2 1=no, 2=yes

ofxn calc*
0.81215
ofxn mod
0.8502

sRun
RunA1
zout
1

Ave
Data
0.00412
Model
0.00398

data/vMax
0.04026

ofxng
369.894

Min d/m
0.88628

ave/min d
87.9408
ave/min m
75.3144

328_Calib_S_A

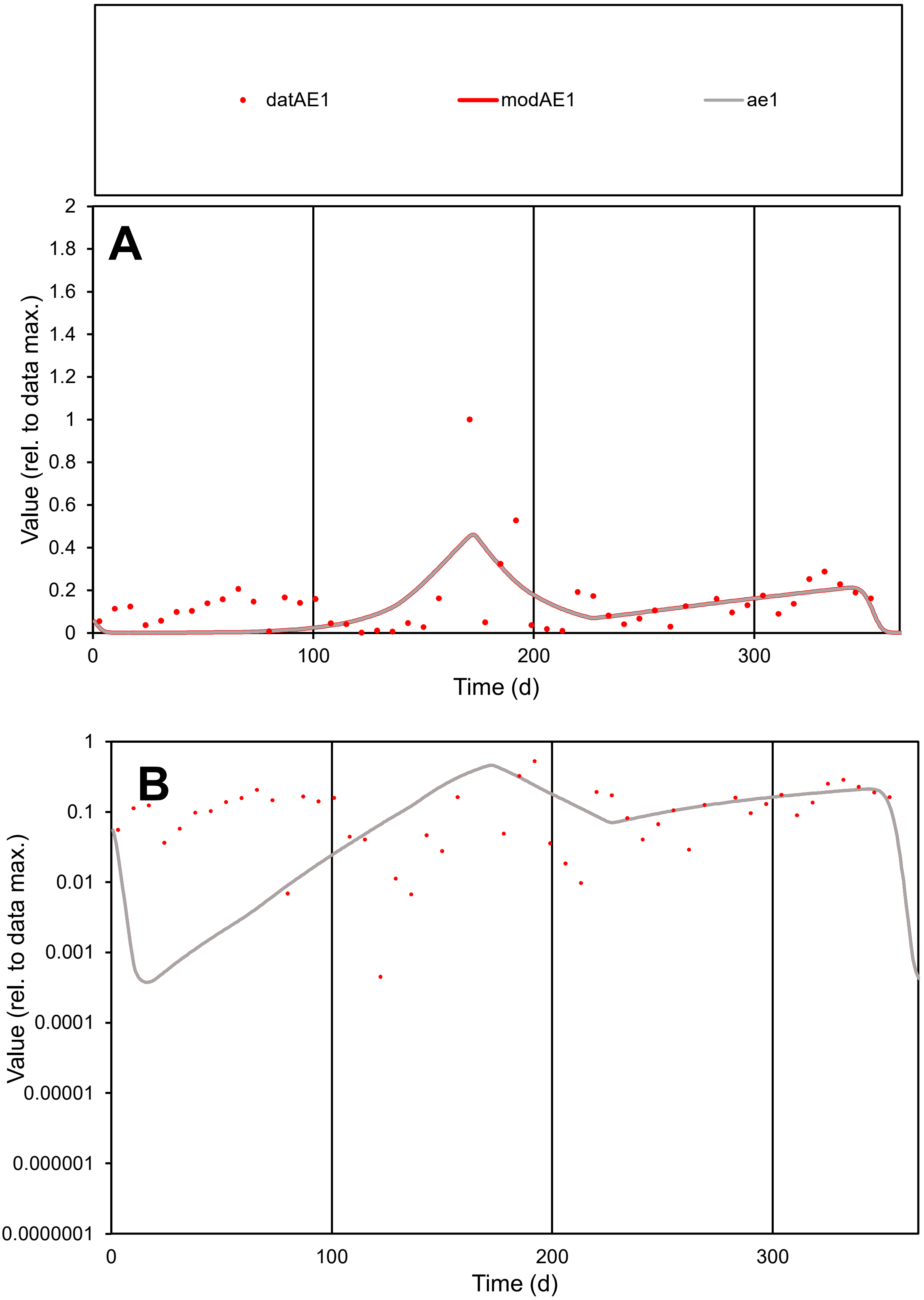


**C1**

Variables
AE1 Provide jtd or Dst
CalibOnly?
2 1 to reproduce of
oNormalize?
2 1=no, 2=yes

ofxn calc*
0.98353
ofxn mod
1.00113

sRun
RunA1
zout
1

Ave
Data
0.00545
Model
0.00456

data/vMax
0.03968

ofxng
369.894

Min d/m
1.16688

ave/min d
303.253
ave/min m
296.418

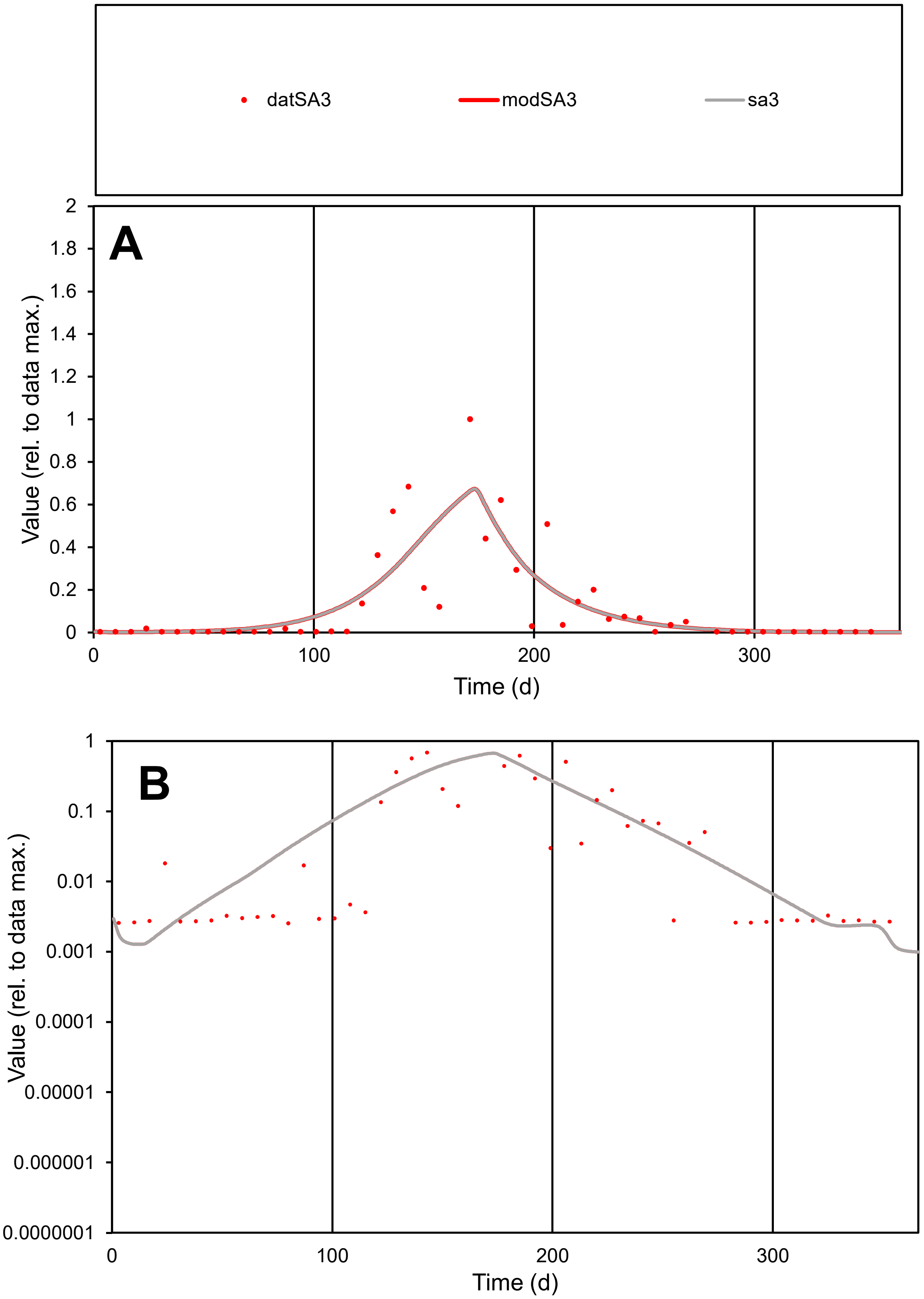


**C1**

Variables
SA3 Provide jtd or Dst
CalibOnly?
2 1 to reproduce of
oNormalize?
2 1=no, 2=yes

ofxn calc*
1.06124
ofxn mod
1.09266

sRun
RunA1
zout
1

Ave
Data
0.00456
Model
0.00463

data/vMax
0.03881

ofxng
369.894

Min d/m
2.00487

ave/min d
46.1029
ave/min m
93.8766

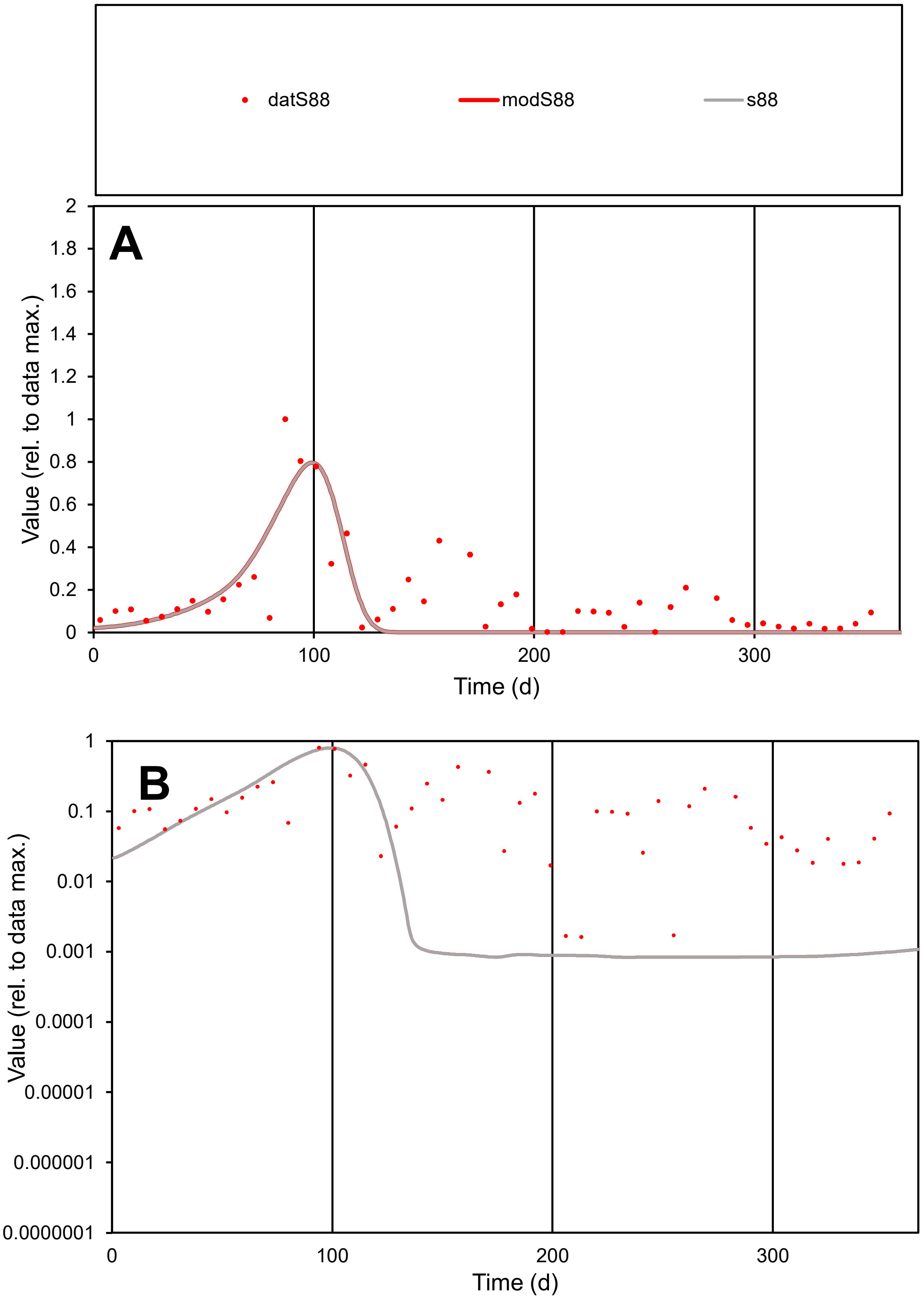


**C1**

Variables
S88 Provide jtd or Dst
CalibOnly?
2 1 to reproduce of
oNormalize?
2 1=no, 2=yes

ofxn calc*
0.90749
ofxn mod
0.92274

sRun
RunA1
zout
1

Ave
Data
0.00599
Model
0.00397

data/vMax
0.03713

ofxng
369.894

Min d/m
1.96042

ave/min d
99.1568
ave/min m
128.857

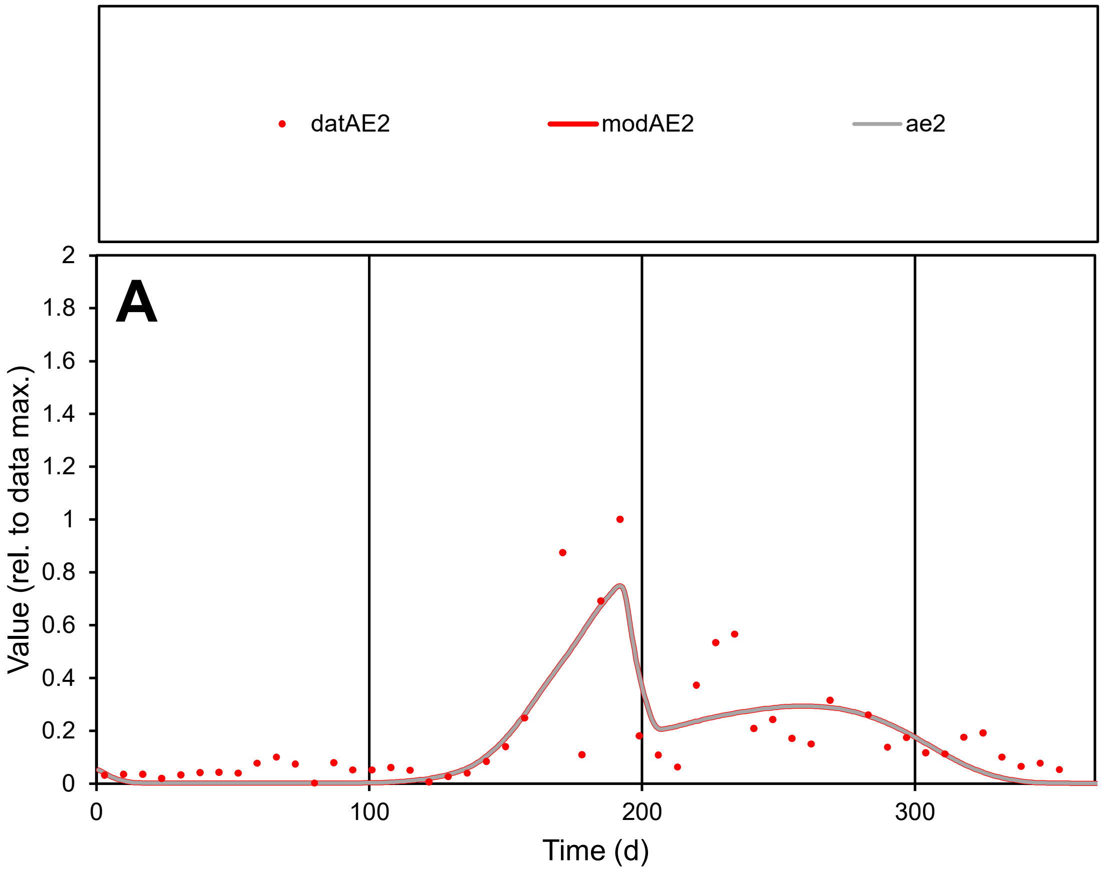


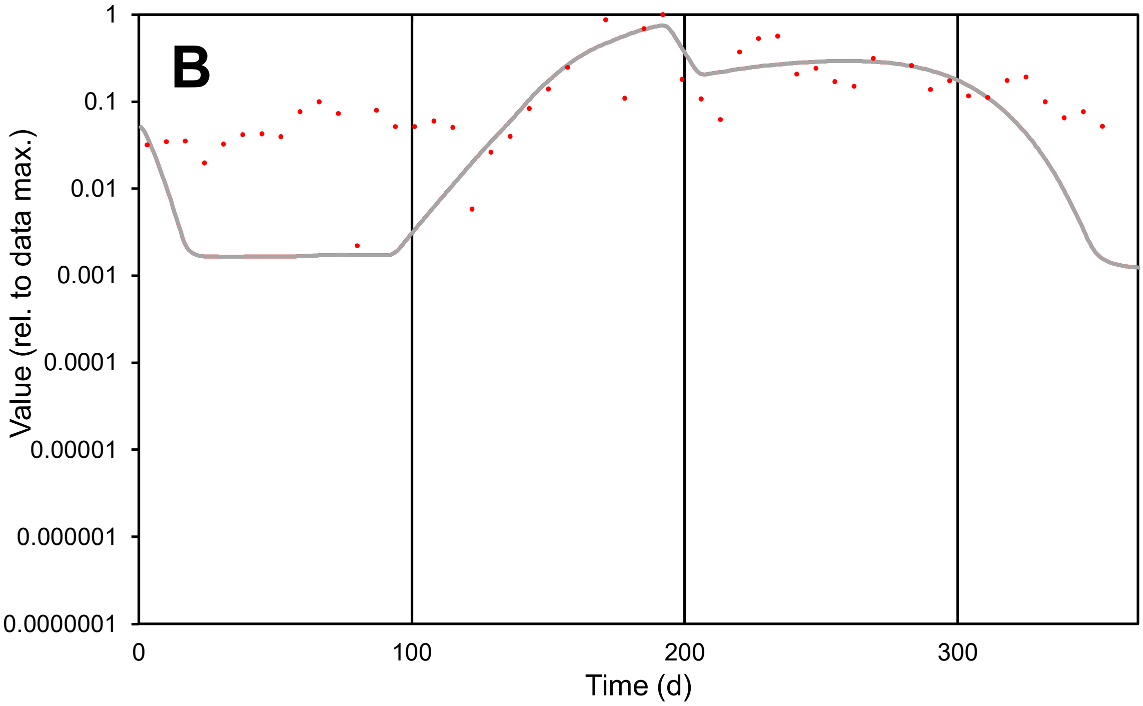


**C1**

Variables
AE2 Provide jtd or Dst
CalibOnly?
2 1 to reproduce of
oNormalize?
2 1=no, 2=yes

ofxn calc*
0.75475
ofxn mod
0.77888

sRun
RunA1
zout
1

Ave
Data
0.00452
Model
0.00392

data/vMax
0.02631

ofxng
369.894

Min d/m
1.47415

ave/min d
77.6879
ave/min m
99.2802

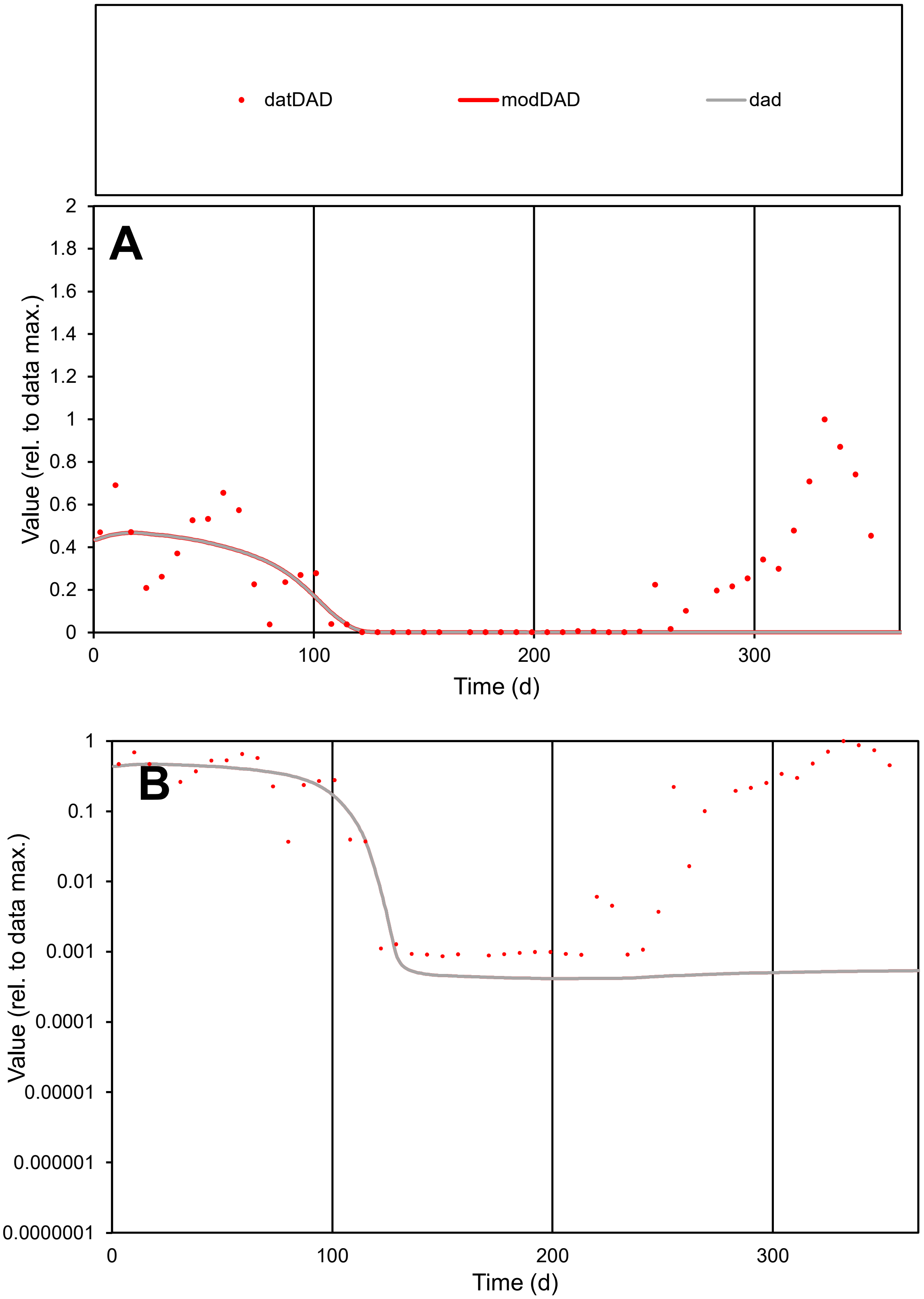


C1

Variables
DAD Provide jtd or Dst
CalibOnly?
2 1 to reproduce of
oNormalize?
2 1=no, 2=yes

ofxn calc*
1.19317
ofxn mod
1.20029

sRun
RunA1
zout
1

Ave
Data
0.0043
Model
0.00212

data/vMax
0.01785

ofxng
369.894

Min d/m
2.09252

ave/min d
278.057
ave/min m
287.034

328_Calib_S_A

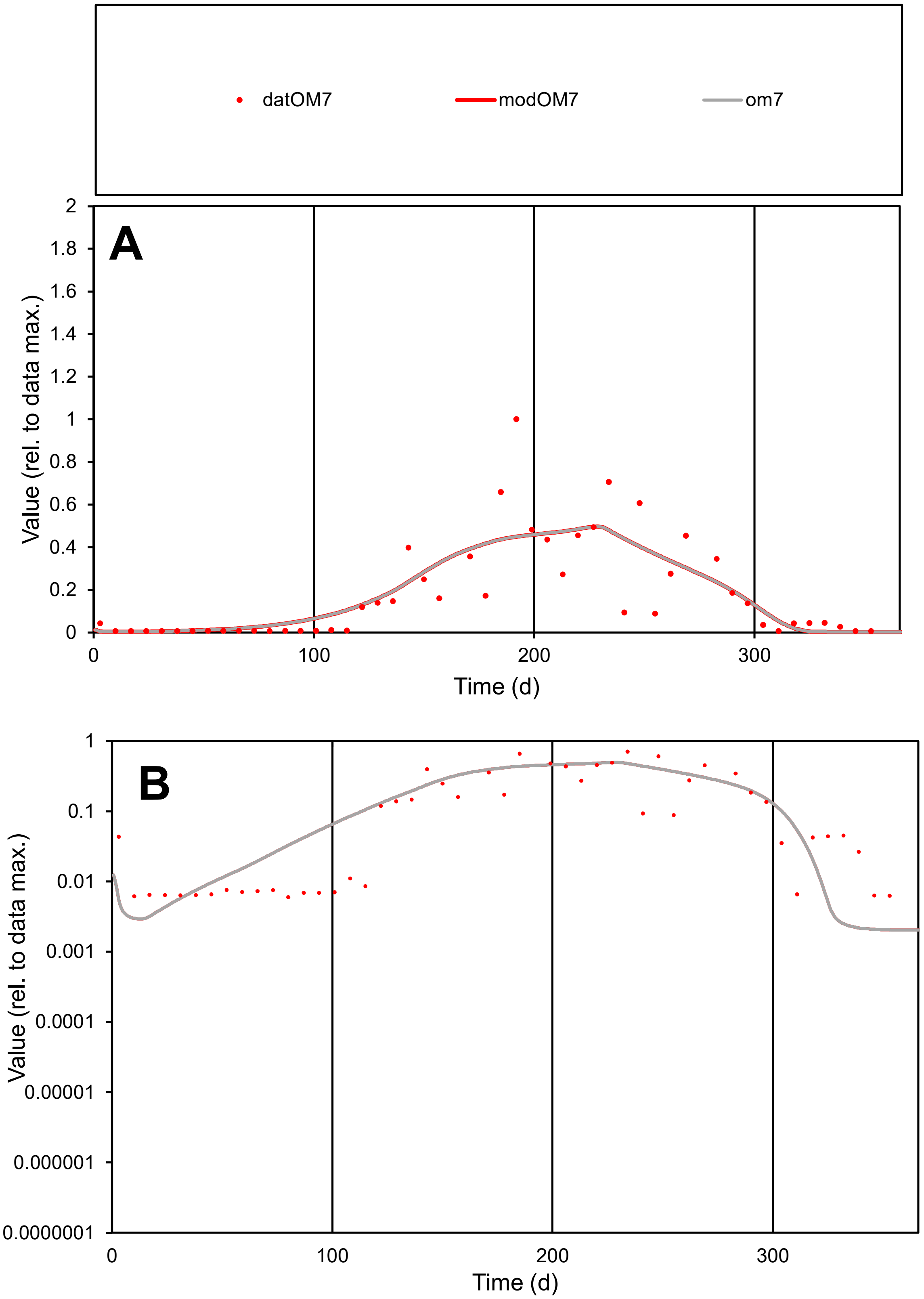


**C1**

Variables
OM7 Provide jtd or Dst
CalibOnly?
2 1 to reproduce of
oNormalize?
2 1=no, 2=yes

ofxn calc*
0.74536
ofxn mod
0.77007

sRun
RunA1
zout
1

Ave
Data
0.00282
Model
0.00288

data/vMax
0.01571

ofxng
369.894

Min d/m
2.92284

ave/min d
30.0292
ave/min m
89.6317

328_Calib_S_A

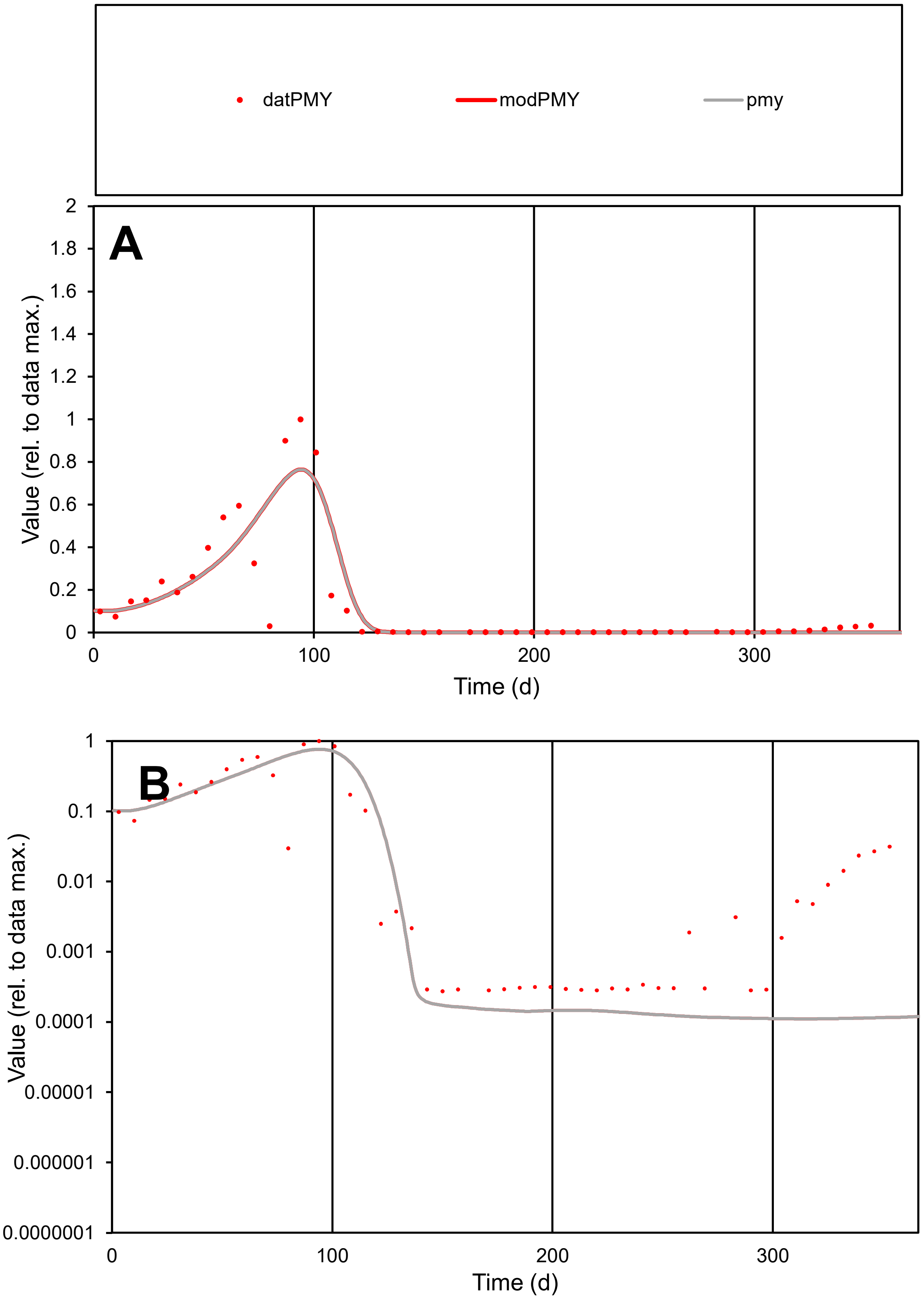


C1

Variables
PMY Provide jtd or Dst
CalibOnly?
2 1 to reproduce of
oNormalize?
2 1=no, 2=yes

ofxn calc*
0.91171
ofxn mod
0.95434

sRun
RunA1
zout
1

Ave
Data
0.00443
Model
0.00442

data/vMax
0.03504

ofxng
369.894

Min d/m
2.47621

ave/min d
459.831
ave/min m
1136.1

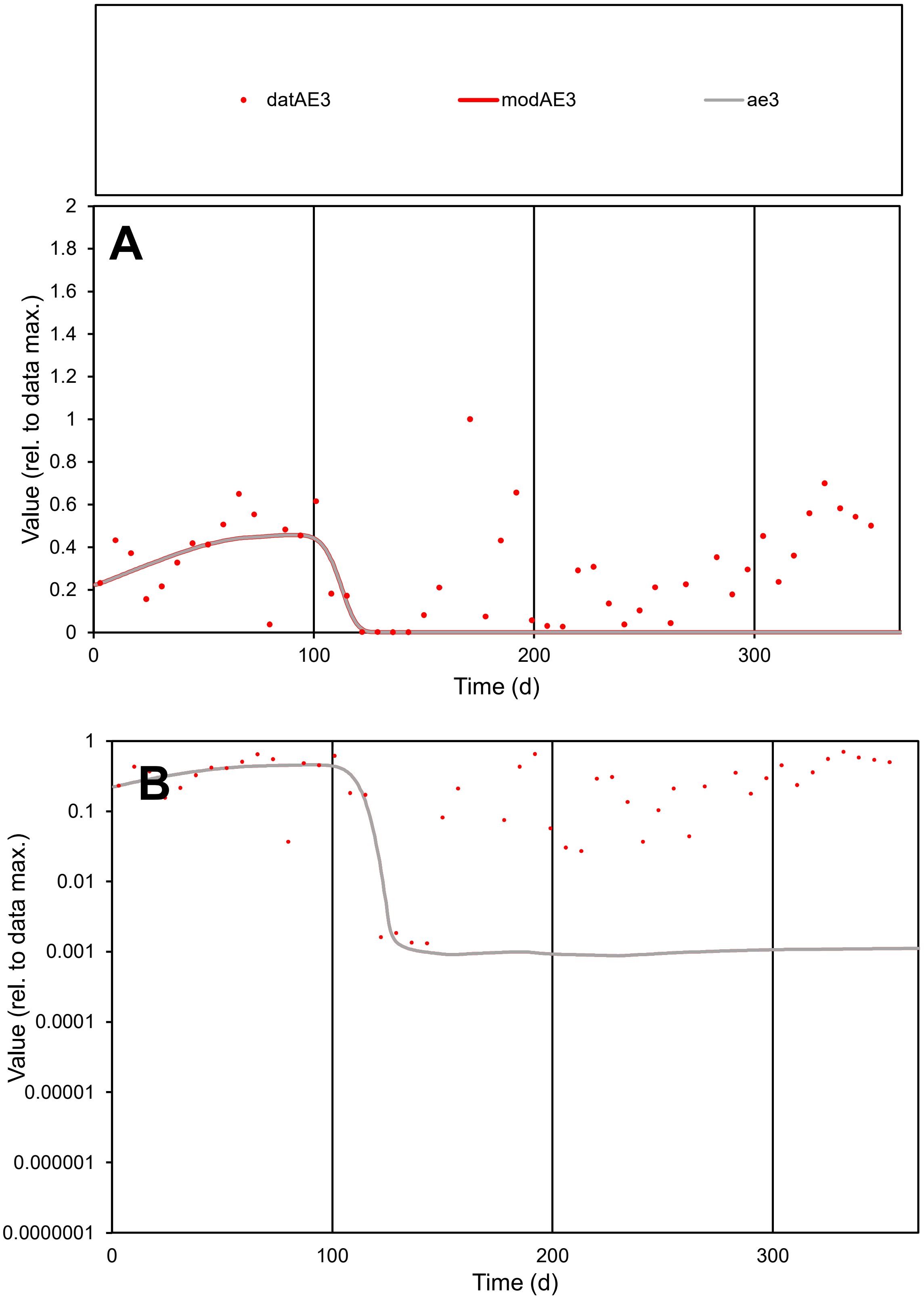


**C1**

Variables
AE3 Provide jtd or Dst
CalibOnly?
2 1 to reproduce of
oNormalize?
2 1=no, 2=yes

ofxn calc*
1.01304
ofxn mod
1.01812

sRun
RunA1
zout
1

Ave
Data
0.00409
Model
0.00171

data/vMax
0.01343

ofxng
369.894

Min d/m
1.50763

ave/min d
229.707
ave/min m
144.547

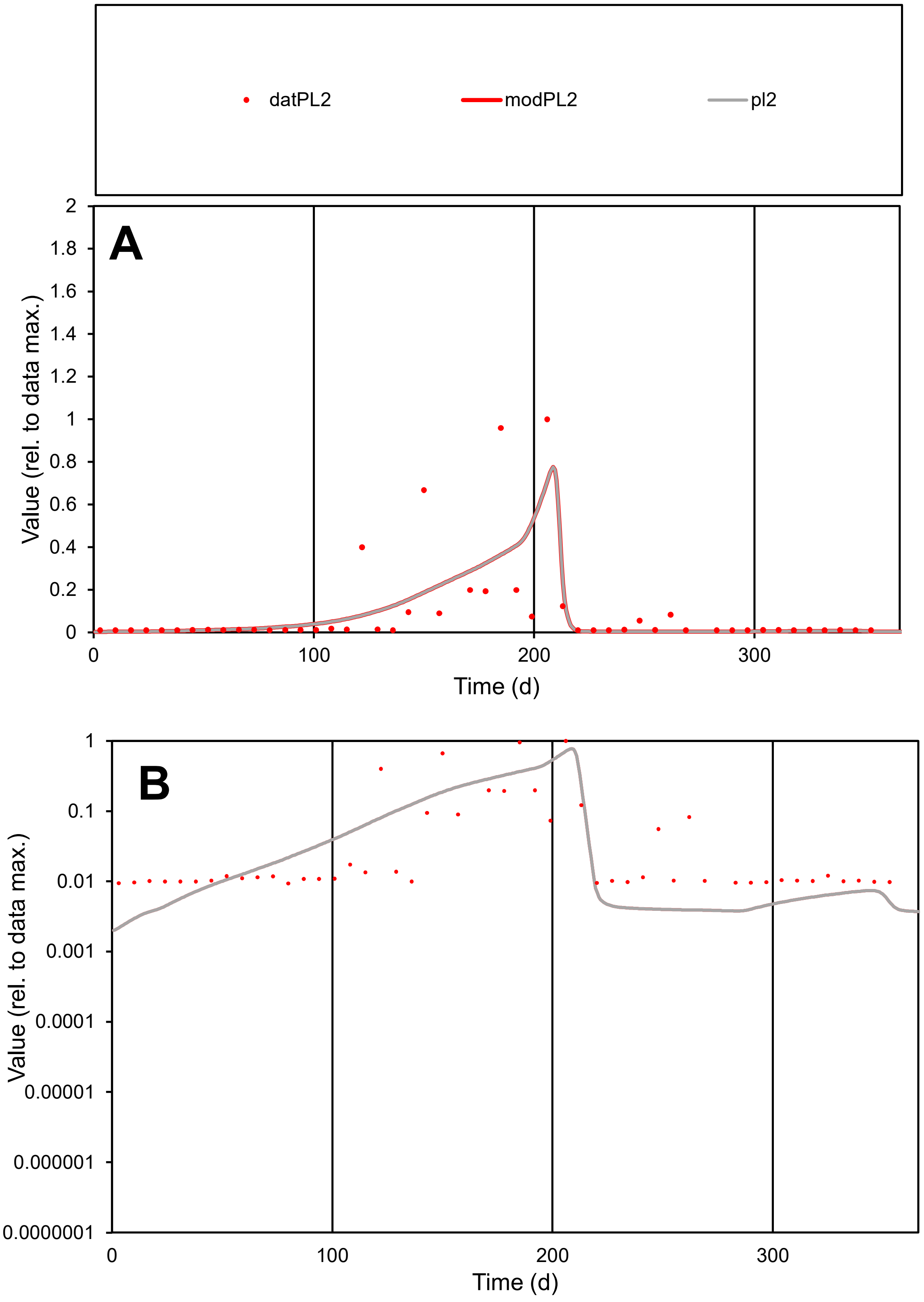


C1

Variables
PL2 Provide jtd or Dst
CalibOnly?
2 1 to reproduce of
oNormalize?
2 1=no, 2=yes

ofxn calc*
1.59648
ofxn mod
1.61532

sRun
RunA1
zout
1

Ave
Data
0.00162
Model
0.00147

data/vMax
0.01756

ofxng
369.894

Min d/m
4.03993

ave/min d
9.87404
ave/min m
36.2412

328_Calib_S_A

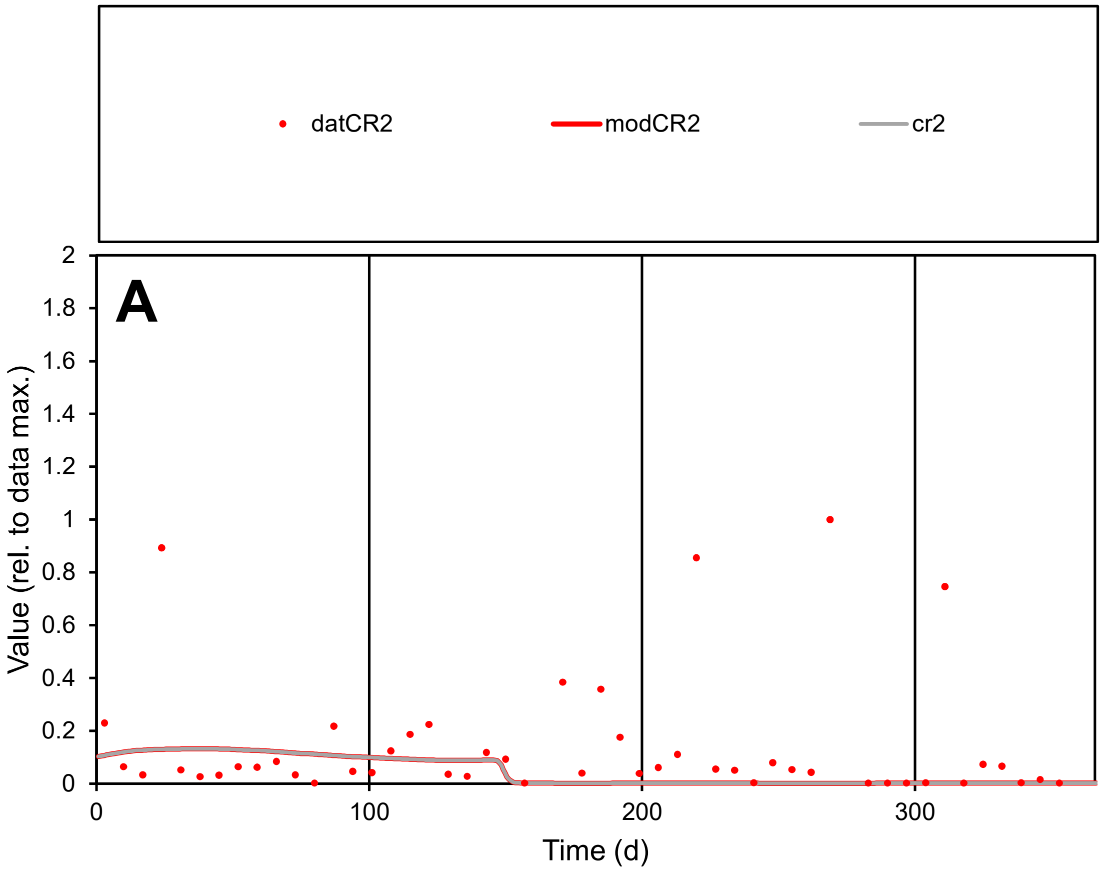


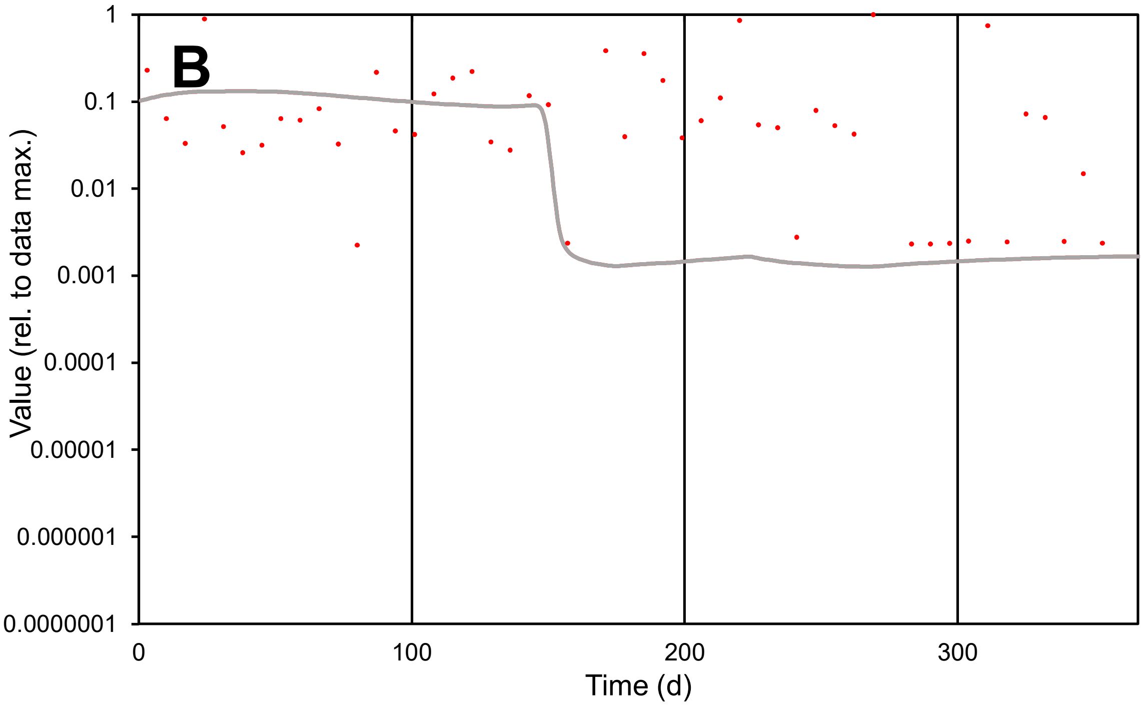


**C1**

Variables
CR2 Provide jtd or Dst
CalibOnly?
2 1 to reproduce of
oNormalize?
2 1=no, 2=yes

ofxn calc*
1.85438
ofxn mod
1.85849

sRun
RunA1
zout
1

Ave
Data
0.00251
Model
0.00088

data/vMax
0.01781

ofxng
369.894

Min d/m
1.77028

ave/min d
62.495
ave/min m
38.936

328_Calib_S_A

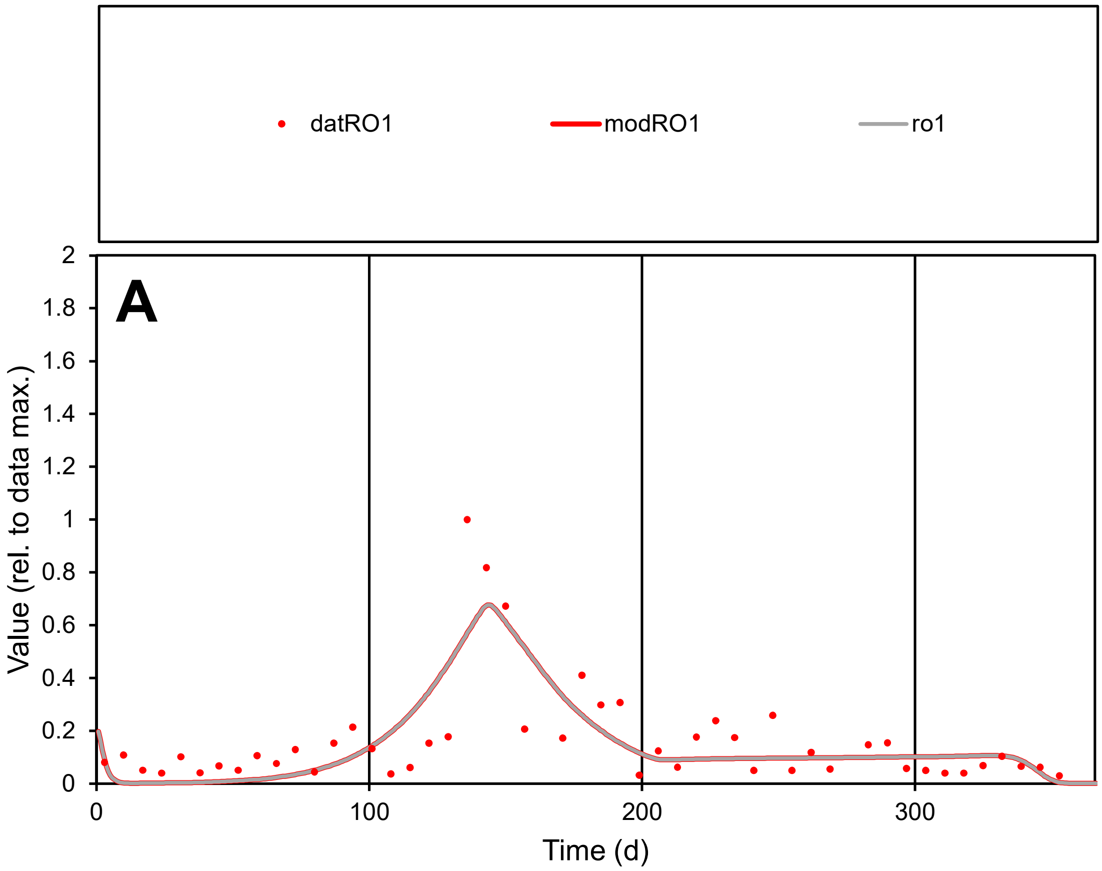


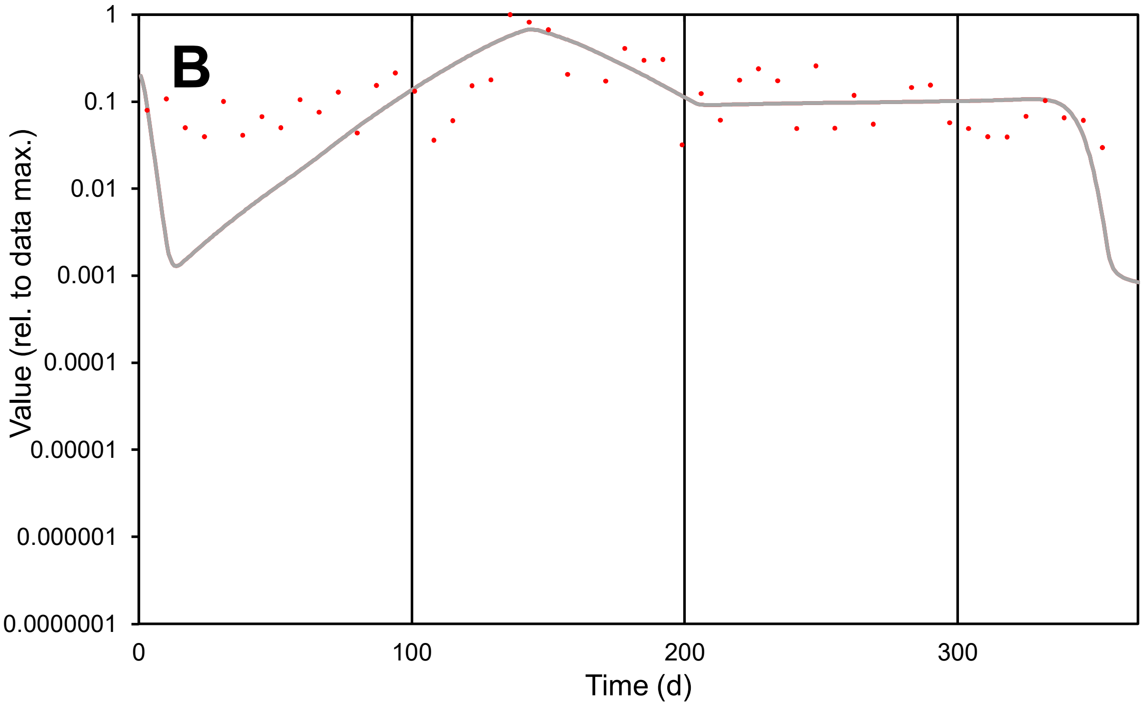


**C1**

Variables
RO1 Provide jtd or Dst
CalibOnly?
2 1 to reproduce of
oNormalize?
2 1=no, 2=yes

ofxn calc*
0.74808
ofxn mod
0.76656

sRun
RunA1
zout
1

Ave
Data
0.00427
Model
0.0039

data/vMax
0.0267

ofxng
369.894

Min d/m
18.2832

ave/min d
5.36581
ave/min m
89.4485

328_Calib_S_A

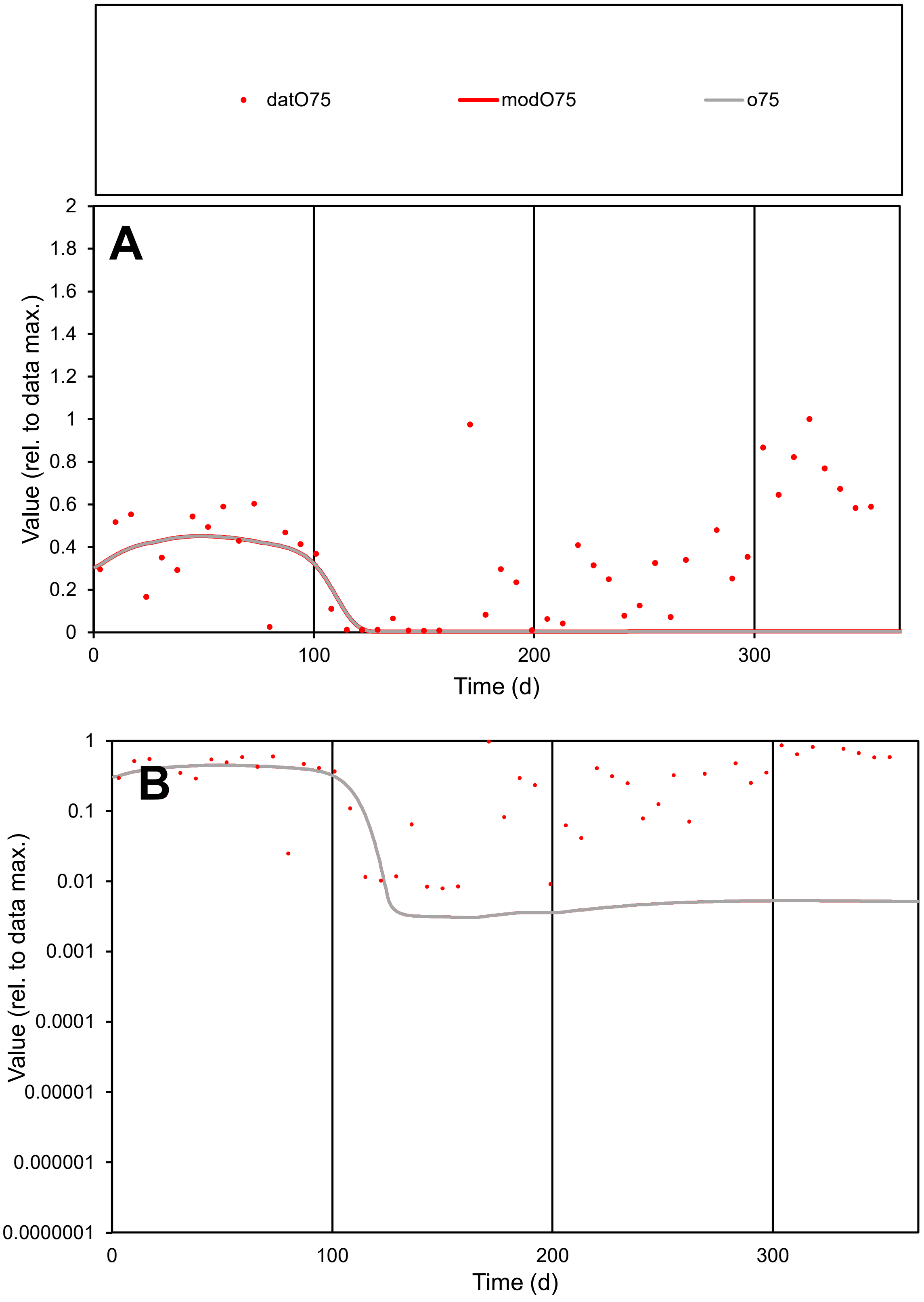


**C1**

Variables

| | |
|---|---|
| O75 | Provide jtd or Dst |
| CalibOnly? | |
| 2 | 1 to reproduce of |
| oNormalize? | |
| 2 | 1=no, 2=yes |

ofxn calc*
1.08604
ofxn mod
1.09024

sRun
RunA1
zout
1

Ave
Data
0.00305
Model
0.00116

data/vMax
0.00881

ofxng
369.894

Min d/m
2.60296

ave/min d
43.4431
ave/min m
42.9243

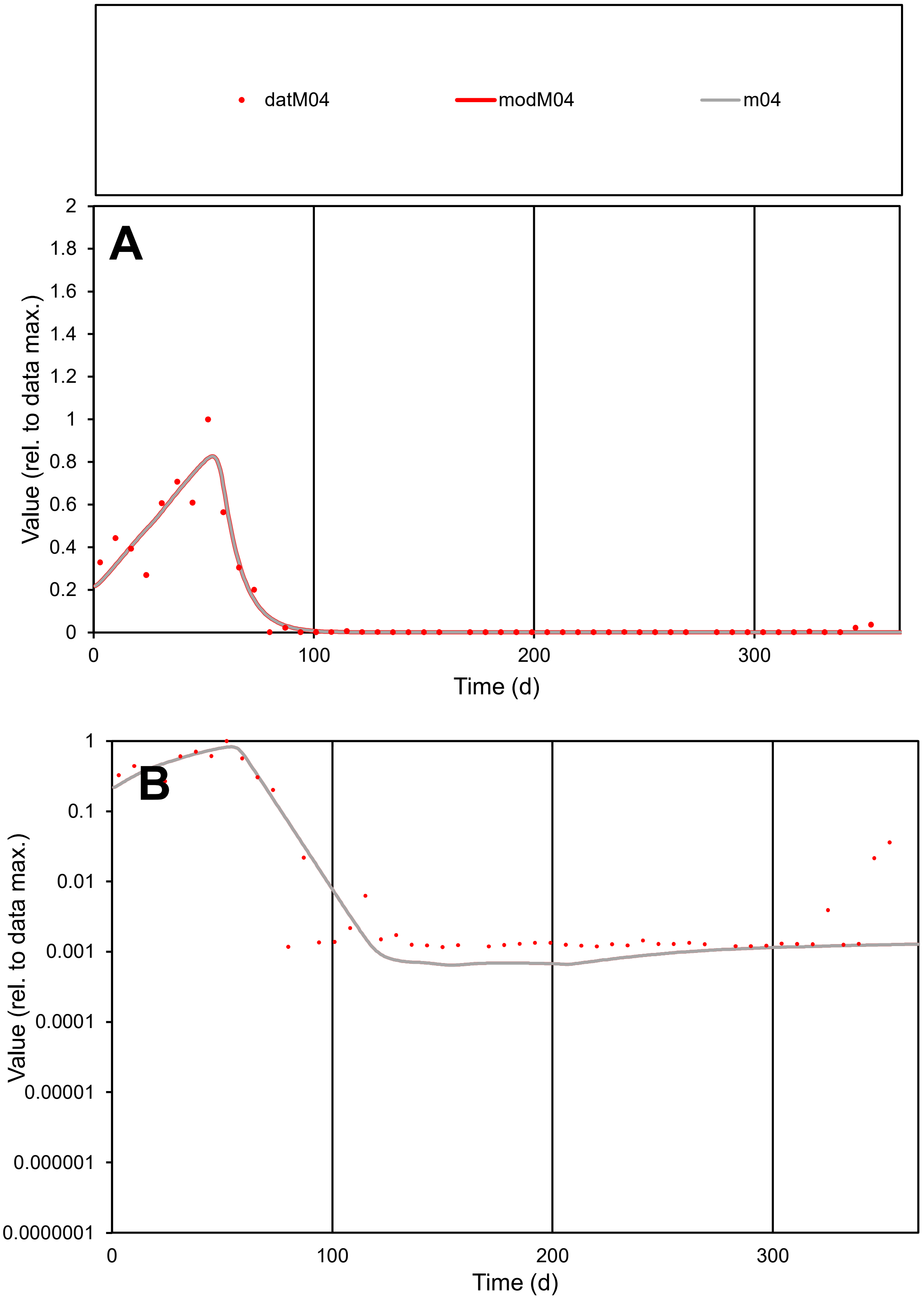


**C1**

Variables
M04 Provide jtd or Dst
CalibOnly?
2 1 to reproduce of
oNormalize?
2 1=no, 2=yes

ofxn calc*
0.45867
ofxn mod
0.52901

sRun
RunA1
zout
1

Ave
Data
0.00343
Model
0.00339

data/vMax
0.03023

ofxng
369.894

Min d/m
1.7926

ave/min d
96.9991
ave/min m
171.914

328_Calib_S_A

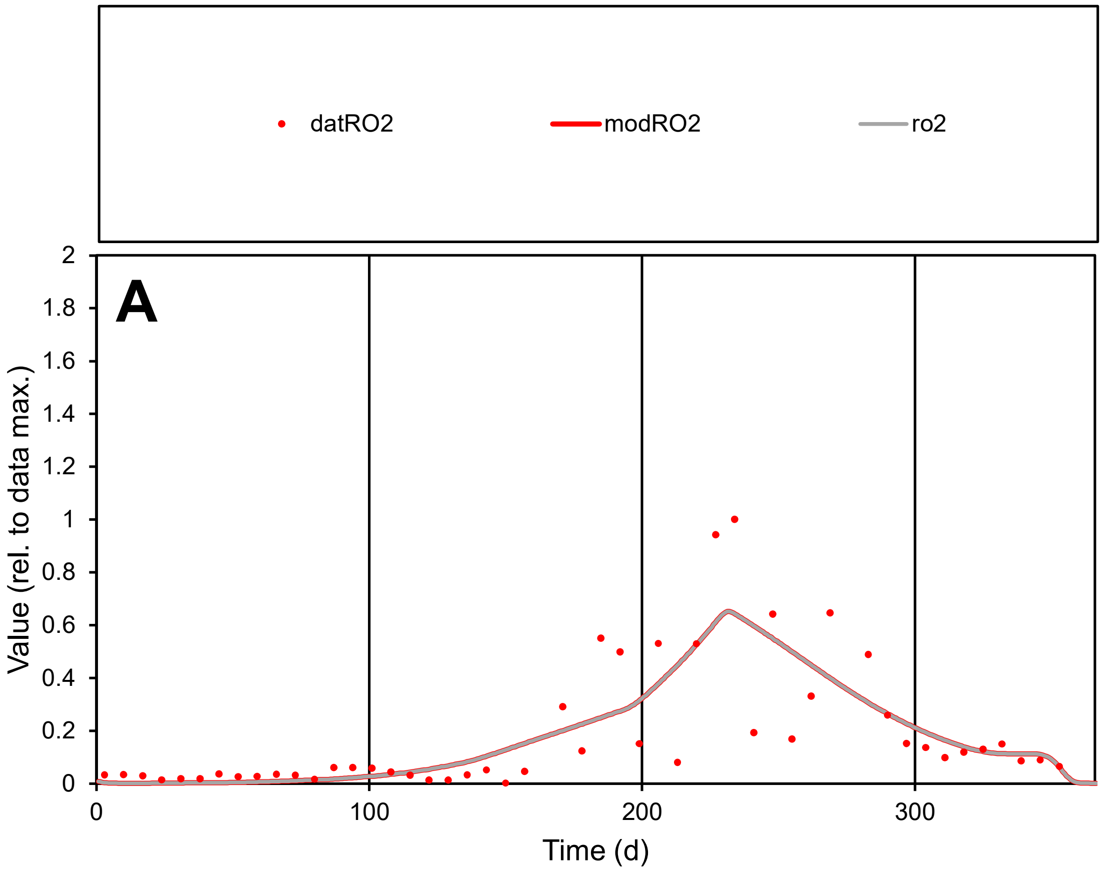


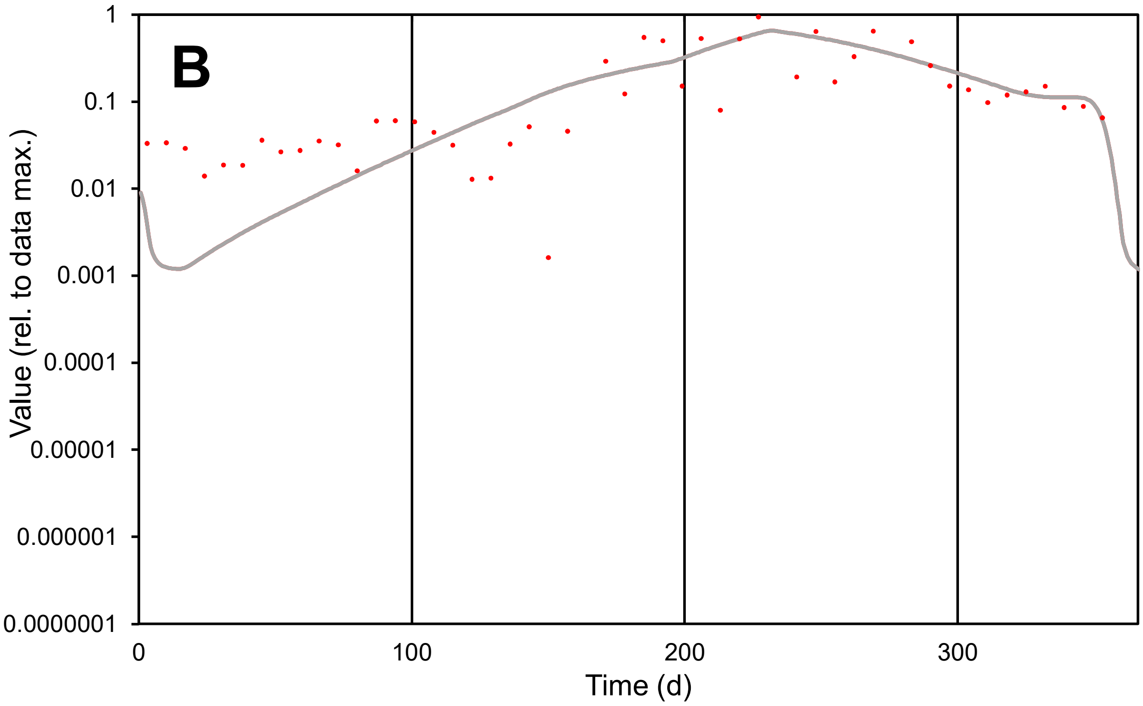


C1

Variables

| | |
|---|---|
| RO2 | Provide jtd or Dst |
| CalibOnly? | |
| 2 | 1 to reproduce of |
| oNormalize? | |
| 2 | 1=no, 2=yes |

ofxn calc*
0.76745
ofxn mod
0.79251

sRun
RunA1
zout
1

Ave
Data
0.00449
Model
0.00438

data/vMax
0.02395

ofxng
369.894

Min d/m
1.32008

ave/min d
116.495
ave/min m
150.066

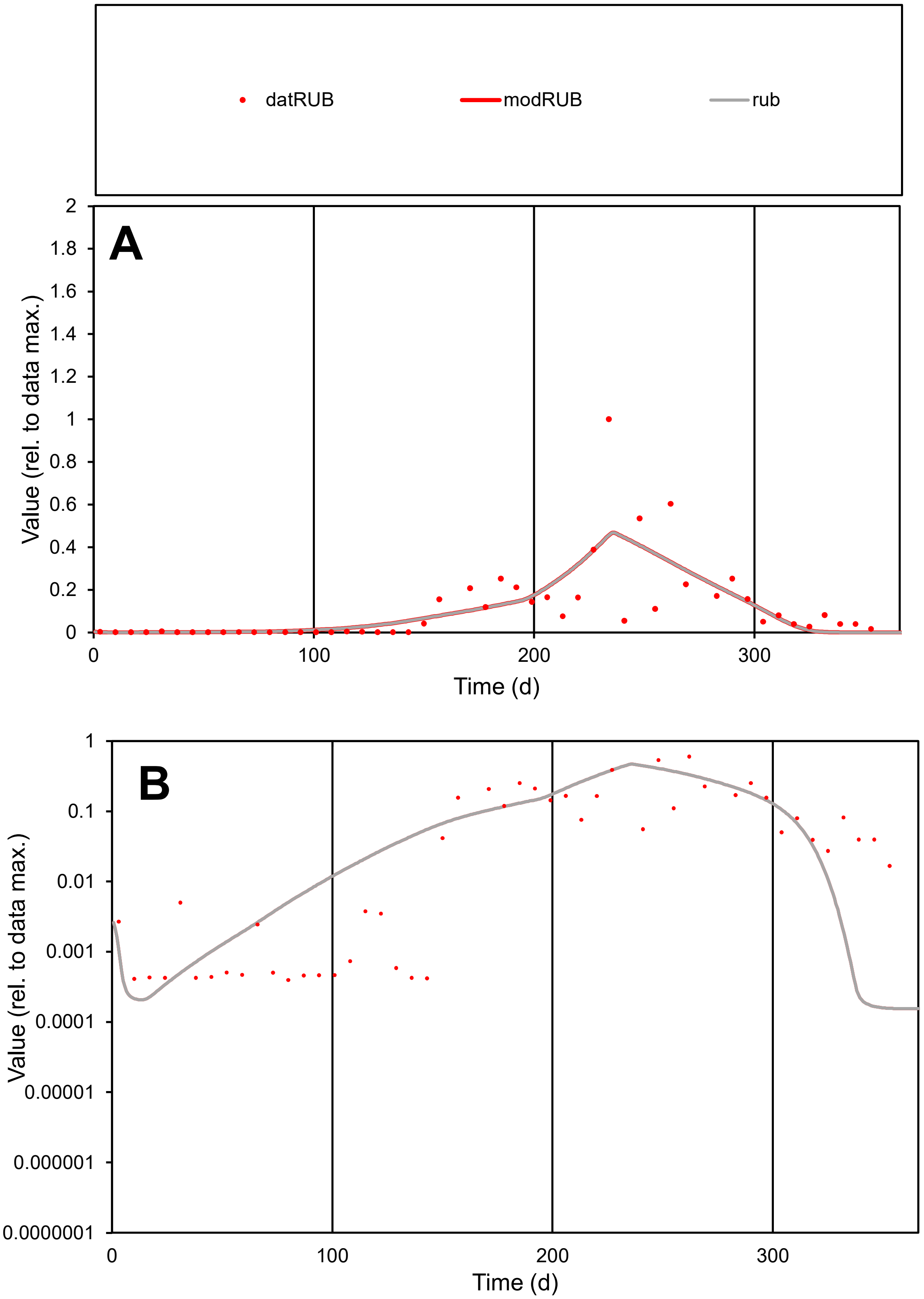


**C1**

Variables
RUB Provide jtd or Dst
CalibOnly?
2 1 to reproduce of
oNormalize?
2 1=no, 2=yes

ofxn calc*
1.09371
ofxn mod
1.11779

sRun
RunA1
zout
1

Ave
Data
0.0028
Model
0.00272

data/vMax
0.02522

ofxng
369.894

Min d/m
2.54892

ave/min d
278.723
ave/min m
691.604

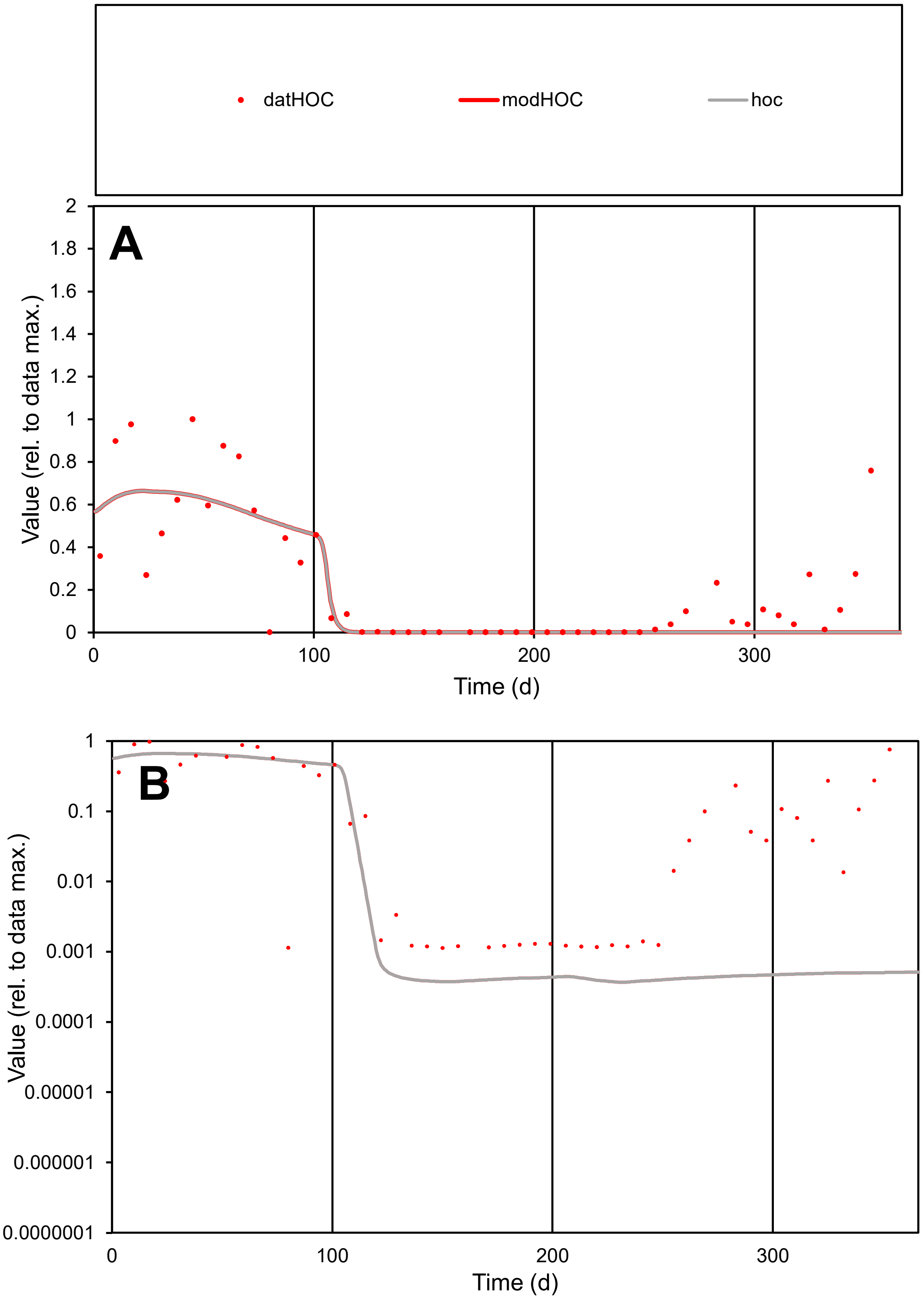


**C1**

Variables

HOC Provide jtd or Dst

CalibOnly?

2 1 to reproduce of

oNormalize?

2 1=no, 2=yes

ofxn calc*

0.85673

ofxn mod

0.87529

sRun

RunA1

zout

1

Ave

Data

0.00249

Model

0.00201

data/vMax

0.01111

ofxng

369.894

Min d/m

3.05031

ave/min d

197.569

ave/min m

487.286

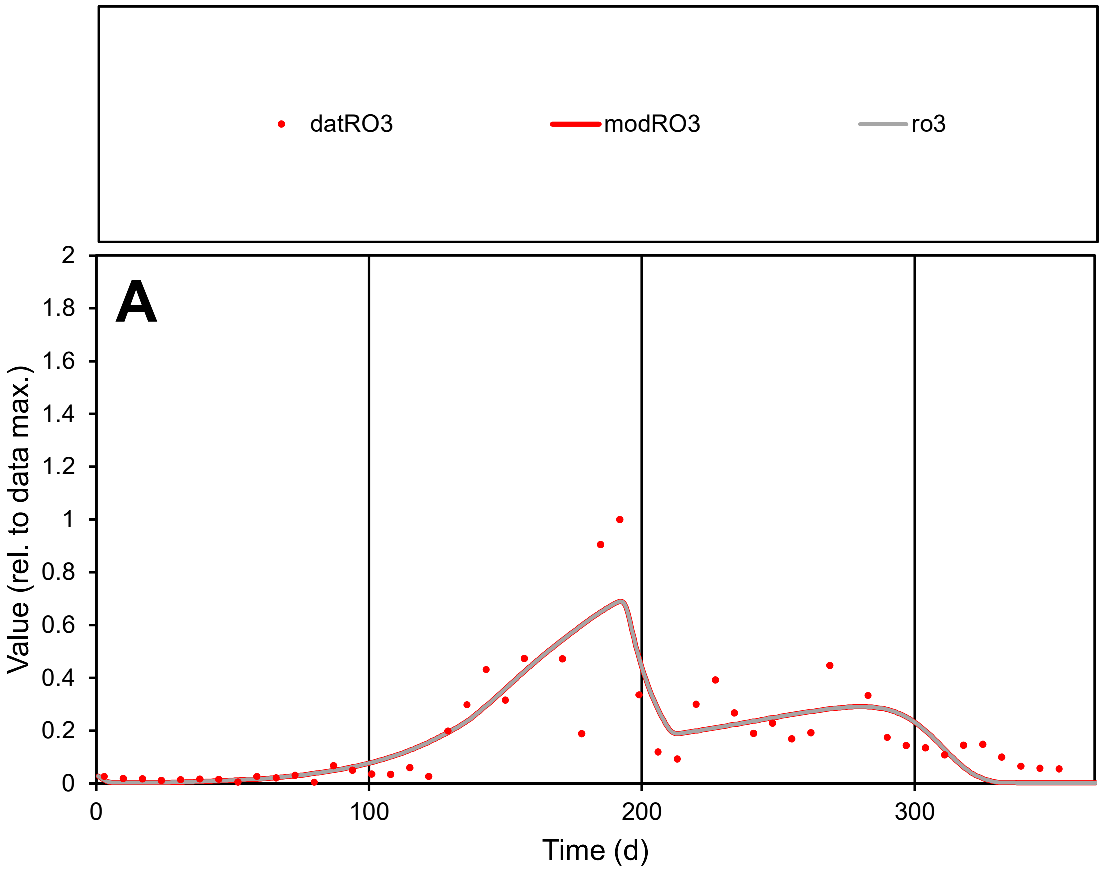


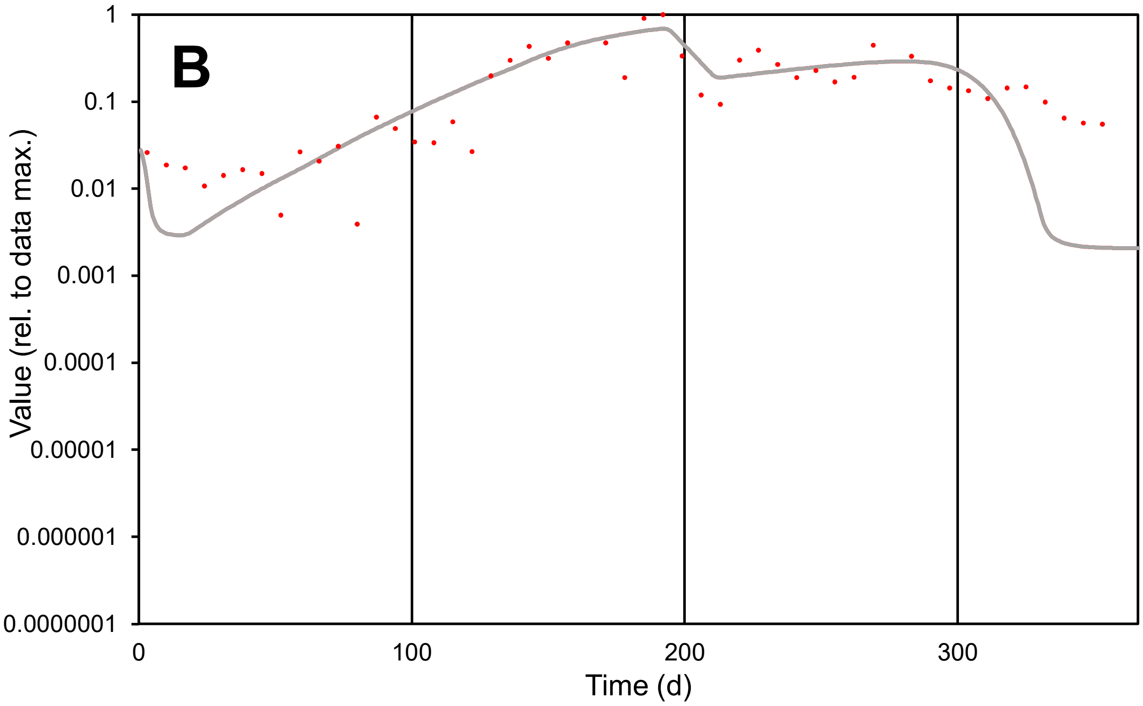


**C1**

Variables
RO3 Provide jtd or Dst
CalibOnly?
2 1 to reproduce of
oNormalize?
2 1=no, 2=yes

ofxn calc*
0.59603
ofxn mod
0.62371

sRun
RunA1
zout
1

Ave
Data
0.00452
Model
0.00446

data/vMax
0.02475

ofxng
369.894

Min d/m
1.87362

ave/min d
46.6717
ave/min m
86.3866

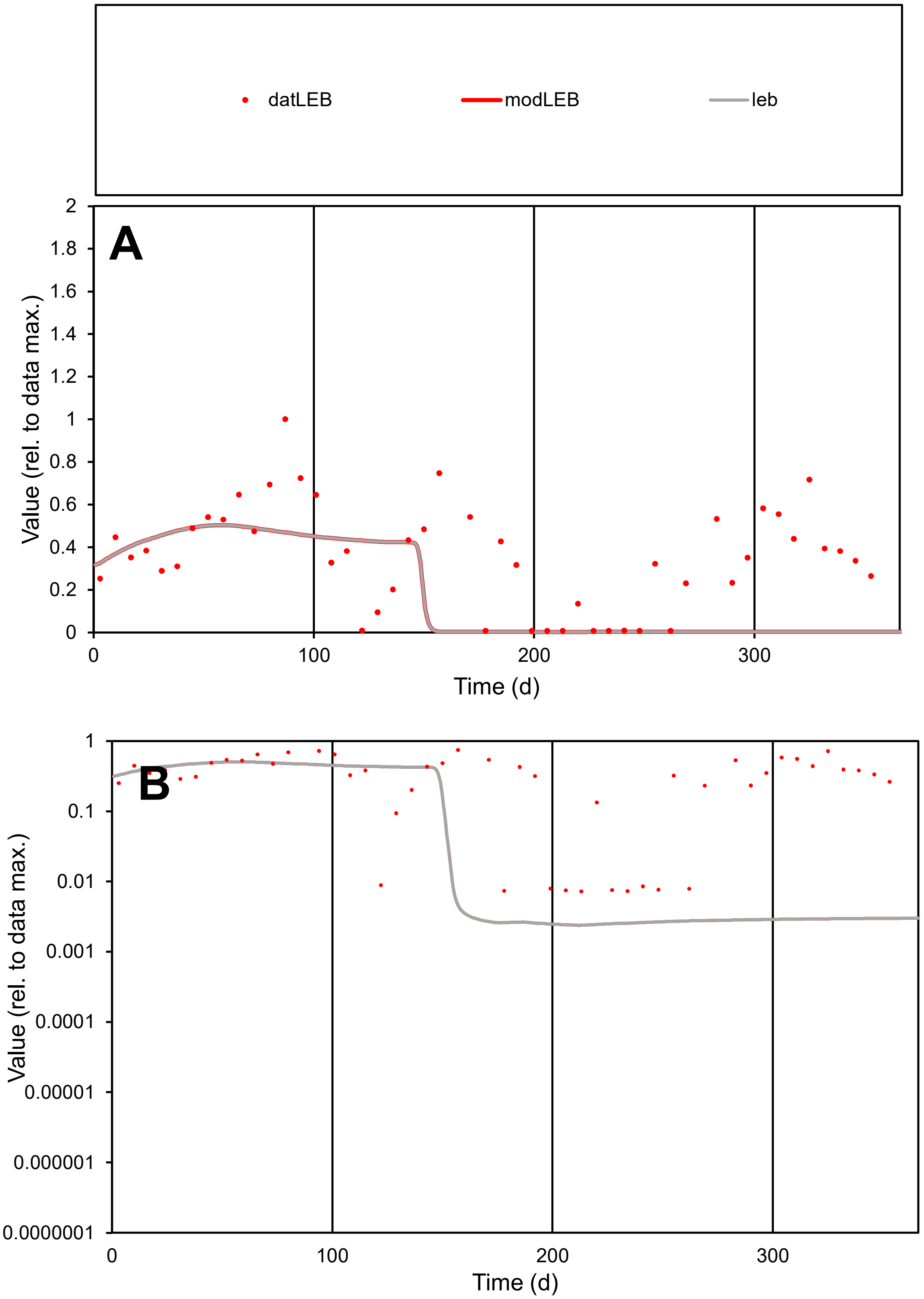


**C1**

Variables
LEB Provide jtd or Dst
CalibOnly?
2 1 to reproduce of
oNormalize?
2 1=no, 2=yes

ofxn calc*
0.8601
ofxn mod
0.86834

sRun
RunA1
zout
1

Ave
Data
0.00356
Model
0.00201

data/vMax
0.01011

ofxng
369.894

Min d/m
3.01195

ave/min d
48.762
ave/min m
82.9018

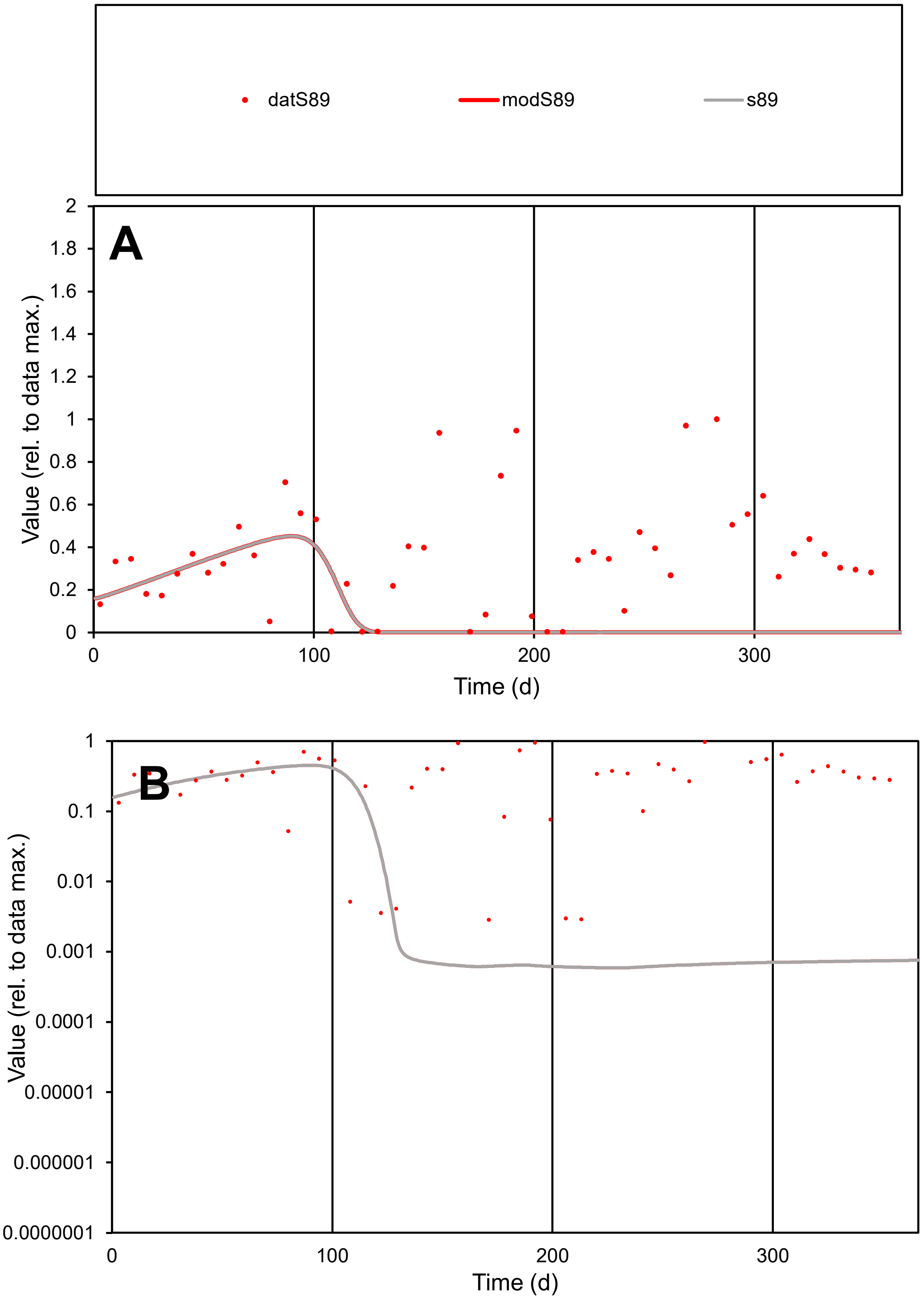


**C1**

Variables
S89 Provide jtd or Dst
CalibOnly?
2 1 to reproduce of
oNormalize?
2 1=no, 2=yes

ofxn calc*
1.10613
ofxn mod
1.11037

sRun
RunA1
zout
1

Ave
Data
0.00463
Model
0.00141

data/vMax
0.013

ofxng
369.894

Min d/m
4.8138

ave/min d
125.146
ave/min m
184.014

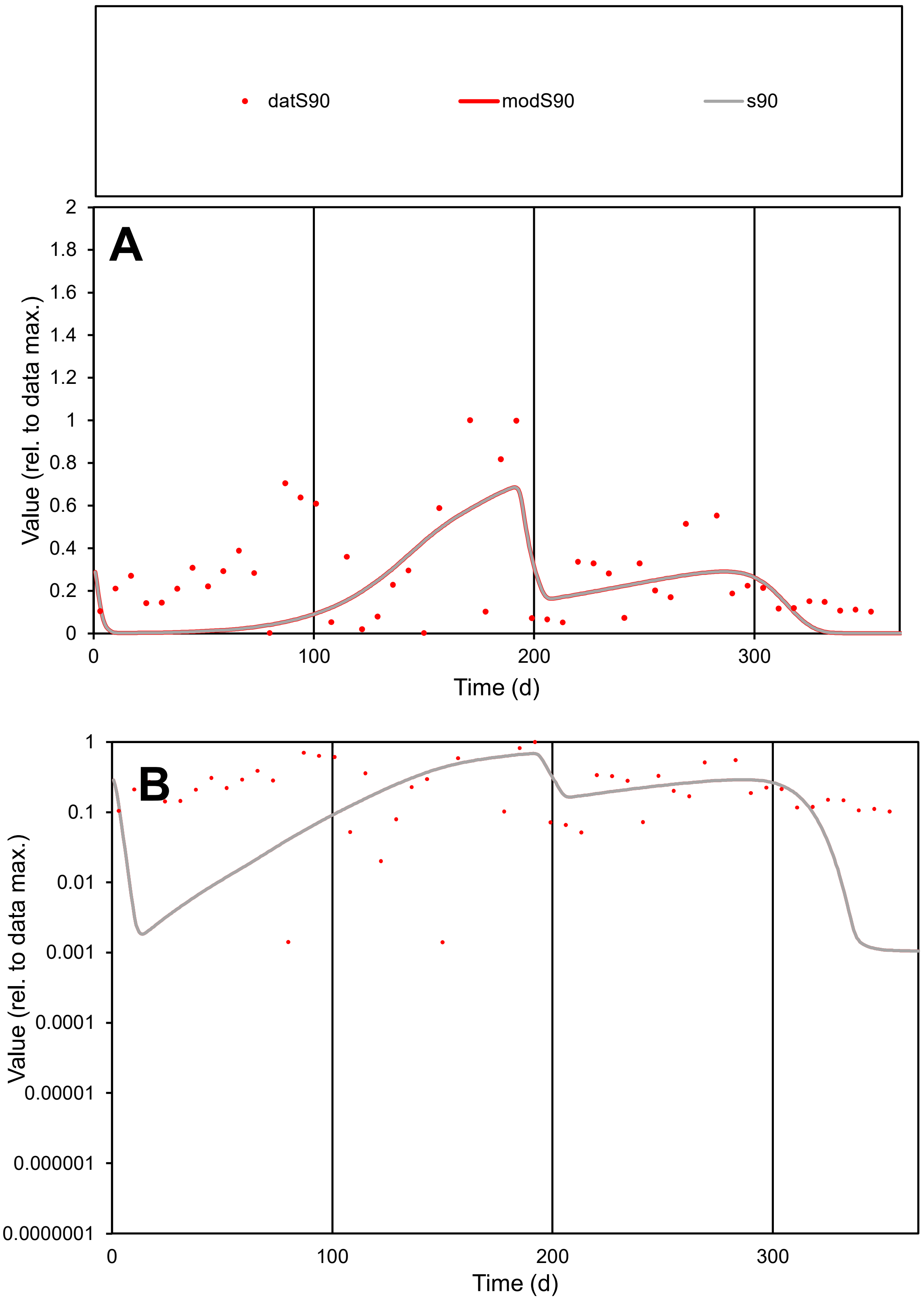


**C1**

Variables
S90 Provide jtd or Dst
CalibOnly?
2 1 to reproduce of
oNormalize?
2 1=no, 2=yes

ofxn calc*
0.87166
ofxn mod
0.88329

sRun
RunA1
zout
1

Ave
Data
0.00391
Model
0.00268

data/vMax
0.01418

ofxng
369.894

Min d/m
1.31209

ave/min d
195.889
ave/min m
176.153

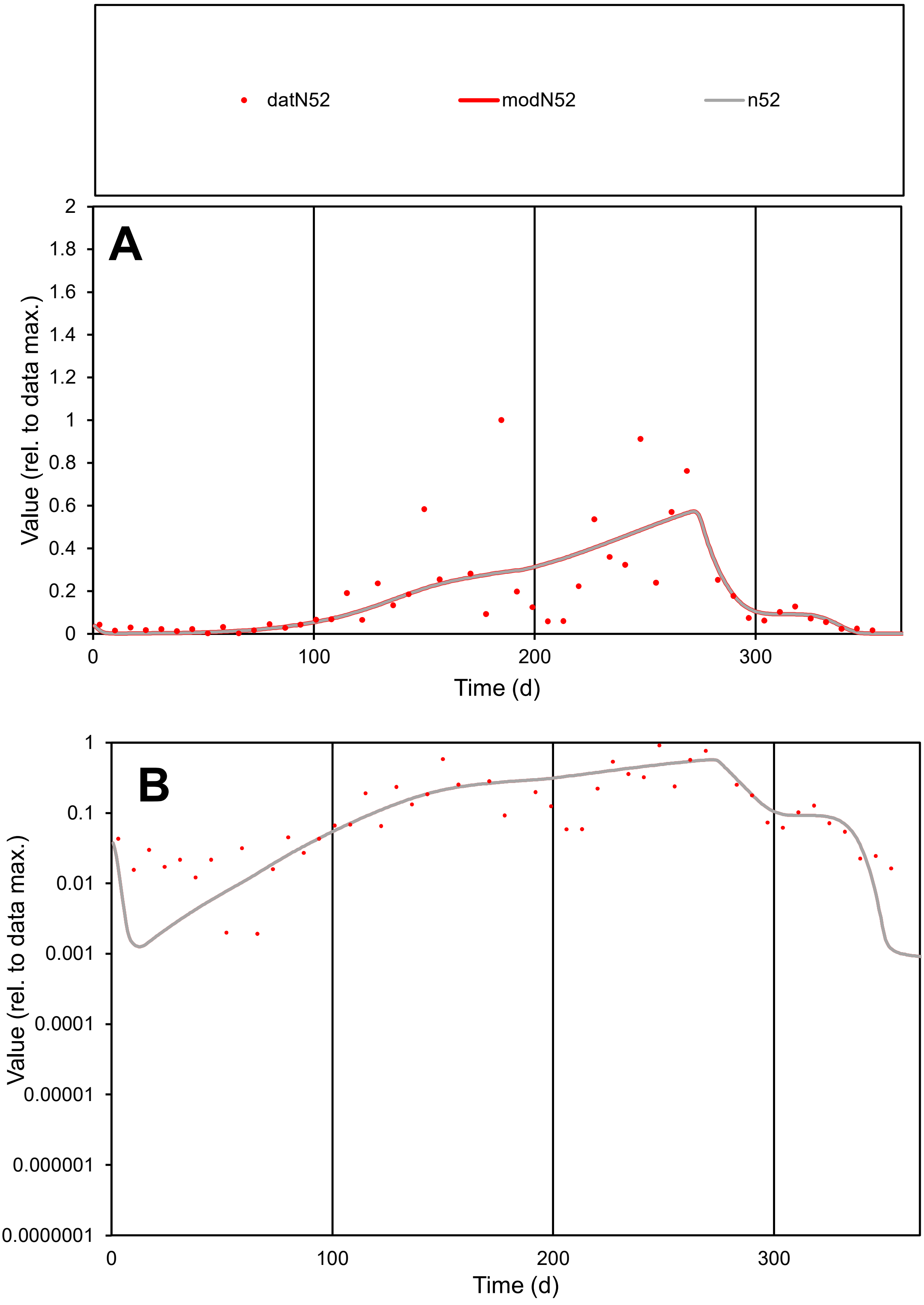


**C1**

Variables
N52 Provide jtd or Dst
CalibOnly?
2 1 to reproduce of
oNormalize?
2 1=no, 2=yes

ofxn calc*
0.89295
ofxn mod
0.91169

sRun
RunA1
zout
1

Ave
Data
0.00279
Model
0.00267

data/vMax
0.01546

ofxng
369.894

Min d/m
1.726

ave/min d
93.9187
ave/min m
155.35

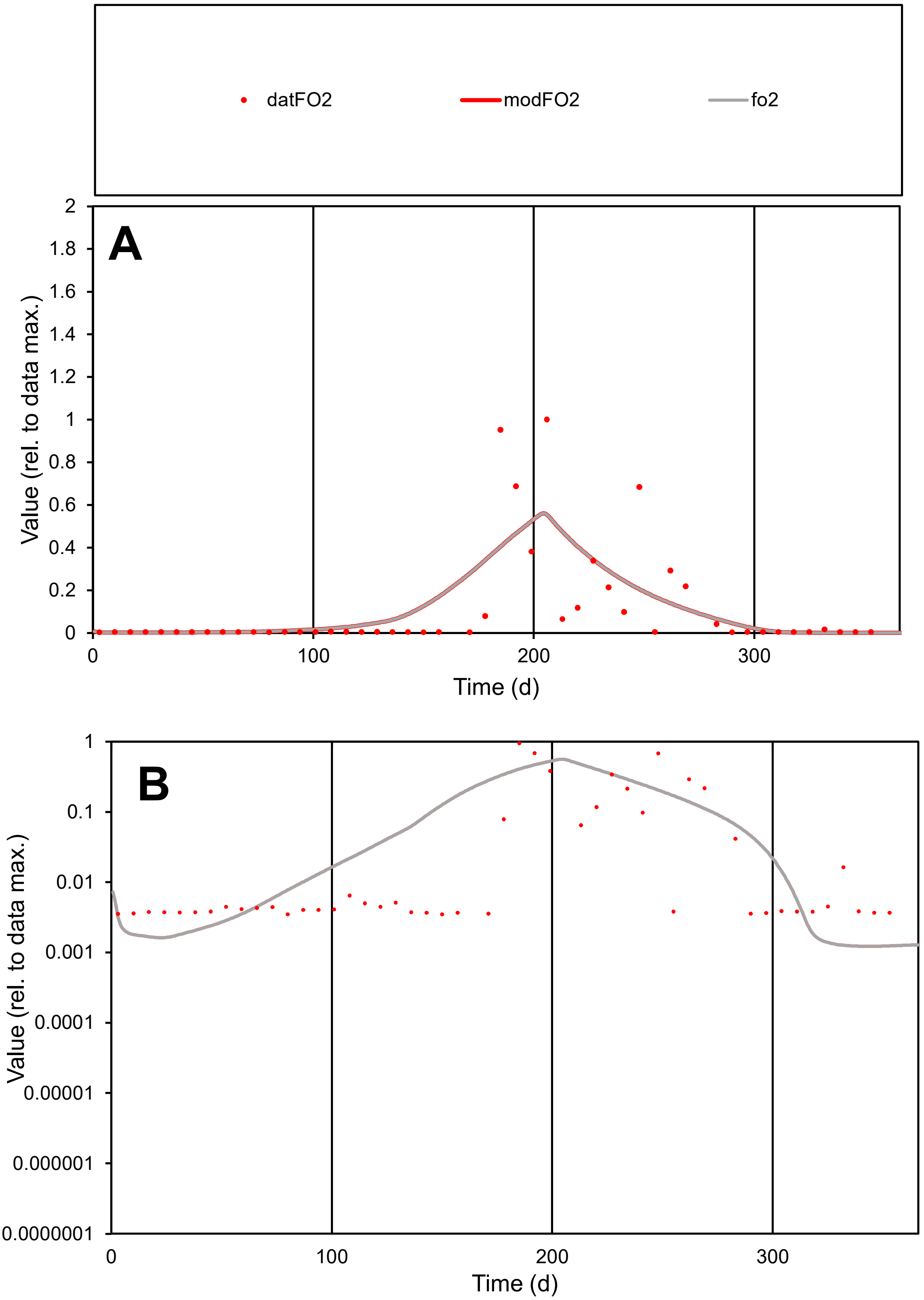


**C1**

Variables
FO2 Provide jtd or Dst
CalibOnly?
2 1 to reproduce of
oNormalize?
2 1=no, 2=yes

ofxn calc*
1.5053
ofxn mod
1.53399

sRun
RunA1
zout
1

Ave
Data
0.0021
Model
0.00224

data/vMax
0.01934

ofxng
369.894

Min d/m
2.83354
ave/min d
31.2399
ave/min m
94.5573

328_Calib_S_A

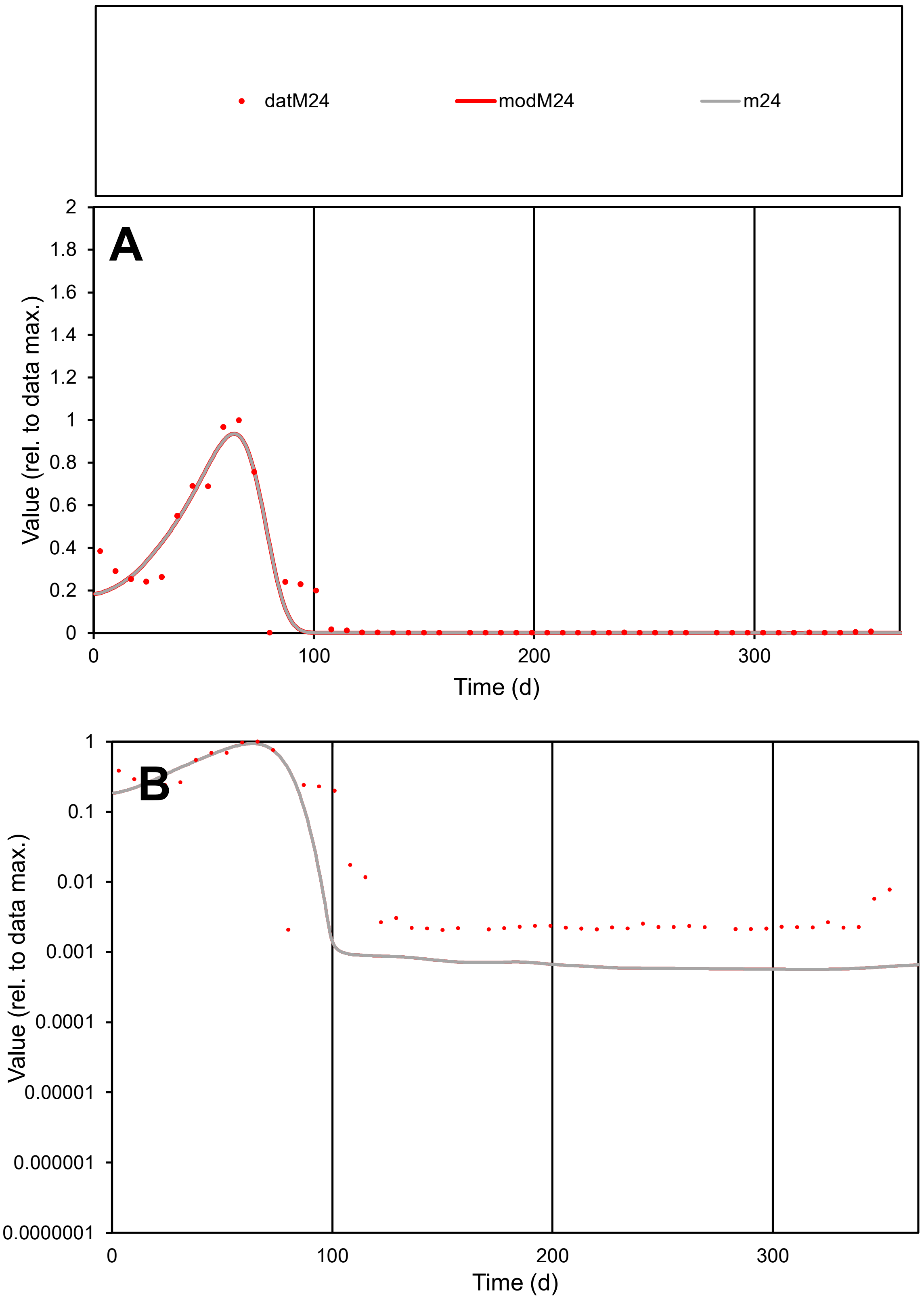


**C1**

Variables

| | |
|---|---|
| M24 | Provide jtd or Dst |

CalibOnly?

| | |
|---|---|
| 2 | 1 to reproduce of |

oNormalize?

| | |
|---|---|
| 2 | 1=no, 2=yes |

ofxn calc*
0.52575
ofxn mod
0.60347

sRun
RunA1
zout
1

Ave
Data
0.00492
Model
0.0046

data/vMax
0.03507

ofxng
369.894

Min d/m
3.63518

ave/min d
67.995
ave/min m
231.047

328_Calib_S_A

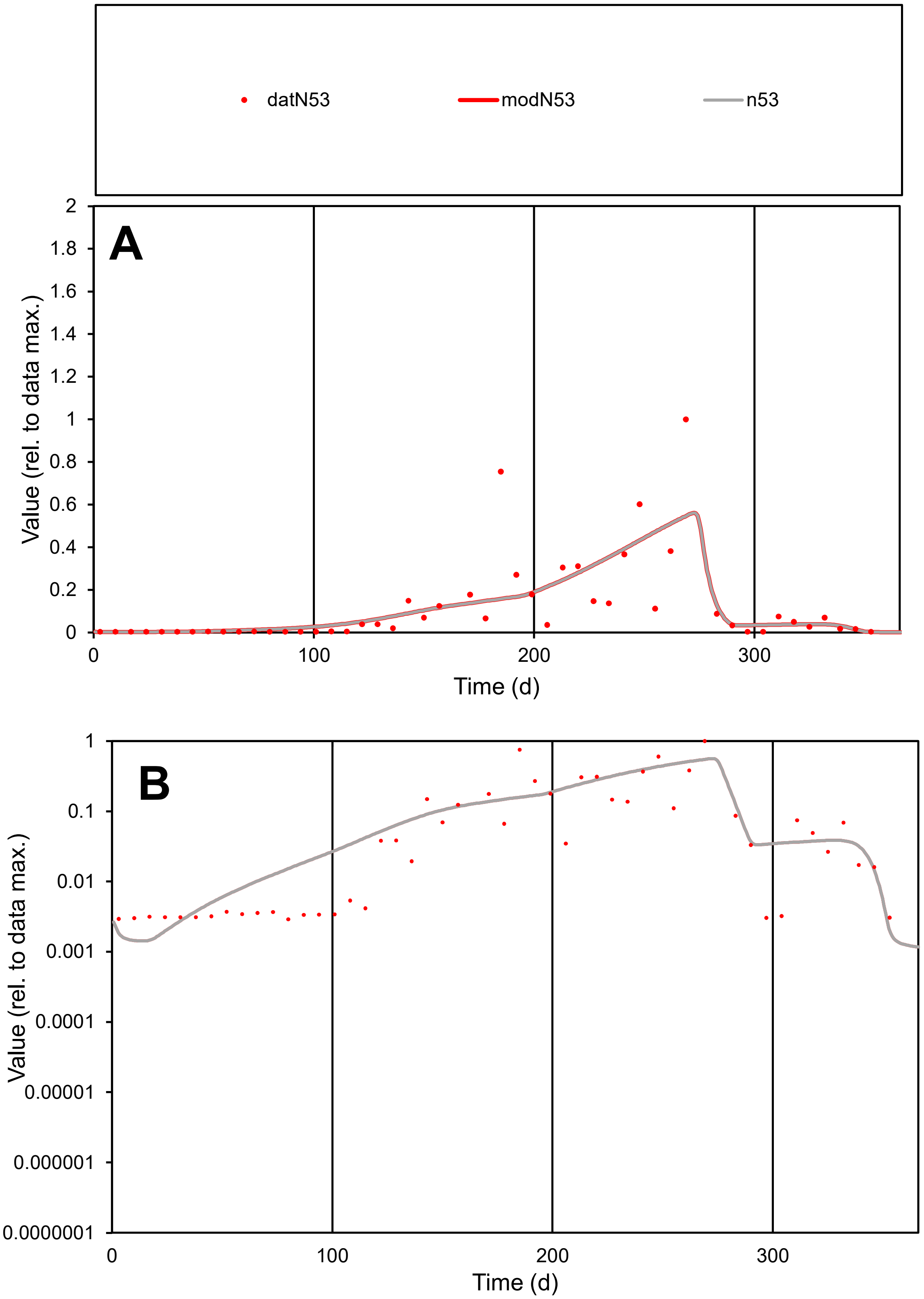


**C1**

Variables
N53 Provide jtd or Dst
CalibOnly?
2 1 to reproduce of
oNormalize?
2 1=no, 2=yes

ofxn calc*
1.14853
ofxn mod
1.1698

sRun
RunA1
zout
1

Ave
Data
0.00148
Model
0.00149

data/vMax
0.01265

ofxng
369.894

Min d/m
2.02978

ave/min d
40.1104
ave/min m
82.0572

328_Calib_S_A

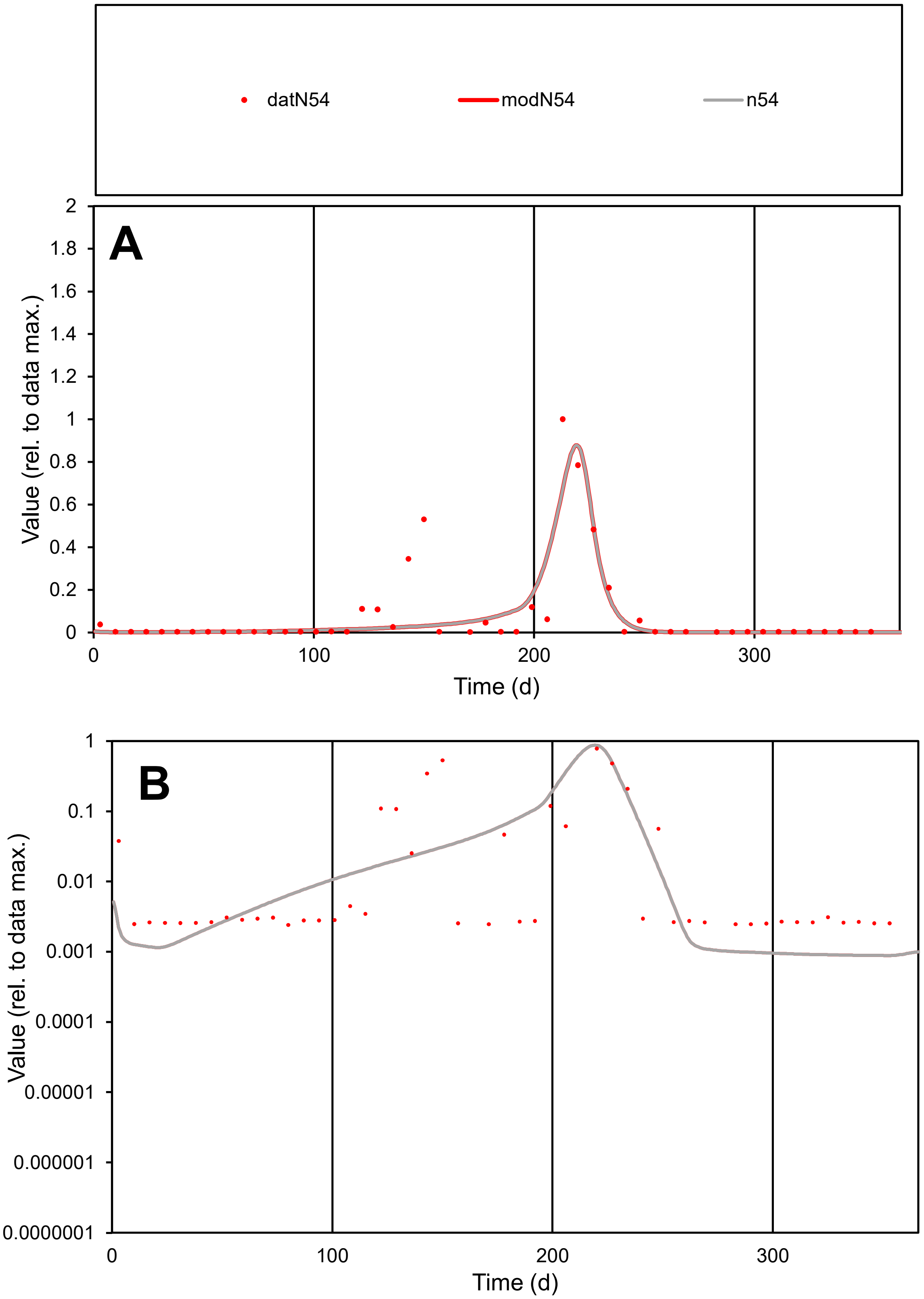


**C1**

Variables

| | |
|---|---|
| N54 | Provide jtd or Dst |
| CalibOnly? | |
| 2 | 1 to reproduce of |
| oNormalize? | |
| 2 | 1=no, 2=yes |

ofxn calc*
1.27876
ofxn mod
1.32385

sRun
RunA1
zout
1

Ave
Data
0.00074
Model
0.00062

data/vMax
0.00902

ofxng
369.894

Min d/m
2.73069

ave/min d
33.9059
ave/min m
77.158

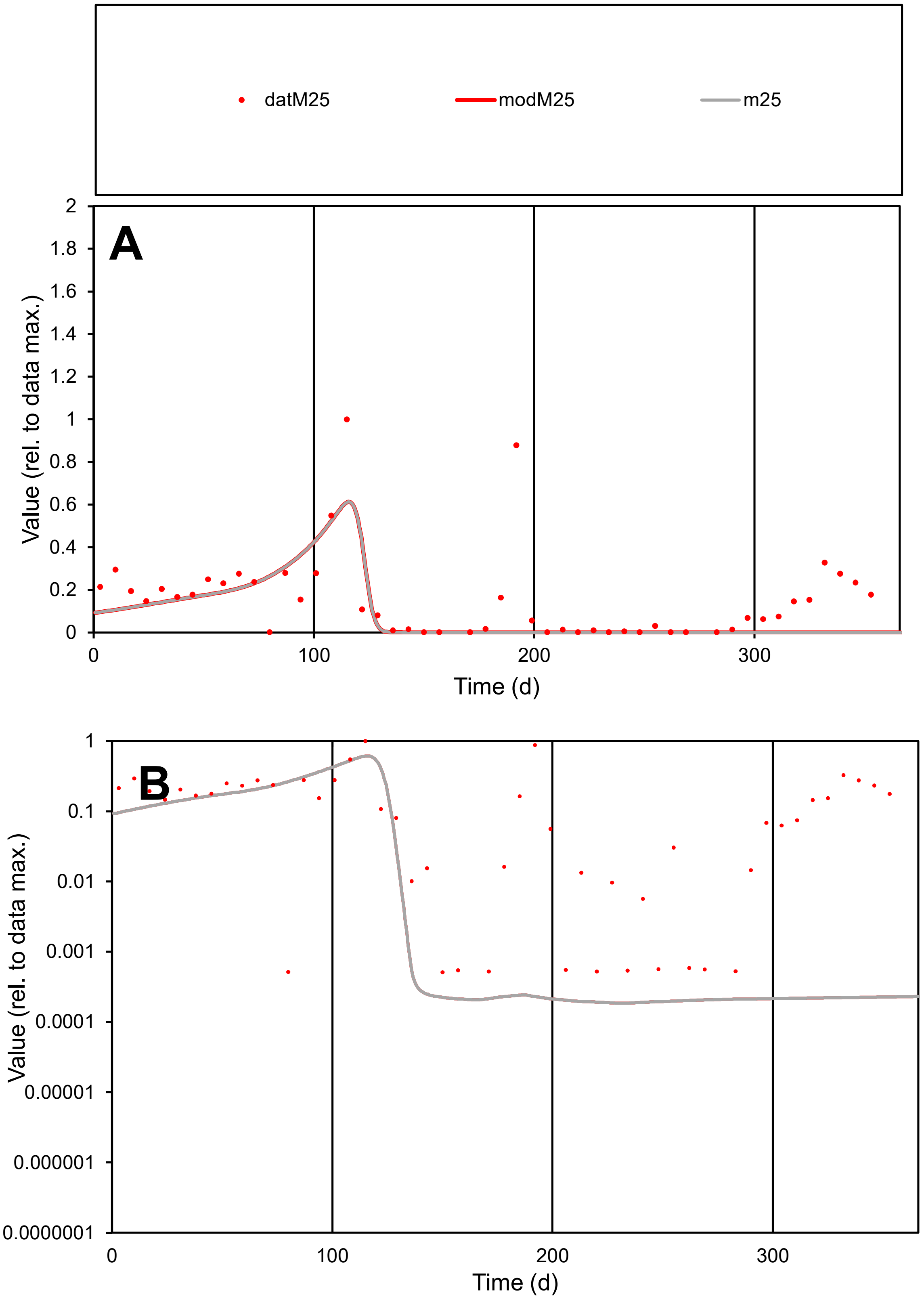


**C1**

Variables
M25 Provide jtd or Dst
CalibOnly?
2 1 to reproduce of
oNormalize?
2 1=no, 2=yes

ofxn calc*
1.13455
ofxn mod
1.15079

sRun
RunA1
zout
1

Ave
Data
0.0029
Model
0.00176

data/vMax
0.01879

ofxng
369.894

Min d/m
2.75221

ave/min d
301.554
ave/min m
502.927

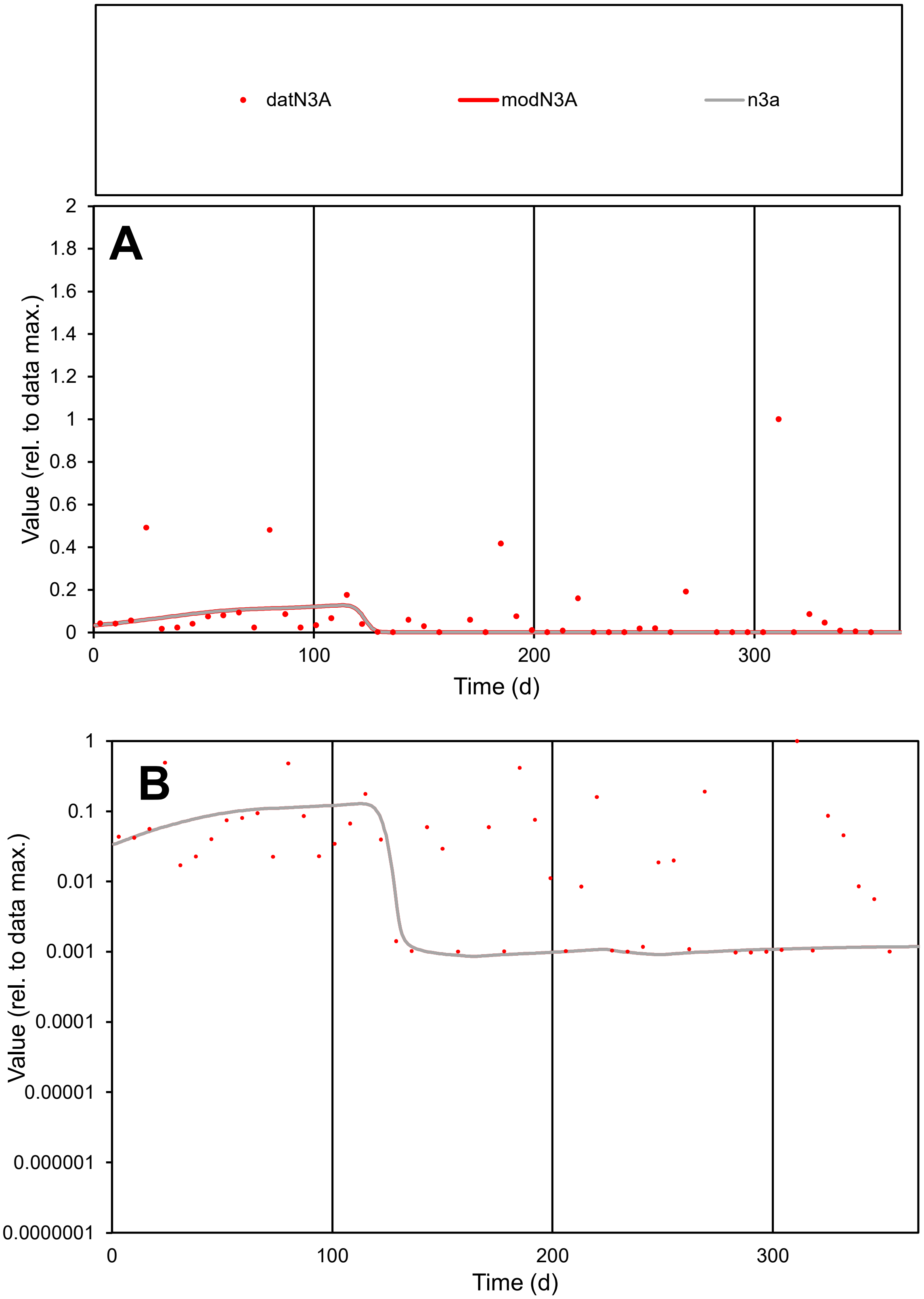


C1

Variables
N3A Provide jtd or Dst
CalibOnly?
2 1 to reproduce of
oNormalize?
2 1=no, 2=yes

ofxn calc*
2.16927
ofxn mod
2.17278

sRun
RunA1
zout
1

Ave
Data
0.00158
Model
0.00064

data/vMax
0.01883

ofxng
369.894

Min d/m
1.11339

ave/min d
85.4093
ave/min m
38.5375

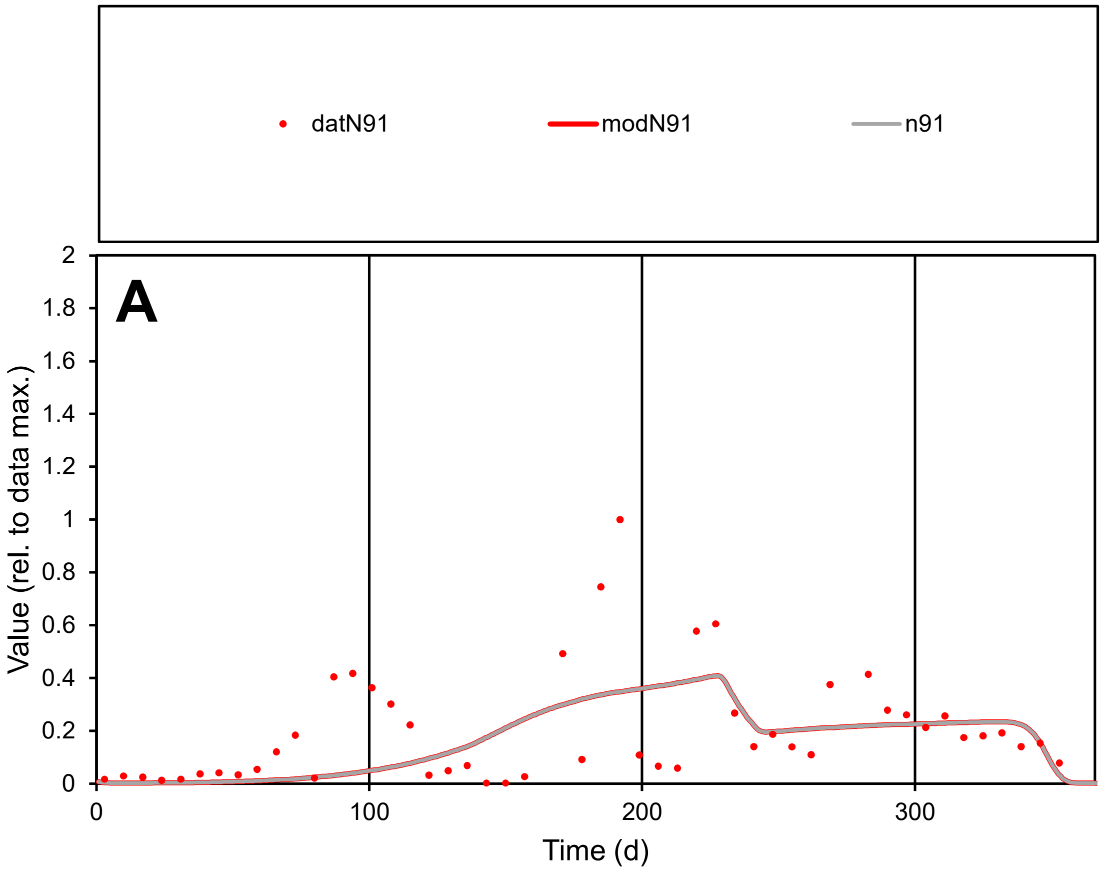


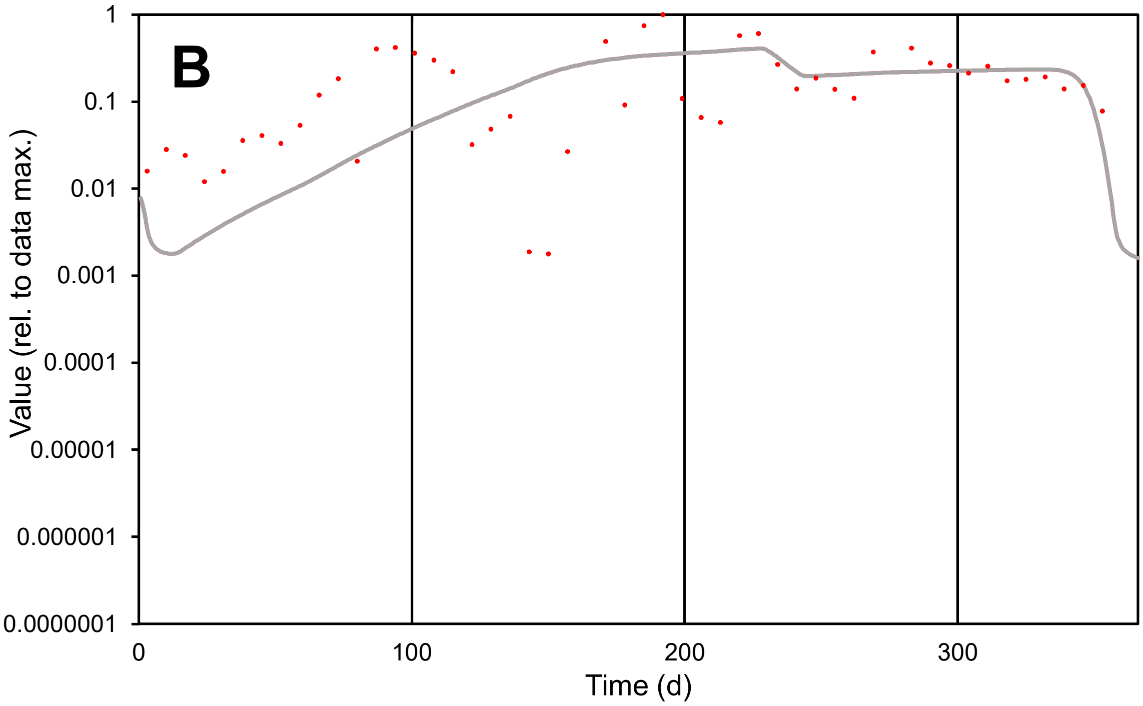


**C1**

Variables

N91 Provide jtd or Dst

CalibOnly?

2 1 to reproduce of

oNormalize?

2 1=no, 2=yes

ofxn calc*

0.94517

ofxn mod

0.9581

sRun

RunA1

zout

1

Ave

Data

0.00353

Model

0.00295

data/vMax

0.01771

ofxng

369.894

Min d/m

1.00381

ave/min d

111.617

ave/min m

93.6006

328_Calib_S_A

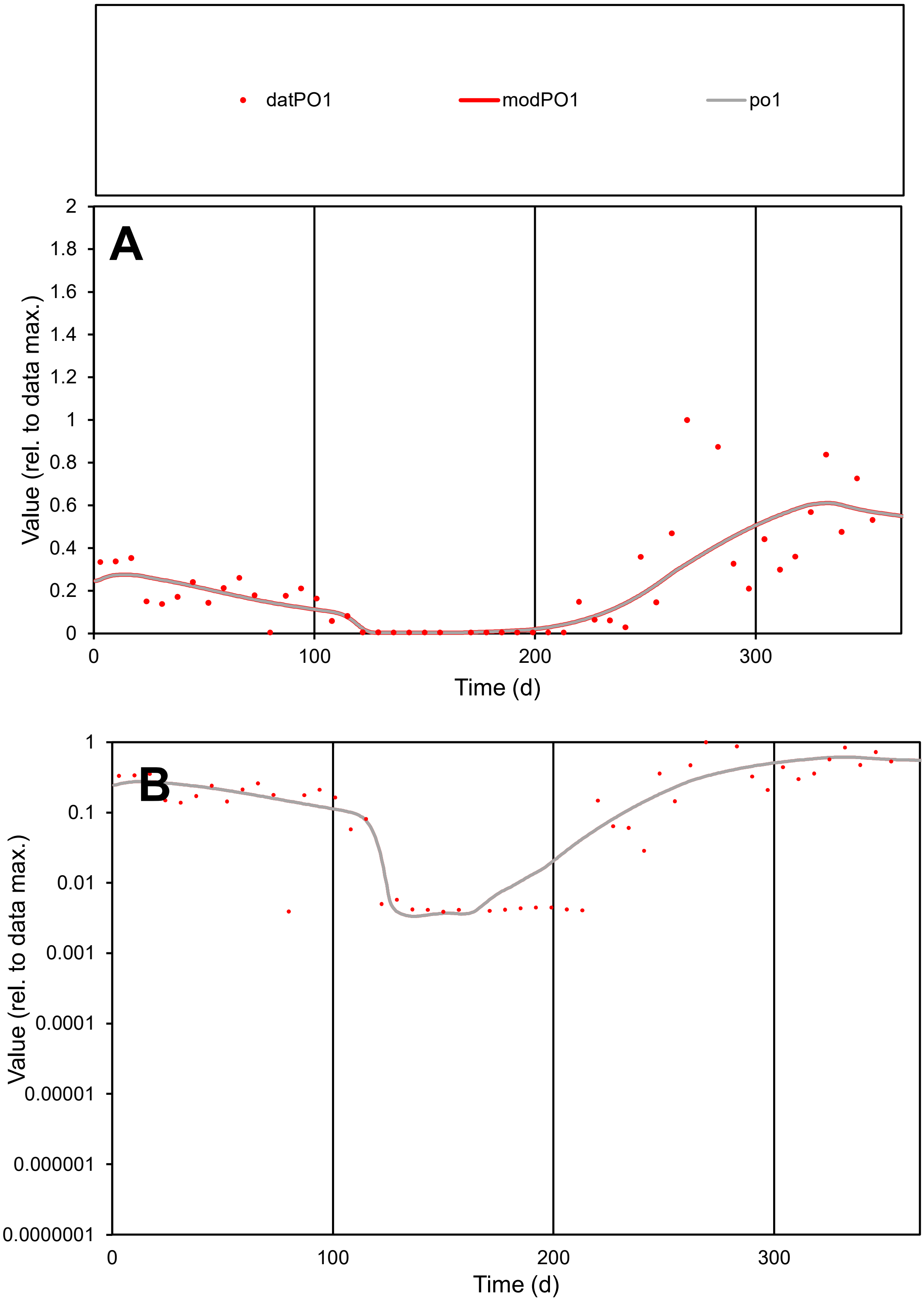


**C1**

Variables

PO1 Provide jtd or Dst

CalibOnly?

2 1 to reproduce of

oNormalize?

2 1=no, 2=yes

ofxn calc*

0.67018

ofxn mod

0.68689

sRun

RunA1

zout

1

Ave

Data

0.00267

Model

0.00258

data/vMax

0.01171

ofxng

369.894

Min d/m

1.16272

ave/min d

58.5451

ave/min m

65.664

328_Calib_S_A

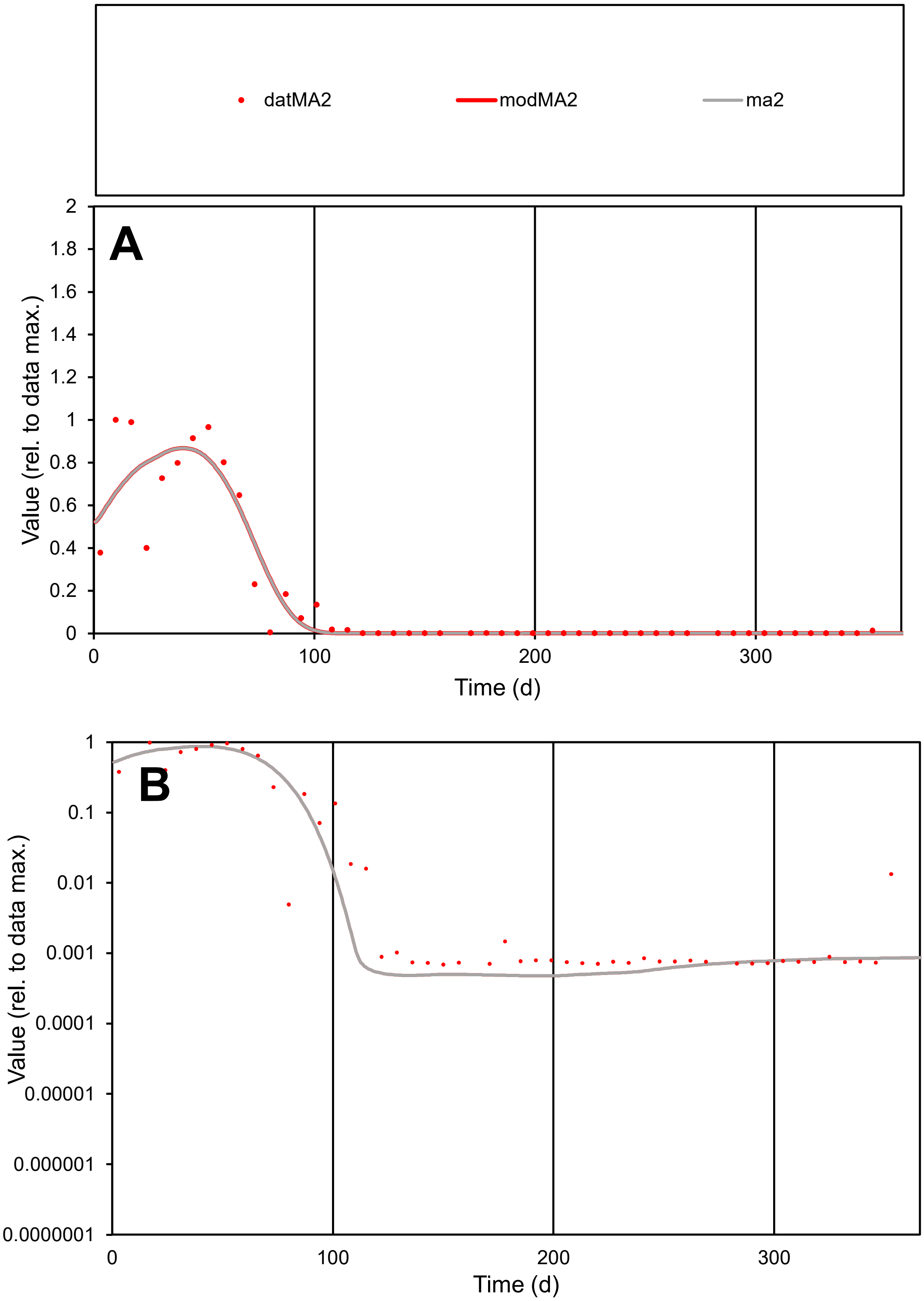


**C1**

Variables

MA2 Provide jtd or Dst

CalibOnly?

2 1 to reproduce of

oNormalize?

2 1=no, 2=yes

ofxn calc*

0.59511

ofxn mod

0.65115

sRun

RunA1

zout

1

Ave

Data

0.00439

Model

0.00433

data/vMax

0.02587

ofxng

369.894

Min d/m

1.44498

ave/min d

245.929

ave/min m

350.41

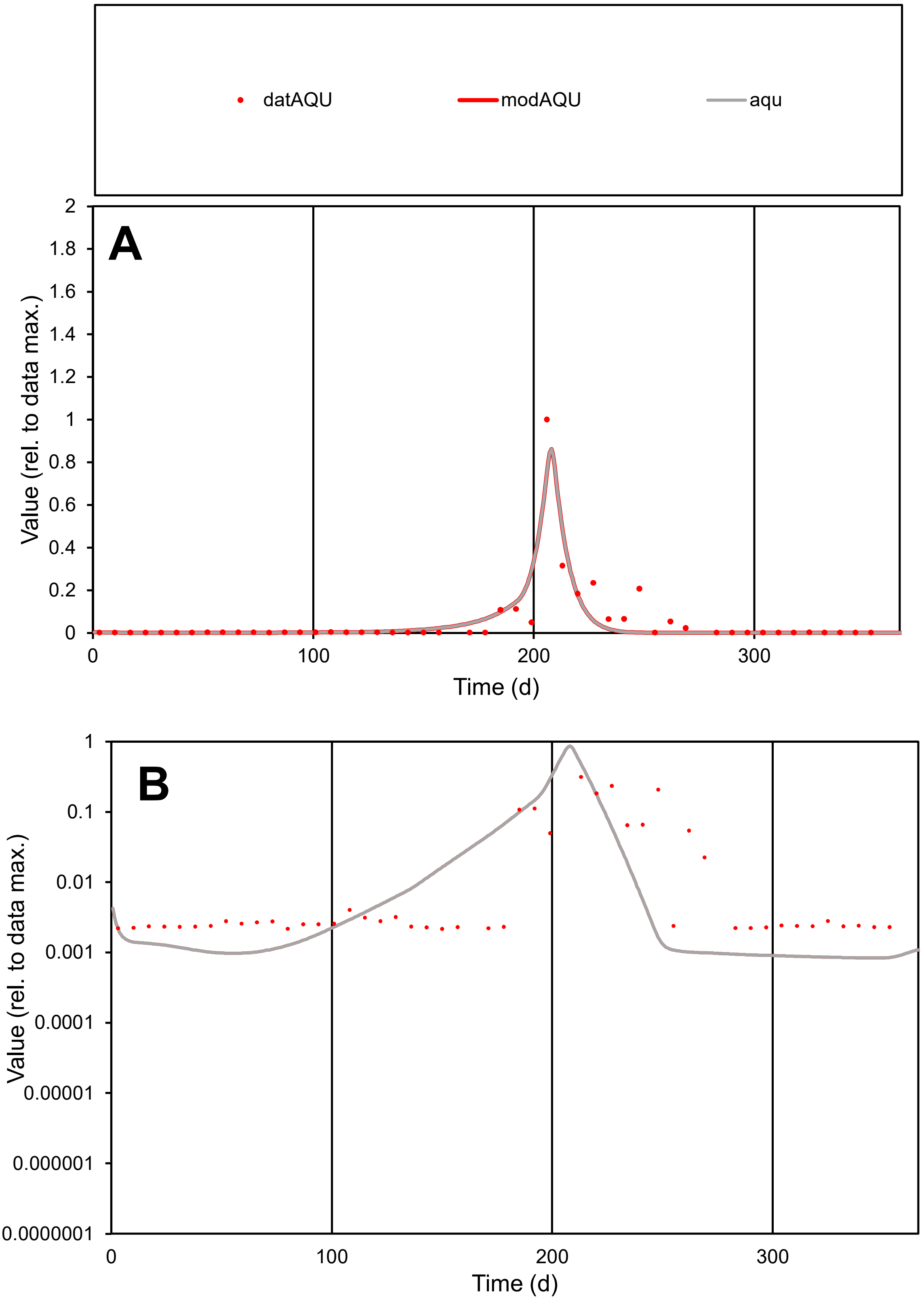


**C1**

Variables
AQU Provide jtd or Dst
CalibOnly?
2 1 to reproduce of
oNormalize?
2 1=no, 2=yes

ofxn calc*
1.07695
ofxn mod
1.21206

sRun
RunA1
zout
1

Ave
Data
0.00308
Model
0.0027

data/vMax
0.06009

ofxng
369.894

Min d/m
2.58679

ave/min d
23.6065
ave/min m
53.493

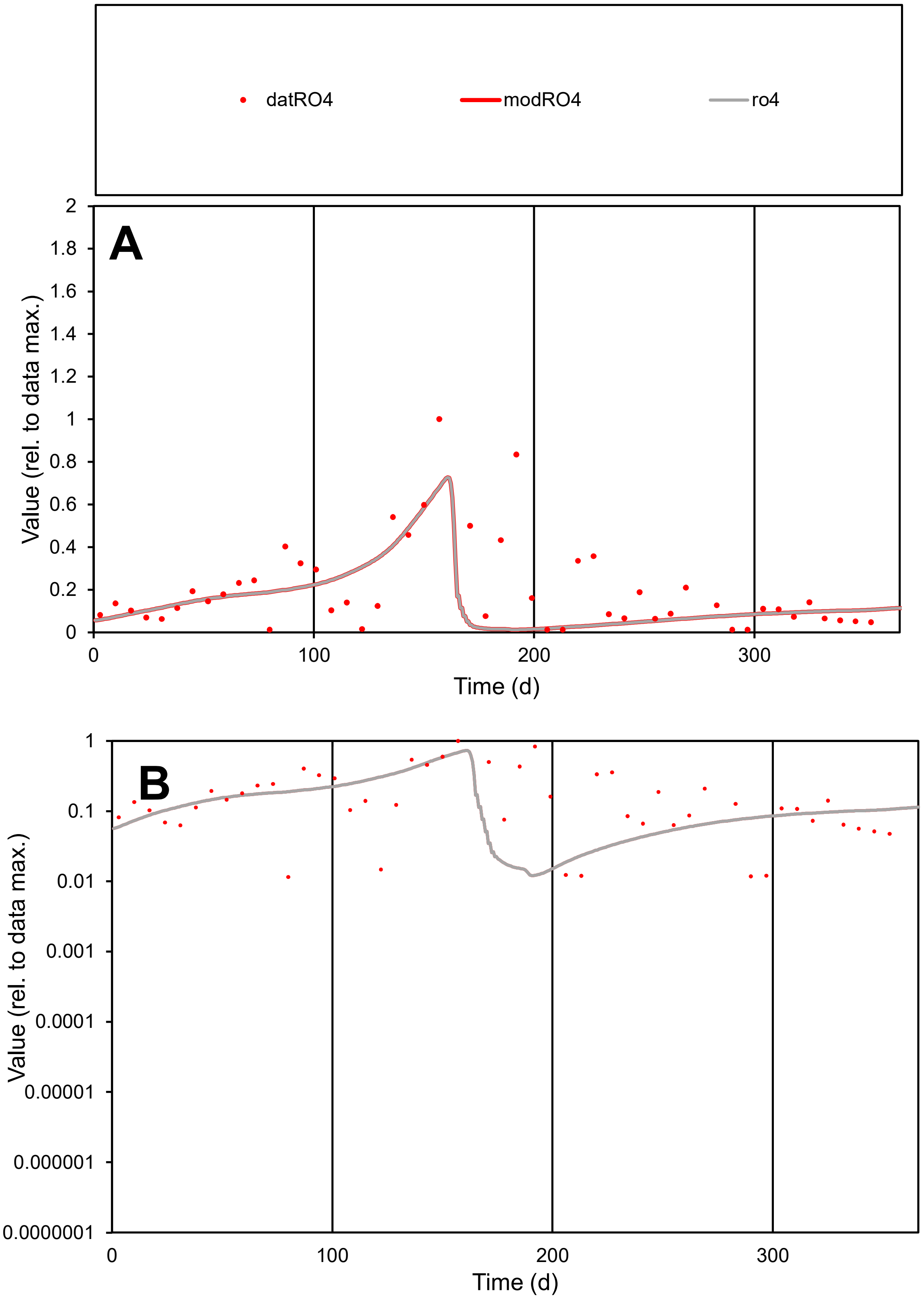


**C1**

Variables
RO4 Provide jtd or Dst
CalibOnly?
2 1 to reproduce of
oNormalize?
2 1=no, 2=yes

ofxn calc*
0.94443
ofxn mod
0.95404

sRun
RunA1
zout
1

Ave
Data
0.00378
Model
0.00279

data/vMax
0.01894

ofxng
369.894

Min d/m
0.93704

ave/min d
17.285
ave/min m
11.9428

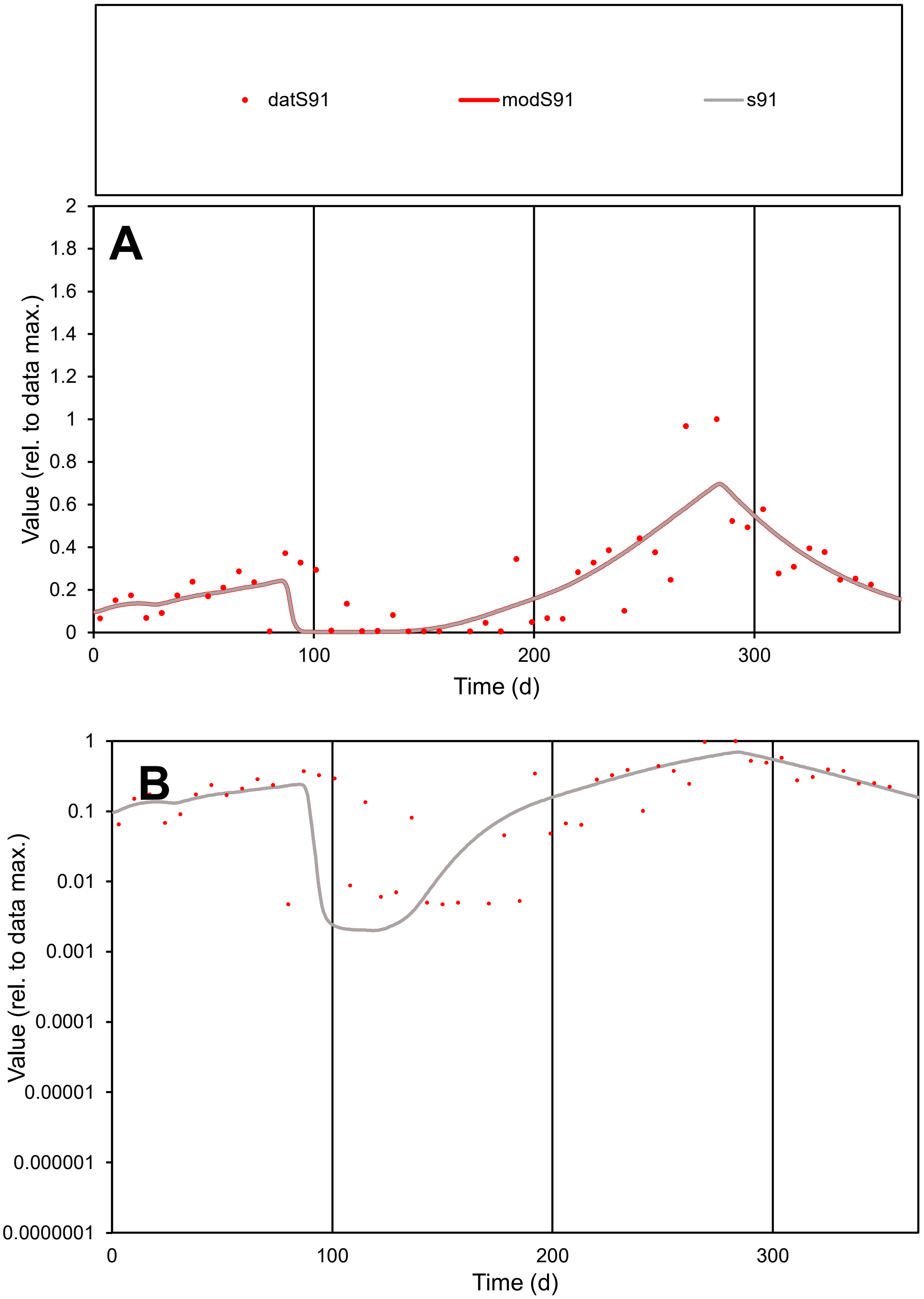


**C1**

Variables

S91 Provide jtd or Dst

CalibOnly?

2 1 to reproduce of

oNormalize?

2 1=no, 2=yes

ofxn calc*

0.5738

ofxn mod

0.59504

sRun

RunA1

zout

1

Ave

Data

0.00325

Model

0.00313

data/vMax

0.01384

ofxng

369.894

Min d/m

2.33999

ave/min d

49.5202

ave/min m

111.841

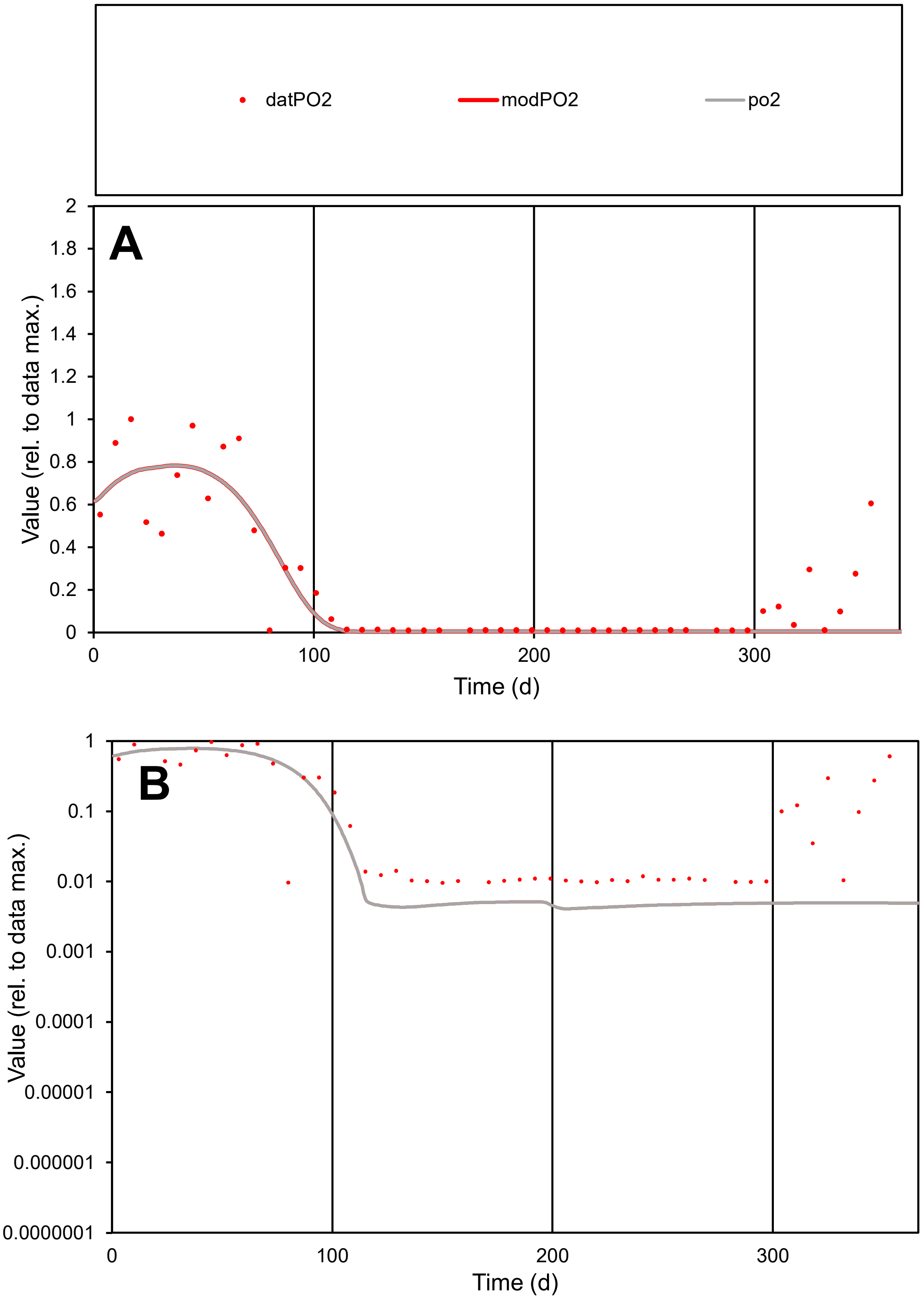


**C1**

Variables

PO2 Provide jtd or Dst

CalibOnly?

2 1 to reproduce of

oNormalize?

2 1=no, 2=yes

ofxn calc*

0.70035

ofxn mod

0.73534

sRun

RunA1

zout

1

Ave

Data

0.00181

Model

0.0015

data/vMax

0.00829

ofxng

369.894

Min d/m

2.3477

ave/min d

22.7295

ave/min m

44.2006

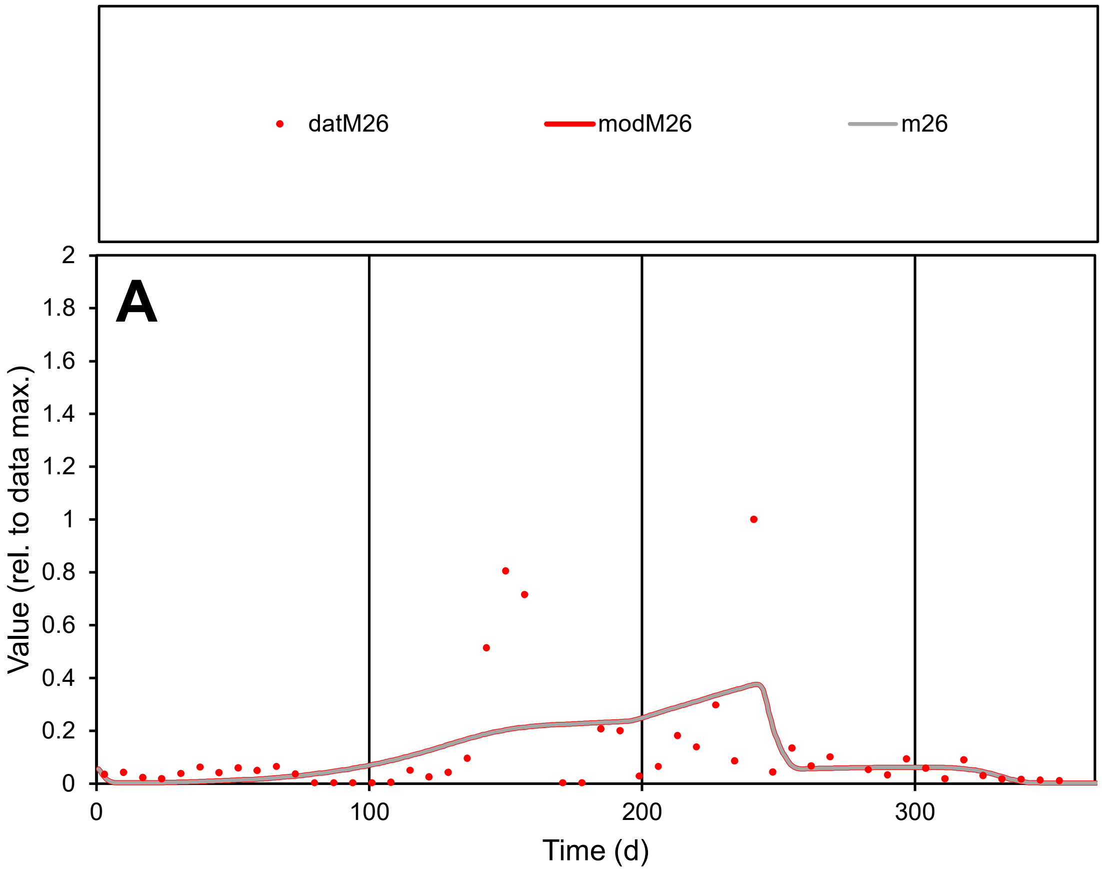


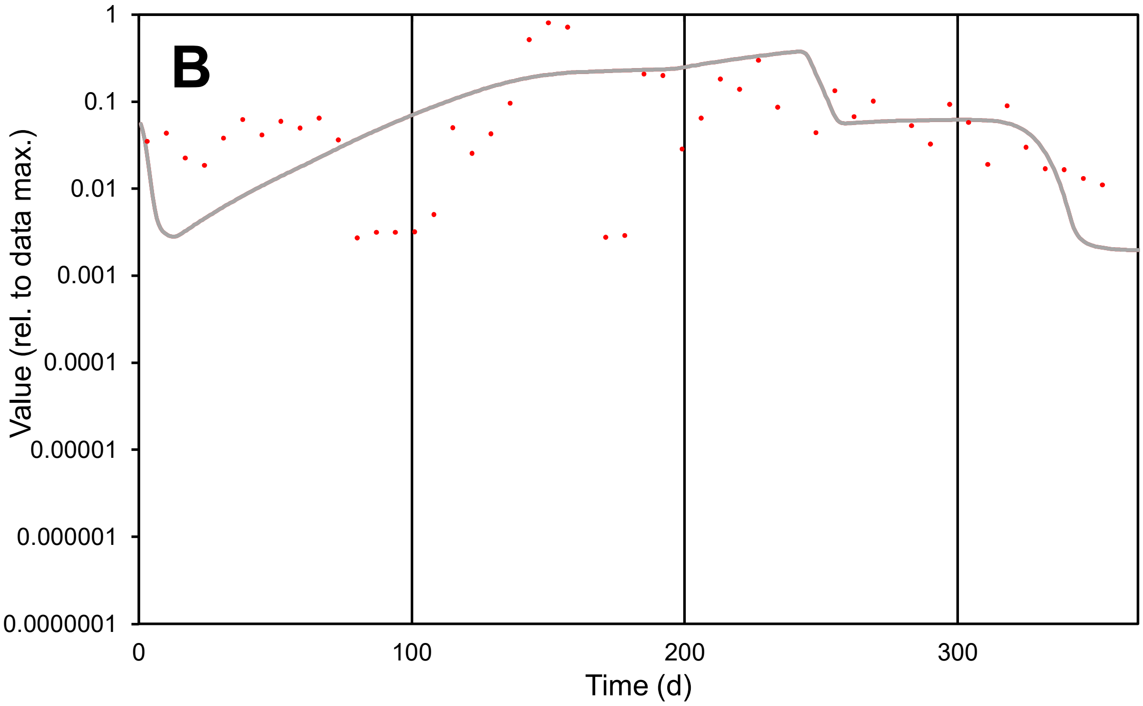


**C1**

Variables
M26 Provide jtd or Dst
CalibOnly?
2 1 to reproduce of
oNormalize?
2 1=no, 2=yes

ofxn calc*
1.49148
ofxn mod
1.50594

sRun
RunA1
zout
1

Ave
Data
0.00353
Model
0.00338

data/vMax
0.03018

ofxng
369.894

Min d/m
1.31938

ave/min d
42.8937
ave/min m
54.2439

328_Calib_S_A

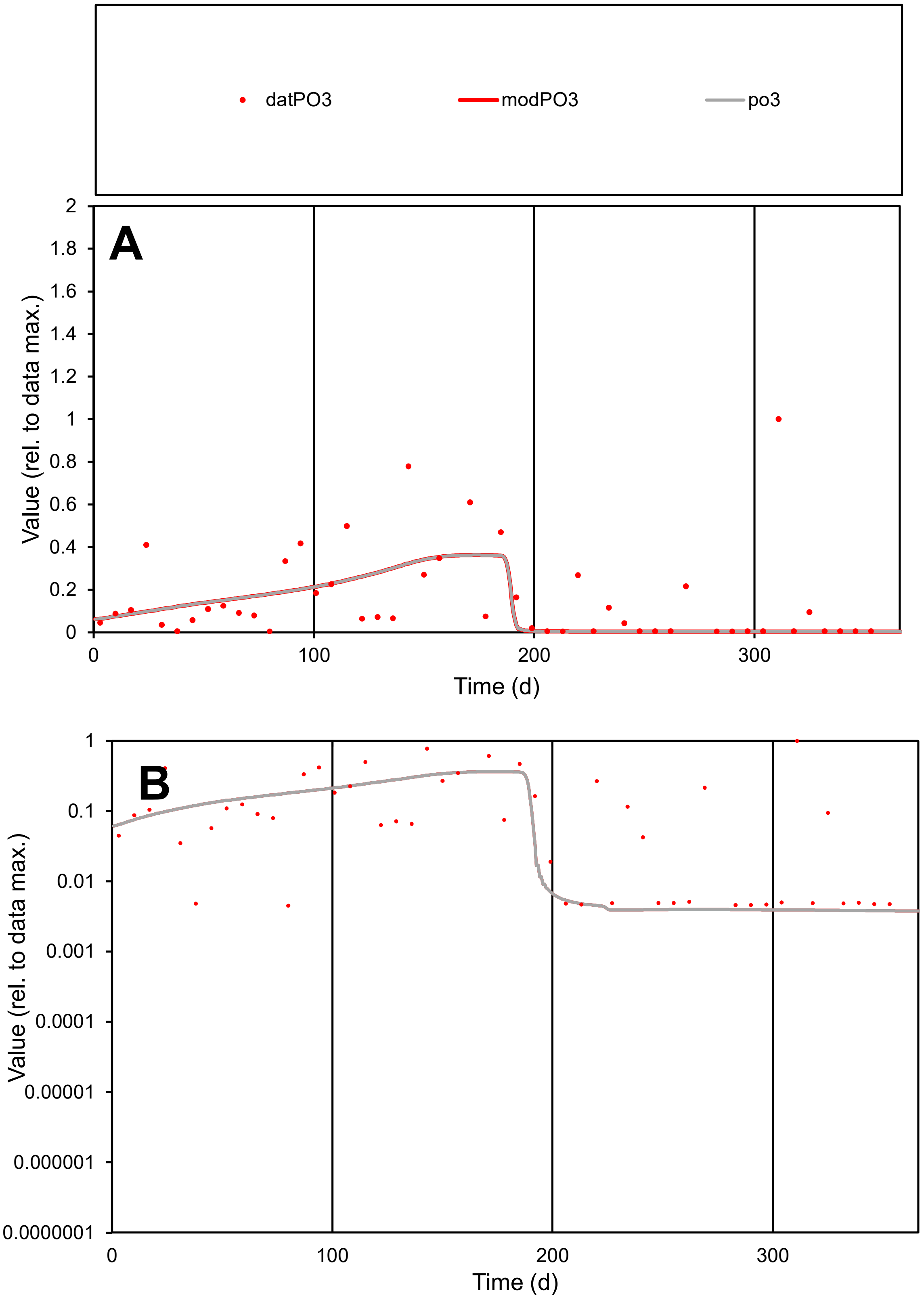


**C1**

Variables
PO3 Provide jtd or Dst
CalibOnly?
2 1 to reproduce of
oNormalize?
2 1=no, 2=yes

ofxn calc*
1.28921
ofxn mod
1.29814

sRun
RunA1
zout
1

Ave
Data
0.0016
Model
0.00122

data/vMax
0.0104

ofxng
369.894

Min d/m
1.184

ave/min d
34.2992
ave/min m
30.9101

328_Calib_S_A

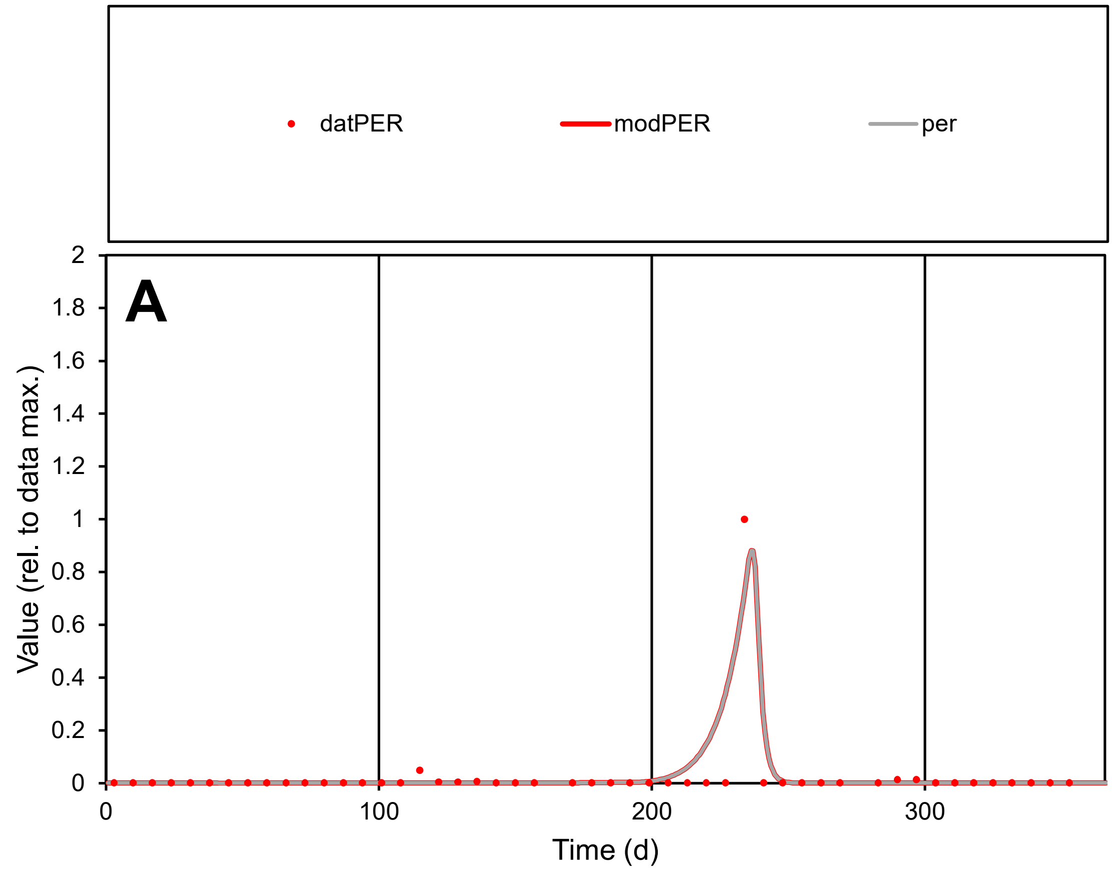


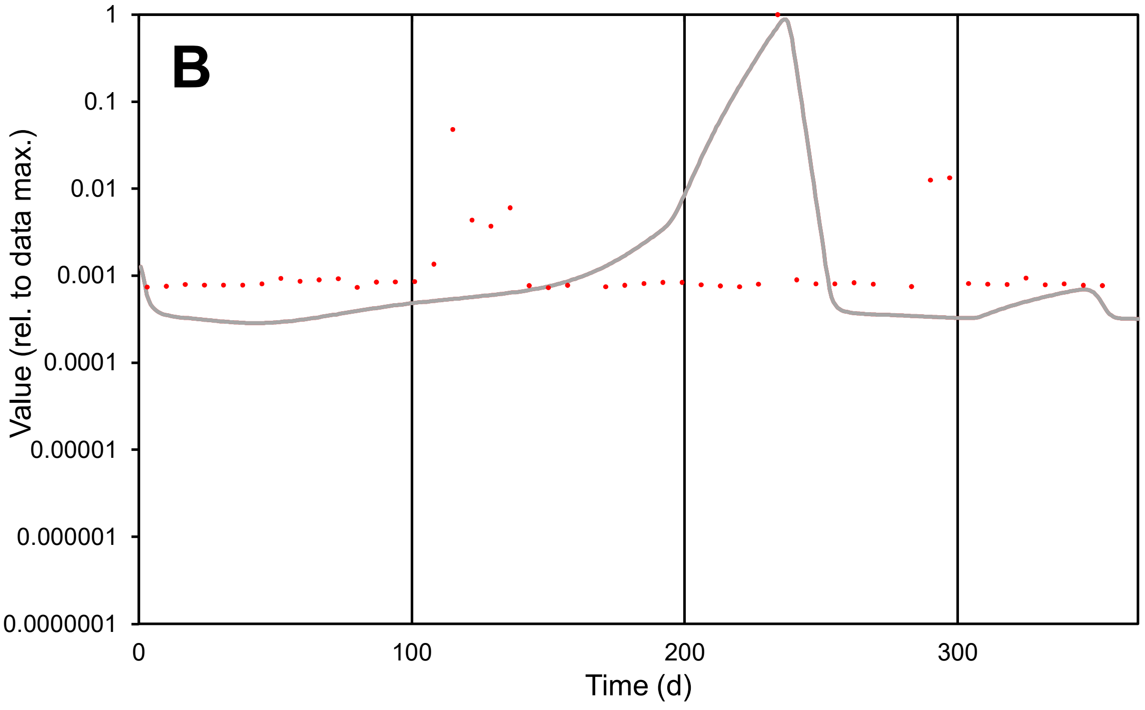


**C1**

Variables

PER Provide jtd or Dst

CalibOnly?

2 1 to reproduce of

oNormalize?

2 1=no, 2=yes

ofxn calc*

2.22784

ofxn mod

2.57097

sRun

RunA1

zout

1

Ave

Data

0.00058

Model

0.00072

data/vMax

0.02513

ofxng

369.894

Min d/m

2.54376

ave/min d

31.4359

ave/min m

100.583

328_Calib_S_A

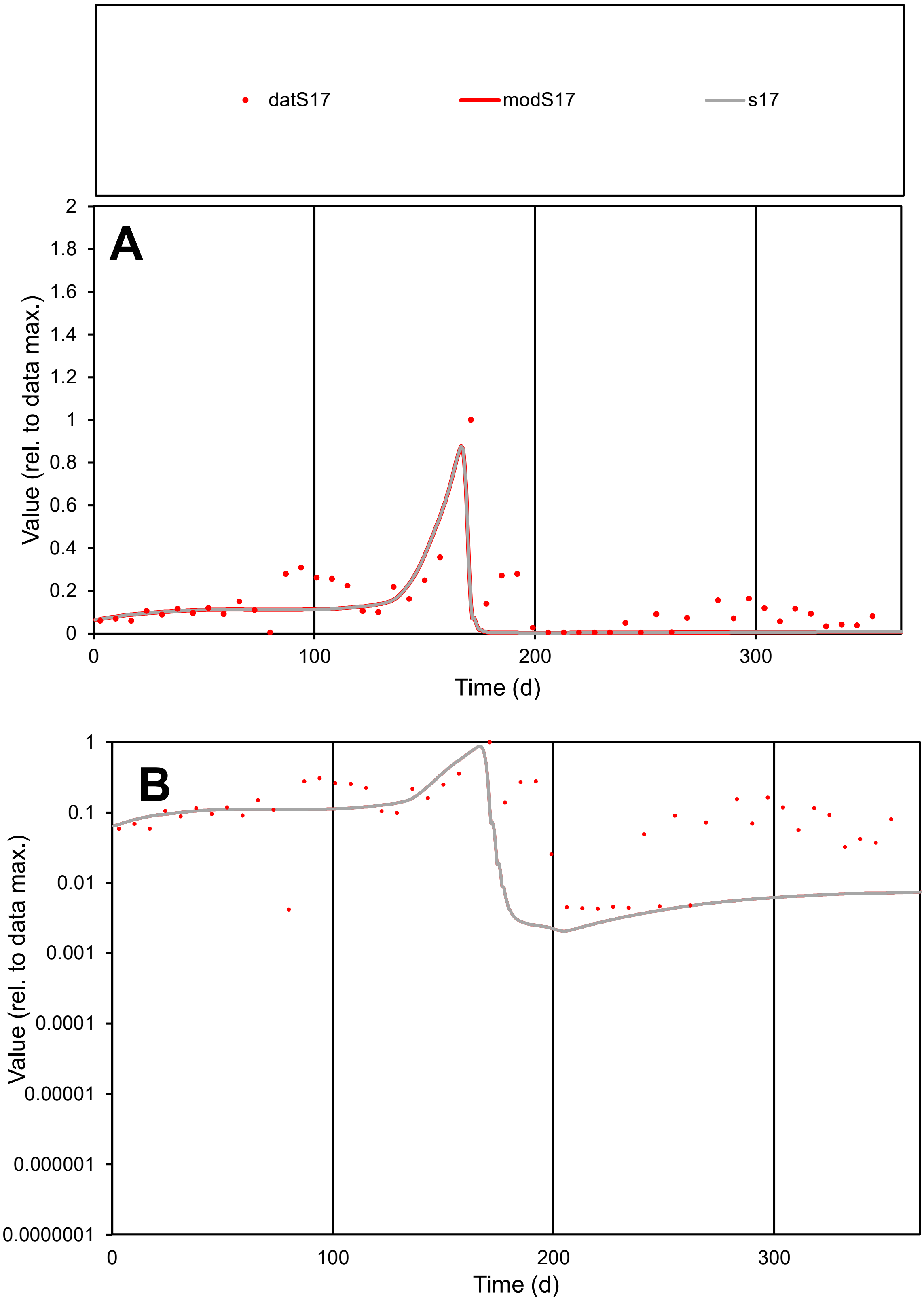


**C1**

Variables
S17 Provide jtd or Dst
CalibOnly?
2 1 to reproduce of
oNormalize?
2 1=no, 2=yes

ofxn calc*
0.77061
ofxn mod
1.02786

sRun
RunA1
zout
1

Ave
Data
0.00345
Model
0.00229

data/vMax
0.026

ofxng
369.894

Min d/m
1.98945
ave/min d
31.544
ave/min m
41.6595

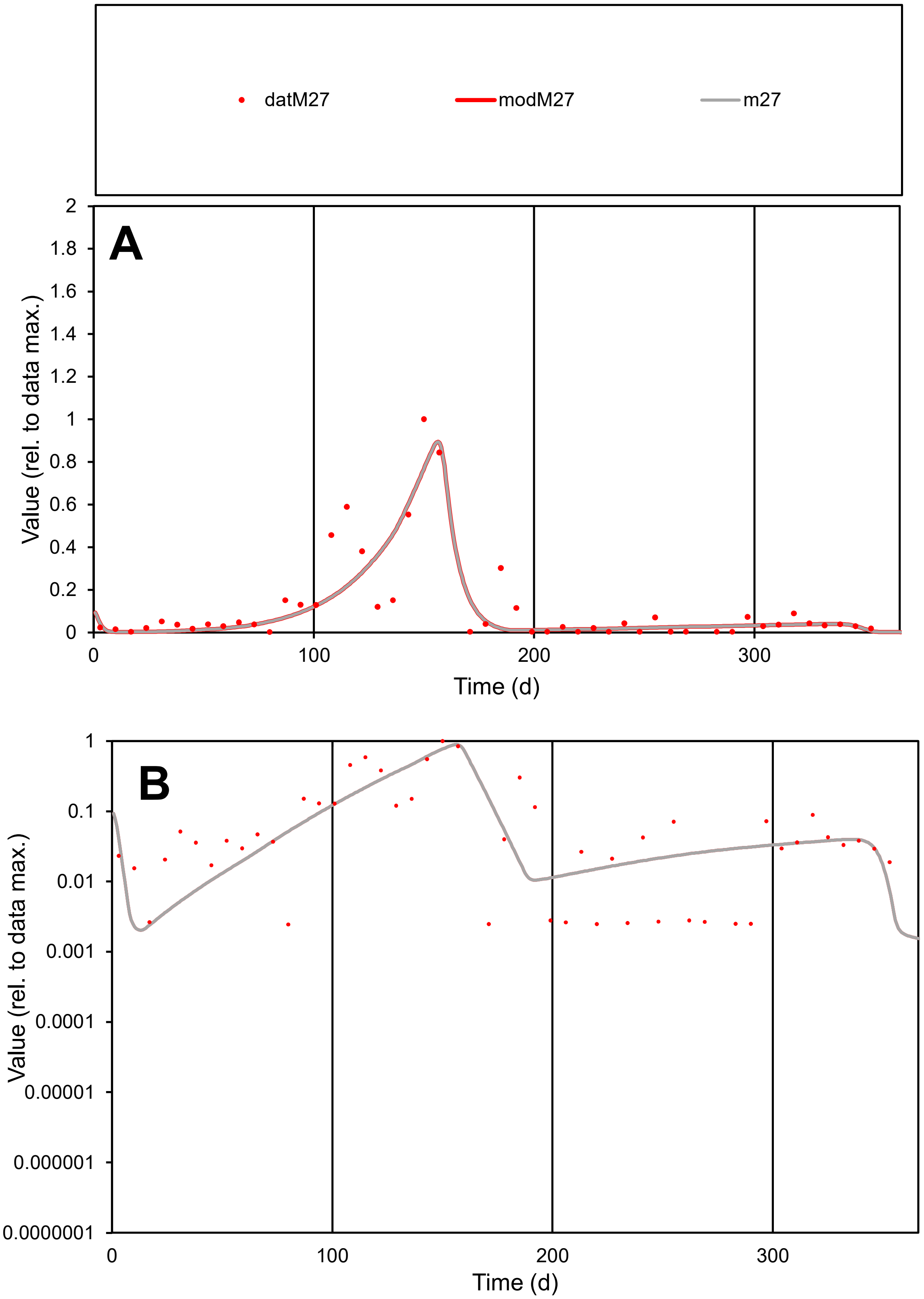


**C1**

Variables
M27 Provide jtd or Dst
CalibOnly?
2 1 to reproduce of
oNormalize?
2 1=no, 2=yes

ofxn calc*
0.88174
ofxn mod
0.91123

sRun
RunA1
zout
1

Ave
Data
0.00319
Model
0.00274

data/vMax
0.02673

ofxng
369.894

Min d/m
1.07289

ave/min d
48.8736
ave/min m
45.0523

328_Calib_S_A

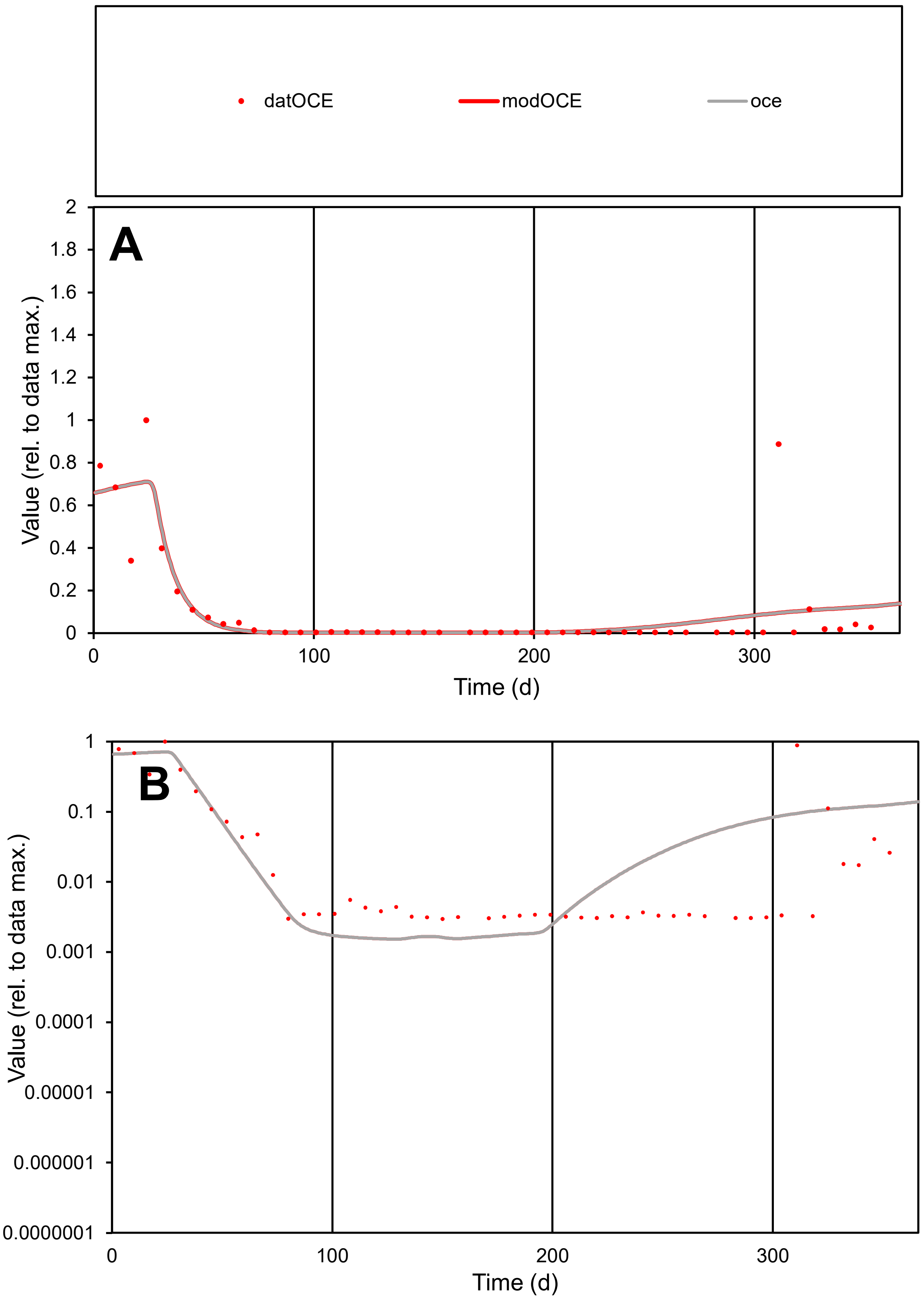


C1

Variables
OCE Provide jtd or Dst
CalibOnly?
2 1 to reproduce of
oNormalize?
2 1=no, 2=yes

ofxn calc*
1.37685
ofxn mod
1.40826

sRun
RunA1
zout
1

Ave
Data
0.00125
Model
0.00125

data/vMax
0.01248

ofxng
369.894

Min d/m
1.93992

ave/min d
33.578
ave/min m
65.2985

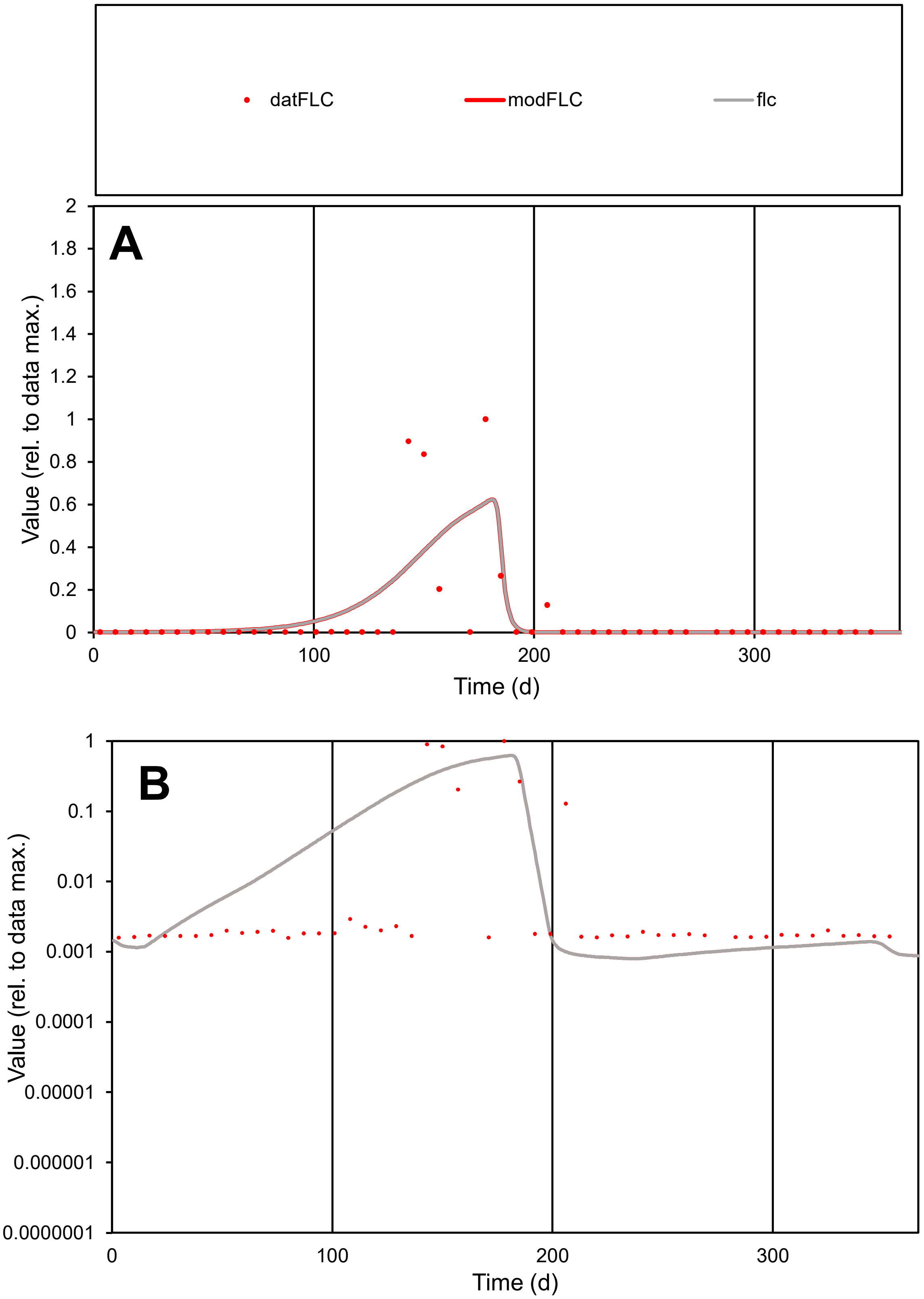


**C1**

Variables
FLC Provide jtd or Dst
CalibOnly?
2 1 to reproduce of
oNormalize?
2 1=no, 2=yes

ofxn calc*
2.24338
ofxn mod
2.27568

sRun
RunA1
zout
1

Ave
Data
0.00054
Model
0.00056

data/vMax
0.0077

ofxng
369.894

Min d/m
1.97133

ave/min d
44.2727
ave/min m
91.5973

328_Calib_S_A

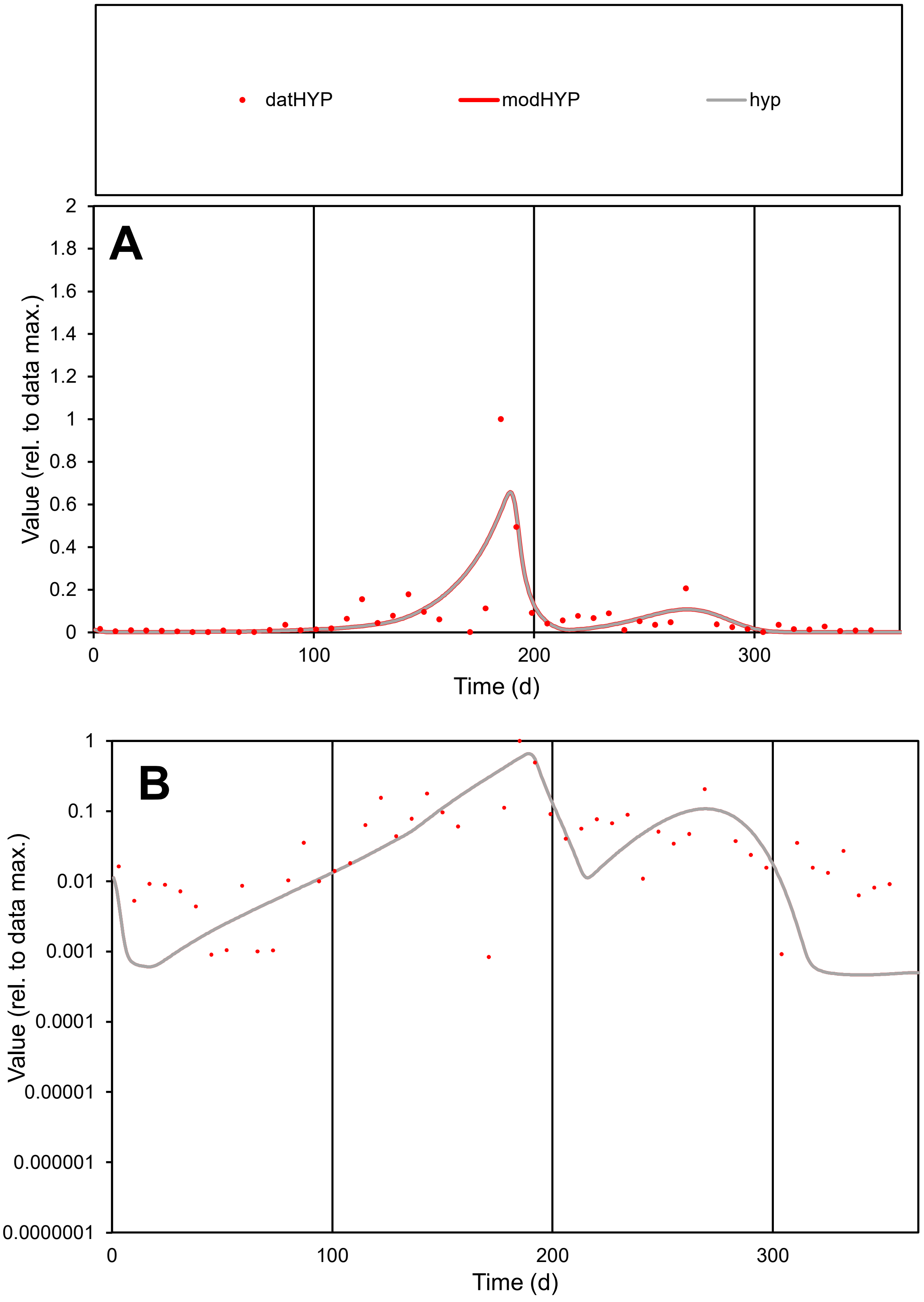


**C1**

Variables
HYP Provide jtd or Dst
CalibOnly?
2 1 to reproduce of
oNormalize?
2 1=no, 2=yes

ofxn calc*
1.31604
ofxn mod
1.35435

sRun
RunA1
zout
1

Ave
Data
0.00135
Model
0.00127

data/vMax
0.01944

ofxng
369.894

Min d/m
1.79326
ave/min d
82.7697
ave/min m
140.211

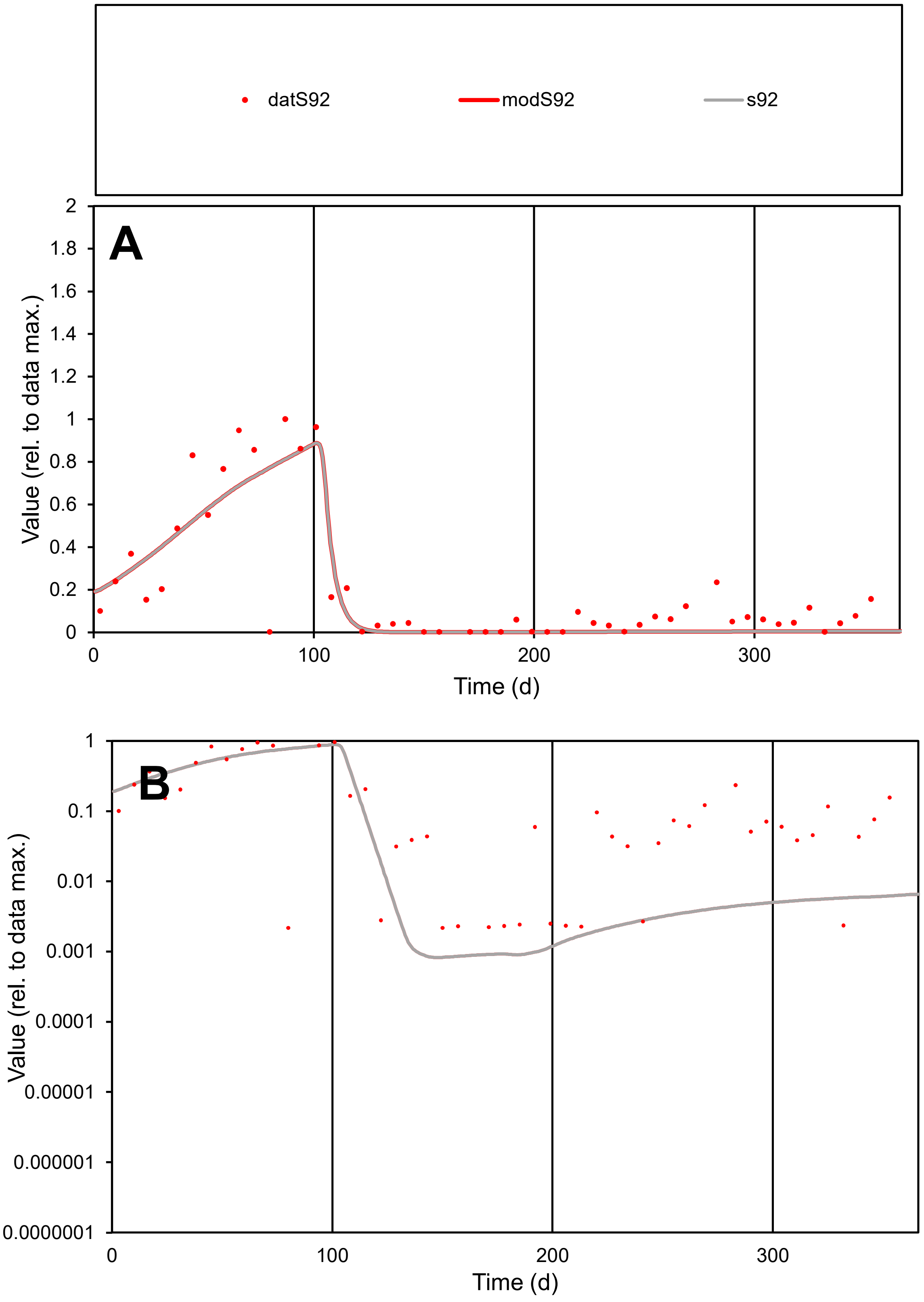


**C1**

Variables
S92 Provide jtd or Dst
CalibOnly?
2 1 to reproduce of
oNormalize?
2 1=no, 2=yes

ofxn calc*
0.70615
ofxn mod
0.73628

sRun
RunA1
zout
1

Ave
Data
0.00224
Model
0.00196

data/vMax
0.01071

ofxng
369.894

Min d/m
2.61343

ave/min d
96.3626
ave/min m
220.342

328_Calib_S_A

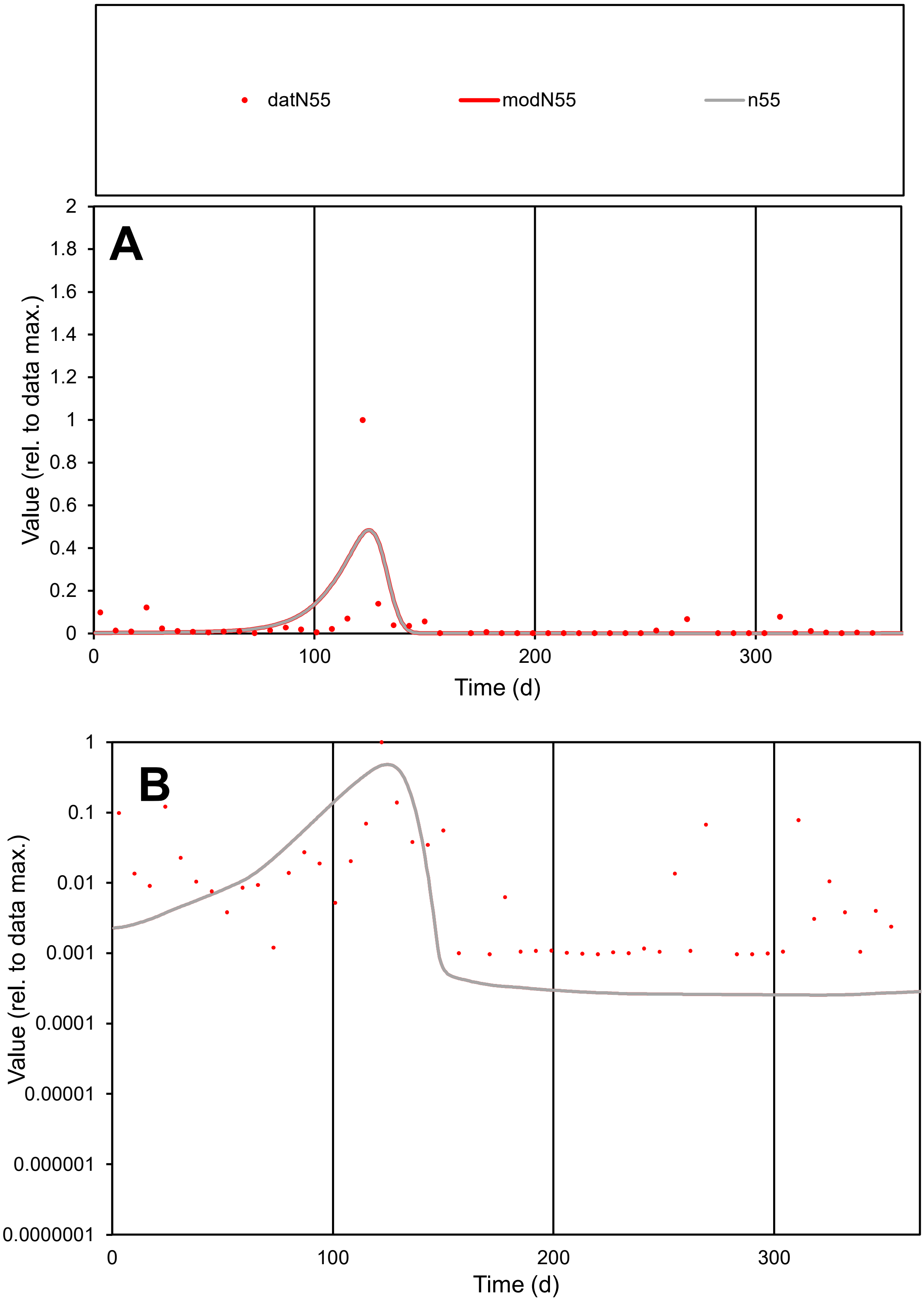


**C1**

Variables

N55 Provide jtd or Dst

CalibOnly?

2 1 to reproduce of

oNormalize?

2 1=no, 2=yes

ofxn calc*

2.49323

ofxn mod

2.62553

sRun

RunA1

zout

1

Ave

Data

0.00156

Model

0.00152

data/vMax

0.03945

ofxng

369.894

Min d/m

3.79047

ave/min d

40.8422

ave/min m

150.51

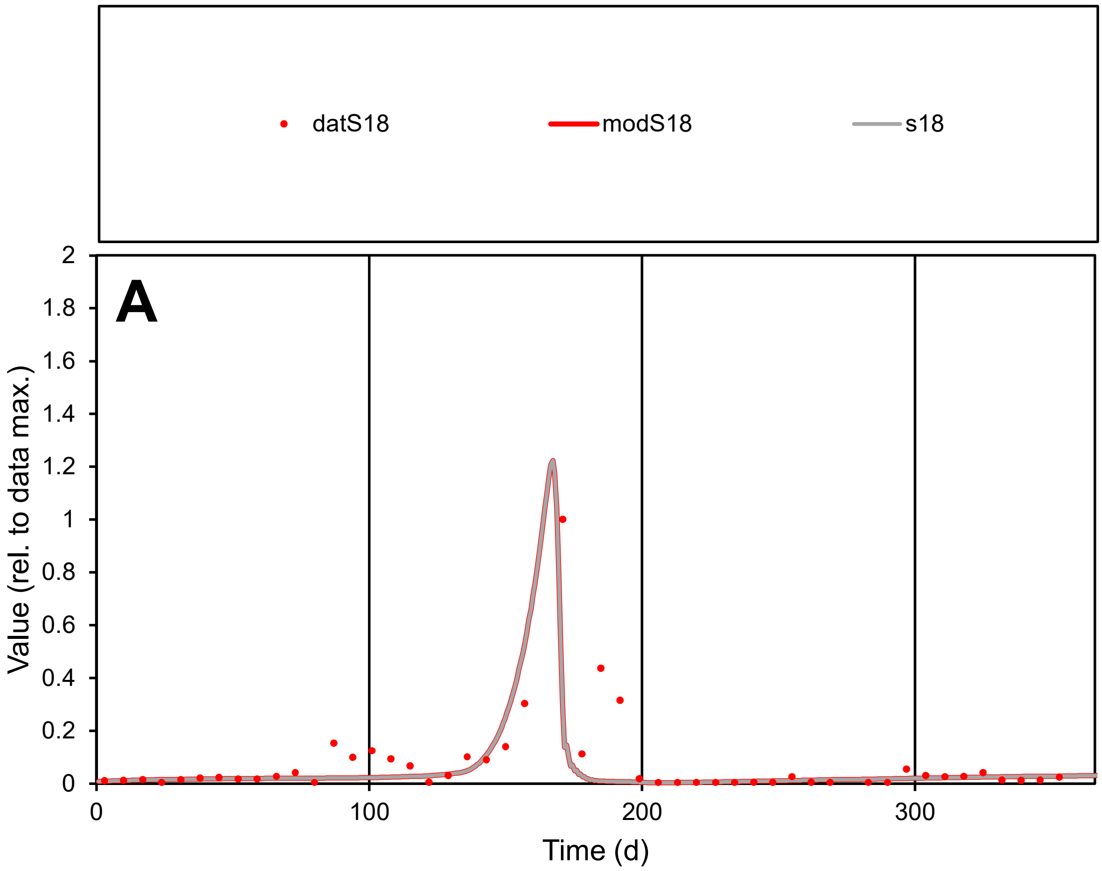


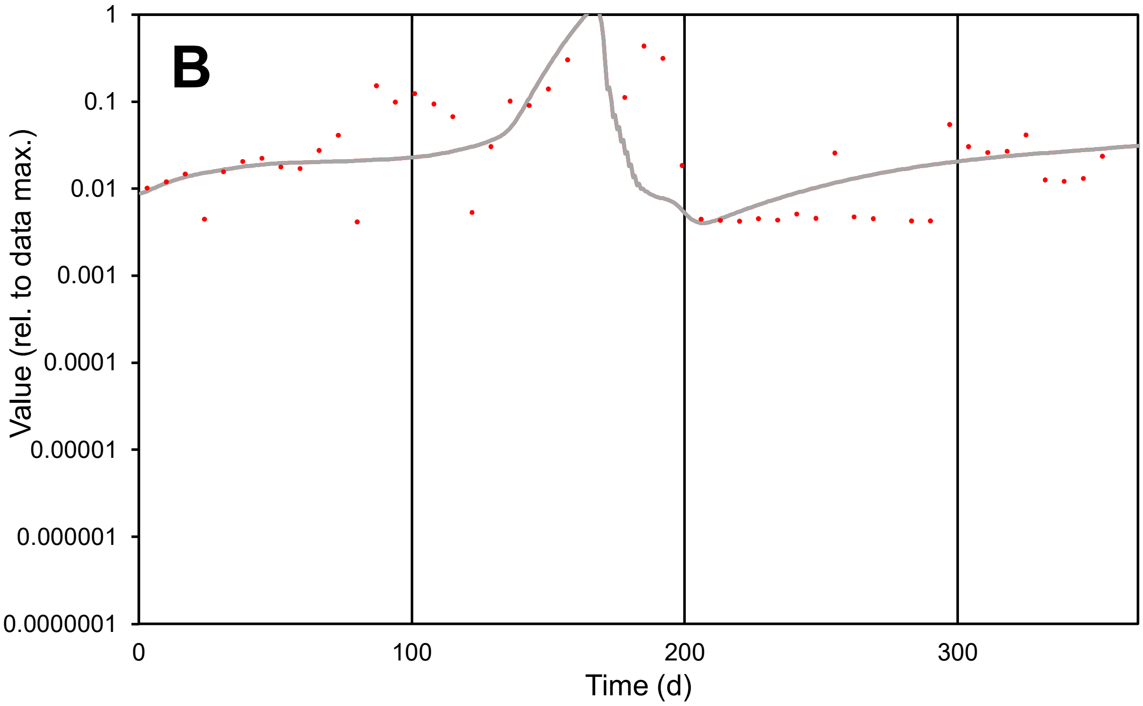


**C1**

Variables
S18 Provide jtd or Dst
CalibOnly?
2 1 to reproduce of
oNormalize?
2 1=no, 2=yes

ofxn calc*
1.28308
ofxn mod
1.57709

sRun
RunA1
zout
1

Ave
Data
0.00474
Model
0.00339

data/vMax
0.06431

ofxng
369.894

Min d/m
1.03398

ave/min d
17.7121
ave/min m
13.0849

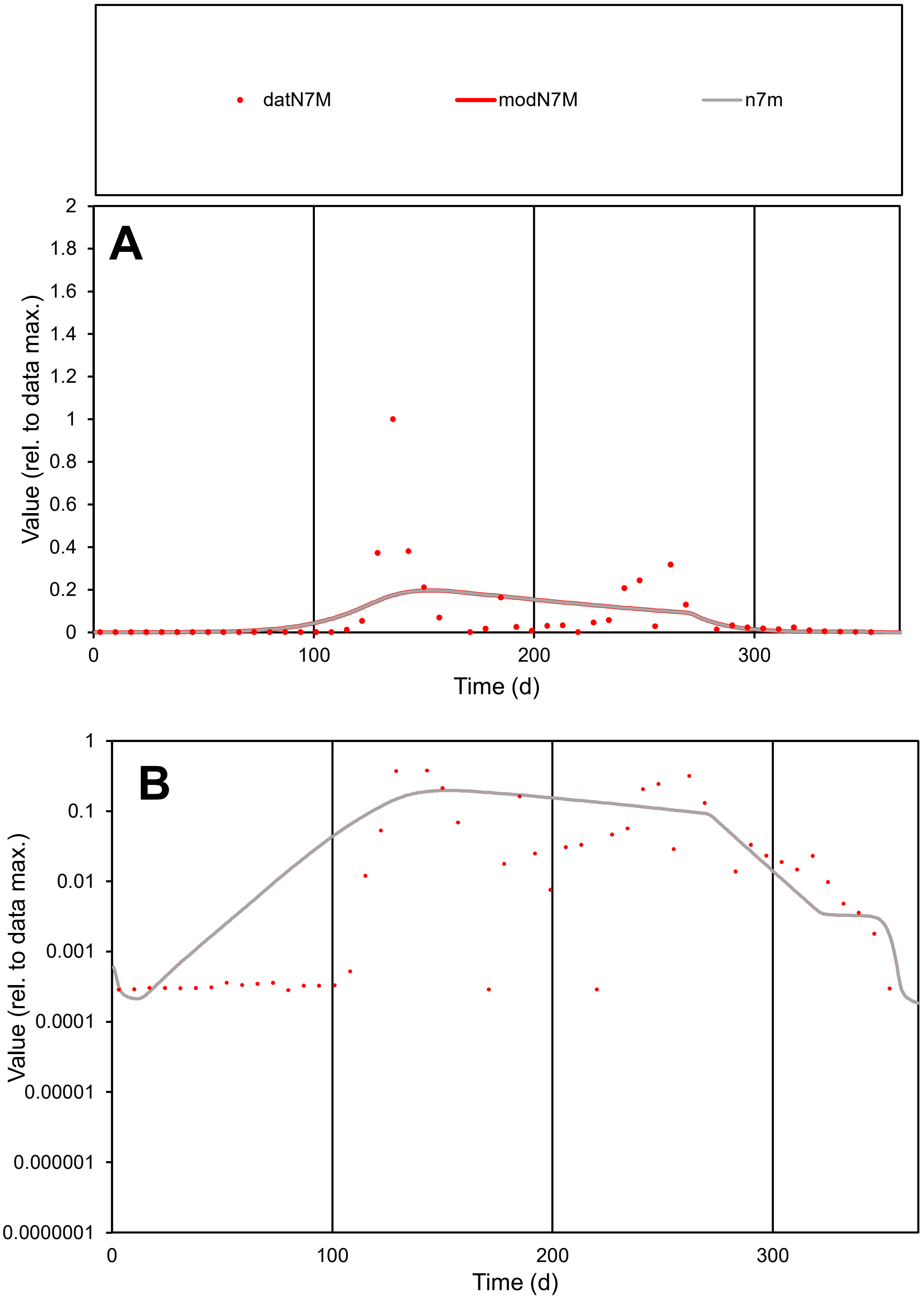


C1

Variables
N7M Provide jtd or Dst
CalibOnly?
2 1 to reproduce of
oNormalize?
2 1=no, 2=yes

ofxn calc*
1.99091
ofxn mod
2.00022

sRun
RunA1
zout
1

Ave
Data
0.00249
Model
0.00241

data/vMax
0.03424

ofxng
369.894

Min d/m
1.3224
ave/min d
256.983
ave/min m
328.843

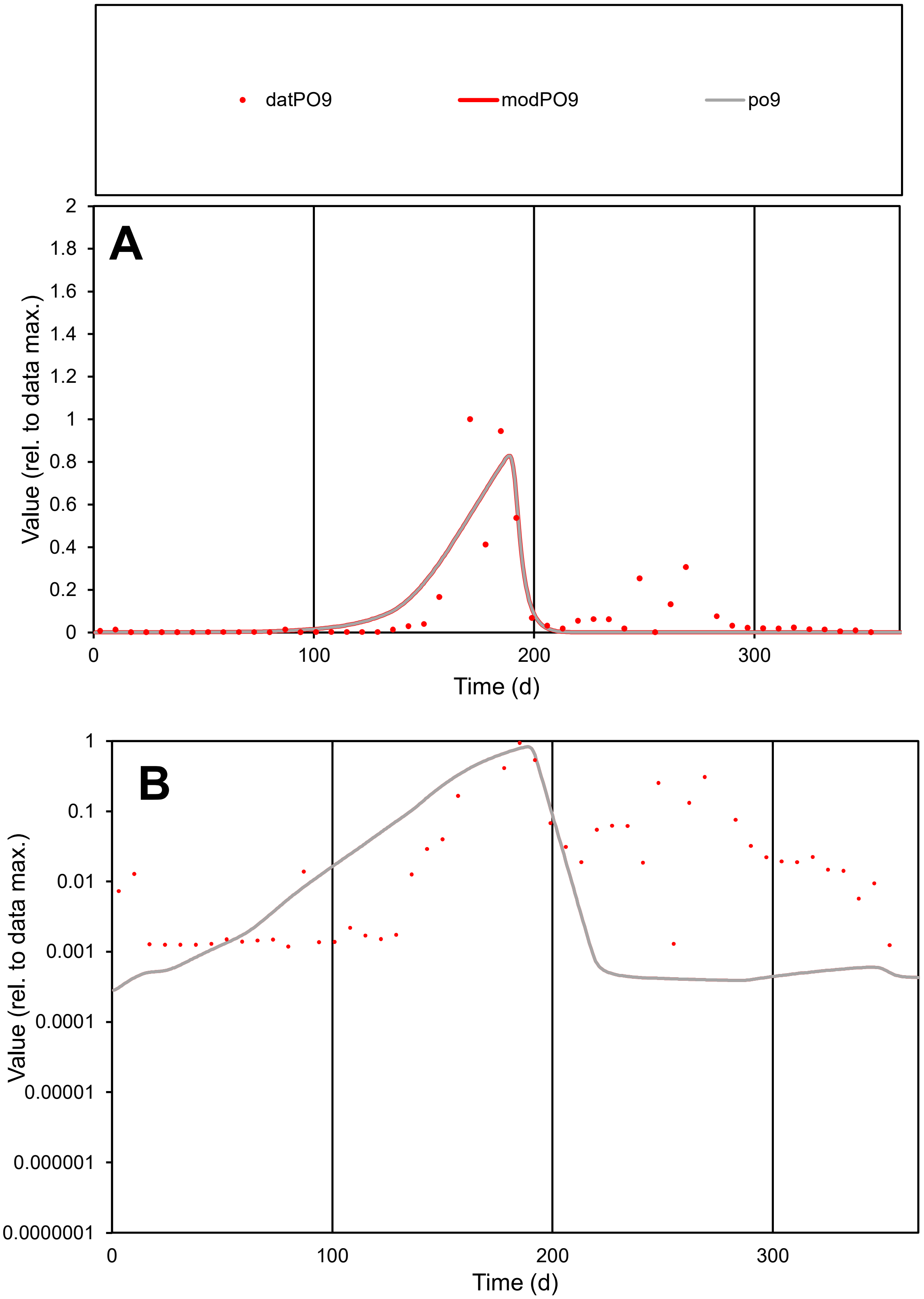


**C1**

Variables
PO9 Provide jtd or Dst
CalibOnly?
2 1 to reproduce of
oNormalize?
2 1=no, 2=yes

ofxn calc*
1.17211
ofxn mod
1.22484

sRun
RunA1
zout
1

Ave
Data
0.0045
Model
0.00377

data/vMax
0.04963

ofxng
369.894

Min d/m
3.58777

ave/min d
76.7193
ave/min m
230.297

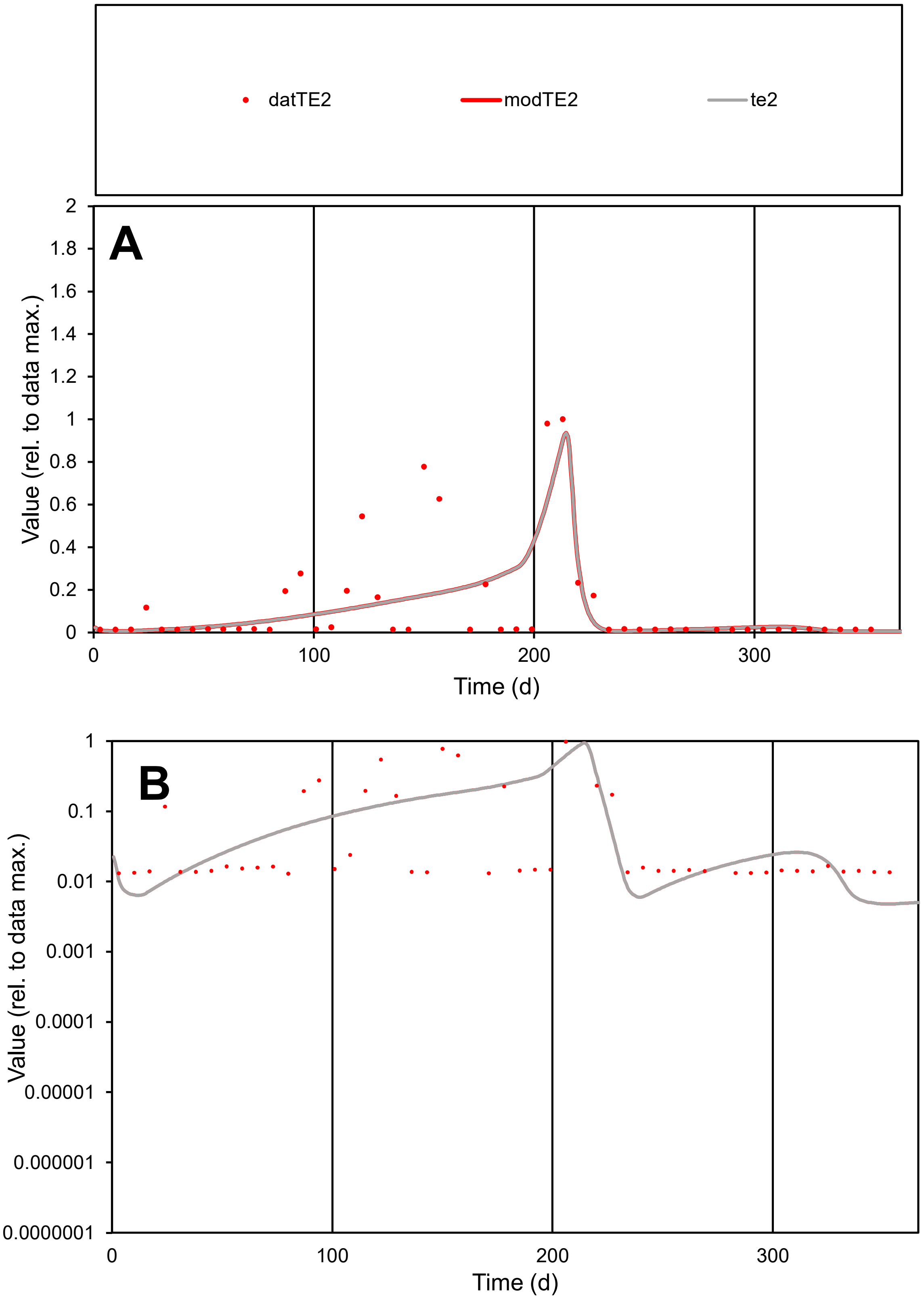


**C1**

Variables
TE2 Provide jtd or Dst
CalibOnly?
2 1 to reproduce of
oNormalize?
2 1=no, 2=yes

ofxn calc*
1.31835
ofxn mod
1.35116

sRun
RunA1
zout
1

Ave
Data
0.00082
Model
0.00071

data/vMax
0.00665

ofxng
369.894

Min d/m
2.68858

ave/min d
9.52788
ave/min m
22.2308

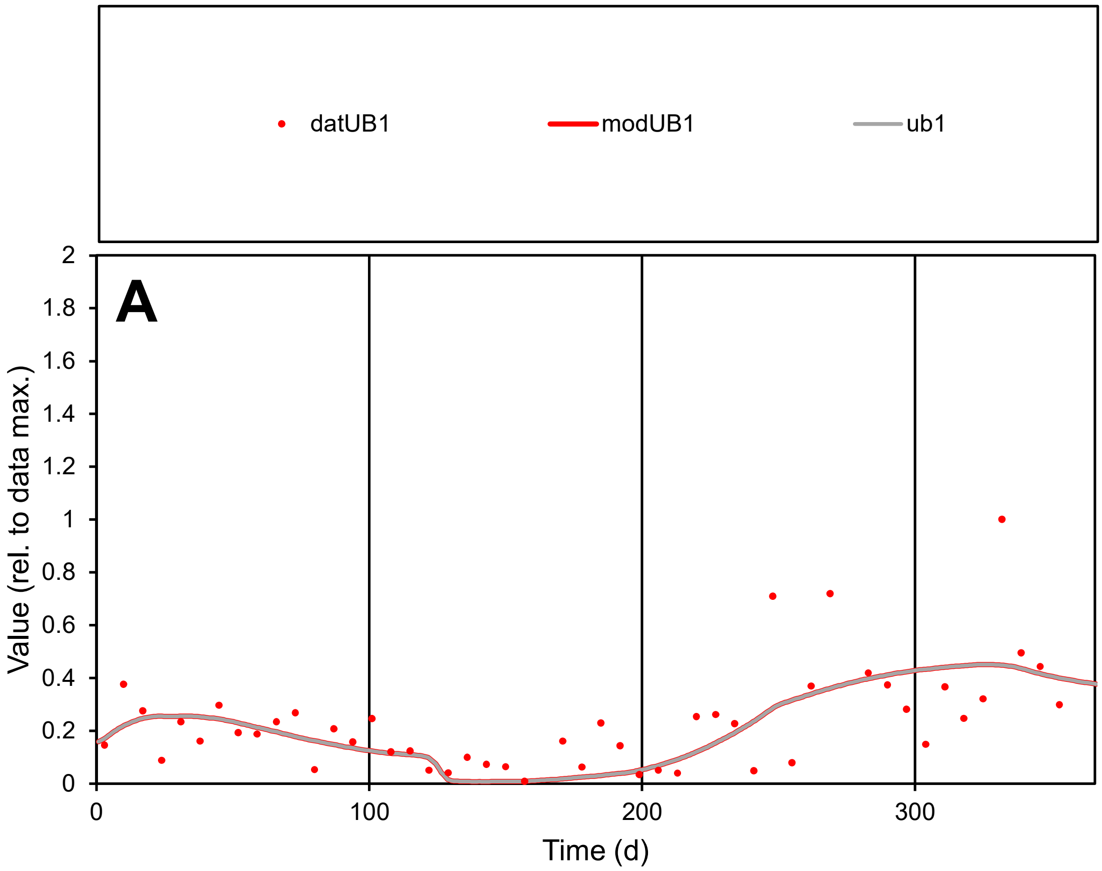


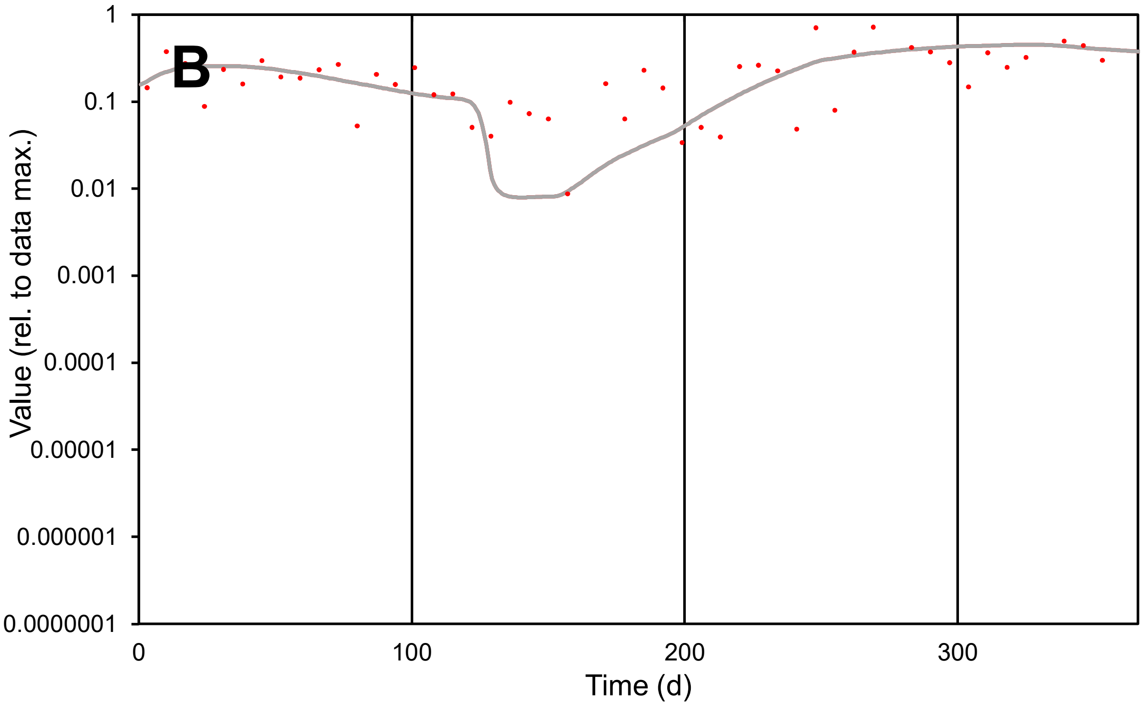


**C1**

Variables
UB1 Provide jtd or Dst
CalibOnly?
2 1 to reproduce of
oNormalize?
2 1=no, 2=yes

ofxn calc*
0.64002
ofxn mod
0.6534

sRun
RunA1
zout
1

Ave
Data
0.00163
Model
0.00149

data/vMax
0.00695

ofxng
369.894

Min d/m
1.09132

ave/min d
26.8619
ave/min m
26.7543

328_Calib_S_A

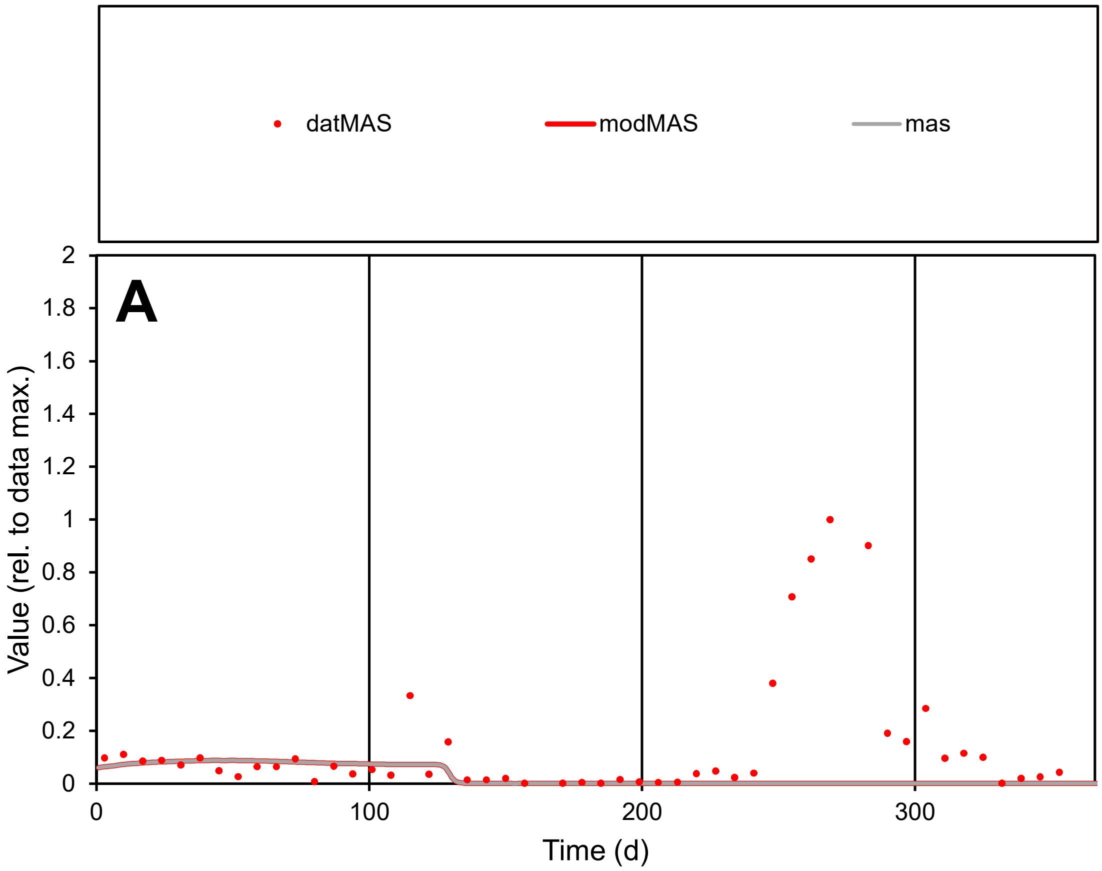


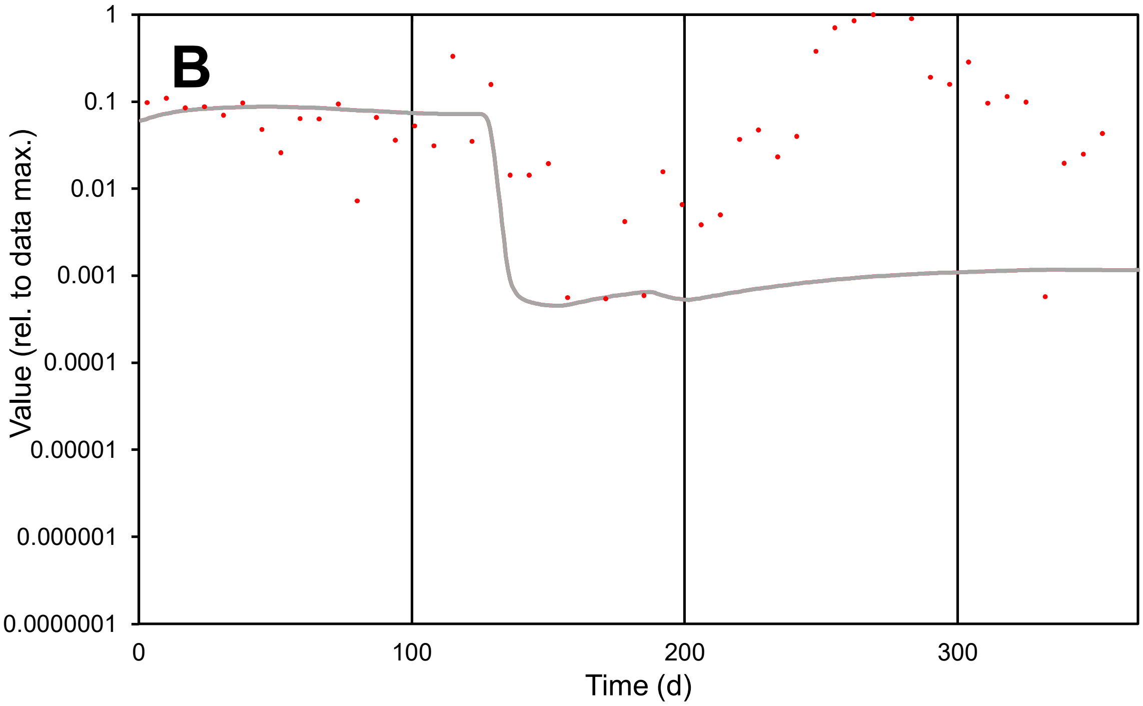


**C1**

Variables
MAS Provide jtd or Dst
CalibOnly?
2 1 to reproduce of
oNormalize?
2 1=no, 2=yes

ofxn calc*
1.95175
ofxn mod
1.95679

sRun
RunA1
zout
1

Ave
Data
0.00308
Model
0.0007

data/vMax
0.02268

ofxng
369.894

Min d/m
1.19961

ave/min d
250.286
ave/min m
68.27

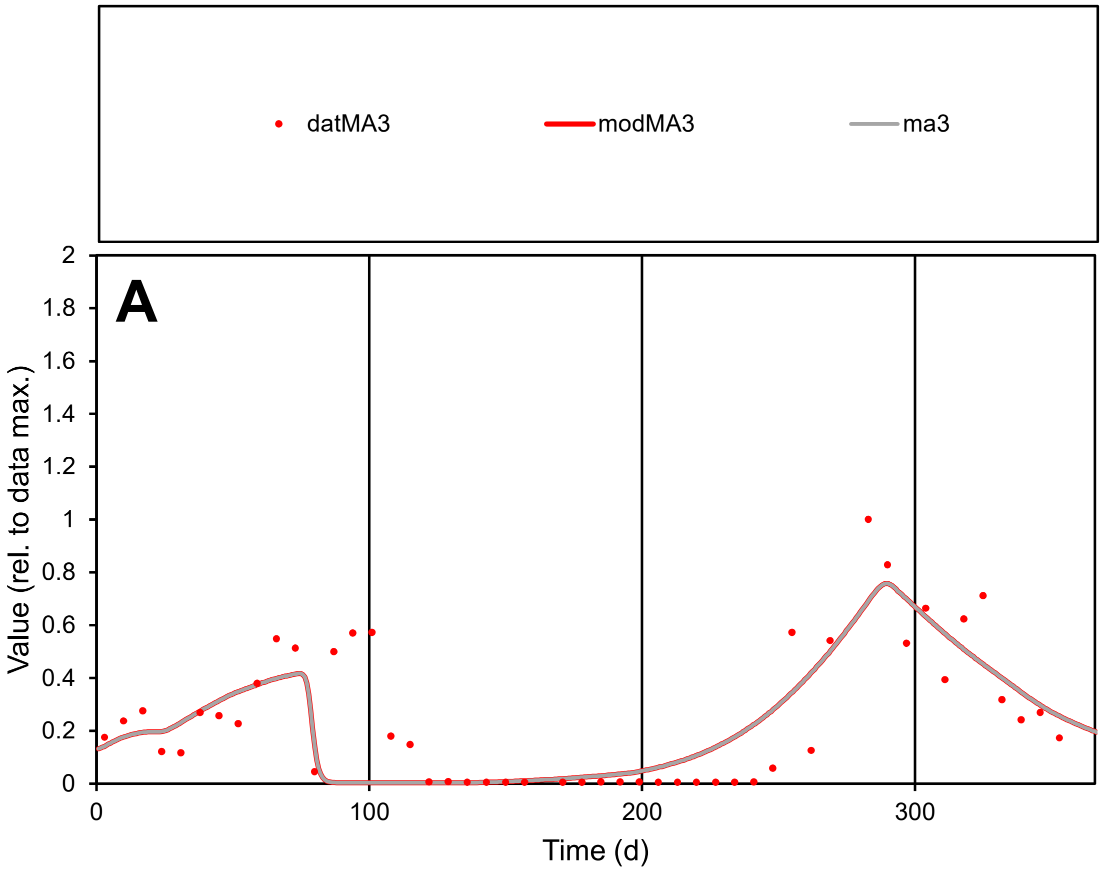


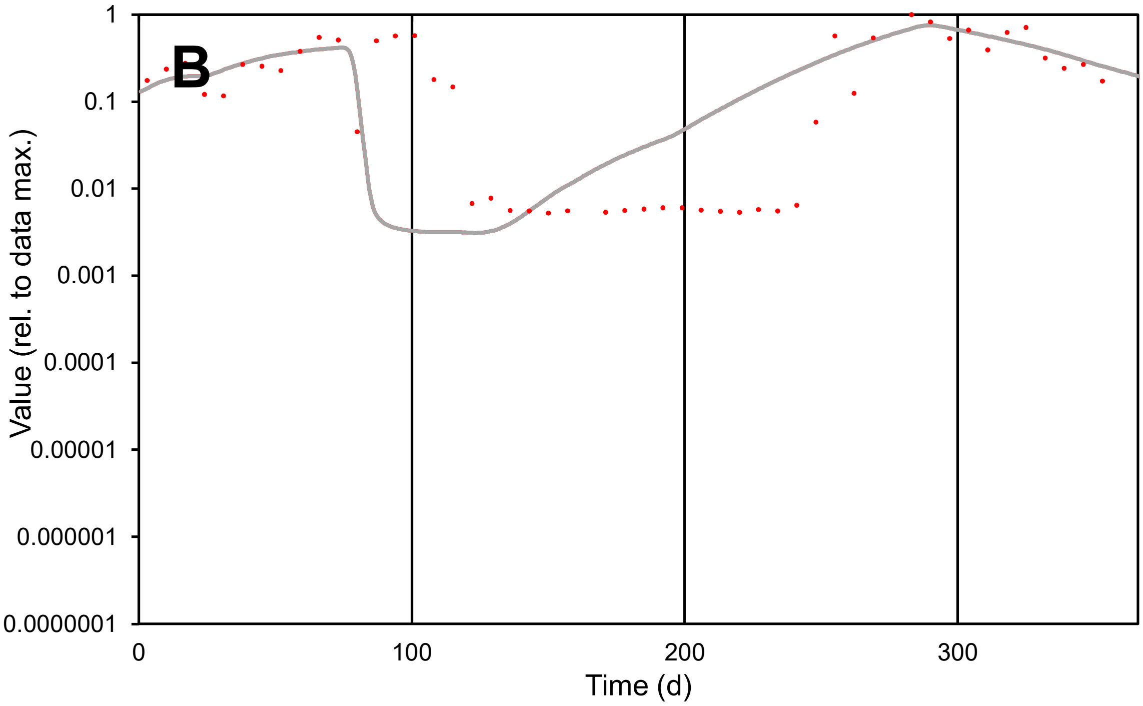


**C1**

Variables
MA3 Provide jtd or Dst
CalibOnly?
2 1 to reproduce of
oNormalize?
2 1=no, 2=yes

ofxn calc*
0.71614
ofxn mod
0.73286

sRun
RunA1
zout
1

Ave
Data
0.00296
Model
0.00272

data/vMax
0.0118

ofxng
369.894

Min d/m
1.69306

ave/min d
47.701
ave/min m
74.2619

328_Calib_S_A

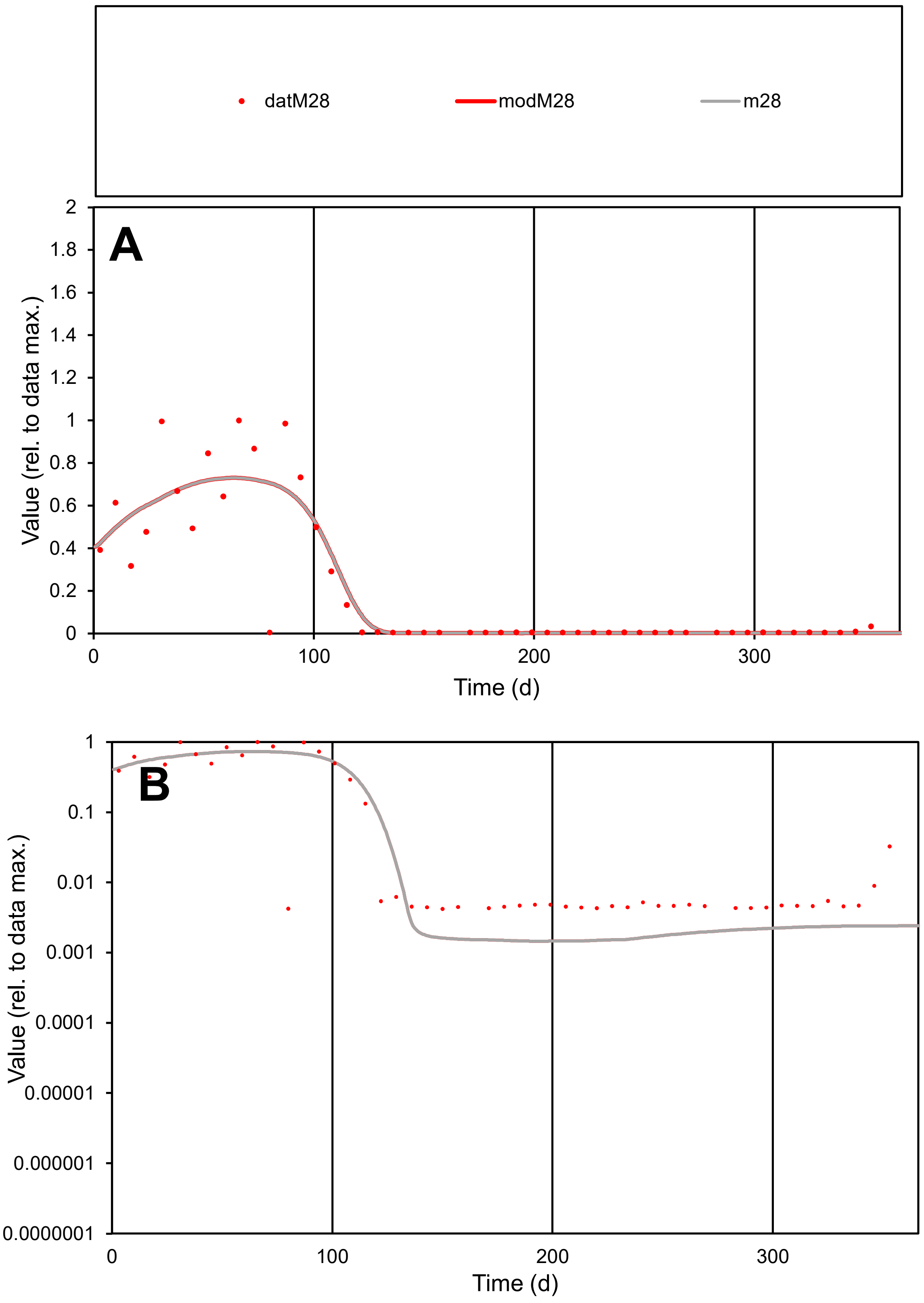


**C1**

Variables
M28 Provide jtd or Dst
CalibOnly?
2 1 to reproduce of
oNormalize?
2 1=no, 2=yes

ofxn calc*
0.68502
ofxn mod
0.71535

sRun
RunA1
zout
1

Ave
Data
0.00265
Model
0.00265

data/vMax
0.01282

ofxng
369.894

Min d/m
2.88806

ave/min d
49.1741
ave/min m
142.003

328_Calib_S_A

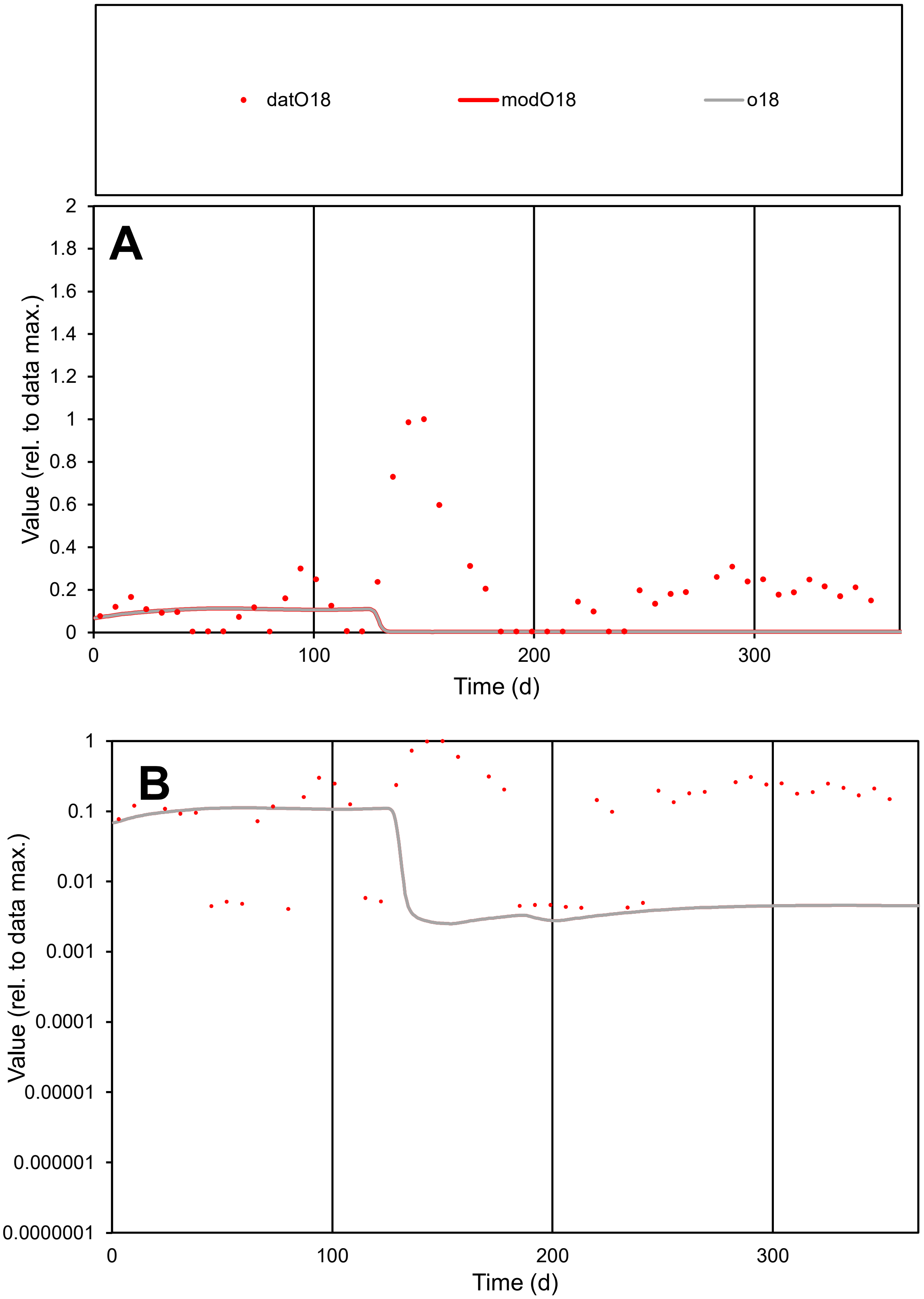


**C1**

Variables

O18 Provide jtd or Dst

CalibOnly?

2 1 to reproduce of

oNormalize?

2 1=no, 2=yes

ofxn calc*

1.49104

ofxn mod

1.4944

sRun

RunA1

zout

1

Ave

Data

0.00239

Model

0.00054

data/vMax

0.01276

ofxng

369.894

Min d/m

1.61608

ave/min d

46.0685

ave/min m

16.8367

328_Calib_S_A

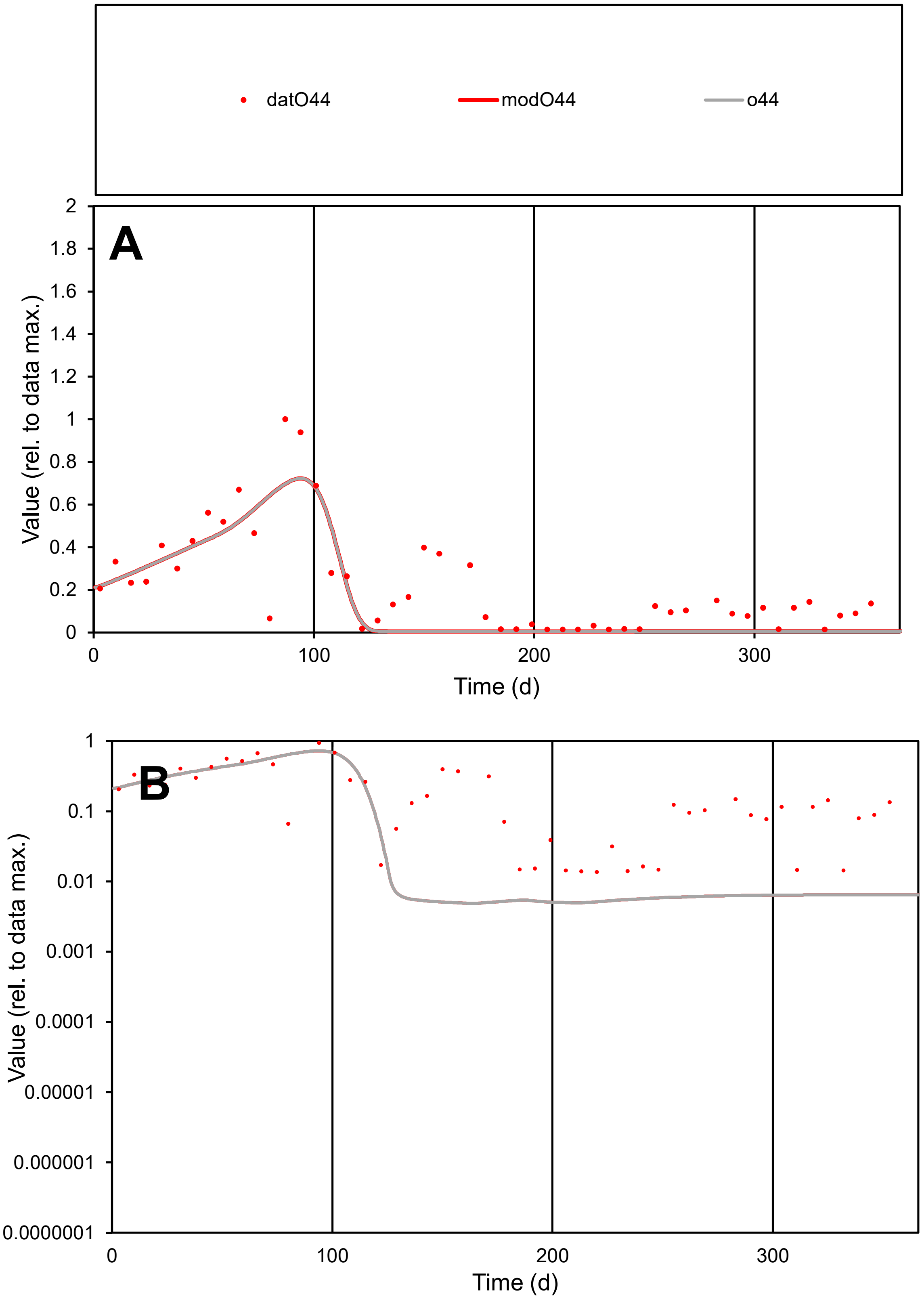


**C1**

Variables
O44 Provide jtd or Dst
CalibOnly?
2 1 to reproduce of
oNormalize?
2 1=no, 2=yes

ofxn calc*
0.7038
ofxn mod
0.72014

sRun
RunA1
zout
1

Ave
Data
0.0018
Model
0.00132

data/vMax
0.00829

ofxng
369.894

Min d/m
2.76502

ave/min d
15.9183
ave/min m
32.3343

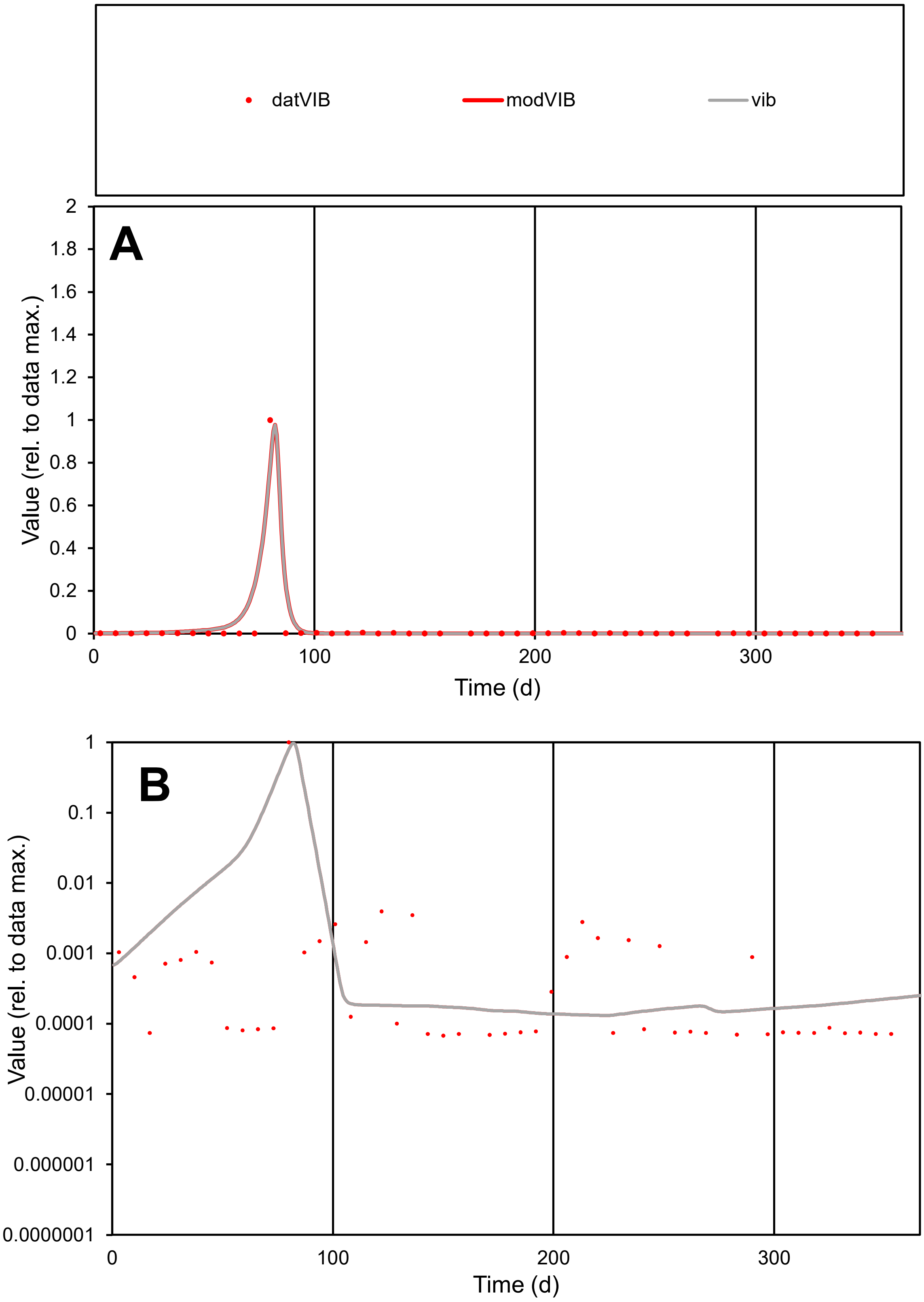


**C1**

Variables
VIB Provide jtd or Dst
CalibOnly?
2 1 to reproduce of
oNormalize?
2 1=no, 2=yes

ofxn calc*
1.35665
ofxn mod
1.71779

sRun
RunA1
zout
1

Ave
Data
0.0055
Model
0.00704

data/vMax
0.2617

ofxng
369.894

Min d/m
0.51512

ave/min d
310.559
ave/min m
204.524

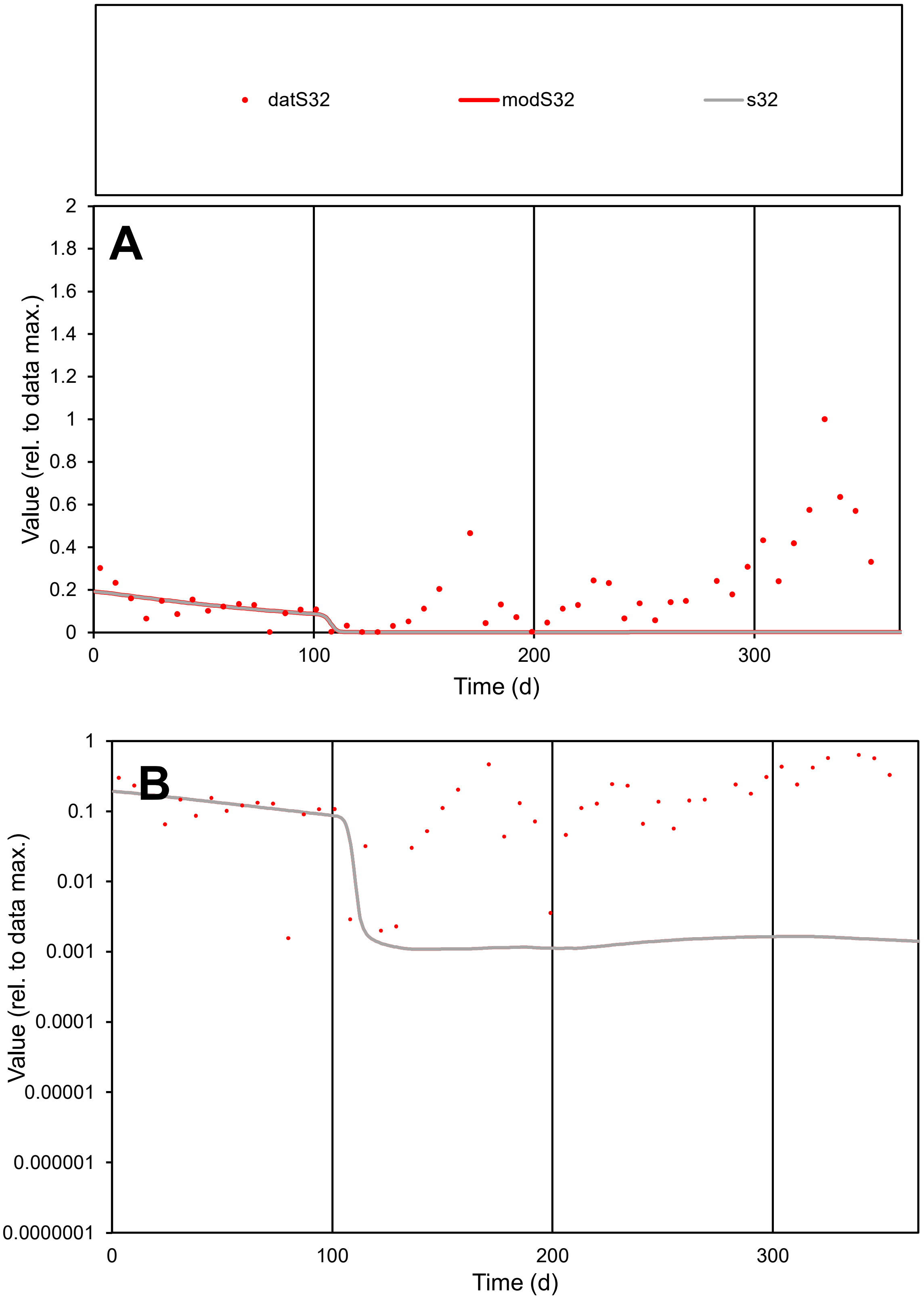


**C1**

Variables
S32 Provide jtd or Dst
CalibOnly?
2 1 to reproduce of
oNormalize?
2 1=no, 2=yes

ofxn calc*
1.36362
ofxn mod
1.3664

sRun
RunA1
zout
1

Ave
Data
0.00132
Model
0.00029

data/vMax
0.00695

ofxng
369.894

Min d/m
1.42669

ave/min d
122.004
ave/min m
37.7948

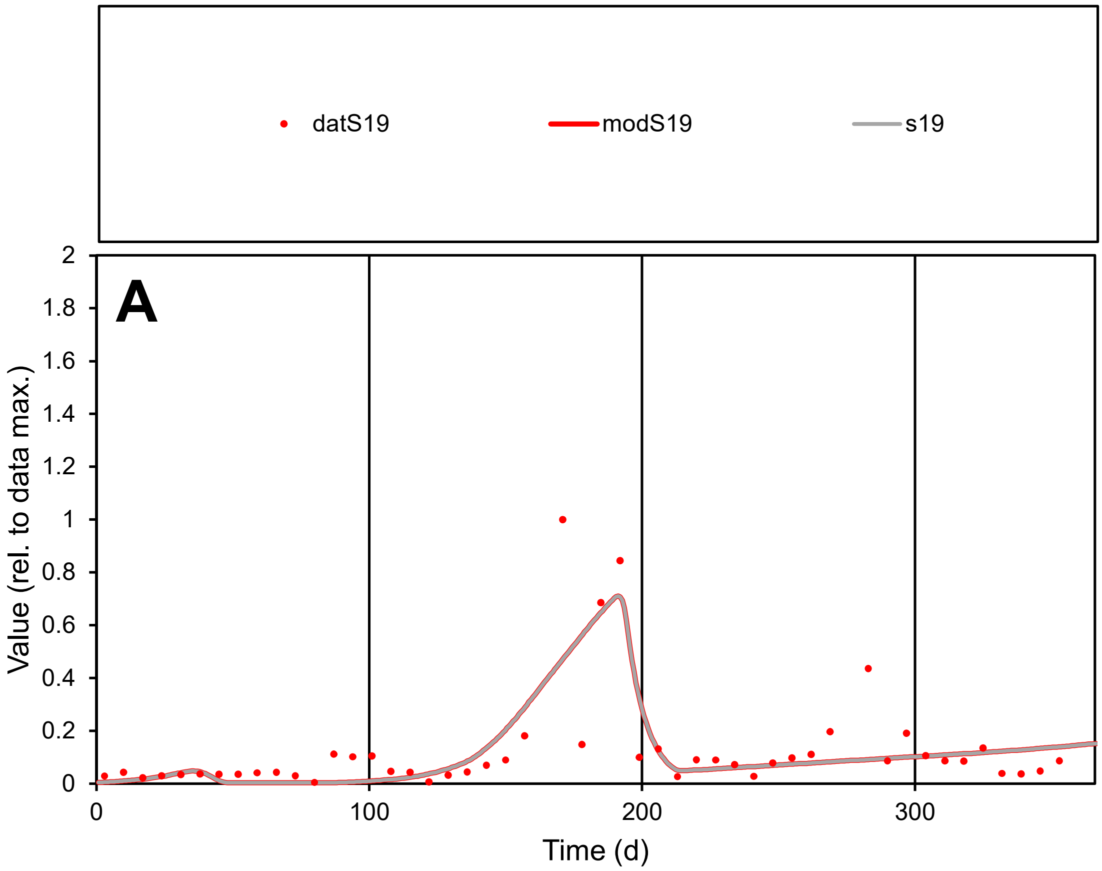


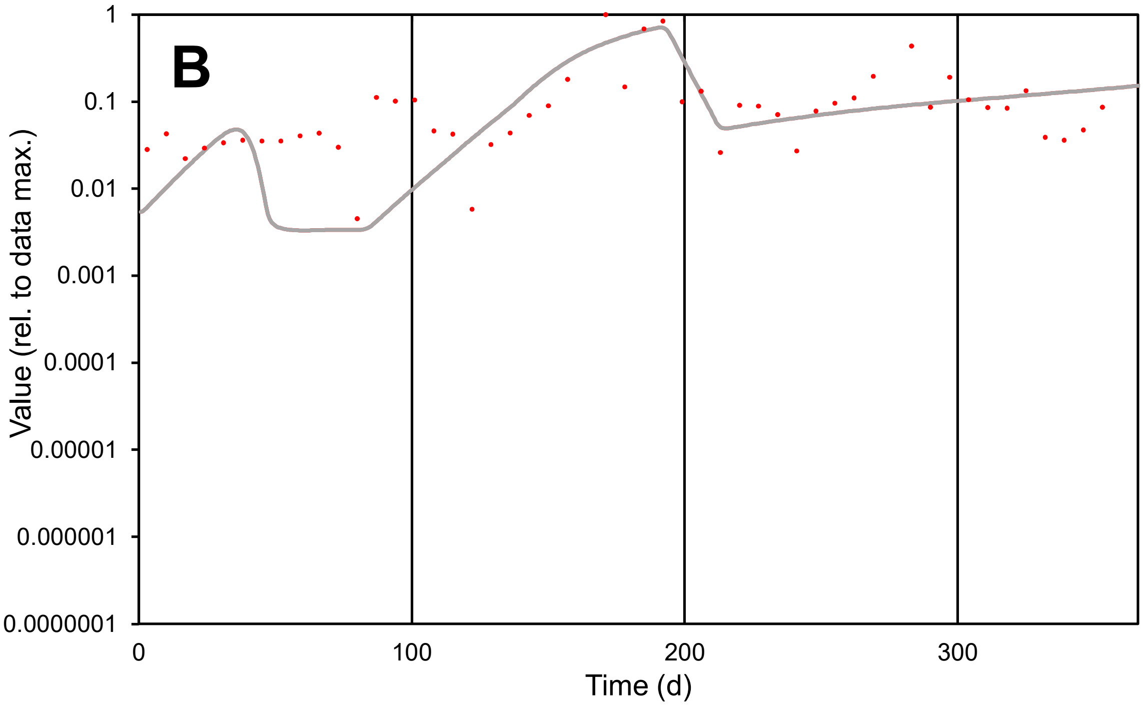


**C1**

Variables
S19 Provide jtd or Dst
CalibOnly?
2 1 to reproduce of
oNormalize?
2 1=no, 2=yes

ofxn calc*
0.95774
ofxn mod
0.98392

sRun
RunA1
zout
1

Ave
Data
0.00133
Model
0.00122

data/vMax
0.01045

ofxng
369.894

Min d/m
1.36557

ave/min d
28.0048
ave/min m
35.0254

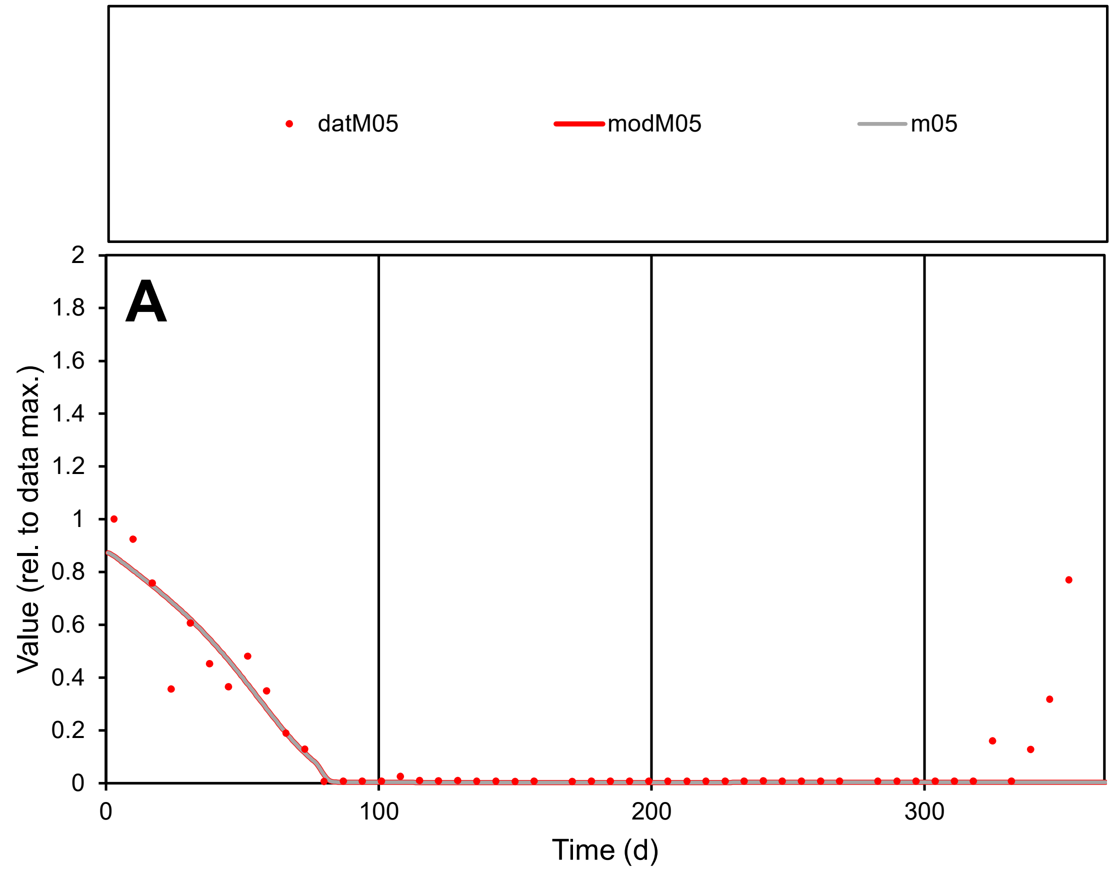


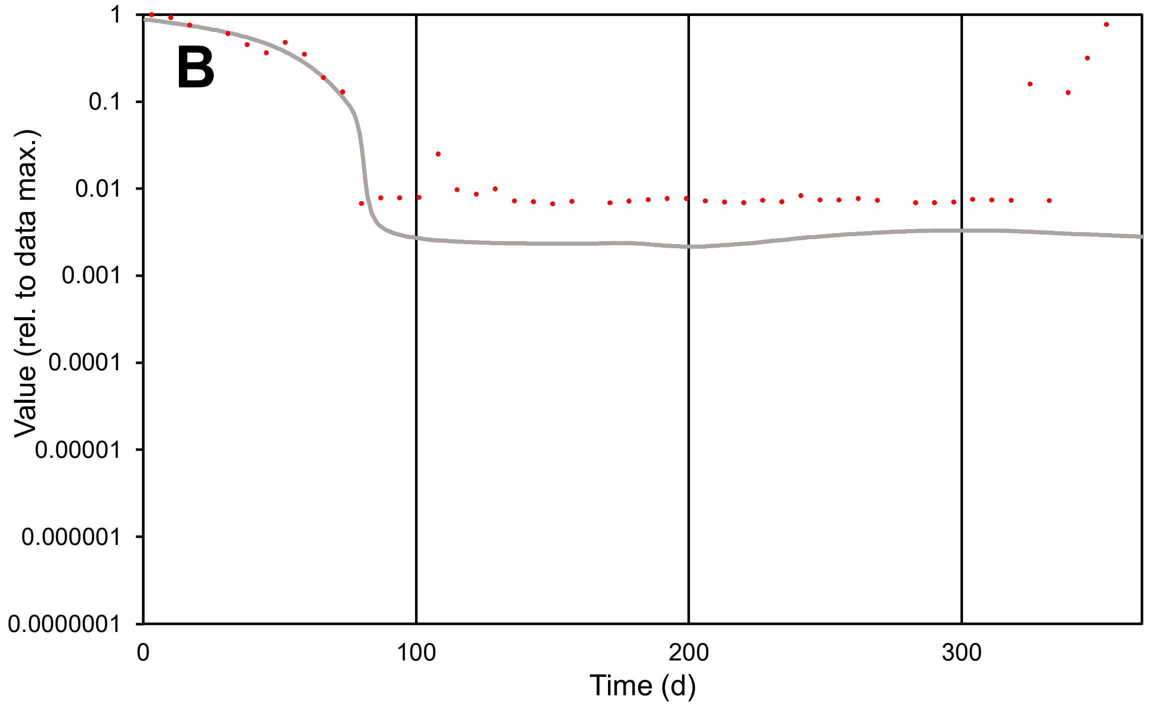


**C1**

Variables
M05 Provide jtd or Dst
CalibOnly?
2 1 to reproduce of
oNormalize?
2 1=no, 2=yes

ofxn calc*
0.91657
ofxn mod
0.94703

sRun
RunA1
zout
1

Ave
Data
0.00095
Model
0.00074

data/vMax
0.00641

ofxng
369.894

Min d/m
3.11175

ave/min d
22.002
ave/min m
53.3927

328_Calib_S_A

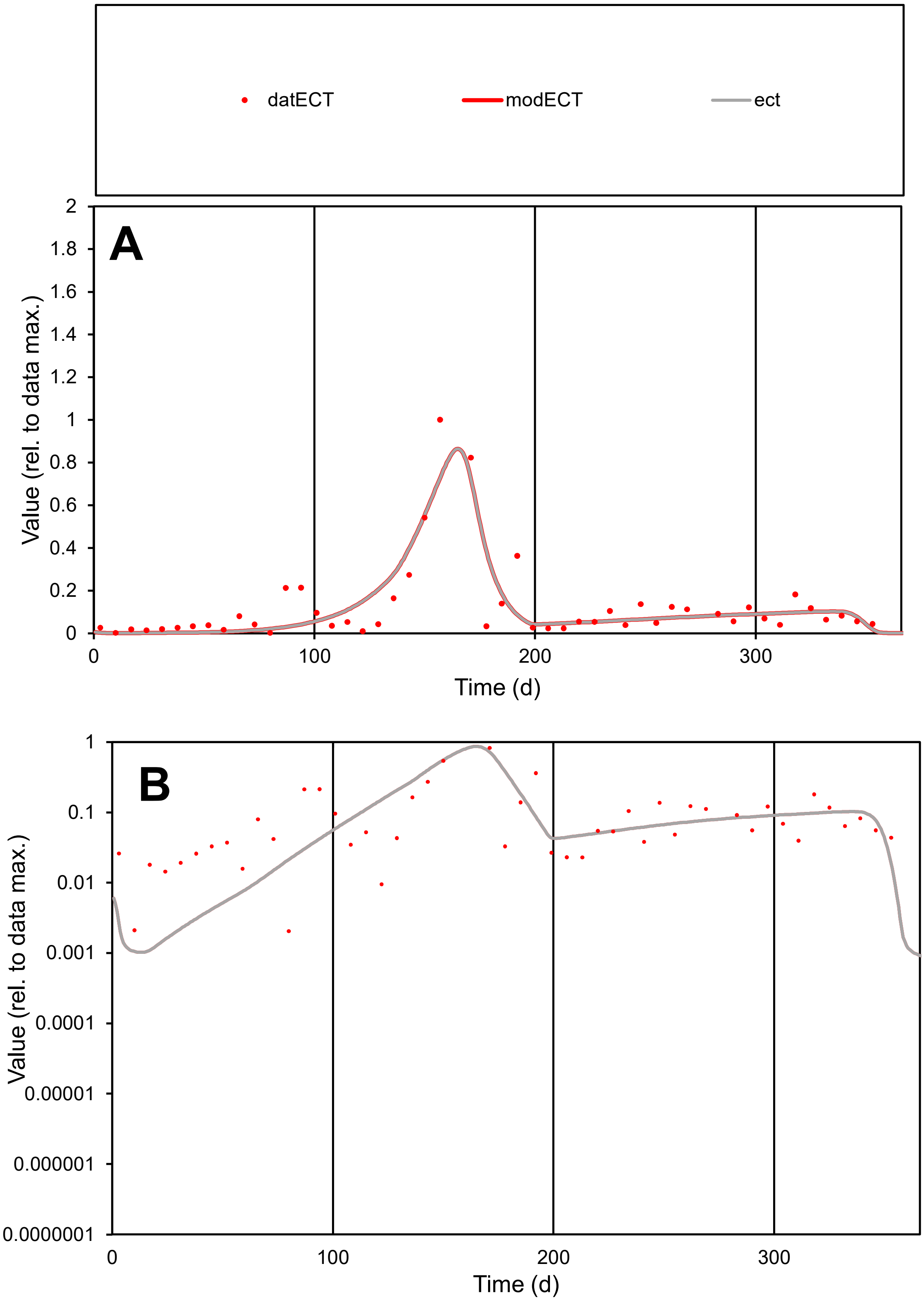


**C1**

Variables
ECT Provide jtd or Dst
CalibOnly?
2 1 to reproduce of
oNormalize?
2 1=no, 2=yes

ofxn calc*
0.75303
ofxn mod
0.79461

sRun
RunA1
zout
1

Ave
Data
0.0017
Model
0.00161

data/vMax
0.01391

ofxng
369.894

Min d/m
1.98536

ave/min d
59.5983
ave/min m
111.936

328_Calib_S_A

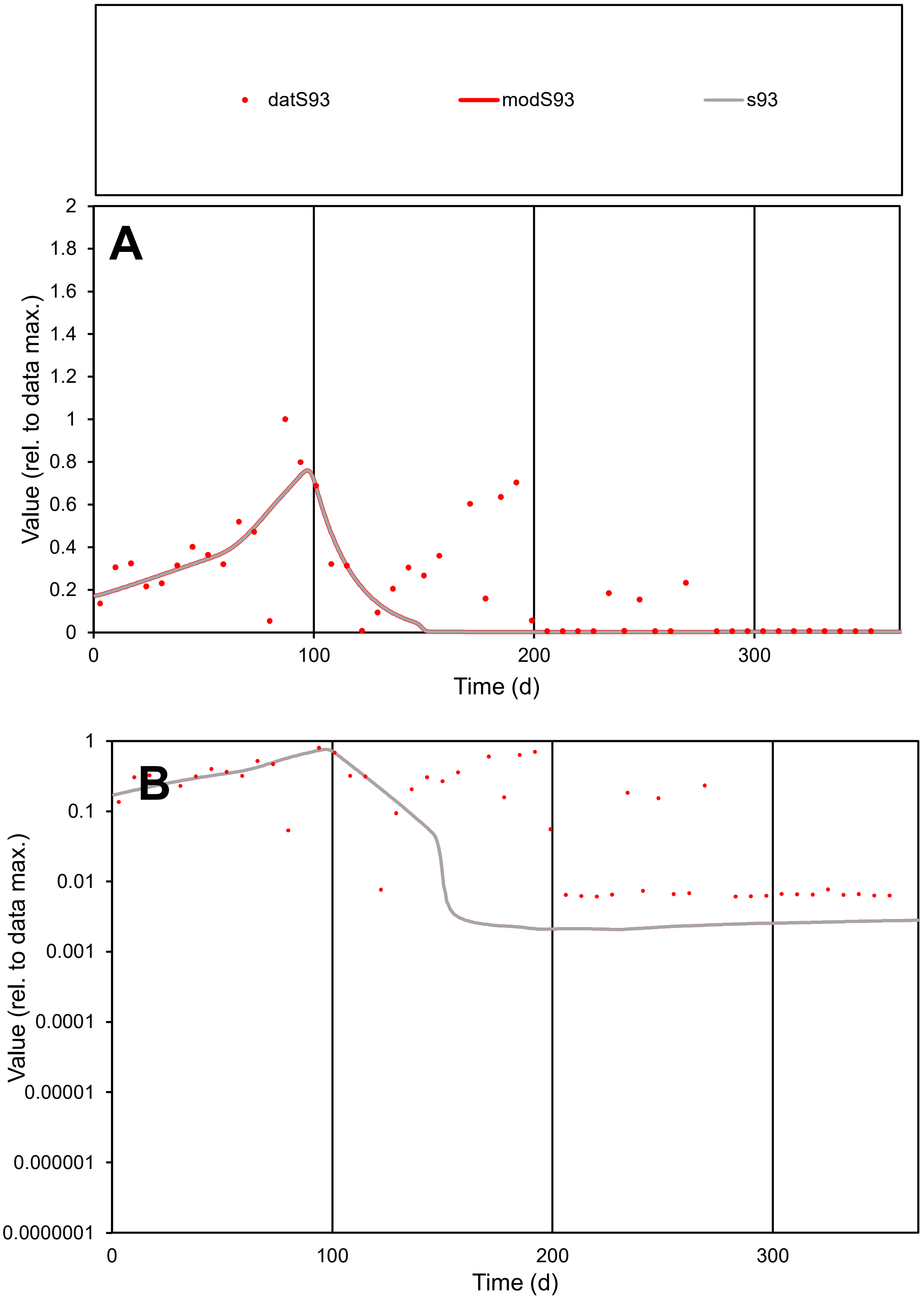


**C1**

Variables
S93 Provide jtd or Dst
CalibOnly?
2 1 to reproduce of
oNormalize?
2 1=no, 2=yes

ofxn calc*
0.9408
ofxn mod
0.95297

sRun
RunA1
zout
1

Ave
Data
0.00242
Model
0.00164

data/vMax
0.01092

ofxng
369.894

Min d/m
2.95619

ave/min d
36.2158
ave/min m
72.3499

328_Calib_S_A

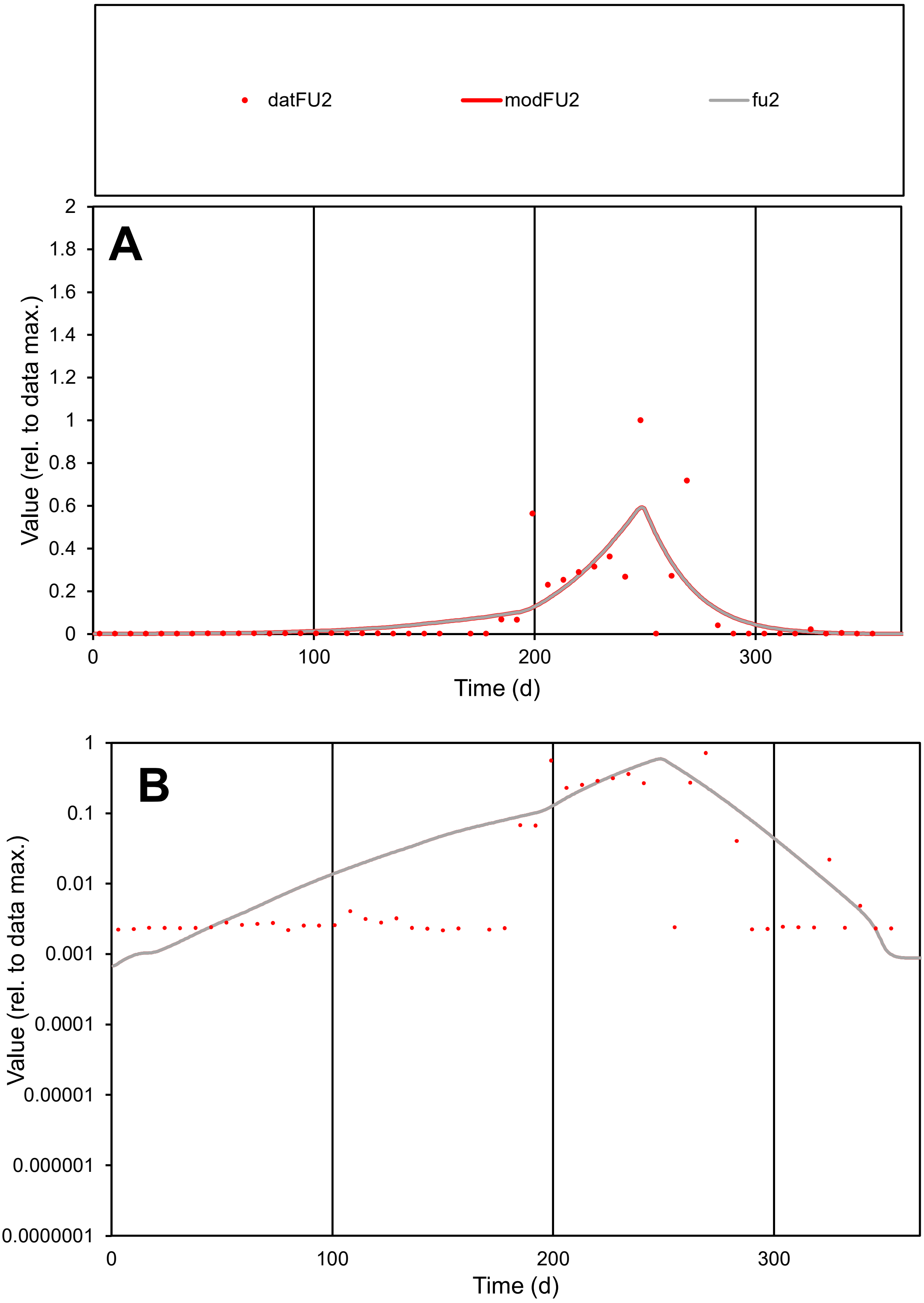


**C1**

Variables

FU2 Provide jtd or Dst

CalibOnly?

2 1 to reproduce of

oNormalize?

2 1=no, 2=yes

ofxn calc*

1.42312

ofxn mod

1.46391

sRun

RunA1

zout

1

Ave

Data

0.00109

Model

0.00112

data/vMax

0.01169

ofxng

369.894

Min d/m

2.78973

ave/min d

42.5975

ave/min m

121.958

328_Calib_S_A

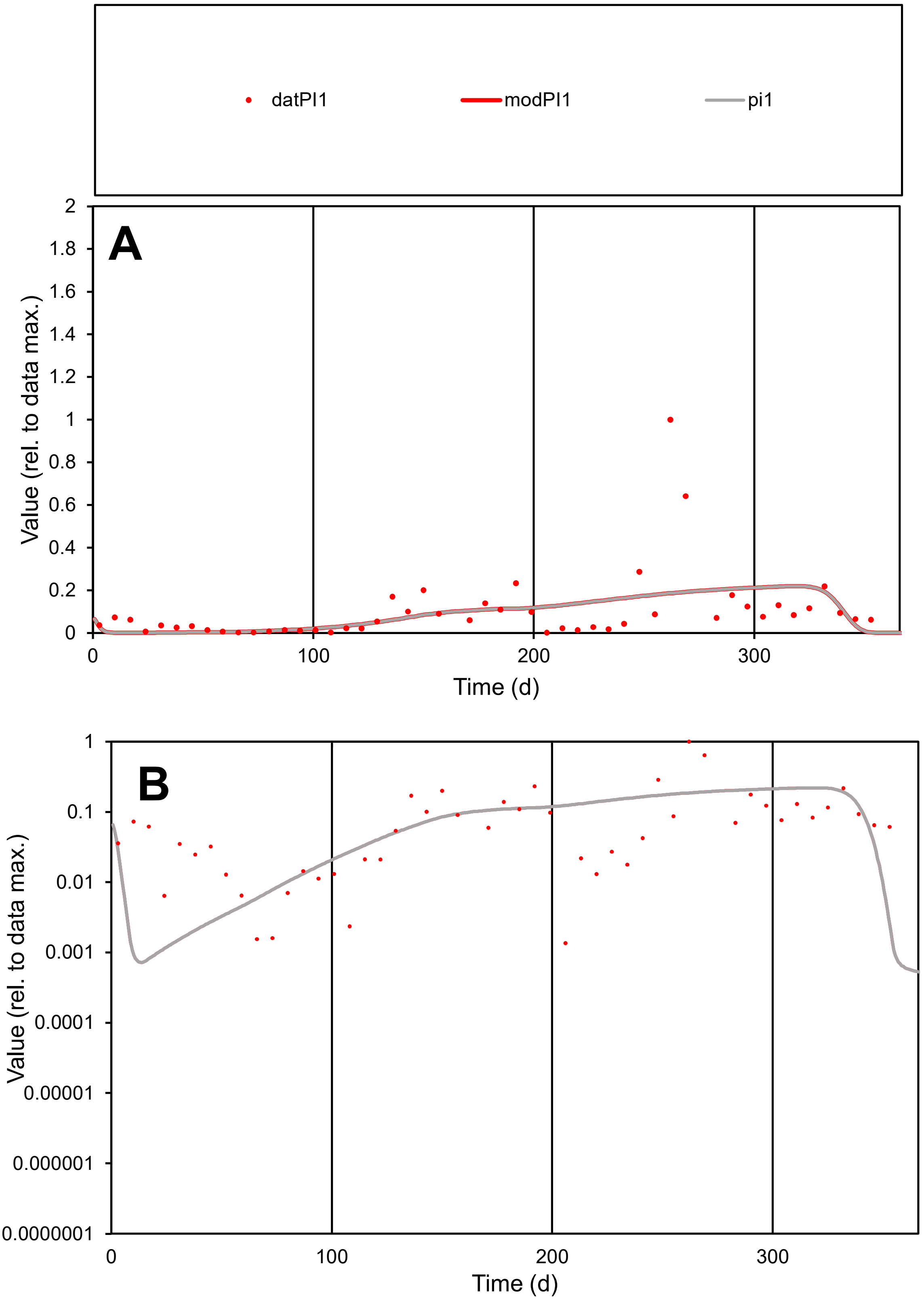


**C1**

Variables

PI1 Provide jtd or Dst

CalibOnly?

2 1 to reproduce of

oNormalize?

2 1=no, 2=yes

ofxn calc*

1.48343

ofxn mod

1.49243

sRun

RunA1

zout

1

Ave

Data

0.00106

Model

0.00098

data/vMax

0.01042

ofxng

369.894

Min d/m

1.76127

ave/min d

74.7279

ave/min m

121.289

328_Calib_S_A

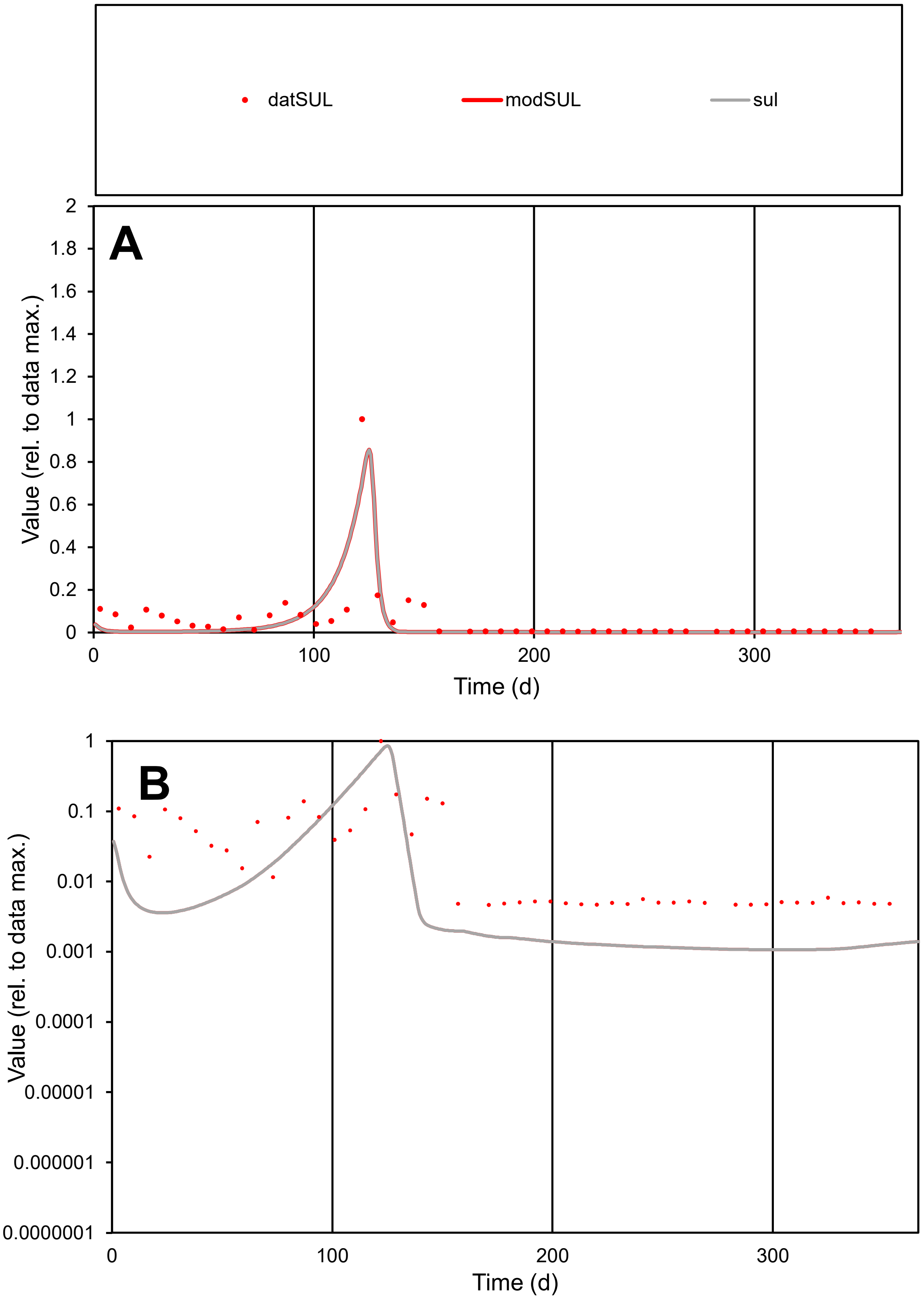


**C1**

Variables

SUL Provide jtd or Dst

CalibOnly?

2 1 to reproduce of

oNormalize?

2 1=no, 2=yes

ofxn calc*

1.21692

ofxn mod

1.31944

sRun

RunA1

zout

1

Ave

Data

0.00098

Model

0.00073

data/vMax

0.01744

ofxng

369.894

Min d/m

4.36686

ave/min d

12.064

ave/min m

39.3474

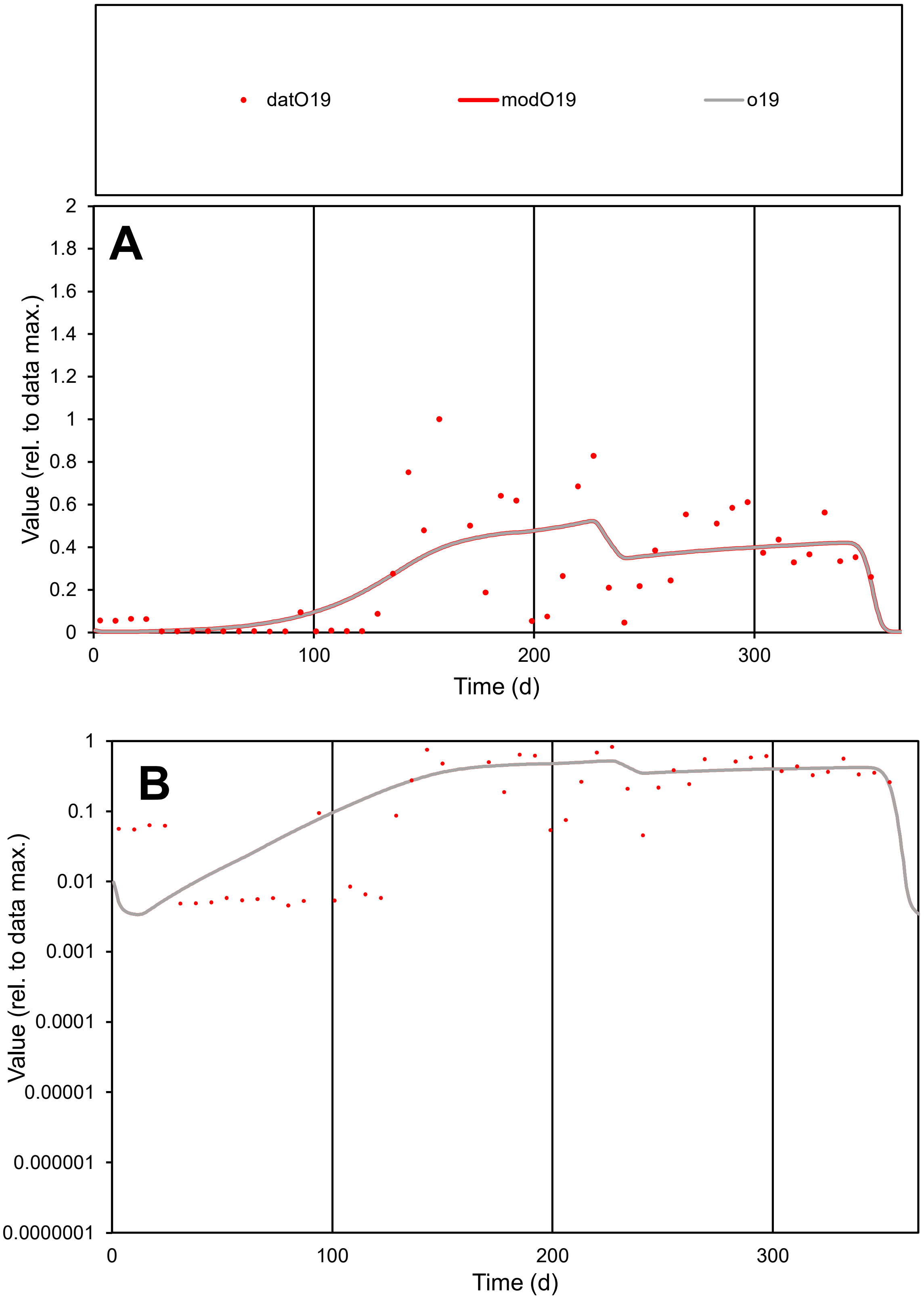


**C1**

Variables
O19 Provide jtd or Dst
CalibOnly?
2 1 to reproduce of
oNormalize?
2 1=no, 2=yes

ofxn calc*
0.68398
ofxn mod
0.69976

sRun
RunA1
zout
1

Ave
Data
0.00168
Model
0.00169

data/vMax
0.00621

ofxng
369.894

Min d/m
1.35493

ave/min d
58.9111
ave/min m
80.4191

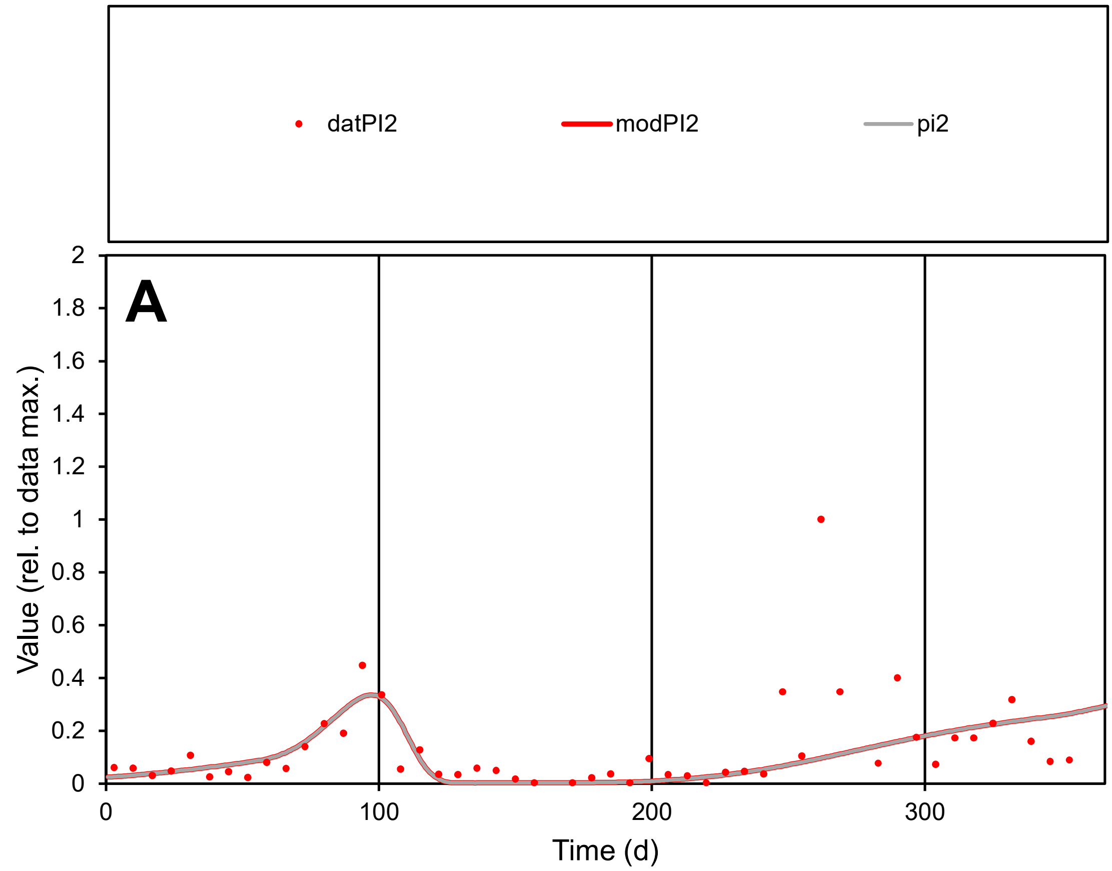


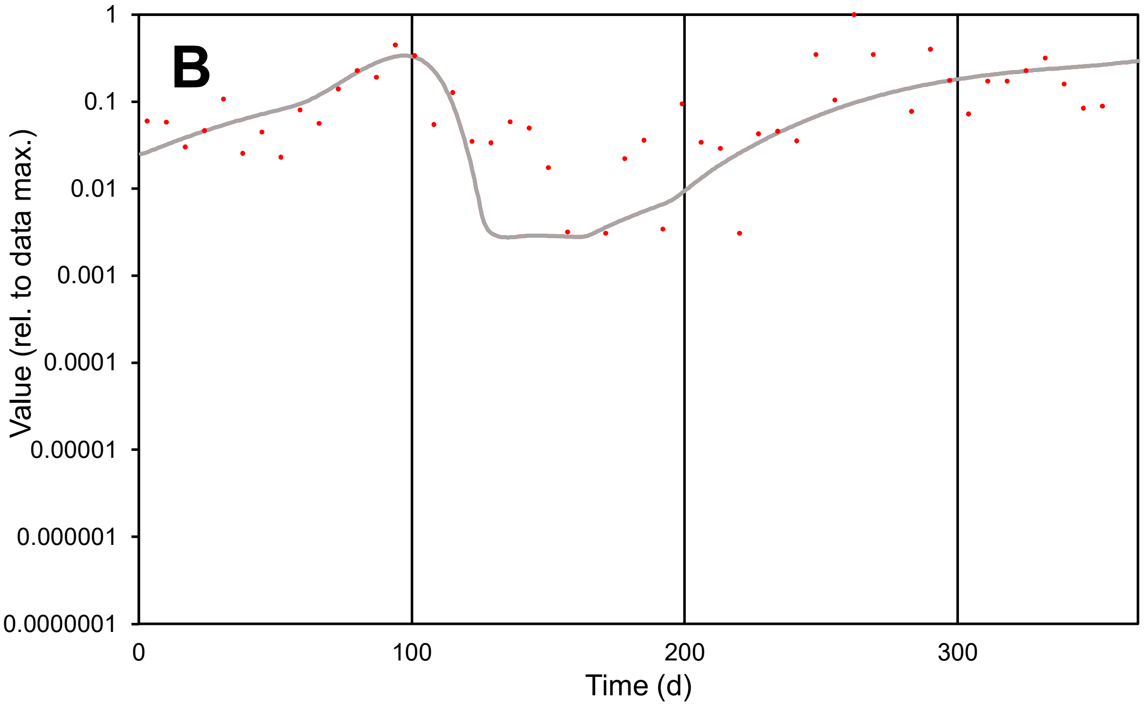


**C1**

Variables
PI2 Provide jtd or Dst
CalibOnly?
2 1 to reproduce of
oNormalize?
2 1=no, 2=yes

ofxn calc*
1.19063
ofxn mod
1.20007

sRun
RunA1
zout
1

Ave
Data
0.00122
Model
0.001

data/vMax
0.00943

ofxng
369.894

Min d/m
1.10526

ave/min d
42.1791
ave/min m
38.0994

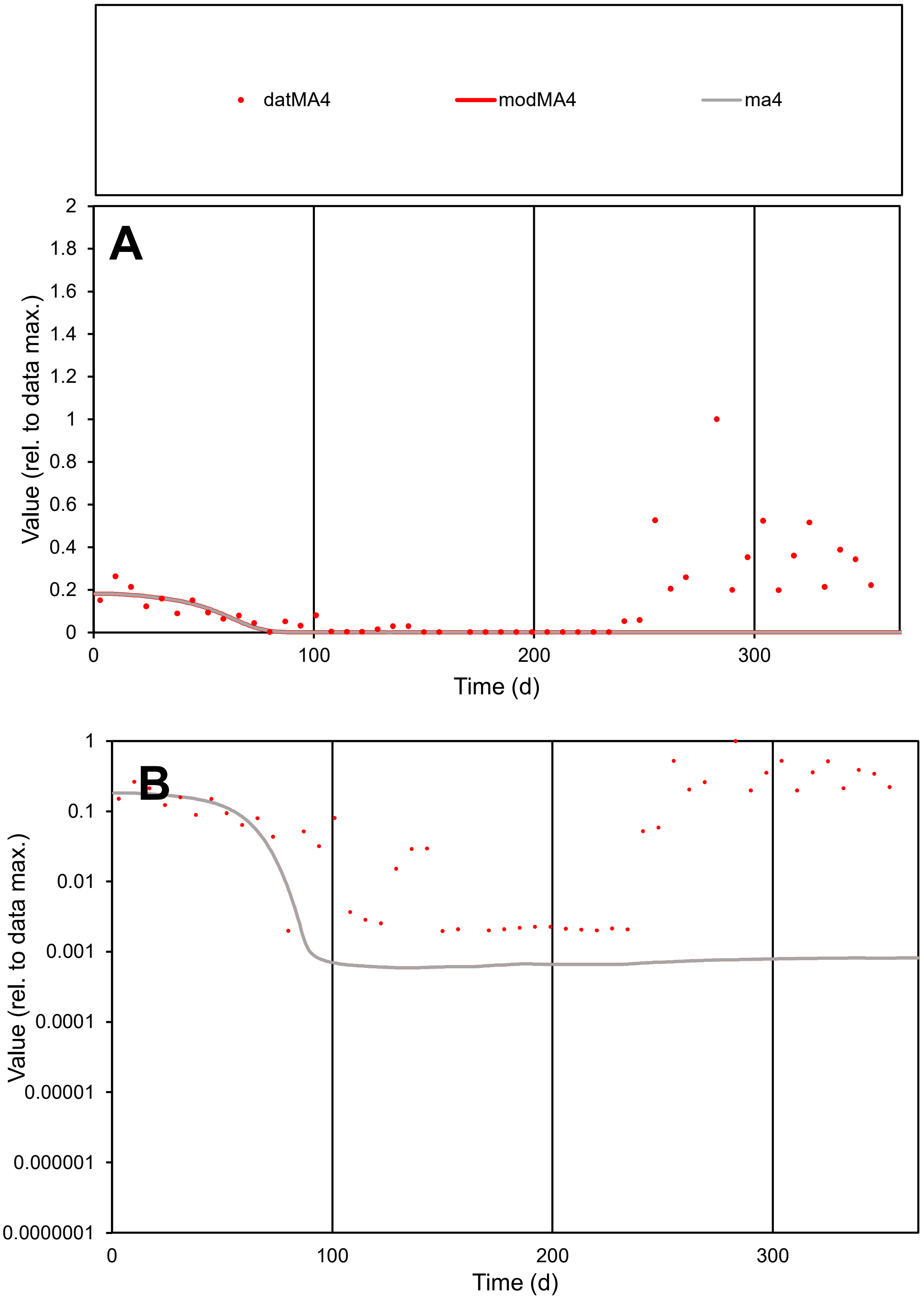


**C1**

Variables
MA4 Provide jtd or Dst
CalibOnly?
2 1 to reproduce of
oNormalize?
2 1=no, 2=yes

ofxn calc*
1.59972
ofxn mod
1.60374

sRun
RunA1
zout
1

Ave
Data
0.00142
Model
0.00029

data/vMax
0.0098

ofxng
369.894

Min d/m
3.31273

ave/min d
73.8067
ave/min m
49.2845

328_Calib_S_A

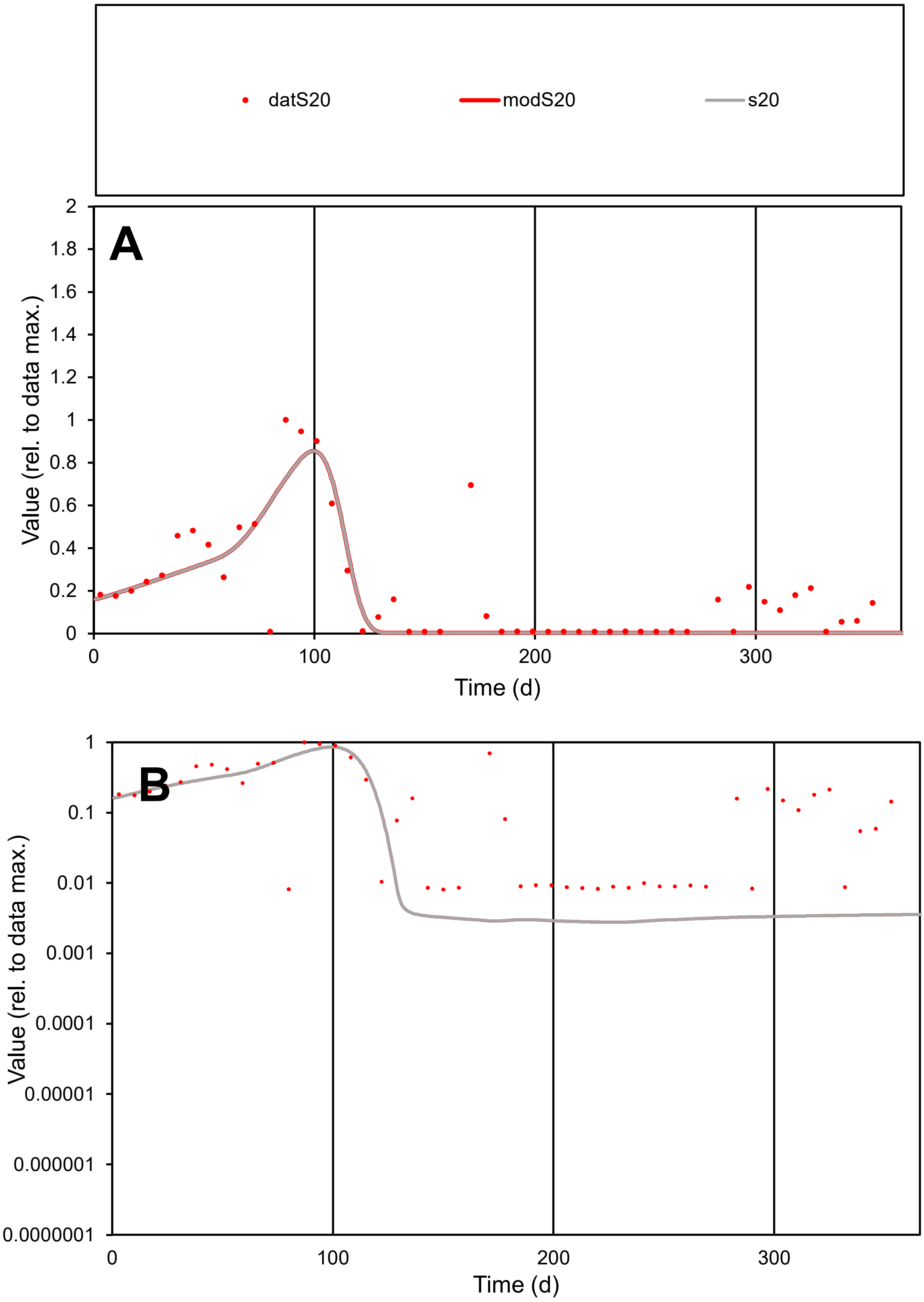


**C1**

Variables

S20 Provide jtd or Dst

CalibOnly?

2 1 to reproduce of

oNormalize?

2 1=no, 2=yes

ofxn calc*

0.79022

ofxn mod

0.80629

sRun

RunA1

zout

1

Ave

Data

0.00196

Model

0.00149

data/vMax

0.00967

ofxng

369.894

Min d/m

2.92106

ave/min d

25.0716

ave/min m

55.7543

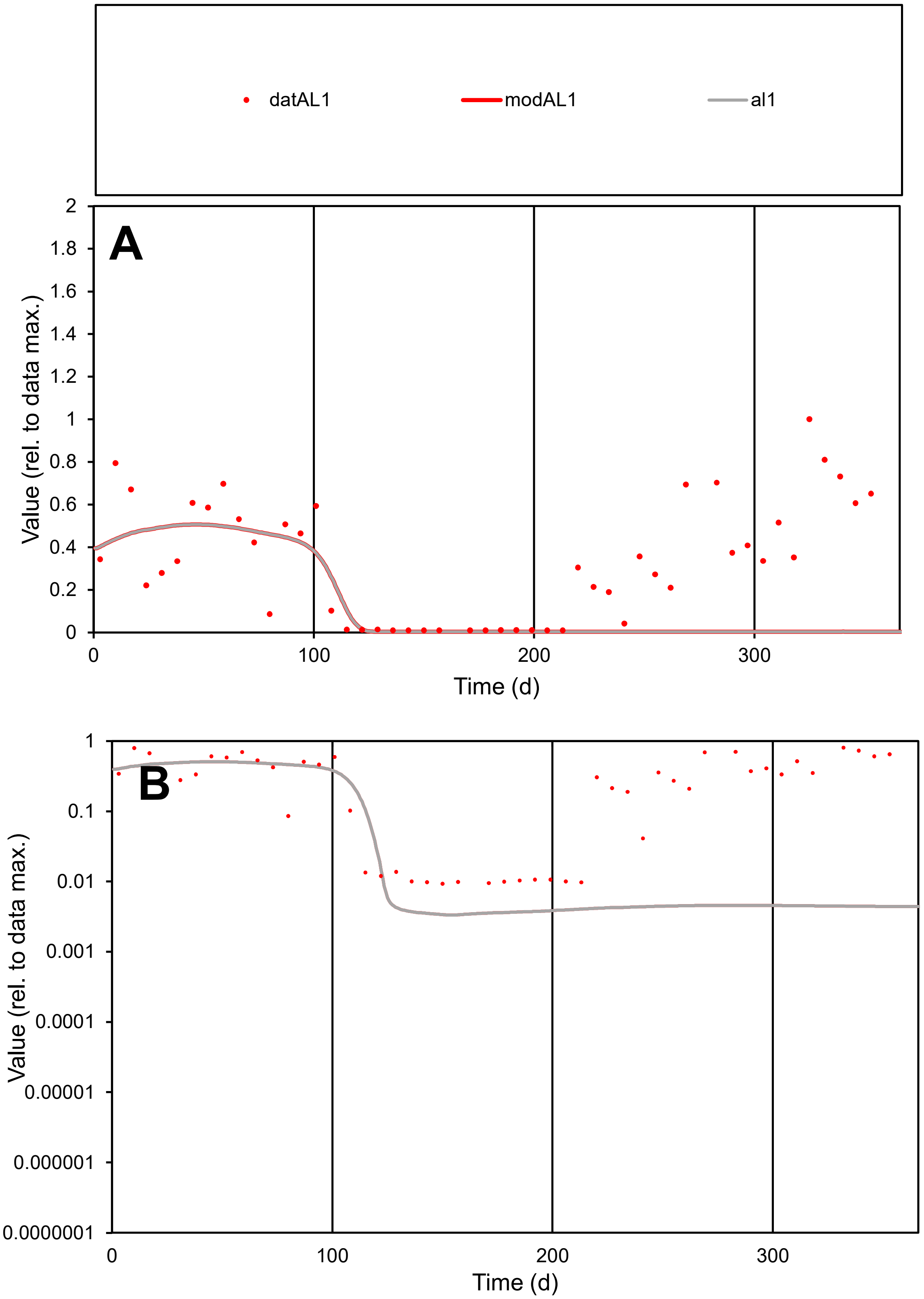


C1

Variables
AL1 Provide jtd or Dst
CalibOnly?
2 1 to reproduce of
oNormalize?
2 1=no, 2=yes

ofxn calc*
1.0342
ofxn mod
1.04041

sRun
RunA1
zout
1

Ave
Data
0.00156
Model
0.00071

data/vMax
0.00472

ofxng
369.894

Min d/m
2.76749

ave/min d
35.4075
ave/min m
44.4926

328_Calib_S_A

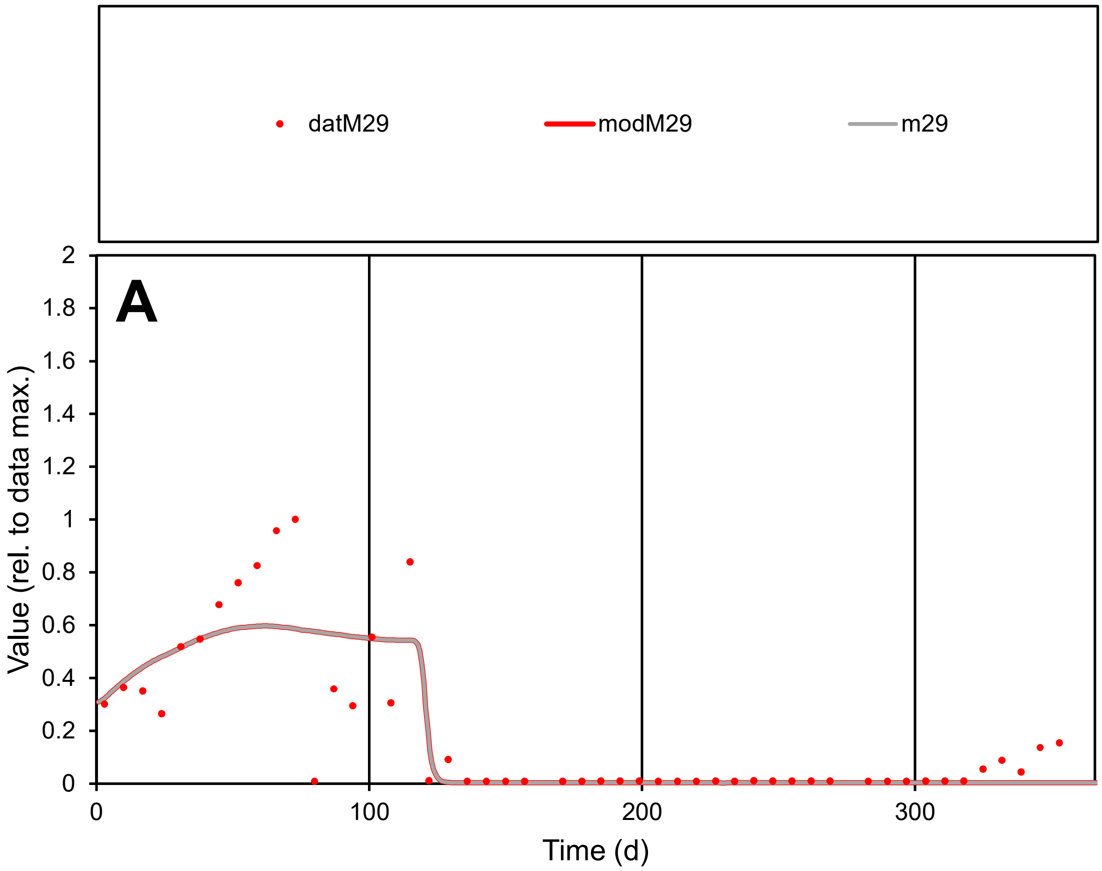


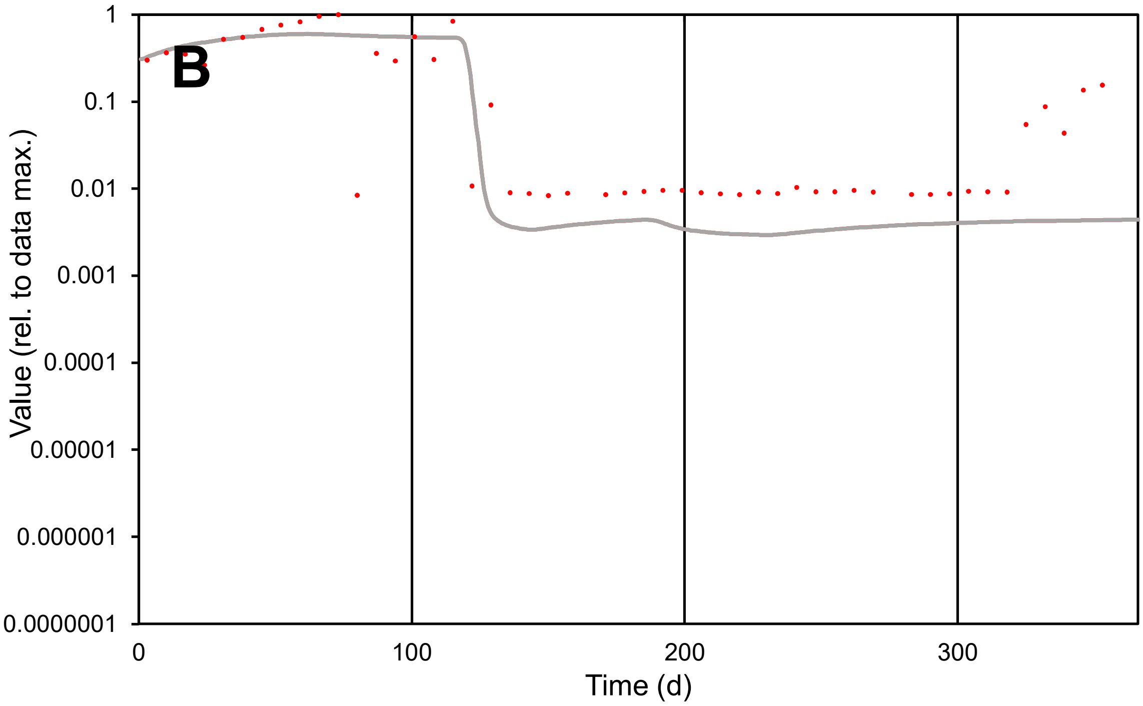


**C1**

Variables
M29 Provide jtd or Dst
CalibOnly?
2 1 to reproduce of
oNormalize?
2 1=no, 2=yes

ofxn calc*
0.75231
ofxn mod
0.77539

sRun
RunA1
zout
1

Ave
Data
0.00129
Model
0.00122

data/vMax
0.00652

ofxng
369.894

Min d/m
2.84439

ave/min d
23.7665
ave/min m
63.6787

328_Calib_S_A

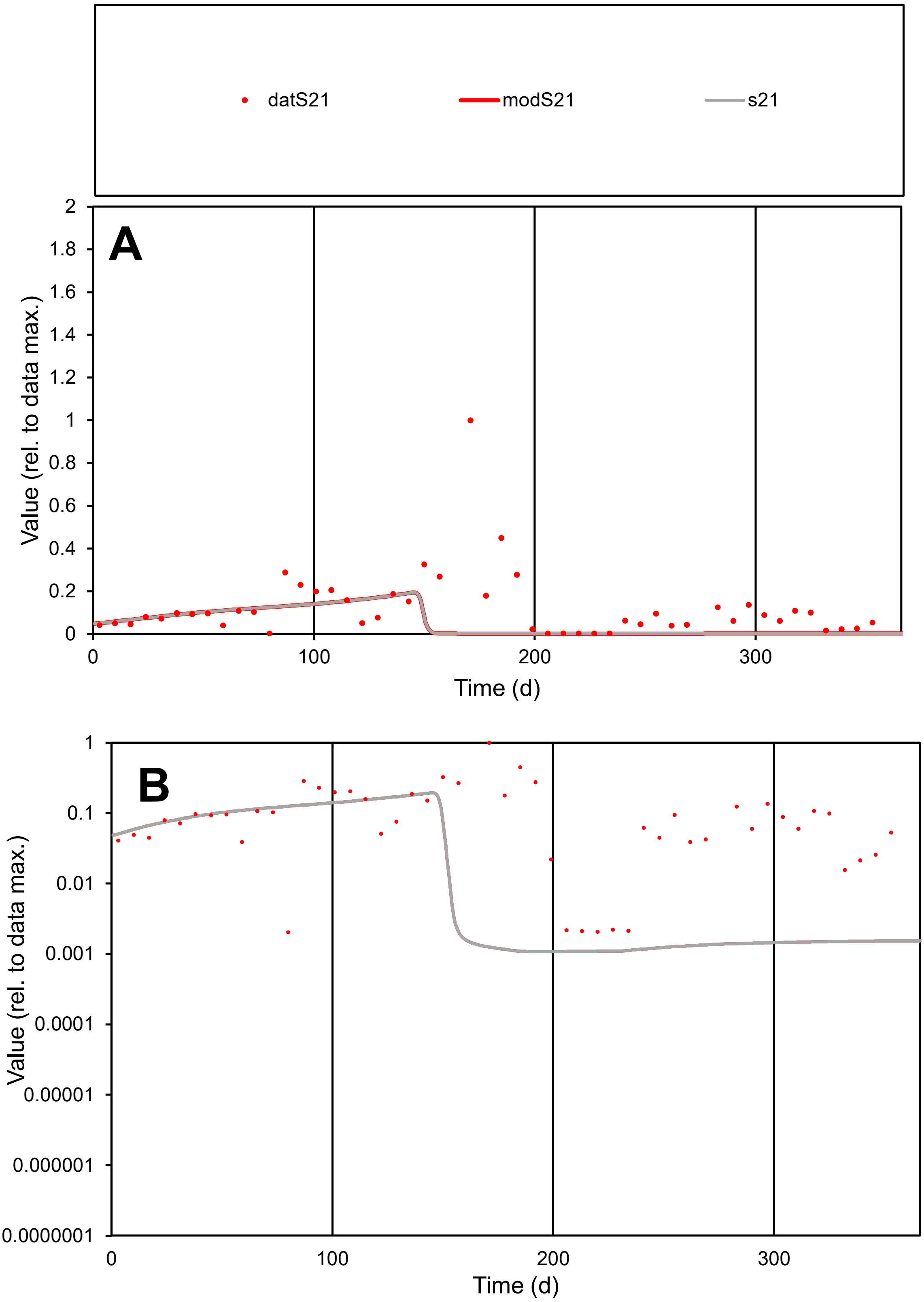


**C1**

Variables

S21 Provide jtd or Dst

CalibOnly?

2 1 to reproduce of

oNormalize?

2 1=no, 2=yes

ofxn calc*

1.47963

ofxn mod

1.48822

sRun

RunA1

zout

1

Ave

Data

0.00247

Model

0.00112

data/vMax

0.02028

ofxng

369.894

Min d/m

1.87768

ave/min d

60.0418

ave/min m

50.9282

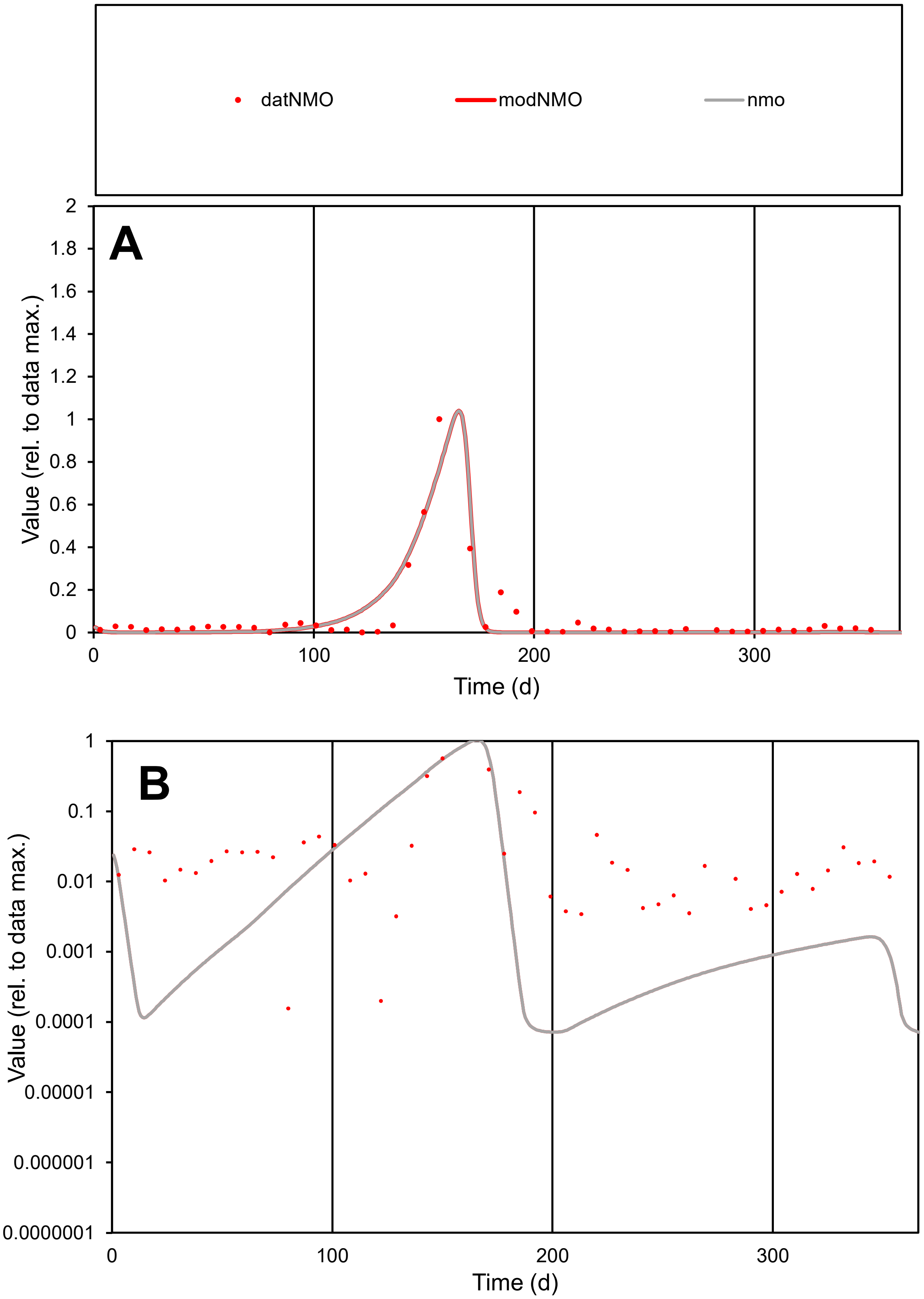


**C1**

Variables
NMO Provide jtd or Dst
CalibOnly?
2 1 to reproduce of
oNormalize?
2 1=no, 2=yes

ofxn calc*
0.87317
ofxn mod
0.94234

sRun
RunA1
zout
1

Ave
Data
0.00309
Model
0.00276

data/vMax
0.04652

ofxng
369.894

Min d/m
2.173

ave/min d
425.404
ave/min m
827.283

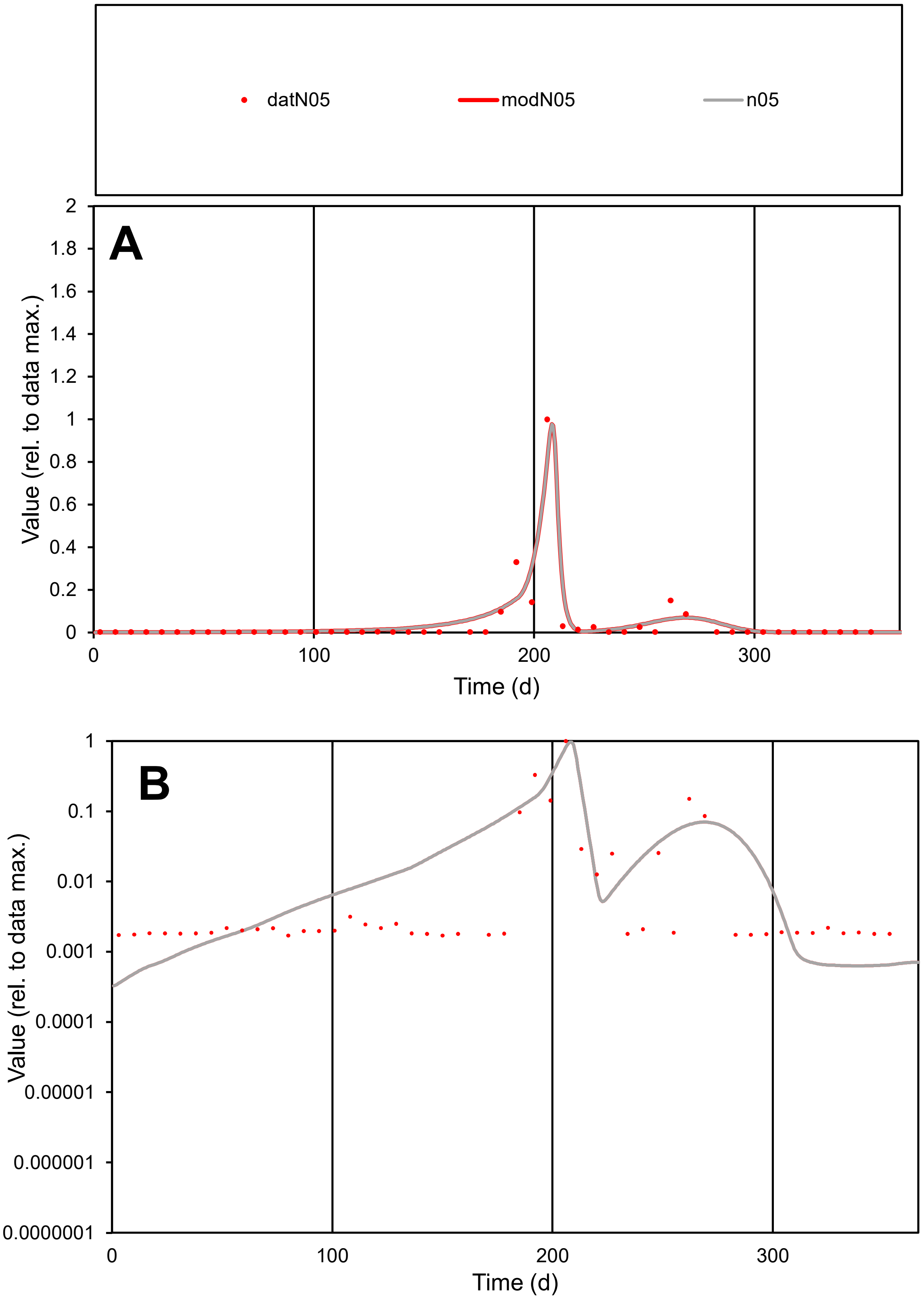


**C1**

Variables
N05 Provide jtd or Dst
CalibOnly?
2 1 to reproduce of
oNormalize?
2 1=no, 2=yes

ofxn calc*
1.05682
ofxn mod
1.1527

sRun
RunA1
zout
1

Ave
Data
0.00095
Model
0.0011

data/vMax
0.02349

ofxng
369.894

Min d/m
4.49431

ave/min d
23.7188
ave/min m
123.49

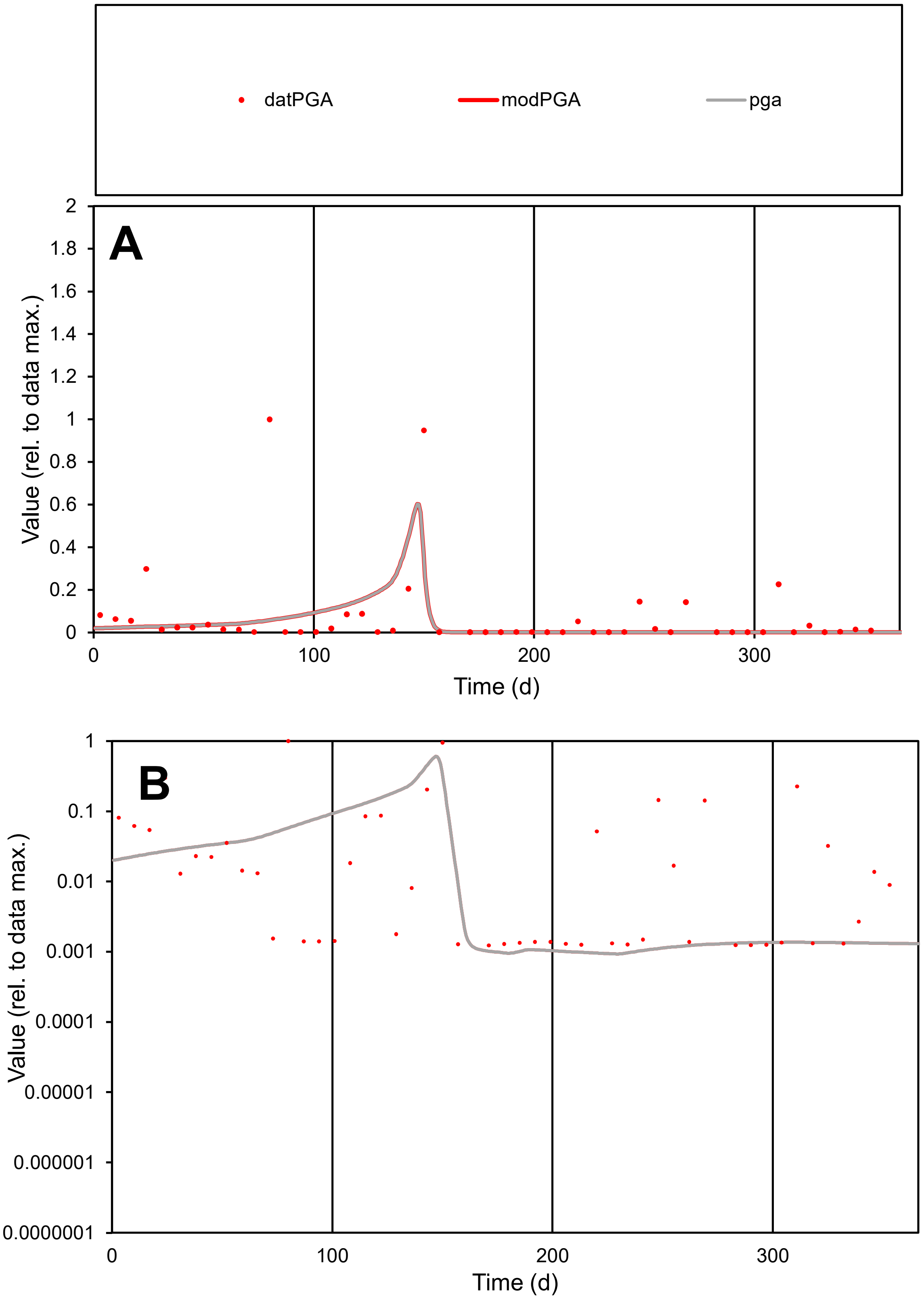


**C1**

Variables
PGA Provide jtd or Dst
CalibOnly?
2 1 to reproduce of
oNormalize?
2 1=no, 2=yes

ofxn calc*
2.32971
ofxn mod
2.4215

sRun
RunA1
zout
1

Ave
Data
0.00105
Model
0.00071

data/vMax
0.01411

ofxng
369.894

Min d/m
1.32346

ave/min d
60.086
ave/min m
53.706

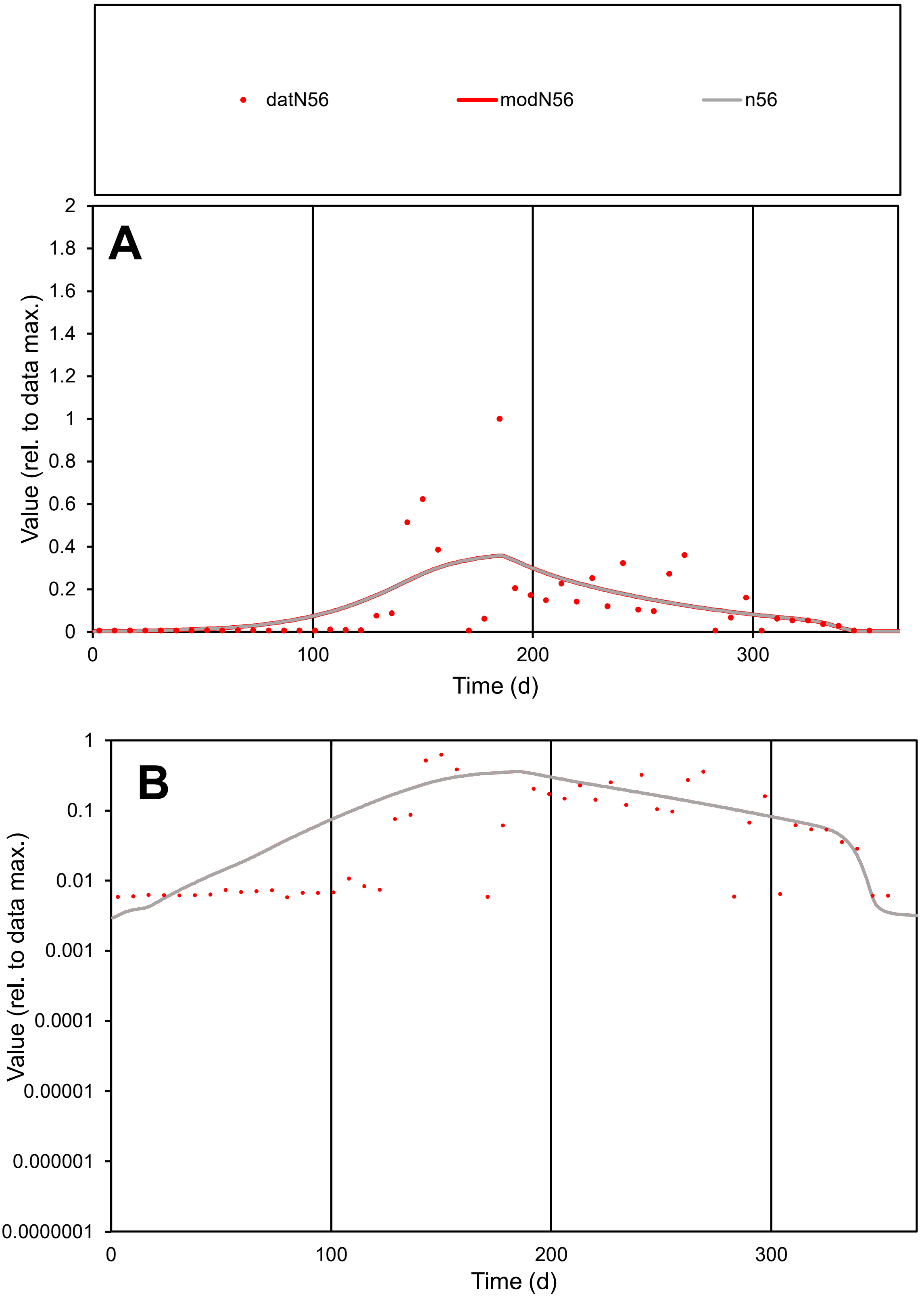


**C1**

Variables
N56 Provide jtd or Dst
CalibOnly?
2 1 to reproduce of
oNormalize?
2 1=no, 2=yes

ofxn calc*
1.24674
ofxn mod
1.26039

sRun
RunA1
zout
1

Ave
Data
0.0007
Model
0.00073

data/vMax
0.00591

ofxng
369.894

Min d/m
1.73695

ave/min d
20.3711
ave/min m
37.205

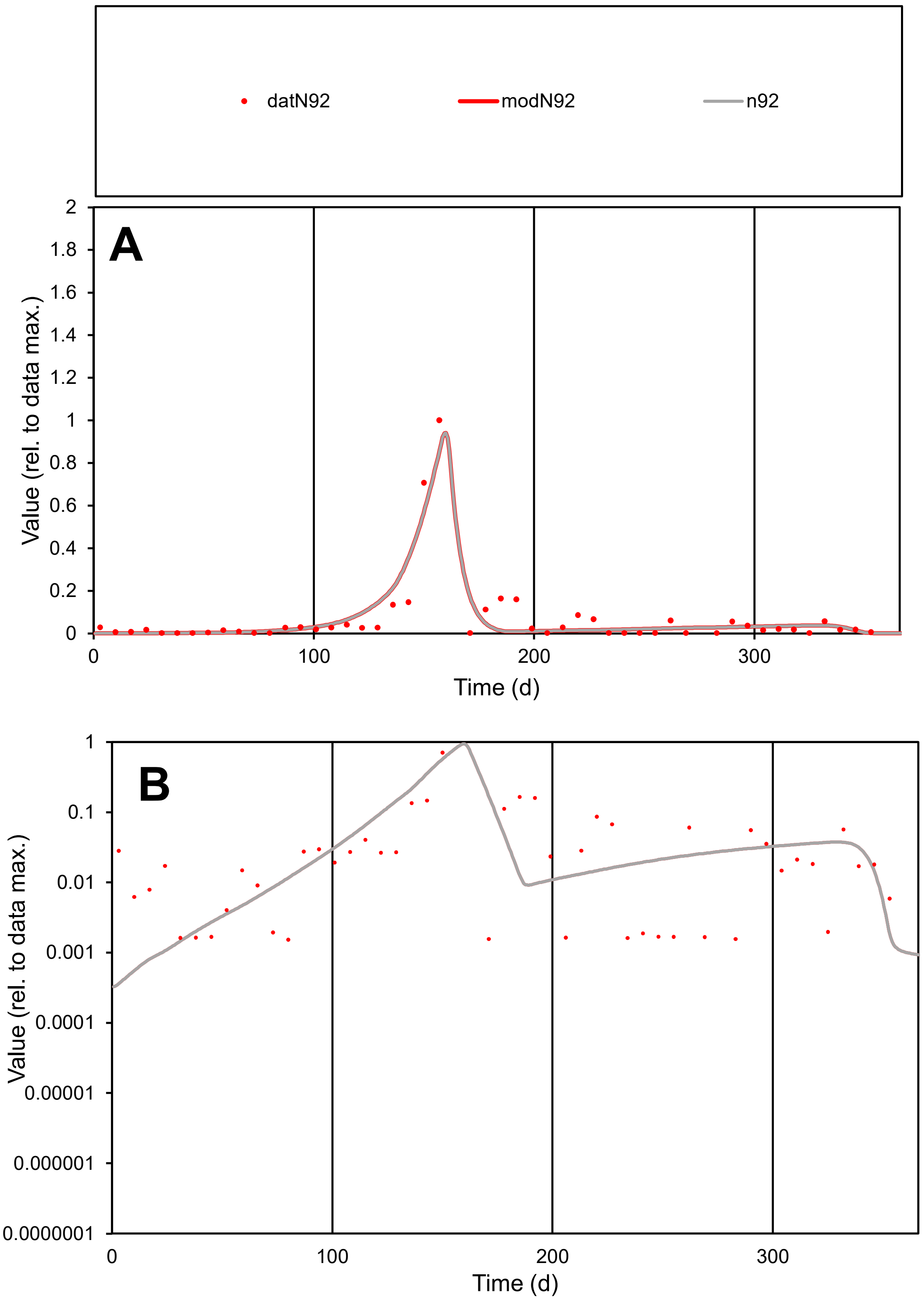


**C1**

Variables
N92 Provide jtd or Dst
CalibOnly?
2 1 to reproduce of
oNormalize?
2 1=no, 2=yes

ofxn calc*
0.8661
ofxn mod
0.93732

sRun
RunA1
zout
1

Ave
Data
0.00165
Model
0.00165

data/vMax
0.02491

ofxng
369.894

Min d/m
3.91402

ave/min d
43.1959
ave/min m
169.782

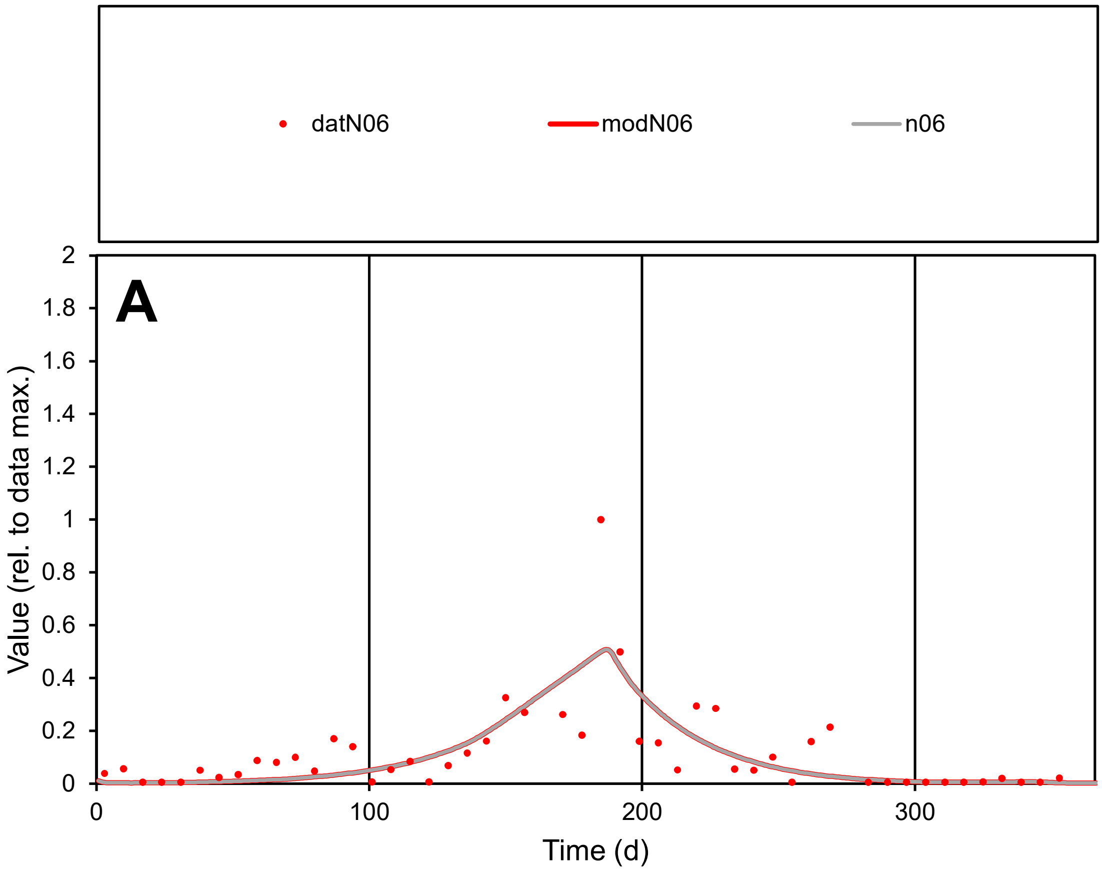


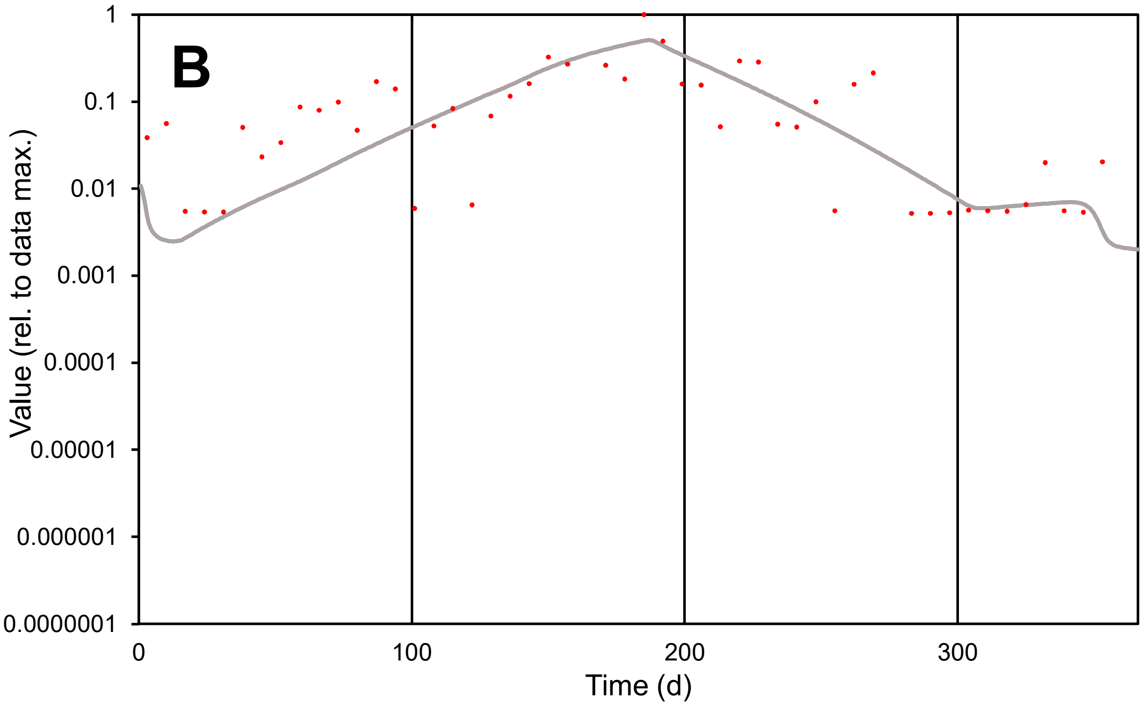


**C1**

Variables
N06 Provide jtd or Dst
CalibOnly?
2 1 to reproduce of
oNormalize?
2 1=no, 2=yes

ofxn calc*
0.96567
ofxn mod
0.98735

sRun
RunA1
zout
1

Ave
Data
0.00144
Model
0.00128

data/vMax
0.01285

ofxng
369.894

Min d/m
2.09302
ave/min d
21.5347
ave/min m
39.9781

328_Calib_S_A

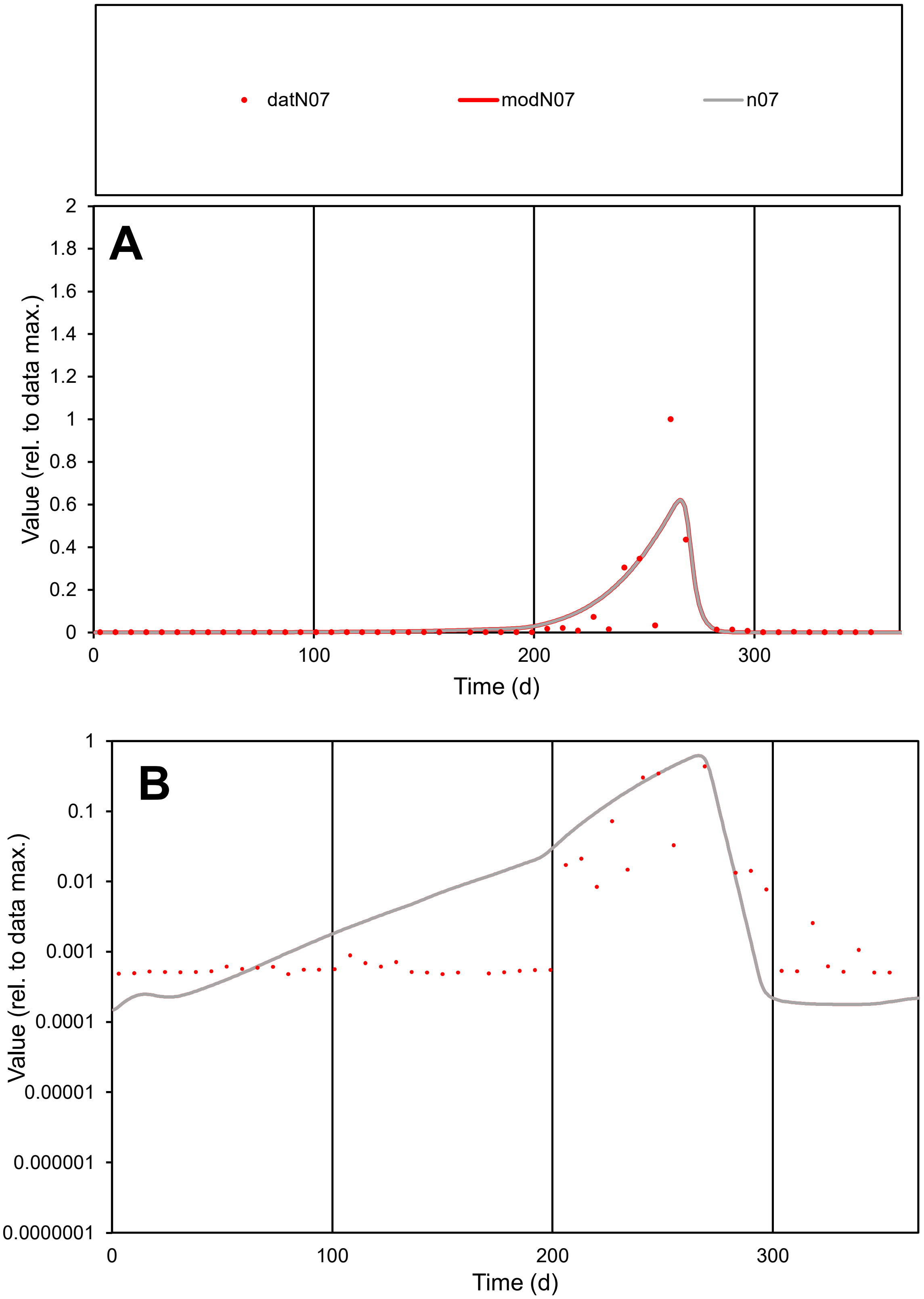


**C1**

Variables
N07 Provide jtd or Dst
CalibOnly?
2 1 to reproduce of
oNormalize?
2 1=no, 2=yes

ofxn calc*
1.90111
ofxn mod
1.94575

sRun
RunA1
zout
1

Ave
Data
0.00167
Model
0.00202

data/vMax
0.03552

ofxng
369.894

Min d/m
2.71351

ave/min d
98.3089
ave/min m
322.408

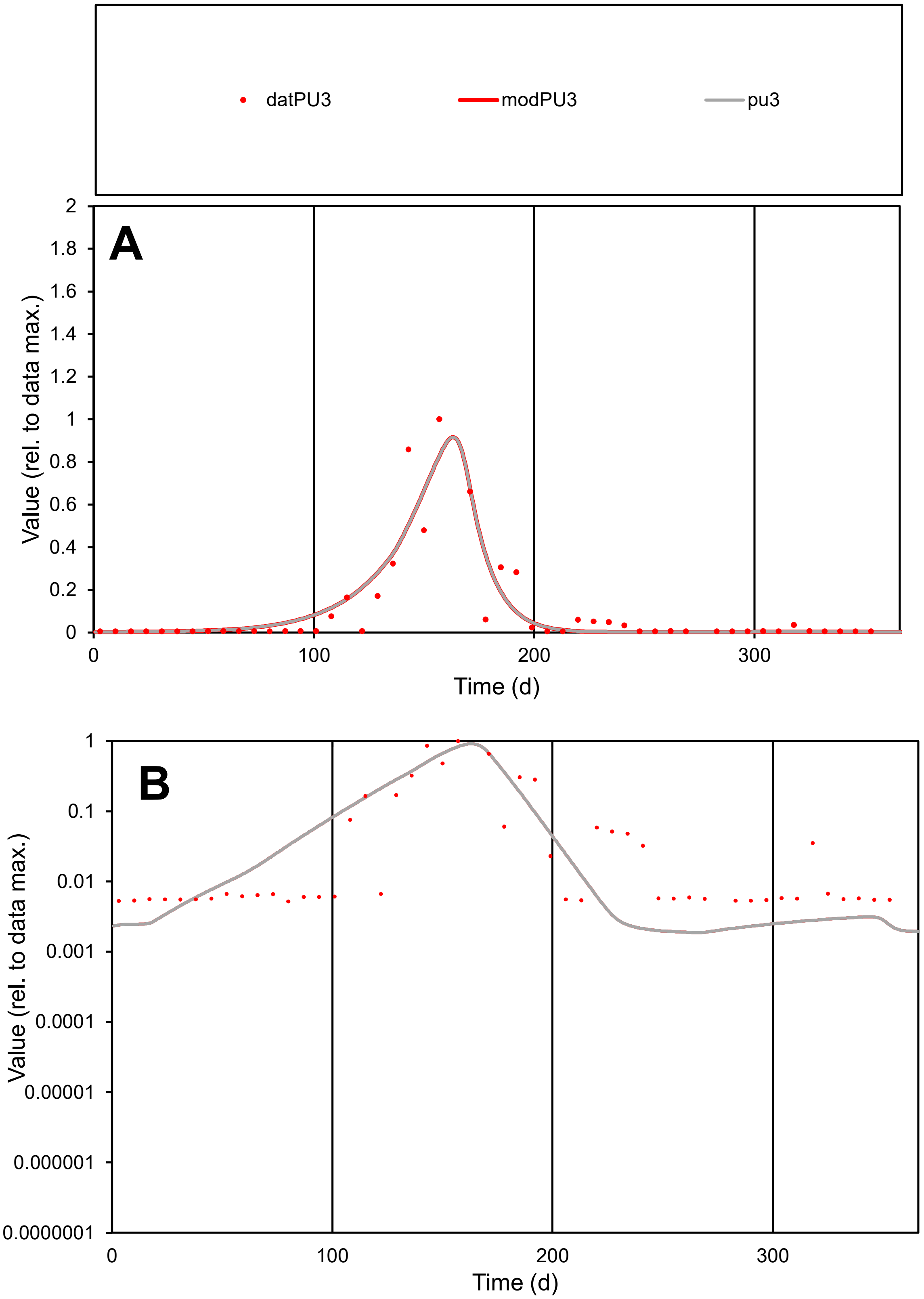


C1

Variables
PU3 Provide jtd or Dst
CalibOnly?
2 1 to reproduce of
oNormalize?
2 1=no, 2=yes

ofxn calc*
0.88106
ofxn mod
0.92822

sRun
RunA1
zout
1

Ave
Data
0.00285
Model
0.00288

data/vMax
0.029

ofxng
369.894

Min d/m
2.77228
ave/min d
18.8463
ave/min m
52.7622

328_Calib_S_A

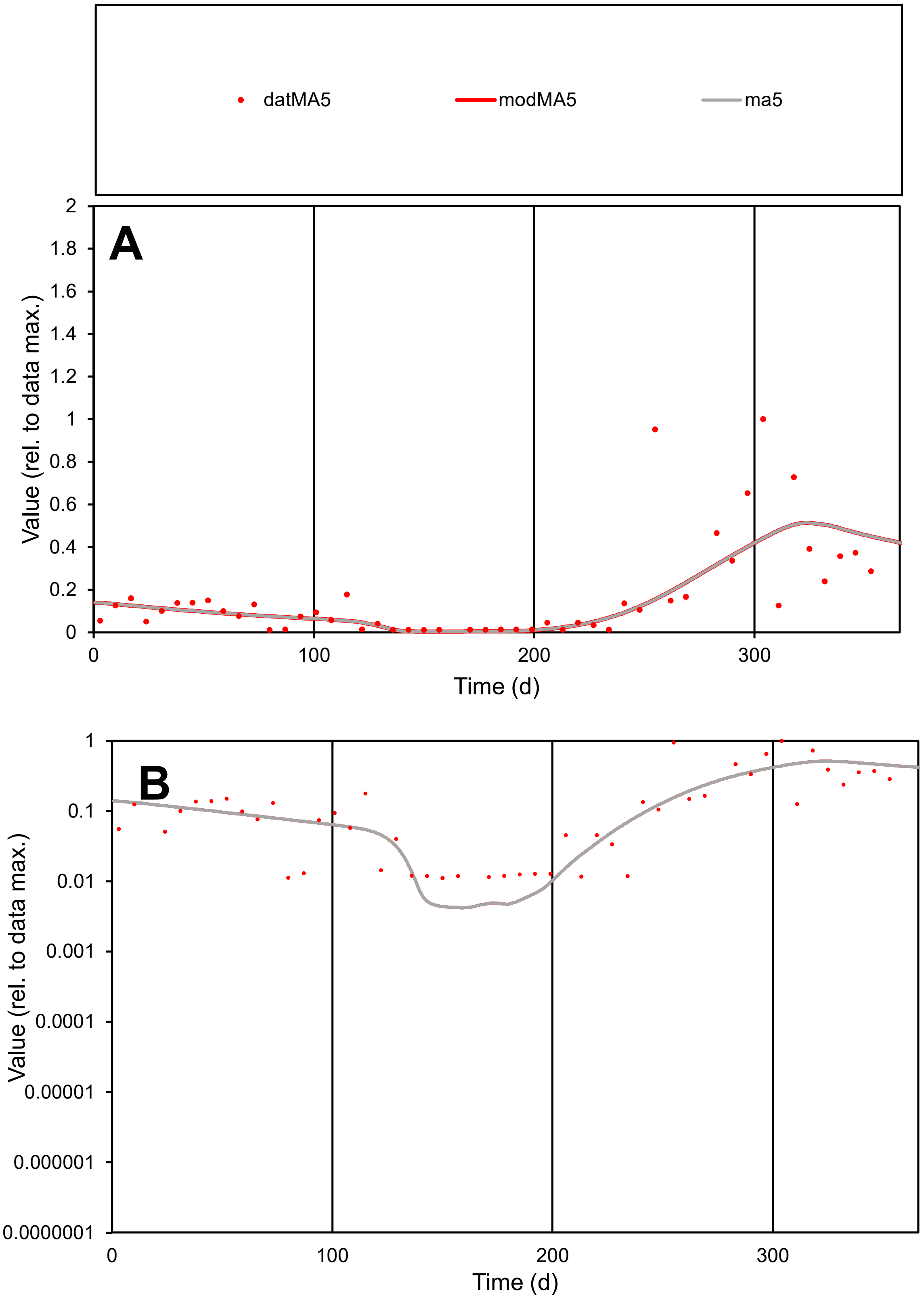


**C1**

Variables
MA5 Provide jtd or Dst
CalibOnly?
2 1 to reproduce of
oNormalize?
2 1=no, 2=yes

ofxn calc*
0.97376
ofxn mod
0.98663

sRun
RunA1
zout
1

Ave
Data
0.00144
Model
0.00131

data/vMax
0.00837

ofxng
369.894

Min d/m
2.67397

ave/min d
15.3331
ave/min m
37.3856

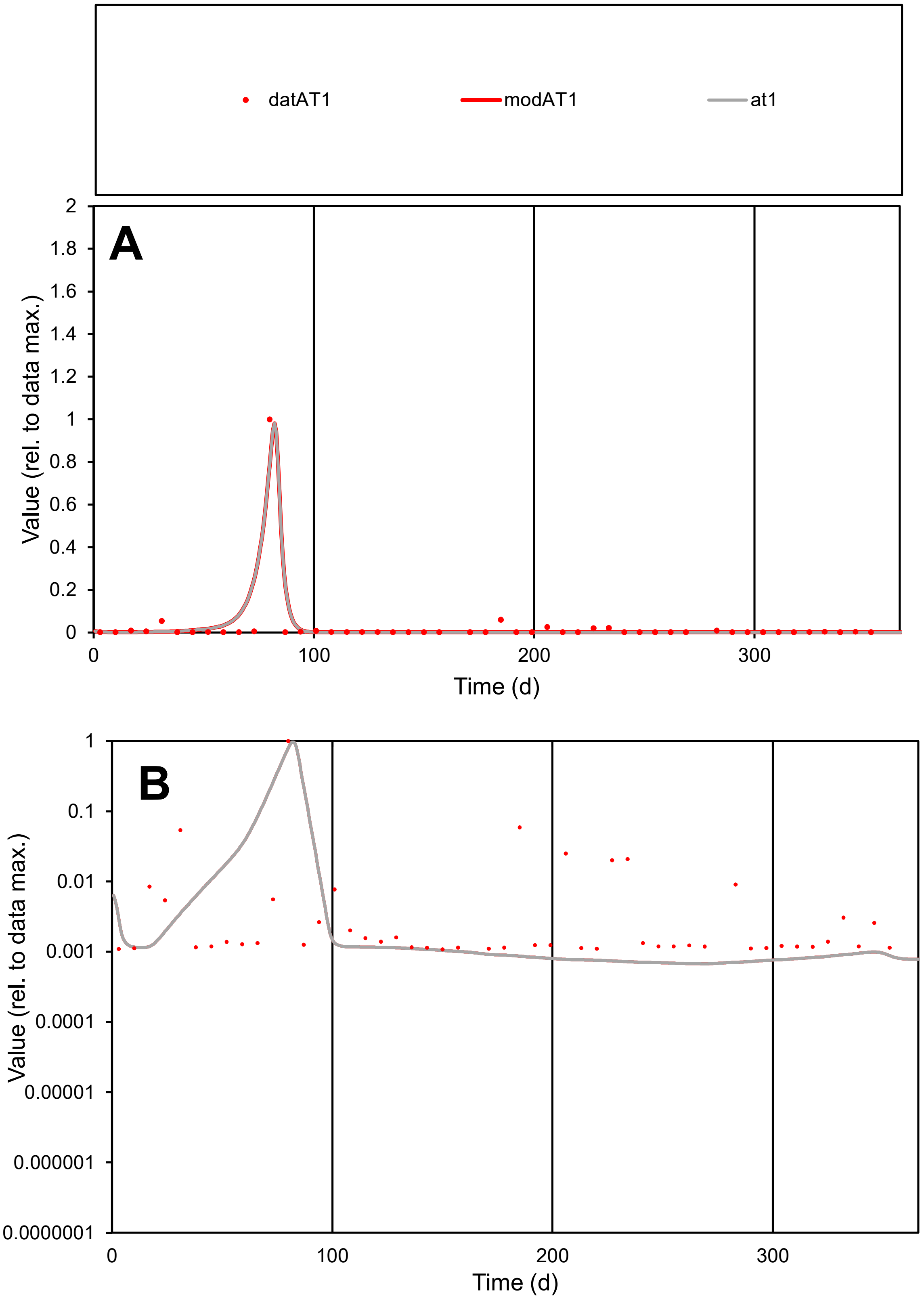


**C1**

Variables
AT1 Provide jtd or Dst
CalibOnly?
2 1 to reproduce of
oNormalize?
2 1=no, 2=yes

ofxn calc*
1.30412
ofxn mod
1.55767

sRun
RunA1
zout
1

Ave
Data
0.00115
Model
0.00126

data/vMax
0.04429

ofxng
369.894

Min d/m
1.60074

ave/min d
23.9431
ave/min m
42.0622

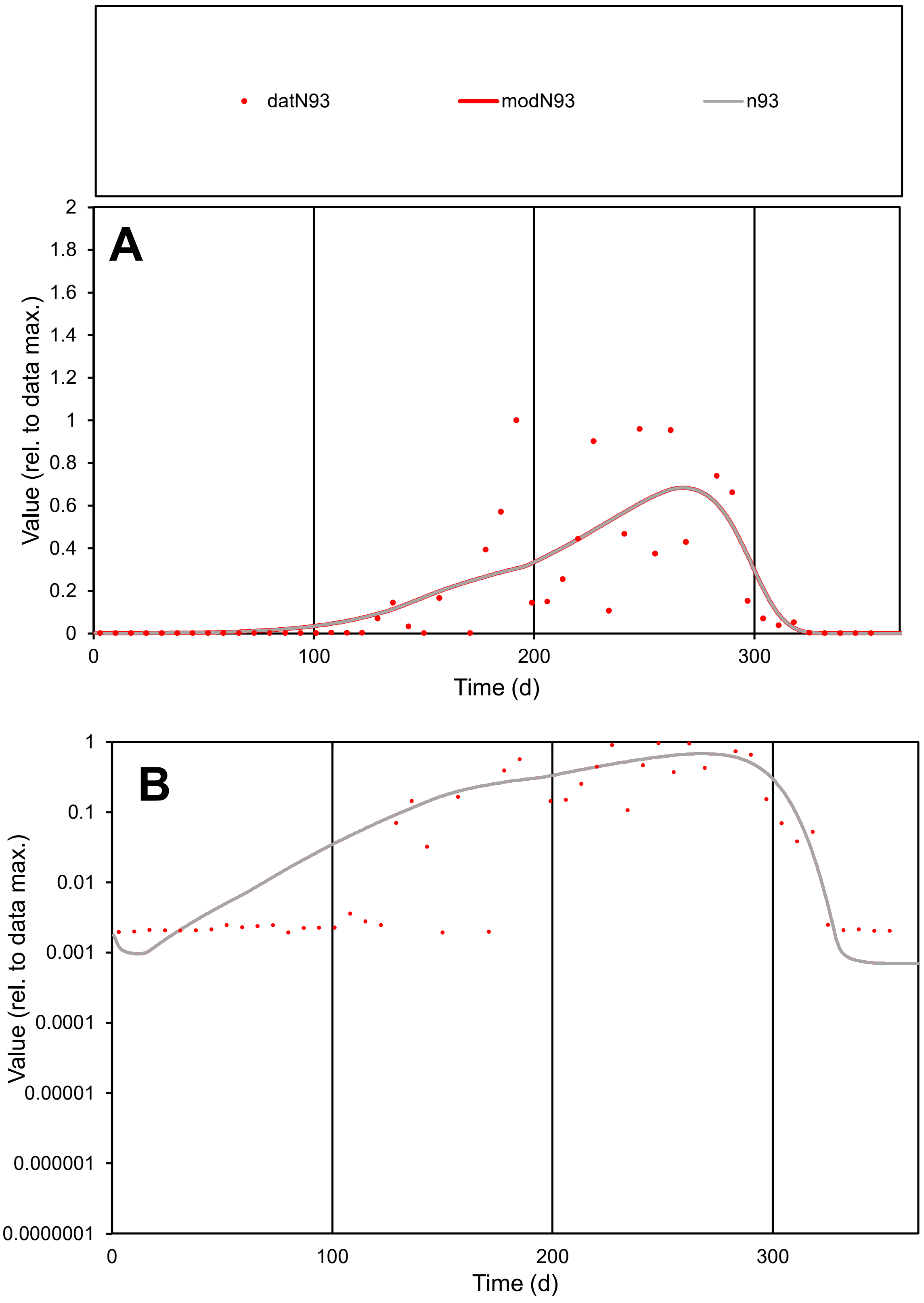


**C1**

Variables
N93 Provide jtd or Dst
CalibOnly?
2 1 to reproduce of
oNormalize?
2 1=no, 2=yes

ofxn calc*
0.94285
ofxn mod
0.96976

sRun
RunA1
zout
1

Ave
Data
0.00171
Model
0.00176

data/vMax
0.00896

ofxng
369.894

Min d/m
2.77422

ave/min d
98.4163
ave/min m
281.677

328_Calib_S_A

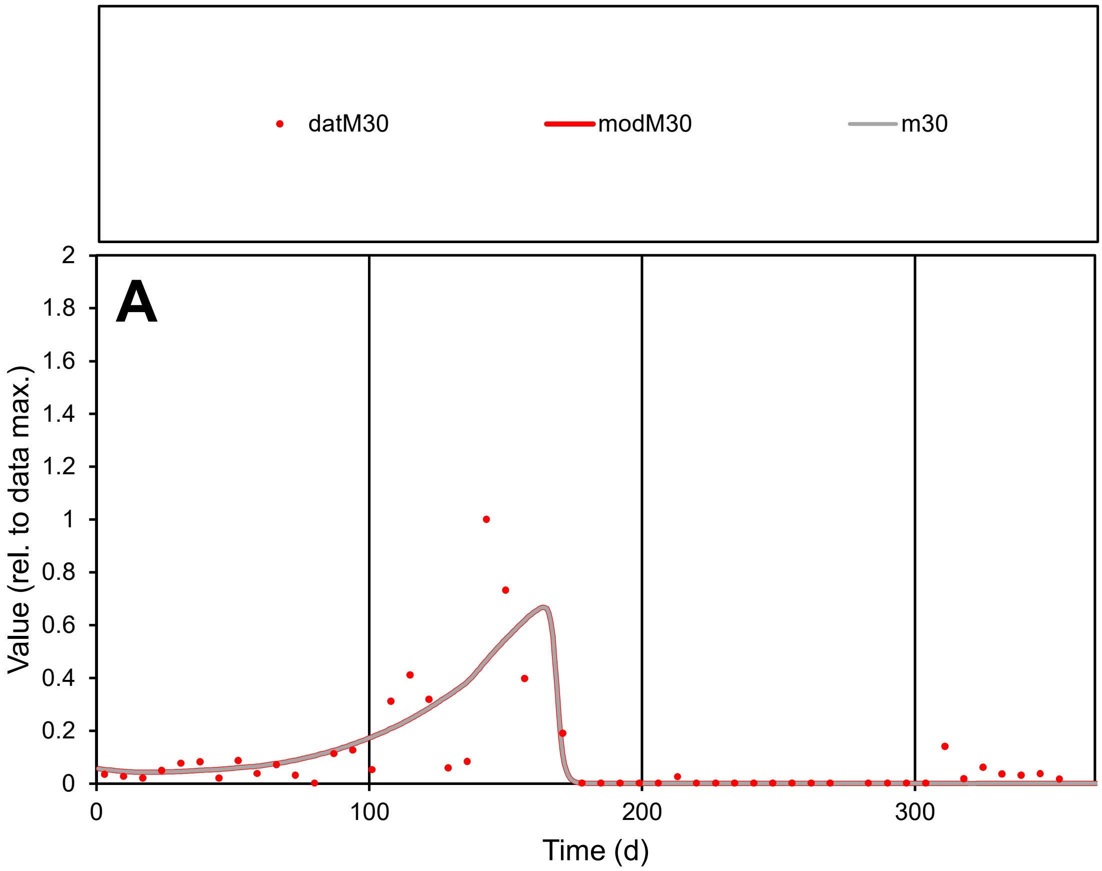


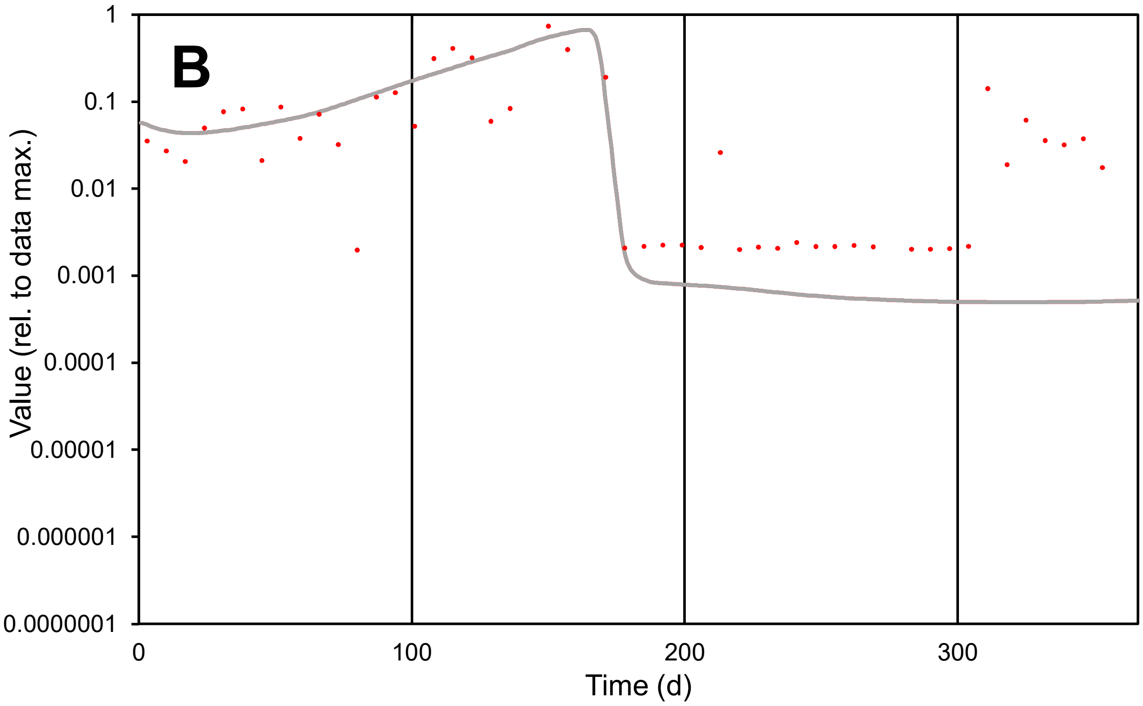


**C1**

Variables

M30 Provide jtd or Dst

CalibOnly?

2 1 to reproduce of

oNormalize?

2 1=no, 2=yes

ofxn calc*

1.16589

ofxn mod

1.18888

sRun

RunA1

zout

1

Ave

Data

0.00203

Model

0.00195

data/vMax

0.02094

ofxng

369.894

Min d/m

3.97872

ave/min d

49.3168

ave/min m

188.382

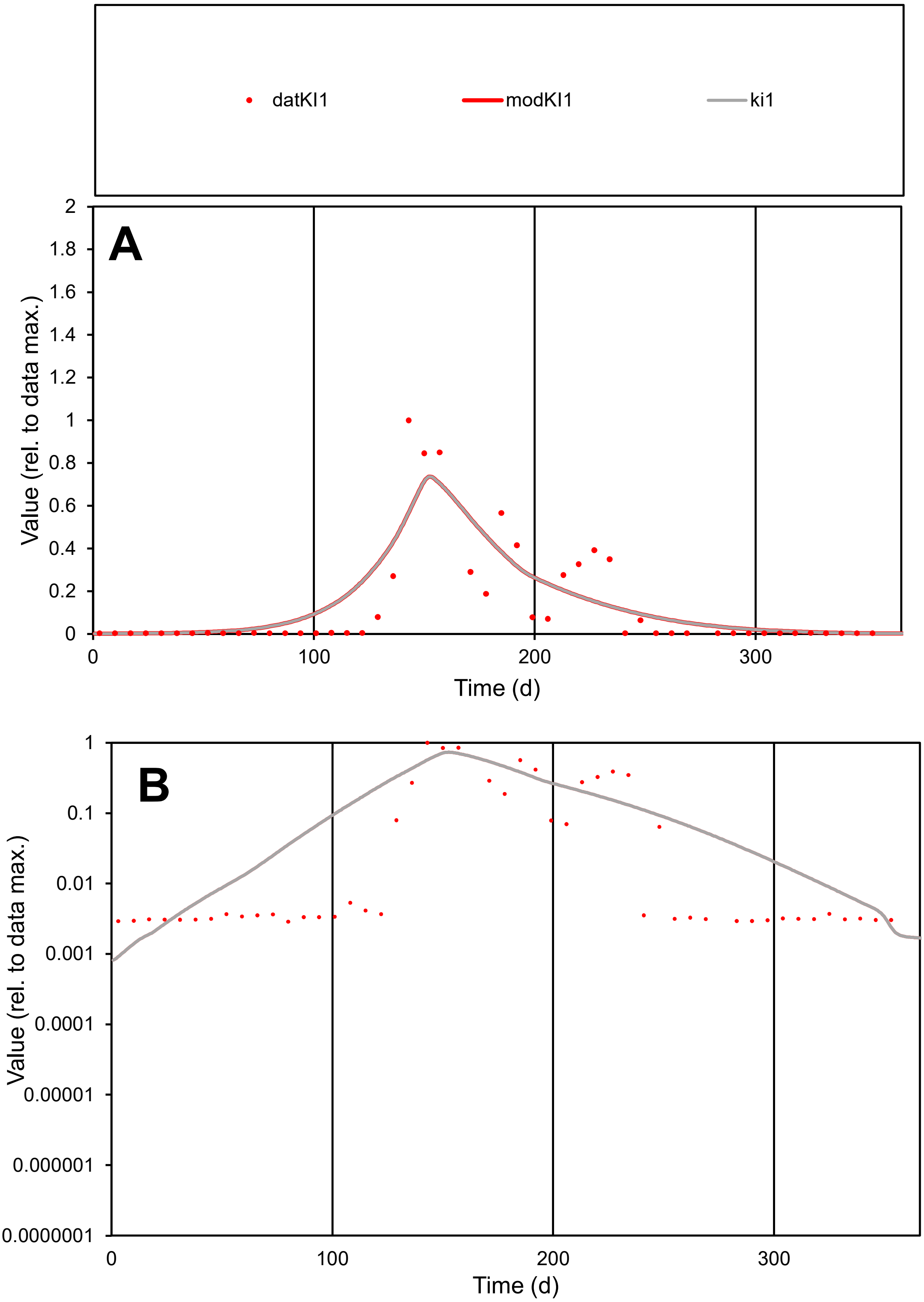


**C1**

Variables

KI1 Provide jtd or Dst

CalibOnly?

2 1 to reproduce of

oNormalize?

2 1=no, 2=yes

ofxn calc*

1.01608

ofxn mod

1.04806

sRun

RunA1

zout

1

Ave

Data

0.00205

Model

0.00232

data/vMax

0.01628

ofxng

369.894

Min d/m

2.90763

ave/min d

43.6743

ave/min m

143.65

328_Calib_S_A

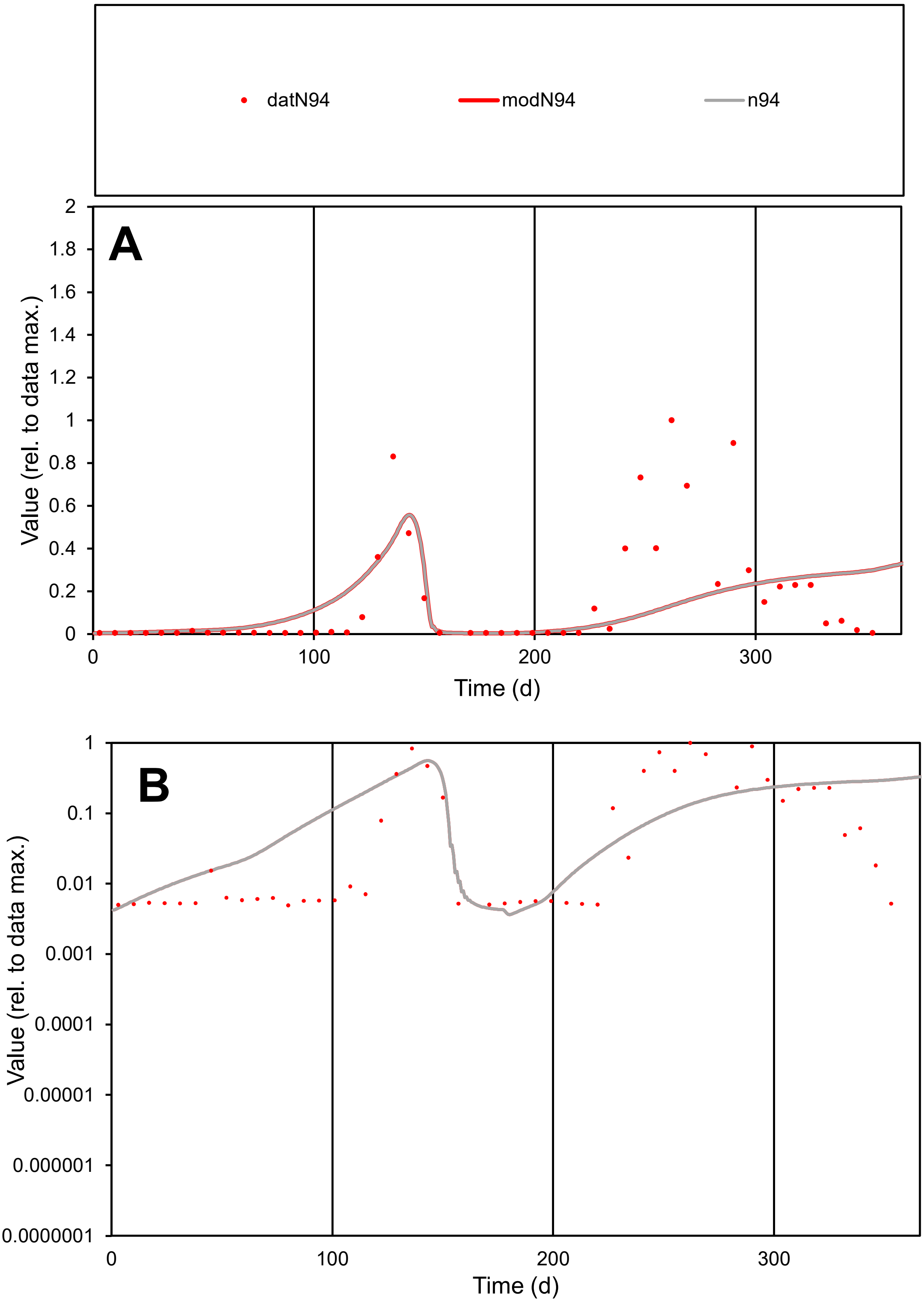


**C1**

Variables
N94 Provide jtd or Dst
CalibOnly?
2 1 to reproduce of
oNormalize?
2 1=no, 2=yes

ofxn calc*
1.44827
ofxn mod
1.45774

sRun
RunA1
zout
1

Ave
Data
0.00147
Model
0.00118

data/vMax
0.0092

ofxng
369.894

Min d/m
1.21928

ave/min d
32.2744
ave/min m
31.5578

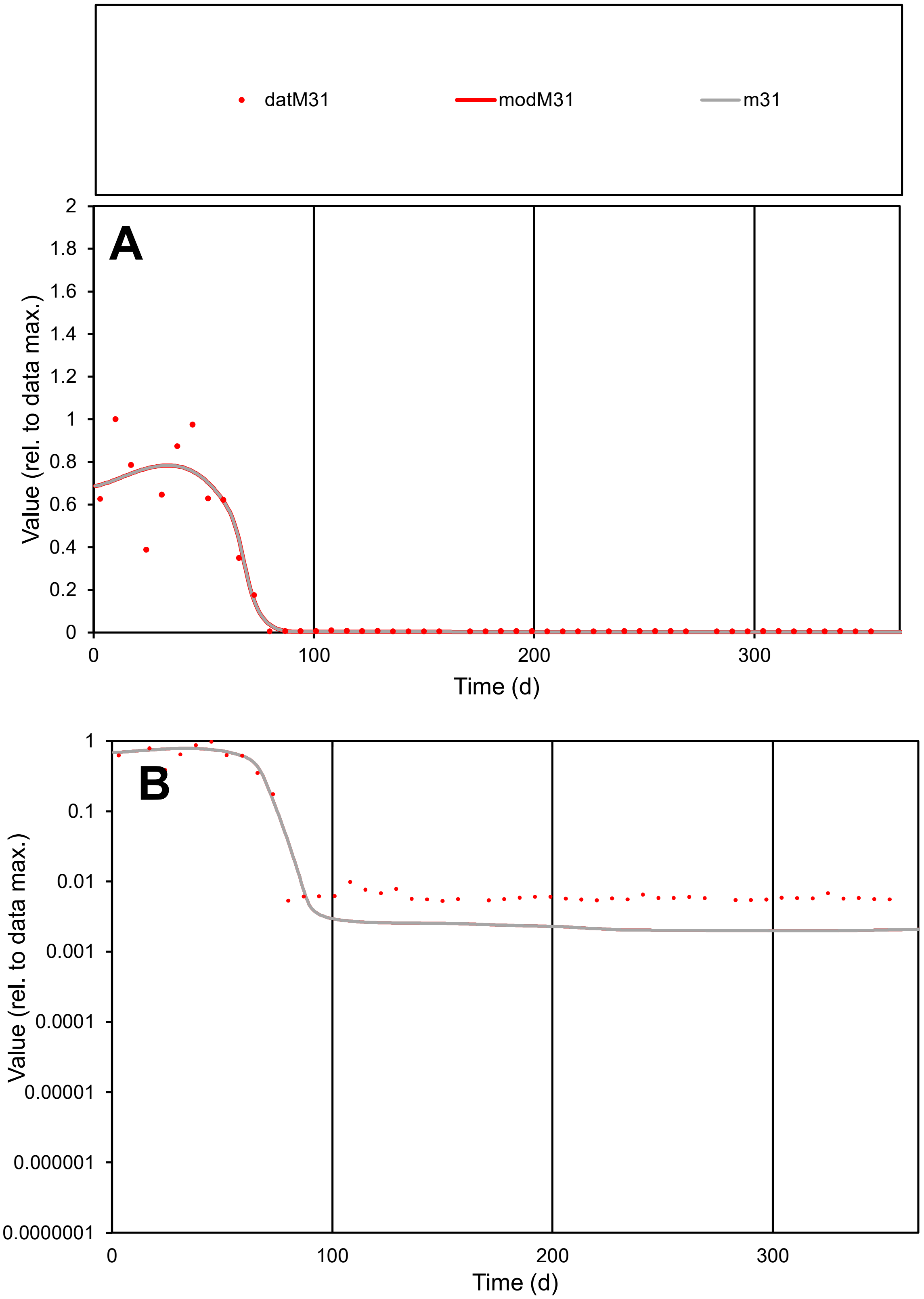


**C1**

Variables
M31 Provide jtd or Dst
CalibOnly?
2 1 to reproduce of
oNormalize?
2 1=no, 2=yes

ofxn calc*
0.53977
ofxn mod
0.60232

sRun
RunA1
zout
1

Ave
Data
0.00229
Model
0.00224

data/vMax
0.01539

ofxng
369.894

Min d/m
2.64821

ave/min d
28.1982
ave/min m
72.9902

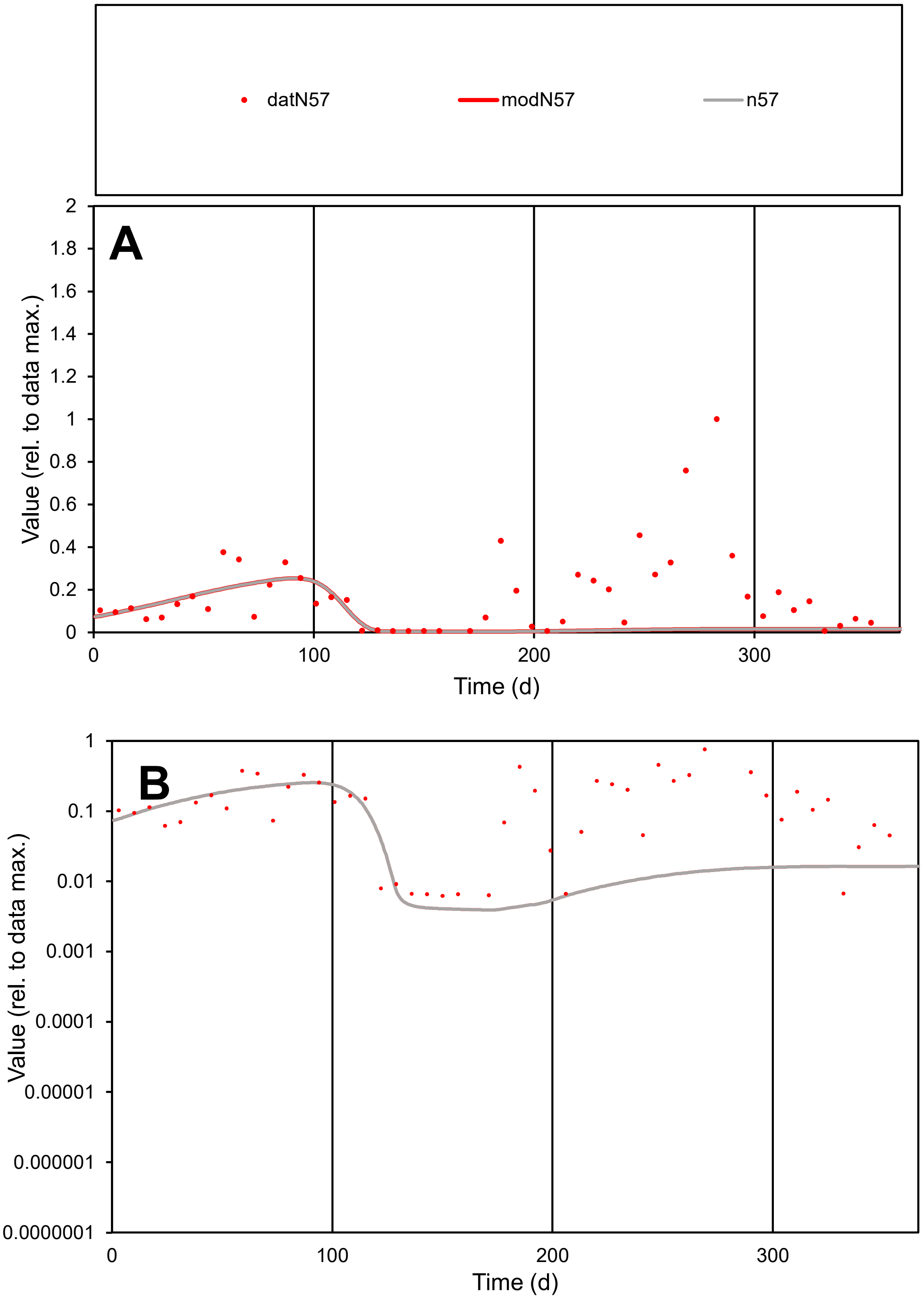


**C1**

Variables
N57 Provide jtd or Dst
CalibOnly?
2 1 to reproduce of
oNormalize?
2 1=no, 2=yes

ofxn calc*
1.32315
ofxn mod
1.32778

sRun
RunA1
zout
1

Ave
Data
0.00101
Model
0.0004

data/vMax
0.00583

ofxng
369.894

Min d/m
1.57479

ave/min d
27.9602
ave/min m
17.3825

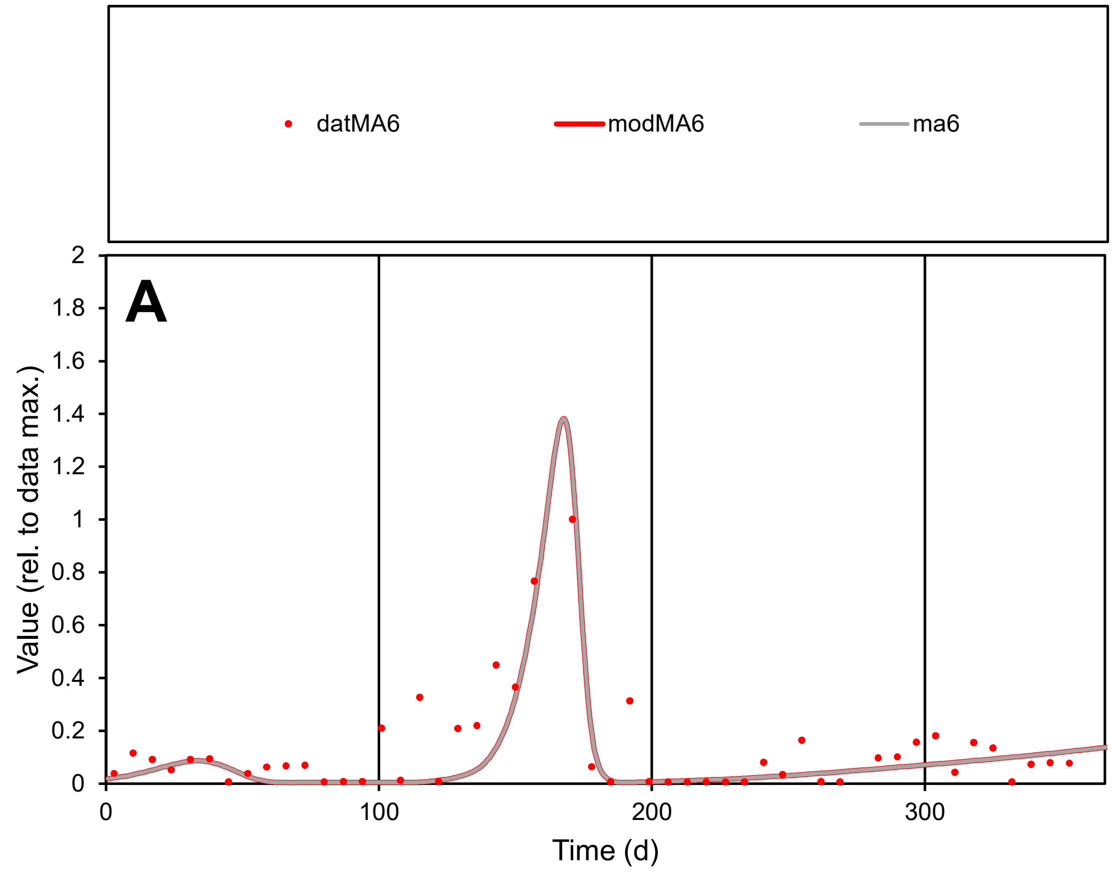


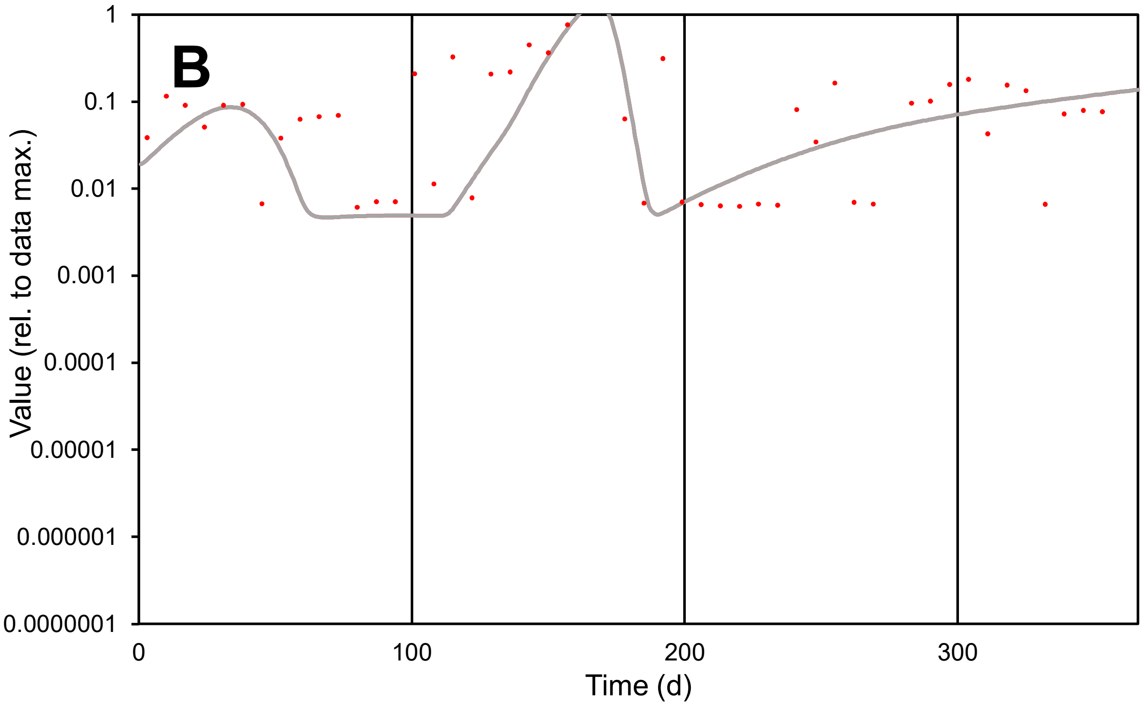


**C1**

Variables
MA6 Provide jtd or Dst
CalibOnly?
2 1 to reproduce of
oNormalize?
2 1=no, 2=yes

ofxn calc*
0.76715
ofxn mod
0.80255

sRun
RunA1
zout
1

Ave
Data
0.00241
Model
0.00168

data/vMax
0.01928

ofxng
369.894

Min d/m
1.30848

ave/min d
20.3713
ave/min m
18.5314

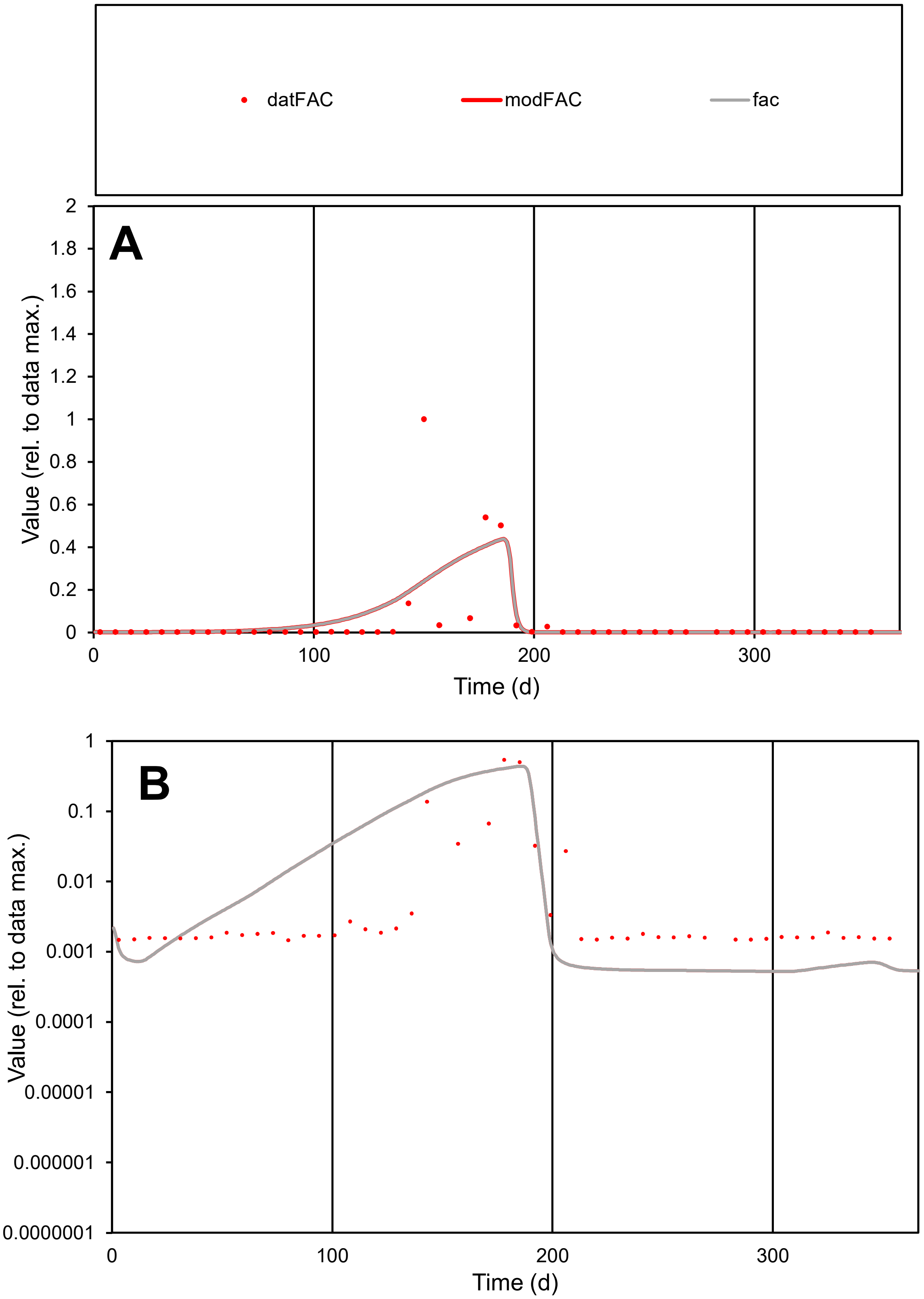


**C1**

Variables
FAC Provide jtd or Dst
CalibOnly?
2 1 to reproduce of
oNormalize?
2 1=no, 2=yes

ofxn calc*
2.58681
ofxn mod
2.61377

sRun
RunA1
zout
1

Ave
Data
0.00292
Model
0.00315

data/vMax
0.05934

ofxng
369.894

Min d/m
2.78631

ave/min d
33.6656
ave/min m
101.08

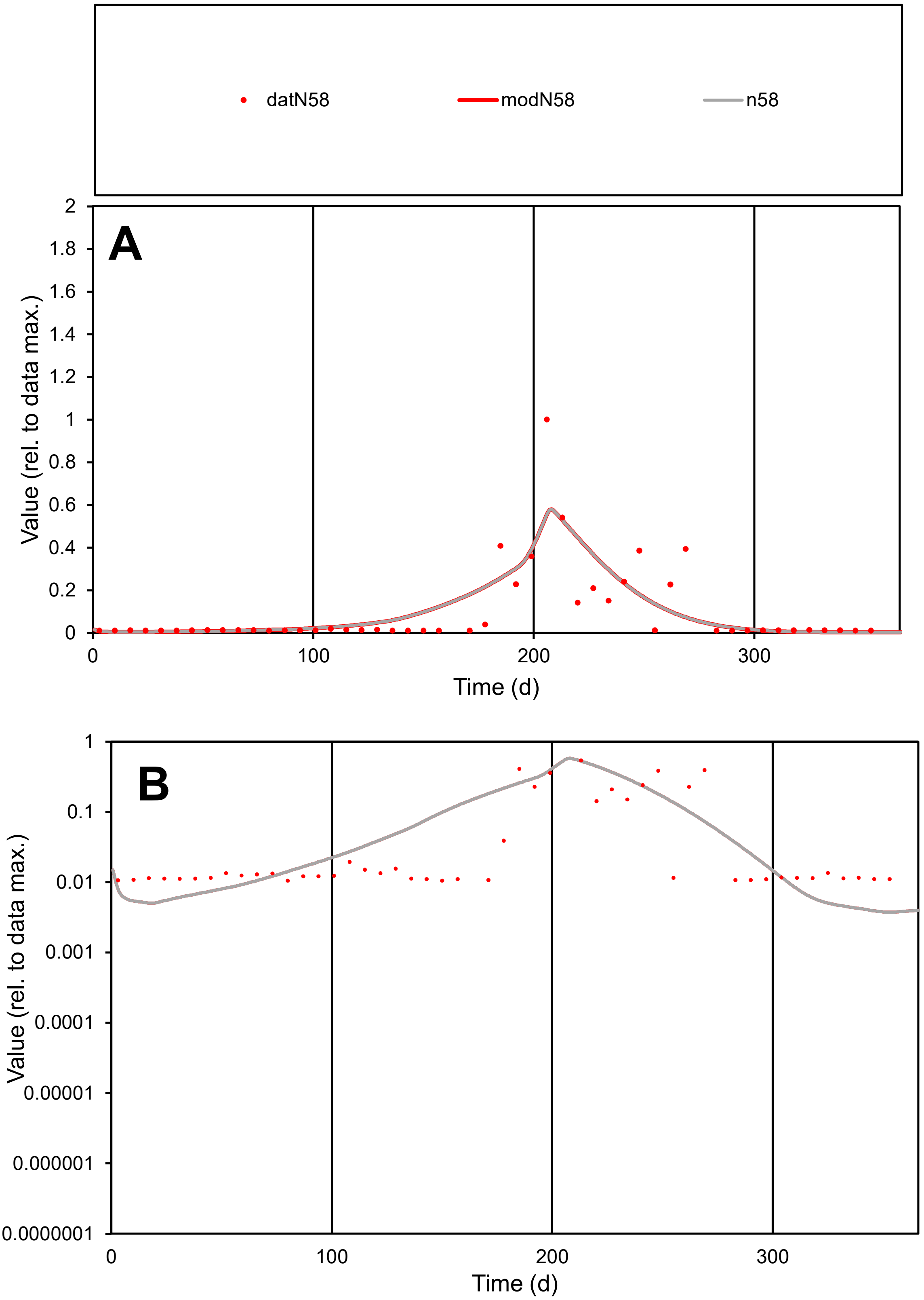


**C1**

Variables

N58 Provide jtd or Dst

CalibOnly?

2 1 to reproduce of

oNormalize?

2 1=no, 2=yes

ofxn calc*

1.14089

ofxn mod

1.169

sRun

RunA1

zout

1

Ave

Data

0.00076

Model

0.0008

data/vMax

0.0078

ofxng

369.894

Min d/m

2.79733

ave/min d

9.26706

ave/min m

27.3425

328_Calib_S_A

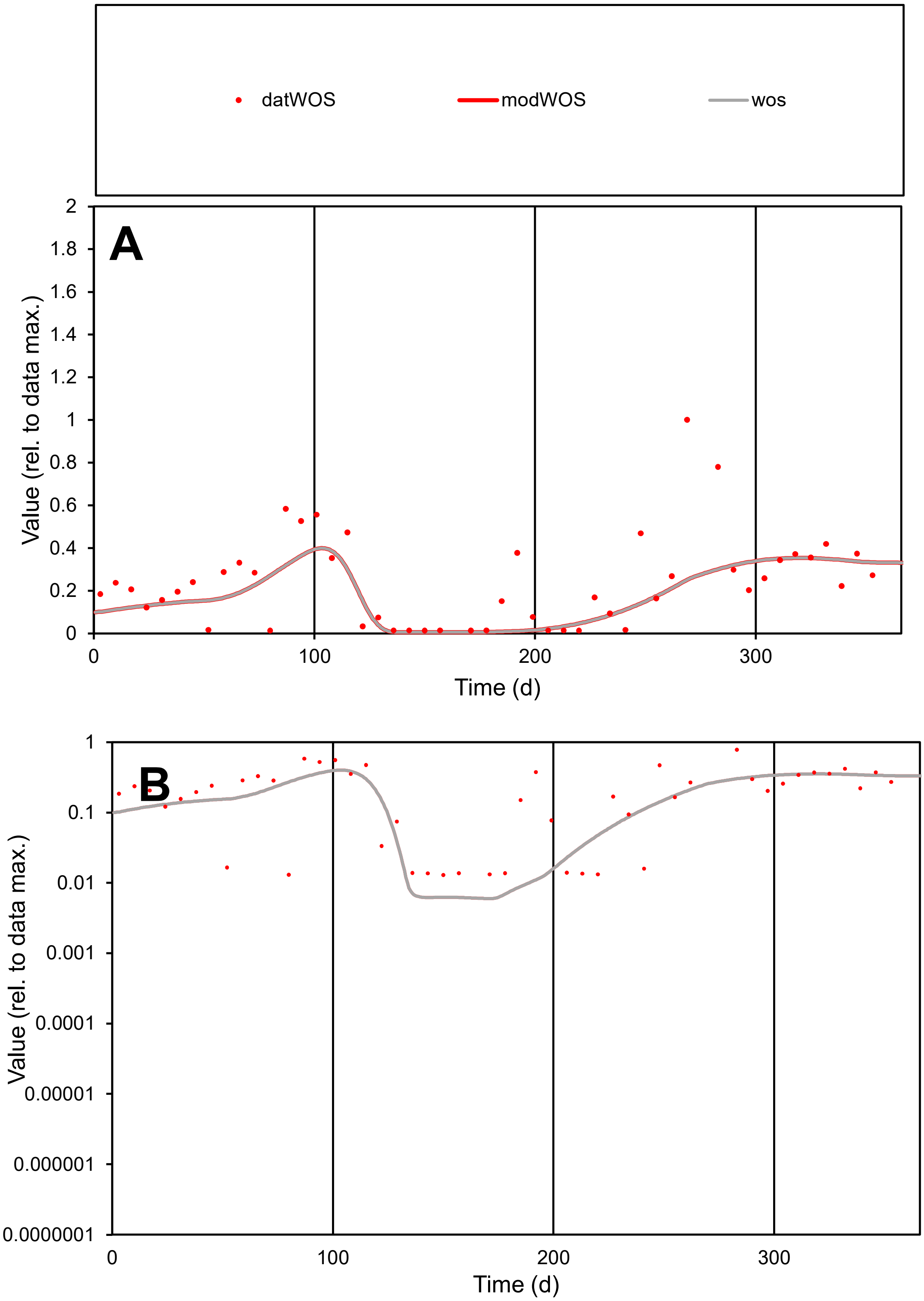


**C1**

Variables
WOS Provide jtd or Dst
CalibOnly?
2 1 to reproduce of
oNormalize?
2 1=no, 2=yes

ofxn calc*
0.71574
ofxn mod
0.72725

sRun
RunA1
zout
1

Ave
Data
0.00126
Model
0.00095

data/vMax
0.00528

ofxng
369.894

Min d/m
2.153

ave/min d
18.4256
ave/min m
30.1046

328_Calib_S_A

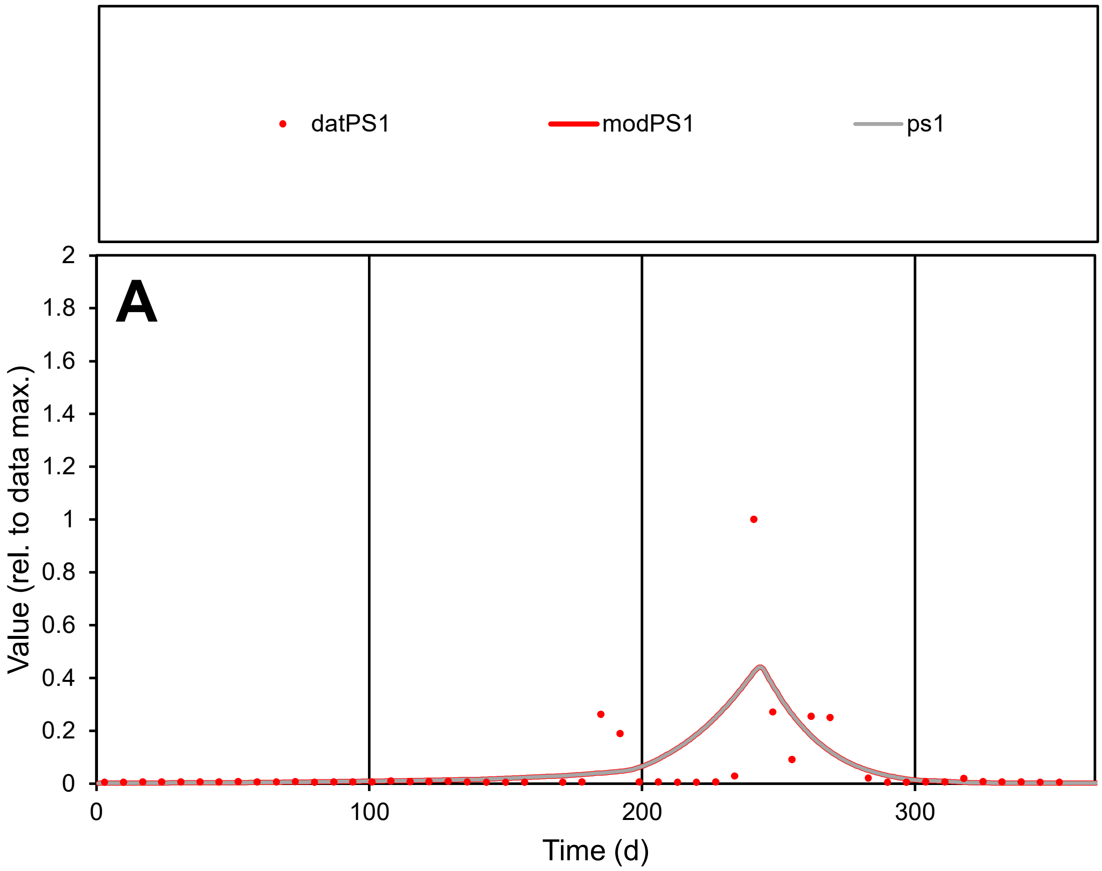


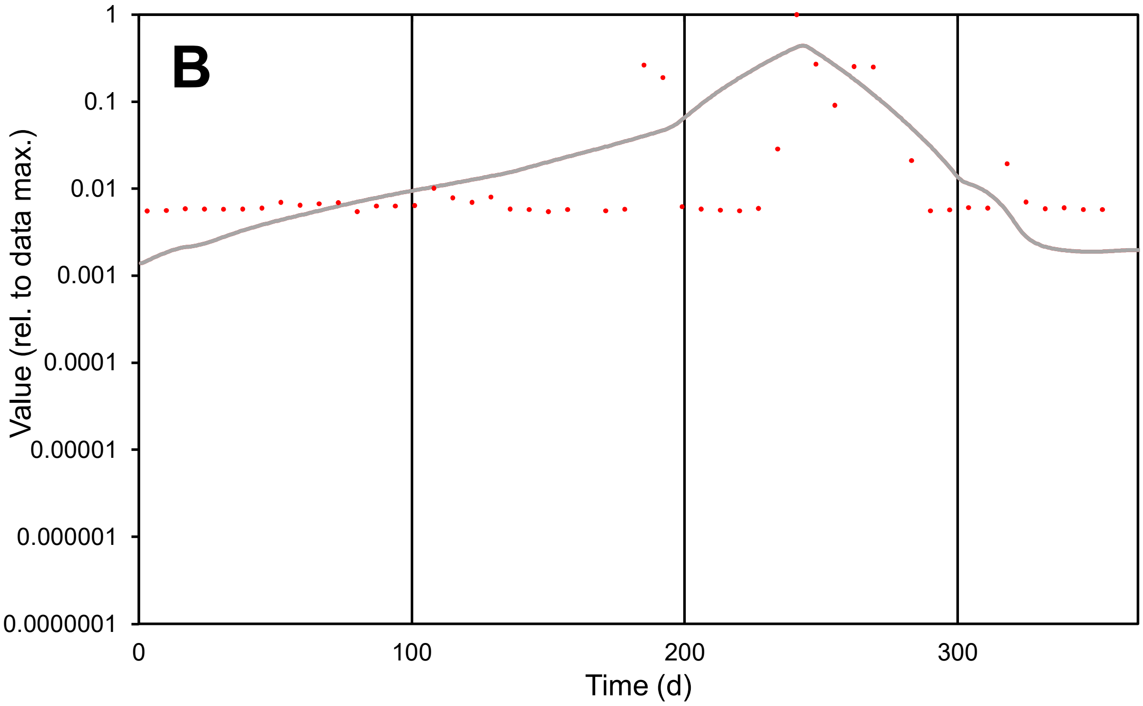


**C1**

Variables
PS1 Provide jtd or Dst
CalibOnly?
2 1 to reproduce of
oNormalize?
2 1=no, 2=yes

ofxn calc*
2.14203
ofxn mod
2.17784

sRun
RunA1
zout
1

Ave
Data
0.00046
Model
0.0005

data/vMax
0.00865

ofxng
369.894

Min d/m
3.50373

ave/min d
9.87783
ave/min m
37.3082

328_Calib_S_A

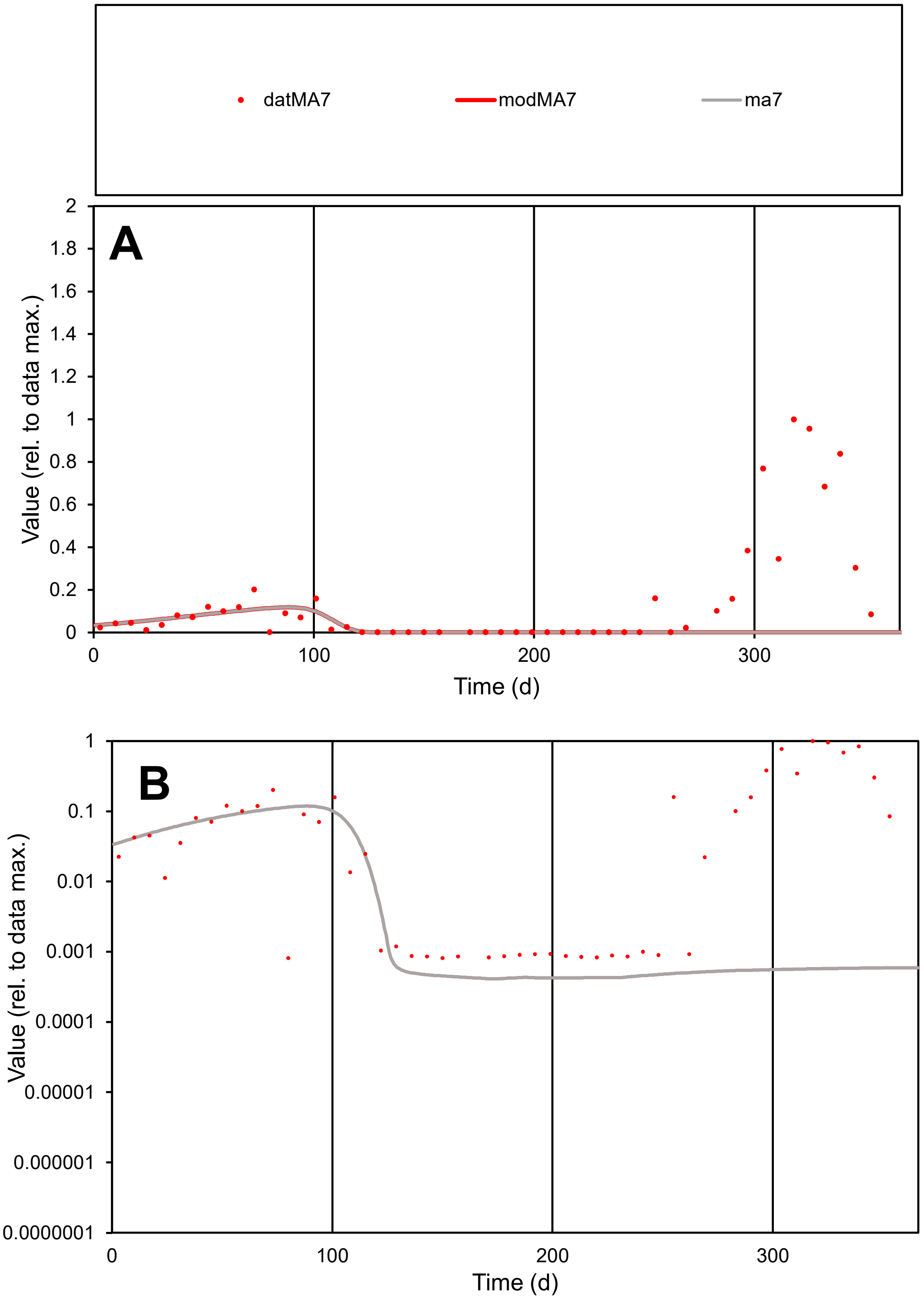


**C1**

Variables
MA7 Provide jtd or Dst
CalibOnly?
2 1 to reproduce of
oNormalize?
2 1=no, 2=yes

ofxn calc*
2.02331
ofxn mod
2.02666

sRun
RunA1
zout
1

Ave
Data
0.00238
Model
0.00045

data/vMax
0.01659

ofxng
369.894

Min d/m
1.97036

ave/min d
177.168
ave/min m
65.8696

328_Calib_S_A

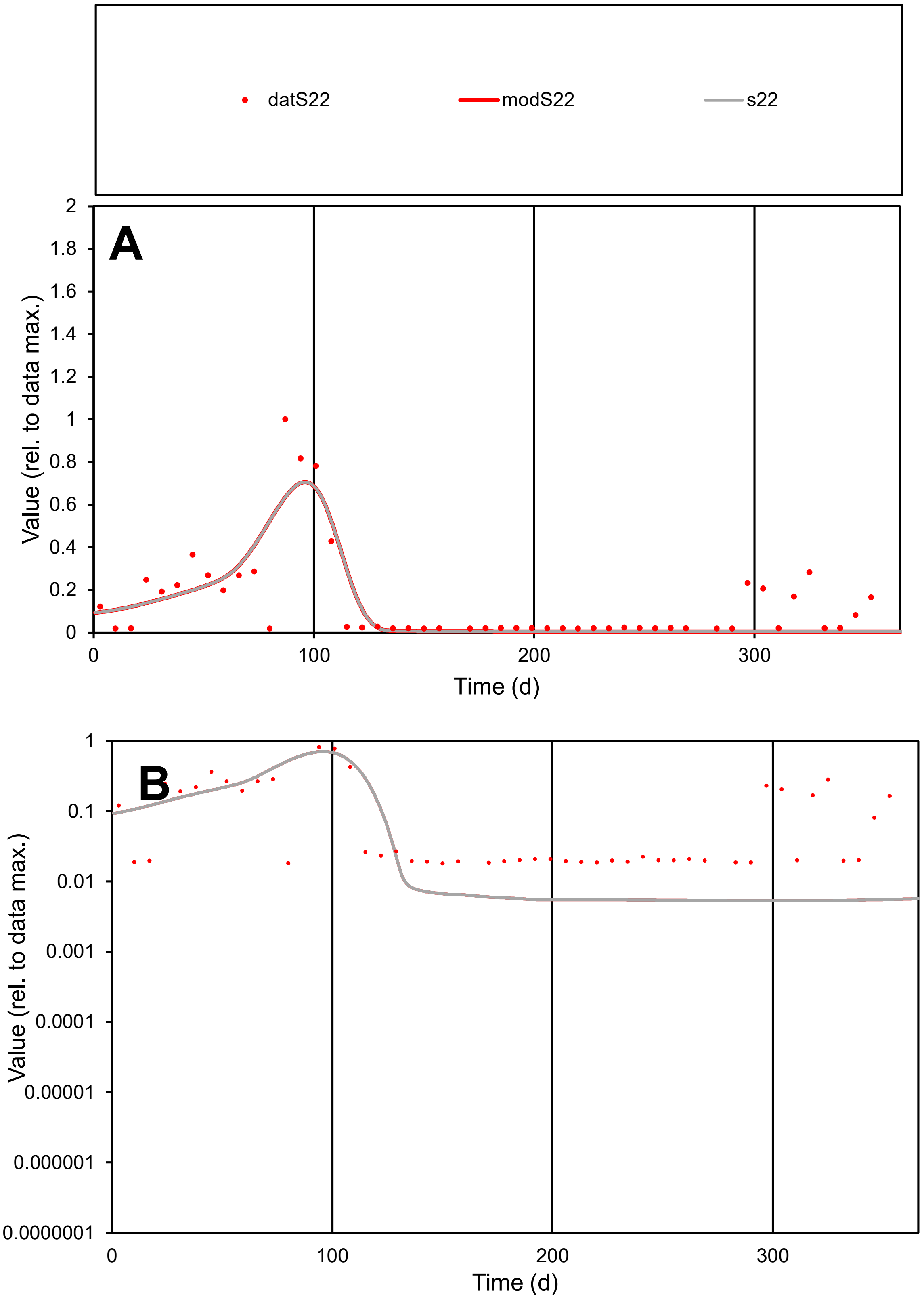


**C1**

Variables
S22 Provide jtd or Dst
CalibOnly?
2 1 to reproduce of
oNormalize?
2 1=no, 2=yes

ofxn calc*
0.87891
ofxn mod
0.9036

sRun
RunA1
zout
1

Ave
Data
0.00114
Model
0.00094

data/vMax
0.00805

ofxng
369.894

Min d/m
3.44293

ave/min d
7.76655
ave/min m
22.0345

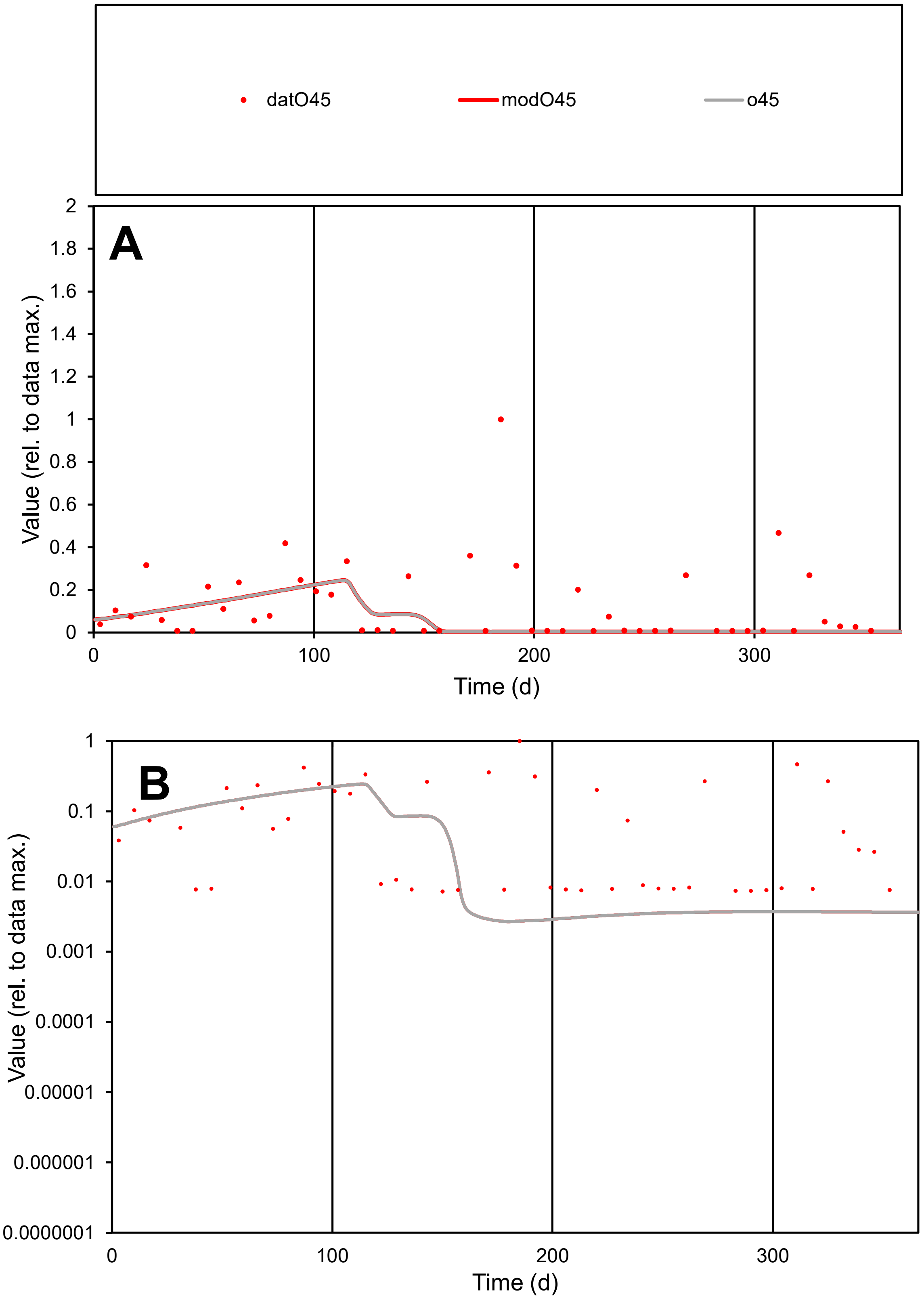


**C1**

Variables
O45 Provide jtd or Dst
CalibOnly?
2 1 to reproduce of
oNormalize?
2 1=no, 2=yes

ofxn calc*
1.55049
ofxn mod
1.5559

sRun
RunA1
zout
1

Ave
Data
0.001
Model
0.00051

data/vMax
0.00796

ofxng
369.894

Min d/m
2.67494

ave/min d
17.4756
ave/min m
23.6625

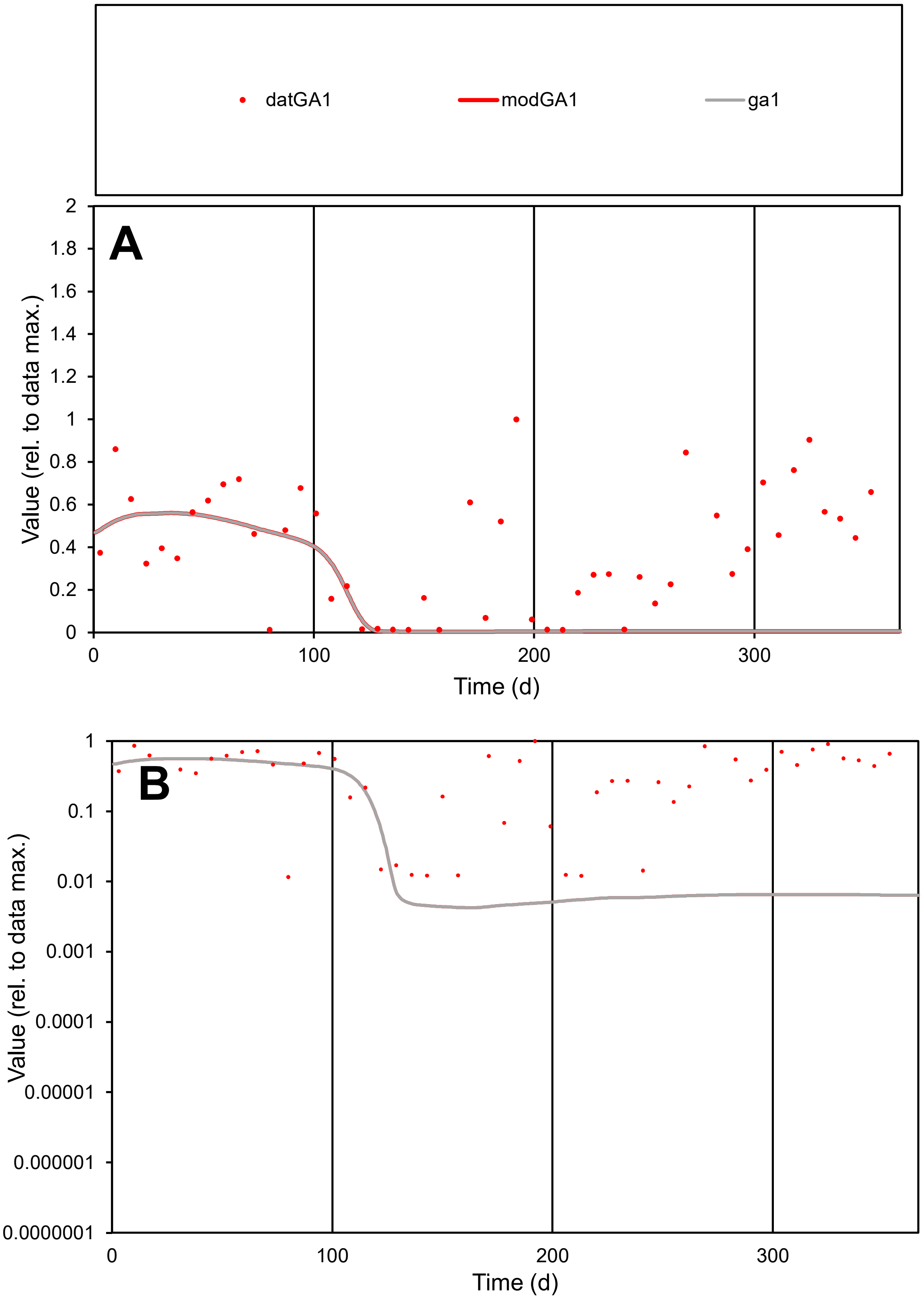


**C1**

Variables
GA1 Provide jtd or Dst
CalibOnly?
2 1 to reproduce of
oNormalize?
2 1=no, 2=yes

ofxn calc*
0.97944
ofxn mod
0.98486

sRun
RunA1
zout
1

Ave
Data
0.00119
Model
0.00052

data/vMax
0.00306

ofxng
369.894

Min d/m
2.73098

ave/min d
33.4499
ave/min m
39.7822

328_Calib_S_A

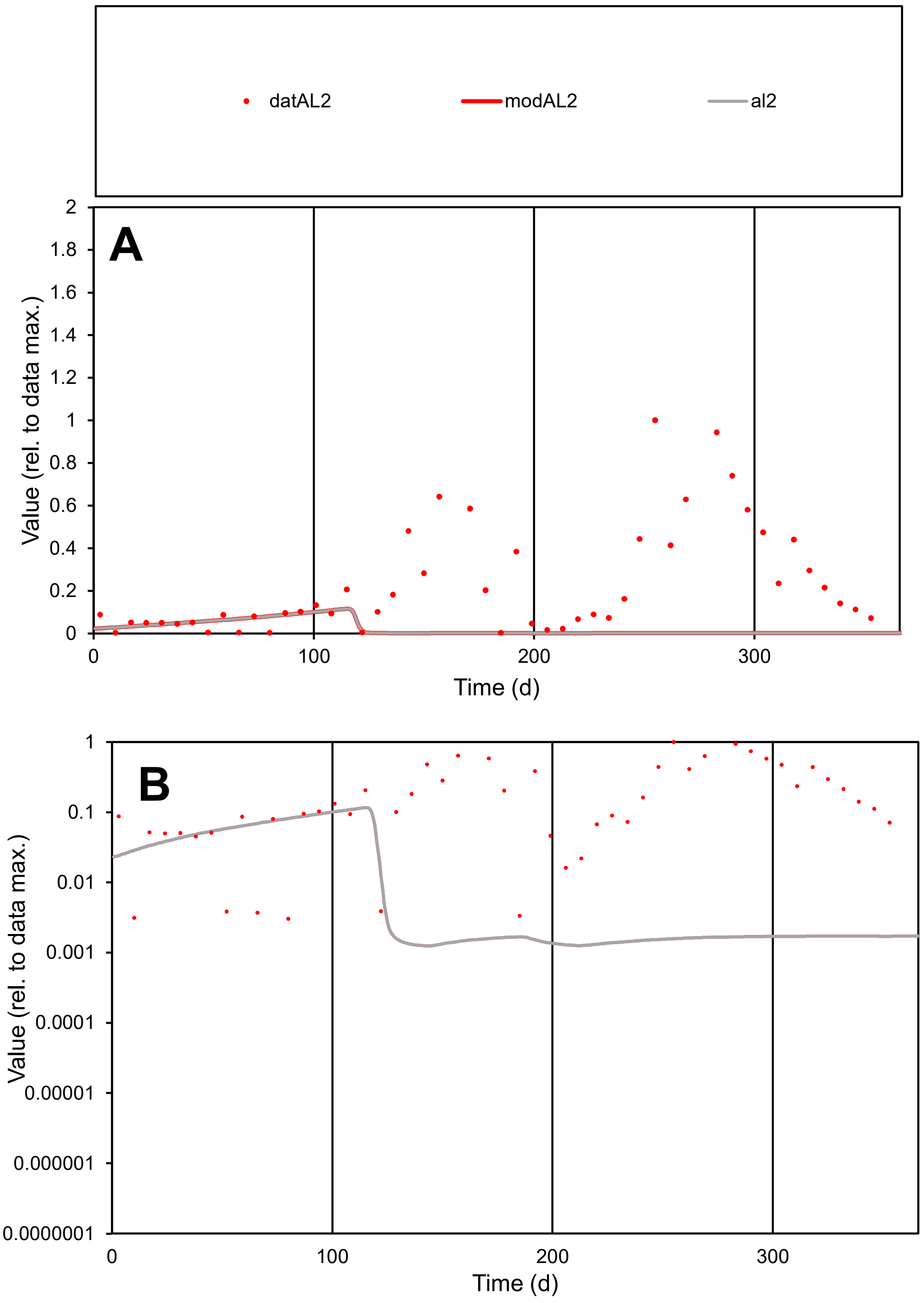


C1

Variables

AL2 Provide jtd or Dst

CalibOnly?

2 1 to reproduce of

oNormalize?

2 1=no, 2=yes

ofxn calc*

1.46582

ofxn mod

1.46884

sRun

RunA1

zout

1

Ave

Data

0.00165

Model

0.00018

data/vMax

0.00722

ofxng

369.894

Min d/m

2.42589

ave/min d

75.7032

ave/min m

19.5063

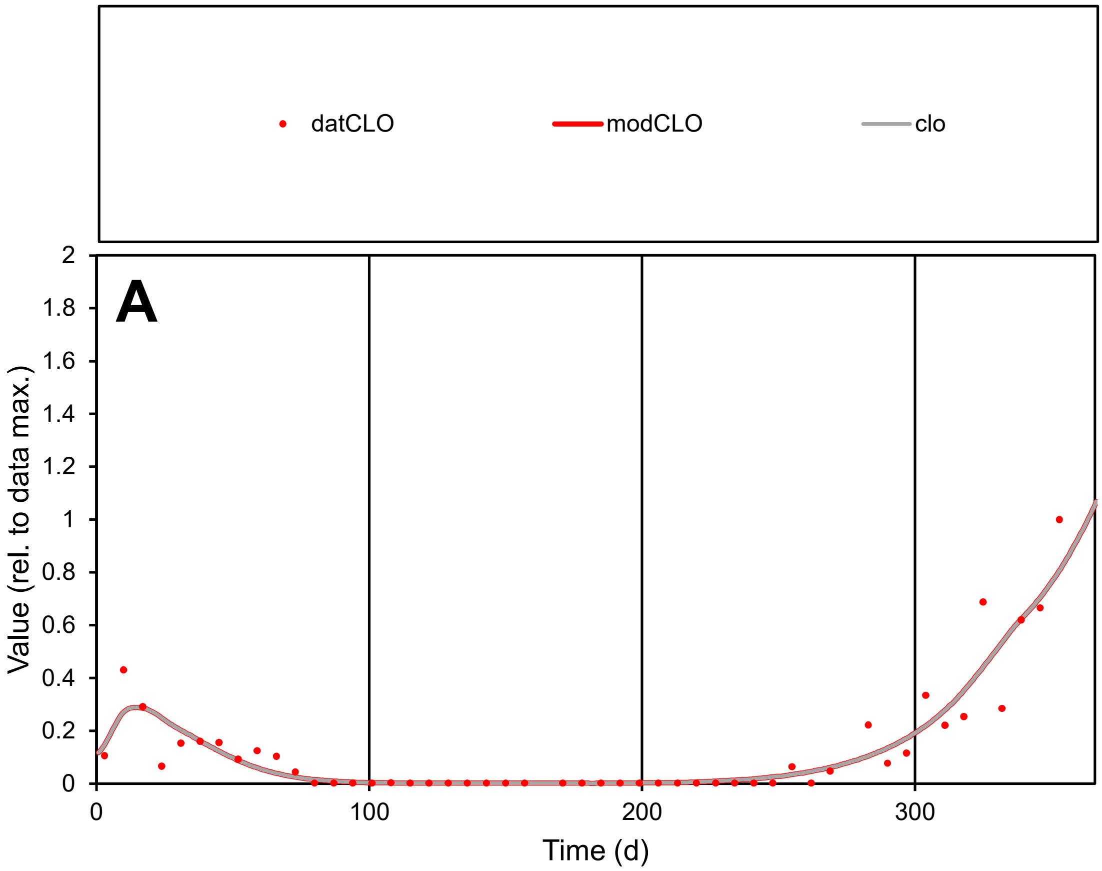


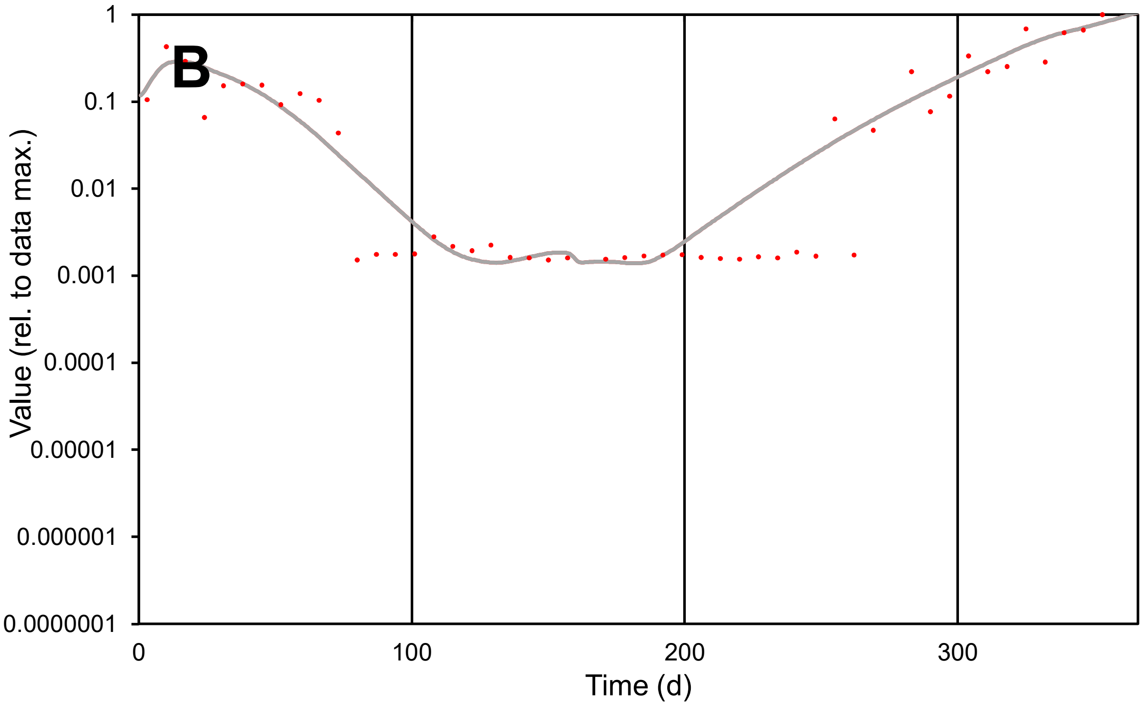


**C1**

Variables
CLO Provide jtd or Dst
CalibOnly?
2 1 to reproduce of
oNormalize?
2 1=no, 2=yes

ofxn calc*
0.57081
ofxn mod
0.60933

sRun
RunA1
zout
1

Ave
Data
0.00113
Model
0.00114

data/vMax
0.00871

ofxng
369.894

Min d/m
1.08217

ave/min d
85.792
ave/min m
93.7189

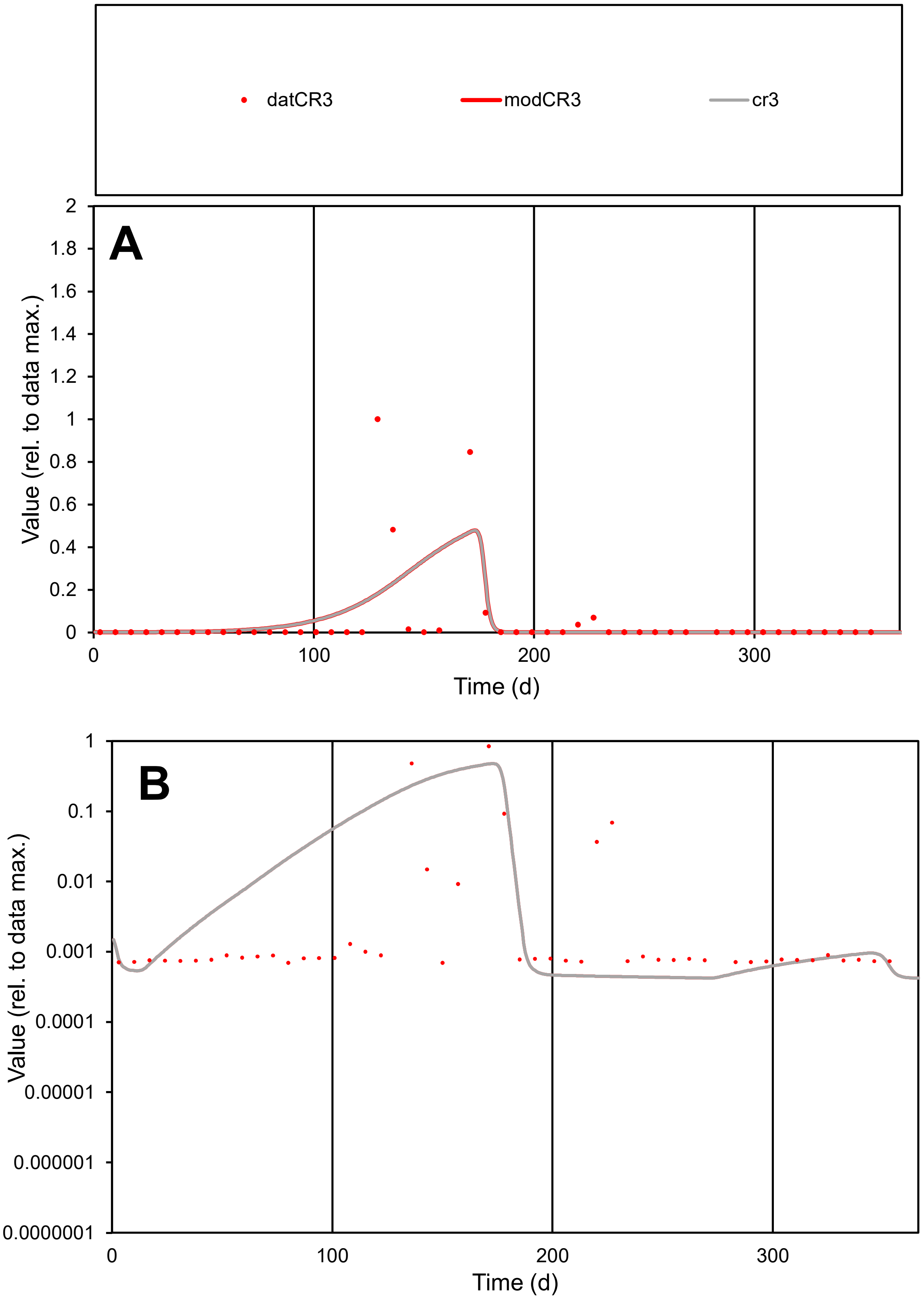


**C1**

Variables
CR3 Provide jtd or Dst
CalibOnly?
2 1 to reproduce of
oNormalize?
2 1=no, 2=yes

ofxn calc*
2.99867
ofxn mod
3.02782

sRun
RunA1
zout
1

Ave
Data
0.00269
Model
0.00265

data/vMax
0.05103

ofxng
369.894

Min d/m
1.64332

ave/min d
75.9162
ave/min m
122.877

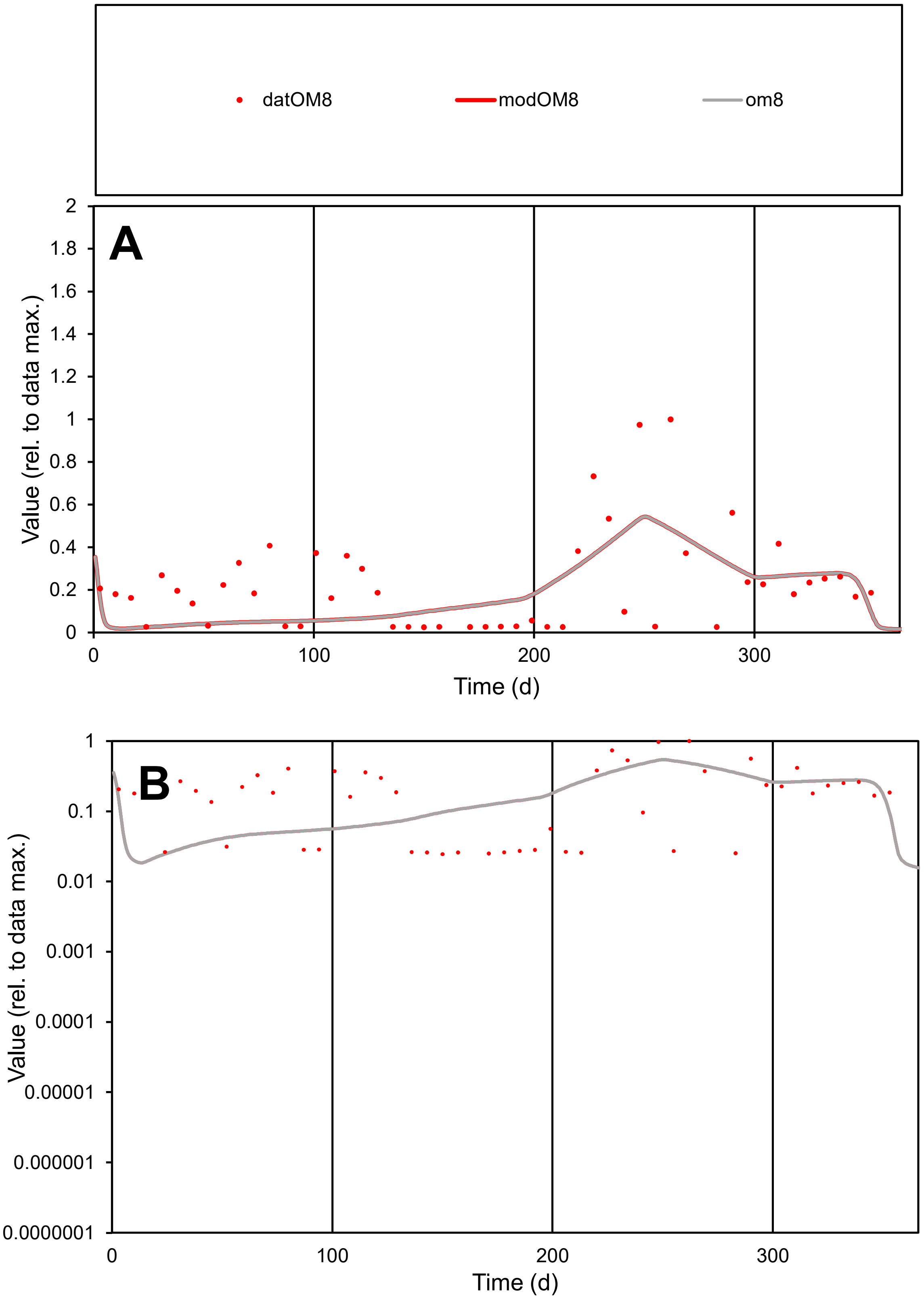


**C1**

Variables
OM8 Provide jtd or Dst
CalibOnly?
2 1 to reproduce of
oNormalize?
2 1=no, 2=yes

ofxn calc*
0.91219
ofxn mod
0.9261

sRun
RunA1
zout
1

Ave
Data
0.00049
Model
0.00041

data/vMax
0.0022

ofxng
369.894

Min d/m
1.29872

ave/min d
9.13191
ave/min m
9.86267

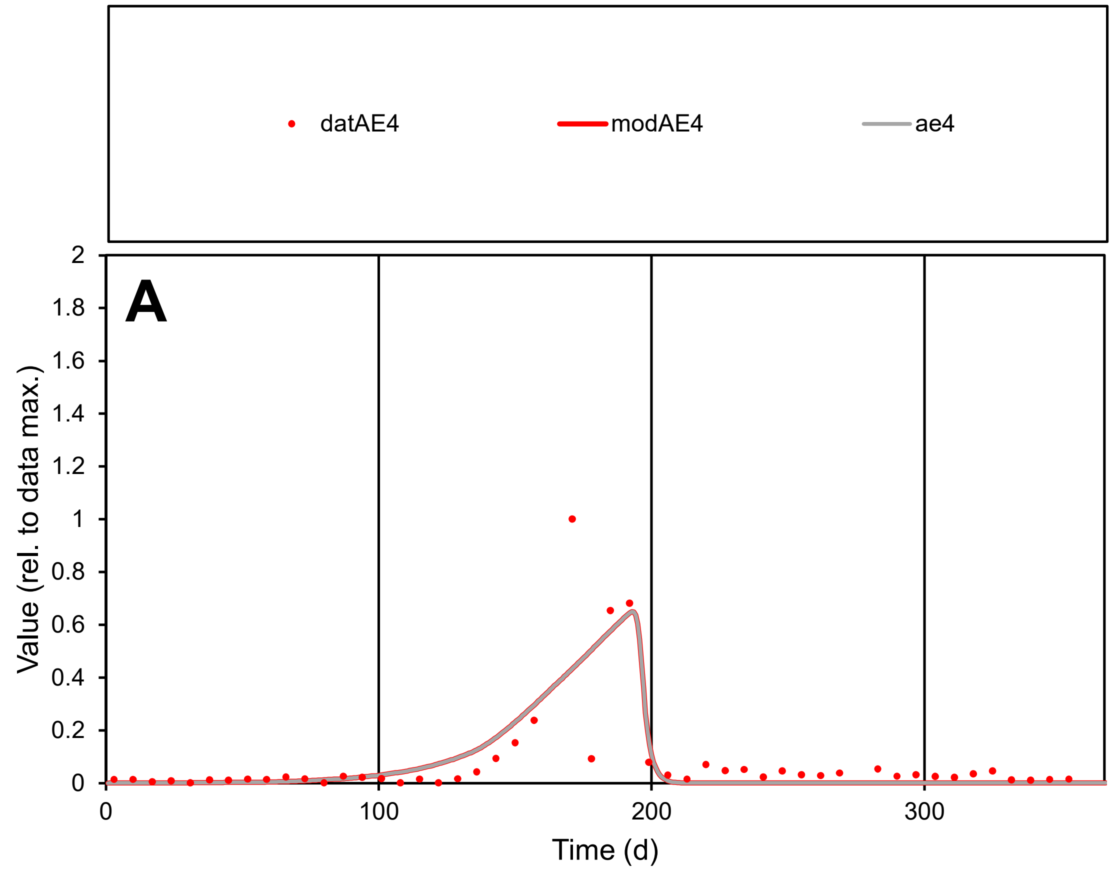


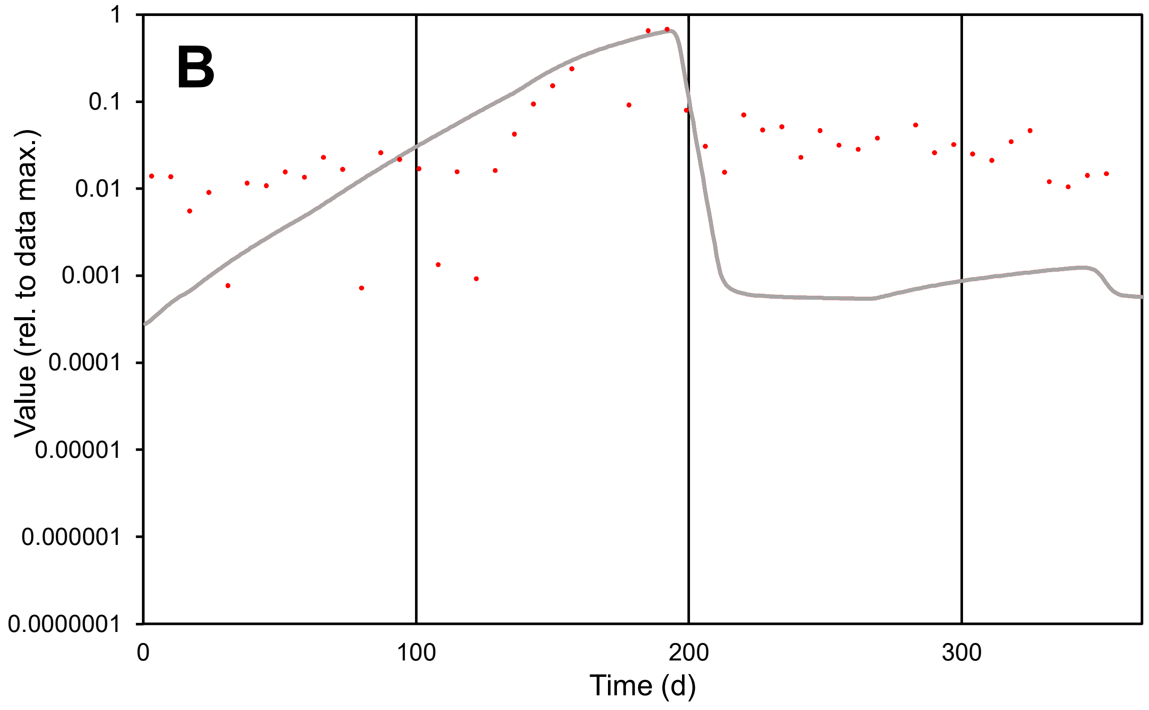


**C1**

Variables
AE4 Provide jtd or Dst
CalibOnly?
2 1 to reproduce of
oNormalize?
2 1=no, 2=yes

ofxn calc*
1.31746
ofxn mod
1.35244

sRun
RunA1
zout
1

Ave
Data
0.0013
Model
0.00117

data/vMax
0.01617

ofxng
369.894

Min d/m
2.12292

ave/min d
111.101
ave/min m
212.349

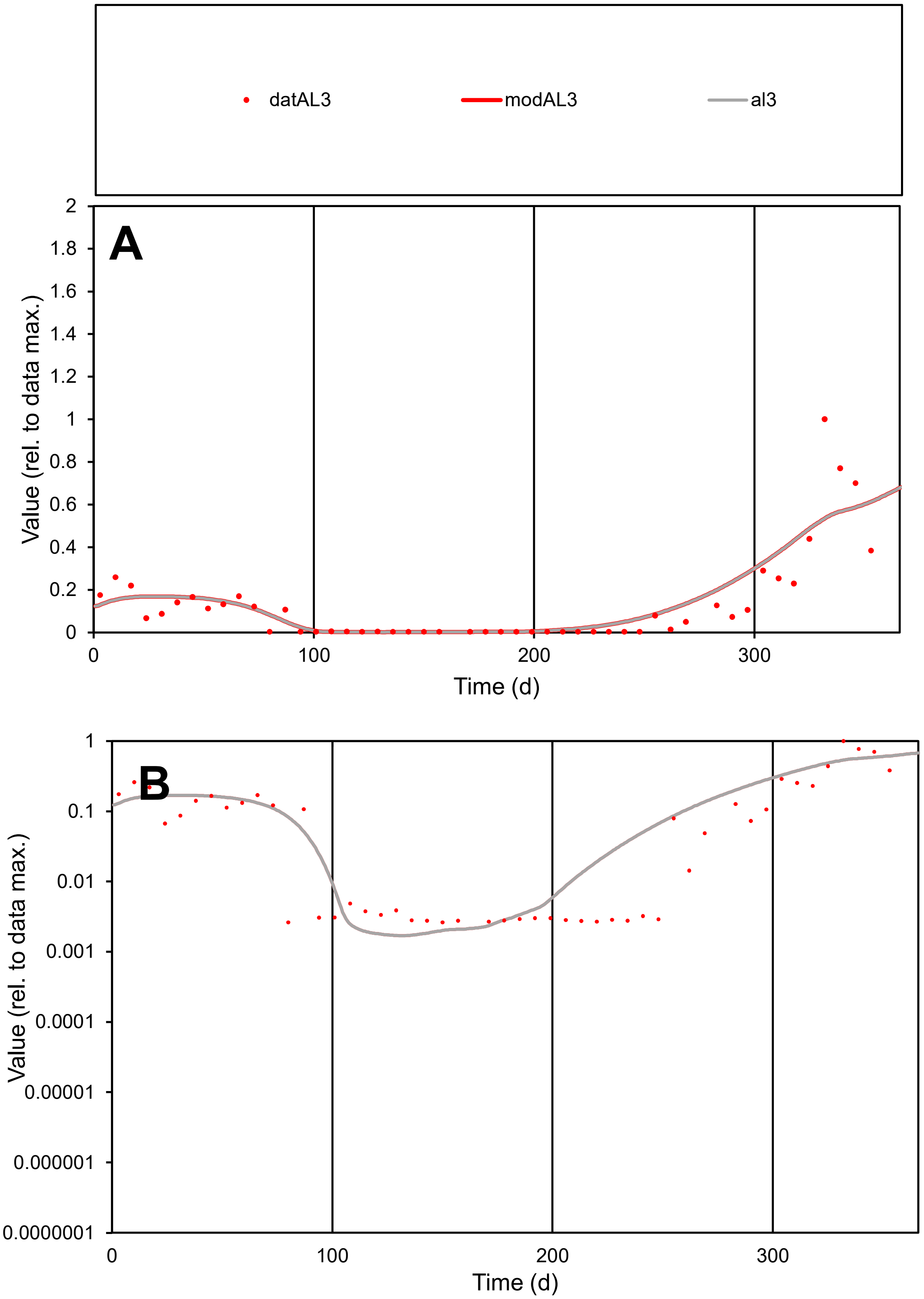


**C1**

Variables
AL3 Provide jtd or Dst
CalibOnly?
2 1 to reproduce of
oNormalize?
2 1=no, 2=yes

ofxn calc*
0.79453
ofxn mod
0.82055

sRun
RunA1
zout
1

Ave
Data
0.00119
Model
0.00134

data/vMax
0.00922

ofxng
369.894

Min d/m
1.53827

ave/min d
49.4302
ave/min m
85.1429

328_Calib_S_A

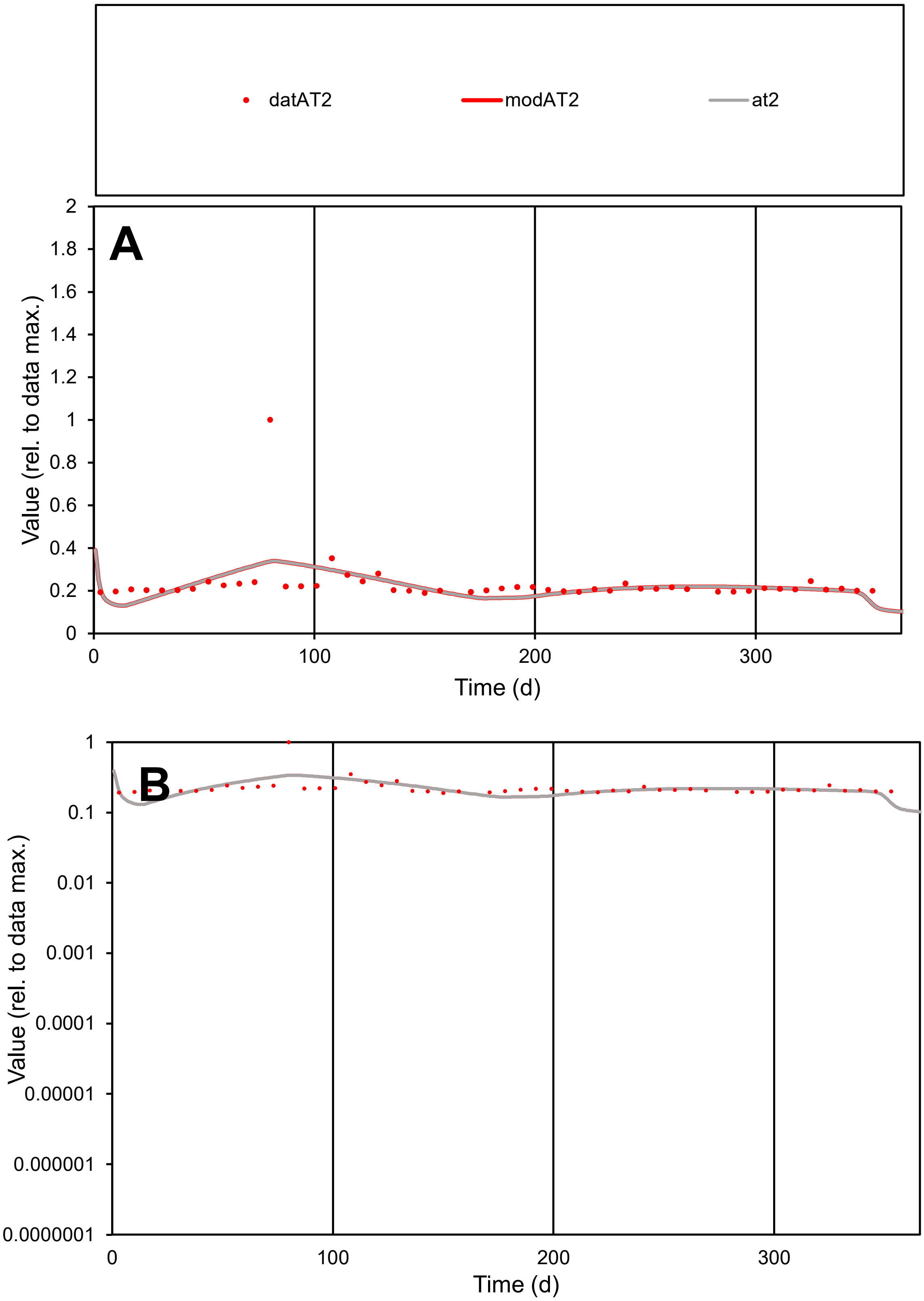


**C1**

Variables

AT2 Provide jtd or Dst

CalibOnly?

2 1 to reproduce of

oNormalize?

2 1=no, 2=yes

ofxn calc*

0.443

ofxn mod

0.45692

sRun

RunA1

zout

1

Ave

Data

9.7E-05

Model

9.3E-05

data/vMax

0.00042

ofxng

369.894

Min d/m

1.48451

ave/min d

1.22475

ave/min m

1.73399

328_Calib_S_A

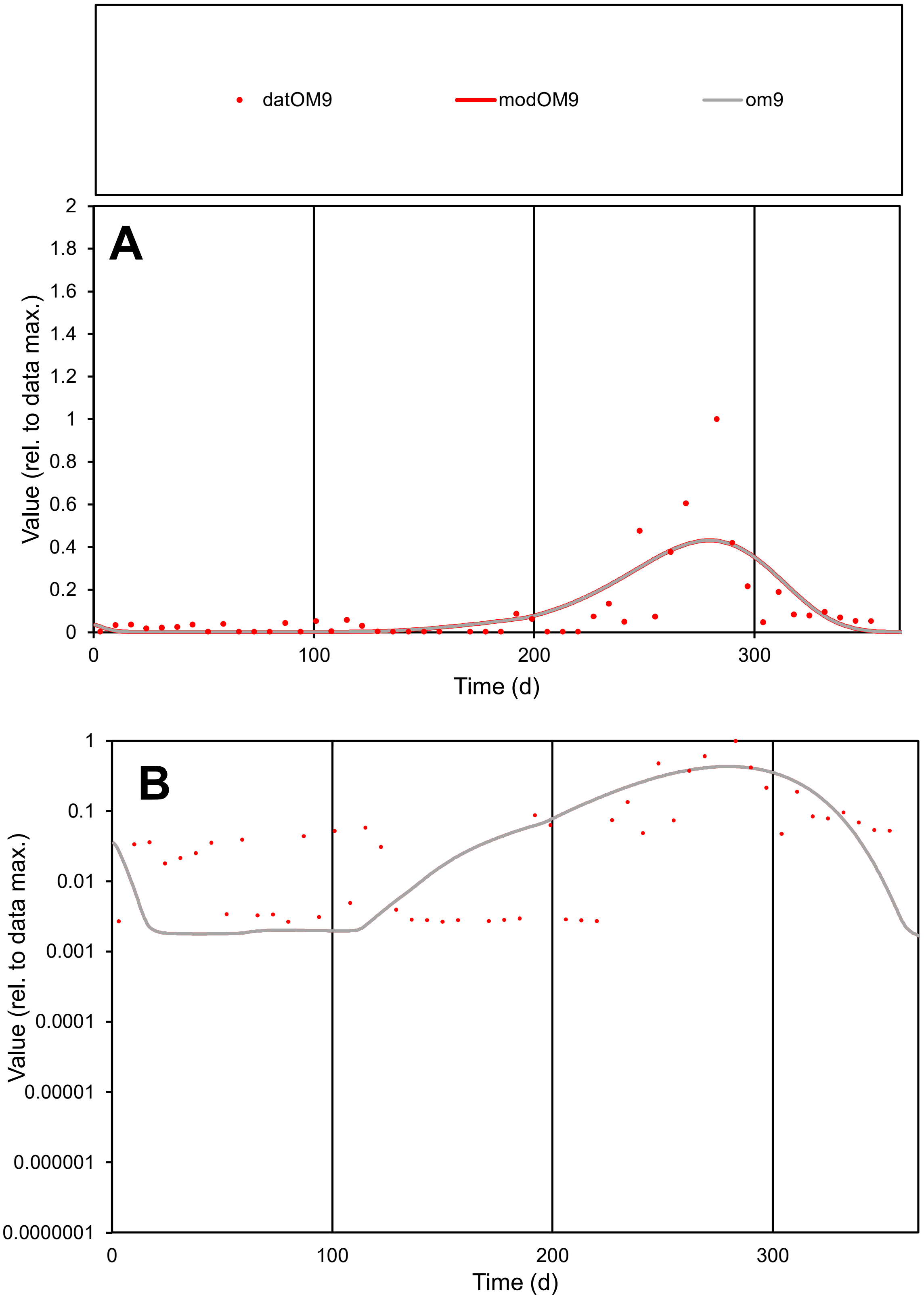


**C1**

Variables
OM9 Provide jtd or Dst
CalibOnly?
2 1 to reproduce of
oNormalize?
2 1=no, 2=yes

ofxn calc*
1.25313
ofxn mod
1.27709

sRun
RunA1
zout
1

Ave
Data
0.00156
Model
0.00169

data/vMax
0.01628

ofxng
369.894

Min d/m
1.49155
ave/min d
36.0989
ave/min m
58.3337

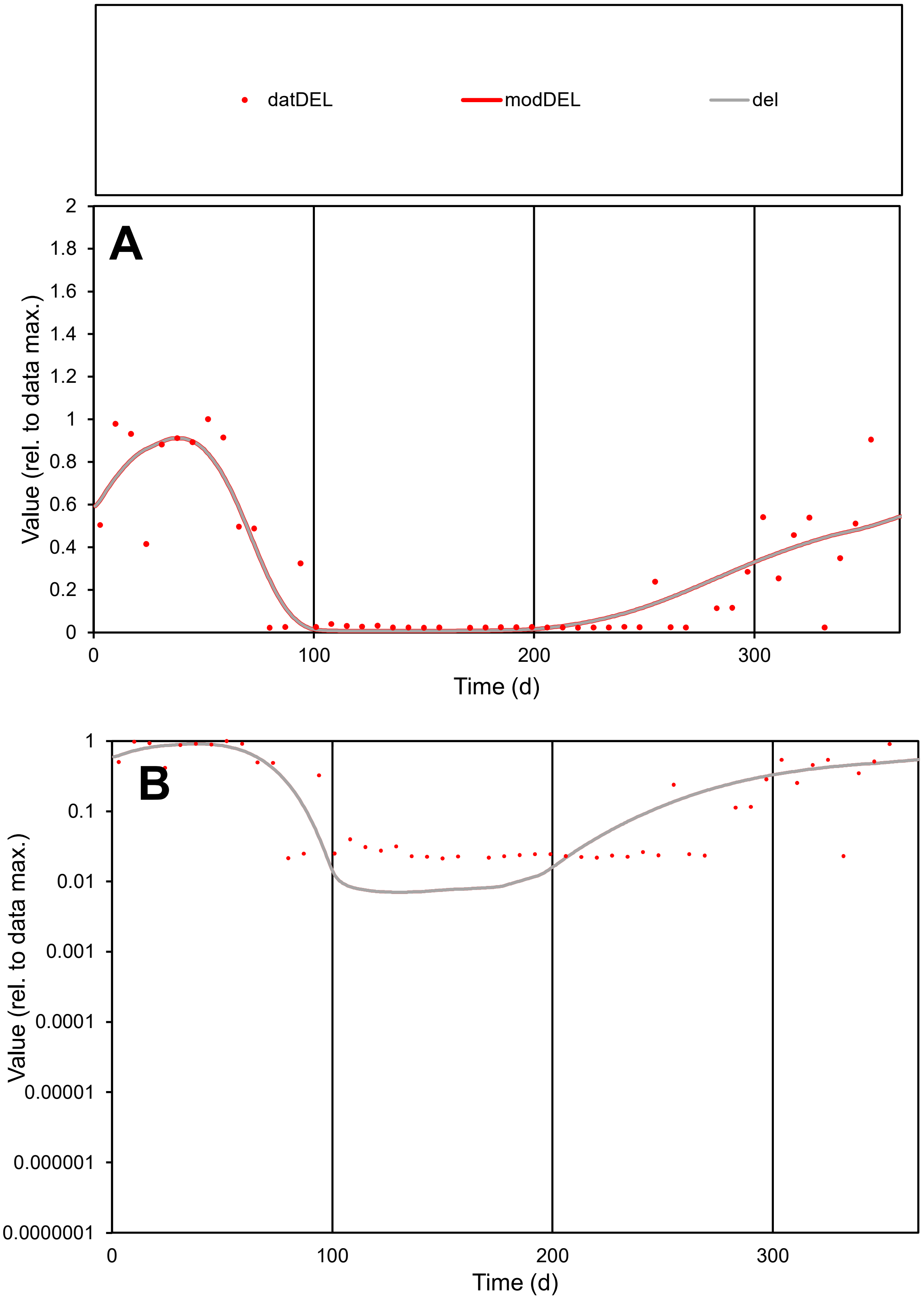


**C1**

Variables
DEL Provide jtd or Dst
CalibOnly?
2 1 to reproduce of
oNormalize?
2 1=no, 2=yes

ofxn calc*
0.51811
ofxn mod
0.55202

sRun
RunA1
zout
1

Ave
Data
0.00081
Model
0.00082

data/vMax
0.0029

ofxng
369.894

Min d/m
3.05372

ave/min d
13.0539
ave/min m
40.6067

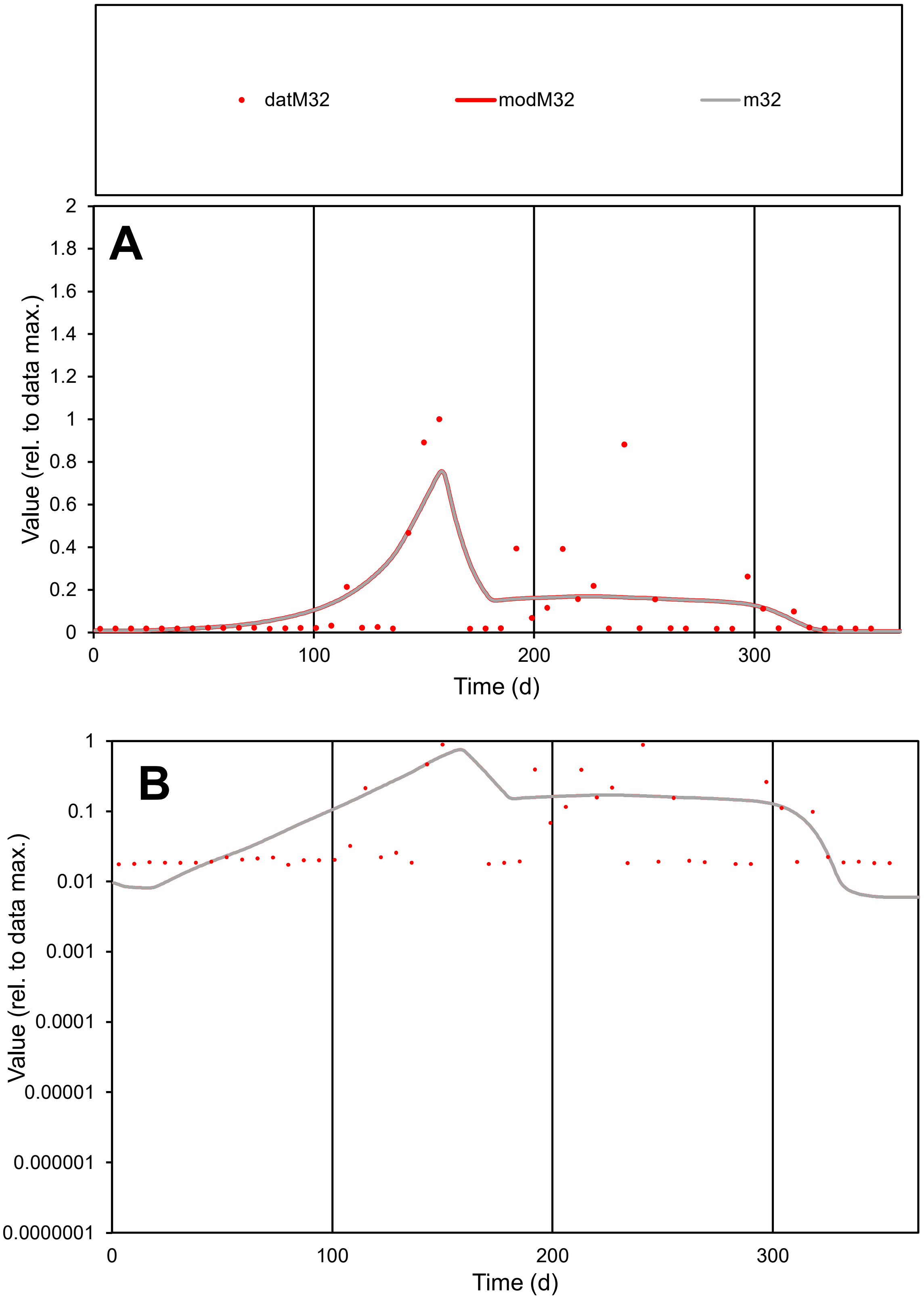


**C1**

Variables
M32 Provide jtd or Dst
CalibOnly?
2 1 to reproduce of
oNormalize?
2 1=no, 2=yes

ofxn calc*
1.27048
ofxn mod
1.29791

sRun
RunA1
zout
1

Ave
Data
0.00151
Model
0.00168

data/vMax
0.0121

ofxng
369.894

Min d/m
2.91194

ave/min d
7.16253
ave/min m
23.2994

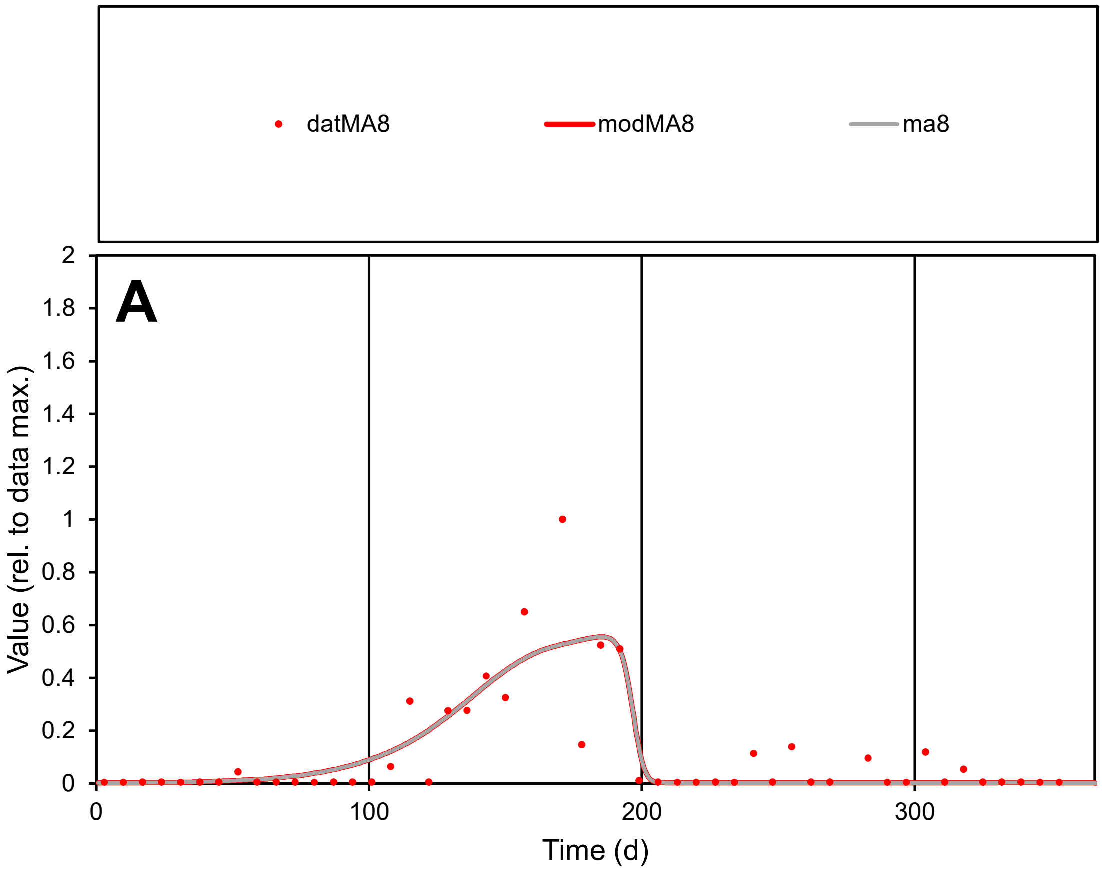


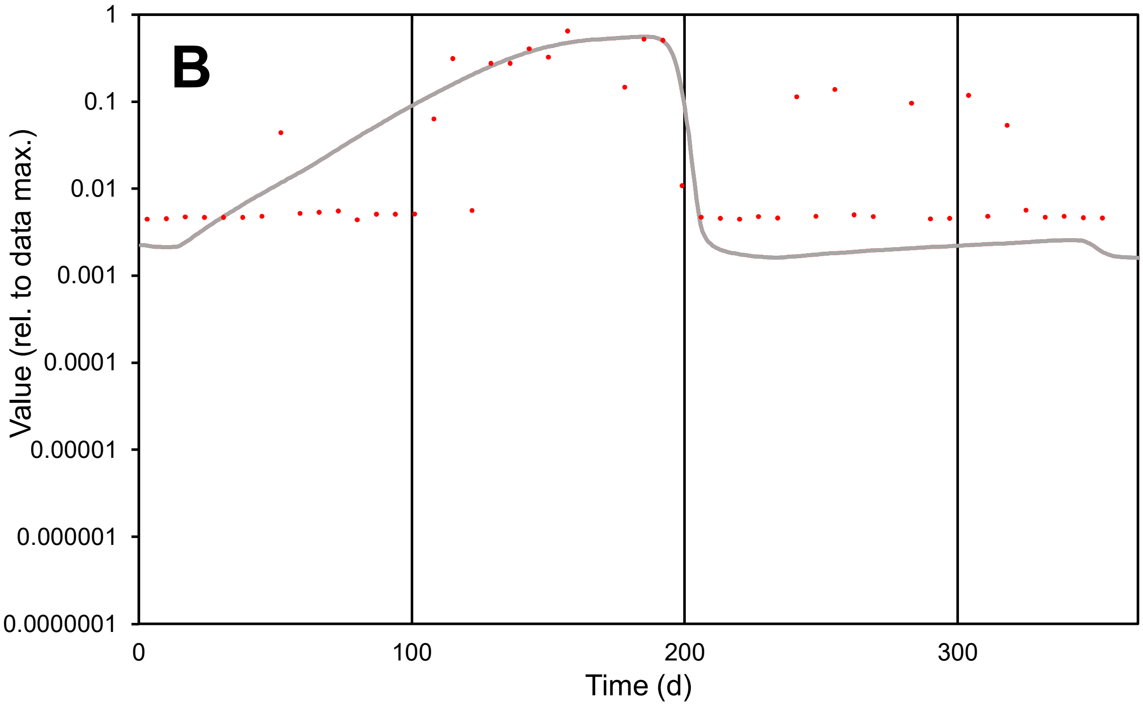


**C1**

Variables
MA8 Provide jtd or Dst
CalibOnly?
2 1 to reproduce of
oNormalize?
2 1=no, 2=yes

ofxn calc*
1.00901
ofxn mod
1.03544

sRun
RunA1
zout
1

Ave
Data
0.00146
Model
0.00138

data/vMax
0.01368

ofxng
369.894

Min d/m
2.71151

ave/min d
24.2353
ave/min m
62.2941

328_Calib_S_A

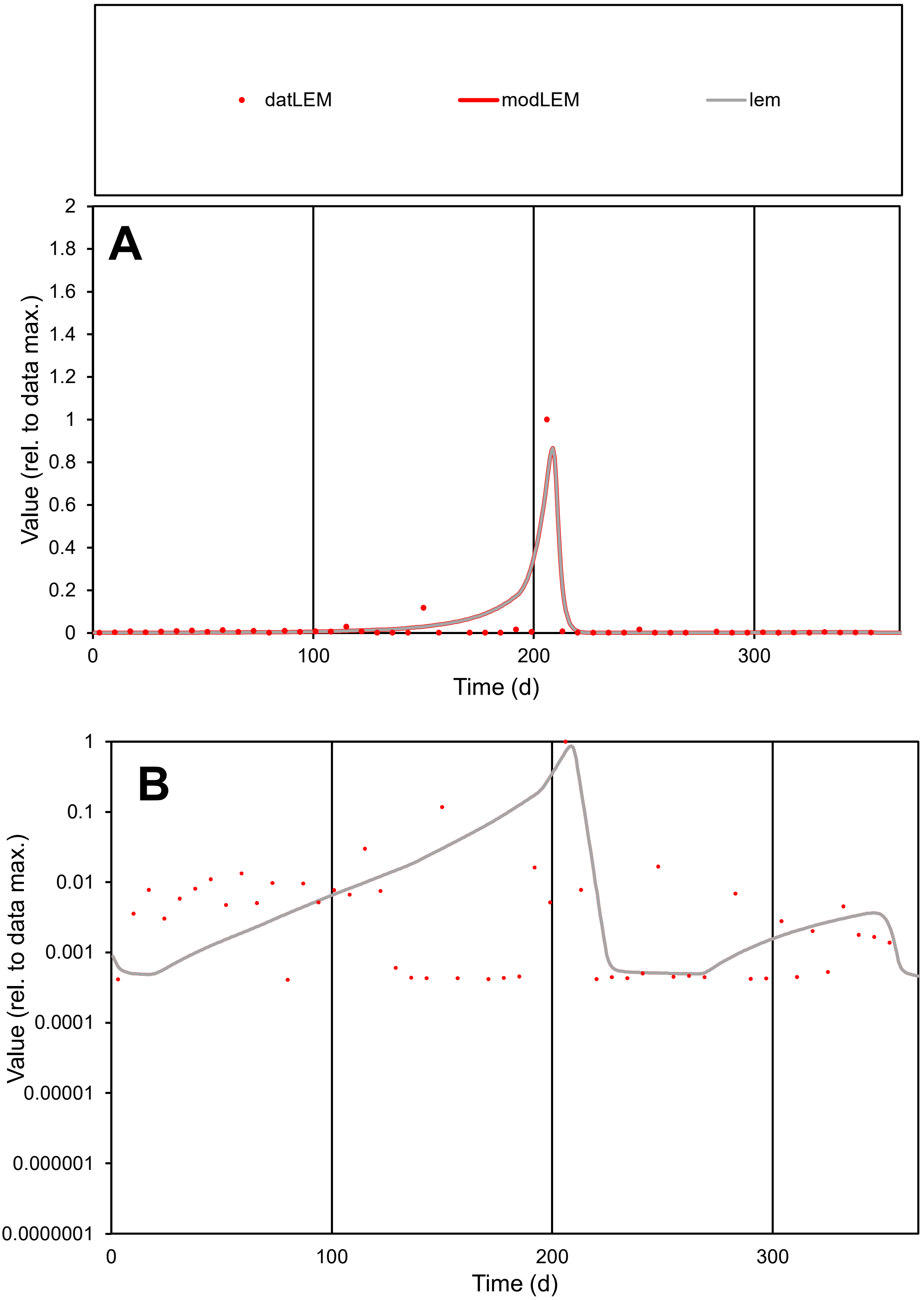


**C1**

Variables
LEM Provide jtd or Dst
CalibOnly?
2 1 to reproduce of
oNormalize?
2 1=no, 2=yes

ofxn calc*
2.10417
ofxn mod
2.31689

sRun
RunA1
zout
1

Ave
Data
0.00144
Model
0.00194

data/vMax
0.05291

ofxng
369.894

Min d/m
0.84017

ave/min d
66.252
ave/min m
75.1992

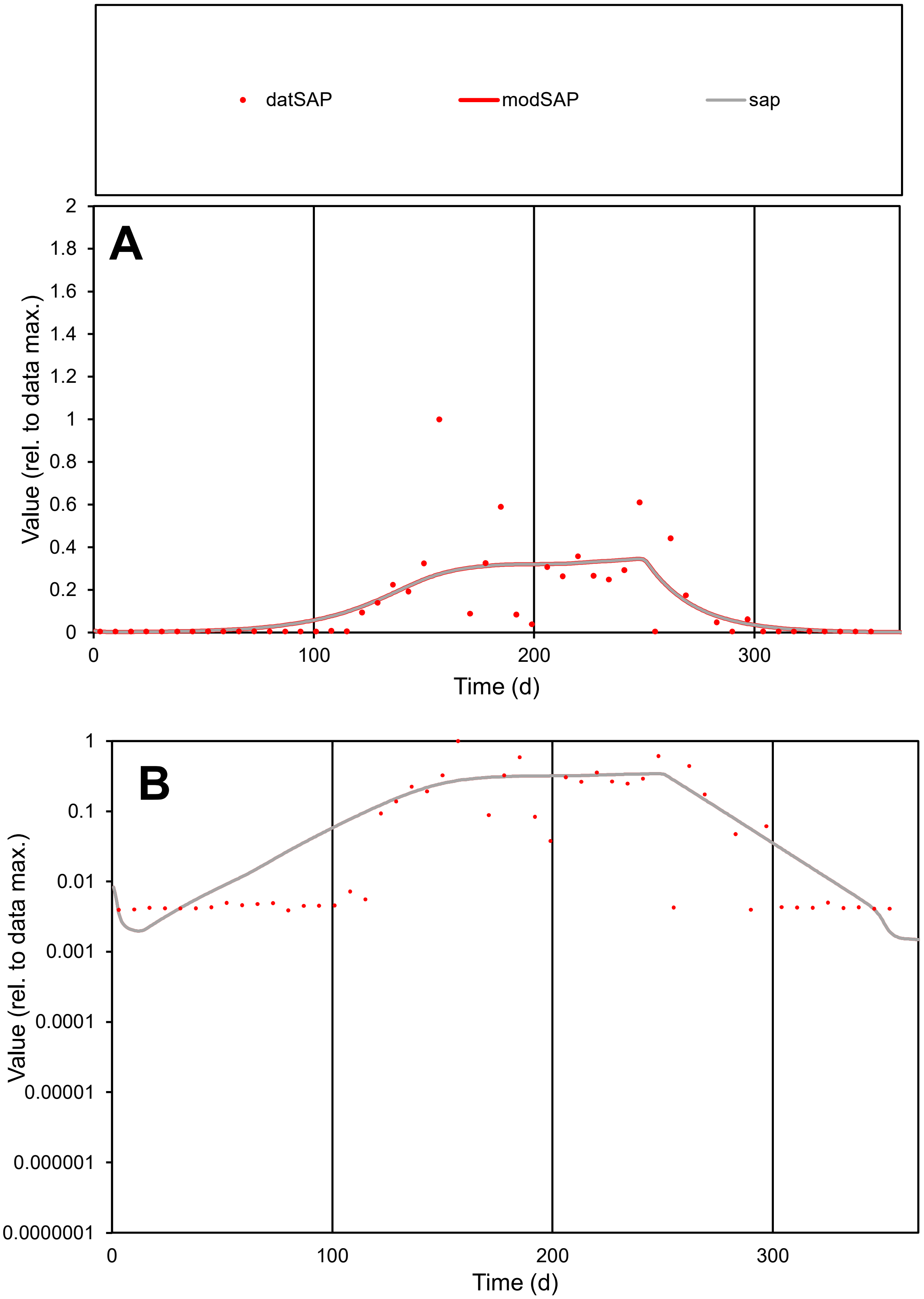


**C1**

Variables
SAP Provide jtd or Dst
CalibOnly?
2 1 to reproduce of
oNormalize?
2 1=no, 2=yes

ofxn calc*
1.125
ofxn mod
1.14373

sRun
RunA1
zout
1

Ave
Data
0.00109
Model
0.00111

data/vMax
0.00849

ofxng
369.894

Min d/m
2.07076

ave/min d
32.9075
ave/min m
69.4075

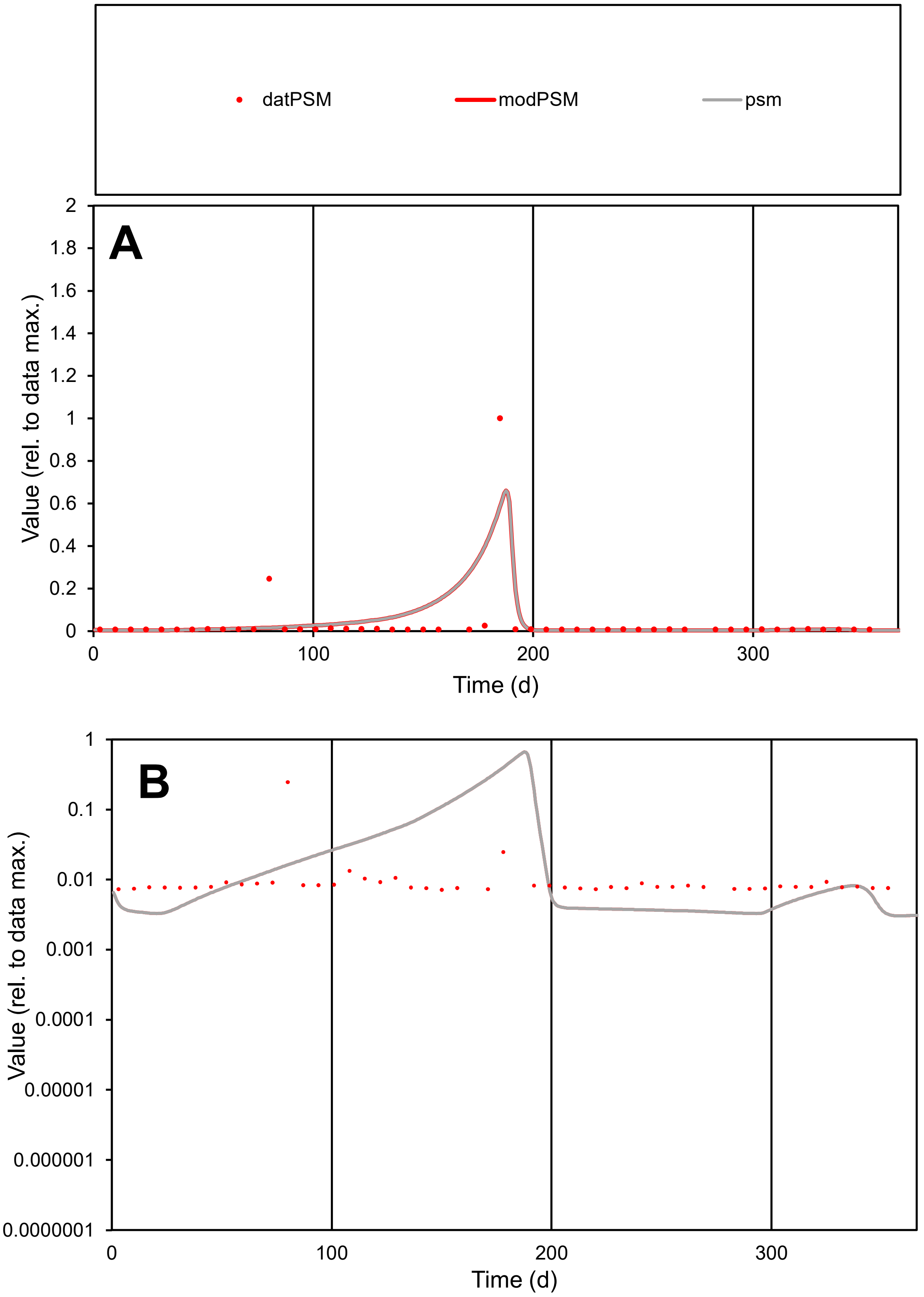


**C1**

Variables

PSM Provide jtd or Dst

CalibOnly?

2 1 to reproduce of

oNormalize?

2 1=no, 2=yes

ofxn calc*

2.79866

ofxn mod

2.89353

sRun

RunA1

zout

1

Ave

Data

0.00013

Model

0.00017

data/vMax

0.00387

ofxng

369.894

Min d/m

2.29246

ave/min d

4.69504

ave/min m

13.9888

328_Calib_S_A

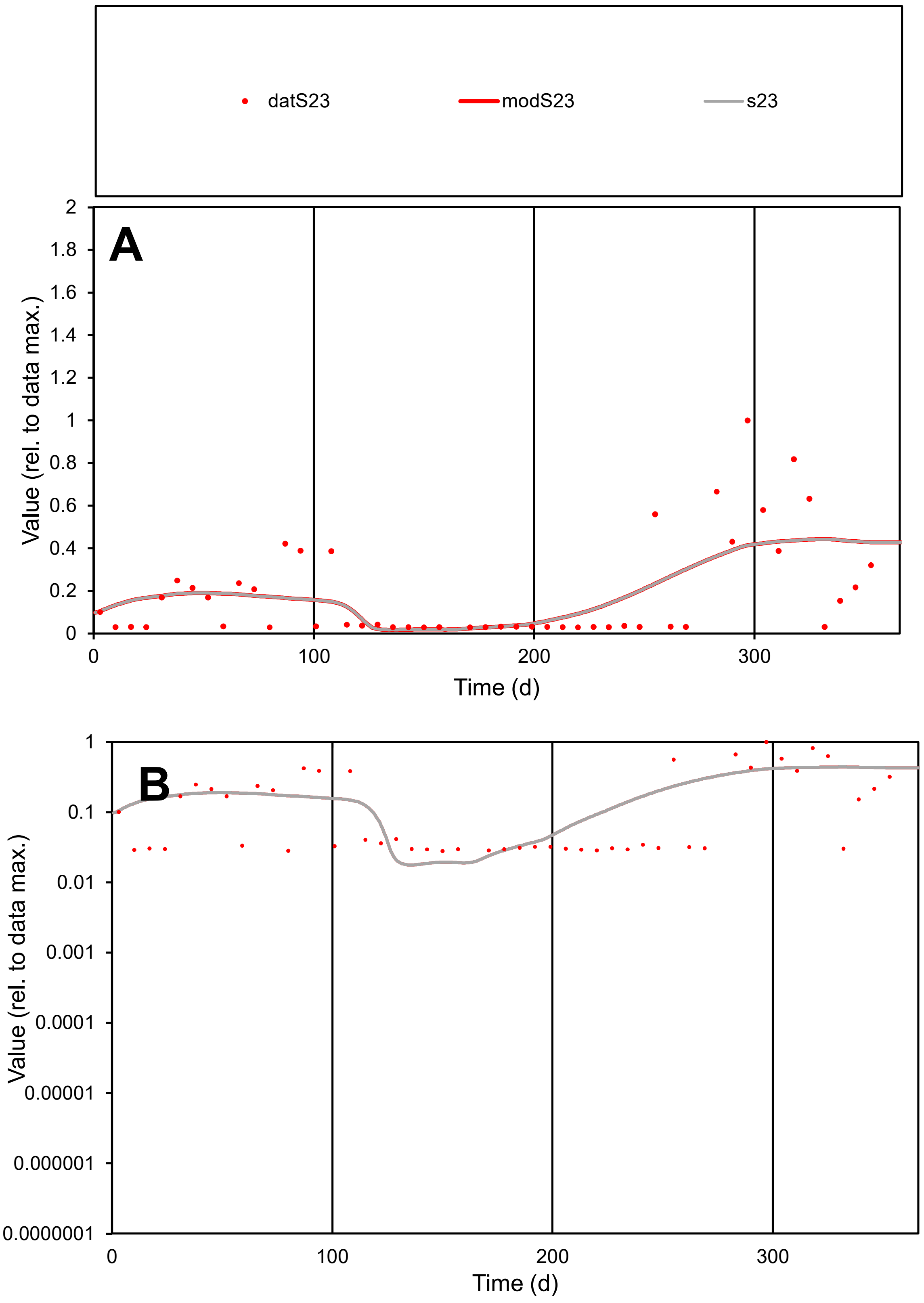


**C1**

Variables
S23 Provide jtd or Dst
CalibOnly?
2 1 to reproduce of
oNormalize?
2 1=no, 2=yes

ofxn calc*
0.94881
ofxn mod
0.96046

sRun
RunA1
zout
1

Ave
Data
0.00108
Model
0.00111

data/vMax
0.00575

ofxng
369.894

Min d/m
1.57493

ave/min d
6.69307
ave/min m
10.832

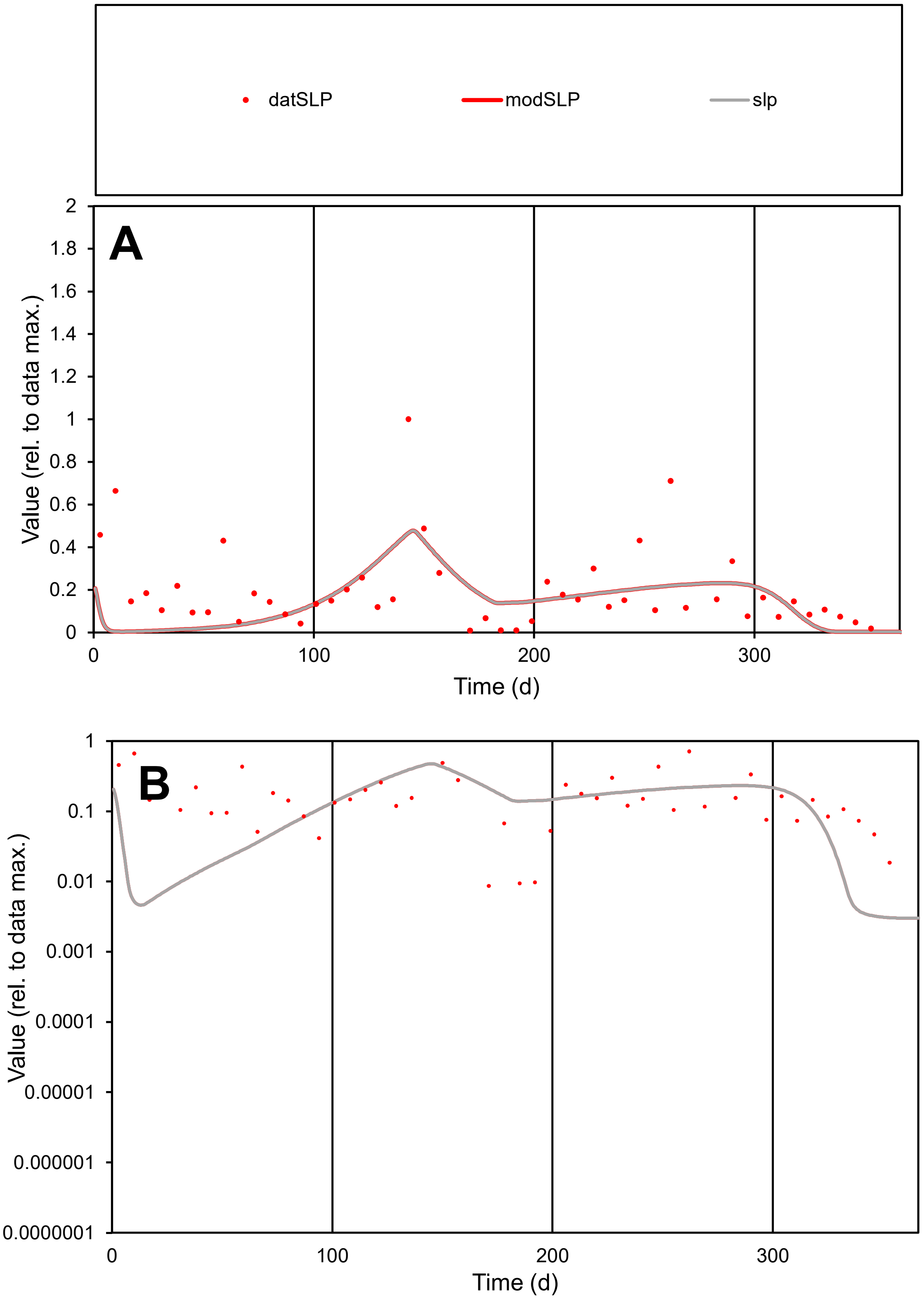


C1

Variables
SLP Provide jtd or Dst
CalibOnly?
2 1 to reproduce of
oNormalize?
2 1=no, 2=yes

ofxn calc*
0.95209
ofxn mod
0.96987

sRun
RunA1
zout
1

Ave
Data
0.00055
Model
0.00042

data/vMax
0.00282

ofxng
369.894

Min d/m
2.85034

ave/min d
22.5403
ave/min m
49.1576

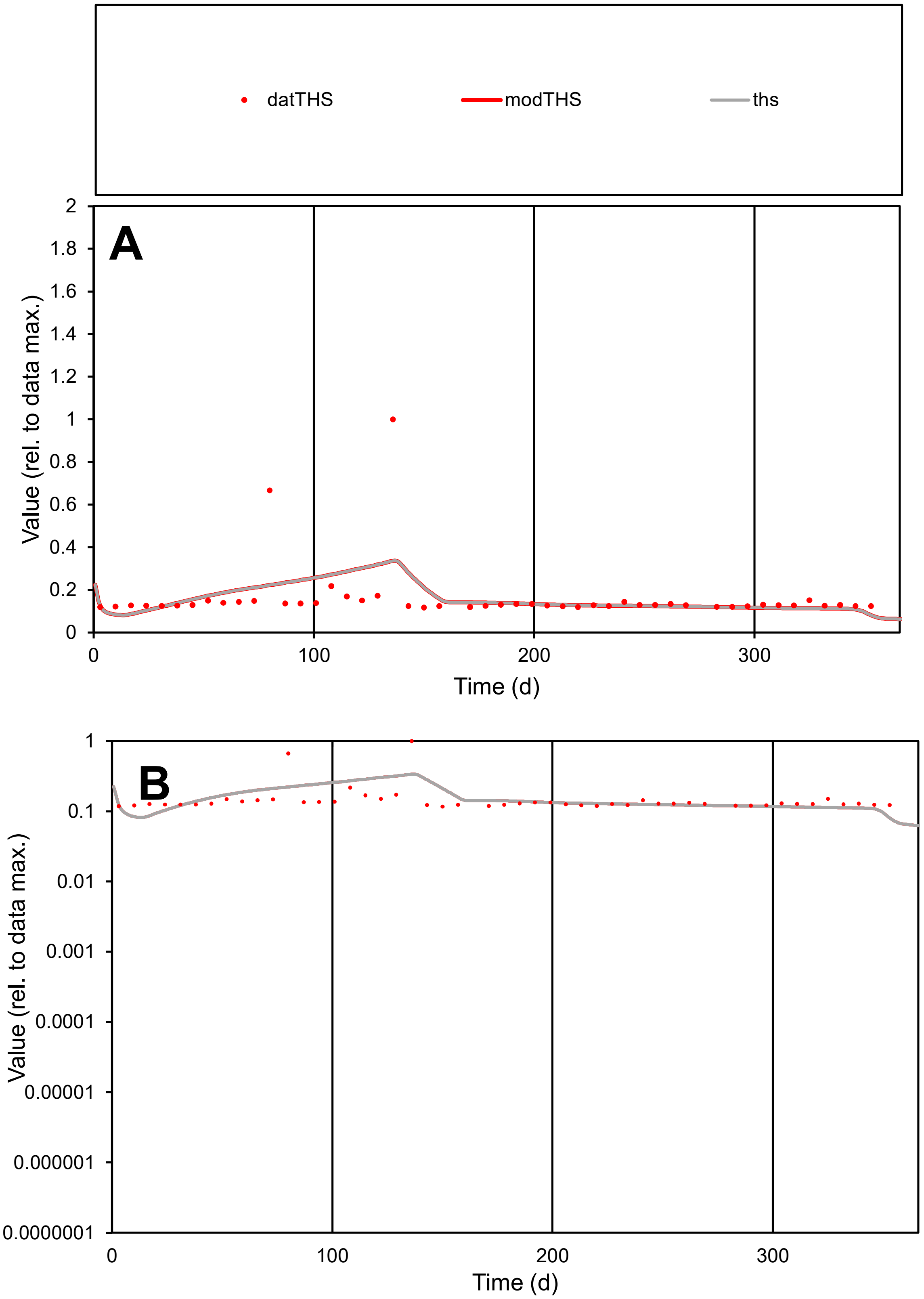


**C1**

Variables
THS Provide jtd or Dst
CalibOnly?
2 1 to reproduce of
oNormalize?
2 1=no, 2=yes

ofxn calc*
0.78695
ofxn mod
0.79596

sRun
RunA1
zout
1

Ave
Data
4E-05
Model
4E-05

data/vMax
0.00025

ofxng
369.894

Min d/m
1.5355

ave/min d
1.3873
ave/min m
2.12281

328_Calib_S_A

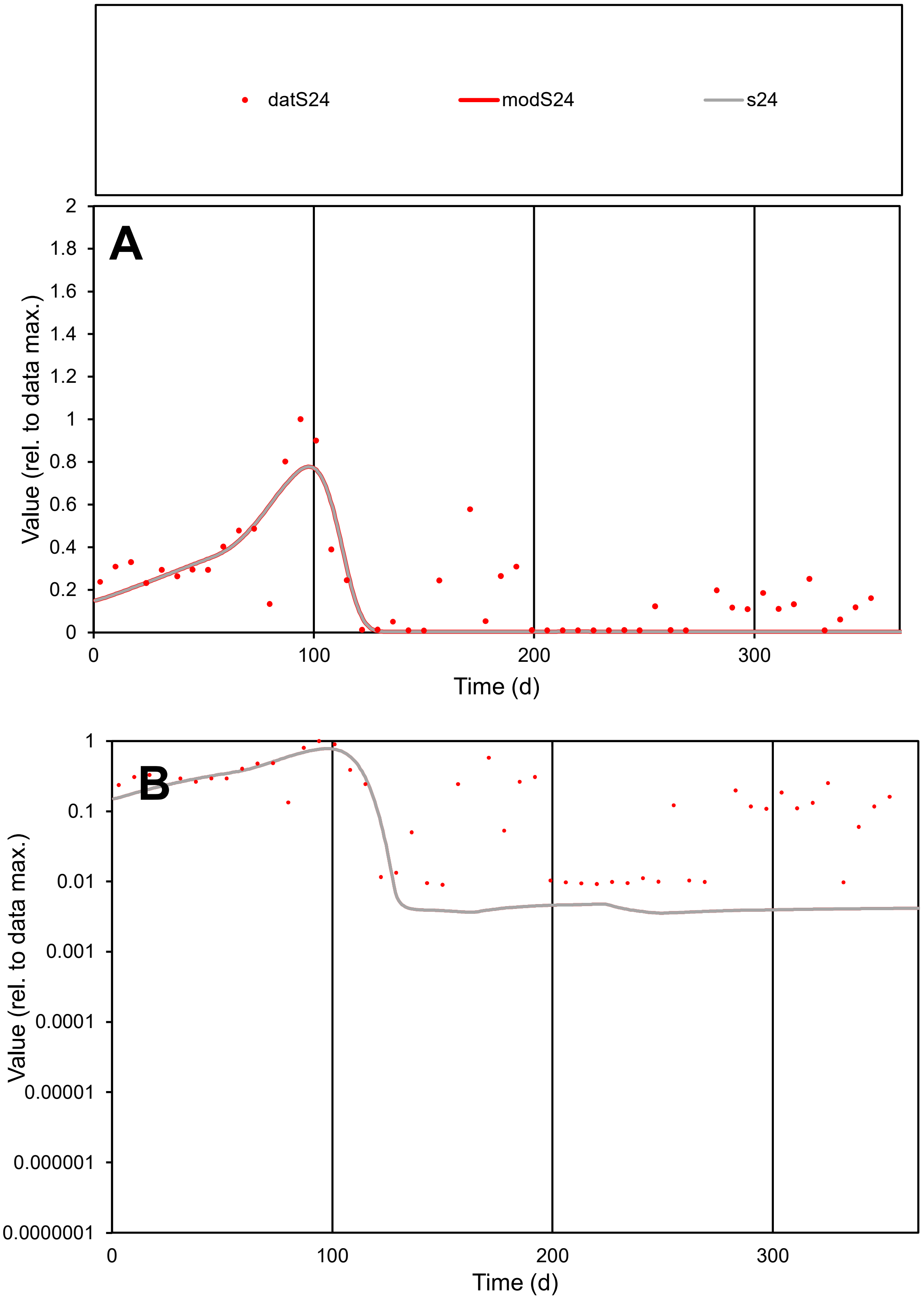


**C1**

Variables
S24 Provide jtd or Dst
CalibOnly?
2 1 to reproduce of
oNormalize?
2 1=no, 2=yes

ofxn calc*
0.73402
ofxn mod
0.74947

sRun
RunA1
zout
1

Ave
Data
0.00136
Model
0.00095

data/vMax
0.00645

ofxng
369.894

Min d/m
2.54661

ave/min d
23.3061
ave/min m
41.3825

328_Calib_S_A

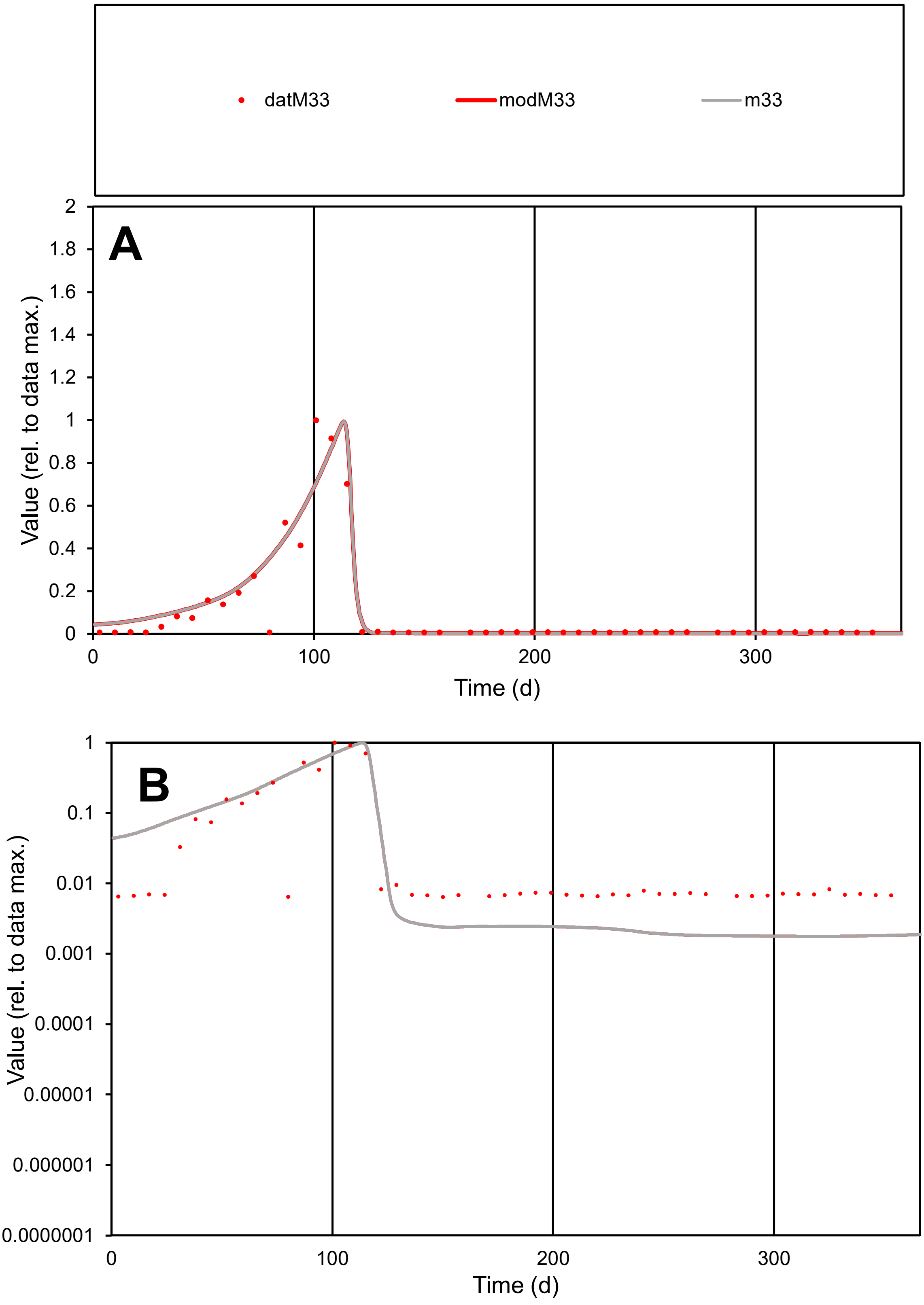


**C1**

Variables
M33 Provide jtd or Dst
CalibOnly?
2 1 to reproduce of
oNormalize?
2 1=no, 2=yes

ofxn calc*
0.72925
ofxn mod
0.77544

sRun
RunA1
zout
1

Ave
Data
0.00104
Model
0.00115

data/vMax
0.01069

ofxng
369.894

Min d/m
3.6315

ave/min d
15.1476
ave/min m
60.8316

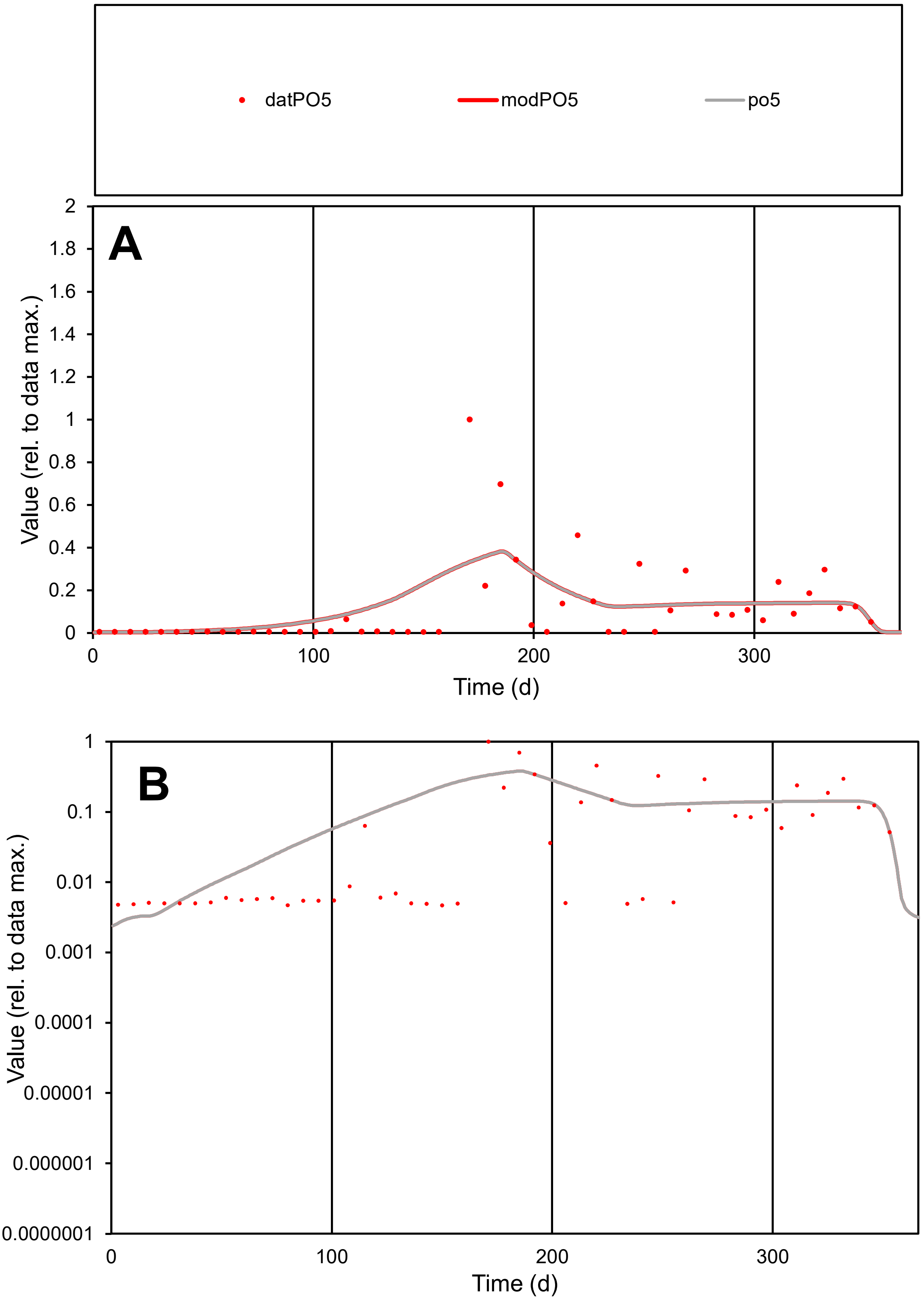


**C1**

Variables
PO5 Provide jtd or Dst
CalibOnly?
2 1 to reproduce of
oNormalize?
2 1=no, 2=yes

ofxn calc*
1.37378
ofxn mod
1.39164

sRun
RunA1
zout
1

Ave
Data
0.0011
Model
0.00125

data/vMax
0.00995

ofxng
369.894

Min d/m
1.70664
ave/min d
23.5985
ave/min m
45.8603

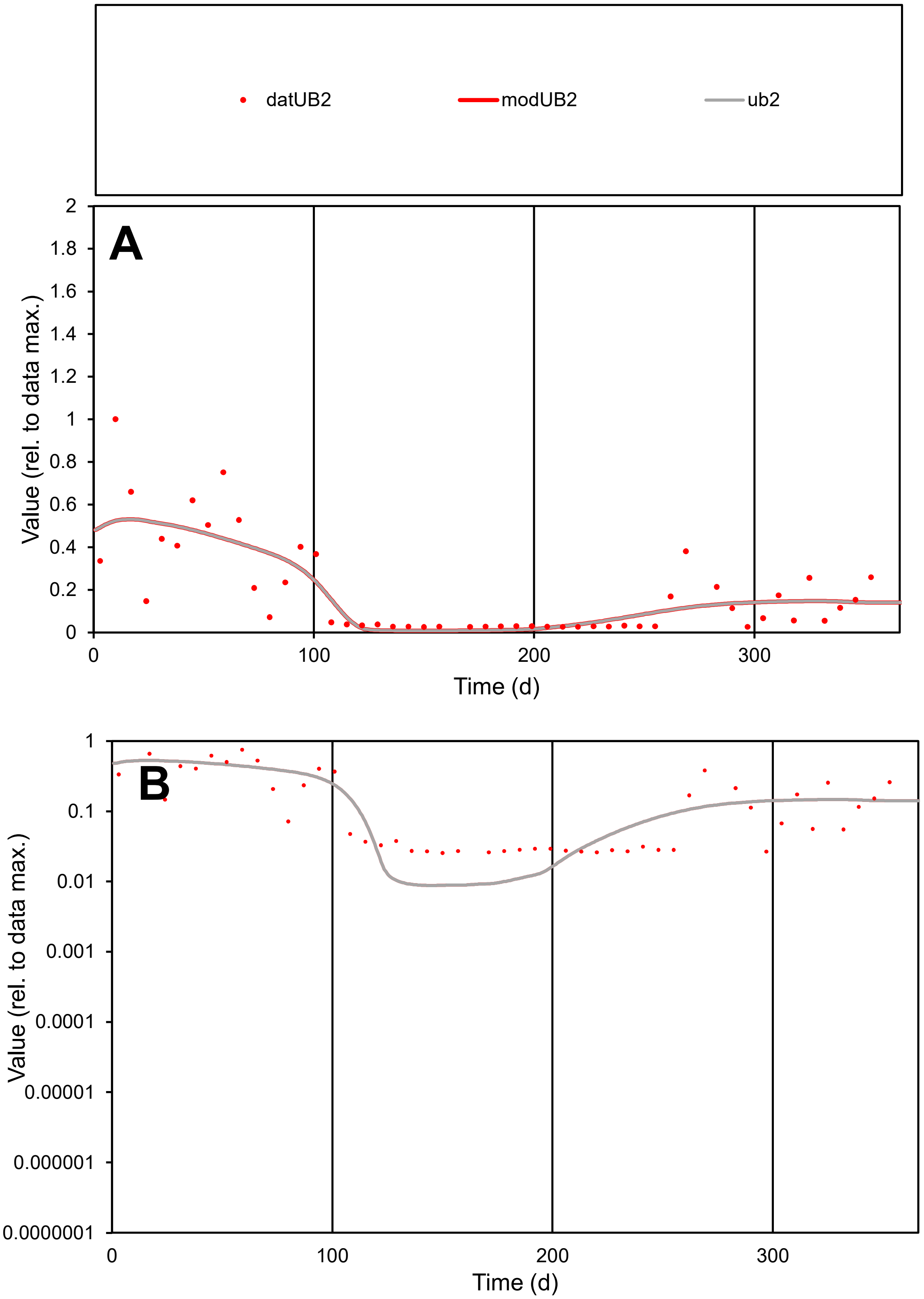


C1

Variables
UB2 Provide jtd or Dst
CalibOnly?
2 1 to reproduce of
oNormalize?
2 1=no, 2=yes

ofxn calc*
0.70178
ofxn mod
0.7241

sRun
RunA1
zout
1

Ave
Data
0.00078
Model
0.00075

data/vMax
0.00409

ofxng
369.894

Min d/m
2.89928

ave/min d
7.47432
ave/min m
20.9792

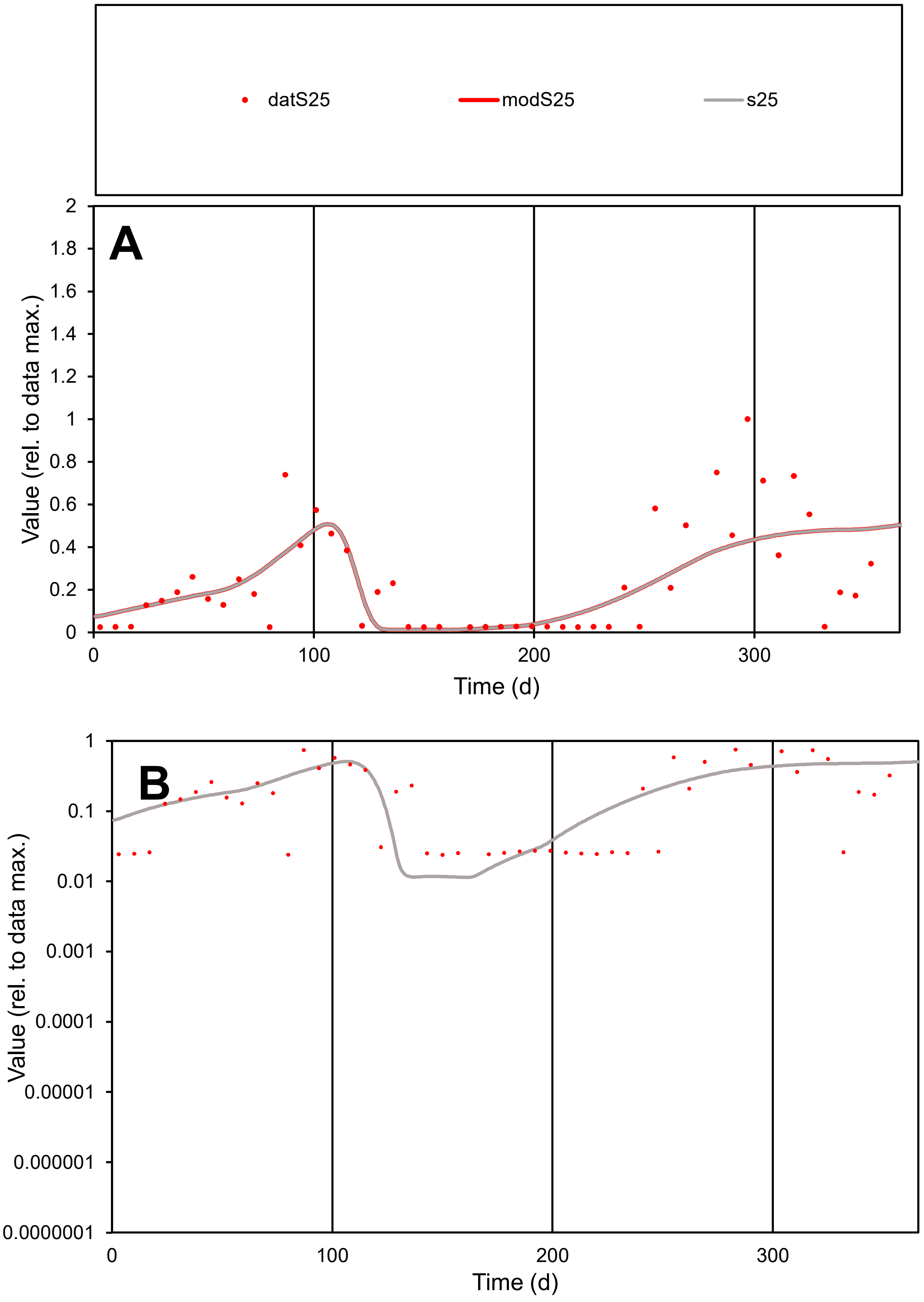


**C1**

Variables
S25 Provide jtd or Dst
CalibOnly?
2 1 to reproduce of
oNormalize?
2 1=no, 2=yes

ofxn calc*
0.75194
ofxn mod
0.76548

sRun
RunA1
zout
1

Ave
Data
0.00169
Model
0.00164

data/vMax
0.00709

ofxng
369.894

Min d/m
2.08652

ave/min d
9.98985
ave/min m
20.2142

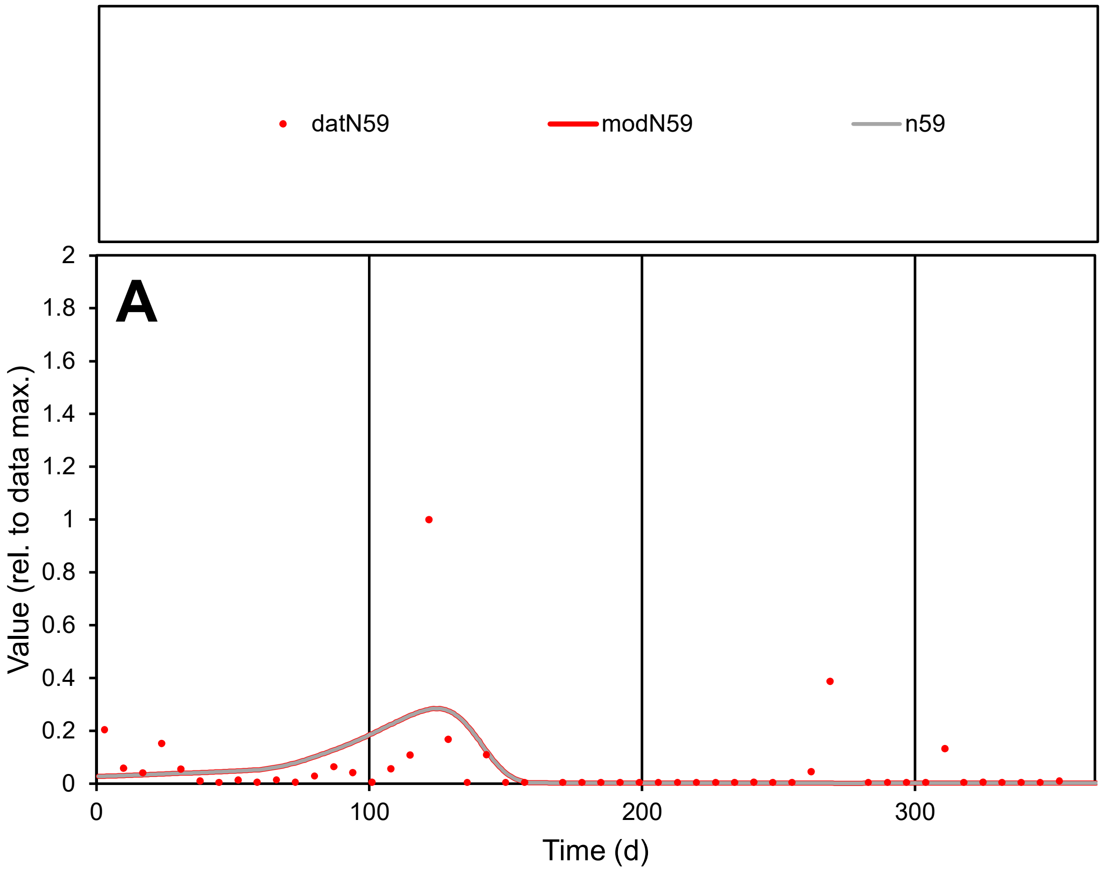


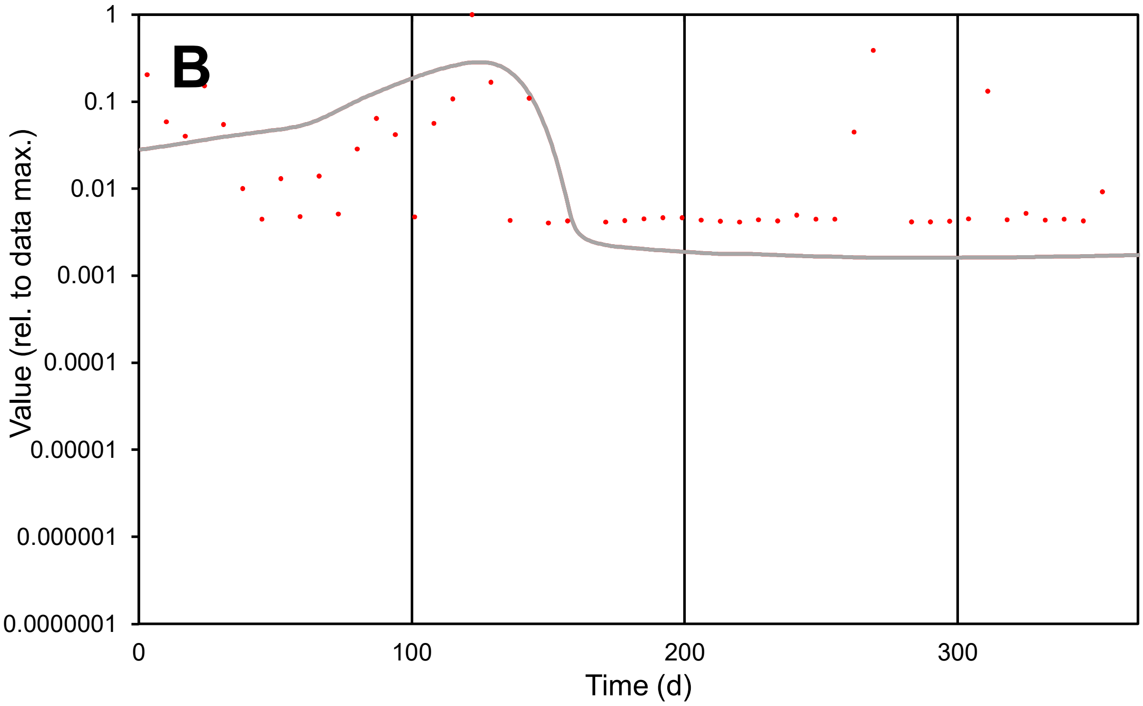


**C1**

Variables
N59 Provide jtd or Dst
CalibOnly?
2 1 to reproduce of
oNormalize?
2 1=no, 2=yes

ofxn calc*
2.33495
ofxn mod
2.35996

sRun
RunA1
zout
1

Ave
Data
0.00059
Model
0.00052

data/vMax
0.01026

ofxng
369.894

Min d/m
2.50697

ave/min d
14.3026
ave/min m
31.3594

328_Calib_S_A

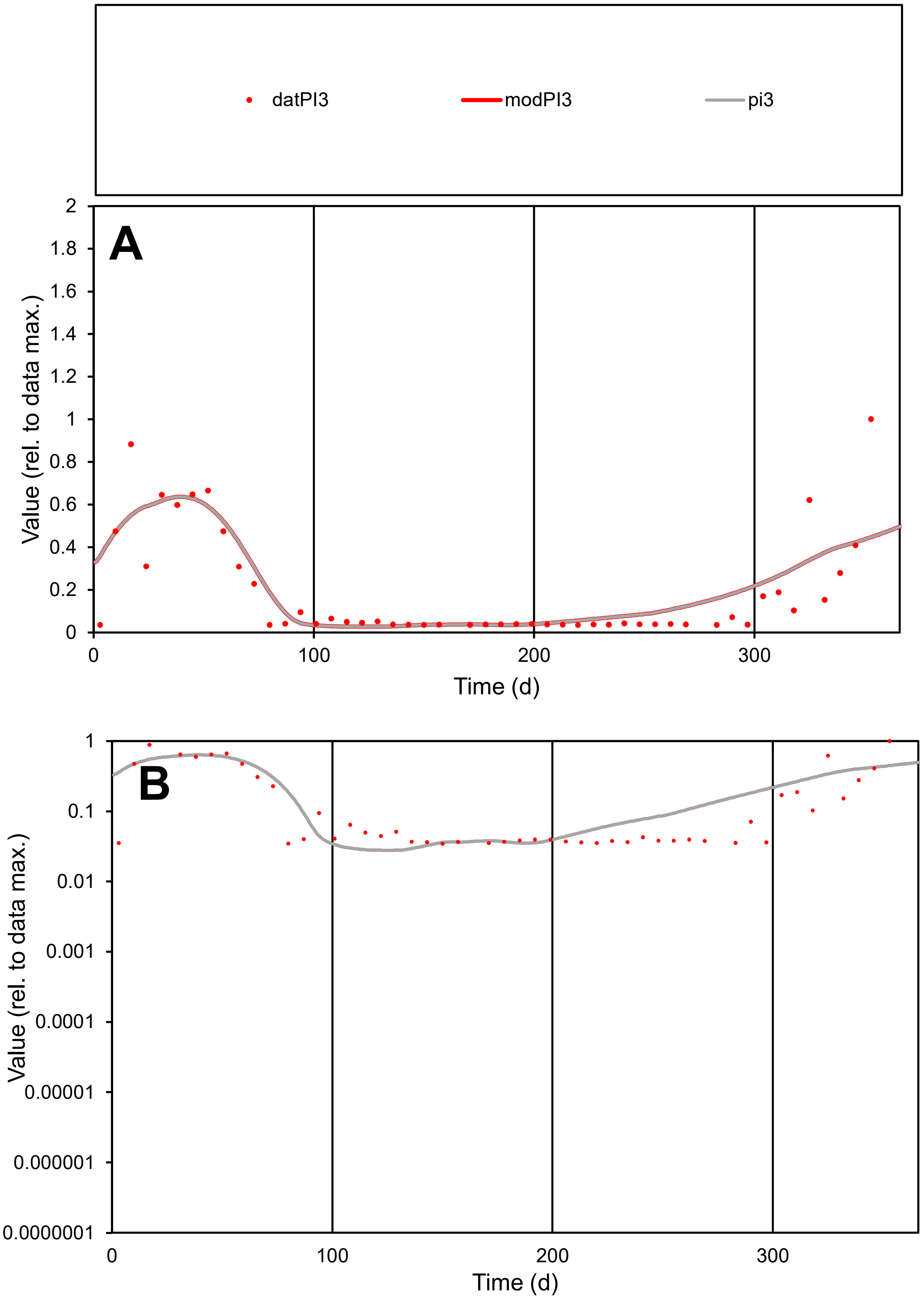


**C1**

Variables

PI3 Provide jtd or Dst

CalibOnly?

2 1 to reproduce of

oNormalize?

2 1=no, 2=yes

ofxn calc*

0.70179

ofxn mod

0.72865

sRun

RunA1

zout

1

Ave

Data

0.0004

Model

0.00044

data/vMax

0.00207

ofxng

369.894

Min d/m

1.24519

ave/min d

5.59873

ave/min m

7.76468

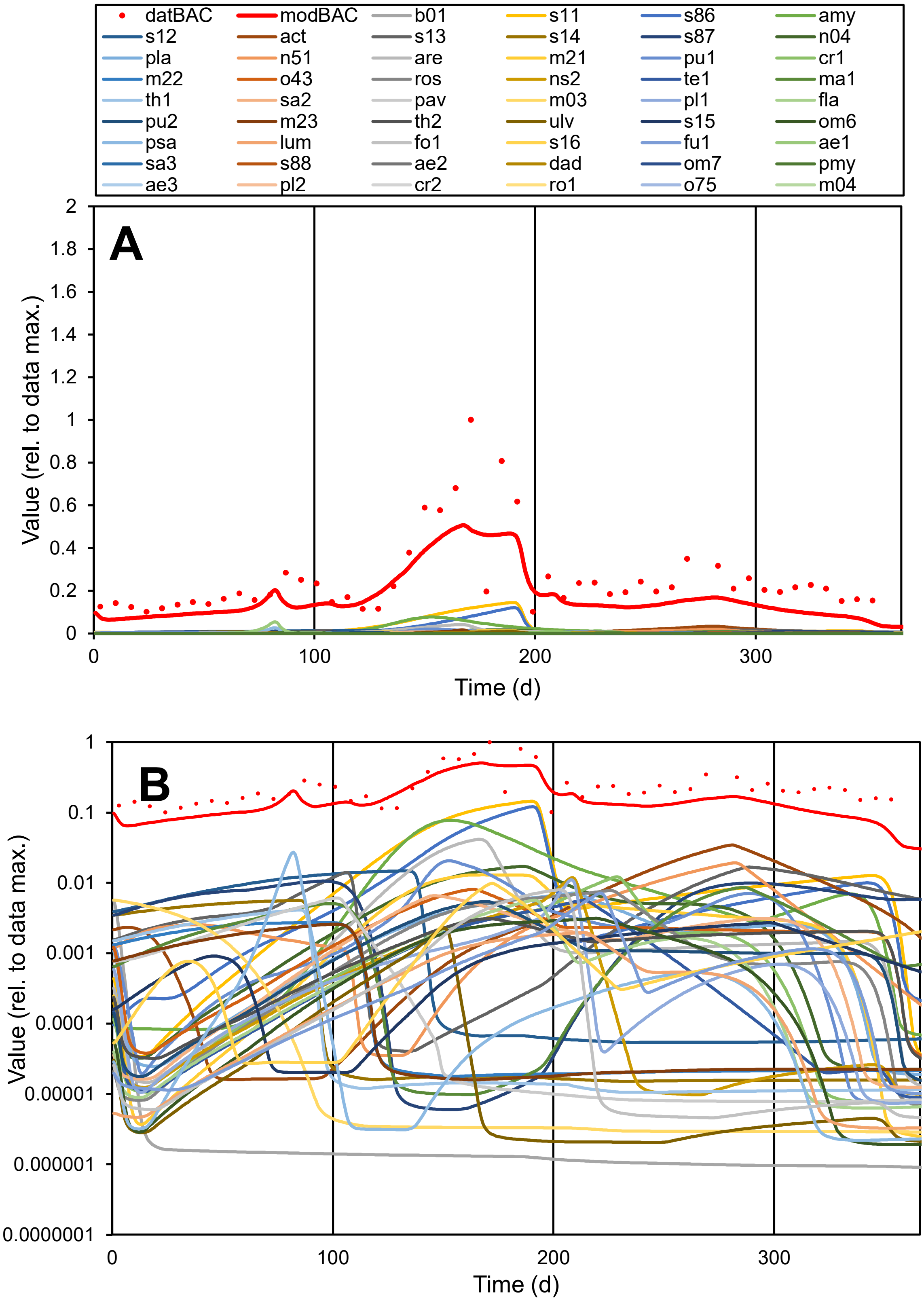


**C1**

Variables

BAC Provide jtd or Dst

CalibOnly?

2 1 to reproduce of

oNormalize?

2 1=no, 2=yes

ofxn calc*

0.52353

ofxn mod

0.53573

sRun

RunA1

zout

1

Ave

Data

1.20369

Model

0.82423

data/vMax

4.709

ofxng

369.894

Min d/m

2.11249

ave/min d

2.53355

ave/min m

3.66488

330_Calib_S_A

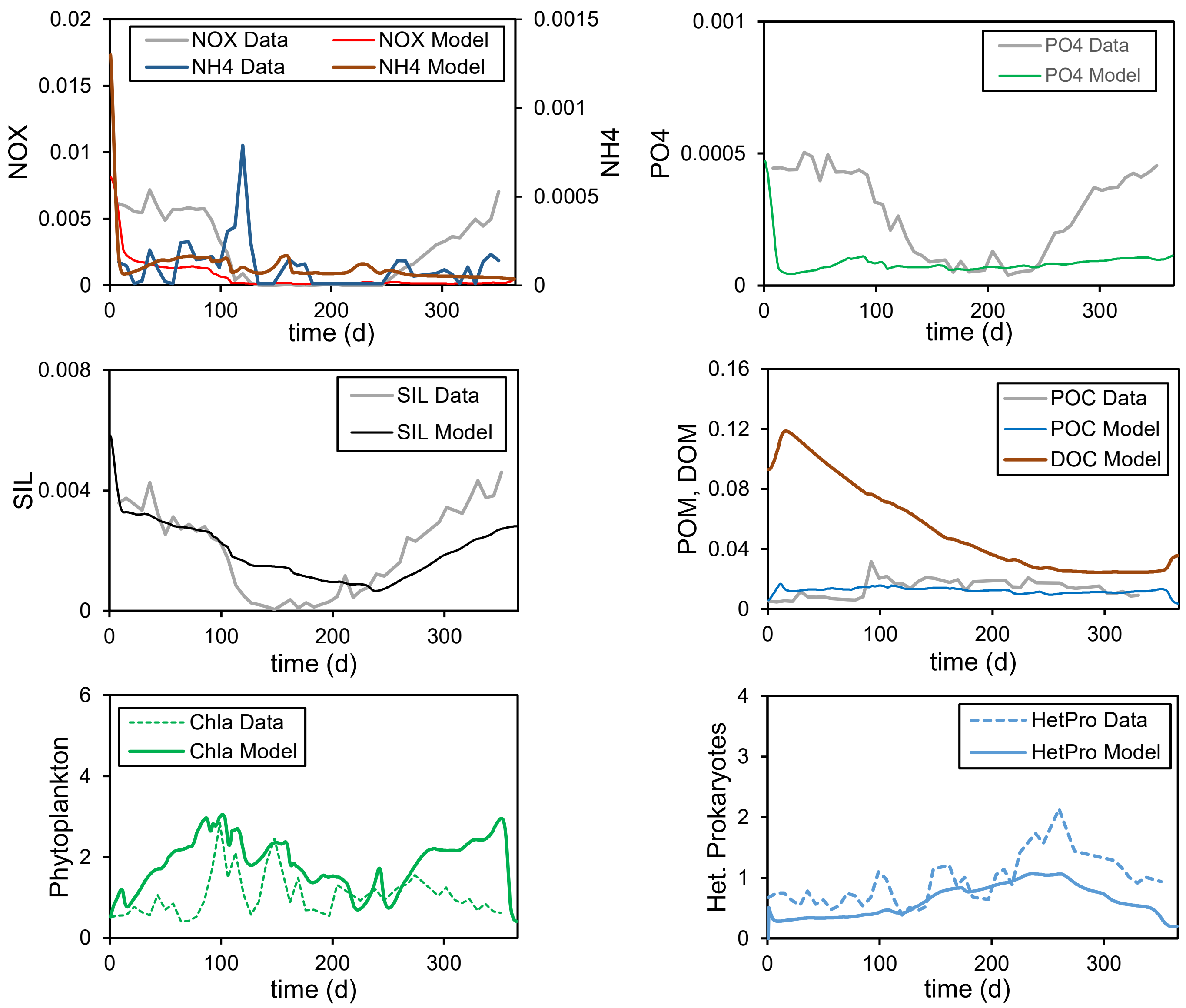

NOX Data
NOX Model
NH4 Data
NH4 Model
NOX
NH4
0.02
0.015
0.01
0.005
0
0.0015
0.001
0.0005
0
100
200
300
time (d)
PO4 Data
PO4 Model
PO4
0.001
0.0005
time (d)
SIL Data
SIL Model
SIL
0.008
0.004
time (d)
POC Data
POC Model
DOC Model
POM, DOM
0.16
0.12
0.08
0.04
time (d)
Chla Data
Chla Model
Phytoplankton
6
4
2
time (d)
HetPro Data
HetPro Model
Het. Prokaryotes
4
3
2
1
time (d)

330_Calib_S_A

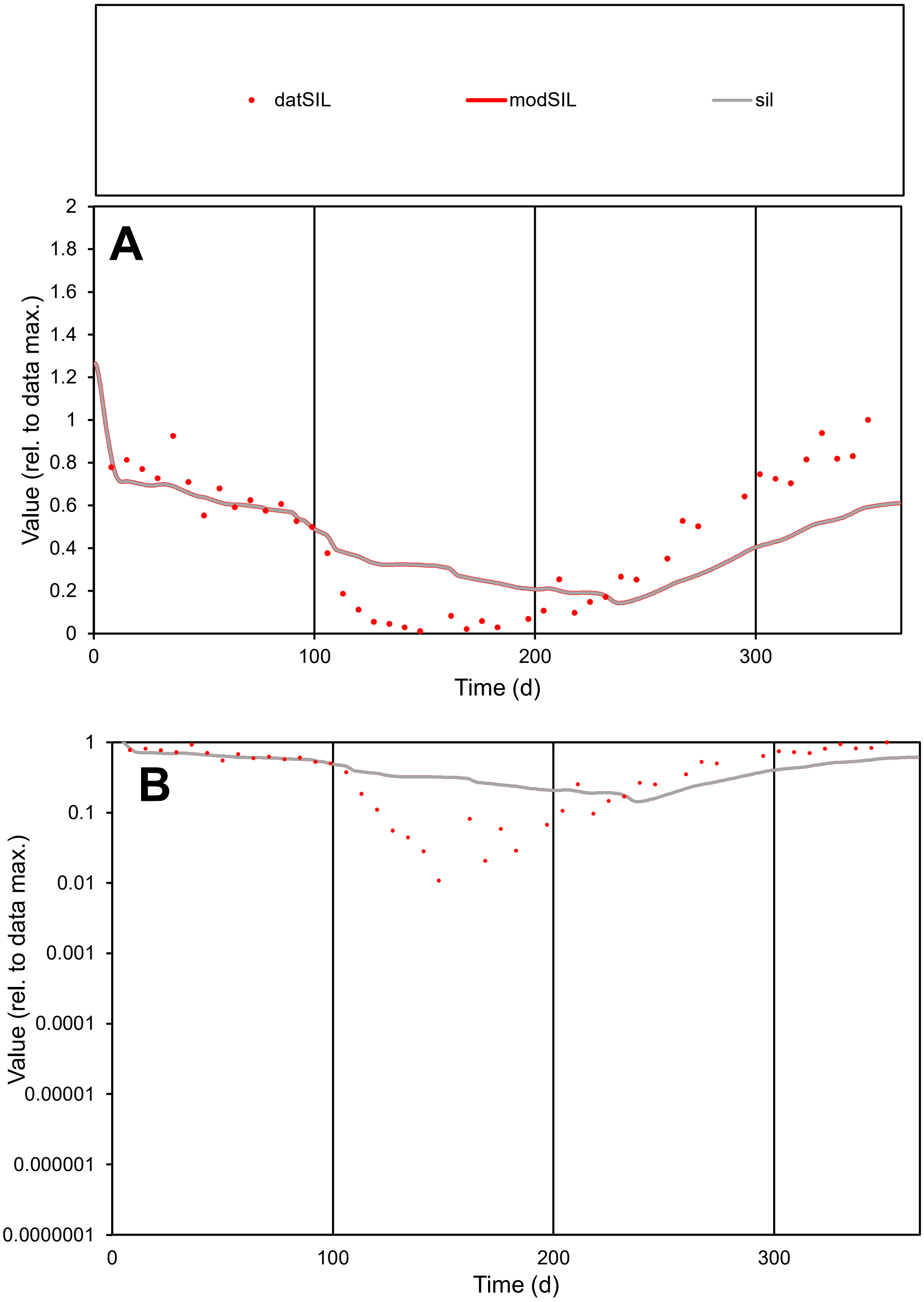


**C1**

Variables

SIL Provide jtd or Dst

CalibOnly?

2 1 to reproduce of

oNormalize?

2 1=no, 2=yes

ofxn calc*

1.3292

ofxn mod

1.37596

sRun

RunA1

zout

1

Ave

Data

0.00208

Model

0.00194

data/vMax

0.00461

ofxng

319.391

Min d/m

0.07452

ave/min d

41.6195

ave/min m

2.89385

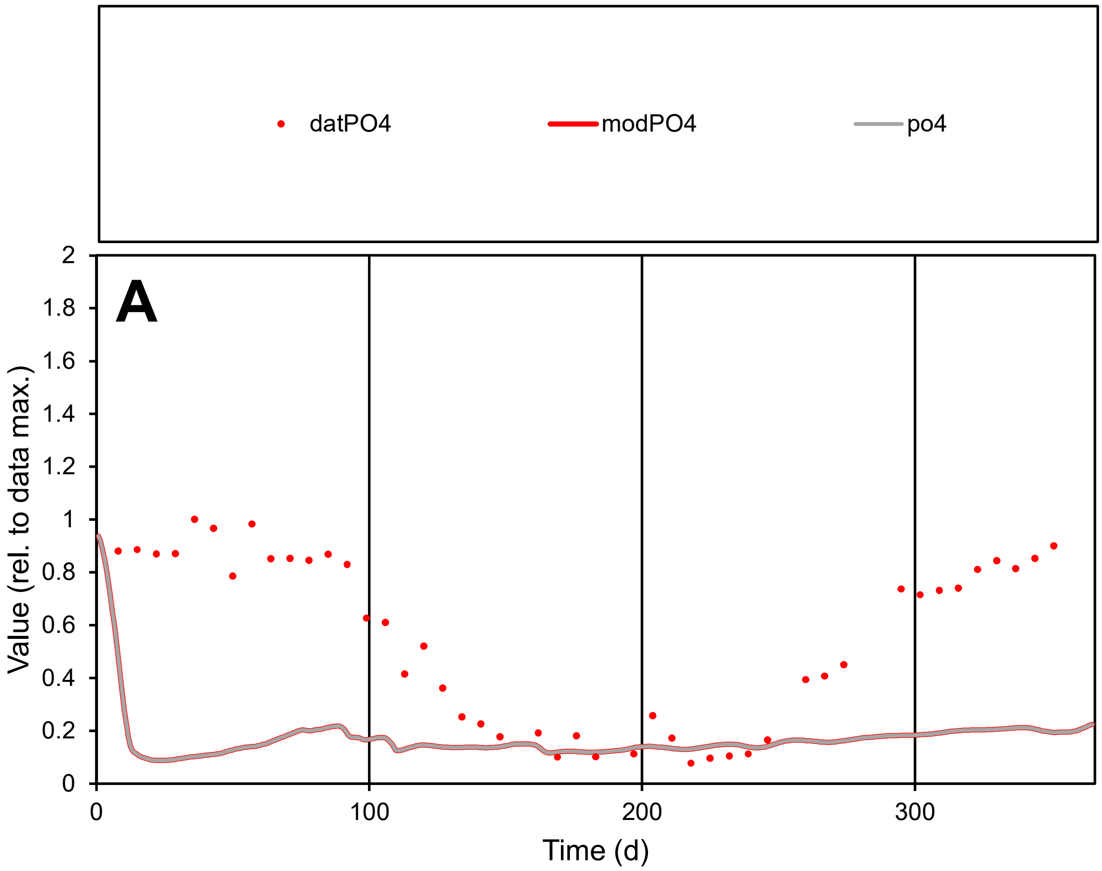


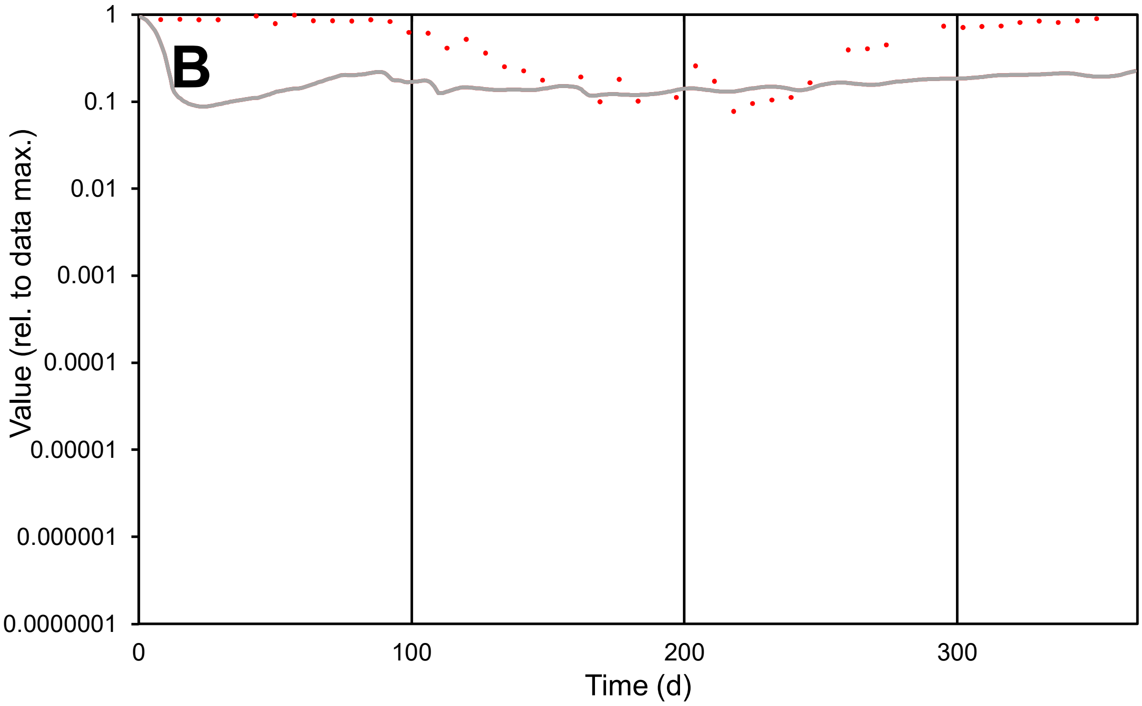


**C1**

Variables
PO4 Provide jtd or Dst
CalibOnly?
2 1 to reproduce of
oNormalize?
2 1=no, 2=yes

ofxn calc*
2.69248
ofxn mod
2.70722

sRun
RunA1
zout
1

Ave
Data
0.00028
Model
8.2E-05

data/vMax
0.0005

ofxng
319.391

Min d/m
0.87903

ave/min d
7.12216
ave/min m
1.84068

330_Calib_S_A

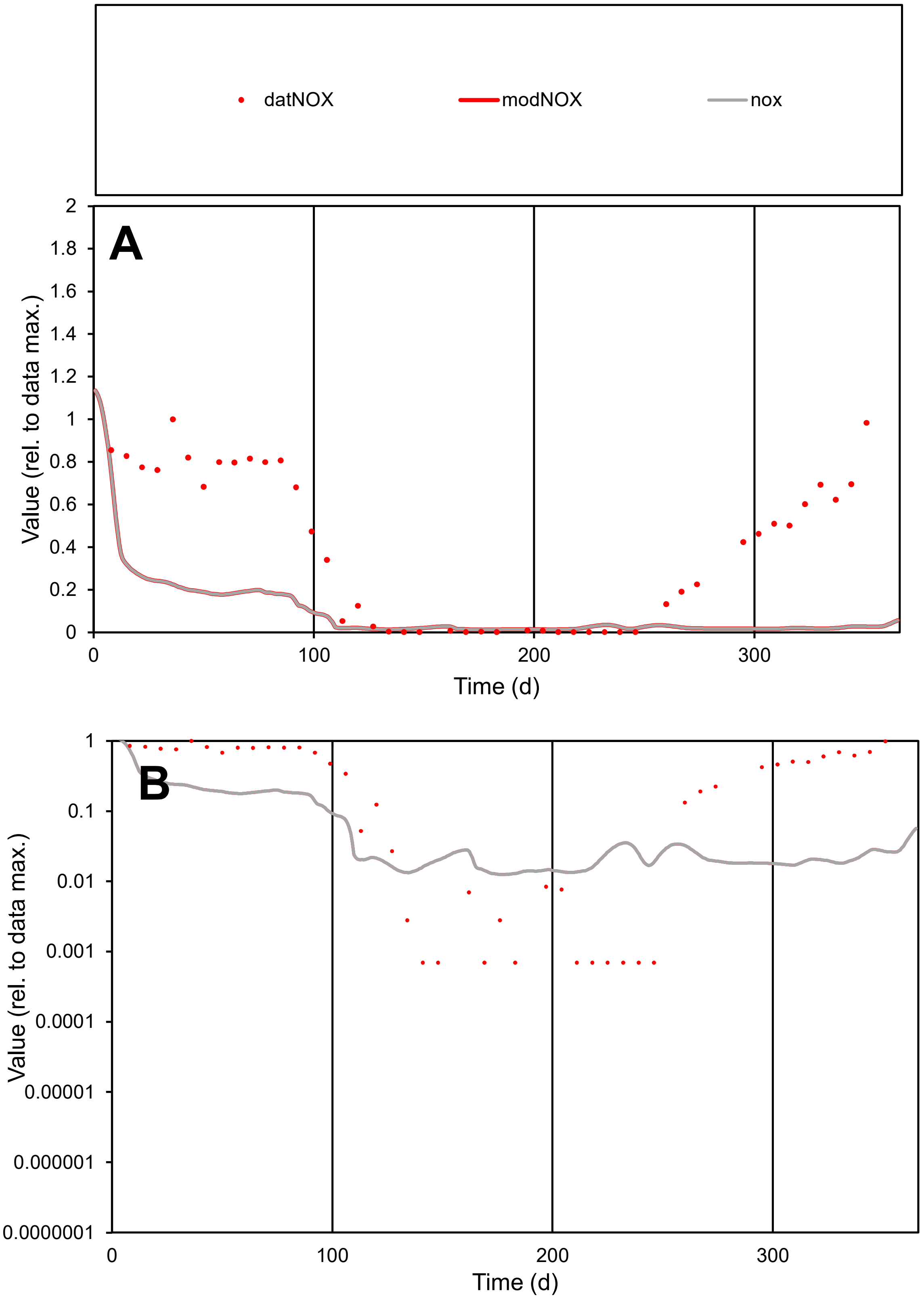


**C1**

Variables
NOX Provide jtd or Dst
CalibOnly?
2 1 to reproduce of
oNormalize?
2 1=no, 2=yes

ofxn calc*
3.235
ofxn mod
3.25667

sRun
RunA1
zout
1

Ave
Data
0.00279
Model
0.00063

data/vMax
0.00717

ofxng
319.391

Min d/m
0.05592

ave/min d
557.673
ave/min m
7.07371

330_Calib_S_A

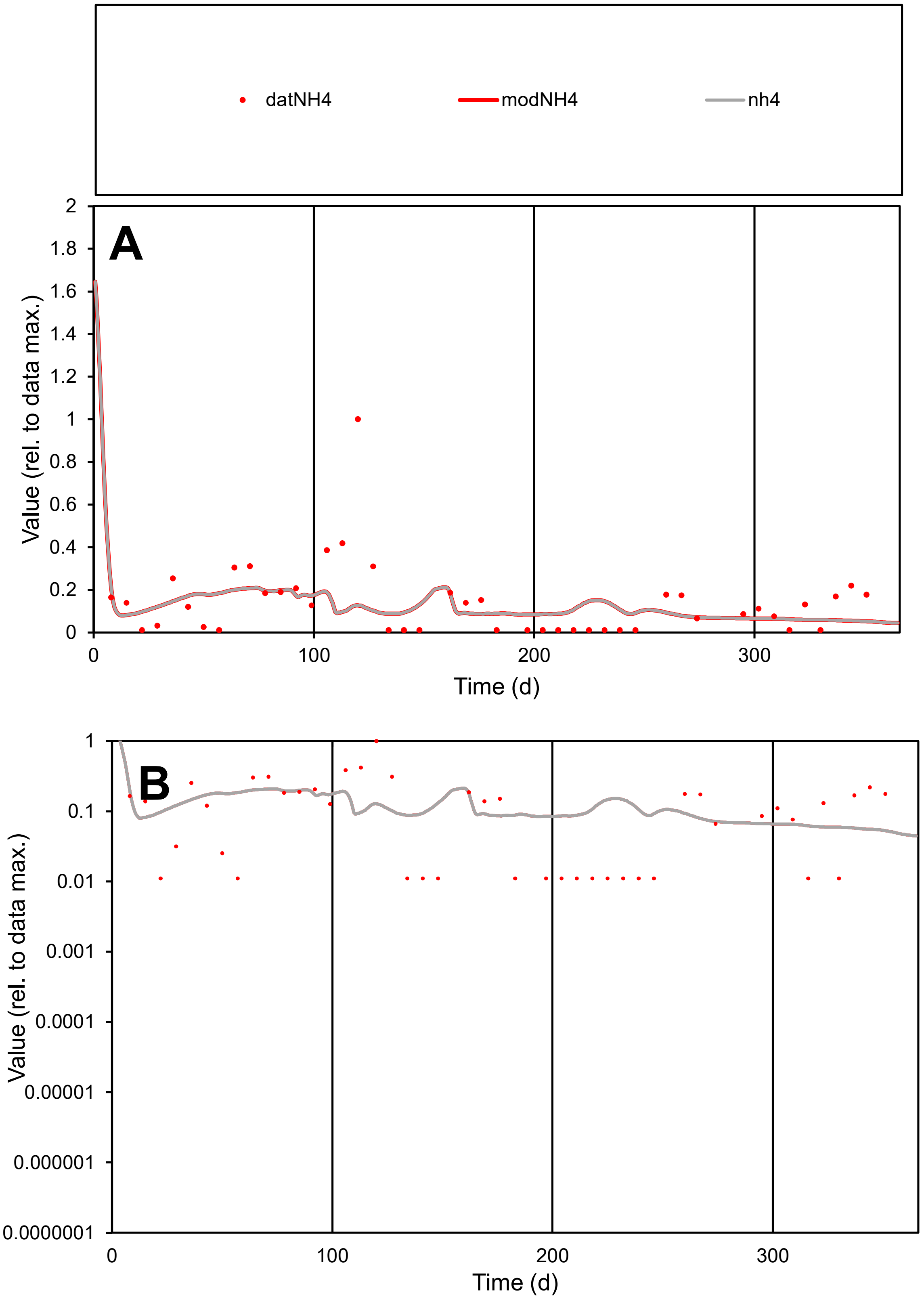


**C1**

Variables
NH4 Provide jtd or Dst
CalibOnly?
2 1 to reproduce of
oNormalize?
2 1=no, 2=yes

ofxn calc*
3.63603
ofxn mod
3.6656

sRun
RunA1
zout
1

Ave
Data
0.00011
Model
9E-05

data/vMax
0.00079

ofxng
319.391

Min d/m
0.21125

ave/min d
12.4612
ave/min m
2.17571

330_Calib_S_A

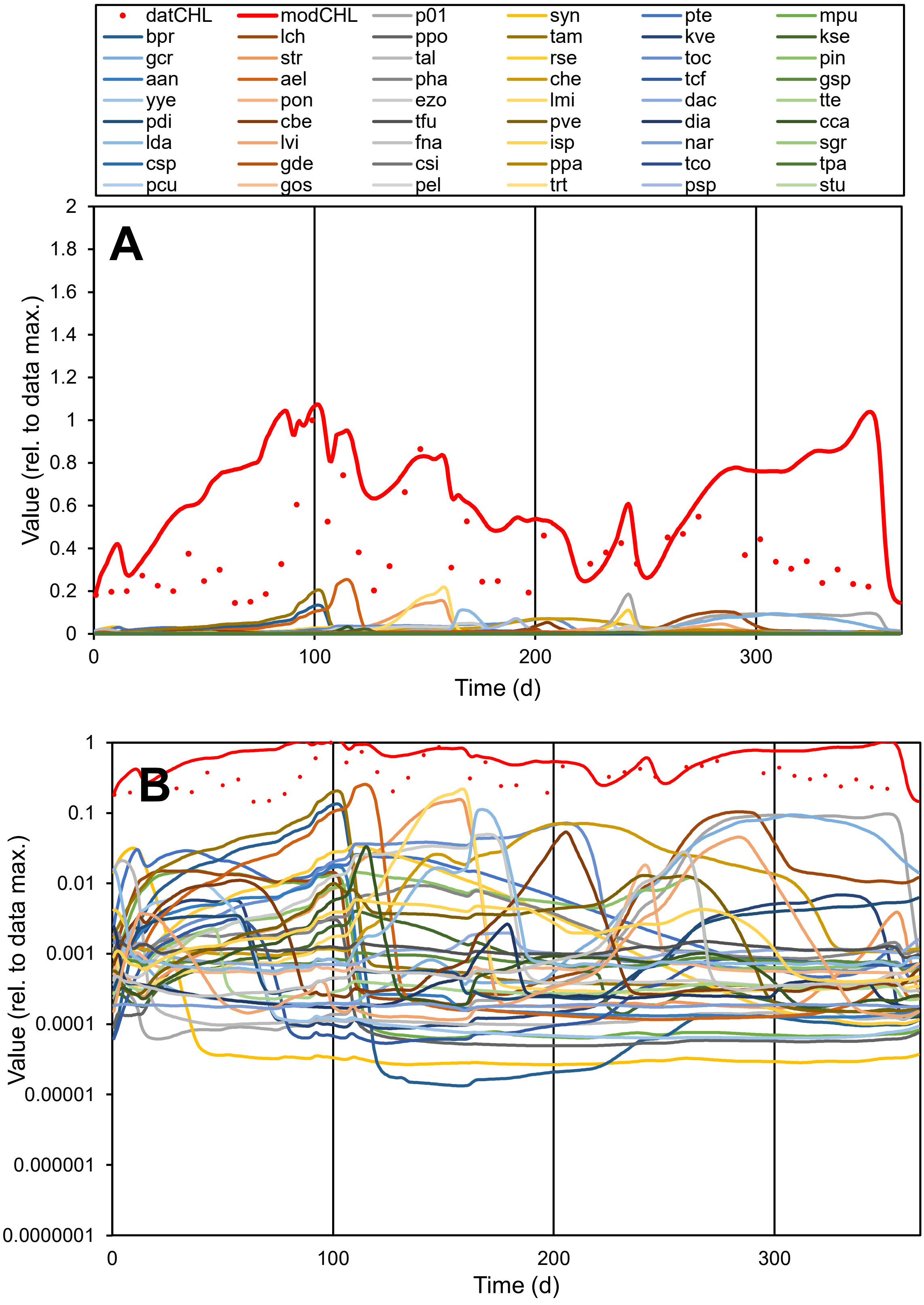


**C1**

Variables

CHL Provide jtd or Dst

CalibOnly?

2 1 to reproduce of

oNormalize?

2 1=no, 2=yes

ofxn calc*

2.7198

ofxn mod

2.98239

sRun

RunA1

zout

1

Ave

Data

1.03269

Model

1.03124

data/vMax

2.841

ofxng

319.391

Min d/m

0.70277

ave/min d

2.51445

ave/min m

1.76461

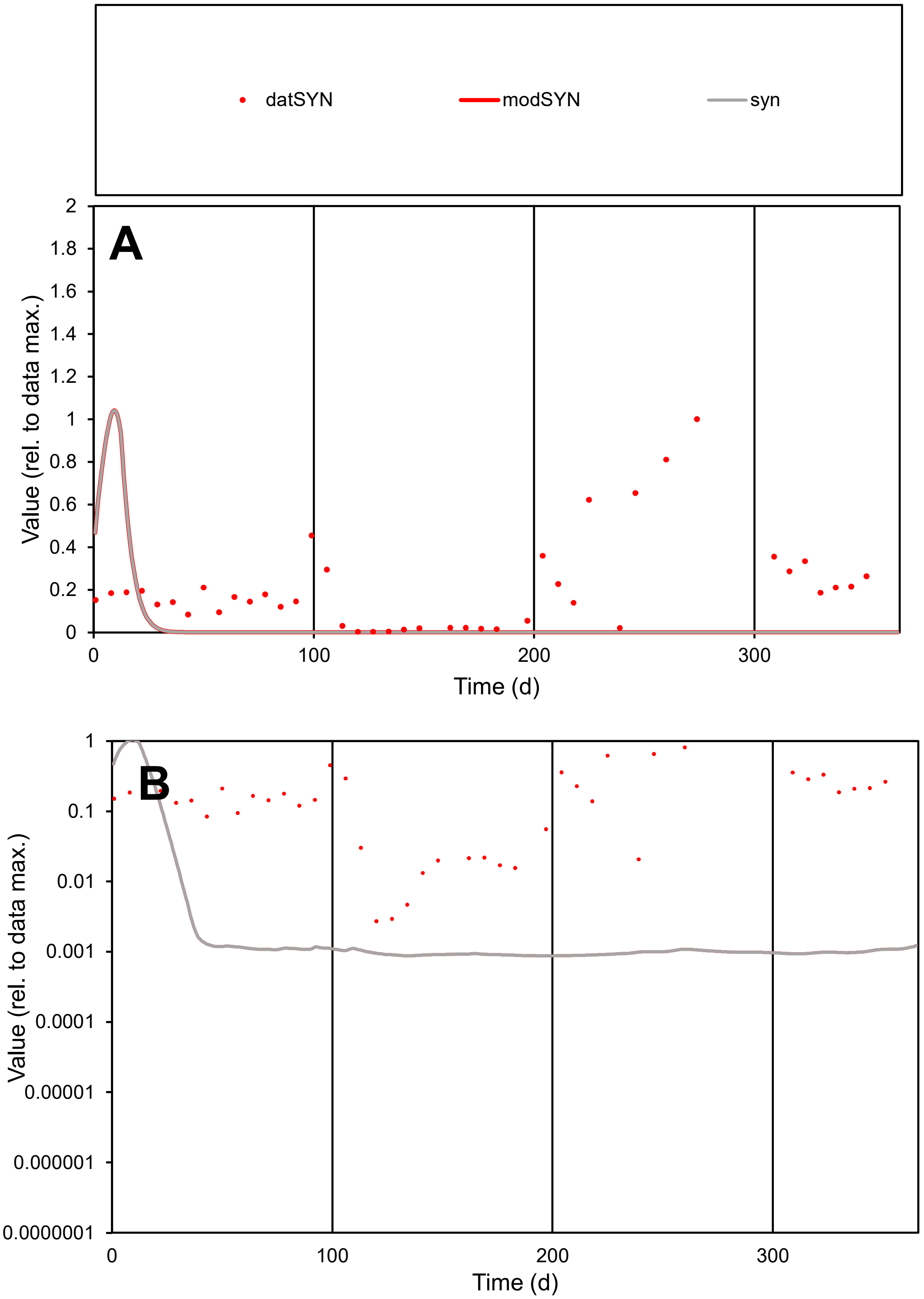


**C1**

Variables

| | |
|---|---|
| SYN | Provide jtd or Dst |
| CalibOnly? | |
| 2 | 1 to reproduce of |
| oNormalize? | |
| 2 | 1=no, 2=yes |

ofxn calc*
1.58195
ofxn mod
1.58985

sRun
RunA1
zout
1

Ave
Data
10.7184
Model
2.67643

data/vMax
51.32

ofxng
319.391

Min d/m
3.10913

ave/min d
76.7796
ave/min m
59.6087

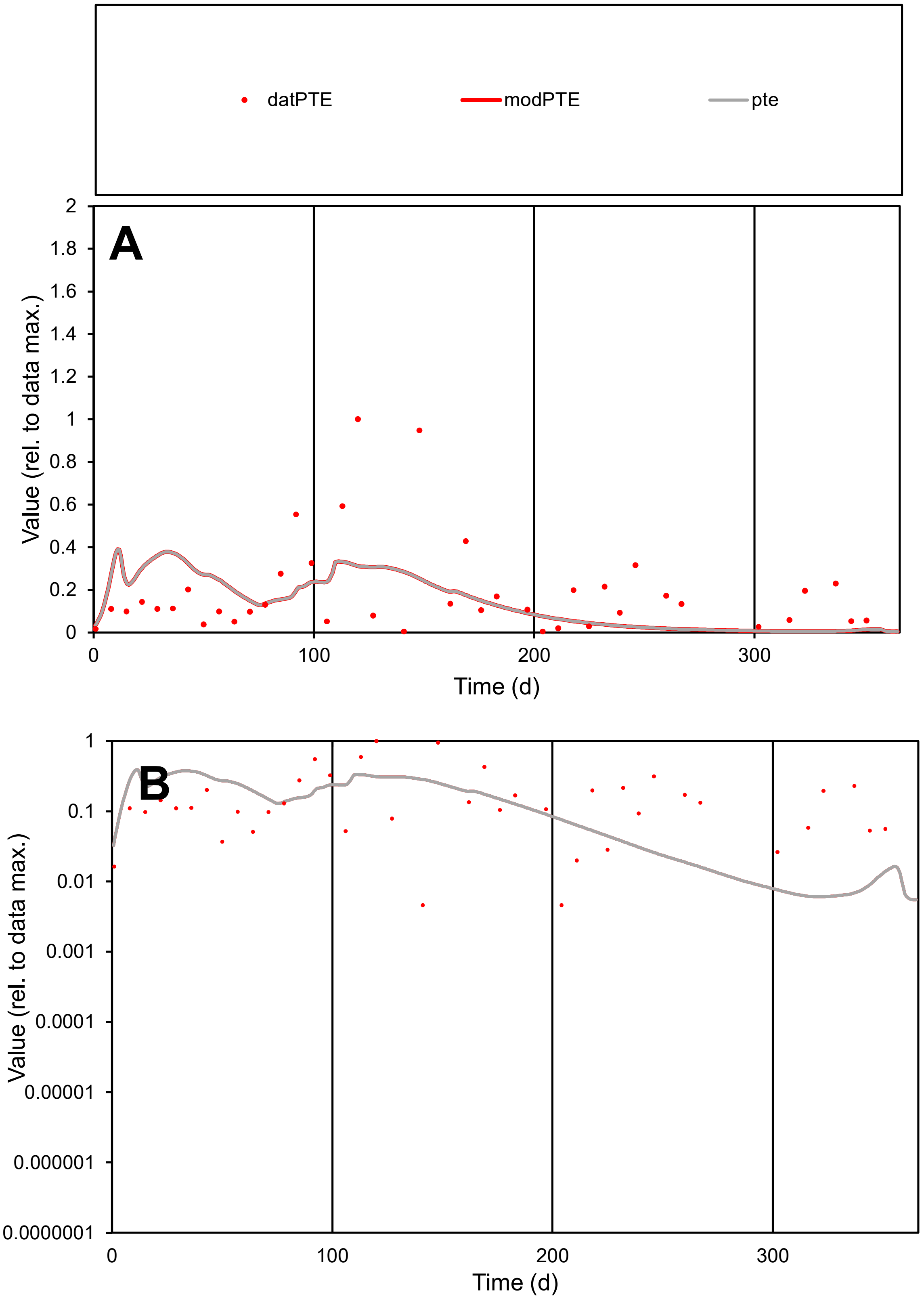


**C1**

Variables
PTE Provide jtd or Dst
CalibOnly?
2 1 to reproduce of
oNormalize?
2 1=no, 2=yes

ofxn calc*
1.16004
ofxn mod
1.16942

sRun
RunA1
zout
1

Ave
Data
0.00035
Model
0.00028

data/vMax
0.00182

ofxng
319.391

Min d/m
0.74843

ave/min d
41.3317
ave/min m
25.0112

330_Calib_S_A

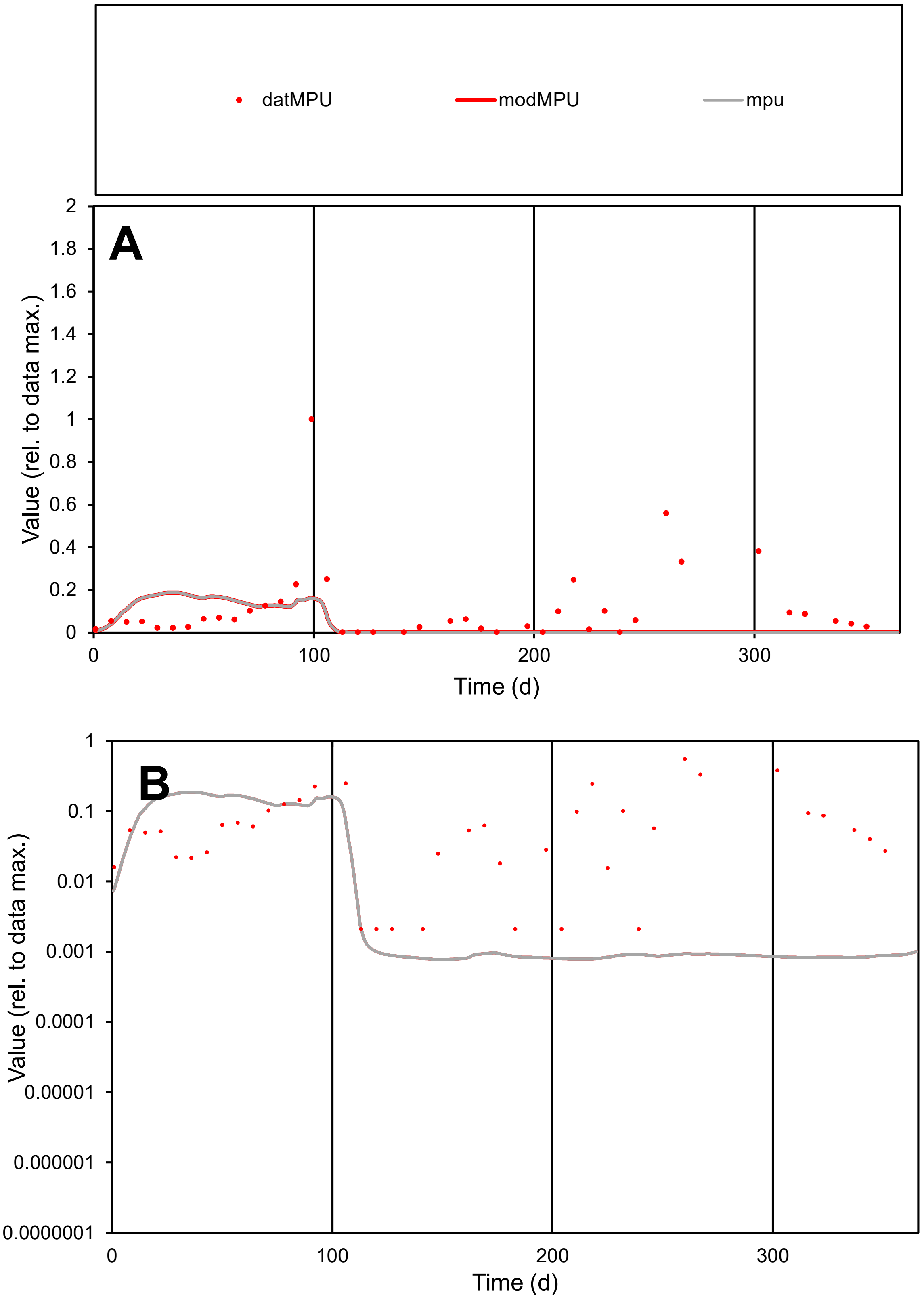


**C1**

Variables

MPU Provide jtd or Dst

CalibOnly?

2 1 to reproduce of

oNormalize?

2 1=no, 2=yes

ofxn calc*

1.73049

ofxn mod

1.73968

sRun

RunA1

zout

1

Ave

Data

0.00022

Model

0.0001

data/vMax

0.00193

ofxng

319.391

Min d/m

2.74412

ave/min d

52.7038

ave/min m

68.248

330_Calib_S_A

• datBPR — modBPR — bpr

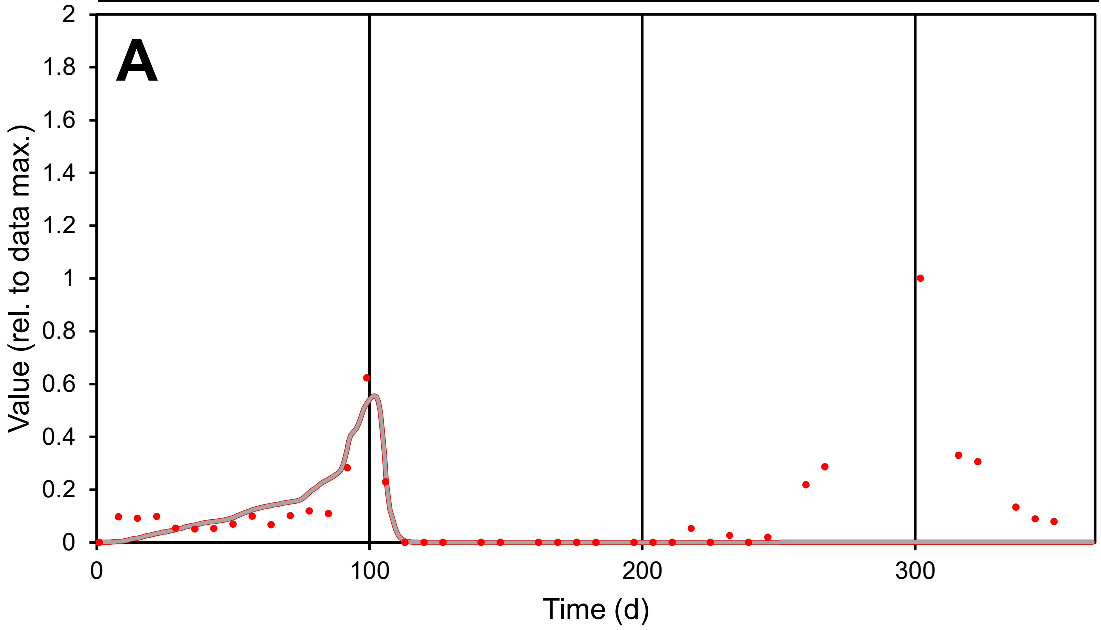


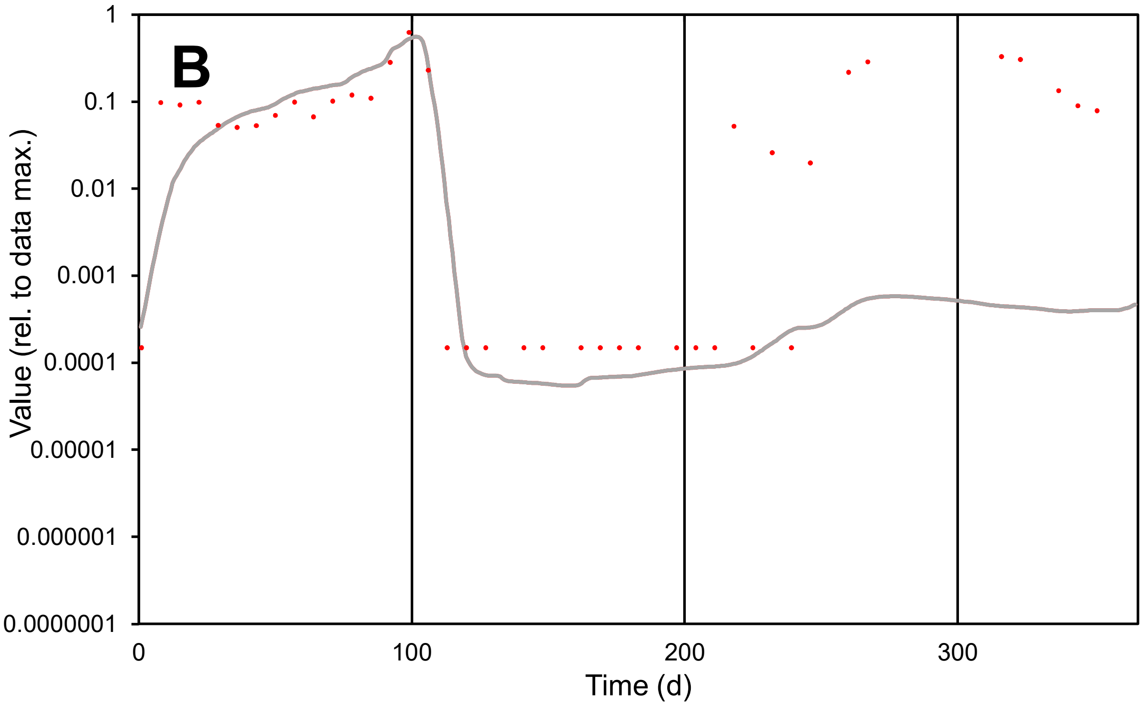


**C1**

Variables
BPR Provide jtd or Dst
CalibOnly?
2 1 to reproduce of
oNormalize?
2 1=no, 2=yes

ofxn calc*
1.6356
ofxn mod
1.64607

sRun
RunA1
zout
1

Ave
Data
0.00014
Model
7.3E-05

data/vMax
0.00125

ofxng
319.391

Min d/m
2.59706

ave/min d
770.152
ave/min m
1021.19

330_Calib_S_A

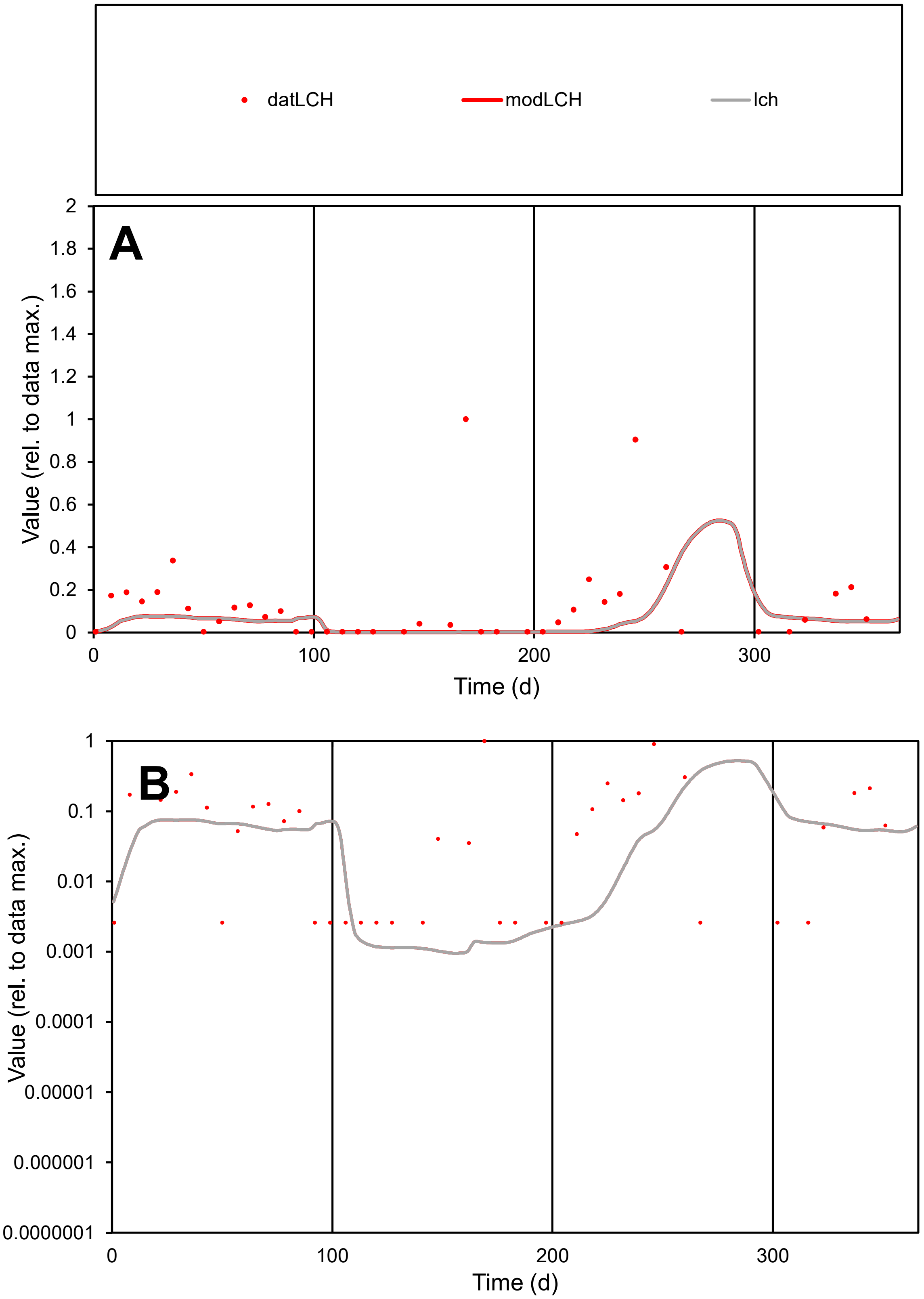


**C1**

Variables
LCH Provide jtd or Dst
CalibOnly?
2 1 to reproduce of
oNormalize?
2 1=no, 2=yes

ofxn calc*
1.80851
ofxn mod
1.81819

sRun
RunA1
zout
1

Ave
Data
0.00012
Model
4.6E-05

data/vMax
0.00094

ofxng
319.391

Min d/m
2.56906

ave/min d
48.7042
ave/min m
48.5516

330_Calib_S_A

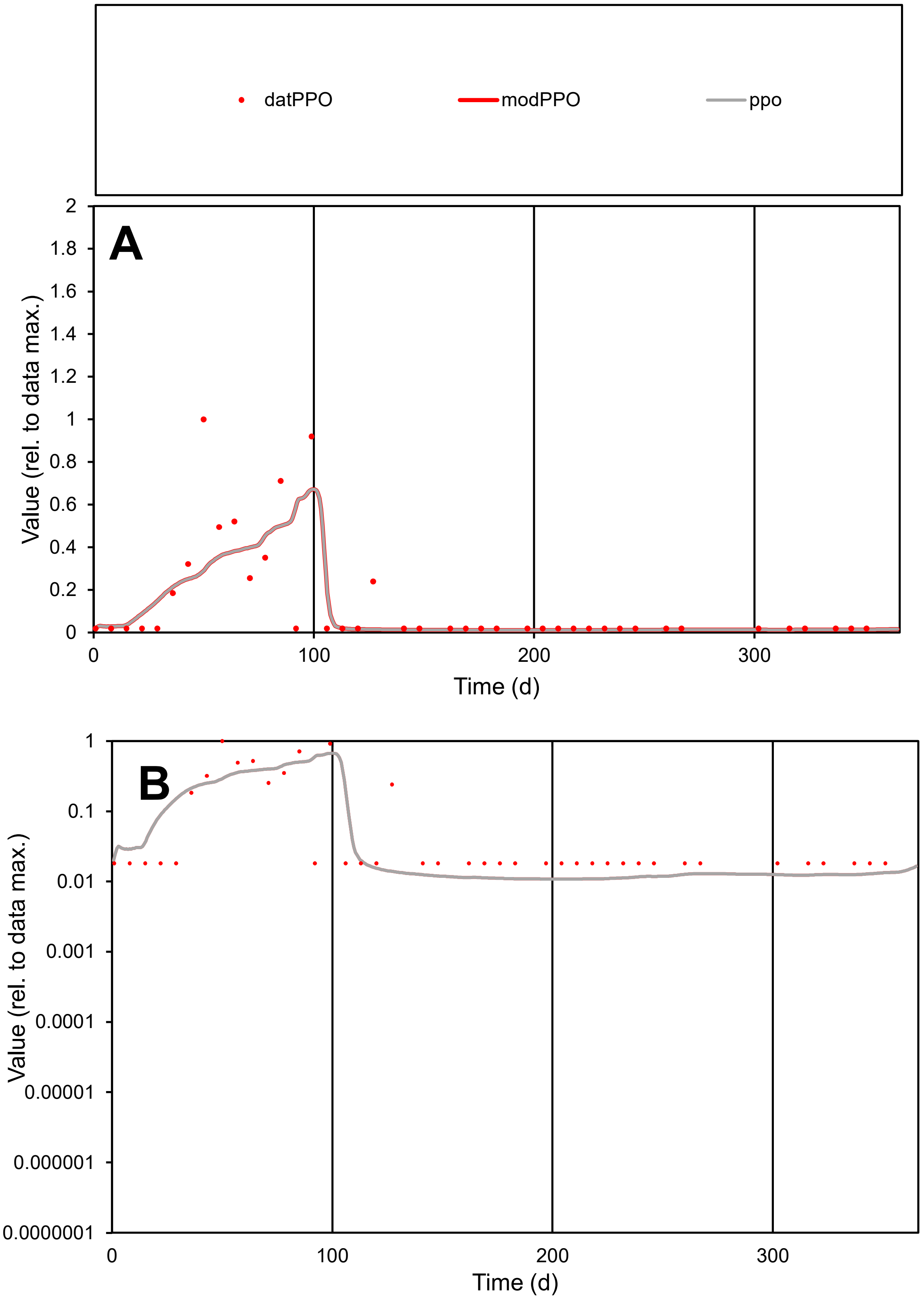


**C1**

Variables
PPO Provide jtd or Dst
CalibOnly?
2 1 to reproduce of
oNormalize?
2 1=no, 2=yes

ofxn calc*
1.14928
ofxn mod
1.20793

sRun
RunA1
zout
1

Ave
Data
1.4E-05
Model
1.2E-05

data/vMax
0.00011

ofxng
319.391

Min d/m
1.67513

ave/min d
7.48554
ave/min m
10.8179

330_Calib_S_A

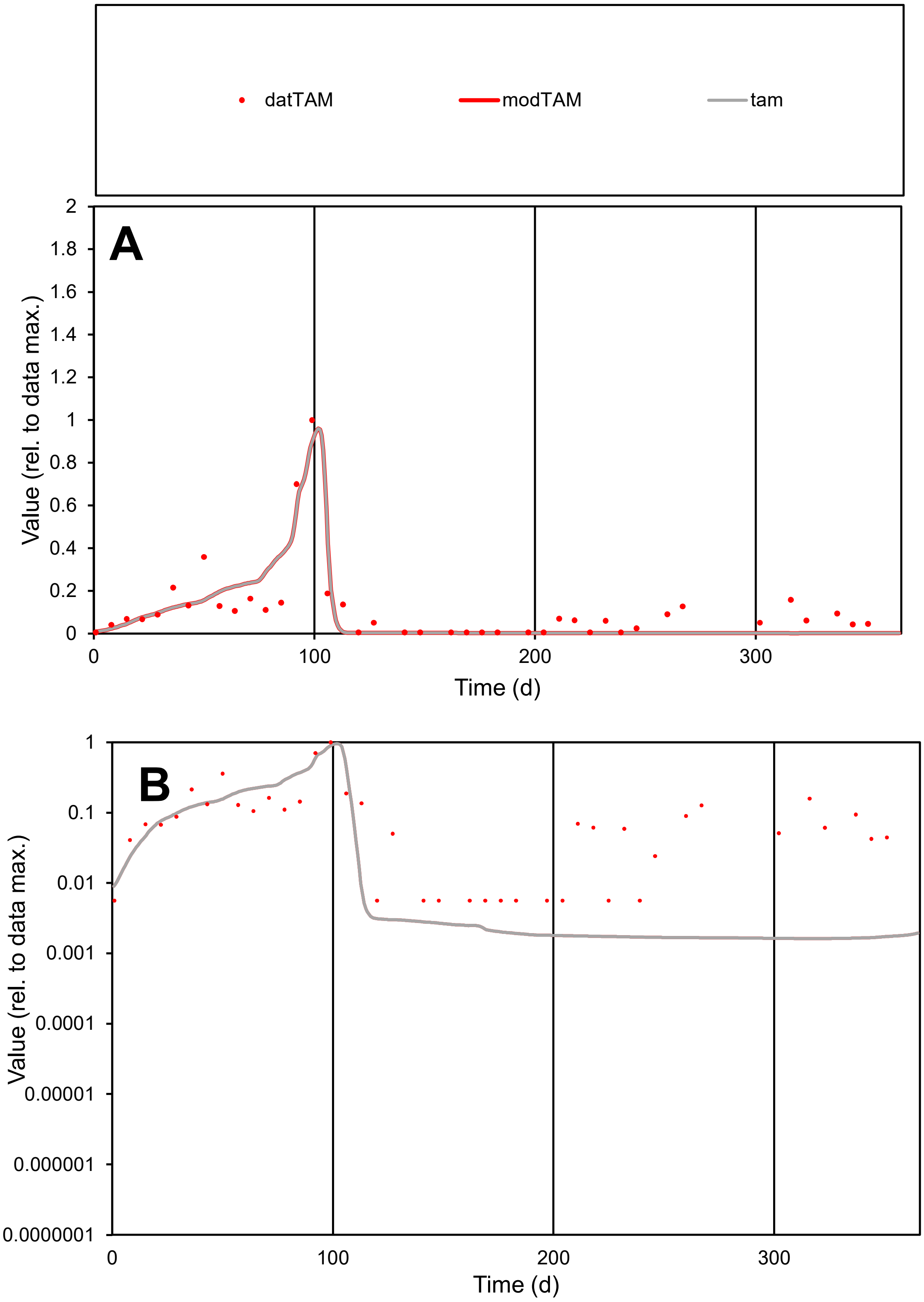


**C1**

Variables

TAM Provide jtd or Dst

CalibOnly?

2 1 to reproduce of

oNormalize?

2 1=no, 2=yes

ofxn calc*

0.6822

ofxn mod

0.7308

sRun

RunA1

zout

1

Ave

Data

0.00012

Model

9.9E-05

data/vMax

0.00102

ofxng

319.391

Min d/m

3.47346

ave/min d

20.115

ave/min m

59.923

330_Calib_S_A

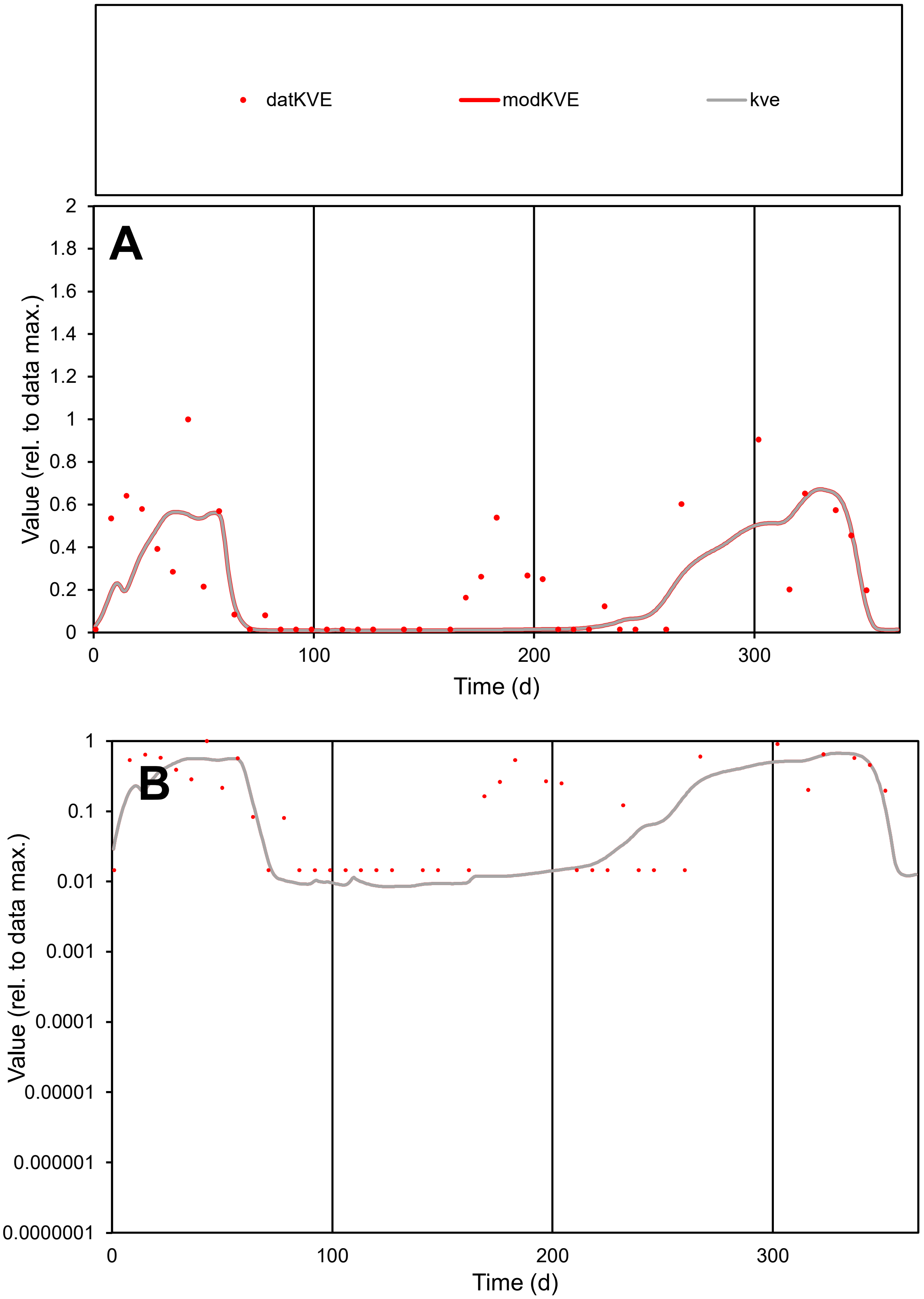


**C1**

Variables
KVE Provide jtd or Dst
CalibOnly?
2 1 to reproduce of
oNormalize?
2 1=no, 2=yes

ofxn calc*
0.82374
ofxn mod
0.84093

sRun
RunA1
zout
1

Ave
Data
5.2E-05
Model
3.9E-05

data/vMax
0.00022

ofxng
319.391

Min d/m
1.70348

ave/min d
16.5004
ave/min m
21.1056

330_Calib_S_A

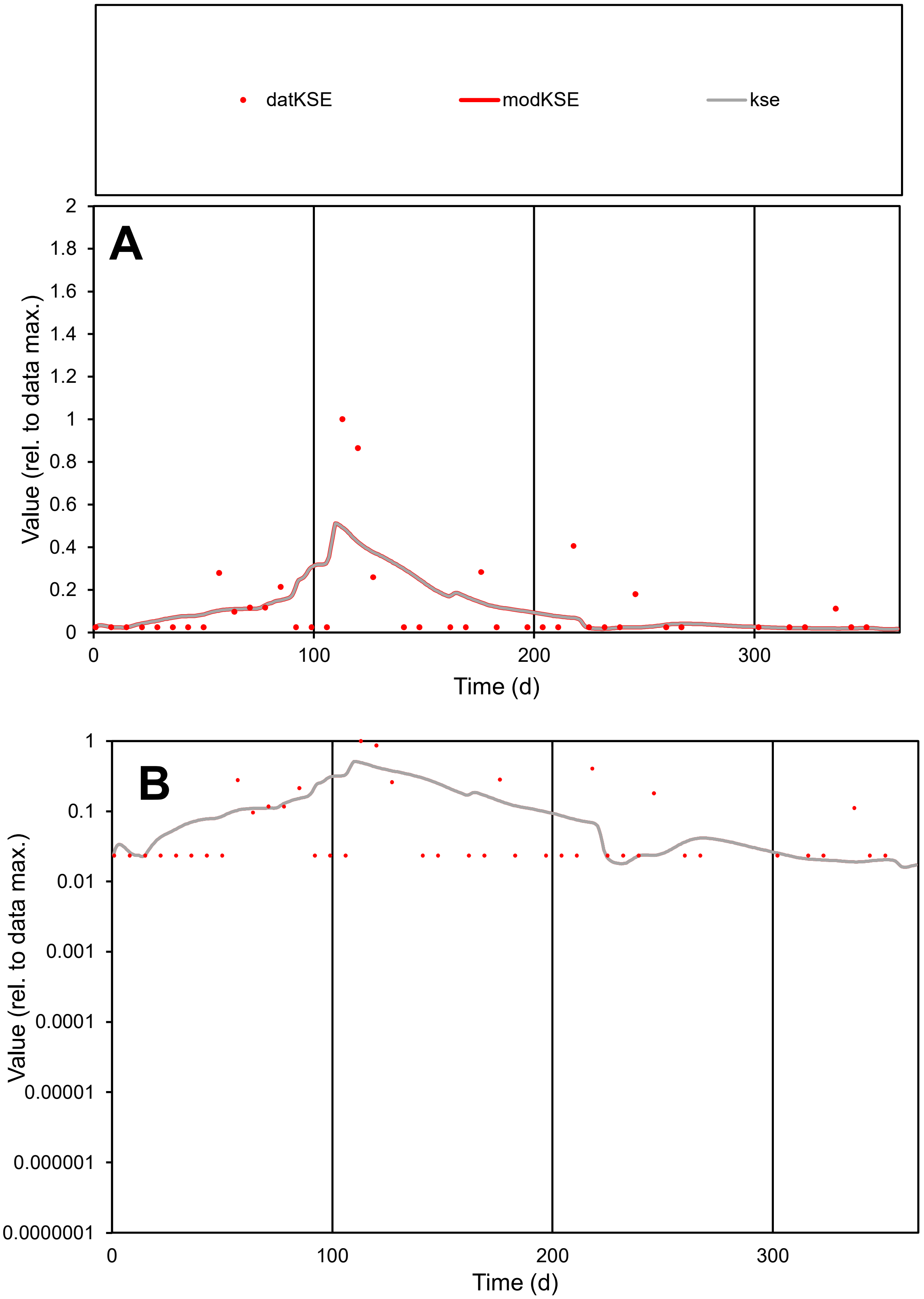


**C1**

Variables

KSE Provide jtd or Dst

CalibOnly?

2 1 to reproduce of

oNormalize?

2 1=no, 2=yes

ofxn calc*

1.43966

ofxn mod

1.45916

sRun

RunA1

zout

1

Ave

Data

1.8E-05

Model

1.9E-05

data/vMax

0.00016

ofxng

319.391

Min d/m

1.2757

ave/min d

4.78084

ave/min m

6.39953

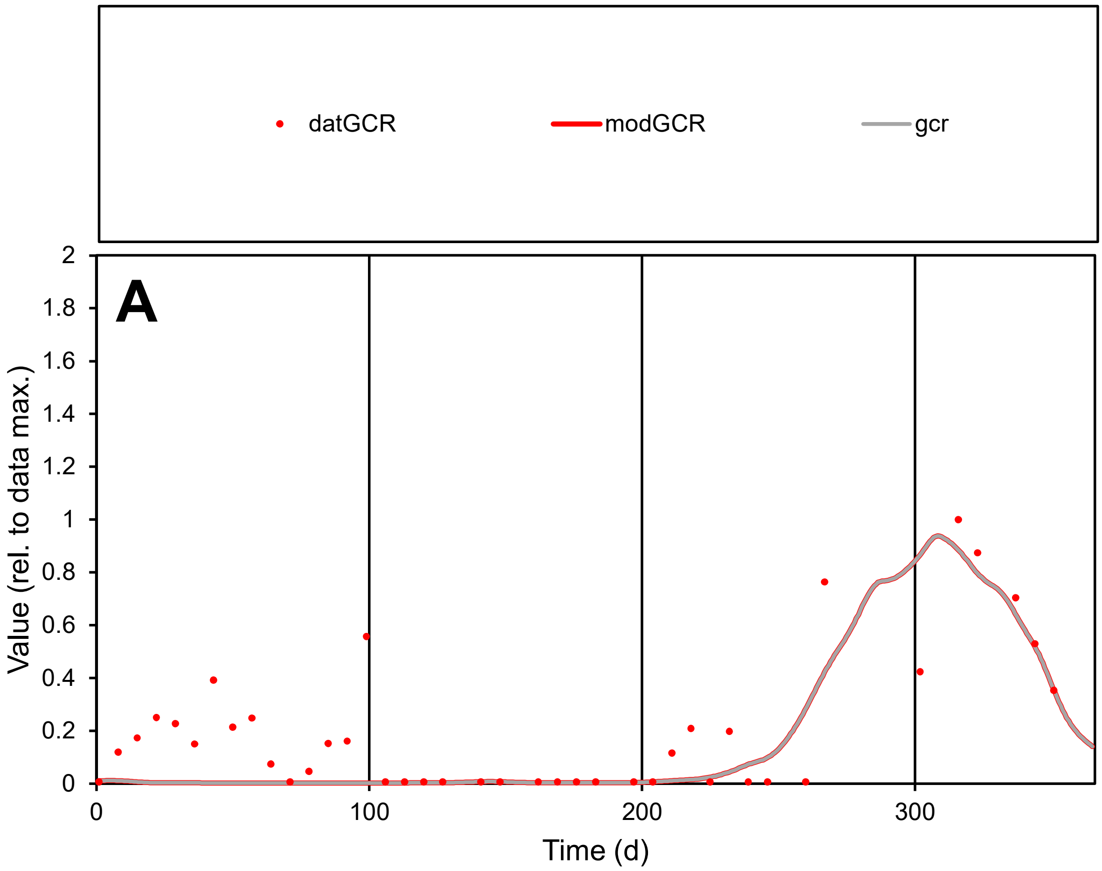


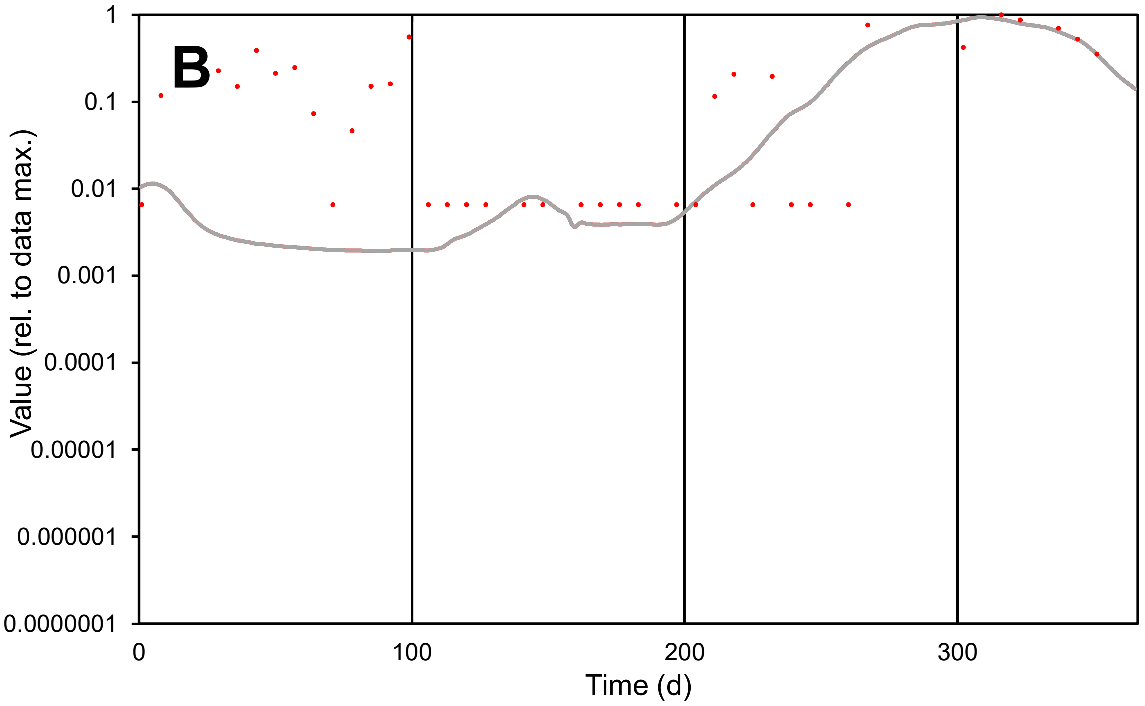


**C1**

Variables
GCR Provide jtd or Dst
CalibOnly?
2 1 to reproduce of
oNormalize?
2 1=no, 2=yes

ofxn calc*
0.89939
ofxn mod
0.9168

sRun
RunA1
zout
1

Ave
Data
9.4E-05
Model
5.9E-05

data/vMax
0.00048

ofxng
319.391

Min d/m
3.41023

ave/min d
29.914
ave/min m
63.9773

330_Calib_S_A

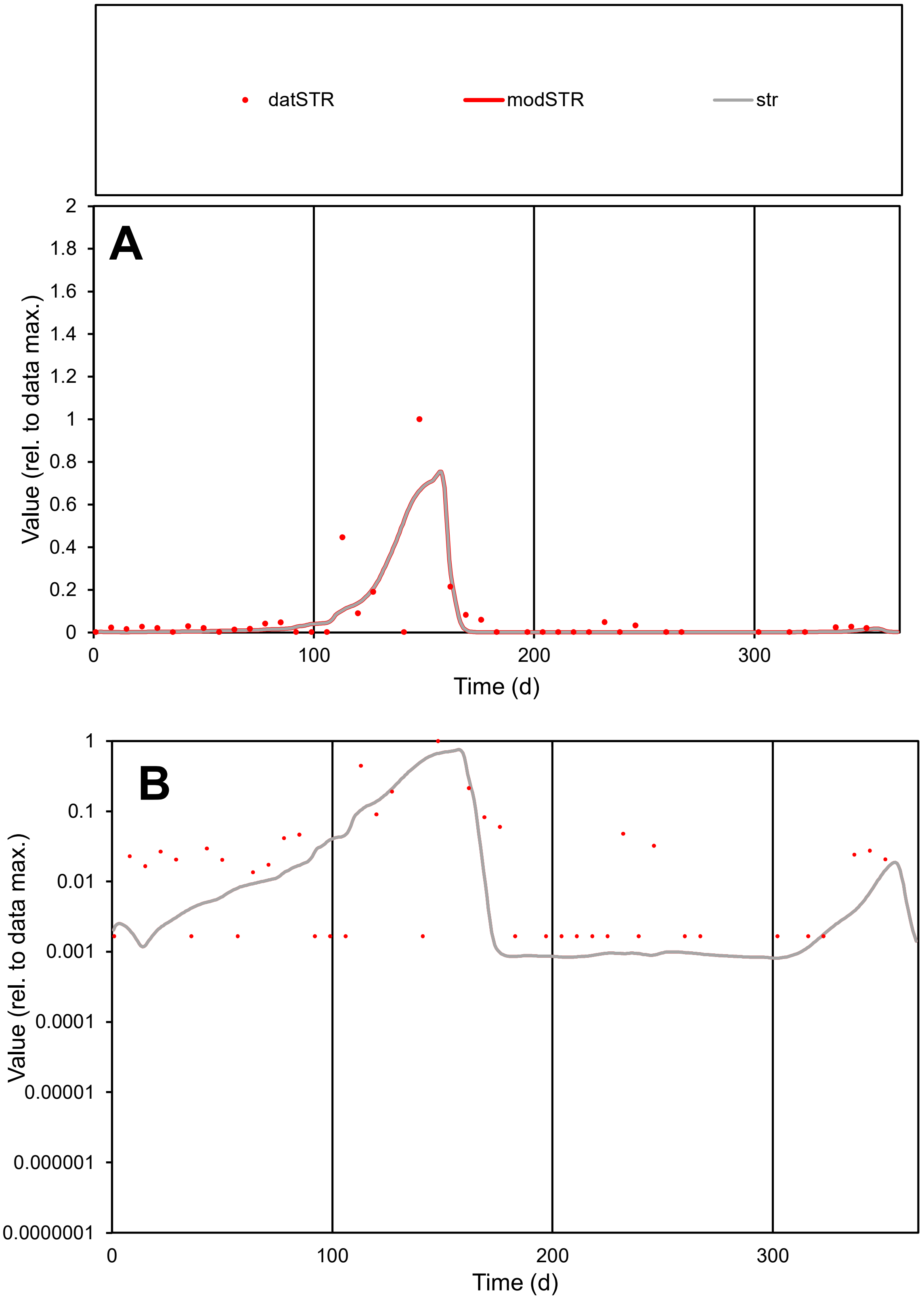


**C1**

Variables
STR Provide jtd or Dst
CalibOnly?
2 1 to reproduce of
oNormalize?
2 1=no, 2=yes

ofxn calc*
1.78298
ofxn mod
1.82809

sRun
RunA1
zout
1

Ave
Data
6E-05
Model
5.1E-05

data/vMax
0.00098

ofxng
319.391

Min d/m
2.036

ave/min d
37.1043
ave/min m
63.7124

330_Calib_S_A

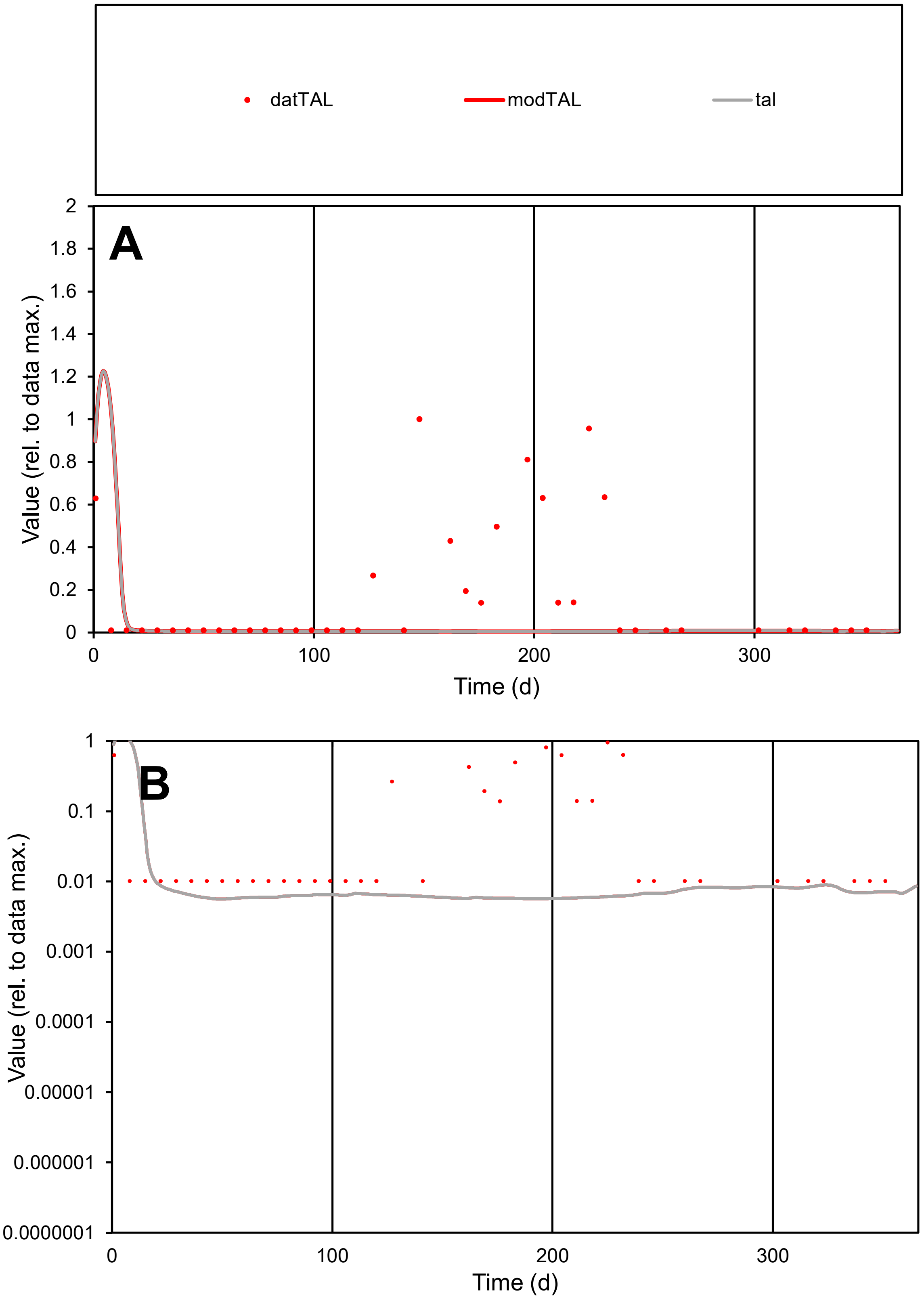

datTAL
modTAL
tal
A
Value (rel. to data max.)
Time (d)
B


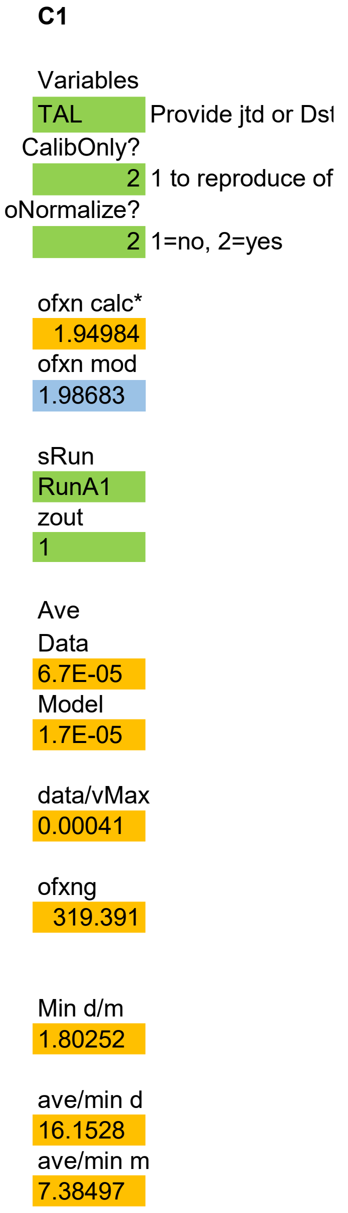

C1
Variables
TAL
Provide jtd or Dst
CalibOnly?
2
1 to reproduce of
oNormalize?
2
1=no, 2=yes
ofxn calc*
1.94984
ofxn mod
1.98683
sRun
RunA1
zout
1
Ave
Data
6.7E-05
Model
1.7E-05
data/vMax
0.00041
ofxng
319.391
Min d/m
1.80252
ave/min d
16.1528
ave/min m
7.38497

330_Calib_S_A

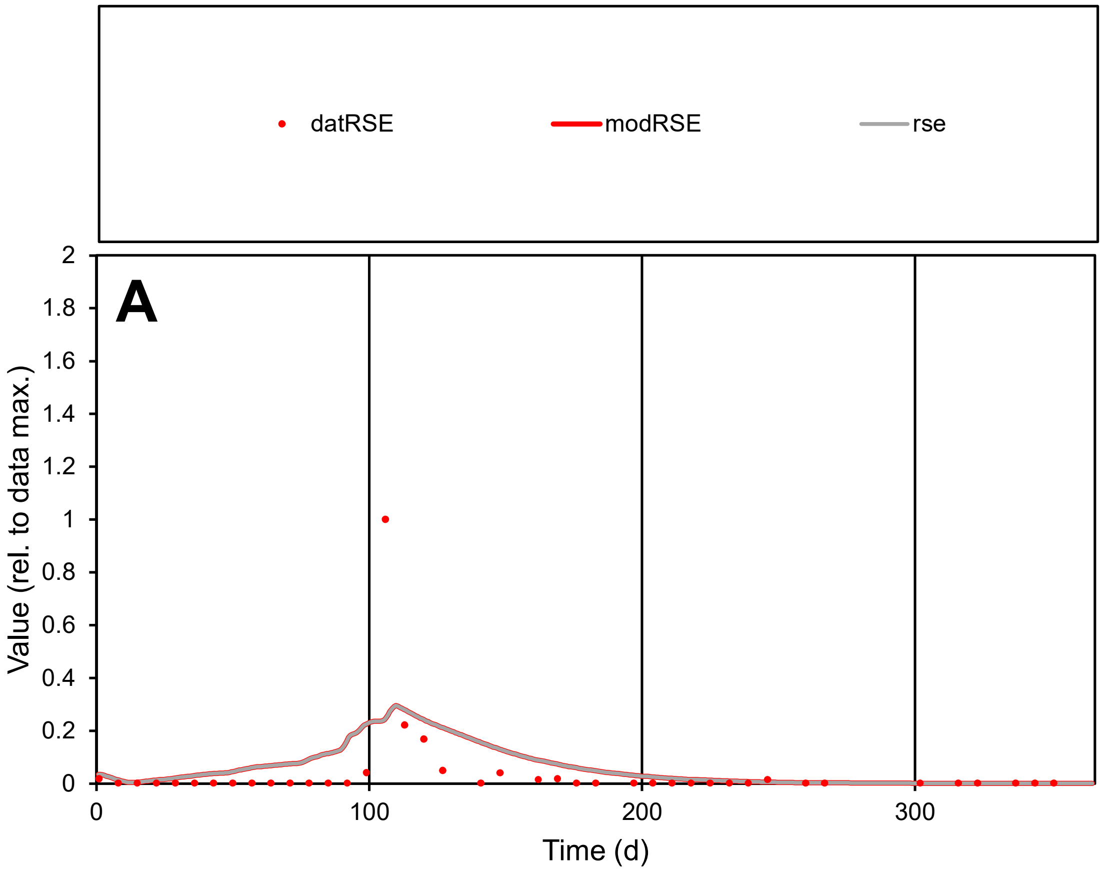


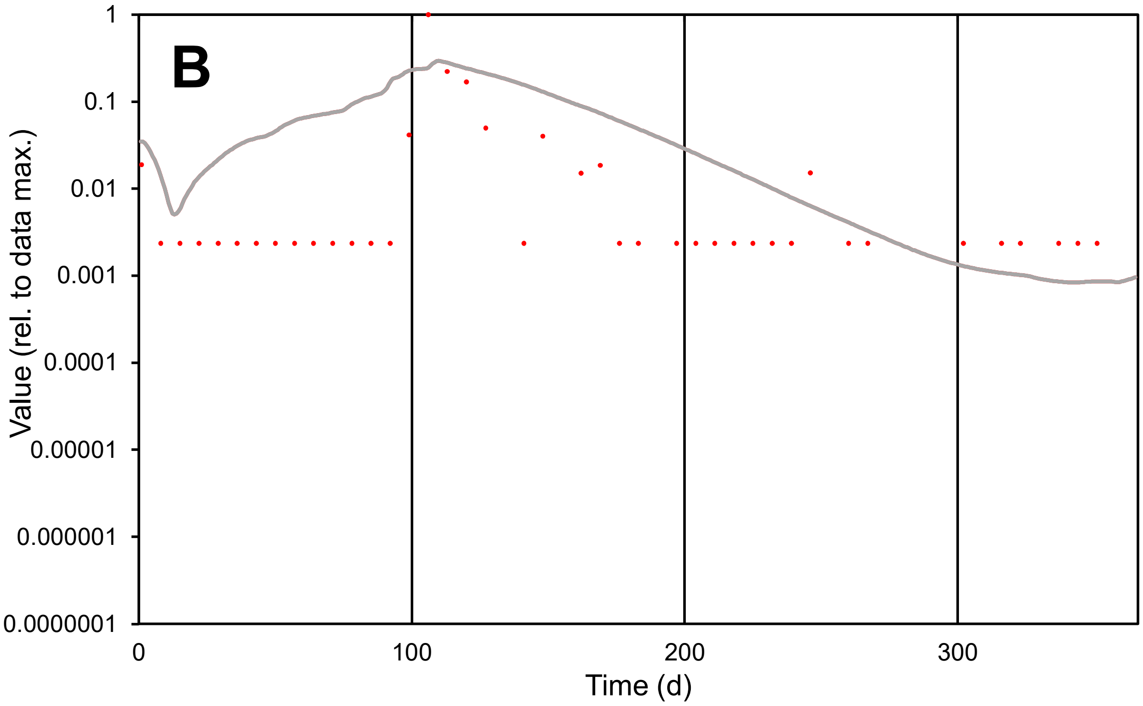


**C1**

Variables
RSE Provide jtd or Dst
CalibOnly?
2 1 to reproduce of
oNormalize?
2 1=no, 2=yes

ofxn calc*
3.20749
ofxn mod
3.24172

sRun
RunA1
zout
1

Ave
Data
8E-05
Model
0.00013

data/vMax
0.00198

ofxng
319.391

Min d/m
2.79054

ave/min d
17.183
ave/min m
75.7627

330_Calib_S_A

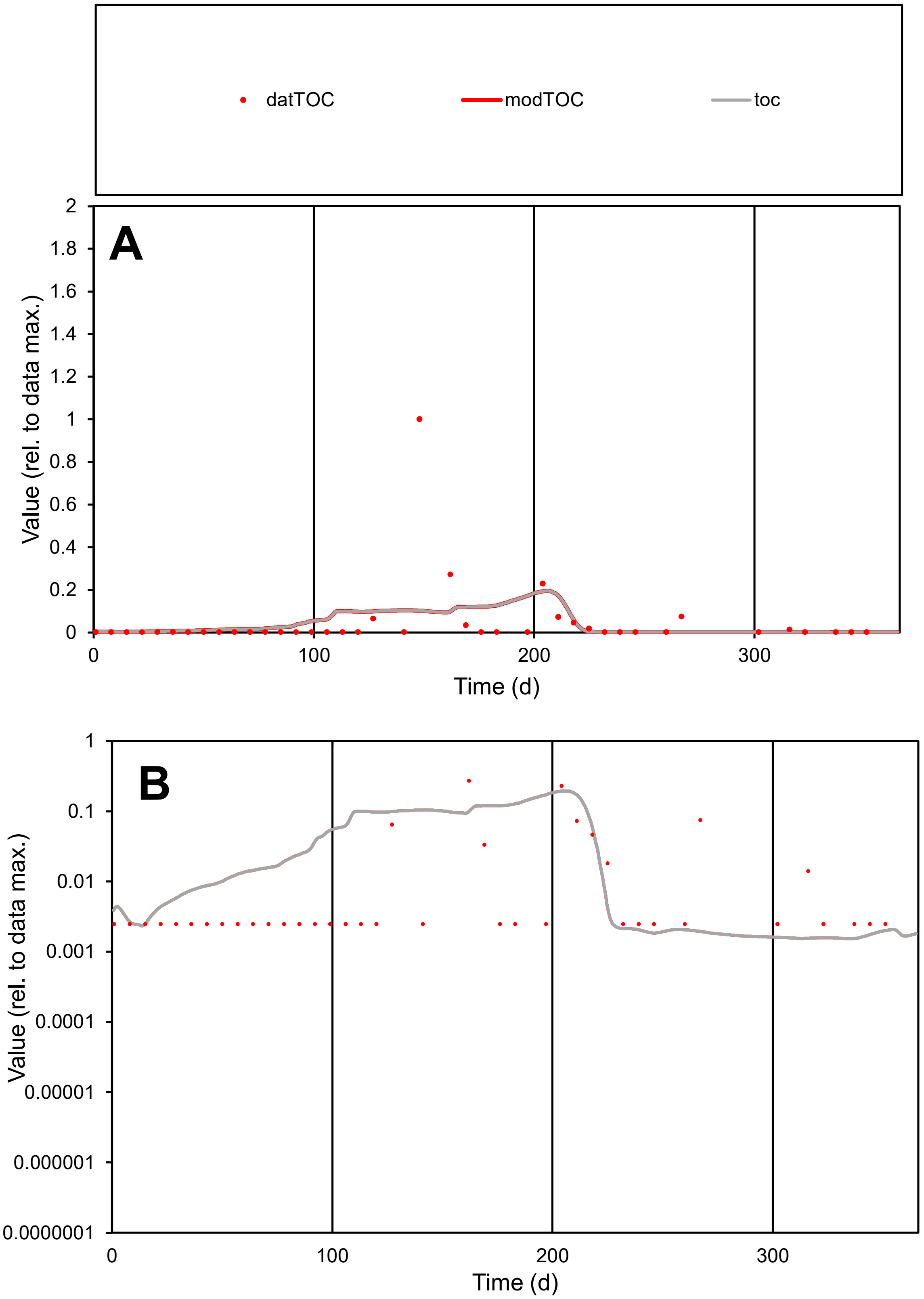


**C1**

Variables
TOC Provide jtd or Dst
CalibOnly?
2 1 to reproduce of
oNormalize?
2 1=no, 2=yes

ofxn calc*
3.28214
ofxn mod
3.28995

sRun
RunA1
zout
1

Ave
Data
8.1E-05
Model
7.9E-05

data/vMax
0.00174

ofxng
319.391

Min d/m
1.58664

ave/min d
18.7205
ave/min m
28.8237

330_Calib_S_A

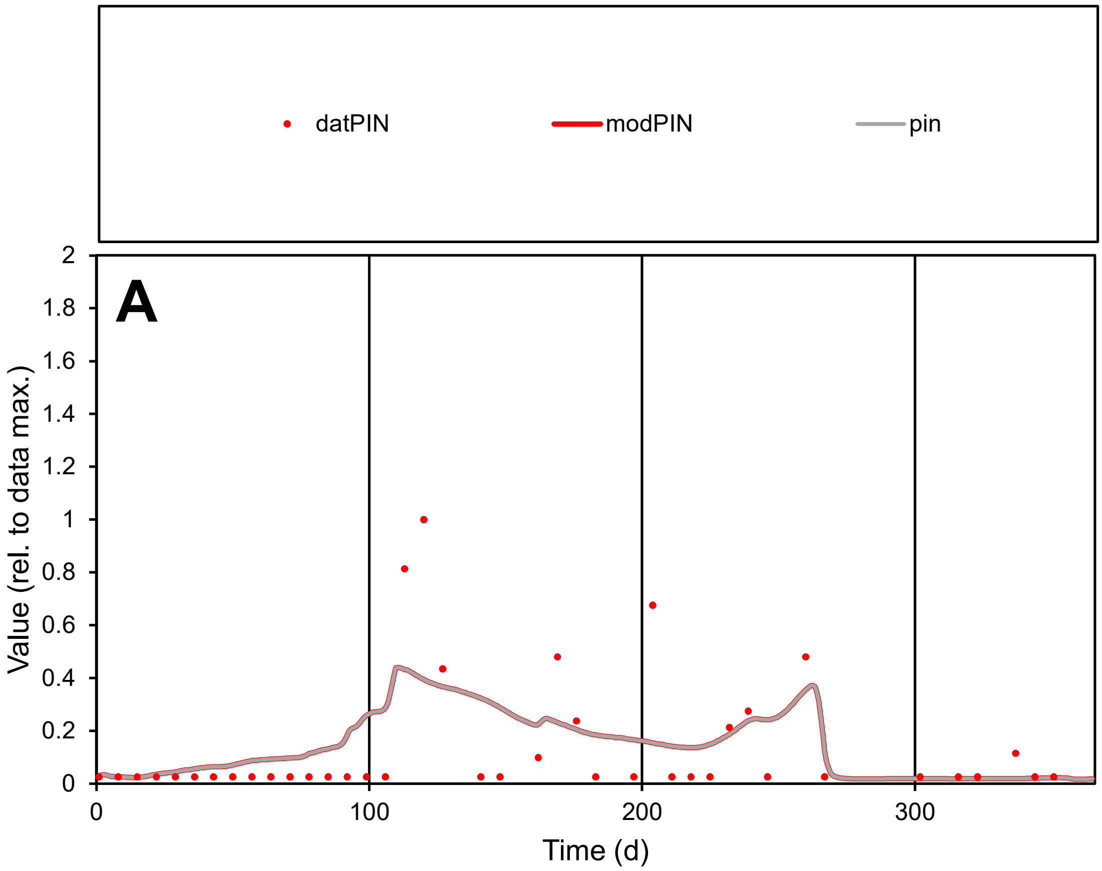


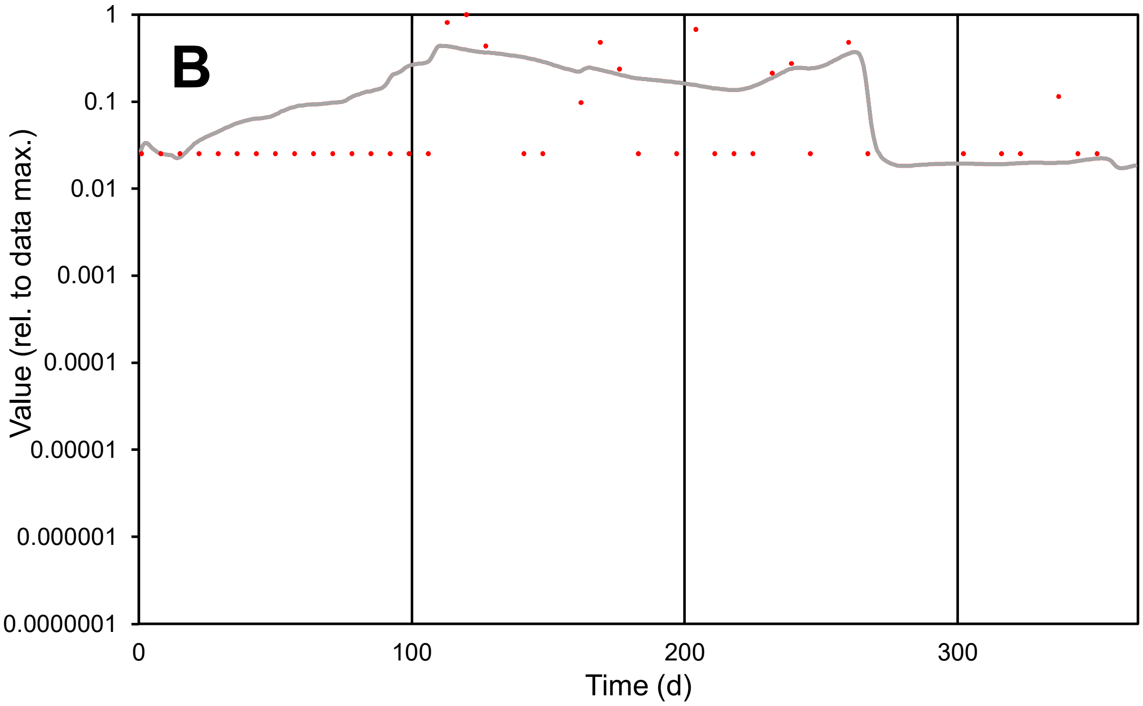


**C1**

Variables
PIN Provide jtd or Dst
CalibOnly?
2 1 to reproduce of
oNormalize?
2 1=no, 2=yes

ofxn calc*
1.34117
ofxn mod
1.3566

sRun
RunA1
zout
1

Ave
Data
2.2E-05
Model
2.4E-05

data/vMax
0.00016

ofxng
319.391

Min d/m
1.323

ave/min d
5.36798
ave/min m
7.82719

330_Calib_S_A

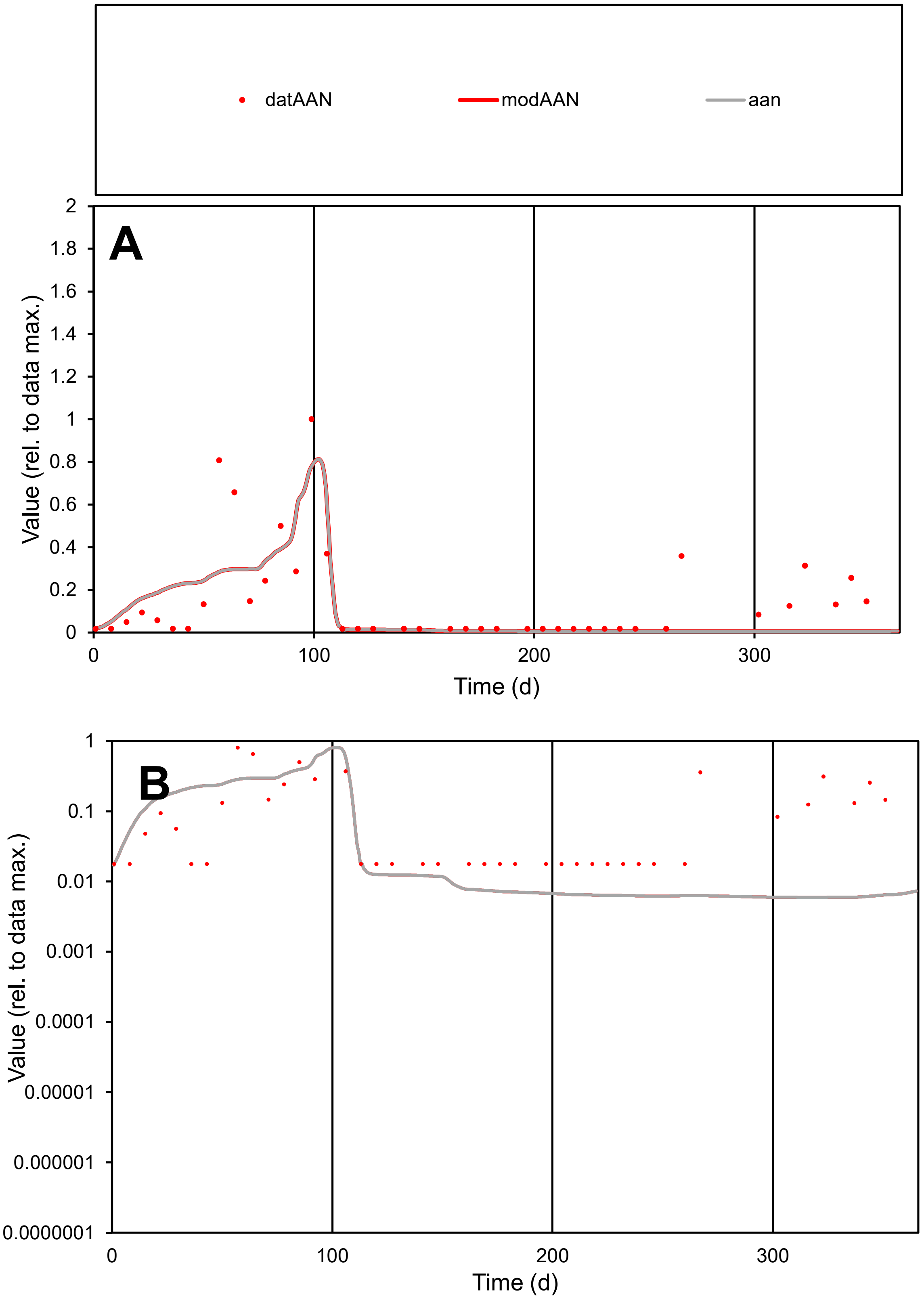


**C1**

Variables
AAN Provide jtd or Dst
CalibOnly?
2 1 to reproduce of
oNormalize?
2 1=no, 2=yes

ofxn calc*
1.0615
ofxn mod
1.08076

sRun
RunA1
zout
1

Ave
Data
4.7E-05
Model
3.6E-05

data/vMax
0.00031

ofxng
319.391

Min d/m
3.00863

ave/min d
8.42026
ave/min m
19.682

330_Calib_S_A

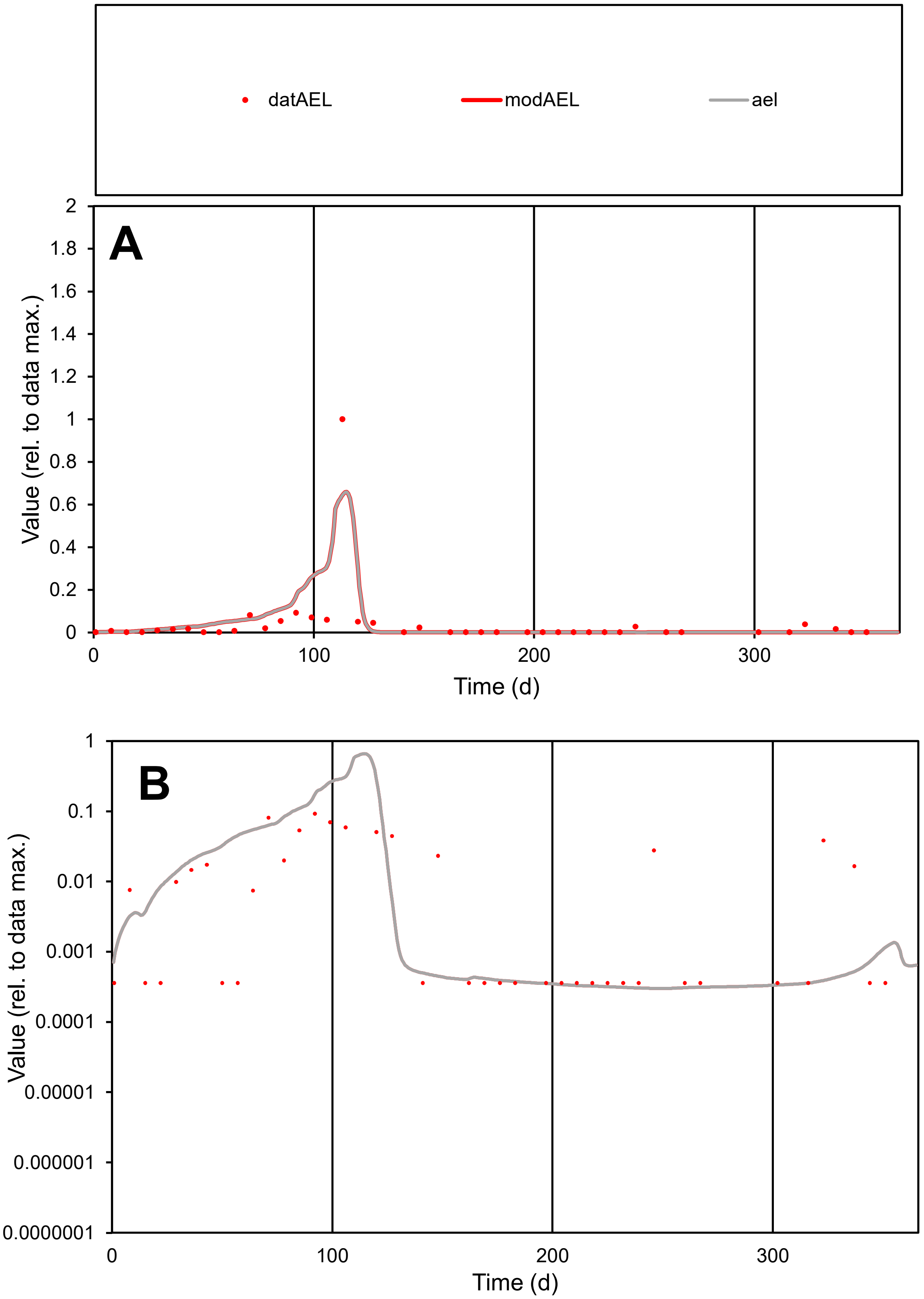


**C1**

Variables

AEL Provide jtd or Dst

CalibOnly?

2 1 to reproduce of

oNormalize?

2 1=no, 2=yes

ofxn calc*

1.93073

ofxn mod

1.98195

sRun

RunA1

zout

1

Ave

Data

7.8E-05

Model

9.5E-05

data/vMax

0.00194

ofxng

319.391

Min d/m

1.19193

ave/min d

111.497

ave/min m

162.837

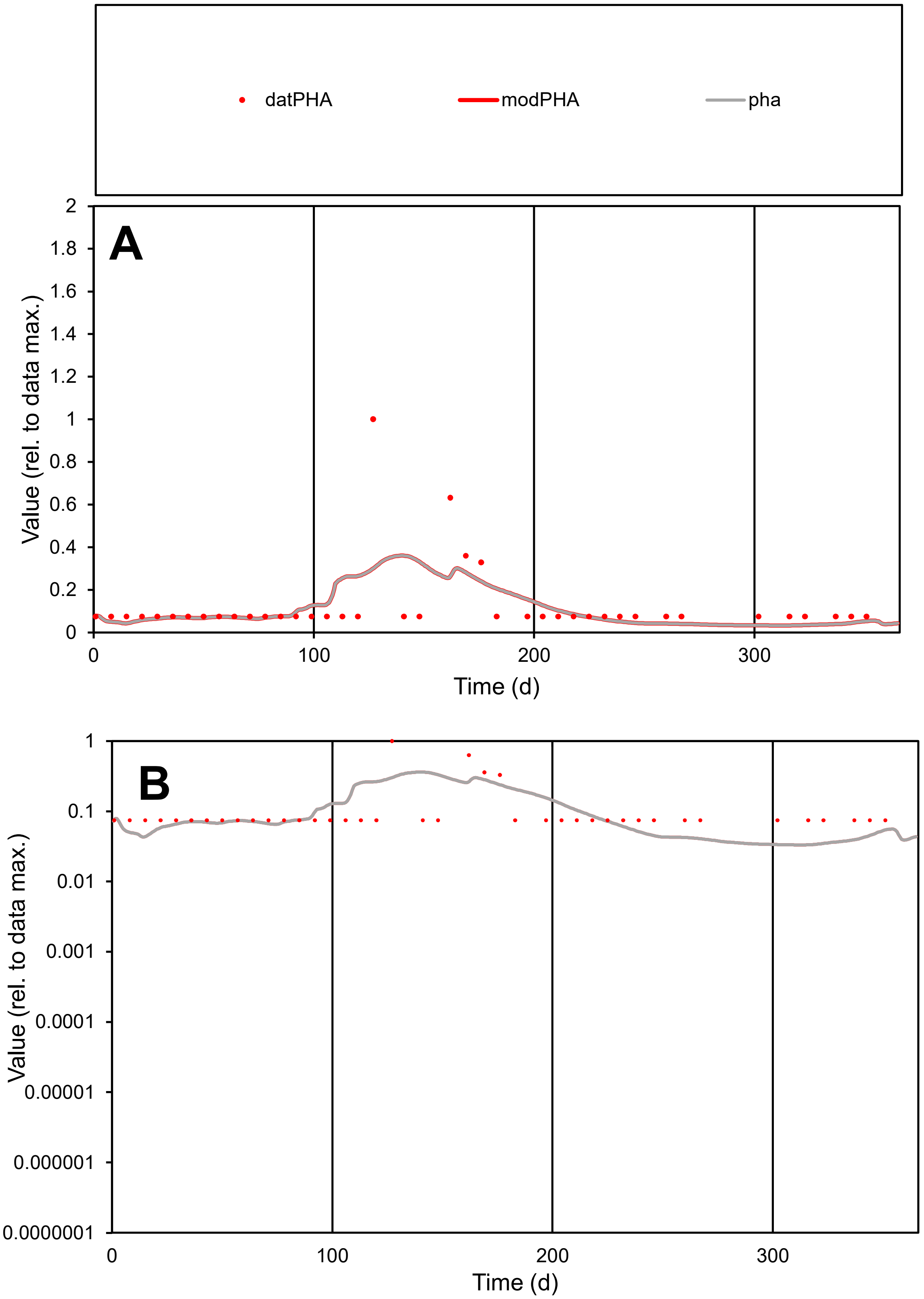


**C1**

Variables
PHA Provide jtd or Dst
CalibOnly?
2 1 to reproduce of
oNormalize?
2 1=no, 2=yes

ofxn calc*
1.17442
ofxn mod
1.18181

sRun
RunA1
zout
1

Ave
Data
1.7E-05
Model
1.6E-05

data/vMax
0.00014

ofxng
319.391

Min d/m
2.23391

ave/min d
1.66097
ave/min m
3.47313

330_Calib_S_A

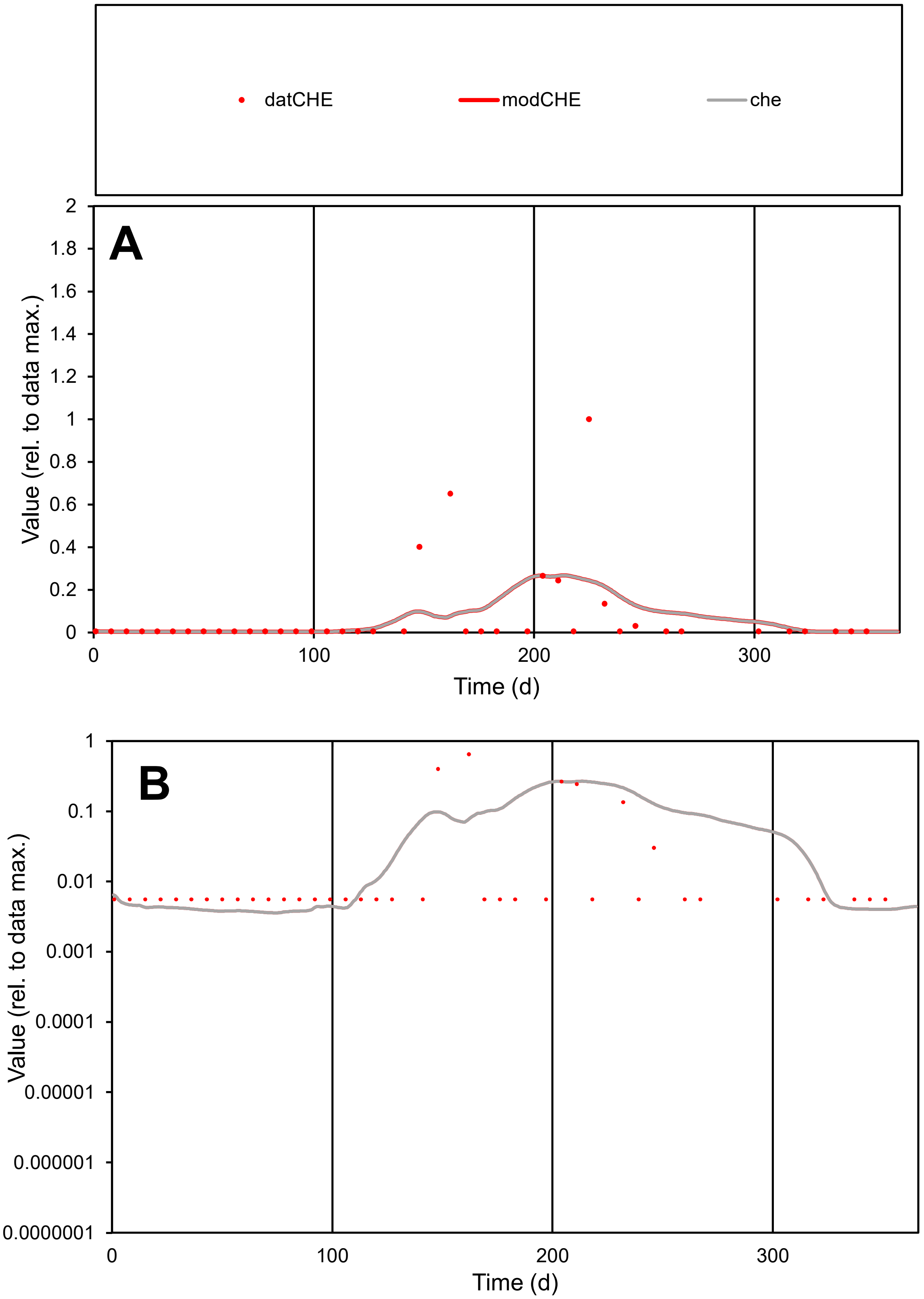


**C1**

Variables
CHE Provide jtd or Dst
CalibOnly?
2 1 to reproduce of
oNormalize?
2 1=no, 2=yes

ofxn calc*
2.41218
ofxn mod
2.42903

sRun
RunA1
zout
1

Ave
Data
8.9E-05
Model
8.3E-05

data/vMax
0.00125

ofxng
319.391

Min d/m
1.55723

ave/min d
12.7071
ave/min m
18.5212

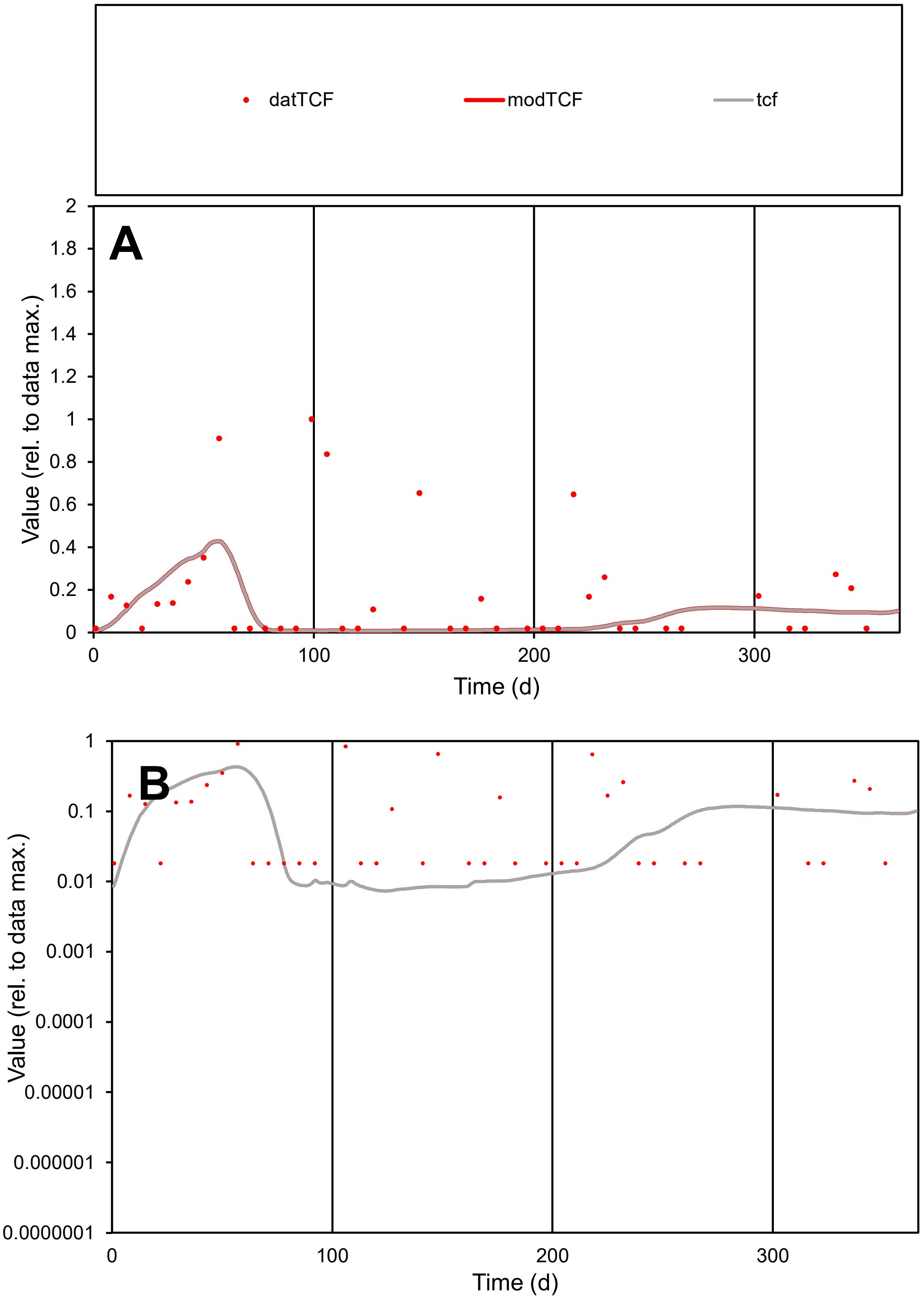


**C1**

Variables
TCF Provide jtd or Dst
CalibOnly?
2 1 to reproduce of
oNormalize?
2 1=no, 2=yes

ofxn calc*
1.60143
ofxn mod
1.61272

sRun
RunA1
zout
1

Ave
Data
2.9E-05
Model
1.5E-05

data/vMax
0.00017

ofxng
319.391

Min d/m
2.44819

ave/min d
9.35452
ave/min m
11.48

330_Calib_S_A

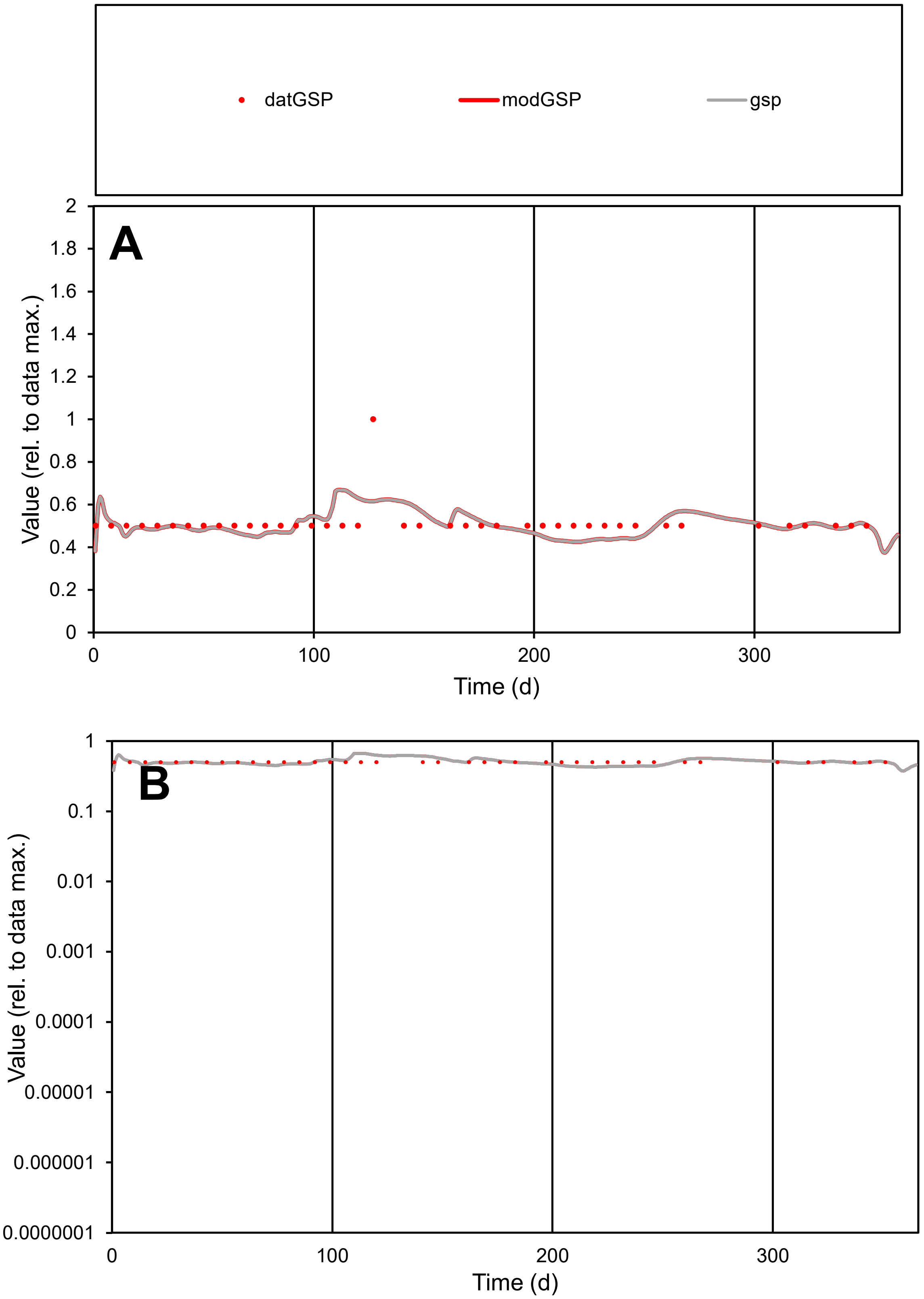


**C1**

Variables

| | |
|---|---|
| GSP | Provide jtd or Dst |
| CalibOnly? | |
| 2 | 1 to reproduce of |
| oNormalize? | |
| 2 | 1=no, 2=yes |

ofxn calc*
0.15587
ofxn mod
0.18888

sRun
RunA1
zout
1

Ave
Data
9.2E-06
Model
9.1E-06

data/vMax
1.8E-05

ofxng
319.391

Min d/m
1.17577

ave/min d
1.02438
ave/min m
1.19144

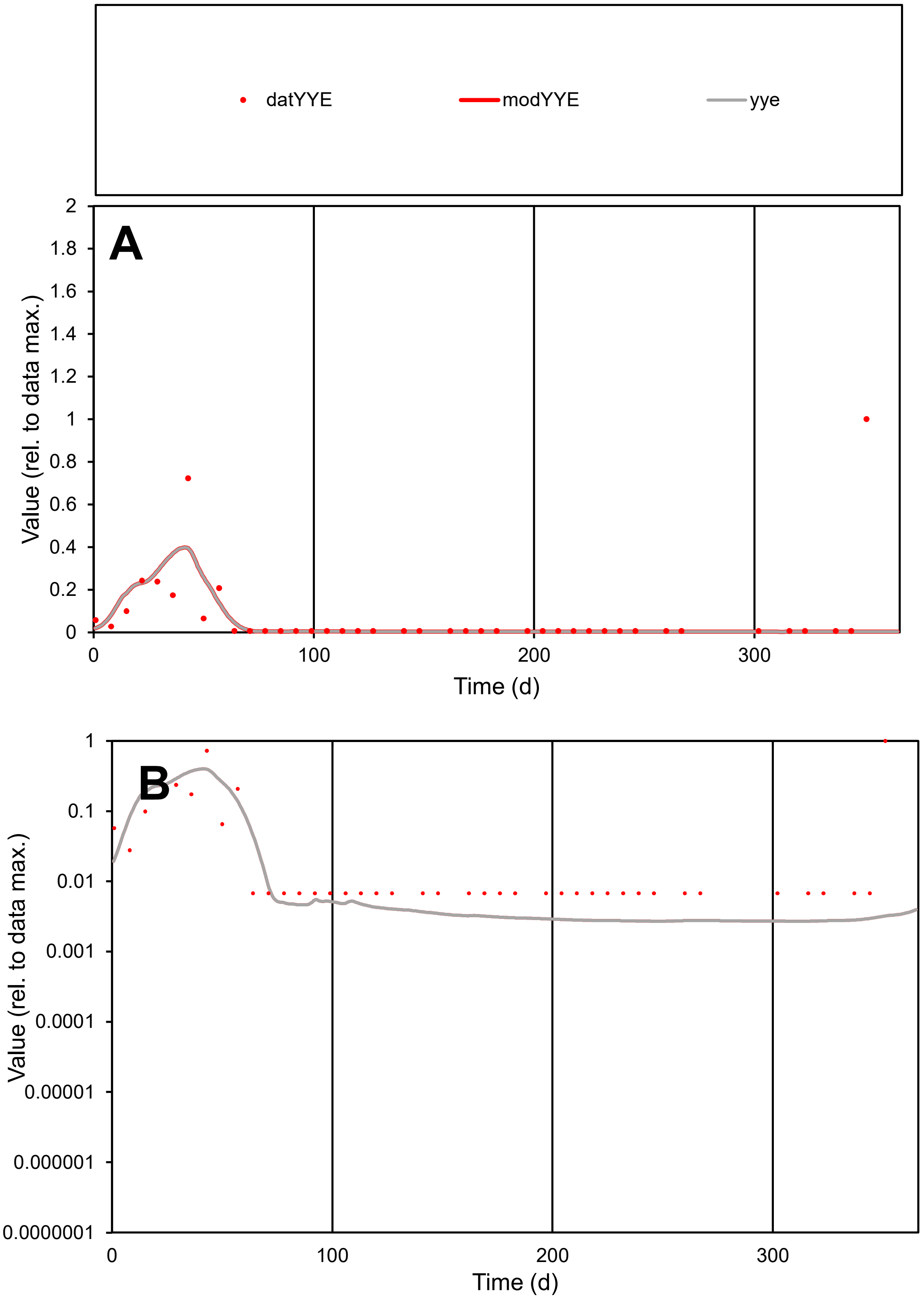


**C1**

Variables
YYE Provide jtd or Dst
CalibOnly?
2 1 to reproduce of
oNormalize?
2 1=no, 2=yes

ofxn calc*
2.29917
ofxn mod
2.31377

sRun
RunA1
zout
1

Ave
Data
4.1E-05
Model
2.8E-05

data/vMax
0.00056

ofxng
319.391

Min d/m
2.51034

ave/min d
10.9517
ave/min m
18.9267

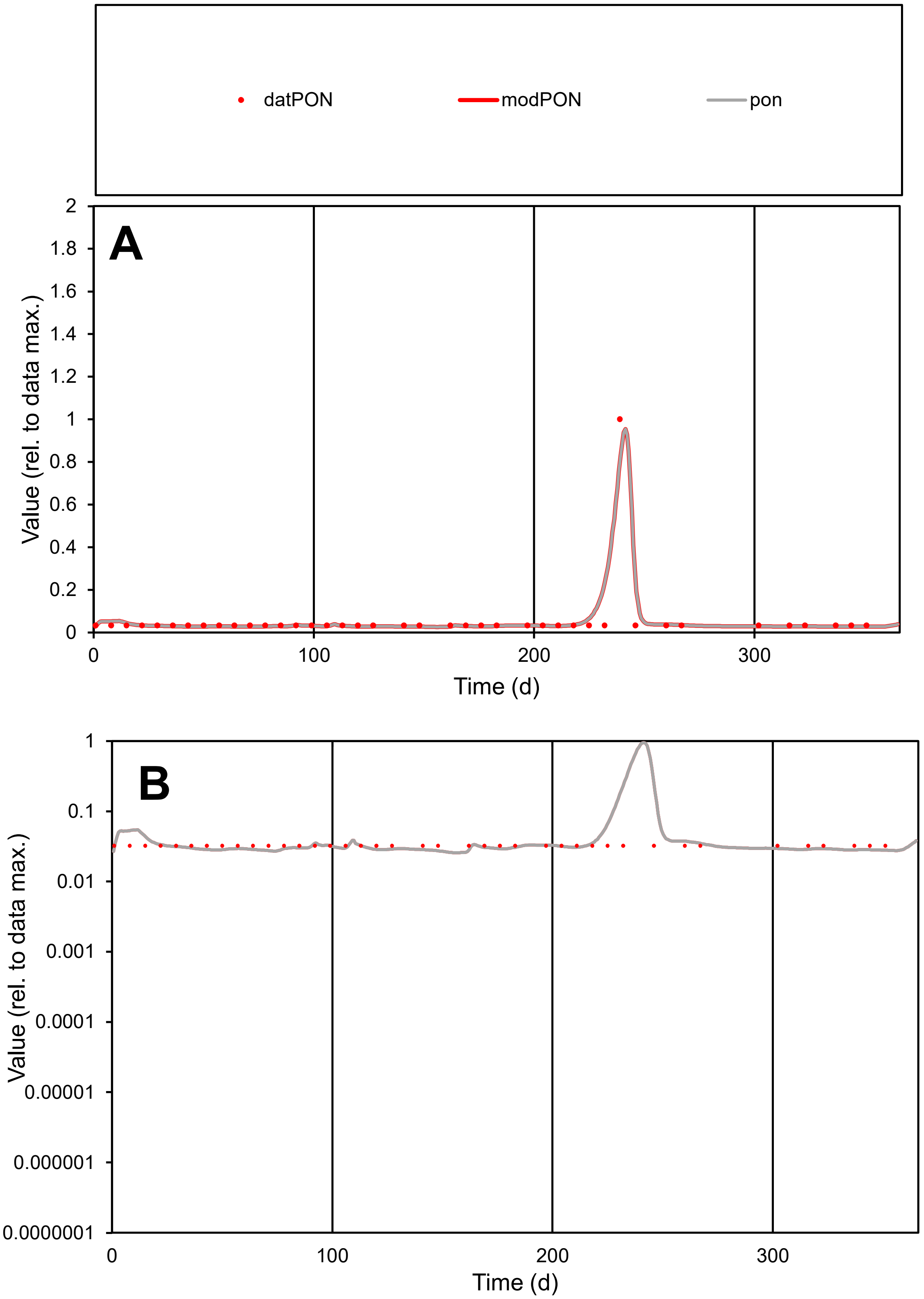


**C1**

Variables
PON Provide jtd or Dst
CalibOnly?
2 1 to reproduce of
oNormalize?
2 1=no, 2=yes

ofxn calc*
0.45509
ofxn mod
0.71149

sRun
RunA1
zout
1

Ave
Data
5.1E-06
Model
5.3E-06

data/vMax
9.1E-05

ofxng
319.391

Min d/m
1.1956

ave/min d
1.72885
ave/min m
2.16982

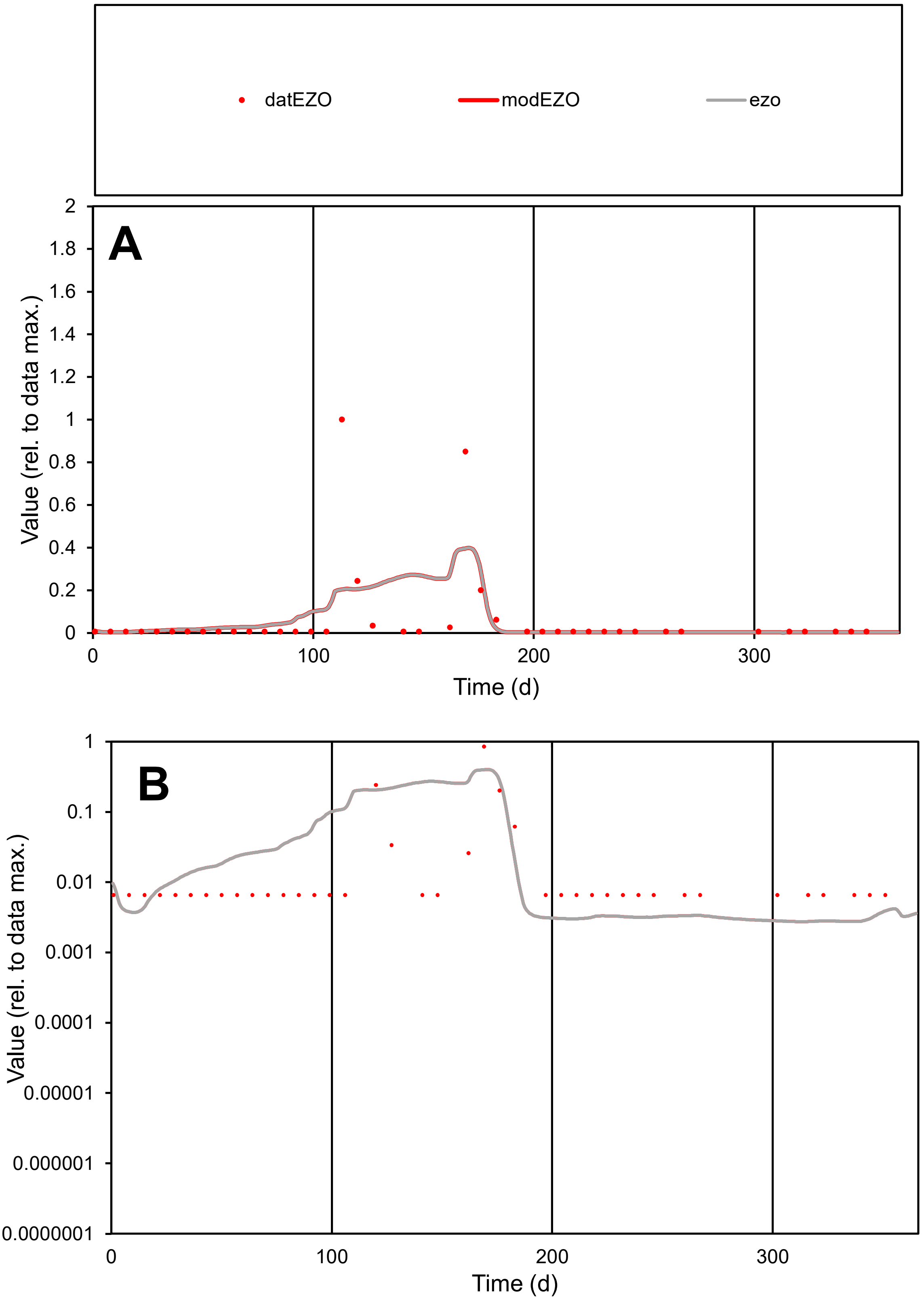


**C1**

Variables
EZO Provide jtd or Dst
CalibOnly?
2 1 to reproduce of
oNormalize?
2 1=no, 2=yes

ofxn calc*
2.52927
ofxn mod
2.549

sRun
RunA1
zout
1

Ave
Data
3.9E-05
Model
3.8E-05

data/vMax
0.0006

ofxng
319.391

Min d/m
2.38104
ave/min d
9.78456
ave/min m
23.1944

330_Calib_S_A

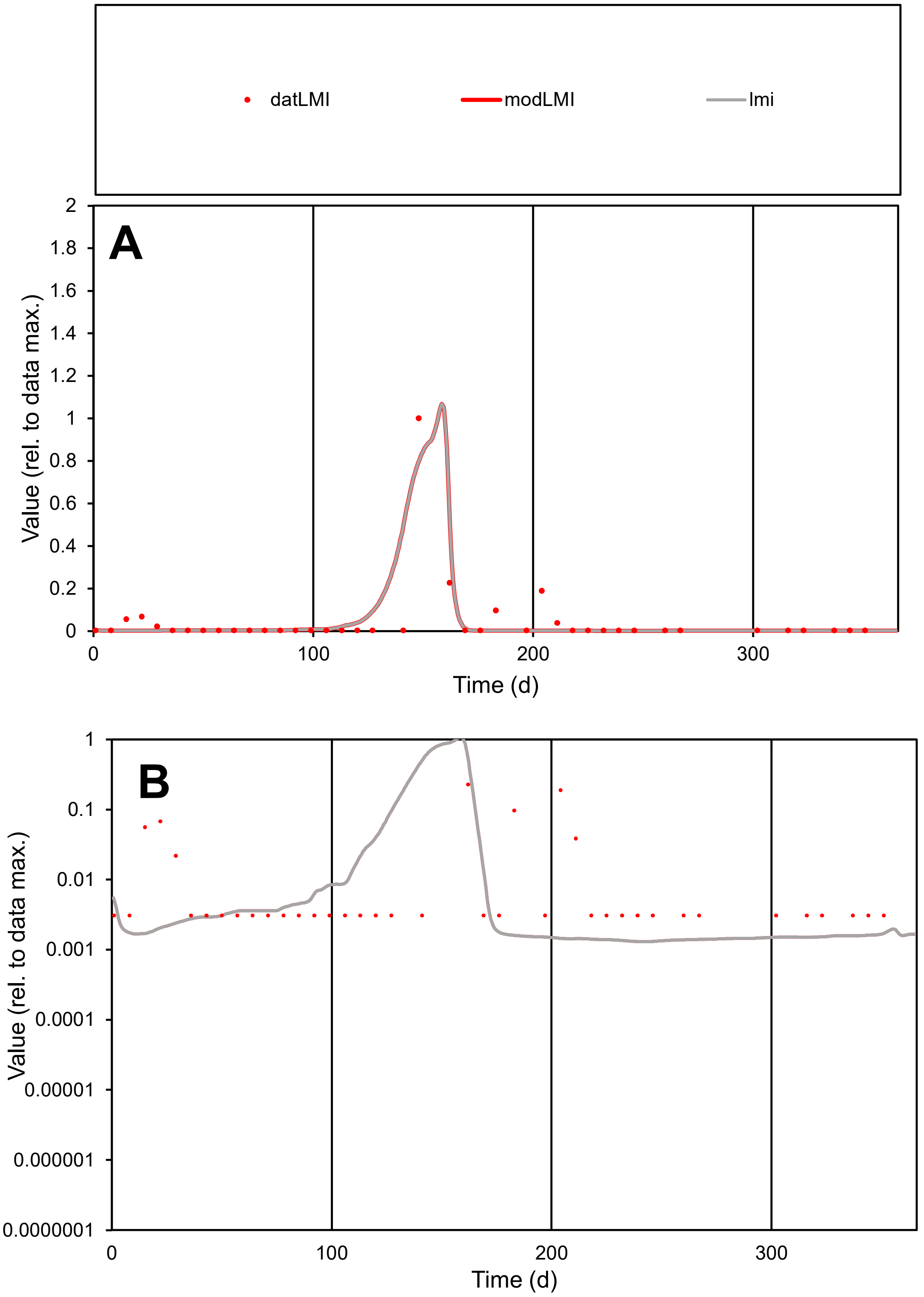


**C1**

Variables
LMI Provide jtd or Dst
CalibOnly?
2 1 to reproduce of
oNormalize?
2 1=no, 2=yes

ofxn calc*
1.66945
ofxn mod
1.97973

sRun
RunA1
zout
1

Ave
Data
4.3E-05
Model
3.9E-05

data/vMax
0.00098

ofxng
319.391

Min d/m
2.36871

ave/min d
14.2257
ave/min m
30.964

330_Calib_S_A

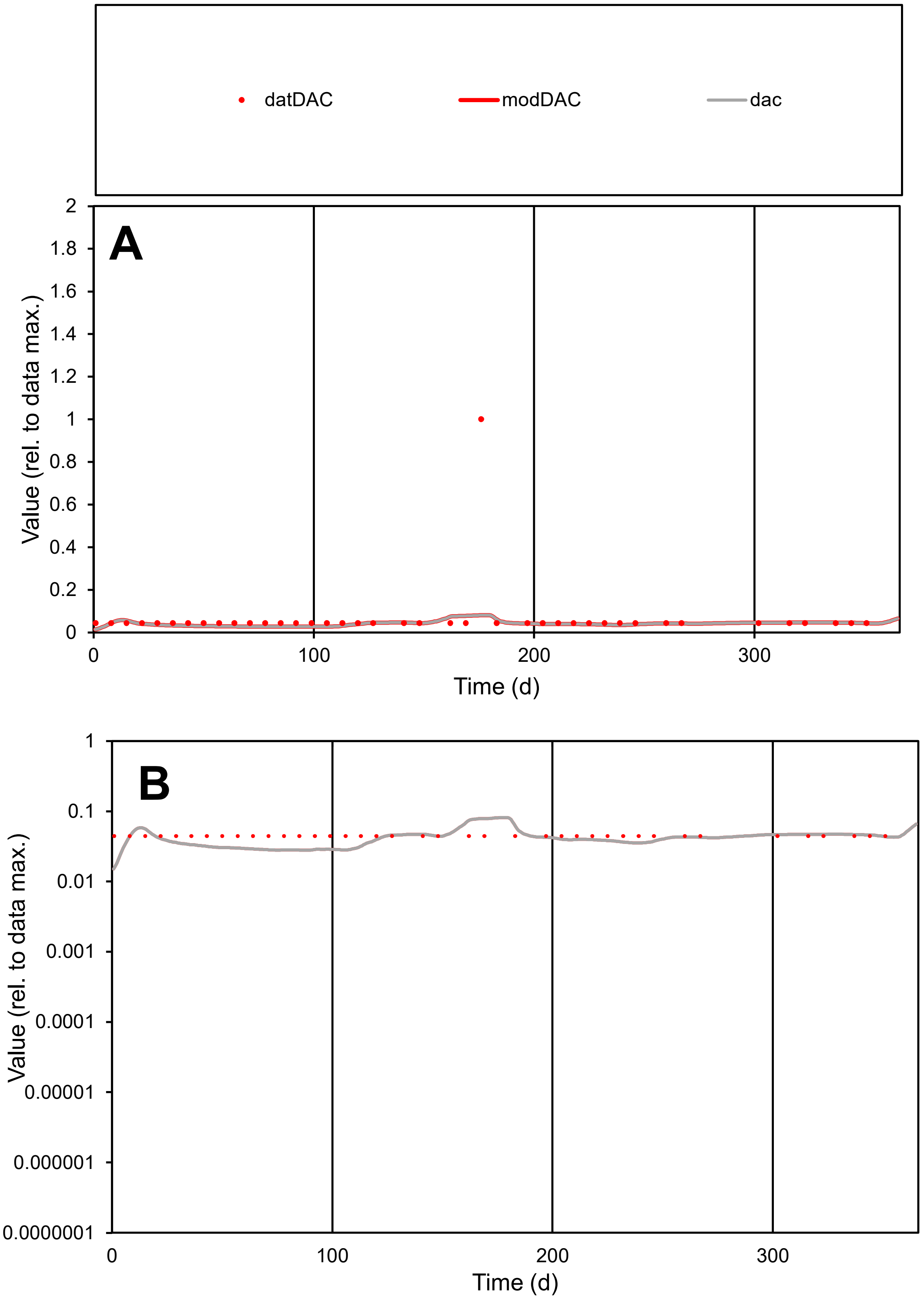


**C1**

Variables
DAC Provide jtd or Dst
CalibOnly?
2 1 to reproduce of
oNormalize?
2 1=no, 2=yes

ofxn calc*
2.12648
ofxn mod
2.1314

sRun
RunA1
zout
1

Ave
Data
1.2E-05
Model
7.2E-06

data/vMax
0.00017

ofxng
319.391

Min d/m
1.88273

ave/min d
1.52507
ave/min m
1.75259

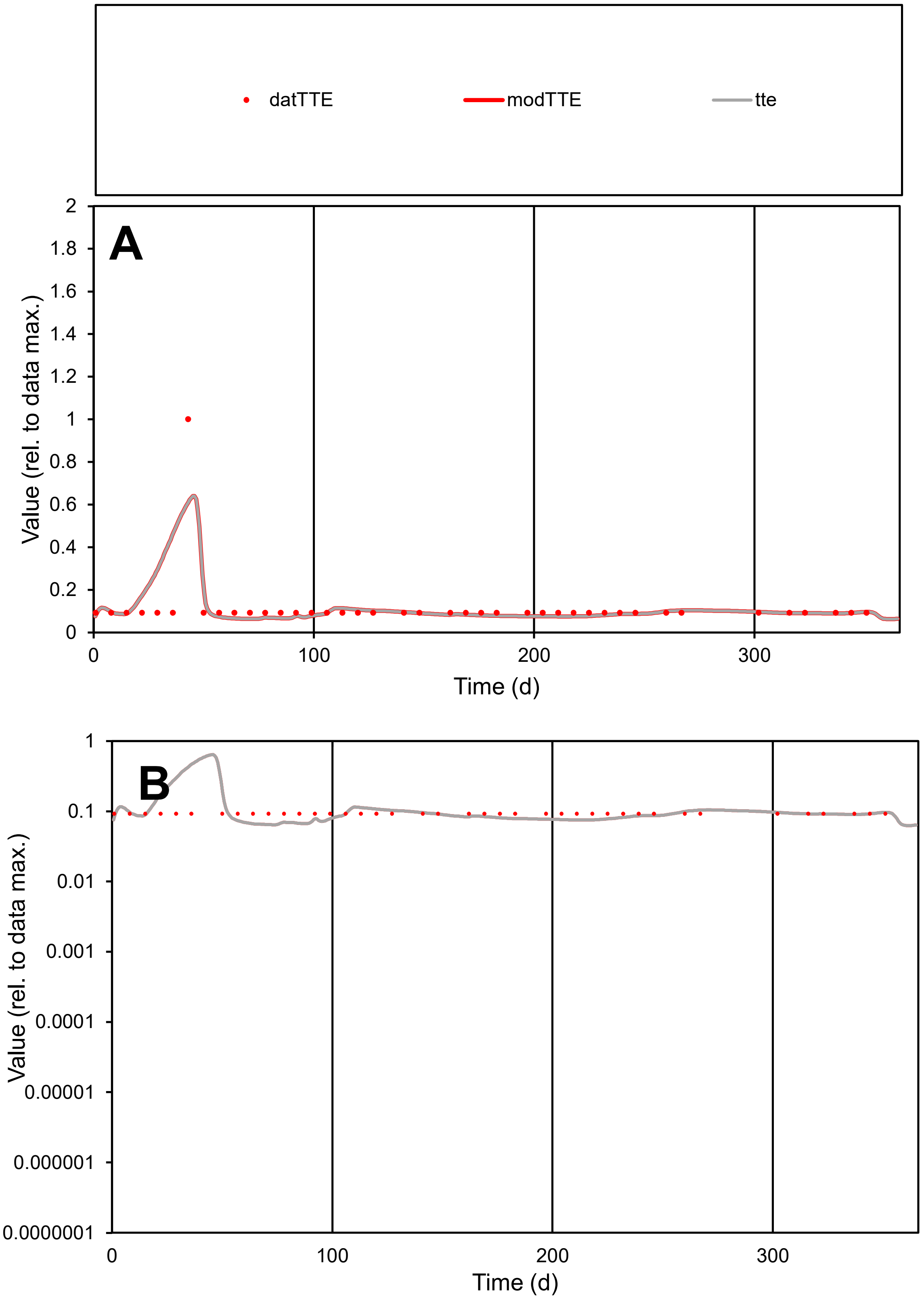


**C1**

Variables
TTE Provide jtd or Dst
CalibOnly?
2 1 to reproduce of
oNormalize?
2 1=no, 2=yes

ofxn calc*
0.78732
ofxn mod
0.80982

sRun
RunA1
zout
1

Ave
Data
4.7E-06
Model
4.8E-06

data/vMax
4.1E-05

ofxng
319.391

Min d/m
1.43934

ave/min d
1.23851
ave/min m
1.82217

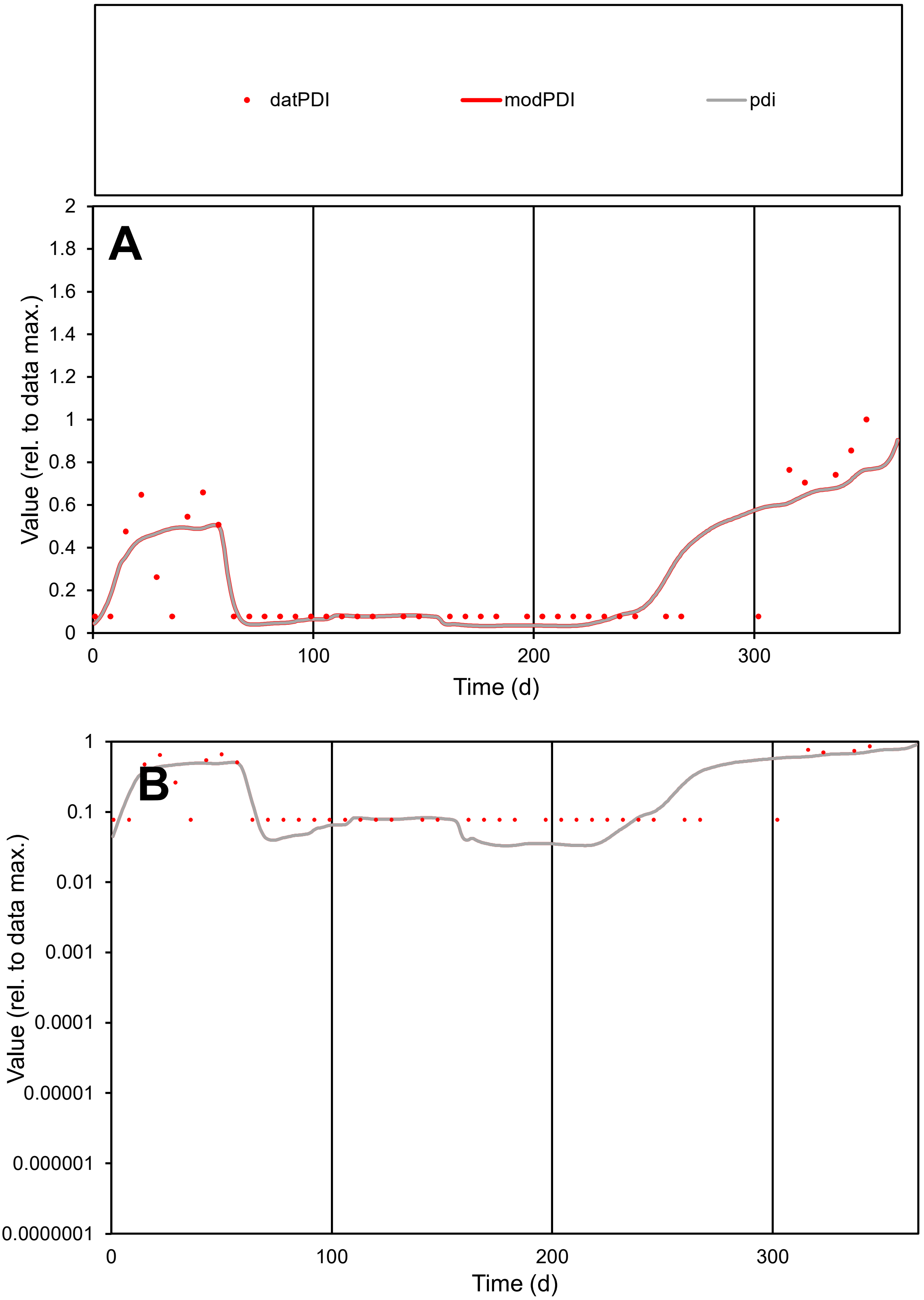


**C1**

Variables
PDI Provide jtd or Dst
CalibOnly?
2 1 to reproduce of
oNormalize?
2 1=no, 2=yes

ofxn calc*
0.60658
ofxn mod
0.62588

sRun
RunA1
zout
1

Ave
Data
1.3E-05
Model
1.3E-05

data/vMax
5.7E-05

ofxng
319.391

Min d/m
2.34704

ave/min d
2.99496
ave/min m
7.0951

330_Calib_S_A

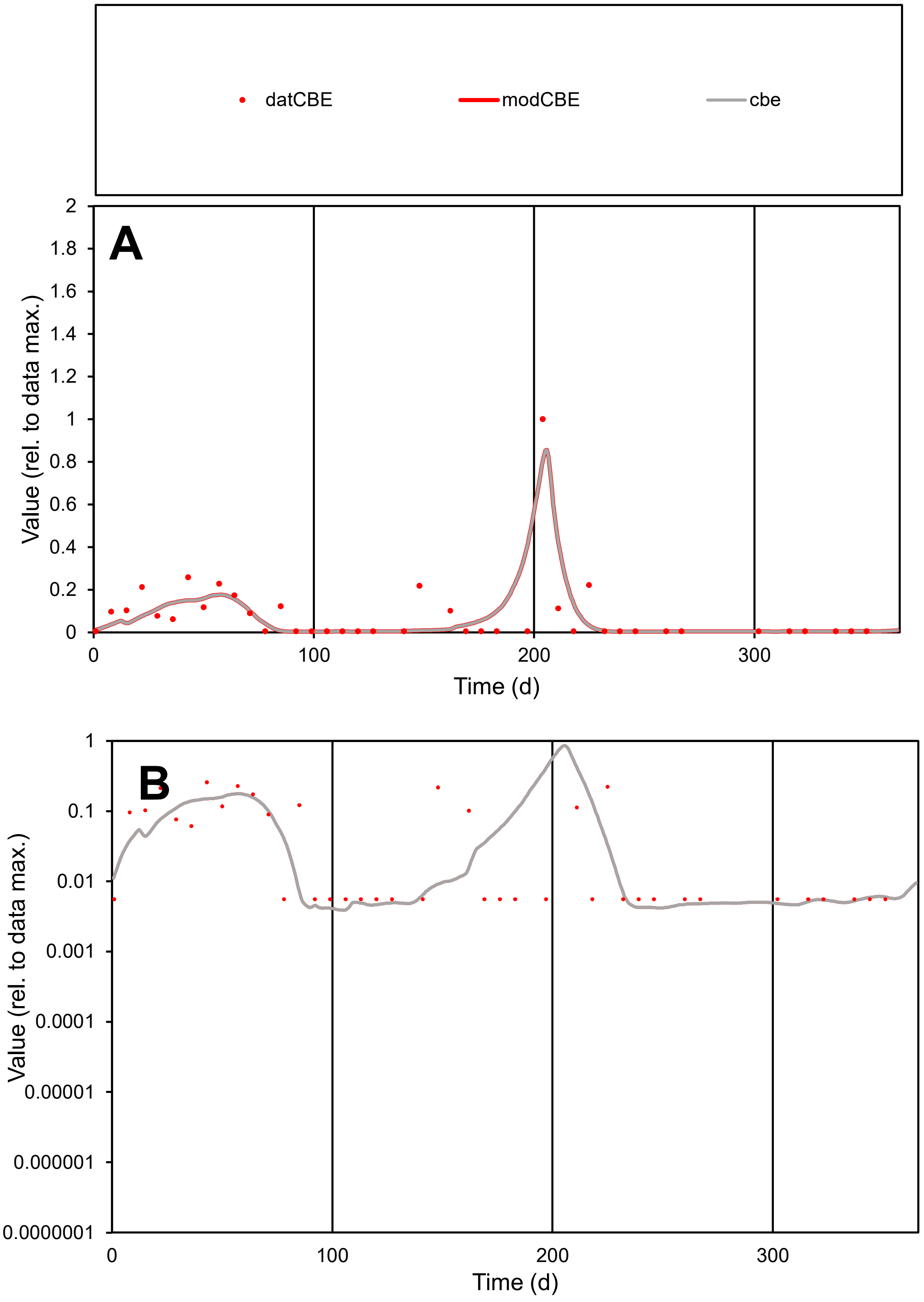


**C1**

Variables

CBE Provide jtd or Dst

CalibOnly?

2 1 to reproduce of

oNormalize?

2 1=no, 2=yes

ofxn calc*

1.06958

ofxn mod

1.19274

sRun

RunA1

zout

1

Ave

Data

2.4E-05

Model

2.2E-05

data/vMax

0.0003

ofxng

319.391

Min d/m

1.36609

ave/min d

14.5773

ave/min m

18.2663

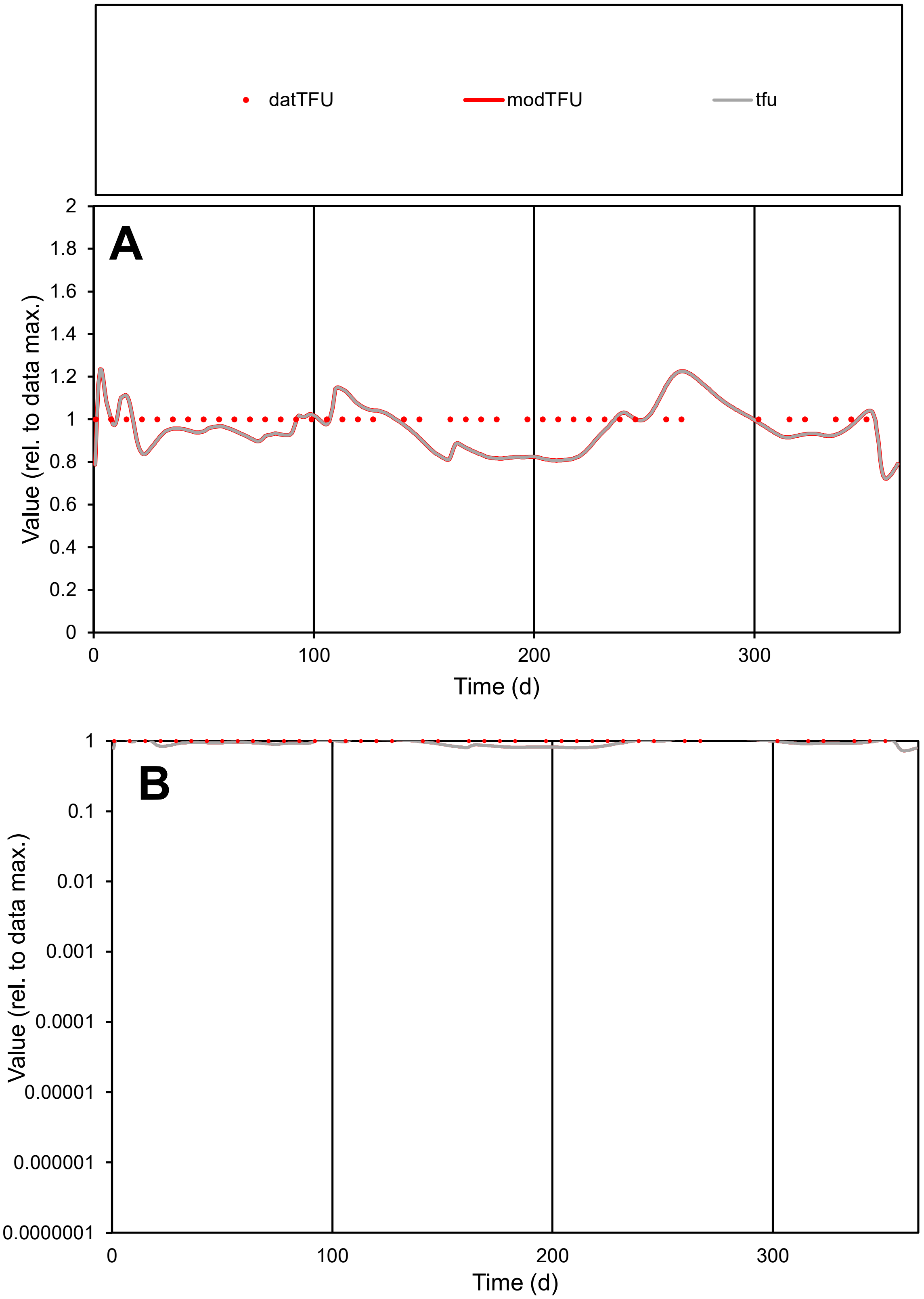


**C1**

Variables
TFU Provide jtd or Dst
CalibOnly?
2 1 to reproduce of
oNormalize?
2 1=no, 2=yes

ofxn calc*
0.10523
ofxn mod
0.14979

sRun
RunA1
zout
1

Ave
Data
1.3E-05
Model
1.2E-05

data/vMax
1.3E-05

ofxng
319.391

Min d/m
1.23786

ave/min d
1
ave/min m
1.18212

330_Calib_S_A

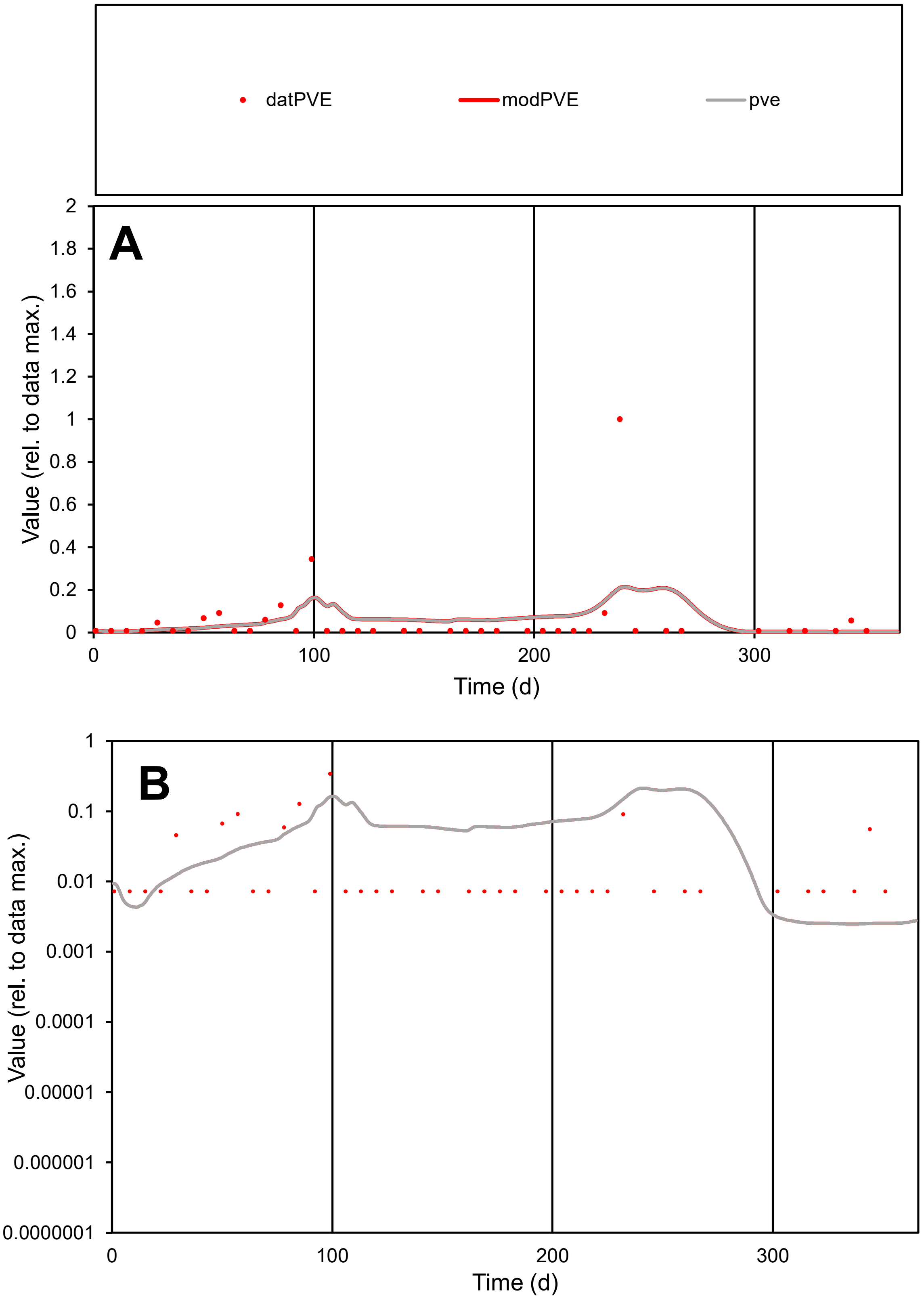


**C1**

Variables
PVE Provide jtd or Dst
CalibOnly?
2 1 to reproduce of
oNormalize?
2 1=no, 2=yes

ofxn calc*
2.7692
ofxn mod
2.80033

sRun
RunA1
zout
1

Ave
Data
1.5E-05
Model
1.8E-05

data/vMax
0.00029

ofxng
319.391

Min d/m
2.91771

ave/min d
7.12079
ave/min m
25.1509

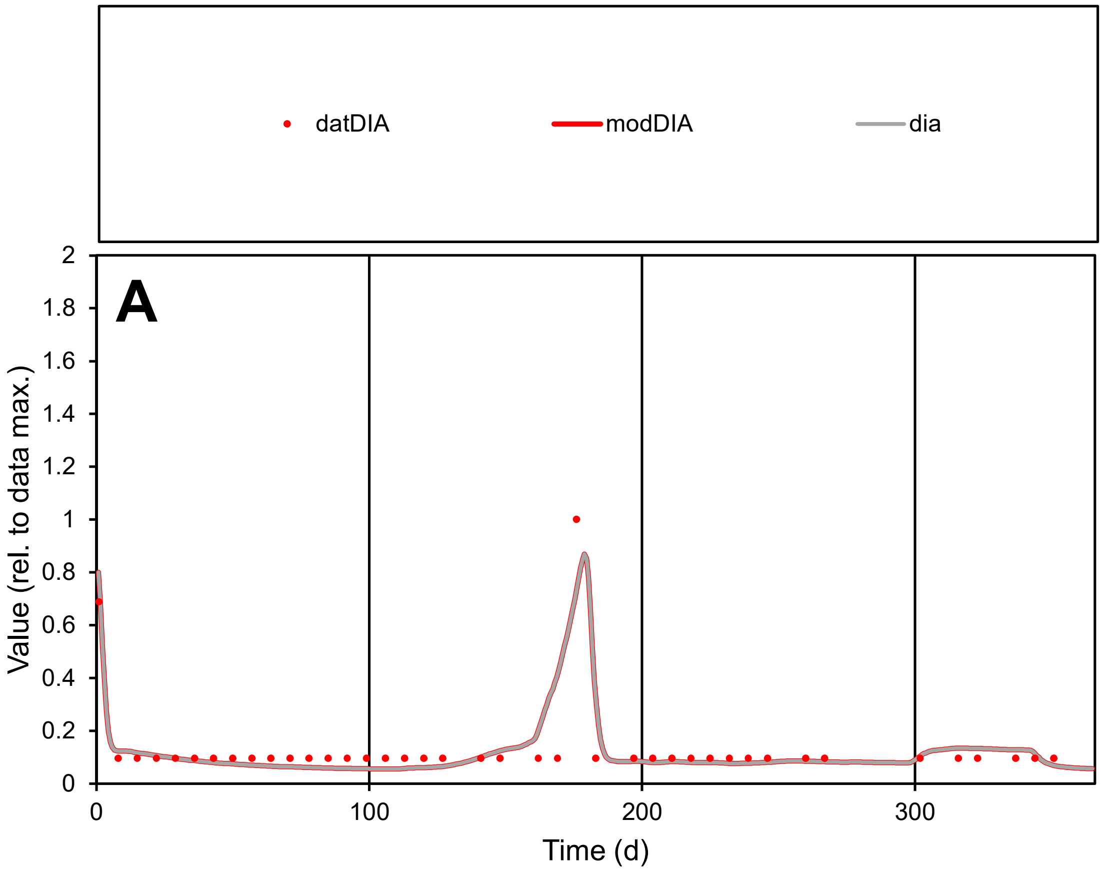


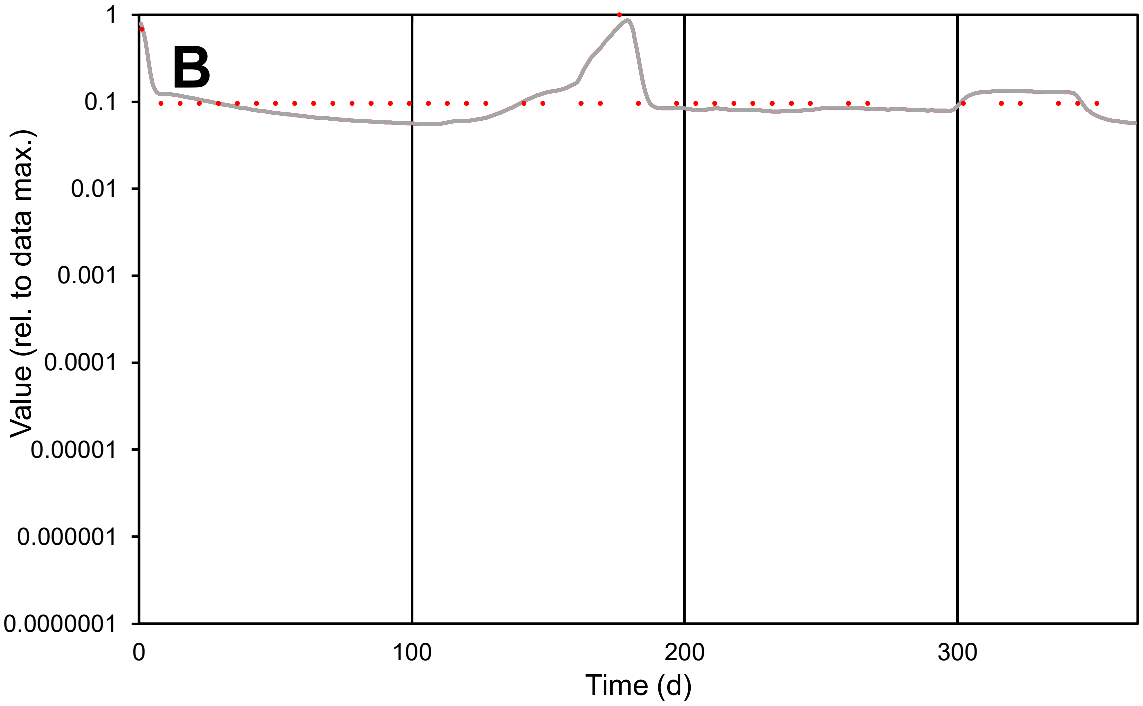


**C1**

Variables
DIA Provide jtd or Dst
CalibOnly?
2 1 to reproduce of
oNormalize?
2 1=no, 2=yes

ofxn calc*
0.46685
ofxn mod
0.51819

sRun
RunA1
zout
1

Ave
Data
8.6E-06
Model
8.4E-06

data/vMax
6.5E-05

ofxng
319.391

Min d/m
1.72684

ave/min d
1.3809
ave/min m
2.34331

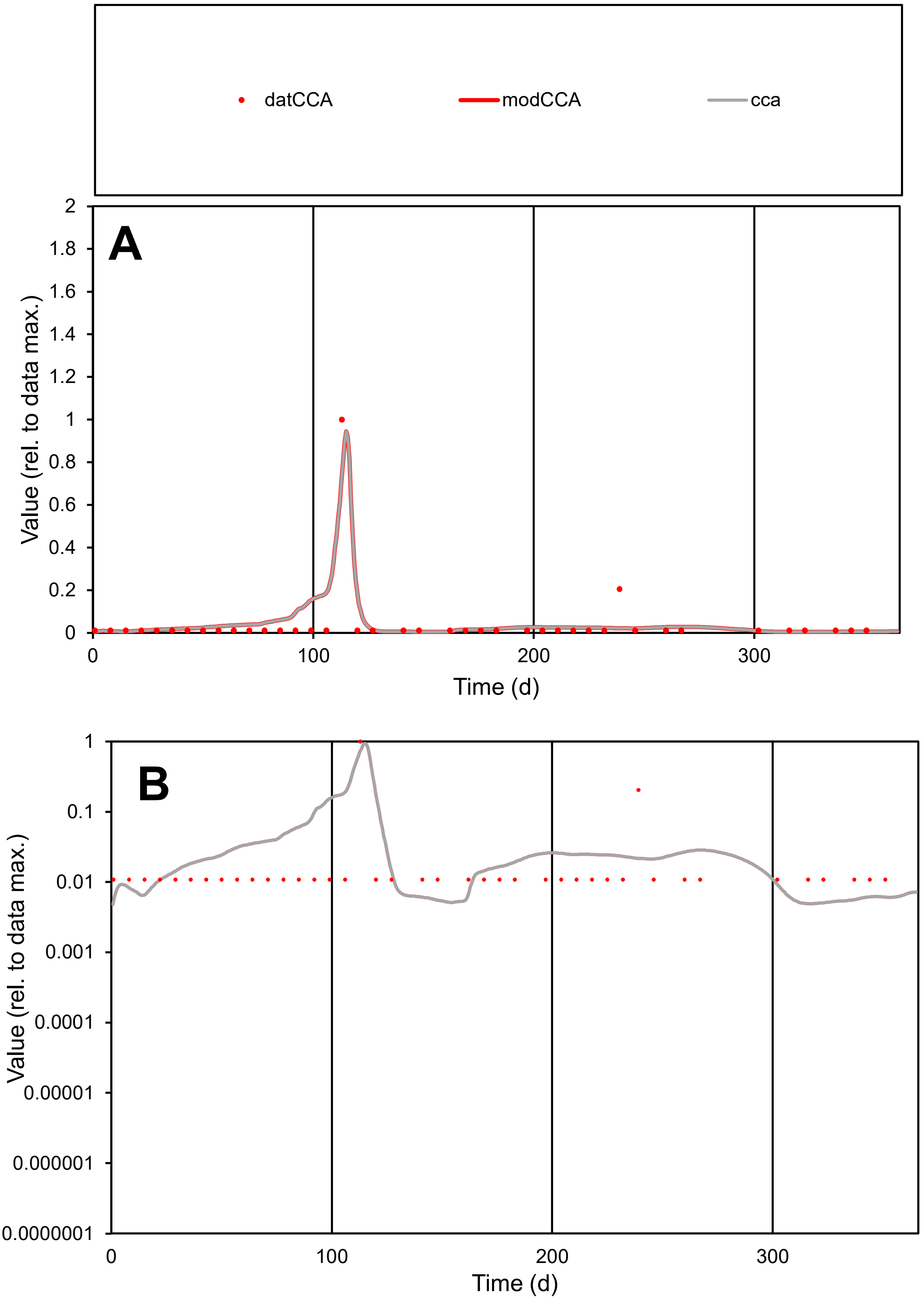


**C1**

Variables
CCA Provide jtd or Dst
CalibOnly?
2 1 to reproduce of
oNormalize?
2 1=no, 2=yes

ofxn calc*
1.30384
ofxn mod
1.38027

sRun
RunA1
zout
1

Ave
Data
6.7E-06
Model
8.5E-06

data/vMax
0.00017

ofxng
319.391

Min d/m
2.20135

ave/min d
3.67071
ave/min m
10.2644

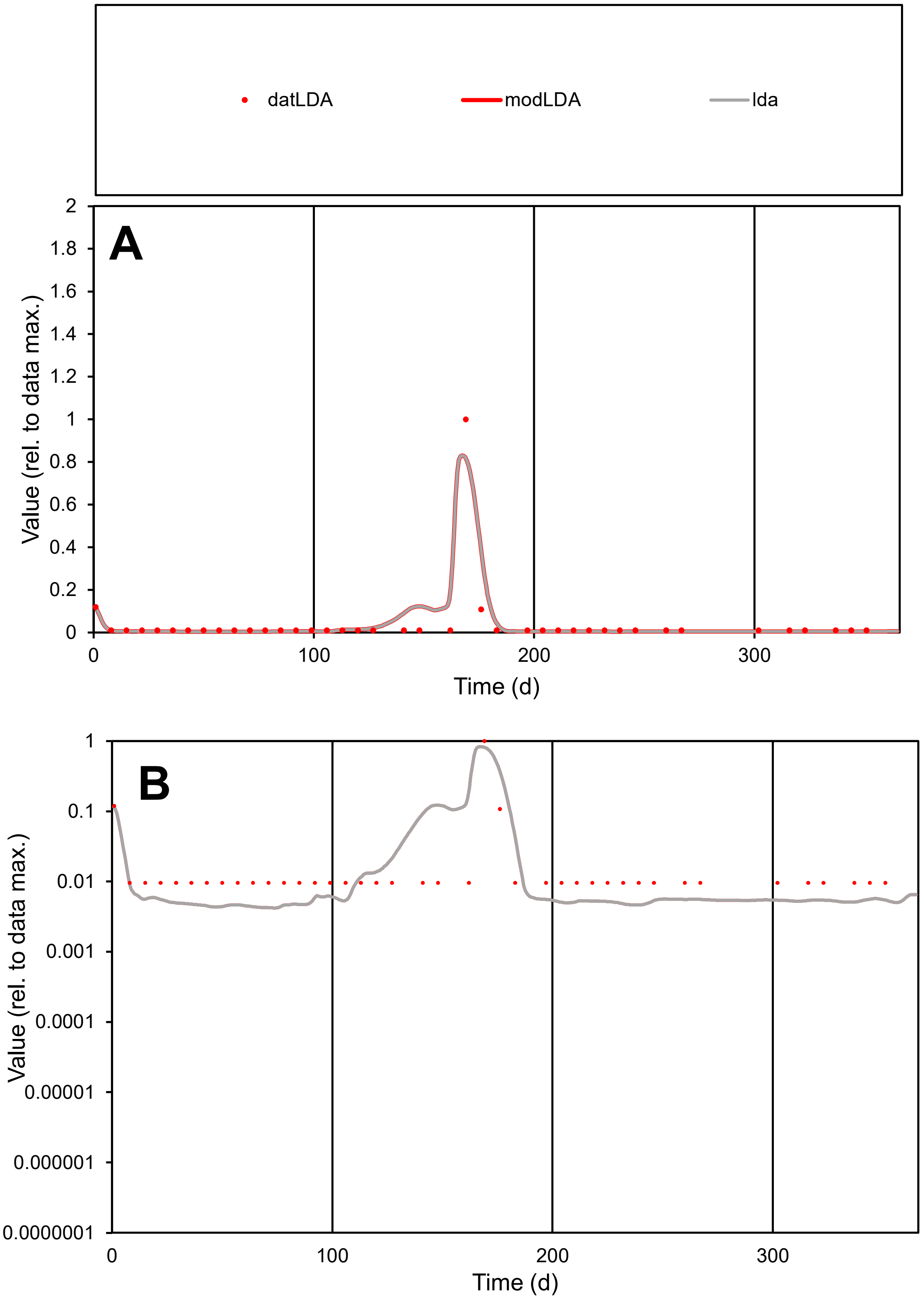


**C1**

Variables
LDA Provide jtd or Dst
CalibOnly?
2 1 to reproduce of
oNormalize?
2 1=no, 2=yes

ofxn calc*
1.20894
ofxn mod
1.35209

sRun
RunA1
zout
1

Ave
Data
2.8E-05
Model
3E-05

data/vMax
0.00072

ofxng
319.391

Min d/m
2.26739

ave/min d
4.05839
ave/min m
9.70536

330_Calib_S_A

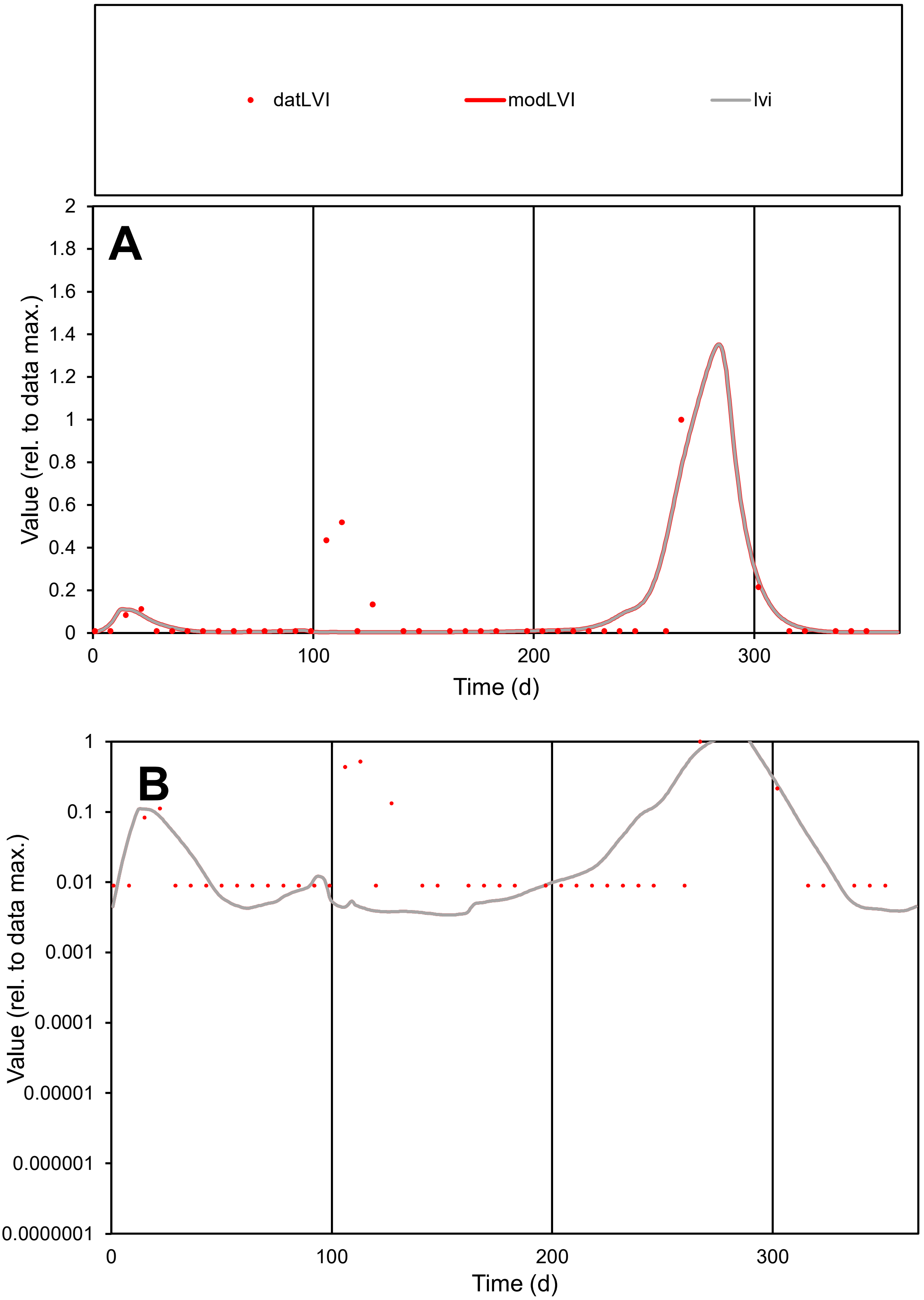


**C1**

Variables

LVI Provide jtd or Dst

CalibOnly?

2 1 to reproduce of

oNormalize?

2 1=no, 2=yes

ofxn calc*

1.82124

ofxn mod

1.8961

sRun

RunA1

zout

1

Ave

Data

1.1E-05

Model

8.4E-06

data/vMax

0.00016

ofxng

319.391

Min d/m

2.58123

ave/min d

7.65356

ave/min m

15.3367

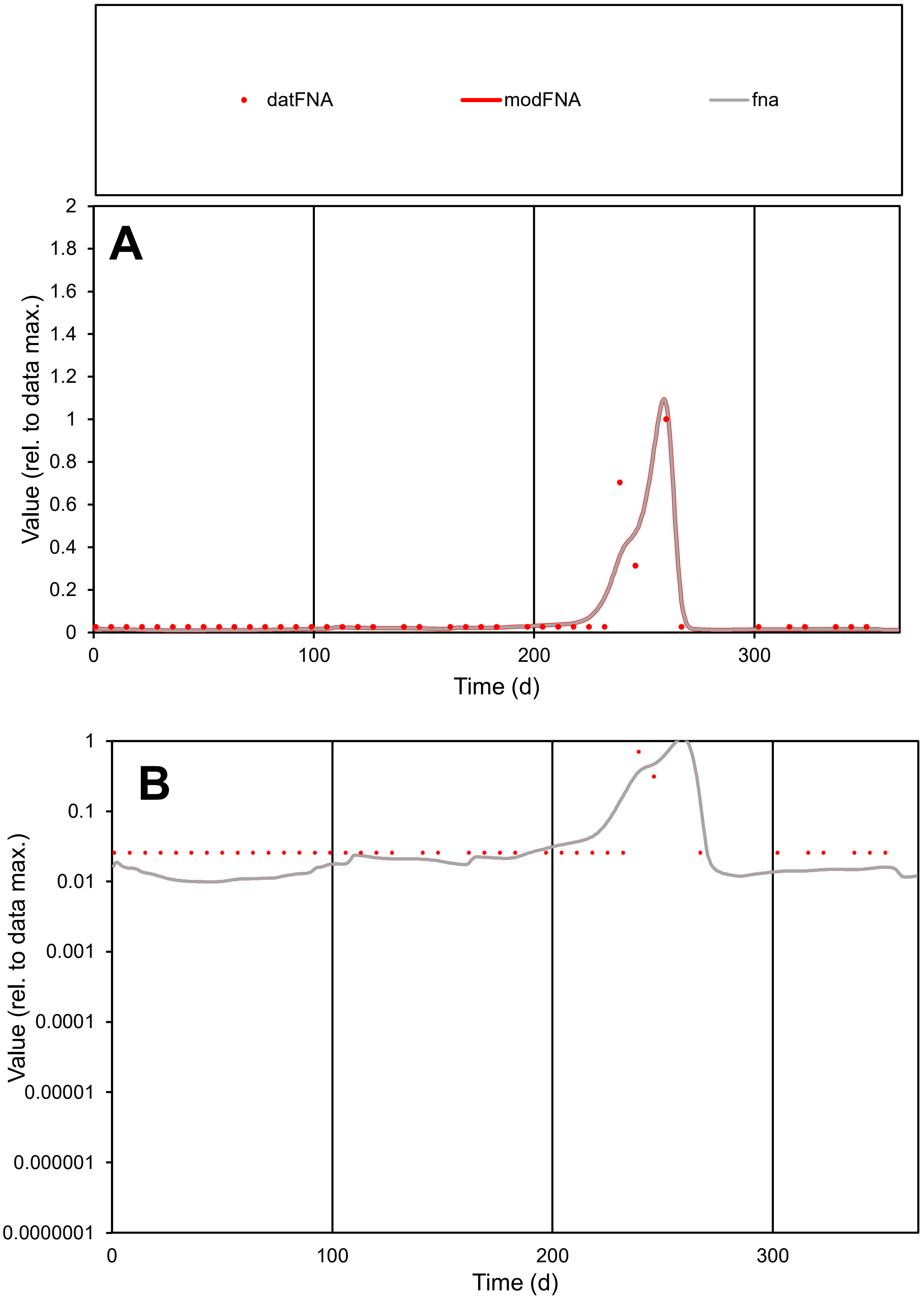


**C1**

Variables

FNA Provide jtd or Dst

CalibOnly?

2 1 to reproduce of

oNormalize?

2 1=no, 2=yes

ofxn calc*

0.80406

ofxn mod

0.86611

sRun

RunA1

zout

1

Ave

Data

8.9E-06

Model

8.2E-06

data/vMax

0.00012

ofxng

319.391

Min d/m

2.60921

ave/min d

2.8434

ave/min m

6.87582

330_Calib_S_A

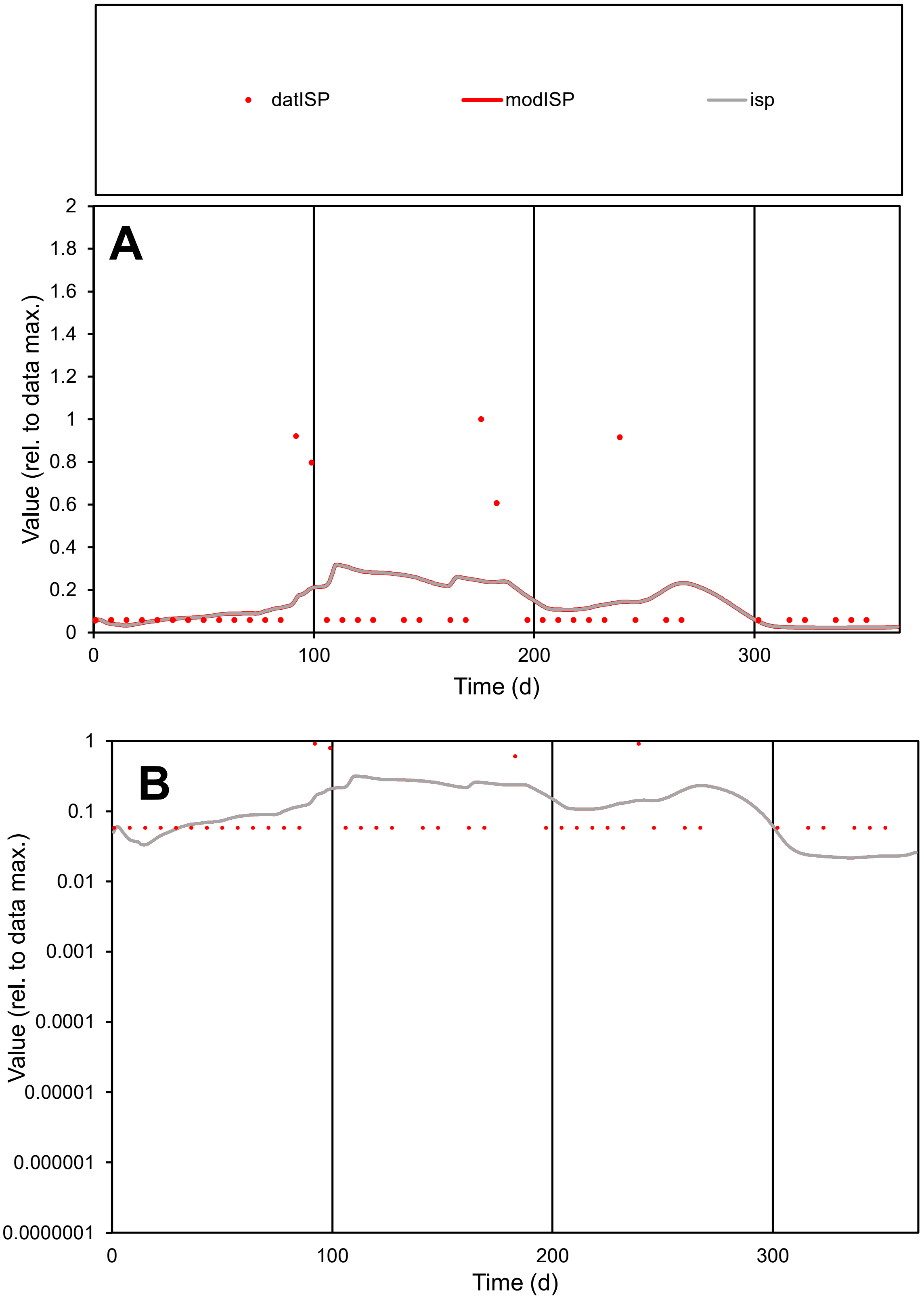


**C1**

Variables
ISP Provide jtd or Dst
CalibOnly?
2 1 to reproduce of
oNormalize?
2 1=no, 2=yes

ofxn calc*
1.64784
ofxn mod
1.65616

sRun
RunA1
zout
1

Ave
Data
1.3E-05
Model
1.2E-05

data/vMax
8.7E-05

ofxng
319.391

Min d/m
2.64394

ave/min d
2.66035
ave/min m
6.14027

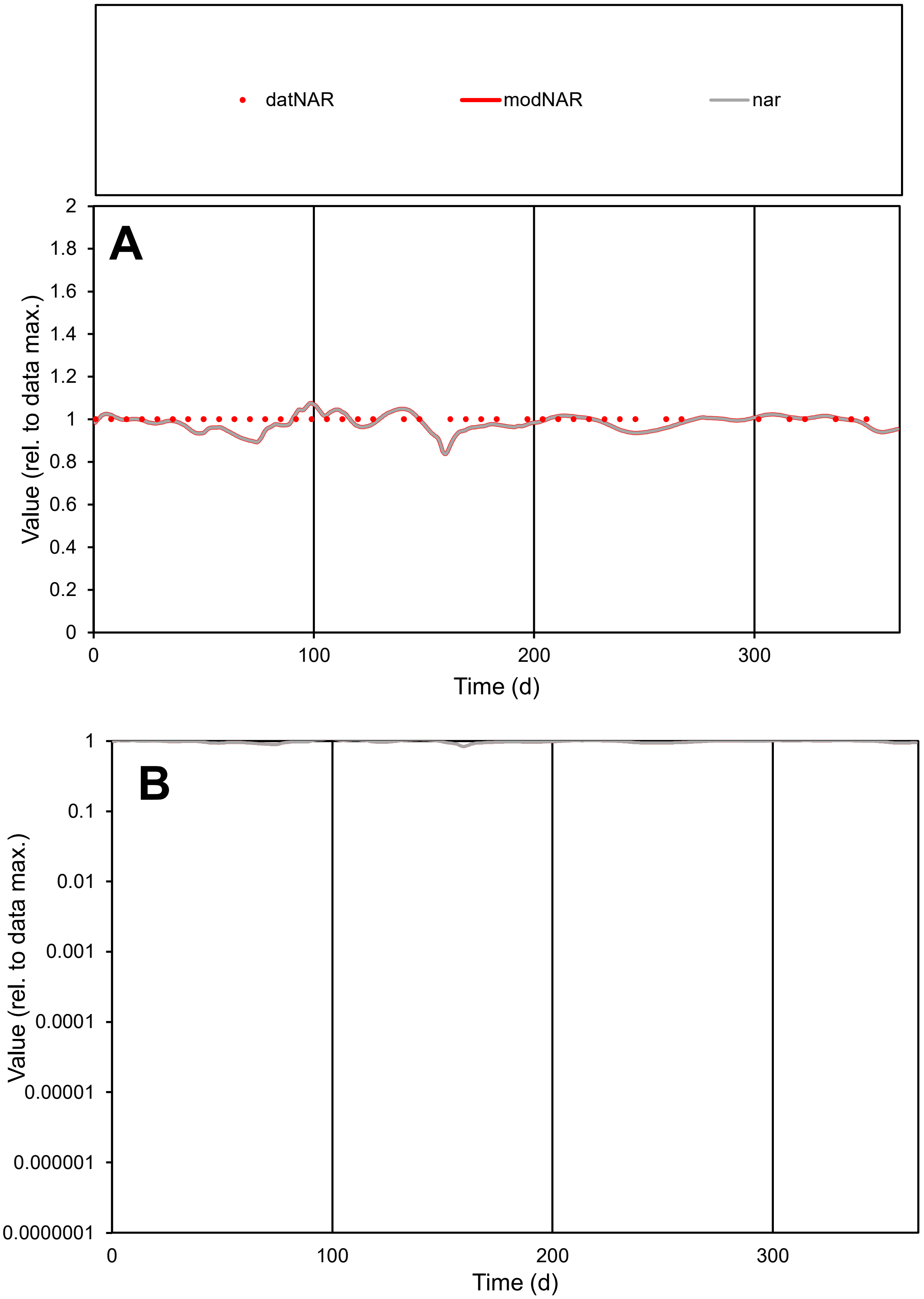


**C1**

Variables

| | |
|---|---|
| NAR | Provide jtd or Dst |
| CalibOnly? | |
| 2 | 1 to reproduce of |
| oNormalize? | |
| 2 | 1=no, 2=yes |

ofxn calc*
0.0386
ofxn mod
0.115

sRun
RunA1
zout
1

Ave
Data
2.8E-06
Model
2.8E-06

data/vMax
2.8E-06

ofxng
319.391

Min d/m
1.11396

ave/min d
1
ave/min m
1.09684

330_Calib_S_A

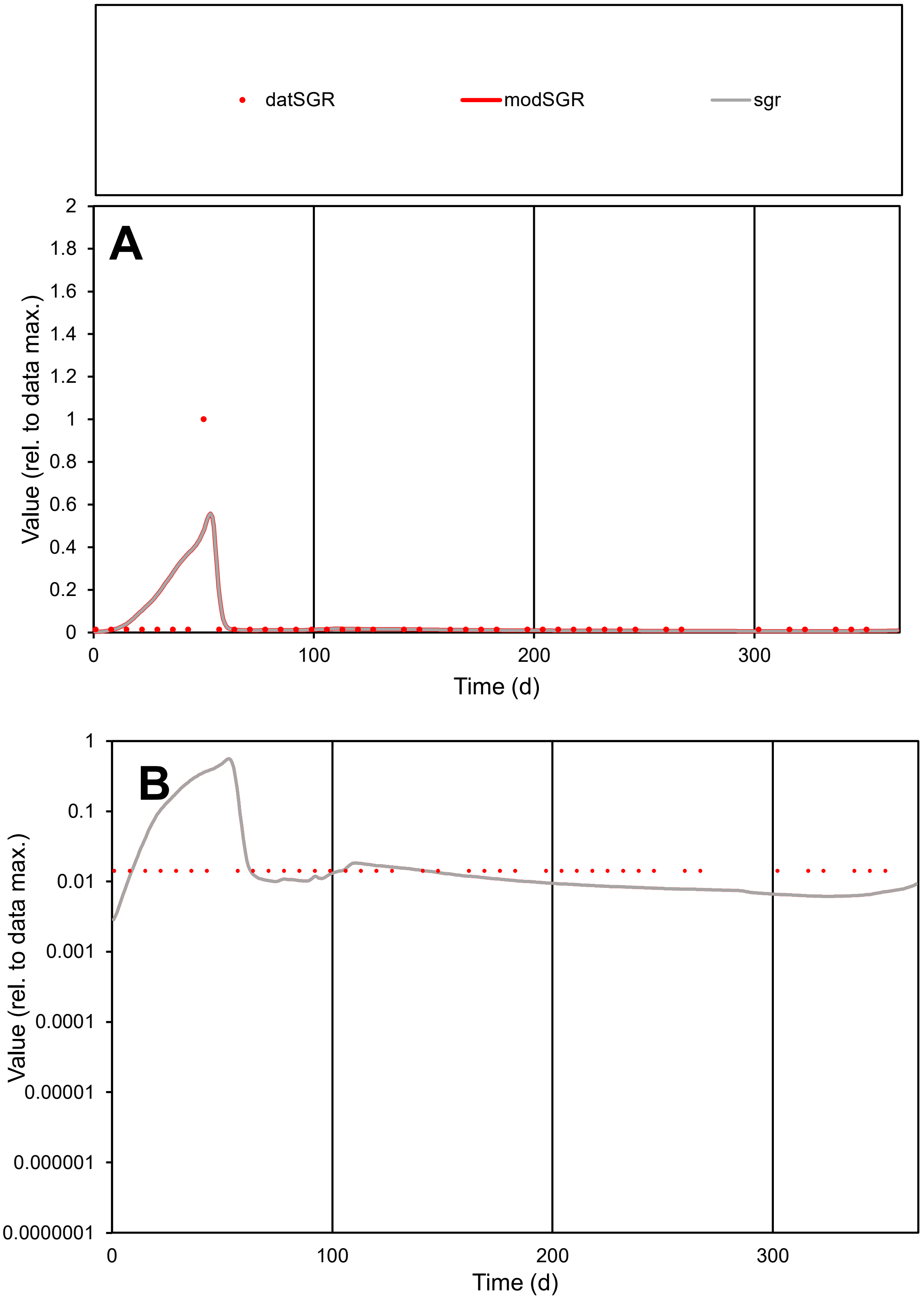


**C1**

Variables
SGR Provide jtd or Dst
CalibOnly?
2 1 to reproduce of
oNormalize?
2 1=no, 2=yes

ofxn calc*
2.69175
ofxn mod
2.81405

sRun
RunA1
zout
1

Ave
Data
1.2E-05
Model
1.4E-05

data/vMax
0.00032

ofxng
319.391

Min d/m
2.91828

ave/min d
2.69933
ave/min m
9.26919

330_Calib_S_A

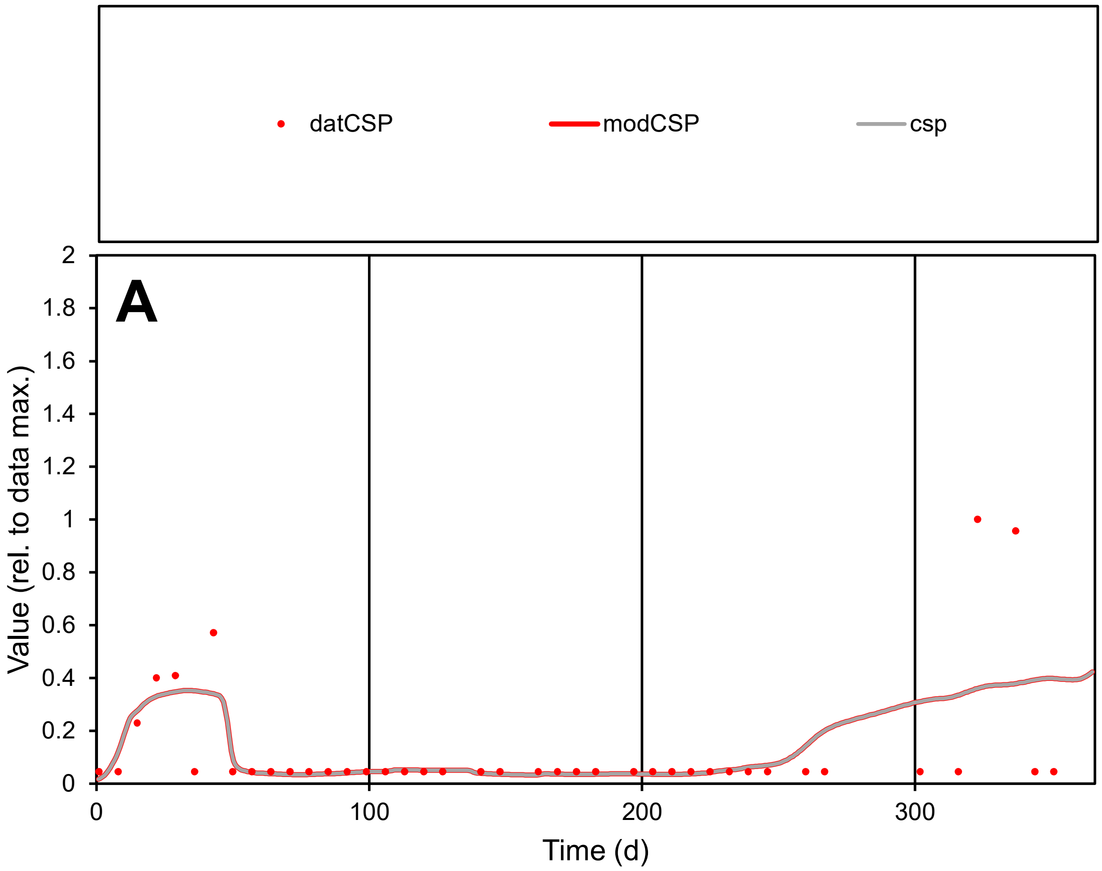


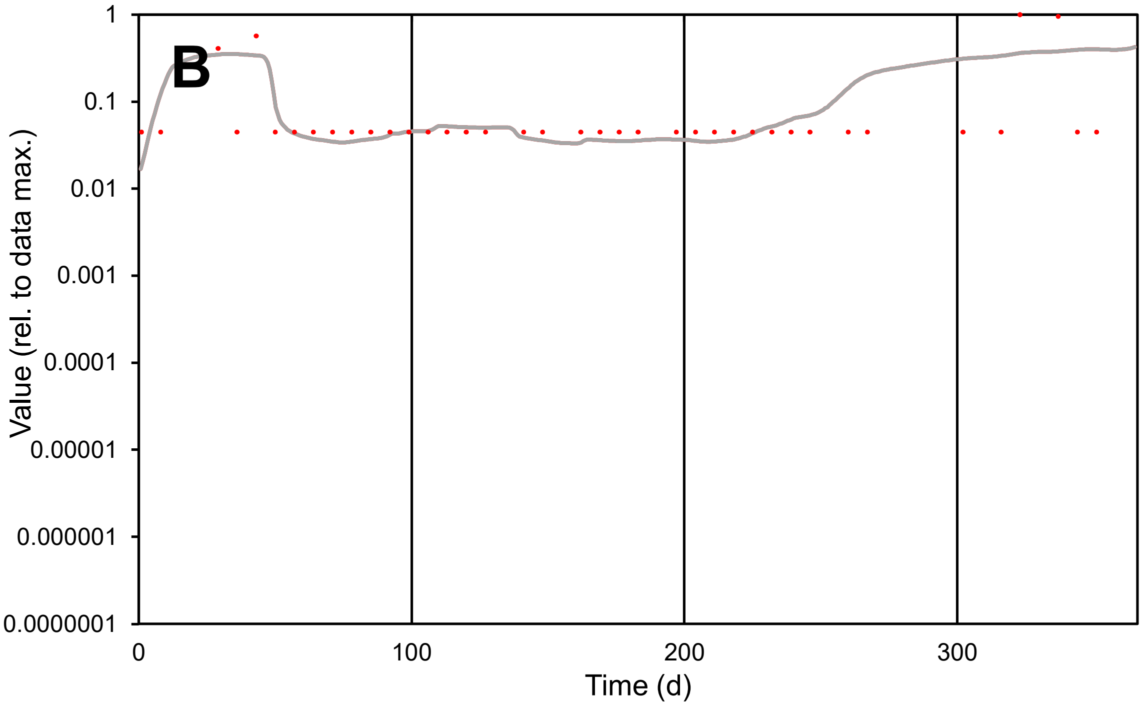


**C1**

Variables

CSP Provide jtd or Dst

CalibOnly?

2 1 to reproduce of

oNormalize?

2 1=no, 2=yes

ofxn calc*

1.44234

ofxn mod

1.45568

sRun

RunA1

zout

1

Ave

Data

6.7E-06

Model

7.1E-06

data/vMax

5.3E-05

ofxng

319.391

Min d/m

1.30295

ave/min d

2.79006

ave/min m

3.86608

330_Calib_S_A

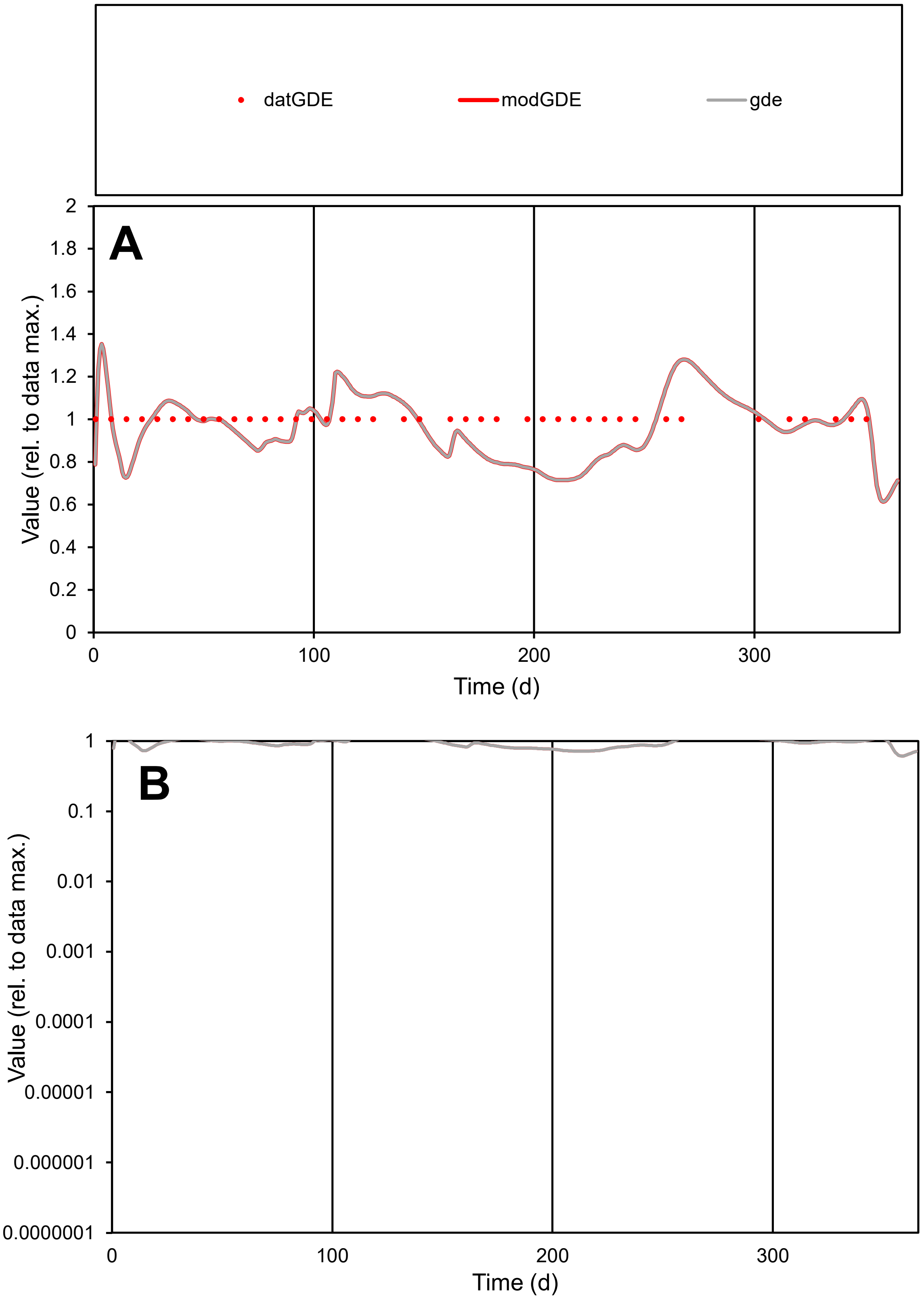


**C1**

Variables
GDE Provide jtd or Dst
CalibOnly?
2 1 to reproduce of
oNormalize?
2 1=no, 2=yes

ofxn calc*
0.13863
ofxn mod
0.17424

sRun
RunA1
zout
1

Ave
Data
2.7E-05
Model
2.6E-05

data/vMax
2.7E-05

ofxng
319.391

Min d/m
1.39776

ave/min d
1
ave/min m
1.33548

330_Calib_S_A

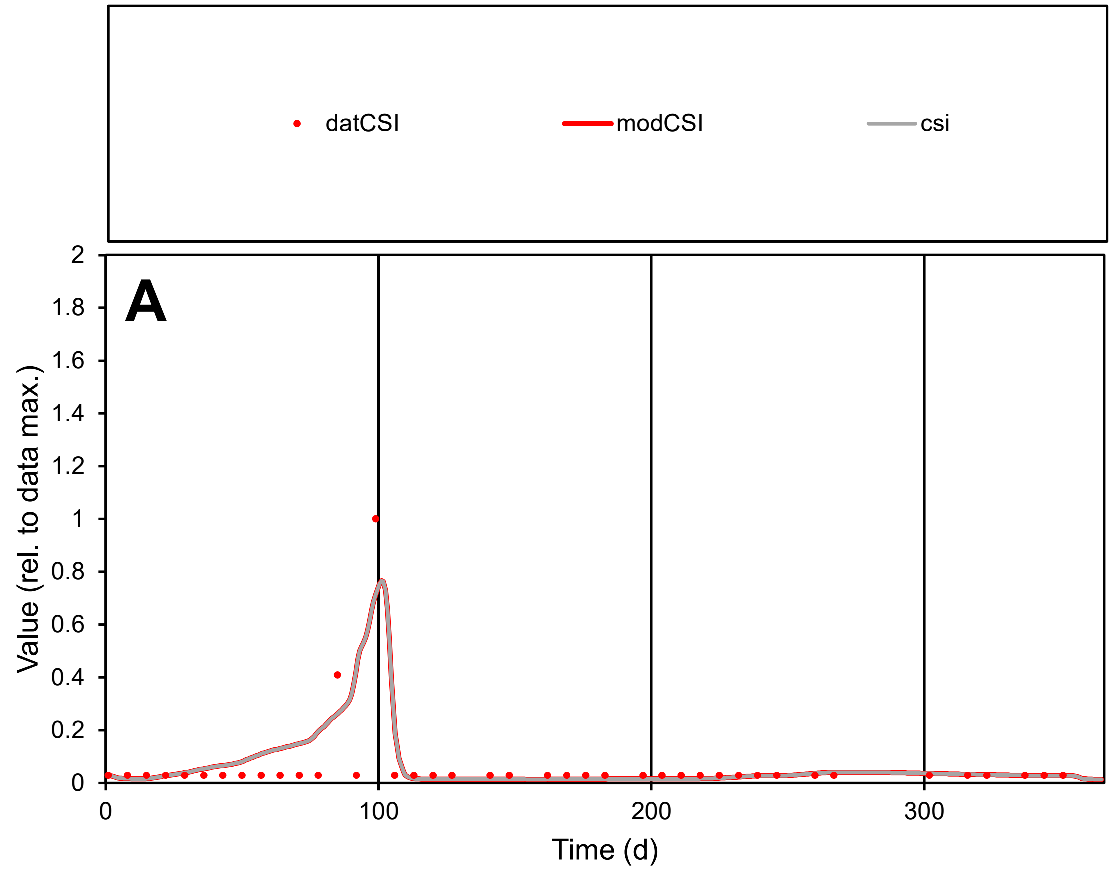


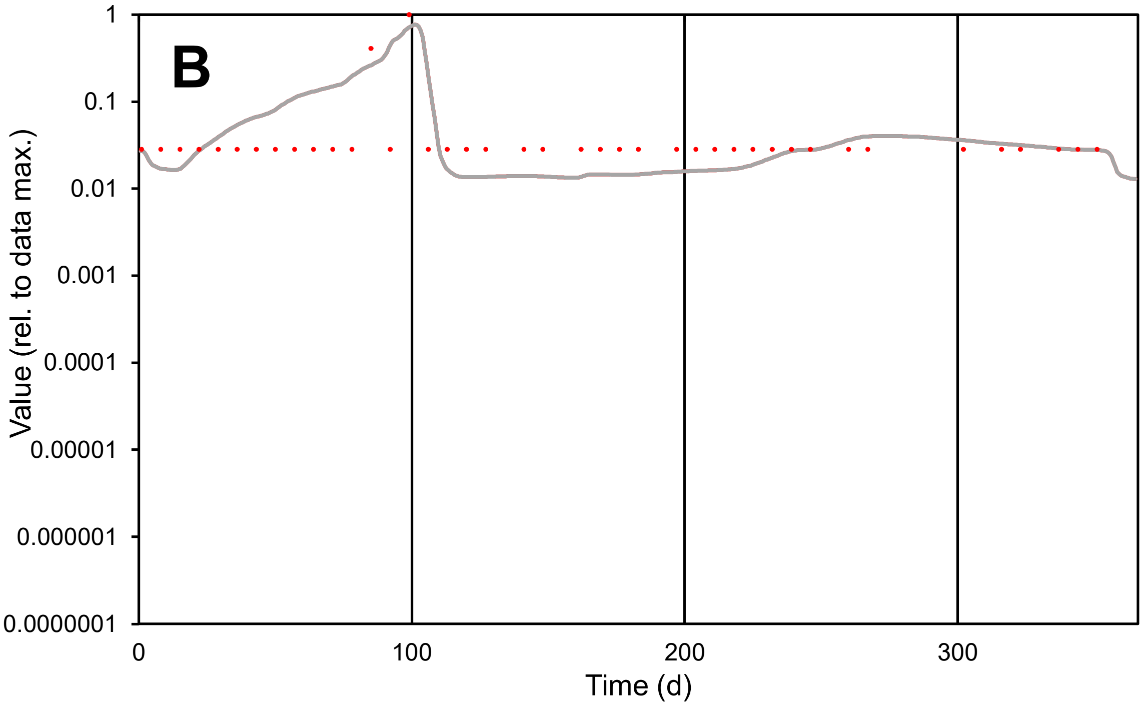


**C1**

Variables
CSI Provide jtd or Dst
CalibOnly?
2 1 to reproduce of
oNormalize?
2 1=no, 2=yes

ofxn calc*
1.24474
ofxn mod
1.39401

sRun
RunA1
zout
1

Ave
Data
5E-06
Model
5.8E-06

data/vMax
8.1E-05

ofxng
319.391

Min d/m
2.09239

ave/min d
2.16271
ave/min m
5.21841

330_Calib_S_A

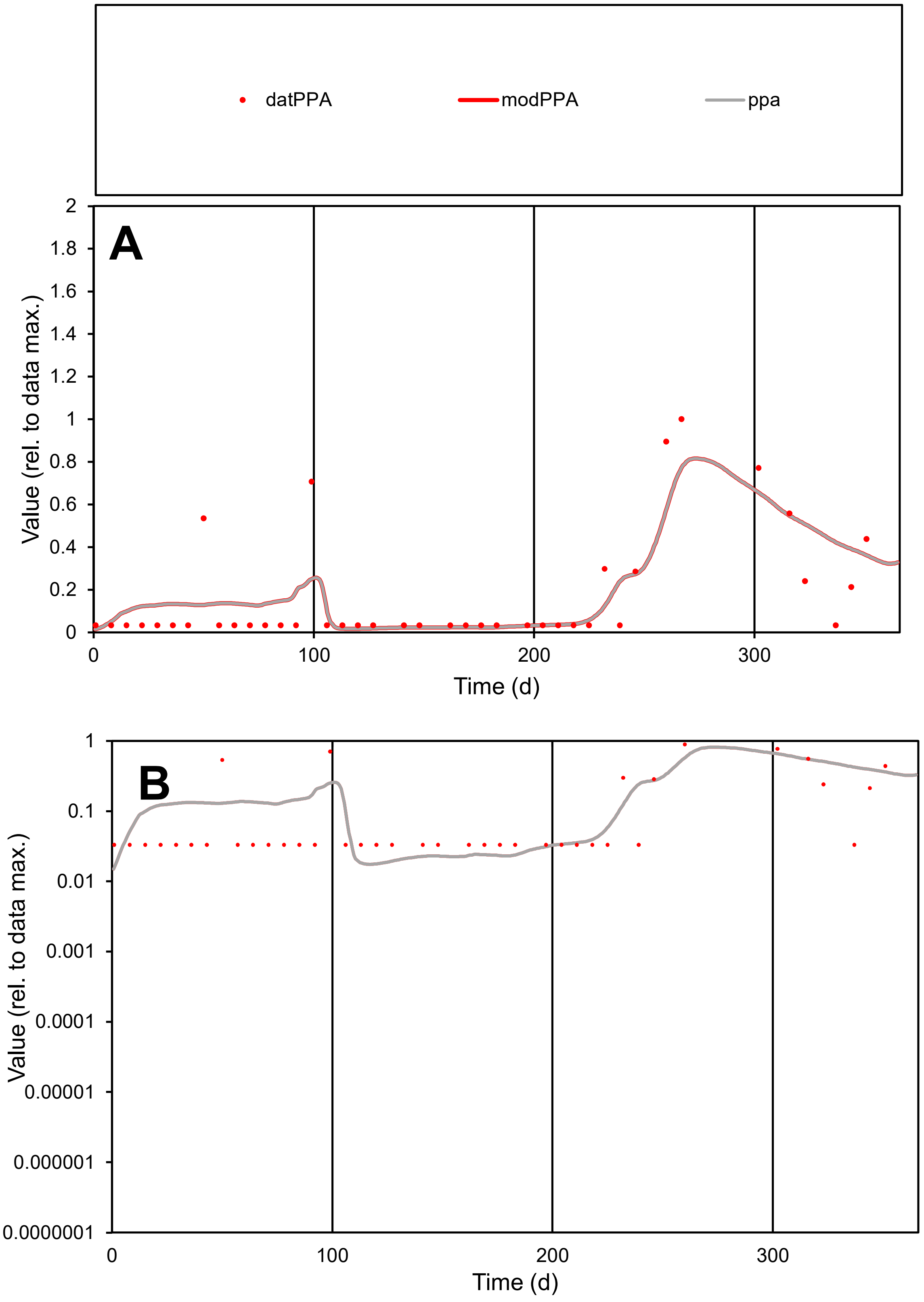


**C1**

Variables

PPA Provide jtd or Dst

CalibOnly?

2 1 to reproduce of

oNormalize?

2 1=no, 2=yes

ofxn calc*

0.88874

ofxn mod

0.91635

sRun

RunA1

zout

1

Ave

Data

1.6E-05

Model

1.7E-05

data/vMax

9.7E-05

ofxng

319.391

Min d/m

1.855

ave/min d

5.10843

ave/min m

9.92541

330_Calib_S_A

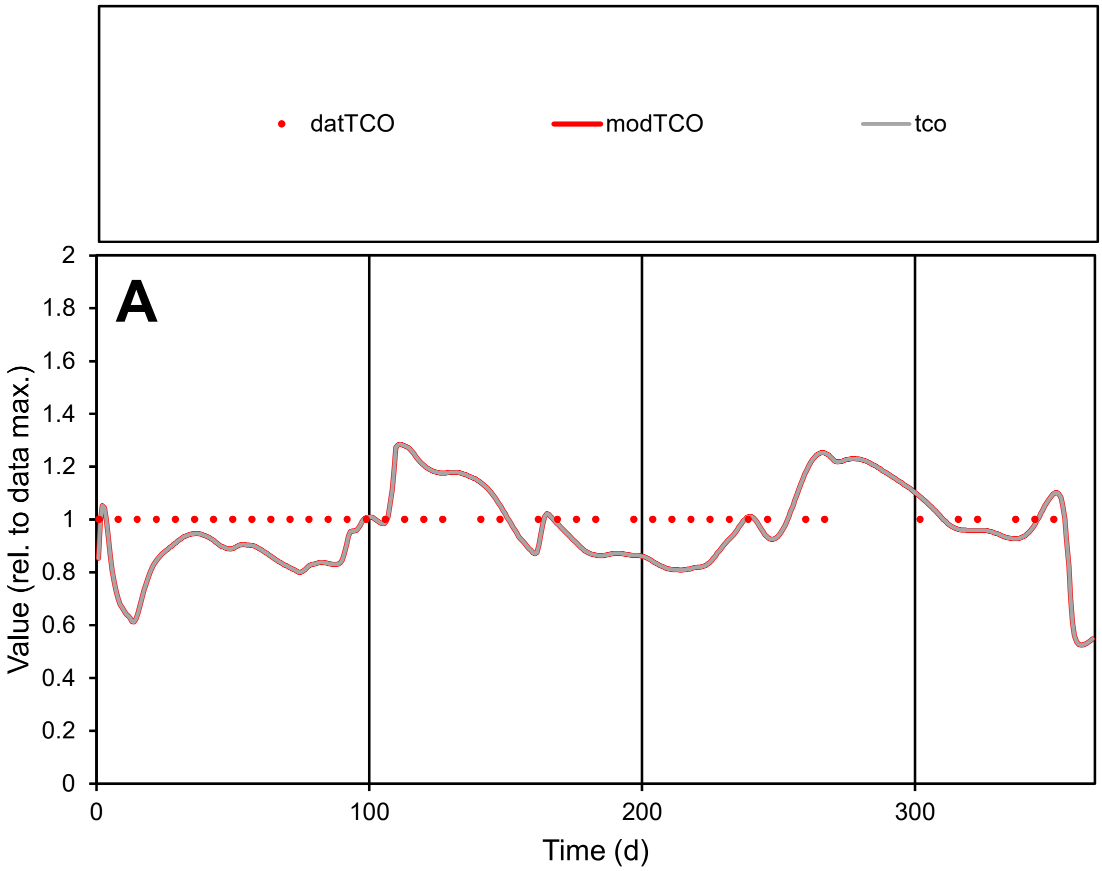


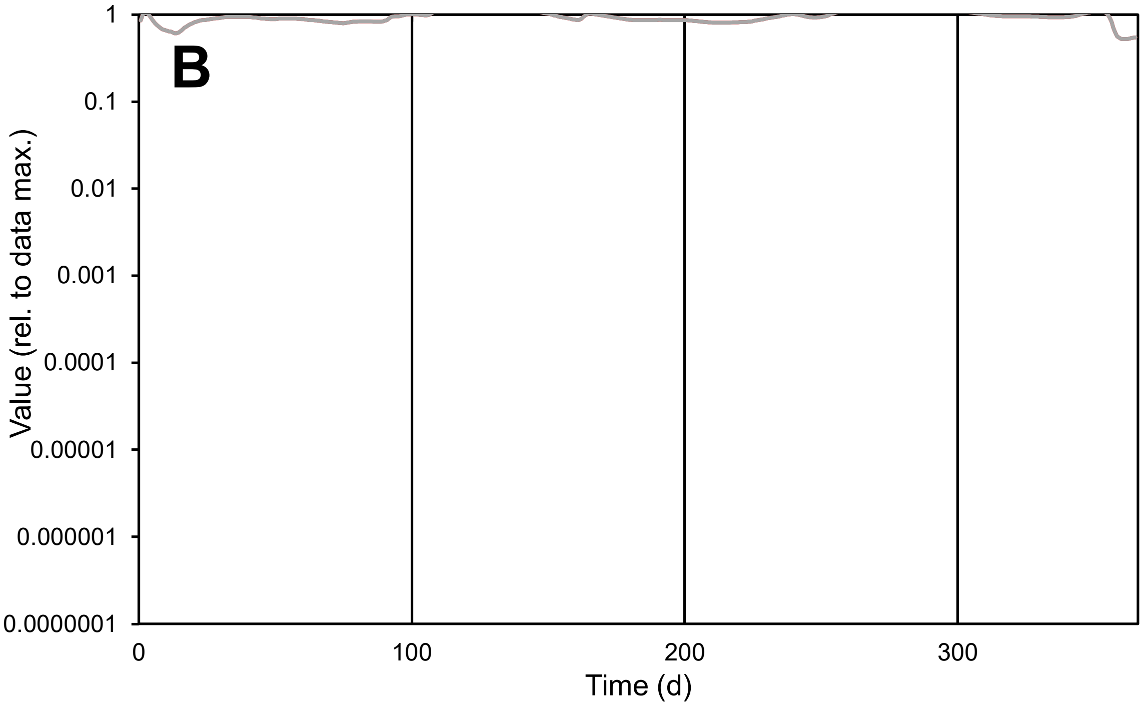


**C1**

Variables
TCO Provide jtd or Dst
CalibOnly?
2 1 to reproduce of
oNormalize?
2 1=no, 2=yes

ofxn calc*
0.14275
ofxn mod
0.17753

sRun
RunA1
zout
1

Ave
Data
8.6E-06
Model
8.2E-06

data/vMax
8.6E-06

ofxng
319.391

Min d/m
1.49835

ave/min d
1
ave/min m
1.43233

330_Calib_S_A

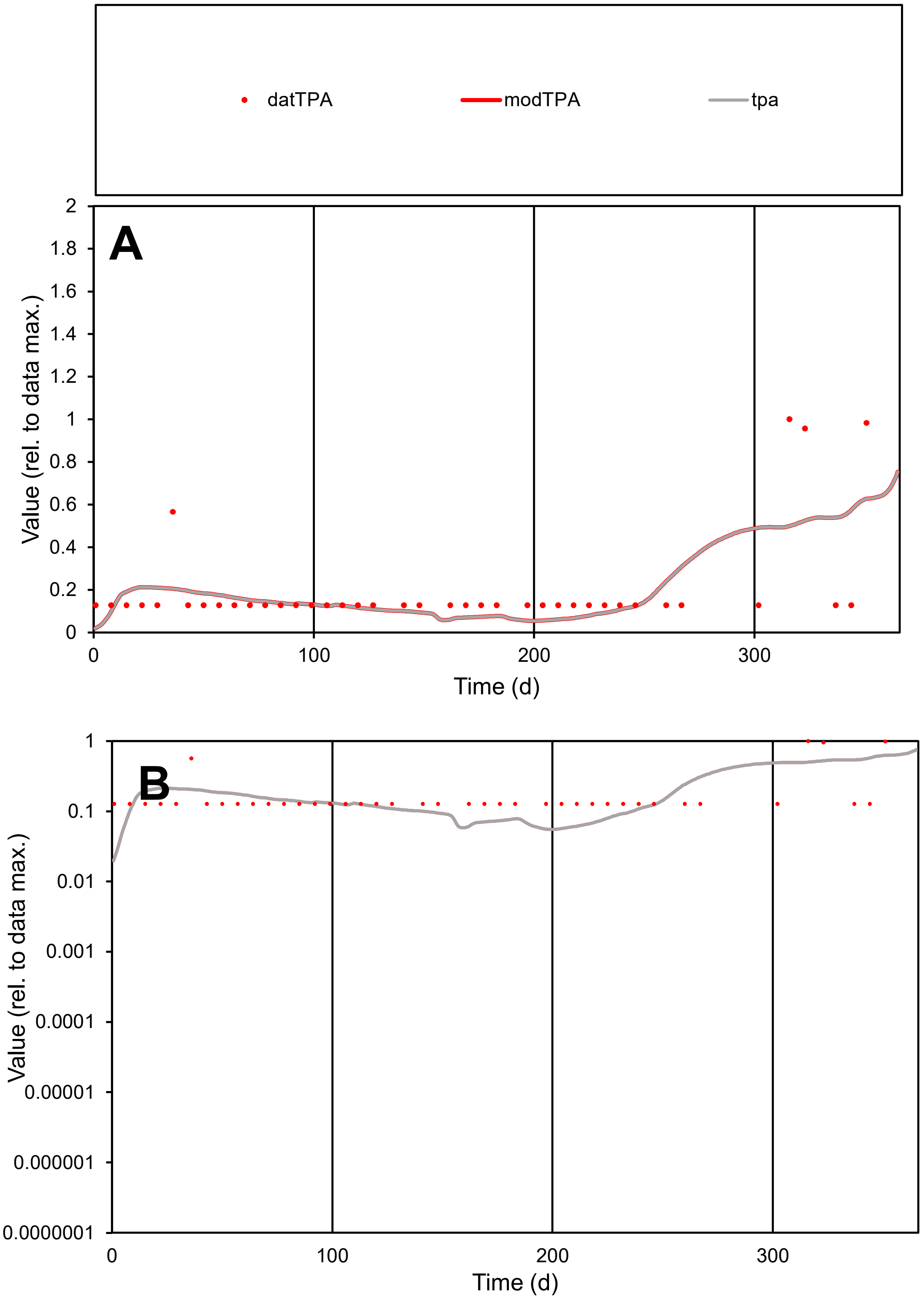


**C1**

Variables
TPA Provide jtd or Dst
CalibOnly?
2 1 to reproduce of
oNormalize?
2 1=no, 2=yes

ofxn calc*
0.89186
ofxn mod
0.90314

sRun
RunA1
zout
1

Ave
Data
7.4E-06
Model
7.1E-06

data/vMax
3.7E-05

ofxng
319.391

Min d/m
3.27828

ave/min d
1.57307
ave/min m
4.91424

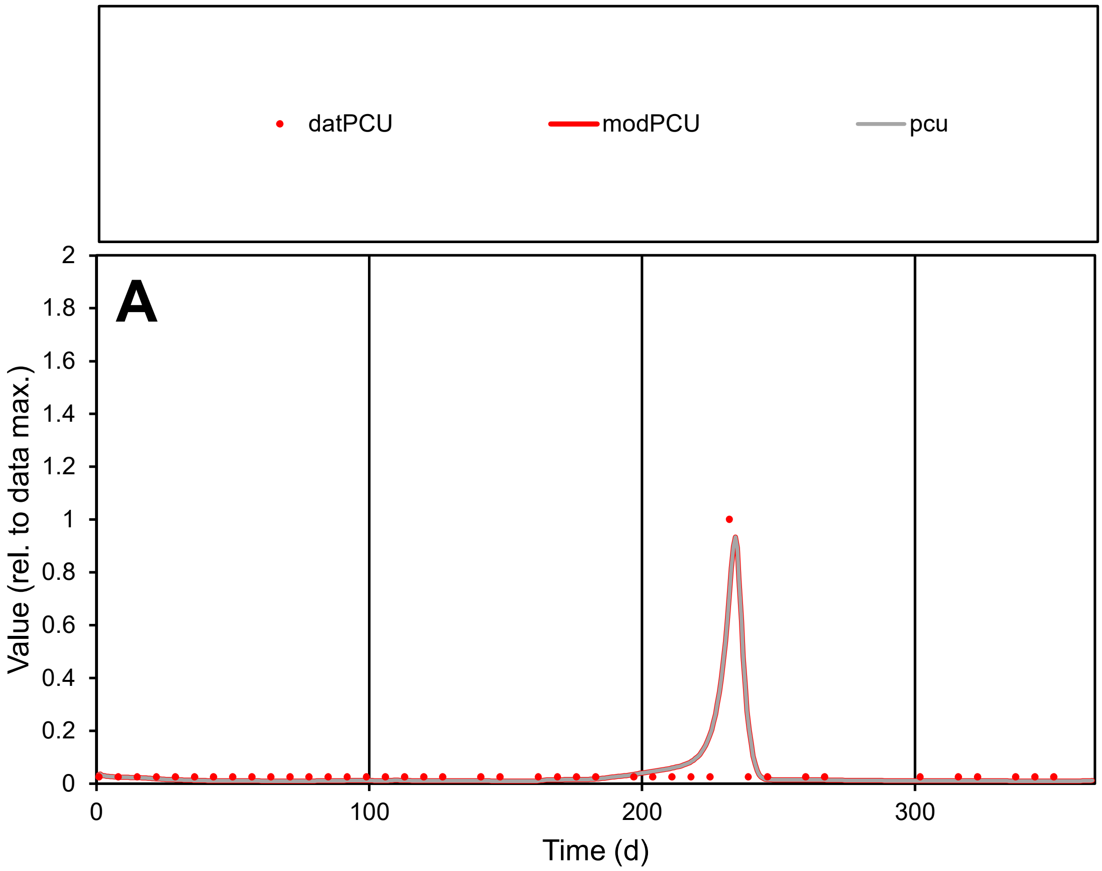


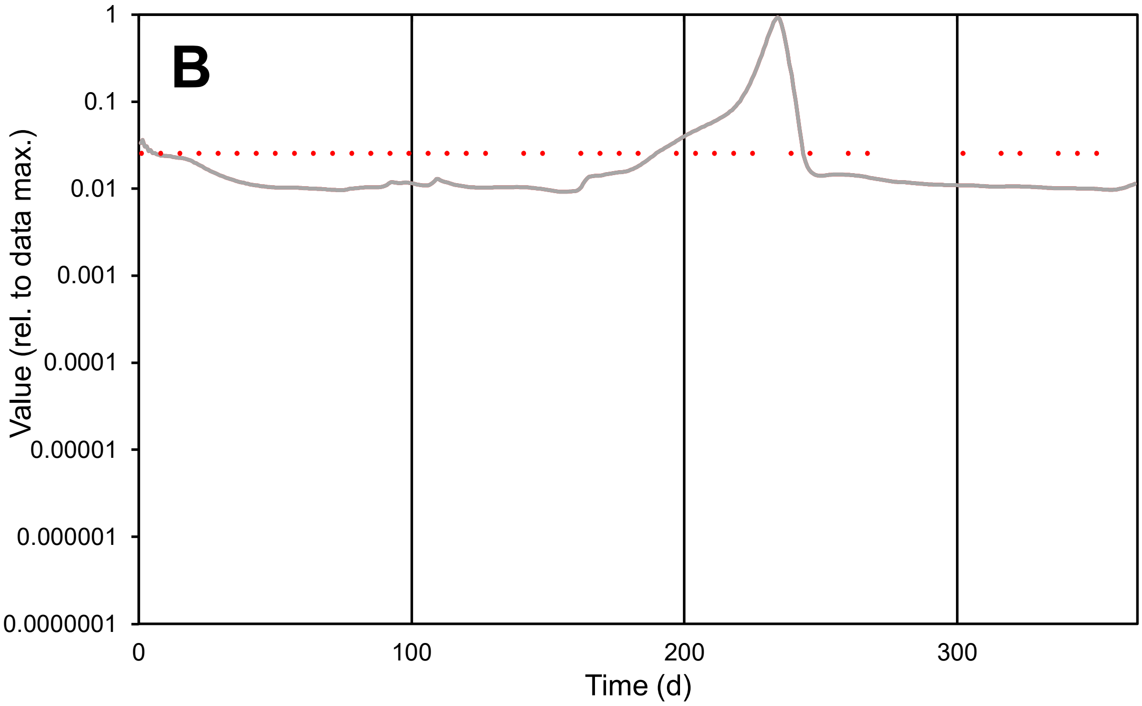


**C1**

Variables

| | |
|---|---|
| PCU | Provide jtd or Dst |
| CalibOnly? | |
| 2 | 1 to reproduce of |
| oNormalize? | |
| 2 | 1=no, 2=yes |
| ofxn calc* | |
| 0.6202 | |
| ofxn mod | |
| 0.82545 | |
| sRun | |
| RunA1 | |
| zout | |
| 1 | |
| Ave | |
| Data | |
| 1.2E-05 | |
| Model | |
| 1.1E-05 | |
| data/vMax | |
| 0.00024 | |
| ofxng | |
| 319.391 | |
| Min d/m | |
| 2.63687 | |
| ave/min d | |
| 1.93326 | |
| ave/min m | |
| 4.51848 | |

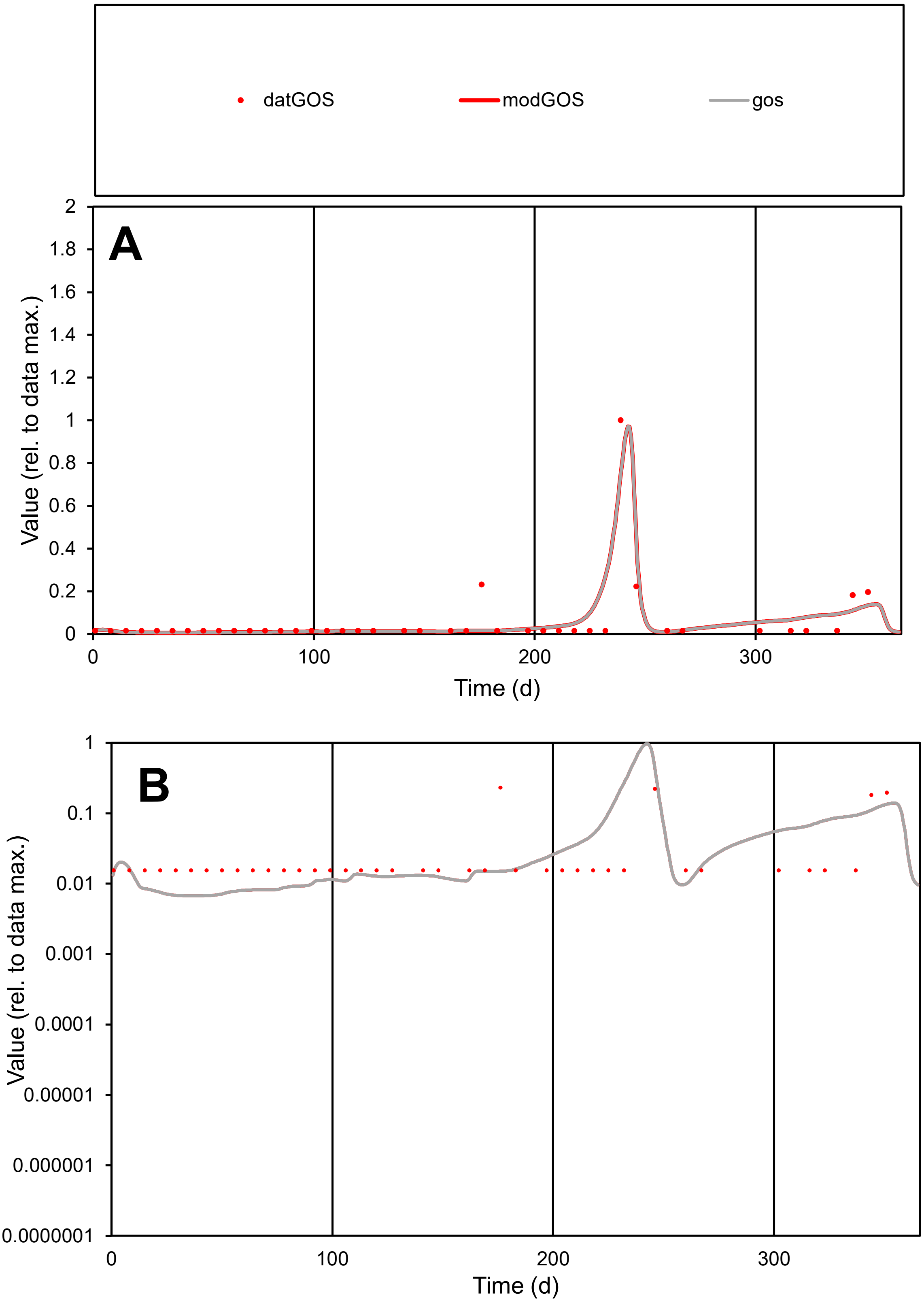


**C1**

Variables

GOS Provide jtd or Dst

CalibOnly?

2 1 to reproduce of

oNormalize?

2 1=no, 2=yes

ofxn calc*

0.959

ofxn mod

1.09533

sRun

RunA1

zout

1

Ave

Data

6.8E-06

Model

7E-06

data/vMax

0.00012

ofxng

319.391

Min d/m

2.30154

ave/min d

3.76477

ave/min m

8.83473

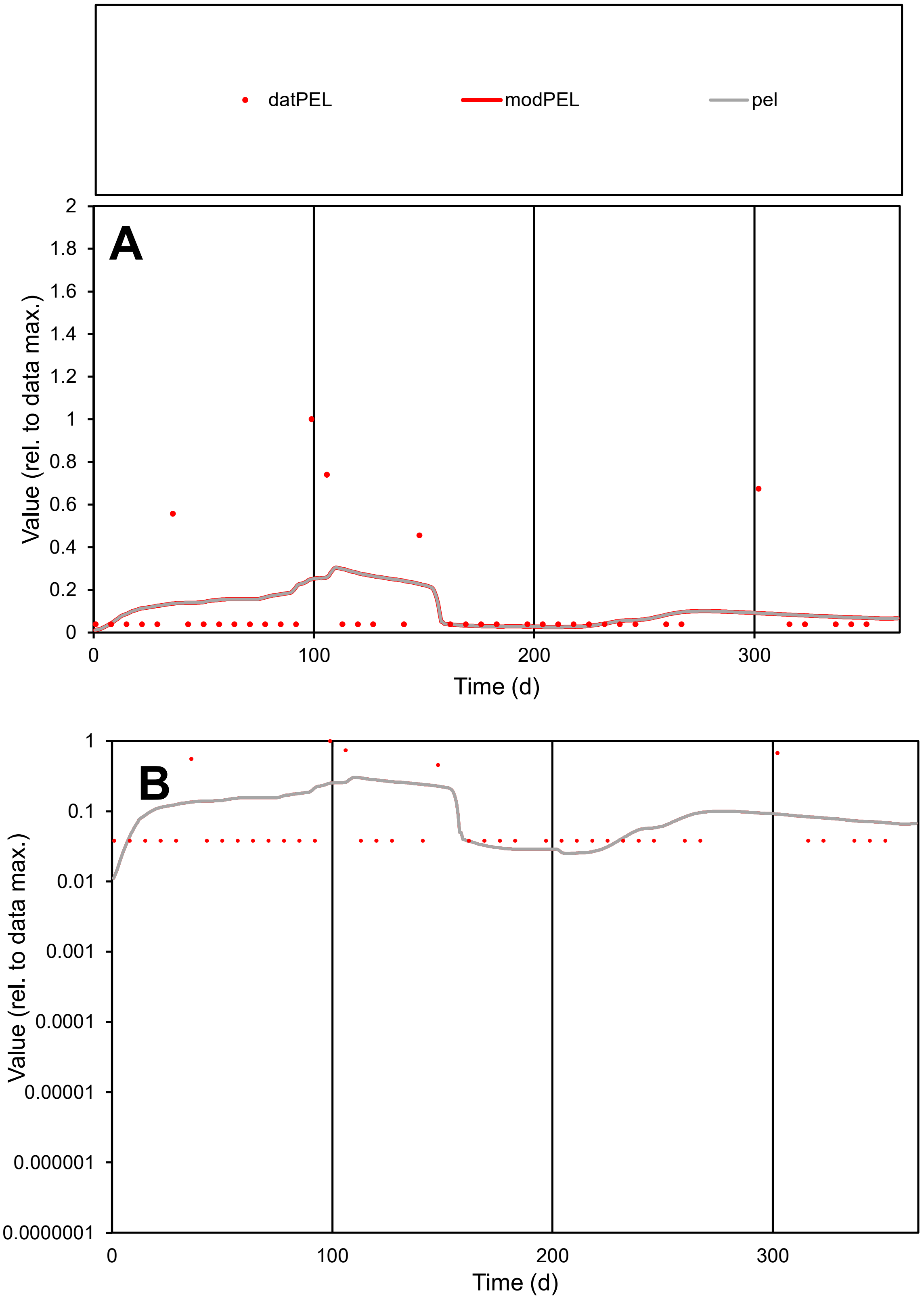


**C1**

Variables

PEL Provide jtd or Dst

CalibOnly?

2 1 to reproduce of

oNormalize?

2 1=no, 2=yes

ofxn calc*

1.74879

ofxn mod

1.75934

sRun

RunA1

zout

1

Ave

Data

1E-05

Model

1E-05

data/vMax

8.6E-05

ofxng

319.391

Min d/m

1.93059

ave/min d

3.08016

ave/min m

5.87301

330_Calib_S_A

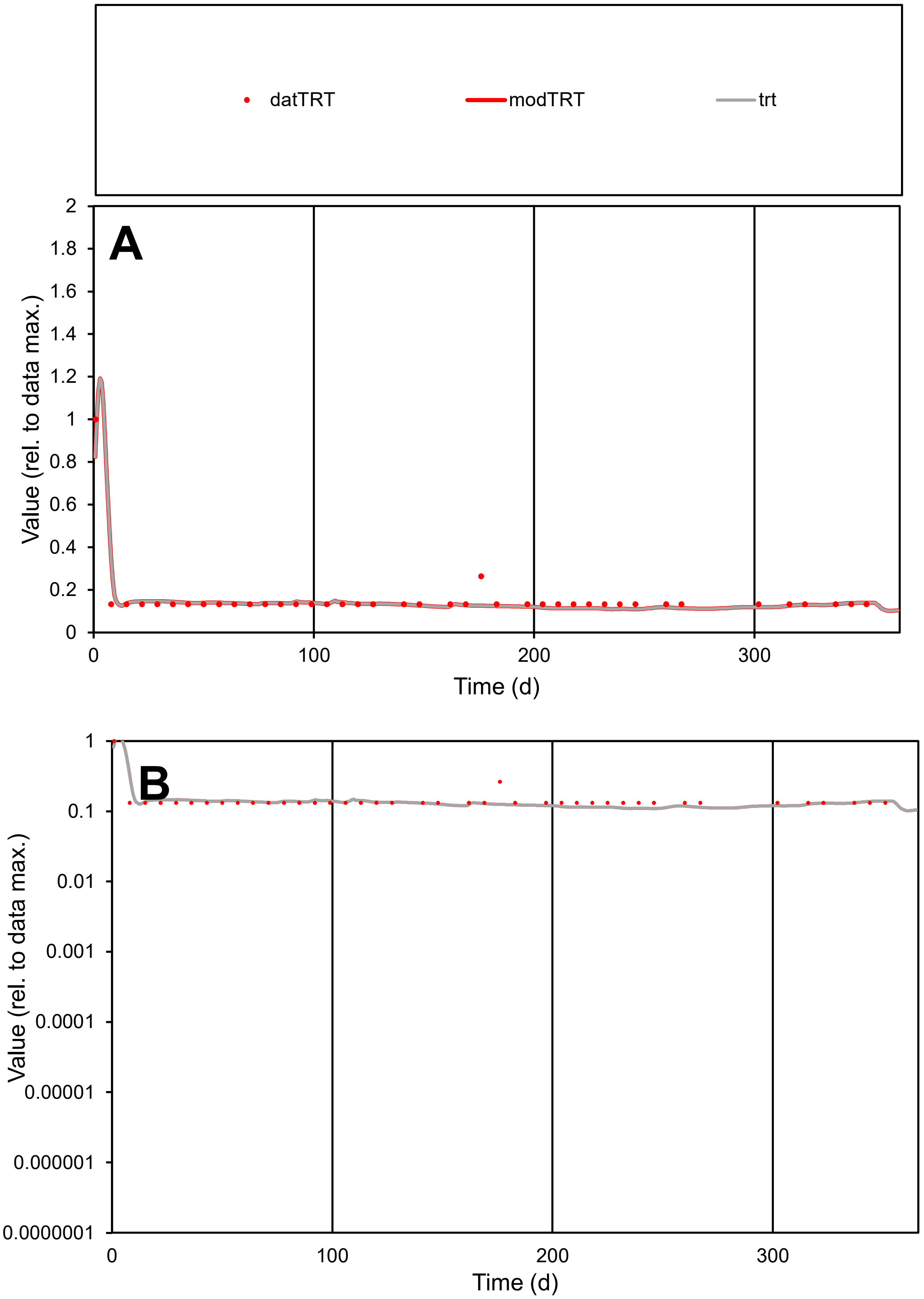


**C1**

Variables
TRT Provide jtd or Dst
CalibOnly?
2 1 to reproduce of
oNormalize?
2 1=no, 2=yes

ofxn calc*
0.16211
ofxn mod
0.20418

sRun
RunA1
zout
1

Ave
Data
2.7E-05
Model
2.7E-05

data/vMax
0.00018

ofxng
319.391

Min d/m
1.20489

ave/min d
1.18506
ave/min m
1.39336

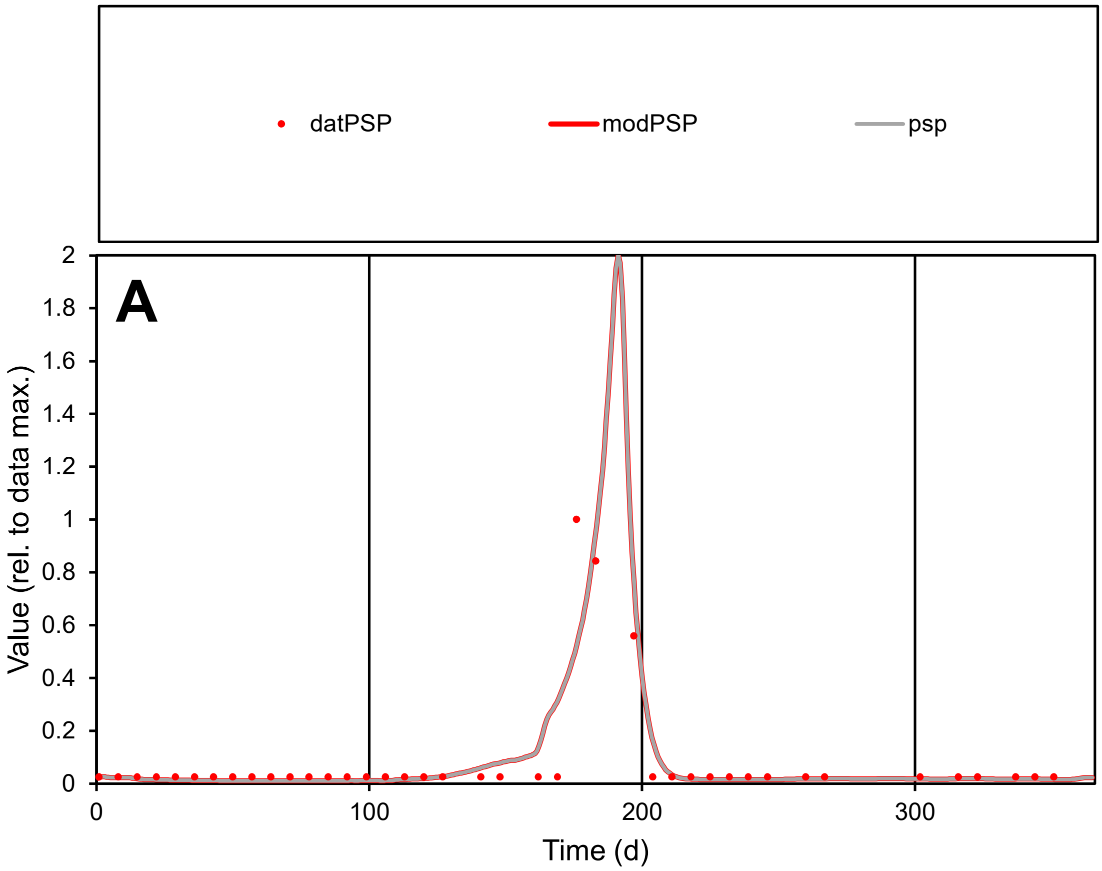


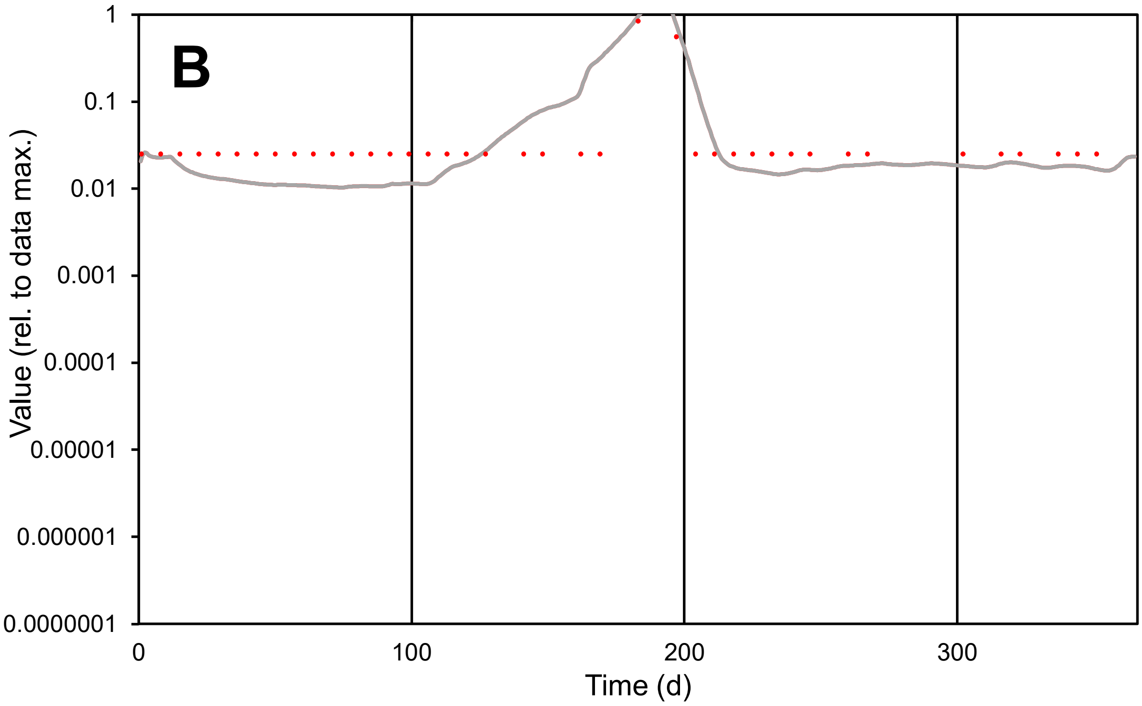


C1

Variables
PSP Provide jtd or Dst
CalibOnly?
2 1 to reproduce of
oNormalize?
2 1=no, 2=yes

ofxn calc*
1.01105
ofxn mod
1.10863

sRun
RunA1
zout
1

Ave
Data
1.4E-05
Model
1.4E-05

data/vMax
0.00017

ofxng
319.391

Min d/m
2.41825

ave/min d
3.25703
ave/min m
7.73063

330_Calib_S_A

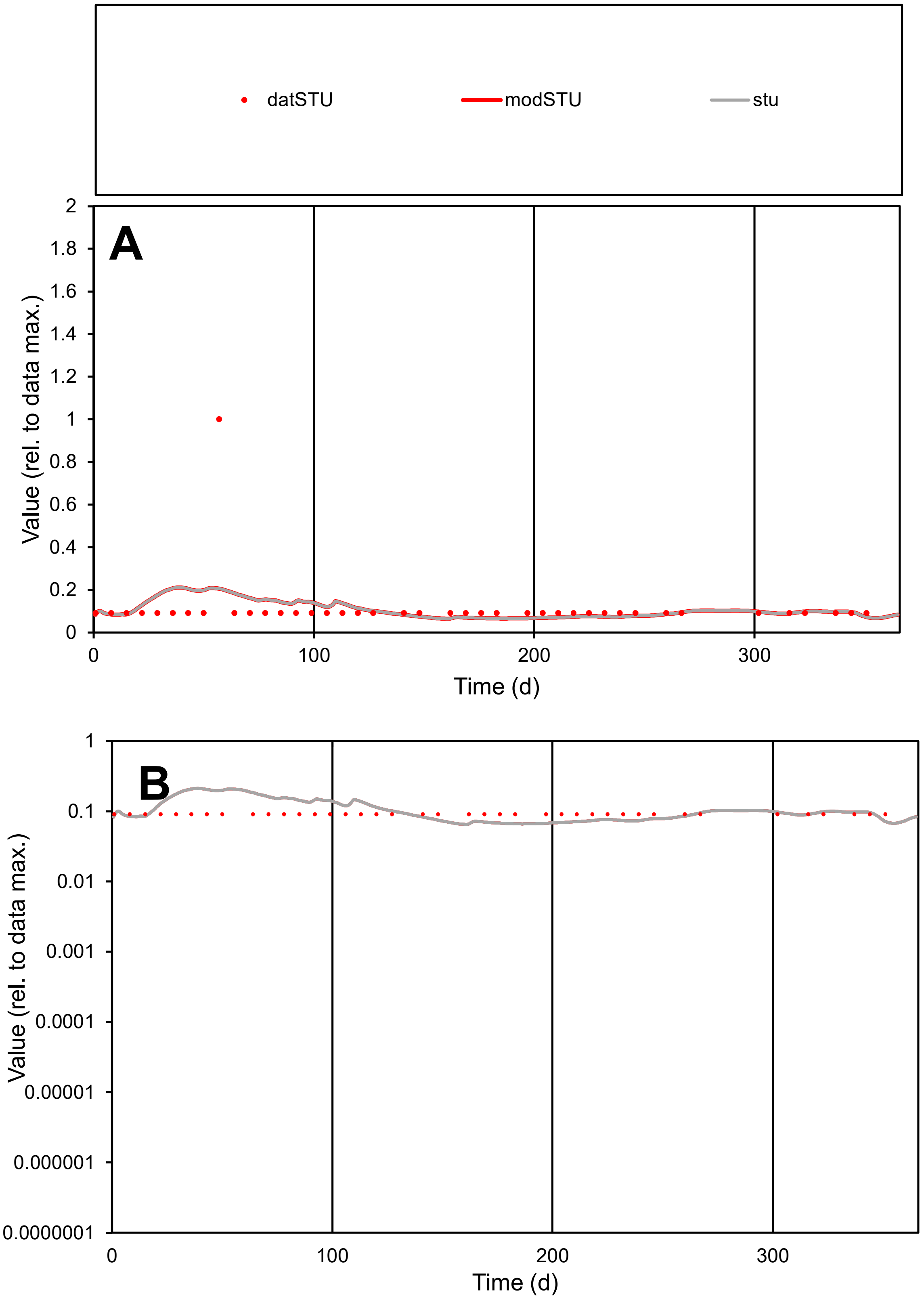


**C1**

Variables
STU Provide jtd or Dst
CalibOnly?
2 1 to reproduce of
oNormalize?
2 1=no, 2=yes

ofxn calc*
1.16839
ofxn mod
1.17657

sRun
RunA1
zout
1

Ave
Data
3.1E-06
Model
3E-06

data/vMax
2.7E-05

ofxng
319.391

Min d/m
1.36678

ave/min d
1.24417
ave/min m
1.66069

330_Calib_S_A

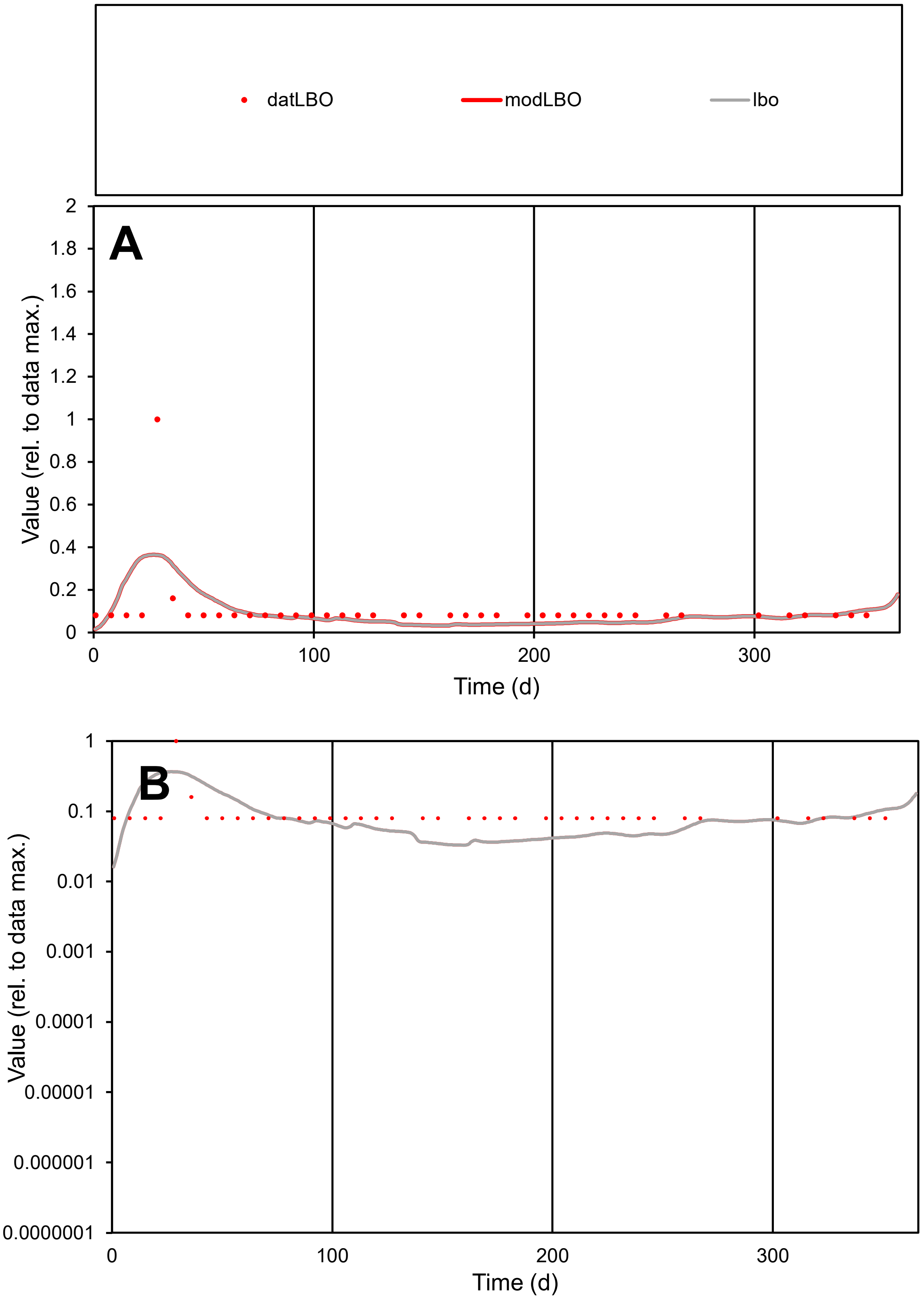


**C1**

Variables
LBO Provide jtd or Dst
CalibOnly?
2 1 to reproduce of
oNormalize?
2 1=no, 2=yes

ofxn calc*
1.15823
ofxn mod
1.17106

sRun
RunA1
zout
1

Ave
Data
1.8E-06
Model
1.6E-06

data/vMax
1.7E-05

ofxng
319.391

Min d/m
2.3163

ave/min d
1.30563
ave/min m
2.75346

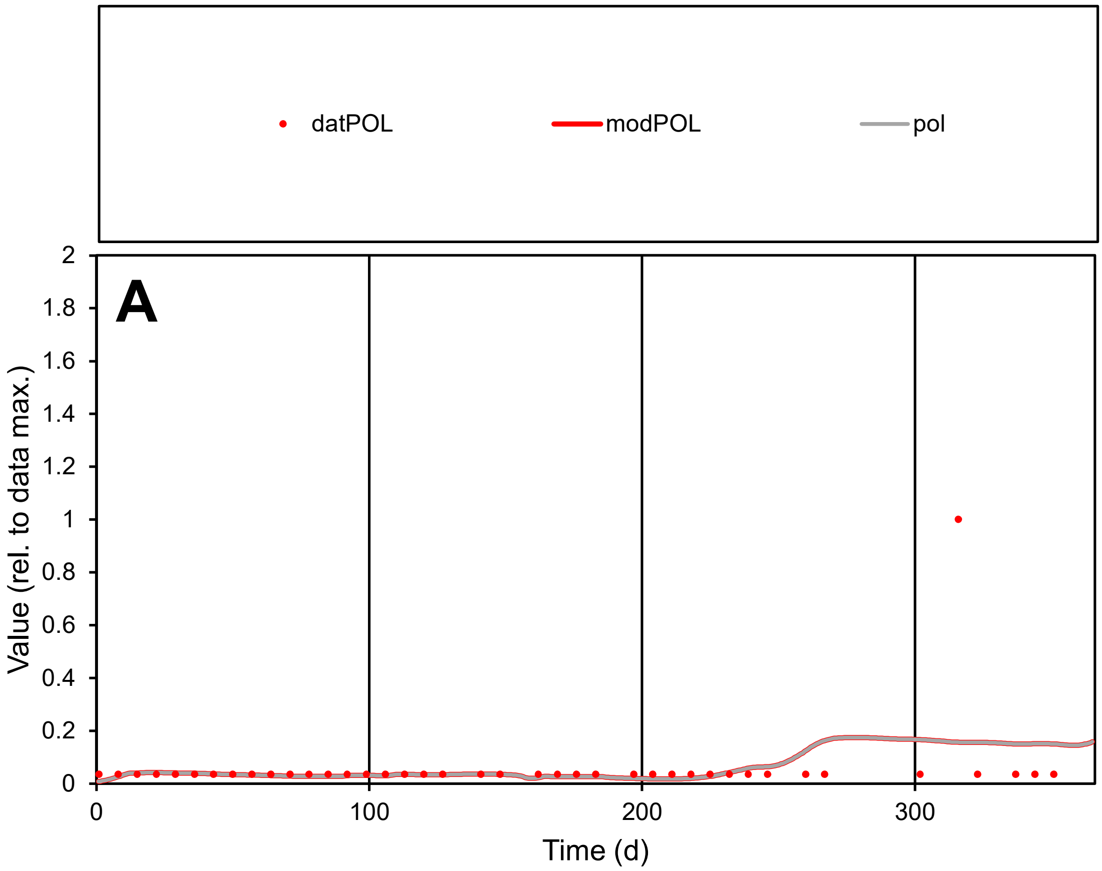


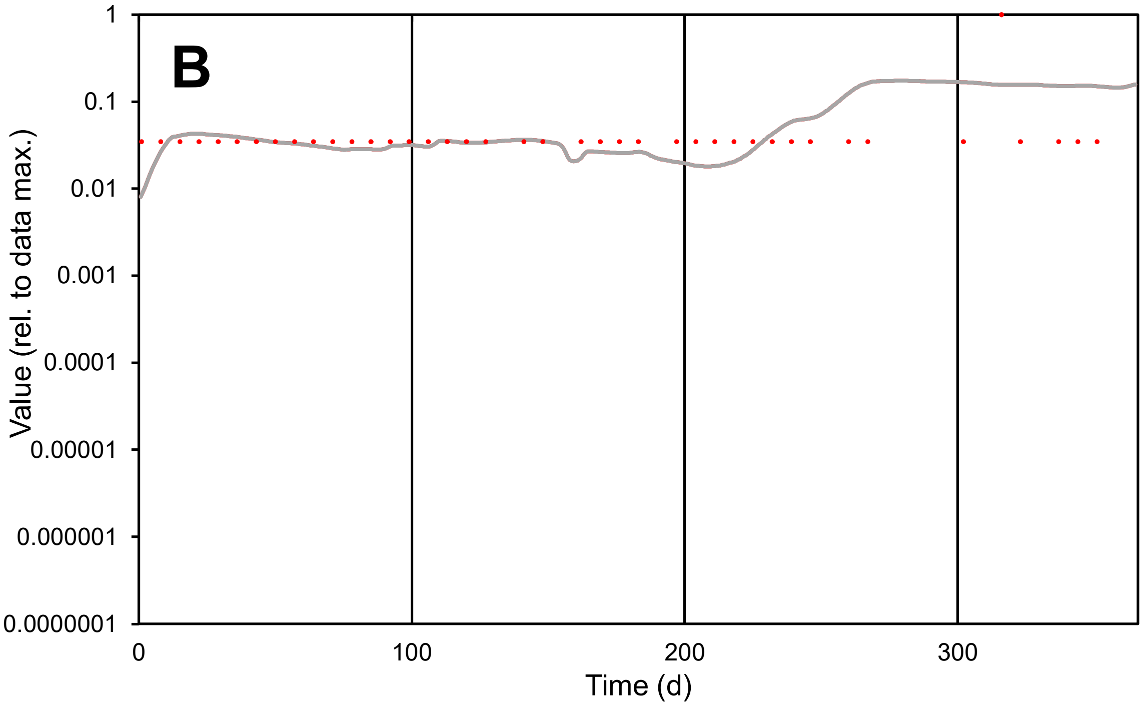


**C1**

| | |
|---|---|
| Variables | |
| POL | Provide jtd or Dst |
| CalibOnly? | |
| 2 | 1 to reproduce of |
| oNormalize? | |
| 2 | 1=no, 2=yes |
| ofxn calc* | |
| 2.41897 | |
| ofxn mod | |
| 2.42371 | |
| sRun | |
| RunA1 | |
| zout | |
| 1 | |
| Ave | |
| Data | |
| 2.7E-06 | |
| Model | |
| 2.6E-06 | |
| data/vMax | |
| 4.6E-05 | |
| ofxng | |
| 319.391 | |
| Min d/m | |
| 2.57752 | |
| ave/min d | |
| 1.6792 | |
| ave/min m | |
| 4.17449 | |

330_Calib_S_A

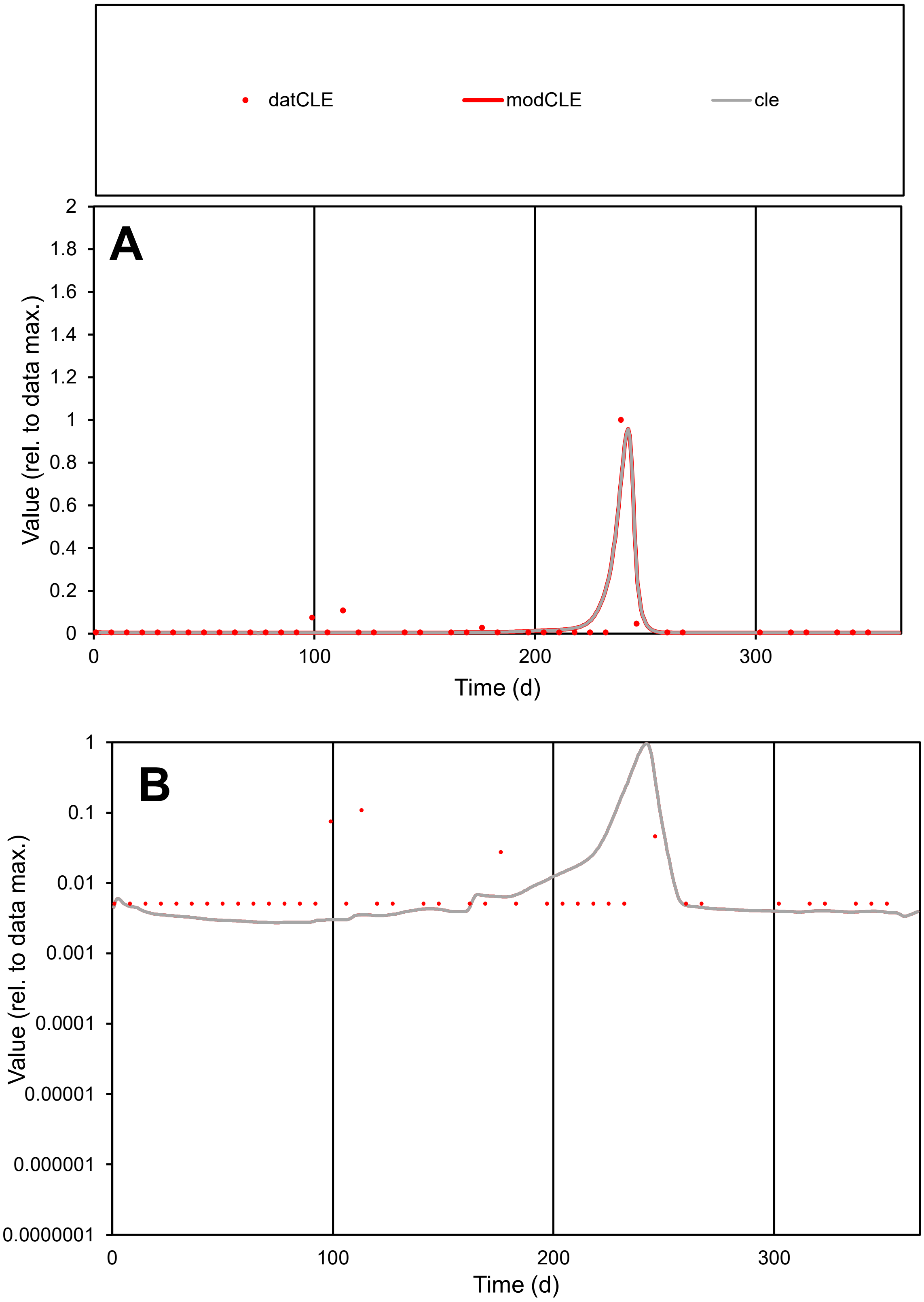


**C1**

Variables
CLE Provide jtd or Dst
CalibOnly?
2 1 to reproduce of
oNormalize?
2 1=no, 2=yes

ofxn calc*
0.96343
ofxn mod
1.32801

sRun
RunA1
zout
1

Ave
Data
3.2E-05
Model
3E-05

data/vMax
0.00092

ofxng
319.391

Min d/m
1.86349

ave/min d
6.90305
ave/min m
12.0053

330_Calib_S_A

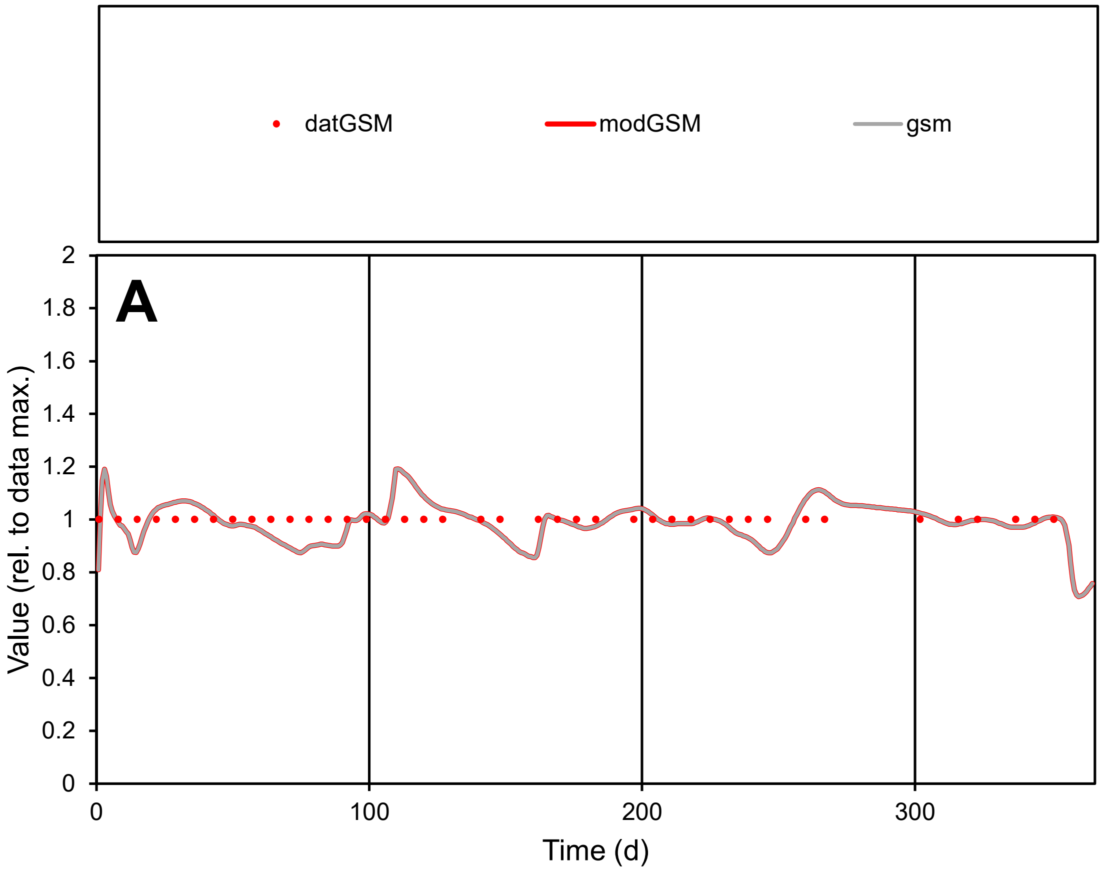


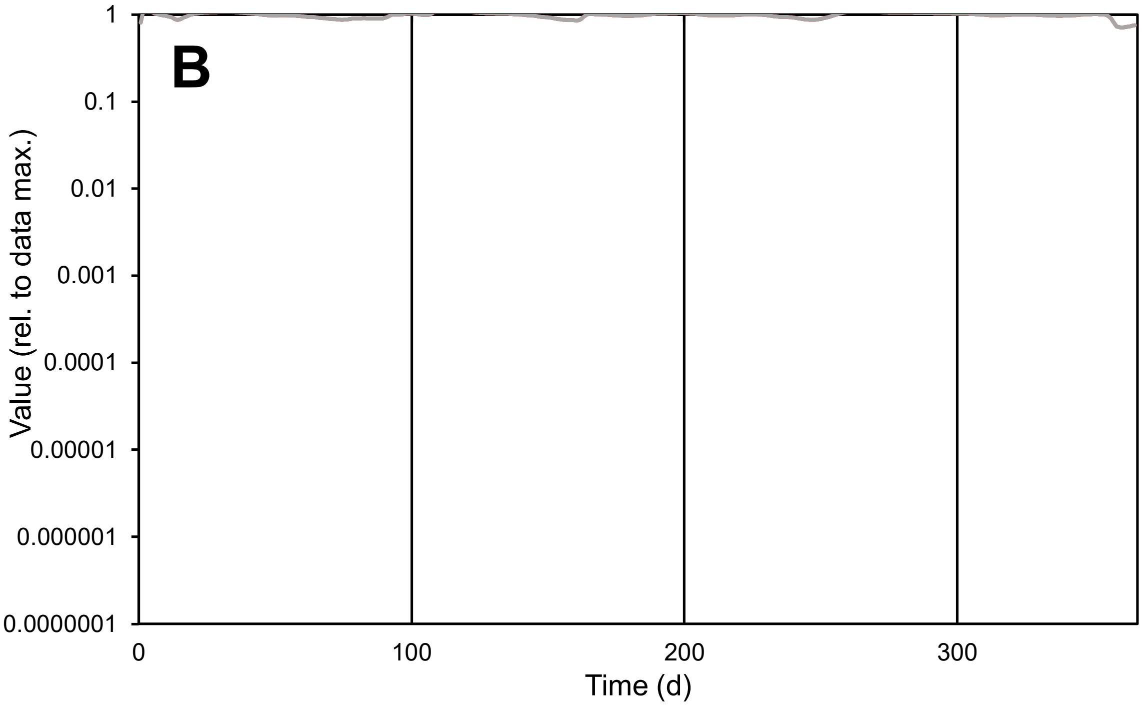


**C1**

Variables
GSM Provide jtd or Dst
CalibOnly?
2 1 to reproduce of
oNormalize?
2 1=no, 2=yes

ofxn calc*
0.05861
ofxn mod
0.1232

sRun
RunA1
zout
1

Ave
Data
1.6E-06
Model
1.6E-06

data/vMax
1.6E-06

ofxng
319.391

Min d/m
1.14255

ave/min d
1
ave/min m
1.13623

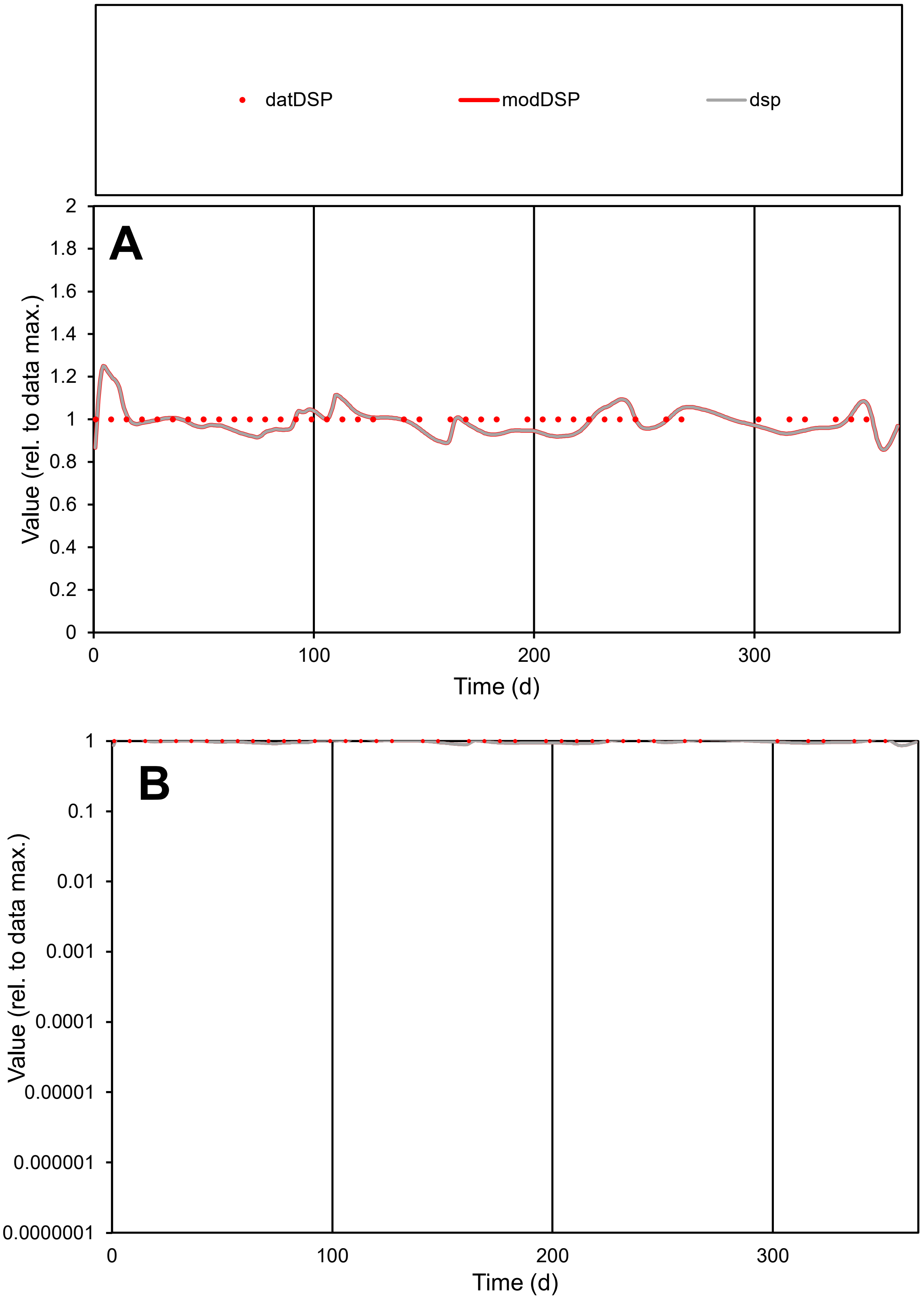


**C1**

Variables
DSP Provide jtd or Dst
CalibOnly?
2 1 to reproduce of
oNormalize?
2 1=no, 2=yes

ofxn calc*
0.05523
ofxn mod
0.12194

sRun
RunA1
zout
1

Ave
Data
5E-06
Model
5E-06

data/vMax
5E-06

ofxng
319.391

Min d/m
1.08663

ave/min d
1
ave/min m
1.07762

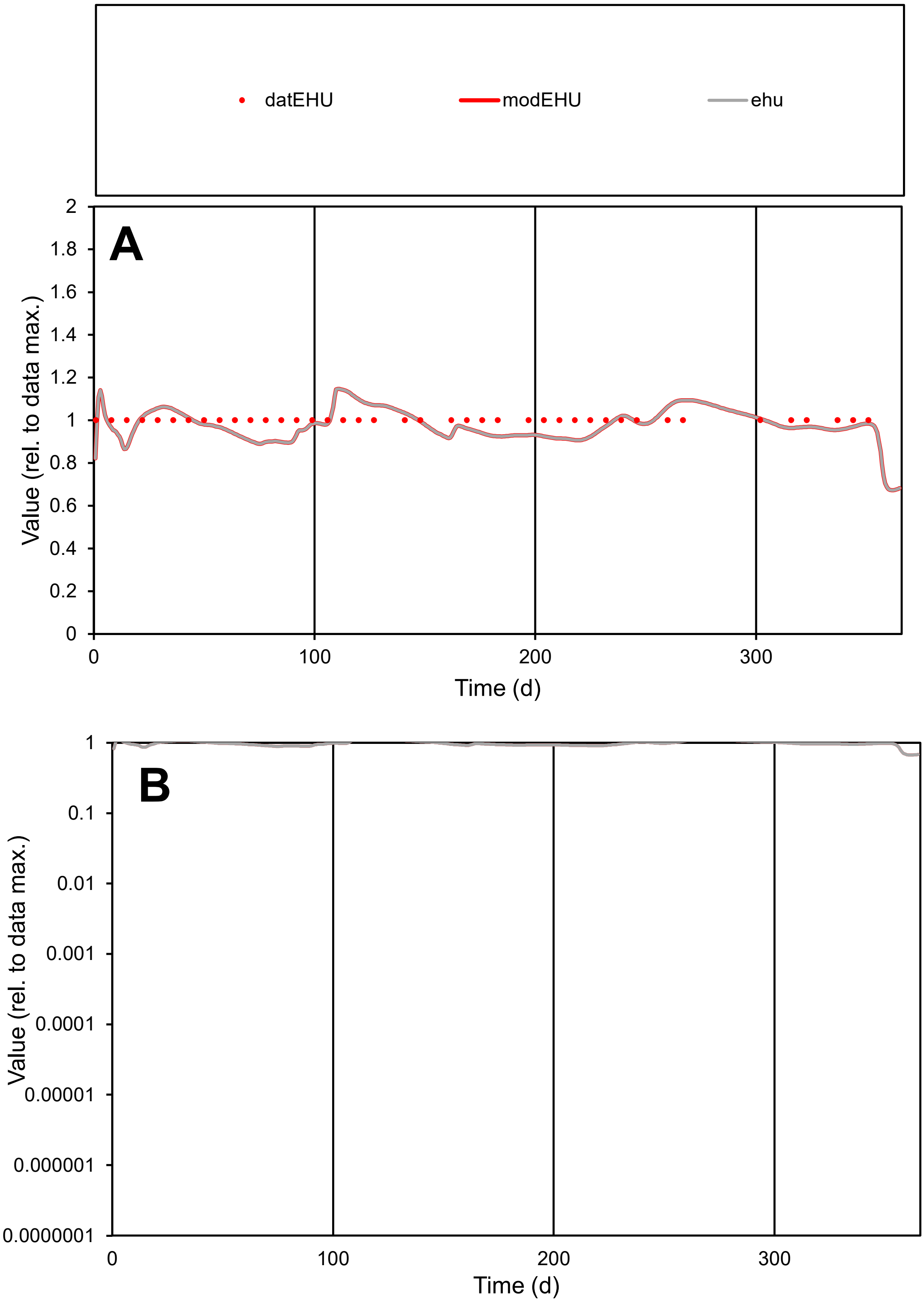


**C1**

Variables
EHU Provide jtd or Dst
CalibOnly?
2 1 to reproduce of
oNormalize?
2 1=no, 2=yes

ofxn calc*
0.06129
ofxn mod
0.12398

sRun
RunA1
zout
1

Ave
Data
5.8E-06
Model
5.6E-06

data/vMax
5.8E-06

ofxng
319.391

Min d/m
1.11596

ave/min d
1
ave/min m
1.09502

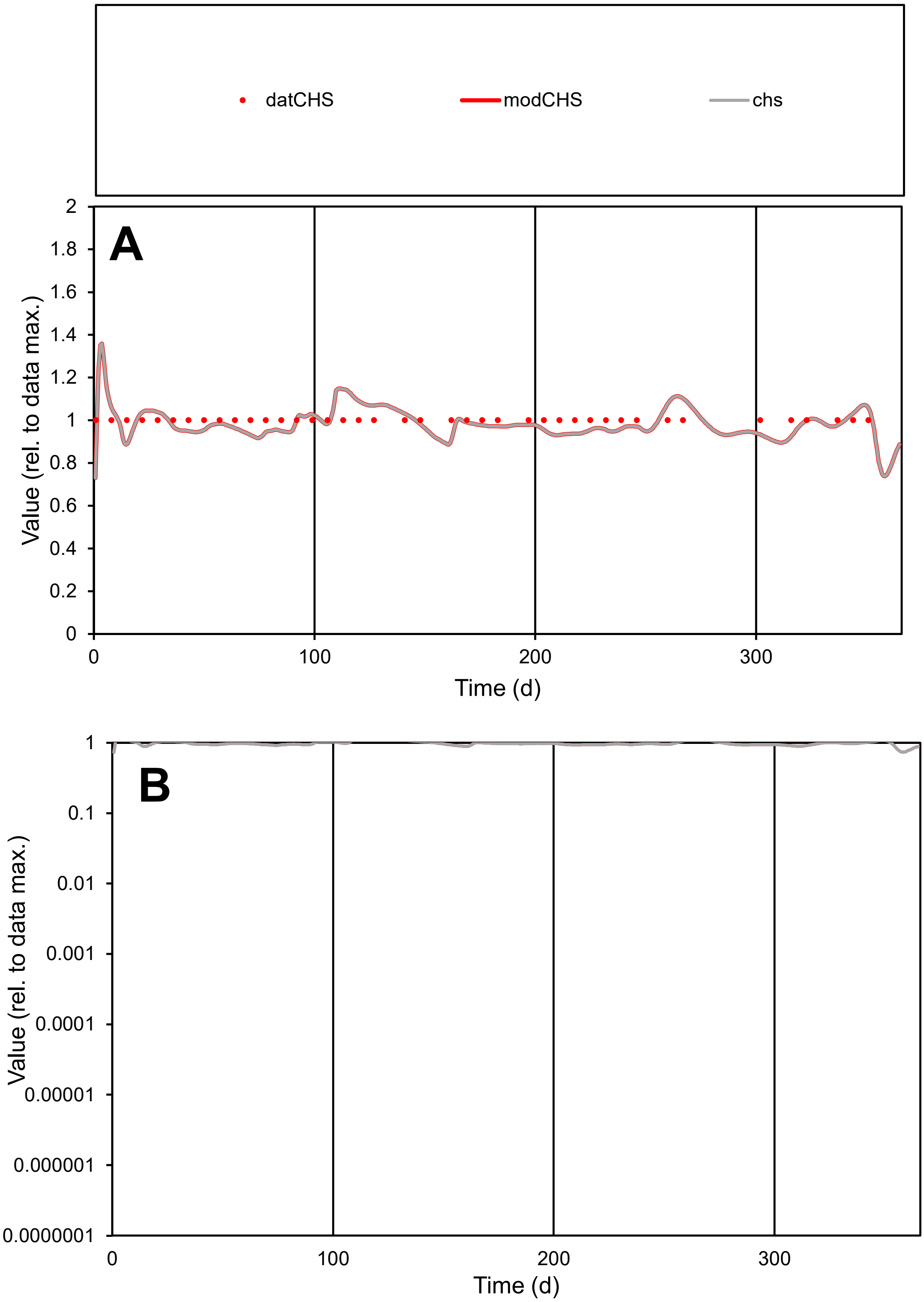


**C1**

Variables
CHS Provide jtd or Dst
CalibOnly?
2 1 to reproduce of
oNormalize?
2 1=no, 2=yes

ofxn calc*
0.05223
ofxn mod
0.12015

sRun
RunA1
zout
1

Ave
Data
2.1E-05
Model
2.1E-05

data/vMax
2.1E-05

ofxng
319.391

Min d/m
1.08085

ave/min d
1
ave/min m
1.07315

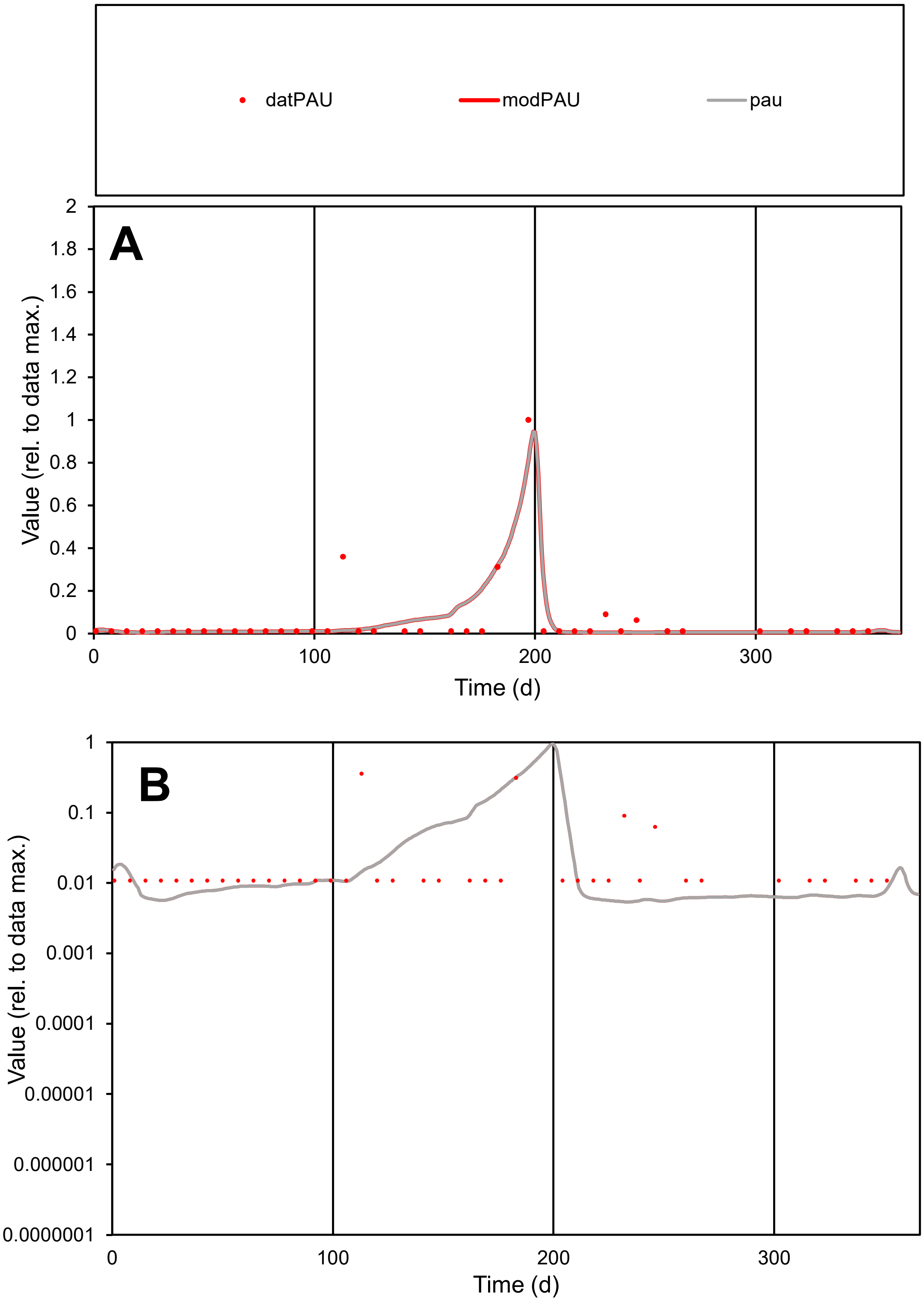


**C1**

Variables

PAU Provide jtd or Dst

CalibOnly?

2 1 to reproduce of

oNormalize?

2 1=no, 2=yes

ofxn calc*

1.31348

ofxn mod

1.38391

sRun

RunA1

zout

1

Ave

Data

2E-05

Model

1.9E-05

data/vMax

0.00036

ofxng

319.391

Min d/m

2.01688

ave/min d

4.98969

ave/min m

9.95065

330_Calib_S_A

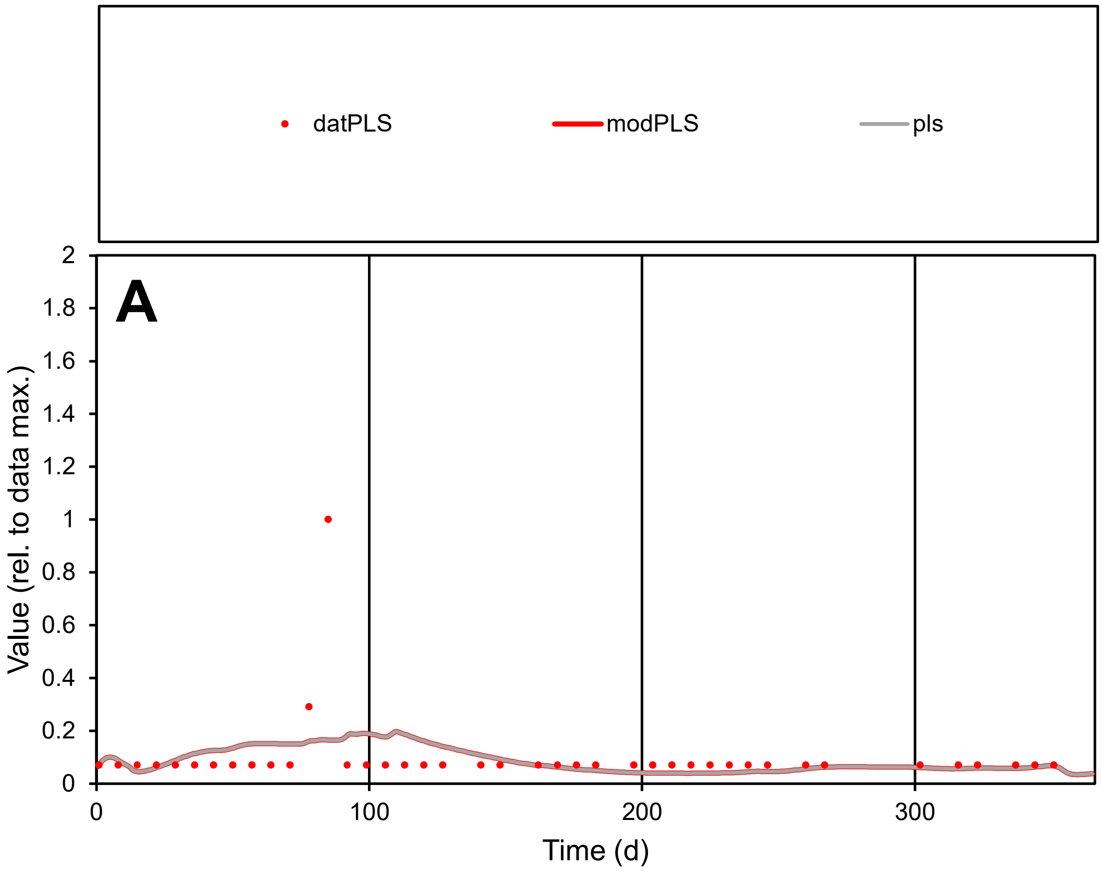


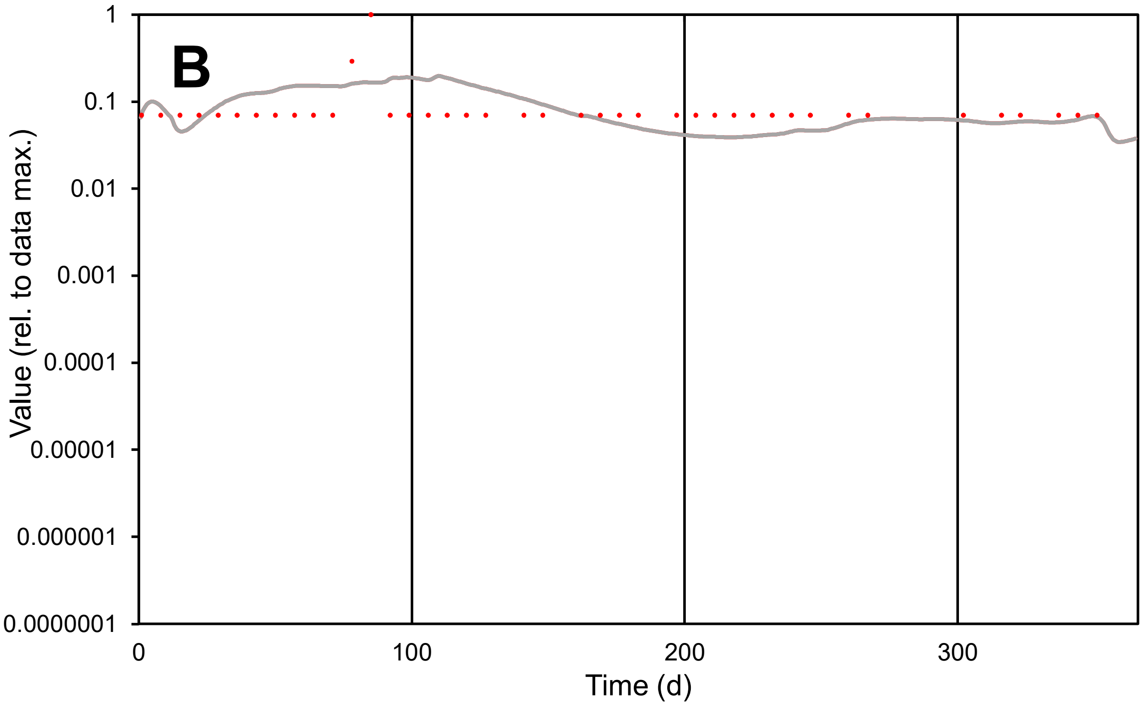


**C1**

Variables

PLS Provide jtd or Dst

CalibOnly?

2 1 to reproduce of

oNormalize?

2 1=no, 2=yes

ofxn calc*

1.4403

ofxn mod

1.44895

sRun

RunA1

zout

1

Ave

Data

5.4E-06

Model

5.1E-06

data/vMax

5.5E-05

ofxng

319.391

Min d/m

1.79151

ave/min d

1.40162

ave/min m

2.37052

330_Calib_S_A

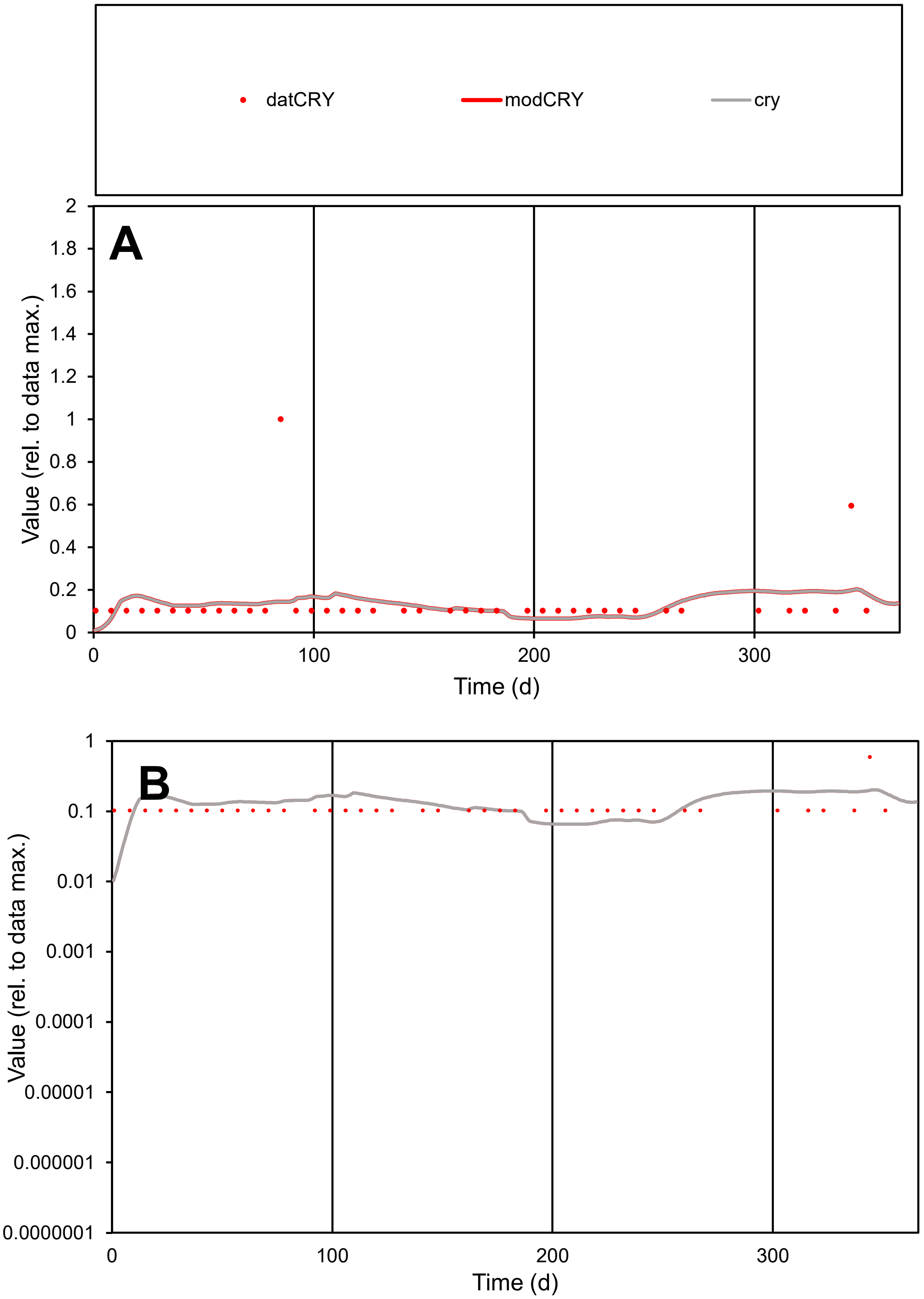


**C1**

Variables
CRY Provide jtd or Dst
CalibOnly?
2 1 to reproduce of
oNormalize?
2 1=no, 2=yes

ofxn calc*
1.13068
ofxn mod
1.13884

sRun
RunA1
zout
1

Ave
Data
4E-06
Model
3.7E-06

data/vMax
2.9E-05

ofxng
319.391

Min d/m
4.66635

ave/min d
1.32908
ave/min m
5.84134

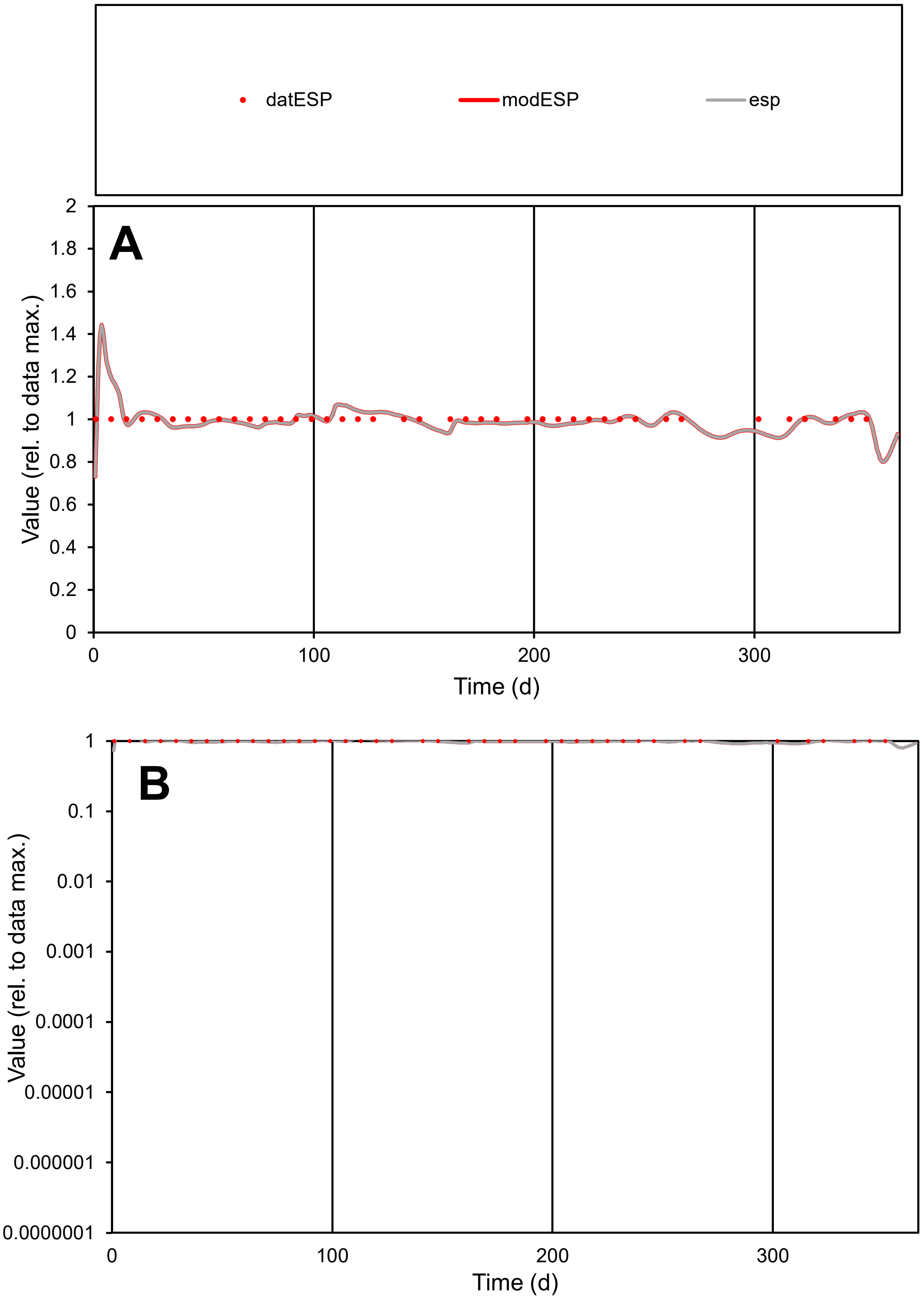


**C1**

Variables
ESP Provide jtd or Dst
CalibOnly?
2 1 to reproduce of
oNormalize?
2 1=no, 2=yes

ofxn calc*
0.03662
ofxn mod
0.11502

sRun
RunA1
zout
1

Ave
Data
8E-06
Model
8E-06

data/vMax
8E-06

ofxng
319.391

Min d/m
1.06875

ave/min d
1
ave/min m
1.06823

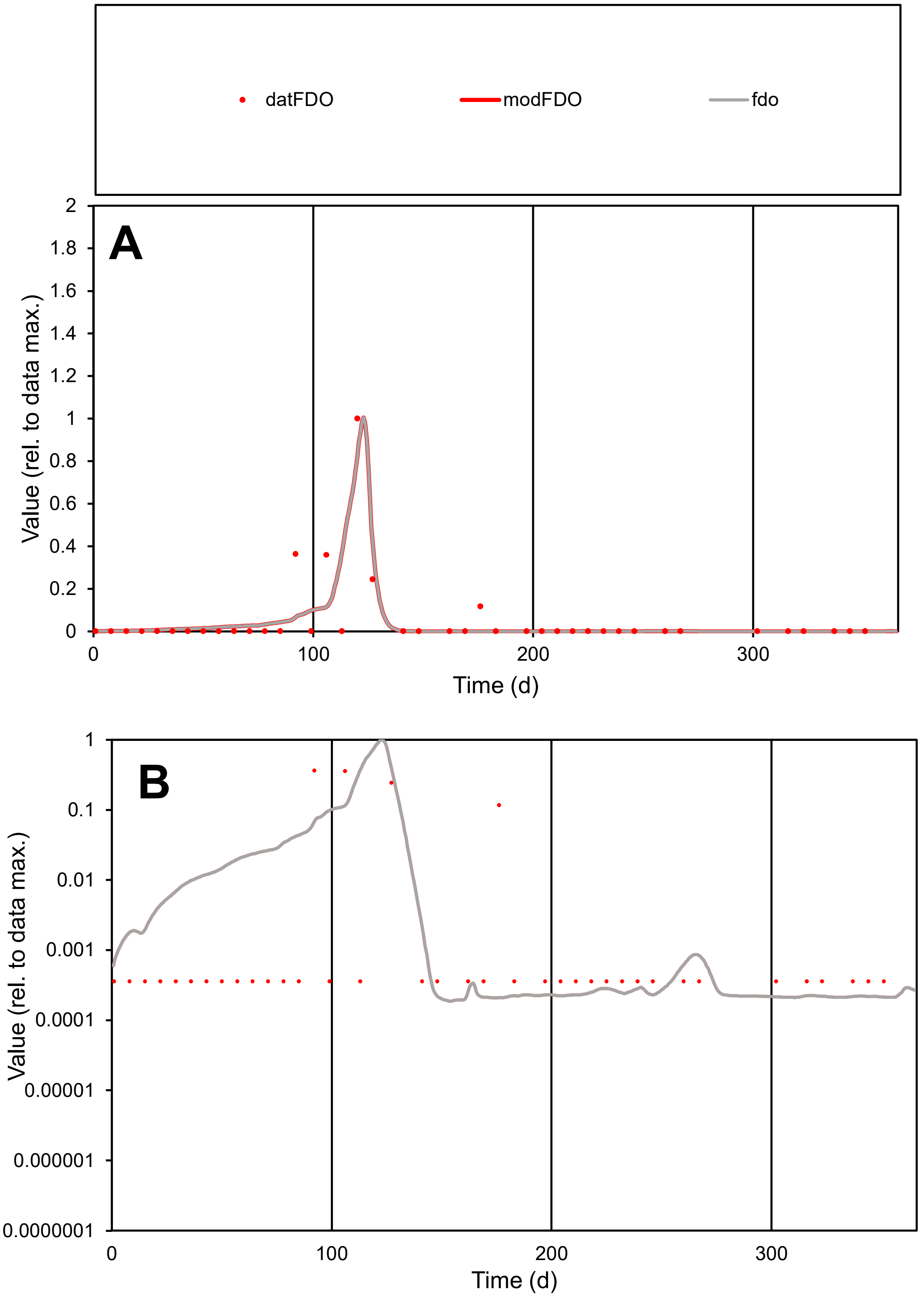


**C1**

Variables

FDO Provide jtd or Dst

CalibOnly?

2 1 to reproduce of

oNormalize?

2 1=no, 2=yes

ofxn calc*

1.50668

ofxn mod

1.61125

sRun

RunA1

zout

1

Ave

Data

1.4E-05

Model

1.4E-05

data/vMax

0.00028

ofxng

319.391

Min d/m

1.71421

ave/min d

143.114

ave/min m

237.136

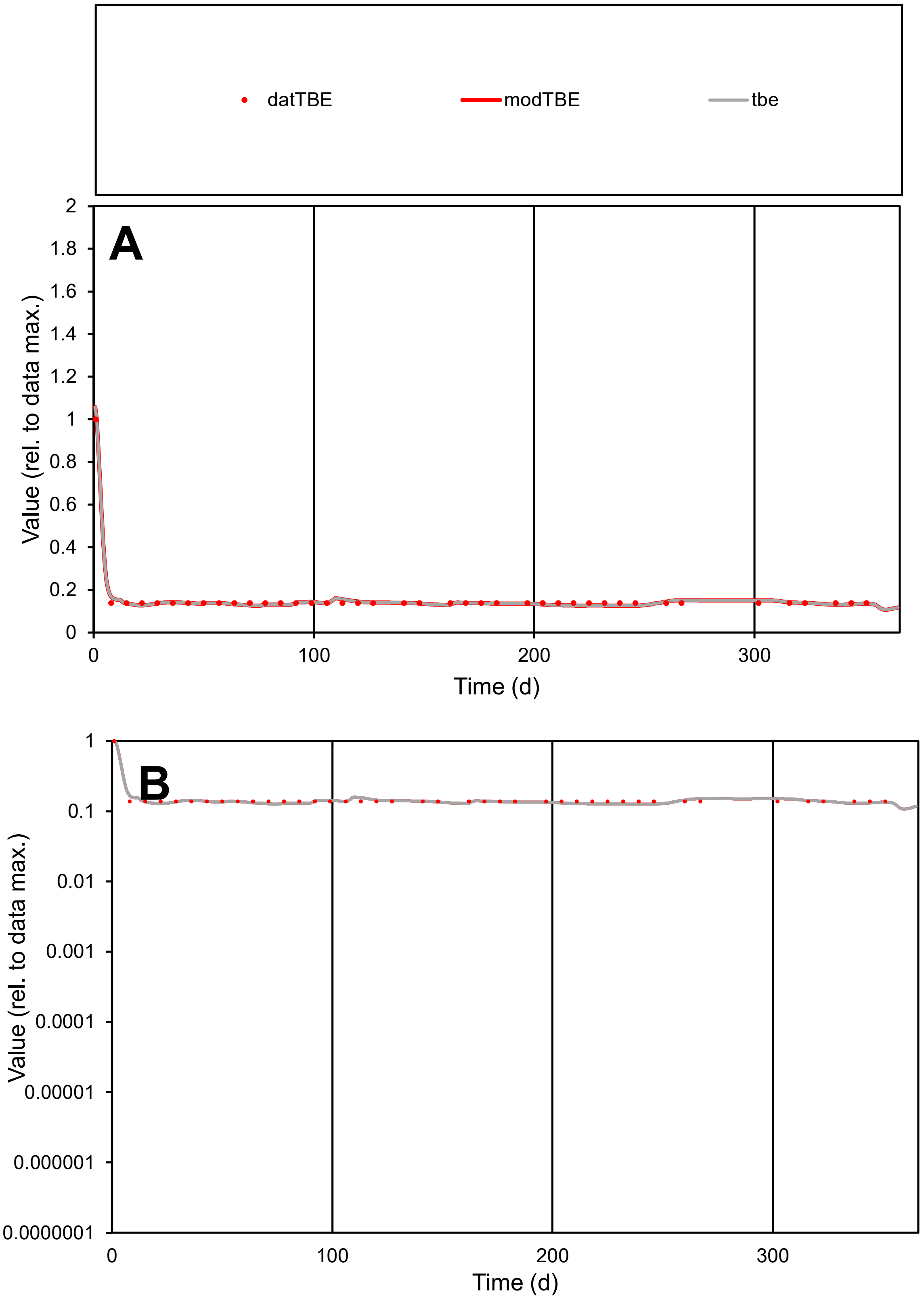


**C1**

Variables
TBE Provide jtd or Dst
CalibOnly?
2 1 to reproduce of
oNormalize?
2 1=no, 2=yes

ofxn calc*
0.0487
ofxn mod
0.13267

sRun
RunA1
zout
1

Ave
Data
7.7E-06
Model
7.7E-06

data/vMax
4.8E-05

ofxng
319.391

Min d/m
1.0882

ave/min d
1.15279
ave/min m
1.24857

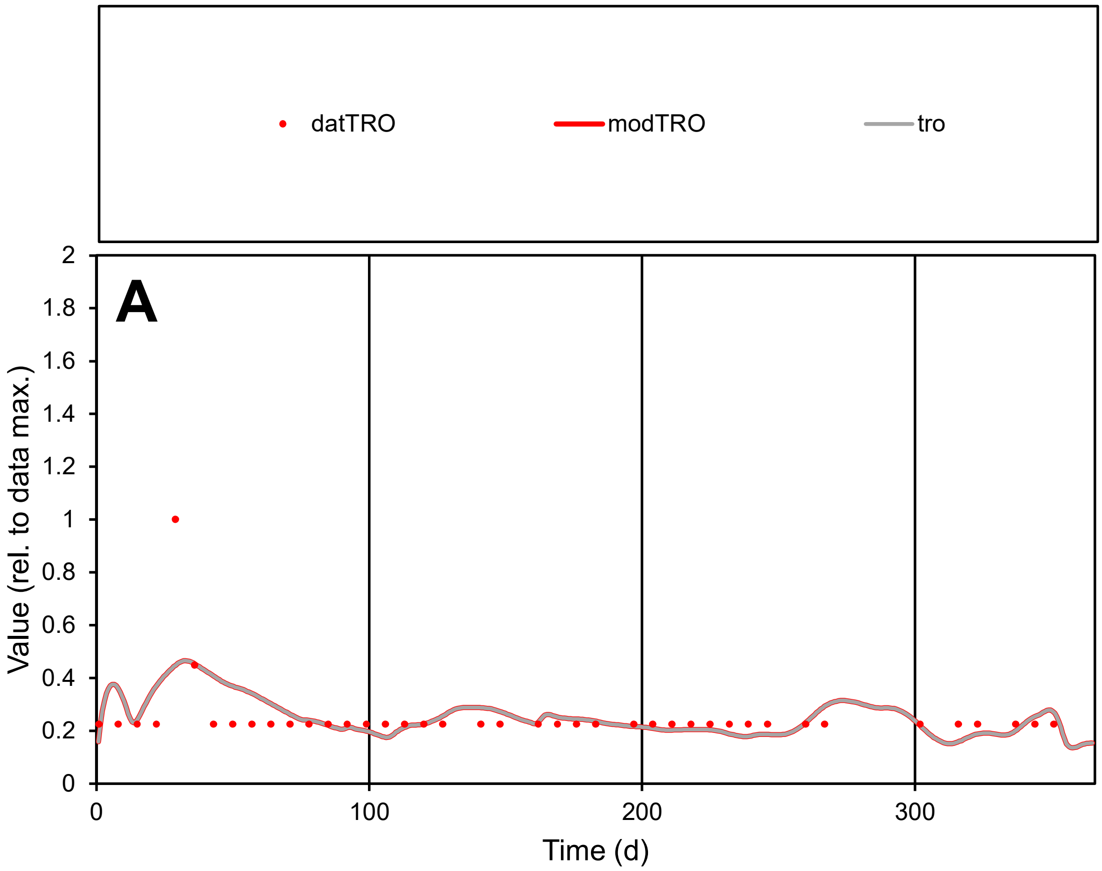


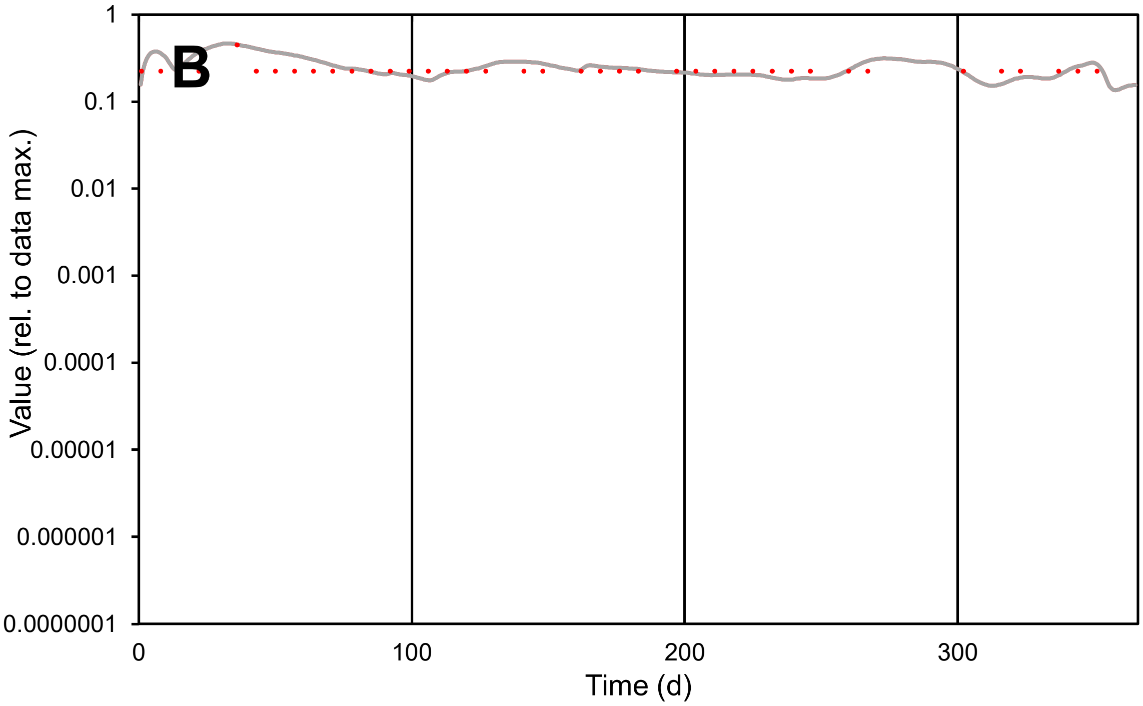


**C1**

Variables
TRO Provide jtd or Dst
CalibOnly?
2 1 to reproduce of
oNormalize?
2 1=no, 2=yes

ofxn calc*
0.40986
ofxn mod
0.42823

sRun
RunA1
zout
1

Ave
Data
5.6E-06
Model
5.7E-06

data/vMax
2.2E-05

ofxng
319.391

Min d/m
1.37377

ave/min d
1.10863
ave/min m
1.55608

330_Calib_S_A

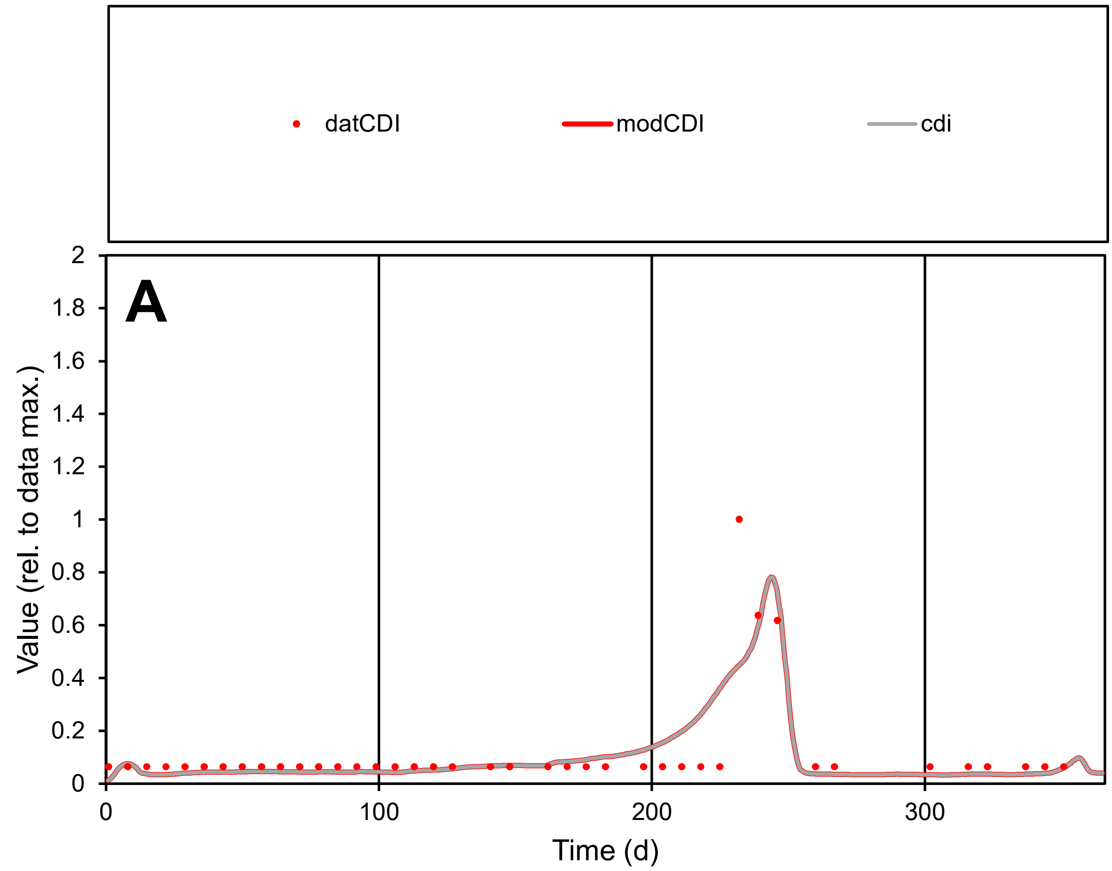


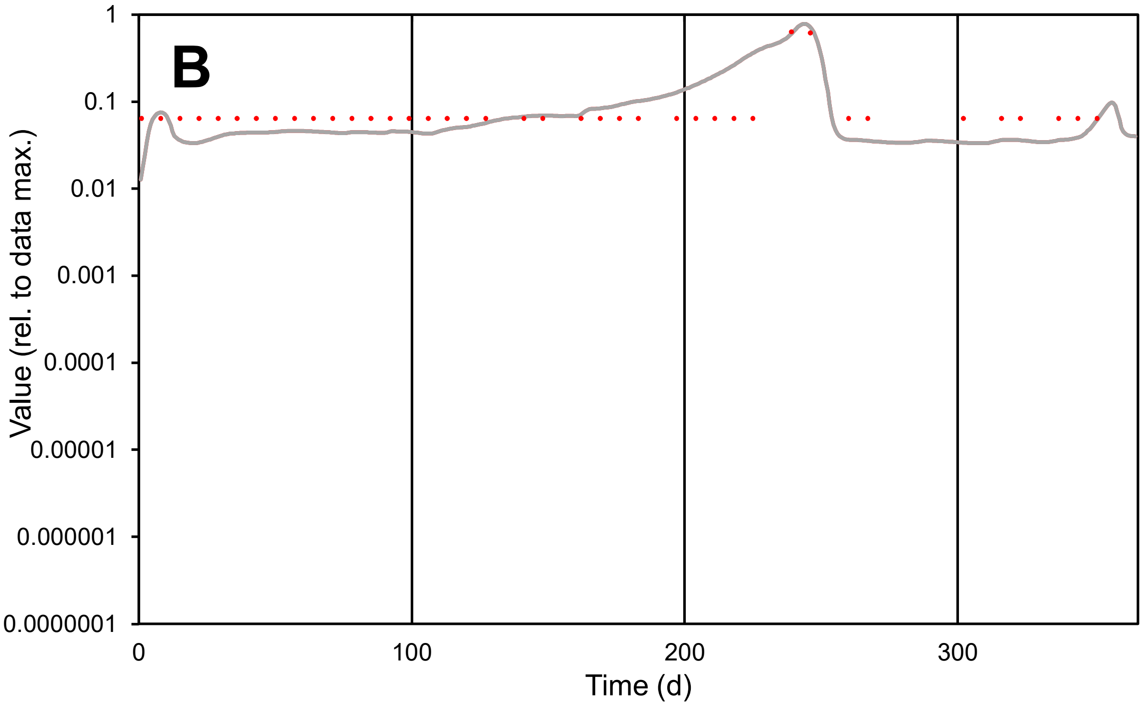


**C1**

Variables

CDI Provide jtd or Dst

CalibOnly?

2 1 to reproduce of

oNormalize?

2 1=no, 2=yes

ofxn calc*

0.93726

ofxn mod

0.9675

sRun

RunA1

zout

1

Ave

Data

3.1E-07

Model

3E-07

data/vMax

2.7E-06

ofxng

319.391

Min d/m

1.90288

ave/min d

1.78443

ave/min m

3.32716

330_Calib_S_A

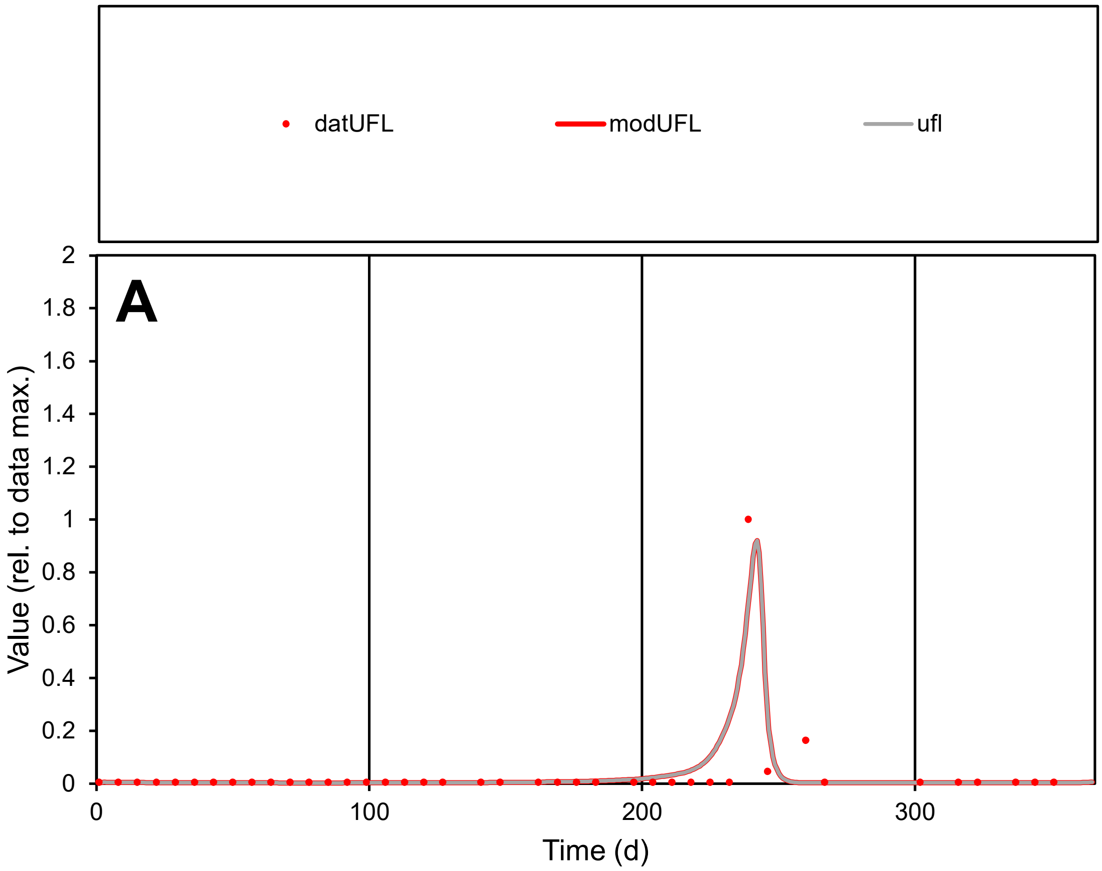


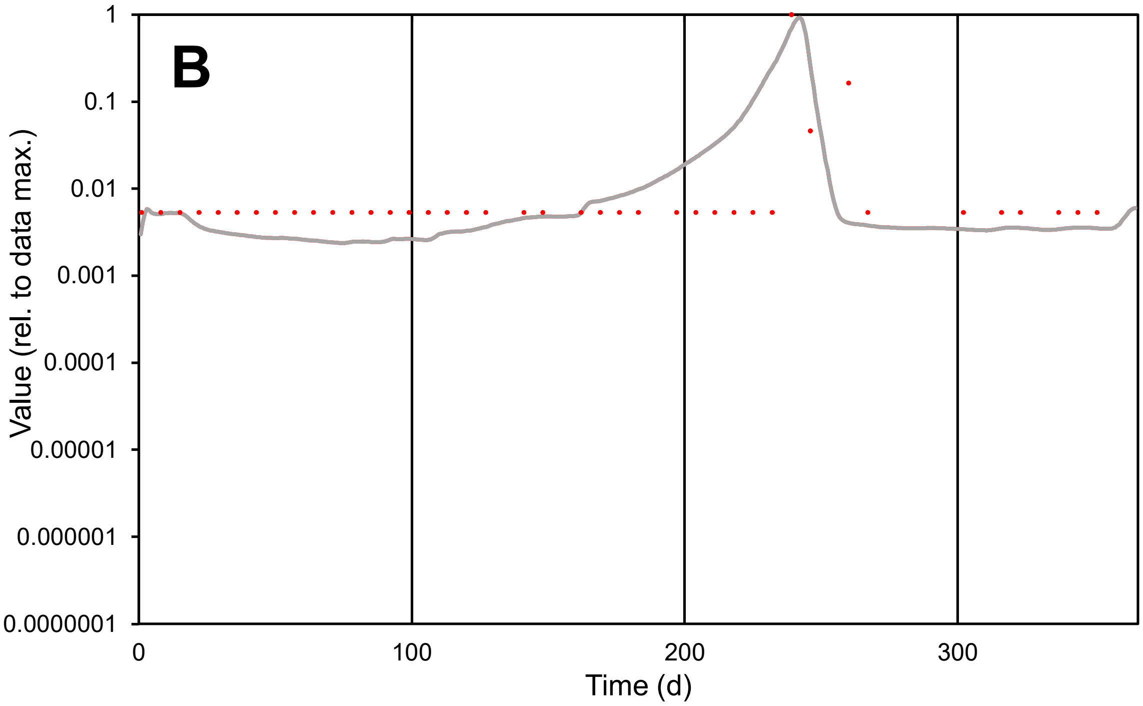


**C1**

Variables
UFL Provide jtd or Dst
CalibOnly?
2 1 to reproduce of
oNormalize?
2 1=no, 2=yes

ofxn calc*
1.31414
ofxn mod
1.64734

sRun
RunA1
zout
1

Ave
Data
2E-05
Model
2E-05

data/vMax
0.00057

ofxng
319.391

Min d/m
2.22045
ave/min d
6.45852
ave/min m
14.6012

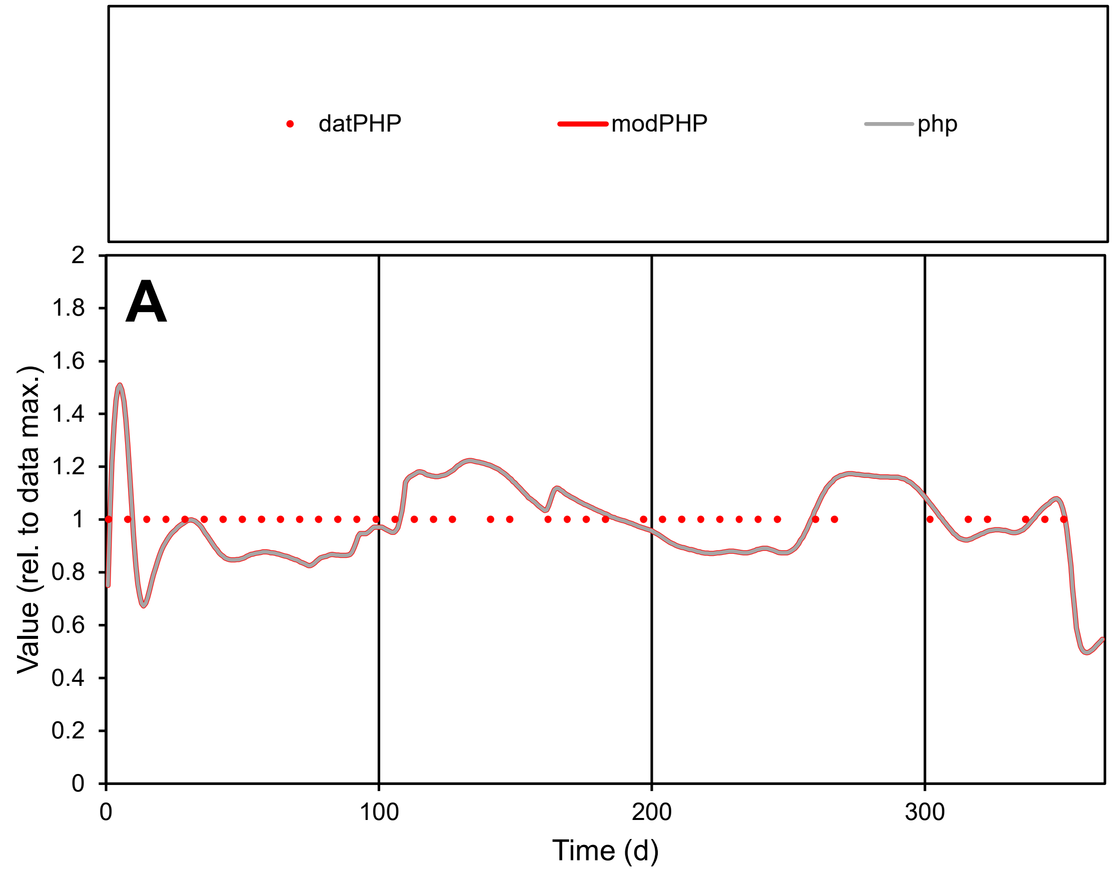


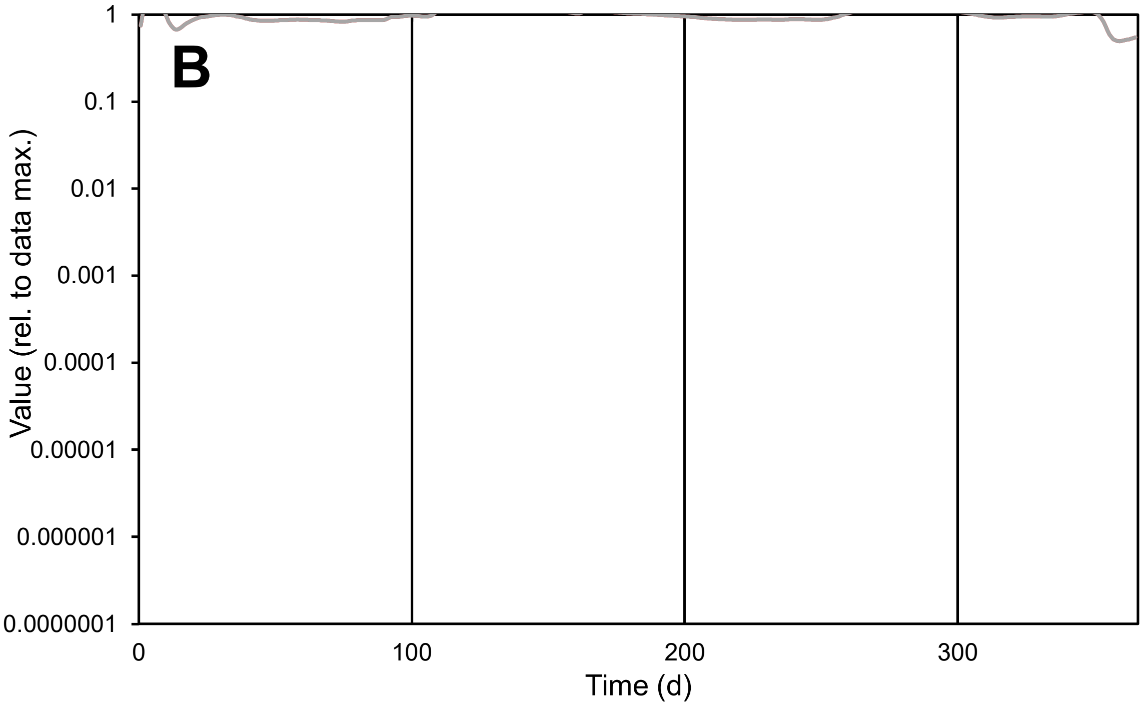


**C1**

Variables
PHP Provide jtd or Dst
CalibOnly?
2 1 to reproduce of
oNormalize?
2 1=no, 2=yes

ofxn calc*
0.1131
ofxn mod
0.15571

sRun
RunA1
zout
1

Ave
Data
6.9E-06
Model
6.8E-06

data/vMax
6.9E-06

ofxng
319.391

Min d/m
1.31036

ave/min d
1
ave/min m
1.2808

330_Calib_S_A

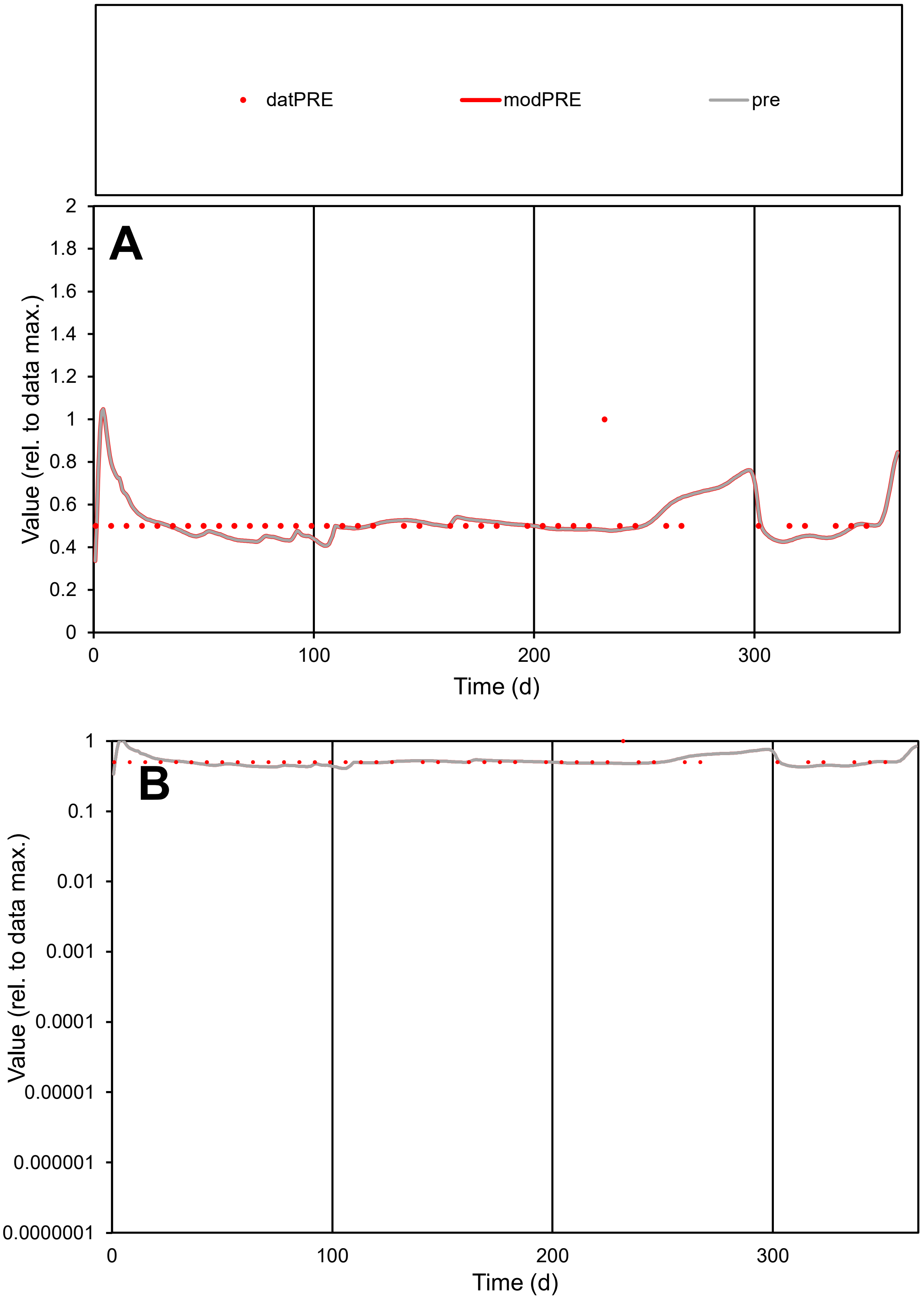


**C1**

Variables
PRE Provide jtd or Dst
CalibOnly?
2 1 to reproduce of
oNormalize?
2 1=no, 2=yes

ofxn calc*
0.19794
ofxn mod
0.22632

sRun
RunA1
zout
1

Ave
Data
1.4E-05
Model
1.3E-05

data/vMax
2.7E-05

ofxng
319.391

Min d/m
1.16884

ave/min d
1.02439
ave/min m
1.16999

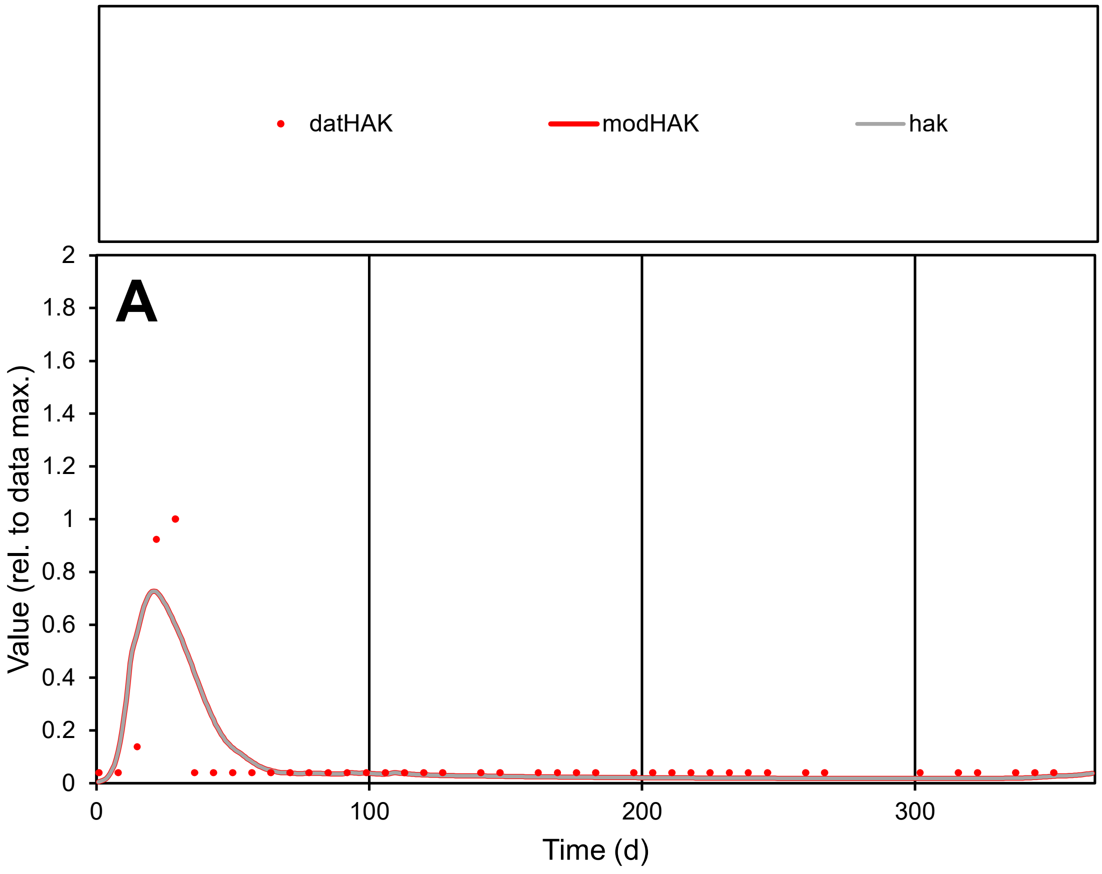


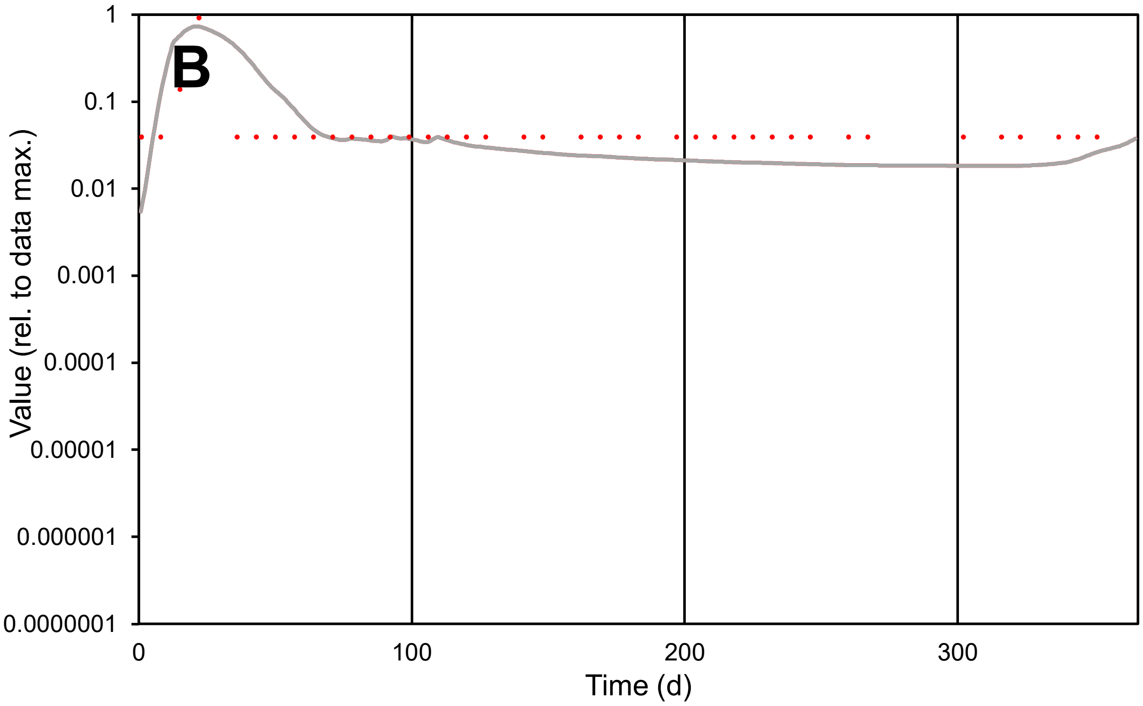


**C1**

Variables
HAK Provide jtd or Dst
CalibOnly?
2 1 to reproduce of
oNormalize?
2 1=no, 2=yes

ofxn calc*
1.32246
ofxn mod
1.39881

sRun
RunA1
zout
1

Ave
Data
1.1E-05
Model
1.1E-05

data/vMax
0.00013

ofxng
319.391

Min d/m
2.1587

ave/min d
2.20592
ave/min m
4.7648

330_Calib_S_A

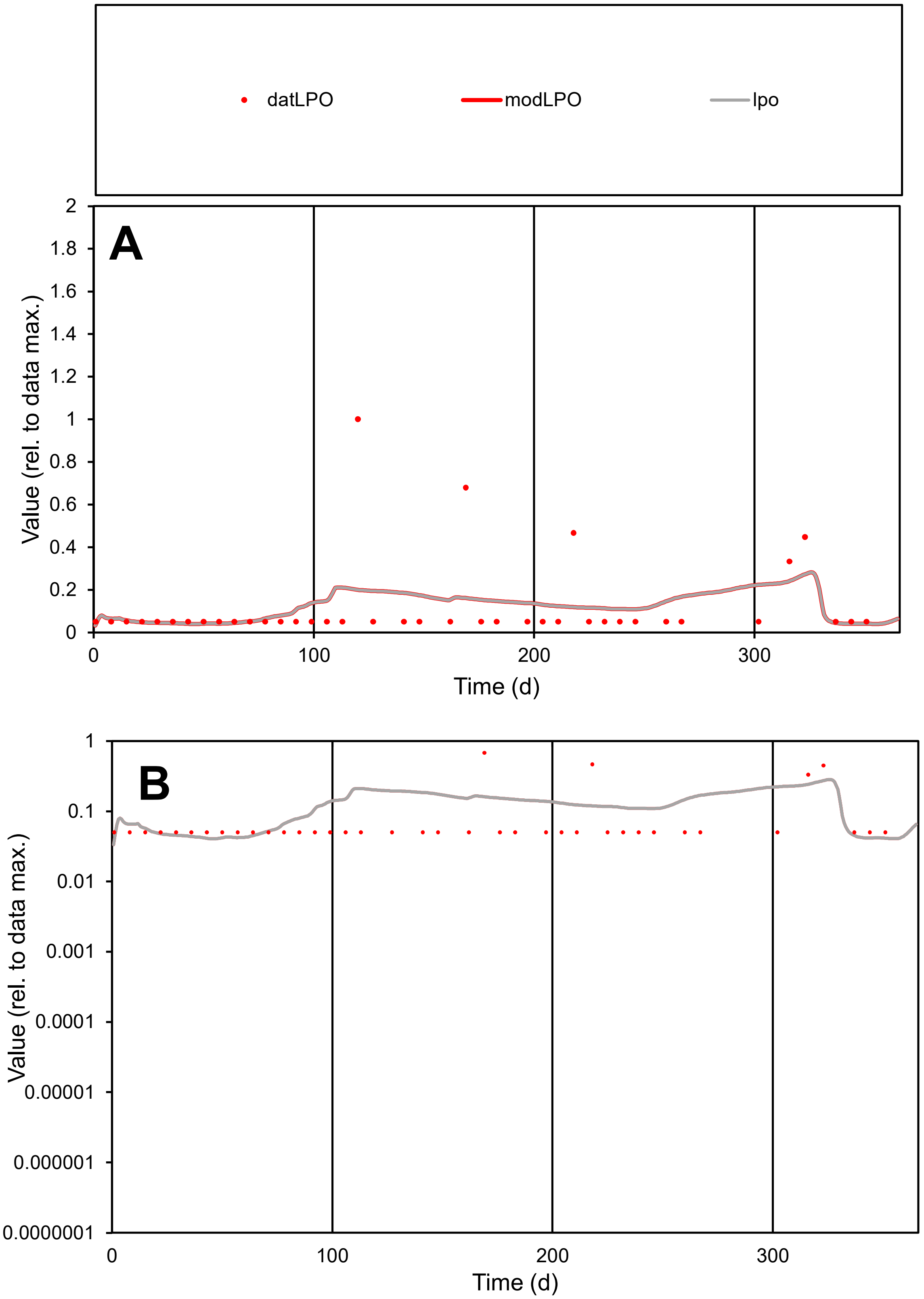


C1

Variables

LPO Provide jtd or Dst

CalibOnly?

2 1 to reproduce of

oNormalize?

2 1=no, 2=yes

ofxn calc*

1.53732

ofxn mod

1.54473

sRun

RunA1

zout

1

Ave

Data

1.3E-05

Model

1.4E-05

data/vMax

0.00012

ofxng

319.391

Min d/m

1.22999

ave/min d

2.30085

ave/min m

2.87259

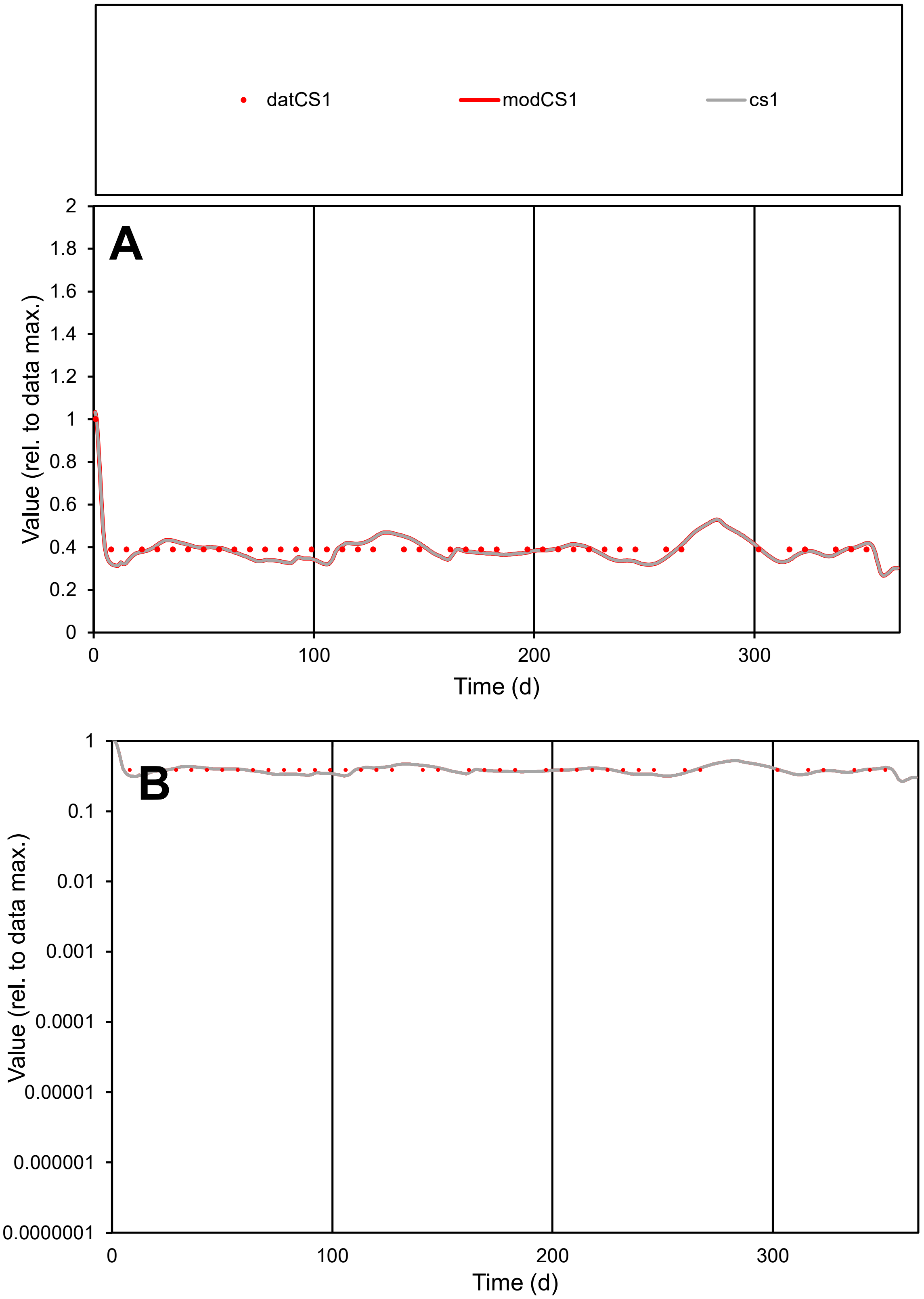


**C1**

Variables
CS1 Provide jtd or Dst
CalibOnly?
2 1 to reproduce of
oNormalize?
2 1=no, 2=yes

ofxn calc*
0.08548
ofxn mod
0.13734

sRun
RunA1
zout
1

Ave
Data
1.8E-05
Model
1.8E-05

data/vMax
4.5E-05

ofxng
319.391

Min d/m
1.23293

ave/min d
1.03828
ave/min m
1.25326

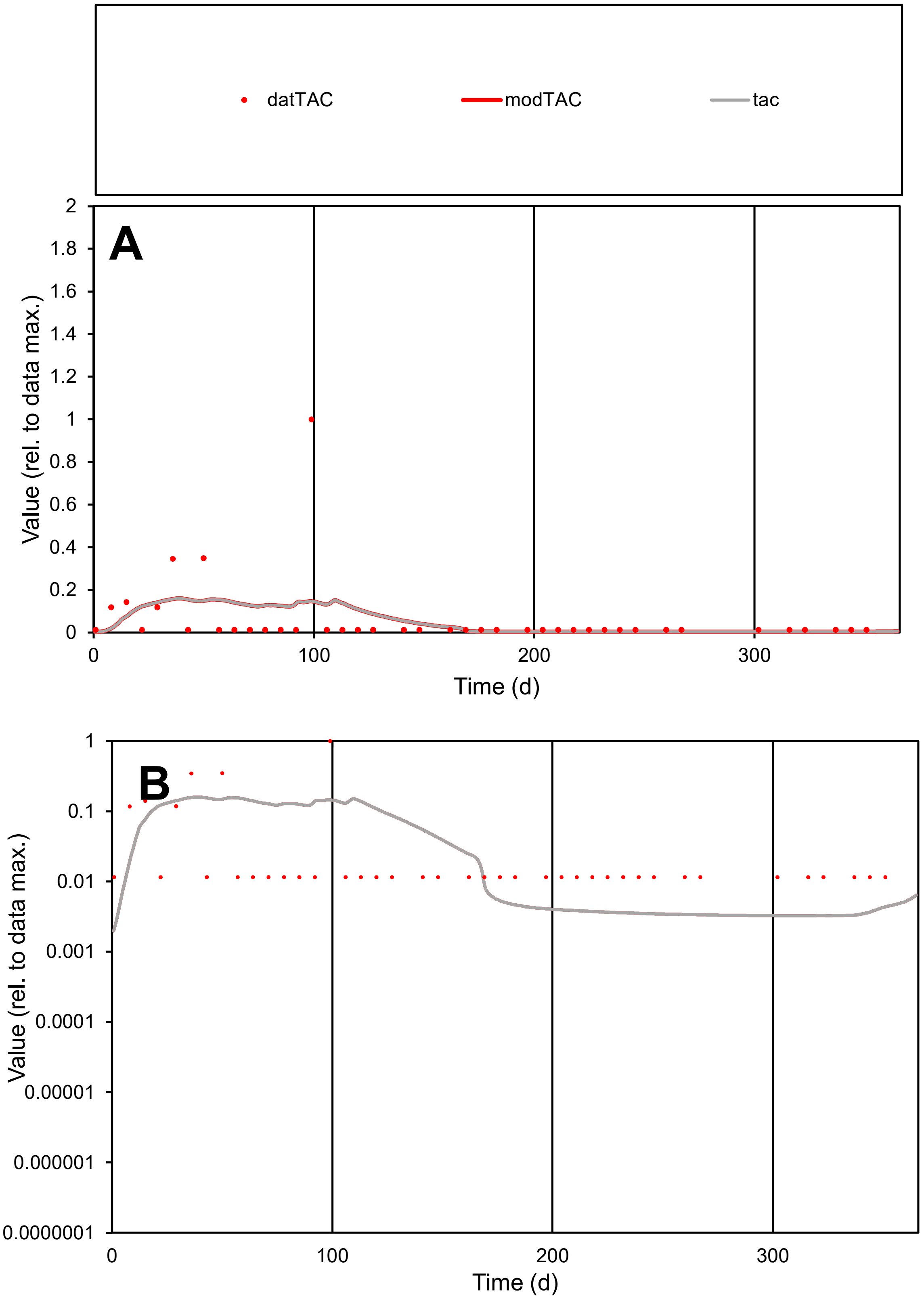


**C1**

Variables
TAC Provide jtd or Dst
CalibOnly?
2 1 to reproduce of
oNormalize?
2 1=no, 2=yes

ofxn calc*
2.56295
ofxn mod
2.58147

sRun
RunA1
zout
1

Ave
Data
6.8E-06
Model
6.7E-06

data/vMax
0.00011

ofxng
319.391

Min d/m
3.54697

ave/min d
5.24786
ave/min m
18.4565

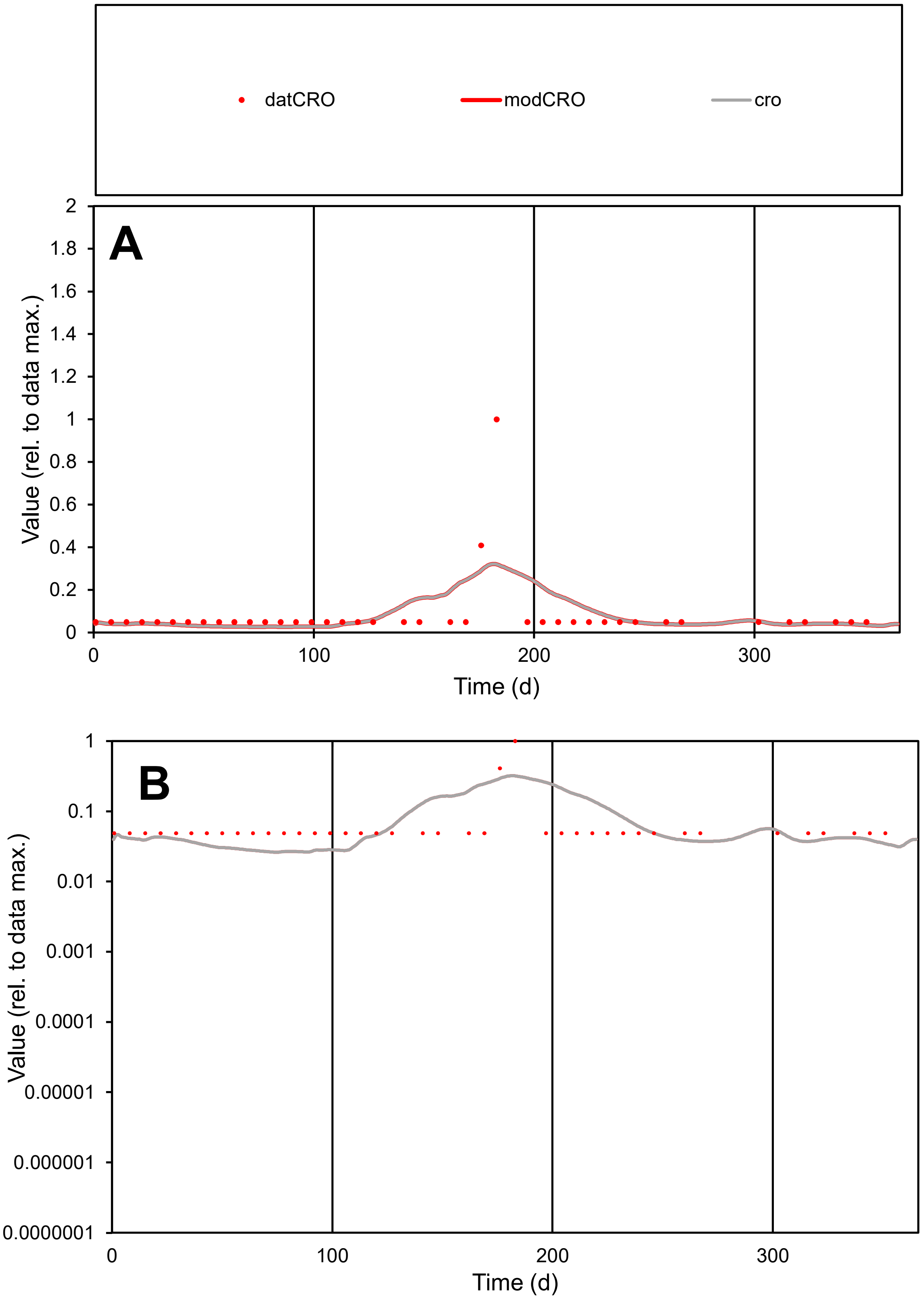


**C1**

Variables
CRO Provide jtd or Dst
CalibOnly?
2 1 to reproduce of
oNormalize?
2 1=no, 2=yes

ofxn calc*
1.55917
ofxn mod
1.58871

sRun
RunA1
zout
1

Ave
Data
1.7E-05
Model
1.7E-05

data/vMax
0.00021

ofxng
319.391

Min d/m
1.84106

ave/min d
1.65604
ave/min m
3.12507

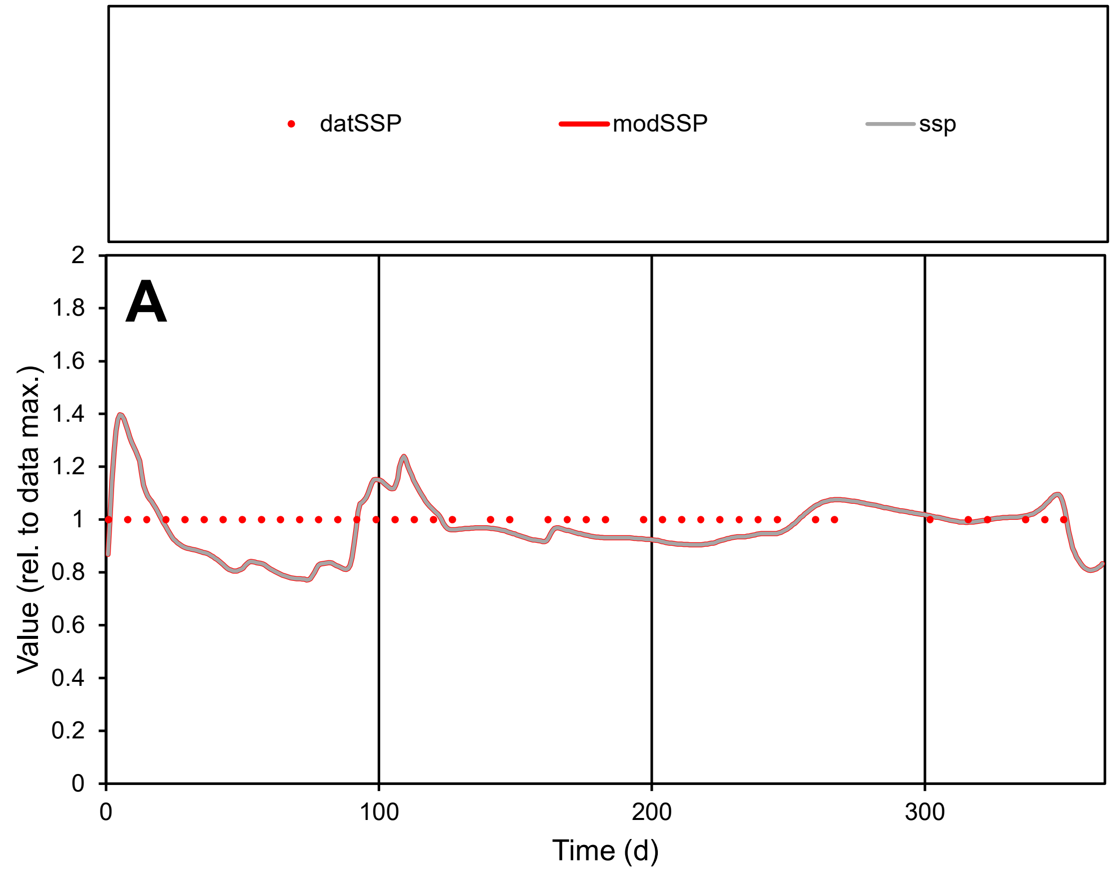


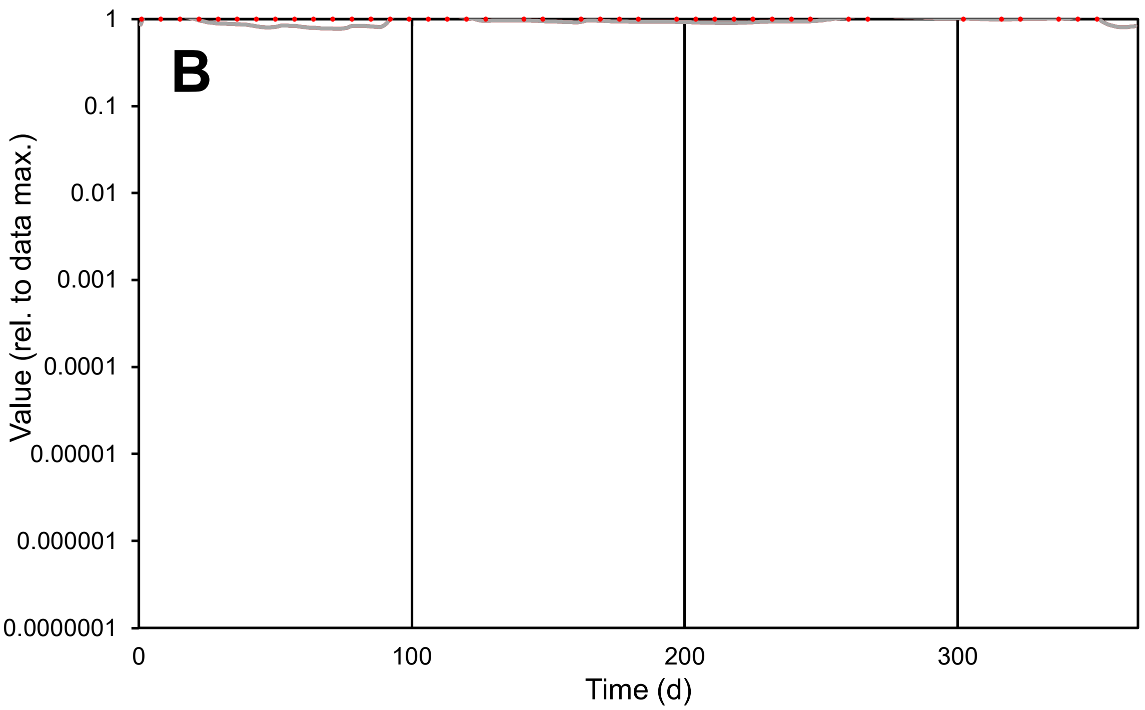


**C1**

Variables
SSP Provide jtd or Dst
CalibOnly?
2 1 to reproduce of
oNormalize?
2 1=no, 2=yes

ofxn calc*
0.11354
ofxn mod
0.15608

sRun
RunA1
zout
1

Ave
Data
1.3E-05
Model
1.3E-05

data/vMax
1.3E-05

ofxng
319.391

Min d/m
1.29073

ave/min d
1
ave/min m
1.24657

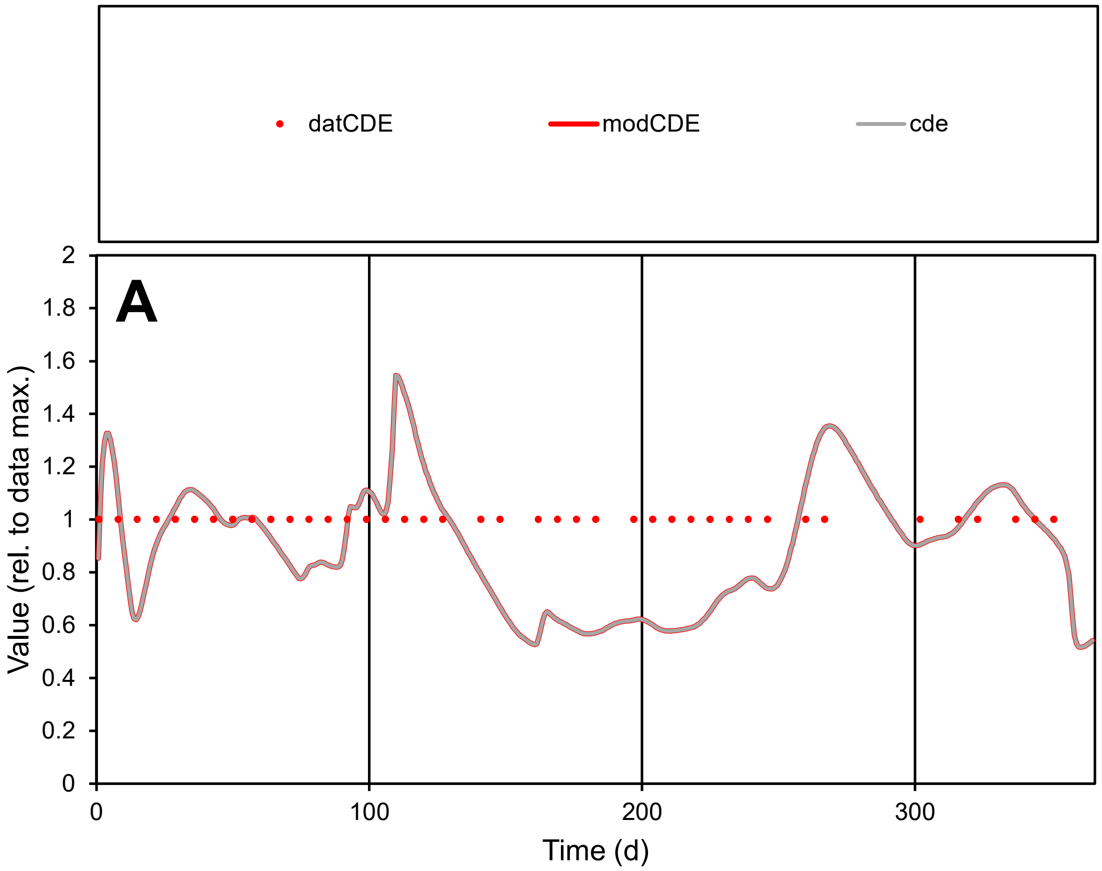


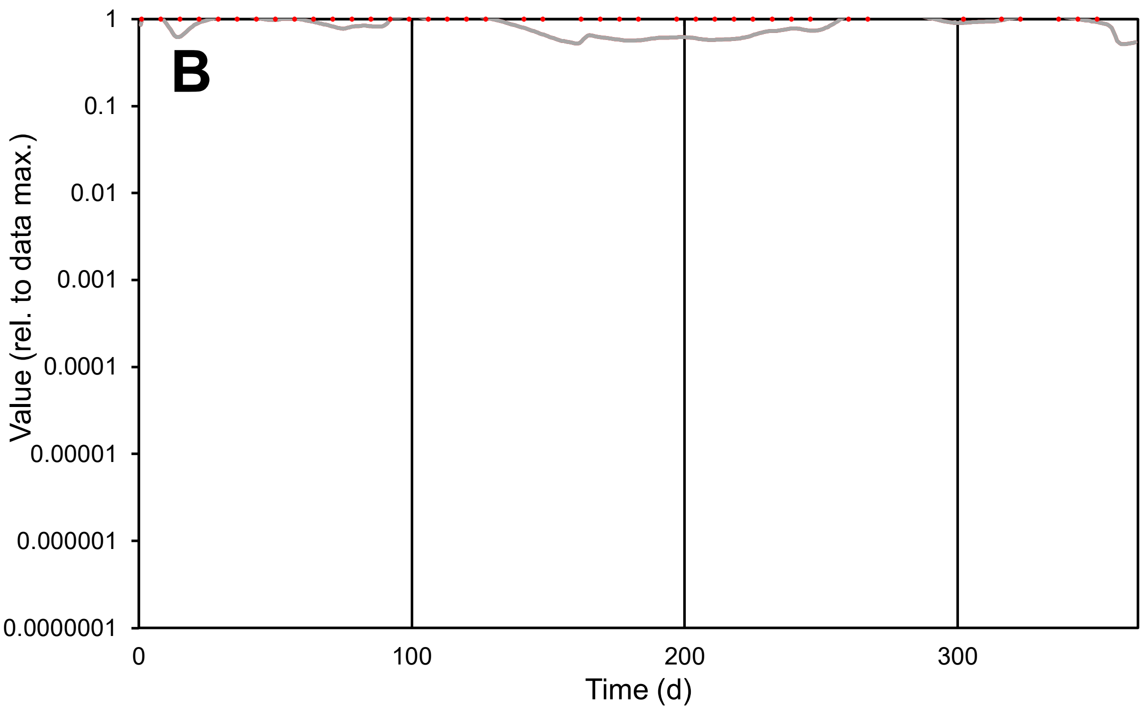


**C1**

Variables
CDE Provide jtd or Dst
CalibOnly?
2 1 to reproduce of
oNormalize?
2 1=no, 2=yes

ofxn calc*
0.24452
ofxn mod
0.26504

sRun
RunA1
zout
1

Ave
Data
1E-05
Model
8.9E-06

data/vMax
1E-05

ofxng
319.391

Min d/m
1.74598

ave/min d
1
ave/min m
1.56331

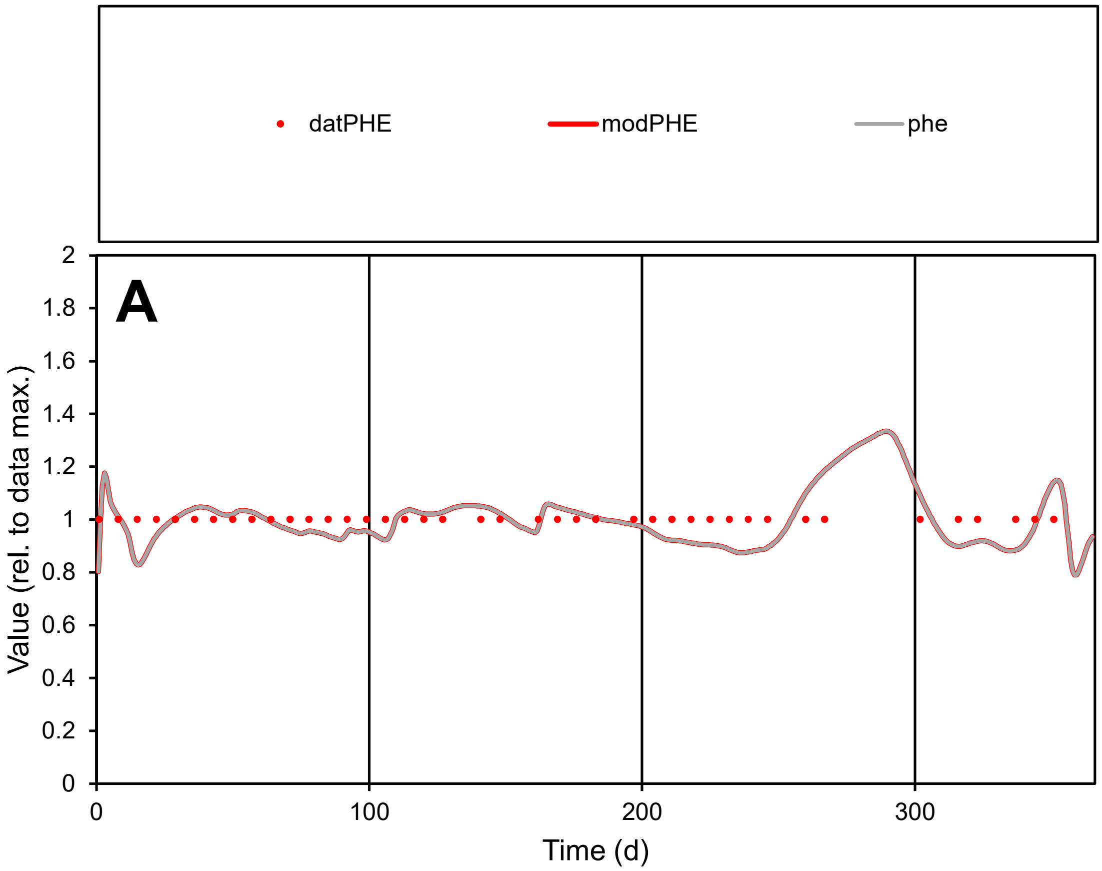


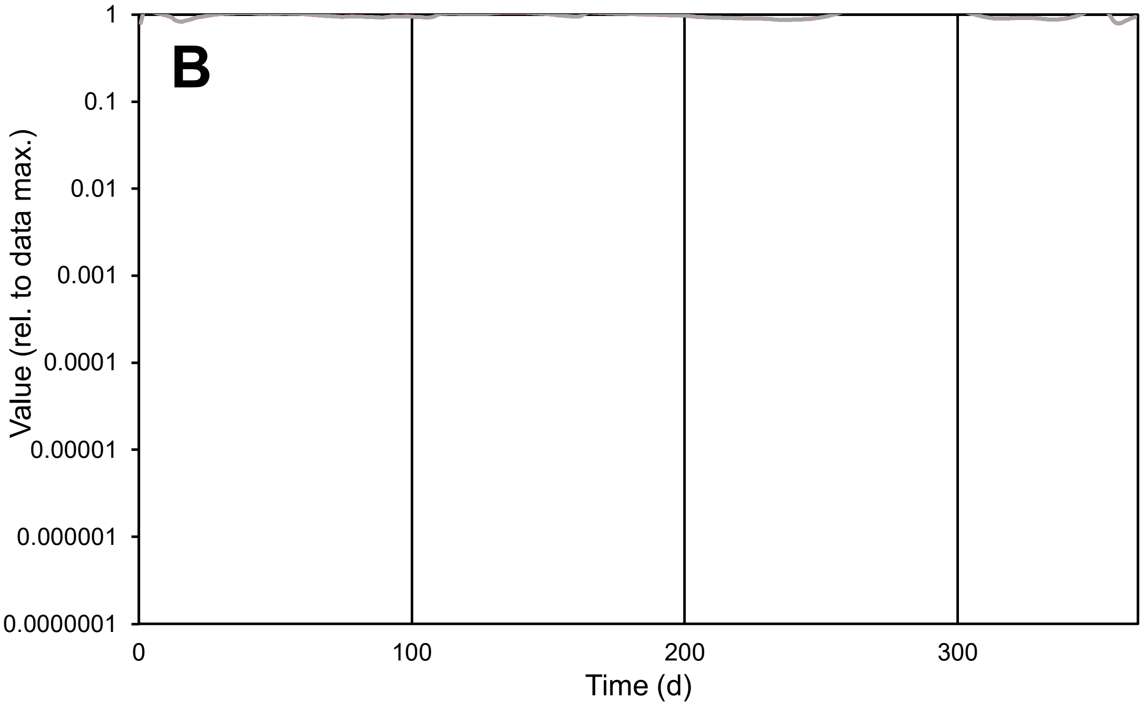


**C1**

Variables
PHE Provide jtd or Dst
CalibOnly?
2 1 to reproduce of
oNormalize?
2 1=no, 2=yes

ofxn calc*
0.07498
ofxn mod
0.13128

sRun
RunA1
zout
1

Ave
Data
4E-06
Model
3.9E-06

data/vMax
4E-06

ofxng
319.391

Min d/m
1.19022

ave/min d
1
ave/min m
1.17348

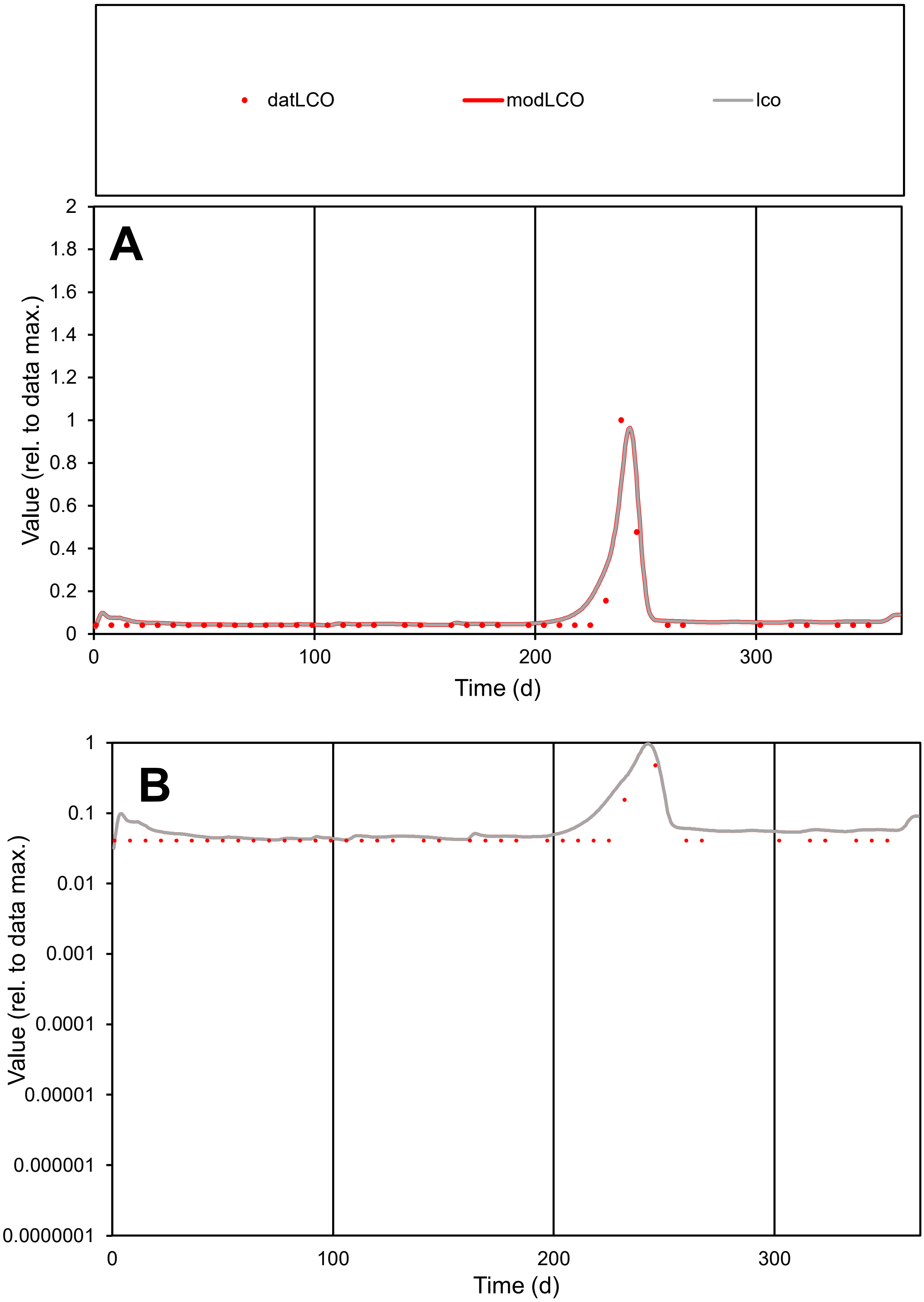


**C1**

Variables
LCO Provide jtd or Dst
CalibOnly?
2 1 to reproduce of
oNormalize?
2 1=no, 2=yes

ofxn calc*
0.54052
ofxn mod
0.63356

sRun
RunA1
zout
1

Ave
Data
1.4E-05
Model
1.7E-05

data/vMax
0.00018

ofxng
319.391

Min d/m
0.97396

ave/min d
1.90326
ave/min m
2.22383

330_Calib_S_A

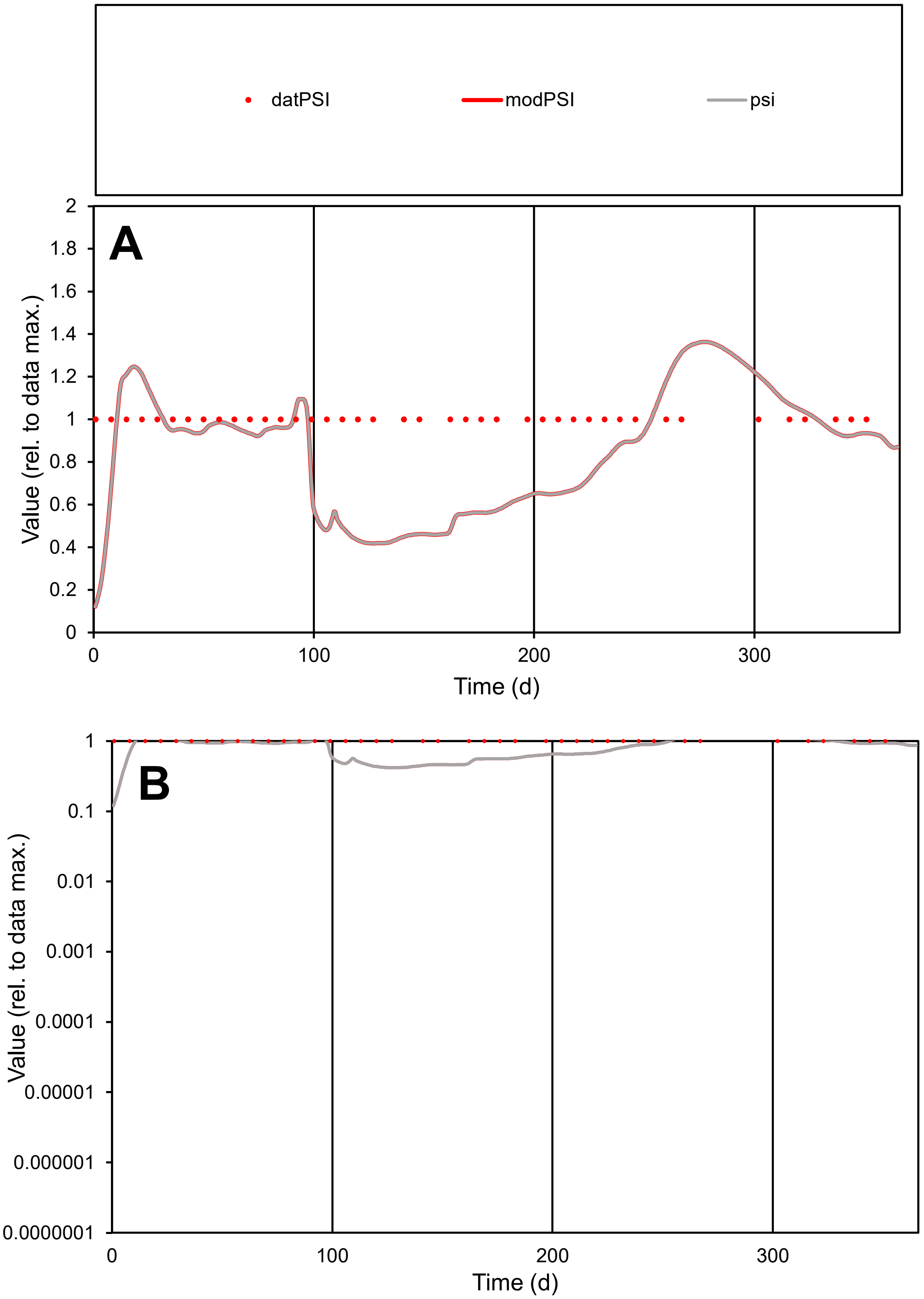


**C1**

Variables
PSI Provide jtd or Dst
CalibOnly?
2 1 to reproduce of
oNormalize?
2 1=no, 2=yes

ofxn calc*
0.32036
ofxn mod
0.3386

sRun
RunA1
zout
1

Ave
Data
3.6E-06
Model
2.9E-06

data/vMax
3.6E-06

ofxng
319.391

Min d/m
3.97608

ave/min d
1
ave/min m
3.24597

330_Calib_S_A

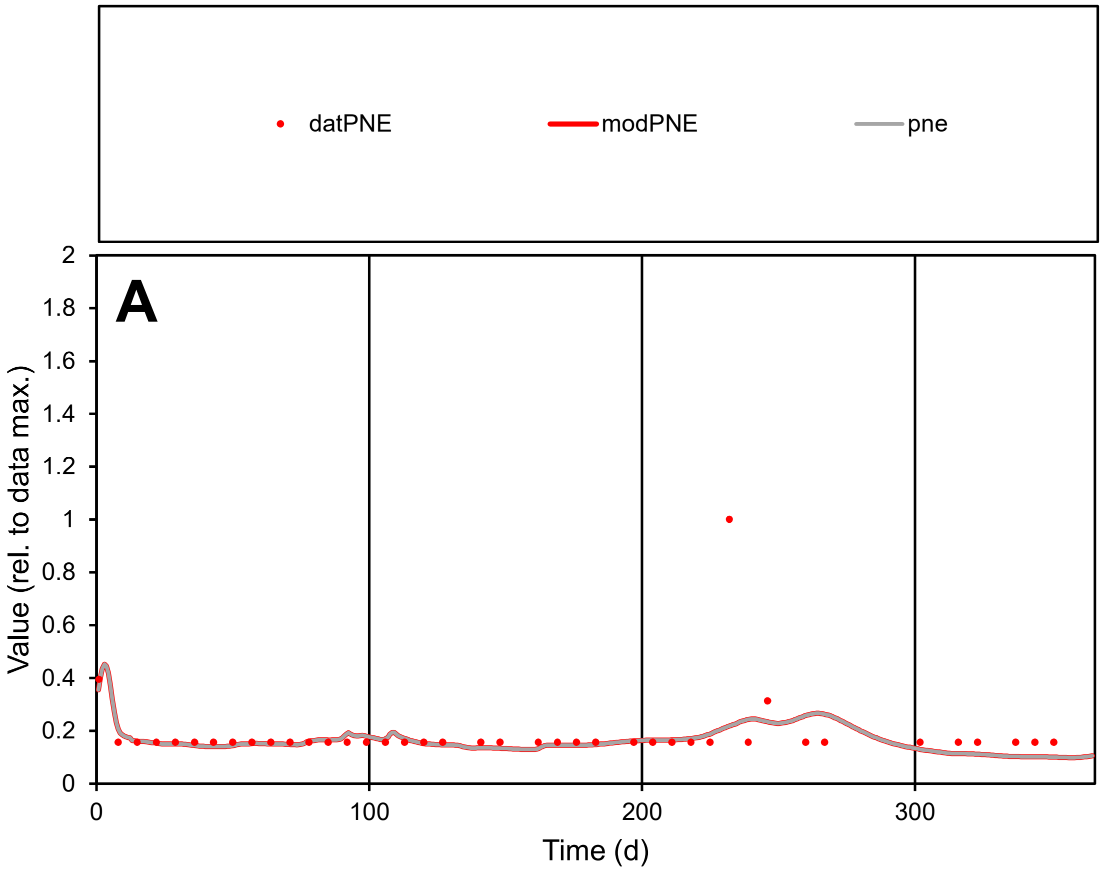


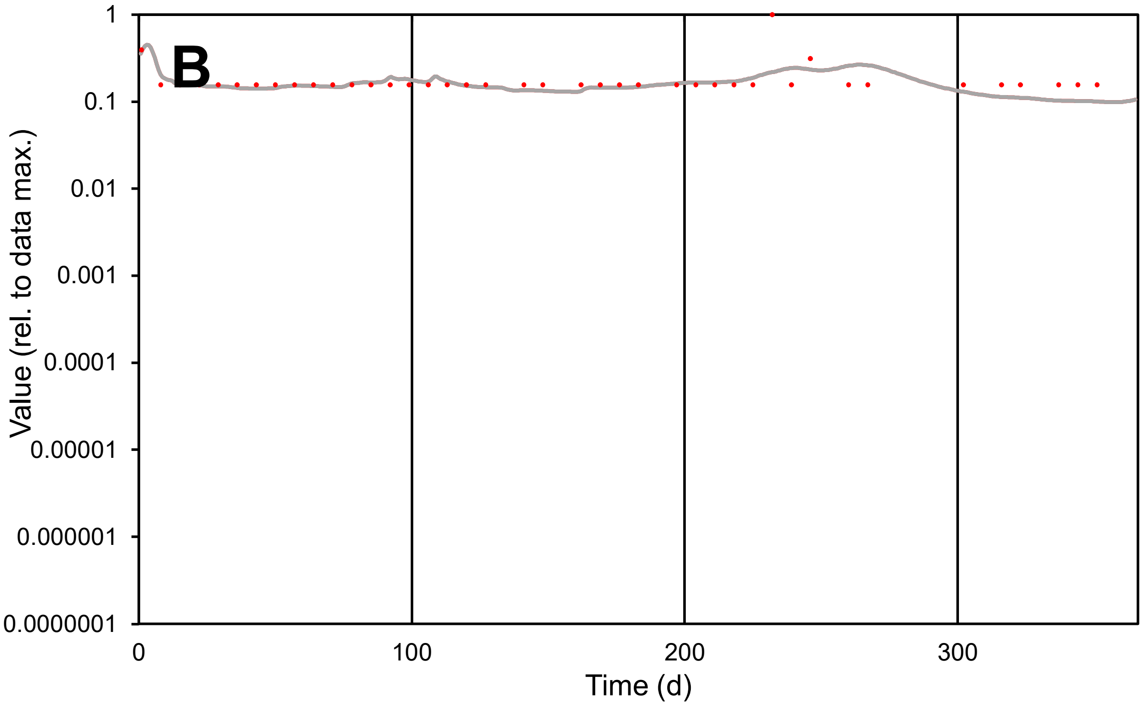


**C1**

Variables
PNE Provide jtd or Dst
CalibOnly?
2 1 to reproduce of
oNormalize?
2 1=no, 2=yes

ofxn calc*
0.67951
ofxn mod
0.69054

sRun
RunA1
zout
1

Ave
Data
1.7E-05
Model
1.6E-05

data/vMax
9.3E-05

ofxng
319.391

Min d/m
1.57024

ave/min d
1.19294
ave/min m
1.67816

330_Calib_S_A

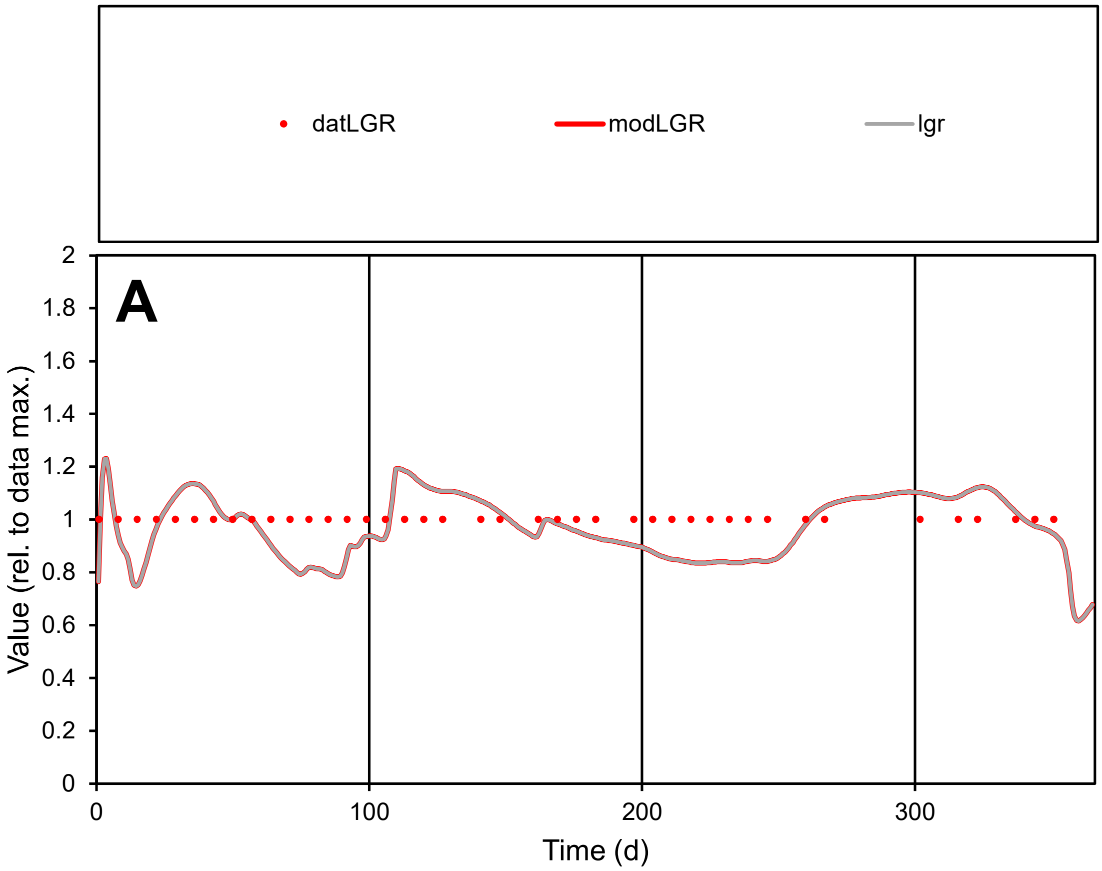


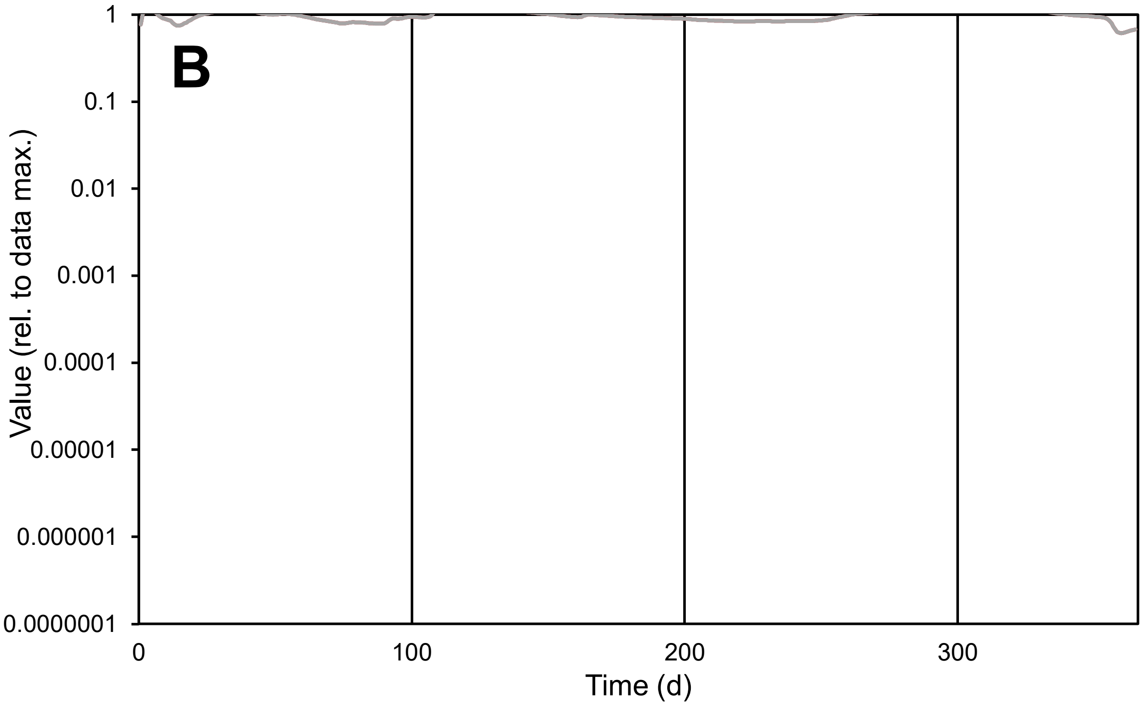


**C1**

Variables

LGR Provide jtd or Dst

CalibOnly?

2 1 to reproduce of

oNormalize?

2 1=no, 2=yes

ofxn calc*

0.11235

ofxn mod

0.15467

sRun

RunA1

zout

1

Ave

Data

5.2E-06

Model

5E-06

data/vMax

5.2E-06

ofxng

319.391

Min d/m

1.26986

ave/min d

1

ave/min m

1.22527

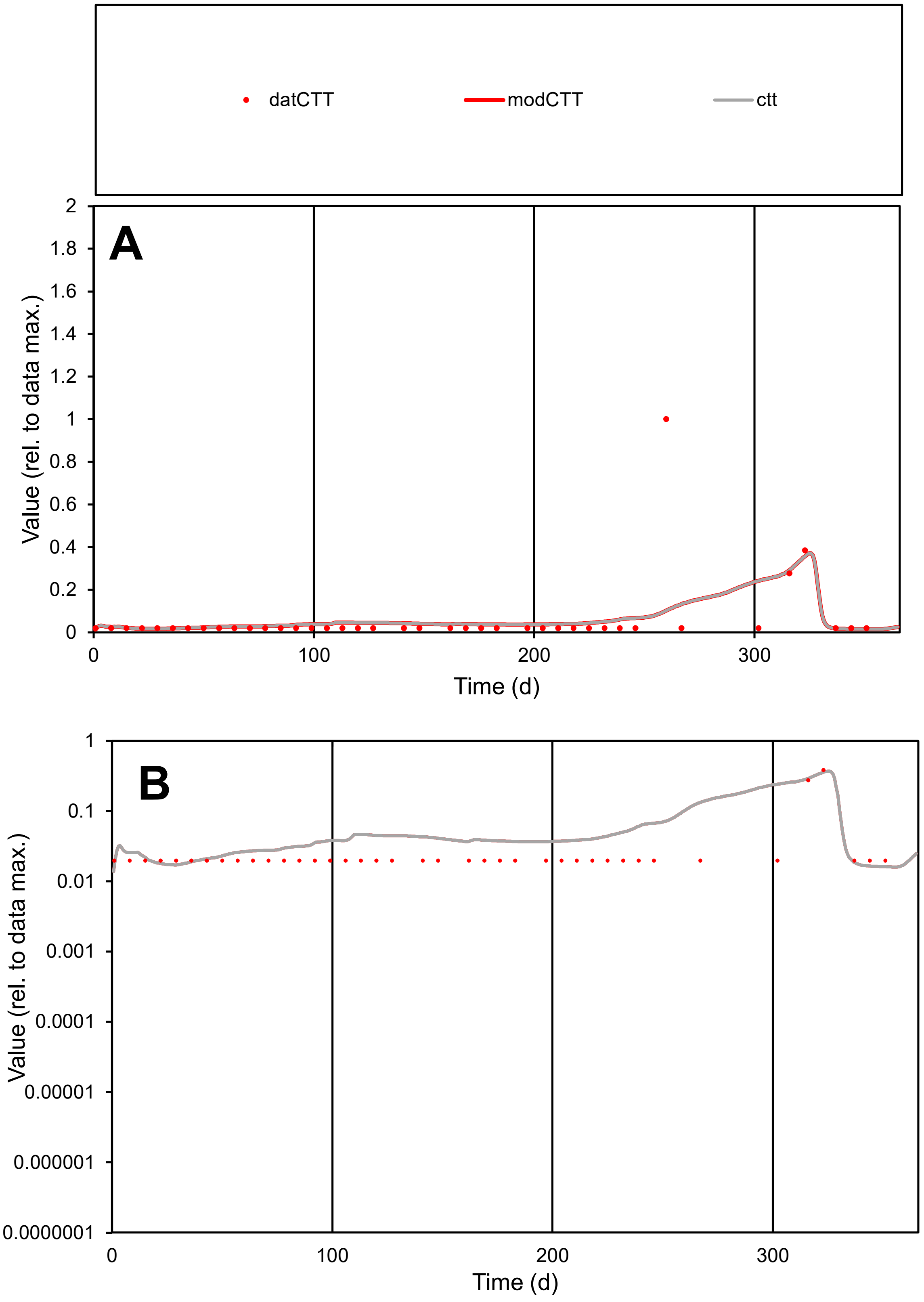


**C1**

Variables
CTT Provide jtd or Dst
CalibOnly?
2 1 to reproduce of
oNormalize?
2 1=no, 2=yes

ofxn calc*
2.48309
ofxn mod
2.49104

sRun
RunA1
zout
1

Ave
Data
4.4E-06
Model
4.4E-06

data/vMax
7.5E-05

ofxng
319.391

Min d/m
1.22158

ave/min d
2.98117
ave/min m
3.59137

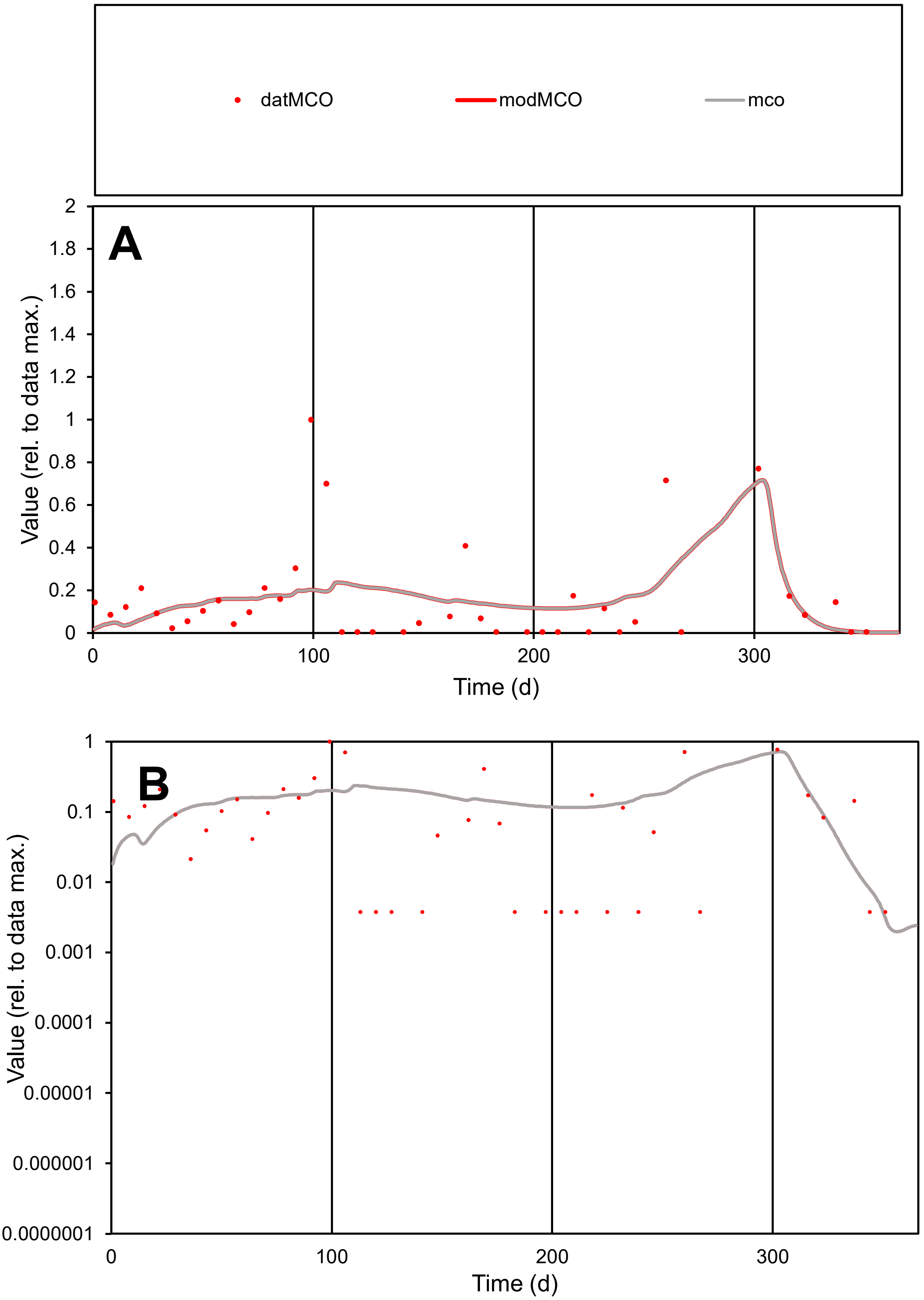


**C1**

Variables

MCO Provide jtd or Dst

CalibOnly?

2 1 to reproduce of

oNormalize?

2 1=no, 2=yes

ofxn calc*

1.30156

ofxn mod

1.31376

sRun

RunA1

zout

1

Ave

Data

2.5E-06

Model

2.5E-06

data/vMax

1.6E-05

ofxng

319.391

Min d/m

1.25694

ave/min d

41.0663

ave/min m

51.6157

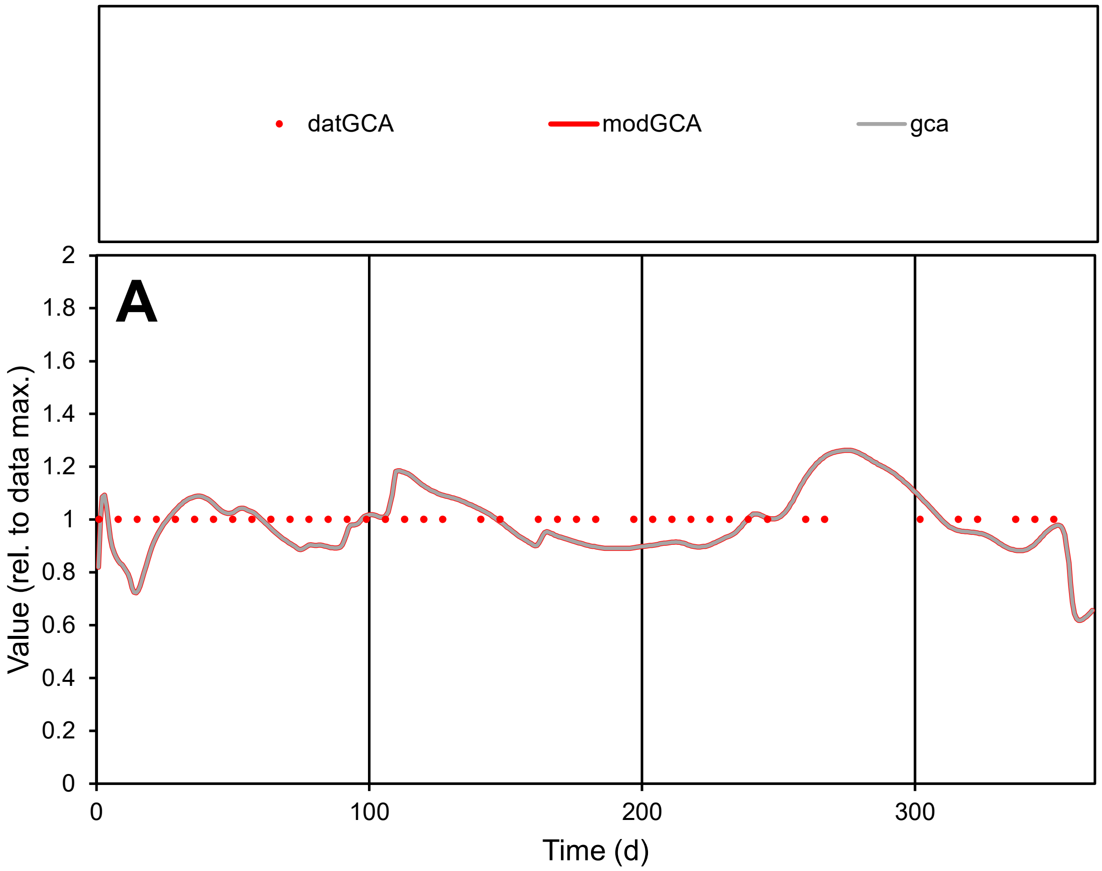


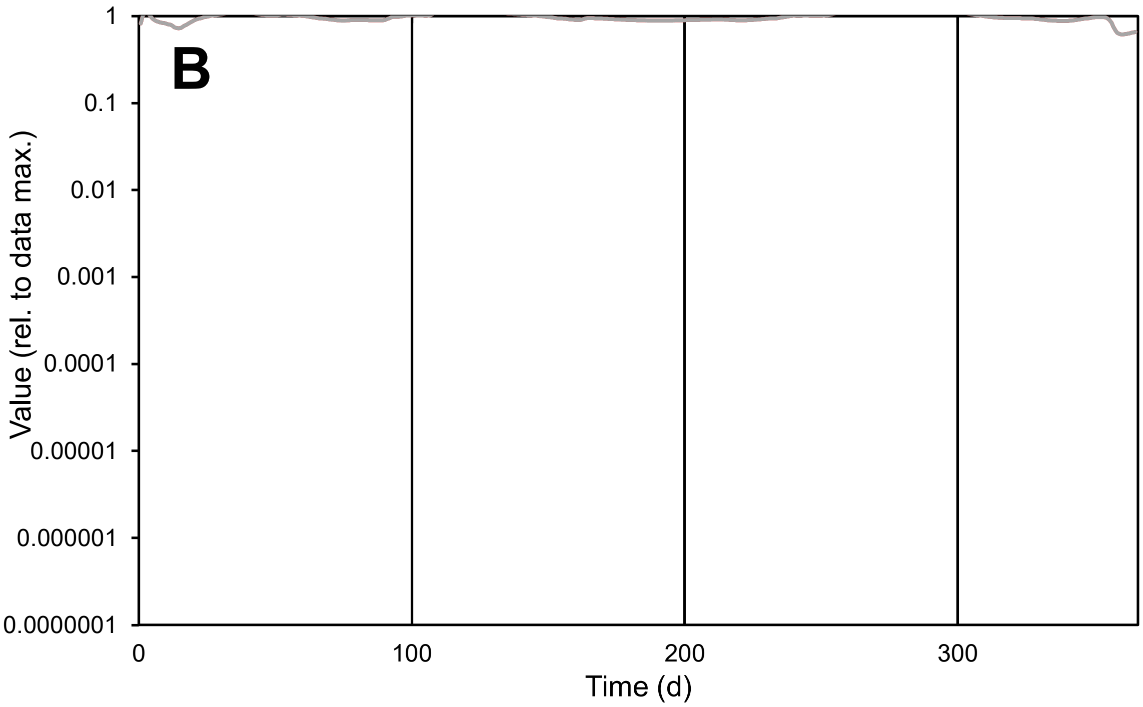


**C1**

Variables
GCA Provide jtd or Dst
CalibOnly?
2 1 to reproduce of
oNormalize?
2 1=no, 2=yes

ofxn calc*
0.09702
ofxn mod
0.1449

sRun
RunA1
zout
1

Ave
Data
8.4E-06
Model
8.2E-06

data/vMax
8.4E-06

ofxng
319.391

Min d/m
1.29584

ave/min d
1
ave/min m
1.27205

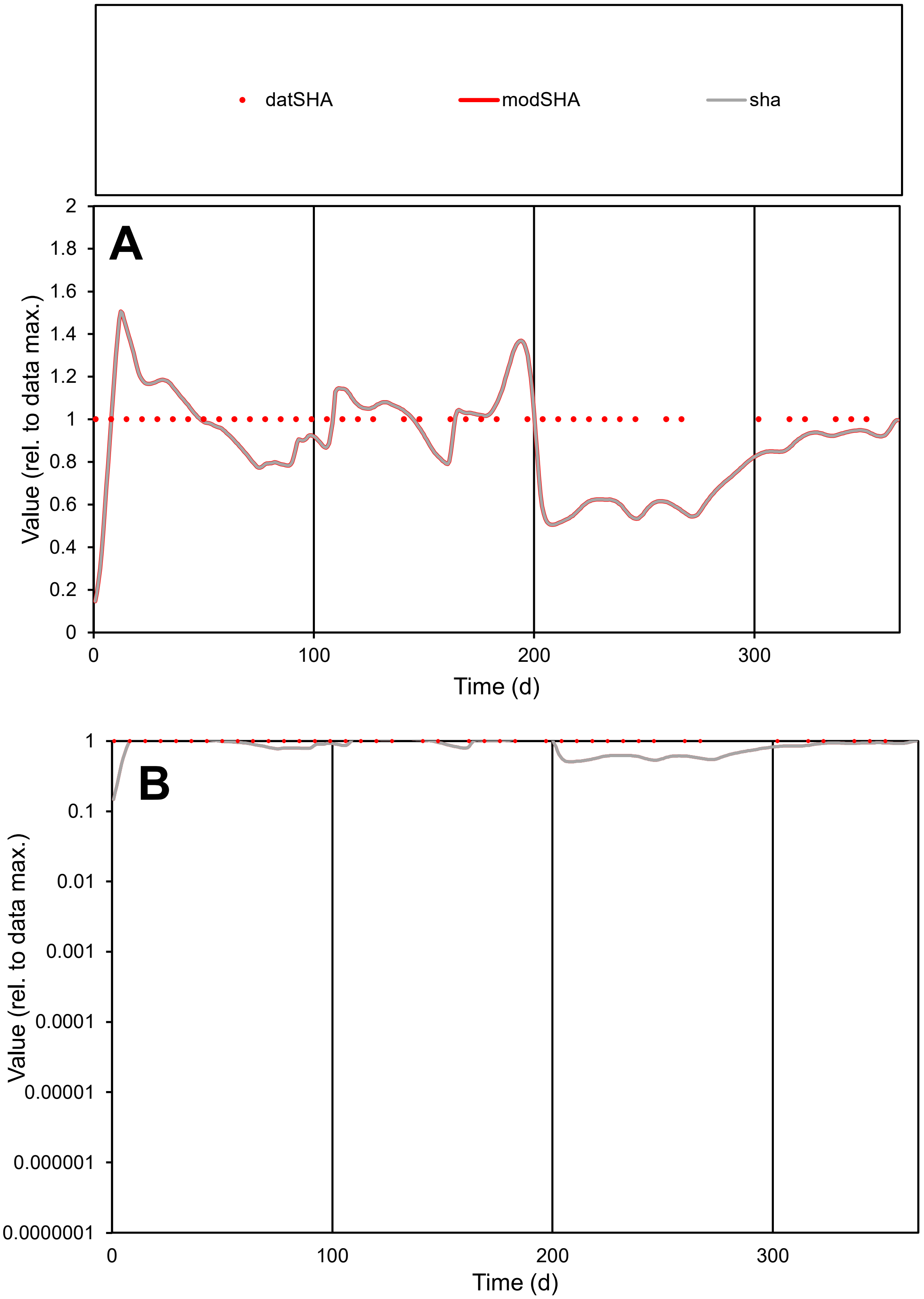


**C1**

Variables
SHA Provide jtd or Dst
CalibOnly?
2 1 to reproduce of
oNormalize?
2 1=no, 2=yes

ofxn calc*
0.25027
ofxn mod
0.27554

sRun
RunA1
zout
1

Ave
Data
7E-07
Model
6.2E-07

data/vMax
7E-07

ofxng
319.391

Min d/m
2.65171

ave/min d
1
ave/min m
2.37104

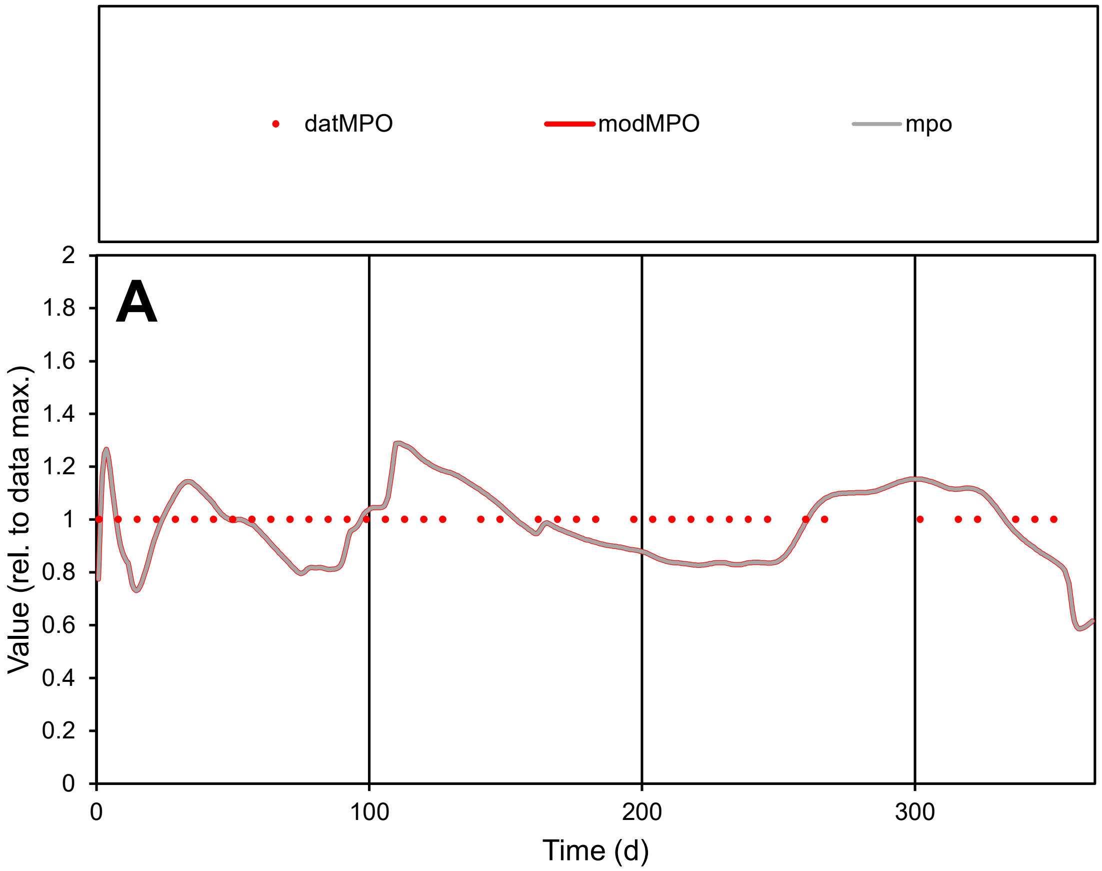


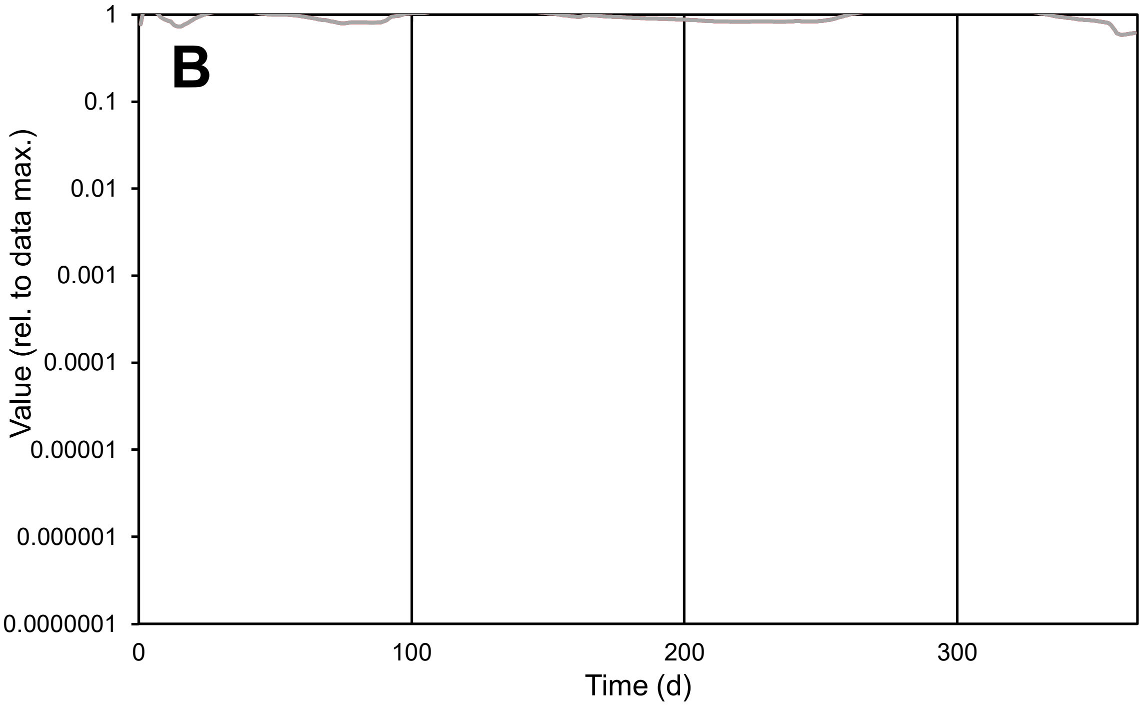


**C1**

Variables
MPO Provide jtd or Dst
CalibOnly?
2 1 to reproduce of
oNormalize?
2 1=no, 2=yes

ofxn calc*
0.13198
ofxn mod
0.16974

sRun
RunA1
zout
1

Ave
Data
1.2E-05
Model
1.2E-05

data/vMax
1.2E-05

ofxng
319.391

Min d/m
1.30811

ave/min d
1
ave/min m
1.27038

330_Calib_S_A

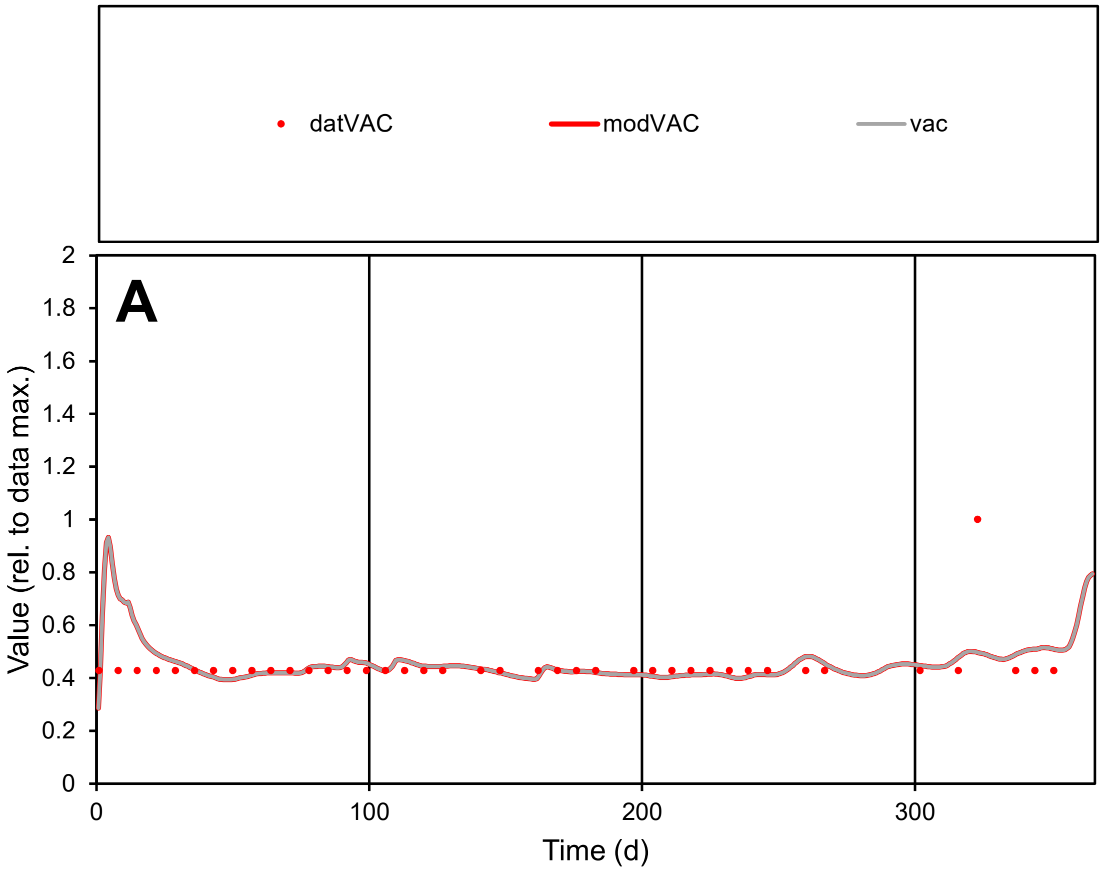


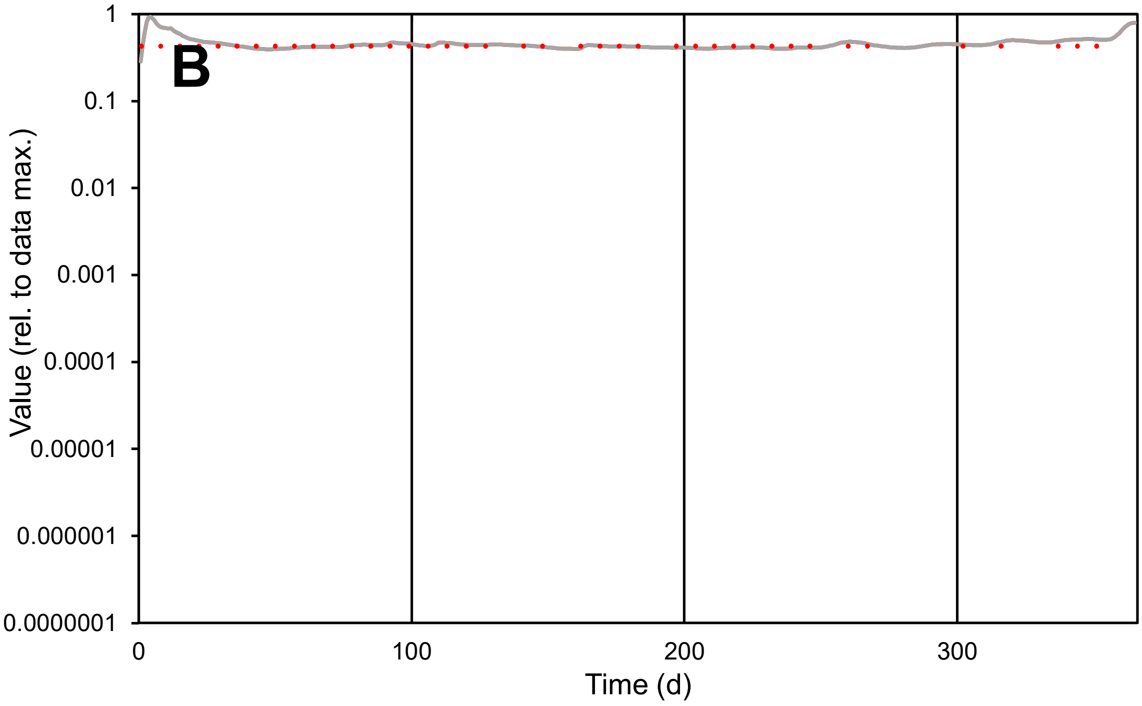


**C1**

Variables

VAC Provide jtd or Dst

CalibOnly?

2 1 to reproduce of

oNormalize?

2 1=no, 2=yes

ofxn calc*

0.21818

ofxn mod

0.24483

sRun

RunA1

zout

1

Ave

Data

9.8E-06

Model

1E-05

data/vMax

2.2E-05

ofxng

319.391

Min d/m

1.0749

ave/min d

1.03261

ave/min m

1.12598

330_Calib_S_A

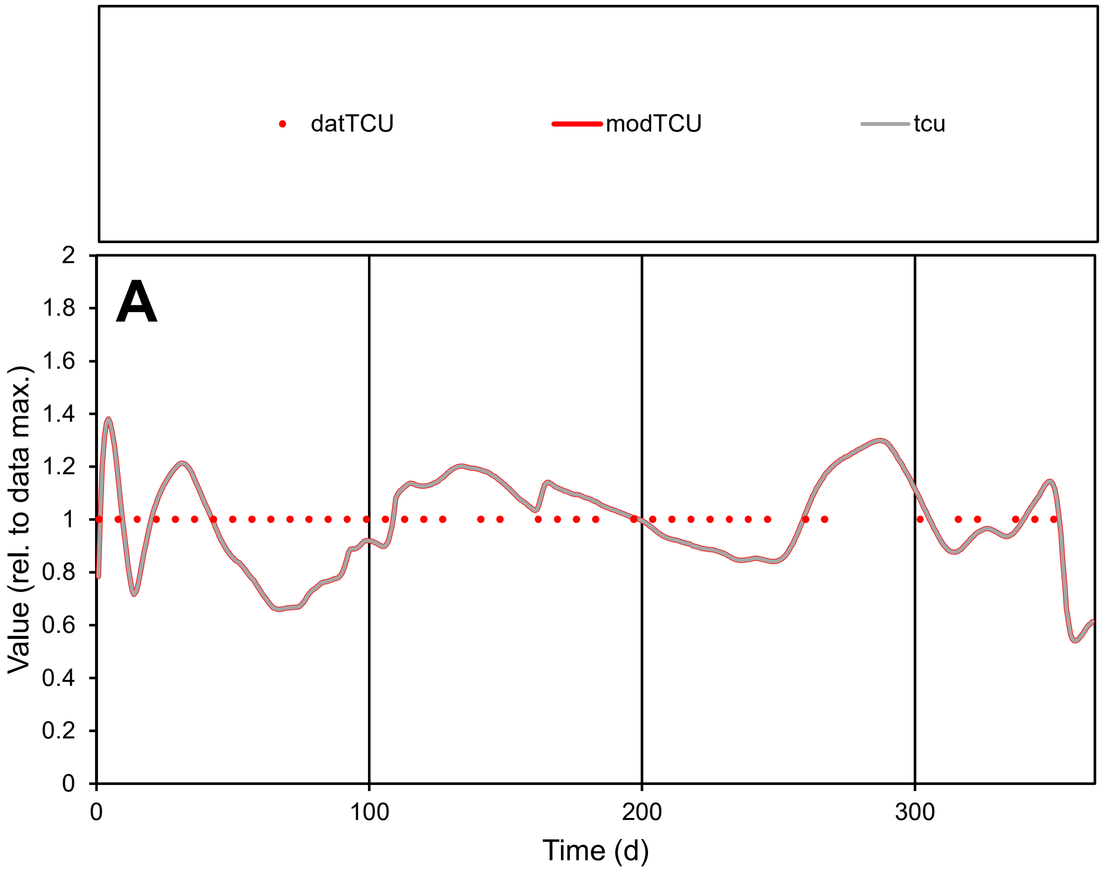


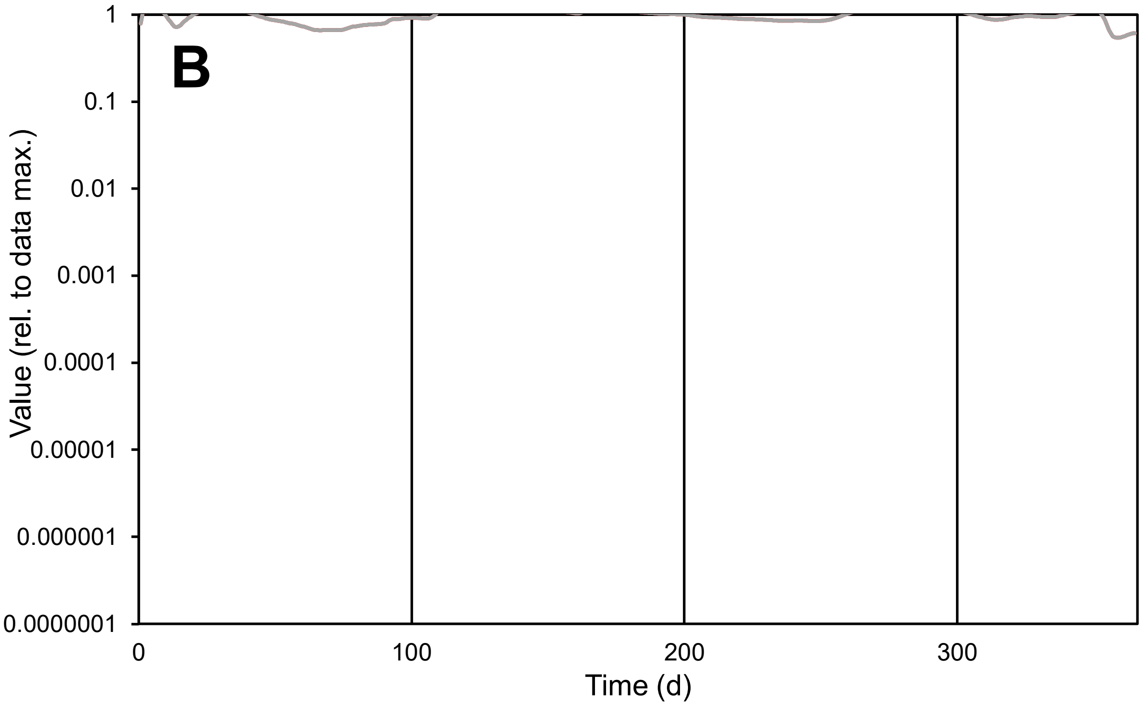


**C1**

Variables

| | |
|---|---|
| TCU | Provide jtd or Dst |
| CalibOnly? | |
| 2 | 1 to reproduce of |
| oNormalize? | |
| 2 | 1=no, 2=yes |
| ofxn calc* | |
| 0.14564 | |
| ofxn mod | |
| 0.18068 | |
| sRun | |
| RunA1 | |
| zout | |
| 1 | |
| Ave | |
| Data | |
| 9.8E-06 | |
| Model | |
| 9.5E-06 | |
| data/vMax | |
| 9.8E-06 | |
| ofxng | |
| 319.391 | |
| Min d/m | |
| 1.49824 | |
| ave/min d | |
| 1 | |
| ave/min m | |
| 1.464 | |

330_Calib_S_A

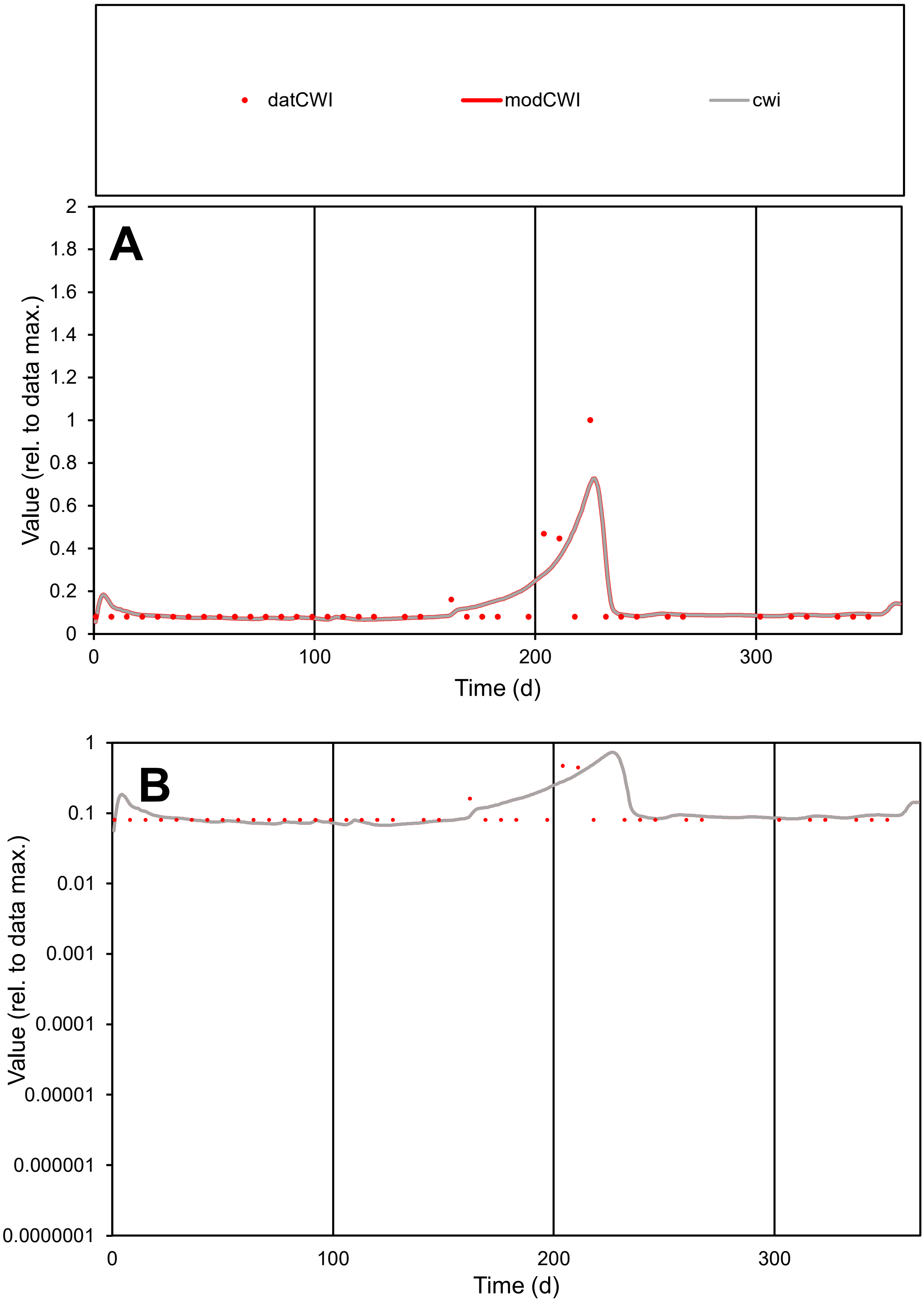


**C1**

Variables
CWI Provide jtd or Dst
CalibOnly?
2 1 to reproduce of
oNormalize?
2 1=no, 2=yes

ofxn calc*
0.71527
ofxn mod
0.75661

sRun
RunA1
zout
1

Ave
Data
2E-05
Model
2.2E-05

data/vMax
0.00017

ofxng
319.391

Min d/m
1.19086

ave/min d
1.53139
ave/min m
1.92862

330_Calib_S_A

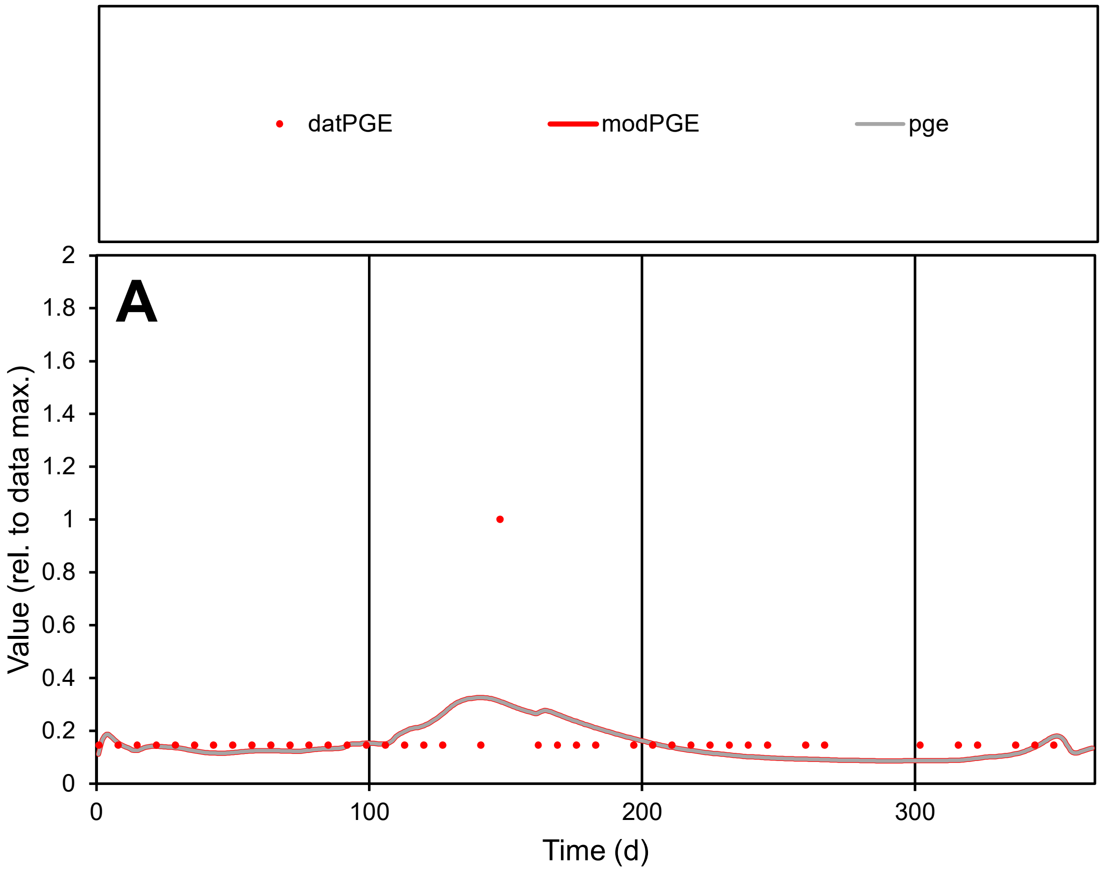


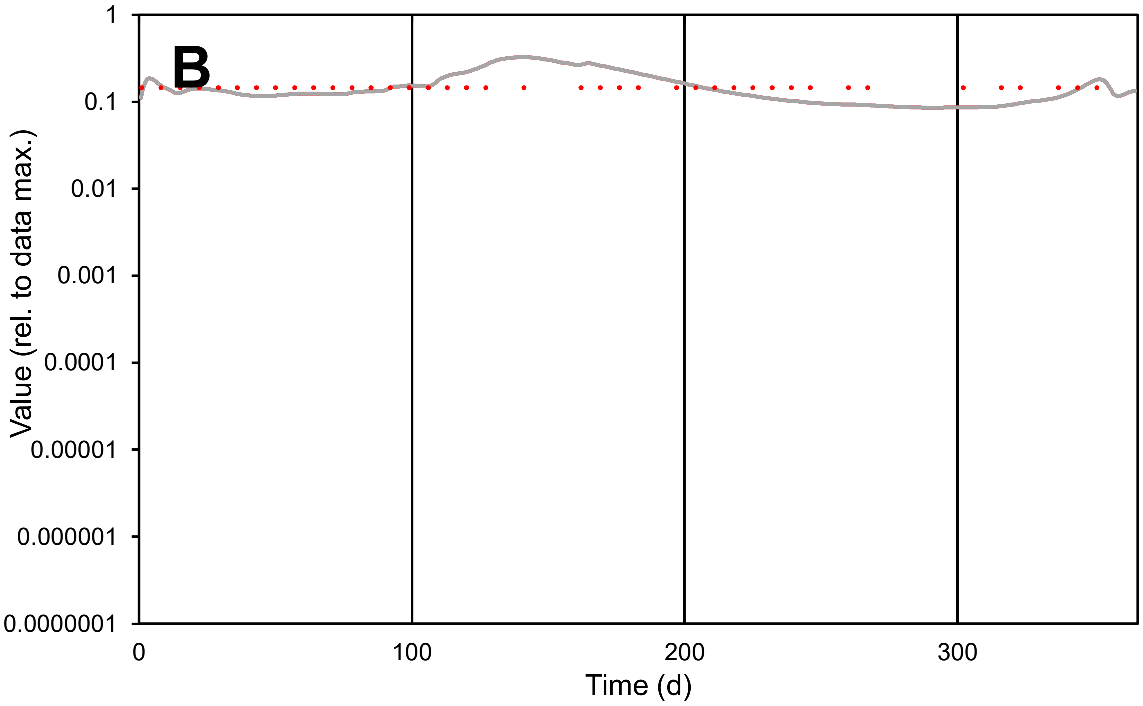


**C1**

Variables
PGE Provide jtd or Dst
CalibOnly?
2 1 to reproduce of
oNormalize?
2 1=no, 2=yes

ofxn calc*
0.73038
ofxn mod
0.74002

sRun
RunA1
zout
1

Ave
Data
6.7E-06
Model
6.2E-06

data/vMax
4E-05

ofxng
319.391

Min d/m
1.67759

ave/min d
1.1433
ave/min m
1.78135

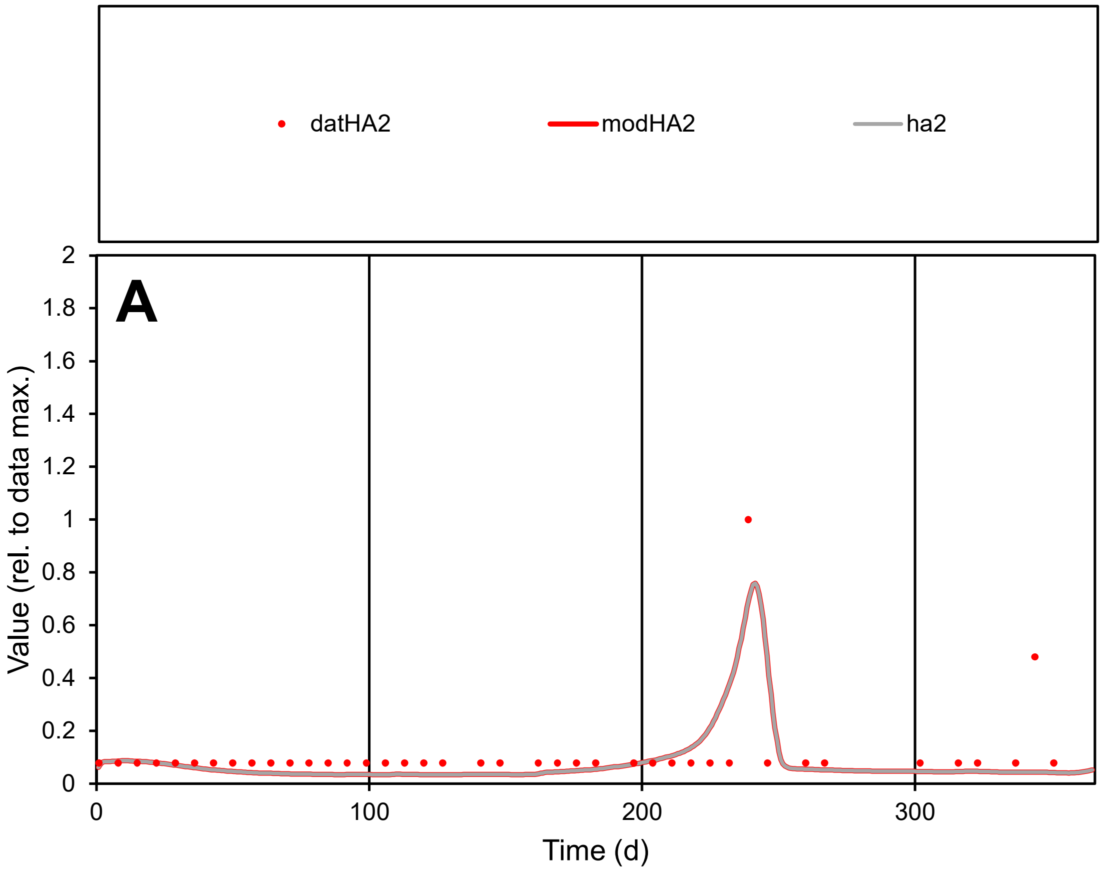


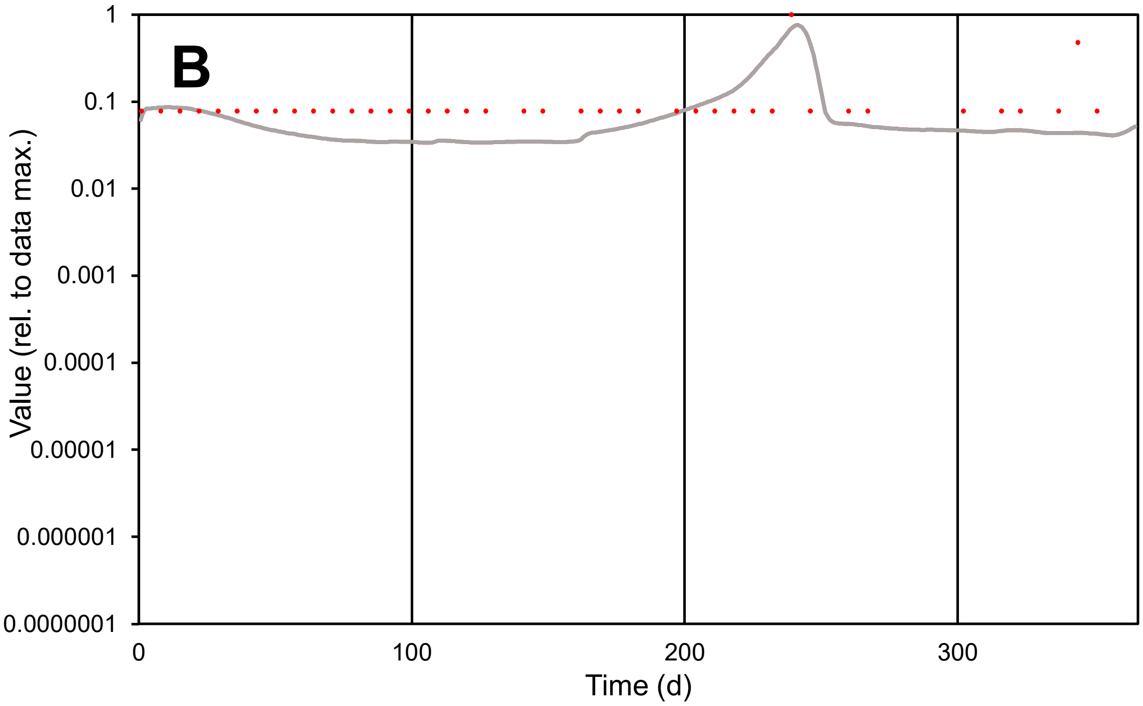


**C1**

Variables
HA2 Provide jtd or Dst
CalibOnly?
2 1 to reproduce of
oNormalize?
2 1=no, 2=yes

ofxn calc*
0.90256
ofxn mod
0.95517

sRun
RunA1
zout
1

Ave
Data
3.7E-06
Model
2.9E-06

data/vMax
3.4E-05

ofxng
319.391

Min d/m
2.28943

ave/min d
1.41401
ave/min m
2.53056

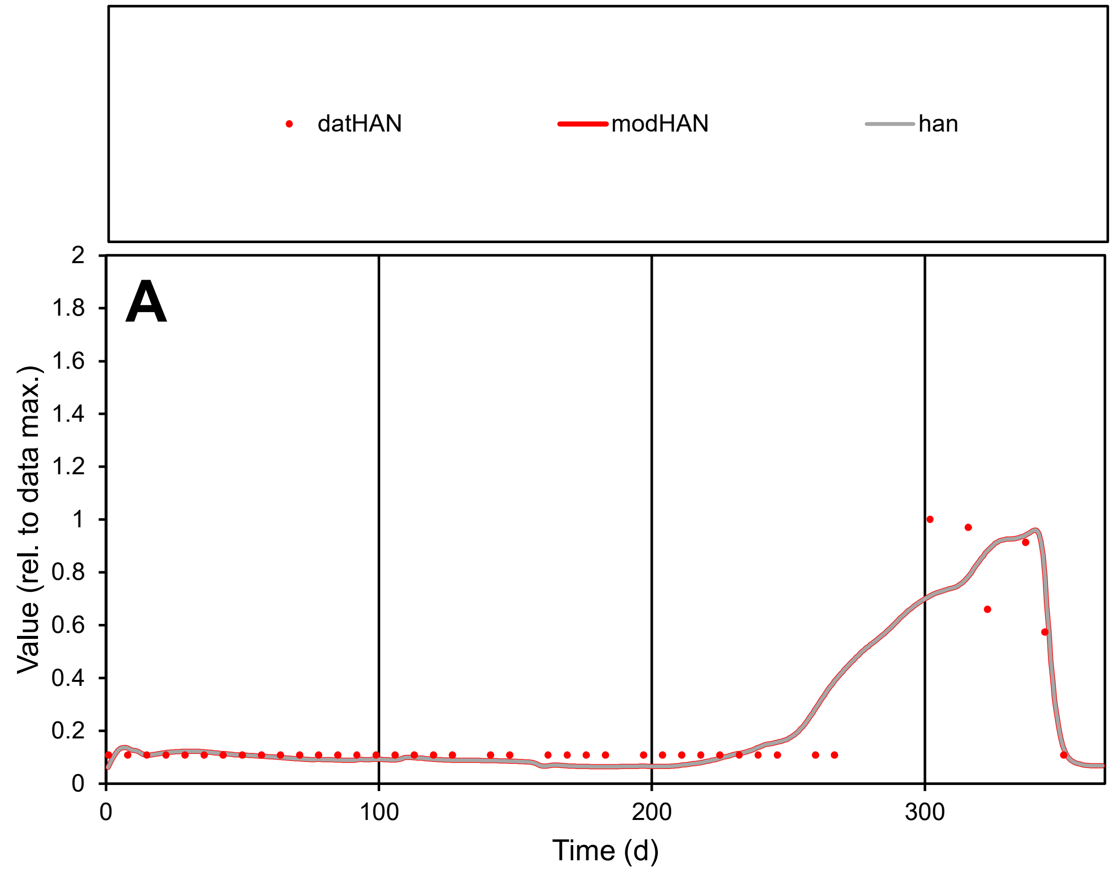


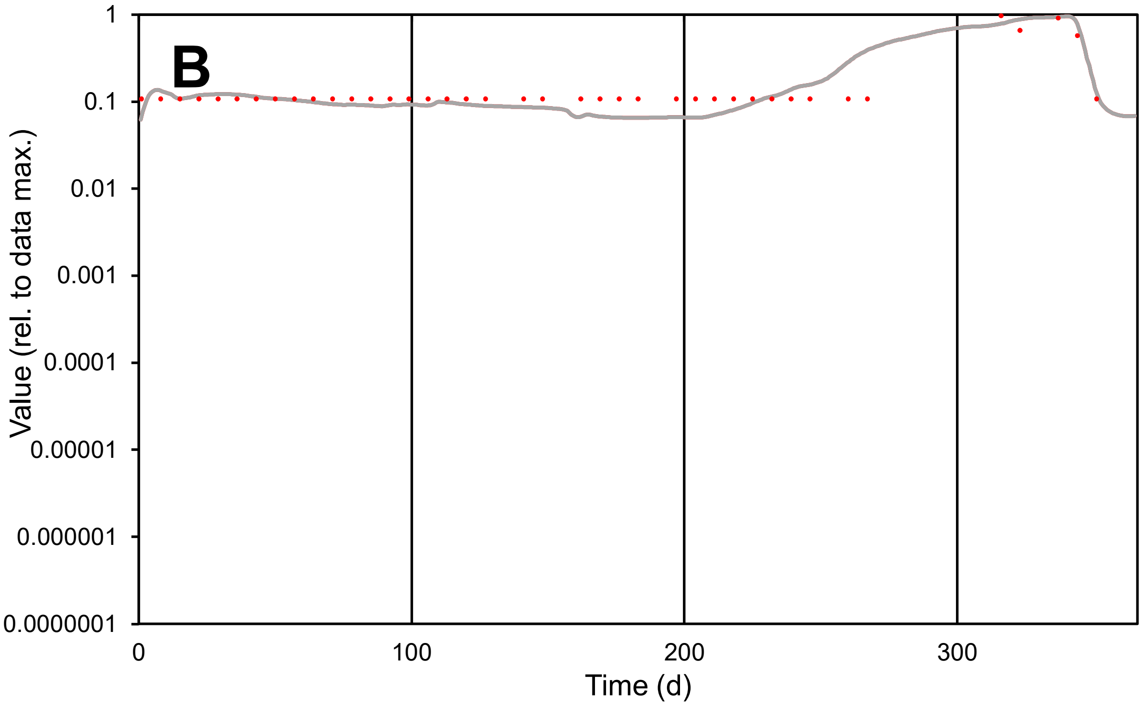


**C1**

Variables
HAN Provide jtd or Dst
CalibOnly?
2 1 to reproduce of
oNormalize?
2 1=no, 2=yes

ofxn calc*
0.4419
ofxn mod
0.4678

sRun
RunA1
zout
1

Ave
Data
9.3E-06
Model
9.2E-06

data/vMax
4.8E-05

ofxng
319.391

Min d/m
1.65664

ave/min d
1.81096
ave/min m
2.9688

330_Calib_S_A

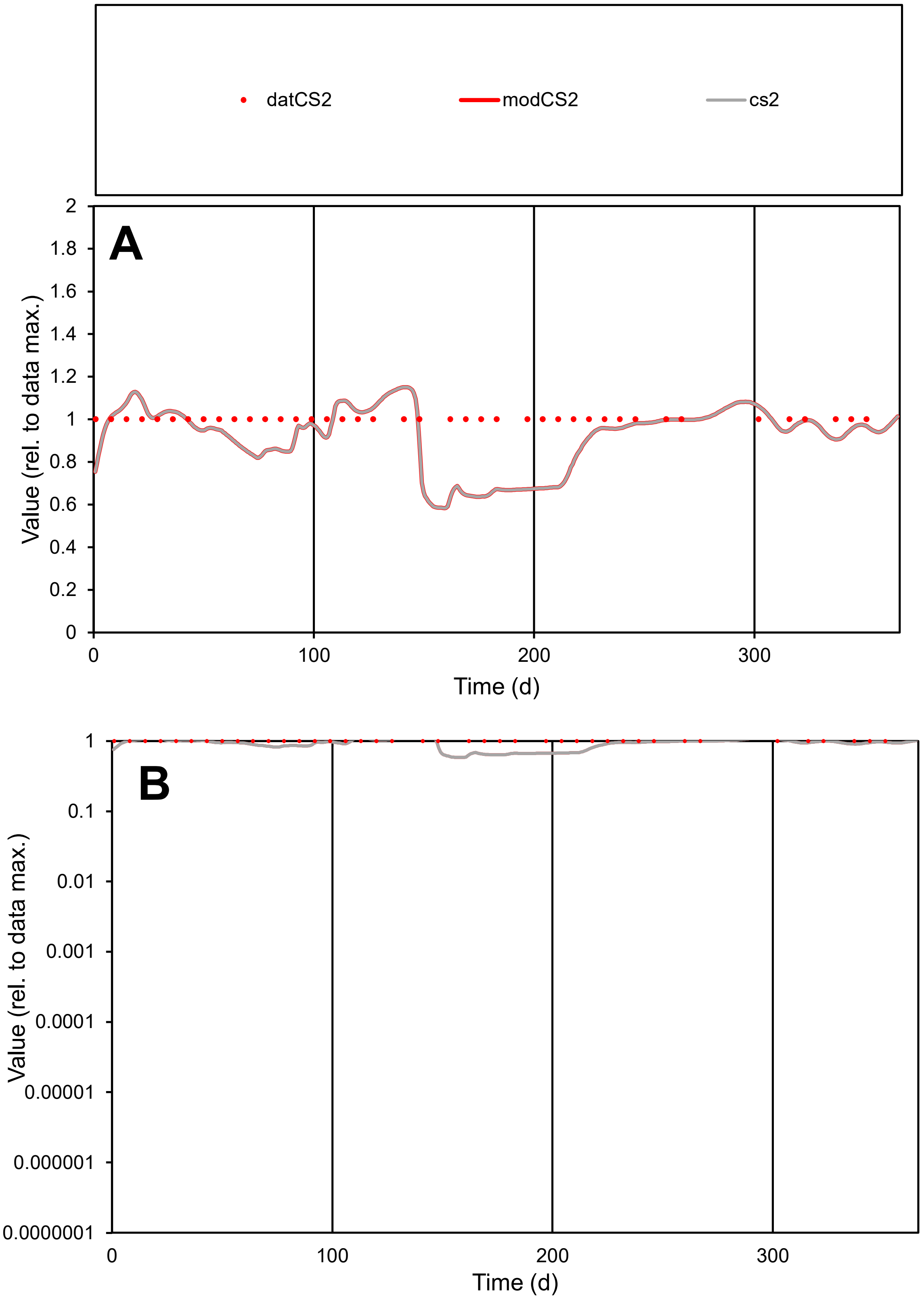


**C1**

Variables
CS2 Provide jtd or Dst
CalibOnly?
2 1 to reproduce of
oNormalize?
2 1=no, 2=yes

ofxn calc*
0.15969
ofxn mod
0.19283

sRun
RunA1
zout
1

Ave
Data
7.6E-06
Model
7E-06

data/vMax
7.6E-06

ofxng
319.391

Min d/m
1.56765

ave/min d
1
ave/min m
1.43791

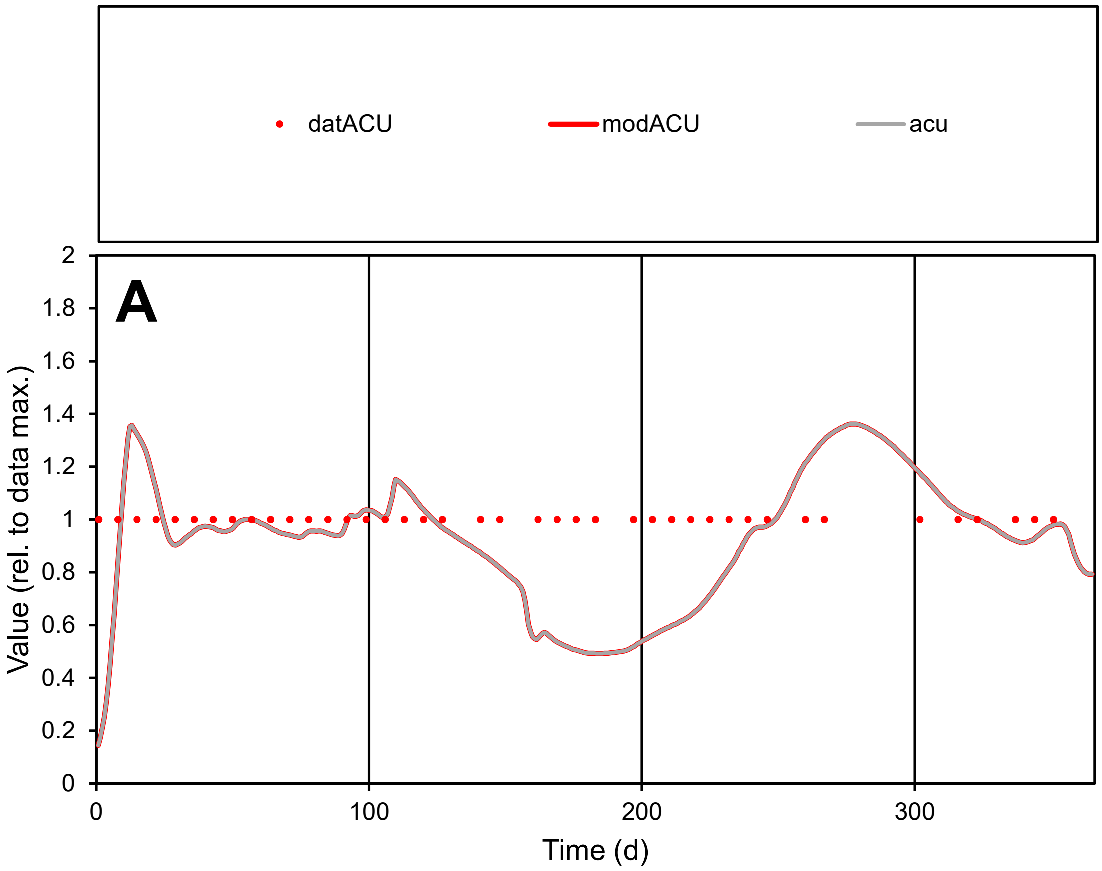


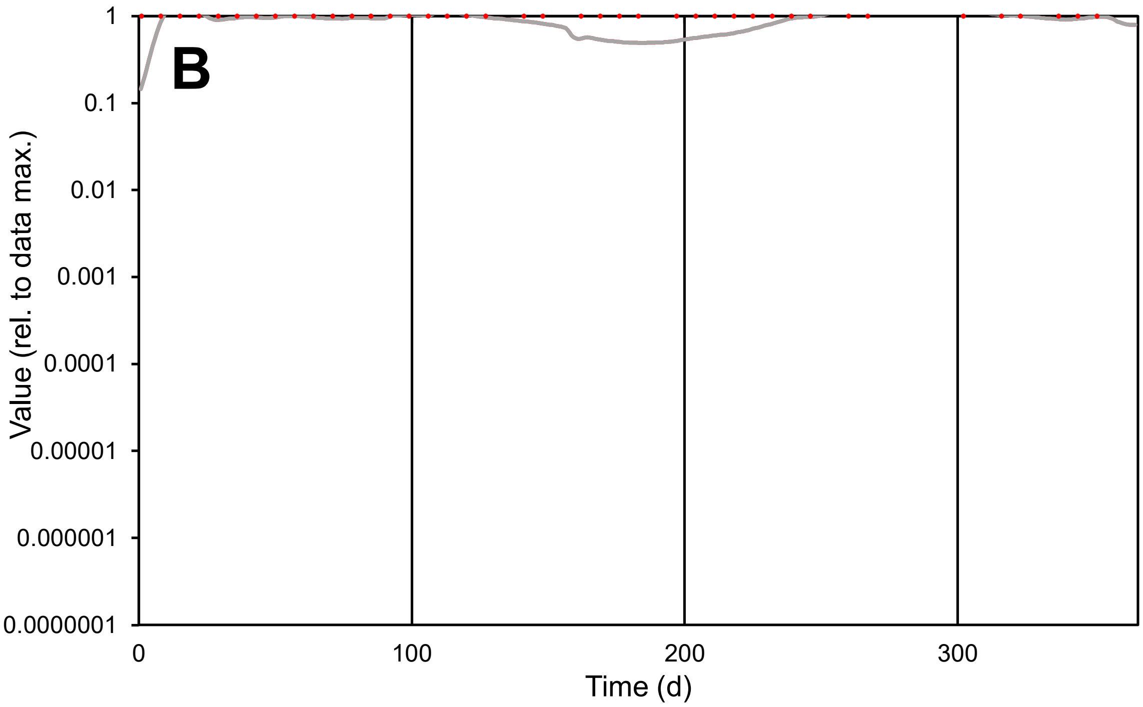


**C1**

Variables
ACU Provide jtd or Dst
CalibOnly?
2 1 to reproduce of
oNormalize?
2 1=no, 2=yes

ofxn calc*
0.24999
ofxn mod
0.27392

sRun
RunA1
zout
1

Ave
Data
1.1E-05
Model
9.9E-06

data/vMax
1.1E-05

ofxng
319.391

Min d/m
3.27246

ave/min d
1
ave/min m
2.92962

330_Calib_S_A

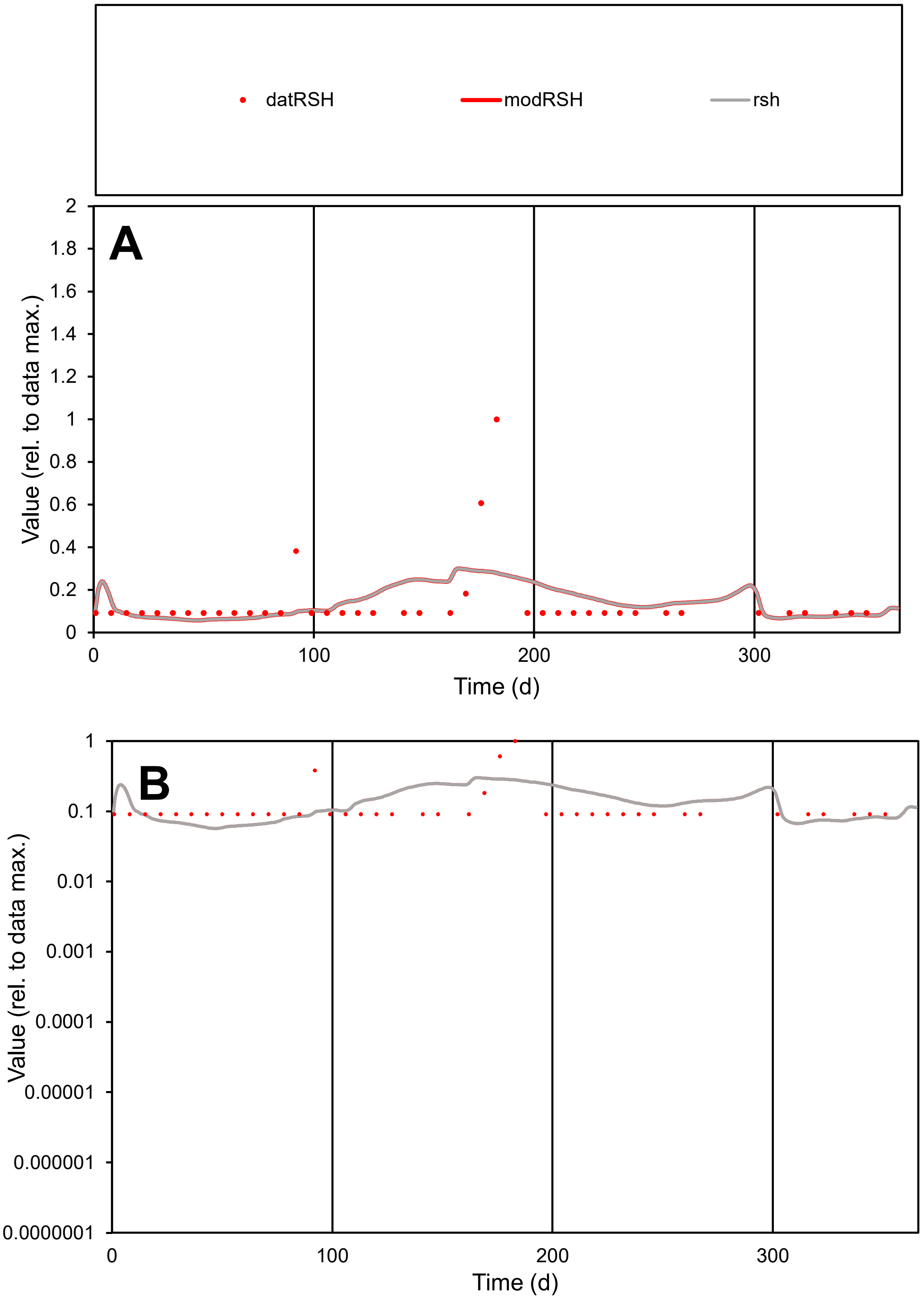


**C1**

Variables
RSH Provide jtd or Dst
CalibOnly?
2 1 to reproduce of
oNormalize?
2 1=no, 2=yes

ofxn calc*
1.09312
ofxn mod
1.10588

sRun
RunA1
zout
1

Ave
Data
2.7E-05
Model
2.7E-05

data/vMax
0.0002

ofxng
319.391

Min d/m
1.57218

ave/min d
1.48557
ave/min m
2.35486

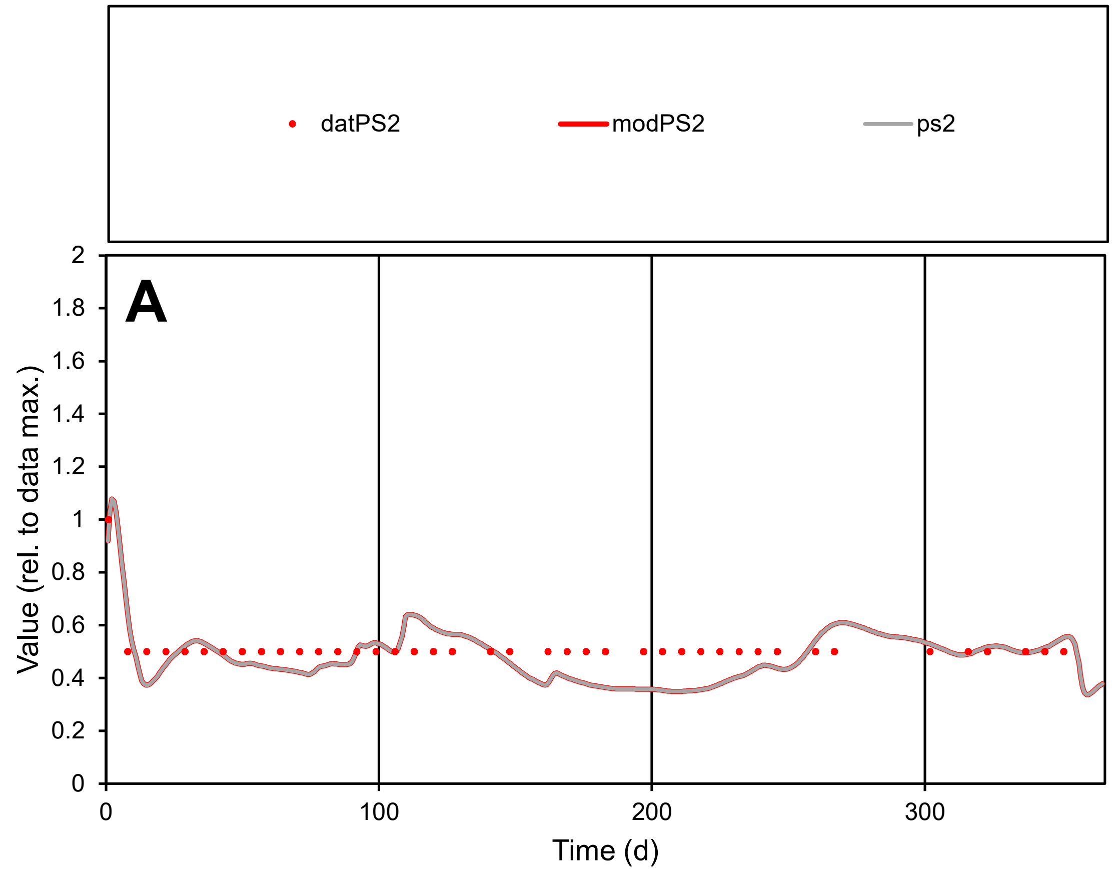


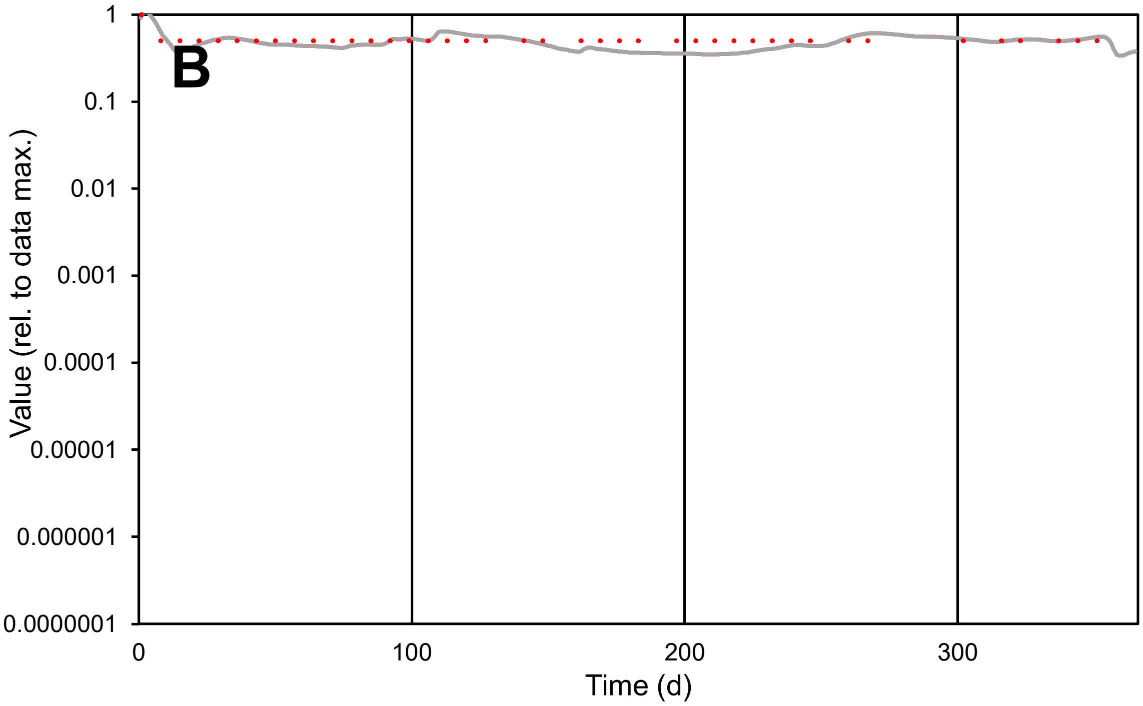


**C1**

Variables
PS2 Provide jtd or Dst
CalibOnly?
2 1 to reproduce of
oNormalize?
2 1=no, 2=yes

ofxn calc*
0.15631
ofxn mod
0.18797

sRun
RunA1
zout
1

Ave
Data
2E-05
Model
1.8E-05

data/vMax
3.8E-05

ofxng
319.391

Min d/m
1.4259

ave/min d
1.02439
ave/min m
1.3784

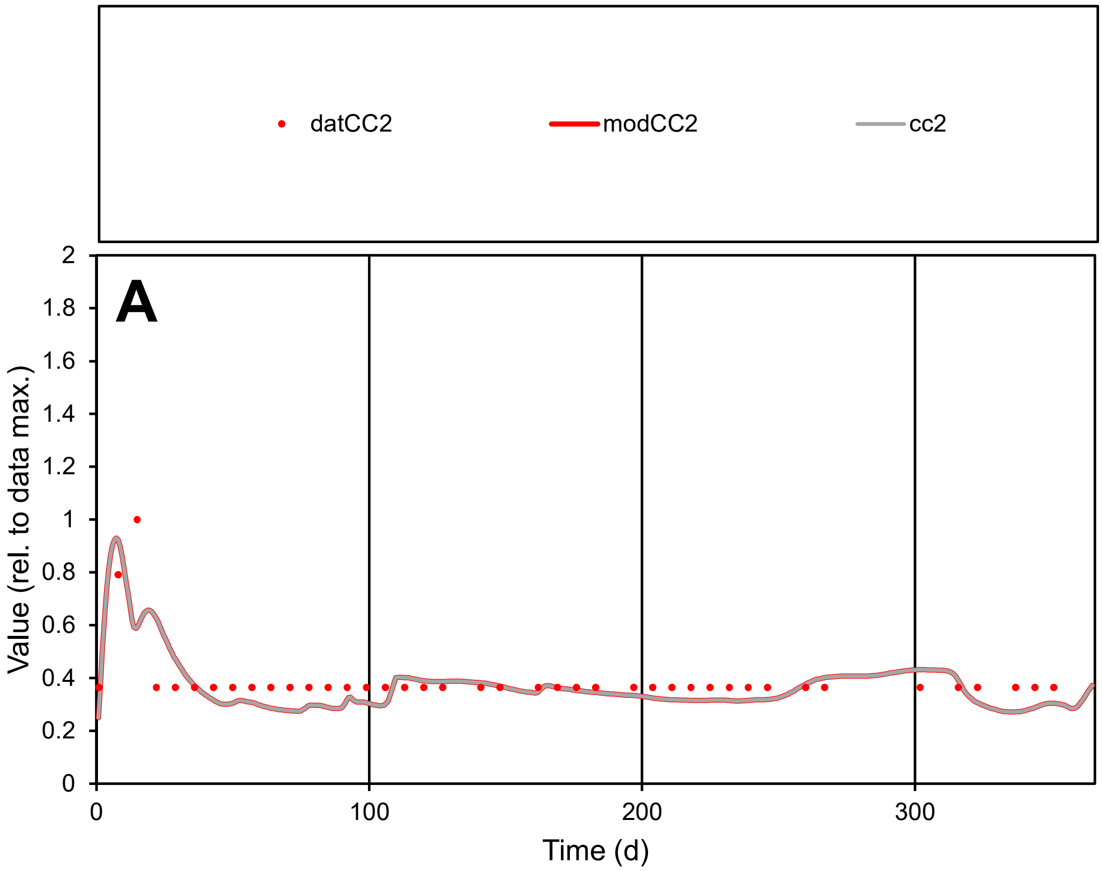


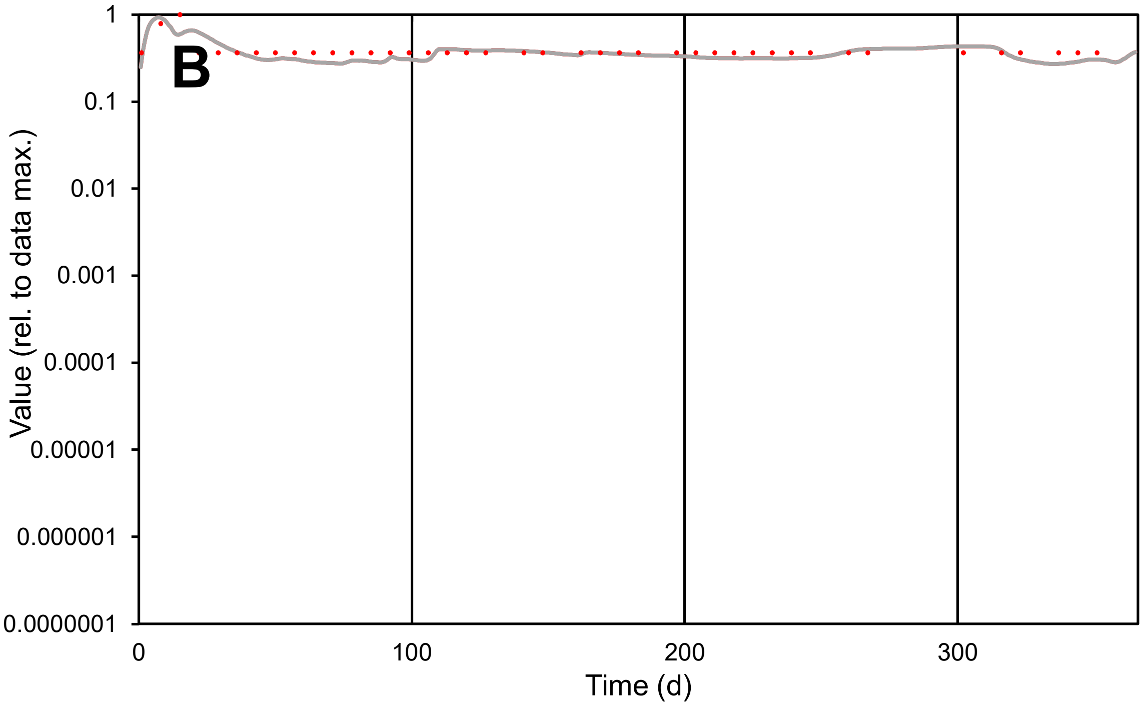


**C1**

Variables
CC2 Provide jtd or Dst
CalibOnly?
2 1 to reproduce of
oNormalize?
2 1=no, 2=yes

ofxn calc*
0.21532
ofxn mod
0.24311

sRun
RunA1
zout
1

Ave
Data
6.3E-06
Model
5.9E-06

data/vMax
1.6E-05

ofxng
319.391

Min d/m
1.3353

ave/min d
1.07113
ave/min m
1.33558

330_Calib_S_A

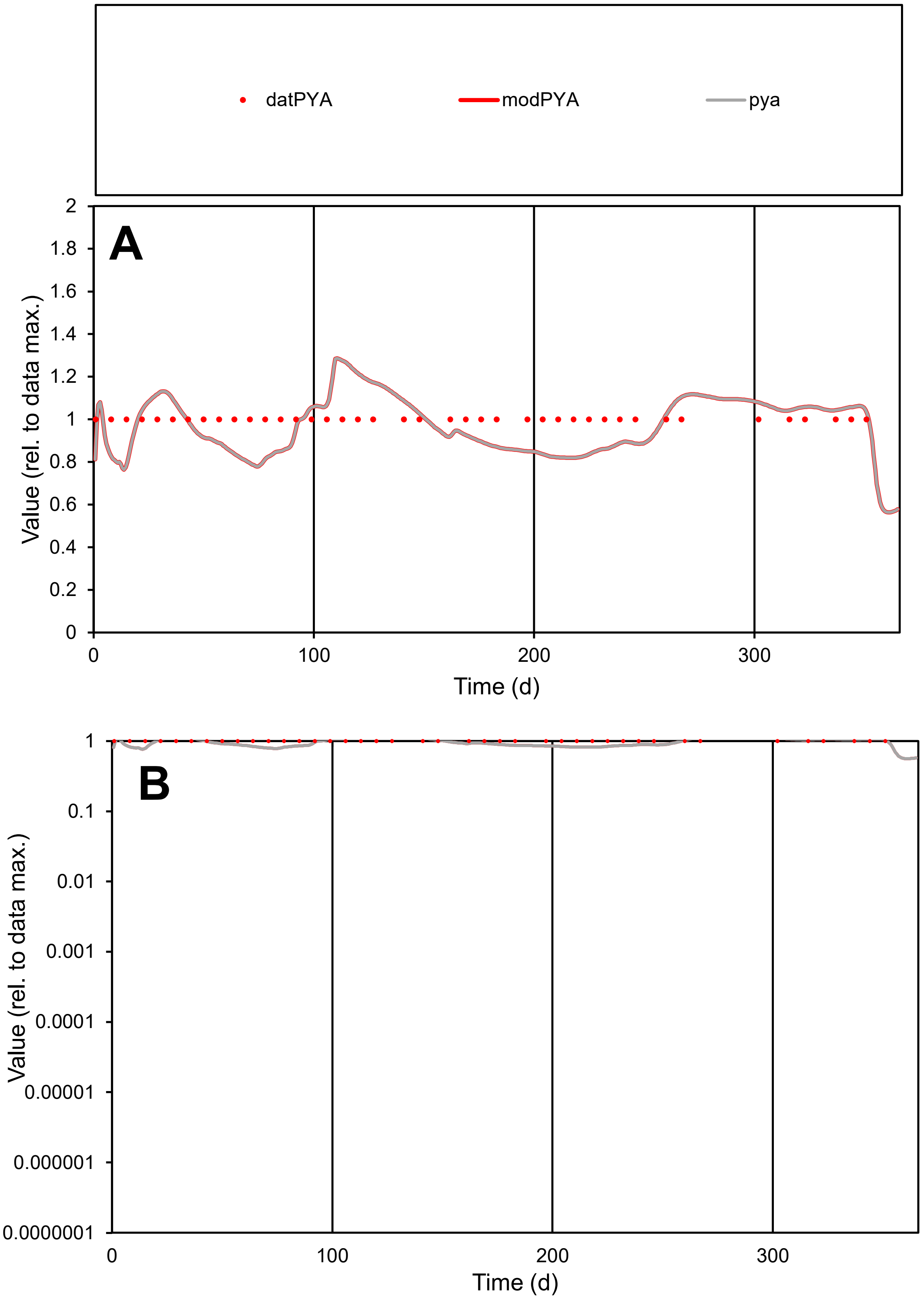


**C1**

Variables
PYA Provide jtd or Dst
CalibOnly?
2 1 to reproduce of
oNormalize?
2 1=no, 2=yes

ofxn calc*
0.1261
ofxn mod
0.16497

sRun
RunA1
zout
1

Ave
Data
1.8E-06
Model
1.8E-06

data/vMax
1.8E-06

ofxng
319.391

Min d/m
1.26713

ave/min d
1
ave/min m
1.2291

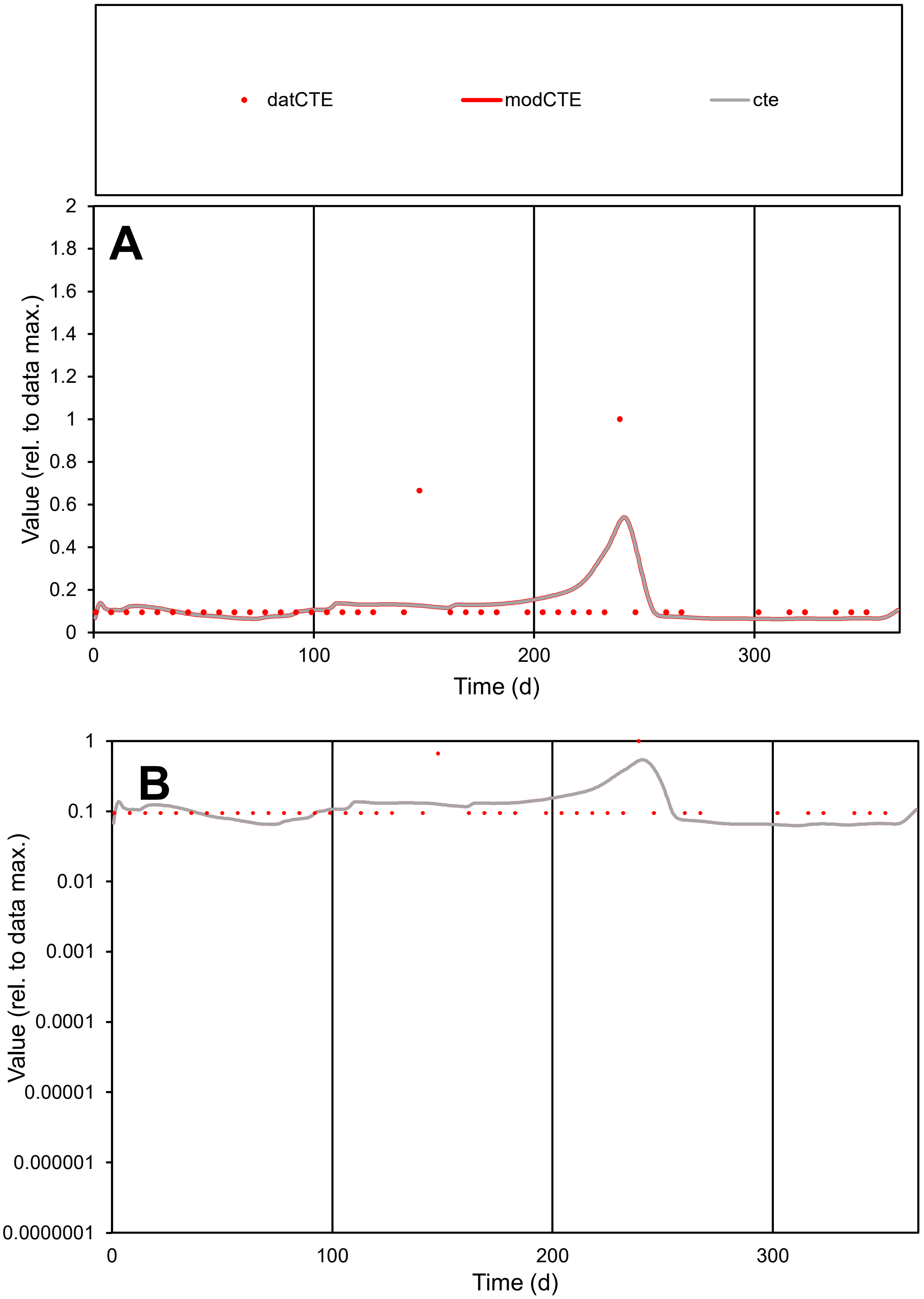


**C1**

Variables

| | |
|---|---|
| CTE | Provide jtd or Dst |
| CalibOnly? | |
| 2 | 1 to reproduce of |
| oNormalize? | |
| 2 | 1=no, 2=yes |

ofxn calc*
1.00704
ofxn mod
1.03189

sRun
RunA1
zout
1

Ave
Data
8.5E-06
Model
8.5E-06

data/vMax
6.5E-05

ofxng
319.391

Min d/m
1.4733

ave/min d
1.3809
ave/min m
2.05423

330_Calib_S_A

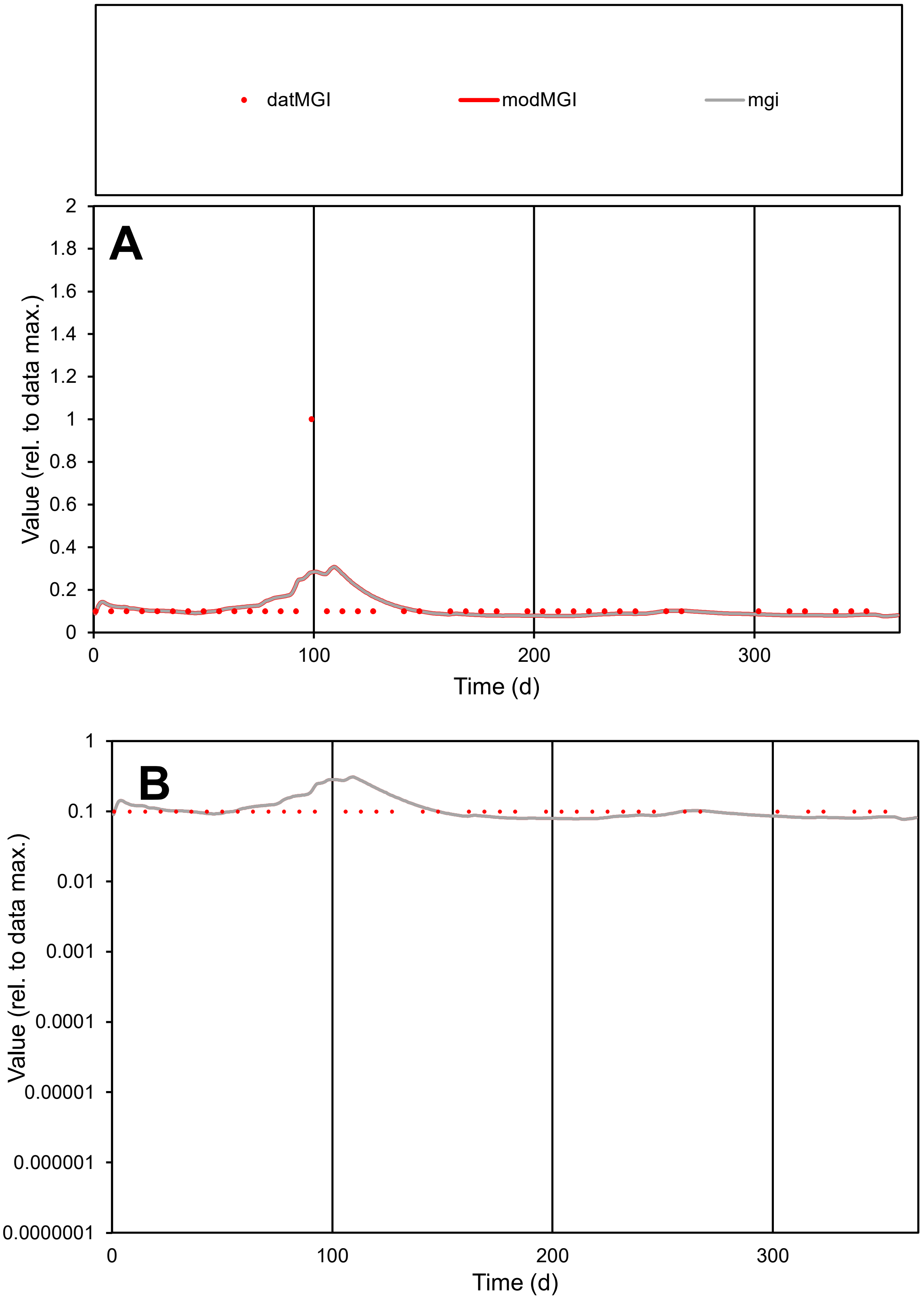


**C1**

Variables

MGI Provide jtd or Dst

CalibOnly?

2 1 to reproduce of

oNormalize?

2 1=no, 2=yes

ofxn calc*

1.01668

ofxn mod

1.02758

sRun

RunA1

zout

1

Ave

Data

1.3E-05

Model

1.2E-05

data/vMax

0.0001

ofxng

319.391

Min d/m

1.26828

ave/min d

1.22211

ave/min m

1.50969

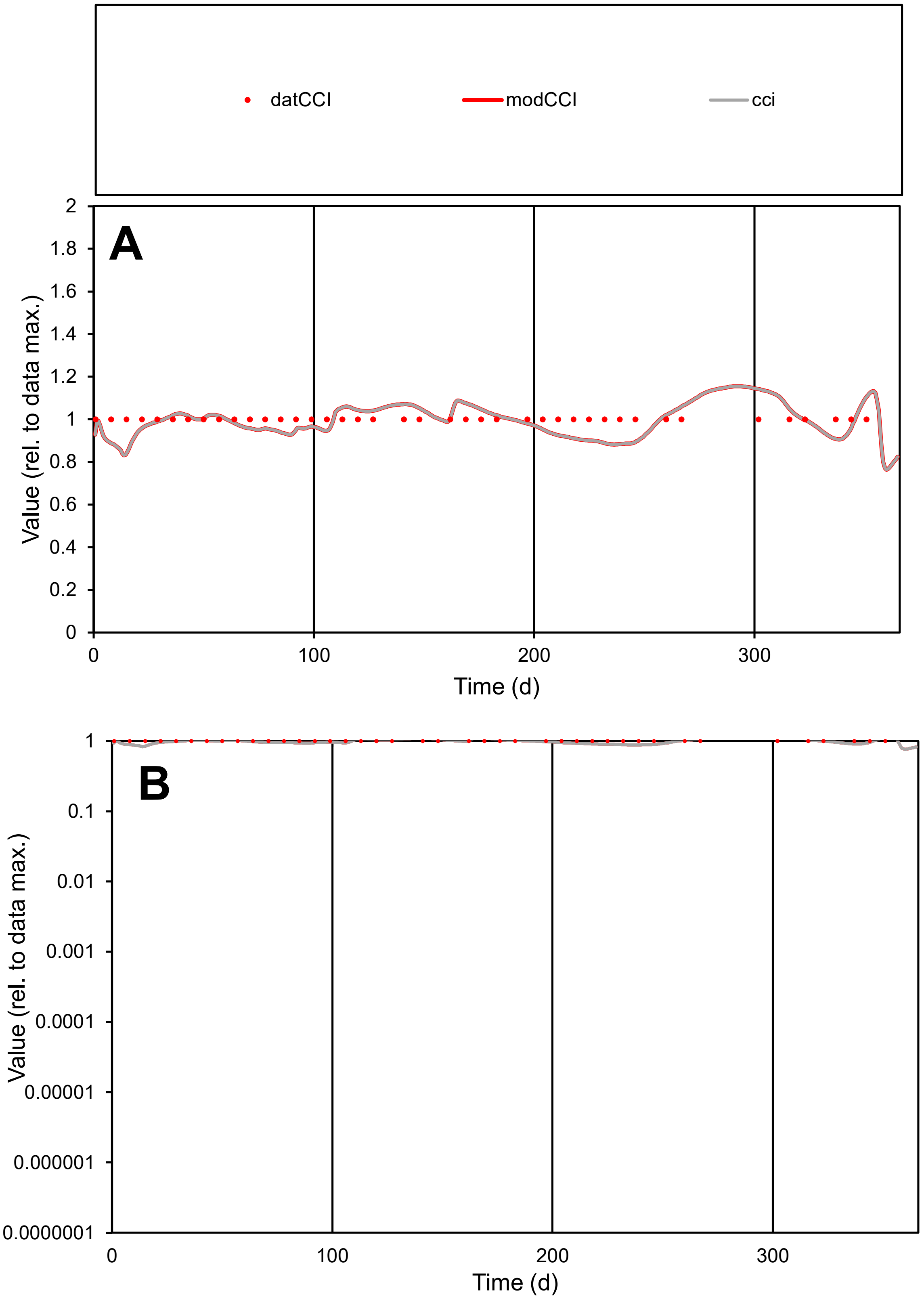


**C1**

Variables
CCI Provide jtd or Dst
CalibOnly?
2 1 to reproduce of
oNormalize?
2 1=no, 2=yes

ofxn calc*
0.0665
ofxn mod
0.12672

sRun
RunA1
zout
1

Ave
Data
1.1E-05
Model
1.1E-05

data/vMax
1.1E-05

ofxng
319.391

Min d/m
1.13776

ave/min d
1
ave/min m
1.12483

330_Calib_S_A

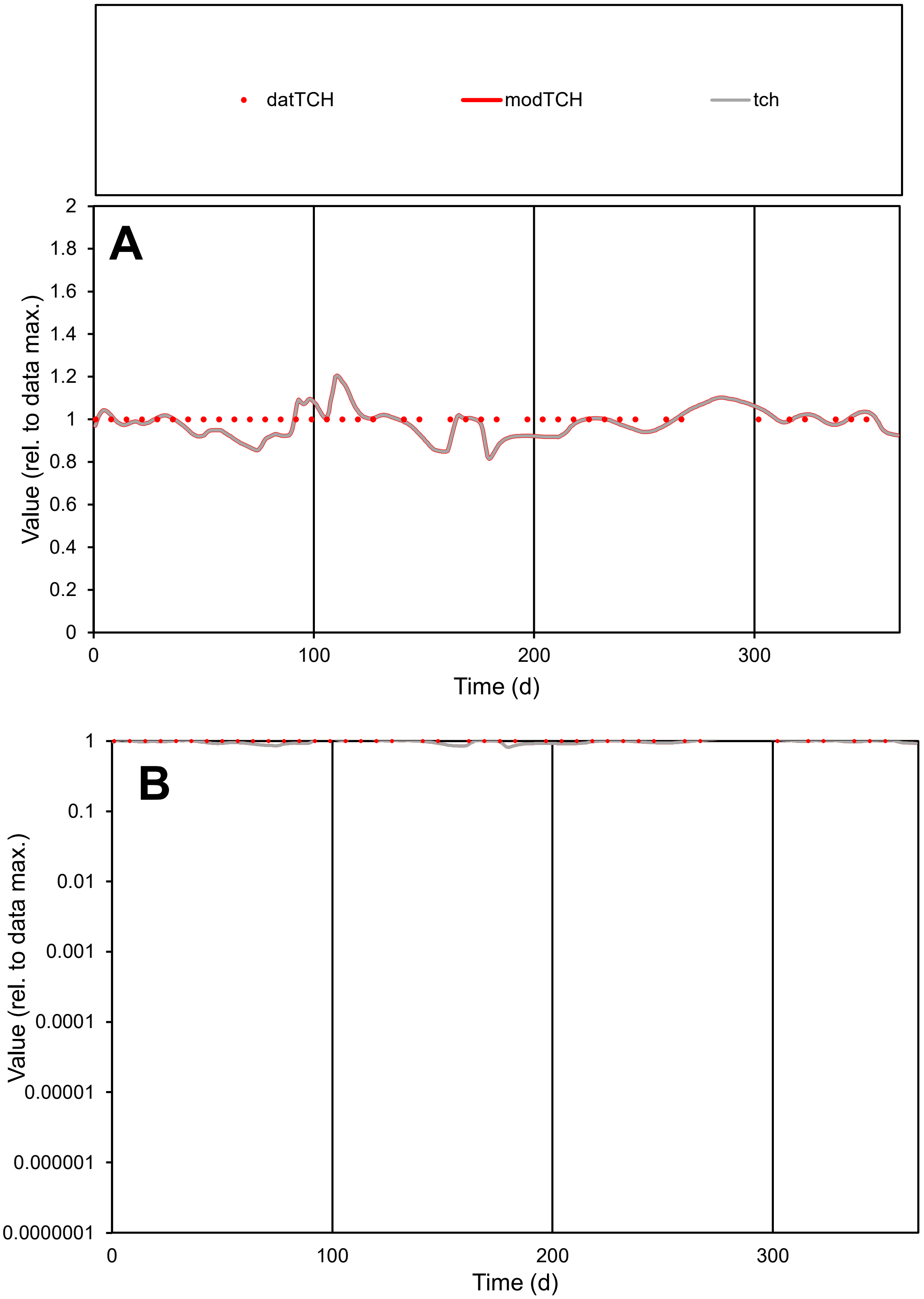


**C1**

Variables
TCH Provide jtd or Dst
CalibOnly?
2 1 to reproduce of
oNormalize?
2 1=no, 2=yes

ofxn calc*
0.06207
ofxn mod
0.12437

sRun
RunA1
zout
1

Ave
Data
1.7E-06
Model
1.7E-06

data/vMax
1.7E-06

ofxng
319.391

Min d/m
1.1584

ave/min d
1
ave/min m
1.13826

330_Calib_S_A

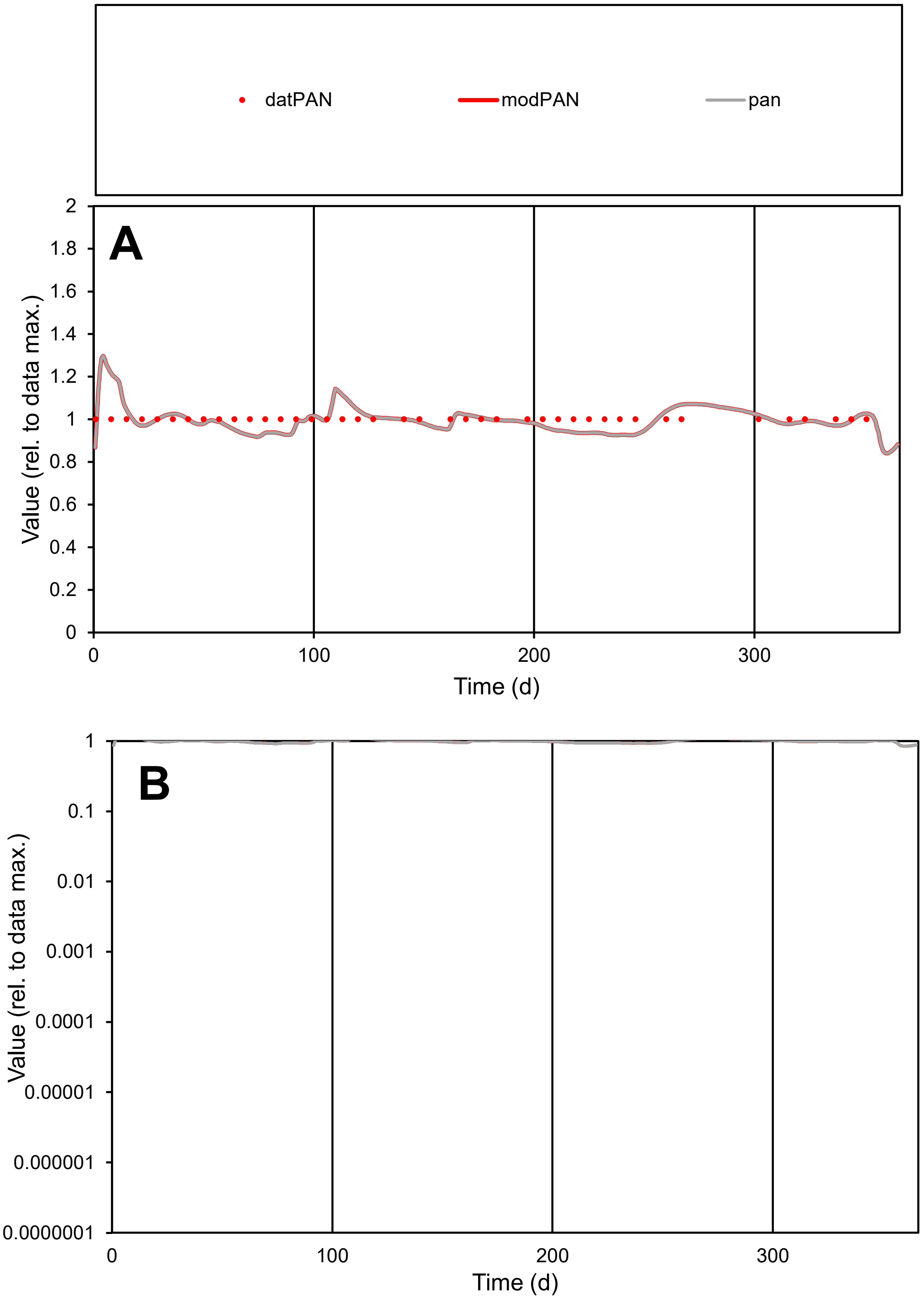


**C1**

Variables
PAN Provide jtd or Dst
CalibOnly?
2 1 to reproduce of
oNormalize?
2 1=no, 2=yes

ofxn calc*
0.05194
ofxn mod
0.12046

sRun
RunA1
zout
1

Ave
Data
9.1E-06
Model
9.1E-06

data/vMax
9.1E-06

ofxng
319.391

Min d/m
1.08326

ave/min d
1
ave/min m
1.07783

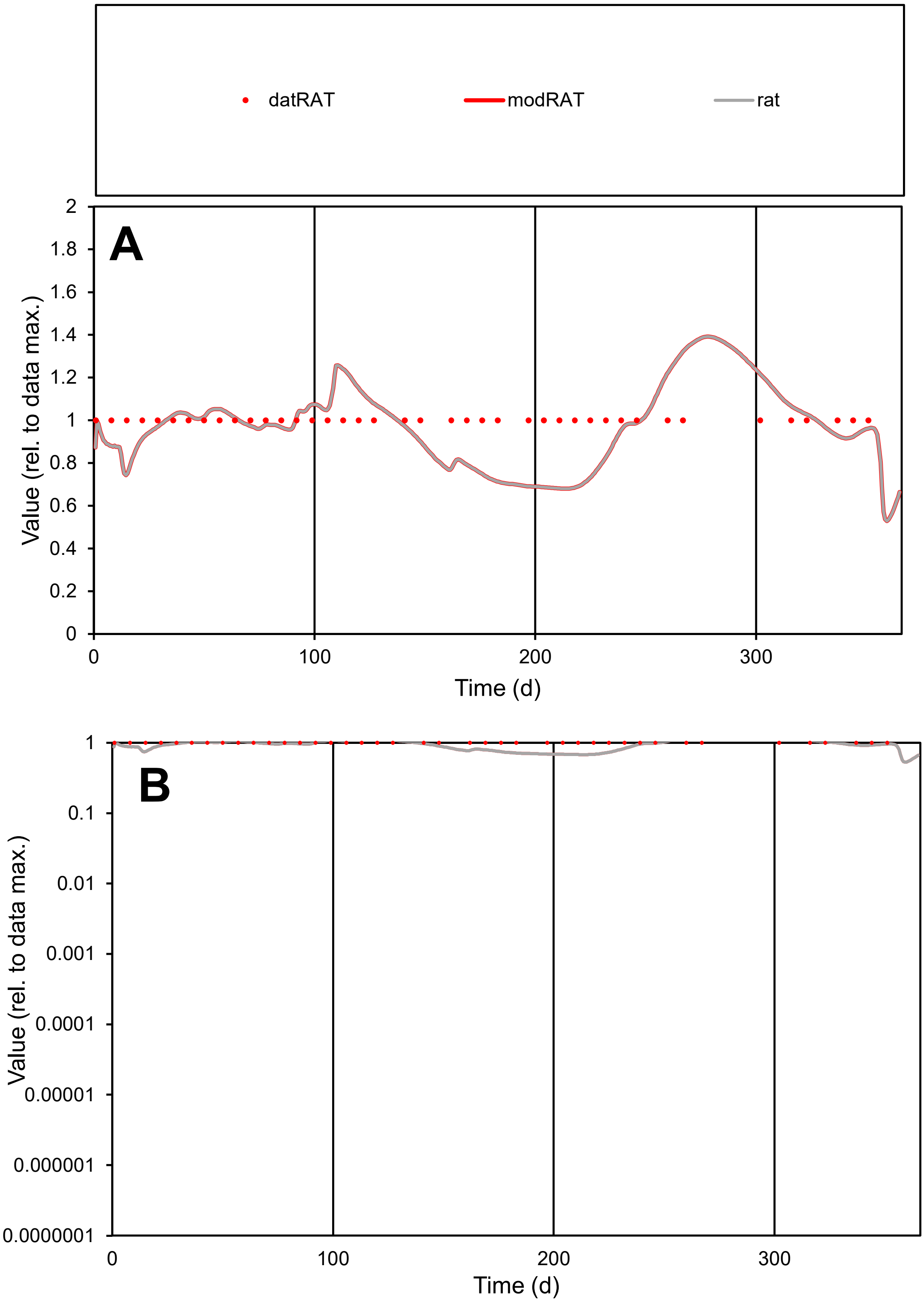


**C1**

Variables
RAT Provide jtd or Dst
CalibOnly?
2 1 to reproduce of
oNormalize?
2 1=no, 2=yes

ofxn calc*
0.16524
ofxn mod
0.1963

sRun
RunA1
zout
1

Ave
Data
1.1E-05
Model
1.1E-05

data/vMax
1.1E-05

ofxng
319.391

Min d/m
1.46956

ave/min d
1
ave/min m
1.40381

330_Calib_S_A

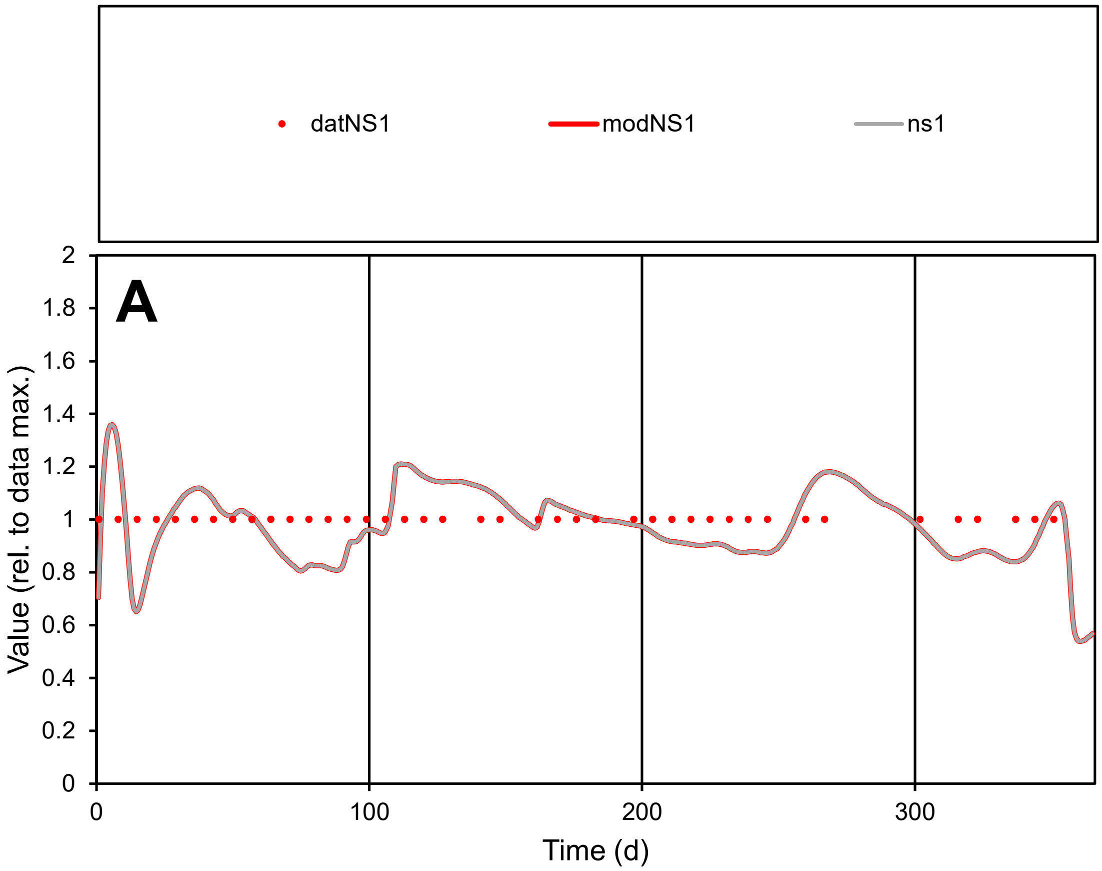


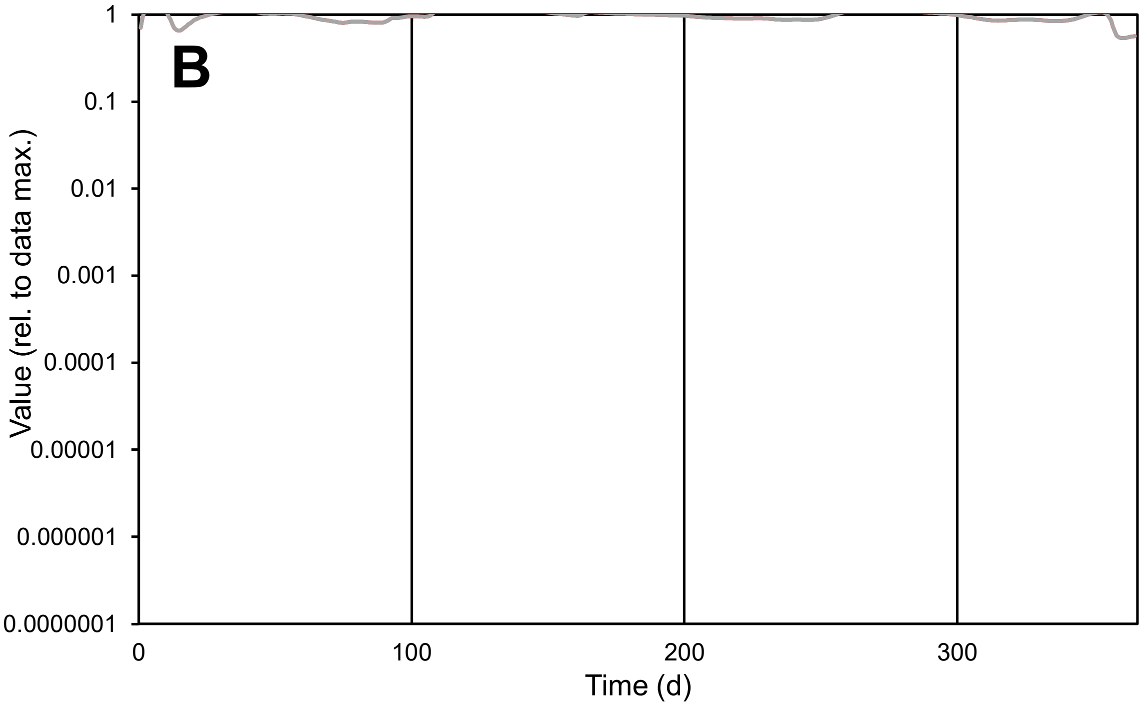


**C1**

Variables
NS1 Provide jtd or Dst
CalibOnly?
2 1 to reproduce of
oNormalize?
2 1=no, 2=yes

ofxn calc*
0.11571
ofxn mod
0.1578

sRun
RunA1
zout
1

Ave
Data
2.7E-05
Model
2.7E-05

data/vMax
2.7E-05

ofxng
319.391

Min d/m
1.41473

ave/min d
1
ave/min m
1.3866

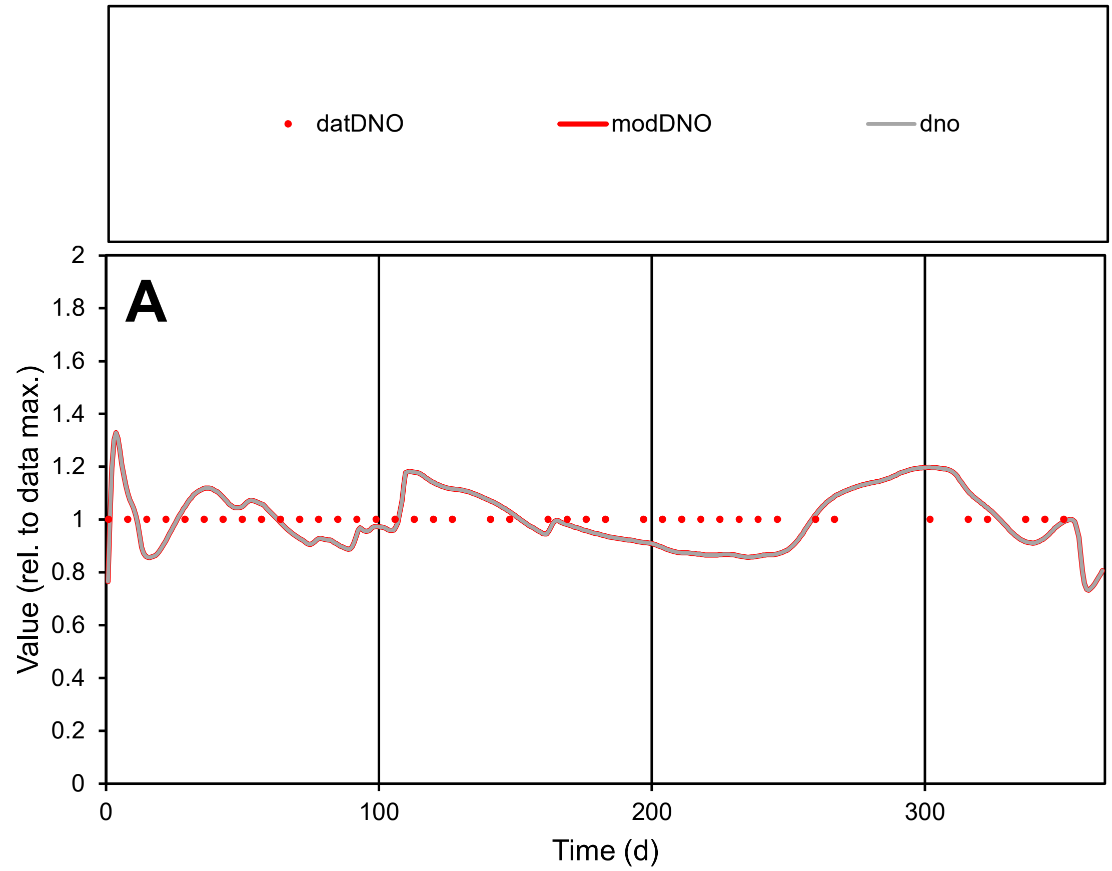


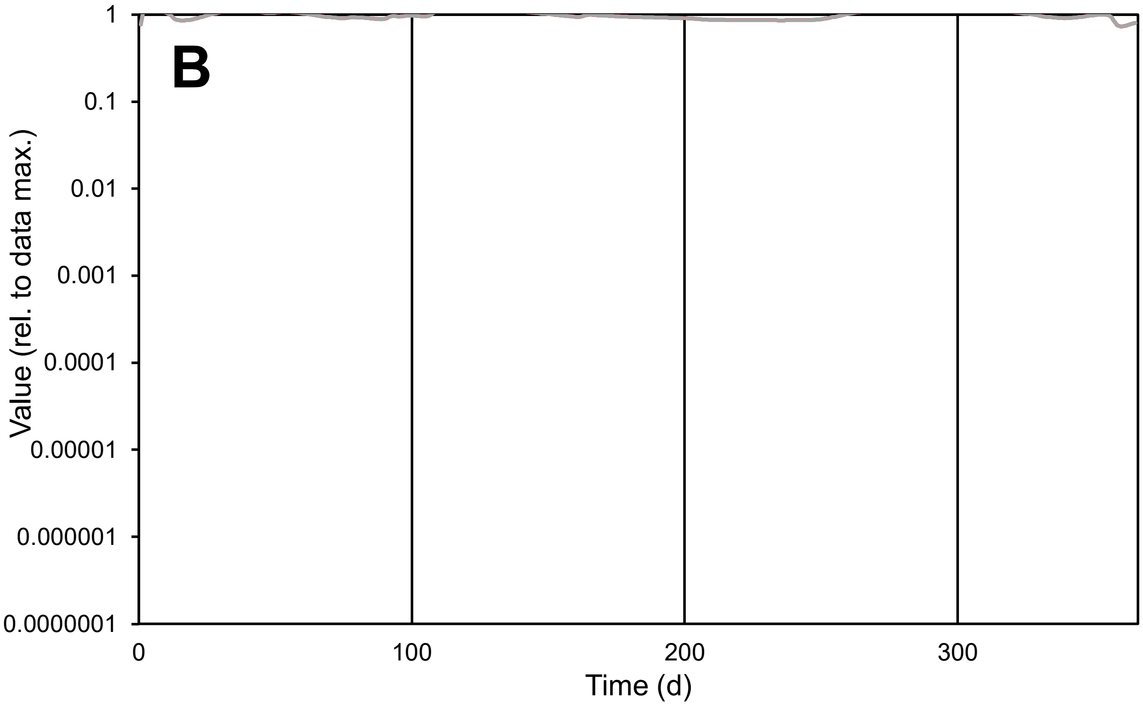


**C1**

Variables

DNO Provide jtd or Dst

CalibOnly?

2 1 to reproduce of

oNormalize?

2 1=no, 2=yes

ofxn calc*

0.09241

ofxn mod

0.14201

sRun

RunA1

zout

1

Ave

Data

1.8E-05

Model

1.8E-05

data/vMax

1.8E-05

ofxng

319.391

Min d/m

1.16549

ave/min d

1

ave/min m

1.15404

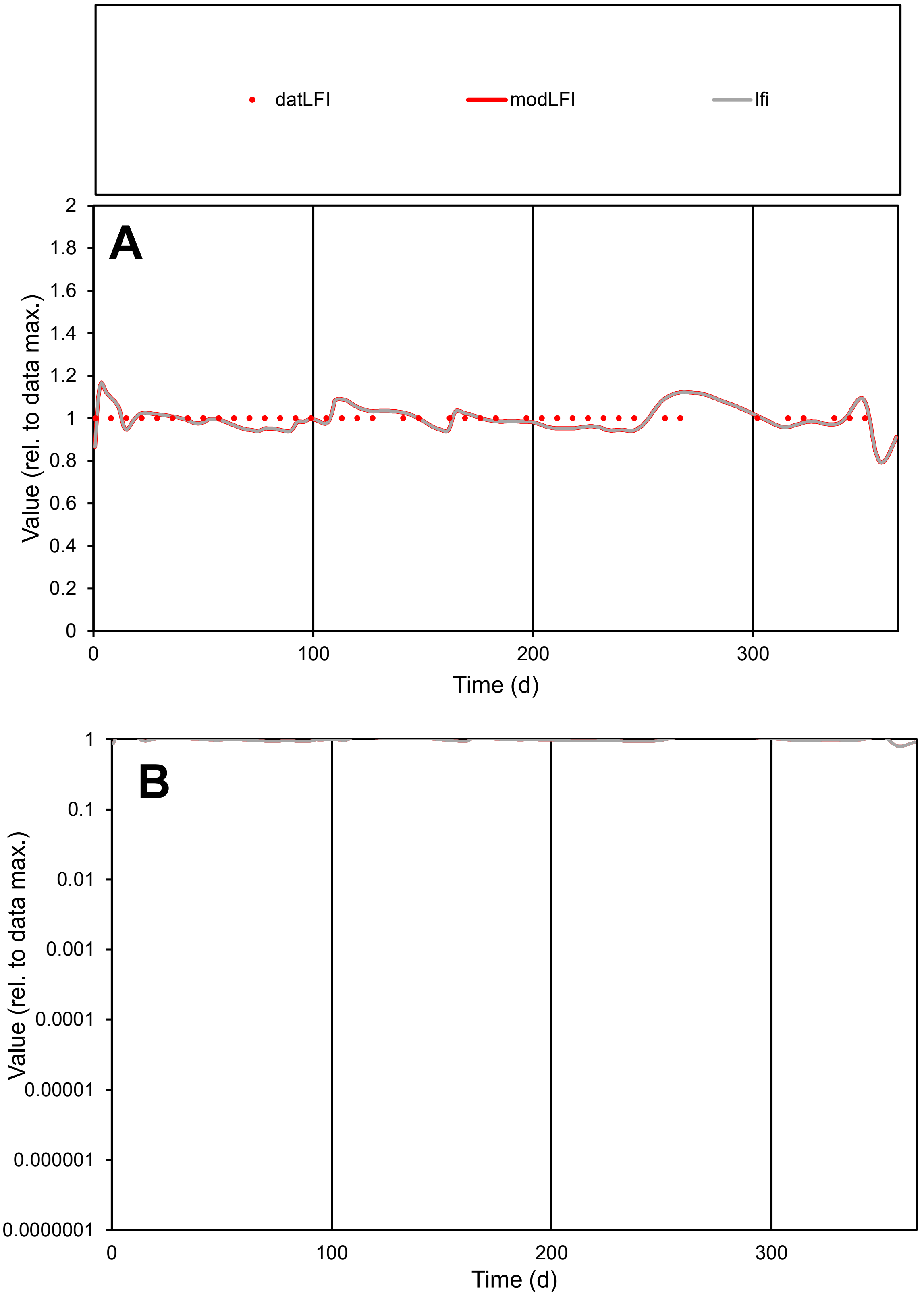


**C1**

Variables
LFI Provide jtd or Dst
CalibOnly?
2 1 to reproduce of
oNormalize?
2 1=no, 2=yes

ofxn calc*
0.04436
ofxn mod
0.11737

sRun
RunA1
zout
1

Ave
Data
8.3E-06
Model
8.3E-06

data/vMax
8.3E-06

ofxng
319.391

Min d/m
1.06212

ave/min d
1
ave/min m
1.06067

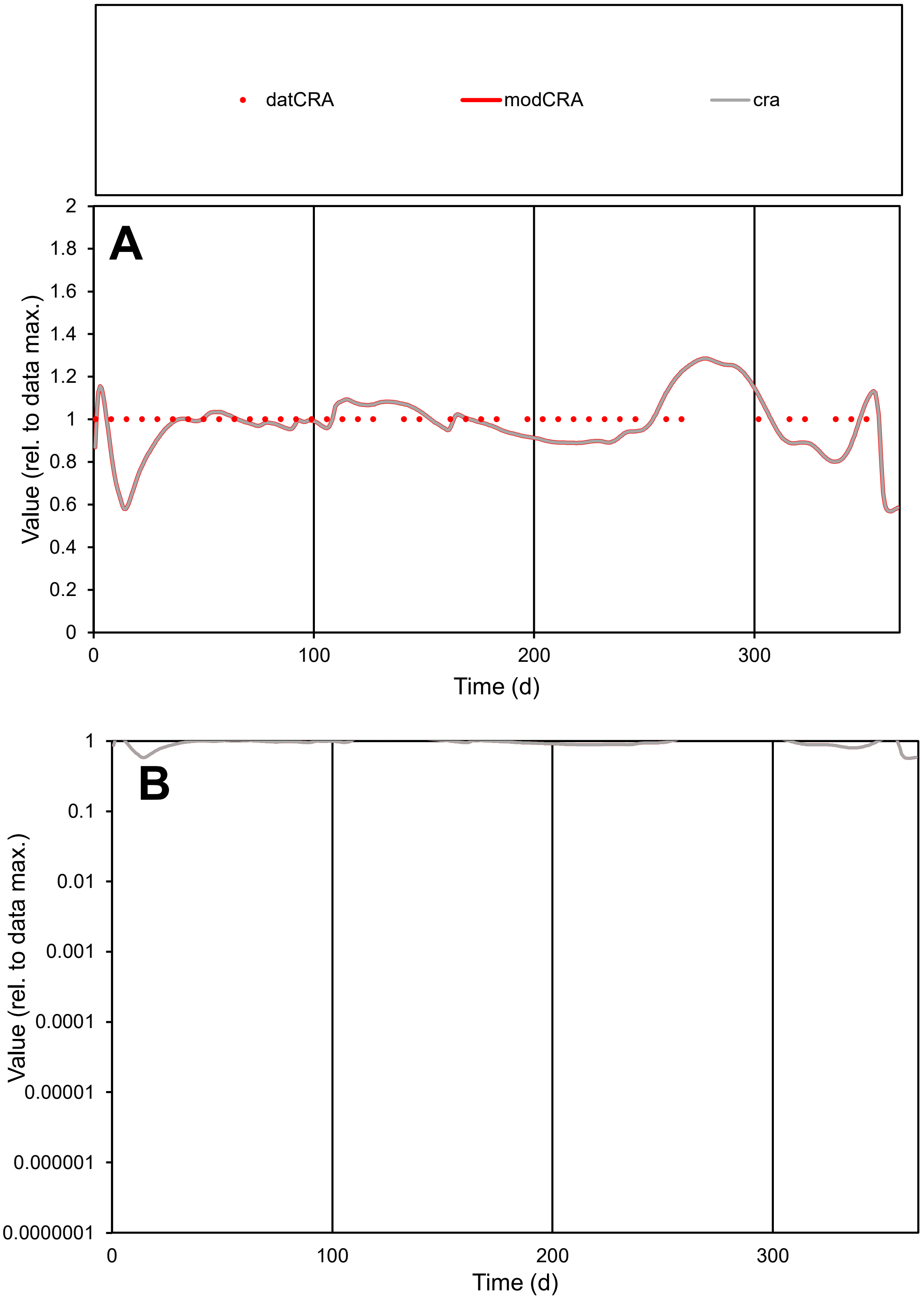


**C1**

Variables
CRA Provide jtd or Dst
CalibOnly?
2 1 to reproduce of
oNormalize?
2 1=no, 2=yes

ofxn calc*
0.11199
ofxn mod
0.15464

sRun
RunA1
zout
1

Ave
Data
4.9E-06
Model
4.7E-06

data/vMax
4.9E-06

ofxng
319.391

Min d/m
1.59371

ave/min d
1
ave/min m
1.54284

330_Calib_S_A

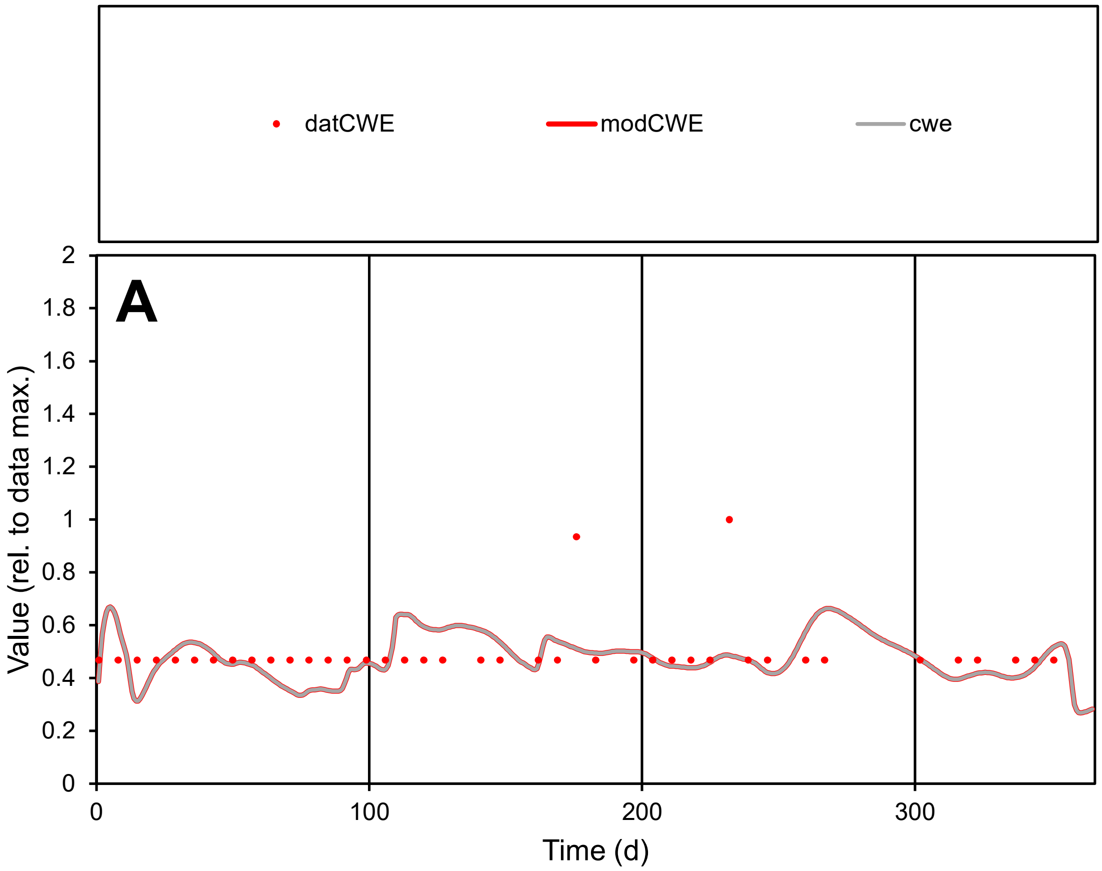


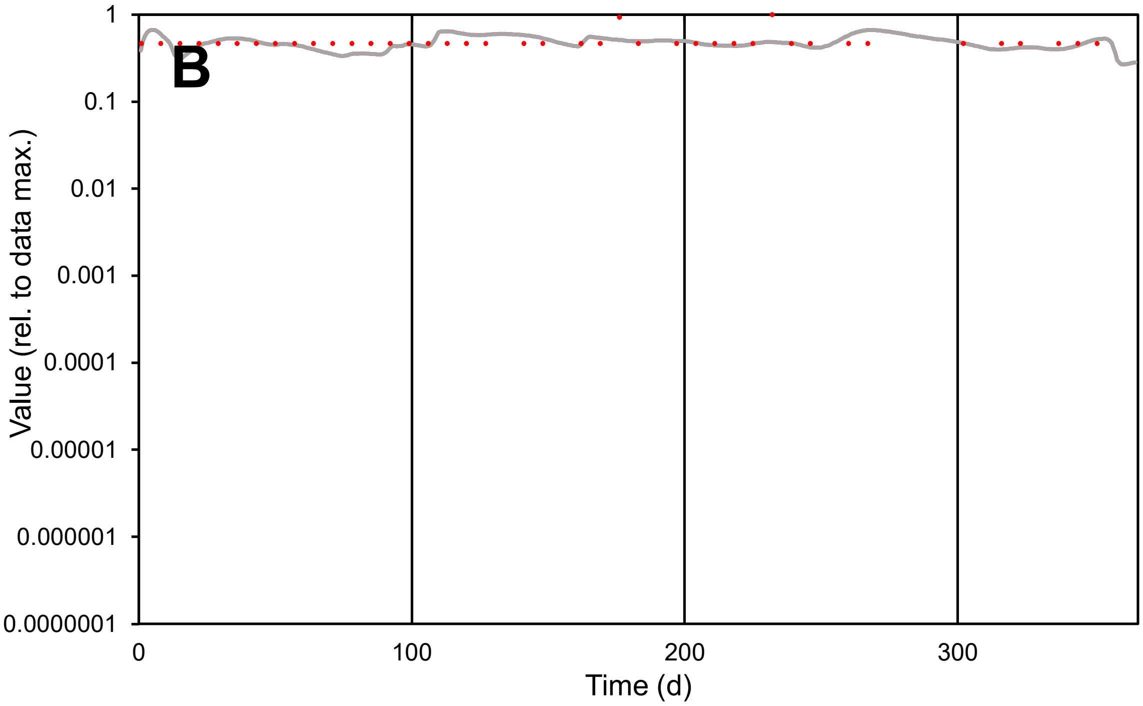


**C1**

Variables
CWE Provide jtd or Dst
CalibOnly?
2 1 to reproduce of
oNormalize?
2 1=no, 2=yes

ofxn calc*
0.26315
ofxn mod
0.28455

sRun
RunA1
zout
1

Ave
Data
2.7E-05
Model
2.6E-05

data/vMax
5.5E-05

ofxng
319.391

Min d/m
1.40341

ave/min d
1.05219
ave/min m
1.43795

330_Calib_S_A

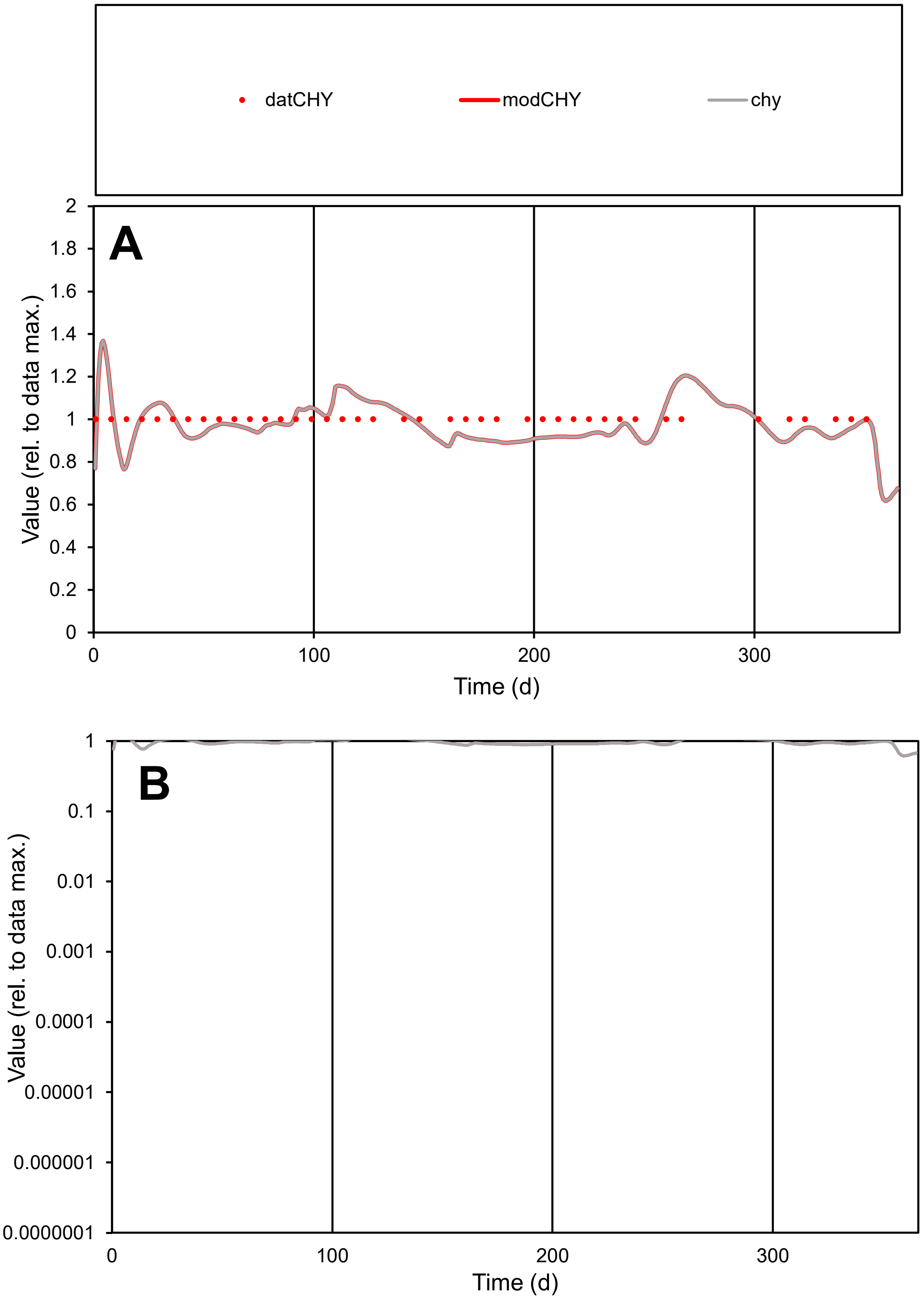


**C1**

Variables

CHY Provide jtd or Dst

CalibOnly?

2 1 to reproduce of

oNormalize?

2 1=no, 2=yes

ofxn calc*

0.07724

ofxn mod

0.13239

sRun

RunA1

zout

1

Ave

Data

6.7E-06

Model

6.6E-06

data/vMax

6.7E-06

ofxng

319.391

Min d/m

1.17409

ave/min d

1

ave/min m

1.15072

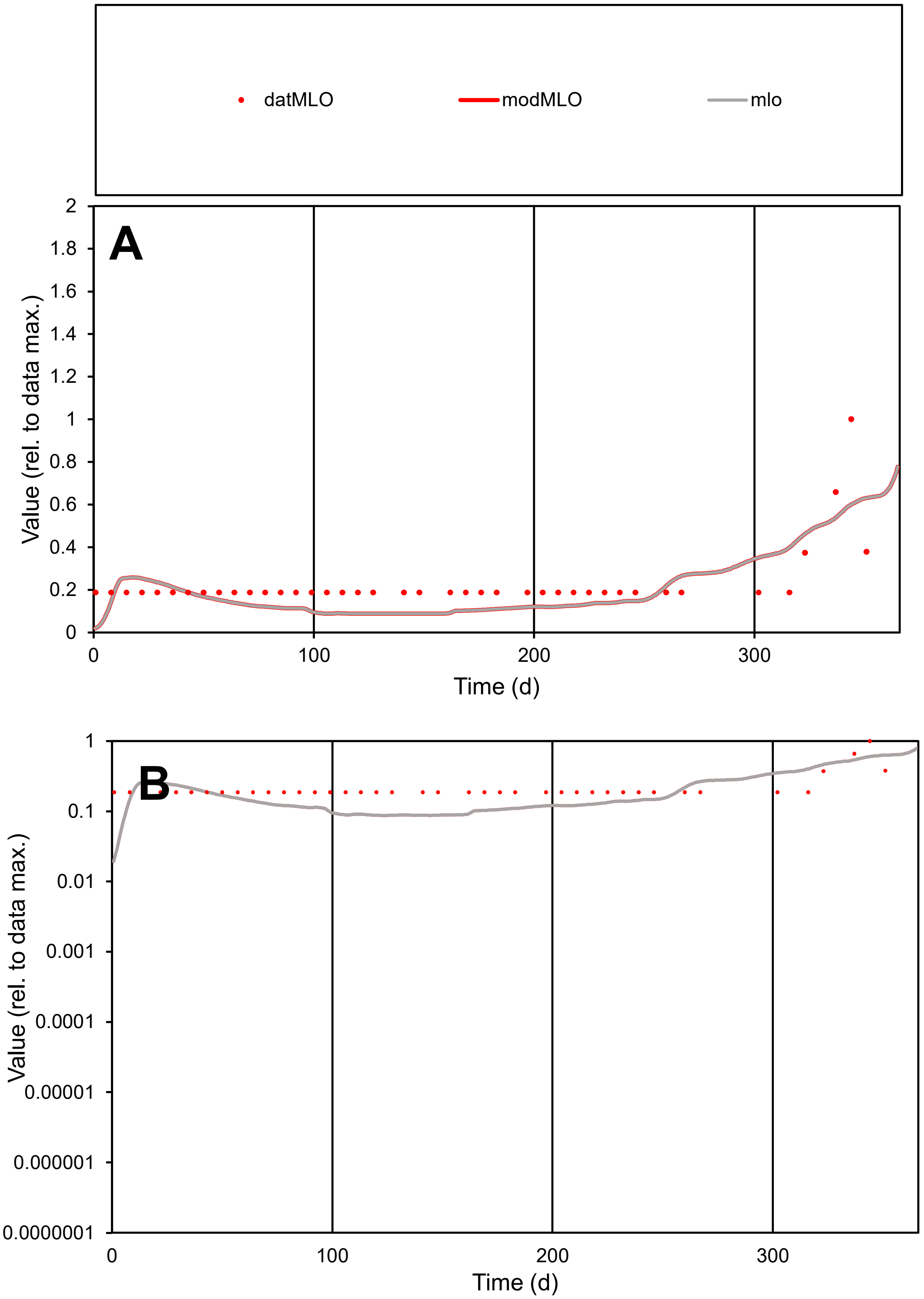


C1

Variables
MLO Provide jtd or Dst
CalibOnly?
2 1 to reproduce of
oNormalize?
2 1=no, 2=yes

ofxn calc*
0.4834
ofxn mod
0.49744

sRun
RunA1
zout
1

Ave
Data
1.3E-05
Model
1.1E-05

data/vMax
5.6E-05

ofxng
319.391

Min d/m
4.09662

ave/min d
1.21671
ave/min m
4.22958

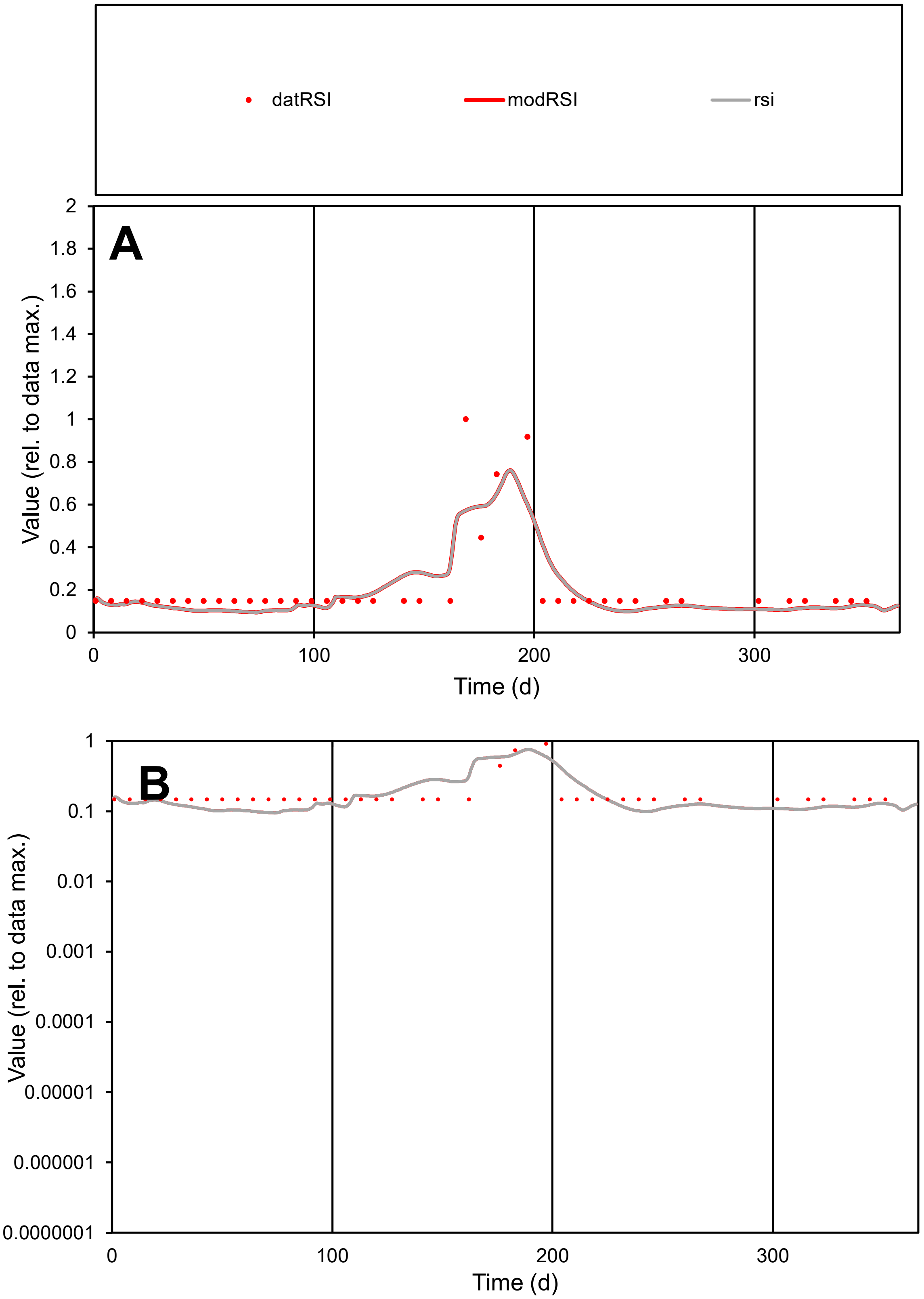


**C1**

Variables
RSI Provide jtd or Dst
CalibOnly?
2 1 to reproduce of
oNormalize?
2 1=no, 2=yes

ofxn calc*
0.49716
ofxn mod
0.54269

sRun
RunA1
zout
1

Ave
Data
1.5E-05
Model
1.3E-05

data/vMax
7E-05

ofxng
319.391

Min d/m
1.54479

ave/min d
1.41501
ave/min m
2.01608

330_Calib_S_A

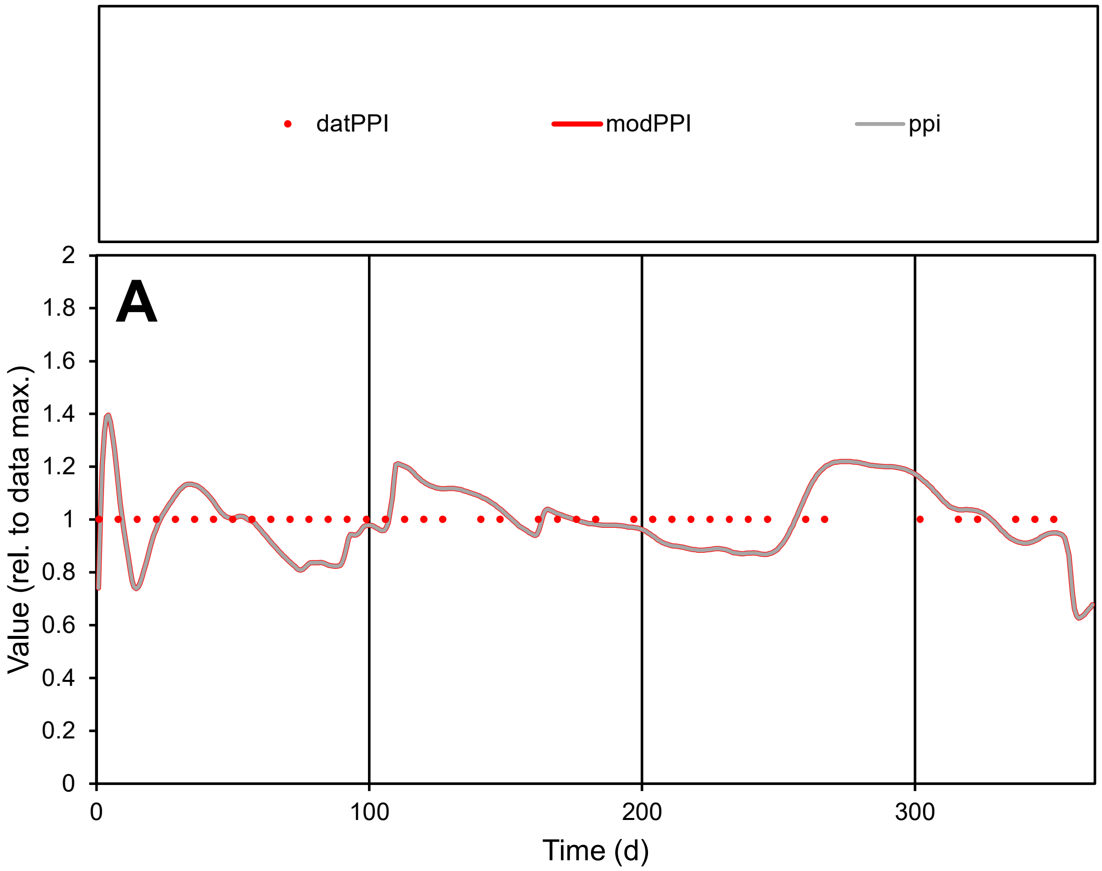


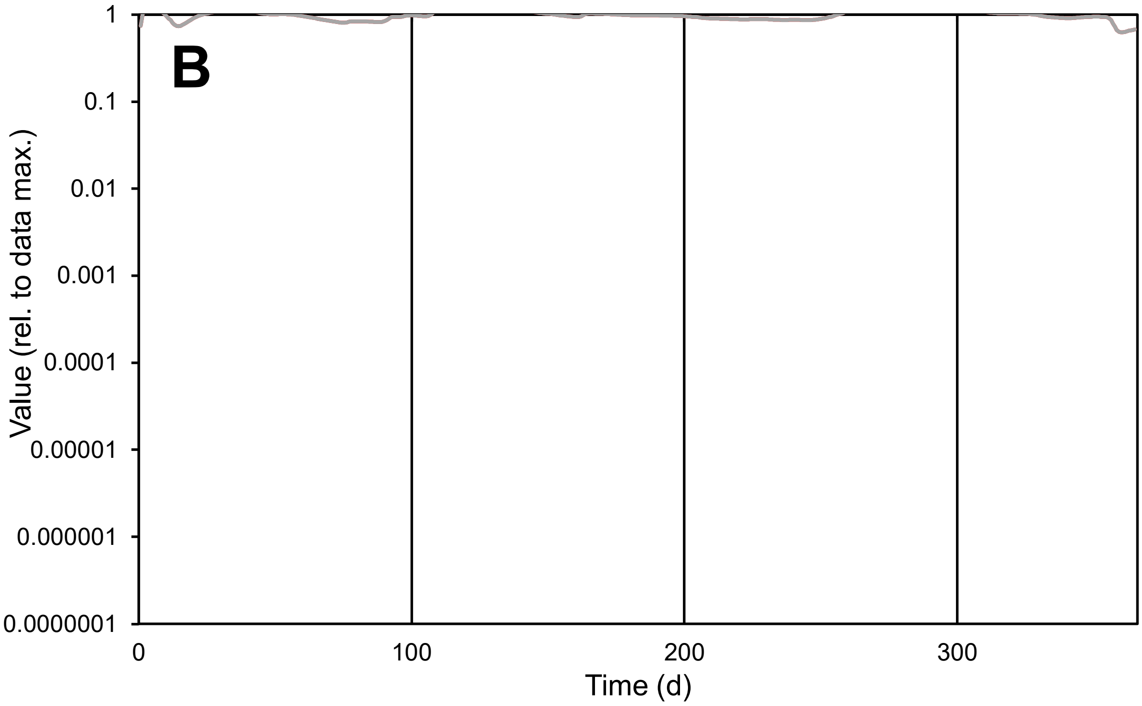


**C1**

Variables
PPI Provide jtd or Dst
CalibOnly?
2 1 to reproduce of
oNormalize?
2 1=no, 2=yes

ofxn calc*
0.10364
ofxn mod
0.14951

sRun
RunA1
zout
1

Ave
Data
1.4E-06
Model
1.4E-06

data/vMax
1.4E-06

ofxng
319.391

Min d/m
1.27965

ave/min d
1
ave/min m
1.26348

330_Calib_S_A

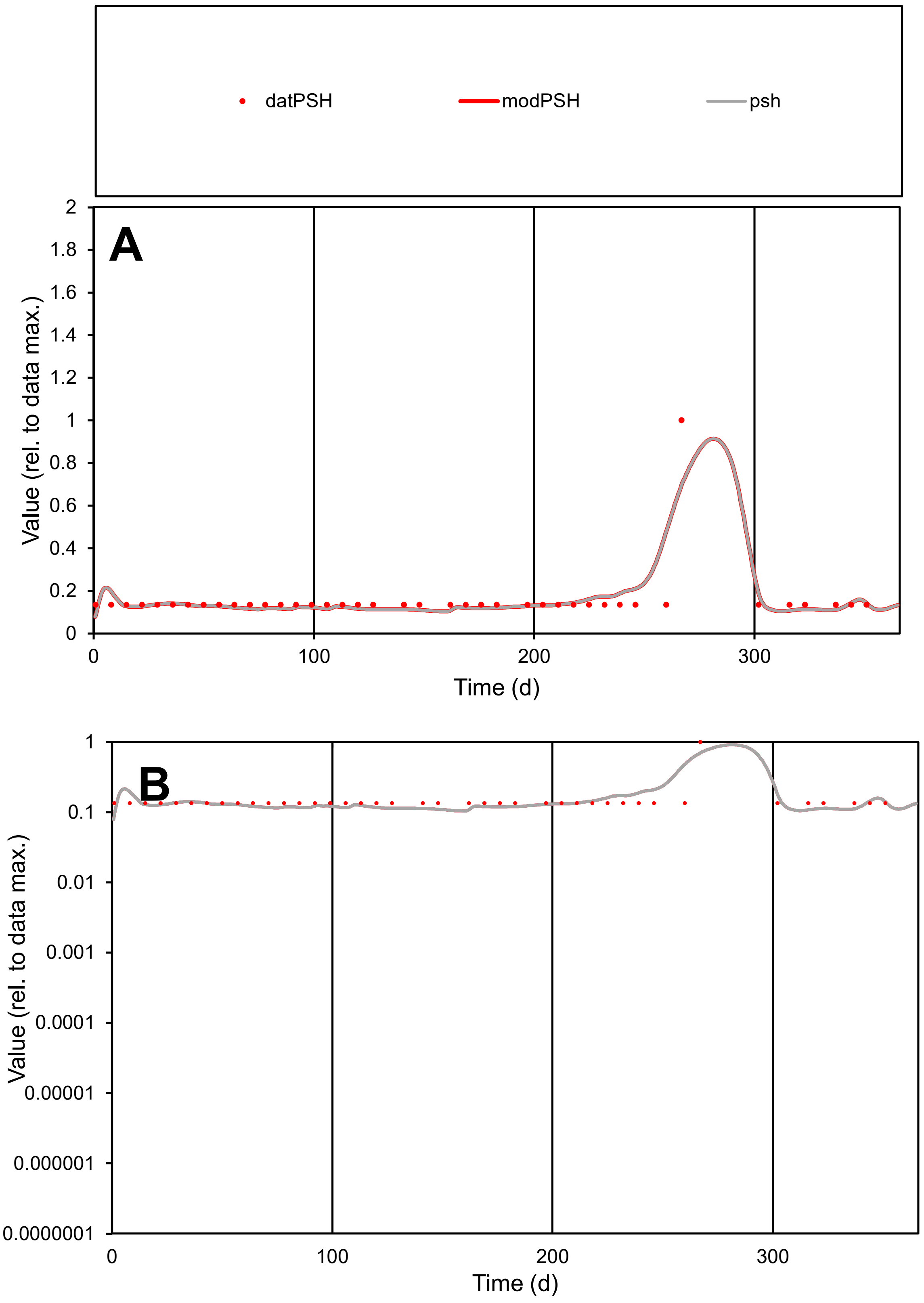


**C1**

Variables
PSH Provide jtd or Dst
CalibOnly?
2 1 to reproduce of
oNormalize?
2 1=no, 2=yes

ofxn calc*
0.47089
ofxn mod
0.49568

sRun
RunA1
zout
1

Ave
Data
1.5E-05
Model
1.5E-05

data/vMax
9.6E-05

ofxng
319.391

Min d/m
1.23563

ave/min d
1.15716
ave/min m
1.43825

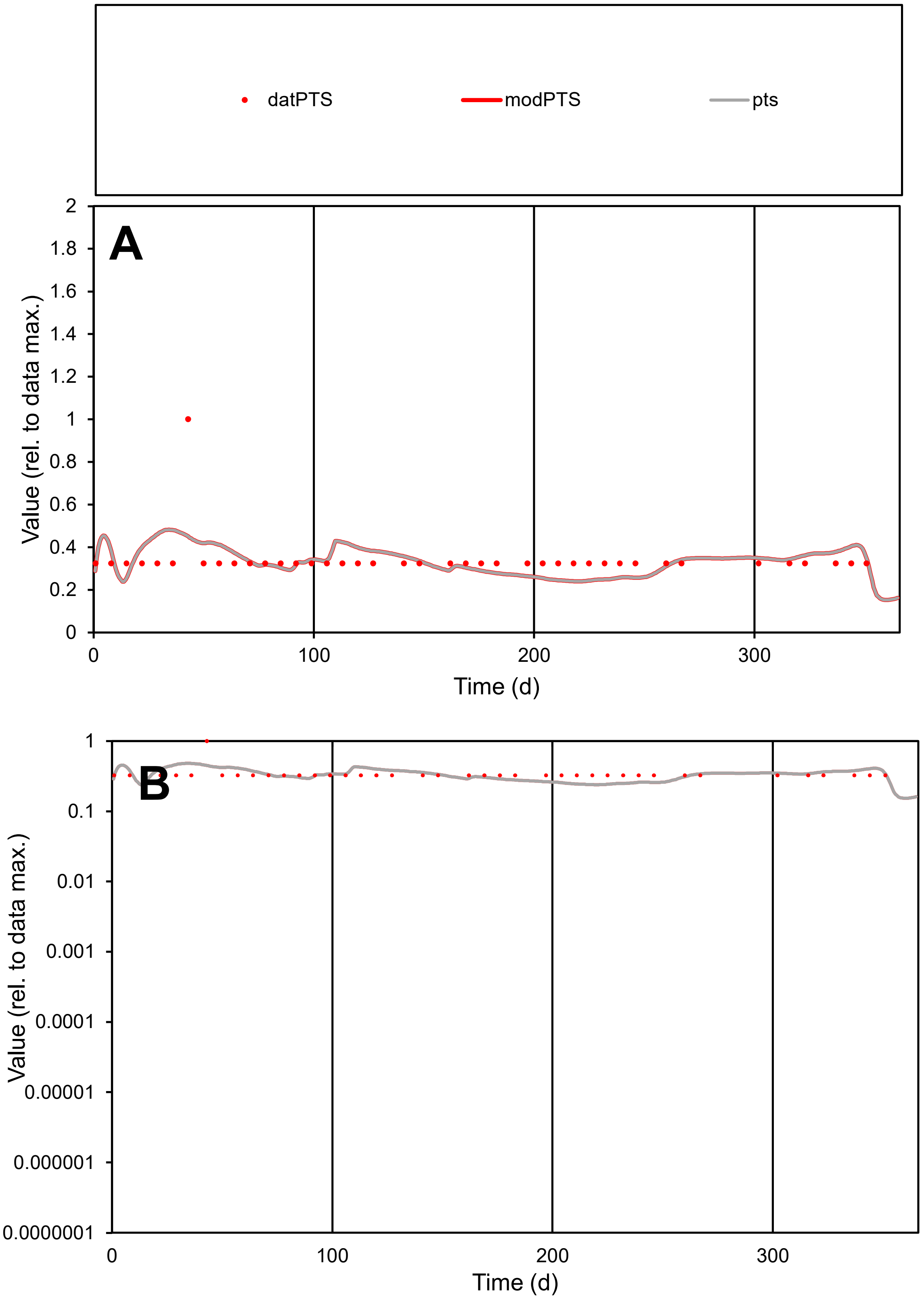


**C1**

Variables

PTS Provide jtd or Dst

CalibOnly?

2 1 to reproduce of

oNormalize?

2 1=no, 2=yes

ofxn calc*

0.31238

ofxn mod

0.33295

sRun

RunA1

zout

1

Ave

Data

1E-05

Model

1E-05

data/vMax

3E-05

ofxng

319.391

Min d/m

1.34804

ave/min d

1.05095

ave/min m

1.40621

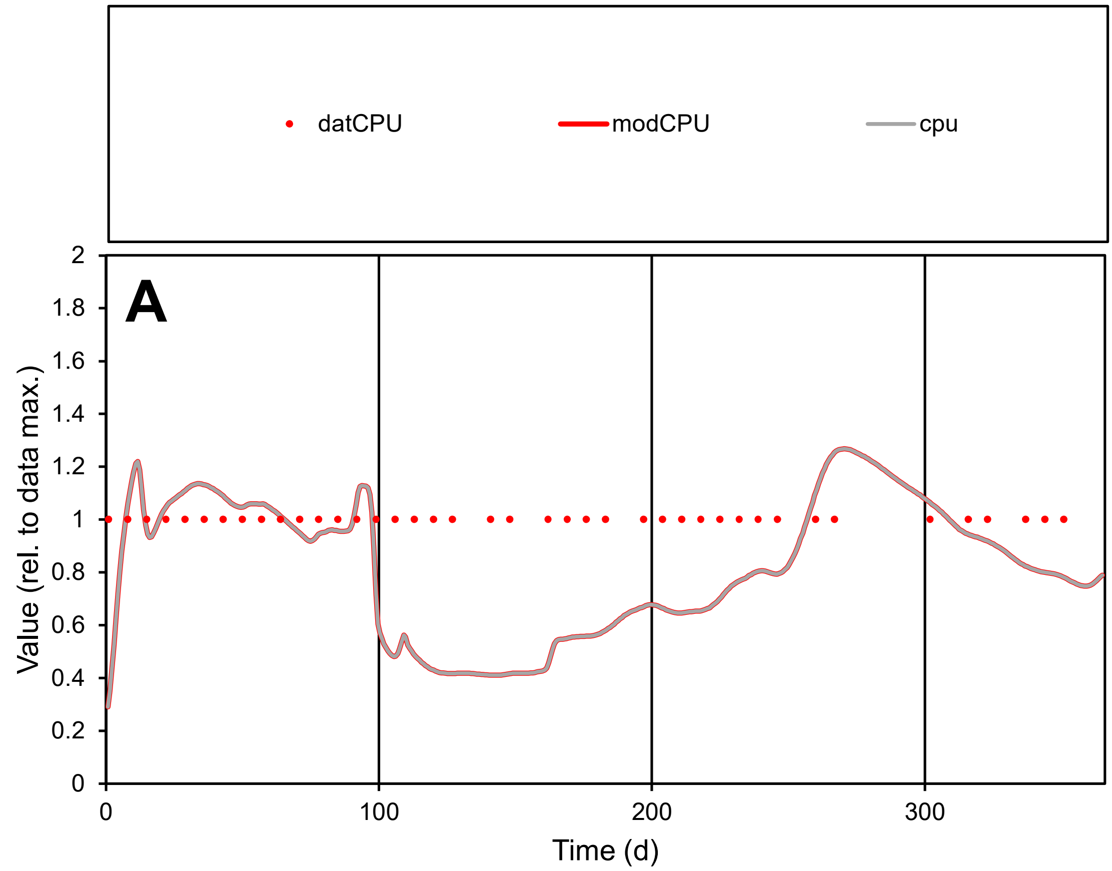


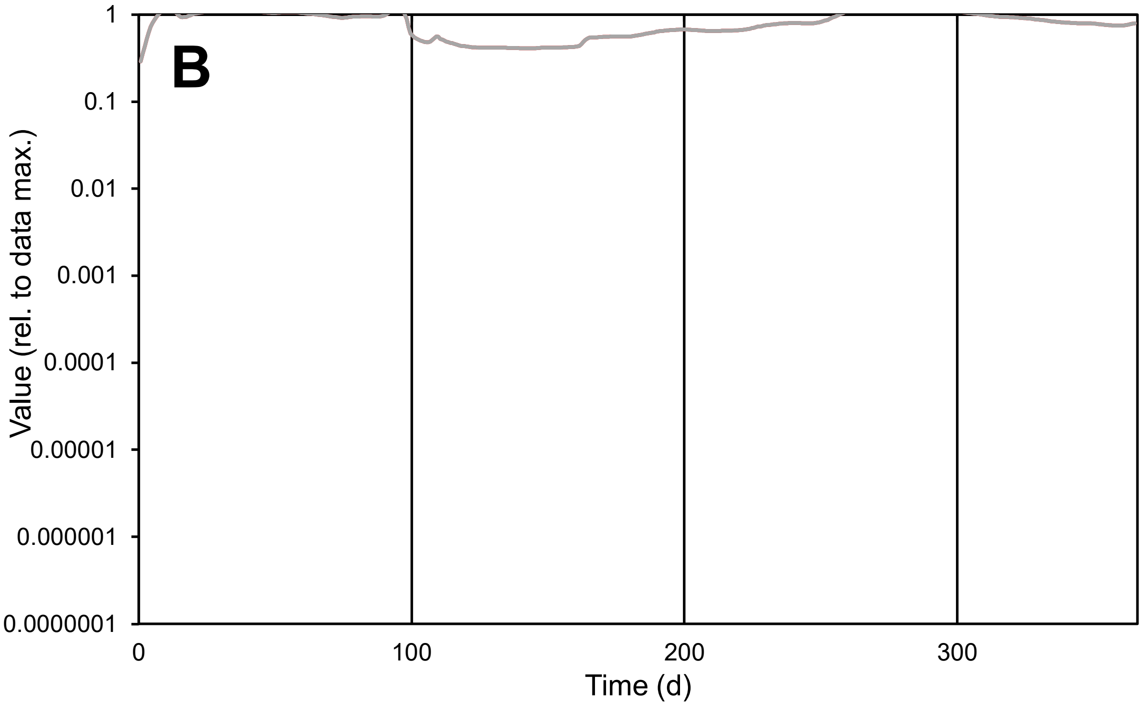


**C1**

Variables
CPU Provide jtd or Dst
CalibOnly?
2 1 to reproduce of
oNormalize?
2 1=no, 2=yes

ofxn calc*
0.31106
ofxn mod
0.33043

sRun
RunA1
zout
1

Ave
Data
4.4E-07
Model
3.6E-07

data/vMax
4.4E-07

ofxng
319.391

Min d/m
2.43234

ave/min d
1
ave/min m
1.96282

330_Calib_S_A

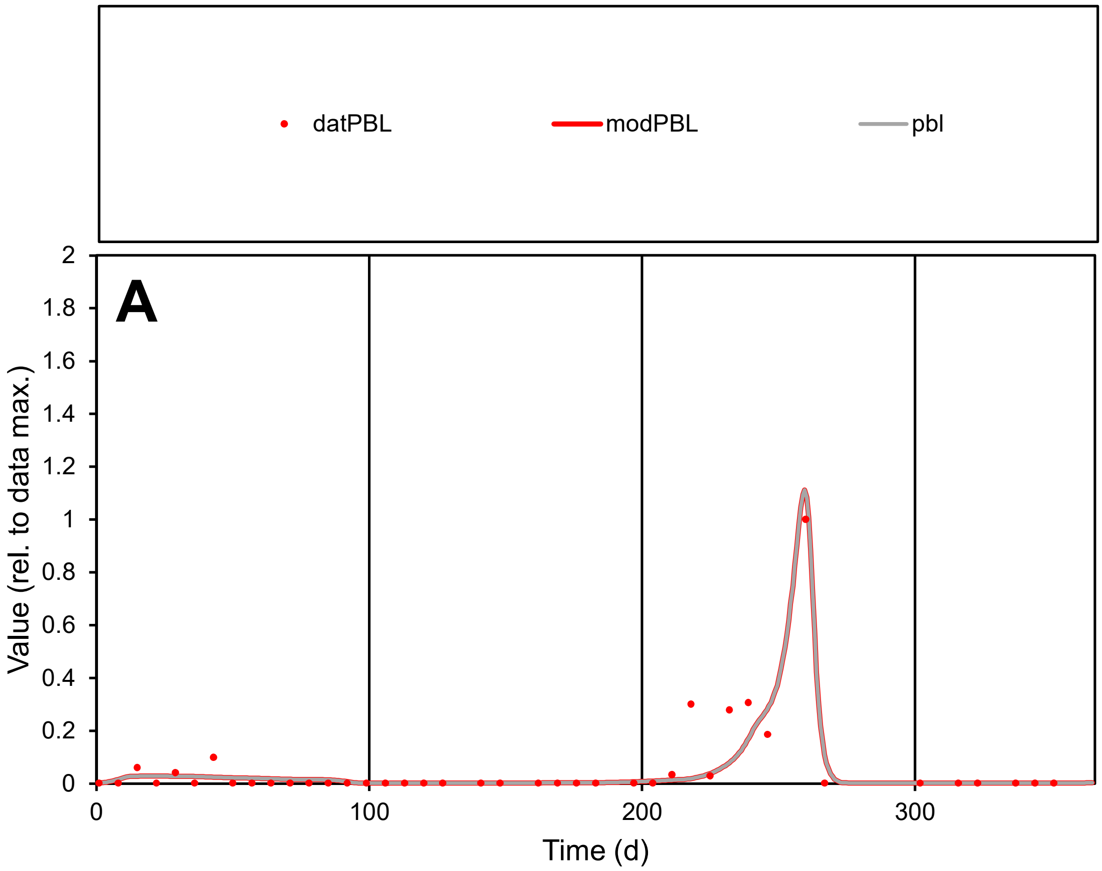


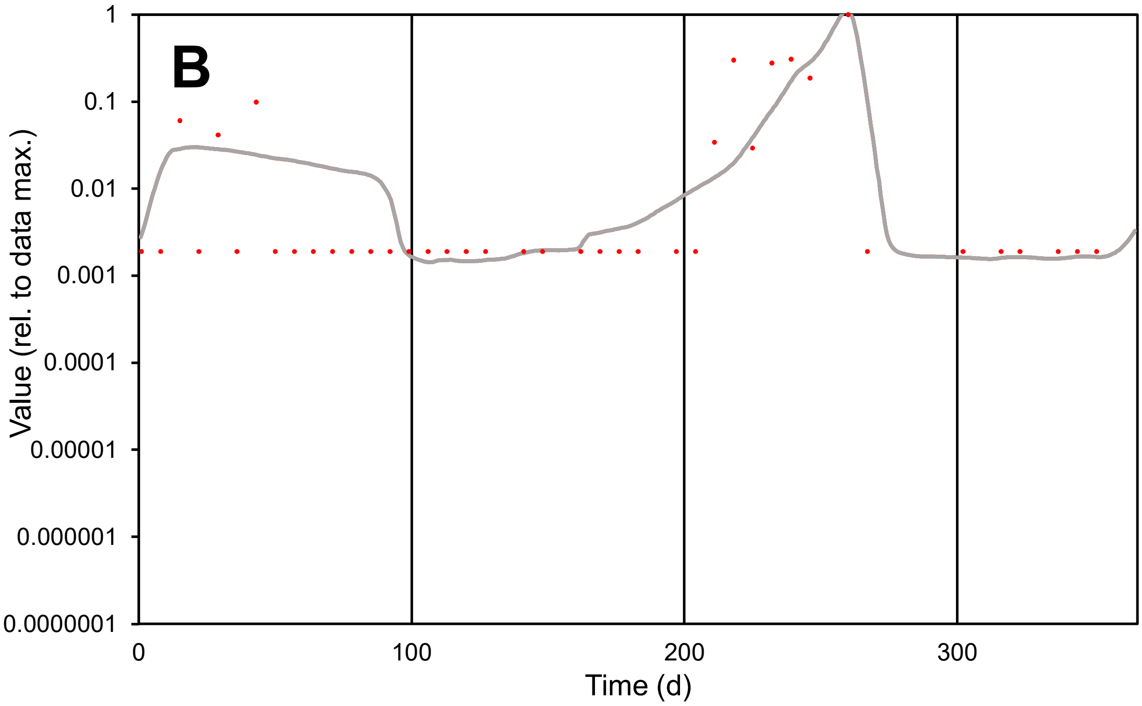


**C1**

Variables
PBL Provide jtd or Dst
CalibOnly?
2 1 to reproduce of
oNormalize?
2 1=no, 2=yes

ofxn calc*
1.01636
ofxn mod
1.0586

sRun
RunA1
zout
1

Ave
Data
1.3E-06
Model
1.1E-06

data/vMax
2.2E-05

ofxng
319.391

Min d/m
1.32462

ave/min d
30.7758
ave/min m
34.1073

330_Calib_S_A

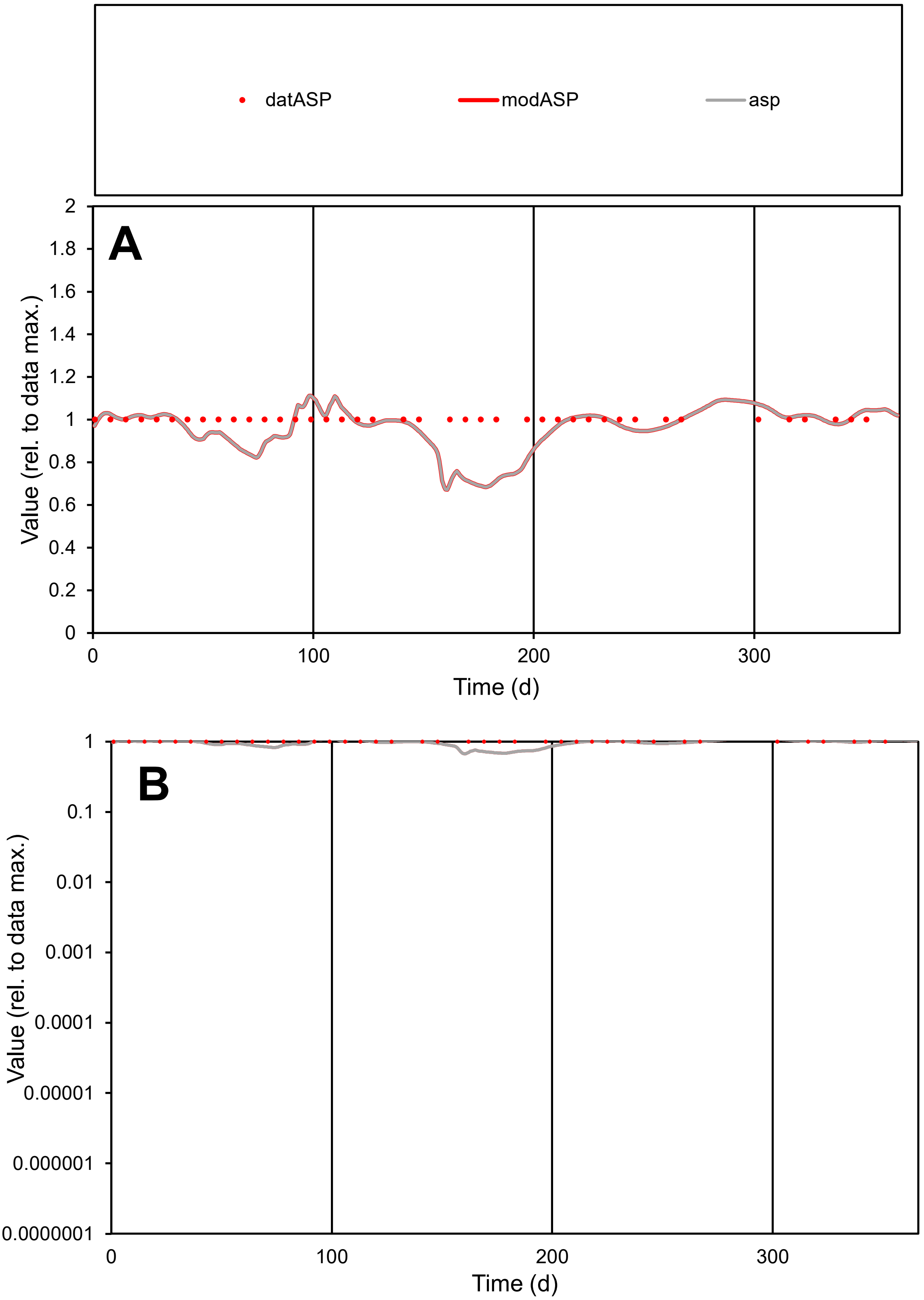


**C1**

Variables
ASP Provide jtd or Dst
CalibOnly?
2 1 to reproduce of
oNormalize?
2 1=no, 2=yes

ofxn calc*
0.10708
ofxn mod
0.15108

sRun
RunA1
zout
1

Ave
Data
5.8E-06
Model
5.5E-06

data/vMax
5.8E-06

ofxng
319.391

Min d/m
1.46045

ave/min d
1
ave/min m
1.39713

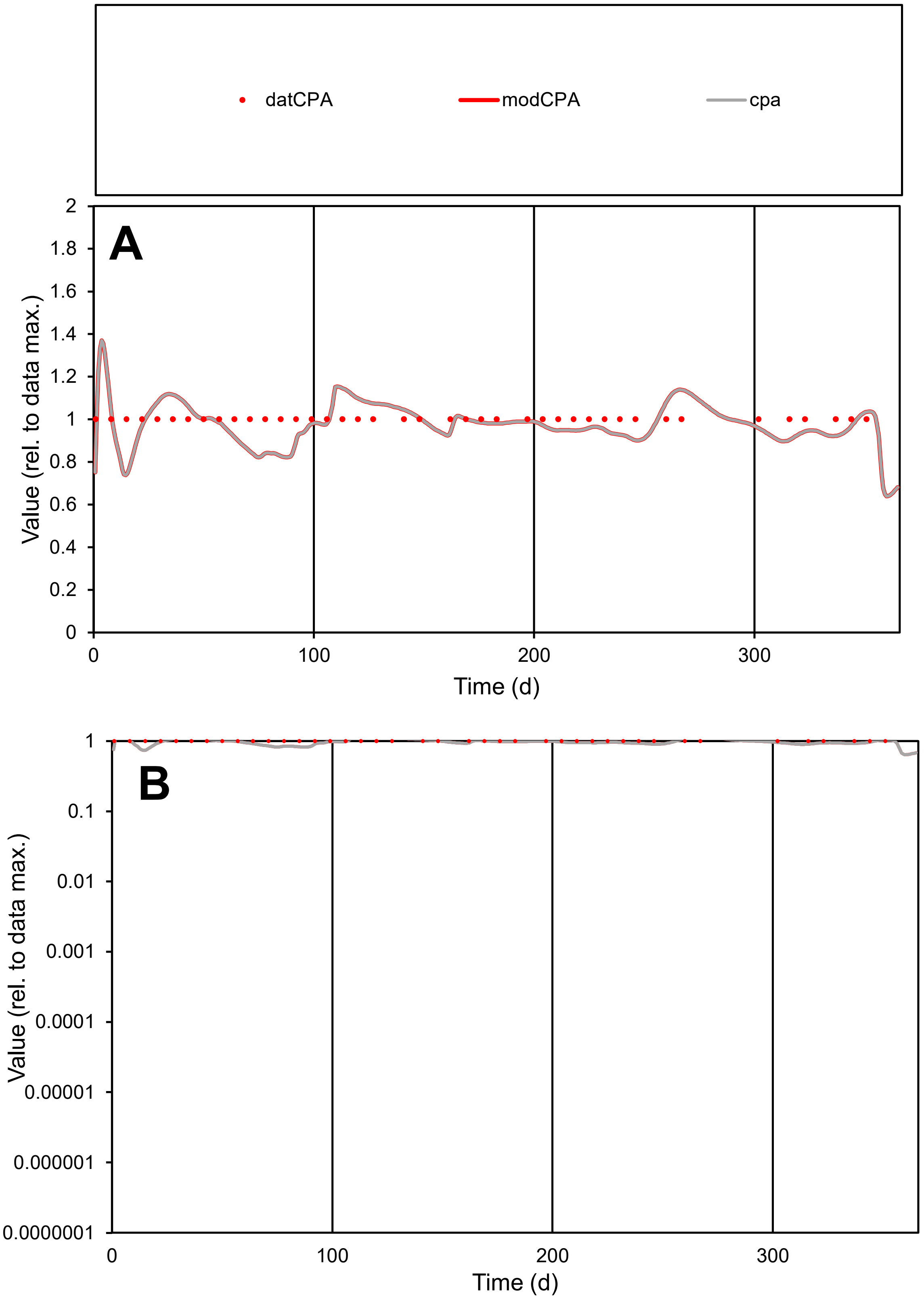


**C1**

Variables
CPA Provide jtd or Dst
CalibOnly?
2 1 to reproduce of
oNormalize?
2 1=no, 2=yes

ofxn calc*
0.08259
ofxn mod
0.13564

sRun
RunA1
zout
1

Ave
Data
6.6E-08
Model
6.5E-08

data/vMax
6.6E-08

ofxng
319.391

Min d/m
1.2695

ave/min d
1
ave/min m
1.24482

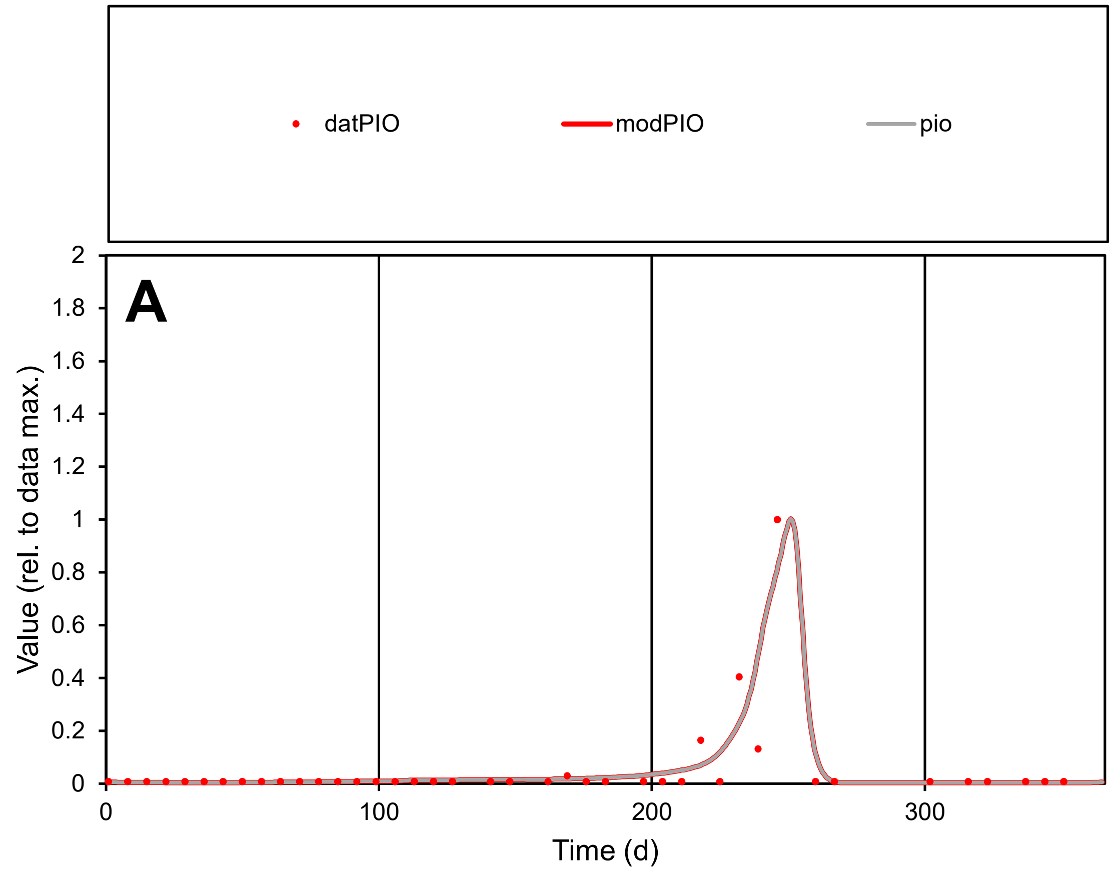


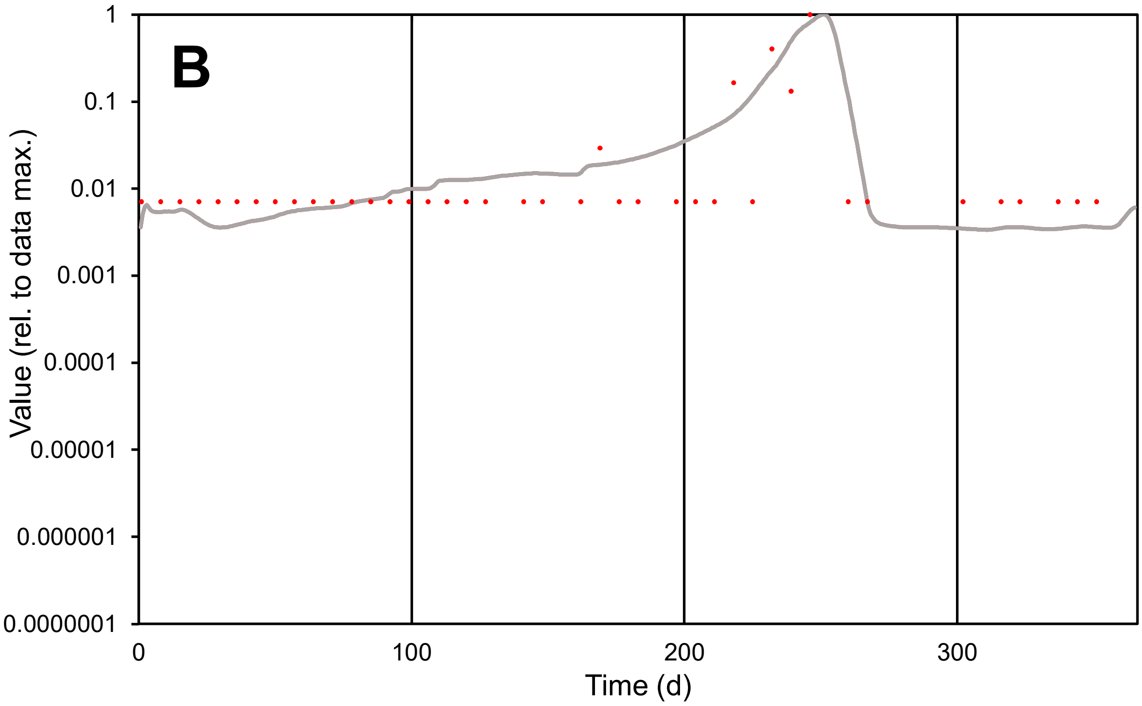


**C1**

Variables
PIO Provide jtd or Dst
CalibOnly?
2 1 to reproduce of
oNormalize?
2 1=no, 2=yes

ofxn calc*
1.33418
ofxn mod
1.47572

sRun
RunA1
zout
1

Ave
Data
9.9E-07
Model
1.1E-06

data/vMax
2.1E-05

ofxng
319.391

Min d/m
2.04421
ave/min d
6.8376
ave/min m
15.6115

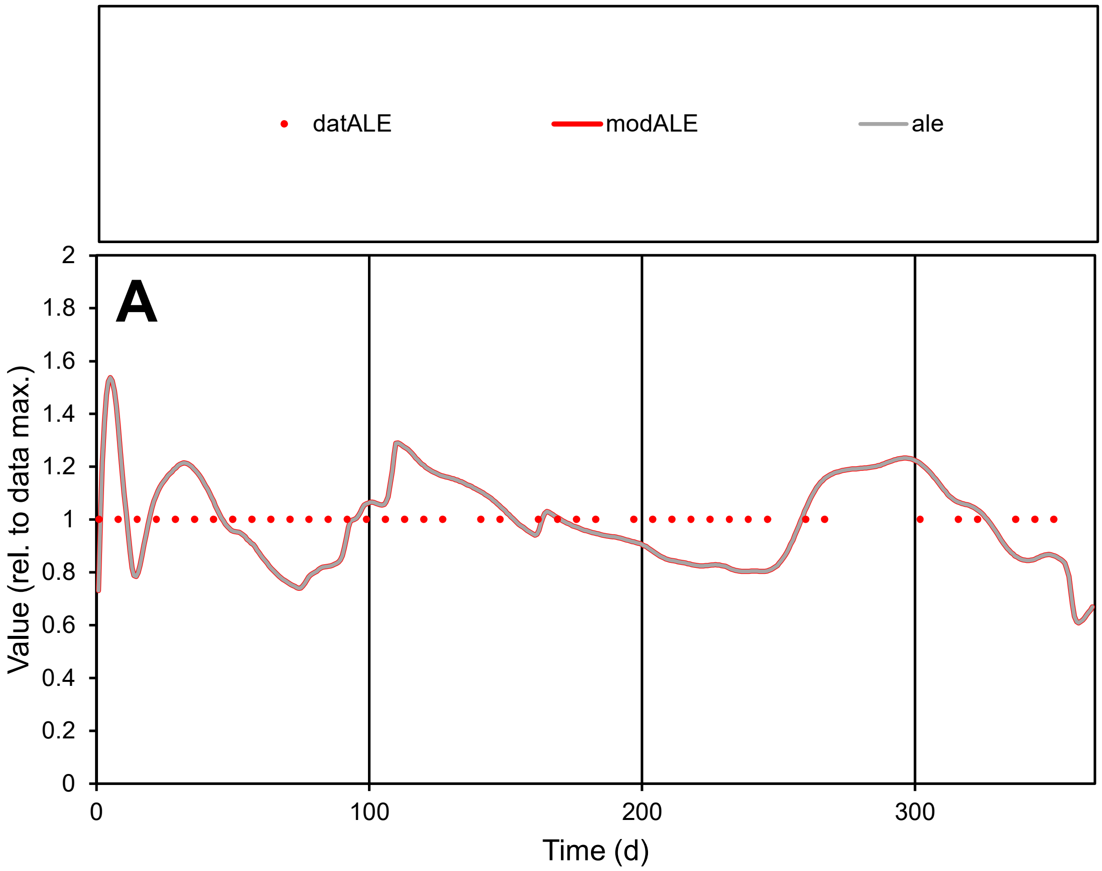


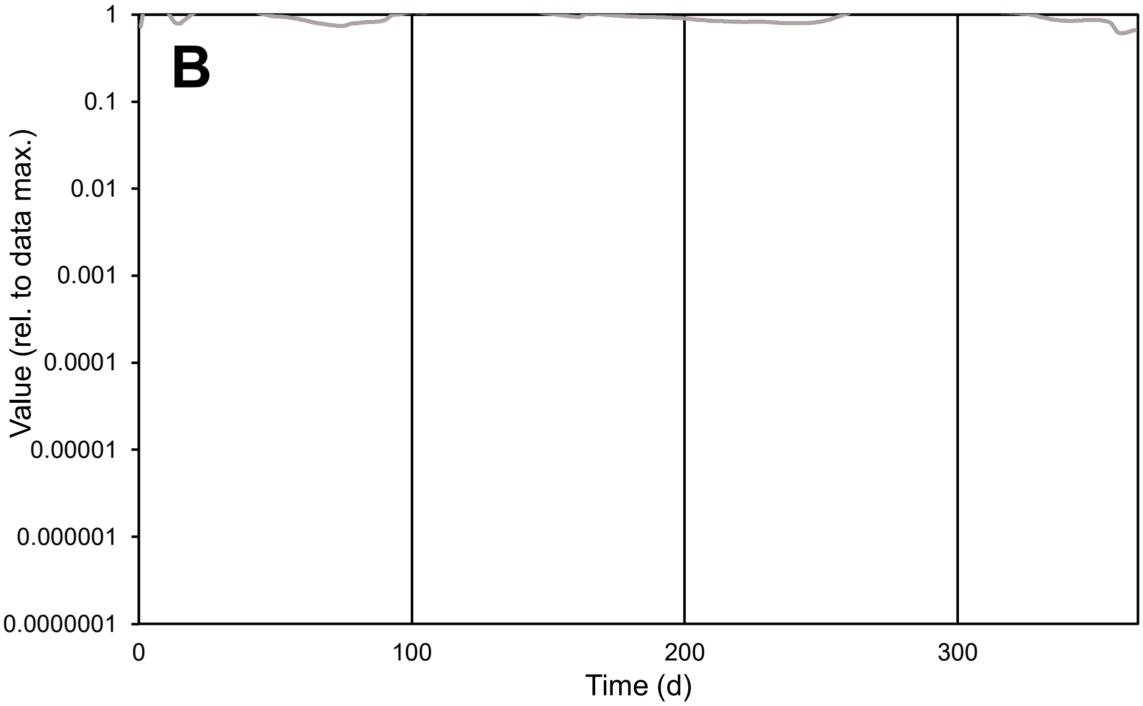


**C1**

Variables
ALE Provide jtd or Dst
CalibOnly?
2 1 to reproduce of
oNormalize?
2 1=no, 2=yes

ofxn calc*
0.14398
ofxn mod
0.17963

sRun
RunA1
zout
1

Ave
Data
1.1E-05
Model
1.1E-05

data/vMax
1.1E-05

ofxng
319.391

Min d/m
1.33333

ave/min d
1
ave/min m
1.30926

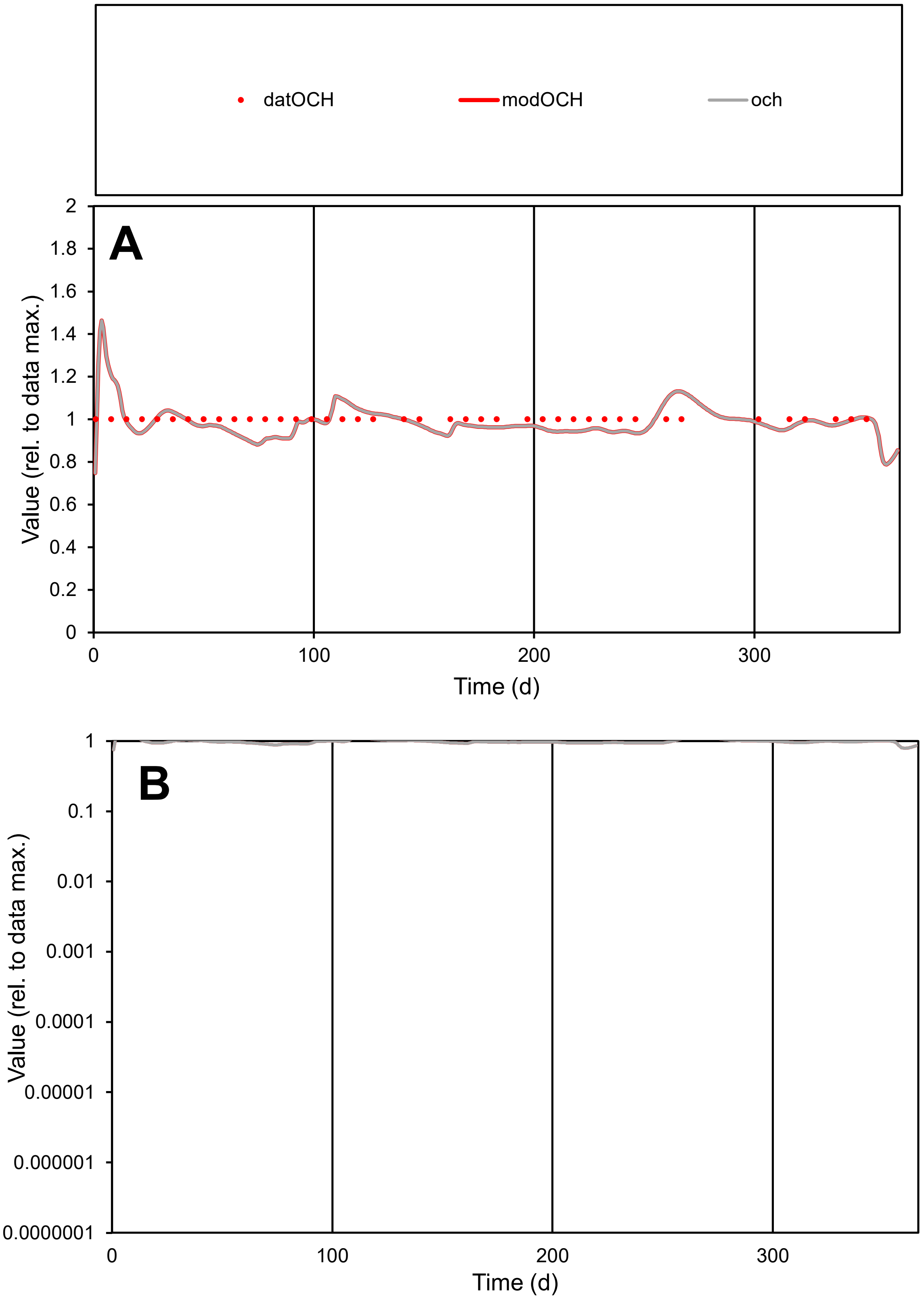


**C1**

Variables
OCH Provide jtd or Dst
CalibOnly?
2 1 to reproduce of
oNormalize?
2 1=no, 2=yes

ofxn calc*
0.05878
ofxn mod
0.12311

sRun
RunA1
zout
1

Ave
Data
4.4E-06
Model
4.3E-06

data/vMax
4.4E-06

ofxng
319.391

Min d/m
1.1214

ave/min d
1
ave/min m
1.10624

330_Calib_S_A

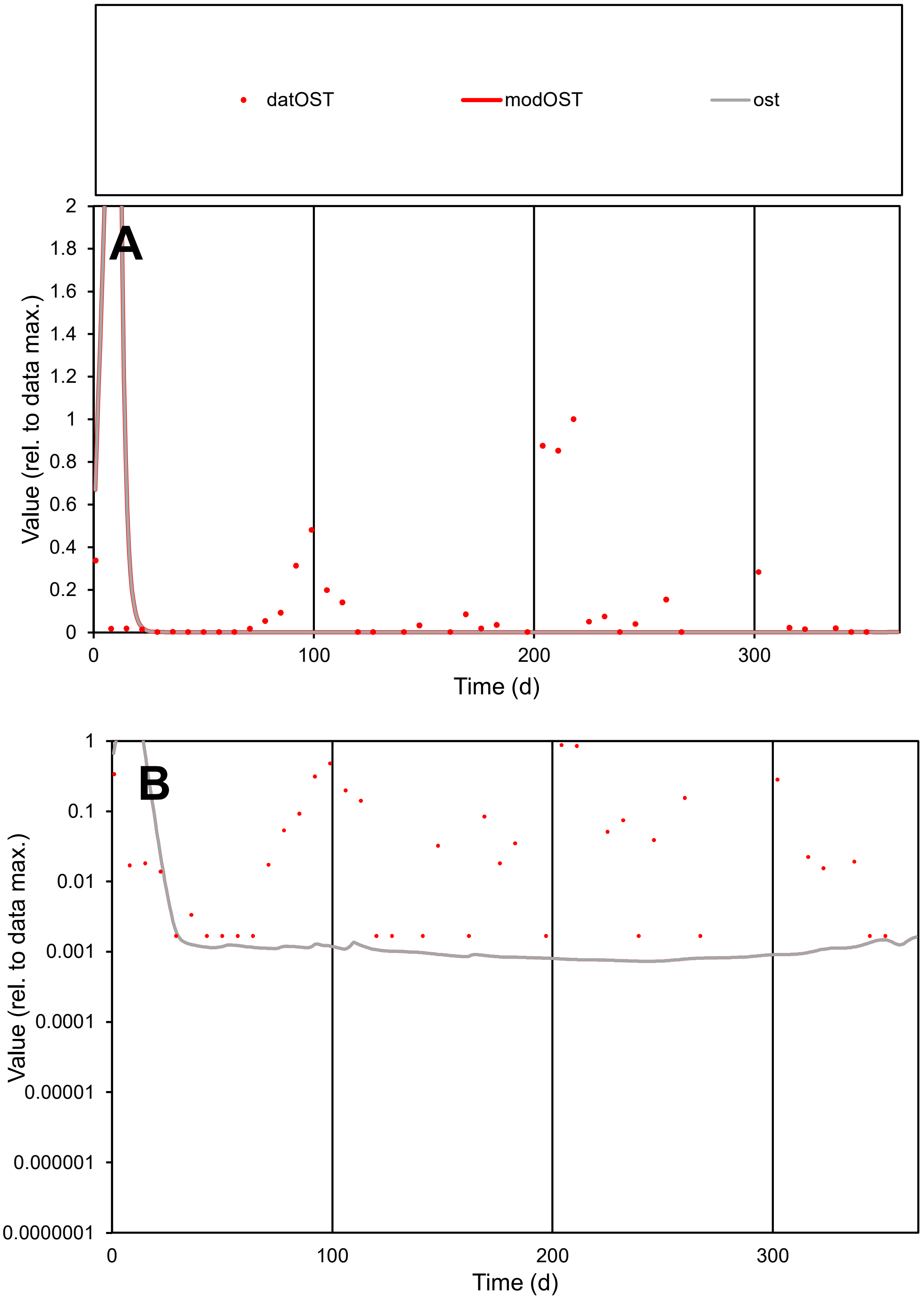


**C1**

Variables
OST Provide jtd or Dst
CalibOnly?
2 1 to reproduce of
oNormalize?
2 1=no, 2=yes

ofxn calc*
3.05106
ofxn mod
3.83584

sRun
RunA1
zout
1

Ave
Data
0.00017
Model
9.1E-05

data/vMax
0.00129

ofxng
319.391

Min d/m
2.2837

ave/min d
76.6297
ave/min m
95.599

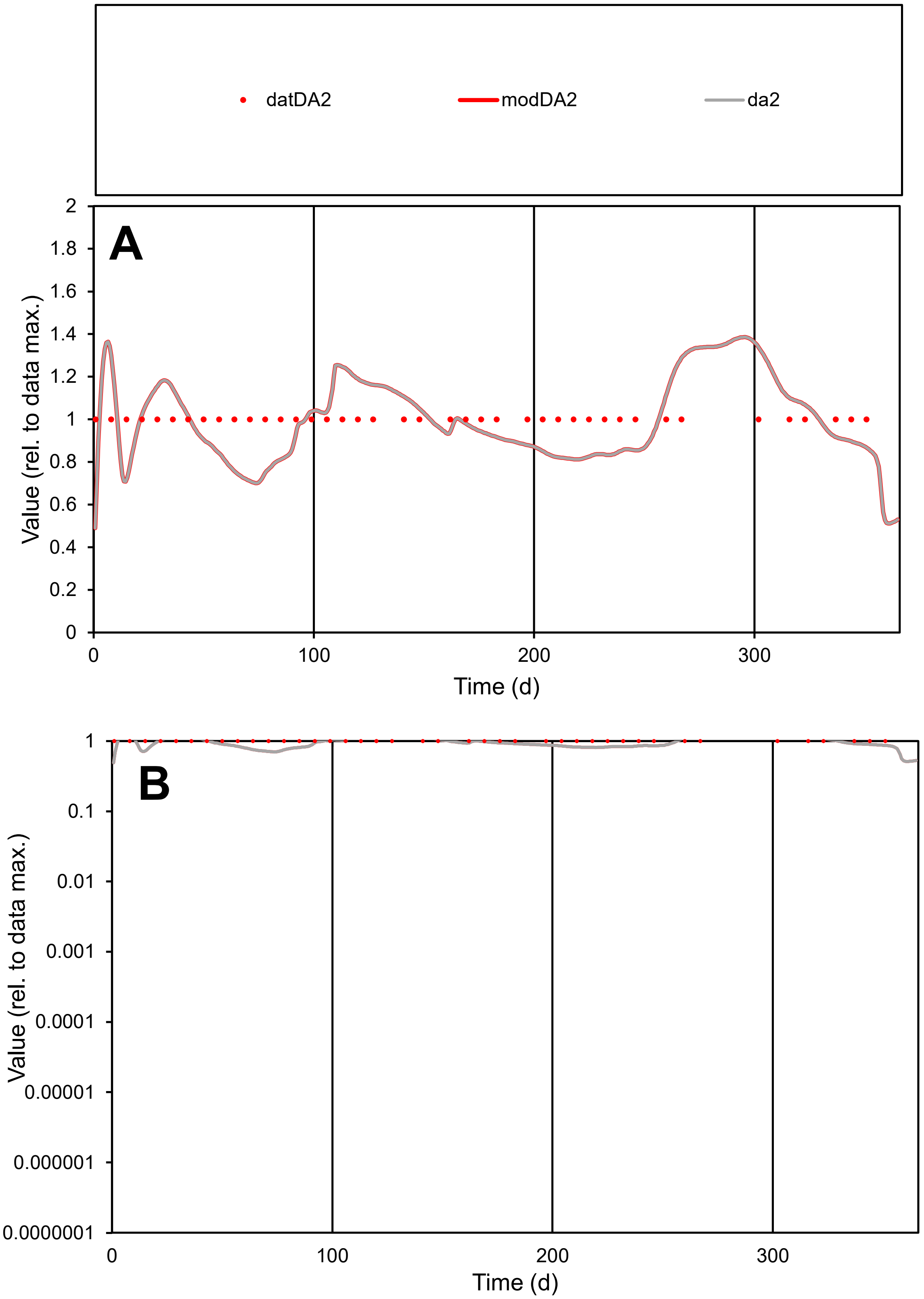


**C1**

Variables
DA2 Provide jtd or Dst
CalibOnly?
2 1 to reproduce of
oNormalize?
2 1=no, 2=yes

ofxn calc*
0.15858
ofxn mod
0.19169

sRun
RunA1
zout
1

Ave
Data
3.1E-06
Model
3E-06

data/vMax
3.1E-06

ofxng
319.391

Min d/m
1.41689

ave/min d
1
ave/min m
1.37901

330_Calib_S_A

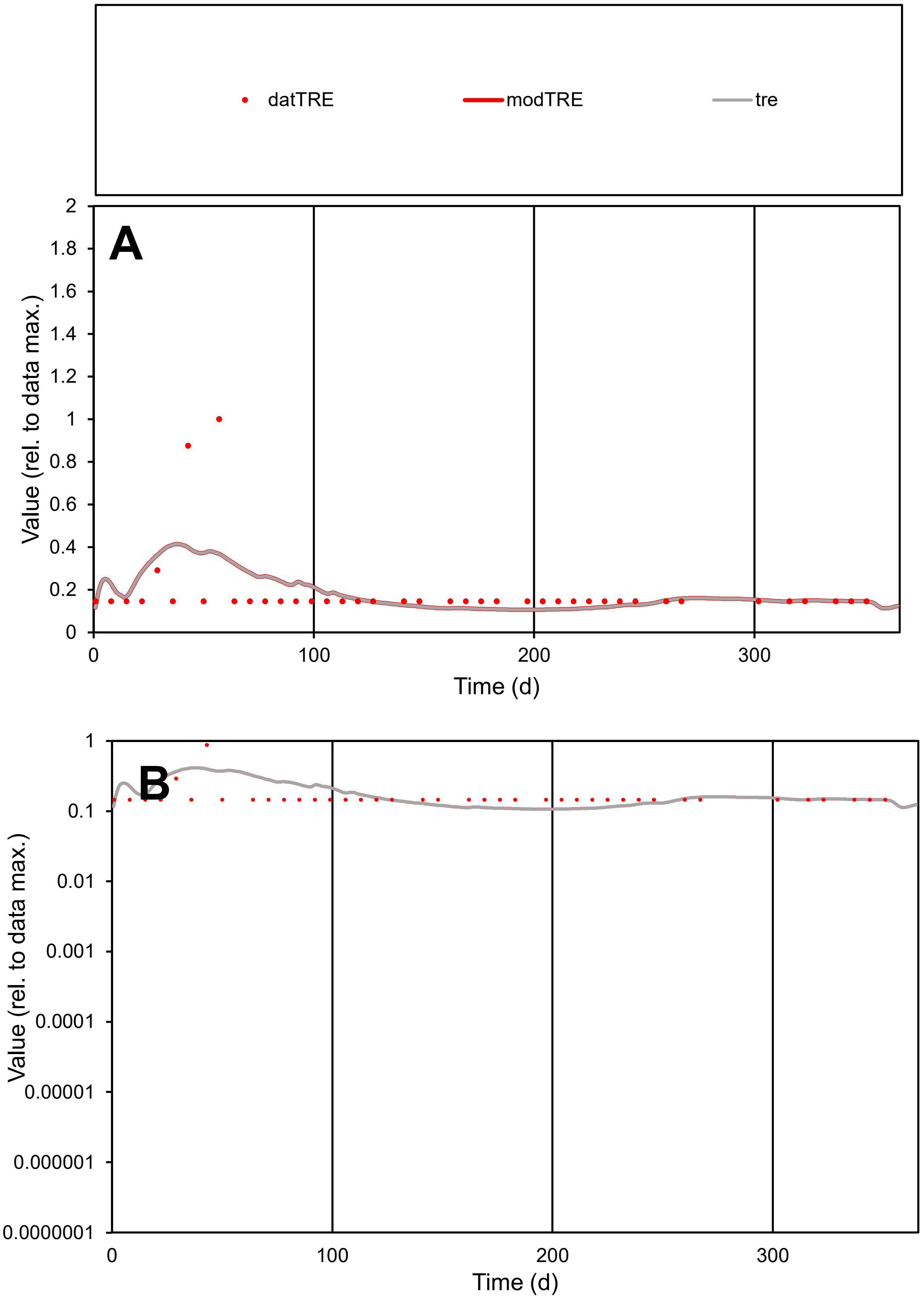


**C1**

Variables
TRE Provide jtd or Dst
CalibOnly?
2 1 to reproduce of
oNormalize?
2 1=no, 2=yes

ofxn calc*
0.78473
ofxn mod
0.79831

sRun
RunA1
zout
1

Ave
Data
8.9E-06
Model
9E-06

data/vMax
4.8E-05

ofxng
319.391

Min d/m
1.3636

ave/min d
1.28988
ave/min m
1.76713

330_Calib_S_A

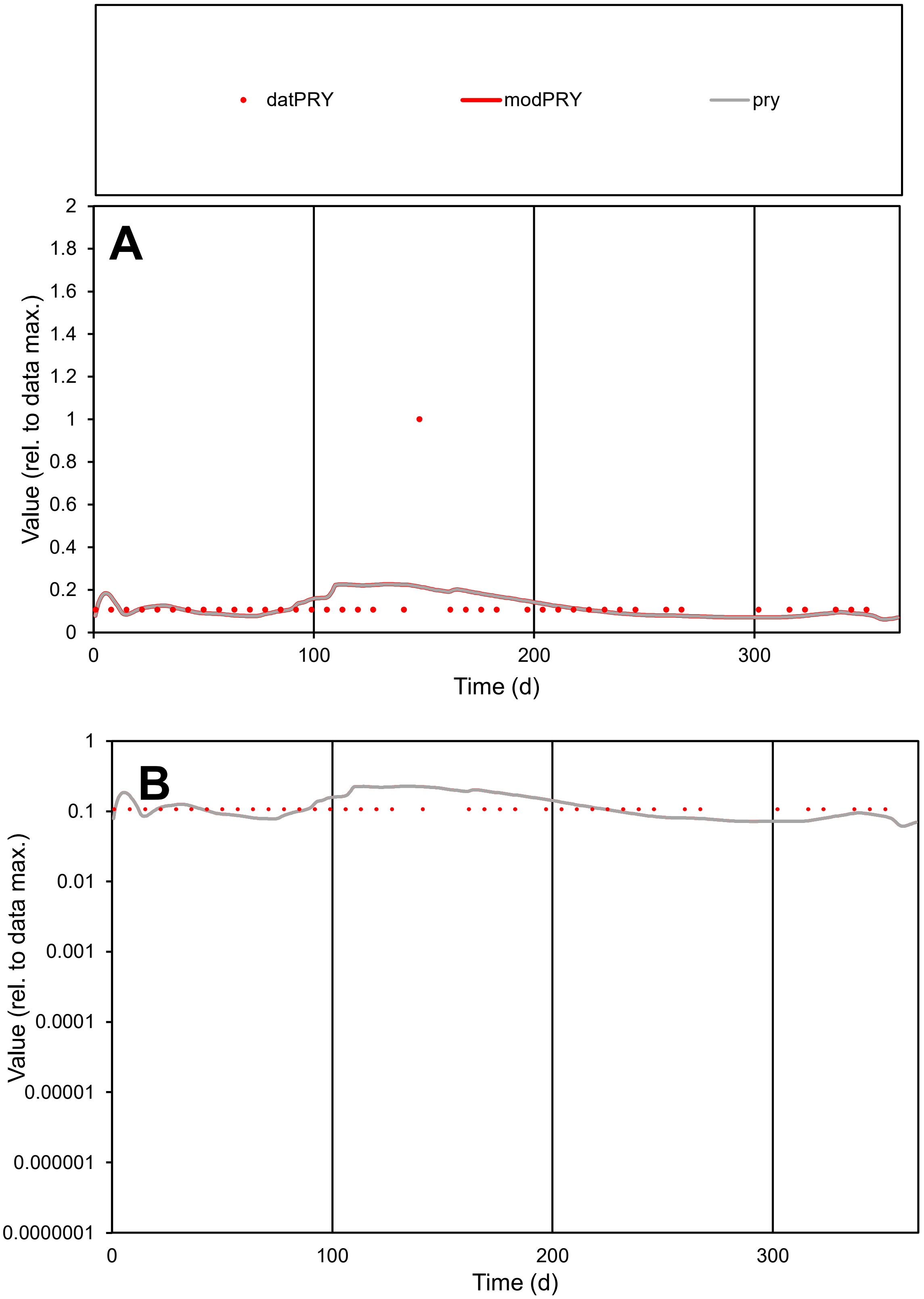


**C1**

Variables
PRY Provide jtd or Dst
CalibOnly?
2 1 to reproduce of
oNormalize?
2 1=no, 2=yes

ofxn calc*
1.02642
ofxn mod
1.03359

sRun
RunA1
zout
1

Ave
Data
1.1E-05
Model
1E-05

data/vMax
8.2E-05

ofxng
319.391

Min d/m
1.49161

ave/min d
1.20312
ave/min m
1.75527

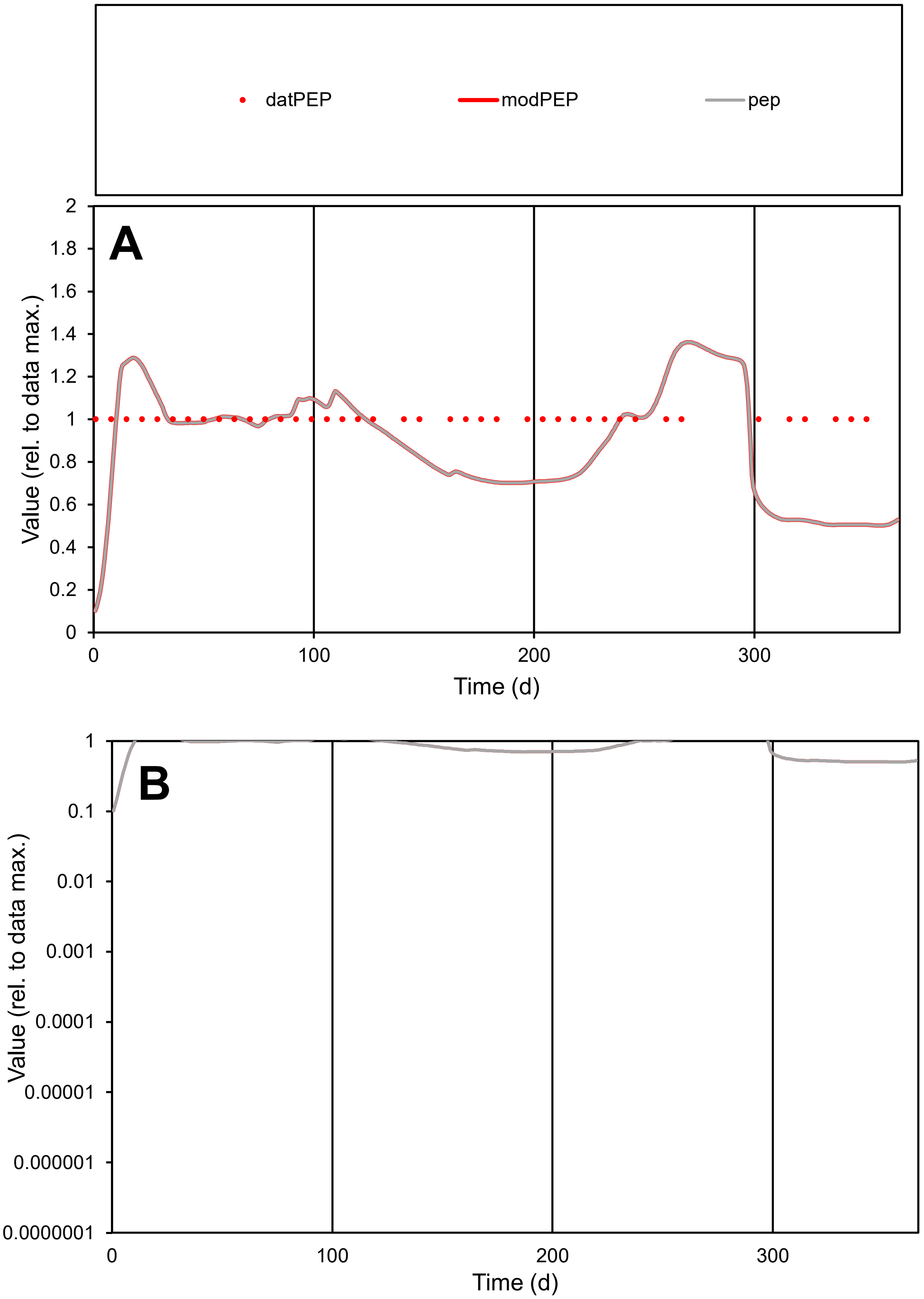


**C1**

Variables
PEP Provide jtd or Dst
CalibOnly?
2 1 to reproduce of
oNormalize?
2 1=no, 2=yes

ofxn calc*
0.27183
ofxn mod
0.29406

sRun
RunA1
zout
1

Ave
Data
4.9E-06
Model
4.3E-06

data/vMax
4.9E-06

ofxng
319.391

Min d/m
4.26448

ave/min d
1
ave/min m
3.74227

330_Calib_S_A

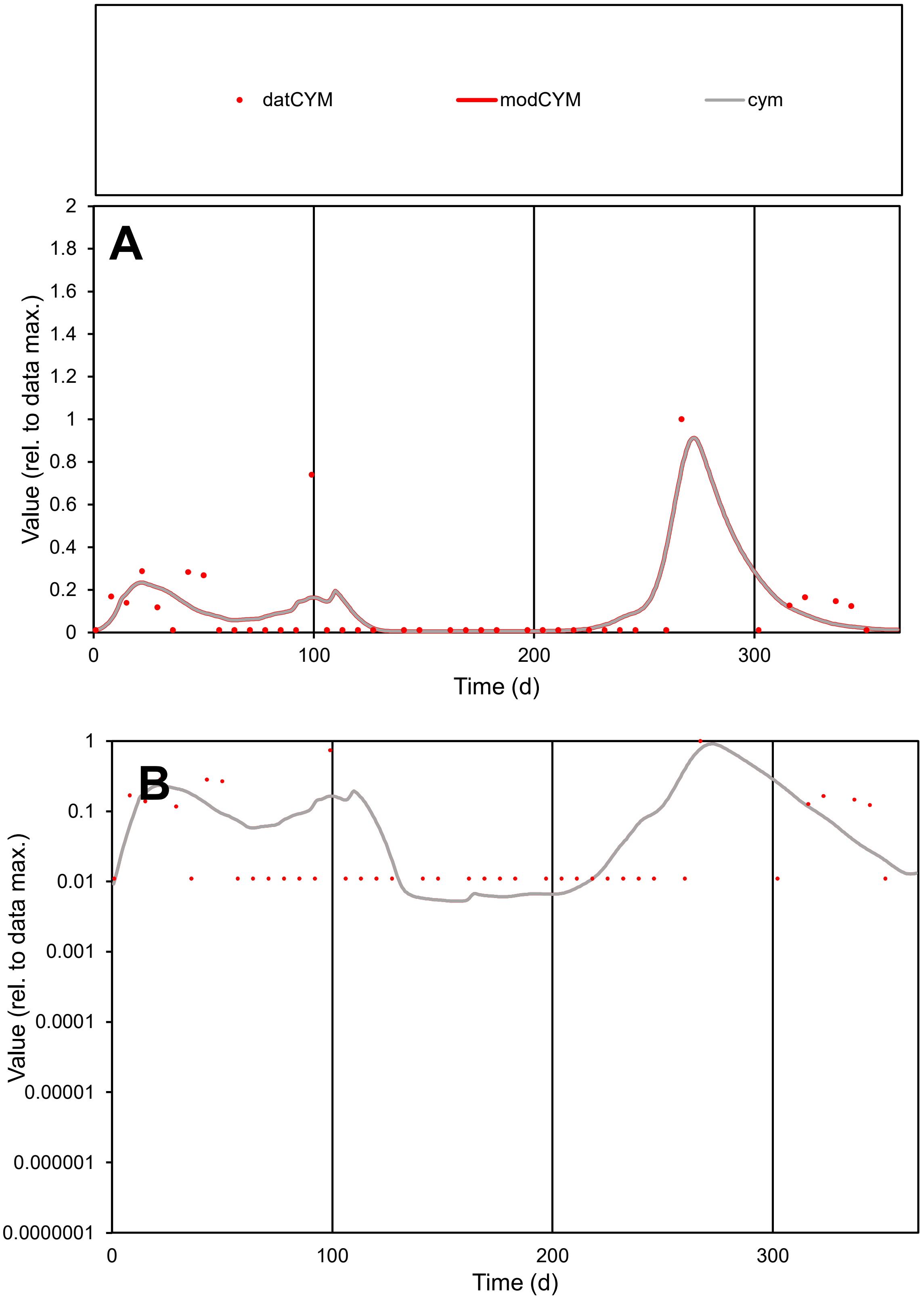


**C1**

Variables
CYM Provide jtd or Dst
CalibOnly?
2 1 to reproduce of
oNormalize?
2 1=no, 2=yes

ofxn calc*
1.36571
ofxn mod
1.43439

sRun
RunA1
zout
1

Ave
Data
1.2E-05
Model
1.2E-05

data/vMax
0.00012

ofxng
319.391

Min d/m
2.03013

ave/min d
8.60485
ave/min m
17.7562

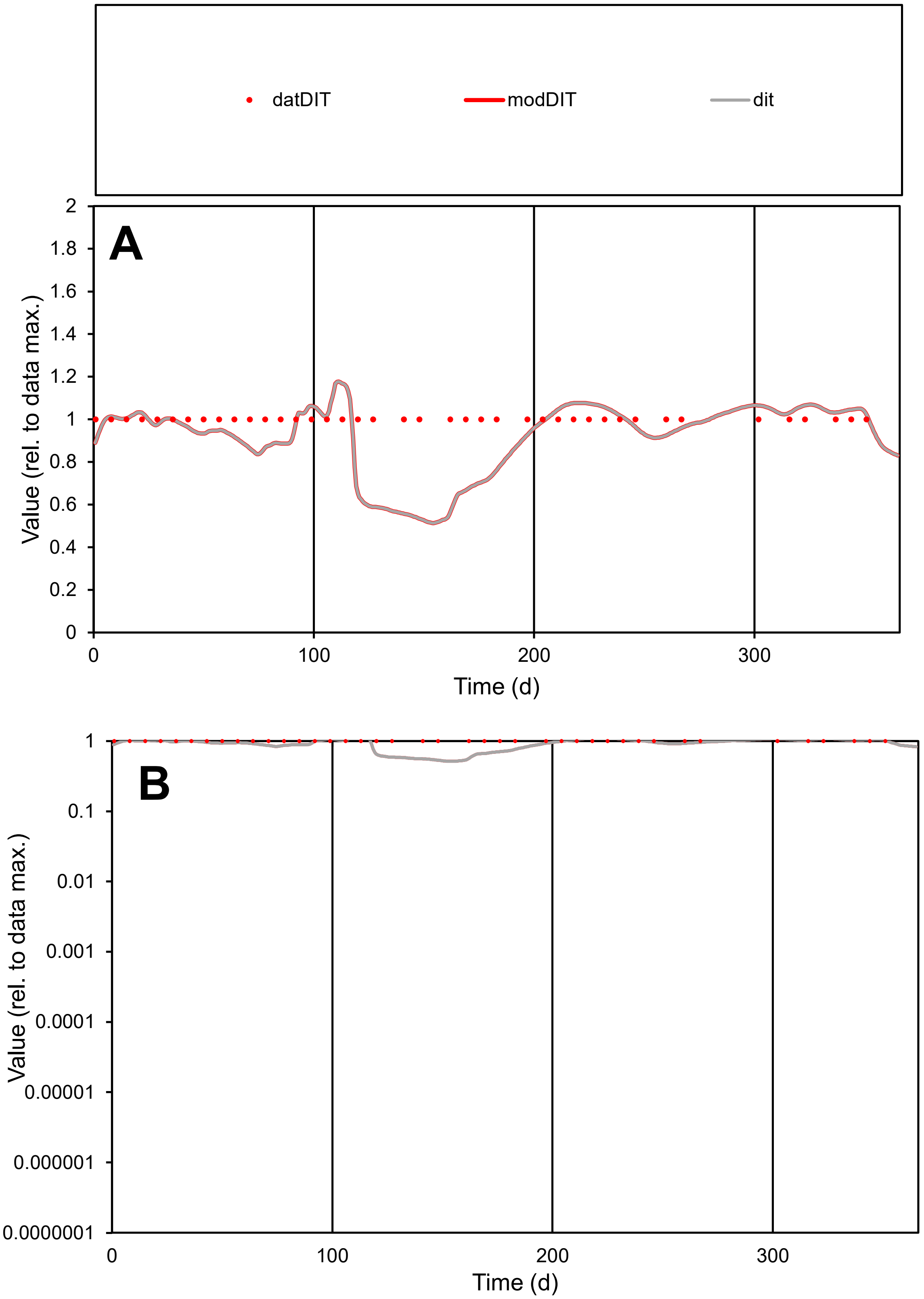


**C1**

Variables
DIT Provide jtd or Dst
CalibOnly?
2 1 to reproduce of
oNormalize?
2 1=no, 2=yes

ofxn calc*
0.17629
ofxn mod
0.20571

sRun
RunA1
zout
1

Ave
Data
5.1E-06
Model
4.7E-06

data/vMax
5.1E-06

ofxng
319.391

Min d/m
1.88844

ave/min d
1
ave/min m
1.75052

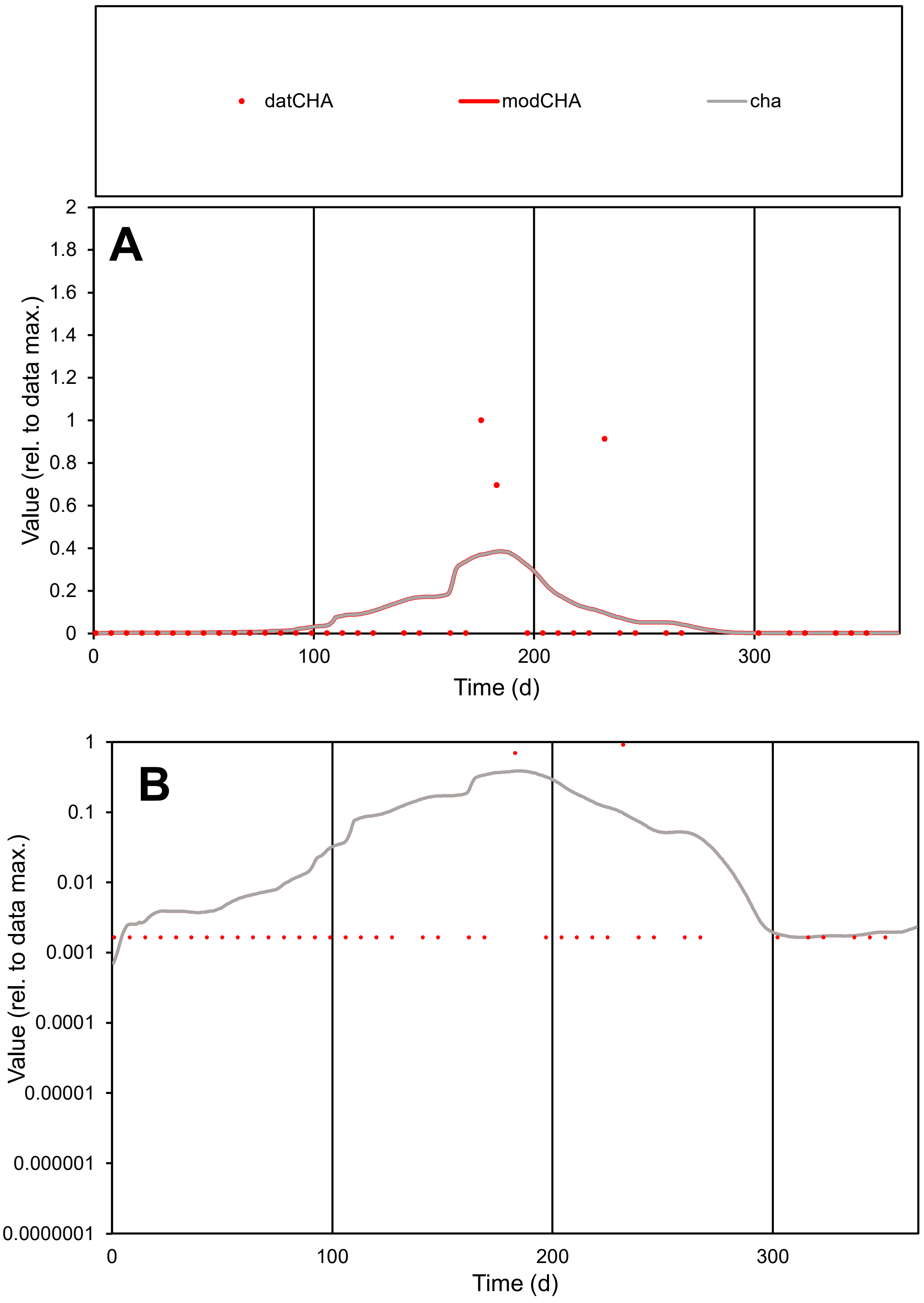


**C1**

Variables
CHA Provide jtd or Dst
CalibOnly?
2 1 to reproduce of
oNormalize?
2 1=no, 2=yes

ofxn calc*
3.02777
ofxn mod
3.05705

sRun
RunA1
zout
1

Ave
Data
3.4E-06
Model
4.2E-06

data/vMax
5.2E-05

ofxng
319.391

Min d/m
1.19462

ave/min d
39.4017
ave/min m
58.021

330_Calib_S_A

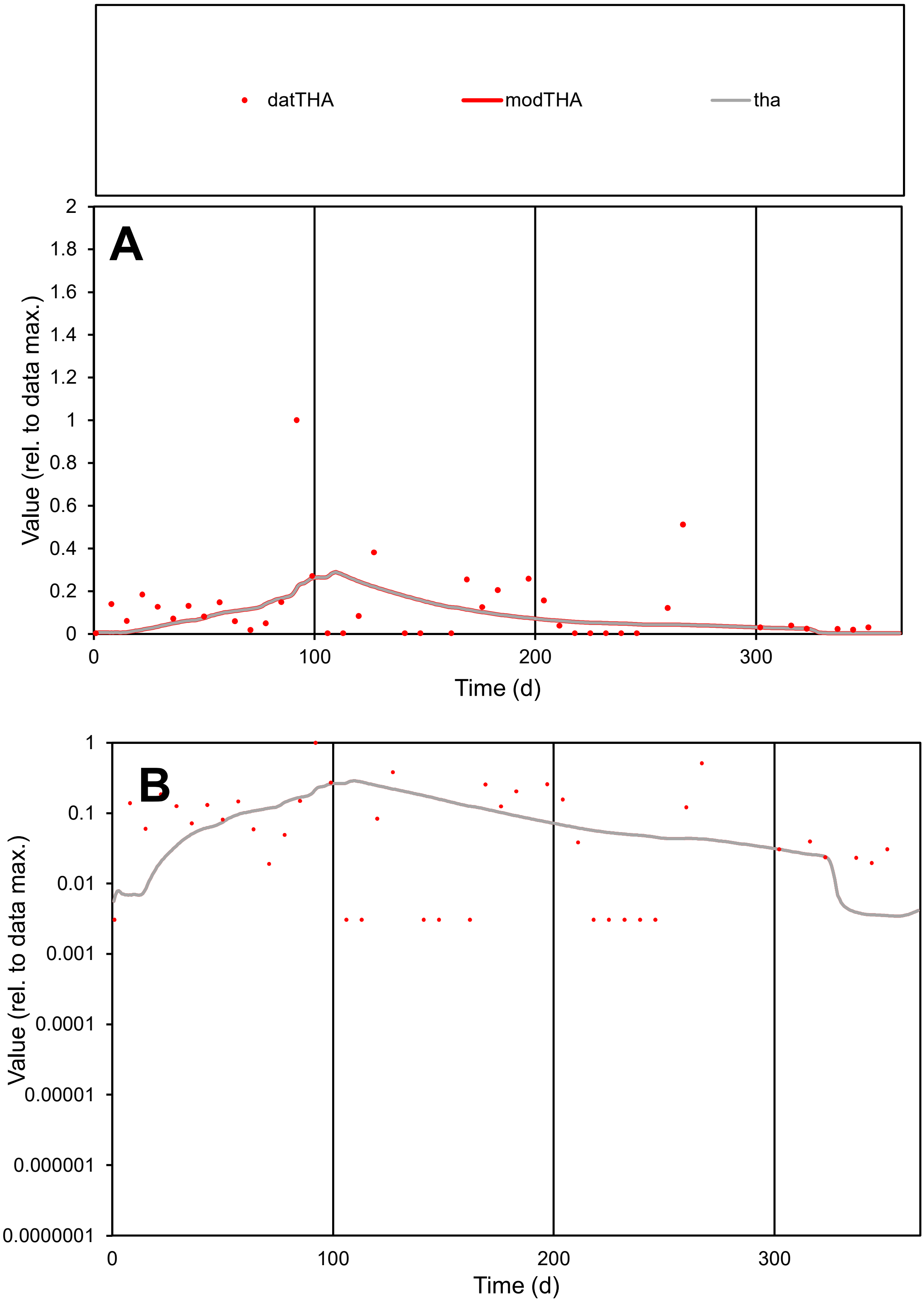


**C1**

Variables

THA Provide jtd or Dst

CalibOnly?

2 1 to reproduce of

oNormalize?

2 1=no, 2=yes

ofxn calc*

1.48424

ofxn mod

1.49053

sRun

RunA1

zout

1

Ave

Data

0.00011

Model

8.3E-05

data/vMax

0.0009

ofxng

319.391

Min d/m

0.87588

ave/min d

38.5364

ave/min m

26.5574

330_Calib_S_A

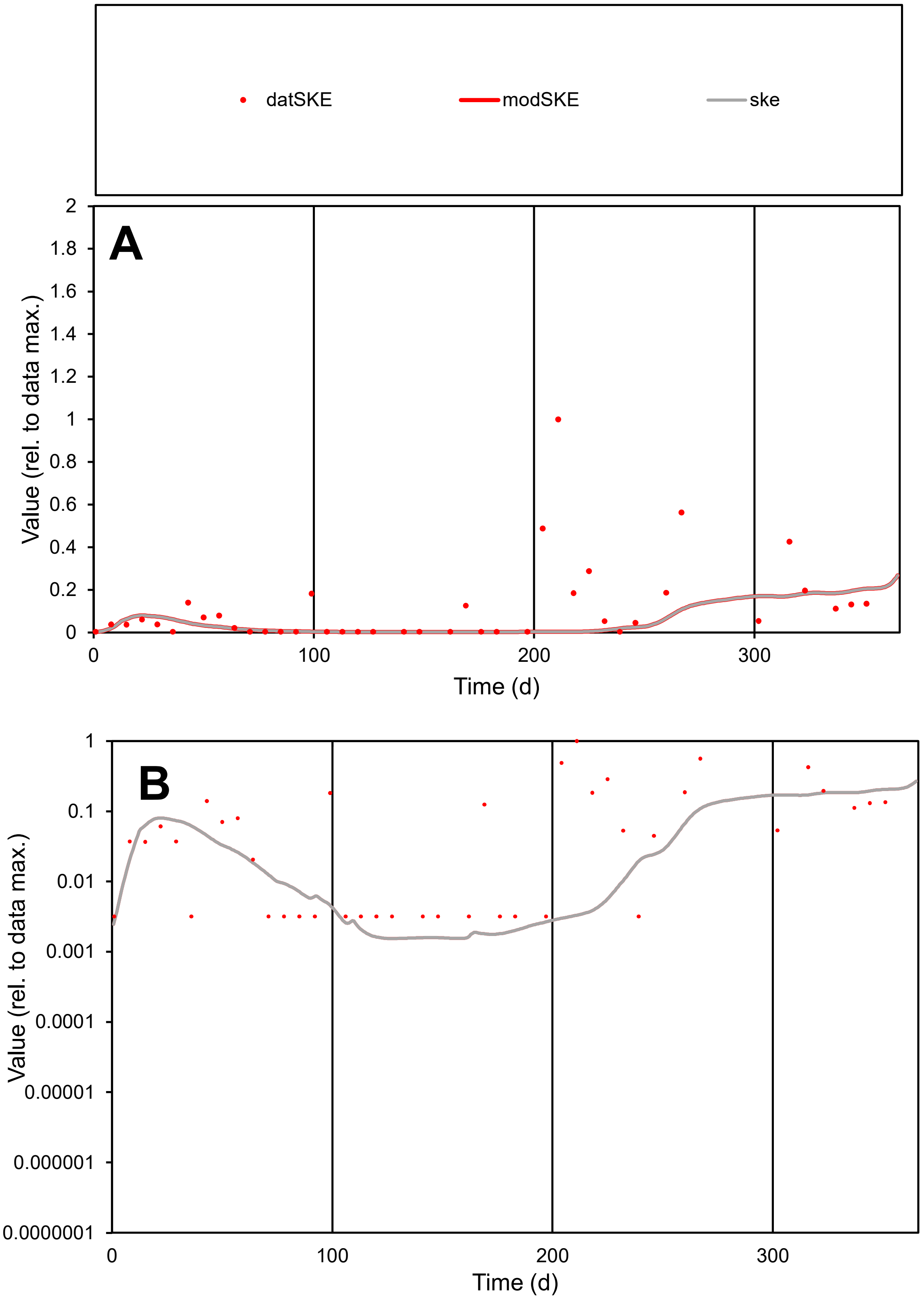


**C1**

| | |
|---|---|
| Variables | |
| SKE | Provide jtd or Dst |
| CalibOnly? | |
| 2 | 1 to reproduce of |
| oNormalize? | |
| 2 | 1=no, 2=yes |
| ofxn calc* | |
| 1.78261 | |
| ofxn mod | |
| 1.78744 | |
| sRun | |
| RunA1 | |
| zout | |
| 1 | |
| Ave | |
| Data | |
| 8.3E-05 | |
| Model | |
| 3.3E-05 | |
| data/vMax | |
| 0.00072 | |
| ofxng | |
| 319.391 | |
| Min d/m | |
| 2.05263 | |
| ave/min d | |
| 36.1063 | |
| ave/min m | |
| 29.2578 | |

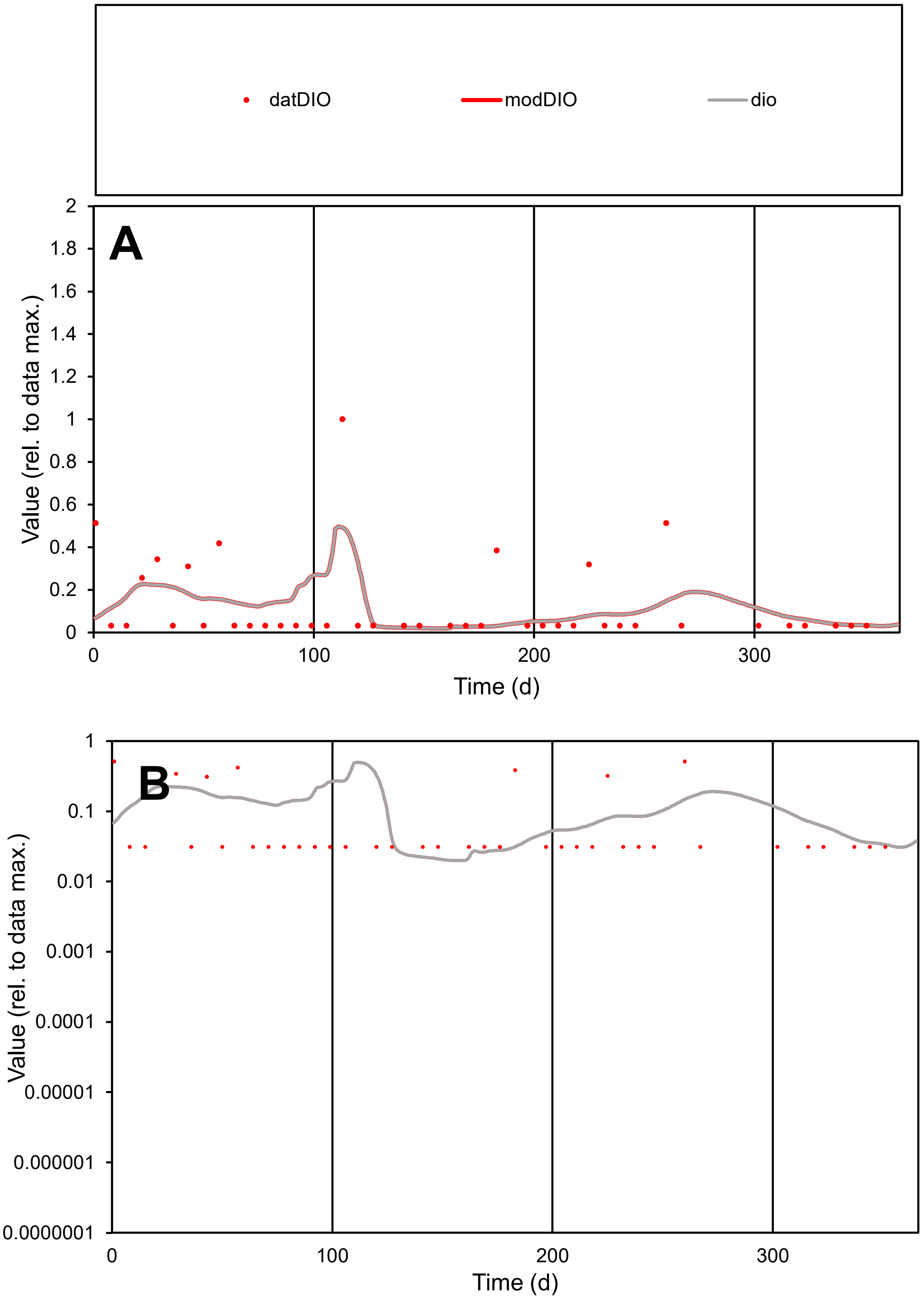


**C1**

Variables
DIO Provide jtd or Dst
CalibOnly?
2 1 to reproduce of
oNormalize?
2 1=no, 2=yes

ofxn calc*
1.37324
ofxn mod
1.39913

sRun
RunA1
zout
1

Ave
Data
1.8E-05
Model
1.8E-05

data/vMax
0.00015

ofxng
319.391

Min d/m
1.4984

ave/min d
3.94934
ave/min m
5.66211

330_Calib_S_A

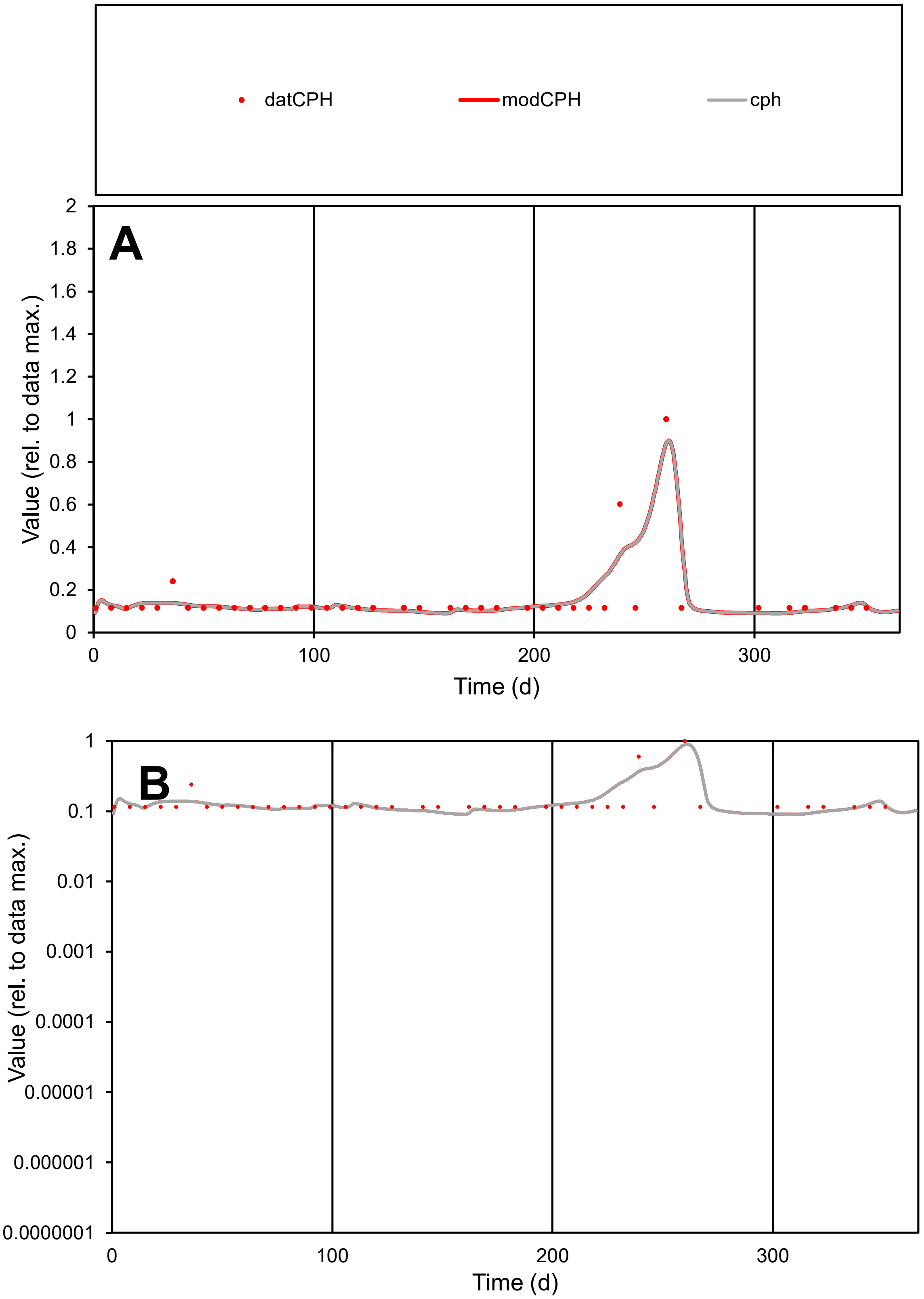


**C1**

Variables
CPH Provide jtd or Dst
CalibOnly?
2 1 to reproduce of
oNormalize?
2 1=no, 2=yes

ofxn calc*
0.46903
ofxn mod
0.5006

sRun
RunA1
zout
1

Ave
Data
1.3E-05
Model
1.3E-05

data/vMax
8.3E-05

ofxng
319.391

Min d/m
1.26295

ave/min d
1.31681
ave/min m
1.72585

330_Calib_S_A

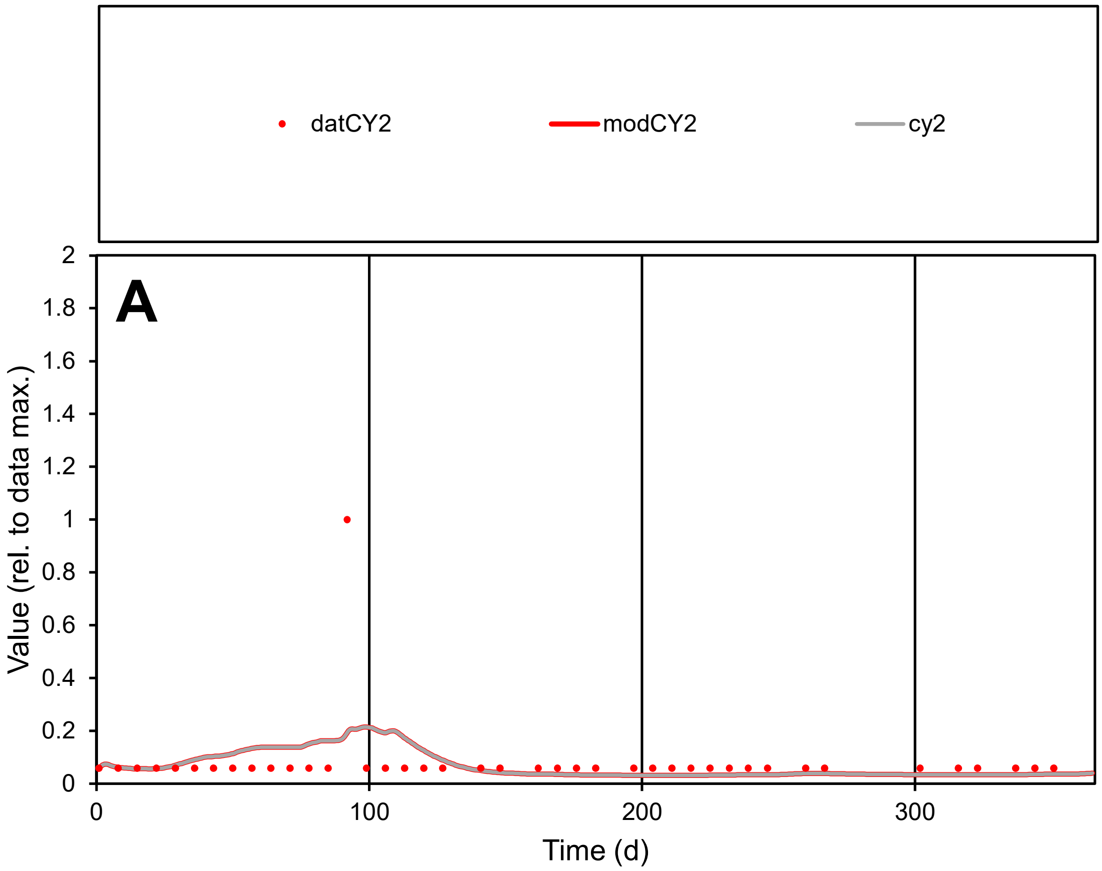


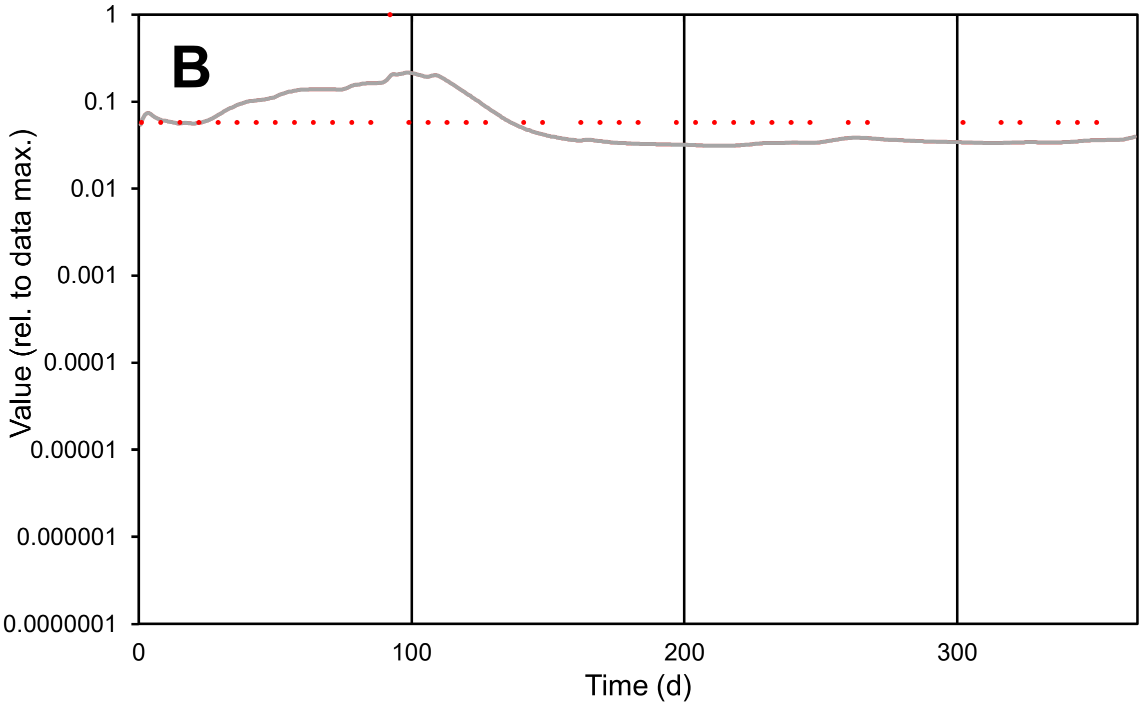


**C1**

Variables
CY2 Provide jtd or Dst
CalibOnly?
2 1 to reproduce of
oNormalize?
2 1=no, 2=yes

ofxn calc*
1.66992
ofxn mod
1.67605

sRun
RunA1
zout
1

Ave
Data
1.6E-05
Model
1.5E-05

data/vMax
0.00019

ofxng
319.391

Min d/m
1.84774

ave/min d
1.39795
ave/min m
2.41954

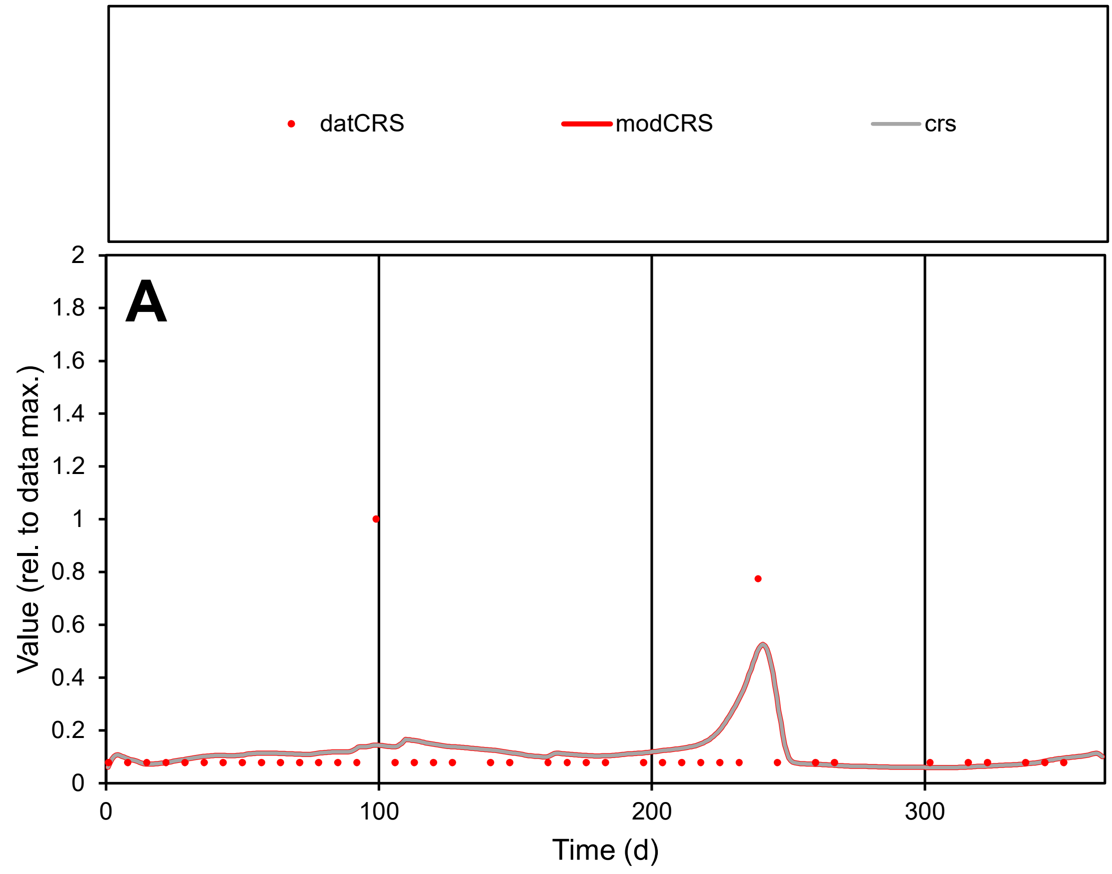


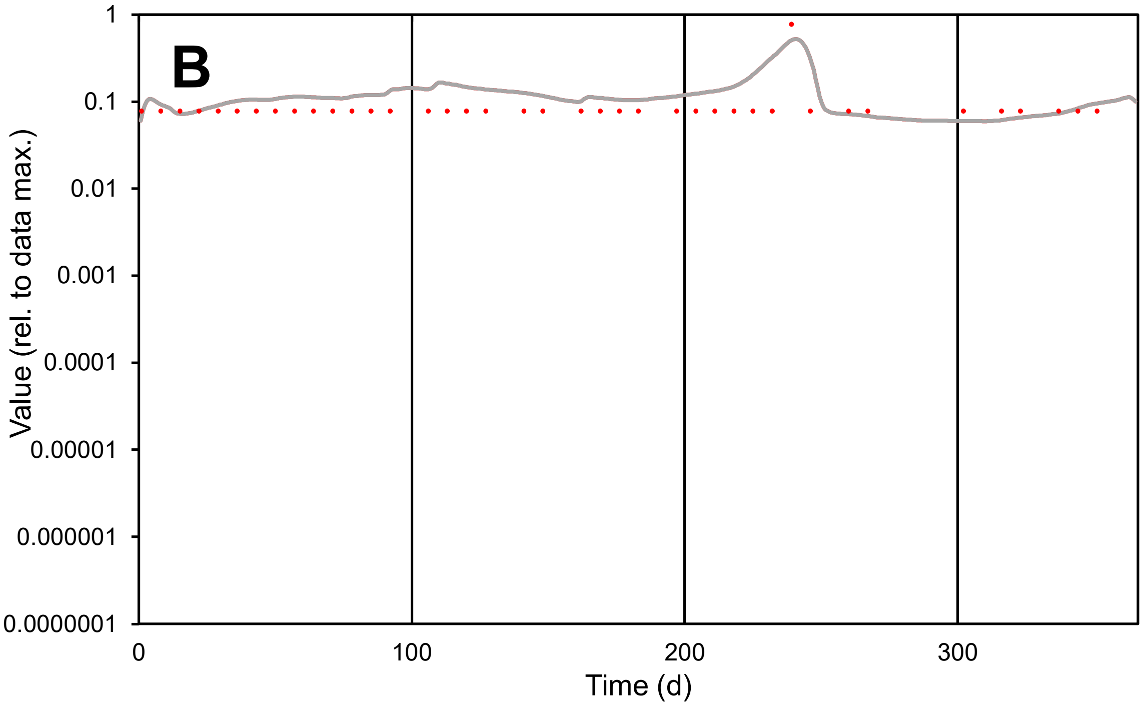


**C1**

Variables
CRS Provide jtd or Dst
CalibOnly?
2 1 to reproduce of
oNormalize?
2 1=no, 2=yes

ofxn calc*
1.27245
ofxn mod
1.29851

sRun
RunA1
zout
1

Ave
Data
8.1E-06
Model
8.6E-06

data/vMax
6.9E-05

ofxng
319.391

Min d/m
1.30898

ave/min d
1.5044
ave/min m
2.08866

330_Calib_S_A

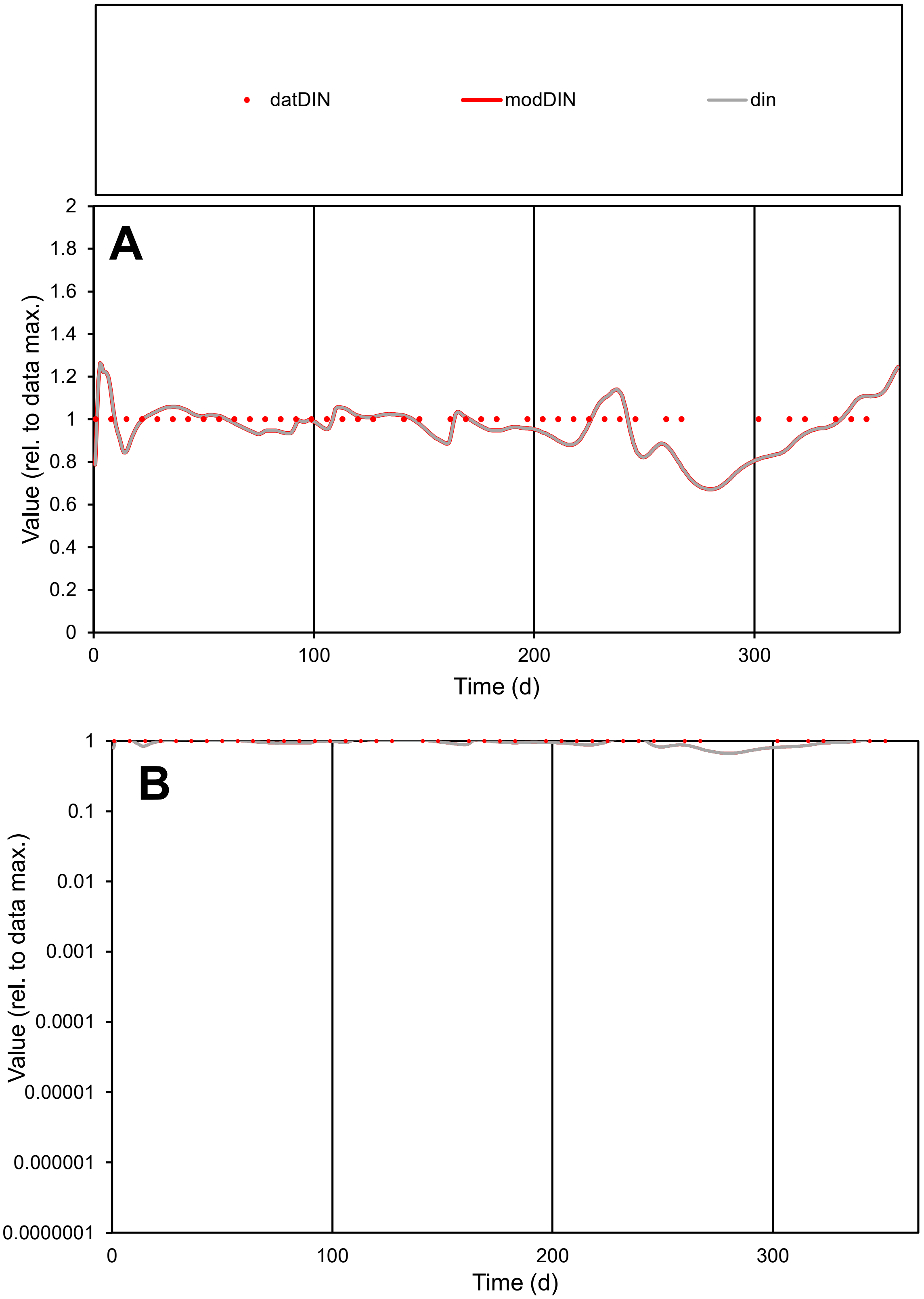


**C1**

Variables
DIN Provide jtd or Dst
CalibOnly?
2 1 to reproduce of
oNormalize?
2 1=no, 2=yes

ofxn calc*
0.07929
ofxn mod
0.13376

sRun
RunA1
zout
1

Ave
Data
2.9E-06
Model
2.8E-06

data/vMax
2.9E-06

ofxng
319.391

Min d/m
1.32294

ave/min d
1
ave/min m
1.28885

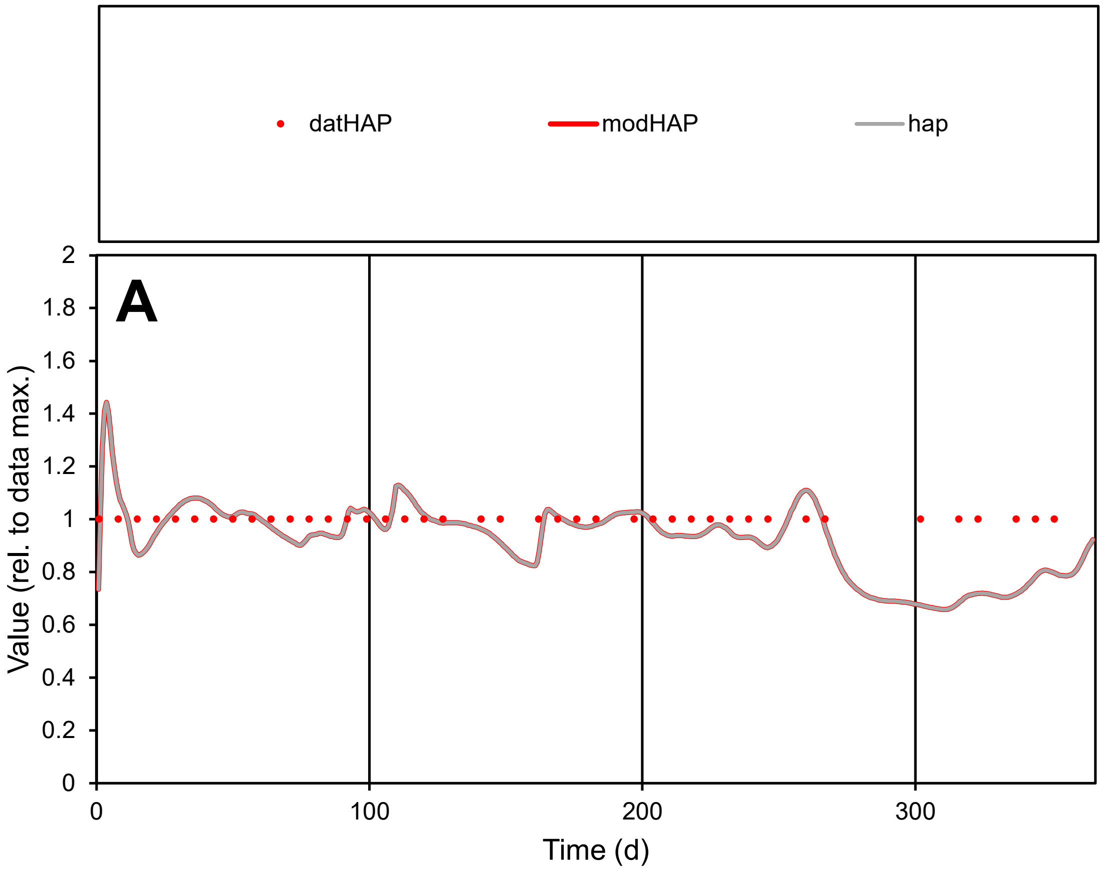


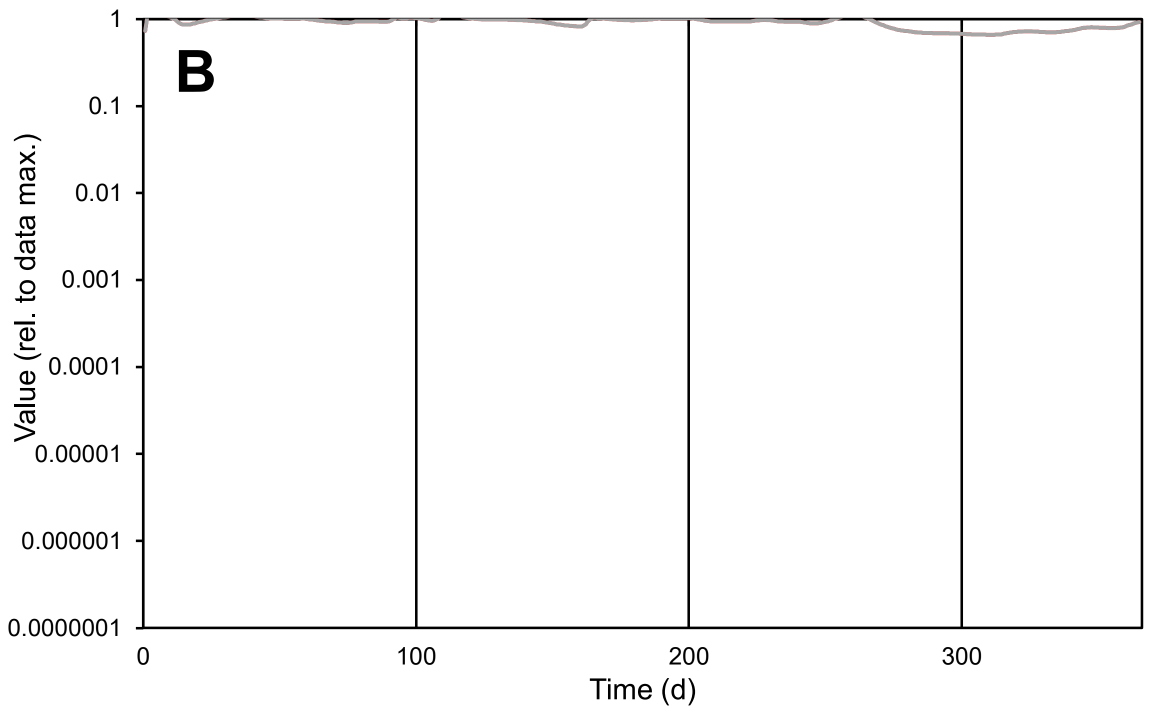


**C1**

Variables
HAP Provide jtd or Dst
CalibOnly?
2 1 to reproduce of
oNormalize?
2 1=no, 2=yes

ofxn calc*
0.11681
ofxn mod
0.15812

sRun
RunA1
zout
1

Ave
Data
1.1E-05
Model
1.1E-05

data/vMax
1.1E-05

ofxng
319.391

Min d/m
1.48931

ave/min d
1
ave/min m
1.41205

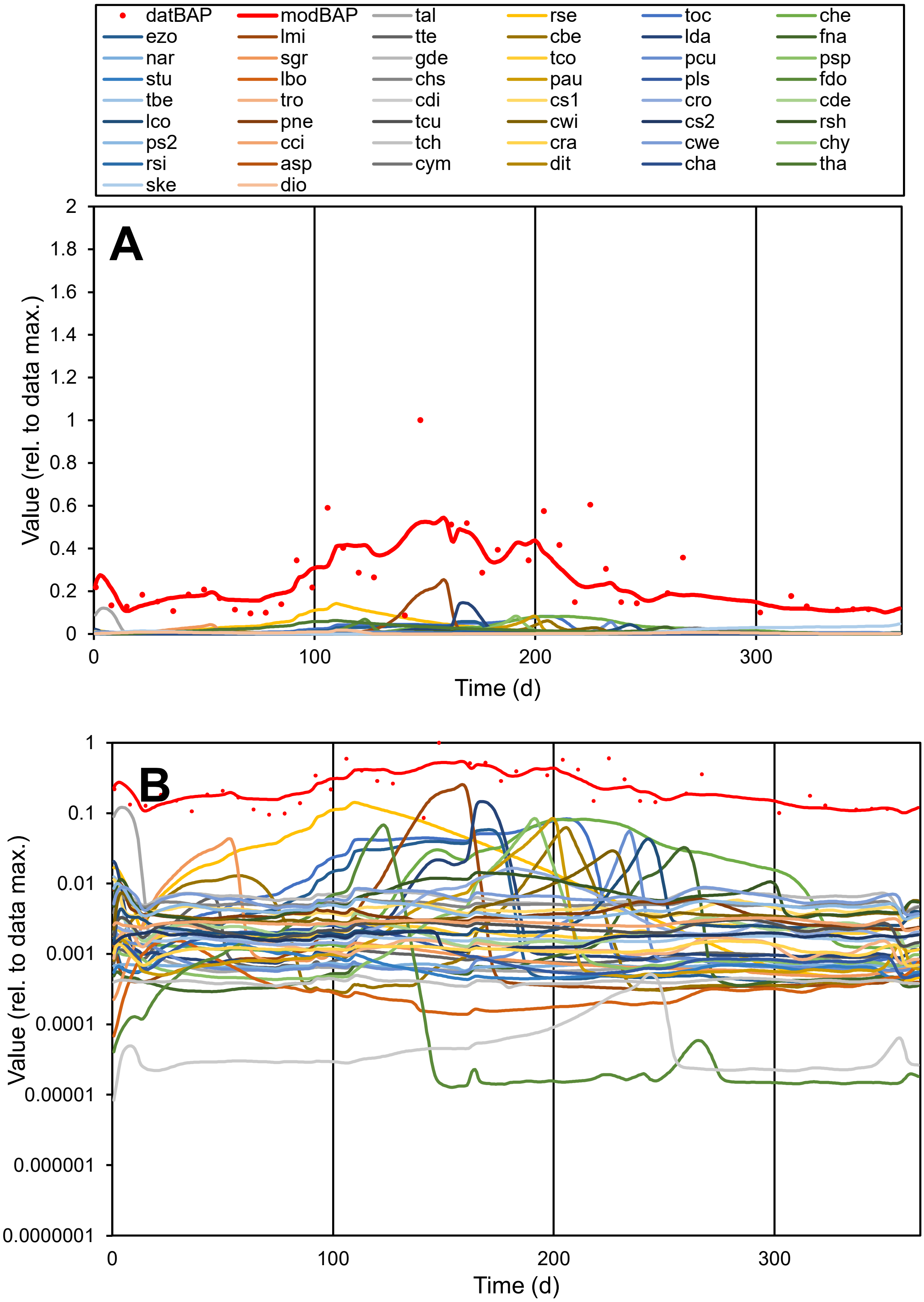


**C1**

Variables

BAP Provide jtd or Dst

CalibOnly?

2 1 to reproduce of

oNormalize?

2 1=no, 2=yes

ofxn calc*

0.51704

ofxn mod

0.5312

sRun

RunA1

zout

1

Ave

Data

0.00108

Model

0.00102

data/vMax

0.0041

ofxng

319.391

Min d/m

0.78613

ave/min d

3.07631

ave/min m

2.29166

330_Calib_S_A

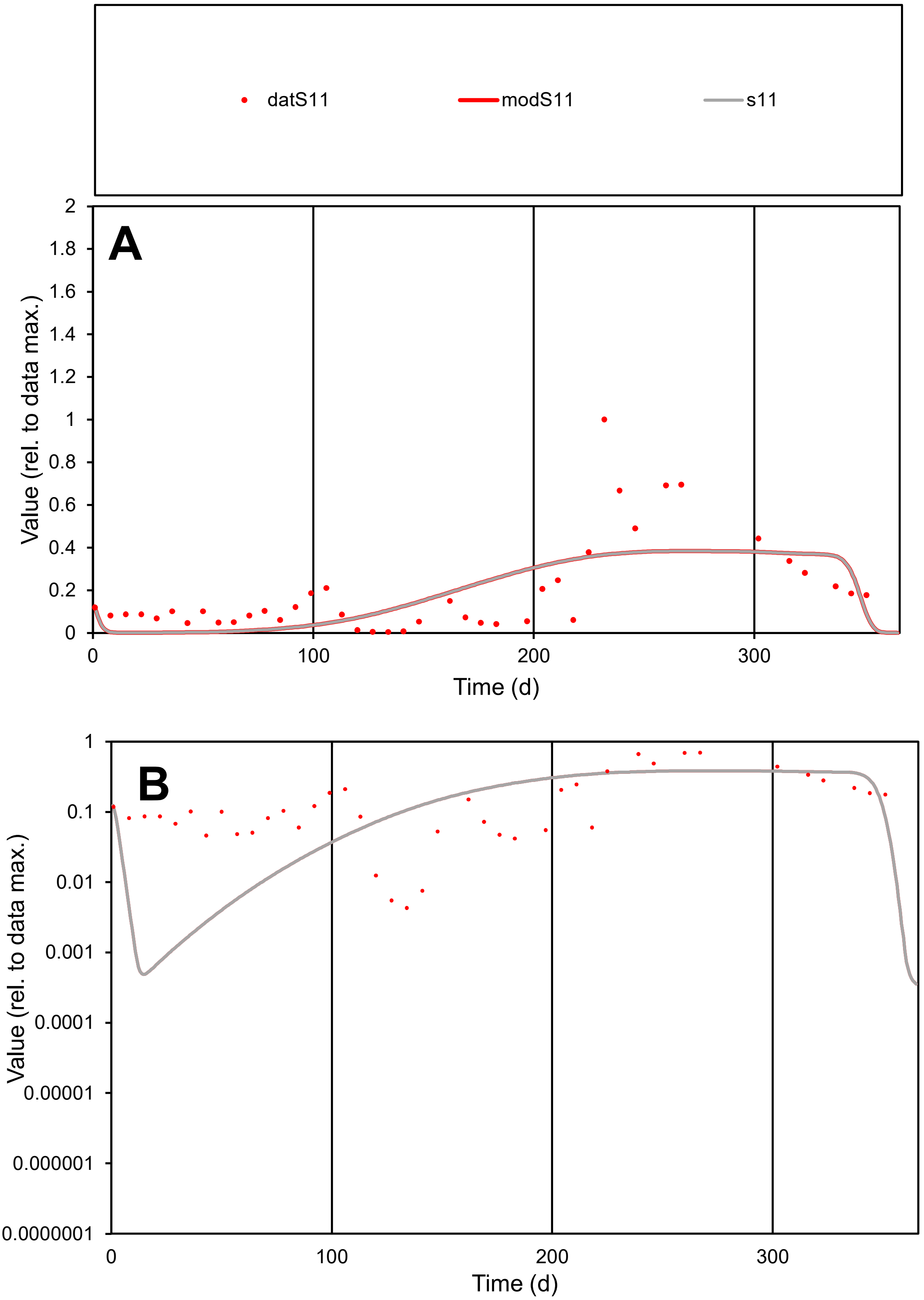


**C1**

Variables
S11 Provide jtd or Dst
CalibOnly?
2 1 to reproduce of
oNormalize?
2 1=no, 2=yes

ofxn calc*
0.85597
ofxn mod
0.8723

sRun
RunA1
zout
1

Ave
Data
0.08708
Model
0.07663

data/vMax
0.448

ofxng
319.391

Min d/m
8.19344

ave/min d
45.2833
ave/min m
326.484

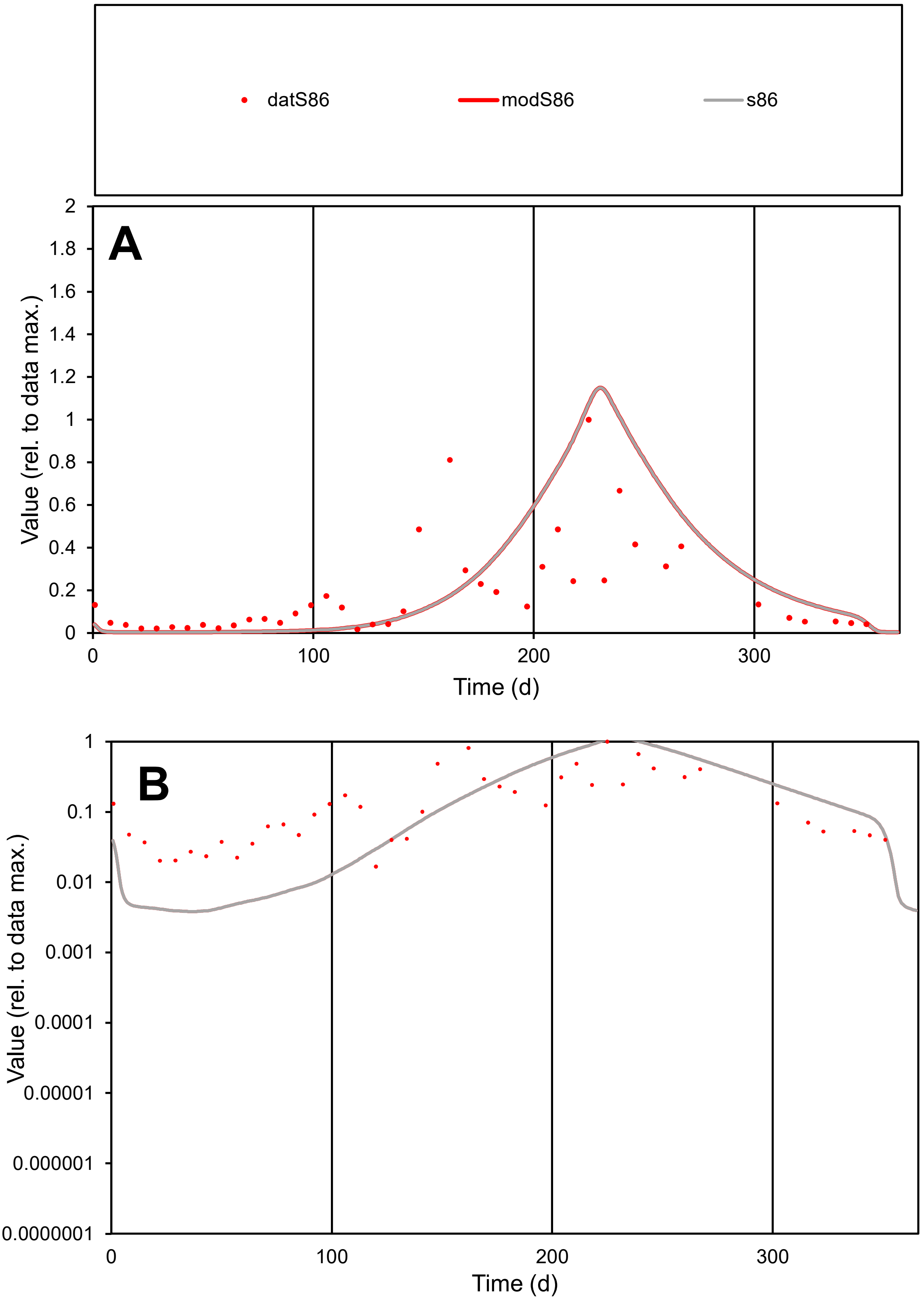


**C1**

Variables
S86 Provide jtd or Dst
CalibOnly?
2 1 to reproduce of
oNormalize?
2 1=no, 2=yes

ofxn calc*
1.35367
ofxn mod
1.37687

sRun
RunA1
zout
1

Ave
Data
0.04823
Model
0.06393

data/vMax
0.2564

ofxng
319.391

Min d/m
4.36617

ave/min d
11.3103
ave/min m
65.4635

330_Calib_S_A

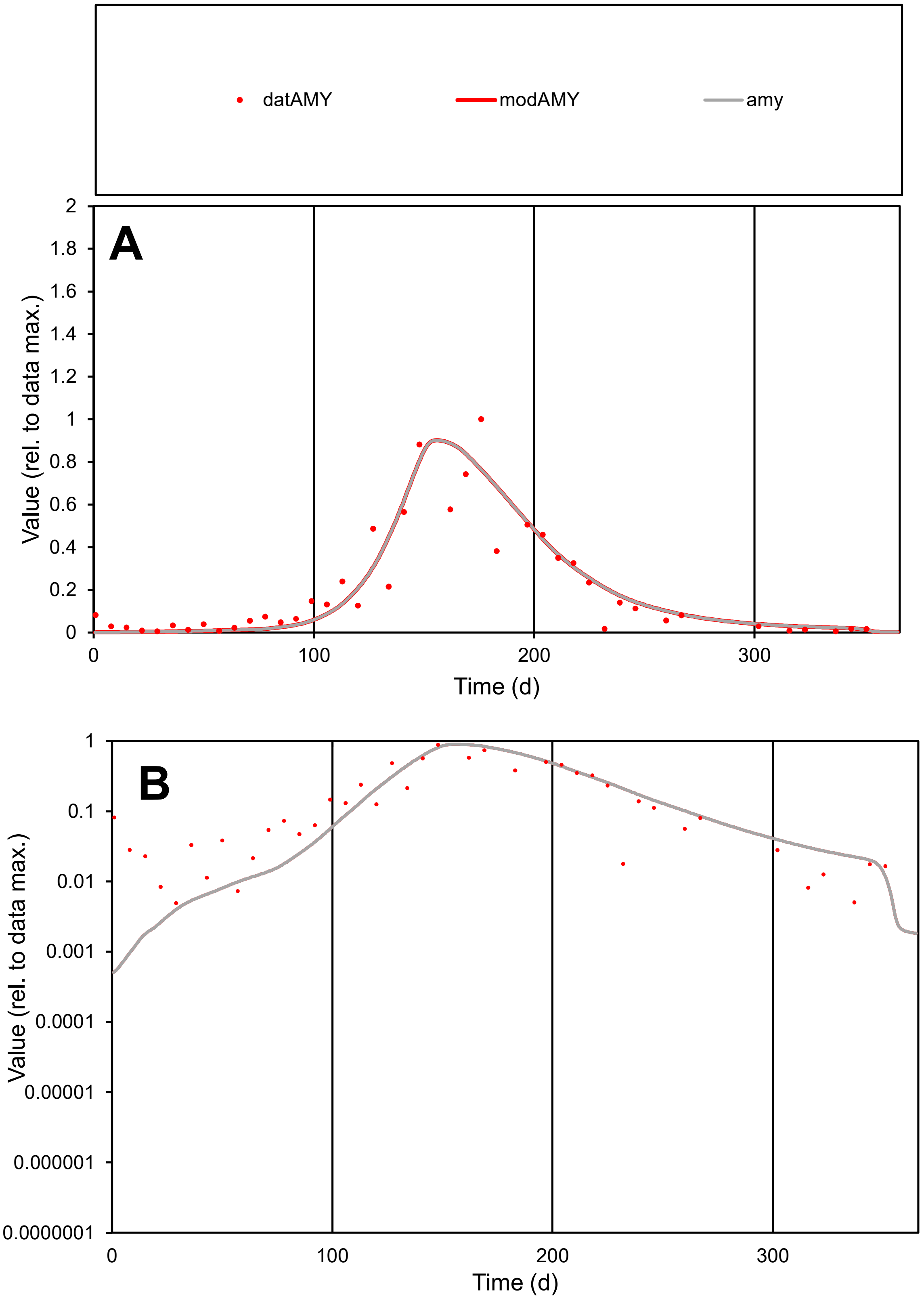


**C1**

Variables
AMY Provide jtd or Dst
CalibOnly?
2 1 to reproduce of
oNormalize?
2 1=no, 2=yes

ofxn calc*
0.50247
ofxn mod
0.53339

sRun
RunA1
zout
1

Ave
Data
0.03608
Model
0.03768

data/vMax
0.1814

ofxng
319.391

Min d/m
9.31491

ave/min d
40.3848
ave/min m
392.863

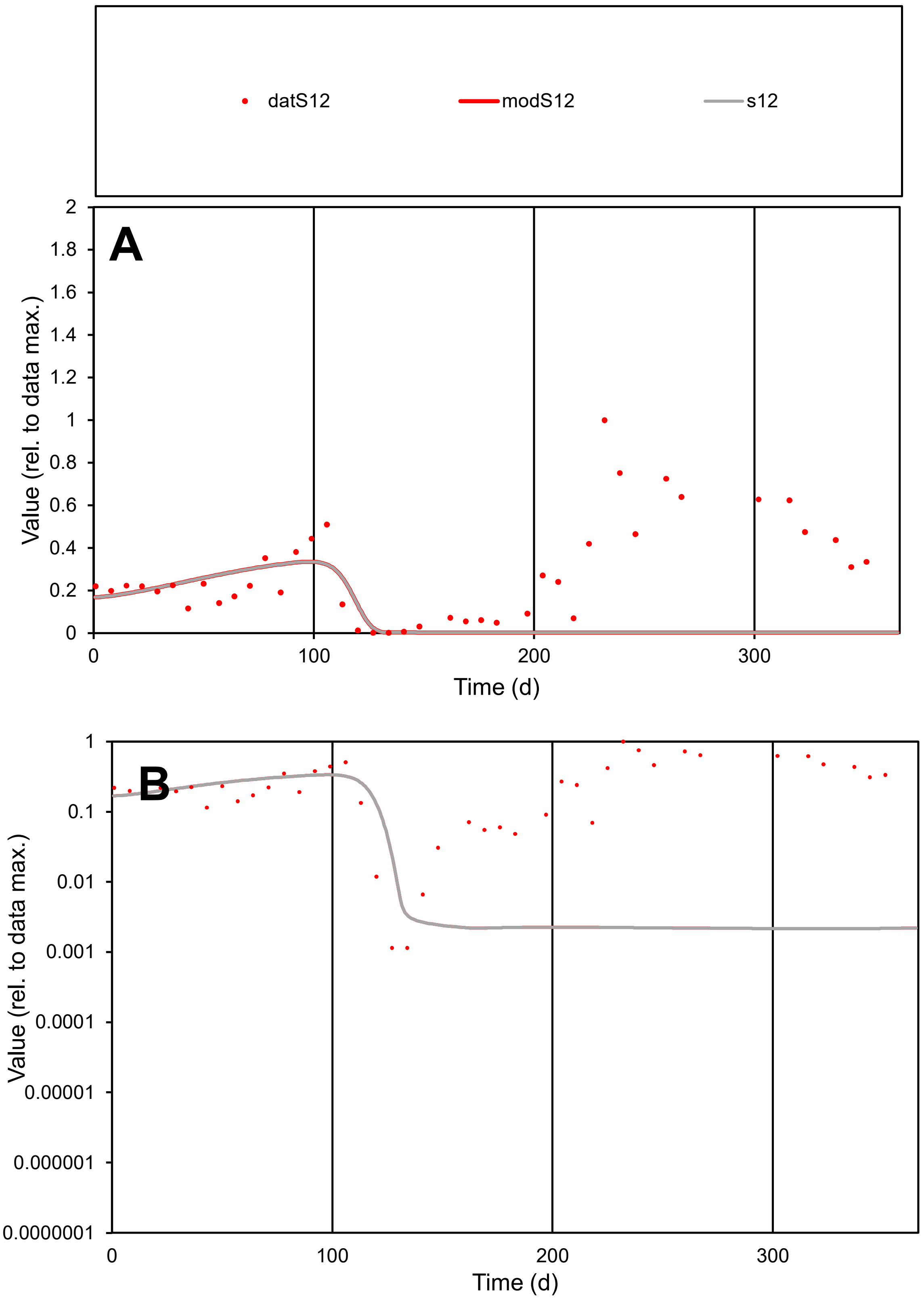


**C1**

Variables
S12 Provide jtd or Dst
CalibOnly?
2 1 to reproduce of
oNormalize?
2 1=no, 2=yes

ofxn calc*
1.16116
ofxn mod
1.16533

sRun
RunA1
zout
1

Ave
Data
0.03686
Model
0.01408

data/vMax
0.1298

ofxng
319.391

Min d/m
0.52901

ave/min d
248.049
ave/min m
50.1086

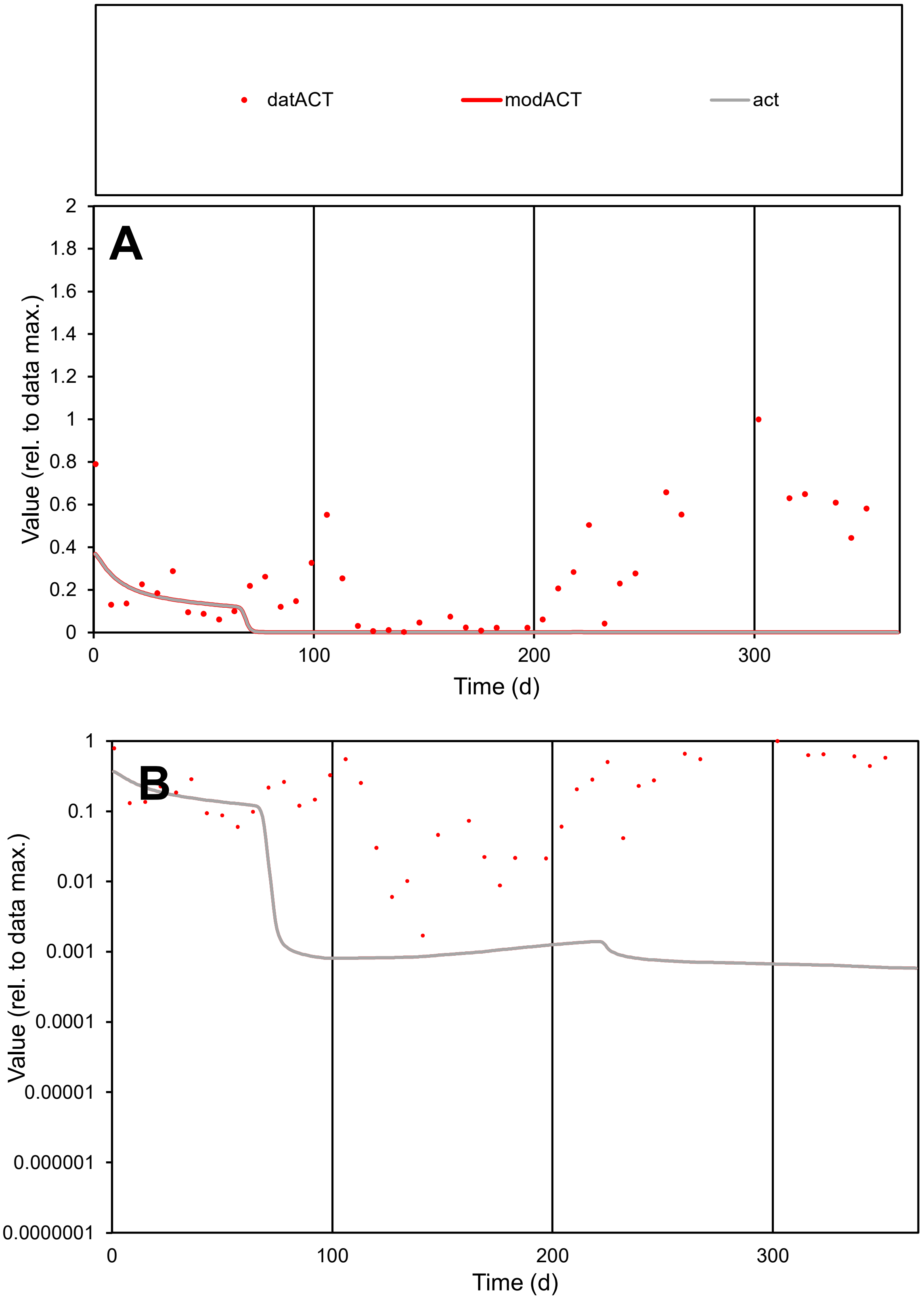


**C1**

Variables
ACT Provide jtd or Dst
CalibOnly?
2 1 to reproduce of
oNormalize?
2 1=no, 2=yes

ofxn calc*
1.29926
ofxn mod
1.3028

sRun
RunA1
zout
1

Ave
Data
0.01784
Model
0.00317

data/vMax
0.06853

ofxng
319.391

Min d/m
2.85819

ave/min d
152.614
ave/min m
77.464

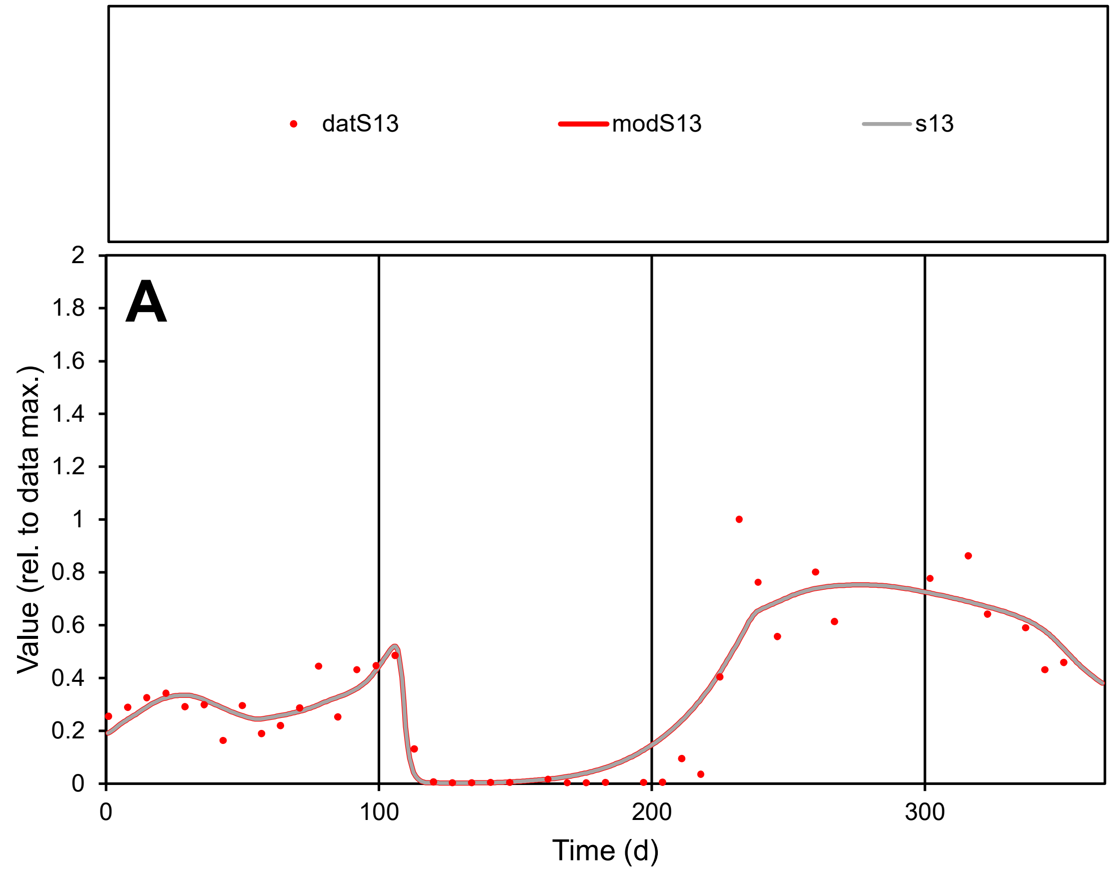


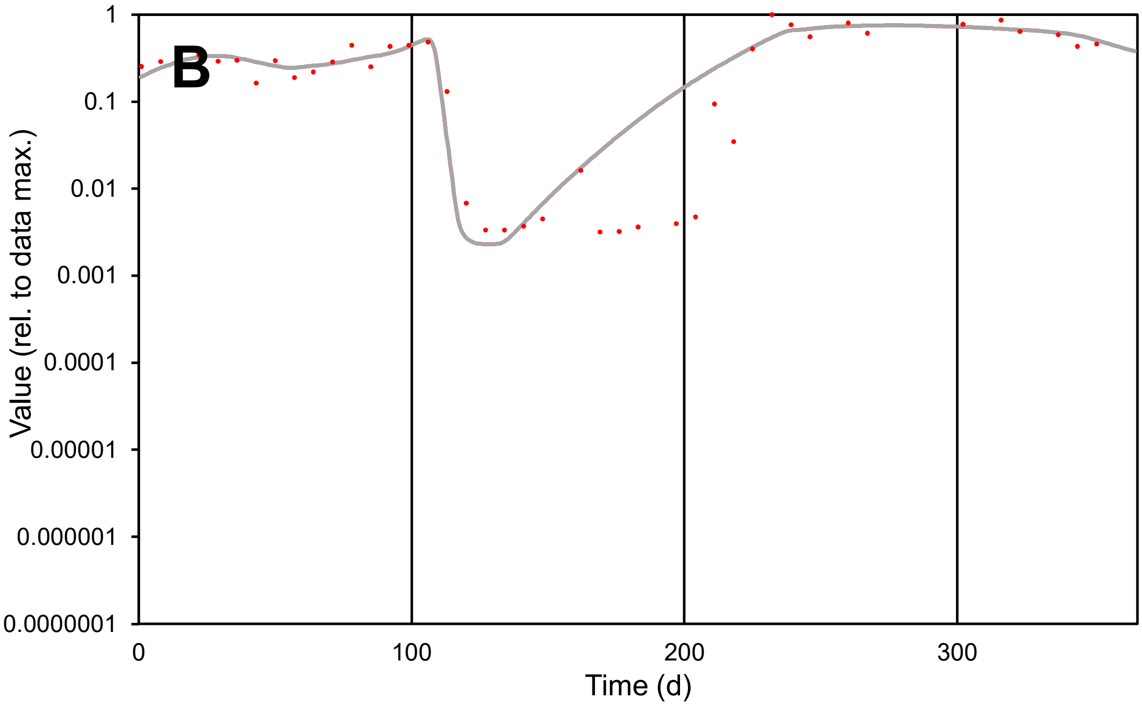


**C1**

Variables
S13 Provide jtd or Dst
CalibOnly?
2 1 to reproduce of
oNormalize?
2 1=no, 2=yes

ofxn calc*
0.34581
ofxn mod
0.37875

sRun
RunA1
zout
1

Ave
Data
0.02779
Model
0.02876

data/vMax
0.08824

ofxng
319.391

Min d/m
1.39286

ave/min d
98.955
ave/min m
142.657

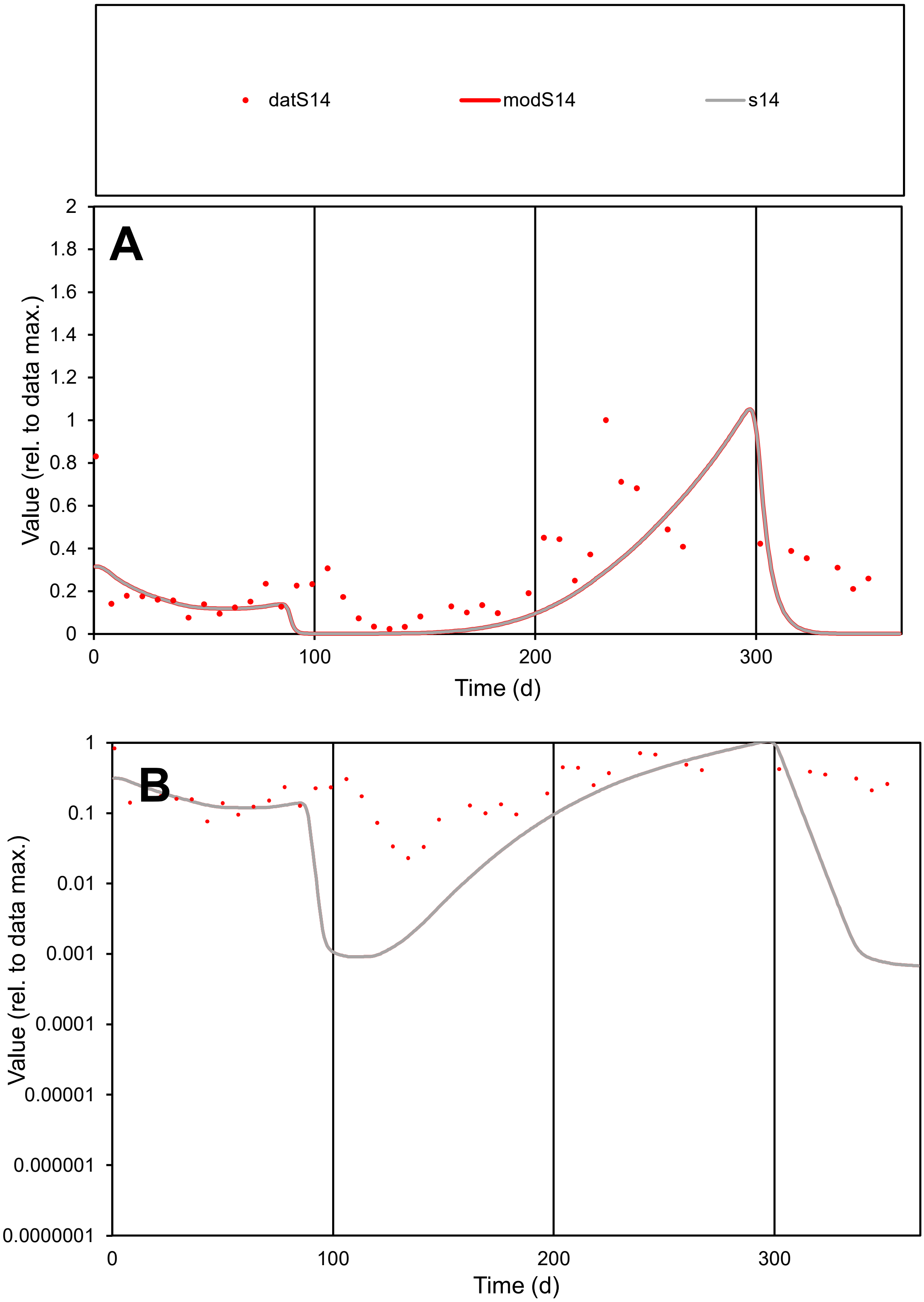


**C1**

Variables
S14 Provide jtd or Dst
CalibOnly?
2 1 to reproduce of
oNormalize?
2 1=no, 2=yes

ofxn calc*
0.81078
ofxn mod
0.8262

sRun
RunA1
zout
1

Ave
Data
0.02659
Model
0.01483

data/vMax
0.1

ofxng
319.391

Min d/m
31.0401

ave/min d
11.5555
ave/min m
200.097

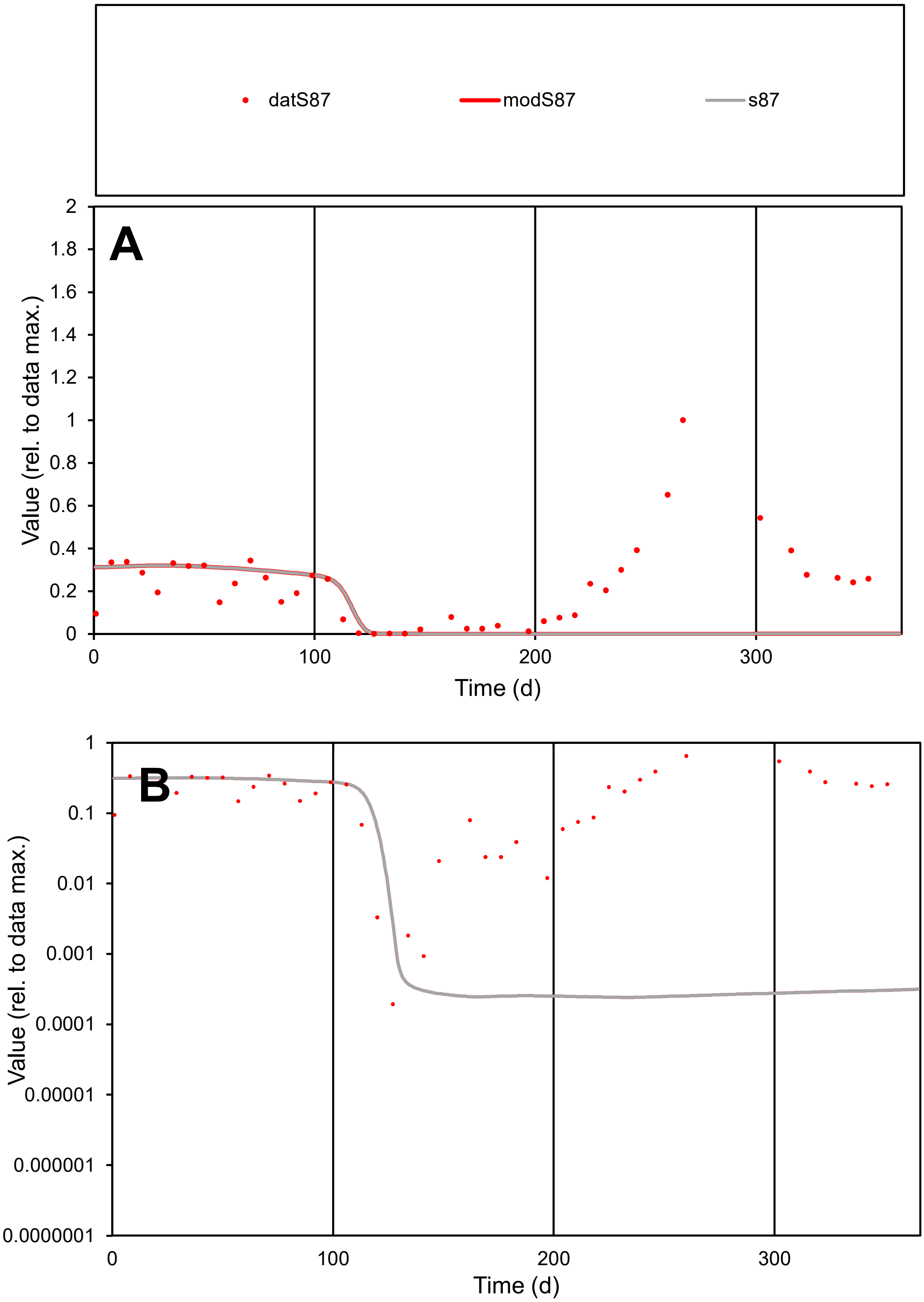


**C1**

Variables
S87 Provide jtd or Dst
CalibOnly?
2 1 to reproduce of
oNormalize?
2 1=no, 2=yes

ofxn calc*
1.12907
ofxn mod
1.13506

sRun
RunA1
zout
1

Ave
Data
0.02158
Model
0.01163

data/vMax
0.09722

ofxng
319.391

Min d/m
0.79957

ave/min d
1148.53
ave/min m
494.803

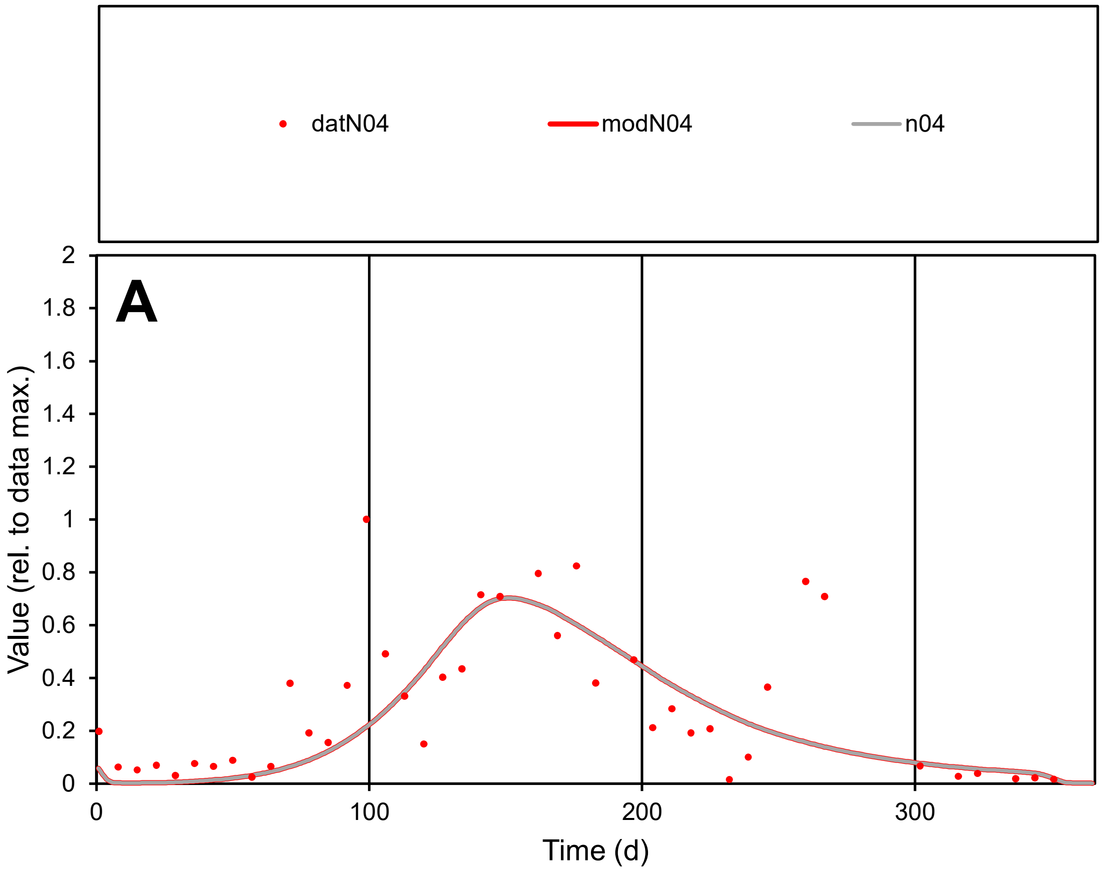


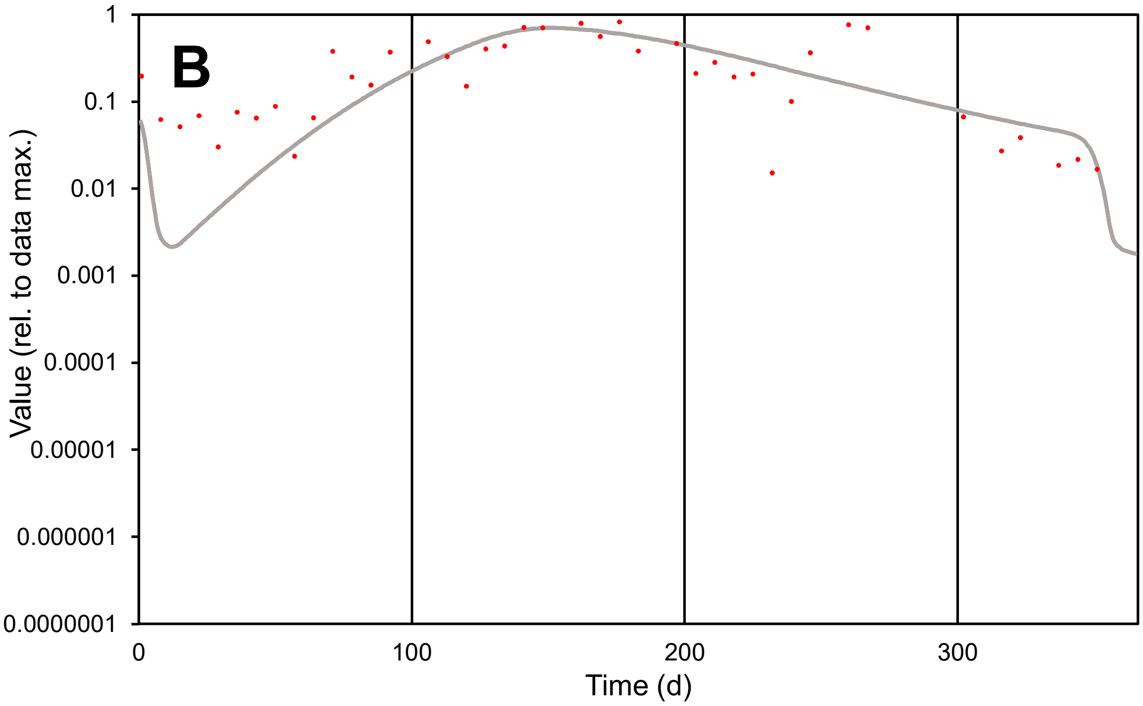


**C1**

Variables
N04 Provide jtd or Dst
CalibOnly?
2 1 to reproduce of
oNormalize?
2 1=no, 2=yes

ofxn calc*
0.74309
ofxn mod
0.75544

sRun
RunA1
zout
1

Ave
Data
0.02153
Model
0.01784

data/vMax
0.07461

ofxng
319.391

Min d/m
6.41605

ave/min d
18.9687
ave/min m
100.848

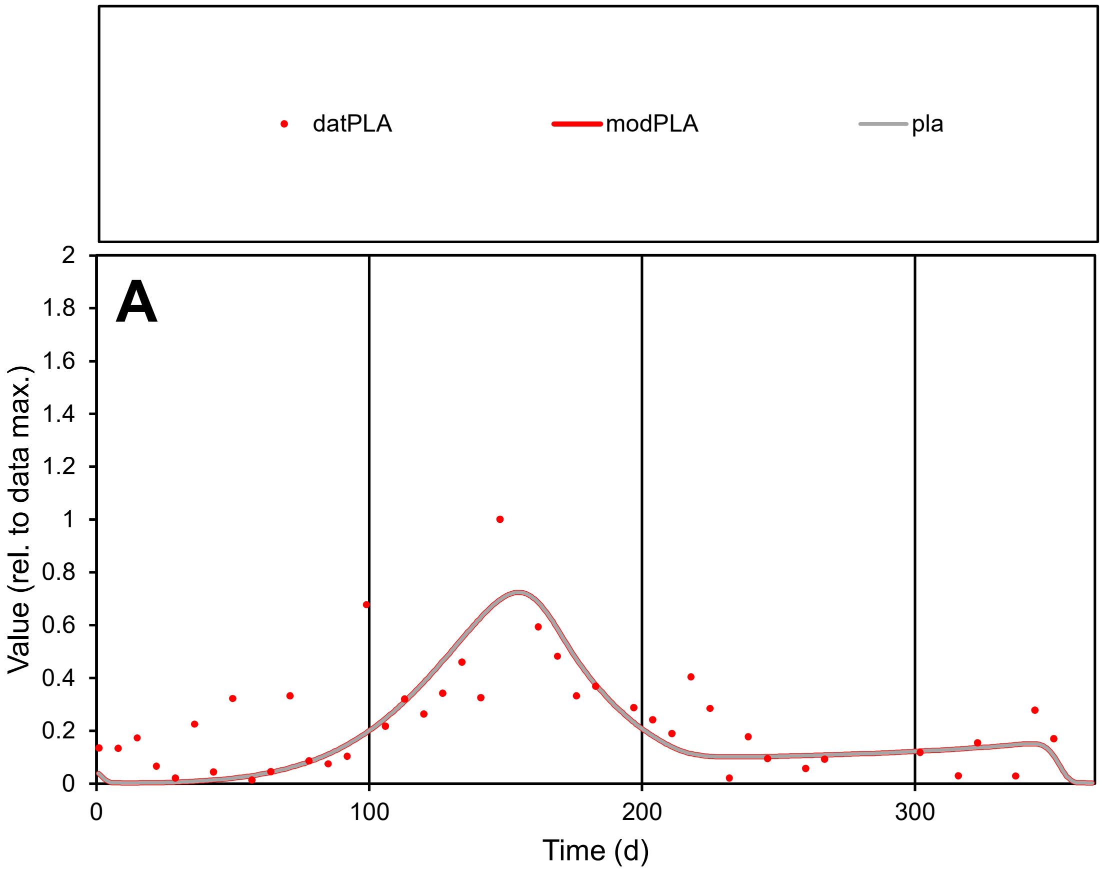


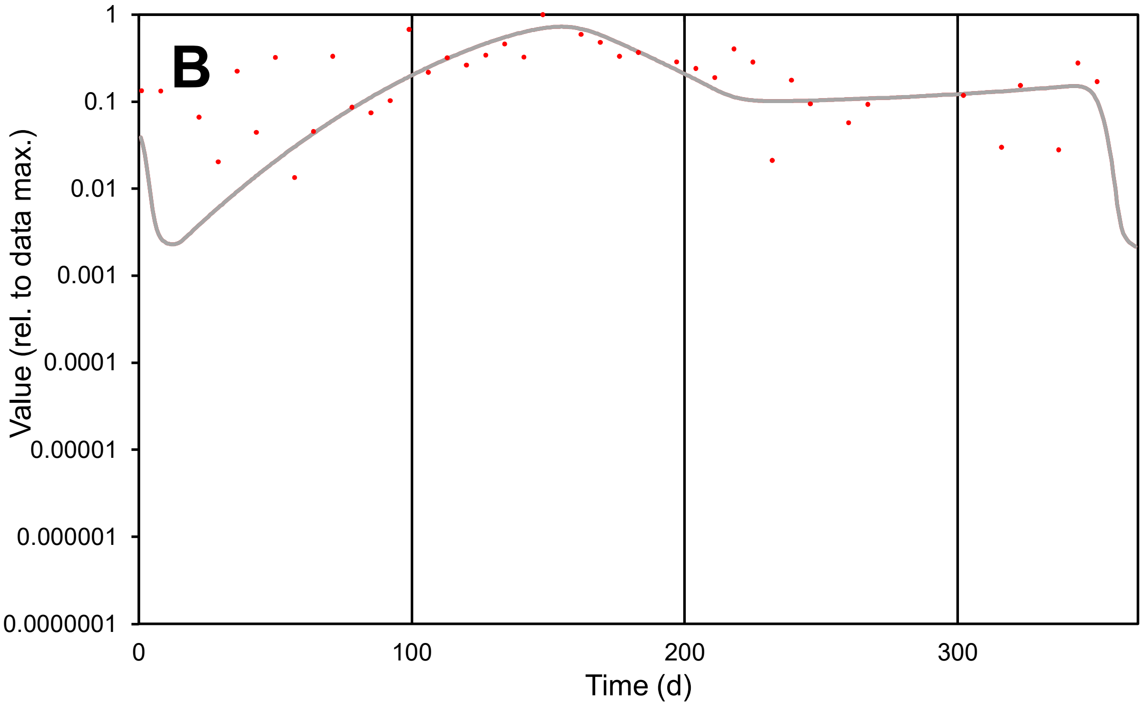


**C1**

Variables

| | |
|---|---|
| PLA | Provide jtd or Dst |
| CalibOnly? | |
| 2 | 1 to reproduce of |
| oNormalize? | |
| 2 | 1=no, 2=yes |
| ofxn calc* | |
| 0.64499 | |
| ofxn mod | |
| 0.66067 | |
| sRun | |
| RunA1 | |
| zout | |
| 1 | |
| Ave | |
| Data | |
| 0.01251 | |
| Model | |
| 0.01056 | |
| data/vMax | |
| 0.05374 | |
| ofxng | |
| 319.391 | |
| Min d/m | |
| 5.48897 | |
| ave/min d | |
| 17.3362 | |
| ave/min m | |
| 80.2677 | |

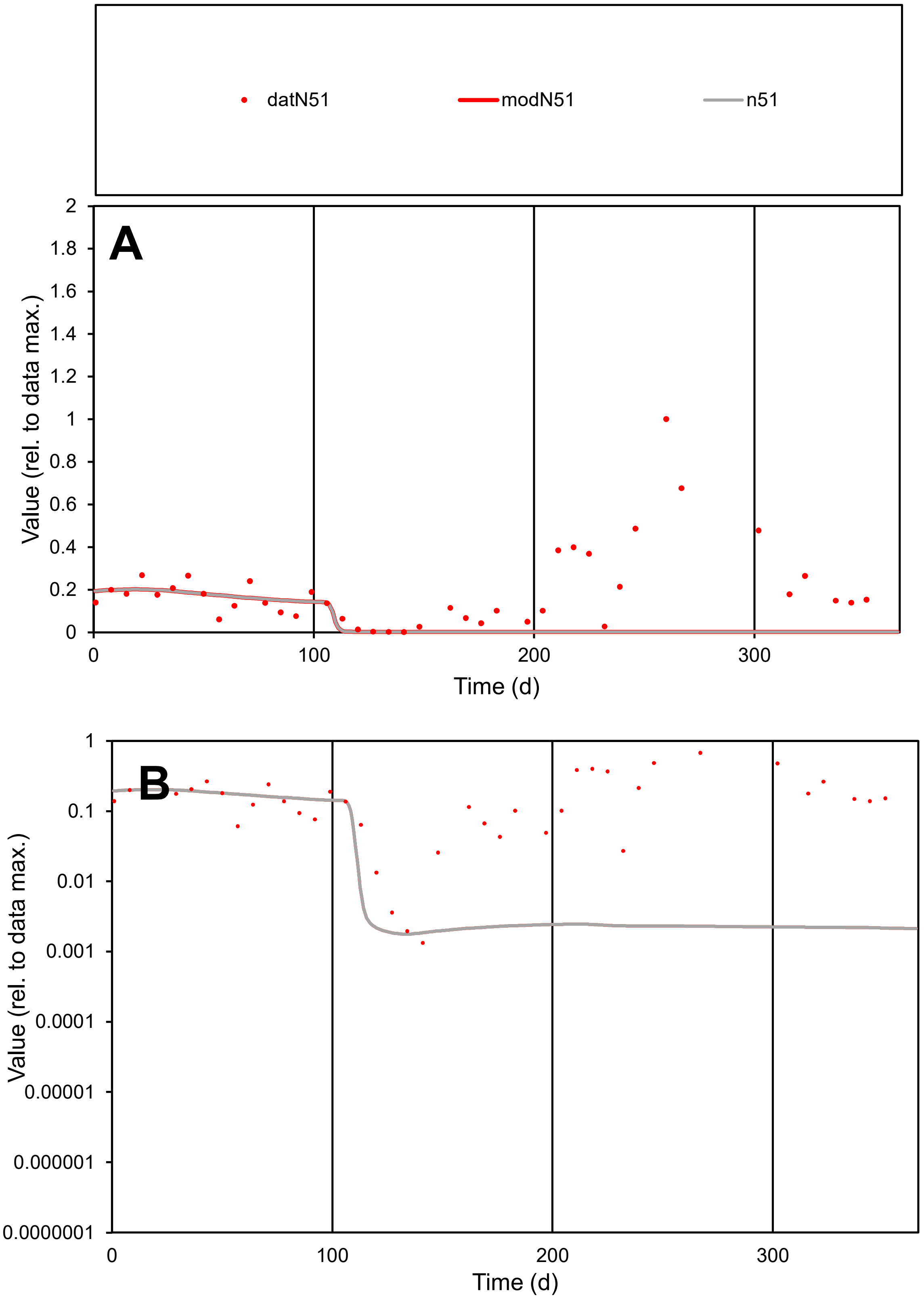


**C1**

Variables
N51 Provide jtd or Dst
CalibOnly?
2 1 to reproduce of
oNormalize?
2 1=no, 2=yes

ofxn calc*
1.28857
ofxn mod
1.2922

sRun
RunA1
zout
1

Ave
Data
0.01328
Model
0.00464

data/vMax
0.06822

ofxng
319.391

Min d/m
0.75319

ave/min d
146.106
ave/min m
38.4474

330_Calib_S_A

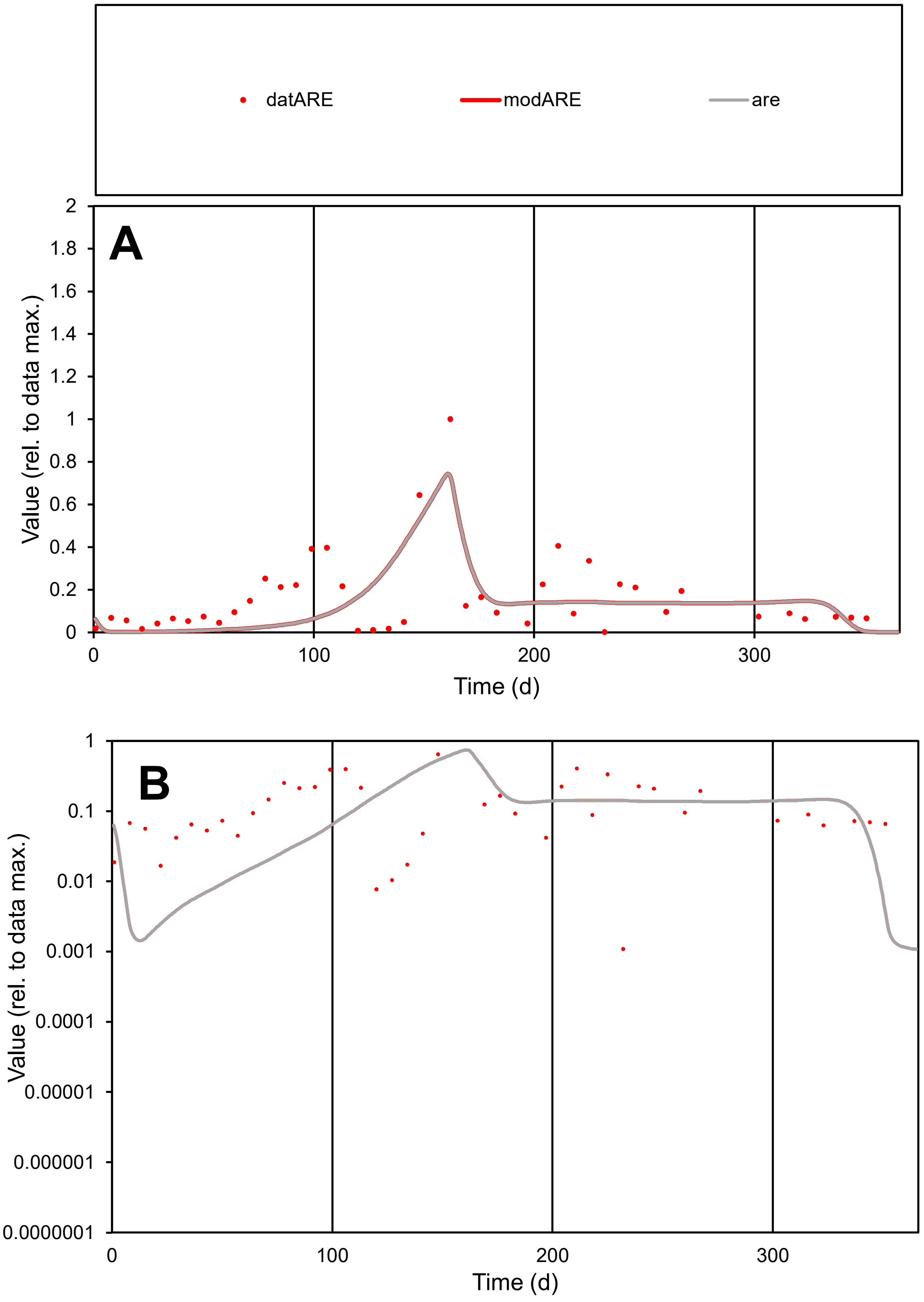


**C1**

Variables

ARE Provide jtd or Dst

CalibOnly?

2 1 to reproduce of

oNormalize?

2 1=no, 2=yes

ofxn calc*

0.93579

ofxn mod

0.97258

sRun

RunA1

zout

1

Ave

Data

0.00853

Model

0.00691

data/vMax

0.05316

ofxng

319.391

Min d/m

0.63068

ave/min d

147.333

ave/min m

75.2974

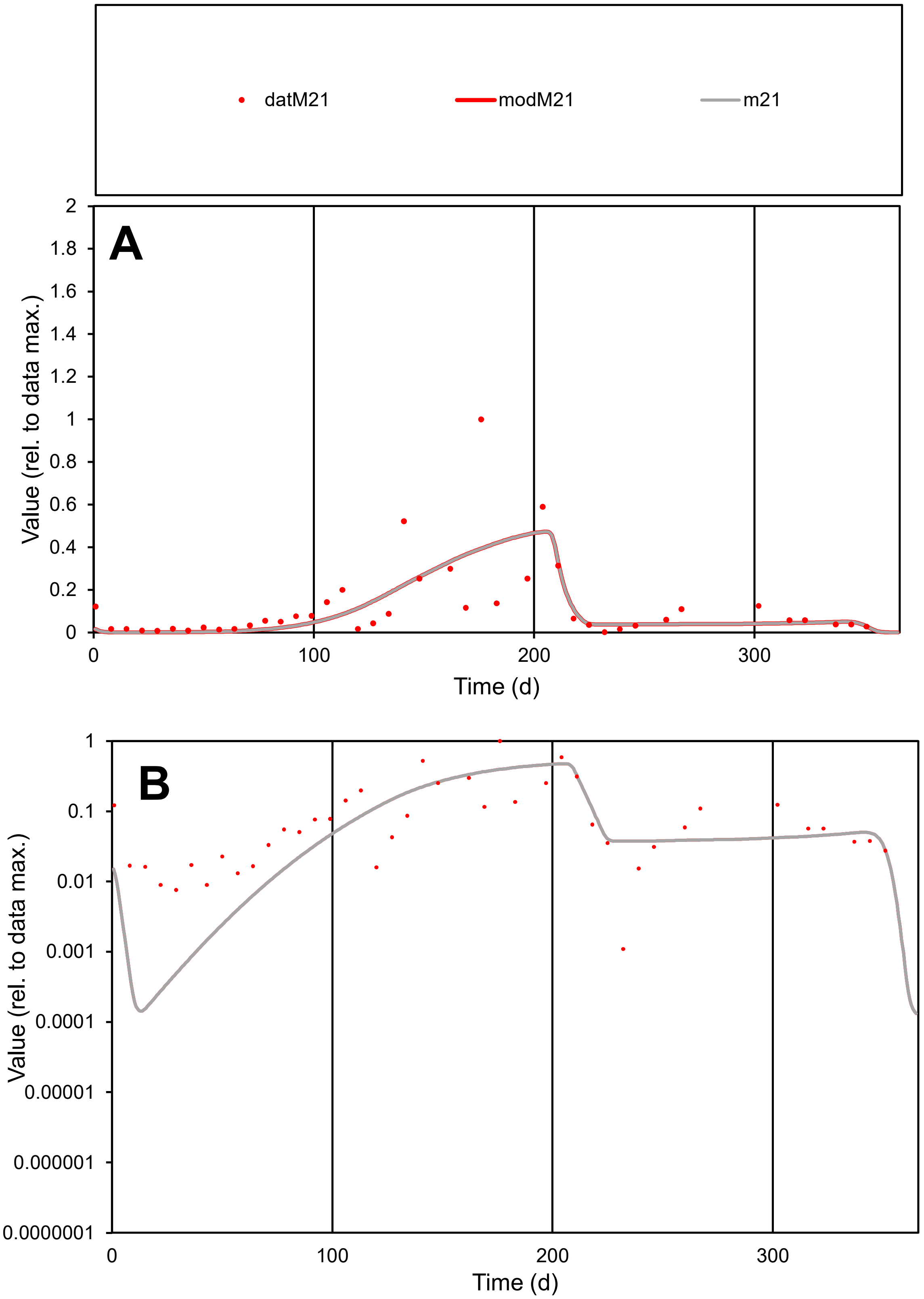


**C1**

Variables
M21 Provide jtd or Dst
CalibOnly?
2 1 to reproduce of
oNormalize?
2 1=no, 2=yes

ofxn calc*
1.07393
ofxn mod
1.08818

sRun
RunA1
zout
1

Ave
Data
0.01593
Model
0.01419

data/vMax
0.1295

ofxng
319.391

Min d/m
6.21007

ave/min d
112.259
ave/min m
621.221

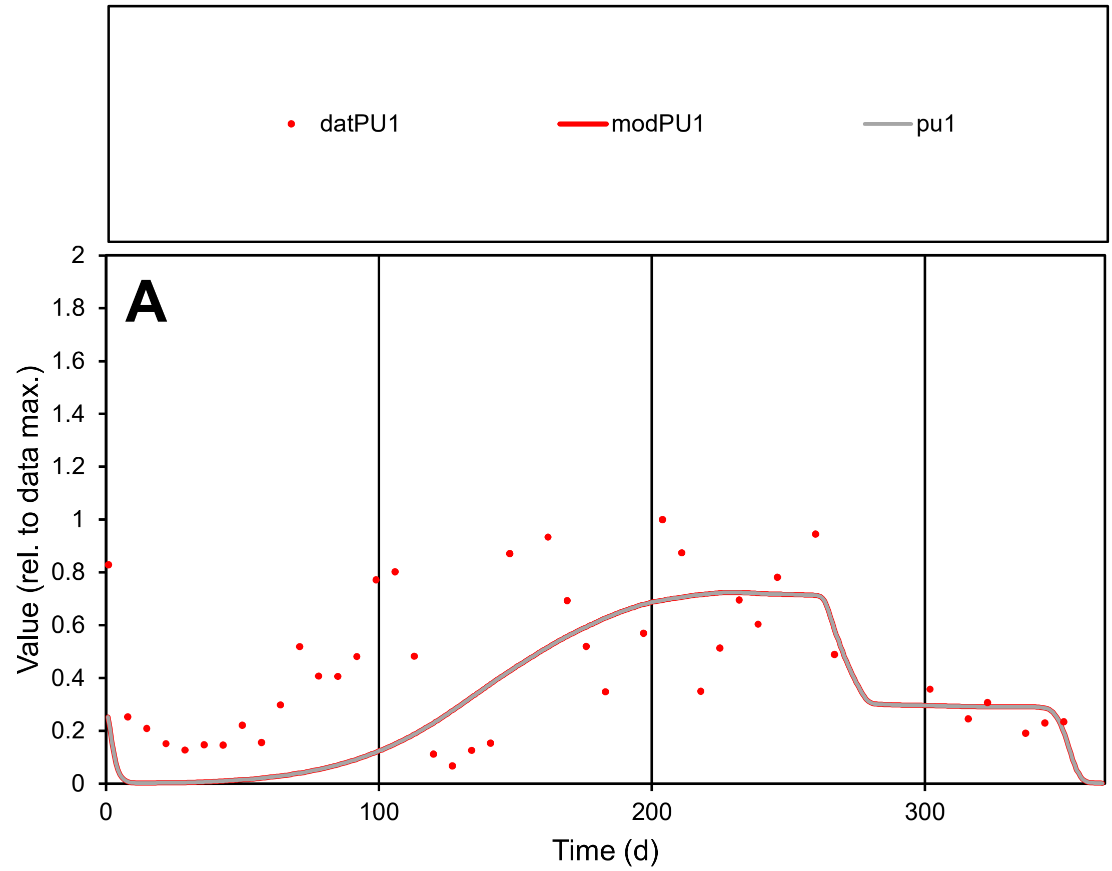


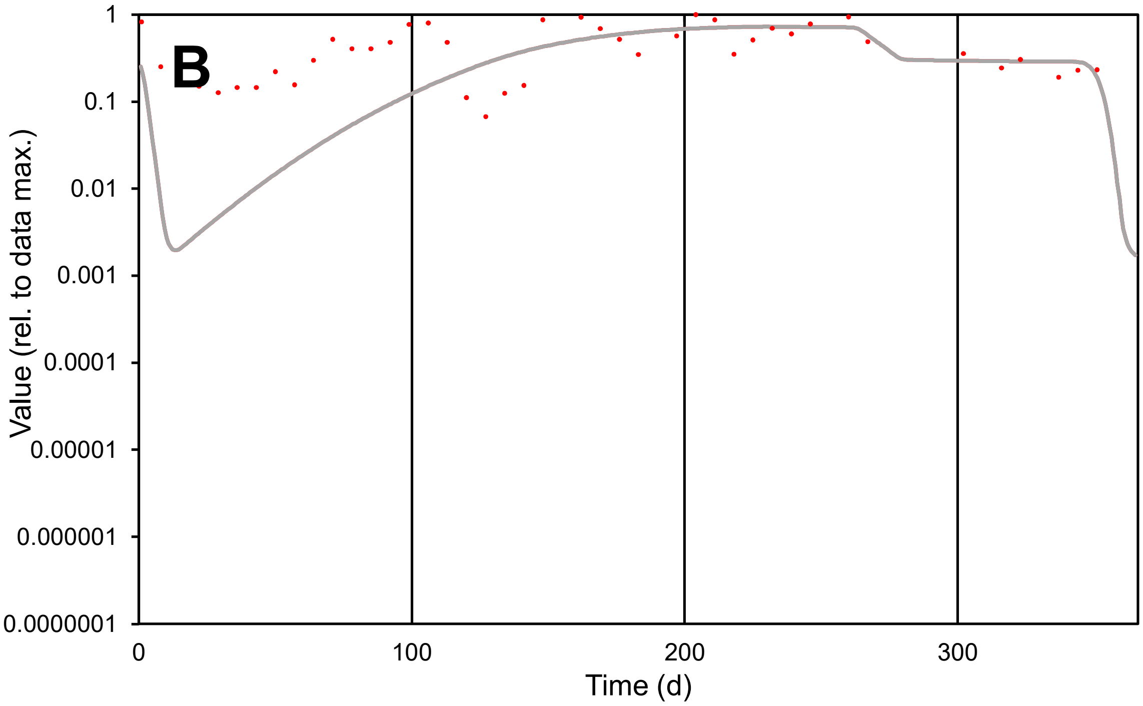


**C1**

Variables
PU1 Provide jtd or Dst
CalibOnly?
2 1 to reproduce of
oNormalize?
2 1=no, 2=yes

ofxn calc*
0.62593
ofxn mod
0.63743

sRun
RunA1
zout
1

Ave
Data
0.01056
Model
0.00773

data/vMax
0.02386

ofxng
319.391

Min d/m
29.9683

ave/min d
6.57794
ave/min m
144.329

330_Calib_S_A

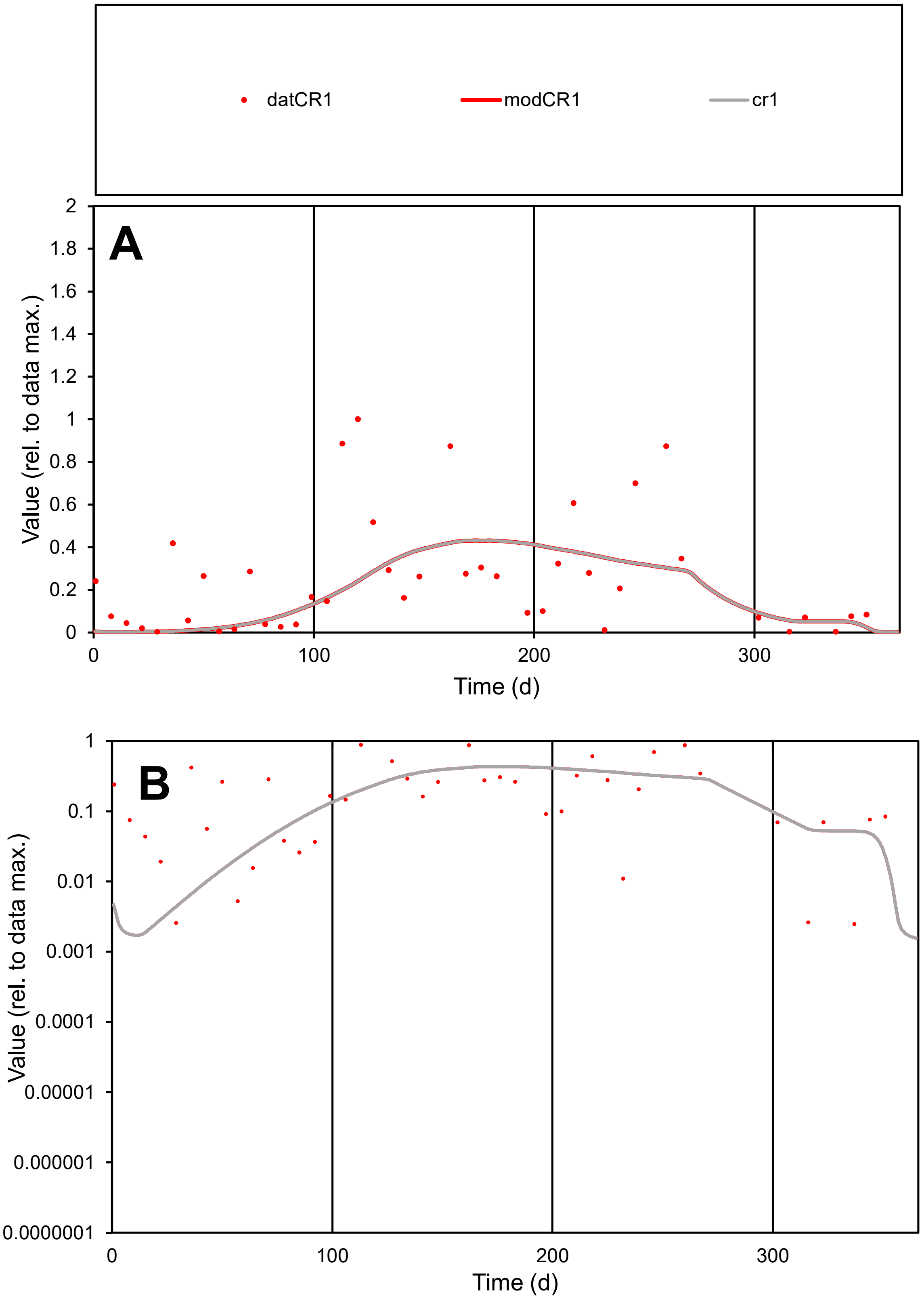


**C1**

Variables
CR1 Provide jtd or Dst
CalibOnly?
2 1 to reproduce of
oNormalize?
2 1=no, 2=yes

ofxn calc*
1.00102
ofxn mod
1.00883

sRun
RunA1
zout
1

Ave
Data
0.00713
Model
0.0055

data/vMax
0.02847

ofxng
319.391

Min d/m
1.43805

ave/min d
100.541
ave/min m
111.571

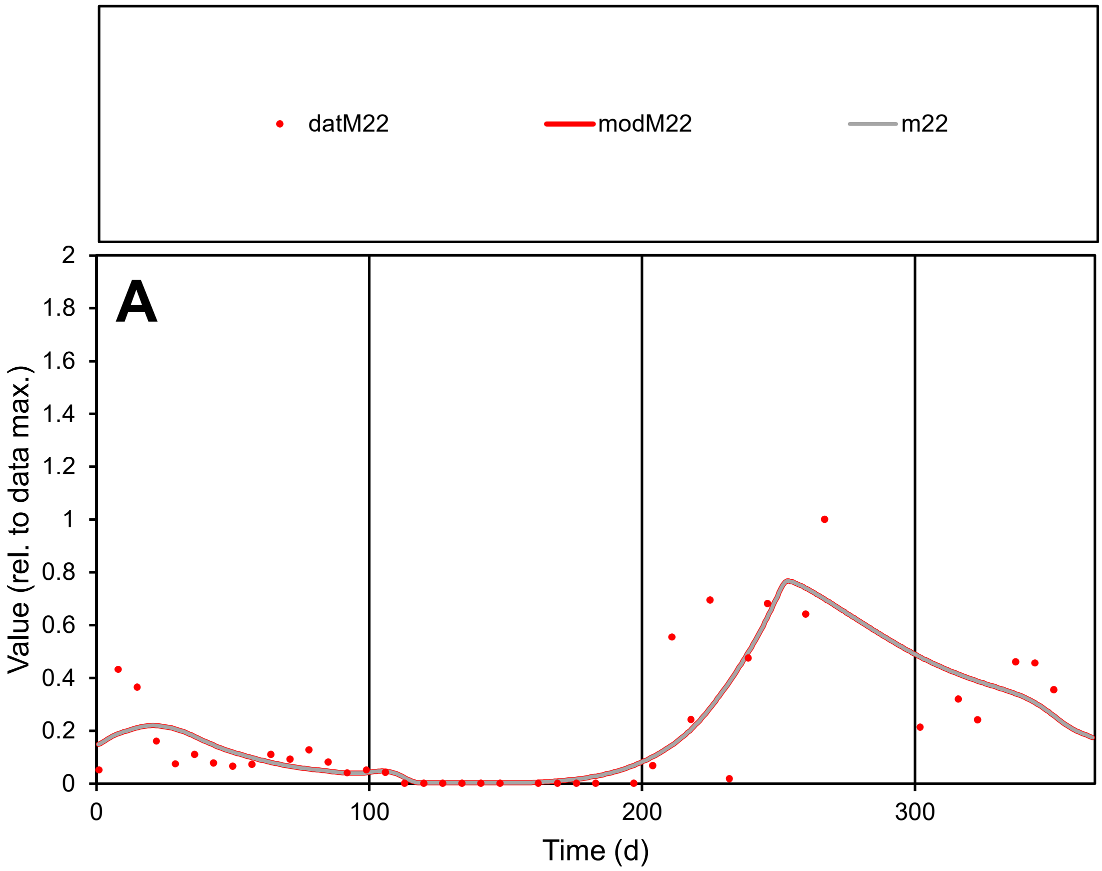


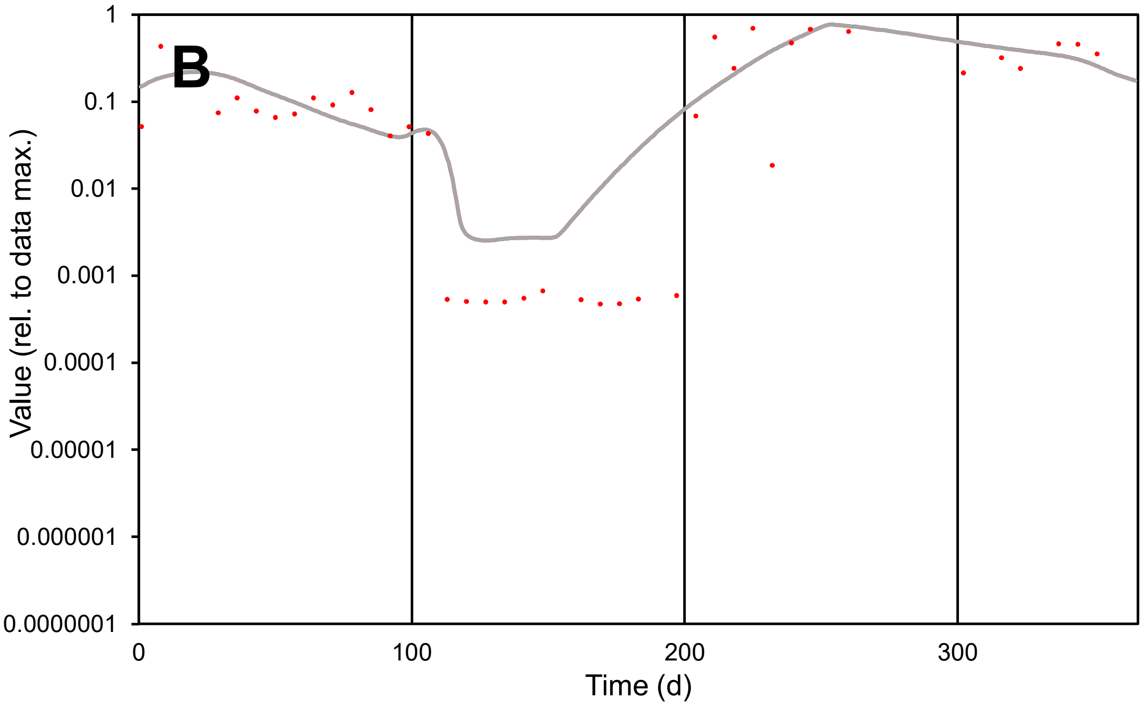


**C1**

Variables
M22 Provide jtd or Dst
CalibOnly?
2 1 to reproduce of
oNormalize?
2 1=no, 2=yes

ofxn calc*
0.70355
ofxn mod
0.72542

sRun
RunA1
zout
1

Ave
Data
0.00914
Model
0.00865

data/vMax
0.04578

ofxng
319.391

Min d/m
0.18554

ave/min d
421.49
ave/min m
74.0086

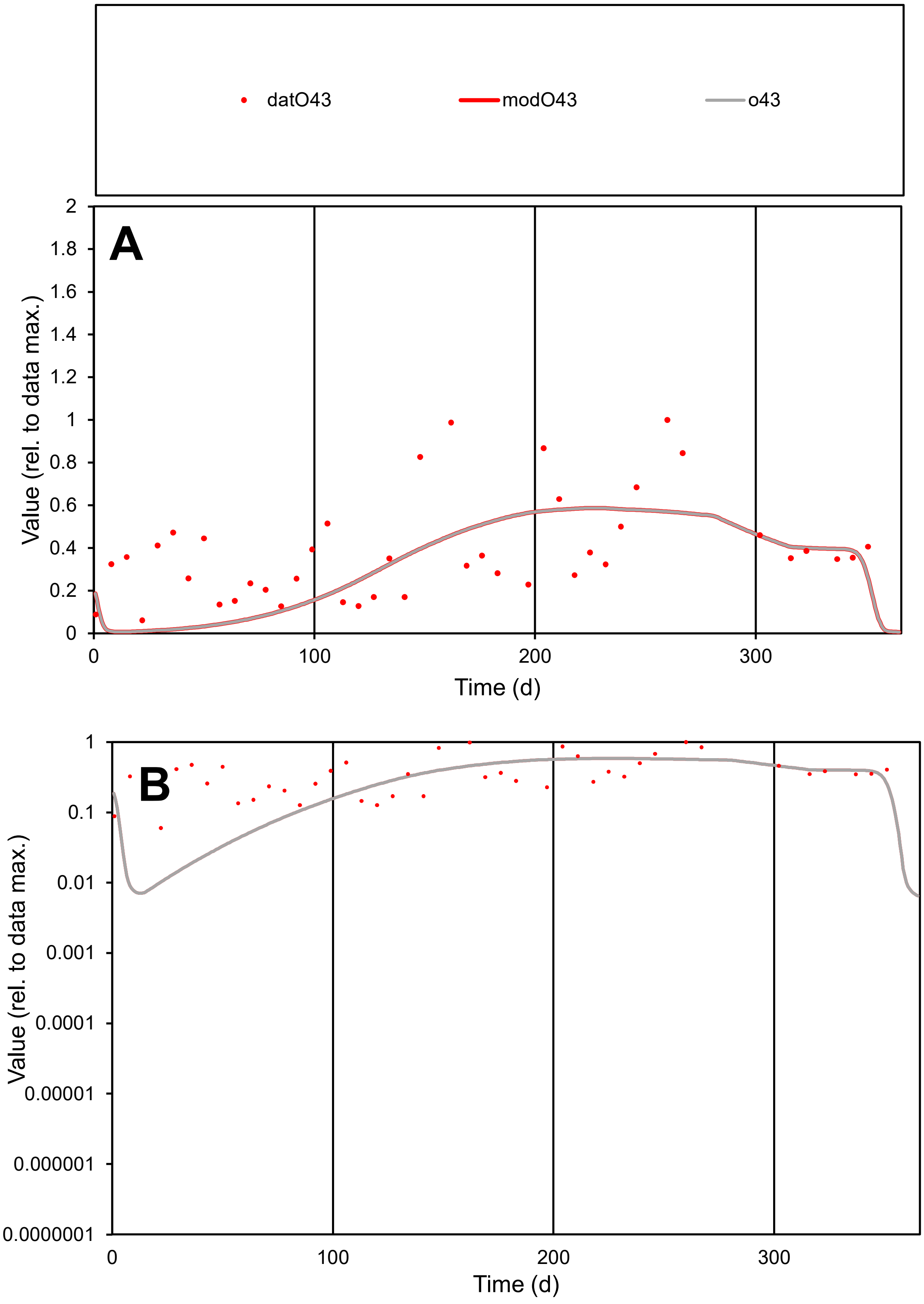


**C1**

Variables

O43 Provide jtd or Dst

CalibOnly?

2 1 to reproduce of

oNormalize?

2 1=no, 2=yes

ofxn calc*

0.62126

ofxn mod

0.63272

sRun

RunA1

zout

1

Ave

Data

0.01006

Model

0.00808

data/vMax

0.0261

ofxng

319.391

Min d/m

7.65251

ave/min d

6.41738

ave/min m

39.4242

330_Calib_S_A

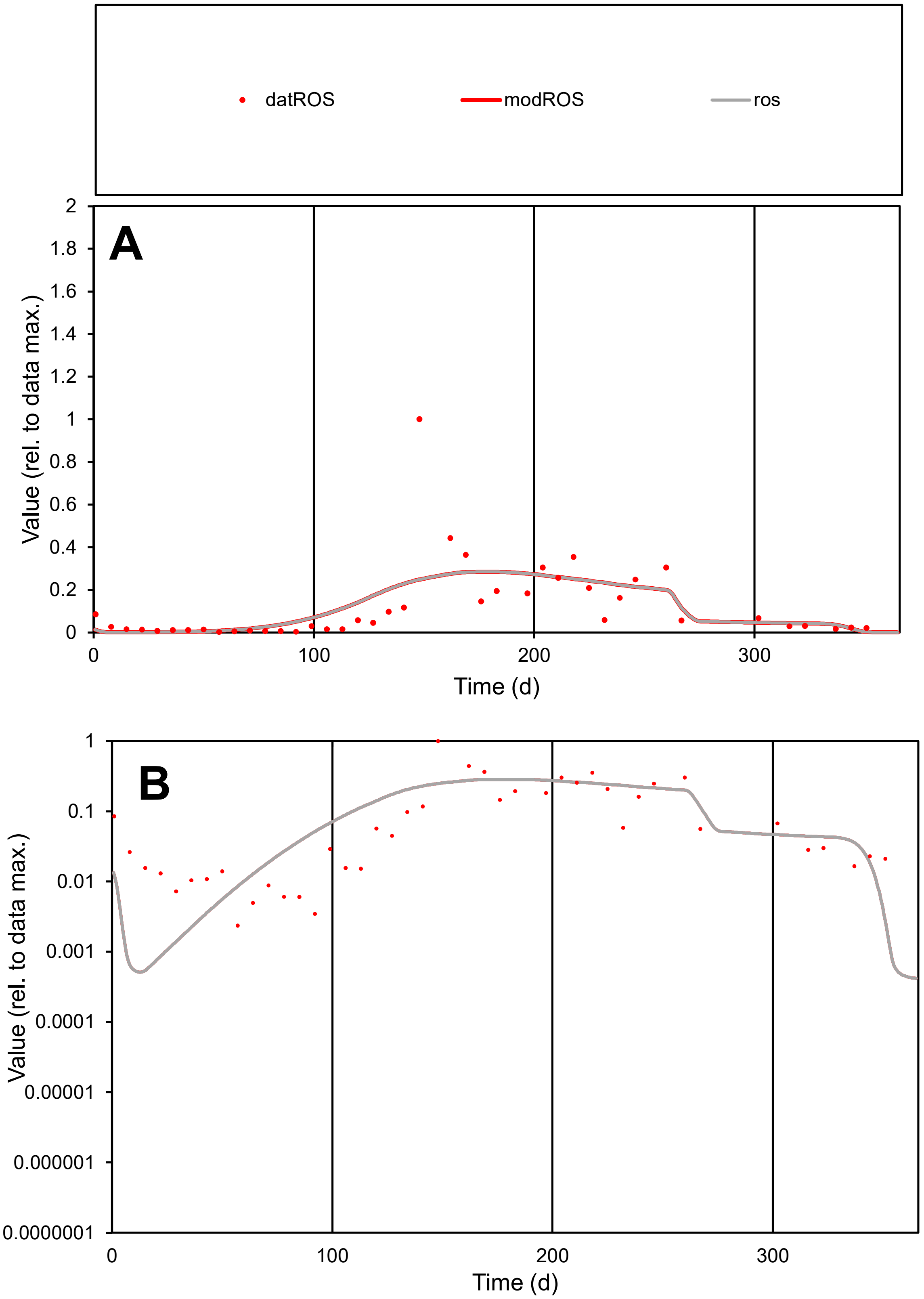


**C1**

Variables
ROS Provide jtd or Dst
CalibOnly?
2 1 to reproduce of
oNormalize?
2 1=no, 2=yes

ofxn calc*
1.12594
ofxn mod
1.13847

sRun
RunA1
zout
1

Ave
Data
0.0094
Model
0.00937

data/vMax
0.07815

ofxng
319.391

Min d/m
4.15082

ave/min d
51.2901
ave/min m
212.2

330_Calib_S_A

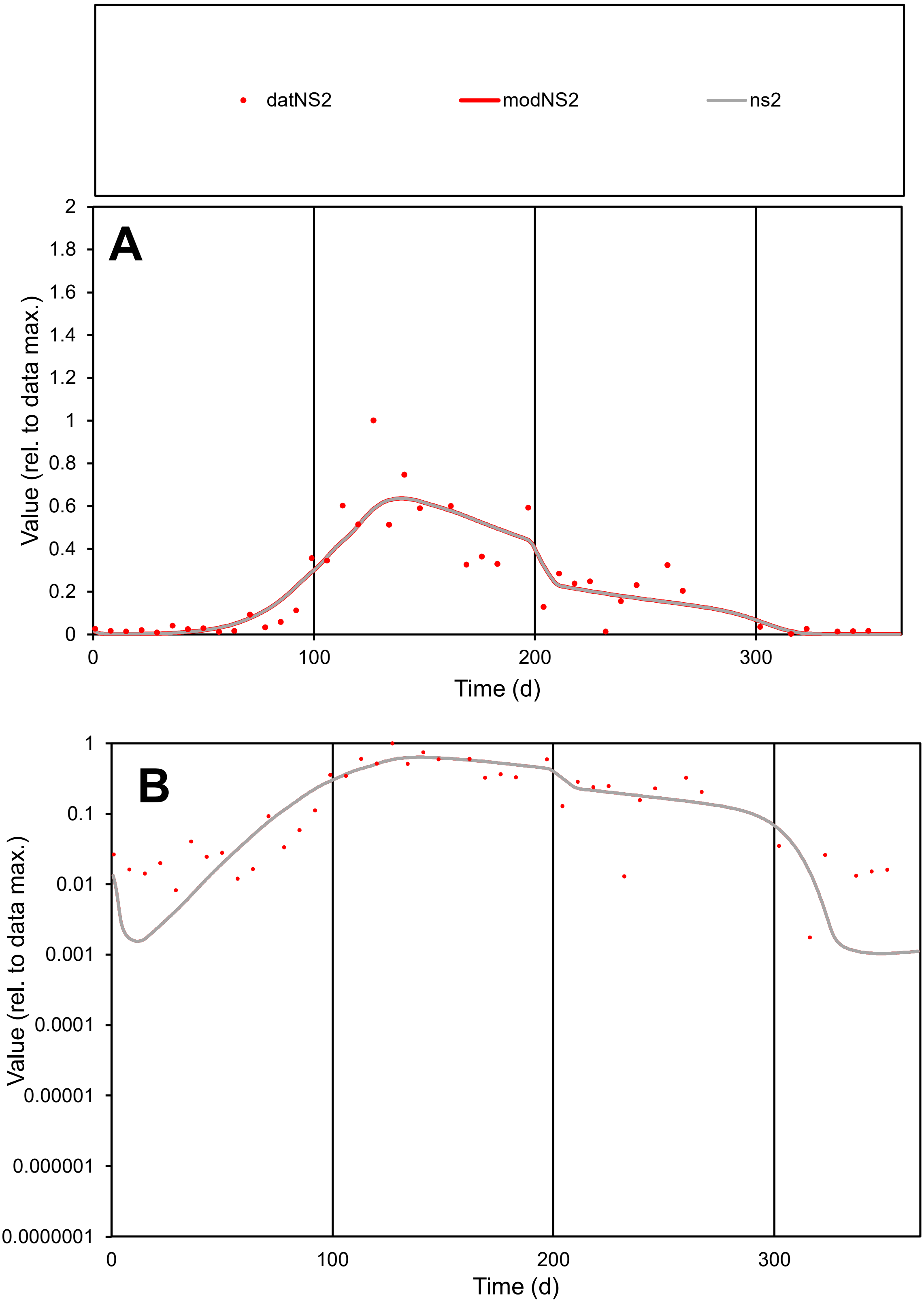


**C1**

Variables
NS2 Provide jtd or Dst
CalibOnly?
2 1 to reproduce of
oNormalize?
2 1=no, 2=yes

ofxn calc*
0.48563
ofxn mod
0.50787

sRun
RunA1
zout
1

Ave
Data
0.00826
Model
0.00822

data/vMax
0.03725

ofxng
319.391

Min d/m
1.68205

ave/min d
126.635
ave/min m
211.916

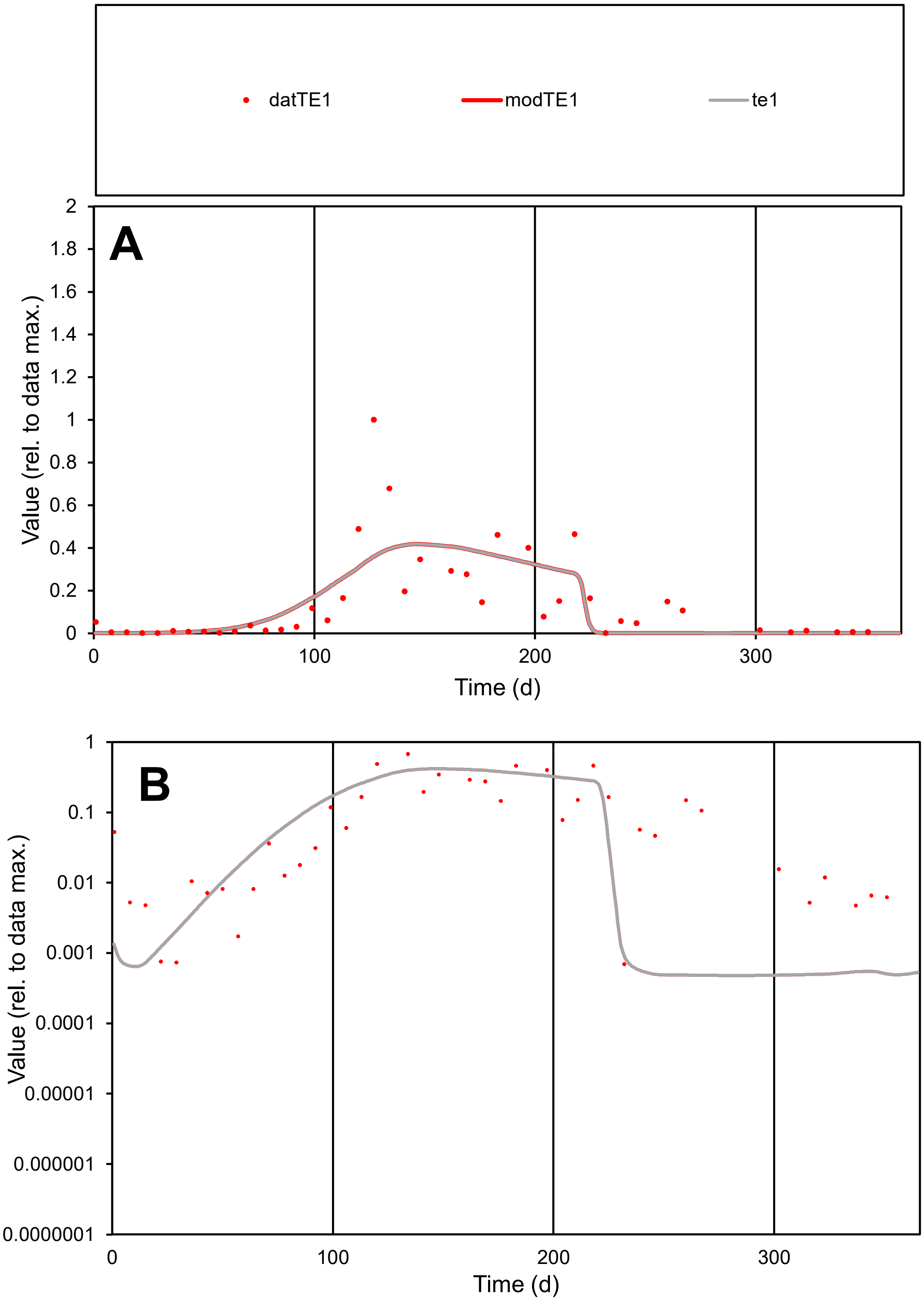


**C1**

Variables
TE1 Provide jtd or Dst
CalibOnly?
2 1 to reproduce of
oNormalize?
2 1=no, 2=yes

ofxn calc*
0.98824
ofxn mod
1.0066

sRun
RunA1
zout
1

Ave
Data
0.01157
Model
0.01103

data/vMax
0.07975

ofxng
319.391

Min d/m
1.44725

ave/min d
207.719
ave/min m
286.693

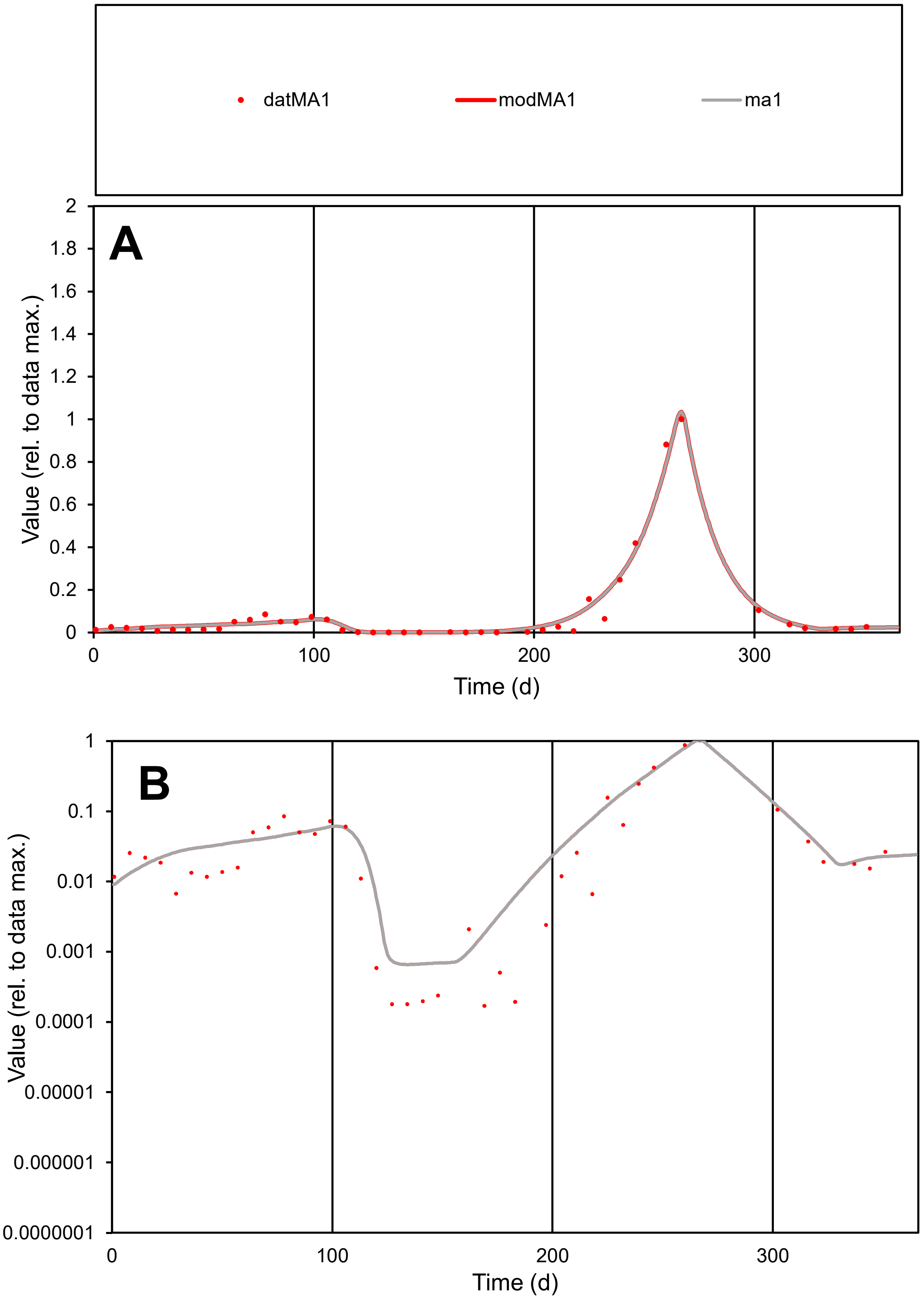


**C1**

Variables
MA1 Provide jtd or Dst
CalibOnly?
2 1 to reproduce of
oNormalize?
2 1=no, 2=yes

ofxn calc*
0.31028
ofxn mod
0.42731

sRun
RunA1
zout
1

Ave
Data
0.00737
Model
0.00788

data/vMax
0.08566

ofxng
319.391

Min d/m
0.25699

ave/min d
507.315
ave/min m
139.429

330_Calib_S_A

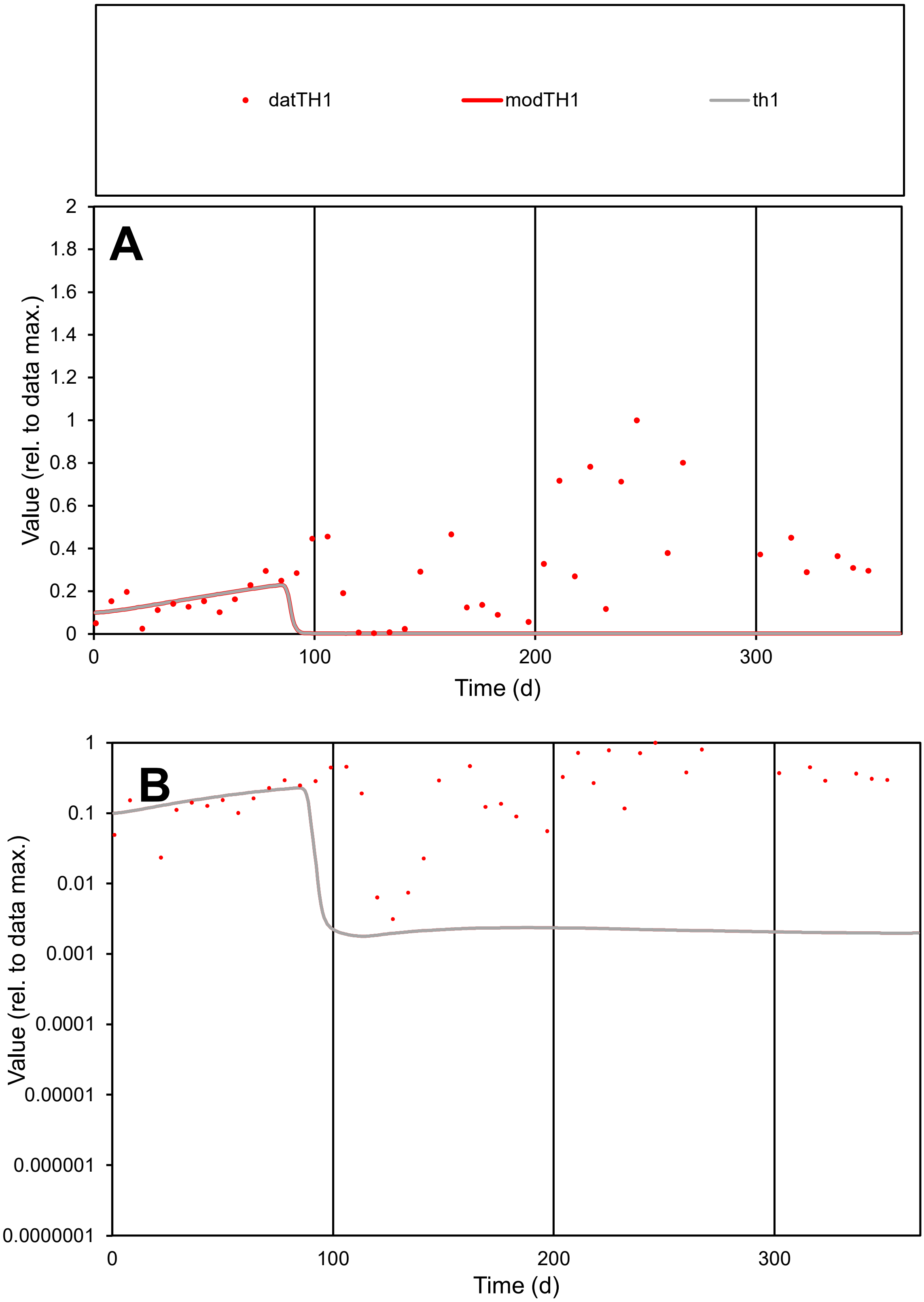


**C1**

Variables
TH1 Provide jtd or Dst
CalibOnly?
2 1 to reproduce of
oNormalize?
2 1=no, 2=yes

ofxn calc*
1.25196
ofxn mod
1.2604

sRun
RunA1
zout
1

Ave
Data
0.00881
Model
0.00179

data/vMax
0.03151

ofxng
319.391

Min d/m
1.76769

ave/min d
89.3031
ave/min m
32.0442

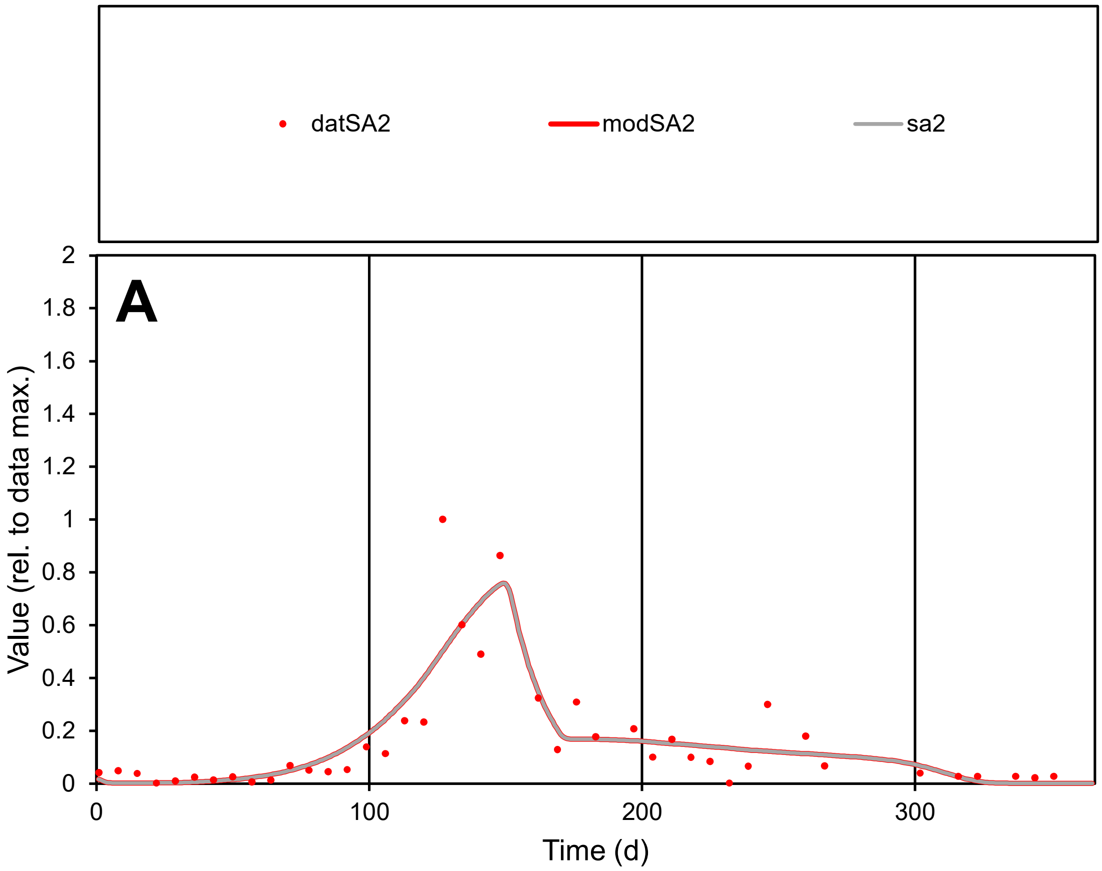


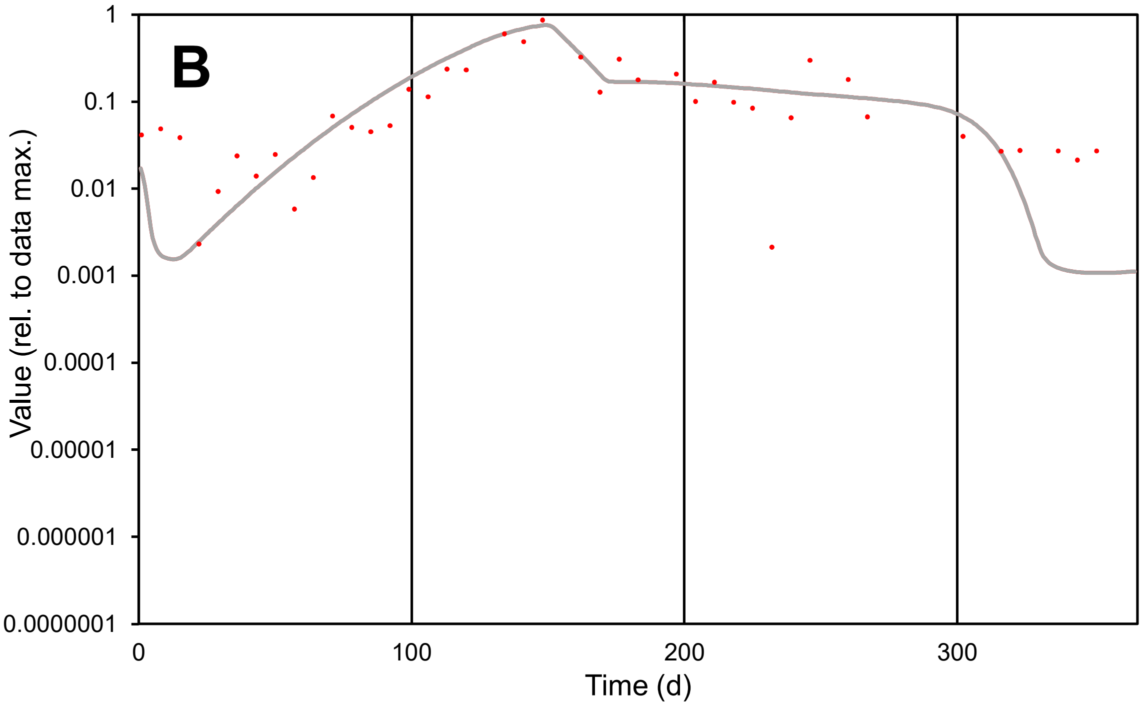


**C1**

Variables
SA2 Provide jtd or Dst
CalibOnly?
2 1 to reproduce of
oNormalize?
2 1=no, 2=yes

ofxn calc*
0.67623
ofxn mod
0.69752

sRun
RunA1
zout
1

Ave
Data
0.0079
Model
0.00796

data/vMax
0.05107

ofxng
319.391

Min d/m
1.97311

ave/min d
72.7358
ave/min m
144.585

330_Calib_S_A

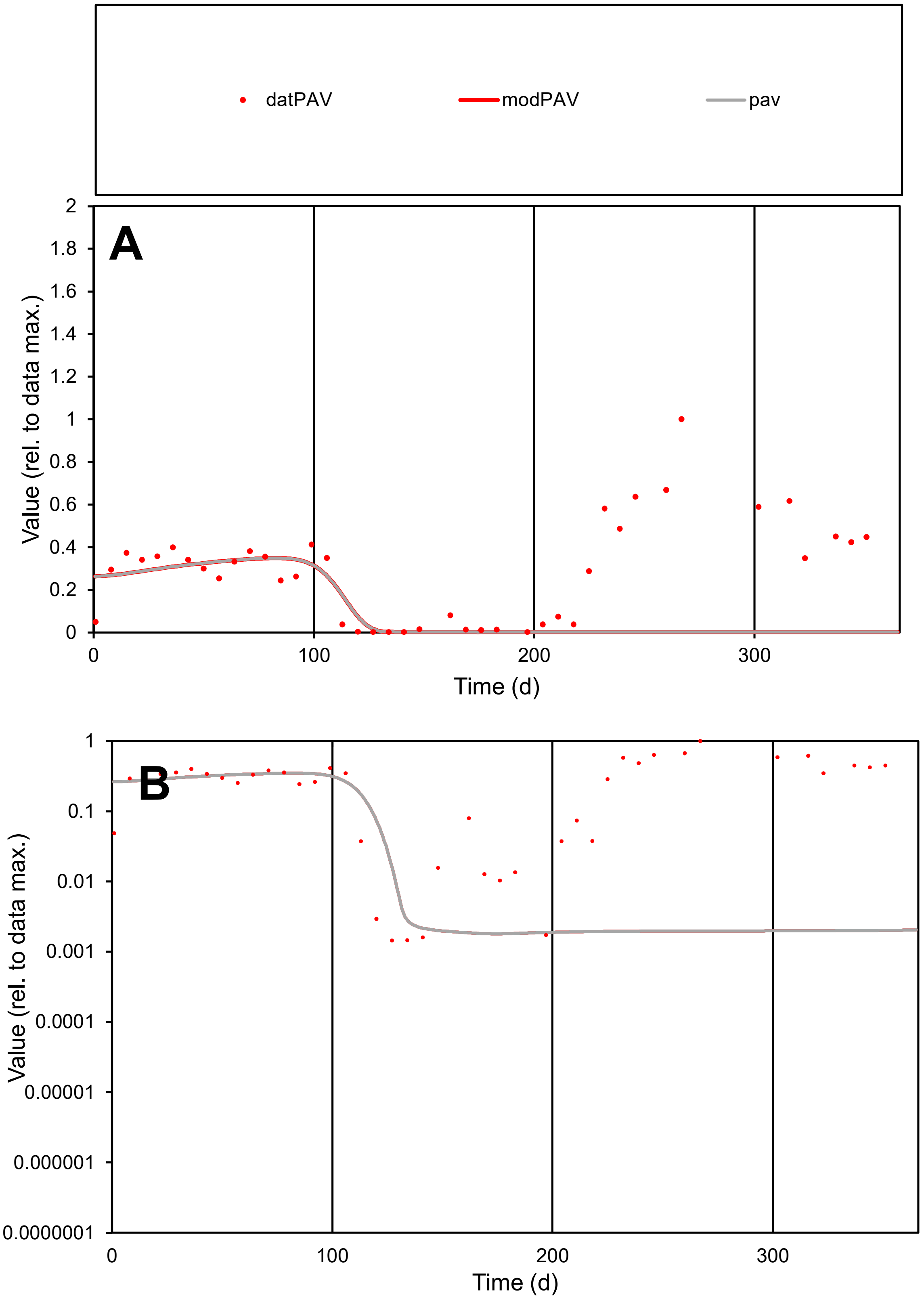


**C1**

Variables
PAV Provide jtd or Dst
CalibOnly?
2 1 to reproduce of
oNormalize?
2 1=no, 2=yes

ofxn calc*
1.096
ofxn mod
1.10081

sRun
RunA1
zout
1

Ave
Data
0.00772
Model
0.00338

data/vMax
0.02724

ofxng
319.391

Min d/m
0.80506

ave/min d
195.586
ave/min m
68.8347

330_Calib_S_A

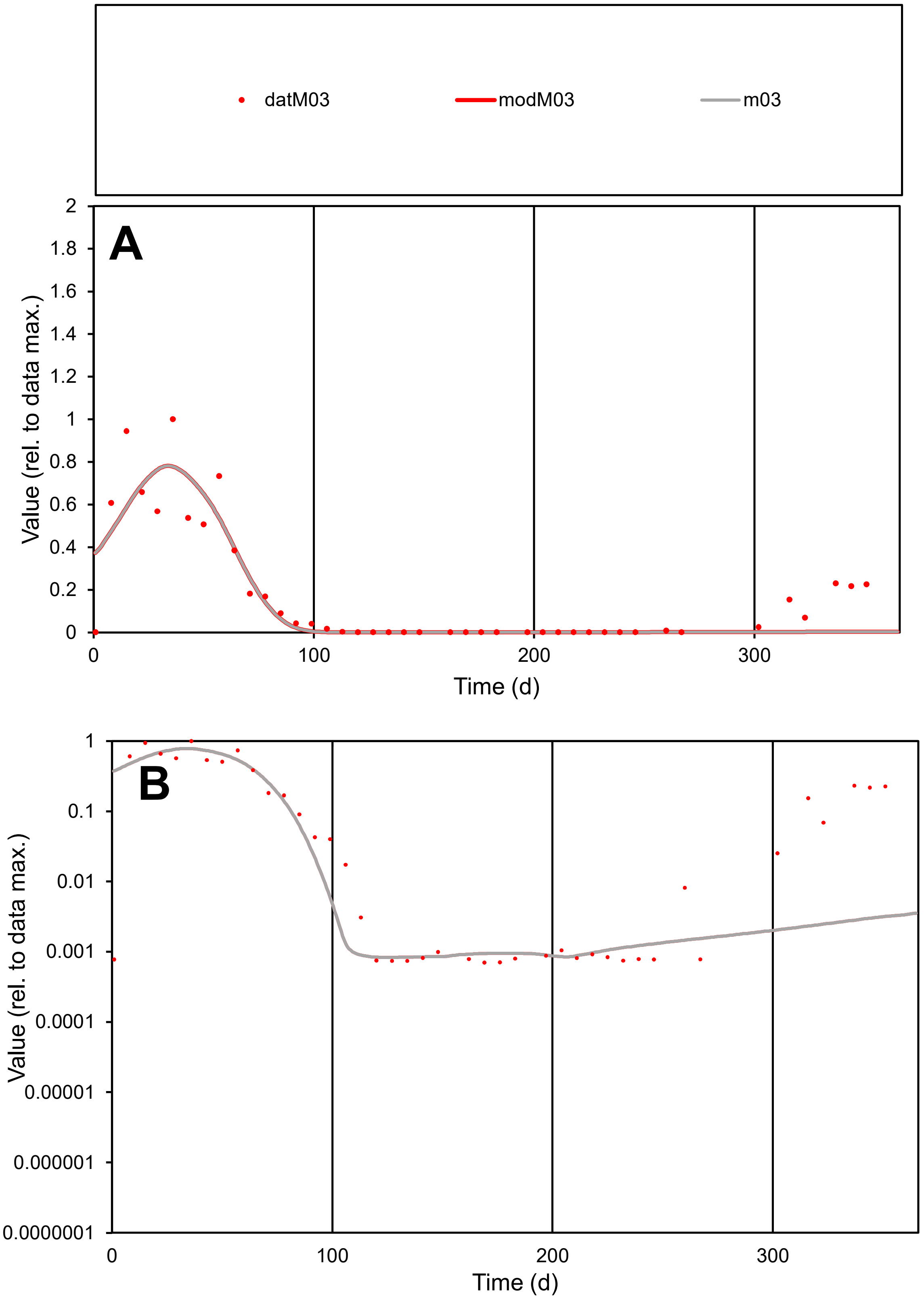


**C1**

Variables
M03 Provide jtd or Dst
CalibOnly?
2 1 to reproduce of
oNormalize?
2 1=no, 2=yes

ofxn calc*
0.69233
ofxn mod
0.72512

sRun
RunA1
zout
1

Ave
Data
0.00446
Model
0.0039

data/vMax
0.02519

ofxng
319.391

Min d/m
0.84715

ave/min d
252.091
ave/min m
186.839

330_Calib_S_A

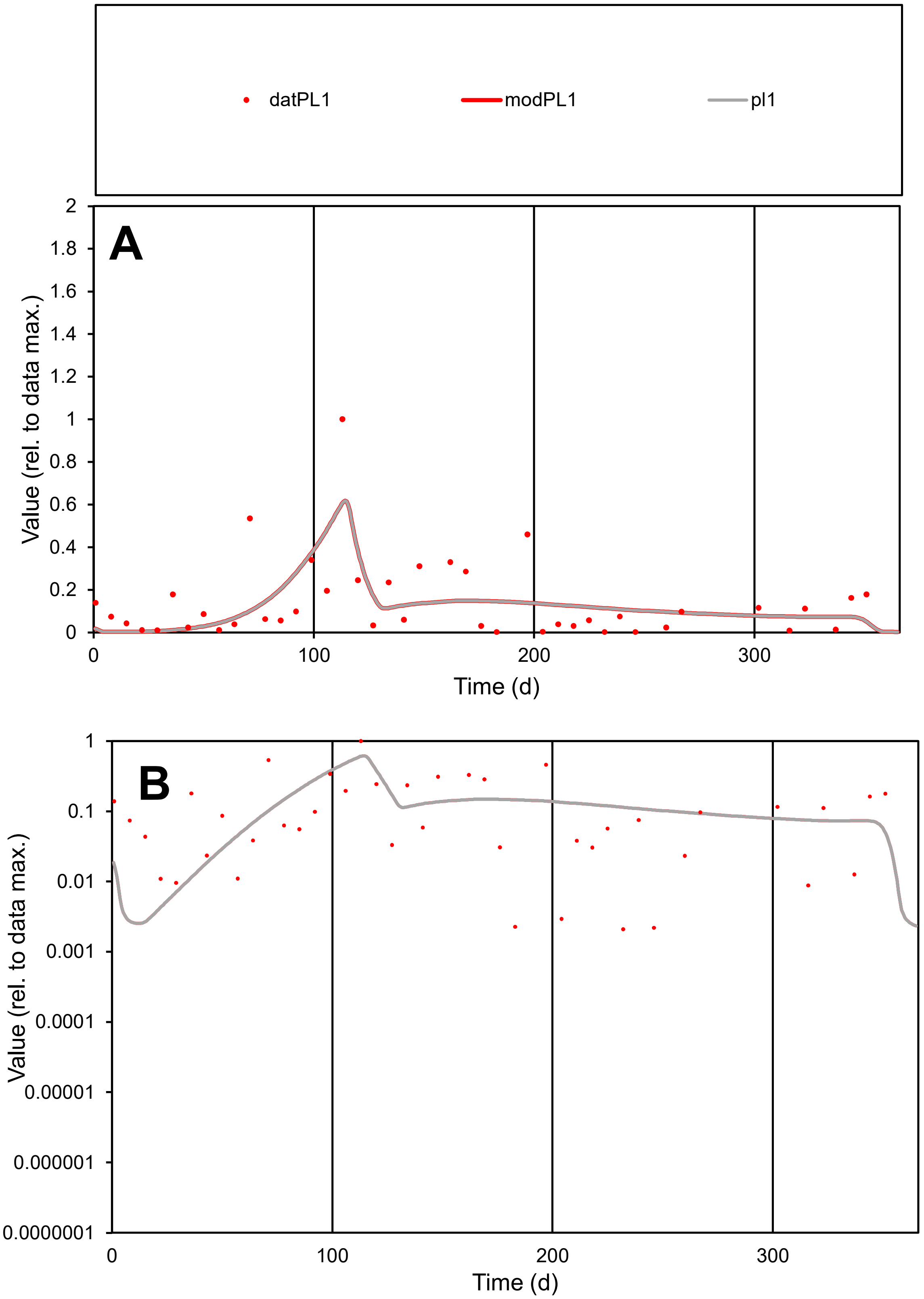


**C1**

Variables
PL1 Provide jtd or Dst
CalibOnly?
2 1 to reproduce of
oNormalize?
2 1=no, 2=yes

ofxn calc*
1.0554
ofxn mod
1.0684

sRun
RunA1
zout
1

Ave
Data
0.00244
Model
0.0023

data/vMax
0.01766

ofxng
319.391

Min d/m
0.8077

ave/min d
65.7738
ave/min m
50.0284

330_Calib_S_A

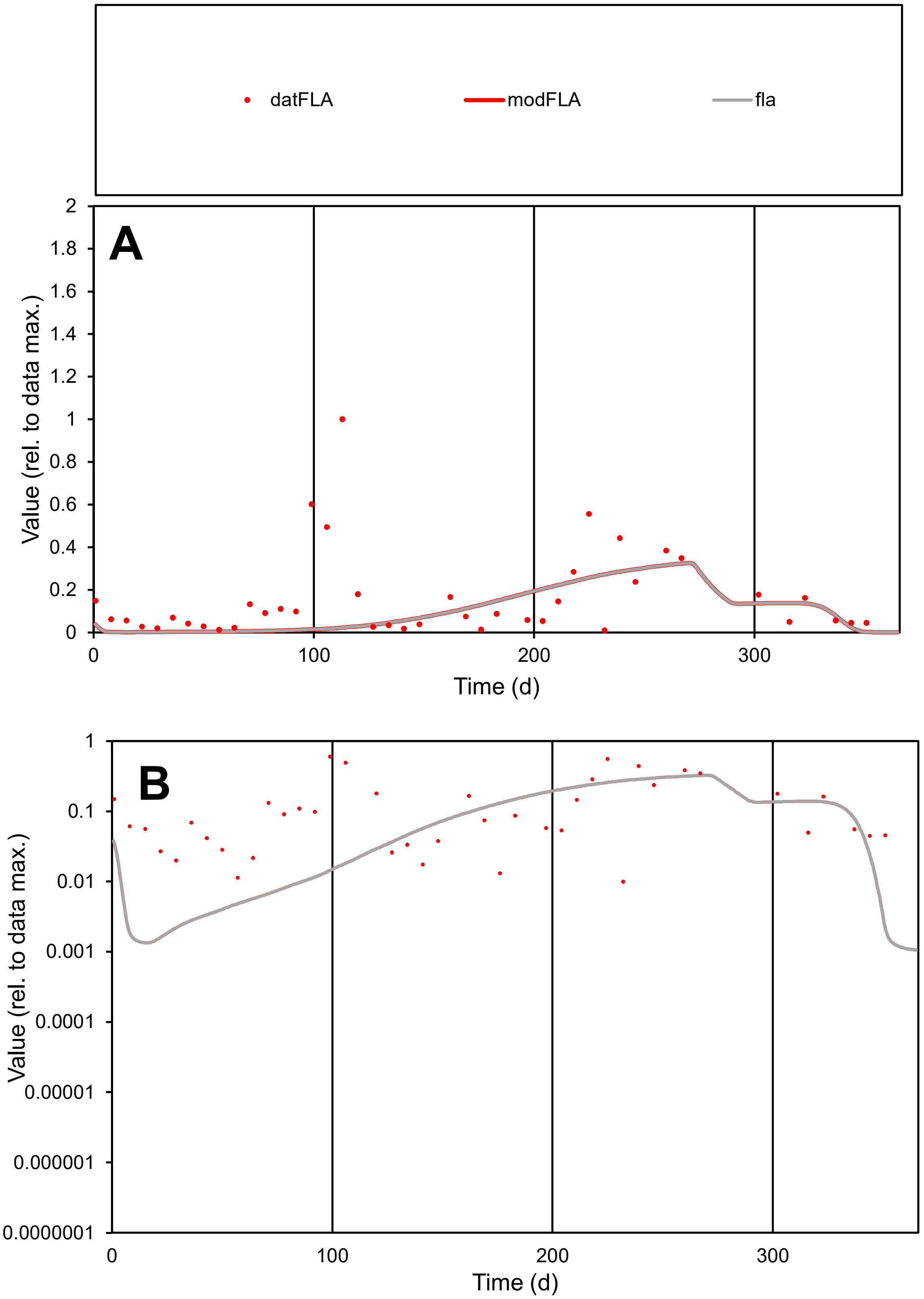


**C1**

Variables
FLA Provide jtd or Dst
CalibOnly?
2 1 to reproduce of
oNormalize?
2 1=no, 2=yes

ofxn calc*
1.32912
ofxn mod
1.33583

sRun
RunA1
zout
1

Ave
Data
0.00499
Model
0.003

data/vMax
0.03126

ofxng
319.391

Min d/m
7.45581

ave/min d
15.9752
ave/min m
71.6033

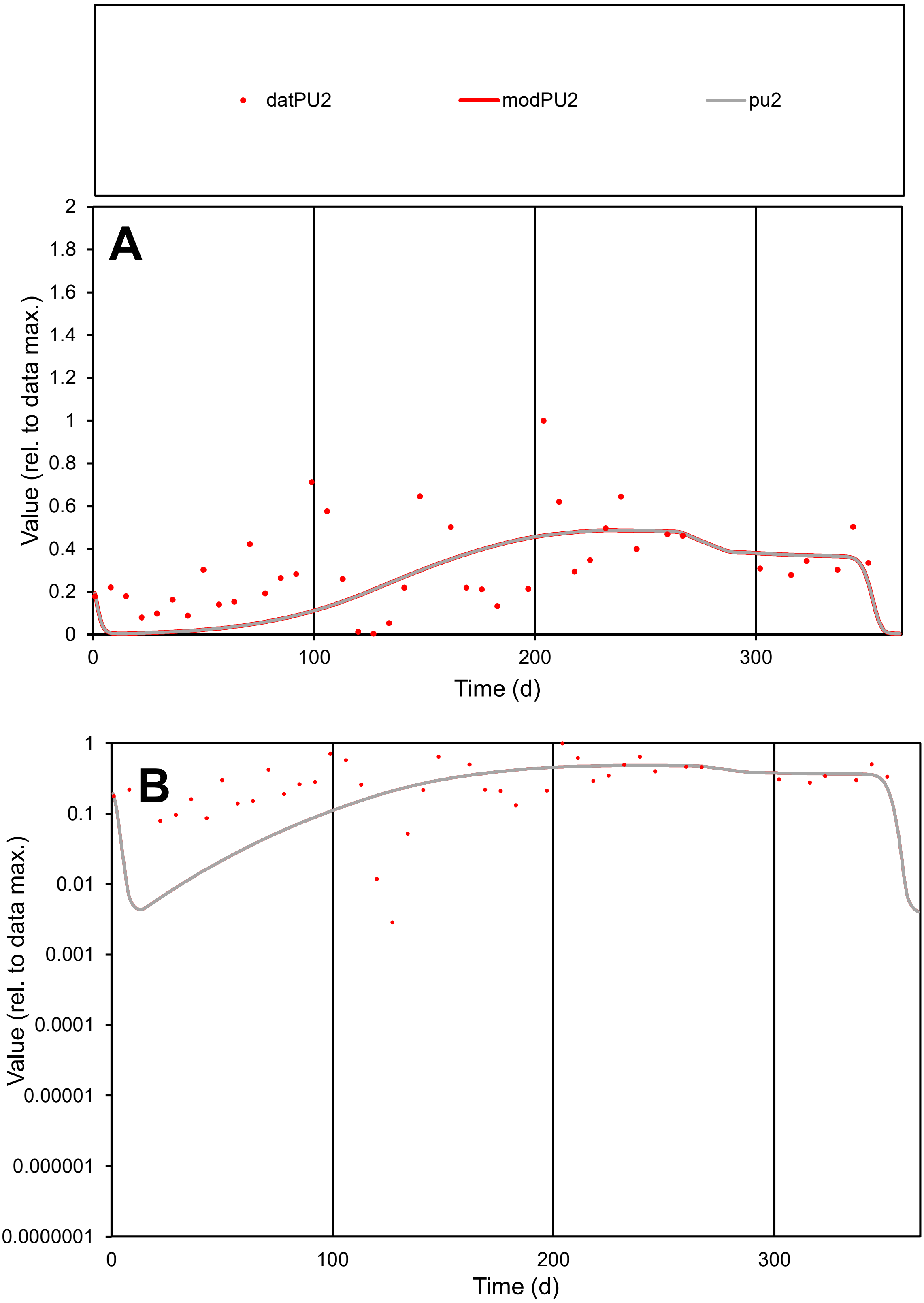


**C1**

Variables
PU2 Provide jtd or Dst
CalibOnly?
2 1 to reproduce of
oNormalize?
2 1=no, 2=yes

ofxn calc*
0.67386
ofxn mod
0.68463

sRun
RunA1
zout
1

Ave
Data
0.0064
Model
0.0051

data/vMax
0.02022

ofxng
319.391

Min d/m
0.58236

ave/min d
109.975
ave/min m
50.9875

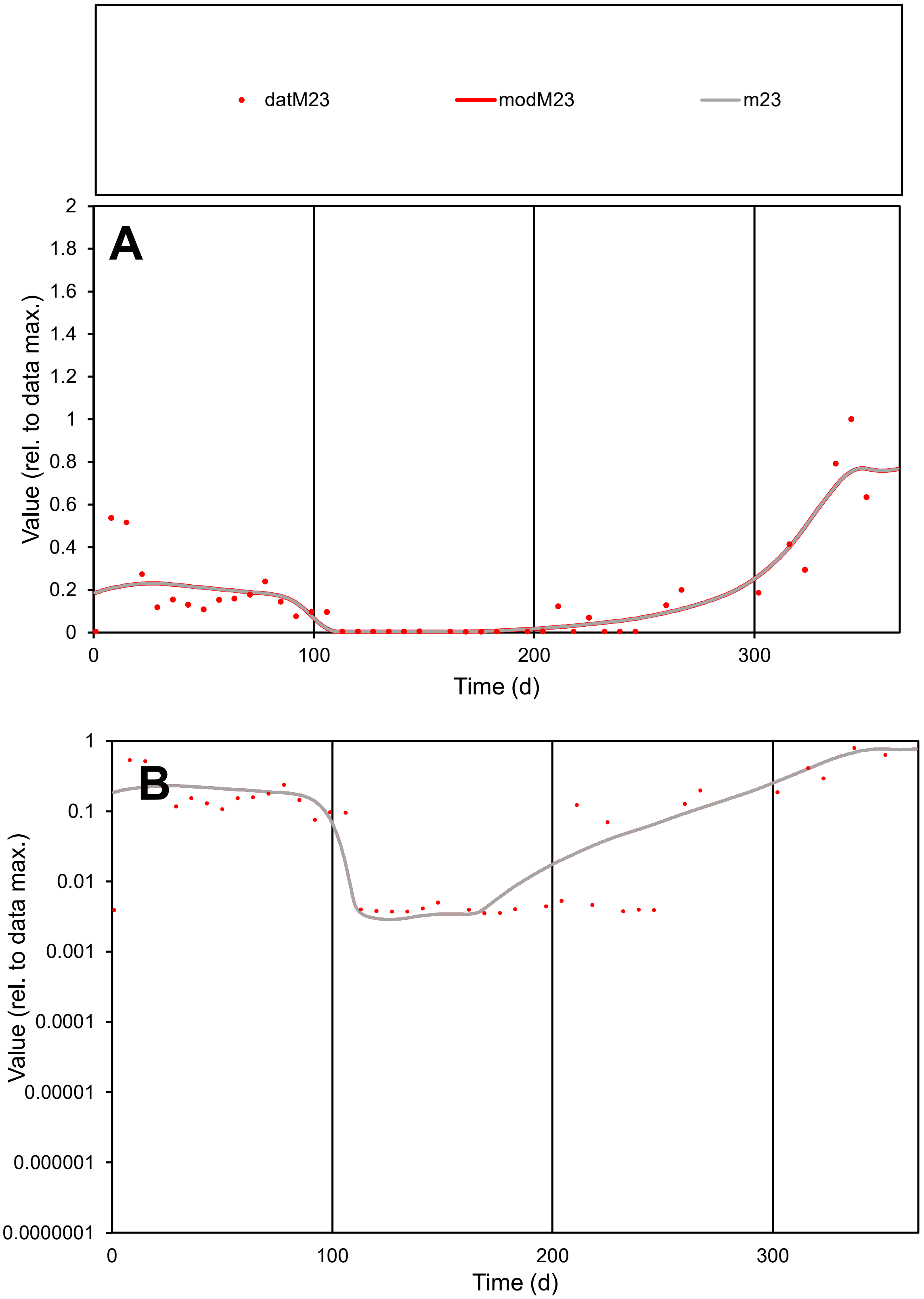


**C1**

| | |
|---|---|
| Variables | |
| M23 | Provide jtd or Dst |
| CalibOnly? | |
| 2 | 1 to reproduce of |
| oNormalize? | |
| 2 | 1=no, 2=yes |
| ofxn calc* | |
| 0.62611 | |
| ofxn mod | |
| 0.64477 | |
| sRun | |
| RunA1 | |
| zout | |
| 1 | |
| Ave | |
| Data | |
| 0.005 | |
| Model | |
| 0.005 | |
| data/vMax | |
| 0.03054 | |
| ofxng | |
| 319.391 | |
| Min d/m | |
| 1.22843 | |
| ave/min d | |
| 46.2506 | |
| ave/min m | |
| 56.7131 | |

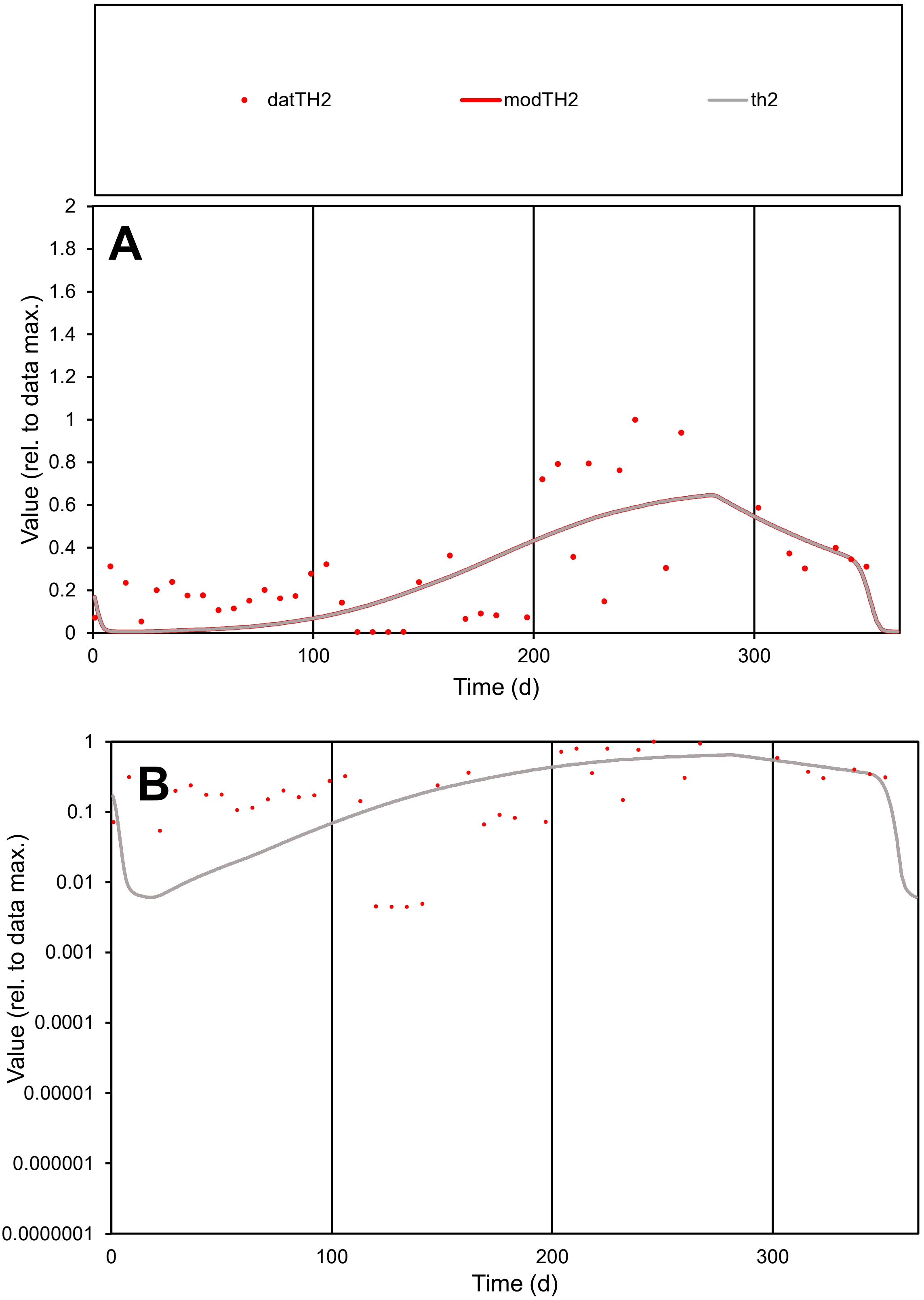


C1

Variables
TH2 Provide jtd or Dst
CalibOnly?
2 1 to reproduce of
oNormalize?
2 1=no, 2=yes

ofxn calc*
0.70034
ofxn mod
0.71521

sRun
RunA1
zout
1

Ave
Data
0.0073
Model
0.00625

data/vMax
0.02519

ofxng
319.391

Min d/m
0.72875

ave/min d
65.0364
ave/min m
40.574

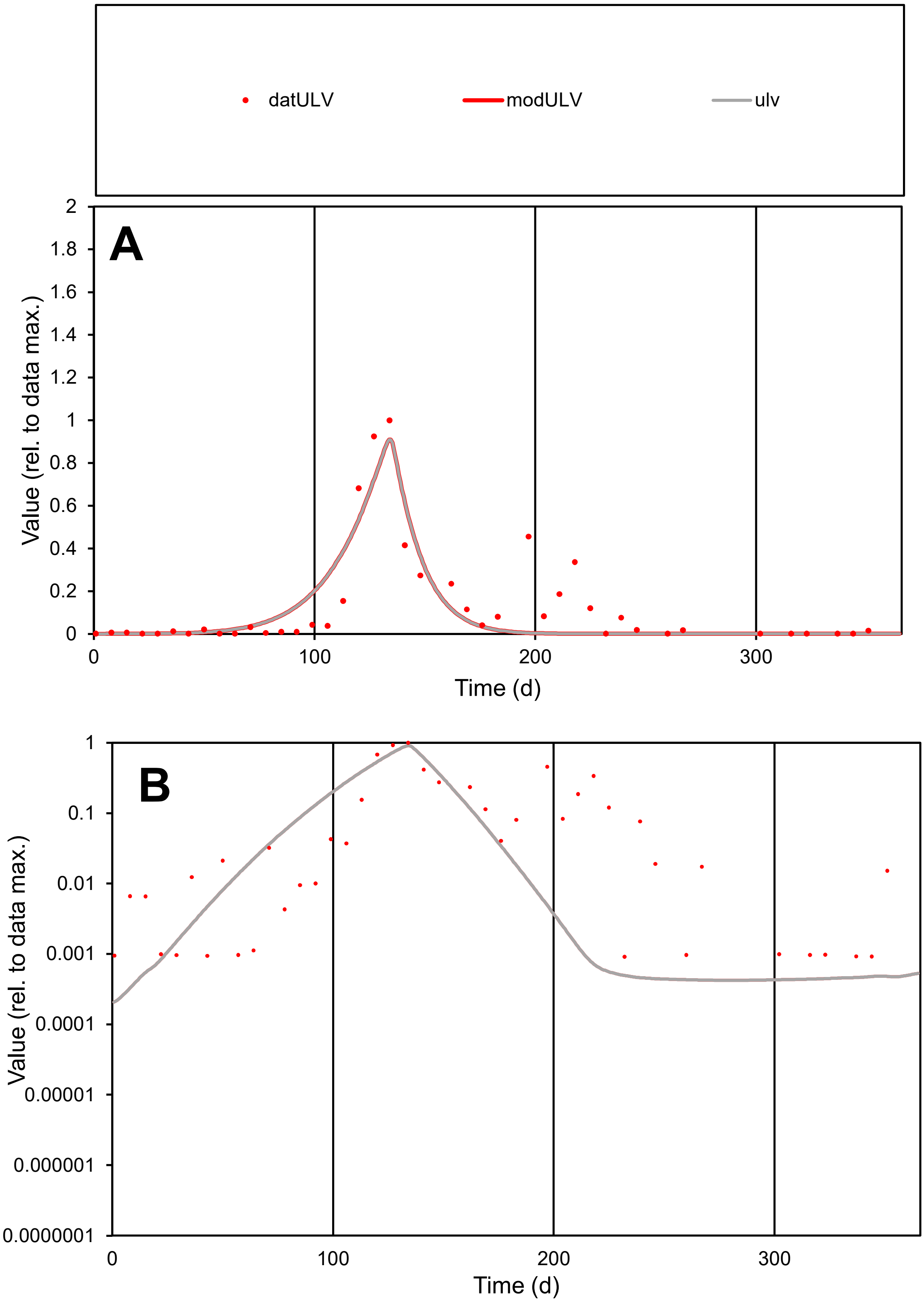


**C1**

Variables
ULV Provide jtd or Dst
CalibOnly?
2 1 to reproduce of
oNormalize?
2 1=no, 2=yes

ofxn calc*
0.95049
ofxn mod
0.97463

sRun
RunA1
zout
1

Ave
Data
0.00261
Model
0.00219

data/vMax
0.02021

ofxng
319.391

Min d/m
4.14937

ave/min d
141.925
ave/min m
494.699

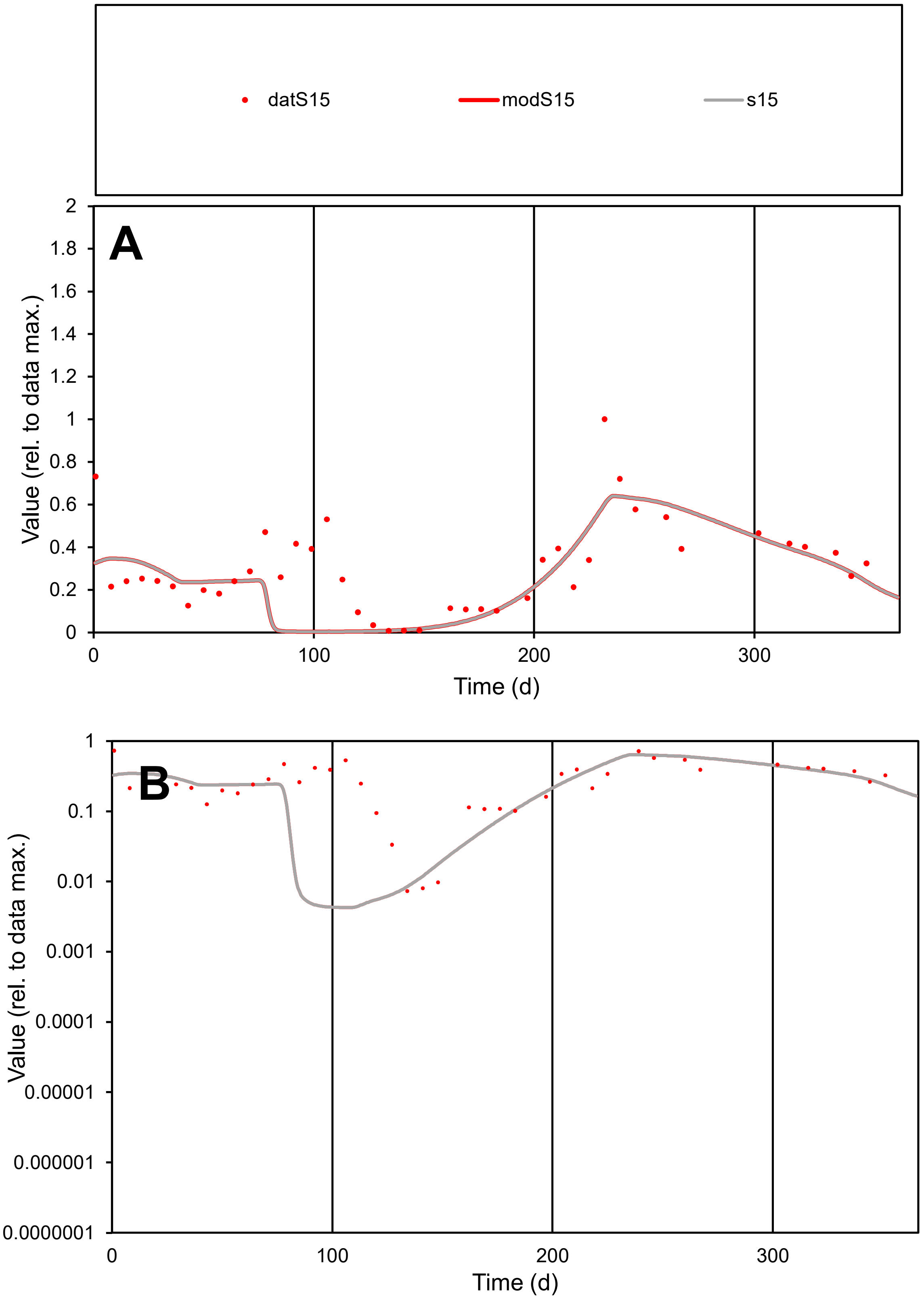


C1

Variables
S15 Provide jtd or Dst
CalibOnly?
2 1 to reproduce of
oNormalize?
2 1=no, 2=yes

ofxn calc*
0.57776
ofxn mod
0.59852

sRun
RunA1
zout
1

Ave
Data
0.0055
Model
0.00451

data/vMax
0.01813

ofxng
319.391

Min d/m
1.71903

ave/min d
41.5623
ave/min m
58.5152

330_Calib_S_A

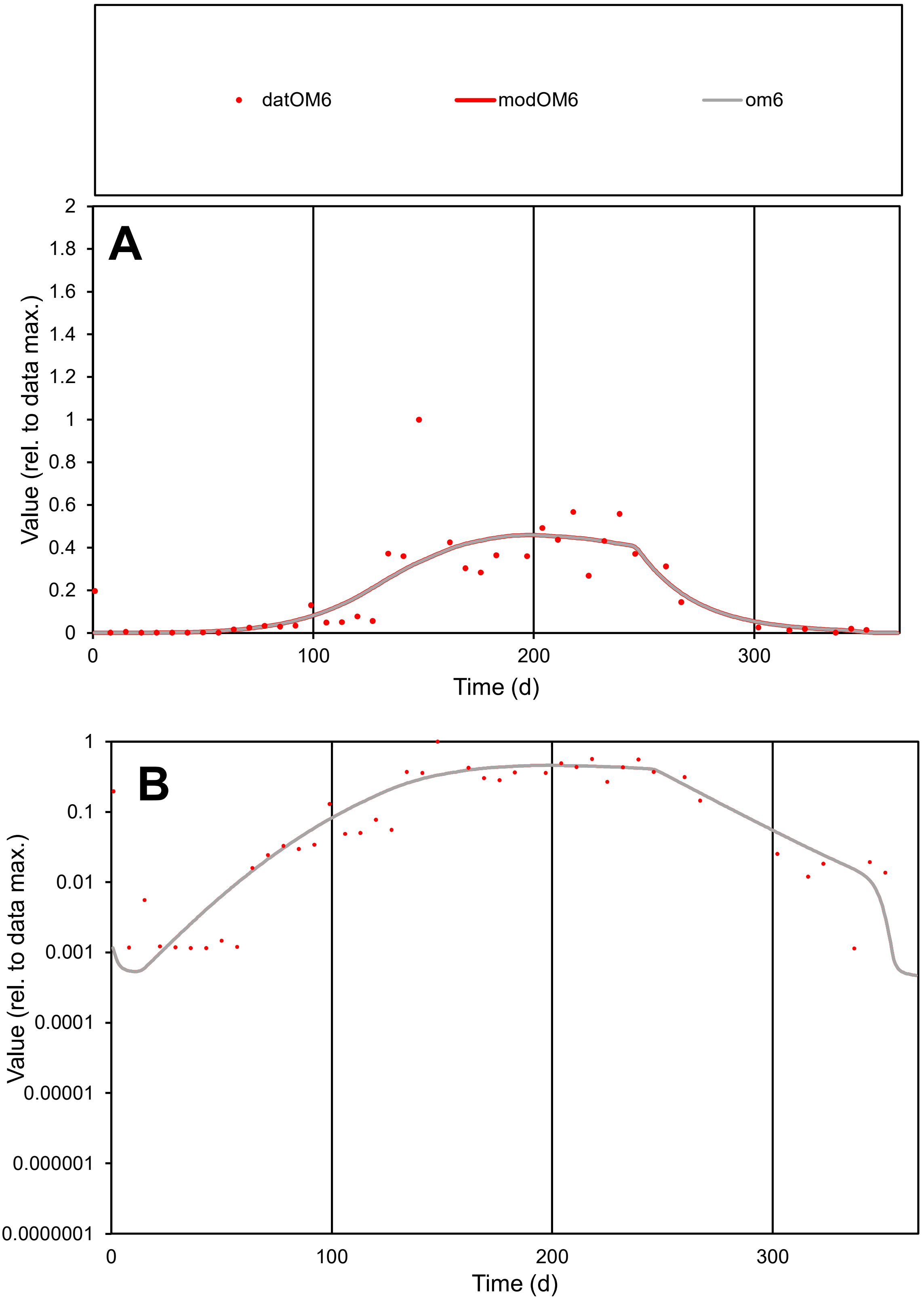


C1

| | |
|---|---|
| Variables | |
| OM6 | Provide jtd or Dst |
| CalibOnly? | |
| 2 | 1 to reproduce of |
| oNormalize? | |
| 2 | 1=no, 2=yes |
| ofxn calc* | |
| 0.67584 | |
| ofxn mod | |
| 0.69701 | |
| sRun | |
| RunA1 | |
| zout | |
| 1 | |
| Ave | |
| Data | |
| 0.00443 | |
| Model | |
| 0.00427 | |
| data/vMax | |
| 0.02375 | |
| ofxng | |
| 319.391 | |
| Min d/m | |
| 2.12363 | |
| ave/min d | |
| 163.321 | |
| ave/min m | |
| 334.007 | |

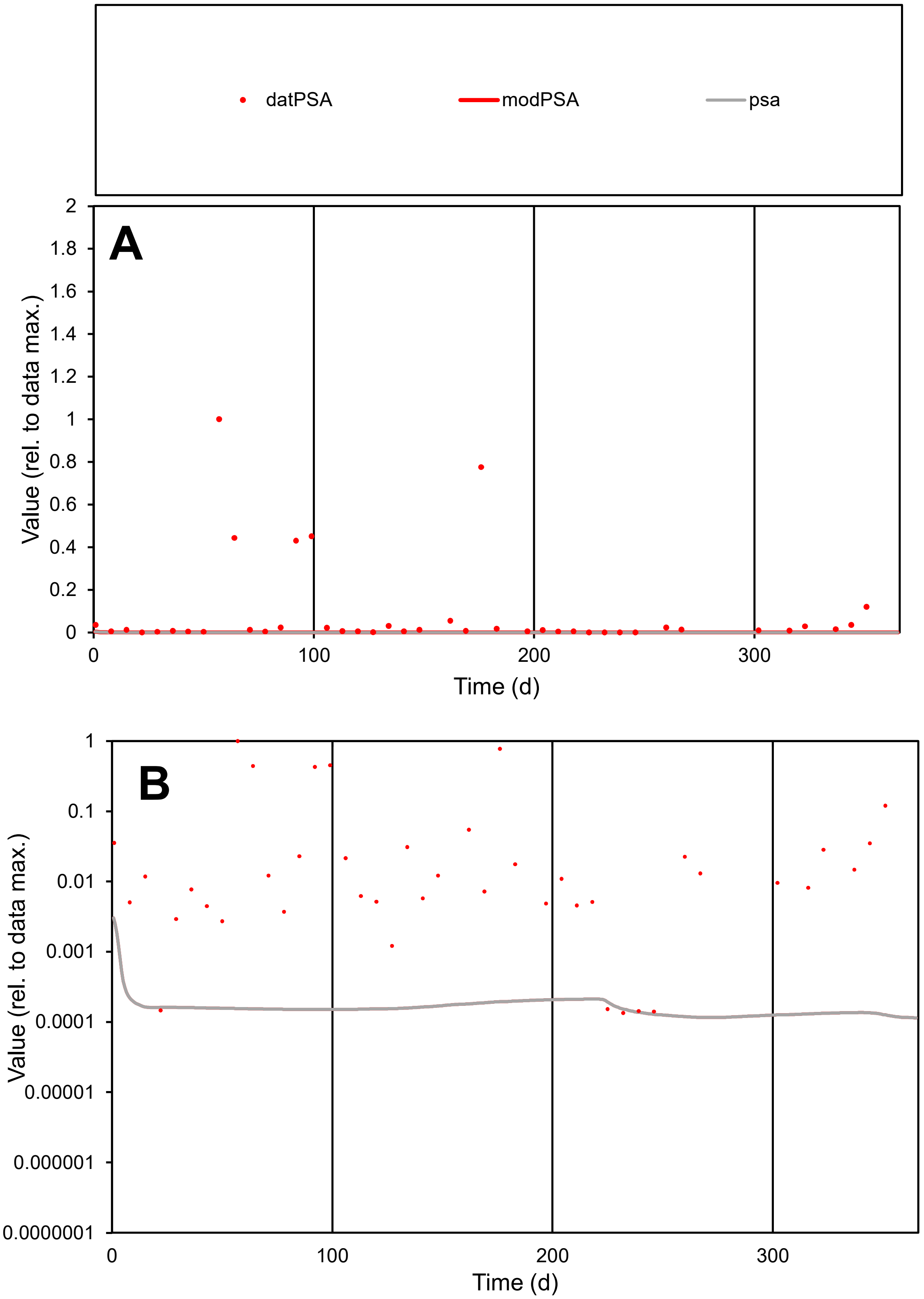


**C1**

Variables
PSA Provide jtd or Dst
CalibOnly?
2 1 to reproduce of
oNormalize?
2 1=no, 2=yes

ofxn calc*
2.63997
ofxn mod
2.64348

sRun
RunA1
zout
1

Ave
Data
0.00607
Model
1.5E-05

data/vMax
0.06991

ofxng
319.391

Min d/m
1.16431

ave/min d
642.432
ave/min m
1.85595

330_Calib_S_A

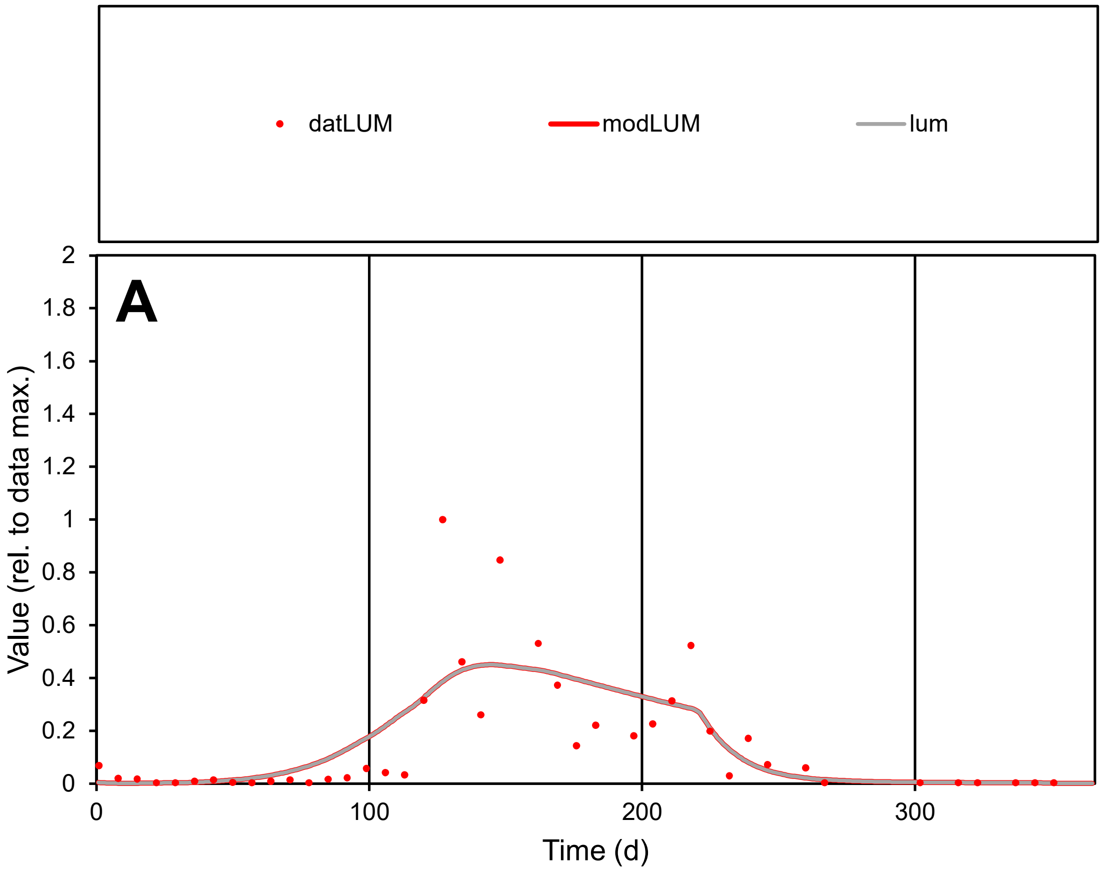


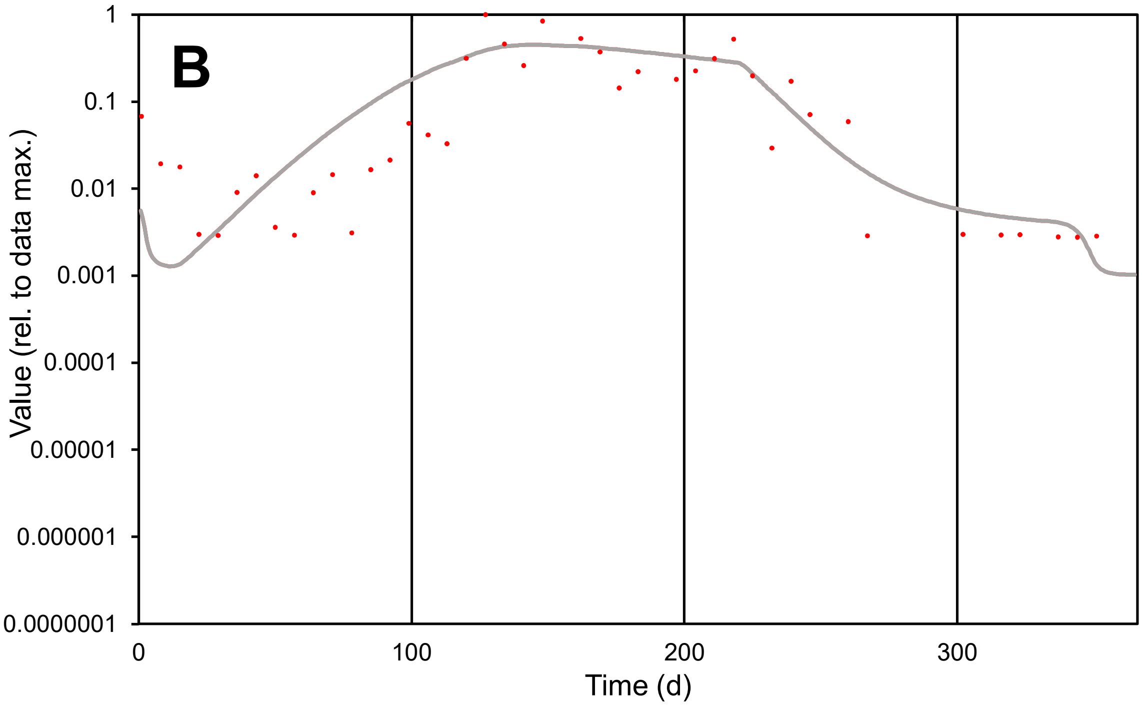


**C1**

Variables

| | |
|---|---|
| LUM | Provide jtd or Dst |
| CalibOnly? | |
| 2 | 1 to reproduce of |
| oNormalize? | |
| 2 | 1=no, 2=yes |

ofxn calc*
0.98006
ofxn mod
0.99487

sRun
RunA1
zout
1

Ave
Data
0.0026
Model
0.00269

data/vMax
0.01739

ofxng
319.391

Min d/m
2.10722

ave/min d
53.9663
ave/min m
117.656

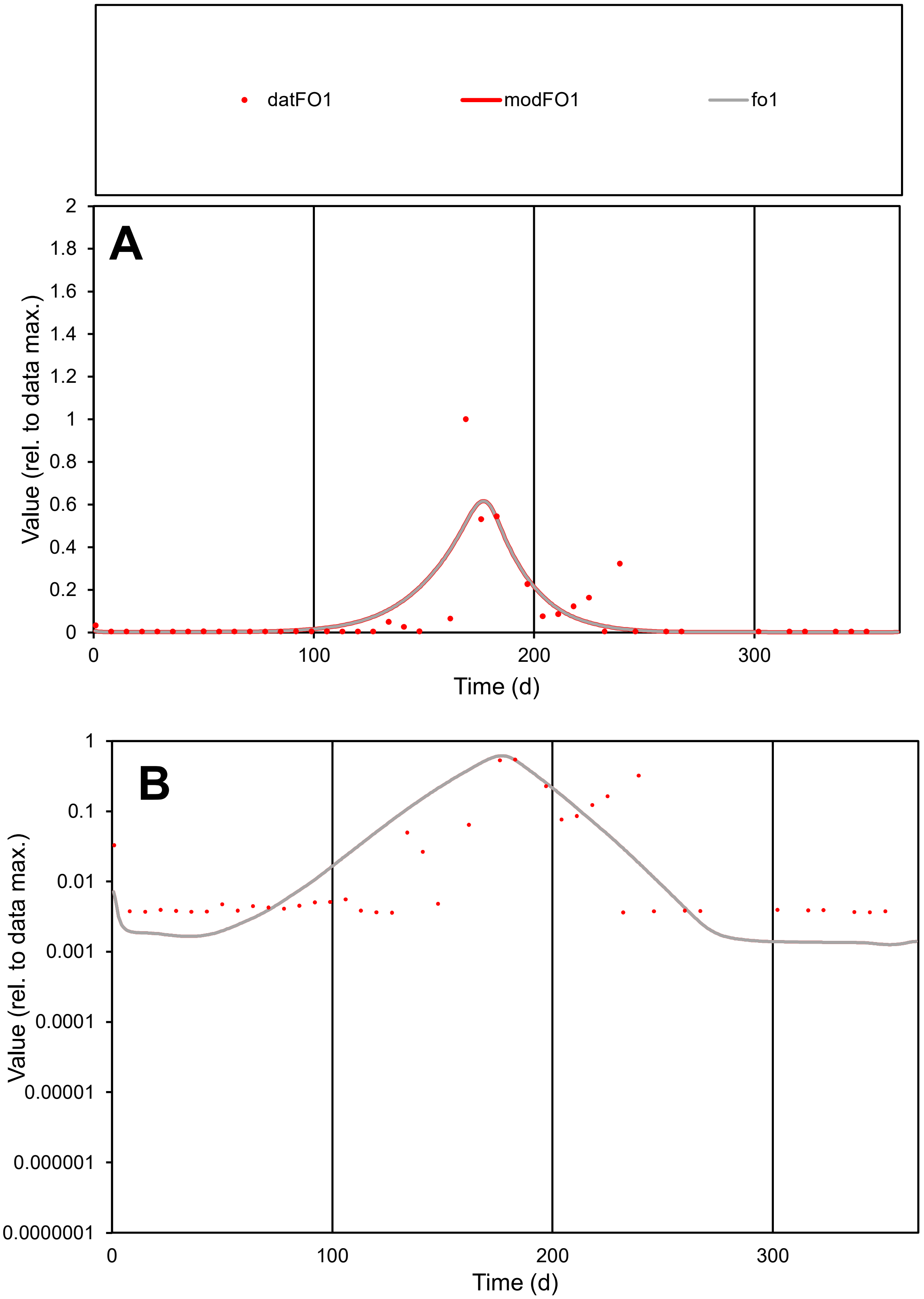


**C1**

Variables
FO1 Provide jtd or Dst
CalibOnly?
2 1 to reproduce of
oNormalize?
2 1=no, 2=yes

ofxn calc*
1.3866
ofxn mod
1.41787

sRun
RunA1
zout
1

Ave
Data
0.00138
Model
0.0014

data/vMax
0.01716

ofxng
319.391

Min d/m
2.85088

ave/min d
22.2704
ave/min m
64.748

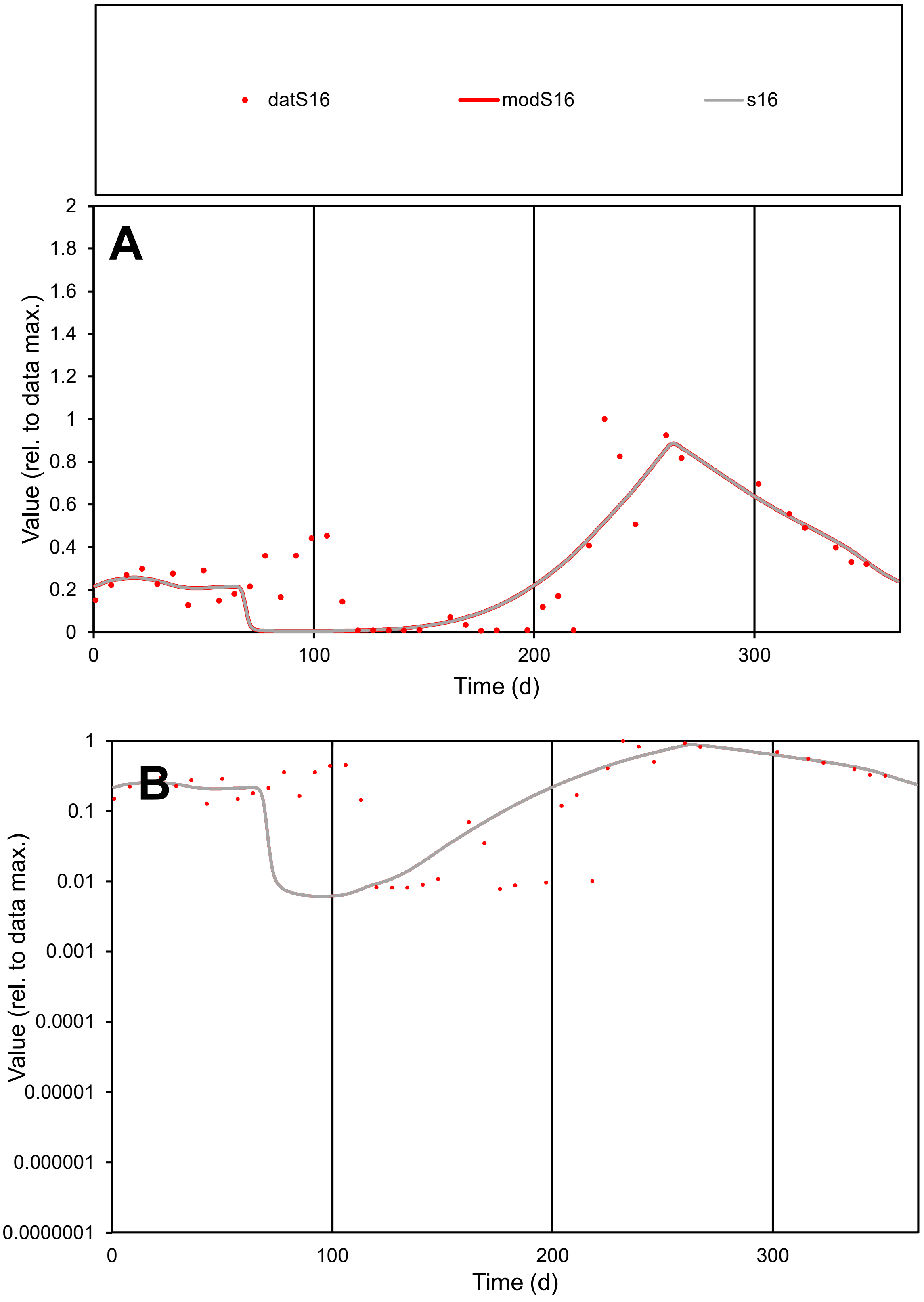


**C1**

Variables
S16 Provide jtd or Dst
CalibOnly?
2 1 to reproduce of
oNormalize?
2 1=no, 2=yes

ofxn calc*
0.60219
ofxn mod
0.62201

sRun
RunA1
zout
1

Ave
Data
0.00466
Model
0.00417

data/vMax
0.01623

ofxng
319.391

Min d/m
1.27703

ave/min d
36.9037
ave/min m
42.1273

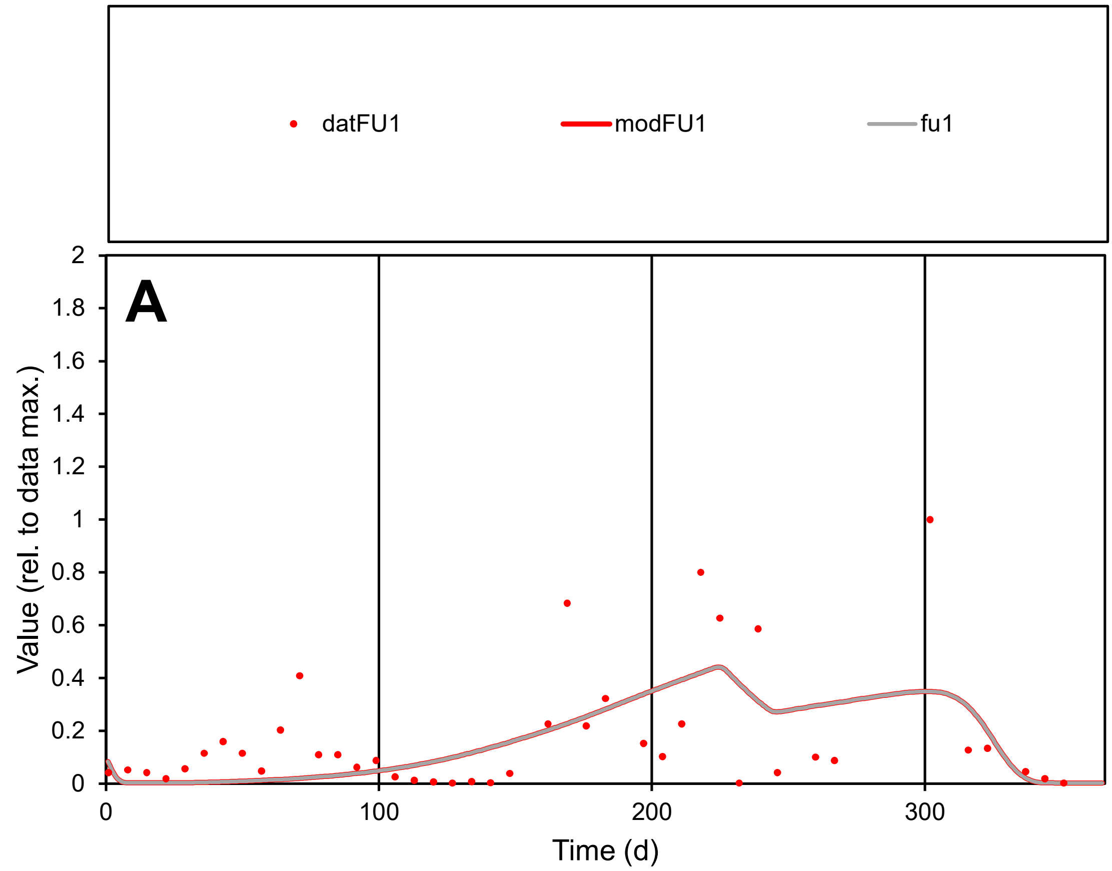


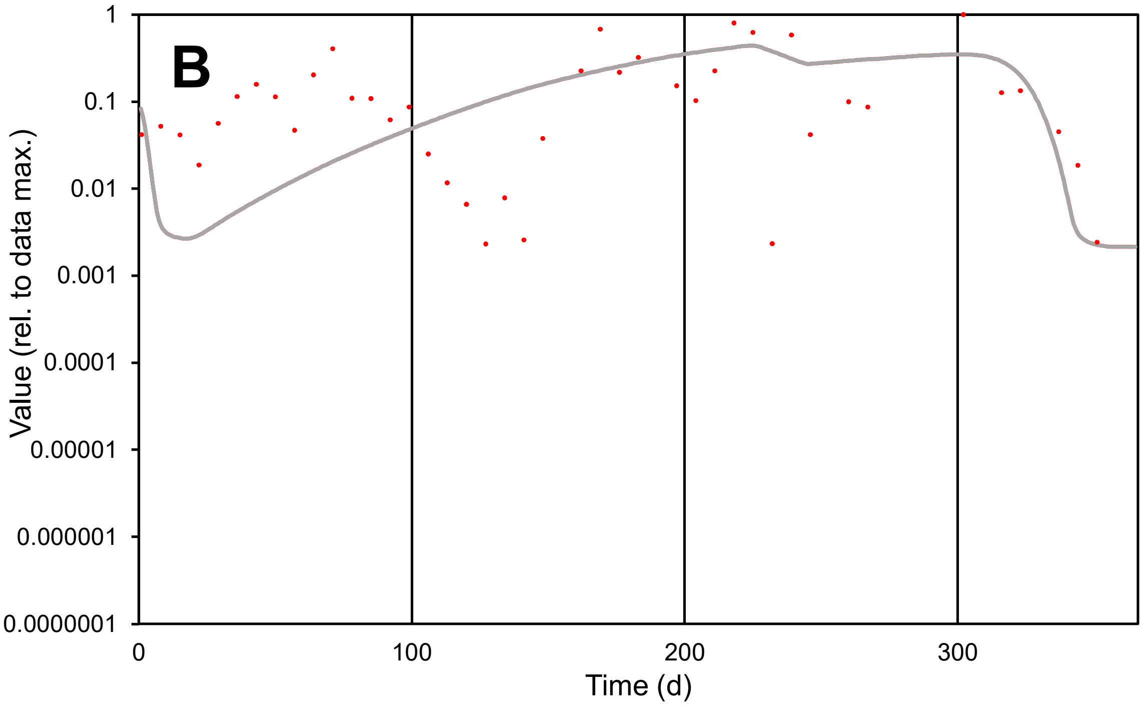


**C1**

Variables
FU1 Provide jtd or Dst
CalibOnly?
2 1 to reproduce of
oNormalize?
2 1=no, 2=yes

ofxn calc*
1.14489
ofxn mod
1.16062

sRun
RunA1
zout
1

Ave
Data
0.00365
Model
0.00321

data/vMax
0.02124

ofxng
319.391

Min d/m
1.03999

ave/min d
73.8558
ave/min m
67.6692

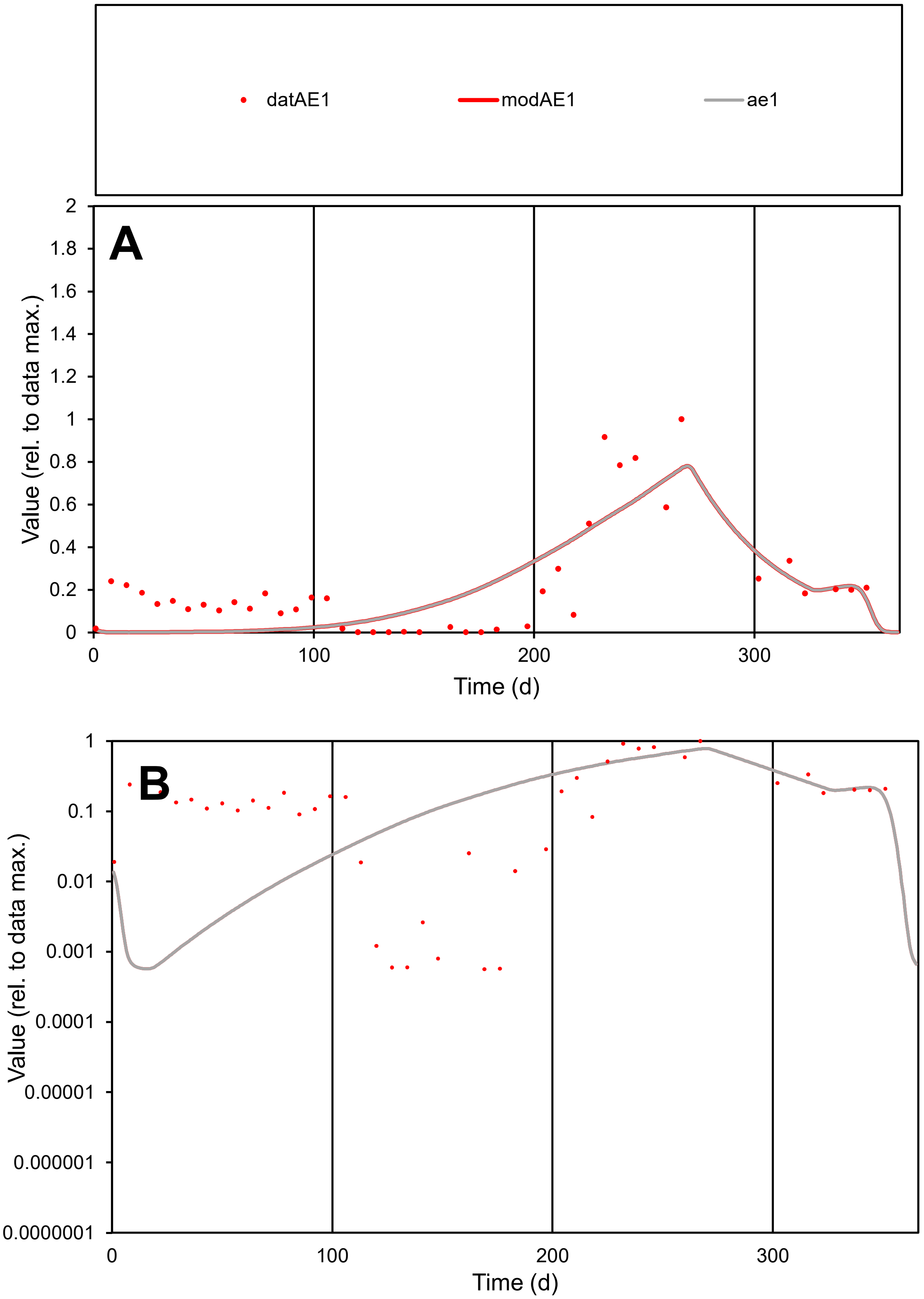


**C1**

Variables
AE1 Provide jtd or Dst
CalibOnly?
2 1 to reproduce of
oNormalize?
2 1=no, 2=yes

ofxn calc*
0.74985
ofxn mod
0.7711

sRun
RunA1
zout
1

Ave
Data
0.00525
Model
0.00474

data/vMax
0.02476

ofxng
319.391

Min d/m
0.99083

ave/min d
374.268
ave/min m
334.284

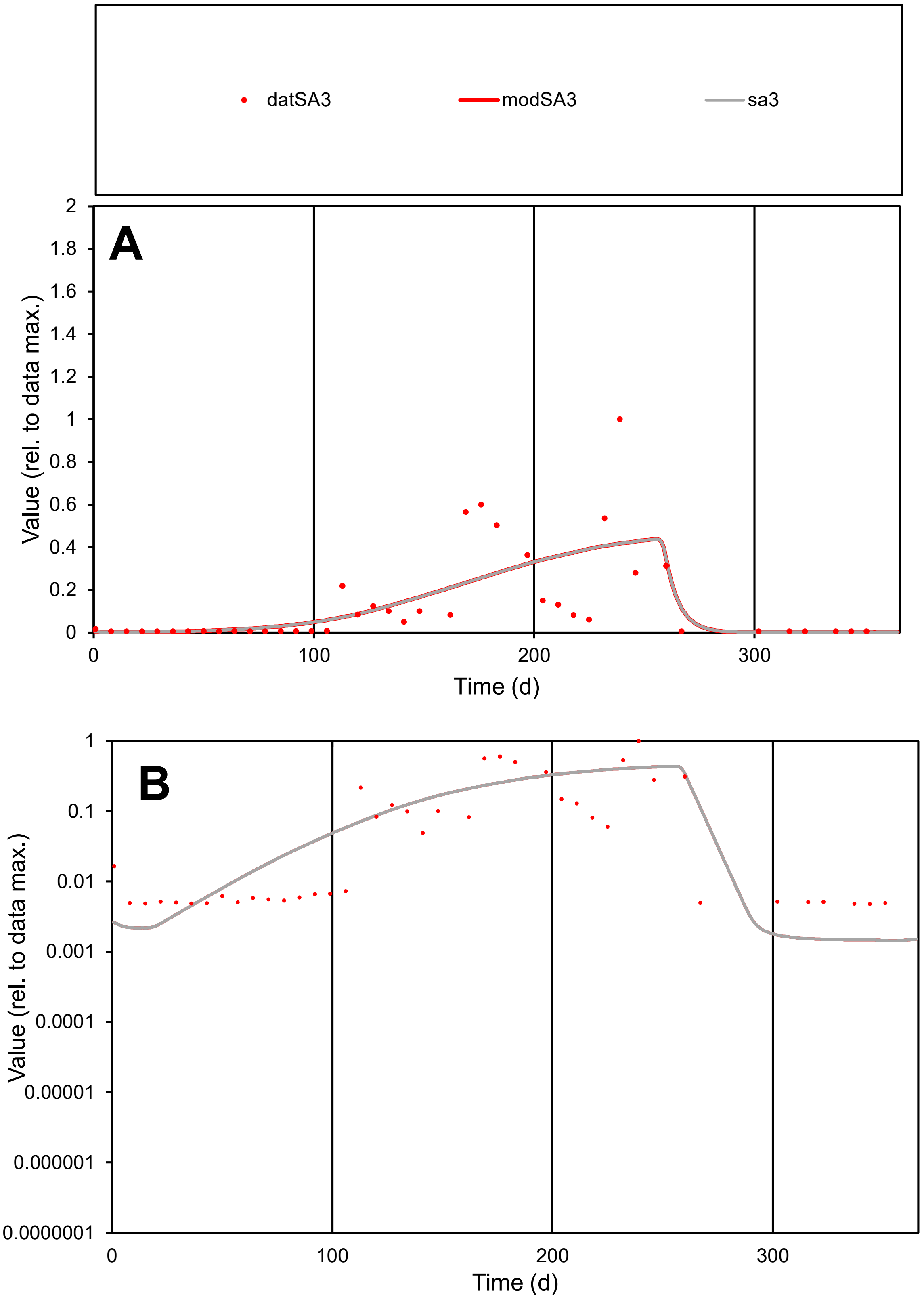


**C1**

Variables
SA3 Provide jtd or Dst
CalibOnly?
2 1 to reproduce of
oNormalize?
2 1=no, 2=yes

ofxn calc*
1.18316
ofxn mod
1.20417

sRun
RunA1
zout
1

Ave
Data
0.00288
Model
0.00285

data/vMax
0.02213

ofxng
319.391

Min d/m
3.31868

ave/min d
27.2778
ave/min m
89.5072

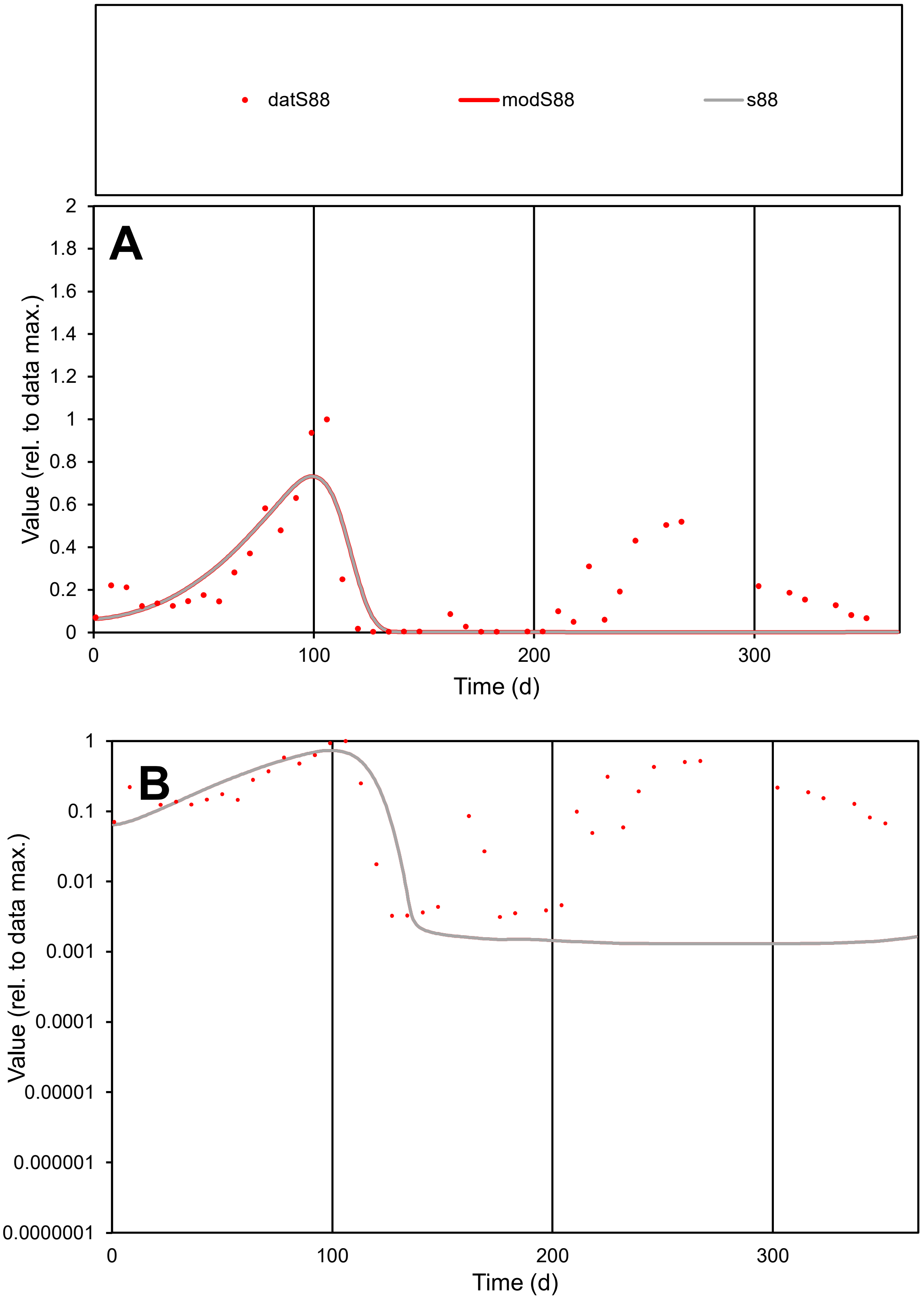


**C1**

Variables
S88 Provide jtd or Dst
CalibOnly?
2 1 to reproduce of
oNormalize?
2 1=no, 2=yes

ofxn calc*
0.83237
ofxn mod
0.84896

sRun
RunA1
zout
1

Ave
Data
0.00404
Model
0.0028

data/vMax
0.01876

ofxng
319.391

Min d/m
2.40188

ave/min d
68.7337
ave/min m
114.667

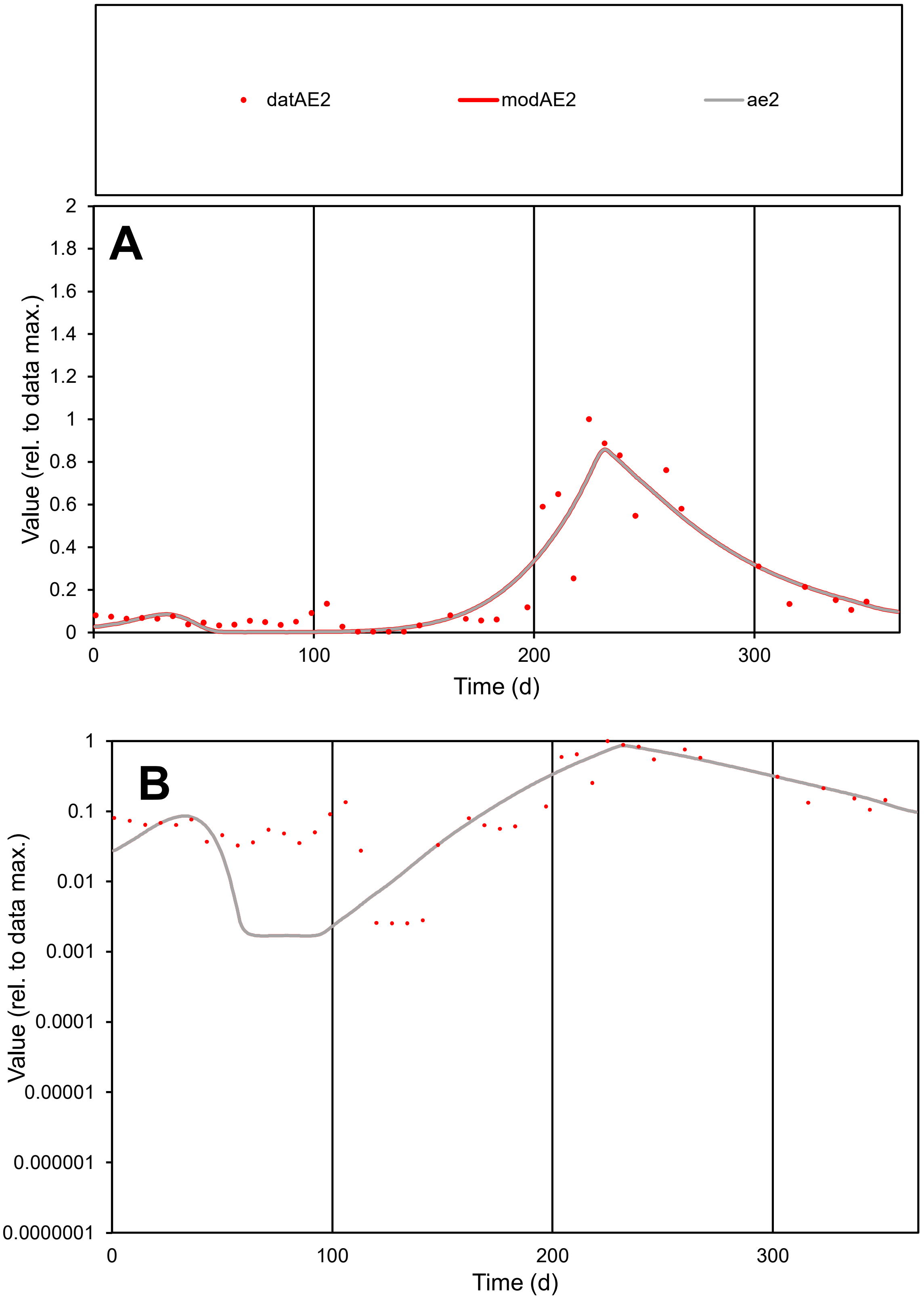


**C1**

Variables
AE2 Provide jtd or Dst
CalibOnly?
2 1 to reproduce of
oNormalize?
2 1=no, 2=yes

ofxn calc*
0.48109
ofxn mod
0.52186

sRun
RunA1
zout
1

Ave
Data
0.00495
Model
0.00477

data/vMax
0.0242

ofxng
319.391

Min d/m
1.50282

ave/min d
80.7435
ave/min m
116.869

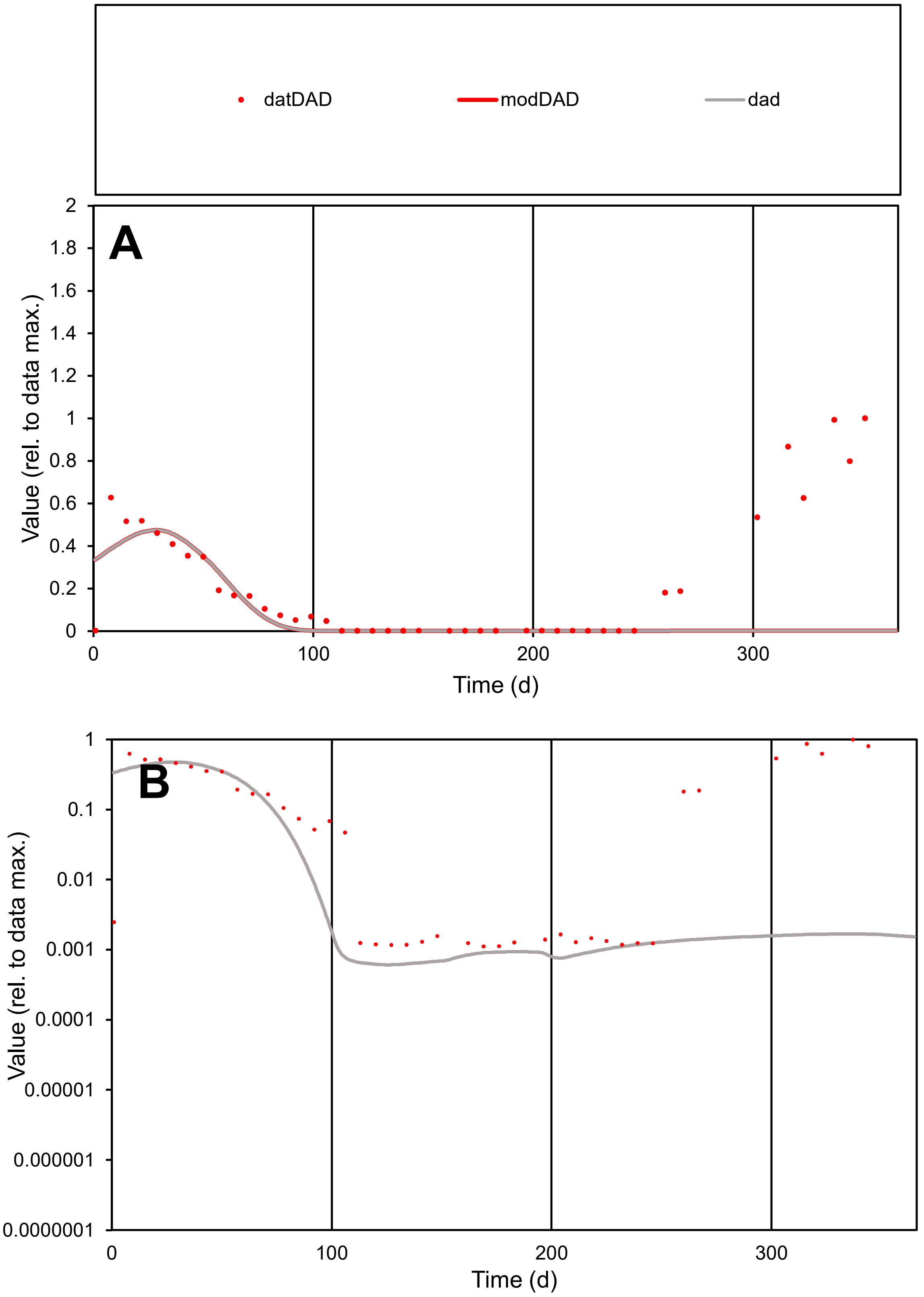


**C1**

Variables

DAD Provide jtd or Dst

CalibOnly?

2 1 to reproduce of

oNormalize?

2 1=no, 2=yes

ofxn calc*

1.44286

ofxn mod

1.44768

sRun

RunA1

zout

1

Ave

Data

0.00311

Model

0.00133

data/vMax

0.01401

ofxng

319.391

Min d/m

1.81648

ave/min d

199.744

ave/min m

155.418

330_Calib_S_A

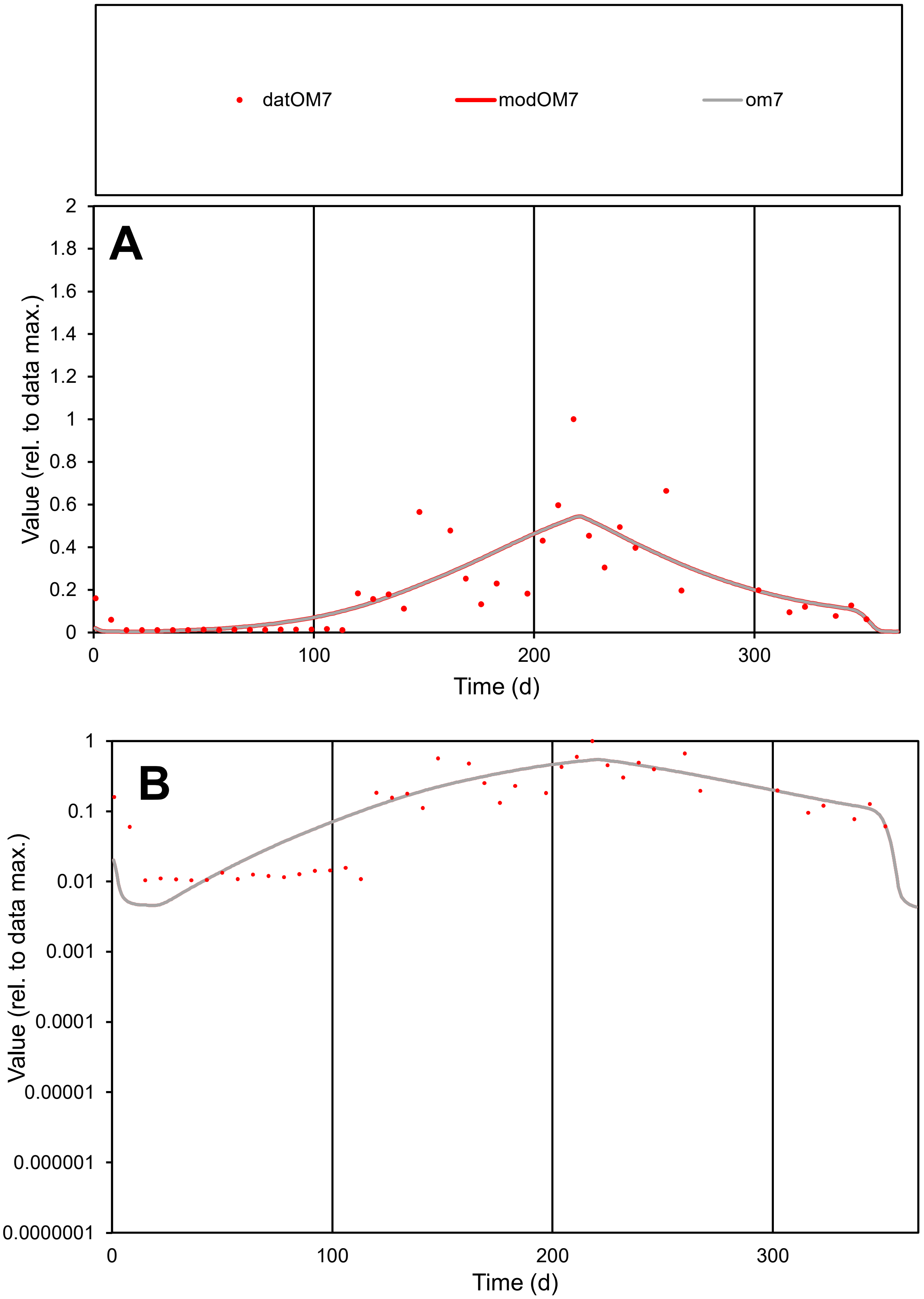


**C1**

Variables
OM7 Provide jtd or Dst
CalibOnly?
2 1 to reproduce of
oNormalize?
2 1=no, 2=yes

ofxn calc*
0.68436
ofxn mod
0.70934

sRun
RunA1
zout
1

Ave
Data
0.00189
Model
0.00188

data/vMax
0.00981

ofxng
319.391

Min d/m
2.28552

ave/min d
18.4419
ave/min m
41.8903

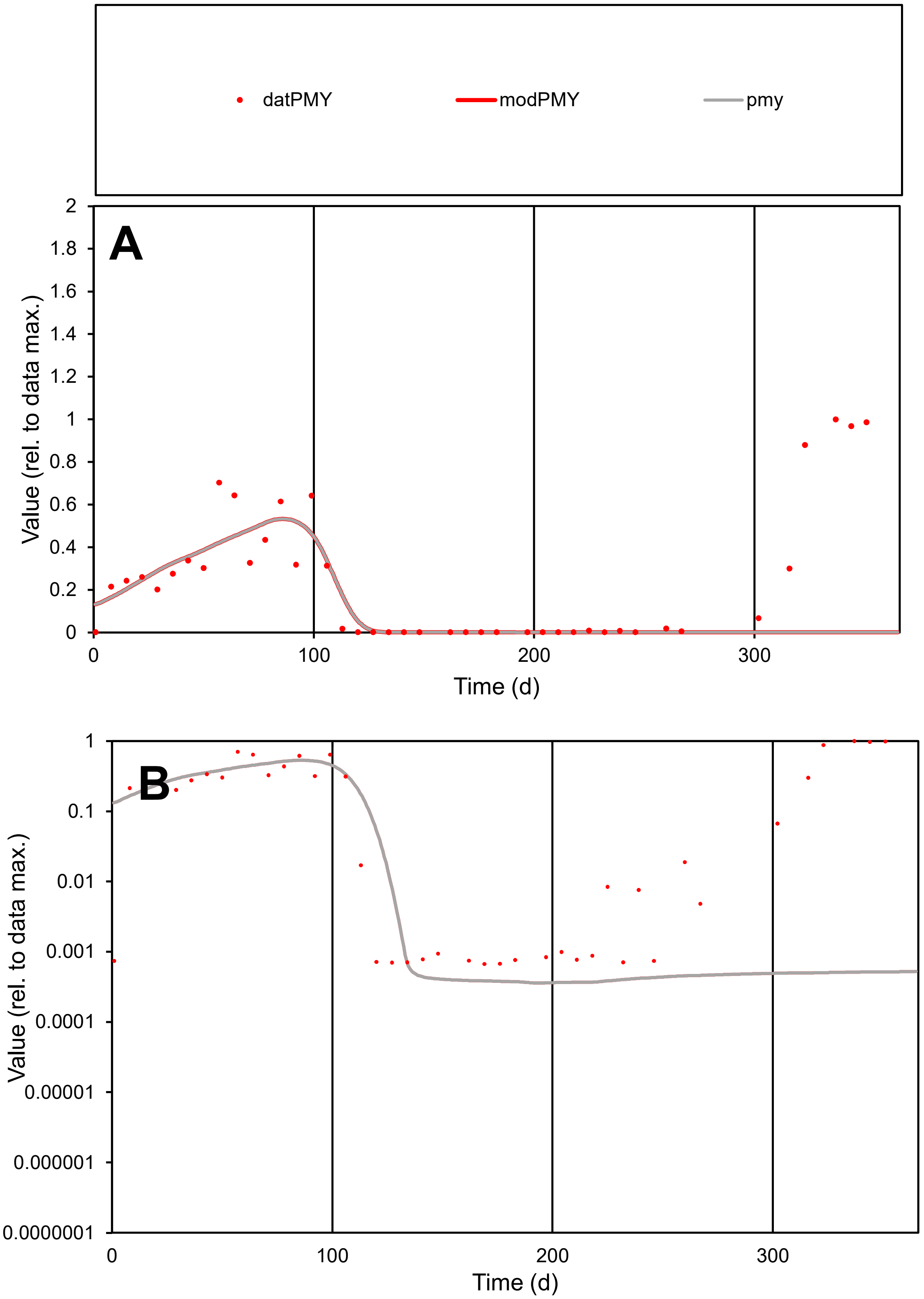


**C1**

Variables
PMY Provide jtd or Dst
CalibOnly?
2 1 to reproduce of
oNormalize?
2 1=no, 2=yes

ofxn calc*
1.29245
ofxn mod
1.29891

sRun
RunA1
zout
1

Ave
Data
0.00349
Model
0.00207

data/vMax
0.01451

ofxng
319.391

Min d/m
1.84021

ave/min d
360.092
ave/min m
394.168

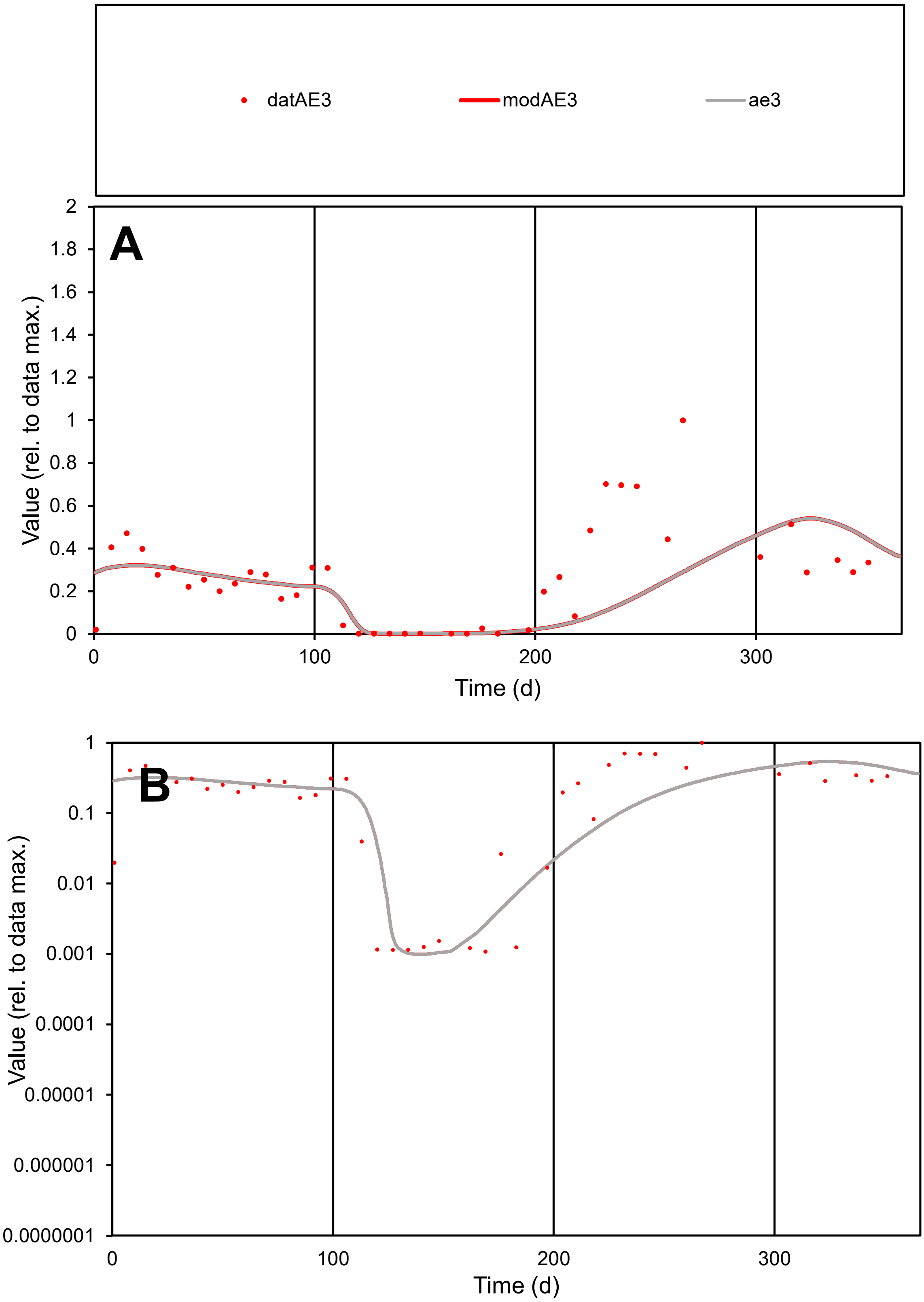


**C1**

Variables
AE3 Provide jtd or Dst
CalibOnly?
2 1 to reproduce of
oNormalize?
2 1=no, 2=yes

ofxn calc*
0.81937
ofxn mod
0.82872

sRun
RunA1
zout
1

Ave
Data
0.00414
Model
0.00323

data/vMax
0.01565

ofxng
319.391

Min d/m
1.08515

ave/min d
244.034
ave/min m
206.706

330_Calib_S_A

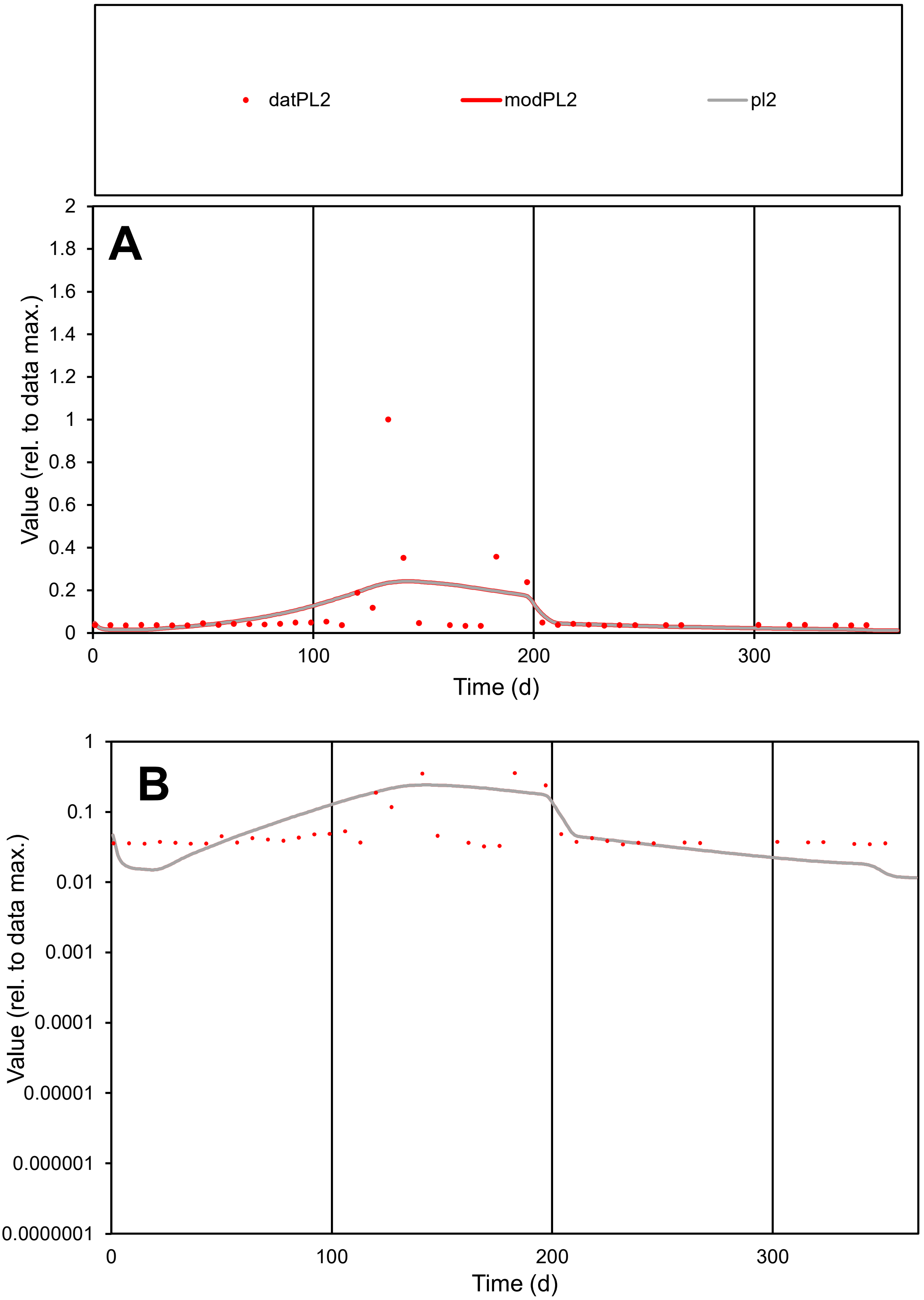


**C1**

Variables
PL2 Provide jtd or Dst
CalibOnly?
2 1 to reproduce of
oNormalize?
2 1=no, 2=yes

ofxn calc*
1.60625
ofxn mod
1.61836

sRun
RunA1
zout
1

Ave
Data
0.00044
Model
0.00045

data/vMax
0.00506

ofxng
319.391

Min d/m
2.47286

ave/min d
2.67844
ave/min m
6.7423

330_Calib_S_A

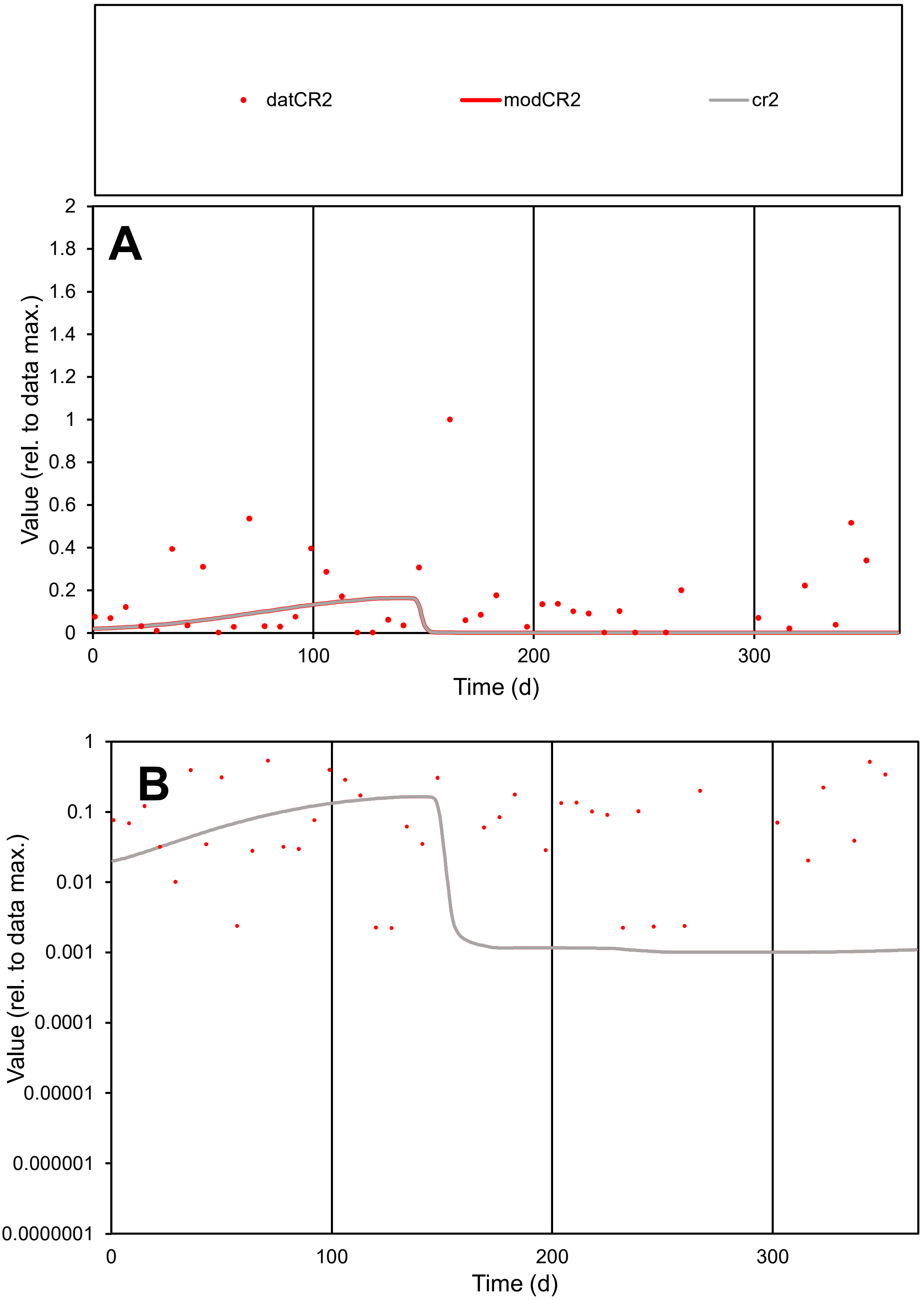


**C1**

Variables
CR2 Provide jtd or Dst
CalibOnly?
2 1 to reproduce of
oNormalize?
2 1=no, 2=yes

ofxn calc*
1.51205
ofxn mod
1.51912

sRun
RunA1
zout
1

Ave
Data
0.00287
Model
0.00095

data/vMax
0.019

ofxng
319.391

Min d/m
2.20229

ave/min d
67.74
ave/min m
49.6082

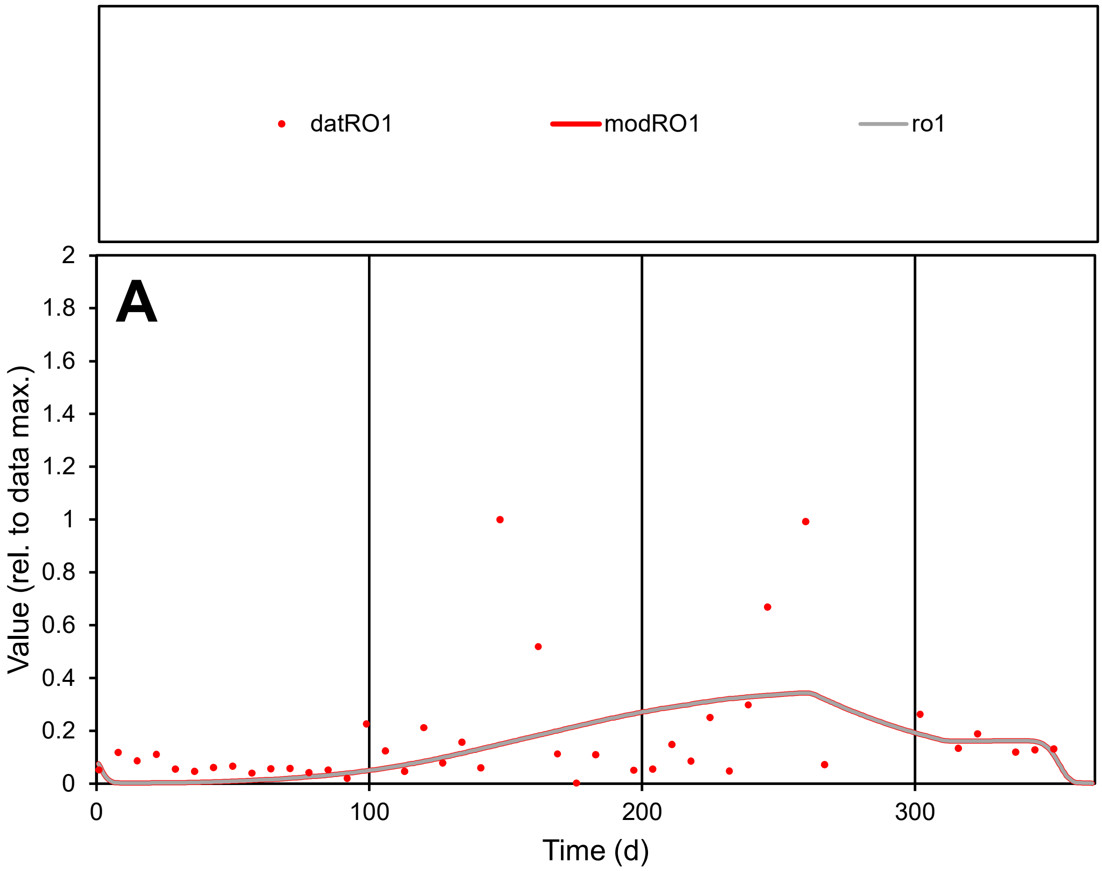


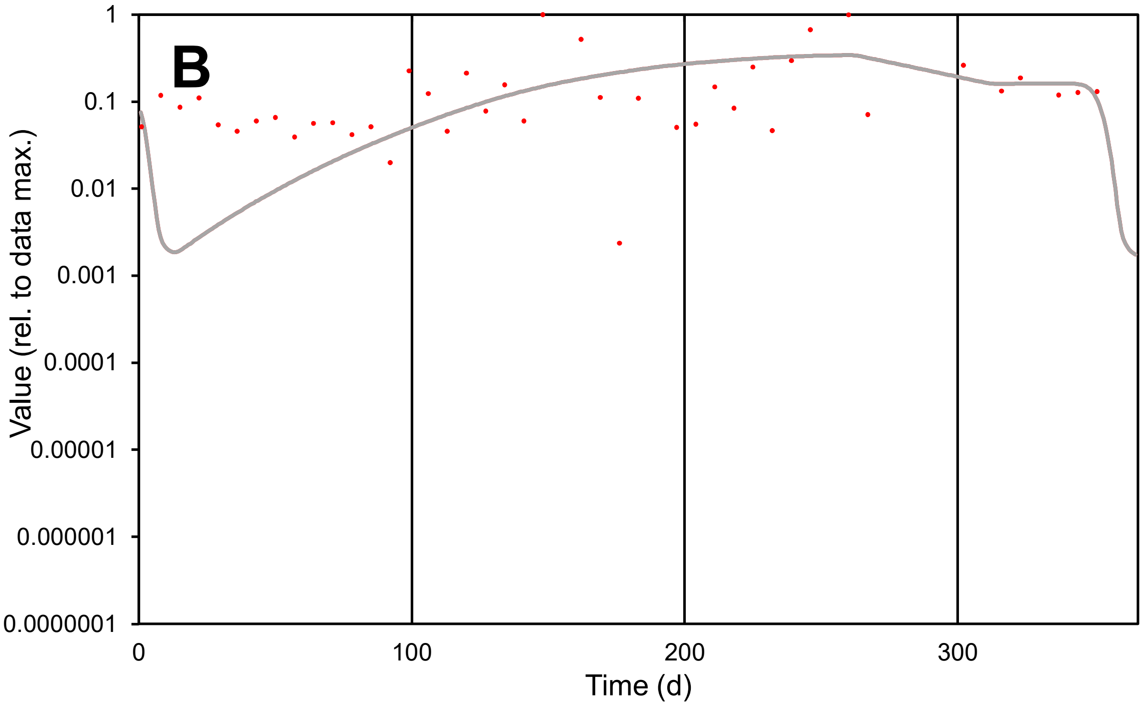


**C1**

Variables
RO1 Provide jtd or Dst
CalibOnly?
2 1 to reproduce of
oNormalize?
2 1=no, 2=yes

ofxn calc*
1.23743
ofxn mod
1.24498

sRun
RunA1
zout
1

Ave
Data
0.00258
Model
0.00213

data/vMax
0.01519

ofxng
319.391

Min d/m
1.12774

ave/min d
71.5522
ave/min m
66.5331

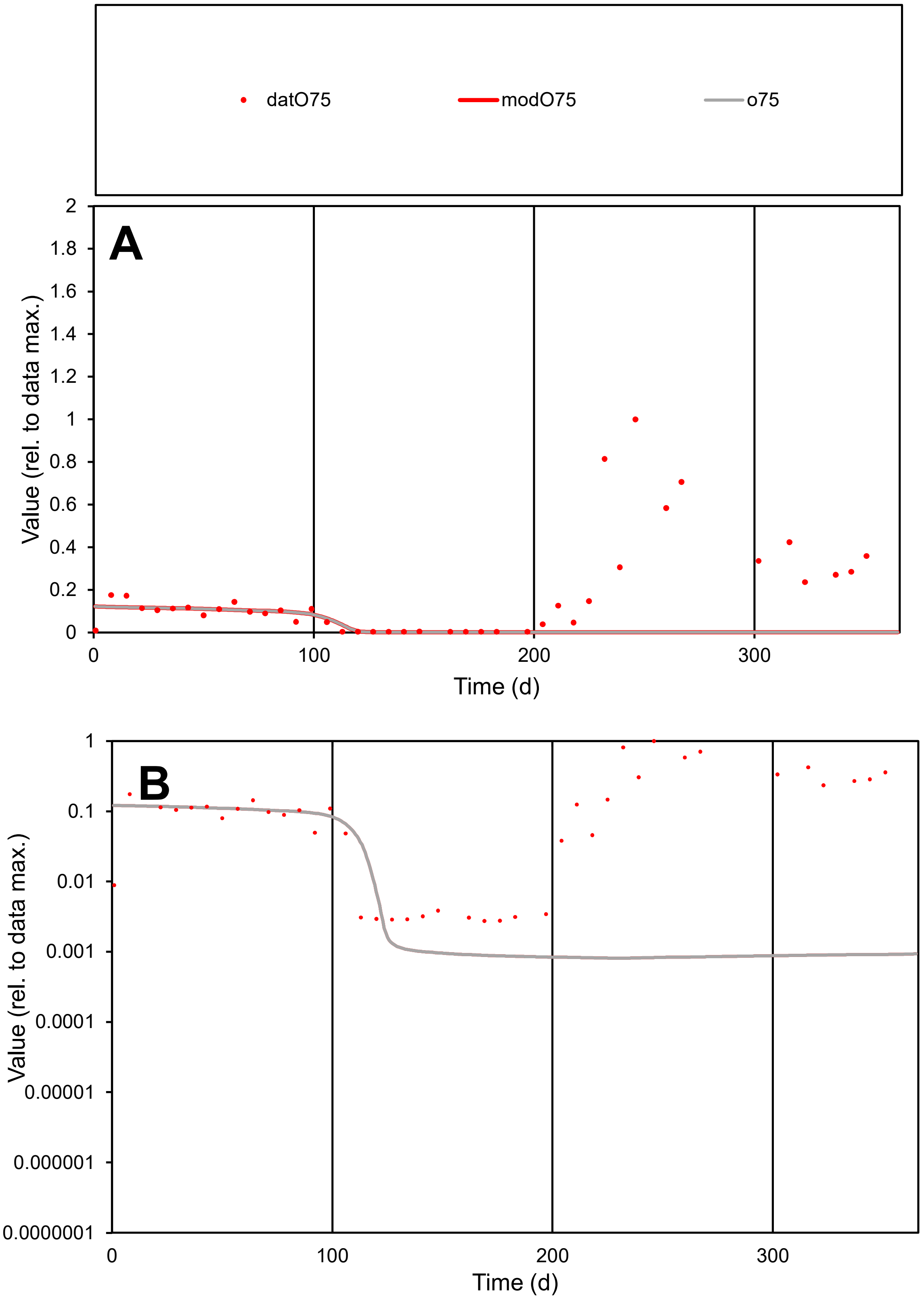


**C1**

Variables
O75 Provide jtd or Dst
CalibOnly?
2 1 to reproduce of
oNormalize?
2 1=no, 2=yes

ofxn calc*
1.59968
ofxn mod
1.60274

sRun
RunA1
zout
1

Ave
Data
0.00451
Model
0.00106

data/vMax
0.02581

ofxng
319.391

Min d/m
3.38506

ave/min d
63.7963
ave/min m
50.764

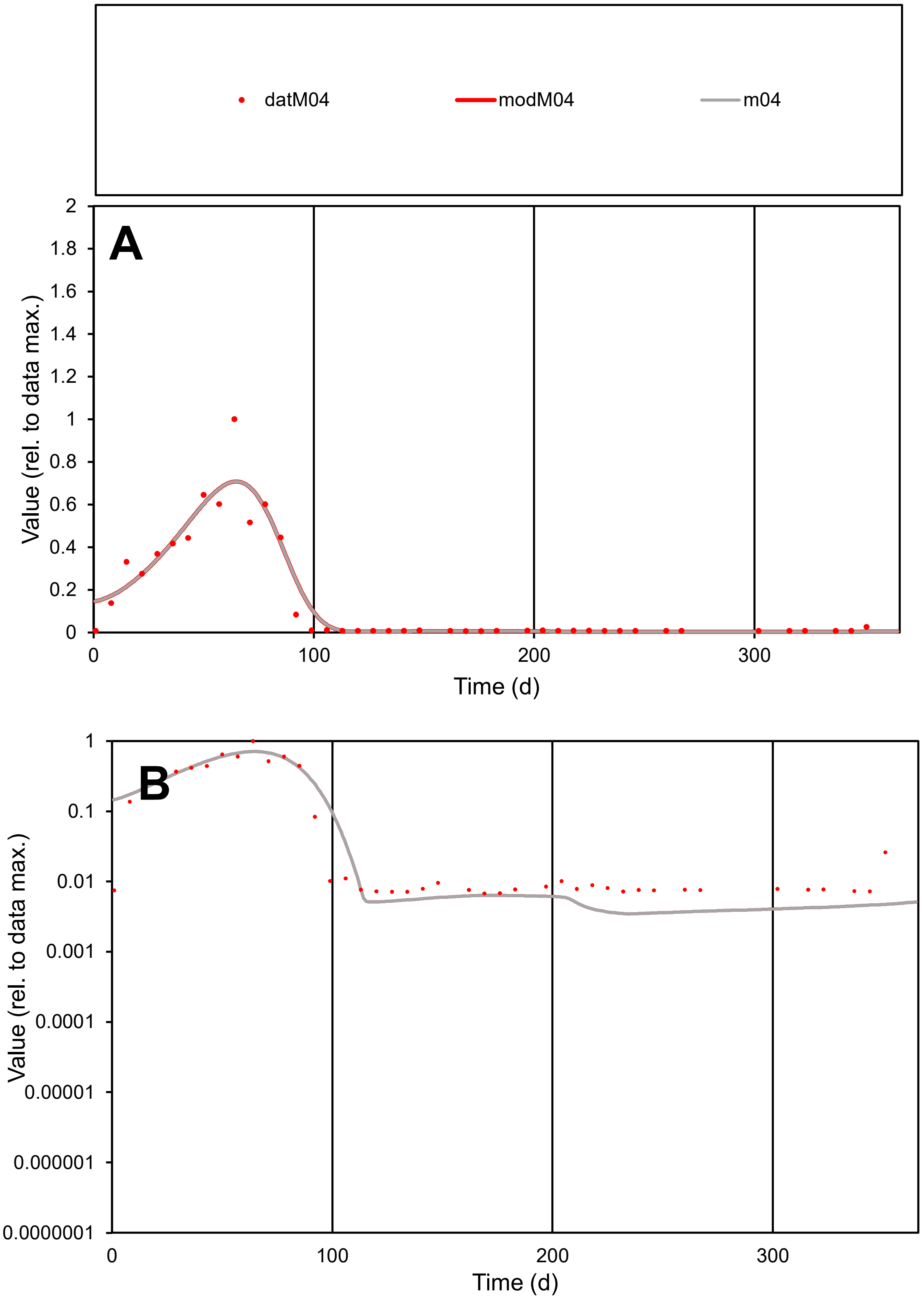


**C1**

Variables
M04 Provide jtd or Dst
CalibOnly?
2 1 to reproduce of
oNormalize?
2 1=no, 2=yes

ofxn calc*
0.44777
ofxn mod
0.50642

sRun
RunA1
zout
1

Ave
Data
0.00076
Model
0.00078

data/vMax
0.00525

ofxng
319.391

Min d/m
1.94688
ave/min d
21.5139
ave/min m
42.5119

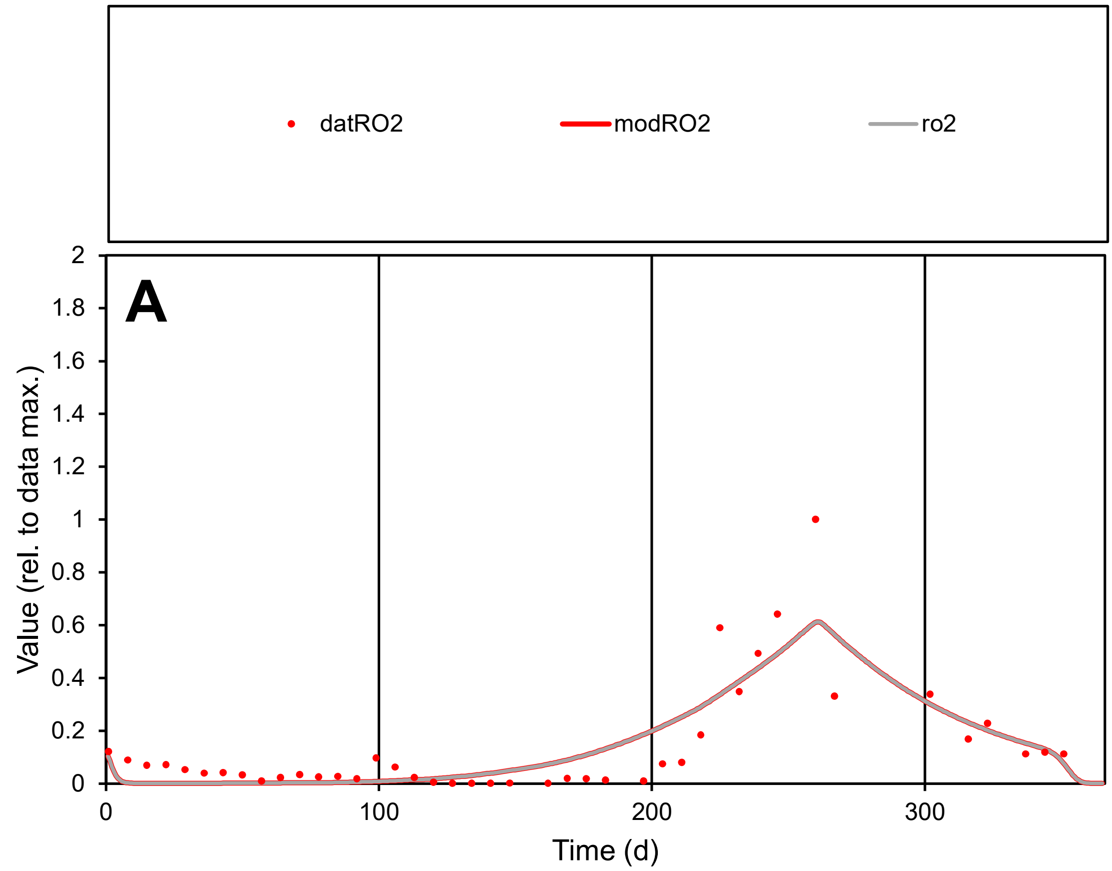


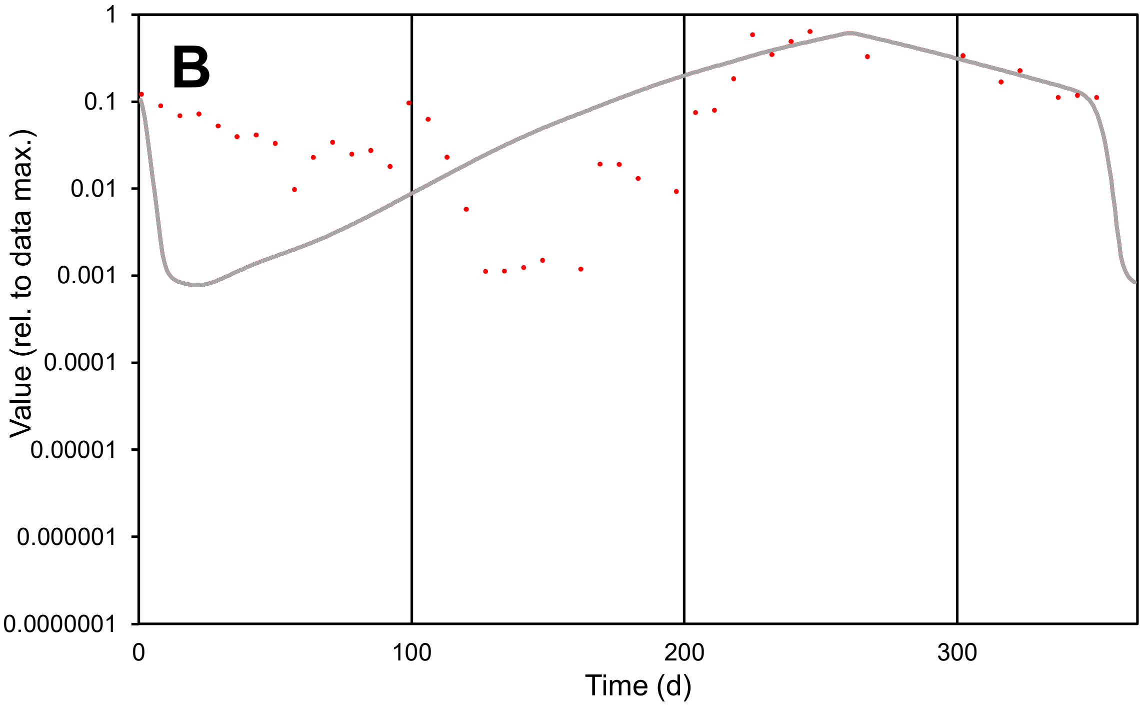


**C1**

Variables

RO2 Provide jtd or Dst

CalibOnly?

2 1 to reproduce of

oNormalize?

2 1=no, 2=yes

ofxn calc*

0.76861

ofxn mod

0.79199

sRun

RunA1

zout

1

Ave

Data

0.00496

Model

0.00485

data/vMax

0.03637

ofxng

319.391

Min d/m

1.42648

ave/min d

121.563

ave/min m

169.377

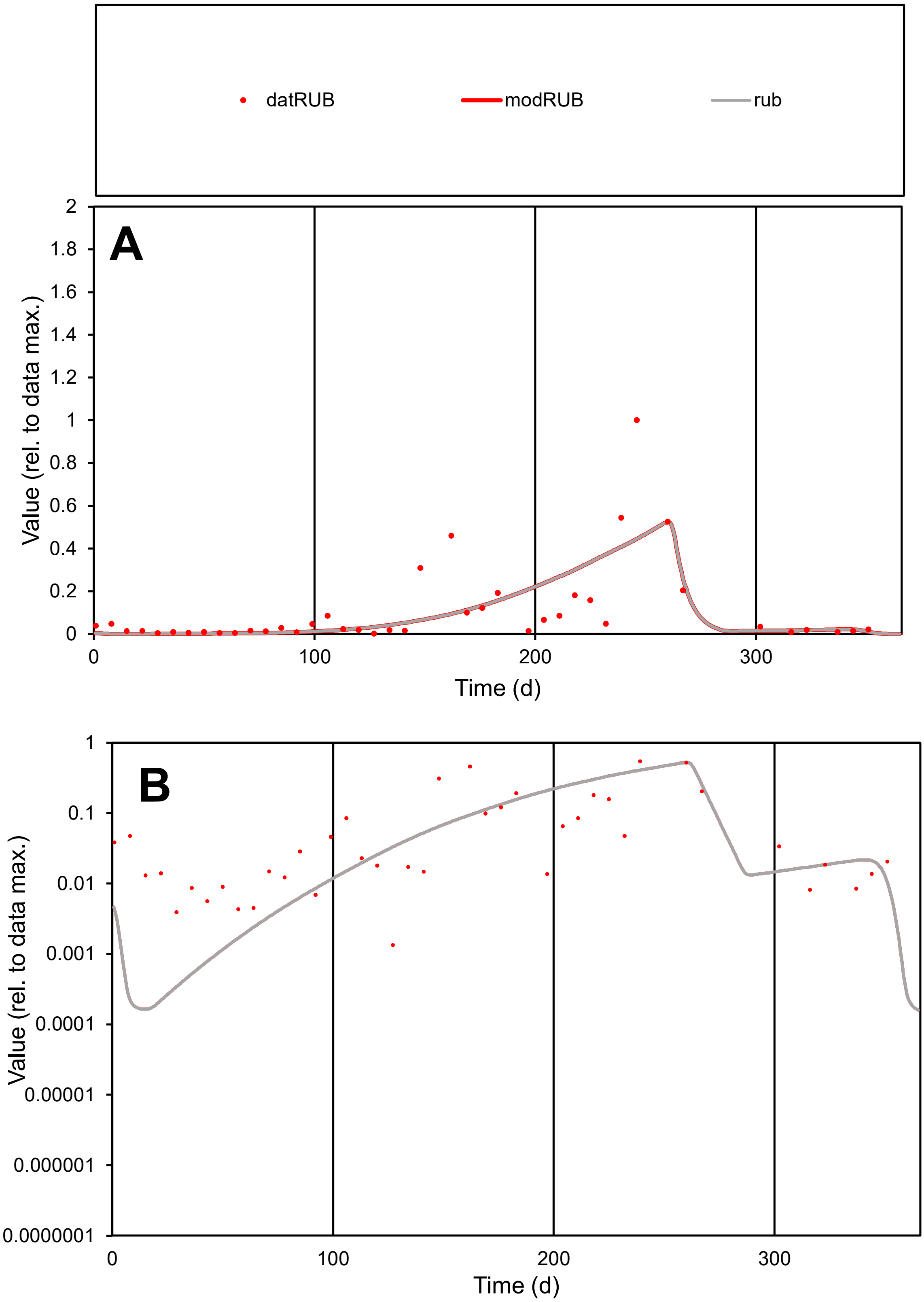


**C1**

Variables
RUB Provide jtd or Dst
CalibOnly?
2 1 to reproduce of
oNormalize?
2 1=no, 2=yes

ofxn calc*
1.26606
ofxn mod
1.28467

sRun
RunA1
zout
1

Ave
Data
0.00341
Model
0.00321

data/vMax
0.03168

ofxng
319.391

Min d/m
8.04883

ave/min d
80.2349
ave/min m
608.426

330_Calib_S_A

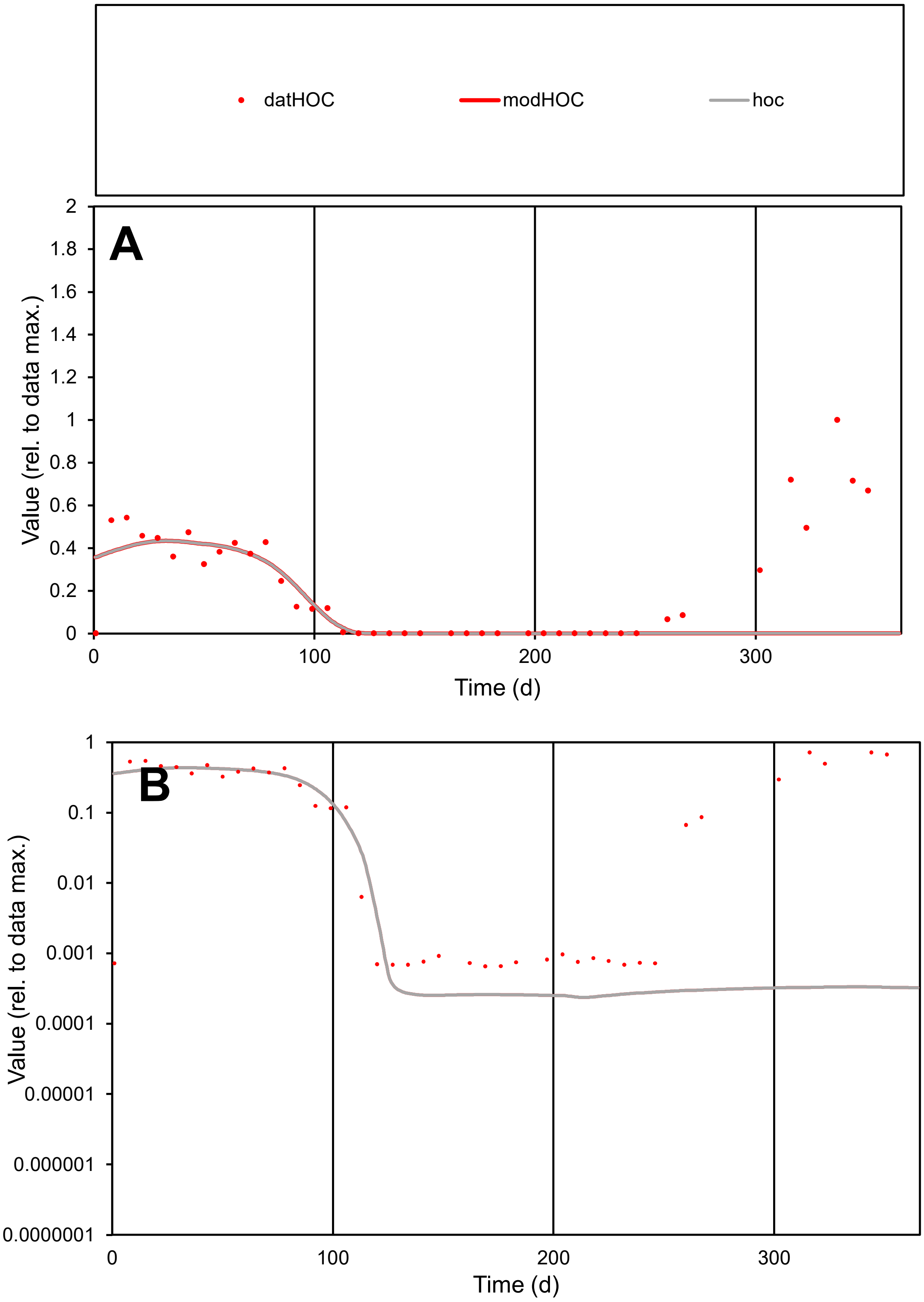


**C1**

Variables

HOC Provide jtd or Dst

CalibOnly?

2 1 to reproduce of

oNormalize?

2 1=no, 2=yes

ofxn calc*

1.19644

ofxn mod

1.20299

sRun

RunA1

zout

1

Ave

Data

0.00436

Model

0.00254

data/vMax

0.01942

ofxng

319.391

Min d/m

2.75054

ave/min d

343.612

ave/min m

551.74

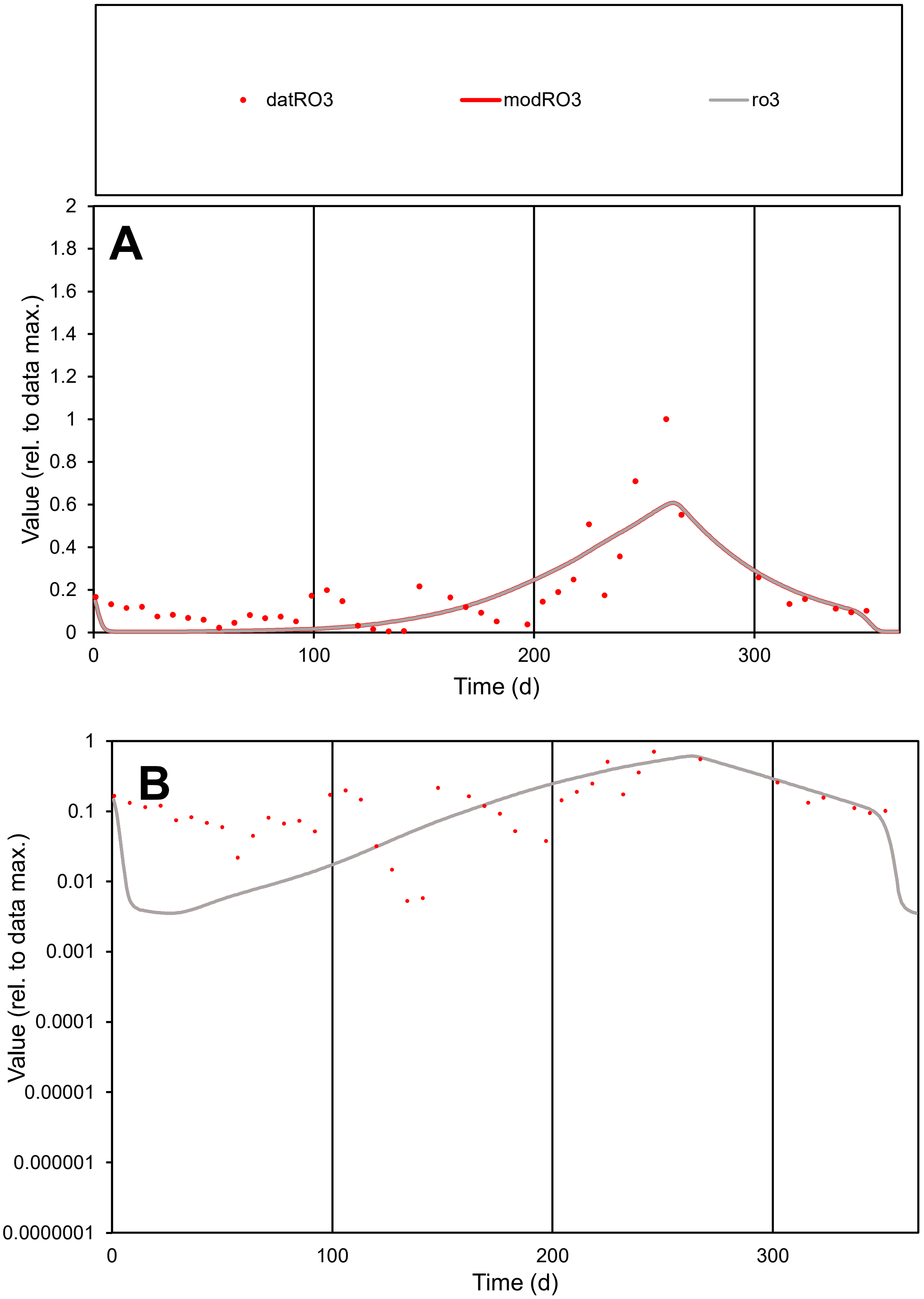


**C1**

Variables
RO3 Provide jtd or Dst
CalibOnly?
2 1 to reproduce of
oNormalize?
2 1=no, 2=yes

ofxn calc*
0.68176
ofxn mod
0.70021

sRun
RunA1
zout
1

Ave
Data
0.00331
Model
0.00285

data/vMax
0.01945

ofxng
319.391

Min d/m
1.48508

ave/min d
32.2849
ave/min m
41.2852

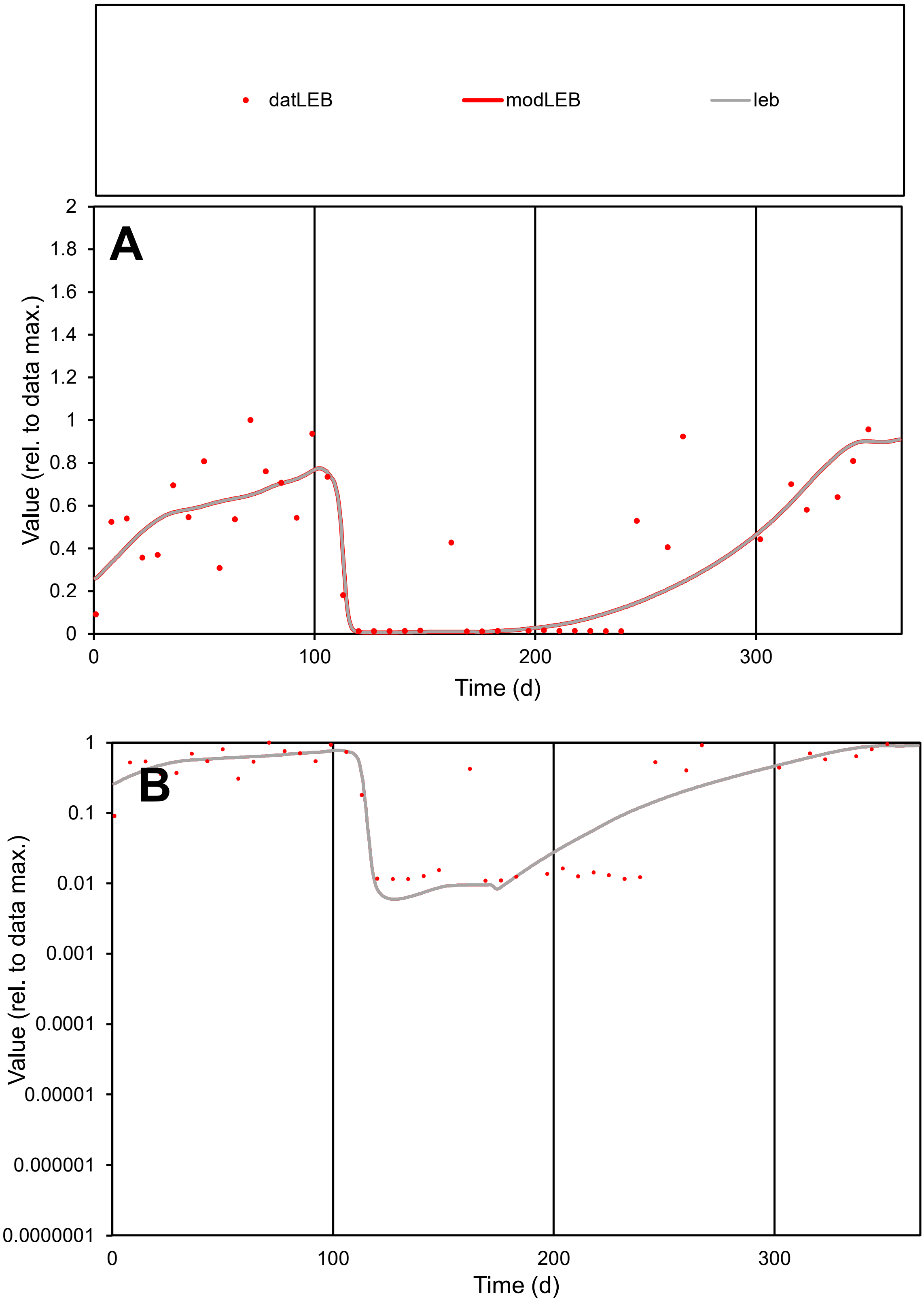


**C1**

Variables
LEB Provide jtd or Dst
CalibOnly?
2 1 to reproduce of
oNormalize?
2 1=no, 2=yes

ofxn calc*
0.47014
ofxn mod
0.48799

sRun
RunA1
zout
1

Ave
Data
0.00249
Model
0.00232

data/vMax
0.00643

ofxng
319.391

Min d/m
1.82692

ave/min d
35.3733
ave/min m
60.3955

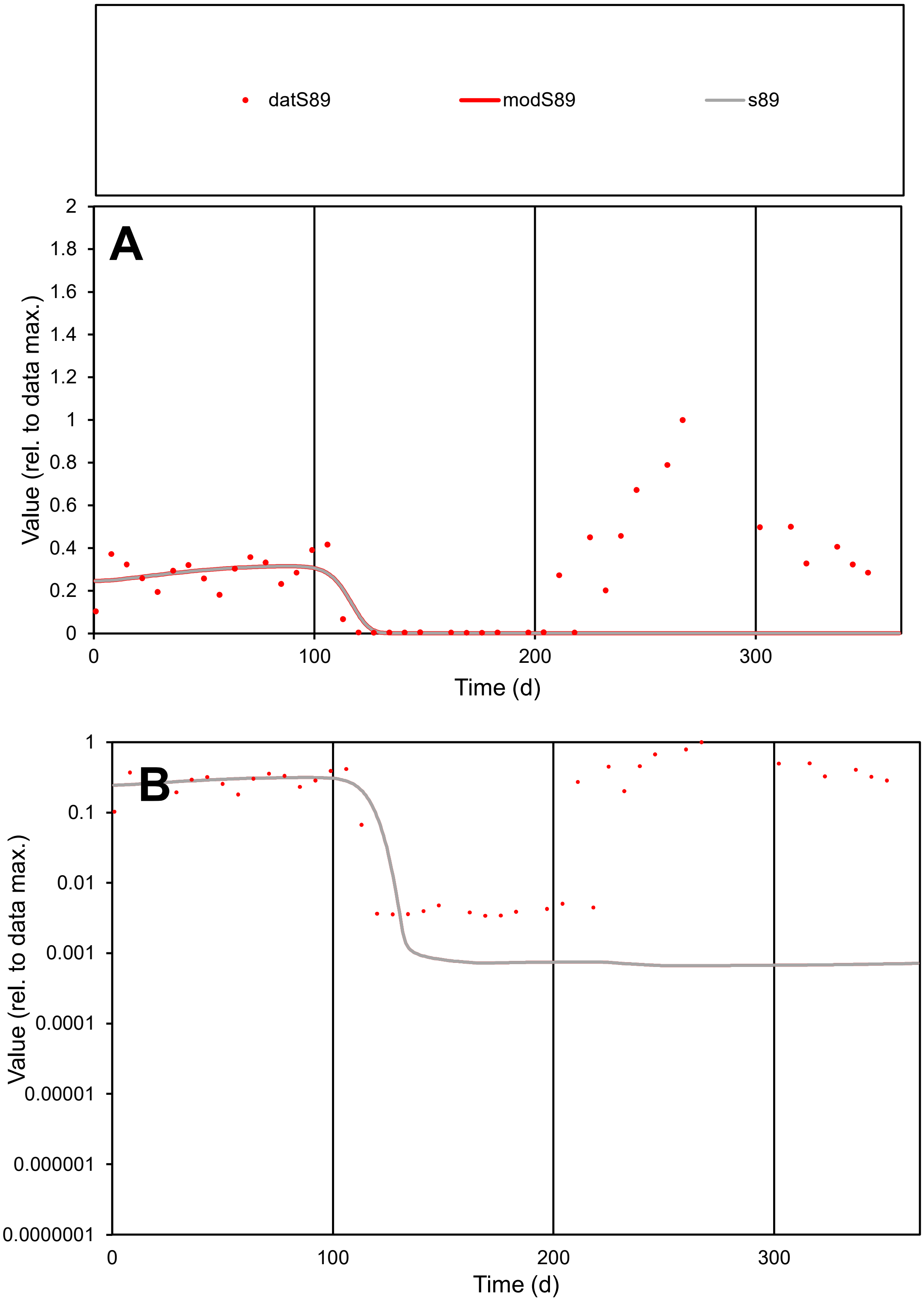


**C1**

| | |
|---|---|
| Variables | |
| S89 | Provide jtd or Dst |
| CalibOnly? | |
| 2 | 1 to reproduce of |
| oNormalize? | |
| 2 | 1=no, 2=yes |
| ofxn calc* | |
| 1.13629 | |
| ofxn mod | |
| 1.14121 | |
| sRun | |
| RunA1 | |
| zout | |
| 1 | |
| Ave | |
| Data | |
| 0.00278 | |
| Model | |
| 0.00123 | |
| data/vMax | |
| 0.01069 | |
| ofxng | |
| 319.391 | |
| Min d/m | |
| 5.09969 | |
| ave/min d | |
| 76.4223 | |
| ave/min m | |
| 172.937 | |

330_Calib_S_A

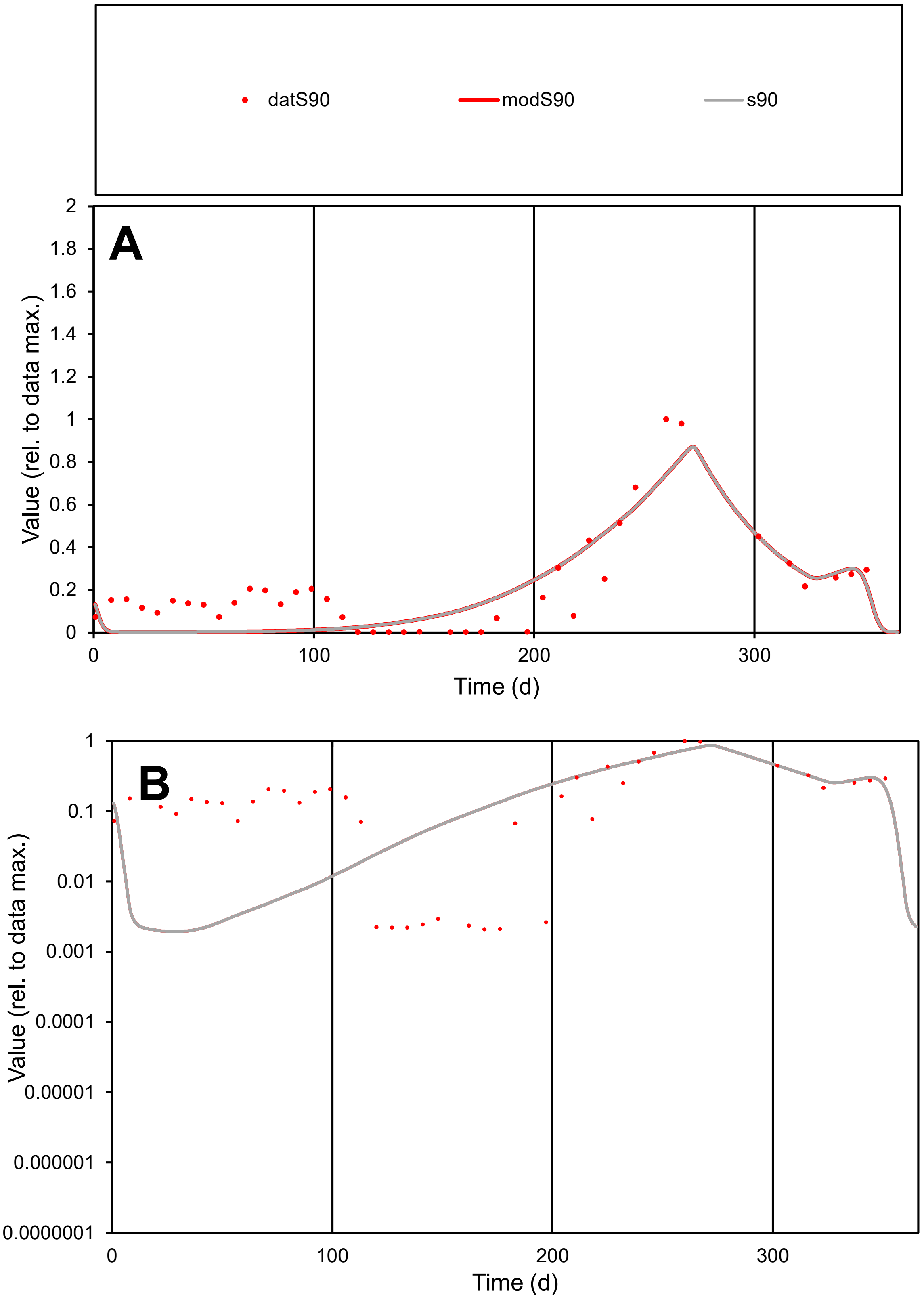


**C1**

Variables
S90 Provide jtd or Dst
CalibOnly?
2 1 to reproduce of
oNormalize?
2 1=no, 2=yes

ofxn calc*
0.6144
ofxn mod
0.63613

sRun
RunA1
zout
1

Ave
Data
0.00198
Model
0.00172

data/vMax
0.00959

ofxng
319.391

Min d/m
1.07953

ave/min d
98.5909
ave/min m
92.1832

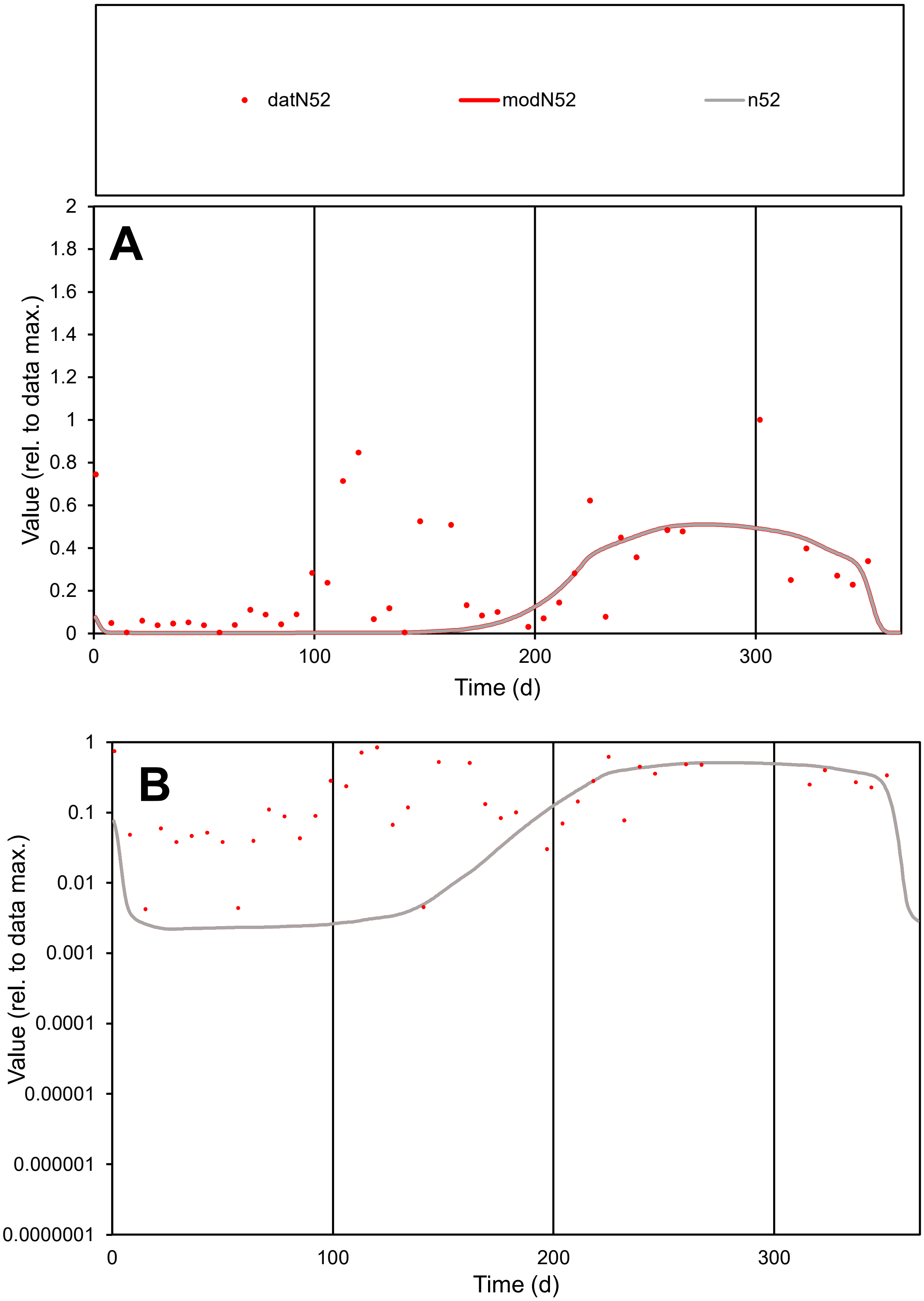


**C1**

Variables
N52 Provide jtd or Dst
CalibOnly?
2 1 to reproduce of
oNormalize?
2 1=no, 2=yes

ofxn calc*
1.05977
ofxn mod
1.06654

sRun
RunA1
zout
1

Ave
Data
0.00155
Model
0.00089

data/vMax
0.00622

ofxng
319.391

Min d/m
1.91232

ave/min d
58.922
ave/min m
64.3855

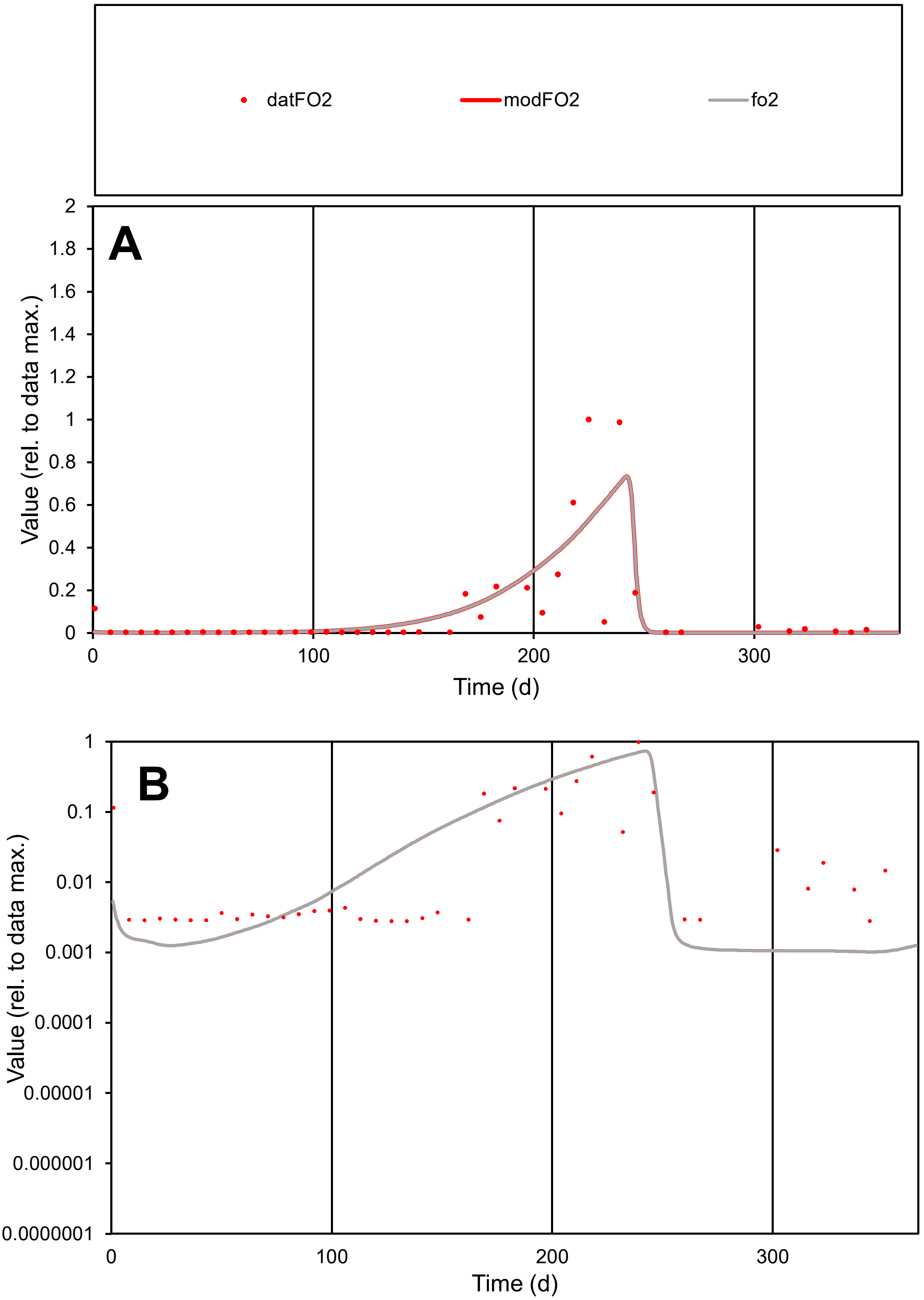


**C1**

Variables
FO2 Provide jtd or Dst
CalibOnly?
2 1 to reproduce of
oNormalize?
2 1=no, 2=yes

ofxn calc*
1.33176
ofxn mod
1.36789

sRun
RunA1
zout
1

Ave
Data
0.00255
Model
0.00264

data/vMax
0.02566

ofxng
319.391

Min d/m
2.72137

ave/min d
35.7057
ave/min m
100.949

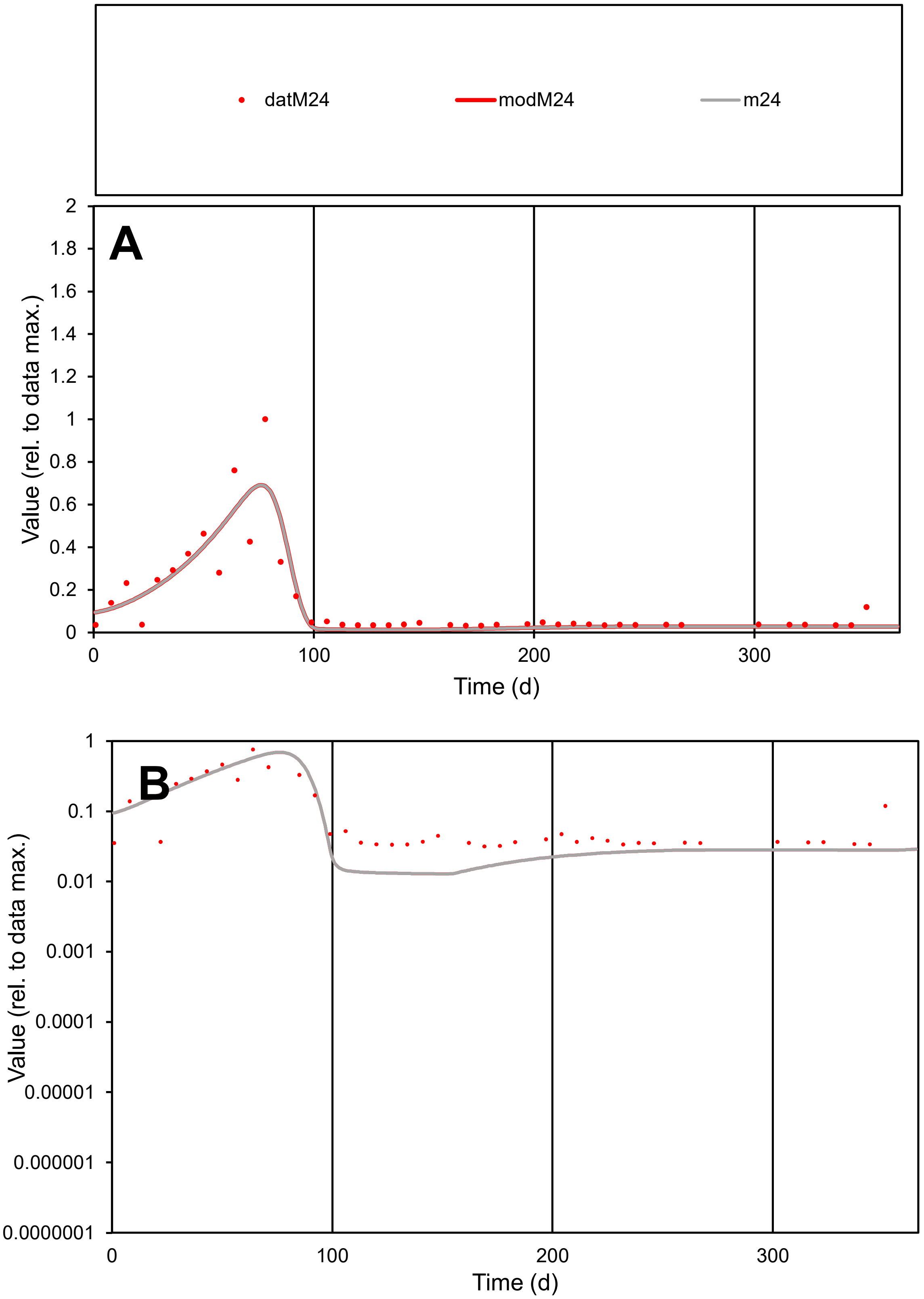


**C1**

Variables
M24 Provide jtd or Dst
CalibOnly?
2 1 to reproduce of
oNormalize?
2 1=no, 2=yes

ofxn calc*
0.60761
ofxn mod
0.63987

sRun
RunA1
zout
1

Ave
Data
0.00032
Model
0.0003

data/vMax
0.00229

ofxng
319.391

Min d/m
2.47955

ave/min d
4.43257
ave/min m
10.2466

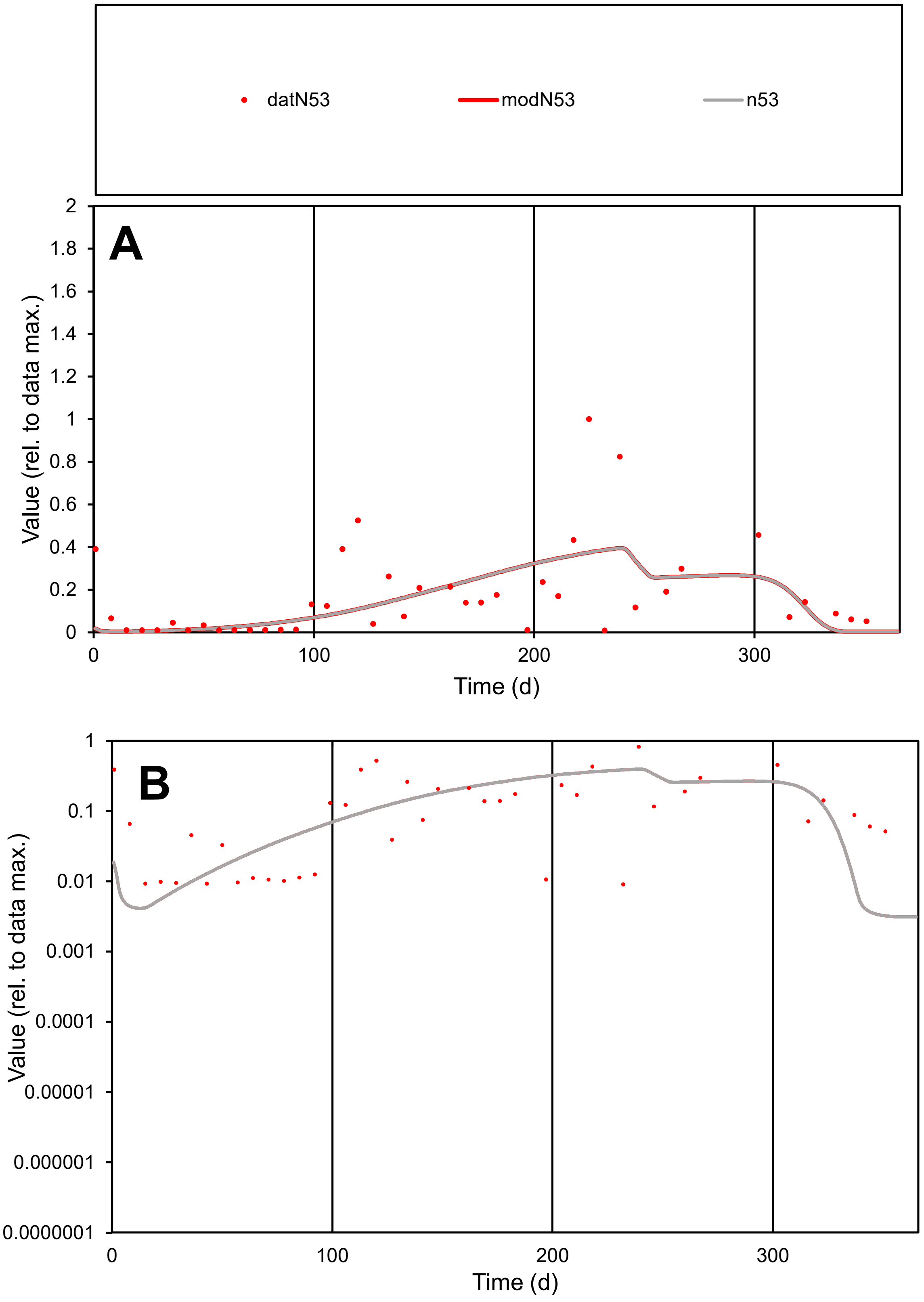


**C1**

Variables
N53 Provide jtd or Dst
CalibOnly?
2 1 to reproduce of
oNormalize?
2 1=no, 2=yes

ofxn calc*
1.07845
ofxn mod
1.09205

sRun
RunA1
zout
1

Ave
Data
0.00074
Model
0.00064

data/vMax
0.00432

ofxng
319.391

Min d/m
2.81962

ave/min d
19.0072
ave/min m
46.241

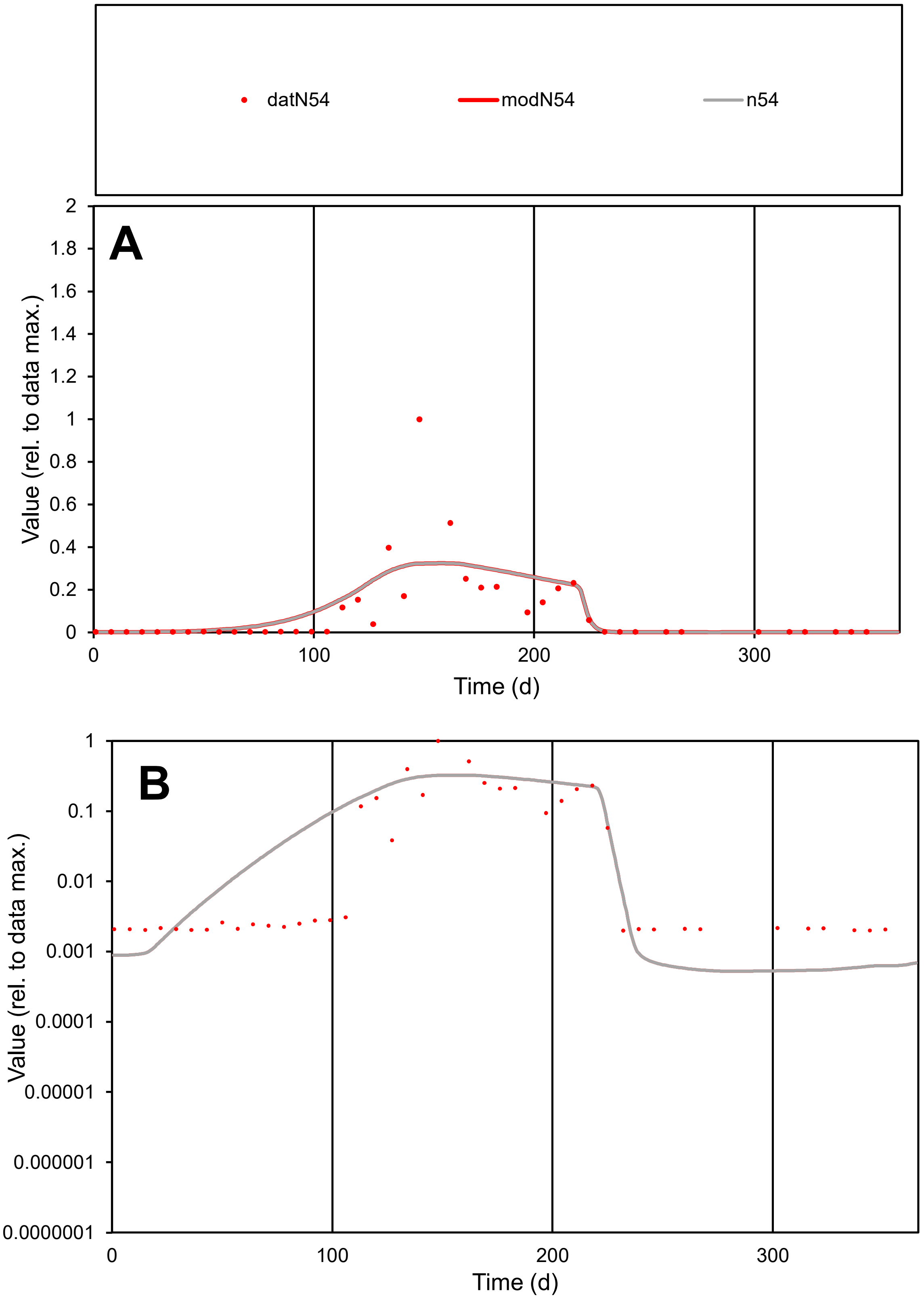


**C1**

Variables
N54 Provide jtd or Dst
CalibOnly?
2 1 to reproduce of
oNormalize?
2 1=no, 2=yes

ofxn calc*
1.3723
ofxn mod
1.38793

sRun
RunA1
zout
1

Ave
Data
0.00107
Model
0.00118

data/vMax
0.01164

ofxng
319.391

Min d/m
3.73486

ave/min d
46.1564
ave/min m
190.405

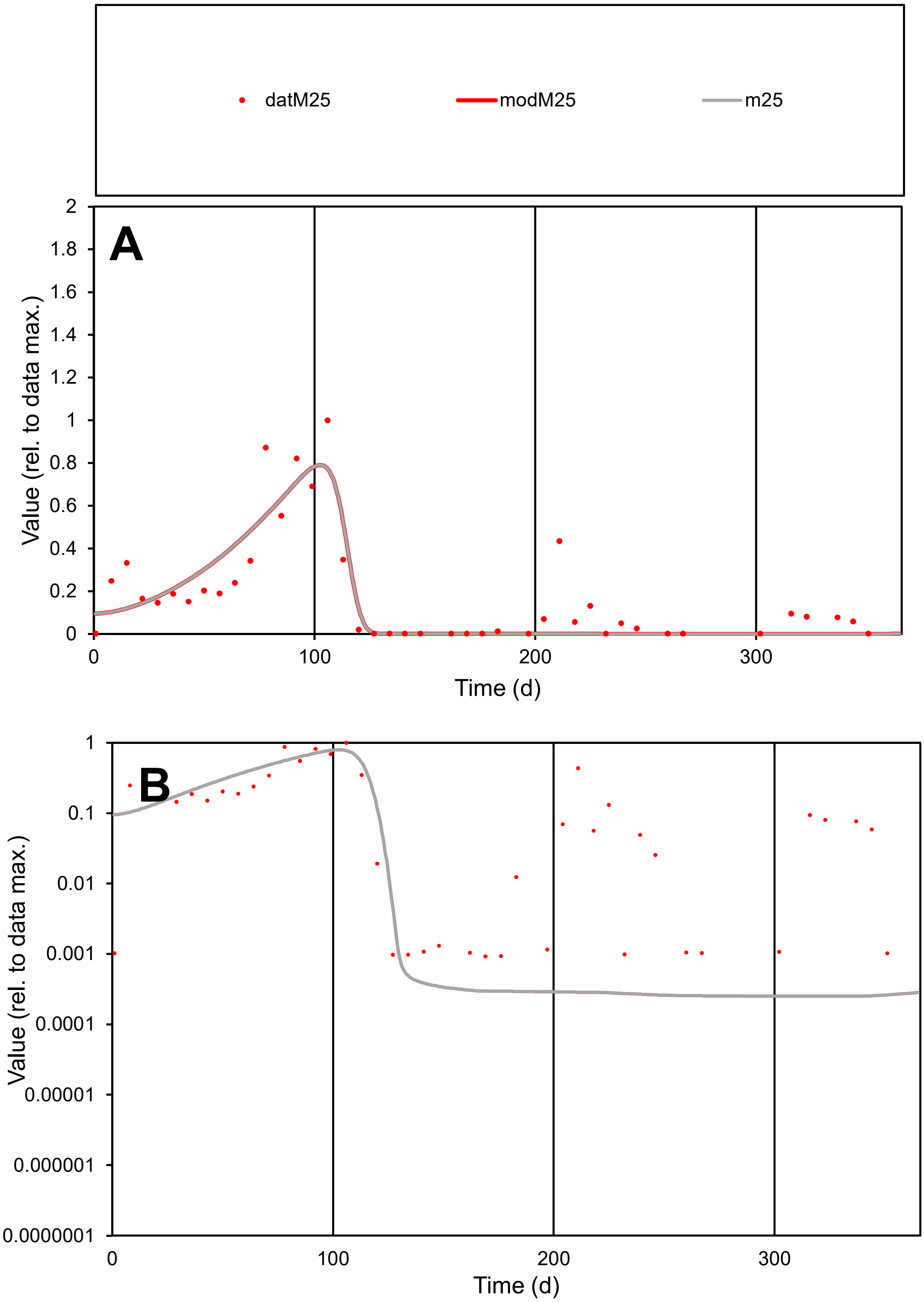


**C1**

Variables
M25 Provide jtd or Dst
CalibOnly?
2 1 to reproduce of
oNormalize?
2 1=no, 2=yes

ofxn calc*
0.67044
ofxn mod
0.6958

sRun
RunA1
zout
1

Ave
Data
0.00189
Model
0.00167

data/vMax
0.01046

ofxng
319.391

Min d/m
3.69656

ave/min d
195.596
ave/min m
637.24

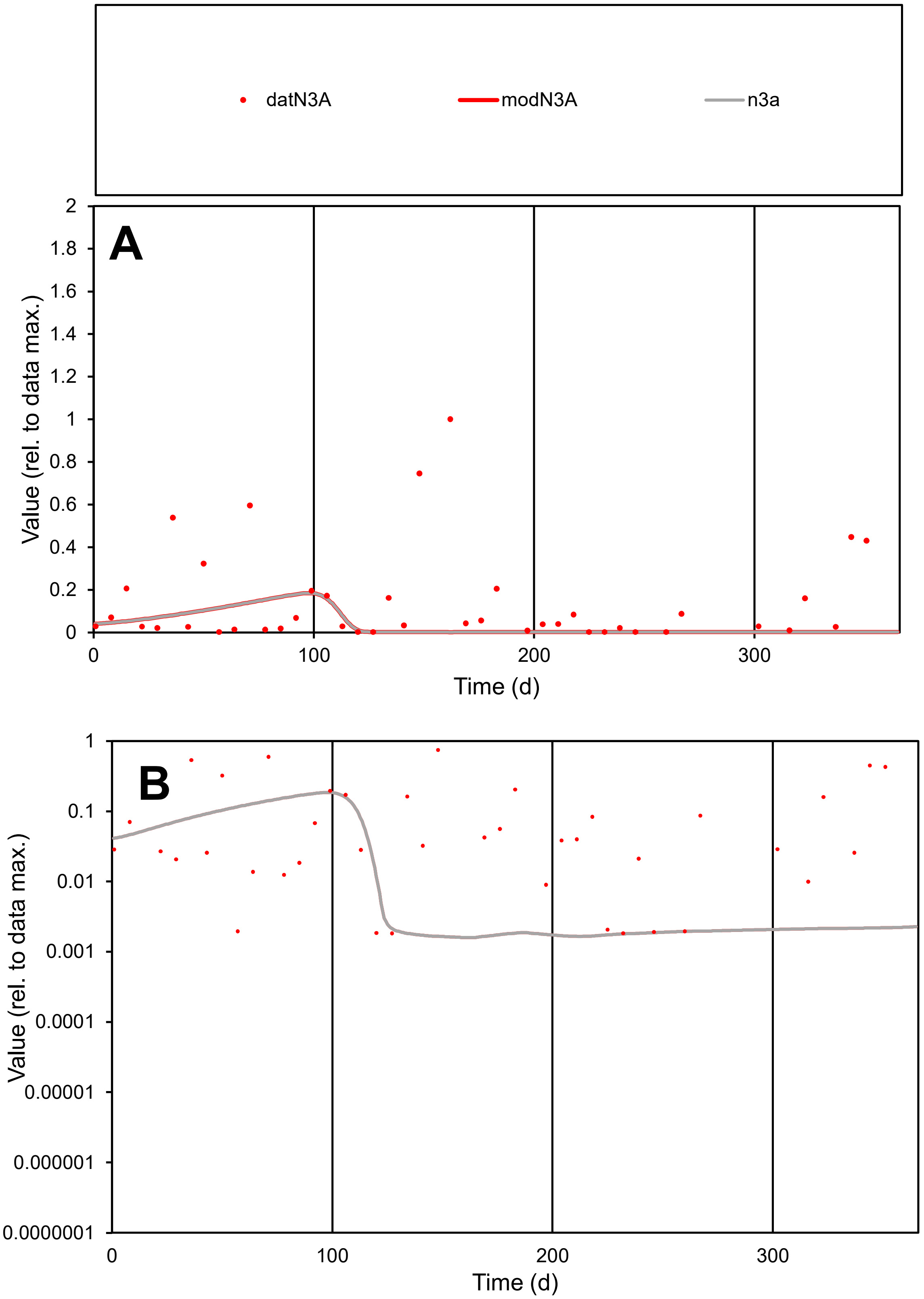


**C1**

Variables

N3A Provide jtd or Dst

CalibOnly?

2 1 to reproduce of

oNormalize?

2 1=no, 2=yes

ofxn calc*

1.75411

ofxn mod

1.75795

sRun

RunA1

zout

1

Ave

Data

0.00148

Model

0.00047

data/vMax

0.01041

ofxng

319.391

Min d/m

1.14864

ave/min d

77.9781

ave/min m

28.1045

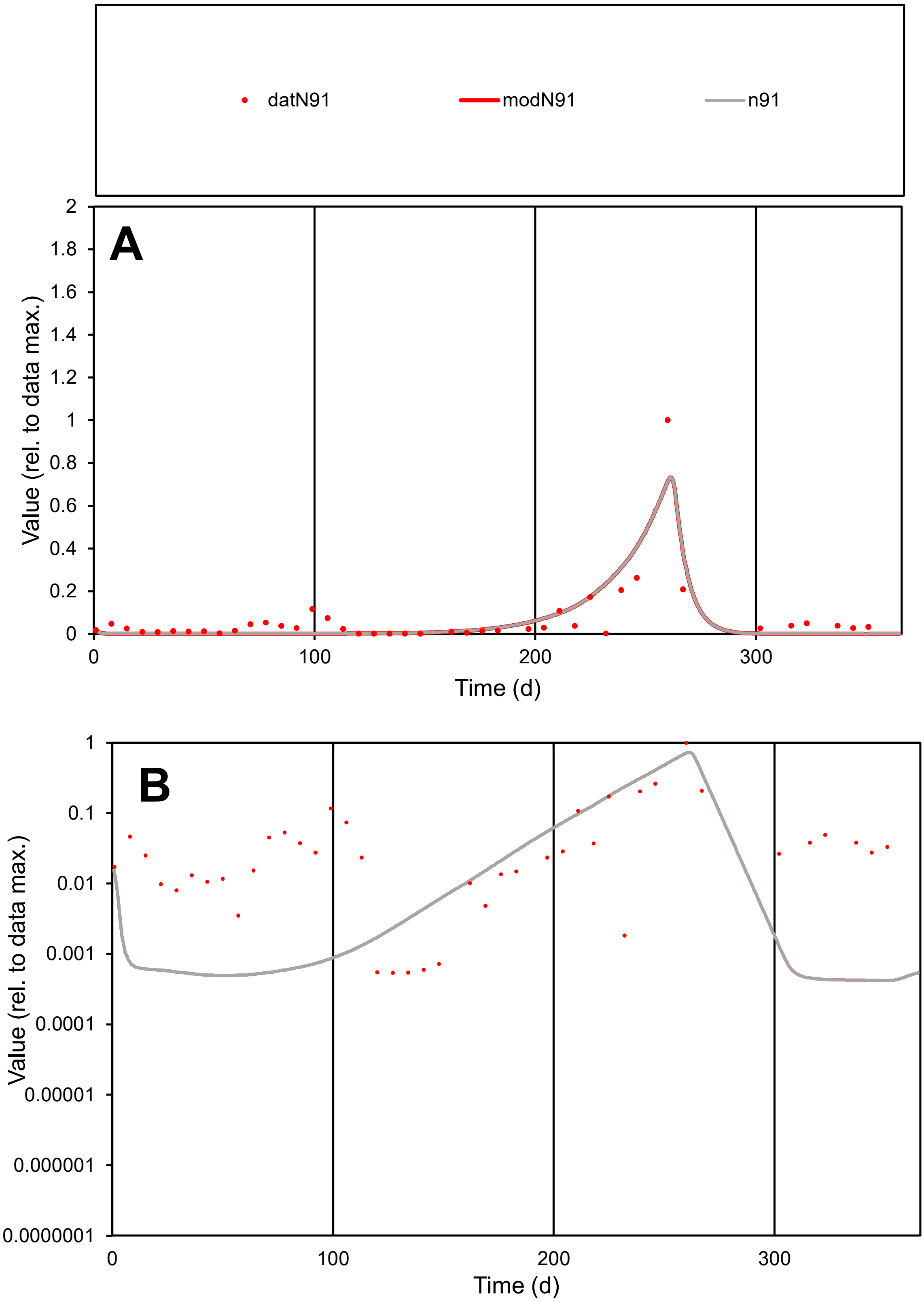


C1

Variables
N91 Provide jtd or Dst
CalibOnly?
2 1 to reproduce of
oNormalize?
2 1=no, 2=yes

ofxn calc*
1.06367
ofxn mod
1.10422

sRun
RunA1
zout
1

Ave
Data
0.0042
Model
0.00394

data/vMax
0.06206

ofxng
319.391

Min d/m
1.28725

ave/min d
125.335
ave/min m
151.35

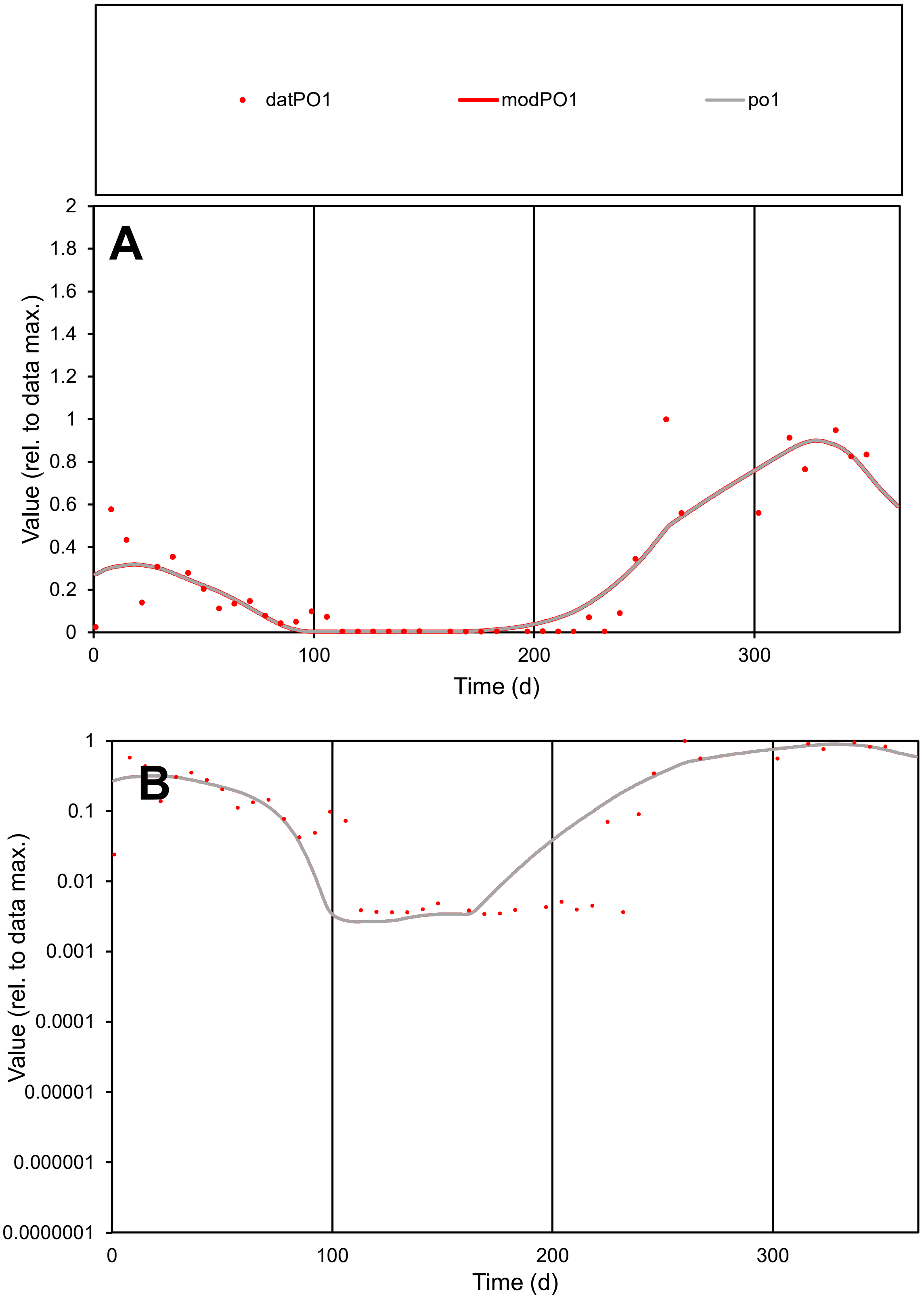


**C1**

Variables
PO1 Provide jtd or Dst
CalibOnly?
2 1 to reproduce of
oNormalize?
2 1=no, 2=yes

ofxn calc*
0.50908
ofxn mod
0.53147

sRun
RunA1
zout
1

Ave
Data
0.00319
Model
0.00318

data/vMax
0.01336

ofxng
319.391

Min d/m
1.28663

ave/min d
69.4367
ave/min m
89.1613

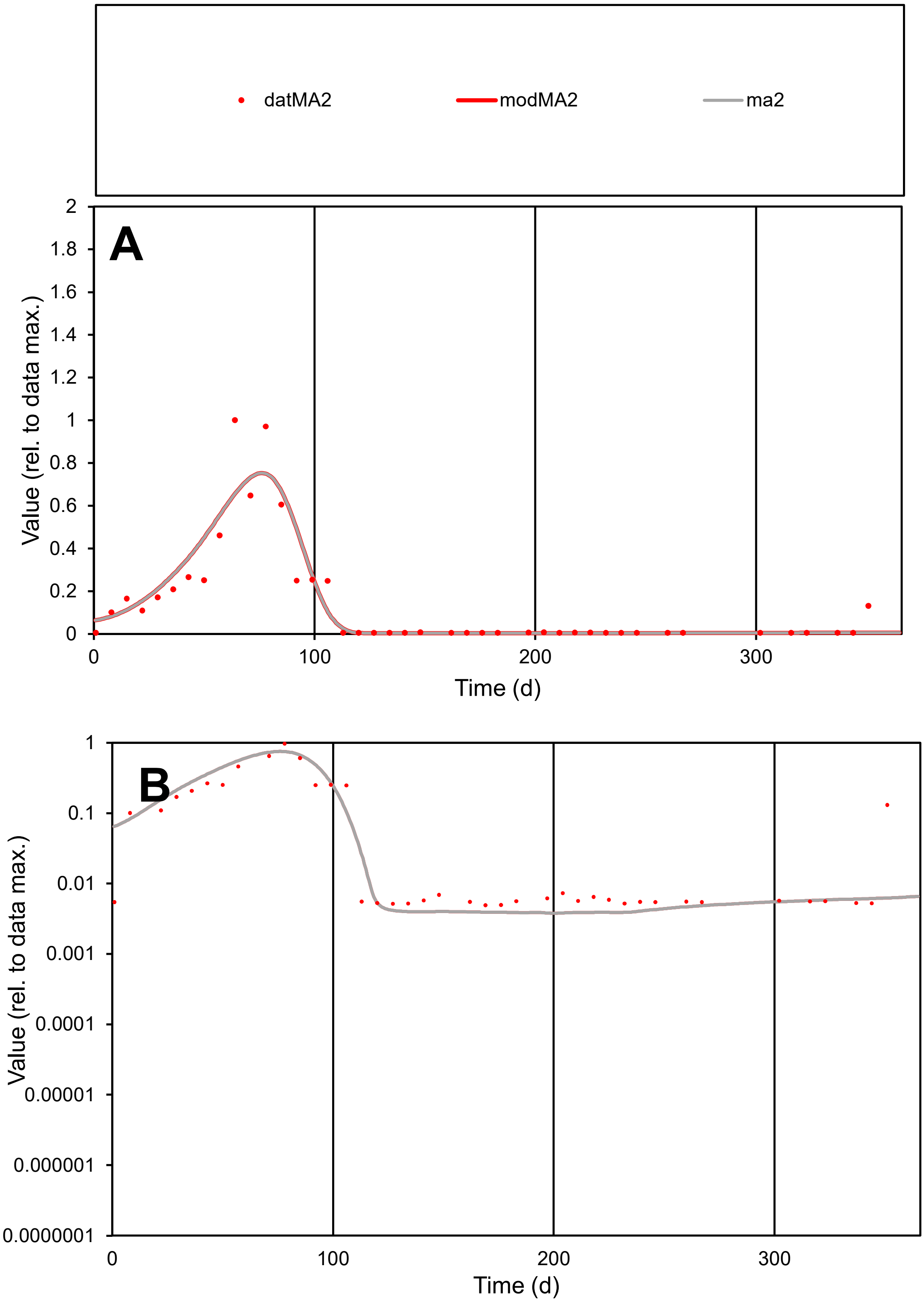


**C1**

Variables
MA2 Provide jtd or Dst
CalibOnly?
2 1 to reproduce of
oNormalize?
2 1=no, 2=yes

ofxn calc*
0.62013
ofxn mod
0.66125

sRun
RunA1
zout
1

Ave
Data
0.00052
Model
0.00053

data/vMax
0.00365

ofxng
319.391

Min d/m
1.29654

ave/min d
28.9702
ave/min m
38.0192

330_Calib_S_A

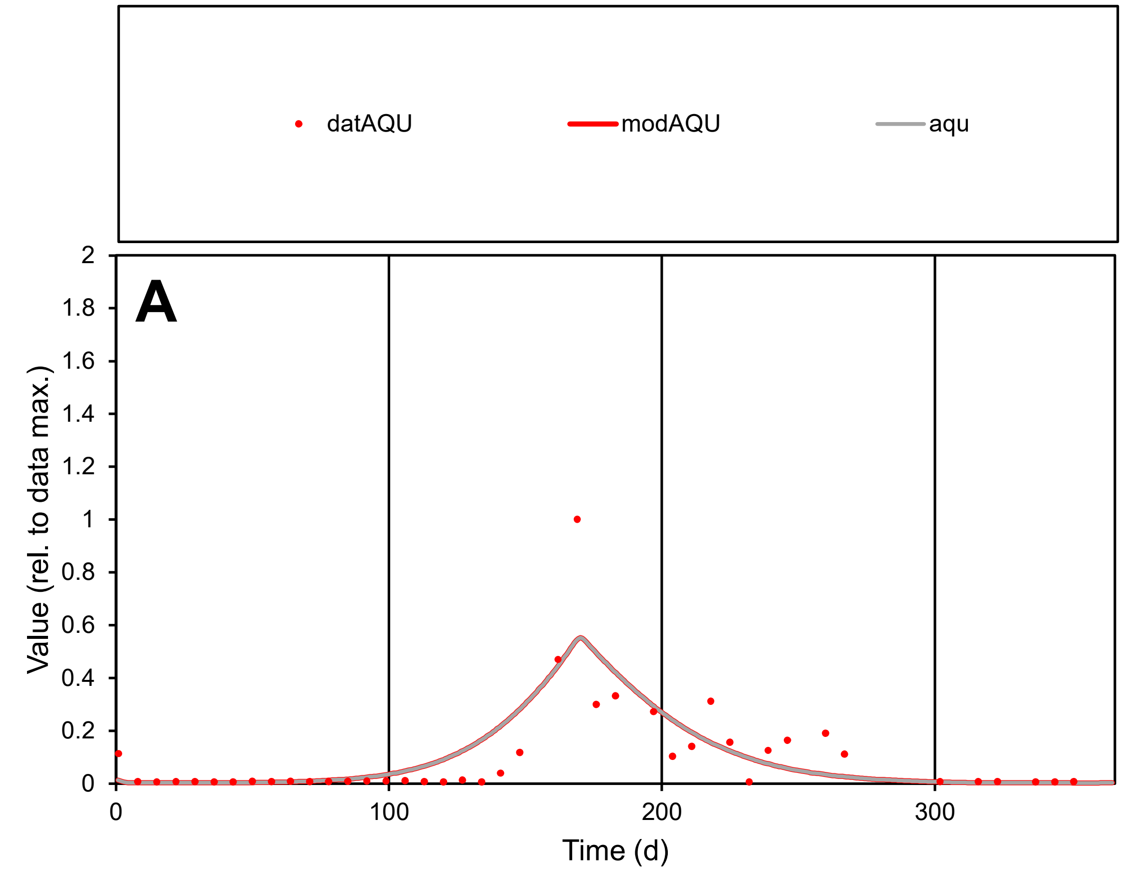


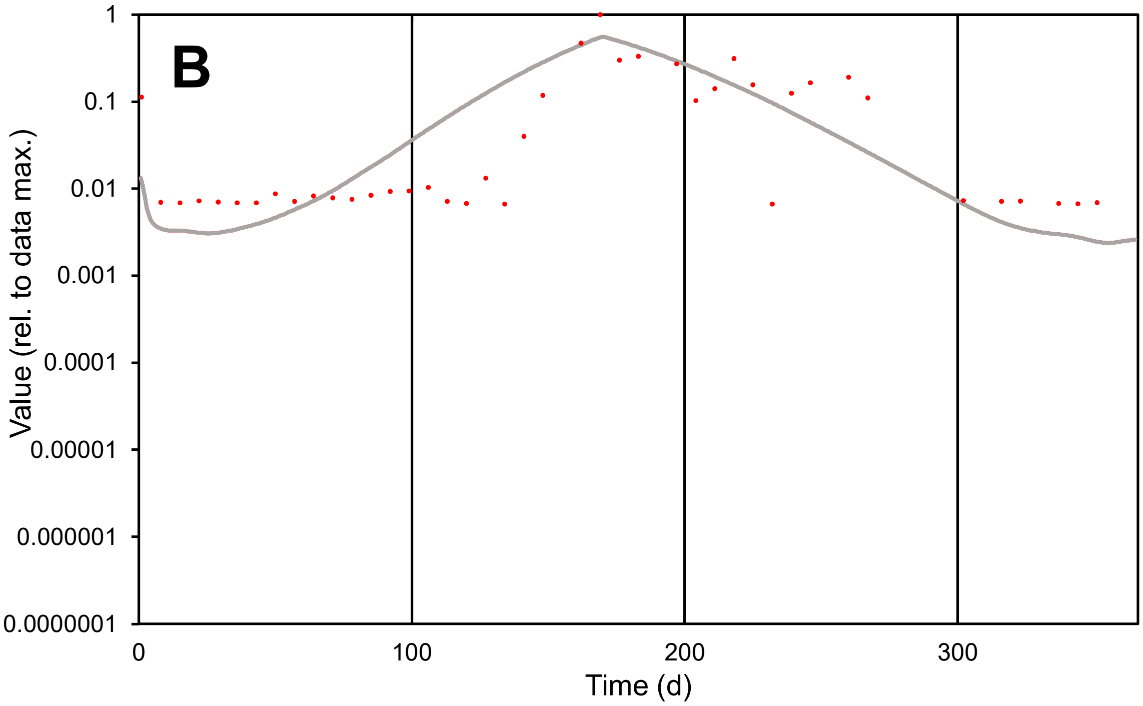


**C1**

Variables
AQU Provide jtd or Dst
CalibOnly?
2 1 to reproduce of
oNormalize?
2 1=no, 2=yes

ofxn calc*
1.0537
ofxn mod
1.08005

sRun
RunA1
zout
1

Ave
Data
0.00206
Model
0.00214

data/vMax
0.02081

ofxng
319.391

Min d/m
2.74525
ave/min d
14.8205
ave/min m
42.3596

330_Calib_S_A

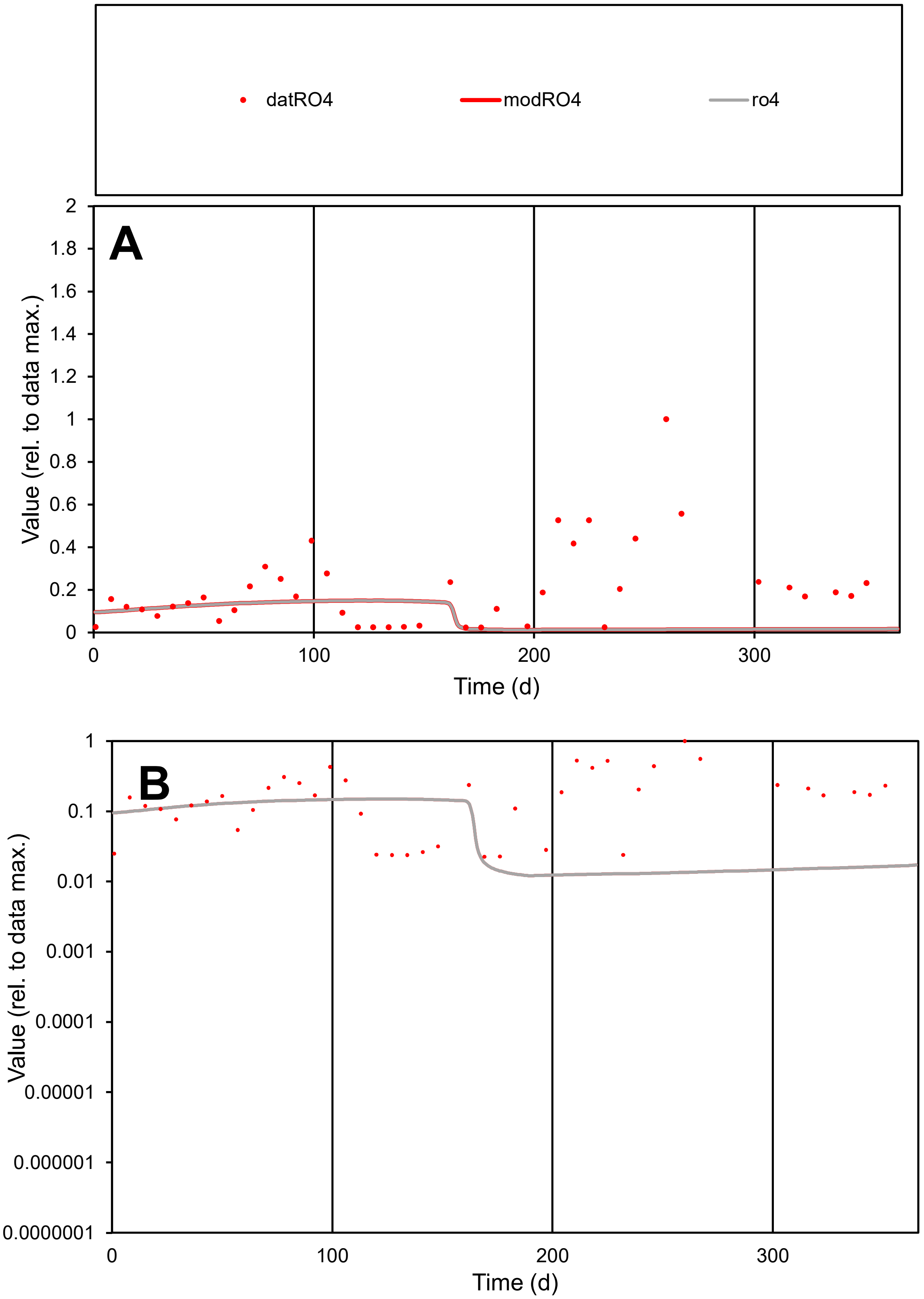


**C1**

Variables
RO4 Provide jtd or Dst
CalibOnly?
2 1 to reproduce of
oNormalize?
2 1=no, 2=yes

ofxn calc*
1.25577
ofxn mod
1.25919

sRun
RunA1
zout
1

Ave
Data
0.00196
Model
0.00077

data/vMax
0.00973

ofxng
319.391

Min d/m
1.82721

ave/min d
8.93566
ave/min m
6.42742

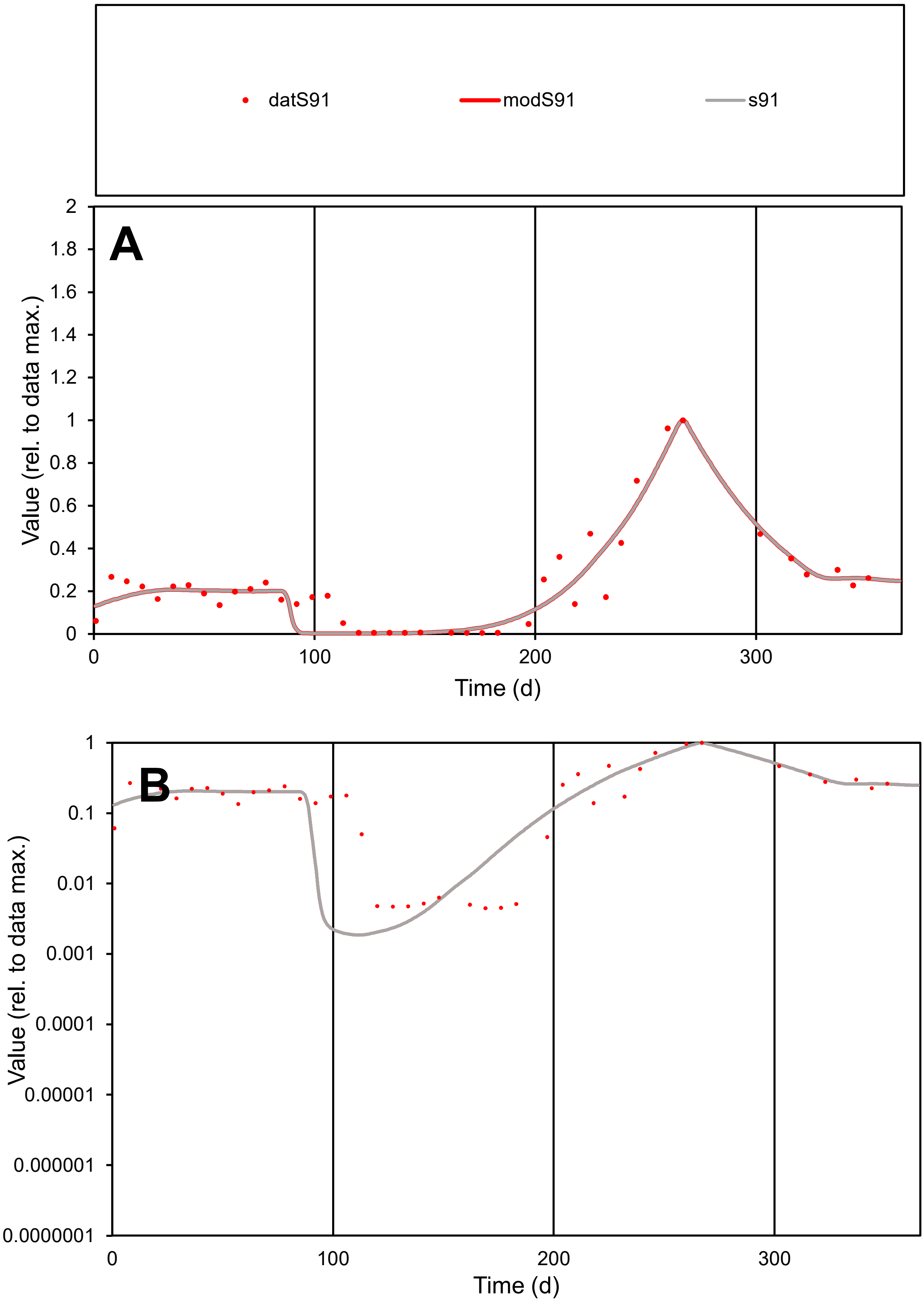


**C1**

Variables
S91 Provide jtd or Dst
CalibOnly?
2 1 to reproduce of
oNormalize?
2 1=no, 2=yes

ofxn calc*
0.34004
ofxn mod
0.37894

sRun
RunA1
zout
1

Ave
Data
0.00335
Model
0.00319

data/vMax
0.01474

ofxng
319.391

Min d/m
2.37987

ave/min d
50.8406
ave/min m
115.239

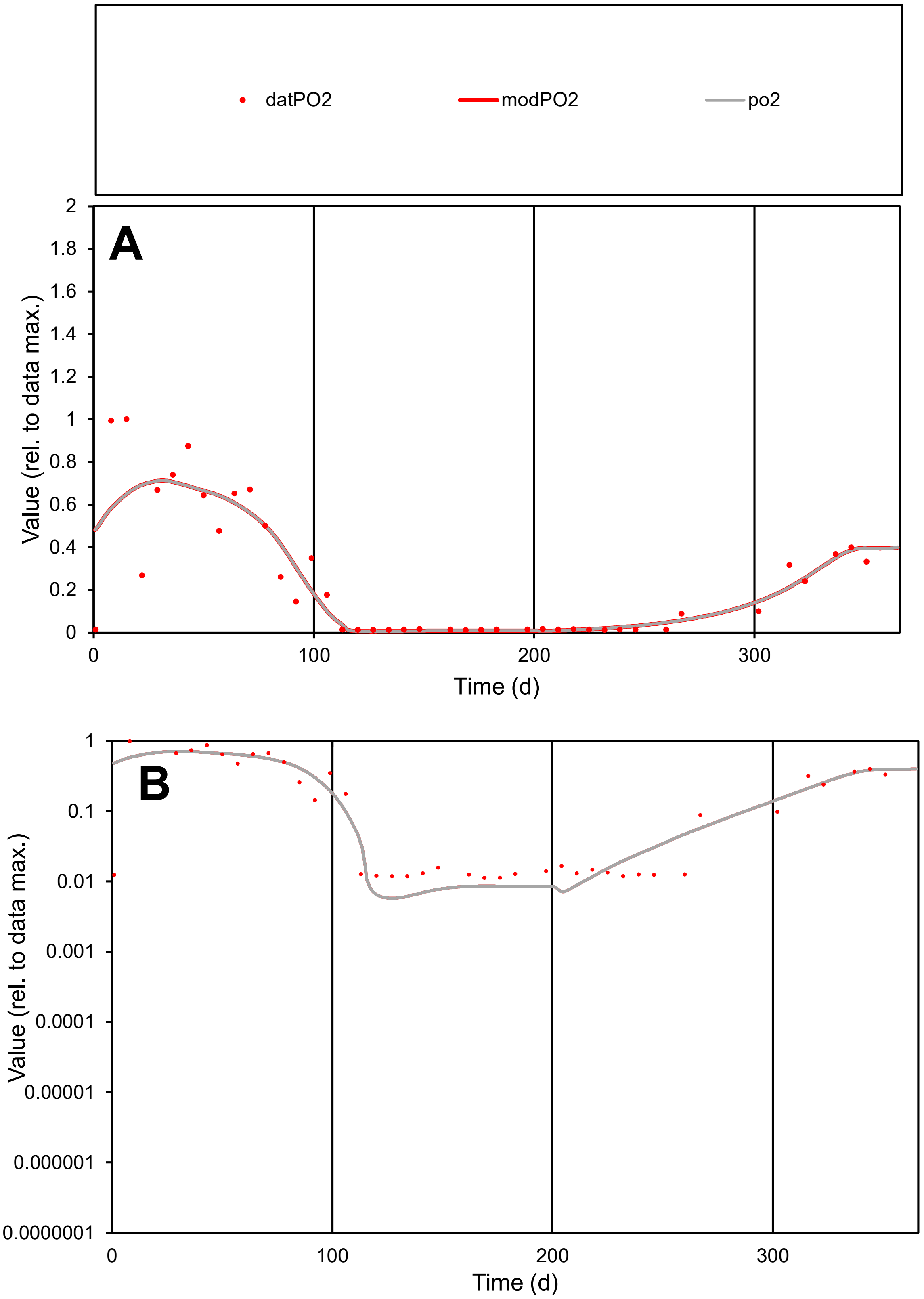


**C1**

Variables
PO2 Provide jtd or Dst
CalibOnly?
2 1 to reproduce of
oNormalize?
2 1=no, 2=yes

ofxn calc*
0.56963
ofxn mod
0.59503

sRun
RunA1
zout
1

Ave
Data
0.00178
Model
0.00179

data/vMax
0.00712

ofxng
319.391

Min d/m
1.94411

ave/min d
22.2766
ave/min m
43.4347

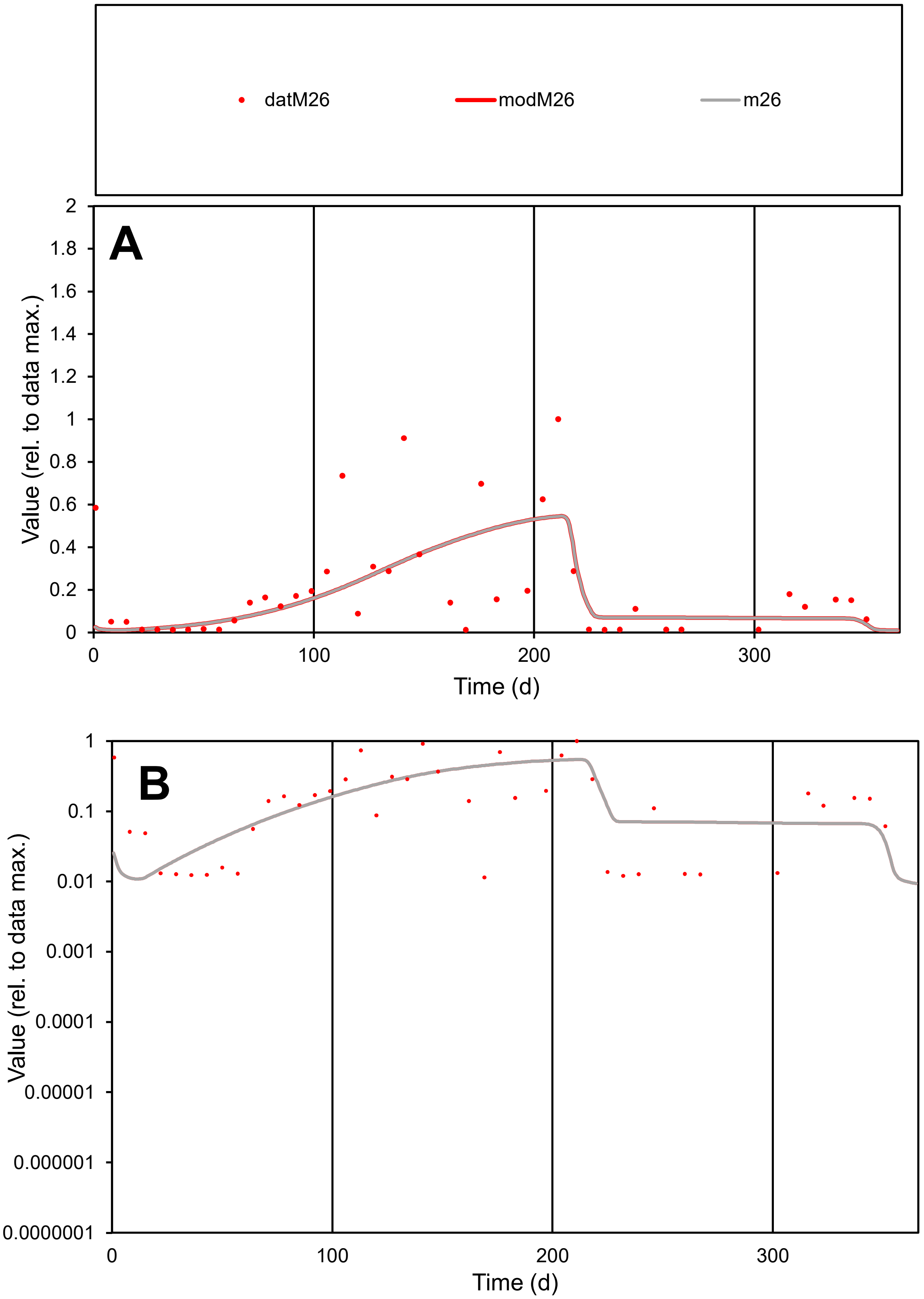


**C1**

Variables
M26 Provide jtd or Dst
CalibOnly?
2 1 to reproduce of
oNormalize?
2 1=no, 2=yes

ofxn calc*
1.00783
ofxn mod
1.02019

sRun
RunA1
zout
1

Ave
Data
0.00147
Model
0.00127

data/vMax
0.00723

ofxng
319.391

Min d/m
1.03495

ave/min d
17.8861
ave/min m
16.0241

330_Calib_S_A

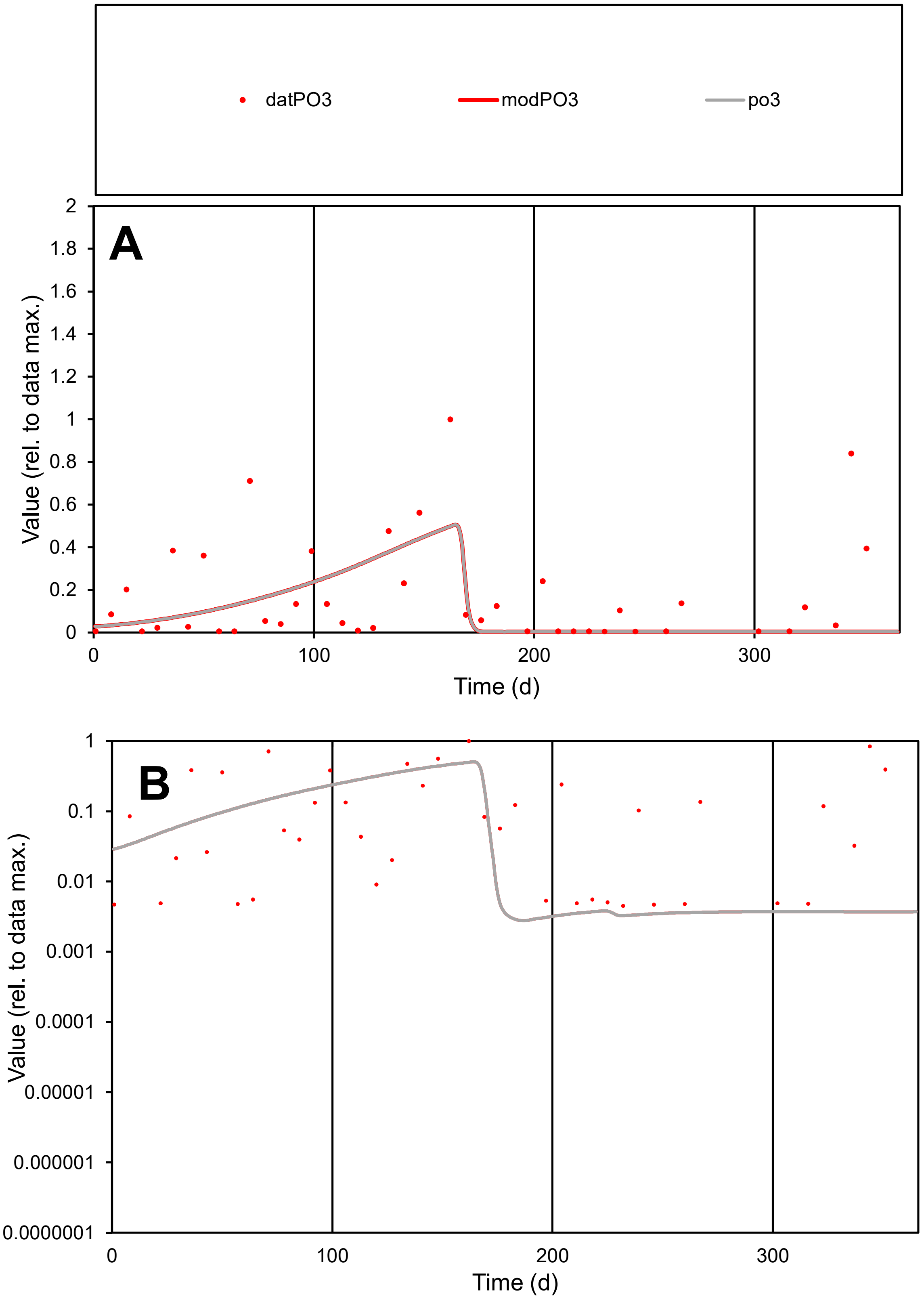


C1

Variables

PO3 Provide jtd or Dst

CalibOnly?

2 1 to reproduce of

oNormalize?

2 1=no, 2=yes

ofxn calc*

1.34526

ofxn mod

1.35458

sRun

RunA1

zout

1

Ave

Data

0.00186

Model

0.00127

data/vMax

0.01104

ofxng

319.391

Min d/m

1.60103

ave/min d

37.3487

ave/min m

41.0162

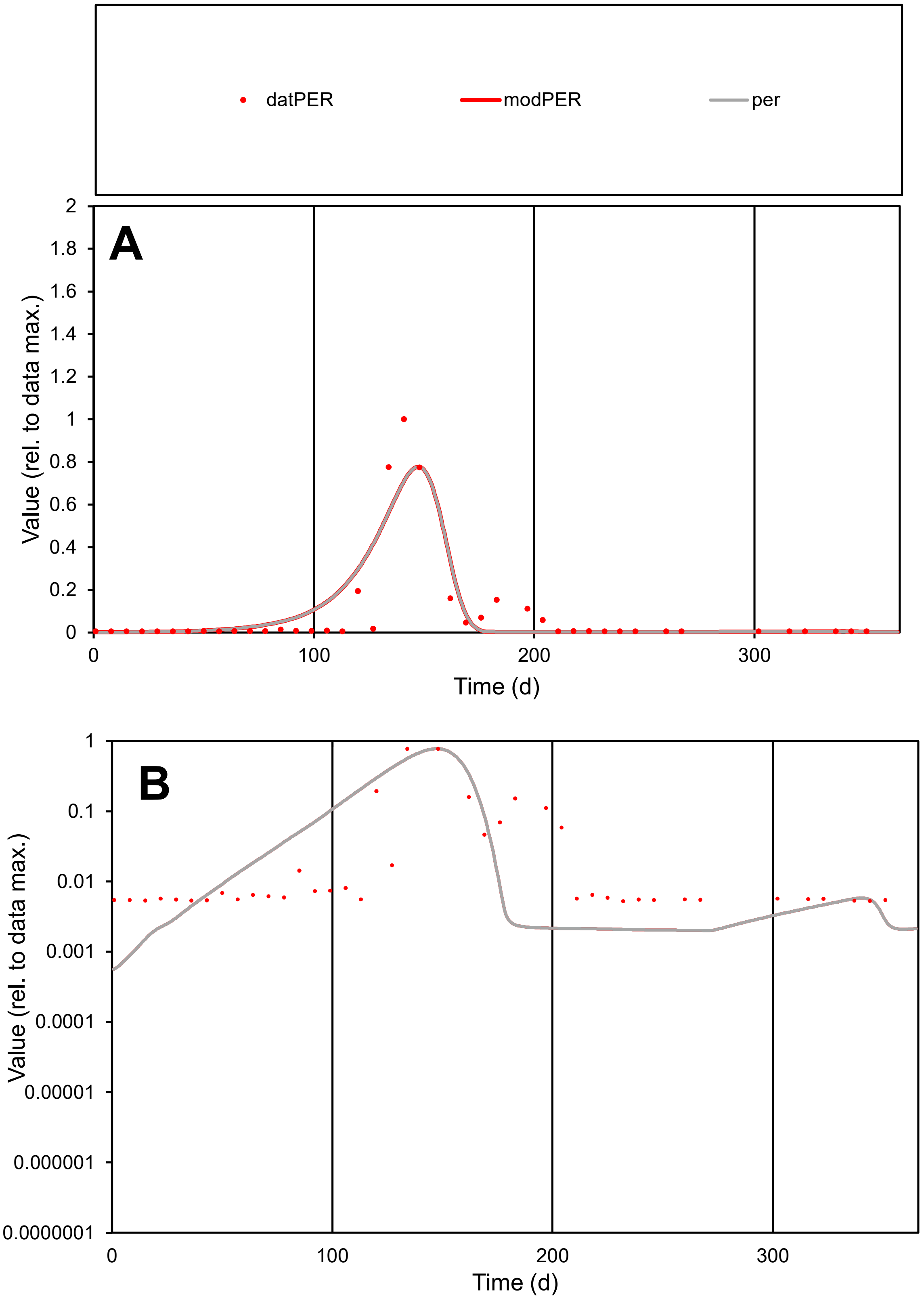


C1

Variables
PER Provide jtd or Dst
CalibOnly?
2 1 to reproduce of
oNormalize?
2 1=no, 2=yes

ofxn calc*
1.13654
ofxn mod
1.19571

sRun
RunA1
zout
1

Ave
Data
0.00032
Model
0.00034

data/vMax
0.00373

ofxng
319.391

Min d/m
8.76682

ave/min d
16.1149
ave/min m
154.013

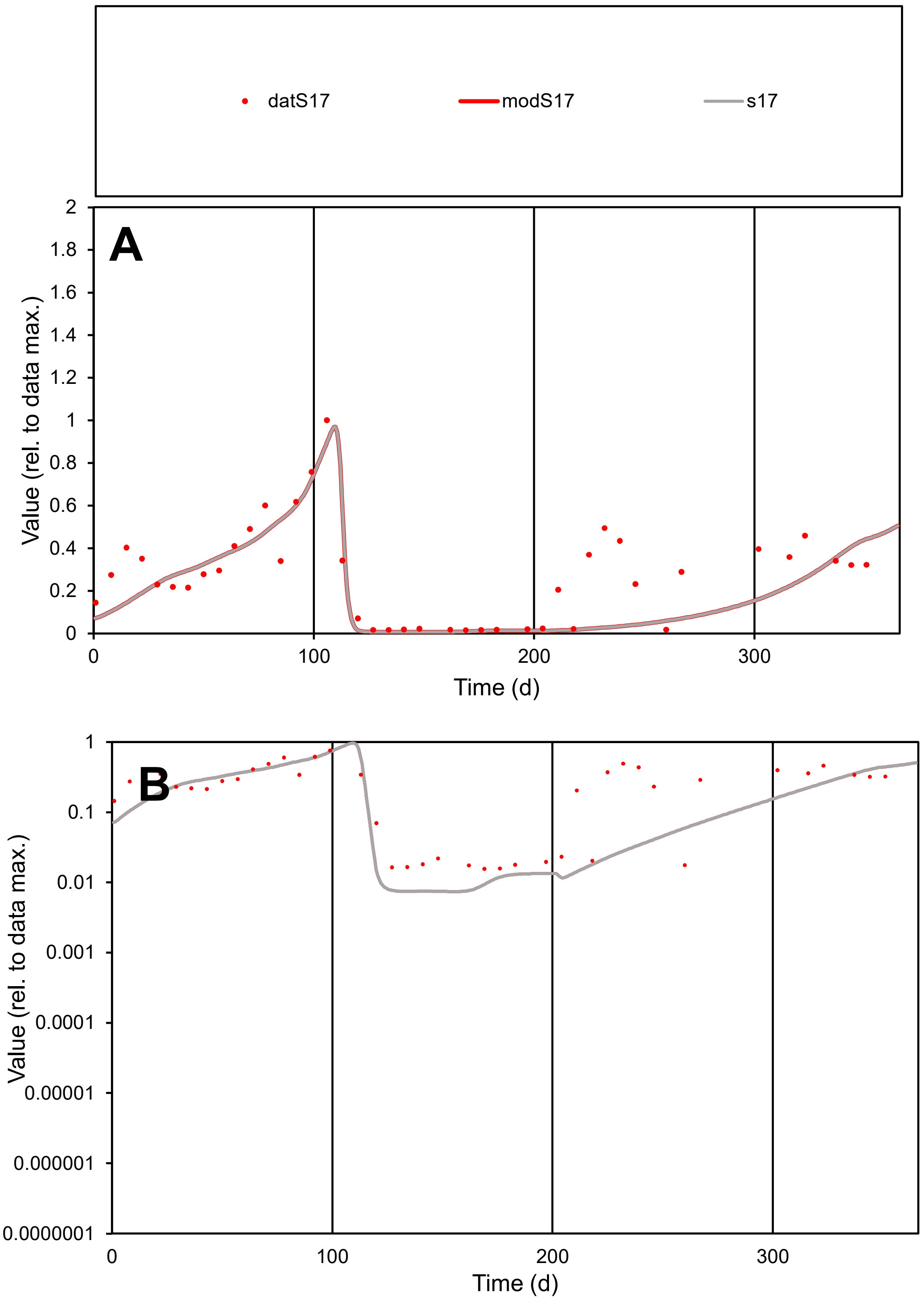


**C1**

Variables
S17 Provide jtd or Dst
CalibOnly?
2 1 to reproduce of
oNormalize?
2 1=no, 2=yes

ofxn calc*
0.5483
ofxn mod
0.56459

sRun
RunA1
zout
1

Ave
Data
0.00192
Model
0.00149

data/vMax
0.00701

ofxng
319.391

Min d/m
2.09489

ave/min d
17.4988
ave/min m
28.5309

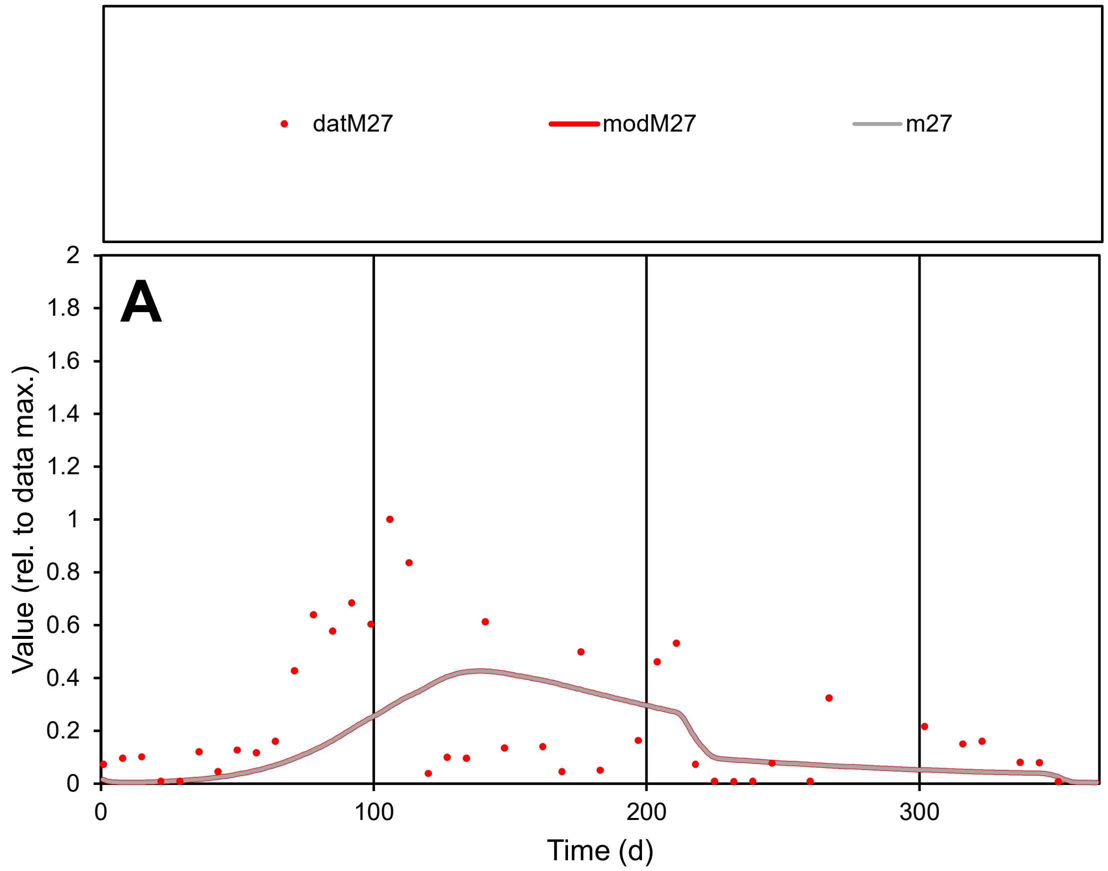


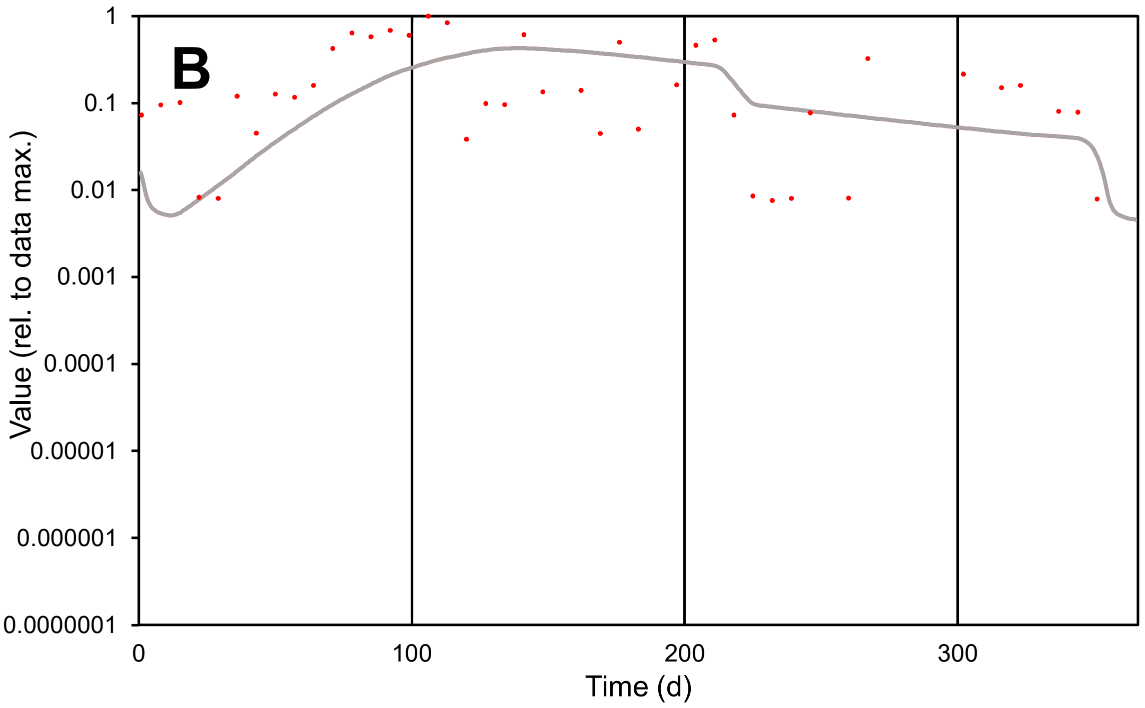


**C1**

Variables
M27 Provide jtd or Dst
CalibOnly?
2 1 to reproduce of
oNormalize?
2 1=no, 2=yes

ofxn calc*
1.08245
ofxn mod
1.08984

sRun
RunA1
zout
1

Ave
Data
0.00211
Model
0.00152

data/vMax
0.00914

ofxng
319.391

Min d/m
1.4531

ave/min d
30.398
ave/min m
31.8391

330_Calib_S_A

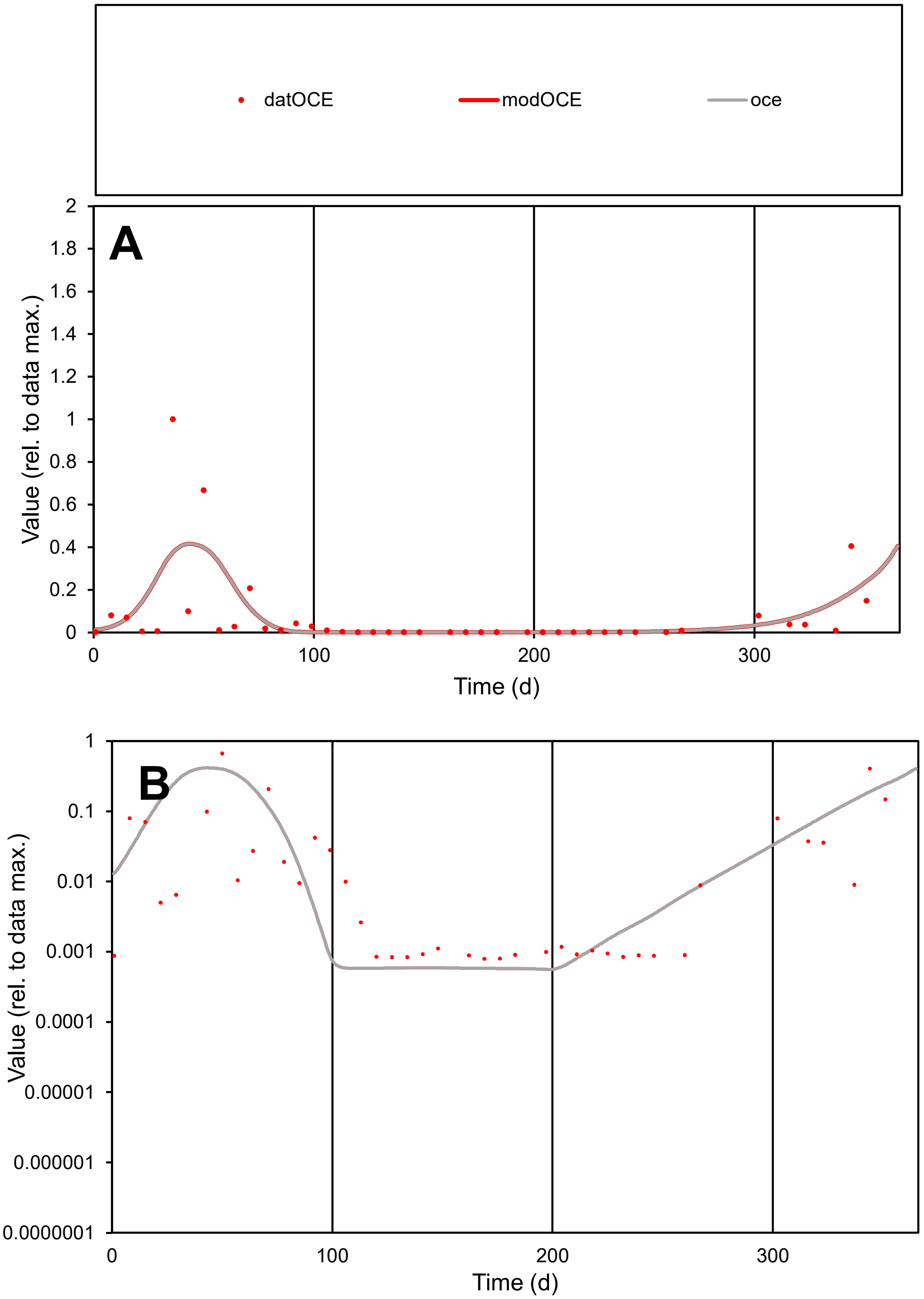


**C1**

Variables
OCE Provide jtd or Dst
CalibOnly?
2 1 to reproduce of
oNormalize?
2 1=no, 2=yes

ofxn calc*
1.91527
ofxn mod
1.97029

sRun
RunA1
zout
1

Ave
Data
0.0034
Model
0.00348

data/vMax
0.04711

ofxng
319.391

Min d/m
1.41692

ave/min d
90.9111
ave/min m
131.996

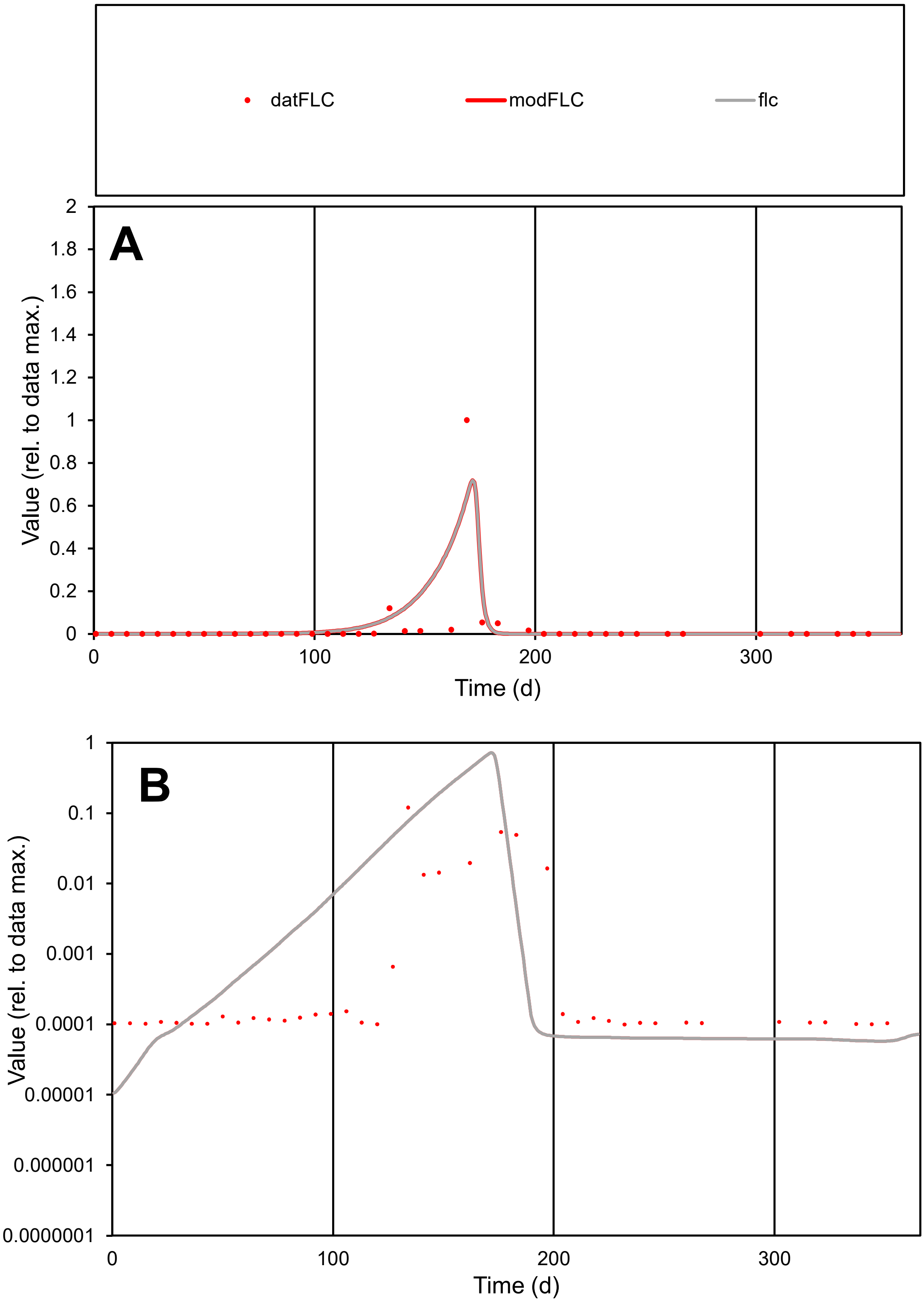


**C1**

Variables

FLC Provide jtd or Dst

CalibOnly?

2 1 to reproduce of

oNormalize?

2 1=no, 2=yes

ofxn calc*

2.62388

ofxn mod

2.8574

sRun

RunA1

zout

1

Ave

Data

0.00397

Model

0.00511

data/vMax

0.1292

ofxng

319.391

Min d/m

8.66487

ave/min d

309.188

ave/min m

3446.66

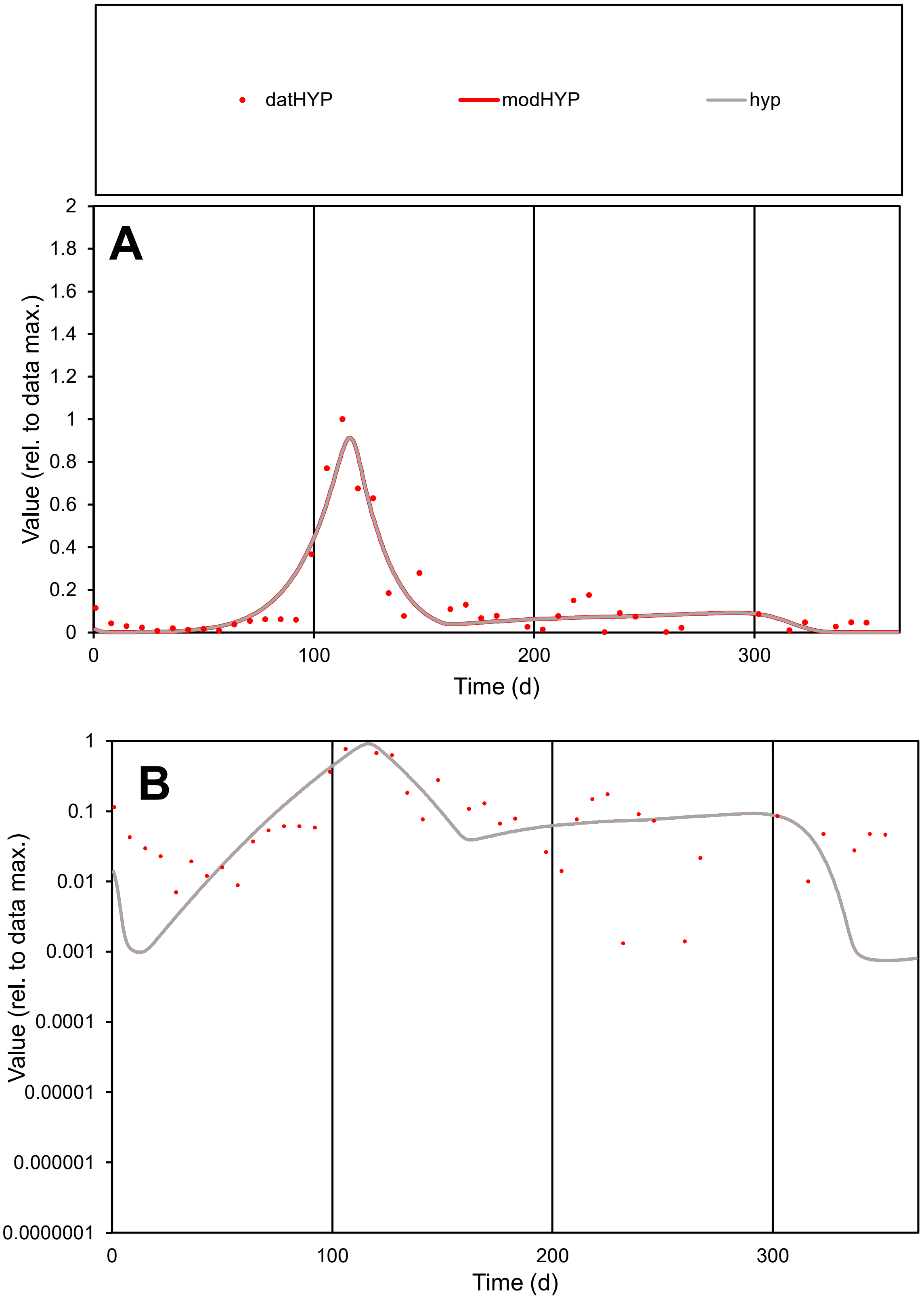


**C1**

Variables
HYP Provide jtd or Dst
CalibOnly?
2 1 to reproduce of
oNormalize?
2 1=no, 2=yes

ofxn calc*
0.55936
ofxn mod
0.59119

sRun
RunA1
zout
1

Ave
Data
0.00178
Model
0.00175

data/vMax
0.01289

ofxng
319.391

Min d/m
1.75916

ave/min d
104.857
ave/min m
181.314

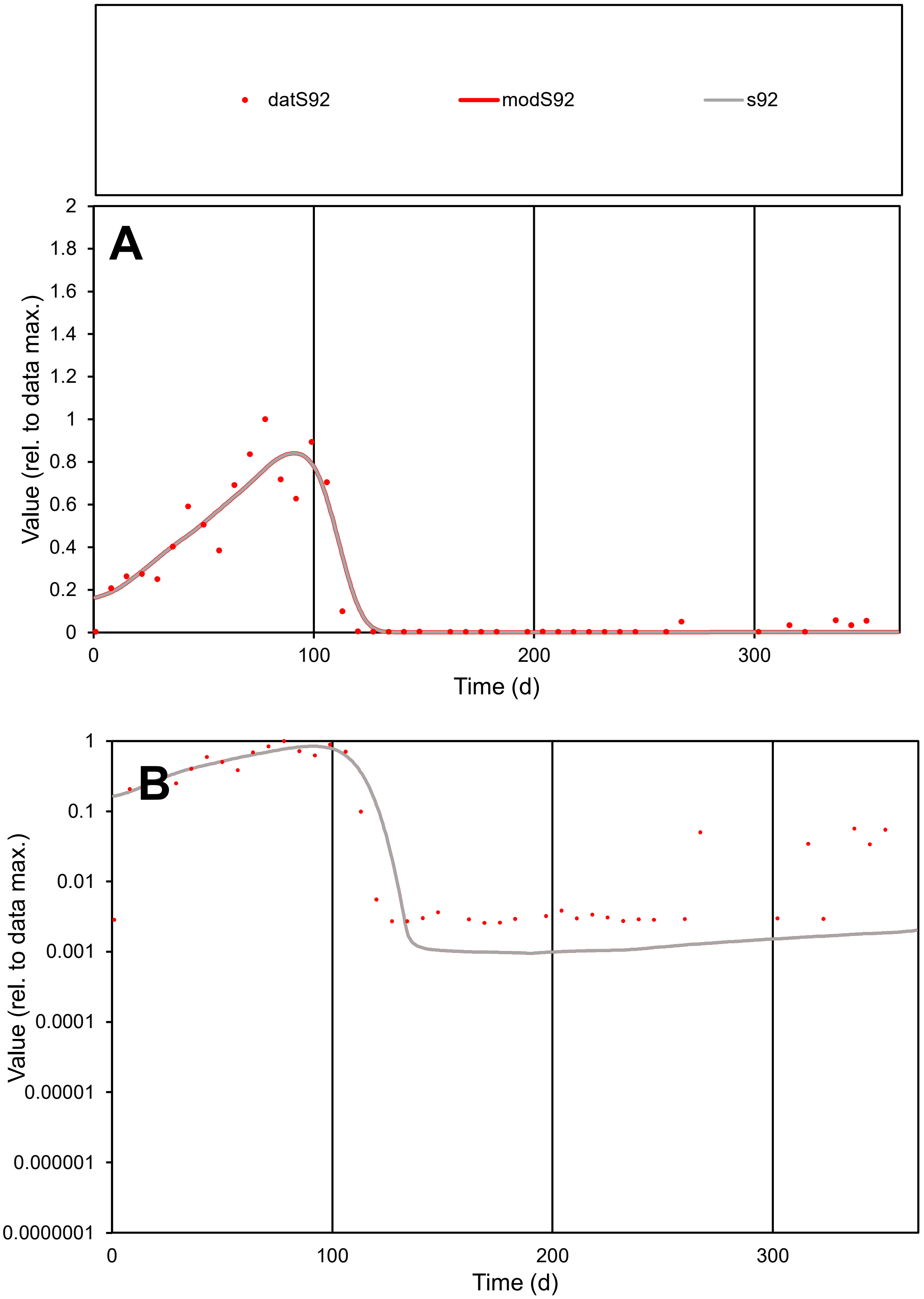


**C1**

Variables
S92 Provide jtd or Dst
CalibOnly?
2 1 to reproduce of
oNormalize?
2 1=no, 2=yes

ofxn calc*
0.40066
ofxn mod
0.45001

sRun
RunA1
zout
1

Ave
Data
0.00189
Model
0.00189

data/vMax
0.00906

ofxng
319.391

Min d/m
2.67673

ave/min d
80.5844
ave/min m
216.178

330_Calib_S_A

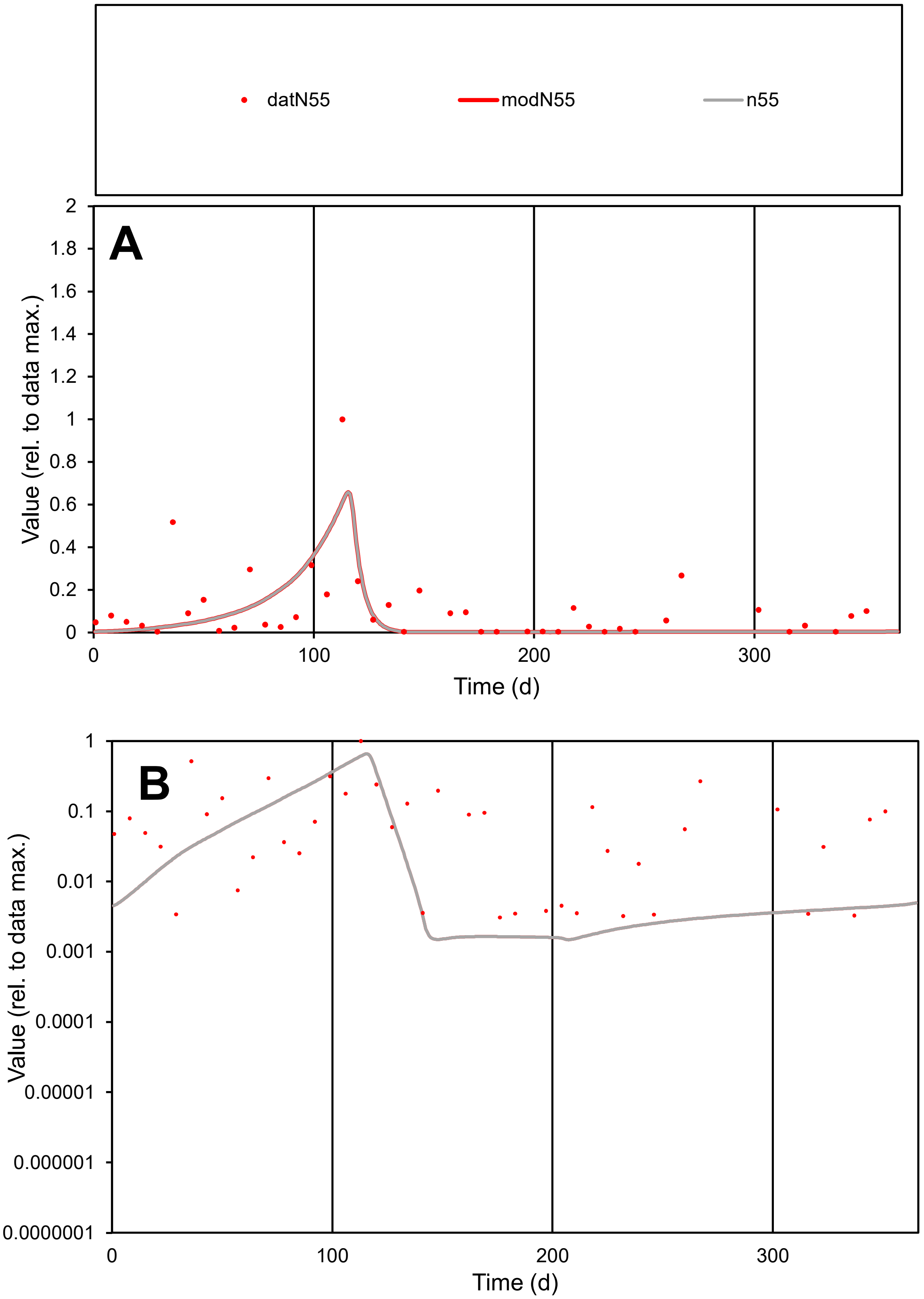


**C1**

Variables
N55 Provide jtd or Dst
CalibOnly?
2 1 to reproduce of
oNormalize?
2 1=no, 2=yes

ofxn calc*
1.24951
ofxn mod
1.26253

sRun
RunA1
zout
1

Ave
Data
0.00134
Model
0.00088

data/vMax
0.01229

ofxng
319.391

Min d/m
2.05643

ave/min d
35.2529
ave/min m
47.7351

330_Calib_S_A

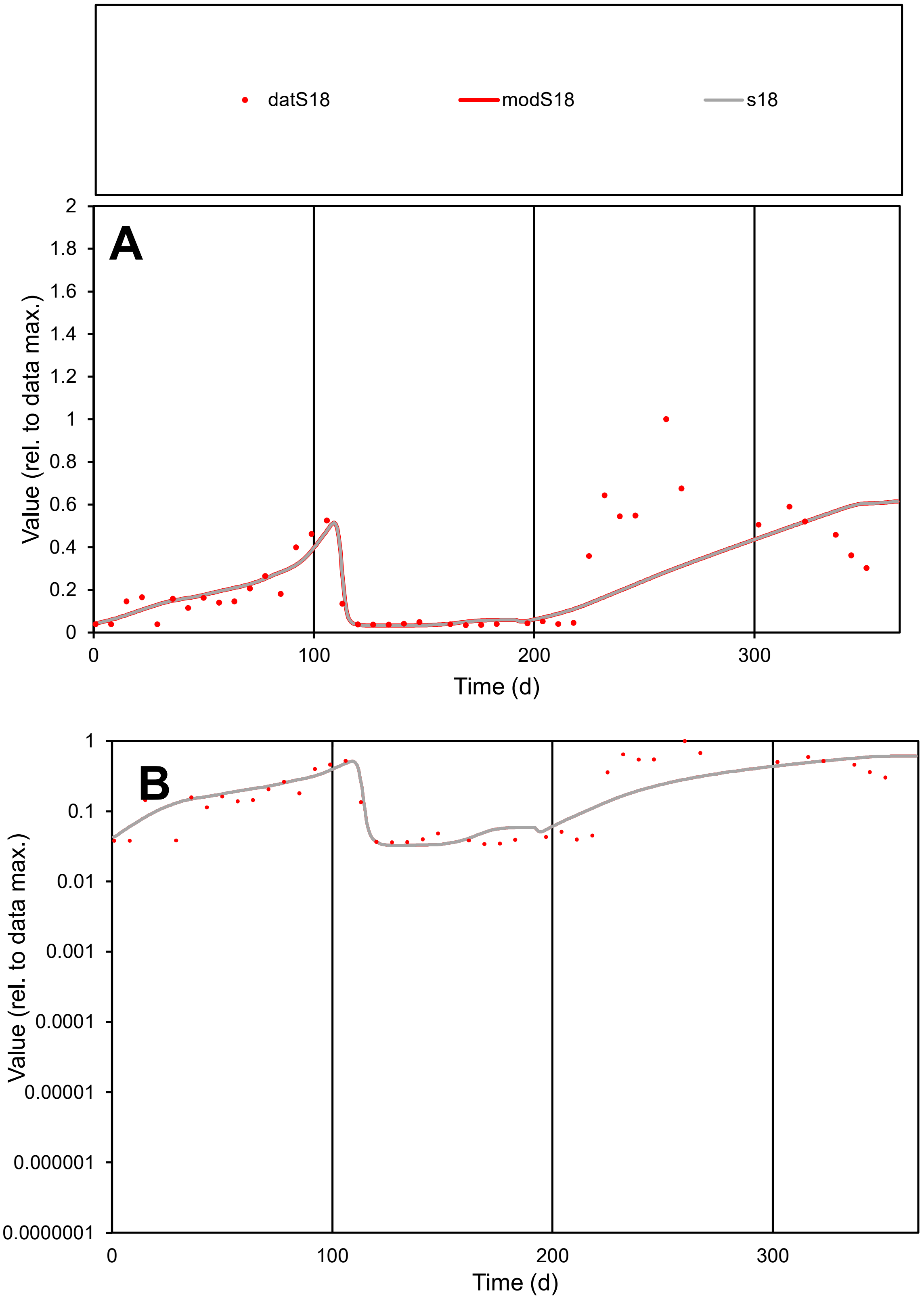


**C1**

Variables

S18 Provide jtd or Dst

CalibOnly?

2 1 to reproduce of

oNormalize?

2 1=no, 2=yes

ofxn calc*

0.72775

ofxn mod

0.73863

sRun

RunA1

zout

1

Ave

Data

0.00192

Model

0.00164

data/vMax

0.00781

ofxng

319.391

Min d/m

1.05722

ave/min d

7.18014

ave/min m

6.46156

330_Calib_S_A

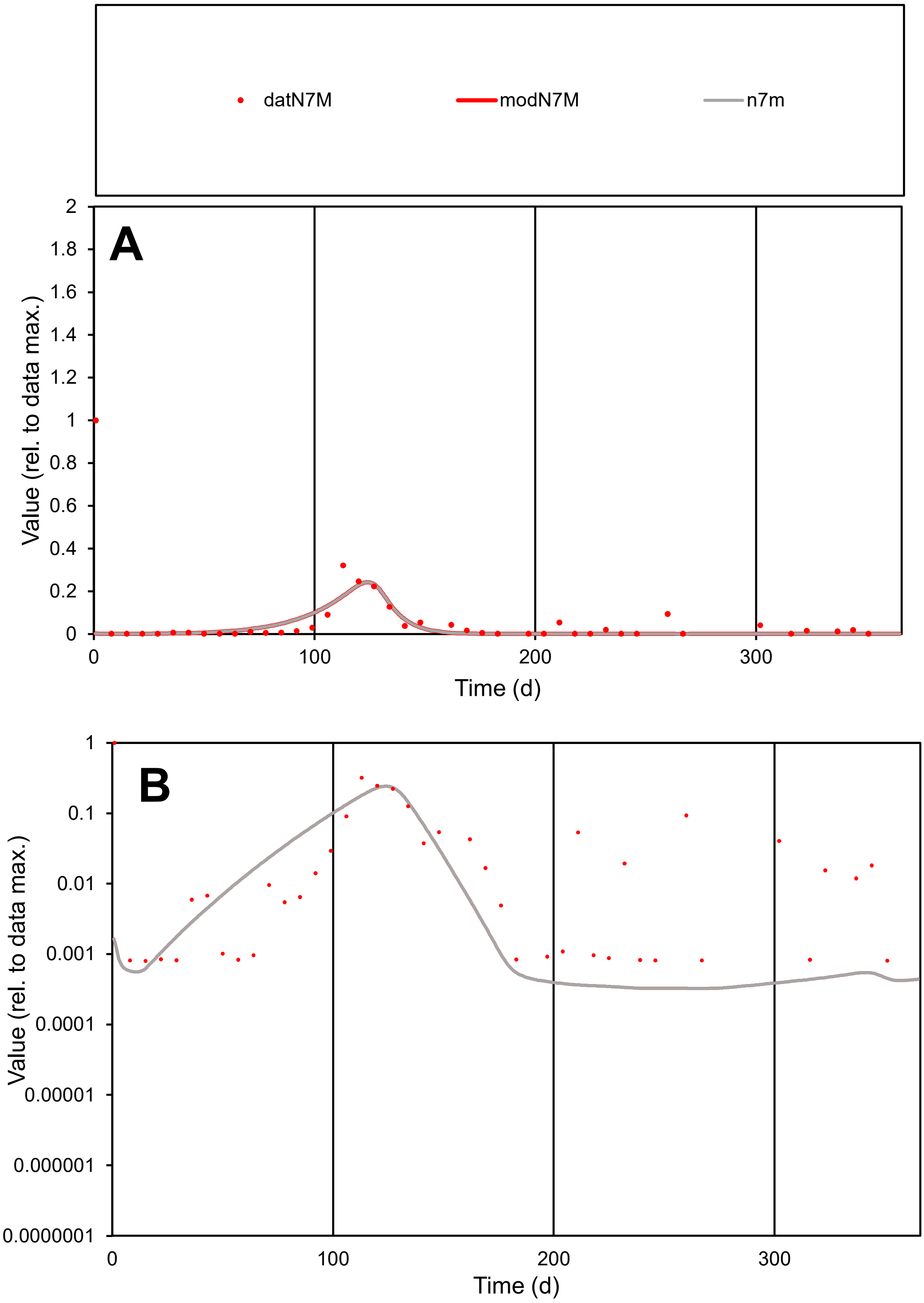


**C1**

Variables

N7M Provide jtd or Dst

CalibOnly?

2 1 to reproduce of

oNormalize?

2 1=no, 2=yes

ofxn calc*

2.6435

ofxn mod

2.64846

sRun

RunA1

zout

1

Ave

Data

0.00079

Model

0.00042

data/vMax

0.01321

ofxng

319.391

Min d/m

2.4696

ave/min d

74.6303

ave/min m

97.9569

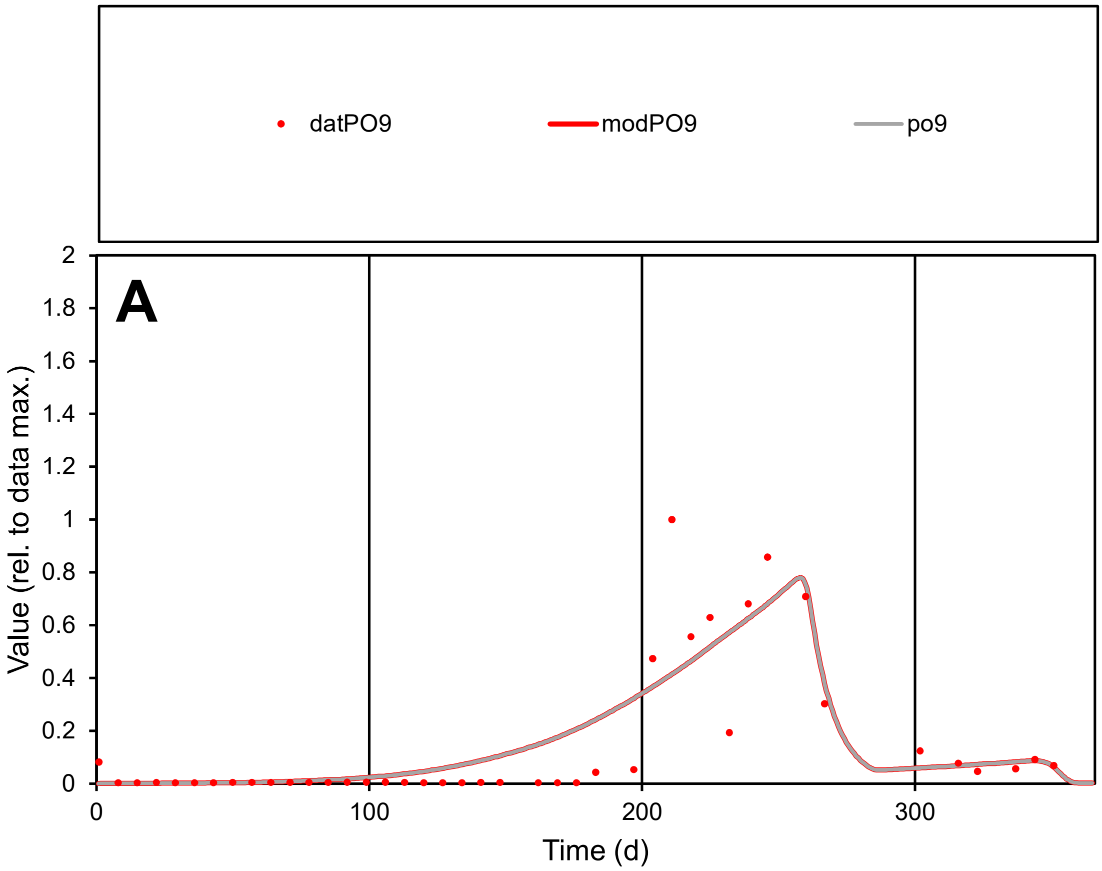


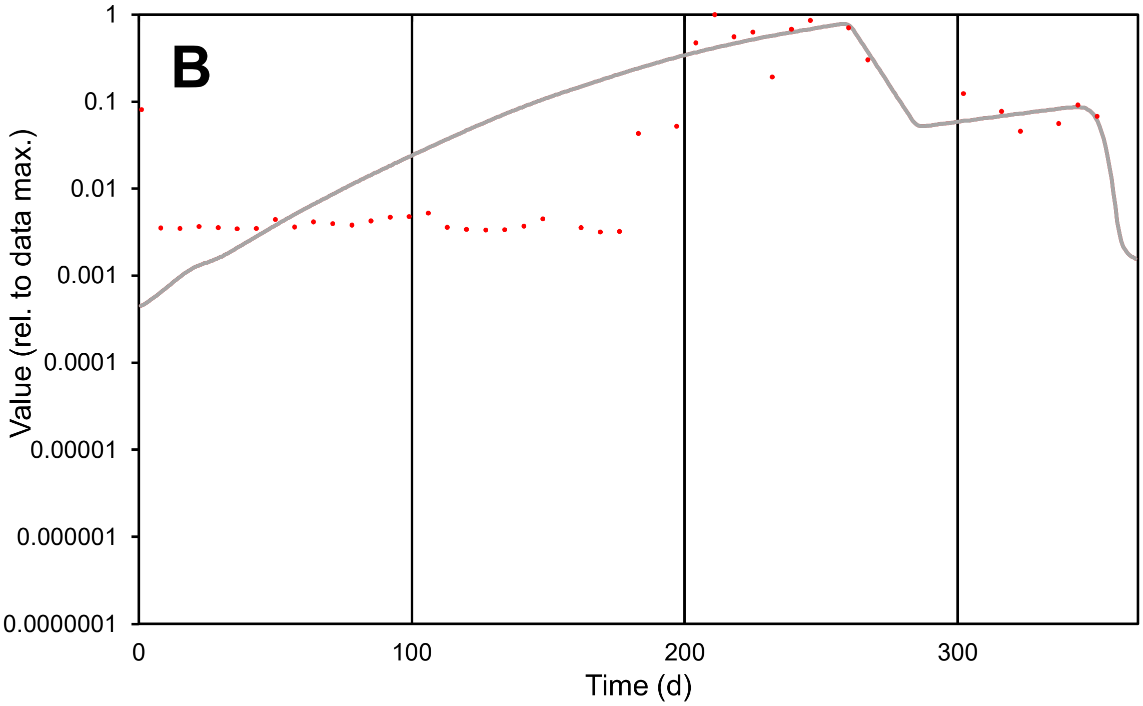


**C1**

Variables
PO9 Provide jtd or Dst
CalibOnly?
2 1 to reproduce of
oNormalize?
2 1=no, 2=yes

ofxn calc*
0.93631
ofxn mod
0.96465

sRun
RunA1
zout
1

Ave
Data
0.00268
Model
0.00298

data/vMax
0.01839

ofxng
319.391

Min d/m
6.92753

ave/min d
45.7217
ave/min m
352.327

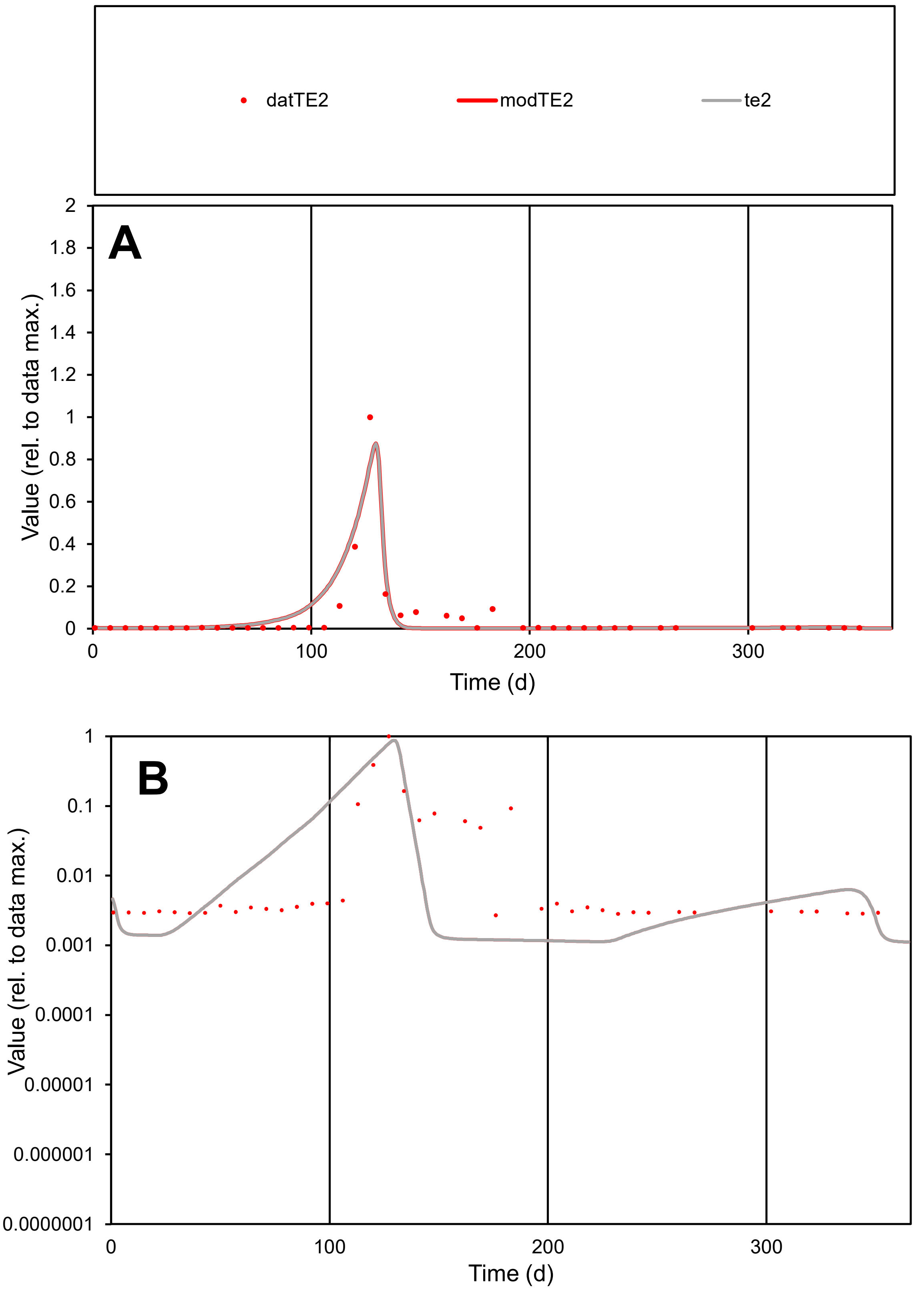


**C1**

Variables
TE2 Provide jtd or Dst
CalibOnly?
2 1 to reproduce of
oNormalize?
2 1=no, 2=yes

ofxn calc*
1.07691
ofxn mod
1.2283

sRun
RunA1
zout
1

Ave
Data
0.00161
Model
0.00175

data/vMax
0.03218

ofxng
319.391

Min d/m
2.38709

ave/min d
18.5528
ave/min m
48.0965

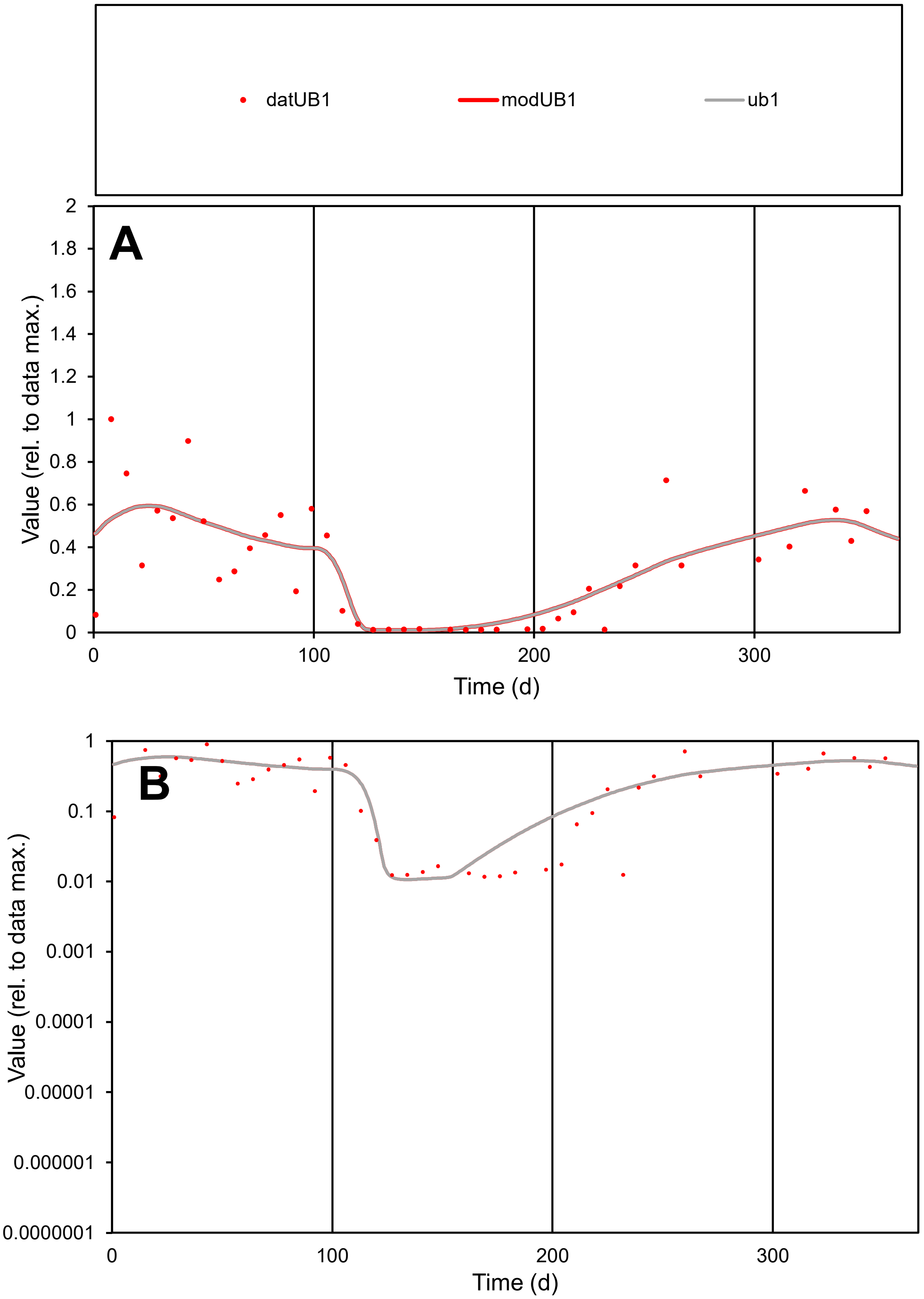


**C1**

Variables
UB1 Provide jtd or Dst
CalibOnly?
2 1 to reproduce of
oNormalize?
2 1=no, 2=yes

ofxn calc*
0.51295
ofxn mod
0.53231

sRun
RunA1
zout
1

Ave
Data
0.00152
Model
0.00154

data/vMax
0.00489

ofxng
319.391

Min d/m
1.10398

ave/min d
26.4227
ave/min m
29.5695

330_Calib_S_A

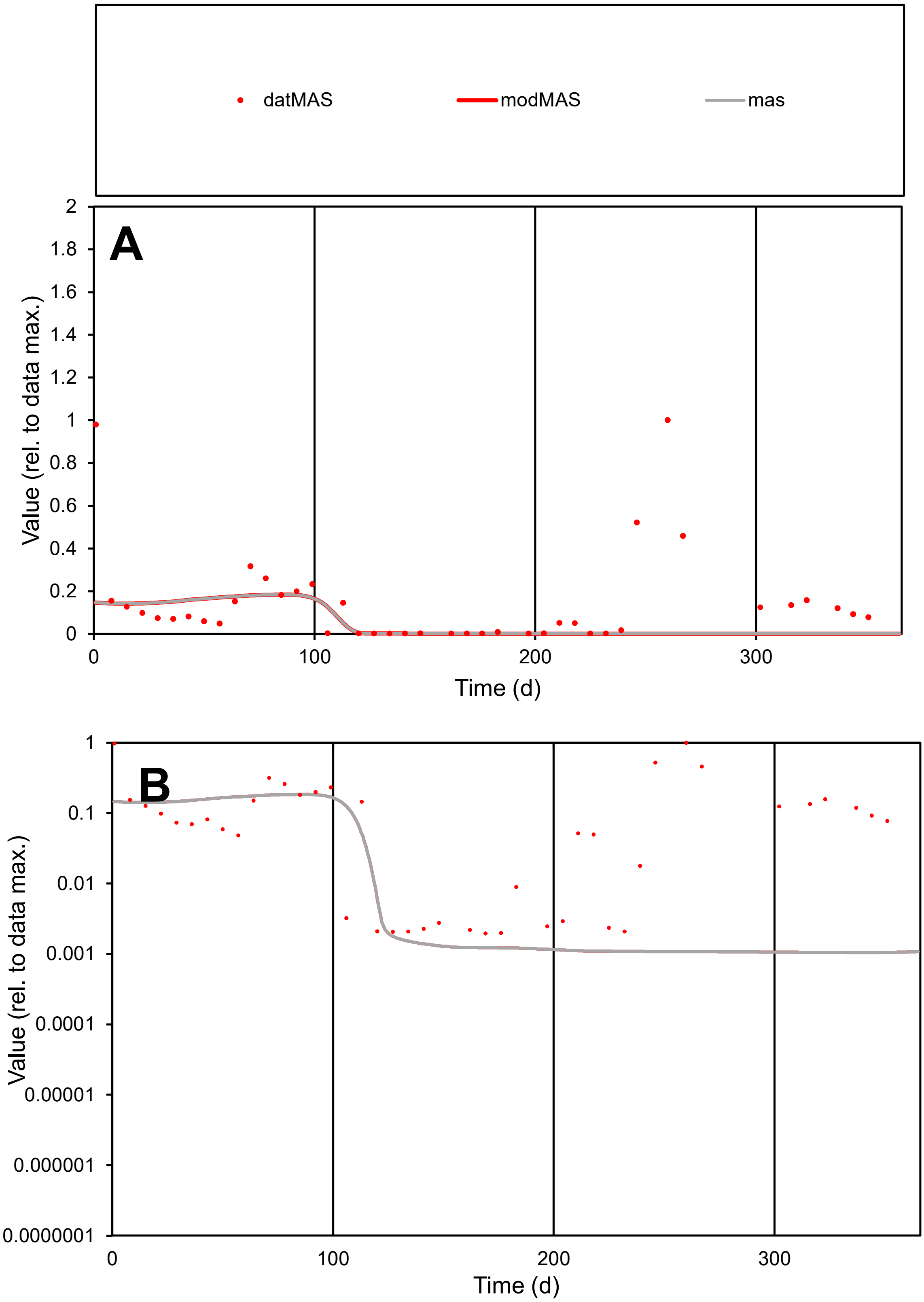


**C1**

Variables
MAS Provide jtd or Dst
CalibOnly?
2 1 to reproduce of
oNormalize?
2 1=no, 2=yes

ofxn calc*
1.65029
ofxn mod
1.6562

sRun
RunA1
zout
1

Ave
Data
0.00088
Model
0.00038

data/vMax
0.00617

ofxng
319.391

Min d/m
1.87781

ave/min d
73.0427
ave/min m
59.4665

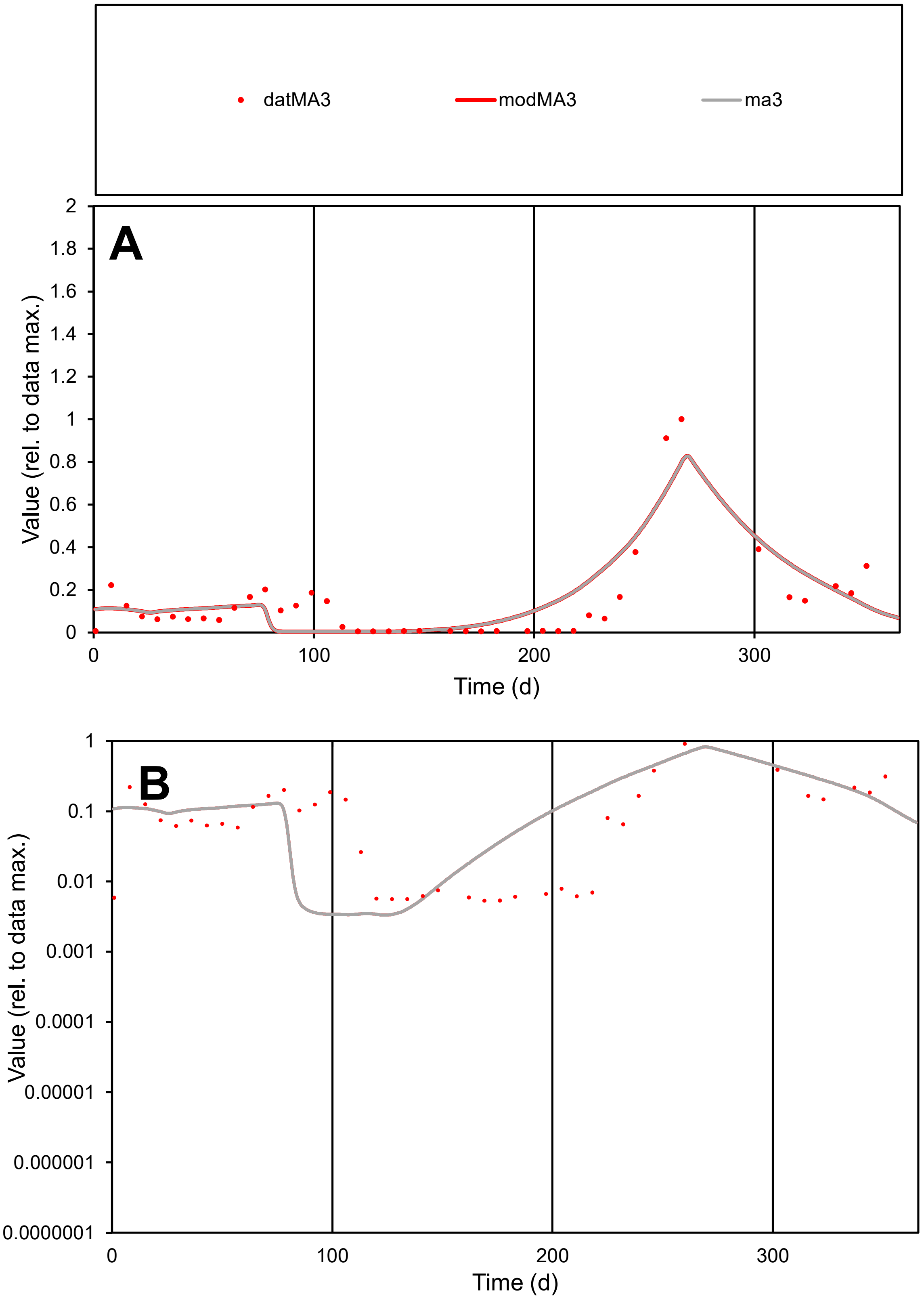


**C1**

Variables
MA3 Provide jtd or Dst
CalibOnly?
2 1 to reproduce of
oNormalize?
2 1=no, 2=yes

ofxn calc*
0.76919
ofxn mod
0.7987

sRun
RunA1
zout
1

Ave
Data
0.00165
Model
0.0018

data/vMax
0.01173

ofxng
319.391

Min d/m
1.57794

ave/min d
26.5125
ave/min m
45.5198

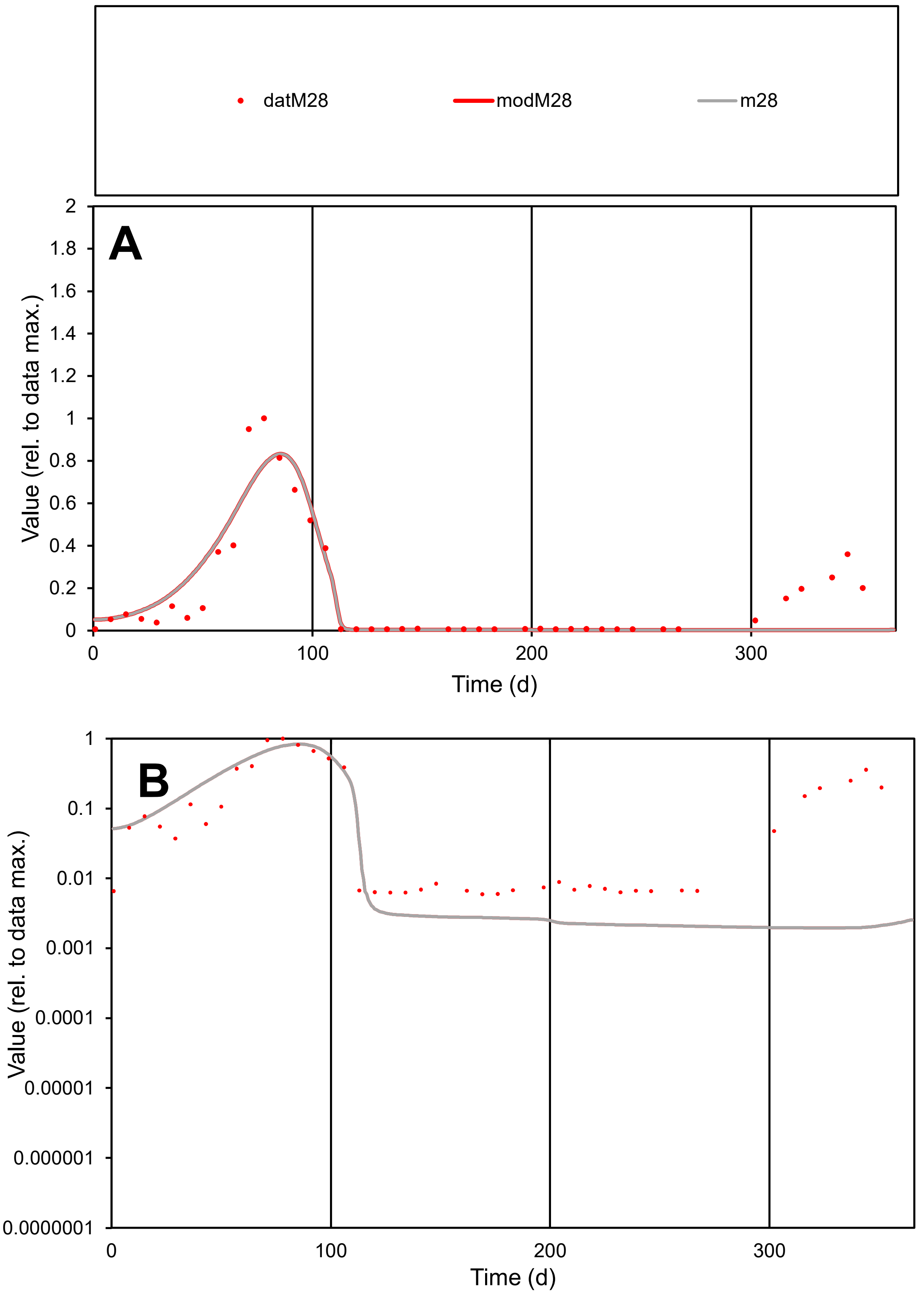


**C1**

Variables
M28 Provide jtd or Dst
CalibOnly?
2 1 to reproduce of
oNormalize?
2 1=no, 2=yes

ofxn calc*
0.68146
ofxn mod
0.71199

sRun
RunA1
zout
1

Ave
Data
0.00151
Model
0.00134

data/vMax
0.00913

ofxng
319.391

Min d/m
3.05177

ave/min d
27.887
ave/min m
75.6492

330_Calib_S_A

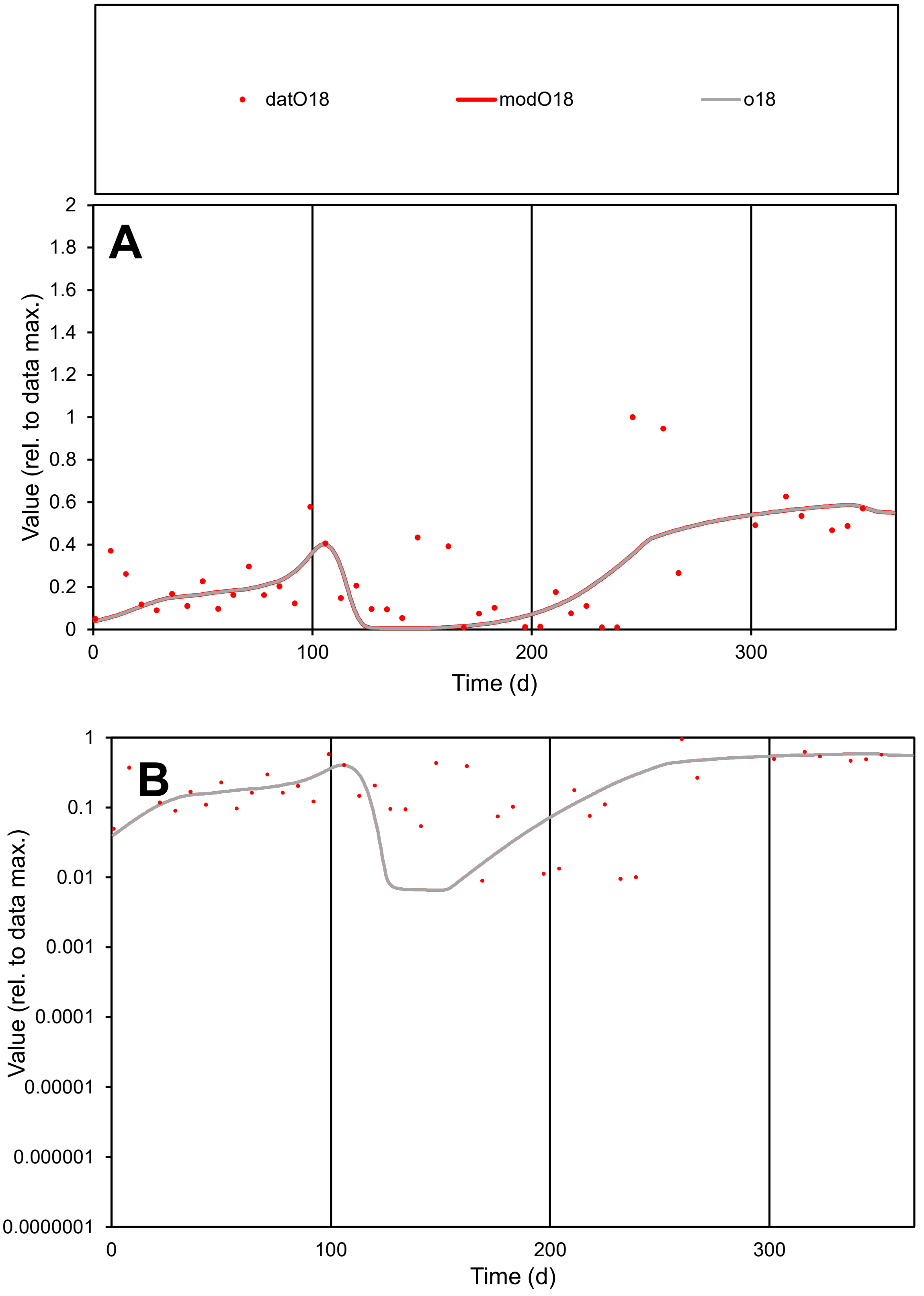


**C1**

Variables
O18 Provide jtd or Dst
CalibOnly?
2 1 to reproduce of
oNormalize?
2 1=no, 2=yes

ofxn calc*
0.72544
ofxn mod
0.73932

sRun
RunA1
zout
1

Ave
Data
0.00149
Model
0.00127

data/vMax
0.00579

ofxng
319.391

Min d/m
1.36481

ave/min d
28.742
ave/min m
33.4478

330_Calib_S_A

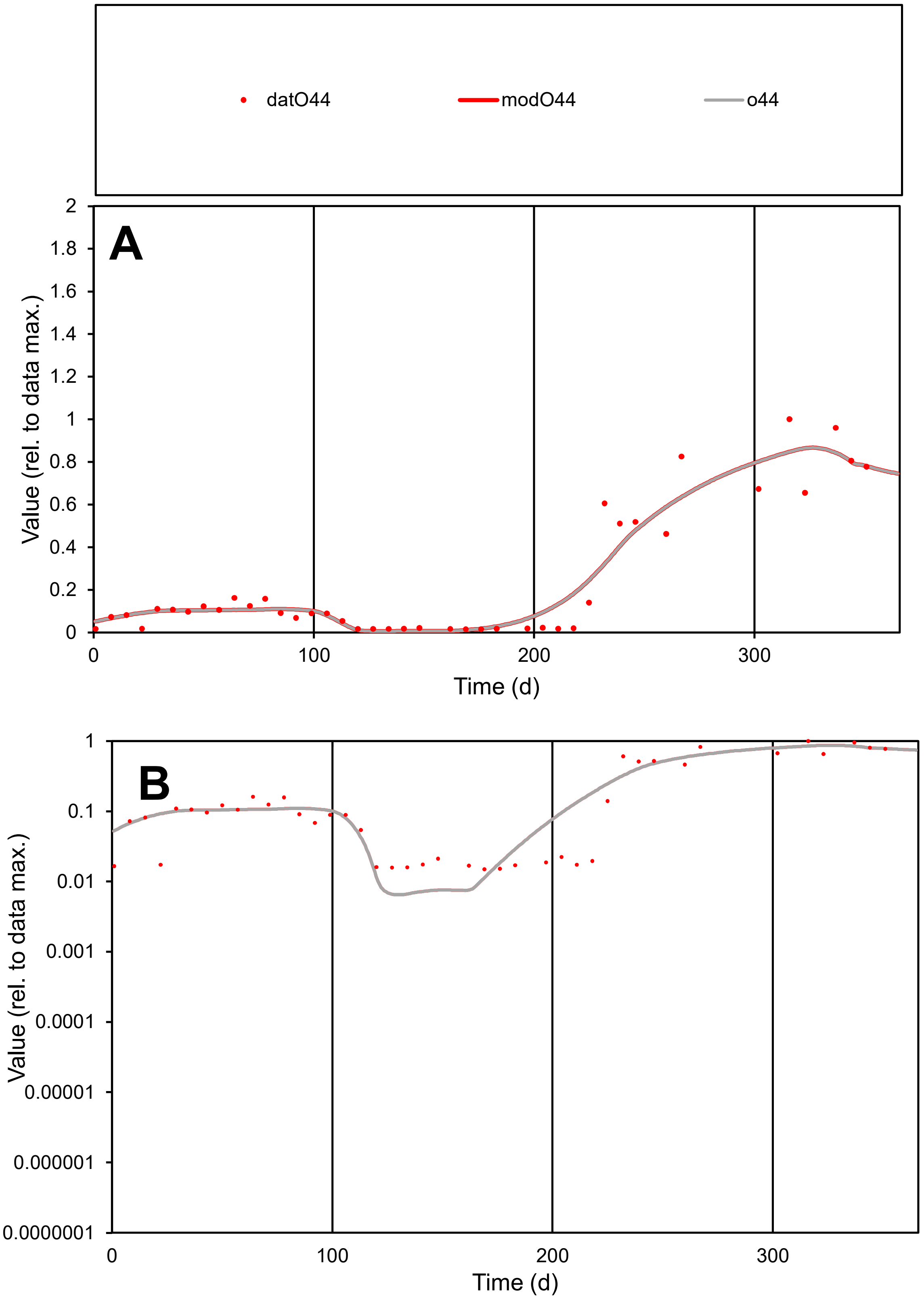


**C1**

Variables
O44 Provide jtd or Dst
CalibOnly?
2 1 to reproduce of
oNormalize?
2 1=no, 2=yes

ofxn calc*
0.36712
ofxn mod
0.40195

sRun
RunA1
zout
1

Ave
Data
0.00171
Model
0.00173

data/vMax
0.00741

ofxng
319.391

Min d/m
2.29261

ave/min d
15.4338
ave/min m
35.7807

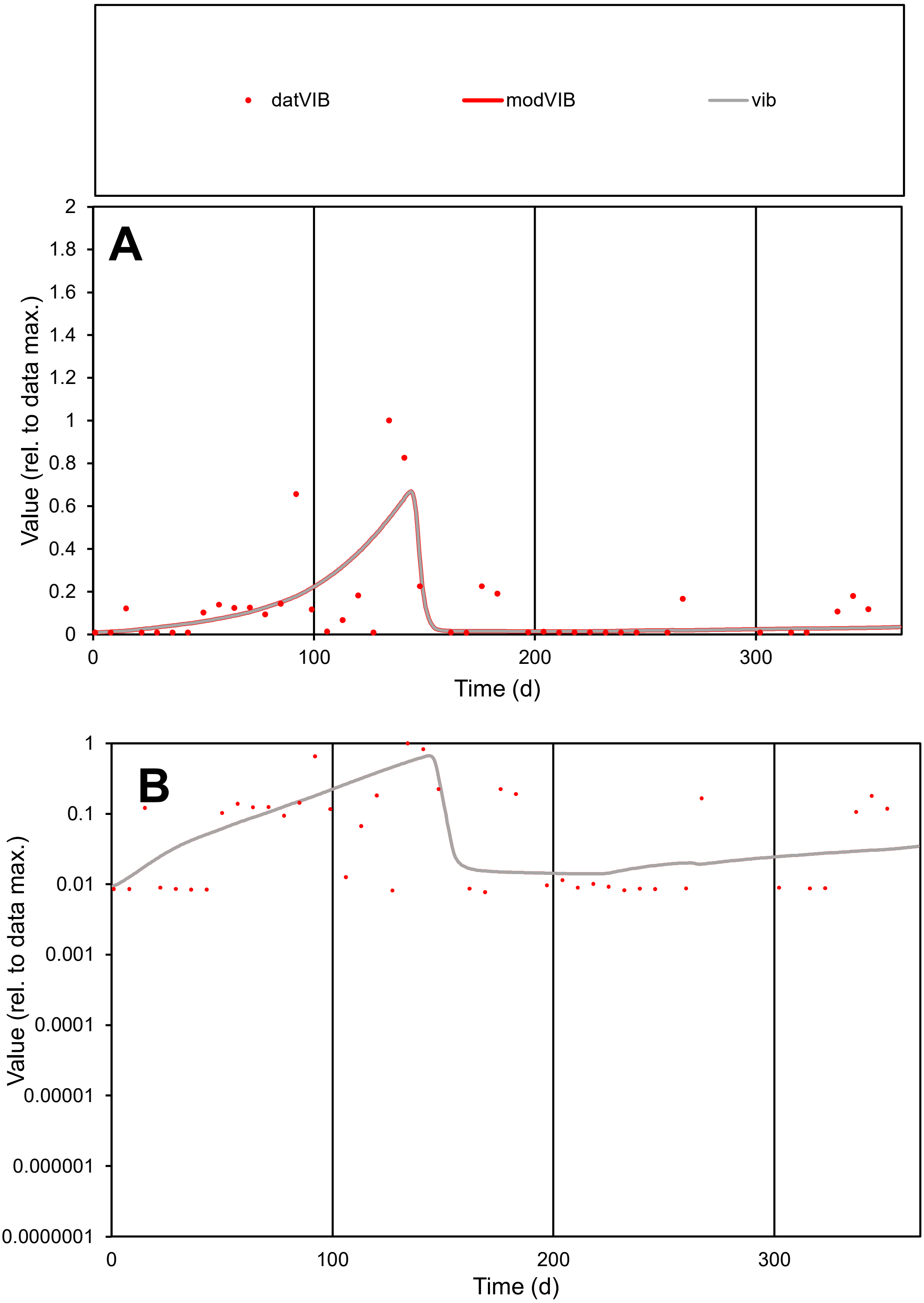


**C1**

Variables

VIB Provide jtd or Dst

CalibOnly?

2 1 to reproduce of

oNormalize?

2 1=no, 2=yes

ofxn calc*

1.26027

ofxn mod

1.27751

sRun

RunA1

zout

1

Ave

Data

0.00028

Model

0.00025

data/vMax

0.00231

ofxng

319.391

Min d/m

0.80415

ave/min d

15.7703

ave/min m

11.1645

330_Calib_S_A

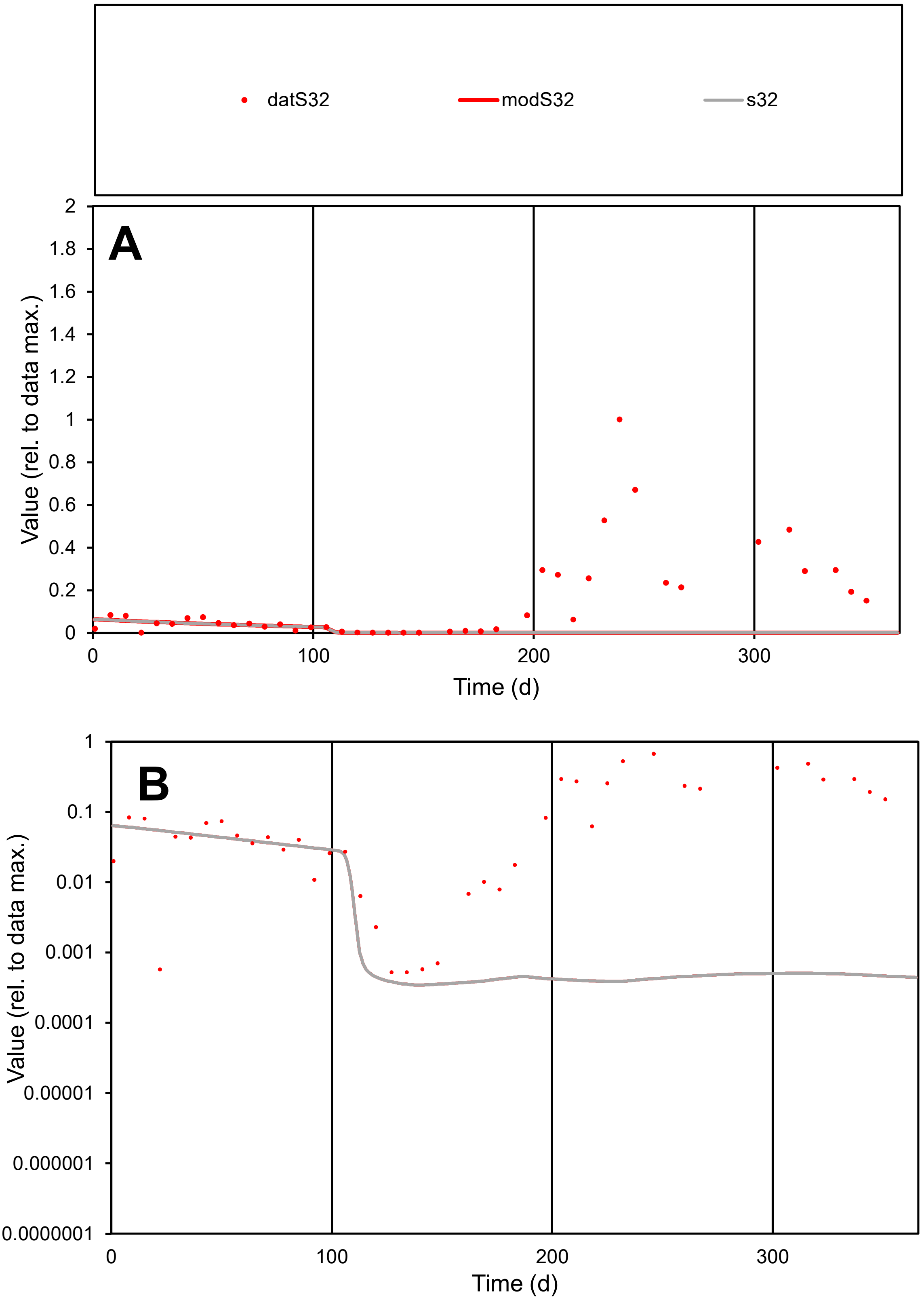


**C1**

Variables
S32 Provide jtd or Dst
CalibOnly?
2 1 to reproduce of
oNormalize?
2 1=no, 2=yes

ofxn calc*
1.72824
ofxn mod
1.73202

sRun
RunA1
zout
1

Ave
Data
0.00322
Model
0.00036

data/vMax
0.02187

ofxng
319.391

Min d/m
1.51363

ave/min d
281.297
ave/min m
48.1304

330_Calib_S_A

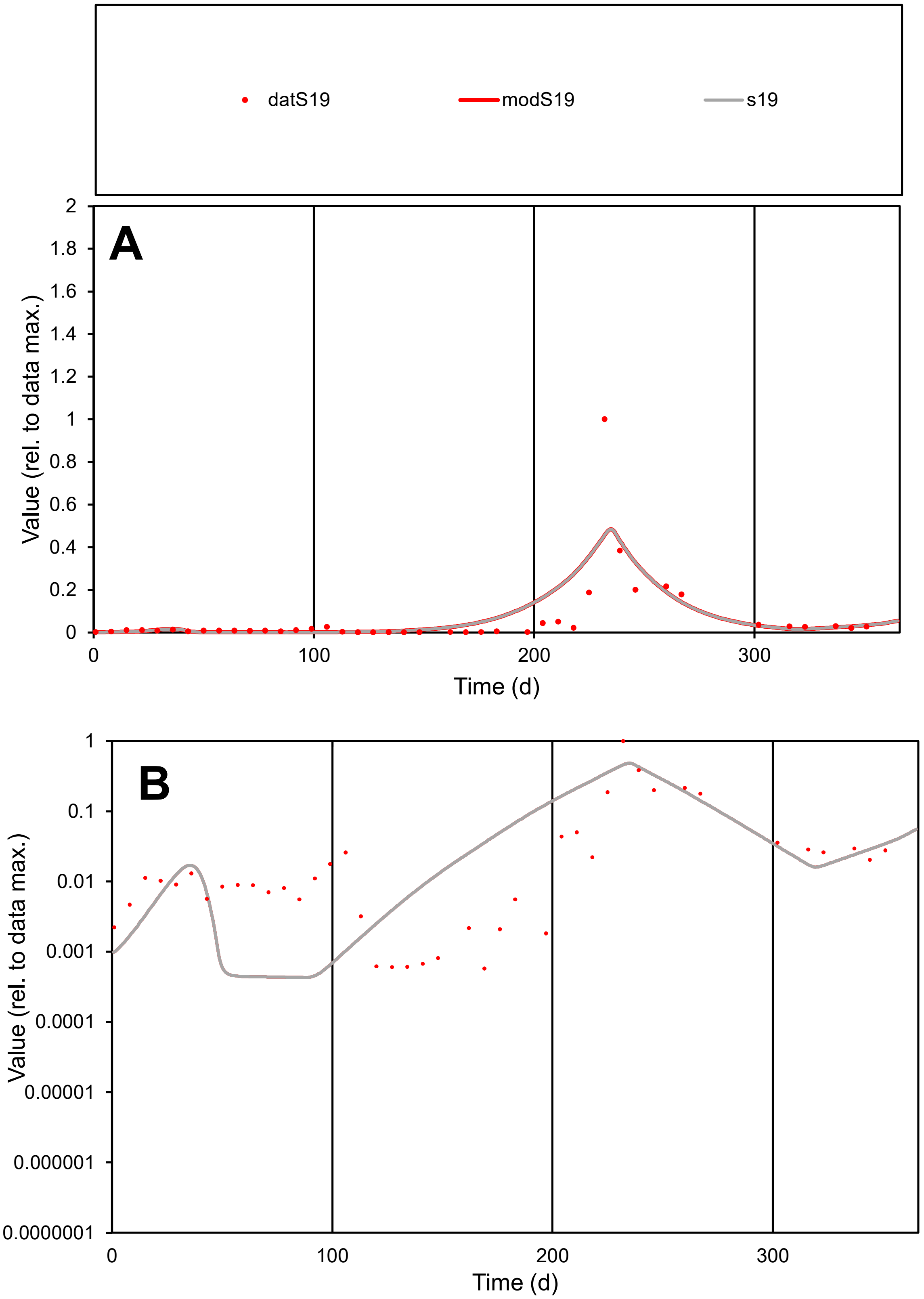


**C1**

Variables

S19 Provide jtd or Dst

CalibOnly?

2 1 to reproduce of

oNormalize?

2 1=no, 2=yes

ofxn calc*

1.65637

ofxn mod

1.70385

sRun

RunA1

zout

1

Ave

Data

0.00515

Model

0.00602

data/vMax

0.08234

ofxng

319.391

Min d/m

1.33568

ave/min d

108.425

ave/min m

169.367

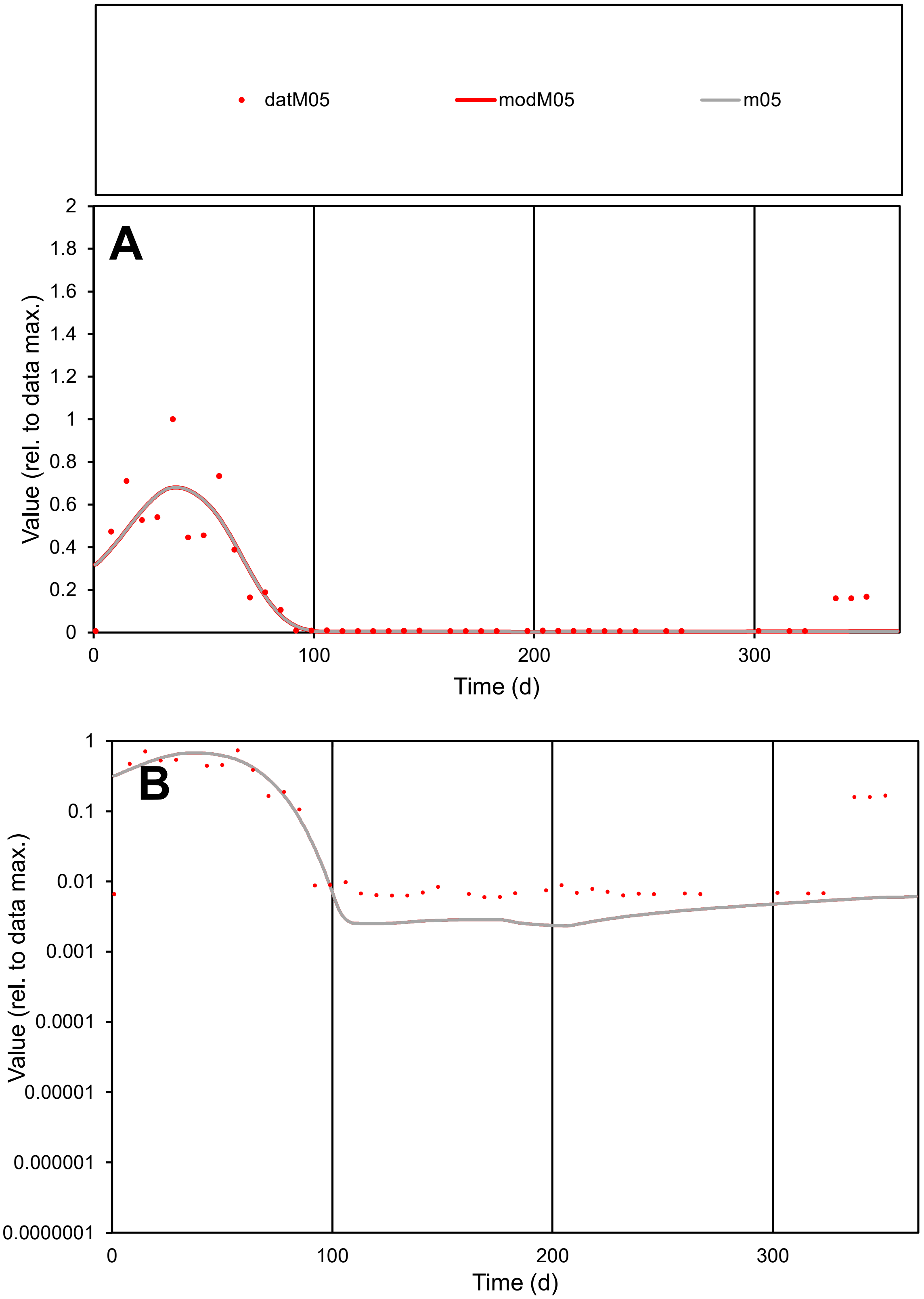


**C1**

Variables
M05 Provide jtd or Dst
CalibOnly?
2 1 to reproduce of
oNormalize?
2 1=no, 2=yes

ofxn calc*
0.69022
ofxn mod
0.72175

sRun
RunA1
zout
1

Ave
Data
0.00111
Model
0.00104

data/vMax
0.00726

ofxng
319.391

Min d/m
2.54157

ave/min d
25.5457
ave/min m
60.6275

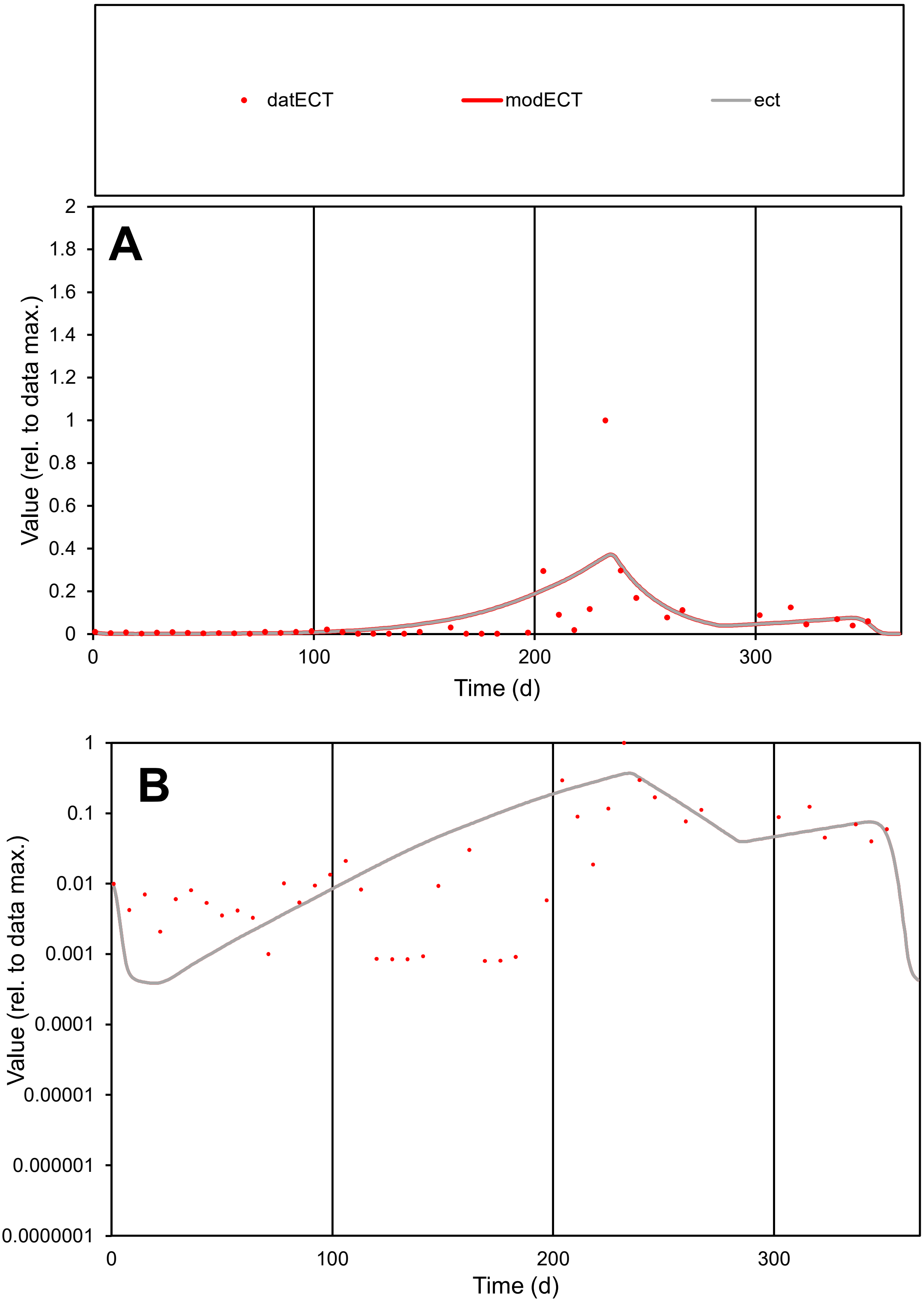


**C1**

Variables
ECT Provide jtd or Dst
CalibOnly?
2 1 to reproduce of
oNormalize?
2 1=no, 2=yes

ofxn calc*
1.83522
ofxn mod
1.86169

sRun
RunA1
zout
1

Ave
Data
0.00235
Model
0.00279

data/vMax
0.03564

ofxng
319.391

Min d/m
2.03574

ave/min d
82.5938
ave/min m
199.43

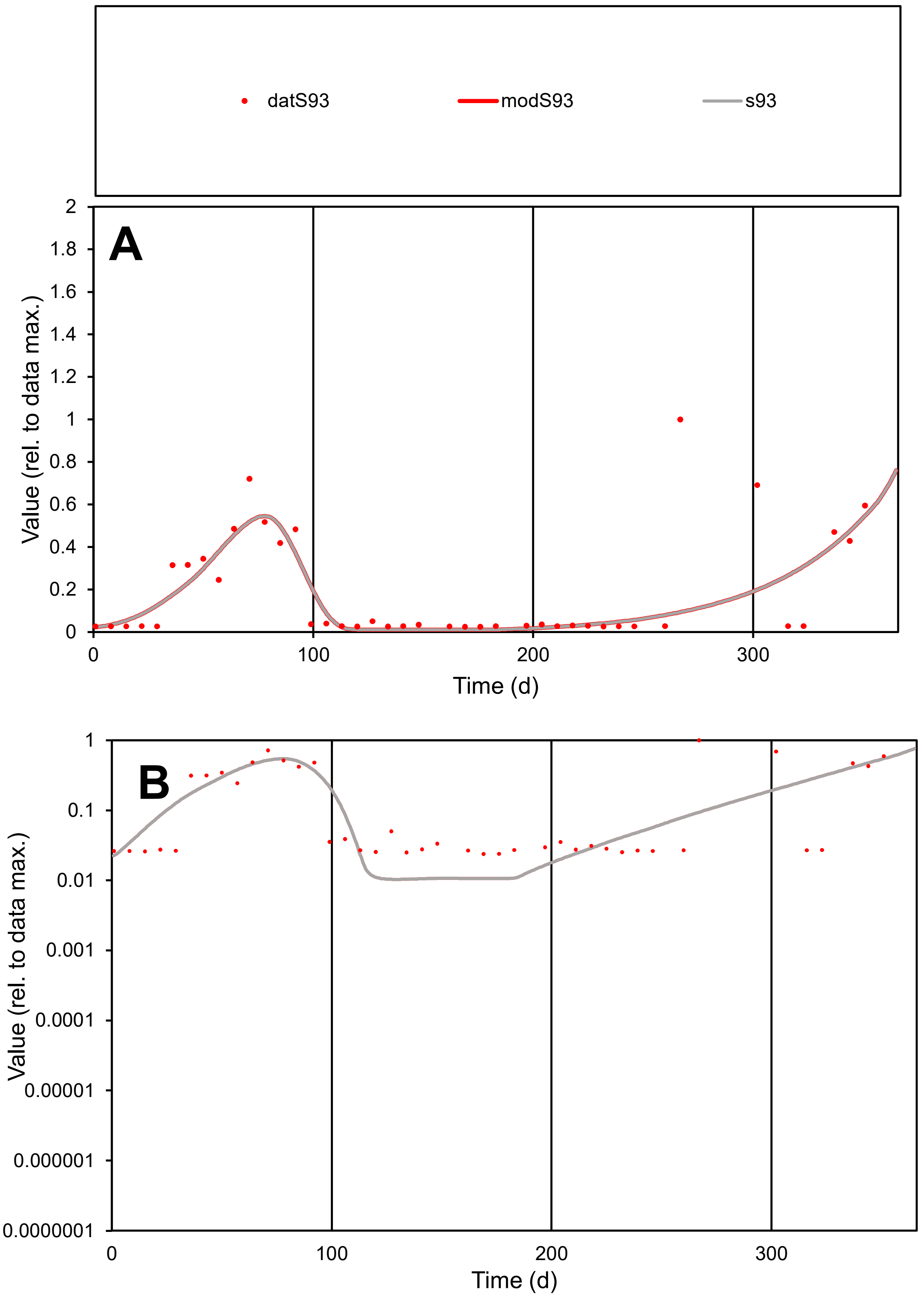


**C1**

Variables

S93 Provide jtd or Dst

CalibOnly?

2 1 to reproduce of

oNormalize?

2 1=no, 2=yes

ofxn calc*

0.95691

ofxn mod

0.97093

sRun

RunA1

zout

1

Ave

Data

0.00051

Model

0.00045

data/vMax

0.00276

ofxng

319.391

Min d/m

2.29713

ave/min d

7.85082

ave/min m

15.734

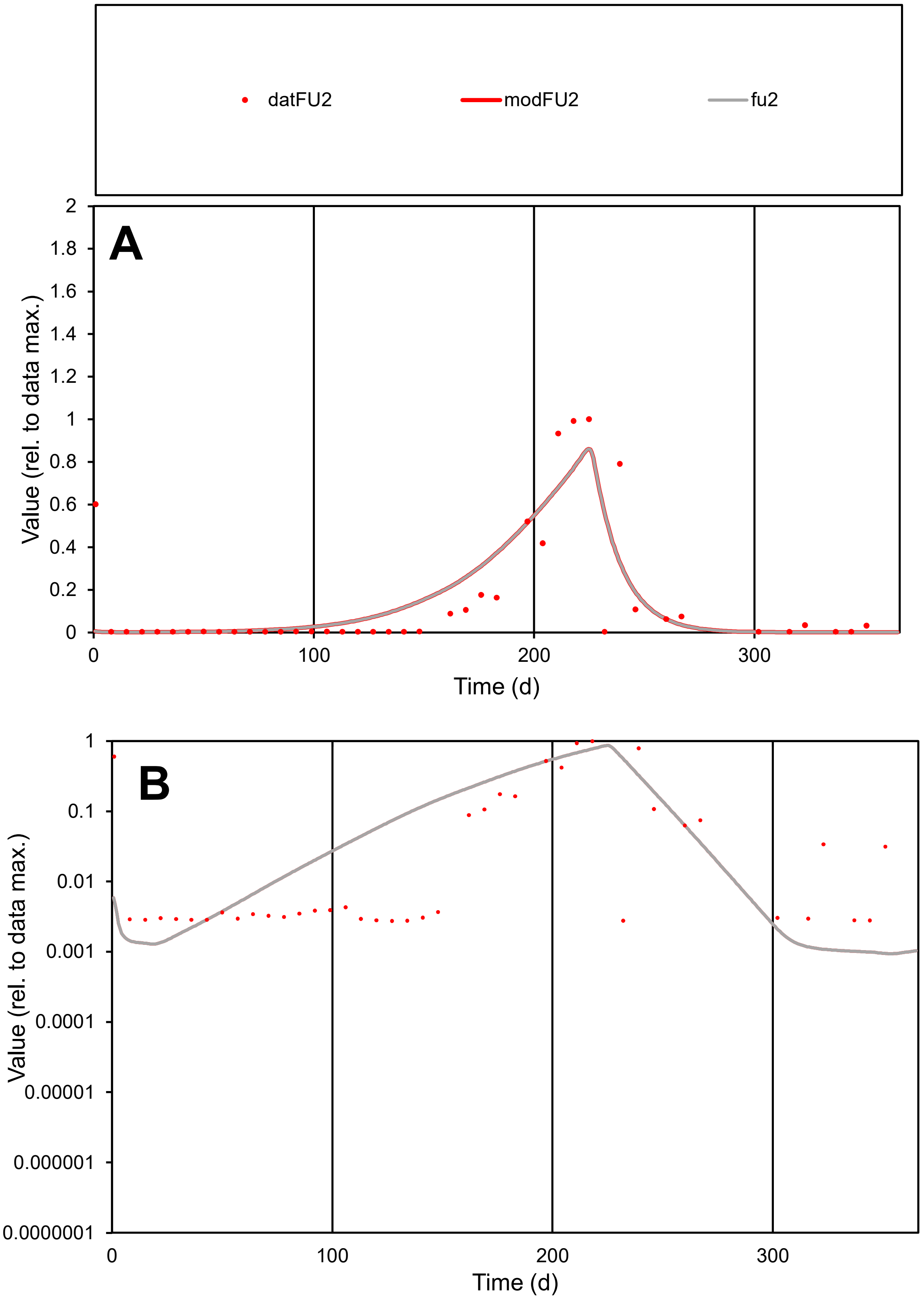


**C1**

Variables

FU2 Provide jtd or Dst

CalibOnly?

2 1 to reproduce of

oNormalize?

2 1=no, 2=yes

ofxn calc*

1.08758

ofxn mod

1.13953

sRun

RunA1

zout

1

Ave

Data

0.00145

Model

0.00148

data/vMax

0.00983

ofxng

319.391

Min d/m

2.92472

ave/min d

53.438

ave/min m

160.302

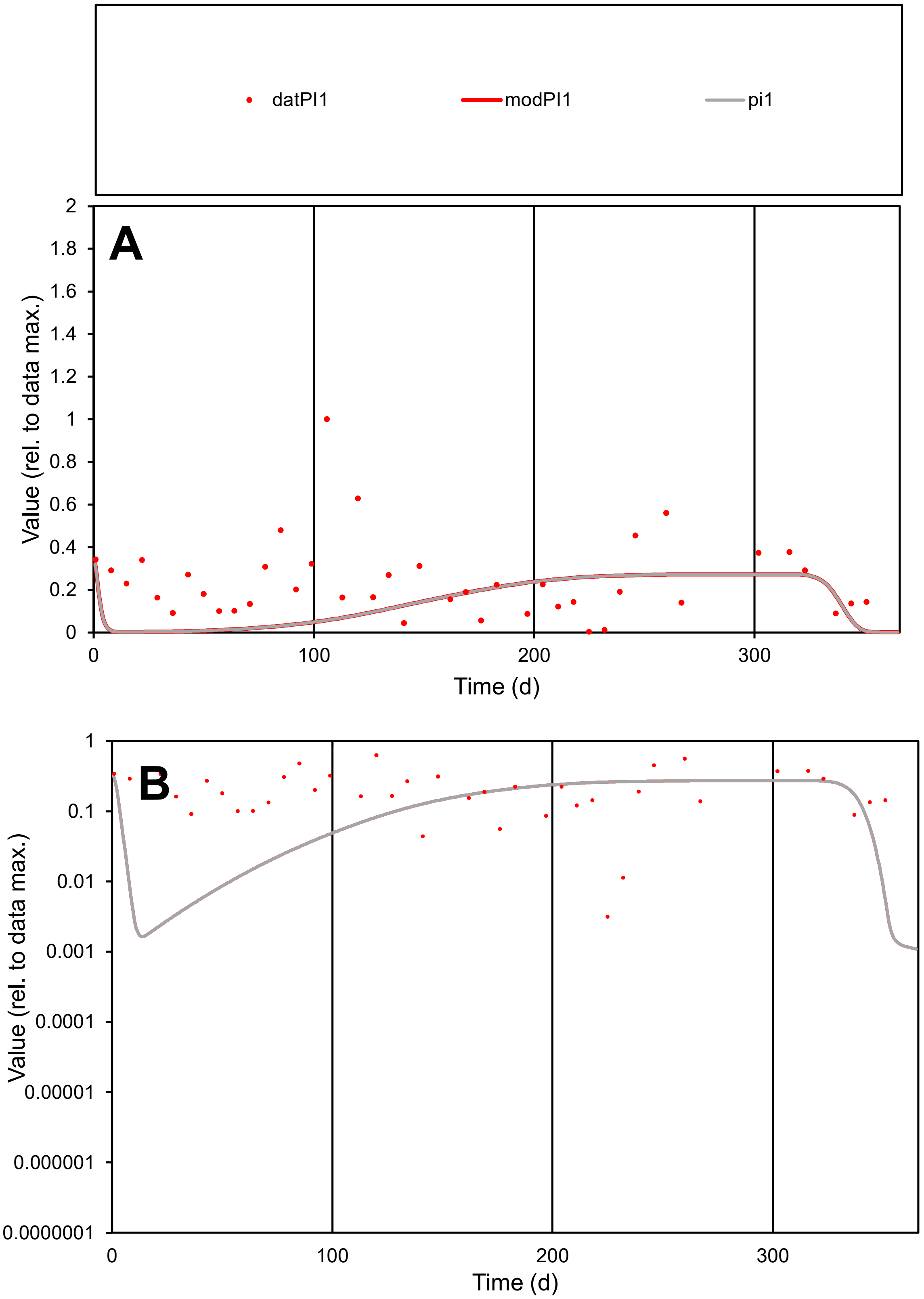


**C1**

Variables
PI1 Provide jtd or Dst
CalibOnly?
2 1 to reproduce of
oNormalize?
2 1=no, 2=yes

ofxn calc*
1.00127
ofxn mod
1.00648

sRun
RunA1
zout
1

Ave
Data
0.00121
Model
0.00067

data/vMax
0.00501

ofxng
319.391

Min d/m
1.72844

ave/min d
76.2386
ave/min m
73.2524

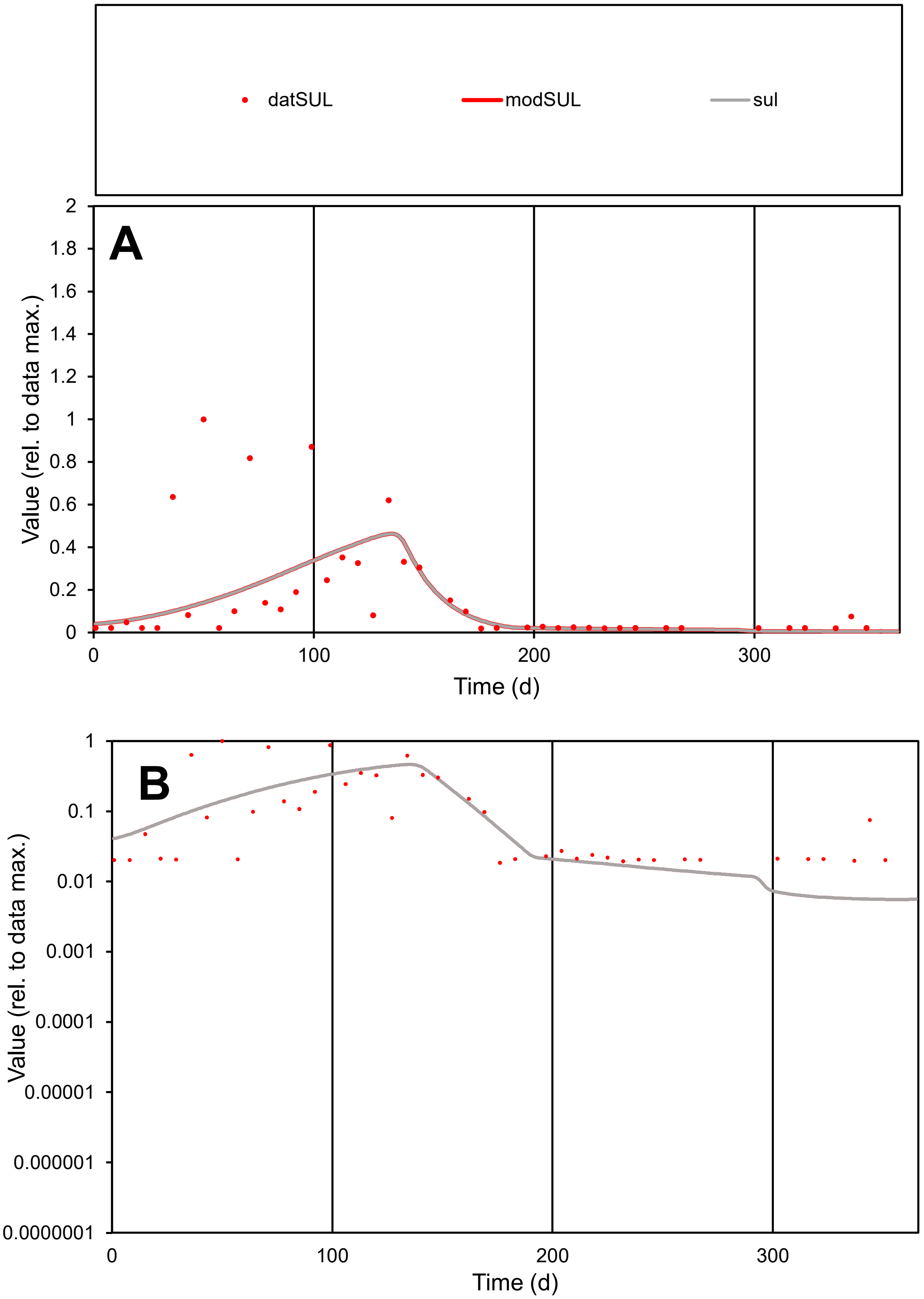


**C1**

Variables
SUL Provide jtd or Dst
CalibOnly?
2 1 to reproduce of
oNormalize?
2 1=no, 2=yes

ofxn calc*
1.2857
ofxn mod
1.29469

sRun
RunA1
zout
1

Ave
Data
0.00073
Model
0.00059

data/vMax
0.00436

ofxng
319.391

Min d/m
3.3177
ave/min d
9.04543
ave/min m
24.2034

330_Calib_S_A

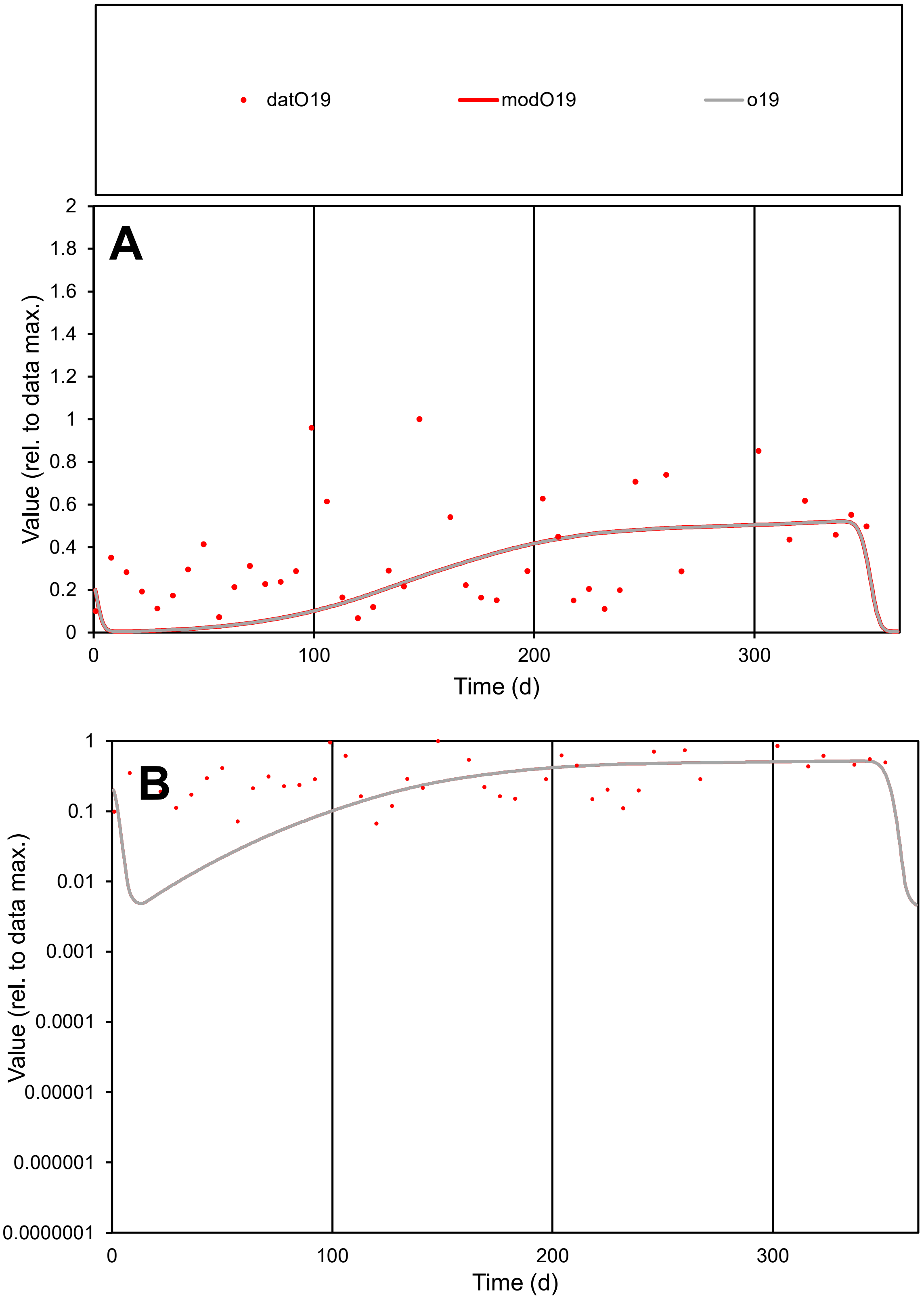


**C1**

Variables
O19 Provide jtd or Dst
CalibOnly?
2 1 to reproduce of
oNormalize?
2 1=no, 2=yes

ofxn calc*
0.76784
ofxn mod
0.77581

sRun
RunA1
zout
1

Ave
Data
0.00165
Model
0.0012

data/vMax
0.00464

ofxng
319.391

Min d/m
12.3168

ave/min d
5.30262
ave/min m
47.4252

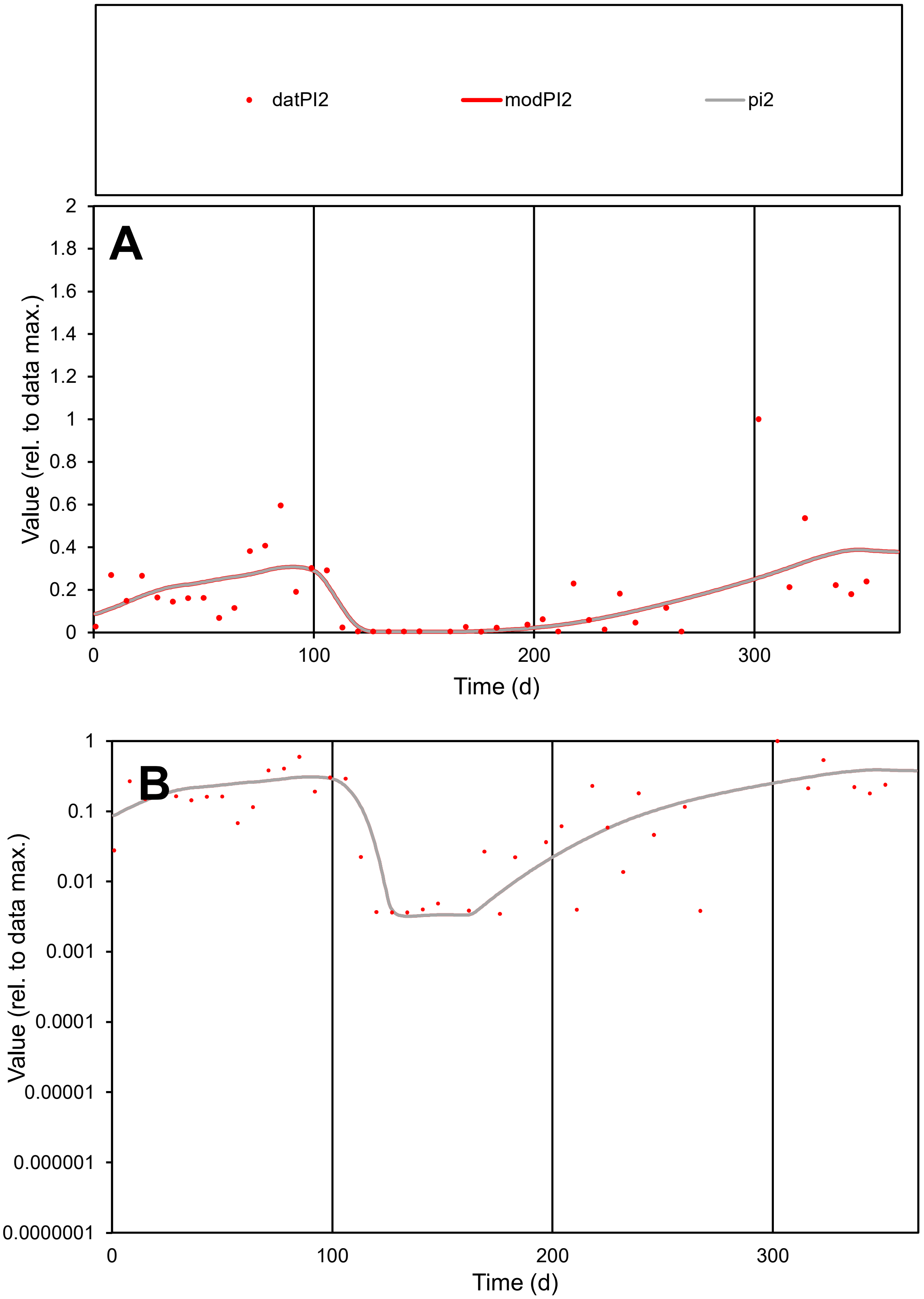


**C1**

Variables
PI2 Provide jtd or Dst
CalibOnly?
2 1 to reproduce of
oNormalize?
2 1=no, 2=yes

ofxn calc*
0.93312
ofxn mod
0.9448

sRun
RunA1
zout
1

Ave
Data
0.00137
Model
0.0013

data/vMax
0.00829

ofxng
319.391

Min d/m
1.08727

ave/min d
47.538
ave/min m
48.9618

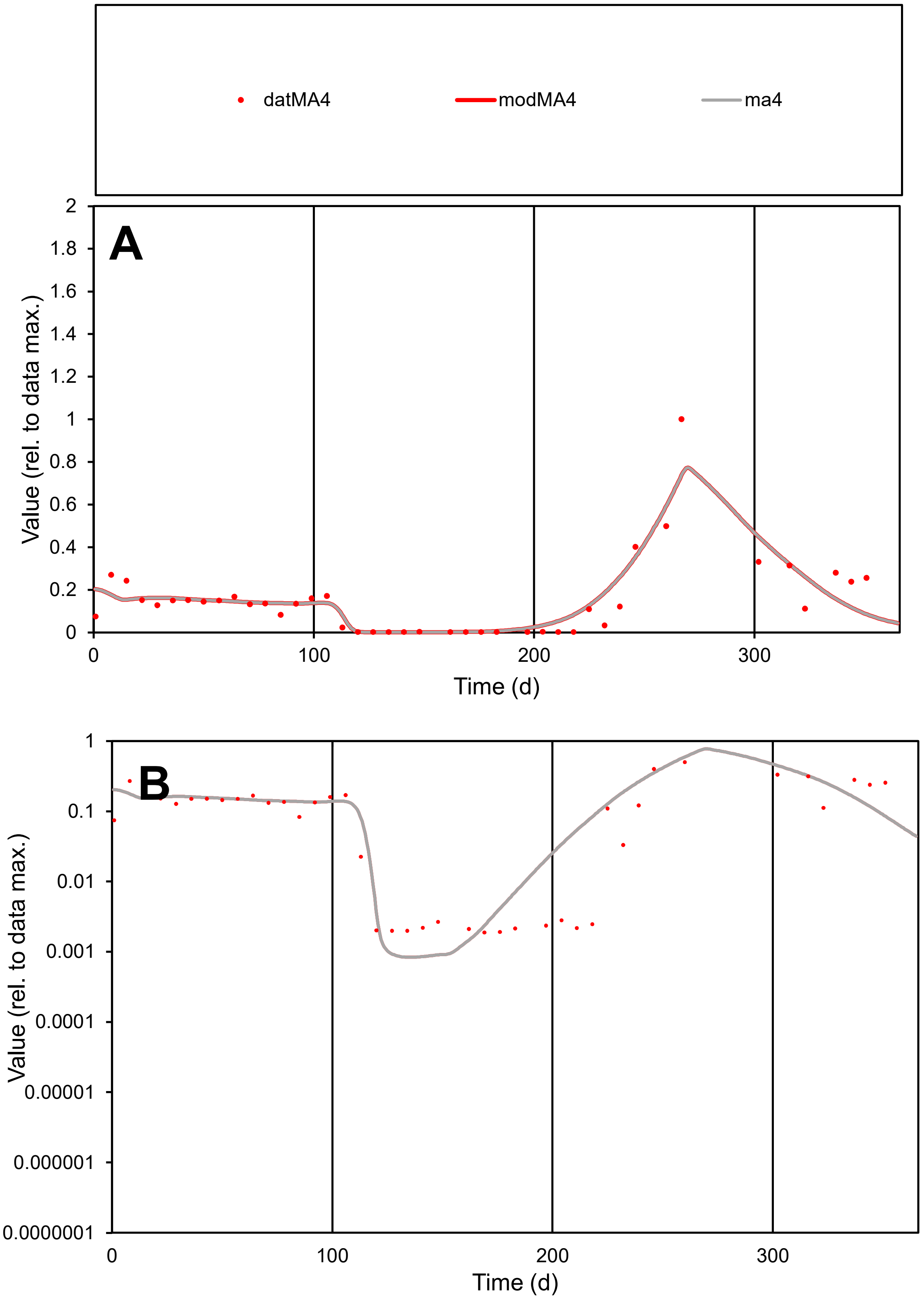


**C1**

Variables
MA4 Provide jtd or Dst
CalibOnly?
2 1 to reproduce of
oNormalize?
2 1=no, 2=yes

ofxn calc*
0.52891
ofxn mod
0.5614

sRun
RunA1
zout
1

Ave
Data
0.00152
Model
0.00156

data/vMax
0.0103

ofxng
319.391

Min d/m
2.24954

ave/min d
78.2604
ave/min m
180.632

330_Calib_S_A

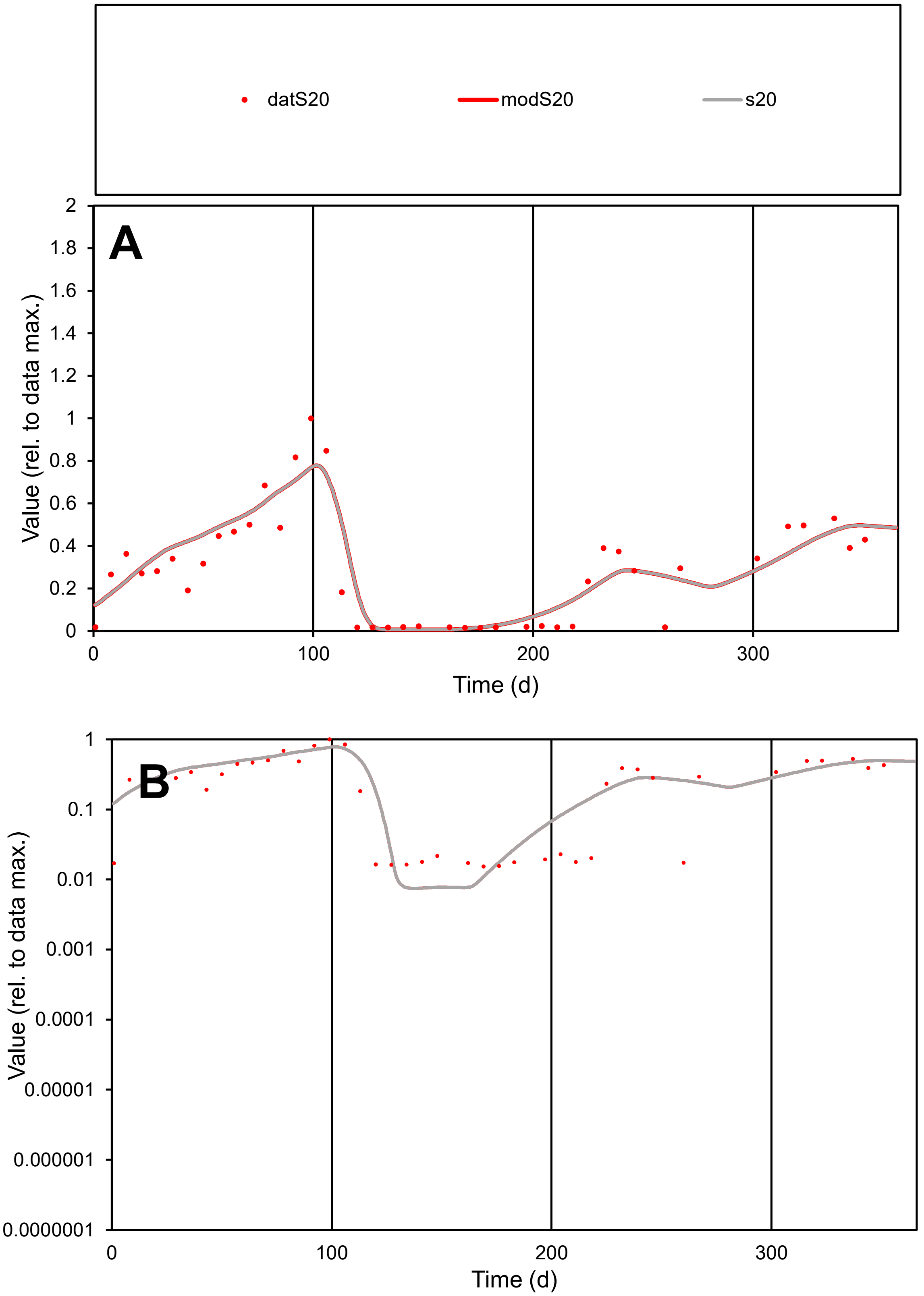


**C1**

Variables
S20 Provide jtd or Dst
CalibOnly?
2 1 to reproduce of
oNormalize?
2 1=no, 2=yes

ofxn calc*
0.3653
ofxn mod
0.40251

sRun
RunA1
zout
1

Ave
Data
0.00145
Model
0.00153

data/vMax
0.0051

ofxng
319.391

Min d/m
2.04186

ave/min d
18.5248
ave/min m
39.8249

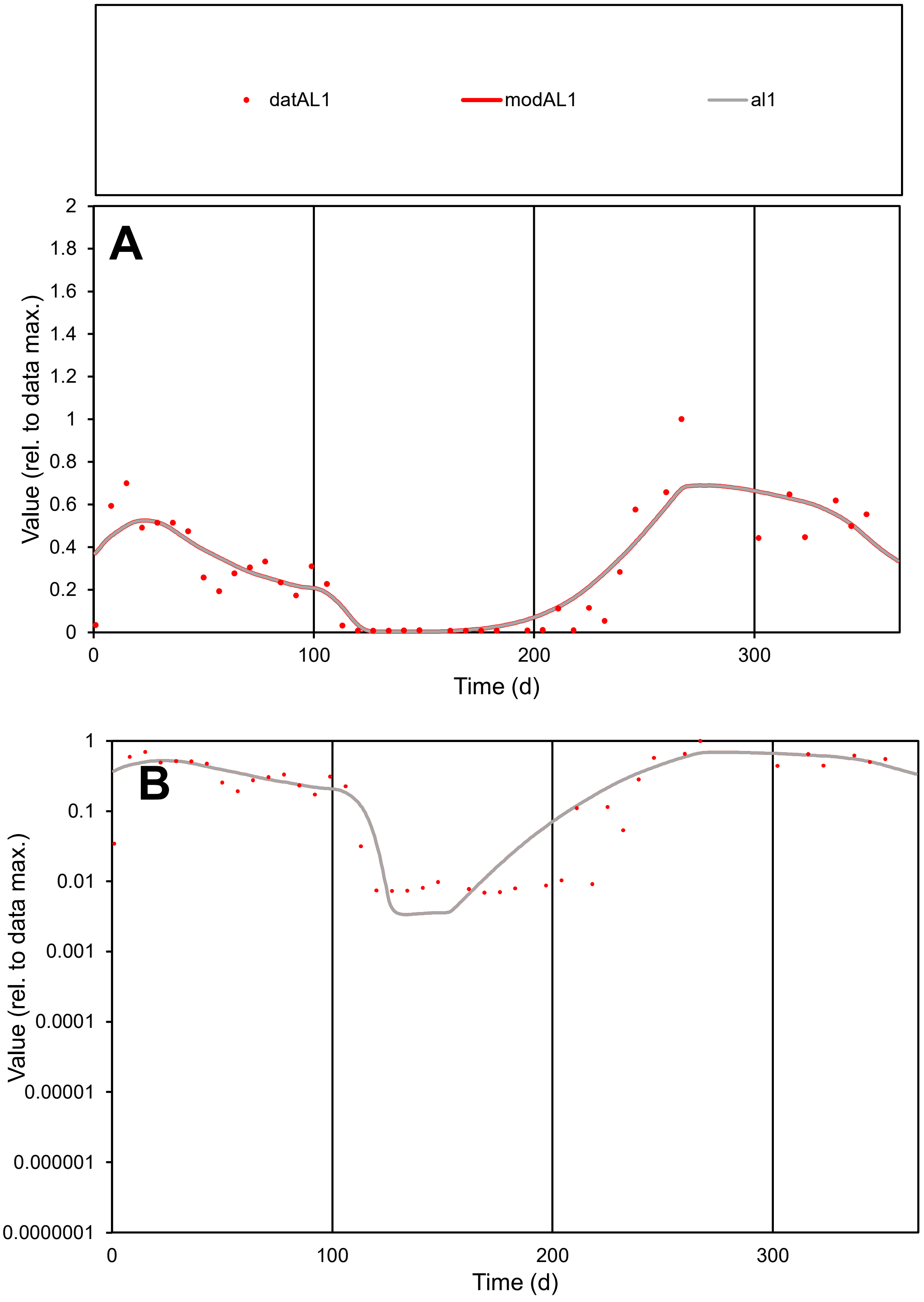


**C1**

Variables
AL1 Provide jtd or Dst
CalibOnly?
2 1 to reproduce of
oNormalize?
2 1=no, 2=yes

ofxn calc*
0.41089
ofxn mod
0.43634

sRun
RunA1
zout
1

Ave
Data
0.00178
Model
0.00185

data/vMax
0.00636

ofxng
319.391

Min d/m
2.05393

ave/min d
40.2678
ave/min m
86.1117

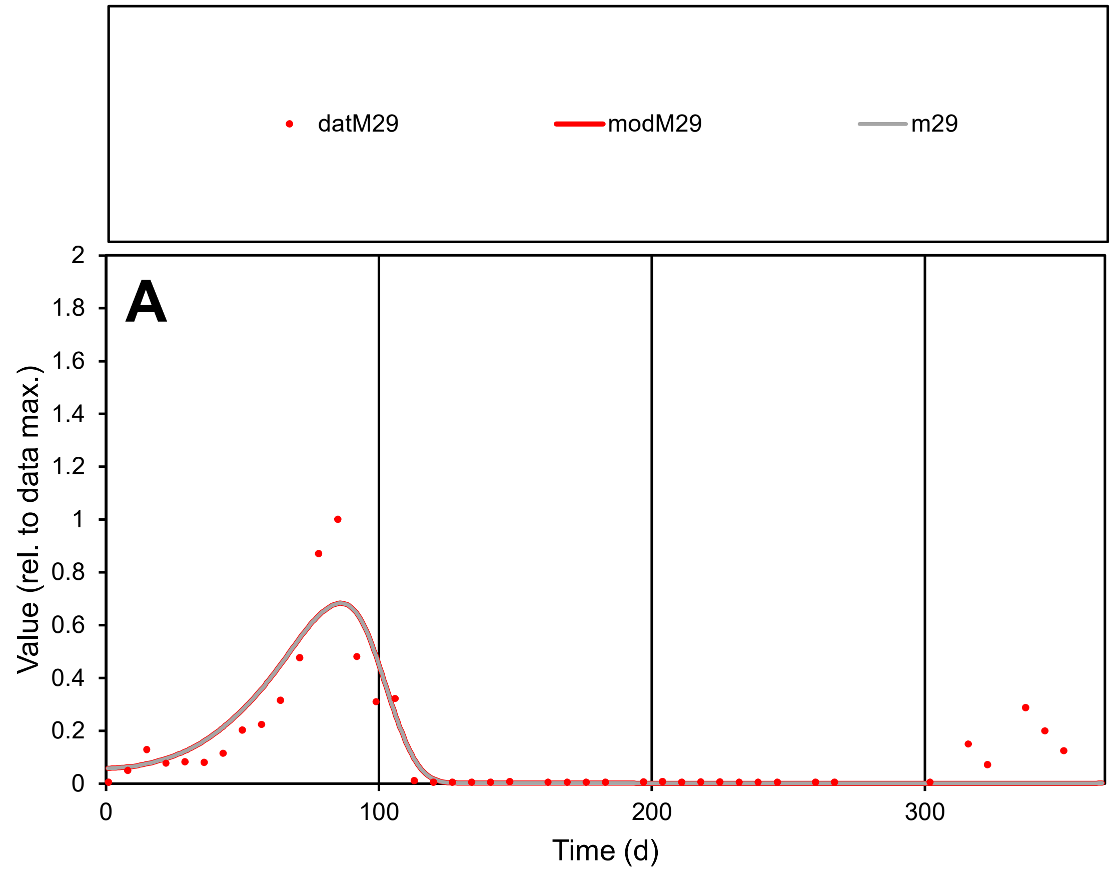


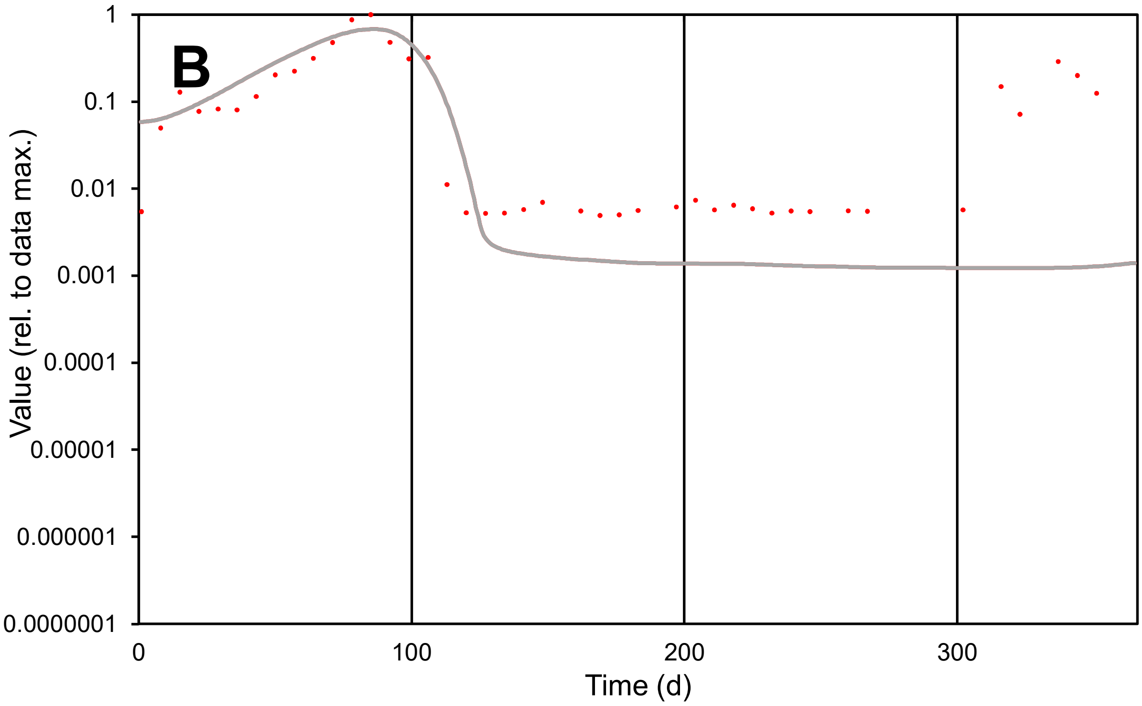


**C1**

Variables
M29 Provide jtd or Dst
CalibOnly?
2 1 to reproduce of
oNormalize?
2 1=no, 2=yes

ofxn calc*
0.75308
ofxn mod
0.7822

sRun
RunA1
zout
1

Ave
Data
0.0015
Model
0.00138

data/vMax
0.01109

ofxng
319.391

Min d/m
4.03461

ave/min d
27.4578
ave/min m
101.826

330_Calib_S_A

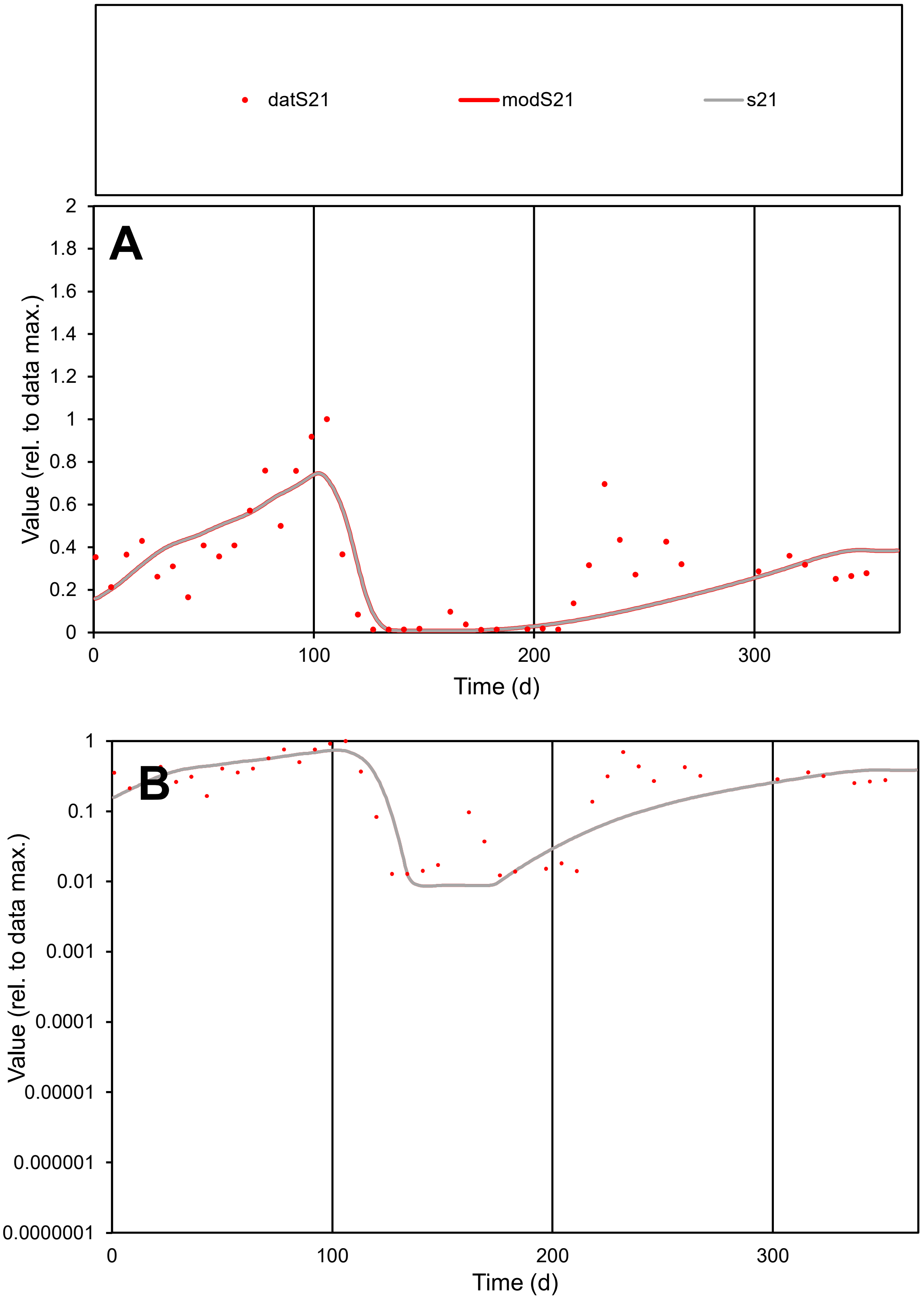


**C1**

Variables
S21 Provide jtd or Dst
CalibOnly?
2 1 to reproduce of
oNormalize?
2 1=no, 2=yes

ofxn calc*
0.54272
ofxn mod
0.56014

sRun
RunA1
zout
1

Ave
Data
0.00103
Model
0.00093

data/vMax
0.00338

ofxng
319.391

Min d/m
1.42695

ave/min d
24.8524
ave/min m
31.8087

330_Calib_S_A

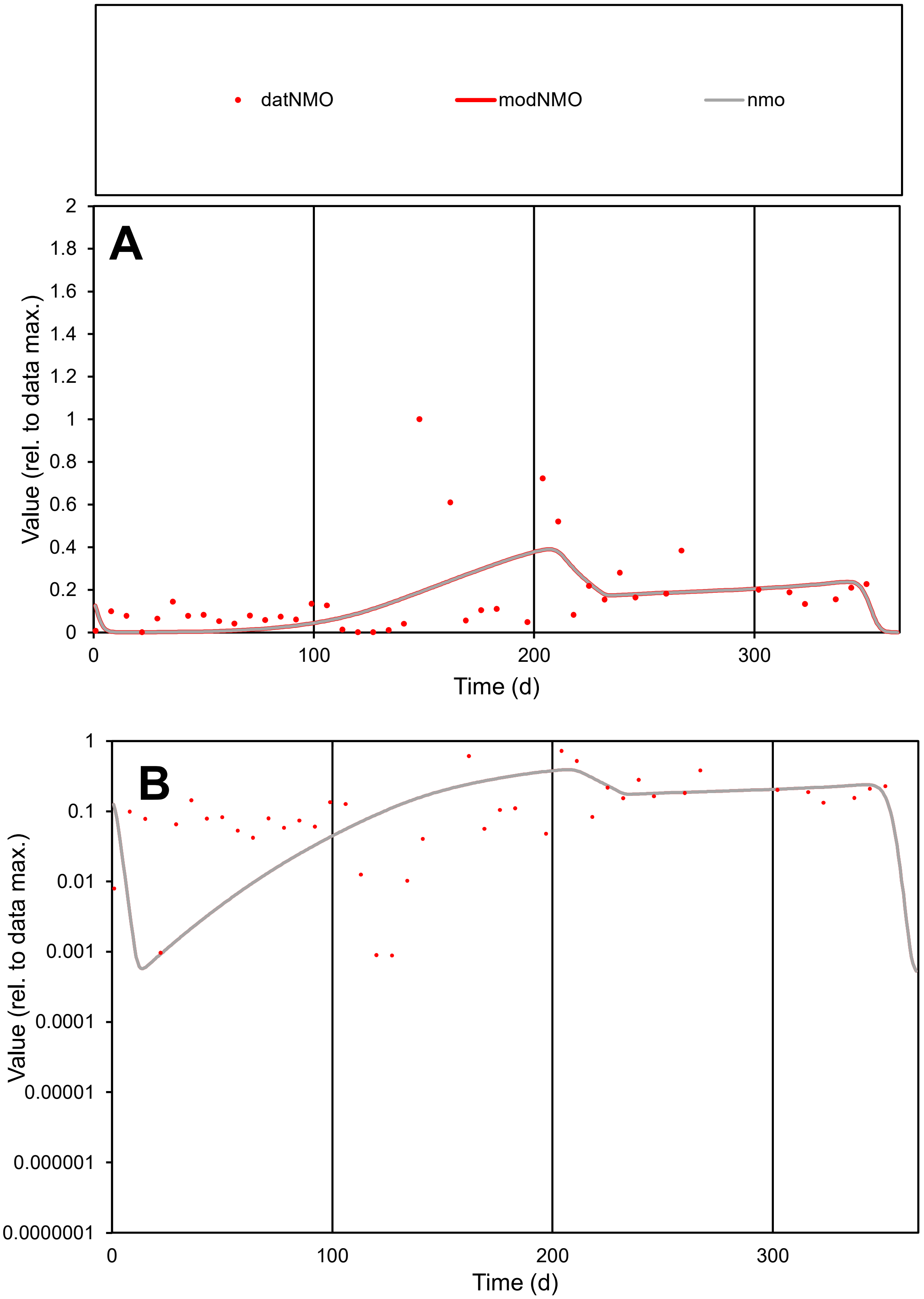


**C1**

Variables
NMO Provide jtd or Dst
CalibOnly?
2 1 to reproduce of
oNormalize?
2 1=no, 2=yes

ofxn calc*
1.09096
ofxn mod
1.10186

sRun
RunA1
zout
1

Ave
Data
0.00144
Model
0.00124

data/vMax
0.00866

ofxng
319.391

Min d/m
1.34233

ave/min d
188.404
ave/min m
216.943

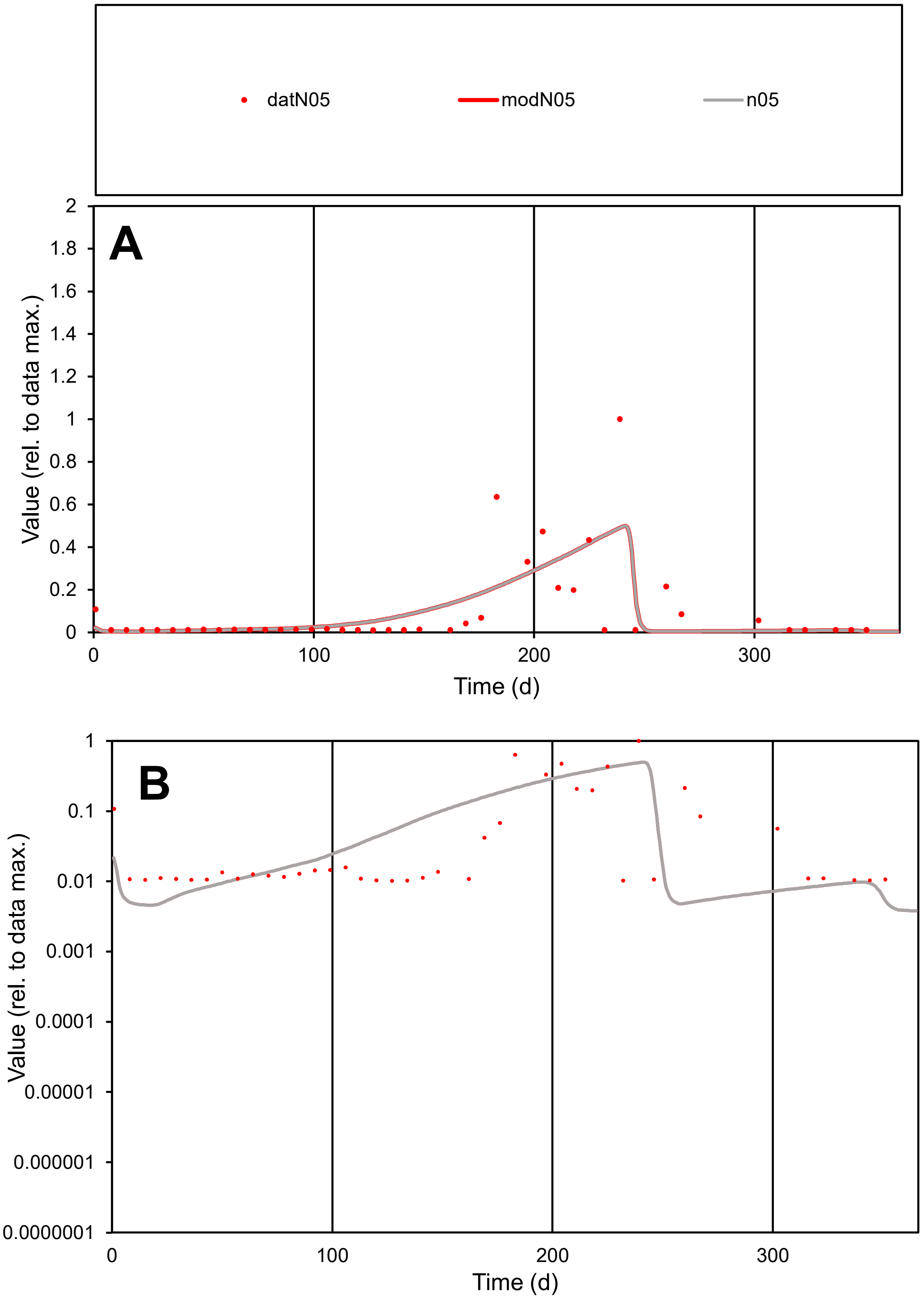


**C1**

Variables
N05 Provide jtd or Dst
CalibOnly?
2 1 to reproduce of
oNormalize?
2 1=no, 2=yes

ofxn calc*
1.40863
ofxn mod
1.4359

sRun
RunA1
zout
1

Ave
Data
0.00041
Model
0.0004

data/vMax
0.00415

ofxng
319.391

Min d/m
2.23835

ave/min d
9.78169
ave/min m
21.3893

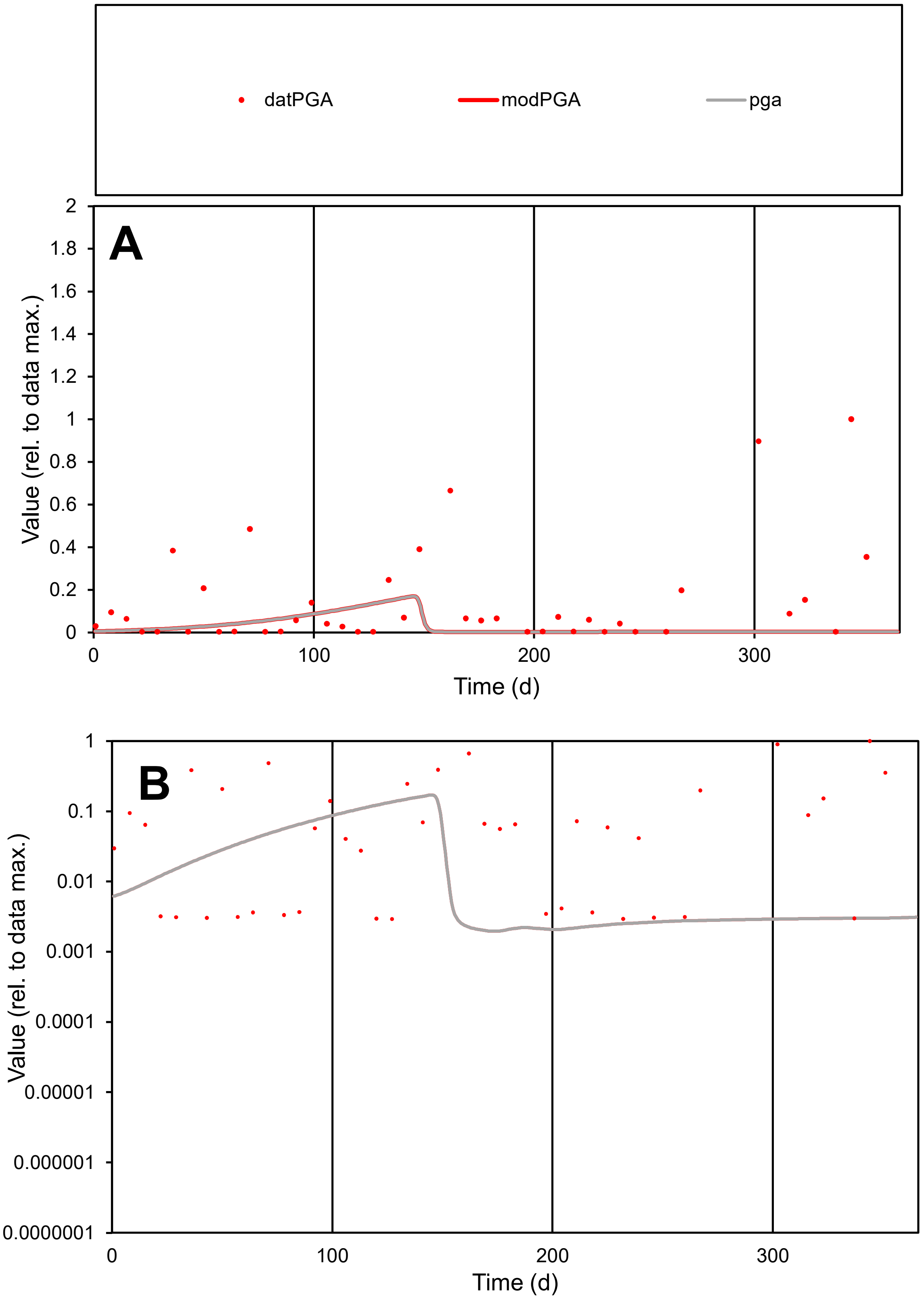


**C1**

Variables
PGA Provide jtd or Dst
CalibOnly?
2 1 to reproduce of
oNormalize?
2 1=no, 2=yes

ofxn calc*
1.85553
ofxn mod
1.85922

sRun
RunA1
zout
1

Ave
Data
0.00088
Model
0.00022

data/vMax
0.00619

ofxng
319.391

Min d/m
1.46272

ave/min d
48.9886
ave/min m
17.7917

330_Calib_S_A

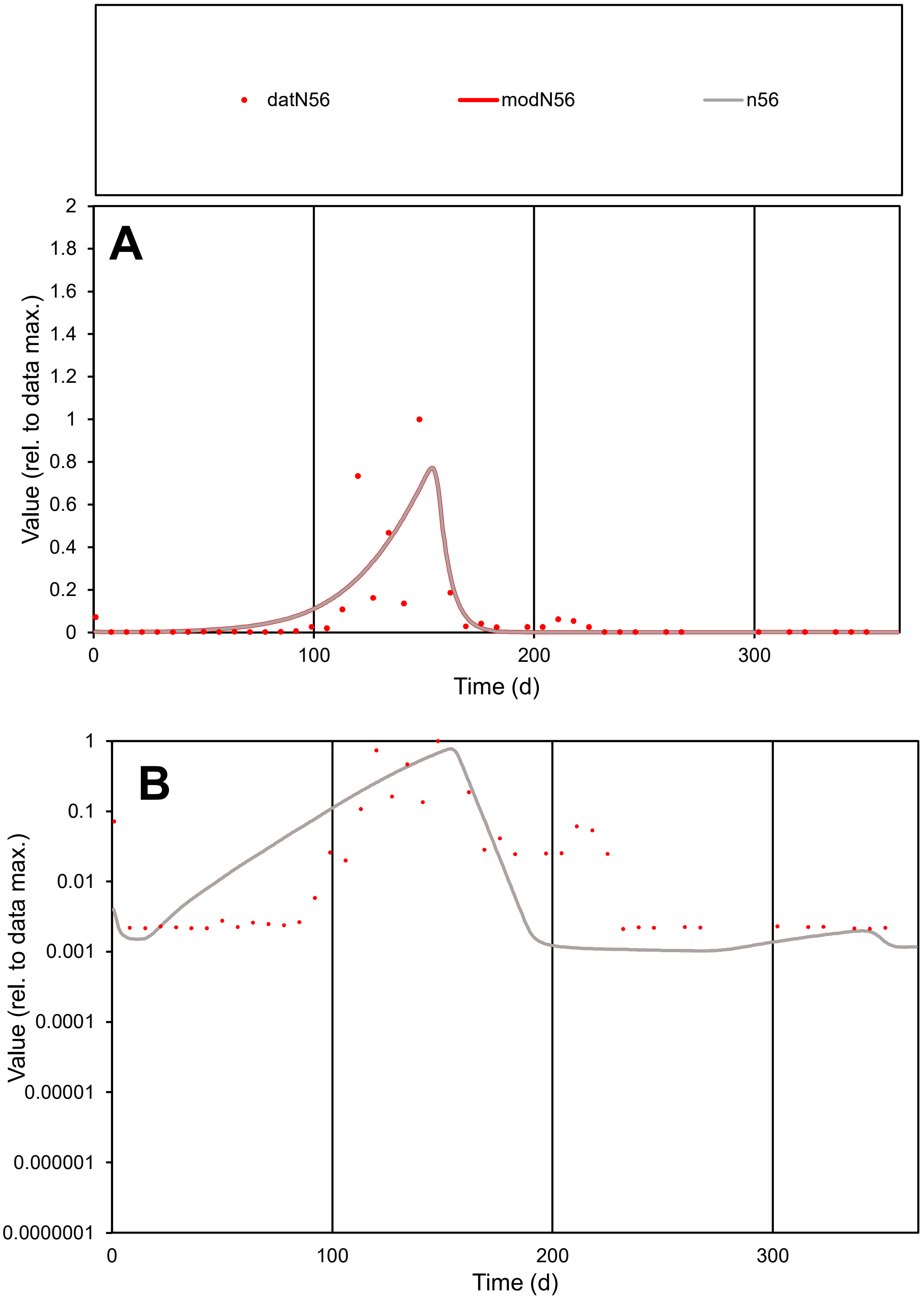


**C1**

Variables
N56 Provide jtd or Dst
CalibOnly?
2 1 to reproduce of
oNormalize?
2 1=no, 2=yes

ofxn calc*
1.4915
ofxn mod
1.51808

sRun
RunA1
zout
1

Ave
Data
0.00134
Model
0.00137

data/vMax
0.01727

ofxng
319.391

Min d/m
2.04724

ave/min d
36.7313
ave/min m
76.808

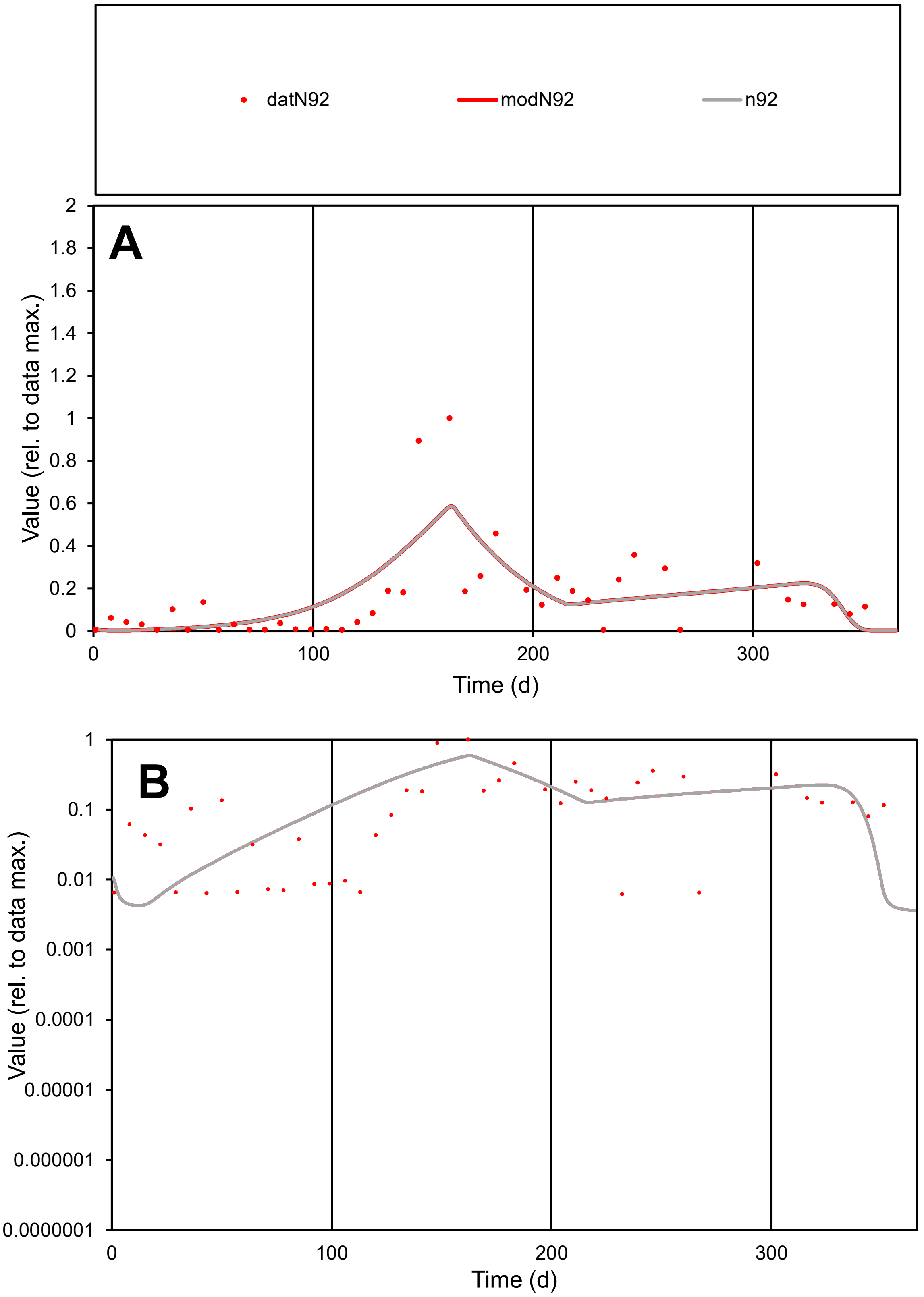


**C1**

Variables
N92 Provide jtd or Dst
CalibOnly?
2 1 to reproduce of
oNormalize?
2 1=no, 2=yes

ofxn calc*
0.92932
ofxn mod
0.9509

sRun
RunA1
zout
1

Ave
Data
0.00101
Model
0.00105

data/vMax
0.0065

ofxng
319.391

Min d/m
1.43094

ave/min d
25.014
ave/min m
36.926

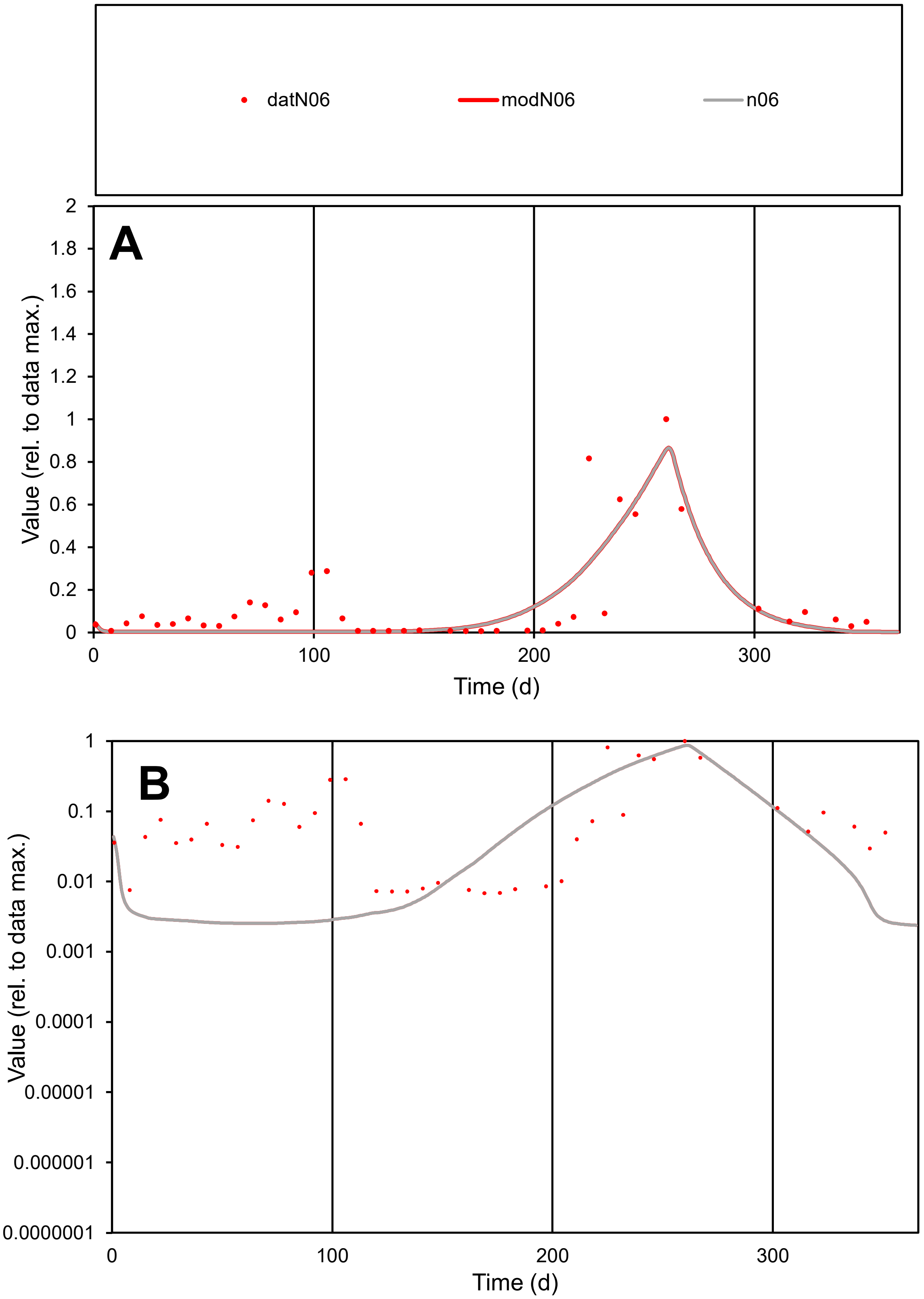


**C1**

Variables
N06 Provide jtd or Dst
CalibOnly?
2 1 to reproduce of
oNormalize?
2 1=no, 2=yes

ofxn calc*
0.9384
ofxn mod
0.96234

sRun
RunA1
zout
1

Ave
Data
0.00132
Model
0.00104

data/vMax
0.00959

ofxng
319.391

Min d/m
2.67939

ave/min d
20.1487
ave/min m
42.6206

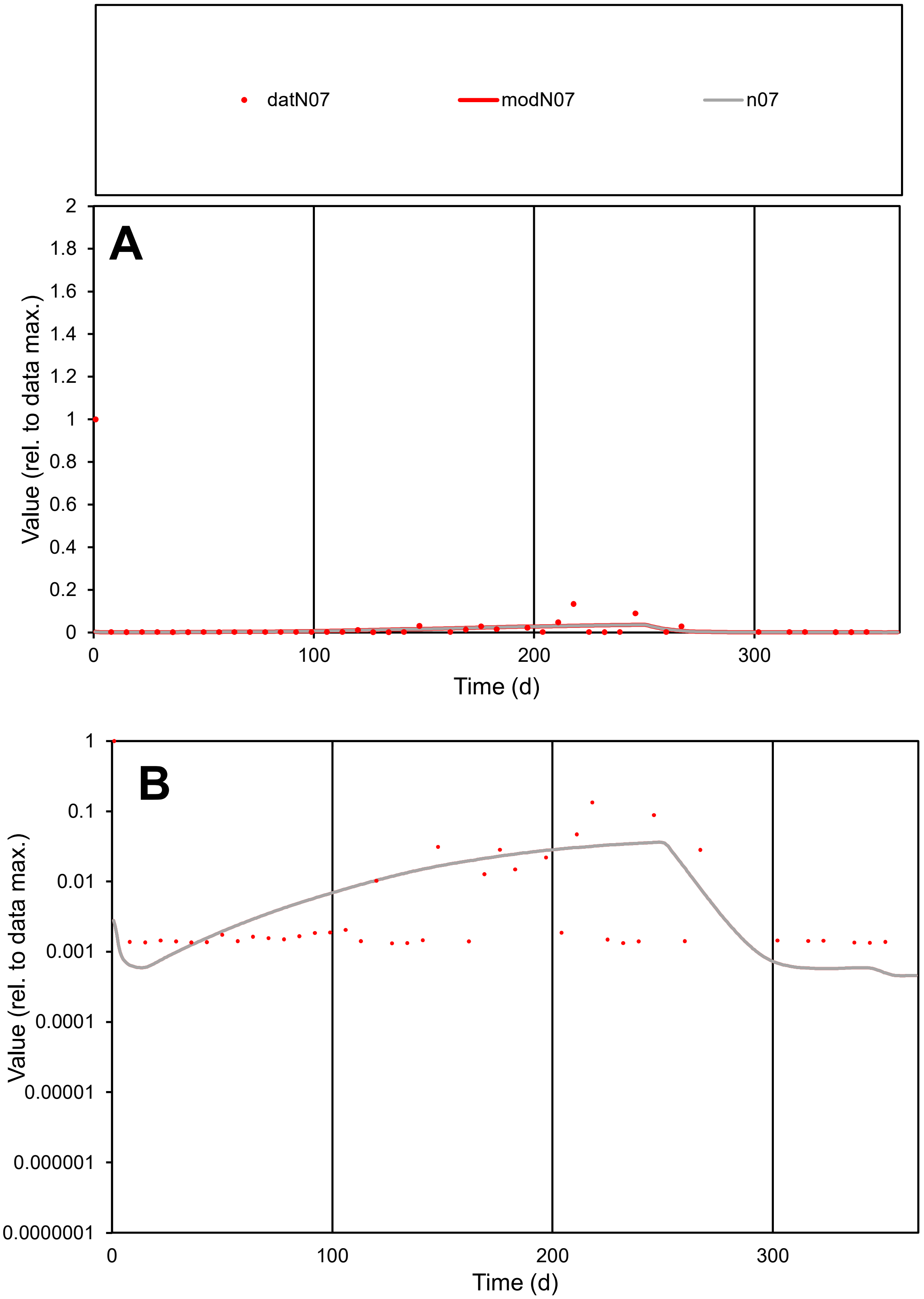


**C1**

Variables
N07 Provide jtd or Dst
CalibOnly?
2 1 to reproduce of
oNormalize?
2 1=no, 2=yes

ofxn calc*
4.46271
ofxn mod
4.46691

sRun
RunA1
zout
1

Ave
Data
0.00048
Model
0.00016

data/vMax
0.01367

ofxng
319.391

Min d/m
2.70088

ave/min d
26.3756
ave/min m
24.2147

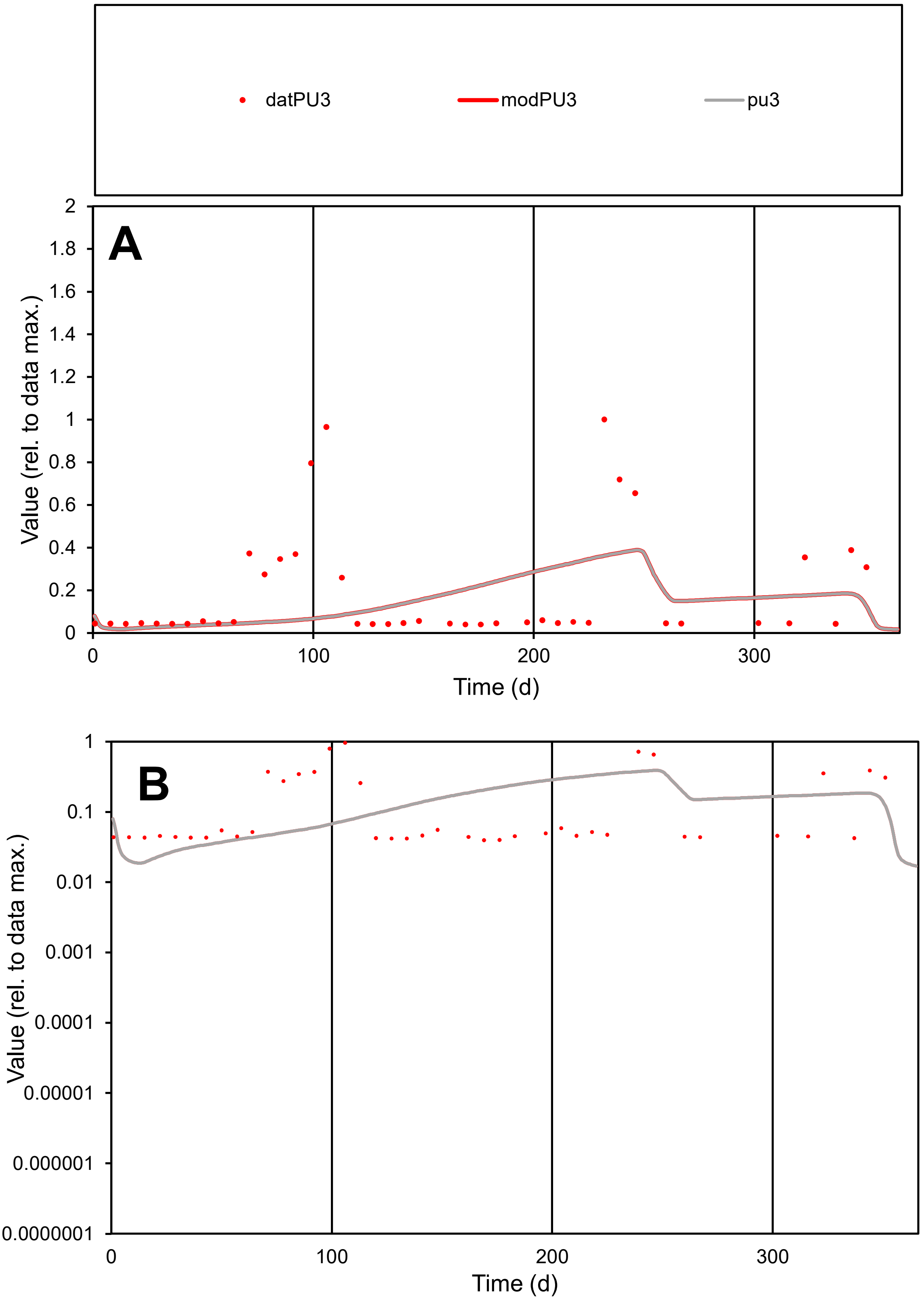


**C1**

Variables
PU3 Provide jtd or Dst
CalibOnly?
2 1 to reproduce of
oNormalize?
2 1=no, 2=yes

ofxn calc*
1.37131
ofxn mod
1.38187

sRun
RunA1
zout
1

Ave
Data
0.00074
Model
0.00057

data/vMax
0.00383

ofxng
319.391

Min d/m
2.03169

ave/min d
4.90243
ave/min m
7.68228

330_Calib_S_A

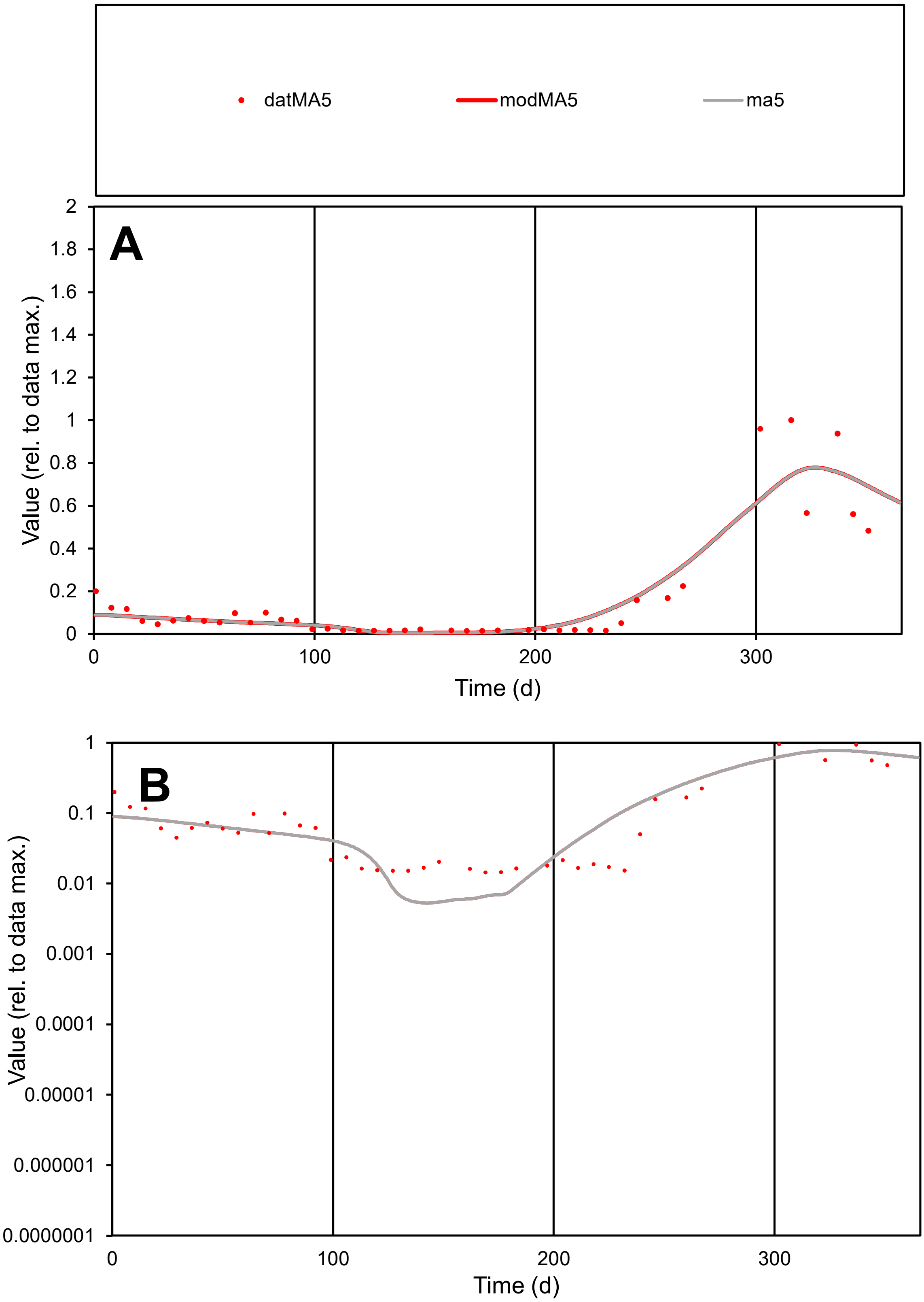


**C1**

Variables
MA5 Provide jtd or Dst
CalibOnly?
2 1 to reproduce of
oNormalize?
2 1=no, 2=yes

ofxn calc*
0.60347
ofxn mod
0.62745

sRun
RunA1
zout
1

Ave
Data
0.00103
Model
0.00104

data/vMax
0.00655

ofxng
319.391

Min d/m
2.72977

ave/min d
10.8851
ave/min m
30.131

330_Calib_S_A

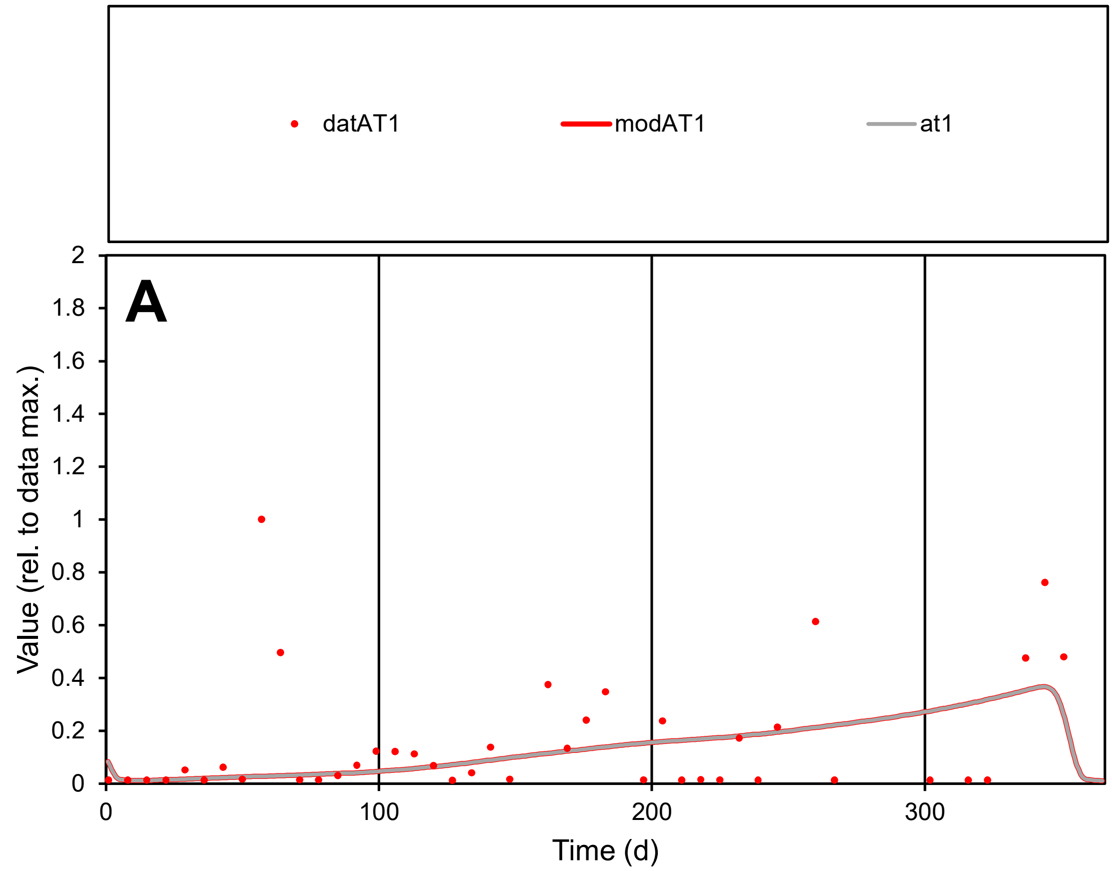


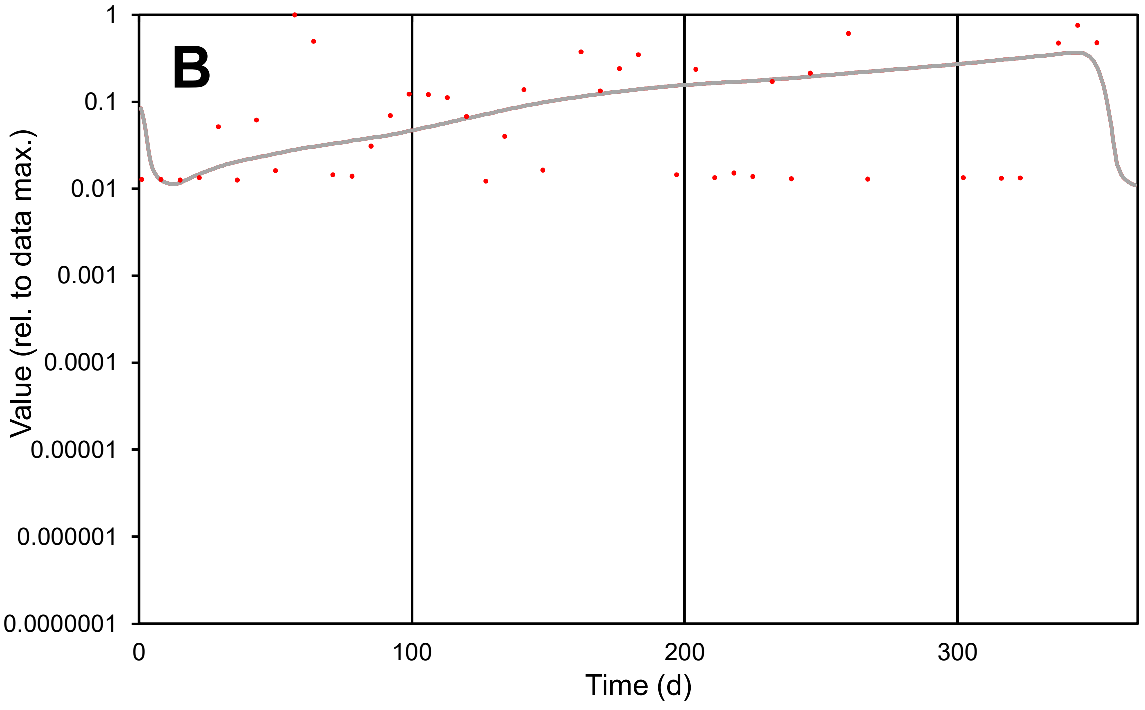


**C1**

Variables
AT1 Provide jtd or Dst
CalibOnly?
2 1 to reproduce of
oNormalize?
2 1=no, 2=yes

ofxn calc*
1.42469
ofxn mod
1.43262

sRun
RunA1
zout
1

Ave
Data
0.00065
Model
0.00051

data/vMax
0.00414

ofxng
319.391

Min d/m
1.03195

ave/min d
12.8605
ave/min m
10.3008

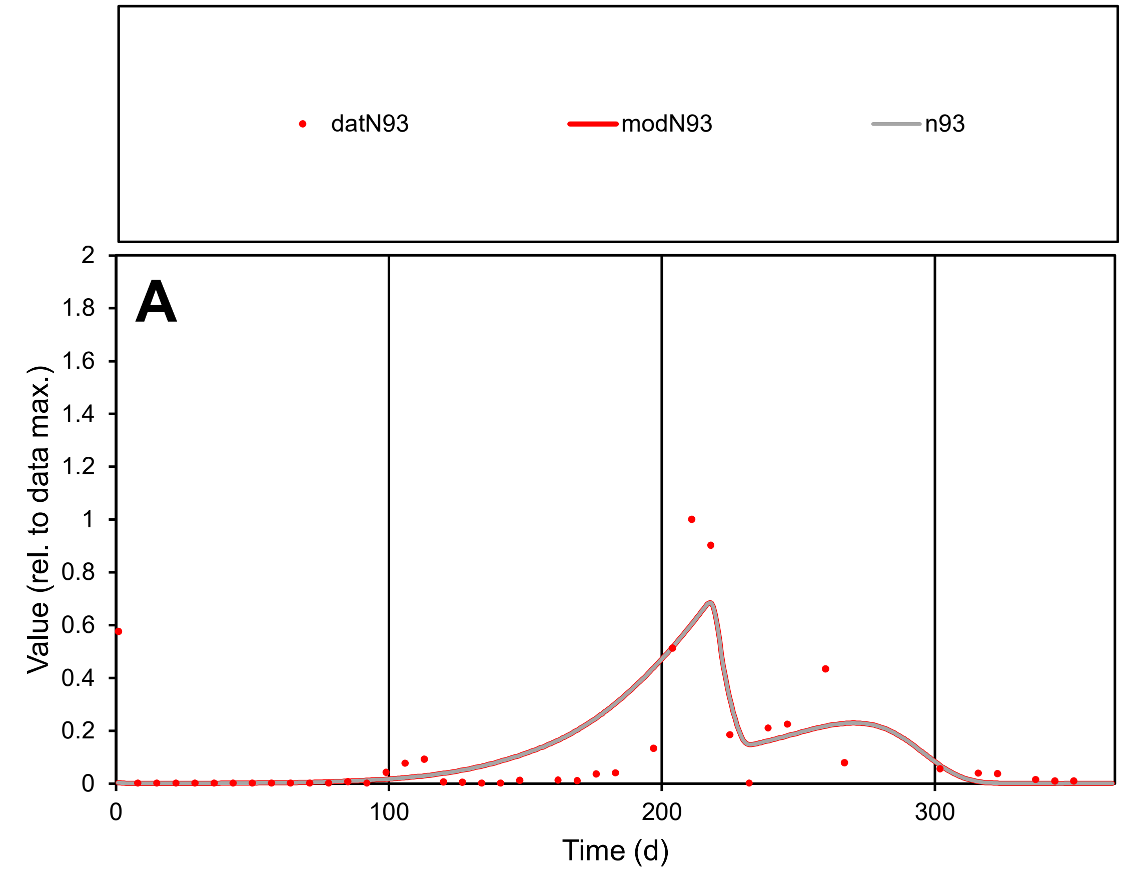


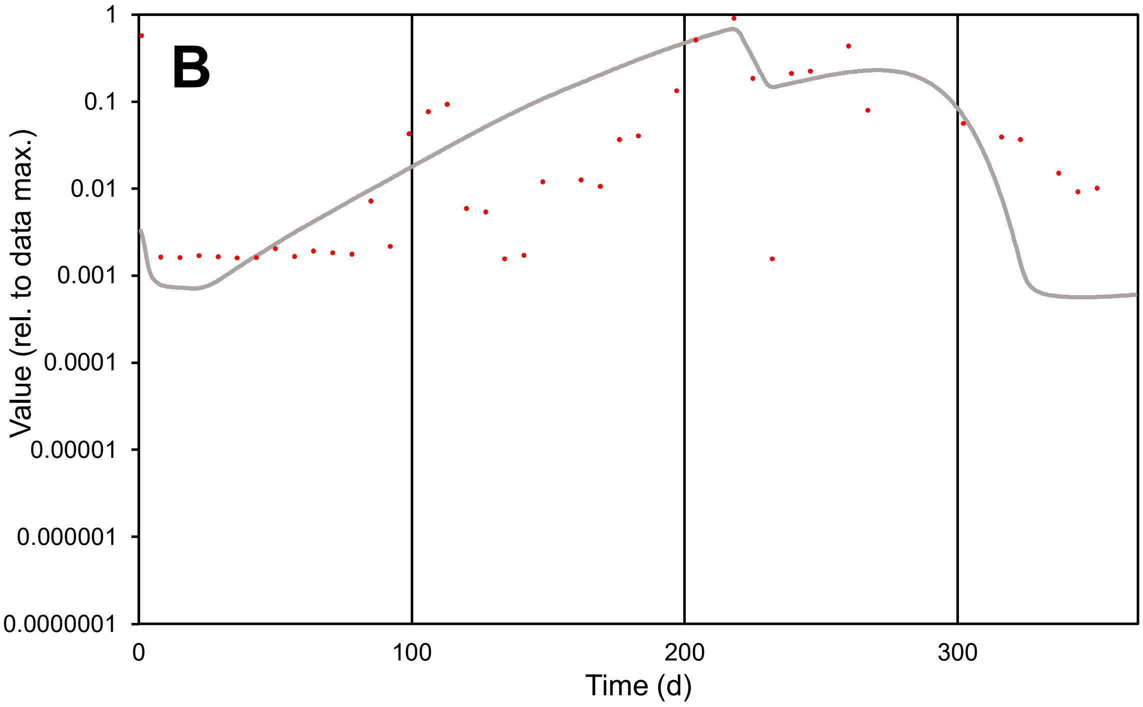


**C1**

Variables

N93 Provide jtd or Dst

CalibOnly?

2 1 to reproduce of

oNormalize?

2 1=no, 2=yes

ofxn calc*

1.29562

ofxn mod

1.33196

sRun

RunA1

zout

1

Ave

Data

0.00135

Model

0.0014

data/vMax

0.01183

ofxng

319.391

Min d/m

2.75713

ave/min d

73.1509

ave/min m

208.526

330_Calib_S_A

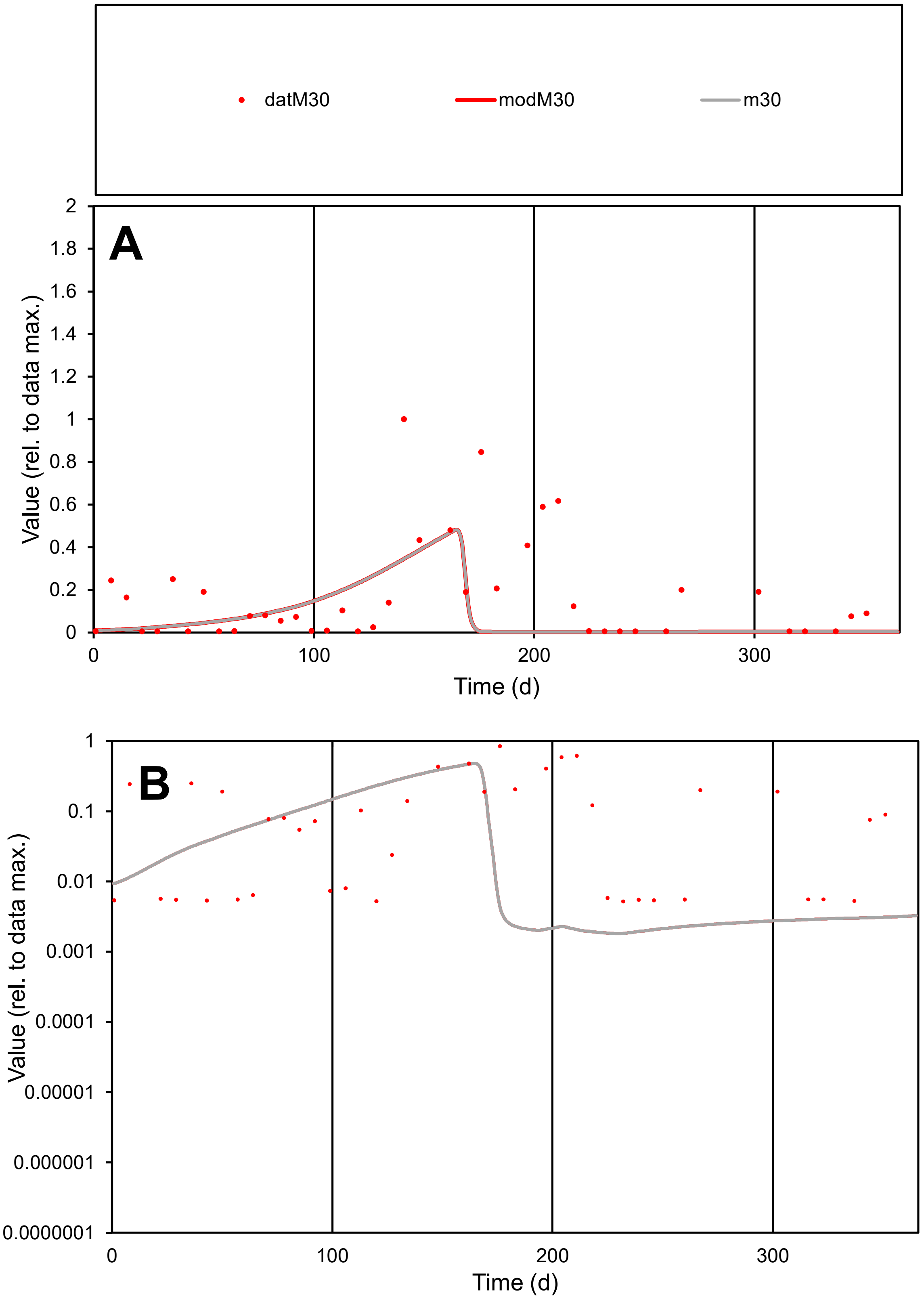


**C1**

Variables
M30 Provide jtd or Dst
CalibOnly?
2 1 to reproduce of
oNormalize?
2 1=no, 2=yes

ofxn calc*
1.47831
ofxn mod
1.4851

sRun
RunA1
zout
1

Ave
Data
0.00139
Model
0.0007

data/vMax
0.0084

ofxng
319.391

Min d/m
2.82783

ave/min d
31.7965
ave/min m
45.4326

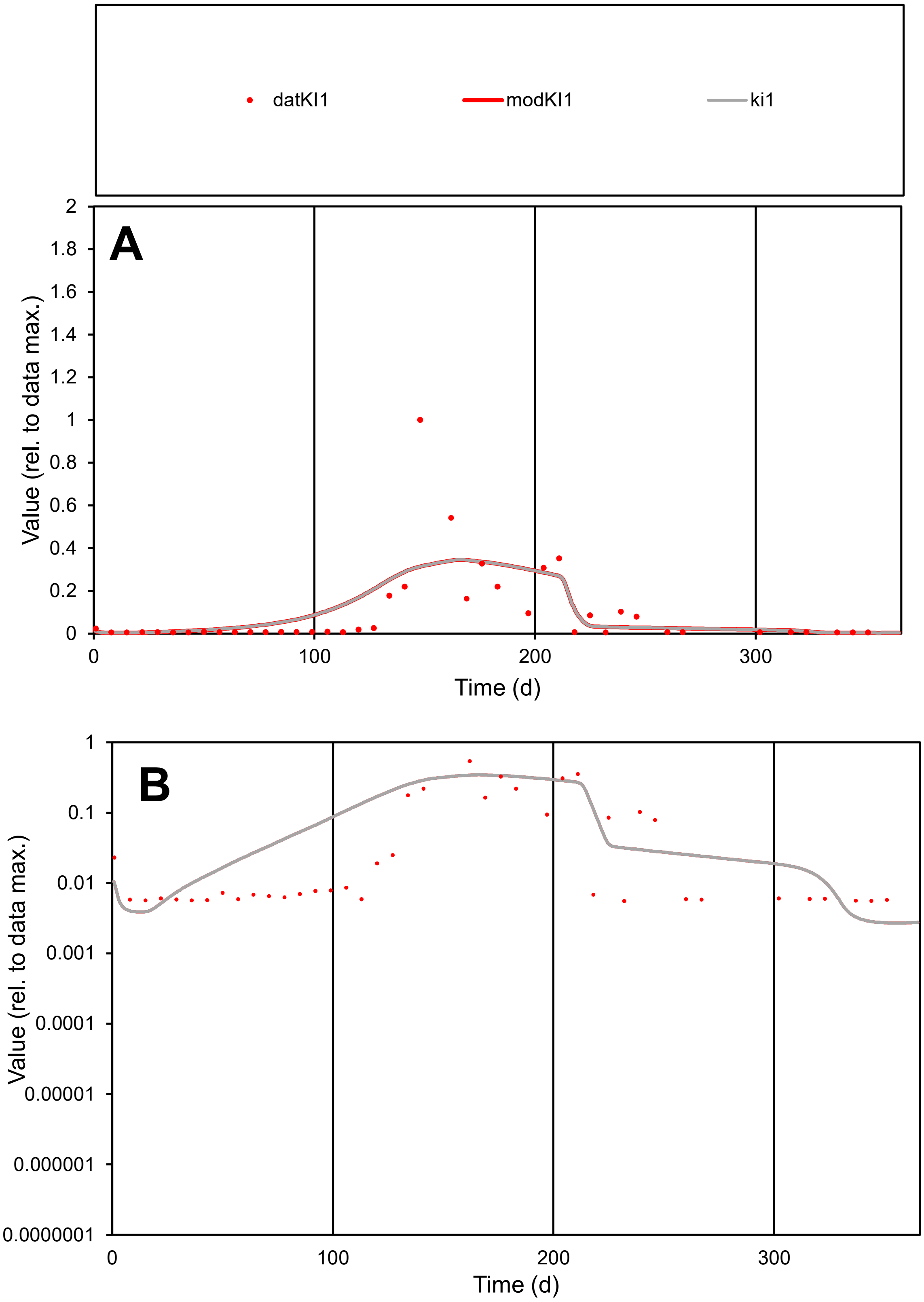


**C1**

Variables
KI1 Provide jtd or Dst
CalibOnly?
2 1 to reproduce of
oNormalize?
2 1=no, 2=yes

ofxn calc*
1.41411
ofxn mod
1.43285

sRun
RunA1
zout
1

Ave
Data
0.00083
Model
0.00093

data/vMax
0.00899

ofxng
319.391

Min d/m
2.04979

ave/min d
16.7421
ave/min m
38.4361

330_Calib_S_A

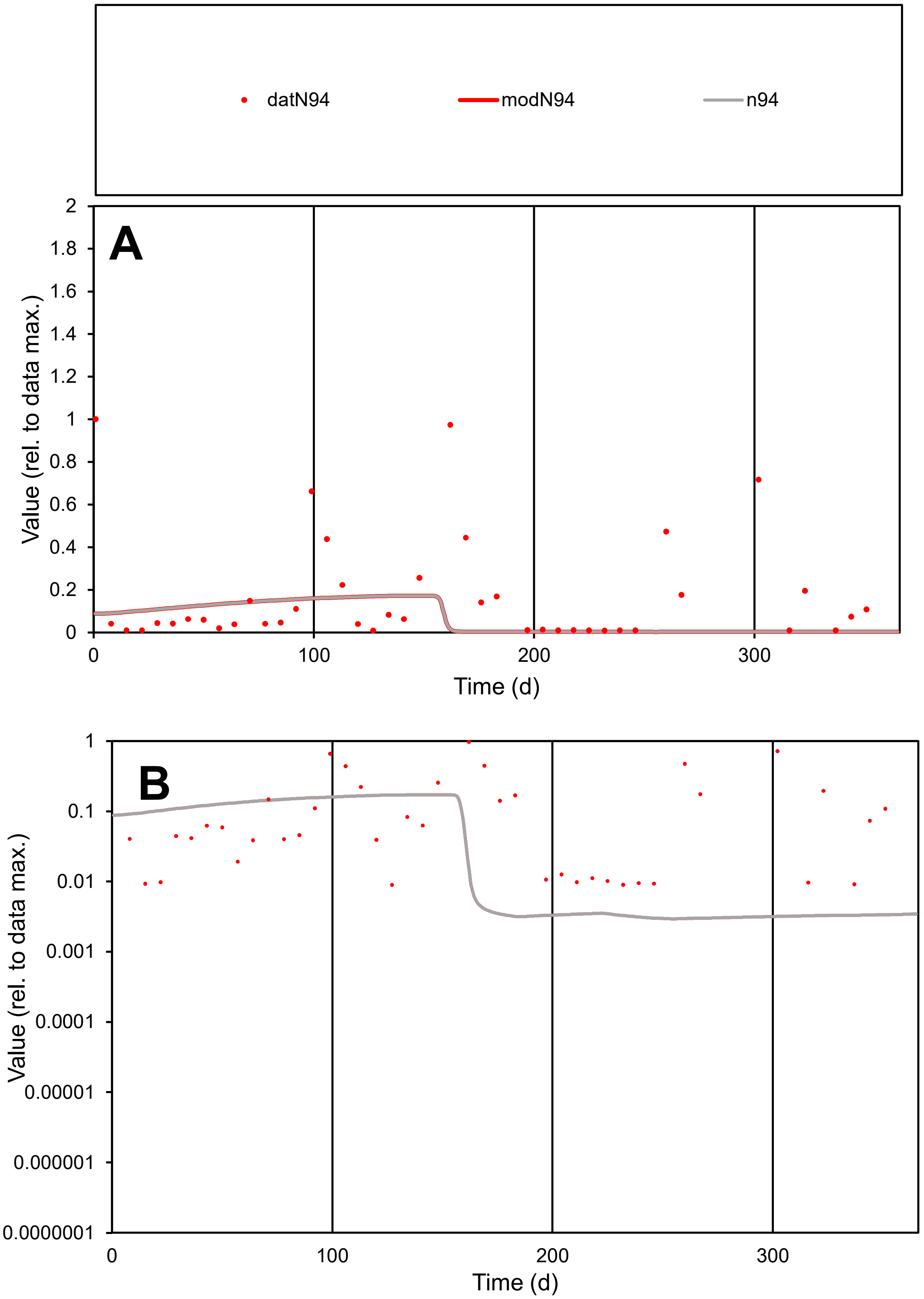


**C1**

Variables
N94 Provide jtd or Dst
CalibOnly?
2 1 to reproduce of
oNormalize?
2 1=no, 2=yes

ofxn calc*
1.67682
ofxn mod
1.68282

sRun
RunA1
zout
1

Ave
Data
0.0009
Model
0.0004

data/vMax
0.00536

ofxng
319.391

Min d/m
3.01759

ave/min d
18.6407
ave/min m
25.2755

330_Calib_S_A

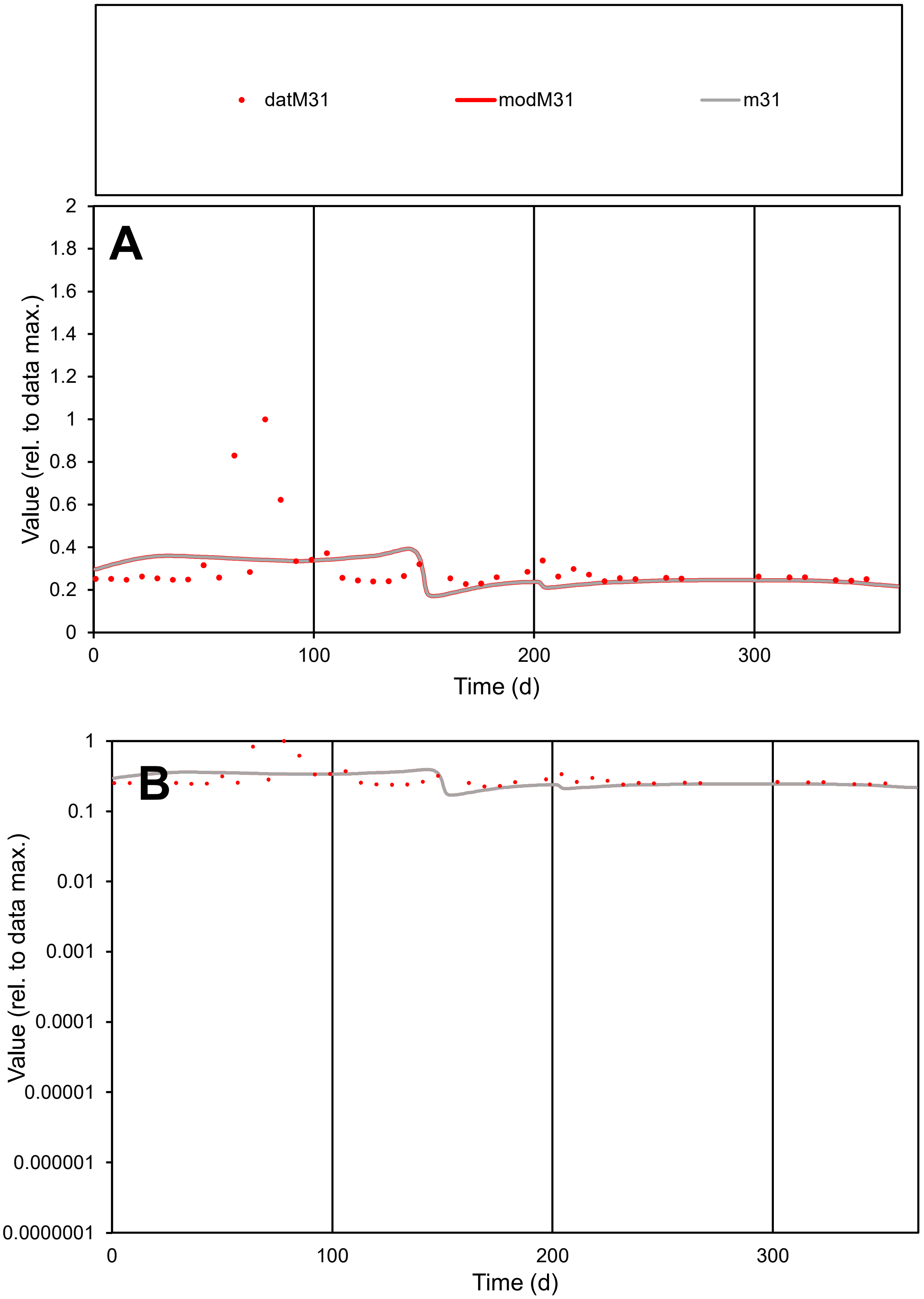


**C1**

Variables
M31 Provide jtd or Dst
CalibOnly?
2 1 to reproduce of
oNormalize?
2 1=no, 2=yes

ofxn calc*
0.48576
ofxn mod
0.49864

sRun
RunA1
zout
1

Ave
Data
0.00011
Model
0.0001

data/vMax
0.00036

ofxng
319.391

Min d/m
1.20962

ave/min d
1.35107
ave/min m
1.54944

330_Calib_S_A

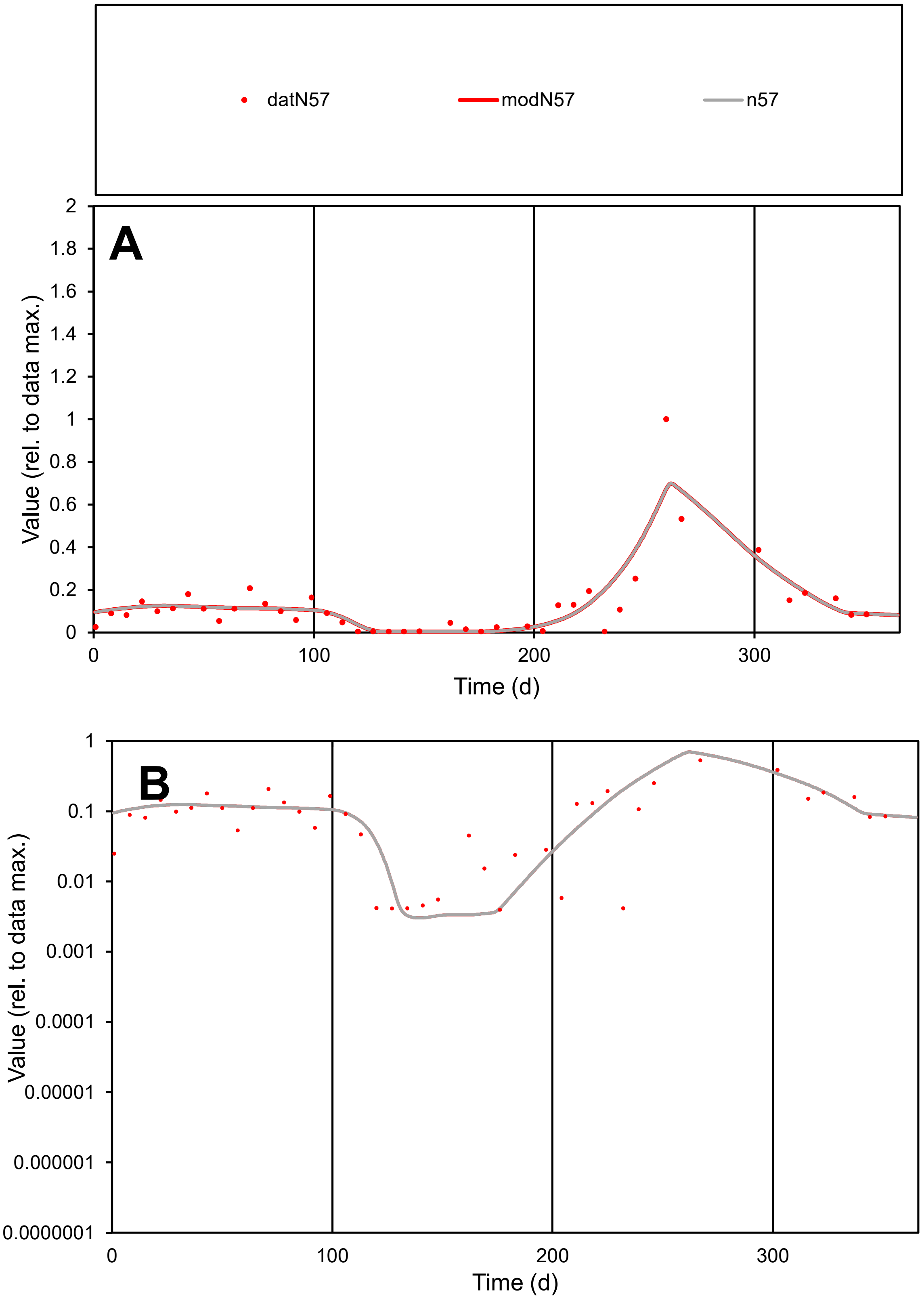


**C1**

Variables
N57 Provide jtd or Dst
CalibOnly?
2 1 to reproduce of
oNormalize?
2 1=no, 2=yes

ofxn calc*
0.61034
ofxn mod
0.63823

sRun
RunA1
zout
1

Ave
Data
0.00118
Model
0.00122

data/vMax
0.00925

ofxng
319.391

Min d/m
1.28887

ave/min d
32.0985
ave/min m
42.8756

330_Calib_S_A

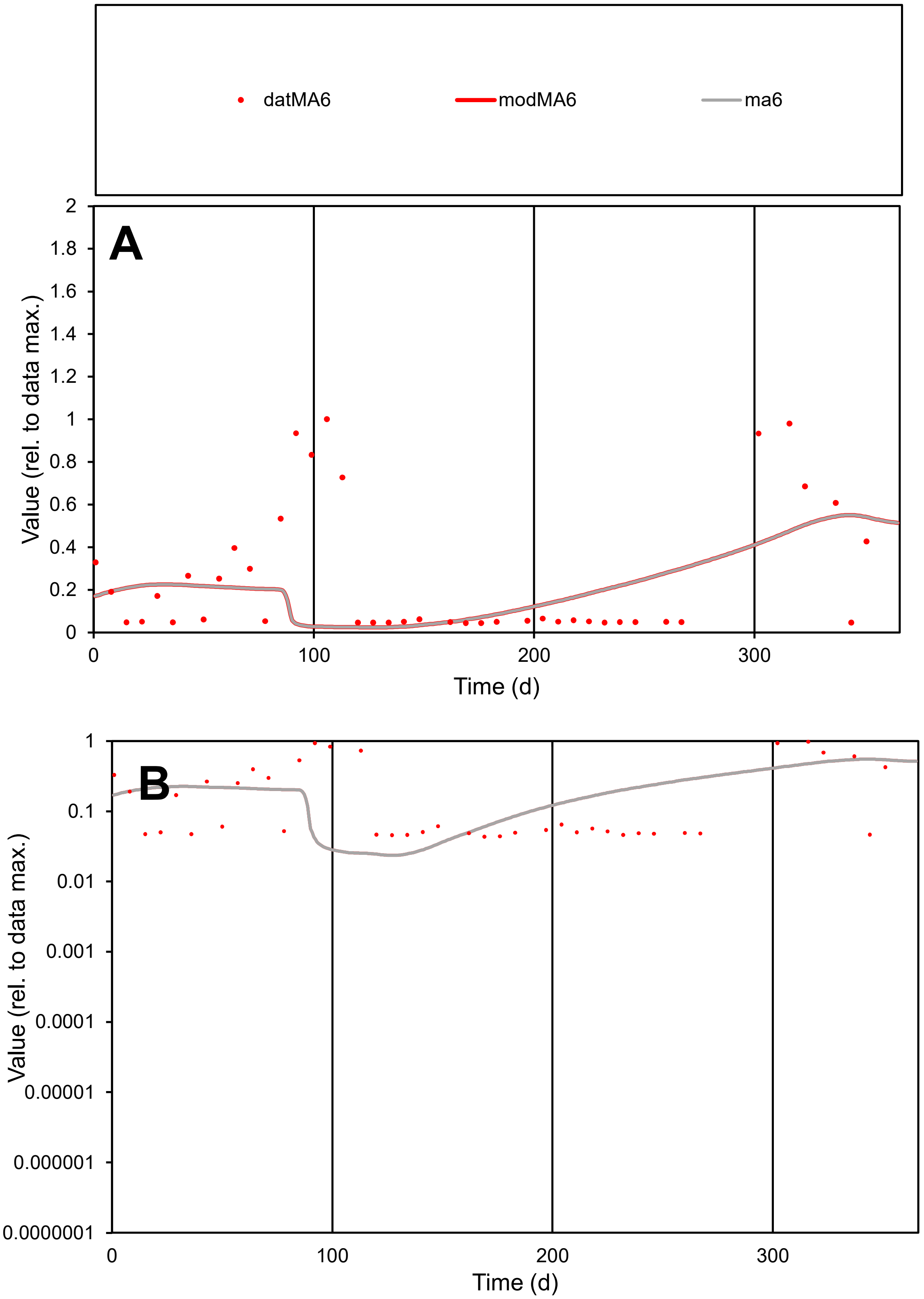


**C1**

Variables
MA6 Provide jtd or Dst
CalibOnly?
2 1 to reproduce of
oNormalize?
2 1=no, 2=yes

ofxn calc*
1.23982
ofxn mod
1.24616

sRun
RunA1
zout
1

Ave
Data
0.0007
Model
0.00054

data/vMax
0.00272

ofxng
319.391

Min d/m
1.83566

ave/min d
5.92749
ave/min m
8.34277

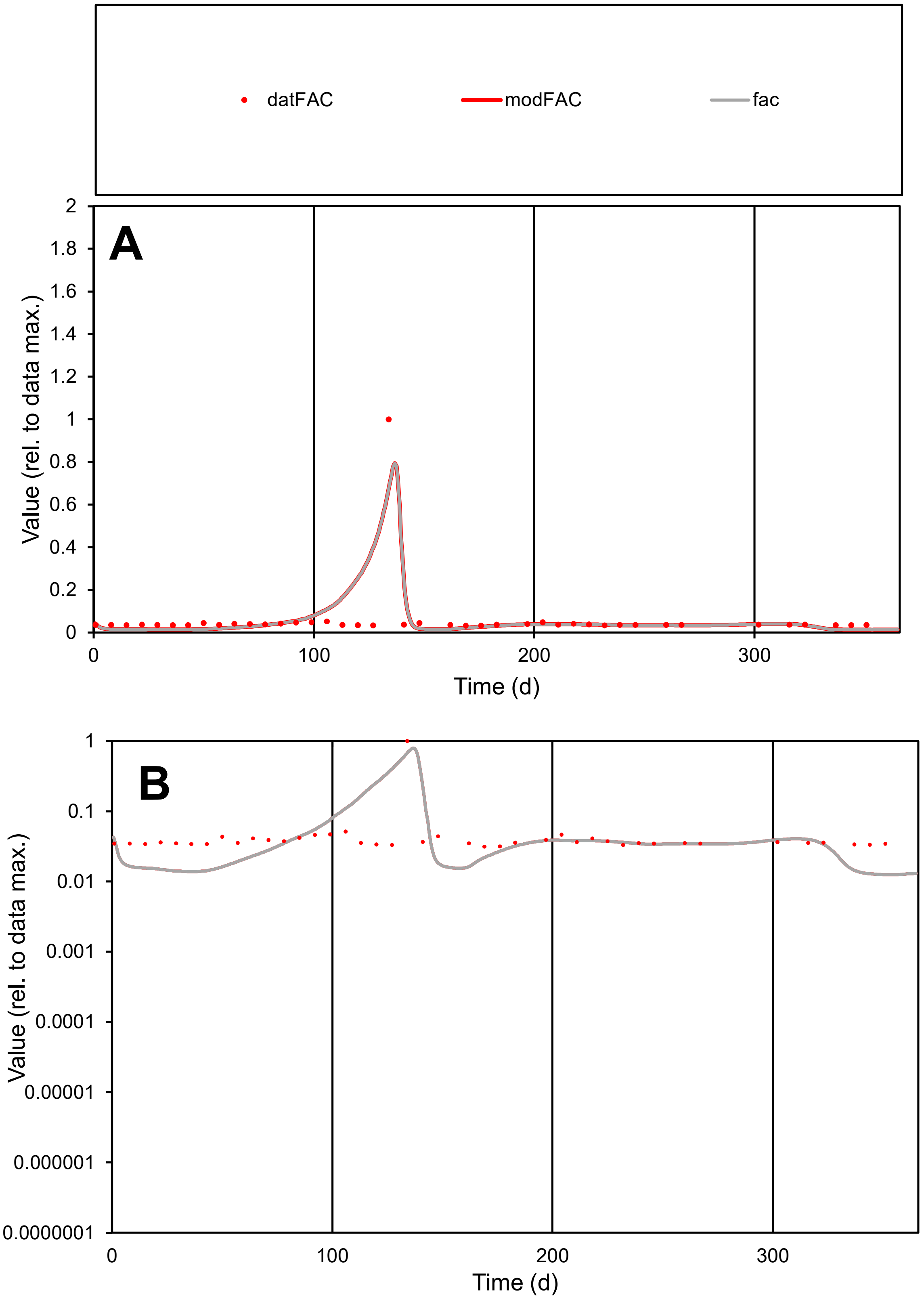


**C1**

Variables
FAC Provide jtd or Dst
CalibOnly?
2 1 to reproduce of
oNormalize?
2 1=no, 2=yes

ofxn calc*
1.25203
ofxn mod
1.38176

sRun
RunA1
zout
1

Ave
Data
0.00017
Model
0.00018

data/vMax
0.00277

ofxng
319.391

Min d/m
2.51405

ave/min d
1.91946
ave/min m
5.23017

330_Calib_S_A

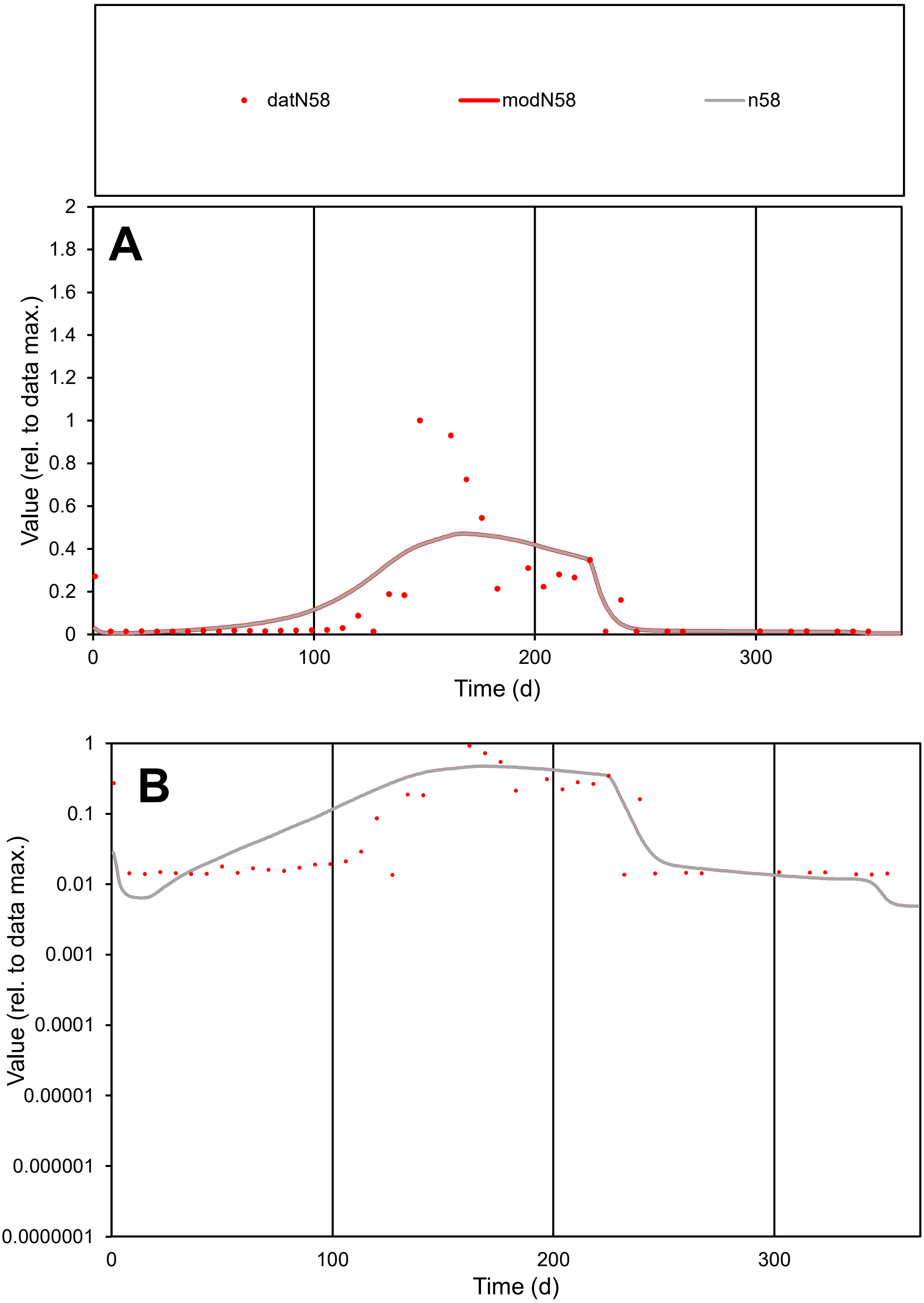


**C1**

Variables
N58 Provide jtd or Dst
CalibOnly?
2 1 to reproduce of
oNormalize?
2 1=no, 2=yes

ofxn calc*
1.06984
ofxn mod
1.0899

sRun
RunA1
zout
1

Ave
Data
0.00094
Model
0.00099

data/vMax
0.00639

ofxng
319.391

Min d/m
2.26129

ave/min d
10.8038
ave/min m
25.9146

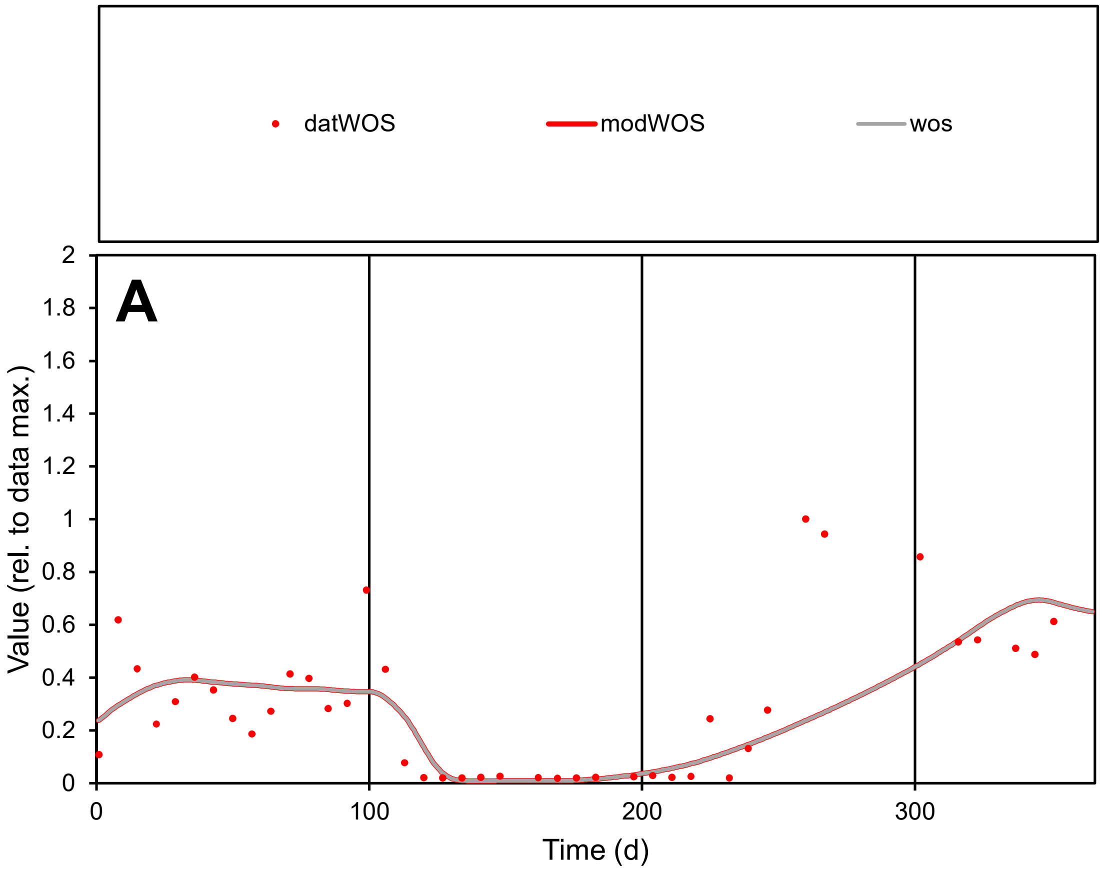


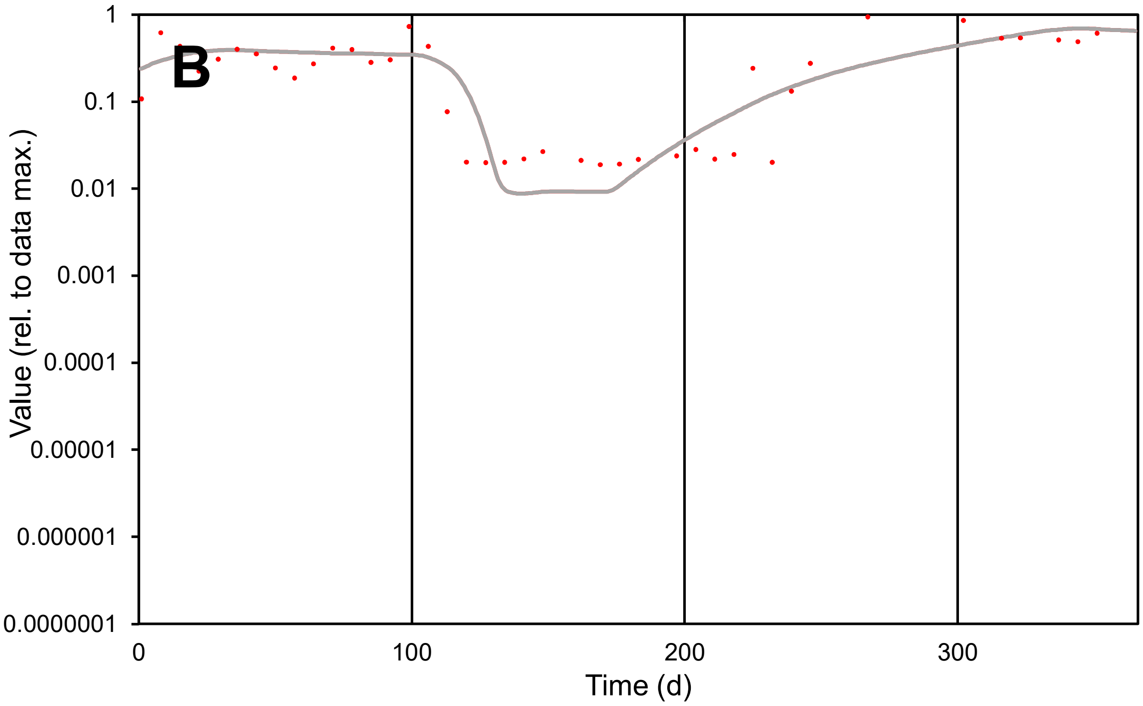


**C1**

Variables
WOS Provide jtd or Dst
CalibOnly?
2 1 to reproduce of
oNormalize?
2 1=no, 2=yes

ofxn calc*
0.69318
ofxn mod
0.70562

sRun
RunA1
zout
1

Ave
Data
0.00106
Model
0.00095

data/vMax
0.00363

ofxng
319.391

Min d/m
2.14063

ave/min d
15.3988
ave/min m
29.5303

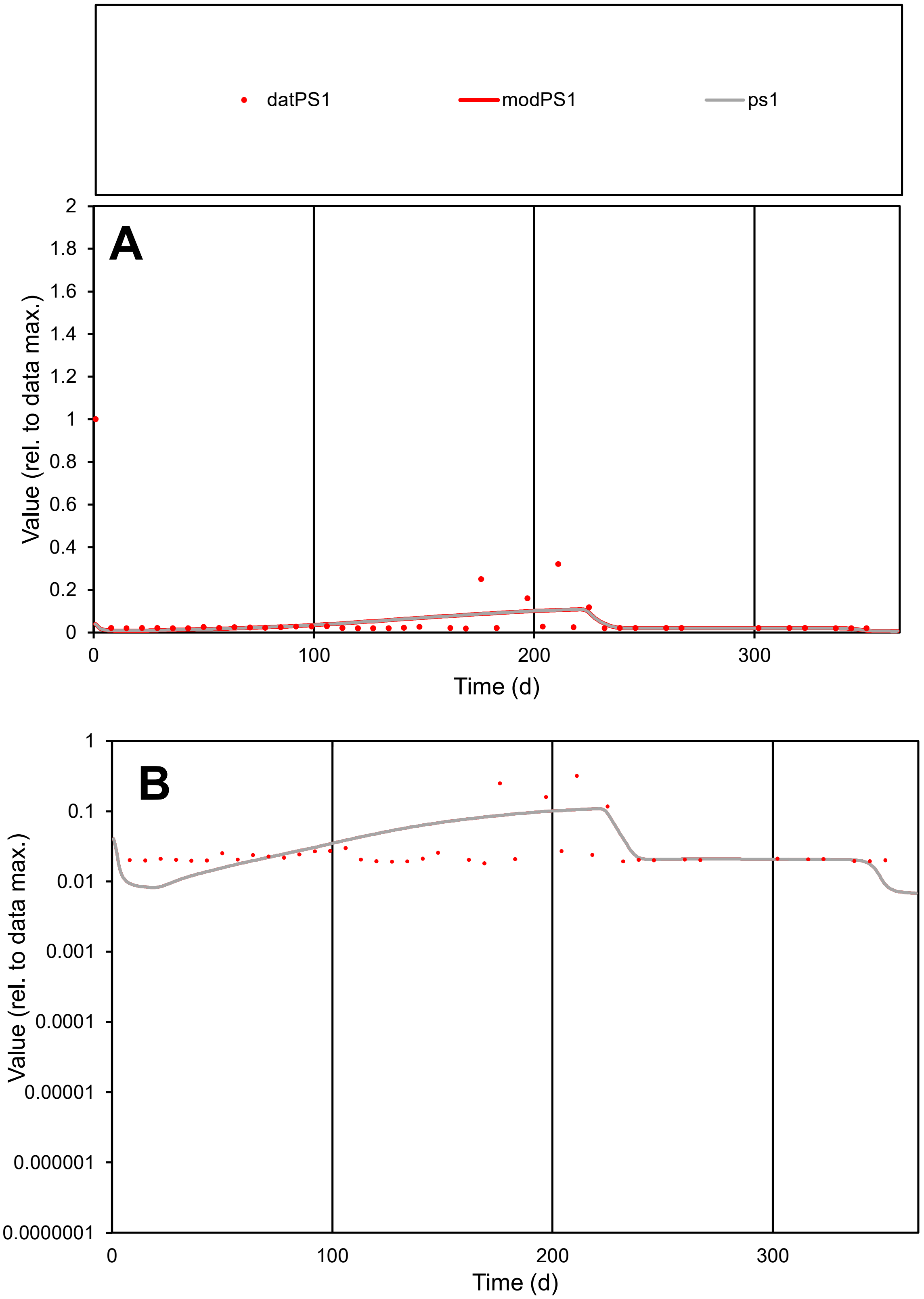


C1

Variables
PS1 Provide jtd or Dst
CalibOnly?
2 1 to reproduce of
oNormalize?
2 1=no, 2=yes

ofxn calc*
2.49966
ofxn mod
2.50475

sRun
RunA1
zout
1

Ave
Data
0.00016
Model
0.00011

data/vMax
0.0026

ofxng
319.391

Min d/m
2.21292

ave/min d
3.46202
ave/min m
5.04151

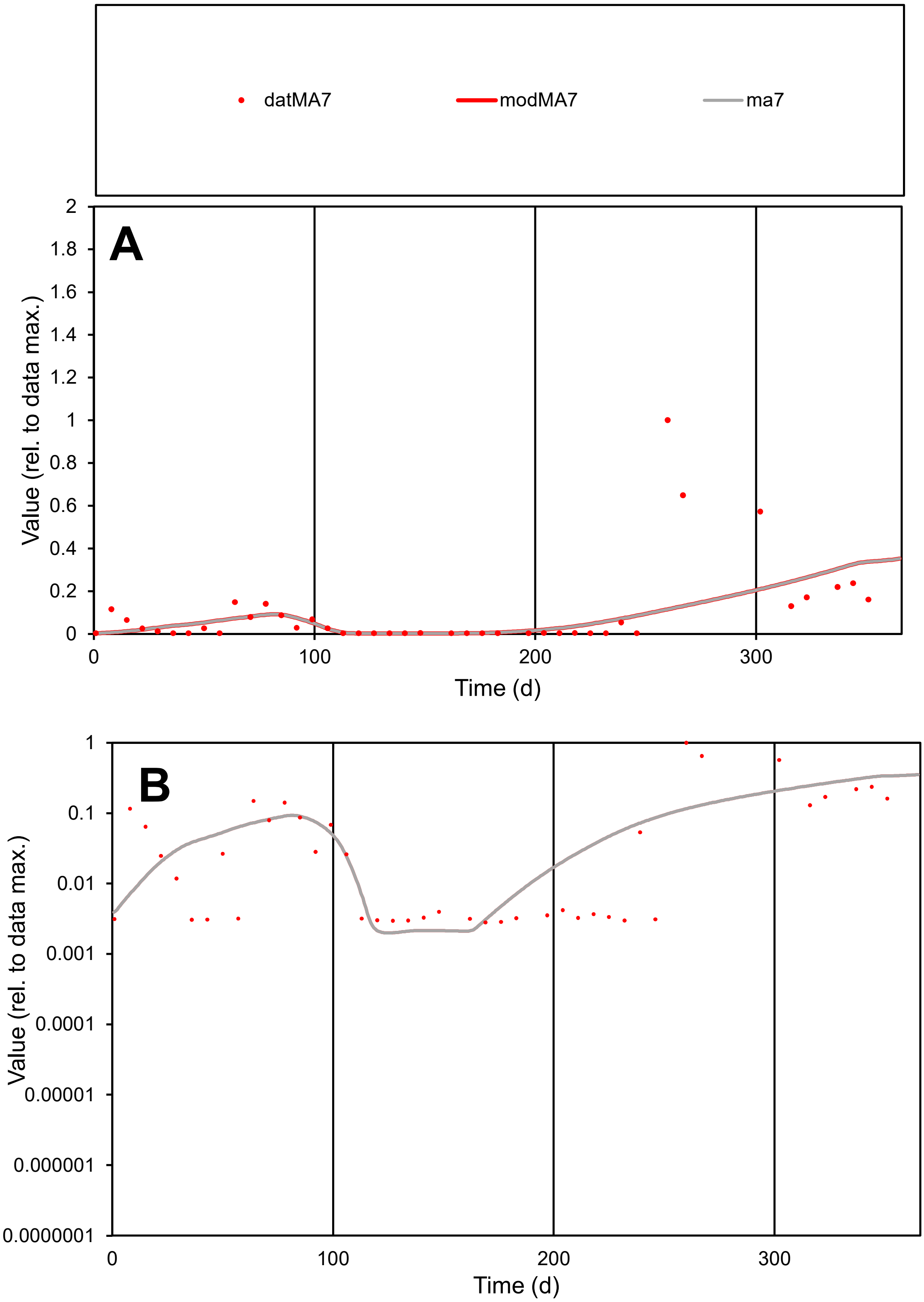


**C1**

Variables
MA7 Provide jtd or Dst
CalibOnly?
2 1 to reproduce of
oNormalize?
2 1=no, 2=yes

ofxn calc*
1.80412
ofxn mod
1.81101

sRun
RunA1
zout
1

Ave
Data
0.00047
Model
0.00036

data/vMax
0.0048

ofxng
319.391

Min d/m
1.4

ave/min d
34.4805
ave/min m
36.8535

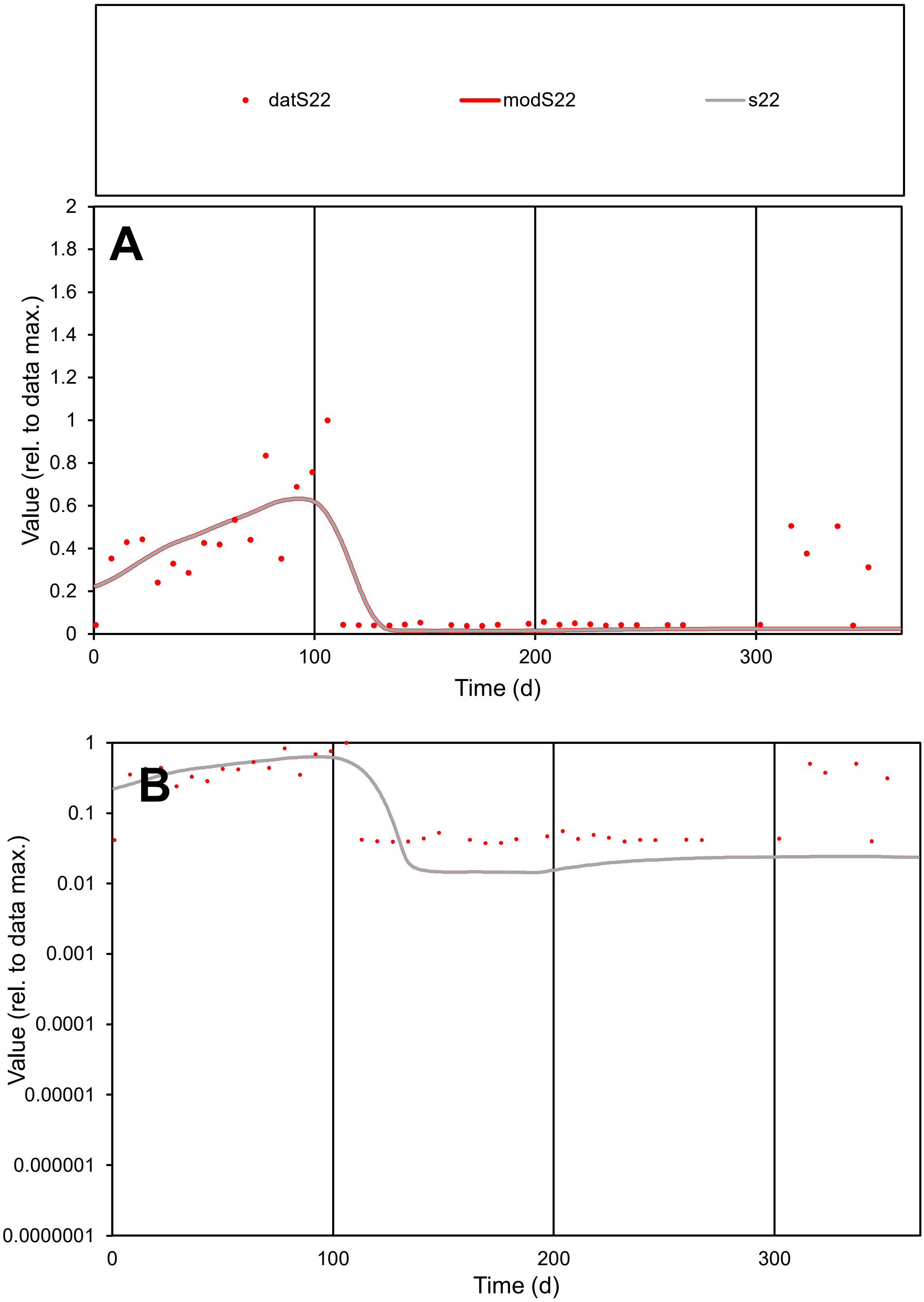


**C1**

Variables

S22 Provide jtd or Dst

CalibOnly?

2 1 to reproduce of

oNormalize?

2 1=no, 2=yes

ofxn calc*

0.72083

ofxn mod

0.74636

sRun

RunA1

zout

1

Ave

Data

0.00096

Model

0.0008

data/vMax

0.00395

ofxng

319.391

Min d/m

2.5846

ave/min d

6.5078

ave/min m

14.0961

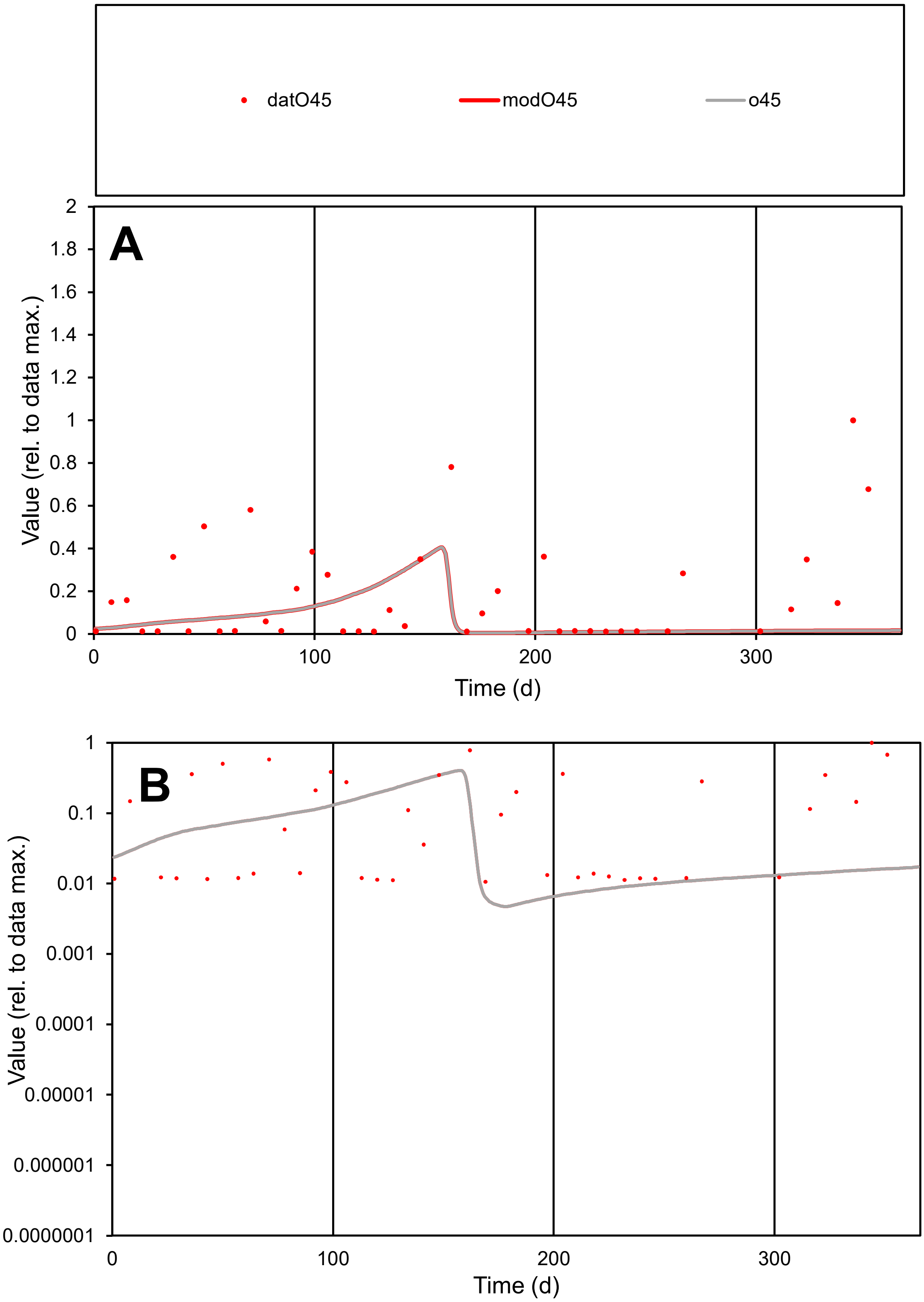


C1

Variables
O45 Provide jtd or Dst
CalibOnly?
2 1 to reproduce of
oNormalize?
2 1=no, 2=yes

ofxn calc*
1.47342
ofxn mod
1.51464

sRun
RunA1
zout
1

Ave
Data
0.00096
Model
0.00042

data/vMax
0.00542

ofxng
319.391

Min d/m
2.23241

ave/min d
16.6956
ave/min m
16.3337

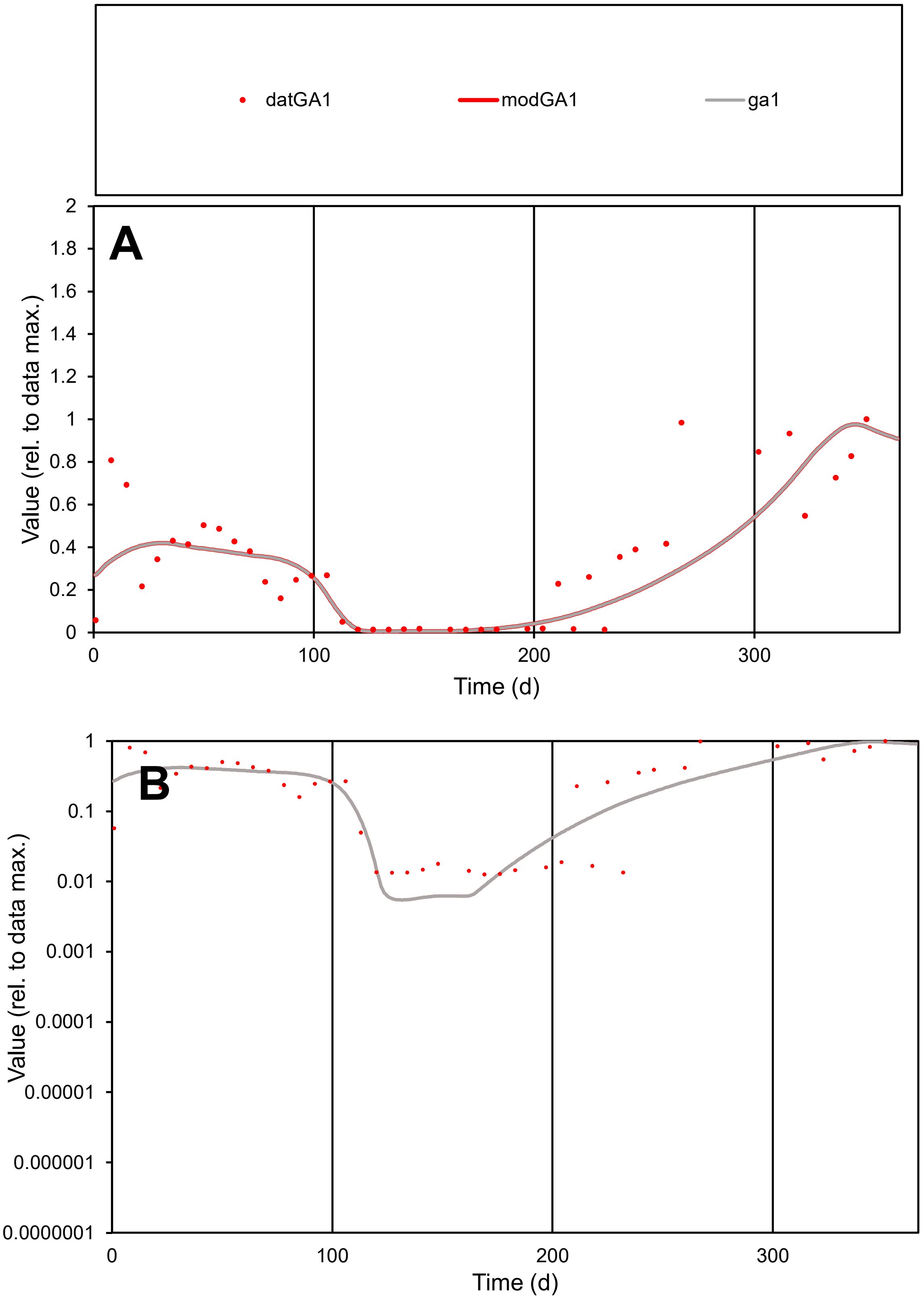


**C1**

Variables
GA1 Provide jtd or Dst
CalibOnly?
2 1 to reproduce of
oNormalize?
2 1=no, 2=yes

ofxn calc*
0.55229
ofxn mod
0.56696

sRun
RunA1
zout
1

Ave
Data
0.00091
Model
0.00081

data/vMax
0.0028

ofxng
319.391

Min d/m
2.27447

ave/min d
25.6945
ave/min m
51.8199

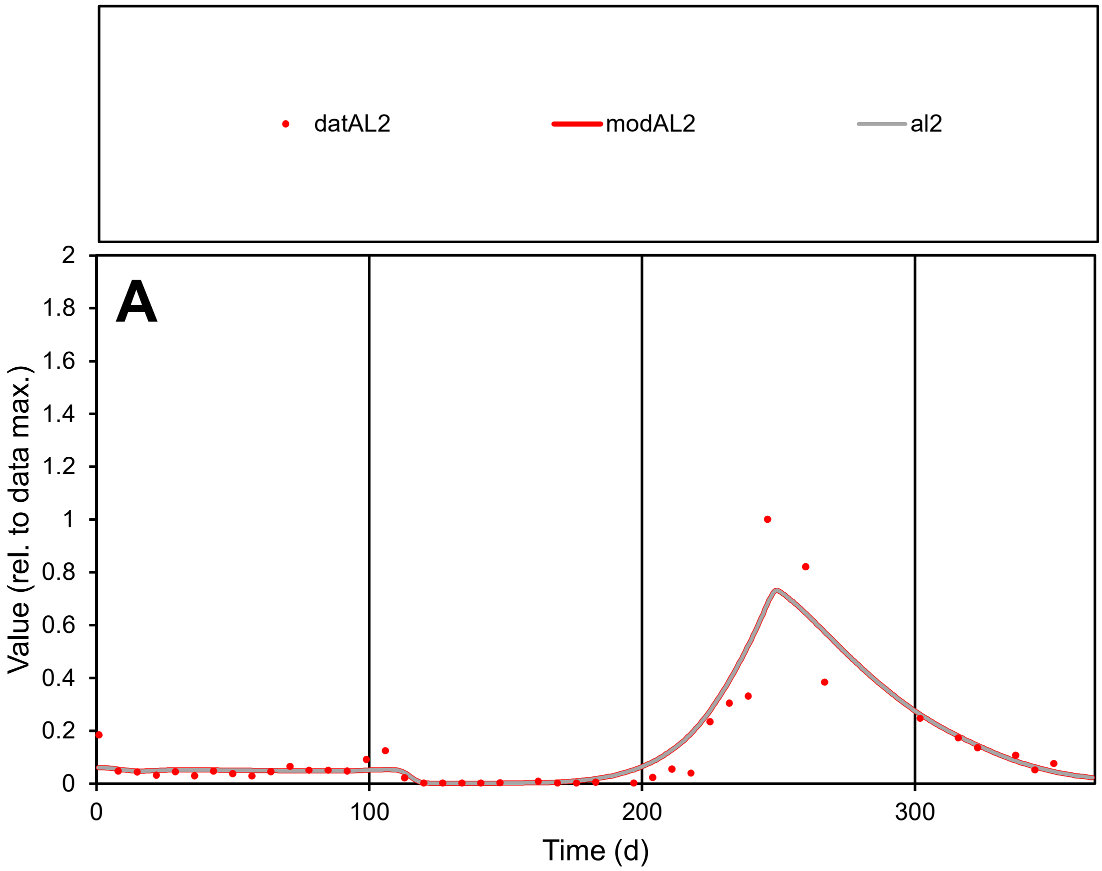


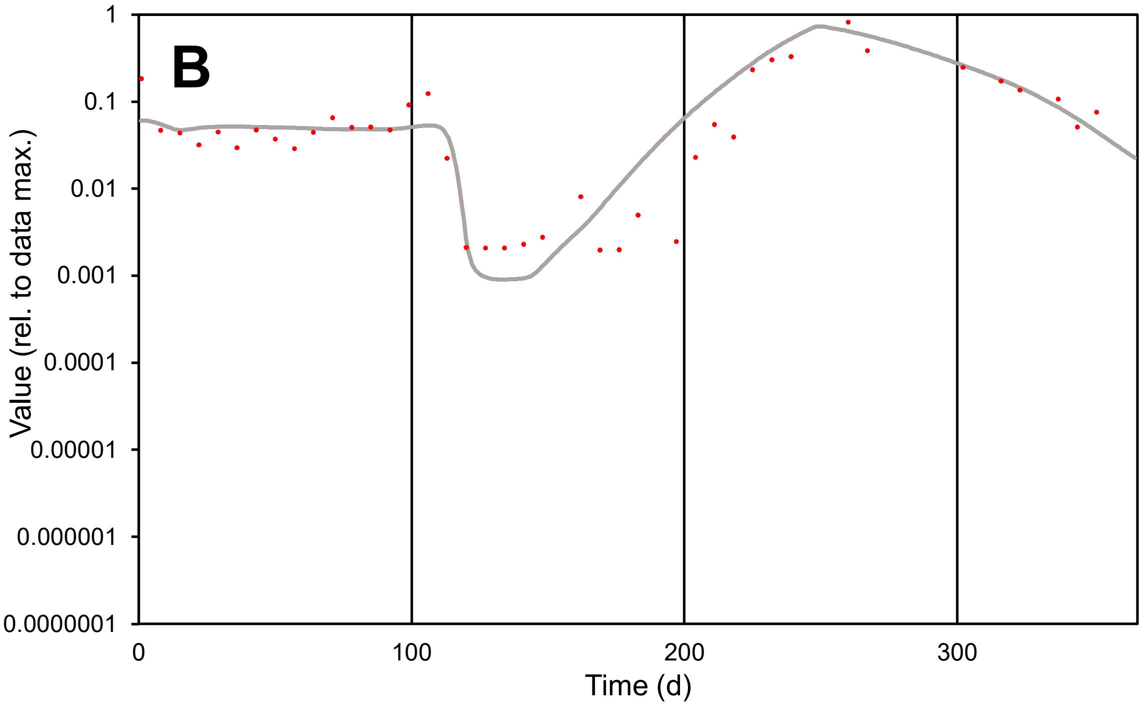


**C1**

Variables
AL2 Provide jtd or Dst
CalibOnly?
2 1 to reproduce of
oNormalize?
2 1=no, 2=yes

ofxn calc*
0.66334
ofxn mod
0.70118

sRun
RunA1
zout
1

Ave
Data
0.00132
Model
0.00138

data/vMax
0.0111

ofxng
319.391

Min d/m
2.17729

ave/min d
60.4456
ave/min m
137.269

330_Calib_S_A

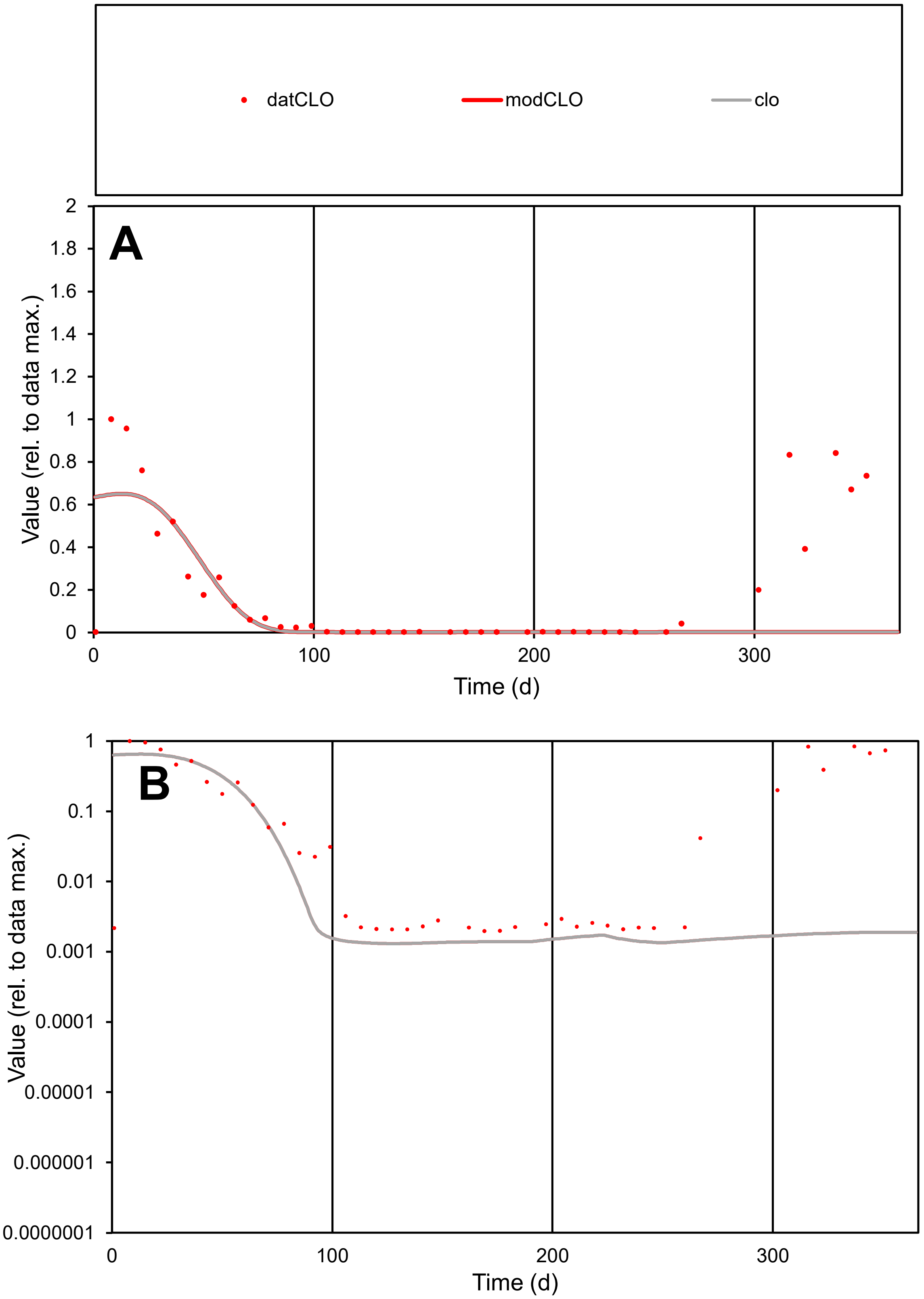


**C1**

Variables
CLO Provide jtd or Dst
CalibOnly?
2 1 to reproduce of
oNormalize?
2 1=no, 2=yes

ofxn calc*
1.37907
ofxn mod
1.38797

sRun
RunA1
zout
1

Ave
Data
0.00136
Model
0.00076

data/vMax
0.00672

ofxng
319.391

Min d/m
1.50722

ave/min d
102.458
ave/min m
86.9552

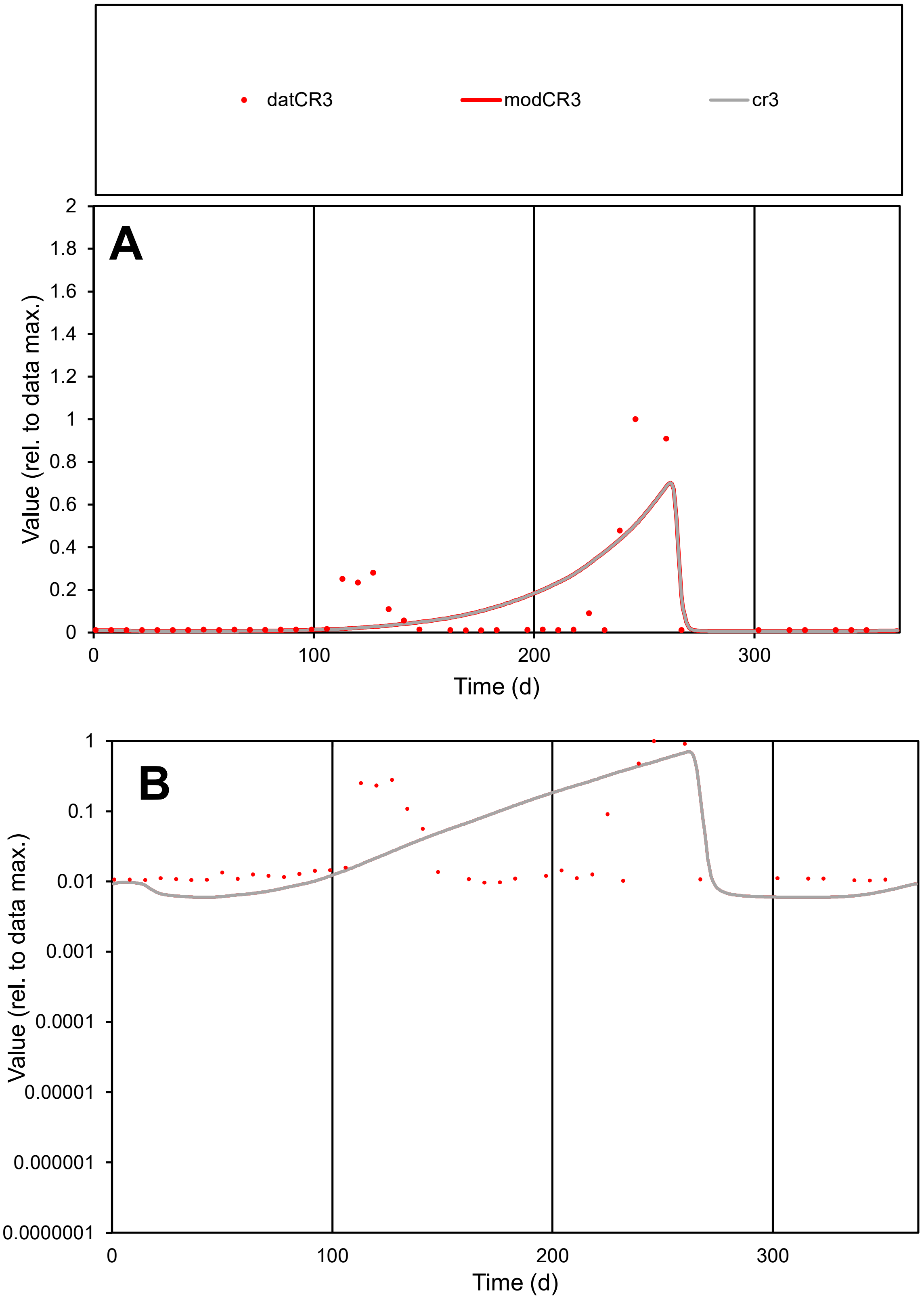


**C1**

Variables
CR3 Provide jtd or Dst
CalibOnly?
2 1 to reproduce of
oNormalize?
2 1=no, 2=yes

ofxn calc*
1.57971
ofxn mod
1.59817

sRun
RunA1
zout
1

Ave
Data
0.00033
Model
0.00036

data/vMax
0.00369

ofxng
319.391

Min d/m
1.61368

ave/min d
9.34884
ave/min m
16.1664

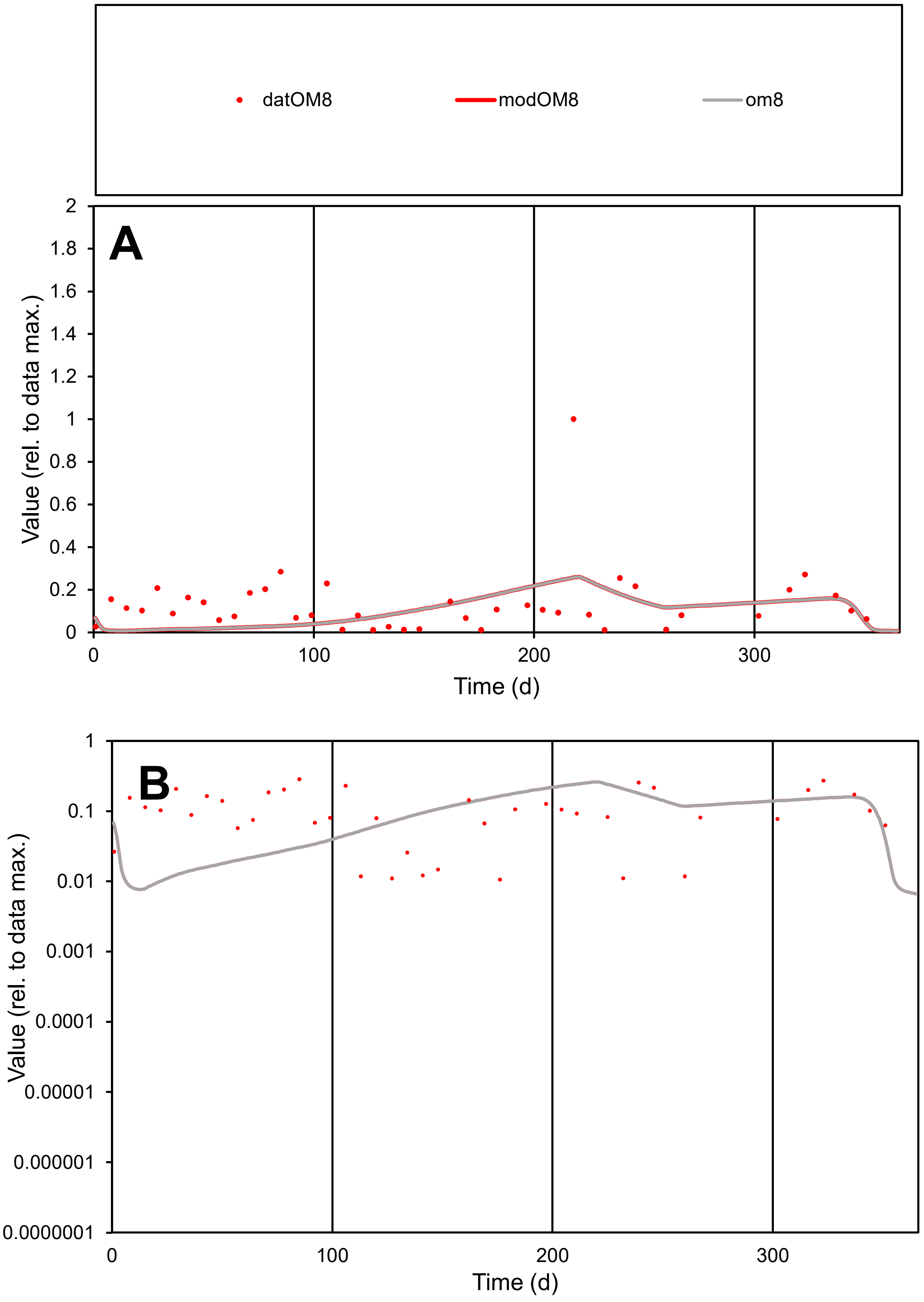


**C1**

Variables

OM8 Provide jtd or Dst

CalibOnly?

2 1 to reproduce of

oNormalize?

2 1=no, 2=yes

ofxn calc*

1.1955

ofxn mod

1.2058

sRun

RunA1

zout

1

Ave

Data

0.00068

Model

0.00052

data/vMax

0.00519

ofxng

319.391

Min d/m

1.30835

ave/min d

12.453

ave/min m

12.3591

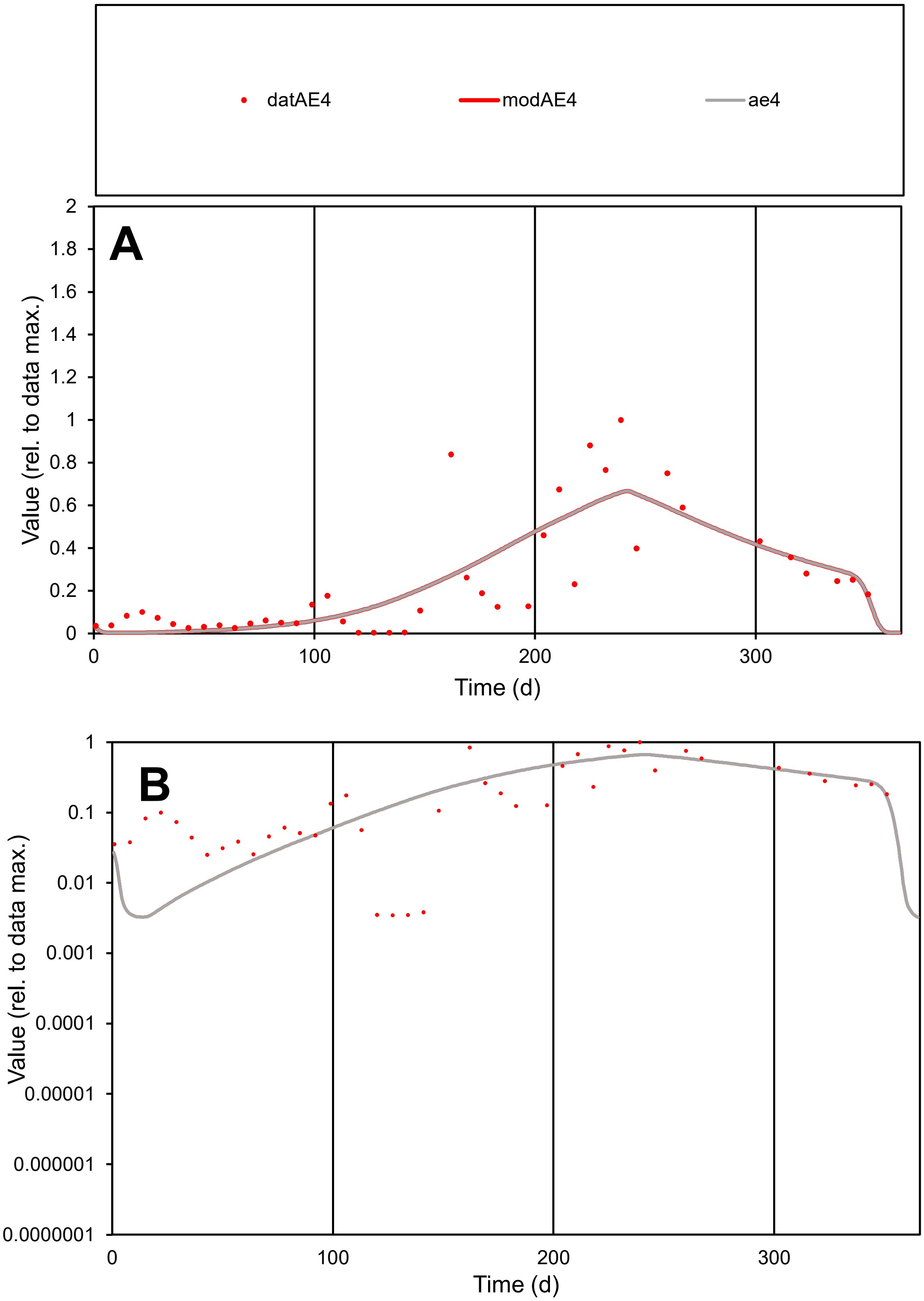


**C1**

Variables
AE4 Provide jtd or Dst
CalibOnly?
2 1 to reproduce of
oNormalize?
2 1=no, 2=yes

ofxn calc*
0.65965
ofxn mod
0.6838

sRun
RunA1
zout
1

Ave
Data
0.00087
Model
0.00086

data/vMax
0.00356

ofxng
319.391

Min d/m
1.02235

ave/min d
70.128
ave/min m
71.3802

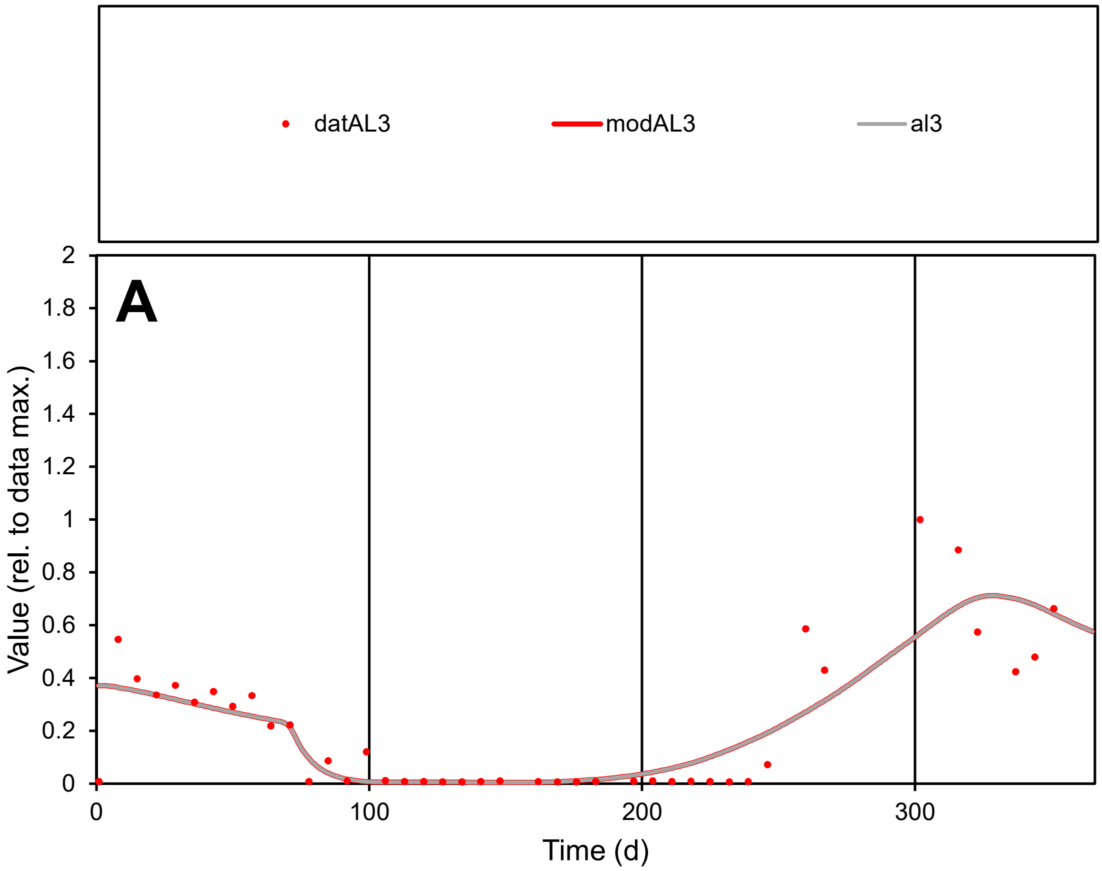


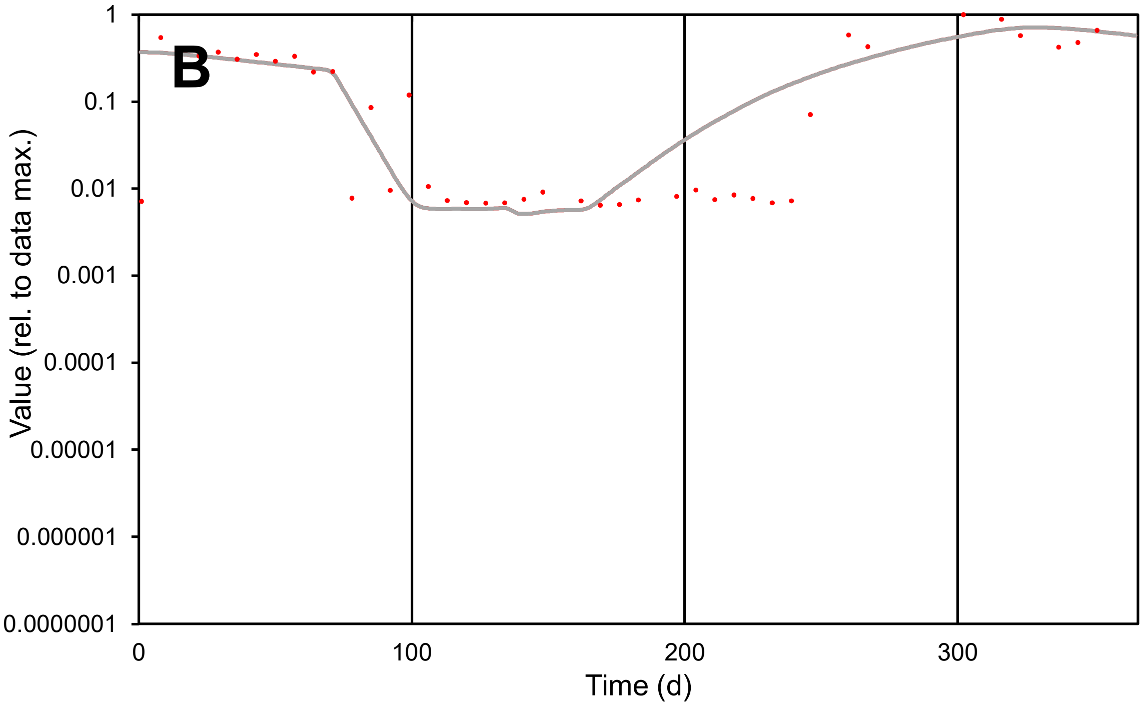


**C1**

Variables

AL3 Provide jtd or Dst

CalibOnly?

2 1 to reproduce of

oNormalize?

2 1=no, 2=yes

ofxn calc*

0.62479

ofxn mod

0.64297

sRun

RunA1

zout

1

Ave

Data

0.00079

Model

0.00079

data/vMax

0.00375

ofxng

319.391

Min d/m

1.25323

ave/min d

32.5352

ave/min m

40.7278

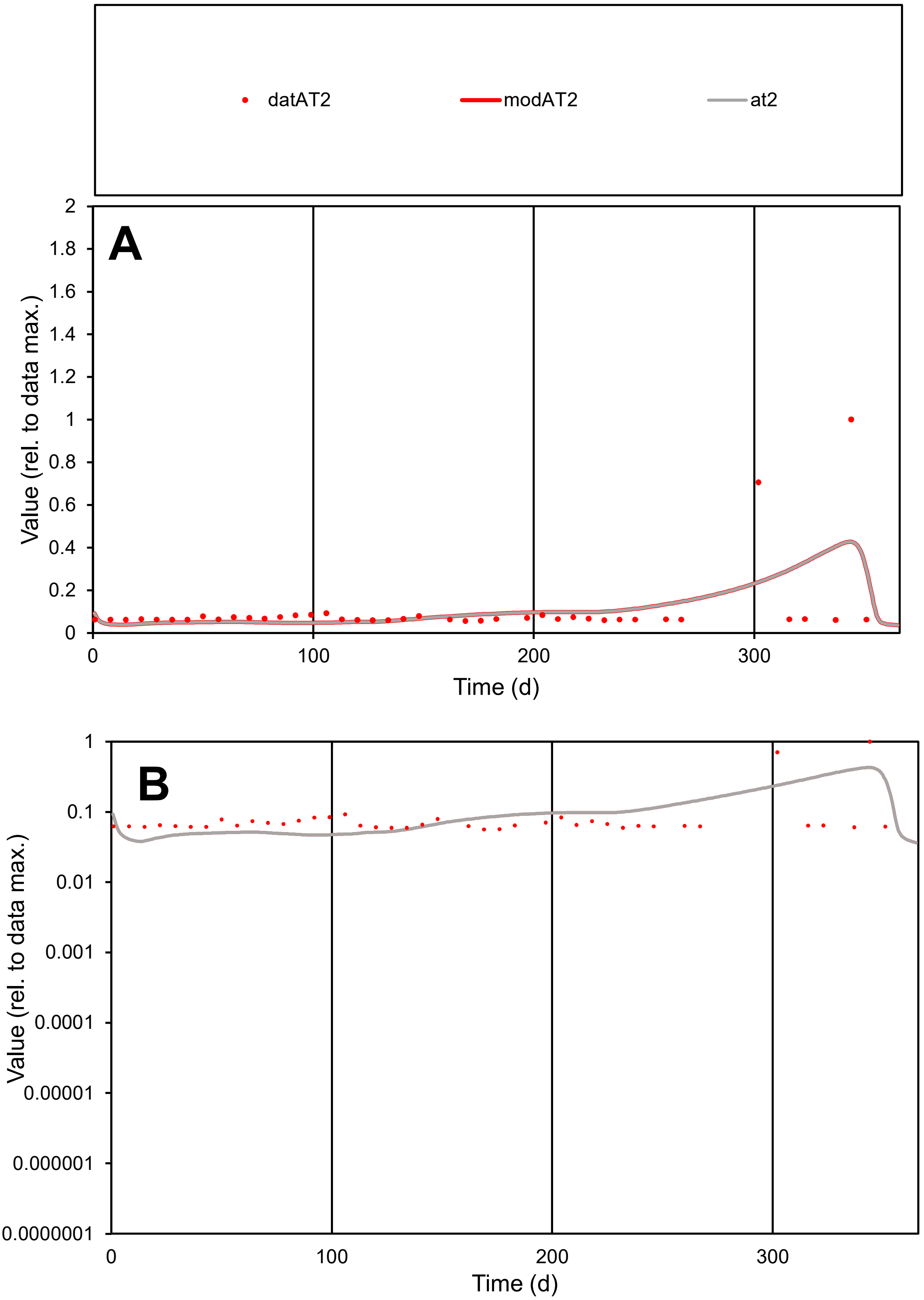


**C1**

Variables
AT2 Provide jtd or Dst
CalibOnly?
2 1 to reproduce of
oNormalize?
2 1=no, 2=yes

ofxn calc*
1.35825
ofxn mod
1.37305

sRun
RunA1
zout
1

Ave
Data
0.00015
Model
0.00015

data/vMax
0.00142

ofxng
319.391

Min d/m
1.42197

ave/min d
1.85343
ave/min m
2.66969

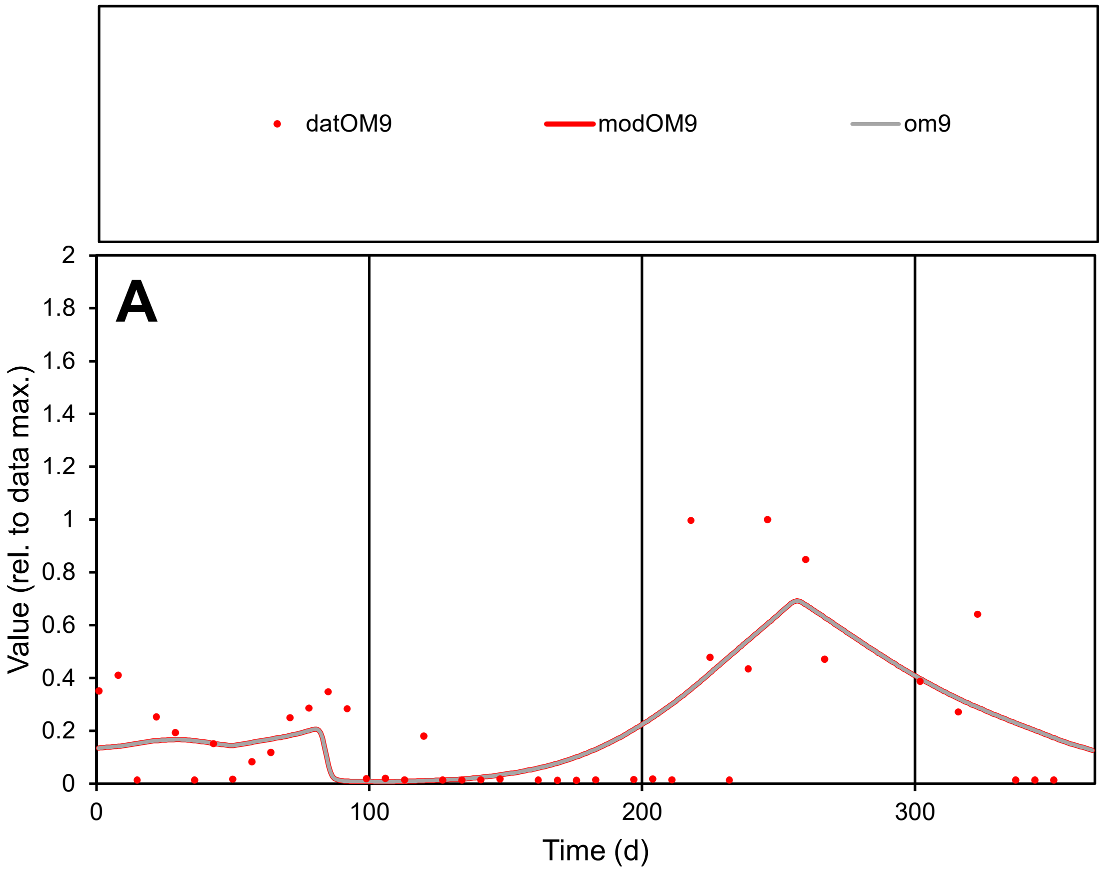


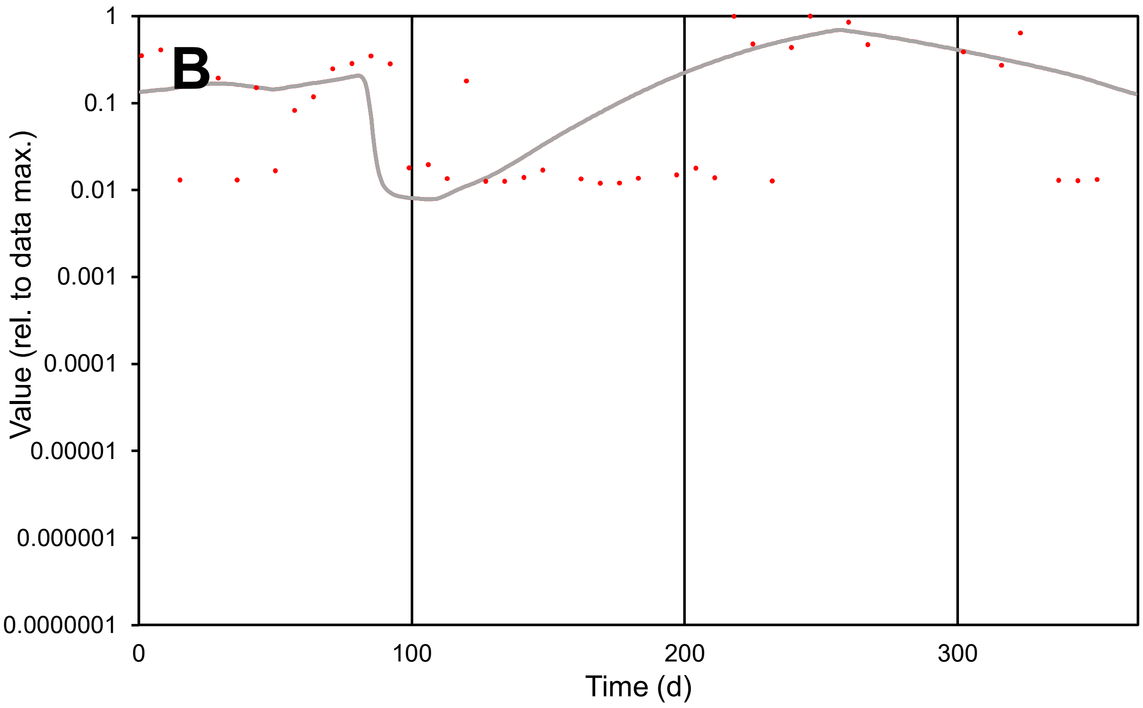


**C1**

Variables

OM9 Provide jtd or Dst

CalibOnly?

2 1 to reproduce of

oNormalize?

2 1=no, 2=yes

ofxn calc*

0.94361

ofxn mod

0.96658

sRun

RunA1

zout

1

Ave

Data

0.00075

Model

0.00075

data/vMax

0.00363

ofxng

319.391

Min d/m

1.52221

ave/min d

17.344

ave/min m

26.1985

330_Calib_S_A

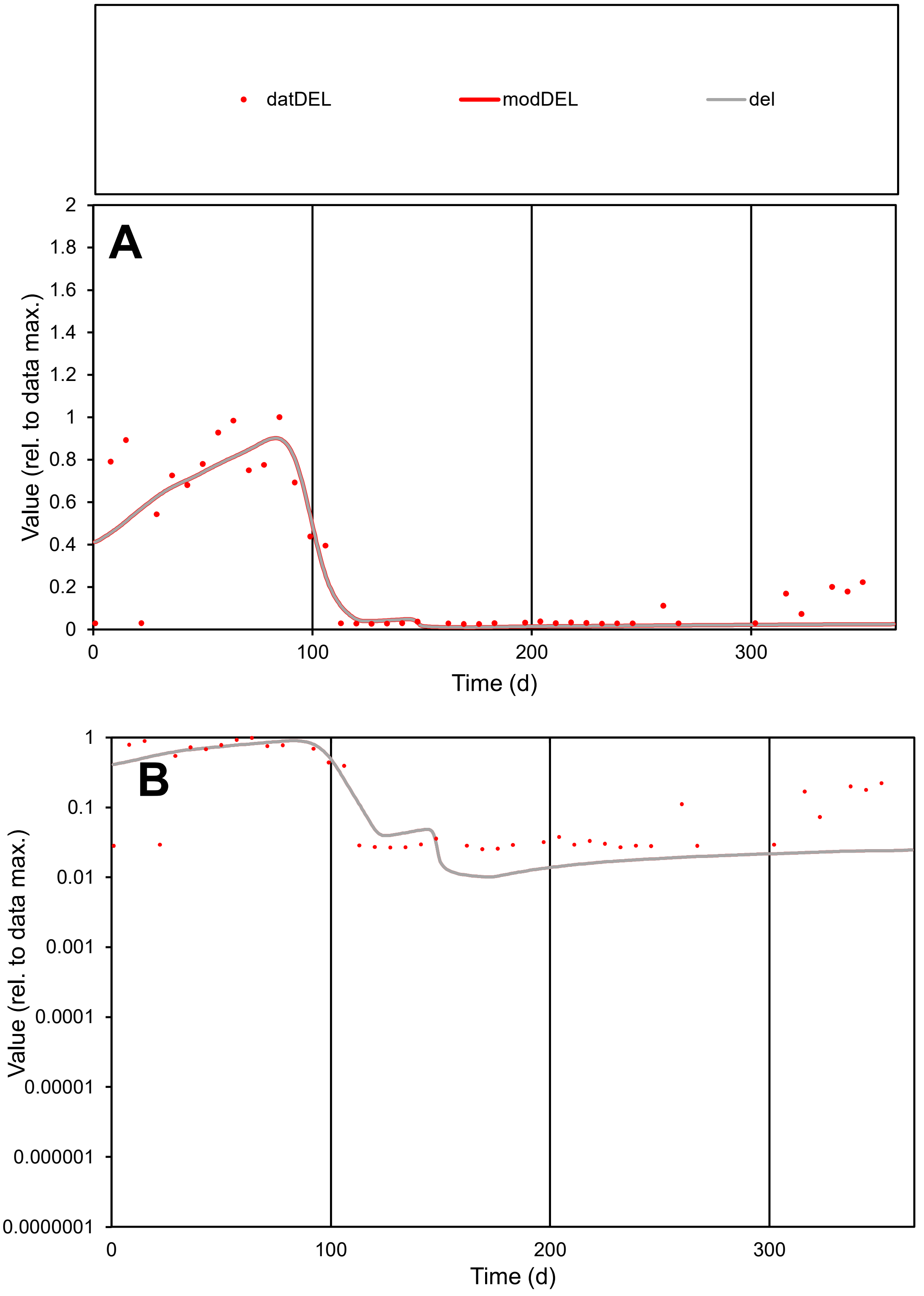


**C1**

Variables

DEL Provide jtd or Dst

CalibOnly?

2 1 to reproduce of

oNormalize?

2 1=no, 2=yes

ofxn calc*

0.52702

ofxn mod

0.55535

sRun

RunA1

zout

1

Ave

Data

0.0007

Model

0.00065

data/vMax

0.00246

ofxng

319.391

Min d/m

2.4938

ave/min d

11.2393

ave/min m

26.0478

330_Calib_S_A

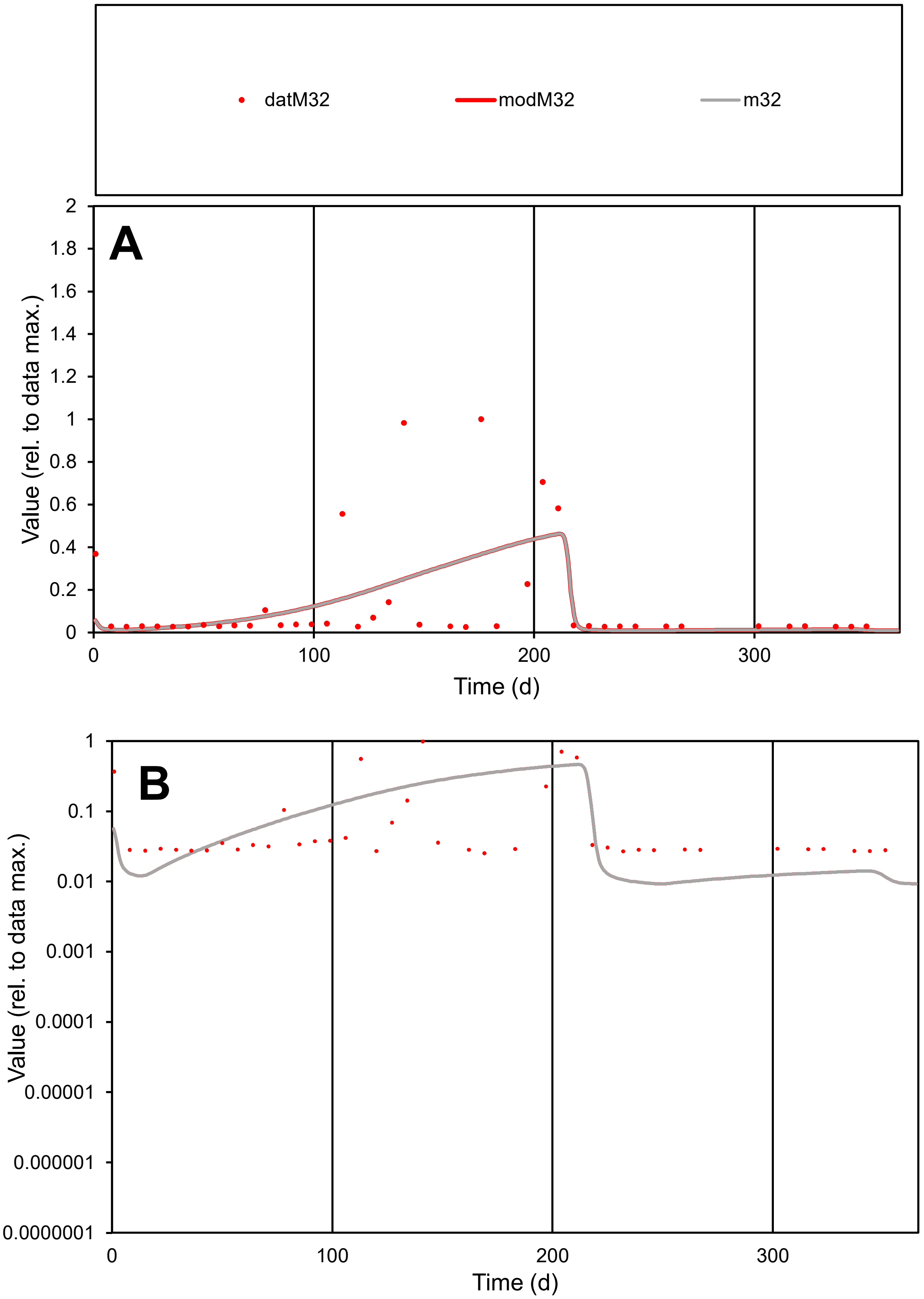


**C1**

Variables
M32 Provide jtd or Dst
CalibOnly?
2 1 to reproduce of
oNormalize?
2 1=no, 2=yes

ofxn calc*
1.50804
ofxn mod
1.51933

sRun
RunA1
zout
1

Ave
Data
0.00113
Model
0.00102

data/vMax
0.0083

ofxng
319.391

Min d/m
2.71894

ave/min d
5.35359
ave/min m
13.2264

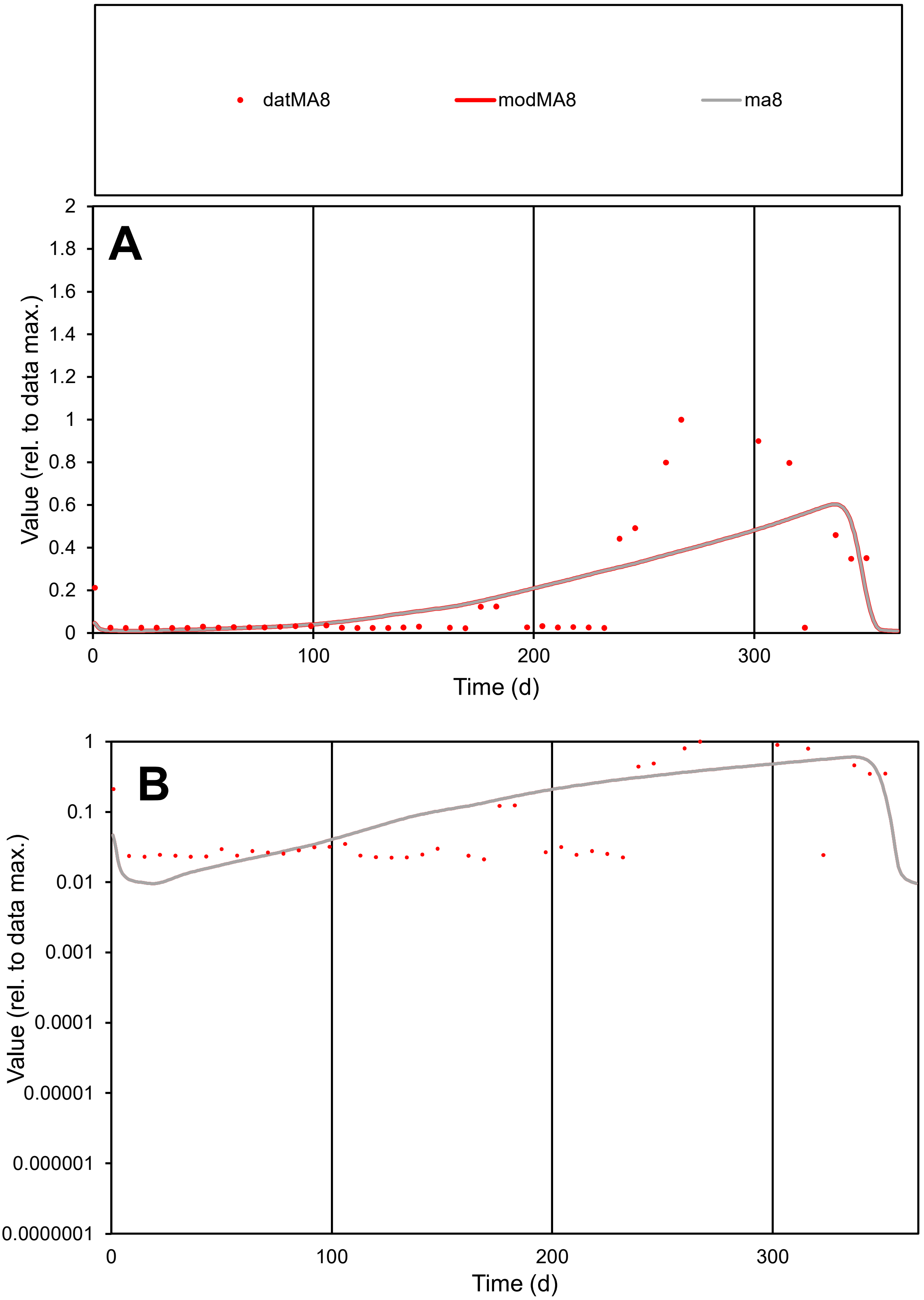


**C1**

Variables
MA8 Provide jtd or Dst
CalibOnly?
2 1 to reproduce of
oNormalize?
2 1=no, 2=yes

ofxn calc*
1.18989
ofxn mod
1.20357

sRun
RunA1
zout
1

Ave
Data
0.00046
Model
0.00048

data/vMax
0.00283

ofxng
319.391

Min d/m
2.20147

ave/min d
7.63703
ave/min m
17.611

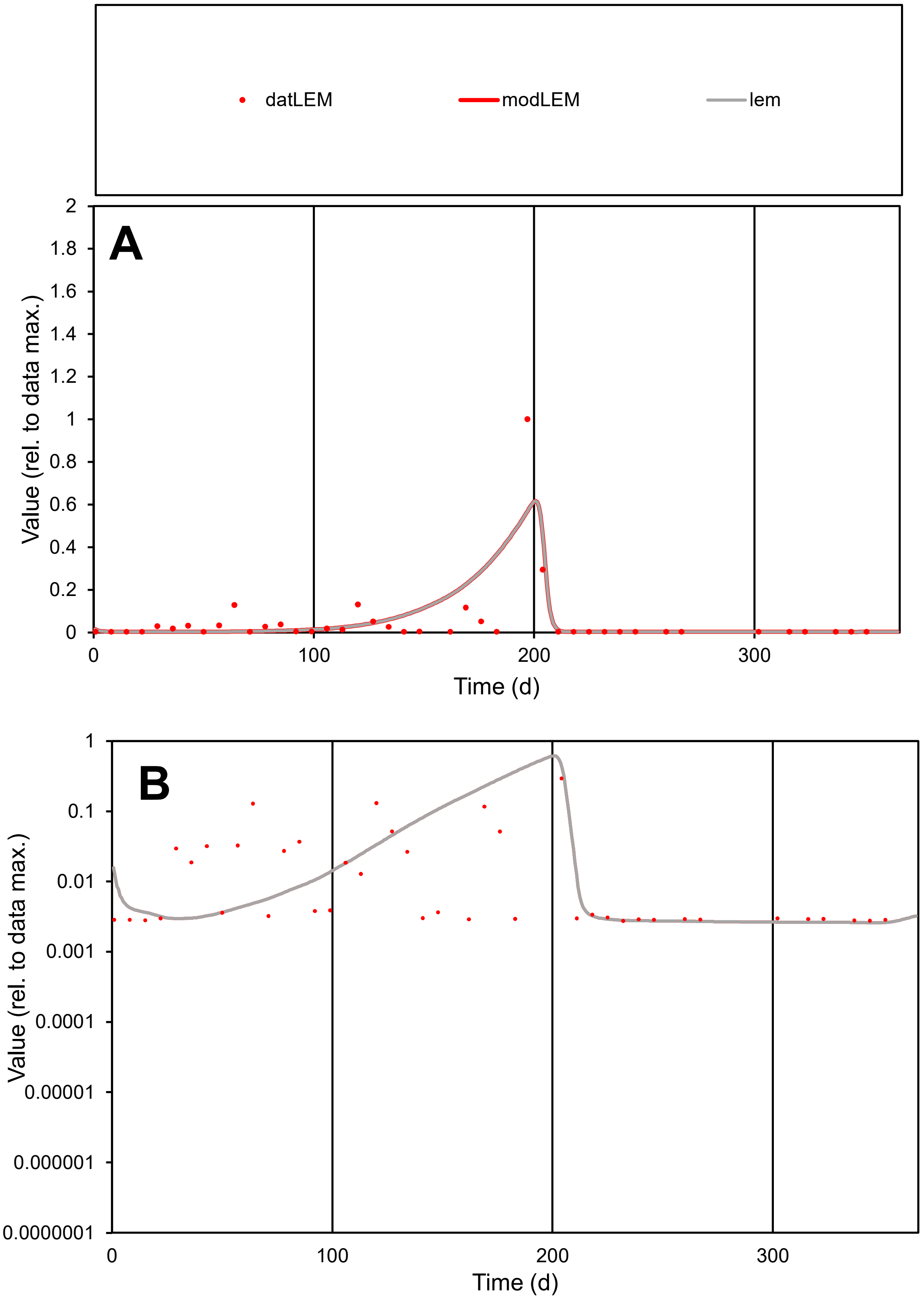


**C1**

| | |
|---|---|
| Variables | |
| LEM | Provide jtd or Dst |
| CalibOnly? | |
| 2 | 1 to reproduce of |
| oNormalize? | |
| 2 | 1=no, 2=yes |
| ofxn calc* | |
| 2.06865 | |
| ofxn mod | |
| 2.10559 | |
| sRun | |
| RunA1 | |
| zout | |
| 1 | |
| Ave | |
| Data | |
| 0.00042 | |
| Model | |
| 0.0005 | |
| data/vMax | |
| 0.00841 | |
| ofxng | |
| 319.391 | |
| Min d/m | |
| 1.05782 | |
| ave/min d | |
| 18.1379 | |
| ave/min m | |
| 22.7646 | |

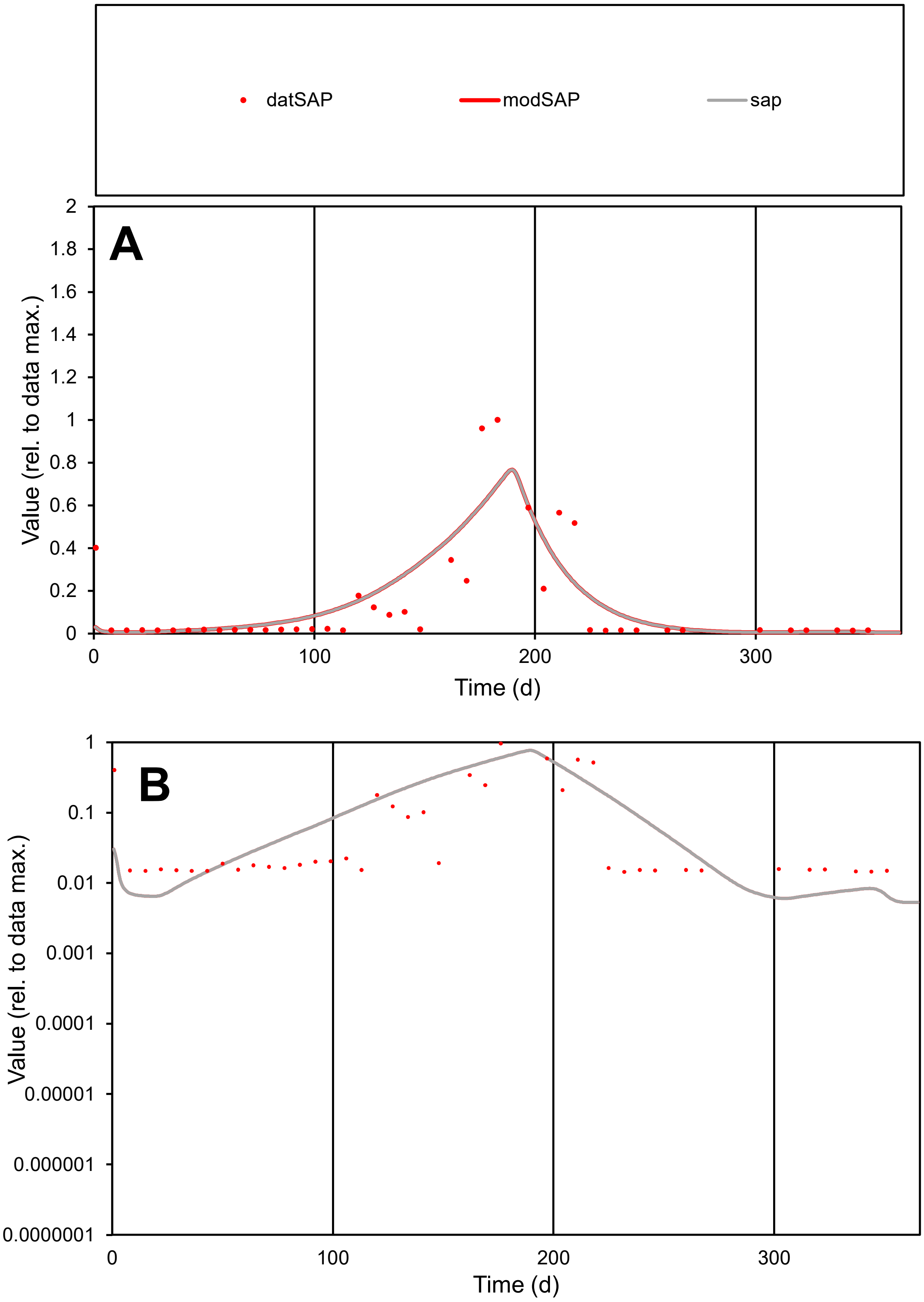


**C1**

Variables
SAP Provide jtd or Dst
CalibOnly?
2 1 to reproduce of
oNormalize?
2 1=no, 2=yes

ofxn calc*
1.02638
ofxn mod
1.05702

sRun
RunA1
zout
1

Ave
Data
0.00034
Model
0.00036

data/vMax
0.00243

ofxng
319.391

Min d/m
2.38425

ave/min d
9.55036
ave/min m
24.4893

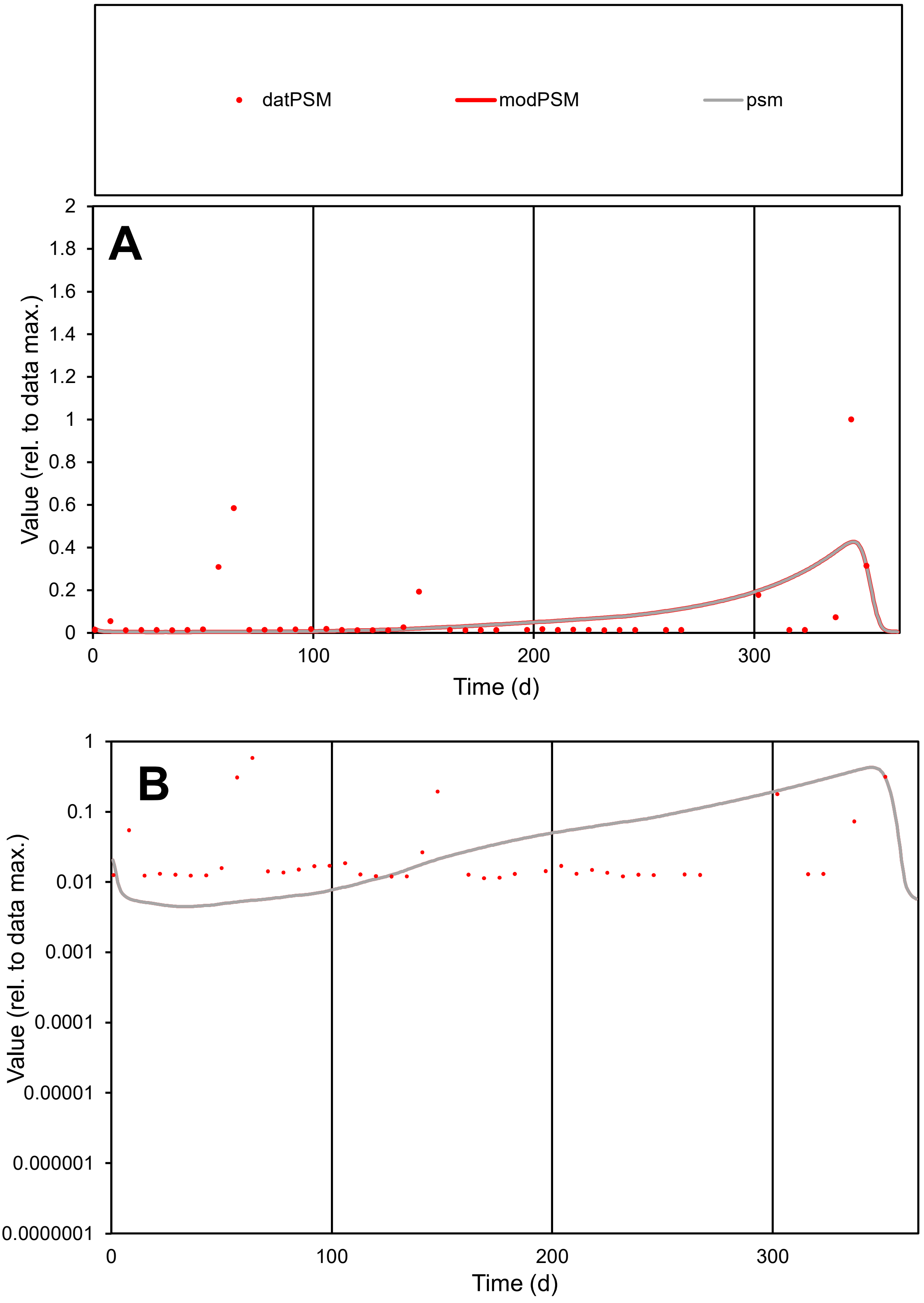


**C1**

Variables
PSM Provide jtd or Dst
CalibOnly?
2 1 to reproduce of
oNormalize?
2 1=no, 2=yes

ofxn calc*
2.10849
ofxn mod
2.11968

sRun
RunA1
zout
1

Ave
Data
0.00019
Model
0.00017

data/vMax
0.00245

ofxng
319.391

Min d/m
2.53461

ave/min d
6.65686
ave/min m
15.3193

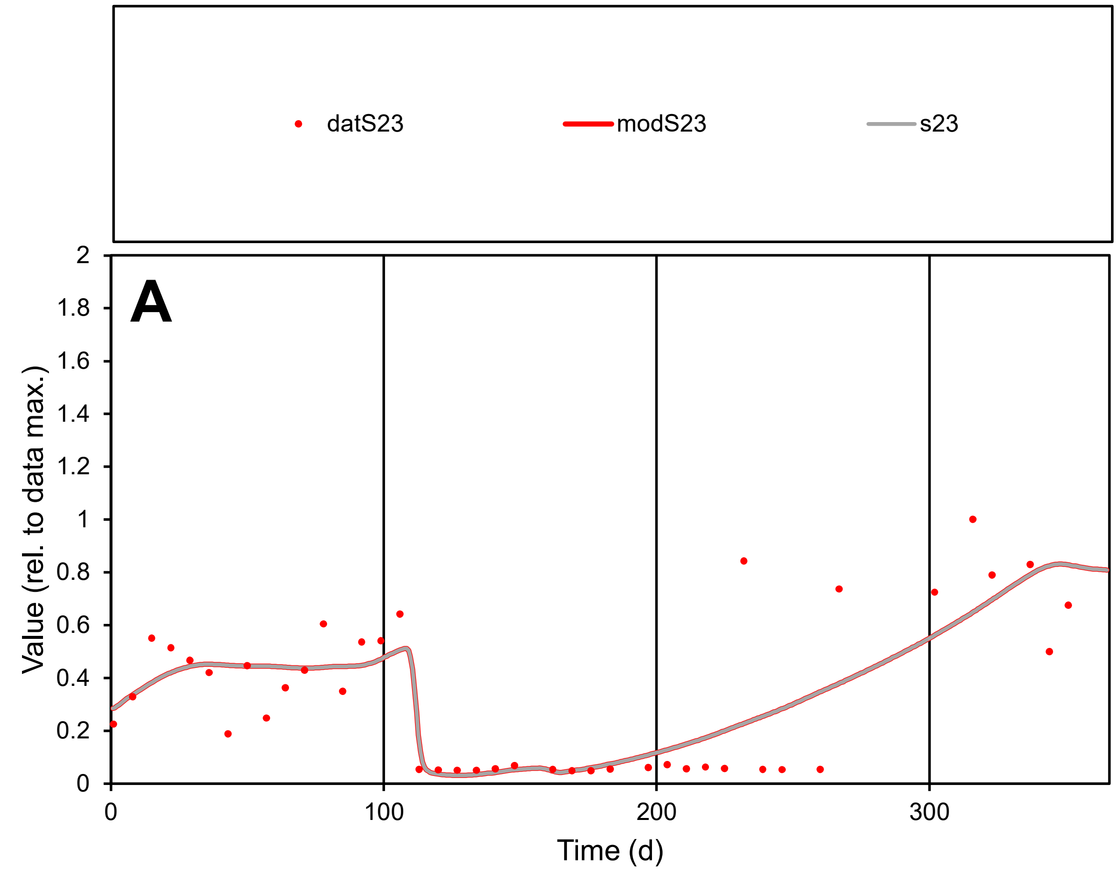


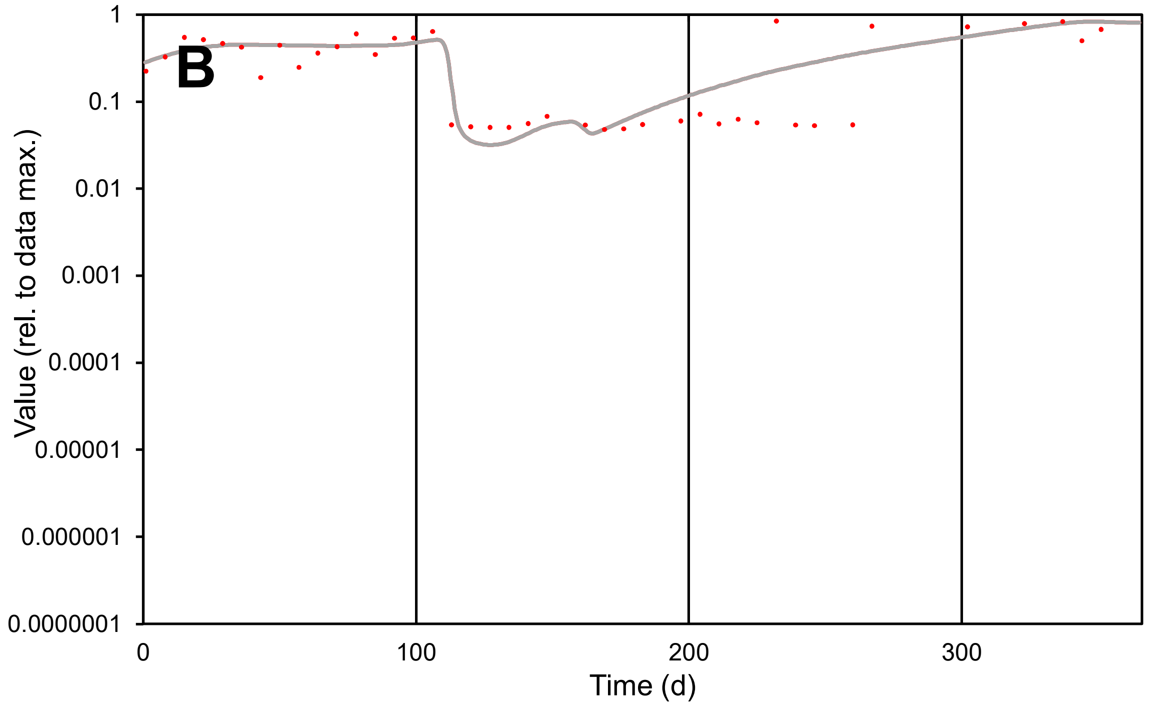


**C1**

Variables
S23 Provide jtd or Dst
CalibOnly?
2 1 to reproduce of
oNormalize?
2 1=no, 2=yes

ofxn calc*
0.513
ofxn mod
0.5299

sRun
RunA1
zout
1

Ave
Data
0.00112
Model
0.00113

data/vMax
0.00337

ofxng
319.391

Min d/m
1.5098

ave/min d
6.91938
ave/min m
10.5383

330_Calib_S_A

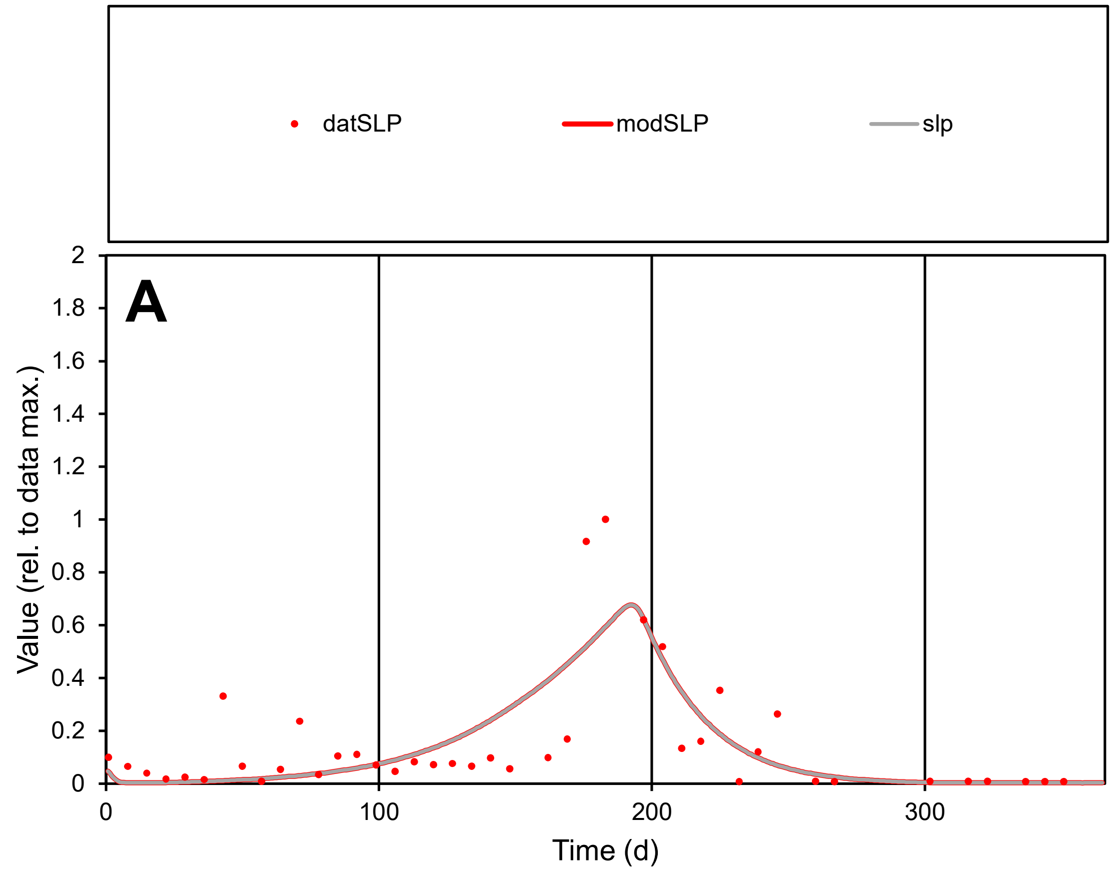


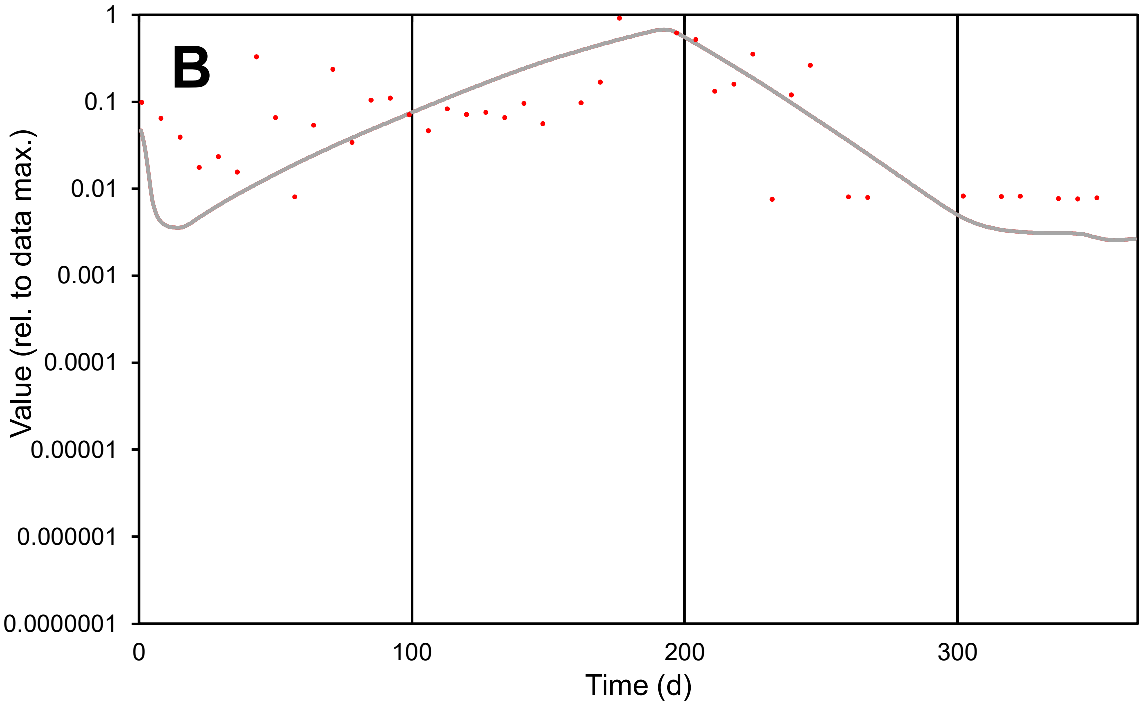


**C1**

Variables

SLP Provide jtd or Dst

CalibOnly?

2 1 to reproduce of

oNormalize?

2 1=no, 2=yes

ofxn calc*

0.97582

ofxn mod

1.00157

sRun

RunA1

zout

1

Ave

Data

0.0005

Model

0.00046

data/vMax

0.00337

ofxng

319.391

Min d/m

2.81284

ave/min d

19.4107

ave/min m

51.0426

330_Calib_S_A

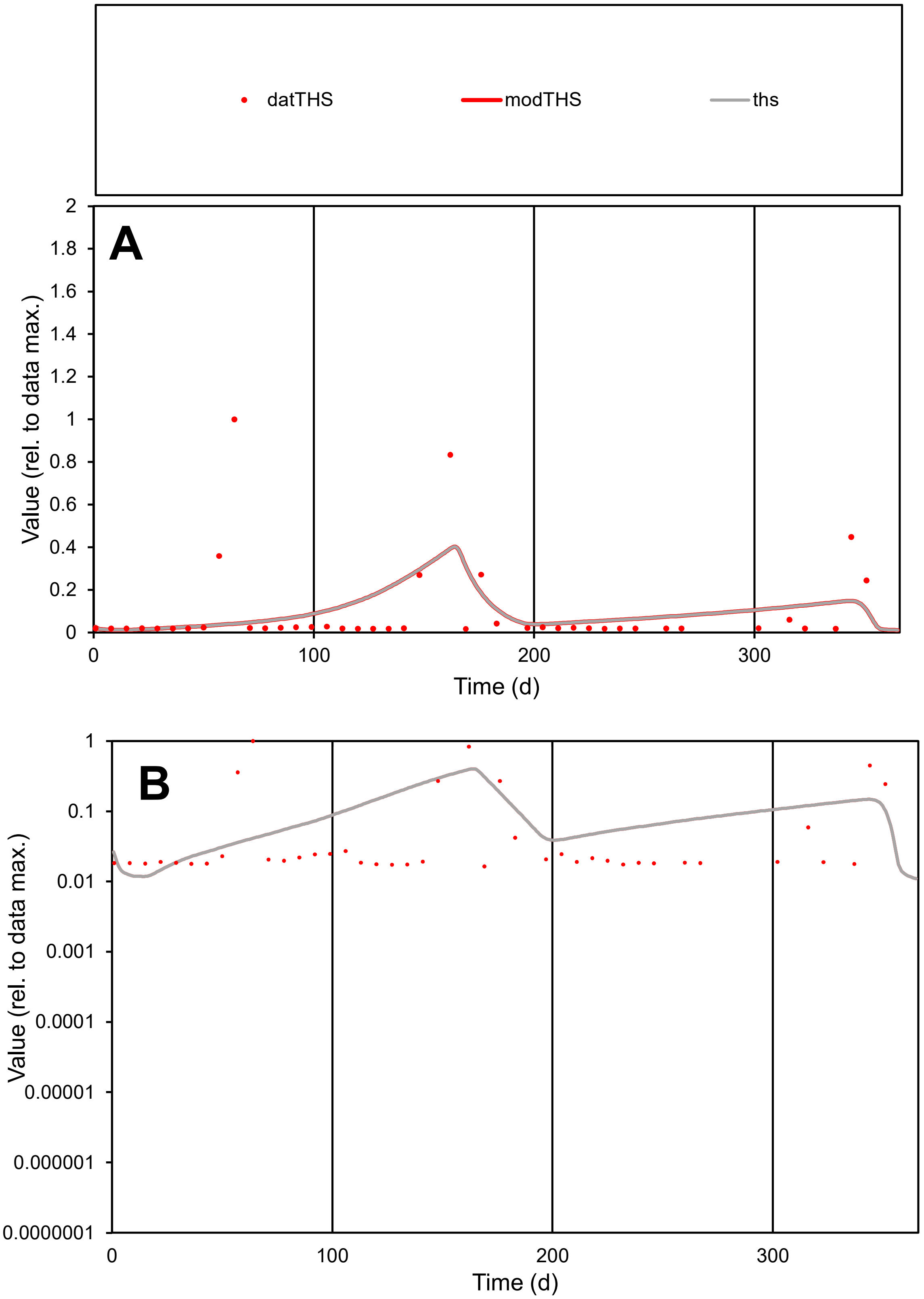


**C1**

Variables
THS Provide jtd or Dst
CalibOnly?
2 1 to reproduce of
oNormalize?
2 1=no, 2=yes

ofxn calc*
1.96694
ofxn mod
1.98339

sRun
RunA1
zout
1

Ave
Data
0.00017
Model
0.00017

data/vMax
0.00176

ofxng
319.391

Min d/m
1.36527

ave/min d
6.02957
ave/min m
8.15305

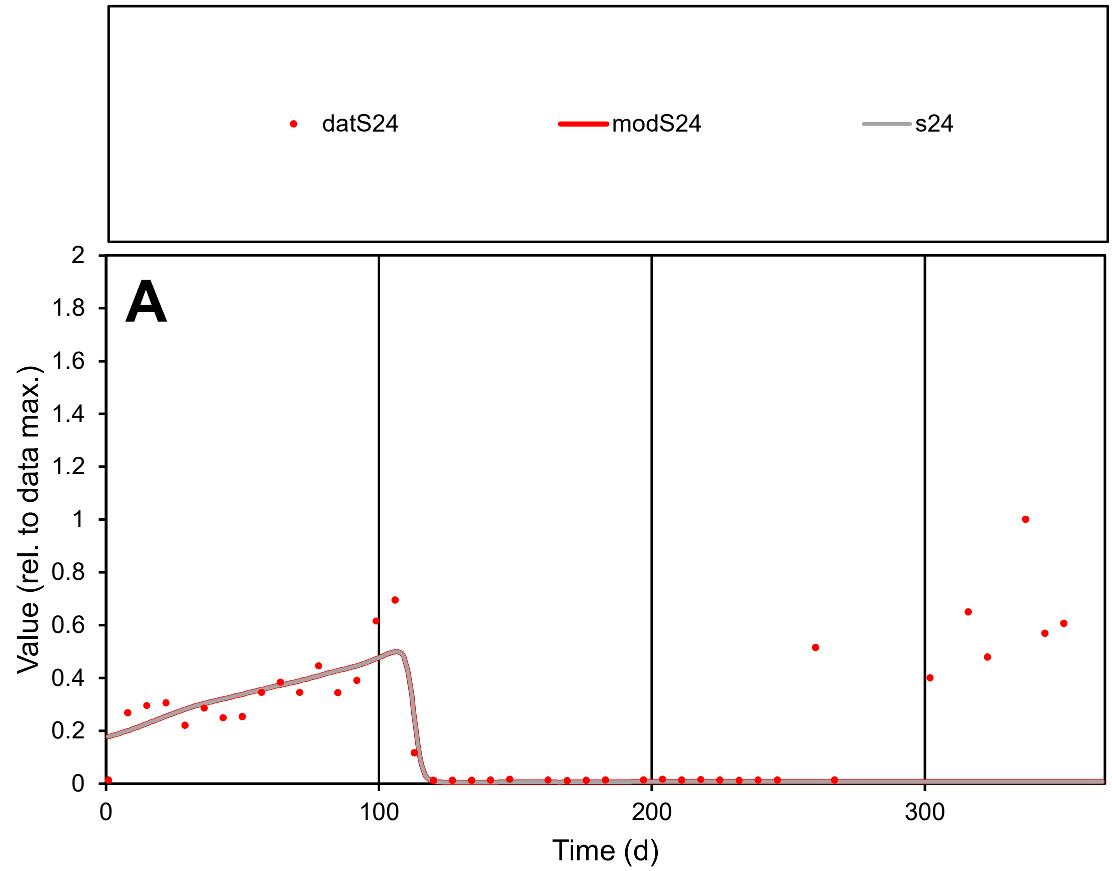


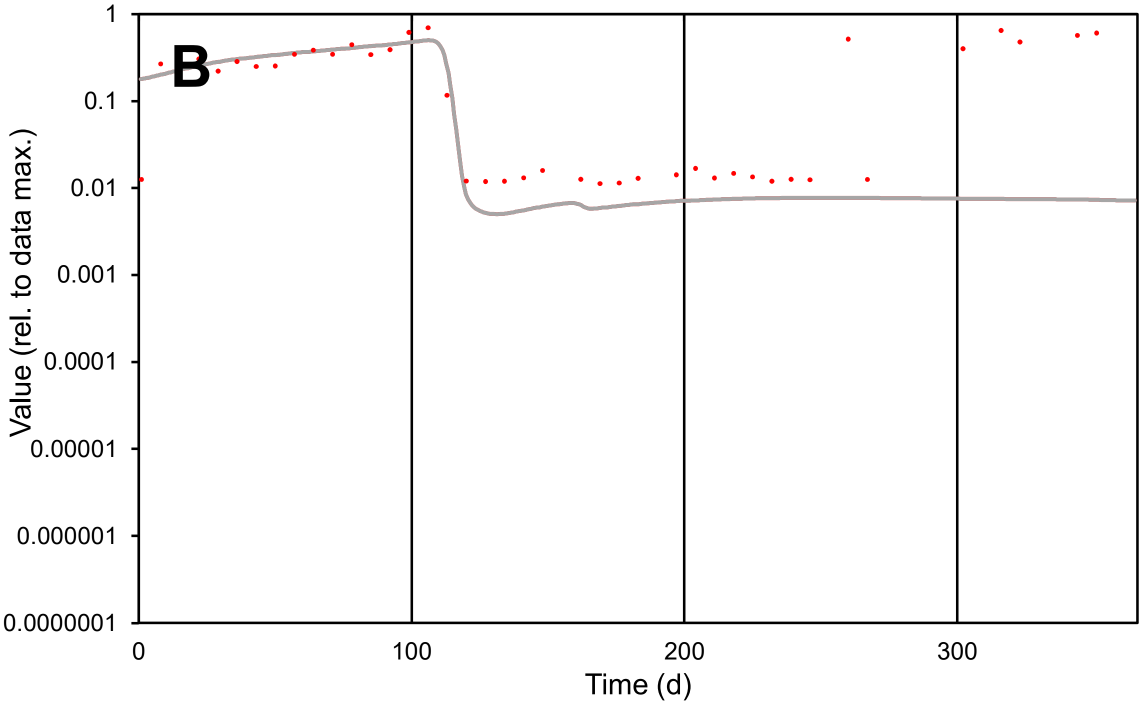


**C1**

Variables

S24 Provide jtd or Dst

CalibOnly?

2 1 to reproduce of

oNormalize?

2 1=no, 2=yes

ofxn calc*

1.08867

ofxn mod

1.09497

sRun

RunA1

zout

1

Ave

Data

0.00124

Model

0.00072

data/vMax

0.00519

ofxng

319.391

Min d/m

2.20618

ave/min d

21.1414

ave/min m

27.3066

330_Calib_S_A

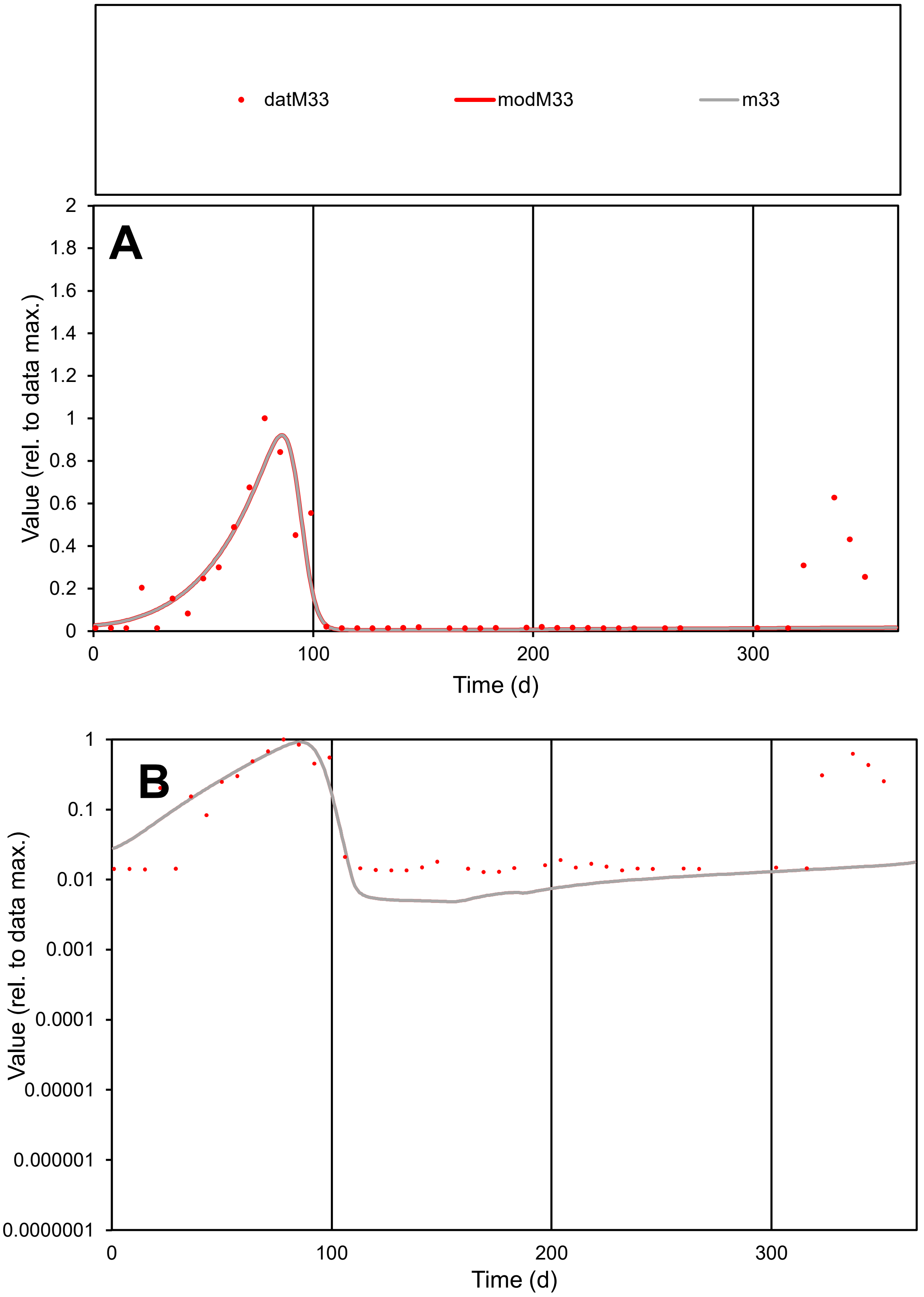


**C1**

Variables
M33 Provide jtd or Dst
CalibOnly?
2 1 to reproduce of
oNormalize?
2 1=no, 2=yes

ofxn calc*
0.87506
ofxn mod
0.90922

sRun
RunA1
zout
1

Ave
Data
0.0009
Model
0.00068

data/vMax
0.00538

ofxng
319.391

Min d/m
2.6174

ave/min d
13.0582
ave/min m
25.9279

330_Calib_S_A

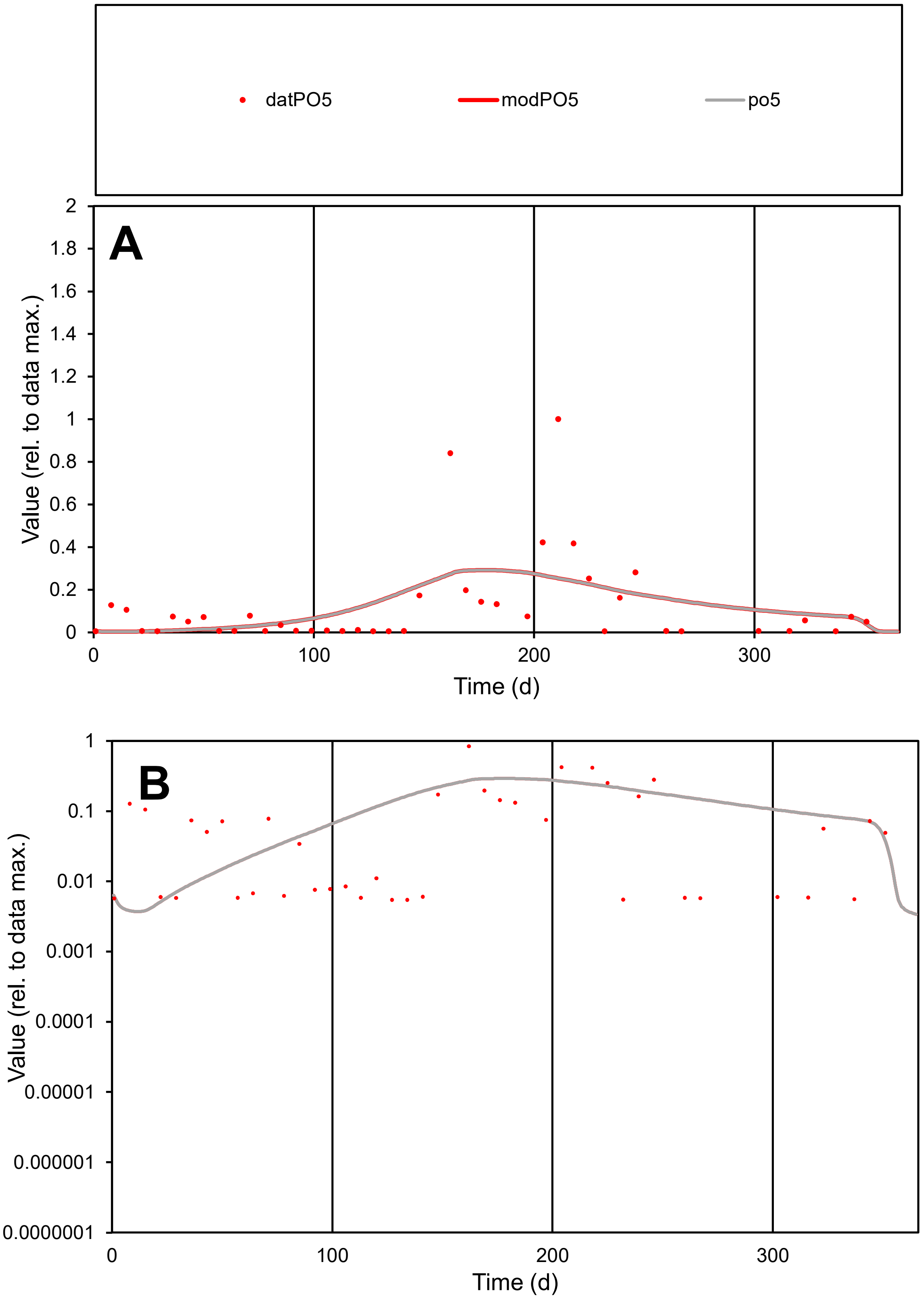


**C1**

Variables
PO5 Provide jtd or Dst
CalibOnly?
2 1 to reproduce of
oNormalize?
2 1=no, 2=yes

ofxn calc*
1.49167
ofxn mod
1.50454

sRun
RunA1
zout
1

Ave
Data
0.00107
Model
0.0011

data/vMax
0.00905

ofxng
319.391

Min d/m
1.45075

ave/min d
21.5851
ave/min m
32.2949

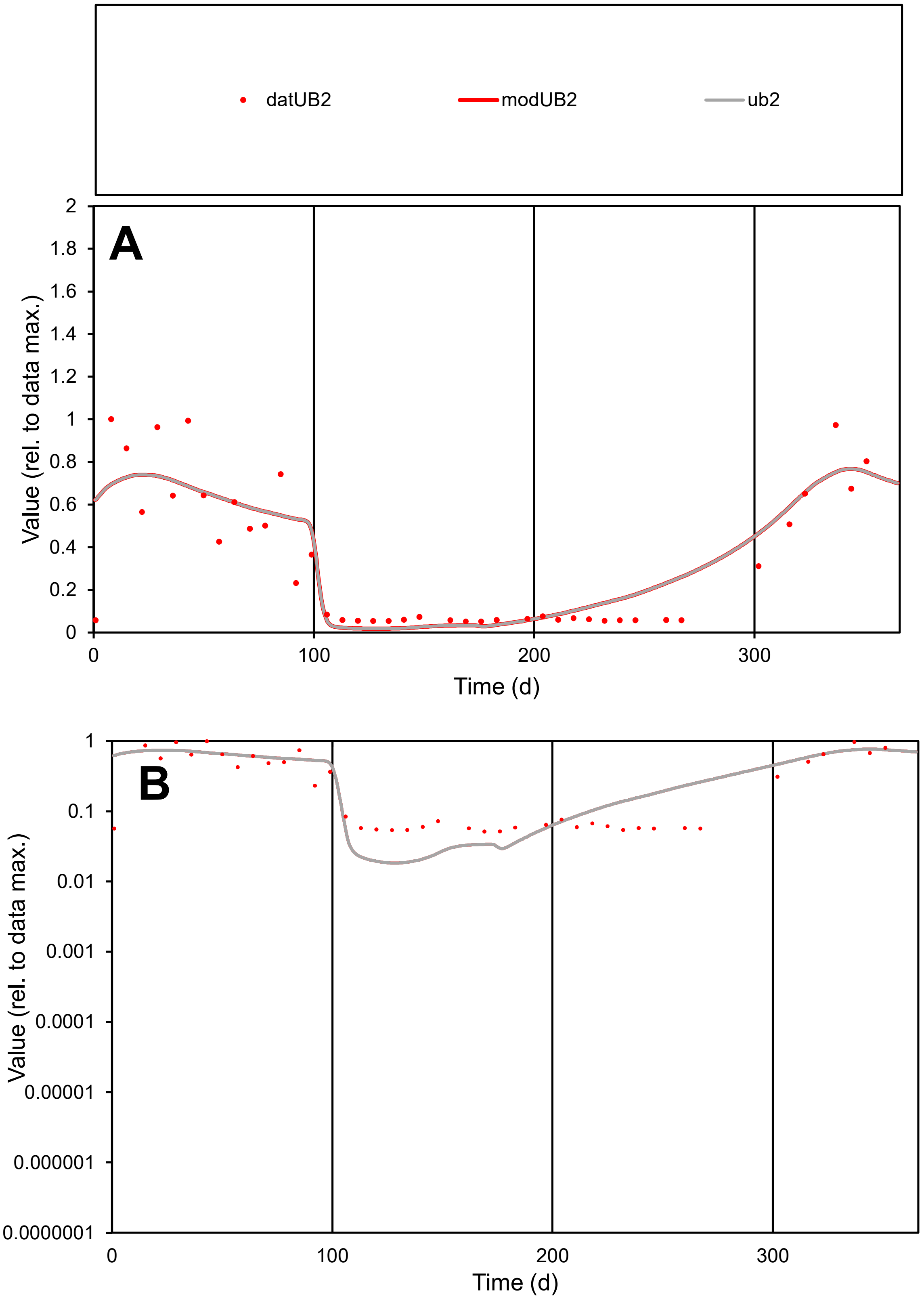


**C1**

Variables
UB2 Provide jtd or Dst
CalibOnly?
2 1 to reproduce of
oNormalize?
2 1=no, 2=yes

ofxn calc*
0.4591
ofxn mod
0.48251

sRun
RunA1
zout
1

Ave
Data
0.00069
Model
0.00074

data/vMax
0.00204

ofxng
319.391

Min d/m
2.78946

ave/min d
6.62836
ave/min m
19.6831

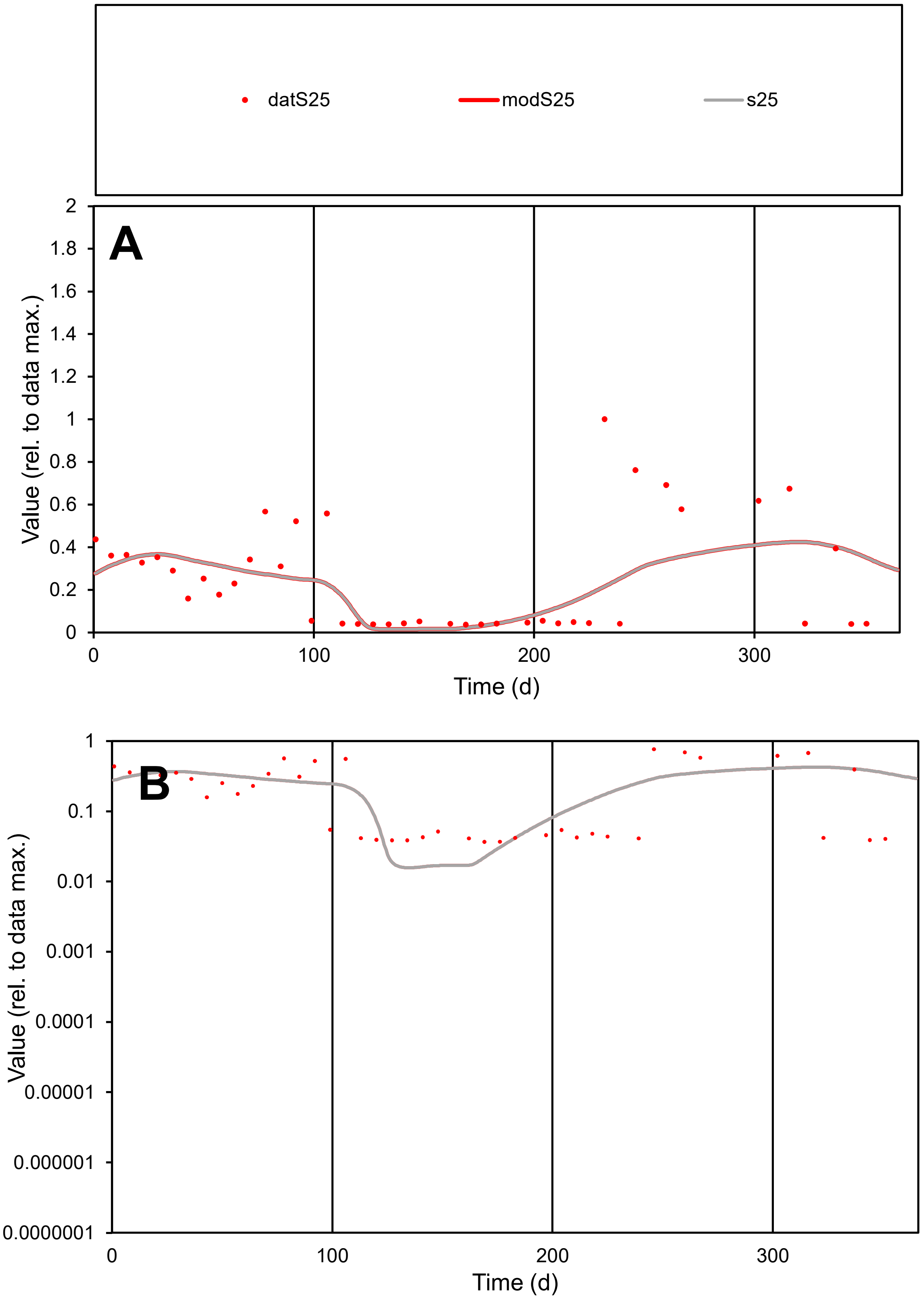


C1

Variables
S25 Provide jtd or Dst
CalibOnly?
2 1 to reproduce of
oNormalize?
2 1=no, 2=yes

ofxn calc*
0.83486
ofxn mod
0.84613

sRun
RunA1
zout
1

Ave
Data
0.0012
Model
0.00107

data/vMax
0.00465

ofxng
319.391

Min d/m
2.32418

ave/min d
7.04491
ave/min m
14.6772

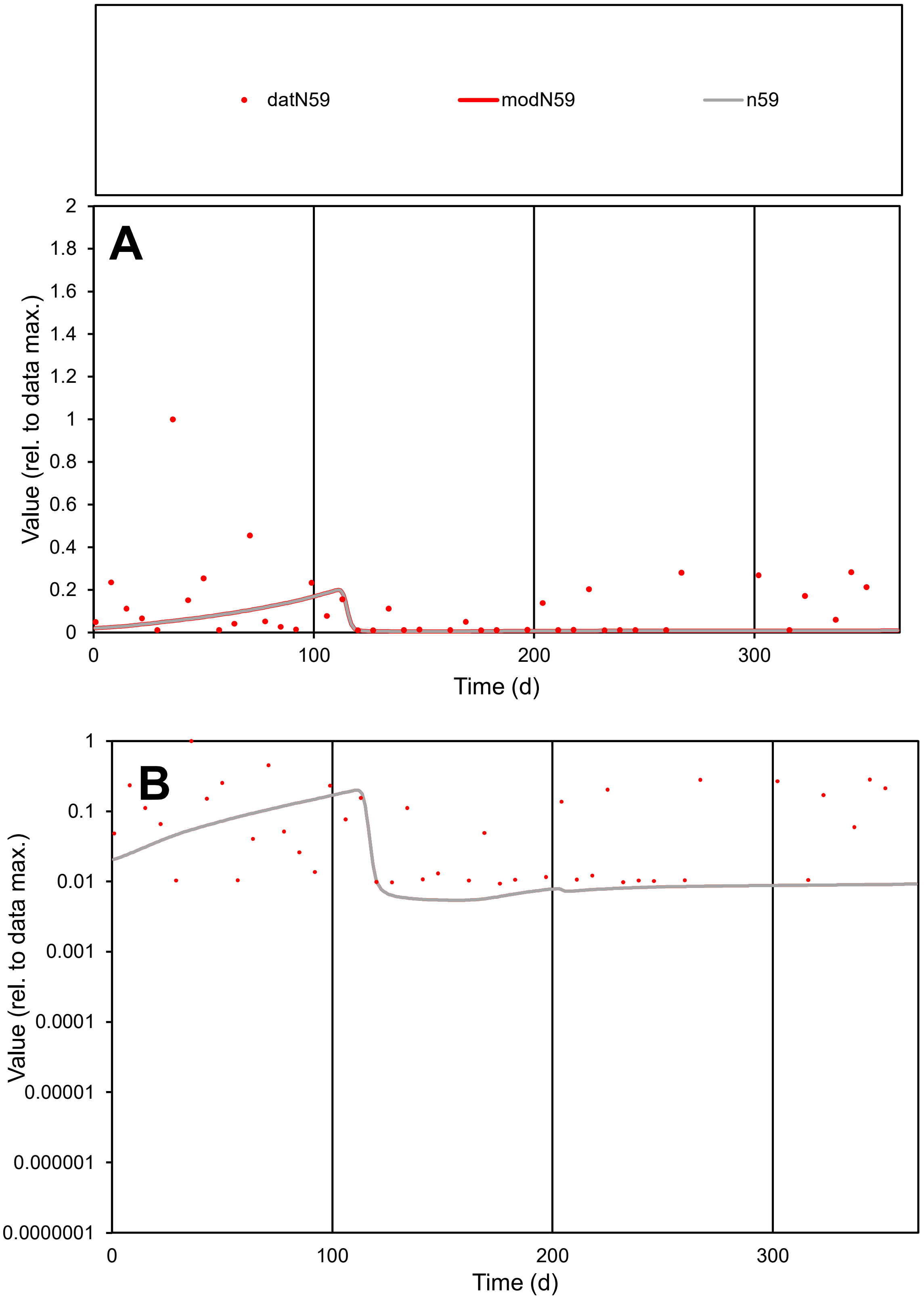


**C1**

Variables
N59 Provide jtd or Dst
CalibOnly?
2 1 to reproduce of
oNormalize?
2 1=no, 2=yes

ofxn calc*
1.63908
ofxn mod
1.64261

sRun
RunA1
zout
1

Ave
Data
0.00052
Model
0.00019

data/vMax
0.0045

ofxng
319.391

Min d/m
1.71621

ave/min d
12.4018
ave/min m
7.64306

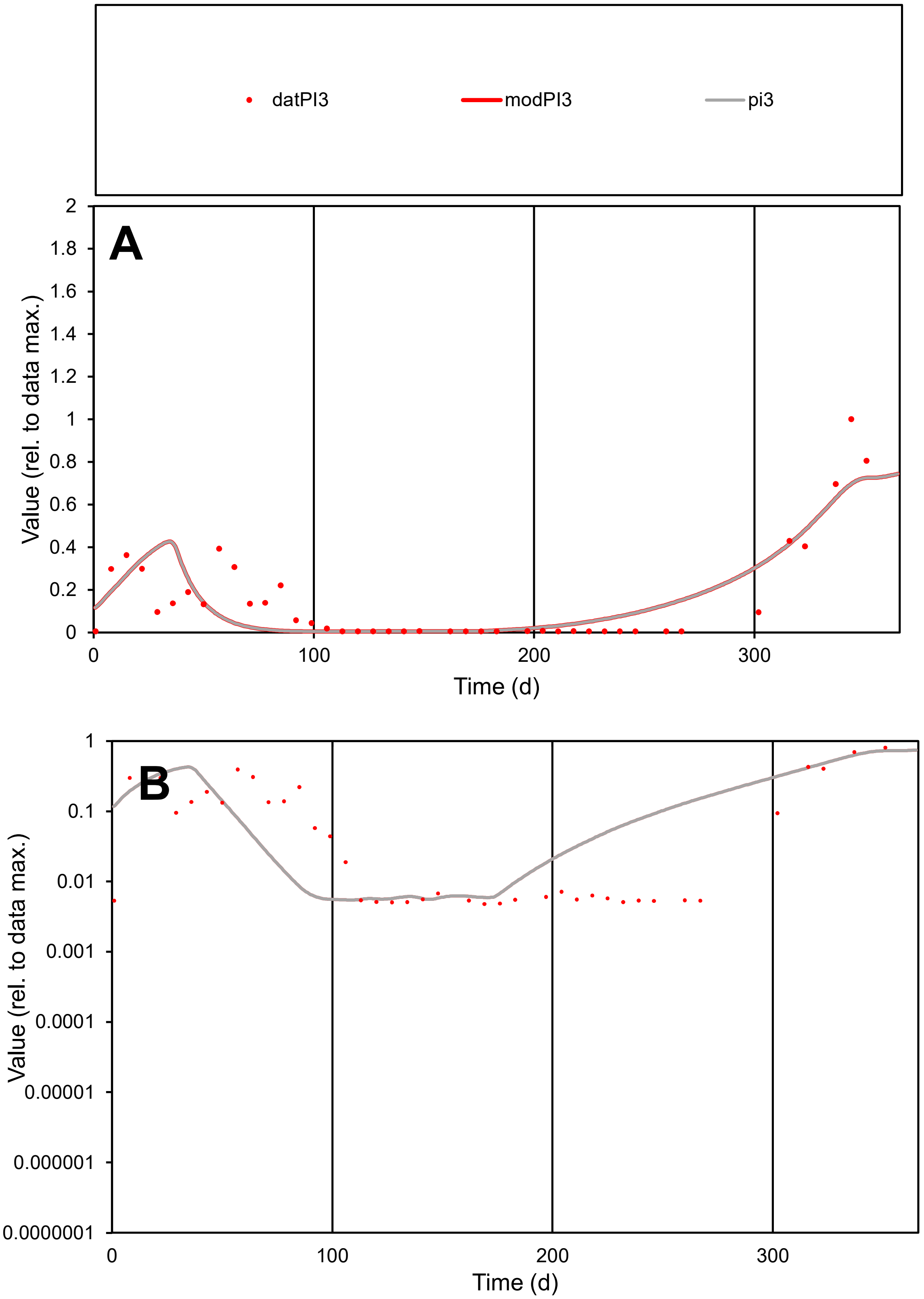


**C1**

Variables
PI3 Provide jtd or Dst
CalibOnly?
2 1 to reproduce of
oNormalize?
2 1=no, 2=yes

ofxn calc*
0.8109
ofxn mod
0.83689

sRun
RunA1
zout
1

Ave
Data
0.00226
Model
0.00225

data/vMax
0.01493

ofxng
319.391

Min d/m
0.88469

ave/min d
31.5937
ave/min m
27.7731

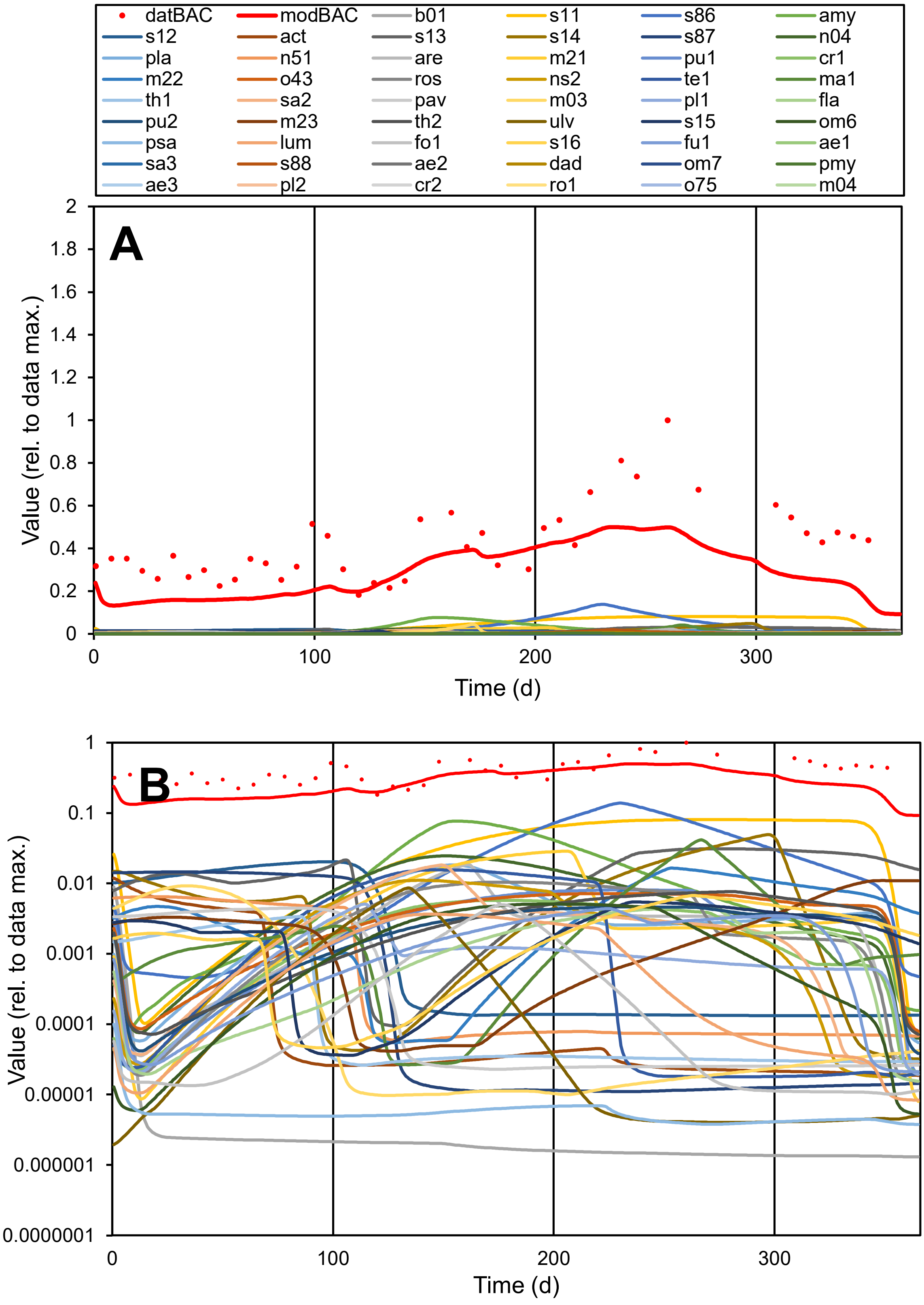


**C1**

Variables

BAC Provide jtd or Dst

CalibOnly?

2 1 to reproduce of

oNormalize?

2 1=no, 2=yes

ofxn calc*

0.45462

ofxn mod

0.46585

sRun

RunA1

zout

1

Ave

Data

0.90039

Model

0.58465

data/vMax

2.134

ofxng

319.391

Min d/m

1.36456

ave/min d

2.32418

ave/min m

2.05937

331_Calib_S_A

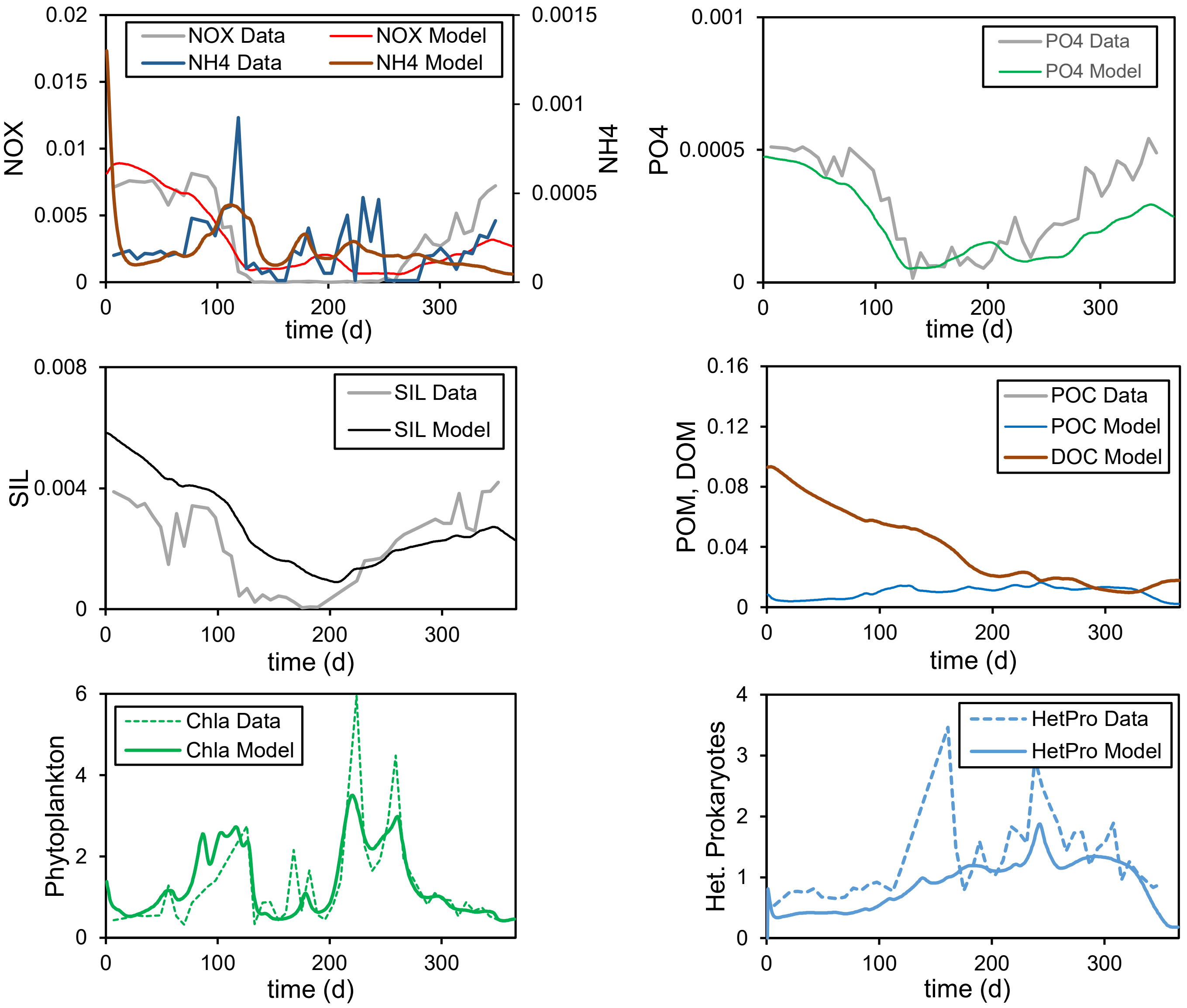

NOX Data
NOX Model
NH4 Data
NH4 Model
NOX
NH4
0.02
0.015
0.01
0.005
0
0.0015
0.001
0.0005
0
100
200
300
time (d)
PO4 Data
PO4 Model
PO4
0.001
0.0005
time (d)
SIL Data
SIL Model
SIL
0.008
0.004
time (d)
POC Data
POC Model
DOC Model
POM, DOM
0.16
0.12
0.08
0.04
time (d)
Chla Data
Chla Model
Phytoplankton
6
4
2
time (d)
HetPro Data
HetPro Model
Het. Prokaryotes
4
3
2
1
time (d)

331_Calib_S_A

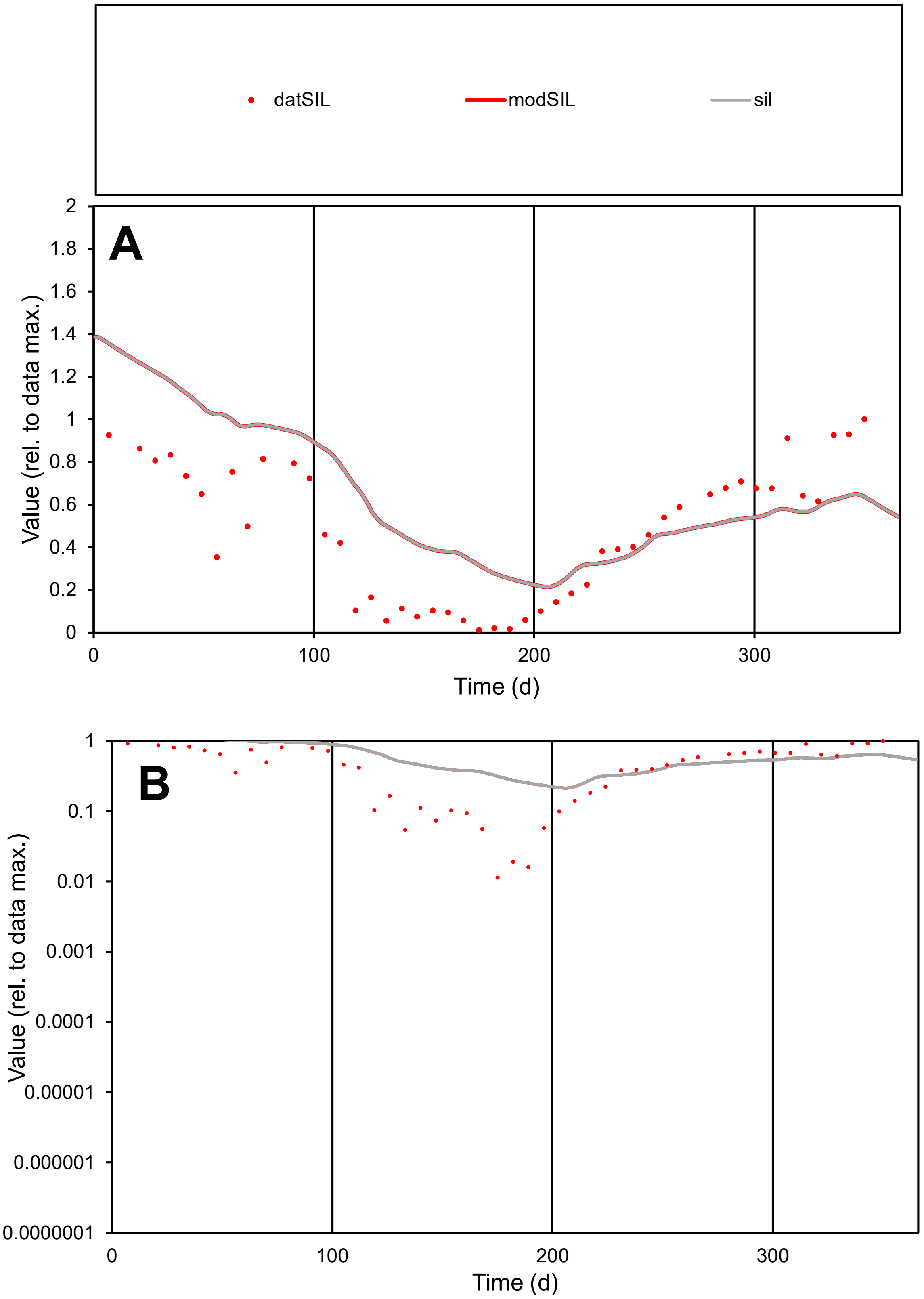


**C1**

Variables
SIL Provide jtd or Dst
CalibOnly?
2 1 to reproduce of
oNormalize?
2 1=no, 2=yes

ofxn calc*
1.8352
ofxn mod
1.88636

sRun
RunA1
zout
1

Ave
Data
0.00199
Model
0.0026

data/vMax
0.0042

ofxng
311.734

Min d/m
0.05269

ave/min d
41.9455
ave/min m
2.88296

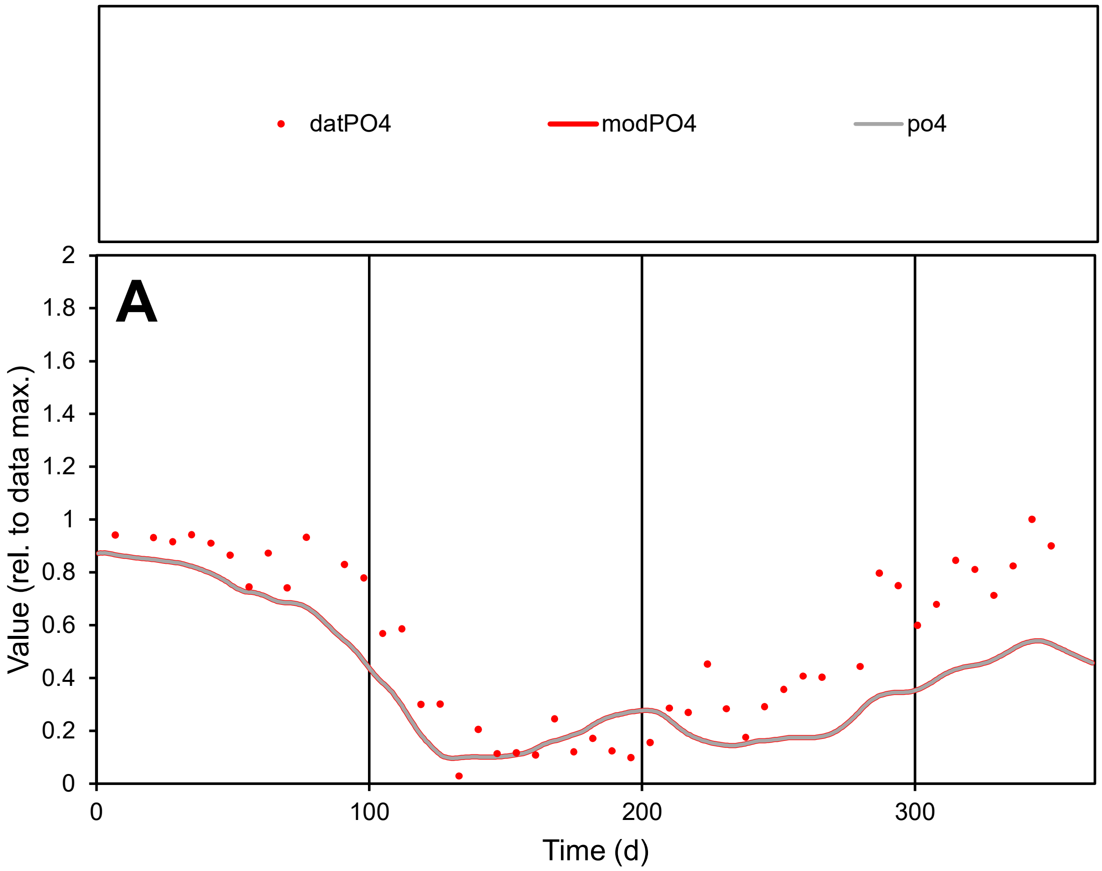


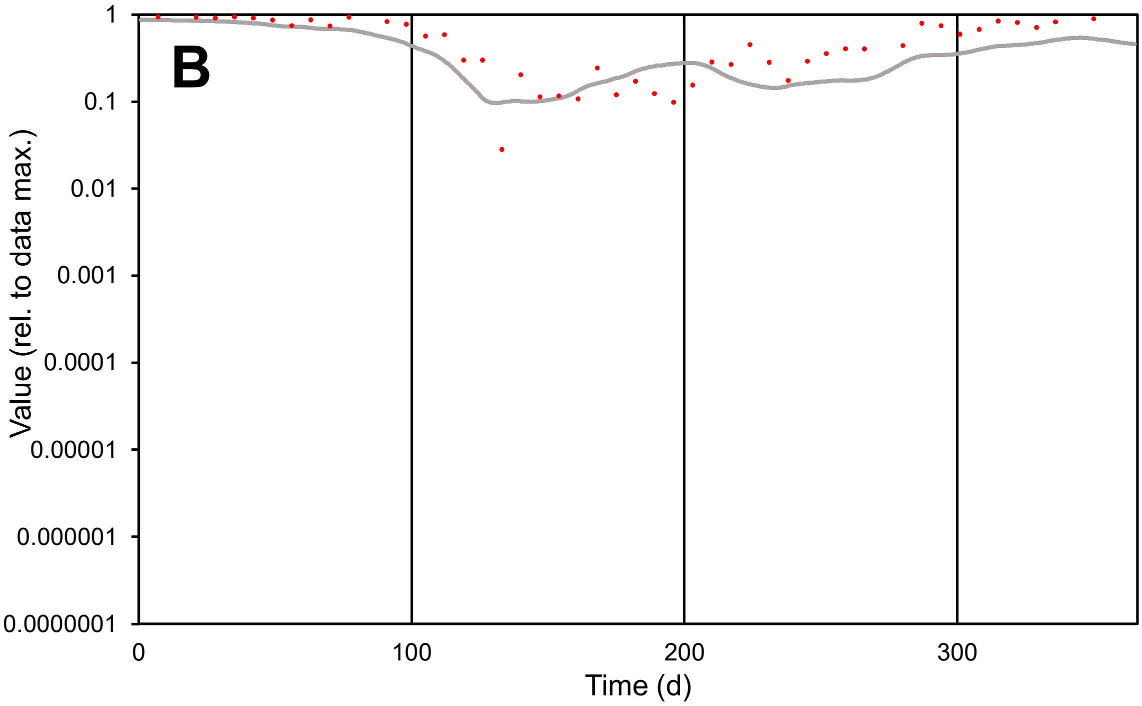


**C1**

Variables
PO4 Provide jtd or Dst
CalibOnly?
2 1 to reproduce of
oNormalize?
2 1=no, 2=yes

ofxn calc*
1.23266
ofxn mod
1.28349

sRun
RunA1
zout
1

Ave
Data
0.00029
Model
0.00021

data/vMax
0.00054

ofxng
311.734

Min d/m
0.28762

ave/min d
18.851
ave/min m
3.87598

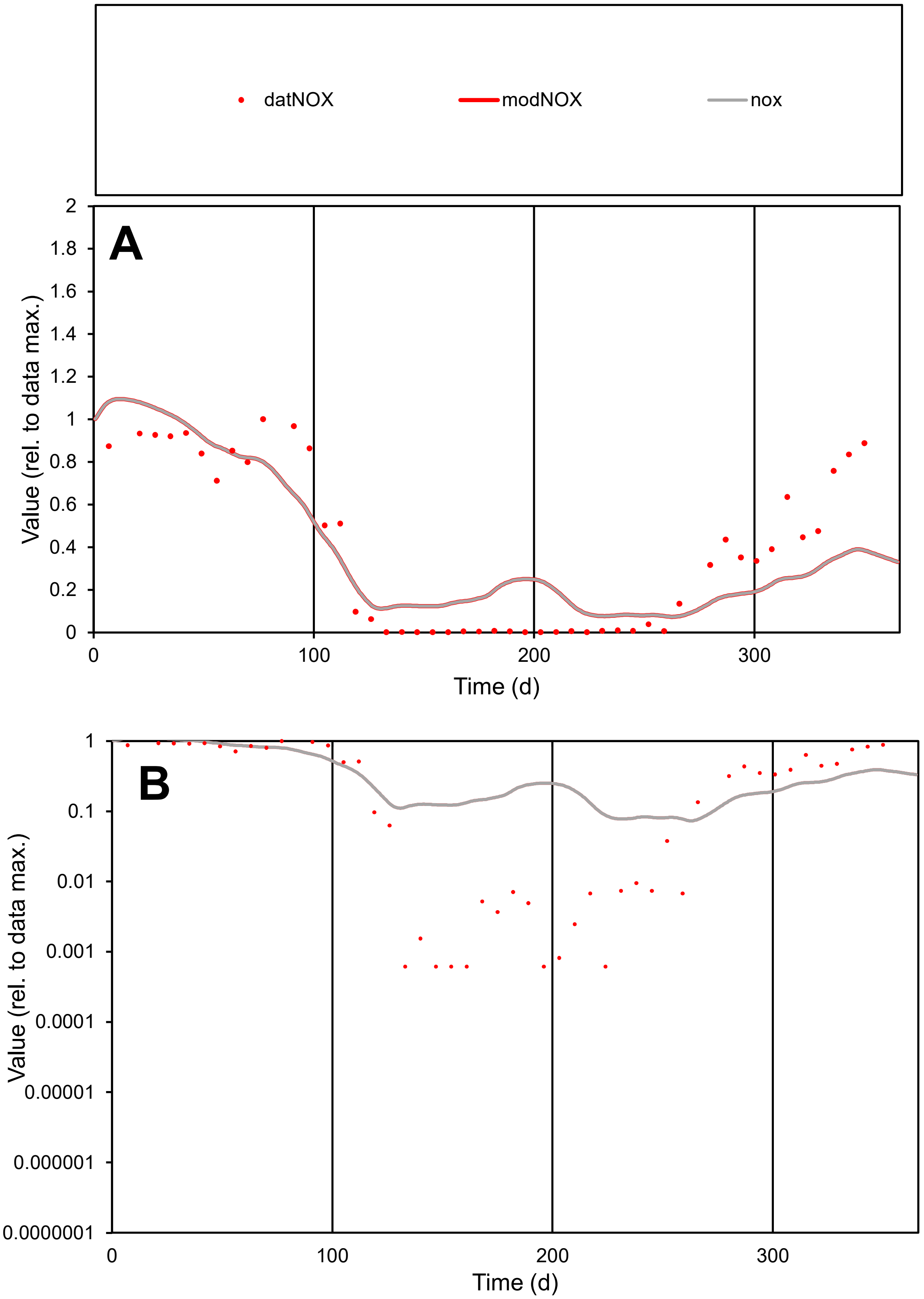


**C1**

Variables
NOX Provide jtd or Dst
CalibOnly?
2 1 to reproduce of
oNormalize?
2 1=no, 2=yes

ofxn calc*
1.55154
ofxn mod
1.62701

sRun
RunA1
zout
1

Ave
Data
0.0031
Model
0.00298

data/vMax
0.00814

ofxng
311.734

Min d/m
0.00818

ave/min d
619.62
ave/min m
4.87733

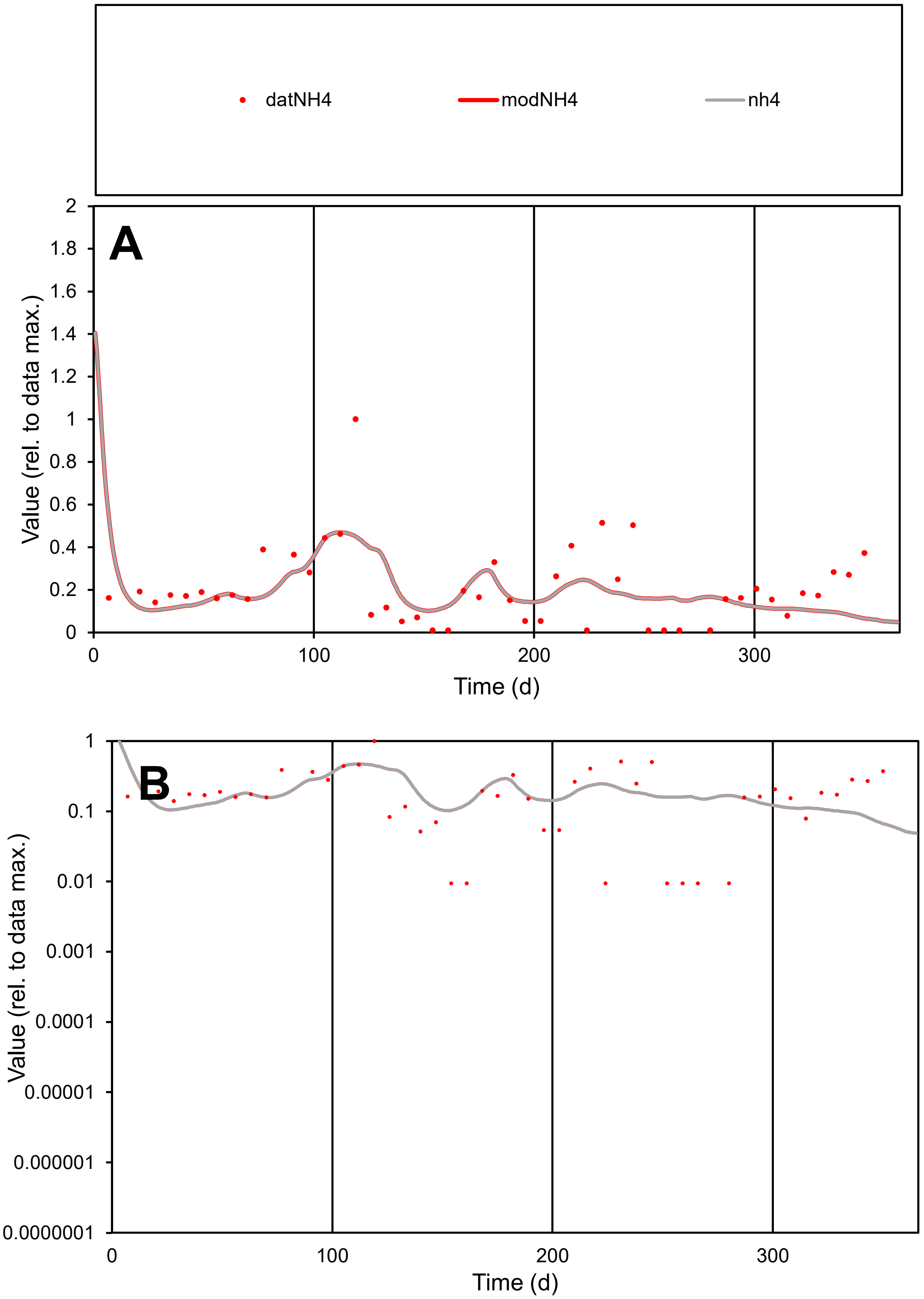


**C1**

Variables
NH4 Provide jtd or Dst
CalibOnly?
2 1 to reproduce of
oNormalize?
2 1=no, 2=yes

ofxn calc*
2.29179
ofxn mod
2.34057

sRun
RunA1
zout
1

Ave
Data
0.00019
Model
0.00018

data/vMax
0.00093

ofxng
311.734

Min d/m
0.1453

ave/min d
21.9847
ave/min m
2.91055

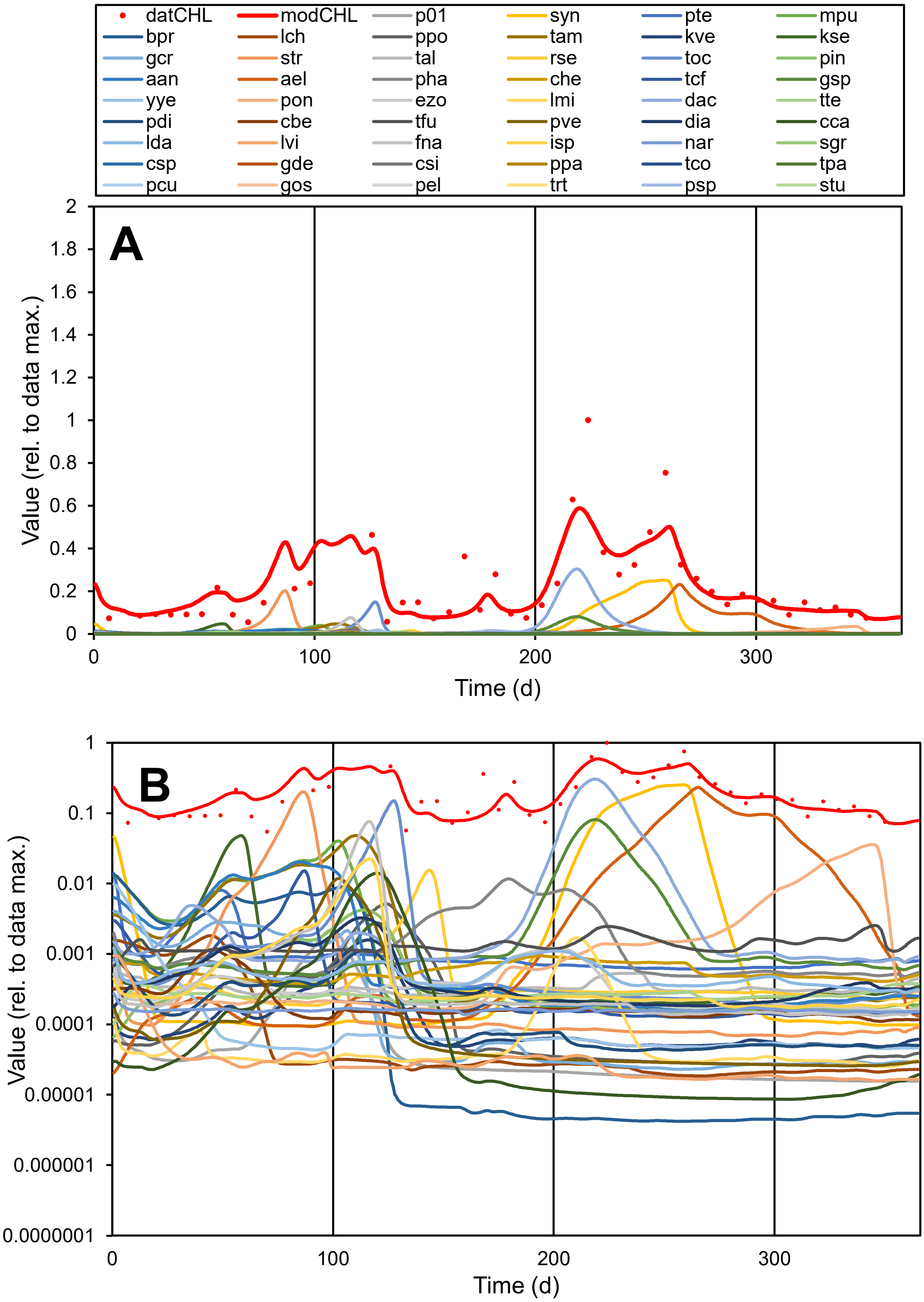


**C1**

Variables

CHL Provide jtd or Dst

CalibOnly?

2 1 to reproduce of

oNormalize?

2 1=no, 2=yes

ofxn calc*

4.74745

ofxn mod

5.00005

sRun

RunA1

zout

1

Ave

Data

1.27698

Model

1.19777

data/vMax

5.949

ofxng

311.734

Min d/m

0.74674

ave/min d

3.91232

ave/min m

2.74027

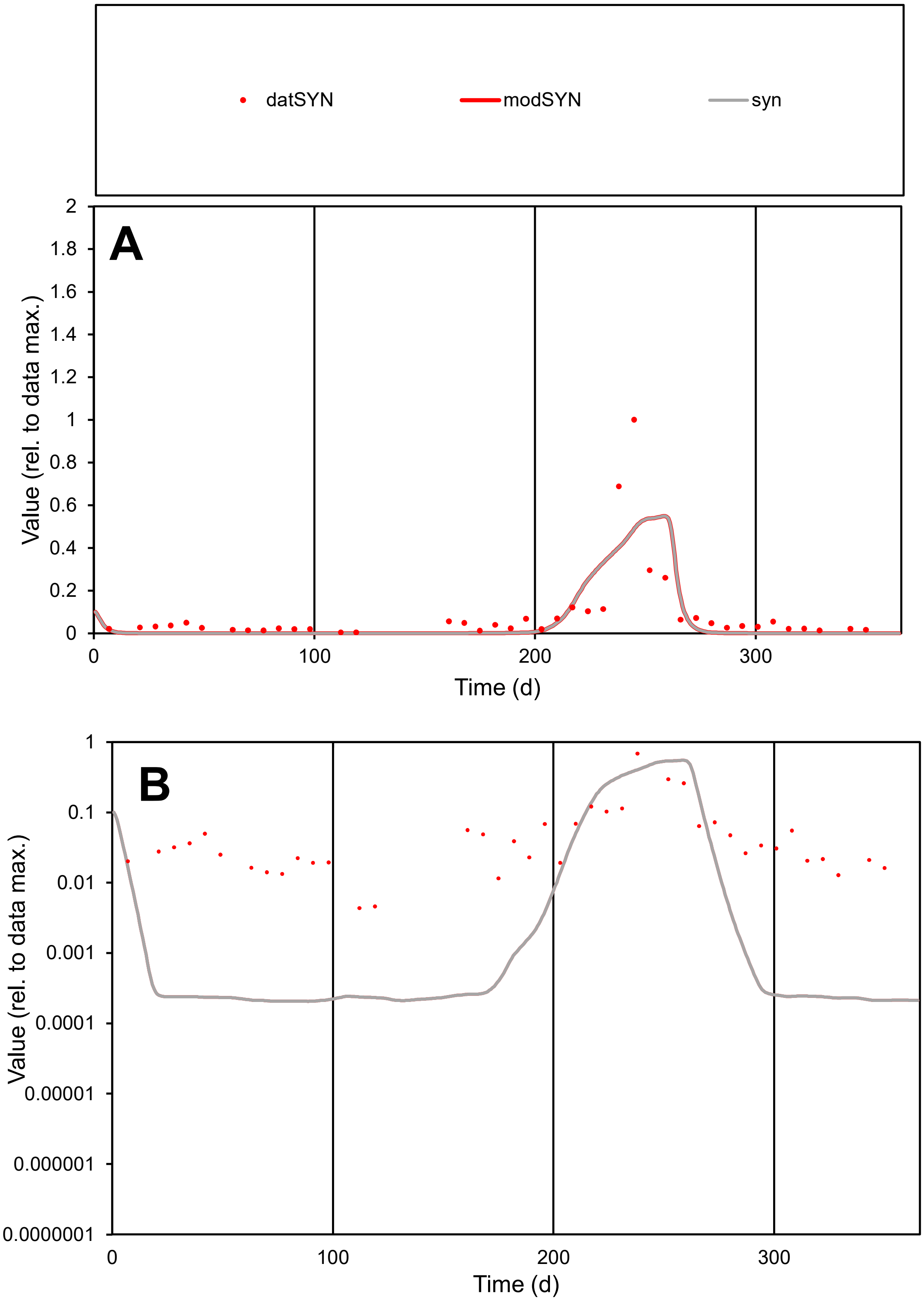


**C1**

Variables
SYN Provide jtd or Dst
CalibOnly?
2 1 to reproduce of
oNormalize?
2 1=no, 2=yes

ofxn calc*
1.31368
ofxn mod
1.33653

sRun
RunA1
zout
1

Ave
Data
28.8899
Model
23.7637

data/vMax
325.2

ofxng
311.734

Min d/m
21.1427

ave/min d
20.3307
ave/min m
353.573

331_Calib_S_A

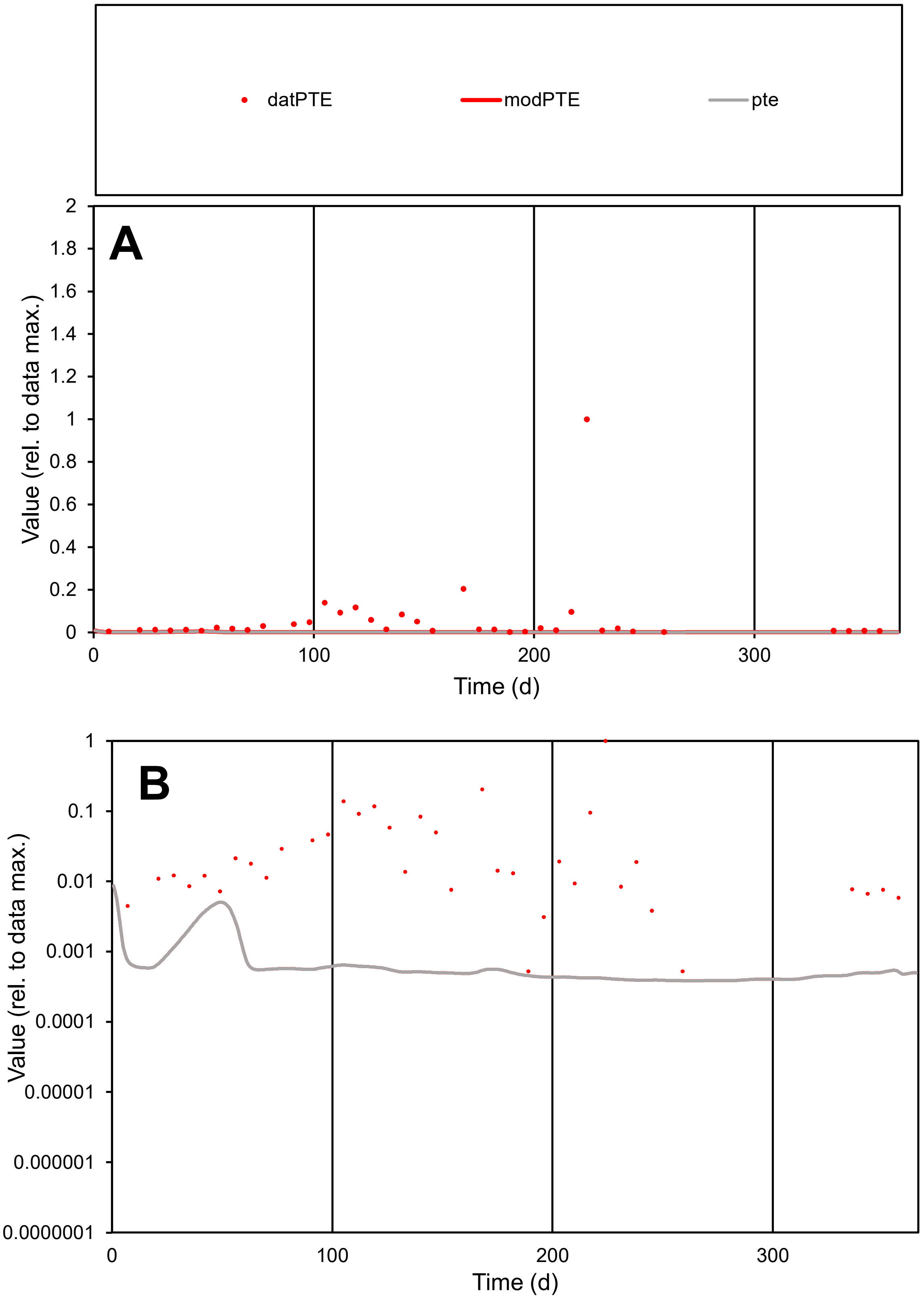


**C1**

Variables
PTE Provide jtd or Dst
CalibOnly?
2 1 to reproduce of
oNormalize?
2 1=no, 2=yes

ofxn calc*
2.91538
ofxn mod
2.92202

sRun
RunA1
zout
1

Ave
Data
0.00095
Model
1.4E-05

data/vMax
0.01592

ofxng
311.734

Min d/m
1.36646

ave/min d
113.15
ave/min m
2.21159

331_Calib_S_A

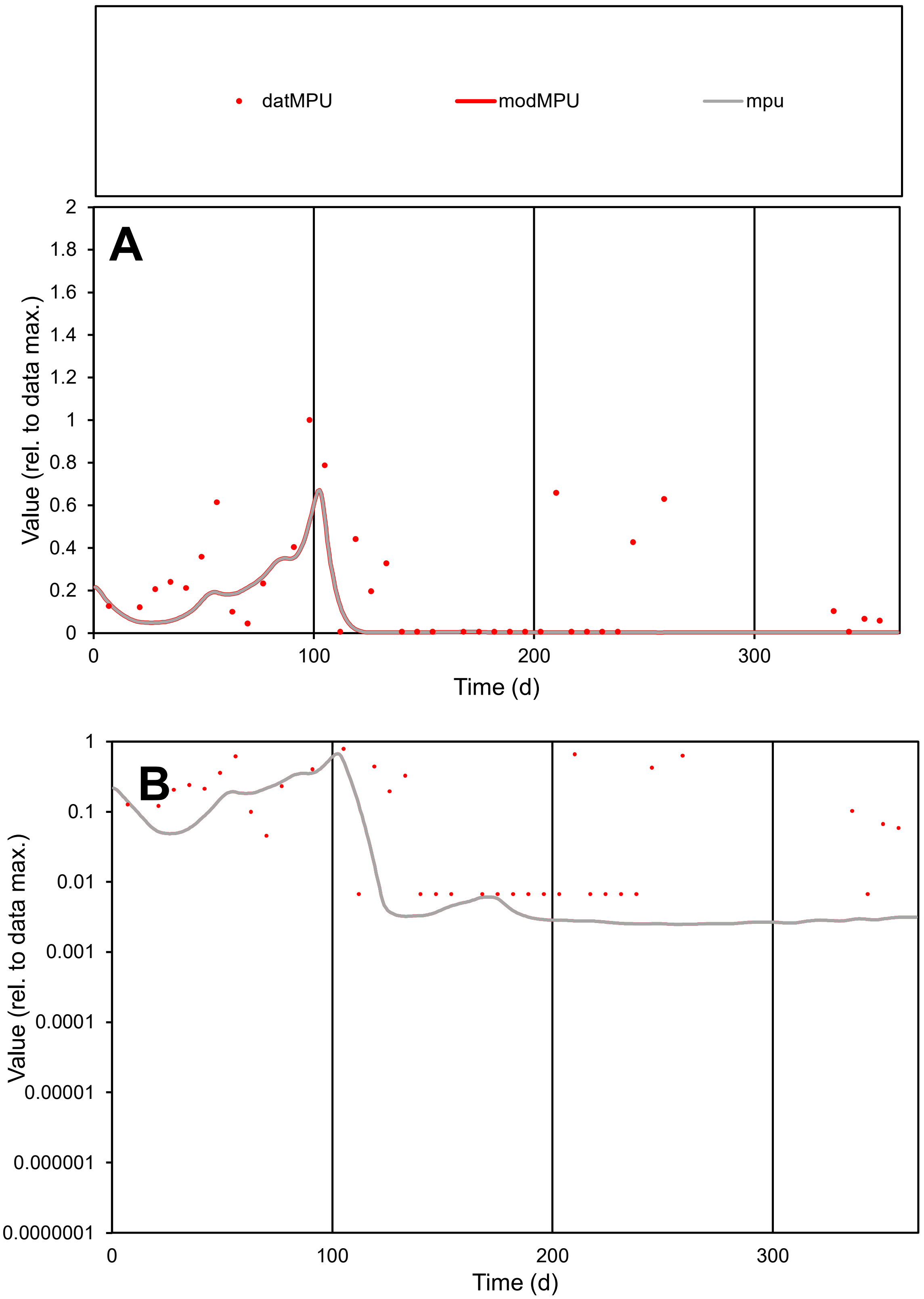


**C1**

Variables
MPU Provide jtd or Dst
CalibOnly?
2 1 to reproduce of
oNormalize?
2 1=no, 2=yes

ofxn calc*
1.13391
ofxn mod
1.1476

sRun
RunA1
zout
1

Ave
Data
0.00012
Model
5E-05

data/vMax
0.00061

ofxng
311.734

Min d/m
2.70596
ave/min d
29.9103
ave/min m
32.8051

331_Calib_S_A

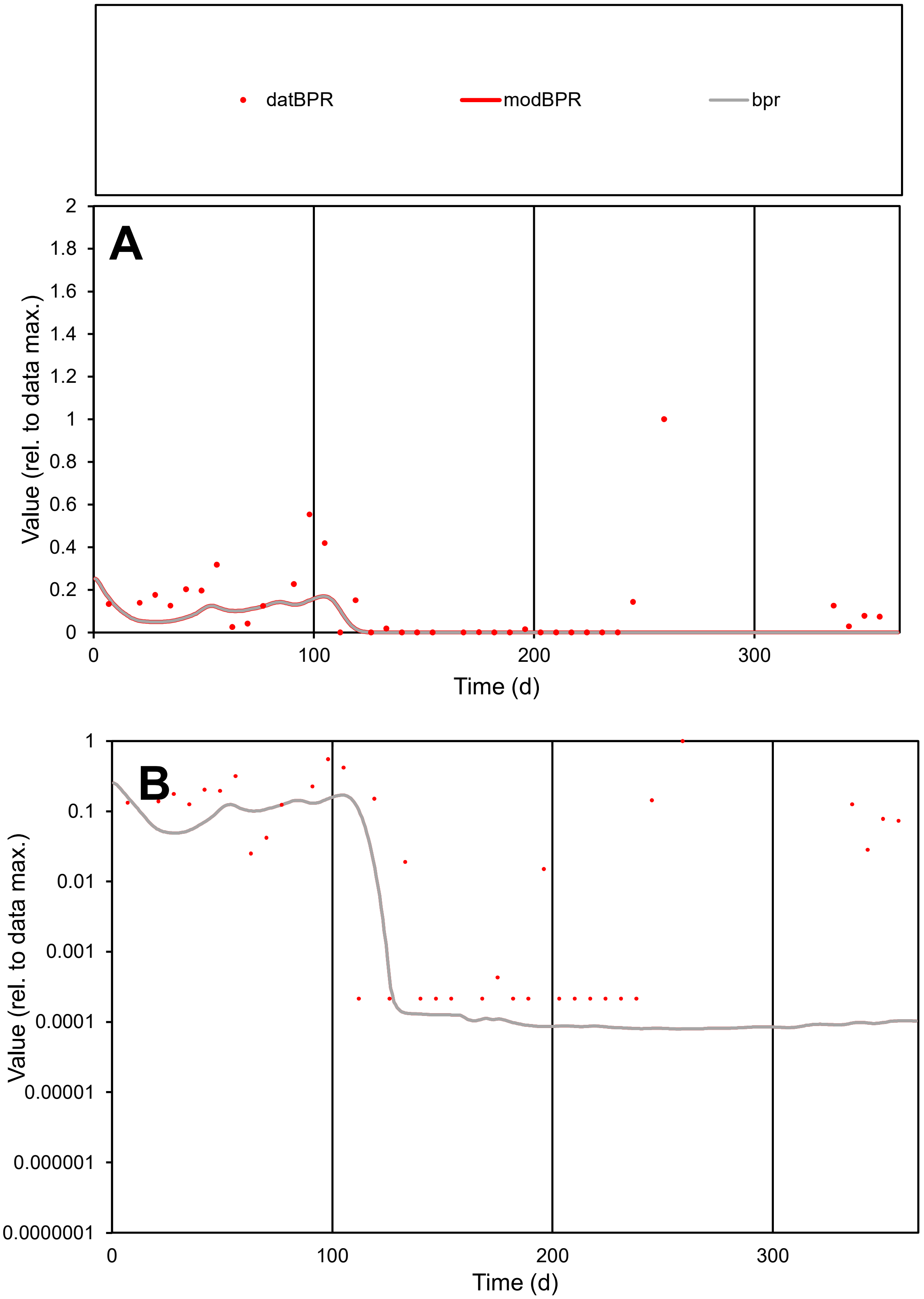


**C1**

Variables

BPR Provide jtd or Dst

CalibOnly?

2 1 to reproduce of

oNormalize?

2 1=no, 2=yes

ofxn calc*

1.66792

ofxn mod

1.67492

sRun

RunA1

zout

1

Ave

Data

0.0001

Model

3.4E-05

data/vMax

0.00086

ofxng

311.734

Min d/m

2.70038

ave/min d

540.902

ave/min m

498.264

331_Calib_S_A

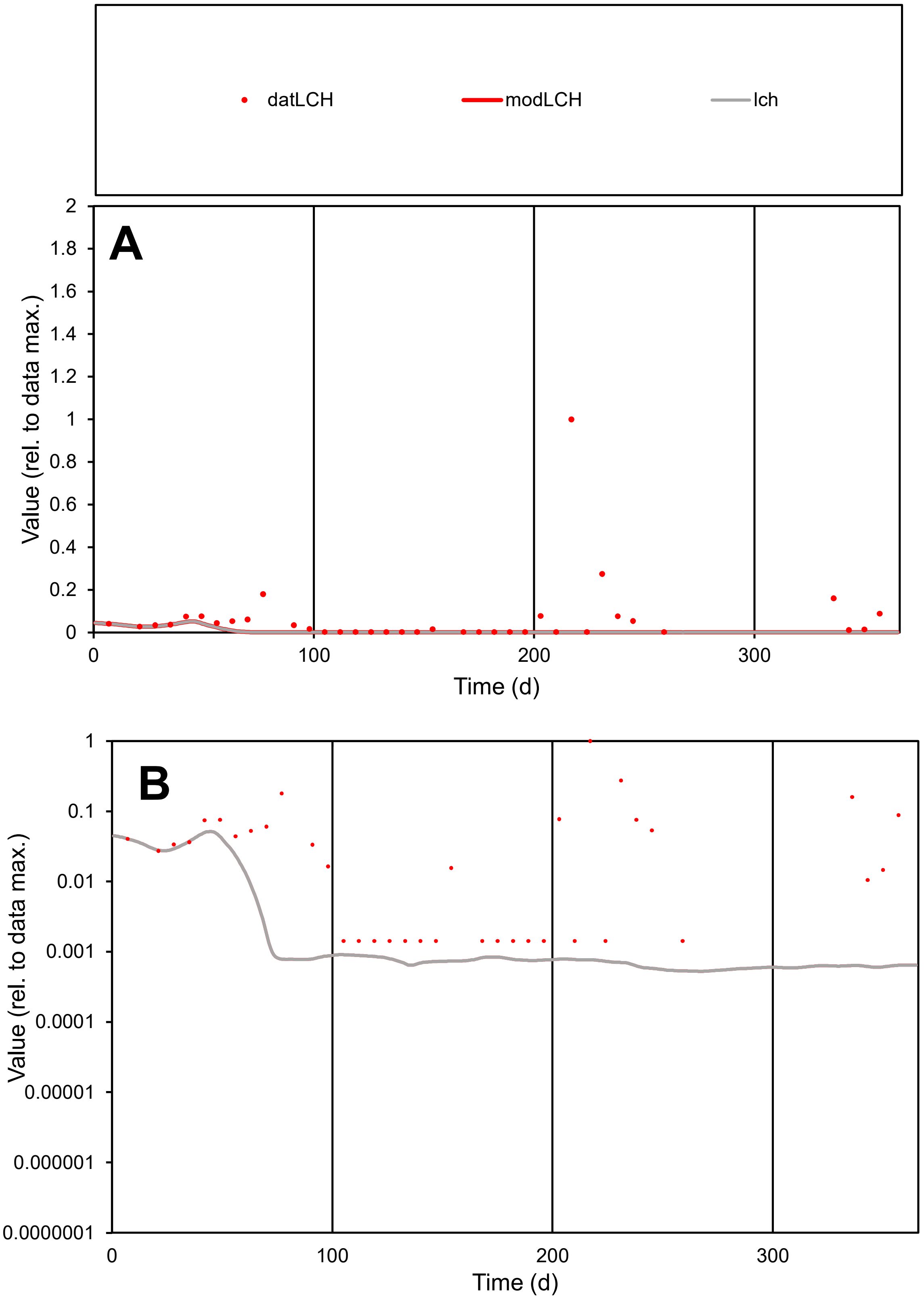


**C1**

Variables
LCH Provide jtd or Dst
CalibOnly?
2 1 to reproduce of
oNormalize?
2 1=no, 2=yes

ofxn calc*
2.65949
ofxn mod
2.66341

sRun
RunA1
zout
1

Ave
Data
0.00011
Model
1.3E-05

data/vMax
0.00171

ofxng
311.734

Min d/m
2.68821

ave/min d
46.585
ave/min m
14.3698

331_Calib_S_A

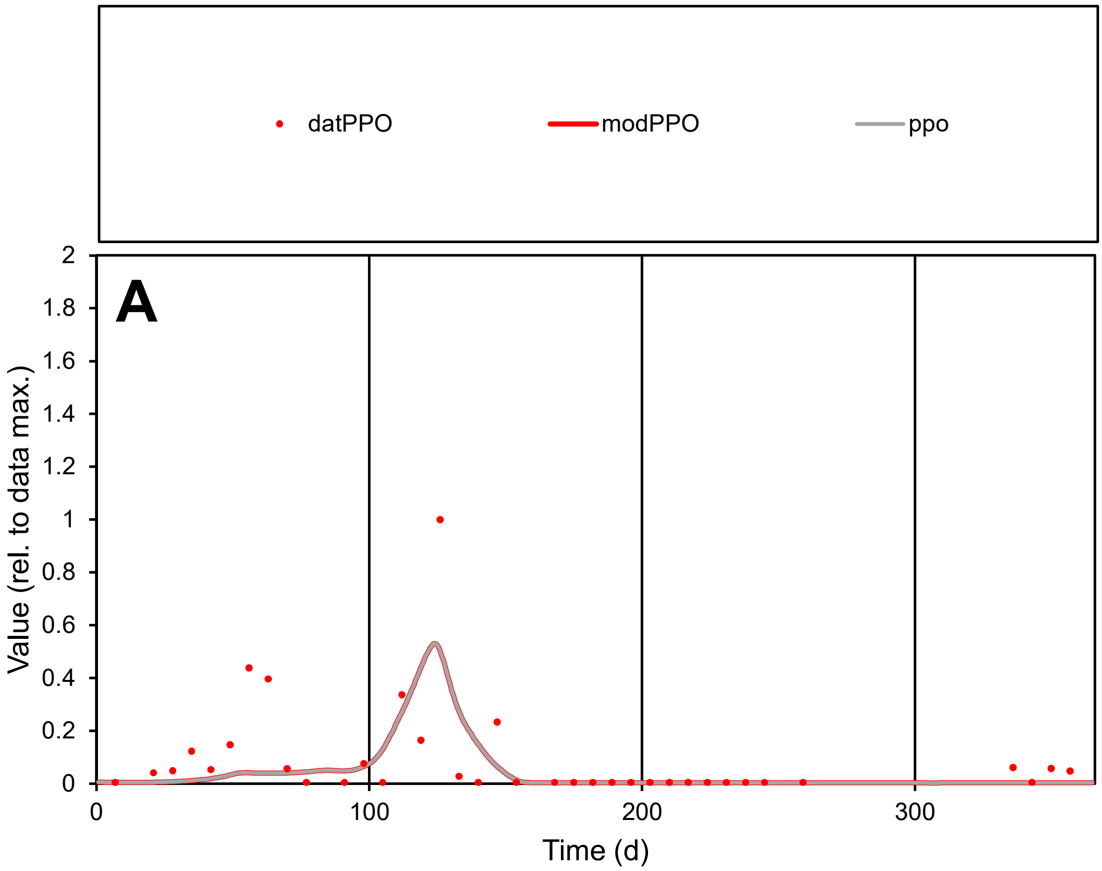


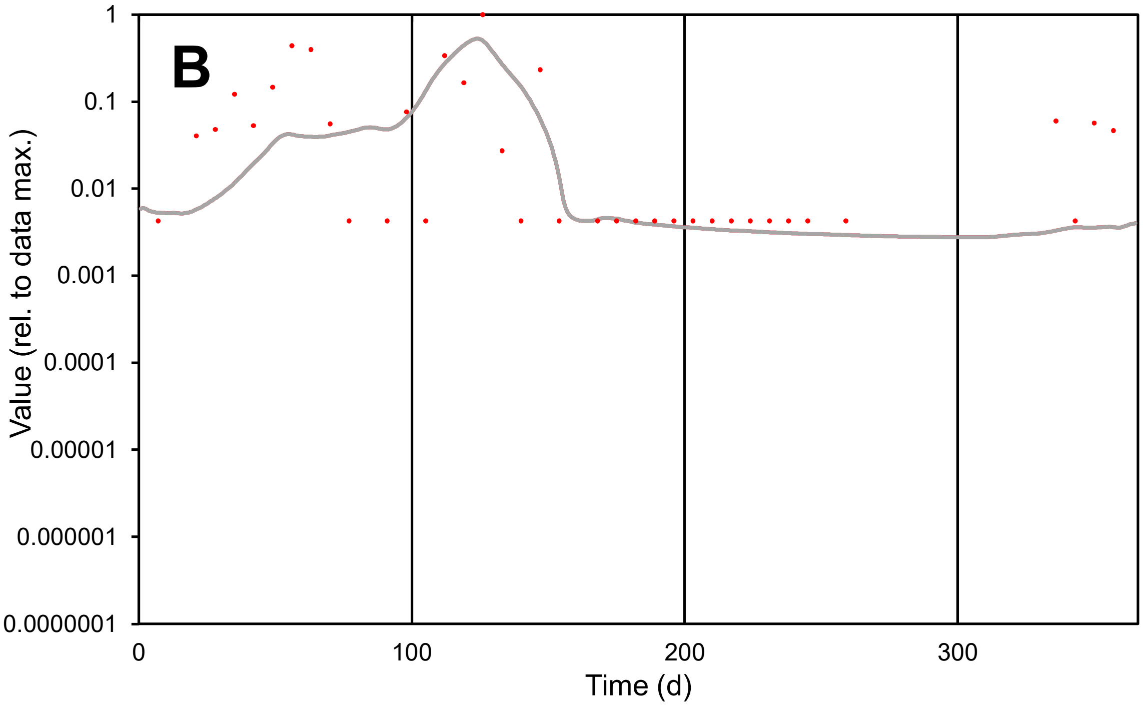


**C1**

Variables
PPO Provide jtd or Dst
CalibOnly?
2 1 to reproduce of
oNormalize?
2 1=no, 2=yes

ofxn calc*
1.56249
ofxn mod
1.59871

sRun
RunA1
zout
1

Ave
Data
4.1E-05
Model
2.6E-05

data/vMax
0.00045

ofxng
311.734

Min d/m
1.45808

ave/min d
21.4693
ave/min m
20.1478

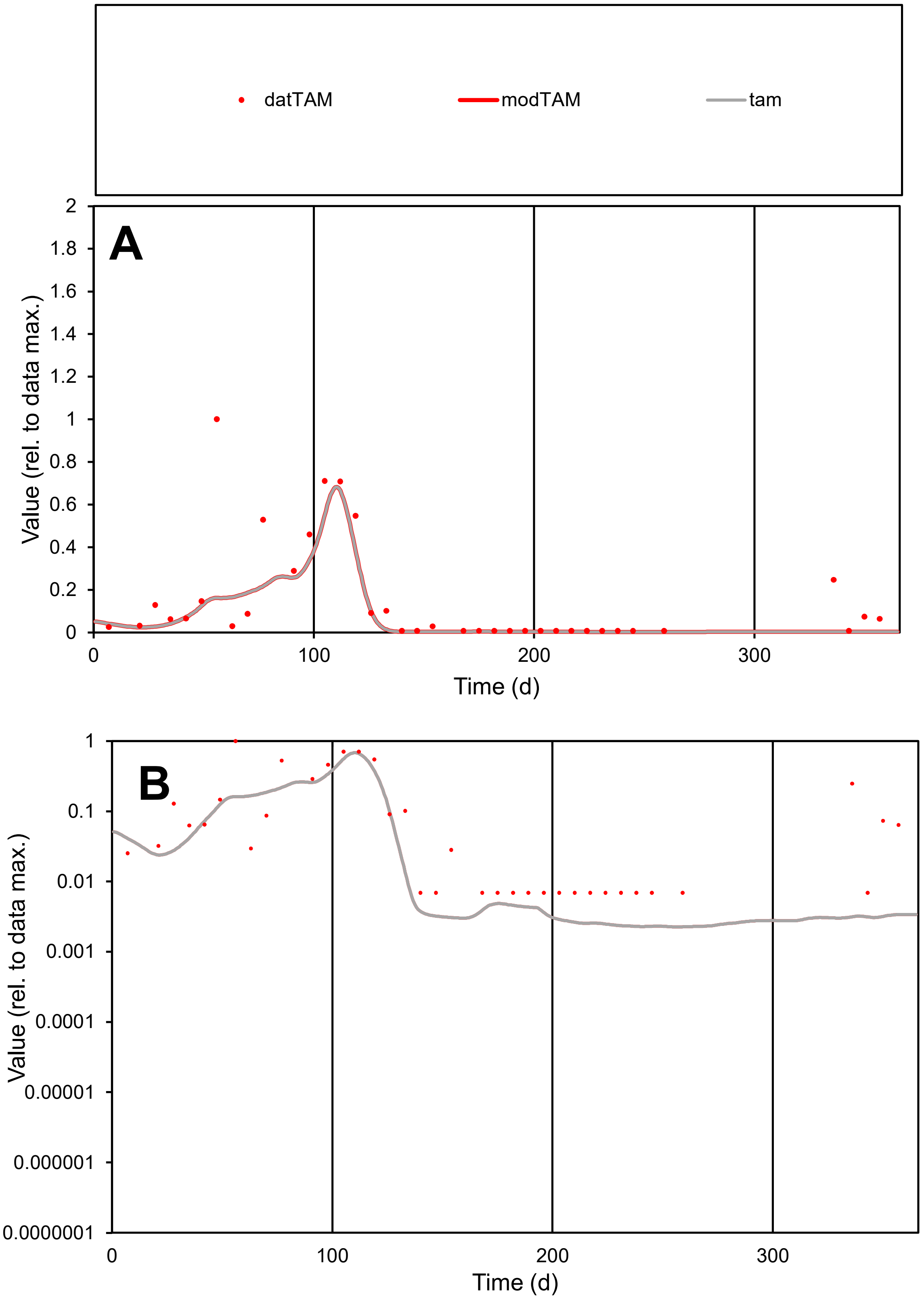


**C1**

Variables
TAM Provide jtd or Dst
CalibOnly?
2 1 to reproduce of
oNormalize?
2 1=no, 2=yes

ofxn calc*
1.08213
ofxn mod
1.09837

sRun
RunA1
zout
1

Ave
Data
0.00012
Model
7.9E-05

data/vMax
0.00083

ofxng
311.734

Min d/m
3.04226

ave/min d
21.6657
ave/min m
41.5068

331_Calib_S_A

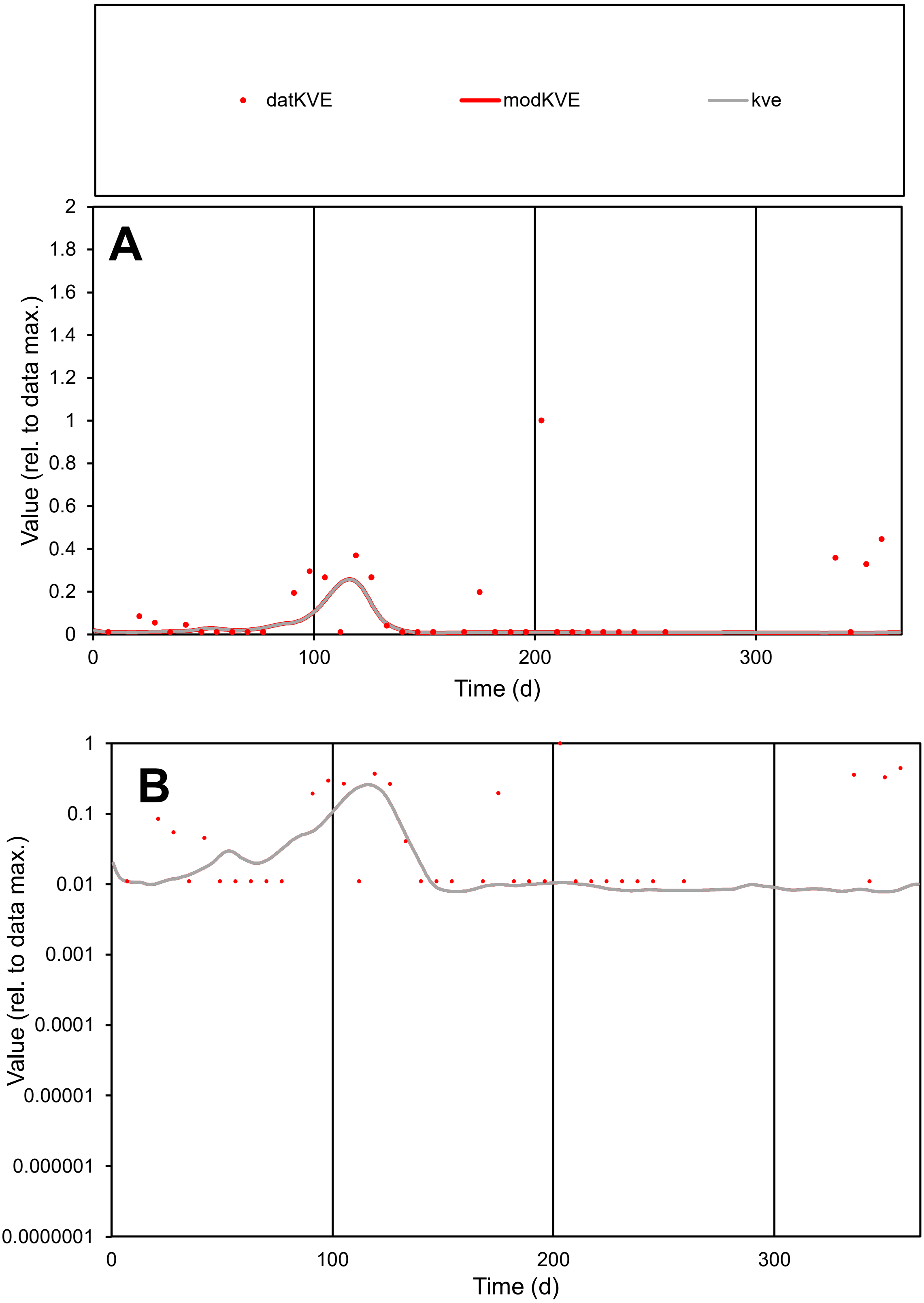


**C1**

Variables
KVE Provide jtd or Dst
CalibOnly?
2 1 to reproduce of
oNormalize?
2 1=no, 2=yes

ofxn calc*
1.82603
ofxn mod
1.83133

sRun
RunA1
zout
1

Ave
Data
3.2E-05
Model
1.1E-05

data/vMax
0.00029

ofxng
311.734

Min d/m
1.39528

ave/min d
10.3372
ave/min m
4.69994

331_Calib_S_A

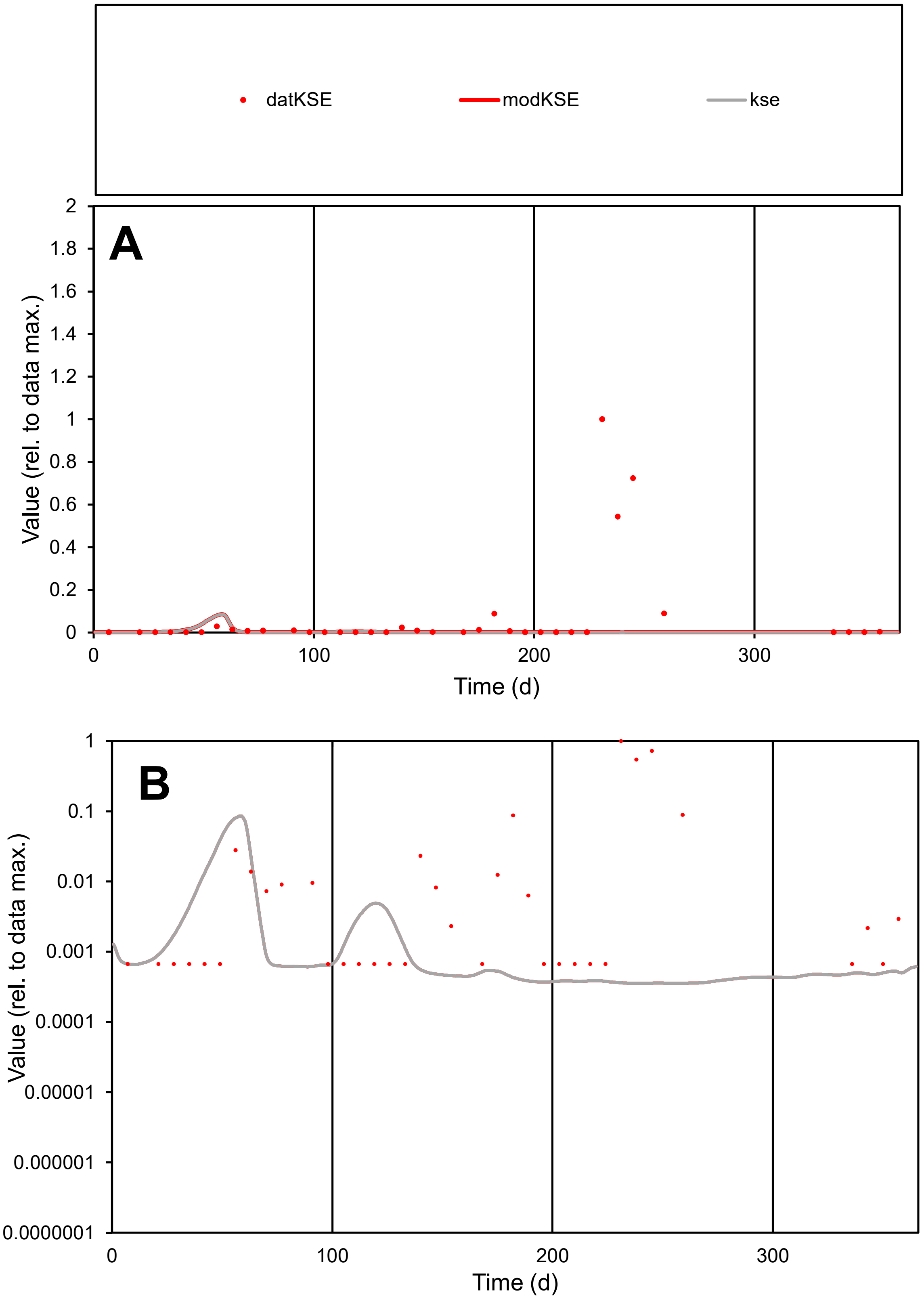


**C1**

Variables

| | |
|---|---|
| KSE | Provide jtd or Dst |

CalibOnly?

| | |
|---|---|
| 2 | 1 to reproduce of |

oNormalize?

| | |
|---|---|
| 2 | 1=no, 2=yes |

ofxn calc*
3.19259
ofxn mod
3.20087

sRun
RunA1
zout
1

Ave
Data
0.00039
Model
2.5E-05

data/vMax
0.00559

ofxng
311.734

Min d/m
1.89965

ave/min d
104.061
ave/min m
12.5844

331_Calib_S_A

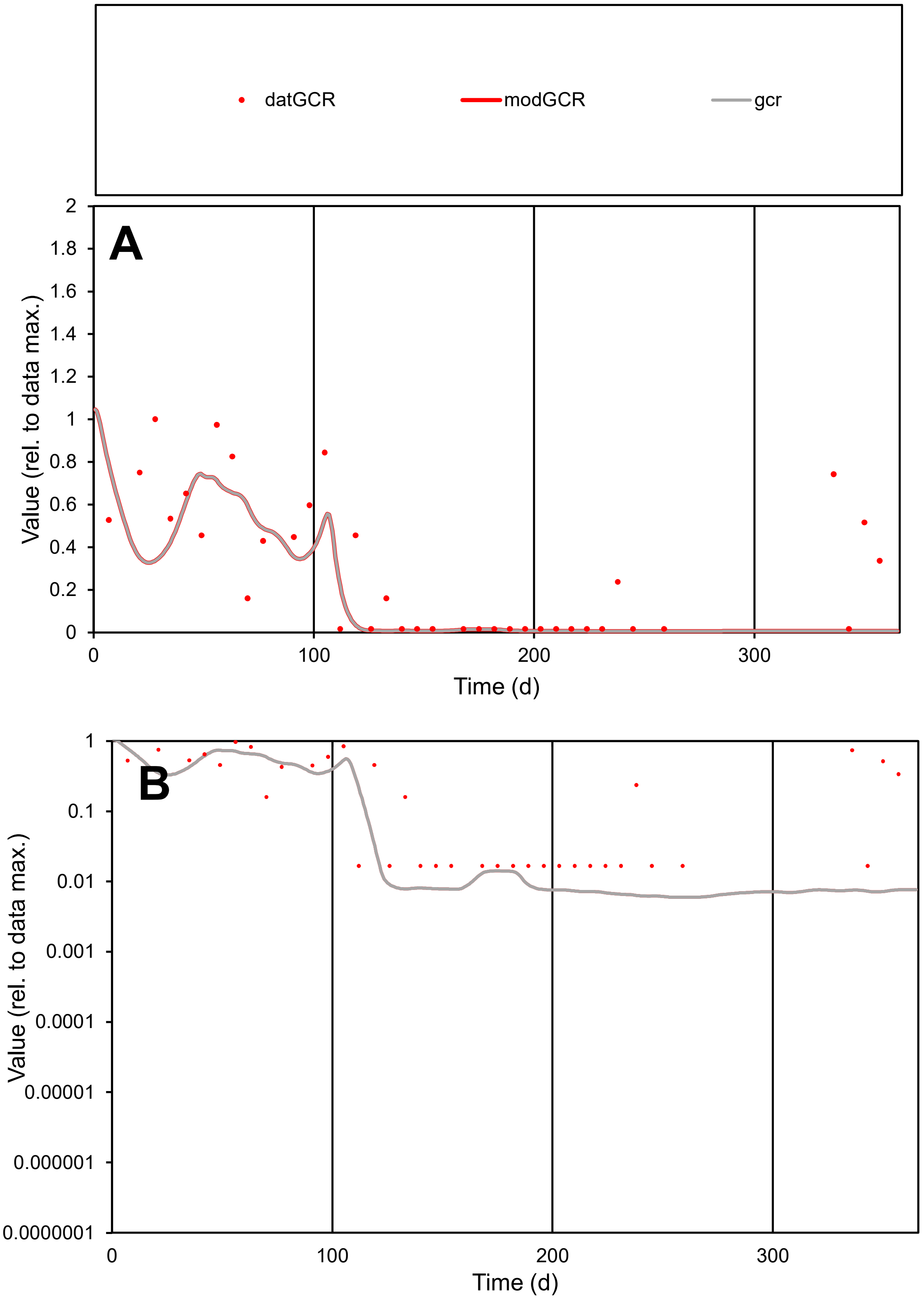


**C1**

Variables
GCR Provide jtd or Dst
CalibOnly?
2 1 to reproduce of
oNormalize?
2 1=no, 2=yes

ofxn calc*
0.8414
ofxn mod
0.85759

sRun
RunA1
zout
1

Ave
Data
5.6E-05
Model
3.7E-05

data/vMax
0.00019

ofxng
311.734

Min d/m
2.78578
ave/min d
17.7323
ave/min m
32.8892

331_Calib_S_A

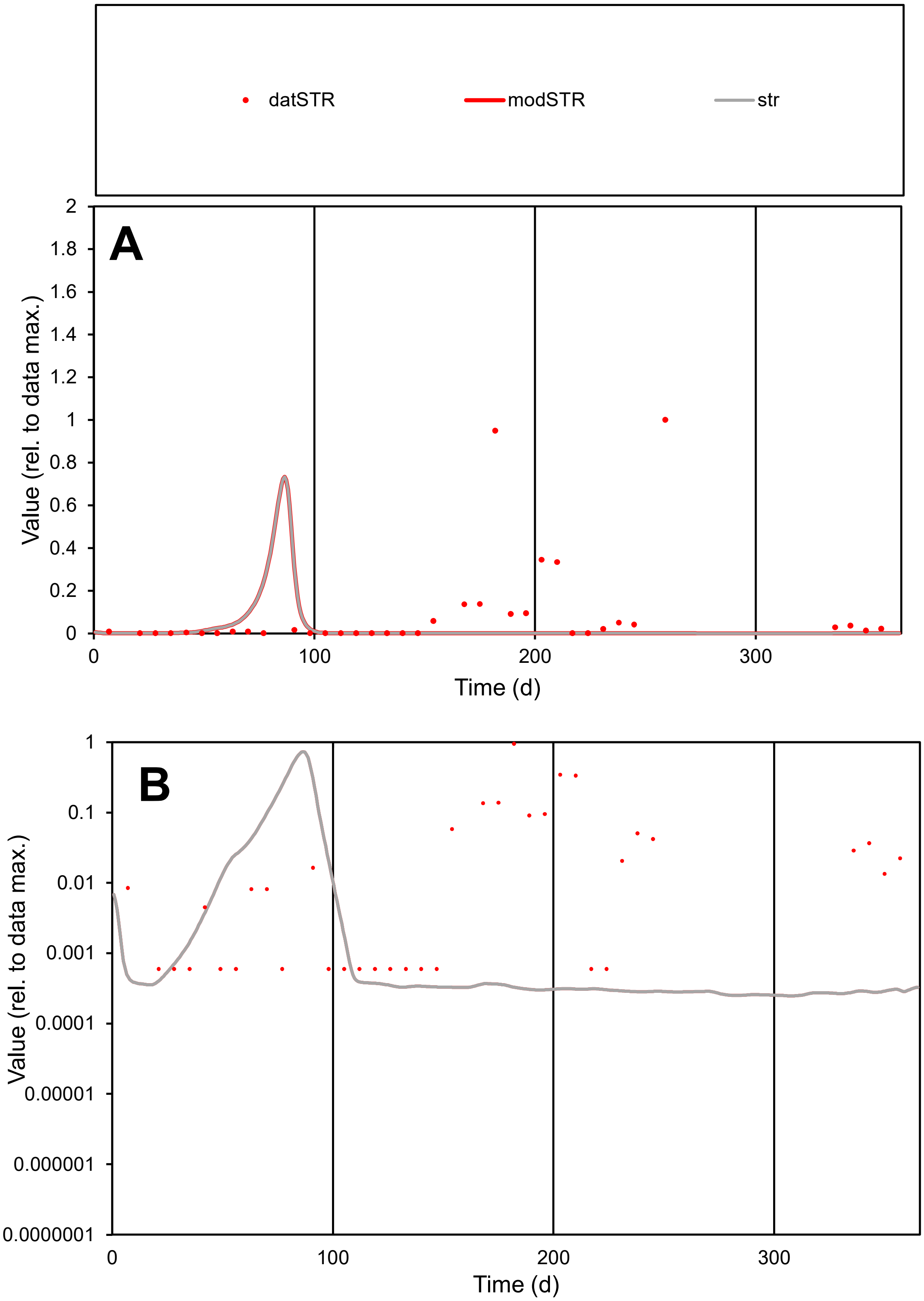


**C1**

Variables
STR Provide jtd or Dst
CalibOnly?
2 1 to reproduce of
oNormalize?
2 1=no, 2=yes

ofxn calc*
2.66084
ofxn mod
2.68302

sRun
RunA1
zout
1

Ave
Data
0.00025
Model
3E-05

data/vMax
0.00273

ofxng
311.734

Min d/m
2.12054

ave/min d
154.411
ave/min m
39.2669

331_Calib_S_A

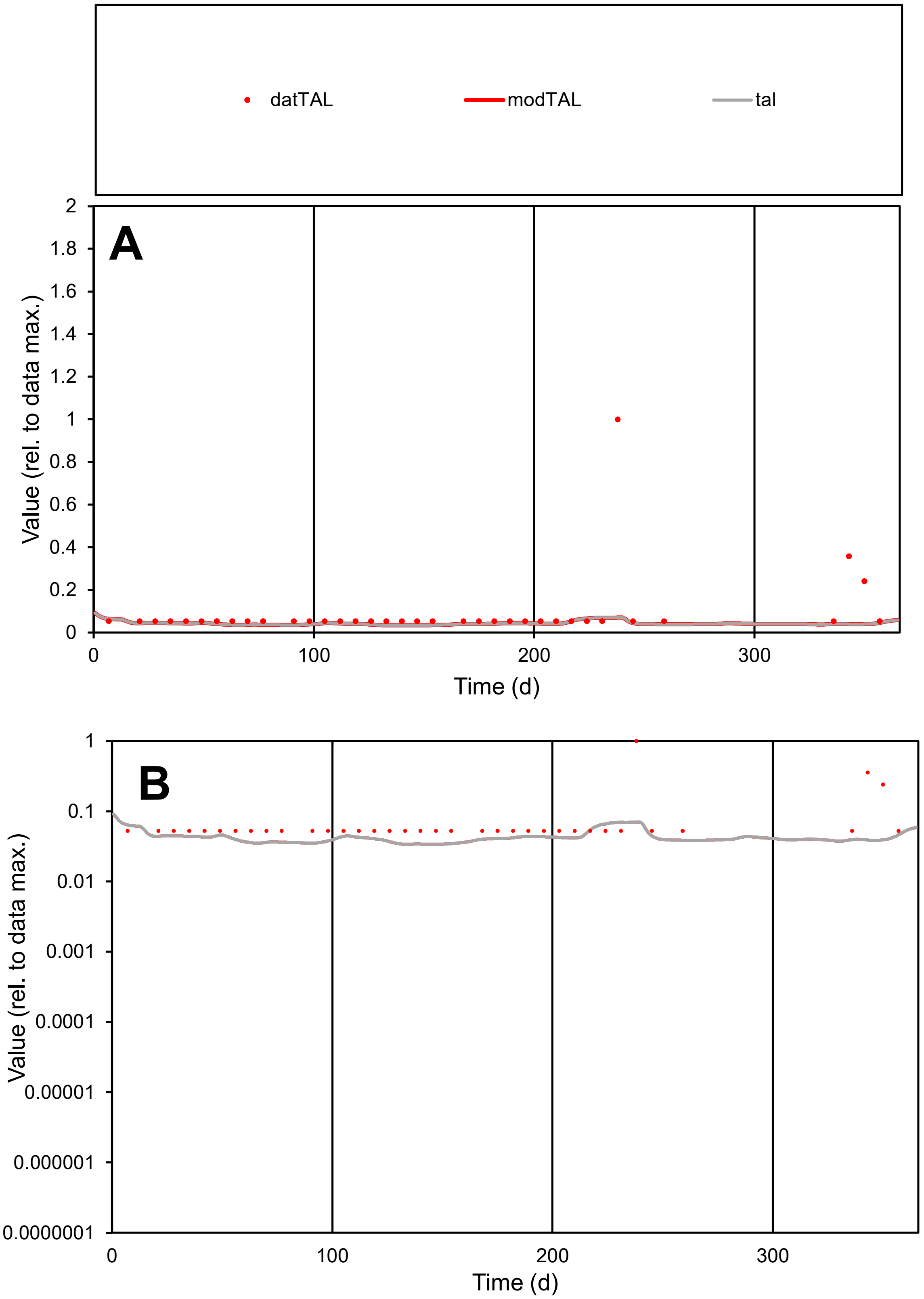


C1

Variables
TAL Provide jtd or Dst
CalibOnly?
2 1 to reproduce of
oNormalize?
2 1=no, 2=yes

ofxn calc*
1.80467
ofxn mod
1.80847

sRun
RunA1
zout
1

Ave
Data
7.2E-06
Model
3.4E-06

data/vMax
7.9E-05

ofxng
311.734

Min d/m
1.55148

ave/min d
1.73749
ave/min m
1.28281

331_Calib_S_A

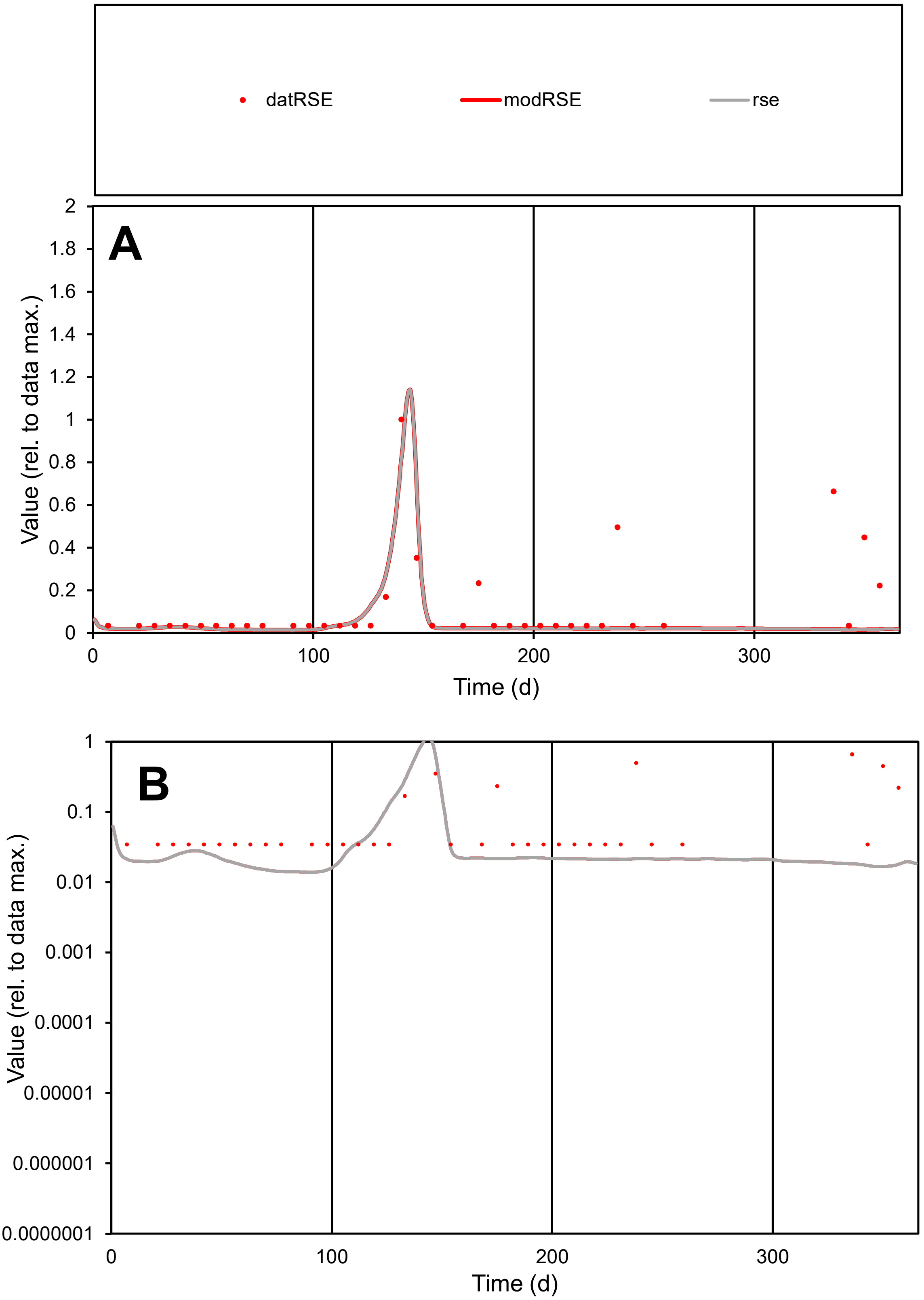


**C1**

Variables
RSE Provide jtd or Dst
CalibOnly?
2 1 to reproduce of
oNormalize?
2 1=no, 2=yes

ofxn calc*
1.28668
ofxn mod
1.29683

sRun
RunA1
zout
1

Ave
Data
1.7E-05
Model
8.9E-06

data/vMax
0.00014

ofxng
311.734

Min d/m
2.48959

ave/min d
3.59087
ave/min m
4.77811

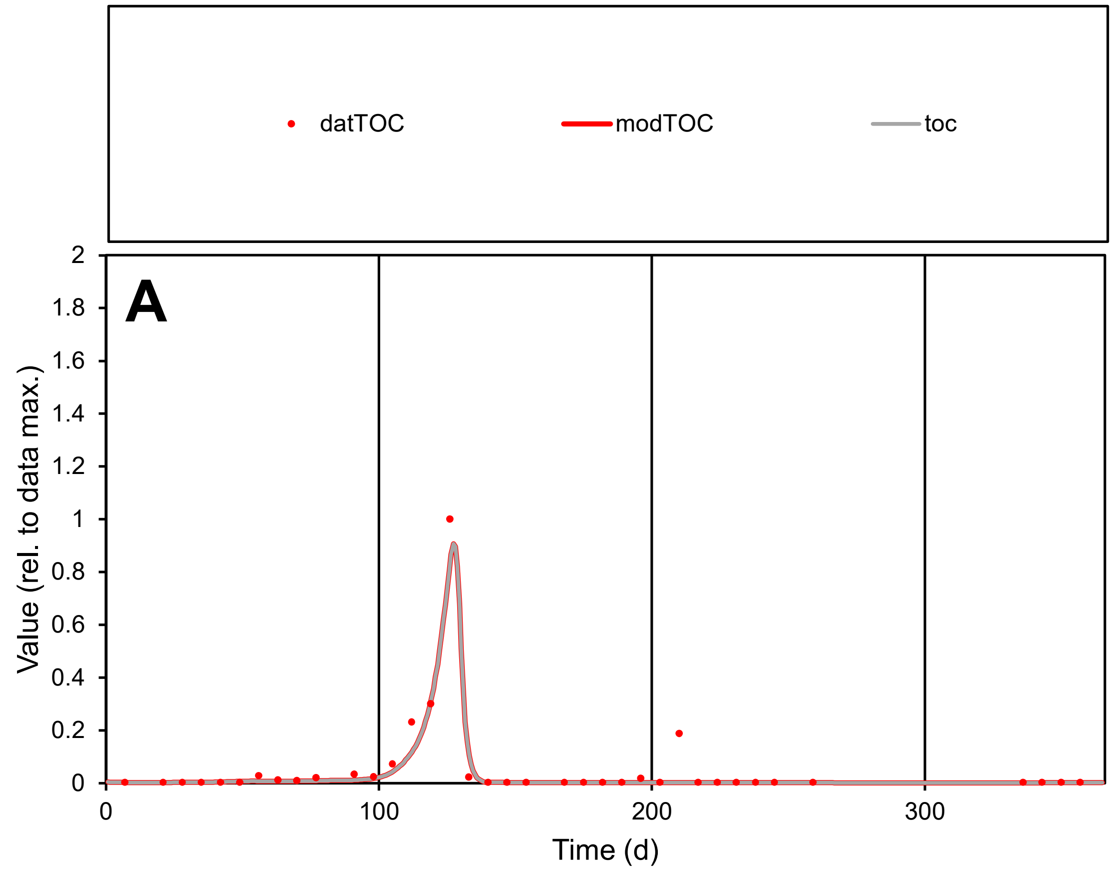


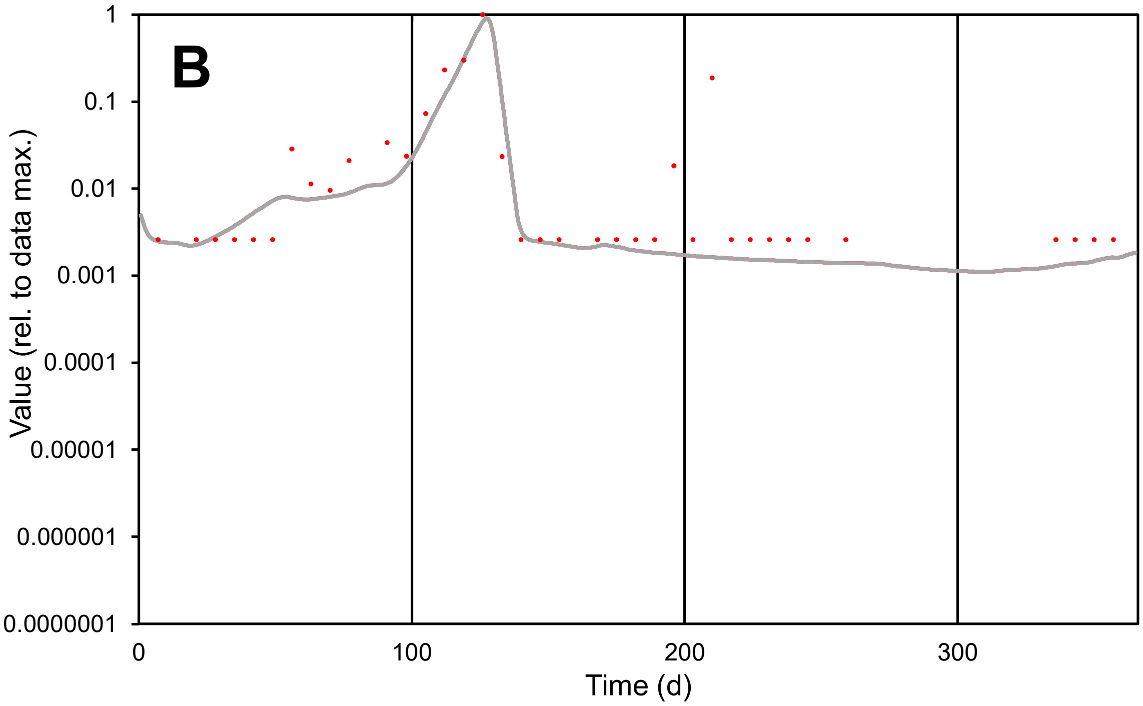


**C1**

Variables
TOC Provide jtd or Dst
CalibOnly?
2 1 to reproduce of
oNormalize?
2 1=no, 2=yes

ofxn calc*
0.71896
ofxn mod
0.77695

sRun
RunA1
zout
1

Ave
Data
9.1E-05
Model
7.3E-05

data/vMax
0.00166

ofxng
311.734

Min d/m
1.96544

ave/min d
21.0813
ave/min m
33.4184

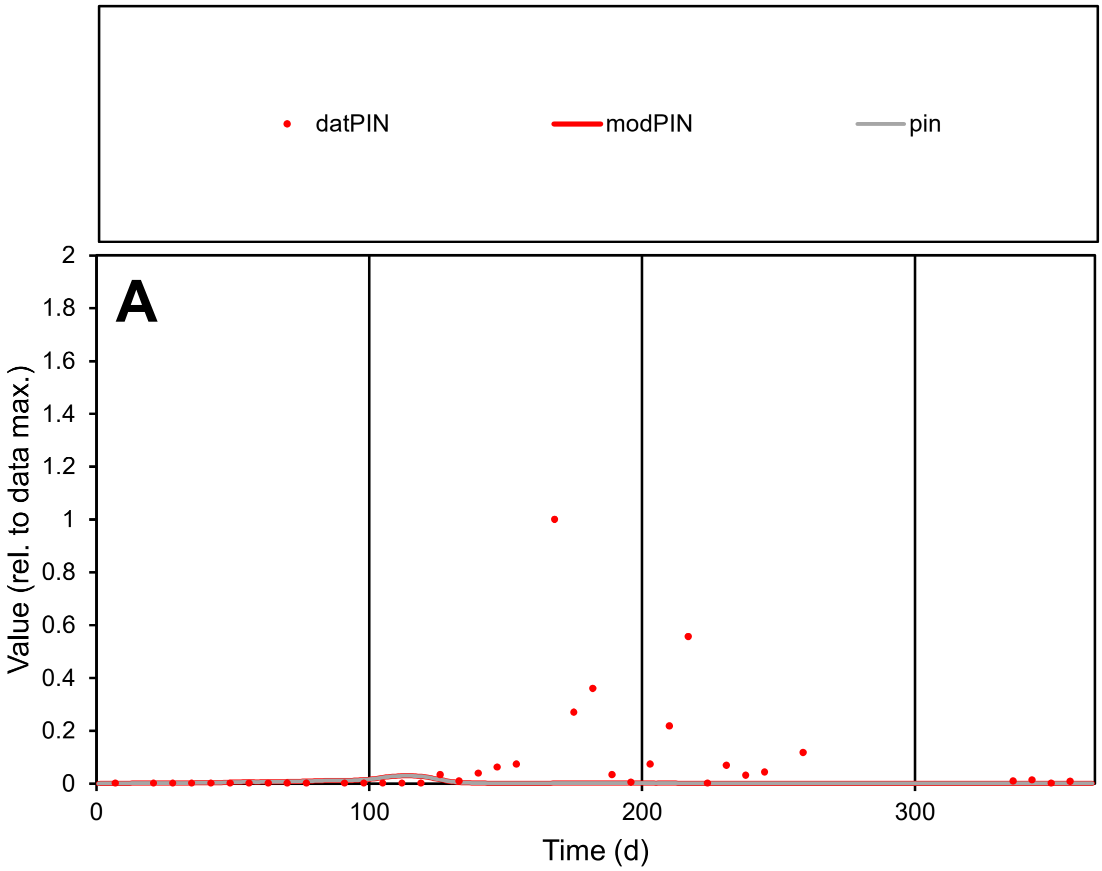


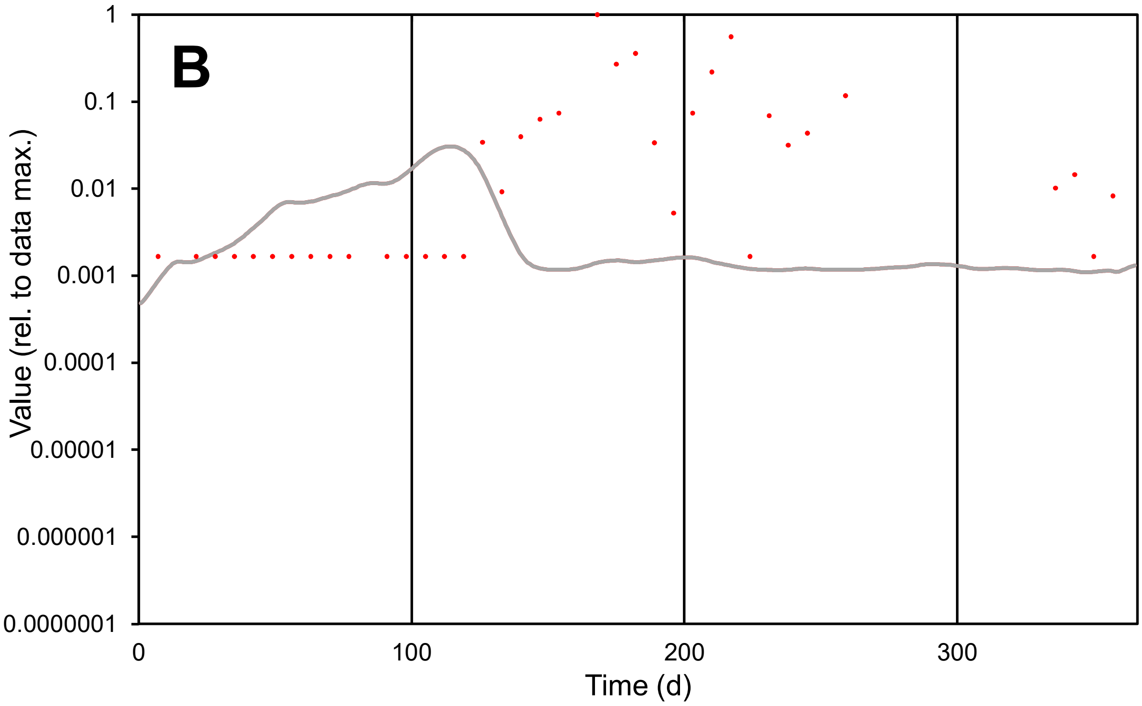


**C1**

Variables

PIN Provide jtd or Dst

CalibOnly?

2 1 to reproduce of

oNormalize?

2 1=no, 2=yes

ofxn calc*

2.50911

ofxn mod

2.51217

sRun

RunA1

zout

1

Ave

Data

0.0002

Model

1.3E-05

data/vMax

0.00241

ofxng

311.734

Min d/m

1.74576

ave/min d

49.6997

ave/min m

5.53046

331_Calib_S_A

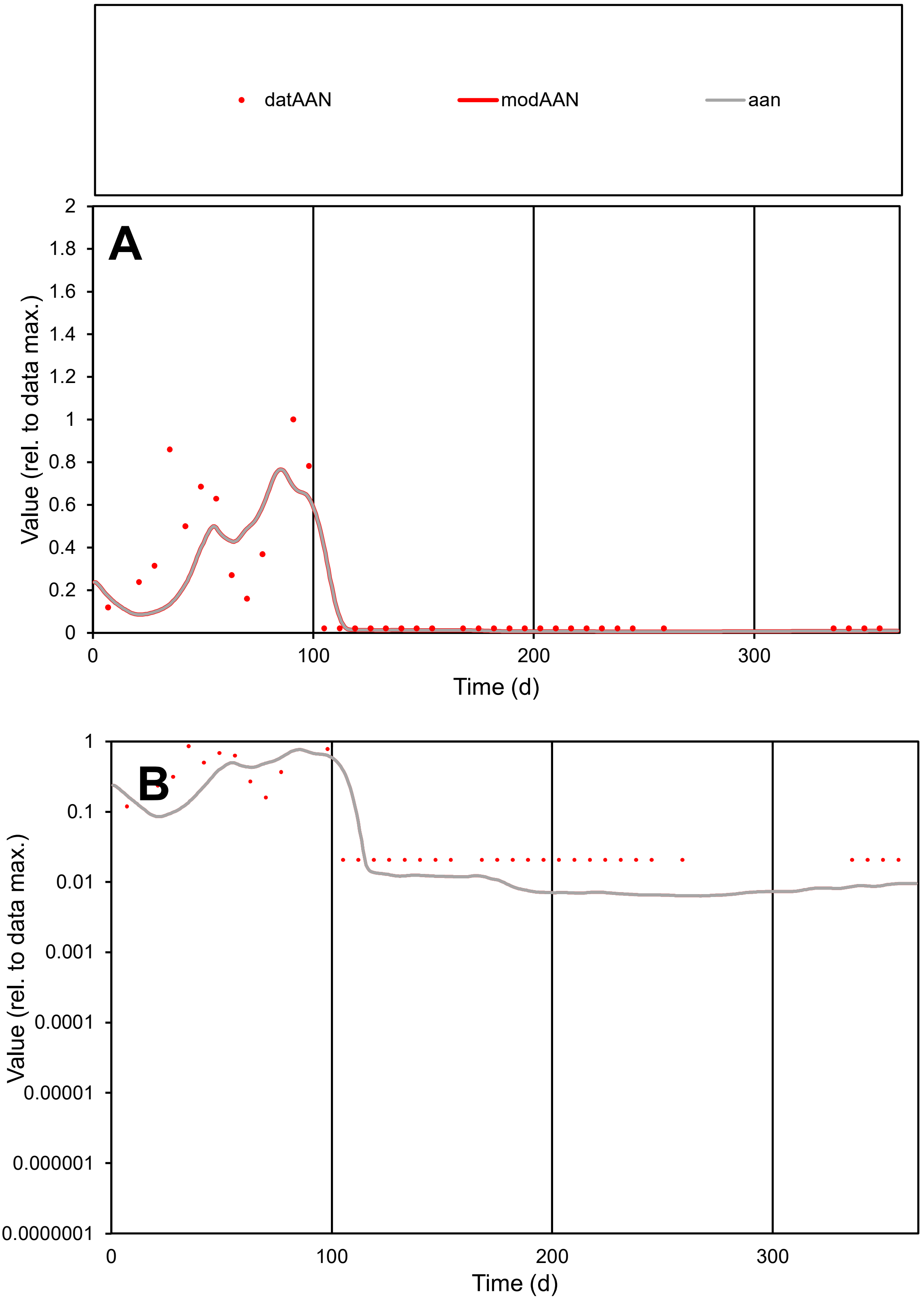


**C1**

Variables
AAN Provide jtd or Dst
CalibOnly?
2 1 to reproduce of
oNormalize?
2 1=no, 2=yes

ofxn calc*
0.98936
ofxn mod
1.02252

sRun
RunA1
zout
1

Ave
Data
4.7E-05
Model
3.6E-05

data/vMax
0.00027

ofxng
311.734

Min d/m
3.23362

ave/min d
8.4217
ave/min m
20.7524

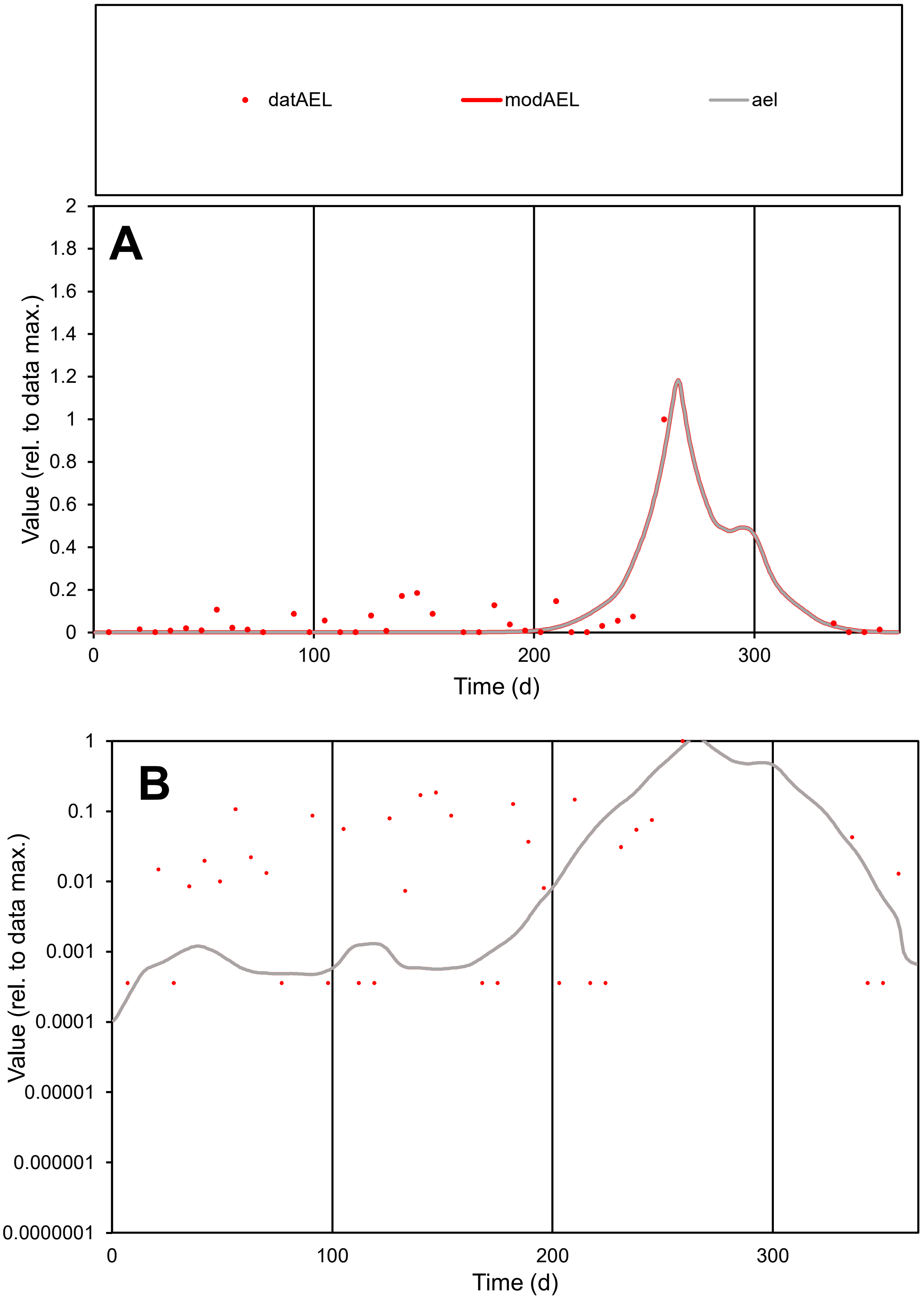


**C1**

Variables
AEL Provide jtd or Dst
CalibOnly?
2 1 to reproduce of
oNormalize?
2 1=no, 2=yes

ofxn calc*
1.20989
ofxn mod
1.28609

sRun
RunA1
zout
1

Ave
Data
0.00013
Model
9.3E-05

data/vMax
0.00195

ofxng
311.734

Min d/m
1.43115

ave/min d
181.702
ave/min m
191.546

331_Calib_S_A

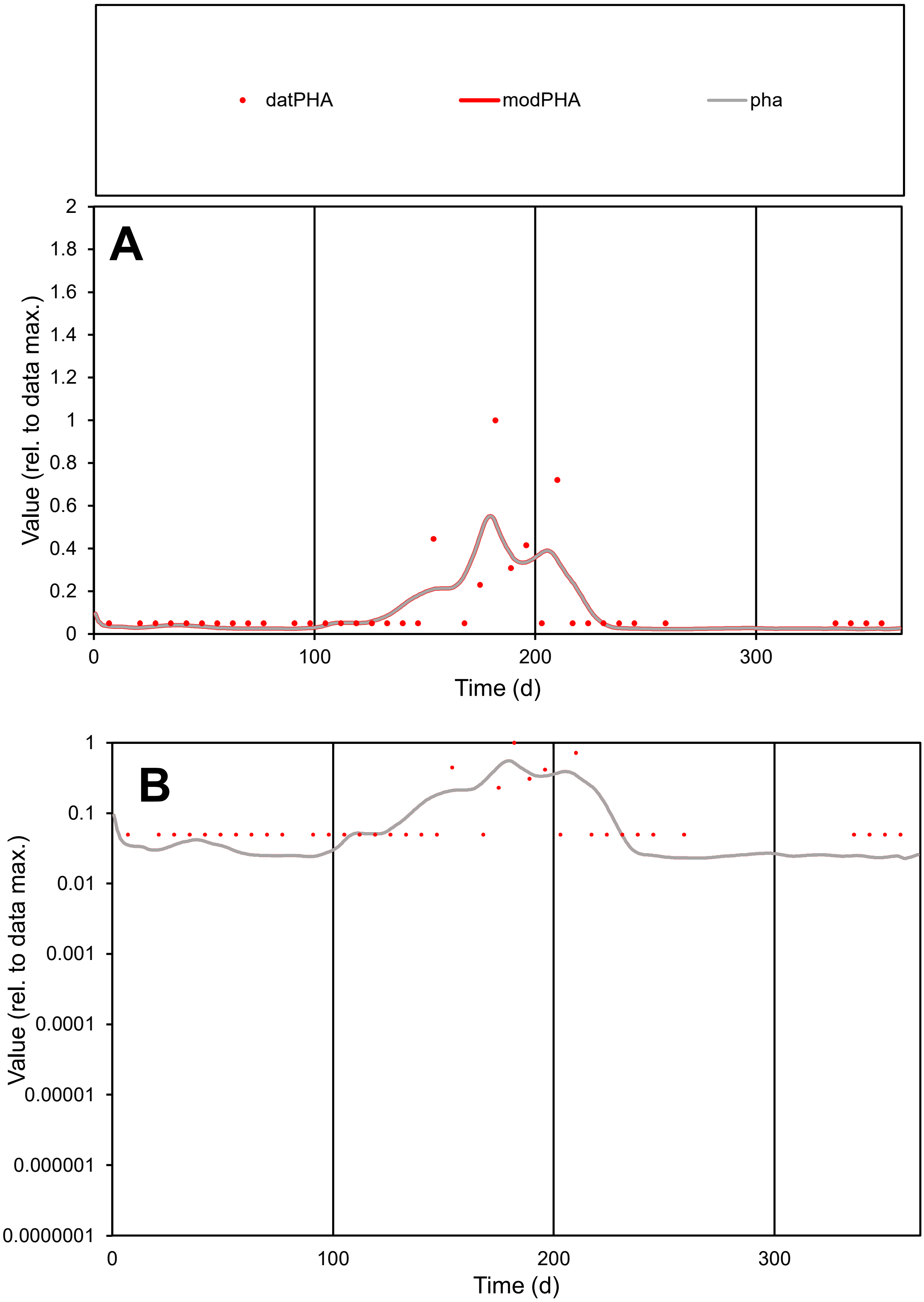


**C1**

Variables
PHA Provide jtd or Dst
CalibOnly?
2 1 to reproduce of
oNormalize?
2 1=no, 2=yes

ofxn calc*
1.11121
ofxn mod
1.15124

sRun
RunA1
zout
1

Ave
Data
2.6E-05
Model
2.5E-05

data/vMax
0.00021

ofxng
311.734

Min d/m
2.14109

ave/min d
2.54168
ave/min m
5.1175

331_Calib_S_A

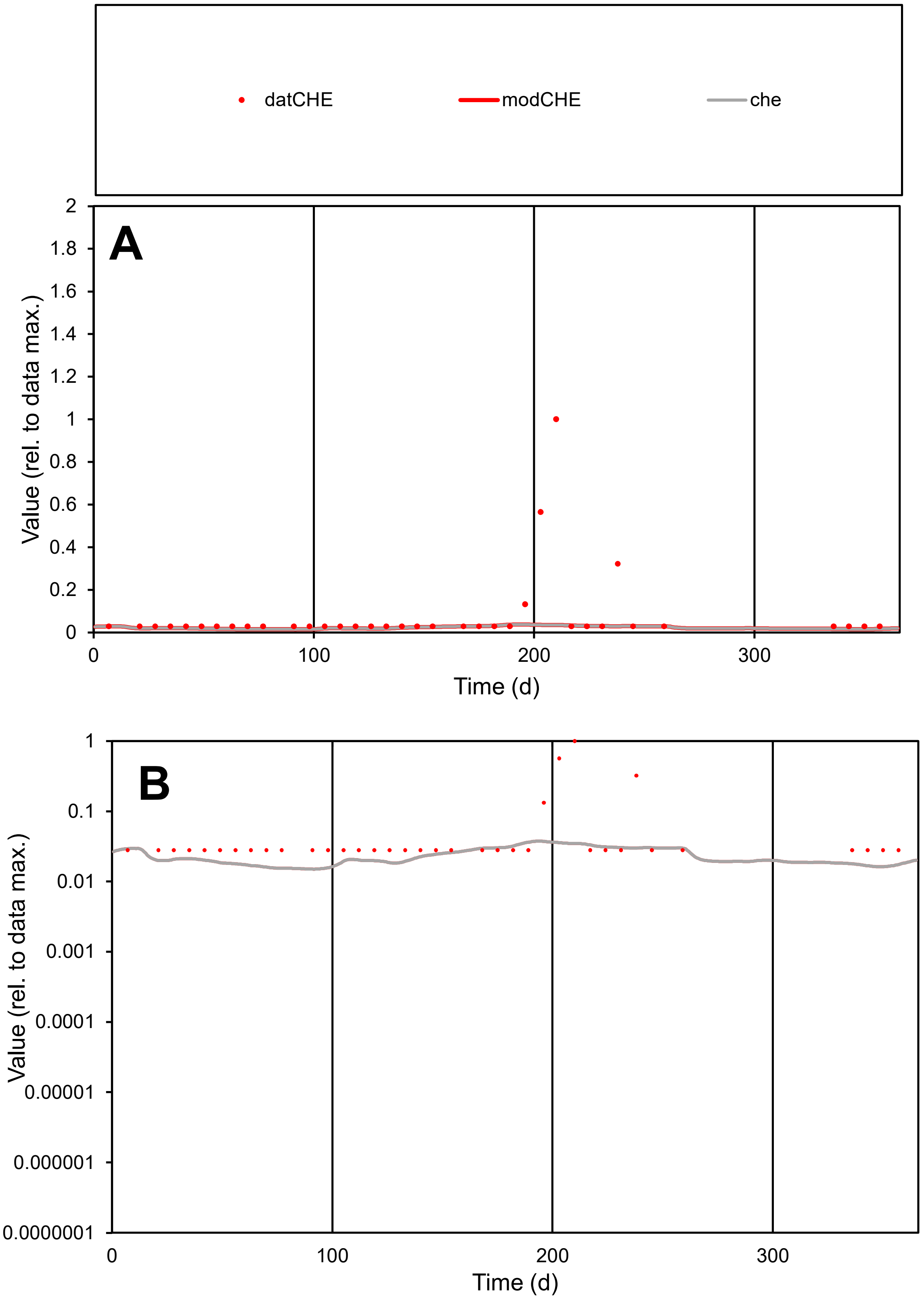


**C1**

Variables

CHE Provide jtd or Dst

CalibOnly?

2 1 to reproduce of

oNormalize?

2 1=no, 2=yes

ofxn calc*

2.36616

ofxn mod

2.37322

sRun

RunA1

zout

1

Ave

Data

2E-05

Model

6E-06

data/vMax

0.00025

ofxng

311.734

Min d/m

1.86256

ave/min d

2.84692

ave/min m

1.59936

331_Calib_S_A

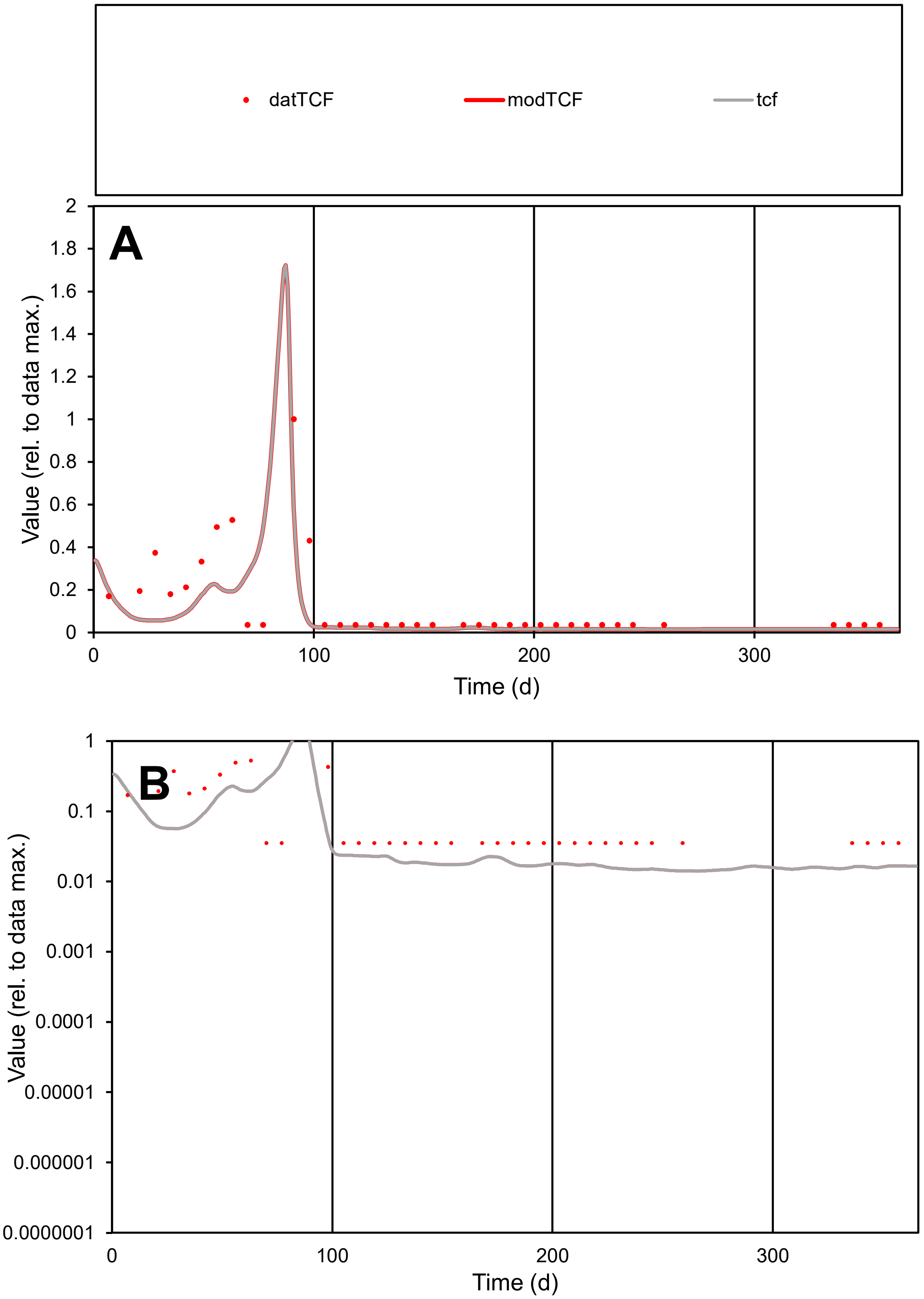


**C1**

Variables
TCF Provide jtd or Dst
CalibOnly?
2 1 to reproduce of
oNormalize?
2 1=no, 2=yes

ofxn calc*
1.0772
ofxn mod
1.21483

sRun
RunA1
zout
1

Ave
Data
1.2E-05
Model
7.3E-06

data/vMax
8.8E-05

ofxng
311.734

Min d/m
2.47933

ave/min d
3.72454
ave/min m
5.78726

331_Calib_S_A

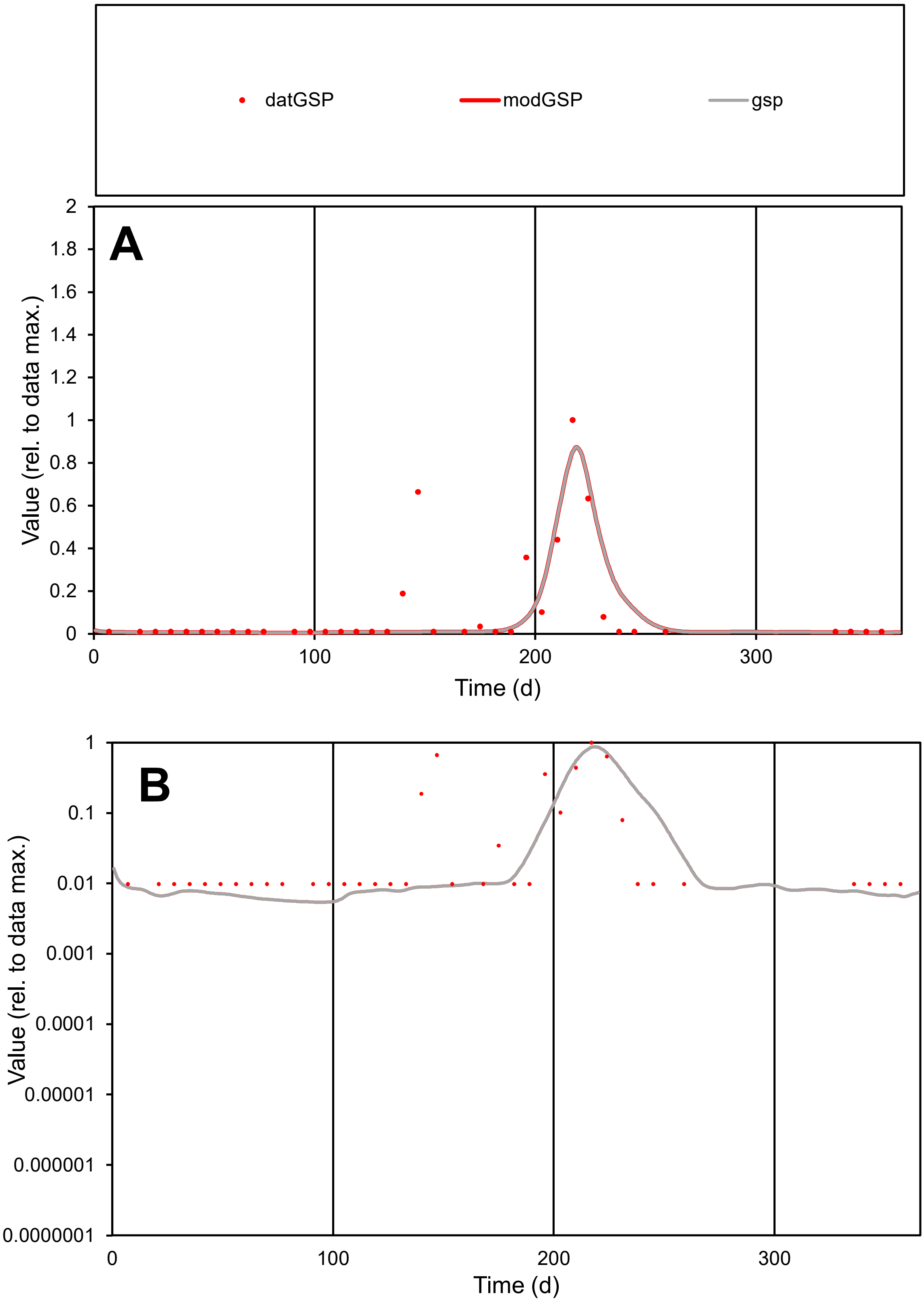


**C1**

Variables

GSP Provide jtd or Dst

CalibOnly?

2 1 to reproduce of

oNormalize?

2 1=no, 2=yes

ofxn calc*

1.2919

ofxn mod

1.3514

sRun

RunA1

zout

1

Ave

Data

9.4E-05

Model

7.9E-05

data/vMax

0.00092

ofxng

311.734

Min d/m

1.8099

ave/min d

10.4277

ave/min m

15.849

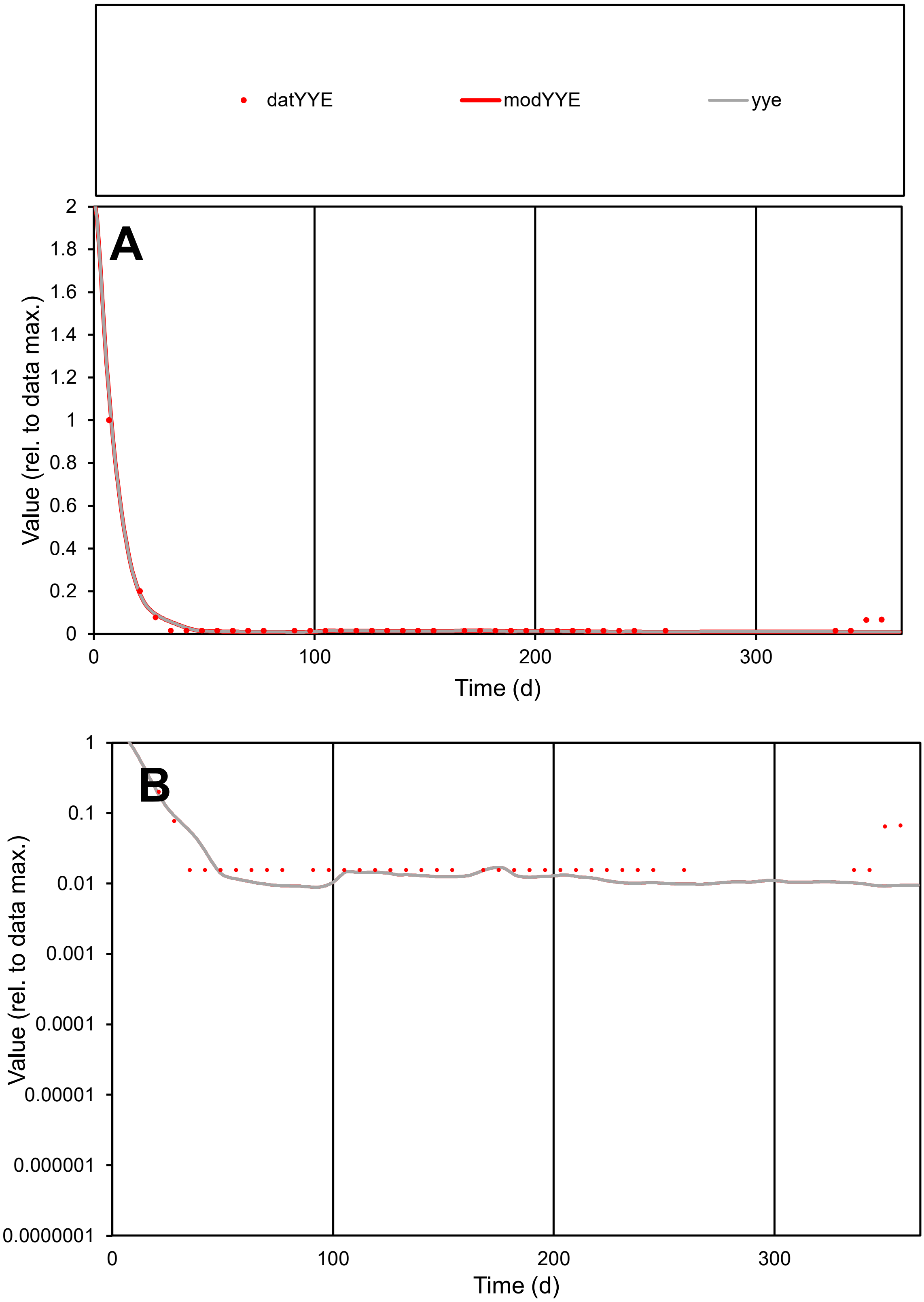


**C1**

Variables
YYE Provide jtd or Dst
CalibOnly?
2 1 to reproduce of
oNormalize?
2 1=no, 2=yes

ofxn calc*
0.2924
ofxn mod
0.36869

sRun
RunA1
zout
1

Ave
Data
1.2E-05
Model
1.1E-05

data/vMax
0.00024

ofxng
311.734

Min d/m
1.76501

ave/min d
3.30534
ave/min m
5.22855

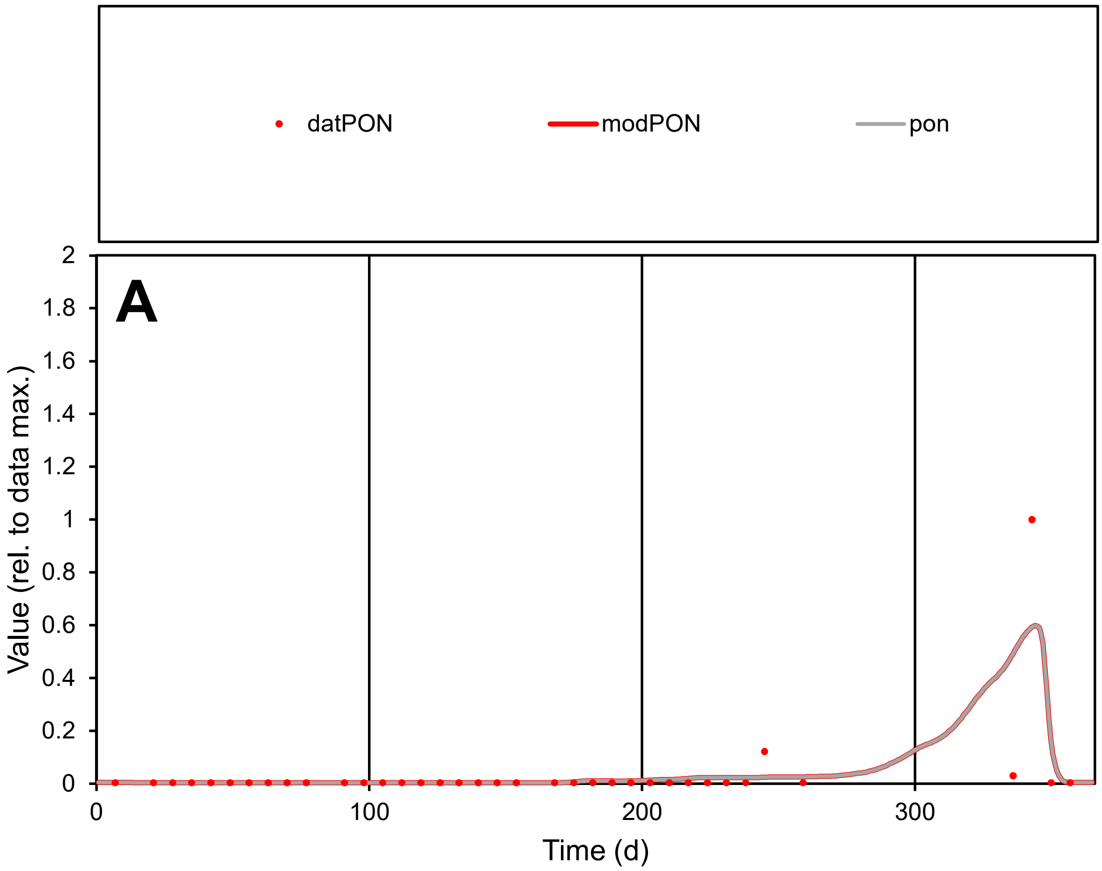


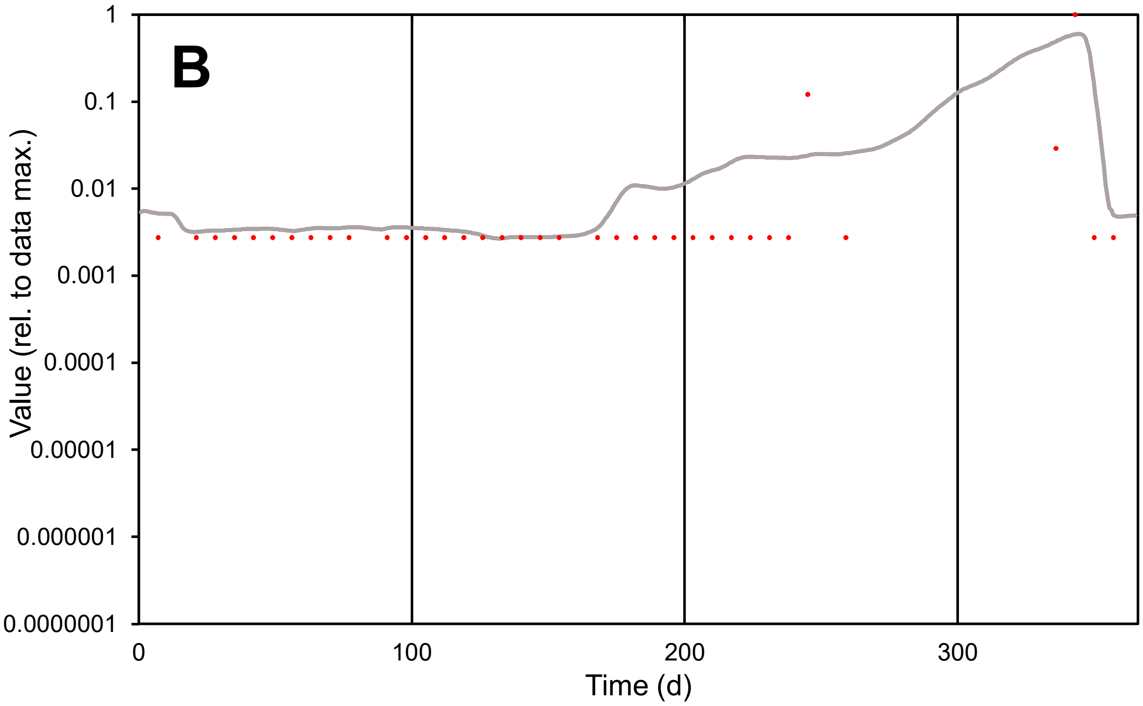


**C1**

Variables
PON Provide jtd or Dst
CalibOnly?
2 1 to reproduce of
oNormalize?
2 1=no, 2=yes

ofxn calc*
3.04595
ofxn mod
3.07578

sRun
RunA1
zout
1

Ave
Data
3.6E-05
Model
4E-05

data/vMax
0.00107

ofxng
311.734

Min d/m
1.00894

ave/min d
12.2562
ave/min m
13.8227

331_Calib_S_A

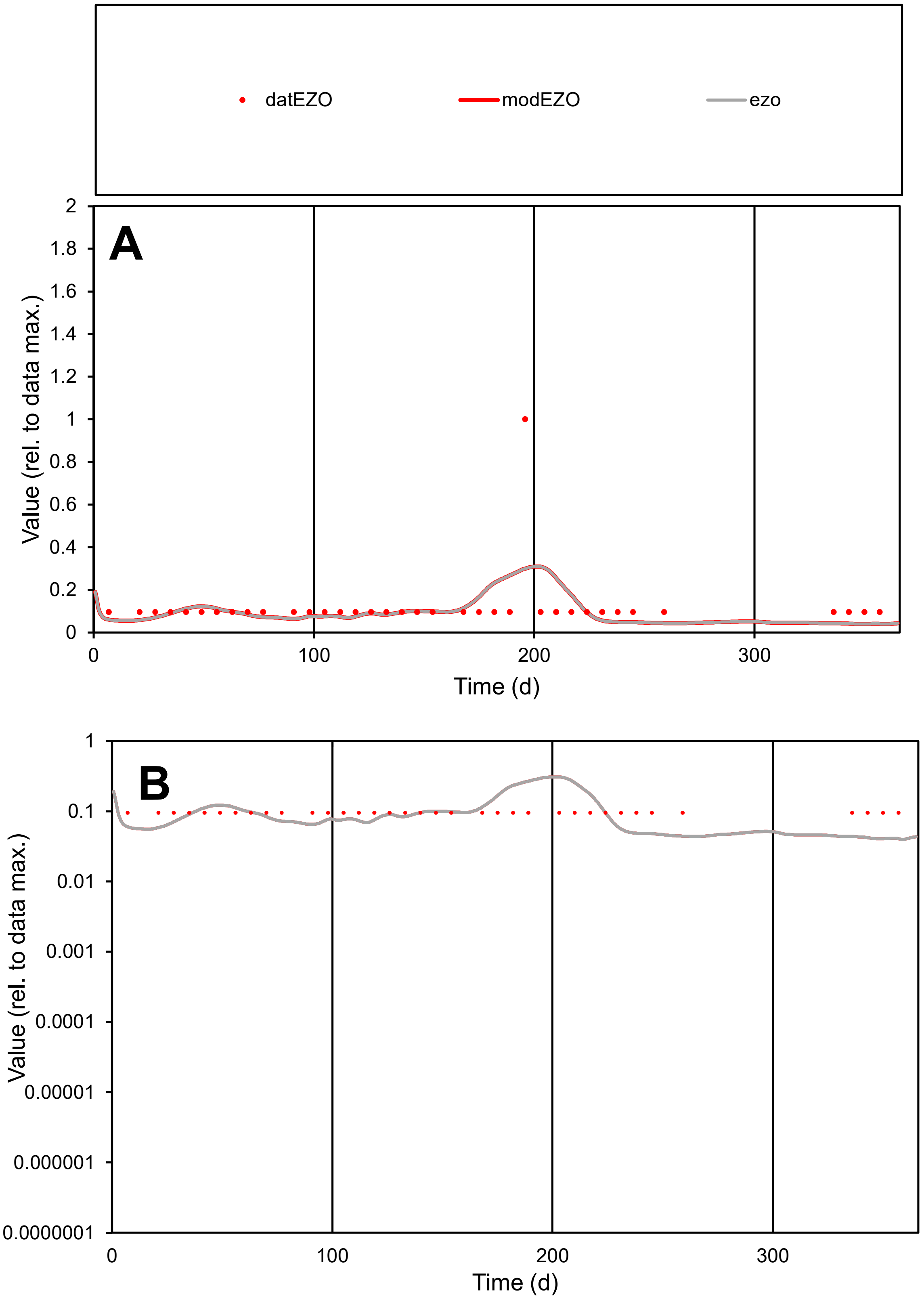


**C1**

Variables
EZO Provide jtd or Dst
CalibOnly?
2 1 to reproduce of
oNormalize?
2 1=no, 2=yes

ofxn calc*
1.09433
ofxn mod
1.10322

sRun
RunA1
zout
1

Ave
Data
5E-06
Model
4.4E-06

data/vMax
4.1E-05

ofxng
311.734

Min d/m
2.40134

ave/min d
1.25606
ave/min m
2.70214

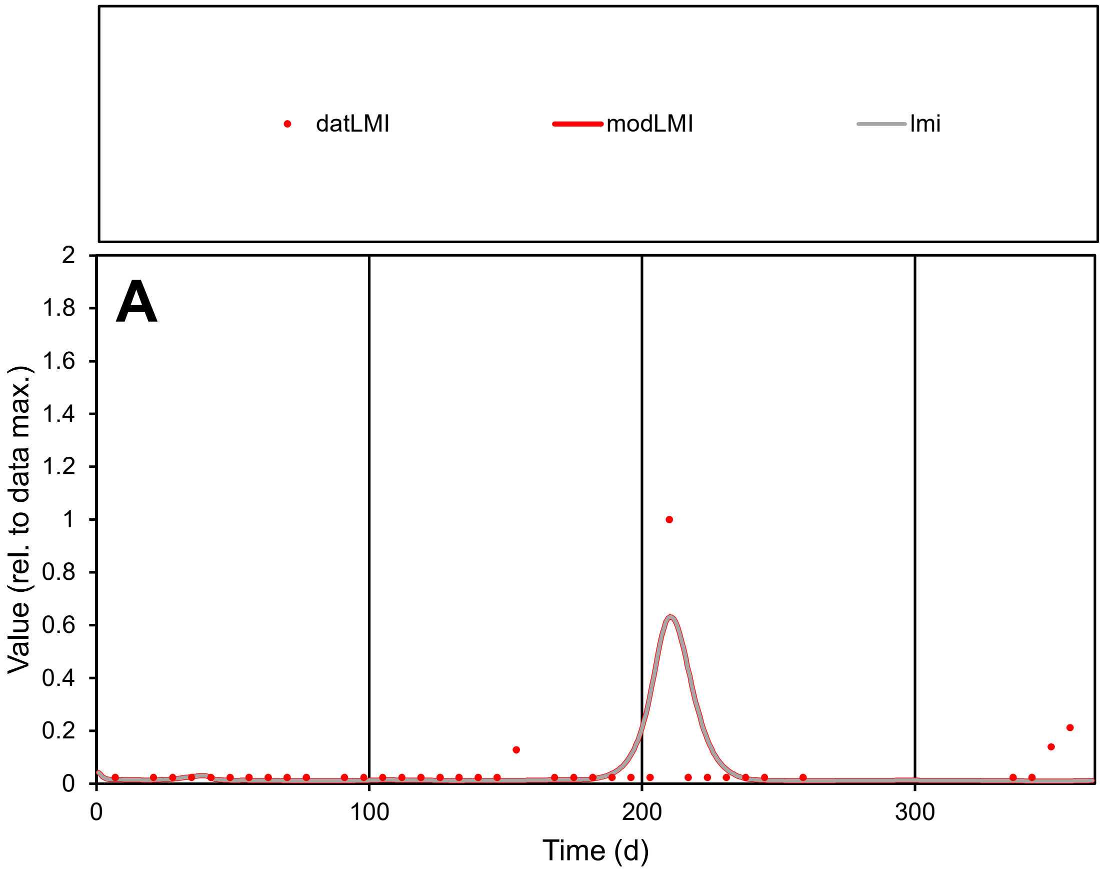


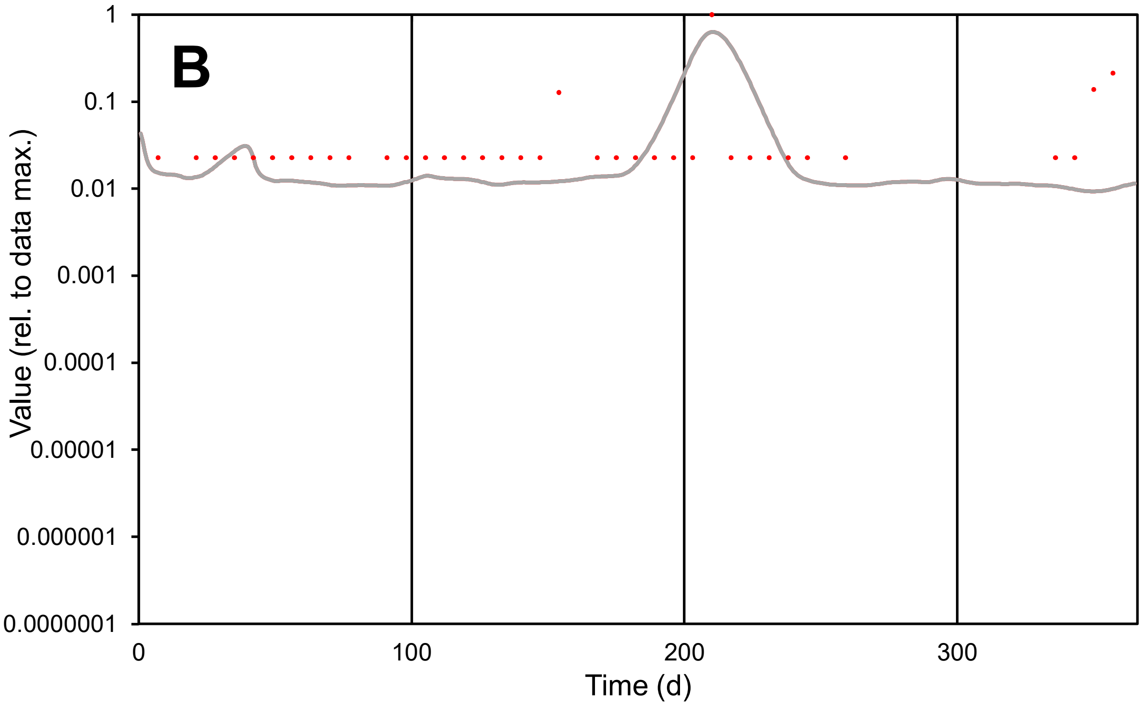


**C1**

Variables
LMI Provide jtd or Dst
CalibOnly?
2 1 to reproduce of
oNormalize?
2 1=no, 2=yes

ofxn calc*
1.61849
ofxn mod
1.76893

sRun
RunA1
zout
1

Ave
Data
8E-06
Model
6.6E-06

data/vMax
0.00013

ofxng
311.734

Min d/m
2.42203

ave/min d
2.65417
ave/min m
5.29819

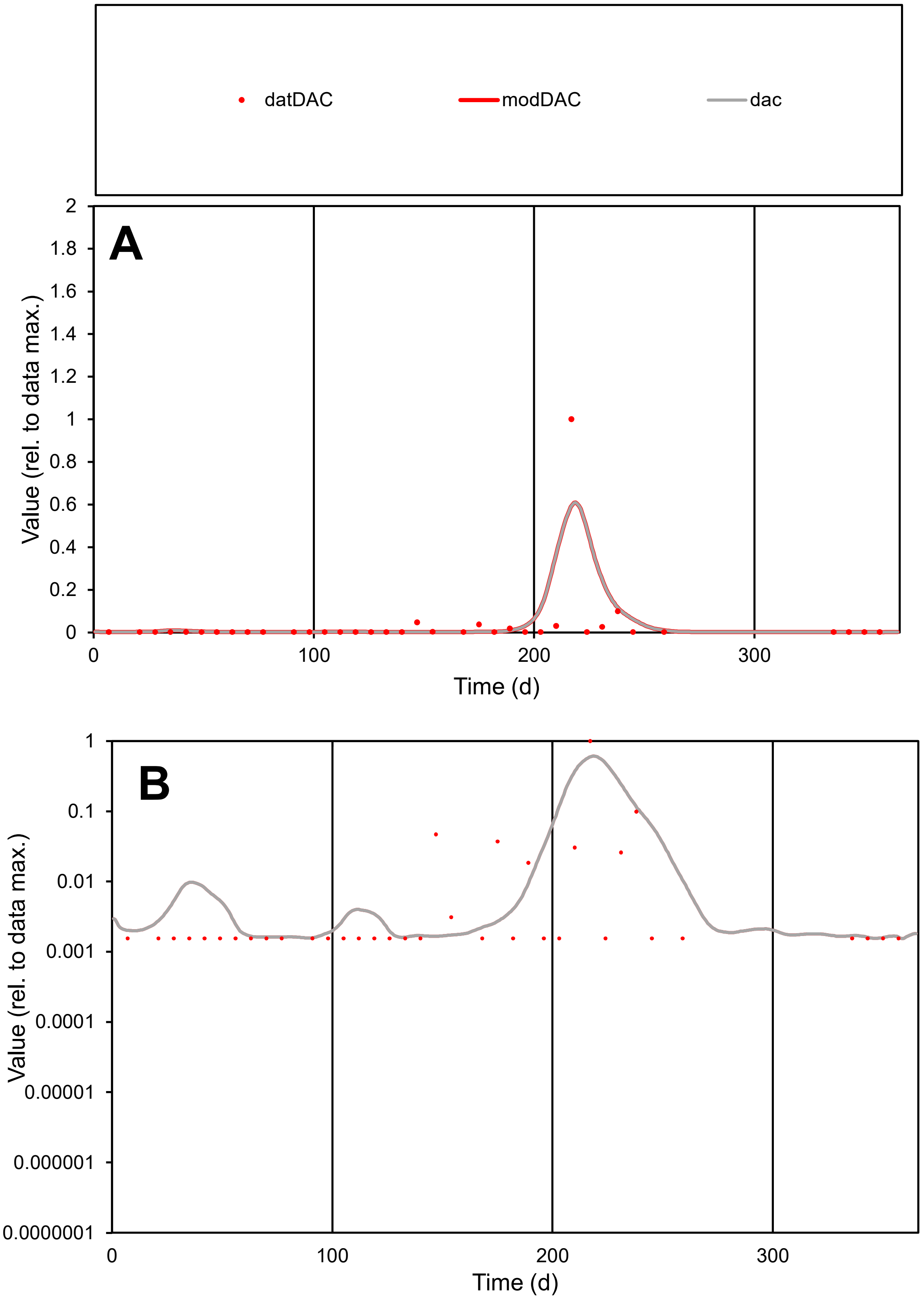


**C1**

Variables
DAC Provide jtd or Dst
CalibOnly?
2 1 to reproduce of
oNormalize?
2 1=no, 2=yes

ofxn calc*
2.80708
ofxn mod
3.14319

sRun
RunA1
zout
1

Ave
Data
0.00017
Model
0.00023

data/vMax
0.00496

ofxng
311.734

Min d/m
1.00903

ave/min d
22.6938
ave/min m
30.6719

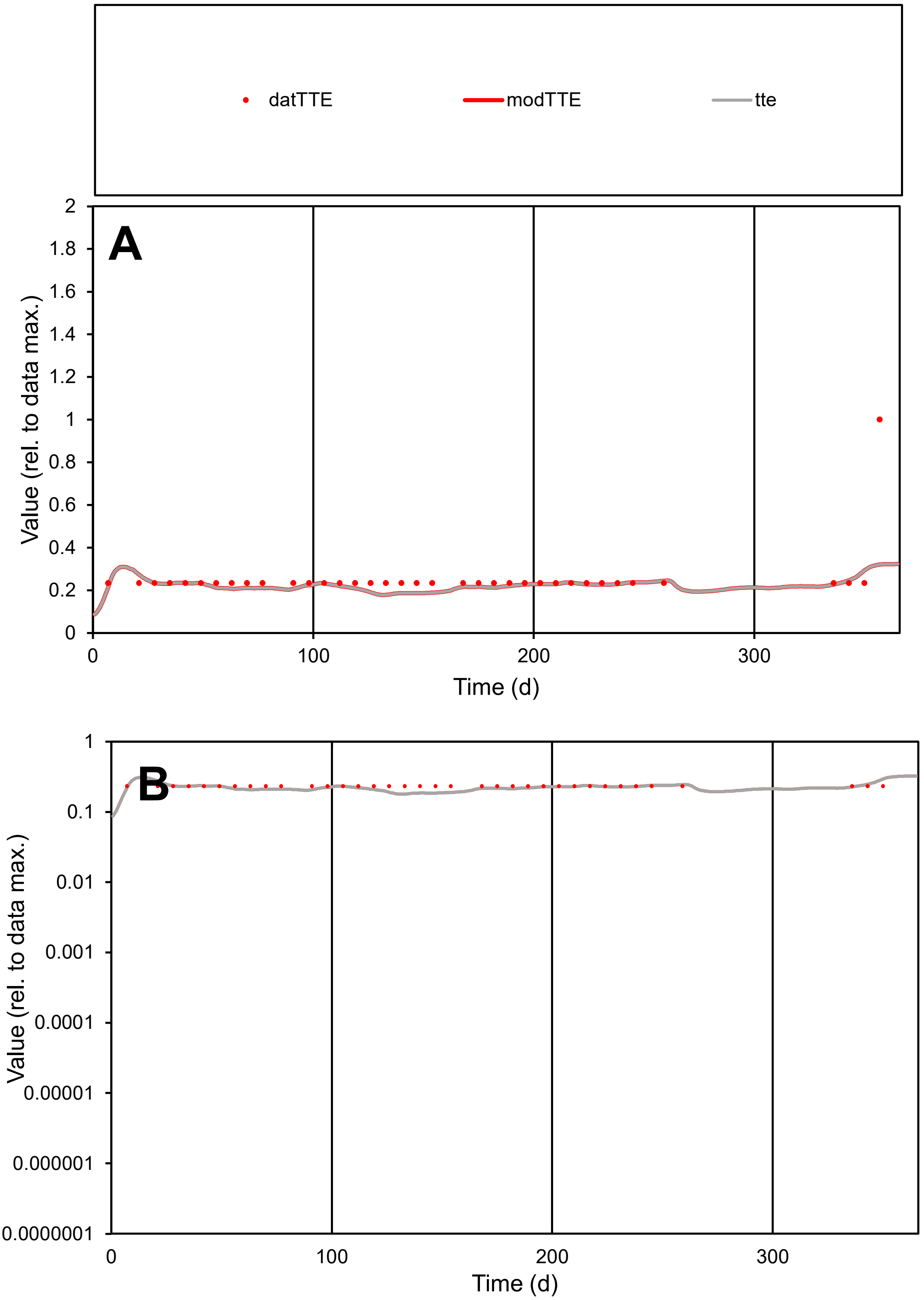


**C1**

Variables

TTE Provide jtd or Dst

CalibOnly?

2 1 to reproduce of

oNormalize?

2 1=no, 2=yes

ofxn calc*

0.4485

ofxn mod

0.46215

sRun

RunA1

zout

1

Ave

Data

4.1E-06

Model

3.6E-06

data/vMax

1.6E-05

ofxng

311.734

Min d/m

1.29511

ave/min d

1.08845

ave/min m

1.25101

331_Calib_S_A

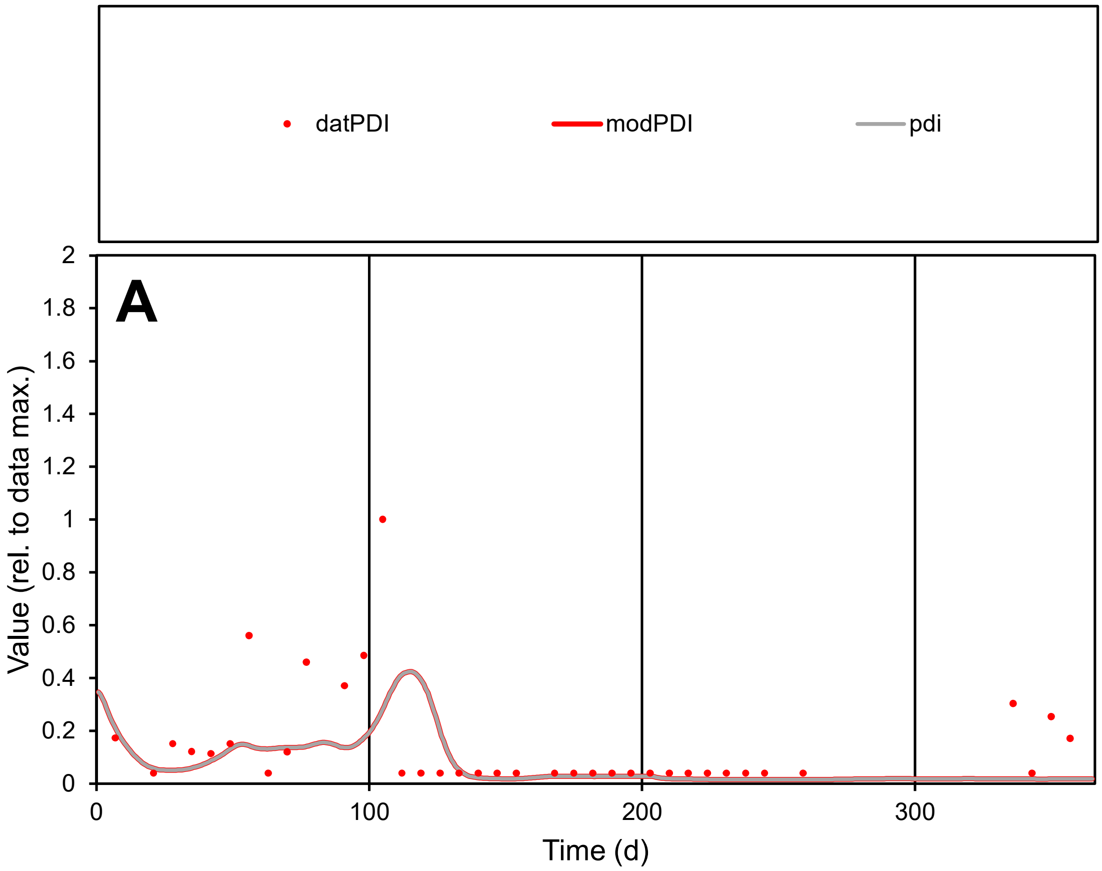


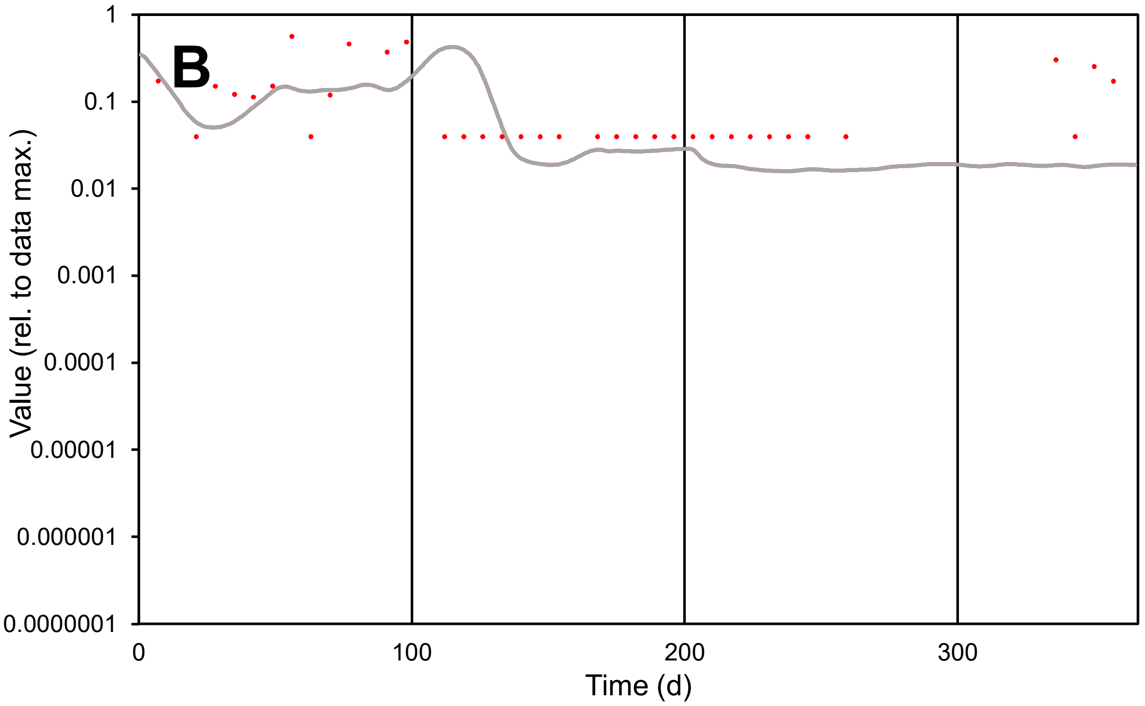


**C1**

Variables
PDI Provide jtd or Dst
CalibOnly?
2 1 to reproduce of
oNormalize?
2 1=no, 2=yes

ofxn calc*
1.30596
ofxn mod
1.31946

sRun
RunA1
zout
1

Ave
Data
1.6E-05
Model
9.3E-06

data/vMax
0.00011

ofxng
311.734

Min d/m
2.48924

ave/min d
3.63769
ave/min m
5.25073

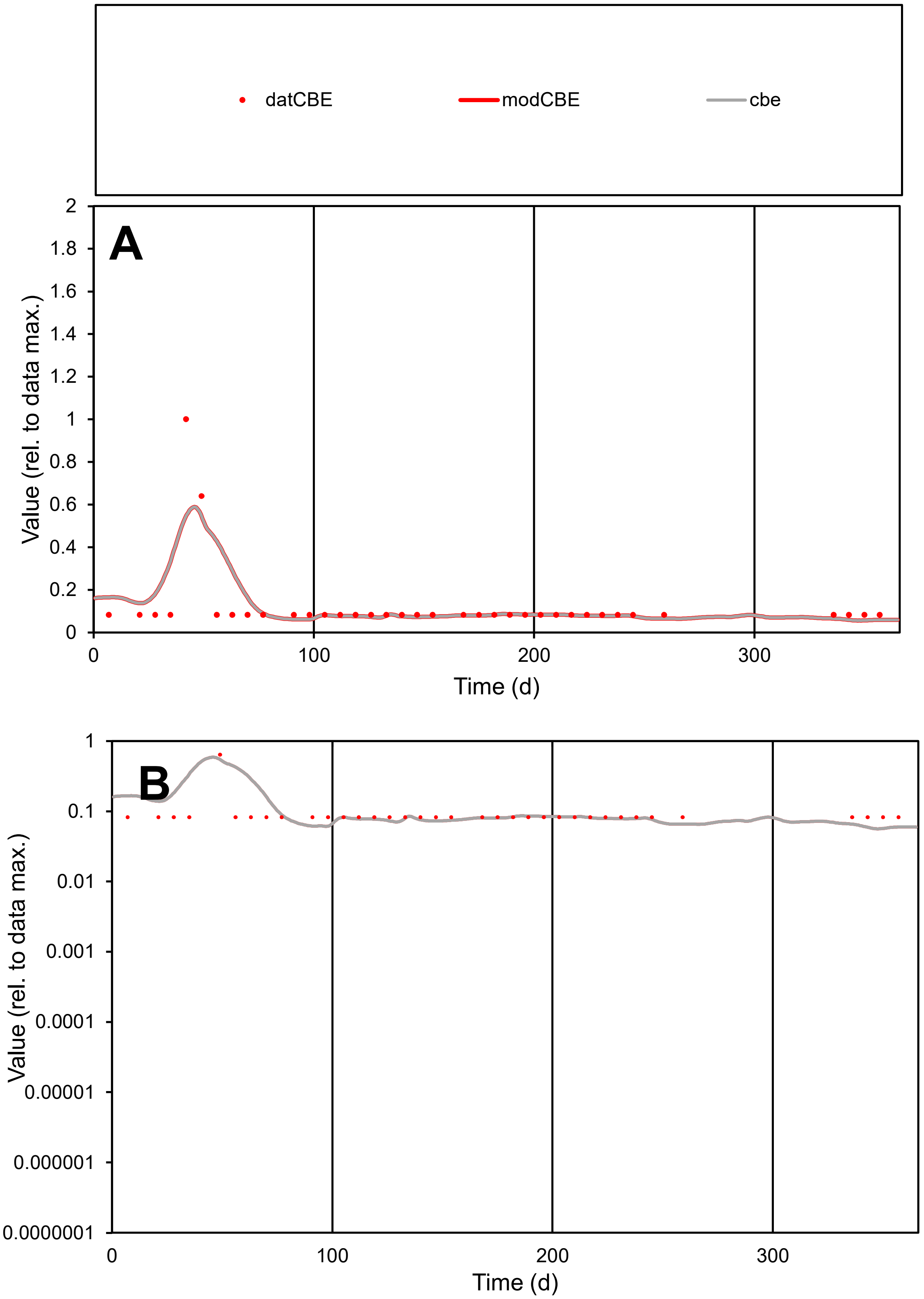


**C1**

Variables
CBE Provide jtd or Dst
CalibOnly?
2 1 to reproduce of
oNormalize?
2 1=no, 2=yes

ofxn calc*
0.86211
ofxn mod
0.8967

sRun
RunA1
zout
1

Ave
Data
2.5E-06
Model
2.6E-06

data/vMax
2E-05

ofxng
311.734

Min d/m
1.43053

ave/min d
1.48438
ave/min m
2.25251

331_Calib_S_A

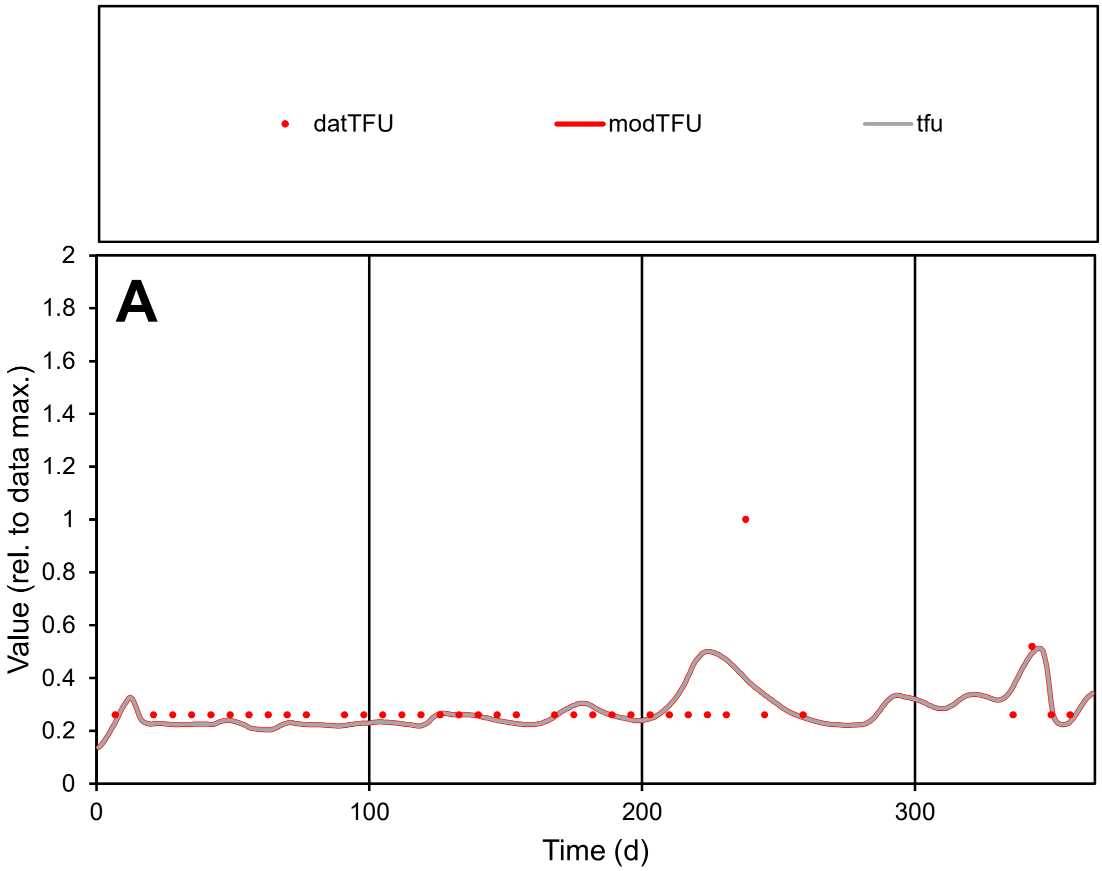


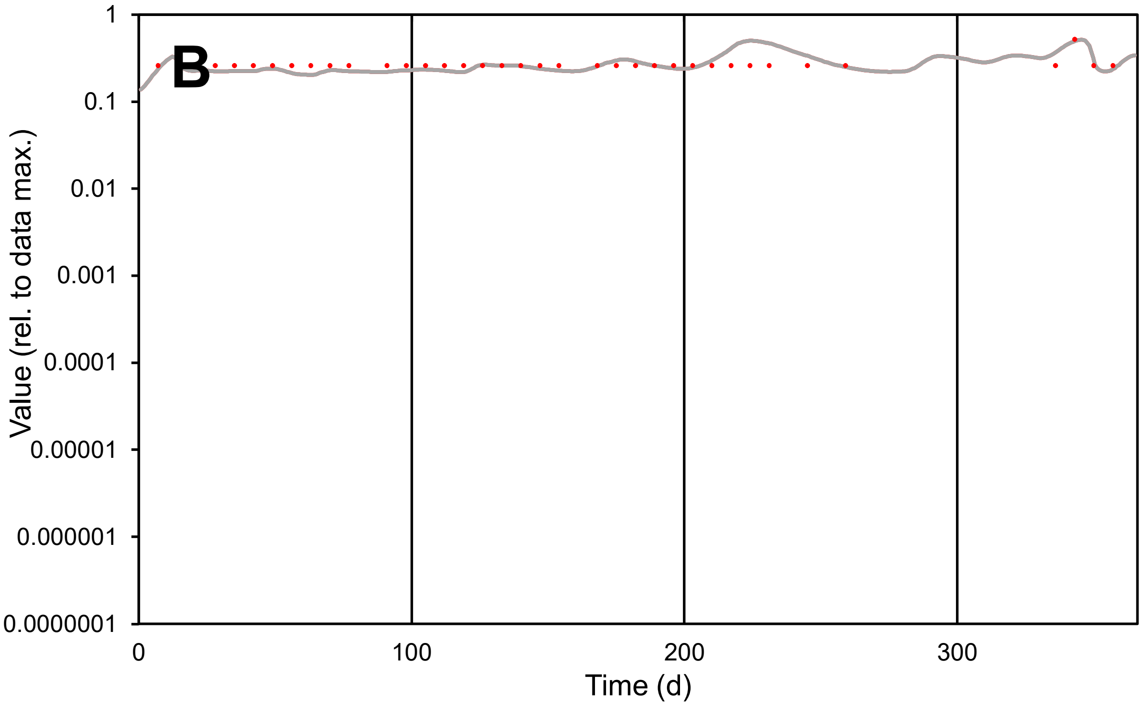


**C1**

Variables
TFU Provide jtd or Dst
CalibOnly?
2 1 to reproduce of
oNormalize?
2 1=no, 2=yes

ofxn calc*
0.42116
ofxn mod
0.4392

sRun
RunA1
zout
1

Ave
Data
1.4E-05
Model
1.4E-05

data/vMax
4.9E-05

ofxng
311.734

Min d/m
1.26238

ave/min d
1.10397
ave/min m
1.3426

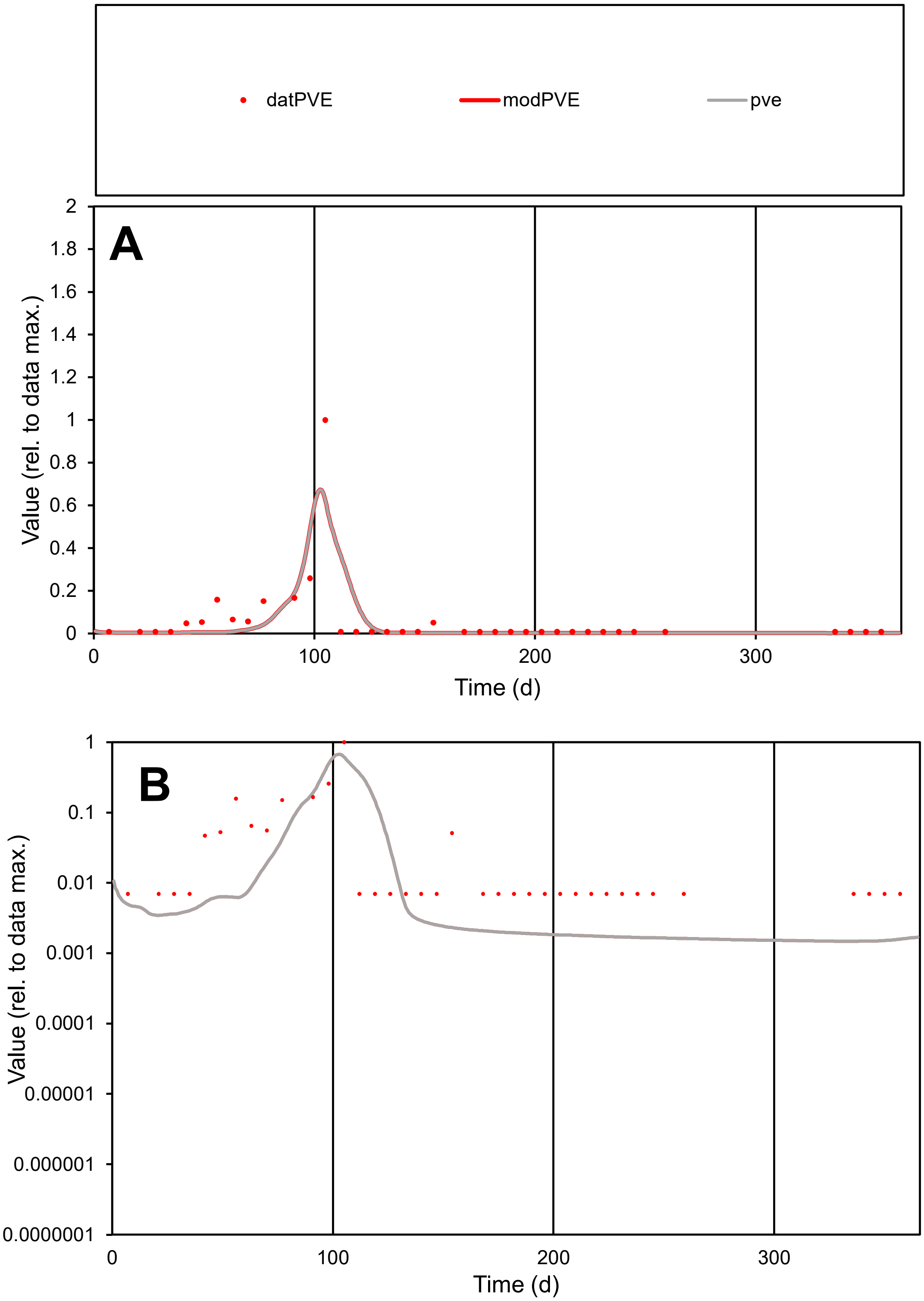


**C1**

Variables
PVE Provide jtd or Dst
CalibOnly?
2 1 to reproduce of
oNormalize?
2 1=no, 2=yes

ofxn calc*
1.58302
ofxn mod
1.71532

sRun
RunA1
zout
1

Ave
Data
1.8E-05
Model
1.5E-05

data/vMax
0.0003

ofxng
311.734

Min d/m
4.72235

ave/min d
8.47416
ave/min m
33.0211

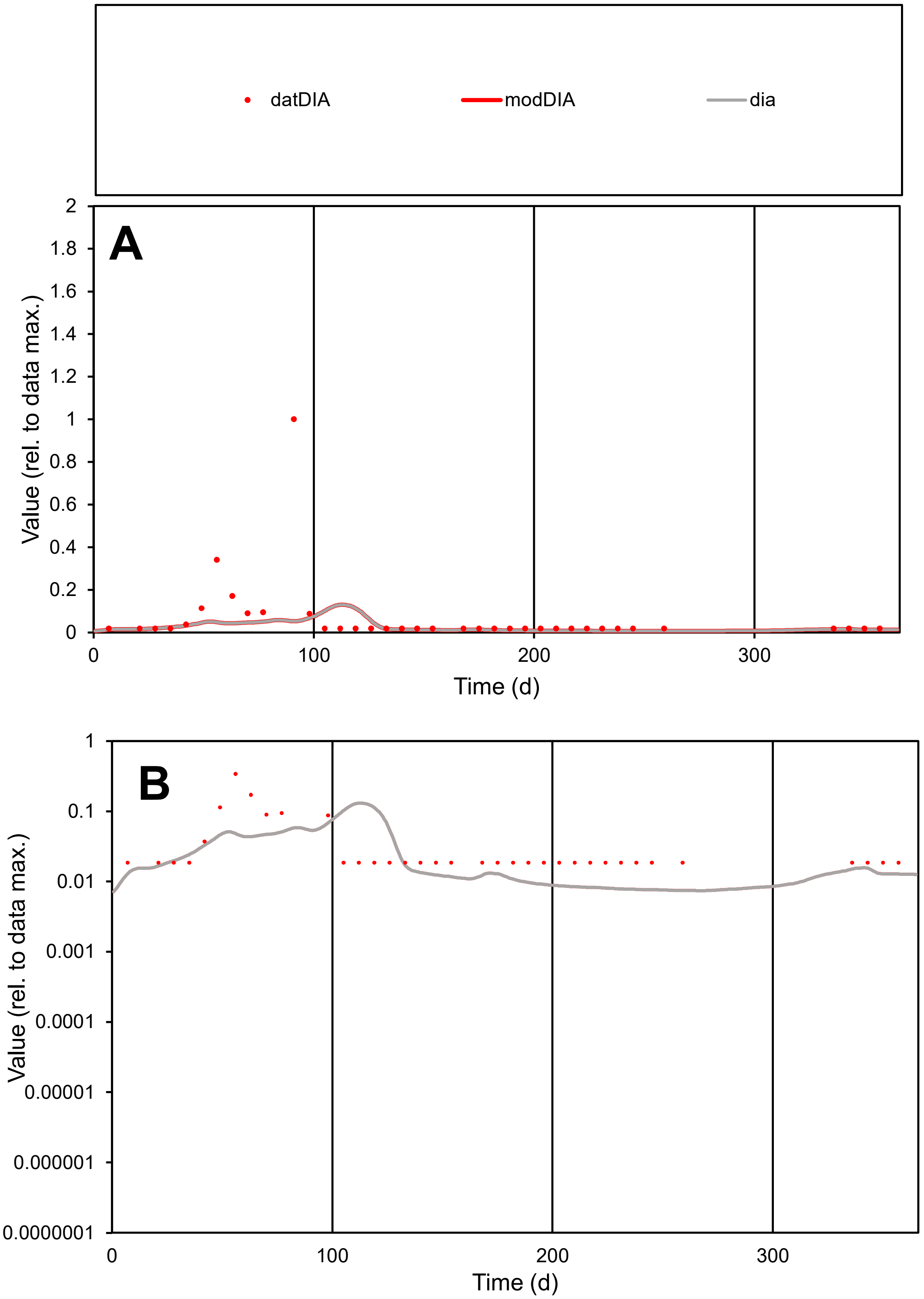


**C1**

Variables

DIA Provide jtd or Dst

CalibOnly?

2 1 to reproduce of

oNormalize?

2 1=no, 2=yes

ofxn calc*

2.49909

ofxn mod

2.50817

sRun

RunA1

zout

1

Ave

Data

2.2E-05

Model

9.7E-06

data/vMax

0.00034

ofxng

311.734

Min d/m

2.4918

ave/min d

3.6032

ave/min m

3.87637

331_Calib_S_A

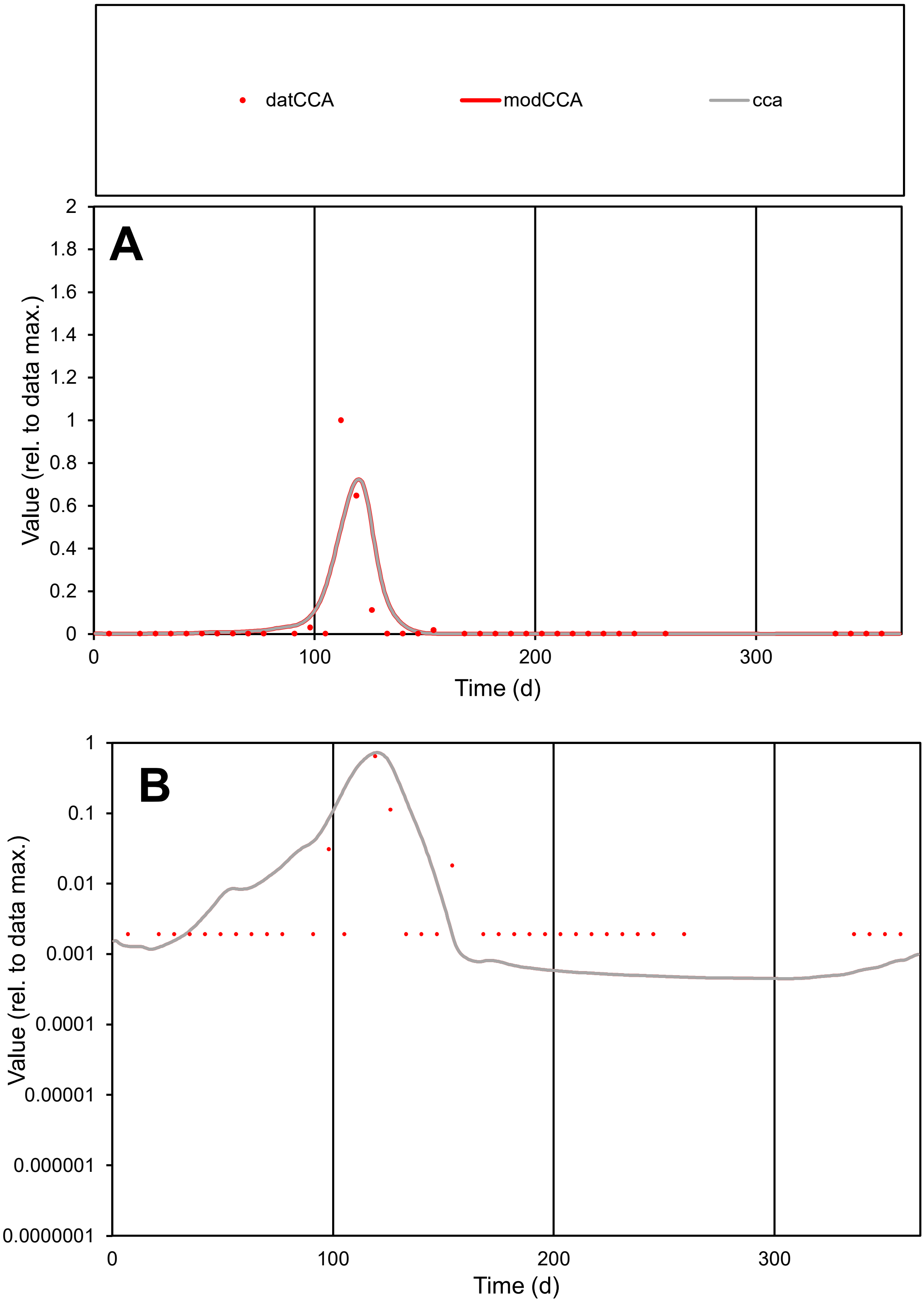


**C1**

Variables
CCA Provide jtd or Dst
CalibOnly?
2 1 to reproduce of
oNormalize?
2 1=no, 2=yes

ofxn calc*
1.70983
ofxn mod
1.88427

sRun
RunA1
zout
1

Ave
Data
4.8E-05
Model
5.3E-05

data/vMax
0.00095

ofxng
311.734

Min d/m
4.09682

ave/min d
26.2346
ave/min m
118.146

331_Calib_S_A

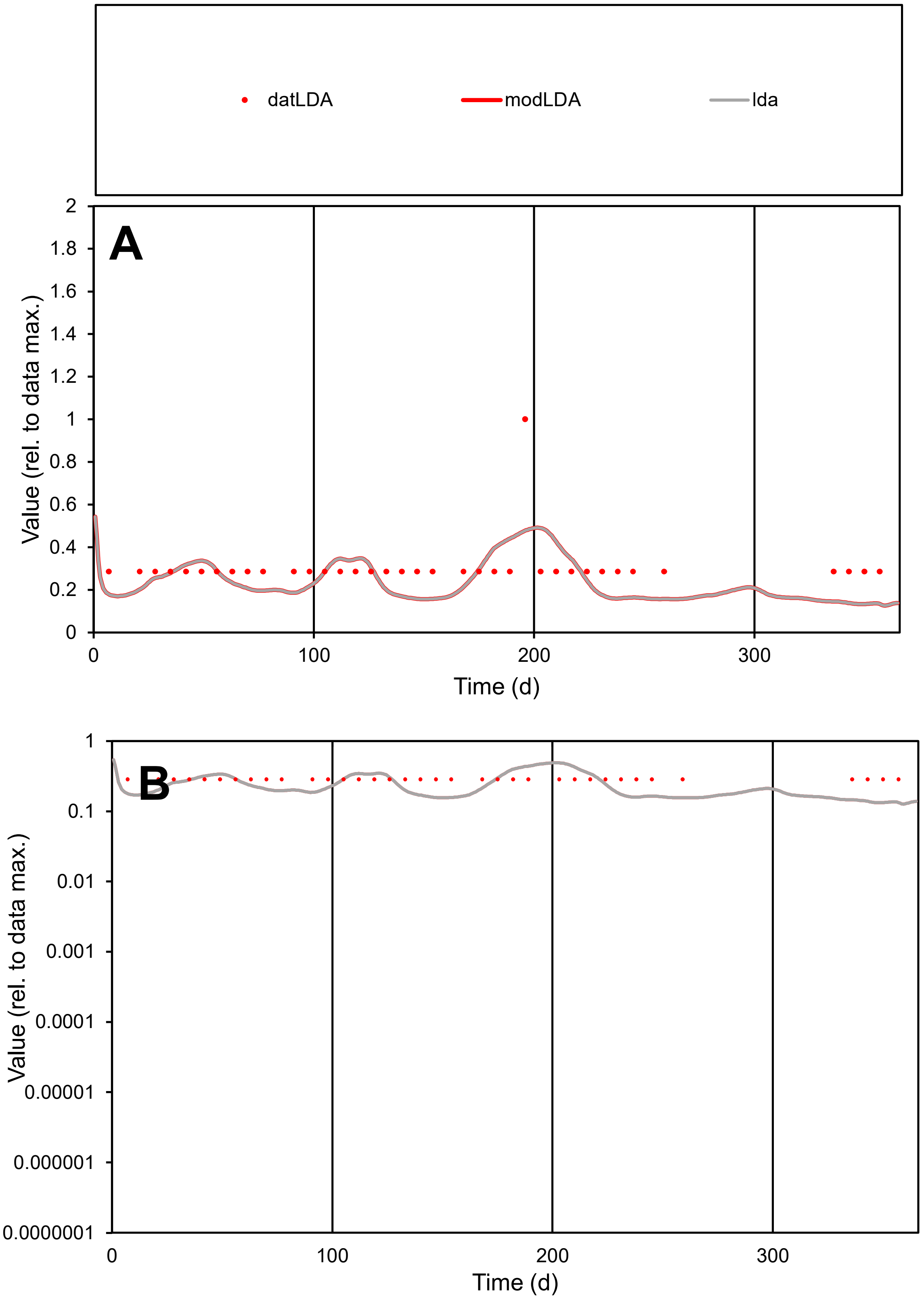


C1

Variables

LDA Provide jtd or Dst

CalibOnly?

2 1 to reproduce of

oNormalize?

2 1=no, 2=yes

ofxn calc*

0.43015

ofxn mod

0.44332

sRun

RunA1

zout

1

Ave

Data

7.4E-06

Model

6.2E-06

data/vMax

2.4E-05

ofxng

311.734

Min d/m

2.14029

ave/min d

1.06777

ave/min m

1.90704

331_Calib_S_A

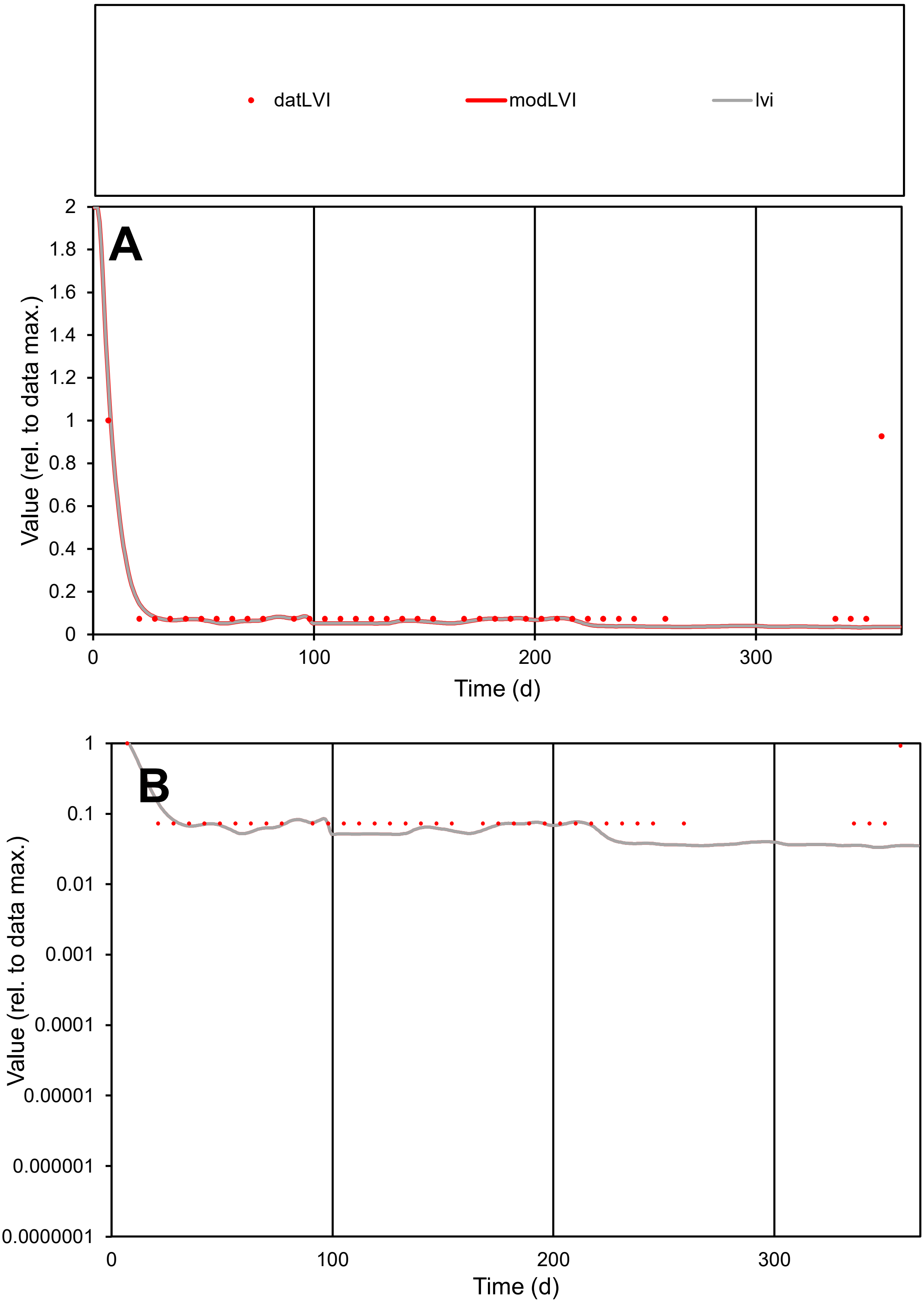


**C1**

Variables
LVI Provide jtd or Dst
CalibOnly?
2 1 to reproduce of
oNormalize?
2 1=no, 2=yes

ofxn calc*
1.22262
ofxn mod
1.22905

sRun
RunA1
zout
1

Ave
Data
2.4E-06
Model
1.6E-06

data/vMax
1.9E-05

ofxng
311.734

Min d/m
2.15161

ave/min d
1.65944
ave/min m
2.49447

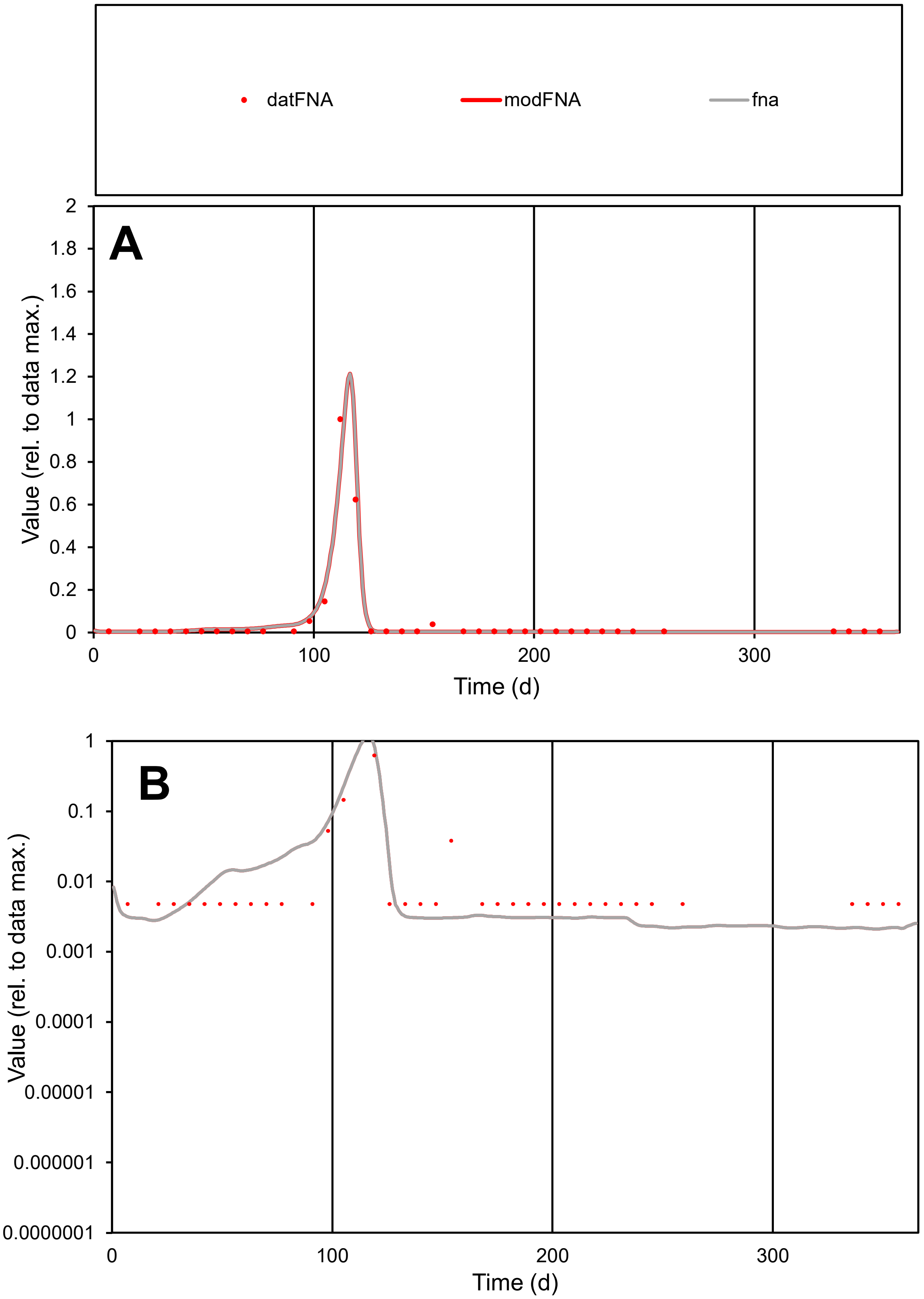


C1

Variables
FNA Provide jtd or Dst
CalibOnly?
2 1 to reproduce of
oNormalize?
2 1=no, 2=yes

ofxn calc*
0.35974
ofxn mod
0.47051

sRun
RunA1
zout
1

Ave
Data
3.5E-05
Model
3.7E-05

data/vMax
0.00065

ofxng
311.734

Min d/m
2.24478

ave/min d
11.3841
ave/min m
26.6431

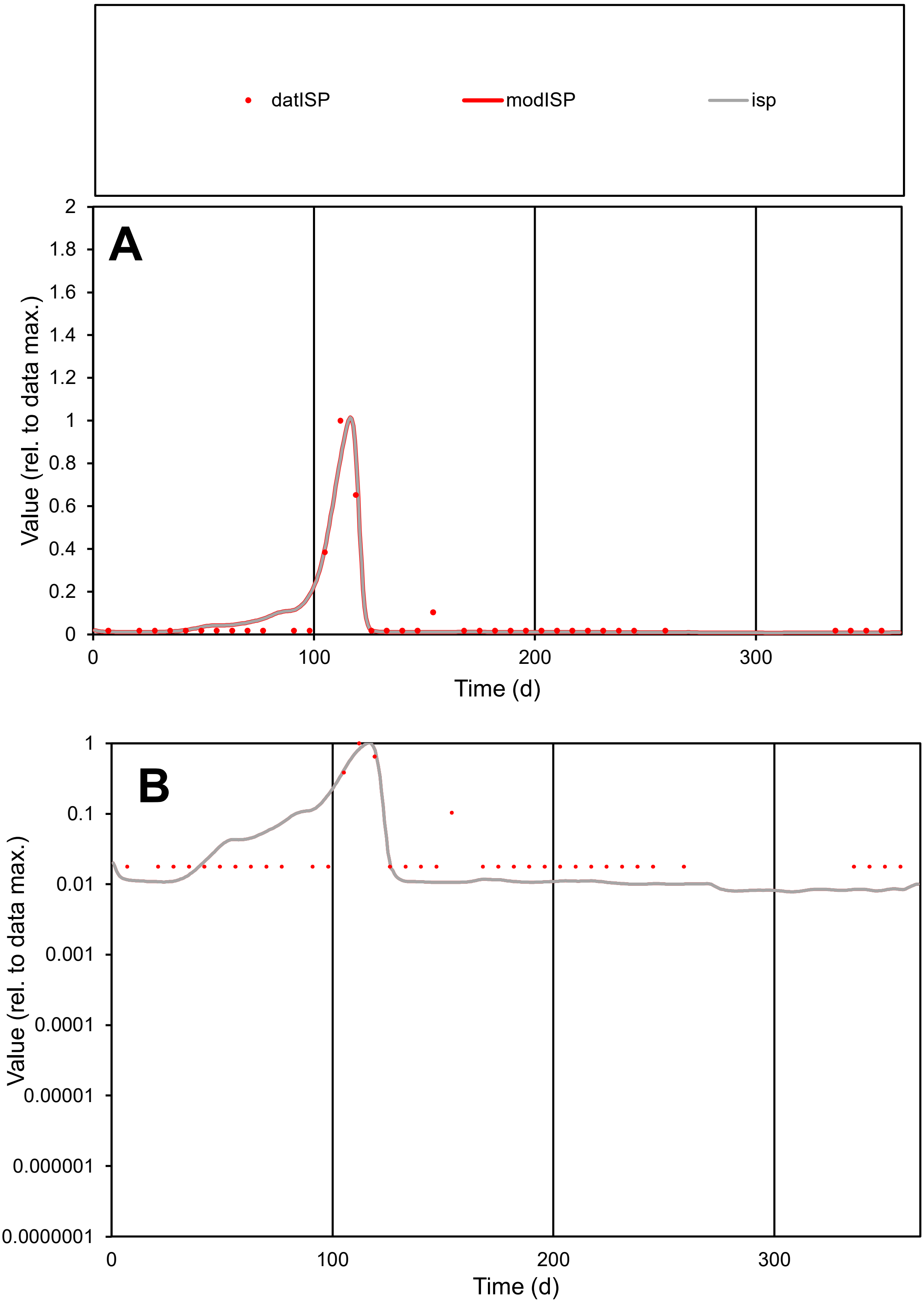


C1

Variables
ISP Provide jtd or Dst
CalibOnly?
2 1 to reproduce of
oNormalize?
2 1=no, 2=yes

ofxn calc*
0.55335
ofxn mod
0.62732

sRun
RunA1
zout
1

Ave
Data
2.1E-05
Model
2.2E-05

data/vMax
0.00028

ofxng
311.734

Min d/m
2.1808

ave/min d
4.13733
ave/min m
9.4118

331_Calib_S_A

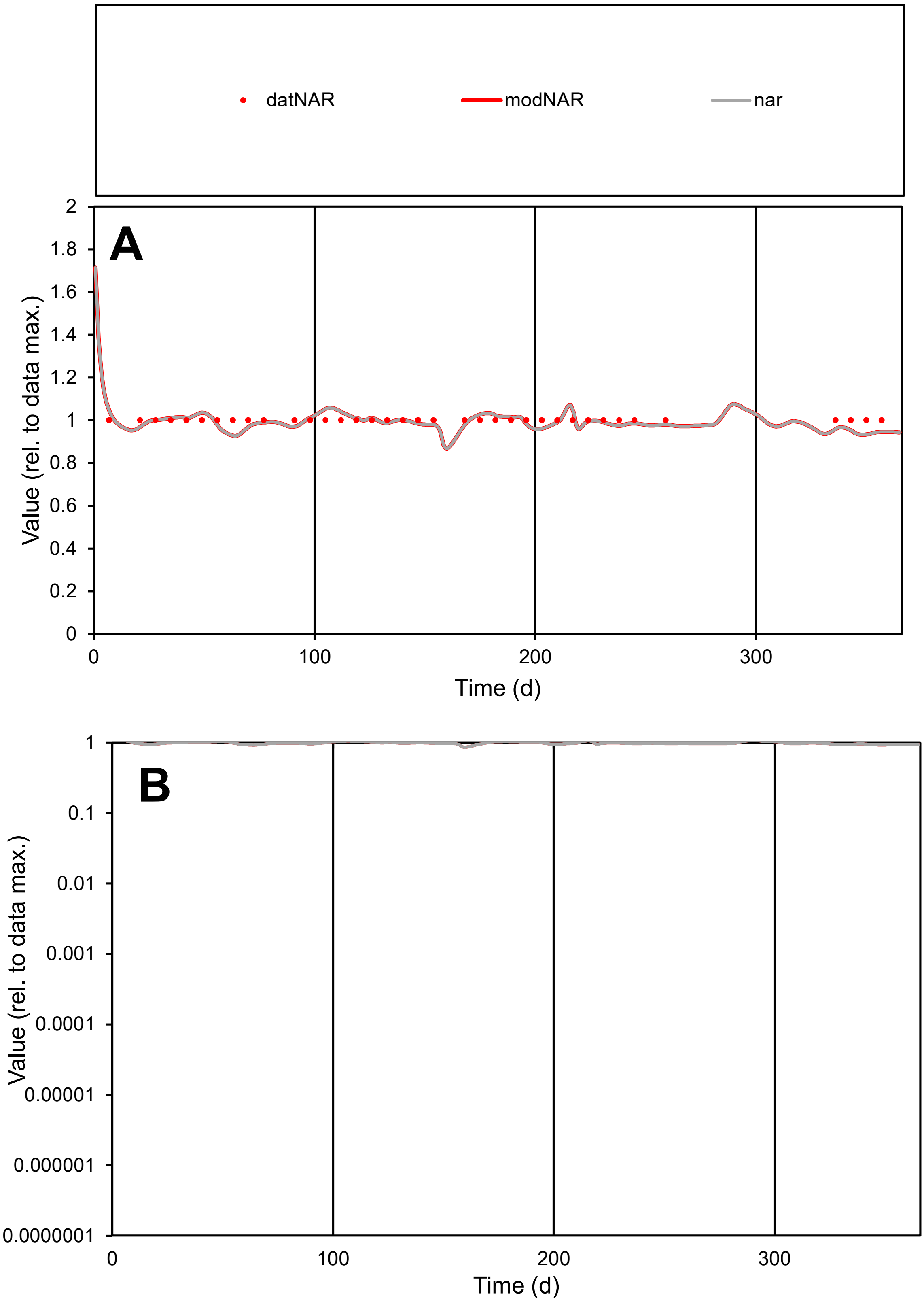


**C1**

Variables
NAR Provide jtd or Dst
CalibOnly?
2 1 to reproduce of
oNormalize?
2 1=no, 2=yes

ofxn calc*
0.02942
ofxn mod
0.11493

sRun
RunA1
zout
1

Ave
Data
2.8E-06
Model
2.8E-06

data/vMax
2.8E-06

ofxng
311.734

Min d/m
1.07742

ave/min d
1
ave/min m
1.06837

331_Calib_S_A

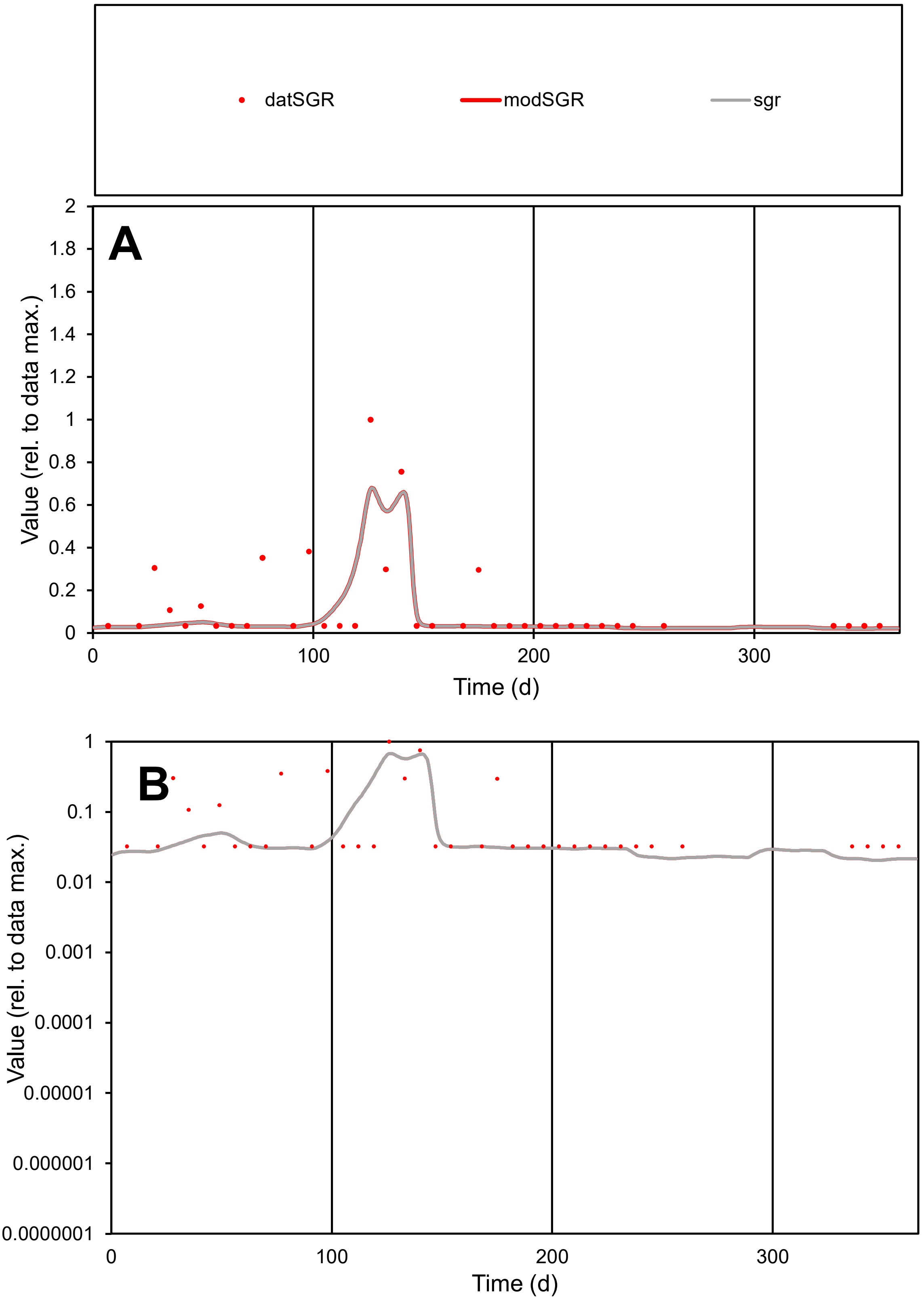


**C1**

Variables
SGR Provide jtd or Dst
CalibOnly?
2 1 to reproduce of
oNormalize?
2 1=no, 2=yes

ofxn calc*
1.06183
ofxn mod
1.0951

sRun
RunA1
zout
1

Ave
Data
1.7E-05
Model
1.3E-05

data/vMax
0.00014

ofxng
311.734

Min d/m
1.55735

ave/min d
3.78319
ave/min m
4.35028

331_Calib_S_A

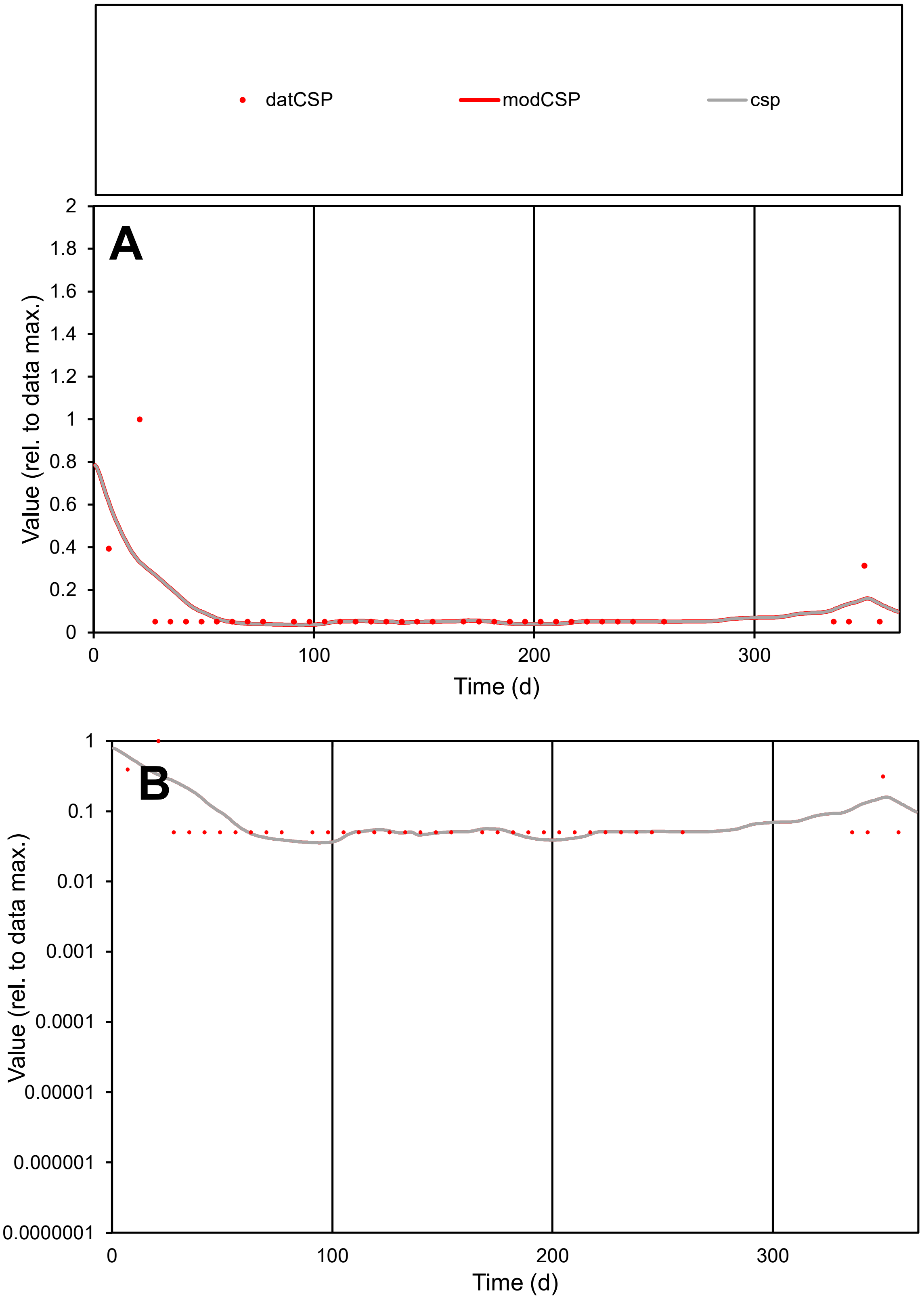


**C1**

Variables

CSP Provide jtd or Dst

CalibOnly?

2 1 to reproduce of

oNormalize?

2 1=no, 2=yes

ofxn calc*

1.36991

ofxn mod

1.4037

sRun

RunA1

zout

1

Ave

Data

4.4E-06

Model

4.3E-06

data/vMax

4.7E-05

ofxng

311.734

Min d/m

1.41494

ave/min d

1.83469

ave/min m

2.57311

331_Calib_S_A

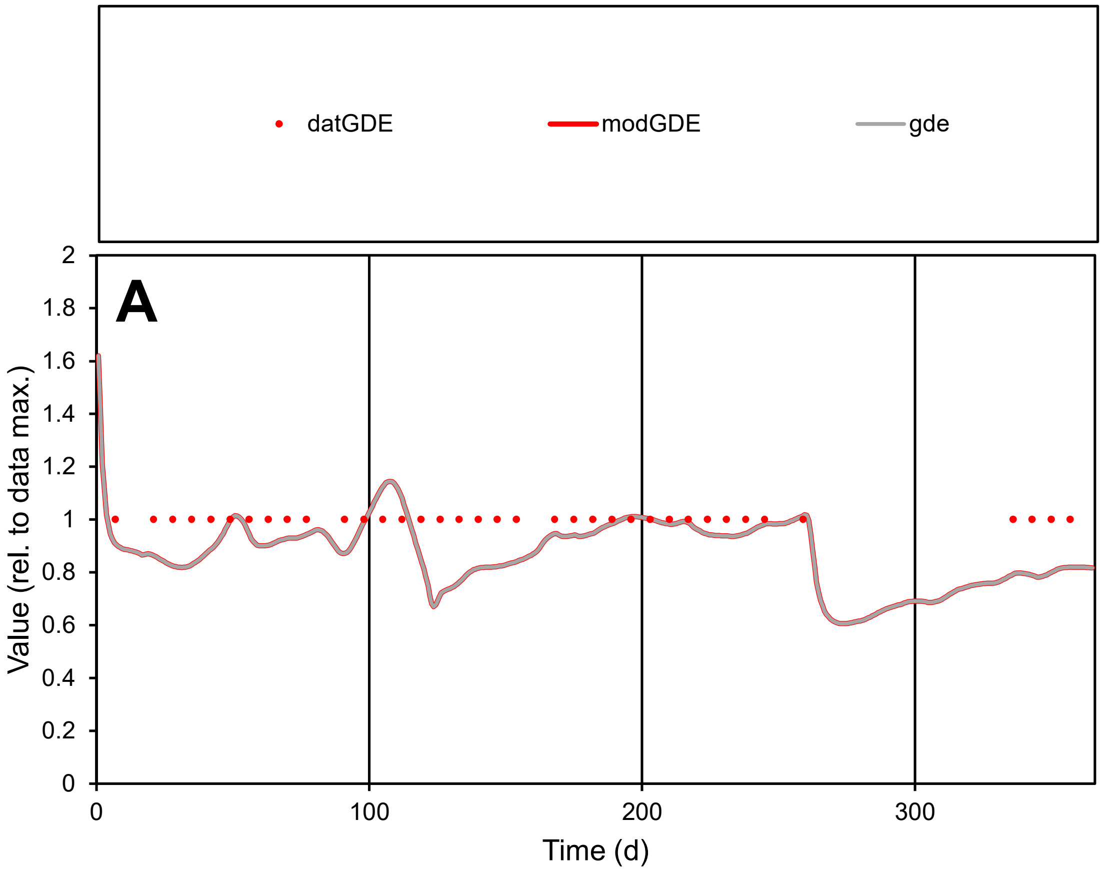


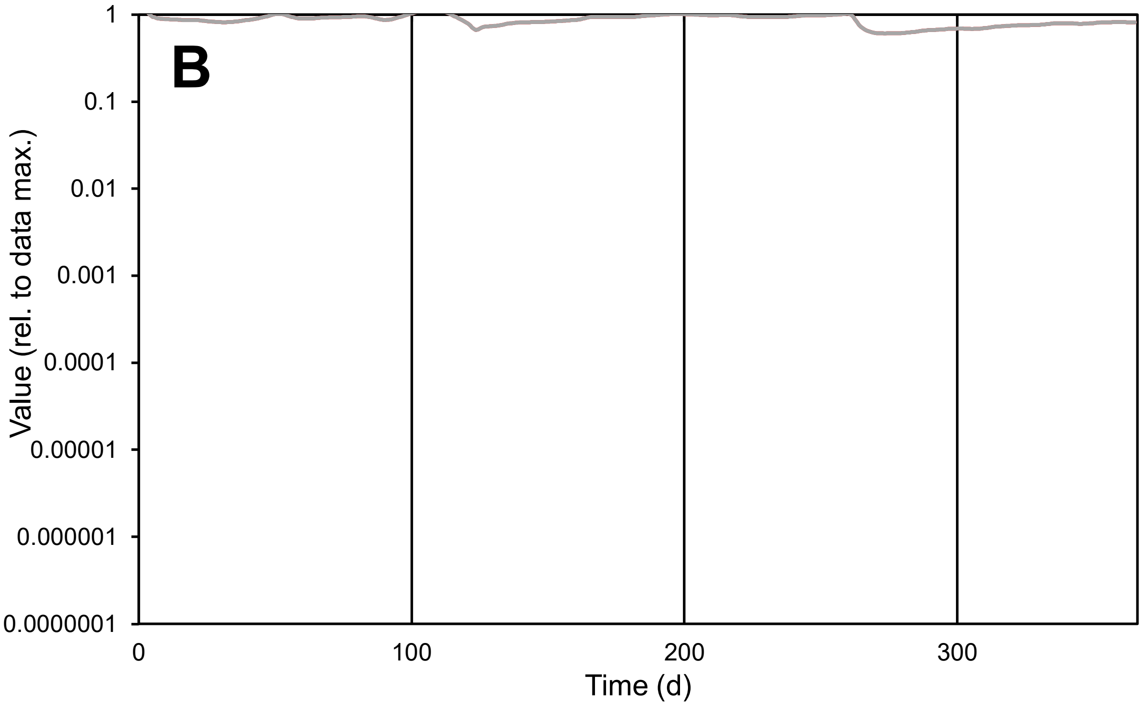


**C1**

Variables
GDE Provide jtd or Dst
CalibOnly?
2 1 to reproduce of
oNormalize?
2 1=no, 2=yes

ofxn calc*
0.12482
ofxn mod
0.16452

sRun
RunA1
zout
1

Ave
Data
2.7E-05
Model
2.5E-05

data/vMax
2.7E-05

ofxng
311.734

Min d/m
1.37194

ave/min d
1
ave/min m
1.2522

331_Calib_S_A

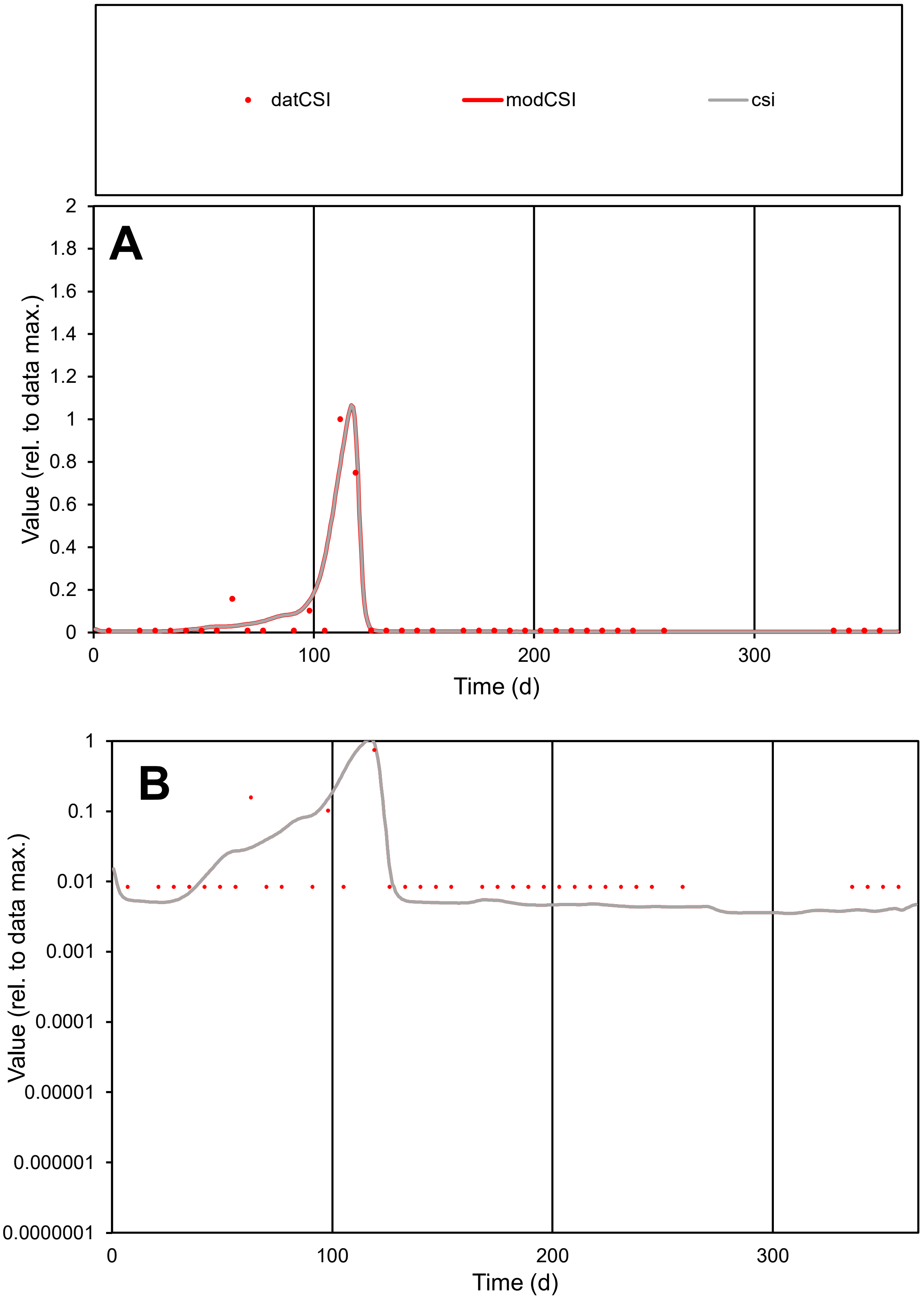


**C1**

Variables
CSI Provide jtd or Dst
CalibOnly?
2 1 to reproduce of
oNormalize?
2 1=no, 2=yes

ofxn calc*
0.90567
ofxn mod
1.03124

sRun
RunA1
zout
1

Ave
Data
1.7E-05
Model
1.8E-05

data/vMax
0.00028

ofxng
311.734

Min d/m
2.19791

ave/min d
7.36843
ave/min m
17.4231

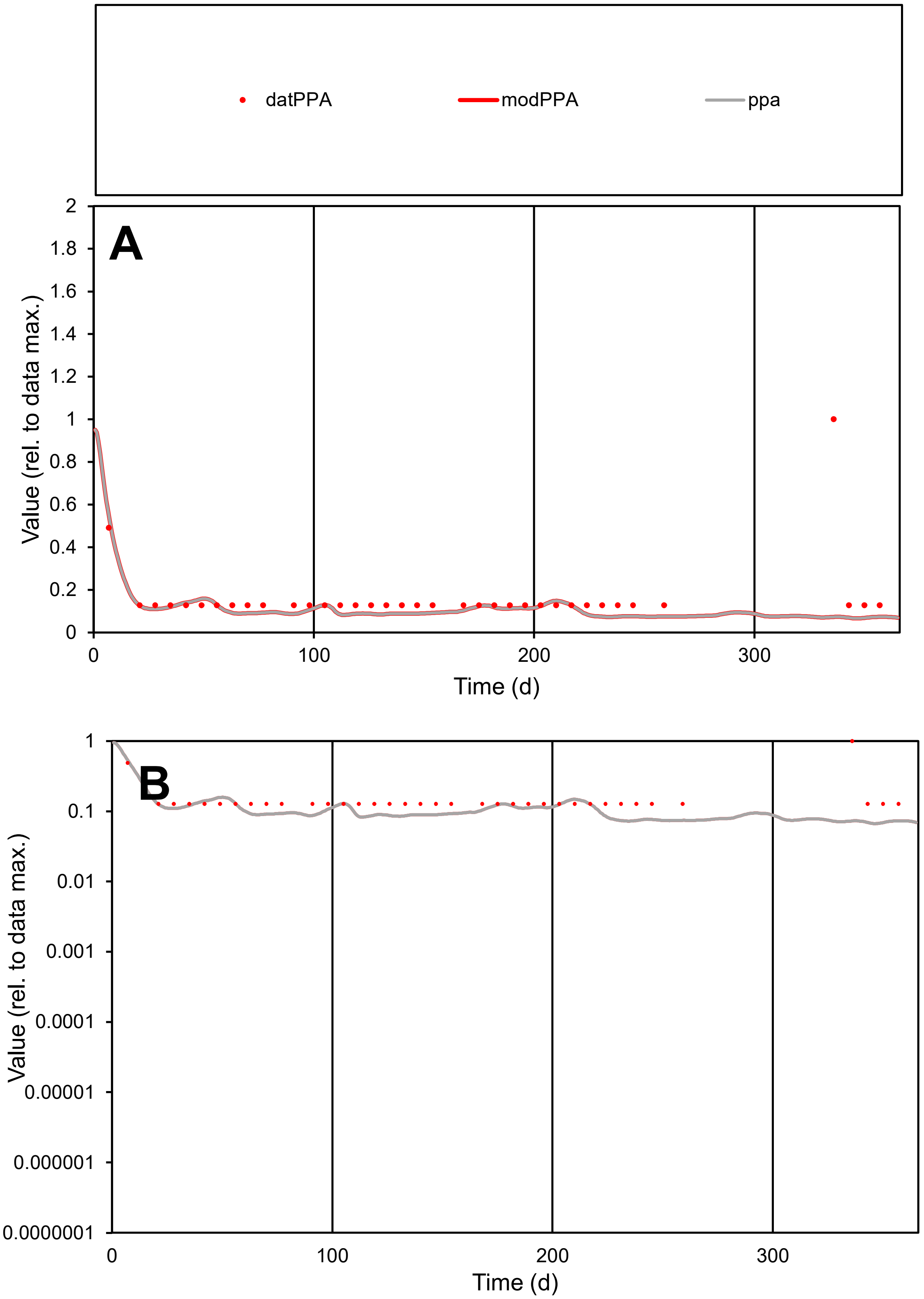


**C1**

Variables
PPA Provide jtd or Dst
CalibOnly?
2 1 to reproduce of
oNormalize?
2 1=no, 2=yes

ofxn calc*
0.96654
ofxn mod
0.9714

sRun
RunA1
zout
1

Ave
Data
4E-06
Model
2.8E-06

data/vMax
2.5E-05

ofxng
311.734

Min d/m
1.87778

ave/min d
1.26015
ave/min m
1.63134

331_Calib_S_A

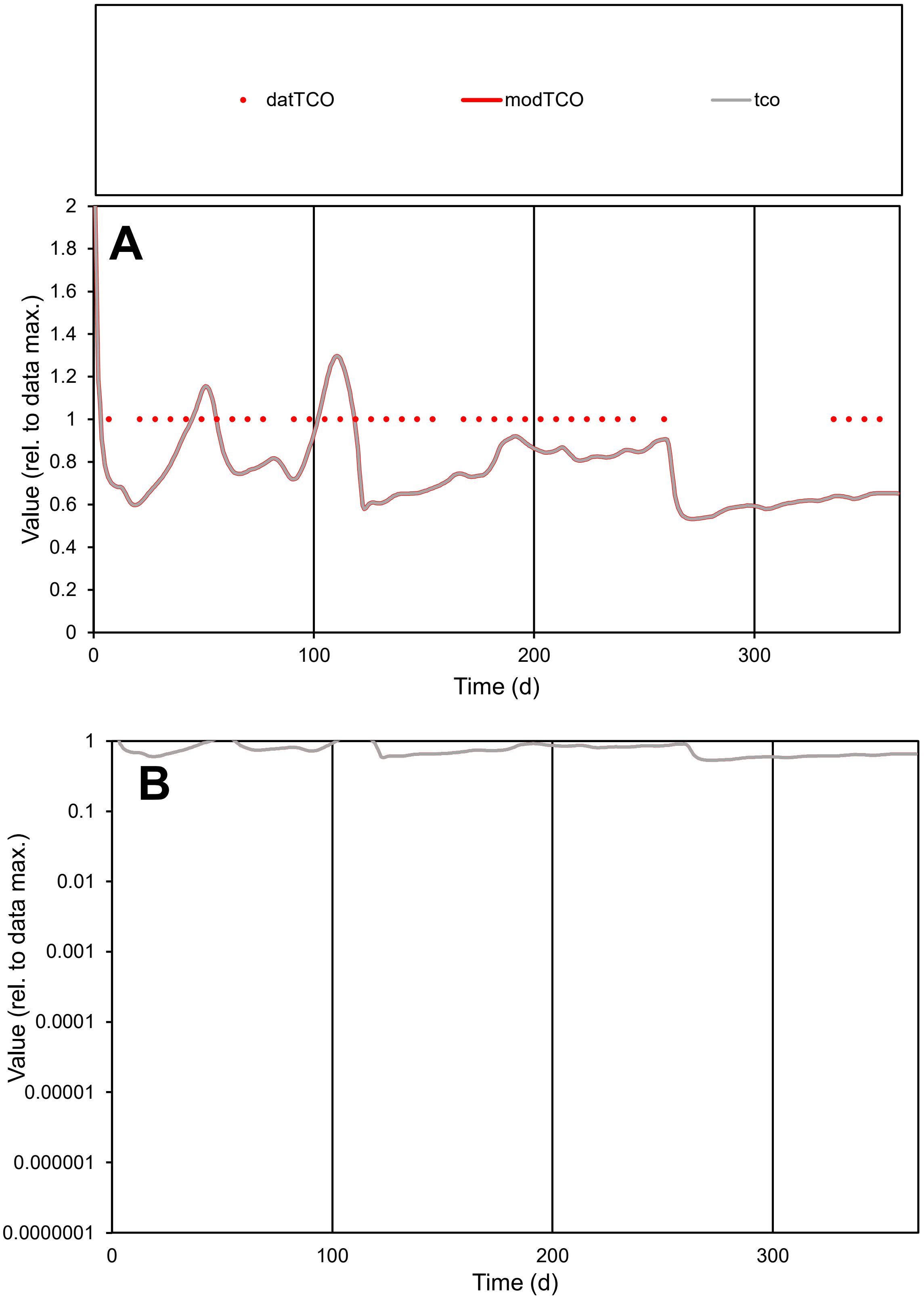


**C1**

Variables
TCO Provide jtd or Dst
CalibOnly?
2 1 to reproduce of
oNormalize?
2 1=no, 2=yes

ofxn calc*
0.24751
ofxn mod
0.26748

sRun
RunA1
zout
1

Ave
Data
8.6E-06
Model
7E-06

data/vMax
8.6E-06

ofxng
311.734

Min d/m
1.64091

ave/min d
1
ave/min m
1.32323

331_Calib_S_A

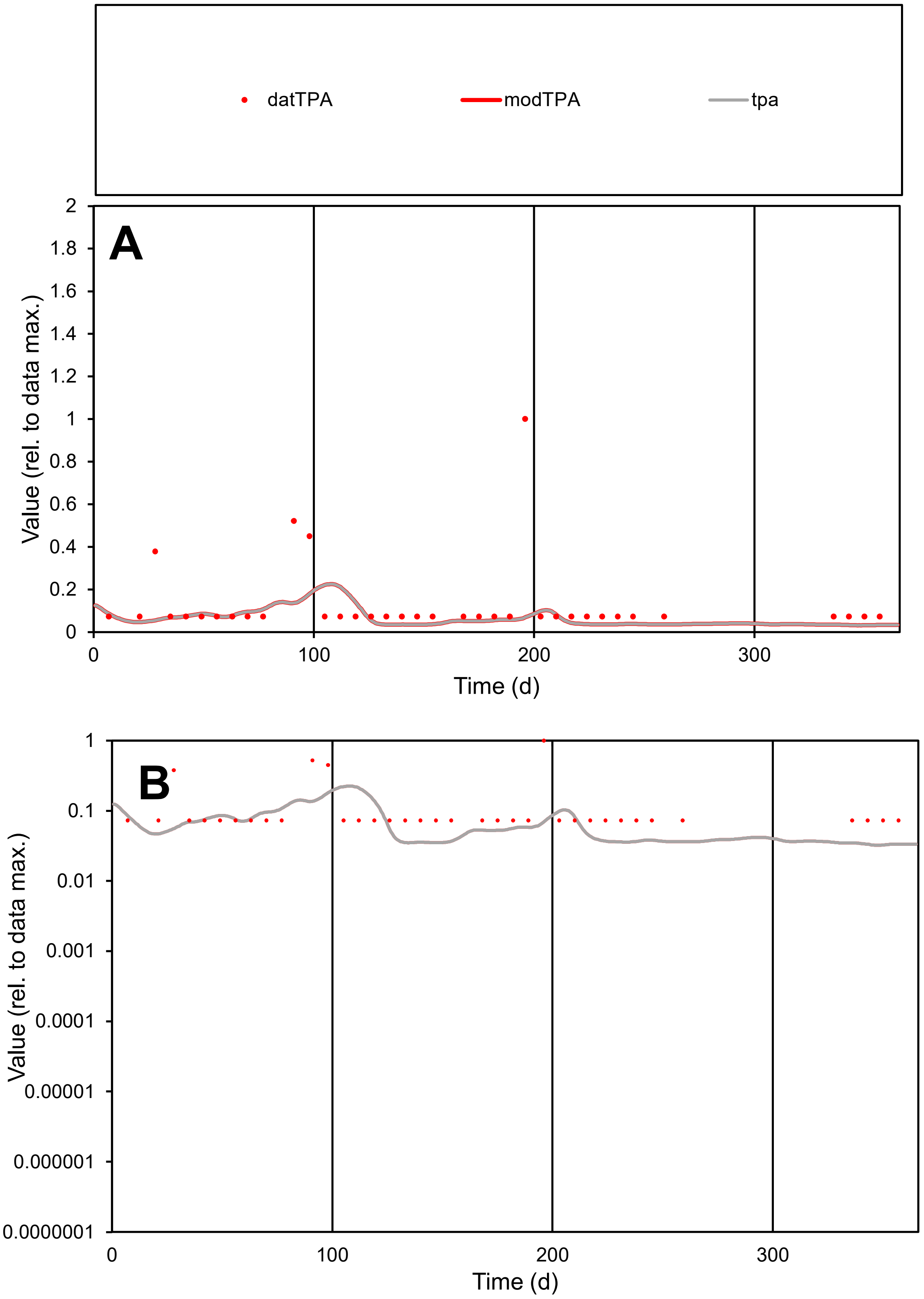


**C1**

Variables

TPA Provide jtd or Dst

CalibOnly?

2 1 to reproduce of

oNormalize?

2 1=no, 2=yes

ofxn calc*

1.42414

ofxn mod

1.42973

sRun

RunA1

zout

1

Ave

Data

8.3E-06

Model

4.7E-06

data/vMax

6.5E-05

ofxng

311.734

Min d/m

2.22515

ave/min d

1.76529

ave/min m

2.19984

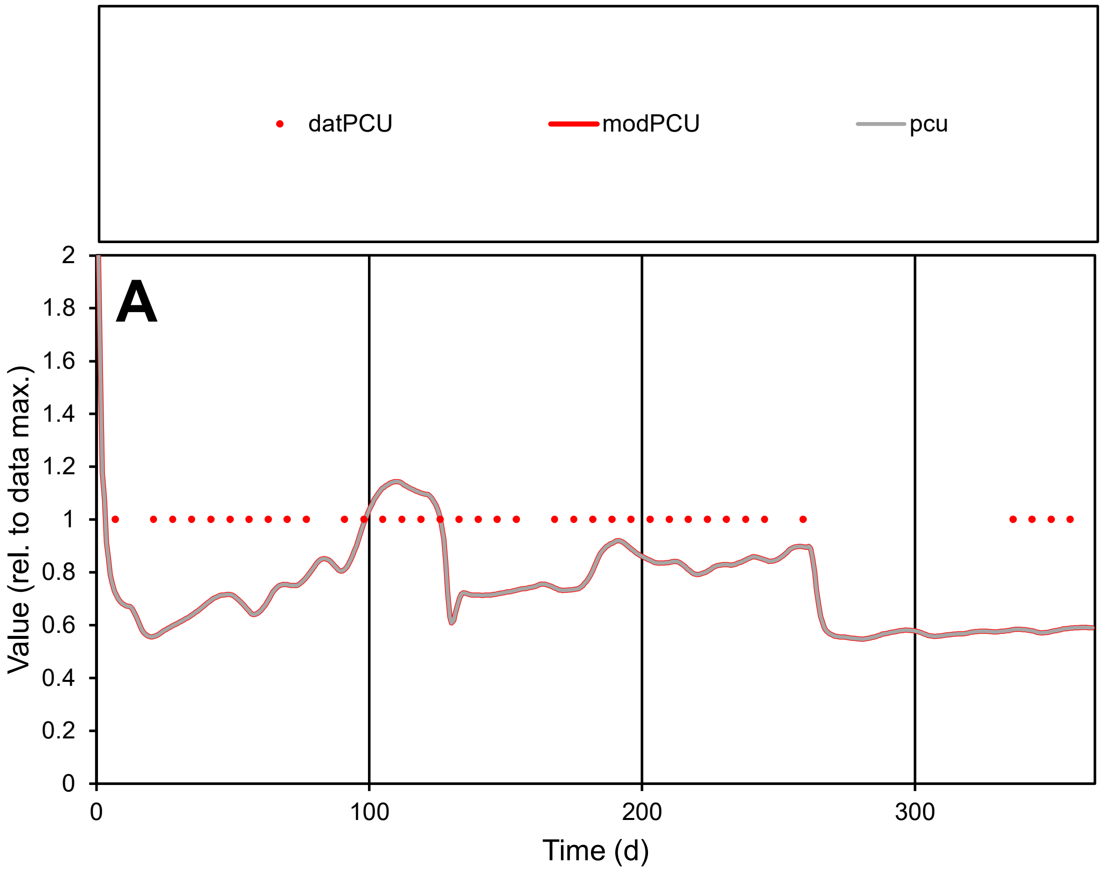


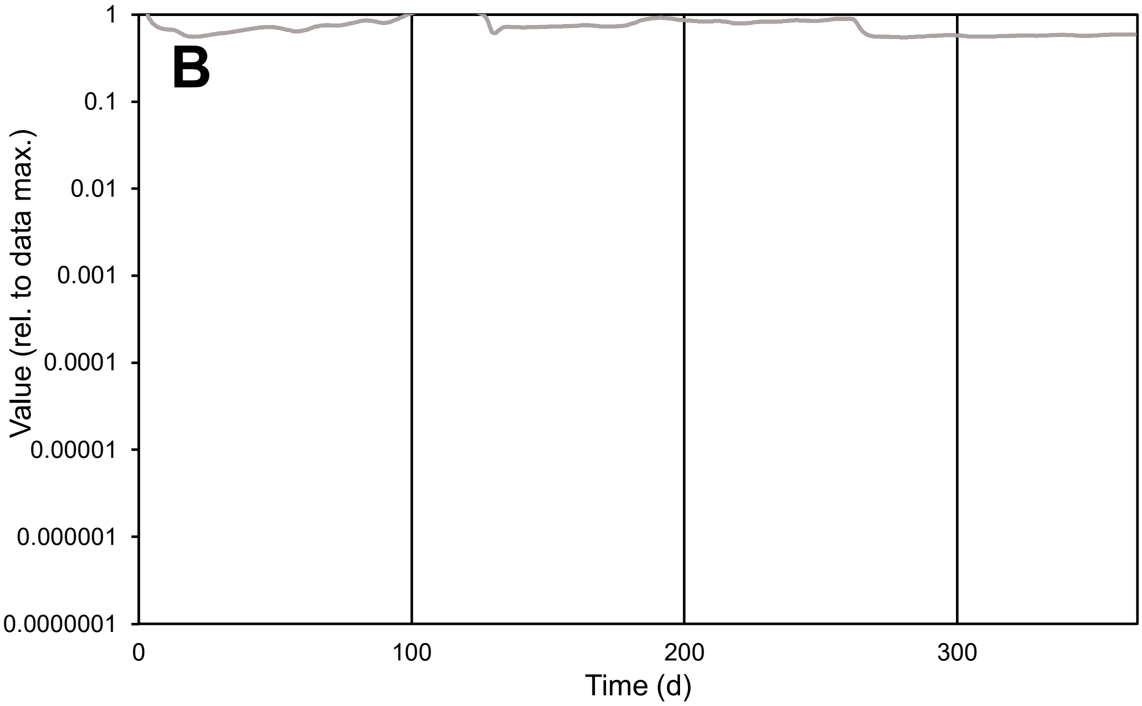


**C1**

Variables
PCU Provide jtd or Dst
CalibOnly?
2 1 to reproduce of
oNormalize?
2 1=no, 2=yes

ofxn calc*
0.26078
ofxn mod
0.27926

sRun
RunA1
zout
1

Ave
Data
6.2E-06
Model
4.9E-06

data/vMax
6.2E-06

ofxng
311.734

Min d/m
1.77275

ave/min d
1
ave/min m
1.39131

331_Calib_S_A

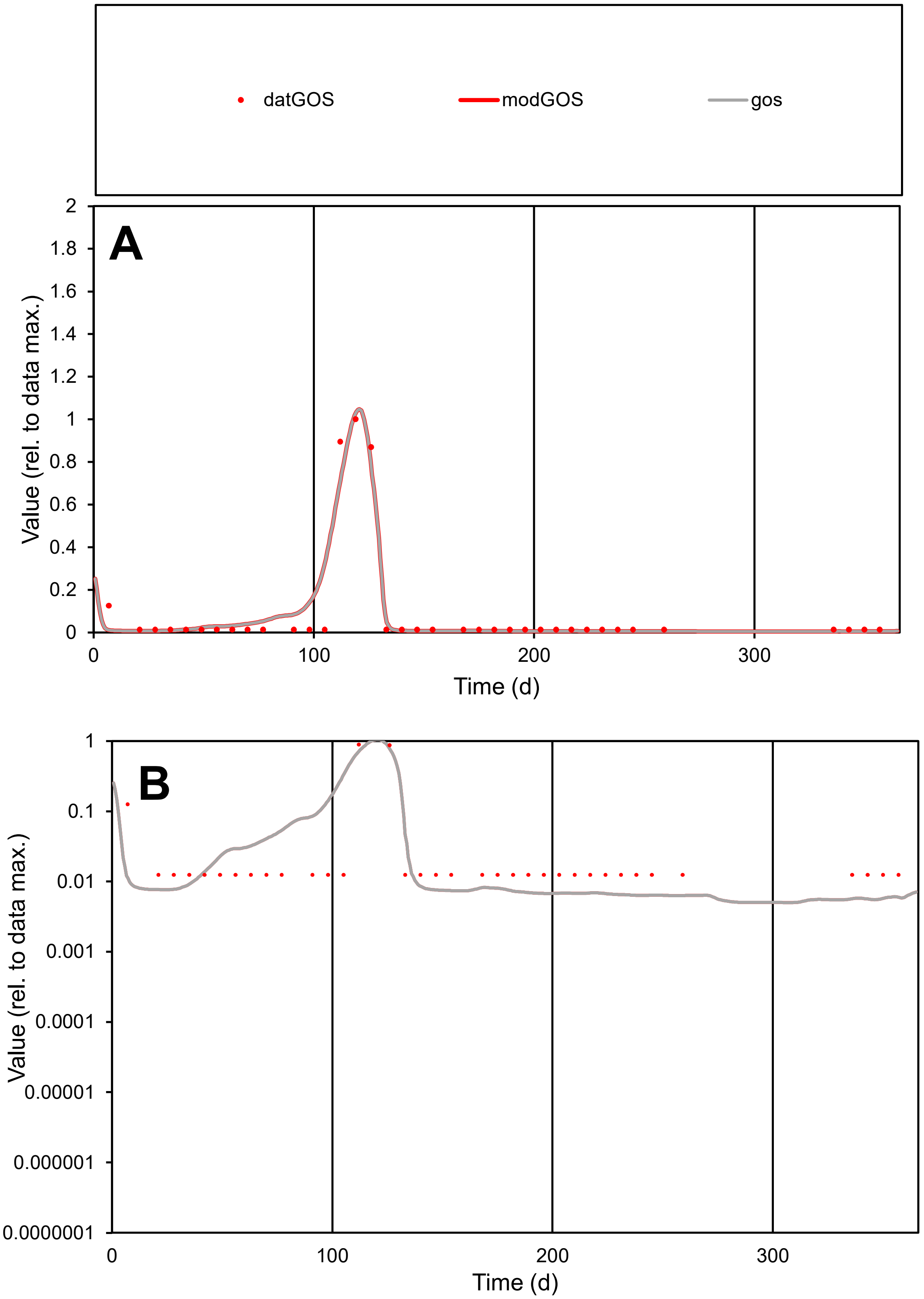


**C1**

Variables
GOS Provide jtd or Dst
CalibOnly?
2 1 to reproduce of
oNormalize?
2 1=no, 2=yes

ofxn calc*
0.62507
ofxn mod
0.77767

sRun
RunA1
zout
1

Ave
Data
1.3E-05
Model
1.4E-05

data/vMax
0.00015

ofxng
311.734

Min d/m
2.21868

ave/min d
7.17851
ave/min m
16.7653

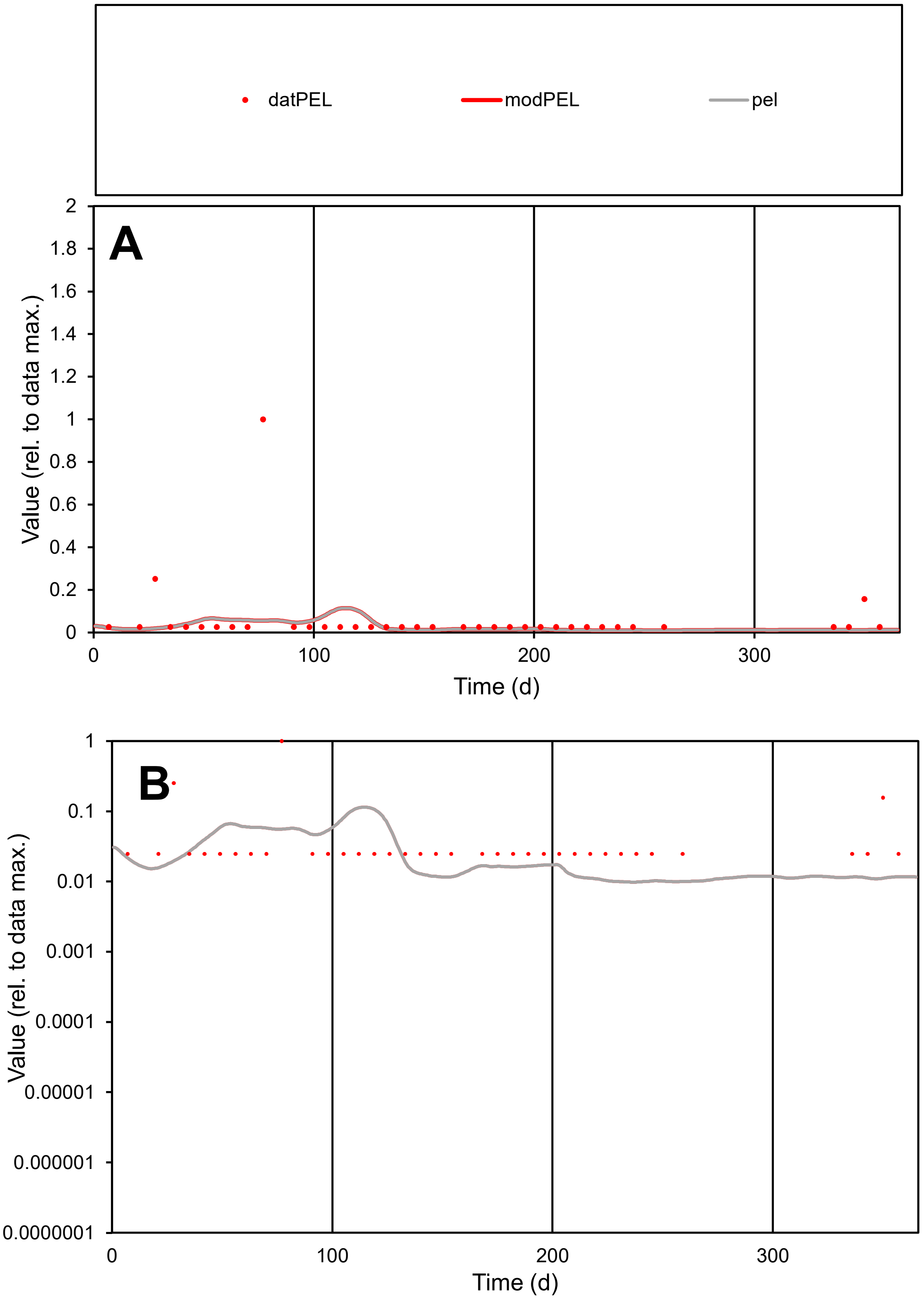


**C1**

Variables
PEL Provide jtd or Dst
CalibOnly?
2 1 to reproduce of
oNormalize?
2 1=no, 2=yes

ofxn calc*
2.68922
ofxn mod
2.69799

sRun
RunA1
zout
1

Ave
Data
8.1E-06
Model
4E-06

data/vMax
0.00013

ofxng
311.734

Min d/m
2.52267

ave/min d
2.45366
ave/min m
3.07905

331_Calib_S_A

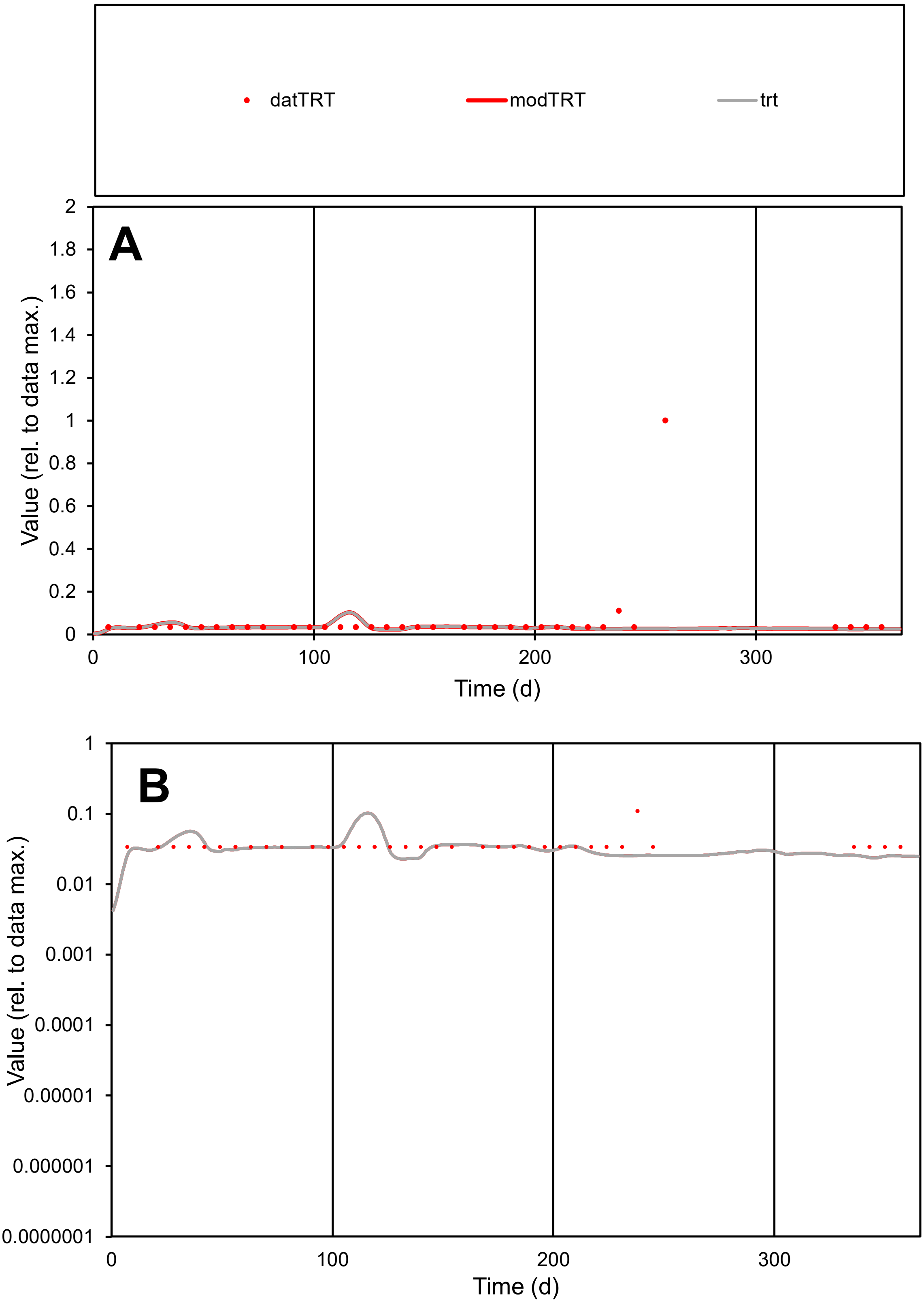


**C1**

Variables
TRT Provide jtd or Dst
CalibOnly?
2 1 to reproduce of
oNormalize?
2 1=no, 2=yes

ofxn calc*
2.59523
ofxn mod
2.6029

sRun
RunA1
zout
1

Ave
Data
4.2E-05
Model
2.3E-05

data/vMax
0.00068

ofxng
311.734

Min d/m
1.47486

ave/min d
1.82866
ave/min m
1.48174

331_Calib_S_A

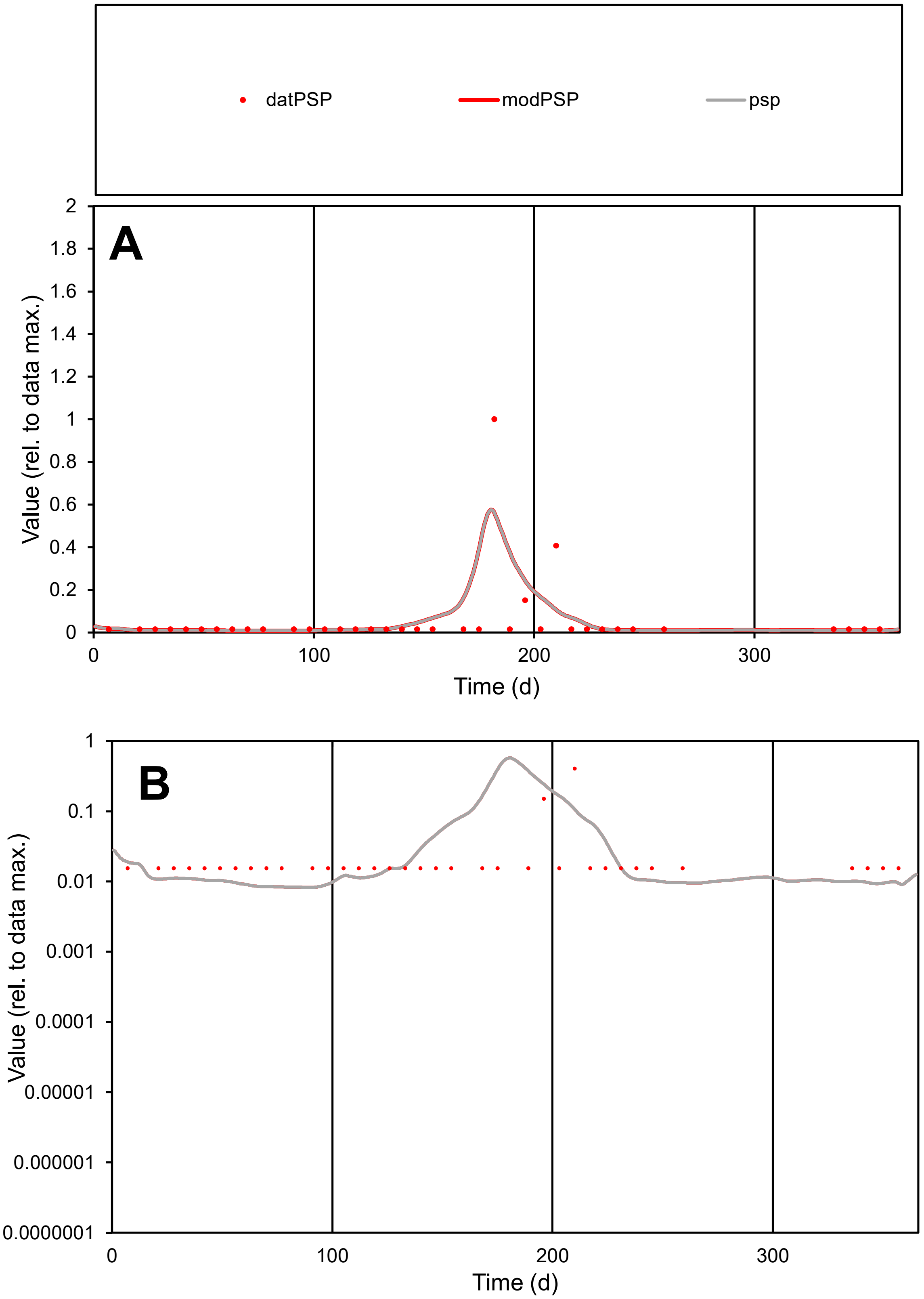


**C1**

Variables
PSP Provide jtd or Dst
CalibOnly?
2 1 to reproduce of
oNormalize?
2 1=no, 2=yes

ofxn calc*
2.06767
ofxn mod
2.23455

sRun
RunA1
zout
1

Ave
Data
1.6E-05
Model
1.7E-05

data/vMax
0.00028

ofxng
311.734

Min d/m
1.8697

ave/min d
3.63946
ave/min m
7.43861

331_Calib_S_A

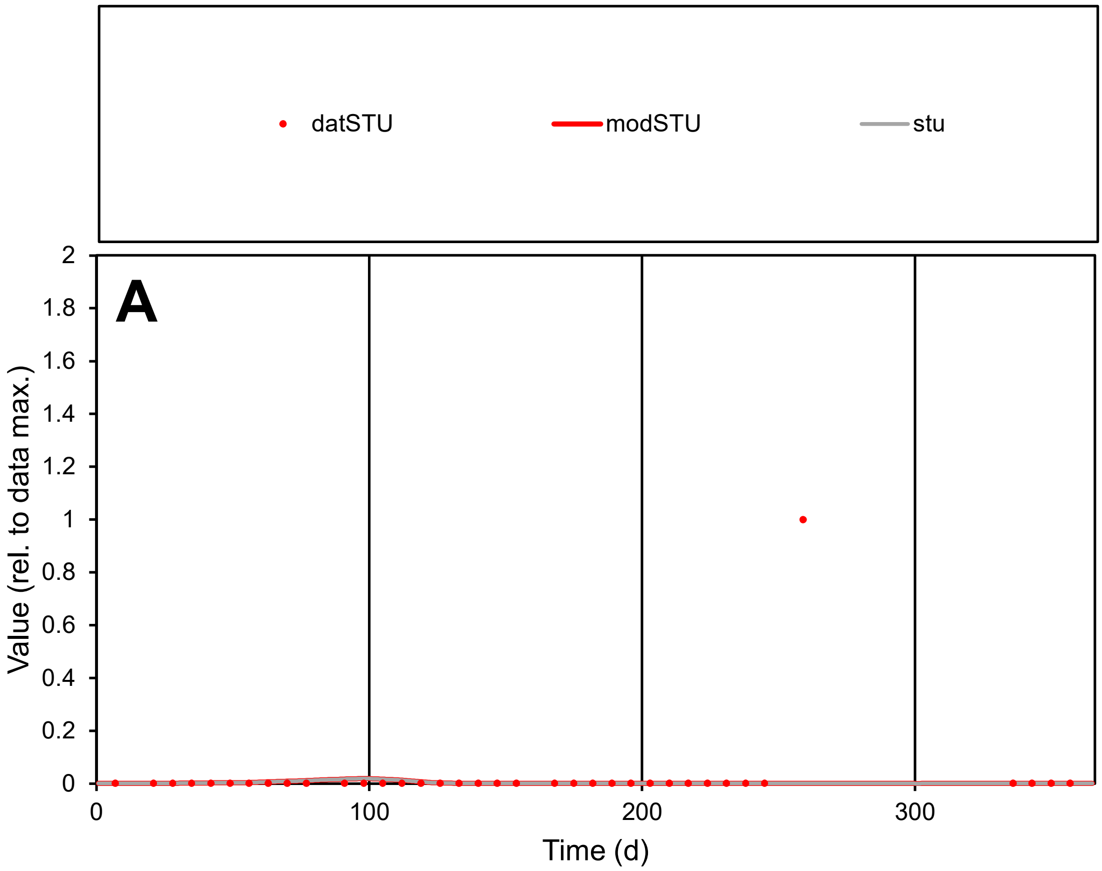


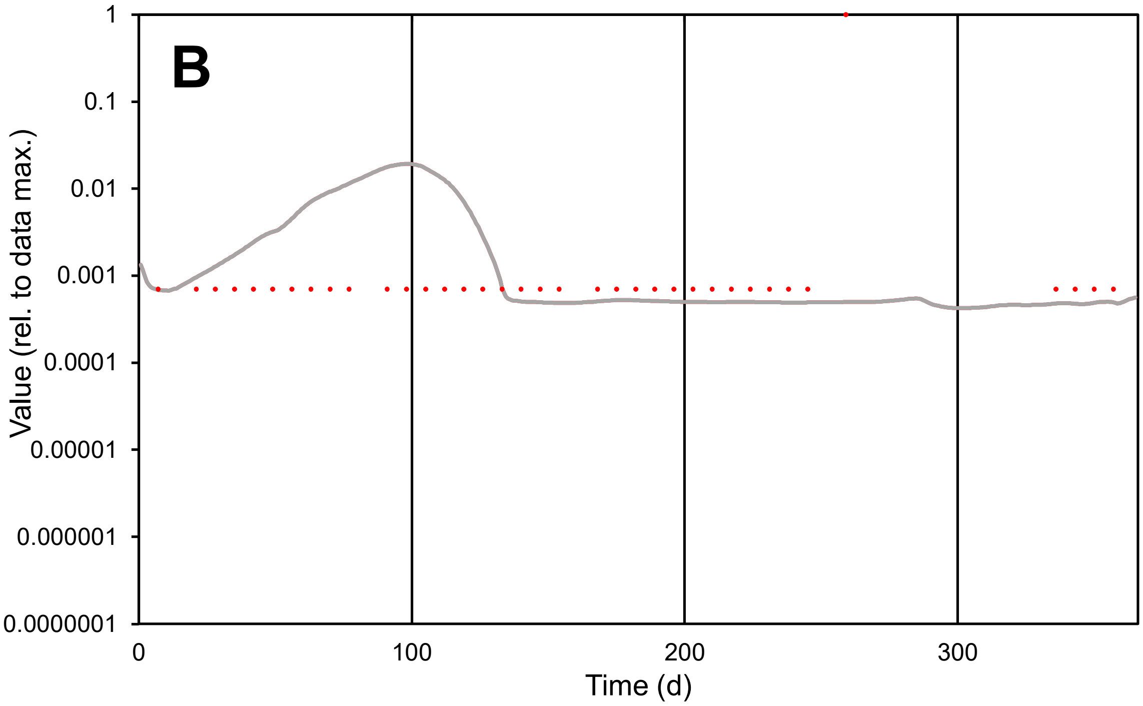


**C1**

Variables
STU Provide jtd or Dst
CalibOnly?
2 1 to reproduce of
oNormalize?
2 1=no, 2=yes

ofxn calc*
5.93337
ofxn mod
5.9456

sRun
RunA1
zout
1

Ave
Data
9.8E-05
Model
1.2E-05

data/vMax
0.00355

ofxng
311.734

Min d/m
1.47865

ave/min d
39.4483
ave/min m
6.93763

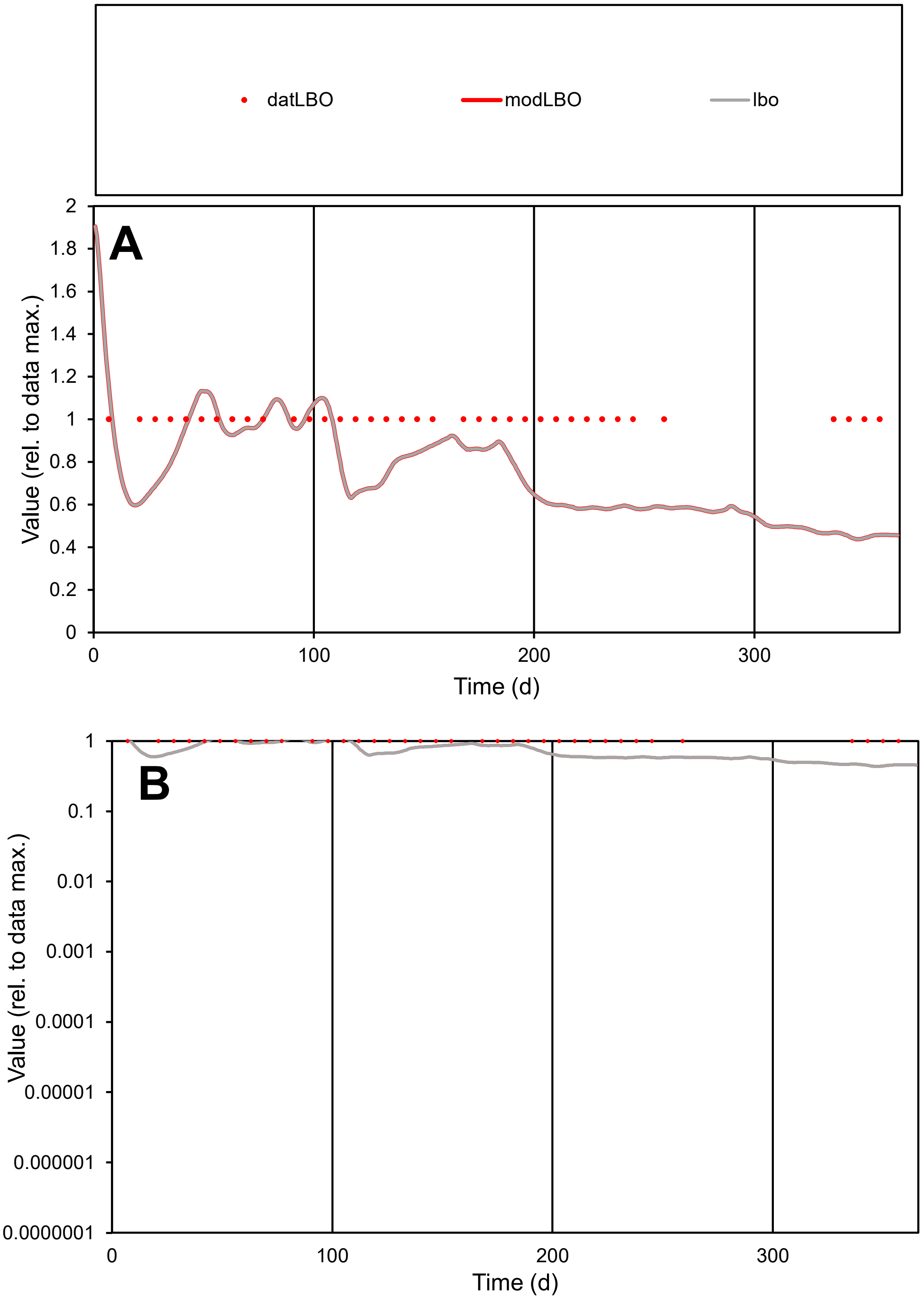


**C1**

Variables

LBO Provide jtd or Dst

CalibOnly?

2 1 to reproduce of

oNormalize?

2 1=no, 2=yes

ofxn calc*

0.30541

ofxn mod

0.32116

sRun

RunA1

zout

1

Ave

Data

1.4E-06

Model

1.1E-06

data/vMax

1.4E-06

ofxng

311.734

Min d/m

2.24434

ave/min d

1

ave/min m

1.72107

331_Calib_S_A

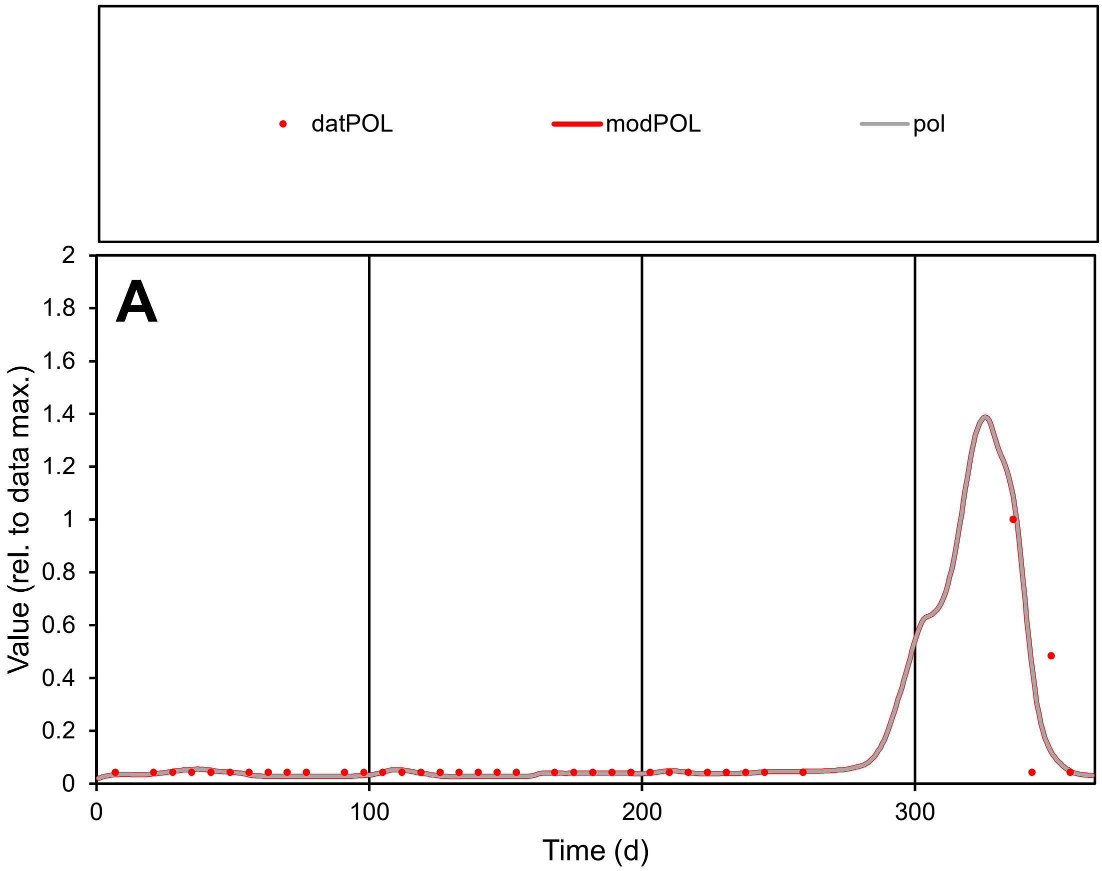


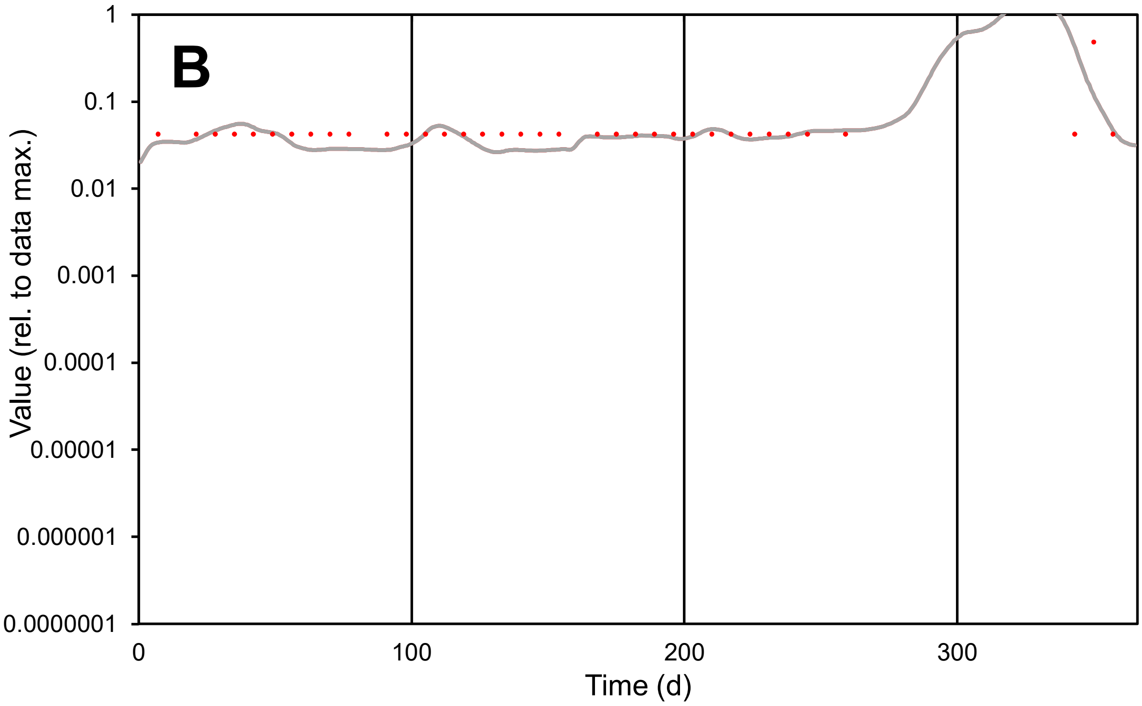


**C1**

| | |
|---|---|
| Variables | |
| POL | Provide jtd or Dst |
| CalibOnly? | |
| 2 | 1 to reproduce of |
| oNormalize? | |
| 2 | 1=no, 2=yes |
| ofxn calc* | |
| 0.89616 | |
| ofxn mod | |
| 0.94443 | |
| sRun | |
| RunA1 | |
| zout | |
| 1 | |
| Ave | |
| Data | |
| 3E-06 | |
| Model | |
| 2.7E-06 | |
| data/vMax | |
| 3.8E-05 | |
| ofxng | |
| 311.734 | |
| Min d/m | |
| 1.55859 | |
| ave/min d | |
| 1.89443 | |
| ave/min m | |
| 2.64493 | |

331_Calib_S_A

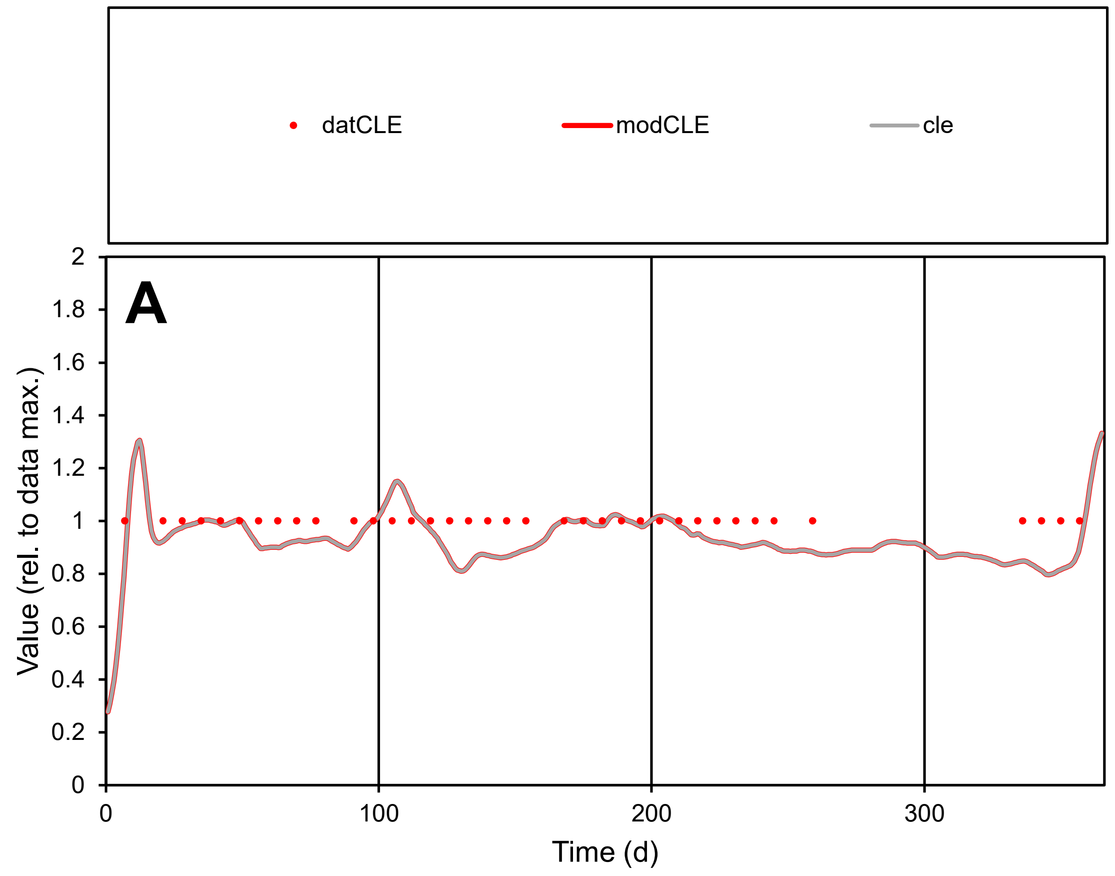


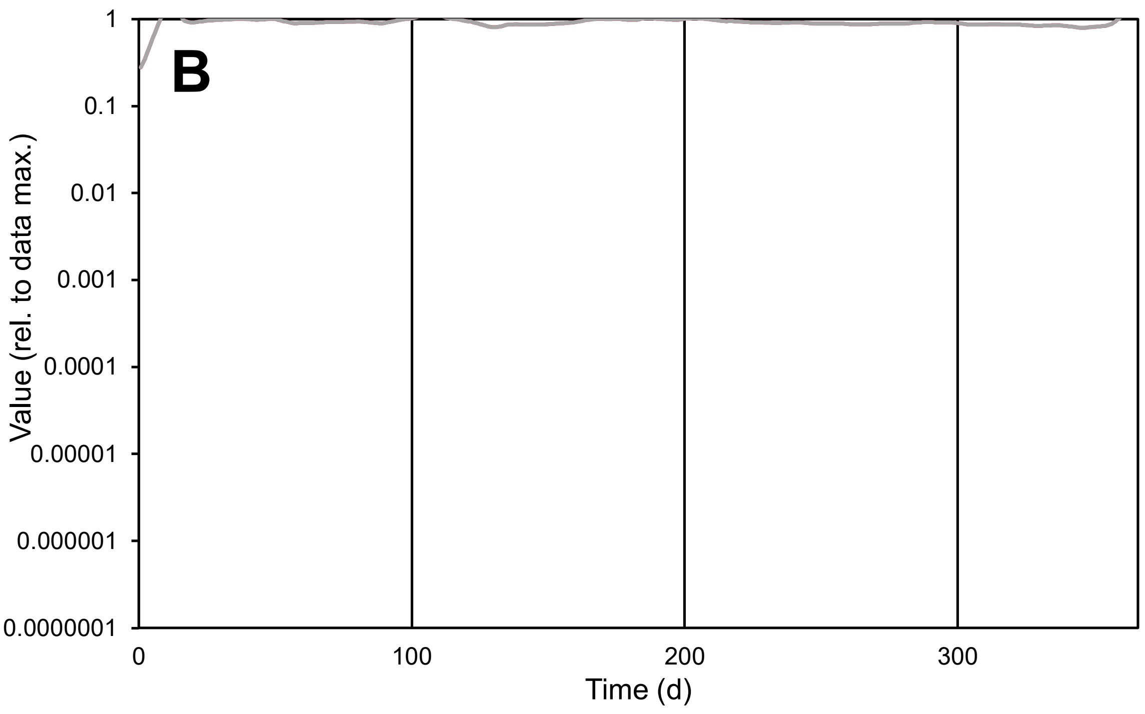


**C1**

Variables
CLE Provide jtd or Dst
CalibOnly?
2 1 to reproduce of
oNormalize?
2 1=no, 2=yes

ofxn calc*
0.0912
ofxn mod
0.1415

sRun
RunA1
zout
1

Ave
Data
4.7E-06
Model
4.4E-06

data/vMax
4.7E-06

ofxng
311.734

Min d/m
1.24463

ave/min d
1
ave/min m
1.17189

331_Calib_S_A

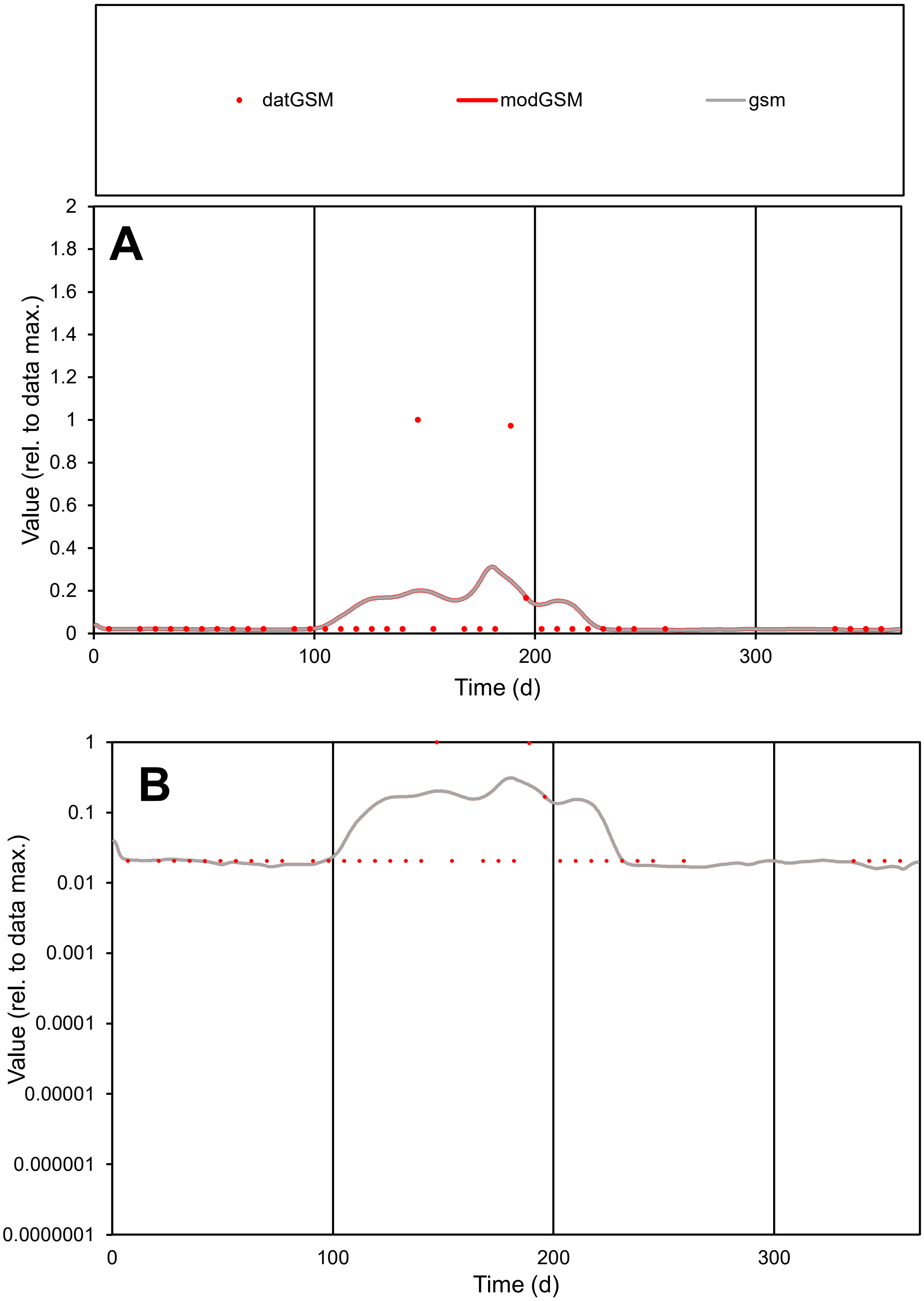


**C1**

Variables
GSM Provide jtd or Dst
CalibOnly?
2 1 to reproduce of
oNormalize?
2 1=no, 2=yes

ofxn calc*
2.57115
ofxn mod
2.61182

sRun
RunA1
zout
1

Ave
Data
5.9E-06
Model
6.3E-06

data/vMax
7.7E-05

ofxng
311.734

Min d/m
1.3082

ave/min d
3.72278
ave/min m
5.19203

331_Calib_S_A

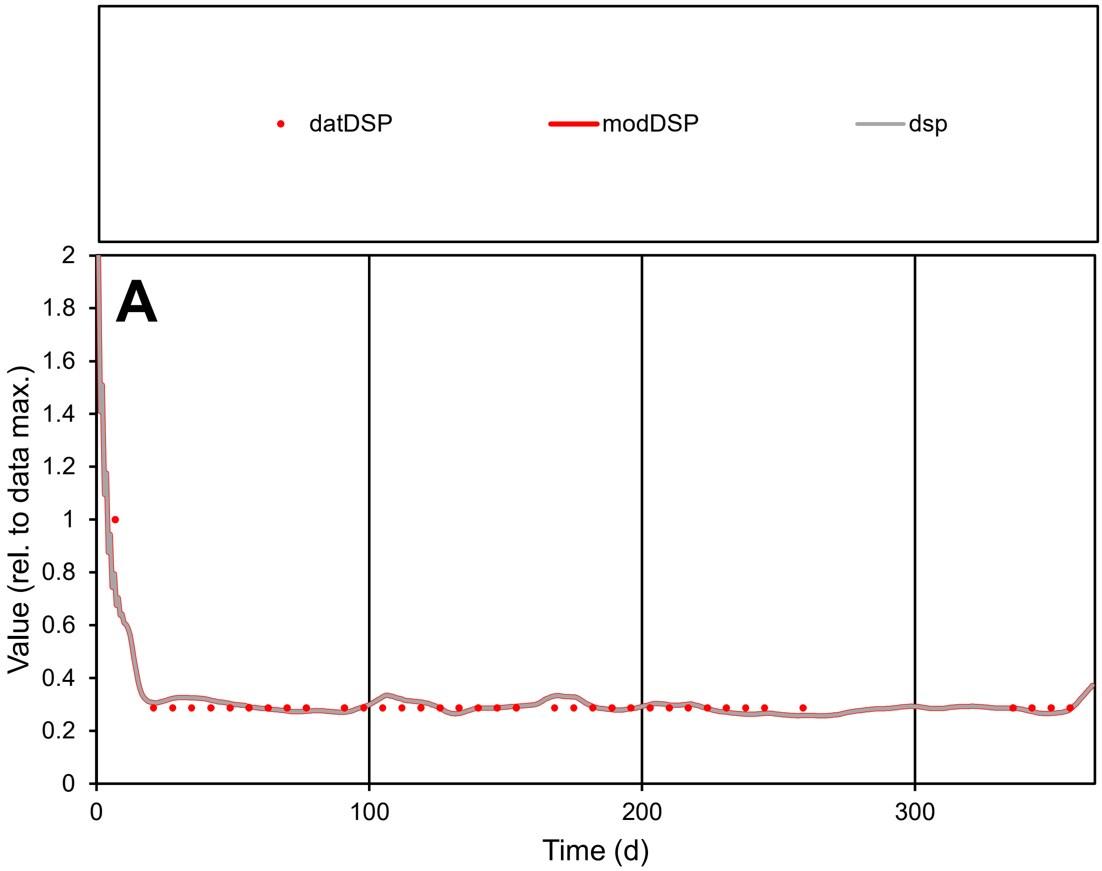


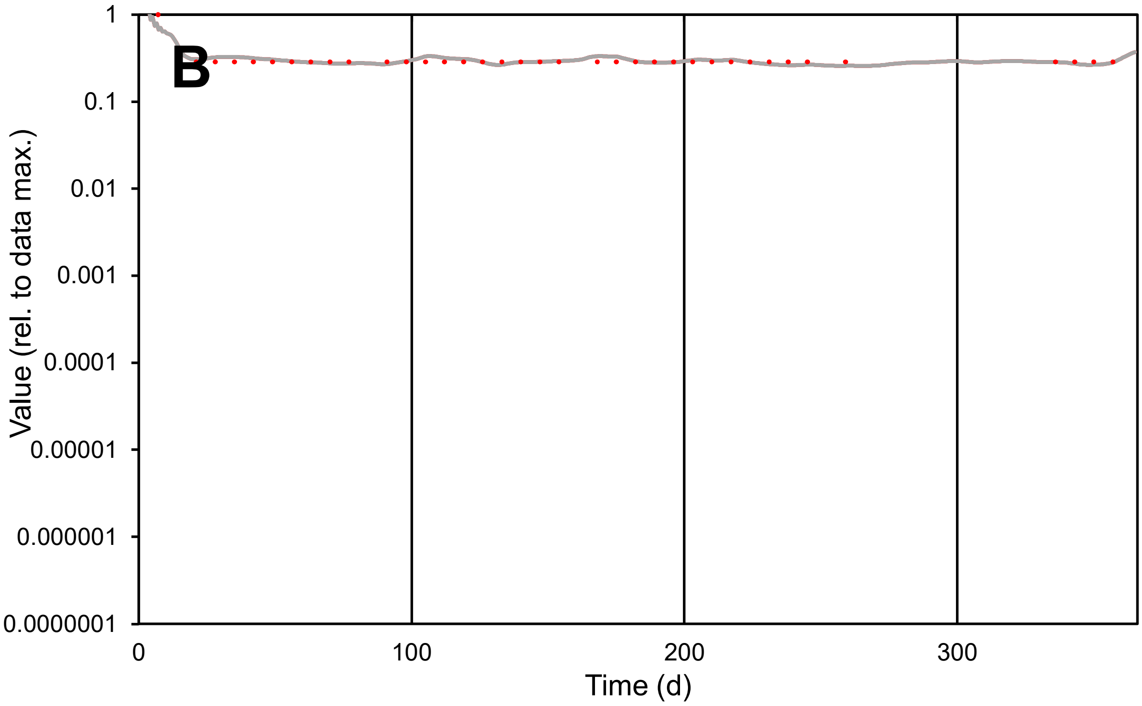


**C1**

Variables
DSP Provide jtd or Dst
CalibOnly?
2 1 to reproduce of
oNormalize?
2 1=no, 2=yes

ofxn calc*
0.17329
ofxn mod
0.20375

sRun
RunA1
zout
1

Ave
Data
5.3E-06
Model
5.3E-06

data/vMax
1.7E-05

ofxng
311.734

Min d/m
1.11198

ave/min d
1.0672
ave/min m
1.17408

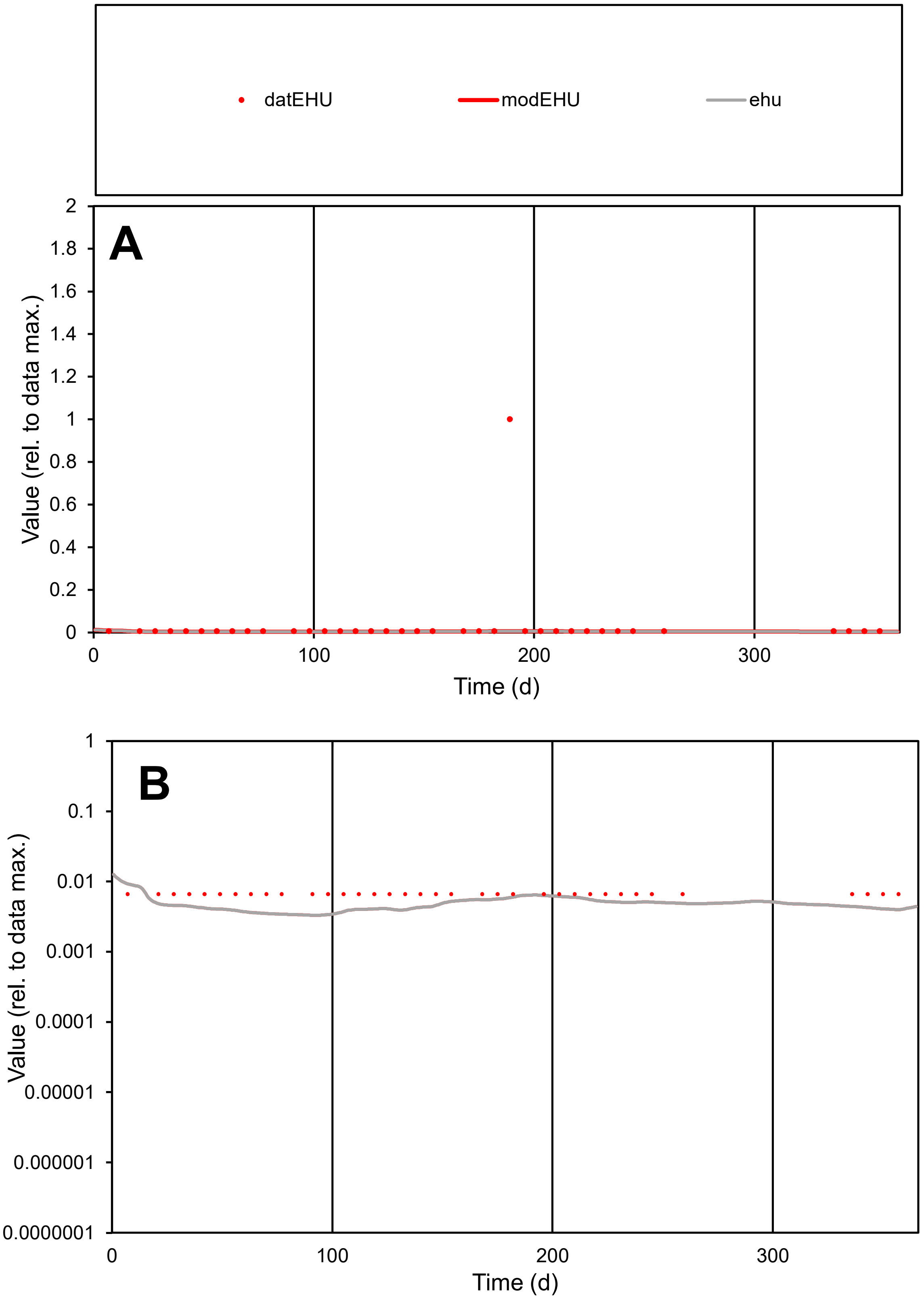


**C1**

Variables
EHU Provide jtd or Dst
CalibOnly?
2 1 to reproduce of
oNormalize?
2 1=no, 2=yes

ofxn calc*
4.88354
ofxn mod
4.89269

sRun
RunA1
zout
1

Ave
Data
2.9E-05
Model
4.1E-06

data/vMax
0.00087

ofxng
311.734

Min d/m
1.98997

ave/min d
5.06624
ave/min m
1.43227

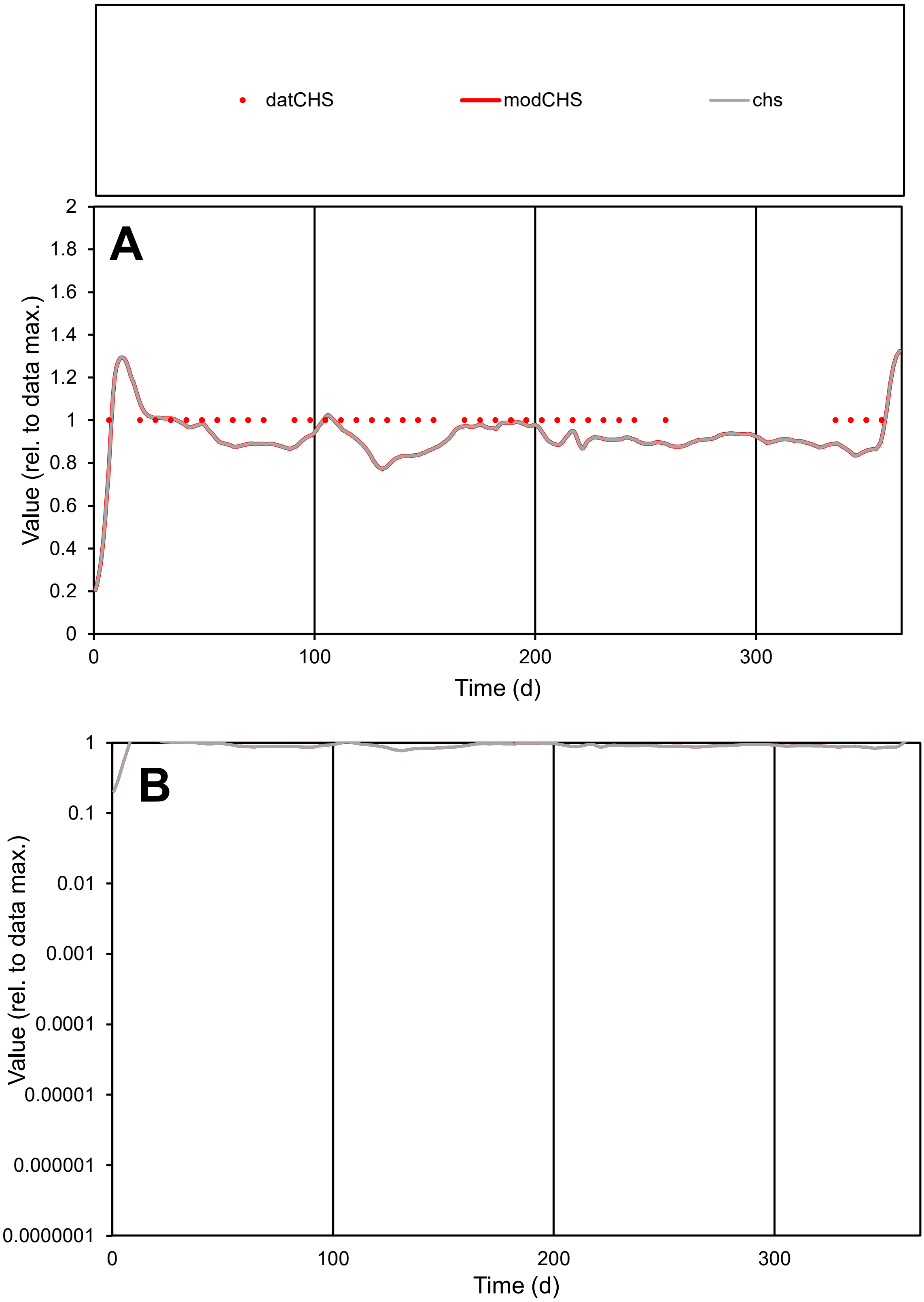


**C1**

Variables

CHS Provide jtd or Dst

CalibOnly?

2 1 to reproduce of

oNormalize?

2 1=no, 2=yes

ofxn calc*

0.09828

ofxn mod

0.14549

sRun

RunA1

zout

1

Ave

Data

2.1E-05

Model

2E-05

data/vMax

2.1E-05

ofxng

311.734

Min d/m

1.24796

ave/min d

1

ave/min m

1.15275

331_Calib_S_A

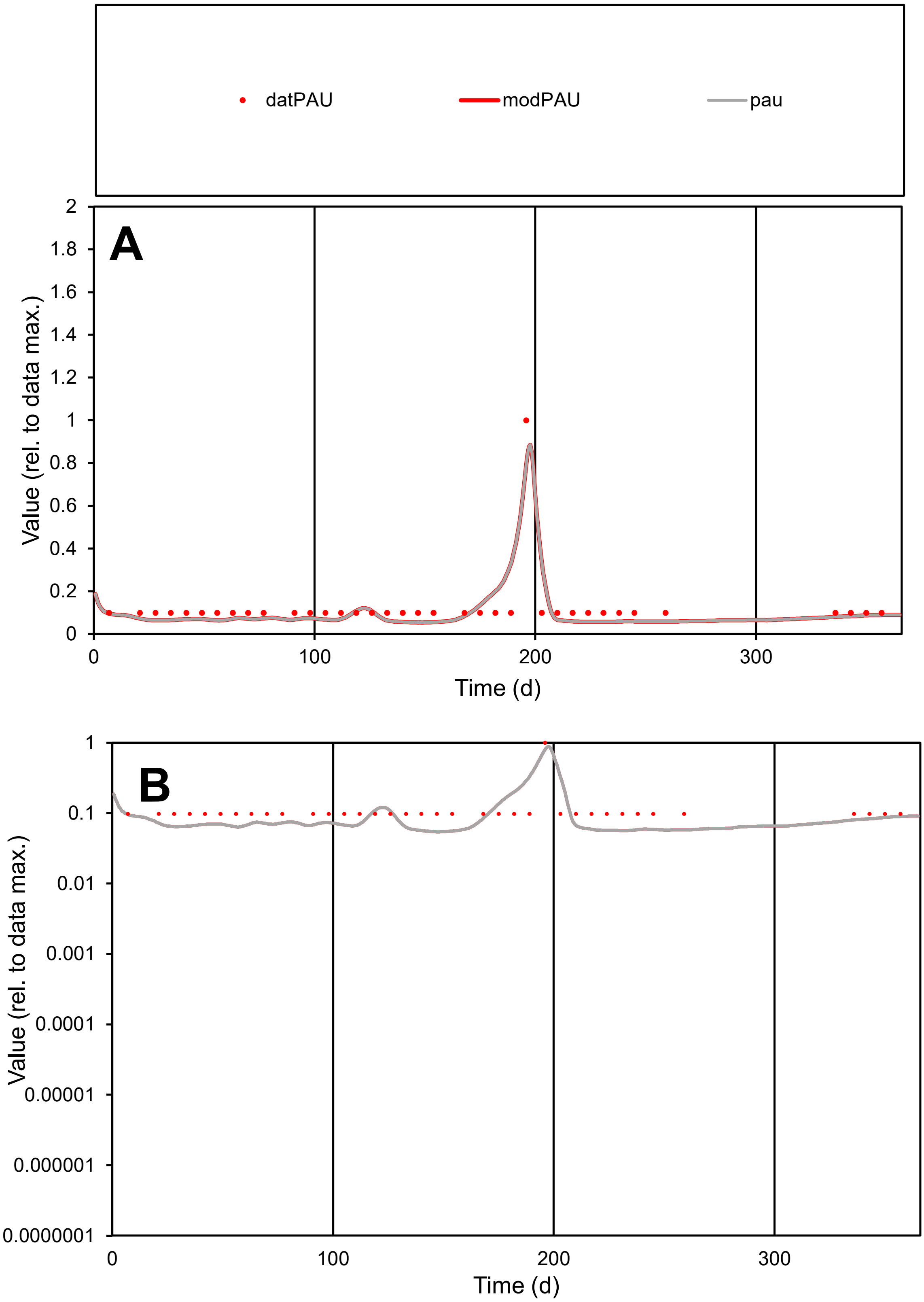


**C1**

Variables
PAU Provide jtd or Dst
CalibOnly?
2 1 to reproduce of
oNormalize?
2 1=no, 2=yes

ofxn calc*
0.45132
ofxn mod
0.4989

sRun
RunA1
zout
1

Ave
Data
4.9E-06
Model
4.4E-06

data/vMax
4E-05

ofxng
311.734

Min d/m
1.80375

ave/min d
1.24914
ave/min m
1.99812

331_Calib_S_A

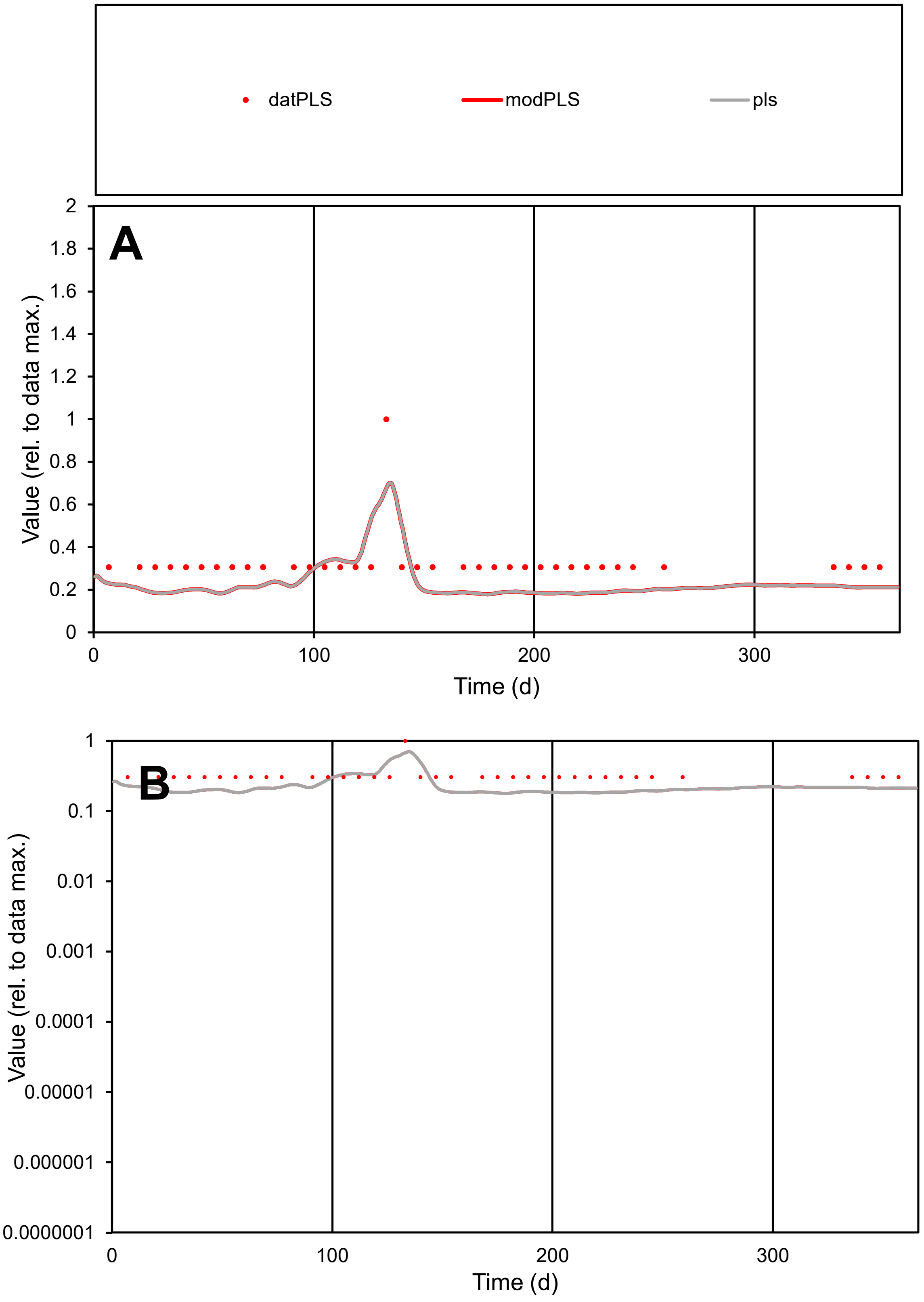


**C1**

Variables
PLS Provide jtd or Dst
CalibOnly?
2 1 to reproduce of
oNormalize?
2 1=no, 2=yes

ofxn calc*
0.36493
ofxn mod
0.38079

sRun
RunA1
zout
1

Ave
Data
4.1E-06
Model
3E-06

data/vMax
1.3E-05

ofxng
311.734

Min d/m
1.69207

ave/min d
1.06142
ave/min m
1.34178

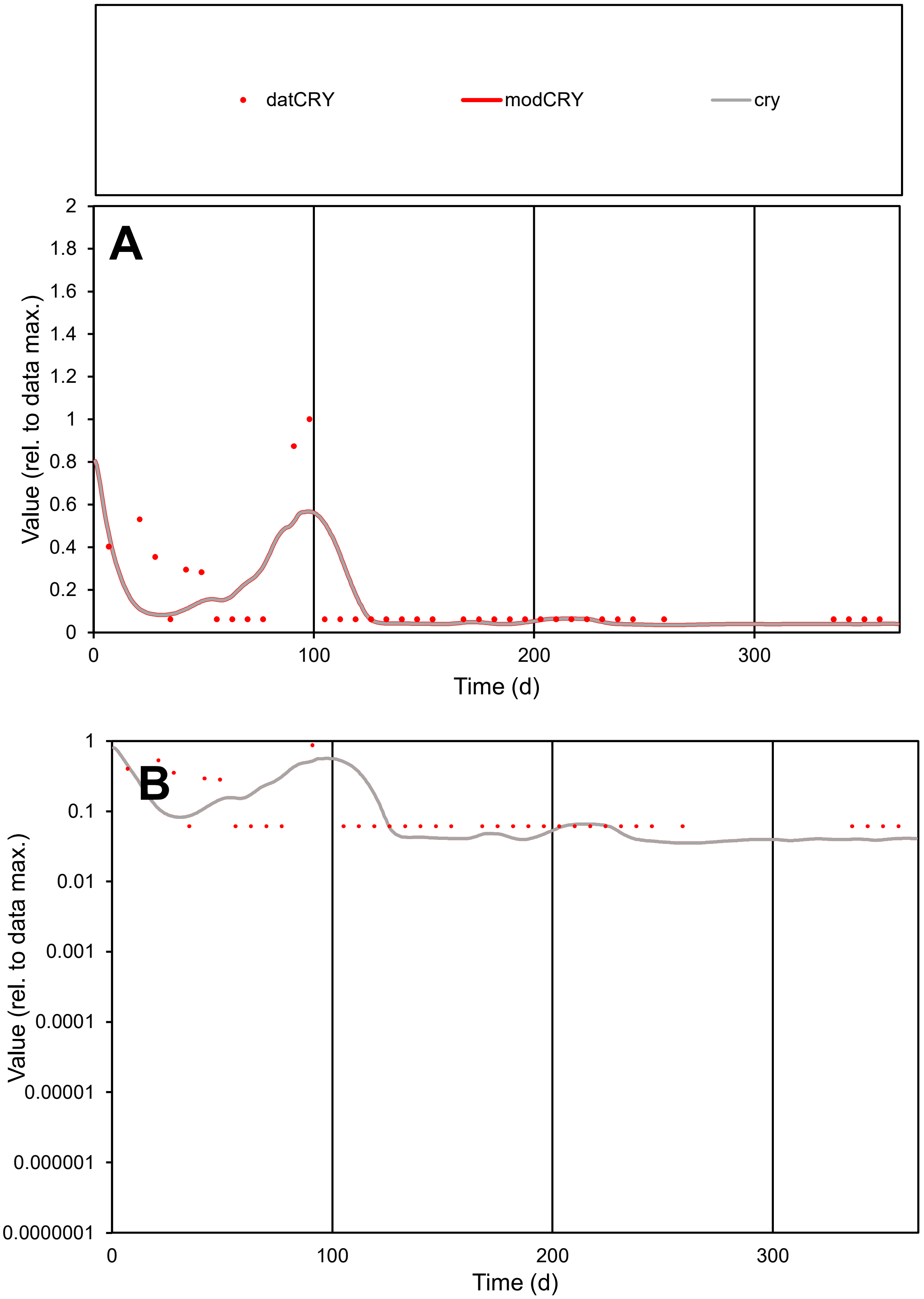


**C1**

Variables
CRY Provide jtd or Dst
CalibOnly?
2 1 to reproduce of
oNormalize?
2 1=no, 2=yes

ofxn calc*
1.06147
ofxn mod
1.09056

sRun
RunA1
zout
1

Ave
Data
7.3E-06
Model
6.4E-06

data/vMax
4.9E-05

ofxng
311.734

Min d/m
1.73706

ave/min d
2.4556
ave/min m
3.71454

331_Calib_S_A

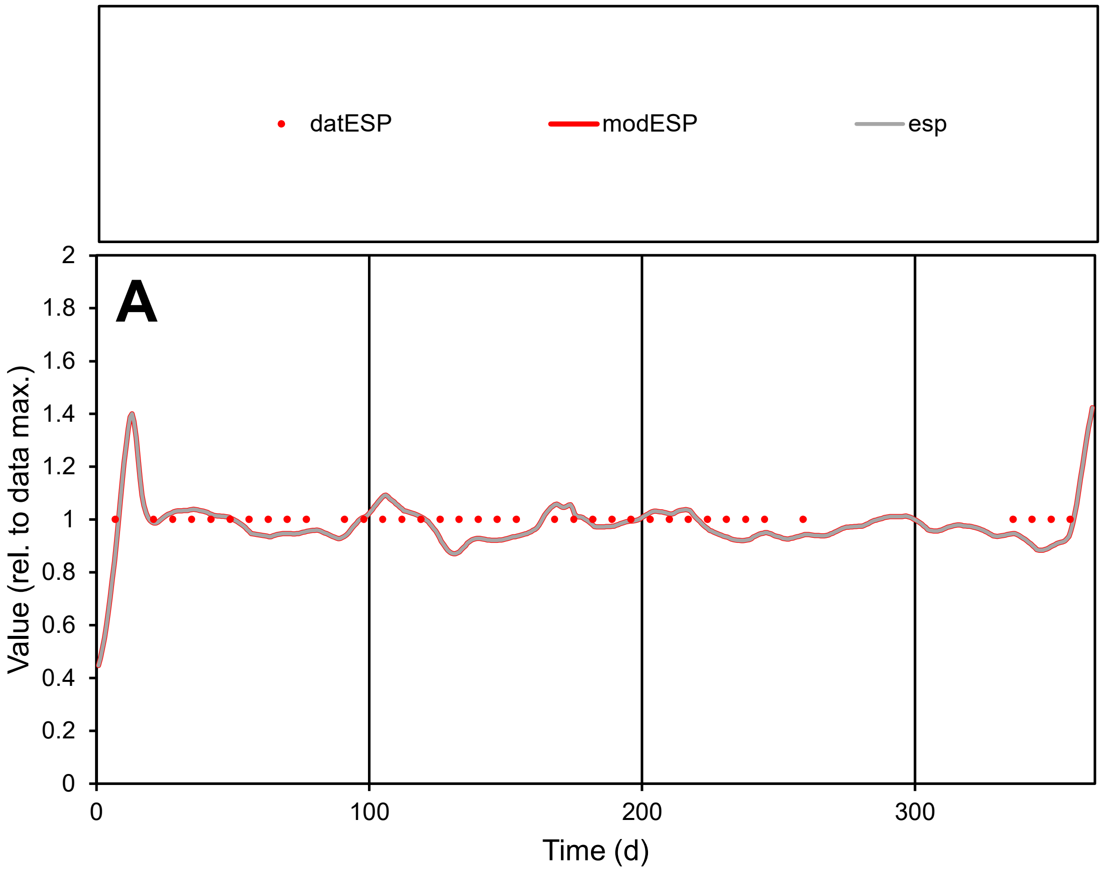


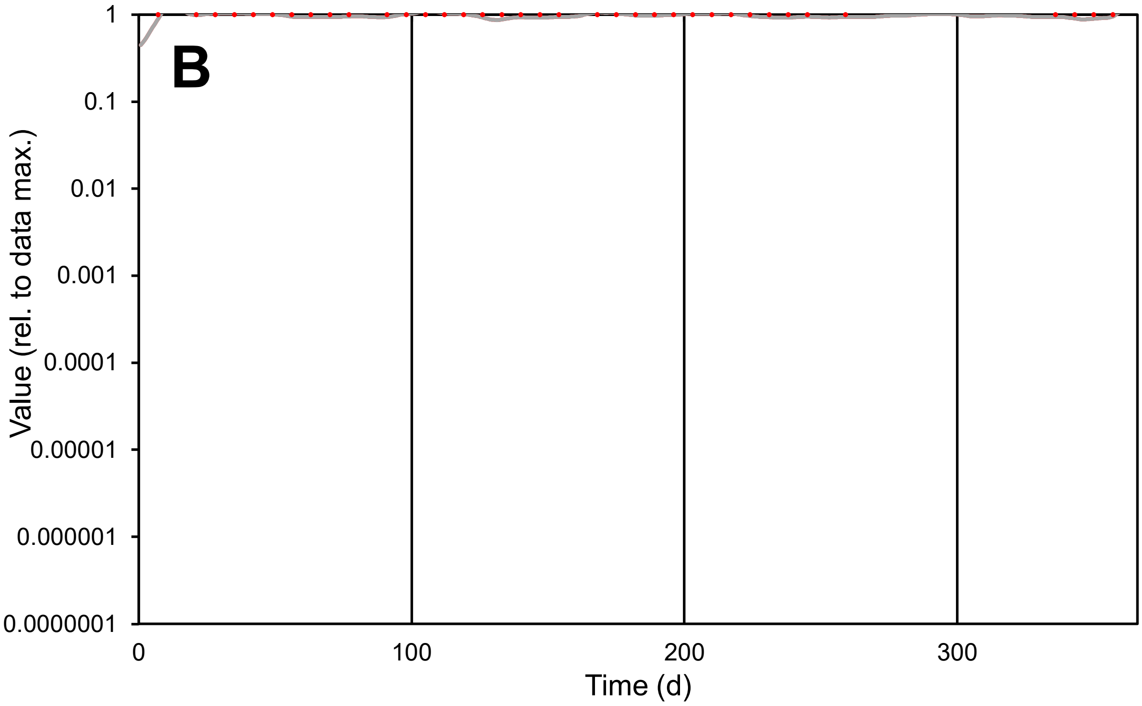


**C1**

Variables
ESP Provide jtd or Dst
CalibOnly?
2 1 to reproduce of
oNormalize?
2 1=no, 2=yes

ofxn calc*
0.05473
ofxn mod
0.12297

sRun
RunA1
zout
1

Ave
Data
8E-06
Model
7.8E-06

data/vMax
8E-06

ofxng
311.734

Min d/m
1.11941

ave/min d
1
ave/min m
1.09201

331_Calib_S_A

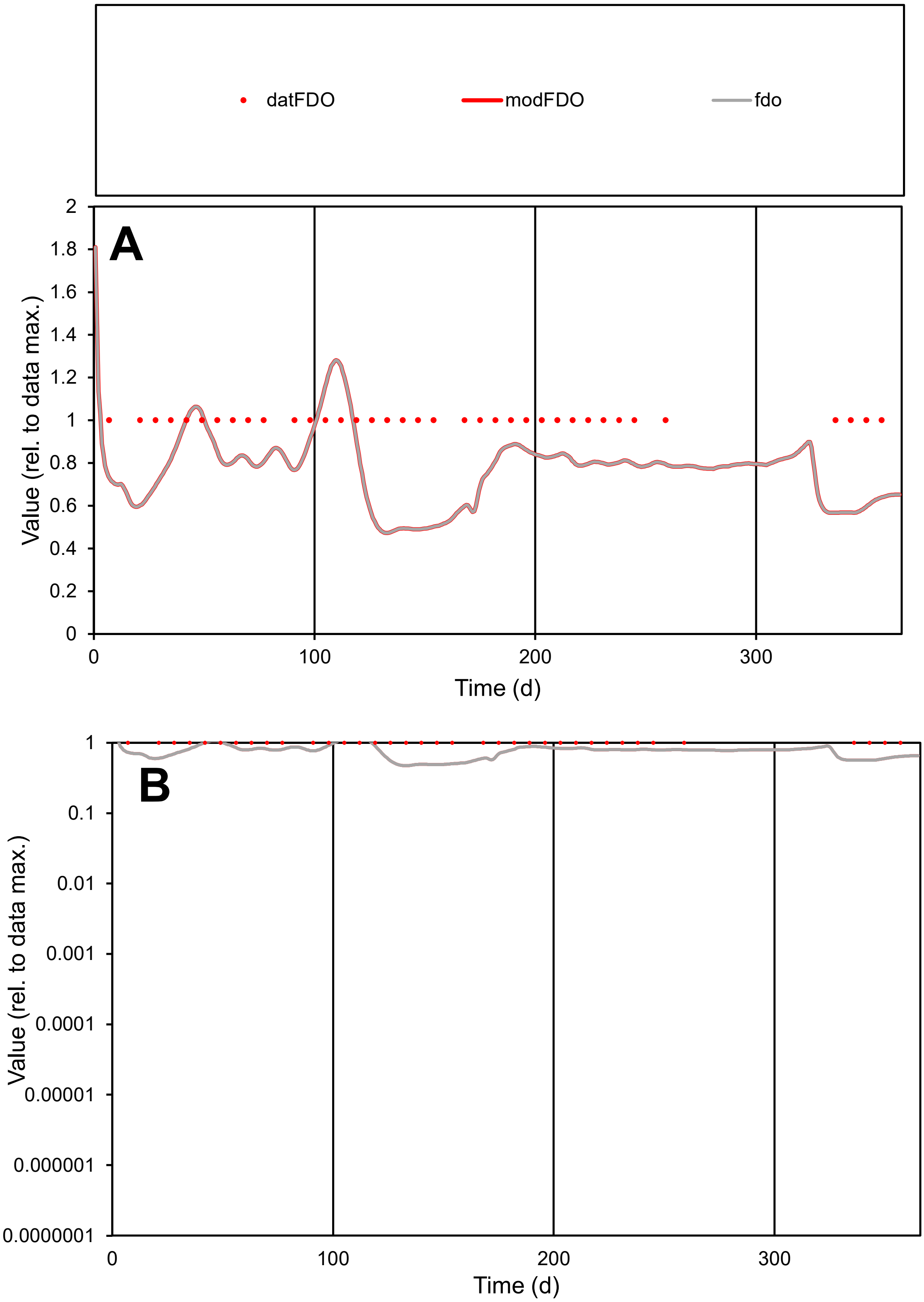


**C1**

Variables
FDO Provide jtd or Dst
CalibOnly?
2 1 to reproduce of
oNormalize?
2 1=no, 2=yes

ofxn calc*
0.29057
ofxn mod
0.30774

sRun
RunA1
zout
1

Ave
Data
1E-07
Model
7.7E-08

data/vMax
1E-07

ofxng
311.734

Min d/m
2.09924

ave/min d
1
ave/min m
1.61414

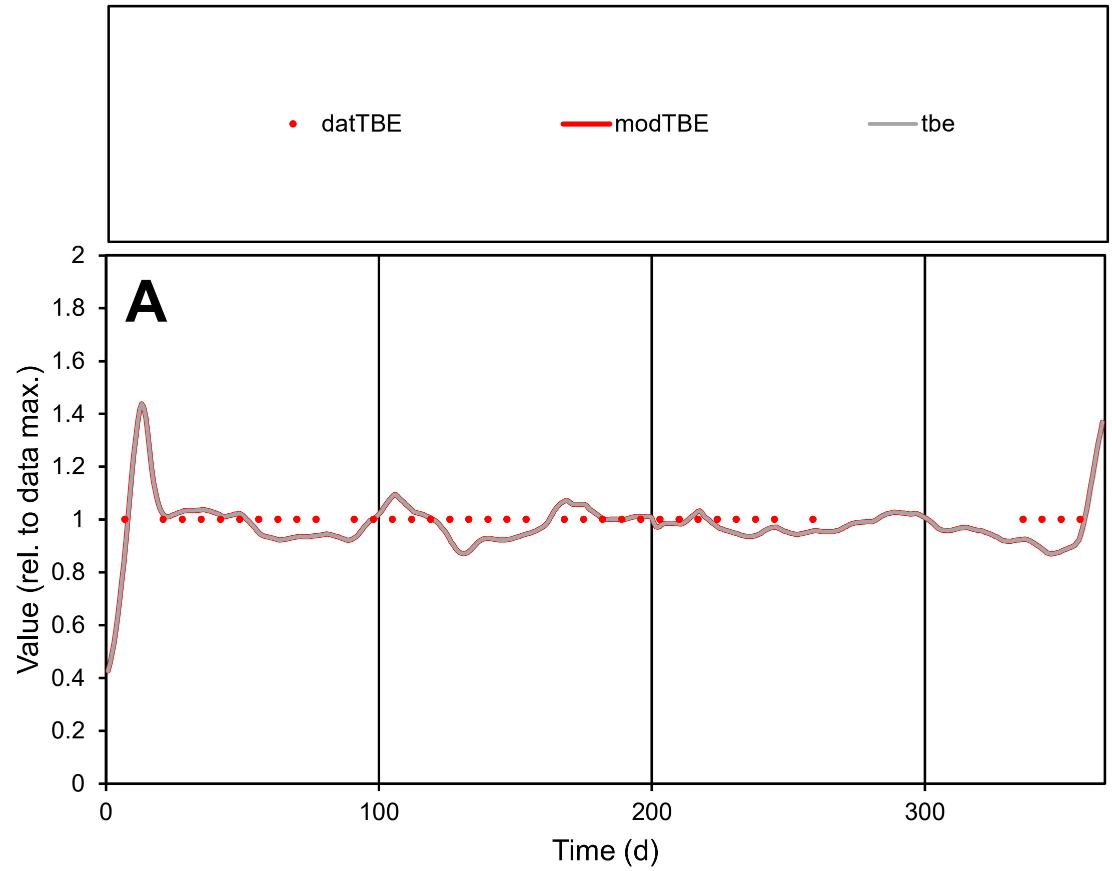


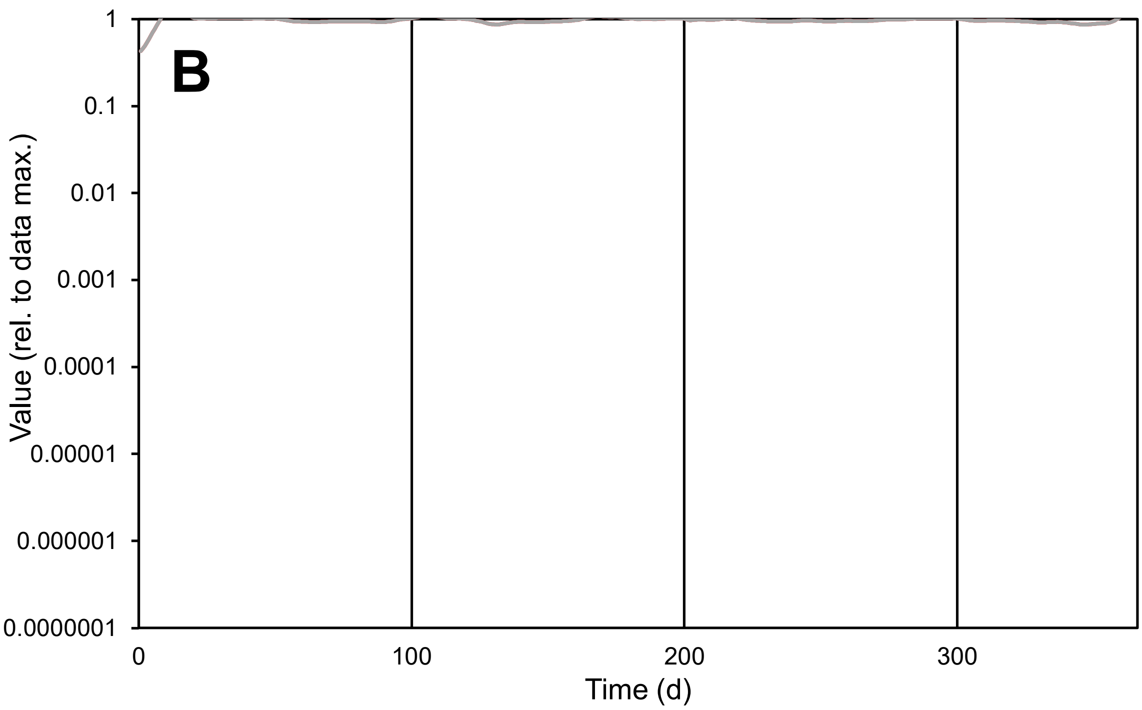


**C1**

Variables
TBE Provide jtd or Dst
CalibOnly?
2 1 to reproduce of
oNormalize?
2 1=no, 2=yes

ofxn calc*
0.05671
ofxn mod
0.12384

sRun
RunA1
zout
1

Ave
Data
6.7E-06
Model
6.5E-06

data/vMax
6.7E-06

ofxng
311.734

Min d/m
1.13598

ave/min d
1
ave/min m
1.10937

331_Calib_S_A

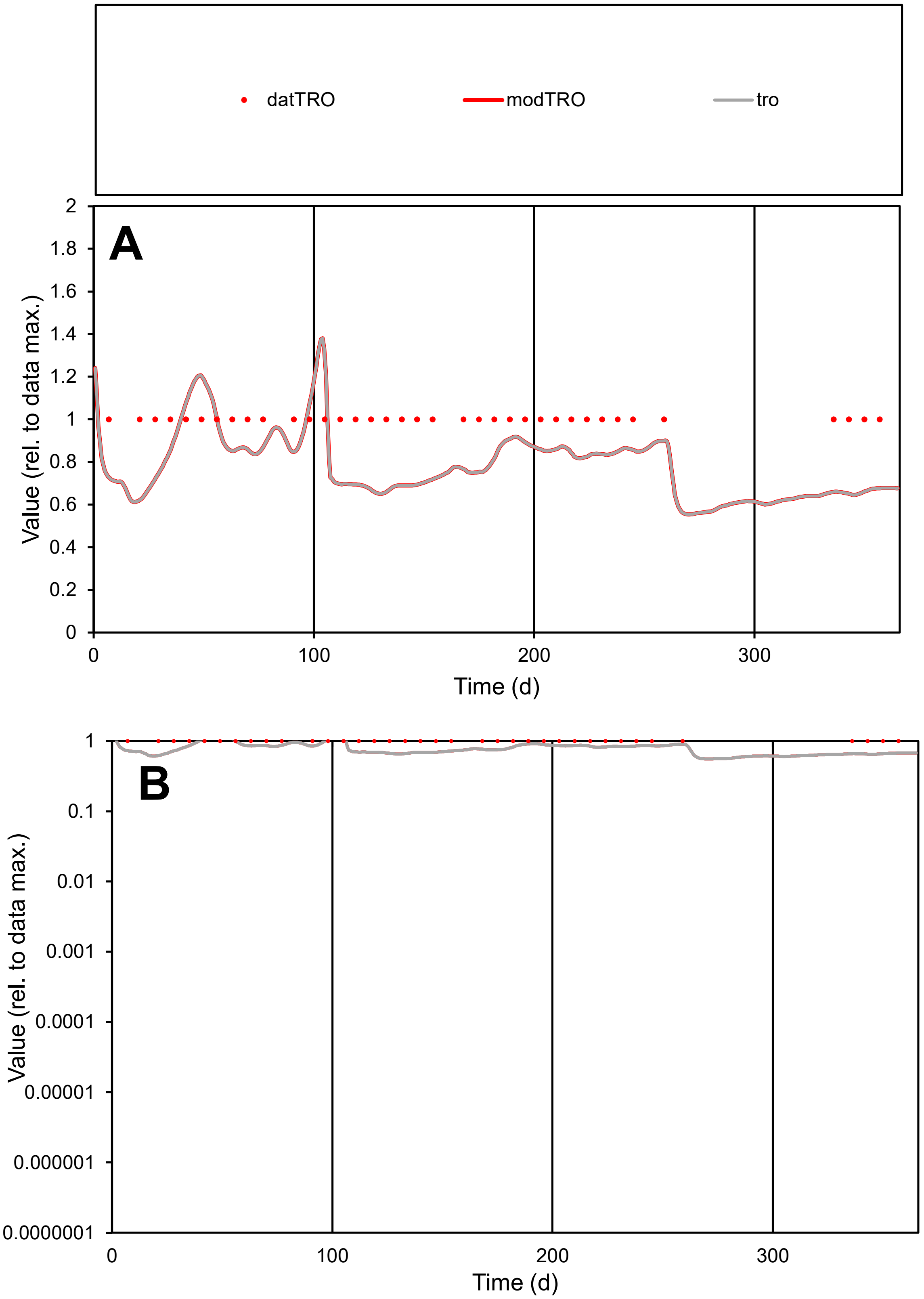


**C1**

Variables
TRO Provide jtd or Dst
CalibOnly?
2 1 to reproduce of
oNormalize?
2 1=no, 2=yes

ofxn calc*
0.22716
ofxn mod
0.24944

sRun
RunA1
zout
1

Ave
Data
5E-06
Model
4.1E-06

data/vMax
5E-06

ofxng
311.734

Min d/m
1.57125

ave/min d
1
ave/min m
1.28938

331_Calib_S_A

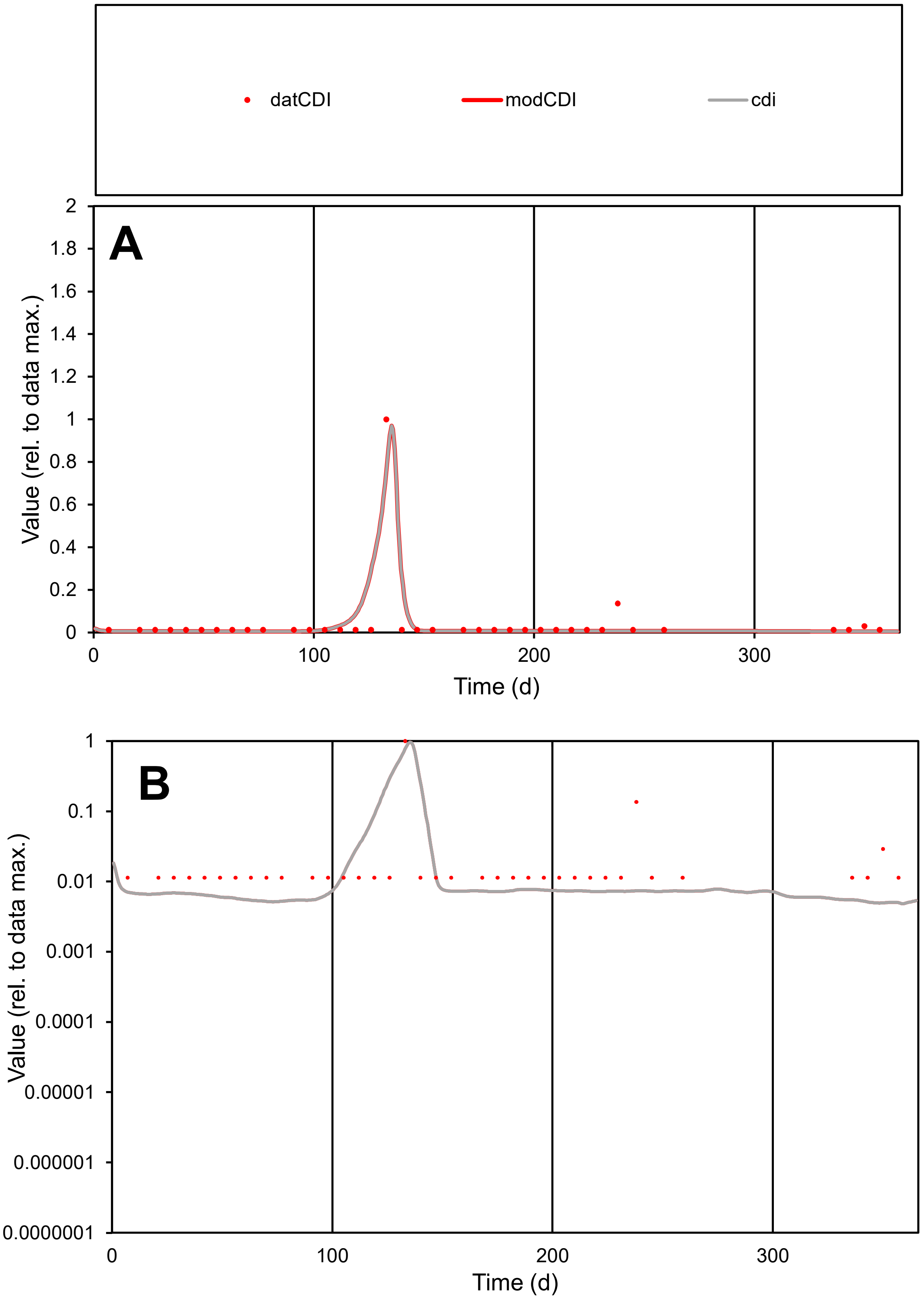


**C1**

Variables
CDI Provide jtd or Dst
CalibOnly?
2 1 to reproduce of
oNormalize?
2 1=no, 2=yes

ofxn calc*
1.08836
ofxn mod
1.29371

sRun
RunA1
zout
1

Ave
Data
6.4E-07
Model
6.4E-07

data/vMax
1.5E-05

ofxng
311.734

Min d/m
2.33549

ave/min d
3.70033
ave/min m
8.65951

331_Calib_S_A

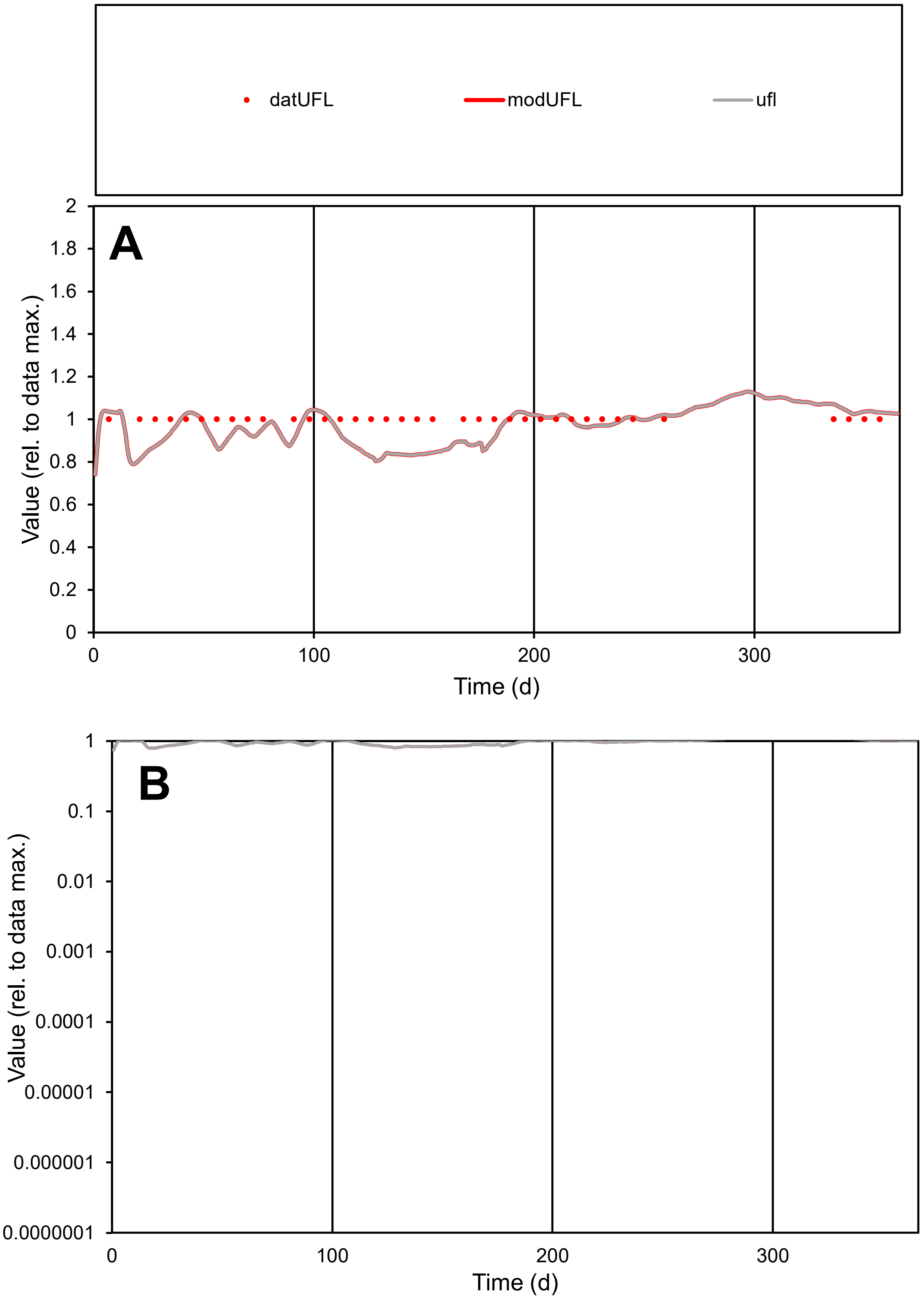


**C1**

Variables
UFL Provide jtd or Dst
CalibOnly?
2 1 to reproduce of
oNormalize?
2 1=no, 2=yes

ofxn calc*
0.08826
ofxn mod
0.14047

sRun
RunA1
zout
1

Ave
Data
3.1E-06
Model
2.9E-06

data/vMax
3.1E-06

ofxng
311.734

Min d/m
1.22333

ave/min d
1
ave/min m
1.16527

331_Calib_S_A

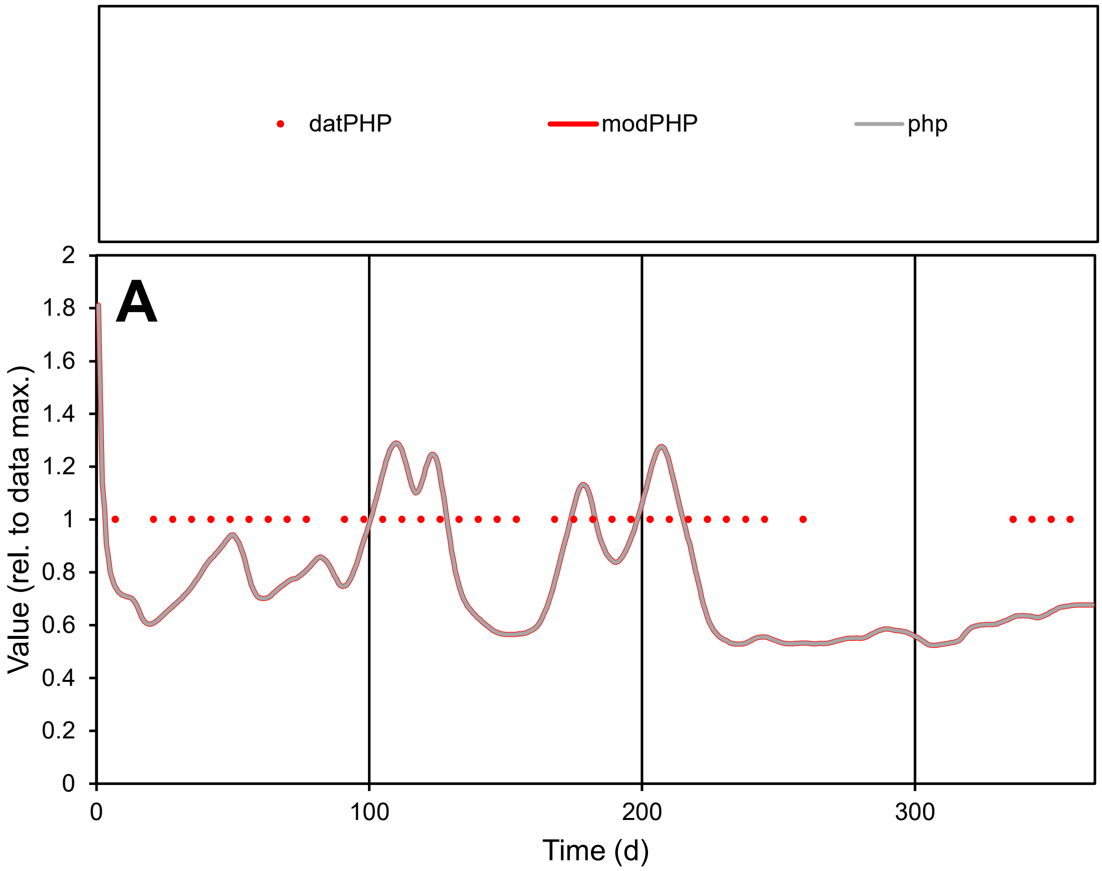


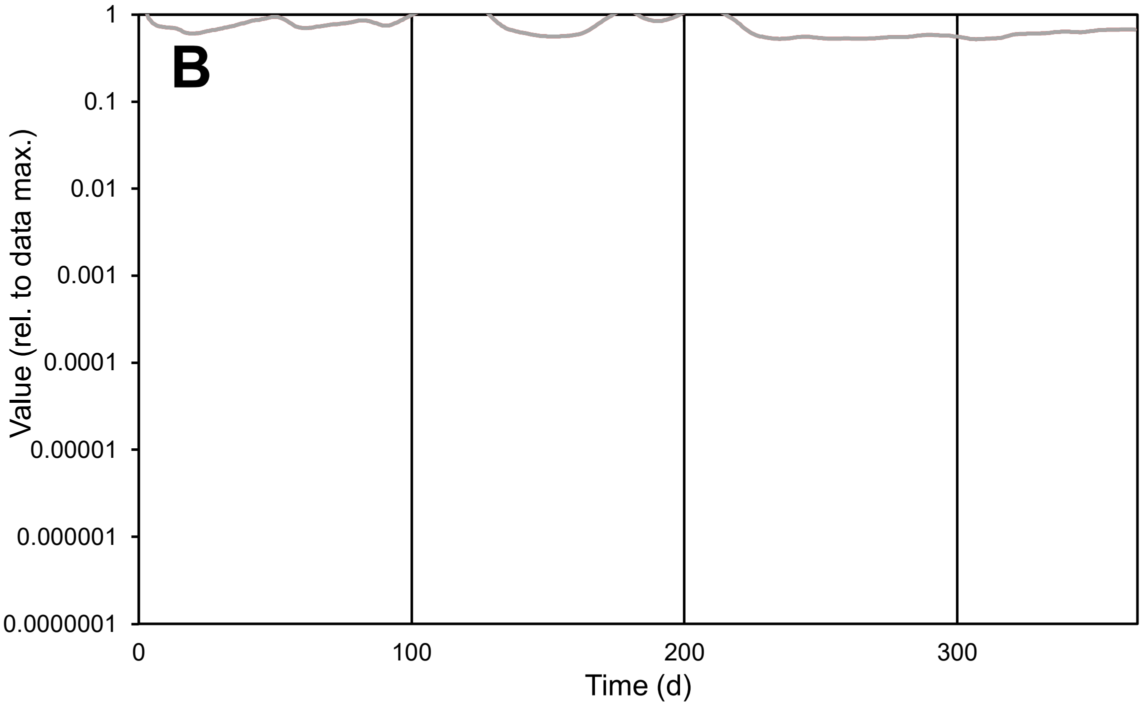


**C1**

Variables

| | |
|---|---|
| PHP | Provide jtd or Dst |
| CalibOnly? | |
| 2 | 1 to reproduce of |
| oNormalize? | |
| 2 | 1=no, 2=yes |
| ofxn calc* | |
| 0.2874 | |
| ofxn mod | |
| 0.30481 | |
| sRun | |
| RunA1 | |
| zout | |
| 1 | |
| Ave | |
| Data | |
| 6.9E-06 | |
| Model | |
| 5.6E-06 | |
| data/vMax | |
| 6.9E-06 | |
| ofxng | |
| 311.734 | |
| Min d/m | |
| 1.88052 | |
| ave/min d | |
| 1 | |
| ave/min m | |
| 1.51586 | |

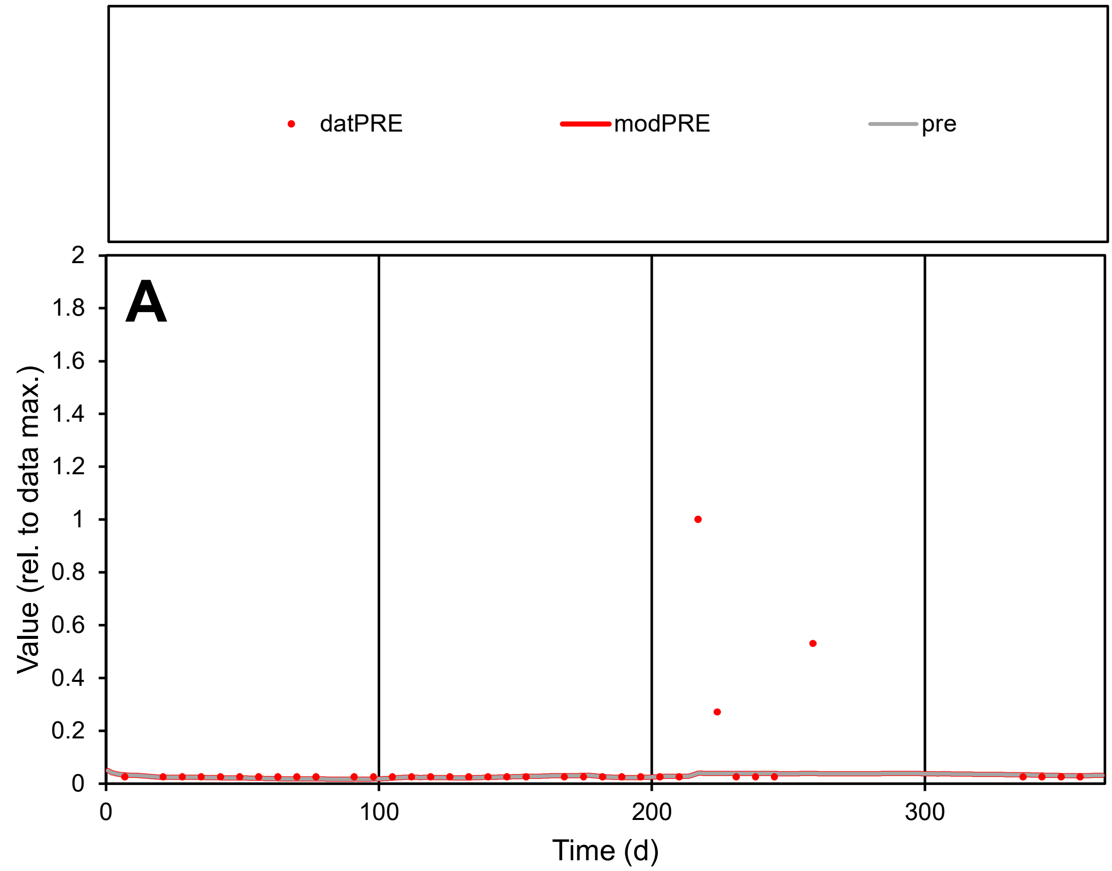


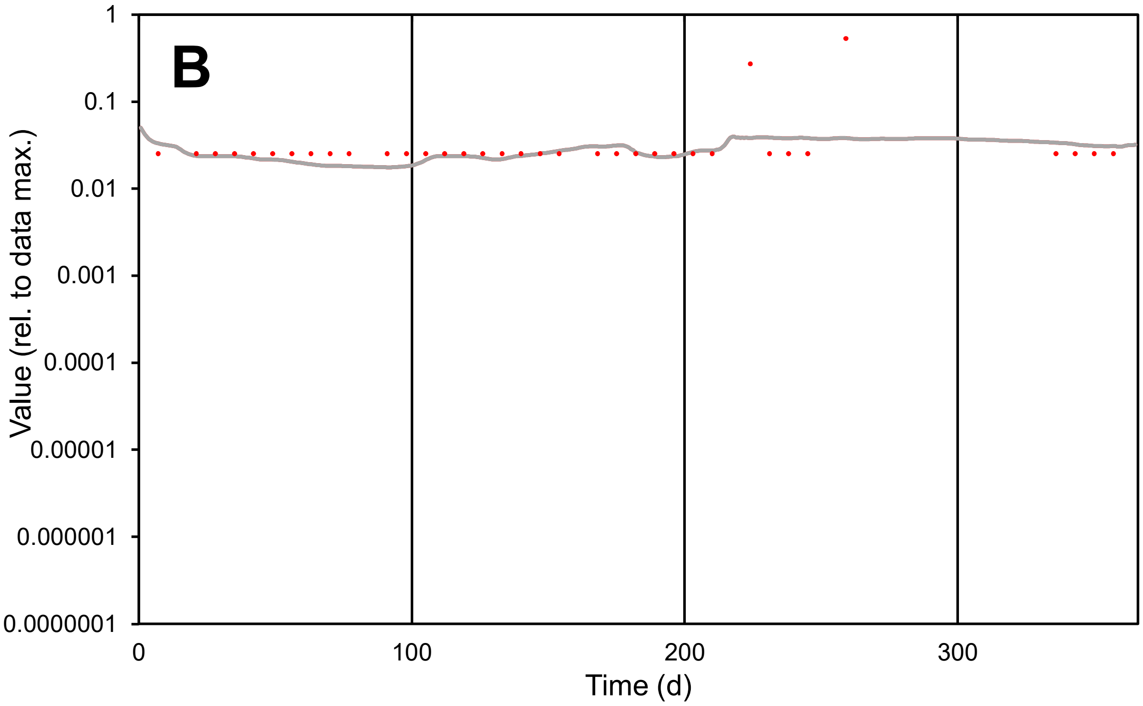


**C1**

Variables
PRE Provide jtd or Dst
CalibOnly?
2 1 to reproduce of
oNormalize?
2 1=no, 2=yes

ofxn calc*
2.52263
ofxn mod
2.52782

sRun
RunA1
zout
1

Ave
Data
3.8E-05
Model
1.4E-05

data/vMax
0.00053

ofxng
311.734

Min d/m
1.44161

ave/min d
2.83798
ave/min m
1.52811

331_Calib_S_A

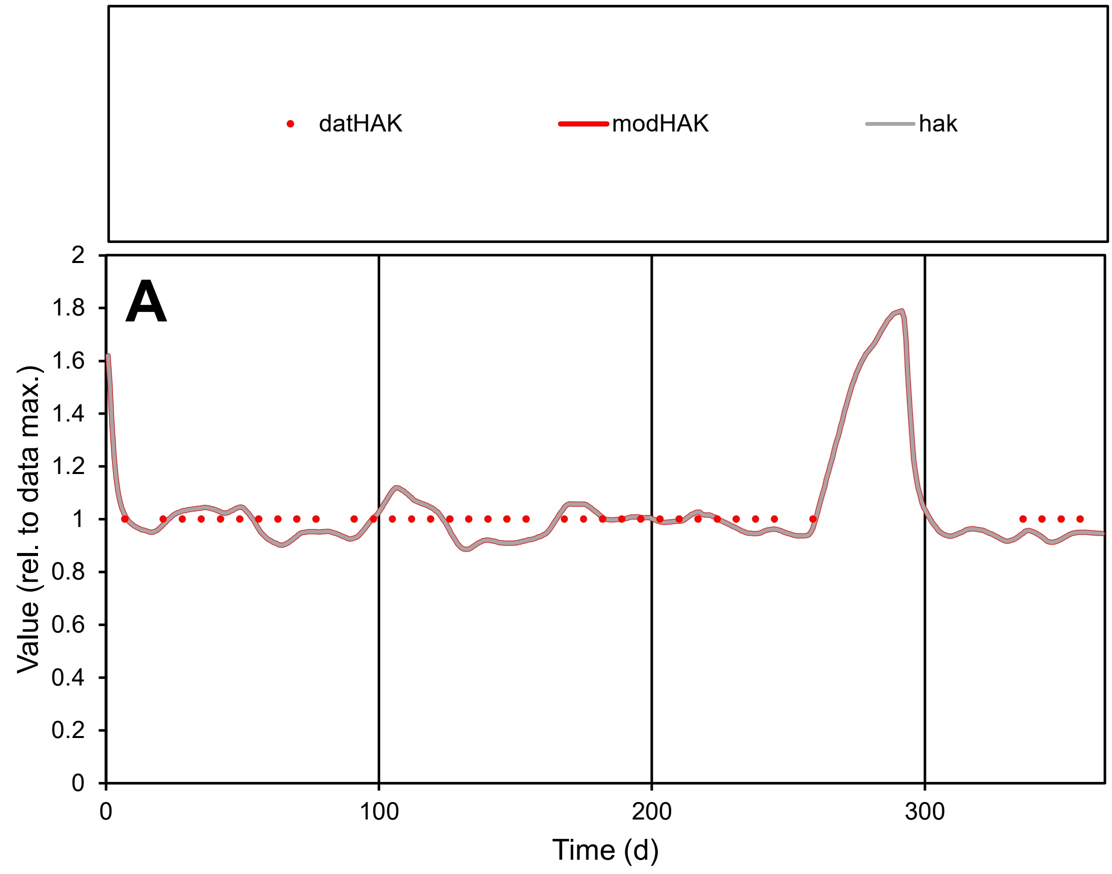


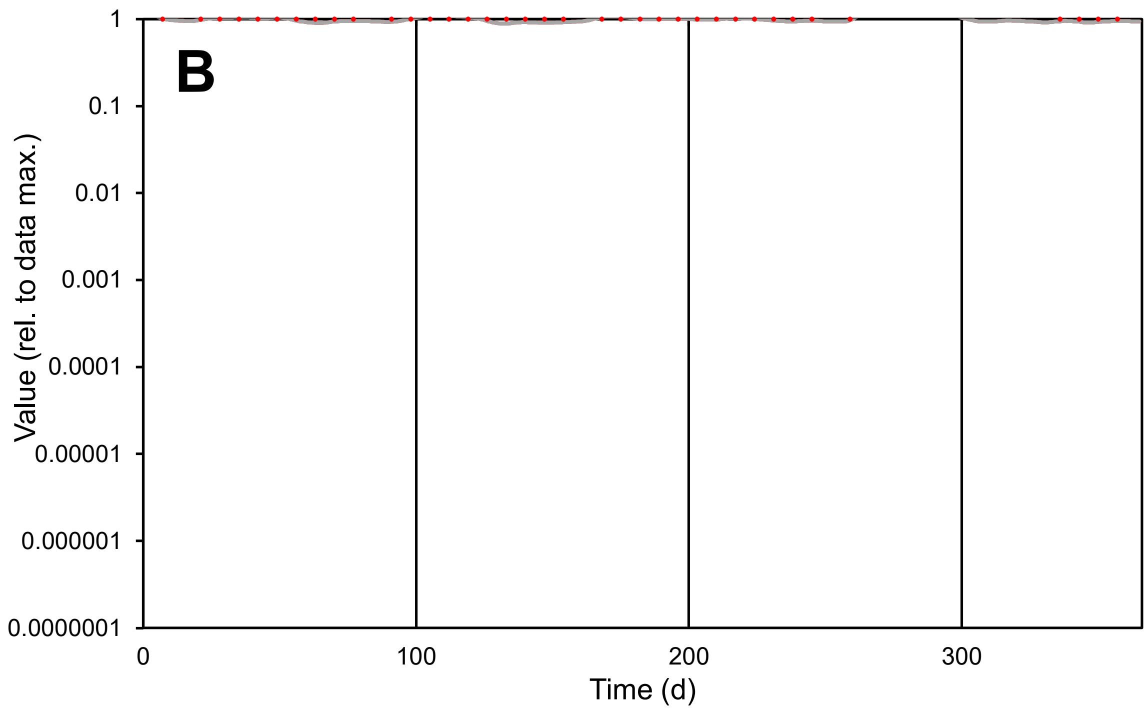


**C1**

Variables
HAK Provide jtd or Dst
CalibOnly?
2 1 to reproduce of
oNormalize?
2 1=no, 2=yes

ofxn calc*
0.055
ofxn mod
0.12338

sRun
RunA1
zout
1

Ave
Data
5.1E-06
Model
5E-06

data/vMax
5.1E-06

ofxng
311.734

Min d/m
1.11546

ave/min d
1
ave/min m
1.09803

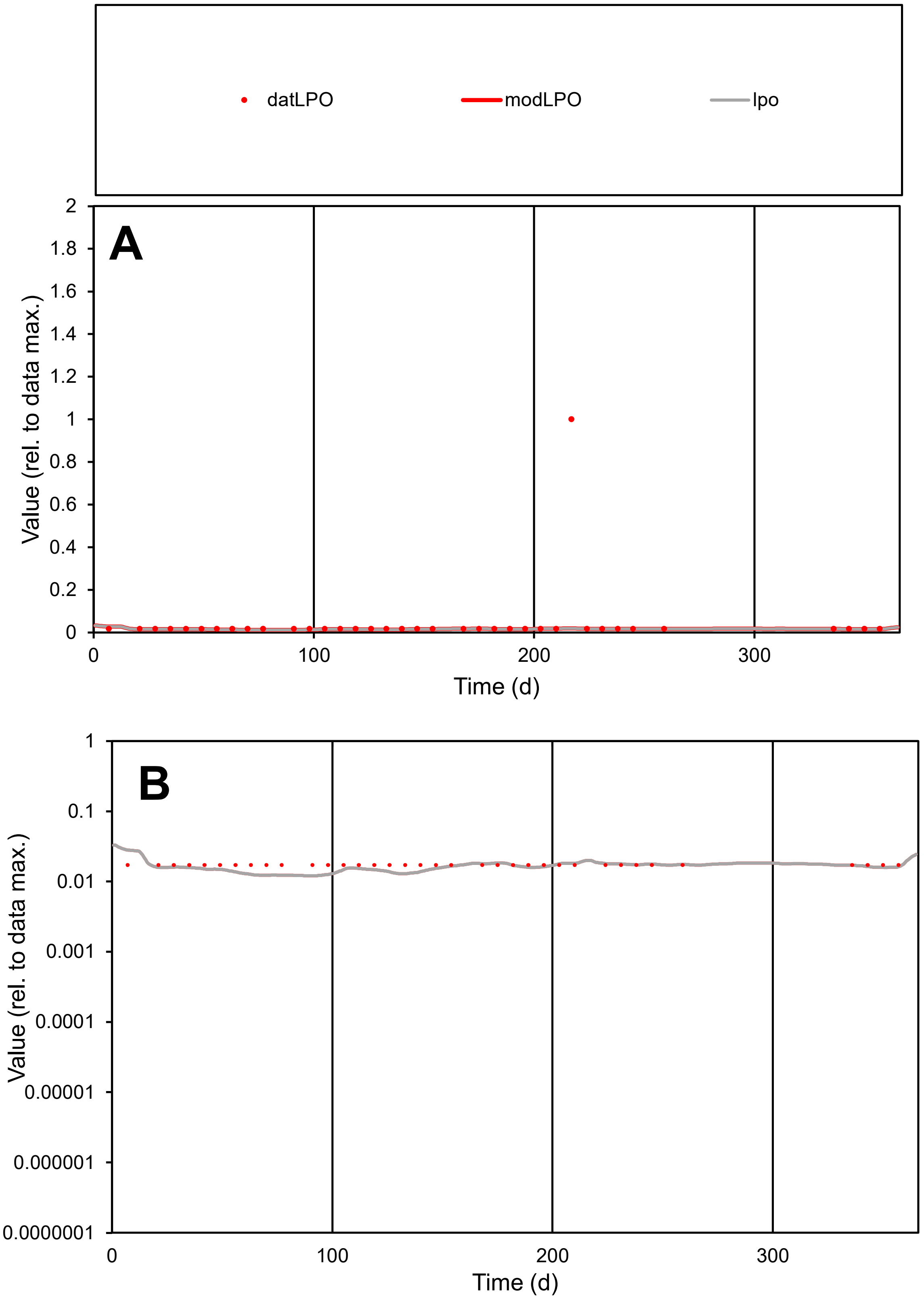


**C1**

Variables
LPO Provide jtd or Dst
CalibOnly?
2 1 to reproduce of
oNormalize?
2 1=no, 2=yes

ofxn calc*
3.68022
ofxn mod
3.68548

sRun
RunA1
zout
1

Ave
Data
1.5E-05
Model
5.5E-06

data/vMax
0.00034

ofxng
311.734

Min d/m
1.42412

ave/min d
2.54032
ave/min m
1.3264

331_Calib_S_A

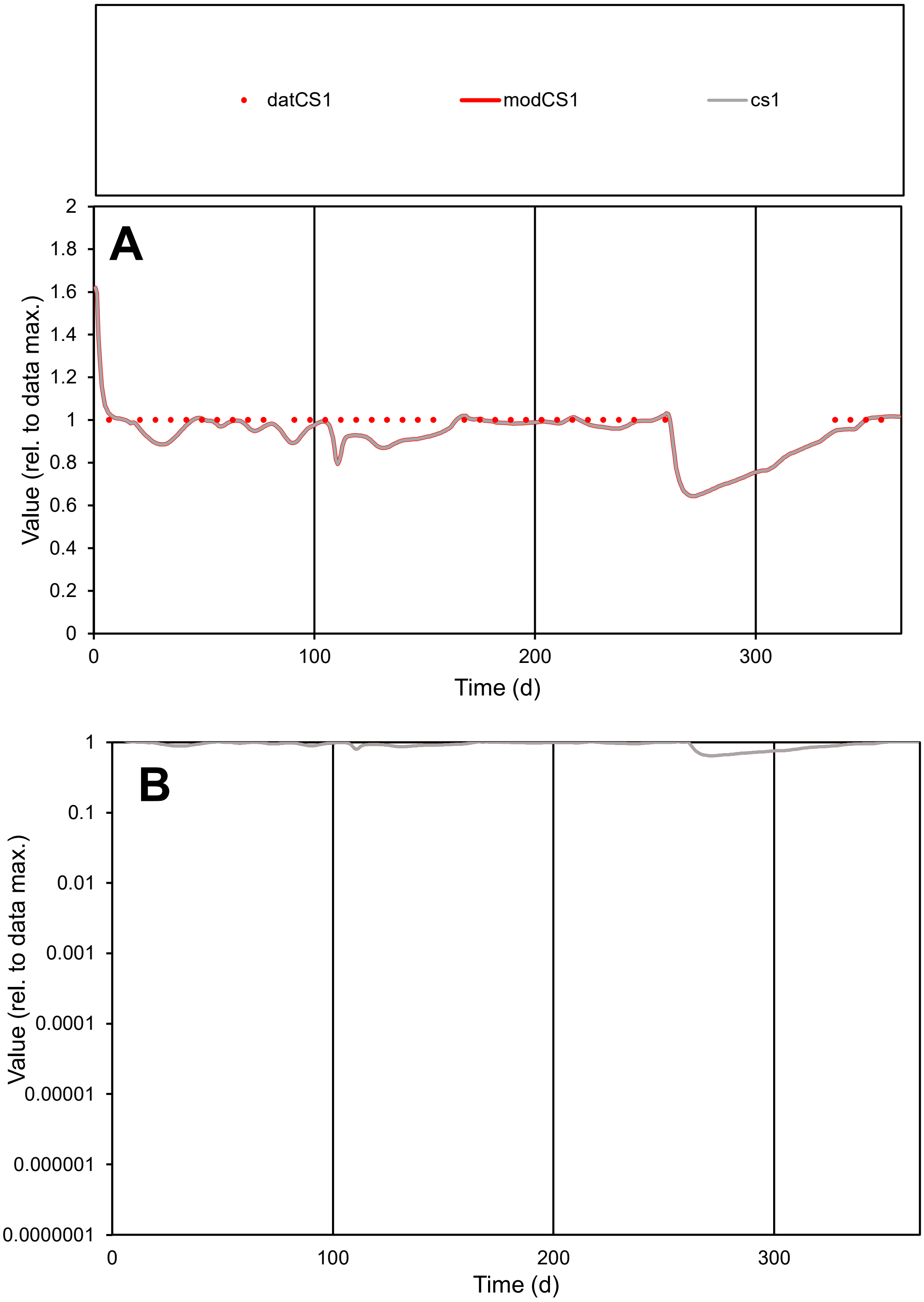


**C1**

Variables
CS1 Provide jtd or Dst
CalibOnly?
2 1 to reproduce of
oNormalize?
2 1=no, 2=yes

ofxn calc*
0.0541
ofxn mod
0.12249

sRun
RunA1
zout
1

Ave
Data
1.8E-05
Model
1.7E-05

data/vMax
1.8E-05

ofxng
311.734

Min d/m
1.13692

ave/min d
1
ave/min m
1.09694

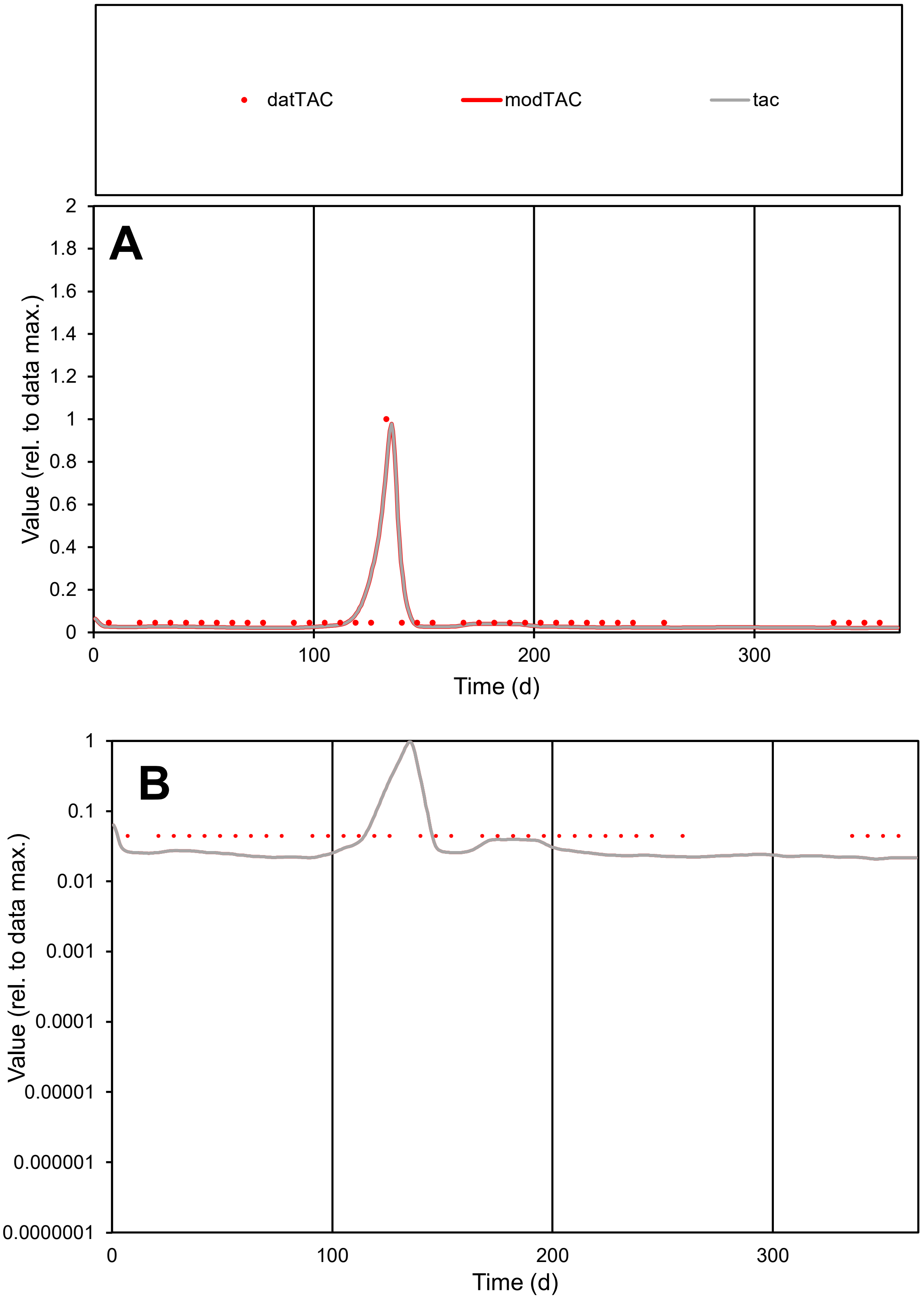


**C1**

Variables
TAC Provide jtd or Dst
CalibOnly?
2 1 to reproduce of
oNormalize?
2 1=no, 2=yes

ofxn calc*
0.5678
ofxn mod
0.67379

sRun
RunA1
zout
1

Ave
Data
2E-06
Model
1.8E-06

data/vMax
2.9E-05

ofxng
311.734

Min d/m
2.10638

ave/min d
1.5803
ave/min m
2.91817

331_Calib_S_A

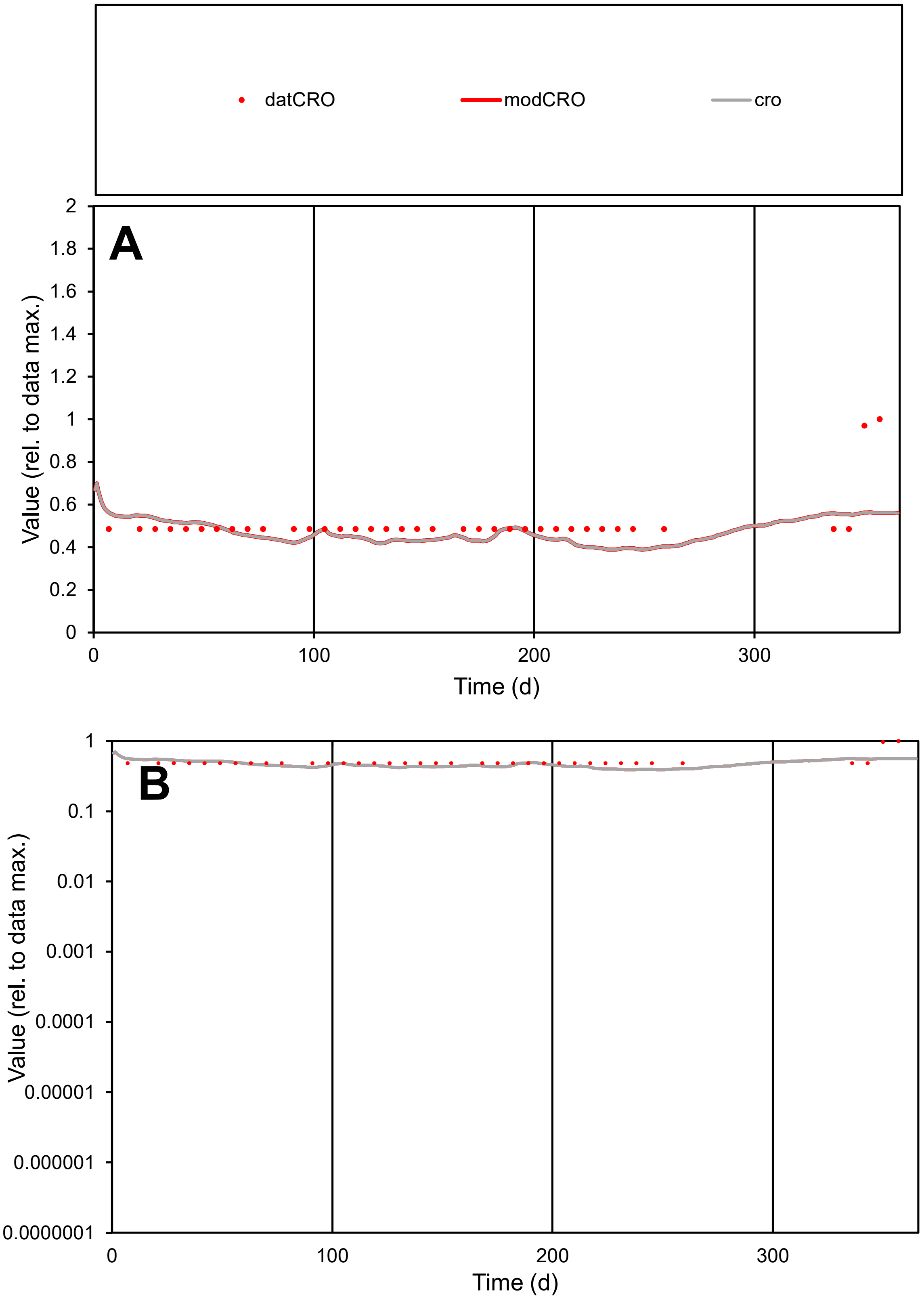


**C1**

Variables
CRO Provide jtd or Dst
CalibOnly?
2 1 to reproduce of
oNormalize?
2 1=no, 2=yes

ofxn calc*
0.21846
ofxn mod
0.24325

sRun
RunA1
zout
1

Ave
Data
1.1E-05
Model
9.8E-06

data/vMax
2.1E-05

ofxng
311.734

Min d/m
1.24121

ave/min d
1.0558
ave/min m
1.19358

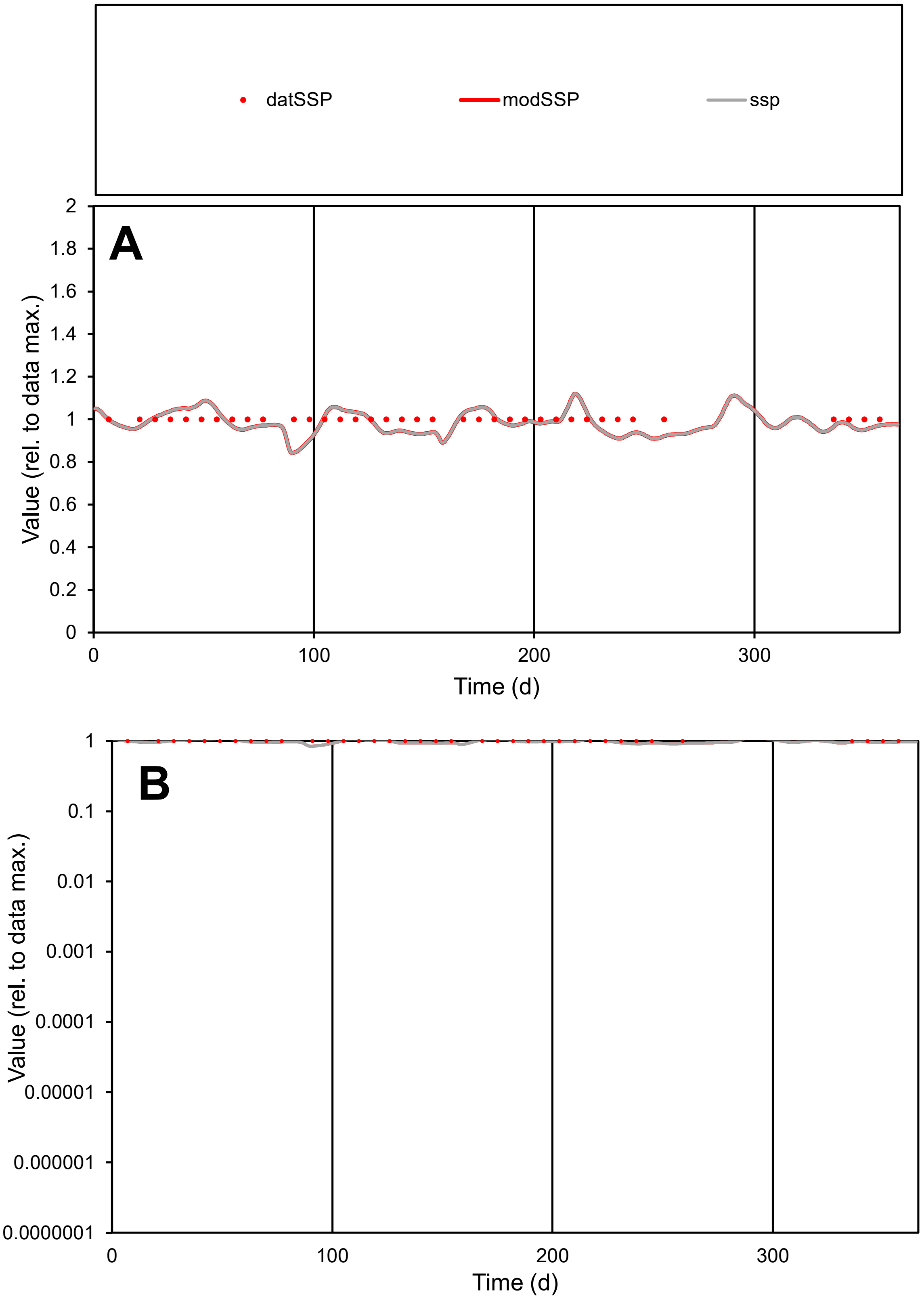


**C1**

Variables
SSP Provide jtd or Dst
CalibOnly?
2 1 to reproduce of
oNormalize?
2 1=no, 2=yes

ofxn calc*
0.0555
ofxn mod
0.12371

sRun
RunA1
zout
1

Ave
Data
1.3E-05
Model
1.3E-05

data/vMax
1.3E-05

ofxng
311.734

Min d/m
1.176

ave/min d
1

ave/min m
1.15863

331_Calib_S_A

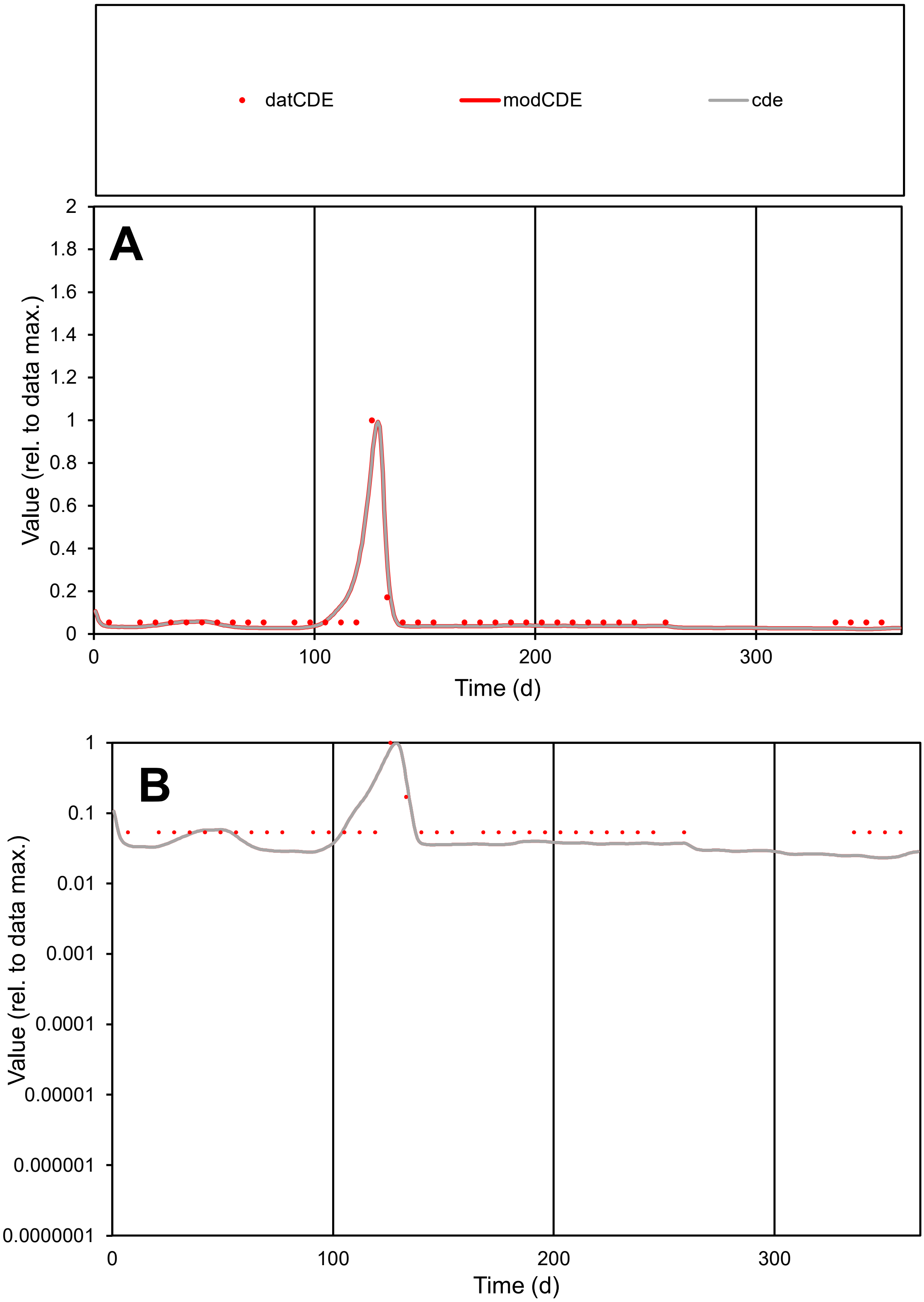


**C1**

Variables
CDE Provide jtd or Dst
CalibOnly?
2 1 to reproduce of
oNormalize?
2 1=no, 2=yes

ofxn calc*
0.45873
ofxn mod
0.58619

sRun
RunA1
zout
1

Ave
Data
1.5E-05
Model
1.4E-05

data/vMax
0.00019

ofxng
311.734

Min d/m
2.30327

ave/min d
1.53815
ave/min m
3.14414

331_Calib_S_A

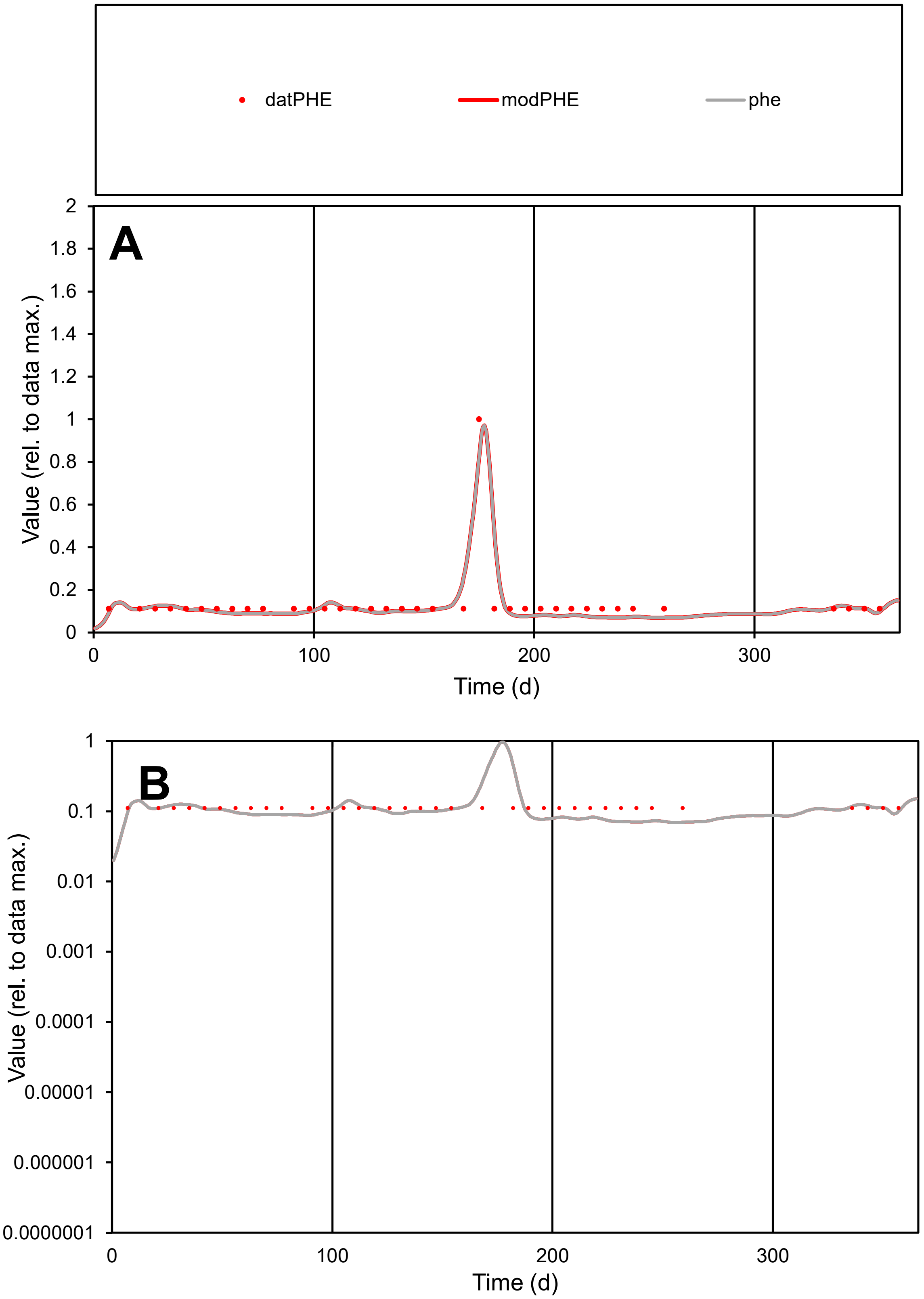


**C1**

Variables
PHE Provide jtd or Dst
CalibOnly?
2 1 to reproduce of
oNormalize?
2 1=no, 2=yes

ofxn calc*
0.25739
ofxn mod
0.33031

sRun
RunA1
zout
1

Ave
Data
4.8E-06
Model
4.5E-06

data/vMax
3.6E-05

ofxng
311.734

Min d/m
1.59189

ave/min d
1.21538
ave/min m
1.80783

331_Calib_S_A

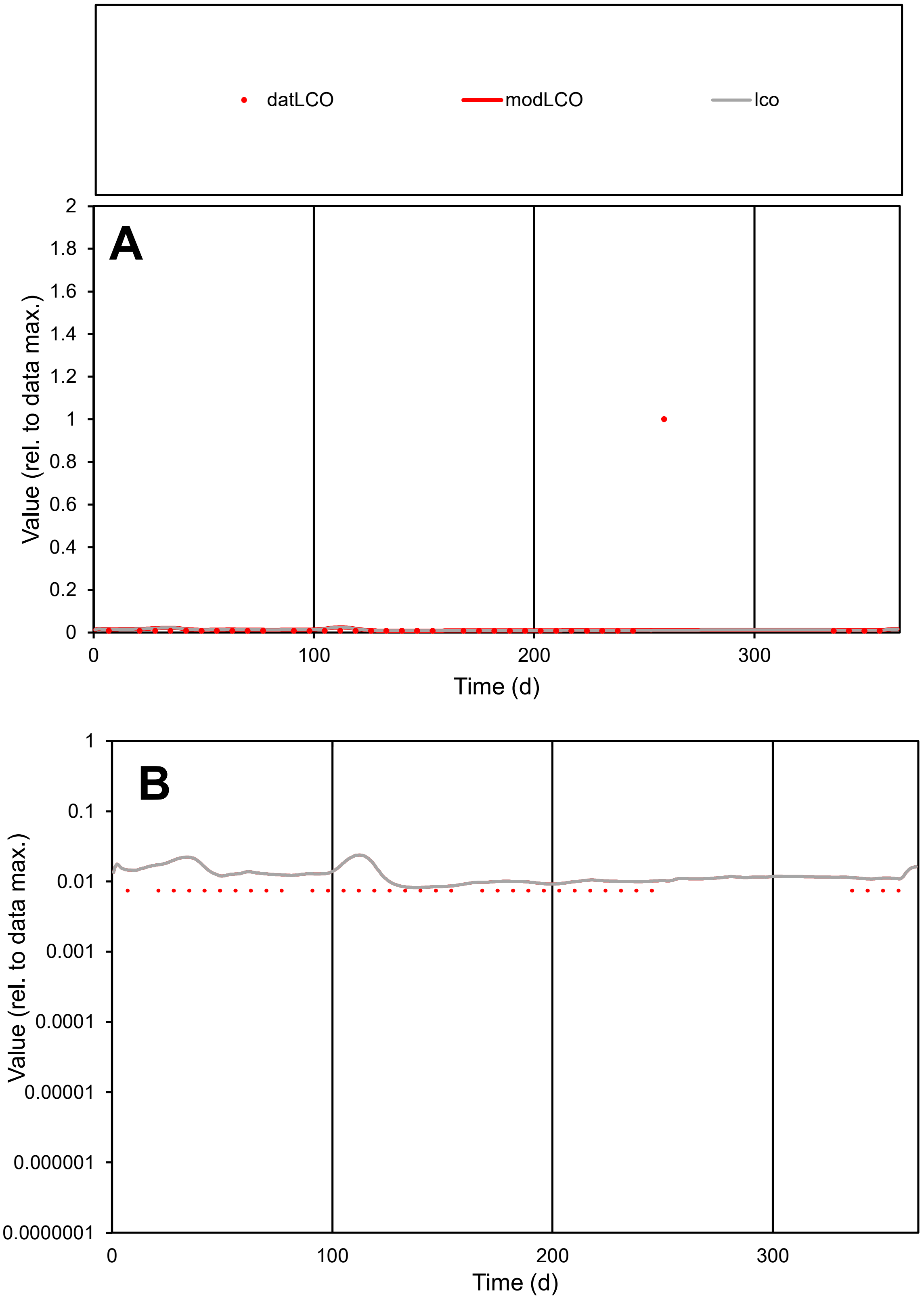


C1

Variables
LCO Provide jtd or Dst
CalibOnly?
2 1 to reproduce of
oNormalize?
2 1=no, 2=yes

ofxn calc*
4.74818
ofxn mod
4.75855

sRun
RunA1
zout
1

Ave
Data
3.4E-05
Model
1.2E-05

data/vMax
0.001

ofxng
311.734

Min d/m
0.90307

ave/min d
4.60613
ave/min m
1.48459

331_Calib_S_A

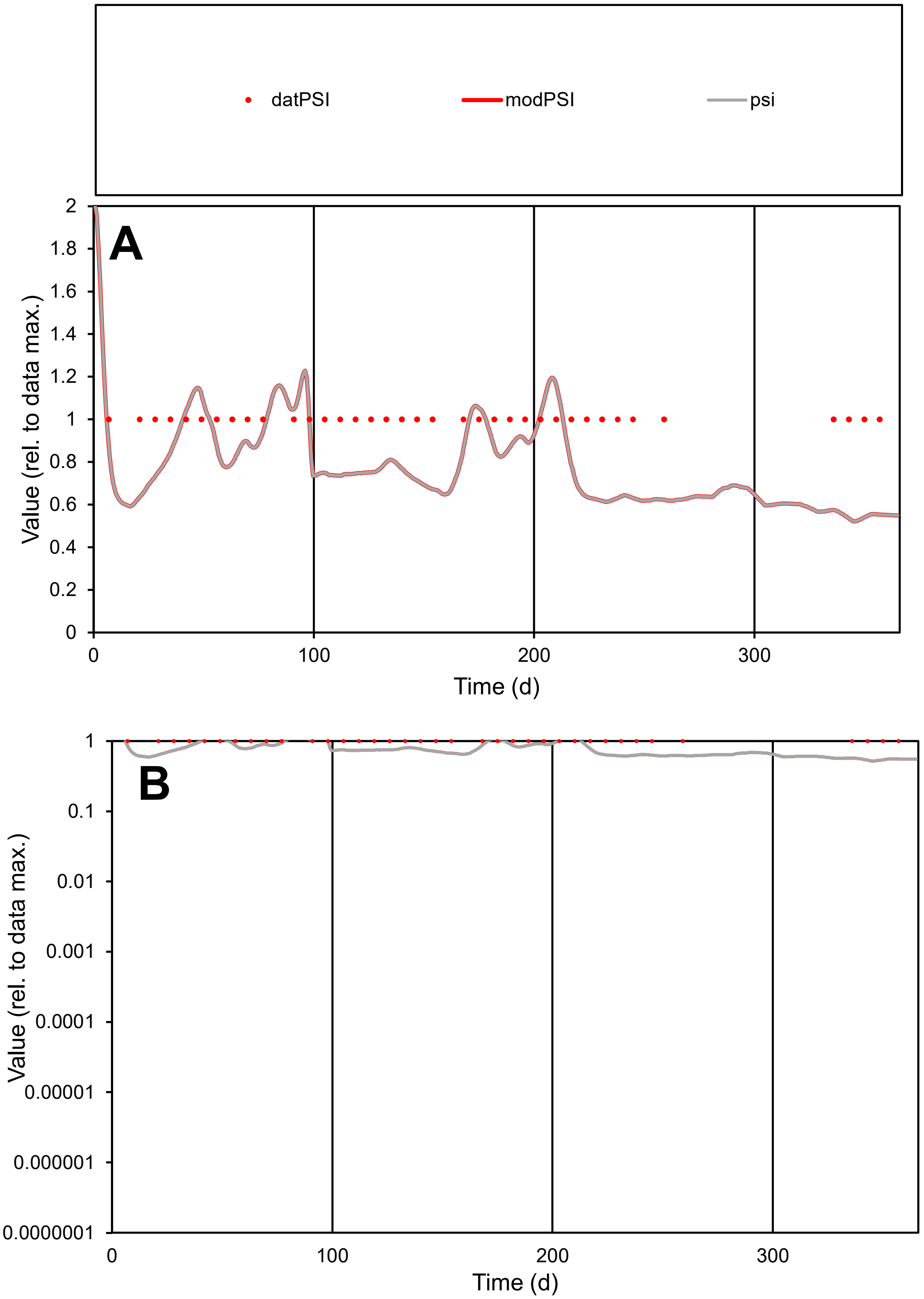


**C1**

Variables
PSI Provide jtd or Dst
CalibOnly?
2 1 to reproduce of
oNormalize?
2 1=no, 2=yes

ofxn calc*
0.26174
ofxn mod
0.2827

sRun
RunA1
zout
1

Ave
Data
3.6E-06
Model
2.8E-06

data/vMax
3.6E-06

ofxng
311.734

Min d/m
1.89505

ave/min d
1
ave/min m
1.51449

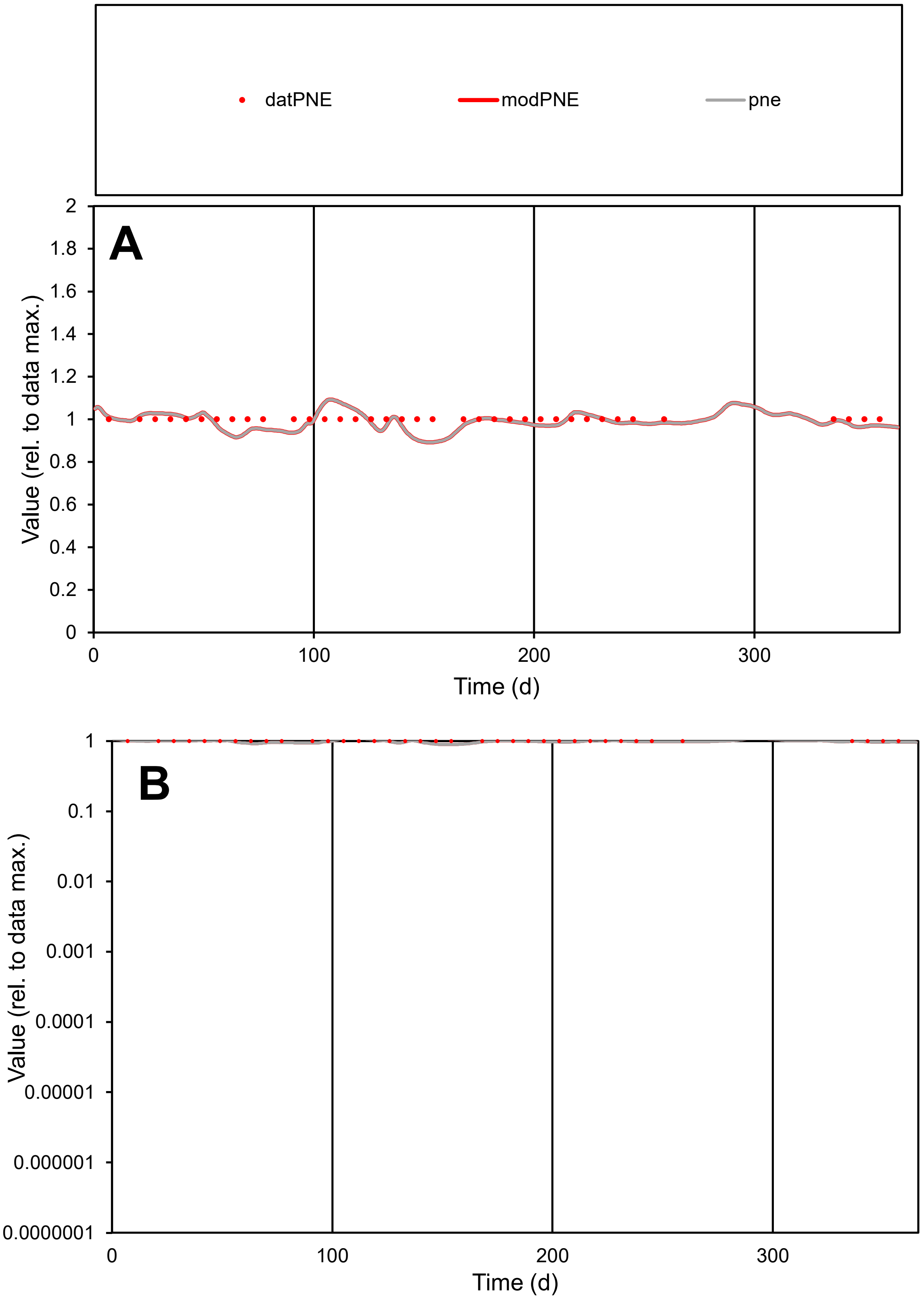


**C1**

Variables

PNE Provide jtd or Dst

CalibOnly?

2 1 to reproduce of

oNormalize?

2 1=no, 2=yes

ofxn calc*

0.04231

ofxn mod

0.11852

sRun

RunA1

zout

1

Ave

Data

1.5E-05

Model

1.4E-05

data/vMax

1.5E-05

ofxng

311.734

Min d/m

1.11778

ave/min d

1

ave/min m

1.10405

331_Calib_S_A

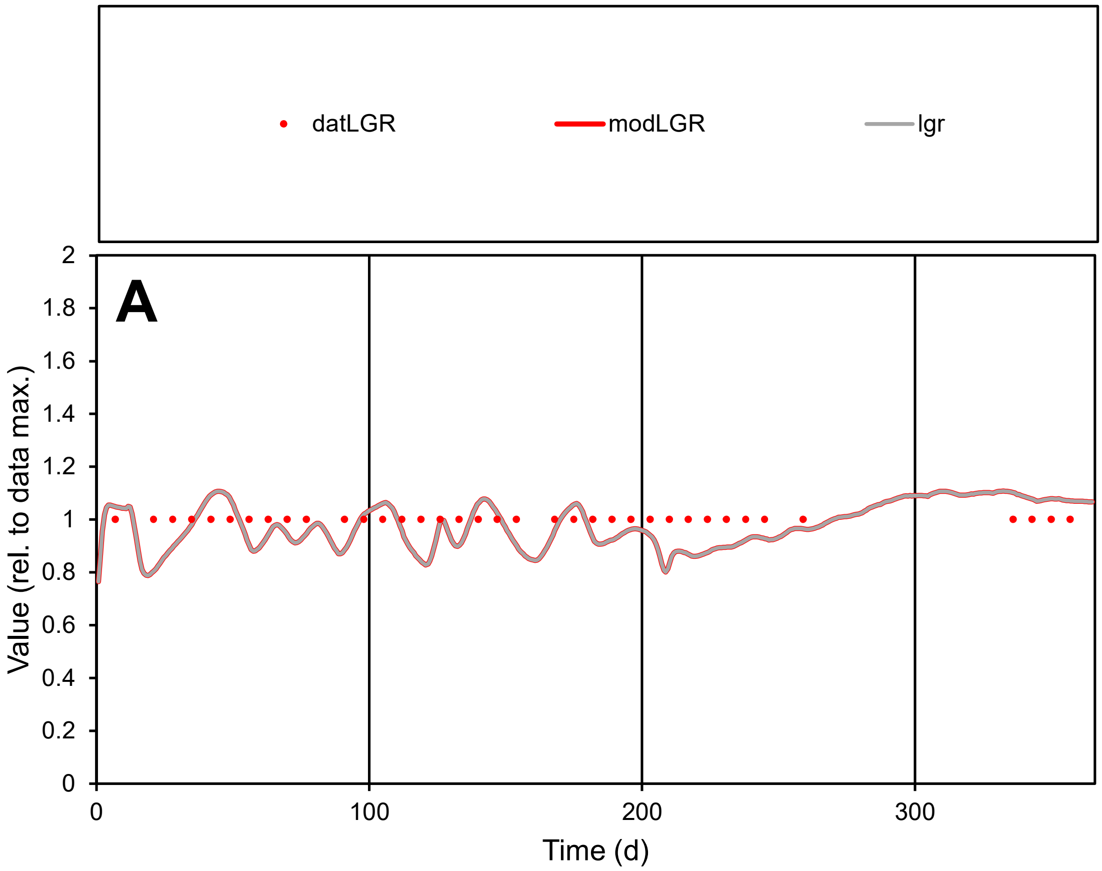


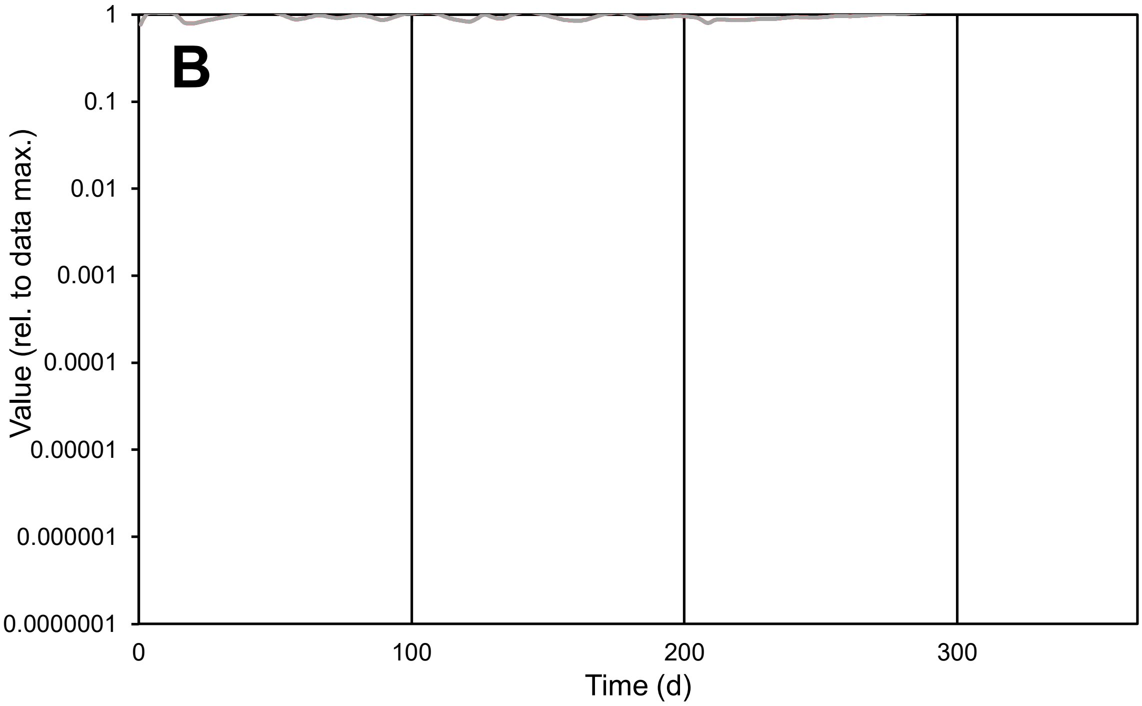


**C1**

Variables
LGR Provide jtd or Dst
CalibOnly?
2 1 to reproduce of
oNormalize?
2 1=no, 2=yes

ofxn calc*
0.08577
ofxn mod
0.13881

sRun
RunA1
zout
1

Ave
Data
5.2E-06
Model
5E-06

data/vMax
5.2E-06

ofxng
311.734

Min d/m
1.2143

ave/min d
1
ave/min m
1.17306

331_Calib_S_A

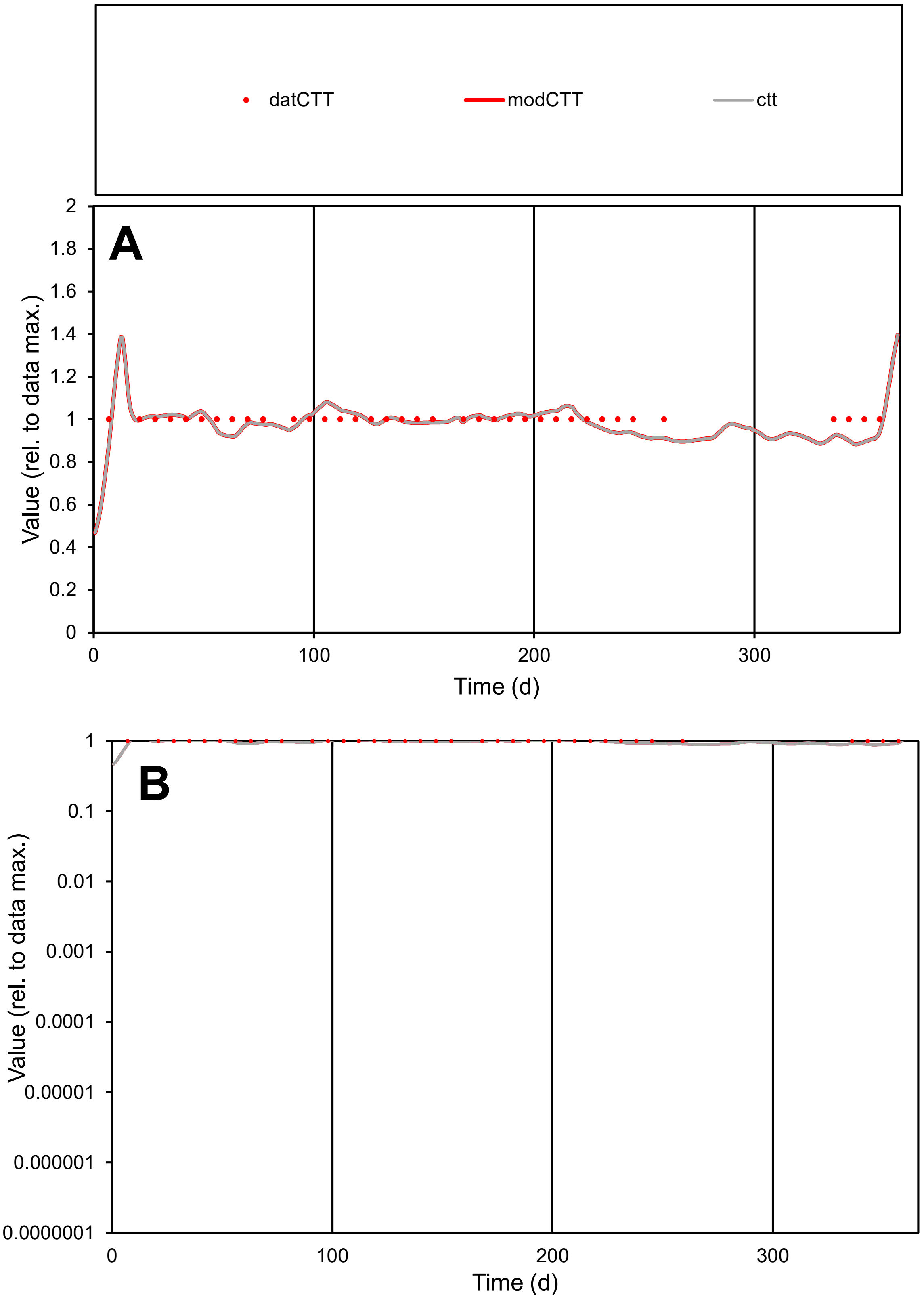


**C1**

Variables
CTT Provide jtd or Dst
CalibOnly?
2 1 to reproduce of
oNormalize?
2 1=no, 2=yes

ofxn calc*
0.04553
ofxn mod
0.11984

sRun
RunA1
zout
1

Ave
Data
1.5E-06
Model
1.5E-06

data/vMax
1.5E-06

ofxng
311.734

Min d/m
1.12094

ave/min d
1
ave/min m
1.1085

331_Calib_S_A

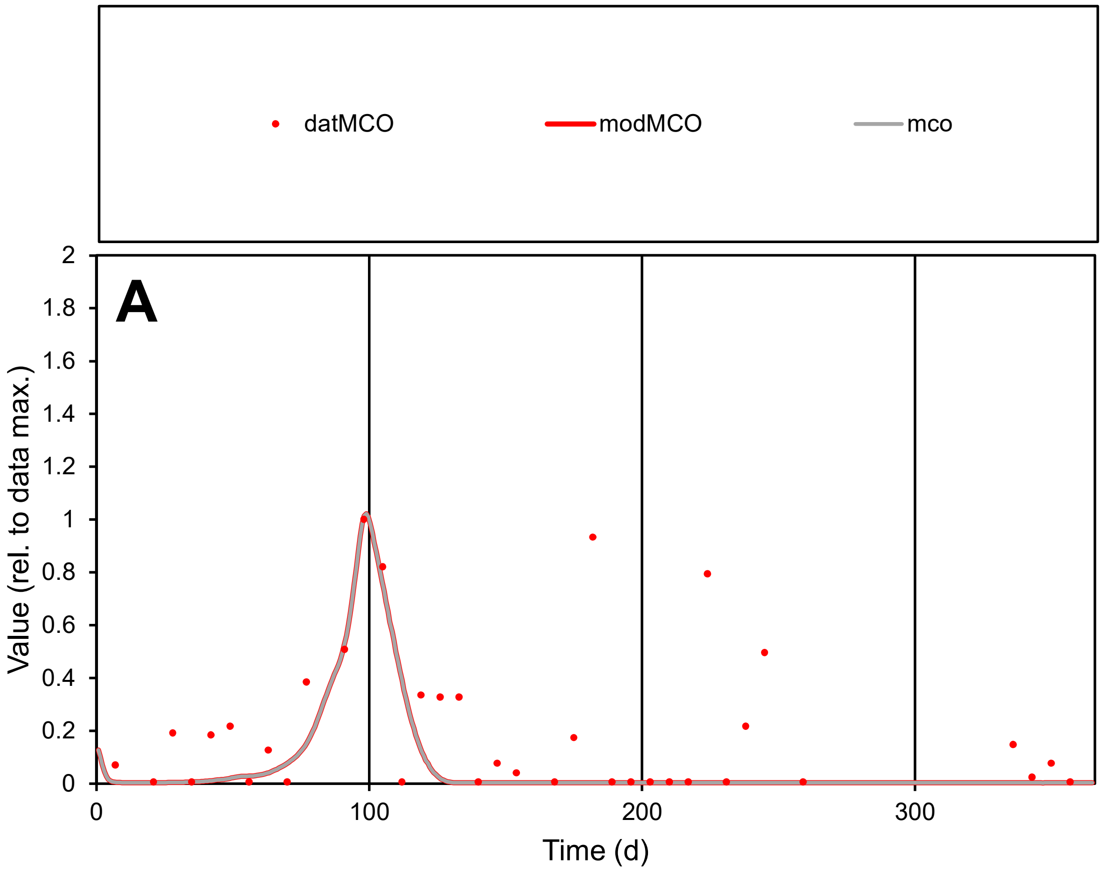


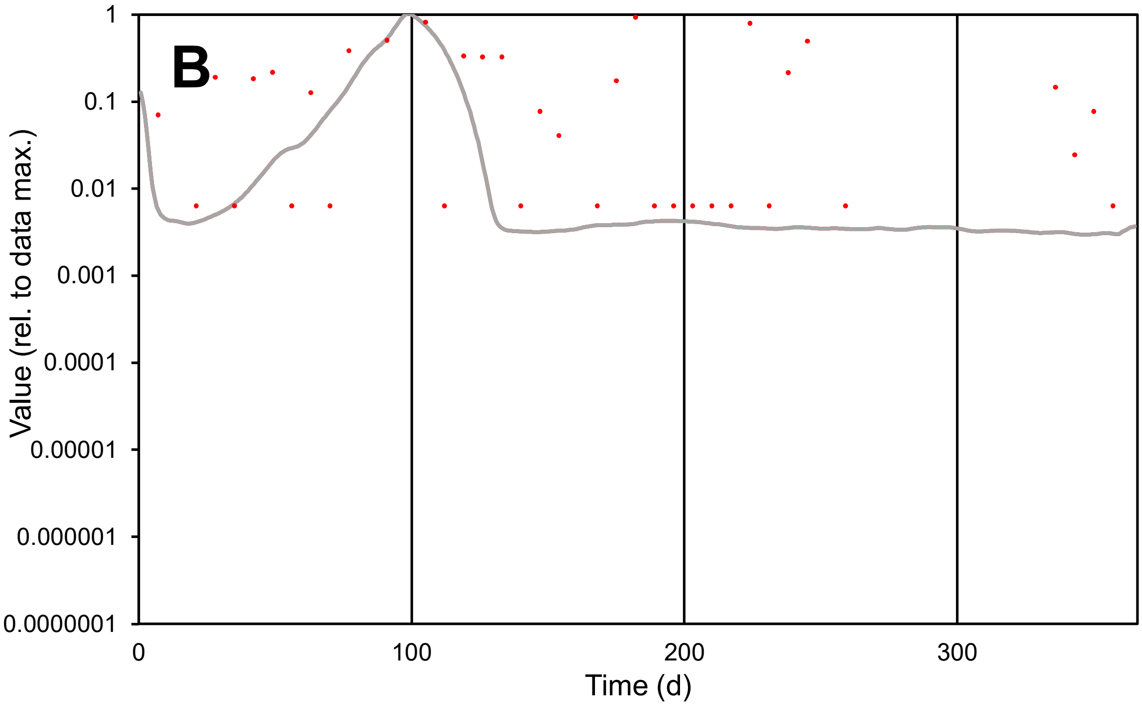


**C1**

Variables

MCO Provide jtd or Dst

CalibOnly?

2 1 to reproduce of

oNormalize?

2 1=no, 2=yes

ofxn calc*

1.22179

ofxn mod

1.23916

sRun

RunA1

zout

1

Ave

Data

2E-06

Model

8.2E-07

data/vMax

9.6E-06

ofxng

311.734

Min d/m

2.12474

ave/min d

32.1084

ave/min m

28.4844

331_Calib_S_A

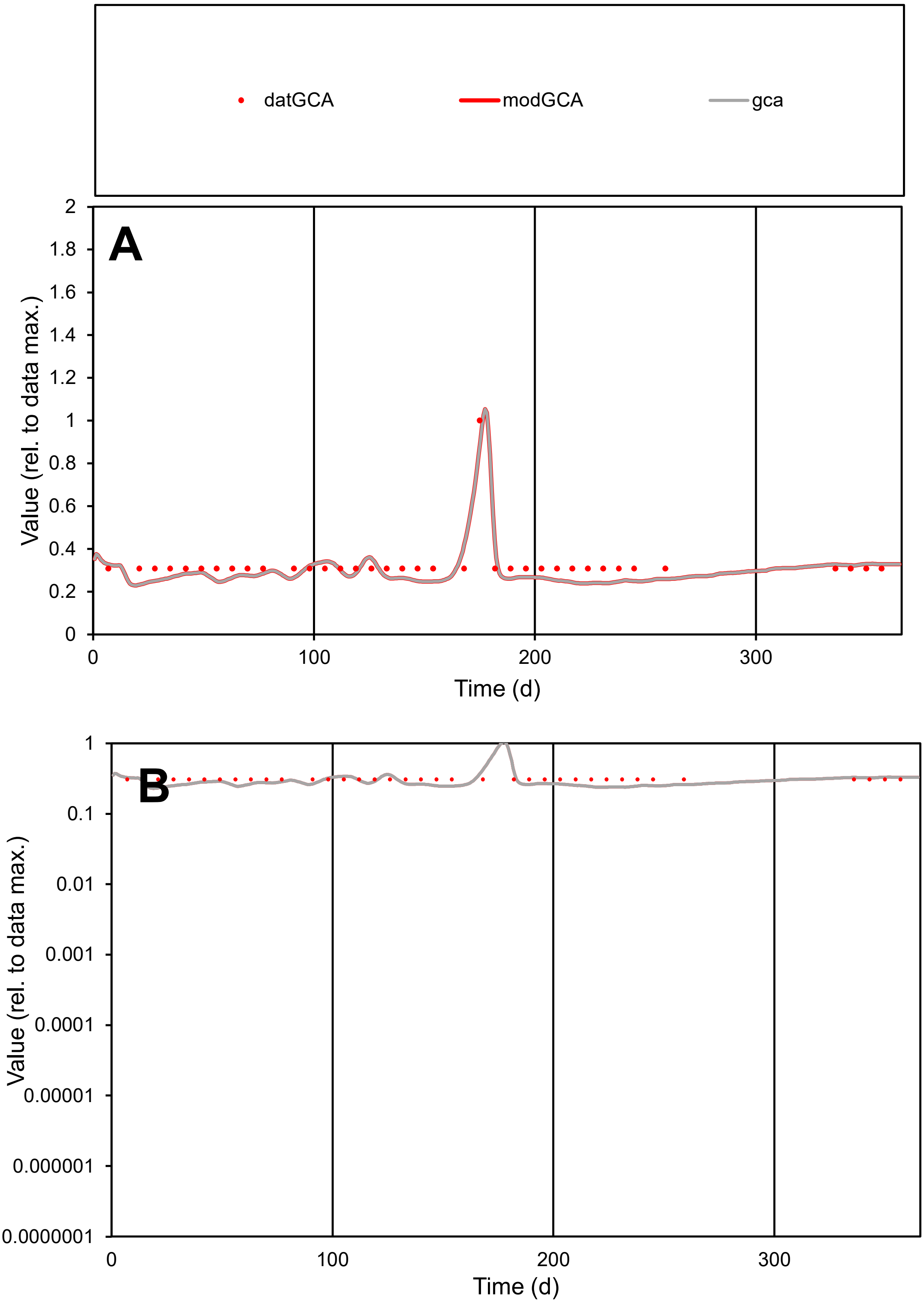


**C1**

Variables

GCA Provide jtd or Dst

CalibOnly?

2 1 to reproduce of

oNormalize?

2 1=no, 2=yes

ofxn calc*

0.14066

ofxn mod

0.17932

sRun

RunA1

zout

1

Ave

Data

8.9E-06

Model

8.2E-06

data/vMax

2.7E-05

ofxng

311.734

Min d/m

1.29624

ave/min d

1.06088

ave/min m

1.2731

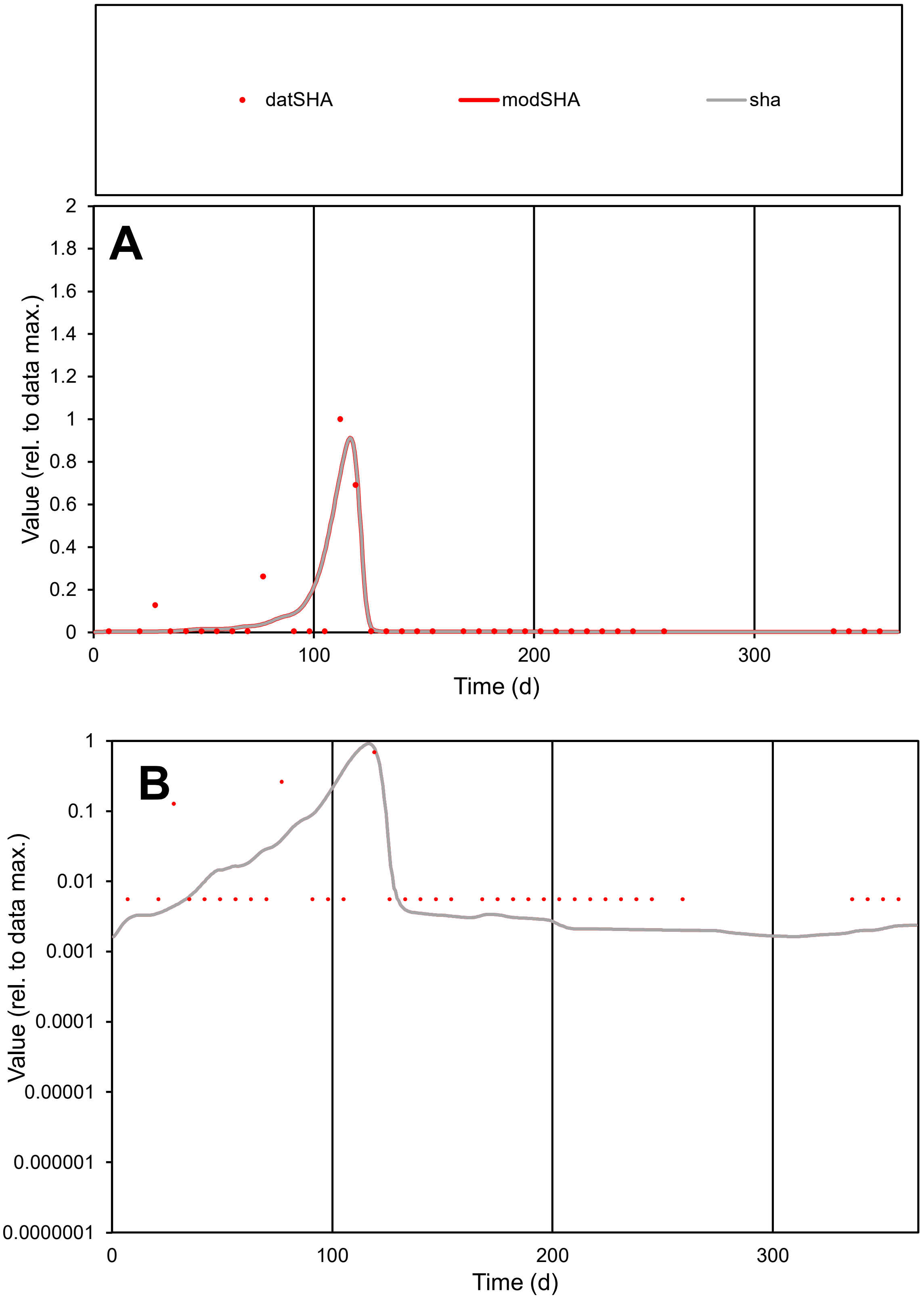


**C1**

Variables
SHA Provide jtd or Dst
CalibOnly?
2 1 to reproduce of
oNormalize?
2 1=no, 2=yes

ofxn calc*
1.24309
ofxn mod
1.35724

sRun
RunA1
zout
1

Ave
Data
7.6E-06
Model
7.6E-06

data/vMax
0.00012

ofxng
311.734

Min d/m
2.87706

ave/min d
10.9412
ave/min m
31.2424

331_Calib_S_A

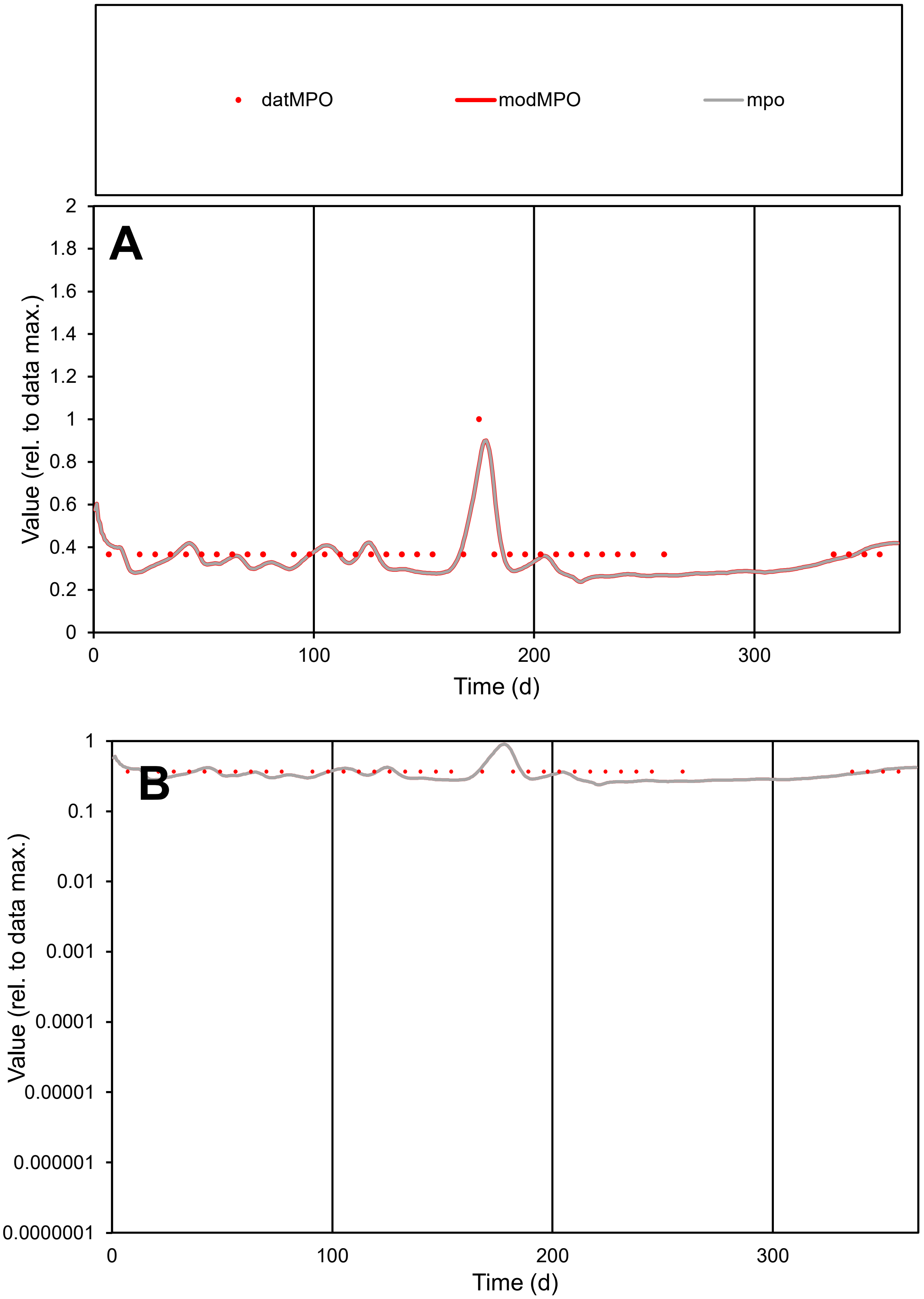


**C1**

Variables
MPO Provide jtd or Dst
CalibOnly?
2 1 to reproduce of
oNormalize?
2 1=no, 2=yes

ofxn calc*
0.18586
ofxn mod
0.21741

sRun
RunA1
zout
1

Ave
Data
1.3E-05
Model
1.2E-05

data/vMax
3.4E-05

ofxng
311.734

Min d/m
1.43154

ave/min d
1.04682
ave/min m
1.36939

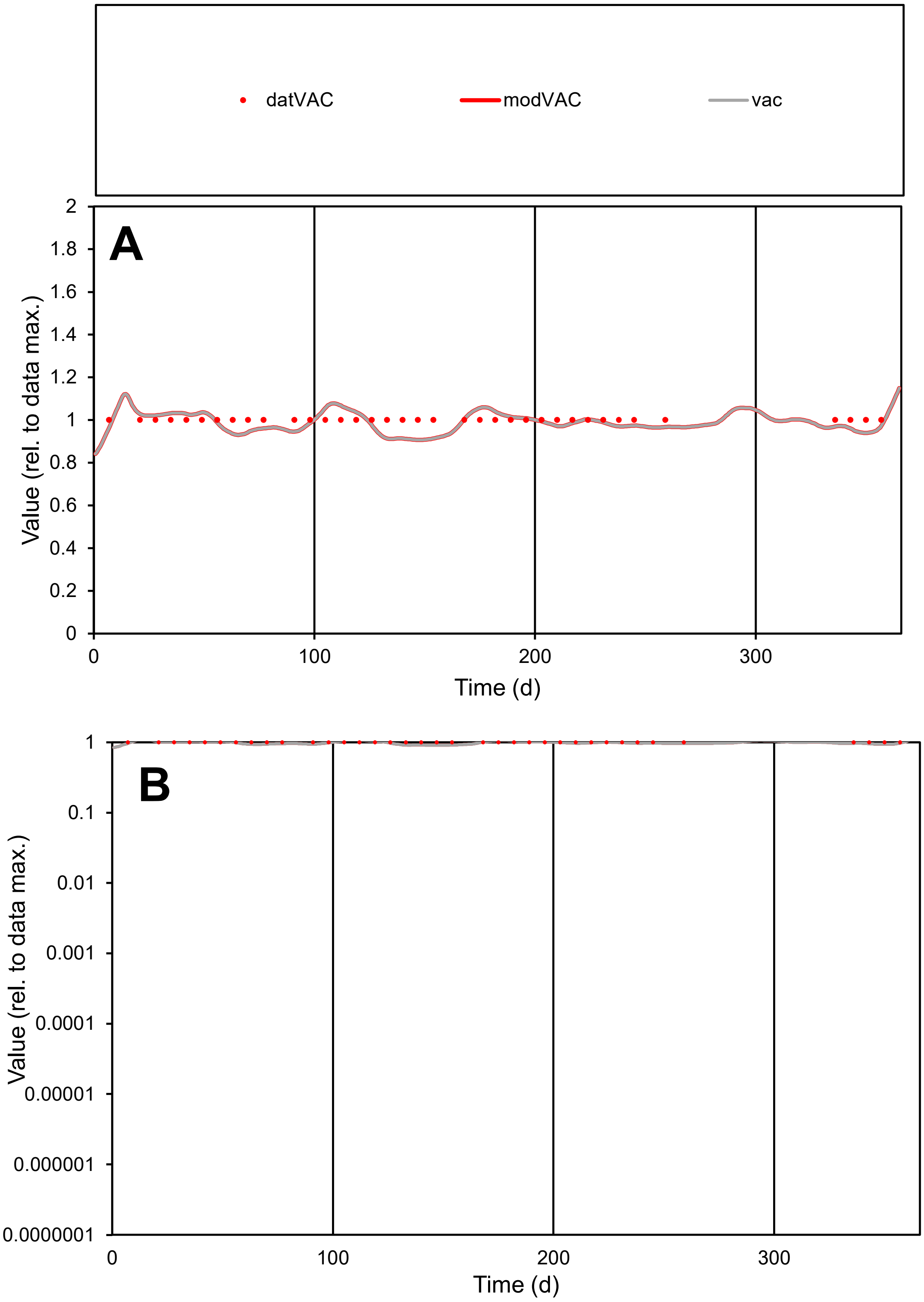


**C1**

Variables
VAC Provide jtd or Dst
CalibOnly?
2 1 to reproduce of
oNormalize?
2 1=no, 2=yes

ofxn calc*
0.0446
ofxn mod
0.11937

sRun
RunA1
zout
1

Ave
Data
9.5E-06
Model
9.4E-06

data/vMax
9.5E-06

ofxng
311.734

Min d/m
1.10311

ave/min d
1
ave/min m
1.08874

331_Calib_S_A

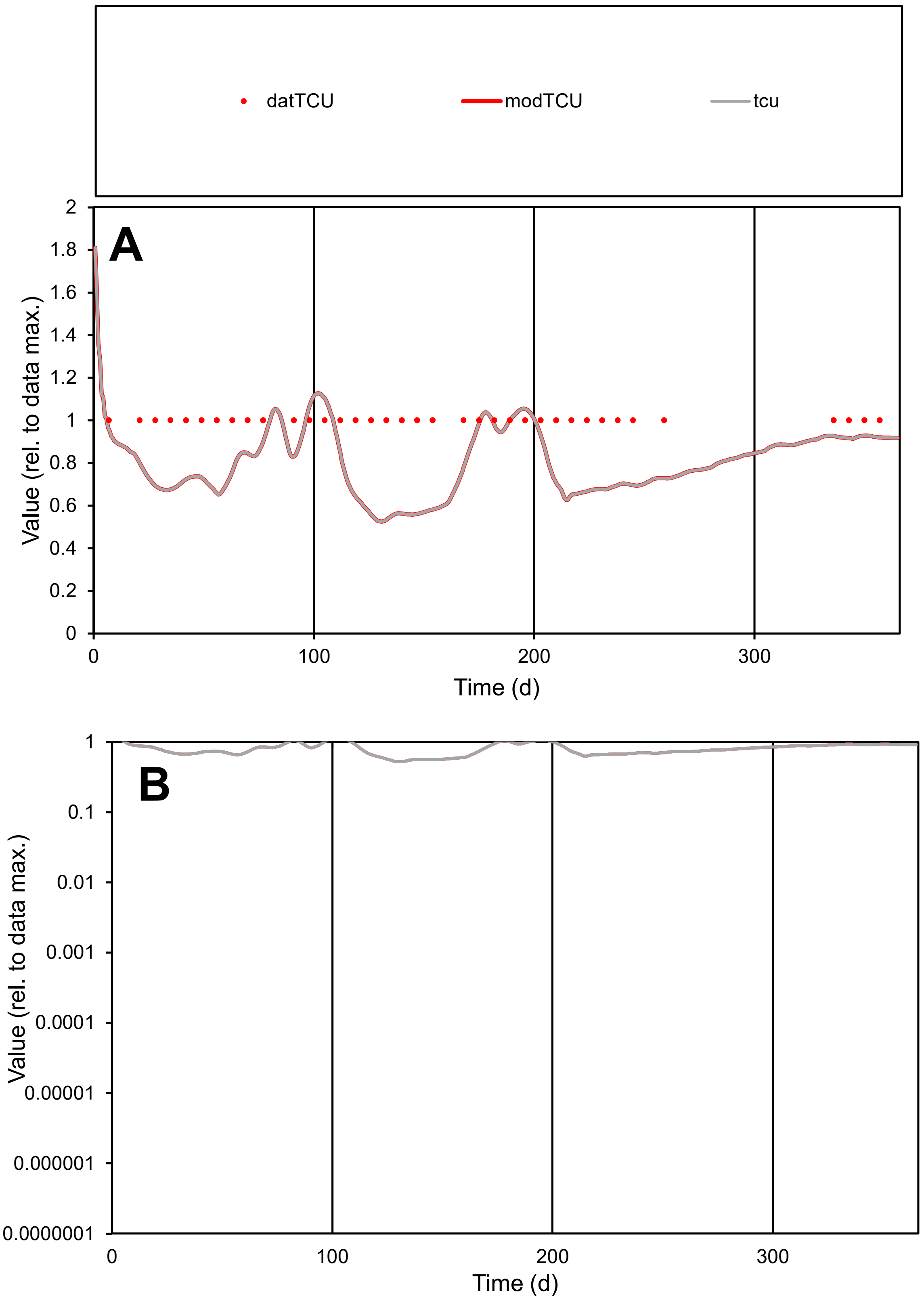


**C1**

Variables
TCU Provide jtd or Dst
CalibOnly?
2 1 to reproduce of
oNormalize?
2 1=no, 2=yes

ofxn calc*
0.26235
ofxn mod
0.28062

sRun
RunA1
zout
1

Ave
Data
9.8E-06
Model
7.7E-06

data/vMax
9.8E-06

ofxng
311.734

Min d/m
1.83983

ave/min d
1
ave/min m
1.45499

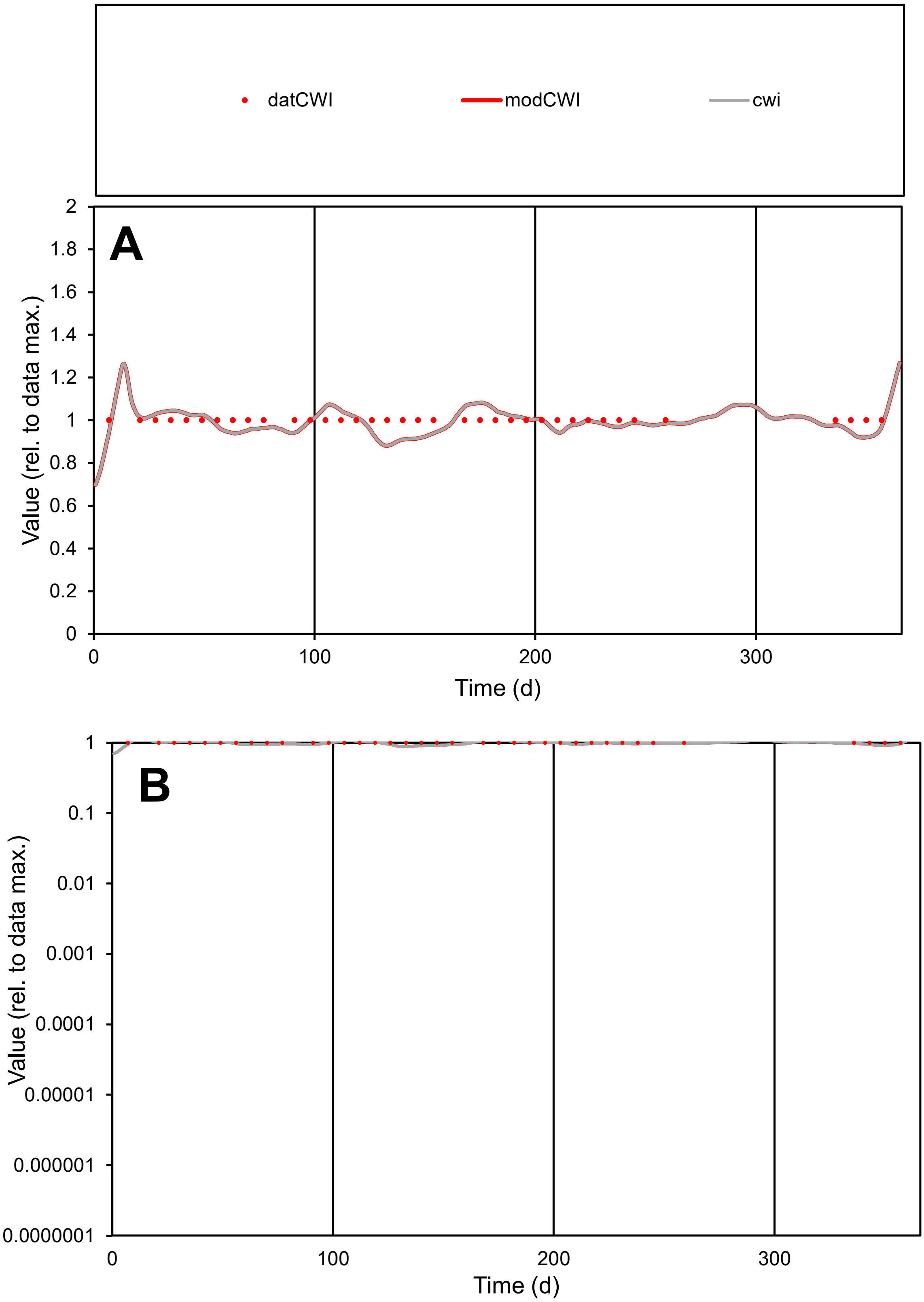


**C1**

Variables
CWI Provide jtd or Dst
CalibOnly?
2 1 to reproduce of
oNormalize?
2 1=no, 2=yes

ofxn calc*
0.04886
ofxn mod
0.12104

sRun
RunA1
zout
1

Ave
Data
1.3E-05
Model
1.3E-05

data/vMax
1.3E-05

ofxng
311.734

Min d/m
1.1301

ave/min d
1
ave/min m
1.11597

331_Calib_S_A

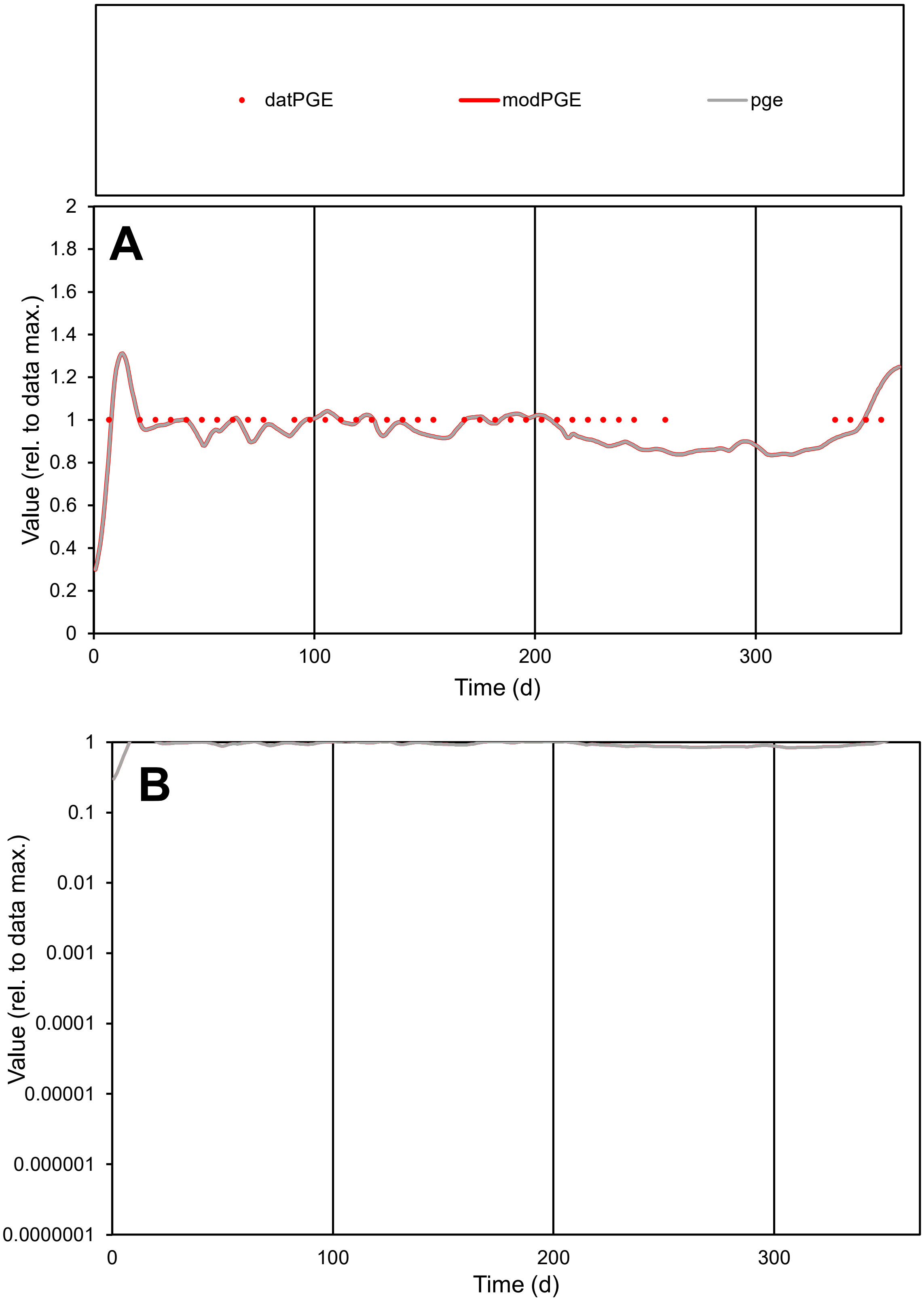


**C1**

Variables
PGE Provide jtd or Dst
CalibOnly?
2 1 to reproduce of
oNormalize?
2 1=no, 2=yes

ofxn calc*
0.06951
ofxn mod
0.12998

sRun
RunA1
zout
1

Ave
Data
5.9E-06
Model
5.7E-06

data/vMax
5.9E-06

ofxng
311.734

Min d/m
1.17818

ave/min d
1
ave/min m
1.14258

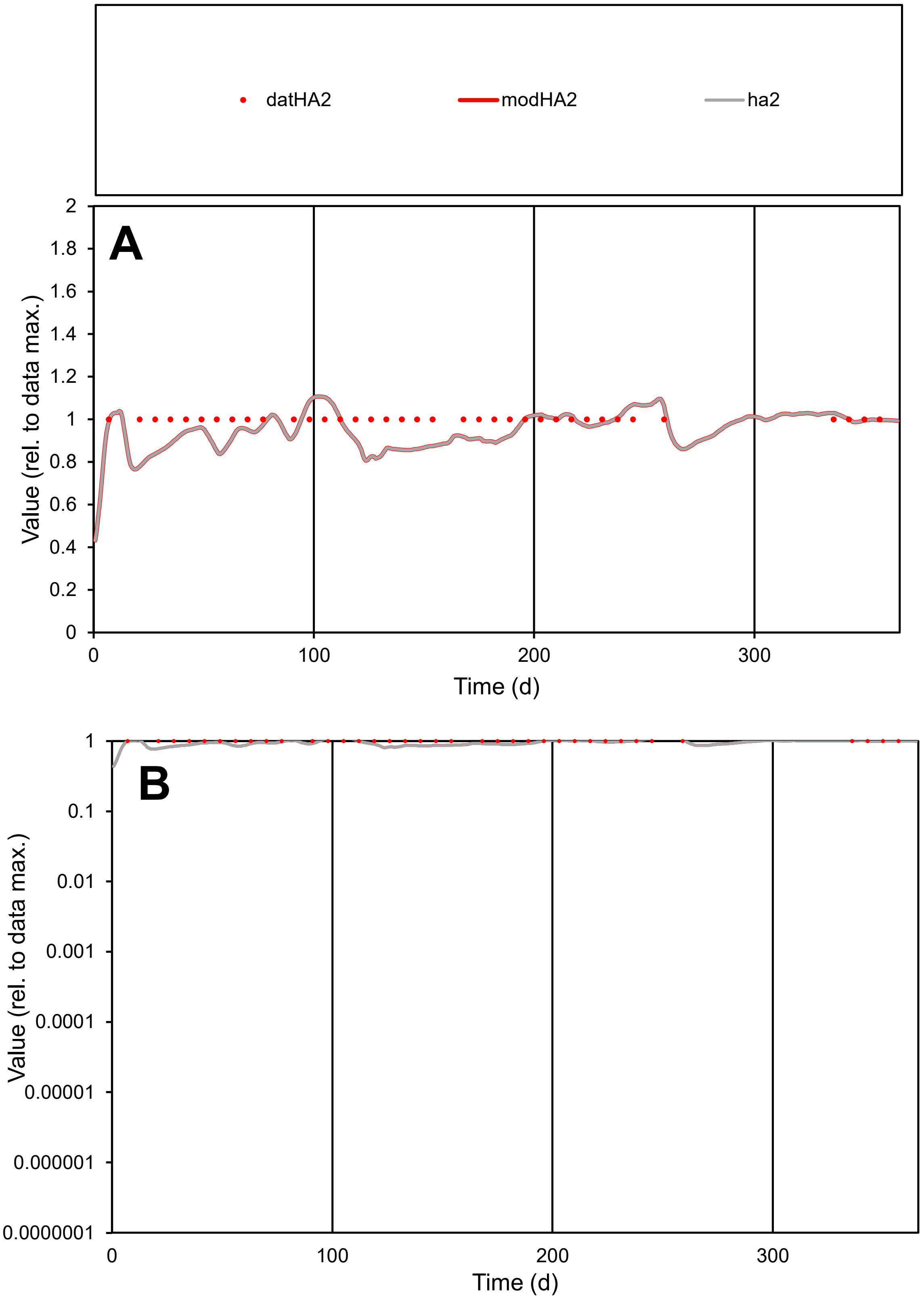


**C1**

Variables
HA2 Provide jtd or Dst
CalibOnly?
2 1 to reproduce of
oNormalize?
2 1=no, 2=yes

ofxn calc*
0.08867
ofxn mod
0.14053

sRun
RunA1
zout
1

Ave
Data
2.6E-06
Model
2.5E-06

data/vMax
2.6E-06

ofxng
311.734

Min d/m
1.2569

ave/min d
1
ave/min m
1.19611

331_Calib_S_A

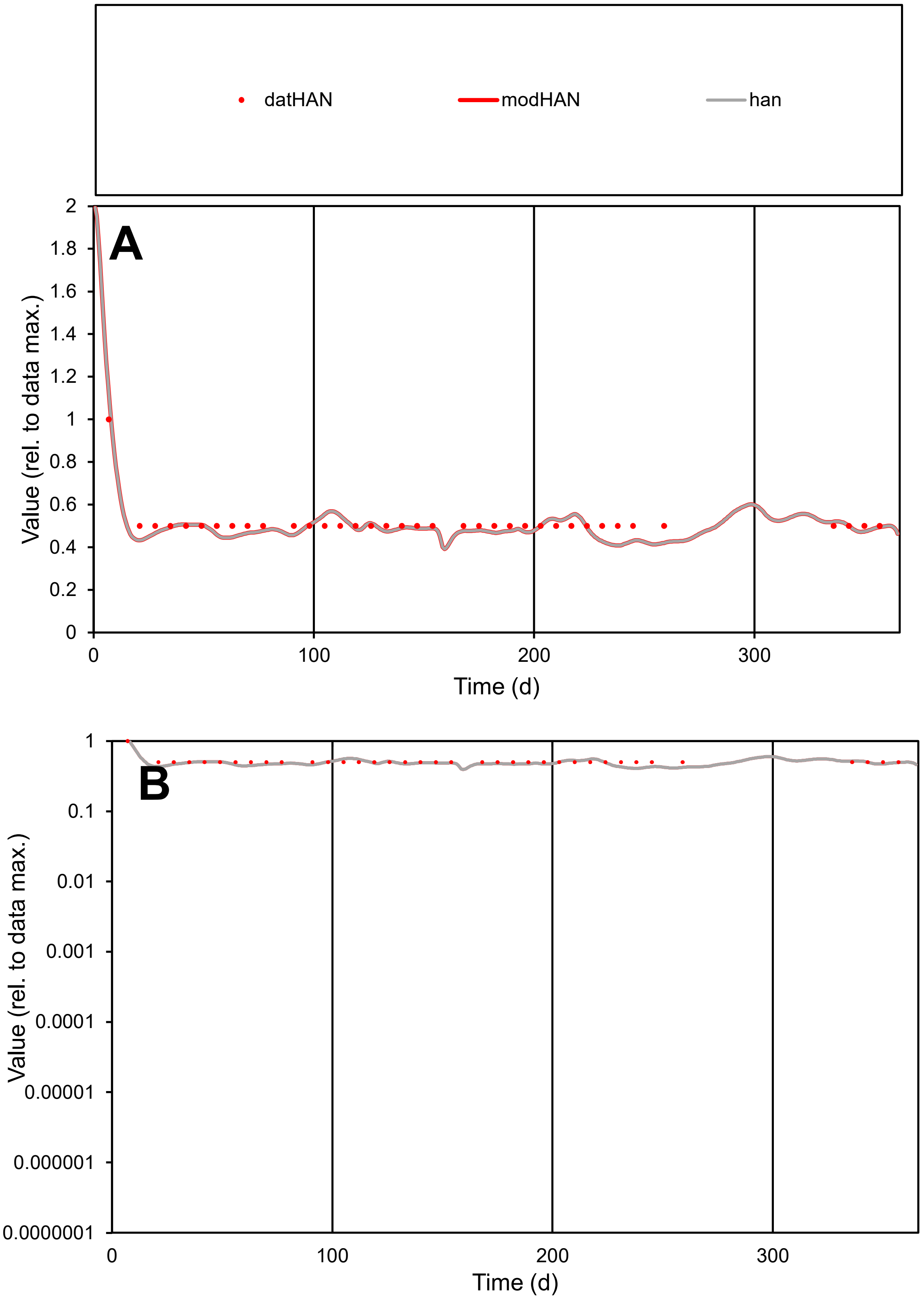


C1

Variables
HAN Provide jtd or Dst
CalibOnly?
2 1 to reproduce of
oNormalize?
2 1=no, 2=yes

ofxn calc*
0.07349
ofxn mod
0.13035

sRun
RunA1
zout
1

Ave
Data
5.3E-06
Model
5.1E-06

data/vMax
1E-05

ofxng
311.734

Min d/m
1.22087

ave/min d
1.02704
ave/min m
1.21371

331_Calib_S_A

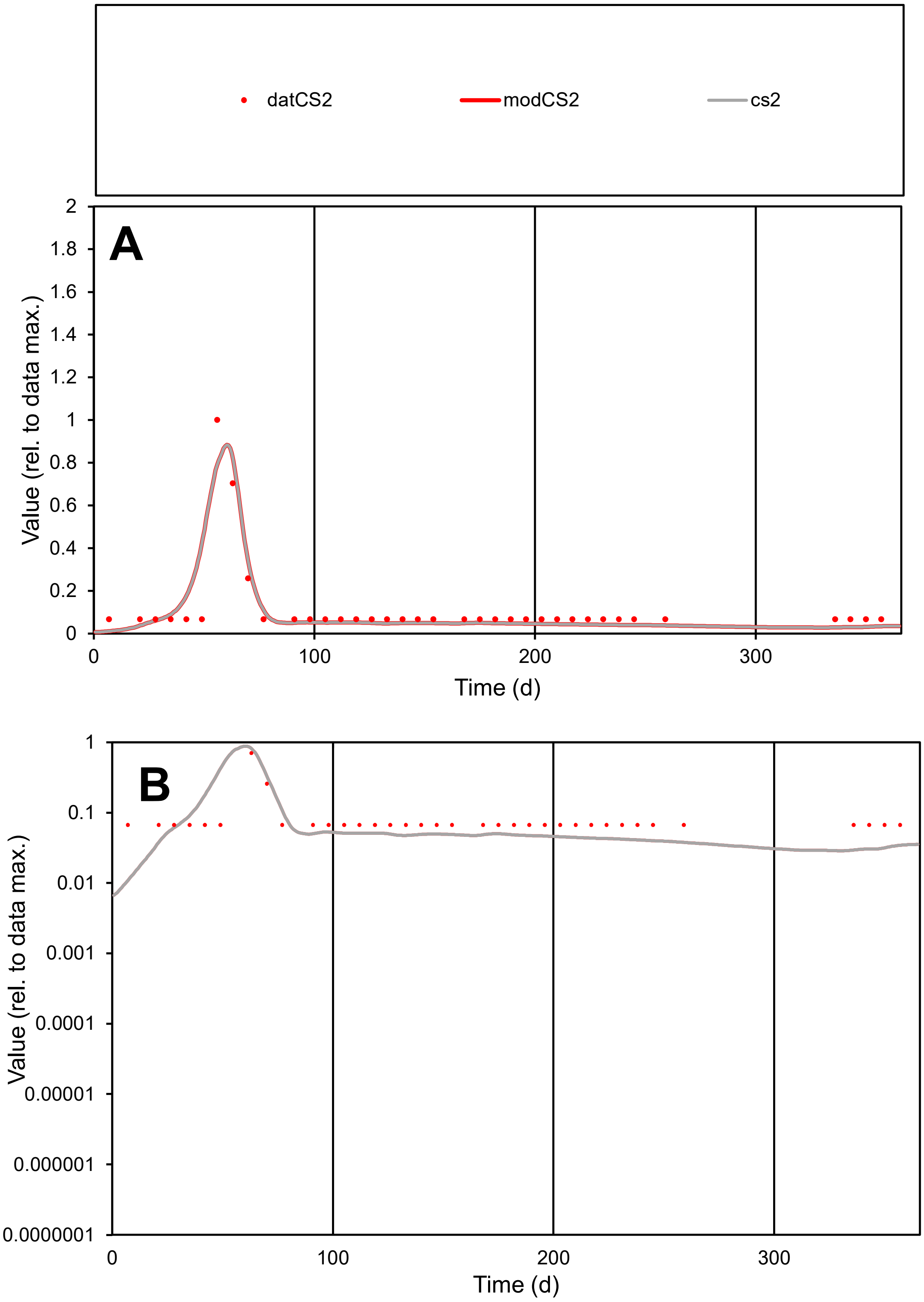


**C1**

Variables
CS2 Provide jtd or Dst
CalibOnly?
2 1 to reproduce of
oNormalize?
2 1=no, 2=yes

ofxn calc*
0.52237
ofxn mod
0.66208

sRun
RunA1
zout
1

Ave
Data
1.3E-05
Model
1.2E-05

data/vMax
0.00011

ofxng
311.734

Min d/m
5.68066

ave/min d
1.71297
ave/min m
8.81523

331_Calib_S_A

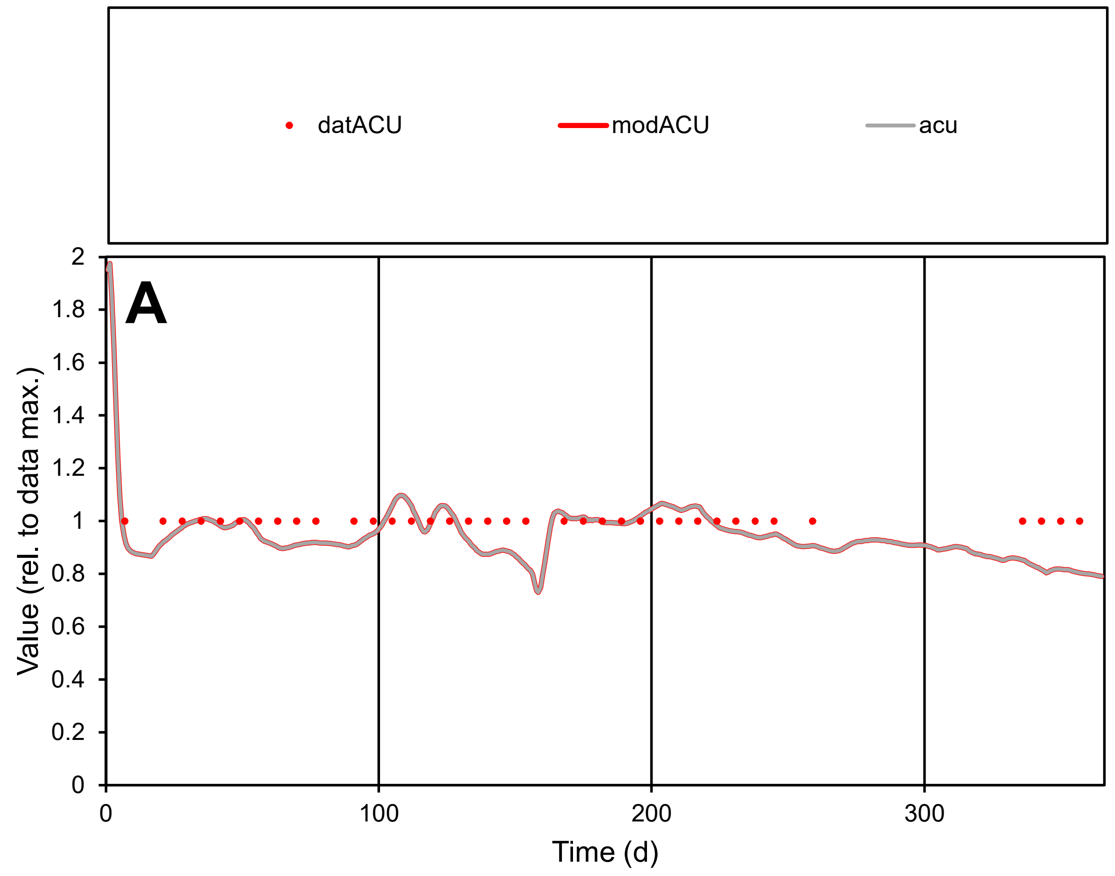


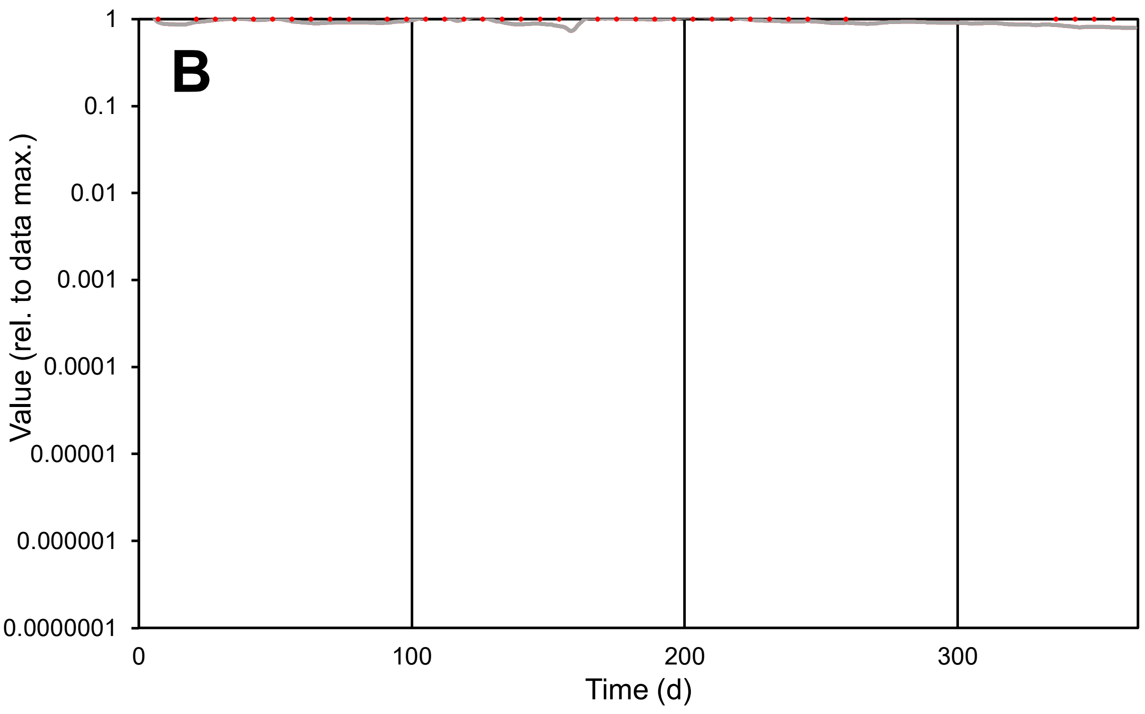


**C1**

Variables

ACU Provide jtd or Dst

CalibOnly?

2 1 to reproduce of

oNormalize?

2 1=no, 2=yes

ofxn calc*

0.08937

ofxn mod

0.1409

sRun

RunA1

zout

1

Ave

Data

1.1E-05

Model

1.1E-05

data/vMax

1.1E-05

ofxng

311.734

Min d/m

1.24705

ave/min d

1

ave/min m

1.18462

331_Calib_S_A

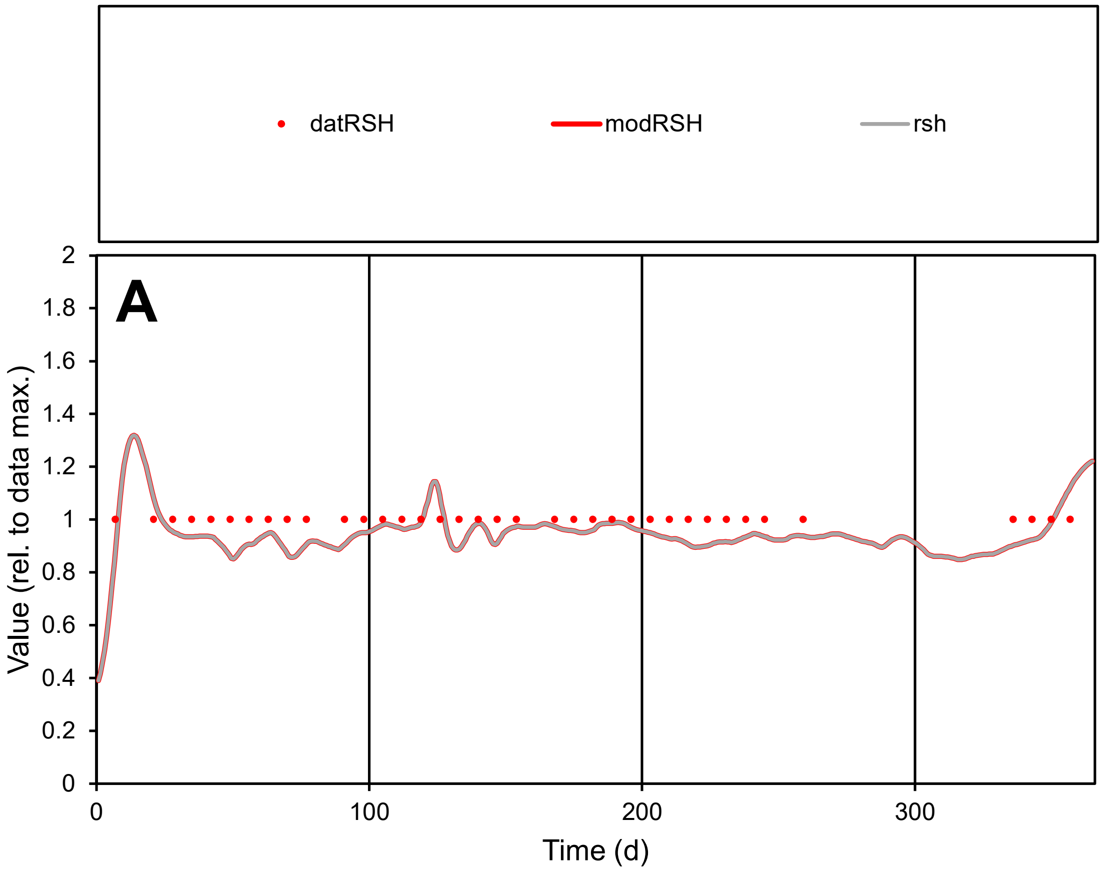


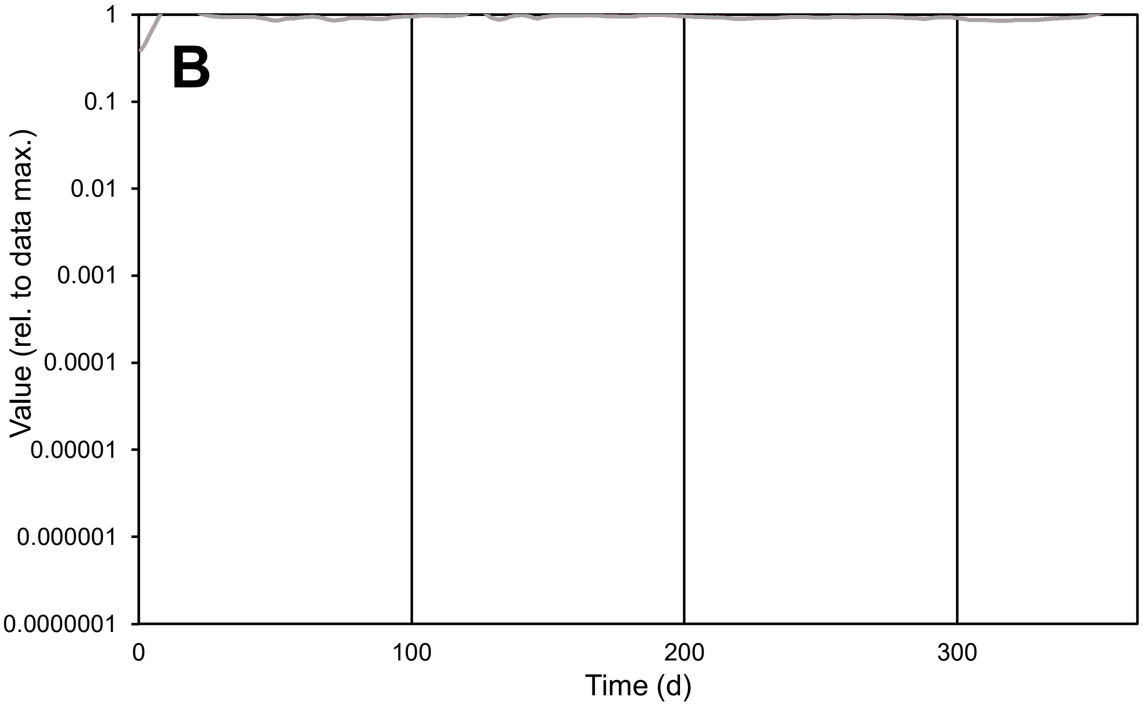


**C1**

Variables

| | |
|---|---|
| RSH | Provide jtd or Dst |
| CalibOnly? | |
| 2 | 1 to reproduce of |
| oNormalize? | |
| 2 | 1=no, 2=yes |

ofxn calc*
0.0715
ofxn mod
0.13025

sRun
RunA1
zout
1

Ave
Data
1.8E-05
Model
1.7E-05

data/vMax
1.8E-05

ofxng
311.734

Min d/m
1.17423

ave/min d
1
ave/min m
1.11735

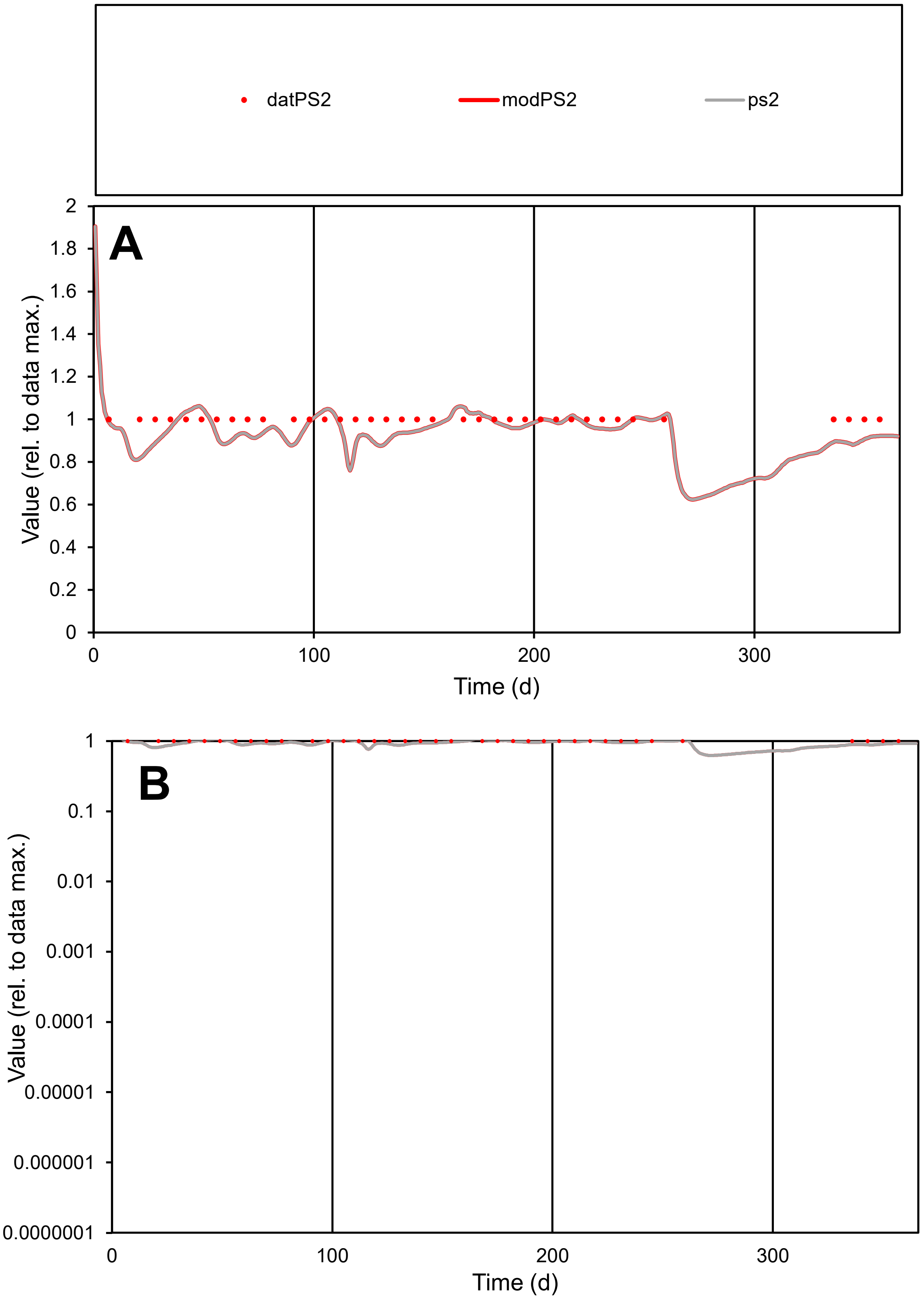


C1

Variables
PS2 Provide jtd or Dst
CalibOnly?
2 1 to reproduce of
oNormalize?
2 1=no, 2=yes

ofxn calc*
0.0682
ofxn mod
0.12889

sRun
RunA1
zout
1

Ave
Data
1.9E-05
Model
1.8E-05

data/vMax
1.9E-05

ofxng
311.734

Min d/m
1.20493

ave/min d
1
ave/min m
1.15327

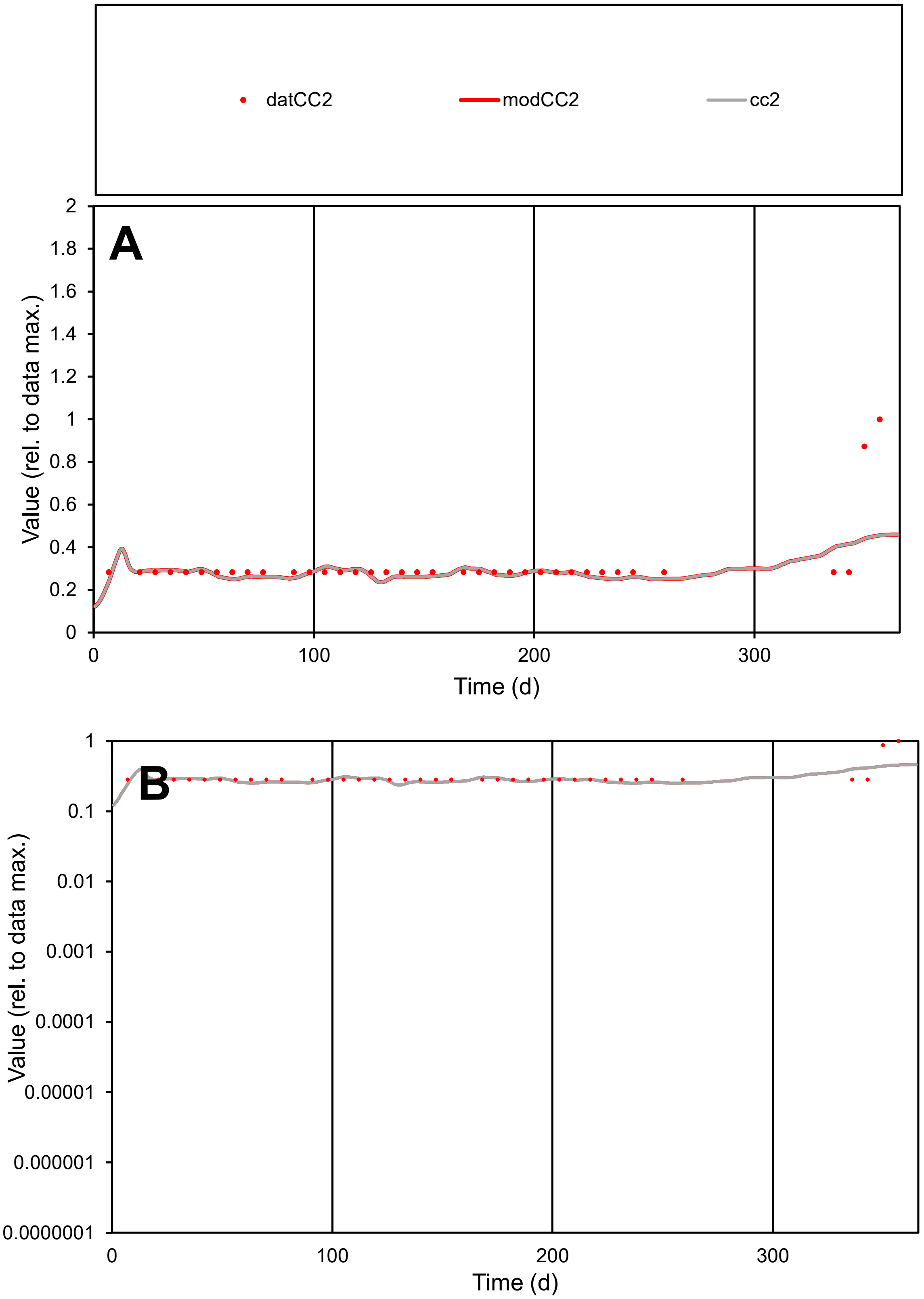


**C1**

Variables
CC2 Provide jtd or Dst
CalibOnly?
2 1 to reproduce of
oNormalize?
2 1=no, 2=yes

ofxn calc*
0.37398
ofxn mod
0.39002

sRun
RunA1
zout
1

Ave
Data
6.6E-06
Model
6E-06

data/vMax
2.1E-05

ofxng
311.734

Min d/m
1.12625

ave/min d
1.12496
ave/min m
1.16059

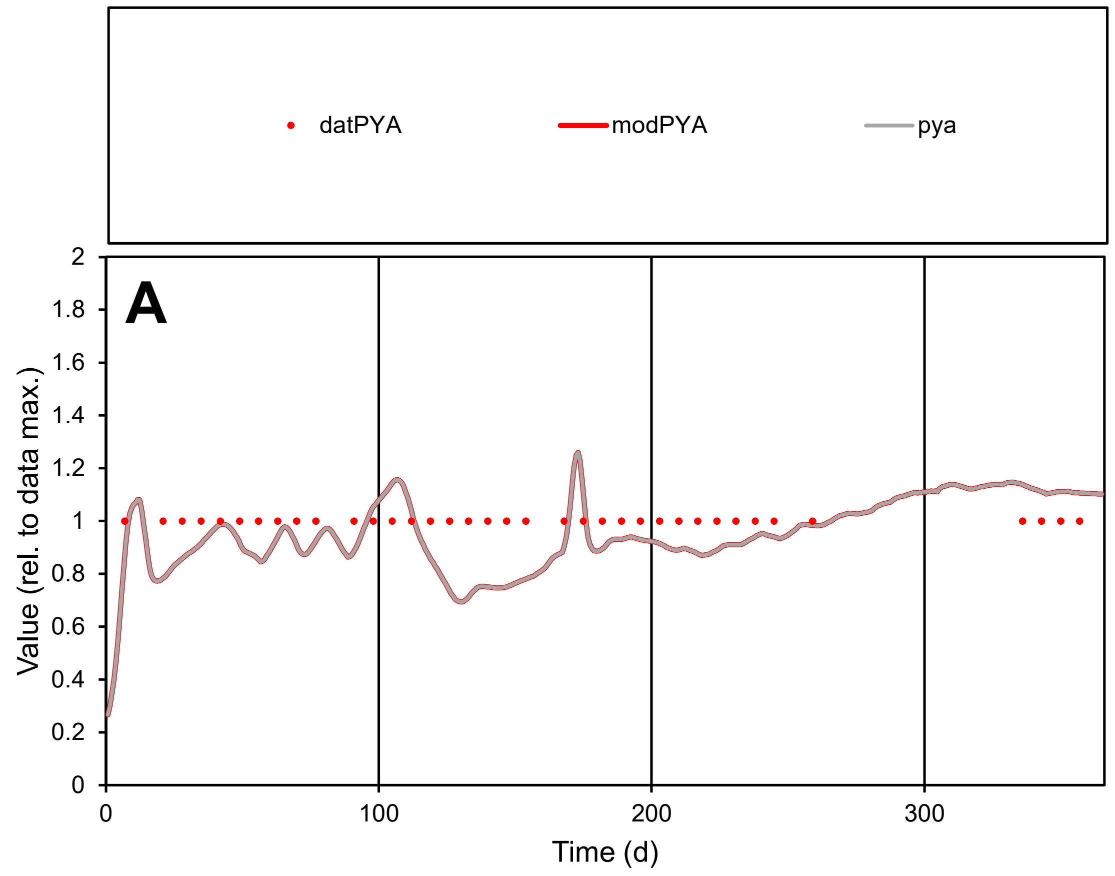


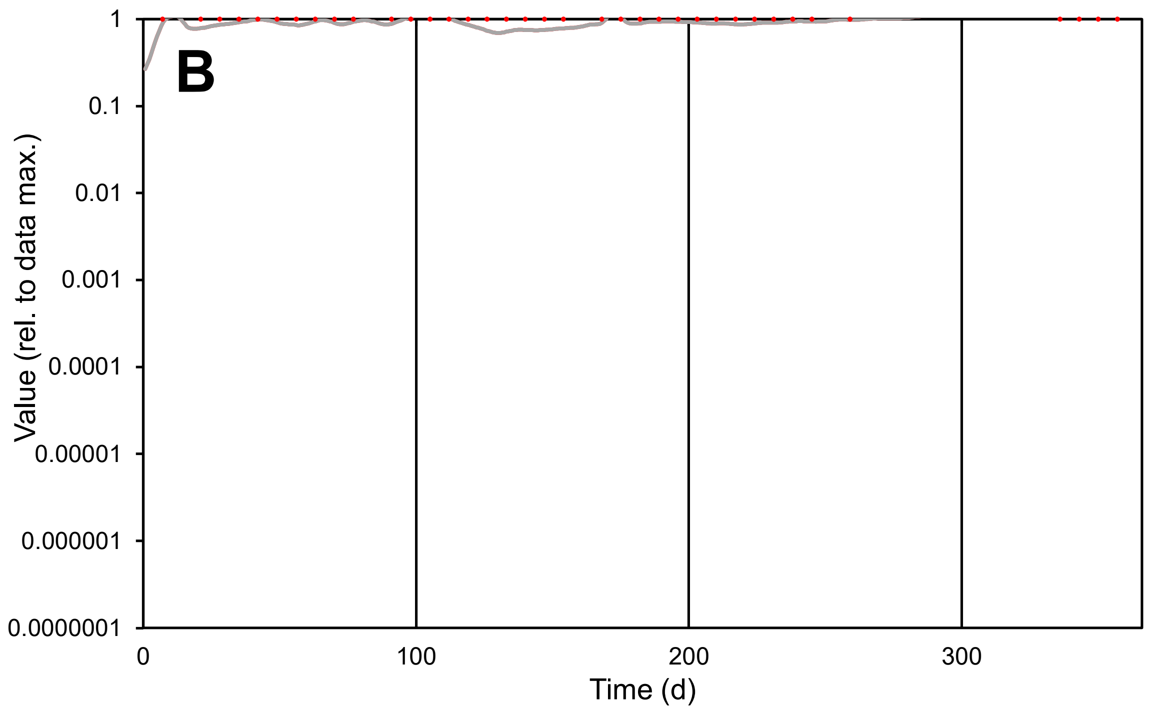


**C1**

Variables
PYA Provide jtd or Dst
CalibOnly?
2 1 to reproduce of
oNormalize?
2 1=no, 2=yes

ofxn calc*
0.13059
ofxn mod
0.16897

sRun
RunA1
zout
1

Ave
Data
1.8E-06
Model
1.7E-06

data/vMax
1.8E-06

ofxng
311.734

Min d/m
1.40031

ave/min d
1
ave/min m
1.30079

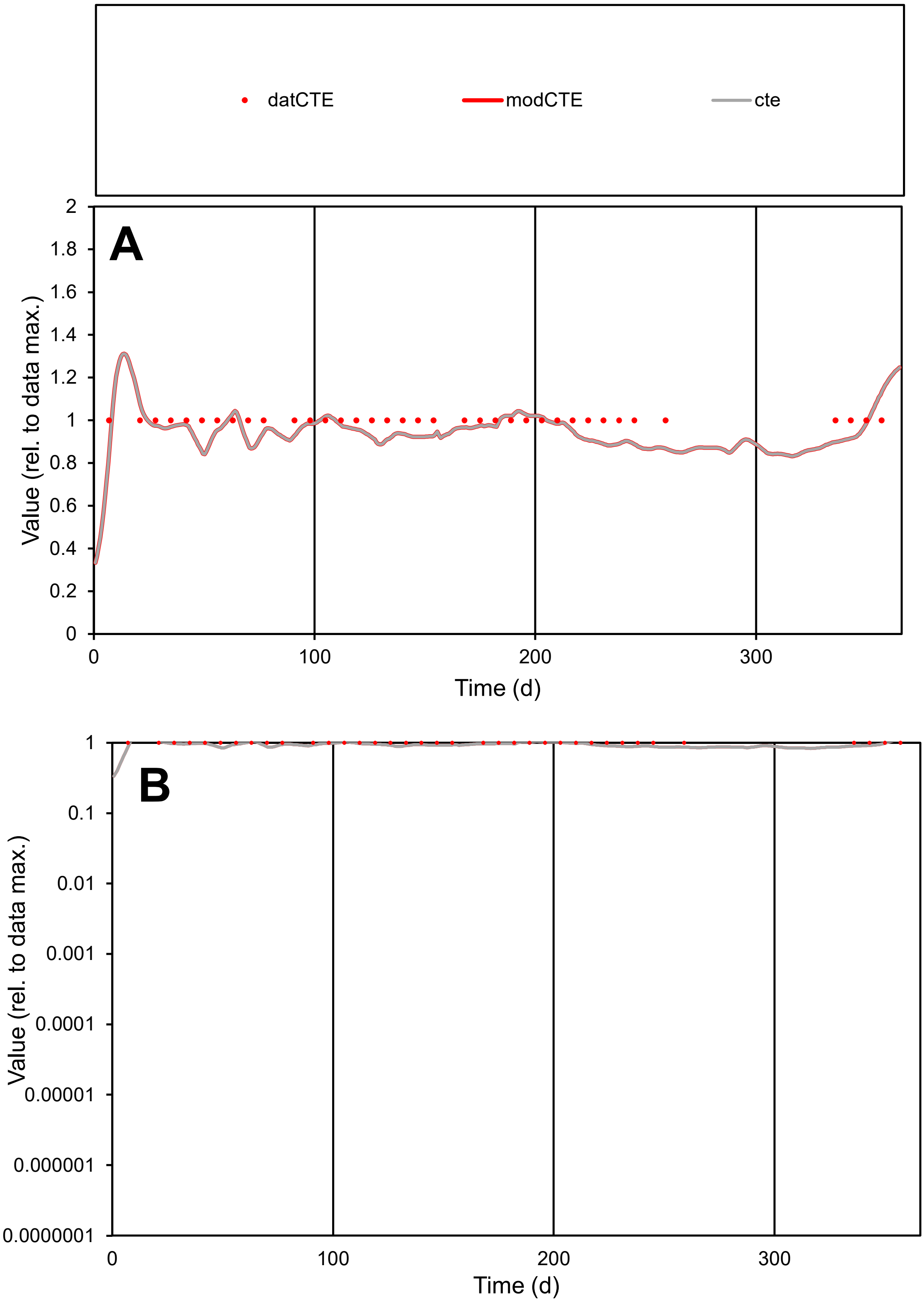


**C1**

Variables
CTE Provide jtd or Dst
CalibOnly?
2 1 to reproduce of
oNormalize?
2 1=no, 2=yes

ofxn calc*
0.07414
ofxn mod
0.1322

sRun
RunA1
zout
1

Ave
Data
6.1E-06
Model
5.9E-06

data/vMax
6.1E-06

ofxng
311.734

Min d/m
1.18744

ave/min d
1
ave/min m
1.1372

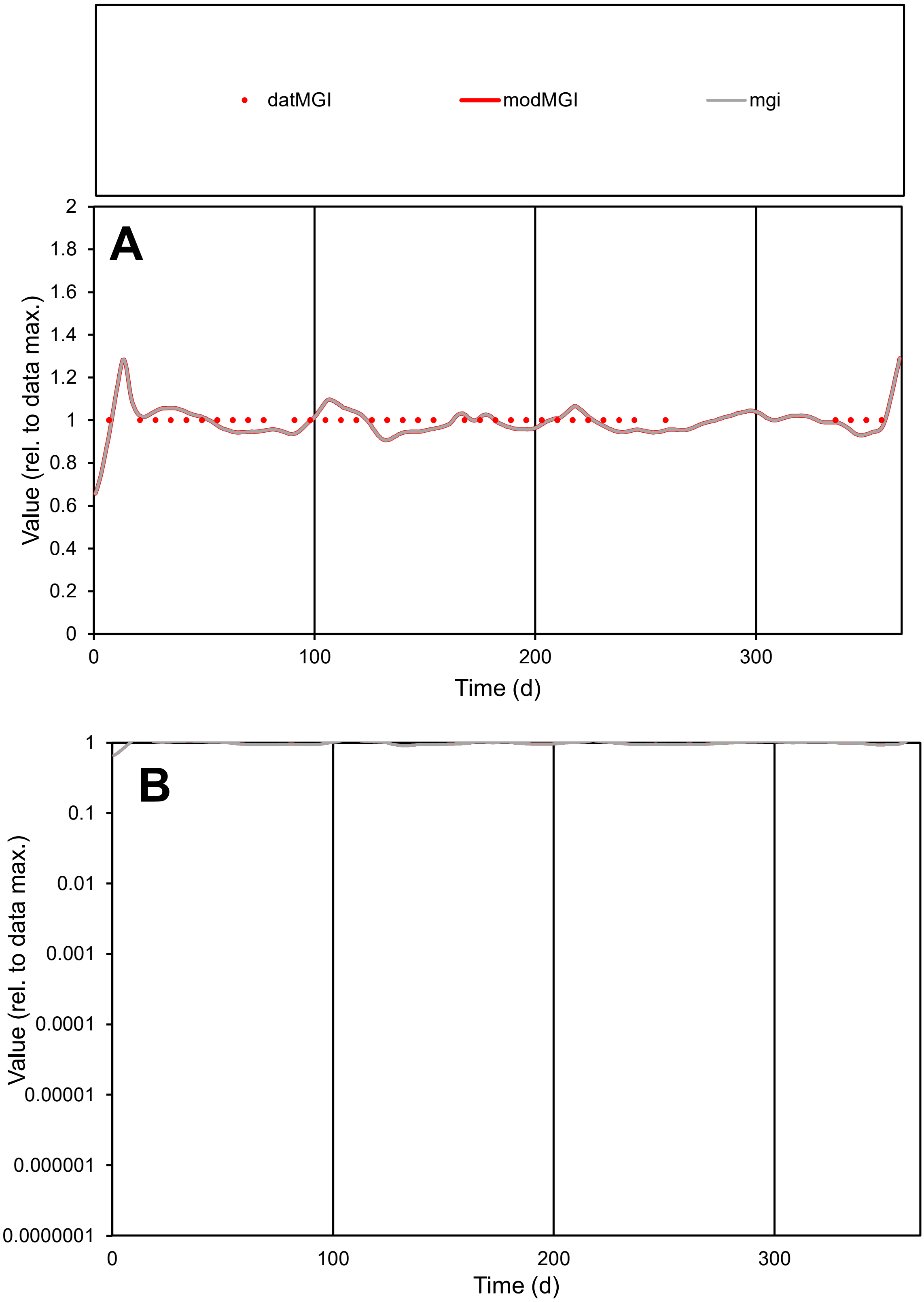


**C1**

Variables
MGI Provide jtd or Dst
CalibOnly?
2 1 to reproduce of
oNormalize?
2 1=no, 2=yes

ofxn calc*
0.04487
ofxn mod
0.11965

sRun
RunA1
zout
1

Ave
Data
1E-05
Model
1E-05

data/vMax
1E-05

ofxng
311.734

Min d/m
1.09589

ave/min d
1
ave/min m
1.08285

331_Calib_S_A

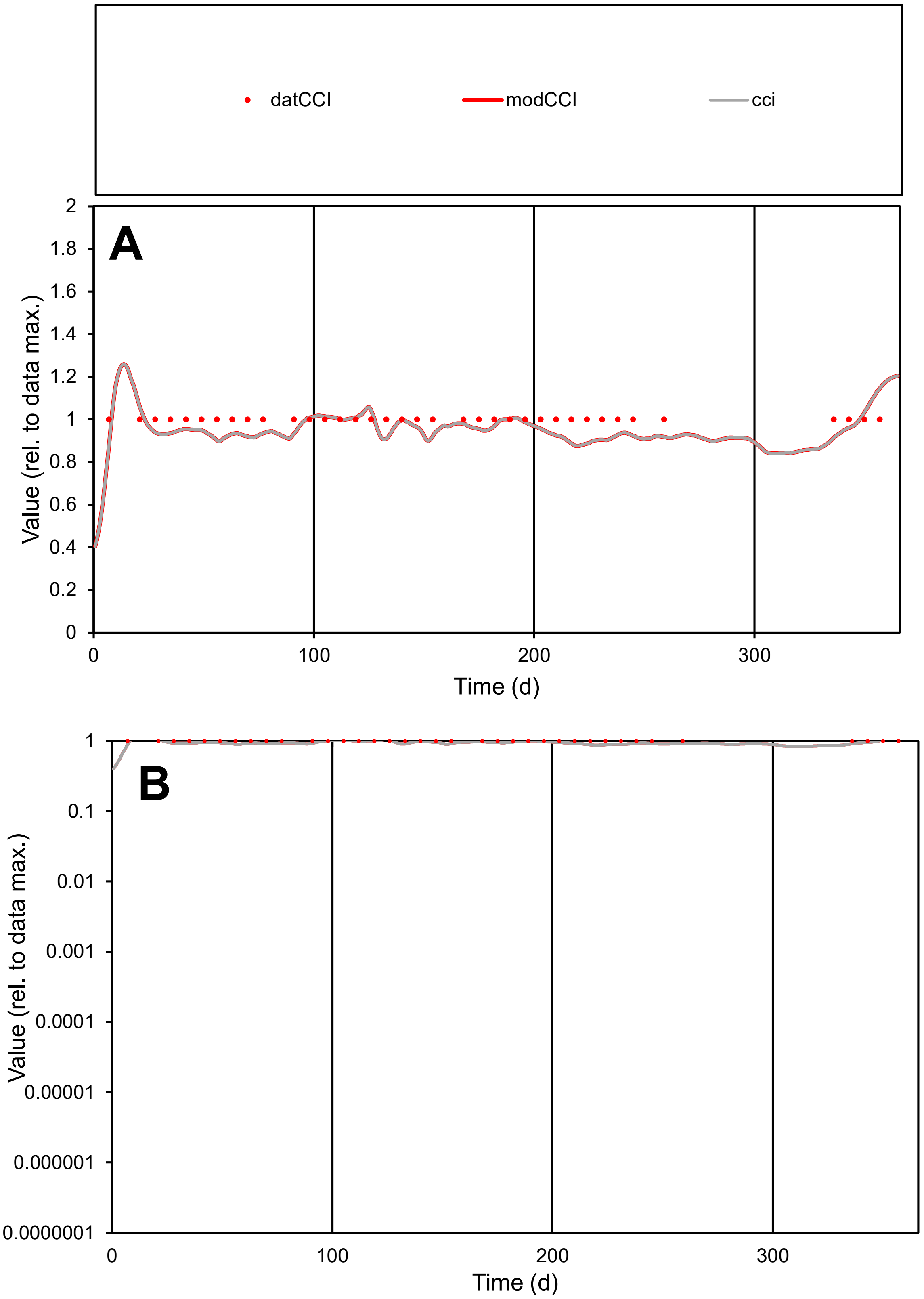


**C1**

Variables
CCI Provide jtd or Dst
CalibOnly?
2 1 to reproduce of
oNormalize?
2 1=no, 2=yes

ofxn calc*
0.06537
ofxn mod
0.12721

sRun
RunA1
zout
1

Ave
Data
1.1E-05
Model
1.1E-05

data/vMax
1.1E-05

ofxng
311.734

Min d/m
1.13776

ave/min d
1
ave/min m
1.09405

331_Calib_S_A

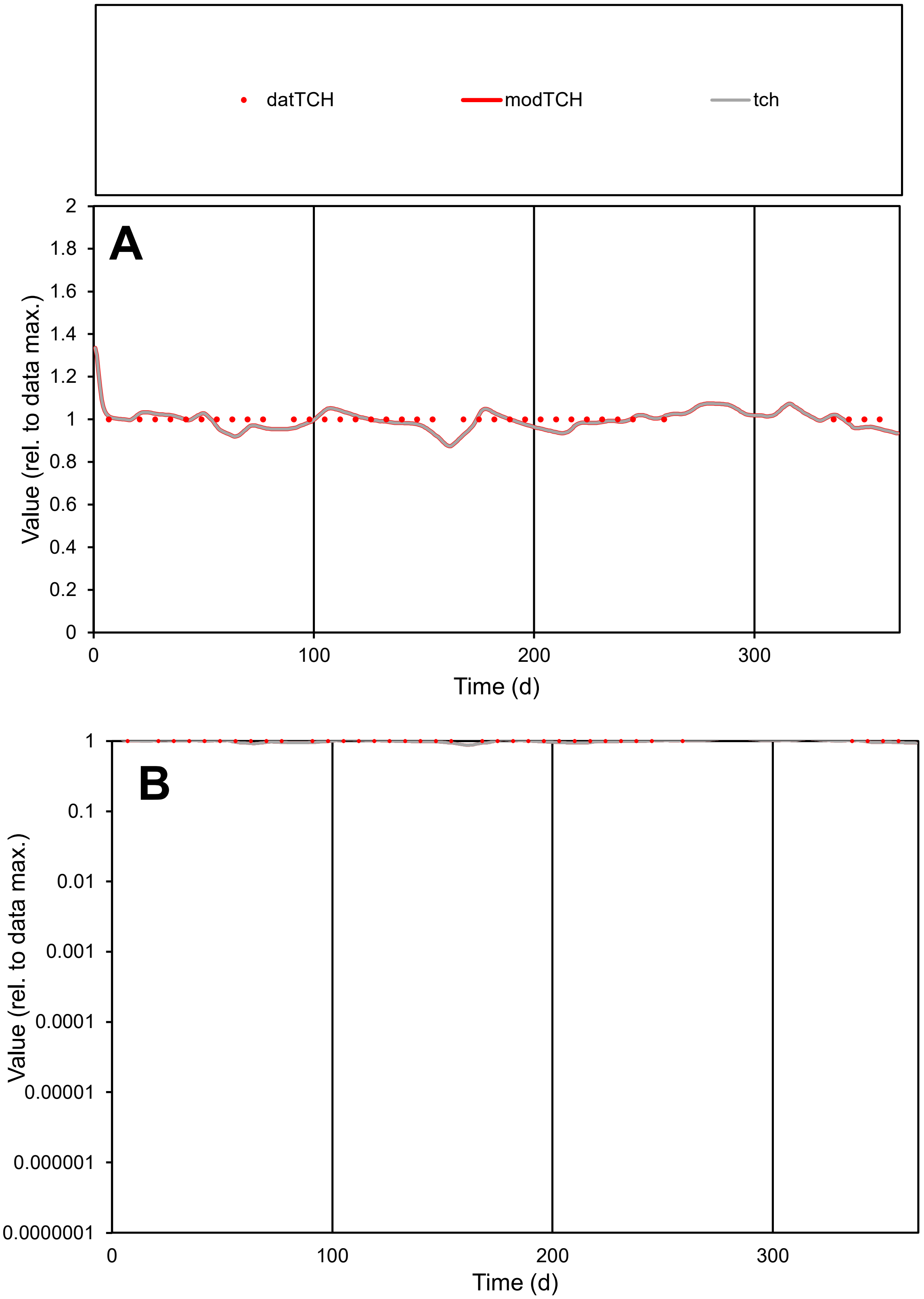


**C1**

Variables

| | |
|---|---|
| TCH | Provide jtd or Dst |
| CalibOnly? | |
| 2 | 1 to reproduce of |
| oNormalize? | |
| 2 | 1=no, 2=yes |

ofxn calc*
0.03543
ofxn mod
0.11628

sRun
RunA1
zout
1

Ave
Data
1.7E-06
Model
1.7E-06

data/vMax
1.7E-06

ofxng
311.734

Min d/m
1.08727

ave/min d
1
ave/min m
1.07434

331_Calib_S_A

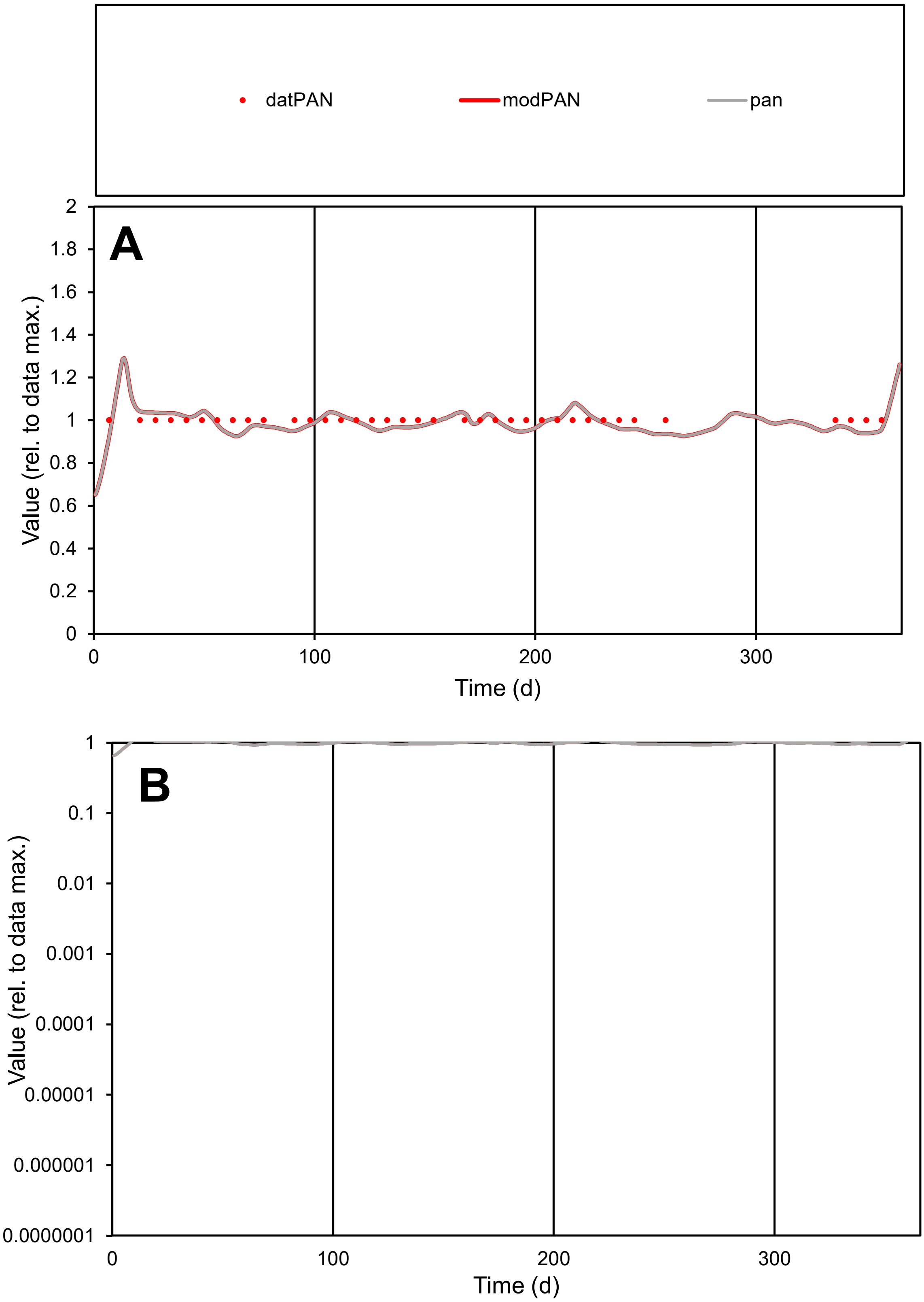


**C1**

Variables
PAN Provide jtd or Dst
CalibOnly?
2 1 to reproduce of
oNormalize?
2 1=no, 2=yes

ofxn calc*
0.03677
ofxn mod
0.11693

sRun
RunA1
zout
1

Ave
Data
9.1E-06
Model
9E-06

data/vMax
9.1E-06

ofxng
311.734

Min d/m
1.08044

ave/min d
1
ave/min m
1.06765

331_Calib_S_A

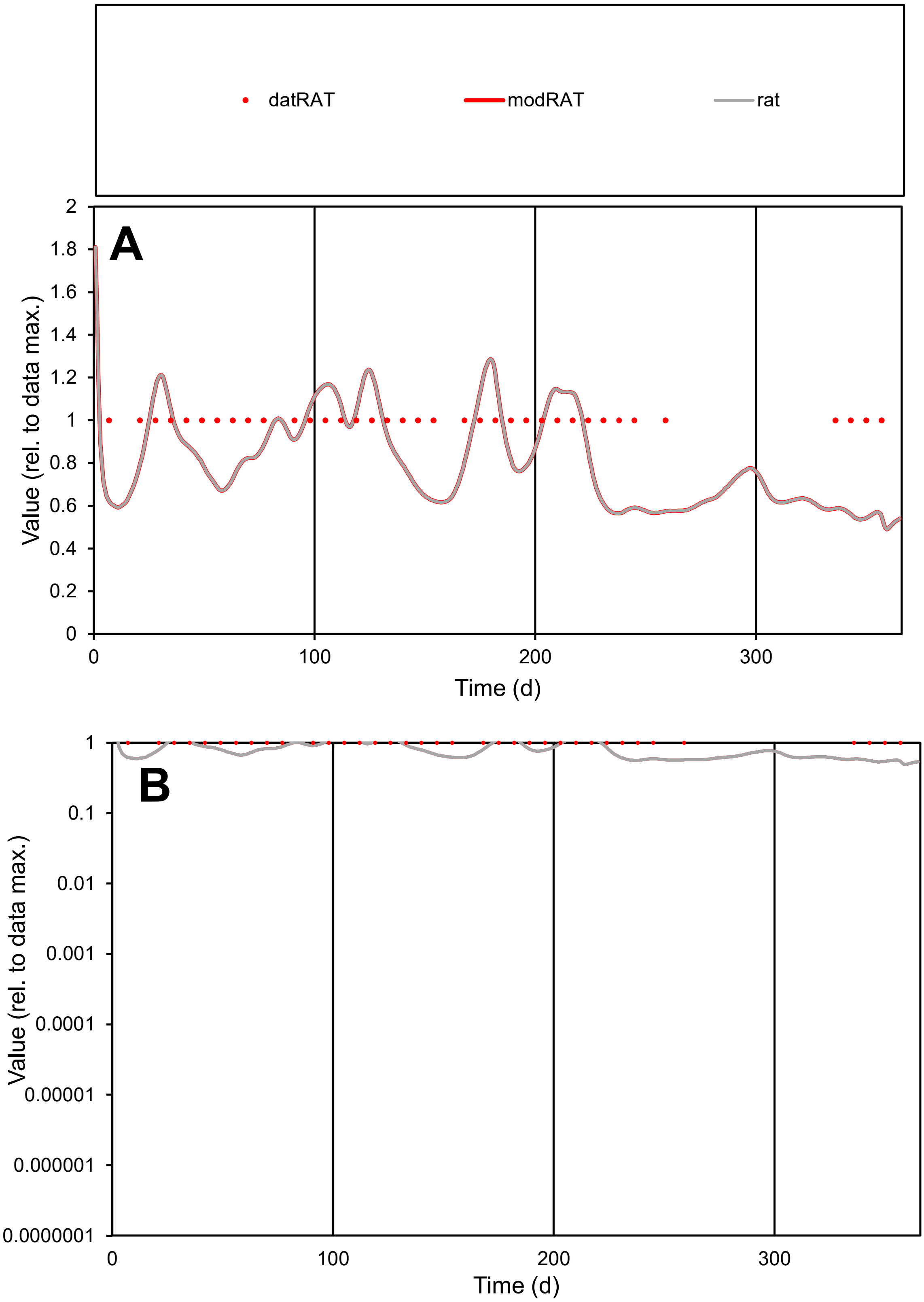


**C1**

Variables

RAT Provide jtd or Dst

CalibOnly?

2 1 to reproduce of

oNormalize?

2 1=no, 2=yes

ofxn calc*

0.26391

ofxn mod

0.28459

sRun

RunA1

zout

1

Ave

Data

1.1E-05

Model

9.6E-06

data/vMax

1.1E-05

ofxng

311.734

Min d/m

1.83594

ave/min d

1

ave/min m

1.55351

331_Calib_S_A

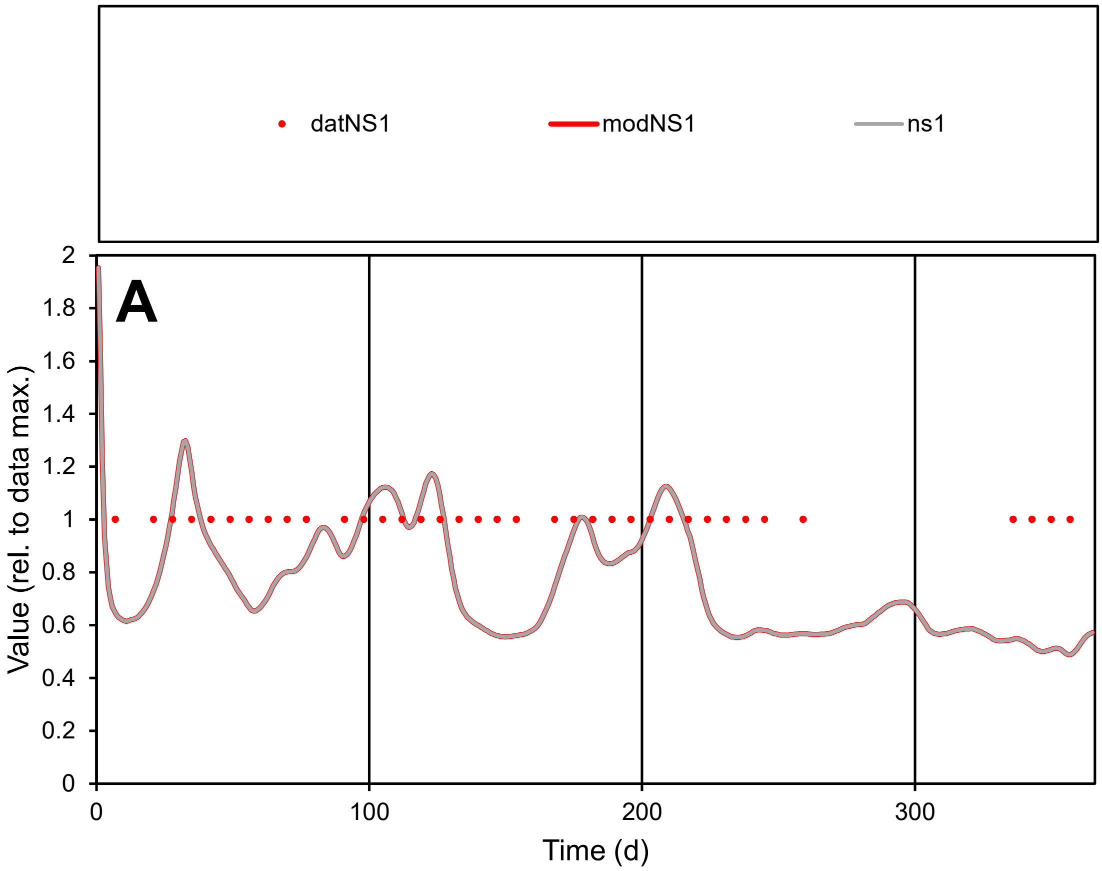


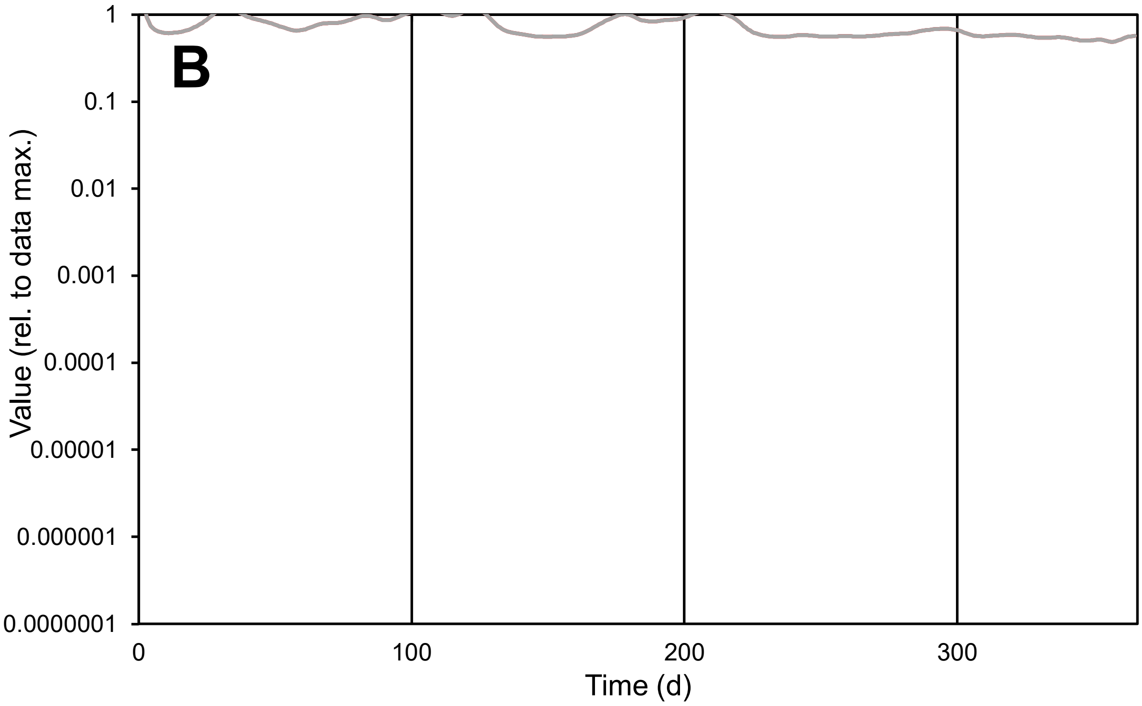


**C1**

Variables

NS1 Provide jtd or Dst

CalibOnly?

2 1 to reproduce of

oNormalize?

2 1=no, 2=yes

ofxn calc*

0.29226

ofxn mod

0.30895

sRun

RunA1

zout

1

Ave

Data

2.7E-05

Model

2.2E-05

data/vMax

2.7E-05

ofxng

311.734

Min d/m

2.00292

ave/min d

1

ave/min m

1.58527

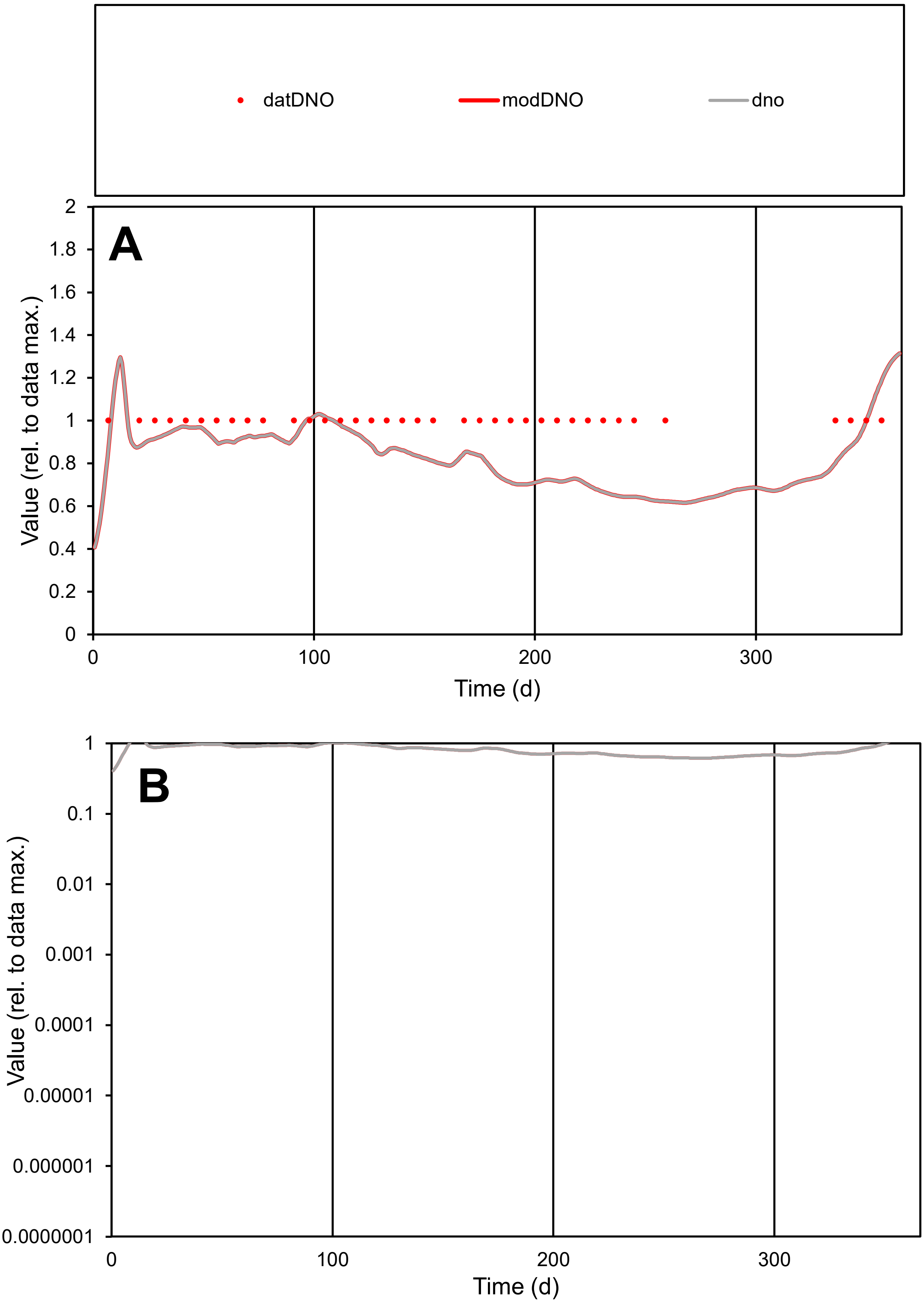


**C1**

Variables

DNO Provide jtd or Dst

CalibOnly?

2 1 to reproduce of

oNormalize?

2 1=no, 2=yes

ofxn calc*

0.19505

ofxn mod

0.22067

sRun

RunA1

zout

1

Ave

Data

1.8E-05

Model

1.5E-05

data/vMax

1.8E-05

ofxng

311.734

Min d/m

1.60889

ave/min d

1

ave/min m

1.37715

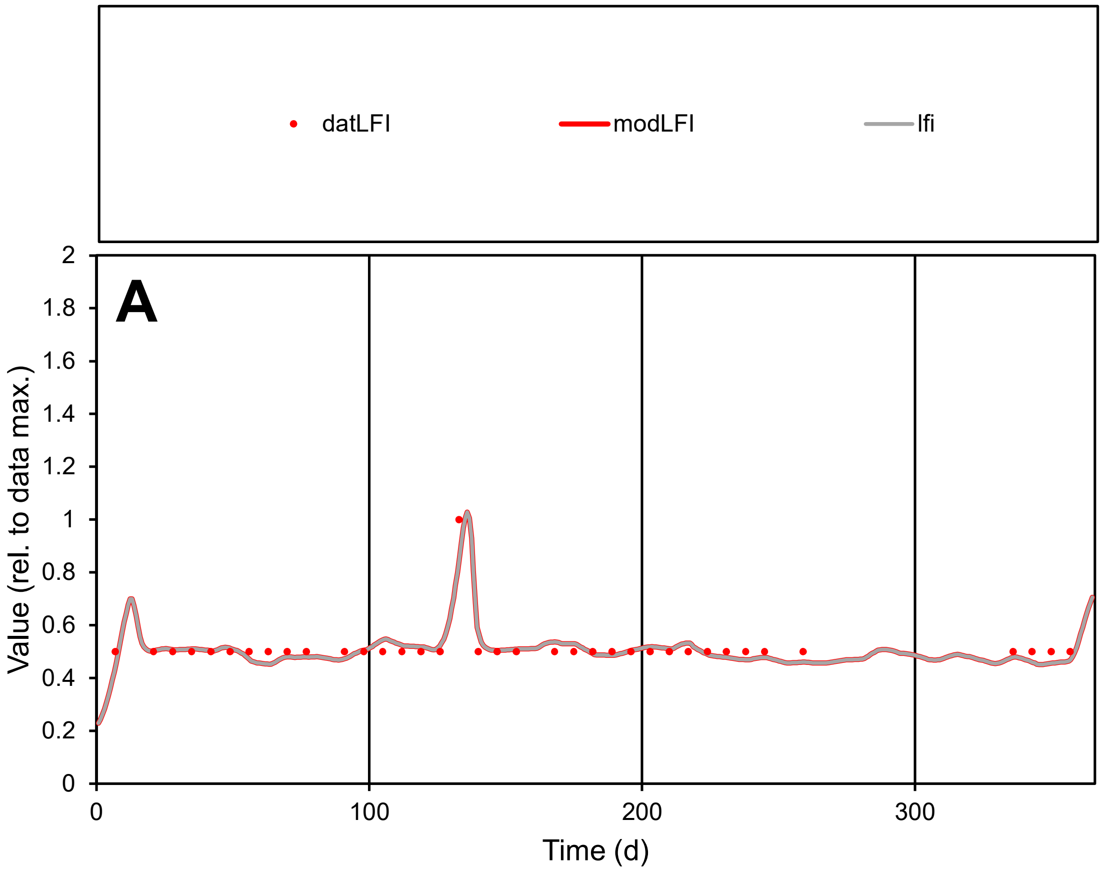


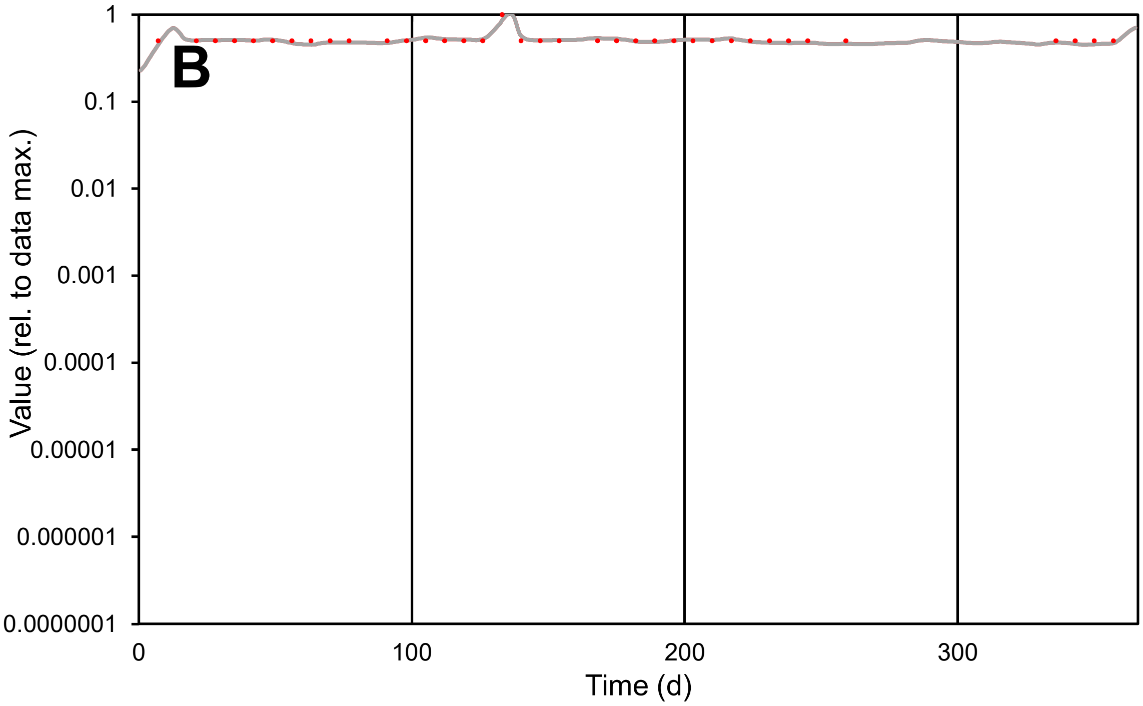


**C1**

Variables

LFI Provide jtd or Dst

CalibOnly?

2 1 to reproduce of

oNormalize?

2 1=no, 2=yes

ofxn calc*

0.04978

ofxn mod

0.12267

sRun

RunA1

zout

1

Ave

Data

8.5E-06

Model

8.5E-06

data/vMax

1.7E-05

ofxng

311.734

Min d/m

1.09661

ave/min d

1.02703

ave/min m

1.12097

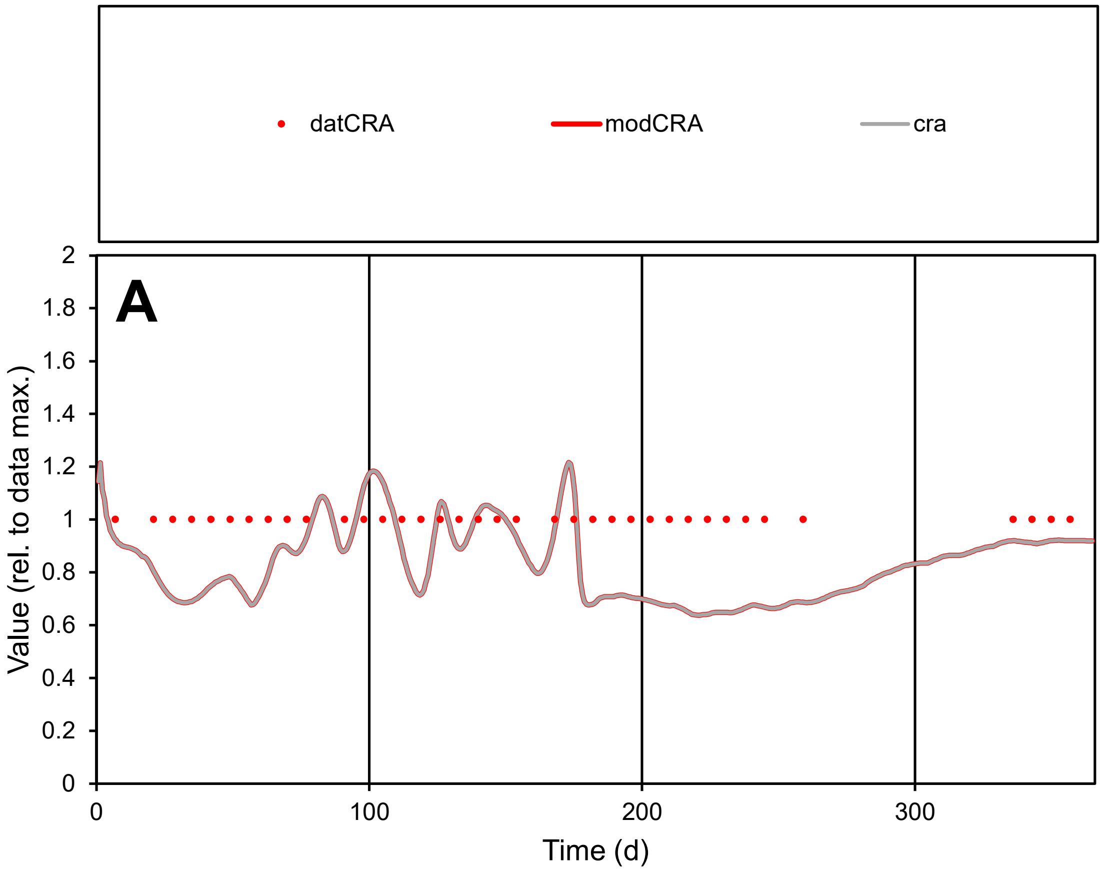


C1

Variables
CRA Provide jtd or Dst
CalibOnly?
2 1 to reproduce of
oNormalize?
2 1=no, 2=yes

ofxn calc*
0.22474
ofxn mod
0.24661

sRun
RunA1
zout
1

Ave
Data
4.9E-06
Model
4E-06

data/vMax
4.9E-06

ofxng
311.734

Min d/m
1.55698

ave/min d
1
ave/min m
1.29586

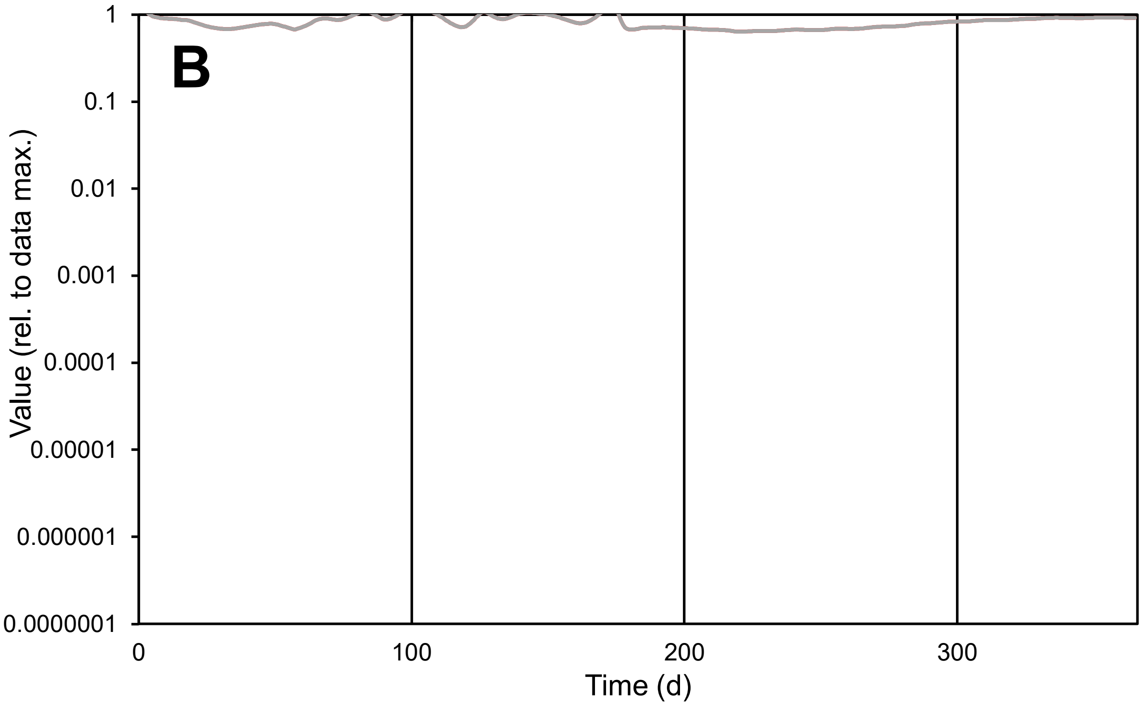

331_Calib_S_A

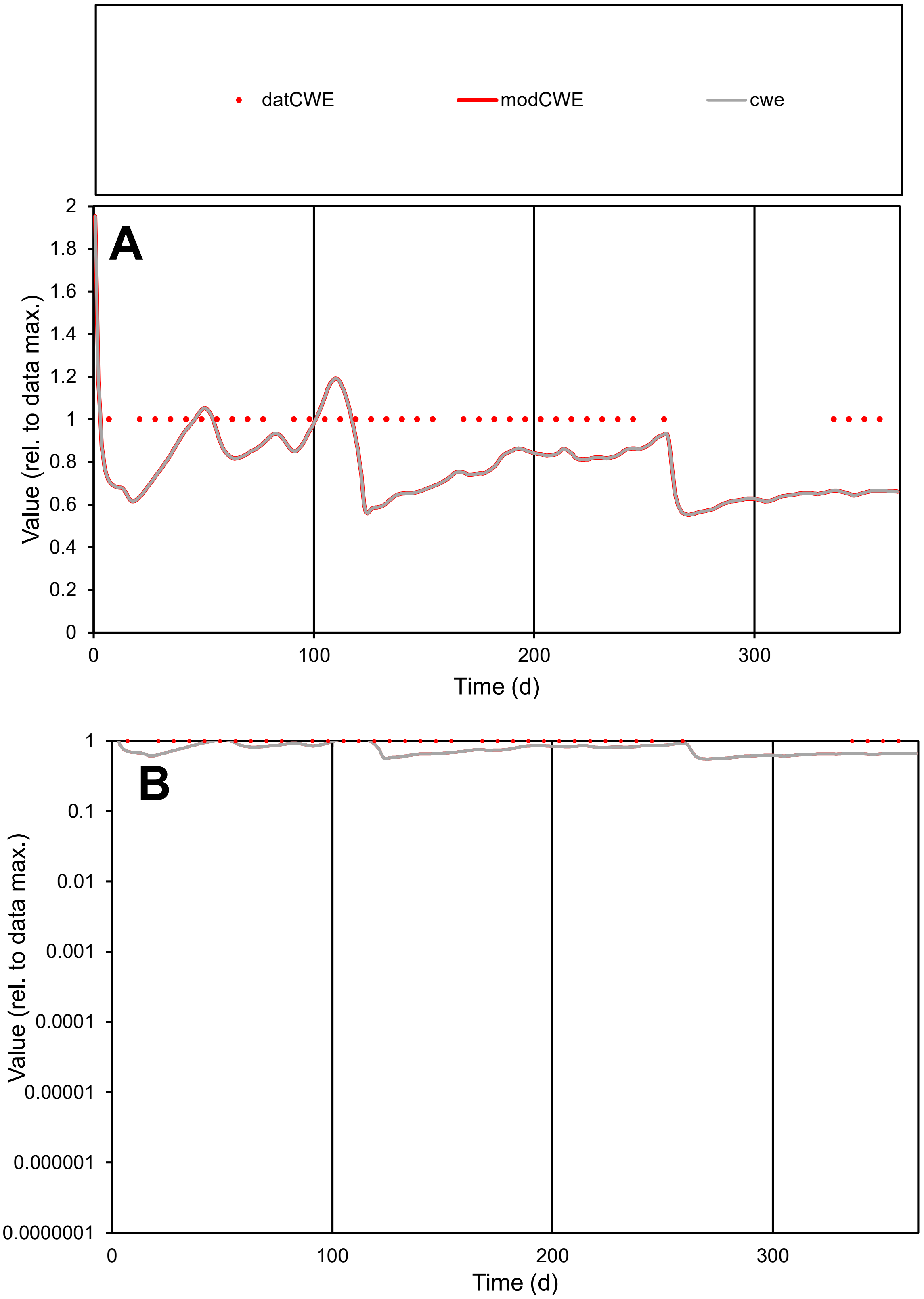


**C1**

Variables

CWE Provide jtd or Dst

CalibOnly?

2 1 to reproduce of

oNormalize?

2 1=no, 2=yes

ofxn calc*

0.23122

ofxn mod

0.25263

sRun

RunA1

zout

1

Ave

Data

2.6E-05

Model

2.1E-05

data/vMax

2.6E-05

ofxng

311.734

Min d/m

1.70912

ave/min d

1

ave/min m

1.38736

331_Calib_S_A

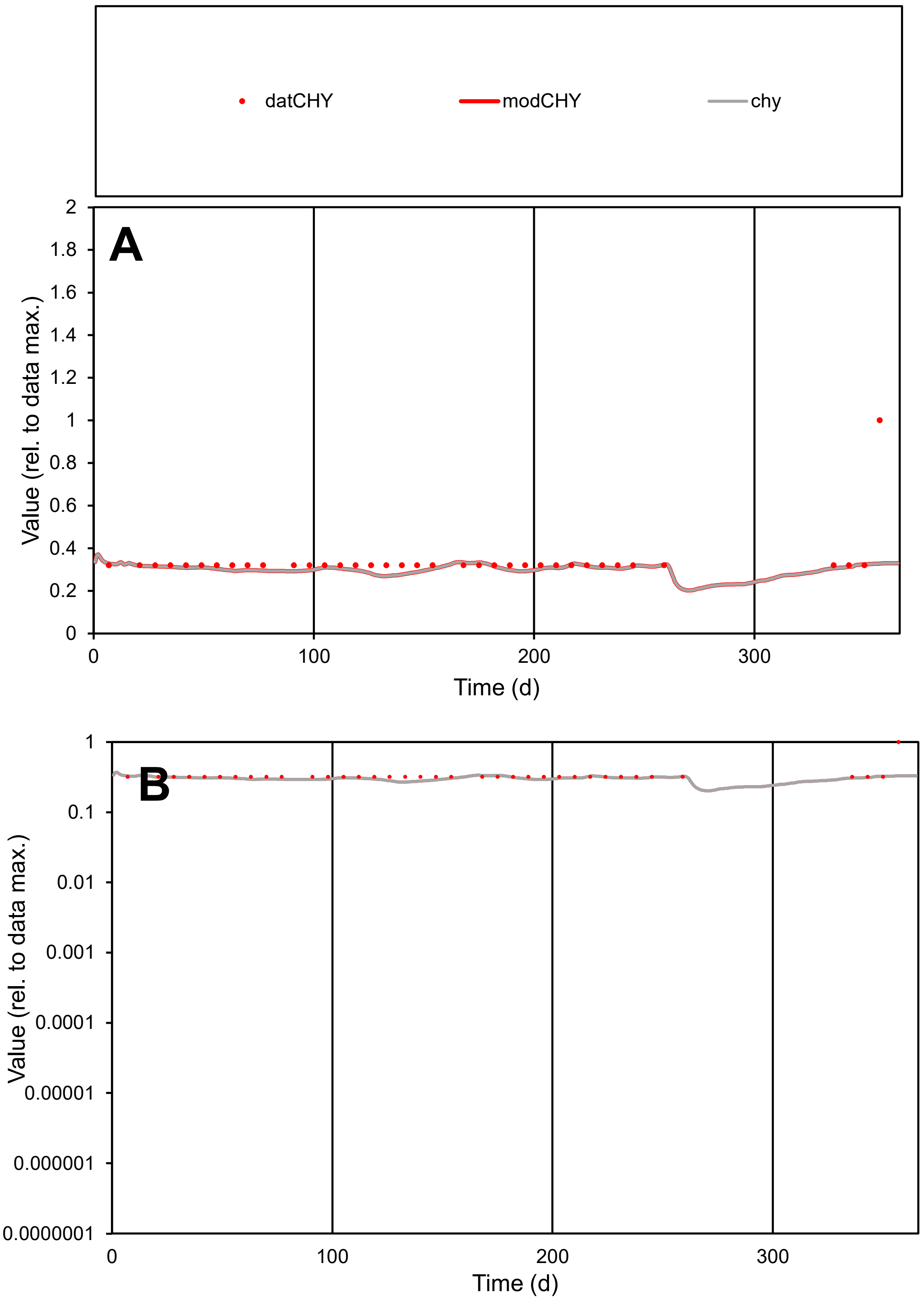


**C1**

Variables

CHY Provide jtd or Dst

CalibOnly?

2 1 to reproduce of

oNormalize?

2 1=no, 2=yes

ofxn calc*

0.3313

ofxn mod

0.34866

sRun

RunA1

zout

1

Ave

Data

7.1E-06

Model

6.4E-06

data/vMax

2.1E-05

ofxng

311.734

Min d/m

1.18217

ave/min d

1.05752

ave/min m

1.13484

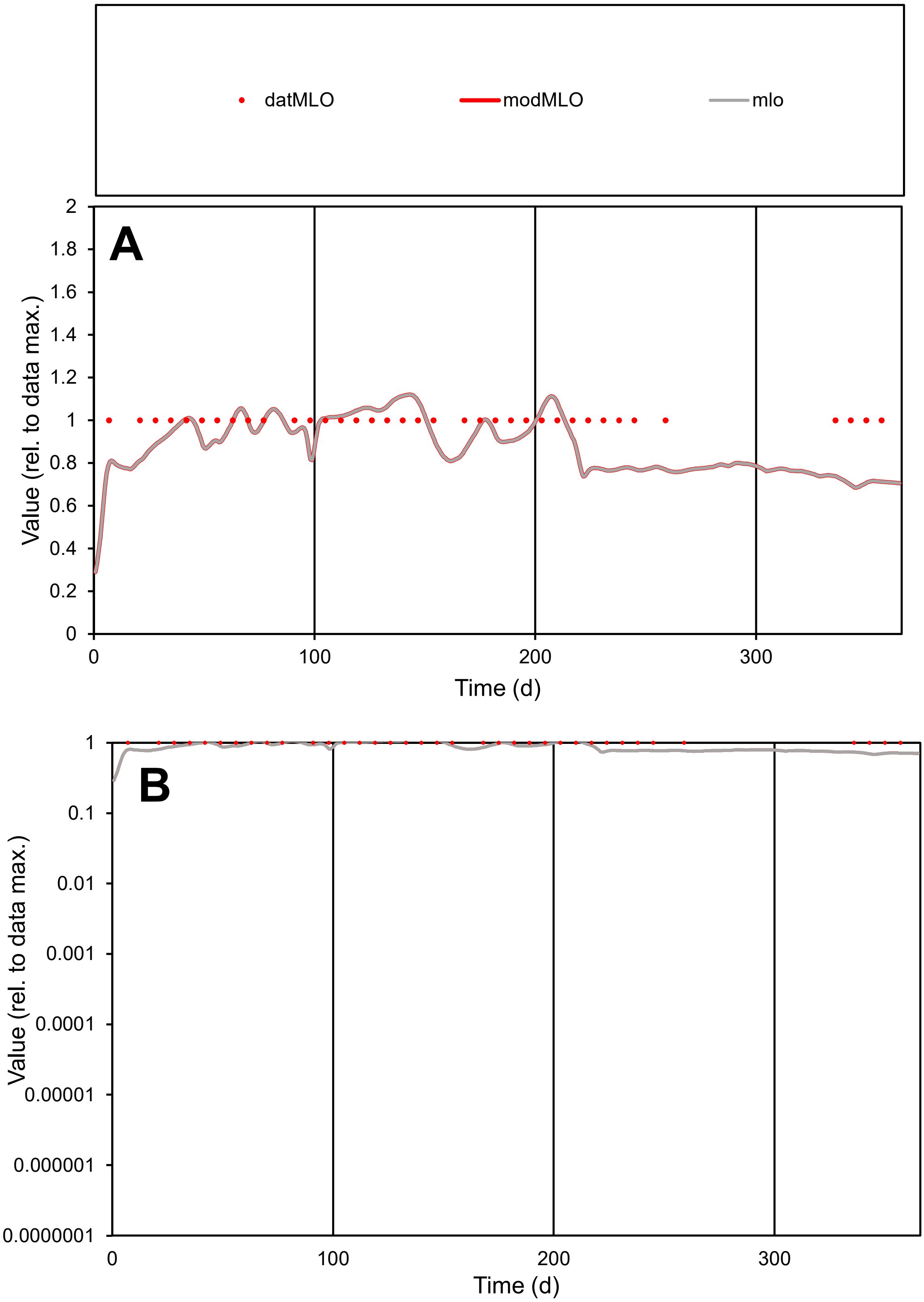


**C1**

Variables
MLO Provide jtd or Dst
CalibOnly?
2 1 to reproduce of
oNormalize?
2 1=no, 2=yes

ofxn calc*
0.15036
ofxn mod
0.18457

sRun
RunA1
zout
1

Ave
Data
1E-05
Model
9.5E-06

data/vMax
1E-05

ofxng
311.734

Min d/m
1.44921

ave/min d
1
ave/min m
1.32

331_Calib_S_A

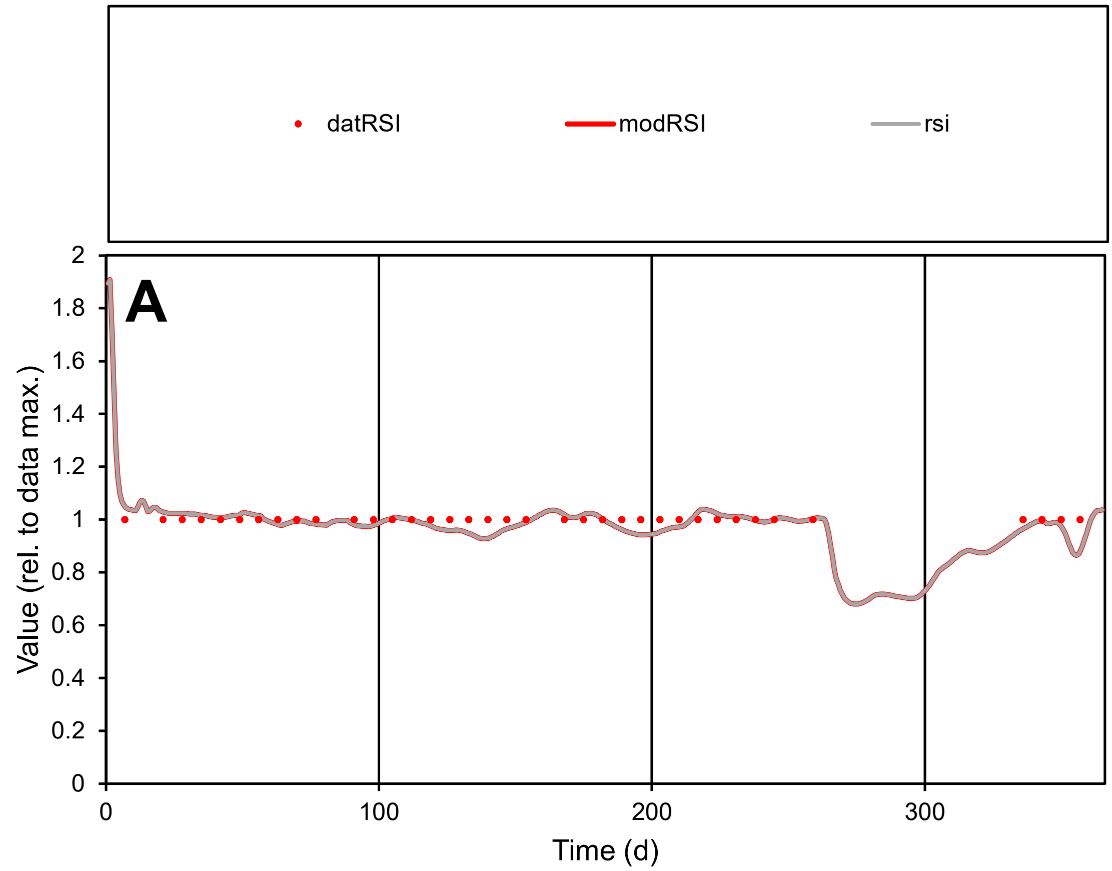


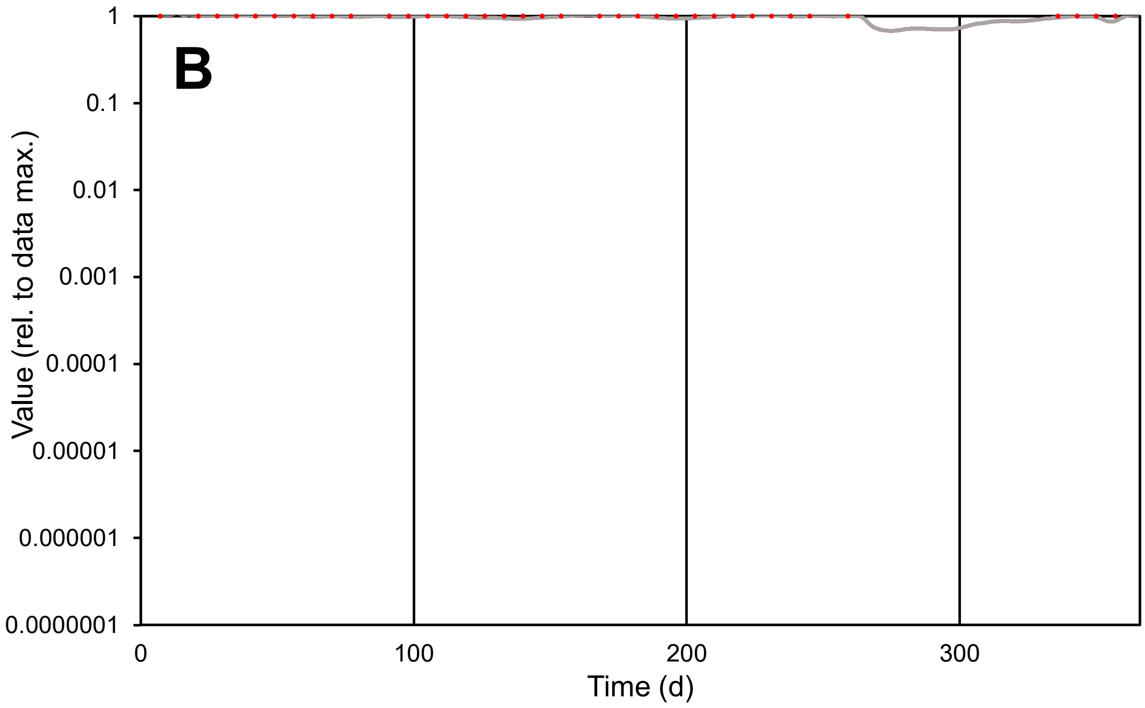


**C1**

Variables
RSI Provide jtd or Dst
CalibOnly?
2 1 to reproduce of
oNormalize?
2 1=no, 2=yes

ofxn calc*
0.03137
ofxn mod
0.11549

sRun
RunA1
zout
1

Ave
Data
1E-05
Model
1E-05

data/vMax
1E-05

ofxng
311.734

Min d/m
1.08213

ave/min d
1
ave/min m
1.07085

331_Calib_S_A

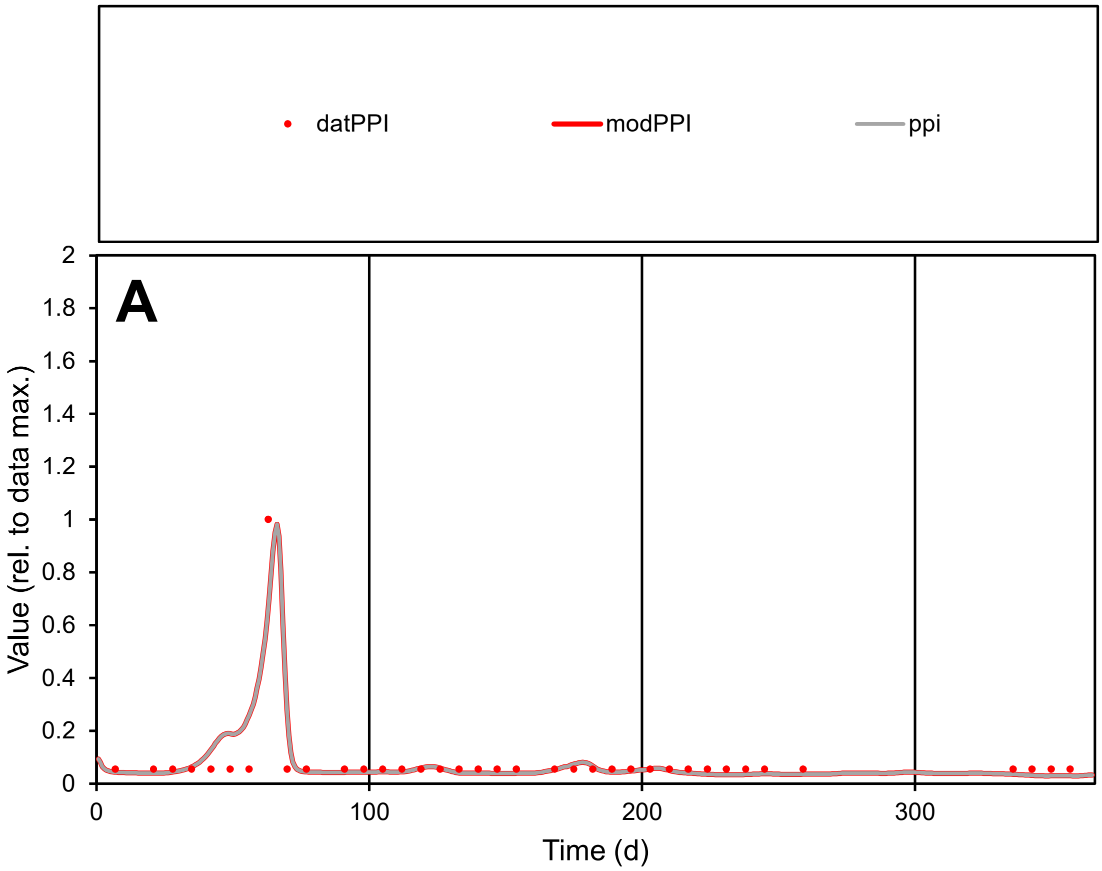


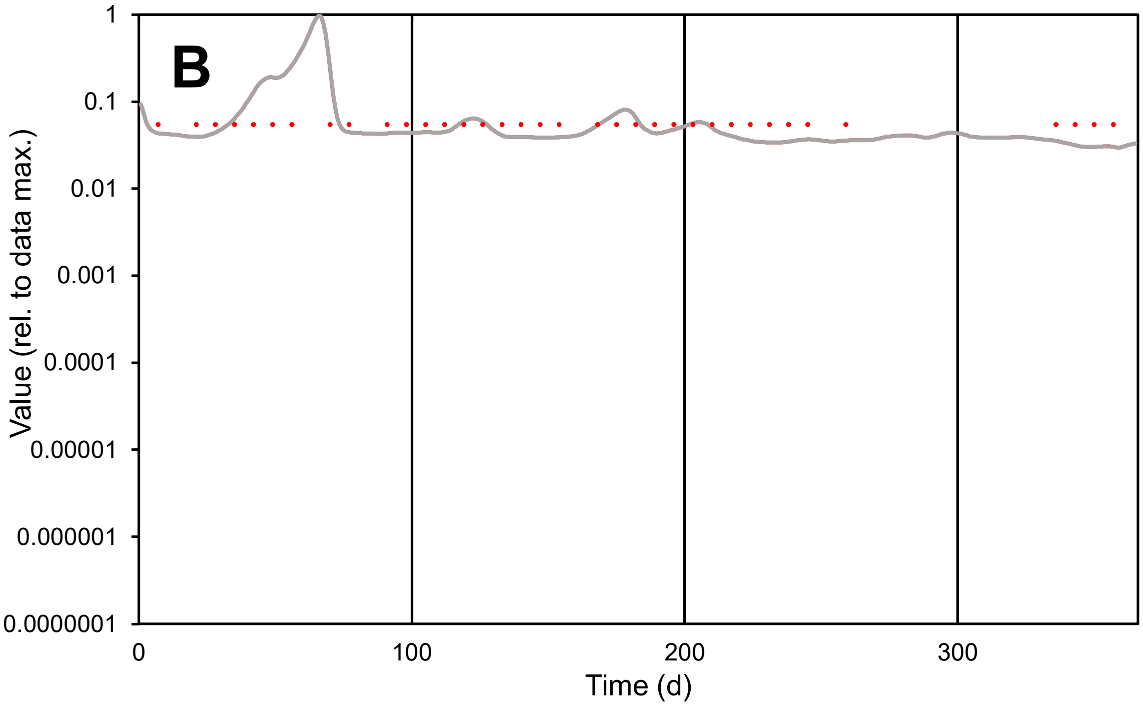


**C1**

Variables

PPI Provide jtd or Dst

CalibOnly?

2 1 to reproduce of

oNormalize?

2 1=no, 2=yes

ofxn calc*

0.57798

ofxn mod

0.67815

sRun

RunA1

zout

1

Ave

Data

2.1E-06

Model

2.1E-06

data/vMax

2.7E-05

ofxng

311.734

Min d/m

1.84533

ave/min d

1.46866

ave/min m

2.68293

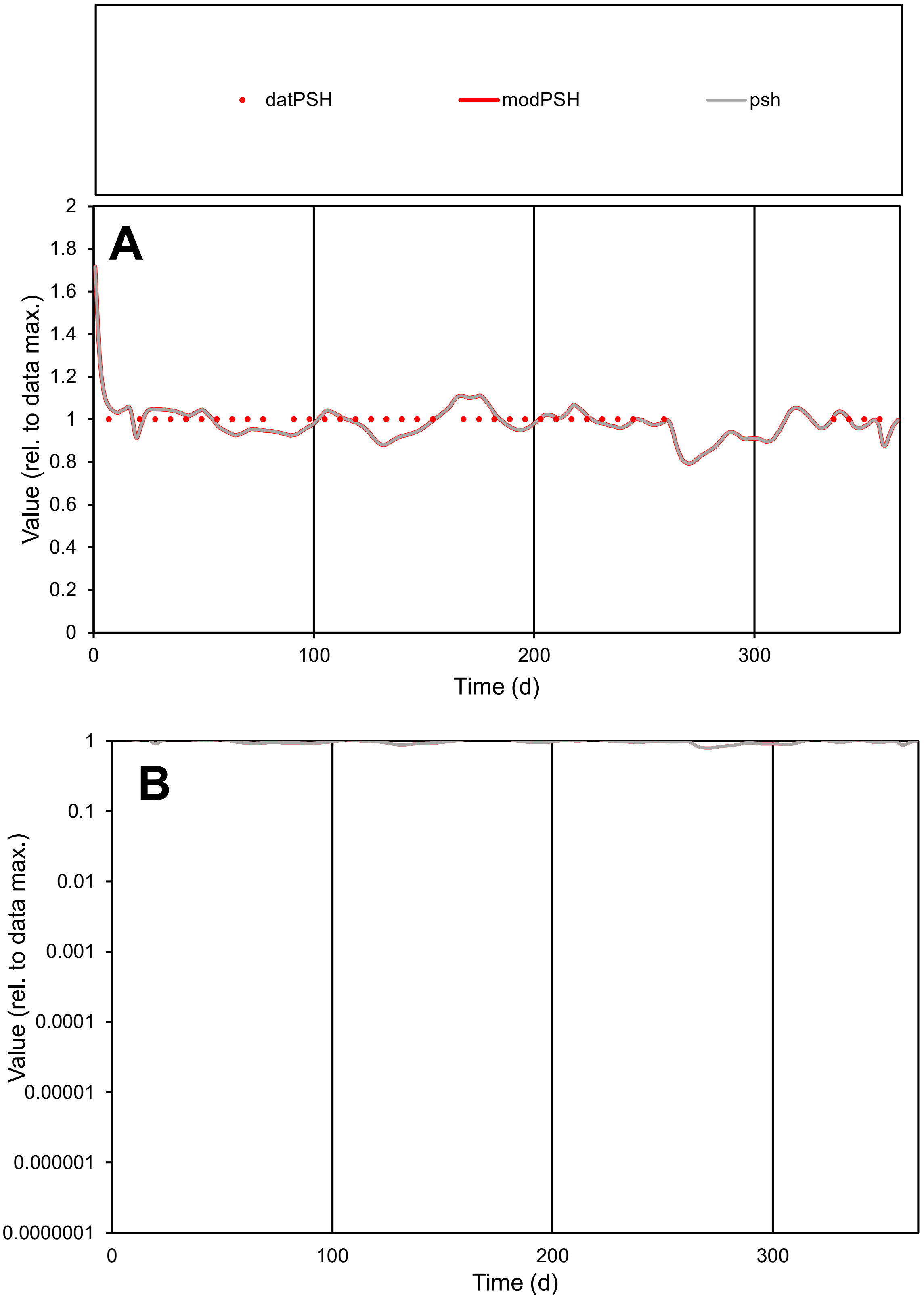


**C1**

Variables
PSH Provide jtd or Dst
CalibOnly?
2 1 to reproduce of
oNormalize?
2 1=no, 2=yes

ofxn calc*
0.05329
ofxn mod
0.12302

sRun
RunA1
zout
1

Ave
Data
1.3E-05
Model
1.3E-05

data/vMax
1.3E-05

ofxng
311.734

Min d/m
1.13757

ave/min d
1
ave/min m
1.12691

331_Calib_S_A

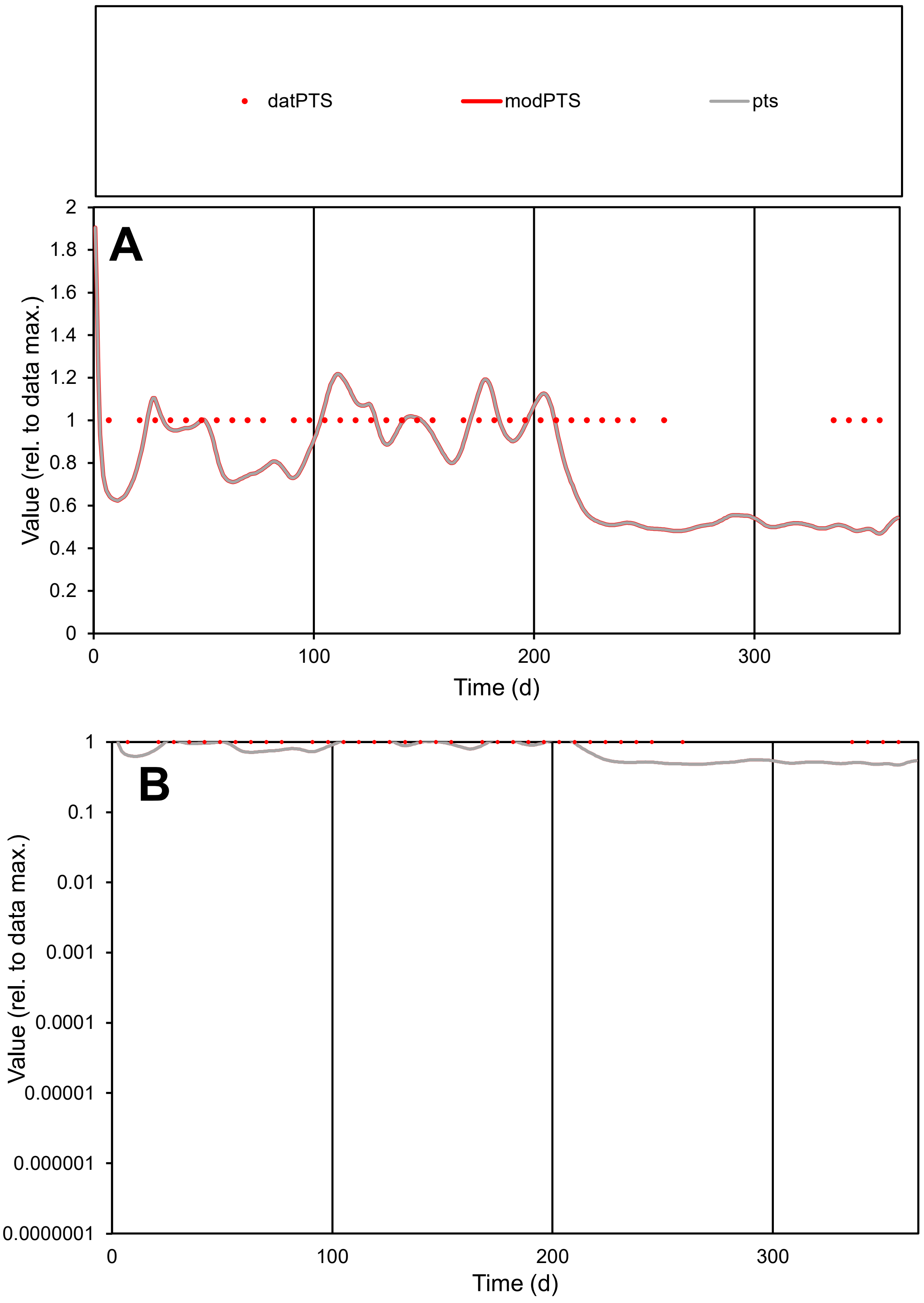


**C1**

Variables
PTS Provide jtd or Dst
CalibOnly?
2 1 to reproduce of
oNormalize?
2 1=no, 2=yes

ofxn calc*
0.28041
ofxn mod
0.29866

sRun
RunA1
zout
1

Ave
Data
9.7E-06
Model
8E-06

data/vMax
9.7E-06

ofxng
311.734

Min d/m
2.09272

ave/min d
1
ave/min m
1.73917

331_Calib_S_A

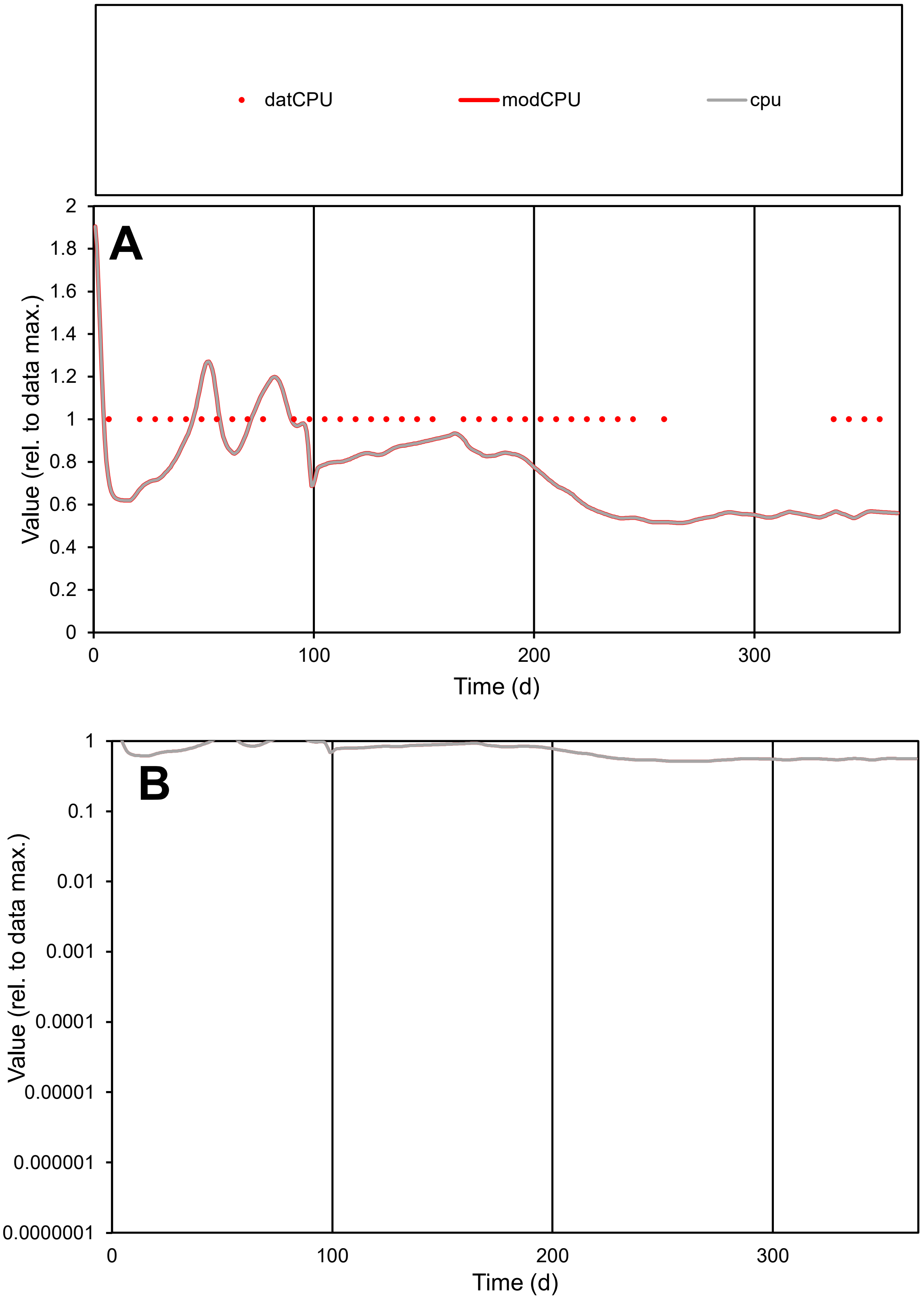


**C1**

Variables

CPU Provide jtd or Dst

CalibOnly?

2 1 to reproduce of

oNormalize?

2 1=no, 2=yes

ofxn calc*

0.28112

ofxn mod

0.29956

sRun

RunA1

zout

1

Ave

Data

4.4E-07

Model

3.4E-07

data/vMax

4.4E-07

ofxng

311.734

Min d/m

1.931

ave/min d

1

ave/min m

1.50577

331_Calib_S_A

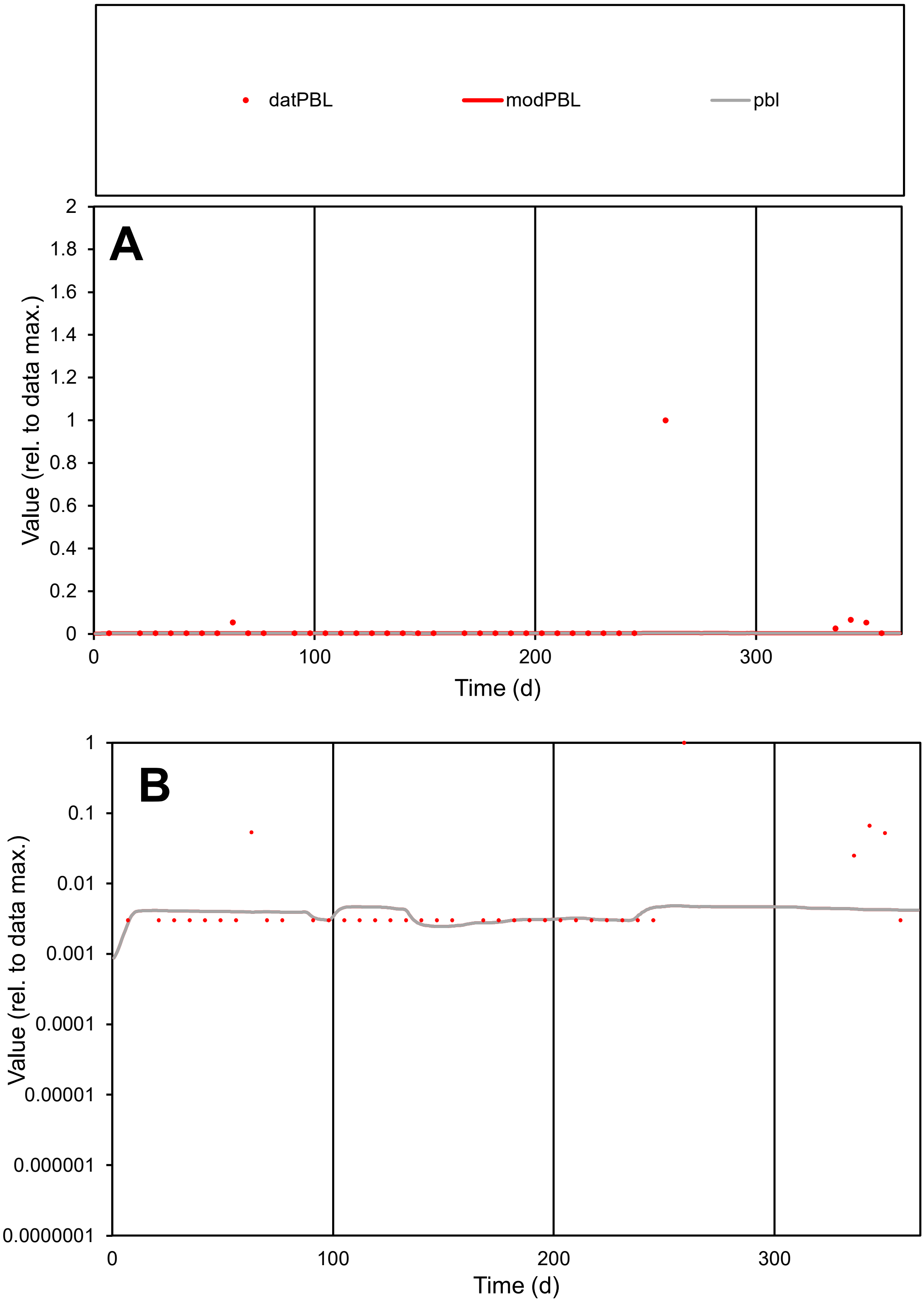


**C1**

Variables

PBL Provide jtd or Dst

CalibOnly?

2 1 to reproduce of

oNormalize?

2 1=no, 2=yes

ofxn calc*

4.70228

ofxn mod

4.71182

sRun

RunA1

zout

1

Ave

Data

4.9E-07

Model

5.2E-08

data/vMax

1.4E-05

ofxng

311.734

Min d/m

1.22687

ave/min d

11.6294

ave/min m

1.49955

331_Calib_S_A

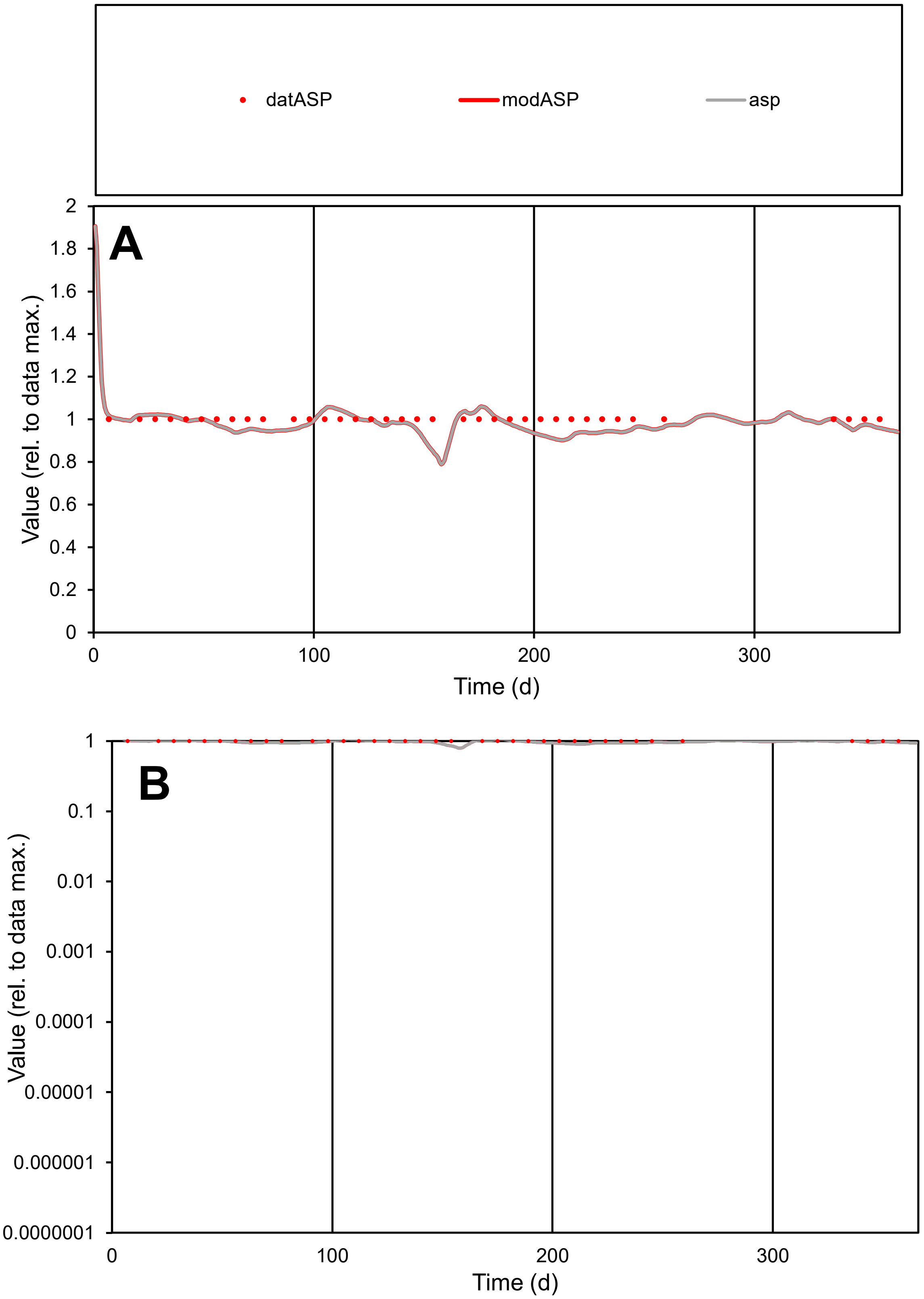


**C1**

Variables
ASP Provide jtd or Dst
CalibOnly?
2 1 to reproduce of
oNormalize?
2 1=no, 2=yes

ofxn calc*
0.05138
ofxn mod
0.12126

sRun
RunA1
zout
1

Ave
Data
5.8E-06
Model
5.6E-06

data/vMax
5.8E-06

ofxng
311.734

Min d/m
1.20241

ave/min d
1
ave/min m
1.17161

331_Calib_S_A

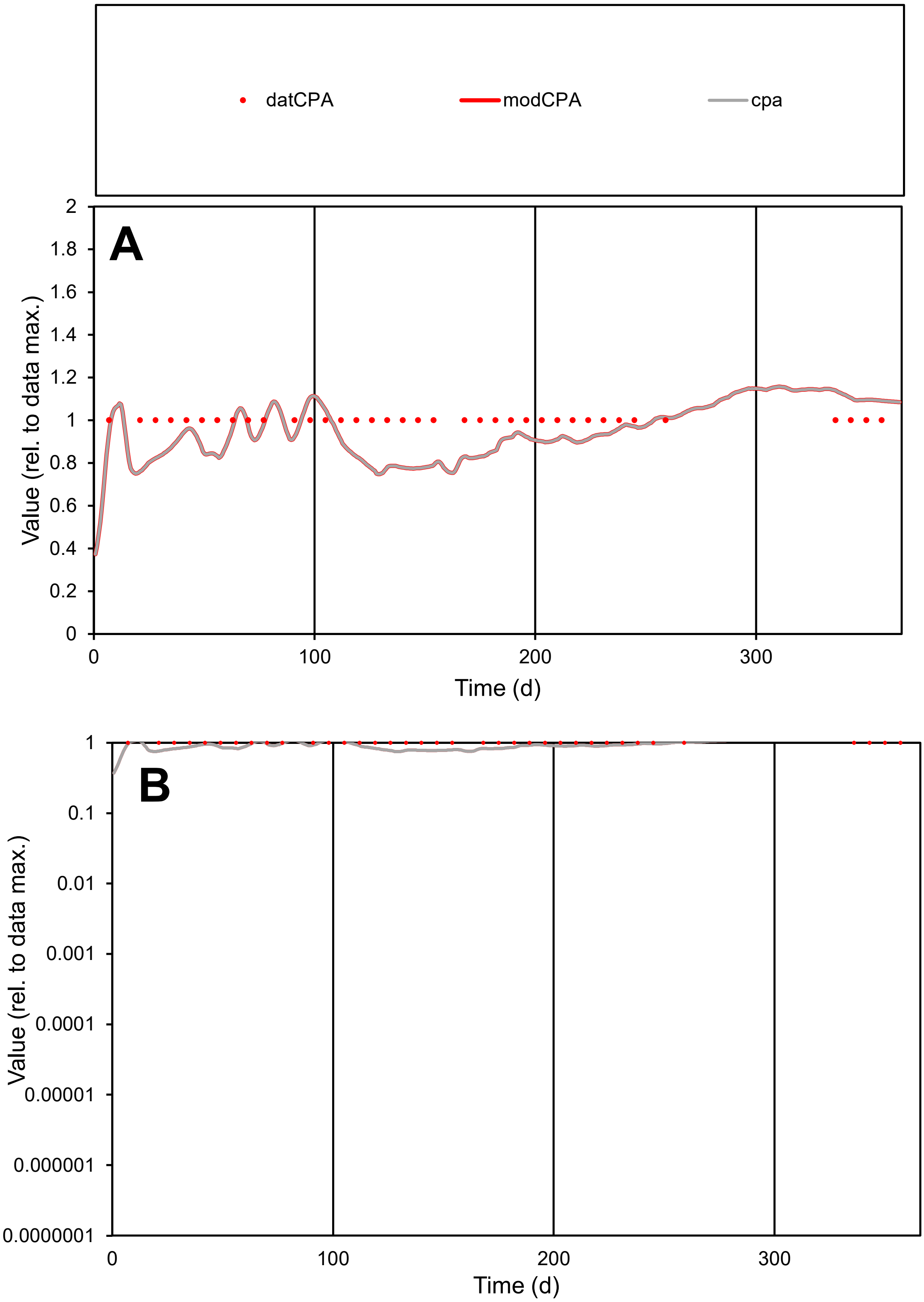


**C1**

Variables
CPA Provide jtd or Dst
CalibOnly?
2 1 to reproduce of
oNormalize?
2 1=no, 2=yes

ofxn calc*
0.12969
ofxn mod
0.16857

sRun
RunA1
zout
1

Ave
Data
6.6E-08
Model
6.1E-08

data/vMax
6.6E-08

ofxng
311.734

Min d/m
1.30851

ave/min d
1
ave/min m
1.2081

331_Calib_S_A

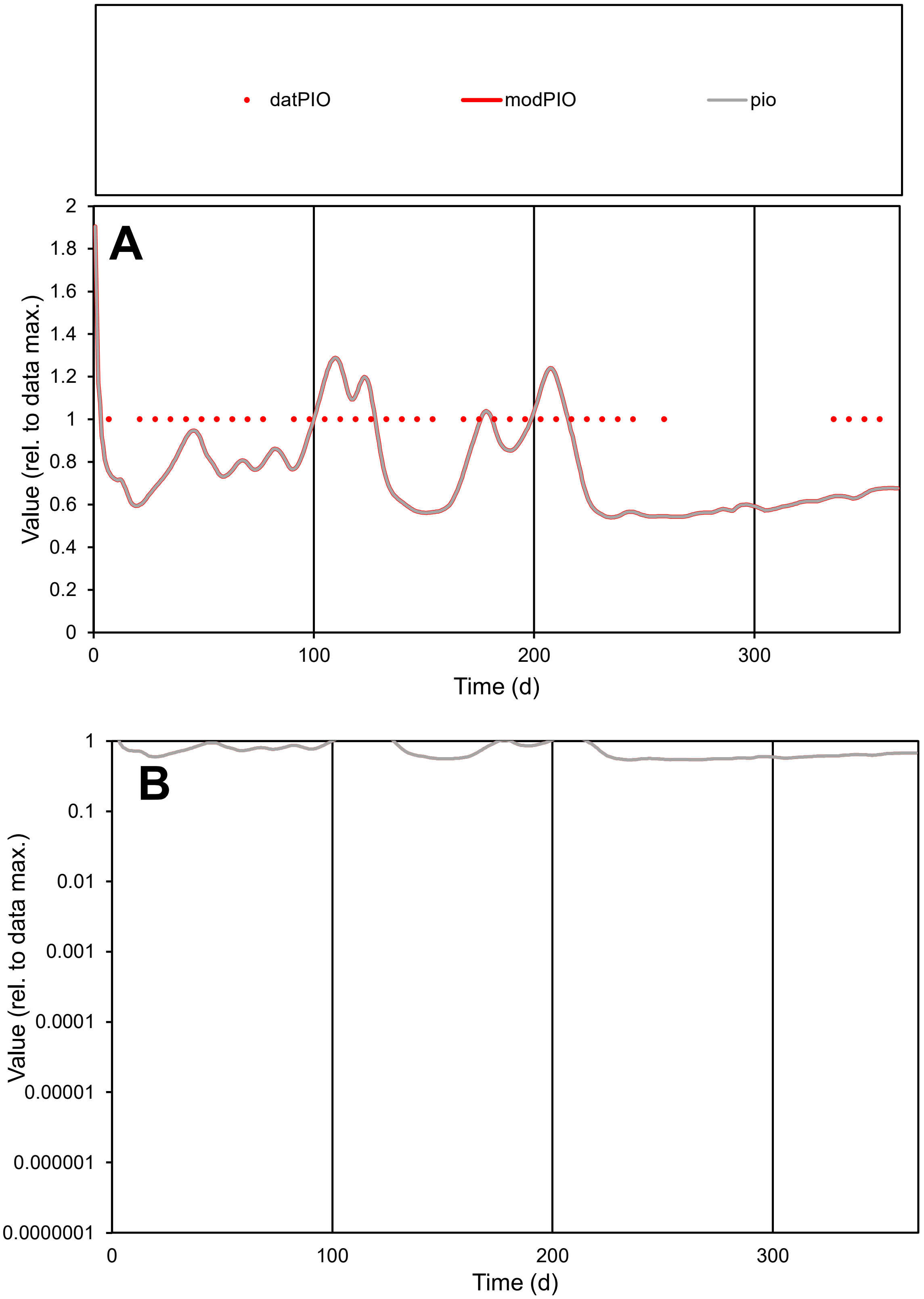


**C1**

Variables
PIO Provide jtd or Dst
CalibOnly?
2 1 to reproduce of
oNormalize?
2 1=no, 2=yes

ofxn calc*
0.28258
ofxn mod
0.3

sRun
RunA1
zout
1

Ave
Data
1.5E-07
Model
1.2E-07

data/vMax
1.5E-07

ofxng
311.734

Min d/m
1.84054

ave/min d
1
ave/min m
1.4801

331_Calib_S_A

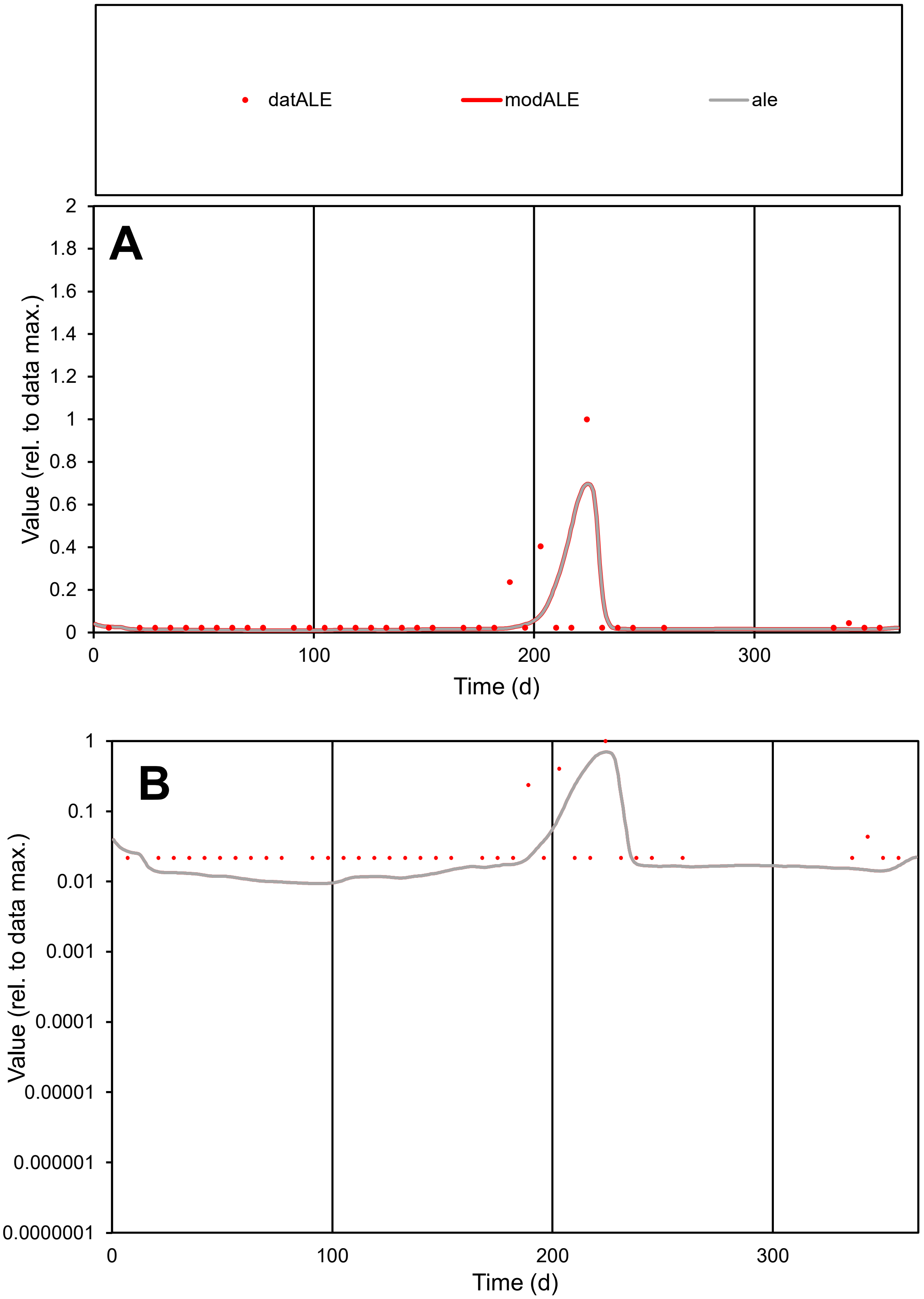


**C1**

Variables

ALE Provide jtd or Dst

CalibOnly?

2 1 to reproduce of

oNormalize?

2 1=no, 2=yes

ofxn calc*

1.62138

ofxn mod

1.77942

sRun

RunA1

zout

1

Ave

Data

3.3E-05

Model

2.6E-05

data/vMax

0.00051

ofxng

311.734

Min d/m

2.3172

ave/min d

2.98668

ave/min m

5.49134

331_Calib_S_A

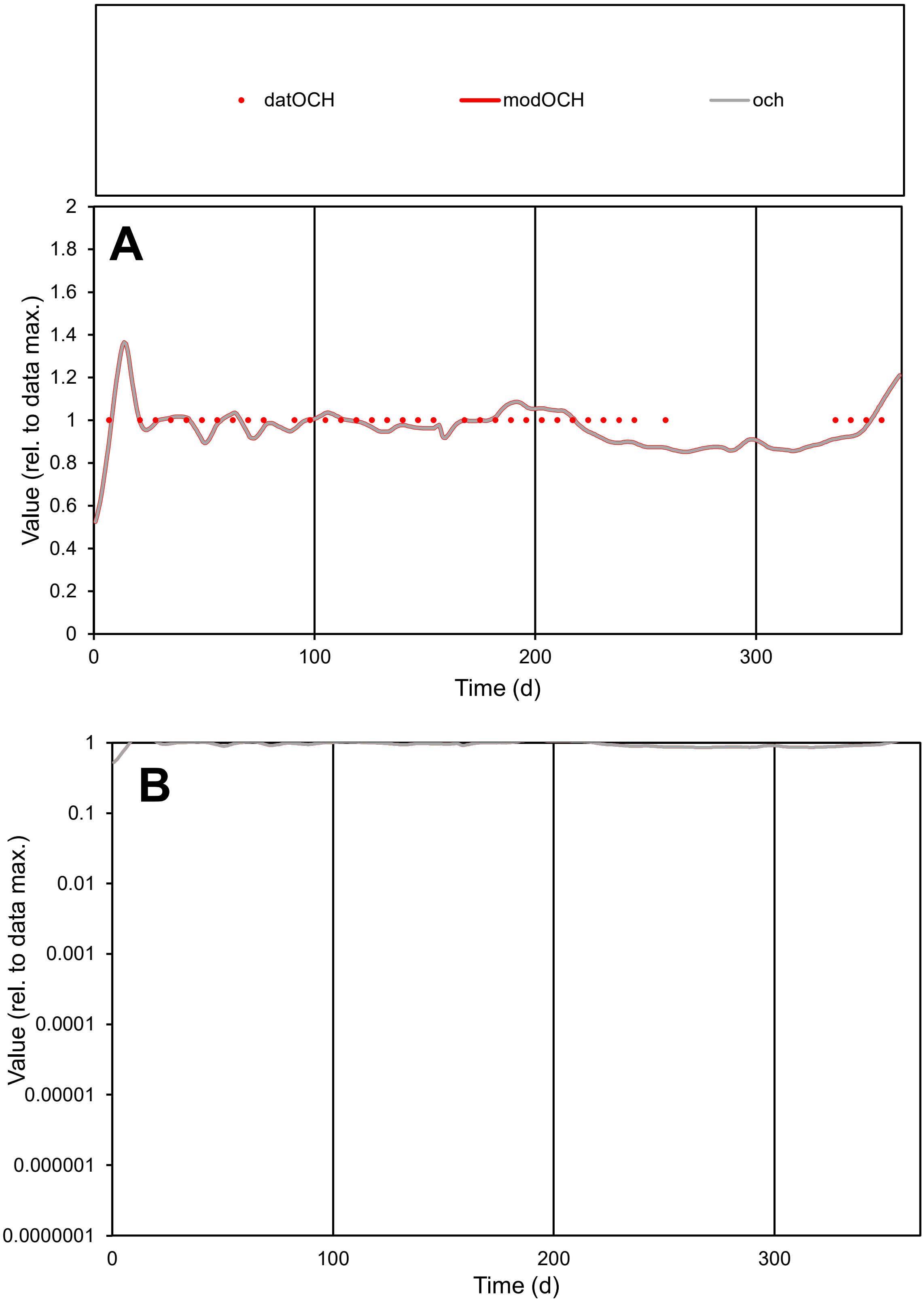


**C1**

Variables
OCH Provide jtd or Dst
CalibOnly?
2 1 to reproduce of
oNormalize?
2 1=no, 2=yes

ofxn calc*
0.05792
ofxn mod
0.12506

sRun
RunA1
zout
1

Ave
Data
4.4E-06
Model
4.3E-06

data/vMax
4.4E-06

ofxng
311.734

Min d/m
1.15135

ave/min d
1
ave/min m
1.13059

331_Calib_S_A

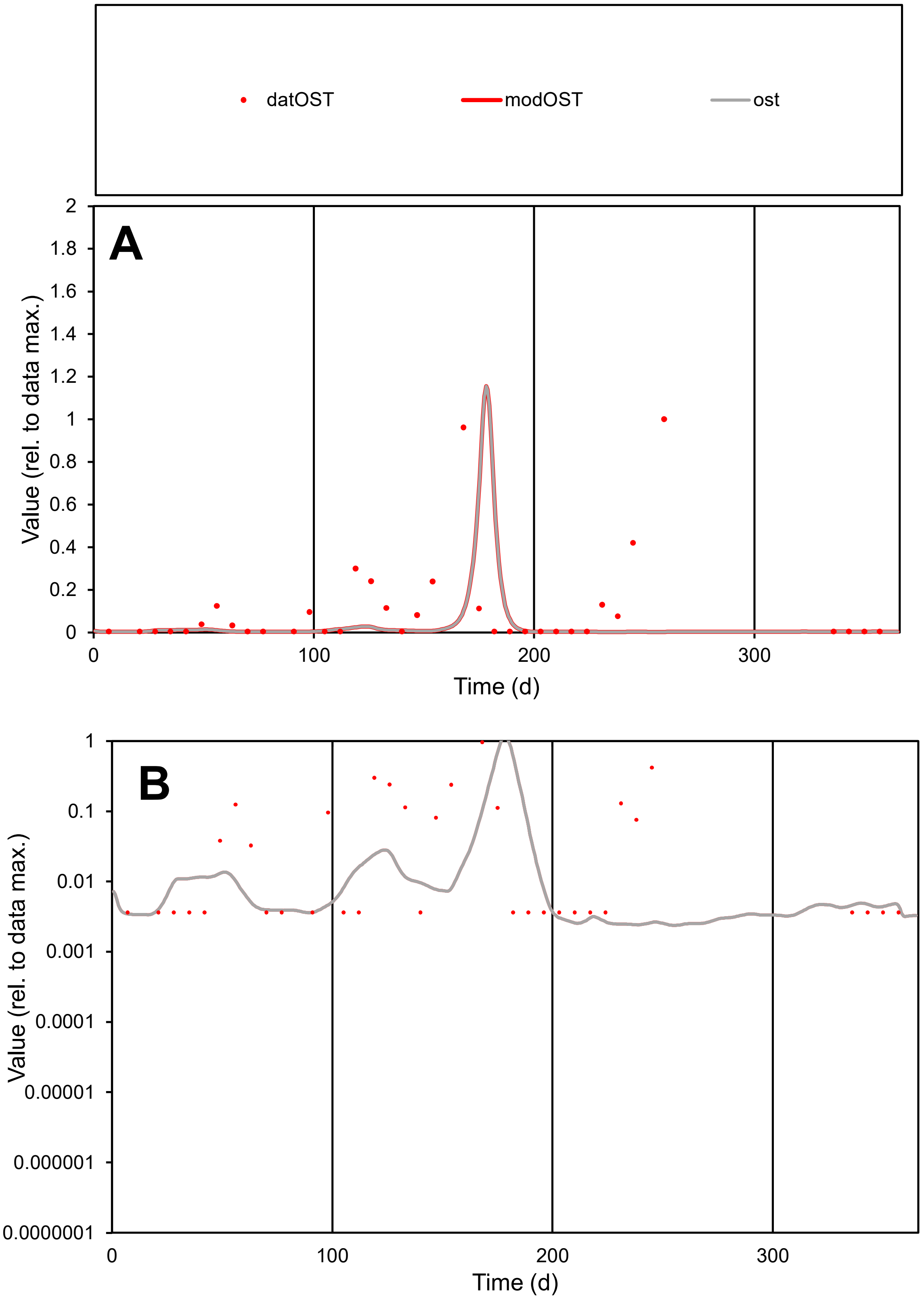


**C1**

Variables
OST Provide jtd or Dst
CalibOnly?
2 1 to reproduce of
oNormalize?
2 1=no, 2=yes

ofxn calc*
2.12948
ofxn mod
2.29212

sRun
RunA1
zout
1

Ave
Data
6.6E-05
Model
1.8E-05

data/vMax
0.0006

ofxng
311.734

Min d/m
1.49345

ave/min d
30.2443
ave/min m
12.7262

331_Calib_S_A

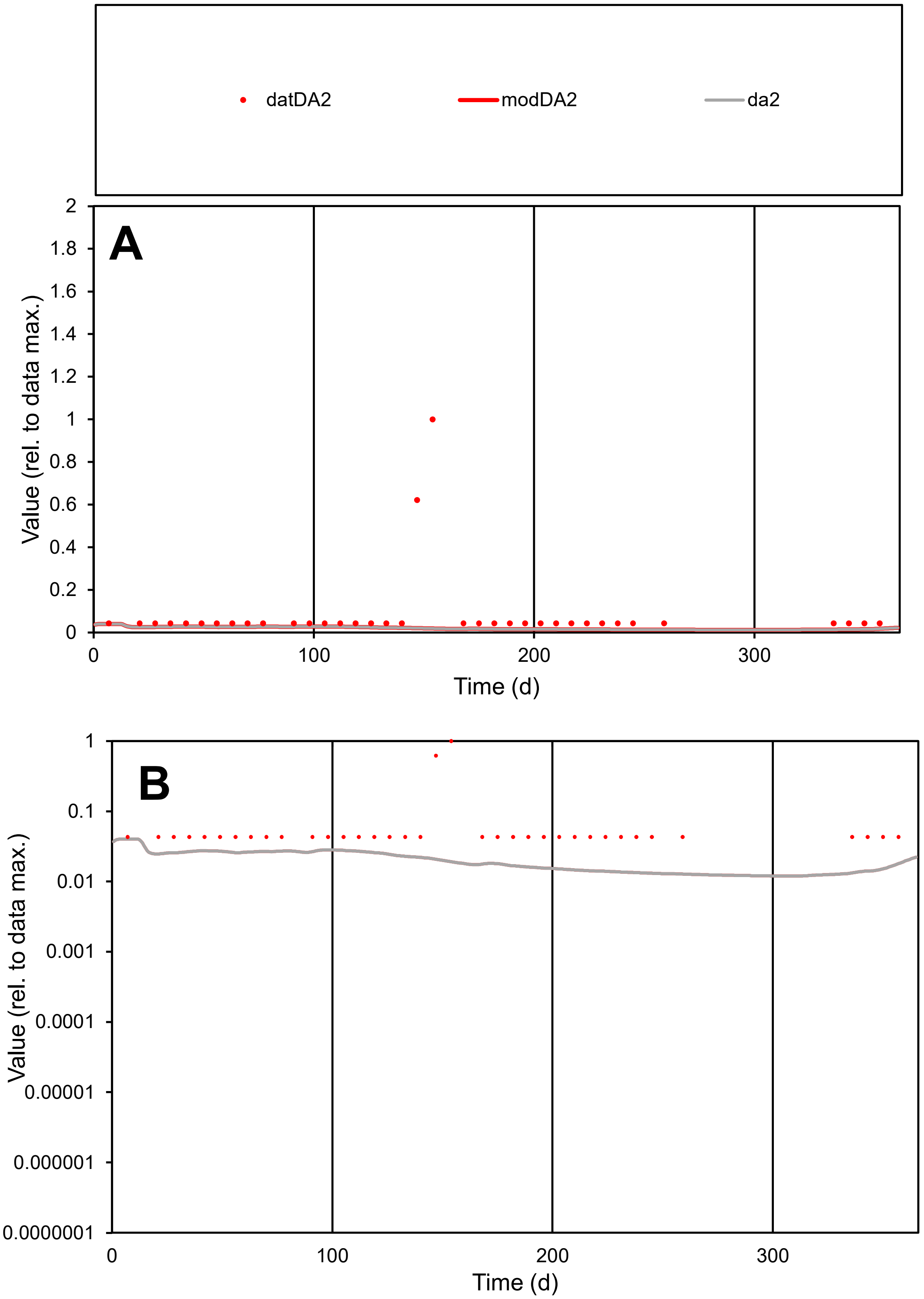


**C1**

Variables
DA2 Provide jtd or Dst
CalibOnly?
2 1 to reproduce of
oNormalize?
2 1=no, 2=yes

ofxn calc*
2.25243
ofxn mod
2.25792

sRun
RunA1
zout
1

Ave
Data
6.1E-06
Model
1.5E-06

data/vMax
7.2E-05

ofxng
311.734

Min d/m
3.38762

ave/min d
1.9631
ave/min m
1.65331

331_Calib_S_A

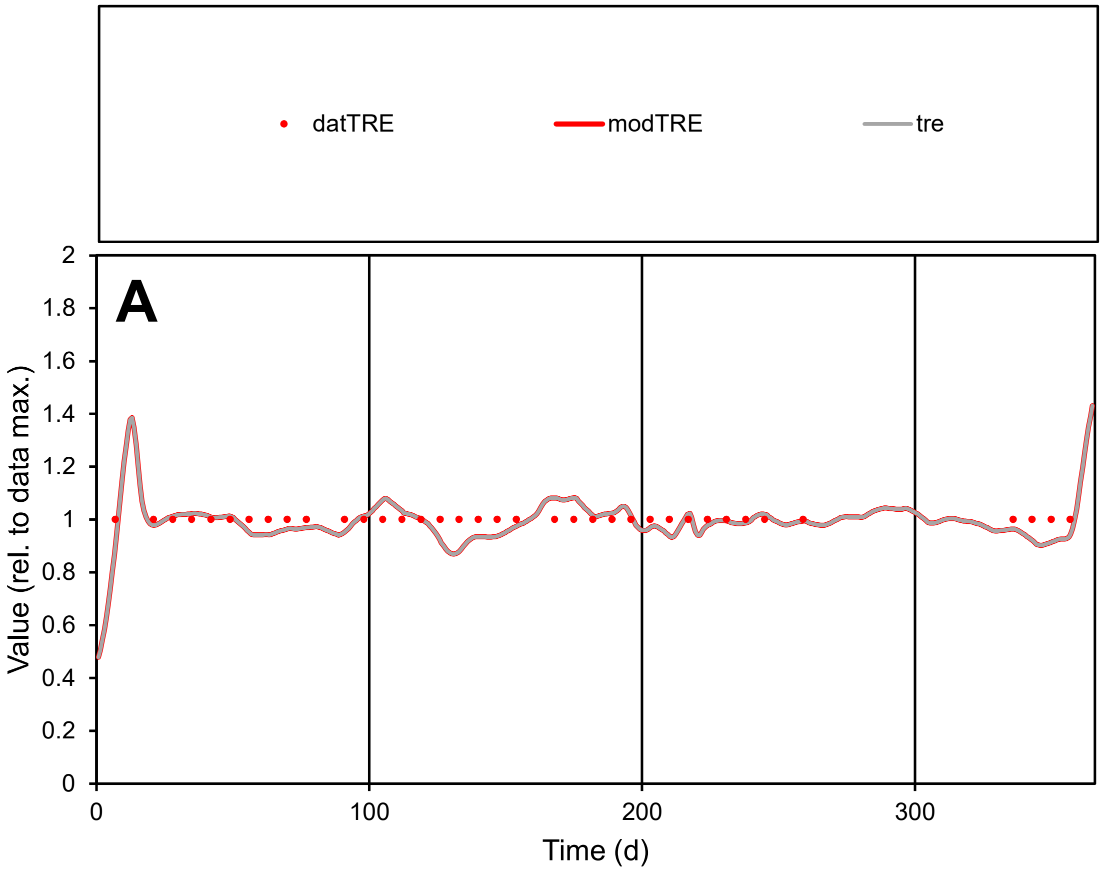


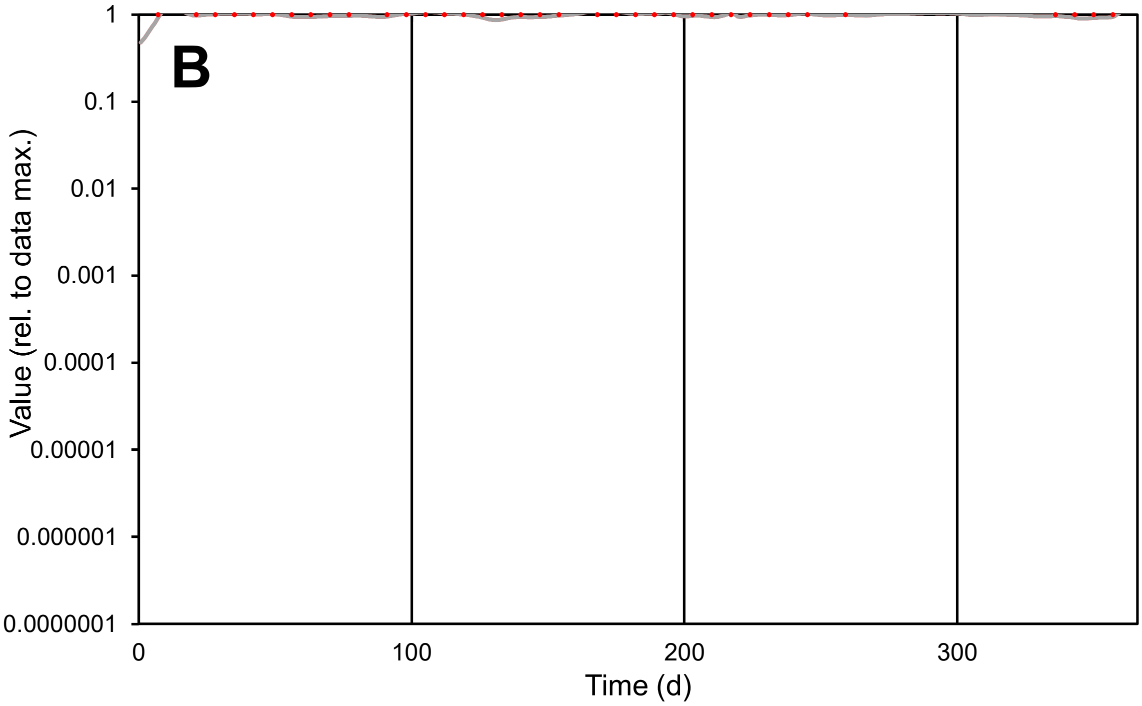


**C1**

Variables
TRE Provide jtd or Dst
CalibOnly?
2 1 to reproduce of
oNormalize?
2 1=no, 2=yes

ofxn calc*
0.04664
ofxn mod
0.12008

sRun
RunA1
zout
1

Ave
Data
6.9E-06
Model
6.8E-06

data/vMax
6.9E-06

ofxng
311.734

Min d/m
1.11338

ave/min d
1
ave/min m
1.09773

331_Calib_S_A

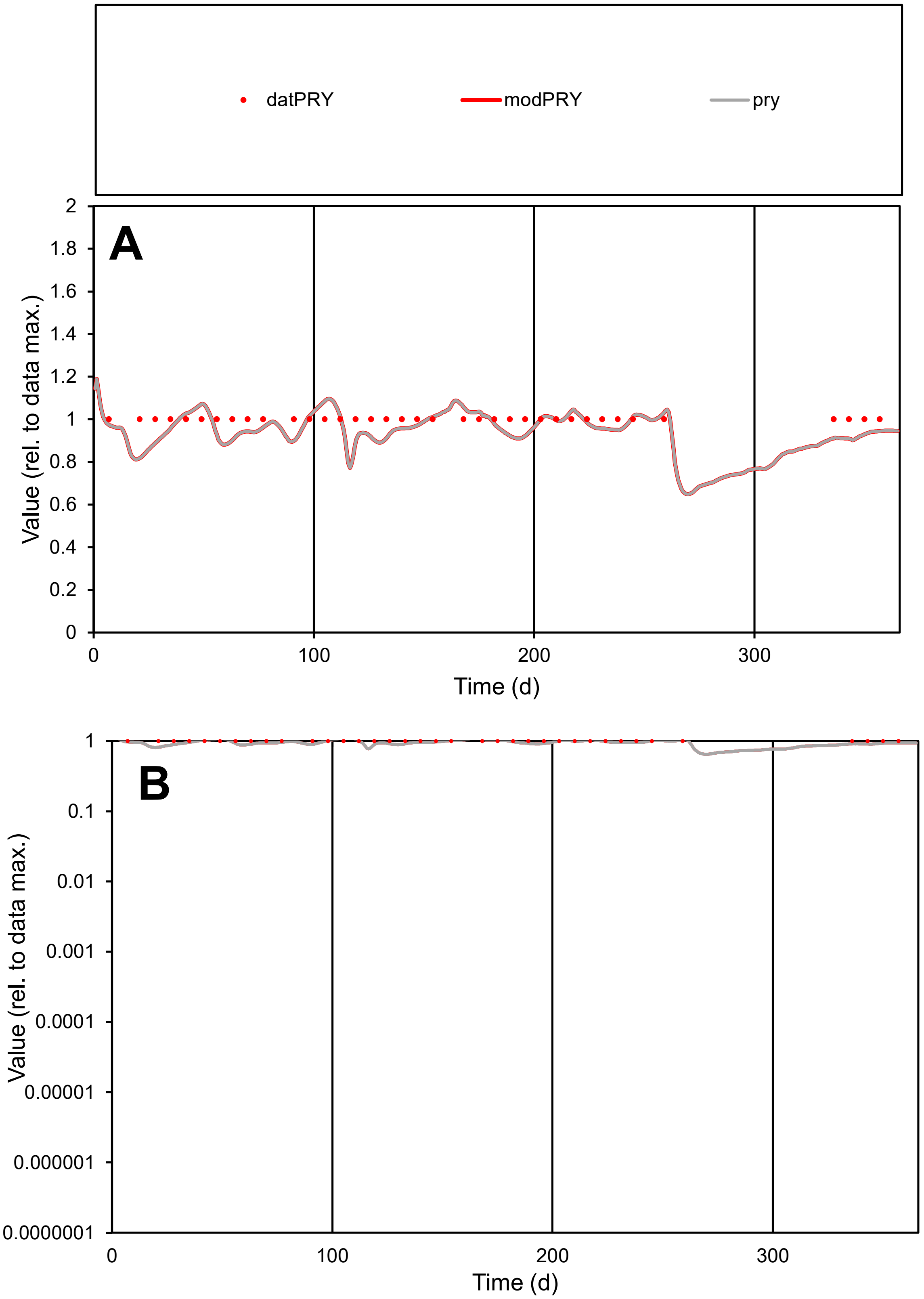


**C1**

Variables
PRY Provide jtd or Dst
CalibOnly?
2 1 to reproduce of
oNormalize?
2 1=no, 2=yes

ofxn calc*
0.06498
ofxn mod
0.12771

sRun
RunA1
zout
1

Ave
Data
8.8E-06
Model
8.5E-06

data/vMax
8.8E-06

ofxng
311.734

Min d/m
1.20295

ave/min d
1
ave/min m
1.16437

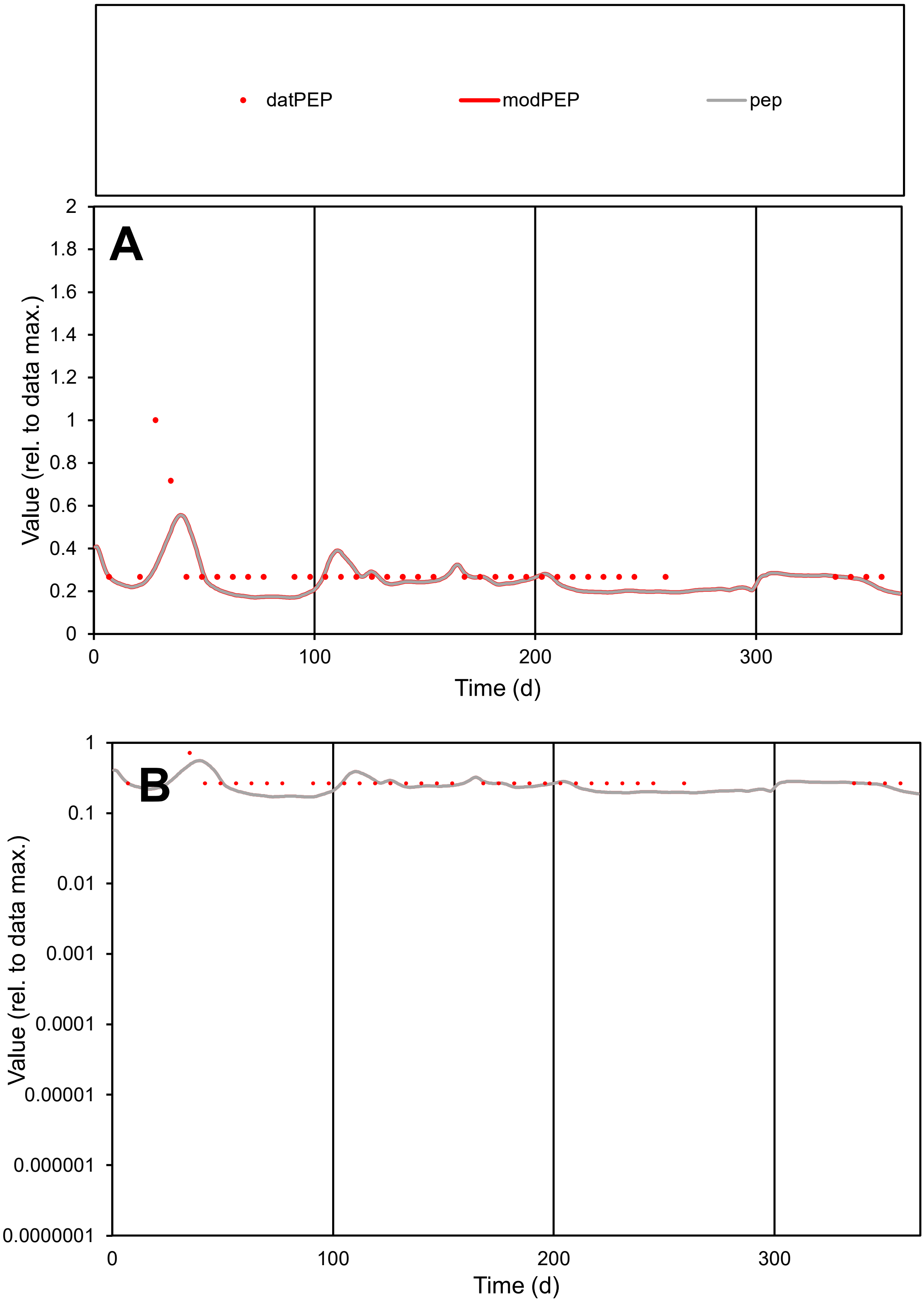


**C1**

Variables
PEP Provide jtd or Dst
CalibOnly?
2 1 to reproduce of
oNormalize?
2 1=no, 2=yes

ofxn calc*
0.43217
ofxn mod
0.44522

sRun
RunA1
zout
1

Ave
Data
5.5E-06
Model
4.7E-06

data/vMax
1.9E-05

ofxng
311.734

Min d/m
1.5506

ave/min d
1.11997
ave/min m
1.47404

331_Calib_S_A

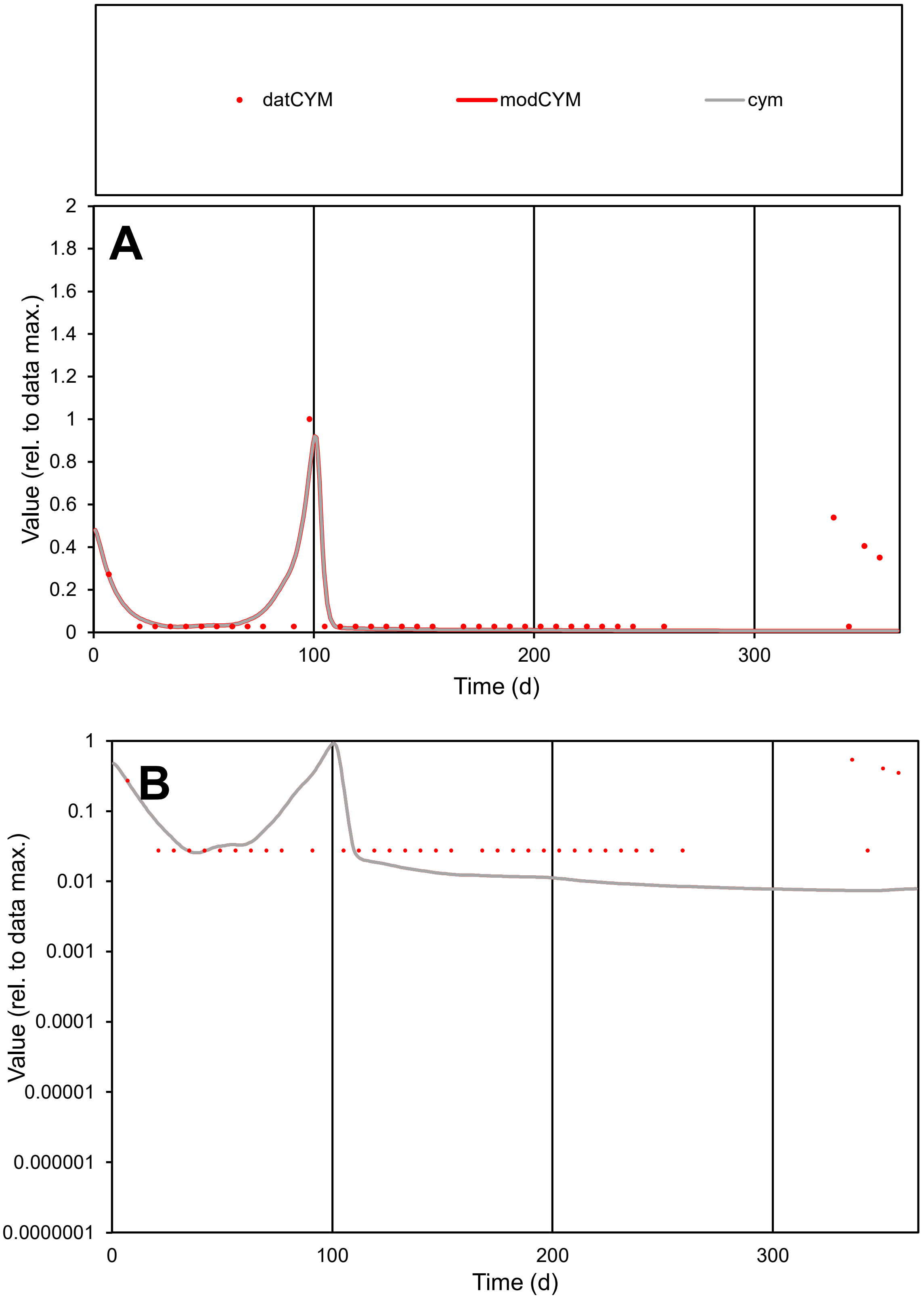


**C1**

Variables
CYM Provide jtd or Dst
CalibOnly?
2 1 to reproduce of
oNormalize?
2 1=no, 2=yes

ofxn calc*
1.4412
ofxn mod
1.47257

sRun
RunA1
zout
1

Ave
Data
4.6E-06
Model
2.9E-06

data/vMax
4.9E-05

ofxng
311.734

Min d/m
3.71756

ave/min d
3.38626
ave/min m
7.9671

331_Calib_S_A

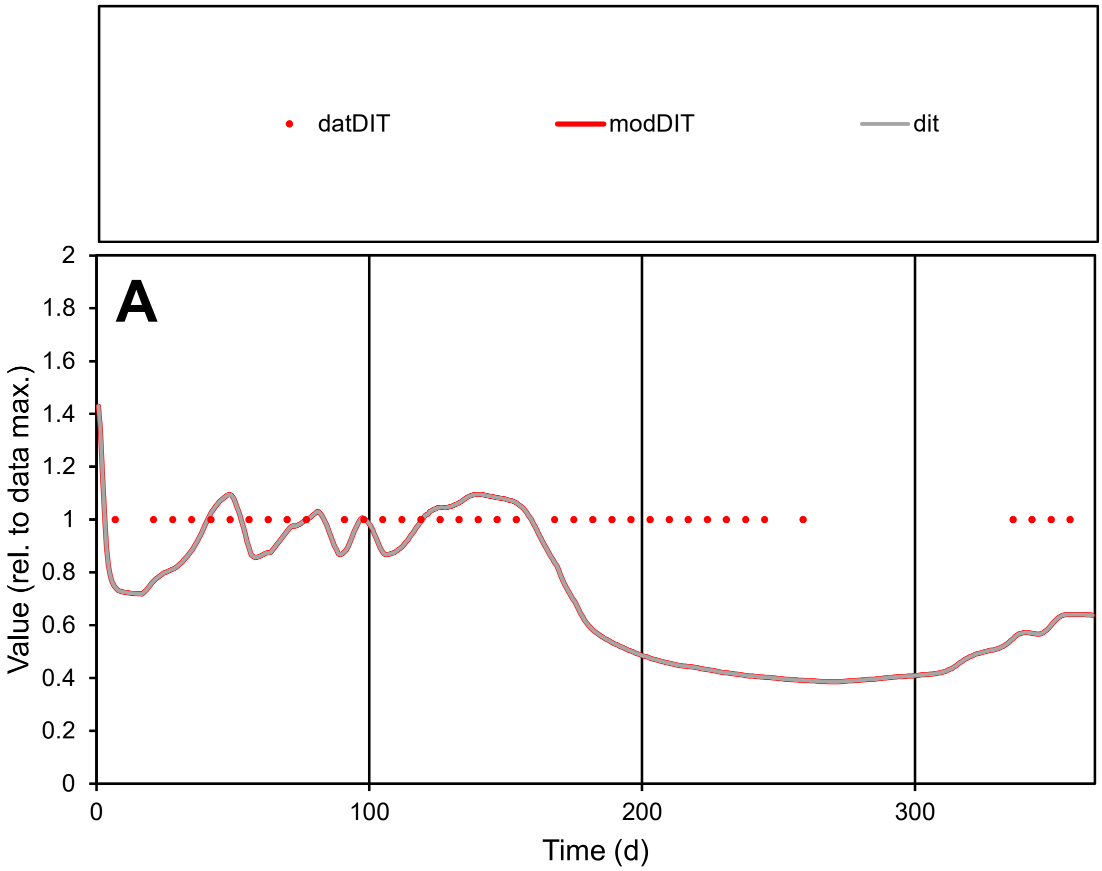


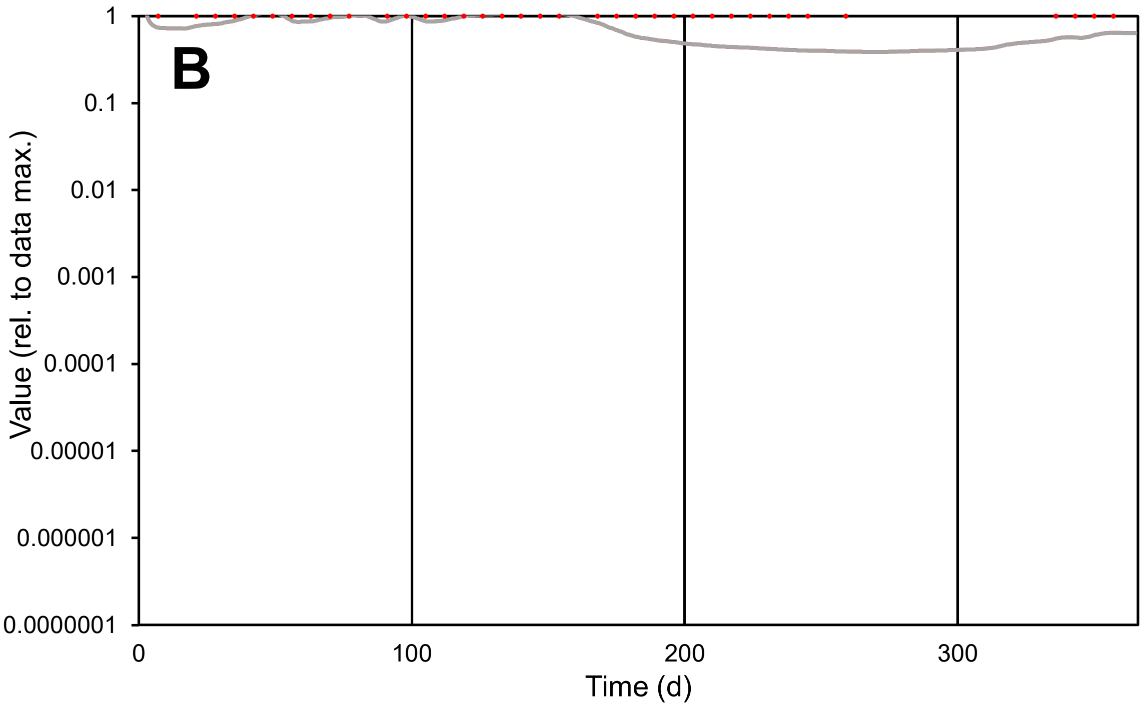


**C1**

Variables
DIT Provide jtd or Dst
CalibOnly?
2 1 to reproduce of
oNormalize?
2 1=no, 2=yes

ofxn calc*
0.34314
ofxn mod
0.35702

sRun
RunA1
zout
1

Ave
Data
5.1E-06
Model
3.9E-06

data/vMax
5.1E-06

ofxng
311.734

Min d/m
2.56241

ave/min d
1
ave/min m
1.93944

331_Calib_S_A

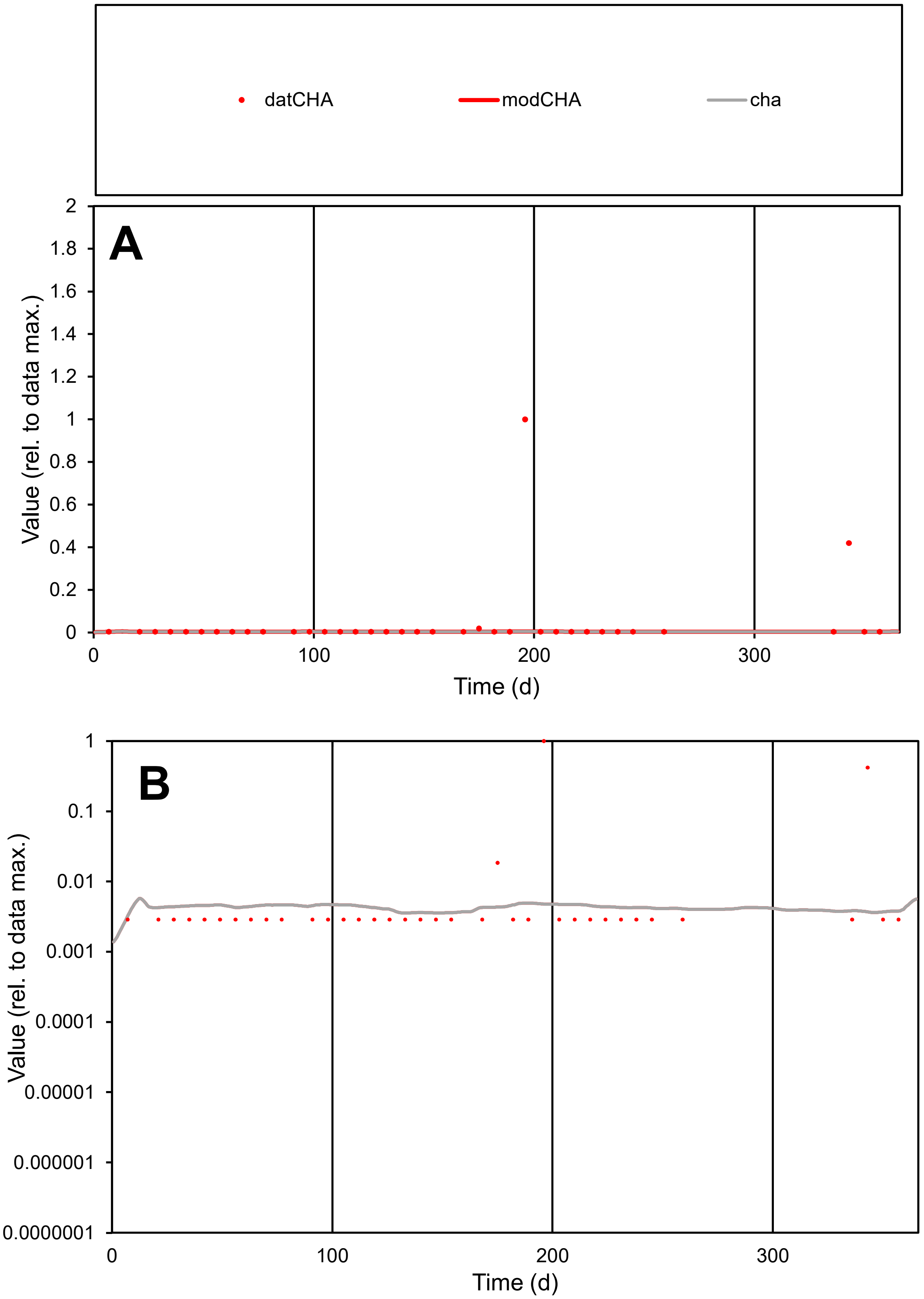


**C1**

Variables

CHA Provide jtd or Dst

CalibOnly?

2 1 to reproduce of

oNormalize?

2 1=no, 2=yes

ofxn calc*

4.27373

ofxn mod

4.27673

sRun

RunA1

zout

1

Ave

Data

1.3E-06

Model

1.3E-07

data/vMax

3E-05

ofxng

311.734

Min d/m

0.87327

ave/min d

14.4796

ave/min m

1.29946

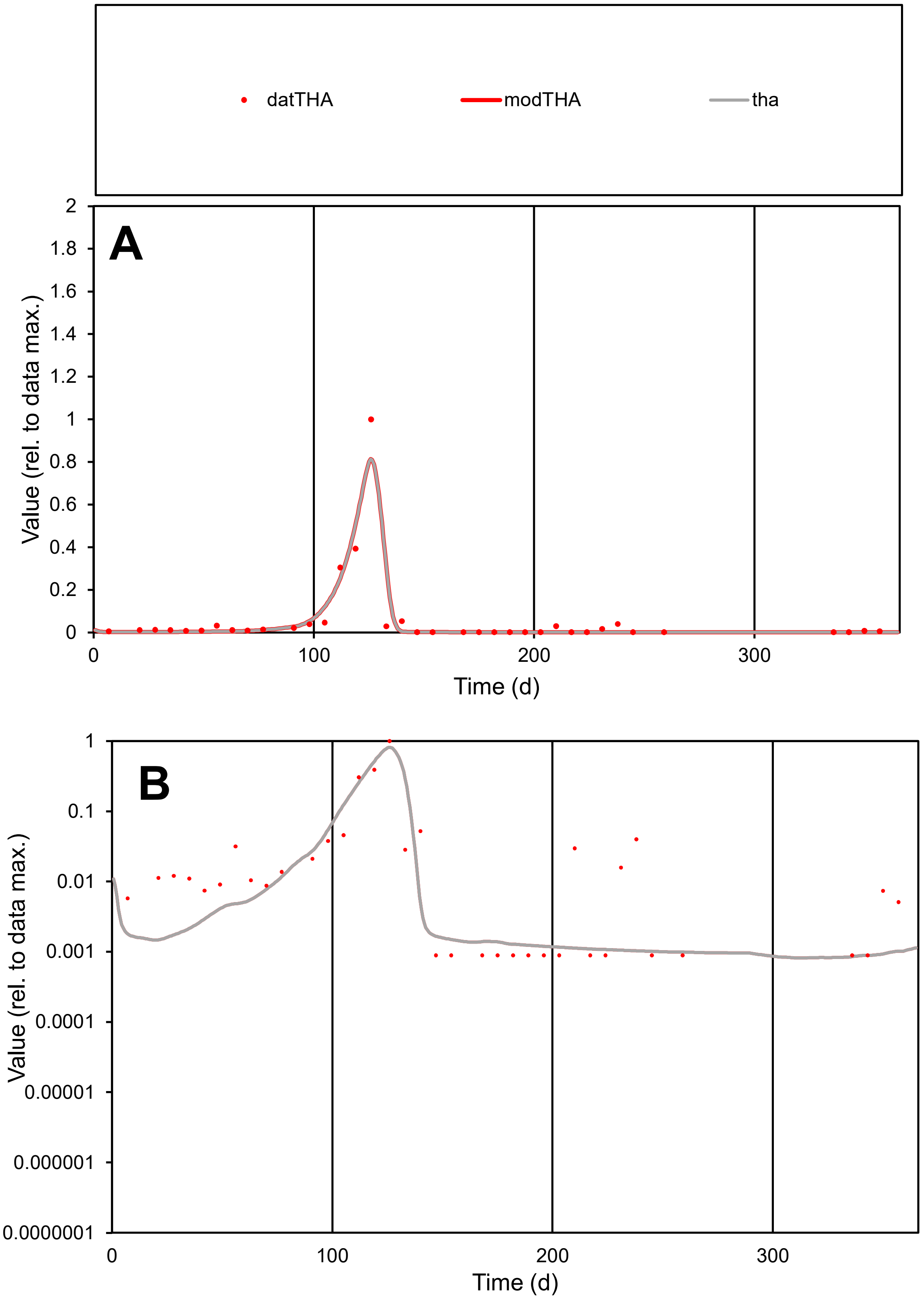


**C1**

Variables
THA Provide jtd or Dst
CalibOnly?
2 1 to reproduce of
oNormalize?
2 1=no, 2=yes

ofxn calc*
0.80221
ofxn mod
0.92106

sRun
RunA1
zout
1

Ave
Data
0.00018
Model
0.00017

data/vMax
0.0031

ofxng
311.734

Min d/m
1.02687

ave/min d
64.3153
ave/min m
62.862

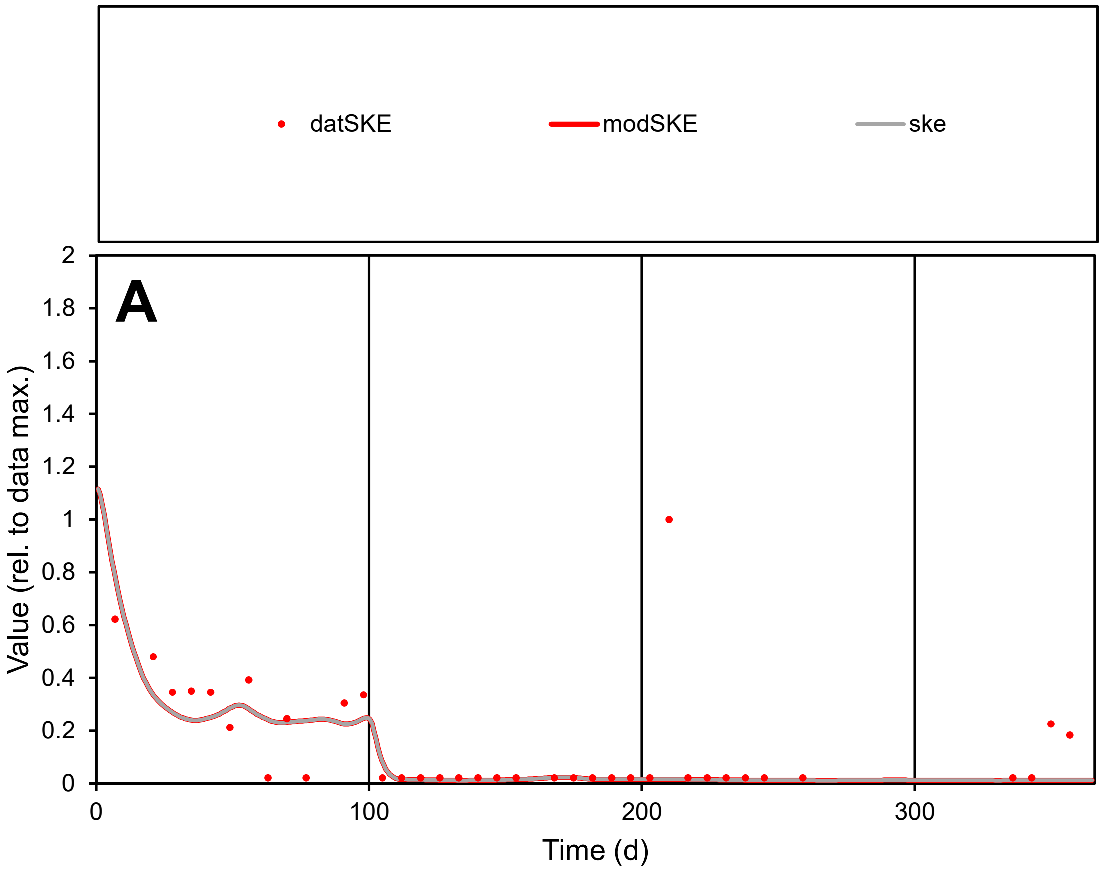


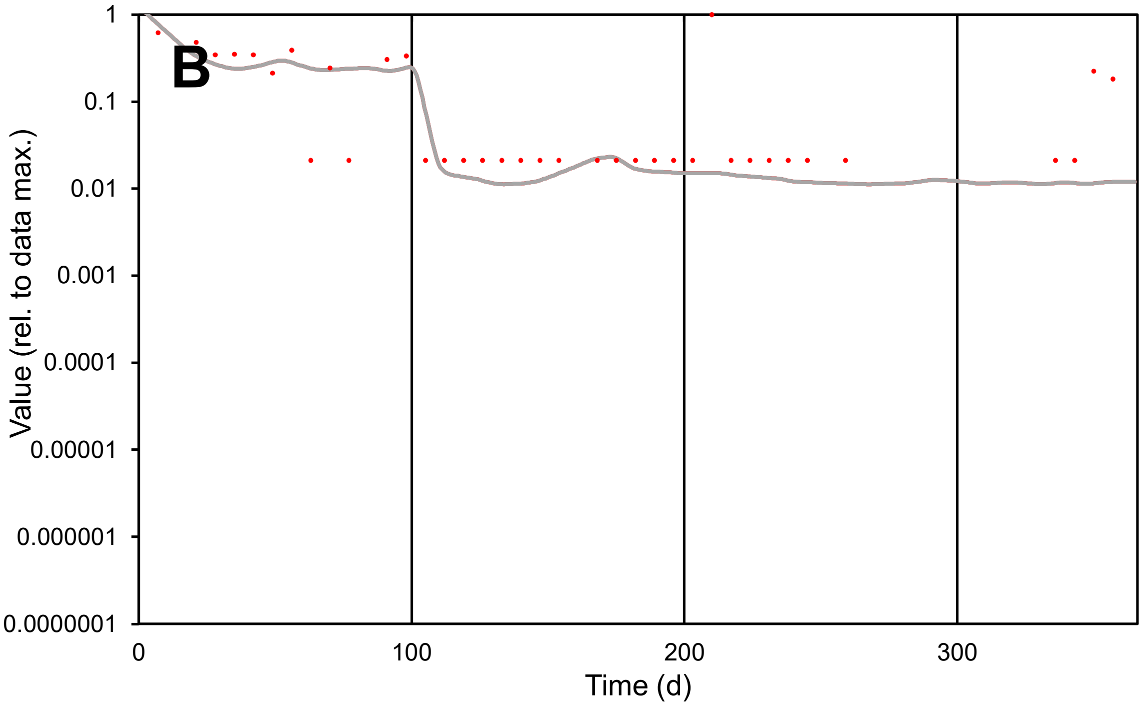


**C1**

Variables
SKE Provide jtd or Dst
CalibOnly?
2 1 to reproduce of
oNormalize?
2 1=no, 2=yes

ofxn calc*
1.22065
ofxn mod
1.23091

sRun
RunA1
zout
1

Ave
Data
1.6E-05
Model
1.2E-05

data/vMax
0.00011

ofxng
311.734

Min d/m
1.87531

ave/min d
7.0905
ave/min m
9.40759

331_Calib_S_A

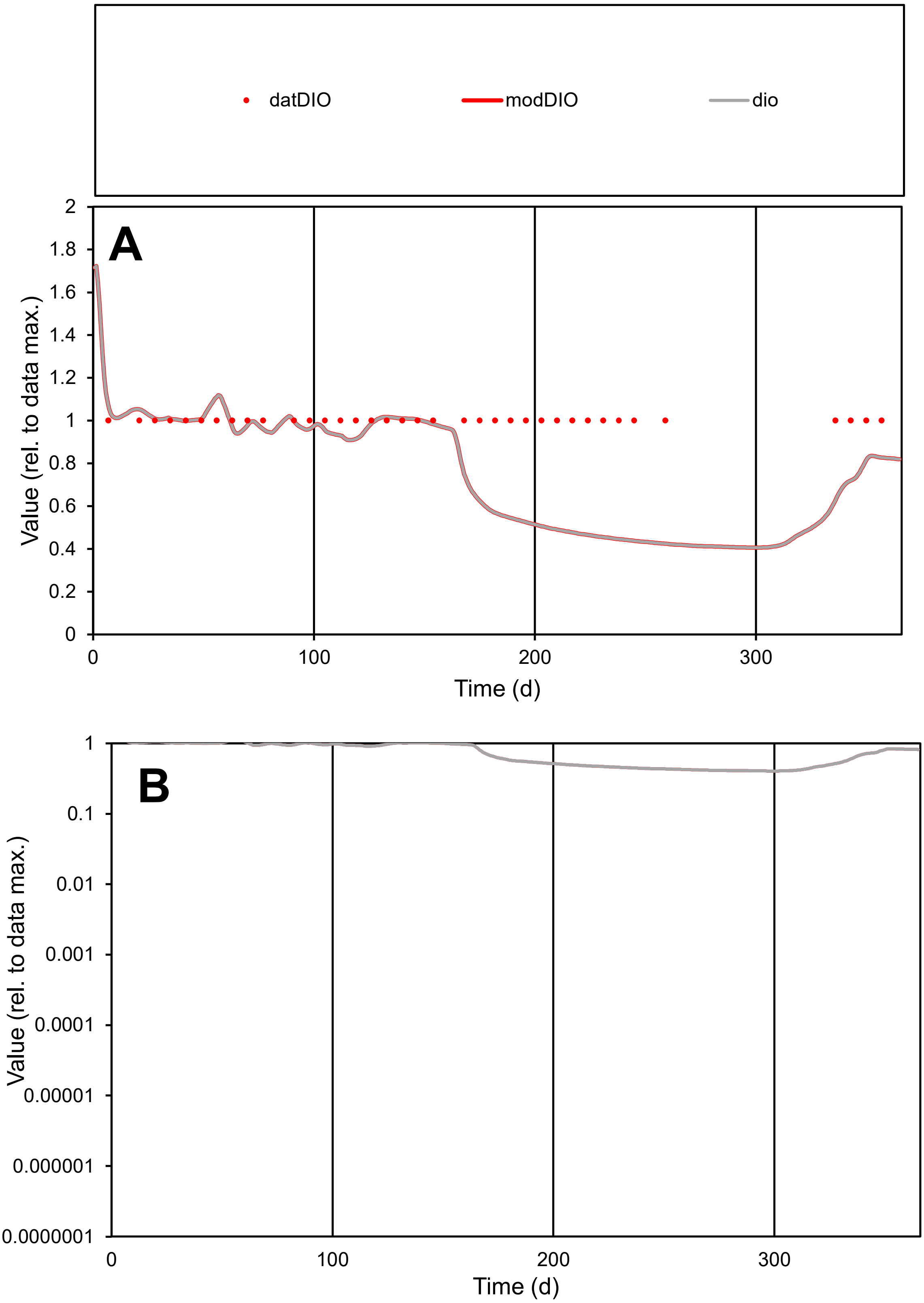


**C1**

Variables

| | |
|---|---|
| DIO | Provide jtd or Dst |
| CalibOnly? | |
| 2 | 1 to reproduce of |
| oNormalize? | |
| 2 | 1=no, 2=yes |
| ofxn calc* | |
| 0.3069 | |
| ofxn mod | |
| 0.32312 | |
| sRun | |
| RunA1 | |
| zout | |
| 1 | |
| Ave | |
| Data | |
| 4.7E-06 | |
| Model | |
| 3.7E-06 | |
| data/vMax | |
| 4.7E-06 | |
| ofxng | |
| 311.734 | |
| Min d/m | |
| 2.36166 | |
| ave/min d | |
| 1 | |
| ave/min m | |
| 1.8854 | |

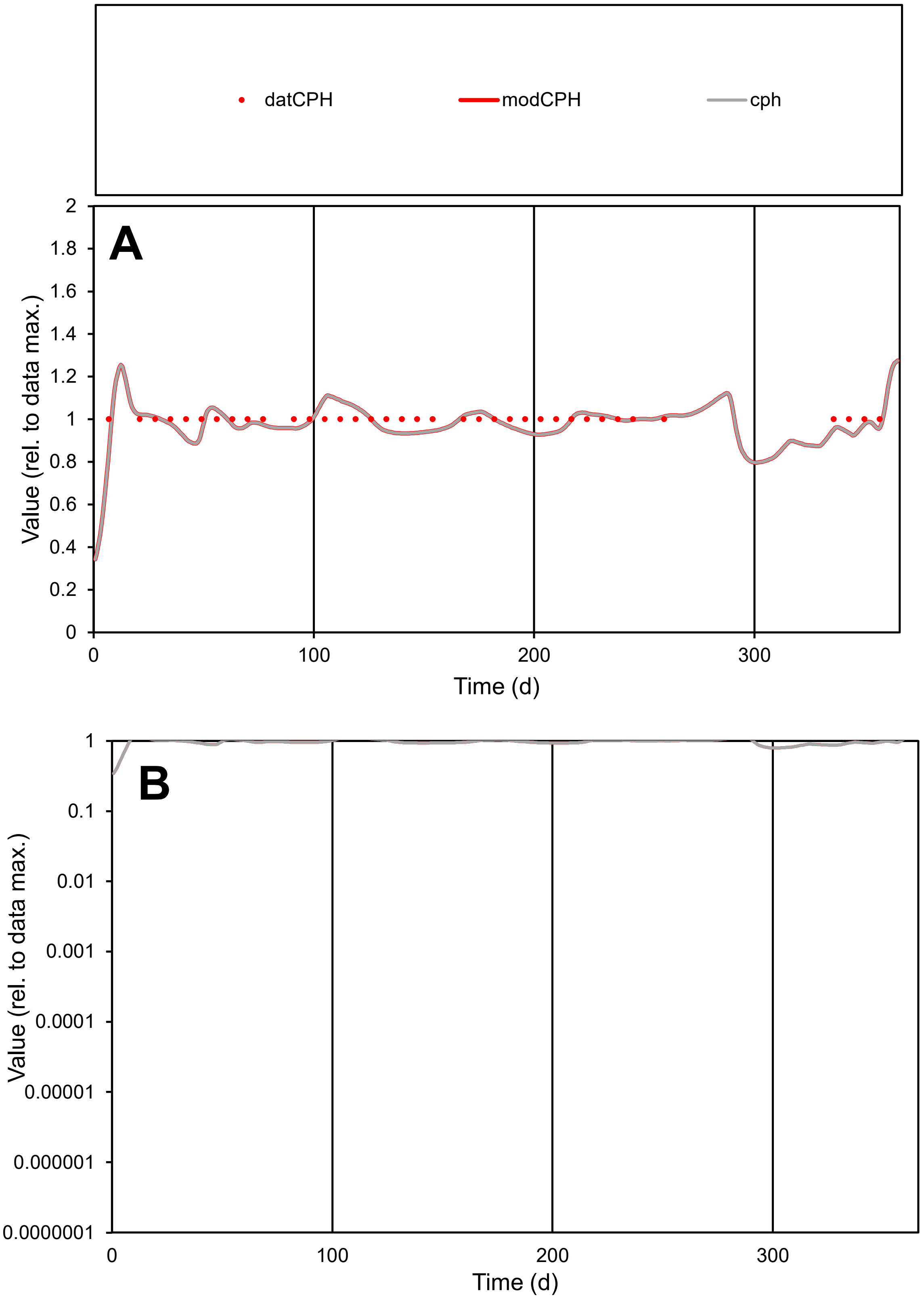


**C1**

Variables

| | |
|---|---|
| CPH | Provide jtd or Dst |

CalibOnly?

| | |
|---|---|
| 2 | 1 to reproduce of |

oNormalize?

| | |
|---|---|
| 2 | 1=no, 2=yes |

ofxn calc*
0.04645
ofxn mod
0.1201

sRun
RunA1
zout
1

Ave
Data
9.6E-06
Model
9.5E-06

data/vMax
9.6E-06

ofxng
311.734

Min d/m
1.11822

ave/min d
1
ave/min m
1.10446

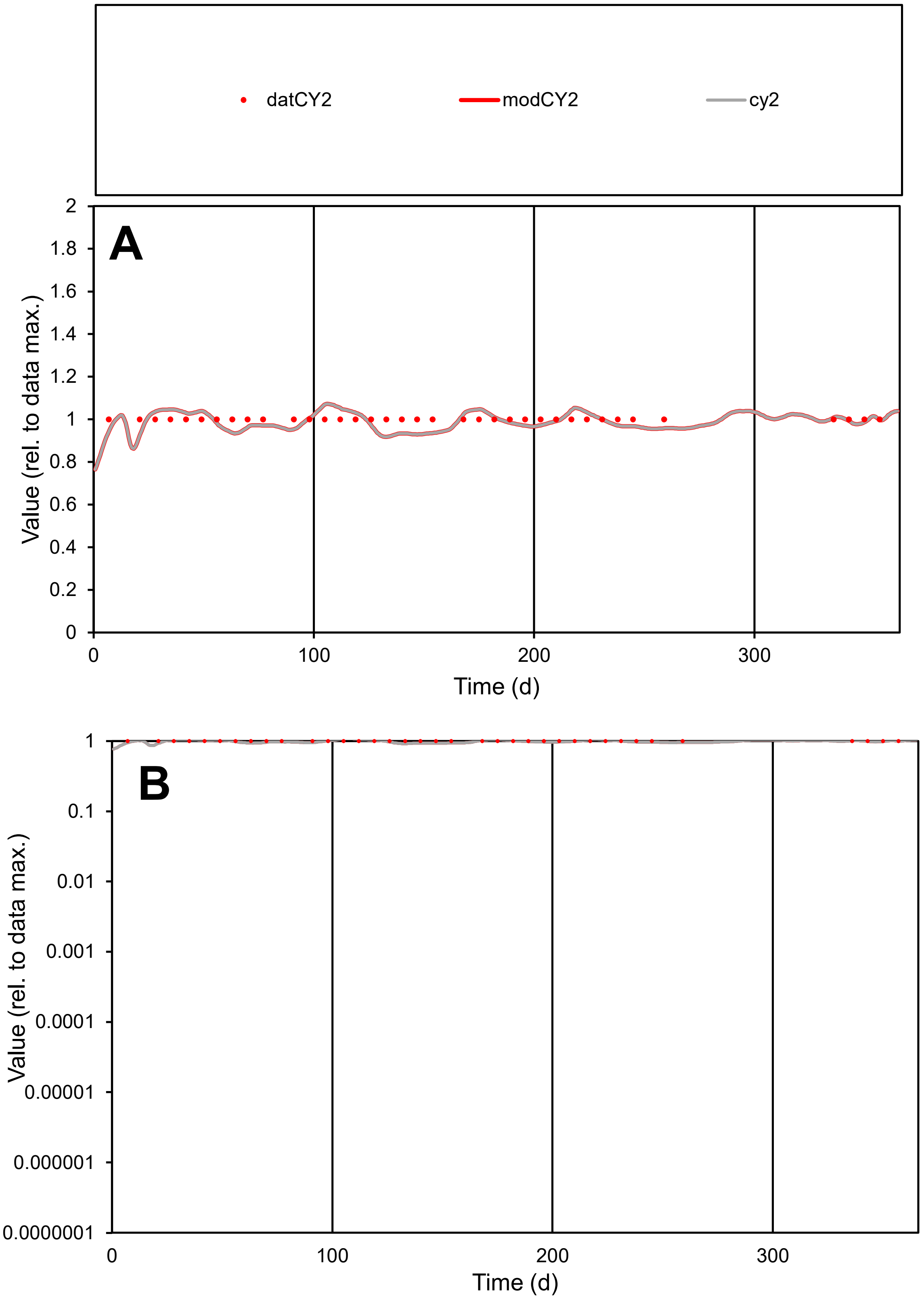


**C1**

| | |
|---|---|
| Variables | |
| CY2 | Provide jtd or Dst |
| CalibOnly? | |
| 2 | 1 to reproduce of |
| oNormalize? | |
| 2 | 1=no, 2=yes |
| ofxn calc* | |
| 0.04022 | |
| ofxn mod | |
| 0.11805 | |
| sRun | |
| RunA1 | |
| zout | |
| 1 | |
| Ave | |
| Data | |
| 1.1E-05 | |
| Model | |
| 1.1E-05 | |
| data/vMax | |
| 1.1E-05 | |
| ofxng | |
| 311.734 | |
| Min d/m | |
| 1.08683 | |
| ave/min d | |
| 1 | |
| ave/min m | |
| 1.07808 | |

331_Calib_S_A

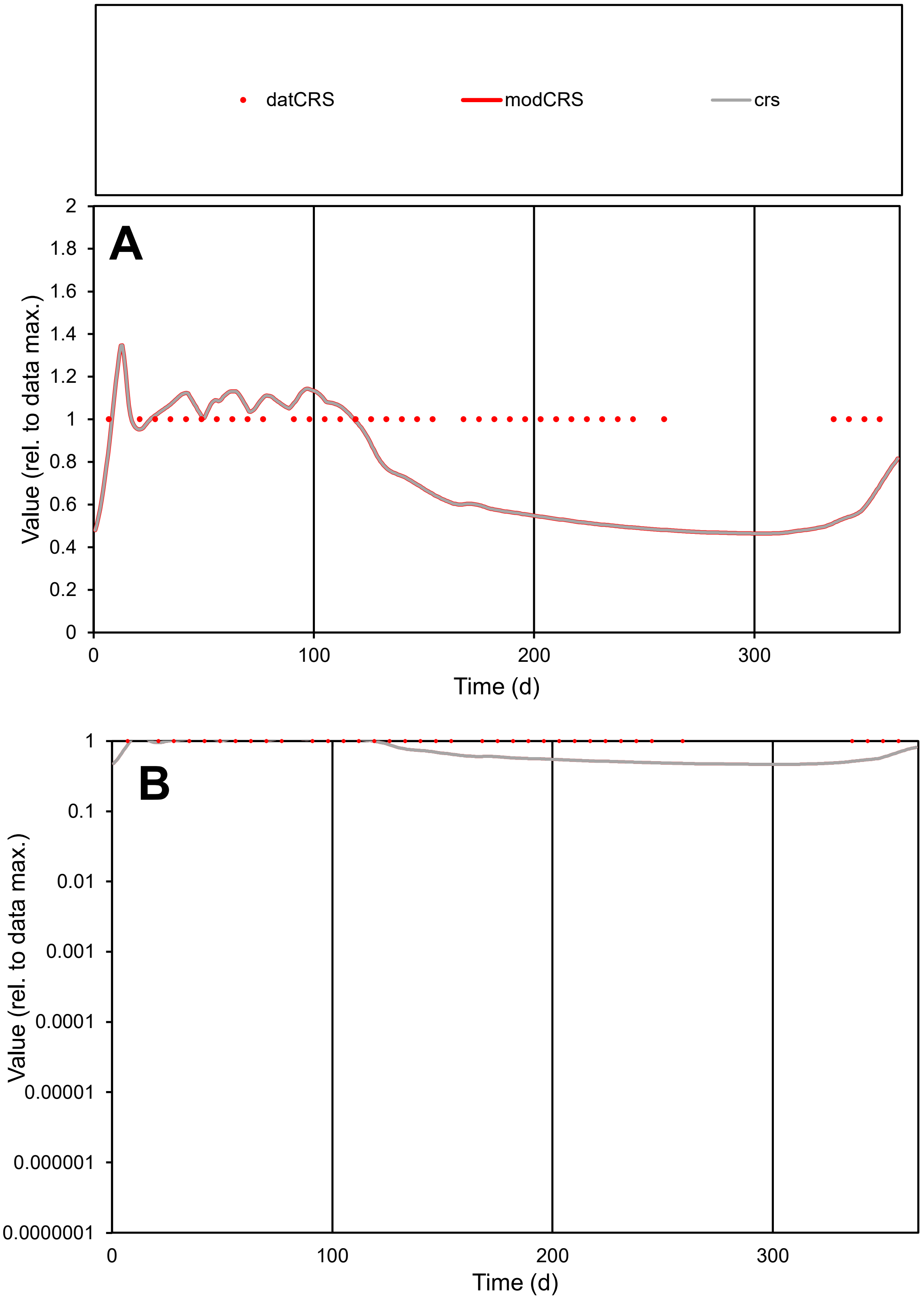


**C1**

Variables

CRS Provide jtd or Dst

CalibOnly?

2 1 to reproduce of

oNormalize?

2 1=no, 2=yes

ofxn calc*

0.32753

ofxn mod

0.3427

sRun

RunA1

zout

1

Ave

Data

5.4E-06

Model

4.2E-06

data/vMax

5.4E-06

ofxng

311.734

Min d/m

2.08157

ave/min d

1

ave/min m

1.62509

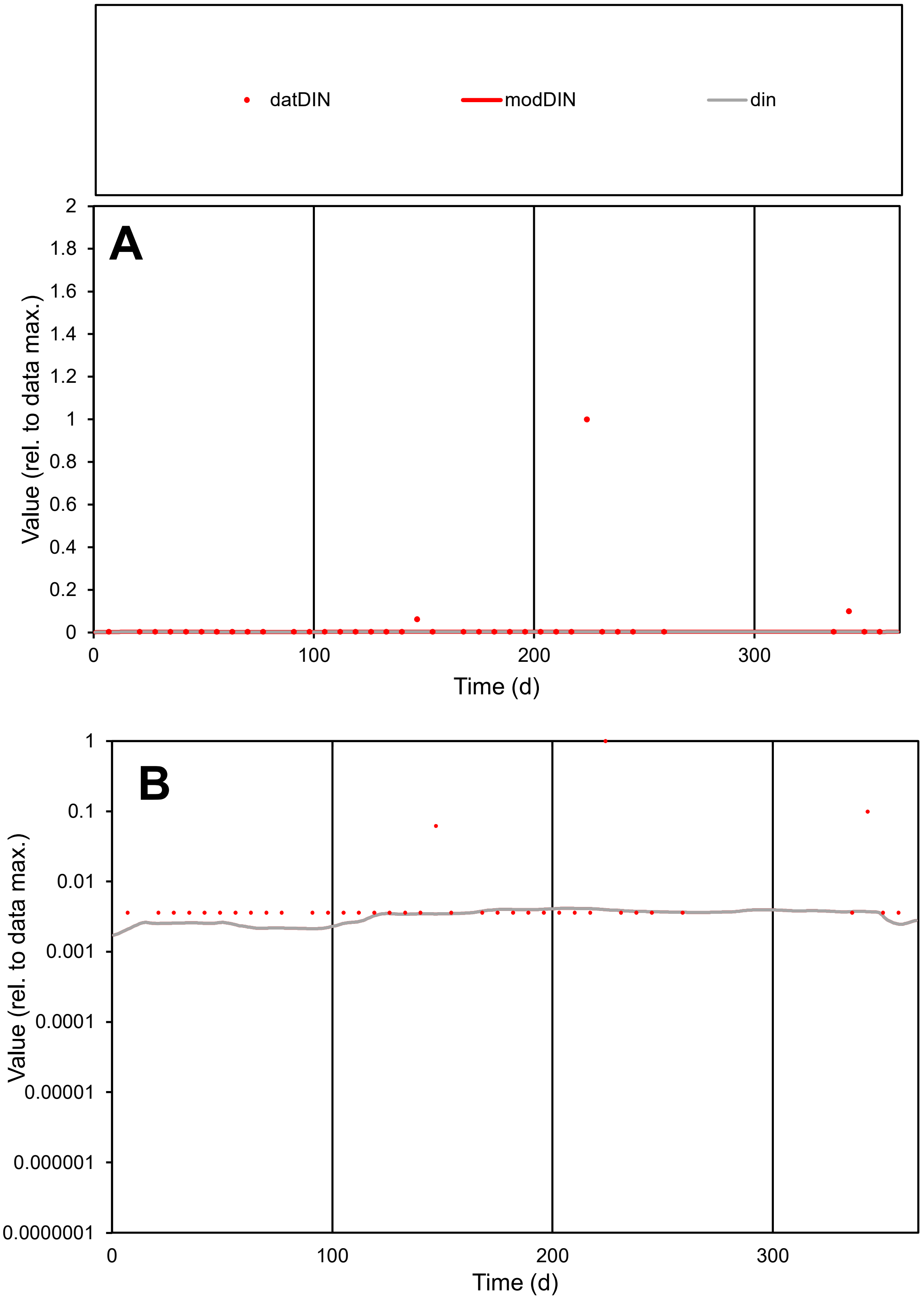


**C1**

Variables
DIN Provide jtd or Dst
CalibOnly?
2 1 to reproduce of
oNormalize?
2 1=no, 2=yes

ofxn calc*
4.75139
ofxn mod
4.76316

sRun
RunA1
zout
1

Ave
Data
2.8E-05
Model
2.6E-06

data/vMax
0.0008

ofxng
311.734

Min d/m
1.6905

ave/min d
9.66943
ave/min m
1.50821

331_Calib_S_A

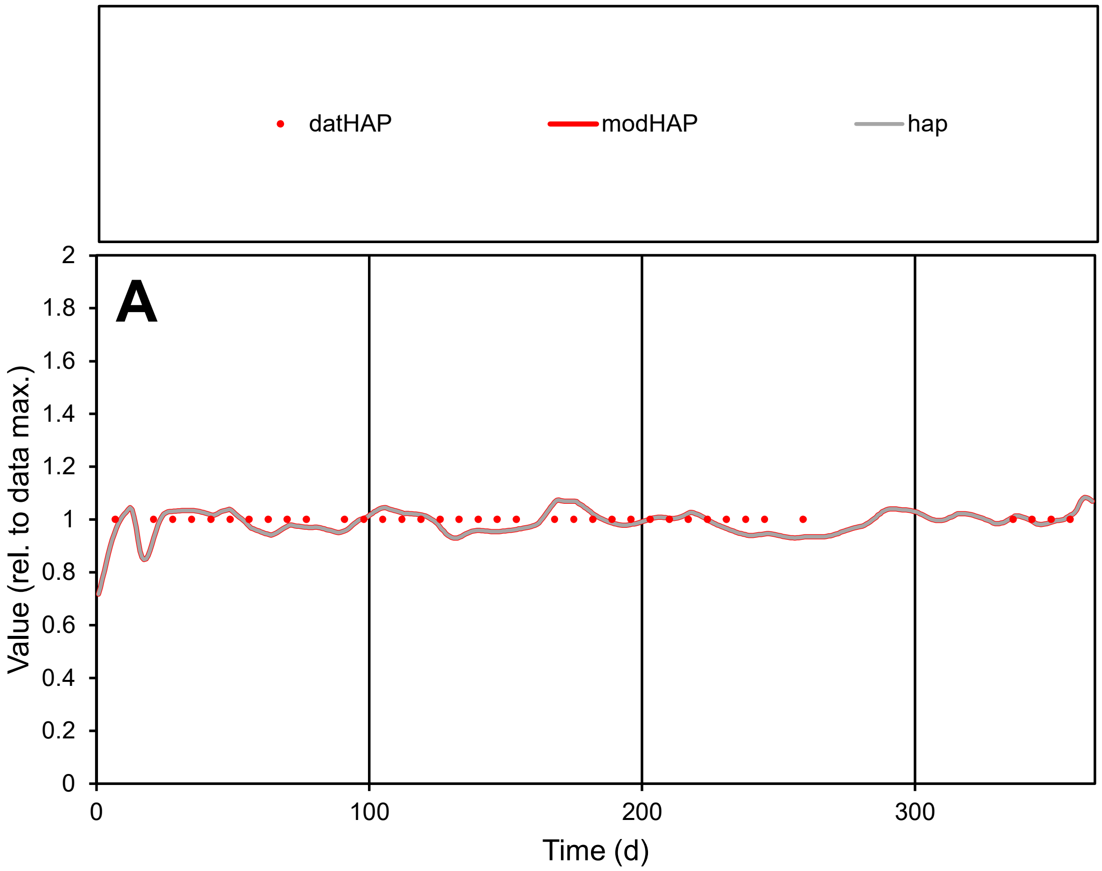


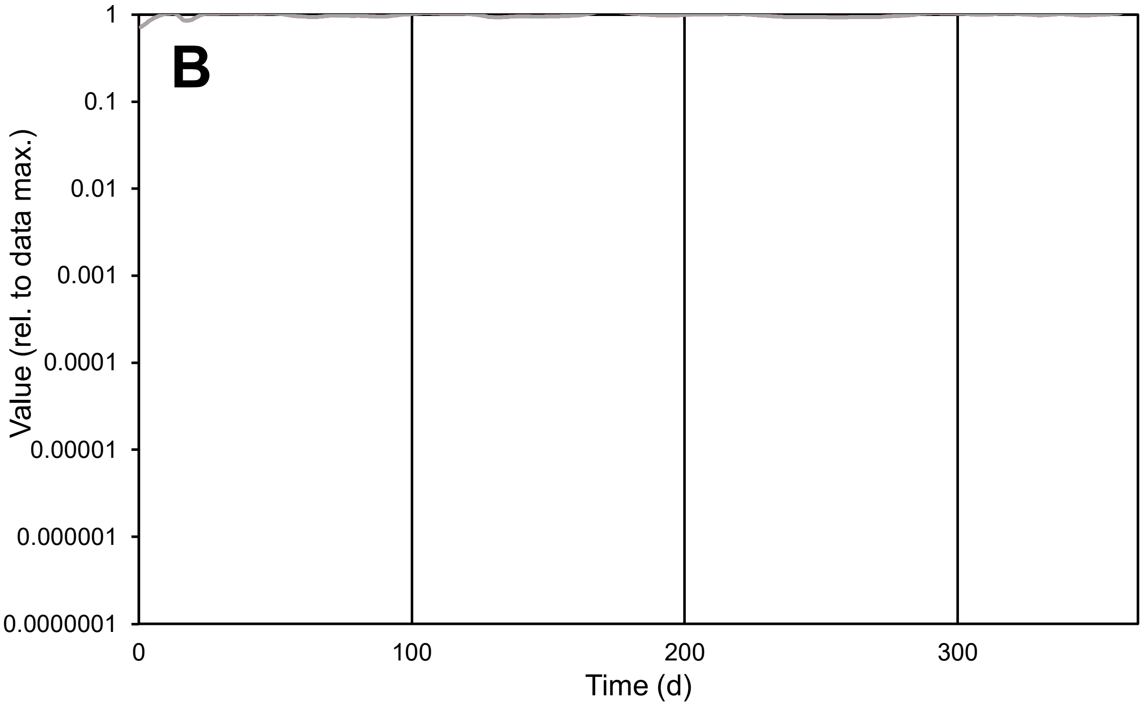


**C1**

Variables
HAP Provide jtd or Dst
CalibOnly?
2 1 to reproduce of
oNormalize?
2 1=no, 2=yes

ofxn calc*
0.03644
ofxn mod
0.11675

sRun
RunA1
zout
1

Ave
Data
1.1E-05
Model
1.1E-05

data/vMax
1.1E-05

ofxng
311.734

Min d/m
1.06975

ave/min d
1
ave/min m
1.06185

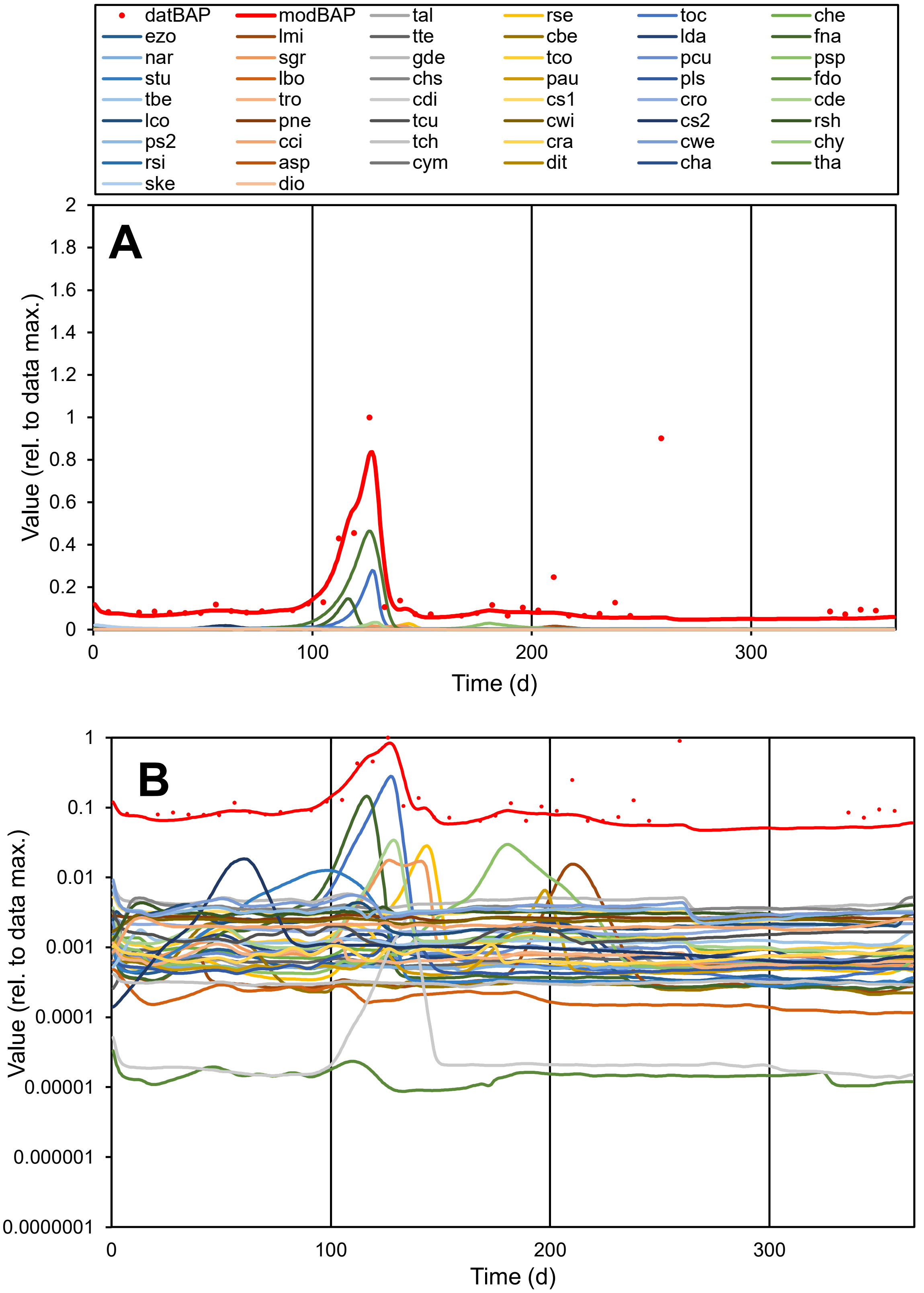


**C1**

Variables

| | |
|---|---|
| BAP | Provide jtd or Dst |

CalibOnly?

| | |
|---|---|
| 2 | 1 to reproduce of |

oNormalize?

| | |
|---|---|
| 2 | 1=no, 2=yes |

ofxn calc*
0.93567

ofxn mod
0.94652

sRun
RunA1

zout
1

Ave
Data
0.00086

Model
0.00067

data/vMax
0.00542

ofxng
311.734

Min d/m
1.24654

ave/min d
2.44435

ave/min m
2.38654

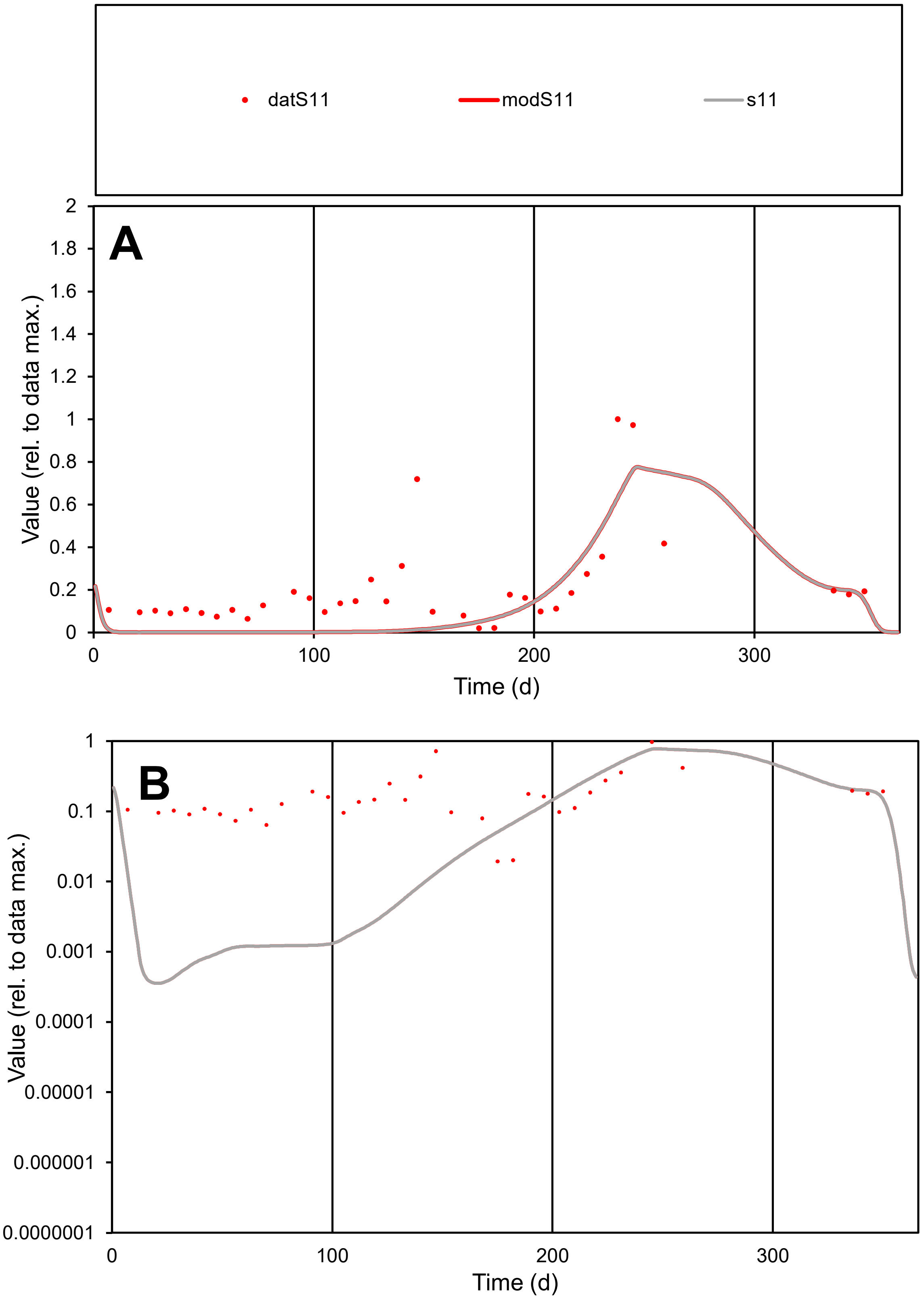


**C1**

Variables
S11 Provide jtd or Dst
CalibOnly?
2 1 to reproduce of
oNormalize?
2 1=no, 2=yes

ofxn calc*
0.78379
ofxn mod
0.80298

sRun
RunA1
zout
1

Ave
Data
0.08793
Model
0.05898

data/vMax
0.4403

ofxng
311.734

Min d/m
53.6008

ave/min d
10.3631
ave/min m
372.57

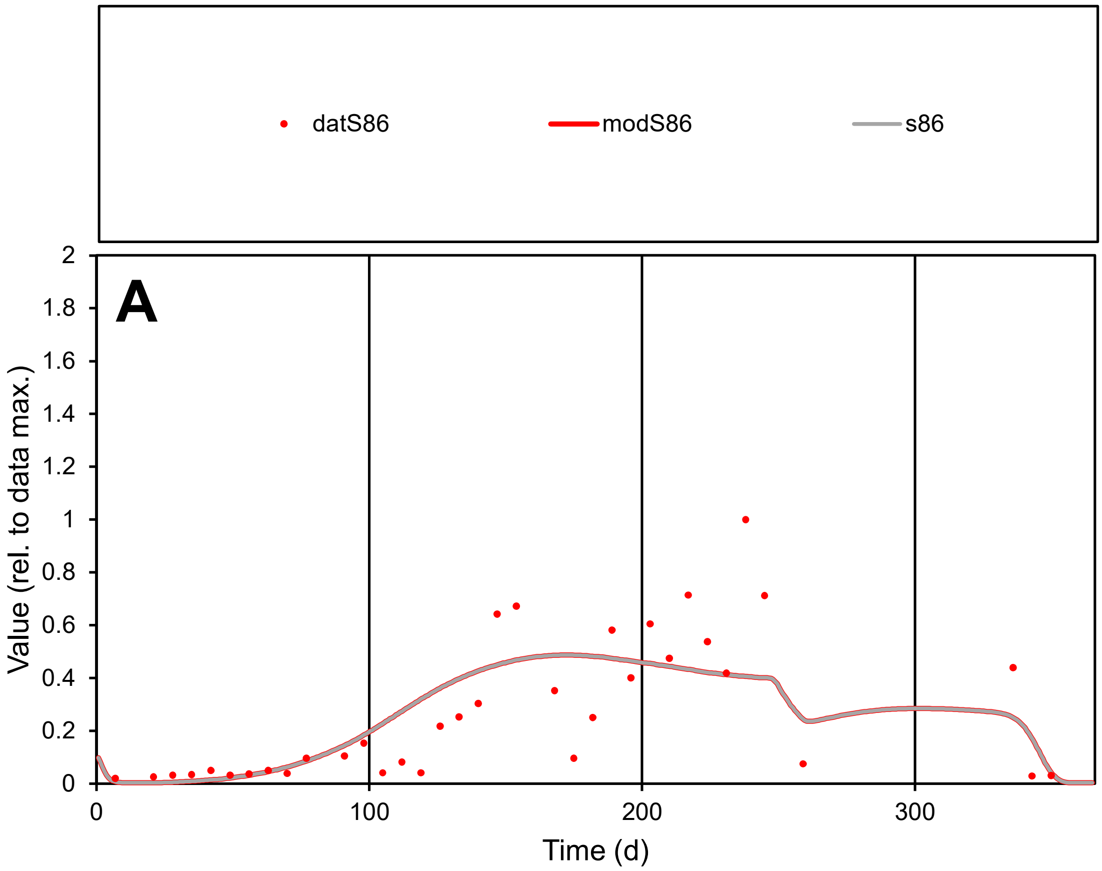


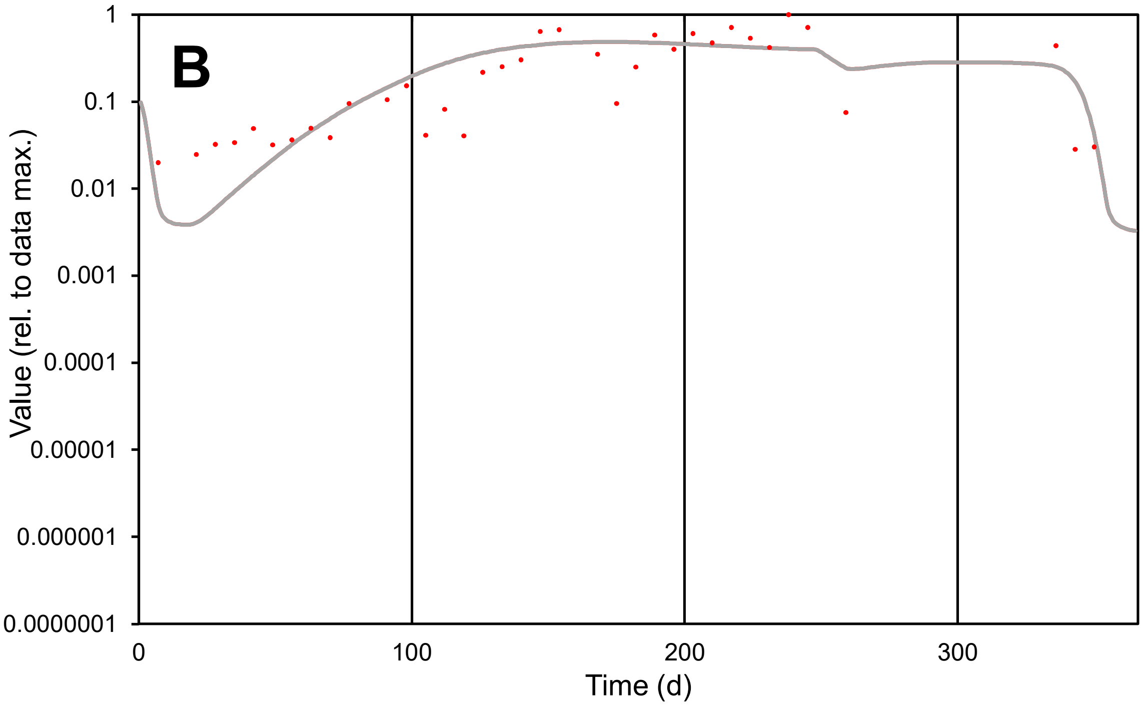


**C1**

Variables
S86 Provide jtd or Dst
CalibOnly?
2 1 to reproduce of
oNormalize?
2 1=no, 2=yes

ofxn calc*
0.73051
ofxn mod
0.7481

sRun
RunA1
zout
1

Ave
Data
0.07221
Model
0.0768

data/vMax
0.2882

ofxng
311.734

Min d/m
4.63636

ave/min d
12.5304
ave/min m
61.7884

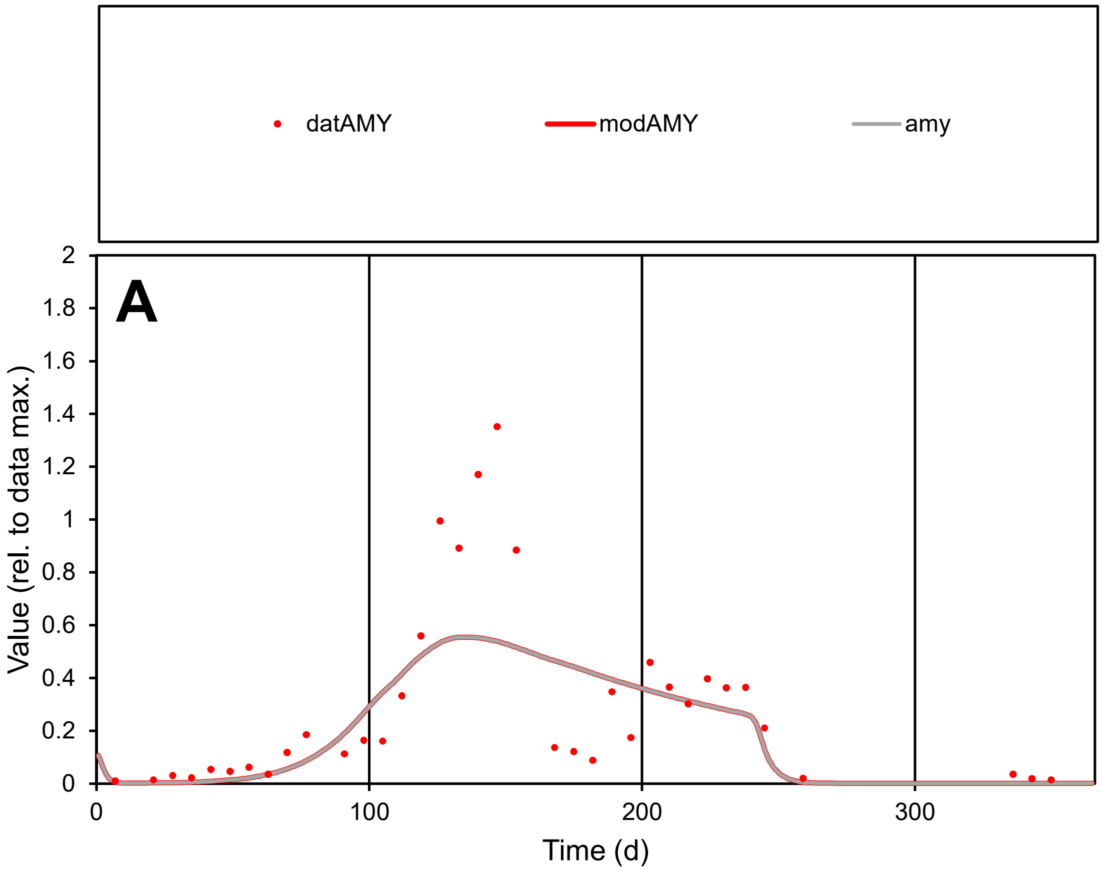


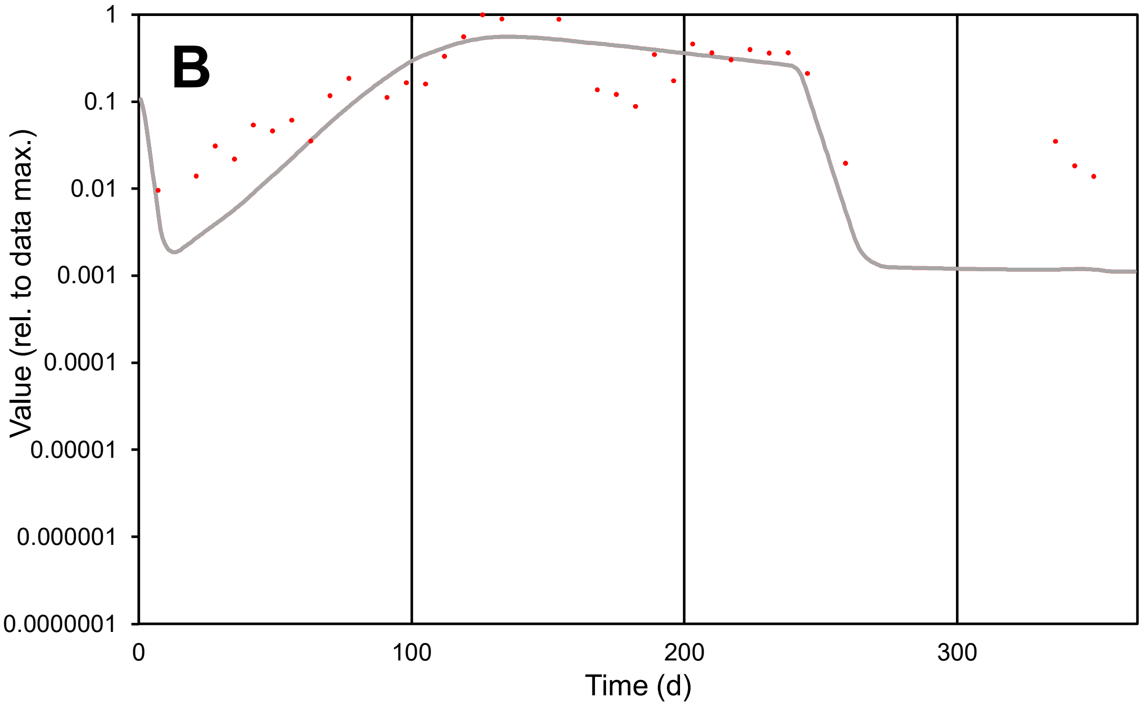


**C1**

| | |
|---|---|
| Variables | |
| AMY | Provide jtd or Dst |
| CalibOnly? | |
| 2 | 1 to reproduce of |
| oNormalize? | |
| 2 | 1=no, 2=yes |
| ofxn calc* | |
| 0.58753 | |
| ofxn mod | |
| 0.60676 | |
| sRun | |
| RunA1 | |
| zout | |
| 1 | |
| Ave | |
| Data | |
| 0.05253 | |
| Model | |
| 0.051 | |
| data/vMax | |
| 0.2106 | |
| ofxng | |
| 311.734 | |
| Min d/m | |
| 8.17628 | |
| ave/min d | |
| 26.0955 | |
| ave/min m | |
| 207.142 | |

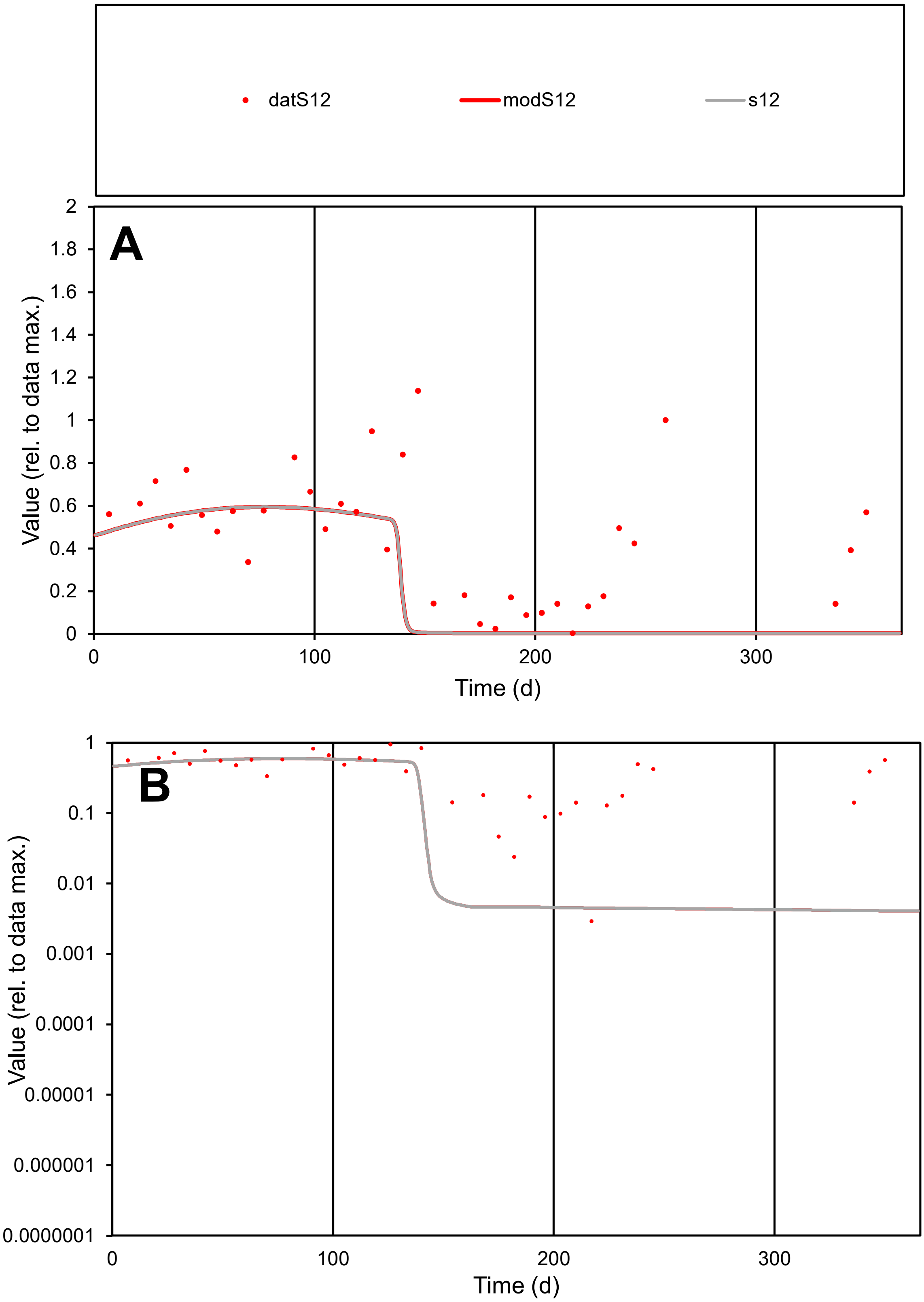


**C1**

Variables

S12 Provide jtd or Dst

CalibOnly?

2 1 to reproduce of

oNormalize?

2 1=no, 2=yes

ofxn calc*

0.68693

ofxn mod

0.70297

sRun

RunA1

zout

1

Ave

Data

0.02883

Model

0.01898

data/vMax

0.06785

ofxng

311.734

Min d/m

0.71552

ave/min d

145.086

ave/min m

68.3348

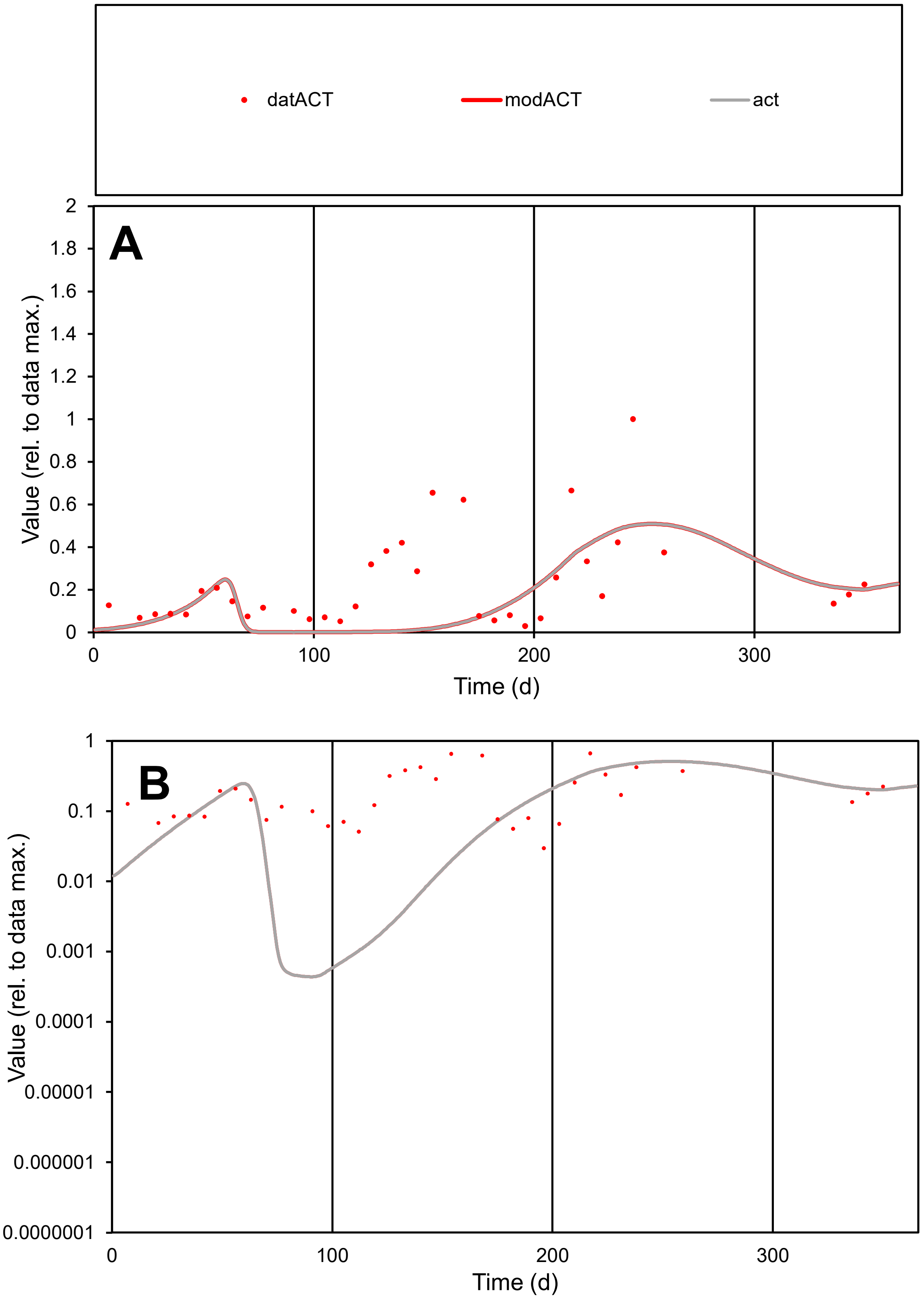


**C1**

Variables
ACT Provide jtd or Dst
CalibOnly?
2 1 to reproduce of
oNormalize?
2 1=no, 2=yes

ofxn calc*
0.89586
ofxn mod
0.90721

sRun
RunA1
zout
1

Ave
Data
0.03595
Model
0.02531

data/vMax
0.1678

ofxng
311.734

Min d/m
67.2428

ave/min d
7.21885
ave/min m
341.745

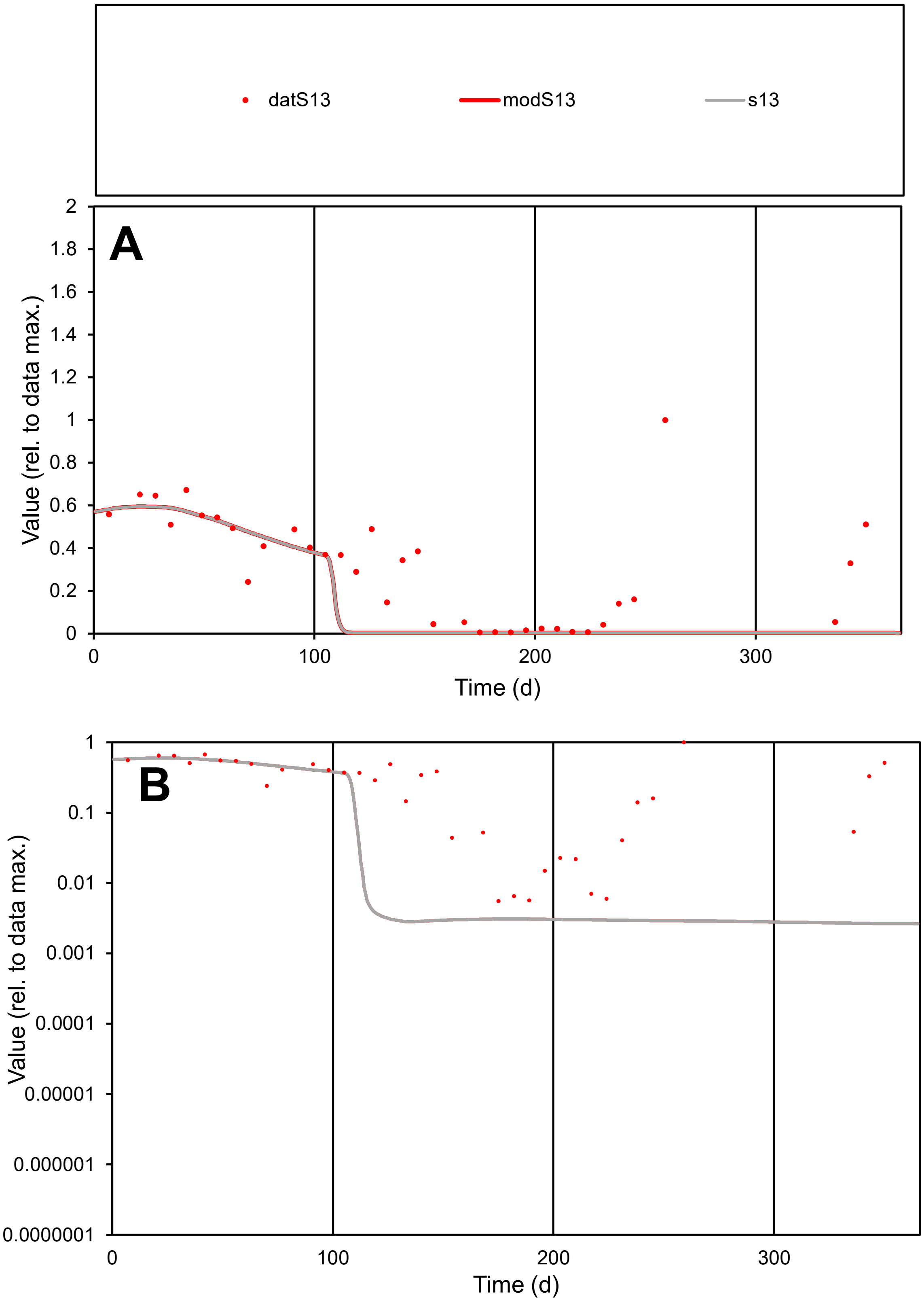


**C1**

Variables
S13 Provide jtd or Dst
CalibOnly?
2 1 to reproduce of
oNormalize?
2 1=no, 2=yes

ofxn calc*
0.77634
ofxn mod
0.79767

sRun
RunA1
zout
1

Ave
Data
0.01647
Model
0.01065

data/vMax
0.05632

ofxng
311.734

Min d/m
2.08166

ave/min d
52.9709
ave/min m
71.2855

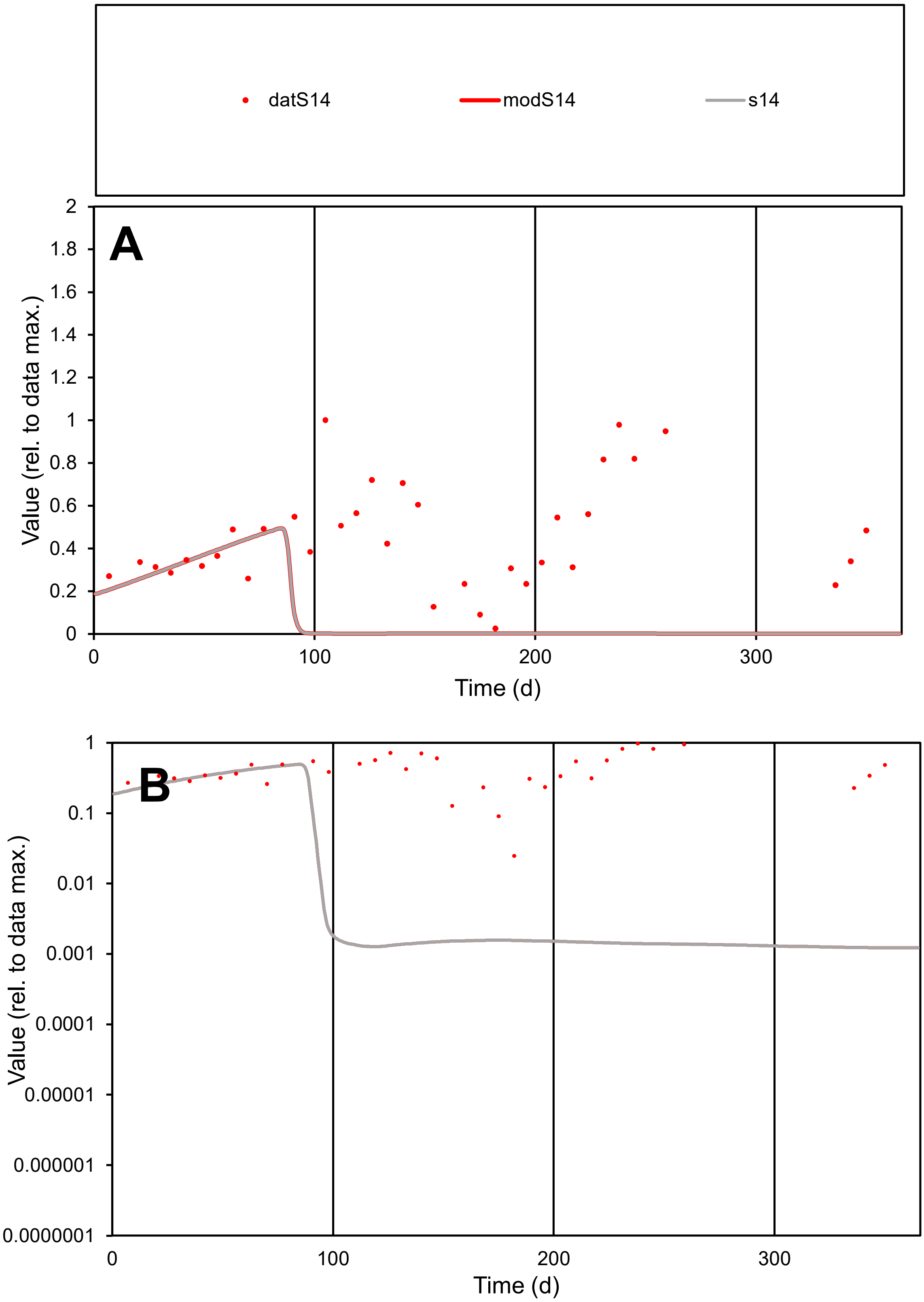


**C1**

Variables

S14 Provide jtd or Dst

CalibOnly?

2 1 to reproduce of

oNormalize?

2 1=no, 2=yes

ofxn calc*

1.03369

ofxn mod

1.04558

sRun

RunA1

zout

1

Ave

Data

0.02647

Model

0.00681

data/vMax

0.06152

ofxng

311.734

Min d/m

20.1058

ave/min d

17.4125

ave/min m

90.0623

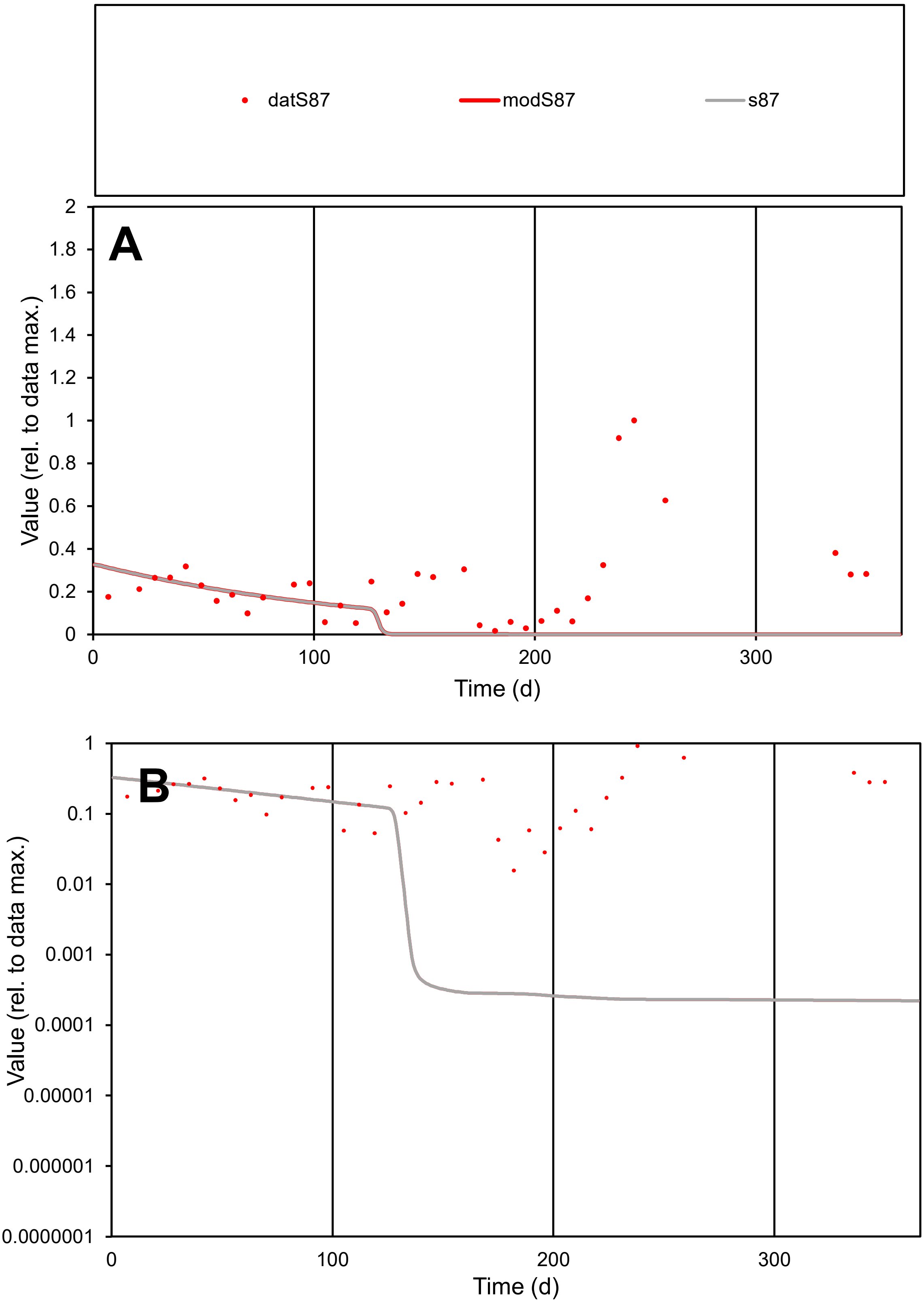


**C1**

Variables
S87 Provide jtd or Dst
CalibOnly?
2 1 to reproduce of
oNormalize?
2 1=no, 2=yes

ofxn calc*
1.25747
ofxn mod
1.26209

sRun
RunA1
zout
1

Ave
Data
0.01942
Model
0.00757

data/vMax
0.08536

ofxng
311.734

Min d/m
70.5728

ave/min d
14.4609
ave/min m
397.851

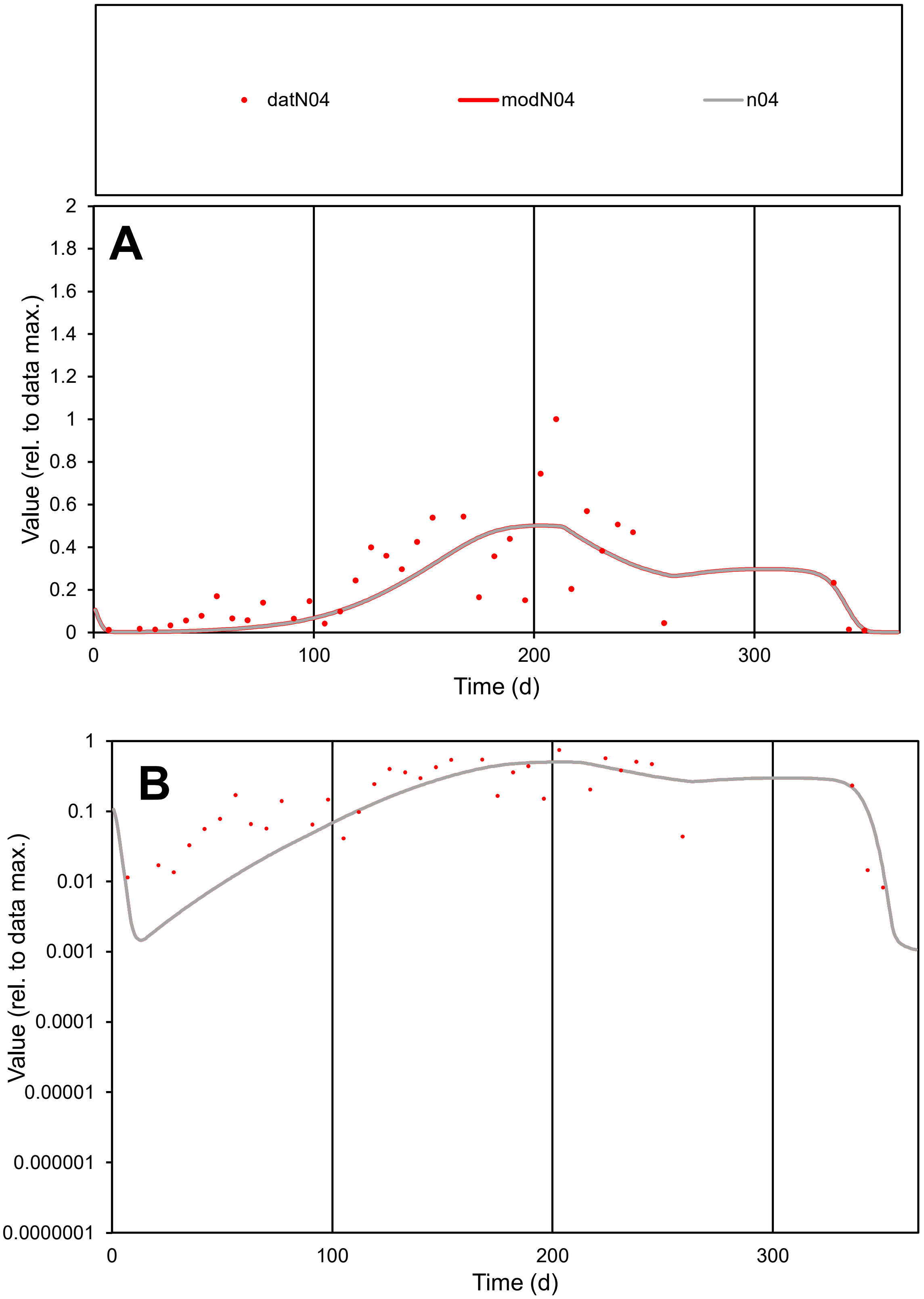


**C1**

Variables
N04 Provide jtd or Dst
CalibOnly?
2 1 to reproduce of
oNormalize?
2 1=no, 2=yes

ofxn calc*
0.64728
ofxn mod
0.66527

sRun
RunA1
zout
1

Ave
Data
0.02782
Model
0.02484

data/vMax
0.118

ofxng
311.734

Min d/m
3.4982

ave/min d
28.673
ave/min m
89.5546

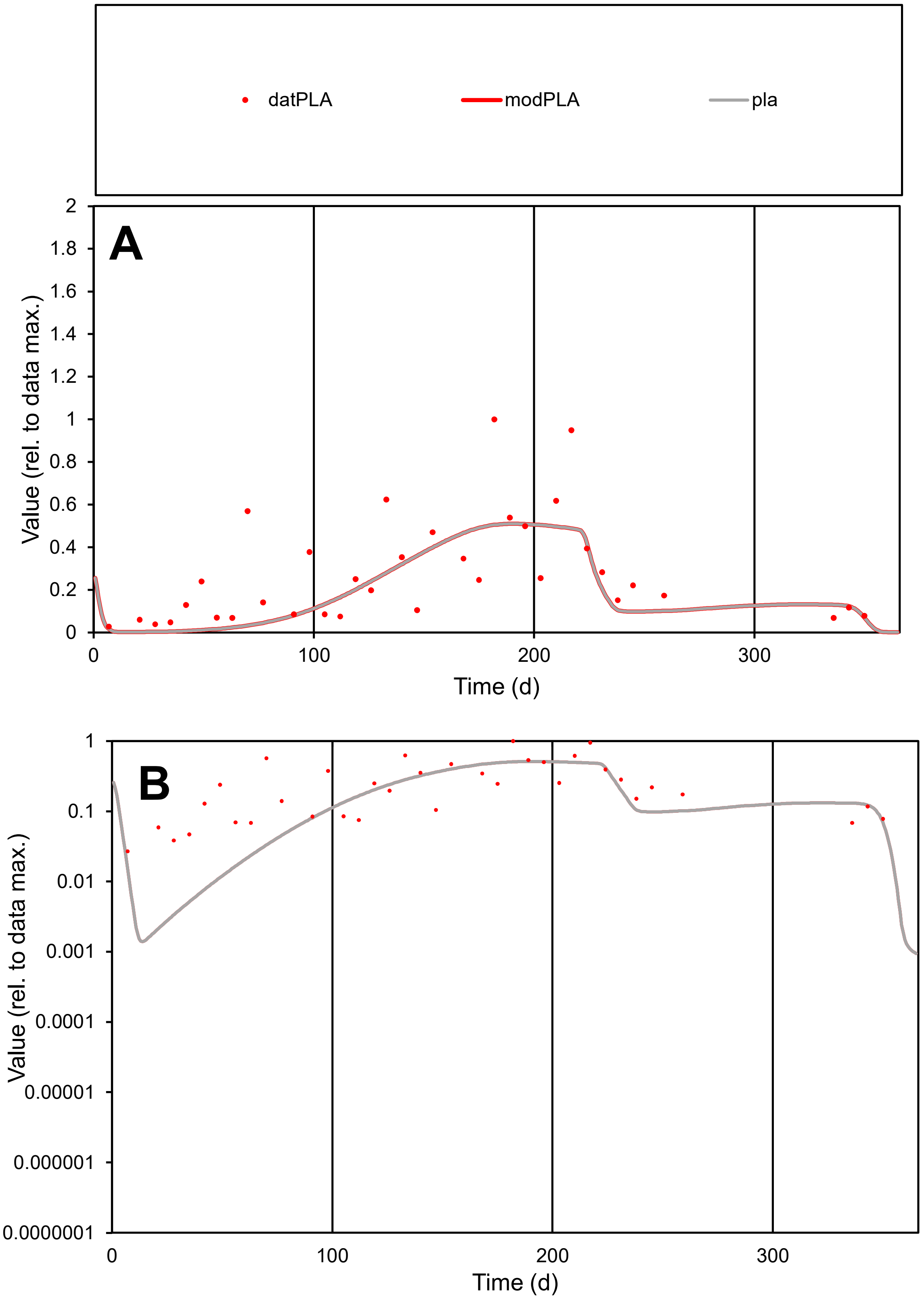


**C1**

Variables
PLA Provide jtd or Dst
CalibOnly?
2 1 to reproduce of
oNormalize?
2 1=no, 2=yes

ofxn calc*
0.70988
ofxn mod
0.72252

sRun
RunA1
zout
1

Ave
Data
0.03125
Model
0.02559

data/vMax
0.1197

ofxng
311.734

Min d/m
11.421

ave/min d
9.67243
ave/min m
90.4426

331_Calib_S_A

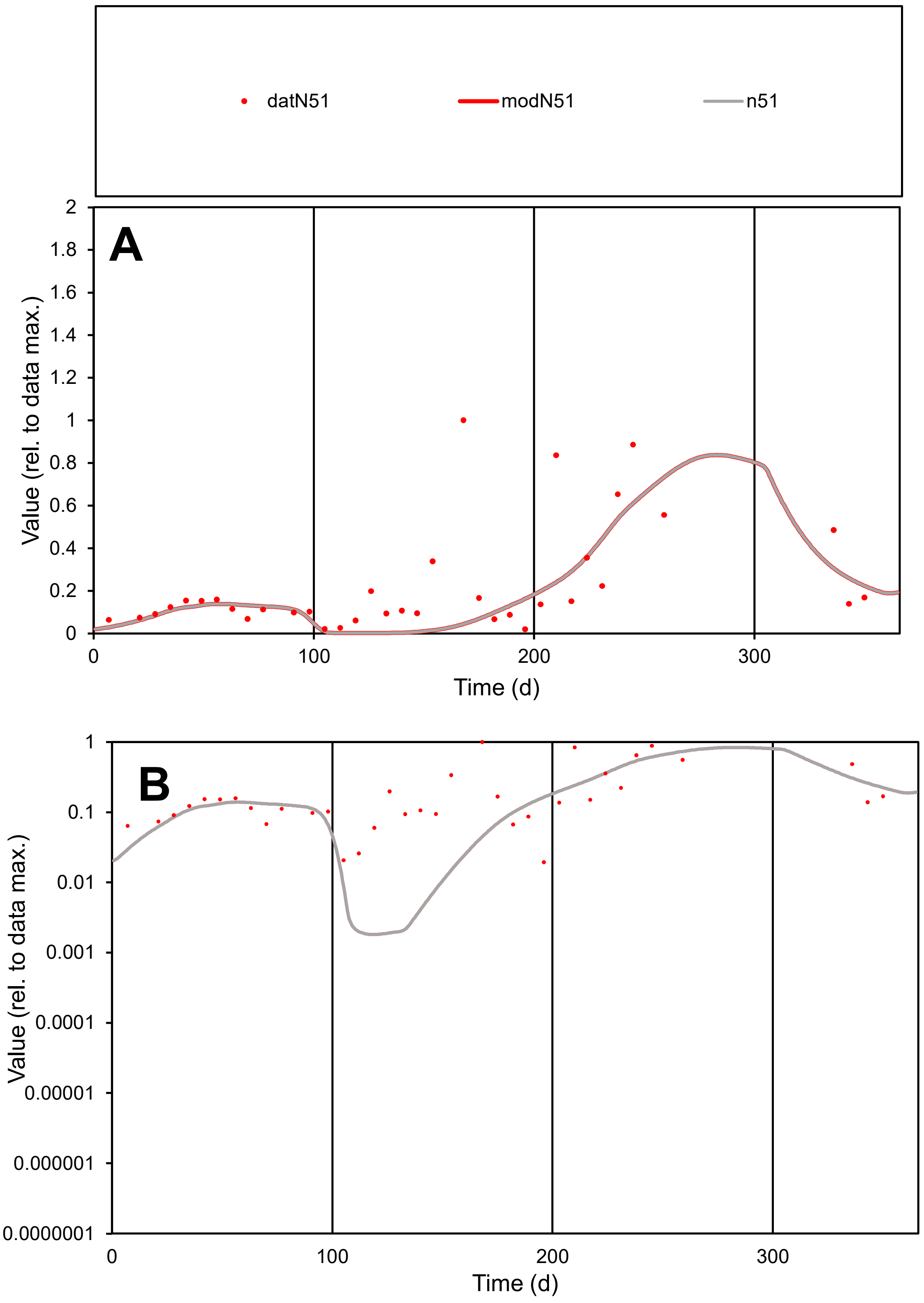


**C1**

Variables
N51 Provide jtd or Dst
CalibOnly?
2 1 to reproduce of
oNormalize?
2 1=no, 2=yes

ofxn calc*
0.96185
ofxn mod
0.97435

sRun
RunA1
zout
1

Ave
Data
0.01746
Model
0.01341

data/vMax
0.07922

ofxng
311.734

Min d/m
10.693

ave/min d
11.3176
ave/min m
92.9395

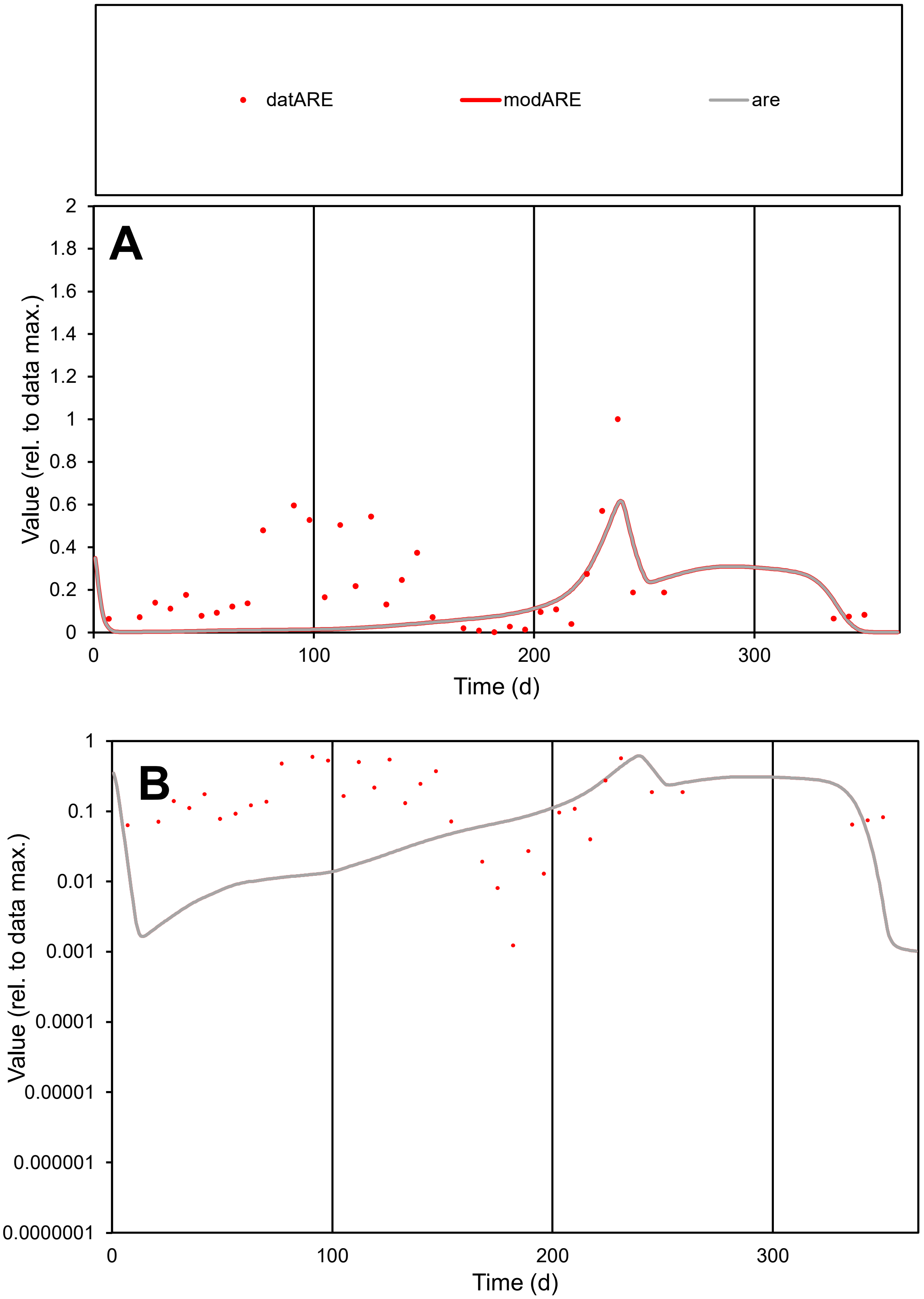


**C1**

Variables

ARE Provide jtd or Dst

CalibOnly?

2 1 to reproduce of

oNormalize?

2 1=no, 2=yes

ofxn calc*

1.08167

ofxn mod

1.0901

sRun

RunA1

zout

1

Ave

Data

0.01153

Model

0.00555

data/vMax

0.05779

ofxng

311.734

Min d/m

0.49179

ave/min d

161.664

ave/min m

38.2608

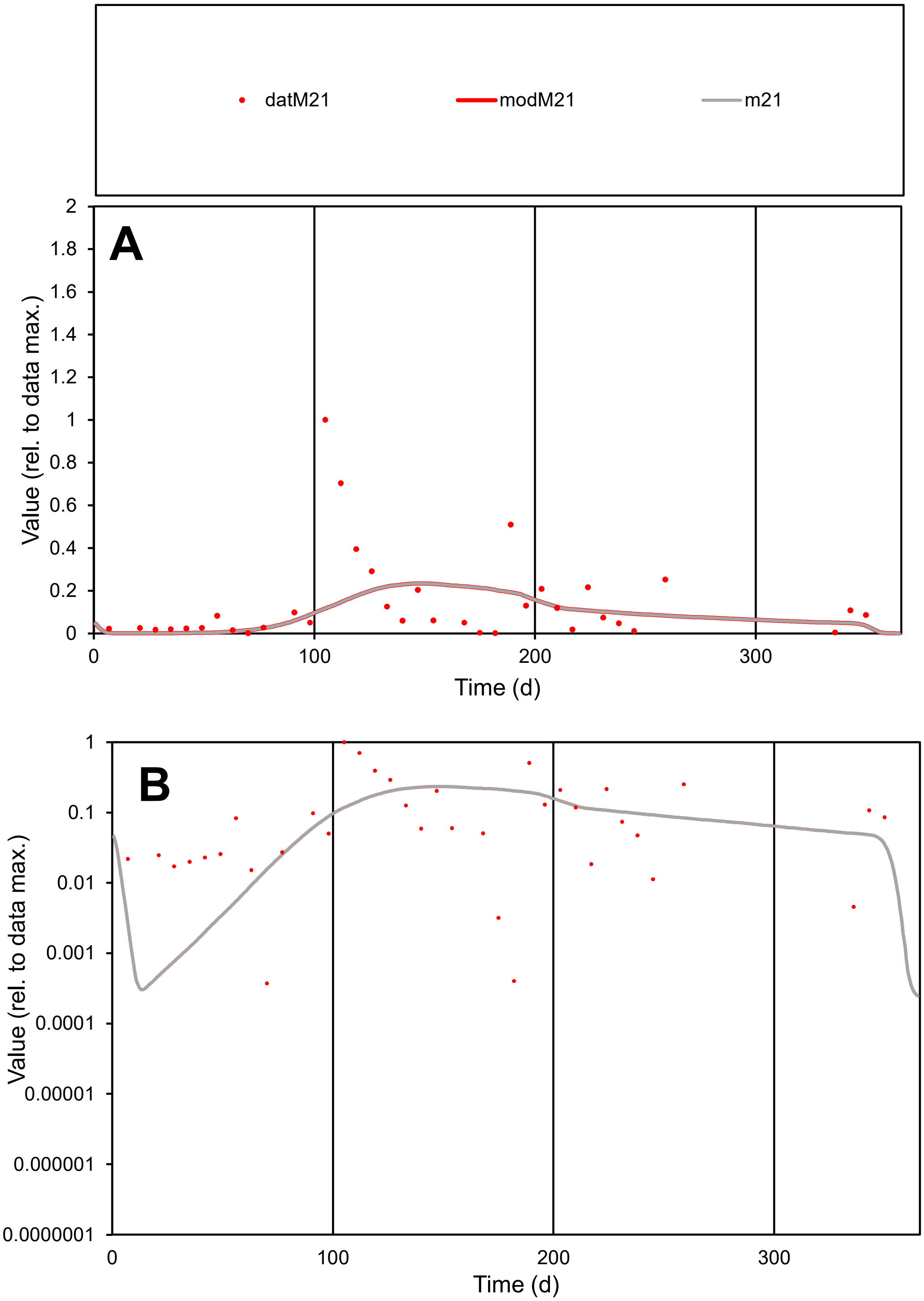


**C1**

Variables
M21 Provide jtd or Dst
CalibOnly?
2 1 to reproduce of
oNormalize?
2 1=no, 2=yes

ofxn calc*
1.51922
ofxn mod
1.52448

sRun
RunA1
zout
1

Ave
Data
0.00881
Model
0.00686

data/vMax
0.06514

ofxng
311.734

Min d/m
0.70345

ave/min d
362.51
ave/min m
198.72

331_Calib_S_A

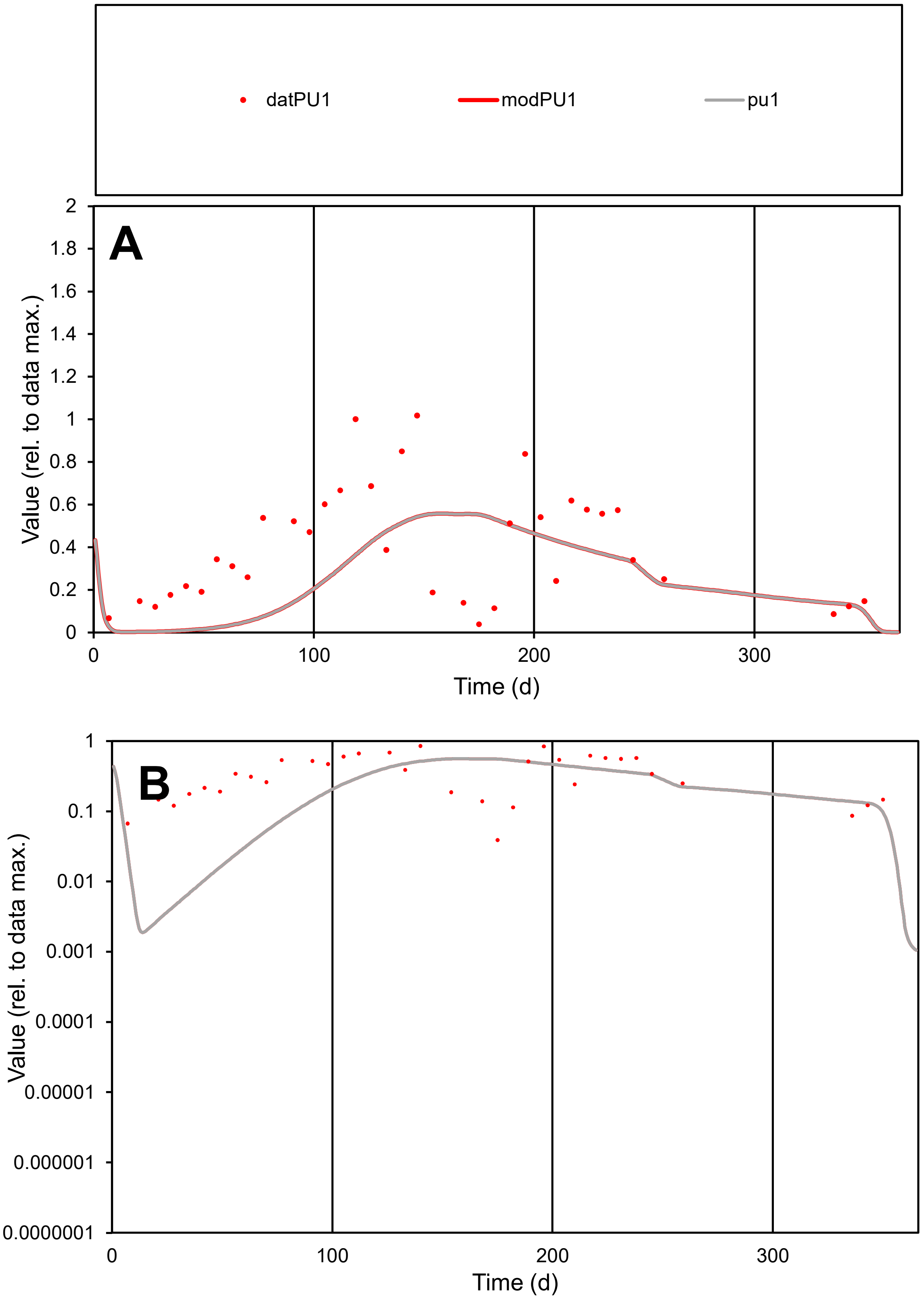


**C1**

Variables
PU1 Provide jtd or Dst
CalibOnly?
2 1 to reproduce of
oNormalize?
2 1=no, 2=yes

ofxn calc*
0.72274
ofxn mod
0.7309

sRun
RunA1
zout
1

Ave
Data
0.014
Model
0.01044

data/vMax
0.03745

ofxng
311.734

Min d/m
12.383

ave/min d
9.61949
ave/min m
88.8743

331_Calib_S_A

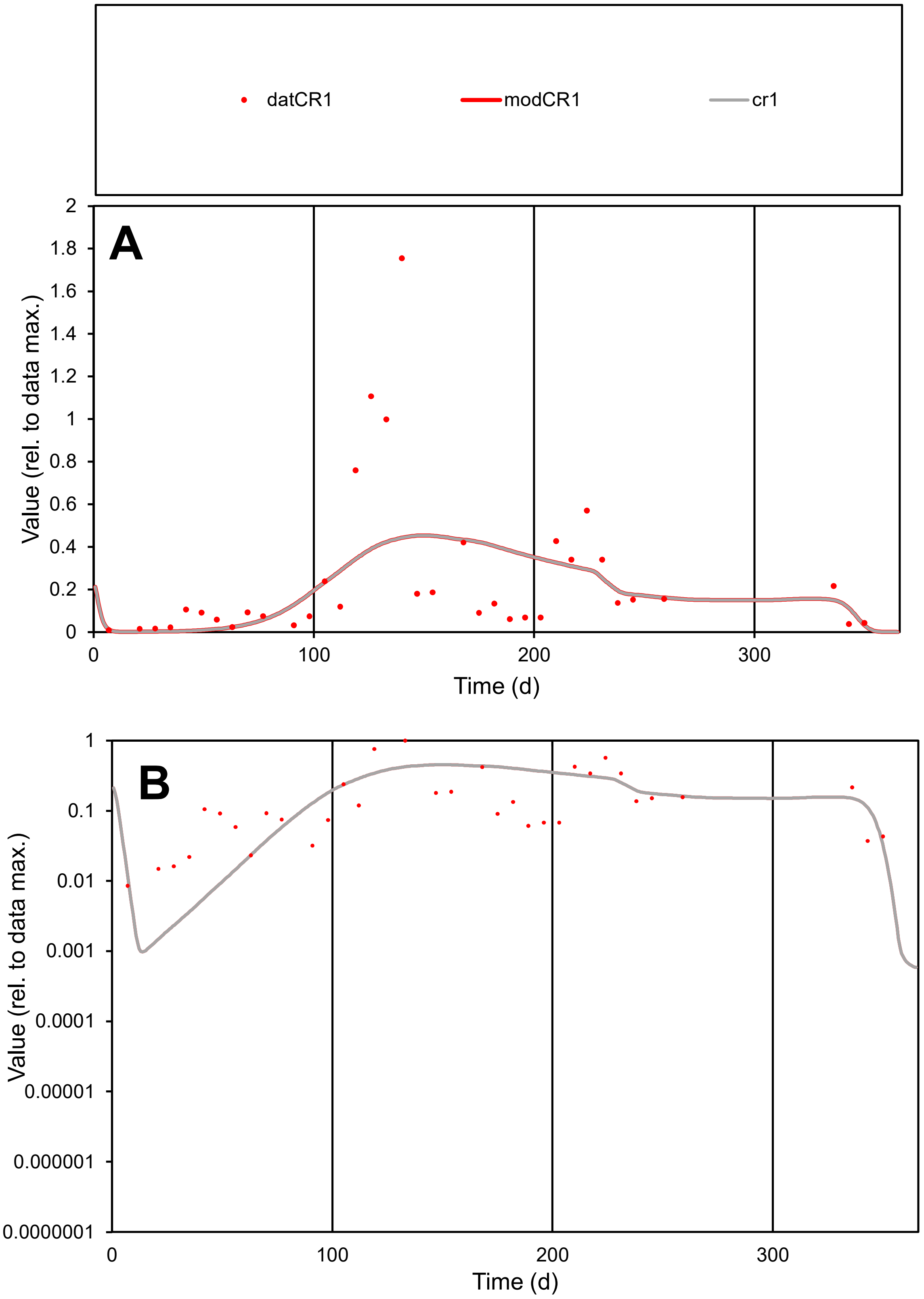


**C1**

Variables
CR1 Provide jtd or Dst
CalibOnly?
2 1 to reproduce of
oNormalize?
2 1=no, 2=yes

ofxn calc*
0.95544
ofxn mod
0.96719

sRun
RunA1
zout
1

Ave
Data
0.01686
Model
0.01701

data/vMax
0.07827

ofxng
311.734

Min d/m
5.19189

ave/min d
25.3234
ave/min m
132.691

331_Calib_S_A

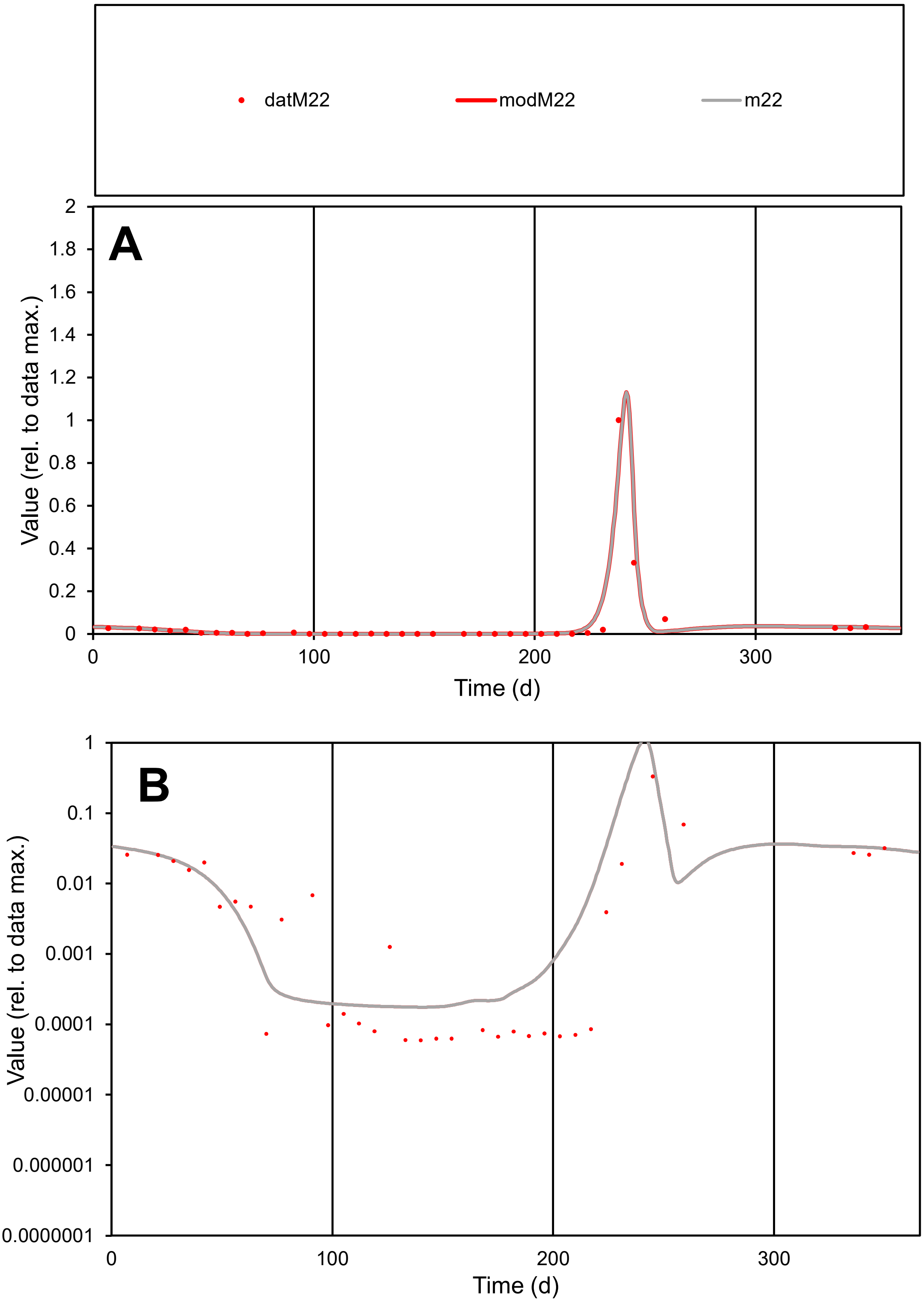


**C1**

| | |
|---|---|
| Variables | |
| M22 | Provide jtd or Dst |
| CalibOnly? | |
| 2 | 1 to reproduce of |
| oNormalize? | |
| 2 | 1=no, 2=yes |
| ofxn calc* | |
| 0.46024 | |
| ofxn mod | |
| 0.68581 | |
| sRun | |
| RunA1 | |
| zout | |
| 1 | |
| Ave | |
| Data | |
| 0.01637 | |
| Model | |
| 0.01639 | |
| data/vMax | |
| 0.3584 | |
| ofxng | |
| 311.734 | |
| Min d/m | |
| 0.34014 | |
| ave/min d | |
| 767.291 | |
| ave/min m | |
| 261.351 | |

331_Calib_S_A

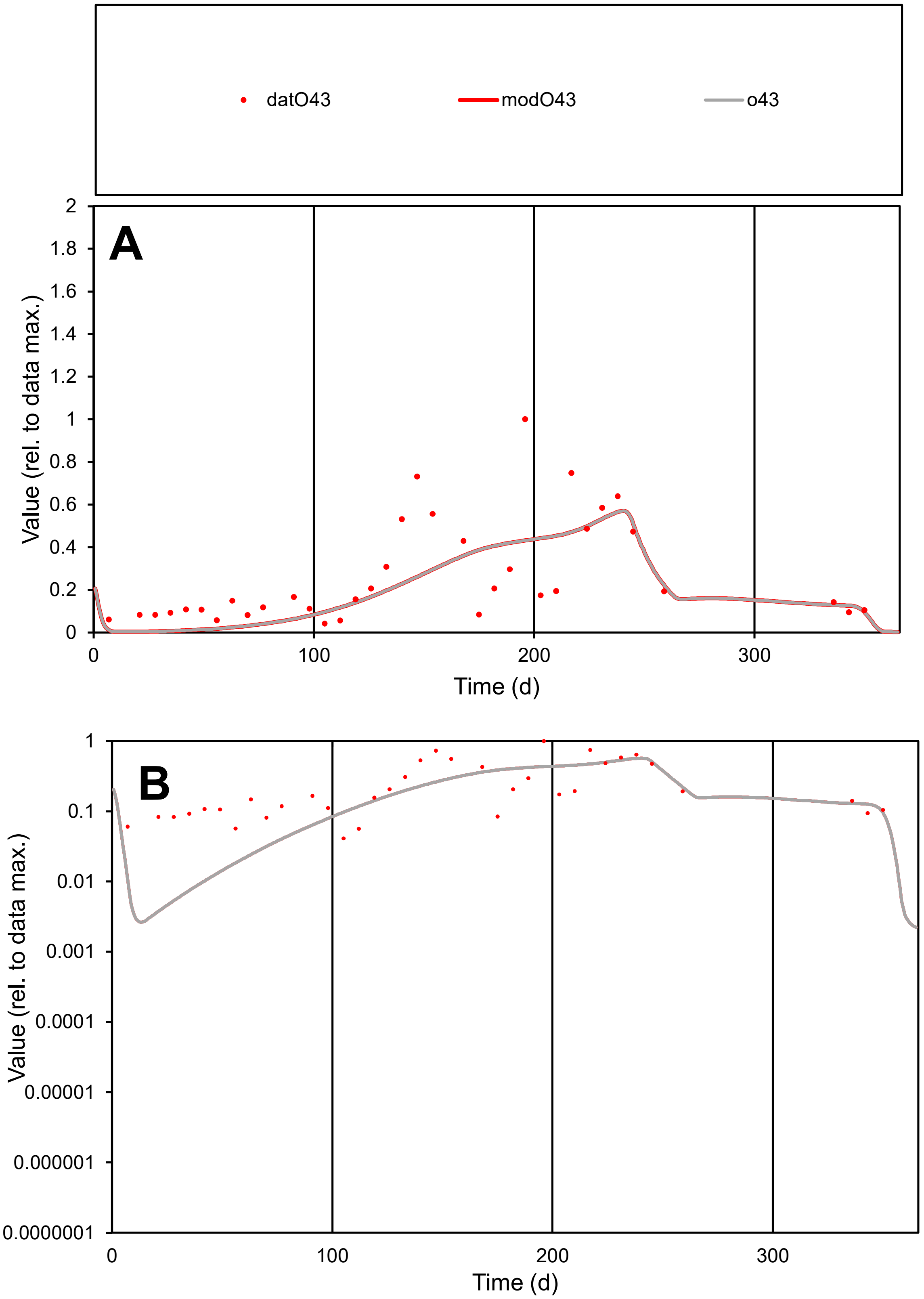


**C1**

Variables
O43 Provide jtd or Dst
CalibOnly?
2 1 to reproduce of
oNormalize?
2 1=no, 2=yes

ofxn calc*
0.62719
ofxn mod
0.64318

sRun
RunA1
zout
1

Ave
Data
0.019
Model
0.01686

data/vMax
0.07654

ofxng
311.734

Min d/m
10.1028

ave/min d
6.03897
ave/min m
54.1374

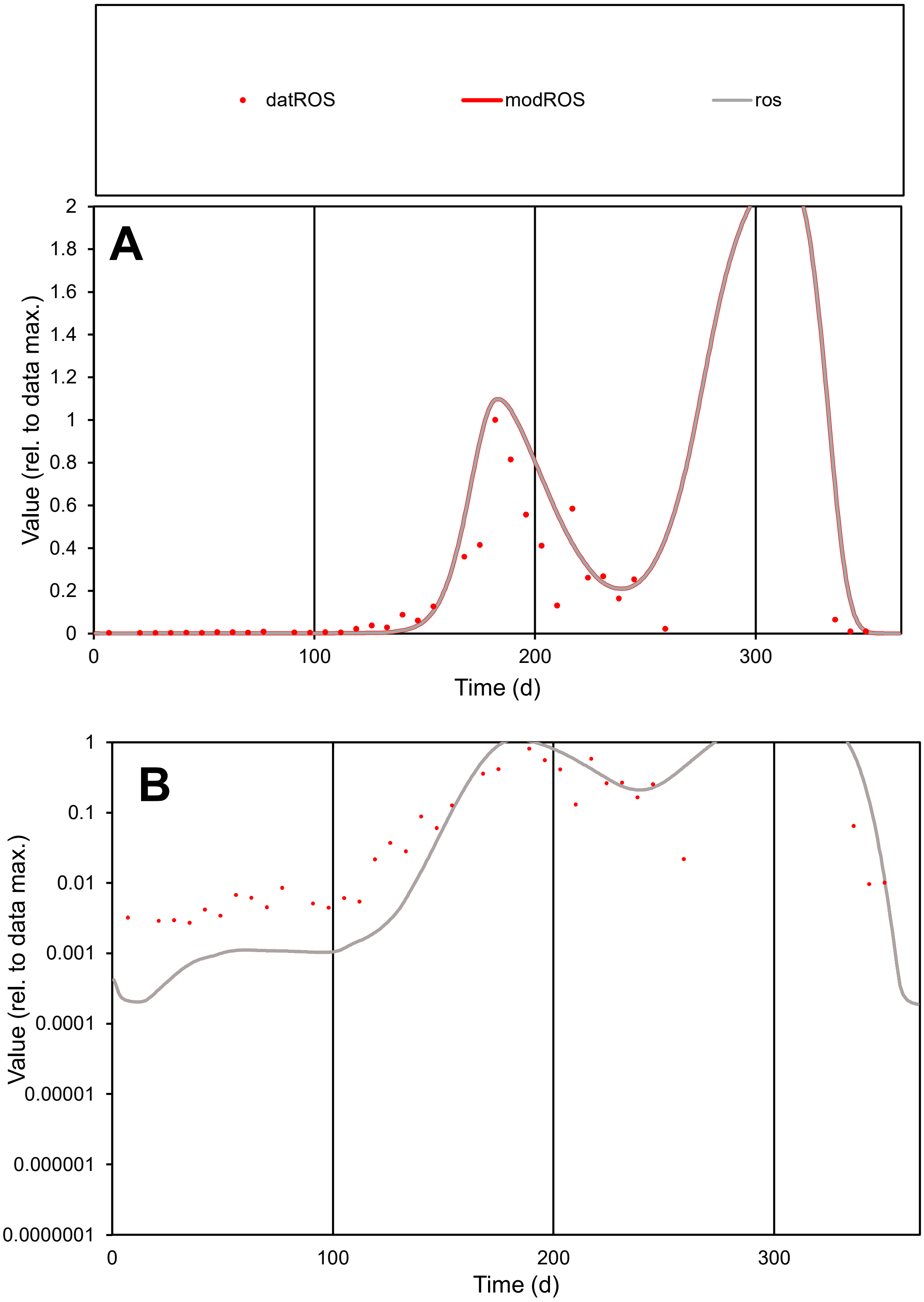


**C1**

Variables
ROS Provide jtd or Dst
CalibOnly?
2 1 to reproduce of
oNormalize?
2 1=no, 2=yes

ofxn calc*
1.03903
ofxn mod
1.12623

sRun
RunA1
zout
1

Ave
Data
0.03007
Model
0.04315

data/vMax
0.1916

ofxng
311.734

Min d/m
13.0702

ave/min d
57.4754
ave/min m
1077.85

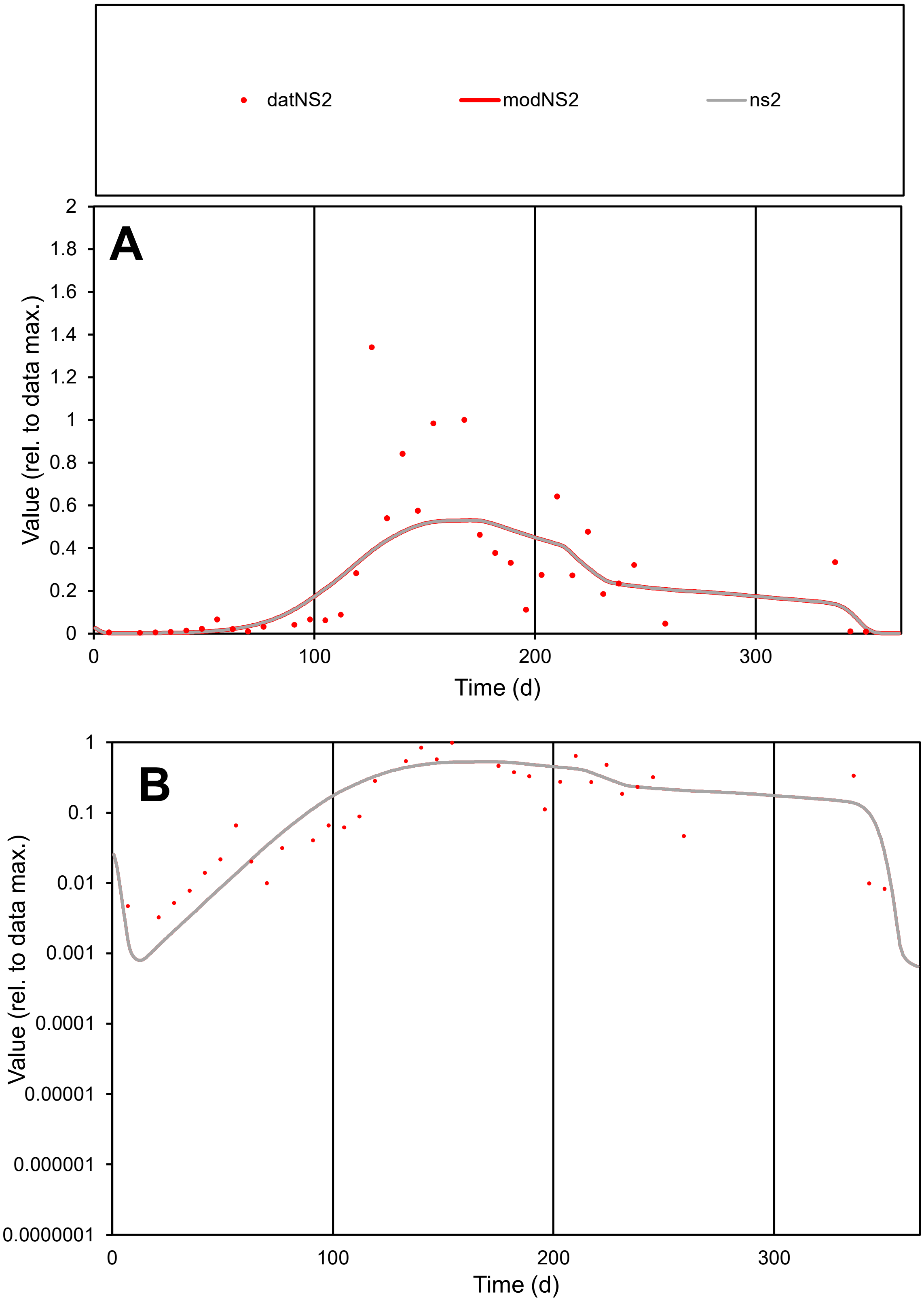


**C1**

Variables
NS2 Provide jtd or Dst
CalibOnly?
2 1 to reproduce of
oNormalize?
2 1=no, 2=yes

ofxn calc*
0.69128
ofxn mod
0.70793

sRun
RunA1
zout
1

Ave
Data
0.0193
Model
0.01922

data/vMax
0.07918

ofxng
311.734

Min d/m
2.26643

ave/min d
74.6426
ave/min m
168.432

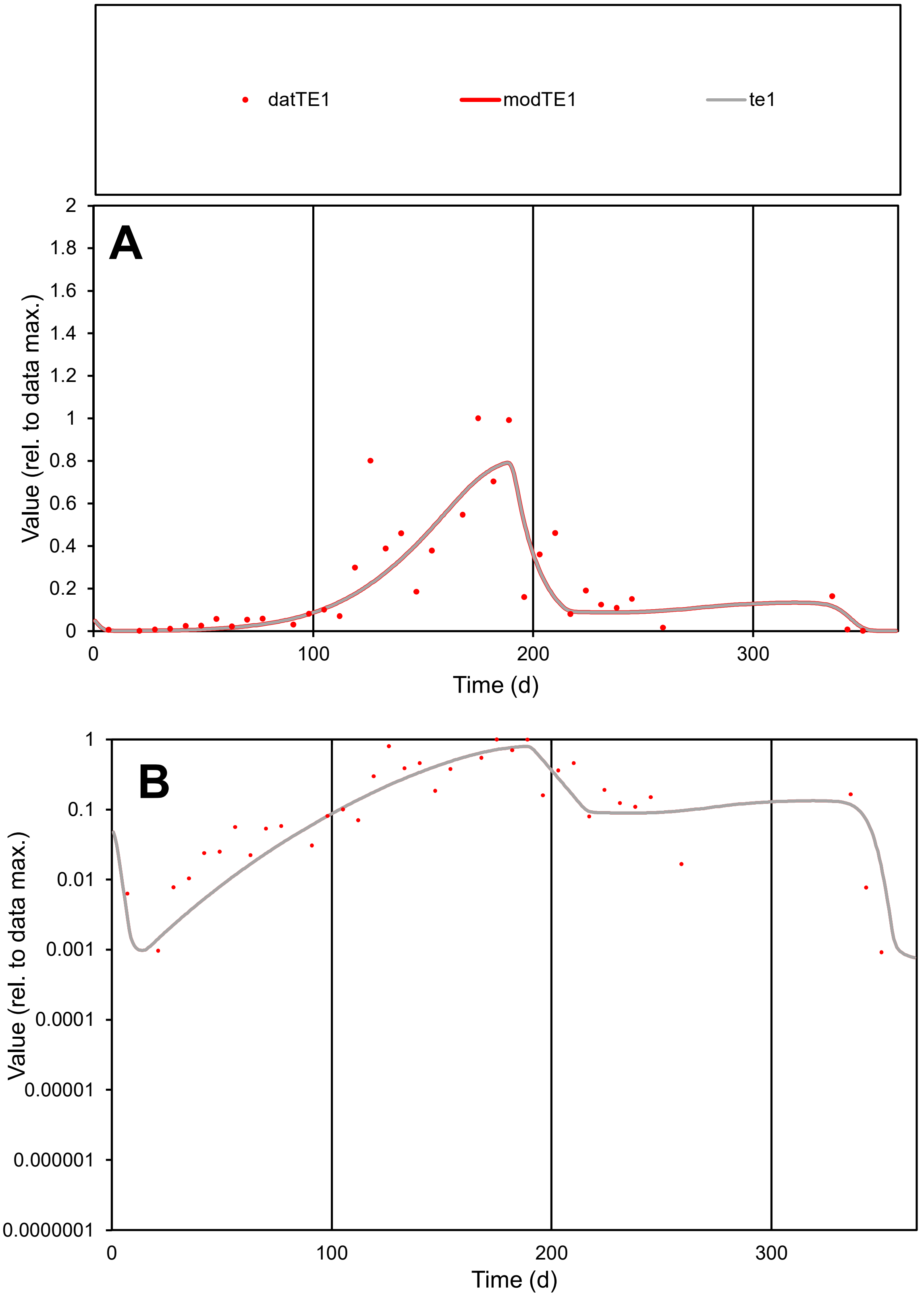


**C1**

Variables
TE1 Provide jtd or Dst
CalibOnly?
2 1 to reproduce of
oNormalize?
2 1=no, 2=yes

ofxn calc*
0.62588
ofxn mod
0.65778

sRun
RunA1
zout
1

Ave
Data
0.01219
Model
0.01136

data/vMax
0.05898

ofxng
311.734

Min d/m
0.63847

ave/min d
225.203
ave/min m
133.992

331_Calib_S_A

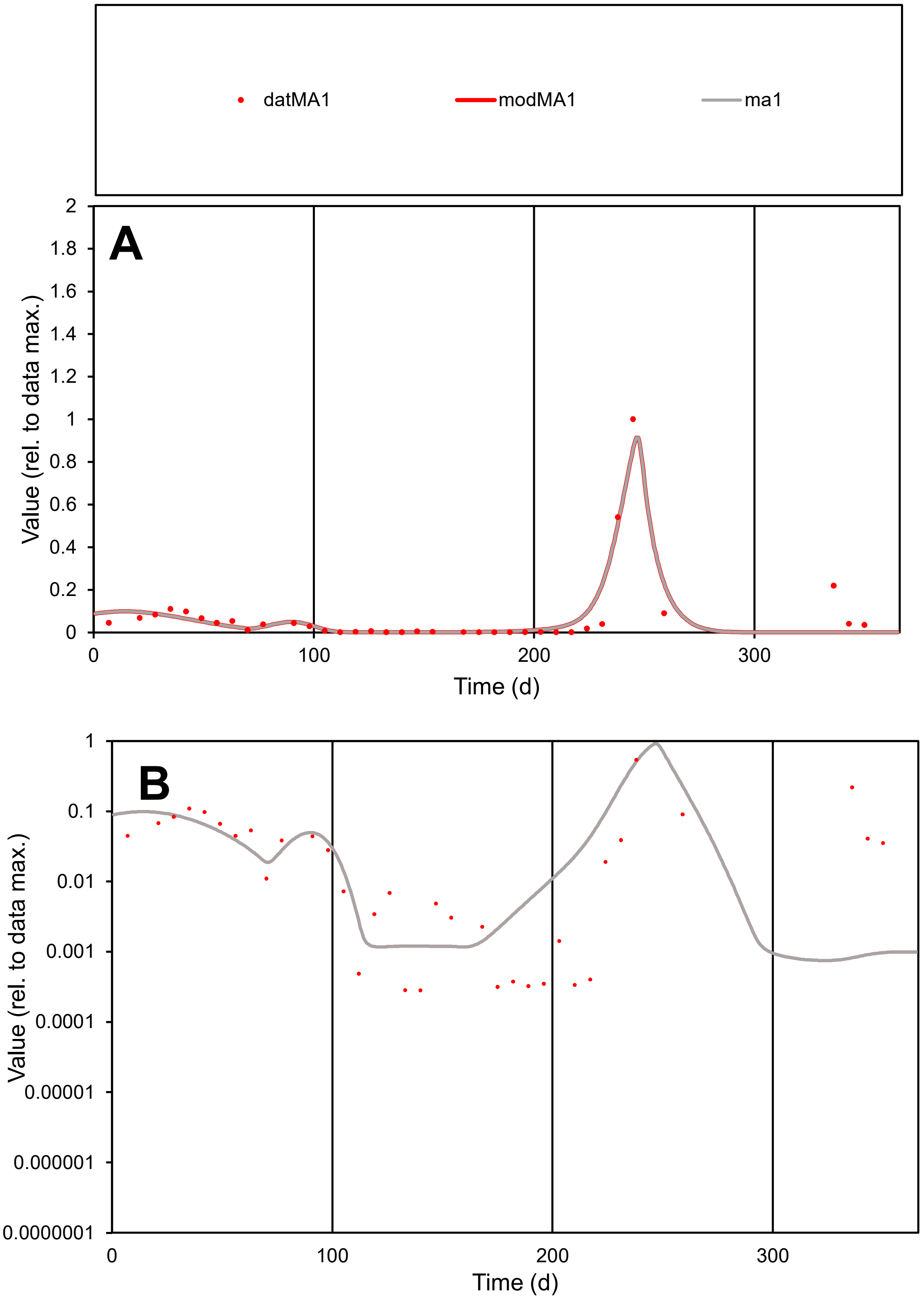


**C1**

Variables
MA1 Provide jtd or Dst
CalibOnly?
2 1 to reproduce of
oNormalize?
2 1=no, 2=yes

ofxn calc*
0.70103
ofxn mod
0.81201

sRun
RunA1
zout
1

Ave
Data
0.00381
Model
0.00381

data/vMax
0.0507

ofxng
311.734

Min d/m
0.33695

ave/min d
266.437
ave/min m
89.8314

331_Calib_S_A

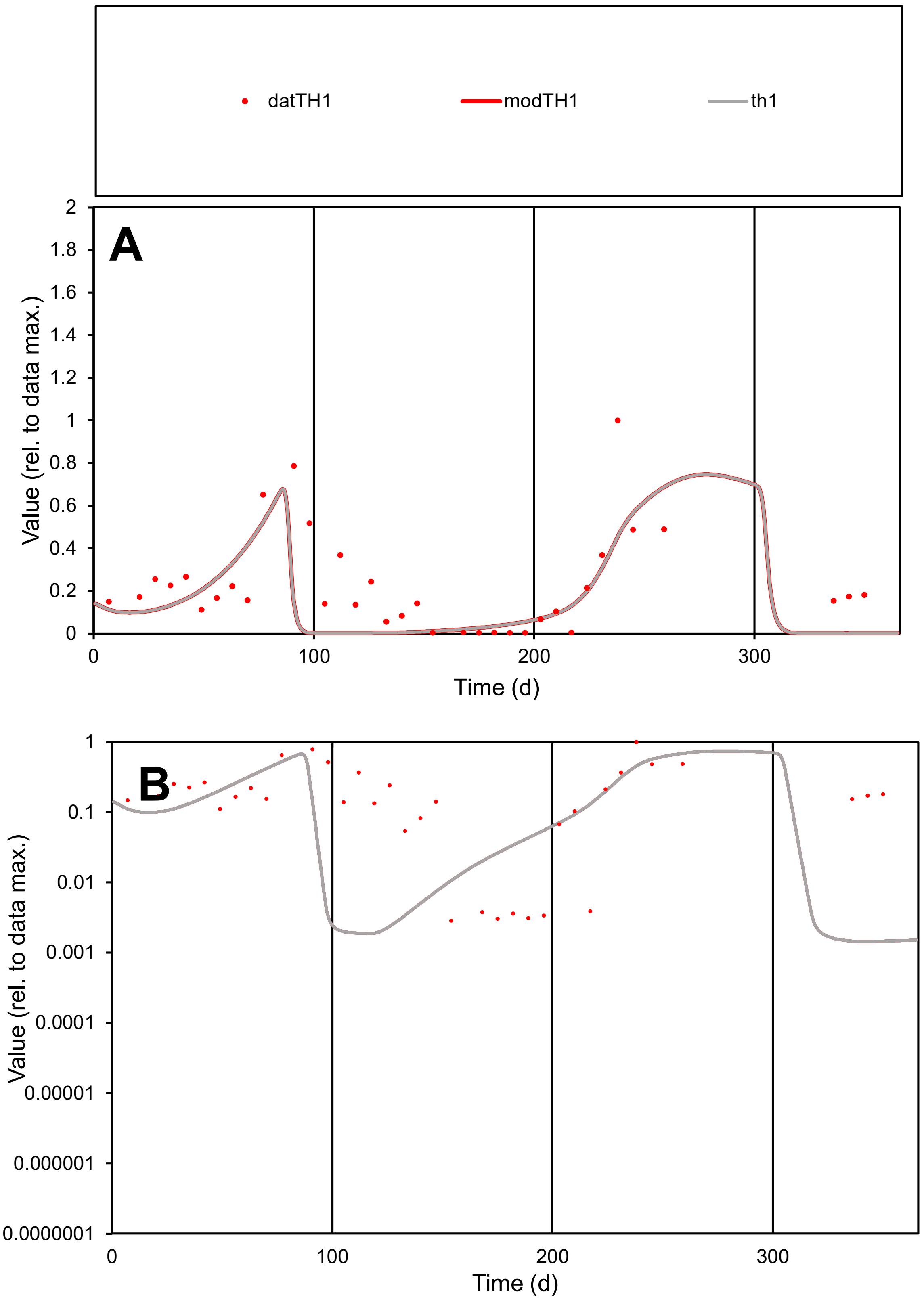


**C1**

Variables

TH1 Provide jtd or Dst

CalibOnly?

2 1 to reproduce of

oNormalize?

2 1=no, 2=yes

ofxn calc*

0.79428

ofxn mod

0.8951

sRun

RunA1

zout

1

Ave

Data

0.0075

Model

0.0055

data/vMax

0.03403

ofxng

311.734

Min d/m

1.97252

ave/min d

77.3733

ave/min m

111.979

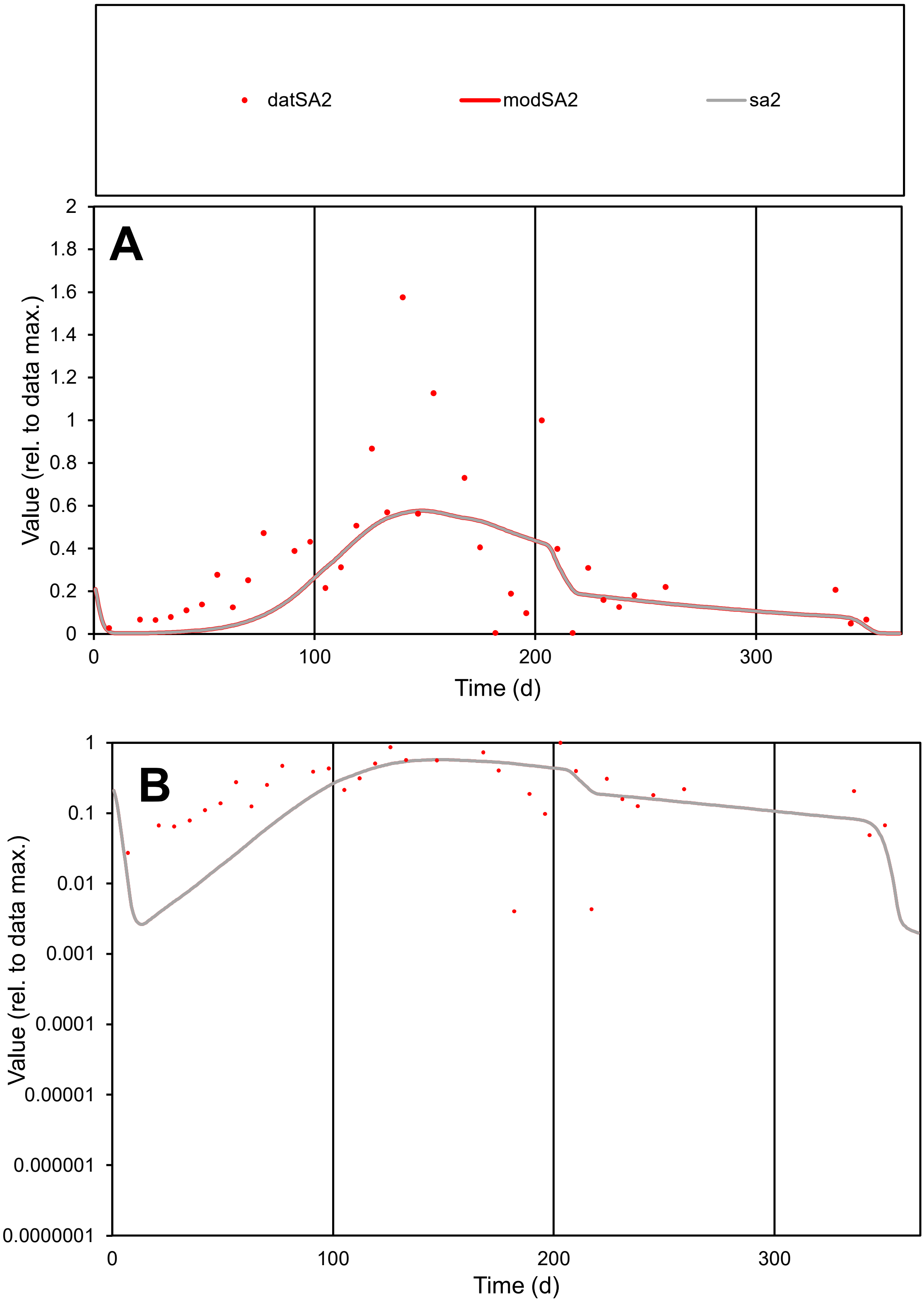


**C1**

Variables

SA2 Provide jtd or Dst

CalibOnly?

2 1 to reproduce of

oNormalize?

2 1=no, 2=yes

ofxn calc*

0.68563

ofxn mod

0.69799

sRun

RunA1

zout

1

Ave

Data

0.00993

Model

0.00841

data/vMax

0.03318

ofxng

311.734

Min d/m

0.97236

ave/min d

74.263

ave/min m

61.1615

331_Calib_S_A

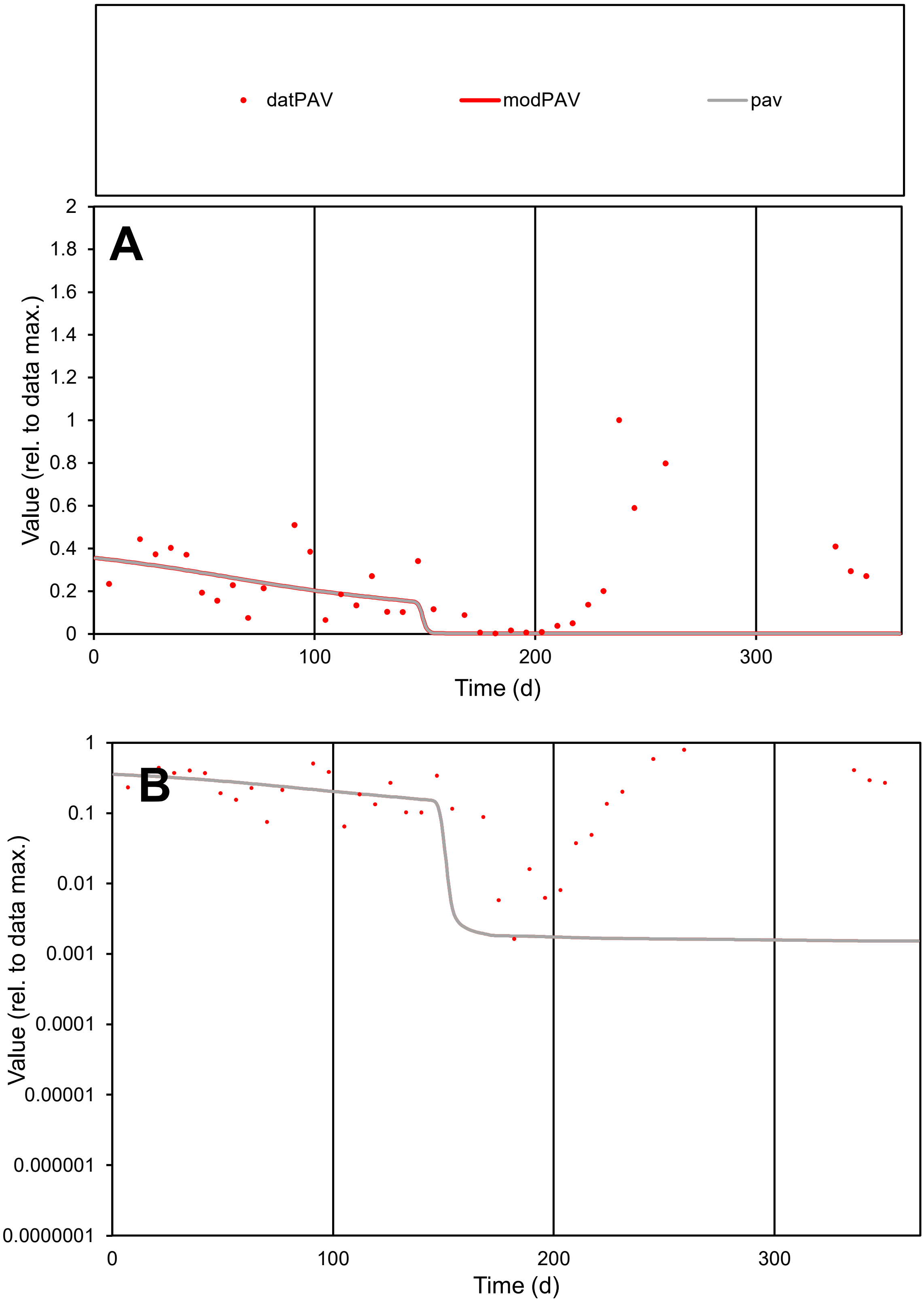


**C1**

Variables
PAV Provide jtd or Dst
CalibOnly?
2 1 to reproduce of
oNormalize?
2 1=no, 2=yes

ofxn calc*
1.14168
ofxn mod
1.14836

sRun
RunA1
zout
1

Ave
Data
0.00708
Model
0.00381

data/vMax
0.02986

ofxng
311.734

Min d/m
1.07273

ave/min d
144.537
ave/min m
83.4051

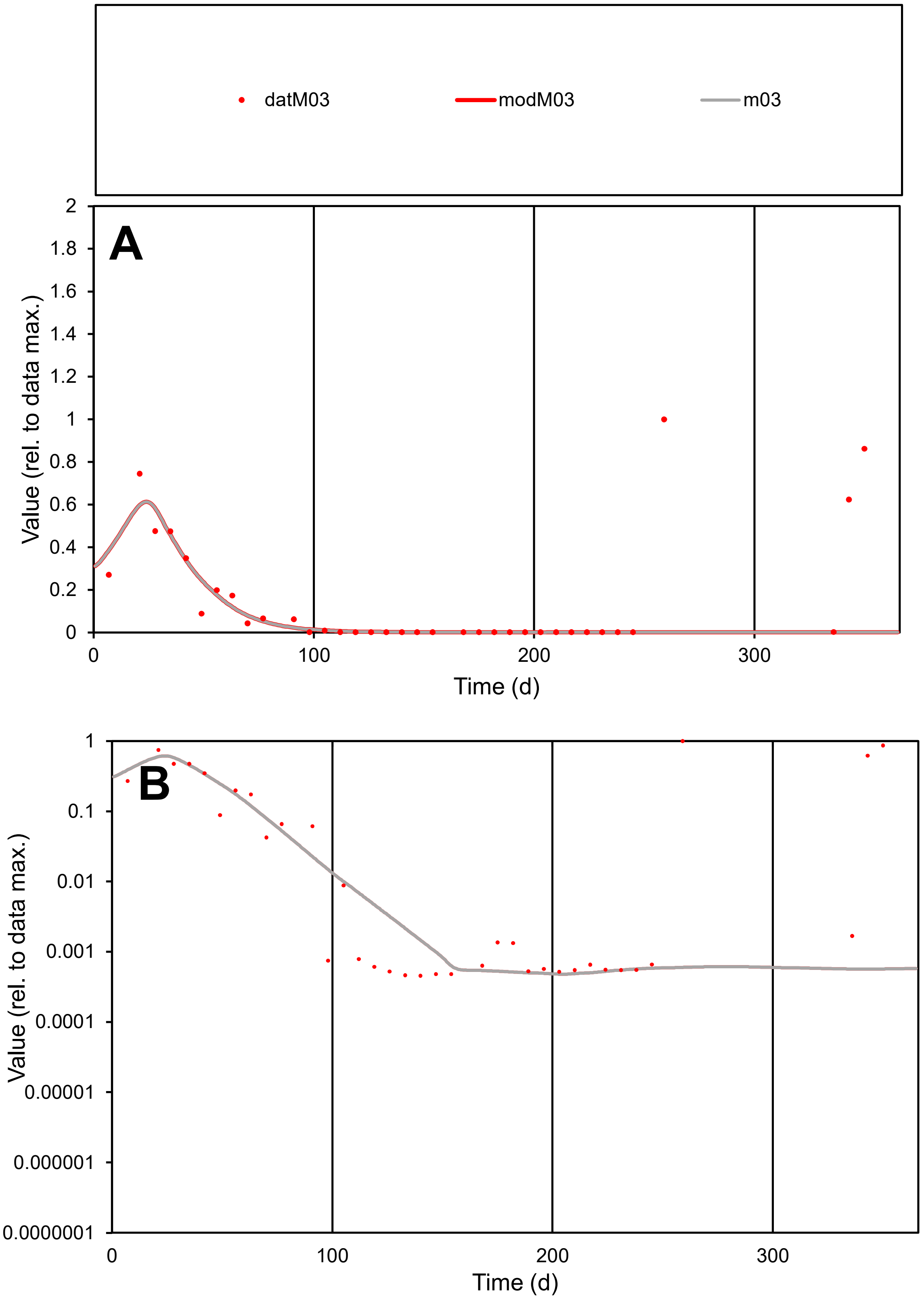


**C1**

Variables
M03 Provide jtd or Dst
CalibOnly?
2 1 to reproduce of
oNormalize?
2 1=no, 2=yes

ofxn calc*
1.63362
ofxn mod
1.64201

sRun
RunA1
zout
1

Ave
Data
0.00577
Model
0.00322

data/vMax
0.03809

ofxng
311.734

Min d/m
0.95392

ave/min d
331.519
ave/min m
176.494

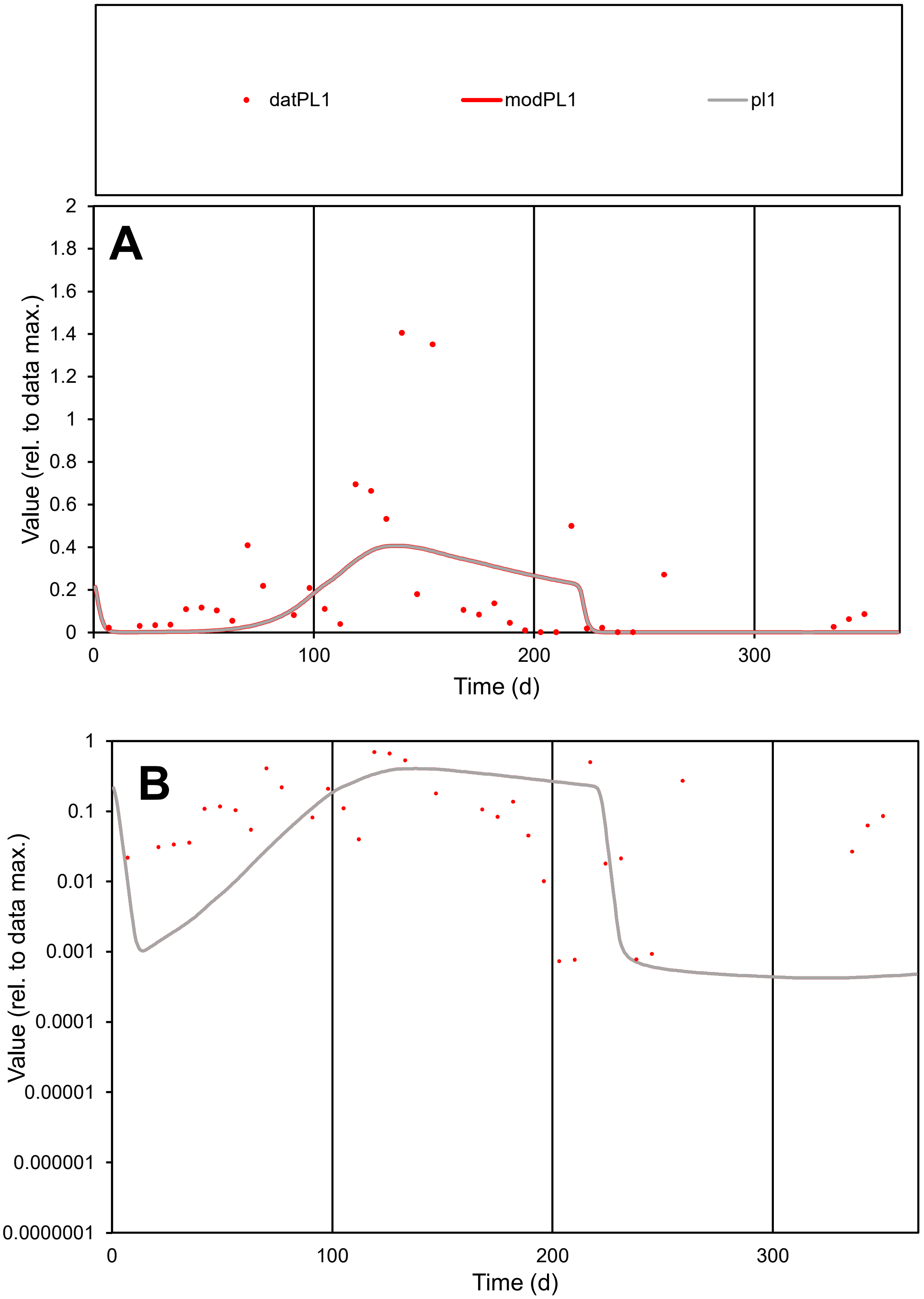


**C1**

Variables
PL1 Provide jtd or Dst
CalibOnly?
2 1 to reproduce of
oNormalize?
2 1=no, 2=yes

ofxn calc*
1.17292
ofxn mod
1.1822

sRun
RunA1
zout
1

Ave
Data
0.00952
Model
0.00826

data/vMax
0.05336

ofxng
311.734

Min d/m
1.70335

ave/min d
243.136
ave/min m
359.223

331_Calib_S_A

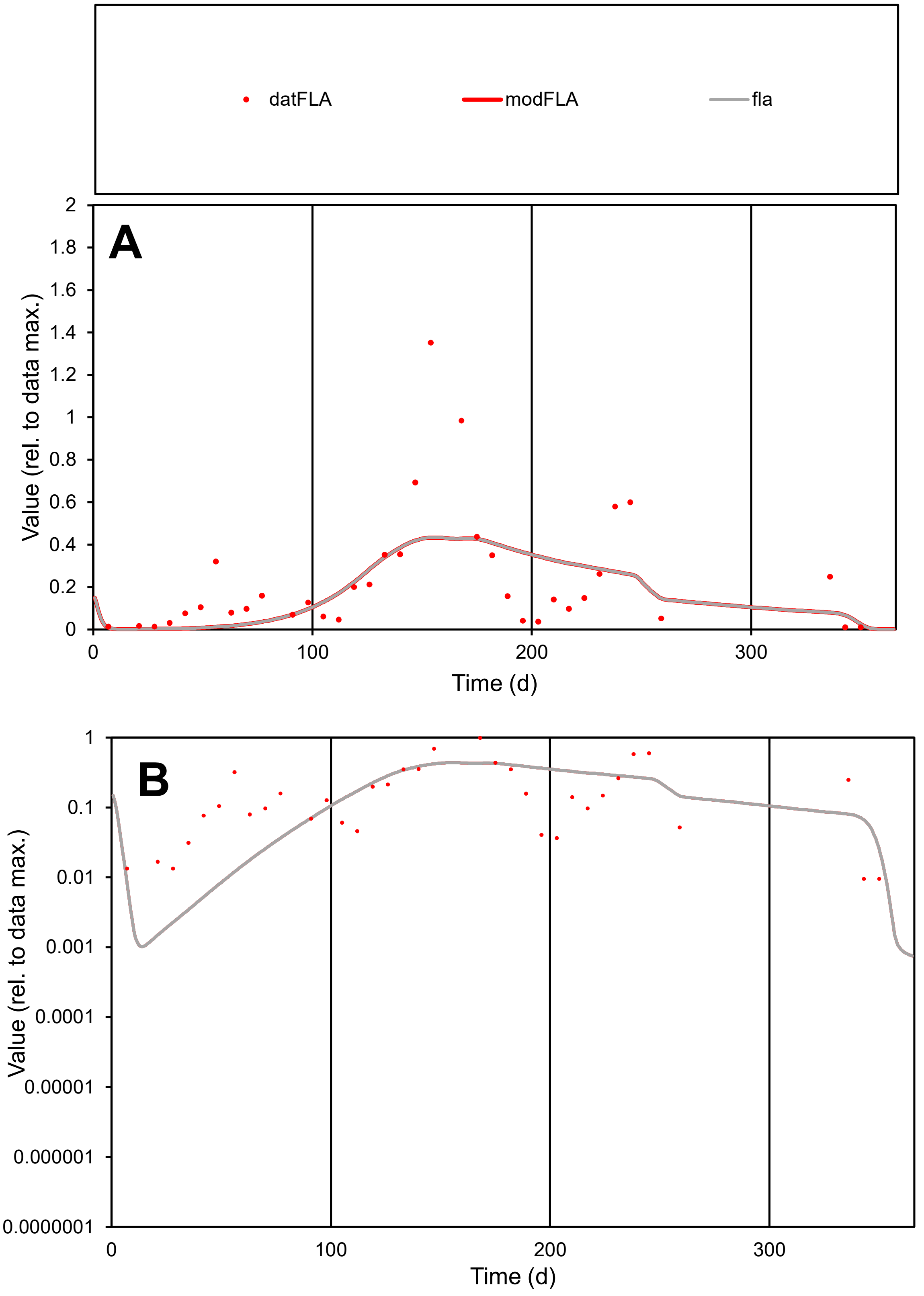


**C1**

Variables
FLA Provide jtd or Dst
CalibOnly?
2 1 to reproduce of
oNormalize?
2 1=no, 2=yes

ofxn calc*
0.94493
ofxn mod
0.9564

sRun
RunA1
zout
1

Ave
Data
0.01061
Model
0.00983

data/vMax
0.04981

ofxng
311.734

Min d/m
5.80118

ave/min d
22.4807
ave/min m
120.766

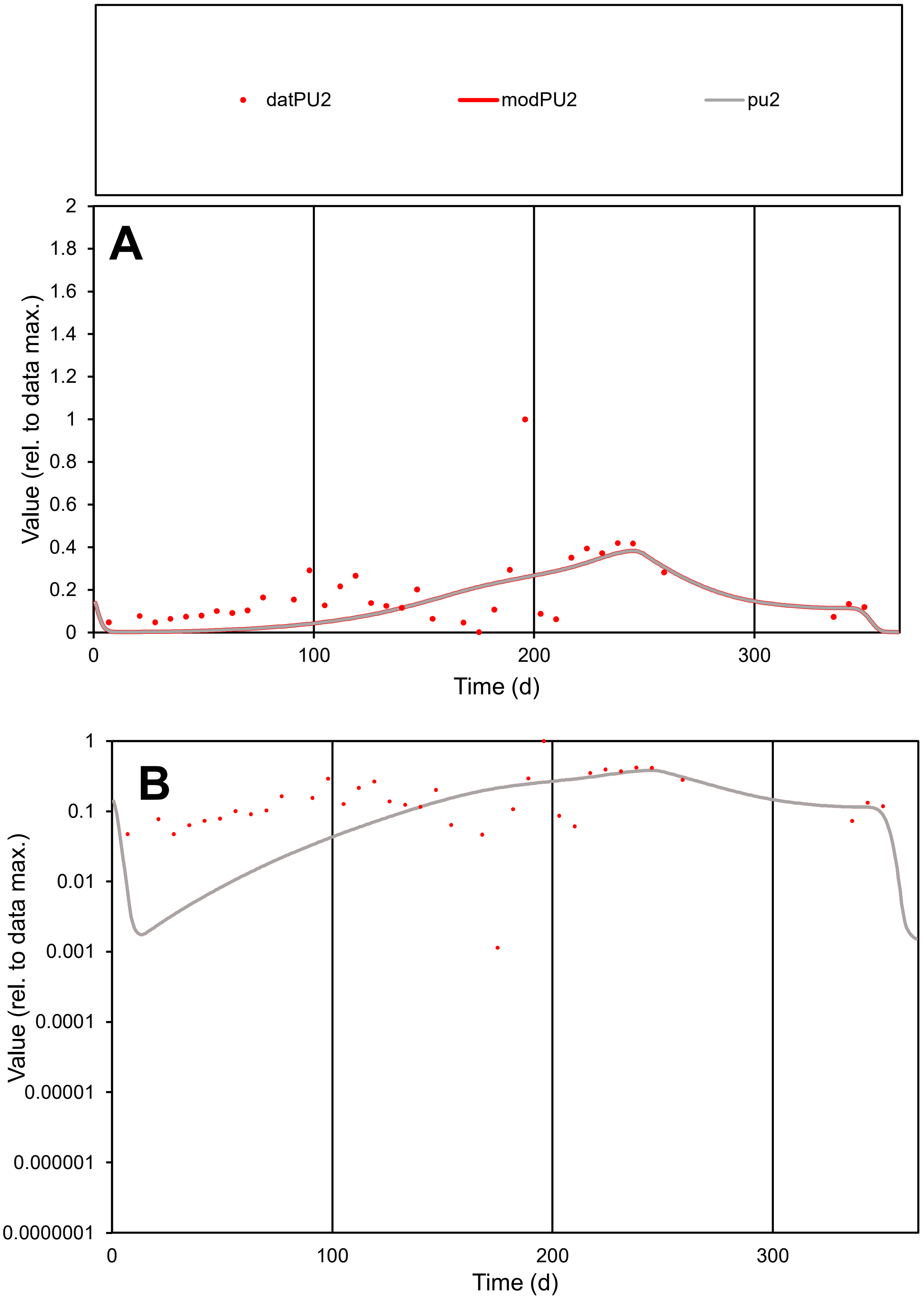


**C1**

Variables
PU2 Provide jtd or Dst
CalibOnly?
2 1 to reproduce of
oNormalize?
2 1=no, 2=yes

ofxn calc*
0.8998
ofxn mod
0.91073

sRun
RunA1
zout
1

Ave
Data
0.00966
Model
0.00753

data/vMax
0.05347

ofxng
311.734

Min d/m
0.43655

ave/min d
157.885
ave/min m
53.7616

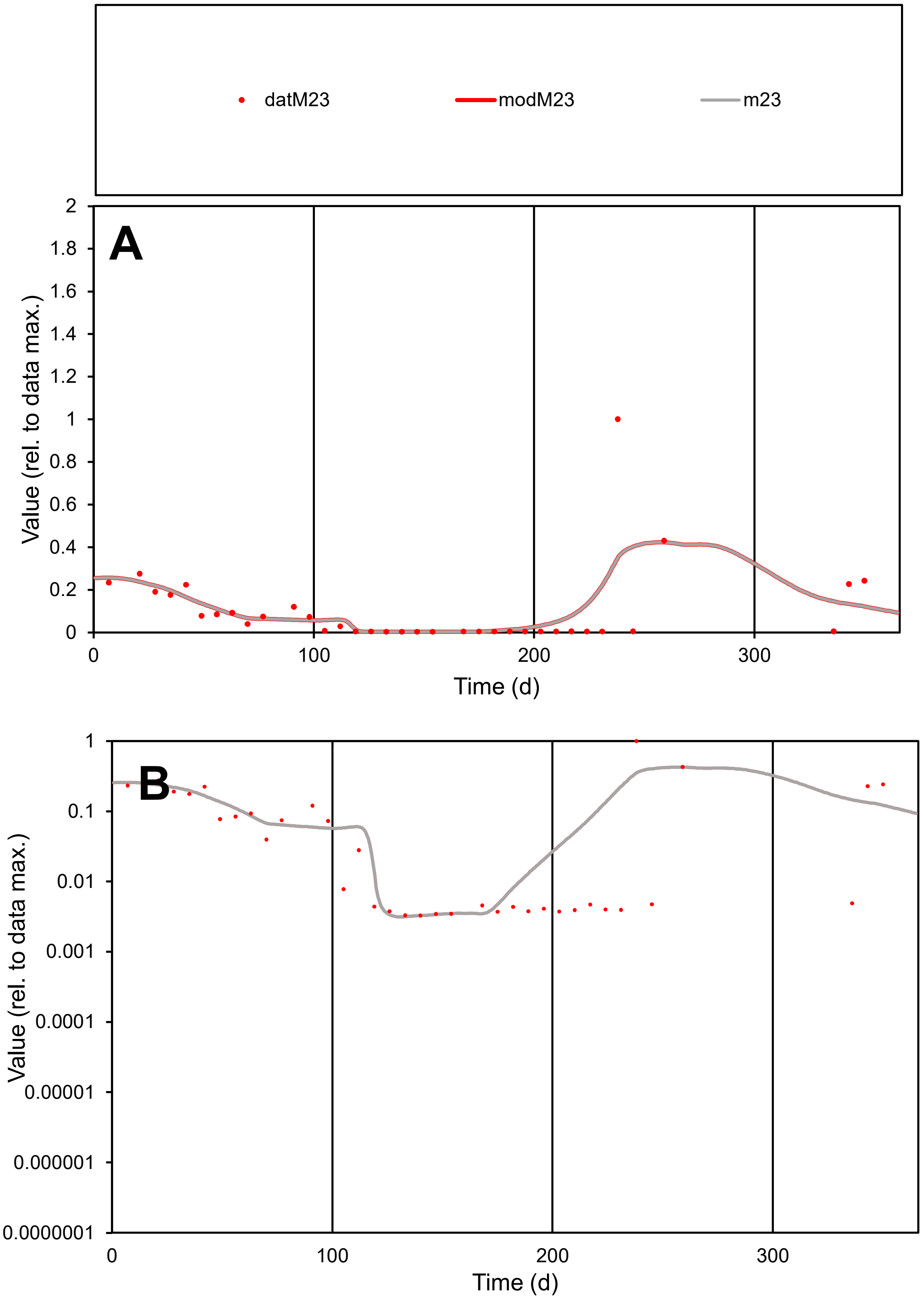


**C1**

Variables
M23 Provide jtd or Dst
CalibOnly?
2 1 to reproduce of
oNormalize?
2 1=no, 2=yes

ofxn calc*
1.34741
ofxn mod
1.37321

sRun
RunA1
zout
1

Ave
Data
0.00331
Model
0.00349

data/vMax
0.03243

ofxng
311.734

Min d/m
1.02899

ave/min d
31.0388
ave/min m
33.7358

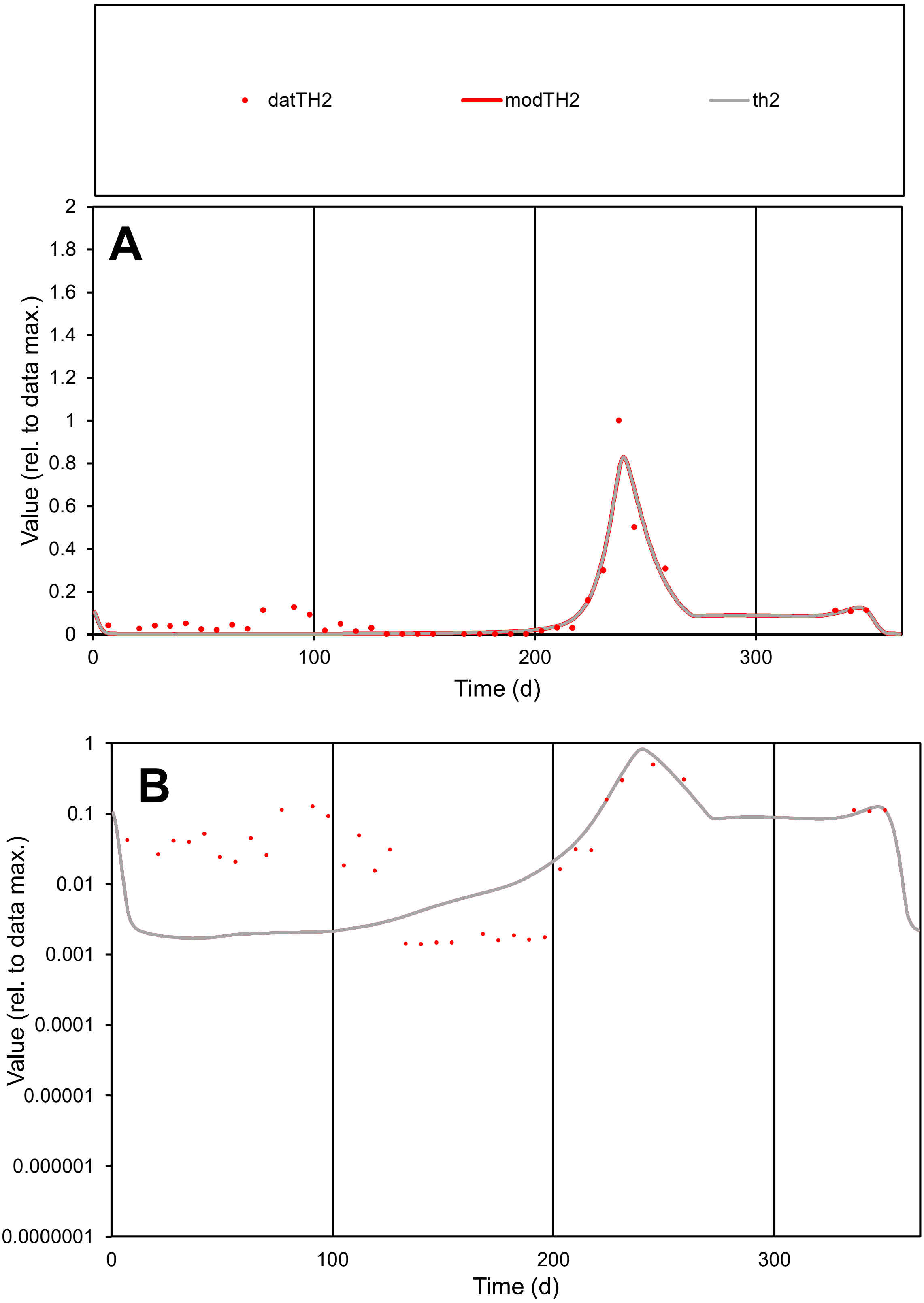


**C1**

Variables
TH2 Provide jtd or Dst
CalibOnly?
2 1 to reproduce of
oNormalize?
2 1=no, 2=yes

ofxn calc*
0.57977
ofxn mod
0.63295

sRun
RunA1
zout
1

Ave
Data
0.00709
Model
0.00577

data/vMax
0.07386

ofxng
311.734

Min d/m
0.82715

ave/min d
67.6315
ave/min m
45.5324

331_Calib_S_A

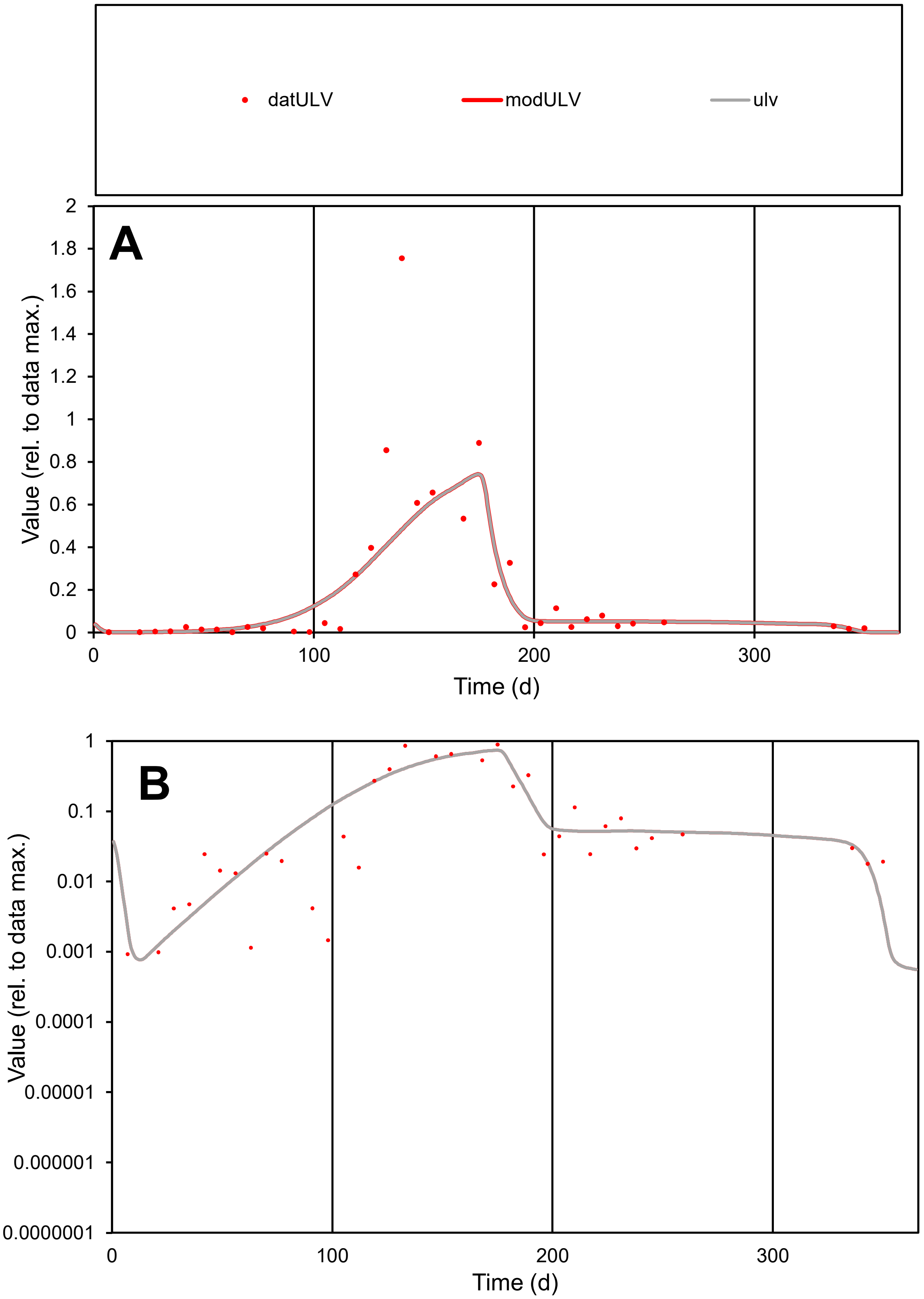


**C1**

Variables
ULV Provide jtd or Dst
CalibOnly?
2 1 to reproduce of
oNormalize?
2 1=no, 2=yes

ofxn calc*
0.73239
ofxn mod
0.7548

sRun
RunA1
zout
1

Ave
Data
0.00308
Model
0.00313

data/vMax
0.01916

ofxng
311.734

Min d/m
0.73915

ave/min d
173.653
ave/min m
130.762

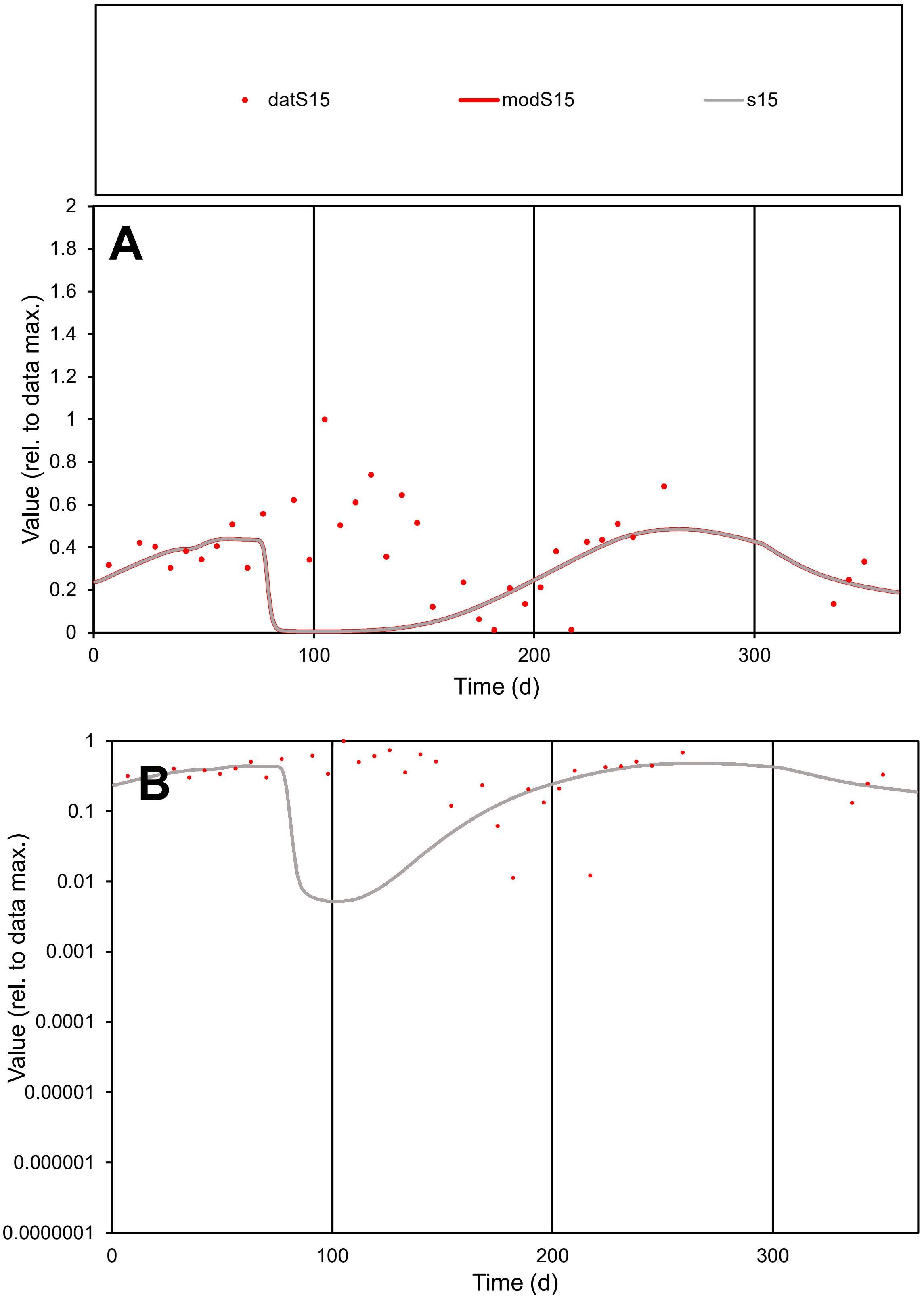


C1

Variables
S15 Provide jtd or Dst
CalibOnly?
2 1 to reproduce of
oNormalize?
2 1=no, 2=yes

ofxn calc*
0.78907
ofxn mod
0.79768

sRun
RunA1
zout
1

Ave
Data
0.00528
Model
0.00347

data/vMax
0.01453

ofxng
311.734

Min d/m
2.16861

ave/min d
32.2859
ave/min m
46.0329

331_Calib_S_A

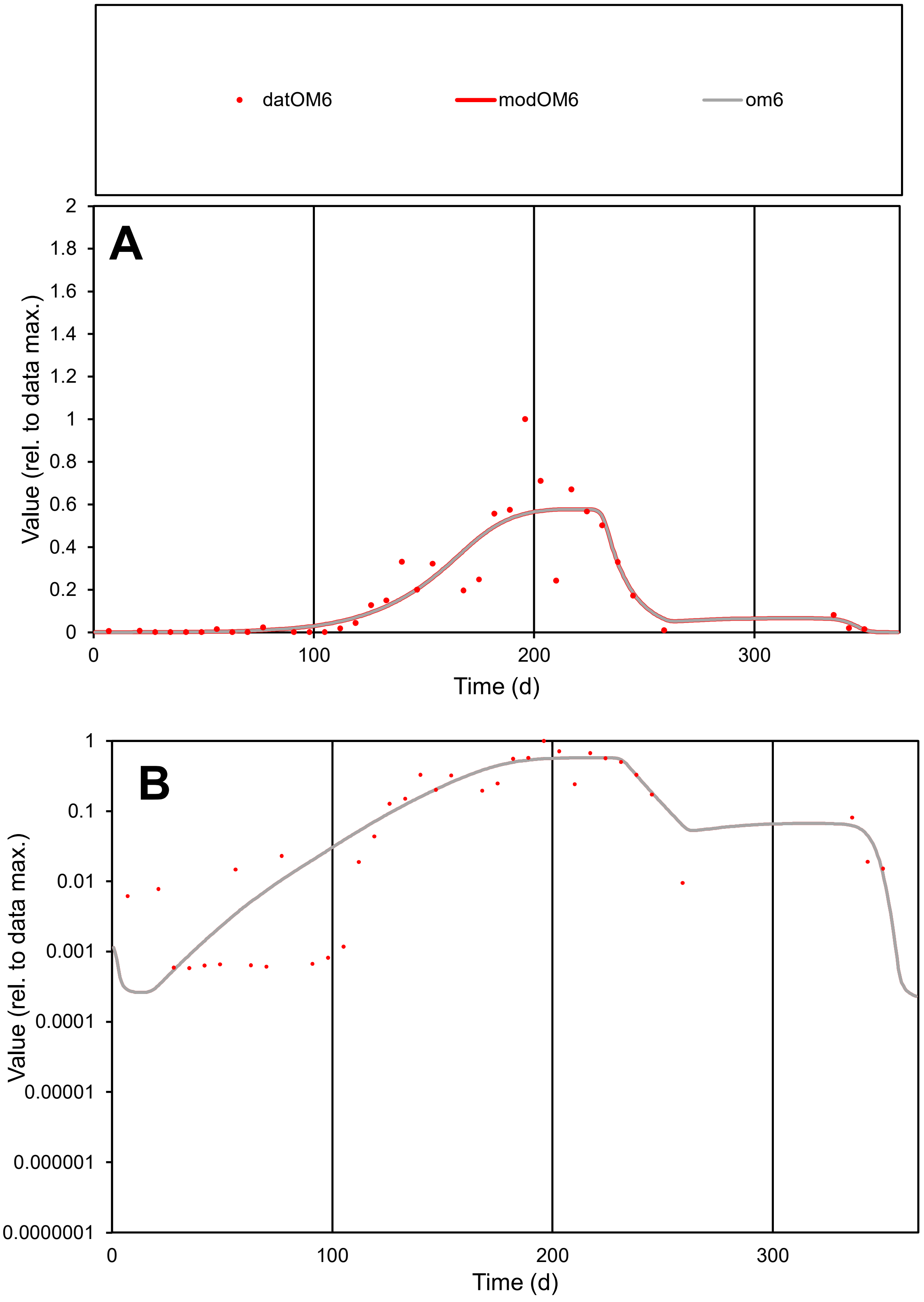


**C1**

Variables
OM6 Provide jtd or Dst
CalibOnly?
2 1 to reproduce of
oNormalize?
2 1=no, 2=yes

ofxn calc*
0.57935
ofxn mod
0.61125

sRun
RunA1
zout
1

Ave
Data
0.00939
Model
0.00964

data/vMax
0.04982

ofxng
311.734

Min d/m
2.10618

ave/min d
324.361
ave/min m
701.44

331_Calib_S_A

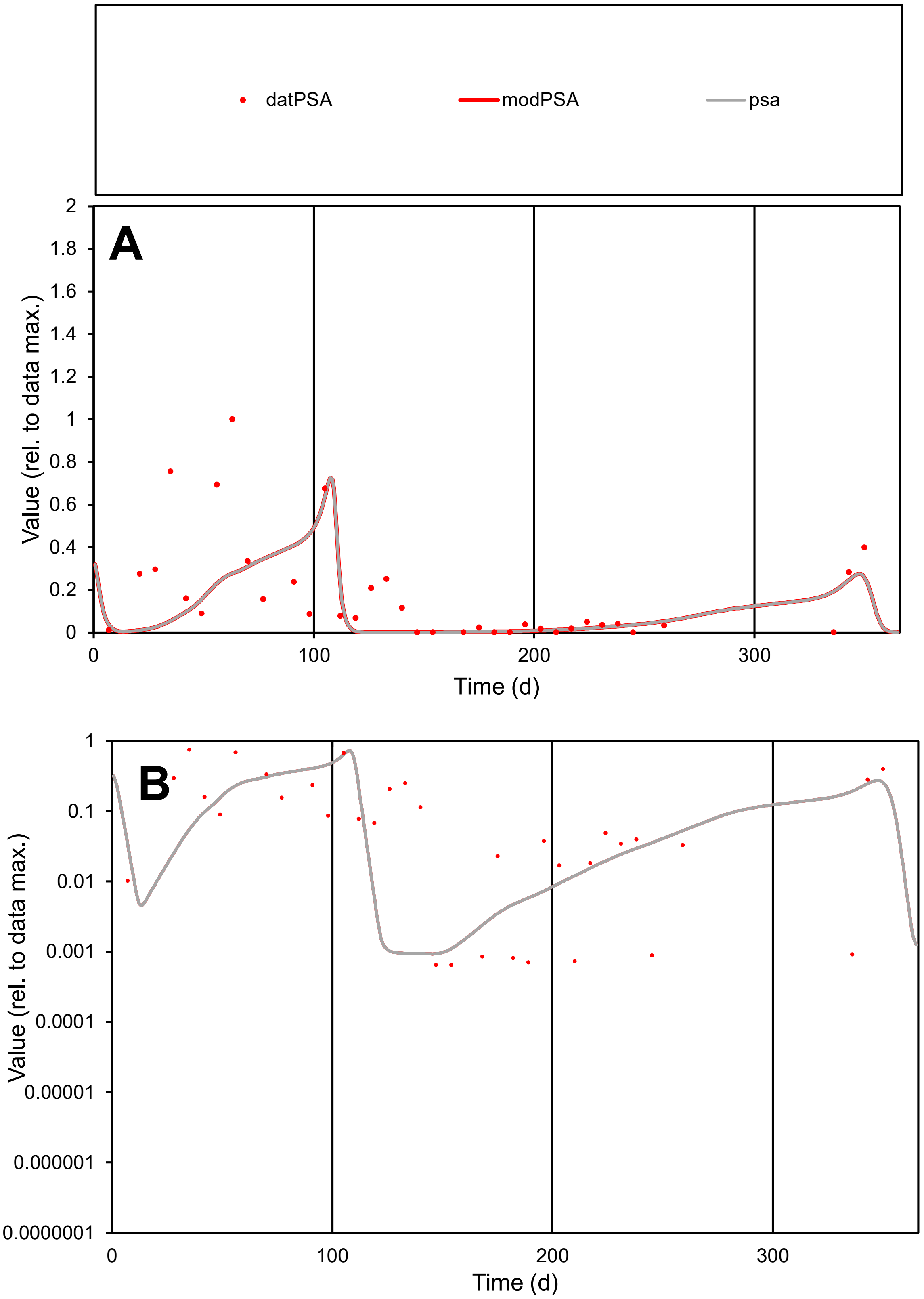


**C1**

Variables
PSA Provide jtd or Dst
CalibOnly?
2 1 to reproduce of
oNormalize?
2 1=no, 2=yes

ofxn calc*
1.24436
ofxn mod
1.25823

sRun
RunA1
zout
1

Ave
Data
0.00247
Model
0.00166

data/vMax
0.01424

ofxng
311.734

Min d/m
0.68484

ave/min d
268.224
ave/min m
123.447

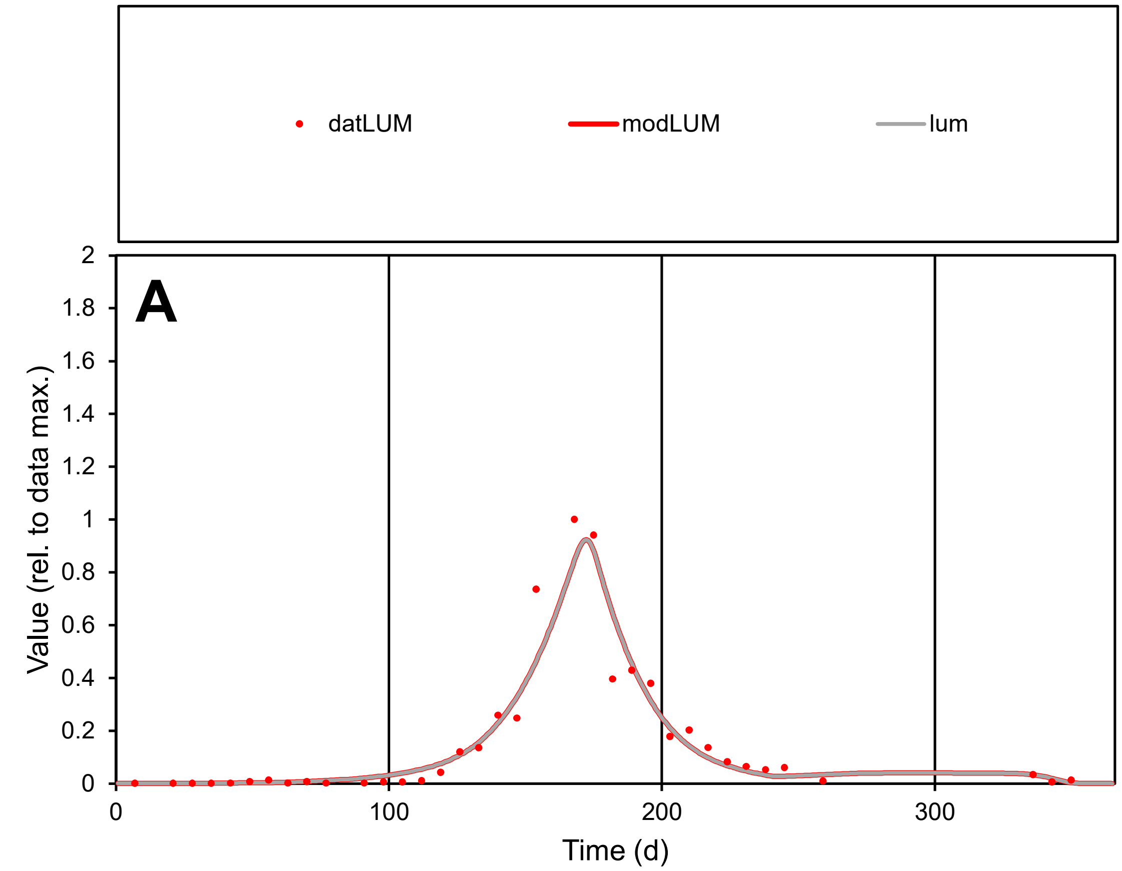


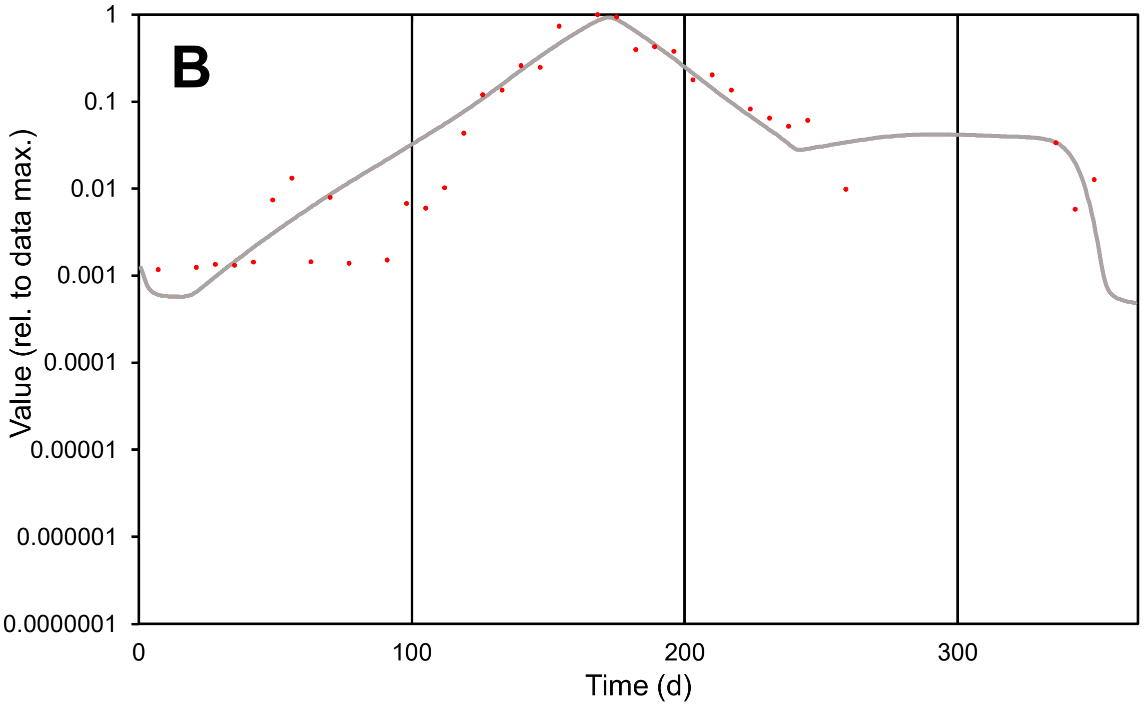


**C1**

Variables
LUM Provide jtd or Dst
CalibOnly?
2 1 to reproduce of
oNormalize?
2 1=no, 2=yes

ofxn calc*
0.41287
ofxn mod
0.45572

sRun
RunA1
zout
1

Ave
Data
0.00561
Model
0.00583

data/vMax
0.03921

ofxng
311.734

Min d/m
1.97258

ave/min d
121.783
ave/min m
249.996

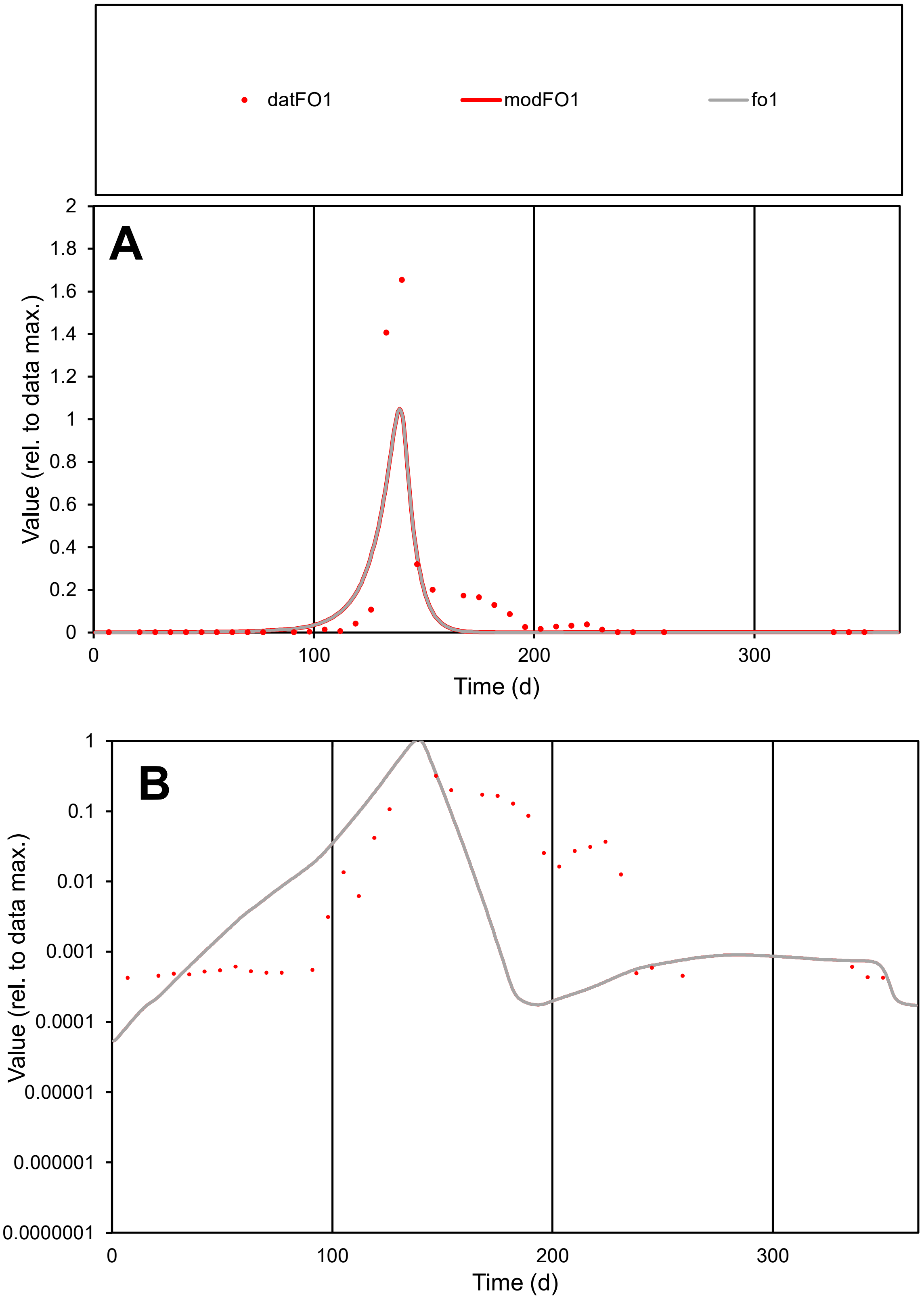


C1

Variables
FO1 Provide jtd or Dst
CalibOnly?
2 1 to reproduce of
oNormalize?
2 1=no, 2=yes

ofxn calc*
0.87486
ofxn mod
0.94627

sRun
RunA1
zout
1

Ave
Data
0.01244
Model
0.01111

data/vMax
0.1408

ofxng
311.734

Min d/m
4.4694

ave/min d
207.651
ave/min m
829.396

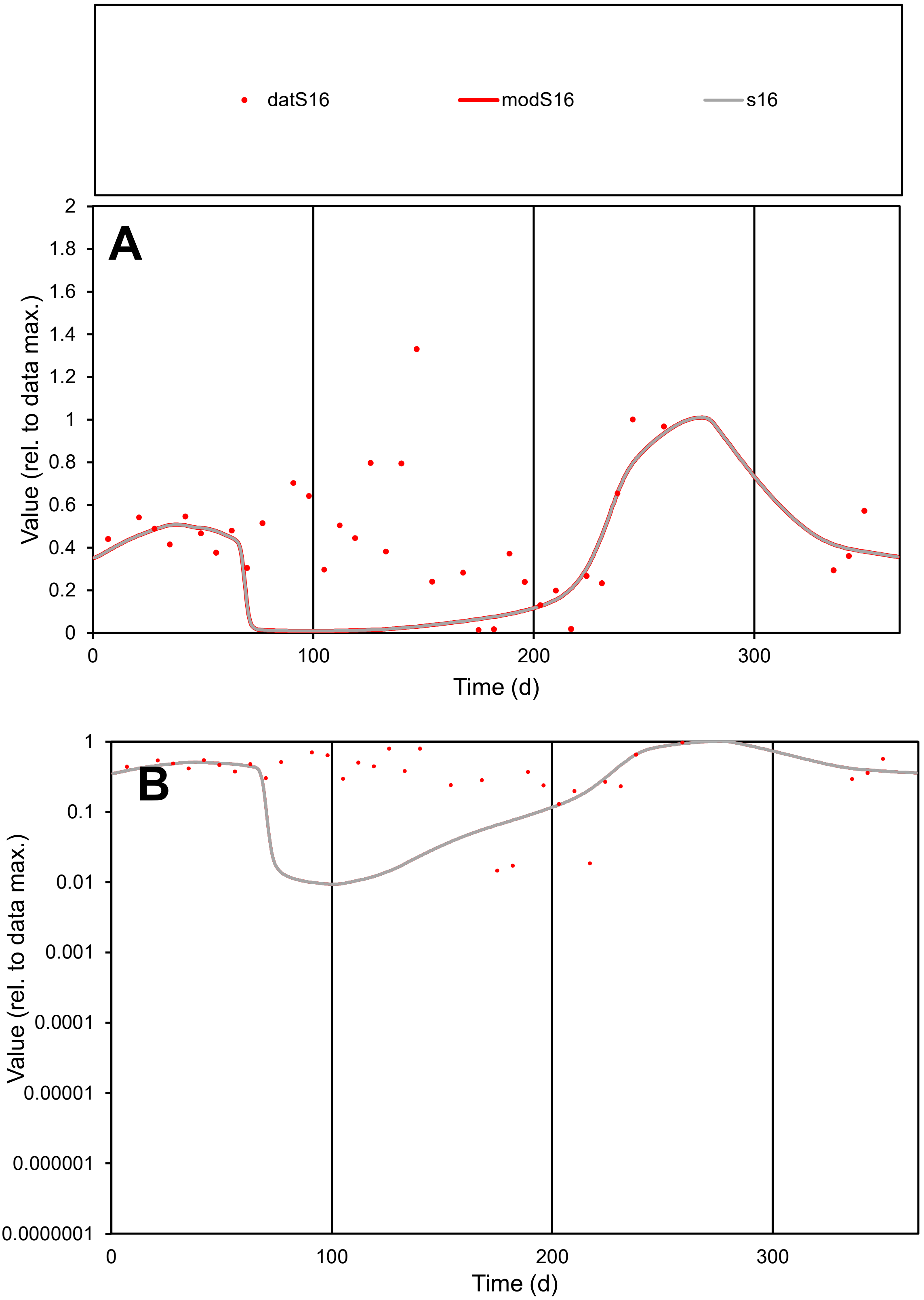


**C1**

Variables
S16 Provide jtd or Dst
CalibOnly?
2 1 to reproduce of
oNormalize?
2 1=no, 2=yes

ofxn calc*
0.73617
ofxn mod
0.74604

sRun
RunA1
zout
1

Ave
Data
0.00401
Model
0.0025

data/vMax
0.00949

ofxng
311.734

Min d/m
1.55374

ave/min d
28.9872
ave/min m
28.035

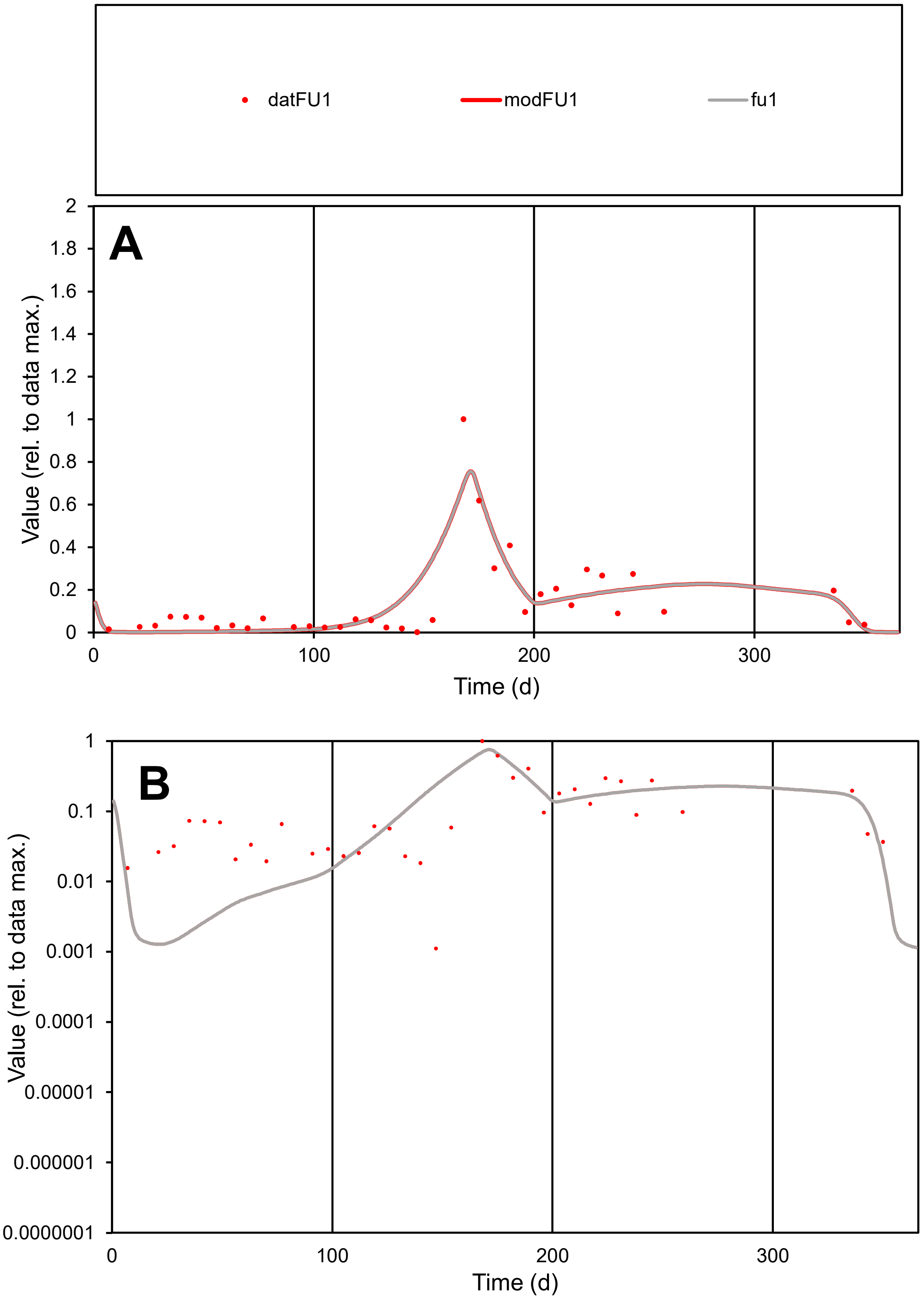


**C1**

Variables
FU1 Provide jtd or Dst
CalibOnly?
2 1 to reproduce of
oNormalize?
2 1=no, 2=yes

ofxn calc*
0.71498
ofxn mod
0.75651

sRun
RunA1
zout
1

Ave
Data
0.00599
Model
0.00604

data/vMax
0.04365

ofxng
311.734

Min d/m
0.86932

ave/min d
123.264
ave/min m
107.997

331_Calib_S_A

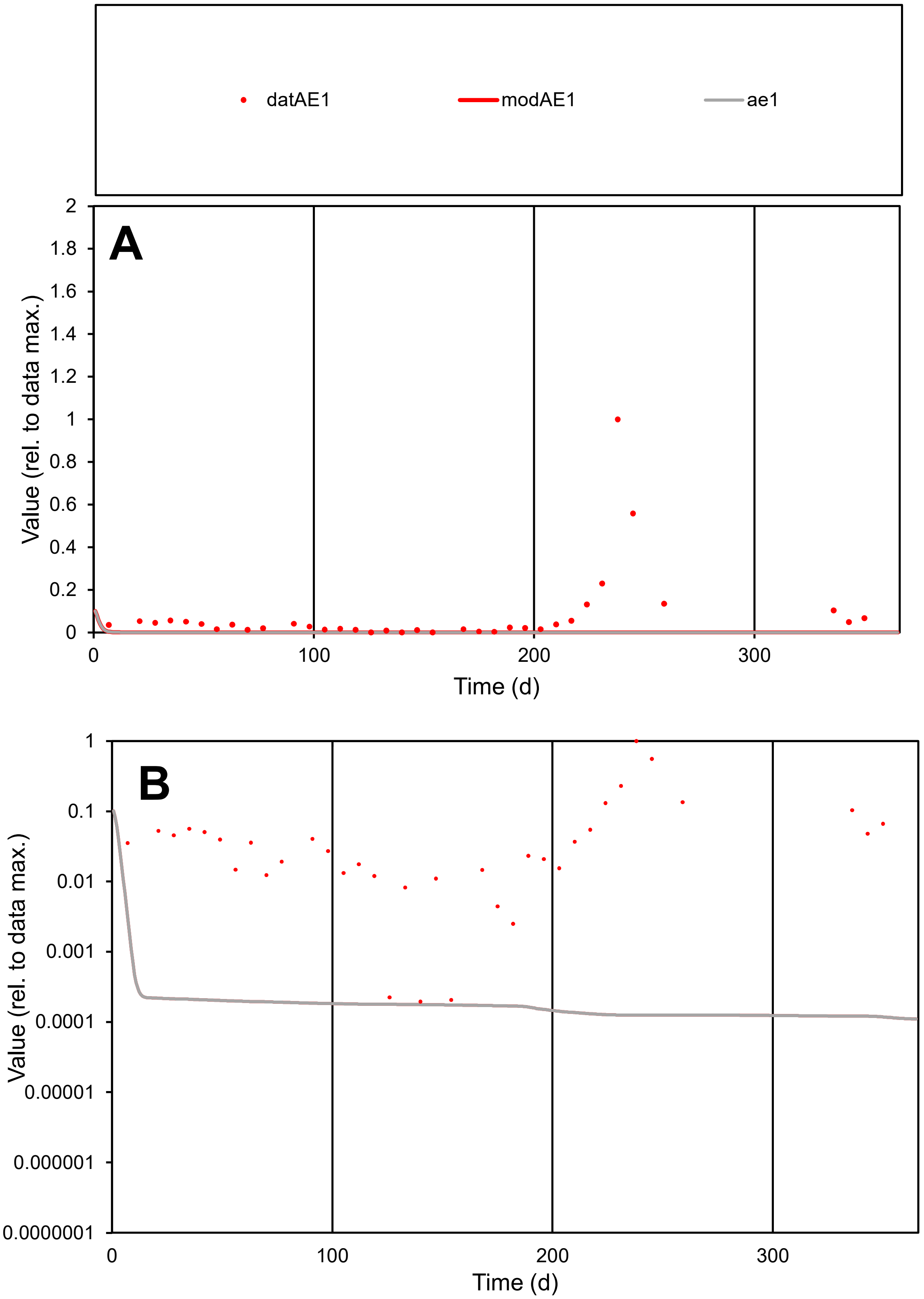


**C1**

Variables

AE1 Provide jtd or Dst

CalibOnly?

2 1 to reproduce of

oNormalize?

2 1=no, 2=yes

ofxn calc*

2.4557

ofxn mod

2.45986

sRun

RunA1

zout

1

Ave

Data

0.00575

Model

6.4E-05

data/vMax

0.07062

ofxng

311.734

Min d/m

1.66446

ave/min d

416.27

ave/min m

7.65969

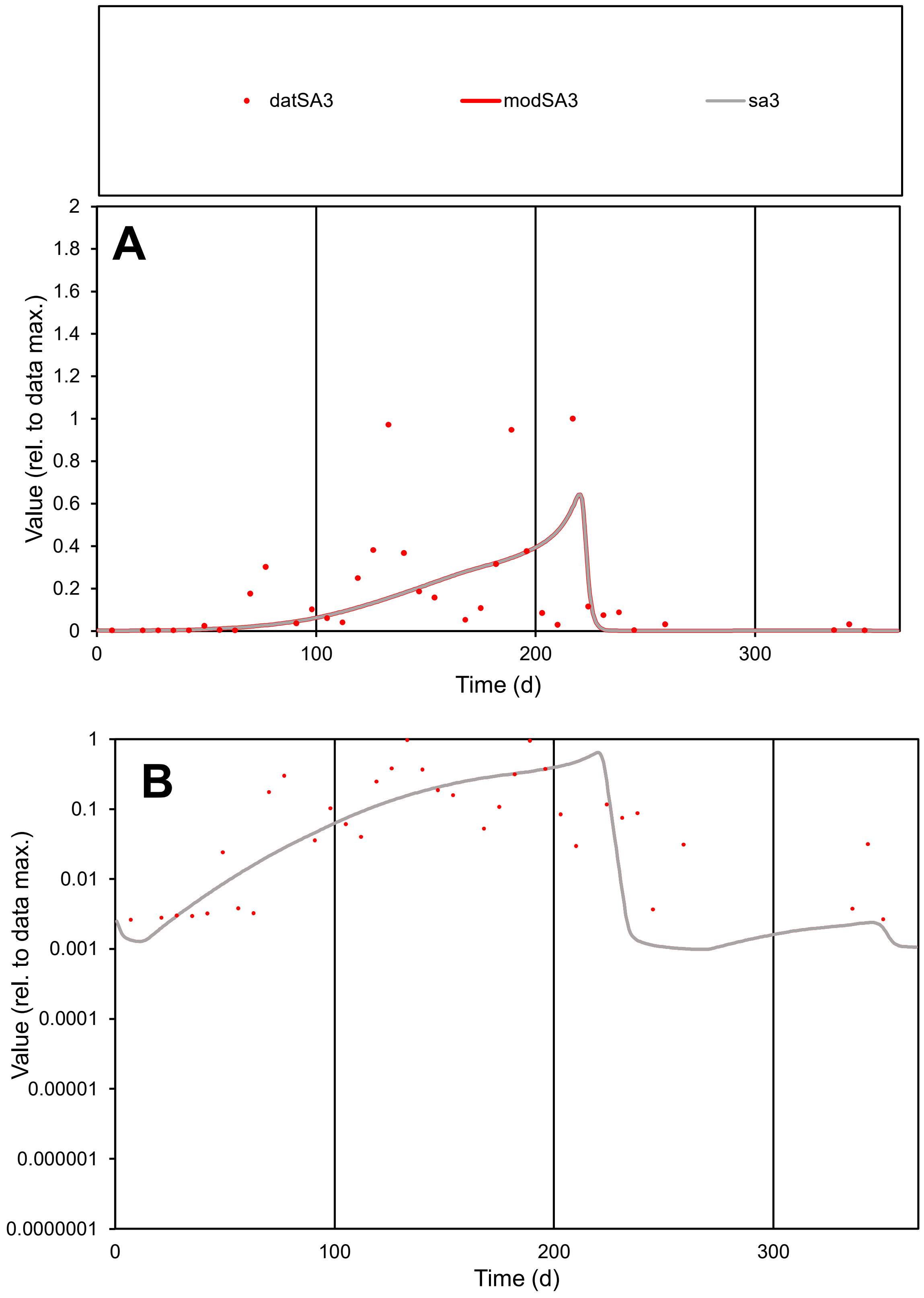


**C1**

Variables
SA3 Provide jtd or Dst
CalibOnly?
2 1 to reproduce of
oNormalize?
2 1=no, 2=yes

ofxn calc*
1.22371
ofxn mod
1.23663

sRun
RunA1
zout
1

Ave
Data
0.00609
Model
0.00504

data/vMax
0.03844

ofxng
311.734

Min d/m
2.61375

ave/min d
60.234
ave/min m
130.277

331_Calib_S_A

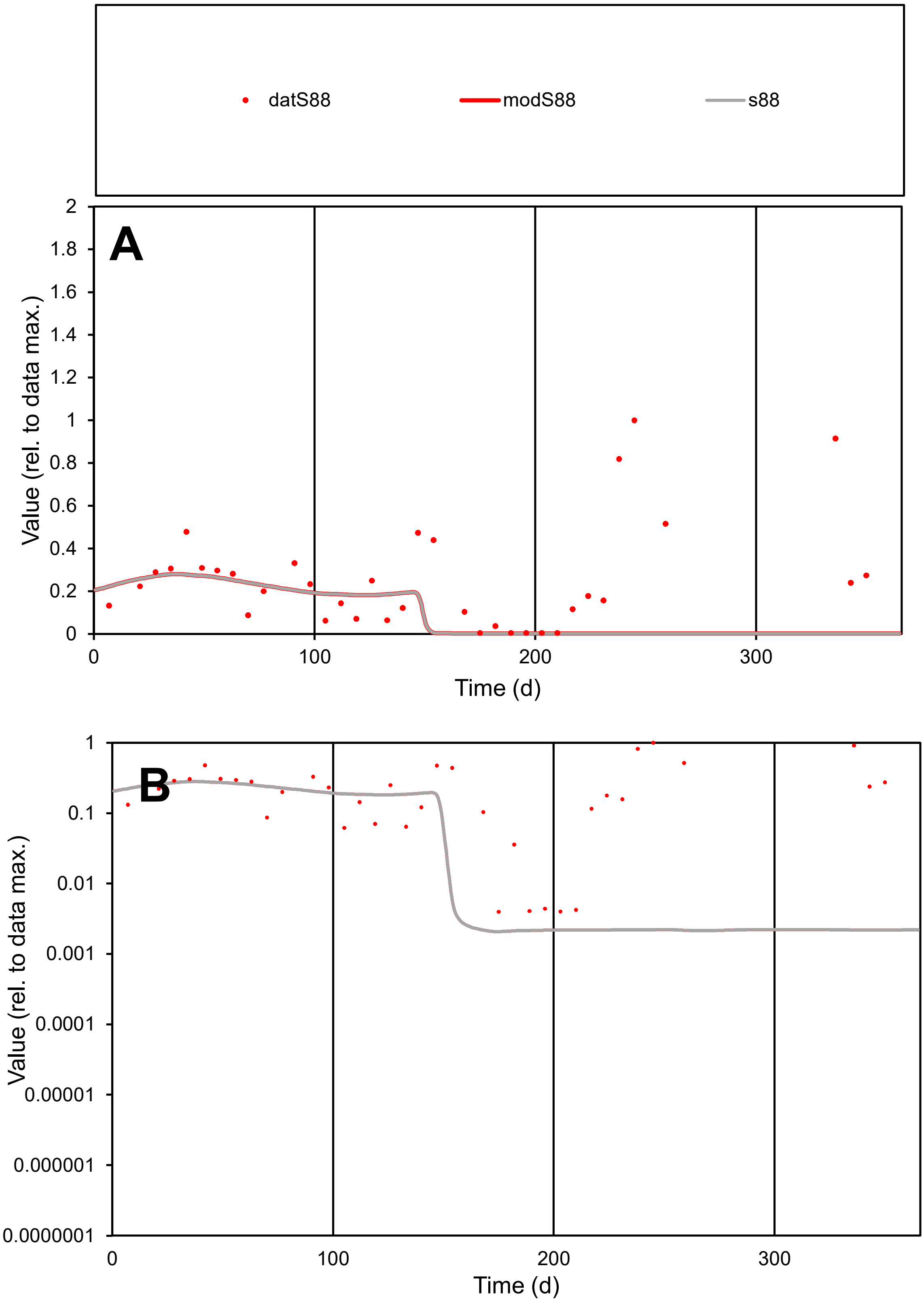


**C1**

Variables
S88 Provide jtd or Dst
CalibOnly?
2 1 to reproduce of
oNormalize?
2 1=no, 2=yes

ofxn calc*
1.23518
ofxn mod
1.23999

sRun
RunA1
zout
1

Ave
Data
0.00396
Model
0.00193

data/vMax
0.01623

ofxng
311.734

Min d/m
1.9095

ave/min d
61.4742
ave/min m
57.3013

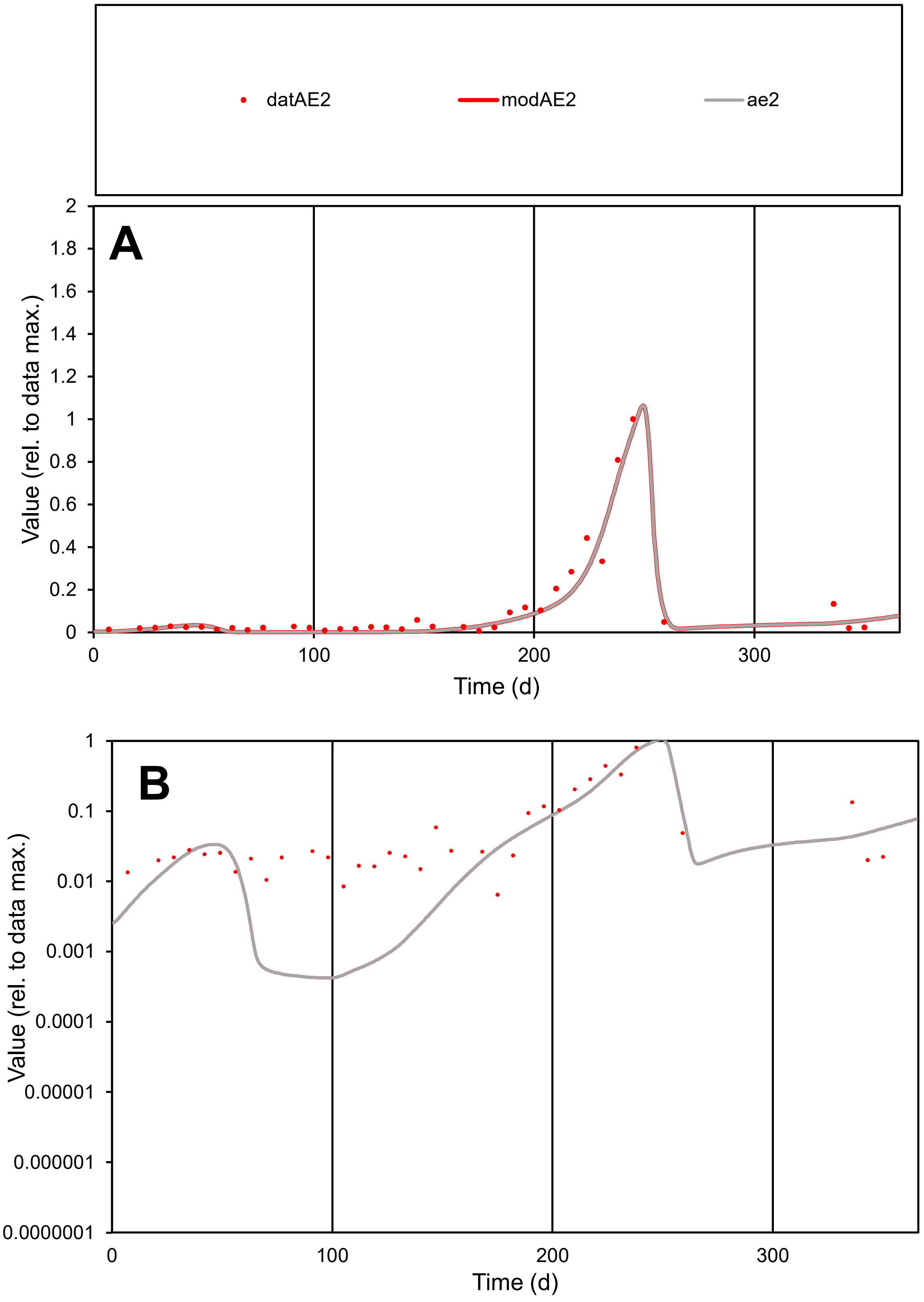


**C1**

Variables
AE2 Provide jtd or Dst
CalibOnly?
2 1 to reproduce of
oNormalize?
2 1=no, 2=yes

ofxn calc*
0.3813
ofxn mod
0.46273

sRun
RunA1
zout
1

Ave
Data
0.00846
Model
0.00757

data/vMax
0.07489

ofxng
311.734

Min d/m
15.3352

ave/min d
17.4387
ave/min m
239.268

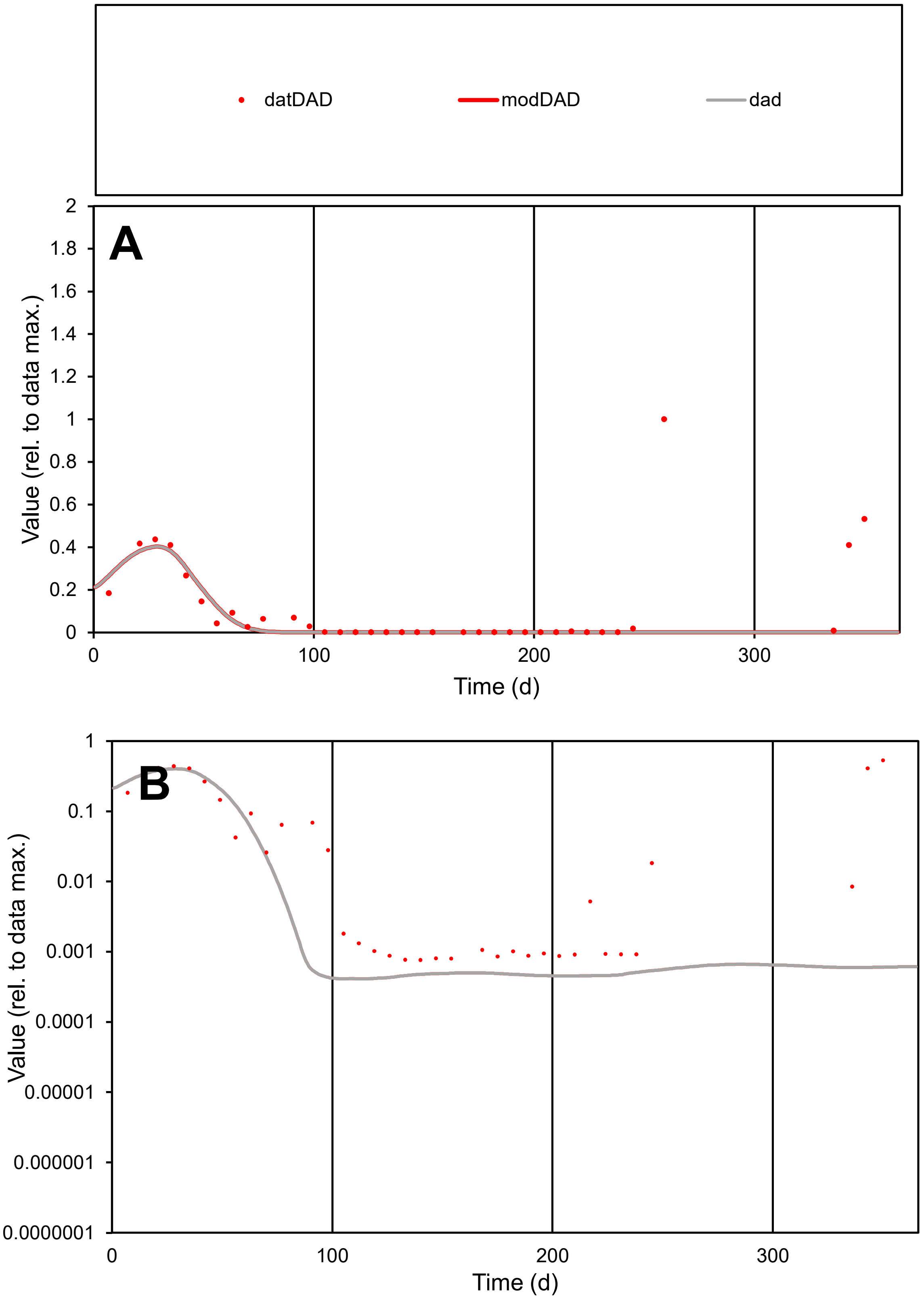


**C1**

Variables
DAD Provide jtd or Dst
CalibOnly?
2 1 to reproduce of
oNormalize?
2 1=no, 2=yes

ofxn calc*
1.74787
ofxn mod
1.75644

sRun
RunA1
zout
1

Ave
Data
0.00233
Model
0.00118

data/vMax
0.02007

ofxng
311.734

Min d/m
1.8559

ave/min d
152.096
ave/min m
142.889

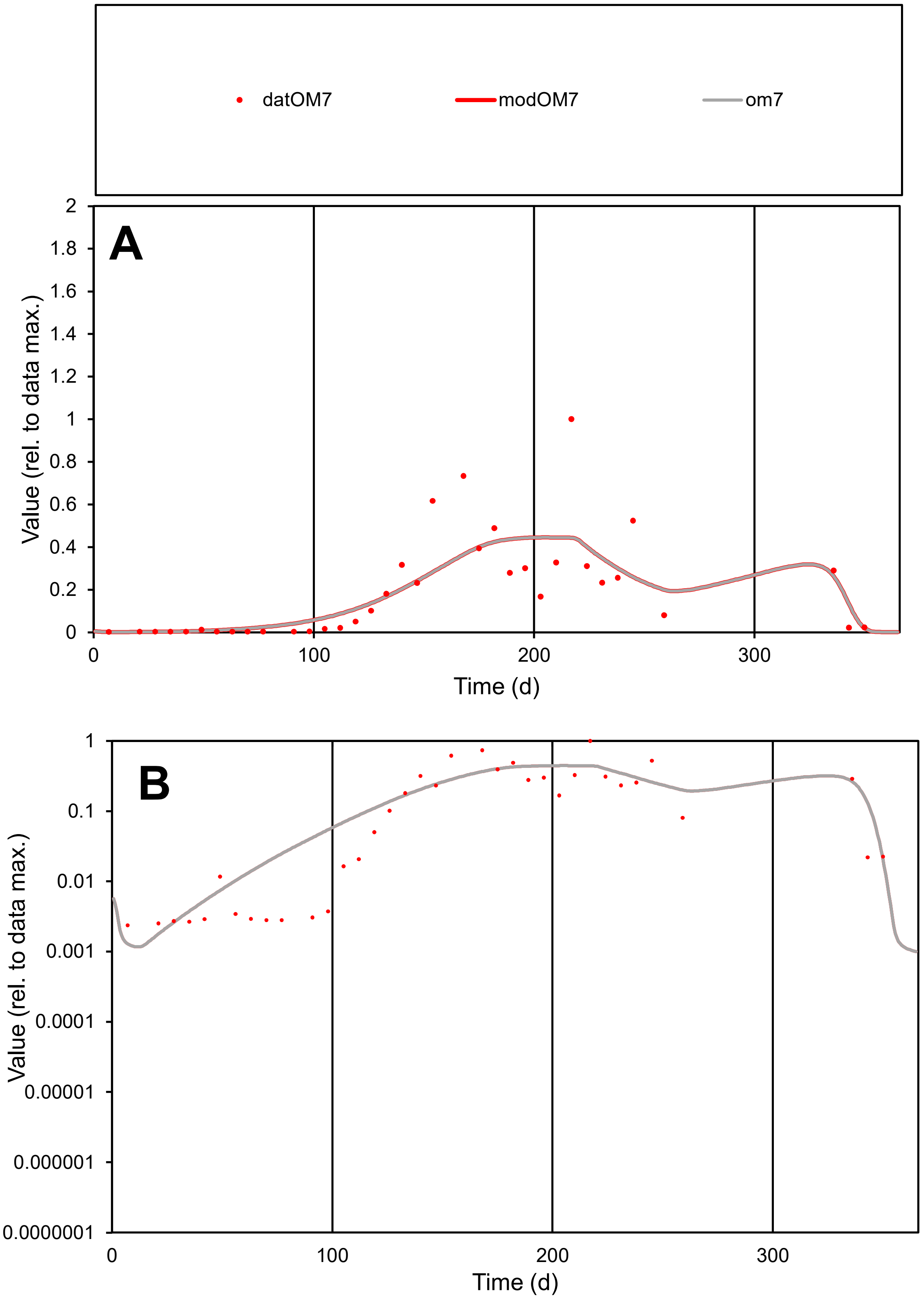


**C1**

Variables

OM7 Provide jtd or Dst

CalibOnly?

2 1 to reproduce of

oNormalize?

2 1=no, 2=yes

ofxn calc*

0.79056

ofxn mod

0.80783

sRun

RunA1

zout

1

Ave

Data

0.00742

Model

0.00762

data/vMax

0.04064

ofxng

311.734

Min d/m

1.89939

ave/min d

77.1919

ave/min m

150.537

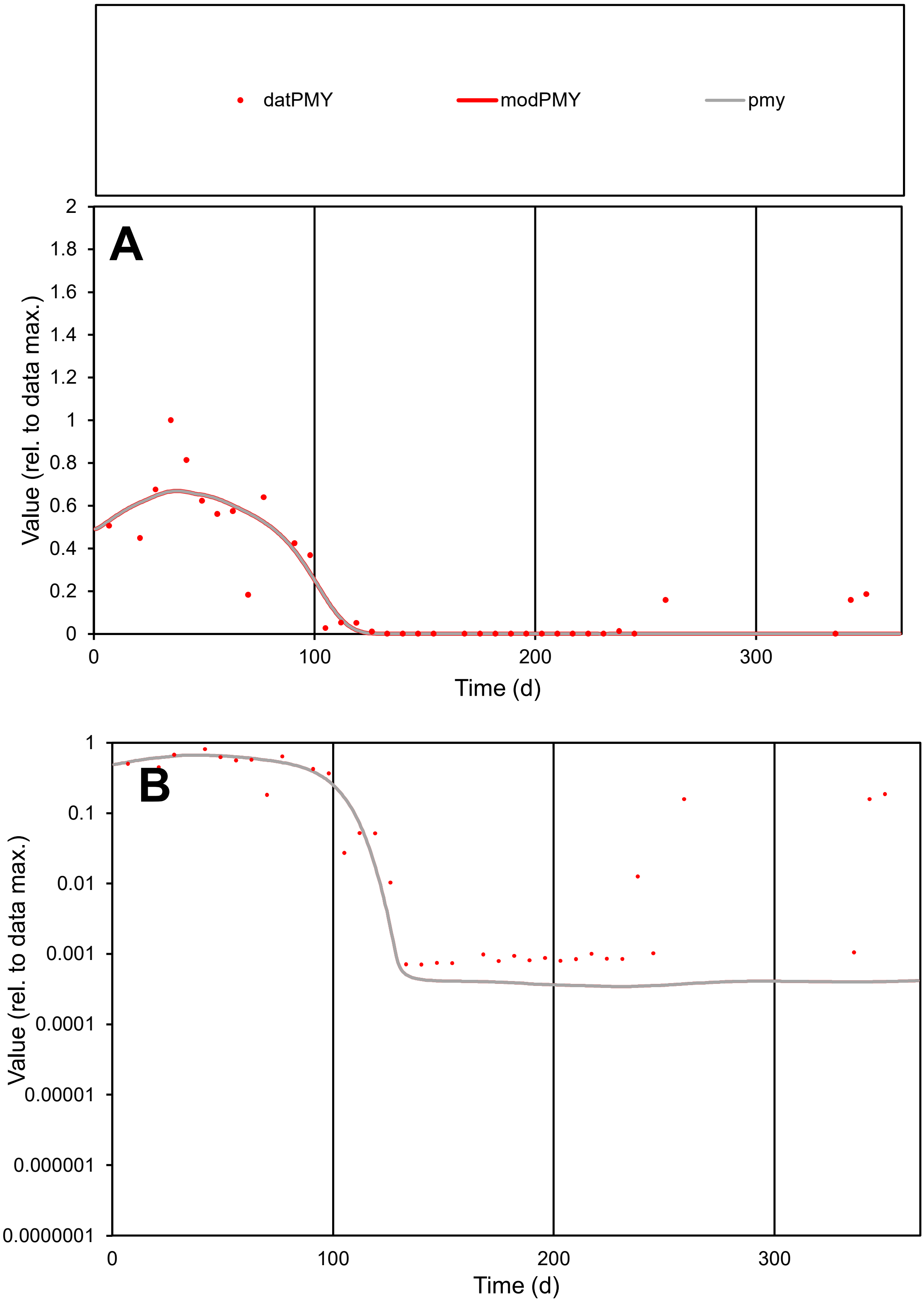


**C1**

Variables
PMY Provide jtd or Dst
CalibOnly?
2 1 to reproduce of
oNormalize?
2 1=no, 2=yes

ofxn calc*
0.52814
ofxn mod
0.56708

sRun
RunA1
zout
1

Ave
Data
0.00281
Model
0.00261

data/vMax
0.01351

ofxng
311.734

Min d/m
2.05279

ave/min d
294.956
ave/min m
562.21

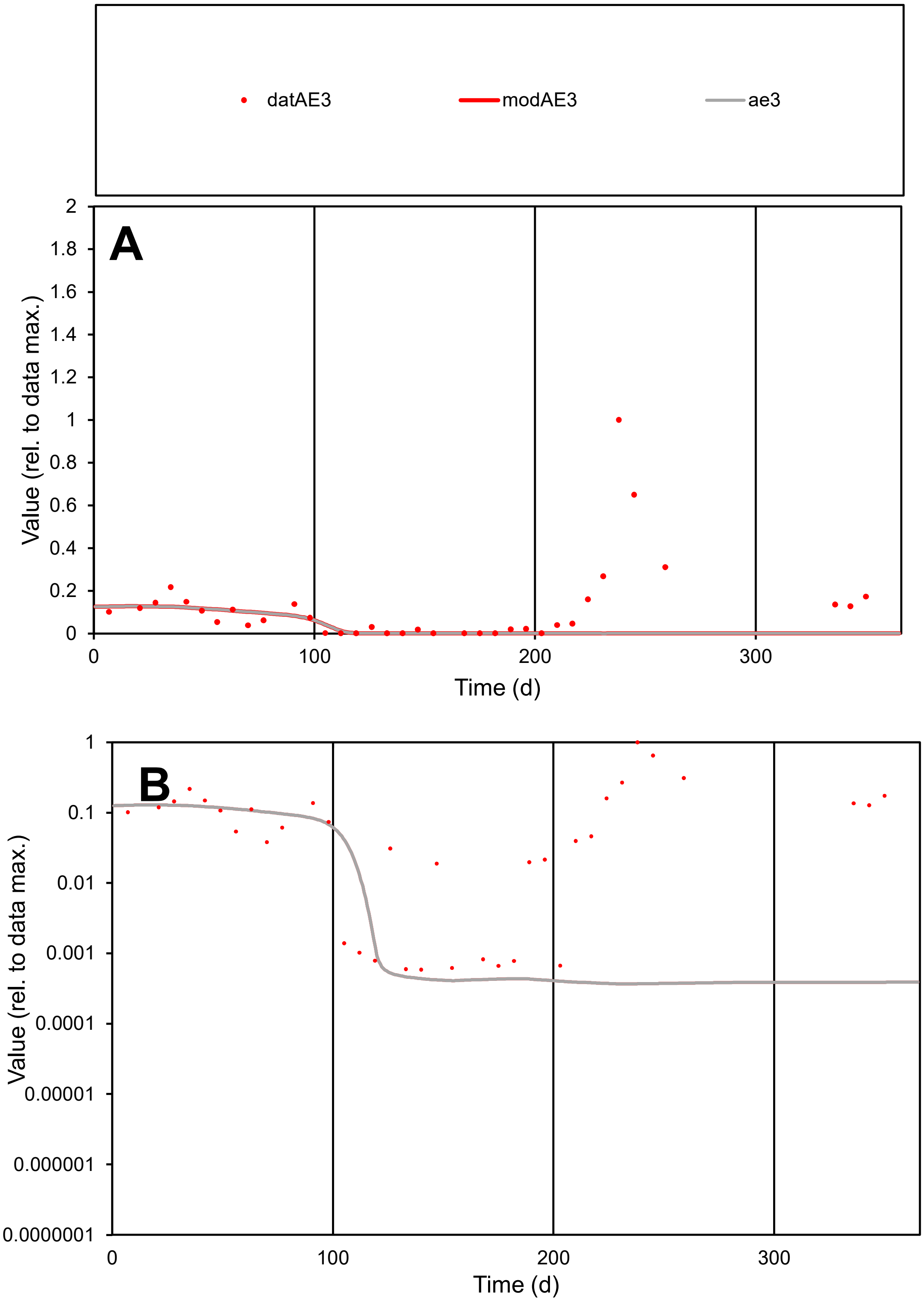


**C1**

Variables
AE3 Provide jtd or Dst
CalibOnly?
2 1 to reproduce of
oNormalize?
2 1=no, 2=yes

ofxn calc*
1.81879
ofxn mod
1.82283

sRun
RunA1
zout
1

Ave
Data
0.00338
Model
0.00106

data/vMax
0.02827

ofxng
311.734

Min d/m
1.60134

ave/min d
203.013
ave/min m
101.447

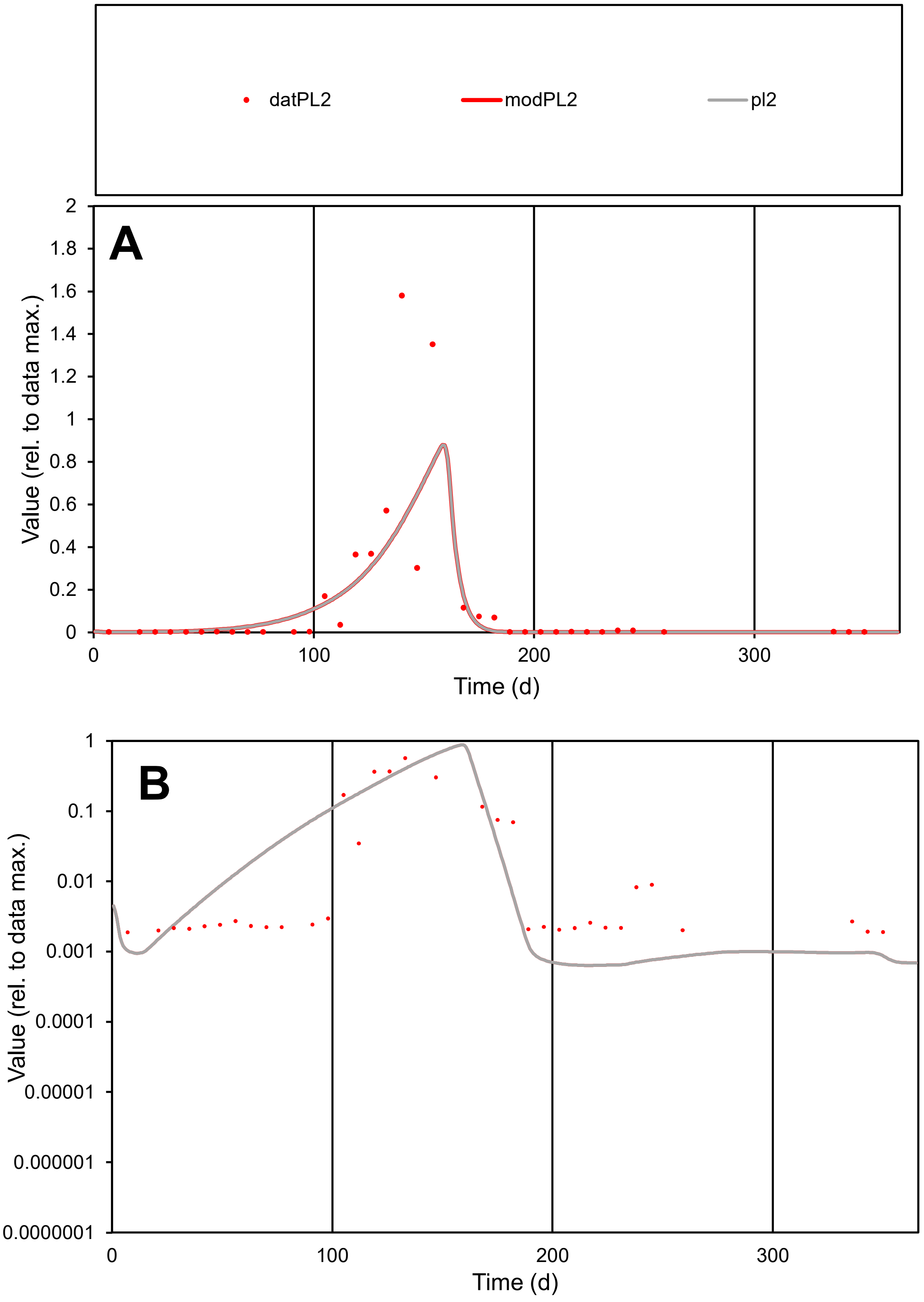


**C1**

Variables
PL2 Provide jtd or Dst
CalibOnly?
2 1 to reproduce of
oNormalize?
2 1=no, 2=yes

ofxn calc*
1.04995
ofxn mod
1.08502

sRun
RunA1
zout
1

Ave
Data
0.00916
Model
0.00944

data/vMax
0.08937

ofxng
311.734

Min d/m
2.95715

ave/min d
54.6326
ave/min m
166.54

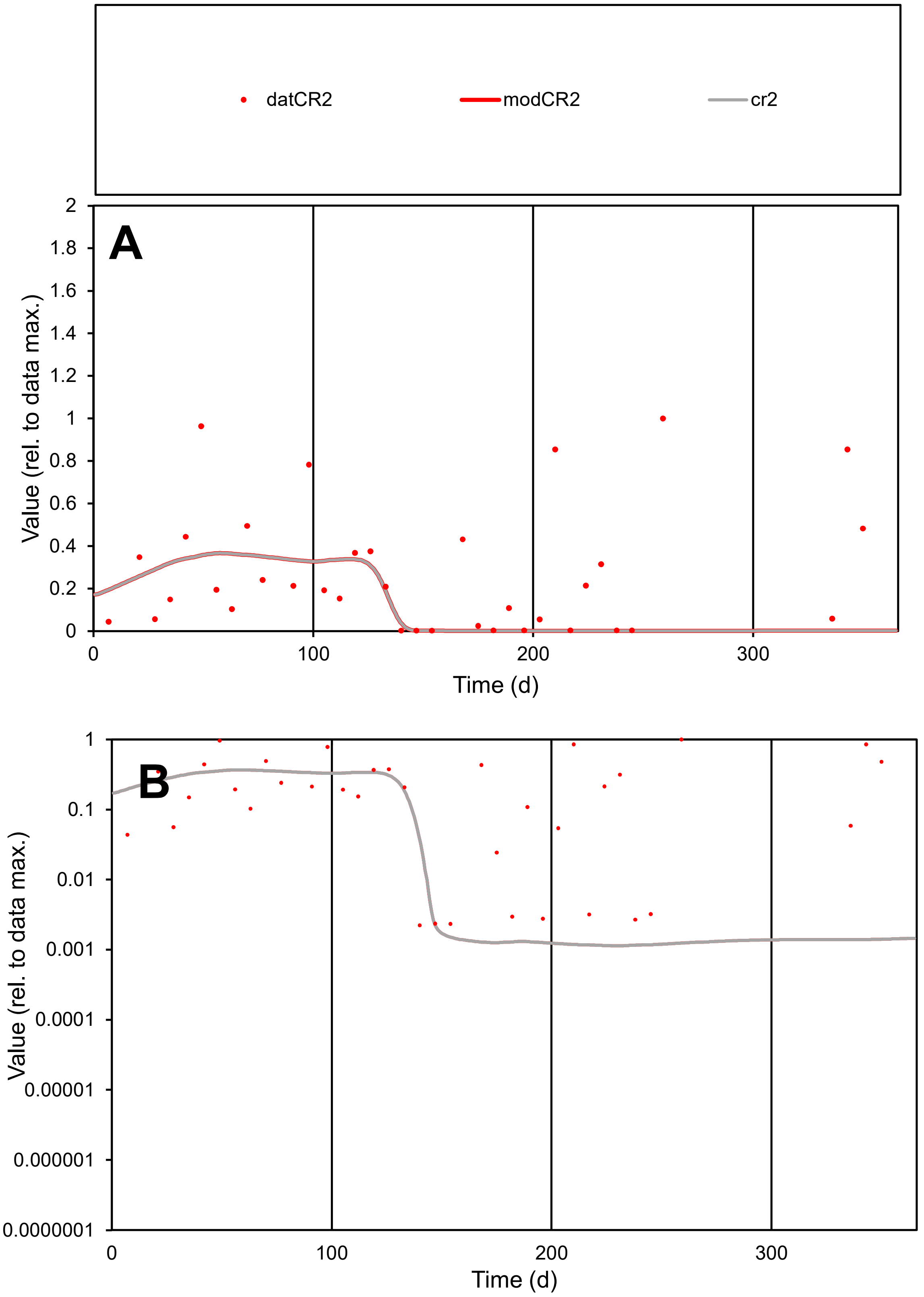


**C1**

Variables

CR2 Provide jtd or Dst

CalibOnly?

2 1 to reproduce of

oNormalize?

2 1=no, 2=yes

ofxn calc*

1.24199

ofxn mod

1.24786

sRun

RunA1

zout

1

Ave

Data

0.00472

Model

0.00265

data/vMax

0.01776

ofxng

311.734

Min d/m

1.95353

ave/min d

119.462

ave/min m

130.771

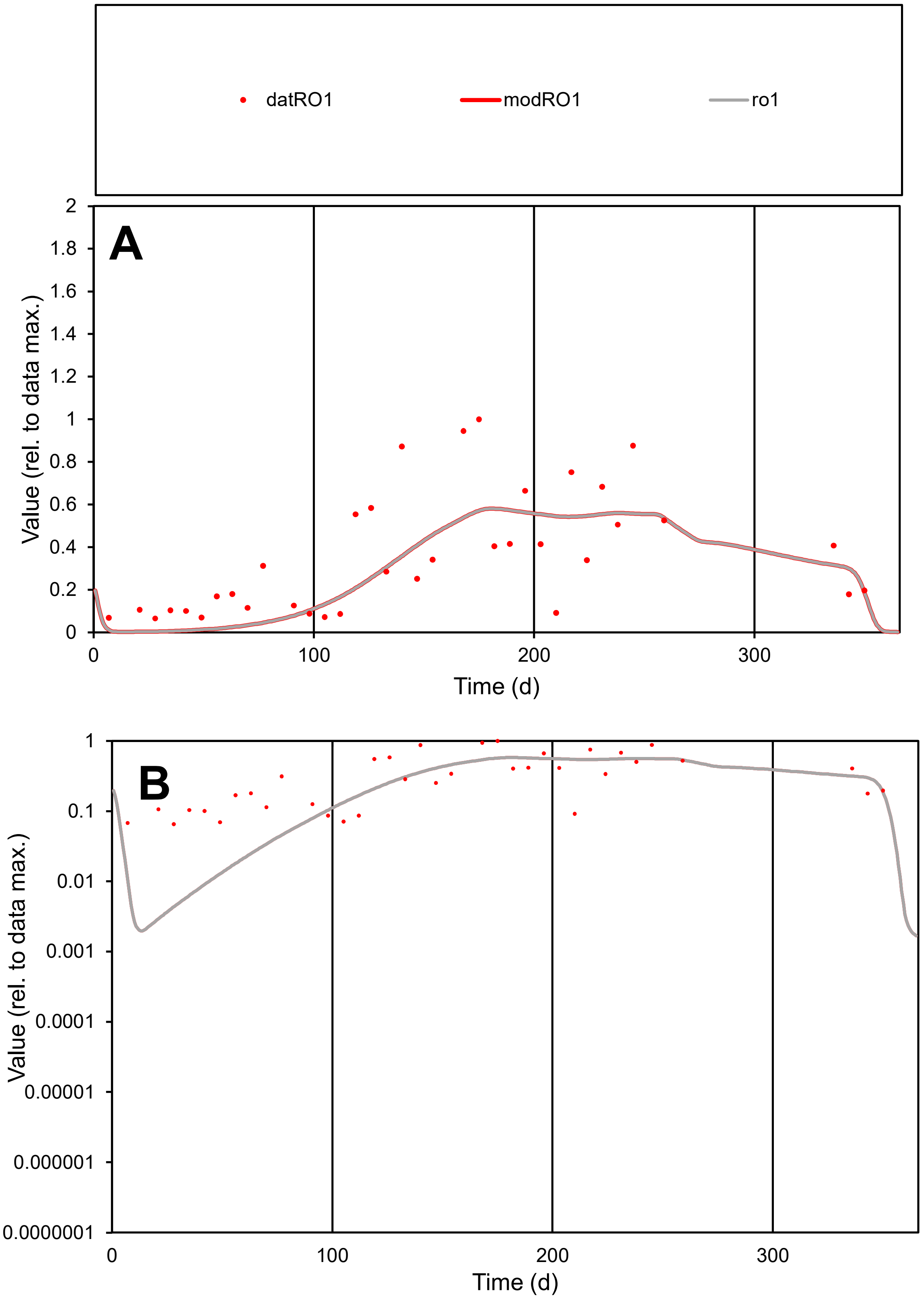


C1

Variables

RO1 Provide jtd or Dst

CalibOnly?

2 1 to reproduce of

oNormalize?

2 1=no, 2=yes

ofxn calc*

0.56187

ofxn mod

0.57657

sRun

RunA1

zout

1

Ave

Data

0.00541

Model

0.00479

data/vMax

0.01602

ofxng

311.734

Min d/m

20.4301

ave/min d

5.17334

ave/min m

93.5531

331_Calib_S_A

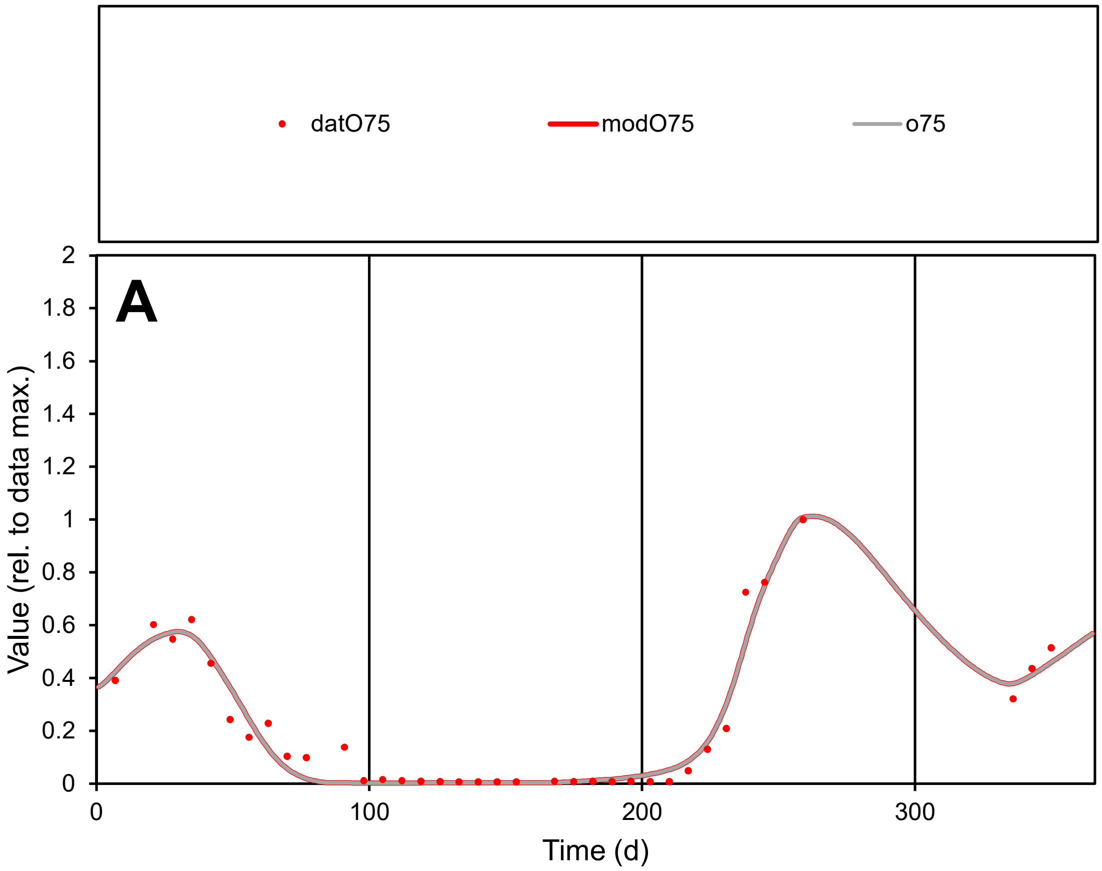


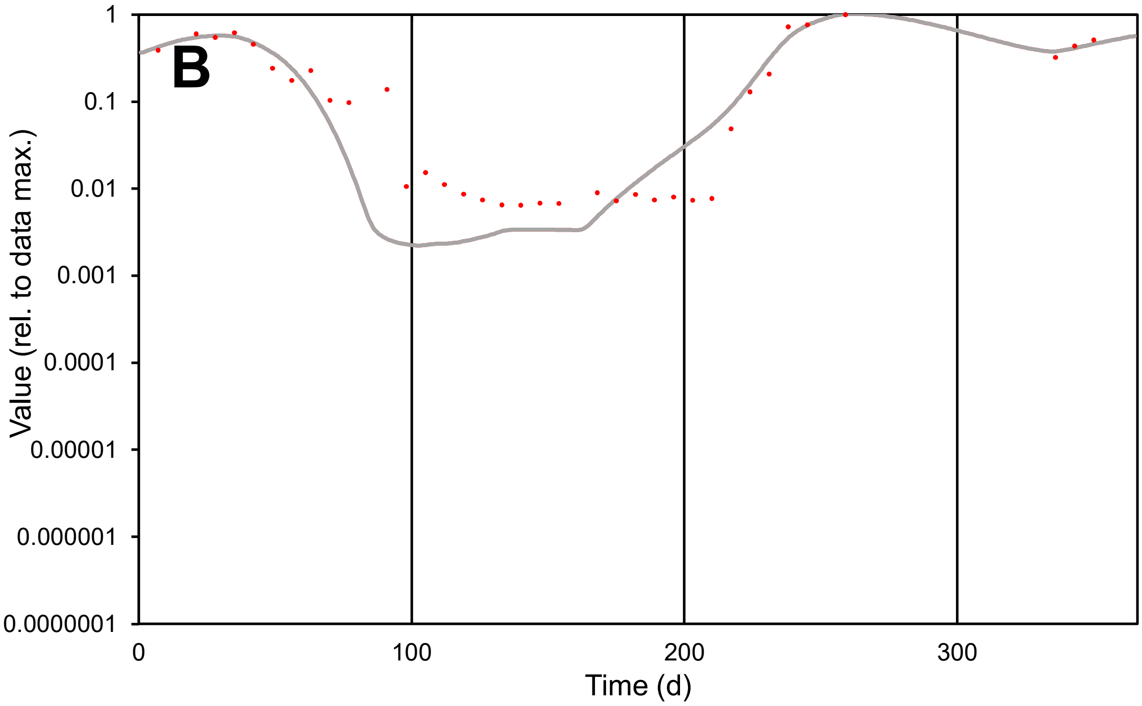


**C1**

Variables

O75 Provide jtd or Dst

CalibOnly?

2 1 to reproduce of

oNormalize?

2 1=no, 2=yes

ofxn calc*

0.24757

ofxn mod

0.30756

sRun

RunA1

zout

1

Ave

Data

0.00236

Model

0.00231

data/vMax

0.01077

ofxng

311.734

Min d/m

2.86249

ave/min d

33.9107

ave/min m

95.2747

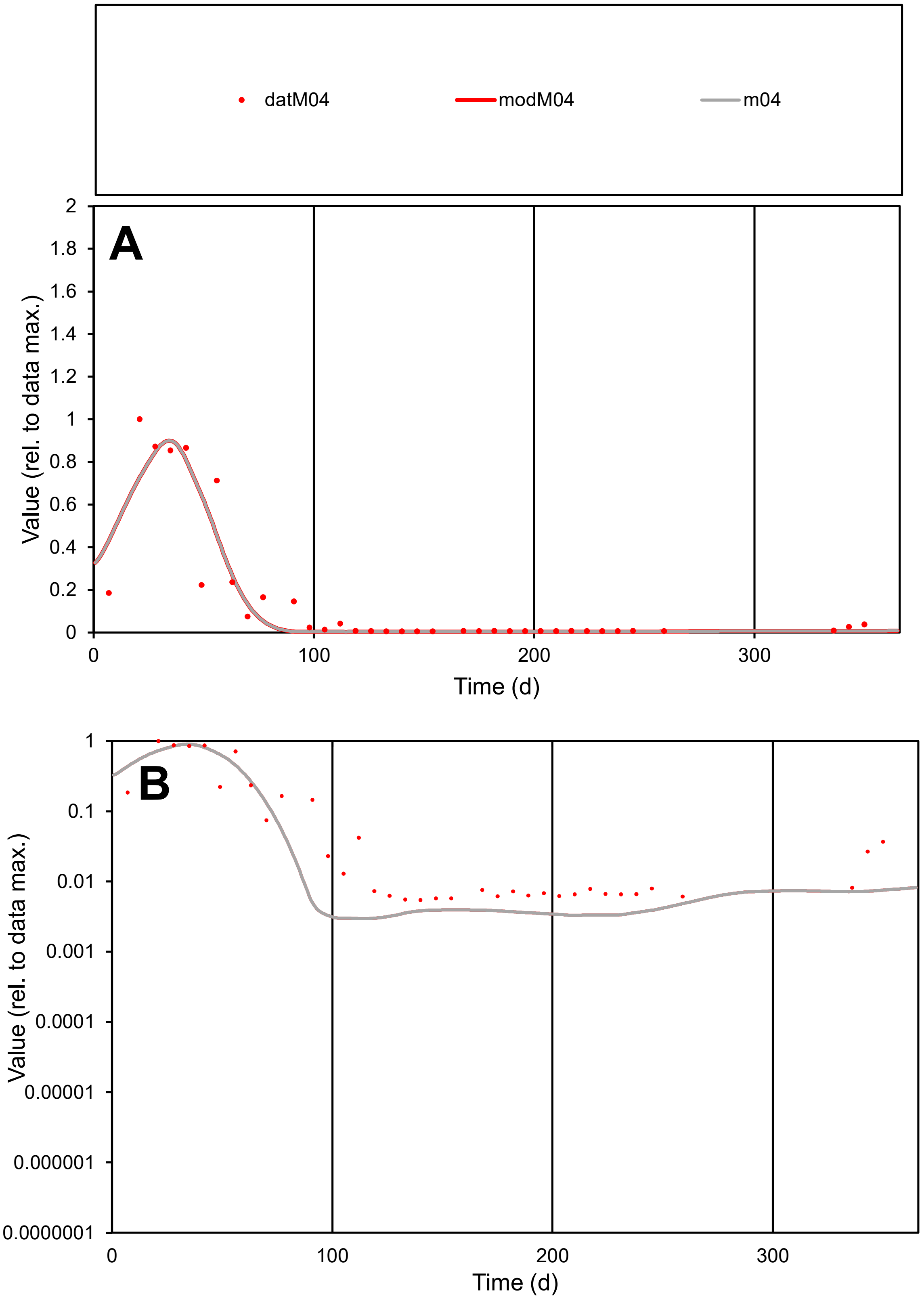


**C1**

Variables

| M04 | Provide jtd or Dst |
|---|---|
| CalibOnly? | |
| 2 | 1 to reproduce of |
| oNormalize? | |
| 2 | 1=no, 2=yes |

ofxn calc*
0.68025
ofxn mod
0.72686

sRun
RunA1
zout
1

Ave
Data
0.001
Model
0.00094

data/vMax
0.00639

ofxng
311.734

Min d/m
1.85321

ave/min d
28.4735
ave/min m
49.6544

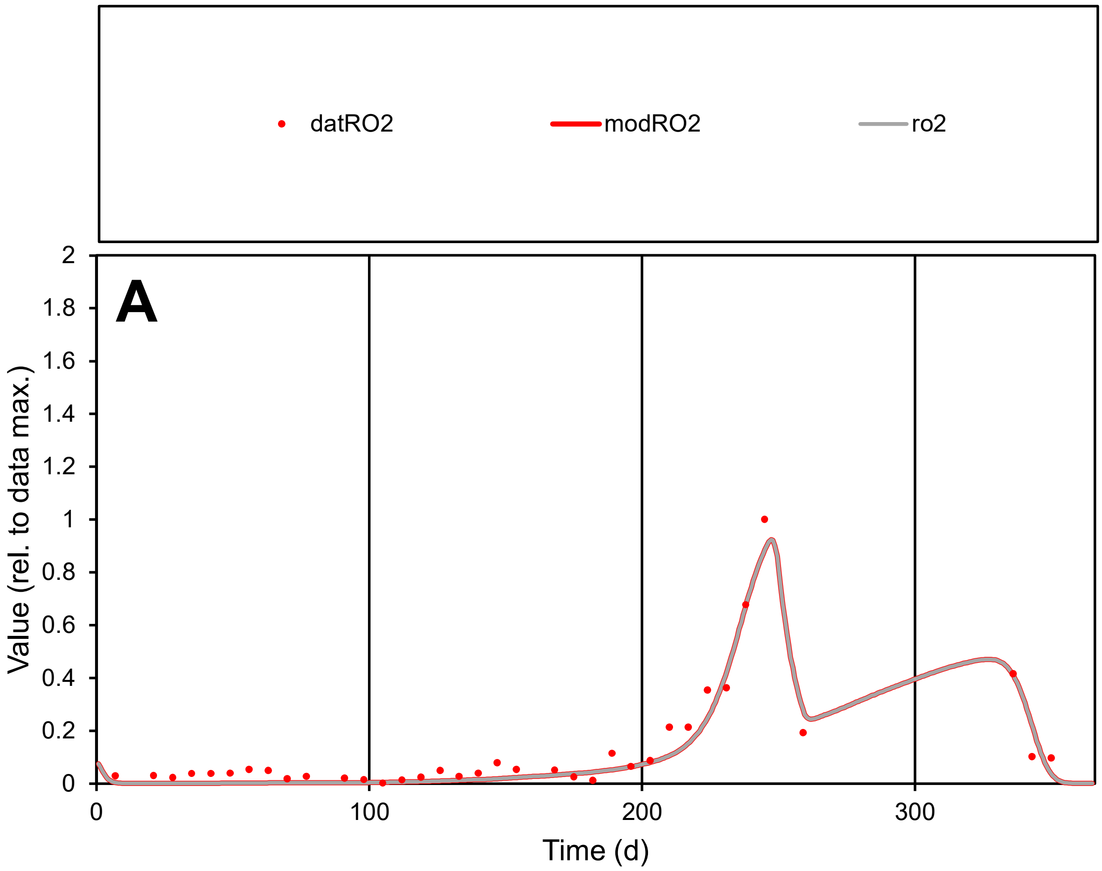


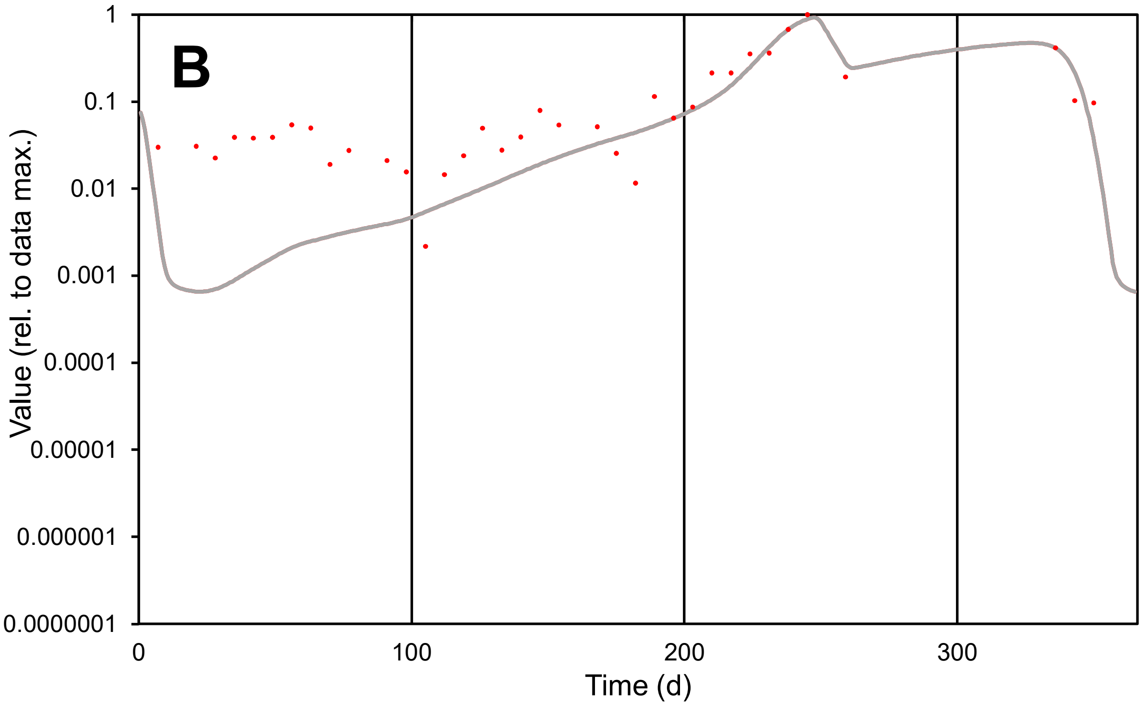


**C1**

Variables
RO2 Provide jtd or Dst
CalibOnly?
2 1 to reproduce of
oNormalize?
2 1=no, 2=yes

ofxn calc*
0.31944
ofxn mod
0.4021

sRun
RunA1
zout
1

Ave
Data
0.0053
Model
0.00452

data/vMax
0.04158

ofxng
311.734

Min d/m
3.30721

ave/min d
58.6695
ave/min m
165.539

331_Calib_S_A

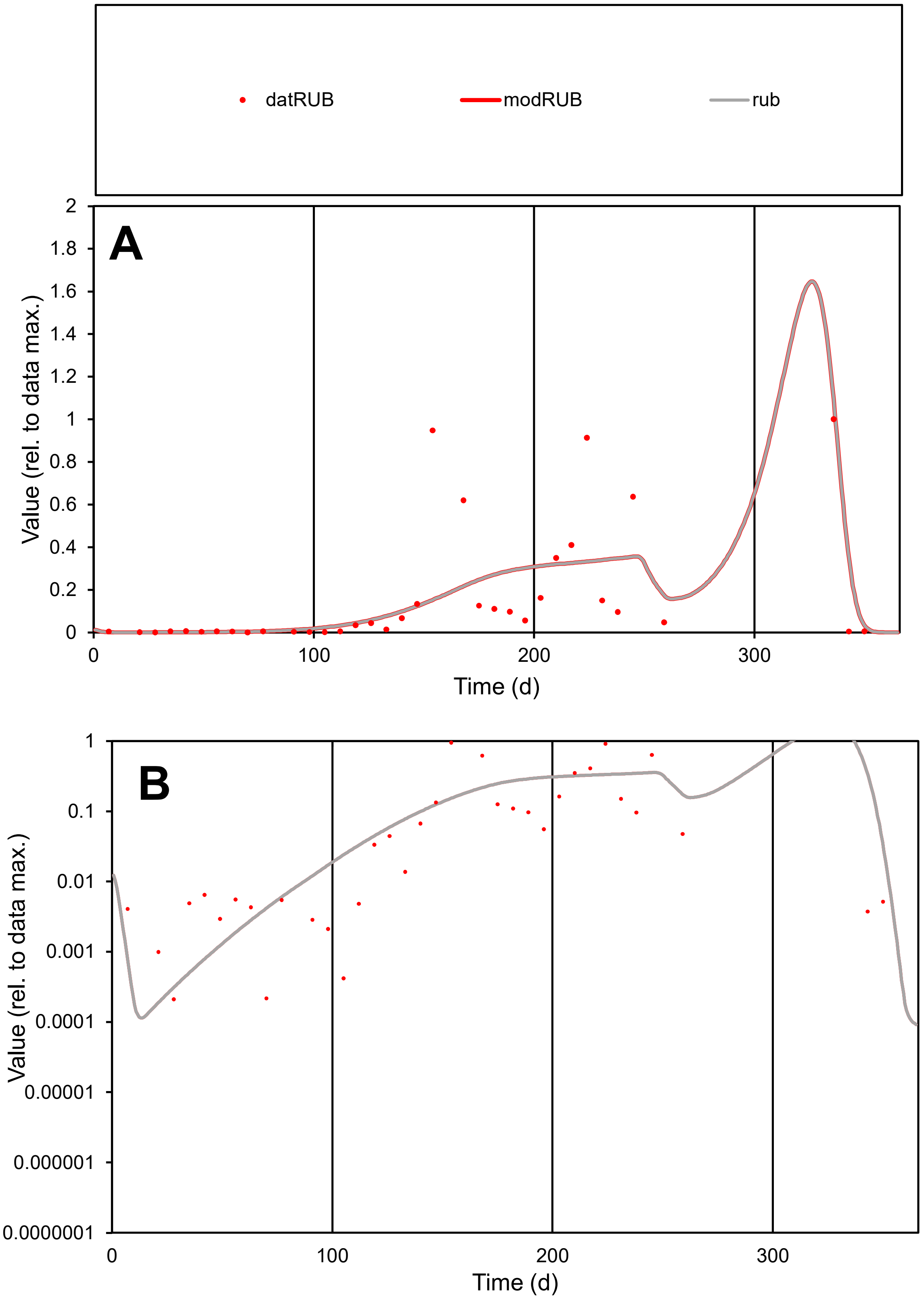


**C1**

Variables
RUB Provide jtd or Dst
CalibOnly?
2 1 to reproduce of
oNormalize?
2 1=no, 2=yes

ofxn calc*
1.13994
ofxn mod
1.16192

sRun
RunA1
zout
1

Ave
Data
0.00891
Model
0.00872

data/vMax
0.05588

ofxng
311.734

Min d/m
1.0034

ave/min d
755.805
ave/min m
742.43

331_Calib_S_A

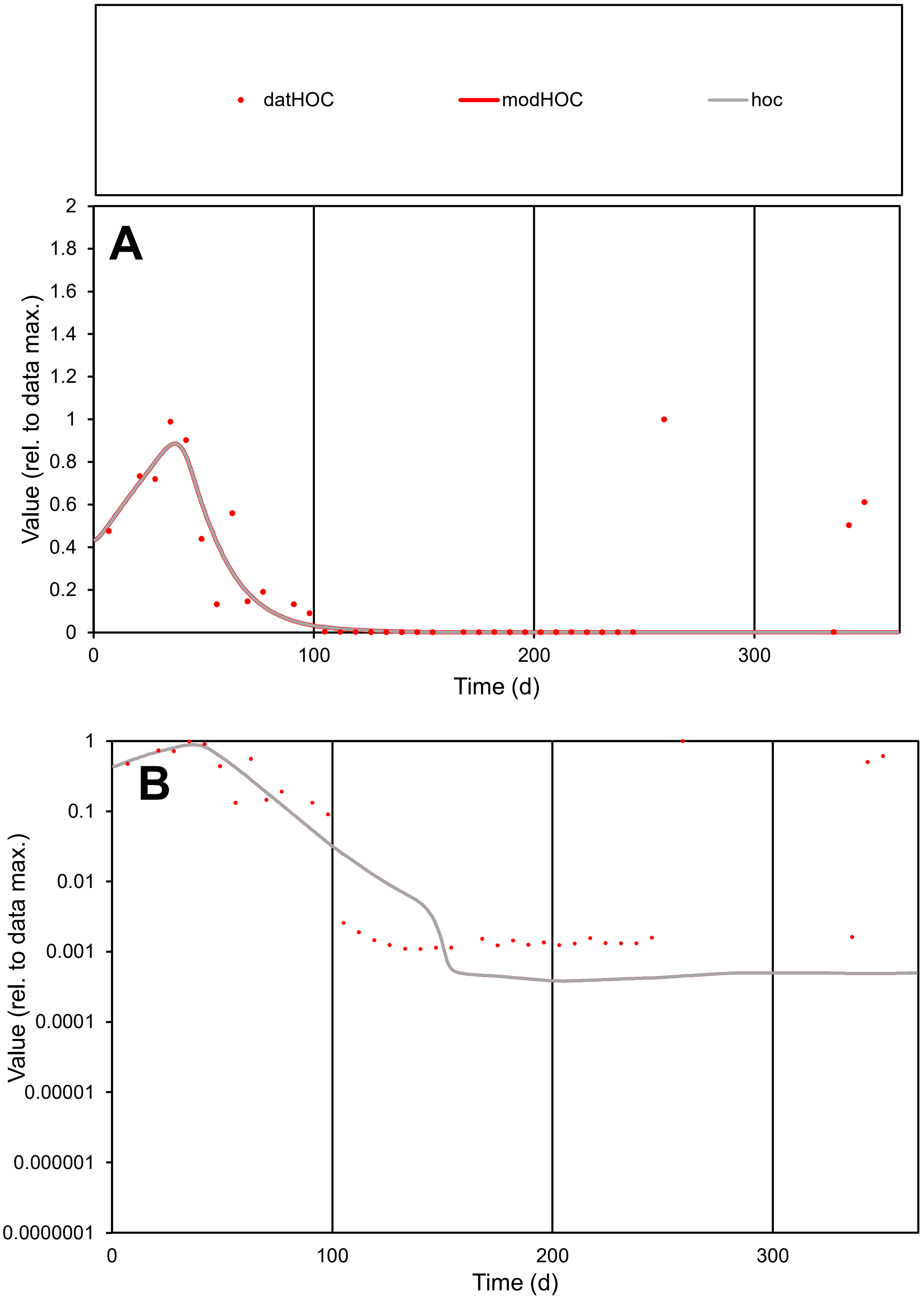


**C1**

Variables
HOC Provide jtd or Dst
CalibOnly?
2 1 to reproduce of
oNormalize?
2 1=no, 2=yes

ofxn calc*
1.06381
ofxn mod
1.08101

sRun
RunA1
zout
1

Ave
Data
0.00243
Model
0.00172

data/vMax
0.01141

ofxng
311.734

Min d/m
2.86535

ave/min d
194.512
ave/min m
395.793

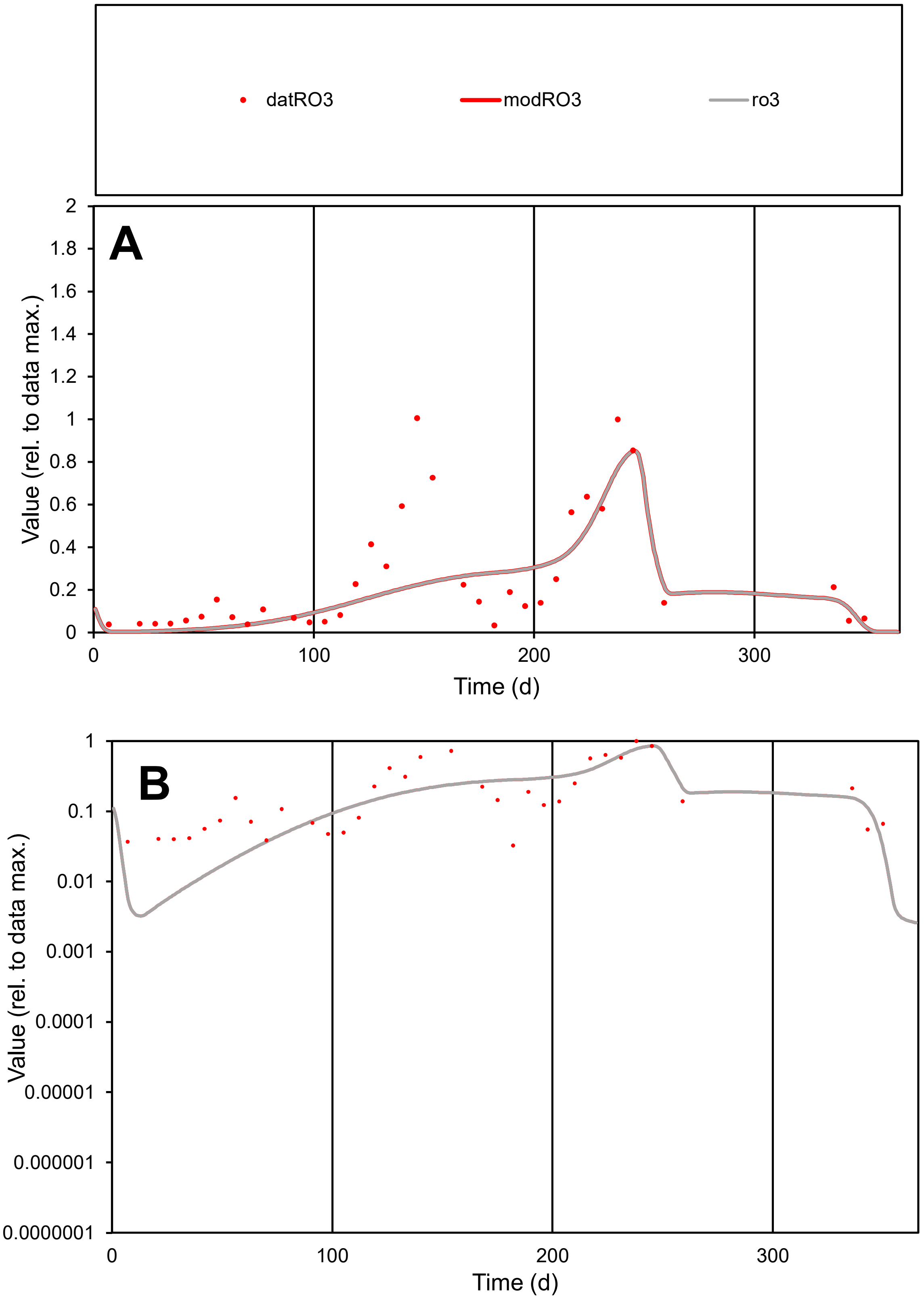


**C1**

Variables
RO3 Provide jtd or Dst
CalibOnly?
2 1 to reproduce of
oNormalize?
2 1=no, 2=yes

ofxn calc*
0.5898
ofxn mod
0.6122

sRun
RunA1
zout
1

Ave
Data
0.00568
Model
0.00507

data/vMax
0.02411

ofxng
311.734

Min d/m
6.57789

ave/min d
7.22844
ave/min m
42.426

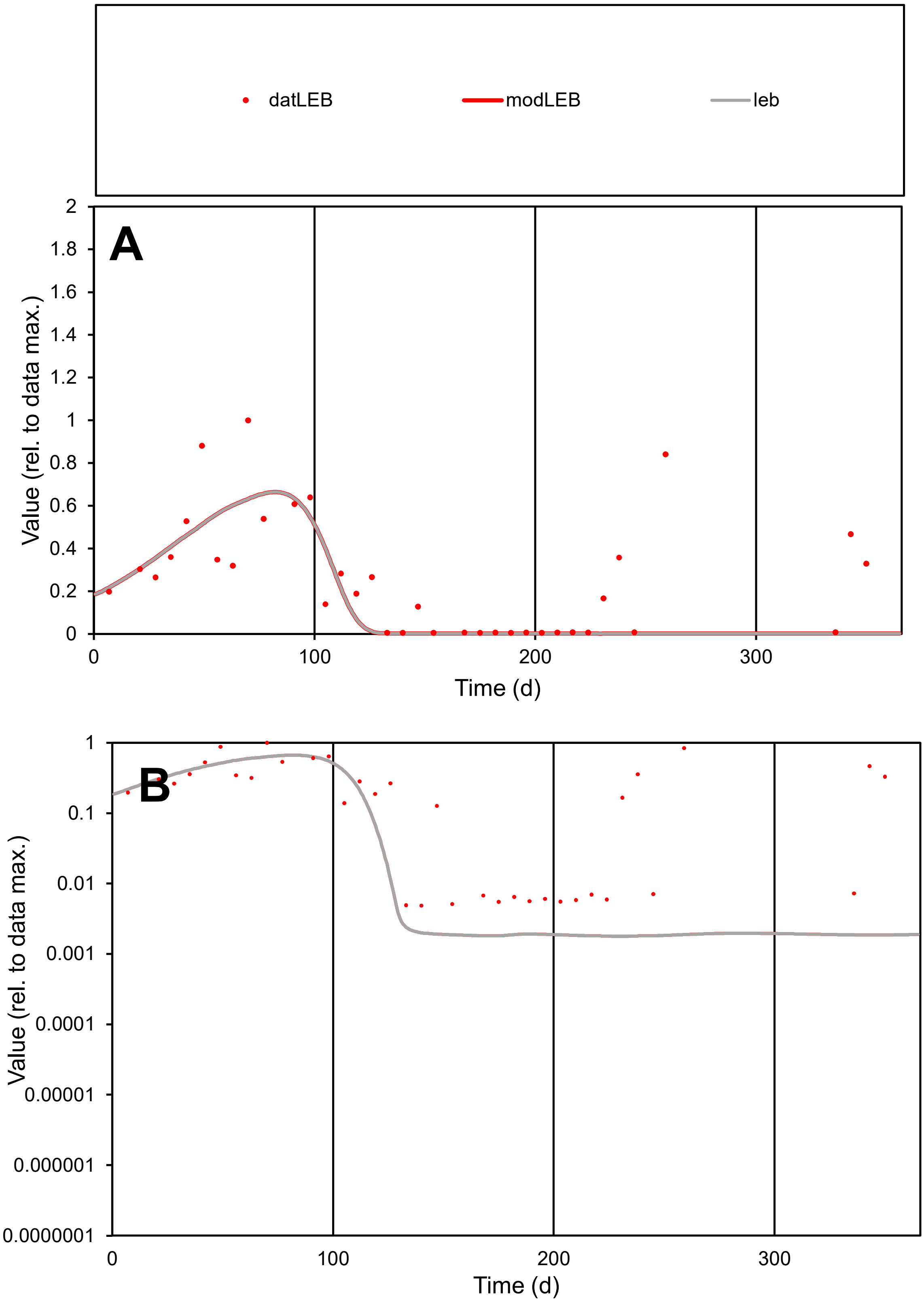


**C1**

Variables
LEB Provide jtd or Dst
CalibOnly?
2 1 to reproduce of
oNormalize?
2 1=no, 2=yes

ofxn calc*
0.8612
ofxn mod
0.87924

sRun
RunA1
zout
1

Ave
Data
0.00359
Model
0.0026

data/vMax
0.01419

ofxng
311.734

Min d/m
2.73753

ave/min d
51.9599
ave/min m
102.783

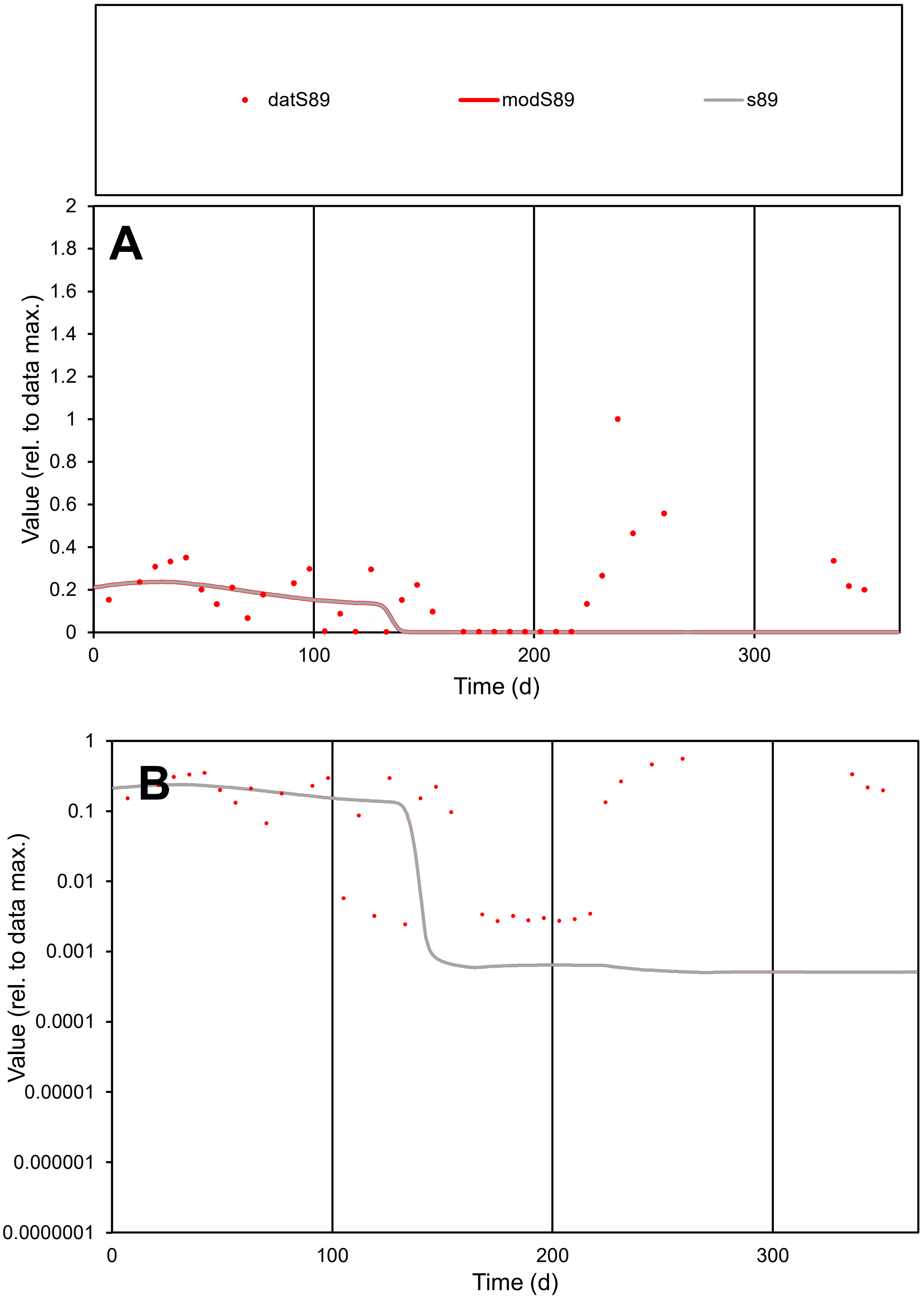


**C1**

Variables
S89 Provide jtd or Dst
CalibOnly?
2 1 to reproduce of
oNormalize?
2 1=no, 2=yes

ofxn calc*
1.29242
ofxn mod
1.29968

sRun
RunA1
zout
1

Ave
Data
0.00267
Model
0.00128

data/vMax
0.01476

ofxng
311.734

Min d/m
4.81912

ave/min d
74.0681
ave/min m
171.433

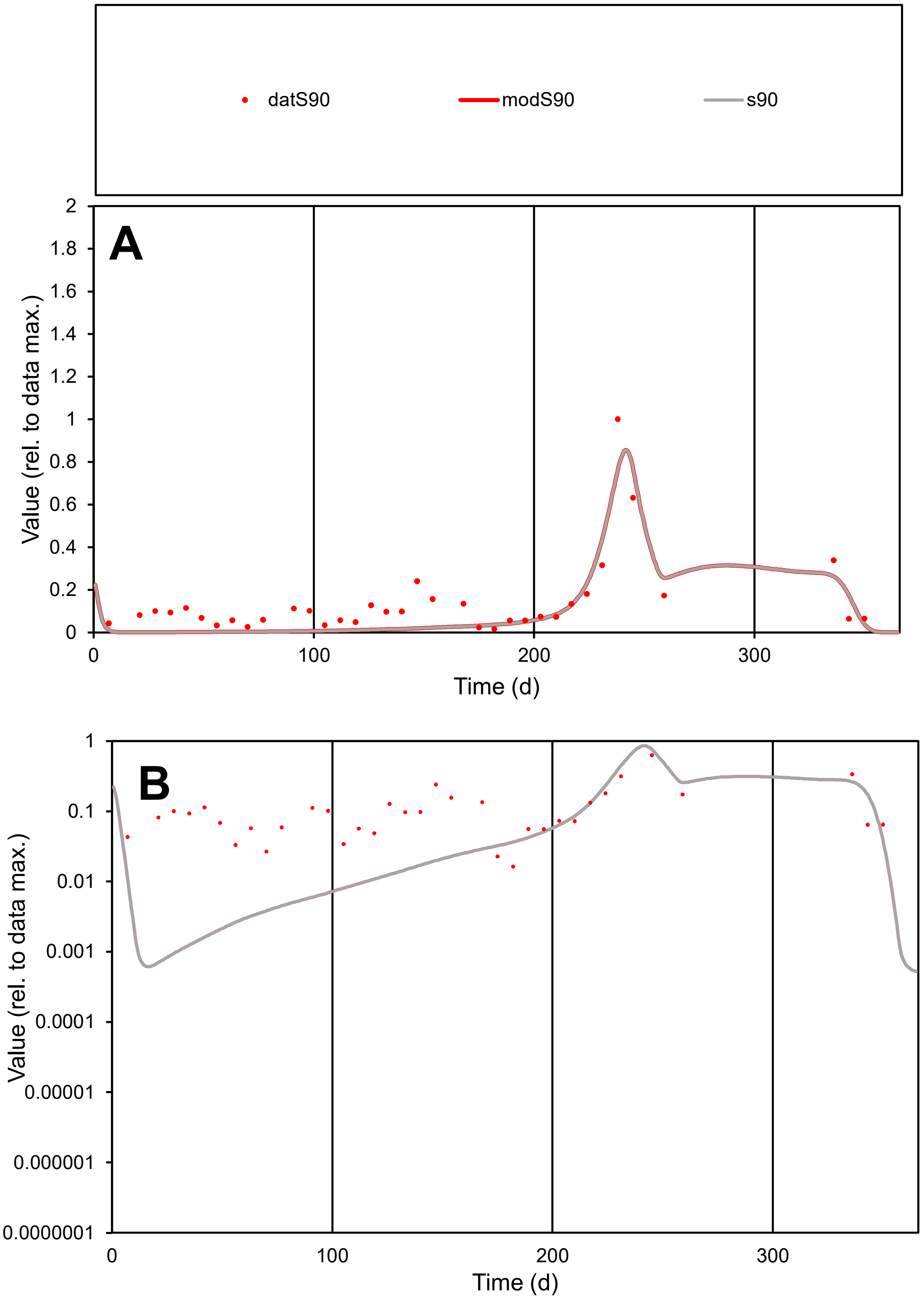


**C1**

Variables
S90 Provide jtd or Dst
CalibOnly?
2 1 to reproduce of
oNormalize?
2 1=no, 2=yes

ofxn calc*
0.56311
ofxn mod
0.59964

sRun
RunA1
zout
1

Ave
Data
0.00505
Model
0.00369

data/vMax
0.03733

ofxng
311.734

Min d/m
21.6869

ave/min d
8.30812
ave/min m
131.447

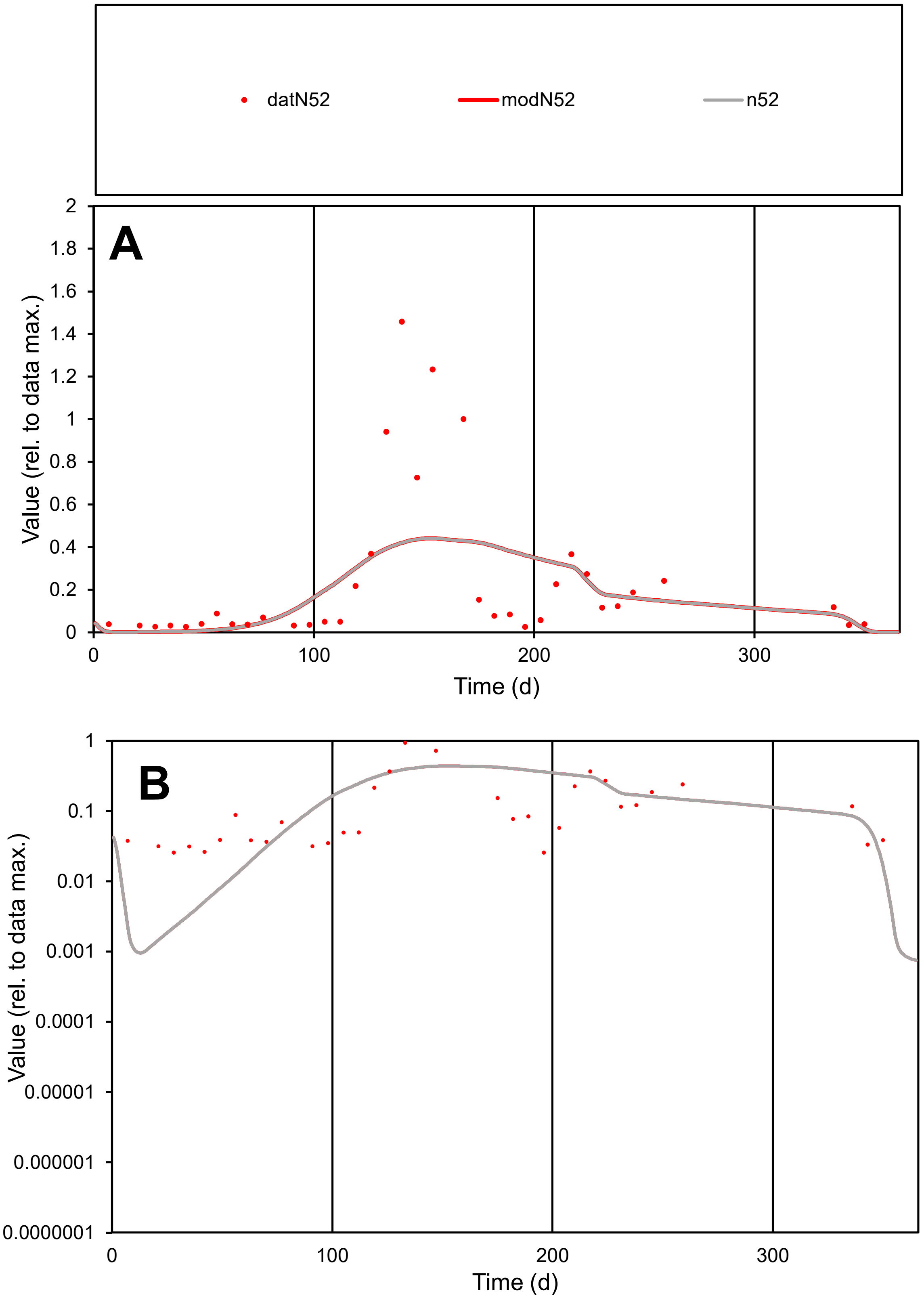


**C1**

Variables
N52 Provide jtd or Dst
CalibOnly?
2 1 to reproduce of
oNormalize?
2 1=no, 2=yes

ofxn calc*
0.99153
ofxn mod
1.00421

sRun
RunA1
zout
1

Ave
Data
0.00416
Model
0.00423

data/vMax
0.02098

ofxng
311.734

Min d/m
16.0173

ave/min d
7.73497
ave/min m
125.882

331_Calib_S_A

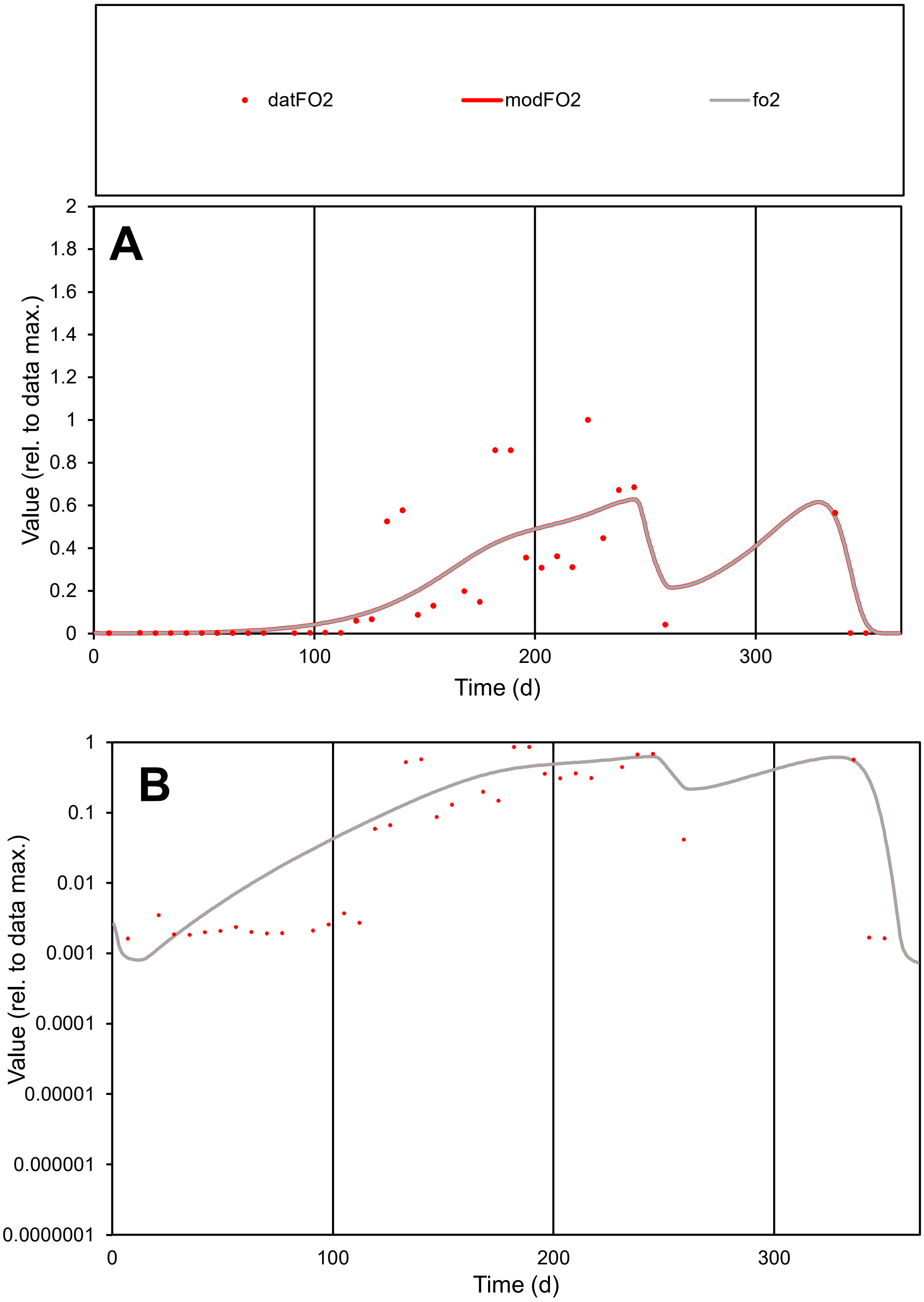


**C1**

Variables
FO2 Provide jtd or Dst
CalibOnly?
2 1 to reproduce of
oNormalize?
2 1=no, 2=yes

ofxn calc*
0.75042
ofxn mod
0.78147

sRun
RunA1
zout
1

Ave
Data
0.0092
Model
0.00957

data/vMax
0.04241

ofxng
311.734

Min d/m
1.94268

ave/min d
133.095
ave/min m
268.865

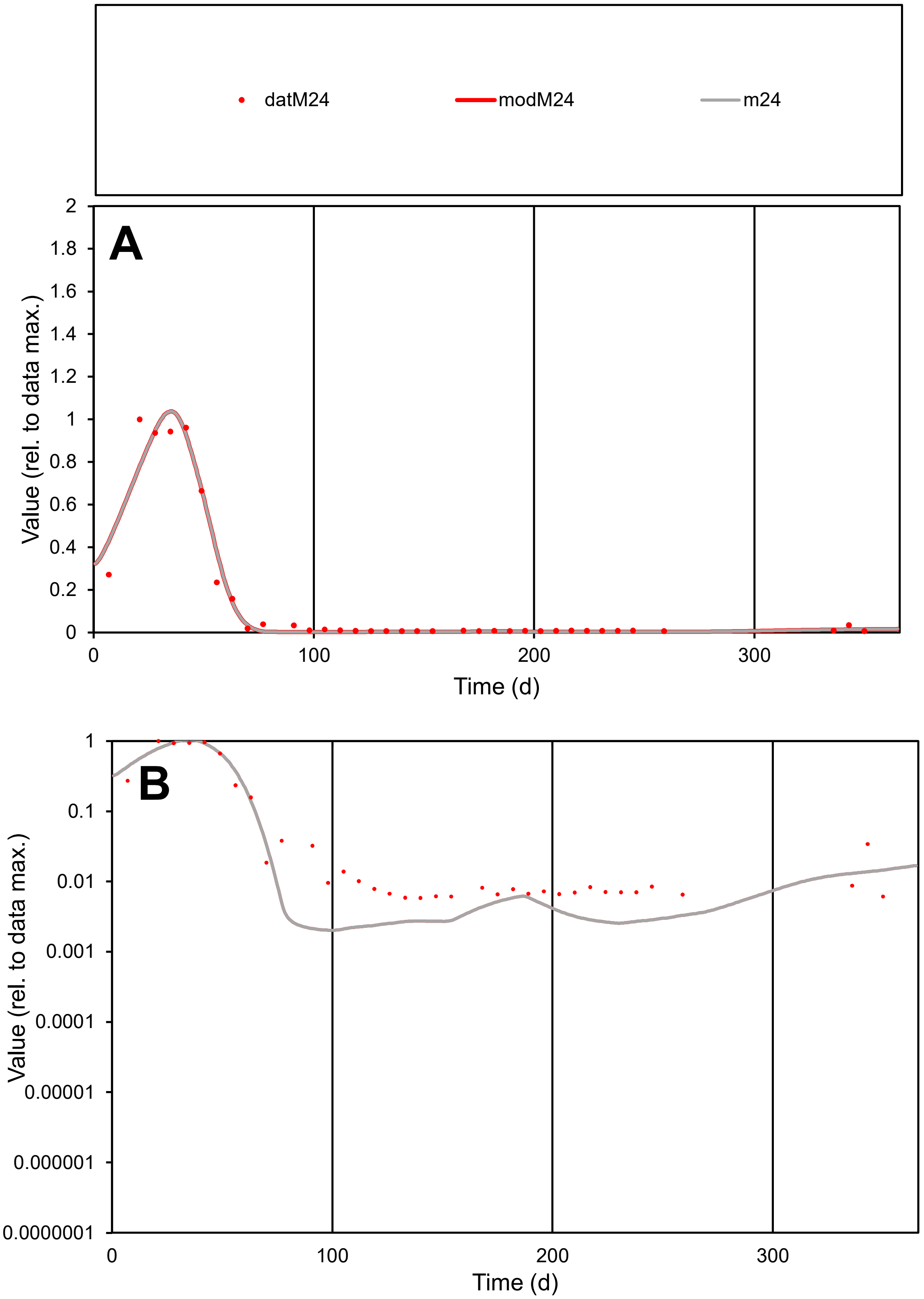


**C1**

Variables
M24 Provide jtd or Dst
CalibOnly?
2 1 to reproduce of
oNormalize?
2 1=no, 2=yes

ofxn calc*
0.32875
ofxn mod
0.43035

sRun
RunA1
zout
1

Ave
Data
0.00186
Model
0.00183

data/vMax
0.01227

ofxng
311.734

Min d/m
2.88084

ave/min d
26.0553
ave/min m
73.5862

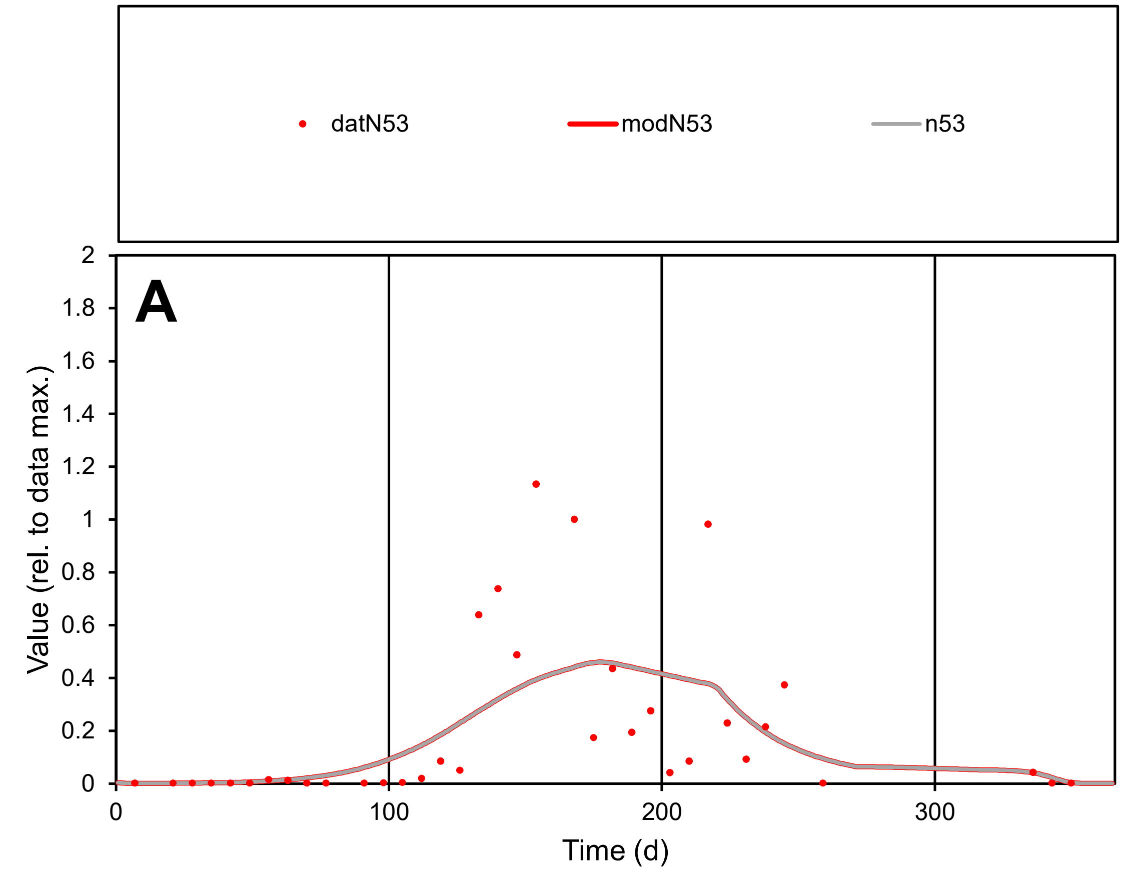


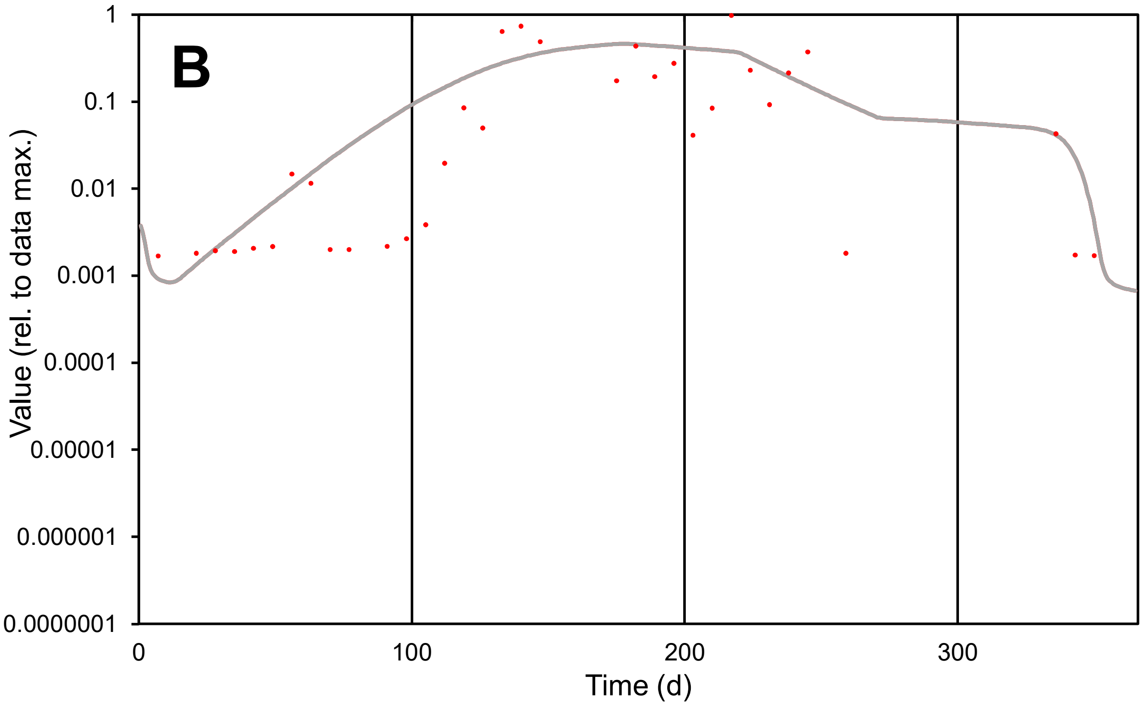


**C1**

Variables

N53 Provide jtd or Dst

CalibOnly?

2 1 to reproduce of

oNormalize?

2 1=no, 2=yes

ofxn calc*

1.13196

ofxn mod

1.14643

sRun

RunA1

zout

1

Ave

Data

0.00397

Model

0.00415

data/vMax

0.02228

ofxng

311.734

Min d/m

1.90293

ave/min d

105.388

ave/min m

209.619

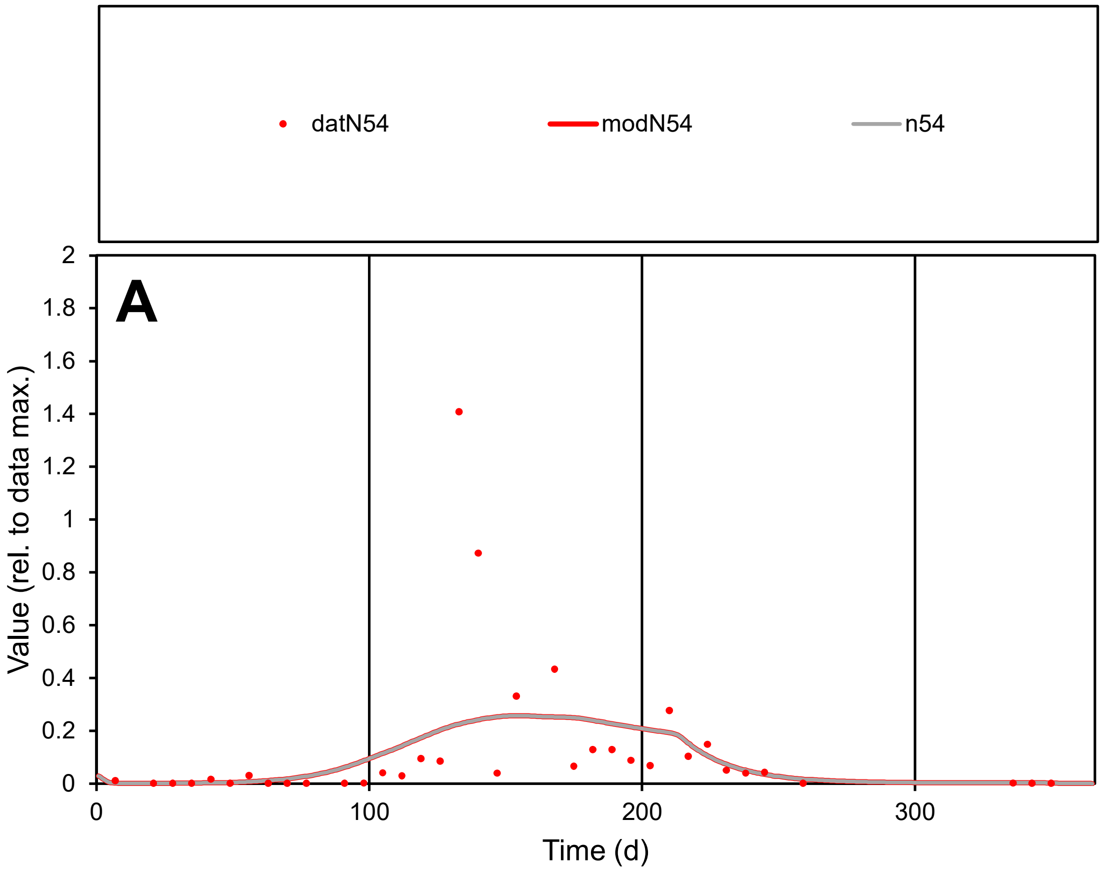


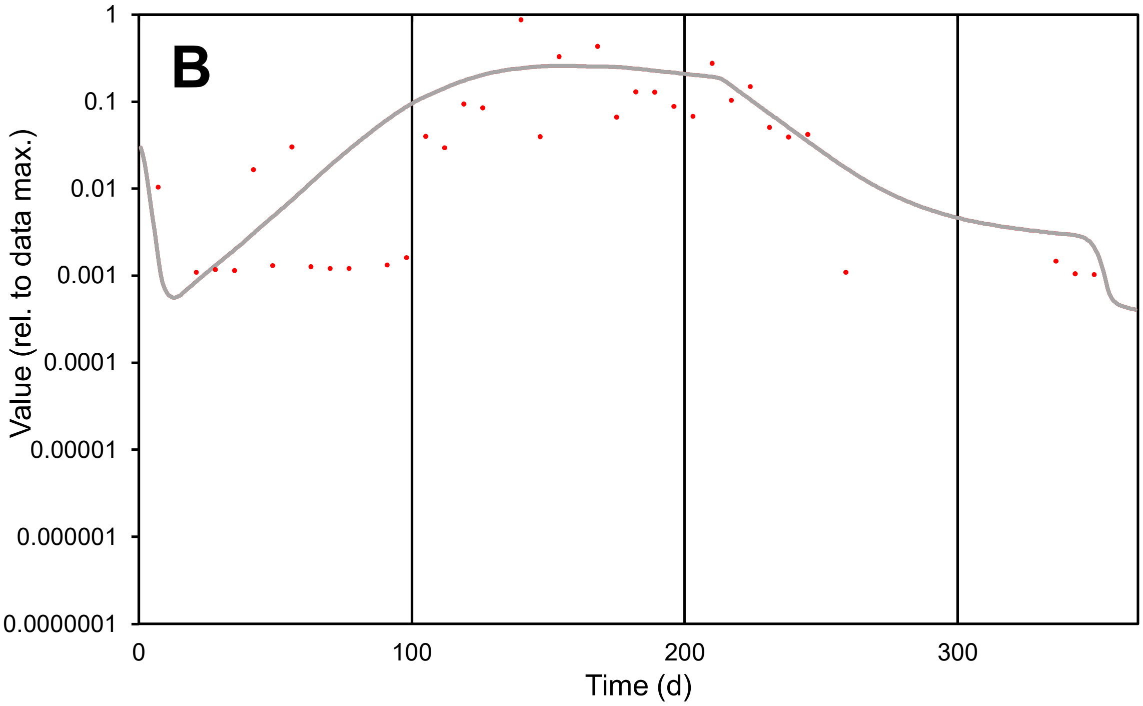


**C1**

Variables
N54 Provide jtd or Dst
CalibOnly?
2 1 to reproduce of
oNormalize?
2 1=no, 2=yes

ofxn calc*
1.56086
ofxn mod
1.57322

sRun
RunA1
zout
1

Ave
Data
0.0022
Model
0.00232

data/vMax
0.02174

ofxng
311.734

Min d/m
1.0977

ave/min d
97.9383
ave/min m
113.21

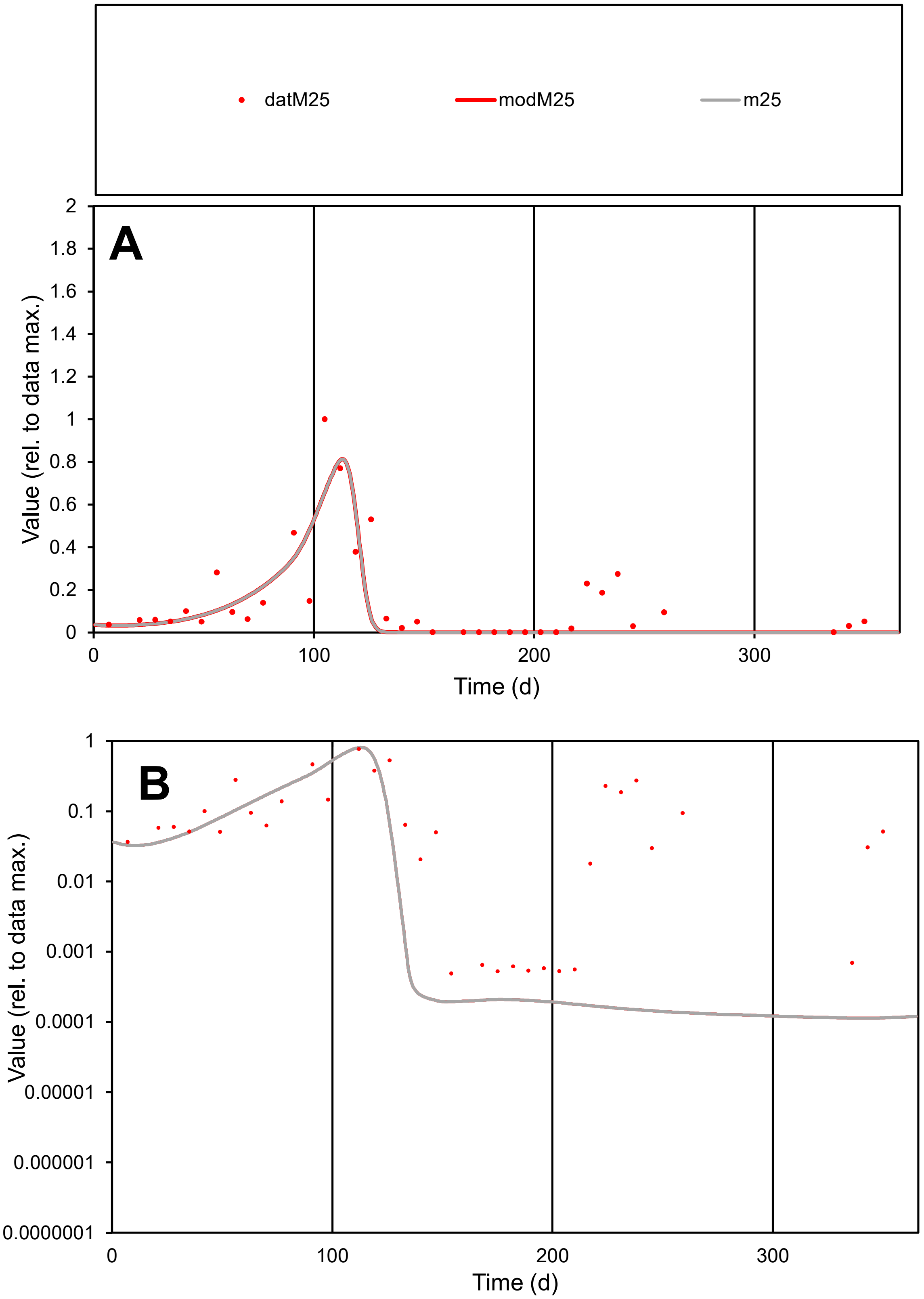


**C1**

Variables
M25 Provide jtd or Dst
CalibOnly?
2 1 to reproduce of
oNormalize?
2 1=no, 2=yes

ofxn calc*
0.85077
ofxn mod
0.89079

sRun
RunA1
zout
1

Ave
Data
0.00288
Model
0.00223

data/vMax
0.02038

ofxng
311.734

Min d/m
4.34952

ave/min d
287.339
ave/min m
965.463

331_Calib_S_A

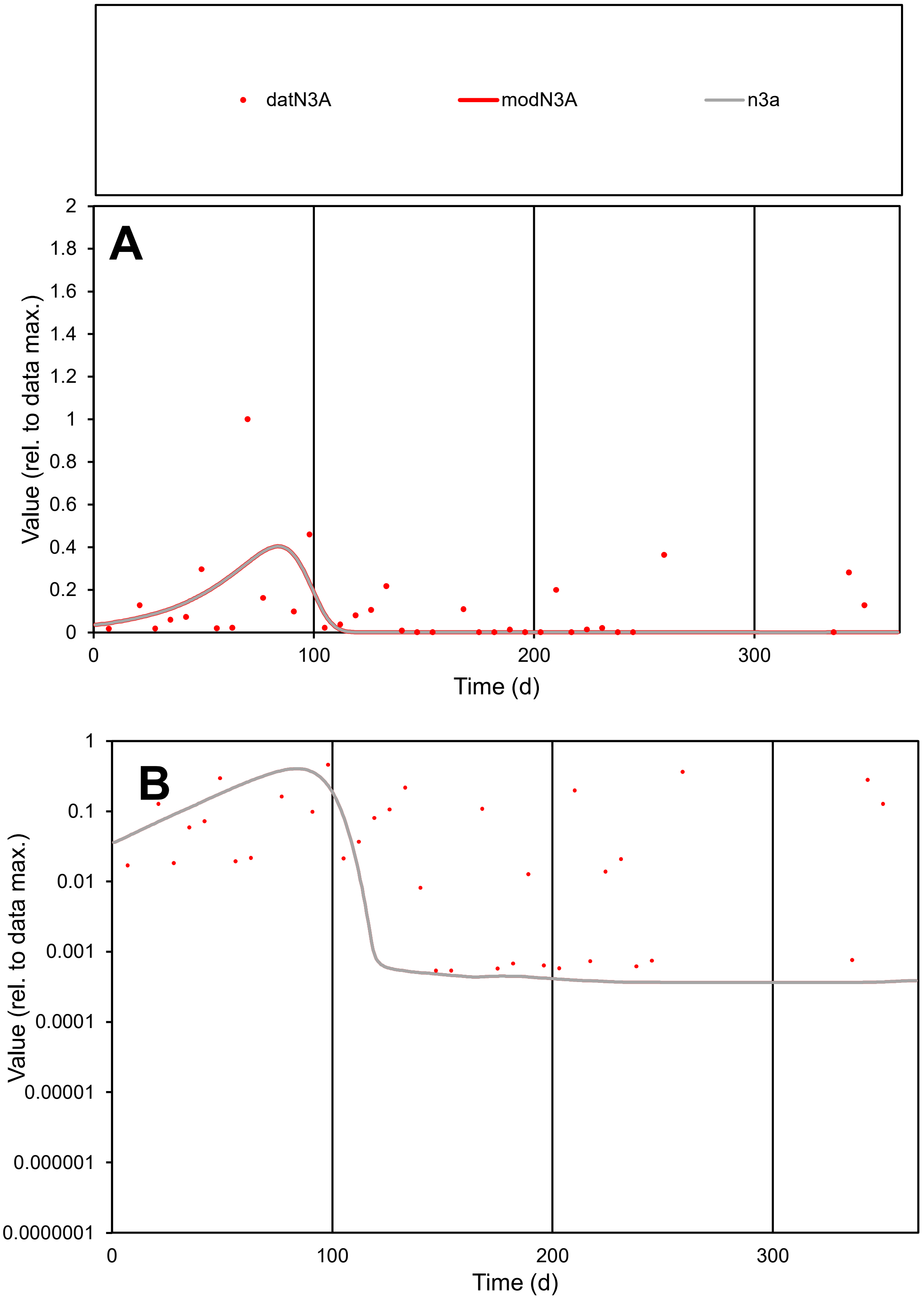


**C1**

Variables

N3A Provide jtd or Dst

CalibOnly?

2 1 to reproduce of

oNormalize?

2 1=no, 2=yes

ofxn calc*

1.5906

ofxn mod

1.60789

sRun

RunA1

zout

1

Ave

Data

0.00371

Model

0.00241

data/vMax

0.03457

ofxng

311.734

Min d/m

1.47393

ave/min d

198.577

ave/min m

190.243

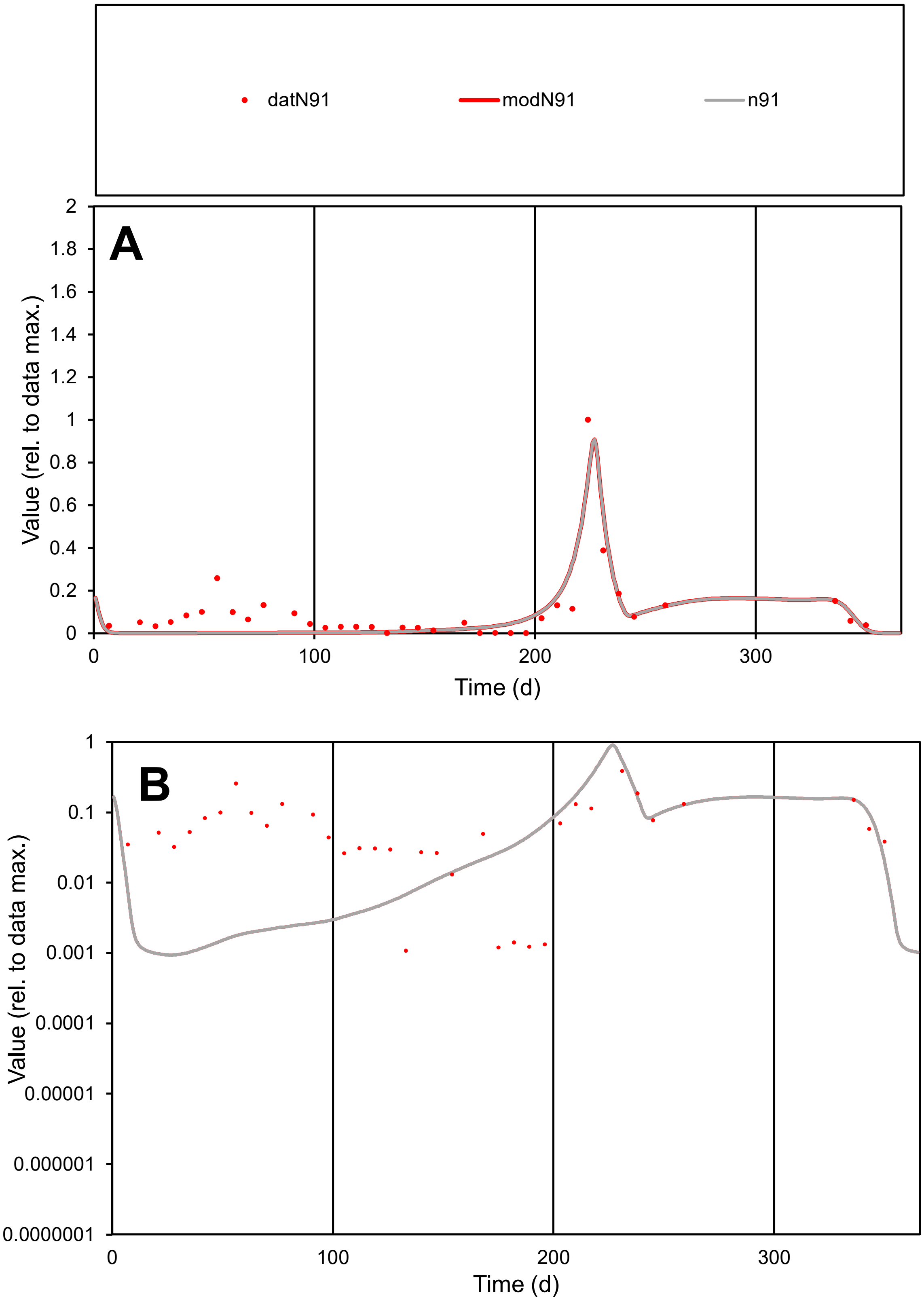


**C1**

Variables
N91 Provide jtd or Dst
CalibOnly?
2 1 to reproduce of
oNormalize?
2 1=no, 2=yes

ofxn calc*
0.76526
ofxn mod
0.83134

sRun
RunA1
zout
1

Ave
Data
0.00294
Model
0.00228

data/vMax
0.02934

ofxng
311.734

Min d/m
1.13669

ave/min d
92.9145
ave/min m
82.0823

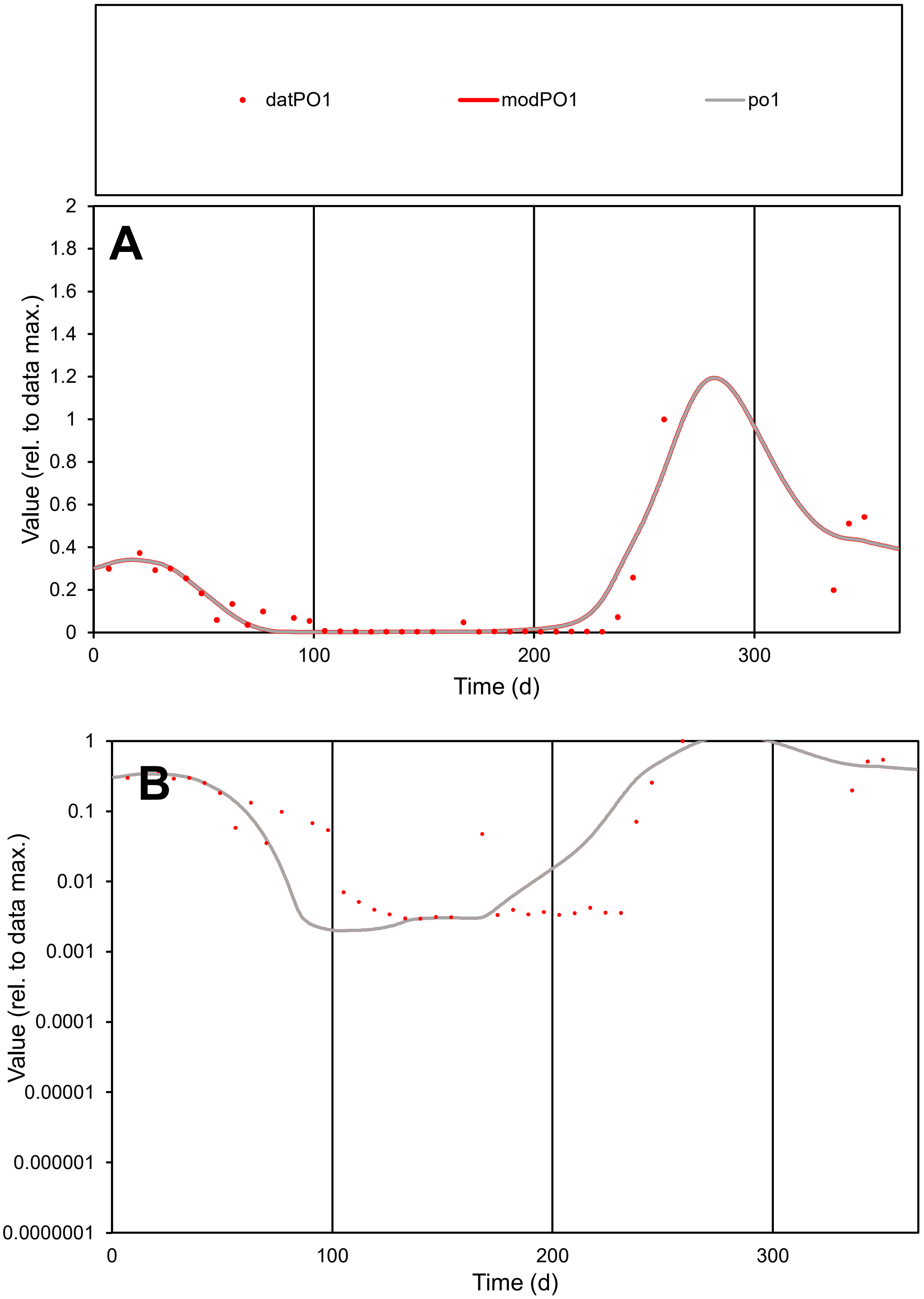


**C1**

Variables
PO1 Provide jtd or Dst
CalibOnly?
2 1 to reproduce of
oNormalize?
2 1=no, 2=yes

ofxn calc*
0.60738
ofxn mod
0.66067

sRun
RunA1
zout
1

Ave
Data
0.00205
Model
0.00217

data/vMax
0.01527

ofxng
311.734

Min d/m
1.4747

ave/min d
45.413
ave/min m
70.807

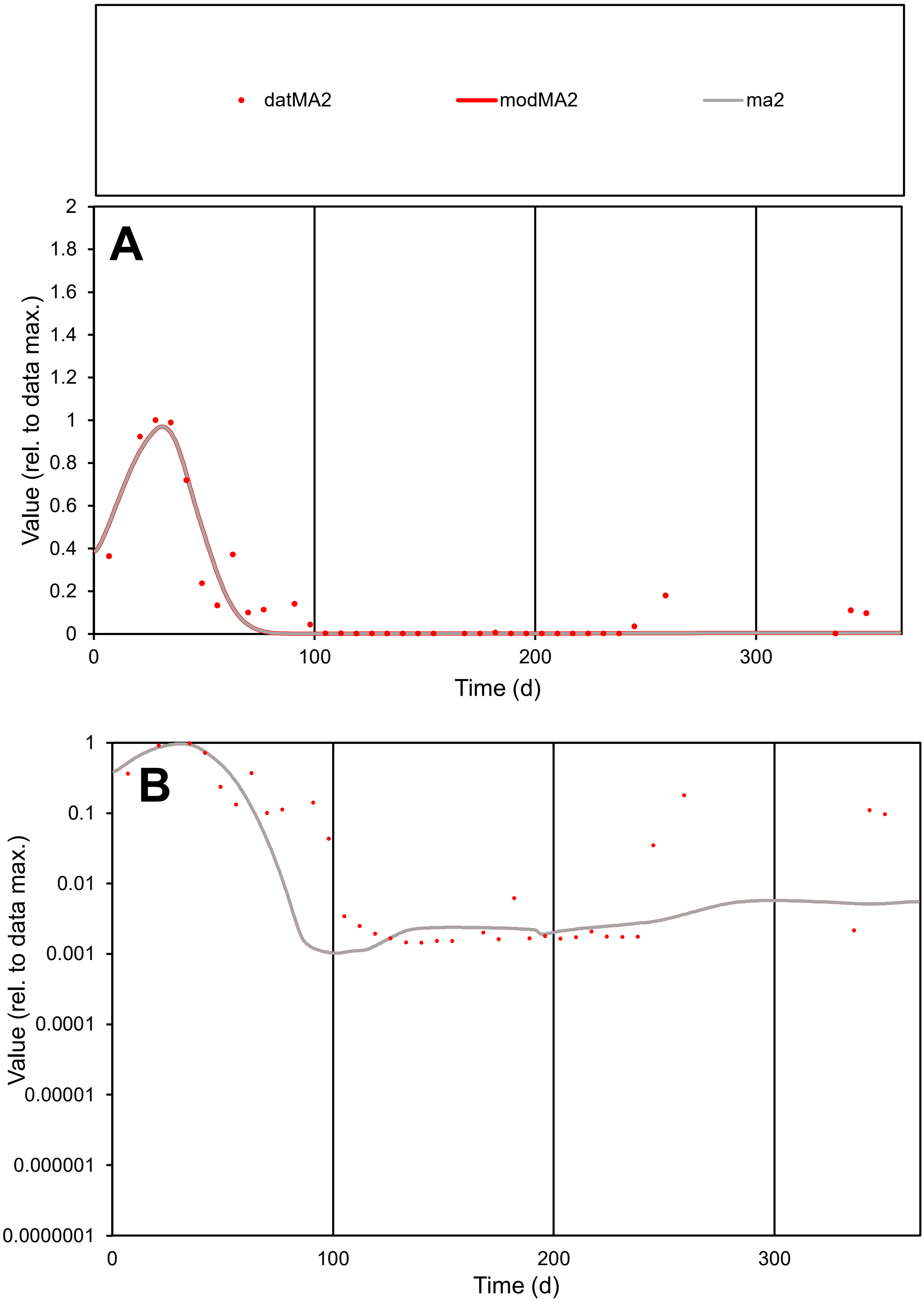


**C1**

Variables
MA2 Provide jtd or Dst
CalibOnly?
2 1 to reproduce of
oNormalize?
2 1=no, 2=yes

ofxn calc*
0.53951
ofxn mod
0.59084

sRun
RunA1
zout
1

Ave
Data
0.0019
Model
0.00168

data/vMax
0.01221

ofxng
311.734

Min d/m
1.39905

ave/min d
107.451
ave/min m
132.803

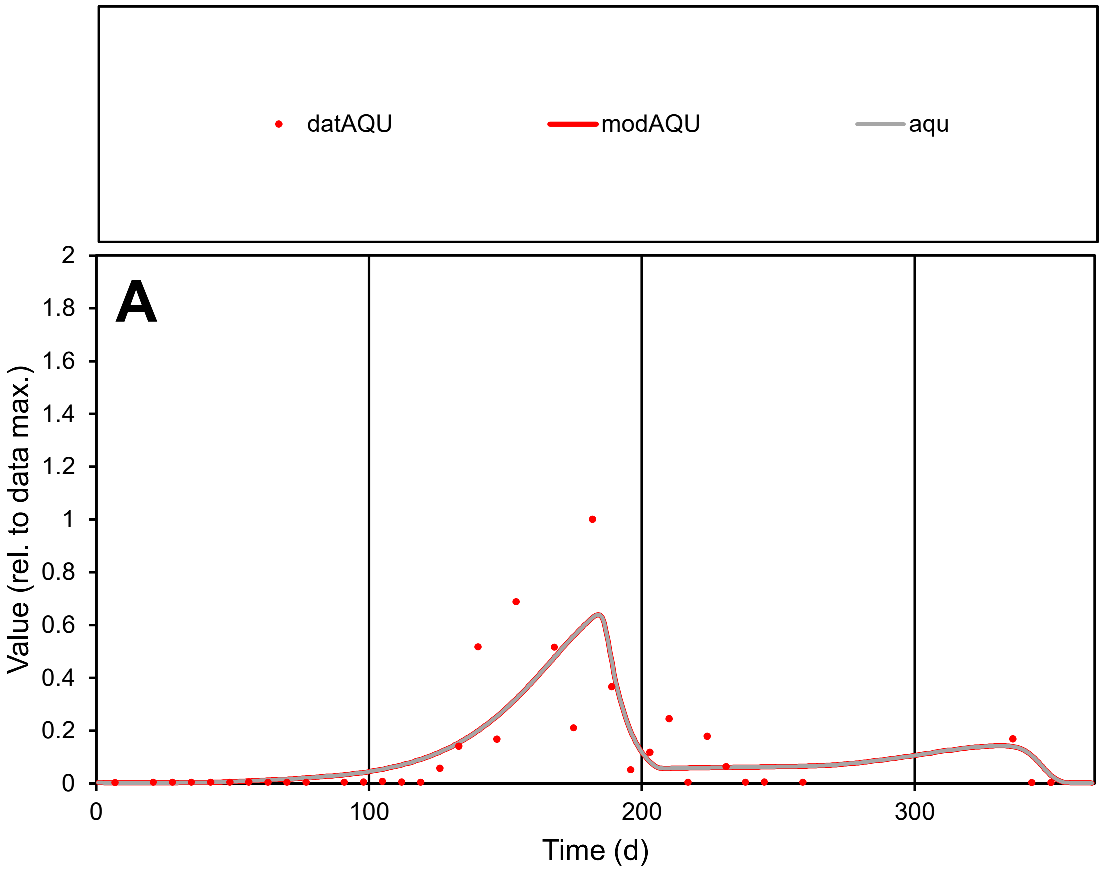


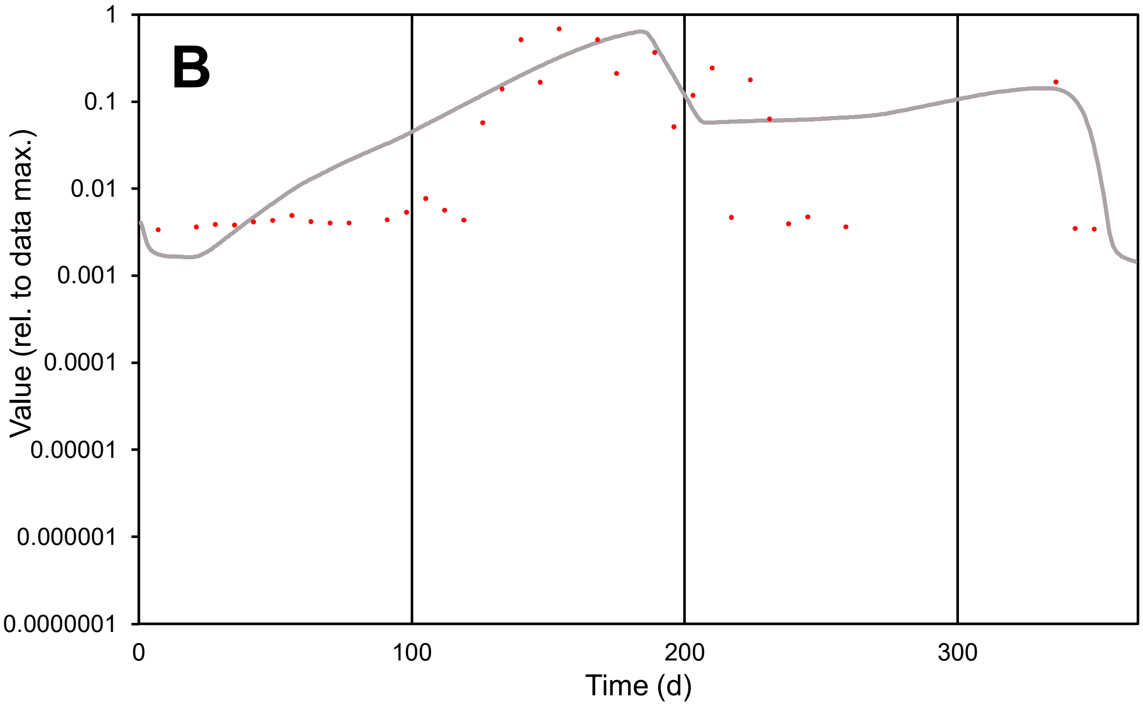


**C1**

Variables
AQU Provide jtd or Dst
CalibOnly?
2 1 to reproduce of
oNormalize?
2 1=no, 2=yes

ofxn calc*
0.95273
ofxn mod
0.98022

sRun
RunA1
zout
1

Ave
Data
0.00447
Model
0.00481

data/vMax
0.0395

ofxng
311.734

Min d/m
1.97117

ave/min d
33.391
ave/min m
70.8263

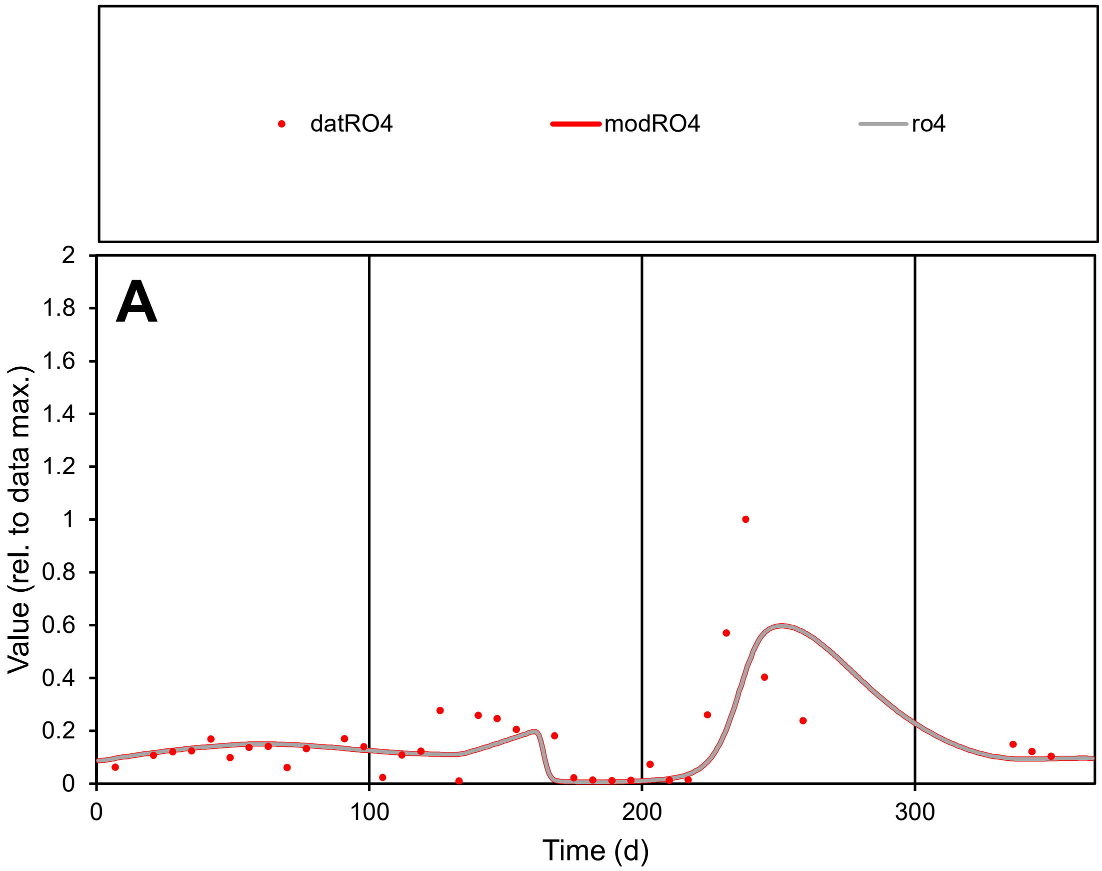


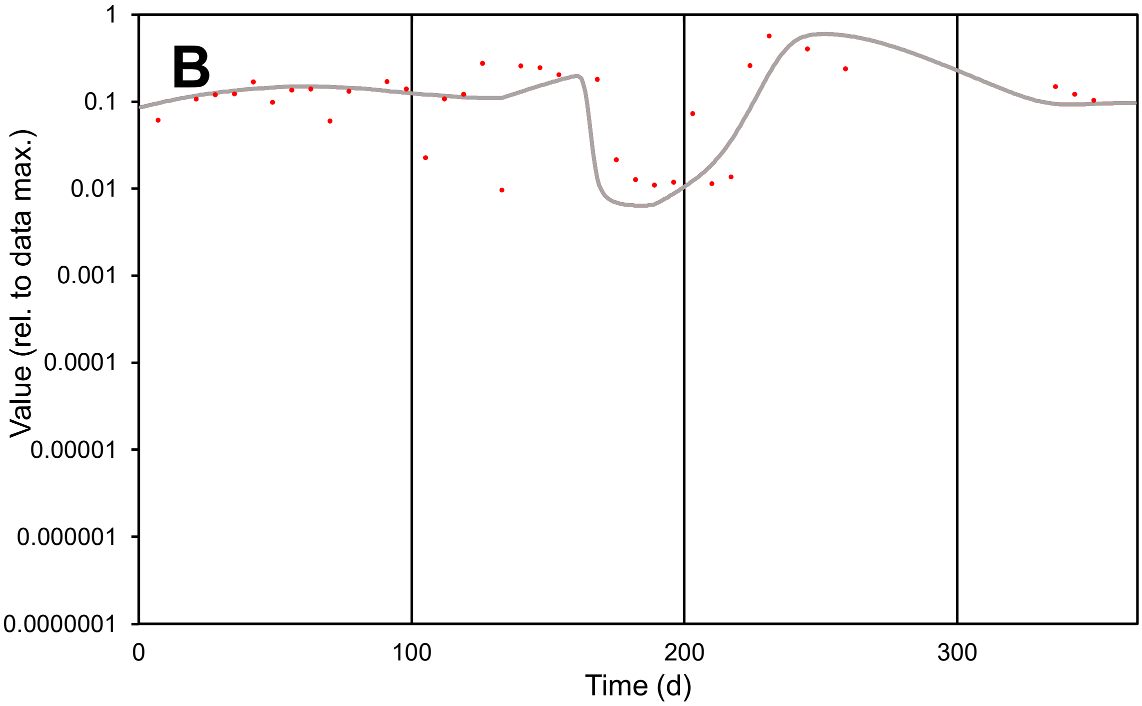


**C1**

Variables
RO4 Provide jtd or Dst
CalibOnly?
2 1 to reproduce of
oNormalize?
2 1=no, 2=yes

ofxn calc*
0.84285
ofxn mod
0.86945

sRun
RunA1
zout
1

Ave
Data
0.00349
Model
0.00319

data/vMax
0.02251

ofxng
311.734

Min d/m
1.51321

ave/min d
16.0267
ave/min m
22.1821

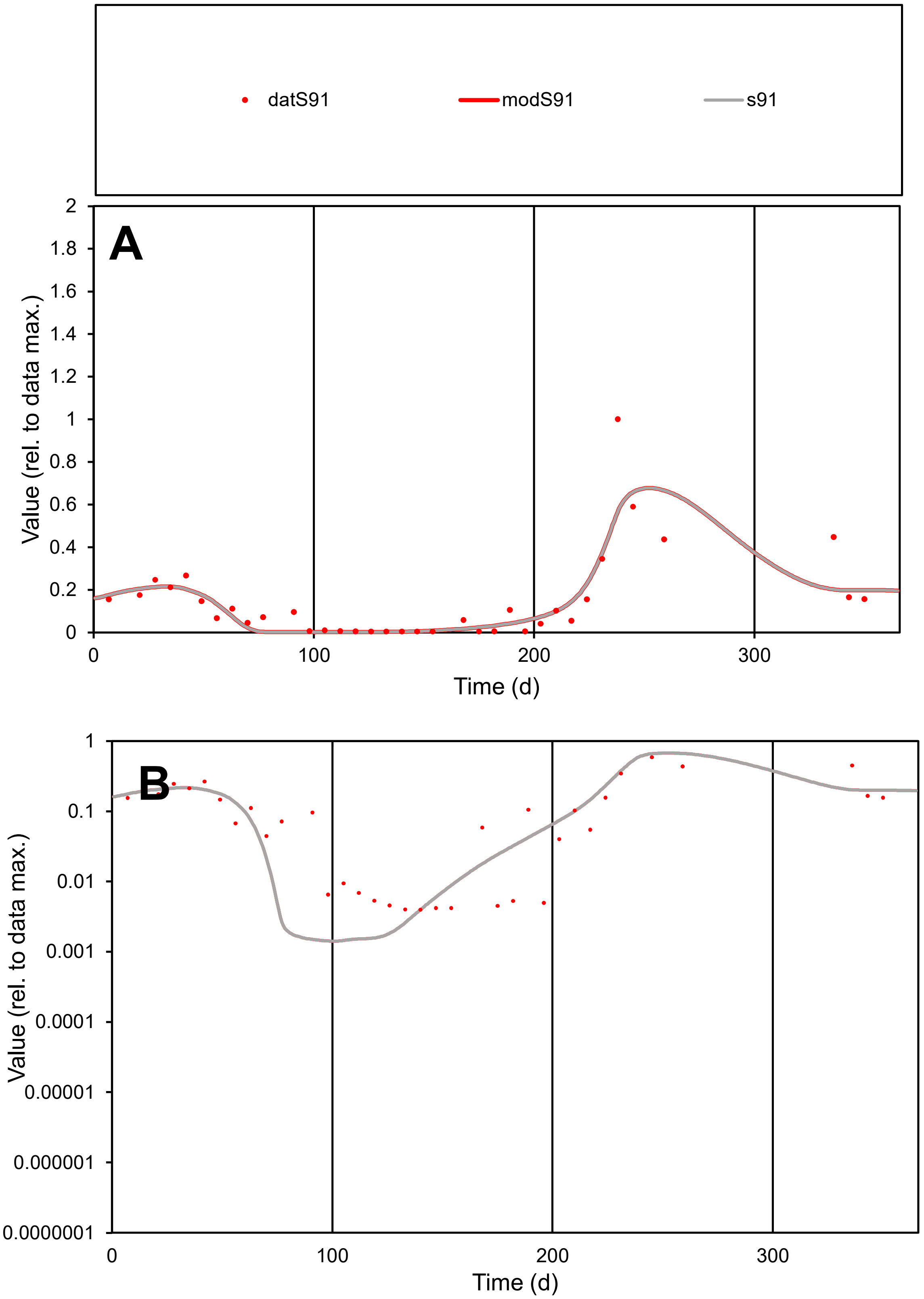


**C1**

Variables
S91 Provide jtd or Dst
CalibOnly?
2 1 to reproduce of
oNormalize?
2 1=no, 2=yes

ofxn calc*
0.65722
ofxn mod
0.68494

sRun
RunA1
zout
1

Ave
Data
0.00241
Model
0.00231

data/vMax
0.0163

ofxng
311.734

Min d/m
2.80302

ave/min d
37.0984
ave/min m
99.7178

331_Calib_S_A

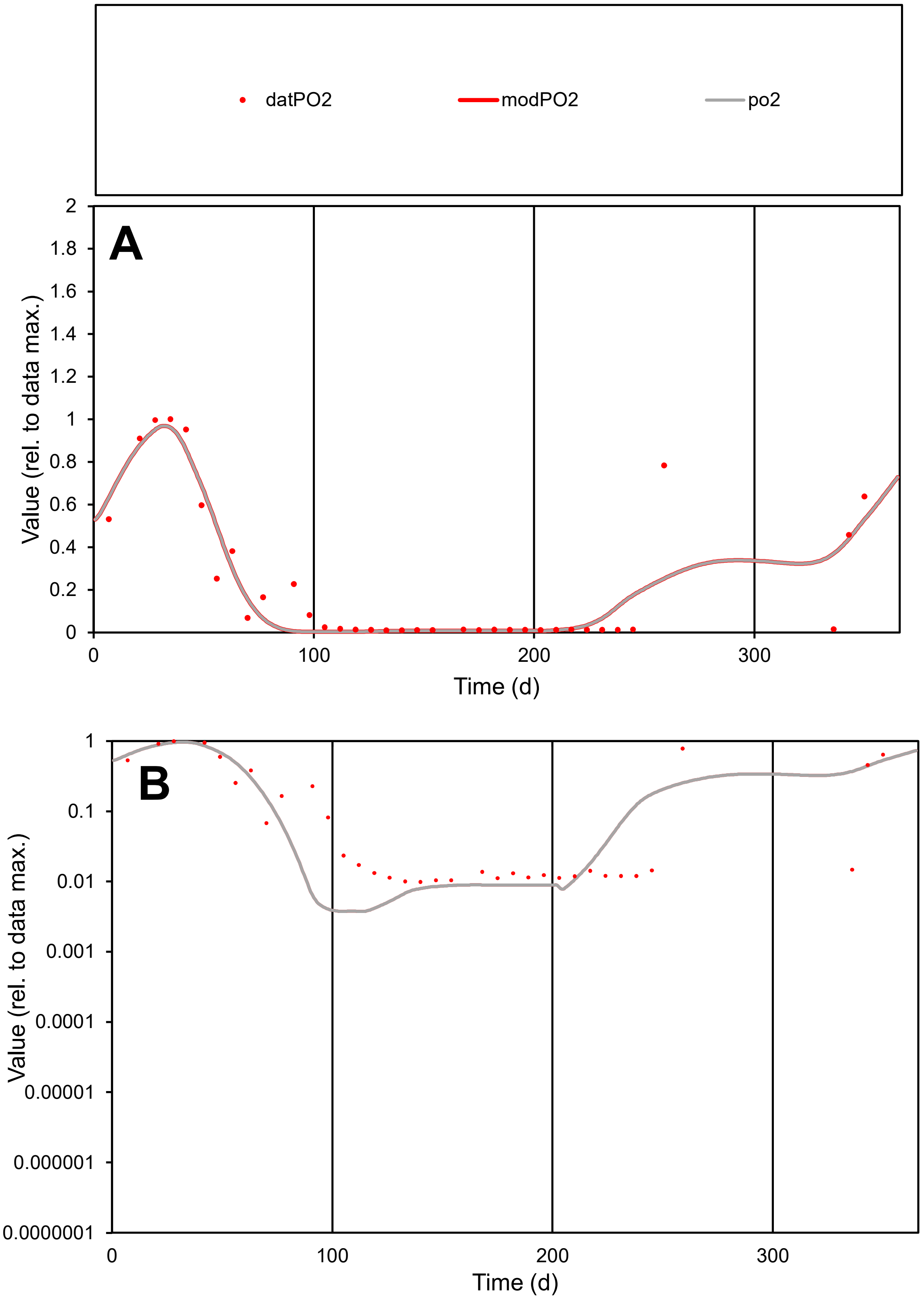


**C1**

Variables
PO2 Provide jtd or Dst
CalibOnly?
2 1 to reproduce of
oNormalize?
2 1=no, 2=yes

ofxn calc*
0.5618
ofxn mod
0.59733

sRun
RunA1
zout
1

Ave
Data
0.00183
Model
0.00177

data/vMax
0.00794

ofxng
311.734

Min d/m
2.6486

ave/min d
23.2917
ave/min m
59.582

331_Calib_S_A

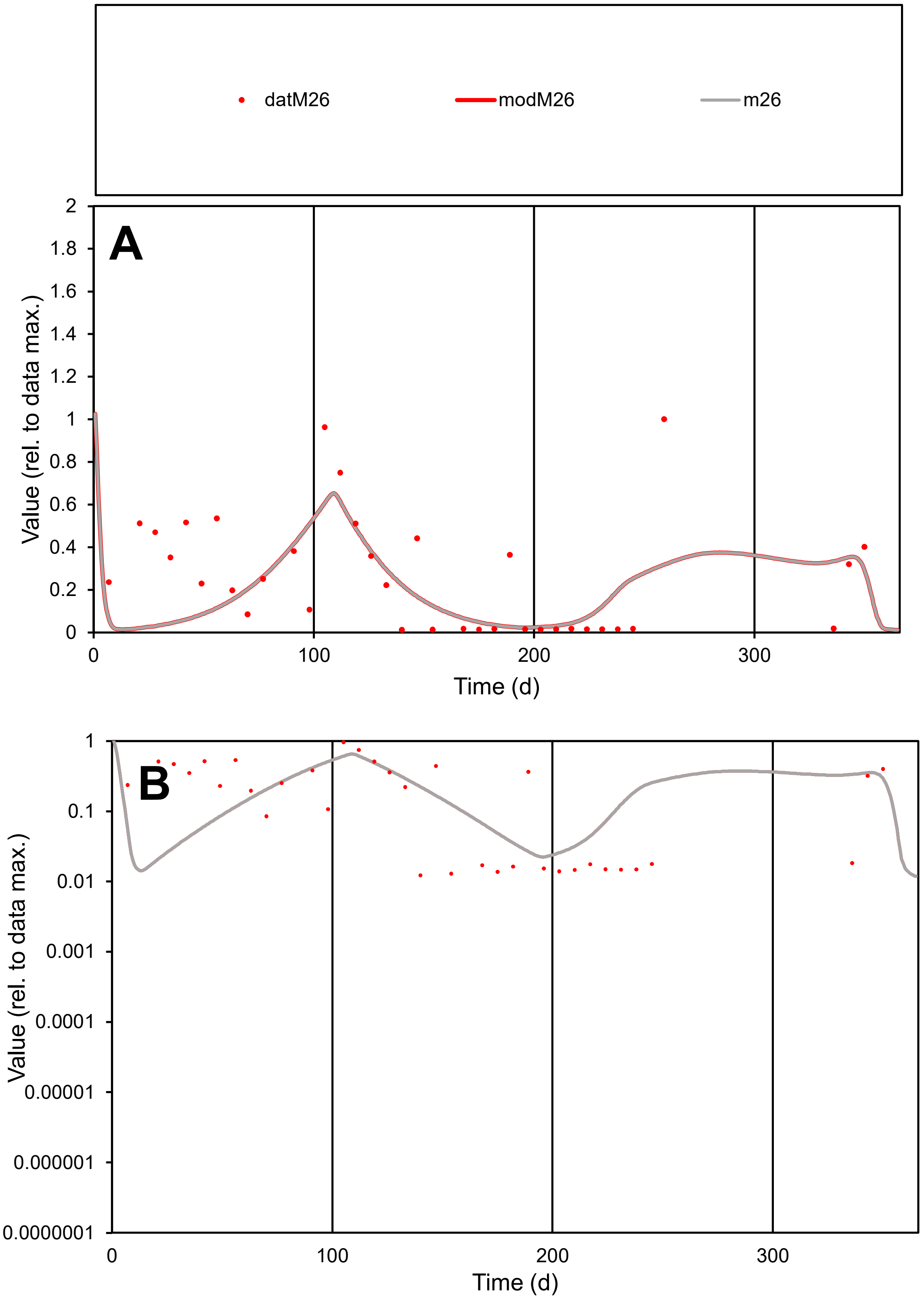


C1

Variables
M26 Provide jtd or Dst
CalibOnly?
2 1 to reproduce of
oNormalize?
2 1=no, 2=yes

ofxn calc*
0.9602
ofxn mod
0.97489

sRun
RunA1
zout
1

Ave
Data
0.00168
Model
0.00131

data/vMax
0.00661

ofxng
311.734

Min d/m
0.54138

ave/min d
20.6839
ave/min m
8.75943

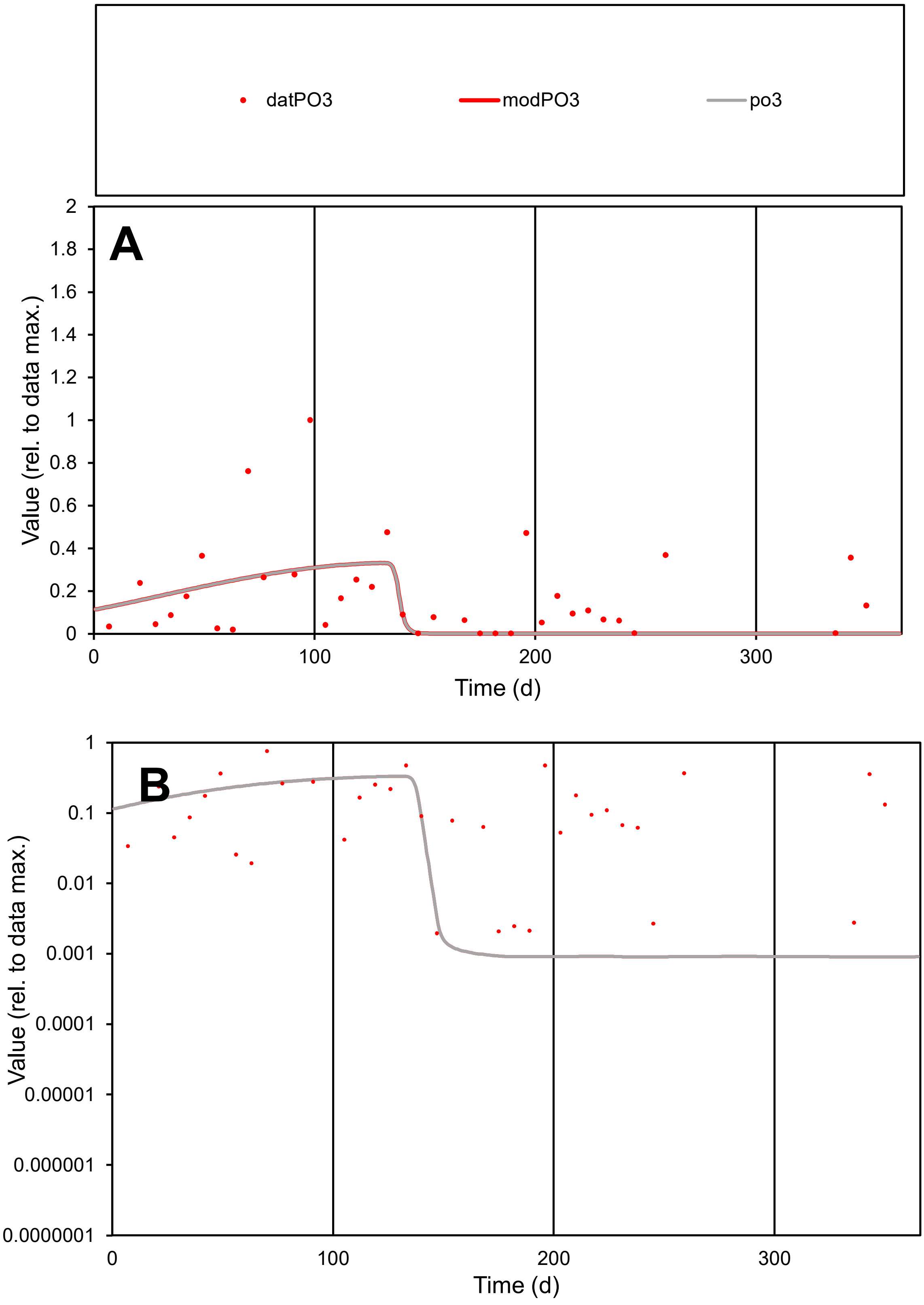


**C1**

Variables
PO3 Provide jtd or Dst
CalibOnly?
2 1 to reproduce of
oNormalize?
2 1=no, 2=yes

ofxn calc*
1.19607
ofxn mod
1.20413

sRun
RunA1
zout
1

Ave
Data
0.00436
Model
0.00305

data/vMax
0.02484

ofxng
311.734

Min d/m
2.17608
ave/min d
89.849
ave/min m
136.727

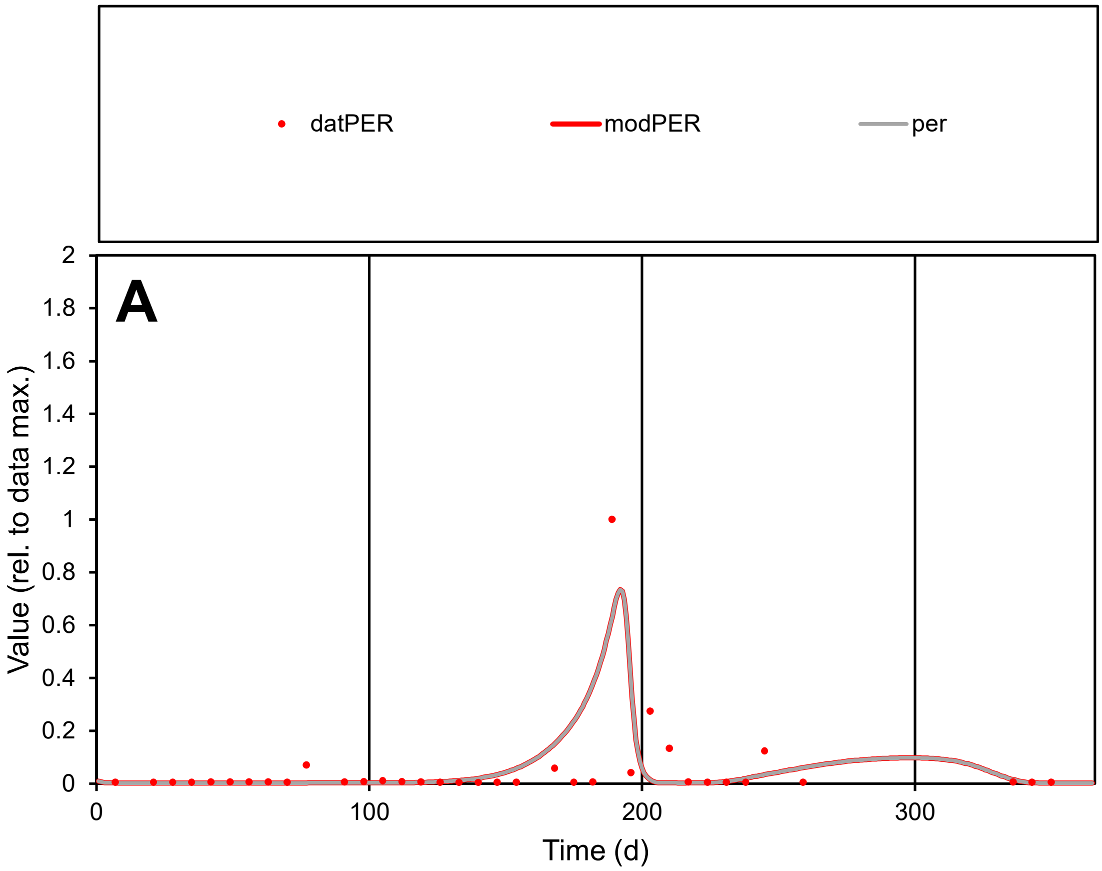


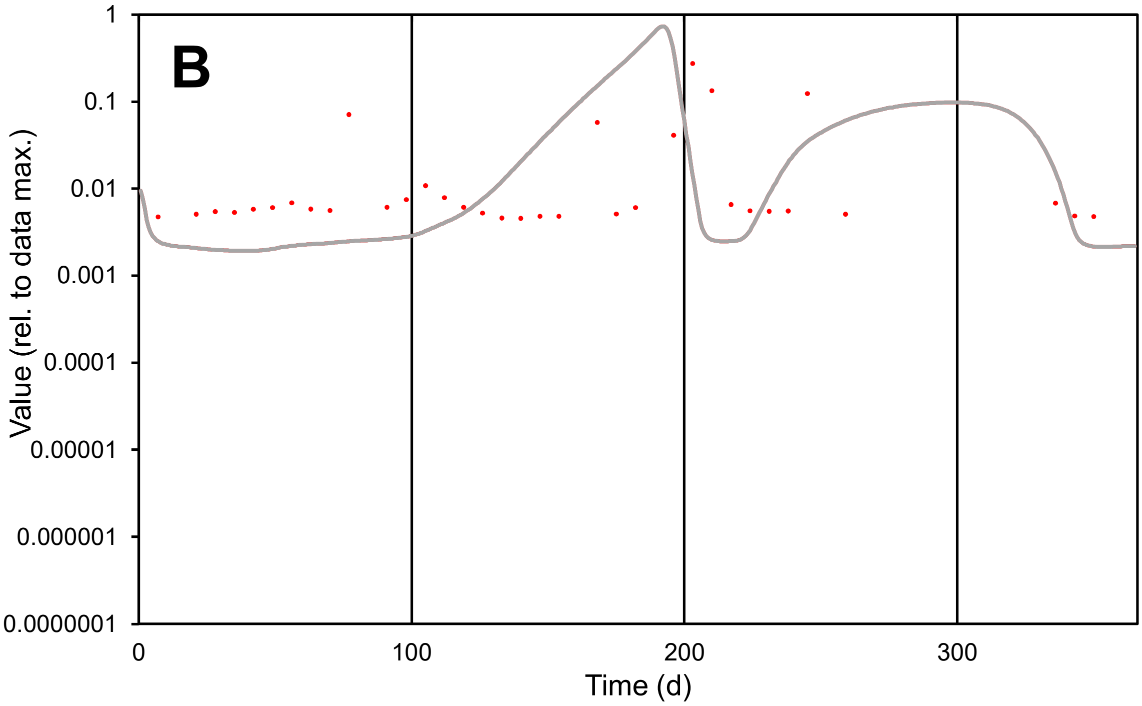


**C1**

| | |
|---|---|
| Variables | |
| PER | Provide jtd or Dst |
| CalibOnly? | |
| 2 | 1 to reproduce of |
| oNormalize? | |
| 2 | 1=no, 2=yes |
| ofxn calc* | |
| 1.94703 | |
| ofxn mod | |
| 2.06039 | |
| sRun | |
| RunA1 | |
| zout | |
| 1 | |
| Ave | |
| Data | |
| 0.00021 | |
| Model | |
| 0.00021 | |
| data/vMax | |
| 0.00397 | |
| ofxng | |
| 311.734 | |
| Min d/m | |
| 2.36546 | |
| ave/min d | |
| 11.3797 | |
| ave/min m | |
| 26.824 | |

331_Calib_S_A

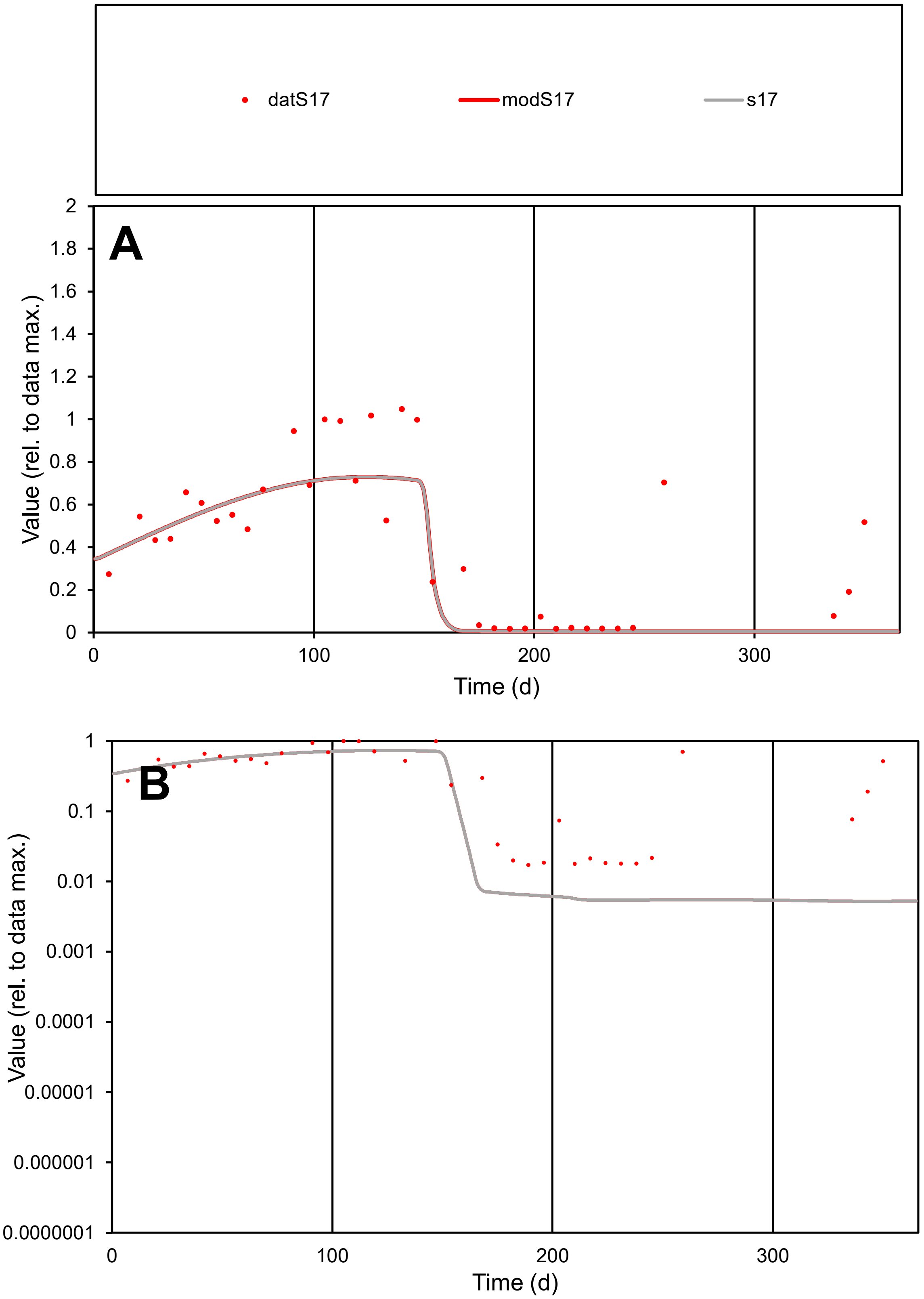


**C1**

Variables
S17 Provide jtd or Dst
CalibOnly?
2 1 to reproduce of
oNormalize?
2 1=no, 2=yes

ofxn calc*
0.48369
ofxn mod
0.50144

sRun
RunA1
zout
1

Ave
Data
0.00284
Model
0.00244

data/vMax
0.0072

ofxng
311.734

Min d/m
3.28738

ave/min d
22.8805
ave/min m
64.5712

331_Calib_S_A

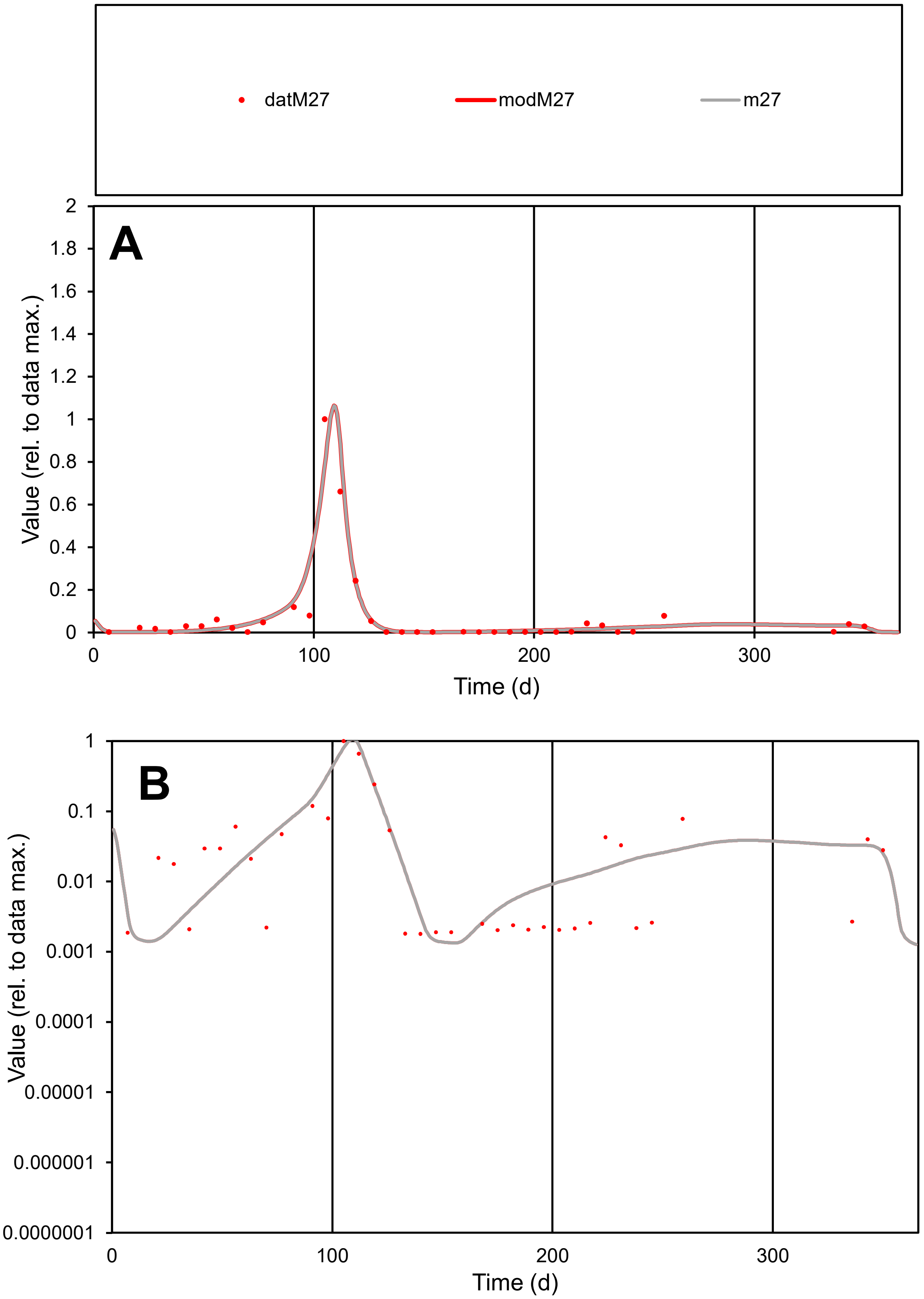


**C1**

Variables
M27 Provide jtd or Dst
CalibOnly?
2 1 to reproduce of
oNormalize?
2 1=no, 2=yes

ofxn calc*
0.56209
ofxn mod
0.7349

sRun
RunA1
zout
1

Ave
Data
0.00261
Model
0.00267

data/vMax
0.03575

ofxng
311.734

Min d/m
1.34956

ave/min d
40.5536
ave/min m
56.0829

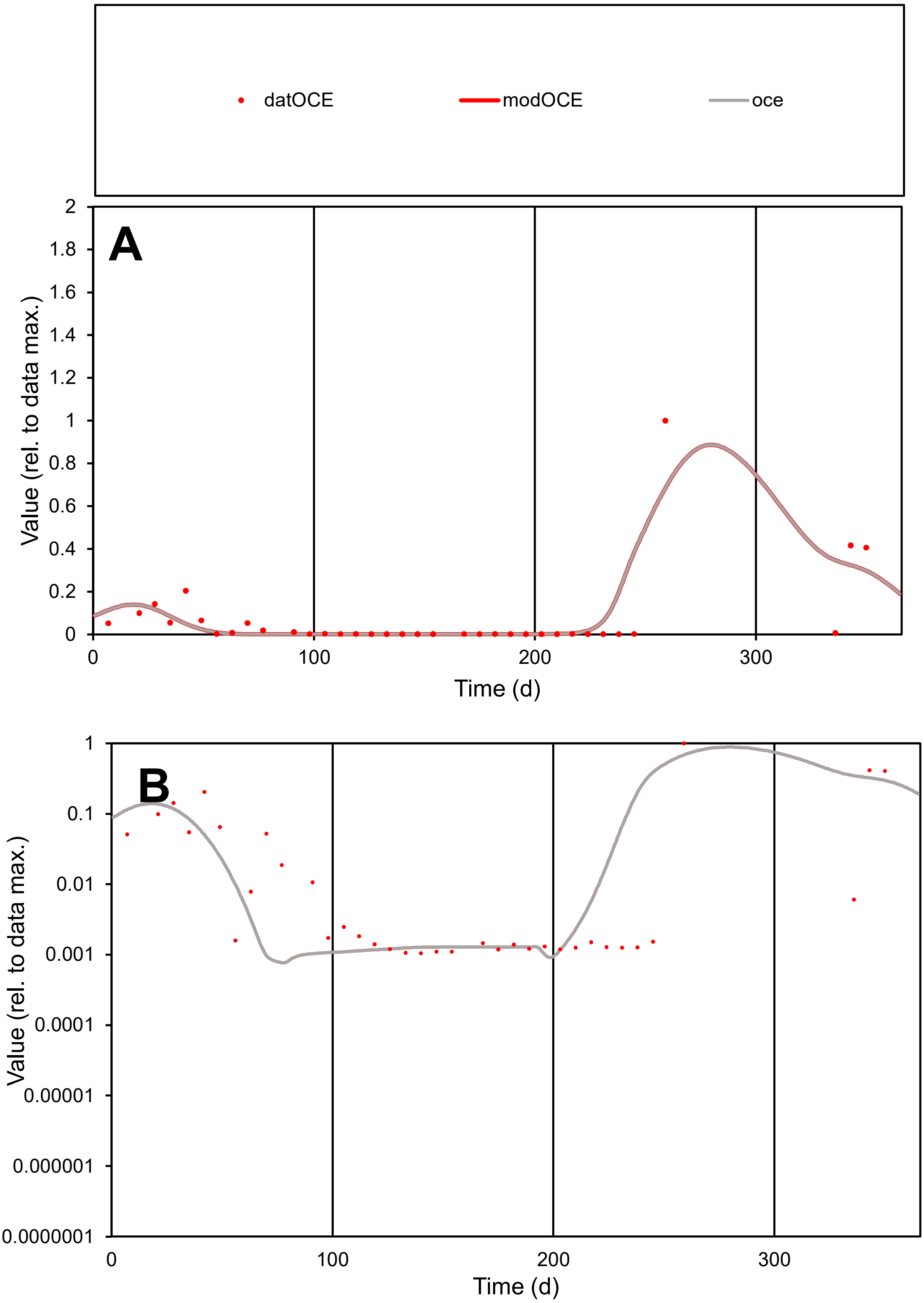


**C1**

Variables
OCE Provide jtd or Dst
CalibOnly?
2 1 to reproduce of
oNormalize?
2 1=no, 2=yes

ofxn calc*
1.44165
ofxn mod
1.56623

sRun
RunA1
zout
1

Ave
Data
0.00249
Model
0.00267

data/vMax
0.03496

ofxng
311.734

Min d/m
1.36024

ave/min d
67.7569
ave/min m
98.7077

331_Calib_S_A

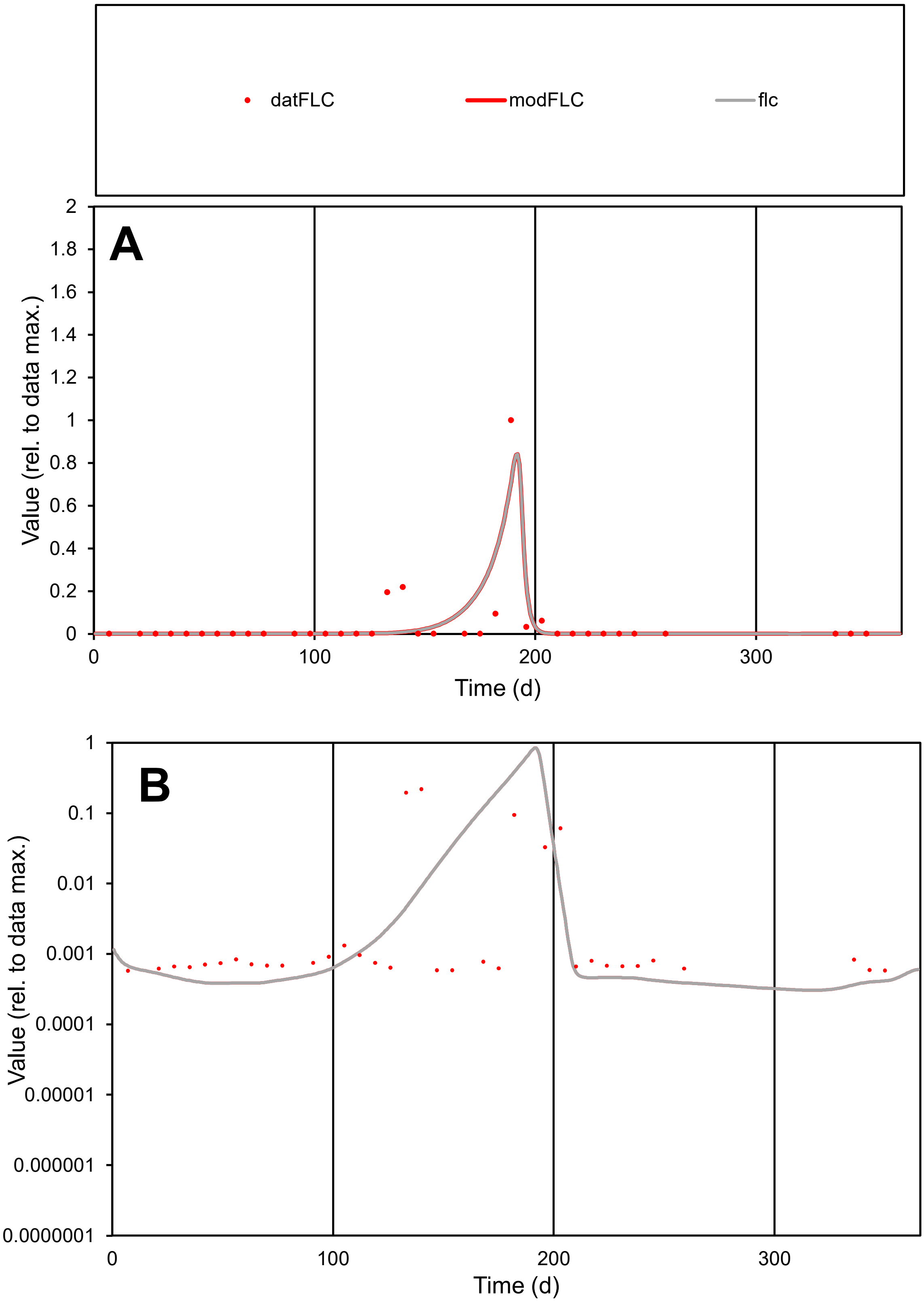


**C1**

Variables
FLC Provide jtd or Dst
CalibOnly?
2 1 to reproduce of
oNormalize?
2 1=no, 2=yes

ofxn calc*
1.71927
ofxn mod
1.9236

sRun
RunA1
zout
1

Ave
Data
0.00088
Model
0.00097

data/vMax
0.02139

ofxng
311.734

Min d/m
1.55002

ave/min d
70.7119
ave/min m
120.847

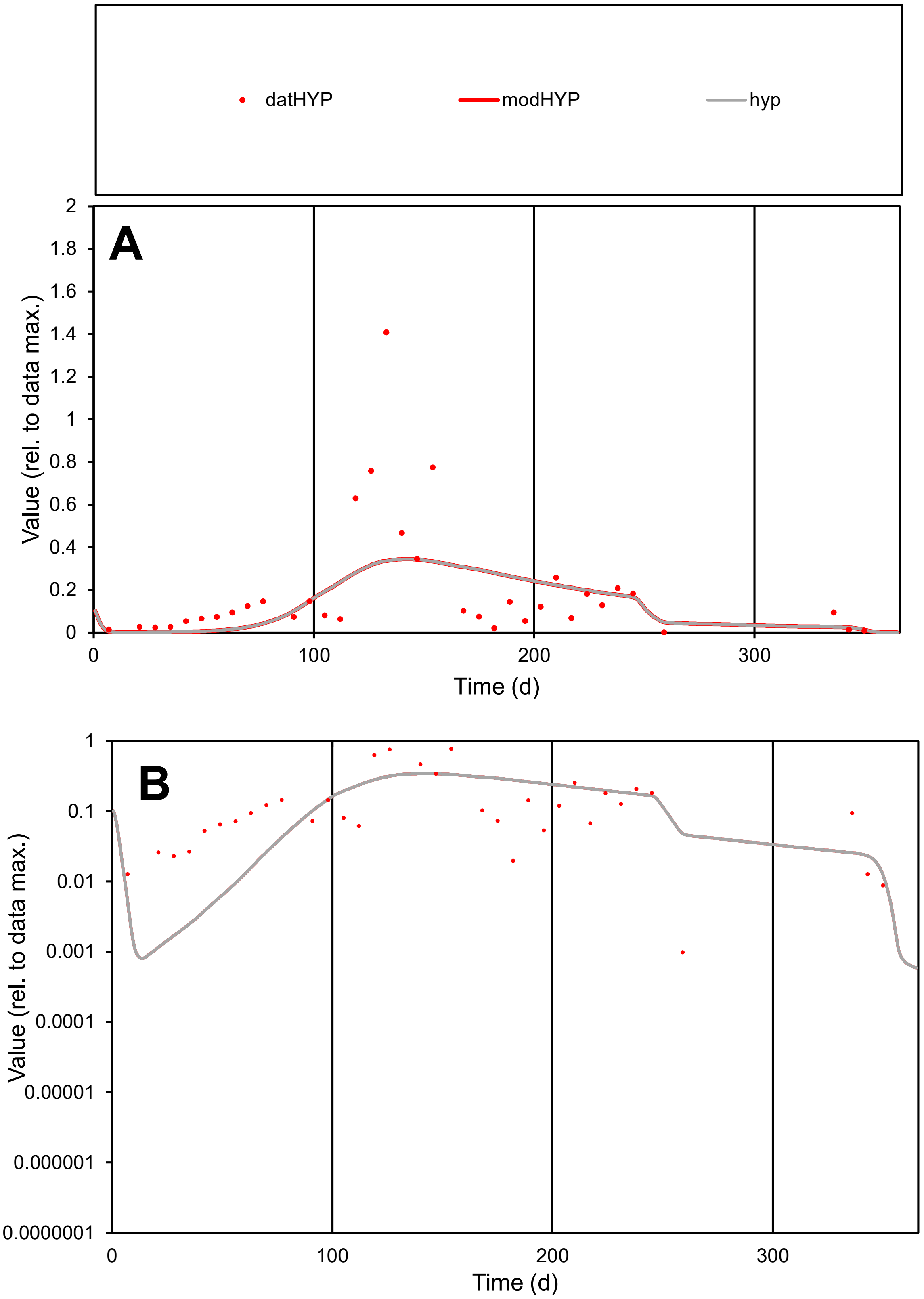


**C1**

Variables
HYP Provide jtd or Dst
CalibOnly?
2 1 to reproduce of
oNormalize?
2 1=no, 2=yes

ofxn calc*
1.01785
ofxn mod
1.02928

sRun
RunA1
zout
1

Ave
Data
0.00294
Model
0.0028

data/vMax
0.01788

ofxng
311.734

Min d/m
0.78183

ave/min d
167.23
ave/min m
124.83

331_Calib_S_A

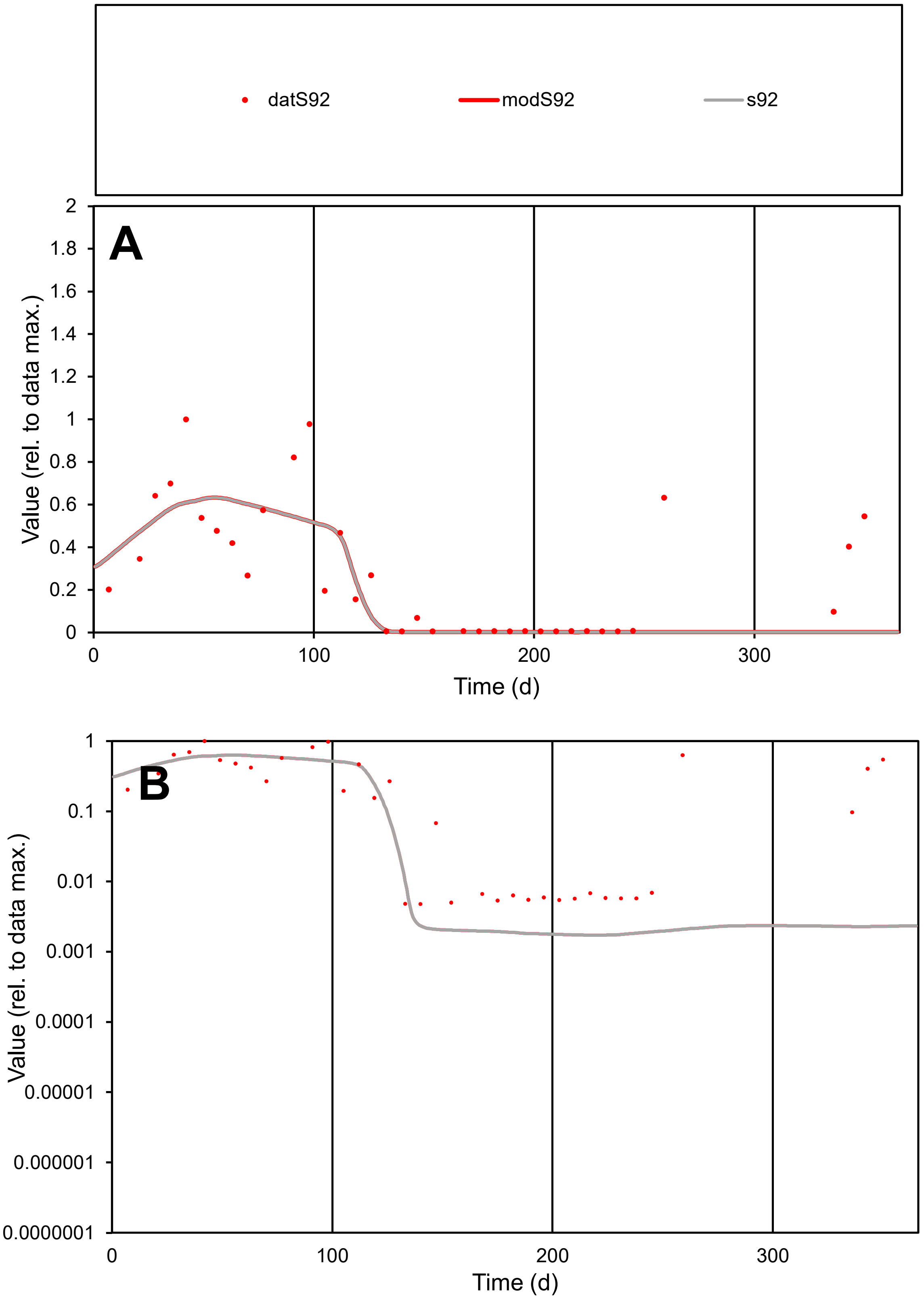


**C1**

Variables
S92 Provide jtd or Dst
CalibOnly?
2 1 to reproduce of
oNormalize?
2 1=no, 2=yes

ofxn calc*
0.79072
ofxn mod
0.80831

sRun
RunA1
zout
1

Ave
Data
0.00131
Model
0.00107

data/vMax
0.00482

ofxng
311.734

Min d/m
2.76882

ave/min d
56.8588
ave/min m
128.358

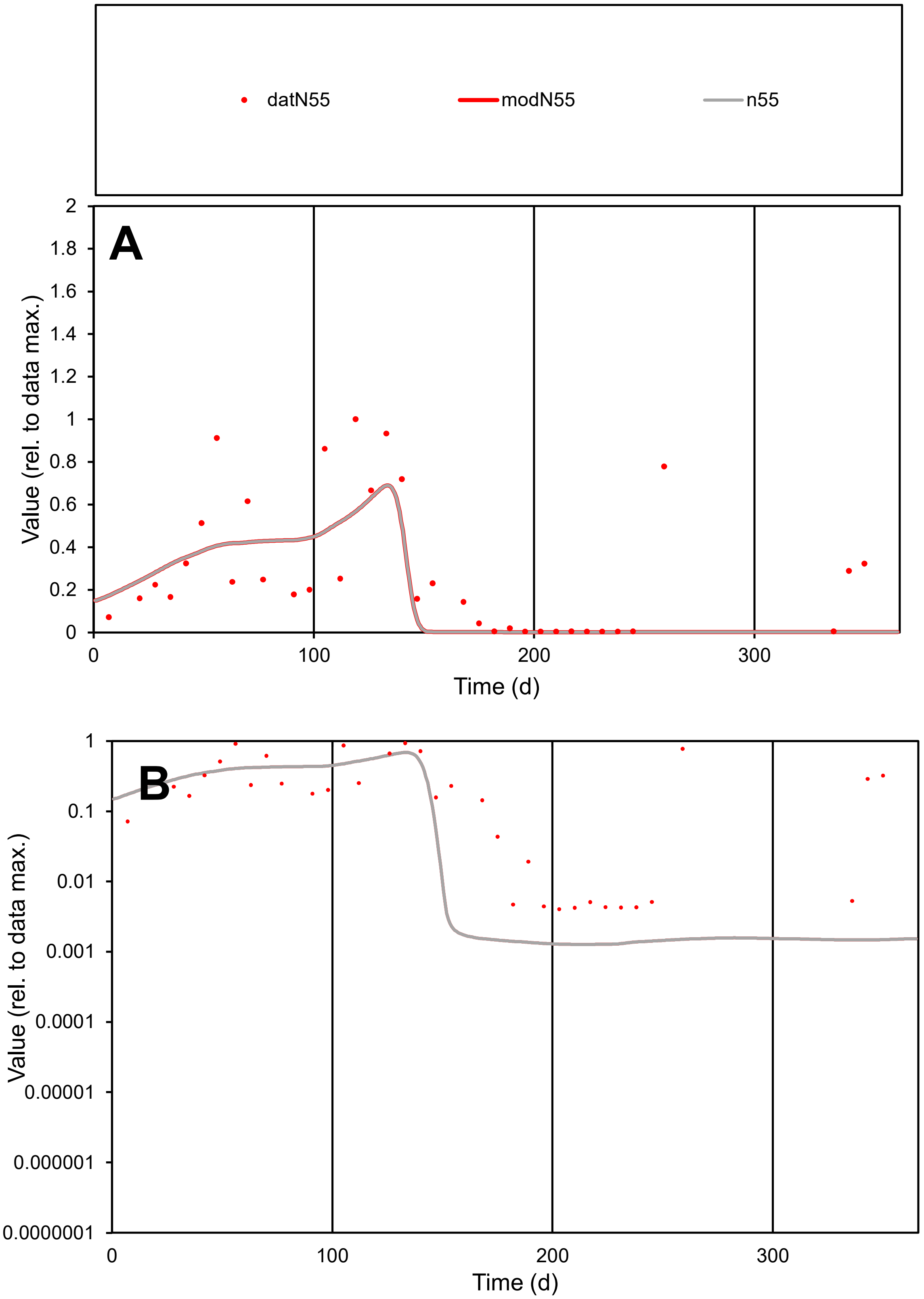


**C1**

Variables
N55 Provide jtd or Dst
CalibOnly?
2 1 to reproduce of
oNormalize?
2 1=no, 2=yes

ofxn calc*
0.85703
ofxn mod
0.8683

sRun
RunA1
zout
1

Ave
Data
0.00273
Model
0.00226

data/vMax
0.01041

ofxng
311.734

Min d/m
3.18196

ave/min d
65.0297
ave/min m
171.061

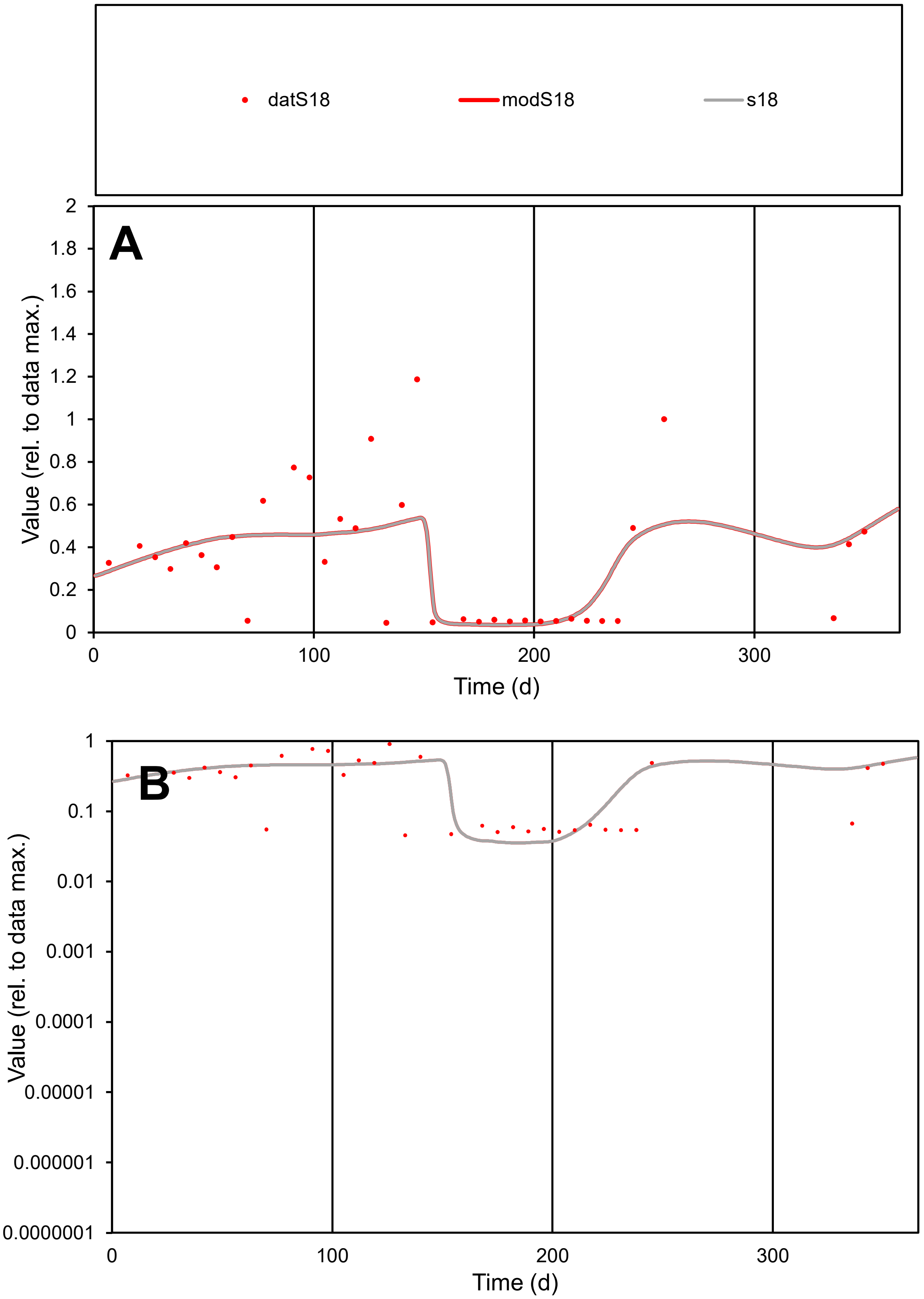


**C1**

Variables
S18 Provide jtd or Dst
CalibOnly?
2 1 to reproduce of
oNormalize?
2 1=no, 2=yes

ofxn calc*
0.58778
ofxn mod
0.60802

sRun
RunA1
zout
1

Ave
Data
0.00187
Model
0.00193

data/vMax
0.00588

ofxng
311.734

Min d/m
1.28214

ave/min d
7.02044
ave/min m
9.2867

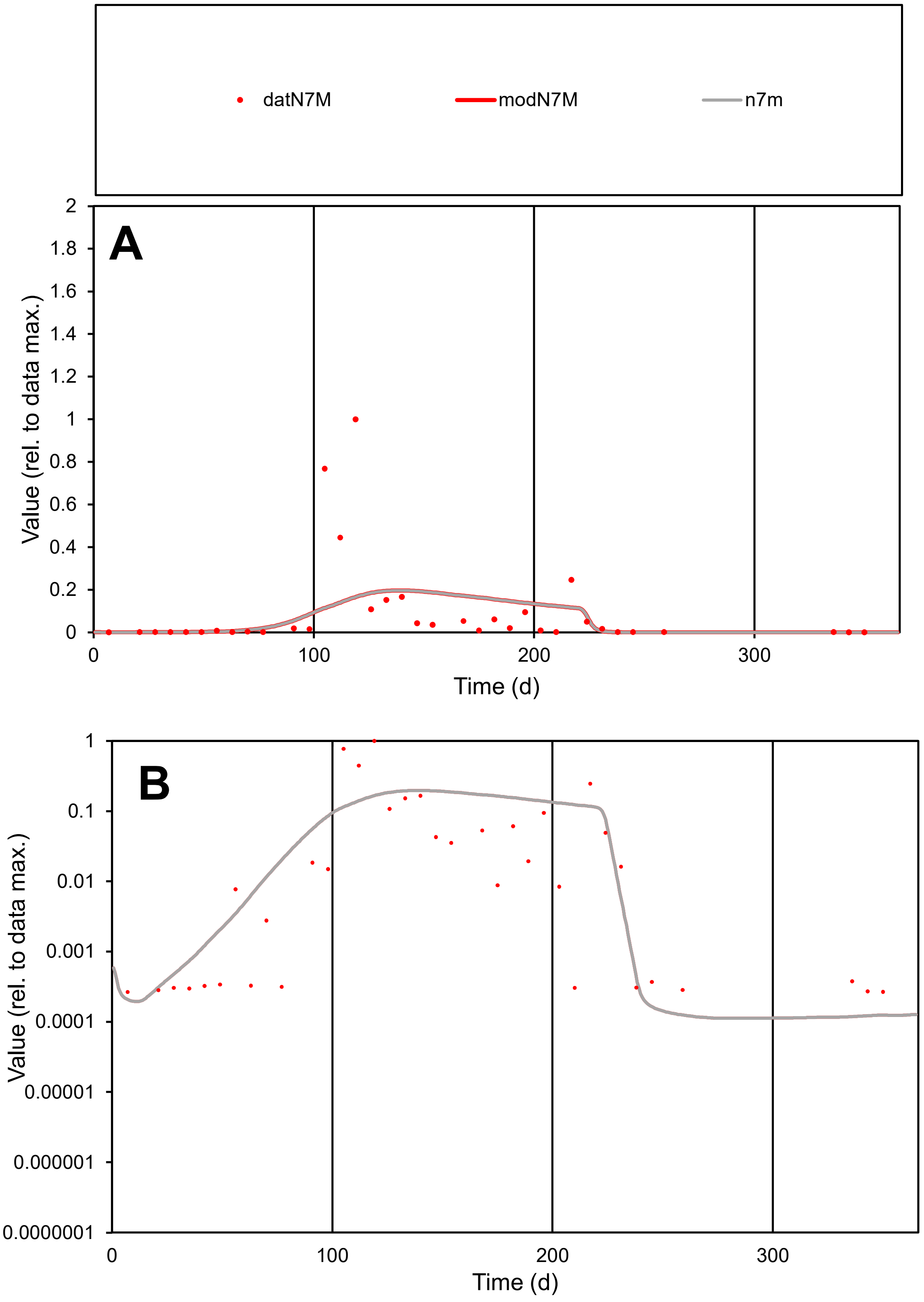


**C1**

Variables
N7M Provide jtd or Dst
CalibOnly?
2 1 to reproduce of
oNormalize?
2 1=no, 2=yes

ofxn calc*
2.24978
ofxn mod
2.25399

sRun
RunA1
zout
1

Ave
Data
0.00328
Model
0.00288

data/vMax
0.03743

ofxng
311.734

Min d/m
2.21301

ave/min d
331.296
ave/min m
644.091

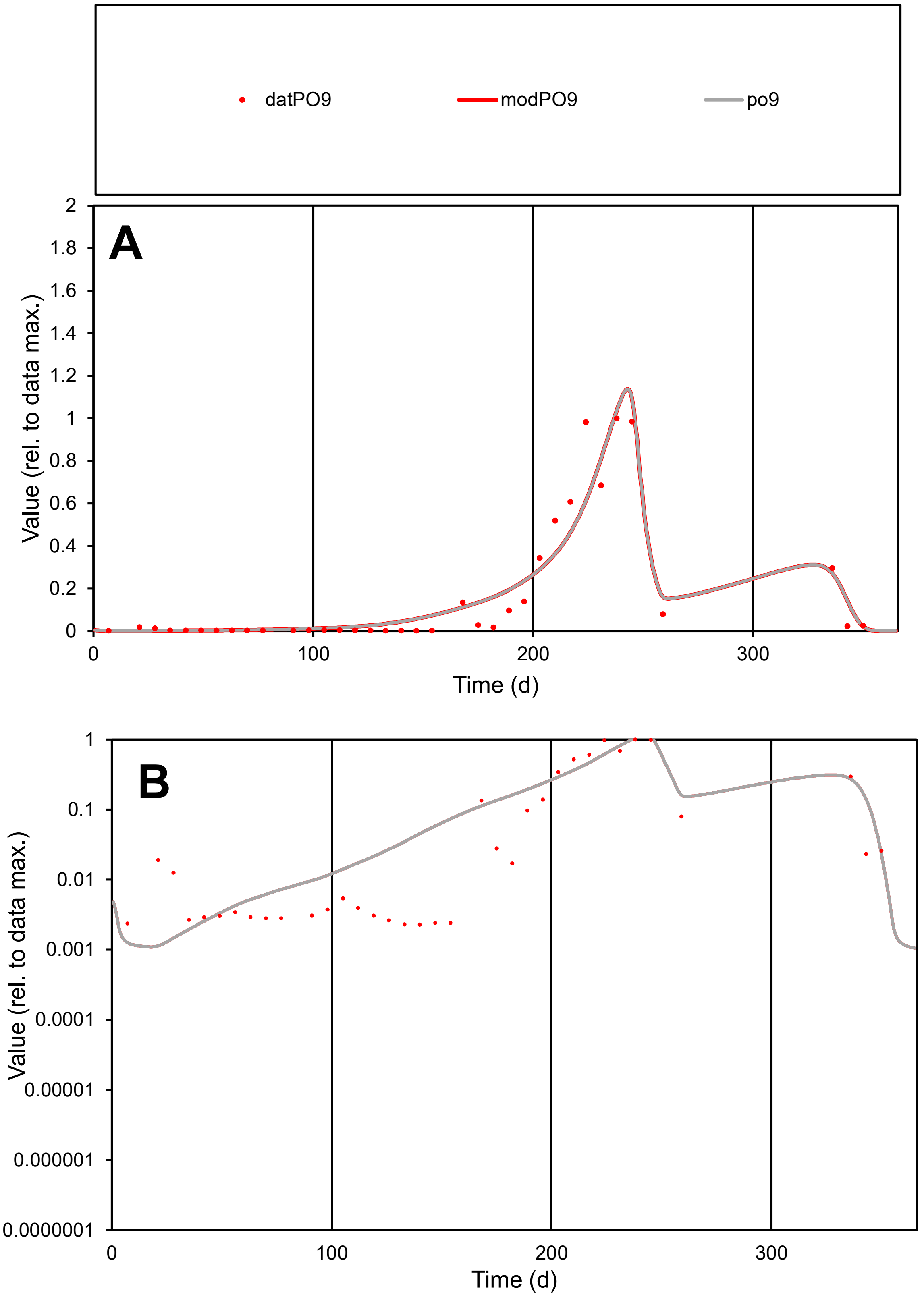


**C1**

Variables

PO9 Provide jtd or Dst

CalibOnly?

2 1 to reproduce of

oNormalize?

2 1=no, 2=yes

ofxn calc*

0.49194

ofxn mod

0.56381

sRun

RunA1

zout

1

Ave

Data

0.00425

Model

0.0045

data/vMax

0.02531

ofxng

311.734

Min d/m

1.93077

ave/min d

73.6246

ave/min m

150.668

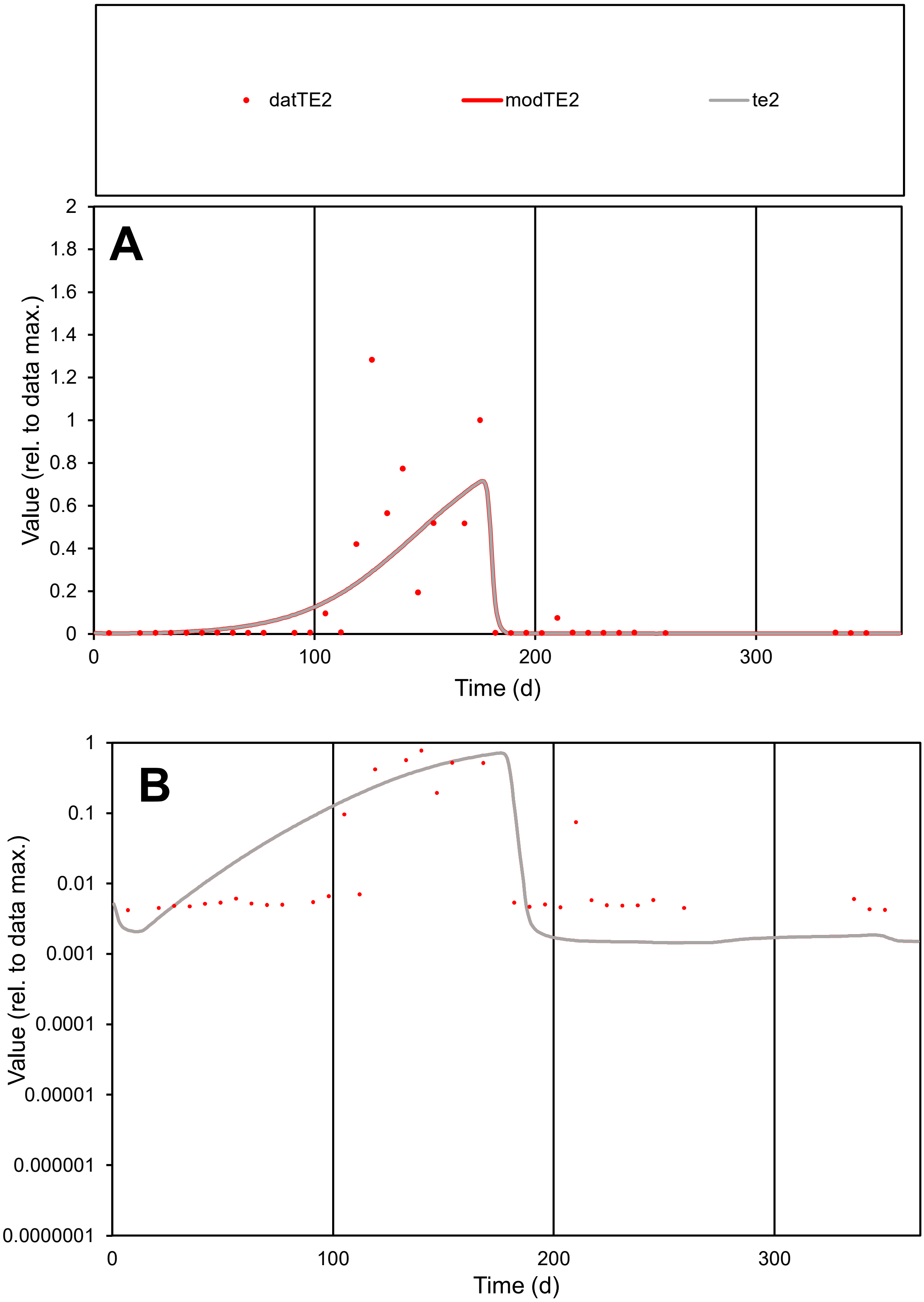


**C1**

Variables
TE2 Provide jtd or Dst
CalibOnly?
2 1 to reproduce of
oNormalize?
2 1=no, 2=yes

ofxn calc*
1.12687
ofxn mod
1.14306

sRun
RunA1
zout
1

Ave
Data
0.00262
Model
0.00263

data/vMax
0.02088

ofxng
311.734

Min d/m
2.90479

ave/min d
29.8619
ave/min m
87.092

331_Calib_S_A

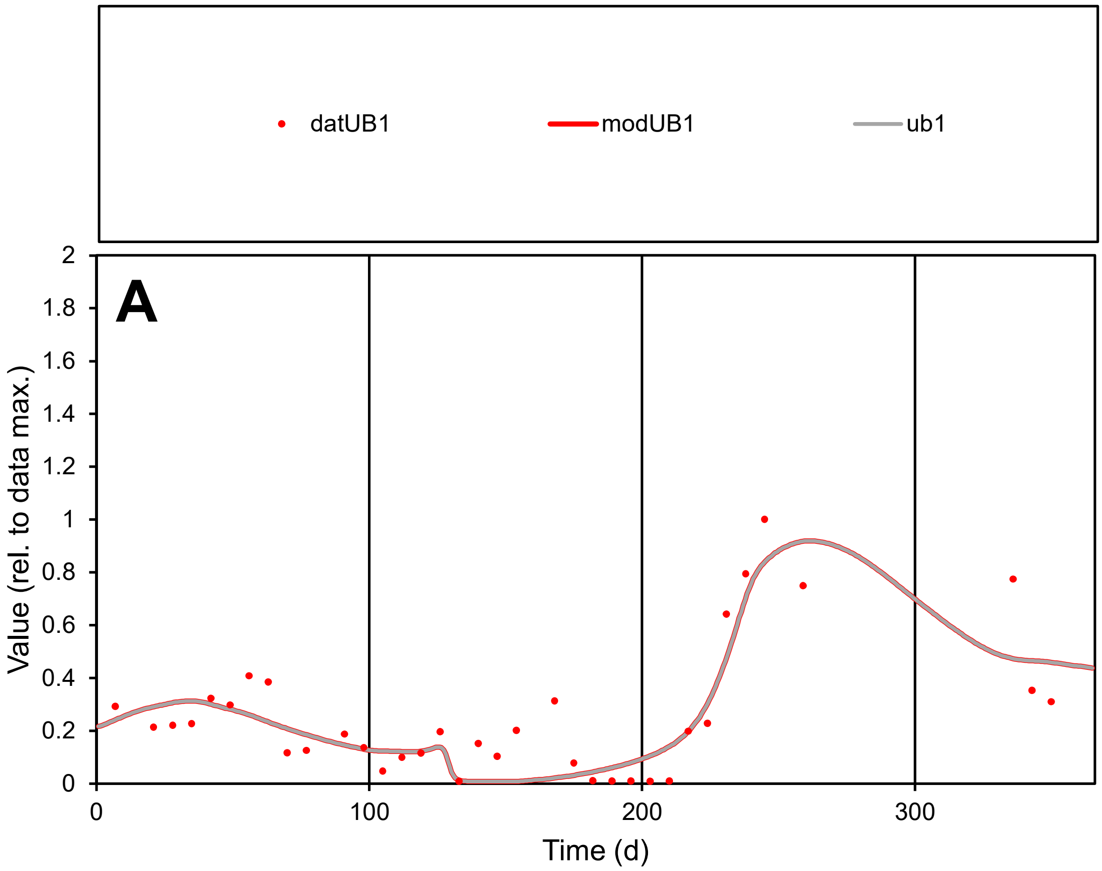


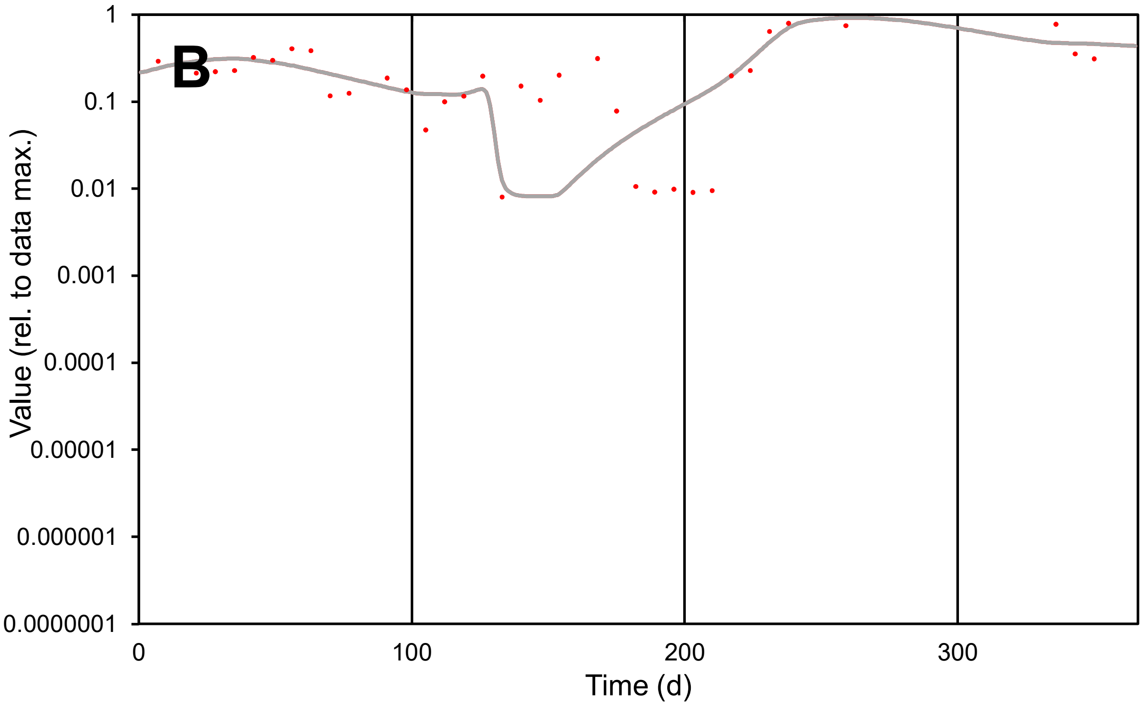


**C1**

| | |
|---|---|
| Variables | |
| UB1 | Provide jtd or Dst |
| CalibOnly? | |
| 2 | 1 to reproduce of |
| oNormalize? | |
| 2 | 1=no, 2=yes |
| ofxn calc* | |
| 0.43471 | |
| ofxn mod | |
| 0.46385 | |
| sRun | |
| RunA1 | |
| zout | |
| 1 | |
| Ave | |
| Data | |
| 0.0018 | |
| Model | |
| 0.00175 | |
| data/vMax | |
| 0.0071 | |
| ofxng | |
| 311.734 | |
| Min d/m | |
| 0.98211 | |
| ave/min d | |
| 31.6009 | |
| ave/min m | |
| 30.0889 | |

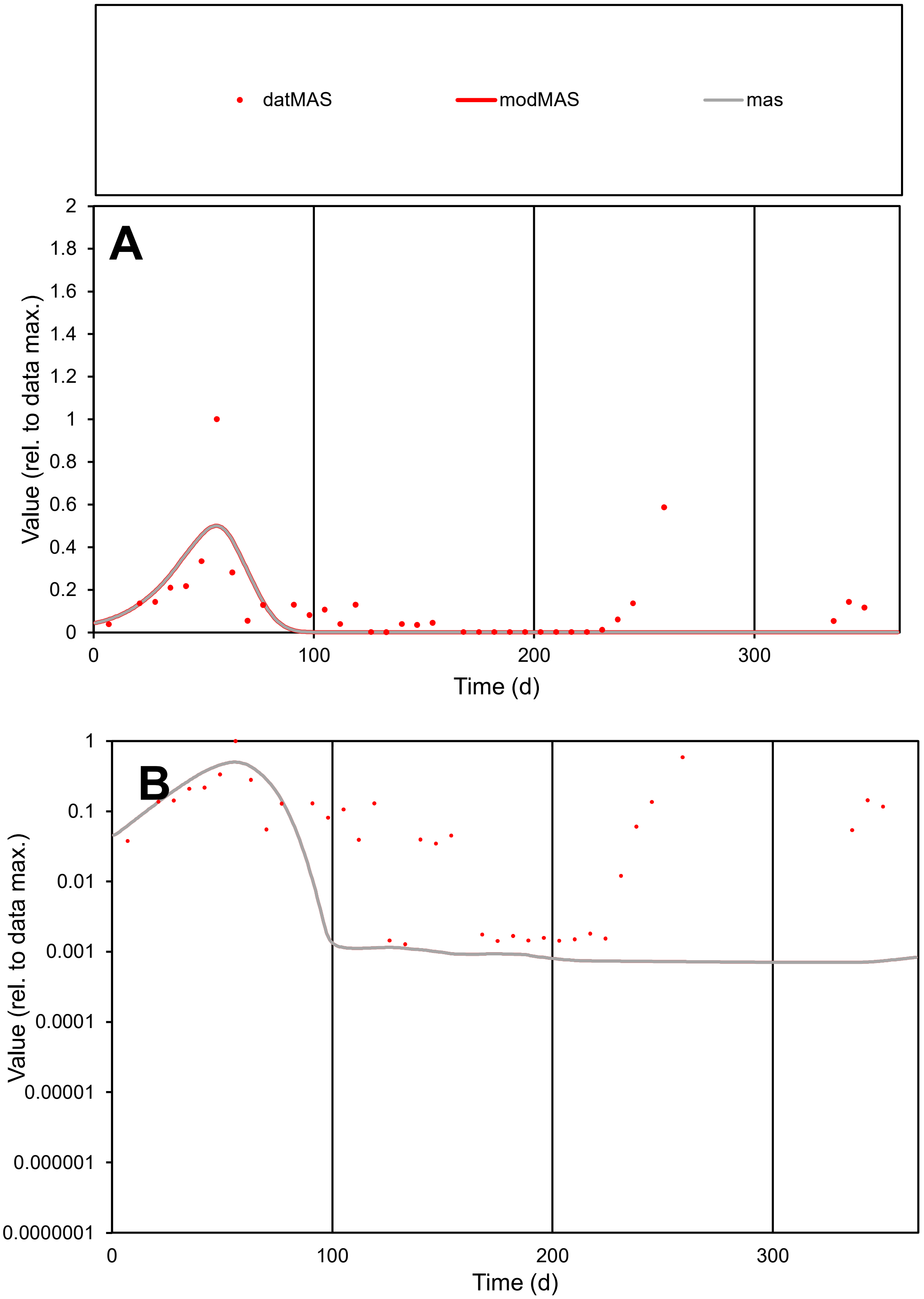


**C1**

Variables
MAS Provide jtd or Dst
CalibOnly?
2 1 to reproduce of
oNormalize?
2 1=no, 2=yes

ofxn calc*
1.26247
ofxn mod
1.29161

sRun
RunA1
zout
1

Ave
Data
0.00111
Model
0.00073

data/vMax
0.00944

ofxng
311.734

Min d/m
1.8009

ave/min d
92.3111
ave/min m
109.575

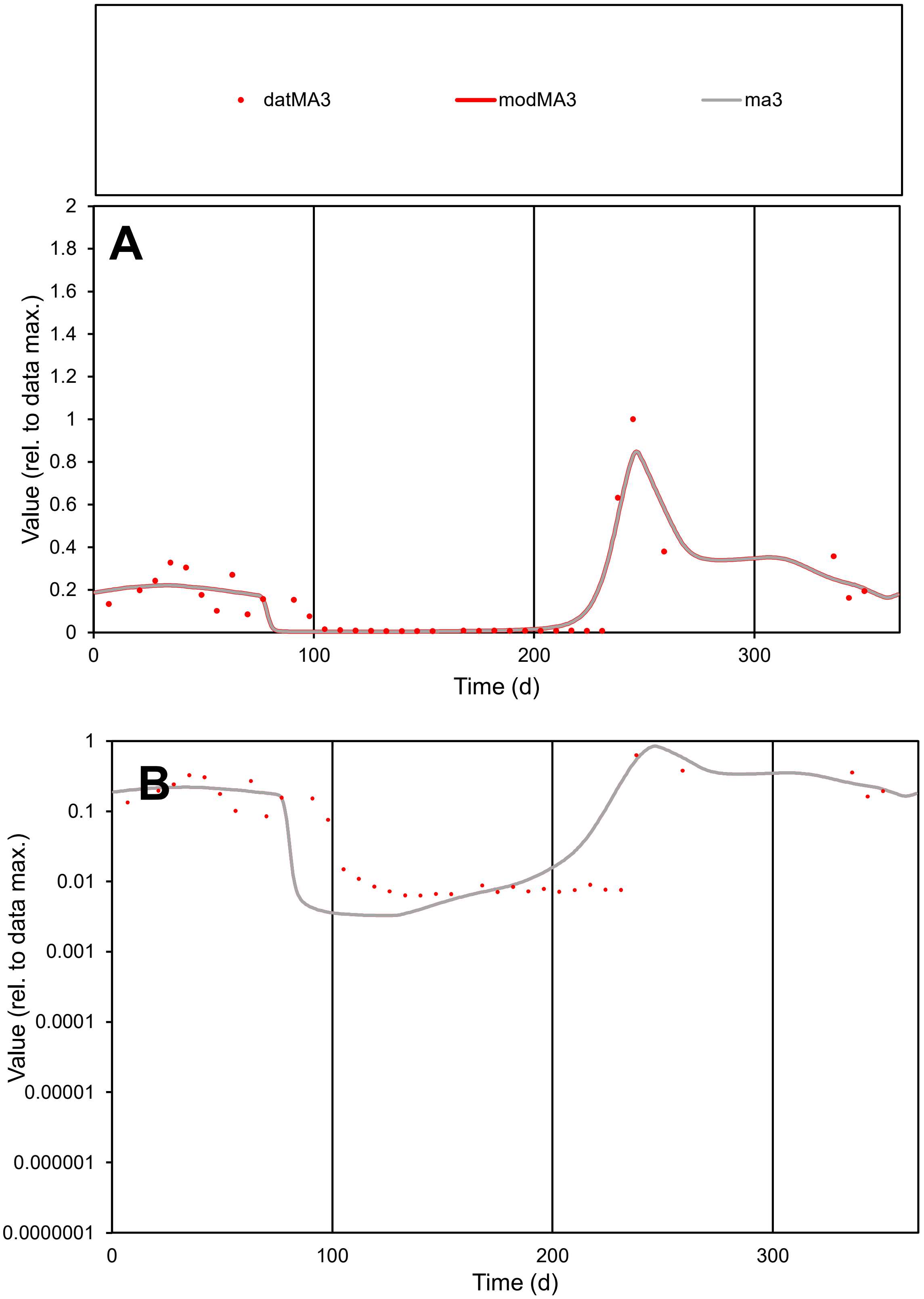


**C1**

Variables
MA3 Provide jtd or Dst
CalibOnly?
2 1 to reproduce of
oNormalize?
2 1=no, 2=yes

ofxn calc*
0.52045
ofxn mod
0.57543

sRun
RunA1
zout
1

Ave
Data
0.00138
Model
0.00139

data/vMax
0.00974

ofxng
311.734

Min d/m
1.92409

ave/min d
22.452
ave/min m
43.5997

331_Calib_S_A

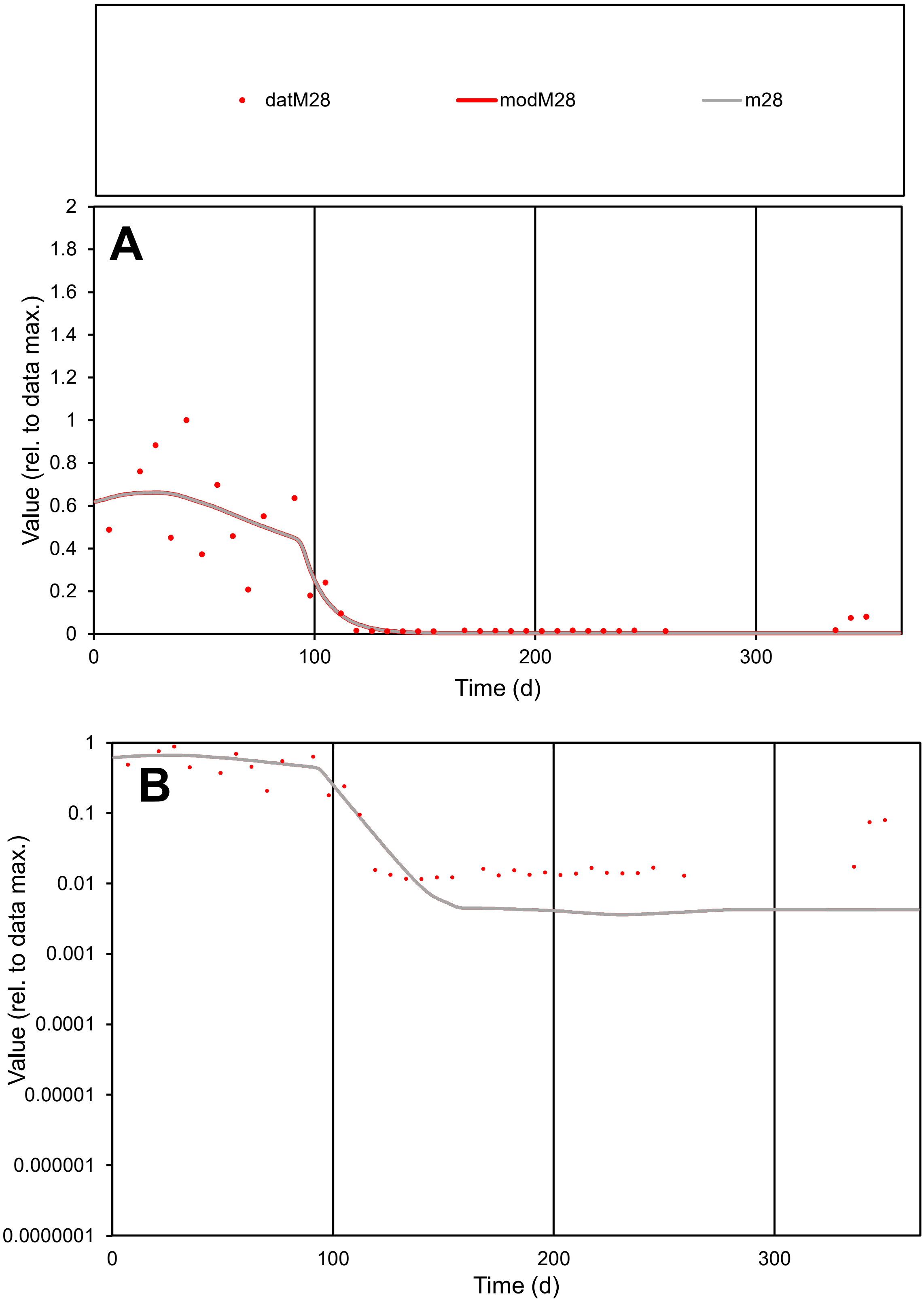


**C1**

Variables
M28 Provide jtd or Dst
CalibOnly?
2 1 to reproduce of
oNormalize?
2 1=no, 2=yes

ofxn calc*
0.57203
ofxn mod
0.60538

sRun
RunA1
zout
1

Ave
Data
0.00095
Model
0.00091

data/vMax
0.0046

ofxng
311.734

Min d/m
3.20806

ave/min d
17.8333
ave/min m
54.6021

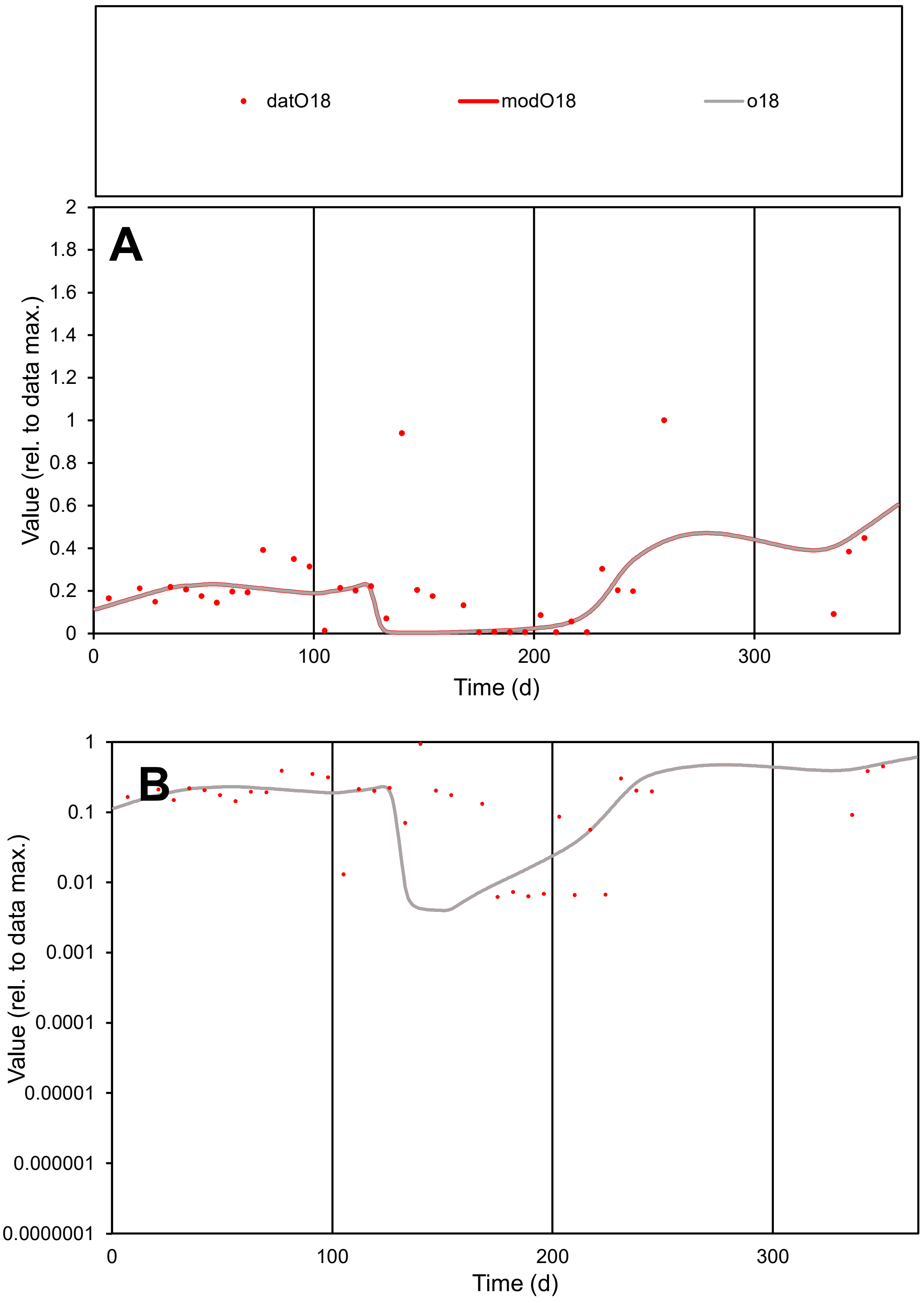


**C1**

Variables
O18 Provide jtd or Dst
CalibOnly?
2 1 to reproduce of
oNormalize?
2 1=no, 2=yes

ofxn calc*
0.81556
ofxn mod
0.83037

sRun
RunA1
zout
1

Ave
Data
0.00183
Model
0.00158

data/vMax
0.00926

ofxng
311.734

Min d/m
1.54462

ave/min d
31.8354
ave/min m
42.5587

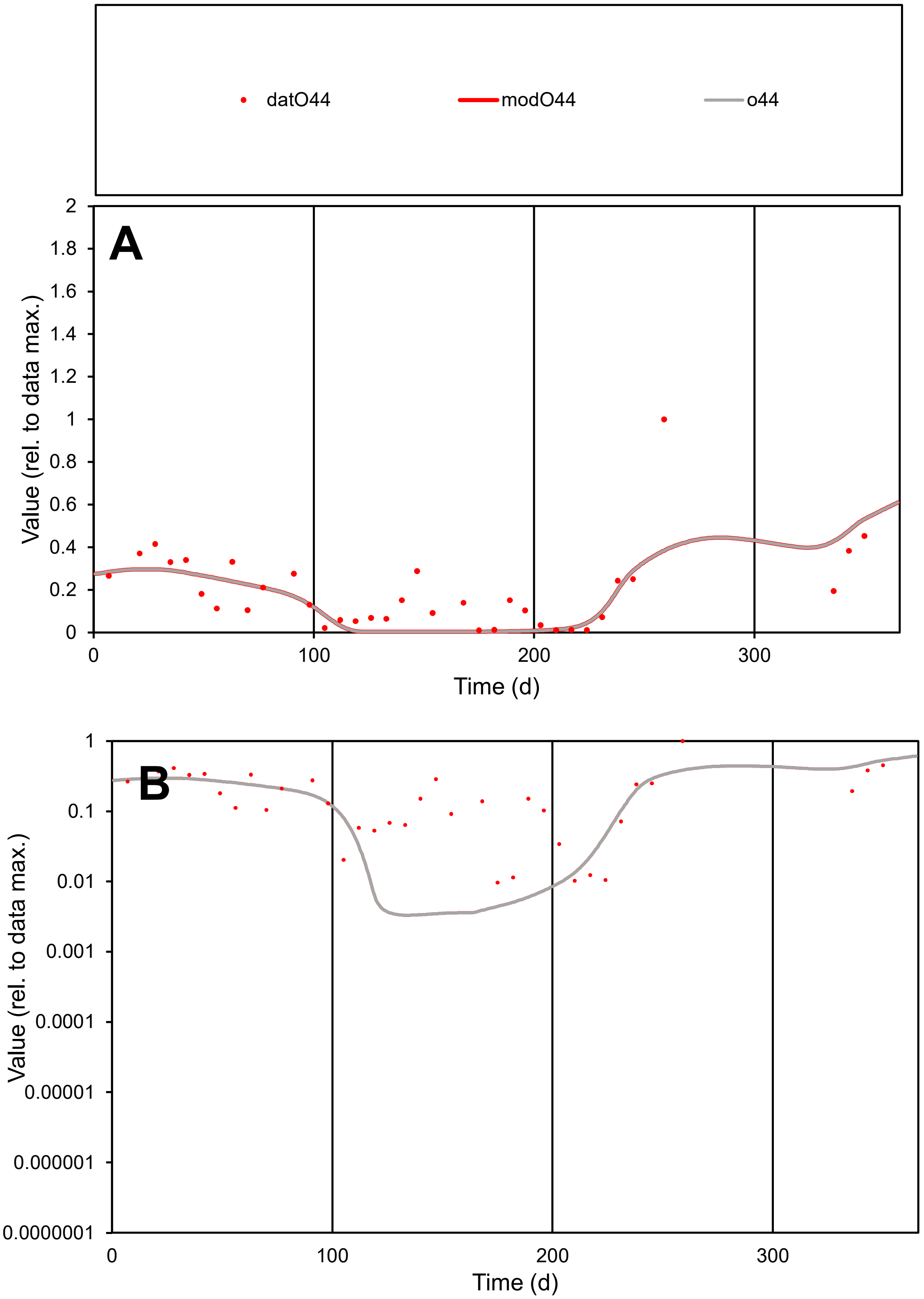


**C1**

Variables
O44 Provide jtd or Dst
CalibOnly?
2 1 to reproduce of
oNormalize?
2 1=no, 2=yes

ofxn calc*
0.72004
ofxn mod
0.73448

sRun
RunA1
zout
1

Ave
Data
0.00238
Model
0.00199

data/vMax
0.0127

ofxng
311.734

Min d/m
2.91331

ave/min d
19.3126
ave/min m
47.0492

331_Calib_S_A

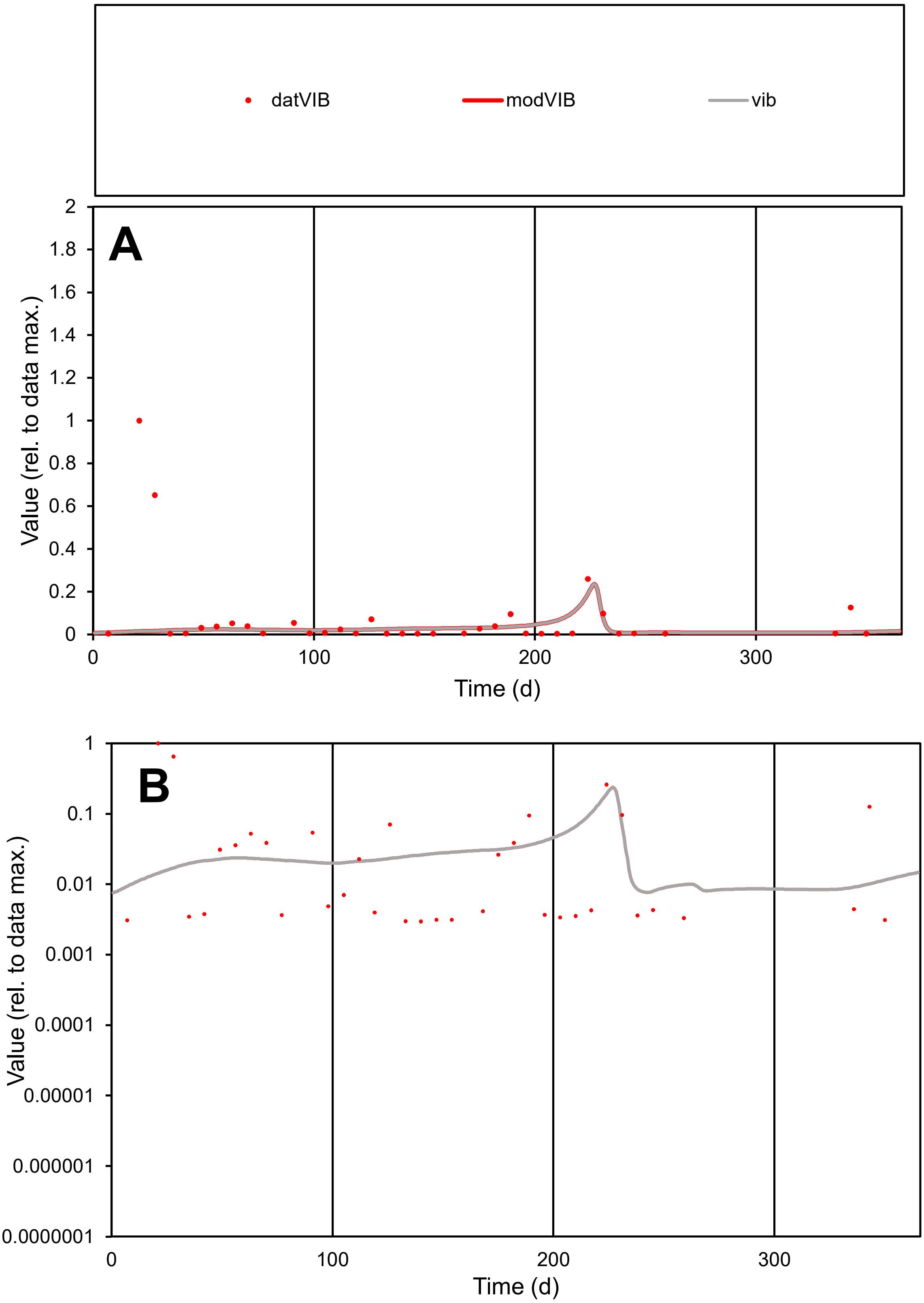


**C1**

Variables
VIB Provide jtd or Dst
CalibOnly?
2 1 to reproduce of
oNormalize?
2 1=no, 2=yes

ofxn calc*
2.6865
ofxn mod
2.6944

sRun
RunA1
zout
1

Ave
Data
0.00044
Model
0.00019

data/vMax
0.00591

ofxng
311.734

Min d/m
0.37234

ave/min d
24.8766
ave/min m
4.00192

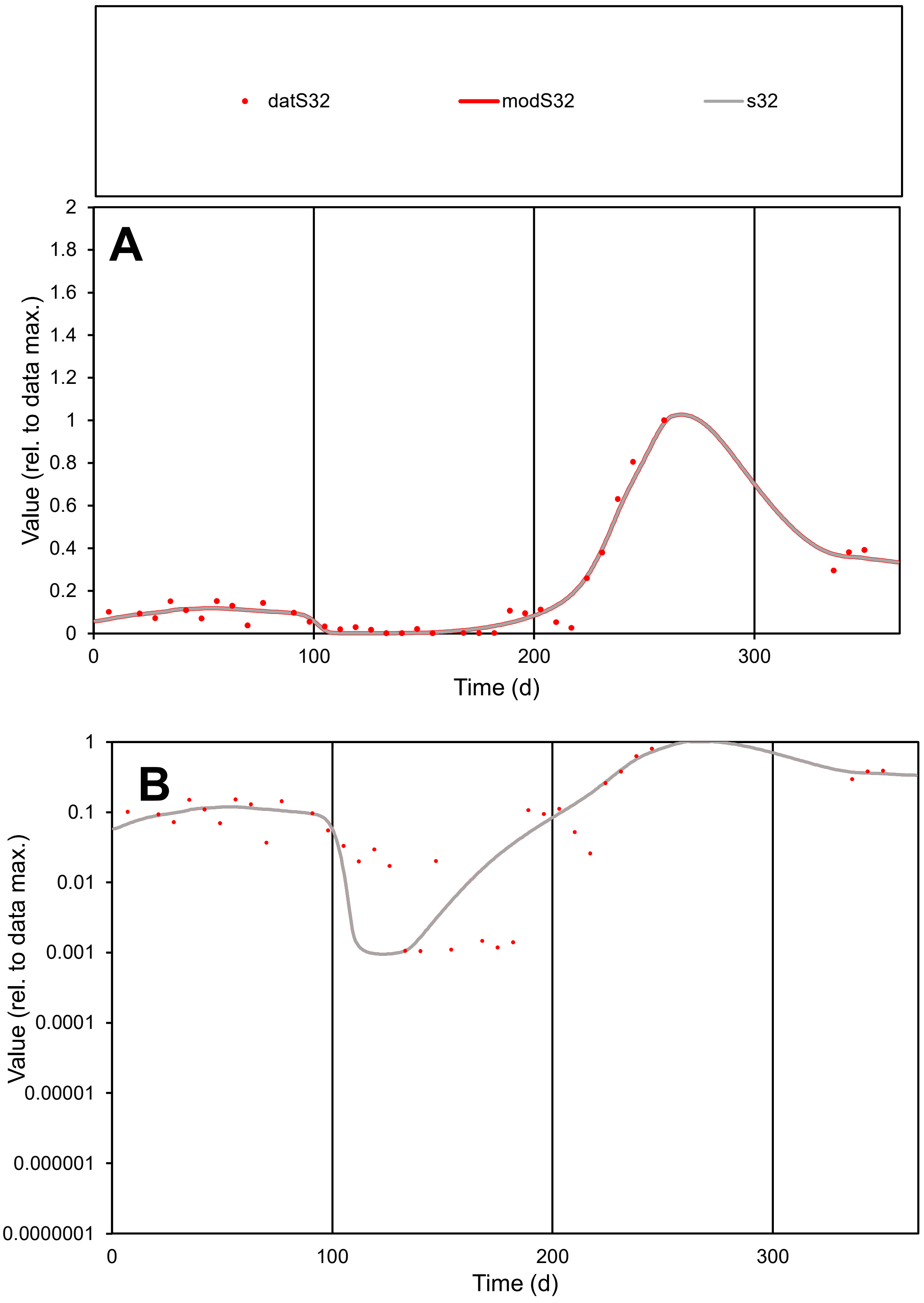


C1

Variables
S32 Provide jtd or Dst
CalibOnly?
2 1 to reproduce of
oNormalize?
2 1=no, 2=yes

ofxn calc*
0.25766
ofxn mod
0.33526

sRun
RunA1
zout
1

Ave
Data
0.00165
Model
0.00168

data/vMax
0.01012

ofxng
311.734

Min d/m
1.09774

ave/min d
154.469
ave/min m
172.566

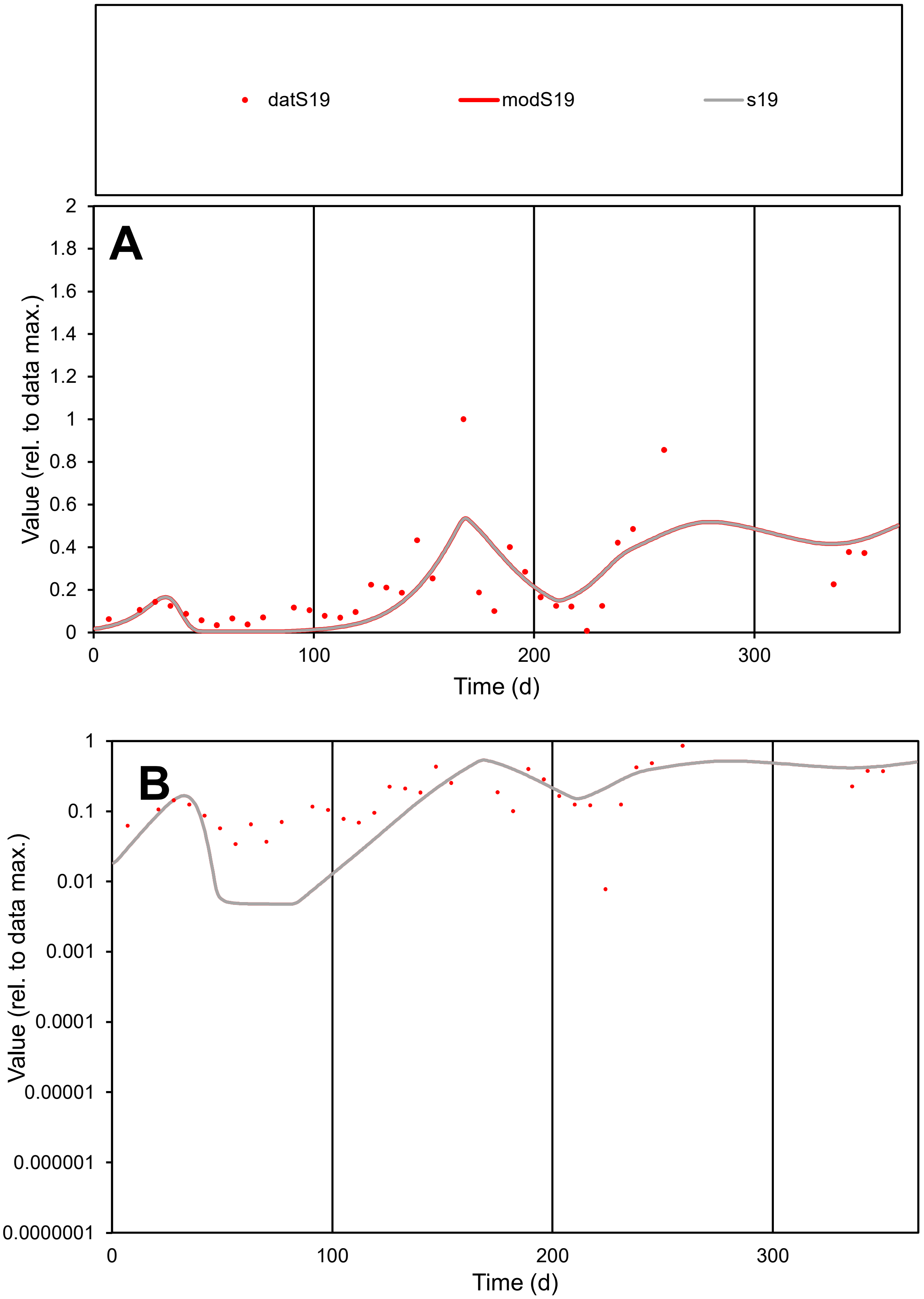


**C1**

Variables
S19 Provide jtd or Dst
CalibOnly?
2 1 to reproduce of
oNormalize?
2 1=no, 2=yes

ofxn calc*
0.6934
ofxn mod
0.70966

sRun
RunA1
zout
1

Ave
Data
0.00151
Model
0.0014

data/vMax
0.00731

ofxng
311.734

Min d/m
1.63537

ave/min d
26.4897
ave/min m
40.2399

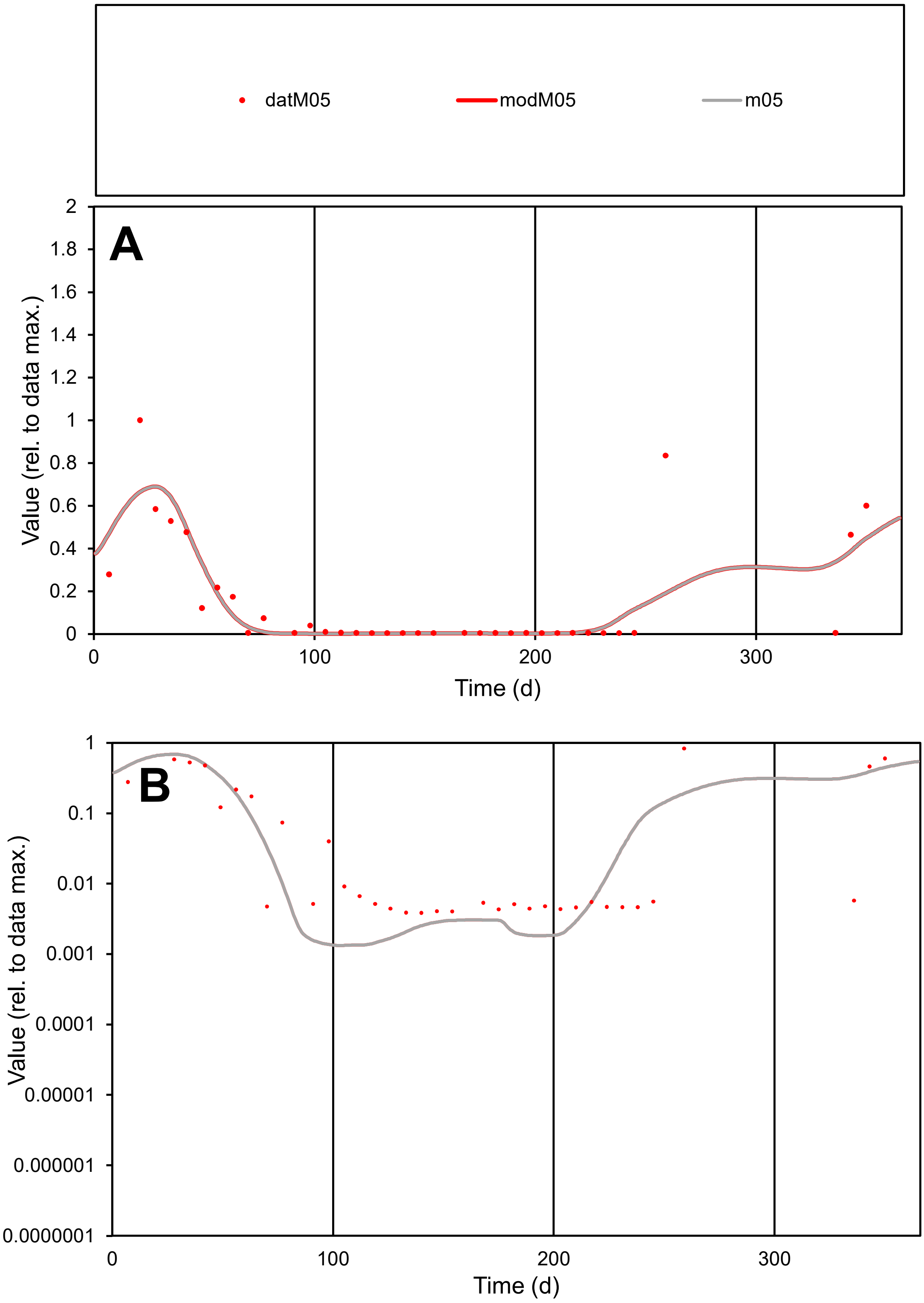


**C1**

Variables
M05 Provide jtd or Dst
CalibOnly?
2 1 to reproduce of
oNormalize?
2 1=no, 2=yes

ofxn calc*
0.94849
ofxn mod
0.98028

sRun
RunA1
zout
1

Ave
Data
0.0017
Model
0.00158

data/vMax
0.0111

ofxng
311.734

Min d/m
2.88319

ave/min d
39.7852
ave/min m
106.7

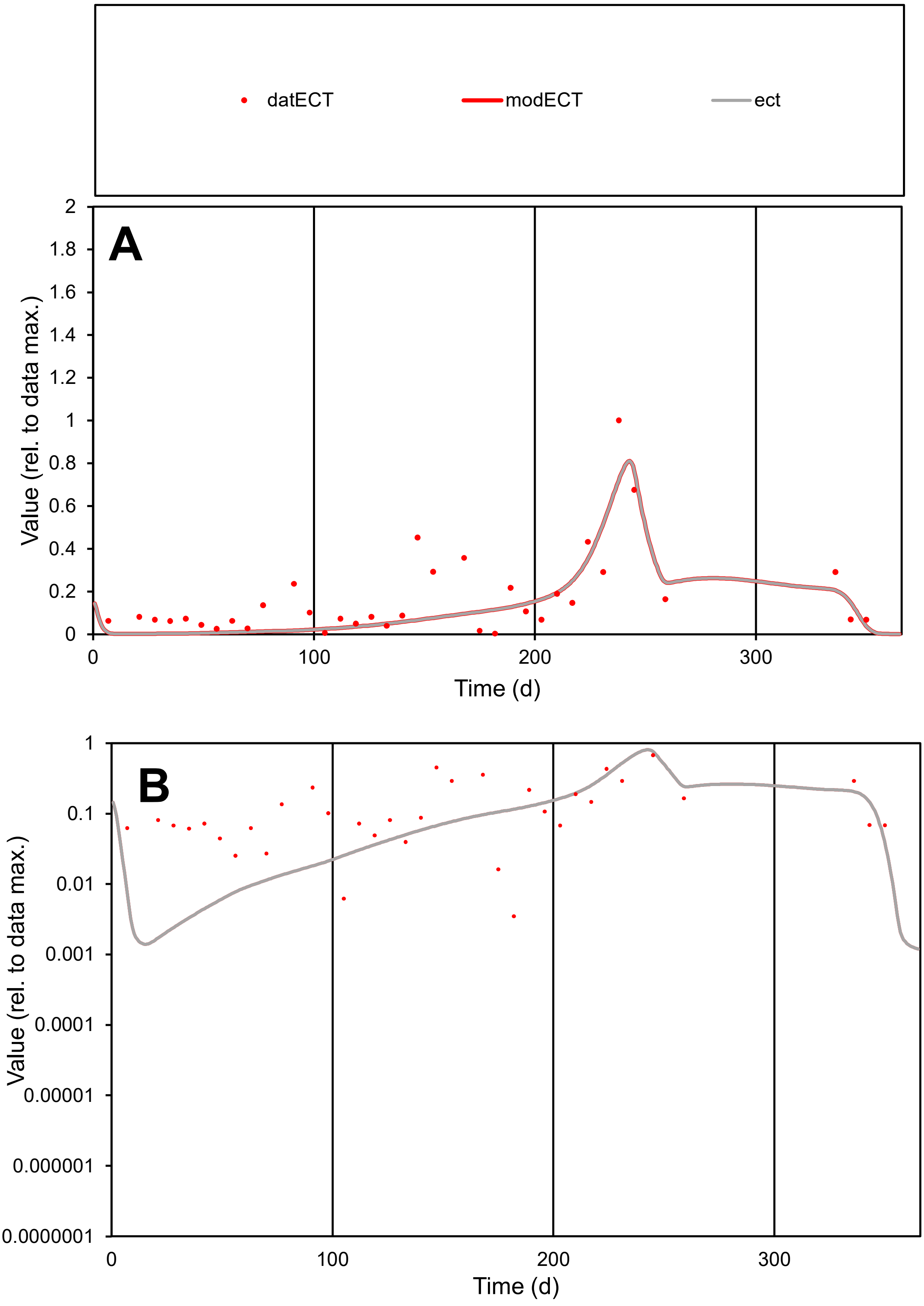


**C1**

Variables

ECT Provide jtd or Dst

CalibOnly?

2 1 to reproduce of

oNormalize?

2 1=no, 2=yes

ofxn calc*

0.67439

ofxn mod

0.70109

sRun

RunA1

zout

1

Ave

Data

0.00175

Model

0.0014

data/vMax

0.01071

ofxng

311.734

Min d/m

1.88322

ave/min d

47.0199

ave/min m

70.7415

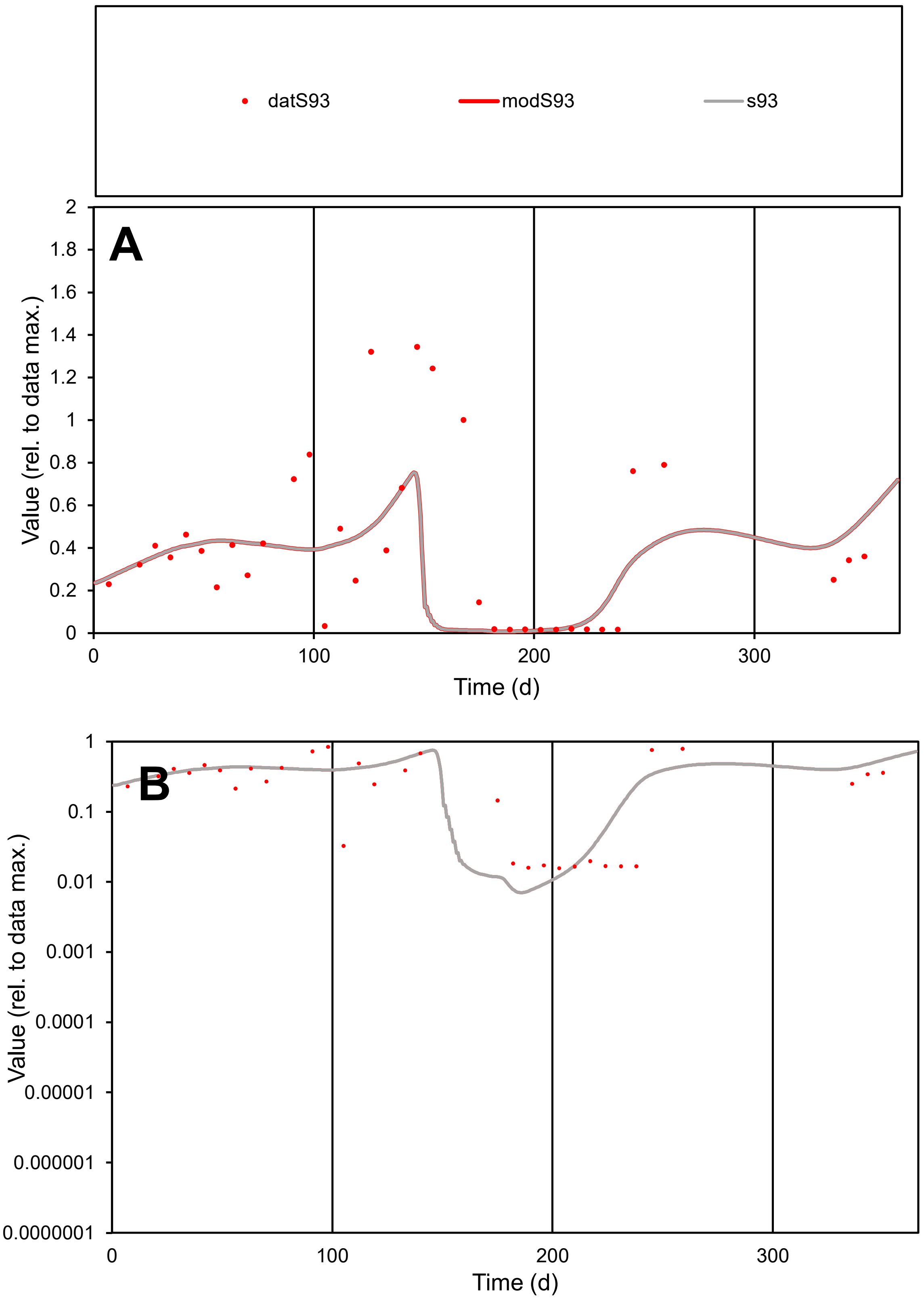


**C1**

Variables

S93 Provide jtd or Dst

CalibOnly?

2 1 to reproduce of

oNormalize?

2 1=no, 2=yes

ofxn calc*

0.81855

ofxn mod

0.83205

sRun

RunA1

zout

1

Ave

Data

0.0017

Model

0.00146

data/vMax

0.00467

ofxng

311.734

Min d/m

2.06528

ave/min d

23.2068

ave/min m

41.0422

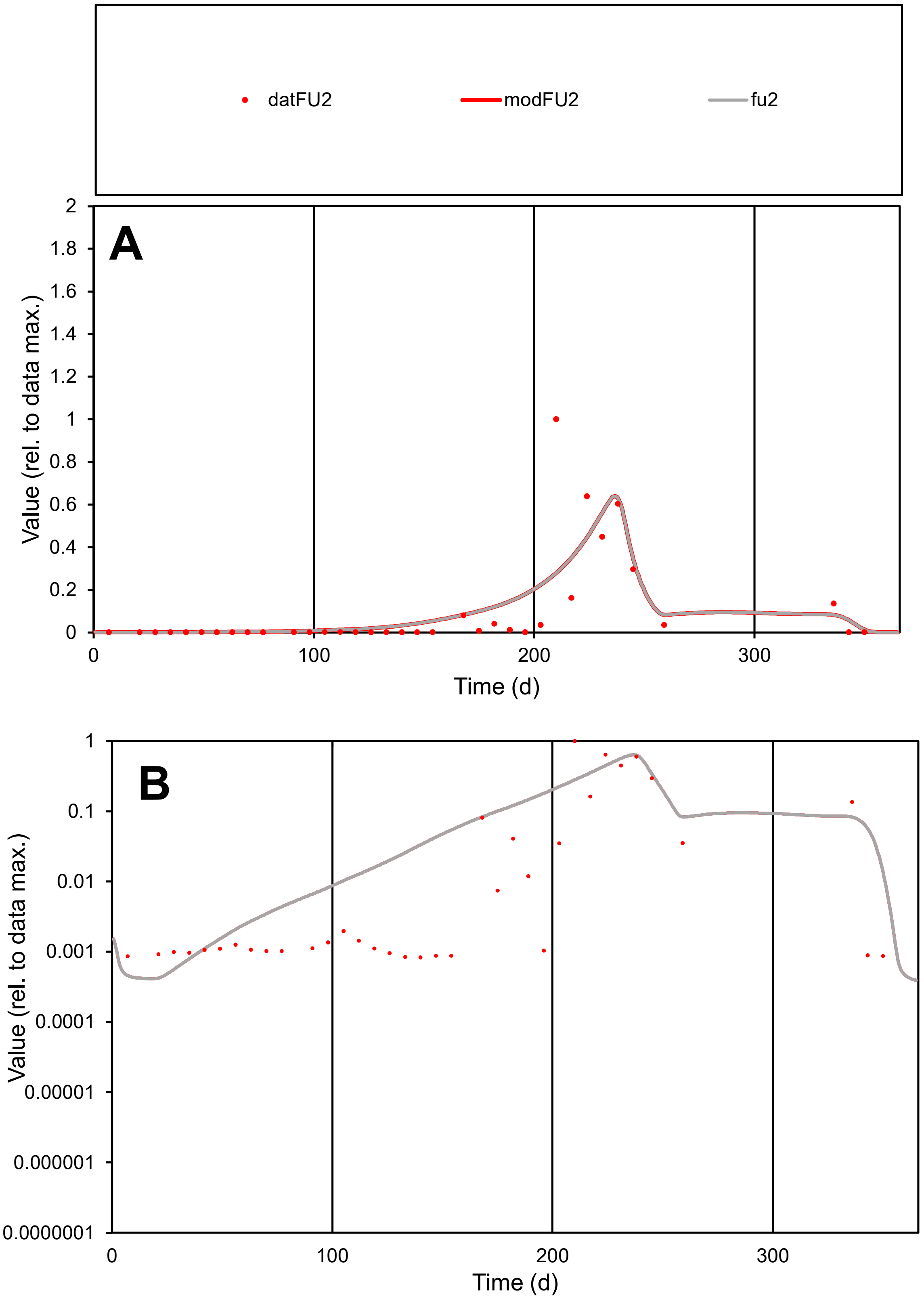


**C1**

Variables
FU2 Provide jtd or Dst
CalibOnly?
2 1 to reproduce of
oNormalize?
2 1=no, 2=yes

ofxn calc*
1.42166
ofxn mod
1.45711

sRun
RunA1
zout
1

Ave
Data
0.00297
Model
0.00328

data/vMax
0.03034

ofxng
311.734

Min d/m
1.89288

ave/min d
117.404
ave/min m
245.428

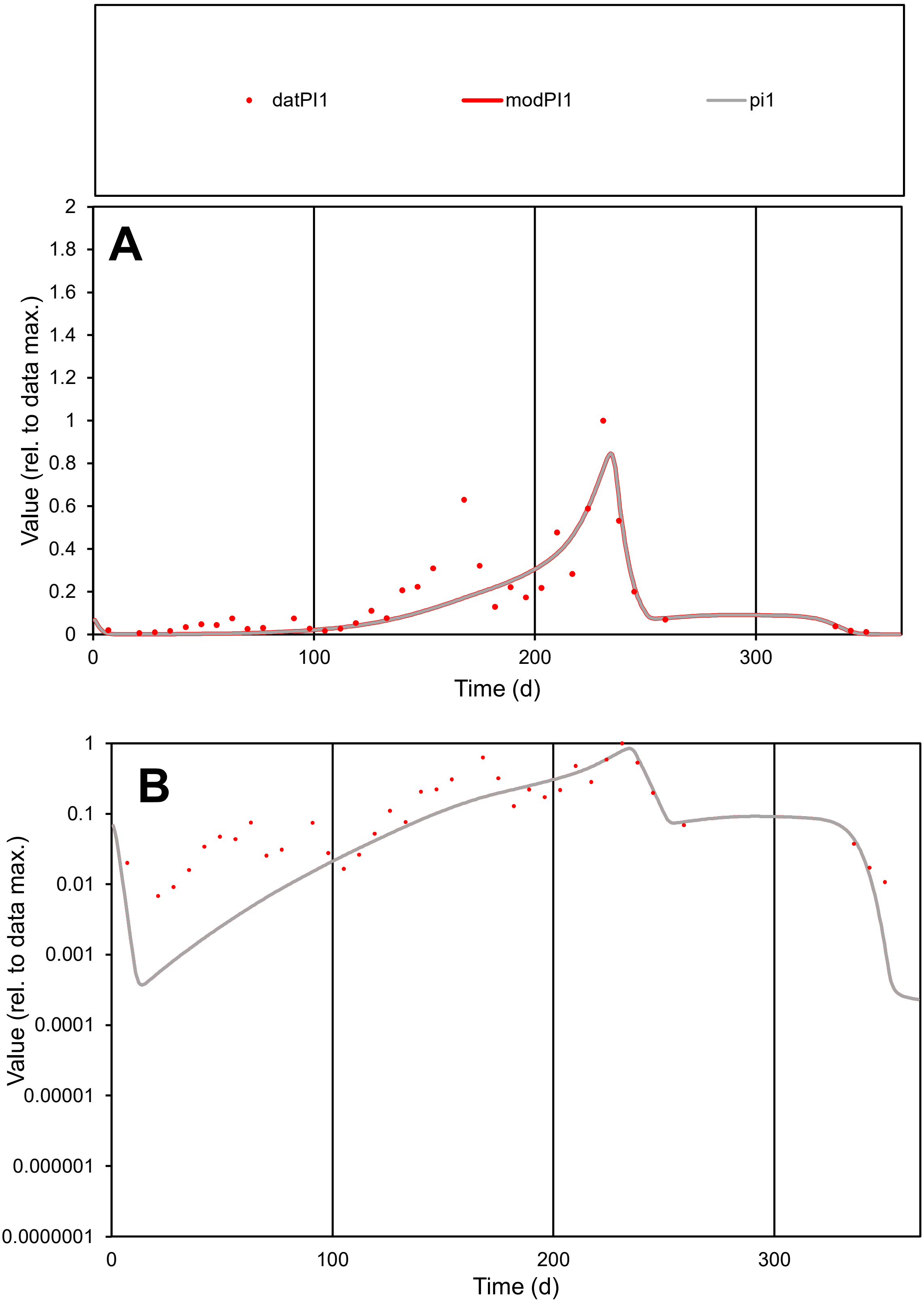


**C1**

Variables
PI1 Provide jtd or Dst
CalibOnly?
2 1 to reproduce of
oNormalize?
2 1=no, 2=yes

ofxn calc*
0.59562
ofxn mod
0.6311

sRun
RunA1
zout
1

Ave
Data
0.00409
Model
0.00354

data/vMax
0.02432

ofxng
311.734

Min d/m
11.4139

ave/min d
24.5763
ave/min m
243.027

331_Calib_S_A

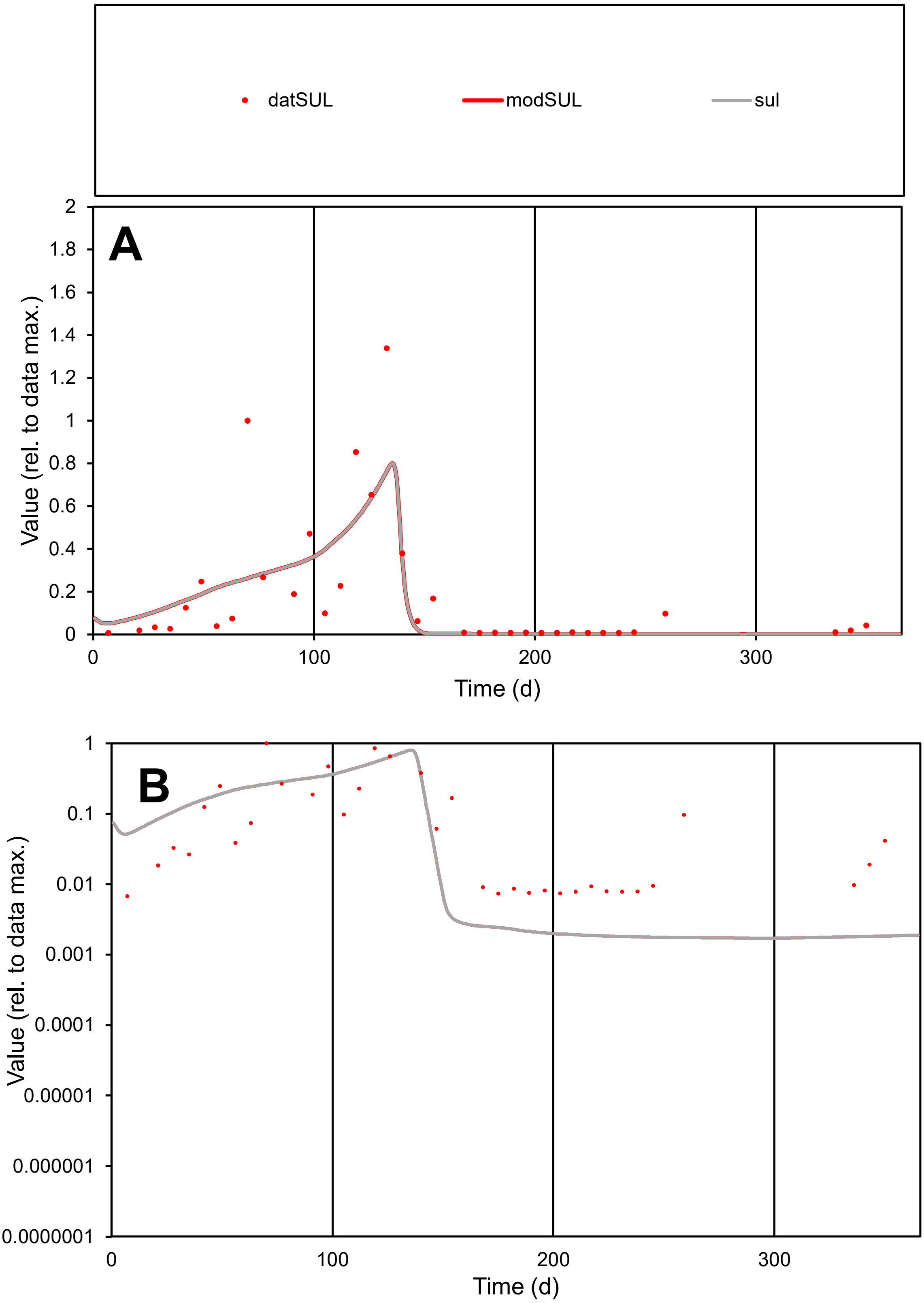


C1

Variables

SUL Provide jtd or Dst

CalibOnly?

2 1 to reproduce of

oNormalize?

2 1=no, 2=yes

ofxn calc*

1.0402

ofxn mod

1.05978

sRun

RunA1

zout

1

Ave

Data

0.00191

Model

0.00187

data/vMax

0.01201

ofxng

311.734

Min d/m

3.88608

ave/min d

23.4749

ave/min m

89.1797

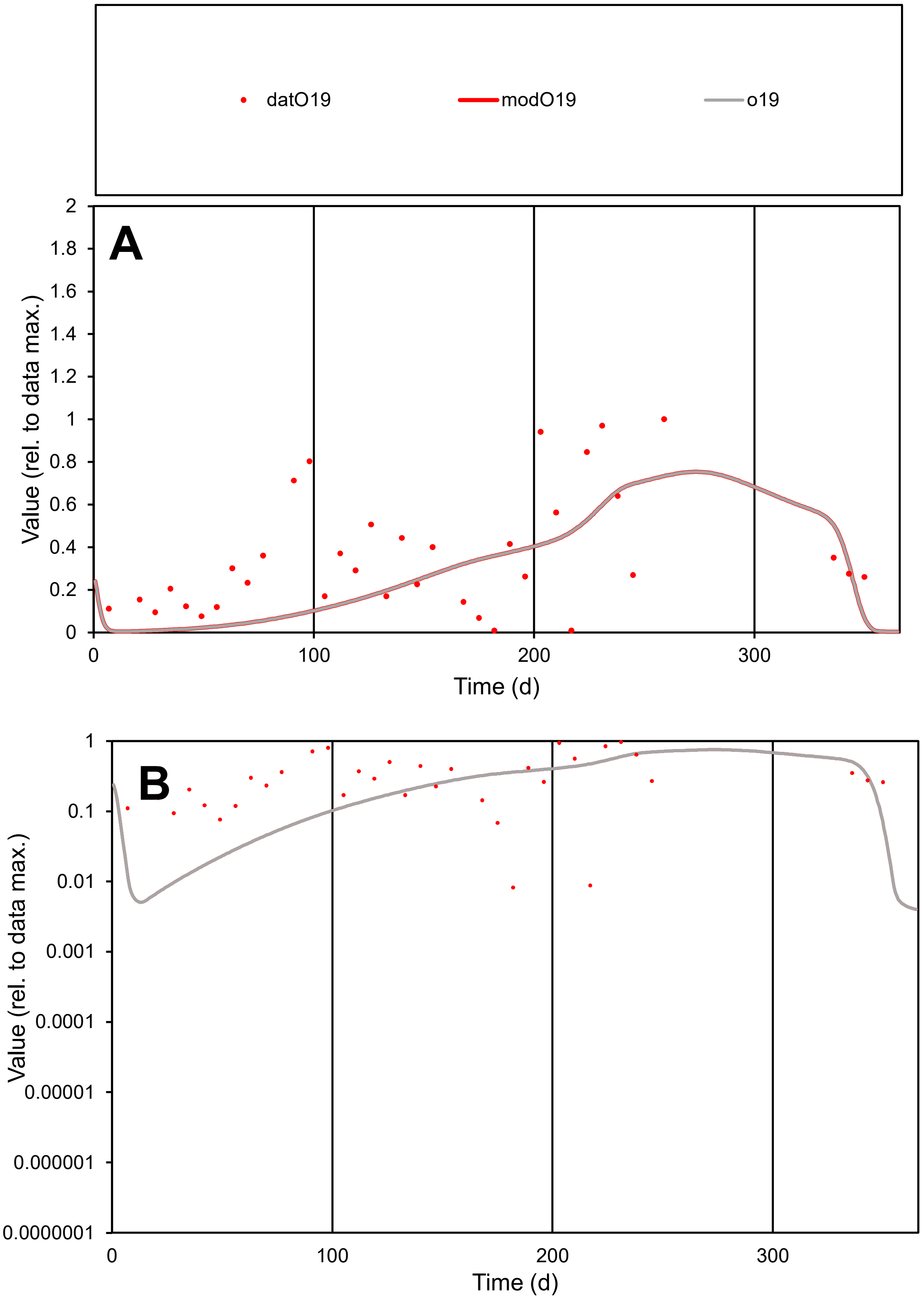


**C1**

Variables
O19 Provide jtd or Dst
CalibOnly?
2 1 to reproduce of
oNormalize?
2 1=no, 2=yes

ofxn calc*
0.78054
ofxn mod
0.79411

sRun
RunA1
zout
1

Ave
Data
0.00156
Model
0.00117

data/vMax
0.00455

ofxng
311.734

Min d/m
1.08137

ave/min d
41.8653
ave/min m
34.1202

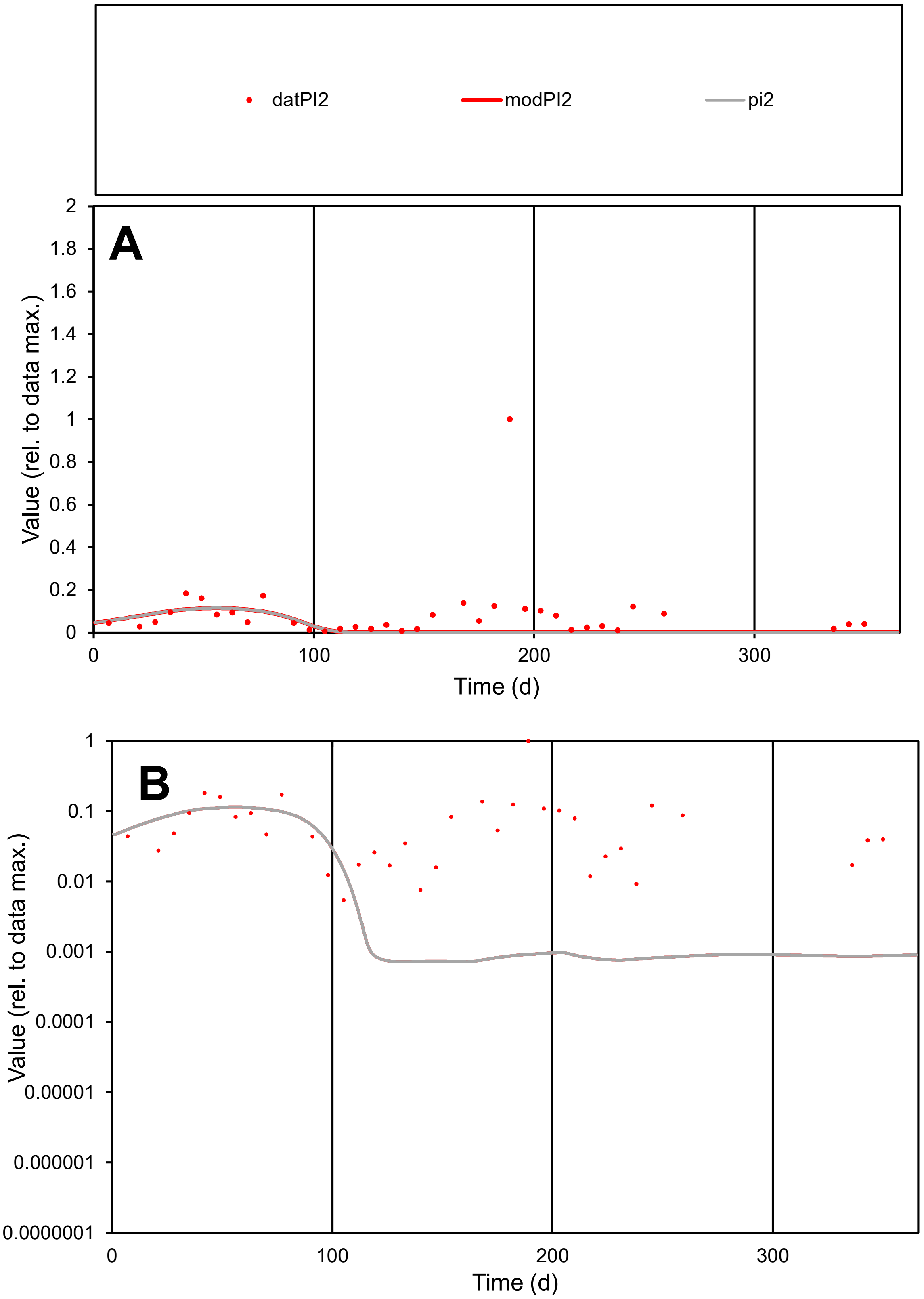


**C1**

Variables
PI2 Provide jtd or Dst
CalibOnly?
2 1 to reproduce of
oNormalize?
2 1=no, 2=yes

ofxn calc*
2.01109
ofxn mod
2.01621

sRun
RunA1
zout
1

Ave
Data
0.00271
Model
0.00095

data/vMax
0.03089

ofxng
311.734

Min d/m
5.99012

ave/min d
20.2961
ave/min m
42.4358

331_Calib_S_A

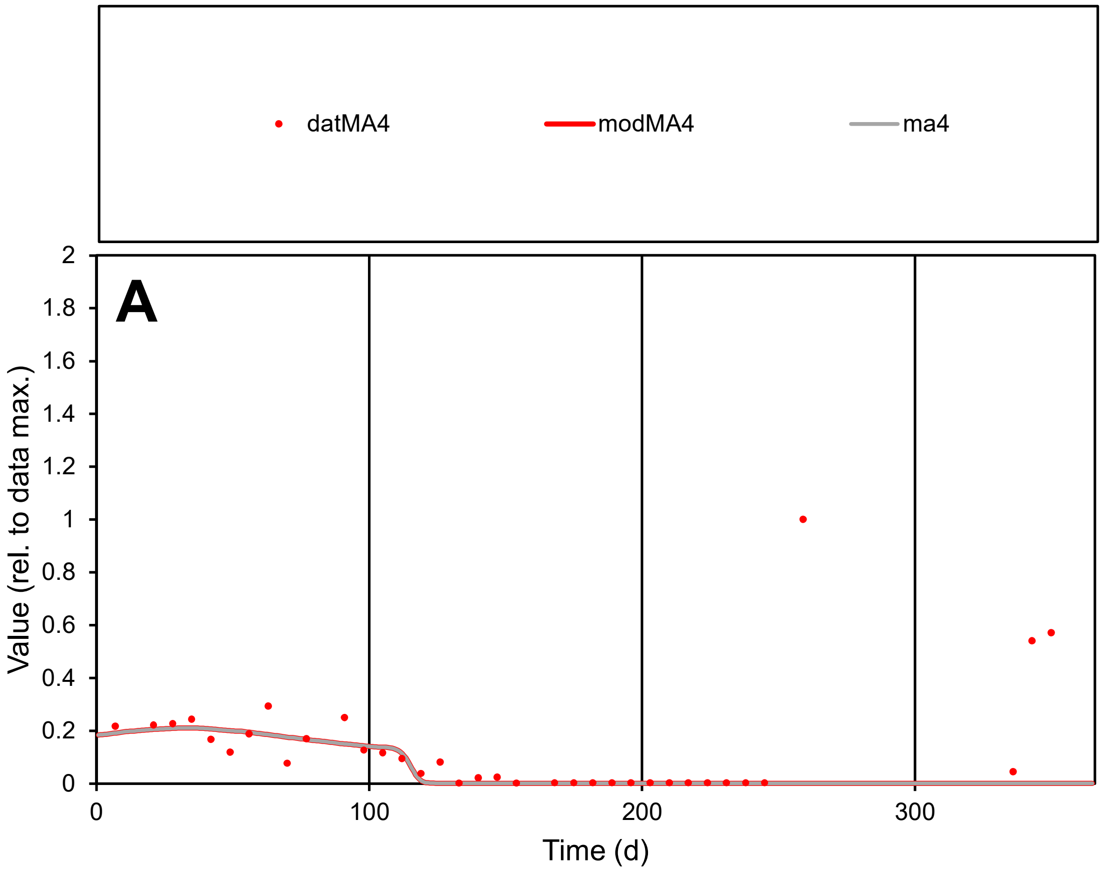


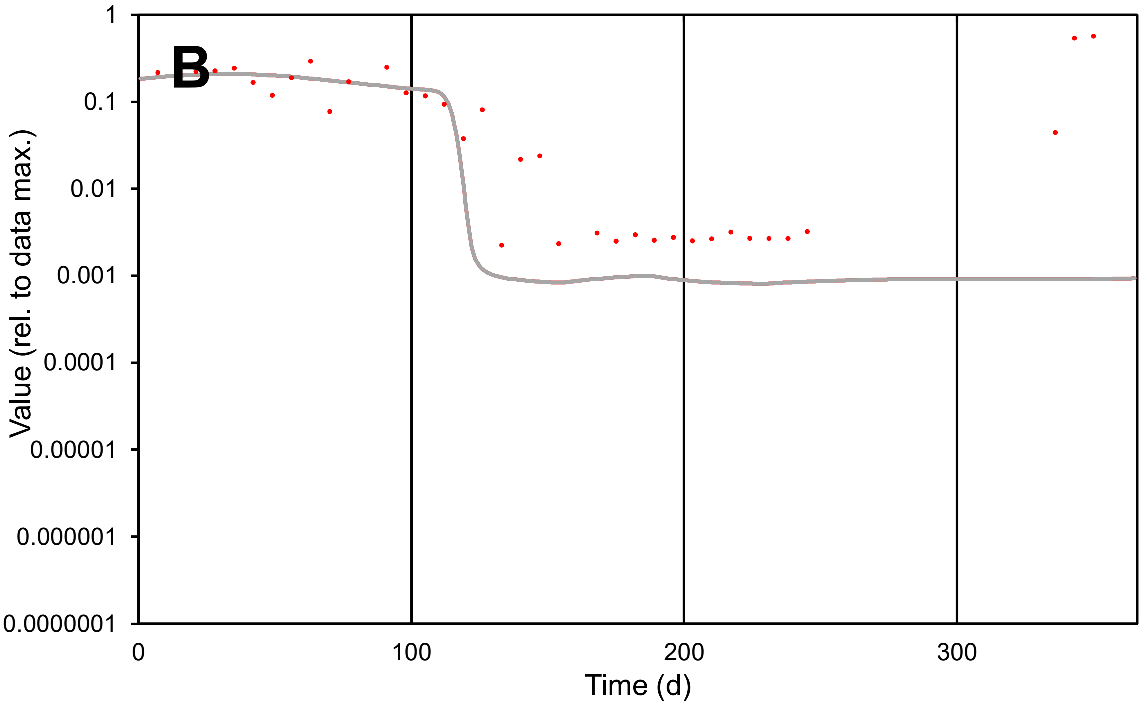


**C1**

Variables

MA4 Provide jtd or Dst

CalibOnly?

2 1 to reproduce of

oNormalize?

2 1=no, 2=yes

ofxn calc*

1.60039

ofxn mod

1.6063

sRun

RunA1

zout

1

Ave

Data

0.00115

Model

0.0006

data/vMax

0.00858

ofxng

311.734

Min d/m

2.76654

ave/min d

59.7654

ave/min m

86.1309

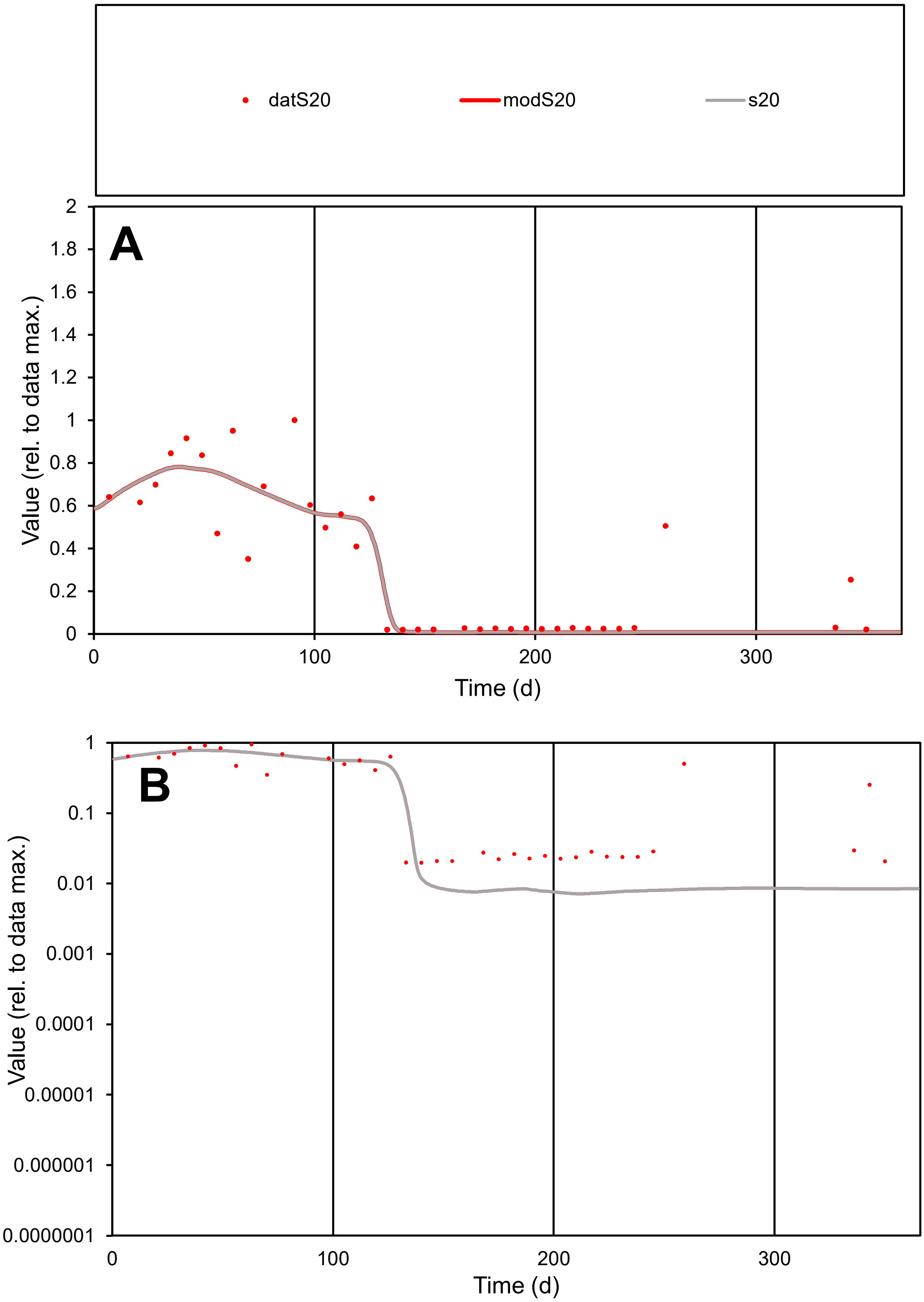


**C1**

Variables
S20 Provide jtd or Dst
CalibOnly?
2 1 to reproduce of
oNormalize?
2 1=no, 2=yes

ofxn calc*
0.45533
ofxn mod
0.48399

sRun
RunA1
zout
1

Ave
Data
0.00127
Model
0.00116

data/vMax
0.0039

ofxng
311.734

Min d/m
2.76189

ave/min d
16.4507
ave/min m
41.4258

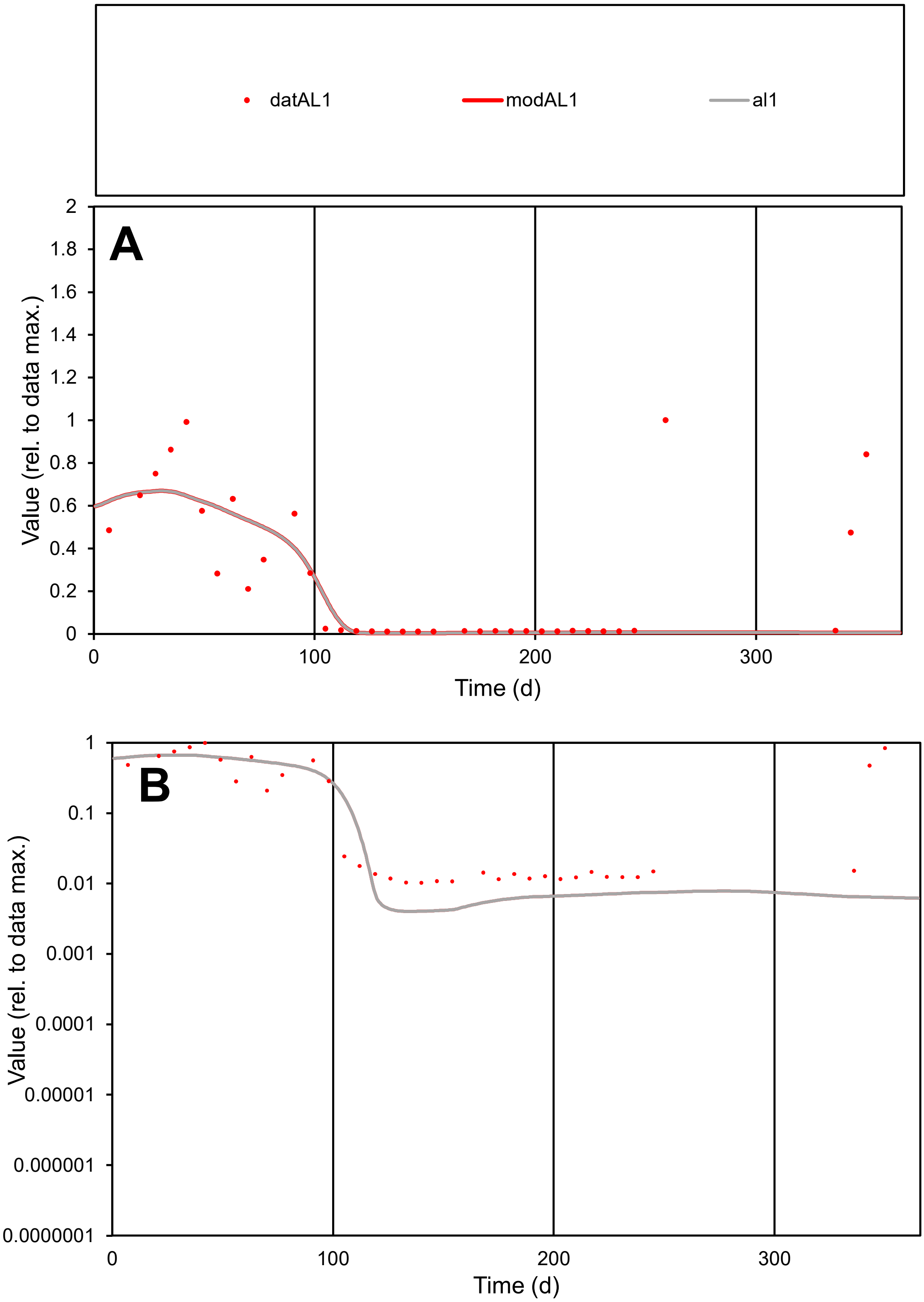


C1

Variables
AL1 Provide jtd or Dst
CalibOnly?
2 1 to reproduce of
oNormalize?
2 1=no, 2=yes

ofxn calc*
0.99717
ofxn mod
1.01145

sRun
RunA1
zout
1

Ave
Data
0.00109
Model
0.00082

data/vMax
0.00425

ofxng
311.734

Min d/m
2.53945
ave/min d
25.0421
ave/min m
48.1426

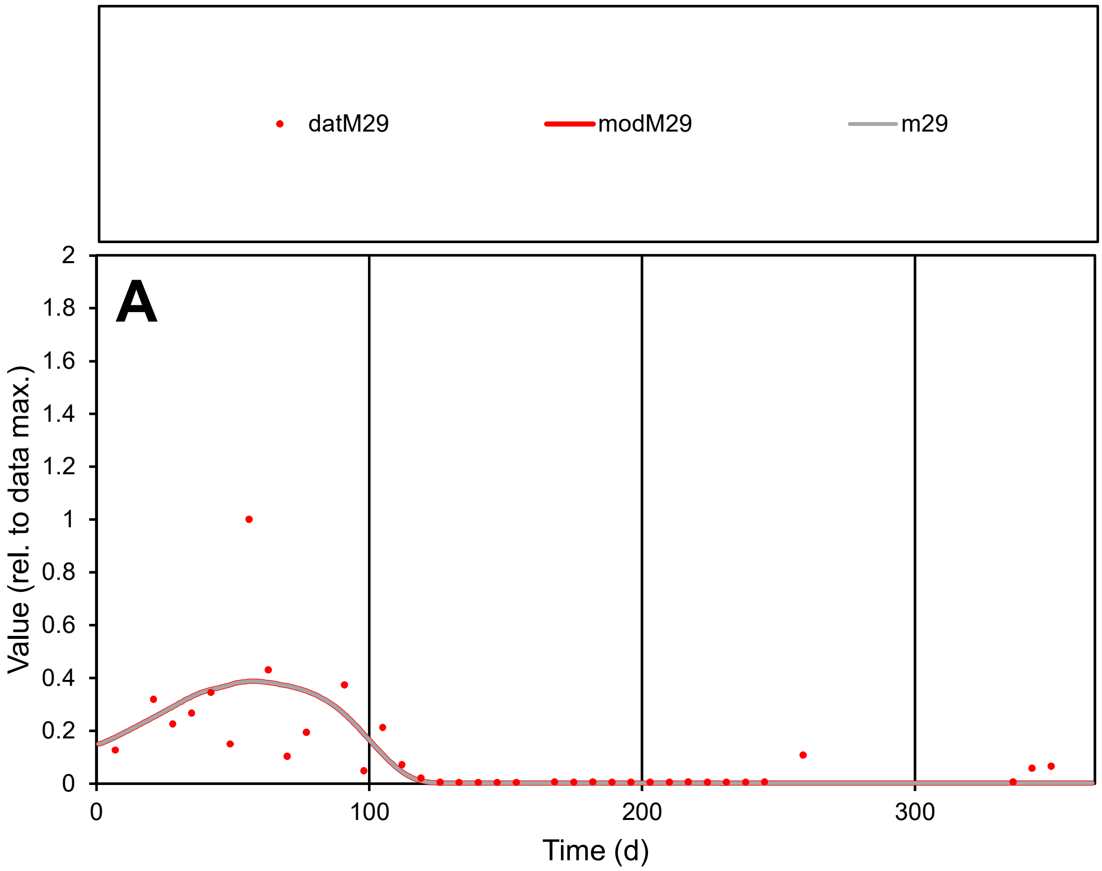


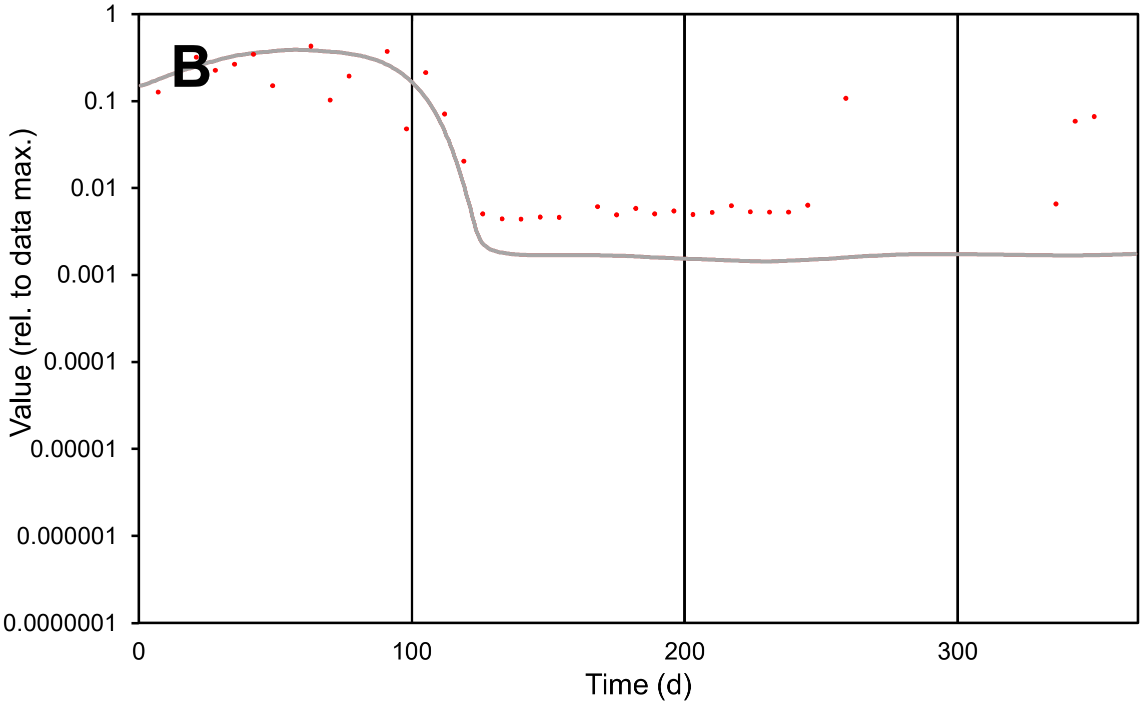


**C1**

Variables

M29 Provide jtd or Dst

CalibOnly?

2 1 to reproduce of

oNormalize?

2 1=no, 2=yes

ofxn calc*

1.10072

ofxn mod

1.12539

sRun

RunA1

zout

1

Ave

Data

0.00144

Model

0.00134

data/vMax

0.01229

ofxng

311.734

Min d/m

3.04291

ave/min d

26.6939

ave/min m

75.4503

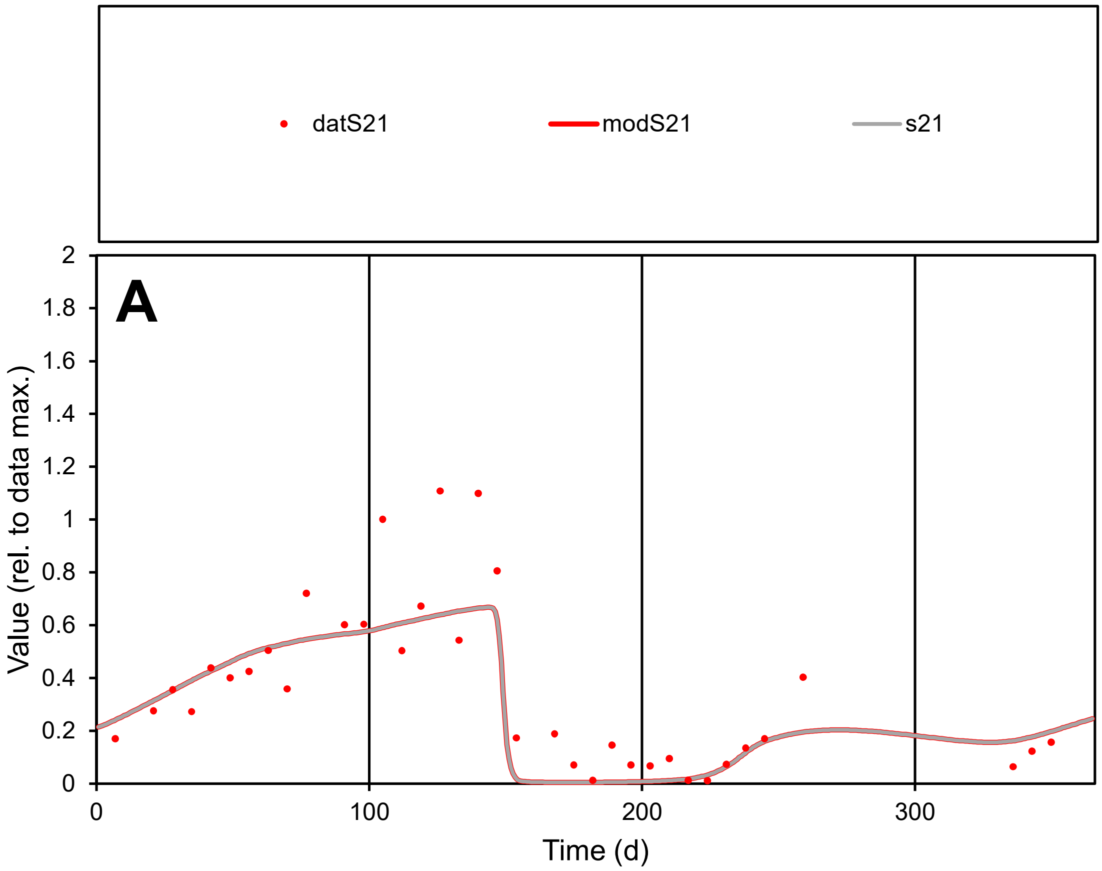


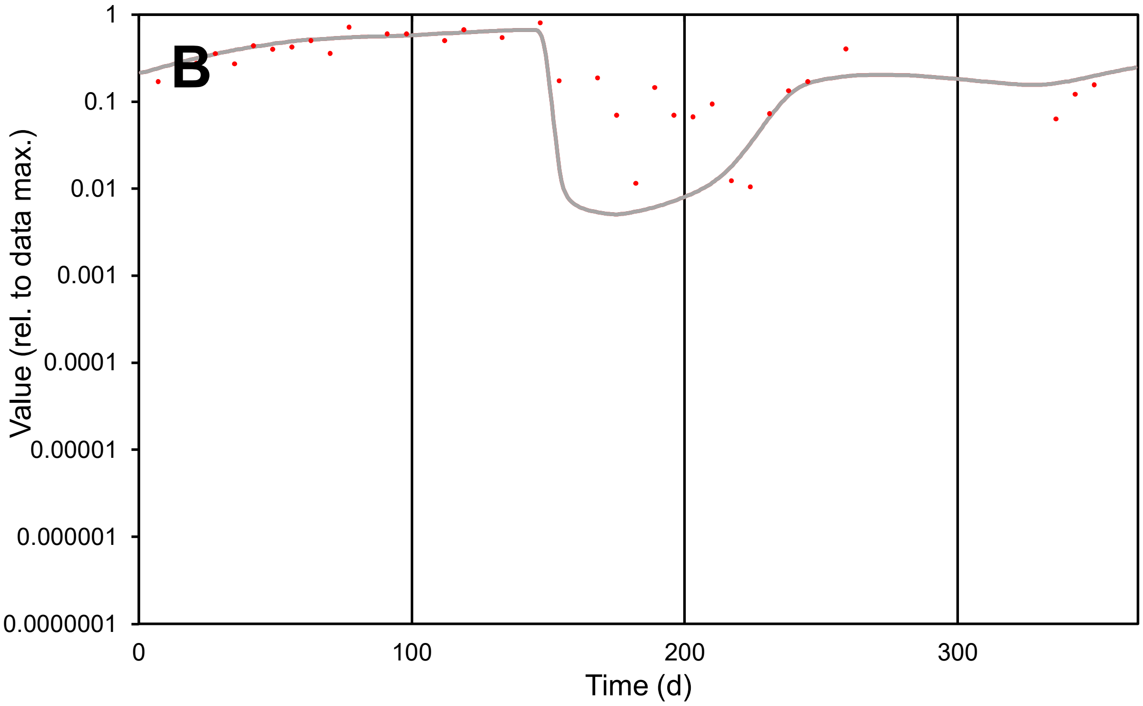


**C1**

| | |
|---|---|
| Variables | |
| S21 | Provide jtd or Dst |
| CalibOnly? | |
| 2 | 1 to reproduce of |
| oNormalize? | |
| 2 | 1=no, 2=yes |
| ofxn calc* | |
| 0.36589 | |
| ofxn mod | |
| 0.39319 | |
| sRun | |
| RunA1 | |
| zout | |
| 1 | |
| Ave | |
| Data | |
| 0.00151 | |
| Model | |
| 0.00145 | |
| data/vMax | |
| 0.00469 | |
| ofxng | |
| 311.734 | |
| Min d/m | |
| 2.0648 | |
| ave/min d | |
| 30.5919 | |
| ave/min m | |
| 60.5826 | |

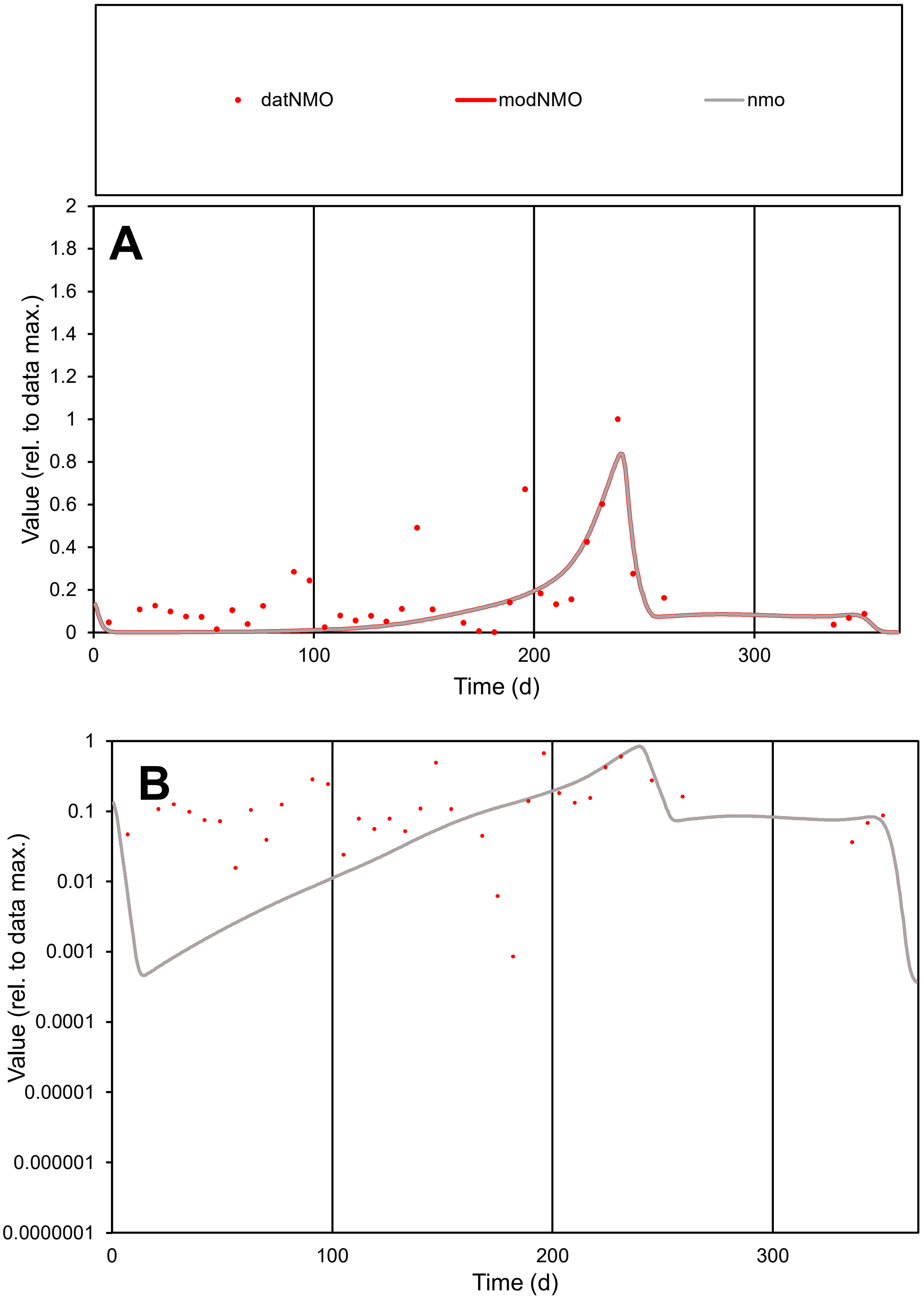


**C1**

Variables
NMO Provide jtd or Dst
CalibOnly?
2 1 to reproduce of
oNormalize?
2 1=no, 2=yes

ofxn calc*
0.8044
ofxn mod
0.8236

sRun
RunA1
zout
1

Ave
Data
0.00188
Model
0.00136

data/vMax
0.01111

ofxng
311.734

Min d/m
1.30862

ave/min d
197.745
ave/min m
187.772

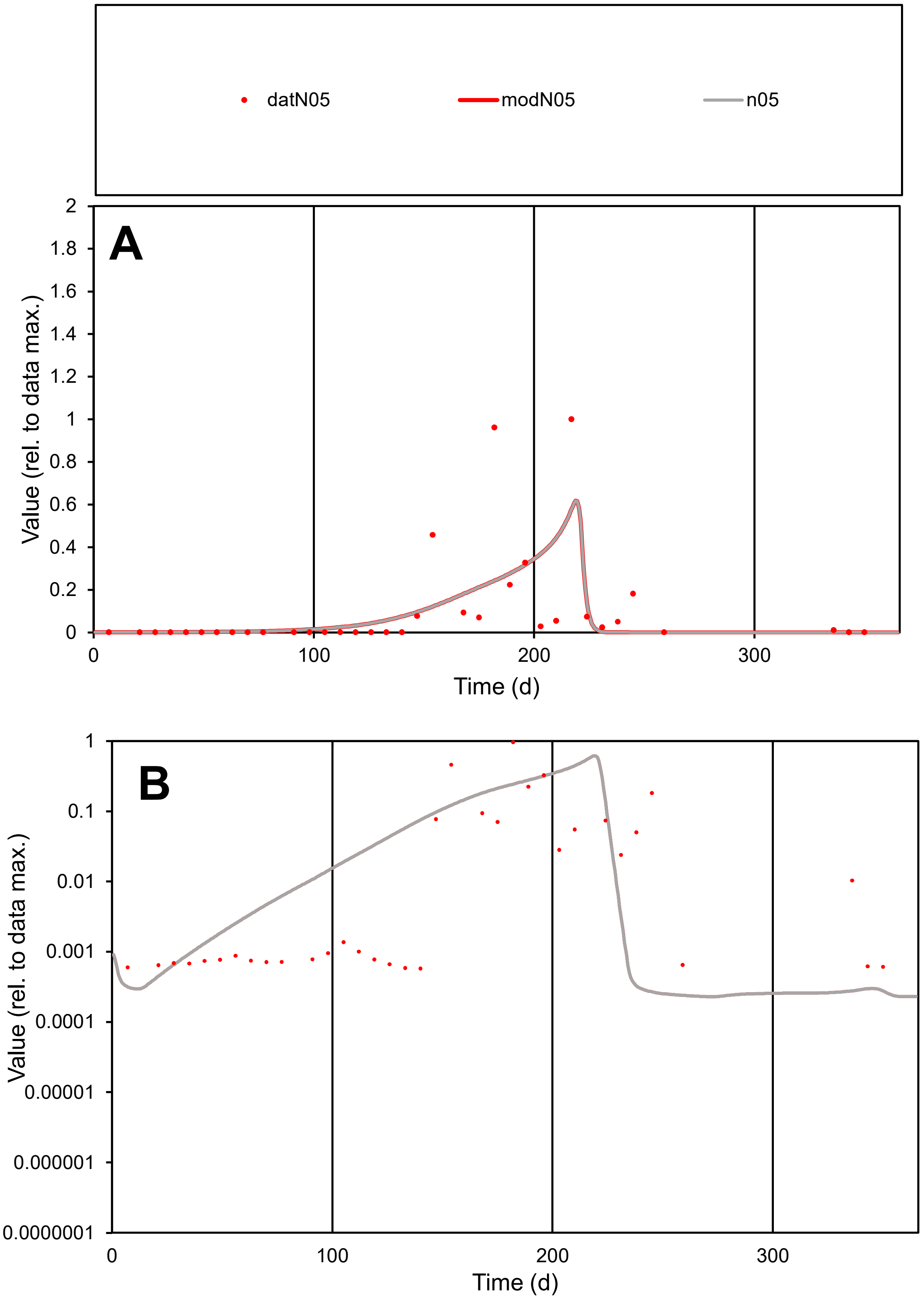


**C1**

Variables
N05 Provide jtd or Dst
CalibOnly?
2 1 to reproduce of
oNormalize?
2 1=no, 2=yes

ofxn calc*
1.76002
ofxn mod
1.77613

sRun
RunA1
zout
1

Ave
Data
0.00663
Model
0.0062

data/vMax
0.06794

ofxng
311.734

Min d/m
2.43457

ave/min d
168.11
ave/min m
382.415

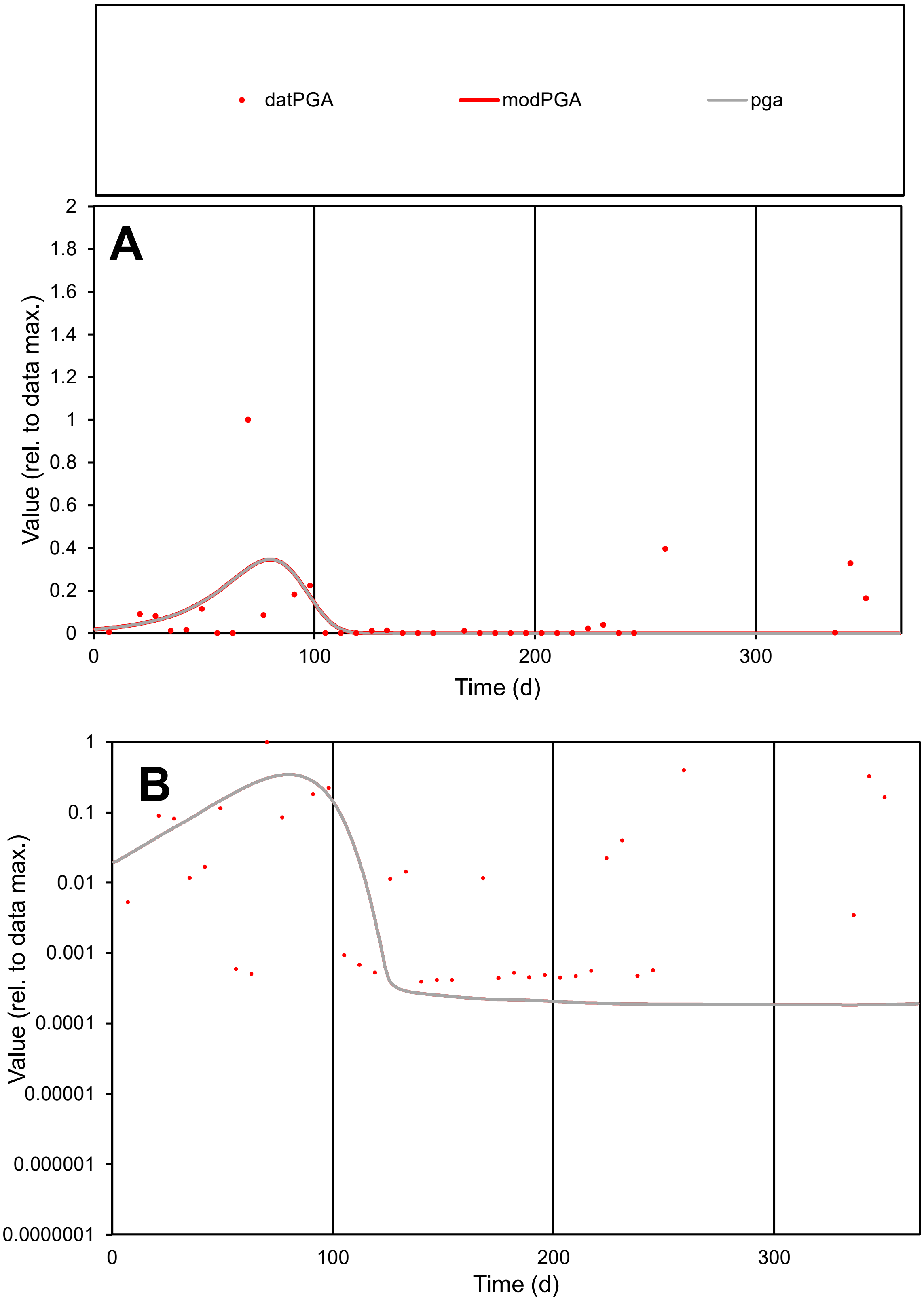


**C1**

Variables
PGA Provide jtd or Dst
CalibOnly?
2 1 to reproduce of
oNormalize?
2 1=no, 2=yes

ofxn calc*
2.09592
ofxn mod
2.11071

sRun
RunA1
zout
1

Ave
Data
0.00334
Model
0.00245

data/vMax
0.04285

ofxng
311.734

Min d/m
2.14431

ave/min d
197.954
ave/min m
312.022

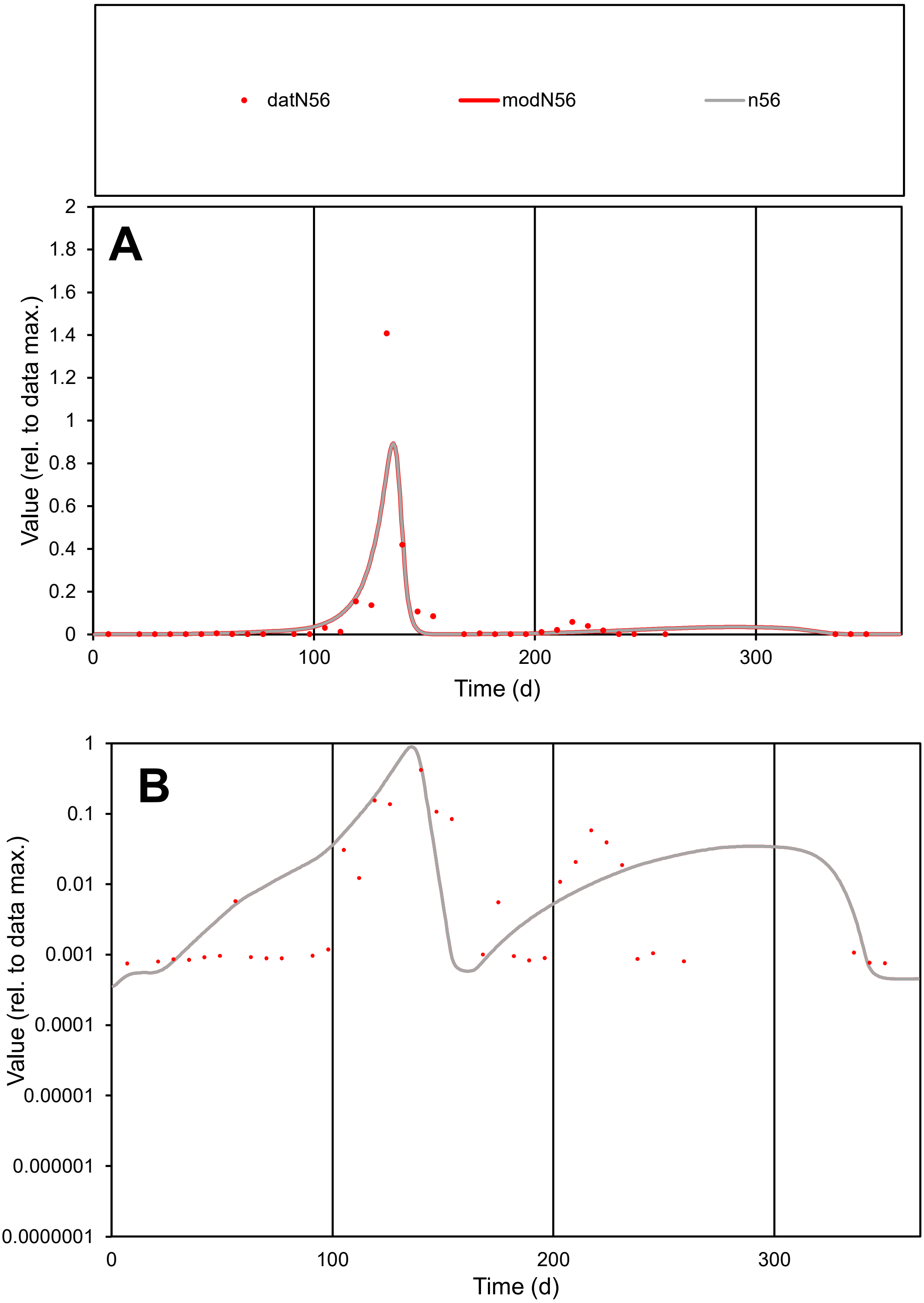


**C1**

Variables
N56 Provide jtd or Dst
CalibOnly?
2 1 to reproduce of
oNormalize?
2 1=no, 2=yes

ofxn calc*
1.01532
ofxn mod
1.16146

sRun
RunA1
zout
1

Ave
Data
0.00241
Model
0.00275

data/vMax
0.0468

ofxng
311.734

Min d/m
1.62843

ave/min d
68.7216
ave/min m
127.636

331_Calib_S_A

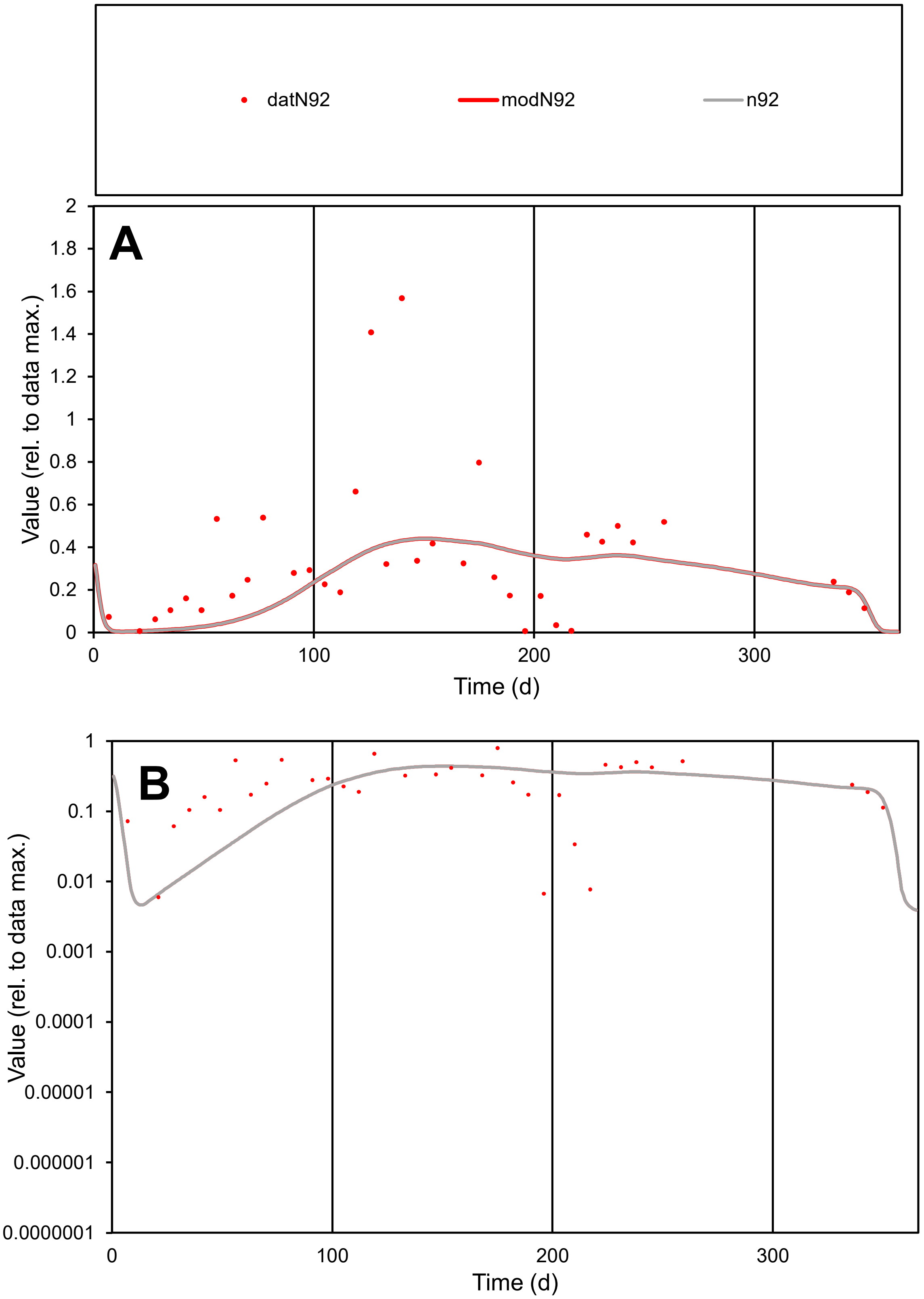


**C1**

Variables
N92 Provide jtd or Dst
CalibOnly?
2 1 to reproduce of
oNormalize?
2 1=no, 2=yes

ofxn calc*
0.76656
ofxn mod
0.77507

sRun
RunA1
zout
1

Ave
Data
0.00212
Model
0.00179

data/vMax
0.00696

ofxng
311.734

Min d/m
0.8357

ave/min d
50.778
ave/min m
35.8213

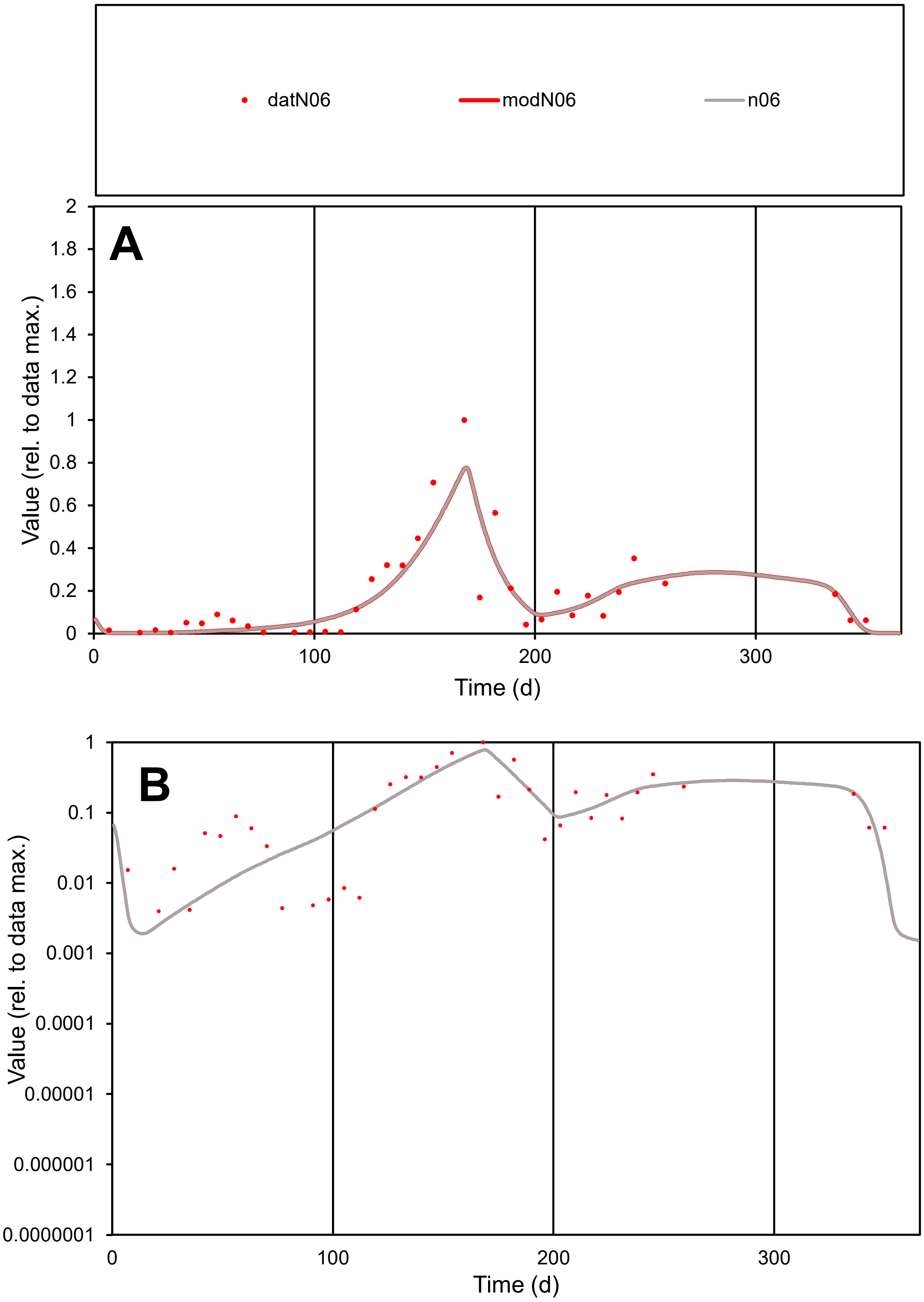


**C1**

Variables
N06 Provide jtd or Dst
CalibOnly?
2 1 to reproduce of
oNormalize?
2 1=no, 2=yes

ofxn calc*
0.57013
ofxn mod
0.6195

sRun
RunA1
zout
1

Ave
Data
0.00278
Model
0.0028

data/vMax
0.01796

ofxng
311.734

Min d/m
1.44753

ave/min d
39.0394
ave/min m
56.794

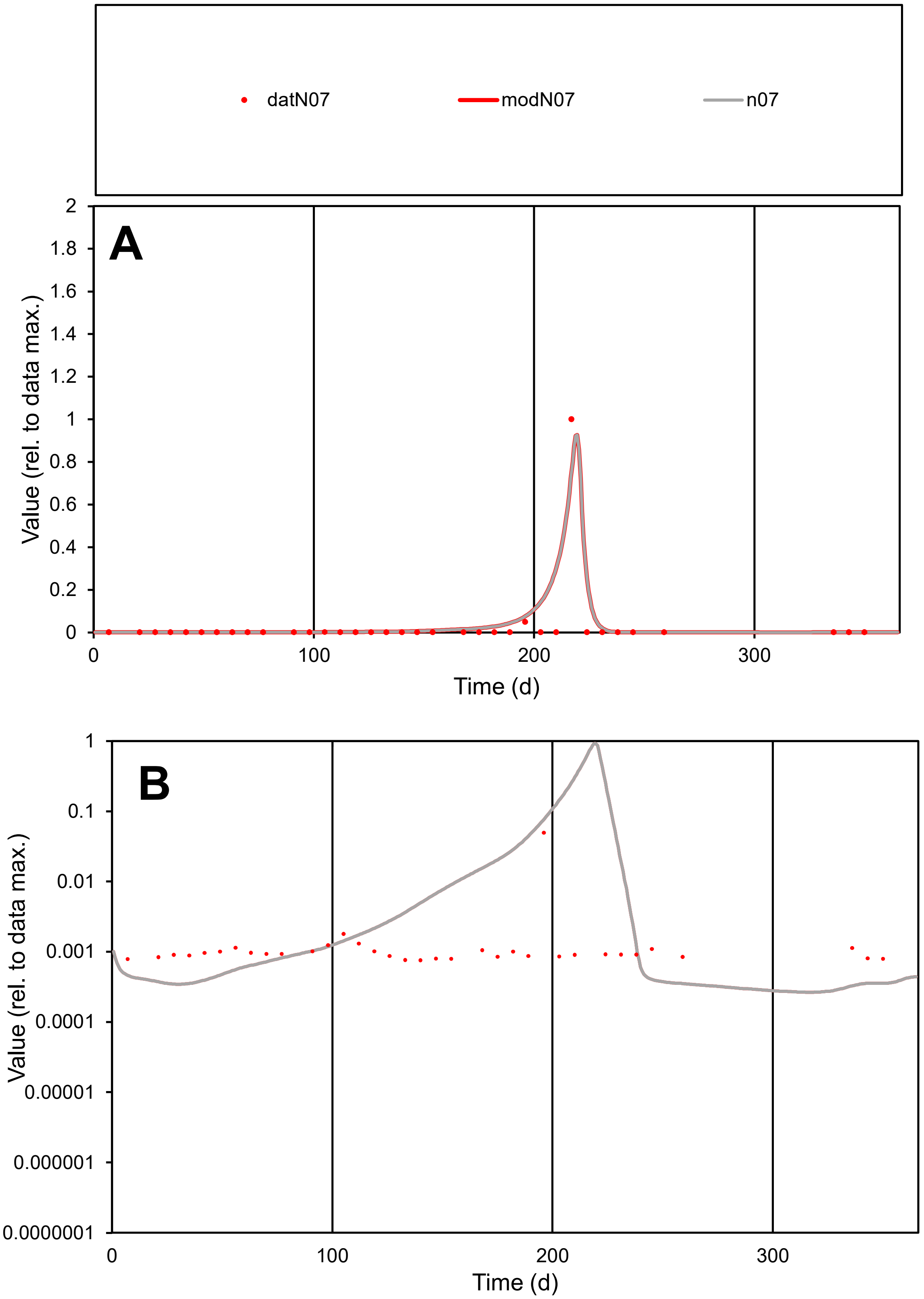


**C1**

Variables
N07 Provide jtd or Dst
CalibOnly?
2 1 to reproduce of
oNormalize?
2 1=no, 2=yes

ofxn calc*
1.64793
ofxn mod
2.02995

sRun
RunA1
zout
1

Ave
Data
0.00067
Model
0.00094

data/vMax
0.02228

ofxng
311.734

Min d/m
2.25902

ave/min d
39.7409
ave/min m
126.003

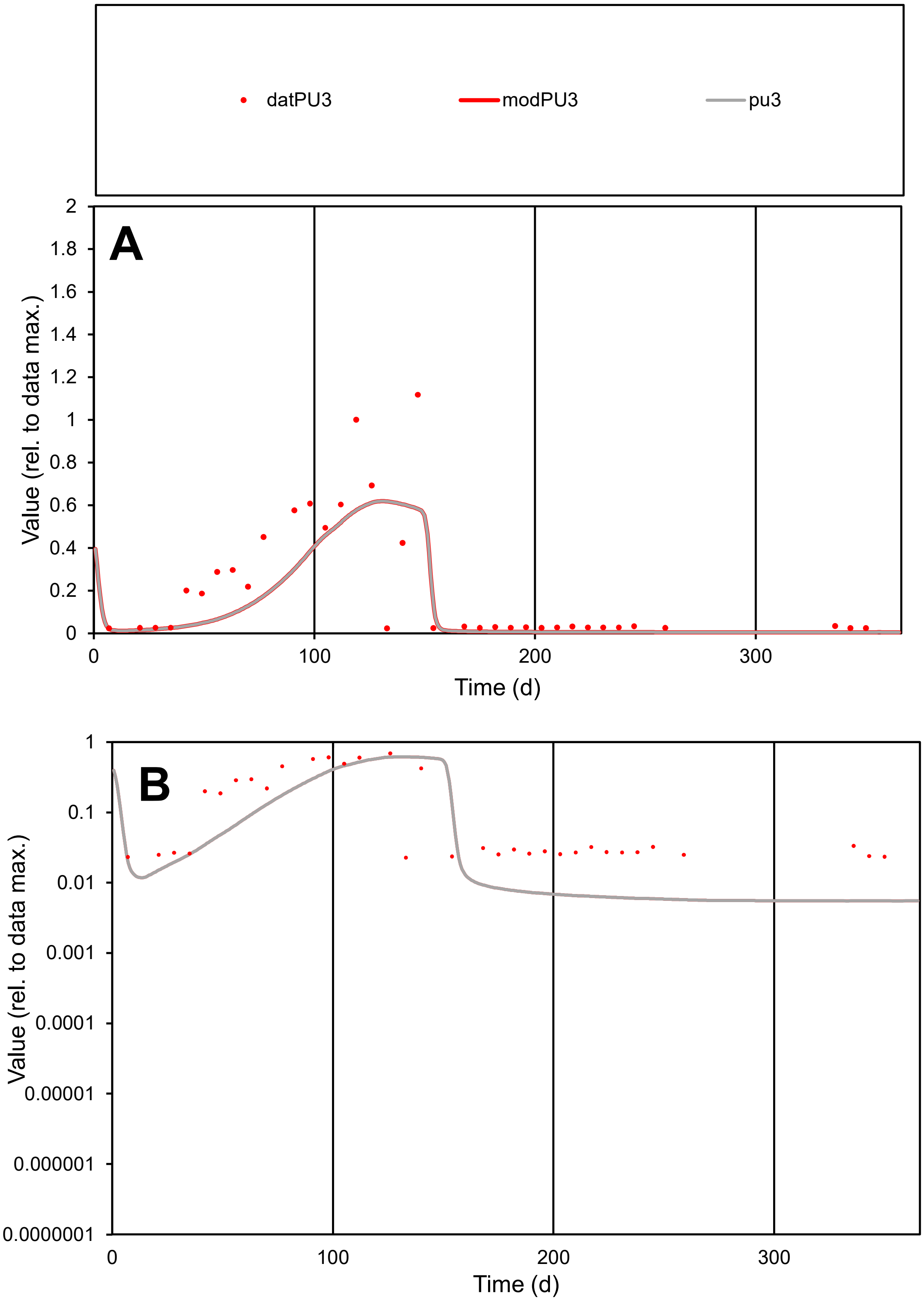


**C1**

Variables
PU3 Provide jtd or Dst
CalibOnly?
2 1 to reproduce of
oNormalize?
2 1=no, 2=yes

ofxn calc*
0.87484
ofxn mod
0.88681

sRun
RunA1
zout
1

Ave
Data
0.0013
Model
0.00102

data/vMax
0.00664

ofxng
311.734

Min d/m
4.08424

ave/min d
8.68023
ave/min m
27.7435

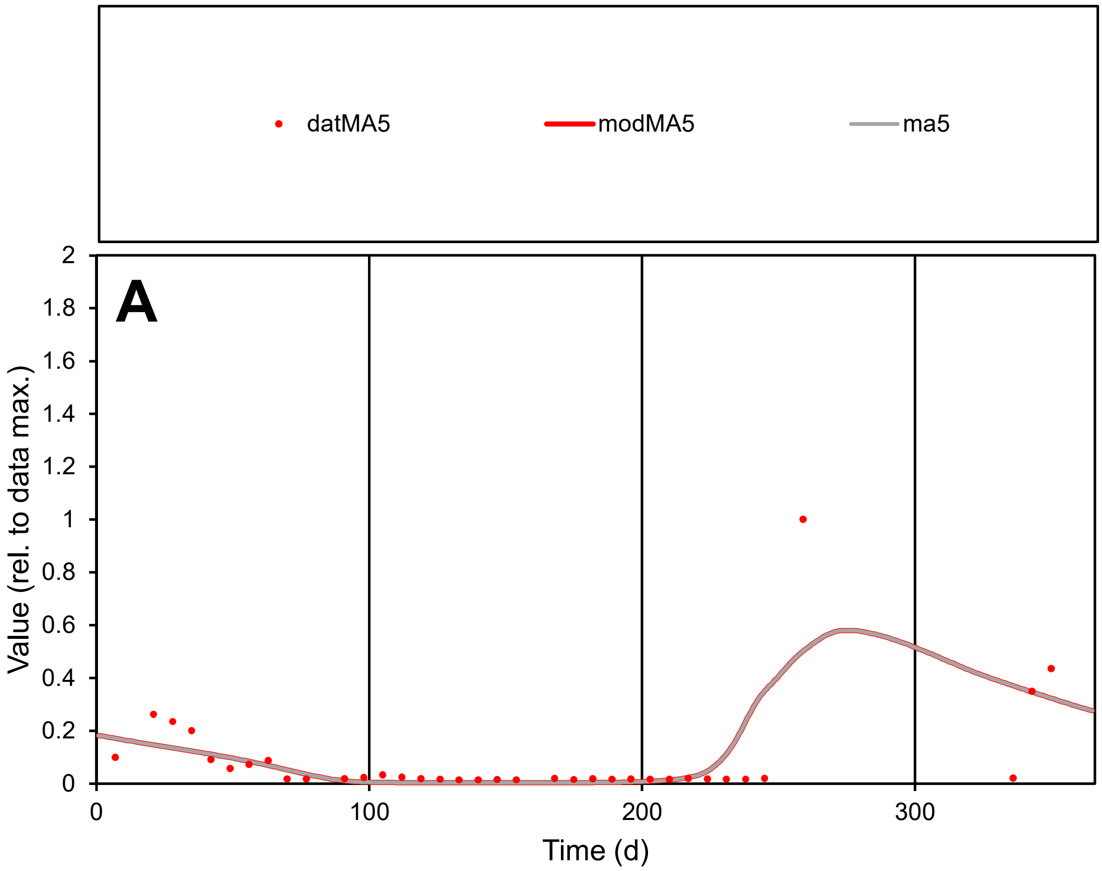


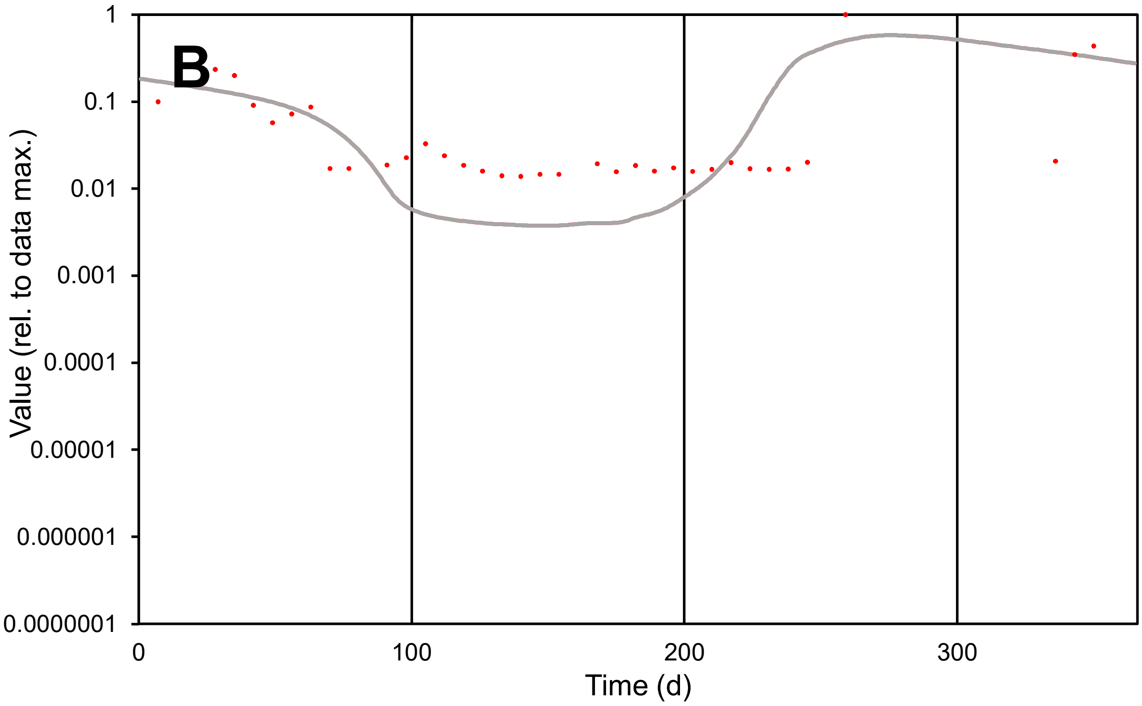


**C1**

| | |
|---|---|
| Variables | |
| MA5 | Provide jtd or Dst |
| CalibOnly? | |
| 2 | 1 to reproduce of |
| oNormalize? | |
| 2 | 1=no, 2=yes |
| ofxn calc* | |
| 1.35283 | |
| ofxn mod | |
| 1.38036 | |
| sRun | |
| RunA1 | |
| zout | |
| 1 | |
| Ave | |
| Data | |
| 0.00062 | |
| Model | |
| 0.00063 | |
| data/vMax | |
| 0.00669 | |
| ofxng | |
| 311.734 | |
| Min d/m | |
| 3.71195 | |
| ave/min d | |
| 6.68737 | |
| ave/min m | |
| 25.0699 | |

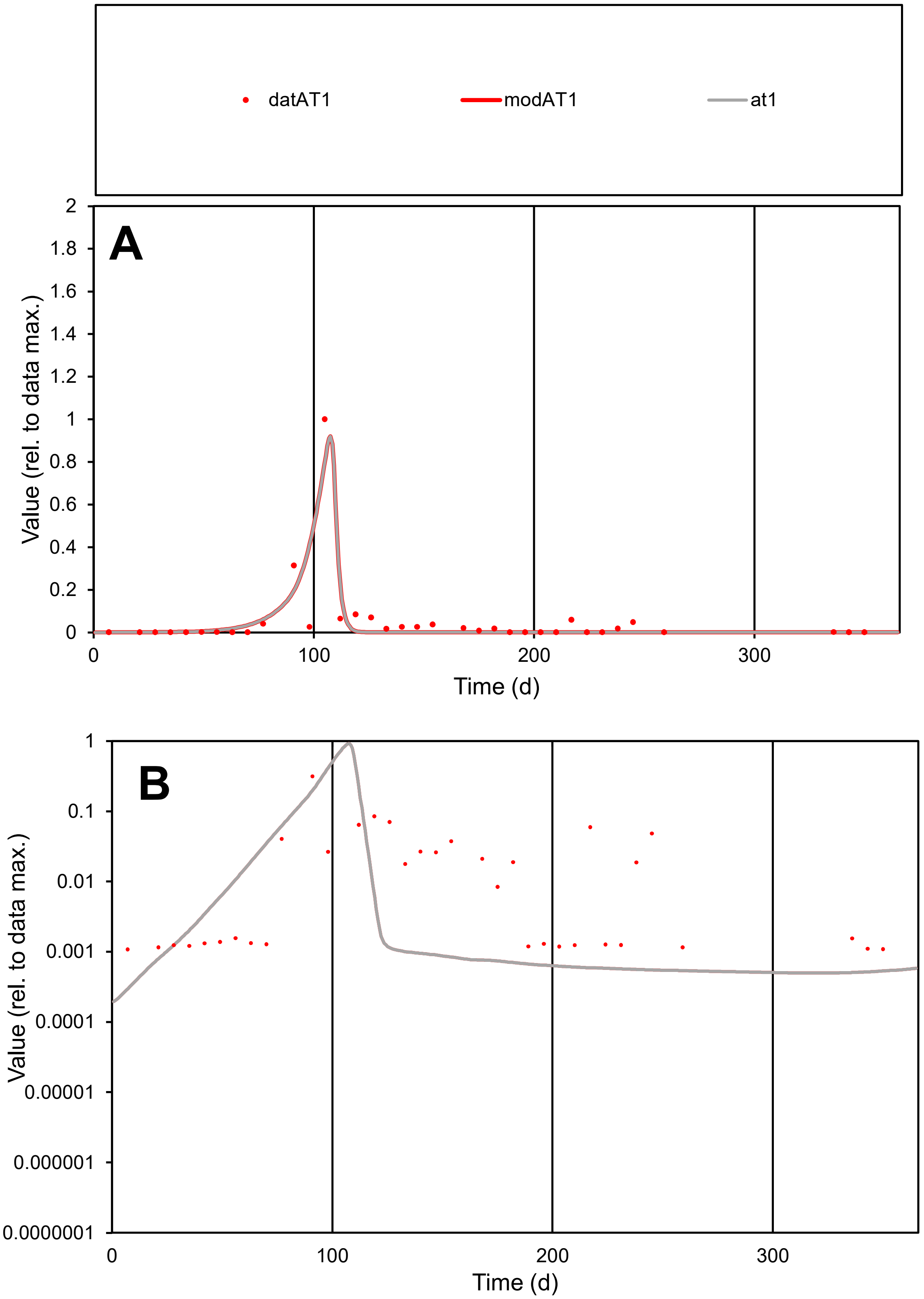


**C1**

Variables

AT1 Provide jtd or Dst

CalibOnly?

2 1 to reproduce of

oNormalize?

2 1=no, 2=yes

ofxn calc*

1.18767

ofxn mod

1.36358

sRun

RunA1

zout

1

Ave

Data

0.00235

Model

0.00216

data/vMax

0.04552

ofxng

311.734

Min d/m

3.41765

ave/min d

47.6938

ave/min m

149.874

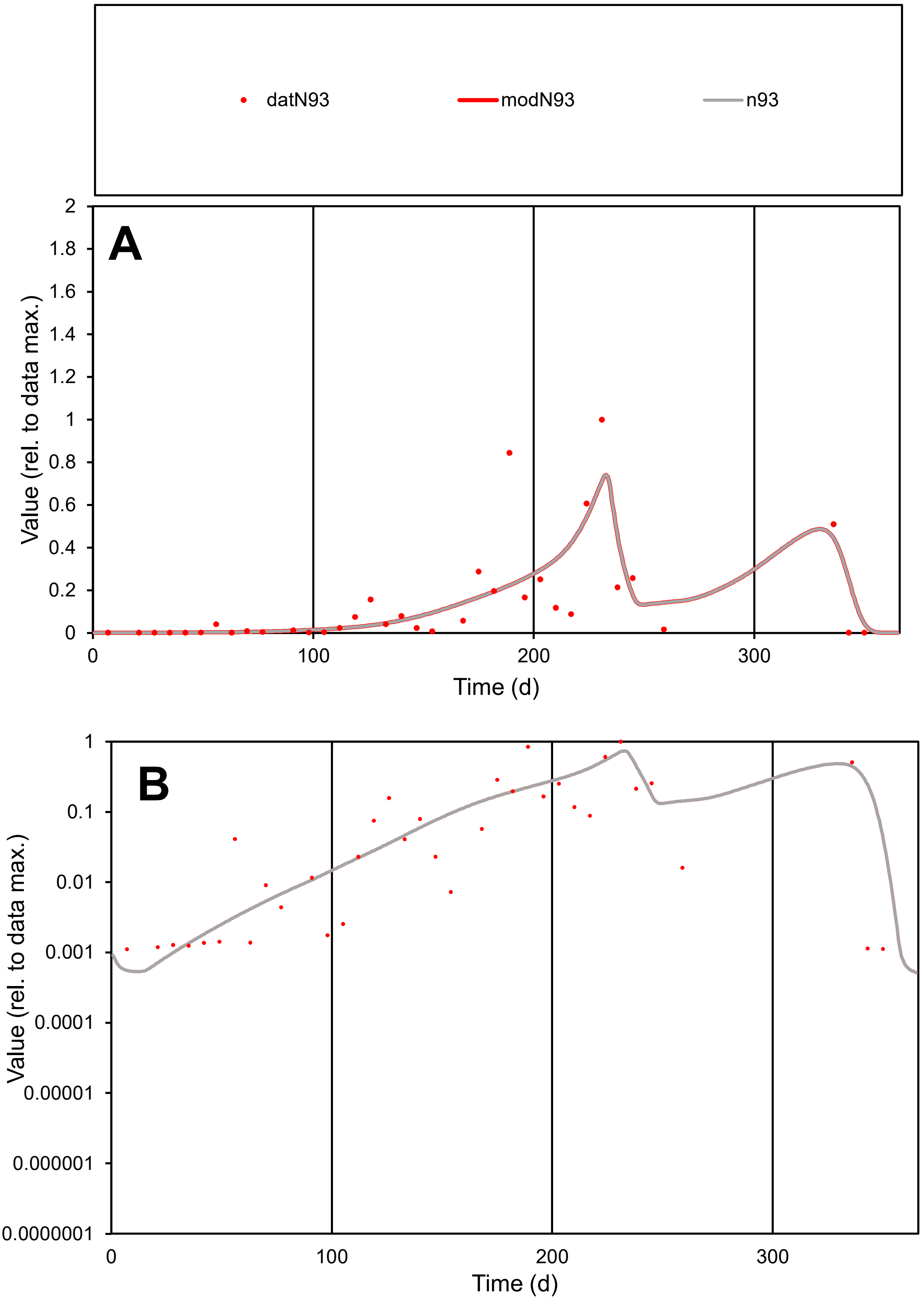


**C1**

Variables
N93 Provide jtd or Dst
CalibOnly?
2 1 to reproduce of
oNormalize?
2 1=no, 2=yes

ofxn calc*
1.03072
ofxn mod
1.07037

sRun
RunA1
zout
1

Ave
Data
0.00222
Model
0.00225

data/vMax
0.016

ofxng
311.734

Min d/m
2.03351

ave/min d
124.532
ave/min m
256.904

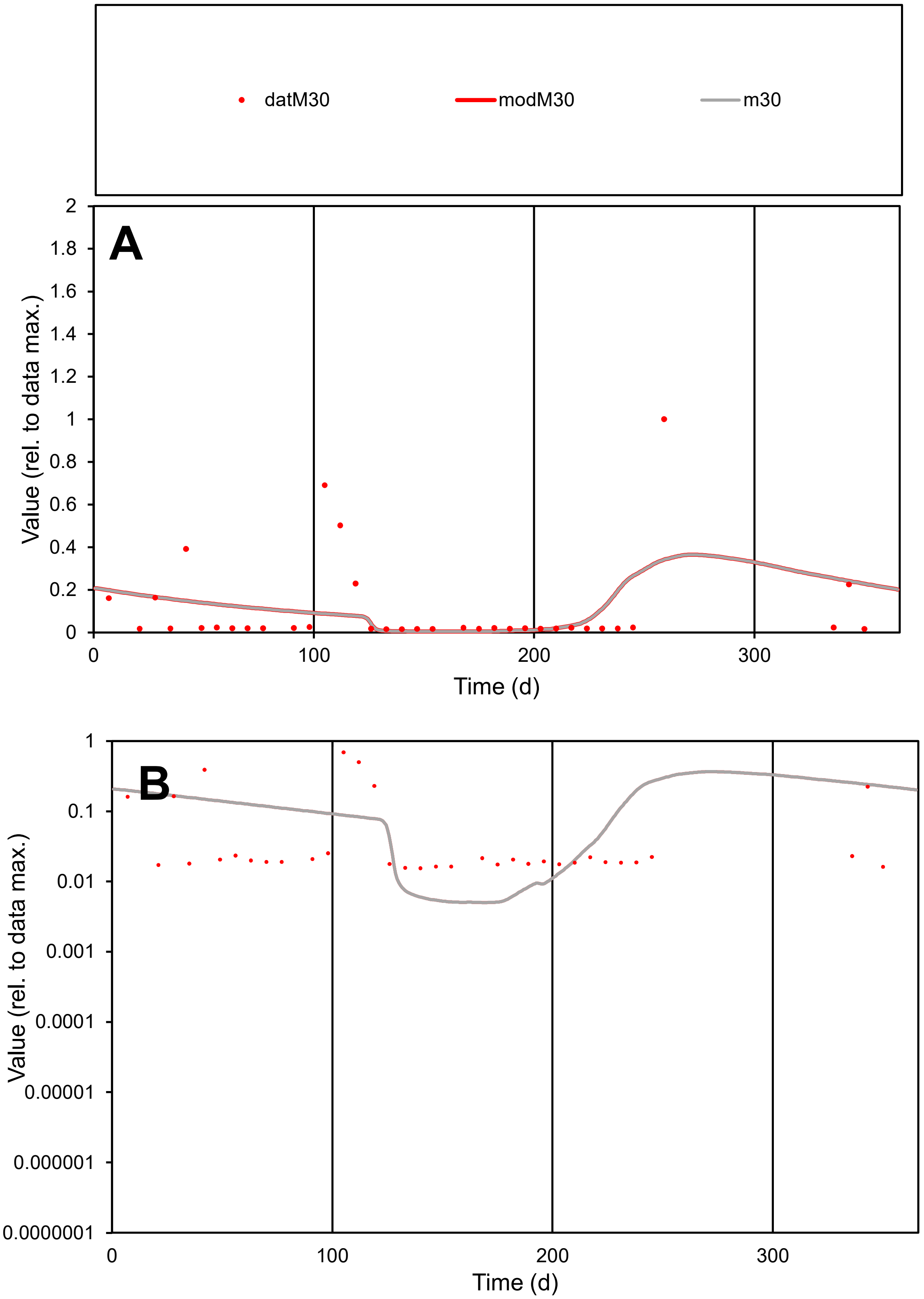


**C1**

Variables
M30 Provide jtd or Dst
CalibOnly?
2 1 to reproduce of
oNormalize?
2 1=no, 2=yes

ofxn calc*
1.78883
ofxn mod
1.80193

sRun
RunA1
zout
1

Ave
Data
0.00028
Model
0.00027

data/vMax
0.00262

ofxng
311.734

Min d/m
3.09793

ave/min d
7.00123
ave/min m
20.6144

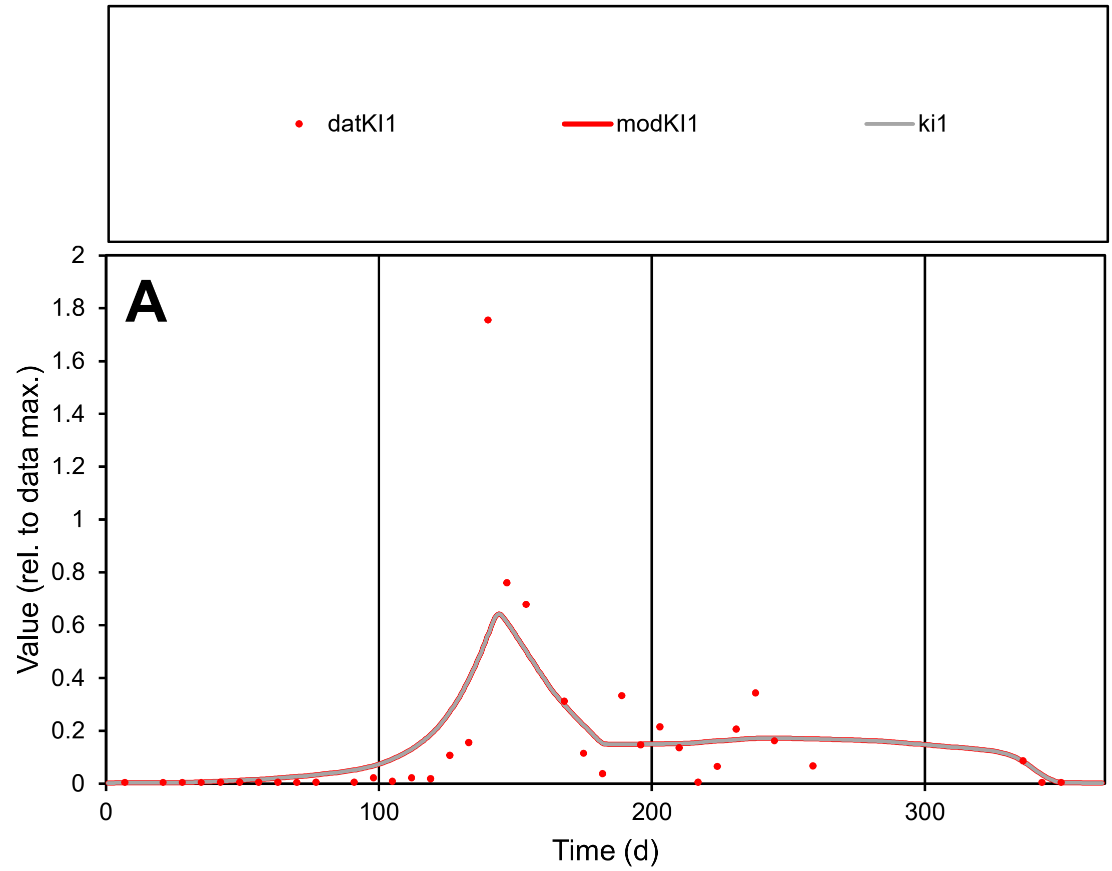


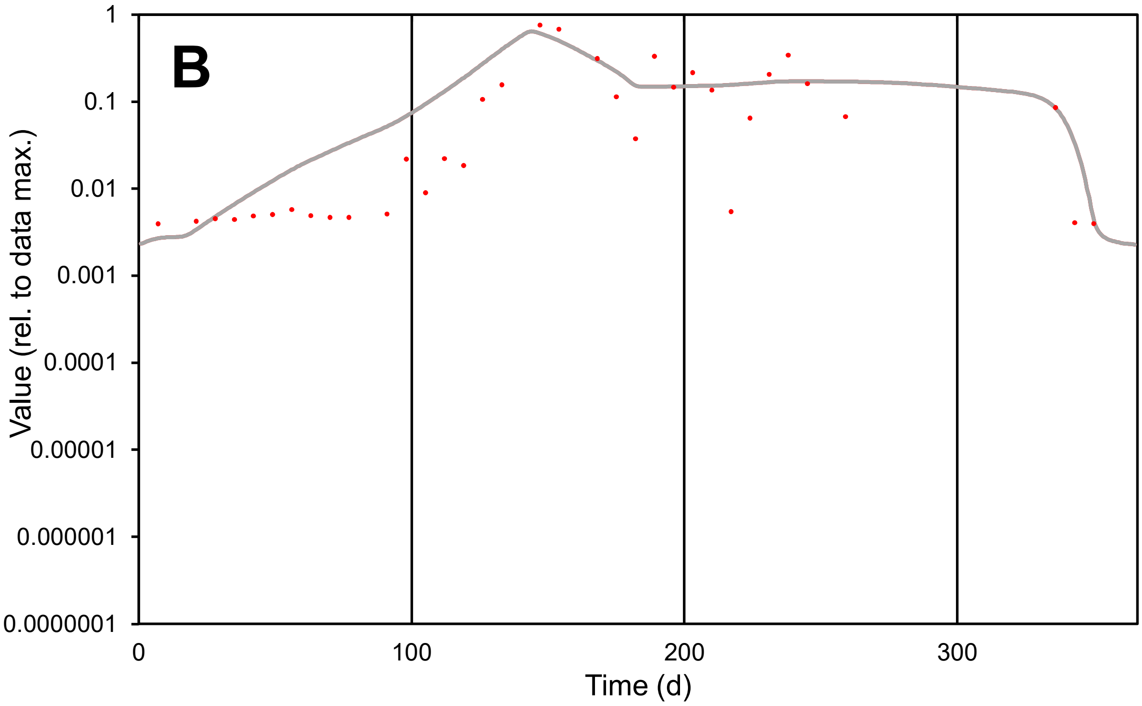


**C1**

Variables

KI1 Provide jtd or Dst

CalibOnly?

2 1 to reproduce of

oNormalize?

2 1=no, 2=yes

ofxn calc*

0.89144

ofxn mod

0.92374

sRun

RunA1

zout

1

Ave

Data

0.00155

Model

0.00179

data/vMax

0.01214

ofxng

311.734

Min d/m

1.45659

ave/min d

32.4057

ave/min m

54.4064

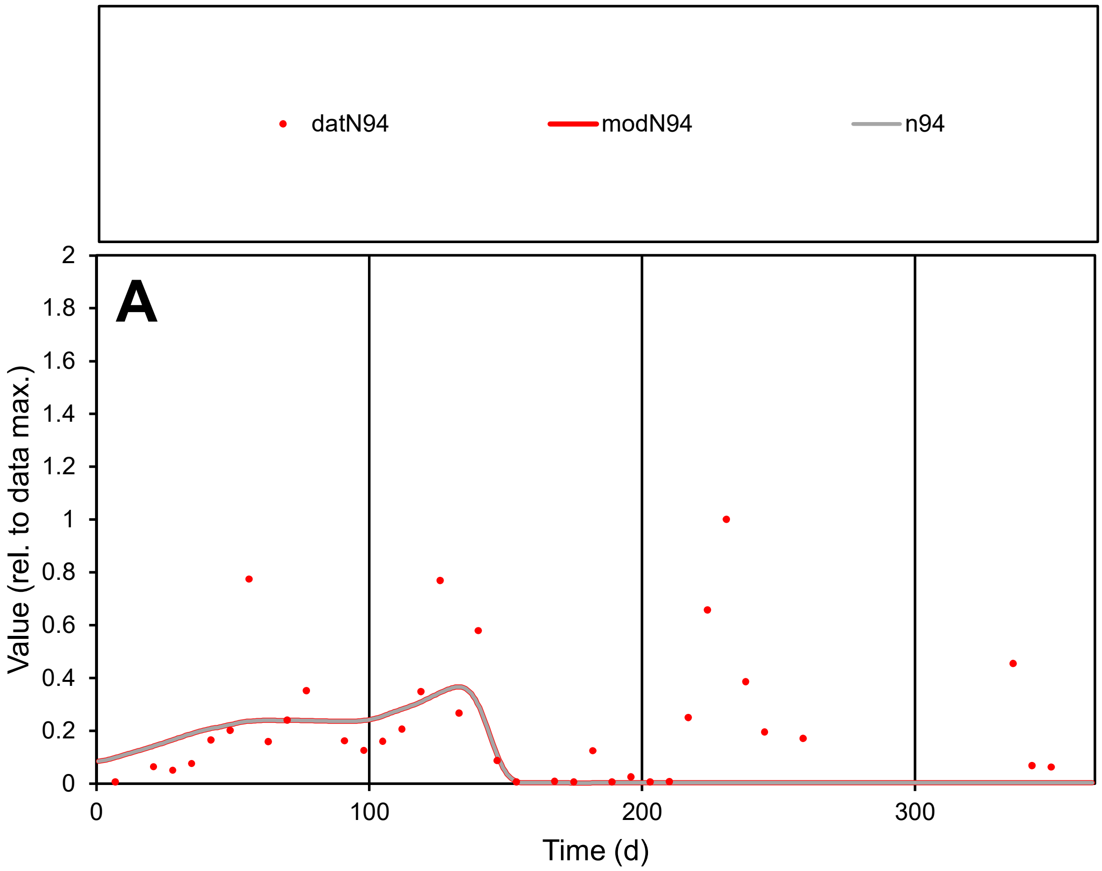


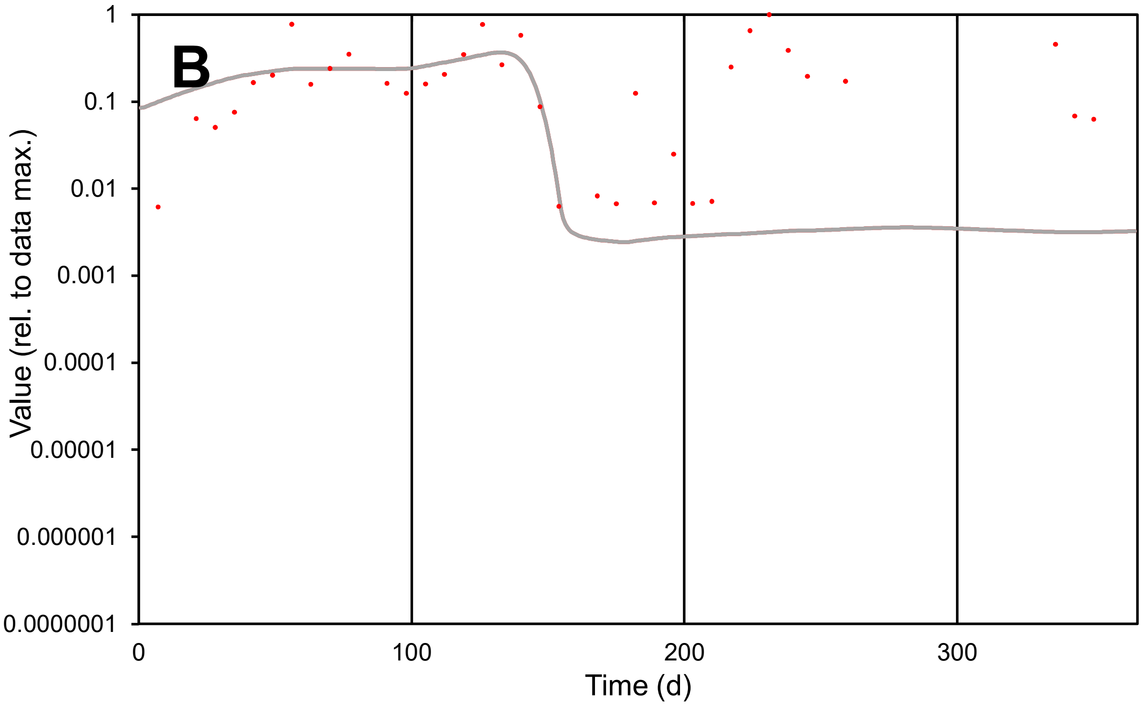


**C1**

Variables
N94 Provide jtd or Dst
CalibOnly?
2 1 to reproduce of
oNormalize?
2 1=no, 2=yes

ofxn calc*
1.20936
ofxn mod
1.21615

sRun
RunA1
zout
1

Ave
Data
0.0016
Model
0.00093

data/vMax
0.00752

ofxng
311.734

Min d/m
2.53428

ave/min d
34.3515
ave/min m
50.585

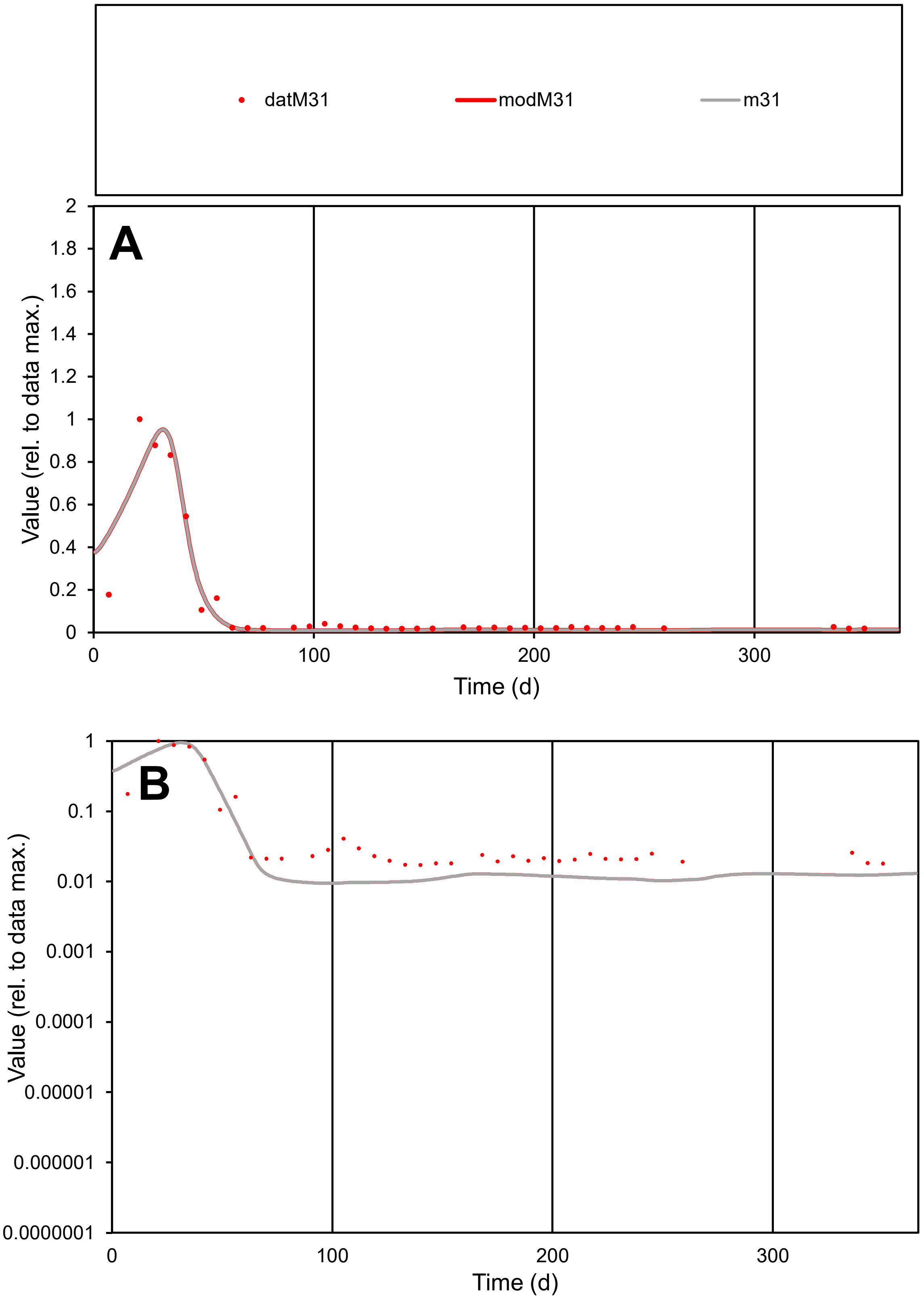


**C1**

Variables
M31 Provide jtd or Dst
CalibOnly?
2 1 to reproduce of
oNormalize?
2 1=no, 2=yes

ofxn calc*
0.53577
ofxn mod
0.6085

sRun
RunA1
zout
1

Ave
Data
0.00056
Model
0.00053

data/vMax
0.00467

ofxng
311.734

Min d/m
1.82362

ave/min d
7.004
ave/min m
12.1235

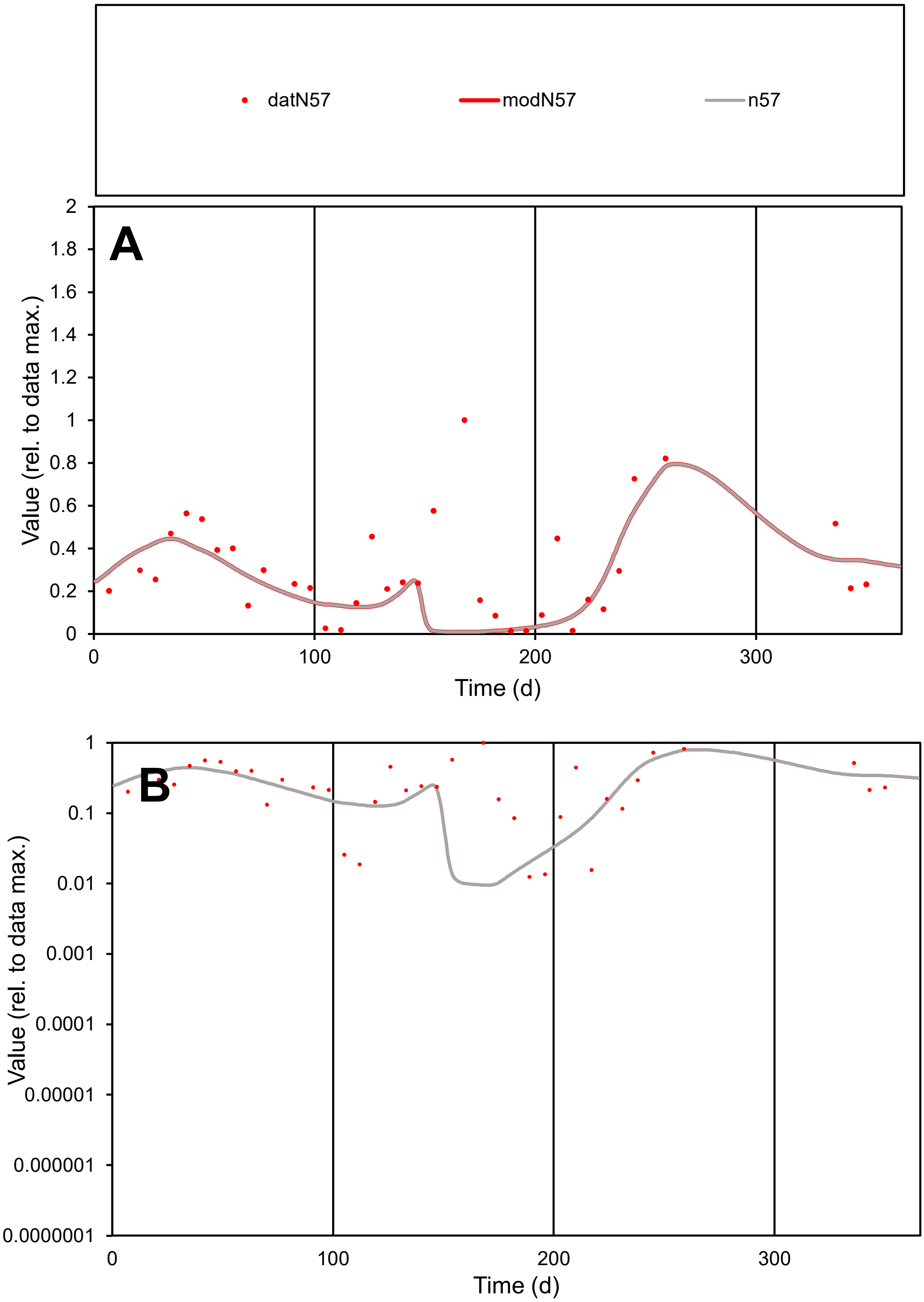


**C1**

Variables
N57 Provide jtd or Dst
CalibOnly?
2 1 to reproduce of
oNormalize?
2 1=no, 2=yes

ofxn calc*
0.75114
ofxn mod
0.76347

sRun
RunA1
zout
1

Ave
Data
0.00094
Model
0.00077

data/vMax
0.00329

ofxng
311.734

Min d/m
1.31386

ave/min d
22.8739
ave/min m
24.7198

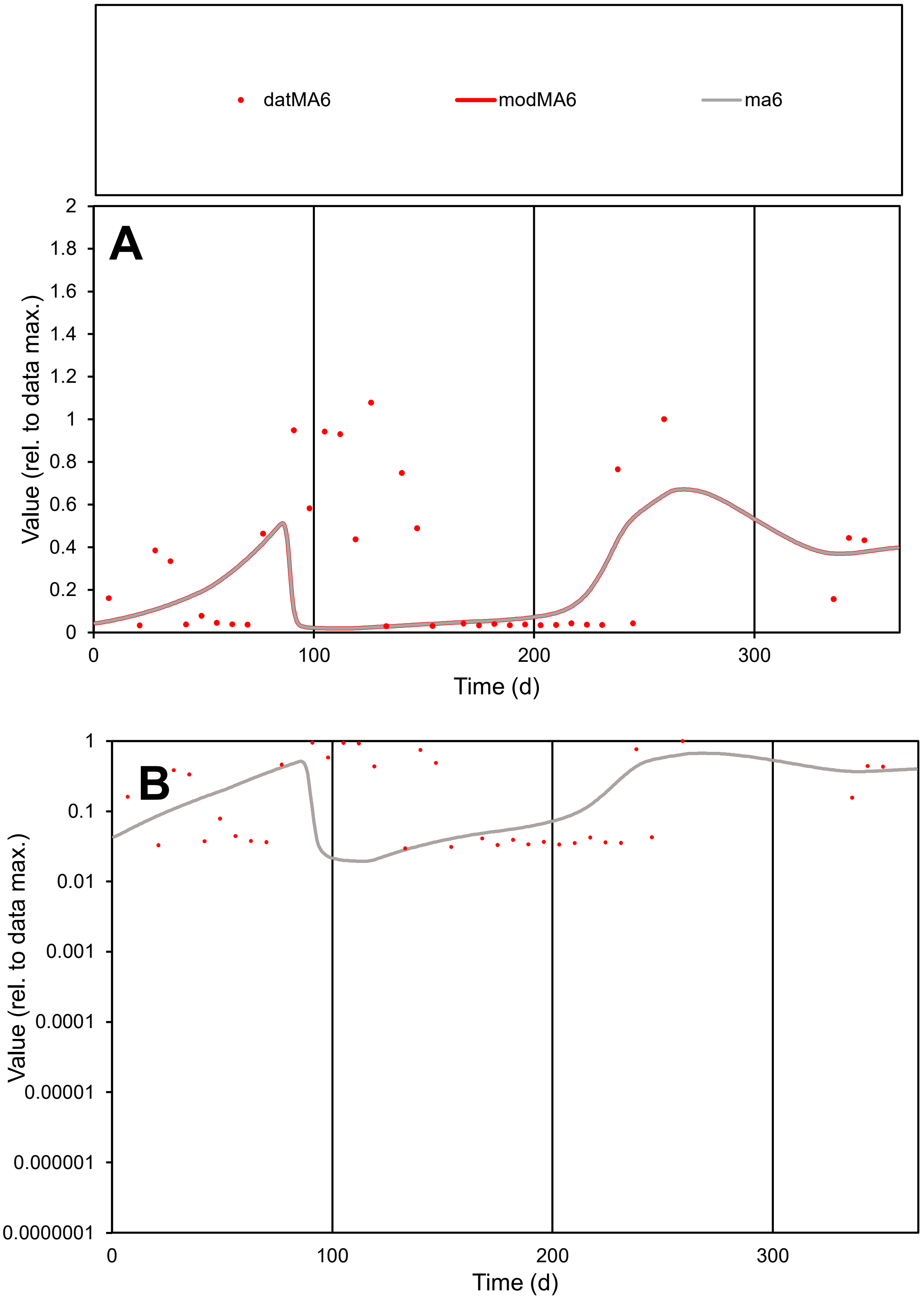


**C1**

Variables
MA6 Provide jtd or Dst
CalibOnly?
2 1 to reproduce of
oNormalize?
2 1=no, 2=yes

ofxn calc*
1.1986
ofxn mod
1.25316

sRun
RunA1
zout
1

Ave
Data
0.00113
Model
0.00072

data/vMax
0.00395

ofxng
311.734

Min d/m
1.53655

ave/min d
9.56384
ave/min m
9.38444

331_Calib_S_A

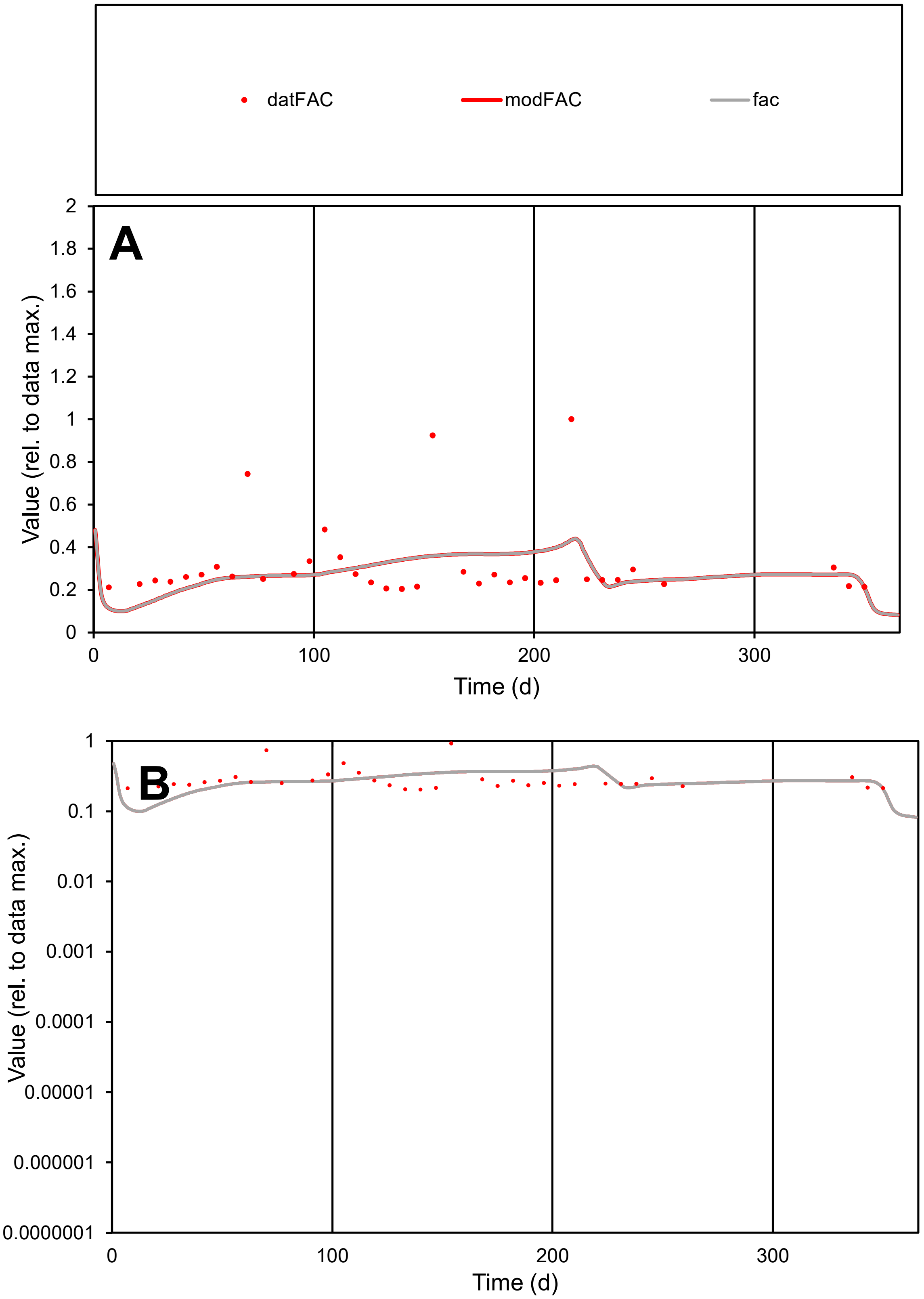


C1

Variables

FAC Provide jtd or Dst

CalibOnly?

2 1 to reproduce of

oNormalize?

2 1=no, 2=yes

ofxn calc*

0.52499

ofxn mod

0.53698

sRun

RunA1

zout

1

Ave

Data

0.00013

Model

0.00012

data/vMax

0.00042

ofxng

311.734

Min d/m

1.8843

ave/min d

1.50204

ave/min m

2.63005

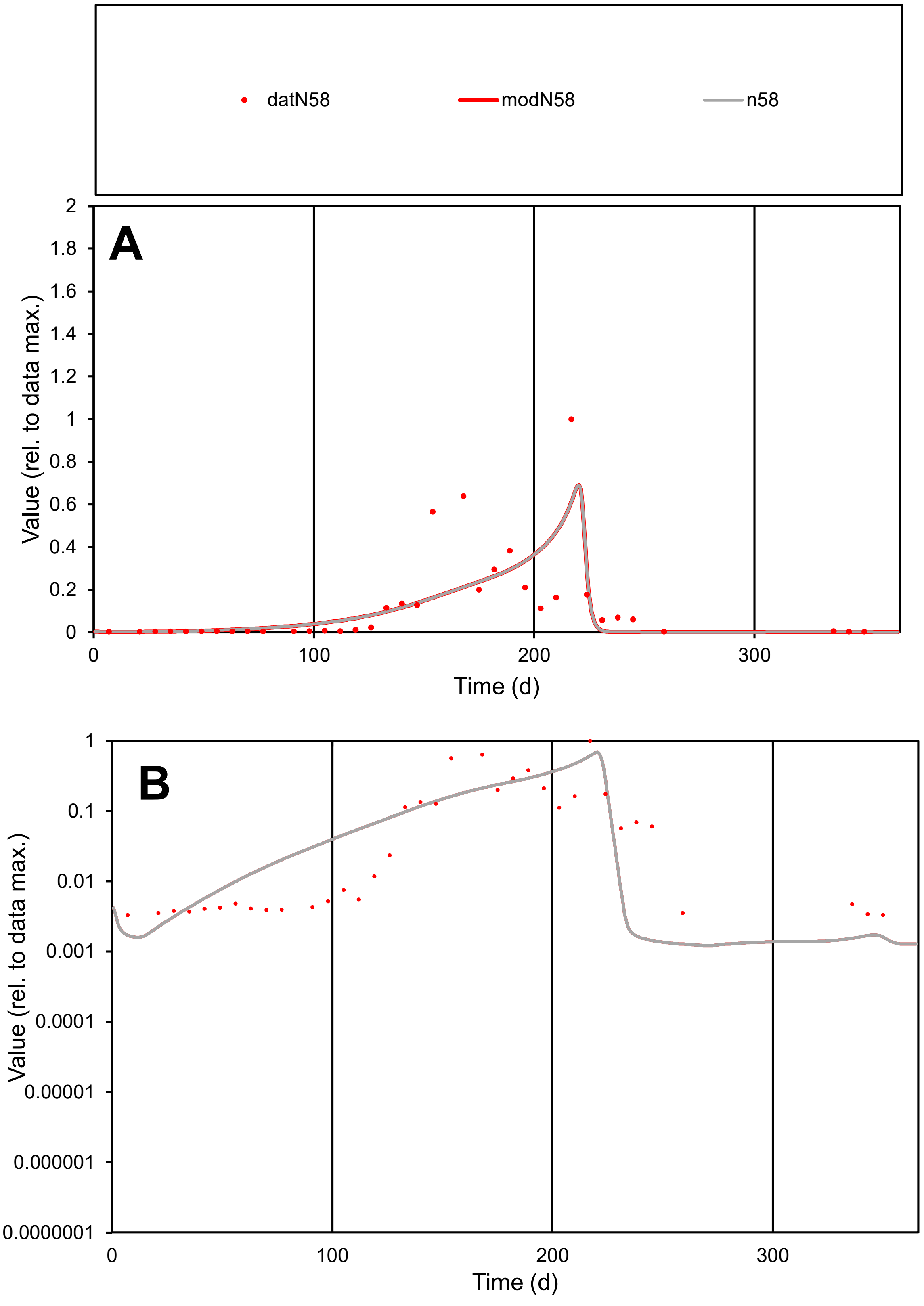


**C1**

Variables

N58 Provide jtd or Dst

CalibOnly?

2 1 to reproduce of

oNormalize?

2 1=no, 2=yes

ofxn calc*

1.11694

ofxn mod

1.13648

sRun

RunA1

zout

1

Ave

Data

0.00292

Model

0.00286

data/vMax

0.02537

ofxng

311.734

Min d/m

2.6155

ave/min d

34.7308

ave/min m

89.0931

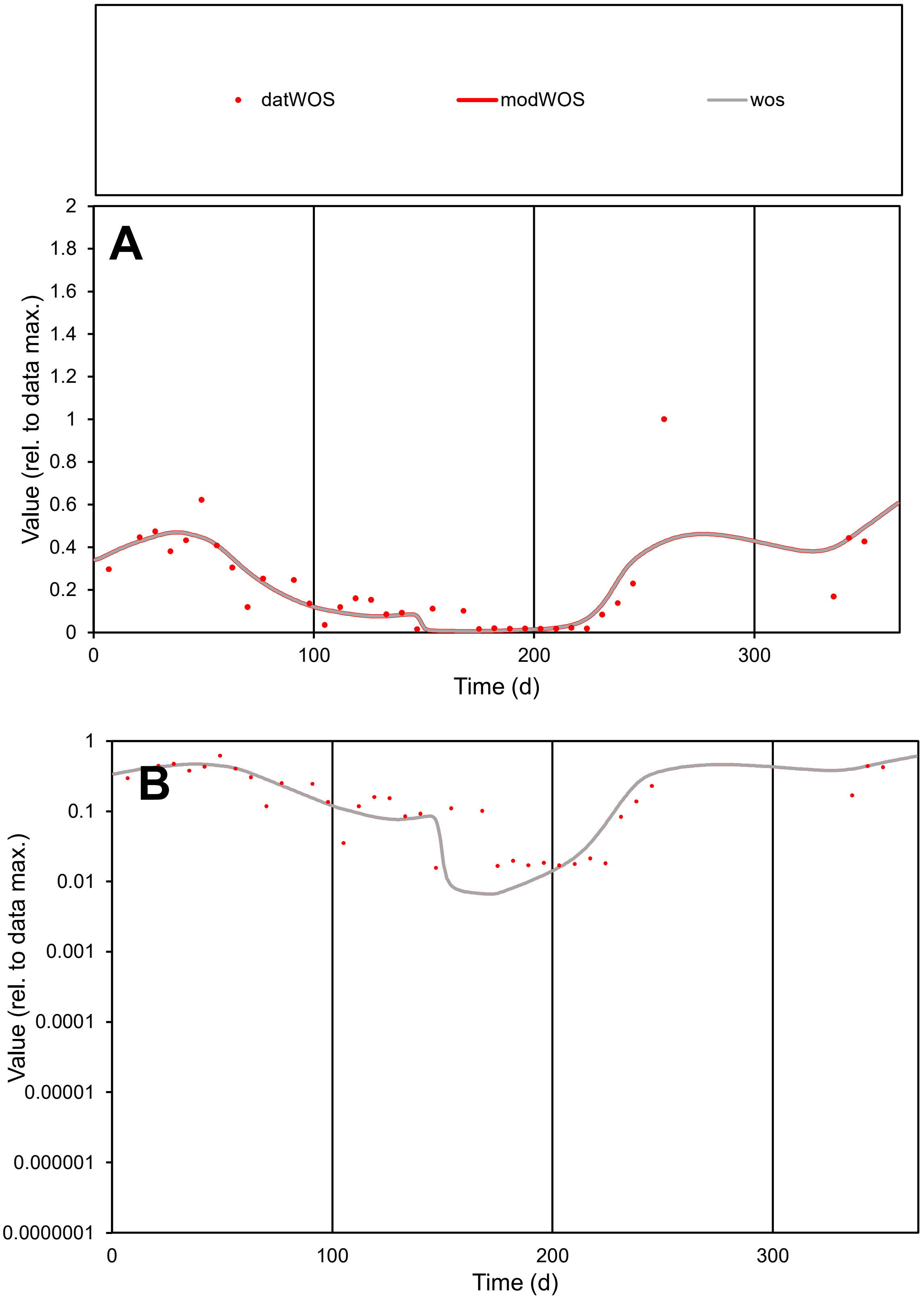


**C1**

Variables
WOS Provide jtd or Dst
CalibOnly?
2 1 to reproduce of
oNormalize?
2 1=no, 2=yes

ofxn calc*
0.5819
ofxn mod
0.60458

sRun
RunA1
zout
1

Ave
Data
0.00095
Model
0.00093

data/vMax
0.00455

ofxng
311.734

Min d/m
2.34618

ave/min d
13.2681
ave/min m
30.7879

331_Calib_S_A

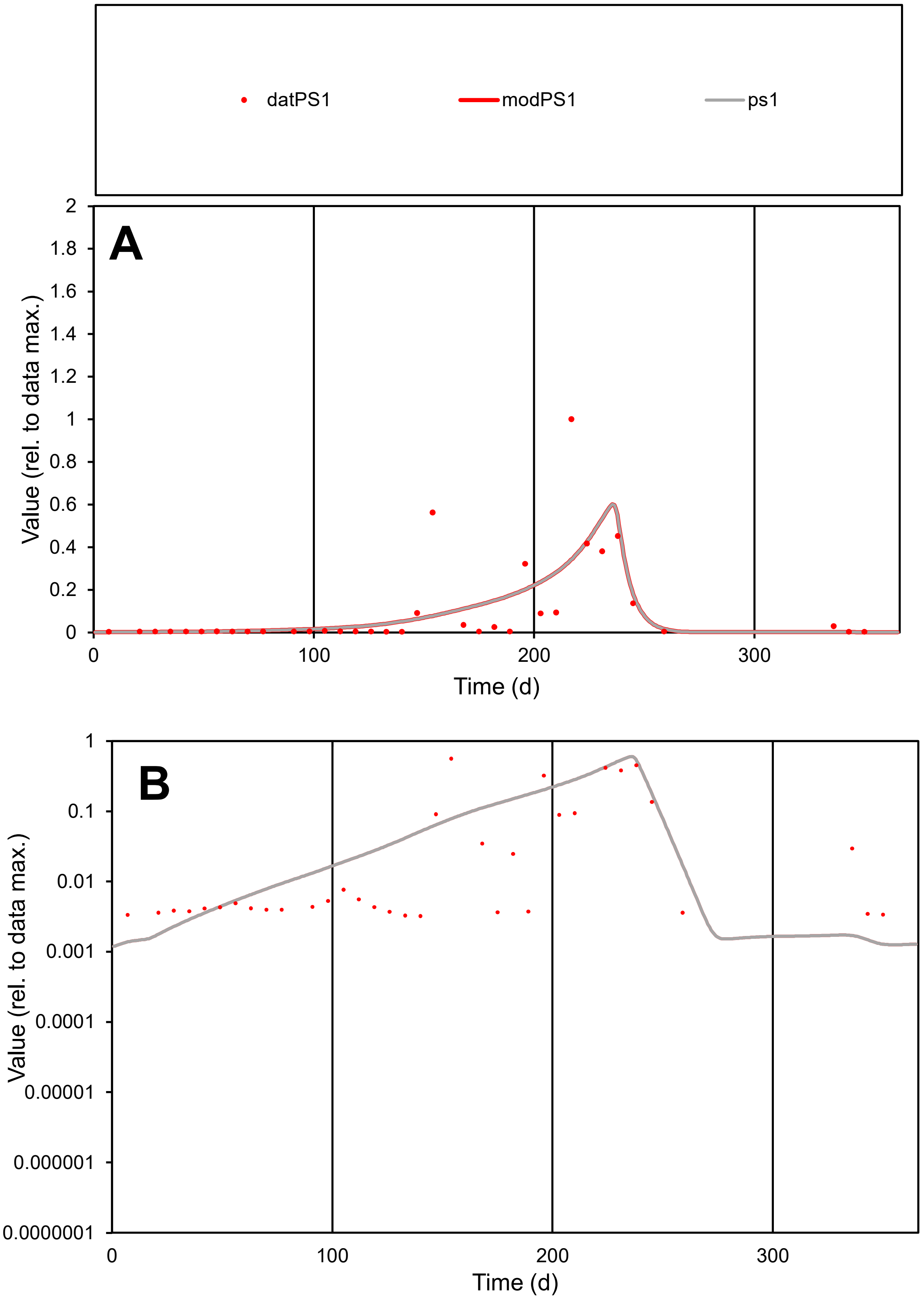


**C1**

Variables
PS1 Provide jtd or Dst
CalibOnly?
2 1 to reproduce of
oNormalize?
2 1=no, 2=yes

ofxn calc*
1.41025
ofxn mod
1.43473

sRun
RunA1
zout
1

Ave
Data
0.00142
Model
0.00146

data/vMax
0.01438

ofxng
311.734

Min d/m
2.55044

ave/min d
30.5509
ave/min m
80.2113

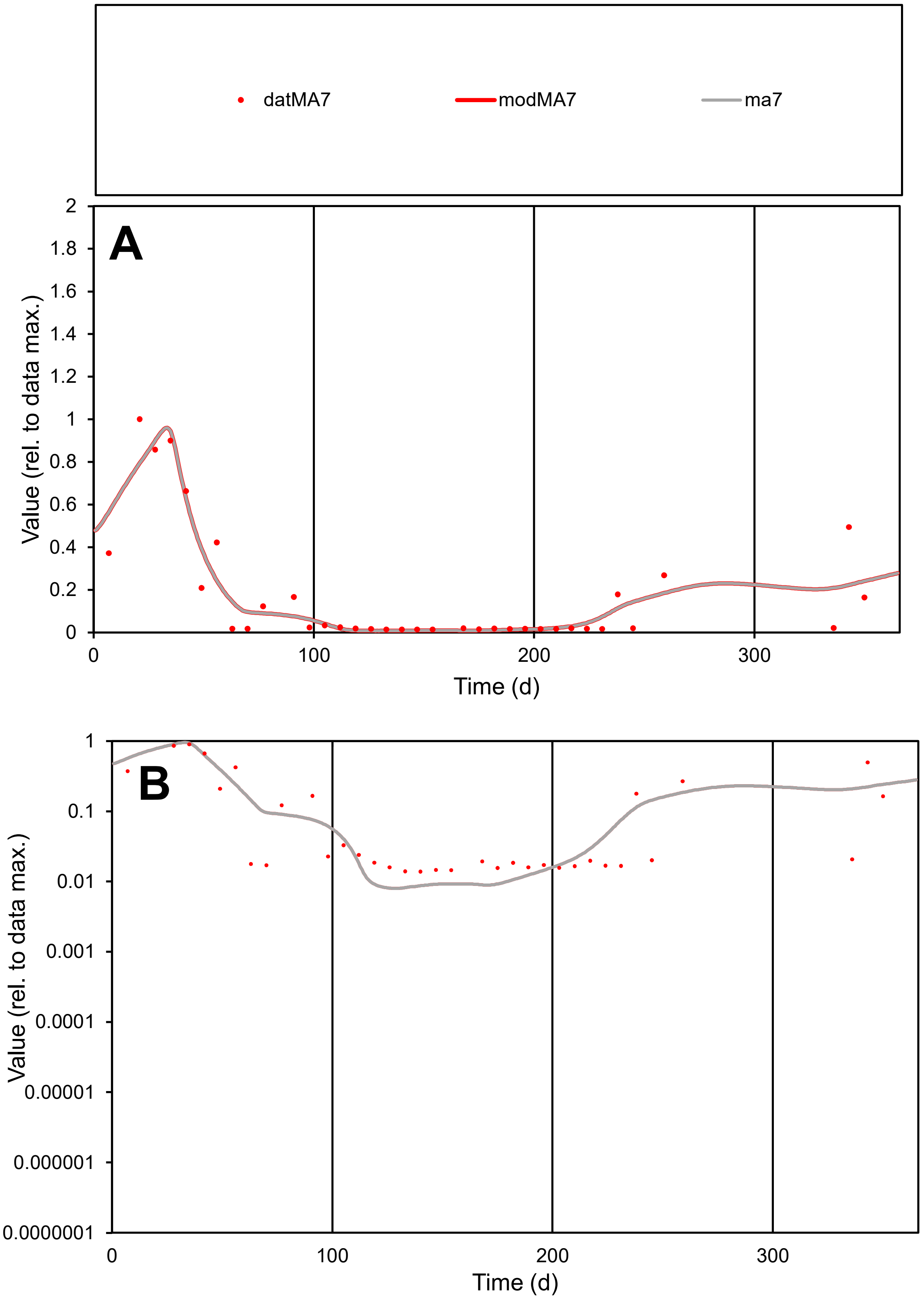


**C1**

Variables
MA7 Provide jtd or Dst
CalibOnly?
2 1 to reproduce of
oNormalize?
2 1=no, 2=yes

ofxn calc*
0.54048
ofxn mod
0.58609

sRun
RunA1
zout
1

Ave
Data
0.00017
Model
0.00017

data/vMax
0.00096

ofxng
311.734

Min d/m
1.72912

ave/min d
12.5141
ave/min m
22.053

331_Calib_S_A

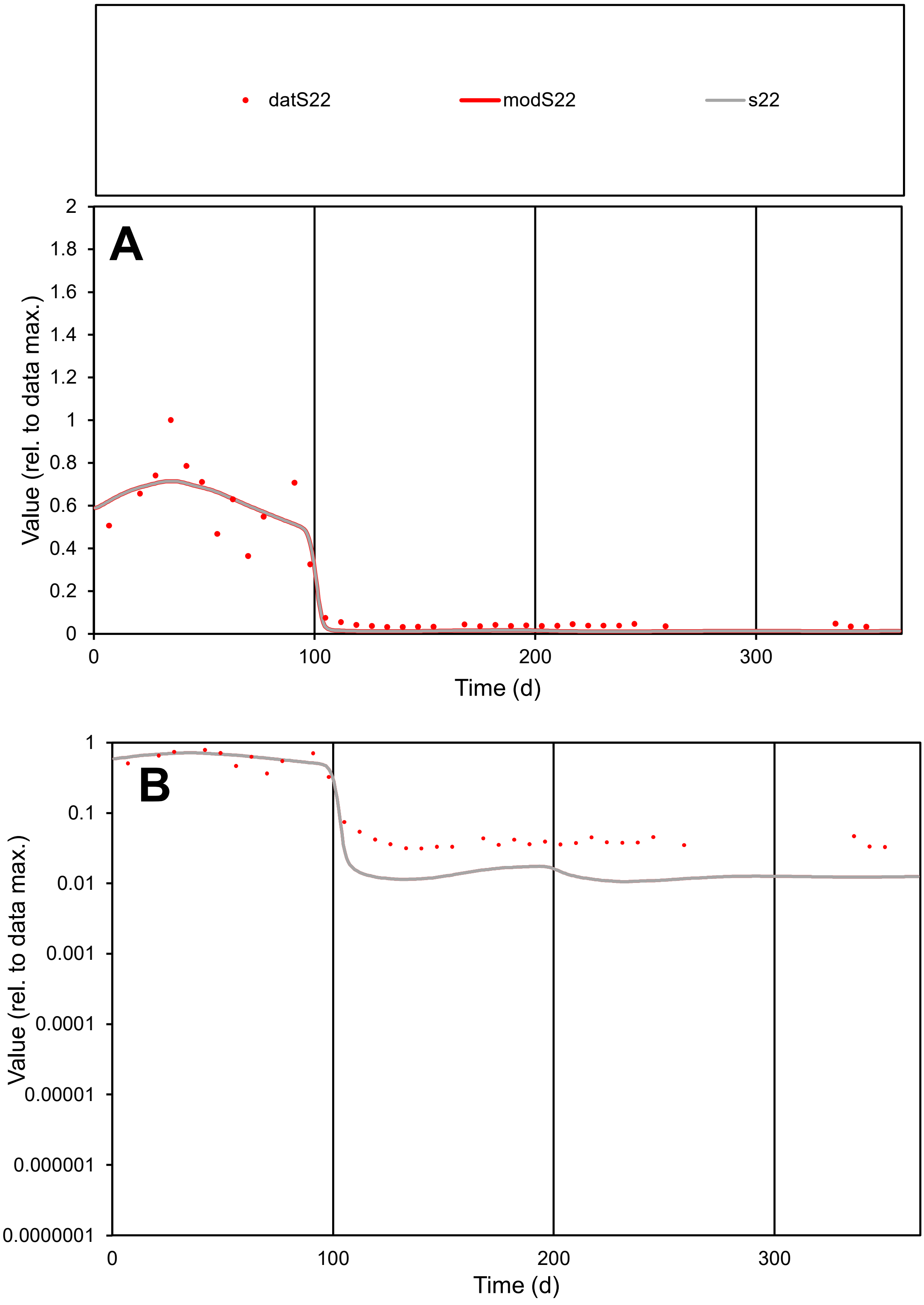


**C1**

Variables
S22 Provide jtd or Dst
CalibOnly?
2 1 to reproduce of
oNormalize?
2 1=no, 2=yes

ofxn calc*
0.3588
ofxn mod
0.40682

sRun
RunA1
zout
1

Ave
Data
0.00108
Model
0.00099

data/vMax
0.00462

ofxng
311.734

Min d/m
2.97192

ave/min d
7.43006
ave/min m
20.3895

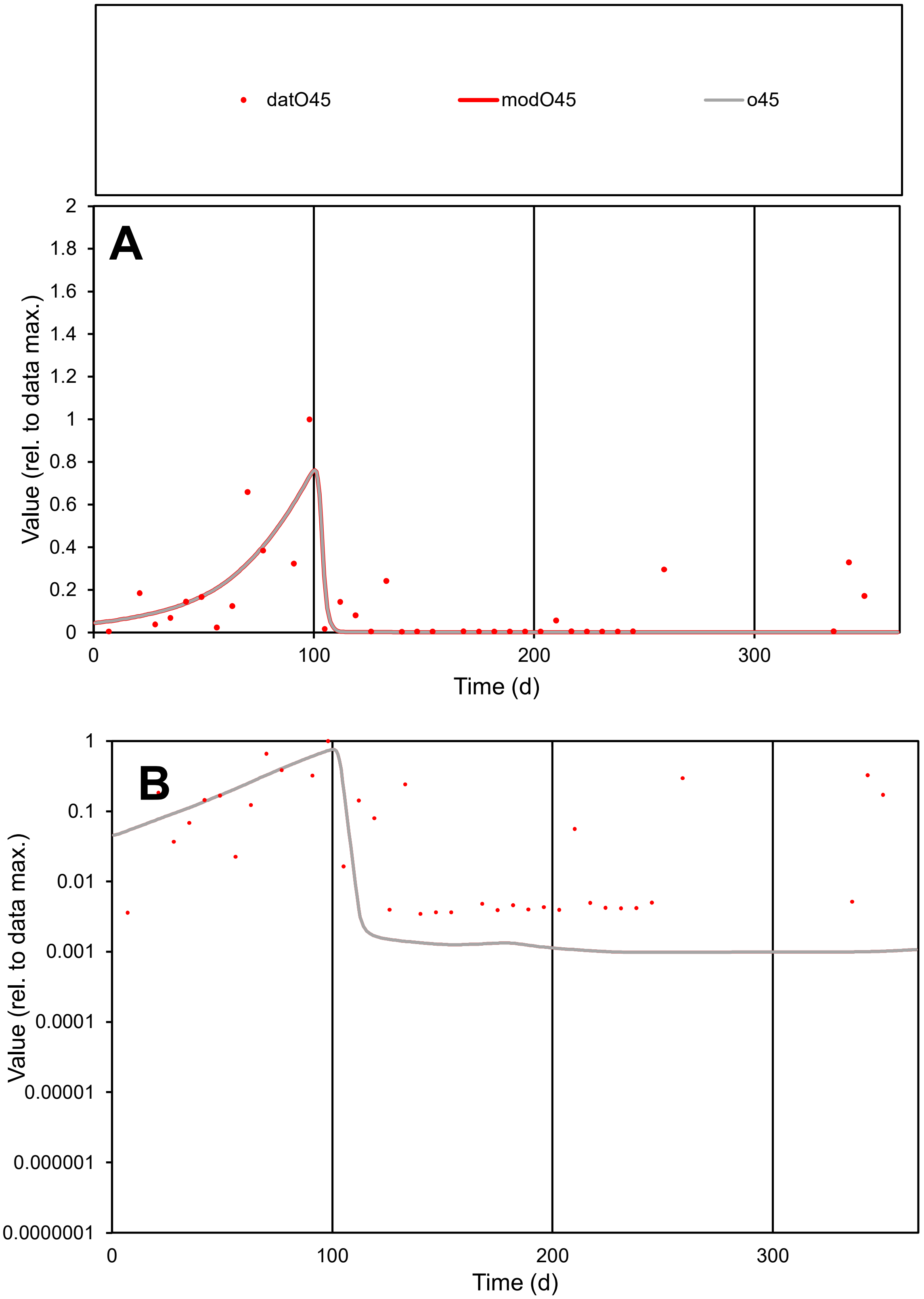


**C1**

Variables
O45 Provide jtd or Dst
CalibOnly?
2 1 to reproduce of
oNormalize?
2 1=no, 2=yes

ofxn calc*
1.04724
ofxn mod
1.07556

sRun
RunA1
zout
1

Ave
Data
0.00201
Model
0.00151

data/vMax
0.01628

ofxng
311.734

Min d/m
3.54899

ave/min d
35.567
ave/min m
95.0833

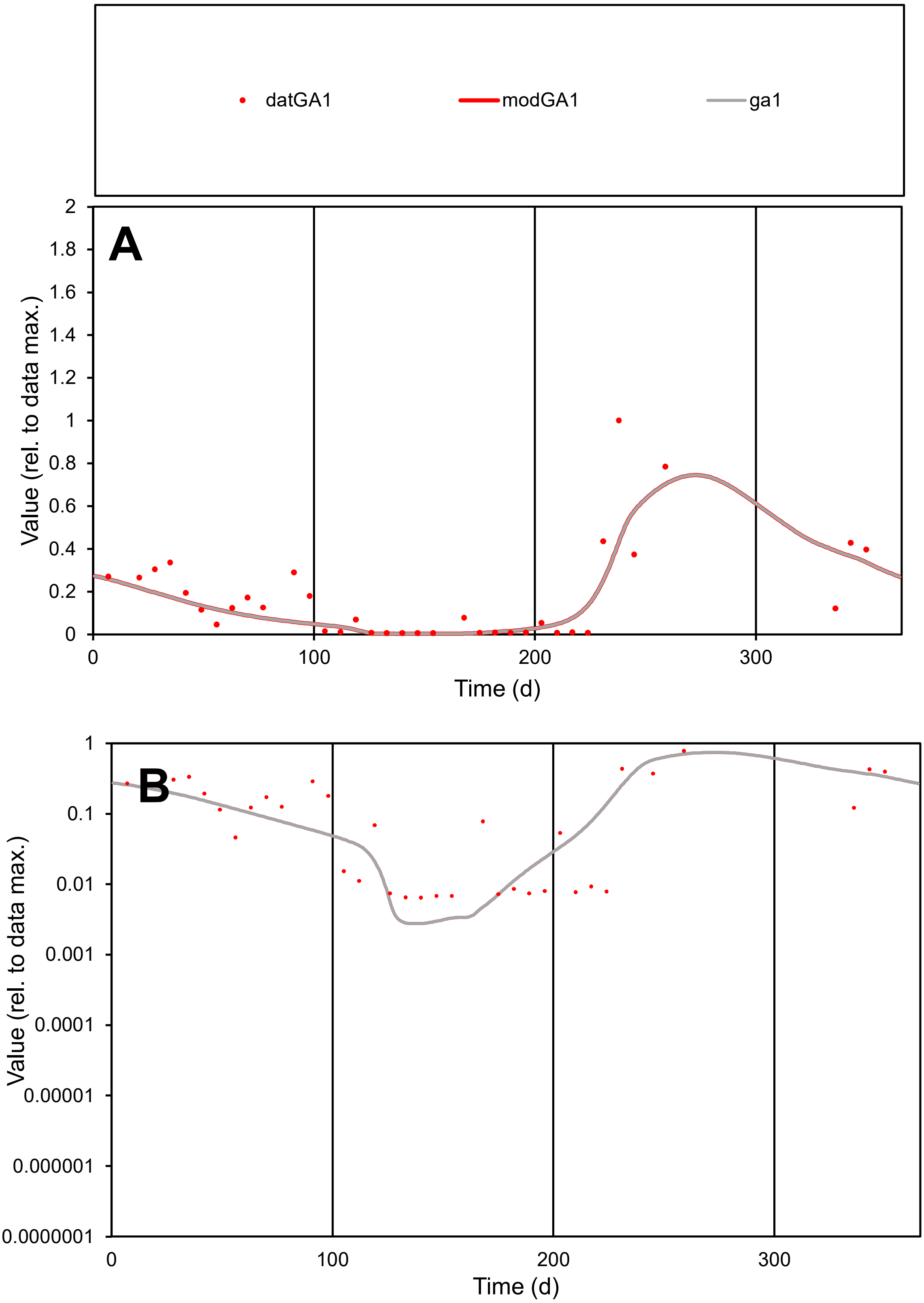


**C1**

Variables
GA1 Provide jtd or Dst
CalibOnly?
2 1 to reproduce of
oNormalize?
2 1=no, 2=yes

ofxn calc*
0.73939
ofxn mod
0.75855

sRun
RunA1
zout
1

Ave
Data
0.00094
Model
0.00078

data/vMax
0.00541

ofxng
311.734

Min d/m
2.32205

ave/min d
27.0003
ave/min m
51.9279

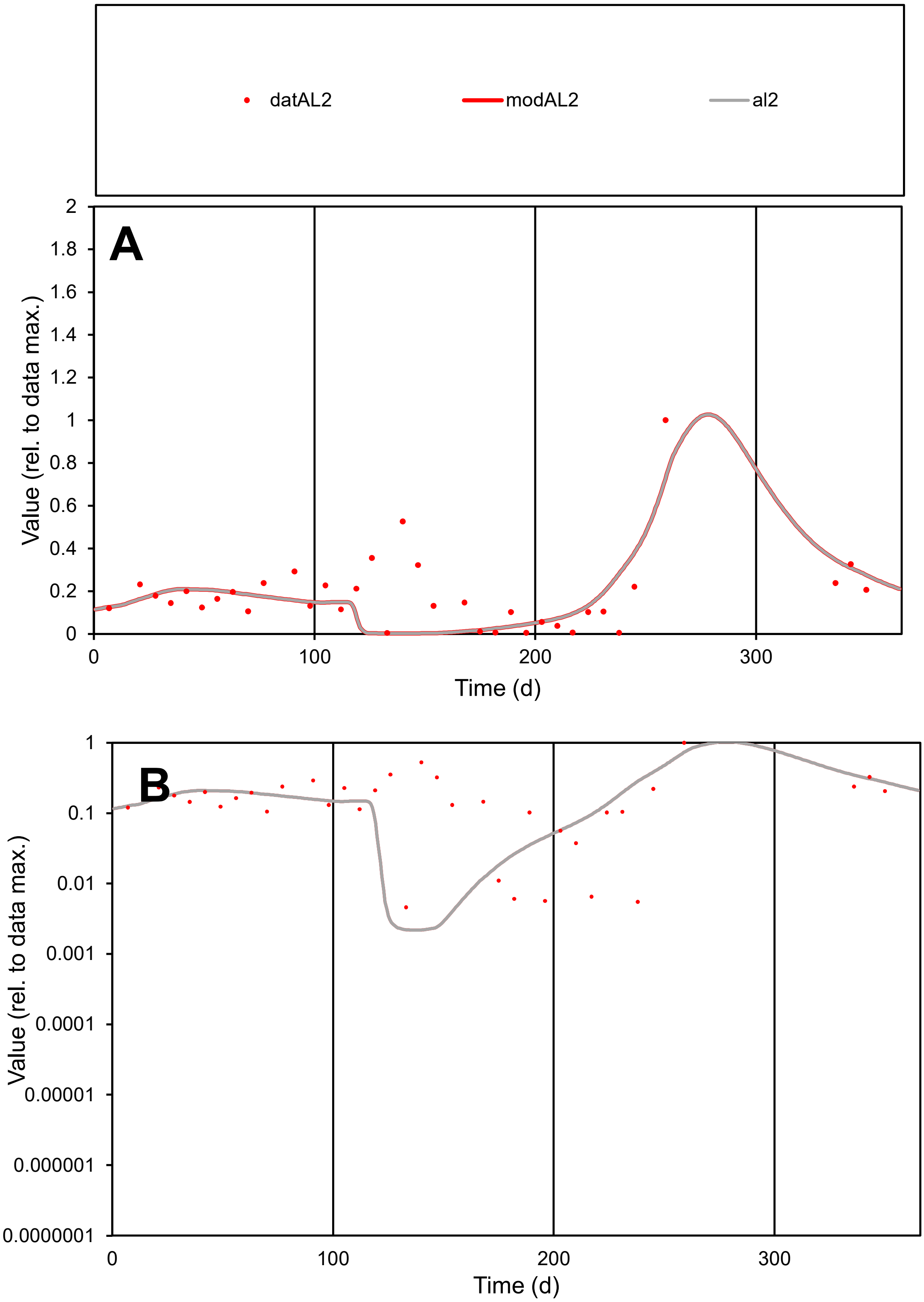


**C1**

Variables

AL2 Provide jtd or Dst

CalibOnly?

2 1 to reproduce of

oNormalize?

2 1=no, 2=yes

ofxn calc*

0.68113

ofxn mod

0.70733

sRun

RunA1

zout

1

Ave

Data

0.0008

Model

0.00074

data/vMax

0.00472

ofxng

311.734

Min d/m

2.09547

ave/min d

37.0452

ave/min m

70.99

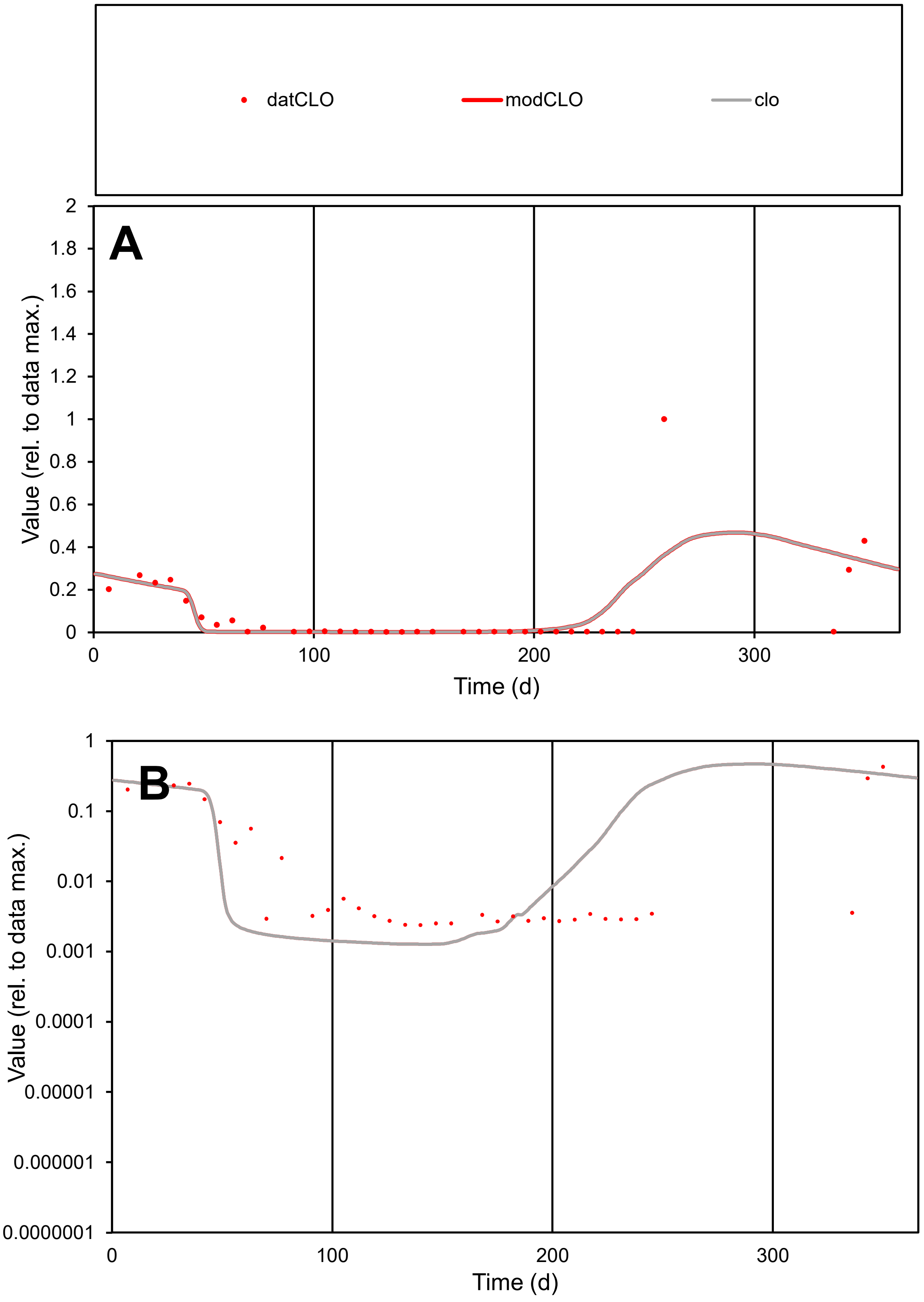


**C1**

Variables
CLO Provide jtd or Dst
CalibOnly?
2 1 to reproduce of
oNormalize?
2 1=no, 2=yes

ofxn calc*
1.55875
ofxn mod
1.58425

sRun
RunA1
zout
1

Ave
Data
0.00047
Model
0.00048

data/vMax
0.00545

ofxng
311.734

Min d/m
1.88595

ave/min d
35.7365
ave/min m
69.3354

331_Calib_S_A

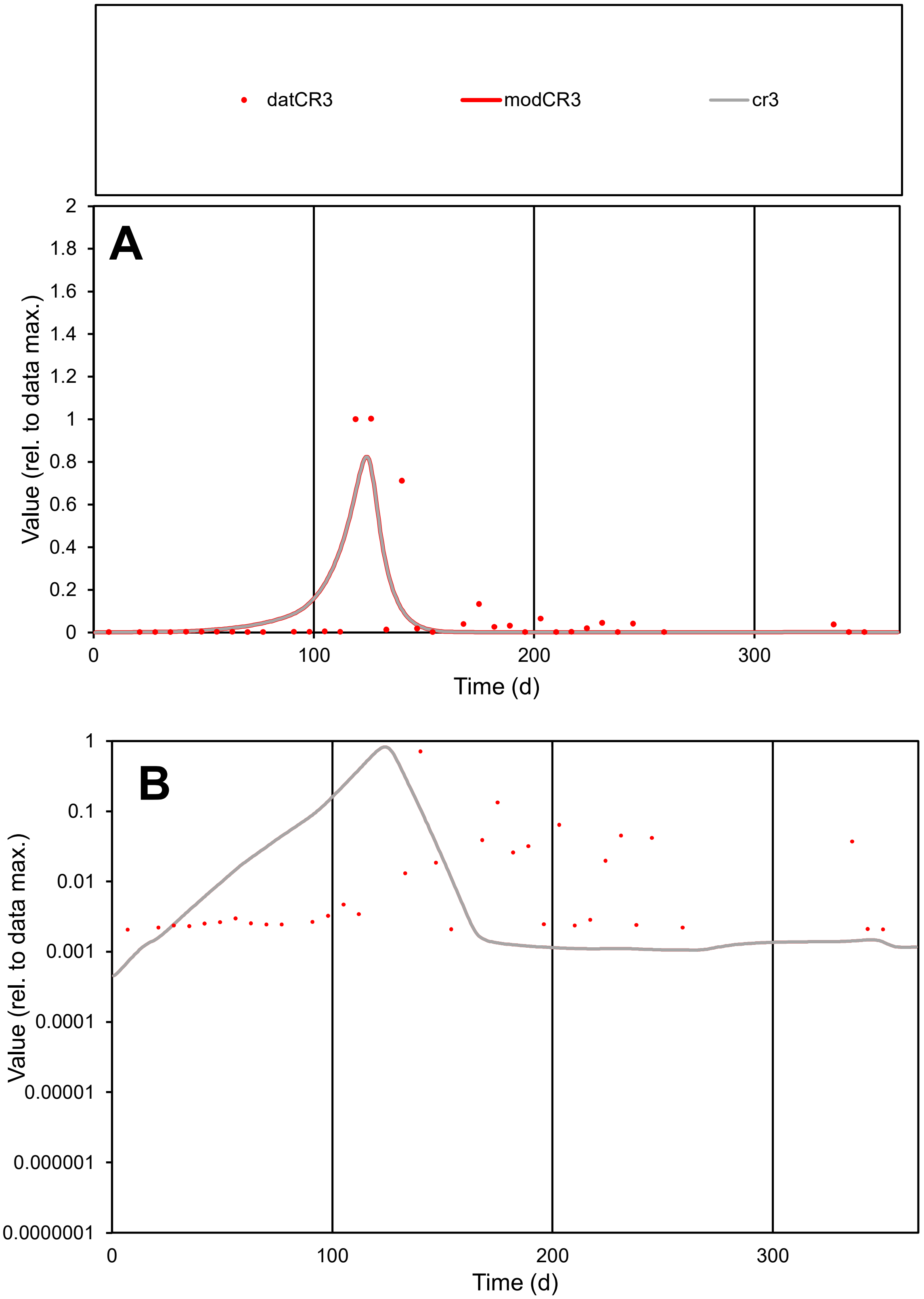


**C1**

Variables

CR3 Provide jtd or Dst

CalibOnly?

2 1 to reproduce of

oNormalize?

2 1=no, 2=yes

ofxn calc*

1.49449

ofxn mod

1.58319

sRun

RunA1

zout

1

Ave

Data

0.00129

Model

0.0013

data/vMax

0.01765

ofxng

311.734

Min d/m

2.69453

ave/min d

35.4846

ave/min m

96.333

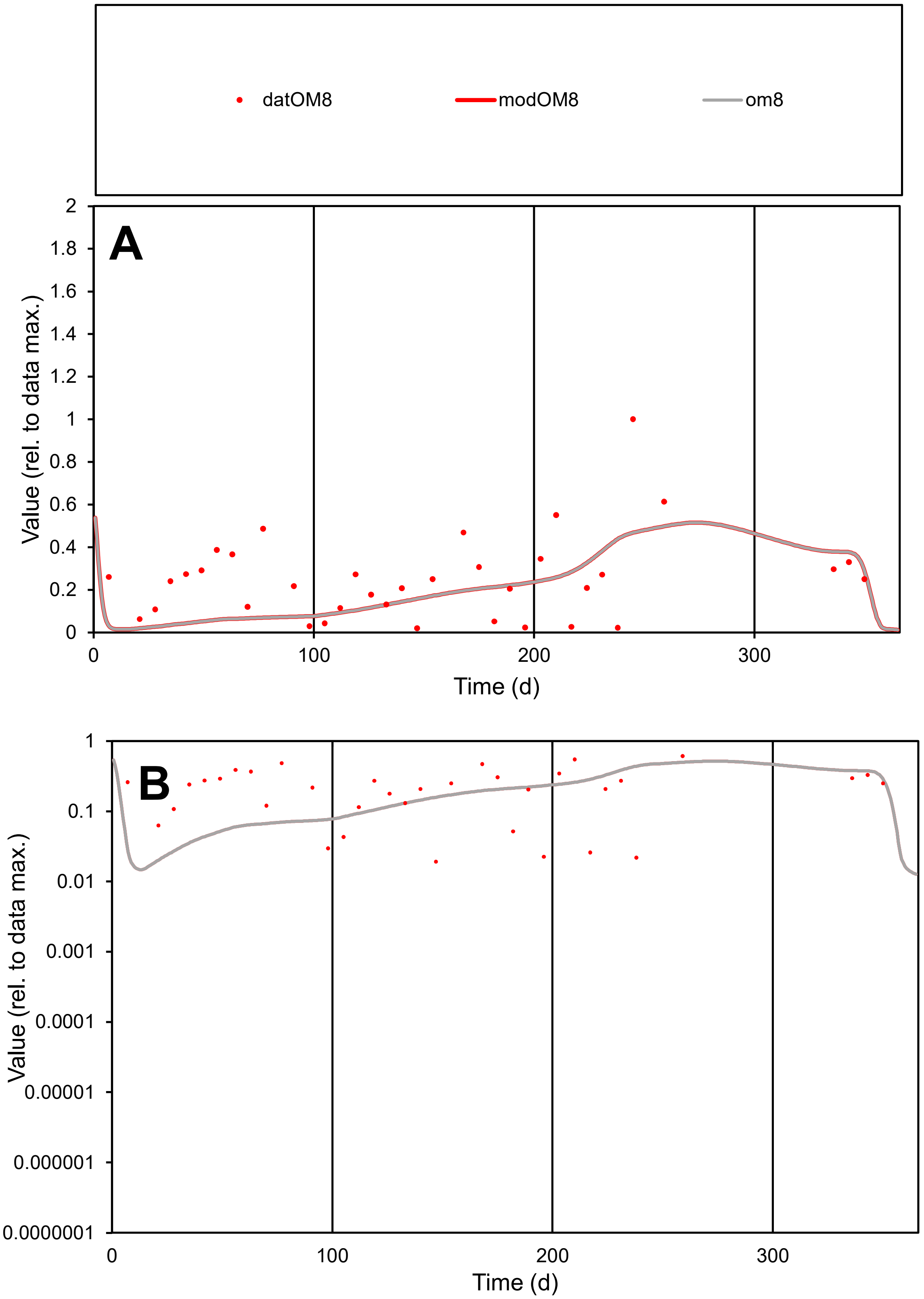


**C1**

Variables
OM8 Provide jtd or Dst
CalibOnly?
2 1 to reproduce of
oNormalize?
2 1=no, 2=yes

ofxn calc*
0.82153
ofxn mod
0.83846

sRun
RunA1
zout
1

Ave
Data
0.00072
Model
0.00055

data/vMax
0.00294

ofxng
311.734

Min d/m
0.88922

ave/min d
12.7363
ave/min m
8.69708

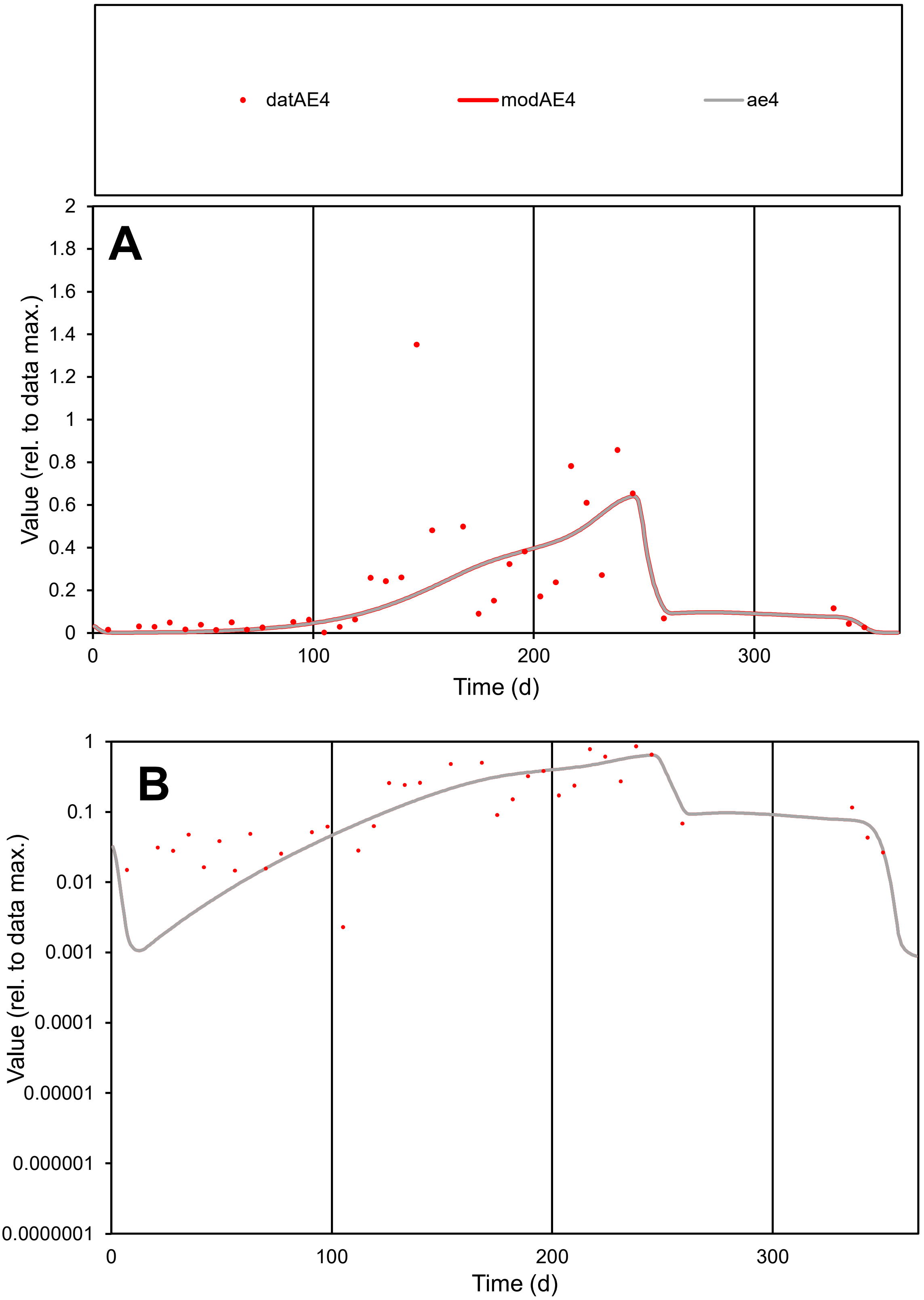


**C1**

Variables

| | |
|---|---|
| AE4 | Provide jtd or Dst |
| CalibOnly? | |
| 2 | 1 to reproduce of |
| oNormalize? | |
| 2 | 1=no, 2=yes |
| ofxn calc* | |
| 0.85312 | |
| ofxn mod | |
| 0.87041 | |
| sRun | |
| RunA1 | |
| zout | |
| 1 | |
| Ave | |
| Data | |
| 0.00252 | |
| Model | |
| 0.00225 | |
| data/vMax | |
| 0.0119 | |
| ofxng | |
| 311.734 | |
| Min d/m | |
| 1.33301 | |
| ave/min d | |
| 92.364 | |
| ave/min m | |
| 109.803 | |

331_Calib_S_A

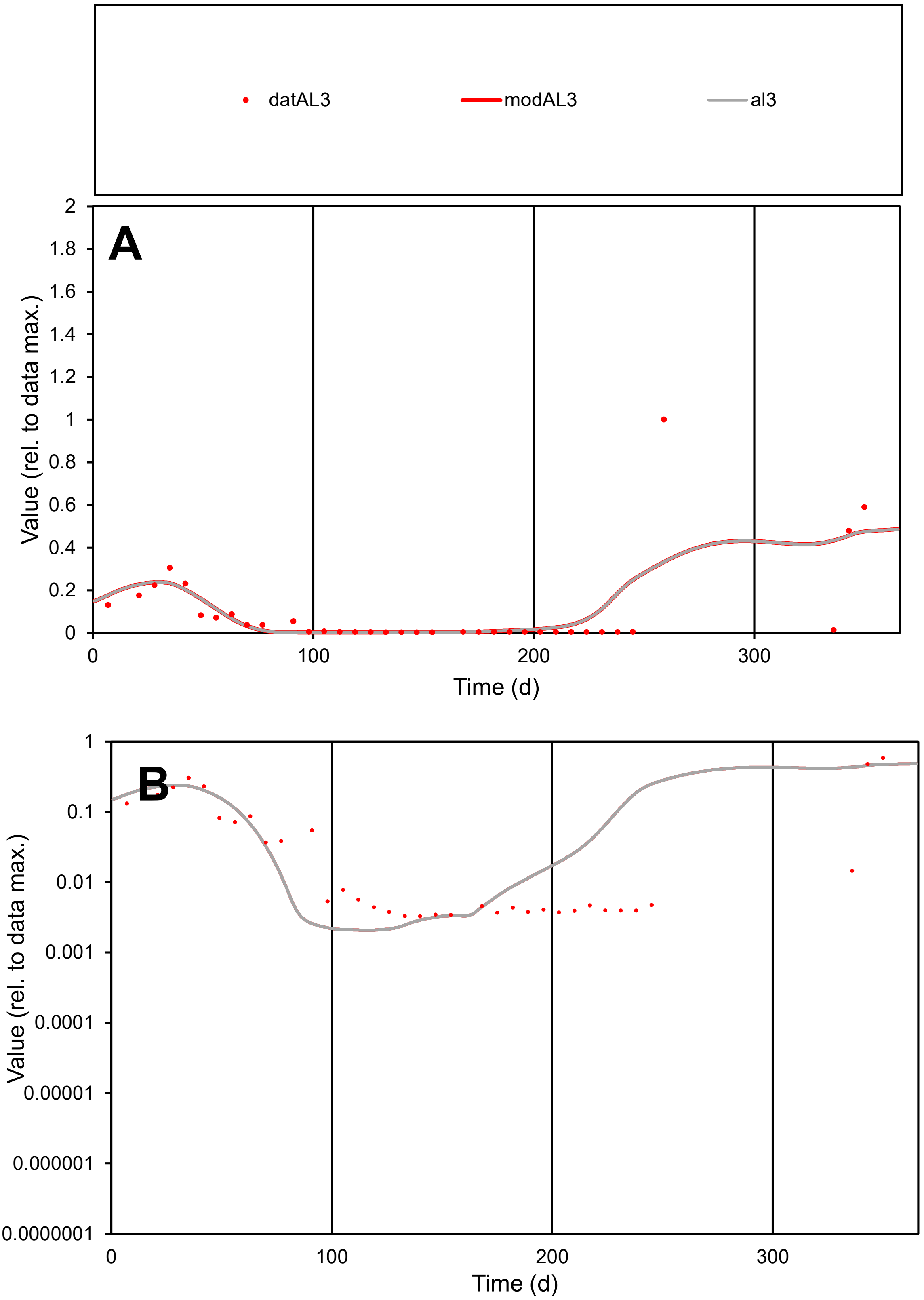


**C1**

Variables
AL3 Provide jtd or Dst
CalibOnly?
2 1 to reproduce of
oNormalize?
2 1=no, 2=yes

ofxn calc*
1.44018
ofxn mod
1.46302

sRun
RunA1
zout
1

Ave
Data
0.00073
Model
0.00079

data/vMax
0.00729

ofxng
311.734

Min d/m
1.57804

ave/min d
30.6591
ave/min m
52.1071

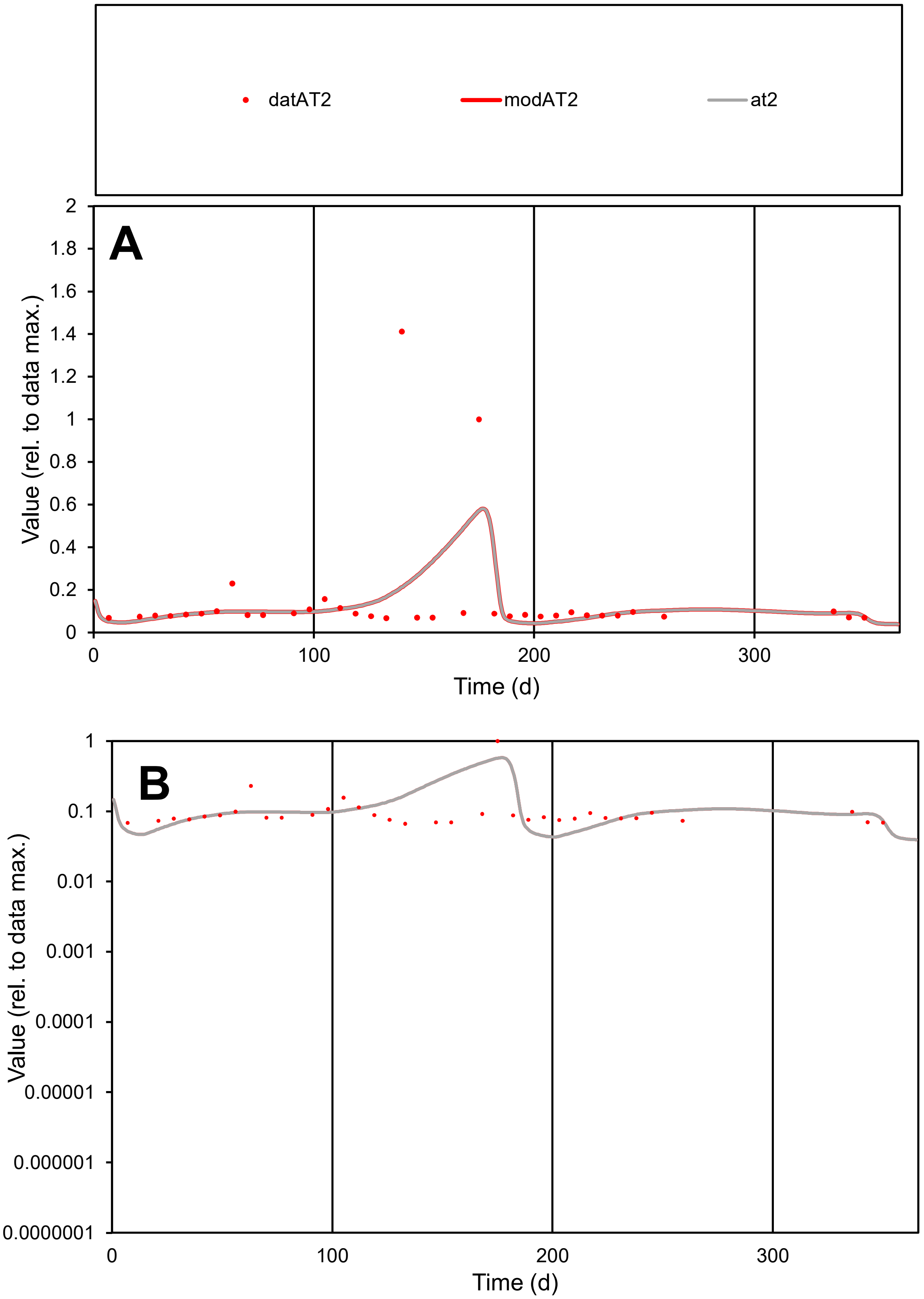


**C1**

Variables
AT2 Provide jtd or Dst
CalibOnly?
2 1 to reproduce of
oNormalize?
2 1=no, 2=yes

ofxn calc*
1.13584
ofxn mod
1.15229

sRun
RunA1
zout
1

Ave
Data
0.00016
Model
0.00015

data/vMax
0.0012

ofxng
311.734

Min d/m
1.50959

ave/min d
2.01642
ave/min m
2.92914

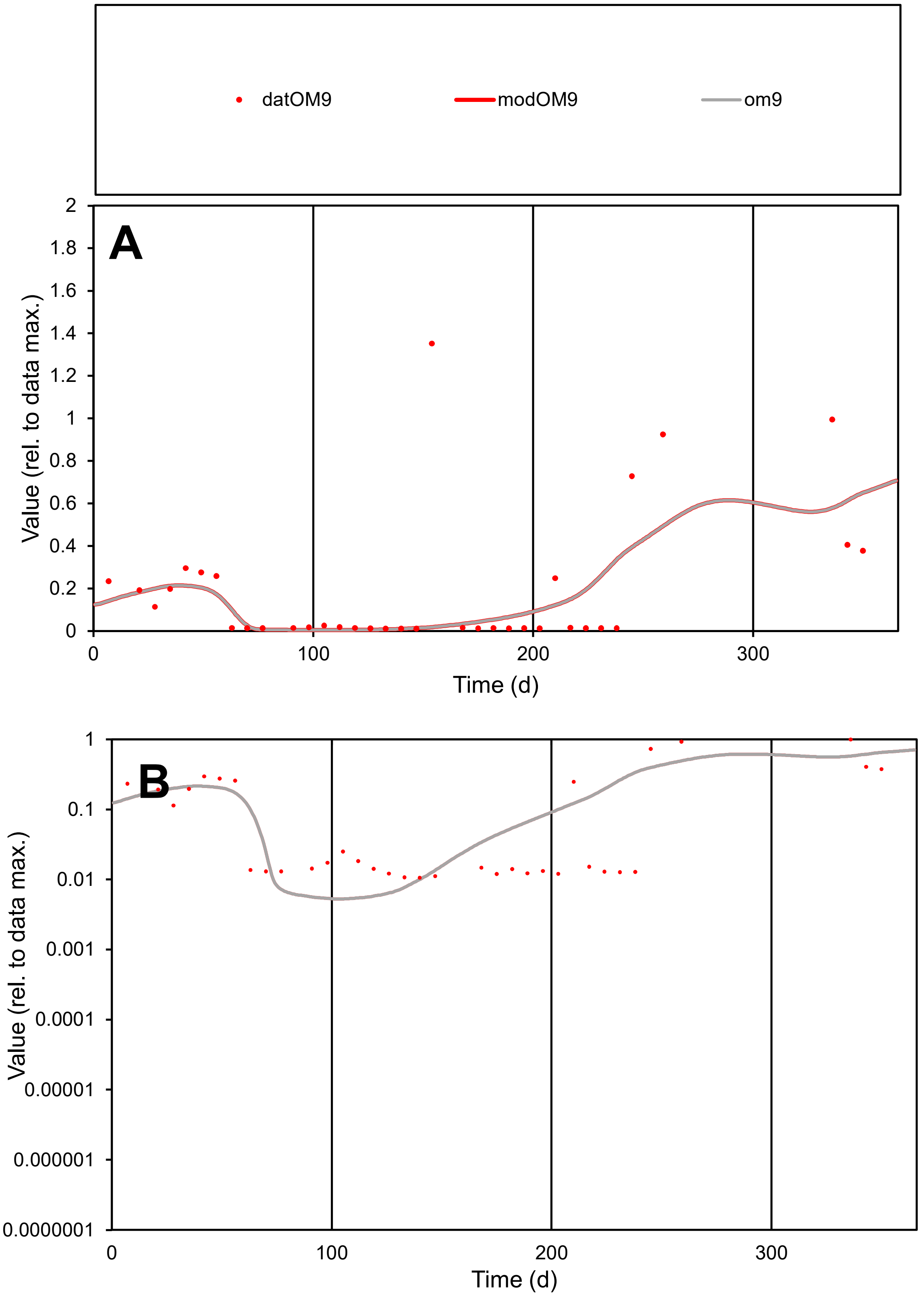


**C1**

Variables

OM9 Provide jtd or Dst

CalibOnly?

2 1 to reproduce of

oNormalize?

2 1=no, 2=yes

ofxn calc*

1.23434

ofxn mod

1.25086

sRun

RunA1

zout

1

Ave

Data

0.00073

Model

0.00064

data/vMax

0.00404

ofxng

311.734

Min d/m

2.00234

ave/min d

17.1455

ave/min m

29.9307

331_Calib_S_A

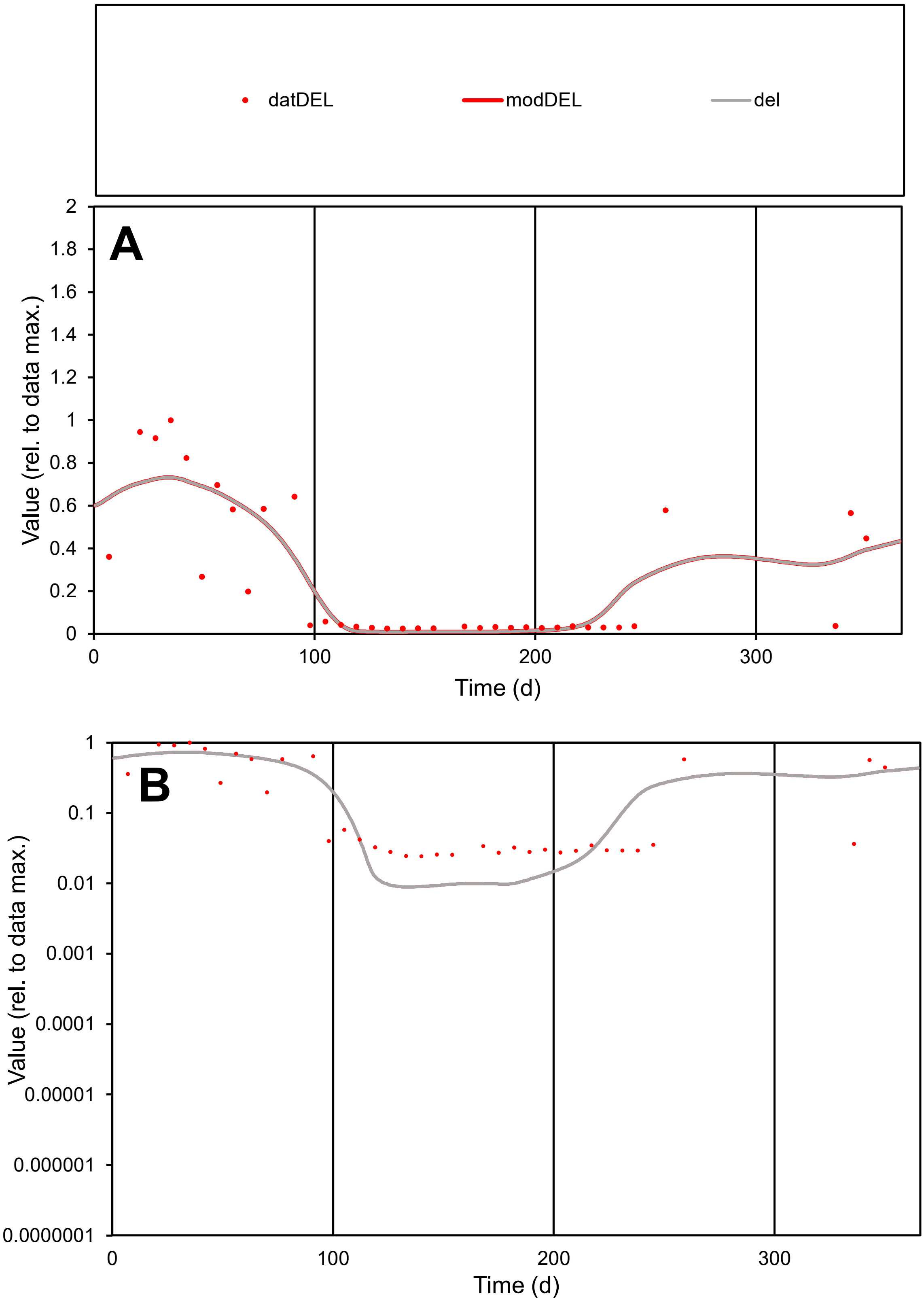


**C1**

Variables
DEL Provide jtd or Dst
CalibOnly?
2 1 to reproduce of
oNormalize?
2 1=no, 2=yes

ofxn calc*
0.63343
ofxn mod
0.66238

sRun
RunA1
zout
1

Ave
Data
0.00065
Model
0.00066

data/vMax
0.00253

ofxng
311.734

Min d/m
2.72444

ave/min d
10.6426
ave/min m
29.2532

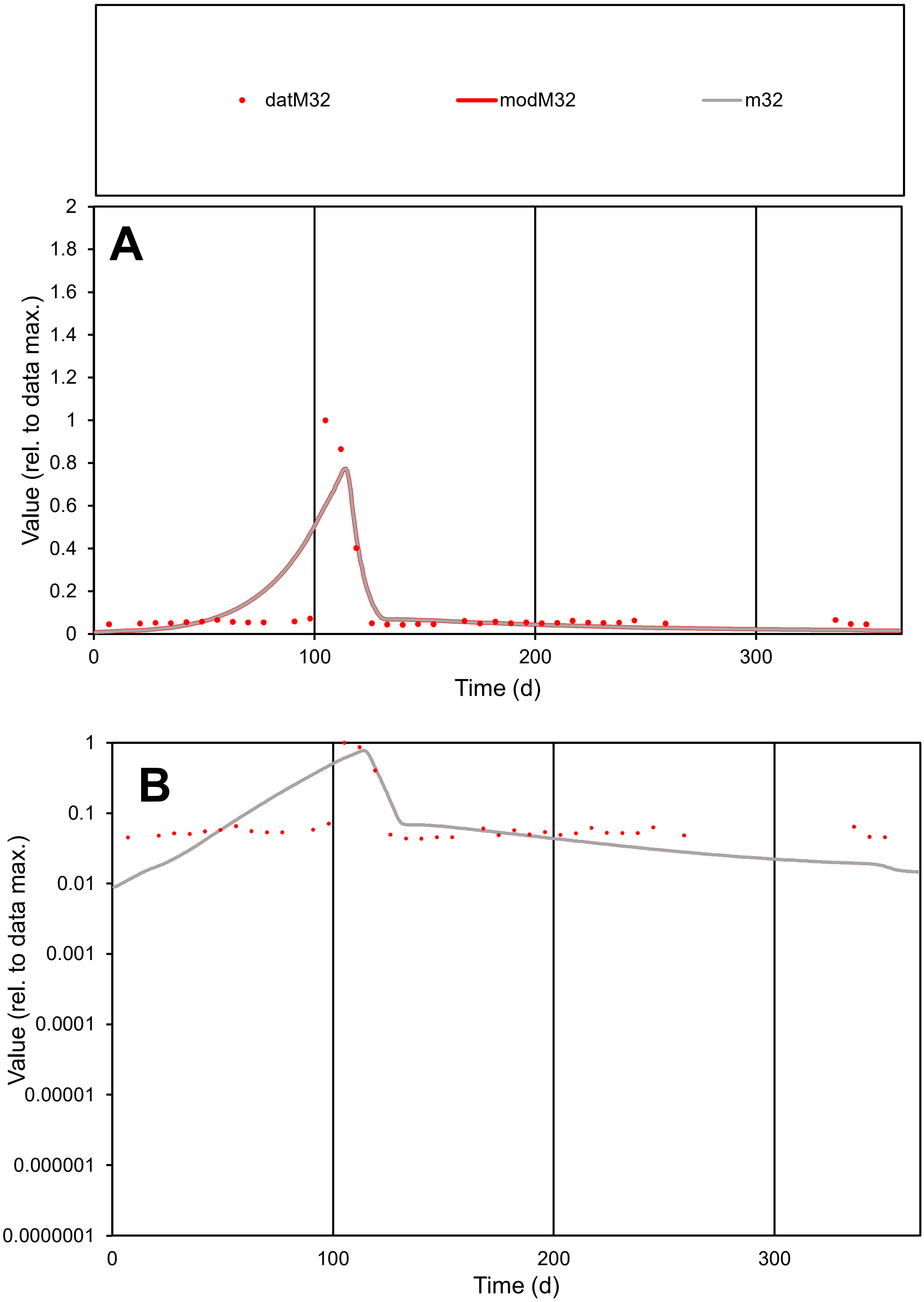


**C1**

Variables
M32 Provide jtd or Dst
CalibOnly?
2 1 to reproduce of
oNormalize?
2 1=no, 2=yes

ofxn calc*
0.99534
ofxn mod
1.02725

sRun
RunA1
zout
1

Ave
Data
0.00053
Model
0.00057

data/vMax
0.00479

ofxng
311.734

Min d/m
3.73175

ave/min d
2.5794
ave/min m
10.2926

331_Calib_S_A

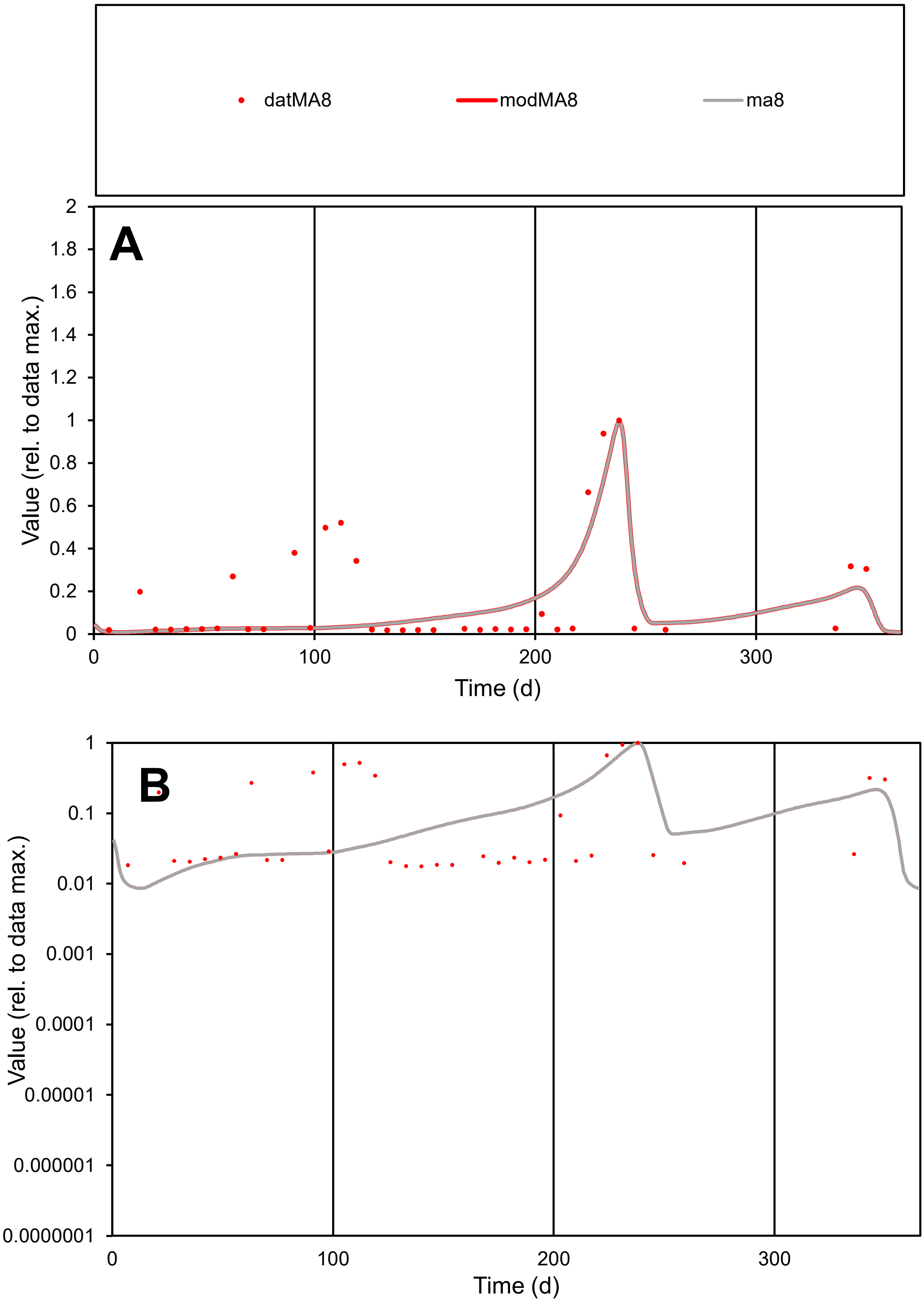


**C1**

Variables

MA8 Provide jtd or Dst

CalibOnly?

2 1 to reproduce of

oNormalize?

2 1=no, 2=yes

ofxn calc*

1.02265

ofxn mod

1.05307

sRun

RunA1

zout

1

Ave

Data

0.00056

Model

0.00047

data/vMax

0.00336

ofxng

311.734

Min d/m

1.87949

ave/min d

9.55304

ave/min m

14.8038

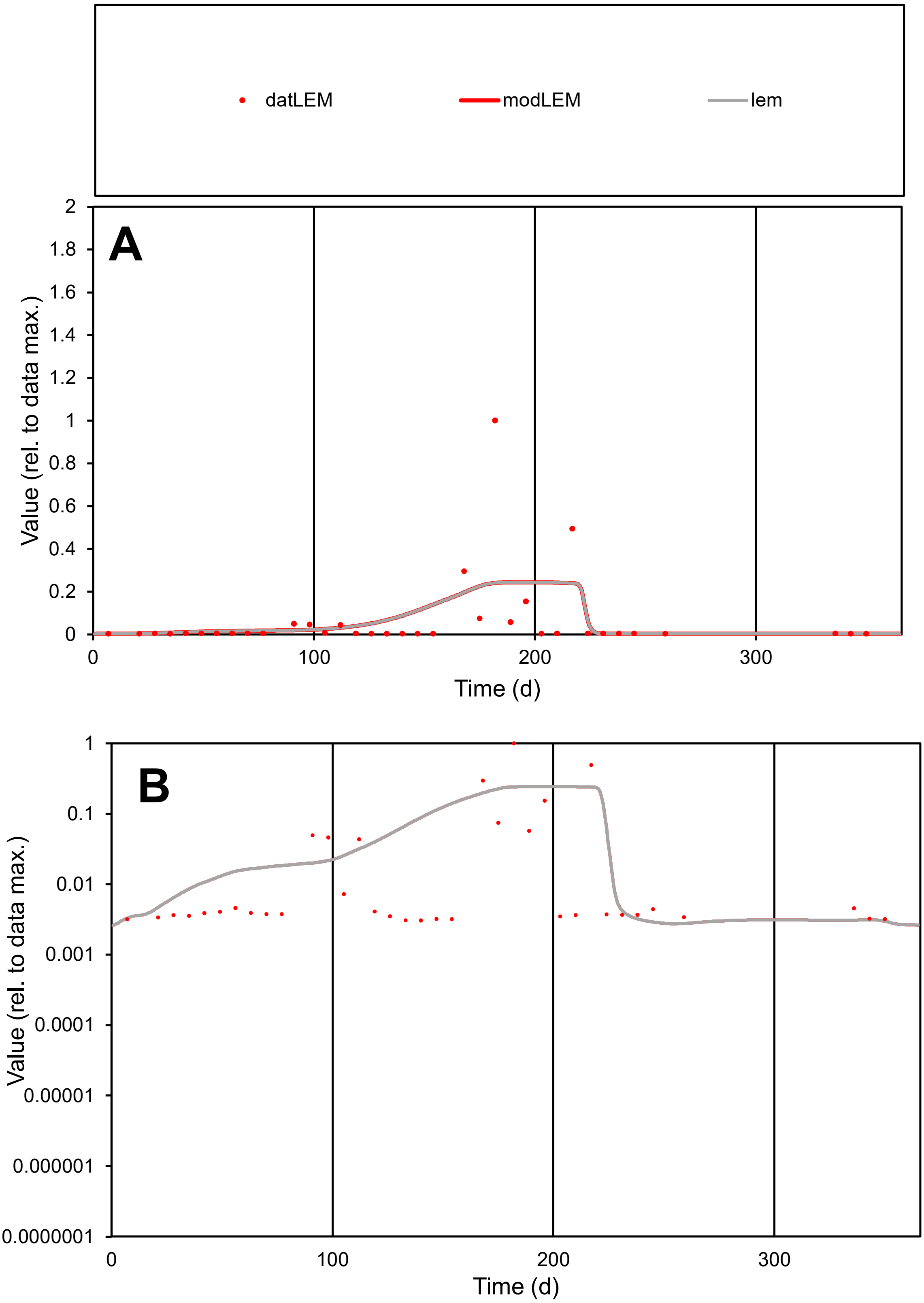


C1

Variables
LEM Provide jtd or Dst
CalibOnly?
2 1 to reproduce of
oNormalize?
2 1=no, 2=yes

ofxn calc*
2.42546
ofxn mod
2.43837

sRun
RunA1
zout
1

Ave
Data
0.00045
Model
0.00051

data/vMax
0.00699

ofxng
311.734

Min d/m
1.09147

ave/min d
21.0616
ave/min m
26.1885

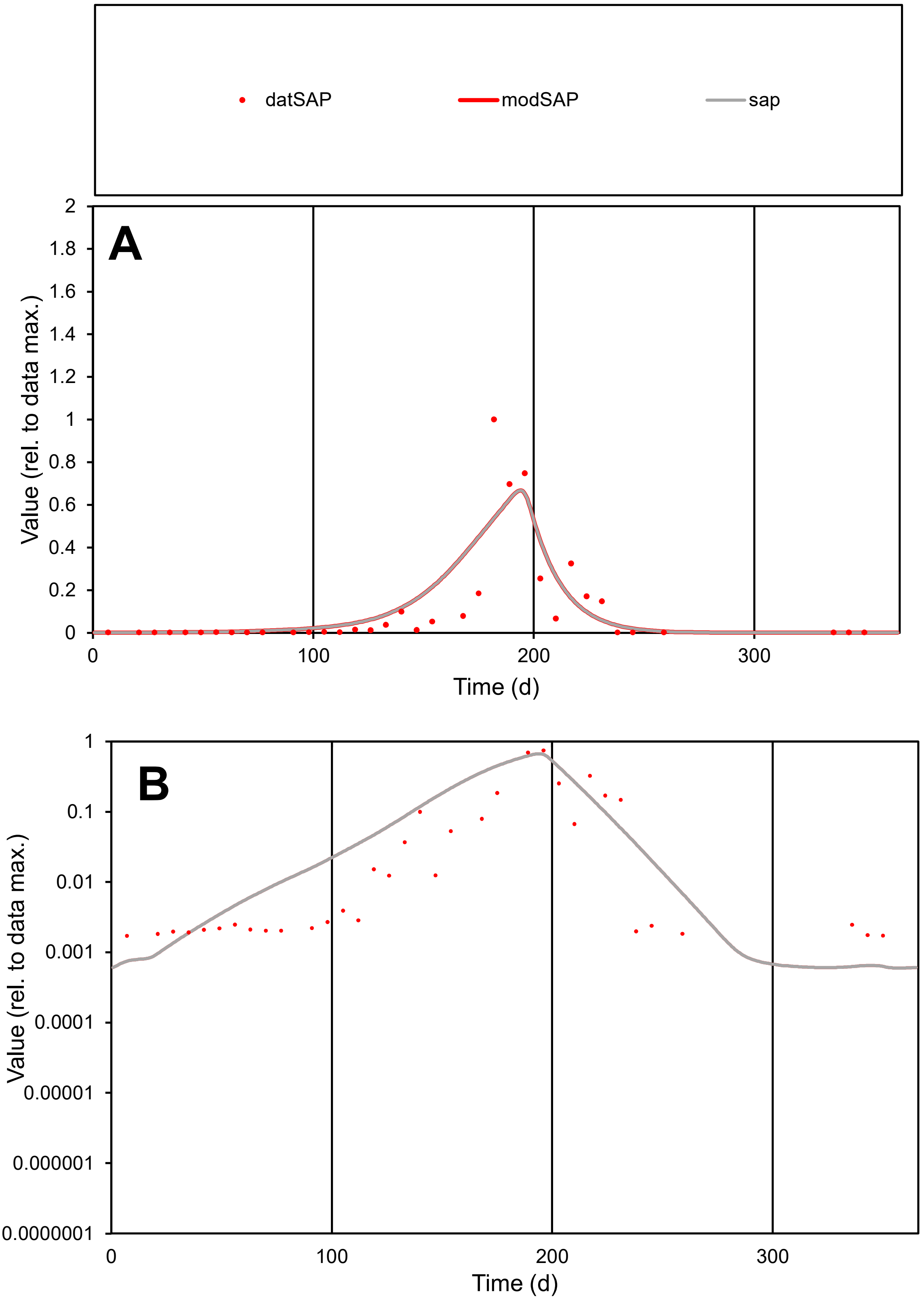


**C1**

Variables
SAP Provide jtd or Dst
CalibOnly?
2 1 to reproduce of
oNormalize?
2 1=no, 2=yes

ofxn calc*
1.12443
ofxn mod
1.15717

sRun
RunA1
zout
1

Ave
Data
0.00213
Model
0.00245

data/vMax
0.01976

ofxng
311.734

Min d/m
2.76163

ave/min d
62.8583
ave/min m
199.95

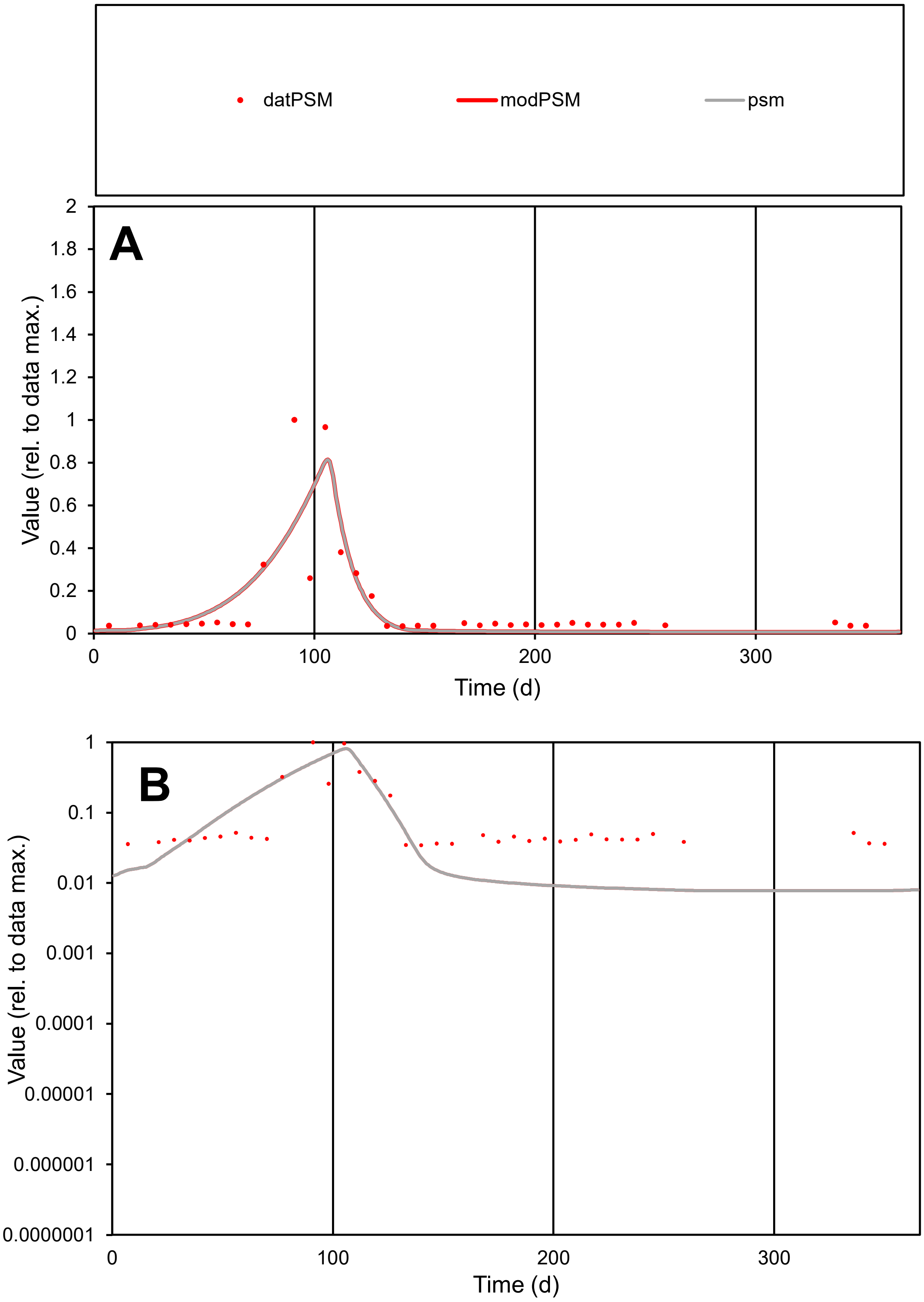


**C1**

Variables

PSM Provide jtd or Dst

CalibOnly?

2 1 to reproduce of

oNormalize?

2 1=no, 2=yes

ofxn calc*

0.91281

ofxn mod

0.94422

sRun

RunA1

zout

1

Ave

Data

0.0001

Model

9.2E-05

data/vMax

0.0008

ofxng

311.734

Min d/m

4.4403

ave/min d

3.67103

ave/min m

14.8843

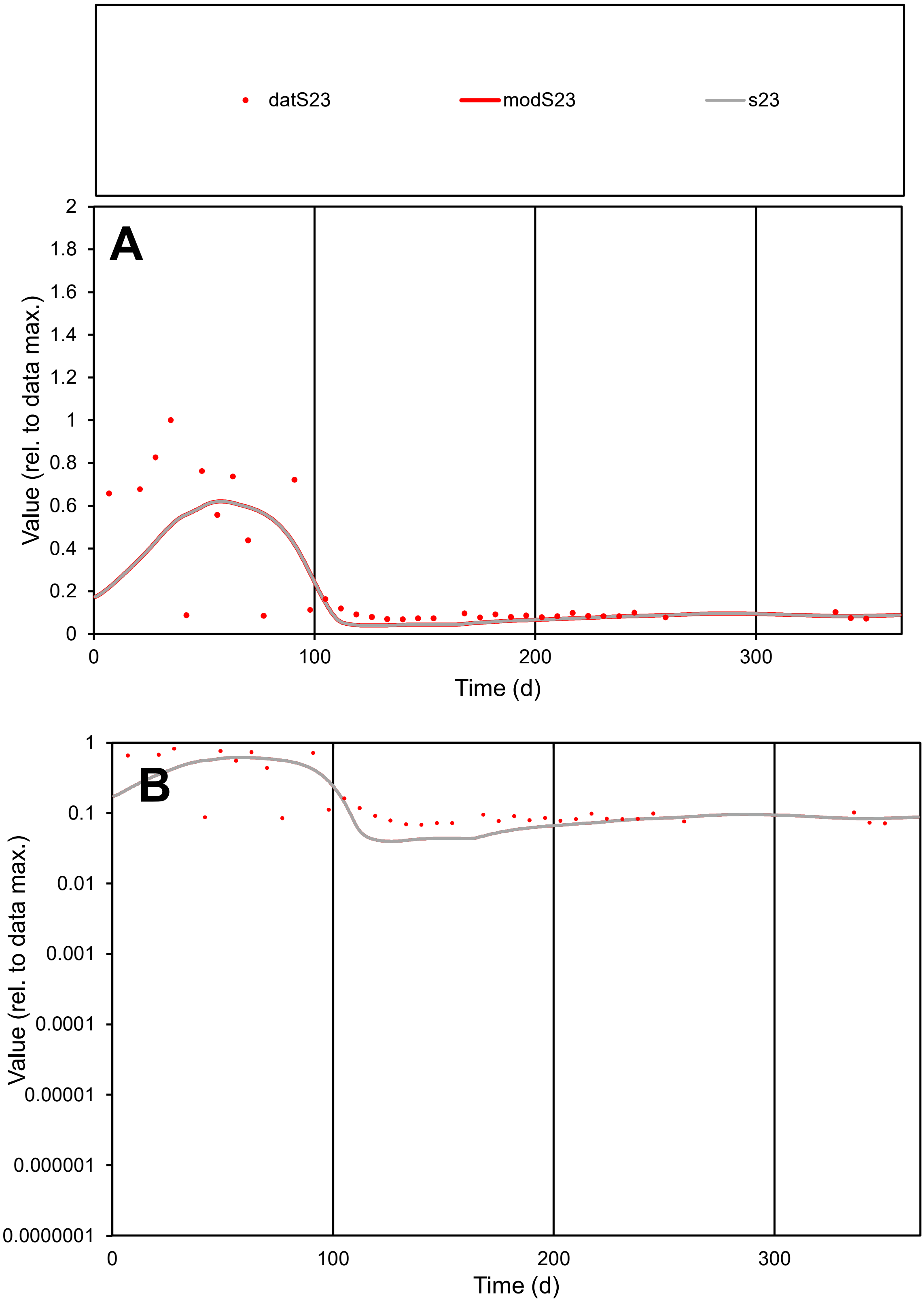


**C1**

Variables
S23 Provide jtd or Dst
CalibOnly?
2 1 to reproduce of
oNormalize?
2 1=no, 2=yes

ofxn calc*
0.78229
ofxn mod
0.79998

sRun
RunA1
zout
1

Ave
Data
0.00056
Model
0.00047

data/vMax
0.00232

ofxng
311.734

Min d/m
1.71907

ave/min d
3.54576
ave/min m
5.1222

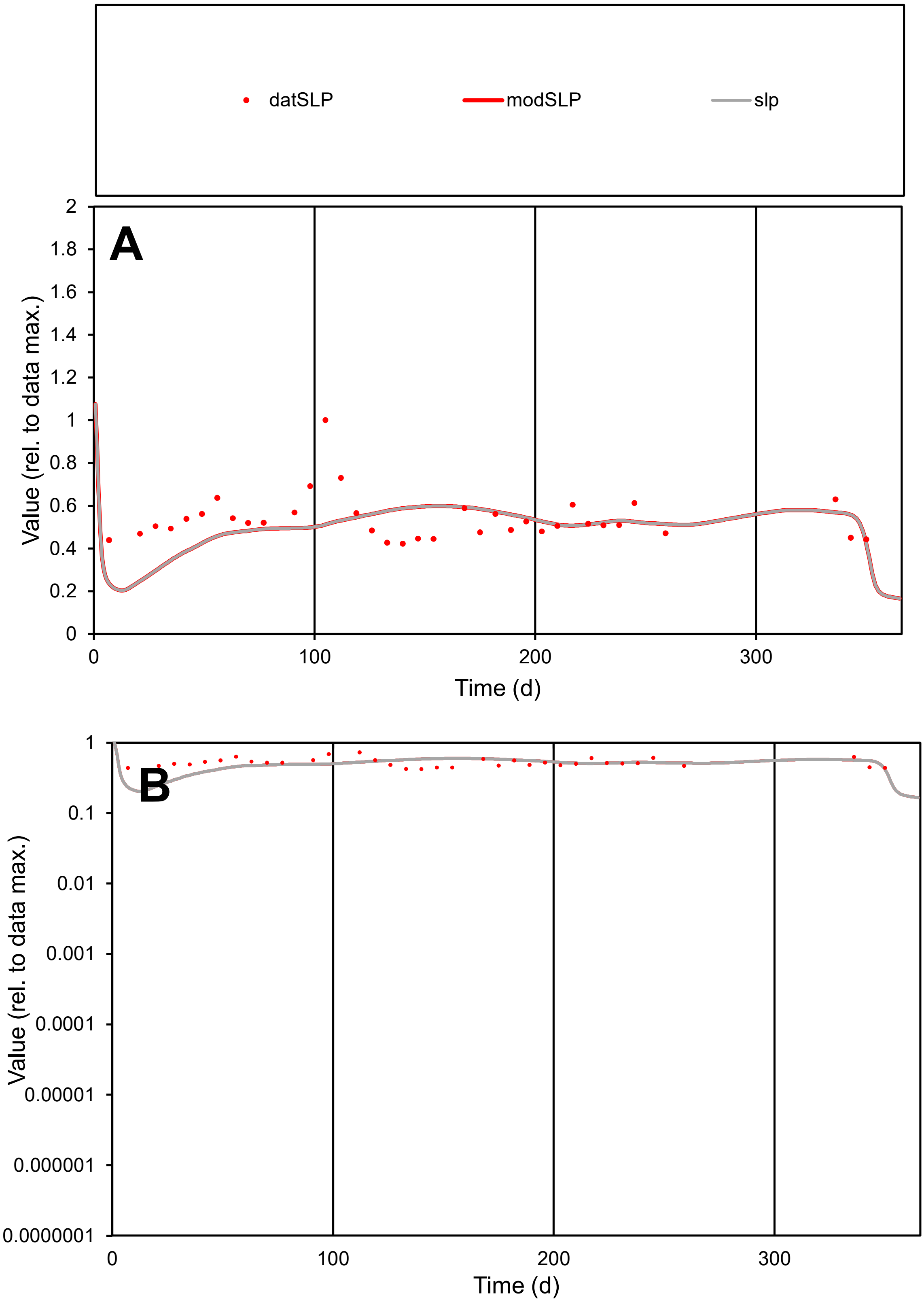


**C1**

Variables
SLP Provide jtd or Dst
CalibOnly?
2 1 to reproduce of
oNormalize?
2 1=no, 2=yes

ofxn calc*
0.25861
ofxn mod
0.28086

sRun
RunA1
zout
1

Ave
Data
3E-05
Model
2.8E-05

data/vMax
5.6E-05

ofxng
311.734

Min d/m
1.88474

ave/min d
1.27437
ave/min m
2.22231

331_Calib_S_A

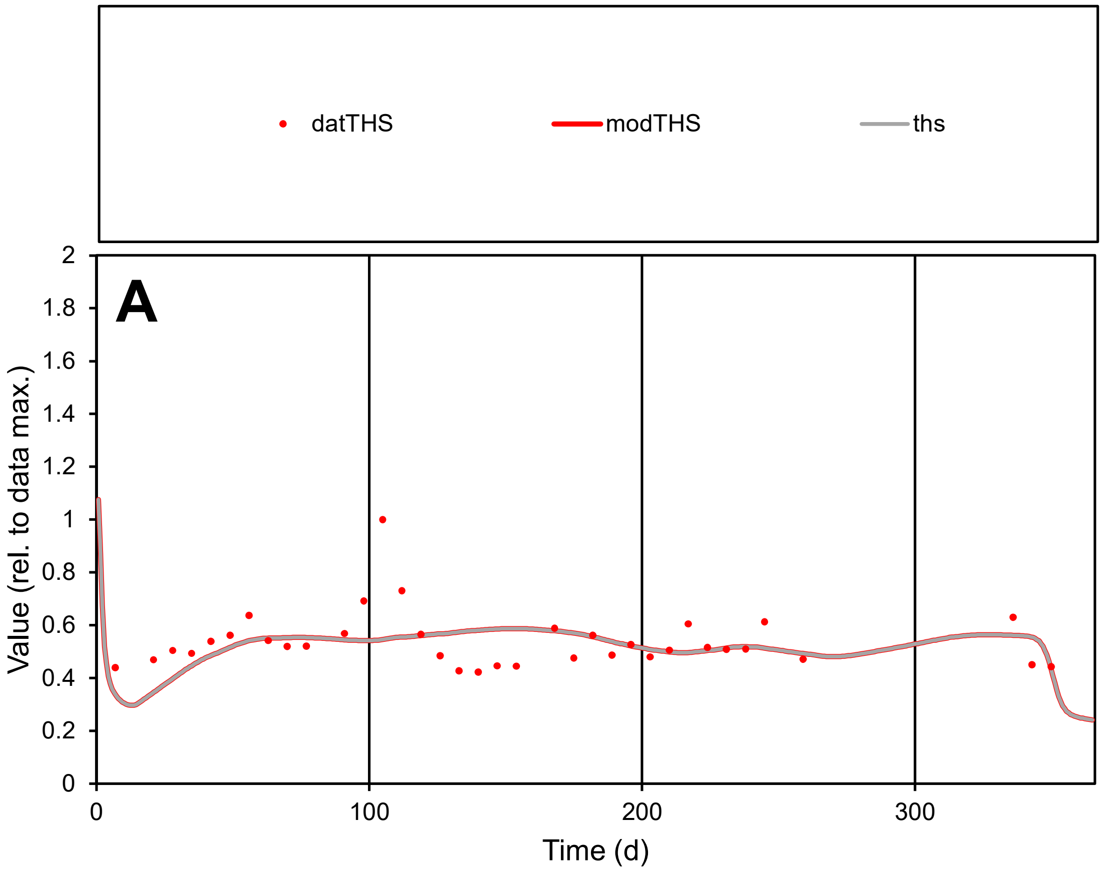


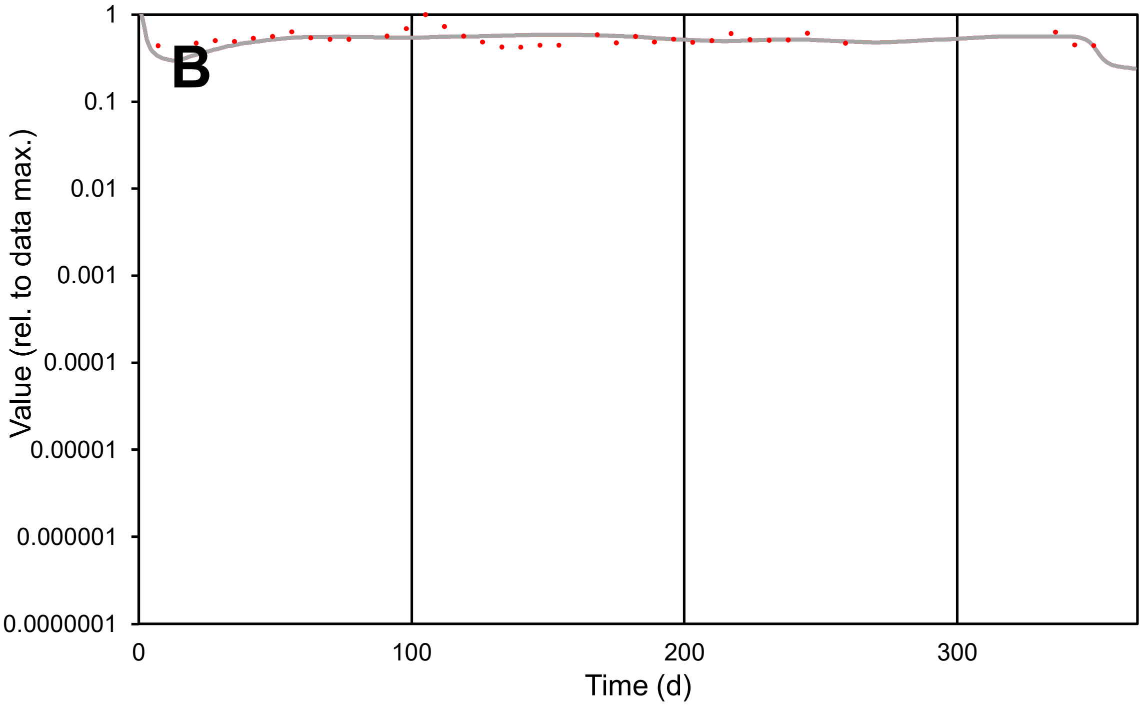


**C1**

Variables
THS Provide jtd or Dst
CalibOnly?
2 1 to reproduce of
oNormalize?
2 1=no, 2=yes

ofxn calc*
0.20951
ofxn mod
0.23659

sRun
RunA1
zout
1

Ave
Data
3.6E-05
Model
3.5E-05

data/vMax
6.7E-05

ofxng
311.734

Min d/m
1.3067

ave/min d
1.27456
ave/min m
1.60827

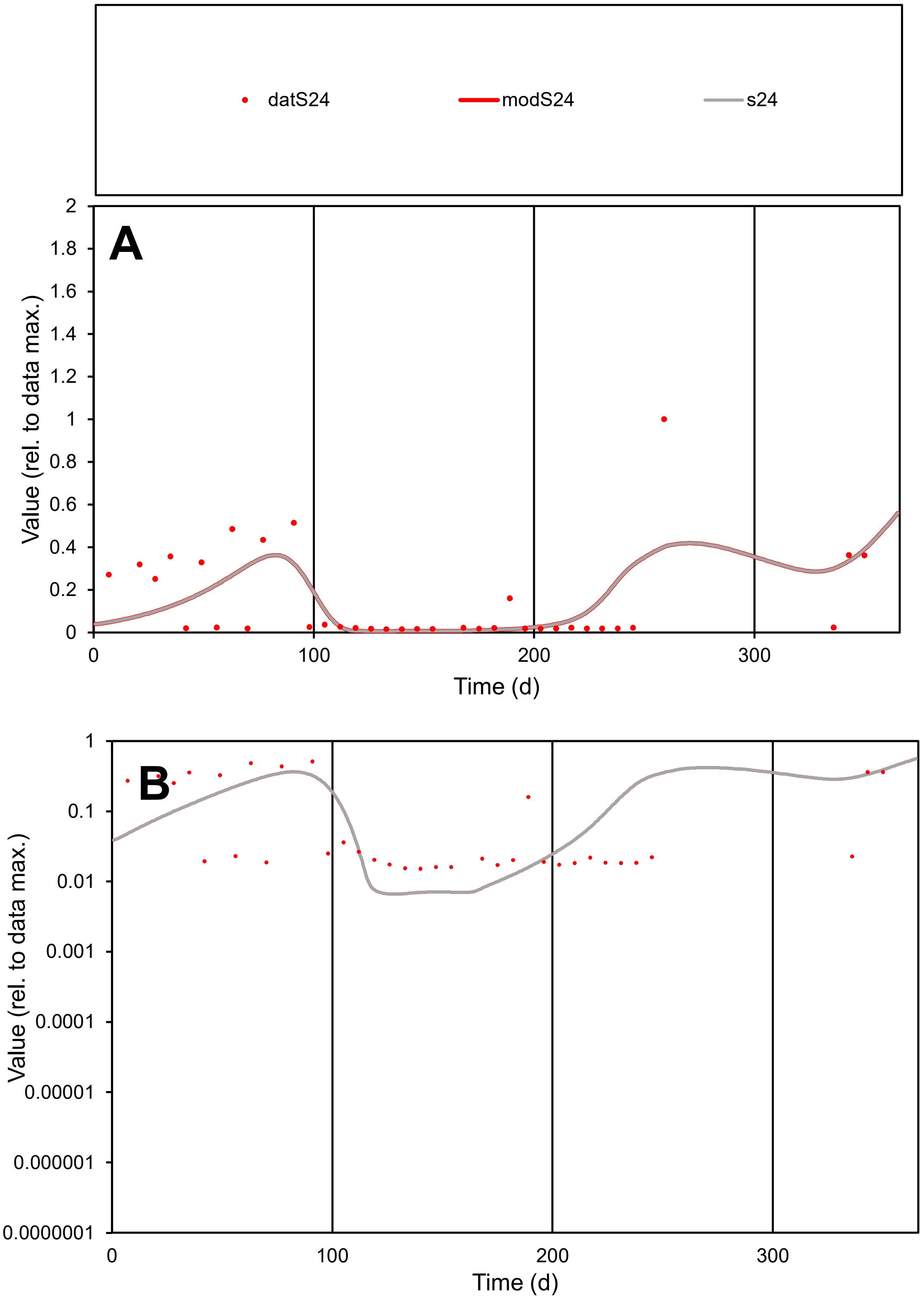


**C1**

Variables
S24 Provide jtd or Dst
CalibOnly?
2 1 to reproduce of
oNormalize?
2 1=no, 2=yes

ofxn calc*
1.16333
ofxn mod
1.18631

sRun
RunA1
zout
1

Ave
Data
0.00056
Model
0.00052

data/vMax
0.00377

ofxng
311.734

Min d/m
2.30188

ave/min d
9.69919
ave/min m
20.7987

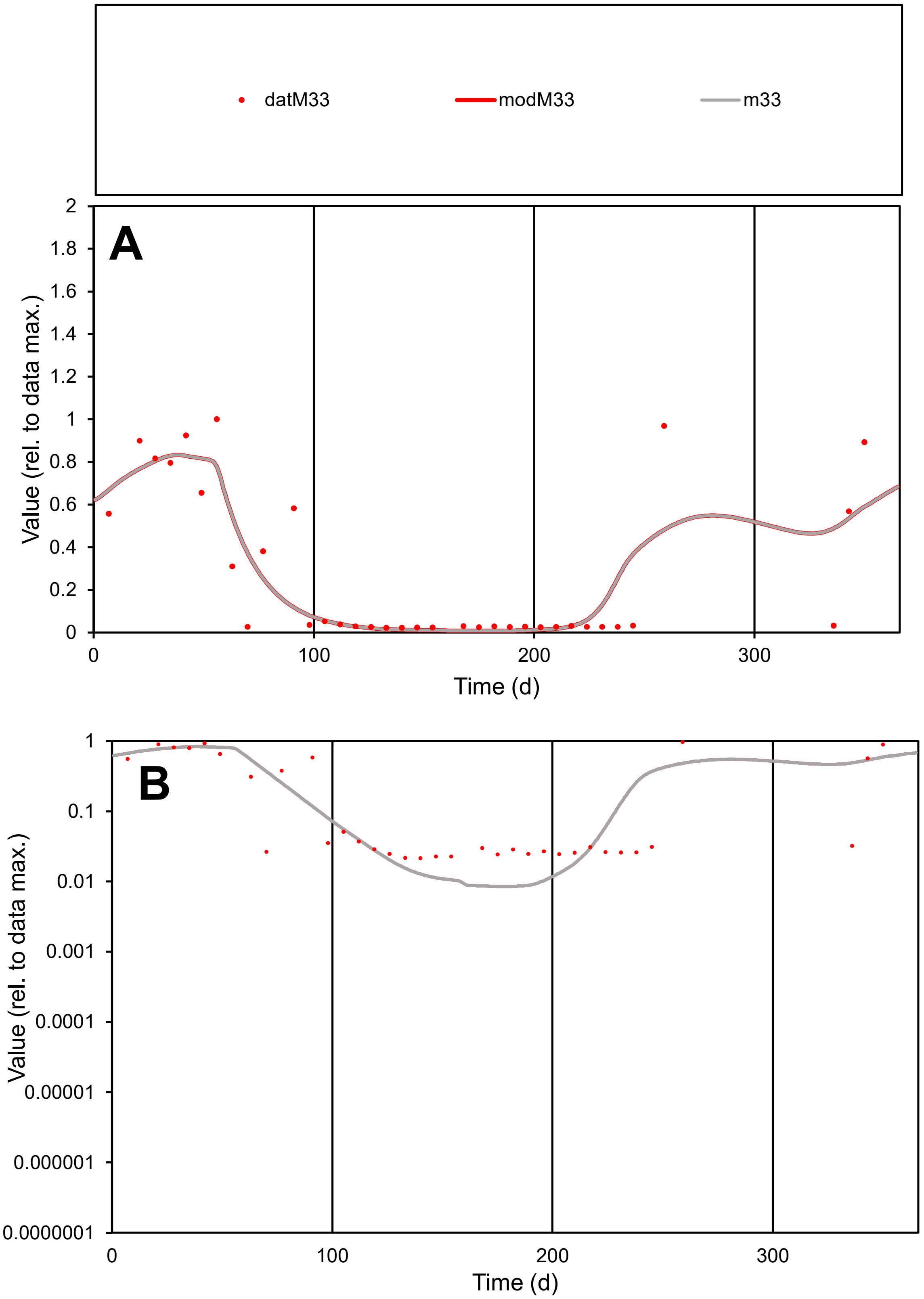


**C1**

Variables
M33 Provide jtd or Dst
CalibOnly?
2 1 to reproduce of
oNormalize?
2 1=no, 2=yes

ofxn calc*
0.64907
ofxn mod
0.68315

sRun
RunA1
zout
1

Ave
Data
0.00087
Model
0.00087

data/vMax
0.00315

ofxng
311.734

Min d/m
2.53403

ave/min d
12.8898
ave/min m
32.4215

331_Calib_S_A

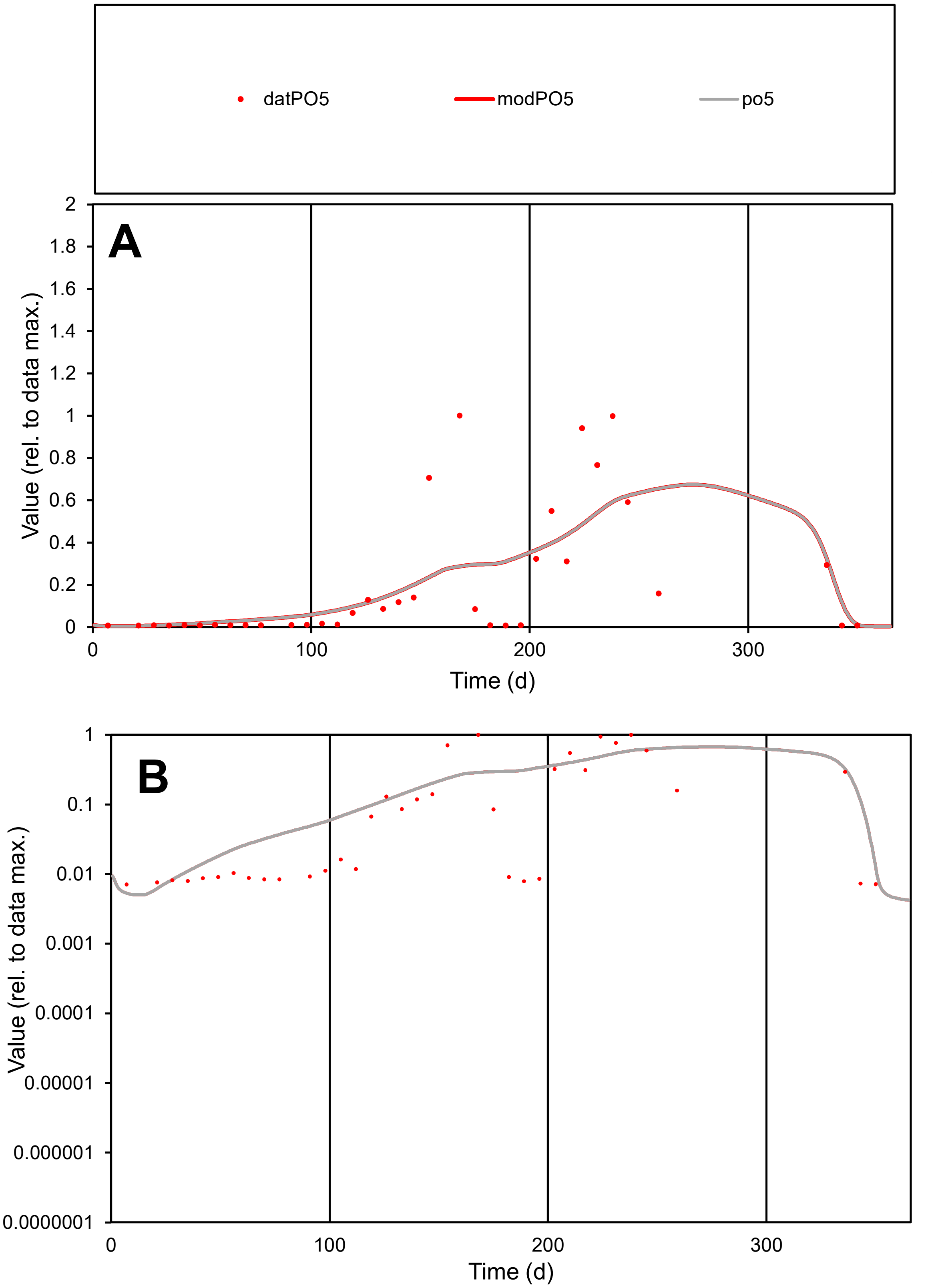


**C1**

Variables
PO5 Provide jtd or Dst
CalibOnly?
2 1 to reproduce of
oNormalize?
2 1=no, 2=yes

ofxn calc*
1.08351
ofxn mod
1.10284

sRun
RunA1
zout
1

Ave
Data
0.00133
Model
0.00139

data/vMax
0.00674

ofxng
311.734

Min d/m
1.36208

ave/min d
27.7821
ave/min m
39.7039

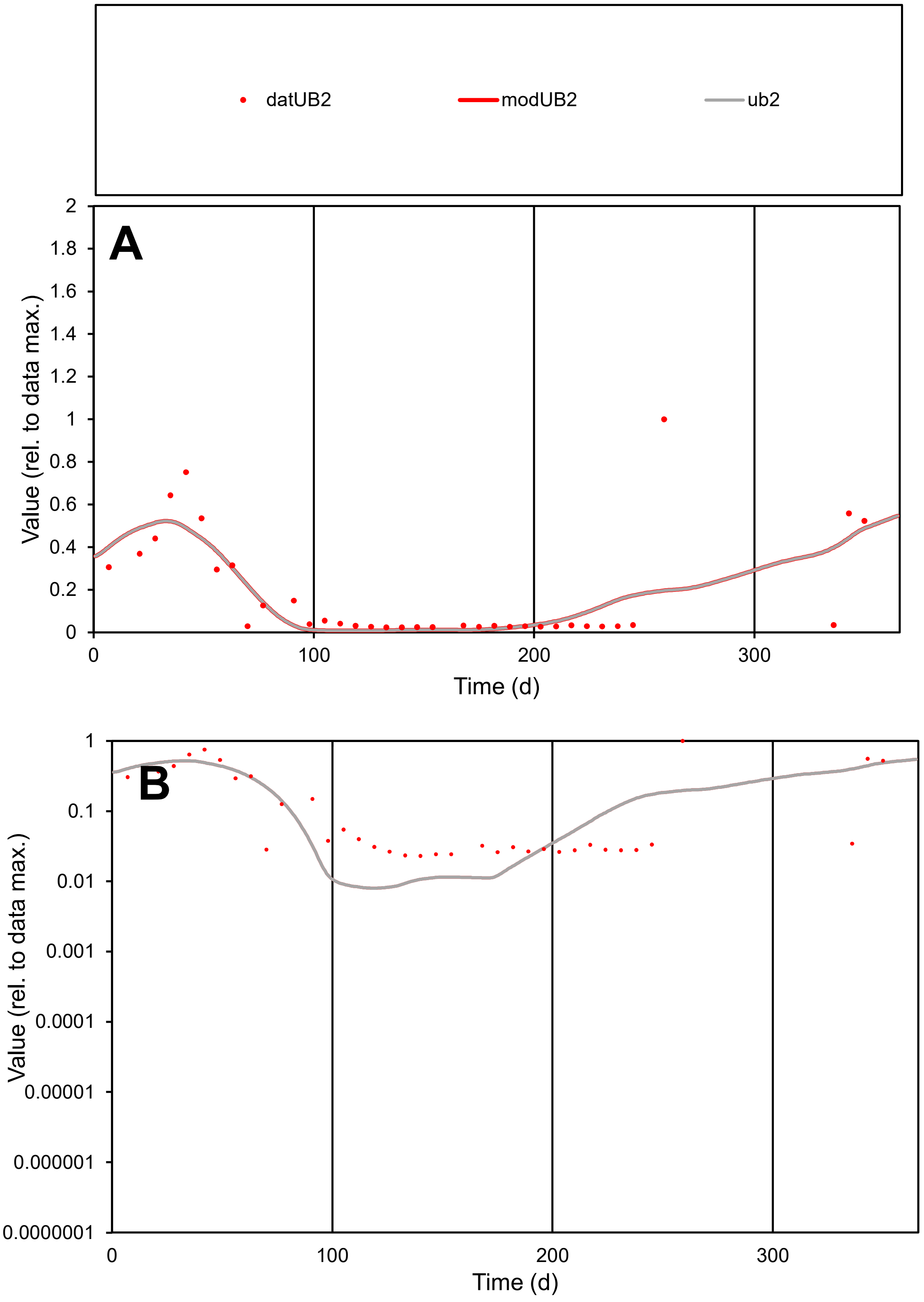


**C1**

Variables
UB2 Provide jtd or Dst
CalibOnly?
2 1 to reproduce of
oNormalize?
2 1=no, 2=yes

ofxn calc*
0.90522
ofxn mod
0.92535

sRun
RunA1
zout
1

Ave
Data
0.00083
Model
0.00078

data/vMax
0.00446

ofxng
311.734

Min d/m
2.88311

ave/min d
8.05857
ave/min m
21.7498

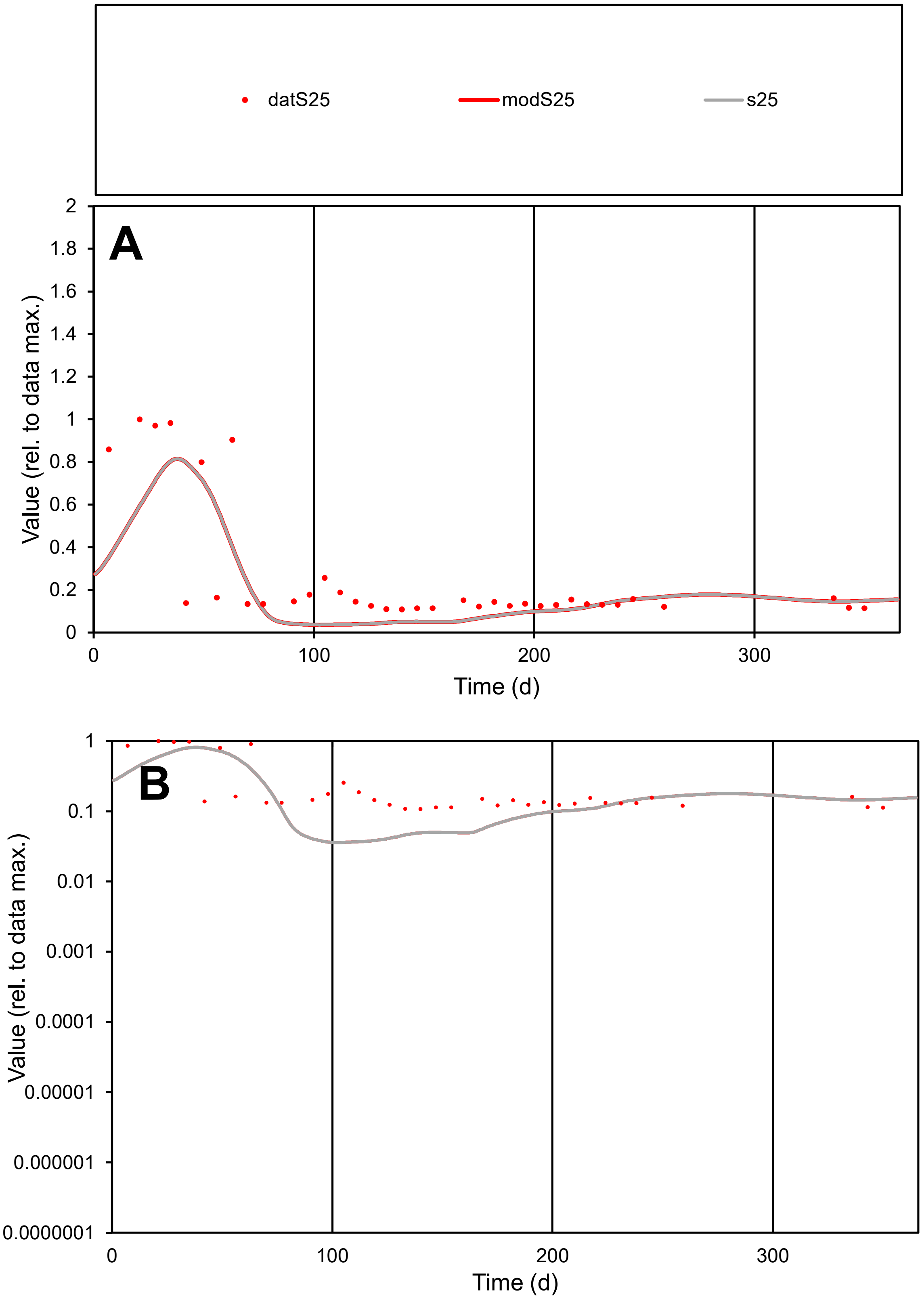


**C1**

Variables

S25 Provide jtd or Dst

CalibOnly?

2 1 to reproduce of

oNormalize?

2 1=no, 2=yes

ofxn calc*

0.74914

ofxn mod

0.76503

sRun

RunA1

zout

1

Ave

Data

0.00042

Model

0.00033

data/vMax

0.00155

ofxng

311.734

Min d/m

2.99428

ave/min d

2.49498

ave/min m

5.83956

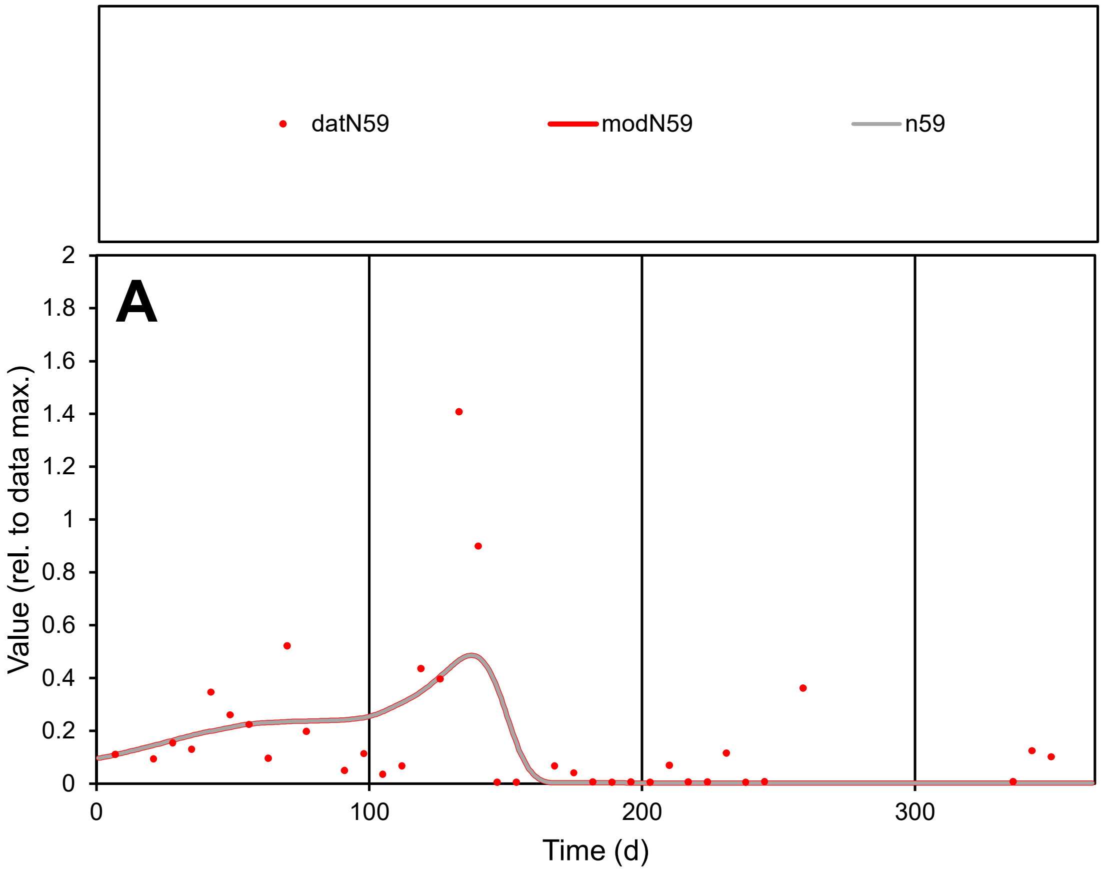


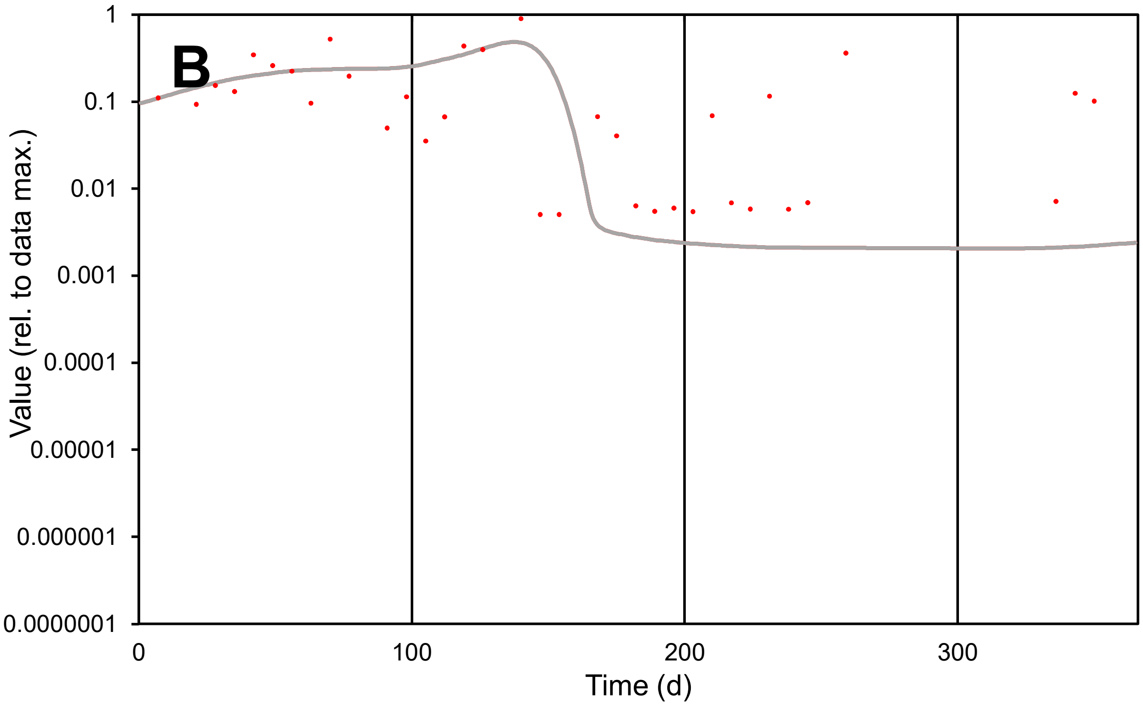


**C1**

| | |
|---|---|
| Variables | |
| N59 | Provide jtd or Dst |
| CalibOnly? | |
| 2 | 1 to reproduce of |
| oNormalize? | |
| 2 | 1=no, 2=yes |
| ofxn calc* | |
| 0.99187 | |
| ofxn mod | |
| 1.01537 | |
| sRun | |
| RunA1 | |
| zout | |
| 1 | |
| Ave | |
| Data | |
| 0.00132 | |
| Model | |
| 0.00122 | |
| data/vMax | |
| 0.00854 | |
| ofxng | |
| 311.734 | |
| Min d/m | |
| 2.41662 | |
| ave/min d | |
| 30.7328 | |
| ave/min m | |
| 68.4321 | |

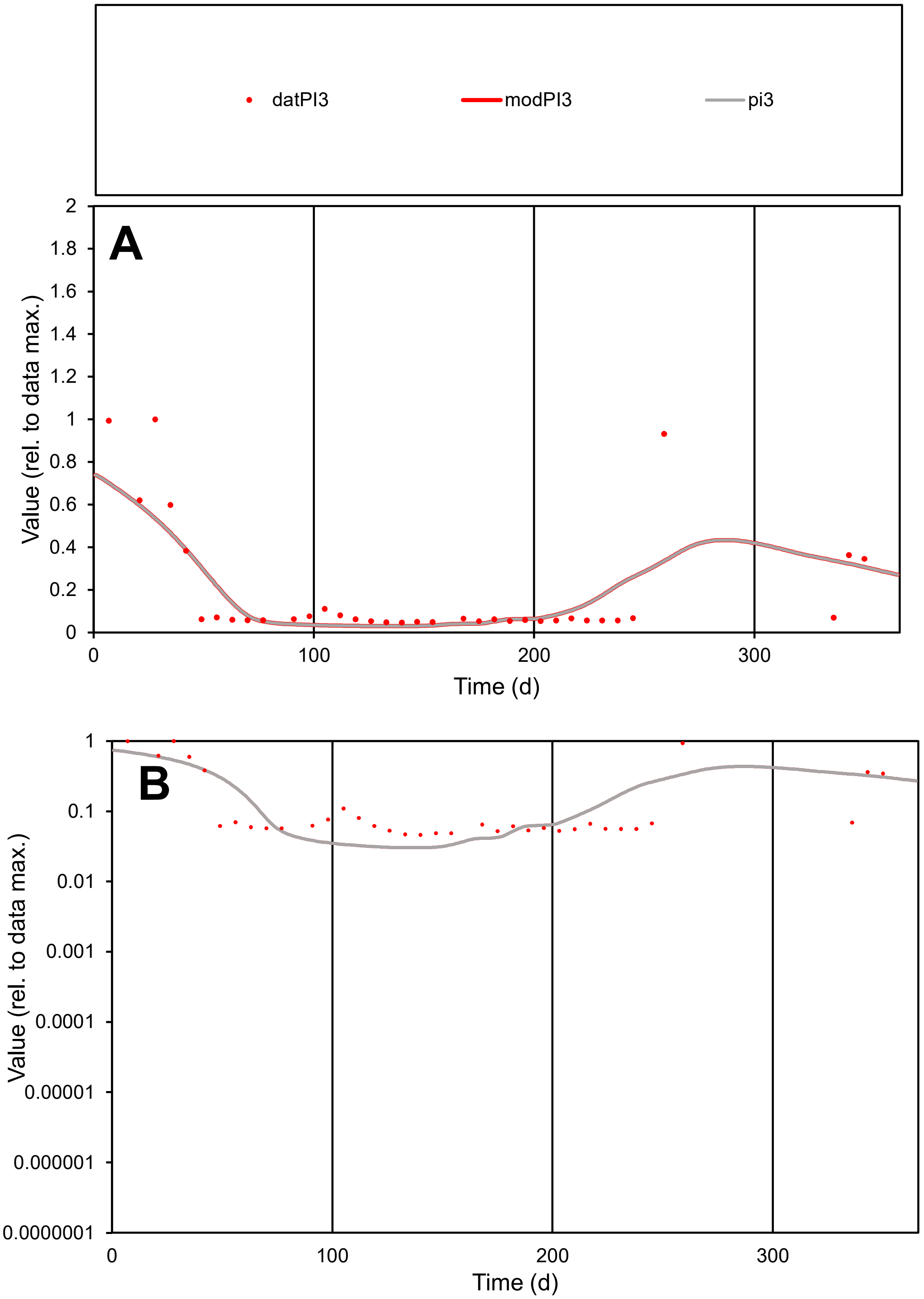


**C1**

Variables
PI3 Provide jtd or Dst
CalibOnly?
2 1 to reproduce of
oNormalize?
2 1=no, 2=yes

ofxn calc*
0.83089
ofxn mod
0.84853

sRun
RunA1
zout
1

Ave
Data
0.00029
Model
0.00027

data/vMax
0.00152

ofxng
311.734

Min d/m
1.5304

ave/min d
4.15905
ave/min m
5.78103

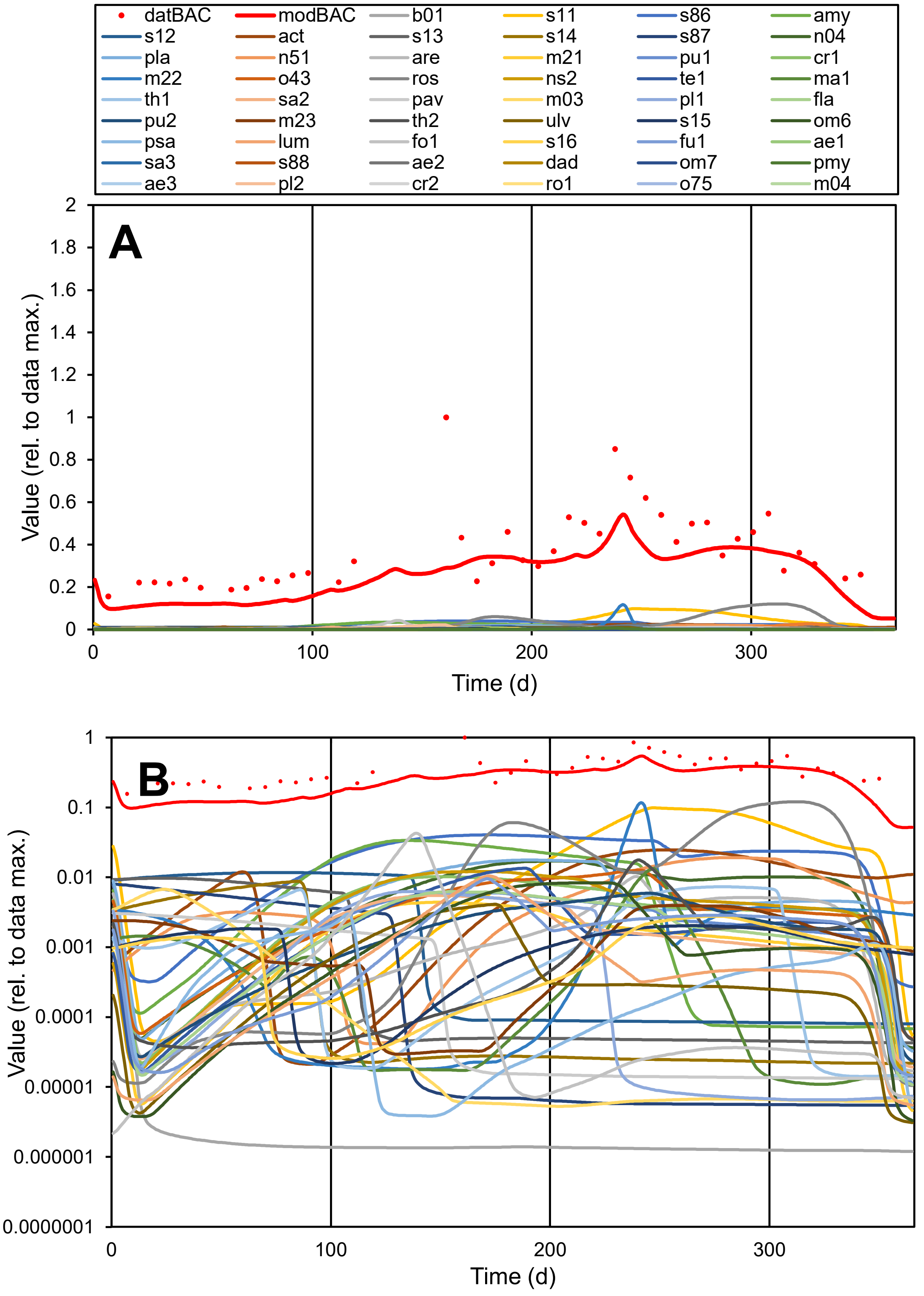


**C1**

Variables

BAC Provide jtd or Dst

CalibOnly?

2 1 to reproduce of

oNormalize?

2 1=no, 2=yes

ofxn calc*

0.44162

ofxn mod

0.45584

sRun

RunA1

zout

1

Ave

Data

1.30276

Model

0.93351

data/vMax

3.463

ofxng

311.734

Min d/m

1.72425

ave/min d

2.41699

ave/min m

2.98628
